# Proteomic analysis of human milk reveals nutritional and immune benefits in the colostrum from mothers with COVID-19

**DOI:** 10.1101/2022.02.25.481966

**Authors:** Juanjuan Guo, Minjie Tan, Jing Zhu, Ye Tian, Huanyu Liu, Fan Luo, Jianbin Wang, Yanyi Huang, Yuanzhen Zhang, Yuexin Yang, Guanbo Wang

## Abstract

The range of benefits breastfeeding provides neonates and infants include nutrition, improved neonatal survival, and reduced morbidity from certain diseases. It also aids maternal health by speeding postpartum recovery. However, due to concern about the risk of SARS-CoV-2 transmission and the lack of evidence of breastmilk’s protective effects against the virus, whether mothers with COVID-19 should be encouraged to breastfeed is under debate. Here, we present the results of proteomic and glycoproteomic studies of breast milk (colostrum and mature milk) from mothers with confirmed COVID-19. All colostrum samples exhibited significantly upregulated immune-related proteins, especially whey proteins with antiviral properties against SARS-CoV-2, and increased glycosylation levels and heterogeneity at those proteins. Such adaptive differences in milk from COVID-19 mothers tend to fade in mature milk from the same mothers one month postpartum. These results suggest the immune benefits of colostrum from mothers with COVID-19 and provide molecular-level insights that aid breastmilk feeding decisions in cases of active infection.

## INTRODUCTION

Severe acute respiratory syndrome coronavirus 2 (SARS-CoV-2) has caused over 171 million confirmed cases of coronavirus disease 2019 (COVID-19), which has resulted in over 3 million deaths globally as of June 2, 2021^1^. Based on data from the United States, approximately 98,948 pregnant women had confirmed COVID-19 infections from January 22 to June 28, 2021, 0.14% of all COVID-19 cases. A systematic evidence review that followed the procedures in the Cochrane handbook for systematic reviews of interventions, identified studies that included mothers with suspected or confirmed COVID-19. As a result, in January 2021 the World Health Organization (WHO) updated their Interim Guidance on Clinical Management of COVID-19 to recommend that mothers with suspected or confirmed COVID-19 be encouraged to initiate and continue breastfeeding^2^. However, concerns over the risk of viral transmission from mother to infant have inevitably hindered the choice to breastfeed. A three-month study in 2020 reported that the breastfeeding rate of infants under one week of age by mothers with COVID-19 was 8.8% in China^3^.

Breastmilk contains not only basic nutrients but also developmental immune system factors (e.g., antibacterial components, immunoglobulins, and immune cells) that help protect newborns from environmental pathogens. Tromp et al. (2017) found that breastfeeding was associated with a 30% reduction in lower respiratory tract infections in young children^4^. A member of the enveloped single-stranded RNA virus family, SARS-CoV-2 infects human bronchial epithelial cells, pneumocytes, and upper respiratory tract cells, and that can then lead to severe, life-threatening respiratory disease and lung injury^5^. Respiratory mucosal immunity with IgA, especially secretory IgA (sIgA), plays a major role in mounting an antiviral defense^6^. As the dominant antibody in milk, sIgA has a stronger protective effect than do IgGs because it can directly neutralize the virus while not activating the complement to cause inflammation^7^. The protection of breastfeeding against respiratory infections in infants is generally recognized^8^, and breast milk is an important sIgA source. However, the immune benefits of feeding breastmilk from mothers with COVID-19 remains elusive because of a lack of molecular-level characterization of that milk.

In this work, we used mass spectrometry (MS)-based proteomic analysis to characterize the immune-related proteins in breast milk collected from both six COVID-19 patients and ten healthy donors and then compared the results from those two groups. Both colostrum and mature milk were examined to monitor dynamic proteomic changes. Most human milk proteins are glycosylated and the glycosylation pattern importantly aids in morbidity reduction^9^ by both affecting proteolytic susceptibility^10^ and serving as immunomodulators^11^ and competitive inhibitors of pathogen binding^11,12^. Therefore, we expanded the scope of this study to the glycoproteomic level by thoroughly characterizing a panel of key immune-related proteins’ glycosylation patterns to reveal the COVID-19-induced dynamic glycoproteome evolvement. We show that combined proteomic and glycoproteomic strategies can provide straightforward nutrition and immunity information that can help professionals recommend appropriate infant feeding procedures for infected mothers.

## RESULTS

### Alteration of proteome profiles in milk from COVID patients

The basic clinical characteristics between the patient group and control group were not significantly different (*p*>0.05, Table 1). Based on the results of nested RT-PCR methods ^13^, all milk samples tested negative for SARS-CoV-2, which ruled out the risk of direct viral transmission through the milk. Also, milk is recognized to be mature after 16 days postpartum and completely mature by 4 to 6 weeks postpartum^14^. Meanwhile, the protein content in human milk gradually decreases throughout lactation, but the rate of decrease is much lower at the fully mature stage^15^ and the composition remains relatively consistent^14^. Therefore, the mature milk samples collected in this study represent a relative stable lactational period (Figure 1A). We conducted proteomic and glycoproteomic analyses of milk compositional changes in four sample groups: colostrum samples from COVID-19 patients (COVID-19 colostrum, n = 6) and from healthy donors (Ctrl colostrum, n = 10), and mature milk samples from COVID-19 patients (COVID-19 mature, n = 6) and from healthy donors (Ctrl mature, n = 10) (Figure 1B).

**Table 1.** Clinical characteristics of sample donors.

|  | Characteristics | Case (n=6) | Control (n=10) | p-value |
| --- | --- | --- | --- | --- |
| **Maternal** | age, median (IQR<sup>a</sup>) [years] | 30.00 (27.75, 30.00) | 33.50 (31.00, 34.75) | 0.08 |
|  | pre-pregnancy BMI<sup>b</sup>, median (IQR) [kg/m<sup>2</sup>] | 19.00 (18.23, 22.03) | 21.55 (20.35, 22.13) | 0.22 |
|  | gestational weight gain, median (IQR) [kg] | 18.50 (15.75, 19.00) | 14.00 (12.25, 15.75) | 0.06 |
|  | multiparous, n [%] | 3 (50) | 8 (80) | 0.30 |
|  | undergraduate, n [%] | 3 (50) | 7 (70) | 0.61 |
|  | postgraduate, n [%] | 3 (50) | 3 (30) | 0.61 |
| **Diseases during pregnancy** | anemia, n [%] | 0 (0) | 0 (0) | NA |
|  | diabetes, n [%] | 1 (17) | 3 (30) | 1.00 |
|  | hypertension, n [%] | 0 (0) | 0 (0) | NA |
|  | pre-eclampsia, n [%] | 1 (17) | 0 (0) | 0.38 |
| **Infant** | male, n [%] | 3 (50) | 6 (60) | 1.00 |
|  | gestational age on admission, median (IQR) [wk] | 37.50 (37.00, 38.75) | 39.00 (38.25, 39.00) | 0.24 |
|  | birth weight, median (IQR) [kg] | 2.99 (2.79, 3.14) | 3.24 (3.10, 3.62) | 0.07 |
|  | birth length, median (IQR) [cm] | 49.50 (48.25, 50) | 49.00 (48.00, 49.75) | 0.82 |
|  | head circumference, median (IQR) [cm] | 34.50 (34.00, 35.00) | 35.00 (35.00, 35.25) | 0.08 |
|  | Apgar 1-min, median (IQR) | 9 (9, 9) | 9 (9, 10) | 0.08 |
|  | Apgar 5-min, median (IQR) | 10 (10, 10) | 10 (10, 10) | 0.84 |

<sup>a</sup>IQR: interquartile range
<sup>b</sup>BMI: body masses index

**Figure 1.**
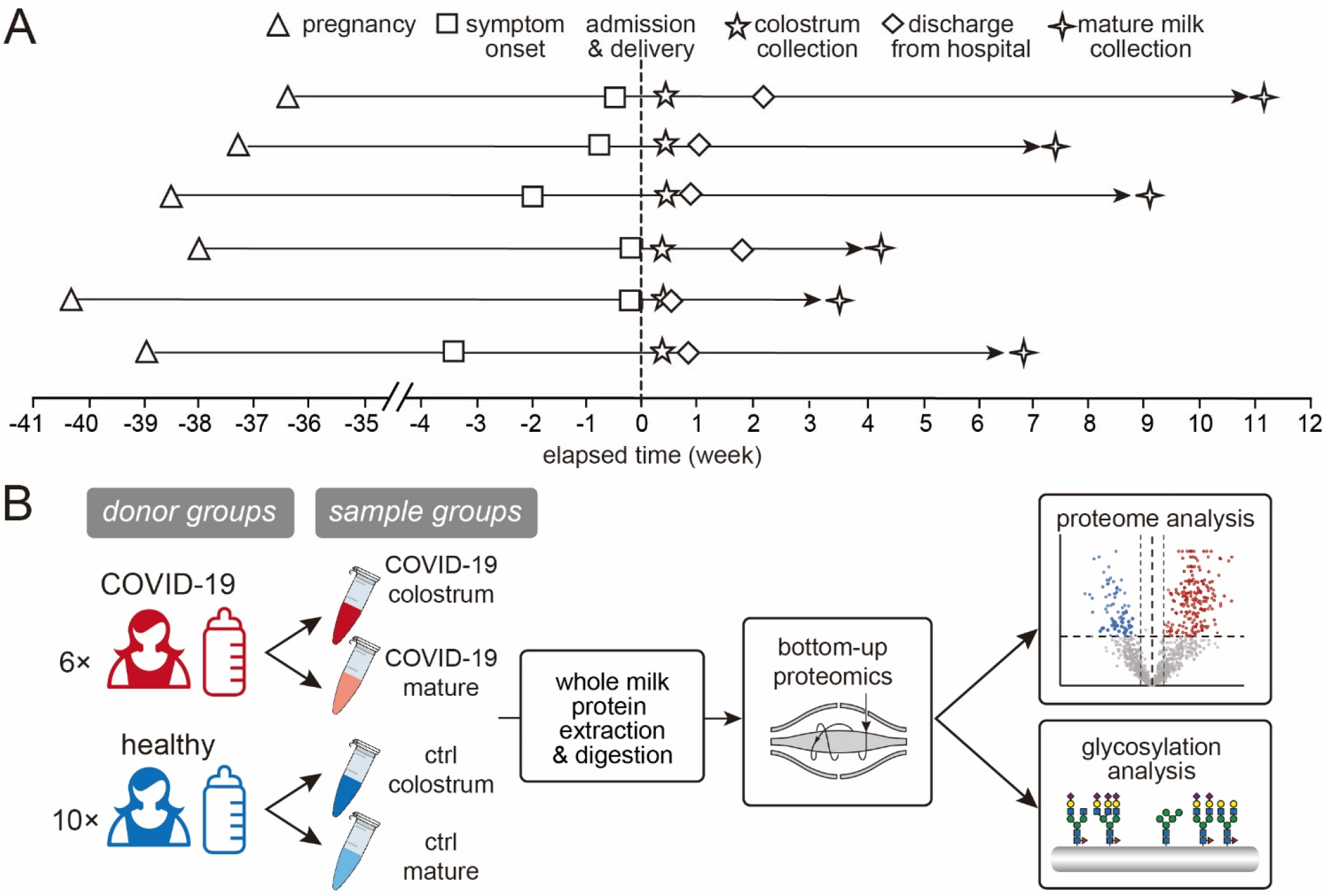
Study information and workflow. (**A**)The timeline of milk sample collection from the 6 COVID-19 patients and (**B**) a schematic workflow of this study.

Of the 1573 proteins identified in the four milk groups, 1439 were sufficiently abundant to be quantitatively monitored, and 1123 of those (78%) were shared by all groups (Figure 2A, Table S1). Principal components analysis (PCA) suggested a distinction between the COVID-19 colostrum group and the other three groups, indicating an alteration in milk proteome upon active SARS-CoV-2 infection (Figure 2B). The total protein content in the COVID-19 colostrum group were 4.1 times higher than that in the control colostrum group; particularly, casein proteins were 3.9 times lower and whey proteins were 7.2 times higher (Figure 2C). As the major components of milk, whey proteins and caseins (comprising about 45%–97% and 3%–55% of human milk proteins, respectively) figure importantly in nutrition and immunity^16^, including being sources of amino acids, enhancing micronutrient bioavailability, providing immunogenic training for innate and adaptive immunity, and promoting intestinal growth and maturation via interactions with the microbiome^17^. The proportion of casein proteins in the COVID-19 colostrum group was lower than that of the other groups (*p* < 0.0001; Figure 2C), likely because of the downregulation of all caseins (α-S1-, β-, κ-caseins) and an increase in the total abundance of whey proteins. In detail, Among the highly abundant milk proteins, beta-casein and immunoglobulins (predominated by sIgA1) were significantly down- and upregulated, respectively (Figure 2D). The differences between COVID-19 milk and Ctrl milk protein contents at the mature stage were not significant (Figure 2C and 2D).

**Figure 2.**
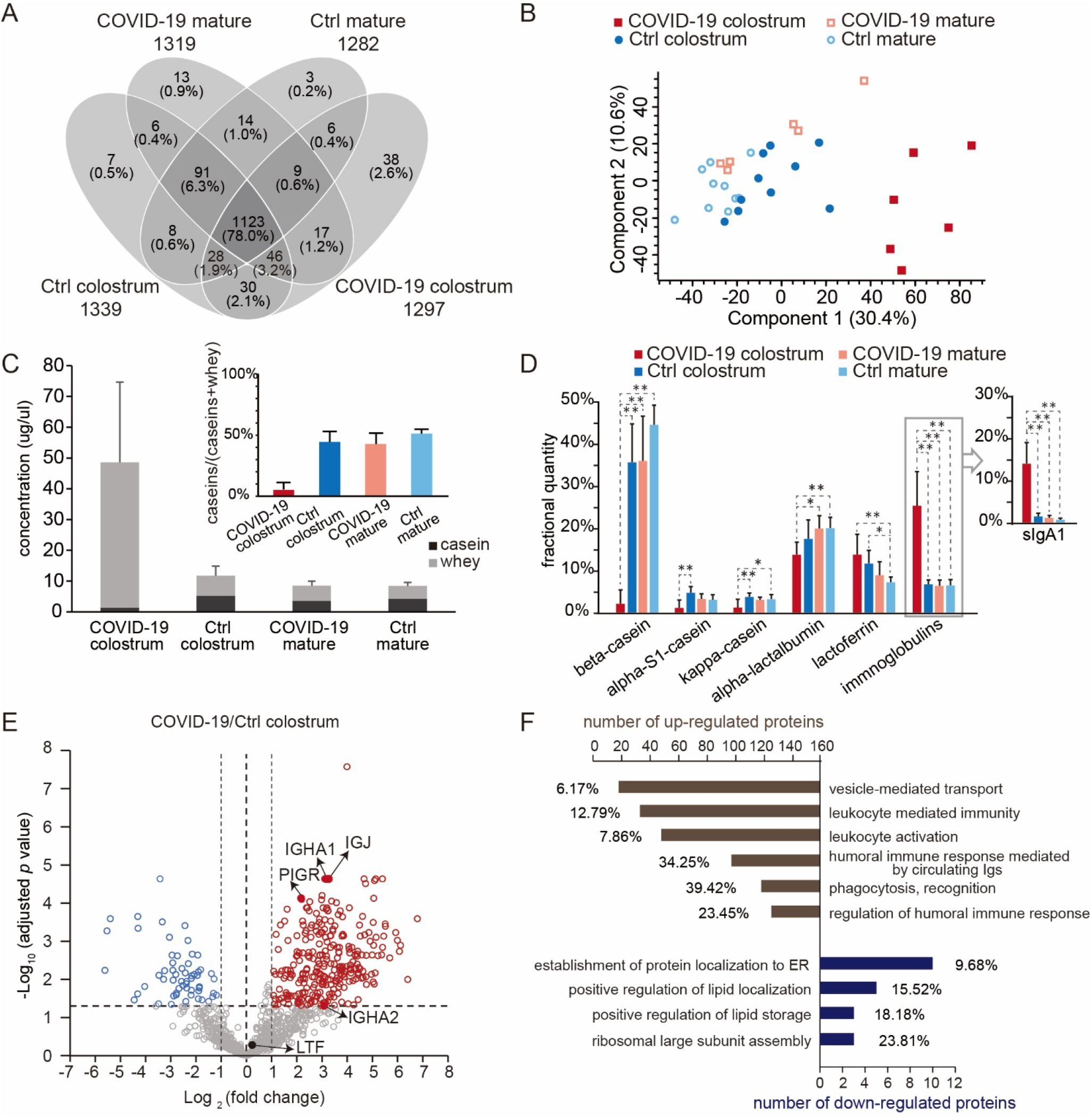
Comparison of human milk proteomes in different sample groups. (**A**) A Venn diagram showing the numbers of proteins that could be quantitatively monitored. The number of proteins for each group are shown below the group name. (**B**) Principal components analysis results of the milk proteome. (**C**) Bar graphs of the quantities of casein (black) and whey (gray) proteins. The inset shows the fractions of caseins. Bars indicate means and whiskers indicate SDs. (**D**) Comparisons of the fractional quantities of major milk proteins. For one-way ANOVA with Tukey HSD test, * *p* < 0.05, ** *p* < 0.01. (**E**) A volcano plot showing proteome changes between COVID-19 colostrum and Ctrl colostrum groups. (**F**) Significantly enriched biological processes of differently expressed proteins (*p* of each Gene Ontology [GO] term < 0.0002). The graph shows both numbers and percentages of differently expressed proteins in each term group.

### Upregulation of immune-related proteins suggest the adaptive immune benefit of colostrum from COVID-19 patients

In comparing the COVID-19 colostrum and Ctrl colostrum groups (cut-off *p* < 0.05 and fold change >2), we found 282 upregulated proteins and 58 downregulated proteins (Figure 2E) in association with 34 and 7 enriched biological processes, respectively, among which we specify the most prominent ones in Figure 2F (see Table S2 for the list of enriched GO terms and their related proteins). The increased expressions of immunoglobulins in response to the viral infection served as the major contributors to those term groups. We did not directly measure sIgA because of the bottom-up approach used in this study. Consider that the intact multimeric form of sIgA incorporates IgA1/IgA2, a joining chain (JC), and a secretory component from the endoproteolytic cleavage of the polymeric immunoglobulin receptor (PIGR)^18^. So, since JC, PIGR, light chains, and variable regions may be used to assemble other forms of immunoglobulins, we instead used the detected quantities of immunoglobulin heavy constant A1 and A2 (IGHA1 and IGHA2) to represent the abundances of sIgA1 and sIgA2. Notably, ribosomal proteins were among the significantly downregulated proteins and they contributed to the GO term for both ribosomal large subunit assembly and protein localization establishment to the endoplasmic reticulum (ER, Figure 2F), both of which are activities associated with viral replication and transport.

Biological pathway analysis revealed 15 differentially expressed proteins that participate in the pathway related to coronavirus disease. Nine ribosomal proteins that might inhibit protein expression in the mammary gland were down-regulated, while some complement proteins (e.g., complement factor B, complement C3, and complement C4B) increased. Altogether, the proteome in COVID-19 colostrum featured a much higher proportion of immunoglobulins than did that of the Ctrl colostrum, thus implying that the former likely provides infants adaptive immune protection against coronavirus.

### The increased microheterogeneity of overall protein glycosylation and the glycosylation levels of immune-related proteins in the COVID-19 colostrum proteome

Glycoprotein microheterogeneity originates from the various glycan structures that may attach to identical glycosylation sites on cell surfaces. Because microheterogeneity leads to differing preferences of glycan-based interactions and reorganization of dynamic epitopes^19^, it thus facilitates virus neutralization. Our glycoproteomic measurements identified 820 unique *N*-glycopeptides from 48 different glycoproteins (Figure 3A), results that mostly agreed with those of a previous study^20^. Most of those glycoproteins were within the top 200 most abundant milk proteins (Figure S1A). Also, most of them had one to six identified glycosites, but tenascin (TNC), which is involved in protection against viral infections^21^, had 12 glycosites (Figure S1B). In the COVID-19 colostrum glycoproteome, we observed a high degree of microheterogeneity (Figure S1C) that was clearly distinct from those of the other three samples groups (Figure 3B). Among 103 glycosites, 66.0% had more than 1 glycoform and 18.4% harbored more than 10 glycoforms. Taking the glycosylation pattern into account, we detected only 178 (21.7%) of the 820 glycopeptides in all sample groups, while 33.2% (272) of the glycopeptides were detected exclusively in the COVID-19 colostrum group (Figure 3A). This was most likely because of a dramatically improved glycosylation pattern diversity and slightly increased number of glycosites in that group (Figure S1D) rather than more protein identities (Figure S1E).

**Figure 3.**
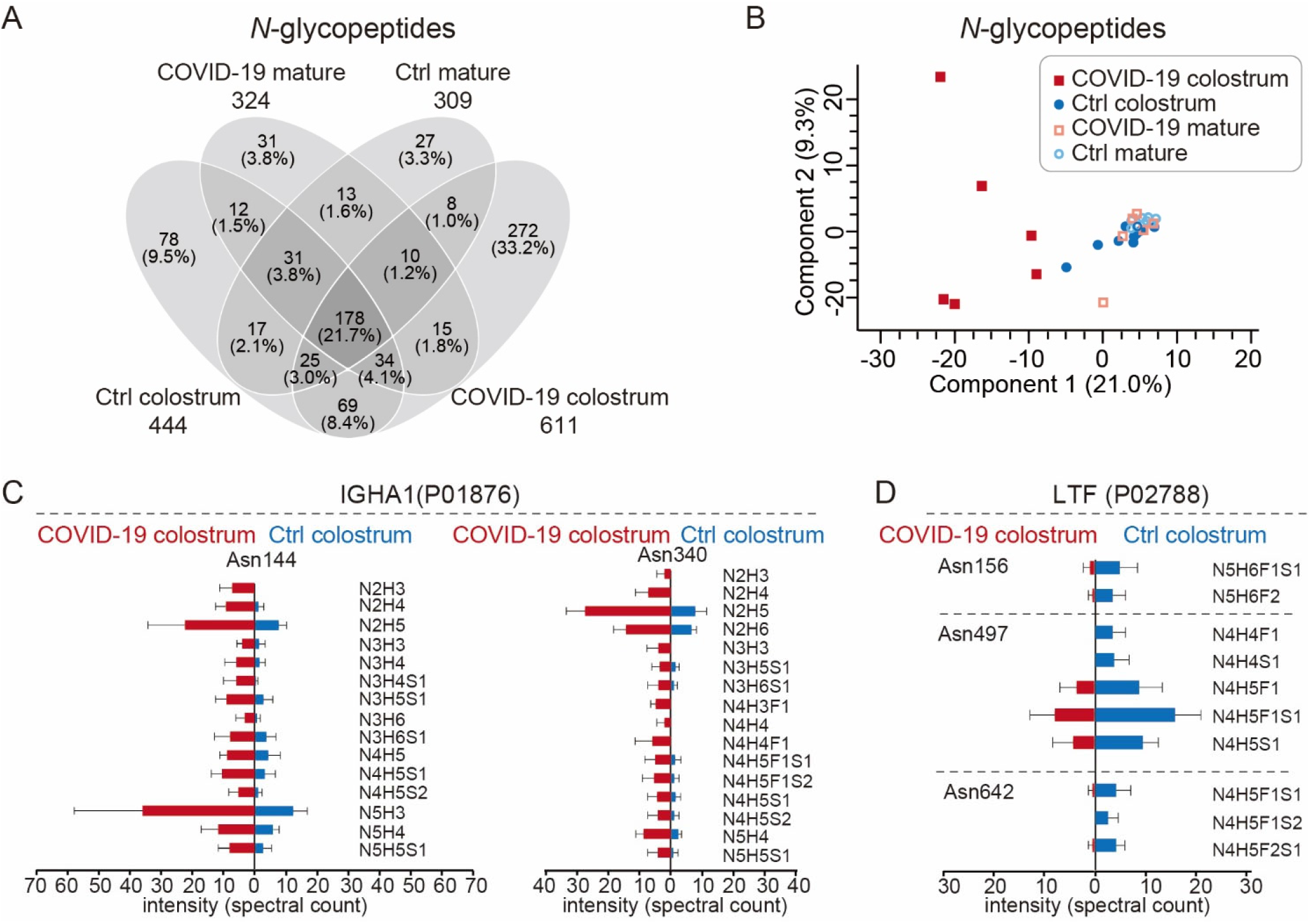
The glycoproteomes of each sample group. (**A**) A Venn diagram showing the numbers of *N*-glycopeptides that could be quantitatively monitored. The number of *N*-glycopeptides for each group are shown below the group name. (**B**) Principal components analysis plot of *N*-glycopeptides. (**C** and **D**) Site-specific changes in *N*-glycopeptides of immunoglobulin heavy constant alpha 1 (IGHA1, **C**) and lactoferrin (LTF, **D**) between COVID-19 colostrum and Ctrl colostrum groups. Bars indicate means and whiskers indicate SDs.

We observed 83 upregulated and 17 downregulated *N*-glycopeptides, as well as site-specific *N*-glycosylation changes, in the COVID-19 colostrum (Figure S2). As the major components in milk, the two most abundant glycoproteins, lactoferrin (LTF) and sIgA1, exhibited different glycosylation pattern alteration trends in response to COVID-19. The elevated level of sIgA, a key component in adaptive immunity, was accompanied by increases in both microheterogeneity at IGHA1’s Asn144, and Asn340 *N*-glycosylation sites and those glycoforms’ expression levels (Figure 3C). In contrast, LTF, which is associated with innate immunity^22^, showed reduced microheterogeneity (Figure 3D).

### COVID-19 colostrum’s immune-related proteome profile changes fade with milk maturation

Compared to levels in the COVID-19 colostrum, the proportions of whey proteins, especially sIgA1, decreased significantly in the COVID-19 mature group (Figure 2C and 2D). Besides the overlapping proteins that were differentially expressed at different lactational stages in the Ctrl groups, 201 proteins were downregulated while 83 proteins were upregulated in COVID-19 milk during its maturation (Figure S3A). Fourteen and eight biological processes changed significantly for down- and up-regulated proteins, respectively (Table S3, Figure S3B). Leukocyte mediated immunity, adaptive immune response, and complement activations involved the largest number of down-regulated proteins, and phagocytosis recognition and regulation of complement activation involved the highest proportion of all associated genes. Up-regulated proteins during maturation were clustered largely in the SRP-dependent co-translational protein targeting membrane and ubiquitin ligase inhibitor activity GO terms. Also, multiple ribosomal proteins contributed to those clusters. The downregulated proteins clearly show that immune-related proteins had obviously decreased during the mature milk phase and with the COVID-19 patients’ recoveries.

The COVID-19 colostrum’s unique glycosylation features also appeared to fade with milk maturation. Seven *N*-glycopeptides were back-upregulated and 55 were back-downregulated in the COVID-19 mature sample group (Figure S4A). The glycosite-specific changes at sIgA and LTF in COVID-19 colostrum, with respect to Ctrl groups, underwent reverse evolvement in the COVID-19 mature group, producing glycosylation patterns that were similar to those of the Ctrl mature milk (Figure S4B). Using the changes between Ctrl colostrum and Ctrl mature as the reference, sIgA’s microheterogeneity and glycosylation levels at Asn144 and Asn340 changed more prominently than other IGHA1 *N*-glycopeptides did (Figure S4C), and so did LTF at Asn156, 497, and 642 (Figure S4D). See Figure S5 for an extended list of *N*-glycoproteome site-specific changes between Ctrl colostrum and Ctrl mature groups. In summary, the proteome and glycoproteome profiles exhibited in the mature milk of the COVID-19 mature and Ctrl mature groups (Figure 2B, 3B, S6A and S6B) were not substantially different.

## DISCUSSION

In healthy mother-infant dyads, human milk provides nutritional and protective components for newborns facing pathogenic challenges^17^. During inflammation or infection in either mother^23^ or infant^24^, the changes in breastmilk components not only reflects the maternal-infant health status^25,26^, but also provides the neonate with protection from diseases ranging from diarrhea to acute respiratory tract infections^27^. However, since SARS-CoV-2 RNA has been detected in human breastmilk, women with COVID-19 are greatly concerned about feeding their infants breastmilk^28-31^. Chinese experts have recommended that COVID-19 infected mothers delay breastfeeding for periods of up to two weeks after delivery^32^, and many patients chose to begin breastfeeding after recovery. Also, some hospitals are implementing practices that conflict with WHO’s policy on the subject^33^. All infected mothers of newborns must understand the nutritional and immune benefits of breastmilk so they may knowledgeably consider the option of feeding it to their infants.

COVID-19 infection clearly leads to significant changes in profiles of both proteome and glycosylation patterns in breastmilk, and they both undergo dynamic evolution as milk maturation proceeds. Two major components of human breastmilk, casein and whey proteins, are down- and up-regulated, respectively, and several studies have found that the composition of human breastmilk changes according to the infant’s demands^34,35^. Fetus oxygen is supplied completely through placental transport, and the most striking features of COVID-19 placentas are decidual arteriopathy and other maternal vascular malperfusion features, either one of which can cause fetal hypoxia^36^. Hypoxia in neonatal rats leads to delayed enzyme maturation in the small intestine, which is the main site for protein absorption. The activities of those enzymes returned to control levels 3 weeks after recovery^37^. Maternal stressors in late pregnancy can also impair the fetal gut, further limiting the ability of the newborn to absorb nutrients, especially proteins from colostrum^38^. Absorption of high-MW casein proteins is more difficult than absorption of relatively low-MW whey proteins. Given that COVID-19 neonates likely suffered fetal hypoxia and maternal stresses due to the disease, the decreased levels of casein proteins and increased whey protein abundance in COVID-19 colostrum benefits the neonatal gut’s ability to effectively absorb milk proteins during the first few days postpartum. As the neonatal gut recovers from hypoxia damage, both casein and whey protein abundances return to normal levels in mature milk (Figure 2C). Since caseins in human milk provide essential amino acids^39^ and transport divalent cations (such as Ca^2+^ and Zn^2+^)^40,41^, the decline of caseins in COVID-19 colostrum might affect absorption and transportation of some essential minerals in the gut of the nursing infant.

Whey proteins show antiviral activity against SARS-CoV-2 and its related pangolin coronavirus (GX_P2V) by blocking viral attachment and viral replication^42^, and immunoglobulins are major elevators of overall whey protein levels (Figure 2D). Immunoglobulins in breast milk trigger antigenic responses to infectious pathogens of maternal mucosa-associated lymphoid tissue, or the bronchomammary axis, and is most likely encountered by the infant^43,44^. Immunoglobulin expression in blood is dramatically increased in the serum proteome of COVID-19 patients who had seroconversion, and most of the immunoglobulin regions are significantly changed in response to SARS-CoV-2 infection^45^. SARS-CoV-2-specific antibodies detected in milk samples collected from COVID-19 patients were capable of neutralizing SARS-CoV-2 in vitro^46,47^. Breastmilk immunoglobulins complement the neonate’s immature adaptive immune system^48^ and protect it by preventing infectious agents on the mucosal membranes from entering tissues, rather than by functioning in the circulation. This contrasts with IgG’s transplacental protection that is most efficient against microbes in the blood and tissues and that functions through a complementary system that induces energy consumption and destructive tissue inflammation^49^. The major immunoglobulin in human milk, sIgA, serves as a first line defense in mucosal areas, using mechanisms that include intracellular neutralization, virus excretion, and immune exclusion^50,51^. When a pathogen enters the mother’s upper airways, the Peyer’s patch acquires the pathogen and its antigens are presented by M cells to circulating B cells that then migrate to the serosal (basolateral) side of the mammary epithelial cell, and it produces IgAs. As the IgAs move from the serosal to the luminal (apical) side of the mammary epithelial cell, they are glycosylated and complexed to form sIgA, which is secreted into milk^17,52^. Since sIgA is more resistant to protein hydrolysis than IgG is, it can function on the mucosal membranes primarily in the gastrointestinal tract and to some extent in the respiratory tract as well^49^. Because the volume of milk ingested increases as a neonate grows, sIgA, which usually occurs in higher concentrations in colostrum than in mature milk, is nevertheless maintained at a relatively constant total amount throughout lactation. On average, the proportion of sIgA1 in the milk proteins of COVID-19 colostrum was 8.6 times greater than in the Ctrl colostrum (Figure 2D), thus indicating that the mothers’ responses to the infection and additional adaptive immunological protection for the infants from viruses may be introduced through the gastrointestinal or respiratory tract. Indeed, it has been reported that a portion of sIgA present in milk from COVID-19 patients were RBD-specific^53,54^. Such protection may be further strengthened by other adaptive immunity-related proteins that appear in higher abundances in COVID-19 colostrum than in Ctrl colostrum (Figure 2F).

Ribosomal proteins are the major class of downregulated whey proteins in COVID-19 colostrum (Figure 2F), even with the increased total amount of whey proteins. Ribosomal proteins are essential for translating genetic information from mRNA into proteins^55^, and they dictate ribosomal large subunit assembly and the establishment of protein localization to the ER. Since SARS-CoV-2 has no cellular structure of its own, it can only replicate by utilizing ribosomes in infected cells to make viral polyprotein and then processing and transporting that polyprotein formed in ER ribosomes^56^. Downregulation of those ribosomal proteins in COVID-19 colostrum may be associated with the milk protein synthesis status that helps inhibit viral replication and viral polyprotein transport in the mammary glands of infected, lactating mothers.

The glycoproteins, especially immunoglobulins in COVID-19 colostrum, generally exhibit much higher degrees of glycan microheterogeneity (Figure 3), a feature that may help recognize more dynamic epitopes and favor different glycan-based interactions^19^. Consider that SARS-CoV-2 assembly and its attachment and entry to a host cell is closely related to its heavily glycosylated spike protein^57^, which has 22 occupied *N*-glycosylated sites^58^ and diverse glycosylation^59^. Glycans at the surface of SARS-CoV-2 are essential for viral entry, as shown in vitro when blocked *N*-glycan biosynthesis at the oligomannose stage dramatically reduced viral entry into HEK293T cells^60^. When SARS-CoV-2 enters the human body, it first recognizes sialic acid and searches for its receptor. Next, very likely the viral envelope glycans interact with sialic acid-binding lectins that are expressed in a host cell-dependent manner^61^. Accordingly, the more that human milk glycoproteins have glycan microheterogeneity, the better their ability might be to interact with SARS-CoV-2 via multiple glycan-based molecular events in the infants’ gastrointestinal tract. That then helps block SARS-CoV-2 binding to its receptor, ACE2, on the cell surface.

The WHO’s breastfeeding recommendations for COVID-19 mothers are largely based on both the significant consequences of not breastfeeding and the separation between mother and child, as well as on the low risk of vertical transmission of COVID-19 through breastfeeding. Here, the proteomic and glycoproteomic data suggest the nutritional and immune benefits of breastmilk feeding at the molecular level. Since all prominent differences between COVID-19 and Ctrl colostrum proteomes, in terms of the casein/whey ratio, differently expressed proteins and the related significantly enriched biological processes, and the microheterogeneity of glycosylation all fade as milk maturation proceeds, those benefits may be attenuated by delaying breastmilk feeding. Based on the premise that contact transmission between COVID-19 mothers and their newborns is prevented, our results provide helpful information for selection of approaches to lactation.

## METHODS

### Sample collection

Human milk samples were collected from six patients with confirmed COVID-19 and from ten healthy donors. First, colostrum samples were collected within 3-4 days post-partum, and then mature milk samples were collected 30-45 days postpartum, upon the COVID-19 patients’ recoveries and release from quarantine or during the healthy donors’ postpartum checkups. The sample sizes and groups were (1) 6 in the group of colostrum from COVID-19 patients (COVID-19 colostrum), (2) 6 in the group of mature milk from COVID-19 patients (COVID-19 mature), (3) 10 in the group of colostrum from healthy donors (Ctrl colostrum), and (4) 10 in the group of mature milk from healthy donors (Ctrl mature) (Figure 1B). This study was approved by the Medical Ethics Committee of Zhongnan Hospital of Wuhan University (approval no. 2020031). Written consent was obtained from each patient.

### Milk sample preparation for proteomic analysis

The in-solution digestion of whole milk was adapted from previous methods^62^. Briefly, a 20 μl milk sample was added to 80 μl of protein lysis buffer (ThermoFisher, USA) with a protease inhibitor (ThermoFisher, USA). The whole milk buffer was sonicated 20 mins in a water bath with cold water and put on ice for another 30 mins. Impurities were removed by methanol/chloroform protein precipitation and the precipitate was air dried. The pellet was further dissolved in an 8 M urea (Sigma, Germany)/50 mM ammonia bicarbonate (Alfa Aesar, Germany) solution, incubated with 10 mM dithiothreitol (Sigma, Canada) at 37 °C for 1 h and followed by alkylation with 20 mM iodoacetamide (Sigma, USA) at 37°C in the dark for 30 mins. For digestion, the samples were incubated with a trypsin/lys-C mixture (w:w = 1:50; Promega, USA) at 37°C for 16 h and then the digest reaction was quenched with 1% formic acid (Fisher Scientific, Czech Republic). The resulting peptides were desalted using a C18 column (ThermoFisher, USA) prior to lyophilization. The same tryptic peptides for proteomic analysis were applied directly to glycoproteomic analysis without further glycopeptide enrichment.

### Liquid chromatography–tandem mass spectrometry

In preparation for the liquid chromatography (LC)–tandem MS (MS/MS) analysis the lyophilized tryptic peptides were reconstituted with 10% formic acid. The peptides were then separated by an UltiMate 3000 RSLCnano LC System (Thermo Fisher Scientific, USA) equipped with an Acclaim PepMap 100-C18 trap column (ThermoFisher) and an Acclaim PepMap RSLC-C18 analytical column (ThermoFisher). Mobile-phase solvents A and B consisted of 0.1% formic acid in water and acetonitrile, respectively. Trapping was performed at a flow rate of 10 μl/min for 5 min with 3% B and separation was performed at a passive split flow of 300 nl/min for 95 min with 8% to 50% B over 51 min, 50% to 99% B over 14 min, 99% B for 15 min, and 99% to 3% B over 10 min. The eluted peptides were electrosprayed at a 2.0 kV spray and analyzed by an Orbitrap Eclipse Tribrid mass spectrometer (Thermo Fisher Scientific, USA) coupled to the LC system. The MS analyses were operated in data-dependent acquisition (DDA) mode using higher-energy collision dissociation (HCD) for MS/MS fragmentation.

### MS settings for proteomic analysis

The term “proteomic analysis” in this work refers to analysis of the proteome regardless of protein modifications. The MS1 scan range was 300–1500 m/z with a 60,000 resolution at 200 m/z, and the automated gain control (AGC) set at 4e5 with a maximum injection time of 50 ms. With charge-states screening enabled, precursor ions in 2+ to 7+ charge states with intensities >5e4 were mass-selected for MS/MS (isolation window: 1.6 m/z). A 45 s dynamic exclusion was set with a 10-ppm exclusion window. The mass spectrometer was operated in DDA mode with a 2 s cycle time. The MS/MS scans were acquired with a 1.4 m/z isolation window, followed by HCD dissociation with a normalized collision energy of 30% and Orbitrap detection with a 100– 2000 m/z scan range, a 30,000 resolution (at 200 m/z), and an AGC of 4e5 with a 54 ms maximum injection time.

### MS settings for glycoproteomic analysis

The term “glycoproteomic analysis” in this work refers to the determination of the positions and identities of the repertoire of glycans attached to the identified peptides. The MS1 scan range was 375–2000 m/z with a 60,000 resolution at 200 m/z, and the AGC was set at 4e5 with a maximum injection time of 50 ms. With charge-states screening enabled, precursor ions in 2+ to 7+ charge states with intensities >5e4 were mass-selected for MS/MS (isolation window: 1.6 m/z). A 60 s dynamic exclusion was set with a 10-ppm exclusion window and a 3 s cycle time. MS/MS spectra were acquired with Orbitrap across a 120–4000 m/z range at 30,000 resolution with the AGC target set to 5e4 and a 75 ms maximum injection time. An HCD was performed with 30% normalized collision energy. Detection of at least three out of 15 oxonium ions of glycopeptides (127.0390+, 145.0495+, 163.0601+, 243.0264+, 405.0793+, 138.055+, 168.0655+, 186.0761+, 204.0867+, 274.0921+, 292.1027+, 366.1395+, 407.1660+, 512.1974+, 657.2349+) triggers electron-transfer/higher-energy collision dissociation (EThcD) for the same precursor (AGC target: 2e5; maximum injection time: 250 ms; supplemental activation for electron-transfer dissociation: 30% normalized collision energy). In addition, the same samples were analyzed in parallel using the same settings except that stepped HCD (i.e., collision energy of 20%, 30%, 40% and a 1.6 Th isolation window) instead of EThcD was used to fragment the glycans and the maximum injection time was set to 75 ms.

### Data processing

Following Zhu et al. (2020)^19^, the raw shotgun LC-MS/MS data were searched with Proteome Discoverer version 2.3 (Thermo Scientific) using the Sequest HT search engine against a UniProt Swiss-Prot database^63^: *Homo sapiens* (canonical and isoform) (October 2020; 26,566 entries). We used fixed Cys carbamidomethylation, variable methionine (Met) oxidation of peptides, and variable Met loss and acetylation of protein N-terminuses as search variables. Cleavage specificity was set for trypsin with an allowed maximum of two missed cleavages.

For database searching, we used a 10-ppm precursor mass tolerance and a fragment mass tolerance of 0.02 Da followed by data filtering using Percolator, thus resulting in a 1% false discovery rate (FDR). We accepted only peptide to spectrum matches (PSMs) with XCorr >2.2, and we then used the full proteome search result as a focused database for glycopeptide identification (1573 entries). For label-free quantification (LFQ), we used the minora feature detector node with high PSM confidence, a minimum of five non-zero points in a chromatographic trace, a minimum of two isotopes, and a 0.2 min maximum retention time difference for isotope peaks. Then Proteome Discoverer consensus workflow was used to both open the search results and enable retention time (RT) alignment with a maximum 5 min RT shift and 10 ppm mass tolerance to match the precursor between runs. We used the extracted ion chromatogram intensities for the LFQ of peptides. The abundance of each identified protein was determined by averaging ion signals representing each protein’s three best ionized unique peptides. Finally, we estimated each protein’s relative abundance as the proportion of its abundance to the sum of protein abundances ^64^.

Byonic version 3.10.2 (Protein Metrics Inc.) was used to search the glycoproteomic data against the previously determined targeted milk protein database (1573 entries). The search was based on the above-mentioned shotgun proteome analysis strategy and used the following search parameters: trypsin digestion with a maximum of three missed cleavages; 10 ppm precursor ion mass tolerance; fragmentation type, HCD for stepped HCD files and EThcD for EThcD files; 20 ppm fragment mass tolerance; cysteine carbamidomethylation as a fixed modification; Met oxidation and acetylation at the peptide N-terminus as variable modifications. For glycan analysis, we used a Byonic database of 132 human glycan entries. The maximum number of precursors per scan was set to one and the FDR to 1%. Only PSMs to the EThcD spectra or more than two PSMs to the HCD spectra, both with non-negligible error probabilities |log Prob|>2.0 and Byonic scores ≥200, were accepted. The total number of spectra counts for each glycopeptide was used as the quantified value for further analysis.

### Statistical Analysis

All statistical analyses were performed using SPSS 21.0 (SPSS Inc., Chicago, IL, USA), Perseus v.1.6.14.0^65^, or under the R Statistical Computing Environment (v. 4.0.2). The Mann-Whitney *U*-test was used to assess differences in skewed data. The Fisher’s exact probability method was used to test the categorical variables, and one-way analysis of variance or Student’s *t*-test were used to compare the differences in normally distributed data. Principal component analysis and plots were used to conduct data reduction and to visualize the differences in all samples. Statistical significance for all tests was *P* < 0.05. Proteins with an adjusted *p*-value<0.05 were considered to have undergone significant protein enrichment. For identification of up and down-regulated proteins, a Cytoscape (v3.6.0) App ClueGOwas^66^ used based on the terms “biological processes” in the human database. Statistical significance was set for a K score of 0.8. The function “GO Term fusion” was selected and GO term restriction levels were set at 1–20, with a minimum of two proteins or 4% proteins in each GO term.

## Data and Software Availabilities

The proteome data (dataset identifier PXD024776) have been deposited to the ProteomeXchange Consortium (http://proteomecentral.proteomexchange.org) via the iProX partner repository PRIDE^67^. Raw and fasta files for glycopeptome data have been deposited to MassIVE (https://massive.ucsd.edu/ProteoSAFe/static/massive.jsp) with the identifier MSV000087229.

## ACKNOWLEDGEMENTS

This study was supported by grants from National Natural Science Foundation of China (NSFC 21974069), Hubei Provincial Science and Technology Department Novel Coronavirus Pneumonia Emergency Science and Technology Project (Grant No. 2020FCA011), Wuhan Novel Coronavirus Pneumonia Emergency Science and Technology Tackling Key Project (Grant No. 2020020201010011), China Postdoctoral Science Foundation (No. 2021T140067), and the Fundamental Research Funds for the Central Universities (Grant No. 2042020kfxg17).

## AUTHOR CONTRIBUTIONS

J.Z., G.W., Y.Z., Y.Y., and Y.H. designed the research. J.G. and H.L. collected samples and informed patient consents. F.L. stored the specimens and performed virus detection. J.Z., M.T., and Y.T. performed proteomic research and data analysis. M.T., J.Z., G.W., and J.G. wrote the manuscript. J.W. and Y.H. revised the manuscript.

