## Supplementary material for "Proteomic analysis of human milk reveals nutritional and immune benefits in the colostrum from mothers with COVID-19": SI

**Table of Contents**

Supplementary Figures……………………………………………………………………………………………………………………….. S2-S7

Supplementary Tables ……………………………………………………………………………………………………………………S8-S165

Address correspondence to:

✉ Yanyi Huang,

✉ Yuanzhen Zhang,

✉ Yuexin Yang,

✉ Guanbo Wang,

**SUPPLEMENTARY FIGURES**

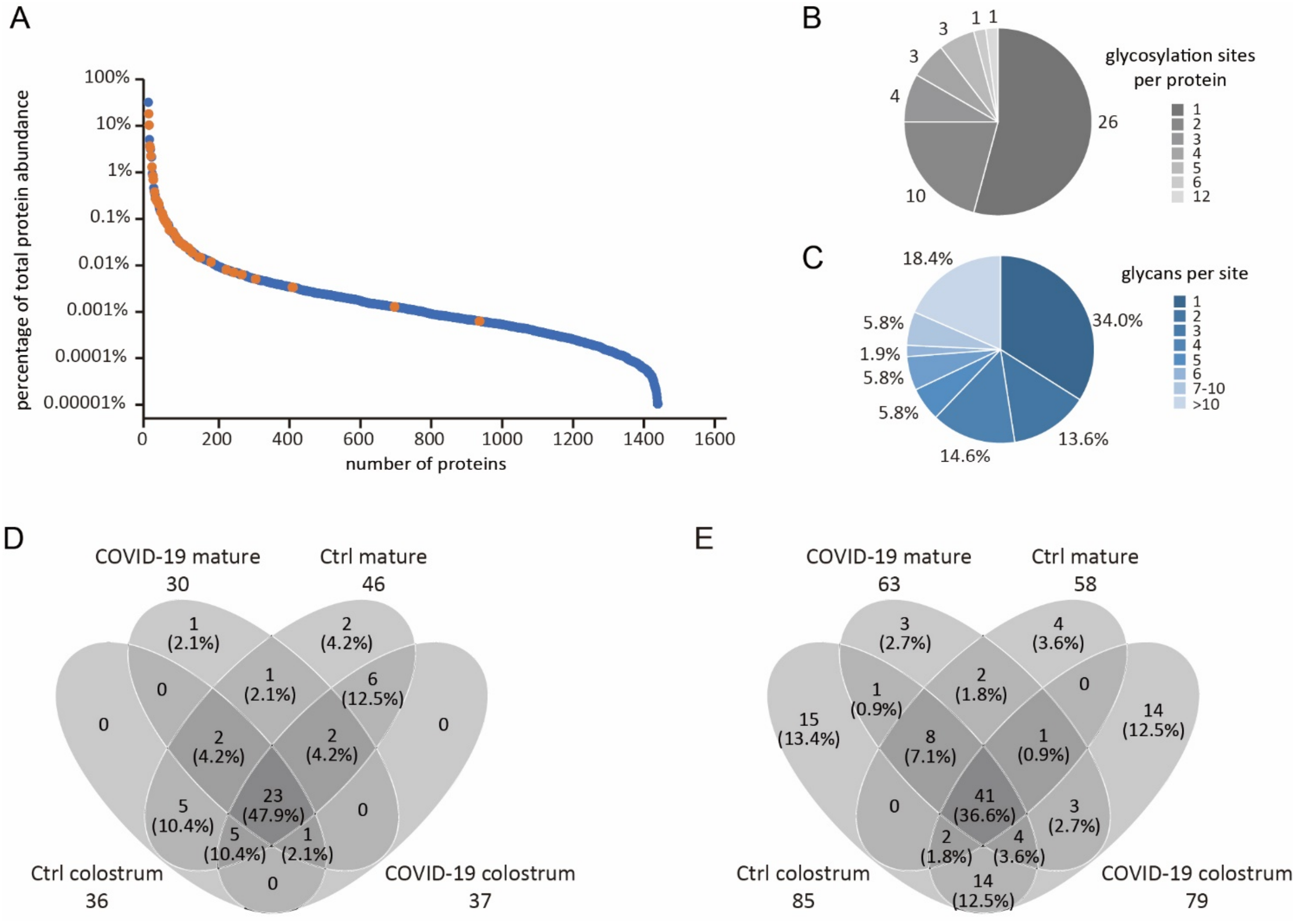


**Figure S1.** Overview of *N*-glycoproteomes in different sample groups. (**A**) Dynamic range of identified proteins. Orange dots indicate identified glycoproteins and blue dots represent proteins without identified glycosylation sites. (**B**) Distributions of the numbers of glycosylation sites detected at each identified glycoprotein. (**C**) Distributions of the glycosylation sites detected with different numbers of glycans. (**D** and **E**) Venn diagrams illustrating (**D**) the numbers of glycoproteins and (**E**) the numbers of glycopeptides identified in each sample group.

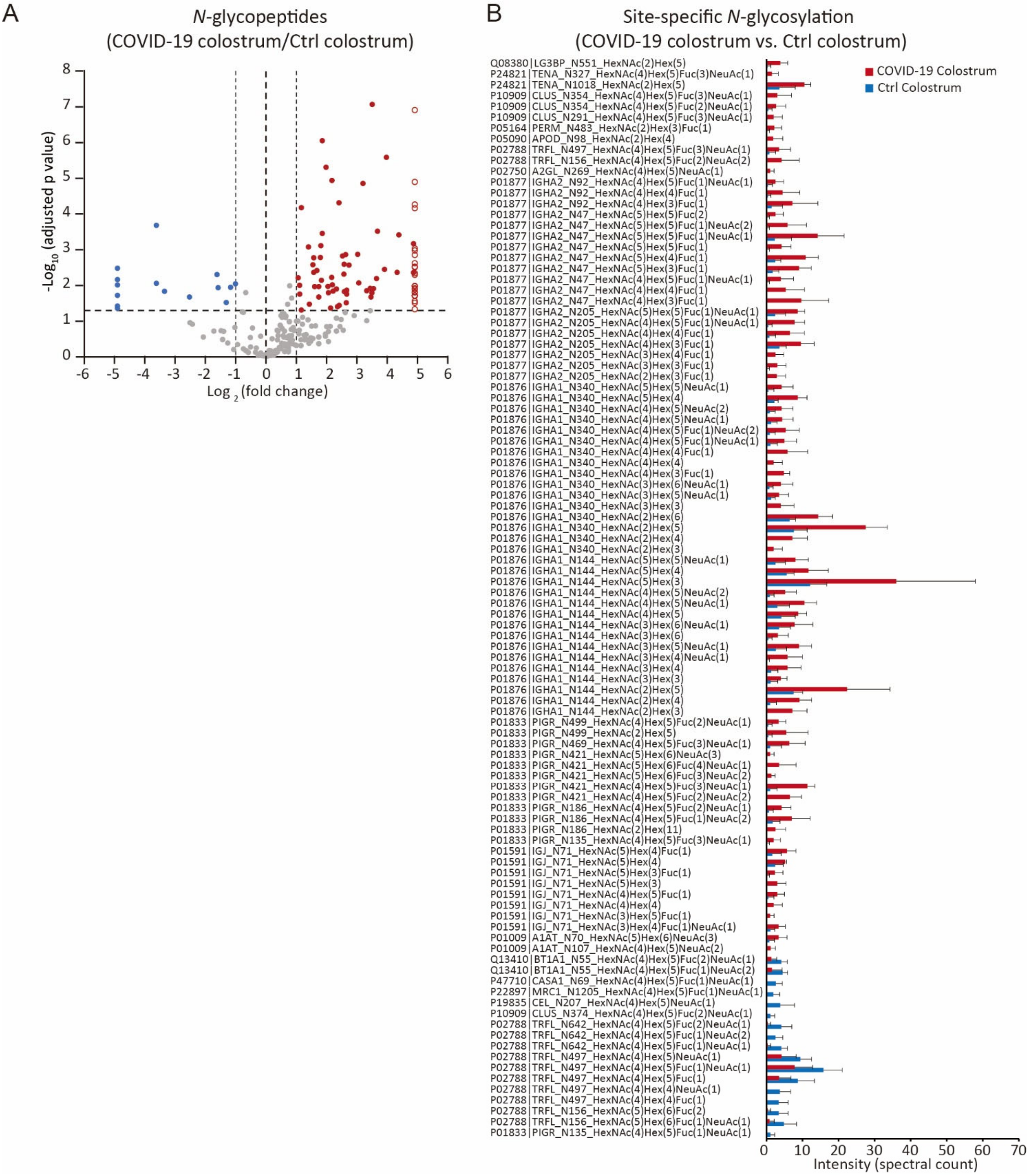


**Figure S2.** Comparison of *N*-glycoproteome profiles in the COVID-19 colostrum and Ctrl colostrum groups. (**A**) A volcano plot showing changes in glycopeptides. The open circles represent glycopeptides that were exclusively detectable in the COVID-19 colostrum group; their abundances in the COVID-19 mature group were manually set as a fixed minimum quantitative value, thereby giving rise to an identical fold-change value in the plot. (**B**) A bar graph showing changes in site-specific *N*-glycosylation. Bars indicate means and whiskers indicate SDs.

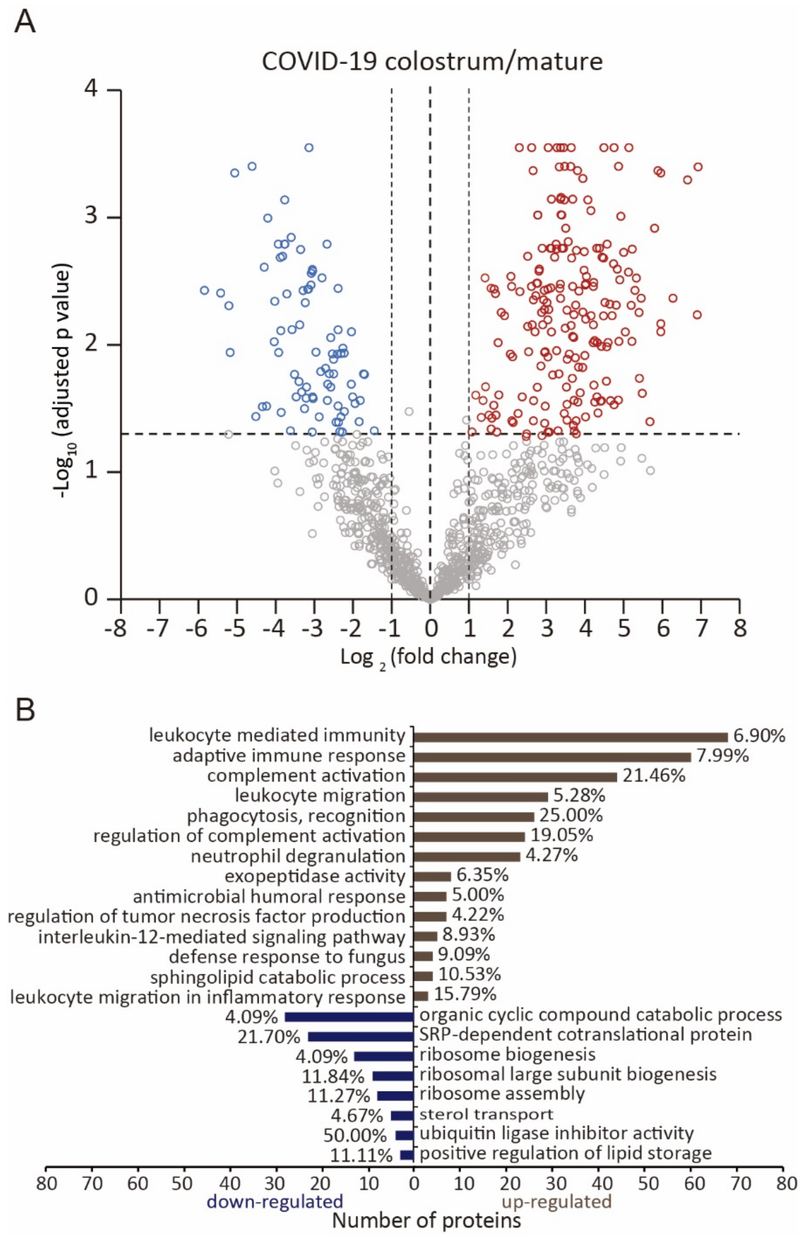


**Figure S3.** Comparison of COVID-19 colostrum and COVID-19 mature sample group proteomes. (**A**) A volcano plot of proteomes in COVID-19 colostrum and COVID-19 mature milk. (**B**) Enriched biological processes of differentially expressed proteins ($p$ of each Gene Ontology [GO] term < 0.0002). The graph shows both numbers and percentages of differently expressed proteins in each term group.

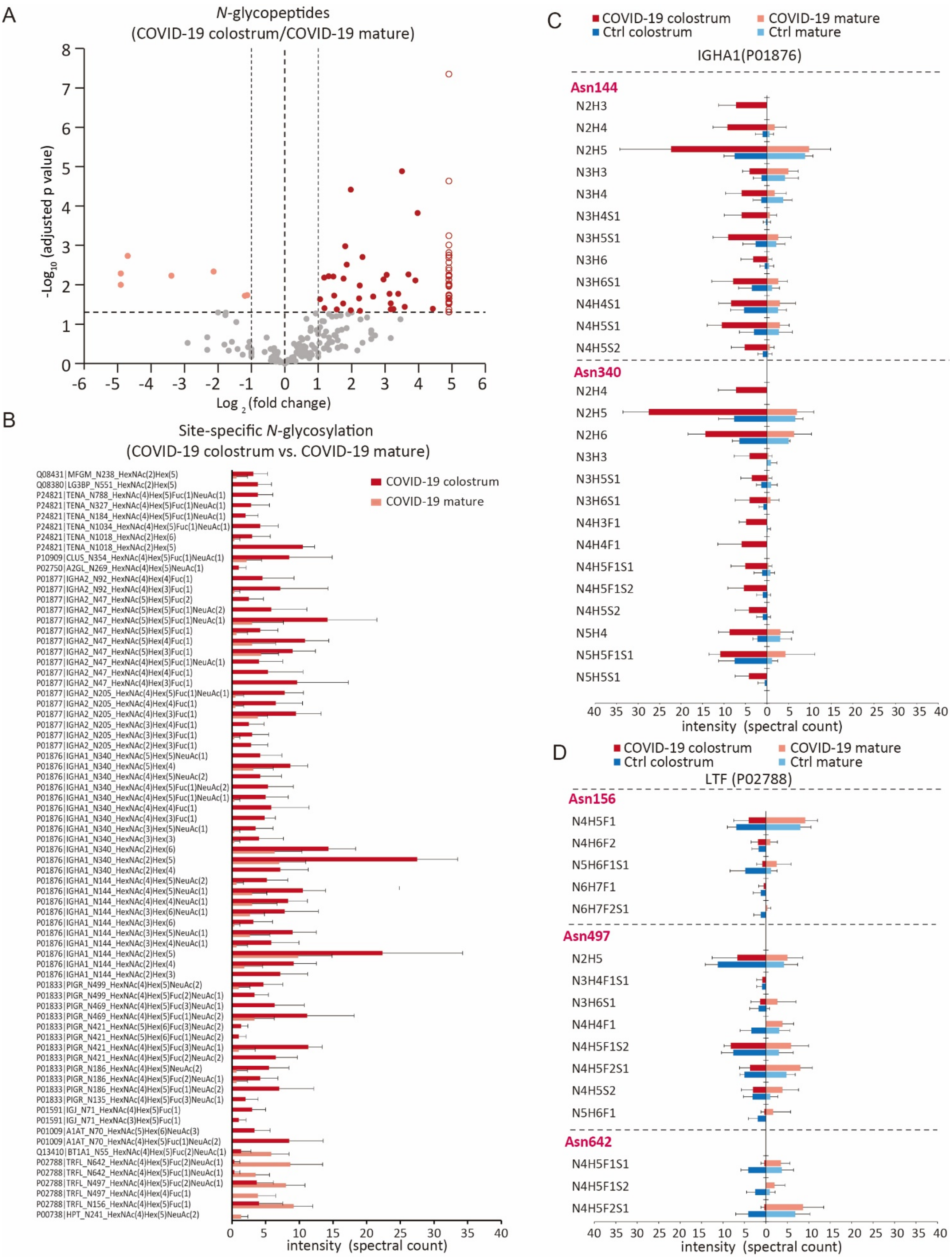


**Figure S4.** Comparisons of the *N*-glycoproteome profiles between the four sample groups. (**A**) A volcano plot showing changes in glycopeptides during milk maturation. The open circles represent glycopeptides that were exclusively detectable in the COVID-19 colostrum group; their abundances in the COVID-19 mature group were manually set as a fixed minimum quantitative value, thereby giving rise to an identical fold-change value in the plot. (**B**) A bar graph showing changes in site-specific *N*-glycosylation during milk maturation. (**C**) Comparisons of site-specific immunoglobulin heavy constant alpha 1 (IGHA1) *N*-glycopeptides in the four sample groups. (**D**) Comparisons of site-specific lactoferrin (LTF) *N*-glycopeptides in the four sample groups. Bars indicate means and whiskers indicate SDs.

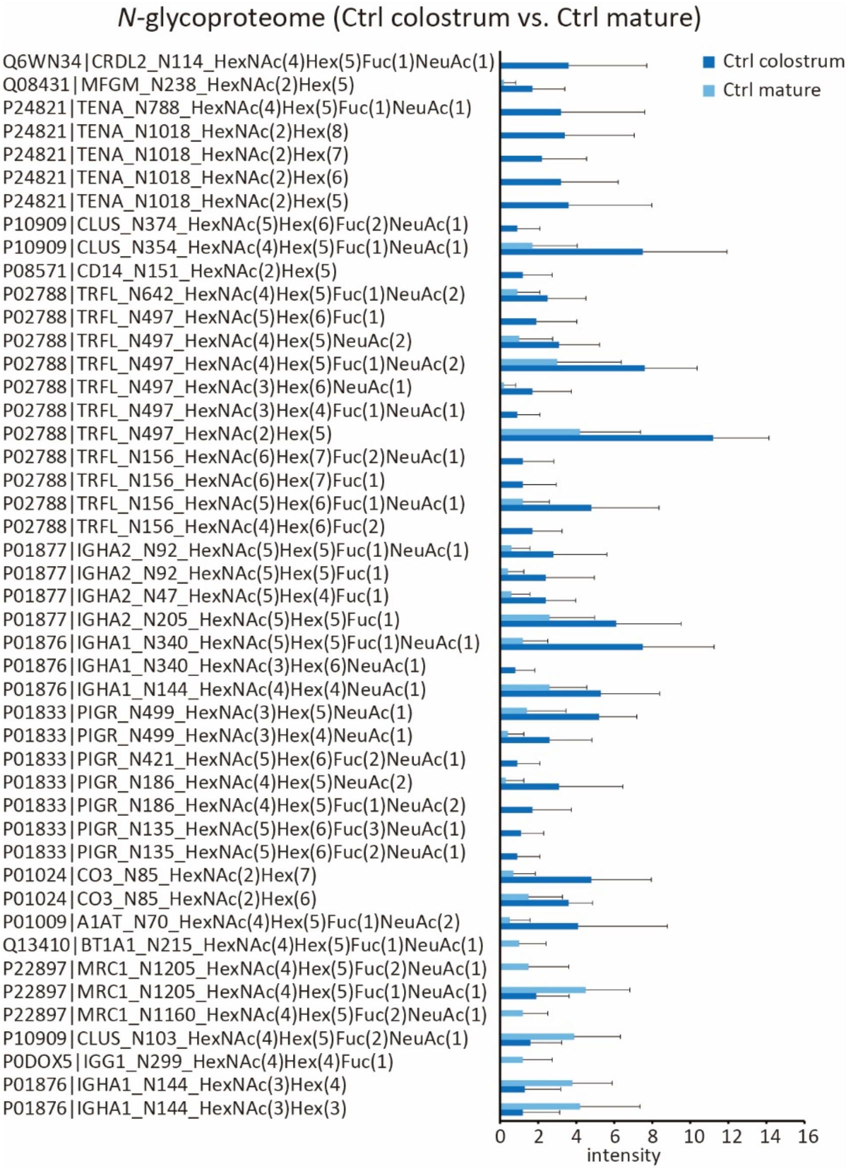


**Figure S5.** Changes in site-specific *N*-glycosylation during maturation of breastmilk from healthy women (Ctrl groups). Bars indicate means and whiskers indicate SDs.

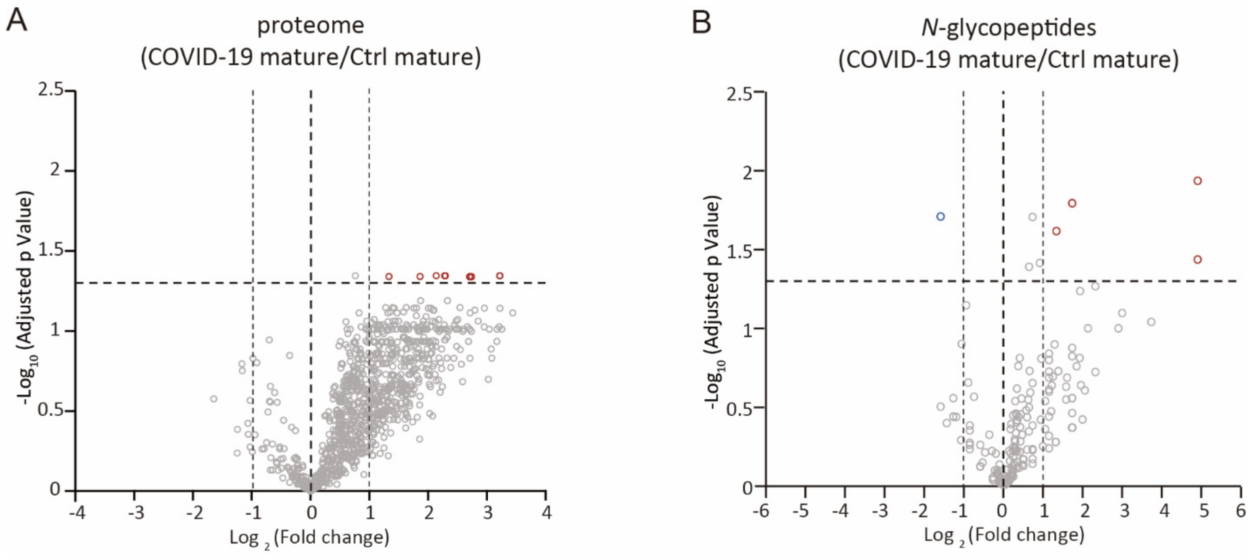


**Figure S6.** Comparisons of the proteome (**A**) and glycoproteome (**B**) profiles between mature milk samples from COVID-19 patients (COVID-19 mature) and healthy women (Ctrl mature). The red dots in the volcano plots indicate up-regulated (**A**) proteins and (**B**) *N*-glycopeptides in COVID-19 milk and the blue dots indicate the down-regulated ones.

**SUPPLEMENTARY FIGURE CAPTIONS**

**Table S1.** List of proteins identified in all sample groups

**Table S2.** Enriched gene ontology (GO) terms associated with the differentially expressed proteins in COVID-19 colostrum and Ctrl colostrum sample groups

**Table S3.** Enriched GO terms associated with the differentially expressed proteins in COVID-19 colostrum and COVID-19 mature sample groups

**Table S4.** Peptide to spectrum matches (PSMs) of identified glycopeptides

**Table S5.** List of unique *N*-glycopeptides

Table S1. List of proteins identified in all sample groups

| Accession | Description | Exp. q-value: Combined | Sum PEP Score | Coverage [%] | # Peptides | # PSMs | # Unique Peptides | # AAs | MW [kDa] | calc. pI | Score Sequest HT: Sequest HT | # Peptides (by Search Engine): Sequest HT | Abundances (Normalized): Normal-Colostrum-1 |
|---|---|---|---|---|---|---|---|---|---|---|---|---|---|
| P49327 | Fatty acid synthase OS=Homo sapiens OX=9606 GN=FASN PE=1 SV=3 | 0 | 2163.927 | 80 | 175 | 4025 | 175 | 2511 | 273.3 | 6.44 | 15093.74 | 175 | 226443261.9 |
| P47989 | Xanthine dehydrogenase/oxidase OS=Homo sapiens OX=9606 GN=XDH PE=1 SV=4 | 0 | 2121.419 | 89 | 131 | 4004 | 130 | 1333 | 146.3 | 7.66 | 16760.81 | 131 | 187185032.7 |
| P24821 | Tenascin OS=Homo sapiens OX=9606 GN=TNC PE=1 SV=3 | 0 | 2098.384 | 76 | 167 | 3646 | 167 | 2201 | 240.7 | 4.89 | 13788.03 | 167 | 448033597.6 |
| P01024 | Complement C3 OS=Homo sapiens OX=9606 GN=C3 PE=1 SV=2 | 0 | 1982.278 | 89 | 184 | 4300 | 184 | 1663 | 187 | 6.4 | 14884.82 | 184 | 81127813.08 |
| P02768 | Albumin OS=Homo sapiens OX=9606 GN=ALB PE=1 SV=2 | 0 | 1789.937 | 90 | 106 | 7241 | 106 | 609 | 69.3 | 6.28 | 24284.37 | 106 | 1287407506 |
| P0C0L5 | Complement C4-B OS=Homo sapiens OX=9606 GN=C4B PE=1 SV=2 | 0 | 1674.28 | 77 | 131 | 3200 | 8 | 1744 | 192.6 | 7.27 | 13113.81 | 131 | 19116449.22 |
| P0C0L4 | Complement C4-A OS=Homo sapiens OX=9606 GN=C4A PE=1 SV=2 | 0 | 1619.104 | 76 | 129 | 3176 | 6 | 1744 | 192.7 | 7.08 | 12856.47 | 129 | 7156730.213 |
| P02788 | Lactotransferrin OS=Homo sapiens OX=9606 GN=LTF PE=1 SV=6 | 0 | 1617.425 | 94 | 123 | 16698 | 123 | 710 | 78.1 | 8.12 | 44799.52 | 123 | 11946022845 |
| P22897 | Macrophage mannose receptor 1 OS=Homo sapiens OX=9606 GN=MRC1 PE=1 SV=1 | 0 | 1288.084 | 63 | 123 | 2885 | 123 | 1456 | 165.9 | 6.54 | 9680.37 | 123 | 33650686.51 |
| P04114 | Apolipoprotein B-100 OS=Homo sapiens OX=9606 GN=APOB PE=1 SV=2 | 0 | 1205.868 | 53 | 190 | 2168 | 190 | 4563 | 515.3 | 7.05 | 5960.28 | 190 | 79286736.56 |
| P01833 | Polymeric immunoglobulin receptor OS=Homo sapiens OX=9606 GN=PIGR PE=1 SV=4 | 0 | 946.876 | 70 | 75 | 4562 | 75 | 764 | 83.2 | 5.74 | 15097.45 | 75 | 3072265173 |
| P02787 | Serotransferrin OS=Homo sapiens OX=9606 GN=TF PE=1 SV=3 | 0 | 870.94 | 80 | 70 | 1620 | 70 | 698 | 77 | 7.12 | 6230.04 | 70 | 66837446.57 |
| P19835 | Bile salt-activated lipase OS=Homo sapiens OX=9606 GN=CEL PE=1 SV=3 | 0 | 724.092 | 58 | 55 | 3218 | 55 | 753 | 79.3 | 5.34 | 10670.03 | 55 | 909001084.9 |
| P35579 | Myosin-9 OS=Homo sapiens OX=9606 GN=MYH9 PE=1 SV=4 | 0 | 697.067 | 56 | 114 | 560 | 105 | 1960 | 226.4 | 5.6 | 1434.78 | 114 | 1890932180 |
| P01009 | Alpha-1-antitrypsin OS=Homo sapiens OX=9606 GN=SERPINA1 PE=1 SV=3 | 0 | 616.334 | 69 | 45 | 1339 | 45 | 418 | 46.7 | 5.59 | 4330.32 | 45 | 97783311.99 |
| Q99541 | Perilipin-2 OS=Homo sapiens OX=9606 GN=PLIN2 PE=1 SV=2 | 0 | 598.801 | 76 | 42 | 1111 | 42 | 437 | 48 | 6.8 | 4149.91 | 42 | 81455181.1 |
| P60709 | Actin, cytoplasmic 1 OS=Homo sapiens OX=9606 GN=ACTB PE=1 SV=1 | 0 | 581.148 | 95 | 40 | 1209 | 1 | 375 | 41.7 | 5.48 | 4117.08 | 40 | 41306163.91 |
| P63261 | Actin, cytoplasmic 2 OS=Homo sapiens OX=9606 GN=ACTG1 PE=1 SV=1 | 0 | 572.474 | 95 | 40 | 1185 | 1 | 375 | 41.8 | 5.48 | 3992.22 | 40 | |
| P01011 | Alpha-1-antichymotrypsin OS=Homo sapiens OX=9606 GN=SERPINA3 PE=1 SV=2 | 0 | 570.318 | 71 | 42 | 1374 | 42 | 423 | 47.6 | 5.52 | 5252.87 | 42 | 396264079.5 |
| Q13410 | Butyrophilin subfamily 1 member A1 OS=Homo sapiens OX=9606 GN=BTN1A1 PE=1 SV=3 | 0 | 540.4 | 60 | 44 | 1398 | 43 | 526 | 58.9 | 5.54 | 4643.85 | 44 | 254999227.8 |
| O60664 | Perilipin-3 OS=Homo sapiens OX=9606 GN=PLIN3 PE=1 SV=3 | 0 | 500.16 | 91 | 34 | 715 | 34 | 434 | 47 | 5.44 | 2967.84 | 34 | 16292801.87 |
| P00450 | Ceruloplasmin OS=Homo sapiens OX=9606 GN=CP PE=1 SV=1 | 0 | 498.335 | 71 | 54 | 819 | 54 | 1065 | 122.1 | 5.72 | 2902.46 | 54 | 1486140.818 |
| P01876 | Immunoglobulin heavy constant alpha 1 OS=Homo sapiens OX=9606 GN=IGHA1 PE=1 SV=2 | 0 | 494.394 | 98 | 34 | 3033 | 21 | 353 | 37.6 | 6.51 | 11072.29 | 34 | 3531401315 |
| P01871 | Immunoglobulin heavy constant mu OS=Homo sapiens OX=9606 GN=IGHM PE=1 SV=4 | 0 | 463.944 | 70 | 39 | 895 | 10 | 453 | 49.4 | 6.77 | 3300.69 | 39 | 106312209.4 |
| Q08431 | Lactadherin OS=Homo sapiens OX=9606 GN=MFGE8 PE=1 SV=3 | 0 | 453.923 | 86 | 36 | 1307 | 36 | 387 | 43.1 | 8.15 | 4714.2 | 36 | 488928114.7 |
| P07996 | Thrombospondin-1 OS=Homo sapiens OX=9606 GN=THBS1 PE=1 SV=2 | 0 | 444.461 | 54 | 61 | 705 | 61 | 1170 | 129.3 | 4.94 | 2141.98 | 61 | 13602275.22 |
| P02774 | Vitamin D-binding protein OS=Homo sapiens OX=9606 GN=GC PE=1 SV=2 | 0 | 439.631 | 78 | 43 | 932 | 43 | 474 | 52.9 | 5.45 | 3684.57 | 43 | 23083092.05 |
| P10909 | Clusterin OS=Homo sapiens OX=9606 GN=CLU PE=1 SV=1 | 0 | 438.115 | 56 | 35 | 1286 | 35 | 449 | 52.5 | 6.27 | 3973.52 | 35 | 855342794.8 |
| Q16851 | UTP--glucose-1-phosphate uridylyltransferase OS=Homo sapiens OX=9606 GN=UGP2 PE=1 SV=5 | 0 | 436.013 | 84 | 52 | 891 | 52 | 508 | 56.9 | 8.15 | 3037.94 | 52 | 4585790.318 |
| P07602 | Prosaposin OS=Homo sapiens OX=9606 GN=PSAP PE=1 SV=2 | 0 | 430.915 | 81 | 39 | 894 | 39 | 524 | 58.1 | 5.17 | 2968.53 | 39 | 57813140.16 |
| P01023 | Alpha-2-macroglobulin OS=Homo sapiens OX=9606 GN=A2M PE=1 SV=3 | 0 | 418.643 | 60 | 63 | 760 | 56 | 1474 | 163.2 | 6.46 | 2122.74 | 63 | 2236921.573 |
| P01877 | Immunoglobulin heavy constant alpha 2 OS=Homo sapiens OX=9606 GN=IGHA2 PE=1 SV=4 | 0 | 395.285 | 85 | 30 | 2163 | 4 | 340 | 36.6 | 6.27 | 7558.47 | 30 | 274256674.9 |
| Q6WN34 | Chordin-like protein 2 OS=Homo sapiens OX=9606 GN=CHRDL2 PE=1 SV=1 | 0 | 390.975 | 70 | 28 | 755 | 28 | 429 | 47.5 | 7.88 | 2924.74 | 28 | 202665493.3 |
| O00391 | Sulfhydryl oxidase 1 OS=Homo sapiens OX=9606 GN=QSOX1 PE=1 SV=3 | 0 | 389.242 | 58 | 40 | 949 | 40 | 747 | 82.5 | 8.92 | 3288.51 | 40 | 45077017.95 |
| P06733 | Alpha-enolase OS=Homo sapiens OX=9606 GN=ENO1 PE=1 SV=2 | 0 | 387.499 | 79 | 38 | 698 | 38 | 434 | 47.1 | 7.39 | 2650.24 | 38 | 14387230.28 |
| P02647 | Apolipoprotein A-I OS=Homo sapiens OX=9606 GN=APOA1 PE=1 SV=1 | 0 | 385.587 | 79 | 36 | 865 | 36 | 267 | 30.8 | 5.76 | 2730.15 | 36 | 57522339.48 |
| P00751 | Complement factor B OS=Homo sapiens OX=9606 GN=CFB PE=1 SV=2 | 0 | 381.117 | 62 | 46 | 684 | 46 | 764 | 85.5 | 7.06 | 2088.58 | 46 | 20332343.17 |
| Q9HC84 | Mucin-5B OS=Homo sapiens OX=9606 GN=MUC5B PE=1 SV=3 | 0 | 378.433 | 19 | 65 | 191 | 65 | 5762 | 596 | 6.64 | 584.43 | 65 | 7282477.882 |
| P07900 | Heat shock protein HSP 90-alpha OS=Homo sapiens OX=9606 GN=HSP90AA1 PE=1 SV=5 | 0 | 378.044 | 56 | 45 | 658 | 28 | 732 | 84.6 | 5.02 | 1917.99 | 45 | 16782001.62 |
| P0DOX2 | Immunoglobulin alpha-2 heavy chain OS=Homo sapiens OX=9606 PE=1 SV=2 | 0 | 372.428 | 83 | 36 | 2029 | 9 | 455 | 48.9 | 6.67 | 6693.65 | 36 | 12460979.61 |
| P04003 | C4b-binding protein alpha chain OS=Homo sapiens OX=9606 GN=C4BPA PE=1 SV=2 | 0 | 356.838 | 70 | 37 | 624 | 37 | 597 | 67 | 7.3 | 2143.67 | 37 | 11380069.54 |
| P02751 | Fibronectin OS=Homo sapiens OX=9606 GN=FN1 PE=1 SV=5 | 0 | 352.395 | 34 | 58 | 672 | 58 | 2477 | 272.2 | 5.5 | 1718.01 | 58 | 179389522.3 |
| P13639 | Elongation factor 2 OS=Homo sapiens OX=9606 GN=EEF2 PE=1 SV=4 | 0 | 352.271 | 70 | 51 | 768 | 51 | 858 | 95.3 | 6.83 | 2185.49 | 51 | 1446907.677 |
| P07237 | Protein disulfide-isomerase OS=Homo sapiens OX=9606 GN=P4HB PE=1 SV=3 | 0 | 347.383 | 79 | 41 | 716 | 41 | 508 | 57.1 | 4.87 | 1951.72 | 41 | 5240754.96 |
| Q96KP4 | Cytosolic non-specific dipeptidase OS=Homo sapiens OX=9606 GN=CNDP2 PE=1 SV=2 | 0 | 346.957 | 84 | 38 | 529 | 38 | 475 | 52.8 | 5.97 | 1796.29 | 38 | 1527402.268 |
| P06396 | Gelsolin OS=Homo sapiens OX=9606 GN=GSN PE=1 SV=1 | 0 | 346.667 | 60 | 35 | 659 | 35 | 782 | 85.6 | 6.28 | 2113.17 | 35 | 2695916.412 |
| P14625 | Endoplasmin OS=Homo sapiens OX=9606 GN=HSP90B1 PE=1 SV=1 | 0 | 343.7 | 59 | 50 | 623 | 41 | 803 | 92.4 | 4.84 | 1523.53 | 50 | 6861886.385 |
| P00709 | Alpha-lactalbumin OS=Homo sapiens OX=9606 GN=LALBA PE=1 SV=1 | 0 | 339.593 | 54 | 20 | 8705 | 20 | 142 | 16.2 | 5 | 20687.09 | 20 | 20046816627 |
| Q99102 | Mucin-4 OS=Homo sapiens OX=9606 GN=MUC4 PE=1 SV=4 | 0 | 339.275 | 26 | 34 | 734 | 34 | 2169 | 231.4 | 6.27 | 2818.99 | 34 | 780083728.8 |
| P08571 | Monocyte differentiation antigen CD14 OS=Homo sapiens OX=9606 GN=CD14 PE=1 SV=2 | 0 | 338.663 | 69 | 22 | 643 | 22 | 375 | 40.1 | 6.23 | 2714 | 22 | 111929031.8 |
| Q08380 | Galectin-3-binding protein OS=Homo sapiens OX=9606 GN=LGALS3BP PE=1 SV=1 | 0 | 337.253 | 53 | 29 | 661 | 29 | 585 | 65.3 | 5.27 | 2647.06 | 29 | 83820288.92 |
| P11021 | Endoplasmic reticulum chaperone BiP OS=Homo sapiens OX=9606 GN=HSPA5 PE=1 SV=2 | 0 | 334.807 | 67 | 44 | 716 | 42 | 654 | 72.3 | 5.16 | 1946.33 | 44 | 19797470.91 |
| Q14697 | Neutral alpha-glucosidase AB OS=Homo sapiens OX=9606 GN=GANAB PE=1 SV=3 | 0 | 329.329 | 73 | 45 | 624 | 45 | 944 | 106.8 | 6.14 | 1943.88 | 45 | 7392670.043 |
| P22079 | Lactoperoxidase OS=Homo sapiens OX=9606 GN=LPO PE=1 SV=2 | 0 | 324.949 | 62 | 41 | 518 | 41 | 712 | 80.2 | 8.62 | 1523.9 | 41 | |
| P04406 | Glyceraldehyde-3-phosphate dehydrogenase OS=Homo sapiens OX=9606 GN=GAPDH PE=1 SV= | 0 | 319.314 | 81 | 27 | 556 | 26 | 335 | 36 | 8.46 | 2344.34 | 27 | 31427076.75 |
| P80303 | Nucleobindin-2 OS=Homo sapiens OX=9606 GN=NUCB2 PE=1 SV=3 | 0 | 316.566 | 63 | 33 | 520 | 31 | 420 | 50.2 | 5.12 | 1730.68 | 33 | 14515930.41 |
| Q02818 | Nucleobindin-1 OS=Homo sapiens OX=9606 GN=NUCB1 PE=1 SV=4 | 0 | 310.241 | 70 | 36 | 580 | 34 | 461 | 53.8 | 5.25 | 1743.14 | 36 | 44012718 |
| P50395 | Rab GDP dissociation inhibitor beta OS=Homo sapiens OX=9606 GN=GDI2 PE=1 SV=2 | 0 | 306.58 | 85 | 37 | 513 | 29 | 445 | 50.6 | 6.47 | 1713.25 | 37 | 2316310.672 |
| P02675 | Fibrinogen beta chain OS=Homo sapiens OX=9606 GN=FGB PE=1 SV=2 | 0 | 306.313 | 76 | 40 | 581 | 40 | 491 | 55.9 | 8.27 | 1812.37 | 40 | 6872547.771 |
| P29401 | Transketolase OS=Homo sapiens OX=9606 GN=TKT PE=1 SV=3 | 0 | 303.728 | 83 | 39 | 476 | 39 | 623 | 67.8 | 7.66 | 1363.25 | 39 | 3196078.107 |
| P0DOX5 | Immunoglobulin gamma-1 heavy chain OS=Homo sapiens OX=9606 PE=1 SV=2 | 0 | 301.149 | 55 | 20 | 686 | 11 | 449 | 49.3 | 8.72 | 2383.11 | 20 | 60756238.58 |
| P05164 | Myeloperoxidase OS=Homo sapiens OX=9606 GN=MPO PE=1 SV=1 | 0 | 295.861 | 55 | 43 | 209 | 38 | 745 | 83.8 | 8.97 | 642.22 | 43 | 267719186.9 |
| P00738 | Haptoglobin OS=Homo sapiens OX=9606 GN=HP PE=1 SV=1 | 0 | 292.745 | 68 | 29 | 713 | 29 | 406 | 45.2 | 6.58 | 2250.5 | 29 | 24179325.37 |
| P0DOX6 | Immunoglobulin mu heavy chain OS=Homo sapiens OX=9606 PE=1 SV=2 | 0 | 292.412 | 49 | 32 | 724 | 2 | 576 | 63.4 | 7.87 | 2351.25 | 32 | 590348.7646 |
| P47710 | Alpha-S1-casein OS=Homo sapiens OX=9606 GN=CSN1S1 PE=1 SV=1 | 0 | 291.134 | 55 | 18 | 1353 | 18 | 185 | 21.7 | 5.38 | 5159.68 | 18 | 5371362580 |
| P04075 | Fructose-bisphosphate aldolase A OS=Homo sapiens OX=9606 GN=ALDOA PE=1 SV=2 | 0 | 291.069 | 89 | 31 | 547 | 29 | 364 | 39.4 | 8.09 | 1707.84 | 31 | 6982554.713 |
| P08238 | Heat shock protein HSP 90-beta OS=Homo sapiens OX=9606 GN=HSP90AB1 PE=1 SV=4 | 0 | 290.161 | 54 | 40 | 608 | 16 | 724 | 83.2 | 5.03 | 1661.93 | 40 | 2329108.362 |
| P13796 | Plastin-2 OS=Homo sapiens OX=9606 GN=LCP1 PE=1 SV=6 | 0 | 287.069 | 85 | 45 | 200 | 45 | 627 | 70.2 | 5.43 | 672.45 | 45 | 13660259.82 |
| P53396 | ATP-citrate synthase OS=Homo sapiens OX=9606 GN=ACLY PE=1 SV=3 | 0 | 282.351 | 50 | 44 | 288 | 44 | 1101 | 120.8 | 7.33 | 716.16 | 44 | 4163064.395 |
| P12814 | Alpha-actinin-1 OS=Homo sapiens OX=9606 GN=ACTN1 PE=1 SV=2 | 0 | 282.323 | 63 | 44 | 412 | 27 | 892 | 103 | 5.41 | 1133.14 | 44 | 1358471.26 |
| P11142 | Heat shock cognate 71 kDa protein OS=Homo sapiens OX=9606 GN=HSPA8 PE=1 SV=1 | 0 | 282.265 | 65 | 39 | 609 | 34 | 646 | 70.9 | 5.52 | 1884.2 | 39 | 48345477.63 |
| P14618 | Pyruvate kinase PKM OS=Homo sapiens OX=9606 GN=PKM PE=1 SV=4 | 0 | 282.255 | 80 | 35 | 311 | 35 | 531 | 57.9 | 7.84 | 958.41 | 35 | 5684276.718 |
| P68032 | Actin, alpha cardiac muscle 1 OS=Homo sapiens OX=9606 GN=ACTC1 PE=1 SV=1 | 0 | 280.459 | 49 | 26 | 788 | 5 | 377 | 42 | 5.39 | 2020.41 | 26 | 28556608.38 |
| Q6UX06 | Olfactomedin-4 OS=Homo sapiens OX=9606 GN=OLFM4 PE=1 SV=1 | 0 | 279.816 | 62 | 35 | 284 | 35 | 510 | 57.2 | 5.69 | 1026.53 | 35 | 38336012.04 |
| P15144 | Aminopeptidase N OS=Homo sapiens OX=9606 GN=ANPEP PE=1 SV=4 | 0 | 276.534 | 54 | 50 | 271 | 50 | 967 | 109.5 | 5.48 | 743.79 | 50 | 2224850.066 |
| P01834 | Immunoglobulin kappa constant OS=Homo sapiens OX=9606 GN=IGKC PE=1 SV=2 | 0 | 275.251 | 93 | 17 | 1067 | 5 | 107 | 11.8 | 6.52 | 3761.63 | 17 | 3677536313 |
| O60488 | Long-chain-fatty-acid--CoA ligase 4 OS=Homo sapiens OX=9606 GN=ACSL4 PE=1 SV=2 | 0 | 272.871 | 72 | 41 | 469 | 38 | 711 | 79.1 | 8.38 | 1333.41 | 41 | 3836891.665 |
| O15232 | Matrilin-3 OS=Homo sapiens OX=9606 GN=MATN3 PE=1 SV=2 | 0 | 266.359 | 77 | 28 | 389 | 28 | 486 | 52.8 | 6.7 | 1228.16 | 28 | 17872472.08 |
| P00747 | Plasminogen OS=Homo sapiens OX=9606 GN=PLG PE=1 SV=2 | 0 | 265.484 | 56 | 39 | 466 | 39 | 810 | 90.5 | 7.24 | 1202.93 | 39 | 11696174.09 |
| P06858 | Lipoprotein lipase OS=Homo sapiens OX=9606 GN=LPL PE=1 SV=1 | 0 | 263.807 | 55 | 25 | 625 | 25 | 475 | 53.1 | 8.15 | 2395.34 | 25 | 100959846.6 |
| P26038 | Moesin OS=Homo sapiens OX=9606 GN=MSN PE=1 SV=3 | 0 | 260.07 | 68 | 46 | 452 | 30 | 577 | 67.8 | 6.4 | 1051.66 | 46 | 402020.4763 |
| P00558 | Phosphoglycerate kinase 1 OS=Homo sapiens OX=9606 GN=PGK1 PE=1 SV=3 | 0 | 259.783 | 88 | 34 | 398 | 34 | 417 | 44.6 | 8.1 | 1170.75 | 34 | 4230968.976 |
| P0DOX7 | Immunoglobulin kappa light chain OS=Homo sapiens OX=9606 PE=1 SV=1 | 0 | 259.207 | 63 | 17 | 962 | 4 | 214 | 23.4 | 7.17 | 3607.19 | 17 | 59895937.02 |
| P25311 | Zinc-alpha-2-glycoprotein OS=Homo sapiens OX=9606 GN=AZGP1 PE=1 SV=2 | 0 | 257.835 | 65 | 29 | 586 | 29 | 298 | 34.2 | 6.05 | 1874.14 | 29 | 41540033.88 |
| P15311 | Ezrin OS=Homo sapiens OX=9606 GN=EZR PE=1 SV=4 | 0 | 256.725 | 63 | 42 | 669 | 27 | 586 | 69.4 | 6.27 | 1579.54 | 42 | 3568884.473 |
| Q14195 | Dihydropyrimidinase-related protein 3 OS=Homo sapiens OX=9606 GN=DPYSL3 PE=1 SV=1 | 0 | 253.038 | 82 | 30 | 332 | 26 | 570 | 61.9 | 6.49 | 949.47 | 30 | 1982122.838 |
| P68104 | Elongation factor 1-alpha 1 OS=Homo sapiens OX=9606 GN=EEF1A1 PE=1 SV=1 | 0 | 251.176 | 67 | 27 | 496 | 27 | 462 | 50.1 | 9.01 | 1941.73 | 27 | 15361091.31 |
| P06727 | Apolipoprotein A-IV OS=Homo sapiens OX=9606 GN=APOA4 PE=1 SV=4 | 0 | 247.82 | 72 | 35 | 556 | 35 | 396 | 45.3 | 5.38 | 1366.24 | 35 | 3165164.396 |
| O43707 | Alpha-actinin-4 OS=Homo sapiens OX=9606 GN=ACTN4 PE=1 SV=2 | 0 | 247.765 | 55 | 39 | 431 | 22 | 911 | 104.8 | 5.44 | 1072.82 | 39 | 437320.5499 |
| P23280 | Carbonic anhydrase 6 OS=Homo sapiens OX=9606 GN=CA6 PE=1 SV=3 | 0 | 246.827 | 67 | 15 | 492 | 15 | 308 | 35.3 | 7.02 | 2276.56 | 15 | 176281156.7 |
| P36871 | Phosphoglucomutase-1 OS=Homo sapiens OX=9606 GN=PGM1 PE=1 SV=3 | 0 | 246.083 | 86 | 37 | 441 | 37 | 562 | 61.4 | 6.76 | 1191.1 | 37 | 3169770.157 |
| P21399 | Cytoplasmic aconitate hydratase OS=Homo sapiens OX=9606 GN=ACO1 PE=1 SV=3 | 0 | 243.618 | 58 | 38 | 341 | 38 | 889 | 98.3 | 6.68 | 754.47 | 38 | 4053448.347 |
| P01008 | Antithrombin-III OS=Homo sapiens OX=9606 GN=SERPINC1 PE=1 SV=1 | 0 | 241.316 | 60 | 26 | 376 | 26 | 464 | 52.6 | 6.71 | 1295.79 | 26 | 4636798.019 |
| P02649 | Apolipoprotein E OS=Homo sapiens OX=9606 GN=APOE PE=1 SV=1 | 0 | 240.763 | 77 | 28 | 611 | 28 | 317 | 36.1 | 5.73 | 1891.13 | 28 | 11123765.58 |
| P02679 | Fibrinogen gamma chain OS=Homo sapiens OX=9606 GN=FGG PE=1 SV=3 | 0 | 240.15 | 65 | 27 | 447 | 27 | 453 | 51.5 | 5.62 | 1441.99 | 27 | 8995843.199 |
| P48449 | Lanosterol synthase OS=Homo sapiens OX=9606 GN=LSS PE=1 SV=1 | 0 | 235.085 | 62 | 41 | 468 | 41 | 732 | 83.3 | 6.61 | 1083.09 | 41 | 1488142.932 |
| Q13228 | Methanethiol oxidase OS=Homo sapiens OX=9606 GN=SELENBP1 PE=1 SV=2 | 0 | 232.392 | 77 | 34 | 448 | 34 | 472 | 52.4 | 6.37 | 1314.67 | 34 | 69264959.57 |
| P02766 | Transthyretin OS=Homo sapiens OX=9606 GN=TTR PE=1 SV=1 | 0 | 231.494 | 73 | 13 | 359 | 13 | 147 | 15.9 | 5.76 | 1708.54 | 13 | 37571956.76 |
| Q00610 | Clathrin heavy chain 1 OS=Homo sapiens OX=9606 GN=CLTC PE=1 SV=5 | 0 | 229.801 | 35 | 40 | 195 | 40 | 1675 | 191.5 | 5.69 | 479.68 | 40 | 4183891.171 |
| P08133 | Annexin A6 OS=Homo sapiens OX=9606 GN=ANXA6 PE=1 SV=3 | 0 | 228.771 | 66 | 42 | 216 | 42 | 673 | 75.8 | 5.6 | 535.56 | 42 | 2128795.697 |
| P10451 | Osteopontin OS=Homo sapiens OX=9606 GN=SPP1 PE=1 SV=1 | 0 | 226.477 | 57 | 15 | 377 | 15 | 314 | 35.4 | 4.58 | 1492.72 | 15 | 73315967.18 |
| Q15833 | Syntaxin-binding protein 2 OS=Homo sapiens OX=9606 GN=STXBP2 PE=1 SV=2 | 0 | 226.265 | 70 | 32 | 356 | 32 | 593 | 66.4 | 6.55 | 953.1 | 32 | 4328875.671 |
| P02765 | Alpha-2-HS-glycoprotein OS=Homo sapiens OX=9606 GN=AHSG PE=1 SV=2 | 0 | 226.23 | 50 | 15 | 338 | 15 | 367 | 39.3 | 5.72 | 1500.51 | 15 | 28653949.1 |
| P01859 | Immunoglobulin heavy constant gamma 2 OS=Homo sapiens OX=9606 GN=IGHG2 PE=1 SV=2 | 0 | 222.565 | 66 | 17 | 574 | 9 | 326 | 35.9 | 7.59 | 1730.62 | 17 | 9933556.903 |
| Q16555 | Dihydropyrimidinase-related protein 2 OS=Homo sapiens OX=9606 GN=DPYSL2 PE=1 SV=1 | 0 | 222.245 | 69 | 30 | 252 | 26 | 572 | 62.3 | 6.38 | 698.12 | 30 | 18015741.2 |
| P15291 | Beta-1,4-galactosyltransferase 1 OS=Homo sapiens OX=9606 GN=B4GALT1 PE=1 SV=5 | 0 | 220.792 | 57 | 18 | 513 | 18 | 398 | 43.9 | 8.65 | 1875.94 | 18 | 123237216.2 |
| P05413 | Fatty acid-binding protein, heart OS=Homo sapiens OX=9606 GN=FABP3 PE=1 SV=4 | 0 | 219.231 | 95 | 19 | 645 | 19 | 133 | 14.8 | 6.8 | 1479.86 | 19 | 97483511.39 |
| P0DTE7 | Alpha-amylase 1B OS=Homo sapiens OX=9606 GN=AMY1B PE=1 SV=1 | 0 | 217.194 | 69 | 25 | 405 | 25 | 511 | 57.7 | 6.93 | 1374.48 | 25 | 13274369.32 |
| P02671 | Fibrinogen alpha chain OS=Homo sapiens OX=9606 GN=FGA PE=1 SV=2 | 0 | 214.703 | 39 | 31 | 509 | 31 | 866 | 94.9 | 6.01 | 1254.92 | 31 | 5628087.553 |
| P36222 | Chitinase-3-like protein 1 OS=Homo sapiens OX=9606 GN=CHI3L1 PE=1 SV=2 | 0 | 212.645 | 75 | 29 | 430 | 29 | 383 | 42.6 | 8.46 | 1391.05 | 29 | 27921177.1 |
| O95573 | Long-chain-fatty-acid--CoA ligase 3 OS=Homo sapiens OX=9606 GN=ACSL3 PE=1 SV=3 | 0 | 211.585 | 66 | 33 | 367 | 30 | 720 | 80.4 | 8.38 | 926.62 | 33 | 2926302.914 |
| P60174 | Triosephosphate isomerase OS=Homo sapiens OX=9606 GN=TPI1 PE=1 SV=4 | 0 | 210.907 | 91 | 22 | 400 | 22 | 249 | 26.7 | 6.9 | 1299.26 | 22 | 4274603.196 |
| P21333 | Filamin-A OS=Homo sapiens OX=9606 GN=FLNA PE=1 SV=4 | 0 | 208.906 | 26 | 47 | 70 | 45 | 2647 | 280.6 | 6.06 | 171.48 | 47 | 408073.3835 |
| P18065 | Insulin-like growth factor-binding protein 2 OS=Homo sapiens OX=9606 GN=IGFBP2 PE=1 SV=2 | 0 | 208.546 | 71 | 21 | 398 | 21 | 325 | 34.8 | 7.5 | 1392.03 | 21 | 8233179.221 |
| P02790 | Hemopexin OS=Homo sapiens OX=9606 GN=HPX PE=1 SV=2 | 0 | 206.99 | 65 | 26 | 486 | 26 | 462 | 51.6 | 7.02 | 1556.85 | 26 | 4234454.055 |
| Q01518 | Adenylyl cyclase-associated protein 1 OS=Homo sapiens OX=9606 GN=CAP1 PE=1 SV=5 | 0 | 206.592 | 69 | 34 | 159 | 34 | 475 | 51.9 | 8.06 | 417.52 | 34 | 3381576.812 |
| Q96S86 | Hyaluronan and proteoglycan link protein 3 OS=Homo sapiens OX=9606 GN=HAPLN3 PE=2 SV=1 | 0 | 206.251 | 81 | 28 | 524 | 28 | 360 | 40.9 | 6.52 | 1464.79 | 28 | 17310952.98 |
| P15328 | Folate receptor alpha OS=Homo sapiens OX=9606 GN=FOLR1 PE=1 SV=3 | 0 | 206.201 | 65 | 17 | 351 | 17 | 257 | 29.8 | 7.97 | 1337.02 | 17 | 6756540.83 |
| P27105 | Stomatin OS=Homo sapiens OX=9606 GN=STOM PE=1 SV=3 | 0 | 203.384 | 75 | 23 | 325 | 23 | 288 | 31.7 | 7.88 | 1094.84 | 23 | 15657228.01 |
| O75891 | Cytosolic 10-formyltetrahydrofolate dehydrogenase OS=Homo sapiens OX=9606 GN=ALDH1L1 PE=1 SV=2 | 0 | 203.105 | 54 | 35 | 157 | 35 | 902 | 98.8 | 5.94 | 391.75 | 35 | 18138757.88 |
| P0DOY3 | Immunoglobulin lambda constant 3 OS=Homo sapiens OX=9606 GN=IGLC3 PE=1 SV=1 | 0 | 201.103 | 96 | 16 | 670 | 10 | 106 | 11.3 | 7.24 | 2243.67 | 16 | 19528724.09 |
| P08670 | Vimentin OS=Homo sapiens OX=9606 GN=VIM PE=1 SV=4 | 0 | 200.134 | 61 | 38 | 152 | 37 | 466 | 53.6 | 5.12 | 458.55 | 38 | 173652305.6 |
| P46940 | Ras GTPase-activating-like protein IQGAP1 OS=Homo sapiens OX=9606 GN=IQGAP1 PE=1 SV=1 | 0 | 197.711 | 33 | 36 | 143 | 36 | 1657 | 189.1 | 6.48 | 361.19 | 36 | 14159502.6 |
| P30101 | Protein disulfide-isomerase A3 OS=Homo sapiens OX=9606 GN=PDIA3 PE=1 SV=4 | 0 | 196.767 | 55 | 28 | 431 | 28 | 505 | 56.7 | 6.35 | 1023.65 | 28 | 6548241.133 |
| P55072 | Transitional endoplasmic reticulum ATPase OS=Homo sapiens OX=9606 GN=VCP PE=1 SV=4 | 0 | 195.153 | 54 | 33 | 267 | 33 | 806 | 89.3 | 5.26 | 749.8 | 33 | 3450394.545 |
| O75874 | Isocitrate dehydrogenase [NADP] cytoplasmic OS=Homo sapiens OX=9606 GN=IDH1 PE=1 SV=2 | 0 | 193.454 | 74 | 28 | 307 | 26 | 414 | 46.6 | 7.01 | 763.04 | 28 | 36765527.42 |
| P30086 | Phosphatidylethanolamine-binding protein 1 OS=Homo sapiens OX=9606 GN=PEBP1 PE=1 SV= | 0 | 193.395 | 88 | 18 | 232 | 18 | 187 | 21 | 7.53 | 899.72 | 18 | 29264386.18 |
| P01133 | Pro-epidermal growth factor OS=Homo sapiens OX=9606 GN=EGF PE=1 SV=2 | 0 | 193.203 | 33 | 33 | 441 | 33 | 1207 | 133.9 | 5.85 | 975.46 | 33 | 111122093.5 |
| P08603 | Complement factor H OS=Homo sapiens OX=9606 GN=CFH PE=1 SV=4 | 0 | 192.821 | 40 | 37 | 354 | 35 | 1231 | 139 | 6.61 | 783.19 | 37 | 2867476.895 |
| O75083 | WD repeat-containing protein 1 OS=Homo sapiens OX=9606 GN=WDR1 PE=1 SV=4 | 0 | 192.435 | 62 | 25 | 300 | 25 | 606 | 66.2 | 6.65 | 791.4 | 25 | 1337472.405 |
| P27797 | Calreticulin OS=Homo sapiens OX=9606 GN=CALR PE=1 SV=1 | 0 | 192.366 | 72 | 23 | 415 | 23 | 417 | 48.1 | 4.44 | 1343.24 | 23 | 4790633.375 |
| Q9P2E9 | Ribosome-binding protein 1 OS=Homo sapiens OX=9606 GN=RRBP1 PE=1 SV=5 | 0 | 190.451 | 27 | 33 | 220 | 33 | 1410 | 152.4 | 8.6 | 460.96 | 33 | 66360002.91 |
| P63104 | 14-3-3 protein zeta/delta OS=Homo sapiens OX=9606 GN=YWHAZ PE=1 SV=1 | 0 | 190.15 | 75 | 22 | 373 | 17 | 245 | 27.7 | 4.79 | 1111.97 | 22 | 10651991.39 |
| Q8WUM4 | Programmed cell death 6-interacting protein OS=Homo sapiens OX=9606 GN=PDCD6IP PE=1 SV=1 | 0 | 188.929 | 51 | 38 | 309 | 38 | 868 | 96 | 6.52 | 659.1 | 38 | 22018013.94 |
| P43490 | Nicotinamide phosphoribosyltransferase OS=Homo sapiens OX=9606 GN=NAMPT PE=1 SV=1 | 0 | 188.724 | 59 | 26 | 397 | 26 | 491 | 55.5 | 7.15 | 1092.6 | 26 | 1016013.926 |
| O60603 | Toll-like receptor 2 OS=Homo sapiens OX=9606 GN=TLR2 PE=1 SV=1 | 0 | 187.386 | 50 | 33 | 352 | 33 | 784 | 89.8 | 6.61 | 888.7 | 33 | 880634.4105 |
| Q14624 | Inter-alpha-trypsin inhibitor heavy chain H4 OS=Homo sapiens OX=9606 GN=ITIH4 PE=1 SV=4 | 0 | 184.336 | 45 | 29 | 432 | 29 | 930 | 103.3 | 6.98 | 1232.79 | 29 | 9302408.248 |
| P07498 | Kappa-casein OS=Homo sapiens OX=9606 GN=CSN3 PE=1 SV=3 | 0 | 180.357 | 68 | 11 | 1345 | 11 | 182 | 20.3 | 8.92 | 4034.07 | 11 | 3325734525 |
| P05814 | Beta-casein OS=Homo sapiens OX=9606 GN=CSN2 PE=1 SV=4 | 0 | 179.058 | 76 | 14 | 5243 | 14 | 226 | 25.4 | 5.78 | 15513.25 | 14 | 25610231676 |
| P02750 | Leucine-rich alpha-2-glycoprotein OS=Homo sapiens OX=9606 GN=LRG1 PE=1 SV=2 | 0 | 178.907 | 63 | 20 | 410 | 20 | 347 | 38.2 | 6.95 | 1535.6 | 20 | 104048851 |
| P16671 | Platelet glycoprotein 4 OS=Homo sapiens OX=9606 GN=CD36 PE=1 SV=2 | 0 | 178.238 | 44 | 20 | 419 | 20 | 472 | 53 | 7.96 | 1449.29 | 20 | 86860928.57 |
| P53634 | Dipeptidyl peptidase 1 OS=Homo sapiens OX=9606 GN=CTSC PE=1 SV=2 | 0 | 177.776 | 64 | 24 | 319 | 24 | 463 | 51.8 | 6.99 | 976.99 | 24 | 14805288.56 |
| P01042 | Kininogen-1 OS=Homo sapiens OX=9606 GN=KNG1 PE=1 SV=2 | 0 | 175.99 | 37 | 22 | 403 | 22 | 644 | 71.9 | 6.81 | 1252.28 | 22 | 3329217.208 |
| O00763 | Acetyl-CoA carboxylase 2 OS=Homo sapiens OX=9606 GN=ACACB PE=1 SV=3 | 0 | 175.167 | 22 | 37 | 105 | 37 | 2458 | 276.4 | 6.49 | 259.7 | 37 | 1404944.059 |
| P09543 | 2',3'-cyclic-nucleotide 3'-phosphodiesterase OS=Homo sapiens OX=9606 GN=CNP PE=1 SV=2 | 0 | 173.904 | 63 | 32 | 366 | 32 | 421 | 47.5 | 9.07 | 790.37 | 32 | 24837695.64 |
| P01034 | Cystatin-C OS=Homo sapiens OX=9606 GN=CST3 PE=1 SV=1 | 0 | 173.878 | 58 | 10 | 386 | 10 | 146 | 15.8 | 8.75 | 1233.91 | 10 | 45822524.29 |
| P07355 | Annexin A2 OS=Homo sapiens OX=9606 GN=ANXA2 PE=1 SV=2 | 0 | 173.535 | 77 | 28 | 376 | 28 | 339 | 38.6 | 7.75 | 1134.14 | 28 | 2518717.997 |
| P0DMV9 | Heat shock 70 kDa protein 1B OS=Homo sapiens OX=9606 GN=HSPA1B PE=1 SV=1 | 0 | 173.104 | 65 | 30 | 182 | 25 | 641 | 70 | 5.66 | 472.19 | 30 | 4645472.093 |
| P06702 | Protein S100-A9 OS=Homo sapiens OX=9606 GN=S100A9 PE=1 SV=1 | 0 | 172.989 | 91 | 15 | 275 | 15 | 114 | 13.2 | 6.13 | 949.75 | 15 | 195781361.1 |
| O00560 | Syntenin-1 OS=Homo sapiens OX=9606 GN=SDCBP PE=1 SV=1 | 0 | 171.471 | 77 | 17 | 356 | 17 | 298 | 32.4 | 7.53 | 1310 | 17 | 25072465.64 |
| P01860 | Immunoglobulin heavy constant gamma 3 OS=Homo sapiens OX=9606 GN=IGHG3 PE=1 SV=2 | 0 | 171.369 | 54 | 16 | 472 | 6 | 377 | 41.3 | 7.9 | 1165.22 | 16 | 1032420.916 |
| P04083 | Annexin A1 OS=Homo sapiens OX=9606 GN=ANXA1 PE=1 SV=2 | 0 | 170.085 | 73 | 25 | 211 | 25 | 346 | 38.7 | 7.02 | 594.03 | 25 | 5923582.714 |
| P0DOX8 | Immunoglobulin lambda-1 light chain OS=Homo sapiens OX=9606 PE=1 SV=1 | 0 | 169.182 | 57 | 19 | 719 | 12 | 216 | 22.8 | 6.76 | 2233.09 | 19 | 98085756.09 |
| P02749 | Beta-2-glycoprotein 1 OS=Homo sapiens OX=9606 GN=APOH PE=1 SV=3 | 0 | 166.617 | 65 | 18 | 360 | 18 | 345 | 38.3 | 7.97 | 1286.3 | 18 | 5995992.808 |
| P23284 | Peptidyl-prolyl cis-trans isomerase B OS=Homo sapiens OX=9606 GN=PPIB PE=1 SV=2 | 0 | 165.789 | 68 | 23 | 421 | 23 | 216 | 23.7 | 9.41 | 1057.55 | 23 | 13221415 |
| P02794 | Ferritin heavy chain OS=Homo sapiens OX=9606 GN=FTH1 PE=1 SV=2 | 0 | 165.612 | 77 | 15 | 300 | 15 | 183 | 21.2 | 5.55 | 826.36 | 15 | 5992475.406 |
| P52209 | 6-phosphogluconate dehydrogenase, decarboxylating OS=Homo sapiens OX=9606 GN=PGD PE=1 SV= | 0 | 164.03 | 73 | 26 | 213 | 26 | 483 | 53.1 | 7.23 | 483.24 | 26 | 5664273.979 |
| P33121 | Long-chain-fatty-acid--CoA ligase 1 OS=Homo sapiens OX=9606 GN=ACSL1 PE=1 SV=1 | 0 | 164.016 | 49 | 32 | 206 | 32 | 698 | 77.9 | 7.15 | 505.5 | 32 | 638155.2783 |
| P62879 | Guanine nucleotide-binding protein G(I)/G(S)/G(T) subunit beta-2 OS=Homo sapiens OX=9606 GN=GNB2 PE=1 SV= | 0 | 163.652 | 86 | 18 | 324 | 10 | 340 | 37.3 | 6 | 1067.03 | 18 | 3732573.837 |
| P07339 | Cathepsin D OS=Homo sapiens OX=9606 GN=CTSD PE=1 SV=1 | 0 | 163.184 | 61 | 22 | 351 | 22 | 412 | 44.5 | 6.54 | 949.87 | 22 | 7059893.796 |
| P01591 | Immunoglobulin J chain OS=Homo sapiens OX=9606 GN=JCHAIN PE=1 SV=4 | 0 | 160.341 | 79 | 20 | 651 | 20 | 159 | 18.1 | 5.24 | 2016.01 | 20 | 644685018.3 |
| P61626 | Lysozyme C OS=Homo sapiens OX=9606 GN=LYZ PE=1 SV=1 | 0 | 159.416 | 69 | 15 | 599 | 15 | 148 | 16.5 | 9.16 | 1722.09 | 15 | 275000072 |
| P23381 | Tryptophan--tRNA ligase, cytoplasmic OS=Homo sapiens OX=9606 GN=WARS1 PE=1 SV=2 | 0 | 158.85 | 55 | 18 | 154 | 18 | 471 | 53.1 | 6.23 | 429.99 | 18 | 1158561.162 |
| Q9Y4L1 | Hypoxia up-regulated protein 1 OS=Homo sapiens OX=9606 GN=HYOU1 PE=1 SV=1 | 0 | 158.848 | 37 | 28 | 209 | 28 | 999 | 111.3 | 5.22 | 422.01 | 28 | 3600857.327 |
| P62937 | Peptidyl-prolyl cis-trans isomerase A OS=Homo sapiens OX=9606 GN=PPIA PE=1 SV=2 | 0 | 157.663 | 86 | 15 | 388 | 15 | 165 | 18 | 7.81 | 1332.39 | 15 | 24752715.52 |
| P04217 | Alpha-1B-glycoprotein OS=Homo sapiens OX=9606 GN=A1BG PE=1 SV=4 | 0 | 156.763 | 60 | 18 | 387 | 18 | 495 | 54.2 | 5.86 | 1271.59 | 18 | 14245053.48 |
| Q6PCB0 | von Willebrand factor A domain-containing protein 1 OS=Homo sapiens OX=9606 GN=VWA1 PE=1 SV=1 | 0 | 156.083 | 59 | 15 | 248 | 15 | 445 | 46.8 | 7.68 | 1028.96 | 15 | 6743744.326 |
| P15941 | Mucin-1 OS=Homo sapiens OX=9606 GN=MUC1 PE=1 SV=3 | 0 | 155.984 | 7 | 11 | 325 | 11 | 1255 | 122 | 7.47 | 1148.02 | 11 | 72922512.94 |
| P23528 | Cofilin-1 OS=Homo sapiens OX=9606 GN=CFL1 PE=1 SV=3 | 0 | 155.791 | 86 | 18 | 270 | 16 | 166 | 18.5 | 8.09 | 1078.86 | 18 | 19591056.25 |
| P49189 | 4-trimethylaminobutyraldehyde dehydrogenase OS=Homo sapiens OX=9606 GN=ALDH9A1 PE=1 SV= | 0 | 155.279 | 57 | 24 | 273 | 24 | 494 | 53.8 | 5.87 | 766.22 | 24 | 19816511.41 |
| P61769 | Beta-2-microglobulin OS=Homo sapiens OX=9606 GN=B2M PE=1 SV=1 | 0 | 155.165 | 78 | 14 | 256 | 14 | 119 | 13.7 | 6.52 | 1250.43 | 14 | 206857642.1 |
| P98160 | Basement membrane-specific heparan sulfate proteoglycan core protein OS=Homo sapiens OX=9606 GN=HSPG2 PE=1 SV=4 | 0 | 154.909 | 12 | 37 | 76 | 37 | 4391 | 468.5 | 6.51 | 187 | 37 | 16355994.01 |
| P12273 | Prolactin-inducible protein OS=Homo sapiens OX=9606 GN=PIP PE=1 SV=1 | 0 | 154.693 | 77 | 16 | 352 | 16 | 146 | 16.6 | 8.05 | 1419.05 | 16 | 99671750.95 |
| Q02809 | Procollagen-lysine,2-oxoglutarate 5-dioxygenase 1 OS=Homo sapiens OX=9606 GN=PLOD1 PE=1 SV=2 | 0 | 154.682 | 54 | 31 | 365 | 31 | 727 | 83.5 | 6.95 | 781.23 | 31 | 1066229.308 |
| P19823 | Inter-alpha-trypsin inhibitor heavy chain H2 OS=Homo sapiens OX=9606 GN=ITIH2 PE=1 SV=2 | 0 | 154.29 | 41 | 26 | 256 | 26 | 946 | 106.4 | 6.86 | 605.92 | 26 | 11154288.07 |
| P05156 | Complement factor I OS=Homo sapiens OX=9606 GN=CFI PE=1 SV=2 | 0 | 153.945 | 41 | 21 | 392 | 21 | 583 | 65.7 | 7.5 | 1067.44 | 21 | 6241603.627 |
| P07195 | L-lactate dehydrogenase B chain OS=Homo sapiens OX=9606 GN=LDHB PE=1 SV=2 | 0 | 153.517 | 71 | 20 | 296 | 19 | 334 | 36.6 | 6.05 | 713.84 | 20 | 3417885.797 |
| P01861 | Immunoglobulin heavy constant gamma 4 OS=Homo sapiens OX=9606 GN=IGHG4 PE=1 SV=1 | 0 | 152.356 | 68 | 14 | 374 | 7 | 327 | 35.9 | 7.36 | 1101.55 | 14 | 18438782.2 |
| P14550 | Aldo-keto reductase family 1 member A1 OS=Homo sapiens OX=9606 GN=AKR1A1 PE=1 SV=3 | 0 | 151.855 | 82 | 22 | 277 | 22 | 325 | 36.6 | 6.79 | 769.44 | 22 | 1837351.458 |
| P62873 | Guanine nucleotide-binding protein G(I)/G(S)/G(T) subunit beta-1 OS=Homo sapiens OX=9606 GN=GNB1 PE=1 SV= | 0 | 148.877 | 72 | 17 | 285 | 10 | 340 | 37.4 | 6 | 1007.41 | 17 | 5342235.891 |
| O00300 | Tumor necrosis factor receptor superfamily member 11B OS=Homo sapiens OX=9606 GN=TNFRSF11B PE=1 SV=3 | 0 | 147.55 | 54 | 20 | 231 | 20 | 401 | 46 | 8.29 | 598.98 | 20 | 2569563.781 |
| P05155 | Plasma protease C1 inhibitor OS=Homo sapiens OX=9606 GN=SERPING1 PE=1 SV=2 | 0 | 146.979 | 44 | 22 | 392 | 22 | 500 | 55.1 | 6.55 | 1102.87 | 22 | 5832392.731 |
| P35241 | Radixin OS=Homo sapiens OX=9606 GN=RDX PE=1 SV=1 | 0 | 145.661 | 45 | 30 | 371 | 12 | 583 | 68.5 | 6.37 | 862.81 | 30 | |
| Q5IS80 | Amyloid-beta A4 protein OS=Pan troglodytes OX=9598 GN=APP PE=2 SV=1 | 0 | 145.569 | 32 | 24 | 329 | 23 | 770 | 86.9 | 4.82 | 783.92 | 24 | 6022839.845 |
| P61586 | Transforming protein RhoA OS=Homo sapiens OX=9606 GN=RHOA PE=1 SV=1 | 0 | 145.344 | 80 | 17 | 362 | 6 | 193 | 21.8 | 6.1 | 1218.01 | 17 | 1515518.656 |
| P00734 | Prothrombin OS=Homo sapiens OX=9606 GN=F2 PE=1 SV=2 | 0 | 145.185 | 49 | 22 | 229 | 22 | 622 | 70 | 5.9 | 574.84 | 22 | 2368080.806 |
| P62258 | 14-3-3 protein epsilon OS=Homo sapiens OX=9606 GN=YWHAE PE=1 SV=1 | 0 | 144.557 | 62 | 19 | 257 | 17 | 255 | 29.2 | 4.74 | 791.96 | 19 | 82482290.77 |
| Q9Y490 | Talin-1 OS=Homo sapiens OX=9606 GN=TLN1 PE=1 SV=3 | 0 | 142.283 | 18 | 29 | 41 | 29 | 2541 | 269.6 | 6.07 | 107.8 | 29 | 351810.3809 |
| Q9NV23 | S-acyl fatty acid synthase thioesterase, medium chain OS=Homo sapiens OX=9606 GN=OLAH PE=1 SV=1 | 0 | 141.918 | 65 | 16 | 149 | 16 | 265 | 29.9 | 6.23 | 495.79 | 16 | 1654375.311 |
| P31946 | 14-3-3 protein beta/alpha OS=Homo sapiens OX=9606 GN=YWHAB PE=1 SV=3 | 0 | 140.734 | 75 | 20 | 239 | 11 | 246 | 28.1 | 4.83 | 640.72 | 20 | 2337982.825 |
| Q92485 | Acid sphingomyelinase-like phosphodiesterase 3b OS=Homo sapiens OX=9606 GN=SMPDL3B PE=1 SV=2 | 0 | 139.693 | 59 | 16 | 212 | 16 | 455 | 50.8 | 5.64 | 594.78 | 16 | 30668970.76 |
| P04004 | Vitronectin OS=Homo sapiens OX=9606 GN=VTN PE=1 SV=1 | 0 | 139.232 | 44 | 18 | 313 | 18 | 478 | 54.3 | 5.8 | 942.75 | 18 | 20862022.72 |
| Q9BRK5 | 45 kDa calcium-binding protein OS=Homo sapiens OX=9606 GN=SDF4 PE=1 SV=1 | 0 | 138.451 | 56 | 14 | 228 | 14 | 362 | 41.8 | 4.86 | 726.57 | 14 | 8437539.81 |
| P08758 | Annexin A5 OS=Homo sapiens OX=9606 GN=ANXA5 PE=1 SV=2 | 0 | 137.201 | 76 | 21 | 246 | 21 | 320 | 35.9 | 5.05 | 693.49 | 21 | 2117966.007 |
| P50148 | Guanine nucleotide-binding protein G(q) subunit alpha OS=Homo sapiens OX=9606 GN=GNAQ PE=1 SV=4 | 0 | 137.046 | 70 | 22 | 220 | 13 | 359 | 42.1 | 5.68 | 603.82 | 22 | 1518054.68 |
| Q562R1 | Beta-actin-like protein 2 OS=Homo sapiens OX=9606 GN=ACTBL2 PE=1 SV=2 | 0 | 136.636 | 33 | 14 | 381 | 3 | 376 | 42 | 5.59 | 884.28 | 14 | |
| Q06210 | Glutamine--fructose-6-phosphate aminotransferase [isomerizing] 1 OS=Homo sapiens OX=9606 GN=GFPT1 PE=1 SV=3 | 0 | 136.448 | 41 | 21 | 135 | 21 | 699 | 78.8 | 7.11 | 388.17 | 21 | 4222684.156 |
| P12821 | Angiotensin-converting enzyme OS=Homo sapiens OX=9606 GN=ACE PE=1 SV=1 | 0 | 135.242 | 29 | 27 | 257 | 27 | 1306 | 149.6 | 6.39 | 637.18 | 27 | 992295.8133 |
| P12429 | Annexin A3 OS=Homo sapiens OX=9606 GN=ANXA3 PE=1 SV=3 | 0 | 133.982 | 74 | 22 | 80 | 22 | 323 | 36.4 | 5.92 | 228.8 | 22 | 2302137.752 |
| P08697 | Alpha-2-antiplasmin OS=Homo sapiens OX=9606 GN=SERPINF2 PE=1 SV=3 | 0 | 130.7 | 54 | 15 | 212 | 15 | 491 | 54.5 | 6.29 | 675.21 | 15 | 2607077.988 |
| P84077 | ADP-ribosylation factor 1 OS=Homo sapiens OX=9606 GN=ARF1 PE=1 SV=2 | 0 | 130.389 | 82 | 15 | 232 | 9 | 181 | 20.7 | 6.8 | 772.57 | 15 | 6049730.037 |
| Q9UNQ0 | Broad substrate specificity ATP-binding cassette transporter ABCG2 OS=Homo sapiens OX=9606 GN=ABCG2 PE=1 SV=3 | 0 | 129.806 | 38 | 19 | 229 | 19 | 655 | 72.3 | 8.69 | 645.84 | 19 | 3415520.451 |
| Q9NP72 | Ras-related protein Rab-18 OS=Homo sapiens OX=9606 GN=RAB18 PE=1 SV=1 | 0 | 128.209 | 76 | 17 | 349 | 17 | 206 | 23 | 5.24 | 1154.58 | 17 | 18252582.88 |
| Q99536 | Synaptic vesicle membrane protein VAT-1 homolog OS=Homo sapiens OX=9606 GN=VAT1 PE=1 SV=2 | 0 | 127.311 | 61 | 17 | 149 | 17 | 393 | 41.9 | 6.29 | 478.88 | 17 | 2384284.128 |
| P31146 | Coronin-1A OS=Homo sapiens OX=9606 GN=CORO1A PE=1 SV=4 | 0 | 127.012 | 57 | 20 | 74 | 19 | 461 | 51 | 6.68 | 250.04 | 20 | 39107930.01 |
| O43490 | Prominin-1 OS=Homo sapiens OX=9606 GN=PROM1 PE=1 SV=1 | 0 | 126.944 | 34 | 24 | 121 | 24 | 865 | 97.1 | 7.27 | 367.33 | 24 | 691546.2776 |
| P10643 | Complement component C7 OS=Homo sapiens OX=9606 GN=C7 PE=1 SV=2 | 0 | 126.697 | 44 | 25 | 139 | 25 | 843 | 93.5 | 6.48 | 292.88 | 25 | 18582587.99 |
| P69905 | Hemoglobin subunit alpha OS=Homo sapiens OX=9606 GN=HBA1 PE=1 SV=2 | 0 | 126.269 | 92 | 11 | 62 | 11 | 142 | 15.2 | 8.68 | 234.93 | 11 | |
| P20061 | Transcobalamin-1 OS=Homo sapiens OX=9606 GN=TCN1 PE=1 SV=2 | 0 | 125.872 | 39 | 14 | 213 | 14 | 433 | 48.2 | 5.03 | 741.93 | 14 | 3141857.072 |
| O00299 | Chloride intracellular channel protein 1 OS=Homo sapiens OX=9606 GN=CLIC1 PE=1 SV=4 | 0 | 125.135 | 85 | 17 | 300 | 17 | 241 | 26.9 | 5.17 | 805.64 | 17 | 3733258.096 |
| P05109 | Protein S100-A8 OS=Homo sapiens OX=9606 GN=S100A8 PE=1 SV=1 | 0 | 123.485 | 91 | 17 | 247 | 17 | 93 | 10.8 | 7.03 | 704.8 | 17 | 95308099.53 |
| P07437 | Tubulin beta chain OS=Homo sapiens OX=9606 GN=TUBB PE=1 SV=2 | 0 | 123.073 | 59 | 19 | 200 | 5 | 444 | 49.6 | 4.89 | 511.34 | 19 | 806625.7836 |
| P68363 | Tubulin alpha-1B chain OS=Homo sapiens OX=9606 GN=TUBA1B PE=1 SV=1 | 0 | 121.812 | 62 | 21 | 185 | 21 | 451 | 50.1 | 5.06 | 473.55 | 21 | 1849687.826 |
| P52565 | Rho GDP-dissociation inhibitor 1 OS=Homo sapiens OX=9606 GN=ARHGDIA PE=1 SV=3 | 0 | 121.112 | 83 | 12 | 150 | 12 | 204 | 23.2 | 5.11 | 516.53 | 12 | 3813341.575 |
| P09960 | Leukotriene A-4 hydrolase OS=Homo sapiens OX=9606 GN=LTA4H PE=1 SV=2 | 0 | 121.086 | 47 | 22 | 54 | 22 | 611 | 69.2 | 6.18 | 143.76 | 22 | 1575354.968 |
| Q92876 | Kallikrein-6 OS=Homo sapiens OX=9606 GN=KLK6 PE=1 SV=1 | 0 | 119.845 | 70 | 11 | 222 | 11 | 244 | 26.8 | 7.44 | 794.45 | 11 | 8549028.076 |
| P00338 | L-lactate dehydrogenase A chain OS=Homo sapiens OX=9606 GN=LDHA PE=1 SV=2 | 0 | 119.456 | 63 | 20 | 125 | 19 | 332 | 36.7 | 8.27 | 320.17 | 20 | 1786964.709 |

Table S1. Cont.

| Accession | Description | Exp. q-value: Combined | Sum PEP Score | Coverage [%] | # Peptides | # PSMs | # Unique Peptides | # AAs | MW [kDa] | calc. pI | Score Sequest HT: Sequest HT | # Peptides (by Search Engine): Sequest HT | Abundances (Normalized): Normal-Colostrum-1 |
|---|---|---|---|---|---|---|---|---|---|---|---|---|---|
| P06744 | Glucose-6-phosphate isomerase OS=Homo sapiens OX=9606 GN=GPI PE=1 SV=4 | 0 | 119.427 | 47 | 20 | 157 | 20 | 558 | 63.1 | 8.32 | 466.6 | 20 | 2044646.689 |
| Q06830 | Peroxiredoxin-1 OS=Homo sapiens OX=9606 GN=PRDX1 PE=1 SV=1 | 0 | 118.723 | 77 | 22 | 213 | 18 | 199 | 22.1 | 8.13 | 514.01 | 22 | 2564460.642 |
| P08134 | Rho-related GTP-binding protein RhoC OS=Homo sapiens OX=9606 GN=RHOC PE=1 SV=1 | 0 | 117.519 | 69 | 13 | 311 | 2 | 193 | 22 | 6.58 | 1039.51 | 13 | |
| P30041 | Peroxiredoxin-6 OS=Homo sapiens OX=9606 GN=PRDX6 PE=1 SV=3 | 0 | 117.483 | 80 | 18 | 172 | 18 | 224 | 25 | 6.38 | 469.59 | 18 | 233000.4187 |
| P04040 | Catalase OS=Homo sapiens OX=9606 GN=CAT PE=1 SV=3 | 0 | 116.84 | 60 | 21 | 67 | 21 | 527 | 59.7 | 7.39 | 171.2 | 21 | 769406.0209 |
| P04792 | Heat shock protein beta-1 OS=Homo sapiens OX=9606 GN=HSPB1 PE=1 SV=2 | 0 | 116.522 | 87 | 14 | 116 | 14 | 205 | 22.8 | 6.4 | 367.81 | 14 | |
| P07737 | Profilin-1 OS=Homo sapiens OX=9606 GN=PFN1 PE=1 SV=2 | 0 | 116.303 | 98 | 16 | 250 | 16 | 140 | 15 | 8.27 | 784.33 | 16 | 11431699.19 |
| P62820 | Ras-related protein Rab-1A OS=Homo sapiens OX=9606 GN=RAB1A PE=1 SV=3 | 0 | 115.355 | 81 | 16 | 213 | 7 | 205 | 22.7 | 6.21 | 637.44 | 16 | 1408571.721 |
| P31949 | Protein S100-A11 OS=Homo sapiens OX=9606 GN=S100A11 PE=1 SV=2 | 0 | 114.497 | 87 | 10 | 152 | 10 | 105 | 11.7 | 7.12 | 433.4 | 10 | 7577944.798 |
| O00468 | Agrin OS=Homo sapiens OX=9606 GN=AGRN PE=1 SV=6 | 0 | 114.456 | 18 | 24 | 69 | 24 | 2068 | 217.2 | 6.39 | 160.59 | 24 | 337893.4974 |
| P63244 | Receptor of activated protein C kinase 1 OS=Homo sapiens OX=9606 GN=RACK1 PE=1 SV=3 | 0 | 114.359 | 66 | 15 | 160 | 15 | 317 | 35.1 | 7.69 | 367.68 | 15 | 3055574.586 |
| P11908 | Ribose-phosphate pyrophosphokinase 2 OS=Homo sapiens OX=9606 GN=PRPS2 PE=1 SV=2 | 0 | 114.018 | 57 | 15 | 85 | 15 | 318 | 34.7 | 6.61 | 263.45 | 15 | |
| P02748 | Complement component C9 OS=Homo sapiens OX=9606 GN=C9 PE=1 SV=2 | 0 | 113.437 | 40 | 19 | 284 | 19 | 559 | 63.1 | 5.59 | 723.59 | 19 | 9721503.489 |
| P61158 | Actin-related protein 3 OS=Homo sapiens OX=9606 GN=ACTR3 PE=1 SV=3 | 0 | 113.303 | 66 | 20 | 106 | 20 | 418 | 47.3 | 5.88 | 233.33 | 20 | 1883760.043 |
| P68371 | Tubulin beta-4B chain OS=Homo sapiens OX=9606 GN=TUBB4B PE=1 SV=1 | 0 | 112.138 | 52 | 16 | 176 | 2 | 445 | 49.8 | 4.89 | 478.34 | 16 | |
| Q9H173 | Nucleotide exchange factor SIL1 OS=Homo sapiens OX=9606 GN=SIL1 PE=1 SV=1 | 0 | 111.809 | 56 | 19 | 322 | 19 | 461 | 52.1 | 5.36 | 797.13 | 19 | 17688086.54 |
| Q9BY76 | Angiopoietin-related protein 4 OS=Homo sapiens OX=9606 GN=ANGPTL4 PE=1 SV=2 | 0 | 111.195 | 45 | 13 | 236 | 13 | 406 | 45.2 | 8.85 | 750.4 | 13 | 8836641.568 |
| P42330 | Aldo-keto reductase family 1 member C3 OS=Homo sapiens OX=9606 GN=AKR1C3 PE=1 SV=4 | 0 | 111.138 | 67 | 19 | 145 | 9 | 323 | 36.8 | 7.94 | 397.97 | 19 | 1093267.218 |
| Q9NR31 | GTP-binding protein SAR1a OS=Homo sapiens OX=9606 GN=SAR1A PE=1 SV=1 | 0 | 110.742 | 75 | 15 | 160 | 10 | 198 | 22.4 | 6.68 | 417.95 | 15 | 755680.8086 |
| P61019 | Ras-related protein Rab-2A OS=Homo sapiens OX=9606 GN=RAB2A PE=1 SV=1 | 0 | 110.543 | 79 | 16 | 173 | 16 | 212 | 23.5 | 6.54 | 493.13 | 16 | 603436.0225 |
| P29992 | Guanine nucleotide-binding protein subunit alpha-11 OS=Homo sapiens OX=9606 GN=GNA11 PE=1 SV=2 | 0 | 109.3 | 58 | 18 | 170 | 7 | 359 | 42.1 | 5.69 | 385.65 | 18 | 1564938.181 |
| P25774 | Cathepsin S OS=Homo sapiens OX=9606 GN=CTSS PE=1 SV=3 | 0 | 109.03 | 56 | 16 | 283 | 16 | 331 | 37.5 | 8.34 | 753.03 | 16 | 4627696.368 |
| Q99497 | Parkinson disease protein 7 OS=Homo sapiens OX=9606 GN=PARK7 PE=1 SV=2 | 0 | 108.654 | 81 | 12 | 138 | 12 | 189 | 19.9 | 6.79 | 431.48 | 12 | 32453909.01 |
| P11678 | Eosinophil peroxidase OS=Homo sapiens OX=9606 GN=EPX PE=1 SV=2 | 0 | 108.081 | 35 | 23 | 46 | 18 | 715 | 81 | 10.29 | 114.4 | 23 | |
| P04843 | Dolichyl-diphosphooligosaccharide--protein glycosyltransferase subunit 1 OS=Homo sapiens OX=9606 GN=RPN1 PE=1 SV=1 | 0 | 107.959 | 40 | 18 | 88 | 18 | 607 | 68.5 | 6.38 | 223.44 | 18 | 35907435.89 |
| P68871 | Hemoglobin subunit beta OS=Homo sapiens OX=9606 GN=HBB PE=1 SV=2 | 0 | 107.881 | 96 | 14 | 93 | 7 | 147 | 16 | 7.28 | 317.38 | 14 | |
| Q04828 | Aldo-keto reductase family 1 member C1 OS=Homo sapiens OX=9606 GN=AKR1C1 PE=1 SV=1 | 0 | 107.817 | 62 | 17 | 128 | 7 | 323 | 36.8 | 7.88 | 352.66 | 17 | 49243172.95 |
| O60543 | Cell death activator CIDE-A OS=Homo sapiens OX=9606 GN=CIDEA PE=1 SV=1 | 0 | 107.135 | 69 | 13 | 221 | 13 | 219 | 24.7 | 9.23 | 646.86 | 13 | 8635007.832 |
| O00584 | Ribonuclease T2 OS=Homo sapiens OX=9606 GN=RNASET2 PE=1 SV=2 | 0 | 107.088 | 50 | 12 | 136 | 12 | 256 | 29.5 | 7.08 | 347.73 | 12 | 4008308.507 |
| P51149 | Ras-related protein Rab-7a OS=Homo sapiens OX=9606 GN=RAB7A PE=1 SV=1 | 0 | 106.73 | 84 | 17 | 235 | 17 | 207 | 23.5 | 6.7 | 645.58 | 17 | 6868349.778 |
| P11234 | Ras-related protein Ral-B OS=Homo sapiens OX=9606 GN=RALB PE=1 SV=1 | 0 | 106.523 | 70 | 14 | 178 | 6 | 206 | 23.4 | 6.62 | 527.58 | 14 | 321382.8284 |
| P07741 | Adenine phosphoribosyltransferase OS=Homo sapiens OX=9606 GN=APRT PE=1 SV=2 | 0 | 106.258 | 91 | 13 | 118 | 13 | 180 | 19.6 | 6.02 | 336.03 | 13 | 4553086.512 |
| O00764 | Pyridoxal kinase OS=Homo sapiens OX=9606 GN=PDXK PE=1 SV=1 | 0 | 106.166 | 70 | 15 | 106 | 15 | 312 | 35.1 | 6.13 | 229.36 | 15 | |
| Q15782 | Chitinase-3-like protein 2 OS=Homo sapiens OX=9606 GN=CHI3L2 PE=1 SV=1 | 0 | 105.56 | 60 | 16 | 101 | 16 | 390 | 43.5 | 7.53 | 337.46 | 16 | 2407031.565 |
| P02652 | Apolipoprotein A-II OS=Homo sapiens OX=9606 GN=APOA2 PE=1 SV=1 | 0 | 105.173 | 69 | 10 | 254 | 10 | 100 | 11.2 | 6.62 | 919.68 | 10 | 13187711.86 |
| P30044 | Peroxiredoxin-5, mitochondrial OS=Homo sapiens OX=9606 GN=PRDX5 PE=1 SV=4 | 0 | 103.711 | 66 | 13 | 137 | 13 | 214 | 22.1 | 8.7 | 431.58 | 13 | |
| P01019 | Angiotensinogen OS=Homo sapiens OX=9606 GN=AGT PE=1 SV=1 | 0 | 103.673 | 36 | 13 | 178 | 13 | 485 | 53.1 | 6.32 | 695.86 | 13 | 2178164.644 |
| P04899 | Guanine nucleotide-binding protein G(i) subunit alpha-2 OS=Homo sapiens OX=9606 GN=GNAI2 PE=1 SV=3 | 0 | 103.646 | 52 | 17 | 142 | 9 | 355 | 40.4 | 5.54 | 340.19 | 17 | 569546.198 |
| Q6UXI9 | Nephronectin OS=Homo sapiens OX=9606 GN=NPNT PE=2 SV=3 | 0 | 103.584 | 44 | 20 | 252 | 20 | 565 | 61.9 | 8.34 | 581.4 | 20 | 5964102.293 |
| P08246 | Neutrophil elastase OS=Homo sapiens OX=9606 GN=ELANE PE=1 SV=1 | 0 | 102.981 | 54 | 11 | 73 | 11 | 267 | 28.5 | 9.35 | 236.57 | 11 | 13711143.06 |
| P63241 | Eukaryotic translation initiation factor 5A-1 OS=Homo sapiens OX=9606 GN=EIF5A PE=1 SV=2 | 0 | 101.645 | 88 | 11 | 114 | 11 | 154 | 16.8 | 5.24 | 395.84 | 11 | 2185011.726 |
| P36578 | 60S ribosomal protein L4 OS=Homo sapiens OX=9606 GN=RPL4 PE=1 SV=5 | 0 | 101.121 | 38 | 17 | 157 | 17 | 427 | 47.7 | 11.06 | 352.09 | 17 | 277646.2033 |
| Q7KZF4 | Staphylococcal nuclease domain-containing protein 1 OS=Homo sapiens OX=9606 GN=SND1 PE=1 SV=1 | 0 | 101.096 | 32 | 19 | 66 | 19 | 910 | 101.9 | 7.17 | 111.18 | 19 | 2761918.329 |
| P29762 | Cellular retinoic acid-binding protein 1 OS=Homo sapiens OX=9606 GN=CRABP1 PE=1 SV=2 | 0 | 101.068 | 85 | 13 | 177 | 12 | 137 | 15.6 | 5.38 | 429.95 | 13 | 635225.901 |
| O60547 | GDP-mannose 4,6 dehydratase OS=Homo sapiens OX=9606 GN=GMDS PE=1 SV=1 | 0 | 100.239 | 59 | 18 | 132 | 18 | 372 | 41.9 | 7.31 | 250.13 | 18 | 1447568.72 |
| Q8N9U0 | Tandem C2 domains nuclear protein OS=Homo sapiens OX=9606 GN=TC2N PE=1 SV=2 | 0 | 99.614 | 54 | 22 | 194 | 22 | 490 | 55.2 | 9.19 | 360.91 | 22 | 5057933.005 |
| P30740 | Leukocyte elastase inhibitor OS=Homo sapiens OX=9606 GN=SERPINB1 PE=1 SV=1 | 0 | 99.177 | 64 | 20 | 61 | 20 | 379 | 42.7 | 6.28 | 166.45 | 20 | 955112.4299 |
| P30085 | UMP-CMP kinase OS=Homo sapiens OX=9606 GN=CMPK1 PE=1 SV=3 | 0 | 98.663 | 64 | 13 | 106 | 13 | 196 | 22.2 | 5.57 | 266.78 | 13 | |
| P36955 | Pigment epithelium-derived factor OS=Homo sapiens OX=9606 GN=SERPINF1 PE=1 SV=4 | 0 | 98.494 | 51 | 17 | 234 | 17 | 418 | 46.3 | 6.38 | 544.95 | 17 | 2532515.903 |
| P62805 | Histone H4 OS=Homo sapiens OX=9606 GN=H4C1 PE=1 SV=2 | 0 | 97.641 | 59 | 13 | 169 | 13 | 103 | 11.4 | 11.36 | 433.63 | 13 | 50018261.83 |
| O14786 | Neuropilin-1 OS=Homo sapiens OX=9606 GN=NRP1 PE=1 SV=3 | 0 | 97.388 | 25 | 15 | 123 | 15 | 923 | 103.1 | 5.88 | 314.47 | 15 | 1359387.691 |
| P05090 | Apolipoprotein D OS=Homo sapiens OX=9606 GN=APOD PE=1 SV=1 | 0 | 97.177 | 41 | 14 | 189 | 14 | 189 | 21.3 | 5.15 | 470.48 | 14 | 1466476.389 |
| P29373 | Cellular retinoic acid-binding protein 2 OS=Homo sapiens OX=9606 GN=CRABP2 PE=1 SV=2 | 0 | 97.067 | 69 | 12 | 178 | 11 | 138 | 15.7 | 5.4 | 516.61 | 12 | 4849341.194 |
| P22314 | Ubiquitin-like modifier-activating enzyme 1 OS=Homo sapiens OX=9606 GN=UBA1 PE=1 SV=3 | 0 | 96.926 | 31 | 21 | 71 | 21 | 1058 | 117.8 | 5.76 | 162.05 | 21 | |
| O00339 | Matrilin-2 OS=Homo sapiens OX=9606 GN=MATN2 PE=1 SV=4 | 0 | 96.865 | 32 | 21 | 99 | 21 | 956 | 106.8 | 6.27 | 179.73 | 21 | 174289.137 |
| P22307 | Non-specific lipid-transfer protein OS=Homo sapiens OX=9606 GN=SCP2 PE=1 SV=2 | 0 | 95.945 | 28 | 17 | 93 | 17 | 547 | 59 | 6.89 | 264.98 | 17 | 2490247.916 |
| P06576 | ATP synthase subunit beta, mitochondrial OS=Homo sapiens OX=9606 GN=ATP5F1B PE=1 SV=3 | 0 | 95.293 | 44 | 15 | 42 | 15 | 529 | 56.5 | 5.4 | 158.99 | 15 | 5308083.687 |
| P32119 | Peroxiredoxin-2 OS=Homo sapiens OX=9606 GN=PRDX2 PE=1 SV=5 | 0 | 95.168 | 87 | 17 | 182 | 16 | 198 | 21.9 | 5.97 | 384.14 | 17 | 528606.9404 |
| P07858 | Cathepsin B OS=Homo sapiens OX=9606 GN=CTSB PE=1 SV=3 | 0 | 94.357 | 55 | 13 | 165 | 13 | 339 | 37.8 | 6.3 | 414.55 | 13 | 2716719.928 |
| P61247 | 40S ribosomal protein S3a OS=Homo sapiens OX=9606 GN=RPS3A PE=1 SV=2 | 0 | 94.179 | 55 | 15 | 182 | 15 | 264 | 29.9 | 9.73 | 295.58 | 15 | 389296.3428 |
| Q99538 | Legumain OS=Homo sapiens OX=9606 GN=LGMN PE=1 SV=1 | 0 | 94.157 | 44 | 14 | 97 | 14 | 433 | 49.4 | 6.55 | 227.41 | 14 | 382720.6608 |
| P61224 | Ras-related protein Rap-1b OS=Homo sapiens OX=9606 GN=RAP1B PE=1 SV=1 | 0 | 94.013 | 79 | 15 | 192 | 4 | 184 | 20.8 | 5.78 | 525.11 | 15 | |
| Q9Y6R7 | IgGFc-binding protein OS=Homo sapiens OX=9606 GN=FCGBP PE=1 SV=3 | 0 | 93.406 | 10 | 20 | 47 | 20 | 5405 | 571.6 | 5.34 | 120.25 | 20 | 1452003.869 |
| P61981 | 14-3-3 protein gamma OS=Homo sapiens OX=9606 GN=YWHAG PE=1 SV=2 | 0 | 92.975 | 66 | 16 | 197 | 11 | 247 | 28.3 | 4.89 | 440.64 | 16 | 3571670.793 |
| P11940 | Polyadenylate-binding protein 1 OS=Homo sapiens OX=9606 GN=PABPC1 PE=1 SV=2 | 0 | 92.806 | 32 | 15 | 104 | 15 | 636 | 70.6 | 9.5 | 267.37 | 15 | 3018505.002 |
| P06737 | Glycogen phosphorylase, liver form OS=Homo sapiens OX=9606 GN=PYGL PE=1 SV=4 | 0 | 92.004 | 30 | 20 | 39 | 15 | 847 | 97.1 | 7.17 | 109.32 | 20 | |
| P00491 | Purine nucleoside phosphorylase OS=Homo sapiens OX=9606 GN=PNP PE=1 SV=2 | 0 | 91.742 | 66 | 14 | 73 | 14 | 289 | 32.1 | 6.95 | 205.32 | 14 | |
| Q12841 | Follistatin-related protein 1 OS=Homo sapiens OX=9606 GN=FSTL1 PE=1 SV=1 | 0 | 91.689 | 59 | 16 | 202 | 16 | 308 | 35 | 5.52 | 443.72 | 16 | 21513847.6 |
| P18669 | Phosphoglycerate mutase 1 OS=Homo sapiens OX=9606 GN=PGAM1 PE=1 SV=2 | 0 | 91.517 | 72 | 14 | 58 | 14 | 254 | 28.8 | 7.18 | 169.67 | 14 | 13771123.72 |
| P39023 | 60S ribosomal protein L3 OS=Homo sapiens OX=9606 GN=RPL3 PE=1 SV=2 | 0 | 91.49 | 42 | 18 | 158 | 18 | 403 | 46.1 | 10.18 | 301.87 | 18 | |
| P11233 | Ras-related protein Ral-A OS=Homo sapiens OX=9606 GN=RALA PE=1 SV=1 | 0 | 91.479 | 65 | 12 | 123 | 4 | 206 | 23.6 | 7.11 | 377.47 | 12 | 396882.3873 |
| P01033 | Metalloproteinase inhibitor 1 OS=Homo sapiens OX=9606 GN=TIMP1 PE=1 SV=1 | 0 | 90.866 | 59 | 9 | 228 | 9 | 207 | 23.2 | 8.1 | 809.83 | 9 | 9706948.824 |
| P00352 | Retinal dehydrogenase 1 OS=Homo sapiens OX=9606 GN=ALDH1A1 PE=1 SV=2 | 0 | 90.853 | 47 | 18 | 57 | 15 | 501 | 54.8 | 6.73 | 137.14 | 18 | 6154454.532 |
| P23297 | Protein S100-A1 OS=Homo sapiens OX=9606 GN=S100A1 PE=1 SV=2 | 0 | 90.686 | 60 | 7 | 94 | 7 | 94 | 10.5 | 4.5 | 535.2 | 7 | 3337694.066 |
| Q15738 | Sterol-4-alpha-carboxylate 3-dehydrogenase, decarboxylating OS=Homo sapiens OX=9606 GN=NSDHL PE=1 SV=2 | 0 | 90.461 | 55 | 14 | 80 | 14 | 373 | 41.9 | 8.06 | 216.68 | 14 | 27834449.56 |
| O43488 | Aflatoxin B1 aldehyde reductase member 2 OS=Homo sapiens OX=9606 GN=AKR7A2 PE=1 SV=3 | 0 | 89.285 | 48 | 12 | 80 | 12 | 359 | 39.6 | 7.17 | 201.2 | 12 | 9637649.615 |
| O75888 | Tumor necrosis factor ligand superfamily member 13 OS=Homo sapiens OX=9606 GN=TNFSF13 PE=1 SV=1 | 0 | 89.056 | 43 | 10 | 117 | 10 | 250 | 27.4 | 9.63 | 437.19 | 10 | 2444859.764 |
| P01903 | HLA class II histocompatibility antigen, DR alpha chain OS=Homo sapiens OX=9606 GN=HLA-DRA PE=1 SV=1 | 0 | 89.034 | 35 | 11 | 223 | 11 | 254 | 28.6 | 5 | 889.18 | 11 | 14683283.08 |
| P40925 | Malate dehydrogenase, cytoplasmic OS=Homo sapiens OX=9606 GN=MDH1 PE=1 SV=4 | 0 | 88.925 | 65 | 15 | 184 | 15 | 334 | 36.4 | 7.36 | 491.64 | 15 | 13978119.85 |
| Q4VNC1 | Probable cation-transporting ATPase 13A4 OS=Homo sapiens OX=9606 GN=ATP13A4 PE=2 SV=3 | 0 | 88.563 | 23 | 18 | 90 | 18 | 1196 | 133.9 | 6.7 | 207.97 | 18 | 2724276.816 |
| P28799 | Progranulin OS=Homo sapiens OX=9606 GN=GRN PE=1 SV=2 | 0 | 88.411 | 29 | 15 | 134 | 15 | 593 | 63.5 | 6.83 | 211.38 | 15 | 1166090.517 |
| P13667 | Protein disulfide-isomerase A4 OS=Homo sapiens OX=9606 GN=PDIA4 PE=1 SV=2 | 0 | 88.199 | 33 | 19 | 98 | 19 | 645 | 72.9 | 5.07 | 192.93 | 19 | 8096858.741 |
| P27348 | 14-3-3 protein theta OS=Homo sapiens OX=9606 GN=YWHAQ PE=1 SV=1 | 0 | 88.031 | 56 | 16 | 166 | 10 | 245 | 27.7 | 4.78 | 355.79 | 16 | 293194.4385 |
| A5A3E0 | POTE ankyrin domain family member F OS=Homo sapiens OX=9606 GN=POTEF PE=1 SV=2 | 0 | 87.924 | 11 | 10 | 252 | 3 | 1075 | 121.4 | 6.2 | 814.16 | 10 | |
| P05154 | Plasma serine protease inhibitor OS=Homo sapiens OX=9606 GN=SERPINA5 PE=1 SV=3 | 0 | 87.552 | 58 | 17 | 118 | 17 | 406 | 45.6 | 9.26 | 338.07 | 17 | 456733.9915 |
| O14745 | Na(+)/H(+) exchange regulatory cofactor NHE-RF1 OS=Homo sapiens OX=9606 GN=SLC9A3R1 PE=1 SV=4 | 0 | 87.297 | 56 | 17 | 235 | 17 | 358 | 38.8 | 5.77 | 485.27 | 17 | 2601030.346 |
| P61764 | Syntaxin-binding protein 1 OS=Homo sapiens OX=9606 GN=STXBP1 PE=1 SV=1 | 0 | 86.32 | 44 | 17 | 73 | 17 | 594 | 67.5 | 6.96 | 183.48 | 17 | 665820.095 |
| Q99574 | Neuroserpin OS=Homo sapiens OX=9606 GN=SERPINI1 PE=1 SV=1 | 0 | 85.924 | 41 | 13 | 104 | 13 | 410 | 46.4 | 4.91 | 338.46 | 13 | 1654347.433 |
| P60953 | Cell division control protein 42 homolog OS=Homo sapiens OX=9606 GN=CDC42 PE=1 SV=2 | 0 | 85.448 | 64 | 10 | 209 | 9 | 191 | 21.2 | 6.55 | 692.89 | 10 | 6161732.663 |
| P80511 | Protein S100-A12 OS=Homo sapiens OX=9606 GN=S100A12 PE=1 SV=2 | 0 | 85.201 | 74 | 9 | 38 | 9 | 92 | 10.6 | 6.25 | 168.78 | 9 | 6383075.795 |
| P52566 | Rho GDP-dissociation inhibitor 2 OS=Homo sapiens OX=9606 GN=ARHGDIB PE=1 SV=3 | 0 | 85.06 | 77 | 11 | 54 | 11 | 201 | 23 | 5.21 | 185.58 | 11 | 1048192.498 |
| P29508 | Serpin B3 OS=Homo sapiens OX=9606 GN=SERPINB3 PE=1 SV=2 | 0 | 85 | 50 | 18 | 20 | 18 | 390 | 44.5 | 6.81 | 63.99 | 18 | |
| Q15181 | Inorganic pyrophosphatase OS=Homo sapiens OX=9606 GN=PPA1 PE=1 SV=2 | 0 | 84.939 | 54 | 11 | 90 | 11 | 289 | 32.6 | 5.86 | 357.32 | 11 | 1456956.787 |
| P60028 | Rab GDP dissociation inhibitor alpha OS=Pan troglodytes OX=9598 GN=GDI1 PE=2 SV=1 | 0 | 84.327 | 27 | 10 | 153 | 2 | 447 | 50.6 | 5.14 | 619.88 | 10 | |
| Q8N474 | Secreted frizzled-related protein 1 OS=Homo sapiens OX=9606 GN=SFRP1 PE=1 SV=1 | 0 | 84.123 | 51 | 12 | 173 | 12 | 314 | 35.4 | 8.85 | 503.47 | 12 | 4584144.229 |
| O75369 | Filamin-B OS=Homo sapiens OX=9606 GN=FLNB PE=1 SV=2 | 0 | 83.747 | 11 | 23 | 83 | 21 | 2602 | 278 | 5.73 | 144.96 | 23 | 2052264.08 |
| Q9H223 | EH domain-containing protein 4 OS=Homo sapiens OX=9606 GN=EHD4 PE=1 SV=1 | 0 | 83.721 | 44 | 21 | 116 | 18 | 541 | 61.1 | 6.76 | 234.76 | 21 | 691500.7927 |
| Q58FF7 | Putative heat shock protein HSP 90-beta-3 OS=Homo sapiens OX=9606 GN=HSP90AB3P PE=5 SV=1 | 0 | 83.489 | 22 | 15 | 170 | 1 | 597 | 68.3 | 4.79 | 435.27 | 15 | |
| Q9H0U4 | Ras-related protein Rab-1B OS=Homo sapiens OX=9606 GN=RAB1B PE=1 SV=1 | 0 | 83.353 | 78 | 13 | 201 | 4 | 201 | 22.2 | 5.73 | 512.88 | 13 | 3176964.848 |
| Q15084 | Protein disulfide-isomerase A6 OS=Homo sapiens OX=9606 GN=PDIA6 PE=1 SV=1 | 0 | 83.146 | 42 | 14 | 125 | 14 | 440 | 48.1 | 5.08 | 357.08 | 14 | |
| P08865 | 40S ribosomal protein SA OS=Homo sapiens OX=9606 GN=RPSA PE=1 SV=4 | 0 | 82.74 | 54 | 11 | 160 | 11 | 295 | 32.8 | 4.87 | 512.45 | 11 | 1620177.88 |
| Q14204 | Cytoplasmic dynein 1 heavy chain 1 OS=Homo sapiens OX=9606 GN=DYNC1H1 PE=1 SV=5 | 0 | 81.928 | 7 | 28 | 52 | 28 | 4646 | 532.1 | 6.4 | 78.26 | 28 | 1760583.526 |
| P41218 | Myeloid cell nuclear differentiation antigen OS=Homo sapiens OX=9606 GN=MNDA PE=1 SV=1 | 0 | 81.427 | 40 | 17 | 47 | 17 | 407 | 45.8 | 9.76 | 126.21 | 17 | 1266299.445 |
| P10412 | Histone H1.4 OS=Homo sapiens OX=9606 GN=H1-4 PE=1 SV=2 | 0 | 81.227 | 33 | 12 | 43 | 3 | 219 | 21.9 | 11.03 | 107.07 | 12 | 750613.3582 |
| P12272 | Parathyroid hormone-related protein OS=Homo sapiens OX=9606 GN=PTHLH PE=1 SV=1 | 0 | 81.208 | 43 | 9 | 91 | 9 | 177 | 20.2 | 10.18 | 269.91 | 9 | |
| P10586 | Receptor-type tyrosine-protein phosphatase F OS=Homo sapiens OX=9606 GN=PTPRF PE=1 SV=2 | 0 | 80.605 | 17 | 24 | 90 | 24 | 1907 | 212.7 | 6.3 | 171.31 | 24 | 2562612.244 |
| P62424 | 60S ribosomal protein L7a OS=Homo sapiens OX=9606 GN=RPL7A PE=1 SV=2 | 0 | 80.564 | 50 | 13 | 173 | 13 | 266 | 30 | 10.61 | 412.19 | 13 | 3085015.69 |
| Q9H4G4 | Golgi-associated plant pathogenesis-related protein 1 OS=Homo sapiens OX=9606 GN=GLIPR2 PE=1 SV=3 | 0 | 80.383 | 75 | 10 | 178 | 10 | 154 | 17.2 | 9.41 | 598.03 | 10 | 13974198.65 |
| A0A0B4J1V0 | Immunoglobulin heavy variable 3-15 OS=Homo sapiens OX=9606 GN=IGHV3-15 PE=3 SV=1 | 0 | 80.363 | 57 | 12 | 135 | 7 | 119 | 12.9 | 8.62 | 347.19 | 12 | 1030139.203 |
| P25398 | 40S ribosomal protein S12 OS=Homo sapiens OX=9606 GN=RPS12 PE=1 SV=3 | 0 | 79.894 | 68 | 8 | 114 | 8 | 132 | 14.5 | 7.21 | 378.84 | 8 | 455217.4526 |
| Q9NZ08 | Endoplasmic reticulum aminopeptidase 1 OS=Homo sapiens OX=9606 GN=ERAP1 PE=1 SV=3 | 0 | 79.65 | 29 | 23 | 107 | 23 | 941 | 107.2 | 6.46 | 235.21 | 23 | 4985960.239 |
| Q9HAV0 | Guanine nucleotide-binding protein subunit beta-4 OS=Homo sapiens OX=9606 GN=GNB4 PE=1 SV=3 | 0 | 78.901 | 50 | 13 | 198 | 7 | 340 | 37.5 | 6 | 535.77 | 13 | 735566.8599 |
| Q02790 | Peptidyl-prolyl cis-trans isomerase FKBP4 OS=Homo sapiens OX=9606 GN=FKBP4 PE=1 SV=3 | 0 | 78.689 | 40 | 17 | 112 | 17 | 459 | 51.8 | 5.43 | 239.86 | 17 | 886550.8197 |
| P60981 | Destrin OS=Homo sapiens OX=9606 GN=DSTN PE=1 SV=3 | 0 | 78.585 | 84 | 15 | 116 | 13 | 165 | 18.5 | 7.85 | 242.61 | 15 | 2037161.394 |
| Q9HB40 | Retinoid-inducible serine carboxypeptidase OS=Homo sapiens OX=9606 GN=SCPEP1 PE=1 SV=1 | 0 | 78.516 | 36 | 13 | 216 | 13 | 452 | 50.8 | 5.81 | 503.44 | 13 | 2784448.094 |
| P09467 | Fructose-1,6-bisphosphatase 1 OS=Homo sapiens OX=9606 GN=FBP1 PE=1 SV=5 | 0 | 78.01 | 46 | 11 | 73 | 10 | 338 | 36.8 | 6.99 | 165.85 | 11 | 653545.7187 |
| P54802 | Alpha-N-acetylglucosaminidase OS=Homo sapiens OX=9606 GN=NAGLU PE=1 SV=2 | 0 | 77.948 | 31 | 15 | 61 | 15 | 743 | 82.2 | 6.65 | 154.88 | 15 | 527701495.8 |
| P06753 | Tropomyosin alpha-3 chain OS=Homo sapiens OX=9606 GN=TPM3 PE=1 SV=2 | 0 | 77.813 | 38 | 10 | 95 | 5 | 285 | 32.9 | 4.72 | 201.8 | 10 | 6984533.87 |
| Q9UL46 | Proteasome activator complex subunit 2 OS=Homo sapiens OX=9606 GN=PSME2 PE=1 SV=4 | 0 | 77.726 | 64 | 13 | 65 | 13 | 239 | 27.4 | 5.73 | 162.34 | 13 | 751688.6413 |
| P34896 | Serine hydroxymethyltransferase, cytosolic OS=Homo sapiens OX=9606 GN=SHMT1 PE=1 SV=1 | 0 | 77.661 | 48 | 17 | 59 | 17 | 483 | 53 | 7.71 | 117.49 | 17 | 463836.2028 |
| Q13162 | Peroxiredoxin-4 OS=Homo sapiens OX=9606 GN=PRDX4 PE=1 SV=1 | 0 | 77.546 | 53 | 11 | 139 | 8 | 271 | 30.5 | 6.29 | 386.86 | 11 | 17139698.96 |
| O00161 | Synaptosomal-associated protein 23 OS=Homo sapiens OX=9606 GN=SNAP23 PE=1 SV=1 | 0 | 77.395 | 72 | 15 | 161 | 15 | 211 | 23.3 | 5.01 | 417.01 | 15 | 683696.9308 |
| P80188 | Neutrophil gelatinase-associated lipocalin OS=Homo sapiens OX=9606 GN=LCN2 PE=1 SV=2 | 0 | 77.306 | 64 | 12 | 94 | 12 | 198 | 22.6 | 8.91 | 310.49 | 12 | 5745481.268 |
| P62834 | Ras-related protein Rap-1A OS=Homo sapiens OX=9606 GN=RAP1A PE=1 SV=1 | 0 | 77.042 | 65 | 13 | 189 | 2 | 184 | 21 | 6.67 | 502.04 | 13 | |
| P18085 | ADP-ribosylation factor 4 OS=Homo sapiens OX=9606 GN=ARF4 PE=1 SV=3 | 0 | 76.764 | 57 | 11 | 141 | 7 | 180 | 20.5 | 7.14 | 386.05 | 11 | |
| Q06323 | Proteasome activator complex subunit 1 OS=Homo sapiens OX=9606 GN=PSME1 PE=1 SV=1 | 0 | 76.586 | 61 | 14 | 110 | 14 | 249 | 28.7 | 6.02 | 262.27 | 14 | 2868745.014 |
| P25705 | ATP synthase subunit alpha, mitochondrial OS=Homo sapiens OX=9606 GN=ATP5F1A PE=1 SV=1 | 0 | 76.422 | 33 | 14 | 30 | 14 | 553 | 59.7 | 9.13 | 90.38 | 14 | 430638.4055 |
| P19440 | Glutathione hydrolase 1 proenzyme OS=Homo sapiens OX=9606 GN=GGT1 PE=1 SV=2 | 0 | 76.305 | 29 | 13 | 262 | 13 | 569 | 61.4 | 7.12 | 778.7 | 13 | 16196774.77 |
| P07686 | Beta-hexosaminidase subunit beta OS=Homo sapiens OX=9606 GN=HEXB PE=1 SV=3 | 0 | 76.103 | 39 | 18 | 79 | 17 | 556 | 63.1 | 6.76 | 157.57 | 18 | 2712918.742 |
| P17900 | Ganglioside GM2 activator OS=Homo sapiens OX=9606 GN=GM2A PE=1 SV=4 | 0 | 75.741 | 63 | 10 | 171 | 10 | 193 | 20.8 | 5.31 | 366.78 | 10 | 2769088.638 |
| P60842 | Eukaryotic initiation factor 4A-I OS=Homo sapiens OX=9606 GN=EIF4A1 PE=1 SV=1 | 0 | 74.893 | 42 | 12 | 126 | 6 | 406 | 46.1 | 5.48 | 323.17 | 12 | 232384.9478 |
| Q14515 | SPARC-like protein 1 OS=Homo sapiens OX=9606 GN=SPARCL1 PE=1 SV=2 | 0 | 74.791 | 28 | 13 | 49 | 13 | 664 | 75.2 | 4.81 | 139.8 | 13 | 2342004.532 |
| Q01459 | Di-N-acetylchitobiase OS=Homo sapiens OX=9606 GN=CTBS PE=1 SV=1 | 0 | 74.721 | 40 | 11 | 137 | 11 | 385 | 43.7 | 6.64 | 337.67 | 11 | 5374230.452 |
| P43652 | Afamin OS=Homo sapiens OX=9606 GN=AFM PE=1 SV=1 | 0 | 74.712 | 35 | 17 | 162 | 17 | 599 | 69 | 5.9 | 244.47 | 17 | 1387380.439 |
| Q99828 | Calcium and integrin-binding protein 1 OS=Homo sapiens OX=9606 GN=CIB1 PE=1 SV=4 | 0 | 74.654 | 77 | 12 | 157 | 12 | 191 | 21.7 | 4.78 | 409.9 | 12 | 1979776.523 |
| P60660 | Myosin light polypeptide 6 OS=Homo sapiens OX=9606 GN=MYL6 PE=1 SV=2 | 0 | 74.612 | 75 | 10 | 135 | 10 | 151 | 16.9 | 4.65 | 386.78 | 10 | 6440208.686 |
| Q66K79 | Carboxypeptidase Z OS=Homo sapiens OX=9606 GN=CPZ PE=1 SV=2 | 0 | 74.456 | 34 | 18 | 126 | 18 | 652 | 73.6 | 7.97 | 221.7 | 18 | |
| P13987 | CD59 glycoprotein OS=Homo sapiens OX=9606 GN=CD59 PE=1 SV=1 | 0 | 74.314 | 41 | 8 | 193 | 8 | 128 | 14.2 | 6.48 | 598.14 | 8 | 37596901.5 |
| P01780 | Immunoglobulin heavy variable 3-7 OS=Homo sapiens OX=9606 GN=IGHV3-7 PE=1 SV=2 | 0 | 74.221 | 68 | 13 | 220 | 7 | 117 | 12.9 | 6.57 | 519.6 | 13 | 10635305.14 |
| O14773 | Tripeptidyl-peptidase 1 OS=Homo sapiens OX=9606 GN=TPP1 PE=1 SV=2 | 0 | 73.817 | 43 | 12 | 61 | 12 | 563 | 61.2 | 6.48 | 208.68 | 12 | 1552759.177 |
| P08195 | 4F2 cell-surface antigen heavy chain OS=Homo sapiens OX=9606 GN=SLC3A2 PE=1 SV=3 | 0 | 73.566 | 28 | 14 | 93 | 14 | 630 | 68 | 5.01 | 215.22 | 14 | 3646444.076 |
| Q9Y240 | C-type lectin domain family 11 member A OS=Homo sapiens OX=9606 GN=CLEC11A PE=1 SV=1 | 0 | 73.565 | 44 | 16 | 63 | 16 | 323 | 35.7 | 5.16 | 186.02 | 16 | 4290244.925 |
| Q12805 | EGF-containing fibulin-like extracellular matrix protein 1 OS=Homo sapiens OX=9606 GN=EFEMP1 PE=1 SV=2 | 0 | 73.498 | 39 | 14 | 79 | 14 | 493 | 54.6 | 5.07 | 203.75 | 14 | 3953095.996 |
| P05107 | Integrin beta-2 OS=Homo sapiens OX=9606 GN=ITGB2 PE=1 SV=2 | 0 | 73.4 | 26 | 18 | 30 | 18 | 769 | 84.7 | 6.95 | 86.52 | 18 | 375130.7594 |
| P01911 | HLA class II histocompatibility antigen, DRB1 beta chain OS=Homo sapiens OX=9606 GN=HLA-DRB1 PE=1 SV=2 | 0 | 73.3 | 54 | 11 | 102 | 6 | 266 | 29.9 | 7.74 | 326.52 | 11 | |
| P23396 | 40S ribosomal protein S3 OS=Homo sapiens OX=9606 GN=RPS3 PE=1 SV=2 | 0 | 73.176 | 60 | 15 | 149 | 15 | 243 | 26.7 | 9.66 | 389.08 | 15 | 1219624.402 |
| P11413 | Glucose-6-phosphate 1-dehydrogenase OS=Homo sapiens OX=9606 GN=G6PD PE=1 SV=4 | 0 | 72.725 | 43 | 18 | 41 | 18 | 515 | 59.2 | 6.84 | 83.11 | 18 | |
| Q96LJ7 | Dehydrogenase/reductase SDR family member 1 OS=Homo sapiens OX=9606 GN=DHRS1 PE=1 SV=1 | 0 | 72.667 | 63 | 15 | 134 | 15 | 313 | 33.9 | 7.83 | 323.7 | 15 | 1015117.001 |
| Q9Y5Y6 | Suppressor of tumorigenicity 14 protein OS=Homo sapiens OX=9606 GN=ST14 PE=1 SV=2 | 0 | 72.119 | 22 | 12 | 105 | 12 | 855 | 94.7 | 6.55 | 244.03 | 12 | 1152931.882 |
| O15143 | Actin-related protein 2/3 complex subunit 1B OS=Homo sapiens OX=9606 GN=ARPC1B PE=1 SV=3 | 0 | 72.069 | 45 | 12 | 29 | 12 | 372 | 40.9 | 8.35 | 97.29 | 12 | 179707.2832 |
| P39656 | Dolichyl-diphosphooligosaccharide--protein glycosyltransferase 48 kDa subunit OS=Homo sapiens OX=9606 GN=DDOST PE=1 SV=4 | 0 | 72.064 | 46 | 12 | 58 | 12 | 456 | 50.8 | 6.55 | 167.05 | 12 | 1961236.358 |
| P40121 | Macrophage-capping protein OS=Homo sapiens OX=9606 GN=CAPG PE=1 SV=2 | 0 | 71.739 | 51 | 10 | 65 | 10 | 348 | 38.5 | 6.19 | 196.29 | 10 | 1667579.323 |
| Q13277 | Syntaxin-3 OS=Homo sapiens OX=9606 GN=STX3 PE=1 SV=3 | 0 | 71.602 | 38 | 12 | 139 | 12 | 289 | 33.1 | 5.44 | 420.76 | 12 | 879301.7102 |
| Q5JWF2 | Guanine nucleotide-binding protein G(s) subunit alpha isoforms XLas OS=Homo sapiens OX=9606 GN=GNAS PE=1 SV=2 | 0 | 71.596 | 19 | 16 | 96 | 15 | 1037 | 111 | 5.03 | 177.8 | 16 | 2019587.201 |
| Q9NZP8 | Complement C1r subcomponent-like protein OS=Homo sapiens OX=9606 GN=C1RL PE=1 SV=2 | 0 | 71.573 | 35 | 13 | 126 | 11 | 487 | 53.5 | 7.2 | 306.92 | 13 | |
| P08311 | Cathepsin G OS=Homo sapiens OX=9606 GN=CTSG PE=1 SV=2 | 0 | 70.294 | 53 | 12 | 49 | 12 | 255 | 28.8 | 11.19 | 158.32 | 12 | 5446087.865 |
| P21964 | Catechol O-methyltransferase OS=Homo sapiens OX=9606 GN=COMT PE=1 SV=2 | 0 | 70.227 | 56 | 10 | 60 | 10 | 271 | 30 | 5.47 | 168.08 | 10 | |
| P18124 | 60S ribosomal protein L7 OS=Homo sapiens OX=9606 GN=RPL7 PE=1 SV=1 | 0 | 70.094 | 44 | 13 | 118 | 13 | 248 | 29.2 | 10.65 | 255.26 | 13 | 1693887.403 |
| P02792 | Ferritin light chain OS=Homo sapiens OX=9606 GN=FTL PE=1 SV=2 | 0 | 70.035 | 56 | 11 | 35 | 11 | 175 | 20 | 5.78 | 131.74 | 11 | 1067407.774 |
| O94760 | N(G),N(G)-dimethylarginine dimethylaminohydrolase 1 OS=Homo sapiens OX=9606 GN=DDAH1 PE=1 SV=3 | 0 | 69.249 | 70 | 13 | 82 | 12 | 285 | 31.1 | 5.81 | 190.13 | 13 | 358873.1705 |
| P62701 | 40S ribosomal protein S4, X isoform OS=Homo sapiens OX=9606 GN=RPS4X PE=1 SV=2 | 0 | 68.963 | 53 | 16 | 155 | 16 | 263 | 29.6 | 10.15 | 353.01 | 16 | 2697700678 |
| P36969 | Phospholipid hydroperoxide glutathione peroxidase OS=Homo sapiens OX=9606 GN=GPX4 PE=1 SV=3 | 0 | 68.646 | 59 | 13 | 154 | 13 | 197 | 22.2 | 8.37 | 315.45 | 13 | |
| P05388 | 60S acidic ribosomal protein P0 OS=Homo sapiens OX=9606 GN=RPLP0 PE=1 SV=1 | 0 | 68.586 | 47 | 11 | 120 | 11 | 317 | 34.3 | 5.97 | 314.05 | 11 | 2310276.408 |
| Q13438 | Protein OS-9 OS=Homo sapiens OX=9606 GN=OS9 PE=1 SV=1 | 0 | 68.545 | 30 | 13 | 93 | 13 | 667 | 75.5 | 4.87 | 256.58 | 13 | 43709305.61 |
| P20742 | Pregnancy zone protein OS=Homo sapiens OX=9606 GN=PZP PE=1 SV=4 | 0 | 68.238 | 12 | 13 | 104 | 6 | 1482 | 163.8 | 6.38 | 263.85 | 13 | |
| Q10471 | Polypeptide N-acetylgalactosaminyltransferase 2 OS=Homo sapiens OX=9606 GN=GALNT2 PE=1 SV=1 | 0 | 68.012 | 39 | 17 | 102 | 17 | 571 | 64.7 | 8.35 | 151.73 | 17 | 4147252.248 |
| Q7Z4W1 | L-xylulose reductase OS=Homo sapiens OX=9606 GN=DCXR PE=1 SV=2 | 0 | 67.775 | 66 | 13 | 90 | 13 | 244 | 25.9 | 8.1 | 221.29 | 13 | 4935811.747 |
| P63000 | Ras-related C3 botulinum toxin substrate 1 OS=Homo sapiens OX=9606 GN=RAC1 PE=1 SV=1 | 0 | 67.686 | 60 | 11 | 266 | 6 | 192 | 21.4 | 8.5 | 609.01 | 11 | 2738310.606 |
| P79483 | HLA class II histocompatibility antigen, DR beta 3 chain OS=Homo sapiens OX=9606 GN=HLA-DRB3 PE=1 SV=1 | 0 | 67.658 | 39 | 7 | 93 | 3 | 266 | 29.9 | 8.56 | 325.48 | 7 | 1635717.29 |
| P50591 | Tumor necrosis factor ligand superfamily member 10 OS=Homo sapiens OX=9606 GN=TNFSF10 PE=1 SV=1 | 0 | 67.238 | 60 | 14 | 210 | 14 | 281 | 32.5 | 7.42 | 427.76 | 14 | 2133395.243 |
| P08754 | Guanine nucleotide-binding protein G(i) subunit alpha OS=Homo sapiens OX=9606 GN=GNAI3 PE=1 SV=3 | 0 | 67.164 | 37 | 11 | 127 | 3 | 354 | 40.5 | 5.69 | 310.95 | 11 | |
| Q04917 | 14-3-3 protein eta OS=Homo sapiens OX=9606 GN=YWHAH PE=1 SV=4 | 0 | 66.614 | 58 | 14 | 140 | 11 | 246 | 28.2 | 4.84 | 264.84 | 14 | 680247.6915 |
| P04350 | Tubulin beta-4A chain OS=Homo sapiens OX=9606 GN=TUBB4A PE=1 SV=2 | 0 | 66.241 | 42 | 12 | 128 | 1 | 444 | 49.6 | 4.88 | 322.66 | 12 | 2026017.539 |
| A0A0B4J1V1 | Immunoglobulin heavy variable 3-21 OS=Homo sapiens OX=9606 GN=IGHV3-21 PE=1 SV=1 | 0 | 65.946 | 63 | 8 | 179 | 2 | 117 | 12.8 | 8.28 | 494.06 | 8 | |
| Q86SQ4 | Adhesion G-protein coupled receptor G6 OS=Homo sapiens OX=9606 GN=ADGRG6 PE=1 SV=3 | 0 | 65.818 | 10 | 11 | 164 | 11 | 1221 | 136.6 | 7.87 | 411.02 | 11 | 13522510.16 |
| P01768 | Immunoglobulin heavy variable 3-30 OS=Homo sapiens OX=9606 GN=IGHV3-30 PE=1 SV=2 | 0 | 65.769 | 63 | 8 | 167 | 3 | 117 | 12.9 | 8.92 | 478.97 | 8 | 3278556.73 |
| P19827 | Inter-alpha-trypsin inhibitor heavy chain H1 OS=Homo sapiens OX=9606 GN=ITIH1 PE=1 SV=3 | 0 | 64.99 | 22 | 16 | 89 | 16 | 911 | 101.3 | 6.79 | 229.43 | 16 | 788532997.4 |
| P04439 | HLA class I histocompatibility antigen, A alpha chain OS=Homo sapiens OX=9606 GN=HLA-A PE=1 SV=2 | 0 | 64.819 | 35 | 9 | 99 | 3 | 365 | 40.8 | 6 | 305.98 | 9 | 3156962.735 |
| P48637 | Glutathione synthetase OS=Homo sapiens OX=9606 GN=GSS PE=1 SV=1 | 0 | 64.797 | 35 | 13 | 33 | 13 | 474 | 52.4 | 5.92 | 75.85 | 13 | 1450707.958 |
| P50990 | T-complex protein 1 subunit theta OS=Homo sapiens OX=9606 GN=CCT8 PE=1 SV=4 | 0 | 64.721 | 31 | 15 | 74 | 15 | 548 | 59.6 | 5.6 | 123.83 | 15 | 374311.2104 |
| P37837 | Transaldolase OS=Homo sapiens OX=9606 GN=TALDO1 PE=1 SV=2 | 0 | 64.689 | 35 | 13 | 41 | 13 | 337 | 37.5 | 6.81 | 106.52 | 13 | 787286.1796 |
| P40227 | T-complex protein 1 subunit zeta OS=Homo sapiens OX=9606 GN=CCT6A PE=1 SV=3 | 0 | 64.632 | 41 | 15 | 69 | 15 | 531 | 58 | 6.68 | 168.48 | 15 | 636491.3664 |
| P27824 | Calnexin OS=Homo sapiens OX=9606 GN=CANX PE=1 SV=2 | 0 | 64.6 | 29 | 12 | 61 | 12 | 592 | 67.5 | 4.6 | 156.38 | 12 | 1870611.527 |
| Q6X4U4 | Sclerostin domain-containing protein 1 OS=Homo sapiens OX=9606 GN=SOSTDC1 PE=1 SV=2 | 0 | 64.545 | 43 | 7 | 158 | 7 | 206 | 23.3 | 9.74 | 474.28 | 7 | 10966447.64 |
| P68431 | Histone H3.1 OS=Homo sapiens OX=9606 GN=H3C1 PE=1 SV=2 | 0 | 64.194 | 48 | 7 | 125 | 1 | 136 | 15.4 | 11.12 | 306.75 | 7 | 2272414.591 |
| P05387 | 60S acidic ribosomal protein P2 OS=Homo sapiens OX=9606 GN=RPLP2 PE=1 SV=1 | 0 | 63.919 | 70 | 5 | 73 | 5 | 115 | 11.7 | 4.54 | 281 | 5 | 1721475.534 |
| Q14118 | Dystroglycan OS=Homo sapiens OX=9606 GN=DAG1 PE=1 SV=2 | 0 | 63.816 | 20 | 11 | 164 | 11 | 895 | 97.4 | 8.56 | 368.64 | 11 | 10317916.01 |
| P16403 | Histone H1.2 OS=Homo sapiens OX=9606 GN=H1-2 PE=1 SV=2 | 0 | 63.677 | 31 | 10 | 43 | 1 | 213 | 21.4 | 10.93 | 102.28 | 10 | 1501226.716 |
| P37802 | Transgelin-2 OS=Homo sapiens OX=9606 GN=TAGLN2 PE=1 SV=3 | 0 | 63.446 | 68 | 10 | 77 | 10 | 199 | 22.4 | 8.25 | 193.03 | 10 | 102395.5494 |
| Q14764 | Major vault protein OS=Homo sapiens OX=9606 GN=MVP PE=1 SV=4 | 0 | 62.845 | 26 | 16 | 38 | 16 | 893 | 99.3 | 5.48 | 79.4 | 16 | |
| Q15365 | Poly(rC)-binding protein 1 OS=Homo sapiens OX=9606 GN=PCBP1 PE=1 SV=2 | 0 | 62.793 | 51 | 11 | 135 | 8 | 356 | 37.5 | 7.09 | 291.65 | 11 | 2316786.408 |
| P24158 | Myeloblastin OS=Homo sapiens OX=9606 GN=PRTN3 PE=1 SV=3 | 0 | 62.755 | 43 | 6 | 32 | 6 | 256 | 27.8 | 8.35 | 102.59 | 6 | 4805921.841 |
| Q9UHL4 | Dipeptidyl peptidase 2 OS=Homo sapiens OX=9606 GN=DPP7 PE=1 SV=3 | 0 | 62.669 | 38 | 12 | 100 | 12 | 492 | 54.3 | 6.32 | 195.06 | 12 | 1248902.488 |
| Q02878 | 60S ribosomal protein L6 OS=Homo sapiens OX=9606 GN=RPL6 PE=1 SV=3 | 0 | 62.074 | 41 | 12 | 155 | 12 | 288 | 32.7 | 10.58 | 225.04 | 12 | 399272.7367 |
| P78371 | T-complex protein 1 subunit beta OS=Homo sapiens OX=9606 GN=CCT2 PE=1 SV=4 | 0 | 61.922 | 33 | 13 | 74 | 13 | 535 | 57.5 | 6.46 | 171.8 | 13 | 4529360.609 |
| P01764 | Immunoglobulin heavy variable 3-23 OS=Homo sapiens OX=9606 GN=IGHV3-23 PE=1 SV=2 | 0 | 61.724 | 63 | 7 | 159 | 2 | 117 | 12.6 | 8.28 | 404.15 | 7 | 30538214.11 |
| P30040 | Endoplasmic reticulum resident protein 29 OS=Homo sapiens OX=9606 GN=ERP29 PE=1 SV=4 | 0 | 61.537 | 53 | 10 | 72 | 10 | 261 | 29 | 7.31 | 176.23 | 10 | |
| Q14376 | UDP-glucose 4-epimerase OS=Homo sapiens OX=9606 GN=GALE PE=1 SV=2 | 0 | 61.501 | 55 | 13 | 24 | 13 | 348 | 38.3 | 6.73 | 68.12 | 13 | |
| Q9HAT2 | Sialate O-acetylesterase OS=Homo sapiens OX=9606 GN=SIAE PE=1 SV=1 | 0 | 61.234 | 35 | 9 | 57 | 9 | 523 | 58.3 | 7.33 | 148.79 | 9 | 1044831.791 |
| Q16777 | Histone H2A type 2-C OS=Homo sapiens OX=9606 GN=H2AC20 PE=1 SV=4 | 0 | 60.852 | 58 | 7 | 98 | 3 | 129 | 14 | 10.9 | 267.75 | 7 | 7433499.817 |
| Q9Y6B6 | GTP-binding protein SAR1b OS=Homo sapiens OX=9606 GN=SAR1B PE=1 SV=1 | 0 | 60.603 | 77 | 11 | 105 | 6 | 198 | 22.4 | 6.11 | 265.39 | 11 | |
| P31151 | Protein S100-A7 OS=Homo sapiens OX=9606 GN=S100A7 PE=1 SV=4 | 0 | 60.377 | 58 | 9 | 31 | 9 | 101 | 11.5 | 6.77 | 97.2 | 9 | 16432517.3 |
| Q08345 | Epithelial discoidin domain-containing receptor 1 OS=Homo sapiens OX=9606 GN=DDR1 PE=1 SV=1 | 0 | 60.181 | 16 | 12 | 197 | 12 | 913 | 101.1 | 6.83 | 407.48 | 12 | 4455742.899 |
| P01782 | Immunoglobulin heavy variable 3-9 OS=Homo sapiens OX=9606 GN=IGHV3-9 PE=1 SV=2 | 0 | 59.881 | 63 | 7 | 171 | 3 | 118 | 12.9 | 7.08 | 572.36 | 7 | 10363978.2 |
| P13693 | Translationally-controlled tumor protein OS=Homo sapiens OX=9606 GN=TPT1 PE=1 SV=1 | 0 | 59.85 | 60 | 7 | 96 | 7 | 172 | 19.6 | 4.93 | 283.25 | 7 | 613168.4202 |
| P20339 | Ras-related protein Rab-5A OS=Homo sapiens OX=9606 GN=RAB5A PE=1 SV=2 | 0 | 59.725 | 48 | 7 | 116 | 4 | 215 | 23.6 | 8.15 | 403.29 | 7 | 598140.6942 |
| P62979 | Ubiquitin-40S ribosomal protein S27a OS=Homo sapiens OX=9606 GN=RPS27A PE=1 SV=2 | 0 | 59.687 | 56 | 9 | 147 | 9 | 156 | 18 | 9.64 | 381.11 | 9 | 21231575.57 |
| Q96JB6 | Lysyl oxidase homolog 4 OS=Homo sapiens OX=9606 GN=LOXL4 PE=1 SV=1 | 0 | 59.469 | 28 | 14 | 23 | 14 | 756 | 84.4 | 7.46 | 52.5 | 14 | 341785.136 |
| P62753 | 40S ribosomal protein S6 OS=Homo sapiens OX=9606 GN=RPS6 PE=1 SV=1 | 0 | 59.436 | 34 | 8 | 75 | 8 | 249 | 28.7 | 10.84 | 216.36 | 8 | 604617.207 |
| P22392 | Nucleoside diphosphate kinase B OS=Homo sapiens OX=9606 GN=NME2 PE=1 SV=1 | 0 | 59.369 | 61 | 10 | 169 | 5 | 152 | 17.3 | 8.41 | 352.82 | 10 | 1075636.579 |
| P02042 | Hemoglobin subunit delta OS=Homo sapiens OX=9606 GN=HBD PE=1 SV=2 | 0 | 59.24 | 65 | 10 | 46 | 3 | 147 | 16 | 8.05 | 149.46 | 10 | |
| P62263 | 40S ribosomal protein S14 OS=Homo sapiens OX=9606 GN=RPS14 PE=1 SV=3 | 0 | 59.179 | 67 | 8 | 66 | 8 | 151 | 16.3 | 10.05 | 128.41 | 8 | 100866.8193 |
| P15880 | 40S ribosomal protein S2 OS=Homo sapiens OX=9606 GN=RPS2 PE=1 SV=2 | 0 | 59.068 | 40 | 9 | 95 | 9 | 293 | 31.3 | 10.24 | 231.36 | 9 | 651102.7584 |
| P25815 | Protein S100-P OS=Homo sapiens OX=9606 GN=S100P PE=1 SV=2 | 0 | 58.953 | 76 | 5 | 28 | 5 | 95 | 10.4 | 4.88 | 101.8 | 5 | 871065.396 |
| P67936 | Tropomyosin alpha-4 chain OS=Homo sapiens OX=9606 GN=TPM4 PE=1 SV=3 | 0 | 58.915 | 44 | 9 | 79 | 4 | 248 | 28.5 | 4.69 | 158.33 | 9 | 1041966.002 |
| P14780 | Matrix metalloproteinase-9 OS=Homo sapiens OX=9606 GN=MMP9 PE=1 SV=3 | 0 | 58.86 | 28 | 14 | 21 | 14 | 707 | 78.4 | 6.06 | 57.7 | 14 | |
| P01824 | Immunoglobulin heavy variable 4-39 OS=Homo sapiens OX=9606 GN=IGHV4-39 PE=1 SV=2 | 0 | 58.535 | 43 | 6 | 139 | 1 | 125 | 13.9 | 9.26 | 363.16 | 6 | |

Table S1. Cont.

| Accession | Description | Exp. q-value: Combined | Sum PEP Score | Coverage [%] | # Peptides | # PSMs | # Unique Peptides | # AAs | MW [kDa] | calc. pI | Score Sequest HT: Sequest HT | # Peptides (by Search Engine): Sequest HT |
|---|---|---|---|---|---|---|---|---|---|---|---|---|
| Q9NRV9 | Heme-binding protein 1 OS=Homo sapiens OX=9606 GN=HEBP1 PE=1 SV=1 | 0 | 58.418 | 59 | 8 | 101 | 8 | 189 | 21.1 | 5.8 | 307.64 | 8 |
| P51148 | Ras-related protein Rab-5C OS=Homo sapiens OX=9606 GN=RAB5C PE=1 SV=2 | 0 | 58.221 | 54 | 9 | 107 | 6 | 216 | 23.5 | 8.41 | 309.17 | 9 |
| Q15907 | Ras-related protein Rab-11B OS=Homo sapiens OX=9606 GN=RAB11B PE=1 SV=4 | 0 | 57.456 | 60 | 12 | 148 | 11 | 218 | 24.5 | 5.94 | 303.64 | 12 |
| Q9BWD1 | Acetyl-CoA acetyltransferase, cytosolic OS=Homo sapiens OX=9606 GN=ACAT2 PE=1 SV=2 | 0 | 57.436 | 39 | 9 | 34 | 9 | 397 | 41.3 | 6.92 | 87.87 | 9 |
| P34096 | Ribonuclease 4 OS=Homo sapiens OX=9606 GN=RNASE4 PE=1 SV=3 | 0 | 56.874 | 40 | 8 | 97 | 8 | 147 | 16.8 | 9.03 | 295.58 | 8 |
| P17987 | T-complex protein 1 subunit alpha OS=Homo sapiens OX=9606 GN=TCP1 PE=1 SV=1 | 0 | 56.735 | 33 | 12 | 47 | 12 | 556 | 60.3 | 6.11 | 89.61 | 12 |
| P46777 | 60S ribosomal protein L5 OS=Homo sapiens OX=9606 GN=RPL5 PE=1 SV=3 | 0 | 56.716 | 41 | 10 | 65 | 10 | 297 | 34.3 | 9.72 | 121.46 | 10 |
| Q9Y646 | Carboxypeptidase Q OS=Homo sapiens OX=9606 GN=CPQ PE=1 SV=1 | 0 | 56.617 | 32 | 11 | 90 | 11 | 472 | 51.9 | 6.18 | 187.47 | 11 |
| A0M8Q6 | Immunoglobulin lambda constant 7 OS=Homo sapiens OX=9606 GN=IGLC7 PE=1 SV=3 | 0 | 56.606 | 56 | 5 | 311 | 1 | 106 | 11.2 | 8.29 | 1002.18 | 5 |
| P01762 | Immunoglobulin heavy variable 3-11 OS=Homo sapiens OX=9606 GN=IGHV3-11 PE=1 SV=2 | 0 | 56.525 | 63 | 8 | 172 | 2 | 117 | 12.9 | 8.6 | 475.24 | 8 |
| O15498 | Synaptobrevin homolog YKT6 OS=Homo sapiens OX=9606 GN=YKT6 PE=1 SV=1 | 0 | 56.35 | 64 | 12 | 80 | 12 | 198 | 22.4 | 6.92 | 148.77 | 12 |
| O75882 | Attractin OS=Homo sapiens OX=9606 GN=ATRN PE=1 SV=2 | 0 | 56.275 | 12 | 14 | 81 | 14 | 1429 | 158.4 | 7.31 | 174.44 | 14 |
| Q96FQ6 | Protein S100-A16 OS=Homo sapiens OX=9606 GN=S100A16 PE=1 SV=1 | 0 | 56.269 | 78 | 8 | 73 | 8 | 103 | 11.8 | 6.79 | 189.7 | 8 |
| Q01814 | Plasma membrane calcium-transporting ATPase 2 OS=Homo sapiens OX=9606 GN=ATP2B2 PE=1 SV=2 | 0 | 56.16 | 13 | 12 | 46 | 12 | 1243 | 136.8 | 5.91 | 75.09 | 12 |
| P18621 | 60S ribosomal protein L17 OS=Homo sapiens OX=9606 GN=RPL17 PE=1 SV=3 | 0 | 56.113 | 50 | 9 | 52 | 9 | 184 | 21.4 | 10.17 | 137.82 | 9 |
| P11216 | Glycogen phosphorylase, brain form OS=Homo sapiens OX=9606 GN=PYGB PE=1 SV=5 | 0 | 55.975 | 17 | 12 | 33 | 7 | 843 | 96.6 | 6.86 | 69.99 | 12 |
| P46782 | 40S ribosomal protein S5 OS=Homo sapiens OX=9606 GN=RPS5 PE=1 SV=4 | 0 | 55.627 | 61 | 11 | 62 | 11 | 204 | 22.9 | 9.72 | 151.74 | 11 |
| P84085 | ADP-ribosylation factor 5 OS=Homo sapiens OX=9606 GN=ARF5 PE=1 SV=2 | 0 | 55.493 | 59 | 10 | 83 | 4 | 180 | 20.5 | 6.79 | 250.61 | 10 |
| P07108 | Acyl-CoA-binding protein OS=Homo sapiens OX=9606 GN=DBI PE=1 SV=2 | 0 | 55.412 | 79 | 6 | 81 | 6 | 87 | 10 | 6.57 | 254.11 | 6 |
| P01763 | Immunoglobulin heavy variable 3-48 OS=Homo sapiens OX=9606 GN=IGHV3-48 PE=1 SV=2 | 0 | 55.228 | 63 | 8 | 165 | 2 | 117 | 12.8 | 5.36 | 448.69 | 8 |
| P11215 | Integrin alpha-M OS=Homo sapiens OX=9606 GN=ITGAM PE=1 SV=2 | 0 | 54.929 | 14 | 13 | 19 | 13 | 1152 | 127.1 | 7.23 | 59.56 | 13 |
| P60033 | CD81 antigen OS=Homo sapiens OX=9606 GN=CD81 PE=1 SV=1 | 0 | 54.82 | 25 | 3 | 98 | 3 | 236 | 25.8 | 5.29 | 555.75 | 3 |
| Q6UWW8 | Carboxylesterase 3 OS=Homo sapiens OX=9606 GN=CES3 PE=1 SV=1 | 0 | 54.529 | 31 | 10 | 48 | 10 | 571 | 62.2 | 5.62 | 111.04 | 10 |
| Q9H8H3 | Methyltransferase-like protein 7A OS=Homo sapiens OX=9606 GN=METTL7A PE=1 SV=1 | 0 | 54.524 | 47 | 8 | 115 | 8 | 244 | 28.3 | 8.38 | 297.37 | 8 |
| P30046 | D-dopachrome decarboxylase OS=Homo sapiens OX=9606 GN=DDT PE=1 SV=3 | 0 | 53.828 | 90 | 8 | 54 | 8 | 118 | 12.7 | 7.3 | 134.95 | 8 |
| P25787 | Proteasome subunit alpha type-2 OS=Homo sapiens OX=9606 GN=PSMA2 PE=1 SV=2 | 0 | 53.462 | 32 | 6 | 42 | 6 | 234 | 25.9 | 7.43 | 115.43 | 6 |
| P36269 | Glutathione hydrolase 5 proenzyme OS=Homo sapiens OX=9606 GN=GGT5 PE=1 SV=2 | 0 | 53.153 | 22 | 9 | 25 | 9 | 586 | 62.2 | 7.55 | 86.65 | 9 |
| Q9H4A4 | Aminopeptidase B OS=Homo sapiens OX=9606 GN=RNPEP PE=1 SV=2 | 0 | 53.124 | 27 | 13 | 32 | 13 | 650 | 72.5 | 5.74 | 55.44 | 13 |
| Q92520 | Protein FAM3C OS=Homo sapiens OX=9606 GN=FAM3C PE=1 SV=1 | 0 | 53.103 | 51 | 10 | 107 | 10 | 227 | 24.7 | 8.29 | 268.79 | 10 |
| P0DP08 | Immunoglobulin heavy variable 4-38-2 OS=Homo sapiens OX=9606 GN=IGHV4-38-2 PE=3 SV=1 | 0 | 53.075 | 46 | 6 | 138 | 1 | 117 | 13 | 9.29 | 356.61 | 6 |
| Q9Y2J8 | Protein-arginine deiminase type-2 OS=Homo sapiens OX=9606 GN=PADI2 PE=1 SV=2 | 0 | 52.98 | 22 | 11 | 95 | 11 | 665 | 75.5 | 5.59 | 228.2 | 11 |
| Q96DA0 | Zymogen granule protein 16 homolog B OS=Homo sapiens OX=9606 GN=ZG16B PE=1 SV=3 | 0 | 52.928 | 50 | 8 | 147 | 8 | 208 | 22.7 | 7.39 | 464.85 | 8 |
| P04179 | Superoxide dismutase [Mn], mitochondrial OS=Homo sapiens OX=9606 GN=SOD2 PE=1 SV=3 | 0 | 52.817 | 61 | 10 | 67 | 10 | 222 | 24.7 | 8.25 | 191.09 | 10 |
| P62917 | 60S ribosomal protein L8 OS=Homo sapiens OX=9606 GN=RPL8 PE=1 SV=2 | 0 | 52.609 | 44 | 10 | 107 | 10 | 257 | 28 | 11.03 | 164.58 | 10 |
| P09972 | Fructose-bisphosphate aldolase C OS=Homo sapiens OX=9606 GN=ALDOC PE=1 SV=2 | 0 | 52.439 | 25 | 6 | 69 | 4 | 364 | 39.4 | 6.87 | 206.36 | 6 |
| A0A0B4J1X5 | Immunoglobulin heavy variable 3-74 OS=Homo sapiens OX=9606 GN=IGHV3-74 PE=3 SV=1 | 0 | 52.397 | 68 | 9 | 208 | 4 | 117 | 12.8 | 8.66 | 520.76 | 9 |
| P35749 | Myosin-11 OS=Homo sapiens OX=9606 GN=MYH11 PE=1 SV=3 | 0 | 52.389 | 6 | 10 | 68 | 1 | 1972 | 227.2 | 5.5 | 169.36 | 10 |
| P31947 | 14-3-3 protein sigma OS=Homo sapiens OX=9606 GN=SFN PE=1 SV=1 | 0 | 52.359 | 42 | 8 | 120 | 5 | 248 | 27.8 | 4.74 | 267.32 | 8 |
| P01825 | Immunoglobulin heavy variable 4-59 OS=Homo sapiens OX=9606 GN=IGHV4-59 PE=1 SV=2 | 0 | 52.231 | 47 | 6 | 137 | 1 | 116 | 12.9 | 9.29 | 352.43 | 6 |
| P61006 | Ras-related protein Rab-8A OS=Homo sapiens OX=9606 GN=RAB8A PE=1 SV=1 | 0 | 52.221 | 53 | 11 | 102 | 3 | 207 | 23.7 | 9.07 | 207.11 | 11 |
| Q9BR76 | Coronin-1B OS=Homo sapiens OX=9606 GN=CORO1B PE=1 SV=1 | 0 | 52.217 | 35 | 13 | 59 | 13 | 489 | 54.2 | 5.88 | 89.65 | 13 |
| P10768 | S-formylglutathione hydrolase OS=Homo sapiens OX=9606 GN=ESD PE=1 SV=2 | 0 | 52.176 | 60 | 10 | 48 | 10 | 282 | 31.4 | 7.02 | 123.29 | 10 |
| P22352 | Glutathione peroxidase 3 OS=Homo sapiens OX=9606 GN=GPX3 PE=1 SV=2 | 0 | 52.036 | 48 | 9 | 139 | 9 | 226 | 25.5 | 8.13 | 351.59 | 9 |
| P62841 | 40S ribosomal protein S15 OS=Homo sapiens OX=9606 GN=RPS15 PE=1 SV=2 | 0 | 52.014 | 56 | 5 | 32 | 5 | 145 | 17 | 10.39 | 119.11 | 5 |
| Q9NZH0 | G-protein coupled receptor family C group 5 member B OS=Homo sapiens OX=9606 GN=GPRC5B PE=2 SV=2 | 0 | 51.948 | 15 | 5 | 113 | 5 | 403 | 44.8 | 8.24 | 324.67 | 5 |
| P09211 | Glutathione S-transferase P OS=Homo sapiens OX=9606 GN=GSTP1 PE=1 SV=2 | 0 | 51.575 | 61 | 9 | 62 | 9 | 210 | 23.3 | 5.64 | 207.2 | 9 |
| P55058 | Phospholipid transfer protein OS=Homo sapiens OX=9606 GN=PLTP PE=1 SV=1 | 0 | 51.422 | 29 | 8 | 55 | 8 | 493 | 54.7 | 7.01 | 146.99 | 8 |
| Q99653 | Calcineurin B homologous protein 1 OS=Homo sapiens OX=9606 GN=CHP1 PE=1 SV=3 | 0 | 51.417 | 58 | 9 | 56 | 9 | 195 | 22.4 | 5.1 | 110.59 | 9 |
| Q9GZM7 | Tubulointerstitial nephritis antigen-like OS=Homo sapiens OX=9606 GN=TINAGL1 PE=1 SV=1 | 0 | 51.395 | 22 | 8 | 42 | 8 | 467 | 52.4 | 6.99 | 108.52 | 8 |
| Q99519 | Sialidase-1 OS=Homo sapiens OX=9606 GN=NEU1 PE=1 SV=1 | 0 | 51.351 | 37 | 11 | 114 | 11 | 415 | 45.4 | 5.88 | 223.14 | 11 |
| O15144 | Actin-related protein 2/3 complex subunit 2 OS=Homo sapiens OX=9606 GN=ARPC2 PE=1 SV=1 | 0 | 51.063 | 52 | 12 | 26 | 12 | 300 | 34.3 | 7.36 | 43.4 | 12 |
| Q02487 | Desmocollin-2 OS=Homo sapiens OX=9606 GN=DSC2 PE=1 SV=1 | 0 | 51.006 | 7 | 7 | 56 | 7 | 901 | 99.9 | 5.34 | 158.62 | 7 |
| Q9Y5P6 | Mannose-1-phosphate guanyltransferase beta OS=Homo sapiens OX=9606 GN=GMPPB PE=1 SV=2 | 0 | 50.933 | 44 | 10 | 54 | 10 | 360 | 39.8 | 6.61 | 129.68 | 10 |
| P01767 | Immunoglobulin heavy variable 3-53 OS=Homo sapiens OX=9606 GN=IGHV3-53 PE=1 SV=2 | 0 | 50.852 | 63 | 7 | 184 | 1 | 116 | 12.8 | 8.28 | 439 | 7 |
| Q9NZM1 | Myoferlin OS=Homo sapiens OX=9606 GN=MYOF PE=1 SV=1 | 0 | 50.778 | 8 | 15 | 49 | 14 | 2061 | 234.6 | 6.18 | 76.23 | 15 |
| Q9H772 | Gremlin-2 OS=Homo sapiens OX=9606 GN=GREM2 PE=1 SV=1 | 0 | 50.641 | 60 | 7 | 73 | 7 | 168 | 19.3 | 9.17 | 223.52 | 7 |
| P02760 | Protein AMBP OS=Homo sapiens OX=9606 GN=AMBP PE=1 SV=1 | 0 | 50.585 | 33 | 12 | 75 | 12 | 352 | 39 | 6.25 | 174.4 | 12 |
| A0A0A0MS15 | Immunoglobulin heavy variable 3-49 OS=Homo sapiens OX=9606 GN=IGHV3-49 PE=3 SV=1 | 0 | 50.549 | 68 | 11 | 138 | 7 | 119 | 13 | 8.62 | 337.06 | 11 |
| P52790 | Hexokinase-3 OS=Homo sapiens OX=9606 GN=HK3 PE=1 SV=2 | 0 | 50.166 | 19 | 11 | 14 | 11 | 923 | 99 | 5.4 | 41.77 | 11 |
| O75355 | Ectonucleoside triphosphate diphosphohydrolase 3 OS=Homo sapiens OX=9606 GN=ENTPD3 PE=1 SV=2 | 0 | 49.953 | 33 | 11 | 81 | 11 | 529 | 59.1 | 6.42 | 169.68 | 11 |
| P61160 | Actin-related protein 2 OS=Homo sapiens OX=9606 GN=ACTR2 PE=1 SV=1 | 0 | 49.895 | 40 | 11 | 25 | 11 | 394 | 44.7 | 6.74 | 61.76 | 11 |
| O60266 | Adenylate cyclase type 3 OS=Homo sapiens OX=9606 GN=ADCY3 PE=1 SV=3 | 0 | 49.829 | 13 | 12 | 41 | 12 | 1144 | 128.9 | 6.58 | 100.56 | 12 |
| P08572 | Collagen alpha-2(IV) chain OS=Homo sapiens OX=9606 GN=COL4A2 PE=1 SV=4 | 0 | 49.582 | 9 | 10 | 51 | 10 | 1712 | 167.4 | 8.66 | 112.68 | 10 |
| P04066 | Tissue alpha-L-fucosidase OS=Homo sapiens OX=9606 GN=FUCA1 PE=1 SV=4 | 0 | 49.352 | 36 | 10 | 86 | 10 | 466 | 53.7 | 6.84 | 227.18 | 10 |
| P49913 | Cathelicidin antimicrobial peptide OS=Homo sapiens OX=9606 GN=CAMP PE=1 SV=1 | 0 | 49.251 | 35 | 9 | 23 | 9 | 170 | 19.3 | 9.41 | 66.68 | 9 |
| P52907 | F-actin-capping protein subunit alpha-1 OS=Homo sapiens OX=9606 GN=CAPZA1 PE=1 SV=3 | 0 | 49.155 | 60 | 9 | 30 | 7 | 286 | 32.9 | 5.69 | 75.31 | 9 |
| P10321 | HLA class I histocompatibility antigen, C alpha chain OS=Homo sapiens OX=9606 GN=HLA-C PE=1 SV=3 | 0 | 48.826 | 34 | 9 | 52 | 3 | 366 | 40.6 | 6.04 | 156.87 | 9 |
| P06703 | Protein S100-A6 OS=Homo sapiens OX=9606 GN=S100A6 PE=1 SV=1 | 0 | 48.786 | 81 | 8 | 60 | 8 | 90 | 10.2 | 5.48 | 158.27 | 8 |
| P61026 | Ras-related protein Rab-10 OS=Homo sapiens OX=9606 GN=RAB10 PE=1 SV=1 | 0 | 48.686 | 52 | 11 | 95 | 7 | 200 | 22.5 | 8.38 | 190.83 | 11 |
| P58499 | Protein FAM3B OS=Homo sapiens OX=9606 GN=FAM3B PE=1 SV=2 | 0 | 48.685 | 54 | 11 | 144 | 11 | 235 | 26 | 8.75 | 295.12 | 11 |
| P13489 | Ribonuclease inhibitor OS=Homo sapiens OX=9606 GN=RNH1 PE=1 SV=2 | 0 | 48.599 | 41 | 10 | 20 | 10 | 461 | 49.9 | 4.82 | 51.91 | 10 |
| O43278 | Kunitz-type protease inhibitor 1 OS=Homo sapiens OX=9606 GN=SPINT1 PE=1 SV=2 | 0 | 48.436 | 26 | 13 | 195 | 13 | 529 | 58.4 | 6.29 | 426.94 | 13 |
| Q8WVQ1 | Soluble calcium-activated nucleotidase 1 OS=Homo sapiens OX=9606 GN=CANT1 PE=1 SV=1 | 0 | 48.424 | 38 | 10 | 44 | 10 | 401 | 44.8 | 6.09 | 111.48 | 10 |
| P20700 | Lamin-B1 OS=Homo sapiens OX=9606 GN=LMNB1 PE=1 SV=2 | 0 | 48.401 | 20 | 11 | 13 | 11 | 586 | 66.4 | 5.16 | 49.11 | 11 |
| O00462 | Beta-mannosidase OS=Homo sapiens OX=9606 GN=MANBA PE=1 SV=3 | 0 | 48.198 | 20 | 14 | 67 | 13 | 879 | 100.8 | 5.52 | 82.19 | 14 |
| P02462 | Collagen alpha-1(IV) chain OS=Homo sapiens OX=9606 GN=COL4A1 PE=1 SV=4 | 0 | 48.054 | 4 | 6 | 31 | 6 | 1669 | 160.5 | 8.28 | 71.36 | 6 |
| Q9BS26 | Endoplasmic reticulum resident protein 44 OS=Homo sapiens OX=9606 GN=ERP44 PE=1 SV=1 | 0 | 47.968 | 30 | 9 | 30 | 8 | 406 | 46.9 | 5.26 | 58.41 | 9 |
| P22676 | Calretinin OS=Homo sapiens OX=9606 GN=CALB2 PE=2 SV=2 | 0 | 47.968 | 44 | 9 | 41 | 9 | 271 | 31.5 | 5.15 | 103.64 | 9 |
| P62241 | 40S ribosomal protein S8 OS=Homo sapiens OX=9606 GN=RPS8 PE=1 SV=2 | 0 | 47.885 | 46 | 9 | 97 | 9 | 208 | 24.2 | 10.32 | 176.71 | 9 |
| P28838 | Cytosol aminopeptidase OS=Homo sapiens OX=9606 GN=LAP3 PE=1 SV=3 | 0 | 47.879 | 22 | 9 | 32 | 9 | 519 | 56.1 | 7.93 | 76.19 | 9 |
| P19105 | Myosin regulatory light chain 12A OS=Homo sapiens OX=9606 GN=MYL12A PE=1 SV=2 | 0 | 47.788 | 64 | 8 | 51 | 1 | 171 | 19.8 | 4.81 | 138.9 | 8 |
| P18428 | Lipopolysaccharide-binding protein OS=Homo sapiens OX=9606 GN=LBP PE=1 SV=3 | 0 | 47.753 | 26 | 8 | 21 | 8 | 481 | 53.4 | 6.7 | 63.48 | 8 |
| P54289 | Voltage-dependent calcium channel subunit alpha-2/delta-1 OS=Homo sapiens OX=9606 GN=CACNA2D1 PE=1 SV=3 | 0 | 47.668 | 15 | 11 | 65 | 11 | 1103 | 124.5 | 5.27 | 111.15 | 11 |
| P14324 | Farnesyl pyrophosphate synthase OS=Homo sapiens OX=9606 GN=FDPS PE=1 SV=4 | 0 | 47.524 | 26 | 8 | 72 | 8 | 419 | 48.2 | 6.15 | 157.36 | 8 |
| P13497 | Bone morphogenetic protein 1 OS=Homo sapiens OX=9606 GN=BMP1 PE=1 SV=2 | 0 | 47.465 | 21 | 16 | 33 | 16 | 986 | 111.2 | 6.9 | 50.74 | 16 |
| P47756 | F-actin-capping protein subunit beta OS=Homo sapiens OX=9606 GN=CAPZB PE=1 SV=4 | 0 | 47.444 | 41 | 10 | 75 | 10 | 277 | 31.3 | 5.59 | 180.28 | 10 |
| Q16778 | Histone H2B type 2-E OS=Homo sapiens OX=9606 GN=H2BC21 PE=1 SV=3 | 0 | 47.376 | 40 | 7 | 112 | 3 | 126 | 13.9 | 10.32 | 349.89 | 7 |
| Q16651 | Prostasin OS=Homo sapiens OX=9606 GN=PRSS8 PE=1 SV=1 | 0 | 47.362 | 41 | 7 | 68 | 7 | 343 | 36.4 | 5.85 | 338.35 | 7 |
| P16401 | Histone H1.5 OS=Homo sapiens OX=9606 GN=H1-5 PE=1 SV=3 | 0 | 47.188 | 30 | 9 | 21 | 8 | 226 | 22.6 | 10.92 | 49.4 | 9 |
| P26641 | Elongation factor 1-gamma OS=Homo sapiens OX=9606 GN=EEF1G PE=1 SV=3 | 0 | 47.175 | 35 | 9 | 66 | 9 | 437 | 50.1 | 6.67 | 171.47 | 9 |
| Q9HDC9 | Adipocyte plasma membrane-associated protein OS=Homo sapiens OX=9606 GN=APMAP PE=1 SV=2 | 0 | 47.101 | 39 | 12 | 35 | 12 | 416 | 46.5 | 6.16 | 72.41 | 12 |
| Q9NR19 | Acetyl-coenzyme A synthetase, cytoplasmic OS=Homo sapiens OX=9606 GN=ACSS2 PE=1 SV=1 | 0 | 46.965 | 25 | 12 | 74 | 12 | 701 | 78.5 | 6.46 | 84.96 | 12 |
| P02753 | Retinol-binding protein 4 OS=Homo sapiens OX=9606 GN=RBP4 PE=1 SV=3 | 0 | 46.953 | 62 | 7 | 46 | 7 | 201 | 23 | 6.07 | 146.44 | 7 |
| P10599 | Thioredoxin OS=Homo sapiens OX=9606 GN=TXN PE=1 SV=3 | 0 | 46.82 | 89 | 8 | 102 | 8 | 105 | 11.7 | 4.92 | 335.58 | 8 |
| P20160 | Azurocidin OS=Homo sapiens OX=9606 GN=AZU1 PE=1 SV=3 | 0 | 46.793 | 53 | 8 | 32 | 8 | 251 | 26.9 | 9.5 | 91.06 | 8 |
| Q03591 | Complement factor H-related protein 1 OS=Homo sapiens OX=9606 GN=CFHR1 PE=1 SV=2 | 0 | 46.784 | 39 | 9 | 116 | 3 | 330 | 37.6 | 7.39 | 249.18 | 9 |
| P27635 | 60S ribosomal protein L10 OS=Homo sapiens OX=9606 GN=RPL10 PE=1 SV=4 | 0 | 46.754 | 38 | 6 | 64 | 6 | 214 | 24.6 | 10.08 | 124.31 | 6 |
| Q8WUH6 | Transmembrane protein 263 OS=Homo sapiens OX=9606 GN=TMEM263 PE=1 SV=1 | 0 | 46.631 | 58 | 5 | 95 | 5 | 116 | 11.7 | 9.32 | 354.82 | 5 |
| P48643 | T-complex protein 1 subunit epsilon OS=Homo sapiens OX=9606 GN=CCT5 PE=1 SV=1 | 0 | 46.571 | 25 | 10 | 43 | 10 | 541 | 59.6 | 5.66 | 143.68 | 10 |
| P01893 | Putative HLA class I histocompatibility antigen, alpha chain H OS=Homo sapiens OX=9606 GN=HLA-H PE=5 SV=3 | 0 | 46.521 | 30 | 7 | 59 | 1 | 362 | 40.9 | 6.3 | 187.31 | 7 |
| P18206 | Vinculin OS=Homo sapiens OX=9606 GN=VCL PE=1 SV=4 | 0 | 46.447 | 14 | 10 | 11 | 10 | 1134 | 123.7 | 5.66 | 34.3 | 10 |
| P15153 | Ras-related C3 botulinum toxin substrate 2 OS=Homo sapiens OX=9606 GN=RAC2 PE=1 SV=1 | 0 | 46.33 | 43 | 9 | 115 | 4 | 192 | 21.4 | 7.61 | 286.23 | 9 |
| P01889 | HLA class I histocompatibility antigen, B alpha chain OS=Homo sapiens OX=9606 GN=HLA-B PE=1 SV=3 | 0 | 46.179 | 29 | 7 | 58 | 1 | 362 | 40.4 | 5.85 | 169.55 | 7 |
| P19875 | C-X-C motif chemokine 2 OS=Homo sapiens OX=9606 GN=CXCL2 PE=1 SV=1 | 0 | 46.179 | 39 | 4 | 95 | 4 | 107 | 11.4 | 10.55 | 333.35 | 4 |
| P01817 | Immunoglobulin heavy variable 2-5 OS=Homo sapiens OX=9606 GN=IGHV2-5 PE=1 SV=2 | 0 | 46.115 | 66 | 6 | 36 | 6 | 119 | 13.2 | 8.29 | 127.58 | 6 |
| P60900 | Proteasome subunit alpha type-6 OS=Homo sapiens OX=9606 GN=PSMA6 PE=1 SV=1 | 0 | 45.926 | 46 | 9 | 57 | 9 | 246 | 27.4 | 6.76 | 121.73 | 9 |
| P62081 | 40S ribosomal protein S7 OS=Homo sapiens OX=9606 GN=RPS7 PE=1 SV=1 | 0 | 45.696 | 58 | 11 | 100 | 11 | 194 | 22.1 | 10.1 | 163.95 | 11 |
| O15540 | Fatty acid-binding protein, brain OS=Homo sapiens OX=9606 GN=FABP7 PE=1 SV=3 | 0 | 45.586 | 85 | 8 | 72 | 8 | 132 | 14.9 | 5.59 | 143.89 | 8 |
| O60568 | Multifunctional procollagen lysine hydroxylase and glycosyltransferase LH3 OS=Homo sapiens OX=9606 GN=PLOD3 PE=1 SV=1 | 0 | 45.56 | 20 | 11 | 17 | 11 | 738 | 84.7 | 6.05 | 36.95 | 11 |
| P09525 | Annexin A4 OS=Homo sapiens OX=9606 GN=ANXA4 PE=1 SV=4 | 0 | 45.309 | 43 | 11 | 57 | 11 | 319 | 35.9 | 6.13 | 133.99 | 11 |
| O14950 | Myosin regulatory light chain 12B OS=Homo sapiens OX=9606 GN=MYL12B PE=1 SV=2 | 0 | 45.158 | 63 | 8 | 51 | 1 | 172 | 19.8 | 4.84 | 137.79 | 8 |
| P29692 | Elongation factor 1-delta OS=Homo sapiens OX=9606 GN=EEF1D PE=1 SV=5 | 0 | 45.11 | 42 | 8 | 63 | 8 | 281 | 31.1 | 5.01 | 108.84 | 8 |
| P01619 | Immunoglobulin kappa variable 3-20 OS=Homo sapiens OX=9606 GN=IGKV3-20 PE=1 SV=2 | 0 | 45.043 | 47 | 4 | 88 | 2 | 116 | 12.5 | 4.96 | 318.91 | 4 |
| Q30154 | HLA class II histocompatibility antigen, DR beta 5 chain OS=Homo sapiens OX=9606 GN=HLA-DRB5 PE=1 SV=1 | 0 | 44.993 | 41 | 7 | 91 | 2 | 266 | 30 | 6.93 | 325.77 | 7 |
| A0A0C4DH42 | Immunoglobulin heavy variable 3-66 OS=Homo sapiens OX=9606 GN=IGHV3-66 PE=3 SV=1 | 0 | 44.925 | 63 | 7 | 184 | 1 | 116 | 12.7 | 8.16 | 435.35 | 7 |
| P28070 | Proteasome subunit beta type-4 OS=Homo sapiens OX=9606 GN=PSMB4 PE=1 SV=4 | 0 | 44.905 | 42 | 7 | 46 | 7 | 264 | 29.2 | 5.97 | 130.62 | 7 |
| O95837 | Guanine nucleotide-binding protein subunit alpha-14 OS=Homo sapiens OX=9606 GN=GNA14 PE=1 SV=1 | 0 | 44.861 | 23 | 7 | 72 | 1 | 355 | 41.5 | 6.07 | 148.29 | 7 |
| Q96MK3 | Pseudokinase FAM20A OS=Homo sapiens OX=9606 GN=FAM20A PE=1 SV=4 | 0 | 44.663 | 28 | 9 | 18 | 9 | 541 | 61.4 | 7.99 | 44.52 | 9 |
| P62745 | Rho-related GTP-binding protein RhoB OS=Homo sapiens OX=9606 GN=RHOB PE=1 SV=1 | 0 | 44.648 | 51 | 7 | 73 | 5 | 196 | 22.1 | 5.24 | 237.64 | 7 |
| P19971 | Thymidine phosphorylase OS=Homo sapiens OX=9606 GN=TYMP PE=1 SV=2 | 0 | 44.634 | 25 | 8 | 13 | 8 | 482 | 49.9 | 5.53 | 47.81 | 8 |
| A0A0B4J1V6 | Immunoglobulin heavy variable 3-73 OS=Homo sapiens OX=9606 GN=IGHV3-73 PE=3 SV=1 | 0 | 44.56 | 50 | 9 | 91 | 4 | 119 | 12.8 | 9.17 | 215.9 | 9 |
| O43692 | Peptidase inhibitor 15 OS=Homo sapiens OX=9606 GN=PI15 PE=1 SV=1 | 0 | 44.342 | 45 | 8 | 17 | 8 | 258 | 29 | 8.03 | 63.83 | 8 |
| P50897 | Palmitoyl-protein thioesterase 1 OS=Homo sapiens OX=9606 GN=PPT1 PE=1 SV=1 | 0 | 44.28 | 28 | 5 | 28 | 5 | 306 | 34.2 | 6.52 | 82.09 | 5 |
| P49908 | Selenoprotein P OS=Homo sapiens OX=9606 GN=SELENOP PE=1 SV=3 | 0 | 44.267 | 22 | 10 | 170 | 10 | 381 | 43.2 | 7.87 | 467.85 | 10 |
| Q9UBI6 | Guanine nucleotide-binding protein G(I)/G(S)/G(O) subunit gamma-12 OS=Homo sapiens OX=9606 GN=GNG12 PE=1 SV=3 | 0 | 44.086 | 71 | 6 | 85 | 6 | 72 | 8 | 8.97 | 277.36 | 6 |
| P84243 | Histone H3.3 OS=Homo sapiens OX=9606 GN=H3-3A PE=1 SV=2 | 0 | 44.056 | 48 | 7 | 109 | 1 | 136 | 15.3 | 11.27 | 209.99 | 7 |
| P07384 | Calpain-1 catalytic subunit OS=Homo sapiens OX=9606 GN=CAPN1 PE=1 SV=1 | 0 | 44.054 | 23 | 11 | 32 | 11 | 714 | 81.8 | 5.67 | 55.26 | 11 |
| P49755 | Transmembrane emp24 domain-containing protein 10 OS=Homo sapiens OX=9606 GN=TMED10 PE=1 SV=2 | 0 | 44.045 | 31 | 8 | 28 | 8 | 219 | 25 | 7.44 | 56.51 | 8 |
| Q15286 | Ras-related protein Rab-35 OS=Homo sapiens OX=9606 GN=RAB35 PE=1 SV=1 | 0 | 43.914 | 62 | 10 | 63 | 8 | 201 | 23 | 8.29 | 72.76 | 10 |
| Q5QNW6 | Histone H2B type 2-F OS=Homo sapiens OX=9606 GN=H2BC18 PE=1 SV=3 | 0 | 43.789 | 40 | 6 | 103 | 2 | 126 | 13.9 | 10.32 | 320.89 | 6 |
| Q71DI3 | Histone H3.2 OS=Homo sapiens OX=9606 GN=H3C15 PE=1 SV=3 | 0 | 43.7 | 48 | 7 | 109 | 1 | 136 | 15.4 | 11.27 | 218.28 | 7 |
| A0A0B4J1Y9 | Immunoglobulin heavy variable 3-72 OS=Homo sapiens OX=9606 GN=IGHV3-72 PE=3 SV=1 | 0 | 43.543 | 66 | 11 | 149 | 6 | 119 | 13.2 | 7.85 | 273.17 | 11 |
| P04062 | Lysosomal acid glucosylceramidase OS=Homo sapiens OX=9606 GN=GBA PE=1 SV=3 | 0 | 43.383 | 26 | 9 | 50 | 9 | 536 | 59.7 | 7.61 | 171.47 | 9 |
| O43175 | D-3-phosphoglycerate dehydrogenase OS=Homo sapiens OX=9606 GN=PHGDH PE=1 SV=4 | 0 | 43.33 | 35 | 11 | 24 | 11 | 533 | 56.6 | 6.71 | 48.38 | 11 |
| P13284 | Gamma-interferon-inducible lysosomal thiol reductase OS=Homo sapiens OX=9606 GN=IFI30 PE=1 SV=3 | 0 | 43.168 | 38 | 5 | 37 | 5 | 250 | 27.9 | 4.88 | 112.64 | 5 |
| O95436 | Sodium-dependent phosphate transport protein 2B OS=Homo sapiens OX=9606 GN=SLC34A2 PE=1 SV=3 | 0 | 42.881 | 19 | 10 | 104 | 10 | 690 | 75.7 | 8.15 | 151.51 | 10 |
| Q7L7L0 | Histone H2A type 3 OS=Homo sapiens OX=9606 GN=H2AW PE=1 SV=3 | 0 | 42.847 | 35 | 6 | 85 | 2 | 130 | 14.1 | 11.05 | 219.42 | 6 |
| P04844 | Dolichyl-diphosphooligosaccharide--protein glycosyltransferase subunit 2 OS=Homo sapiens OX=9606 GN=RPN2 PE=1 SV=3 | 0 | 42.42 | 26 | 8 | 28 | 8 | 631 | 69.2 | 5.69 | 90.82 | 8 |
| Q13217 | DnaJ homolog subfamily C member 3 OS=Homo sapiens OX=9606 GN=DNAJC3 PE=1 SV=1 | 0 | 42.42 | 23 | 9 | 51 | 9 | 504 | 57.5 | 6.15 | 63.91 | 9 |
| Q06481 | Amyloid-like protein 2 OS=Homo sapiens OX=9606 GN=APLP2 PE=1 SV=2 | 0 | 42.307 | 16 | 10 | 42 | 9 | 763 | 86.9 | 4.79 | 94.29 | 10 |
| Q8NBS9 | Thioredoxin domain-containing protein 5 OS=Homo sapiens OX=9606 GN=TXNDC5 PE=1 SV=2 | 0 | 42.296 | 25 | 8 | 61 | 8 | 432 | 47.6 | 5.97 | 127.58 | 8 |
| P51884 | Lumican OS=Homo sapiens OX=9606 GN=LUM PE=1 SV=2 | 0 | 42.246 | 28 | 7 | 41 | 7 | 338 | 38.4 | 6.61 | 81.8 | 7 |
| Q86X29 | Lipolysis-stimulated lipoprotein receptor OS=Homo sapiens OX=9606 GN=LSR PE=1 SV=4 | 0 | 42.142 | 19 | 9 | 107 | 9 | 649 | 71.4 | 7.97 | 167.11 | 9 |
| Q92930 | Ras-related protein Rab-8B OS=Homo sapiens OX=9606 GN=RAB8B PE=1 SV=2 | 0 | 42.14 | 46 | 10 | 93 | 2 | 207 | 23.6 | 9.07 | 171.06 | 10 |
| P01814 | Immunoglobulin heavy variable 2-70 OS=Homo sapiens OX=9606 GN=IGHV2-70 PE=1 SV=2 | 0 | 41.962 | 39 | 4 | 24 | 2 | 119 | 13.3 | 6.52 | 90.29 | 4 |
| P98164 | Low-density lipoprotein receptor-related protein 2 OS=Homo sapiens OX=9606 GN=LRP2 PE=1 SV=3 | 0 | 41.796 | 3 | 10 | 16 | 10 | 4655 | 521.6 | 5.08 | 46.66 | 10 |
| P62269 | 40S ribosomal protein S18 OS=Homo sapiens OX=9606 GN=RPS18 PE=1 SV=3 | 0 | 41.66 | 62 | 13 | 150 | 13 | 152 | 17.7 | 10.99 | 240.57 | 13 |
| P06310 | Immunoglobulin kappa variable 2-30 OS=Homo sapiens OX=9606 GN=IGKV2-30 PE=3 SV=2 | 0 | 41.18 | 44 | 5 | 63 | 3 | 120 | 13.2 | 8.51 | 189.41 | 5 |
| Q9BXJ1 | Complement C1q tumor necrosis factor-related protein 1 OS=Homo sapiens OX=9606 GN=C1QTNF1 PE=1 SV=1 | 0 | 41.062 | 28 | 6 | 90 | 6 | 281 | 31.7 | 6.9 | 265.72 | 6 |
| Q07020 | 60S ribosomal protein L18 OS=Homo sapiens OX=9606 GN=RPL18 PE=1 SV=2 | 0 | 41.04 | 43 | 8 | 115 | 8 | 188 | 21.6 | 11.72 | 264.55 | 8 |
| P30050 | 60S ribosomal protein L12 OS=Homo sapiens OX=9606 GN=RPL12 PE=1 SV=1 | 0 | 40.708 | 45 | 5 | 74 | 5 | 165 | 17.8 | 9.42 | 221.94 | 5 |
| P52823 | Stanniocalcin-1 OS=Homo sapiens OX=9606 GN=STC1 PE=1 SV=1 | 0 | 40.462 | 32 | 7 | 48 | 7 | 247 | 27.6 | 7.99 | 98.59 | 7 |
| P61020 | Ras-related protein Rab-5B OS=Homo sapiens OX=9606 GN=RAB5B PE=1 SV=1 | 0 | 40.424 | 42 | 7 | 85 | 4 | 215 | 23.7 | 8.13 | 212.87 | 7 |
| Q14512 | Fibroblast growth factor-binding protein 1 OS=Homo sapiens OX=9606 GN=FGFBP1 PE=1 SV=1 | 0 | 40.349 | 31 | 6 | 44 | 6 | 234 | 26.2 | 9.1 | 88.16 | 6 |
| P33908 | Mannosyl-oligosaccharide 1,2-alpha-mannosidase IA OS=Homo sapiens OX=9606 GN=MAN1A1 PE=1 SV=3 | 0 | 40.339 | 14 | 8 | 39 | 8 | 653 | 72.9 | 6.47 | 60.27 | 8 |
| A0A0C4DH31 | Immunoglobulin heavy variable 1-18 OS=Homo sapiens OX=9606 GN=IGHV1-18 PE=3 SV=1 | 0 | 40.329 | 61 | 5 | 107 | 4 | 117 | 12.8 | 8.84 | 322.76 | 5 |
| P15531 | Nucleoside diphosphate kinase A OS=Homo sapiens OX=9606 GN=NME1 PE=1 SV=1 | 0 | 40.321 | 74 | 9 | 123 | 4 | 152 | 17.1 | 6.19 | 249.44 | 9 |
| P01701 | Immunoglobulin lambda variable 1-51 OS=Homo sapiens OX=9606 GN=IGLV1-51 PE=1 SV=2 | 0 | 39.726 | 43 | 6 | 73 | 6 | 117 | 12.2 | 7.03 | 199.3 | 6 |
| Q8NBJ4 | Golgi membrane protein 1 OS=Homo sapiens OX=9606 GN=GOLM1 PE=1 SV=1 | 0 | 39.714 | 27 | 11 | 72 | 11 | 401 | 45.3 | 4.97 | 99.47 | 11 |
| P0DJI8 | Serum amyloid A-1 protein OS=Homo sapiens OX=9606 GN=SAA1 PE=1 SV=1 | 0 | 39.681 | 70 | 7 | 29 | 5 | 122 | 13.5 | 6.79 | 71.08 | 7 |
| Q9UHD0 | Interleukin-19 OS=Homo sapiens OX=9606 GN=IL19 PE=1 SV=2 | 0 | 39.436 | 37 | 6 | 49 | 6 | 177 | 20.4 | 7.69 | 173.32 | 6 |
| P07948 | Tyrosine-protein kinase Lyn OS=Homo sapiens OX=9606 GN=LYN PE=1 SV=3 | 0 | 39.435 | 26 | 11 | 44 | 9 | 512 | 58.5 | 7.11 | 69.94 | 11 |
| P21926 | CD9 antigen OS=Homo sapiens OX=9606 GN=CD9 PE=1 SV=4 | 0 | 39.352 | 25 | 6 | 119 | 6 | 228 | 25.4 | 7.15 | 415.09 | 6 |
| A0A075B6Q5 | Immunoglobulin heavy variable 3-64 OS=Homo sapiens OX=9606 GN=IGHV3-64 PE=3 SV=1 | 0 | 39.34 | 41 | 4 | 55 | 3 | 118 | 12.9 | 7.85 | 166.69 | 4 |
| P62280 | 40S ribosomal protein S11 OS=Homo sapiens OX=9606 GN=RPS11 PE=1 SV=3 | 0 | 39.306 | 53 | 11 | 105 | 11 | 158 | 18.4 | 10.3 | 136.11 | 11 |
| P04632 | Calpain small subunit 1 OS=Homo sapiens OX=9606 GN=CAPNS1 PE=1 SV=1 | 0 | 39.242 | 41 | 9 | 48 | 9 | 268 | 28.3 | 5.2 | 97.69 | 9 |
| Q01469 | Fatty acid-binding protein 5 OS=Homo sapiens OX=9606 GN=FABP5 PE=1 SV=3 | 0 | 39.19 | 60 | 7 | 21 | 7 | 135 | 15.2 | 7.01 | 58.02 | 7 |
| P28676 | Grancalcin OS=Homo sapiens OX=9606 GN=GCA PE=1 SV=2 | 0 | 39.133 | 58 | 10 | 22 | 10 | 217 | 24 | 5.21 | 55.57 | 10 |
| P46781 | 40S ribosomal protein S9 OS=Homo sapiens OX=9606 GN=RPS9 PE=1 SV=3 | 0 | 38.937 | 45 | 13 | 114 | 13 | 194 | 22.6 | 10.65 | 156.76 | 13 |
| Q14240 | Eukaryotic initiation factor 4A-II OS=Homo sapiens OX=9606 GN=EIF4A2 PE=1 SV=2 | 0 | 38.829 | 20 | 7 | 94 | 1 | 407 | 46.4 | 5.48 | 252.78 | 7 |
| Q96IU4 | Protein ABHD14B OS=Homo sapiens OX=9606 GN=ABHD14B PE=1 SV=1 | 0 | 38.677 | 34 | 6 | 38 | 6 | 210 | 22.3 | 6.4 | 66.72 | 6 |
| P63220 | 40S ribosomal protein S21 OS=Homo sapiens OX=9606 GN=RPS21 PE=1 SV=1 | 0 | 38.591 | 64 | 5 | 58 | 5 | 83 | 9.1 | 8.5 | 167.17 | 5 |
| O60506 | Heterogeneous nuclear ribonucleoprotein Q OS=Homo sapiens OX=9606 GN=SYNCRIP PE=1 SV=2 | 0 | 38.515 | 17 | 8 | 27 | 8 | 623 | 69.6 | 8.59 | 54.58 | 8 |
| P21281 | V-type proton ATPase subunit B, brain isoform OS=Homo sapiens OX=9606 GN=ATP6V1B2 PE=1 SV=3 | 0 | 38.456 | 24 | 8 | 13 | 8 | 511 | 56.5 | 5.81 | 35.94 | 8 |
| Q9NRJ3 | C-C motif chemokine 28 OS=Homo sapiens OX=9606 GN=CCL28 PE=1 SV=1 | 0 | 38.34 | 22 | 4 | 51 | 4 | 127 | 14.3 | 10.2 | 181.86 | 4 |
| O60888 | Protein CutA OS=Homo sapiens OX=9606 GN=CUTA PE=1 SV=2 | 0 | 38.293 | 41 | 4 | 51 | 4 | 179 | 19.1 | 5.5 | 192.52 | 4 |
| P39019 | 40S ribosomal protein S19 OS=Homo sapiens OX=9606 GN=RPS19 PE=1 SV=2 | 0 | 38.203 | 48 | 7 | 33 | 7 | 145 | 16.1 | 10.32 | 68.73 | 7 |
| P35268 | 60S ribosomal protein L22 OS=Homo sapiens OX=9606 GN=RPL22 PE=1 SV=2 | 0 | 38.185 | 46 | 6 | 44 | 6 | 128 | 14.8 | 9.19 | 172.16 | 6 |
| P08708 | 40S ribosomal protein S17 OS=Homo sapiens OX=9606 GN=RPS17 PE=1 SV=2 | 0 | 38.098 | 52 | 5 | 39 | 5 | 135 | 15.5 | 9.85 | 77.11 | 5 |
| P62277 | 40S ribosomal protein S13 OS=Homo sapiens OX=9606 GN=RPS13 PE=1 SV=2 | 0 | 37.926 | 38 | 6 | 55 | 6 | 151 | 17.2 | 10.54 | 106.38 | 6 |
| P49368 | T-complex protein 1 subunit gamma OS=Homo sapiens OX=9606 GN=CCT3 PE=1 SV=4 | 0 | 37.861 | 27 | 11 | 35 | 11 | 545 | 60.5 | 6.49 | 71.34 | 11 |
| A0A087WW87 | Immunoglobulin kappa variable 2-40 OS=Homo sapiens OX=9606 GN=IGKV2-40 PE=3 SV=2 | 0 | 37.825 | 31 | 5 | 76 | 3 | 121 | 13.3 | 4.61 | 262.44 | 5 |
| Q99727 | Metalloproteinase inhibitor 4 OS=Homo sapiens OX=9606 GN=TIMP4 PE=2 SV=1 | 0 | 37.619 | 40 | 7 | 62 | 7 | 224 | 25.5 | 7.8 | 181.7 | 7 |
| P62906 | 60S ribosomal protein L10a OS=Homo sapiens OX=9606 GN=RPL10A PE=1 SV=2 | 0 | 37.527 | 36 | 7 | 48 | 7 | 217 | 24.8 | 9.94 | 81.19 | 7 |
| P50991 | T-complex protein 1 subunit delta OS=Homo sapiens OX=9606 GN=CCT4 PE=1 SV=4 | 0 | 37.415 | 16 | 8 | 30 | 8 | 539 | 57.9 | 7.83 | 59.5 | 8 |
| P32320 | Cytidine deaminase OS=Homo sapiens OX=9606 GN=CDA PE=1 SV=2 | 0 | 37.254 | 80 | 5 | 5 | 5 | 146 | 16.2 | 6.92 | 24.42 | 5 |
| P47895 | Aldehyde dehydrogenase family 1 member A3 OS=Homo sapiens OX=9606 GN=ALDH1A3 PE=1 SV=2 | 0 | 37.209 | 19 | 9 | 31 | 7 | 512 | 56.1 | 7.25 | 66.66 | 9 |
| P25789 | Proteasome subunit alpha type-4 OS=Homo sapiens OX=9606 GN=PSMA4 PE=1 SV=1 | 0 | 37.187 | 40 | 6 | 22 | 6 | 261 | 29.5 | 7.72 | 45.71 | 6 |
| P08185 | Corticosteroid-binding globulin OS=Homo sapiens OX=9606 GN=SERPINA6 PE=1 SV=1 | 0 | 37.126 | 28 | 8 | 81 | 8 | 405 | 45.1 | 6.04 | 177.47 | 8 |
| Q8NBJ7 | Inactive C-alpha-formylglycine-generating enzyme 2 OS=Homo sapiens OX=9606 GN=SUMF2 PE=1 SV=2 | 0 | 36.948 | 37 | 9 | 53 | 9 | 301 | 33.8 | 8 | 81.88 | 9 |
| P17858 | ATP-dependent 6-phosphofructokinase, liver type OS=Homo sapiens OX=9606 GN=PFKL PE=1 SV=6 | 0 | 36.876 | 13 | 7 | 17 | 7 | 780 | 85 | 7.5 | 39.78 | 7 |
| P35754 | Glutaredoxin-1 OS=Homo sapiens OX=9606 GN=GLRX PE=1 SV=2 | 0 | 36.727 | 64 | 7 | 62 | 7 | 106 | 11.8 | 8.09 | 99.38 | 7 |
| P00918 | Carbonic anhydrase 2 OS=Homo sapiens OX=9606 GN=CA2 PE=1 SV=2 | 0 | 36.713 | 34 | 7 | 35 | 7 | 260 | 29.2 | 7.4 | 83.79 | 7 |
| A0A0C4DH38 | Immunoglobulin heavy variable 5-51 OS=Homo sapiens OX=9606 GN=IGHV5-51 PE=3 SV=1 | 0 | 36.699 | 60 | 5 | 134 | 3 | 117 | 12.7 | 8.27 | 424.44 | 5 |
| P43251 | Biotinidase OS=Homo sapiens OX=9606 GN=BTD PE=1 SV=2 | 0 | 36.668 | 19 | 8 | 99 | 8 | 543 | 61.1 | 6.25 | 255.6 | 8 |
| P49419 | Alpha-aminoadipic semialdehyde dehydrogenase OS=Homo sapiens OX=9606 GN=ALDH7A1 PE=1 SV=5 | 0 | 36.554 | 27 | 9 | 19 | 9 | 539 | 58.5 | 7.99 | 35.81 | 9 |
| P00167 | Cytochrome b5 OS=Homo sapiens OX=9606 GN=CYB5A PE=1 SV=2 | 0 | 36.467 | 49 | 6 | 56 | 6 | 134 | 15.3 | 4.96 | 122.48 | 6 |
| P13762 | HLA class II histocompatibility antigen, DR beta 4 chain OS=Homo sapiens OX=9606 GN=HLA-DRB4 PE=1 SV=2 | 0 | 36.435 | 54 | 10 | 46 | 8 | 266 | 29.9 | 7.33 | 90.8 | 10 |
| P02763 | Alpha-1-acid glycoprotein 1 OS=Homo sapiens OX=9606 GN=ORM1 PE=1 SV=1 | 0 | 36.292 | 37 | 6 | 91 | 5 | 201 | 23.5 | 5.02 | 255.75 | 6 |
| O00116 | Alkyldihydroxyacetonephosphate synthase, peroxisomal OS=Homo sapiens OX=9606 GN=AGPS PE=1 SV=1 | 0 | 36.272 | 20 | 11 | 18 | 11 | 658 | 72.9 | 7.34 | 31.39 | 11 |
| Q8WTS1 | 1-acylglycerol-3-phosphate O-acyltransferase ABHD5 OS=Homo sapiens OX=9606 GN=ABHD5 PE=1 SV=1 | 0 | 35.976 | 29 | 6 | 31 | 6 | 349 | 39.1 | 6.61 | 96.32 | 6 |
| Q9NYQ8 | Protocadherin Fat 2 OS=Homo sapiens OX=9606 GN=FAT2 PE=1 SV=2 | 0 | 35.916 | 4 | 13 | 13 | 13 | 4349 | 479 | 5.16 | 18.9 | 13 |
| Q24JP5 | Transmembrane protein 132A OS=Homo sapiens OX=9606 GN=TMEM132A PE=1 SV=1 | 0 | 35.807 | 15 | 9 | 12 | 9 | 1023 | 110 | 5.62 | 29.25 | 9 |
| Q99832 | T-complex protein 1 subunit eta OS=Homo sapiens OX=9606 GN=CCT7 PE=1 SV=2 | 0 | 35.792 | 20 | 8 | 26 | 8 | 543 | 59.3 | 7.65 | 42.59 | 8 |
| A0A0J9YXX1 | Immunoglobulin heavy variable 5-10-1 OS=Homo sapiens OX=9606 GN=IGHV5-10-1 PE=3 SV=1 | 0 | 35.666 | 51 | 5 | 101 | 3 | 117 | 12.8 | 8.28 | 264.75 | 5 |
| A0A0C4DH32 | Immunoglobulin heavy variable 3-20 OS=Homo sapiens OX=9606 GN=IGHV3-20 PE=3 SV=2 | 0 | 35.572 | 44 | 6 | 77 | 2 | 117 | 12.7 | 7.87 | 221.31 | 6 |
| A0A0C4DH43 | Immunoglobulin heavy variable 2-70D OS=Homo sapiens OX=9606 GN=IGHV2-70D PE=3 SV=1 | 0 | 35.549 | 39 | 4 | 41 | 1 | 119 | 13.3 | 8.29 | 115.61 | 4 |
| P02741 | C-reactive protein OS=Homo sapiens OX=9606 GN=CRP PE=1 SV=1 | 0 | 35.286 | 29 | 8 | 61 | 8 | 224 | 25 | 5.63 | 167.28 | 8 |
| P20618 | Proteasome subunit beta type-1 OS=Homo sapiens OX=9606 GN=PSMB1 PE=1 SV=2 | 0 | 35.26 | 38 | 7 | 29 | 7 | 241 | 26.5 | 8.13 | 35.68 | 7 |
| P51571 | Translocon-associated protein subunit delta OS=Homo sapiens OX=9606 GN=SSR4 PE=1 SV=1 | 0 | 35.152 | 31 | 5 | 35 | 5 | 173 | 19 | 6.15 | 88.97 | 5 |
| P62829 | 60S ribosomal protein L23 OS=Homo sapiens OX=9606 GN=RPL23 PE=1 SV=1 | 0 | 35.145 | 53 | 7 | 71 | 7 | 140 | 14.9 | 10.51 | 144.11 | 7 |
| Q9BWS9 | Chitinase domain-containing protein 1 OS=Homo sapiens OX=9606 GN=CHID1 PE=1 SV=1 | 0 | 35.117 | 26 | 9 | 43 | 9 | 393 | 44.9 | 8.63 | 75.05 | 9 |
| A0A0B4J1X8 | Immunoglobulin heavy variable 3-43 OS=Homo sapiens OX=9606 GN=IGHV3-43 PE=3 SV=1 | 0 | 34.982 | 44 | 5 | 111 | 3 | 118 | 13.1 | 5.41 | 310.76 | 5 |
| A0A0C4DH34 | Immunoglobulin heavy variable 4-28 OS=Homo sapiens OX=9606 GN=IGHV4-28 PE=3 SV=1 | 0 | 34.959 | 46 | 6 | 114 | 6 | 117 | 13.1 | 9.29 | 207.63 | 6 |
| P28066 | Proteasome subunit alpha type-5 OS=Homo sapiens OX=9606 GN=PSMA5 PE=1 SV=3 | 0 | 34.956 | 37 | 6 | 46 | 6 | 241 | 26.4 | 4.79 | 110.61 | 6 |
| O14818 | Proteasome subunit alpha type-7 OS=Homo sapiens OX=9606 GN=PSMA7 PE=1 SV=1 | 0 | 34.761 | 45 | 8 | 14 | 8 | 248 | 27.9 | 8.46 | 32.29 | 8 |
| P29122 | Proprotein convertase subtilisin/kexin type 6 OS=Homo sapiens OX=9606 GN=PCSK6 PE=1 SV=1 | 0 | 34.661 | 11 | 9 | 39 | 9 | 969 | 106.4 | 7.68 | 80.79 | 9 |

Table S1. Cont.

| Accession | Description | Exp. q-value: Combined | Sum PEP Score | Coverage [%] | # Peptides | # PSMs | # Unique Peptides | # AAs | MW [kDa] | calc. pI | Score Sequest HT: Sequest HT |
|---|---|---|---|---|---|---|---|---|---|---|---|
| P04271 | Protein S100-B OS=Homo sapiens OX=9606 GN=S100B PE=1 SV=2 | 0 | 34.638 | 40 | 2 | 23 | 2 | 92 | 10.7 | 4.59 | 85.42 |
| P07998 | Ribonuclease pancreatic OS=Homo sapiens OX=9606 GN=RNASE1 PE=1 SV=4 | 0 | 34.627 | 55 | 5 | 23 | 5 | 156 | 17.6 | 8.79 | 54.52 |
| P07814 | Bifunctional glutamate/proline--tRNA ligase OS=Homo sapiens OX=9606 GN=EPRS1 PE=1 SV=5 | 0 | 34.54 | 10 | 12 | 30 | 12 | 1512 | 170.5 | 7.33 | 30.69 |
| P13688 | Carcinoembryonic antigen-related cell adhesion molecule 1 OS=Homo sapiens OX=9606 GN=CEACAM1 PE=1 SV=2 | 0 | 34.495 | 16 | 5 | 33 | 5 | 526 | 57.5 | 5.97 | 116.98 |
| Q9NYU2 | UDP-glucose:glycoprotein glucosyltransferase 1 OS=Homo sapiens OX=9606 GN=UGGT1 PE=1 SV=3 | 0 | 34.461 | 10 | 10 | 14 | 10 | 1555 | 177.1 | 5.63 | 22.81 |
| Q13630 | GDP-L-fucose synthase OS=Homo sapiens OX=9606 GN=GFUS PE=1 SV=1 | 0 | 34.438 | 39 | 9 | 51 | 9 | 321 | 35.9 | 6.6 | 69.04 |
| Q9NZ20 | Group 3 secretory phospholipase A2 OS=Homo sapiens OX=9606 GN=PLA2G3 PE=1 SV=2 | 0 | 34.435 | 17 | 6 | 25 | 6 | 509 | 57.1 | 9.07 | 60.09 |
| P04080 | Cystatin-B OS=Homo sapiens OX=9606 GN=CSTB PE=1 SV=2 | 0 | 34.25 | 77 | 5 | 58 | 5 | 98 | 11.1 | 7.56 | 187.62 |
| A0A0B4J2H0 | Immunoglobulin heavy variable 1-69D OS=Homo sapiens OX=9606 GN=IGHV1-69D PE=1 SV=1 | 0 | 34.151 | 61 | 4 | 120 | 3 | 117 | 12.7 | 8.47 | 258.75 |
| P17931 | Galectin-3 OS=Homo sapiens OX=9606 GN=LGALS3 PE=1 SV=5 | 0 | 34.103 | 28 | 6 | 27 | 6 | 250 | 26.1 | 8.56 | 66.57 |
| Q14019 | Coactosin-like protein OS=Homo sapiens OX=9606 GN=COTL1 PE=1 SV=3 | 0 | 34.087 | 65 | 8 | 13 | 8 | 142 | 15.9 | 5.67 | 35.23 |
| P04839 | Cytochrome b-245 heavy chain OS=Homo sapiens OX=9606 GN=CYBB PE=1 SV=2 | 0 | 33.775 | 18 | 9 | 12 | 9 | 570 | 65.3 | 8.63 | 30.05 |
| P17066 | Heat shock 70 kDa protein 6 OS=Homo sapiens OX=9606 GN=HSPA6 PE=1 SV=2 | 0 | 33.725 | 15 | 7 | 102 | 1 | 643 | 71 | 6.14 | 237.93 |
| P05452 | Tetranectin OS=Homo sapiens OX=9606 GN=CLEC3B PE=1 SV=3 | 0 | 33.531 | 46 | 8 | 66 | 8 | 202 | 22.5 | 5.67 | 157.81 |
| O75594 | Peptidoglycan recognition protein 1 OS=Homo sapiens OX=9606 GN=PGLYRP1 PE=1 SV=1 | 0 | 33.519 | 49 | 5 | 8 | 5 | 196 | 21.7 | 8.59 | 32.01 |
| A0A0B4J1U7 | Immunoglobulin heavy variable 6-1 OS=Homo sapiens OX=9606 GN=IGHV6-1 PE=3 SV=1 | 0 | 33.431 | 41 | 6 | 78 | 6 | 121 | 13.5 | 9.2 | 148.41 |
| Q92820 | Gamma-glutamyl hydrolase OS=Homo sapiens OX=9606 GN=GGH PE=1 SV=2 | 0 | 33.431 | 23 | 5 | 31 | 5 | 318 | 35.9 | 7.11 | 89.21 |
| P51993 | 4-galactosyl-N-acetylglucosaminide 3-alpha-L-fucosyltransferase FUT6 OS=Homo sapiens OX=9606 GN=FUT6 PE=1 SV=1 | 0 | 33.359 | 24 | 7 | 77 | 2 | 359 | 41.8 | 8.68 | 141.6 |
| Q15582 | Transforming growth factor-beta-induced protein ig-h3 OS=Homo sapiens OX=9606 GN=TGFBI PE=1 SV=1 | 0 | 33.357 | 19 | 7 | 10 | 7 | 683 | 74.6 | 7.71 | 36.72 |
| Q6W4X9 | Mucin-6 OS=Homo sapiens OX=9606 GN=MUC6 PE=1 SV=3 | 0 | 33.292 | 5 | 8 | 11 | 8 | 2439 | 256.9 | 7.39 | 25.97 |
| P06865 | Beta-hexosaminidase subunit alpha OS=Homo sapiens OX=9606 GN=HEXA PE=1 SV=2 | 0 | 33.179 | 22 | 8 | 38 | 7 | 529 | 60.7 | 5.16 | 87.24 |
| P62750 | 60S ribosomal protein L23a OS=Homo sapiens OX=9606 GN=RPL23A PE=1 SV=1 | 0 | 33.172 | 28 | 7 | 47 | 7 | 156 | 17.7 | 10.45 | 86.97 |
| Q6UXS0 | C-type lectin domain family 19 member A OS=Homo sapiens OX=9606 GN=CLEC19A PE=1 SV=1 | 0 | 33.085 | 39 | 3 | 7 | 3 | 136 | 15.4 | 6.67 | 35.02 |
| Q71UI9 | Histone H2A.V OS=Homo sapiens OX=9606 GN=H2AZ2 PE=1 SV=3 | 0 | 33.064 | 54 | 6 | 53 | 4 | 128 | 13.5 | 10.58 | 115.07 |
| P01766 | Immunoglobulin heavy variable 3-13 OS=Homo sapiens OX=9606 GN=IGHV3-13 PE=1 SV=2 | 0 | 32.713 | 41 | 4 | 104 | 2 | 116 | 12.5 | 7.08 | 286.95 |
| Q15435 | Protein phosphatase 1 regulatory subunit 7 OS=Homo sapiens OX=9606 GN=PPP1R7 PE=1 SV=1 | 0 | 32.71 | 19 | 5 | 24 | 5 | 360 | 41.5 | 4.91 | 79.72 |
| P61313 | 60S ribosomal protein L15 OS=Homo sapiens OX=9606 GN=RPL15 PE=1 SV=2 | 0 | 32.707 | 35 | 7 | 67 | 7 | 204 | 24.1 | 11.62 | 111.62 |
| P23083 | Immunoglobulin heavy variable 1-2 OS=Homo sapiens OX=9606 GN=IGHV1-2 PE=1 SV=2 | 0 | 32.333 | 43 | 5 | 77 | 4 | 117 | 13.1 | 9.13 | 194.16 |
| Q8IWU5 | Extracellular sulfatase Sulf-2 OS=Homo sapiens OX=9606 GN=SULF2 PE=1 SV=1 | 0 | 32.287 | 12 | 8 | 31 | 8 | 870 | 100.4 | 9.17 | 37.46 |
| O94985 | Calsyntenin-1 OS=Homo sapiens OX=9606 GN=CLSTN1 PE=1 SV=1 | 0 | 32.281 | 14 | 10 | 31 | 10 | 981 | 109.7 | 4.91 | 58.97 |
| A0A075B6K4 | Immunoglobulin lambda variable 3-10 OS=Homo sapiens OX=9606 GN=IGLV3-10 PE=3 SV=2 | 0 | 31.824 | 36 | 7 | 84 | 5 | 115 | 12.4 | 4.83 | 168.07 |
| Q8WTV0 | Scavenger receptor class B member 1 OS=Homo sapiens OX=9606 GN=SCARB1 PE=1 SV=1 | 0 | 31.702 | 15 | 7 | 60 | 7 | 552 | 60.8 | 8.24 | 126.29 |
| P01602 | Immunoglobulin kappa variable 1-5 OS=Homo sapiens OX=9606 GN=IGKV1-5 PE=1 SV=2 | 0 | 31.645 | 27 | 3 | 58 | 2 | 117 | 12.8 | 8.28 | 217.18 |
| P14314 | Glucosidase 2 subunit beta OS=Homo sapiens OX=9606 GN=PRKCSH PE=1 SV=2 | 0 | 31.559 | 19 | 8 | 44 | 8 | 528 | 59.4 | 4.41 | 47.87 |
| Q9Y394 | Dehydrogenase/reductase SDR family member 7 OS=Homo sapiens OX=9606 GN=DHRS7 PE=1 SV=1 | 0 | 31.497 | 24 | 6 | 30 | 6 | 339 | 38.3 | 8.32 | 62.23 |
| P46776 | 60S ribosomal protein L27a OS=Homo sapiens OX=9606 GN=RPL27A PE=1 SV=2 | 0 | 31.488 | 32 | 5 | 19 | 5 | 148 | 16.6 | 11 | 43.13 |
| P01031 | Complement C5 OS=Homo sapiens OX=9606 GN=C5 PE=1 SV=4 | 0 | 31.43 | 7 | 9 | 23 | 9 | 1676 | 188.2 | 6.52 | 40.91 |
| P07947 | Tyrosine-protein kinase Yes OS=Homo sapiens OX=9606 GN=YES1 PE=1 SV=3 | 0 | 31.406 | 19 | 9 | 68 | 7 | 543 | 60.8 | 6.74 | 96.54 |
| P40926 | Malate dehydrogenase, mitochondrial OS=Homo sapiens OX=9606 GN=MDH2 PE=1 SV=3 | 0 | 31.4 | 25 | 6 | 11 | 6 | 338 | 35.5 | 8.68 | 24.84 |
| P01599 | Immunoglobulin kappa variable 1-17 OS=Homo sapiens OX=9606 GN=IGKV1-17 PE=1 SV=2 | 0 | 31.392 | 26 | 4 | 38 | 4 | 117 | 12.8 | 8.68 | 111.21 |
| P02743 | Serum amyloid P-component OS=Homo sapiens OX=9606 GN=APCS PE=1 SV=2 | 0 | 31.348 | 33 | 7 | 112 | 7 | 223 | 25.4 | 6.54 | 276.25 |
| P26583 | High mobility group protein B2 OS=Homo sapiens OX=9606 GN=HMGB2 PE=1 SV=2 | 0 | 31.253 | 32 | 6 | 16 | 4 | 209 | 24 | 7.81 | 40.04 |
| P02654 | Apolipoprotein C-I OS=Homo sapiens OX=9606 GN=APOC1 PE=1 SV=1 | 0 | 31.105 | 29 | 5 | 66 | 5 | 83 | 9.3 | 8.47 | 110.34 |
| P12109 | Collagen alpha-1(VI) chain OS=Homo sapiens OX=9606 GN=COL6A1 PE=1 SV=3 | 0 | 31.015 | 11 | 8 | 18 | 8 | 1028 | 108.5 | 5.43 | 42.43 |
| Q14914 | Prostaglandin reductase 1 OS=Homo sapiens OX=9606 GN=PTGR1 PE=1 SV=2 | 0 | 30.886 | 29 | 6 | 15 | 6 | 329 | 35.8 | 8.29 | 43.68 |
| P11717 | Cation-independent mannose-6-phosphate receptor OS=Homo sapiens OX=9606 GN=IGF2R PE=1 SV=3 | 0 | 30.786 | 6 | 11 | 23 | 11 | 2491 | 274.2 | 5.94 | 24.86 |
| P35555 | Fibrillin-1 OS=Homo sapiens OX=9606 GN=FBN1 PE=1 SV=4 | 0 | 30.782 | 3 | 8 | 10 | 8 | 2871 | 312.1 | 4.93 | 17.87 |
| P31944 | Caspase-14 OS=Homo sapiens OX=9606 GN=CASP14 PE=1 SV=2 | 0 | 30.747 | 37 | 9 | 12 | 9 | 242 | 27.7 | 5.58 | 28.7 |
| P12724 | Eosinophil cationic protein OS=Homo sapiens OX=9606 GN=RNASE3 PE=1 SV=2 | 0 | 30.737 | 35 | 6 | 15 | 6 | 160 | 18.4 | 10.02 | 46.79 |
| O95969 | Secretoglobin family 1D member 2 OS=Homo sapiens OX=9606 GN=SCGB1D2 PE=2 SV=1 | 0 | 30.665 | 40 | 8 | 29 | 8 | 90 | 9.9 | 8.25 | 74.28 |
| Q6IBS0 | Twinfilin-2 OS=Homo sapiens OX=9606 GN=TWF2 PE=1 SV=2 | 0 | 30.654 | 25 | 5 | 10 | 5 | 349 | 39.5 | 6.84 | 33.83 |
| P04433 | Immunoglobulin kappa variable 3-11 OS=Homo sapiens OX=9606 GN=IGKV3-11 PE=1 SV=1 | 0 | 30.472 | 52 | 3 | 84 | 3 | 115 | 12.6 | 4.96 | 284 |
| A0A0C4DH29 | Immunoglobulin heavy variable 1-3 OS=Homo sapiens OX=9606 GN=IGHV1-3 PE=3 SV=1 | 0 | 30.408 | 35 | 4 | 111 | 3 | 117 | 13 | 9.55 | 216.52 |
| O75396 | Vesicle-trafficking protein SEC22b OS=Homo sapiens OX=9606 GN=SEC22B PE=1 SV=4 | 0 | 30.307 | 35 | 6 | 23 | 6 | 215 | 24.6 | 6.92 | 49.01 |
| A0A075B6H9 | Immunoglobulin lambda variable 4-69 OS=Homo sapiens OX=9606 GN=IGLV4-69 PE=1 SV=1 | 0 | 30.288 | 44 | 5 | 15 | 5 | 119 | 12.8 | 6.51 | 42.19 |
| P62913 | 60S ribosomal protein L11 OS=Homo sapiens OX=9606 GN=RPL11 PE=1 SV=2 | 0 | 30.228 | 39 | 8 | 60 | 8 | 178 | 20.2 | 9.6 | 143.96 |
| P61353 | 60S ribosomal protein L27 OS=Homo sapiens OX=9606 GN=RPL27 PE=1 SV=2 | 0 | 29.882 | 49 | 6 | 54 | 6 | 136 | 15.8 | 10.56 | 86.35 |
| P49788 | Retinoic acid receptor responder protein 1 OS=Homo sapiens OX=9606 GN=RARRES1 PE=1 SV=2 | 0 | 29.871 | 28 | 9 | 43 | 9 | 294 | 33.3 | 8.51 | 82.62 |
| P05386 | 60S acidic ribosomal protein P1 OS=Homo sapiens OX=9606 GN=RPLP1 PE=1 SV=1 | 0 | 29.869 | 81 | 4 | 39 | 4 | 114 | 11.5 | 4.32 | 149.59 |
| P63173 | 60S ribosomal protein L38 OS=Homo sapiens OX=9606 GN=RPL38 PE=1 SV=2 | 0 | 29.856 | 36 | 3 | 20 | 3 | 70 | 8.2 | 10.1 | 74.52 |
| Q58FF3 | Putative endoplasmin-like protein OS=Homo sapiens OX=9606 GN=HSP90B2P PE=5 SV=1 | 0 | 29.853 | 13 | 8 | 114 | 1 | 399 | 45.8 | 5.26 | 134.16 |
| Q14847 | LIM and SH3 domain protein 1 OS=Homo sapiens OX=9606 GN=LASP1 PE=1 SV=2 | 0 | 29.688 | 32 | 9 | 46 | 9 | 261 | 29.7 | 7.05 | 73.18 |
| P03950 | Angiogenin OS=Homo sapiens OX=9606 GN=ANG PE=1 SV=1 | 0 | 29.602 | 37 | 5 | 46 | 5 | 147 | 16.5 | 9.64 | 124.06 |
| P09382 | Galectin-1 OS=Homo sapiens OX=9606 GN=LGALS1 PE=1 SV=2 | 0 | 29.6 | 58 | 6 | 21 | 6 | 135 | 14.7 | 5.5 | 45.1 |
| P10301 | Ras-related protein R-Ras OS=Homo sapiens OX=9606 GN=RRAS PE=1 SV=1 | 0 | 29.539 | 39 | 6 | 30 | 5 | 218 | 23.5 | 6.93 | 64.98 |
| A2NJV5 | Immunoglobulin kappa variable 2-29 OS=Homo sapiens OX=9606 GN=IGKV2-29 PE=3 SV=2 | 0 | 29.454 | 28 | 5 | 75 | 2 | 120 | 13.1 | 7.28 | 198.58 |
| P62328 | Thymosin beta-4 OS=Homo sapiens OX=9606 GN=TMSB4X PE=1 SV=2 | 0 | 29.45 | 77 | 4 | 5 | 4 | 44 | 5.1 | 5.06 | 17.59 |
| P30153 | Serine/threonine-protein phosphatase 2A 65 kDa regulatory subunit A alpha isoform OS=Homo sapiens OX=9606 GN=PPP2R1A PE=1 SV=4 | 0 | 29.433 | 17 | 7 | 19 | 7 | 589 | 65.3 | 5.11 | 41.13 |
| P40429 | 60S ribosomal protein L13a OS=Homo sapiens OX=9606 GN=RPL13A PE=1 SV=2 | 0 | 29.42 | 31 | 7 | 89 | 7 | 203 | 23.6 | 10.93 | 130.34 |
| Q8IWA5 | Choline transporter-like protein 2 OS=Homo sapiens OX=9606 GN=SLC44A2 PE=1 SV=3 | 0 | 29.252 | 12 | 8 | 56 | 8 | 706 | 80.1 | 8.57 | 79.32 |
| P61978 | Heterogeneous nuclear ribonucleoprotein K OS=Homo sapiens OX=9606 GN=HNRNPK PE=1 SV=1 | 0 | 29.197 | 22 | 7 | 11 | 7 | 463 | 50.9 | 5.54 | 30 |
| P09237 | Matrilysin OS=Homo sapiens OX=9606 GN=MMP7 PE=1 SV=1 | 0 | 29.156 | 33 | 8 | 27 | 8 | 267 | 29.7 | 7.91 | 63.62 |
| P28062 | Proteasome subunit beta type-8 OS=Homo sapiens OX=9606 GN=PSMB8 PE=1 SV=3 | 0 | 29.03 | 22 | 5 | 12 | 5 | 276 | 30.3 | 7.43 | 38.94 |
| Q15904 | V-type proton ATPase subunit S1 OS=Homo sapiens OX=9606 GN=ATP6AP1 PE=1 SV=2 | 0 | 29.006 | 17 | 6 | 17 | 6 | 470 | 52 | 6.14 | 47.02 |
| P11279 | Lysosome-associated membrane glycoprotein 1 OS=Homo sapiens OX=9606 GN=LAMP1 PE=1 SV=3 | 0 | 28.837 | 12 | 5 | 64 | 5 | 417 | 44.9 | 8.75 | 155.97 |
| A0A0C4DH67 | Immunoglobulin kappa variable 1-8 OS=Homo sapiens OX=9606 GN=IGKV1-8 PE=3 SV=1 | 0 | 28.792 | 50 | 5 | 53 | 4 | 115 | 12.5 | 9.01 | 196.99 |
| Q9BQI0 | Allograft inflammatory factor 1-like OS=Homo sapiens OX=9606 GN=AIF1L PE=1 SV=1 | 0 | 28.763 | 51 | 8 | 50 | 8 | 150 | 17.1 | 7.2 | 72.21 |
| O95716 | Ras-related protein Rab-3D OS=Homo sapiens OX=9606 GN=RAB3D PE=1 SV=1 | 0 | 28.686 | 32 | 7 | 54 | 4 | 219 | 24.3 | 4.93 | 114.53 |
| P60866 | 40S ribosomal protein S20 OS=Homo sapiens OX=9606 GN=RPS20 PE=1 SV=1 | 0 | 28.673 | 32 | 5 | 59 | 5 | 119 | 13.4 | 9.94 | 112.72 |
| Q13347 | Eukaryotic translation initiation factor 3 subunit I OS=Homo sapiens OX=9606 GN=EIF3I PE=1 SV=1 | 0 | 28.661 | 33 | 9 | 21 | 9 | 325 | 36.5 | 5.64 | 26.38 |
| P05543 | Thyroxine-binding globulin OS=Homo sapiens OX=9606 GN=SERPINA7 PE=1 SV=2 | 0 | 28.611 | 22 | 7 | 35 | 7 | 415 | 46.3 | 6.3 | 52.17 |
| P61163 | Alpha-centractin OS=Homo sapiens OX=9606 GN=ACTR1A PE=1 SV=1 | 0 | 28.461 | 22 | 5 | 21 | 5 | 376 | 42.6 | 6.64 | 41.89 |
| P54819 | Adenylate kinase 2, mitochondrial OS=Homo sapiens OX=9606 GN=AK2 PE=1 SV=2 | 0 | 28.421 | 36 | 5 | 6 | 5 | 239 | 26.5 | 7.81 | 23.21 |
| P42766 | 60S ribosomal protein L35 OS=Homo sapiens OX=9606 GN=RPL35 PE=1 SV=2 | 0 | 28.294 | 31 | 5 | 45 | 5 | 123 | 14.5 | 11.05 | 80.23 |
| P62266 | 40S ribosomal protein S23 OS=Homo sapiens OX=9606 GN=RPS23 PE=1 SV=3 | 0 | 28.293 | 49 | 7 | 57 | 7 | 143 | 15.8 | 10.49 | 121.86 |
| Q58FF6 | Putative heat shock protein HSP 90-beta 4 OS=Homo sapiens OX=9606 GN=HSP90AB4P PE=5 SV=1 | 0 | 28.275 | 11 | 5 | 58 | 1 | 505 | 58.2 | 4.73 | 173.81 |
| O95336 | 6-phosphogluconolactonase OS=Homo sapiens OX=9606 GN=PGLS PE=1 SV=2 | 0 | 28.212 | 33 | 6 | 49 | 6 | 258 | 27.5 | 6.05 | 166.47 |
| A0A075B6R2 | Immunoglobulin heavy variable 4-4 OS=Homo sapiens OX=9606 GN=IGHV4-4 PE=3 SV=2 | 0 | 28.207 | 46 | 7 | 81 | 5 | 117 | 12.8 | 9.6 | 208.18 |
| P06312 | Immunoglobulin kappa variable 4-1 OS=Homo sapiens OX=9606 GN=IGKV4-1 PE=1 SV=1 | 0 | 28.186 | 30 | 4 | 99 | 4 | 121 | 13.4 | 5.25 | 267.18 |
| P56937 | 3-keto-steroid reductase/17-beta-hydroxysteroid dehydrogenase 7 OS=Homo sapiens OX=9606 GN=HSD17B7 PE=1 SV=1 | 0 | 28.129 | 30 | 7 | 30 | 7 | 341 | 38.2 | 8.1 | 36.59 |
| P14868 | Aspartate--tRNA ligase, cytoplasmic OS=Homo sapiens OX=9606 GN=DARS1 PE=1 SV=2 | 0 | 28.106 | 18 | 7 | 11 | 7 | 501 | 57.1 | 6.55 | 19.16 |
| P22061 | Protein-L-isoaspartate(D-aspartate) O-methyltransferase OS=Homo sapiens OX=9606 GN=PCMT1 PE=1 SV=4 | 0 | 28.067 | 42 | 5 | 28 | 5 | 227 | 24.6 | 7.21 | 69.27 |
| P49257 | Protein ERGIC-53 OS=Homo sapiens OX=9606 GN=LMAN1 PE=1 SV=2 | 0 | 28.059 | 14 | 6 | 21 | 6 | 510 | 57.5 | 6.77 | 36.95 |
| Q9H336 | Cysteine-rich secretory protein LCCL domain-containing 1 OS=Homo sapiens OX=9606 GN=CRISPLD1 PE=2 SV=1 | 0 | 27.863 | 14 | 6 | 20 | 6 | 500 | 56.9 | 8.12 | 35.8 |
| Q9UM07 | Protein-arginine deiminase type-4 OS=Homo sapiens OX=9606 GN=PADI4 PE=1 SV=2 | 0 | 27.747 | 11 | 6 | 7 | 6 | 663 | 74 | 6.58 | 22.68 |
| P51659 | Peroxisomal multifunctional enzyme type 2 OS=Homo sapiens OX=9606 GN=HSD17B4 PE=1 SV=3 | 0 | 27.643 | 12 | 6 | 8 | 6 | 736 | 79.6 | 8.84 | 22.85 |
| Q96TA1 | Protein Niban 2 OS=Homo sapiens OX=9606 GN=NIBAN2 PE=1 SV=3 | 0 | 27.55 | 16 | 10 | 24 | 10 | 746 | 84.1 | 6.19 | 26.28 |
| P62888 | 60S ribosomal protein L30 OS=Homo sapiens OX=9606 GN=RPL30 PE=1 SV=2 | 0 | 27.51 | 57 | 5 | 35 | 5 | 115 | 12.8 | 9.63 | 81.28 |
| Q9Y696 | Chloride intracellular channel protein 4 OS=Homo sapiens OX=9606 GN=CLIC4 PE=1 SV=4 | 0 | 27.483 | 27 | 5 | 37 | 5 | 253 | 28.8 | 5.59 | 53.41 |
| P30043 | Flavin reductase (NADPH) OS=Homo sapiens OX=9606 GN=BLVRB PE=1 SV=3 | 0 | 27.435 | 33 | 4 | 40 | 4 | 206 | 22.1 | 7.65 | 89.14 |
| P01714 | Immunoglobulin lambda variable 3-19 OS=Homo sapiens OX=9606 GN=IGLV3-19 PE=1 SV=2 | 0 | 27.338 | 44 | 4 | 64 | 4 | 112 | 12 | 4.96 | 162.26 |
| A0A0C4DH25 | Immunoglobulin kappa variable 3D-20 OS=Homo sapiens OX=9606 GN=IGKV3D-20 PE=3 SV=1 | 0 | 27.271 | 52 | 4 | 79 | 2 | 116 | 12.5 | 4.59 | 208.56 |
| Q96AG4 | Leucine-rich repeat-containing protein 59 OS=Homo sapiens OX=9606 GN=LRRC59 PE=1 SV=1 | 0 | 27.268 | 24 | 6 | 31 | 6 | 307 | 34.9 | 9.57 | 63.59 |
| P26373 | 60S ribosomal protein L13 OS=Homo sapiens OX=9606 GN=RPL13 PE=1 SV=4 | 0 | 27.176 | 30 | 7 | 98 | 7 | 211 | 24.2 | 11.65 | 227.78 |
| A0A075B6R9 | Probable non-functional immunoglobulin kappa variable 2D-24 OS=Homo sapiens OX=9606 GN=IGKV2D-24 PE=1 SV=1 | 0 | 27.168 | 38 | 4 | 48 | 3 | 120 | 13.1 | 8.87 | 132.94 |
| P00568 | Adenylate kinase isoenzyme 1 OS=Homo sapiens OX=9606 GN=AK1 PE=1 SV=3 | 0 | 27.13 | 36 | 6 | 34 | 6 | 194 | 21.6 | 8.63 | 69.61 |
| P55001 | Microfibrillar-associated protein 2 OS=Homo sapiens OX=9606 GN=MFAP2 PE=1 SV=1 | 0 | 27.109 | 27 | 4 | 31 | 4 | 183 | 20.8 | 4.97 | 62.68 |
| P25786 | Proteasome subunit alpha type-1 OS=Homo sapiens OX=9606 GN=PSMA1 PE=1 SV=1 | 0 | 27.101 | 32 | 8 | 43 | 8 | 263 | 29.5 | 6.61 | 34.92 |
| Q53GQ0 | Very-long-chain 3-oxoacyl-CoA reductase OS=Homo sapiens OX=9606 GN=HSD17B12 PE=1 SV=2 | 0 | 26.963 | 20 | 4 | 18 | 4 | 312 | 34.3 | 9.32 | 33.94 |
| O95274 | Ly6/PLAUR domain-containing protein 3 OS=Homo sapiens OX=9606 GN=LYPD3 PE=1 SV=2 | 0 | 26.859 | 22 | 4 | 10 | 4 | 346 | 35.9 | 7.75 | 45.4 |
| Q9UBX7 | Kallikrein-11 OS=Homo sapiens OX=9606 GN=KLK11 PE=1 SV=2 | 0 | 26.79 | 28 | 5 | 10 | 5 | 282 | 31 | 8.94 | 28.84 |
| P61254 | 60S ribosomal protein L26 OS=Homo sapiens OX=9606 GN=RPL26 PE=1 SV=1 | 0 | 26.533 | 37 | 8 | 60 | 8 | 145 | 17.2 | 10.55 | 91.17 |
| P36980 | Complement factor H-related protein 2 OS=Homo sapiens OX=9606 GN=CFHR2 PE=1 SV=1 | 0 | 26.525 | 36 | 5 | 40 | 1 | 270 | 30.6 | 6.38 | 73.55 |
| P12830 | Cadherin-1 OS=Homo sapiens OX=9606 GN=CDH1 PE=1 SV=3 | 0 | 26.501 | 8 | 7 | 81 | 7 | 882 | 97.4 | 4.73 | 147.29 |
| P04432 | Immunoglobulin kappa variable 1D-39 OS=Homo sapiens OX=9606 GN=IGKV1D-39 PE=3 SV=2 | 0 | 26.467 | 32 | 3 | 75 | 2 | 117 | 12.7 | 8.66 | 228.46 |
| Q13510 | Acid ceramidase OS=Homo sapiens OX=9606 GN=ASAH1 PE=1 SV=5 | 0 | 26.466 | 19 | 6 | 19 | 6 | 395 | 44.6 | 7.62 | 46.82 |
| A0A0J9YX35 | Immunoglobulin heavy variable 3-64D OS=Homo sapiens OX=9606 GN=IGHV3-64D PE=3 SV=1 | 0 | 26.368 | 44 | 5 | 86 | 4 | 117 | 12.8 | 7.85 | 244.26 |
| P21217 | 3-galactosyl-N-acetylglucosaminide 4-alpha-L-fucosyltransferase FUT3 OS=Homo sapiens OX=9606 GN=FUT3 PE=1 SV=1 | 0 | 26.349 | 19 | 6 | 64 | 1 | 361 | 42.1 | 9.01 | 120.01 |
| Q9H3G5 | Probable serine carboxypeptidase CPVL OS=Homo sapiens OX=9606 GN=CPVL PE=1 SV=2 | 0 | 26.219 | 27 | 8 | 11 | 8 | 476 | 54.1 | 5.62 | 21.89 |
| P22626 | Heterogeneous nuclear ribonucleoproteins A2/B1 OS=Homo sapiens OX=9606 GN=HNRNPA2B1 PE=1 SV=2 | 0 | 26.215 | 23 | 6 | 10 | 6 | 353 | 37.4 | 8.95 | 26.01 |
| O00592 | Podocalyxin OS=Homo sapiens OX=9606 GN=PODXL PE=1 SV=2 | 0 | 26.154 | 10 | 6 | 116 | 6 | 558 | 58.6 | 5.49 | 248.05 |
| P01700 | Immunoglobulin lambda variable 1-47 OS=Homo sapiens OX=9606 GN=IGLV1-47 PE=1 SV=2 | 0 | 26.056 | 54 | 4 | 75 | 2 | 117 | 12.3 | 5.91 | 137.57 |
| Q9H444 | Charged multivesicular body protein 4b OS=Homo sapiens OX=9606 GN=CHMP4B PE=1 SV=1 | 0 | 26.036 | 30 | 6 | 51 | 6 | 224 | 24.9 | 4.82 | 92.35 |
| Q14344 | Guanine nucleotide-binding protein subunit alpha-13 OS=Homo sapiens OX=9606 GN=GNA13 PE=1 SV=2 | 0 | 25.941 | 32 | 9 | 53 | 7 | 377 | 44 | 8 | 71.28 |
| P20036 | HLA class II histocompatibility antigen, DP alpha 1 chain OS=Homo sapiens OX=9606 GN=HLA-DPA1 PE=1 SV=1 | 0 | 25.85 | 18 | 4 | 16 | 4 | 260 | 29.4 | 5.21 | 63.53 |
| O43852 | Calumenin OS=Homo sapiens OX=9606 GN=CALU PE=1 SV=2 | 0 | 25.815 | 14 | 3 | 23 | 3 | 315 | 37.1 | 4.64 | 78.43 |
| P10809 | 60 kDa heat shock protein, mitochondrial OS=Homo sapiens OX=9606 GN=HSPD1 PE=1 SV=2 | 0 | 25.812 | 13 | 5 | 13 | 5 | 573 | 61 | 5.87 | 36.25 |
| P61088 | Ubiquitin-conjugating enzyme E2 N OS=Homo sapiens OX=9606 GN=UBE2N PE=1 SV=1 | 0 | 25.699 | 53 | 7 | 19 | 7 | 152 | 17.1 | 6.57 | 26.71 |
| A0A0B4J1V2 | Immunoglobulin heavy variable 2-26 OS=Homo sapiens OX=9606 GN=IGHV2-26 PE=3 SV=1 | 0 | 25.695 | 60 | 4 | 19 | 4 | 119 | 13.2 | 8.29 | 53.68 |
| O75629 | Protein CREG1 OS=Homo sapiens OX=9606 GN=CREG1 PE=1 SV=1 | 0 | 25.589 | 19 | 2 | 16 | 2 | 220 | 24.1 | 7.59 | 54.88 |
| Q9Y653 | Adhesion G-protein coupled receptor G1 OS=Homo sapiens OX=9606 GN=ADGRG1 PE=1 SV=2 | 0 | 25.581 | 9 | 4 | 9 | 4 | 693 | 77.7 | 8.48 | 34.93 |
| O00159 | Unconventional myosin-Ic OS=Homo sapiens OX=9606 GN=MYO1C PE=1 SV=4 | 0 | 25.528 | 9 | 8 | 17 | 8 | 1063 | 121.6 | 9.41 | 18.39 |
| Q13200 | 26S proteasome non-ATPase regulatory subunit 2 OS=Homo sapiens OX=9606 GN=PSMD2 PE=1 SV=3 | 0 | 25.474 | 11 | 6 | 12 | 6 | 908 | 100.1 | 5.2 | 20.84 |
| P62249 | 40S ribosomal protein S16 OS=Homo sapiens OX=9606 GN=RPS16 PE=1 SV=2 | 0 | 25.451 | 40 | 7 | 63 | 7 | 146 | 16.4 | 10.21 | 104.58 |
| P09429 | High mobility group protein B1 OS=Homo sapiens OX=9606 GN=HMGB1 PE=1 SV=3 | 0 | 25.379 | 33 | 6 | 16 | 4 | 215 | 24.9 | 5.74 | 39.77 |
| P61106 | Ras-related protein Rab-14 OS=Homo sapiens OX=9606 GN=RAB14 PE=1 SV=4 | 0 | 25.248 | 22 | 5 | 20 | 3 | 215 | 23.9 | 6.21 | 56.4 |
| P54727 | UV excision repair protein RAD23 homolog B OS=Homo sapiens OX=9606 GN=RAD23B PE=1 SV=1 | 0 | 25.21 | 21 | 7 | 36 | 7 | 409 | 43.1 | 4.84 | 72.56 |
| P61916 | NPC intracellular cholesterol transporter 2 OS=Homo sapiens OX=9606 GN=NPC2 PE=1 SV=1 | 0 | 25.173 | 49 | 6 | 79 | 6 | 151 | 16.6 | 7.65 | 160.59 |
| Q15366 | Poly(rC)-binding protein 2 OS=Homo sapiens OX=9606 GN=PCBP2 PE=1 SV=1 | 0 | 25.173 | 19 | 5 | 44 | 2 | 365 | 38.6 | 6.79 | 97.5 |
| O75695 | Protein XRP2 OS=Homo sapiens OX=9606 GN=RP2 PE=1 SV=4 | 0 | 24.916 | 20 | 6 | 42 | 6 | 350 | 39.6 | 5.12 | 54.83 |
| P13866 | Sodium/glucose cotransporter 1 OS=Homo sapiens OX=9606 GN=SLC5A1 PE=1 SV=1 | 0 | 24.895 | 10 | 6 | 40 | 6 | 664 | 73.4 | 7.68 | 110.87 |
| O14579 | Coatomer subunit epsilon OS=Homo sapiens OX=9606 GN=COPE PE=1 SV=3 | 0 | 24.838 | 24 | 3 | 9 | 3 | 308 | 34.5 | 5.12 | 20.25 |
| O60259 | Kallikrein-8 OS=Homo sapiens OX=9606 GN=KLK8 PE=1 SV=1 | 0 | 24.763 | 21 | 4 | 30 | 4 | 260 | 28 | 7.58 | 34.04 |
| P39687 | Acidic leucine-rich nuclear phosphoprotein 32 family member A OS=Homo sapiens OX=9606 GN=ANP32A PE=1 SV=1 | 0 | 24.761 | 24 | 5 | 6 | 4 | 249 | 28.6 | 4.09 | 15.44 |
| P58107 | Epiplakin OS=Homo sapiens OX=9606 GN=EPPK1 PE=1 SV=3 | 0 | 24.704 | 9 | 4 | 9 | 3 | 5088 | 555.3 | 5.62 | 18.2 |
| O76061 | Stanniocalcin-2 OS=Homo sapiens OX=9606 GN=STC2 PE=1 SV=1 | 0 | 24.663 | 28 | 5 | 12 | 5 | 302 | 33.2 | 7.3 | 24.89 |
| Q9NR45 | Sialic acid synthase OS=Homo sapiens OX=9606 GN=NANS PE=1 SV=2 | 0 | 24.654 | 23 | 5 | 21 | 5 | 359 | 40.3 | 6.74 | 50.77 |
| O75915 | PRA1 family protein 3 OS=Homo sapiens OX=9606 GN=ARL6IP5 PE=1 SV=1 | 0 | 24.648 | 29 | 6 | 41 | 6 | 188 | 21.6 | 9.77 | 97.52 |
| P04233 | HLA class II histocompatibility antigen gamma chain OS=Homo sapiens OX=9606 GN=CD74 PE=1 SV=3 | 0 | 24.528 | 16 | 4 | 18 | 4 | 296 | 33.5 | 8.44 | 40.21 |
| Q9UL25 | Ras-related protein Rab-21 OS=Homo sapiens OX=9606 GN=RAB21 PE=1 SV=3 | 0 | 24.523 | 34 | 6 | 24 | 6 | 225 | 24.3 | 7.94 | 52.61 |
| Q14210 | Lymphocyte antigen 6D OS=Homo sapiens OX=9606 GN=LY6D PE=1 SV=1 | 0 | 24.494 | 46 | 5 | 37 | 5 | 128 | 13.3 | 8.21 | 93.08 |
| P24298 | Alanine aminotransferase 1 OS=Homo sapiens OX=9606 GN=GPT PE=1 SV=3 | 0 | 24.43 | 23 | 7 | 8 | 7 | 496 | 54.6 | 7.18 | 19 |
| P04440 | HLA class II histocompatibility antigen, DP beta 1 chain OS=Homo sapiens OX=9606 GN=HLA-DPB1 PE=1 SV=1 | 0 | 24.361 | 29 | 5 | 31 | 4 | 258 | 29.1 | 8.57 | 65.07 |
| Q9BRA2 | Thioredoxin domain-containing protein 17 OS=Homo sapiens OX=9606 GN=TXNDC17 PE=1 SV=1 | 0 | 24.341 | 54 | 5 | 29 | 5 | 123 | 13.9 | 5.52 | 48.54 |
| P02656 | Apolipoprotein C-III OS=Homo sapiens OX=9606 GN=APOC3 PE=1 SV=1 | 0 | 24.323 | 34 | 3 | 43 | 3 | 99 | 10.8 | 5.41 | 144.01 |
| P29966 | Myristoylated alanine-rich C-kinase substrate OS=Homo sapiens OX=9606 GN=MARCKS PE=1 SV=4 | 0 | 24.303 | 27 | 5 | 59 | 5 | 332 | 31.5 | 4.45 | 73.23 |
| Q9NRA1 | Platelet-derived growth factor C OS=Homo sapiens OX=9606 GN=PDGFC PE=1 SV=2 | 0 | 24.294 | 20 | 6 | 11 | 6 | 345 | 39 | 6.14 | 25.28 |
| P0DJI9 | Serum amyloid A-2 protein OS=Homo sapiens OX=9606 GN=SAA2 PE=1 SV=1 | 0 | 24.22 | 47 | 4 | 15 | 2 | 122 | 13.5 | 9.14 | 50.6 |
| Q99674 | Cell growth regulator with EF hand domain protein 1 OS=Homo sapiens OX=9606 GN=CGREF1 PE=2 SV=3 | 0 | 24.199 | 29 | 4 | 7 | 4 | 318 | 33.4 | 4.39 | 19.67 |
| Q14CN2 | Calcium-activated chloride channel regulator 4 OS=Homo sapiens OX=9606 GN=CLCA4 PE=1 SV=2 | 0 | 24.095 | 12 | 7 | 12 | 7 | 919 | 101.2 | 5.47 | 29.54 |
| Q9NZ01 | Very-long-chain enoyl-CoA reductase OS=Homo sapiens OX=9606 GN=TECR PE=1 SV=1 | 0 | 24.079 | 16 | 5 | 11 | 5 | 308 | 36 | 9.45 | 26.48 |
| Q05315 | Galectin-10 OS=Homo sapiens OX=9606 GN=CLC PE=1 SV=3 | 0 | 24.068 | 35 | 3 | 7 | 3 | 142 | 16.4 | 7.37 | 16.92 |
| O75131 | Copine-3 OS=Homo sapiens OX=9606 GN=CPNE3 PE=1 SV=1 | 0 | 24.056 | 16 | 7 | 24 | 7 | 537 | 60.1 | 5.85 | 53.1 |
| P62070 | Ras-related protein R-Ras2 OS=Homo sapiens OX=9606 GN=RRAS2 PE=1 SV=1 | 0 | 24.017 | 26 | 4 | 25 | 3 | 204 | 23.4 | 6.01 | 54.95 |
| P46779 | 60S ribosomal protein L28 OS=Homo sapiens OX=9606 GN=RPL28 PE=1 SV=3 | 0 | 23.964 | 31 | 5 | 60 | 5 | 137 | 15.7 | 12.02 | 127.36 |
| P42677 | 40S ribosomal protein S27 OS=Homo sapiens OX=9606 GN=RPS27 PE=1 SV=3 | 0 | 23.926 | 42 | 4 | 31 | 1 | 84 | 9.5 | 9.45 | 66.72 |
| P22894 | Neutrophil collagenase OS=Homo sapiens OX=9606 GN=MMP8 PE=1 SV=1 | 0 | 23.903 | 15 | 6 | 9 | 6 | 467 | 53.4 | 6.87 | 22.41 |
| Q96IJ6 | Mannose-1-phosphate guanyltransferase alpha OS=Homo sapiens OX=9606 GN=GMPPA PE=1 SV=1 | 0 | 23.813 | 16 | 6 | 23 | 6 | 420 | 46.3 | 7.21 | 43.05 |
| Q14974 | Importin subunit beta-1 OS=Homo sapiens OX=9606 GN=KPNB1 PE=1 SV=2 | 0 | 23.779 | 8 | 5 | 20 | 5 | 876 | 97.1 | 4.78 | 43.13 |
| P01703 | Immunoglobulin lambda variable 1-40 OS=Homo sapiens OX=9606 GN=IGLV1-40 PE=1 SV=2 | 0 | 23.594 | 43 | 3 | 47 | 3 | 118 | 12.3 | 5.74 | 173.31 |
| P48723 | Heat shock 70 kDa protein 13 OS=Homo sapiens OX=9606 GN=HSPA13 PE=1 SV=1 | 0 | 23.59 | 21 | 7 | 15 | 7 | 471 | 51.9 | 5.76 | 31.53 |
| O75787 | Renin receptor OS=Homo sapiens OX=9606 GN=ATP6AP2 PE=1 SV=2 | 0 | 23.466 | 21 | 5 | 9 | 5 | 350 | 39 | 6.1 | 29.9 |
| P27169 | Serum paraoxonase/arylesterase 1 OS=Homo sapiens OX=9606 GN=PON1 PE=1 SV=3 | 0 | 23.316 | 25 | 6 | 24 | 6 | 355 | 39.7 | 5.22 | 33.53 |
| Q07065 | Cytoskeleton-associated protein 4 OS=Homo sapiens OX=9606 GN=CKAP4 PE=1 SV=2 | 0 | 23.28 | 9 | 4 | 12 | 4 | 602 | 66 | 5.92 | 17.62 |
| Q9BUP3 | Oxidoreductase HTATIP2 OS=Homo sapiens OX=9606 GN=HTATIP2 PE=1 SV=2 | 0 | 22.981 | 26 | 7 | 22 | 7 | 242 | 27 | 8.38 | 33.49 |
| P59665 | Neutrophil defensin 1 OS=Homo sapiens OX=9606 GN=DEFA1 PE=1 SV=1 | 0 | 22.975 | 20 | 4 | 119 | 4 | 94 | 10.2 | 6.99 | 293.53 |
| P35542 | Serum amyloid A-4 protein OS=Homo sapiens OX=9606 GN=SAA4 PE=1 SV=2 | 0 | 22.956 | 27 | 3 | 35 | 3 | 130 | 14.7 | 9.07 | 62.52 |
| P20851 | C4b-binding protein beta chain OS=Homo sapiens OX=9606 GN=C4BPB PE=1 SV=1 | 0 | 22.903 | 26 | 6 | 24 | 6 | 252 | 28.3 | 5.14 | 54.39 |
| P00736 | Complement C1r subcomponent OS=Homo sapiens OX=9606 GN=C1R PE=1 SV=2 | 0 | 22.891 | 13 | 7 | 36 | 5 | 705 | 80.1 | 6.21 | 81.54 |
| P14384 | Carboxypeptidase M OS=Homo sapiens OX=9606 GN=CPM PE=1 SV=2 | 0 | 22.782 | 17 | 5 | 8 | 5 | 443 | 50.5 | 7.36 | 22.19 |
| P84098 | 60S ribosomal protein L19 OS=Homo sapiens OX=9606 GN=RPL19 PE=1 SV=1 | 0 | 22.78 | 26 | 5 | 21 | 5 | 196 | 23.5 | 11.47 | 50.66 |
| P61513 | 60S ribosomal protein L37a OS=Homo sapiens OX=9606 GN=RPL37A PE=1 SV=2 | 0 | 22.737 | 41 | 3 | 21 | 3 | 92 | 10.3 | 10.43 | 65.99 |
| O43657 | Tetraspanin-6 OS=Homo sapiens OX=9606 GN=TSPAN6 PE=1 SV=1 | 0 | 22.729 | 20 | 4 | 8 | 4 | 245 | 27.5 | 8.1 | 15.45 |
| Q99805 | Transmembrane 9 superfamily member 2 OS=Homo sapiens OX=9606 GN=TM9SF2 PE=1 SV=1 | 0 | 22.717 | 11 | 6 | 18 | 6 | 663 | 75.7 | 7.44 | 41.12 |
| Q6UX73 | UPF0764 protein C16orf89 OS=Homo sapiens OX=9606 GN=C16orf89 PE=1 SV=2 | 0 | 22.57 | 16 | 6 | 54 | 6 | 402 | 45.4 | 6.19 | 139.04 |
| Q969H8 | Myeloid-derived growth factor OS=Homo sapiens OX=9606 GN=MYDGF PE=1 SV=1 | 0 | 22.567 | 27 | 4 | 45 | 4 | 173 | 18.8 | 6.68 | 87.9 |
| O60234 | Glia maturation factor gamma OS=Homo sapiens OX=9606 GN=GMFG PE=1 SV=1 | 0 | 22.499 | 58 | 6 | 11 | 6 | 142 | 16.8 | 5.26 | 29.92 |
| A0A0C4DH33 | Immunoglobulin heavy variable 1-24 OS=Homo sapiens OX=9606 GN=IGHV1-24 PE=3 SV=1 | 0 | 22.442 | 47 | 3 | 5 | 3 | 117 | 12.8 | 5.16 | 22.83 |
| O15145 | Actin-related protein 2/3 complex subunit 3 OS=Homo sapiens OX=9606 GN=ARPC3 PE=1 SV=3 | 0 | 22.312 | 30 | 5 | 31 | 5 | 178 | 20.5 | 8.59 | 47.97 |
| Q04609 | Glutamate carboxypeptidase 2 OS=Homo sapiens OX=9606 GN=FOLH1 PE=1 SV=1 | 0 | 22.283 | 8 | 4 | 30 | 4 | 750 | 84.3 | 6.98 | 72.64 |
| P28072 | Proteasome subunit beta type-6 OS=Homo sapiens OX=9606 GN=PSMB6 PE=1 SV=4 | 0 | 22.256 | 34 | 5 | 27 | 5 | 239 | 25.3 | 4.92 | 53.7 |
| P41091 | Eukaryotic translation initiation factor 2 subunit 3 OS=Homo sapiens OX=9606 GN=EIF2S3 PE=1 SV=3 | 0 | 22.173 | 17 | 4 | 8 | 4 | 472 | 51.1 | 8.4 | 19.53 |
| A0A075B6S5 | Immunoglobulin kappa variable 1-27 OS=Homo sapiens OX=9606 GN=IGKV1-27 PE=3 SV=1 | 0 | 22.135 | 56 | 4 | 52 | 3 | 117 | 12.7 | 8.29 | 204.54 |
| P15814 | Immunoglobulin lambda-like polypeptide 1 OS=Homo sapiens OX=9606 GN=IGLL1 PE=1 SV=1 | 0 | 22.118 | 18 | 3 | 46 | 3 | 213 | 22.9 | 10.07 | 171.2 |
| O60613 | Selenoprotein F OS=Homo sapiens OX=9606 GN=SELENOF PE=1 SV=4 | 0 | 22.11 | 30 | 4 | 25 | 4 | 165 | 18.1 | 5.03 | 48.68 |
| P53602 | Diphosphomevalonate decarboxylase OS=Homo sapiens OX=9606 GN=MVD PE=1 SV=1 | 0 | 22.075 | 27 | 6 | 21 | 6 | 400 | 43.4 | 7.23 | 18.03 |
| Q92882 | Osteoclast-stimulating factor 1 OS=Homo sapiens OX=9606 GN=OSTF1 PE=1 SV=2 | 0 | 22.016 | 34 | 5 | 10 | 5 | 214 | 23.8 | 5.68 | 18.97 |
| Q92597 | Protein NDRG1 OS=Homo sapiens OX=9606 GN=NDRG1 PE=1 SV=1 | 0 | 21.998 | 27 | 5 | 13 | 5 | 394 | 42.8 | 5.82 | 25.84 |
| P04278 | Sex hormone-binding globulin OS=Homo sapiens OX=9606 GN=SHBG PE=1 SV=2 | 0 | 21.907 | 21 | 4 | 8 | 4 | 402 | 43.8 | 6.71 | 22.57 |
| P47897 | Glutamine--tRNA ligase OS=Homo sapiens OX=9606 GN=QARS1 PE=1 SV=1 | 0 | 21.9 | 8 | 5 | 25 | 5 | 775 | 87.7 | 7.15 | 84.04 |
| P09651 | Heterogeneous nuclear ribonucleoprotein A1 OS=Homo sapiens OX=9606 GN=HNRNPA1 PE=1 SV=5 | 0 | 21.87 | 17 | 4 | 8 | 4 | 372 | 38.7 | 9.13 | 24.03 |
| P47755 | F-actin-capping protein subunit alpha-2 OS=Homo sapiens OX=9606 GN=CAPZA2 PE=1 SV=3 | 0 | 21.71 | 29 | 6 | 12 | 4 | 286 | 32.9 | 5.85 | 29.26 |
| Q13765 | Nascent polypeptide-associated complex subunit alpha OS=Homo sapiens OX=9606 GN=NACA PE=1 SV=1 | 0 | 21.636 | 27 | 4 | 26 | 4 | 215 | 23.4 | 4.56 | 78.76 |
| P78324 | Tyrosine-protein phosphatase non-receptor type substrate 1 OS=Homo sapiens OX=9606 GN=SIRPA PE=1 SV=2 | 0 | 21.597 | 18 | 7 | 46 | 7 | 504 | 54.9 | 6.98 | 47.83 |
| Q16348 | Solute carrier family 15 member 2 OS=Homo sapiens OX=9606 GN=SLC15A2 PE=1 SV=2 | 0 | 21.57 | 9 | 8 | 72 | 8 | 729 | 81.7 | 8.16 | 62.79 |
| Q99584 | Protein S100-A13 OS=Homo sapiens OX=9606 GN=S100A13 PE=1 SV=1 | 0 | 21.549 | 65 | 5 | 16 | 5 | 98 | 11.5 | 6.16 | 33.22 |
| P62910 | 60S ribosomal protein L32 OS=Homo sapiens OX=9606 GN=RPL32 PE=1 SV=2 | 0 | 21.524 | 17 | 2 | 16 | 2 | 135 | 15.9 | 11.33 | 29.19 |
| Q687X5 | Metalloreductase STEAP4 OS=Homo sapiens OX=9606 GN=STEAP4 PE=1 SV=1 | 0 | 21.477 | 17 | 5 | 23 | 5 | 459 | 51.9 | 9.29 | 37.35 |
| O75367 | Core histone macro-H2A.1 OS=Homo sapiens OX=9606 GN=MACROH2A1 PE=1 SV=4 | 0 | 21.454 | 30 | 7 | 8 | 7 | 372 | 39.6 | 9.79 | 17 |
| P16152 | Carbonyl reductase [NADPH] 1 OS=Homo sapiens OX=9606 GN=CBR1 PE=1 SV=3 | 0 | 21.451 | 28 | 6 | 10 | 6 | 277 | 30.4 | 8.32 | 18.1 |
| P03973 | Antileukoproteinase OS=Homo sapiens OX=9606 GN=SLPI PE=1 SV=2 | 0 | 21.449 | 28 | 5 | 24 | 5 | 132 | 14.3 | 8.75 | 51.9 |
| Q2I0M4 | Leucine-rich repeat-containing protein 26 OS=Homo sapiens OX=9606 GN=LRRC26 PE=1 SV=2 | 0 | 21.43 | 15 | 3 | 11 | 3 | 334 | 34.8 | 9.17 | 33.8 |
| P62854 | 40S ribosomal protein S26 OS=Homo sapiens OX=9606 GN=RPS26 PE=1 SV=3 | 0 | 21.391 | 31 | 3 | 32 | 3 | 115 | 13 | 11 | 64.13 |
| O75347 | Tubulin-specific chaperone A OS=Homo sapiens OX=9606 GN=TBCA PE=1 SV=3 | 0 | 21.37 | 37 | 3 | 8 | 3 | 108 | 12.8 | 5.29 | 5.71 |
| P08962 | CD63 antigen OS=Homo sapiens OX=9606 GN=CD63 PE=1 SV=2 | 0 | 21.355 | 17 | 3 | 37 | 3 | 238 | 25.6 | 7.81 | 106.53 |
| P24534 | Elongation factor 1-beta OS=Homo sapiens OX=9606 GN=EEF1B2 PE=1 SV=3 | 0 | 21.337 | 24 | 4 | 17 | 4 | 225 | 24.7 | 4.67 | 36.07 |
| Q5D862 | Filaggrin-2 OS=Homo sapiens OX=9606 GN=FLG2 PE=1 SV=1 | 0 | 21.303 | 2 | 4 | 5 | 4 | 2391 | 247.9 | 8.31 | 15.23 |

Table S1. Cont.

| Accession | Description | Exp. q-value: Combined | Sum PEP Score | Coverage [%] | # Peptides | # PSMs | # Unique Peptides | # AAs | MW [kDa] | calc. pI | Score Sequest HT: Sequest HT | # Peptides (by Search Engine): Sequest HT | Abundances (Normalized): Normal-Colostrum-1 | Abundances (Normalized): Normal-Colostrum-2 | Abundances (Normalized): Normal-Colostrum-3 | Abundances (Normalized): Normal-Colostrum-4 | Abundances (Normalized): Normal-Colostrum-5 | Abundances (Normalized): Normal-Colostrum-6 | Abundances (Normalized): Normal-Colostrum-7 | Abundances (Normalized): Normal-Colostrum-8 | Abundances (Normalized): Normal-Colostrum-9 | Abundances (Normalized): Normal-Colostrum-10 | Abundances (Normalized): Covid-colostrum-1 | Abundances (Normalized): Covid-colostrum-2 | Abundances (Normalized): Covid-colostrum-3 | Abundances (Normalized): Covid-colostrum-4 | Abundances (Normalized): Covid-colostrum-5 | Abundances (Normalized): Covid-colostrum-6 | Abundances (Normalized): Normal-mature-1 | Abundances (Normalized): Normal-mature-2 | Abundances (Normalized): Normal-mature-3 | Abundances (Normalized): Normal-mature-4 | Abundances (Normalized): Normal-mature-5 | Abundances (Normalized): Normal-mature-6 | Abundances (Normalized): Normal-mature-7 | Abundances (Normalized): Normal-mature-8 | Abundances (Normalized): Normal-mature-9 | Abundances (Normalized): Normal-mature-10 | Abundances (Normalized): Covid-mature-1 | Abundances (Normalized): Covid-mature-2 | Abundances (Normalized): Covid-mature-3 | Abundances (Normalized): Covid-mature-4 | Abundances (Normalized): Covid-mature-5 | Abundances (Normalized): Covid-mature-6 |
|---|---|---|---|---|---|---|---|---|---|---|---|---|---|---|---|---|---|---|---|---|---|---|---|---|---|---|---|---|---|---|---|---|---|---|---|---|---|---|---|---|---|---|---|---|
| Q9UN76 | Sodium- and chloride-dependent neutral and basic amino acid transporter B(0+) OS=Homo sapiens OX=9606 GN=SLC6A14 PE=2 SV=1 | 0 | 21.239 | 7 | 5 | 85 | 5 | 642 | 72.1 | 8.19 | 213.93 | 5 | 1816669.357 | 1810586.144 | 2309024.446 | 1578238.071 | 1340773.33 | 1524603.084 | 2264077.492 | 1234725.665 | 491806.7084 | 1280352.757 | 686402.5833 | 2024902.586 | 493811.8316 | 381861.8072 | 1995150.087 | 539265.2433 | 456156.3694 | 785070.9646 | 417165.7909 | 966771.3797 | 482629.0934 | 957791.8908 | 1585799.174 | 1119785.777 | 275906.8335 | 770121.0513 | 1657144.813 | 1886651.986 | 1490526.593 | 1229812.145 | 1578176.465 | 1602301.132 |
| O43598 | 2'-deoxynucleoside 5'-phosphate N-hydrolase 1 OS=Homo sapiens OX=9606 GN=DNPH1 PE=1 SV=1 | 0 | 21.143 | 33 | 3 | 12 | 3 | 174 | 19.1 | 5.05 | 36.18 | 3 | 247192.2347 | 447654.5258 | | 780741.0162 | 275816.1258 | 437778.521 | 503167.1013 | 330073.2039 | 247113.8361 | 346106.281 | | 246076.5766 | | | 472678.8115 | | 455651.6972 | 326668.3026 | 1079517.273 | 451257.6077 | 1027776.557 | 428921.4987 | 305494.7052 | 270441.3021 | 467641.1027 | 798538.3355 | 888050.8684 | 694320.848 | 1223765.347 | 1111891.469 | 413668.8389 | 1210608.527 |
| P05198 | Eukaryotic translation initiation factor 2 subunit 1 OS=Homo sapiens OX=9606 GN=EIF2S1 PE=1 SV=3 | 0 | 21.032 | 15 | 4 | 10 | 4 | 315 | 36.1 | 5.08 | 16.33 | 4 | | | | | | | 1886837.755 | | 679409.1877 | | | | | | | | | | | | | | | | | | | 1032545.758 | 1657470.746 | 951866.4679 | | |
| P01717 | Immunoglobulin lambda variable 3-25 OS=Homo sapiens OX=9606 GN=IGLV3-25 PE=1 SV=2 | 0 | 20.865 | 37 | 3 | 63 | 1 | 112 | 12 | 4.5 | 120.57 | 3 | | | | | | | | | | | 9181828 | | | | | | | | | | | | | | | | | | | | | |
| P07357 | Complement component C8 alpha chain OS=Homo sapiens OX=9606 GN=C8A PE=1 SV=2 | 0 | 20.818 | 12 | 5 | 37 | 5 | 584 | 65.1 | 6.47 | 84.53 | 5 | 159418.2564 | 125546.1977 | 708876.9815 | 111164.1974 | 400342.6467 | | 75946.92064 | 203221.2545 | 48319.38452 | 95957.37893 | 220889.4167 | 212106.8466 | 140630.5112 | | 389220.4191 | 132916.1054 | 239705.4084 | | 366194.0606 | 215714.4876 | 123066.9436 | 270470.8803 | 202521.3346 | 283360.1669 | 93851.68818 | 60252.2593 | 156532.1331 | 522892.1409 | | 317967.43 | 133030.7018 | 407865.0602 |
| Q9BTY2 | Plasma alpha-L-fucosidase OS=Homo sapiens OX=9606 GN=FUCA2 PE=1 SV=2 | 0 | 20.798 | 14 | 6 | 28 | 6 | 467 | 54 | 6.25 | 49.22 | 6 | 447151.0812 | 533311.2281 | | 783996.8476 | 1151596.817 | 568957.201 | | 551192.0005 | 387094.7914 | 1112060.337 | 2668547.583 | | 1619384.552 | | 733895.5371 | | | | | 1142221.498 | | | | | 1079038.653 | 2867757.525 | 853703.4768 | | 501386.0851 | | 491449.4798 | |
| Q8WUA8 | Tsukushin OS=Homo sapiens OX=9606 GN=TSKU PE=2 SV=3 | 0 | 20.789 | 20 | 4 | 5 | 4 | 353 | 37.8 | 6.87 | 15.21 | 4 | 1040663.79 | 428600.511 | 245055.0889 | 1108180.371 | | | 1102317.266 | 1210822.699 | 653022.9587 | 683130.8568 | 1780332.875 | | | | 891574.826 | | 1053745.774 | 562877.5669 | 445043.5259 | 1057745.59 | 1105465.723 | 288011.5411 | 146641.2629 | 843216.8417 | 269626.1427 | 84662.67852 | | 1127334.412 | 1564389.806 | | 1307938.56 | 2290254.145 |
| O15162 | Phospholipid scramblase 1 OS=Homo sapiens OX=9606 GN=PLSCR1 PE=1 SV=1 | 0 | 20.78 | 23 | 5 | 24 | 5 | 318 | 35 | 4.94 | 25.79 | 5 | | | 2850745.863 | 266286.3342 | 184431.9973 | | 407549.309 | | 504273.6251 | | 479138.9583 | 2434949.405 | 256048.9402 | 381864.6822 | | | 210613.8651 | | 438596.1012 | 183456.077 | | | 293163.0555 | | 569992.3895 | | | 258913.3469 | | 233265.2758 | 150417.299 | 2500566.375 |
| A0A087WSY4 | Immunoglobulin heavy variable 4-30-2 OS=Homo sapiens OX=9606 GN=IGHV4-30-2 PE=3 SV=1 | 0 | 20.715 | 28 | 4 | 85 | 2 | 118 | 13 | 9.67 | 196.48 | 4 | 1899158.036 | 750594.7075 | 1430262.401 | 1472433.527 | 1661695.302 | 702744.15 | 1394963.122 | 4325397.479 | 707932.3817 | 699310.7324 | 13377802 | 1201108.043 | 5381267.067 | 6126759.367 | 4577681.758 | 6180612.796 | 653725.859 | 564248.2678 | 596847.8127 | 1072449.005 | 822820.9785 | | 534121.3929 | 836207.8872 | 540615.5823 | | 777262.2076 | 467537.7845 | 700765.3083 | 458586.2214 | 737885.3161 | 835153.7596 |
| Q68D85 | Natural cytotoxicity triggering receptor 3 ligand 1 OS=Homo sapiens OX=9606 GN=NCR3LG1 PE=1 SV=1 | 0 | 20.655 | 12 | 4 | 15 | 4 | 454 | 50.8 | 5.59 | 38.21 | 4 | | | | | | | | | | | 519324.5417 | | | | | | 147282.808 | 293106.0635 | 339085.447 | | | 257540.3385 | 702912.6426 | | 789569.2707 | | 1771191.006 | 329217.3394 | | 699052.3449 | | 418131.616 |
| Q53TN4 | Cytochrome b reductase 1 OS=Homo sapiens OX=9606 GN=CYBRD1 PE=1 SV=1 | 0 | 20.629 | 19 | 4 | 33 | 4 | 286 | 31.6 | 8.76 | 66.18 | 4 | | 1393122.537 | | 537903.6984 | 1445490.343 | 415282.821 | 1130392.52 | 305668.9514 | 127404.0002 | 408921.699 | | 1329420.807 | 361841.2037 | 650064.8243 | | 388793.1564 | 964692.4348 | 891921.6477 | | 408457.9902 | 564852.0521 | | 745881.8749 | 351634.3118 | | 111276.7685 | 1848124.877 | 811562.7112 | | 306358.498 | 144447.9533 | 1218885.848 |
| P26992 | Ciliary neurotrophic factor receptor subunit alpha OS=Homo sapiens OX=9606 GN=CNTFR PE=1 SV=2 | 0 | 20.612 | 19 | 4 | 15 | 4 | 372 | 40.6 | 6.76 | 37.71 | 4 | | | | 545451.5616 | 385258.8913 | 285716.2268 | 1288769.844 | 790130.8598 | | 519052.4534 | | 1413267.111 | | | 1421140.028 | | 355137.016 | 950986.9729 | | 605937.1489 | 474103.3283 | 514470.3681 | 821838.8417 | 721417.7364 | | 414851.7233 | 429161.9716 | 627297.6051 | 779876.7649 | | 1041812.794 | 710861.7025 |
| P62244 | 40S ribosomal protein S15a OS=Homo sapiens OX=9606 GN=RPS15A PE=1 SV=2 | 0 | 20.453 | 36 | 5 | 56 | 5 | 130 | 14.8 | 10.13 | 138.43 | 5 | | 253002.6777 | | 1411395.268 | 1356581.956 | 1176608.949 | 1135575.369 | 507592.7556 | 1212418.145 | 1430201.519 | 624132.2917 | 187951.7547 | | | 2160171.393 | | 256529.8002 | | 225270.6747 | 1919623.022 | 2033291.494 | 1251928.224 | | 178181.4276 | 287679.8424 | 1403034.936 | 1493150.374 | 2075309.434 | 2560325.342 | 290940.8636 | 1460889.563 | 4829164.339 |
| P53621 | Coatomer subunit alpha OS=Homo sapiens OX=9606 GN=COPA PE=1 SV=2 | 0 | 20.442 | 8 | 6 | 6 | 6 | 1224 | 138.3 | 7.66 | 7.39 | 6 | | | 1860332.841 | 6140601.537 | 1437335.323 | 1346560.175 | | 1574671.935 | 2269441.042 | 625111.9053 | 12897286.67 | 6306873.667 | | 3045019.569 | 5460194.203 | 270755.9662 | 241643.1027 | | 8925924.676 | 1113765.204 | 634747.7054 | 829041.6807 | 4605118.301 | 6613381.671 | 3898112.309 | 1603095.939 | | 2053829.234 | 178480.1639 | | 8646330.176 | 5803178.994 |
| P49721 | Proteasome subunit beta type-2 OS=Homo sapiens OX=9606 GN=PSMB2 PE=1 SV=1 | 0 | 20.334 | 39 | 5 | 8 | 5 | 201 | 22.8 | 7.02 | 18.72 | 5 | | 171631.5964 | | | | 991166.859 | | | | 489566.2558 | | | | | | | | | | 1084649.855 | 294686.6641 | | | | 1210841.445 | 1082132.987 | | | 547386.5753 | 303277.1818 | 607415.6685 | 742218.0844 |
| Q9UNW1 | Multiple inositol polyphosphate phosphatase 1 OS=Homo sapiens OX=9606 GN=MINPP1 PE=1 SV=1 | 0 | 20.324 | 20 | 6 | 18 | 6 | 487 | 55 | 7.81 | 22.89 | 6 | 18199894.38 | 32851059.54 | 10889699.58 | 76955047.52 | 25516070.29 | 75817123.17 | 80281277.58 | 30420284.31 | 38672167.86 | 228159979.1 | 5708140.667 | 29021093.38 | 1187778.943 | 471693.6233 | 71139491.92 | 542266.6409 | 42133645.05 | 13641293.9 | 76301970.24 | 93022490.91 | 15988569.78 | 590877016.9 | 30236448.1 | 46236713.01 | 27509793.79 | 77164576.41 | 54716475.83 | 938401080.1 | 58055644.54 | 134413330.8 | 168884924.4 | 76210127.68 |

[illegible]

Table S1. Cont.

| Accession | Description | Exp. q-value: Combined | Sum PEP Score | Coverage [%] | # Peptides | # PSMs | # Unique Peptides | # AAs | MW [kDa] | calc. pI | Score Sequest HT: Sequest HT | # Peptides (by Search Engine): Sequest HT |
|---|---|---|---|---|---|---|---|---|---|---|---|---|
| P53004 | Biliverdin reductase A OS=Homo sapiens OX=9606 GN=BLVRA PE=1 SV=2 | 0 | 11.622 | 8 | 1 | 2 | 1 | 296 | 33.4 | 6.44 | 16.76 | 1 |
| Q9H008 | Phospholysine phosphohistidine inorganic pyrophosphate phosphatase OS=Homo sapiens OX=9606 GN=LHPP PE=1 SV=2 | 0 | 11.617 | 12 | 2 | 4 | 2 | 270 | 29.1 | 6.15 | 13.46 | 2 |
| Q96SA4 | Serine incorporator 2 OS=Homo sapiens OX=9606 GN=SERINC2 PE=2 SV=3 | 0 | 11.595 | 6 | 1 | 1 | 1 | 455 | 50.7 | 6.19 | 5.65 | 1 |
| P55735 | Protein SEC13 homolog OS=Homo sapiens OX=9606 GN=SEC13 PE=1 SV=3 | 0 | 11.575 | 12 | 2 | 4 | 2 | 322 | 35.5 | 5.48 | 13.95 | 2 |
| Q15080 | Neutrophil cytosol factor 4 OS=Homo sapiens OX=9606 GN=NCF4 PE=1 SV=2 | 0 | 11.486 | 8 | 2 | 2 | 2 | 339 | 39 | 6.83 | 7.51 | 2 |
| P13671 | Complement component C6 OS=Homo sapiens OX=9606 GN=C6 PE=1 SV=3 | 0 | 11.401 | 4 | 3 | 5 | 3 | 934 | 104.7 | 6.76 | 6.91 | 3 |
| Q96DG6 | Carboxymethylenebutenolidase homolog OS=Homo sapiens OX=9606 GN=CMBL PE=1 SV=1 | 0 | 11.376 | 10 | 2 | 12 | 2 | 245 | 28 | 7.18 | 23.3 | 2 |
| P21796 | Voltage-dependent anion-selective channel protein 1 OS=Homo sapiens OX=9606 GN=VDAC1 PE=1 SV=2 | 0 | 11.369 | 7 | 1 | 3 | 1 | 283 | 30.8 | 8.54 | 10.1 | 1 |
| Q99460 | 26S proteasome non-ATPase regulatory subunit 1 OS=Homo sapiens OX=9606 GN=PSMD1 PE=1 SV=2 | 0 | 11.344 | 3 | 3 | 7 | 3 | 953 | 105.8 | 5.39 | 13.9 | 3 |
| O00160 | Unconventional myosin-If OS=Homo sapiens OX=9606 GN=MYO1F PE=1 SV=3 | 0 | 11.341 | 6 | 3 | 3 | 3 | 1098 | 124.8 | 9.11 | 8.25 | 3 |
| Q8NFU4 | Follicular dendritic cell secreted peptide OS=Homo sapiens OX=9606 GN=FDCSP PE=1 SV=1 | 0 | 11.237 | 36 | 1 | 14 | 1 | 85 | 9.7 | 9.32 | 70.15 | 1 |
| Q9NY33 | Dipeptidyl peptidase 3 OS=Homo sapiens OX=9606 GN=DPP3 PE=1 SV=2 | 0 | 11.218 | 7 | 4 | 5 | 4 | 737 | 82.5 | 5.1 | 9.27 | 4 |
| P55884 | Eukaryotic translation initiation factor 3 subunit B OS=Homo sapiens OX=9606 GN=EIF3B PE=1 SV=3 | 0 | 11.19 | 4 | 3 | 8 | 3 | 814 | 92.4 | 5 | 18.41 | 3 |
| Q9NTK5 | Obg-like ATPase 1 OS=Homo sapiens OX=9606 GN=OLA1 PE=1 SV=2 | 0 | 11.175 | 6 | 2 | 16 | 2 | 396 | 44.7 | 7.81 | 28.08 | 2 |
| Q9UGM3 | Deleted in malignant brain tumors 1 protein OS=Homo sapiens OX=9606 GN=DMBT1 PE=1 SV=2 | 0 | 11.158 | 14 | 4 | 12 | 4 | 2413 | 260.6 | 5.44 | 34.3 | 4 |
| P0DP25 | Calmodulin-3 OS=Homo sapiens OX=9606 GN=CALM3 PE=1 SV=1 | 0 | 11.126 | 21 | 2 | 5 | 2 | 149 | 16.8 | 4.22 | 18.03 | 2 |
| P47914 | 60S ribosomal protein L29 OS=Homo sapiens OX=9606 GN=RPL29 PE=1 SV=2 | 0 | 11.123 | 14 | 2 | 29 | 2 | 159 | 17.7 | 11.66 | 80.08 | 2 |
| P62847 | 40S ribosomal protein S24 OS=Homo sapiens OX=9606 GN=RPS24 PE=1 SV=1 | 0 | 11.104 | 18 | 3 | 29 | 3 | 133 | 15.4 | 10.78 | 74.31 | 3 |
| P55259 | Pancreatic secretory granule membrane major glycoprotein GP2 OS=Homo sapiens OX=9606 GN=GP2 PE=1 SV=3 | 0 | 11.101 | 9 | 4 | 5 | 4 | 537 | 59.4 | 5.24 | 6.98 | 4 |
| Q9Y262 | Eukaryotic translation initiation factor 3 subunit L OS=Homo sapiens OX=9606 GN=EIF3L PE=1 SV=1 | 0 | 11.057 | 7 | 3 | 8 | 3 | 564 | 66.7 | 6.34 | 14.98 | 3 |
| Q96QK1 | Vacuolar protein sorting-associated protein 35 OS=Homo sapiens OX=9606 GN=VPS35 PE=1 SV=2 | 0 | 10.943 | 4 | 3 | 5 | 3 | 796 | 91.6 | 5.49 | 9.45 | 3 |
| Q8IZA0 | Dyslexia-associated protein KIAA0319-like protein OS=Homo sapiens OX=9606 GN=KIAA0319L PE=1 SV=2 | 0 | 10.84 | 2 | 2 | 12 | 2 | 1049 | 115.6 | 6.05 | 26.6 | 2 |
| Q7Z404 | Transmembrane channel-like protein 4 OS=Homo sapiens OX=9606 GN=TMC4 PE=1 SV=3 | 0 | 10.825 | 7 | 3 | 11 | 3 | 712 | 79.2 | 8.94 | 9.71 | 3 |
| P80723 | Brain acid soluble protein 1 OS=Homo sapiens OX=9606 GN=BASP1 PE=1 SV=2 | 0 | 10.779 | 17 | 2 | 2 | 2 | 227 | 22.7 | 4.63 | 7.7 | 2 |
| P09769 | Tyrosine-protein kinase Fgr OS=Homo sapiens OX=9606 GN=FGR PE=1 SV=2 | 0 | 10.655 | 5 | 2 | 13 | 1 | 529 | 59.4 | 5.59 | 25.24 | 2 |
| P14866 | Heterogeneous nuclear ribonucleoprotein L OS=Homo sapiens OX=9606 GN=HNRNPL PE=1 SV=2 | 0 | 10.644 | 5 | 1 | 1 | 1 | 589 | 64.1 | 8.22 | 7.54 | 1 |
| P01721 | Immunoglobulin lambda variable 6-57 OS=Homo sapiens OX=9606 GN=IGLV6-57 PE=1 SV=2 | 0 | 10.62 | 17 | 1 | 18 | 1 | 117 | 12.6 | 4.78 | 79.56 | 1 |
| P35475 | Alpha-L-iduronidase OS=Homo sapiens OX=9606 GN=IDUA PE=1 SV=2 | 0 | 10.616 | 6 | 3 | 4 | 3 | 653 | 72.6 | 9.14 | 8.36 | 3 |
| P61619 | Protein transport protein Sec61 subunit alpha isoform 1 OS=Homo sapiens OX=9606 GN=SEC61A1 PE=1 SV=2 | 0 | 10.57 | 8 | 3 | 12 | 3 | 476 | 52.2 | 8.06 | 16.06 | 3 |
| P27449 | V-type proton ATPase 16 kDa proteolipid subunit OS=Homo sapiens OX=9606 GN=ATP6V0C PE=1 SV=1 | 0 | 10.54 | 20 | 1 | 1 | 1 | 155 | 15.7 | 8.44 | 6.79 | 1 |
| P10253 | Lysosomal alpha-glucosidase OS=Homo sapiens OX=9606 GN=GAA PE=1 SV=4 | 0 | 10.534 | 5 | 3 | 8 | 3 | 952 | 105.3 | 6 | 18.56 | 3 |
| O43813 | Glutathione S-transferase LANCL1 OS=Homo sapiens OX=9606 GN=LANCL1 PE=1 SV=1 | 0 | 10.532 | 9 | 2 | 5 | 2 | 399 | 45.3 | 7.75 | 11.37 | 2 |
| O76003 | Glutaredoxin-3 OS=Homo sapiens OX=9606 GN=GLRX3 PE=1 SV=2 | 0 | 10.521 | 8 | 1 | 23 | 1 | 335 | 37.4 | 5.39 | 111.14 | 1 |
| P35237 | Serpin B6 OS=Homo sapiens OX=9606 GN=SERPINB6 PE=1 SV=3 | 0 | 10.516 | 11 | 3 | 3 | 3 | 376 | 42.6 | 5.27 | 5.88 | 3 |
| Q9UJJ9 | N-acetylglucosamine-1-phosphotransferase subunit gamma OS=Homo sapiens OX=9606 GN=GNPTG PE=1 SV=1 | 0 | 10.469 | 11 | 2 | 3 | 2 | 305 | 34 | 6.95 | 5.34 | 2 |
| Q15063 | Periostin OS=Homo sapiens OX=9606 GN=POSTN PE=1 SV=2 | 0 | 10.439 | 7 | 3 | 4 | 3 | 836 | 93.3 | 7.53 | 12.3 | 3 |
| P21757 | Macrophage scavenger receptor types I and II OS=Homo sapiens OX=9606 GN=MSR1 PE=1 SV=1 | 0 | 10.427 | 6 | 1 | 1 | 1 | 451 | 49.7 | 5.91 | 7.79 | 1 |
| Q6GMV3 | Putative peptidyl-tRNA hydrolase PTRHD1 OS=Homo sapiens OX=9606 GN=PTRHD1 PE=1 SV=1 | 0 | 10.426 | 21 | 1 | 1 | 1 | 140 | 15.8 | 9.1 | 6.85 | 1 |
| P0DTE1 | Probable non-functional immunoglobulin heavy variable 3-38-3 OS=Homo sapiens OX=9606 GN=IGHV3-38-3 PE=1 SV=1 | 0 | 10.424 | 19 | 3 | 38 | 2 | 115 | 12.7 | 9.25 | 77.4 | 3 |
| P61970 | Nuclear transport factor 2 OS=Homo sapiens OX=9606 GN=NUTF2 PE=1 SV=1 | 0 | 10.38 | 46 | 4 | 17 | 4 | 127 | 14.5 | 5.38 | 8.5 | 4 |
| P21291 | Cysteine and glycine-rich protein 1 OS=Homo sapiens OX=9606 GN=CSRP1 PE=1 SV=3 | 0 | 10.371 | 17 | 2 | 3 | 2 | 193 | 20.6 | 8.57 | 6.03 | 2 |
| Q6UXY8 | Transmembrane channel-like protein 5 OS=Homo sapiens OX=9606 GN=TMC5 PE=2 SV=3 | 0 | 10.362 | 4 | 4 | 16 | 4 | 1006 | 114.7 | 8.21 | 14.72 | 4 |
| O14672 | Disintegrin and metalloproteinase domain-containing protein 10 OS=Homo sapiens OX=9606 GN=ADAM10 PE=1 SV=1 | 0 | 10.348 | 5 | 3 | 24 | 3 | 748 | 84.1 | 7.77 | 33.65 | 3 |
| P01715 | Immunoglobulin lambda variable 3-1 OS=Homo sapiens OX=9606 GN=IGLV3-1 PE=1 SV=2 | 0 | 10.27 | 26 | 2 | 19 | 1 | 115 | 12.3 | 4.94 | 39.29 | 2 |
| P06748 | Nucleophosmin OS=Homo sapiens OX=9606 GN=NPM1 PE=1 SV=2 | 0 | 10.254 | 10 | 2 | 2 | 2 | 294 | 32.6 | 4.78 | 7.29 | 2 |
| Q9P2B2 | Prostaglandin F2 receptor negative regulator OS=Homo sapiens OX=9606 GN=PTGFRN PE=1 SV=2 | 0 | 10.252 | 4 | 4 | 6 | 4 | 879 | 98.5 | 6.61 | 10.05 | 4 |
| P19878 | Neutrophil cytosol factor 2 OS=Homo sapiens OX=9606 GN=NCF2 PE=1 SV=2 | 0 | 10.249 | 11 | 3 | 3 | 3 | 526 | 59.7 | 6.16 | 8.67 | 3 |
| P25788 | Proteasome subunit alpha type-3 OS=Homo sapiens OX=9606 GN=PSMA3 PE=1 SV=2 | 0 | 10.243 | 14 | 3 | 23 | 3 | 255 | 28.4 | 5.33 | 53.47 | 3 |
| Q13046 | Putative pregnancy-specific beta-1-glycoprotein 7 OS=Homo sapiens OX=9606 GN=PSG7 PE=5 SV=3 | 0 | 10.23 | 9 | 3 | 5 | 3 | 419 | 47 | 8.43 | 8.39 | 3 |
| P60022 | Beta-defensin 1 OS=Homo sapiens OX=9606 GN=DEFB1 PE=1 SV=1 | 0 | 10.222 | 34 | 1 | 20 | 1 | 68 | 7.4 | 8.6 | 112.71 | 1 |
| Q13404 | Ubiquitin-conjugating enzyme E2 variant 1 OS=Homo sapiens OX=9606 GN=UBE2V1 PE=1 SV=2 | 0 | 10.203 | 26 | 4 | 6 | 4 | 147 | 16.5 | 7.93 | 5.27 | 4 |
| B2RUZ4 | Small integral membrane protein 1 OS=Homo sapiens OX=9606 GN=SMIM1 PE=1 SV=1 | 0 | 10.155 | 36 | 2 | 10 | 2 | 78 | 8.7 | 9.03 | 13.12 | 2 |
| Q9UM54 | Unconventional myosin-VI OS=Homo sapiens OX=9606 GN=MYO6 PE=1 SV=4 | 0 | 10.13 | 4 | 4 | 4 | 4 | 1294 | 149.6 | 8.53 | 3.94 | 4 |
| P99999 | Cytochrome c OS=Homo sapiens OX=9606 GN=CYCS PE=1 SV=2 | 0 | 10.079 | 24 | 2 | 2 | 2 | 105 | 11.7 | 9.57 | 5.73 | 2 |
| O43927 | C-X-C motif chemokine 13 OS=Homo sapiens OX=9606 GN=CXCL13 PE=1 SV=1 | 0 | 10.014 | 23 | 2 | 7 | 2 | 109 | 12.7 | 10.33 | 22.54 | 2 |
| Q9Y678 | Coatomer subunit gamma-1 OS=Homo sapiens OX=9606 GN=COPG1 PE=1 SV=1 | 0 | 10.002 | 6 | 4 | 5 | 4 | 874 | 97.7 | 5.47 | 10.61 | 4 |
| P22059 | Oxysterol-binding protein 1 OS=Homo sapiens OX=9606 GN=OSBP PE=1 SV=1 | 0 | 10.001 | 5 | 3 | 6 | 2 | 807 | 89.4 | 7.3 | 11.98 | 3 |
| P49407 | Beta-arrestin-1 OS=Homo sapiens OX=9606 GN=ARRB1 PE=1 SV=2 | 0 | 9.948 | 14 | 4 | 6 | 4 | 418 | 47 | 6.2 | 5.63 | 4 |
| O00264 | Membrane-associated progesterone receptor component 1 OS=Homo sapiens OX=9606 GN=PGRMC1 PE=1 SV=3 | 0 | 9.93 | 32 | 3 | 4 | 3 | 195 | 21.7 | 4.7 | 9.58 | 3 |
| P60228 | Eukaryotic translation initiation factor 3 subunit E OS=Homo sapiens OX=9606 GN=EIF3E PE=1 SV=1 | 0 | 9.872 | 9 | 4 | 8 | 4 | 445 | 52.2 | 6.04 | 12.34 | 4 |
| P62330 | ADP-ribosylation factor 6 OS=Homo sapiens OX=9606 GN=ARF6 PE=1 SV=2 | 0 | 9.75 | 21 | 3 | 11 | 2 | 175 | 20.1 | 8.95 | 30.17 | 3 |
| O94804 | Serine/threonine-protein kinase 10 OS=Homo sapiens OX=9606 GN=STK10 PE=1 SV=1 | 0 | 9.711 | 2 | 1 | 1 | 1 | 968 | 112.1 | 6.95 | 4.02 | 1 |
| Q99598 | Translin-associated protein X OS=Homo sapiens OX=9606 GN=TSNAX PE=1 SV=1 | 0 | 9.675 | 7 | 1 | 5 | 1 | 290 | 33.1 | 6.55 | 16.65 | 1 |
| O00410 | Importin-5 OS=Homo sapiens OX=9606 GN=IPO5 PE=1 SV=4 | 0 | 9.639 | 2 | 1 | 20 | 1 | 1097 | 123.6 | 4.94 | 62.92 | 1 |
| Q15843 | NEDD8 OS=Homo sapiens OX=9606 GN=NEDD8 PE=1 SV=1 | 0 | 9.596 | 17 | 1 | 1 | 1 | 81 | 9.1 | 8.43 | 3.1 | 1 |
| Q9UHG3 | Prenylcysteine oxidase 1 OS=Homo sapiens OX=9606 GN=PCYOX1 PE=1 SV=3 | 0 | 9.591 | 9 | 4 | 8 | 4 | 505 | 56.6 | 6.18 | 5.74 | 4 |
| P01718 | Immunoglobulin lambda variable 3-27 OS=Homo sapiens OX=9606 GN=IGLV3-27 PE=1 SV=2 | 0 | 9.553 | 19 | 2 | 30 | 1 | 113 | 12.2 | 5.01 | 45.97 | 2 |
| A0A075B6H7 | Probable non-functional immunoglobulin kappa variable 3-7 OS=Homo sapiens OX=9606 GN=IGKV3-7 PE=1 SV=1 | 0 | 9.55 | 42 | 3 | 61 | 2 | 116 | 12.8 | 5.25 | 128 | 3 |
| Q9Y5Z4 | Heme-binding protein 2 OS=Homo sapiens OX=9606 GN=HEBP2 PE=1 SV=1 | 0 | 9.542 | 15 | 3 | 6 | 3 | 205 | 22.9 | 4.63 | 6.2 | 3 |
| Q15382 | GTP-binding protein Rheb OS=Homo sapiens OX=9606 GN=RHEB PE=1 SV=1 | 0 | 9.527 | 14 | 2 | 8 | 2 | 184 | 20.5 | 5.92 | 17.7 | 2 |
| O43776 | Asparagine--tRNA ligase, cytoplasmic OS=Homo sapiens OX=9606 GN=NARS1 PE=1 SV=1 | 0 | 9.509 | 6 | 3 | 4 | 3 | 548 | 62.9 | 6.25 | 4.61 | 3 |
| Q5IS43 | Platelet-activating factor acetylhydrolase IB subunit alpha OS=Pan troglodytes OX=9598 GN=PAFAH1B1 PE=2 SV=3 | 0 | 9.496 | 11 | 4 | 6 | 3 | 410 | 46.7 | 7.37 | 1.87 | 4 |
| Q9Y2Y8 | Proteoglycan 3 OS=Homo sapiens OX=9606 GN=PRG3 PE=1 SV=2 | 0 | 9.442 | 12 | 2 | 4 | 2 | 225 | 25.4 | 4.81 | 7.04 | 2 |
| Q5XG92 | Carboxylesterase 4A OS=Homo sapiens OX=9606 GN=CES4A PE=2 SV=2 | 0 | 9.44 | 5 | 2 | 6 | 2 | 561 | 63.5 | 9.31 | 9.68 | 2 |
| Q15121 | Astrocytic phosphoprotein PEA-15 OS=Homo sapiens OX=9606 GN=PEA15 PE=1 SV=2 | 0 | 9.384 | 18 | 1 | 6 | 1 | 130 | 15 | 5.02 | 11.4 | 1 |
| P00492 | Hypoxanthine-guanine phosphoribosyltransferase OS=Homo sapiens OX=9606 GN=HPRT1 PE=1 SV=2 | 0 | 9.379 | 16 | 3 | 4 | 3 | 218 | 24.6 | 6.68 | 4.65 | 3 |
| Q96FJ2 | Dynein light chain 2, cytoplasmic OS=Homo sapiens OX=9606 GN=DYNLL2 PE=1 SV=1 | 0 | 9.375 | 13 | 2 | 4 | 2 | 89 | 10.3 | 7.37 | 6.3 | 2 |
| O15305 | Phosphomannomutase 2 OS=Homo sapiens OX=9606 GN=PMM2 PE=1 SV=1 | 0 | 9.338 | 13 | 2 | 5 | 2 | 246 | 28.1 | 6.77 | 12.92 | 2 |
| A0A075B6K0 | Immunoglobulin lambda variable 3-16 OS=Homo sapiens OX=9606 GN=IGLV3-16 PE=3 SV=2 | 0 | 9.309 | 18 | 2 | 33 | 1 | 115 | 12.5 | 4.65 | 47.74 | 2 |
| Q9P2X0 | Dolichol-phosphate mannosyltransferase subunit 3 OS=Homo sapiens OX=9606 GN=DPM3 PE=1 SV=2 | 0 | 9.255 | 24 | 2 | 49 | 2 | 92 | 10.1 | 5.94 | 113.69 | 2 |
| Q8N752 | Casein kinase I isoform alpha-like OS=Homo sapiens OX=9606 GN=CSNK1A1L PE=2 SV=2 | 0 | 9.234 | 10 | 3 | 16 | 3 | 337 | 39.1 | 9.45 | 15.44 | 3 |
| Q12907 | Vesicular integral-membrane protein VIP36 OS=Homo sapiens OX=9606 GN=LMAN2 PE=1 SV=1 | 0 | 9.223 | 10 | 2 | 4 | 2 | 356 | 40.2 | 6.95 | 8.37 | 2 |
| O15511 | Actin-related protein 2/3 complex subunit 5 OS=Homo sapiens OX=9606 GN=ARPC5 PE=1 SV=3 | 0 | 9.158 | 17 | 2 | 5 | 2 | 151 | 16.3 | 5.67 | 6.41 | 2 |
| Q08722 | Leukocyte surface antigen CD47 OS=Homo sapiens OX=9606 GN=CD47 PE=1 SV=1 | 0 | 9.154 | 9 | 3 | 45 | 3 | 323 | 35.2 | 7.21 | 75.1 | 3 |
| Q9UBQ7 | Glyoxylate reductase/hydroxypyruvate reductase OS=Homo sapiens OX=9606 GN=GRHPR PE=1 SV=1 | 0 | 9.131 | 15 | 2 | 3 | 2 | 328 | 35.6 | 7.39 | 9.02 | 2 |
| O00481 | Butyrophilin subfamily 3 member A1 OS=Homo sapiens OX=9606 GN=BTN3A1 PE=1 SV=3 | 0 | 9.098 | 3 | 2 | 28 | 1 | 513 | 57.6 | 8.03 | 65.8 | 2 |
| P09603 | Macrophage colony-stimulating factor 1 OS=Homo sapiens OX=9606 GN=CSF1 PE=1 SV=2 | 0 | 9.084 | 6 | 2 | 4 | 2 | 554 | 60.1 | 5.29 | 14.69 | 2 |
| P40763 | Signal transducer and activator of transcription 3 OS=Homo sapiens OX=9606 GN=STAT3 PE=1 SV=2 | 0 | 9.063 | 6 | 2 | 4 | 2 | 770 | 88 | 6.3 | 7.88 | 2 |
| P15289 | Arylsulfatase A OS=Homo sapiens OX=9606 GN=ARSA PE=1 SV=3 | 0 | 9.056 | 6 | 2 | 8 | 2 | 507 | 53.6 | 6.07 | 19.75 | 2 |
| P40617 | ADP-ribosylation factor-like protein 4A OS=Homo sapiens OX=9606 GN=ARL4A PE=1 SV=2 | 0 | 9.051 | 15 | 2 | 6 | 2 | 200 | 22.6 | 9.17 | 13.14 | 2 |
| Q6UWX4 | HHIP-like protein 2 OS=Homo sapiens OX=9606 GN=HHIPL2 PE=1 SV=1 | 0 | 9.031 | 6 | 3 | 3 | 3 | 724 | 80.7 | 9.01 | 6.1 | 3 |
| Q9ULP9 | TBC1 domain family member 24 OS=Homo sapiens OX=9606 GN=TBC1D24 PE=1 SV=2 | 0 | 9.021 | 7 | 4 | 6 | 4 | 559 | 62.9 | 7.36 | 5.89 | 4 |
| A0A0C4DH30 | Probable non-functional immunoglobulin heavy variable 3-16 OS=Homo sapiens OX=9606 GN=IGHV3-16 PE=1 SV=1 | 0 | 9.005 | 47 | 4 | 7 | 4 | 117 | 13.1 | 9.73 | 12.69 | 4 |
| Q96HE7 | ERO1-like protein alpha OS=Homo sapiens OX=9606 GN=ERO1A PE=1 SV=2 | 0 | 8.953 | 5 | 2 | 11 | 2 | 468 | 54.4 | 5.68 | 19.4 | 2 |
| Q04323 | UBX domain-containing protein 1 OS=Homo sapiens OX=9606 GN=UBXN1 PE=1 SV=2 | 0 | 8.942 | 6 | 1 | 2 | 1 | 297 | 33.3 | 5.25 | 7.38 | 1 |
| P18077 | 60S ribosomal protein L35a OS=Homo sapiens OX=9606 GN=RPL35A PE=1 SV=2 | 0 | 8.884 | 23 | 4 | 12 | 4 | 110 | 12.5 | 11.06 | 7.21 | 4 |
| O60784 | Target of Myb protein 1 OS=Homo sapiens OX=9606 GN=TOM1 PE=1 SV=2 | 0 | 8.748 | 12 | 3 | 4 | 3 | 492 | 53.8 | 4.7 | 3.35 | 3 |
| P55957 | BH3-interacting domain death agonist OS=Homo sapiens OX=9606 GN=BID PE=1 SV=1 | 0 | 8.691 | 8 | 1 | 2 | 1 | 195 | 22 | 5.44 | 10.14 | 1 |
| Q96D31 | Calcium release-activated calcium channel protein 1 OS=Homo sapiens OX=9606 GN=ORAI1 PE=1 SV=2 | 0 | 8.678 | 4 | 1 | 13 | 1 | 301 | 32.6 | 8.32 | 47.06 | 1 |
| P20930 | Filaggrin OS=Homo sapiens OX=9606 GN=FLG PE=1 SV=3 | 0 | 8.676 | 1 | 3 | 3 | 3 | 4061 | 434.9 | 9.25 | 5 | 3 |
| P20827 | Ephrin-A1 OS=Homo sapiens OX=9606 GN=EFNA1 PE=1 SV=2 | 0 | 8.672 | 13 | 2 | 33 | 2 | 205 | 23.8 | 6.83 | 63.97 | 2 |
| P38606 | V-type proton ATPase catalytic subunit A OS=Homo sapiens OX=9606 GN=ATP6V1A PE=1 SV=2 | 0 | 8.652 | 5 | 3 | 3 | 3 | 617 | 68.3 | 5.52 | 2.74 | 3 |
| Q96G03 | Phosphoglucomutase-2 OS=Homo sapiens OX=9606 GN=PGM2 PE=1 SV=4 | 0 | 8.618 | 3 | 2 | 4 | 2 | 612 | 68.2 | 6.73 | 5.4 | 2 |
| Q15008 | 26S proteasome non-ATPase regulatory subunit 6 OS=Homo sapiens OX=9606 GN=PSMD6 PE=1 SV=1 | 0 | 8.604 | 5 | 1 | 1 | 1 | 389 | 45.5 | 5.62 | 5.56 | 1 |
| Q96RF0 | Sorting nexin-18 OS=Homo sapiens OX=9606 GN=SNX18 PE=1 SV=2 | 0 | 8.594 | 5 | 2 | 3 | 2 | 628 | 68.9 | 5.68 | 5.39 | 2 |
| P62993 | Growth factor receptor-bound protein 2 OS=Homo sapiens OX=9606 GN=GRB2 PE=1 SV=1 | 0 | 8.544 | 18 | 4 | 4 | 4 | 217 | 25.2 | 6.32 | 9.77 | 4 |
| P25686 | DnaJ homolog subfamily B member 2 OS=Homo sapiens OX=9606 GN=DNAJB2 PE=1 SV=3 | 0 | 8.52 | 10 | 3 | 15 | 3 | 324 | 35.6 | 5.95 | 9.13 | 3 |
| P08236 | Beta-glucuronidase OS=Homo sapiens OX=9606 GN=GUSB PE=1 SV=2 | 0 | 8.484 | 4 | 2 | 5 | 2 | 651 | 74.7 | 7.02 | 9.2 | 2 |
| P62333 | 26S proteasome regulatory subunit 10B OS=Homo sapiens OX=9606 GN=PSMC6 PE=1 SV=1 | 0 | 8.482 | 7 | 2 | 7 | 2 | 389 | 44.1 | 7.49 | 13.39 | 2 |
| P31948 | Stress-induced-phosphoprotein 1 OS=Homo sapiens OX=9606 GN=STIP1 PE=1 SV=1 | 0 | 8.456 | 8 | 3 | 6 | 3 | 543 | 62.6 | 6.8 | 8.65 | 3 |
| P01601 | Immunoglobulin kappa variable 1D-16 OS=Homo sapiens OX=9606 GN=IGKV1D-16 PE=3 SV=2 | 0 | 8.425 | 14 | 1 | 5 | 1 | 117 | 12.7 | 7.74 | 22.11 | 1 |
| P01920 | HLA class II histocompatibility antigen, DQ beta 1 chain OS=Homo sapiens OX=9606 GN=HLA-DQB1 PE=1 SV=2 | 0 | 8.416 | 12 | 3 | 3 | 3 | 261 | 30 | 7.25 | 4.99 | 3 |
| Q99735 | Microsomal glutathione S-transferase 2 OS=Homo sapiens OX=9606 GN=MGST2 PE=1 SV=1 | 0 | 8.408 | 10 | 1 | 11 | 1 | 147 | 16.6 | 9.55 | 22.9 | 1 |
| Q96AD5 | Patatin-like phospholipase domain-containing protein 2 OS=Homo sapiens OX=9606 GN=PNPLA2 PE=1 SV=1 | 0 | 8.405 | 5 | 2 | 2 | 2 | 504 | 55.3 | 7.08 | 5.19 | 2 |
| Q5W0Z9 | Palmitoyltransferase ZDHHC20 OS=Homo sapiens OX=9606 GN=ZDHHC20 PE=1 SV=1 | 0 | 8.387 | 14 | 4 | 15 | 4 | 365 | 42.3 | 7.71 | 10.78 | 4 |
| Q96S97 | Myeloid-associated differentiation marker OS=Homo sapiens OX=9606 GN=MYADM PE=1 SV=2 | 0 | 8.339 | 10 | 2 | 5 | 2 | 322 | 35.3 | 8.15 | 7.7 | 2 |
| Q9C0H2 | Protein tweety homolog 3 OS=Homo sapiens OX=9606 GN=TTYH3 PE=1 SV=3 | 0 | 8.333 | 6 | 2 | 4 | 2 | 523 | 57.5 | 5.39 | 10.88 | 2 |
| Q9UHY7 | Enolase-phosphatase E1 OS=Homo sapiens OX=9606 GN=ENOPH1 PE=1 SV=1 | 0 | 8.318 | 19 | 2 | 2 | 2 | 261 | 28.9 | 4.78 | 4.23 | 2 |
| P61960 | Ubiquitin-fold modifier 1 OS=Homo sapiens OX=9606 GN=UFM1 PE=1 SV=1 | 0 | 8.304 | 27 | 2 | 25 | 2 | 85 | 9.1 | 9.31 | 48.67 | 2 |
| P12838 | Neutrophil defensin 4 OS=Homo sapiens OX=9606 GN=DEFA4 PE=1 SV=2 | 0 | 8.295 | 18 | 1 | 7 | 1 | 97 | 10.5 | 8.02 | 30.01 | 1 |
| Q07654 | Trefoil factor 3 OS=Homo sapiens OX=9606 GN=TFF3 PE=1 SV=2 | 0 | 8.276 | 33 | 2 | 7 | 2 | 94 | 10.2 | 6.92 | 16.22 | 2 |
| P04211 | Immunoglobulin lambda variable 7-43 OS=Homo sapiens OX=9606 GN=IGLV7-43 PE=3 SV=2 | 0 | 8.225 | 21 | 3 | 68 | 1 | 117 | 12.4 | 7.03 | 126.23 | 3 |
| P62136 | Serine/threonine-protein phosphatase PP1-alpha catalytic subunit OS=Homo sapiens OX=9606 GN=PPP1CA PE=1 SV=1 | 0 | 8.192 | 11 | 2 | 6 | 2 | 330 | 37.5 | 6.33 | 14.93 | 2 |
| Q13393 | Phospholipase D1 OS=Homo sapiens OX=9606 GN=PLD1 PE=1 SV=1 | 0 | 8.164 | 3 | 3 | 7 | 3 | 1074 | 124.1 | 8.78 | 6.51 | 3 |
| Q9Y3D6 | Mitochondrial fission 1 protein OS=Homo sapiens OX=9606 GN=FIS1 PE=1 SV=2 | 0 | 8.109 | 11 | 1 | 10 | 1 | 152 | 16.9 | 8.79 | 25.13 | 1 |
| P55786 | Puromycin-sensitive aminopeptidase OS=Homo sapiens OX=9606 GN=NPEPPS PE=1 SV=2 | 0 | 8.036 | 5 | 3 | 3 | 3 | 919 | 103.2 | 5.72 | 2.45 | 3 |
| P55036 | 26S proteasome non-ATPase regulatory subunit 4 OS=Homo sapiens OX=9606 GN=PSMD4 PE=1 SV=1 | 0 | 7.998 | 4 | 1 | 3 | 1 | 377 | 40.7 | 4.79 | 7.47 | 1 |
| Q7LBR1 | Charged multivesicular body protein 1b OS=Homo sapiens OX=9606 GN=CHMP1B PE=1 SV=1 | 0 | 7.982 | 13 | 3 | 32 | 3 | 199 | 22.1 | 8.1 | 48.04 | 3 |
| Q9H9H4 | Vacuolar protein sorting-associated protein 37B OS=Homo sapiens OX=9606 GN=VPS37B PE=1 SV=1 | 0 | 7.967 | 9 | 1 | 2 | 1 | 285 | 31.3 | 7.34 | 8.62 | 1 |
| P81605 | Dermcidin OS=Homo sapiens OX=9606 GN=DCD PE=1 SV=2 | 0 | 7.947 | 22 | 2 | 14 | 2 | 110 | 11.3 | 6.54 | 19.58 | 2 |
| Q13011 | Delta(3,5)-Delta(2,4)-dienoyl-CoA isomerase, mitochondrial OS=Homo sapiens OX=9606 GN=ECH1 PE=1 SV=2 | 0 | 7.916 | 8 | 2 | 2 | 2 | 328 | 35.8 | 8 | 5.13 | 2 |
| O00602 | Ficolin-1 OS=Homo sapiens OX=9606 GN=FCN1 PE=1 SV=2 | 0 | 7.915 | 17 | 2 | 2 | 2 | 326 | 35.1 | 6.86 | 6.42 | 2 |
| P21980 | Protein-glutamine gamma-glutamyltransferase 2 OS=Homo sapiens OX=9606 GN=TGM2 PE=1 SV=2 | 0 | 7.914 | 5 | 3 | 3 | 3 | 687 | 77.3 | 5.22 | 4.73 | 3 |
| P18887 | DNA repair protein XRCC1 OS=Homo sapiens OX=9606 GN=XRCC1 PE=1 SV=2 | 0 | 7.903 | 6 | 3 | 3 | 3 | 633 | 69.4 | 6.39 | 4.39 | 3 |
| Q9UBX5 | Fibulin-5 OS=Homo sapiens OX=9606 GN=FBLN5 PE=1 SV=1 | 0 | 7.84 | 5 | 2 | 3 | 2 | 448 | 50.1 | 4.73 | 8.05 | 2 |
| B5ME19 | Eukaryotic translation initiation factor 3 subunit C-like protein OS=Homo sapiens OX=9606 GN=EIF3CL PE=1 SV=1 | 0 | 7.83 | 2 | 2 | 15 | 2 | 914 | 105.4 | 5.64 | 32.93 | 2 |
| O75923 | Dysferlin OS=Homo sapiens OX=9606 GN=DYSF PE=1 SV=1 | 0 | 7.828 | 2 | 2 | 4 | 1 | 2080 | 237.1 | 5.64 | 11.72 | 2 |
| P63313 | Thymosin beta-10 OS=Homo sapiens OX=9606 GN=TMSB10 PE=1 SV=2 | 0 | 7.828 | 66 | 2 | 4 | 2 | 44 | 5 | 5.36 | 7.39 | 2 |
| Q7L2H7 | Eukaryotic translation initiation factor 3 subunit M OS=Homo sapiens OX=9606 GN=EIF3M PE=1 SV=1 | 0 | 7.711 | 10 | 2 | 3 | 2 | 374 | 42.5 | 5.63 | 7.51 | 2 |
| Q9H3Z4 | DnaJ homolog subfamily C member 5 OS=Homo sapiens OX=9606 GN=DNAJC5 PE=1 SV=1 | 0 | 7.709 | 16 | 2 | 10 | 2 | 198 | 22.1 | 5.07 | 9.72 | 2 |
| P0DOX4 | Immunoglobulin epsilon heavy chain OS=Homo sapiens OX=9606 PE=1 SV=1 | 0 | 7.702 | 4 | 2 | 6 | 2 | 547 | 60.3 | 7.99 | 10.83 | 2 |
| Q9UBS4 | DnaJ homolog subfamily B member 11 OS=Homo sapiens OX=9606 GN=DNAJB11 PE=1 SV=1 | 0 | 7.678 | 6 | 2 | 11 | 2 | 358 | 40.5 | 6.18 | 9.63 | 2 |
| O00754 | Lysosomal alpha-mannosidase OS=Homo sapiens OX=9606 GN=MAN2B1 PE=1 SV=3 | 0 | 7.668 | 4 | 2 | 3 | 2 | 1011 | 113.7 | 7.28 | 5.86 | 2 |
| Q9P0L0 | Vesicle-associated membrane protein-associated protein A OS=Homo sapiens OX=9606 GN=VAPA PE=1 SV=3 | 0 | 7.63 | 11 | 2 | 4 | 2 | 249 | 27.9 | 8.62 | 4.41 | 2 |
| O14556 | Glyceraldehyde-3-phosphate dehydrogenase, testis-specific OS=Homo sapiens OX=9606 GN=GAPDHS PE=1 SV=2 | 0 | 7.629 | 4 | 2 | 45 | 1 | 408 | 44.5 | 8.19 | 70.83 | 2 |
| Q969P0 | Immunoglobulin superfamily member 8 OS=Homo sapiens OX=9606 GN=IGSF8 PE=1 SV=1 | 0 | 7.621 | 3 | 2 | 13 | 2 | 613 | 65 | 8 | 16.49 | 2 |
| Q92484 | Acid sphingomyelinase-like phosphodiesterase 3a OS=Homo sapiens OX=9606 GN=SMPDL3A PE=1 SV=2 | 0 | 7.594 | 7 | 3 | 11 | 3 | 453 | 51.2 | 6.33 | 1.96 | 3 |
| A0A0C4DH35 | Probable non-functional immunoglobulin heavy variable 3-35 OS=Homo sapiens OX=9606 GN=IGHV3-35 PE=1 SV=1 | 0 | 7.572 | 15 | 2 | 44 | 2 | 117 | 12.8 | 7.88 | 129.1 | 2 |
| P48595 | Serpin B10 OS=Homo sapiens OX=9606 GN=SERPINB10 PE=1 SV=1 | 0 | 7.536 | 9 | 3 | 3 | 3 | 397 | 45.4 | 6.16 | 6.28 | 3 |
| P21741 | Midkine OS=Homo sapiens OX=9606 GN=MDK PE=1 SV=1 | 0 | 7.452 | 12 | 2 | 33 | 2 | 143 | 15.6 | 9.79 | 60.86 | 2 |
| P55008 | Allograft inflammatory factor 1 OS=Homo sapiens OX=9606 GN=AIF1 PE=1 SV=1 | 0 | 7.448 | 14 | 1 | 1 | 1 | 147 | 16.7 | 6.24 | 4 | 1 |
| Q969Q0 | 60S ribosomal protein L36a-like OS=Homo sapiens OX=9606 GN=RPL36AL PE=1 SV=3 | 0 | 7.429 | 16 | 2 | 26 | 2 | 106 | 12.5 | 10.65 | 45.53 | 2 |
| P10153 | Non-secretory ribonuclease OS=Homo sapiens OX=9606 GN=RNASE2 PE=1 SV=2 | 0 | 7.408 | 11 | 1 | 1 | 1 | 161 | 18.3 | 8.73 | 4.61 | 1 |
| Q00765 | Receptor expression-enhancing protein 5 OS=Homo sapiens OX=9606 GN=REEP5 PE=1 SV=3 | 0 | 7.39 | 11 | 2 | 3 | 2 | 189 | 21.5 | 8.1 | 4.66 | 2 |
| P10644 | cAMP-dependent protein kinase type I-alpha regulatory subunit OS=Homo sapiens OX=9606 GN=PRKAR1A PE=1 SV=1 | 0 | 7.379 | 6 | 1 | 1 | 1 | 381 | 43 | 5.35 | 5.09 | 1 |
| P61768 | Major prion protein OS=Pan troglodytes OX=9598 GN=PRNP PE=2 SV=1 | 0 | 7.368 | 5 | 1 | 5 | 1 | 253 | 27.6 | 9.14 | 16.08 | 1 |
| Q9NR50 | Translation initiation factor eIF-2B subunit gamma OS=Homo sapiens OX=9606 GN=EIF2B3 PE=1 SV=1 | 0 | 7.355 | 6 | 2 | 2 | 2 | 452 | 50.2 | 6.47 | 3.29 | 2 |
| Q6JBY9 | CapZ-interacting protein OS=Homo sapiens OX=9606 GN=RCSD1 PE=1 SV=1 | 0 | 7.33 | 4 | 1 | 1 | 1 | 416 | 44.5 | 5.4 | 2.63 | 1 |
| Q96BM9 | ADP-ribosylation factor-like protein 8A OS=Homo sapiens OX=9606 GN=ARL8A PE=1 SV=1 | 0 | 7.33 | 19 | 3 | 7 | 3 | 186 | 21.4 | 7.77 | 11.06 | 3 |
| Q08211 | ATP-dependent RNA helicase A OS=Homo sapiens OX=9606 GN=DHX9 PE=1 SV=4 | 0 | 7.306 | 1 | 1 | 1 | 1 | 1270 | 140.9 | 6.84 | 4.13 | 1 |
| Q01629 | Interferon-induced transmembrane protein 2 OS=Homo sapiens OX=9606 GN=IFITM2 PE=1 SV=2 | 0 | 7.296 | 12 | 1 | 2 | 1 | 132 | 14.6 | 7.42 | 5.51 | 1 |
| P60468 | Protein transport protein Sec61 subunit beta OS=Homo sapiens OX=9606 GN=SEC61B PE=1 SV=2 | 0 | 7.262 | 27 | 2 | 23 | 2 | 96 | 10 | 11.56 | 23.92 | 2 |
| Q9UGM5 | Fetuin-B OS=Homo sapiens OX=9606 GN=FETUB PE=1 SV=2 | 0 | 7.223 | 10 | 2 | 3 | 2 | 382 | 42 | 6.83 | 7.31 | 2 |
| P22735 | Protein-glutamine gamma-glutamyltransferase K OS=Homo sapiens OX=9606 GN=TGM1 PE=1 SV=4 | 0 | 7.22 | 4 | 2 | 2 | 2 | 817 | 89.7 | 6.04 | 5.63 | 2 |
| Q14508 | WAP four-disulfide core domain protein 2 OS=Homo sapiens OX=9606 GN=WFDC2 PE=1 SV=2 | 0 | 7.216 | 11 | 1 | 4 | 1 | 124 | 13 | 4.84 | 13.64 | 1 |
| Q13113 | PDZK1-interacting protein 1 OS=Homo sapiens OX=9606 GN=PDZK1IP1 PE=1 SV=1 | 0 | 7.187 | 18 | 2 | 9 | 2 | 114 | 12.2 | 4.83 | 15.03 | 2 |
| A0A075B6J1 | Immunoglobulin lambda variable 5-37 OS=Homo sapiens OX=9606 GN=IGLV5-37 PE=3 SV=1 | 0 | 7.174 | 16 | 1 | 1 | 1 | 123 | 13.3 | 4.94 | 3.15 | 1 |
| Q99747 | Gamma-soluble NSF attachment protein OS=Homo sapiens OX=9606 GN=NAPG PE=1 SV=1 | 0 | 7.166 | 7 | 2 | 4 | 2 | 312 | 34.7 | 5.41 | 0 | 2 |
| O00629 | Importin subunit alpha-3 OS=Homo sapiens OX=9606 GN=KPNA4 PE=1 SV=1 | 0 | 7.158 | 3 | 1 | 1 | 1 | 521 | 57.9 | 4.96 | 4.47 | 1 |
| Q9BW60 | Elongation of very long chain fatty acids protein 1 OS=Homo sapiens OX=9606 GN=ELOVL1 PE=1 SV=1 | 0 | 7.126 | 5 | 1 | 2 | 1 | 279 | 32.6 | 9.6 | 6.49 | 1 |
| Q9Y6N5 | Sulfide:quinone oxidoreductase, mitochondrial OS=Homo sapiens OX=9606 GN=SQOR PE=1 SV=1 | 0 | 7.103 | 3 | 1 | 2 | 1 | 450 | 49.9 | 9.11 | 7.14 | 1 |
| Q9P2J5 | Leucine--tRNA ligase, cytoplasmic OS=Homo sapiens OX=9606 GN=LARS1 PE=1 SV=2 | 0 | 7.031 | 3 | 2 | 2 | 2 | 1176 | 134.4 | 7.3 | 3.4 | 2 |
| P28482 | Mitogen-activated protein kinase 1 OS=Homo sapiens OX=9606 GN=MAPK1 PE=1 SV=3 | 0 | 7.027 | 9 | 3 | 4 | 3 | 360 | 41.4 | 6.98 | 3.57 | 3 |
| Q8IZV5 | Retinol dehydrogenase 10 OS=Homo sapiens OX=9606 GN=RDH10 PE=1 SV=1 | 0 | 6.971 | 7 | 2 | 2 | 2 | 341 | 38.1 | 7.4 | 4.15 | 2 |
| P45974 | Ubiquitin carboxyl-terminal hydrolase 5 OS=Homo sapiens OX=9606 GN=USP5 PE=1 SV=2 | 0 | 6.955 | 2 | 1 | 1 | 1 | 858 | 95.7 | 5.03 | 4.71 | 1 |
| P19652 | Alpha-1-acid glycoprotein 2 OS=Homo sapiens OX=9606 GN=ORM2 PE=1 SV=2 | 0 | 6.928 | 9 | 2 | 37 | 2 | 201 | 23.6 | 5.11 | 81.68 | 2 |
| P22528 | Cornifin-B OS=Homo sapiens OX=9606 GN=SPRR1B PE=1 SV=2 | 0 | 6.927 | 44 | 3 | 9 | 3 | 89 | 9.9 | 8.48 | 4.52 | 3 |
| O15067 | Phosphoribosylformylglycinamidine synthase OS=Homo sapiens OX=9606 GN=PFAS PE=1 SV=4 | 0 | 6.916 | 1 | 1 | 1 | 1 | 1338 | 144.6 | 5.76 | 4.05 | 1 |
| Q96HC4 | PDZ and LIM domain protein 5 OS=Homo sapiens OX=9606 GN=PDLIM5 PE=1 SV=5 | 0 | 6.902 | 6 | 3 | 7 | 3 | 596 | 63.9 | 8.21 | 5.62 | 3 |
| P50454 | Serpin H1 OS=Homo sapiens OX=9606 GN=SERPINH1 PE=1 SV=2 | 0 | 6.895 | 10 | 2 | 3 | 2 | 418 | 46.4 | 8.69 | 6.93 | 2 |
| Q8N307 | Mucin-20 OS=Homo sapiens OX=9606 GN=MUC20 PE=1 SV=3 | 0 | 6.863 | 3 | 2 | 13 | 2 | 709 | 71.9 | 5.07 | 19.89 | 2 |
| P62273 | 40S ribosomal protein S29 OS=Homo sapiens OX=9606 GN=RPS29 PE=1 SV=2 | 0 | 6.816 | 34 | 2 | 18 | 2 | 56 | 6.7 | 10.13 | 16.21 | 2 |
| A0A075B6J9 | Immunoglobulin lambda variable 2-18 OS=Homo sapiens OX=9606 GN=IGLV2-18 PE=3 SV=2 | 0 | 6.78 | 25 | 2 | 11 | 2 | 118 | 12.4 | 4.82 | 16.66 | 2 |
| O75914 | Serine/threonine-protein kinase PAK 3 OS=Homo sapiens OX=9606 GN=PAK3 PE=1 SV=2 | 0 | 6.729 | 5 | 1 | 1 | 1 | 559 | 62.3 | 5.45 | 4.07 | 1 |
| P05204 | Non-histone chromosomal protein HMG-17 OS=Homo sapiens OX=9606 GN=HMGN2 PE=1 SV=3 | 0 | 6.725 | 17 | 1 | 4 | 1 | 90 | 9.4 | 9.99 | 6.22 | 1 |
| O00232 | 26S proteasome non-ATPase regulatory subunit 12 OS=Homo sapiens OX=9606 GN=PSMD12 PE=1 SV=3 | 0 | 6.699 | 3 | 1 | 1 | 1 | 456 | 52.9 | 7.65 | 3.03 | 1 |
| P26640 | Valine--tRNA ligase OS=Homo sapiens OX=9606 GN=VARS1 PE=1 SV=4 | 0 | 6.665 | 3 | 2 | 2 | 2 | 1264 | 140.4 | 7.59 | 2.42 | 2 |
| Q99436 | Proteasome subunit beta type-7 OS=Homo sapiens OX=9606 GN=PSMB7 PE=1 SV=1 | 0 | 6.661 | 12 | 3 | 16 | 3 | 277 | 29.9 | 7.68 | 4.14 | 3 |
| Q13501 | Sequestosome-1 OS=Homo sapiens OX=9606 GN=SQSTM1 PE=1 SV=1 | 0 | 6.645 | 3 | 1 | 2 | 1 | 440 | 47.7 | 5.22 | 1.87 | 1 |
| P01040 | Cystatin-A OS=Homo sapiens OX=9606 GN=CSTA PE=1 SV=1 | 0 | 6.639 | 28 | 1 | 1 | 1 | 98 | 11 | 5.5 | 4.24 | 1 |
| Q9Y376 | Calcium-binding protein 39 OS=Homo sapiens OX=9606 GN=CAB39 PE=1 SV=1 | 0 | 6.633 | 6 | 2 | 2 | 2 | 341 | 39.8 | 6.89 | 5.08 | 2 |
| A0A087WSX0 | Immunoglobulin lambda variable 5-45 OS=Homo sapiens OX=9606 GN=IGLV5-45 PE=3 SV=1 | 0 | 6.62 | 14 | 1 | 2 | 1 | 123 | 13.2 | 7.2 | 8.15 | 1 |
| Q9Y6C9 | Mitochondrial carrier homolog 2 OS=Homo sapiens OX=9606 GN=MTCH2 PE=1 SV=1 | 0 | 6.612 | 8 | 1 | 1 | 1 | 303 | 33.3 | 7.97 | 4.92 | 1 |
| Q13275 | Semaphorin-3F OS=Homo sapiens OX=9606 GN=SEMA3F PE=2 SV=2 | 0 | 6.606 | 4 | 3 | 3 | 3 | 785 | 88.3 | 8.27 | 4.21 | 3 |
| Q00169 | Phosphatidylinositol transfer protein alpha isoform OS=Homo sapiens OX=9606 GN=PITPNA PE=1 SV=2 | 0 | 6.6 | 7 | 2 | 12 | 2 | 270 | 31.8 | 6.55 | 14.64 | 2 |
| P14543 | Nidogen-1 OS=Homo sapiens OX=9606 GN=NID1 PE=1 SV=3 | 0 | 6.591 | 2 | 2 | 4 | 2 | 1247 | 136.3 | 5.29 | 10.66 | 2 |
| P16989 | Y-box-binding protein 3 OS=Homo sapiens OX=9606 GN=YBX3 PE=1 SV=4 | 0 | 6.586 | 5 | 1 | 5 | 1 | 372 | 40.1 | 9.77 | 13.53 | 1 |
| Q9BQ13 | BTB/POZ domain-containing protein KCTD14 OS=Homo sapiens OX=9606 GN=KCTD14 PE=1 SV=2 | 0 | 6.556 | 11 | 2 | 4 | 2 | 255 | 29.6 | 8.59 | 2.03 | 2 |
| P0CG30 | Glutathione S-transferase theta-2B OS=Homo sapiens OX=9606 GN=GSTT2B PE=1 SV=1 | 0 | 6.553 | 16 | 2 | 2 | 2 | 244 | 27.5 | 6.4 | 4.39 | 2 |
| P12236 | ADP/ATP translocase 3 OS=Homo sapiens OX=9606 GN=SLC25A6 PE=1 SV=4 | 0 | 6.538 | 10 | 3 | 3 | 3 | 298 | 32.8 | 9.74 | 2.38 | 3 |
| Q16674 | Melanoma-derived growth regulatory protein OS=Homo sapiens OX=9606 GN=MIA PE=1 SV=1 | 0 | 6.536 | 19 | 3 | 7 | 3 | 131 | 14.5 | 8.79 | 9.72 | 3 |
| P55327 | Tumor protein D52 OS=Homo sapiens OX=9606 GN=TPD52 PE=1 SV=2 | 0 | 6.53 | 6 | 1 | 7 | 1 | 224 | 24.3 | 4.83 | 18.4 | 1 |
| O75822 | Eukaryotic translation initiation factor 3 subunit J OS=Homo sapiens OX=9606 GN=EIF3J PE=1 SV=2 | 0 | 6.495 | 9 | 2 | 2 | 2 | 258 | 29 | 4.83 | 5.59 | 2 |
| P60059 | Protein transport protein Sec61 subunit gamma OS=Homo sapiens OX=9606 GN=SEC61G PE=1 SV=1 | 0 | 6.492 | 18 | 1 | 2 | 1 | 68 | 7.7 | 9.99 | 7.04 | 1 |
| Q15233 | Non-POU domain-containing octamer-binding protein OS=Homo sapiens OX=9606 GN=NONO PE=1 SV=4 | 0 | 6.491 | 5 | 1 | 2 | 1 | 471 | 54.2 | 8.95 | 7.52 | 1 |
| P36952 | Serpin B5 OS=Homo sapiens OX=9606 GN=SERPINB5 PE=1 SV=2 | 0 | 6.468 | 5 | 1 | 1 | 1 | 375 | 42.1 | 6.05 | 3.82 | 1 |
| Q969T9 | WW domain-binding protein 2 OS=Homo sapiens OX=9606 GN=WBP2 PE=1 SV=1 | 0 | 6.464 | 9 | 2 | 2 | 2 | 261 | 28.1 | 5.91 | 4.88 | 2 |
| P02746 | Complement C1q subcomponent subunit B OS=Homo sapiens OX=9606 GN=C1QB PE=1 SV=3 | 0 | 6.46 | 8 | 2 | 4 | 2 | 253 | 26.7 | 8.63 | 9.67 | 2 |
| Q9HAS3 | Solute carrier family 28 member 3 OS=Homo sapiens OX=9606 GN=SLC28A3 PE=1 SV=1 | 0 | 6.458 | 2 | 1 | 5 | 1 | 691 | 76.9 | 7.62 | 15.57 | 1 |
| P62316 | Small nuclear ribonucleoprotein Sm D2 OS=Homo sapiens OX=9606 GN=SNRPD2 PE=1 SV=1 | 0 | 6.428 | 8 | 1 | 1 | 1 | 118 | 13.5 | 9.91 | 1.86 | 1 |
| Q13561 | Dynactin subunit 2 OS=Homo sapiens OX=9606 GN=DCTN2 PE=1 SV=4 | 0 | 6.417 | 7 | 3 | 5 | 3 | 401 | 44.2 | 5.21 | 6.89 | 3 |
| P17213 | Bactericidal permeability-increasing protein OS=Homo sapiens OX=9606 GN=BPI PE=1 SV=4 | 0 | 6.338 | 5 | 1 | 1 | 1 | 487 | 53.9 | 9.38 | 4.11 | 1 |
| Q8IUE6 | Histone H2A type 2-B OS=Homo sapiens OX=9606 GN=H2AC21 PE=1 SV=3 | 0 | 6.315 | 12 | 2 | 24 | 1 | 130 | 14 | 10.89 | 39.1 | 2 |
| O75340 | Programmed cell death protein 6 OS=Homo sapiens OX=9606 GN=PDCD6 PE=1 SV=1 | 0 | 6.31 | 13 | 2 | 5 | 2 | 191 | 21.9 | 5.4 | 6.52 | 2 |
| Q8NC51 | Plasminogen activator inhibitor 1 RNA-binding protein OS=Homo sapiens OX=9606 GN=SERBP1 PE=1 SV=2 | 0 | 6.27 | 8 | 3 | 12 | 3 | 408 | 44.9 | 8.65 | 4.73 | 3 |
| O43633 | Charged multivesicular body protein 2a OS=Homo sapiens OX=9606 GN=CHMP2A PE=1 SV=1 | 0 | 6.214 | 11 | 3 | 4 | 3 | 222 | 25.1 | 5.97 | 5.03 | 3 |
| Q9Y2S2 | Lambda-crystallin homolog OS=Homo sapiens OX=9606 GN=CRYL1 PE=1 SV=3 | 0 | 6.179 | 16 | 3 | 3 | 3 | 319 | 35.4 | 6.18 | 1.91 | 3 |

Table S1. Cont.

| Accession | Description | Exp. q-value: Combined | Sum PEP Score | Coverage [%] | # Peptides | # PSMs | # Unique Peptides | # AAs | MW [kDa] | calc. pI | Score Sequest HT: Sequest HT | # Peptides (by Search Engine): Sequest HT |
|---|---|---|---|---|---|---|---|---|---|---|---|---|
| Q9Y2V2 | Calcium-regulated heat-stable protein 1 OS=Homo sapiens OX=9606 GN=CARHSP1 PE=1 SV=2 | 0 | 6.162 | 11 | 1 | 5 | 1 | 147 | 15.9 | 8.21 | 12.66 | 1 |
| Q01955 | Collagen alpha-3(IV) chain OS=Homo sapiens OX=9606 GN=COL4A3 PE=1 SV=3 | 0 | 6.148 | 1 | 1 | 6 | 1 | 1670 | 161.7 | 9.16 | 10.03 | 1 |
| O00499 | Myc box-dependent-interacting protein 1 OS=Homo sapiens OX=9606 GN=BIN1 PE=1 SV=1 | 0 | 6.132 | 6 | 3 | 4 | 2 | 593 | 64.7 | 5.06 | 4.75 | 3 |
| O15372 | Eukaryotic translation initiation factor 3 subunit H OS=Homo sapiens OX=9606 GN=EIF3H PE=1 SV=1 | 0 | 6.125 | 9 | 3 | 8 | 3 | 352 | 39.9 | 6.54 | 2.54 | 3 |
| P54577 | Tyrosine--tRNA ligase, cytoplasmic OS=Homo sapiens OX=9606 GN=YARS1 PE=1 SV=4 | 0 | 6.096 | 4 | 2 | 2 | 2 | 528 | 59.1 | 7.05 | 3.55 | 2 |
| Q9H0U3 | Magnesium transporter protein 1 OS=Homo sapiens OX=9606 GN=MAGT1 PE=1 SV=1 | 0 | 6.041 | 5 | 2 | 6 | 2 | 335 | 38 | 9.63 | 4.34 | 2 |
| Q9BVK6 | Transmembrane emp24 domain-containing protein 9 OS=Homo sapiens OX=9606 GN=TMED9 PE=1 SV=2 | 0 | 6.036 | 11 | 2 | 5 | 2 | 235 | 27.3 | 8.02 | 4.7 | 2 |
| P61923 | Coatomer subunit zeta-1 OS=Homo sapiens OX=9606 GN=COPZ1 PE=1 SV=1 | 0 | 6.018 | 8 | 1 | 1 | 1 | 177 | 20.2 | 4.81 | 2.93 | 1 |
| Q9Y3E7 | Charged multivesicular body protein 3 OS=Homo sapiens OX=9606 GN=CHMP3 PE=1 SV=3 | 0 | 6.007 | 9 | 2 | 9 | 2 | 222 | 25.1 | 5.12 | 15.19 | 2 |
| Q5JPE7 | Nodal modulator 2 OS=Homo sapiens OX=9606 GN=NOMO2 PE=1 SV=1 | 0 | 5.986 | 2 | 2 | 4 | 2 | 1267 | 139.4 | 5.76 | 6.24 | 2 |
| Q969R2 | Oxysterol-binding protein 2 OS=Homo sapiens OX=9606 GN=OSBP2 PE=1 SV=2 | 0 | 5.984 | 3 | 2 | 3 | 1 | 916 | 101.2 | 6.54 | 4.81 | 2 |
| O75436 | Vacuolar protein sorting-associated protein 26A OS=Homo sapiens OX=9606 GN=VPS26A PE=1 SV=2 | 0 | 5.976 | 7 | 1 | 4 | 1 | 327 | 38.1 | 6.57 | 6.83 | 1 |
| A0A0B4J1U3 | Immunoglobulin lambda variable 1-36 OS=Homo sapiens OX=9606 GN=IGLV1-36 PE=1 SV=5 | 0 | 5.967 | 38 | 2 | 3 | 2 | 117 | 12.5 | 4.78 | 8.57 | 2 |
| P30048 | Thioredoxin-dependent peroxide reductase, mitochondrial OS=Homo sapiens OX=9606 GN=PRDX3 PE=1 SV=3 | 0 | 5.961 | 10 | 2 | 6 | 2 | 256 | 27.7 | 7.78 | 9.2 | 2 |
| P54578 | Ubiquitin carboxyl-terminal hydrolase 14 OS=Homo sapiens OX=9606 GN=USP14 PE=1 SV=3 | 0 | 5.935 | 4 | 2 | 7 | 2 | 494 | 56 | 5.3 | 8.61 | 2 |
| P15151 | Poliovirus receptor OS=Homo sapiens OX=9606 GN=PVR PE=1 SV=2 | 0 | 5.923 | 5 | 2 | 4 | 2 | 417 | 45.3 | 6.52 | 5.01 | 2 |
| P01704 | Immunoglobulin lambda variable 2-14 OS=Homo sapiens OX=9606 GN=IGLV2-14 PE=1 SV=2 | 0 | 5.917 | 13 | 1 | 7 | 1 | 120 | 12.6 | 6.49 | 12.49 | 1 |
| A0A075B6S9 | Probable non-functional immunoglobulinn kappa variable 1-37 OS=Homo sapiens OX=9606 GN=IGKV1-37 PE=1 SV=7 | 0 | 5.889 | 14 | 1 | 7 | 1 | 117 | 12.7 | 9.25 | 16.91 | 1 |
| O75531 | Barrier-to-autointegration factor OS=Homo sapiens OX=9606 GN=BANF1 PE=1 SV=1 | 0 | 5.884 | 13 | 1 | 5 | 1 | 89 | 10.1 | 6.09 | 7.3 | 1 |
| O00487 | 26S proteasome non-ATPase regulatory subunit 14 OS=Homo sapiens OX=9606 GN=PSMD14 PE=1 SV=1 | 0 | 5.859 | 7 | 1 | 2 | 1 | 310 | 34.6 | 6.52 | 6.71 | 1 |
| P49207 | 60S ribosomal protein L34 OS=Homo sapiens OX=9606 GN=RPL34 PE=1 SV=3 | 0 | 5.803 | 13 | 2 | 31 | 2 | 117 | 13.3 | 11.47 | 30.86 | 2 |
| Q96FV2 | Secernin-2 OS=Homo sapiens OX=9606 GN=SCRN2 PE=1 SV=3 | 0 | 5.792 | 3 | 1 | 1 | 1 | 425 | 46.6 | 5.67 | 3.57 | 1 |
| P51858 | Hepatoma-derived growth factor OS=Homo sapiens OX=9606 GN=HDGF PE=1 SV=1 | 0 | 5.777 | 7 | 1 | 1 | 1 | 240 | 26.8 | 4.73 | 4.65 | 1 |
| P16070 | CD44 antigen OS=Homo sapiens OX=9606 GN=CD44 PE=1 SV=3 | 0 | 5.774 | 2 | 1 | 1 | 1 | 742 | 81.5 | 5.33 | 3.48 | 1 |
| P30533 | Alpha-2-macroglobulin receptor-associated protein OS=Homo sapiens OX=9606 GN=LRPAP1 PE=1 SV=1 | 0 | 5.752 | 5 | 2 | 3 | 2 | 357 | 41.4 | 8.78 | 1.87 | 2 |
| O43505 | Beta-1,4-glucuronyltransferase 1 OS=Homo sapiens OX=9606 GN=B4GAT1 PE=1 SV=1 | 0 | 5.751 | 11 | 2 | 3 | 2 | 415 | 47.1 | 7.2 | 7.18 | 2 |
| Q8WWX9 | Selenoprotein M OS=Homo sapiens OX=9606 GN=SELENOM PE=1 SV=3 | 0 | 5.749 | 21 | 2 | 2 | 2 | 145 | 16.2 | 5.54 | 2.25 | 2 |
| Q9NRP0 | Oligosaccharyltransferase complex subunit OSTC OS=Homo sapiens OX=9606 GN=OSTC PE=1 SV=1 | 0 | 5.729 | 8 | 1 | 5 | 1 | 149 | 16.8 | 9.13 | 15.54 | 1 |
| P13498 | Cytochrome b-245 light chain OS=Homo sapiens OX=9606 GN=CYBA PE=1 SV=3 | 0 | 5.702 | 16 | 3 | 5 | 3 | 195 | 21 | 9.54 | 5.52 | 3 |
| P01909 | HLA class II histocompatibility antigen, DQ alpha 1 chain OS=Homo sapiens OX=9606 GN=HLA-DQA1 PE=1 SV=1 | 0 | 5.687 | 6 | 1 | 1 | 1 | 254 | 27.8 | 5.72 | 3.06 | 1 |
| Q86YZ3 | Hornerin OS=Homo sapiens OX=9606 GN=HRNR PE=1 SV=2 | 0 | 5.666 | 2 | 1 | 3 | 1 | 2850 | 282.2 | 10.04 | 6.79 | 1 |
| Q16890 | Tumor protein D53 OS=Homo sapiens OX=9606 GN=TPD52L1 PE=1 SV=1 | 0 | 5.661 | 7 | 1 | 5 | 1 | 204 | 22.4 | 5.62 | 11.07 | 1 |
| P62861 | 40S ribosomal protein S30 OS=Homo sapiens OX=9606 GN=FAU PE=1 SV=1 | 0 | 5.65 | 17 | 1 | 13 | 1 | 59 | 6.6 | 12.15 | 29.14 | 1 |
| Q99714 | 3-hydroxyacyl-CoA dehydrogenase type-2 OS=Homo sapiens OX=9606 GN=HSD17B10 PE=1 SV=3 | 0 | 5.632 | 10 | 1 | 1 | 1 | 261 | 26.9 | 7.78 | 4.37 | 1 |
| Q8IZ83 | Aldehyde dehydrogenase family 16 member A1 OS=Homo sapiens OX=9606 GN=ALDH16A1 PE=1 SV=2 | 0 | 5.601 | 3 | 2 | 2 | 2 | 802 | 85.1 | 6.79 | 2.52 | 2 |
| Q9H098 | Protein FAM107B OS=Homo sapiens OX=9606 GN=FAM107B PE=1 SV=1 | 0 | 5.549 | 16 | 2 | 5 | 2 | 131 | 15.5 | 8.29 | 4.05 | 2 |
| Q96P63 | Serpin B12 OS=Homo sapiens OX=9606 GN=SERPINB12 PE=1 SV=1 | 0 | 5.49 | 4 | 1 | 1 | 1 | 405 | 46.2 | 5.53 | 3.39 | 1 |
| O14498 | Immunoglobulin superfamily containing leucine-rich repeat protein OS=Homo sapiens OX=9606 GN=ISLR PE=2 SV=1 | 0 | 5.464 | 6 | 2 | 2 | 2 | 428 | 46 | 5.15 | 2.3 | 2 |
| P61009 | Signal peptidase complex subunit 3 OS=Homo sapiens OX=9606 GN=SPCS3 PE=1 SV=1 | 0 | 5.4 | 12 | 2 | 9 | 2 | 180 | 20.3 | 8.62 | 12.15 | 2 |
| P0DOX3 | Immunoglobulin delta heavy chain OS=Homo sapiens OX=9606 PE=1 SV=1 | 0 | 5.36 | 3 | 2 | 49 | 2 | 512 | 56.2 | 8.02 | 101.29 | 2 |
| Q92499 | ATP-dependent RNA helicase DDX1 OS=Homo sapiens OX=9606 GN=DDX1 PE=1 SV=2 | 0 | 5.359 | 3 | 1 | 1 | 1 | 740 | 82.4 | 7.23 | 4.29 | 1 |
| P17980 | 26S proteasome regulatory subunit 6A OS=Homo sapiens OX=9606 GN=PSMC3 PE=1 SV=3 | 0 | 5.352 | 6 | 2 | 2 | 2 | 439 | 49.2 | 5.24 | 4.82 | 2 |
| Q13444 | Disintegrin and metalloproteinase domain-containing protein 15 OS=Homo sapiens OX=9606 GN=ADAM15 PE=1 SV=4 | 0 | 5.352 | 2 | 1 | 4 | 1 | 863 | 92.9 | 6.73 | 7.49 | 1 |
| Q01082 | Spectrin beta chain, non-erythrocytic 1 OS=Homo sapiens OX=9606 GN=SPTBN1 PE=1 SV=2 | 0 | 5.328 | 1 | 1 | 1 | 1 | 2364 | 274.4 | 5.57 | 2.97 | 1 |
| Q8N271 | Prominin-2 OS=Homo sapiens OX=9606 GN=PROM2 PE=1 SV=1 | 0 | 5.323 | 2 | 2 | 8 | 2 | 834 | 91.8 | 6.15 | 6.77 | 2 |
| P13010 | X-ray repair cross-complementing protein 5 OS=Homo sapiens OX=9606 GN=XRCC5 PE=1 SV=3 | 0 | 5.316 | 7 | 2 | 2 | 2 | 732 | 82.7 | 5.81 | 4.91 | 2 |
| Q02543 | 60S ribosomal protein L18a OS=Homo sapiens OX=9606 GN=RPL18A PE=1 SV=2 | 0 | 5.262 | 9 | 2 | 13 | 2 | 176 | 20.7 | 10.71 | 20.25 | 2 |
| Q8IZP2 | Putative protein FAM10A4 OS=Homo sapiens OX=9606 GN=ST13P4 PE=5 SV=1 | 0 | 5.249 | 7 | 2 | 6 | 2 | 240 | 27.4 | 5.08 | 8.57 | 2 |
| Q9H074 | Polyadenylate-binding protein-interacting protein 1 OS=Homo sapiens OX=9606 GN=PAIP1 PE=1 SV=1 | 0 | 5.214 | 4 | 2 | 2 | 2 | 479 | 53.5 | 4.81 | 4.13 | 2 |
| Q9UKS6 | Protein kinase C and casein kinase substrate in neurons protein 3 OS=Homo sapiens OX=9606 GN=PACSIN3 PE=1 SV=2 | 0 | 5.199 | 5 | 2 | 4 | 2 | 424 | 48.5 | 6.18 | 6.39 | 2 |
| Q15262 | Receptor-type tyrosine-protein phosphatase kappa OS=Homo sapiens OX=9606 GN=PTPRK PE=1 SV=2 | 0 | 5.18 | 1 | 2 | 2 | 2 | 1439 | 162 | 5.9 | 2.32 | 2 |
| O75368 | SH3 domain-binding glutamic acid-rich-like protein OS=Homo sapiens OX=9606 GN=SH3BGRL PE=1 SV=1 | 0 | 5.144 | 12 | 2 | 4 | 2 | 114 | 12.8 | 5.25 | 5.8 | 2 |
| Q06278 | Aldehyde oxidase OS=Homo sapiens OX=9606 GN=AOX1 PE=1 SV=2 | 0 | 5.126 | 1 | 2 | 49 | 1 | 1338 | 147.8 | 7.17 | 104.24 | 2 |
| Q6EEV6 | Small ubiquitin-related modifier 4 OS=Homo sapiens OX=9606 GN=SUMO4 PE=1 SV=2 | 0 | 5.126 | 13 | 1 | 5 | 1 | 95 | 10.7 | 7.18 | 12.32 | 1 |
| P62191 | 26S proteasome regulatory subunit 4 OS=Homo sapiens OX=9606 GN=PSMC1 PE=1 SV=1 | 0 | 5.126 | 3 | 1 | 3 | 1 | 440 | 49.2 | 6.21 | 3.86 | 1 |
| P80297 | Metallothionein-1X OS=Homo sapiens OX=9606 GN=MT1X PE=1 SV=1 | 0 | 5.097 | 20 | 1 | 2 | 1 | 61 | 6.1 | 7.96 | 3.13 | 1 |
| A0A0C4DH55 | Immunoglobulin kappa variable 3D-7 OS=Homo sapiens OX=9606 GN=IGKV3D-7 PE=3 SV=5 | 0 | 5.081 | 26 | 2 | 37 | 1 | 119 | 13.1 | 5.94 | 69.49 | 2 |
| P62891 | 60S ribosomal protein L39 OS=Homo sapiens OX=9606 GN=RPL39 PE=1 SV=2 | 0 | 5.037 | 20 | 1 | 16 | 1 | 51 | 6.4 | 12.56 | 30.12 | 1 |
| Q03426 | Mevalonate kinase OS=Homo sapiens OX=9606 GN=MVK PE=1 SV=1 | 0 | 4.979 | 3 | 1 | 1 | 1 | 396 | 42.4 | 6.46 | 3.22 | 1 |
| Q3KQV9 | UDP-N-acetylhexosamine pyrophosphorylase-like protein 1 OS=Homo sapiens OX=9606 GN=UAP1L1 PE=1 SV=2 | 0 | 4.979 | 4 | 2 | 2 | 2 | 507 | 57 | 6.32 | 2.36 | 2 |
| Q8N387 | Mucin-15 OS=Homo sapiens OX=9606 GN=MUC15 PE=1 SV=2 | 0 | 4.948 | 4 | 2 | 3 | 2 | 334 | 36.3 | 5.08 | 2.28 | 2 |
| P15374 | Ubiquitin carboxyl-terminal hydrolase isozyme L3 OS=Homo sapiens OX=9606 GN=UCHL3 PE=1 SV=1 | 0 | 4.923 | 9 | 2 | 4 | 2 | 230 | 26.2 | 4.92 | 1.89 | 2 |
| P50135 | Histamine N-methyltransferase OS=Homo sapiens OX=9606 GN=HNMT PE=1 SV=1 | 0 | 4.922 | 6 | 1 | 1 | 1 | 292 | 33.3 | 5.34 | 2.33 | 1 |
| Q3LXA3 | Triokinase/FMN cyclase OS=Homo sapiens OX=9606 GN=TKFC PE=1 SV=2 | 0 | 4.909 | 3 | 1 | 1 | 1 | 575 | 58.9 | 7.49 | 2.36 | 1 |
| Q9NZA1 | Chloride intracellular channel protein 5 OS=Homo sapiens OX=9606 GN=CLIC5 PE=1 SV=3 | 0 | 4.908 | 4 | 1 | 5 | 1 | 410 | 46.5 | 4.82 | 16.99 | 1 |
| P49591 | Serine--tRNA ligase, cytoplasmic OS=Homo sapiens OX=9606 GN=SARS1 PE=1 SV=3 | 0 | 4.906 | 2 | 1 | 2 | 1 | 514 | 58.7 | 6.43 | 1.75 | 1 |
| Q68D91 | Metallo-beta-lactamase domain-containing protein 2 OS=Homo sapiens OX=9606 GN=MBLAC2 PE=1 SV=3 | 0 | 4.844 | 9 | 2 | 4 | 2 | 279 | 31.4 | 6.92 | 0 | 2 |
| P60604 | Ubiquitin-conjugating enzyme E2 G2 OS=Homo sapiens OX=9606 GN=UBE2G2 PE=1 SV=1 | 0 | 4.822 | 9 | 1 | 1 | 1 | 165 | 18.6 | 4.7 | 2.74 | 1 |
| Q6UXA7 | Uncharacterized protein C6orf15 OS=Homo sapiens OX=9606 GN=C6orf15 PE=1 SV=3 | 0 | 4.815 | 7 | 1 | 2 | 1 | 325 | 34.2 | 8.07 | 1.66 | 1 |
| Q9H1E1 | Ribonuclease 7 OS=Homo sapiens OX=9606 GN=RNASE7 PE=1 SV=2 | 0 | 4.783 | 8 | 1 | 1 | 1 | 156 | 17.4 | 9.69 | 3.55 | 1 |
| Q9GZZ9 | Ubiquitin-like modifier-activating enzyme 5 OS=Homo sapiens OX=9606 GN=UBA5 PE=1 SV=1 | 0 | 4.777 | 6 | 1 | 1 | 1 | 404 | 44.8 | 4.84 | 3.14 | 1 |
| Q6NUK1 | Calcium-binding mitochondrial carrier protein SCaMC-1 OS=Homo sapiens OX=9606 GN=SLC25A24 PE=1 SV=2 | 0 | 4.682 | 5 | 1 | 1 | 1 | 477 | 53.3 | 6.33 | 2.4 | 1 |
| P98171 | Rho GTPase-activating protein 4 OS=Homo sapiens OX=9606 GN=ARHGAP4 PE=1 SV=2 | 0 | 4.681 | 2 | 1 | 1 | 1 | 946 | 105 | 6.34 | 3.11 | 1 |
| P53618 | Coatomer subunit beta OS=Homo sapiens OX=9606 GN=COPB1 PE=1 SV=3 | 0 | 4.678 | 1 | 1 | 2 | 1 | 953 | 107.1 | 6.05 | 3.23 | 1 |
| P06454 | Prothymosin alpha OS=Homo sapiens OX=9606 GN=PTMA PE=1 SV=2 | 0 | 4.642 | 14 | 1 | 2 | 1 | 111 | 12.2 | 3.78 | 5.15 | 1 |
| P55145 | Mesencephalic astrocyte-derived neurotrophic factor OS=Homo sapiens OX=9606 GN=MANF PE=1 SV=3 | 0 | 4.633 | 11 | 2 | 5 | 2 | 182 | 20.7 | 8.69 | 4.3 | 2 |
| Q9NPR2 | Semaphorin-4B OS=Homo sapiens OX=9606 GN=SEMA4B PE=1 SV=4 | 0 | 4.631 | 3 | 2 | 5 | 2 | 837 | 92.7 | 7.05 | 5.79 | 2 |
| O75493 | Carbonic anhydrase-related protein 11 OS=Homo sapiens OX=9606 GN=CA11 PE=1 SV=2 | 0 | 4.616 | 7 | 1 | 1 | 1 | 328 | 36.2 | 9.74 | 2.8 | 1 |
| Q9NVS9 | Pyridoxine-5'-phosphate oxidase OS=Homo sapiens OX=9606 GN=PNPO PE=1 SV=1 | 0 | 4.58 | 7 | 1 | 1 | 1 | 261 | 30 | 7.06 | 3.51 | 1 |
| P41567 | Eukaryotic translation initiation factor 1 OS=Homo sapiens OX=9606 GN=EIF1 PE=1 SV=1 | 0 | 4.575 | 16 | 1 | 1 | 1 | 113 | 12.7 | 7.44 | 3.45 | 1 |
| P17174 | Aspartate aminotransferase, cytoplasmic OS=Homo sapiens OX=9606 GN=GOT1 PE=1 SV=3 | 0 | 4.547 | 5 | 2 | 3 | 2 | 413 | 46.2 | 7.01 | 1.96 | 2 |
| Q10472 | Polypeptide N-acetylgalactosaminyltransferase 1 OS=Homo sapiens OX=9606 GN=GALNT1 PE=1 SV=1 | 0 | 4.534 | 4 | 2 | 7 | 2 | 559 | 64.2 | 7.72 | 4.03 | 2 |
| P29622 | Kallistatin OS=Homo sapiens OX=9606 GN=SERPINA4 PE=1 SV=3 | 0 | 4.53 | 5 | 2 | 2 | 2 | 427 | 48.5 | 7.75 | 3.68 | 2 |
| Q9BT67 | NEDD4 family-interacting protein 1 OS=Homo sapiens OX=9606 GN=NDFIP1 PE=1 SV=1 | 0 | 4.513 | 8 | 1 | 1 | 1 | 221 | 24.9 | 4.68 | 3.28 | 1 |
| P04424 | Argininosuccinate lyase OS=Homo sapiens OX=9606 GN=ASL PE=1 SV=4 | 0 | 4.508 | 4 | 2 | 2 | 2 | 464 | 51.6 | 6.48 | 2.35 | 2 |
| Q02750 | Dual specificity mitogen-activated protein kinase kinase 1 OS=Homo sapiens OX=9606 GN=MAP2K1 PE=1 SV=2 | 0 | 4.489 | 7 | 1 | 1 | 1 | 393 | 43.4 | 6.62 | 0 | 1 |
| Q9BYD5 | Cornifelin OS=Homo sapiens OX=9606 GN=CNFN PE=1 SV=2 | 0 | 4.447 | 24 | 1 | 1 | 1 | 112 | 12.4 | 6.1 | 3.25 | 1 |
| O75955 | Flotillin-1 OS=Homo sapiens OX=9606 GN=FLOT1 PE=1 SV=3 | 0 | 4.411 | 8 | 2 | 2 | 2 | 427 | 47.3 | 7.49 | 4.37 | 2 |
| Q86X52 | Chondroitin sulfate synthase 1 OS=Homo sapiens OX=9606 GN=CHSY1 PE=1 SV=3 | 0 | 4.382 | 2 | 1 | 1 | 1 | 802 | 91.7 | 9.23 | 3.76 | 1 |
| Q15599 | Na(+)/H(+) exchange regulatory cofactor NHE-RF2 OS=Homo sapiens OX=9606 GN=SLC9A3R2 PE=1 SV=2 | 0 | 4.379 | 3 | 1 | 3 | 1 | 337 | 37.4 | 7.93 | 4.34 | 1 |
| O95793 | Double-stranded RNA-binding protein Staufen homolog 1 OS=Homo sapiens OX=9606 GN=STAU1 PE=1 SV=2 | 0 | 4.371 | 4 | 2 | 5 | 2 | 577 | 63.1 | 9.44 | 7.98 | 2 |
| Q92734 | Protein TFG OS=Homo sapiens OX=9606 GN=TFG PE=1 SV=2 | 0 | 4.37 | 5 | 2 | 11 | 2 | 400 | 43.4 | 5.1 | 7.25 | 2 |
| Q9BPX5 | Actin-related protein 2/3 complex subunit 5-like protein OS=Homo sapiens OX=9606 GN=ARPC5L PE=1 SV=1 | 0 | 4.364 | 12 | 2 | 3 | 2 | 153 | 16.9 | 6.6 | 1.6 | 2 |
| Q9UI12 | V-type proton ATPase subunit H OS=Homo sapiens OX=9606 GN=ATP6V1H PE=1 SV=1 | 0 | 4.353 | 5 | 1 | 1 | 1 | 483 | 55.8 | 6.48 | 3.37 | 1 |
| Q9NS98 | Semaphorin-3G OS=Homo sapiens OX=9606 GN=SEMA3G PE=1 SV=1 | 0 | 4.342 | 2 | 2 | 3 | 2 | 782 | 86.6 | 7.78 | 3.63 | 2 |
| P52943 | Cysteine-rich protein 2 OS=Homo sapiens OX=9606 GN=CRIP2 PE=1 SV=1 | 0 | 4.332 | 13 | 2 | 5 | 2 | 208 | 22.5 | 8.72 | 0 | 2 |
| P46783 | 40S ribosomal protein S10 OS=Homo sapiens OX=9606 GN=RPS10 PE=1 SV=1 | 0 | 4.328 | 5 | 1 | 22 | 1 | 165 | 18.9 | 10.15 | 67.89 | 1 |
| P16422 | Epithelial cell adhesion molecule OS=Homo sapiens OX=9606 GN=EPCAM PE=1 SV=2 | 0 | 4.32 | 4 | 1 | 3 | 1 | 314 | 34.9 | 7.46 | 8.03 | 1 |
| Q15149 | Plectin OS=Homo sapiens OX=9606 GN=PLEC PE=1 SV=3 | 0 | 4.296 | 1 | 2 | 4 | 1 | 4684 | 531.5 | 5.96 | 2.01 | 2 |
| Q8WUY1 | Protein THEM6 OS=Homo sapiens OX=9606 GN=THEM6 PE=1 SV=2 | 0 | 4.294 | 9 | 2 | 3 | 2 | 208 | 23.9 | 9.55 | 3.62 | 2 |
| Q9UNL2 | Translocon-associated protein subunit gamma OS=Homo sapiens OX=9606 GN=SSR3 PE=1 SV=1 | 0 | 4.278 | 8 | 1 | 2 | 1 | 185 | 21.1 | 9.61 | 5.53 | 1 |
| Q04637 | Eukaryotic translation initiation factor 4 gamma 1 OS=Homo sapiens OX=9606 GN=EIF4G1 PE=1 SV=4 | 0 | 4.254 | 1 | 1 | 1 | 1 | 1599 | 175.4 | 5.33 | 2.09 | 1 |
| O15400 | Syntaxin-7 OS=Homo sapiens OX=9606 GN=STX7 PE=1 SV=4 | 0 | 4.245 | 4 | 1 | 2 | 1 | 261 | 29.8 | 5.55 | 4.99 | 1 |
| P00367 | Glutamate dehydrogenase 1, mitochondrial OS=Homo sapiens OX=9606 GN=GLUD1 PE=1 SV=2 | 0 | 4.232 | 2 | 1 | 2 | 1 | 558 | 61.4 | 7.8 | 3.96 | 1 |
| P28300 | Protein-lysine 6-oxidase OS=Homo sapiens OX=9606 GN=LOX PE=1 SV=2 | 0 | 4.231 | 5 | 1 | 1 | 1 | 417 | 46.9 | 8.09 | 2.73 | 1 |
| P37268 | Squalene synthase OS=Homo sapiens OX=9606 GN=FDFT1 PE=1 SV=1 | 0 | 4.224 | 4 | 1 | 1 | 1 | 417 | 48.1 | 6.54 | 2.86 | 1 |
| P02747 | Complement C1q subcomponent subunit C OS=Homo sapiens OX=9606 GN=C1QC PE=1 SV=3 | 0 | 4.222 | 9 | 2 | 2 | 2 | 245 | 25.8 | 8.41 | 4.5 | 2 |
| O75954 | Tetraspanin-9 OS=Homo sapiens OX=9606 GN=TSPAN9 PE=1 SV=1 | 0 | 4.211 | 5 | 1 | 9 | 1 | 239 | 26.8 | 7.68 | 21.59 | 1 |
| P31689 | DnaJ homolog subfamily A member 1 OS=Homo sapiens OX=9606 GN=DNAJA1 PE=1 SV=2 | 0 | 4.187 | 6 | 1 | 1 | 1 | 397 | 44.8 | 7.08 | 2.52 | 1 |
| O95372 | Acyl-protein thioesterase 2 OS=Homo sapiens OX=9606 GN=LYPLA2 PE=1 SV=1 | 0 | 4.139 | 7 | 2 | 15 | 2 | 231 | 24.7 | 7.23 | 12.63 | 2 |
| Q86VB7 | Scavenger receptor cysteine-rich type 1 protein M130 OS=Homo sapiens OX=9606 GN=CD163 PE=1 SV=2 | 0 | 4.068 | 1 | 1 | 1 | 1 | 1156 | 125.4 | 5.95 | 3.11 | 1 |
| Q9BVC6 | Transmembrane protein 109 OS=Homo sapiens OX=9606 GN=TMEM109 PE=1 SV=1 | 0 | 4.056 | 5 | 1 | 2 | 1 | 243 | 26.2 | 10.48 | 4.28 | 1 |
| O95721 | Synaptosomal-associated protein 29 OS=Homo sapiens OX=9606 GN=SNAP29 PE=1 SV=1 | 0 | 4.052 | 5 | 1 | 1 | 1 | 258 | 29 | 5.81 | 2.2 | 1 |
| Q9NP79 | Vacuolar protein sorting-associated protein VTA1 homolog OS=Homo sapiens OX=9606 GN=VTA1 PE=1 SV=1 | 0 | 4.047 | 8 | 2 | 2 | 2 | 307 | 33.9 | 6.29 | 2.2 | 2 |
| P10114 | Ras-related protein Rap-2a OS=Homo sapiens OX=9606 GN=RAP2A PE=1 SV=1 | 0 | 4.02 | 20 | 2 | 2 | 2 | 183 | 20.6 | 4.82 | 2.01 | 2 |
| P13716 | Delta-aminolevulinic acid dehydratase OS=Homo sapiens OX=9606 GN=ALAD PE=1 SV=1 | 0 | 3.997 | 3 | 1 | 1 | 1 | 330 | 36.3 | 6.79 | 1.81 | 1 |
| Q9UQB8 | Brain-specific angiogenesis inhibitor 1-associated protein 2 OS=Homo sapiens OX=9606 GN=BAIAP2 PE=1 SV=1 | 0 | 3.895 | 4 | 2 | 6 | 2 | 552 | 60.8 | 8.9 | 2.25 | 2 |
| P46459 | Vesicle-fusing ATPase OS=Homo sapiens OX=9606 GN=NSF PE=1 SV=3 | 0 | 3.89 | 2 | 1 | 2 | 1 | 744 | 82.5 | 6.95 | 6.11 | 1 |
| P98187 | Cytochrome P450 4F8 OS=Homo sapiens OX=9606 GN=CYP4F8 PE=1 SV=1 | 0 | 3.878 | 3 | 1 | 1 | 1 | 520 | 60 | 8.43 | 2.62 | 1 |
| P17813 | Endoglin OS=Homo sapiens OX=9606 GN=ENG PE=1 SV=2 | 0 | 3.875 | 2 | 1 | 2 | 1 | 658 | 70.5 | 6.61 | 3.69 | 1 |
| Q14435 | Polypeptide N-acetylgalactosaminyltransferase 3 OS=Homo sapiens OX=9606 GN=GALNT3 PE=1 SV=2 | 0 | 3.874 | 2 | 1 | 5 | 1 | 633 | 72.6 | 7.99 | 2.17 | 1 |
| Q9HB63 | Netrin-4 OS=Homo sapiens OX=9606 GN=NTN4 PE=1 SV=2 | 0 | 3.823 | 5 | 2 | 4 | 2 | 628 | 70 | 8.05 | 0 | 2 |
| P35443 | Thrombospondin-4 OS=Homo sapiens OX=9606 GN=THBS4 PE=1 SV=2 | 0 | 3.823 | 2 | 2 | 3 | 2 | 961 | 105.8 | 4.68 | 6.9 | 2 |
| P55084 | Trifunctional enzyme subunit beta, mitochondrial OS=Homo sapiens OX=9606 GN=HADHB PE=1 SV=3 | 0 | 3.811 | 3 | 1 | 1 | 1 | 474 | 51.3 | 9.41 | 1.66 | 1 |
| A0A075B6I7 | Probable non-functional immunoglobulin lambda variable 5-48 OS=Homo sapiens OX=9606 GN=IGLV5-48 PE=1 SV=2 | 0 | 3.799 | 12 | 1 | 2 | 1 | 105 | 11.2 | 9.51 | 3.8 | 1 |
| P11766 | Alcohol dehydrogenase class-3 OS=Homo sapiens OX=9606 GN=ADH5 PE=1 SV=4 | 0 | 3.789 | 4 | 2 | 7 | 2 | 374 | 39.7 | 7.49 | 1.61 | 2 |
| A0A0G2JS06 | Immunoglobulin lambda variable 5-39 OS=Homo sapiens OX=9606 GN=IGLV5-39 PE=3 SV=1 | 0 | 3.752 | 12 | 2 | 5 | 2 | 123 | 13.4 | 8.63 | 6 | 2 |
| P63208 | S-phase kinase-associated protein 1 OS=Homo sapiens OX=9606 GN=SKP1 PE=1 SV=2 | 0 | 3.749 | 9 | 2 | 5 | 2 | 163 | 18.6 | 4.54 | 1.65 | 2 |
| O00115 | Deoxyribonuclease-2-alpha OS=Homo sapiens OX=9606 GN=DNASE2 PE=1 SV=2 | 0 | 3.737 | 3 | 1 | 3 | 1 | 360 | 39.6 | 8.05 | 6.89 | 1 |
| Q9NQR4 | Omega-amidase NIT2 OS=Homo sapiens OX=9606 GN=NIT2 PE=1 SV=1 | 0 | 3.736 | 4 | 1 | 2 | 1 | 276 | 30.6 | 7.21 | 6.66 | 1 |
| P01236 | Prolactin OS=Homo sapiens OX=9606 GN=PRL PE=1 SV=1 | 0 | 3.723 | 5 | 1 | 11 | 1 | 227 | 25.9 | 6.98 | 15.71 | 1 |
| P31025 | Lipocalin-1 OS=Homo sapiens OX=9606 GN=LCN1 PE=1 SV=1 | 0 | 3.691 | 6 | 1 | 1 | 1 | 176 | 19.2 | 5.58 | 2.6 | 1 |
| Q9UD71 | Protein phosphatase 1 regulatory subunit 1B OS=Homo sapiens OX=9606 GN=PPP1R1B PE=1 SV=2 | 0 | 3.687 | 10 | 1 | 1 | 1 | 204 | 22.9 | 4.53 | 3.54 | 1 |
| A0A075B6I1 | Immunoglobulin lambda variable 4-60 OS=Homo sapiens OX=9606 GN=IGLV4-60 PE=3 SV=1 | 0 | 3.673 | 14 | 1 | 2 | 1 | 120 | 13 | 6.25 | 2.3 | 1 |
| Q5IS99 | Caspase-3 OS=Saimiri boliviensis boliviensis OX=39432 GN=CASP3 PE=2 SV=1 | 0 | 3.656 | 5 | 1 | 2 | 1 | 277 | 31.4 | 6.92 | 0 | 1 |
| P54108 | Cysteine-rich secretory protein 3 OS=Homo sapiens OX=9606 GN=CRISP3 PE=1 SV=1 | 0 | 3.637 | 8 | 2 | 2 | 2 | 245 | 27.6 | 7.8 | 1.66 | 2 |
| Q9H0Q0 | CYFIP-related Rac1 interactor A OS=Homo sapiens OX=9606 GN=CYRIA PE=2 SV=1 | 0 | 3.617 | 4 | 1 | 2 | 1 | 323 | 37.3 | 6.01 | 4.85 | 1 |
| Q8IXL6 | Extracellular serine/threonine protein kinase FAM20C OS=Homo sapiens OX=9606 GN=FAM20C PE=1 SV=2 | 0 | 3.598 | 3 | 1 | 1 | 1 | 584 | 66.2 | 7.74 | 2.23 | 1 |
| P28065 | Proteasome subunit beta type-9 OS=Homo sapiens OX=9606 GN=PSMB9 PE=1 SV=2 | 0 | 3.587 | 5 | 1 | 1 | 1 | 219 | 23.3 | 5.03 | 1.74 | 1 |
| P42229 | Signal transducer and activator of transcription 5A OS=Homo sapiens OX=9606 GN=STAT5A PE=1 SV=1 | 0 | 3.576 | 4 | 2 | 4 | 2 | 794 | 90.6 | 6.39 | 2.26 | 2 |
| O14531 | Dihydropyrimidinase-related protein 4 OS=Homo sapiens OX=9606 GN=DPYSL4 PE=1 SV=2 | 0 | 3.562 | 3 | 1 | 1 | 1 | 572 | 61.8 | 7.09 | 2.33 | 1 |
| P61812 | Transforming growth factor beta-2 proprotein OS=Homo sapiens OX=9606 GN=TGFB2 PE=1 SV=1 | 0 | 3.543 | 4 | 1 | 2 | 1 | 414 | 47.7 | 8.53 | 3.74 | 1 |
| P51857 | Aldo-keto reductase family 1 member D1 OS=Homo sapiens OX=9606 GN=AKR1D1 PE=1 SV=1 | 0 | 3.537 | 3 | 2 | 5 | 1 | 326 | 37.4 | 7.5 | 4.32 | 2 |
| Q9UNF0 | Protein kinase C and casein kinase substrate in neurons protein 2 OS=Homo sapiens OX=9606 GN=PACSIN2 PE=1 SV=2 | 0 | 3.526 | 2 | 1 | 1 | 1 | 486 | 55.7 | 5.2 | 3.46 | 1 |
| Q9BY67 | Cell adhesion molecule 1 OS=Homo sapiens OX=9606 GN=CADM1 PE=1 SV=2 | 0 | 3.509 | 4 | 2 | 5 | 2 | 442 | 48.5 | 5.07 | 0 | 2 |
| P34913 | Bifunctional epoxide hydrolase 2 OS=Homo sapiens OX=9606 GN=EPHX2 PE=1 SV=2 | 0 | 3.49 | 4 | 1 | 1 | 1 | 555 | 62.6 | 6.28 | 2.96 | 1 |
| Q6UVY6 | DBH-like monooxygenase protein 1 OS=Homo sapiens OX=9606 GN=MOXD1 PE=1 SV=1 | 0 | 3.474 | 3 | 2 | 3 | 2 | 613 | 69.6 | 6.43 | 3.84 | 2 |
| P43121 | Cell surface glycoprotein MUC18 OS=Homo sapiens OX=9606 GN=MCAM PE=1 SV=2 | 0 | 3.461 | 2 | 1 | 1 | 1 | 646 | 71.6 | 5.76 | 2.15 | 1 |
| Q15257 | Serine/threonine-protein phosphatase 2A activator OS=Homo sapiens OX=9606 GN=PTPA PE=1 SV=3 | 0 | 3.413 | 8 | 2 | 2 | 2 | 358 | 40.6 | 5.94 | 2.13 | 2 |
| P62195 | 26S proteasome regulatory subunit 8 OS=Homo sapiens OX=9606 GN=PSMC5 PE=1 SV=1 | 0 | 3.4 | 2 | 1 | 2 | 1 | 406 | 45.6 | 7.55 | 4.34 | 1 |
| Q9UKK9 | ADP-sugar pyrophosphatase OS=Homo sapiens OX=9606 GN=NUDT5 PE=1 SV=1 | 0 | 3.39 | 7 | 1 | 1 | 1 | 219 | 24.3 | 4.94 | 0 | 1 |
| Q00688 | Peptidyl-prolyl cis-trans isomerase FKBP3 OS=Homo sapiens OX=9606 GN=FKBP3 PE=1 SV=1 | 0 | 3.375 | 5 | 1 | 1 | 1 | 224 | 25.2 | 9.28 | 2.24 | 1 |
| O60884 | DnaJ homolog subfamily A member 2 OS=Homo sapiens OX=9606 GN=DNAJA2 PE=1 SV=1 | 0 | 3.372 | 3 | 1 | 7 | 1 | 412 | 45.7 | 6.48 | 6.21 | 1 |
| P61421 | V-type proton ATPase subunit d 1 OS=Homo sapiens OX=9606 GN=ATP6V0D1 PE=1 SV=1 | 0 | 3.322 | 3 | 1 | 1 | 1 | 351 | 40.3 | 5 | 2.3 | 1 |
| P40189 | Interleukin-6 receptor subunit beta OS=Homo sapiens OX=9606 GN=IL6ST PE=1 SV=2 | 0 | 3.322 | 1 | 1 | 2 | 1 | 918 | 103.5 | 5.95 | 4.58 | 1 |
| Q92692 | Nectin-2 OS=Homo sapiens OX=9606 GN=NECTIN2 PE=1 SV=1 | 0 | 3.318 | 2 | 1 | 1 | 1 | 538 | 57.7 | 4.82 | 0 | 1 |
| P17612 | cAMP-dependent protein kinase catalytic subunit alpha OS=Homo sapiens OX=9606 GN=PRKACA PE=1 SV=2 | 0 | 3.312 | 5 | 2 | 3 | 2 | 351 | 40.6 | 8.79 | 3.41 | 2 |
| Q92688 | Acidic leucine-rich nuclear phosphoprotein 32 family member B OS=Homo sapiens OX=9606 GN=ANP32B PE=1 SV=1 | 0 | 3.308 | 6 | 2 | 3 | 1 | 251 | 28.8 | 4.06 | 2.51 | 2 |
| P20138 | Myeloid cell surface antigen CD33 OS=Homo sapiens OX=9606 GN=CD33 PE=1 SV=2 | 0 | 3.303 | 3 | 1 | 4 | 1 | 364 | 39.8 | 8.38 | 2.3 | 1 |
| P28074 | Proteasome subunit beta type-5 OS=Homo sapiens OX=9606 GN=PSMB5 PE=1 SV=3 | 0 | 3.301 | 4 | 1 | 1 | 1 | 263 | 28.5 | 6.92 | 2.5 | 1 |
| Q9Y4K3 | TNF receptor-associated factor 6 OS=Homo sapiens OX=9606 GN=TRAF6 PE=1 SV=1 | 0 | 3.271 | 4 | 1 | 3 | 1 | 522 | 59.5 | 6.44 | 6.3 | 1 |
| A6NI72 | Putative neutrophil cytosol factor 1B OS=Homo sapiens OX=9606 GN=NCF1B PE=5 SV=2 | 0 | 3.271 | 2 | 1 | 1 | 1 | 391 | 44.8 | 9.13 | 1.79 | 1 |
| P00403 | Cytochrome c oxidase subunit 2 OS=Homo sapiens OX=9606 GN=MT-CO2 PE=1 SV=1 | 0 | 3.25 | 7 | 2 | 2 | 2 | 227 | 25.5 | 4.82 | 1.97 | 2 |
| Q96M27 | Protein PRRC1 OS=Homo sapiens OX=9606 GN=PRRC1 PE=1 SV=1 | 0 | 3.249 | 7 | 1 | 1 | 1 | 445 | 46.7 | 5.83 | 2.67 | 1 |
| Q9UBC9 | Small proline-rich protein 3 OS=Homo sapiens OX=9606 GN=SPRR3 PE=1 SV=2 | 0 | 3.242 | 9 | 2 | 2 | 2 | 169 | 18.1 | 8.57 | 0 | 2 |
| Q8IX19 | Mast cell-expressed membrane protein 1 OS=Homo sapiens OX=9606 GN=MCEMP1 PE=1 SV=1 | 0 | 3.225 | 7 | 1 | 1 | 1 | 187 | 21.2 | 8.87 | 0 | 1 |
| Q8TF09 | Dynein light chain roadblock-type 2 OS=Homo sapiens OX=9606 GN=DYNLRB2 PE=1 SV=1 | 0 | 3.212 | 13 | 1 | 1 | 1 | 96 | 10.8 | 7.5 | 2.37 | 1 |
| P21695 | Glycerol-3-phosphate dehydrogenase [NAD(+)], cytoplasmic OS=Homo sapiens OX=9606 GN=GPD1 PE=1 SV=4 | 0 | 3.209 | 12 | 2 | 2 | 2 | 349 | 37.5 | 6.18 | 3.38 | 2 |
| Q04900 | Sialomucin core protein 24 OS=Homo sapiens OX=9606 GN=CD164 PE=1 SV=2 | 0 | 3.179 | 15 | 1 | 1 | 1 | 197 | 20.9 | 7.97 | 2.15 | 1 |
| P20042 | Eukaryotic translation initiation factor 2 subunit 2 OS=Homo sapiens OX=9606 GN=EIF2S2 PE=1 SV=2 | 0 | 3.167 | 3 | 1 | 2 | 1 | 333 | 38.4 | 5.8 | 4.75 | 1 |
| Q9Y6X5 | Bis(5'-adenosyl)-triphosphatase ENPP4 OS=Homo sapiens OX=9606 GN=ENPP4 PE=1 SV=3 | 0 | 3.163 | 2 | 1 | 1 | 1 | 453 | 51.6 | 6.15 | 2.65 | 1 |
| Q99623 | Prohibitin-2 OS=Homo sapiens OX=9606 GN=PHB2 PE=1 SV=2 | 0 | 3.16 | 3 | 1 | 2 | 1 | 299 | 33.3 | 9.83 | 5 | 1 |
| A0A075B6K2 | Immunoglobulin lambda variable 3-12 OS=Homo sapiens OX=9606 GN=IGLV3-12 PE=3 SV=2 | 0 | 3.15 | 14 | 1 | 3 | 1 | 115 | 12.4 | 5.67 | 5.02 | 1 |
| Q15027 | Arf-GAP with coiled-coil, ANK repeat and PH domain-containing protein 1 OS=Homo sapiens OX=9606 GN=ACAP1 PE=1 SV=1 | 0 | 3.145 | 2 | 1 | 1 | 1 | 740 | 81.5 | 7.66 | 2.16 | 1 |
| Q96KN1 | Protein LRATD2 OS=Homo sapiens OX=9606 GN=LRATD2 PE=1 SV=1 | 0 | 3.127 | 3 | 1 | 2 | 1 | 310 | 34.5 | 5.54 | 1.95 | 1 |
| Q15274 | Nicotinate-nucleotide pyrophosphorylase [carboxylating] OS=Homo sapiens OX=9606 GN=QPRT PE=1 SV=3 | 0 | 3.122 | 4 | 1 | 1 | 1 | 297 | 30.8 | 6.21 | 0 | 1 |
| Q86VE9 | Serine incorporator 5 OS=Homo sapiens OX=9606 GN=SERINC5 PE=2 SV=1 | 0 | 3.119 | 2 | 1 | 2 | 1 | 423 | 47 | 7.62 | 4.24 | 1 |
| Q14031 | Collagen alpha-6(IV) chain OS=Homo sapiens OX=9606 GN=COL4A6 PE=1 SV=3 | 0 | 3.108 | 2 | 1 | 1 | 1 | 1691 | 163.7 | 9.2 | 2.32 | 1 |
| Q00325 | Phosphate carrier protein, mitochondrial OS=Homo sapiens OX=9606 GN=SLC25A3 PE=1 SV=2 | 0 | 3.106 | 3 | 1 | 1 | 1 | 362 | 40.1 | 9.38 | 2.27 | 1 |
| Q10588 | ADP-ribosyl cyclase/cyclic ADP-ribose hydrolase 2 OS=Homo sapiens OX=9606 GN=BST1 PE=1 SV=2 | 0 | 3.102 | 4 | 1 | 1 | 1 | 318 | 35.7 | 7.8 | 2.29 | 1 |
| P09669 | Cytochrome c oxidase subunit 6C OS=Homo sapiens OX=9606 GN=COX6C PE=1 SV=2 | 0 | 3.101 | 11 | 1 | 1 | 1 | 75 | 8.8 | 10.39 | 2.38 | 1 |
| P53597 | Succinate--CoA ligase [ADP/GDP-forming] subunit alpha, mitochondrial OS=Homo sapiens OX=9606 GN=SUCLG1 PE=1 SV=4 | 0 | 3.094 | 3 | 1 | 4 | 1 | 346 | 36.2 | 8.79 | 5.71 | 1 |
| P11047 | Laminin subunit gamma-1 OS=Homo sapiens OX=9606 GN=LAMC1 PE=1 SV=3 | 0 | 3.092 | 1 | 1 | 1 | 1 | 1609 | 177.5 | 5.12 | 2.37 | 1 |
| Q9BUL8 | Programmed cell death protein 10 OS=Homo sapiens OX=9606 GN=PDCD10 PE=1 SV=1 | 0 | 3.088 | 5 | 1 | 8 | 1 | 212 | 24.7 | 8.19 | 3.32 | 1 |
| Q9H4A3 | Serine/threonine-protein kinase WNK1 OS=Homo sapiens OX=9606 GN=WNK1 PE=1 SV=2 | 0 | 3.079 | 0 | 1 | 7 | 1 | 2382 | 250.6 | 6.34 | 5.28 | 1 |
| A0A0C4DH39 | Immunoglobulin heavy variable 1-58 OS=Homo sapiens OX=9606 GN=IGHV1-58 PE=3 SV=1 | 0 | 3.057 | 13 | 1 | 1 | 1 | 117 | 13 | 9.25 | 2.66 | 1 |
| Q6NVY1 | 3-hydroxyisobutyryl-CoA hydrolase, mitochondrial OS=Homo sapiens OX=9606 GN=HIBCH PE=1 SV=2 | 0 | 3.052 | 3 | 1 | 1 | 1 | 386 | 43.5 | 8.19 | 2.61 | 1 |
| Q9UMX5 | Neudesin OS=Homo sapiens OX=9606 GN=NENF PE=1 SV=1 | 0 | 3.051 | 9 | 1 | 1 | 1 | 172 | 18.8 | 5.69 | 1.89 | 1 |
| Q9UJ68 | Mitochondrial peptide methionine sulfoxide reductase OS=Homo sapiens OX=9606 GN=MSRA PE=1 SV=1 | 0 | 3.05 | 10 | 2 | 5 | 2 | 235 | 26.1 | 8.09 | 2.03 | 2 |
| Q9GZP4 | PITH domain-containing protein 1 OS=Homo sapiens OX=9606 GN=PITHD1 PE=1 SV=1 | 0 | 3.014 | 4 | 1 | 1 | 1 | 211 | 24.2 | 5.74 | 0 | 1 |
| Q15369 | Elongin-C OS=Homo sapiens OX=9606 GN=ELOC PE=1 SV=1 | 0 | 3.004 | 9 | 1 | 5 | 1 | 112 | 12.5 | 4.78 | 7.94 | 1 |
| P0C6X5 | Replicase polyprotein 1ab OS=Human coronavirus NL63 OX=277944 GN=rep PE=1 SV=1 | 0 | 2.99 | 0 | 1 | 29 | 1 | 6729 | 752.2 | 7.06 | 10.11 | 1 |
| O15533 | Tapasin OS=Homo sapiens OX=9606 GN=TAPBP PE=1 SV=1 | 0 | 2.987 | 2 | 1 | 1 | 1 | 448 | 47.6 | 7.15 | 0 | 1 |
| P27701 | CD82 antigen OS=Homo sapiens OX=9606 GN=CD82 PE=1 SV=1 | 0 | 2.97 | 4 | 1 | 8 | 1 | 267 | 29.6 | 5.24 | 4.23 | 1 |
| P05161 | Ubiquitin-like protein ISG15 OS=Homo sapiens OX=9606 GN=ISG15 PE=1 SV=5 | 0 | 2.967 | 5 | 1 | 1 | 1 | 165 | 17.9 | 7.44 | 2.1 | 1 |
| Q99523 | Sortilin OS=Homo sapiens OX=9606 GN=SORT1 PE=1 SV=3 | 0 | 2.964 | 1 | 1 | 3 | 1 | 831 | 92 | 5.74 | 5.99 | 1 |
| Q96A33 | Coiled-coil domain-containing protein 47 OS=Homo sapiens OX=9606 GN=CCDC47 PE=1 SV=1 | 0 | 2.964 | 4 | 1 | 2 | 1 | 483 | 55.8 | 4.87 | 2.7 | 1 |
| Q12904 | Aminoacyl tRNA synthase complex-interacting multifunctional protein 1 OS=Homo sapiens OX=9606 GN=AIMP1 PE=1 SV=2 | 0 | 2.951 | 3 | 1 | 3 | 1 | 312 | 34.3 | 8.43 | 5.88 | 1 |
| Q16706 | Alpha-mannosidase 2 OS=Homo sapiens OX=9606 GN=MAN2A1 PE=1 SV=2 | 0 | 2.948 | 1 | 1 | 1 | 1 | 1144 | 131.1 | 7.58 | 1.74 | 1 |
| Q13155 | Aminoacyl tRNA synthase complex-interacting multifunctional protein 2 OS=Homo sapiens OX=9606 GN=AIMP2 PE=1 SV=2 | 0 | 2.945 | 4 | 1 | 1 | 1 | 320 | 35.3 | 8.22 | 2.34 | 1 |
| O60762 | Dolichol-phosphate mannosyltransferase subunit 1 OS=Homo sapiens OX=9606 GN=DPM1 PE=1 SV=1 | 0 | 2.936 | 8 | 1 | 1 | 1 | 260 | 29.6 | 9.57 | 0 | 1 |
| Q93084 | Sarcoplasmic/endoplasmic reticulum calcium ATPase 3 OS=Homo sapiens OX=9606 GN=ATP2A3 PE=1 SV=2 | 0 | 2.918 | 1 | 1 | 3 | 1 | 1043 | 113.9 | 5.57 | 6.06 | 1 |
| P12081 | Histidine--tRNA ligase, cytoplasmic OS=Homo sapiens OX=9606 GN=HARS1 PE=1 SV=2 | 0 | 2.909 | 3 | 1 | 1 | 1 | 509 | 57.4 | 5.88 | 0 | 1 |
| P35590 | Tyrosine-protein kinase receptor Tie-1 OS=Homo sapiens OX=9606 GN=TIE1 PE=1 SV=1 | 0.001 | 2.902 | 1 | 1 | 1 | 1 | 1138 | 125 | 7.03 | 2.28 | 1 |
| P15086 | Carboxypeptidase B OS=Homo sapiens OX=9606 GN=CPB1 PE=1 SV=4 | 0.001 | 2.89 | 2 | 1 | 1 | 1 | 417 | 47.3 | 6.61 | 2.54 | 1 |
| O43396 | Thioredoxin-like protein 1 OS=Homo sapiens OX=9606 GN=TXNL1 PE=1 SV=3 | 0.001 | 2.879 | 2 | 1 | 2 | 1 | 289 | 32.2 | 4.96 | 1.67 | 1 |
| Q8IZ81 | ELMO domain-containing protein 2 OS=Homo sapiens OX=9606 GN=ELMOD2 PE=1 SV=1 | 0.001 | 2.864 | 3 | 1 | 10 | 1 | 293 | 34.9 | 8.05 | 5.32 | 1 |
| Q9H6S3 | Epidermal growth factor receptor kinase substrate 8-like protein 2 OS=Homo sapiens OX=9606 GN=EPS8L2 PE=1 SV=2 | 0.002 | 2.821 | 1 | 1 | 1 | 1 | 715 | 80.6 | 6.84 | 1.88 | 1 |
| P48509 | CD151 antigen OS=Homo sapiens OX=9606 GN=CD151 PE=1 SV=3 | 0.002 | 2.807 | 4 | 1 | 11 | 1 | 253 | 28.3 | 7.47 | 15.89 | 1 |
| P14406 | Cytochrome c oxidase subunit 7A2, mitochondrial OS=Homo sapiens OX=9606 GN=COX7A2 PE=1 SV=1 | 0.002 | 2.802 | 16 | 1 | 1 | 1 | 83 | 9.4 | 9.76 | 1.87 | 1 |
| Q9H190 | Syntenin-2 OS=Homo sapiens OX=9606 GN=SDCBP2 PE=1 SV=2 | 0.002 | 2.801 | 4 | 1 | 1 | 1 | 292 | 31.6 | 9.07 | 1.84 | 1 |
| Q86UD1 | Out at first protein homolog OS=Homo sapiens OX=9606 GN=OAF PE=2 SV=1 | 0.002 | 2.797 | 3 | 1 | 1 | 1 | 273 | 30.7 | 6.84 | 2.18 | 1 |
| O95834 | Echinoderm microtubule-associated protein-like 2 OS=Homo sapiens OX=9606 GN=EML2 PE=1 SV=1 | 0.002 | 2.789 | 2 | 1 | 1 | 1 | 649 | 70.6 | 6.32 | 2.71 | 1 |
| P61077 | Ubiquitin-conjugating enzyme E2 D3 OS=Homo sapiens OX=9606 GN=UBE2D3 PE=1 SV=1 | 0.002 | 2.789 | 7 | 1 | 3 | 1 | 147 | 16.7 | 7.8 | 2.24 | 1 |
| P62308 | Small nuclear ribonucleoprotein G OS=Homo sapiens OX=9606 GN=SNRPG PE=1 SV=1 | 0.002 | 2.751 | 17 | 1 | 1 | 1 | 76 | 8.5 | 8.88 | 3.22 | 1 |
| P28068 | HLA class II histocompatibility antigen, DM beta chain OS=Homo sapiens OX=9606 GN=HLA-DMB PE=1 SV=1 | 0.002 | 2.746 | 4 | 1 | 6 | 1 | 263 | 28.9 | 7.44 | 12.46 | 1 |
| Q8NFU3 | Thiosulfate:glutathione sulfurtransferase OS=Homo sapiens OX=9606 GN=TSTD1 PE=1 SV=3 | 0.002 | 2.738 | 8 | 1 | 2 | 1 | 115 | 12.5 | 6.07 | 2.23 | 1 |

Table S1. Cont.

| Accession | Description | Exp. q-value: Combined | Sum PEP Score | Coverage [%] | # Peptides | # PSMs | # Unique Peptides | # AAs | MW [kDa] | calc. pI | Score Sequest HT: Sequest HT | # Peptides (by Search Engine): Sequest HT |
|---|---|---|---|---|---|---|---|---|---|---|---|---|
| Q4JQV9 | DNA polymerase processivity factor OS=Varicella-zoster virus (strain Oka vaccine) OX=341980 GN=ORF16 PE=3 SV=1 | 0.002 | 2.736 | 4 | 1 | 5 | 1 | 408 | 46 | 8.03 | 2.84 | 1 |
| P22234 | Multifunctional protein ADE2 OS=Homo sapiens OX=9606 GN=PAICS PE=1 SV=3 | 0.002 | 2.732 | 8 | 1 | 1 | 1 | 425 | 47 | 7.23 | 2.38 | 1 |
| O95394 | Phosphoacetylglucosamine mutase OS=Homo sapiens OX=9606 GN=PGM3 PE=1 SV=1 | 0.002 | 2.727 | 3 | 1 | 2 | 1 | 542 | 59.8 | 6.25 | 3.47 | 1 |
| P68402 | Platelet-activating factor acetylhydrolase IB subunit beta OS=Homo sapiens OX=9606 GN=PAFAH1B2 PE=1 SV=1 | 0.002 | 2.724 | 4 | 1 | 1 | 1 | 229 | 25.6 | 5.92 | 2.59 | 1 |
| P48163 | NADP-dependent malic enzyme OS=Homo sapiens OX=9606 GN=ME1 PE=1 SV=1 | 0.002 | 2.722 | 1 | 1 | 2 | 1 | 572 | 64.1 | 6.13 | 0 | 1 |
| Q01581 | Hydroxymethylglutaryl-CoA synthase, cytoplasmic OS=Homo sapiens OX=9606 GN=HMGCS1 PE=1 SV=2 | 0.002 | 2.685 | 2 | 1 | 1 | 1 | 520 | 57.3 | 5.41 | 0 | 1 |
| O95164 | Ubiquitin-like protein 3 OS=Homo sapiens OX=9606 GN=UBL3 PE=1 SV=1 | 0.002 | 2.676 | 8 | 1 | 4 | 1 | 117 | 13.1 | 6.77 | 4.64 | 1 |
| P10619 | Lysosomal protective protein OS=Homo sapiens OX=9606 GN=CTSA PE=1 SV=2 | 0.002 | 2.668 | 4 | 1 | 2 | 1 | 480 | 54.4 | 6.61 | 5.23 | 1 |
| P05997 | Collagen alpha-2(V) chain OS=Homo sapiens OX=9606 GN=COL5A2 PE=1 SV=3 | 0.002 | 2.663 | 1 | 2 | 2 | 2 | 1499 | 144.8 | 6.46 | 0 | 2 |
| Q86V21 | Acetoacetyl-CoA synthetase OS=Homo sapiens OX=9606 GN=AACS PE=1 SV=1 | 0.002 | 2.657 | 3 | 2 | 2 | 2 | 672 | 75.1 | 6.24 | 0 | 2 |
| Q9NX62 | Golgi-resident adenosine 3',5'-bisphosphate 3'-phosphatase OS=Homo sapiens OX=9606 GN=BPNT2 PE=1 SV=1 | 0.002 | 2.647 | 2 | 1 | 10 | 1 | 359 | 38.7 | 6.86 | 5.27 | 1 |
| O94903 | Pyridoxal phosphate homeostasis protein OS=Homo sapiens OX=9606 GN=PLPBP PE=1 SV=1 | 0.002 | 2.638 | 5 | 1 | 2 | 1 | 275 | 30.3 | 7.5 | 4.74 | 1 |
| A7J390 | RNA-directed RNA polymerase OS=Rotavirus A (strain RVA/Human/Japan/S2/1980/G2P1B[4]) OX=10959 PE=3 SV=1 | 0.002 | 2.586 | 1 | 1 | 2 | 1 | 1088 | 125 | 8.13 | 3.87 | 1 |
| Q7L523 | Ras-related GTP-binding protein A OS=Homo sapiens OX=9606 GN=RRAGA PE=1 SV=1 | 0.002 | 2.581 | 5 | 1 | 1 | 1 | 313 | 36.5 | 7.72 | 2.58 | 1 |
| Q9Y6V0 | Protein piccolo OS=Homo sapiens OX=9606 GN=PCLO PE=1 SV=5 | 0.002 | 2.572 | 0 | 2 | 9 | 2 | 5142 | 560.4 | 6.47 | 3.62 | 2 |
| O95445 | Apolipoprotein M OS=Homo sapiens OX=9606 GN=APOM PE=1 SV=2 | 0.002 | 2.568 | 5 | 1 | 5 | 1 | 188 | 21.2 | 6.01 | 5.34 | 1 |
| Q9Y2A9 | N-acetyllactosaminide beta-1,3-N-acetylglucosaminyltransferase 3 OS=Homo sapiens OX=9606 GN=B3GNT3 PE=1 SV=2 | 0.002 | 2.562 | 3 | 1 | 1 | 1 | 372 | 42.5 | 8.68 | 2.21 | 1 |
| Q9UBW5 | Bridging integrator 2 OS=Homo sapiens OX=9606 GN=BIN2 PE=1 SV=3 | 0.002 | 2.561 | 5 | 2 | 2 | 1 | 565 | 61.8 | 5.15 | 4.81 | 2 |
| Q15293 | Reticulocalbin-1 OS=Homo sapiens OX=9606 GN=RCN1 PE=1 SV=1 | 0.002 | 2.559 | 4 | 1 | 1 | 1 | 331 | 38.9 | 5 | 2.01 | 1 |
| Q96HU8 | GTP-binding protein Di-Ras2 OS=Homo sapiens OX=9606 GN=DIRAS2 PE=1 SV=1 | 0.003 | 2.549 | 6 | 1 | 3 | 1 | 199 | 22.5 | 8.76 | 1.92 | 1 |
| Q13296 | Mammaglobin-A OS=Homo sapiens OX=9606 GN=SCGB2A2 PE=1 SV=1 | 0.003 | 2.549 | 13 | 1 | 1 | 1 | 93 | 10.5 | 4.3 | 3.31 | 1 |
| Q92973 | Transportin-1 OS=Homo sapiens OX=9606 GN=TNPO1 PE=1 SV=2 | 0.004 | 2.548 | 3 | 1 | 1 | 1 | 898 | 102.3 | 4.98 | 2.32 | 1 |
| Q9P0K7 | Ankycorbin OS=Homo sapiens OX=9606 GN=RAI14 PE=1 SV=2 | 0.004 | 2.547 | 1 | 1 | 1 | 1 | 980 | 110 | 6.21 | 2.34 | 1 |
| P62314 | Small nuclear ribonucleoprotein Sm D1 OS=Homo sapiens OX=9606 GN=SNRPD1 PE=1 SV=1 | 0.004 | 2.539 | 17 | 1 | 1 | 1 | 119 | 13.3 | 11.56 | 1.86 | 1 |
| P23246 | Splicing factor, proline- and glutamine-rich OS=Homo sapiens OX=9606 GN=SFPQ PE=1 SV=2 | 0.004 | 2.535 | 2 | 1 | 1 | 1 | 707 | 76.1 | 9.44 | 0 | 1 |
| P22532 | Small proline-rich protein 2D OS=Homo sapiens OX=9606 GN=SPRR2D PE=2 SV=2 | 0.004 | 2.526 | 13 | 1 | 1 | 1 | 72 | 7.9 | 8.37 | 0 | 1 |
| Q08554 | Desmocollin-1 OS=Homo sapiens OX=9606 GN=DSC1 PE=1 SV=2 | 0.005 | 2.507 | 2 | 1 | 1 | 1 | 894 | 99.9 | 5.43 | 0 | 1 |
| Q9UI42 | Carboxypeptidase A4 OS=Homo sapiens OX=9606 GN=CPA4 PE=1 SV=2 | 0.005 | 2.503 | 5 | 1 | 1 | 1 | 421 | 47.3 | 6.7 | 2.2 | 1 |
| Q9HD42 | Charged multivesicular body protein 1a OS=Homo sapiens OX=9606 GN=CHMP1A PE=1 SV=1 | 0.005 | 2.492 | 4 | 1 | 6 | 1 | 196 | 21.7 | 8.06 | 3.4 | 1 |
| Q8IYJ3 | Synaptotagmin-like protein 1 OS=Homo sapiens OX=9606 GN=SYTL1 PE=1 SV=1 | 0.005 | 2.489 | 4 | 1 | 1 | 1 | 562 | 61.8 | 5.48 | 2.99 | 1 |
| Q86WV6 | Stimulator of interferon genes protein OS=Homo sapiens OX=9606 GN=STING1 PE=1 SV=1 | 0.005 | 2.47 | 7 | 1 | 1 | 1 | 379 | 42.2 | 7.05 | 2.25 | 1 |
| P61011 | Signal recognition particle 54 kDa protein OS=Homo sapiens OX=9606 GN=SRP54 PE=1 SV=1 | 0.005 | 2.452 | 2 | 1 | 3 | 1 | 504 | 55.7 | 8.75 | 3.68 | 1 |
| P43686 | 26S proteasome regulatory subunit 6B OS=Homo sapiens OX=9606 GN=PSMC4 PE=1 SV=2 | 0.005 | 2.44 | 2 | 1 | 1 | 1 | 418 | 47.3 | 5.21 | 1.92 | 1 |
| Q12840 | Kinesin heavy chain isoform 5A OS=Homo sapiens OX=9606 GN=KIF5A PE=1 SV=2 | 0.005 | 2.44 | 1 | 1 | 1 | 1 | 1032 | 117.3 | 5.9 | 1.68 | 1 |
| O75352 | Mannose-P-dolichol utilization defect 1 protein OS=Homo sapiens OX=9606 GN=MPDU1 PE=1 SV=2 | 0.005 | 2.415 | 4 | 1 | 6 | 1 | 247 | 26.6 | 8.94 | 3.41 | 1 |
| Q8IY18 | Structural maintenance of chromosomes protein 5 OS=Homo sapiens OX=9606 GN=SMC5 PE=1 SV=2 | 0.005 | 2.413 | 1 | 1 | 3 | 1 | 1101 | 128.7 | 8.38 | 5.85 | 1 |
| Q00577 | Transcriptional activator protein Pur-alpha OS=Homo sapiens OX=9606 GN=PURA PE=1 SV=2 | 0.005 | 2.4 | 2 | 1 | 1 | 1 | 322 | 34.9 | 6.44 | 1.72 | 1 |
| P00750 | Tissue-type plasminogen activator OS=Homo sapiens OX=9606 GN=PLAT PE=1 SV=1 | 0.005 | 2.4 | 2 | 1 | 1 | 1 | 562 | 62.9 | 7.8 | 0 | 1 |
| Q96C24 | Synaptotagmin-like protein 4 OS=Homo sapiens OX=9606 GN=SYTL4 PE=1 SV=2 | 0.005 | 2.399 | 1 | 1 | 3 | 1 | 671 | 76 | 8.98 | 8.59 | 1 |
| Q16836 | Hydroxyacyl-coenzyme A dehydrogenase, mitochondrial OS=Homo sapiens OX=9606 GN=HADH PE=1 SV=3 | 0.005 | 2.393 | 10 | 1 | 1 | 1 | 314 | 34.3 | 8.85 | 2.26 | 1 |
| A0A075B6T7 | T cell receptor alpha variable 6 OS=Homo sapiens OX=9606 GN=TRAV6 PE=3 SV=2 | 0.005 | 2.383 | 8 | 1 | 10 | 1 | 132 | 15.4 | 9.1 | 14.42 | 1 |
| P35998 | 26S proteasome regulatory subunit 7 OS=Homo sapiens OX=9606 GN=PSMC2 PE=1 SV=3 | 0.005 | 2.373 | 3 | 1 | 1 | 1 | 433 | 48.6 | 5.95 | 0 | 1 |
| Q86UL3 | Glycerol-3-phosphate acyltransferase 4 OS=Homo sapiens OX=9606 GN=GPAT4 PE=1 SV=1 | 0.005 | 2.366 | 4 | 1 | 1 | 1 | 456 | 52 | 9.19 | 2.2 | 1 |
| O00182 | Galectin-9 OS=Homo sapiens OX=9606 GN=LGALS9 PE=1 SV=2 | 0.005 | 2.365 | 2 | 1 | 1 | 1 | 355 | 39.5 | 9.17 | 1.61 | 1 |
| Q86VP1 | Tax1-binding protein 1 OS=Homo sapiens OX=9606 GN=TAX1BP1 PE=1 SV=2 | 0.005 | 2.356 | 1 | 1 | 1 | 1 | 789 | 90.8 | 5.43 | 1.86 | 1 |
| Q9UBM7 | 7-dehydrocholesterol reductase OS=Homo sapiens OX=9606 GN=DHCR7 PE=1 SV=1 | 0.005 | 2.354 | 2 | 1 | 6 | 1 | 475 | 54.5 | 8.7 | 1.84 | 1 |
| O75390 | Citrate synthase, mitochondrial OS=Homo sapiens OX=9606 GN=CS PE=1 SV=2 | 0.005 | 2.343 | 2 | 1 | 1 | 1 | 466 | 51.7 | 8.32 | 0 | 1 |
| P61601 | Neurocalcin-delta OS=Homo sapiens OX=9606 GN=NCALD PE=1 SV=2 | 0.005 | 2.338 | 4 | 1 | 4 | 1 | 193 | 22.2 | 5.35 | 0 | 1 |
| Q96AC1 | Fermitin family homolog 2 OS=Homo sapiens OX=9606 GN=FERMT2 PE=1 SV=1 | 0.008 | 2.271 | 3 | 1 | 1 | 1 | 680 | 77.8 | 6.7 | 0 | 1 |
| P12956 | X-ray repair cross-complementing protein 6 OS=Homo sapiens OX=9606 GN=XRCC6 PE=1 SV=2 | 0.008 | 2.264 | 5 | 1 | 1 | 1 | 609 | 69.8 | 6.64 | 0 | 1 |
| O75503 | Ceroid-lipofuscinosis neuronal protein 5 OS=Homo sapiens OX=9606 GN=CLN5 PE=1 SV=2 | 0.009 | 2.251 | 3 | 1 | 1 | 1 | 358 | 41.5 | 7.4 | 1.8 | 1 |
| Q96HP4 | Oxidoreductase NAD-binding domain-containing protein 1 OS=Homo sapiens OX=9606 GN=OXNAD1 PE=1 SV=1 | 0.009 | 2.247 | 2 | 1 | 3 | 1 | 312 | 34.8 | 8.37 | 5.75 | 1 |
| O94886 | CSC1-like protein 1 OS=Homo sapiens OX=9606 GN=TMEM63A PE=1 SV=3 | 0.01 | 2.243 | 1 | 1 | 1 | 1 | 807 | 92.1 | 7.27 | 0 | 1 |
| P15170 | Eukaryotic peptide chain release factor GTP-binding subunit ERF3A OS=Homo sapiens OX=9606 GN=GSPT1 PE=1 SV=1 | 0.01 | 2.241 | 2 | 1 | 1 | 1 | 499 | 55.7 | 5.62 | 0 | 1 |

| Accession | Abundances (Normalized): Normal-Colostrum-1 | Abundances (Normalized): Normal-Colostrum-2 | Abundances (Normalized): Normal-Colostrum-3 | Abundances (Normalized): Normal-Colostrum-4 | Abundances (Normalized): Normal-Colostrum-5 | Abundances (Normalized): Normal-Colostrum-6 | Abundances (Normalized): Normal-Colostrum-7 | Abundances (Normalized): Normal-Colostrum-8 | Abundances (Normalized): Normal-Colostrum-9 | Abundances (Normalized): Normal-Colostrum-10 |
|---|---|---|---|---|---|---|---|---|---|---|
| Q4JQV9 | 12947939.48 | 671093.2219 | 2251554.948 | | | | 1403517.674 | | 748072.4755 | |
| P22234 | | | | | | | | | | |
| O95394 | | | | | | | | | | |
| P68402 | | 603463.5278 | | 1180369.056 | | | | | | |
| P48163 | | | | | | | | | | |
| Q01581 | | 752364.357 | 412758.8241 | 516105.4897 | 303666.2047 | 329996.2855 | | 626176.2979 | | 209371.8425 |
| O95164 | | | | | | | | | 619672.1697 | |
| P10619 | | | | | | | | | | |
| P05997 | | 819486.8559 | | | 699234.0167 | 532792.8199 | 890477.8777 | 770081.9843 | 352816.0845 | 646094.8393 |
| Q86V21 | | 571937.8156 | | 1275621.322 | 948400.8783 | 2047603.644 | | 1278669.4 | 1422629.347 | 1636515.168 |
| Q9NX62 | | | | | | | | | | |
| O94903 | | | 1938201.646 | | | | | | | 1490366.058 |
| A7J390 | 3188844.274 | 1575111.998 | 4294850.802 | 3481733.256 | 3414085.554 | 3907719.896 | 4786893.932 | 1977831.931 | 2380847.396 | 4764650.899 |
| Q7L523 | | | | | | 1457663.344 | | | | 1494531.695 |
| Q9Y6V0 | 1492907.34 | 1682402.063 | 857290.4105 | 619942.4672 | 1764646.834 | 1356670.247 | 2741747.903 | 3100041.482 | 1103396.607 | 605618.4154 |
| O95445 | 373148.9384 | 314748.5509 | | | 1033384.677 | | | | | |
| Q9Y2A9 | | | | | | | | | | |
| Q9UBW5 | | | | | | | | | | |
| Q15293 | | | | | | | | | | |
| Q96HU8 | | | 3182450.369 | | 3497185.187 | | 5751098.663 | | 1586964.021 | |
| Q13296 | | | | | | | | | | |
| Q92973 | | | | | | | | | | |
| Q9P0K7 | | | | | | | | | | |
| P62314 | | | | | | | | | | |
| P23246 | | | | | | | | | | |
| P22532 | | 112886.1798 | | | | | | | | 63013.5946 |
| Q08554 | | | | | | | | | | |
| Q9UI42 | | | | | | | | | | |
| Q9HD42 | 10366838.4 | 2737578.06 | 3096676.684 | 4577033.915 | 4002835.422 | 3806128.261 | 4443659.991 | 2658906.96 | 1861121.69 | 3124688.626 |
| Q8IYJ3 | | | | | | | | | | |
| Q86WV6 | | | | | | | | | | |
| P61011 | 4934551.072 | 932623.6066 | 965546.7607 | 2186935.211 | 1512062.693 | 3968582.527 | 2319701.693 | 4581798.078 | 5069232.872 | 5053951.229 |
| P43686 | | | | 559837.9366 | | | | | | |
| Q12840 | | | | | | | | | | |
| O75352 | | | | | | 381182.898 | | | | |
| Q8IY18 | | 290139.8316 | | | | | 557186.9305 | | | |
| Q00577 | 95825034.23 | | | | | | | | | |
| P00750 | | | | | | | | | | |
| Q96C24 | | | | 1284232.646 | | | | | | 516564.4953 |
| Q16836 | 4163575.13 | 6314370.882 | 4909063.402 | | | | | | | 12532762.53 |
| A0A075B6T7 | | | 561545.602 | | | | | 656862.5804 | 620837.1087 | |
| P35998 | | | | | | | | | | |
| Q86UL3 | | | | | | | | | | |
| O00182 | | | | 447934.0241 | | 433548.7269 | 393512.2761 | | | 629215.162 |
| Q86VP1 | | | | | | | | | 6611378.369 | 4857224.908 |
| Q9UBM7 | | | | | | | | 694729.9697 | | |
| O75390 | | | | | | | | | | |
| P61601 | | | 823415.1289 | 1044375.699 | | | | | | 2207527.56 |
| Q96AC1 | | | | | | | | | | |
| P12956 | | | | | | | | | | |
| O75503 | | | 480606.1233 | 453562.8439 | 798017.8768 | 813628.5004 | | | | 970177.6371 |
| Q96HP4 | | | 1962287.456 | | | | | | | |
| O94886 | 2731822.571 | 1548827.242 | 2249932.853 | 2213428.169 | 2367235.819 | 816170.3306 | 1392995.302 | 1497653.767 | 1314008.858 | 925416.3708 |
| P15170 | | | | | | | 1007604.862 | | 475624.4048 | |

| Accession | Abundances (Normalized): Covid-colostrum-1 | Abundances (Normalized): Covid-colostrum-2 | Abundances (Normalized): Covid-colostrum-3 | Abundances (Normalized): Covid-colostrum-4 | Abundances (Normalized): Covid-colostrum-5 | Abundances (Normalized): Covid-colostrum-6 |
|---|---|---|---|---|---|---|
| Q4JQV9 | | 3812925.666 | 32991257.22 | 246468114.2 | | 2427006.346 |
| P22234 | | | | | | |
| O95394 | | | | | | |
| P68402 | | | | | | |
| P48163 | | | | | | |
| Q01581 | 8249181 | | | | | |
| O95164 | | | | 2111884.484 | 1099423.25 | 868404.5302 |
| P10619 | | | | | | |
| P05997 | 1301610 | 1105358.906 | | | 706504.3743 | |
| Q86V21 | | | 4353937.919 | | 1189021.339 | |
| Q9NX62 | | 1136952.113 | | | | |
| O94903 | 2471585.5 | | | 555319.6777 | 1212455.128 | |
| A7J390 | | 2947870.437 | | | 3079026.659 | |
| Q7L523 | 1791608.5 | | | | 3406596.473 | |
| Q9Y6V0 | 1410633.75 | 2744626.265 | 400357.1527 | 214160.0646 | 1522640.88 | 634239.2328 |
| O95445 | | | | | | |
| Q9Y2A9 | | | | | | |
| Q9UBW5 | | | | | | |
| Q15293 | | | | | | |
| Q96HU8 | | | 1471798.329 | | | 1055332.87 |
| Q13296 | | | | 11681501.34 | | |
| Q92973 | | | | | | |
| Q9P0K7 | | | | | | |
| P62314 | | | | | | |
| P23246 | | | | | | |
| P22532 | | | 439567.455 | | | 288333.563 |
| Q08554 | | | | | | |
| Q9UI42 | | | | | | |
| Q9HD42 | 5119961.5 | 12341742.26 | 6525801.852 | 10085485.41 | 26266682.07 | 2089884.377 |
| Q8IYJ3 | | | | | | |
| Q86WV6 | 2797889.5 | | | | | |
| P61011 | 43874849 | 3898426.927 | 10915964.01 | 8509863.37 | 10317365.86 | 52406492.08 |
| P43686 | | | | | | |
| Q12840 | | | | | | |
| O75352 | | | | | | |
| Q8IY18 | | 229852.3513 | | | | |
| Q00577 | | | 131422648.3 | 928028338.9 | | |
| P00750 | | | | | | |
| Q96C24 | | | | | | |
| Q16836 | 5452674 | 4248353.668 | | | | |
| A0A075B6T7 | | | | | | |
| P35998 | 1005436.625 | | | 934293.038 | | 6726936.329 |
| Q86UL3 | | | | | | |
| O00182 | | | 686292.5168 | | 1303681.055 | 520102.8021 |
| Q86VP1 | | | | | | 8494116.924 |
| Q9UBM7 | | | 596461.4168 | 3005007.072 | | |
| O75390 | 3460762.25 | | | | 1220096.98 | |
| P61601 | 1224429.375 | | | | 1520966.236 | |
| Q96AC1 | | | | | | |
| P12956 | 3181965 | | | | | |
| O75503 | | 4935120.03 | | 6564019.711 | | 1170411.532 |
| Q96HP4 | | | | | | |
| O94886 | | 4126742.26 | 1161690.984 | | 3416835.752 | 1509162.508 |
| P15170 | | | | | | |

| Accession | Abundances (Normalized): Normal-mature-1 | Abundances (Normalized): Normal-mature-2 | Abundances (Normalized): Normal-mature-3 | Abundances (Normalized): Normal-mature-4 | Abundances (Normalized): Normal-mature-5 | Abundances (Normalized): Normal-mature-6 | Abundances (Normalized): Normal-mature-7 | Abundances (Normalized): Normal-mature-8 | Abundances (Normalized): Normal-mature-9 | Abundances (Normalized): Normal-mature-10 |
|---|---|---|---|---|---|---|---|---|---|---|
| Q4JQV9 | 3982274.327 | 1397745.007 | 50766886.06 | 1339052.824 | 16139930.06 | 832576.2473 | 1080840.636 | 2946560.608 | 598372.1028 | |
| P22234 | | | | | | | | | | |
| O95394 | | | | | | | | | | |
| P68402 | | | | | 767843.6096 | 651755.1304 | | | | 2077299.063 |
| P48163 | | | | | | | | | | |
| Q01581 | | 601695.0925 | | 836438.3779 | 1334916.574 | | 360276.738 | | | 230320.8545 |
| O95164 | 1340285.907 | | | 586457.4228 | | 634166.2962 | | 822871.8242 | 706240.5227 | |
| P10619 | | | | | | | | | | |
| P05997 | 1407341.227 | 1234403.708 | 1406097.147 | 1221645.884 | 913735.1637 | 768831.2021 | 998308.08 | | 508613.6473 | 294249.6699 |
| Q86V21 | | 877377.7687 | 438655.0882 | 869287.7261 | | 524364.34 | 633158.5821 | | 886087.5213 | 1051758.653 |
| Q9NX62 | 552821.5078 | 563261.7146 | | | | 639609.7825 | 257231.8802 | | 412706.2854 | 189167.8097 |
| O94903 | 1138313.968 | 2826541.338 | | | | | | 1389771.922 | | |
| A7J390 | 3228872.583 | 1819339.917 | 1821629.113 | 4302399.859 | 1641643.122 | 2403381.68 | 3193997.926 | 1836167.313 | 1256103.61 | 2057011.691 |
| Q7L523 | | | | | | | | | | 2411024.395 |
| Q9Y6V0 | 3341288.324 | 1220318.841 | 2673618.17 | 3136306.721 | 3928688.958 | 2999596.722 | 730210.6213 | 473598.3352 | 1175173.836 | 222471.0036 |
| O95445 | 665123.1538 | 558407.199 | 1095626.139 | 733779.4135 | 598241.7994 | 945582.86 | 701119.3342 | 447926.899 | 477978.6372 | 314585.5751 |
| Q9Y2A9 | | | | | | | | | | |
| Q9UBW5 | | | | | | | | | | |
| Q15293 | | | | | | | | | | |
| Q96HU8 | | | 1365187.753 | 2478175.89 | | | 3735185.204 | | | 1659425.953 |
| Q13296 | | | | | | | | | | |
| Q92973 | | | | | | | | | | |
| Q9P0K7 | | | | | | | | | | |
| P62314 | | | | | | | | | | |
| P23246 | | | | | | | | | | |
| P22532 | | | | | | | | | | |
| Q08554 | | | | | | | | | | |
| Q9UI42 | | | | | | | | | | |
| Q9HD42 | 2037851.766 | 1952372.787 | 2375636.647 | 3355088.586 | 1782461.498 | 2344454.768 | 2065700.281 | 3660728.375 | 833507.6366 | 1010816.065 |
| Q8IYJ3 | | | | | | | | | | |
| Q86WV6 | | | | | | | | | | |
| P61011 | 3608592.023 | 2216547.342 | | 2759699.431 | | 1644846.288 | 1211416.016 | 3229927.379 | 1090803.07 | 395581.3426 |
| P43686 | | | | | | | | | | |
| Q12840 | | | | | | | | | | |
| O75352 | | | | | 590346.9407 | 350433.428 | | | | 275085.3401 |
| Q8IY18 | 113221.3876 | 397002.2587 | | 231058.4515 | 544321.5604 | 307155.2833 | | | | |
| Q00577 | | | 242317249.9 | | 43602743.49 | | | | | |
| P00750 | | | | | | | | | | |
| Q96C24 | | | | 936819.1115 | 344626.2288 | | 411251.8558 | | 846905.6766 | 676498.7622 |
| Q16836 | 5002645.418 | 7205173.111 | 5292230.233 | | 6616026.564 | 2853190.469 | 5410835.729 | | | |
| A0A075B6T7 | | | | 623473.5827 | | | | | 1309438.877 | |
| P35998 | | | | | | | | | | |
| Q86UL3 | | | | | | | | | | |
| O00182 | | | 323419.6039 | | 518738.0186 | | | | | |
| Q86VP1 | | | 24191065.5 | | | | | | | |
| Q9UBM7 | | | 832634.3275 | | 757121.7487 | 409196.5241 | | | | 384548.9971 |
| O75390 | | | | | | | 1743820.152 | | | |
| P61601 | | | | | | | | | | 1350516.466 |
| Q96AC1 | | | | | | | | | | |
| P12956 | | | | | | | | | | |
| O75503 | 920961.0889 | | | | | | 764321.6538 | 507468.4605 | | 733883.3861 |
| Q96HP4 | | | | | | | 662080.1074 | | | 1626169.227 |
| O94886 | | 434044.079 | 758733.1325 | 1502120.581 | 919683.8153 | 1144820.45 | 900271.513 | 907477.0566 | 742766.7131 | 811834.4186 |
| P15170 | | | | | | | | | | |

| Accession | Abundances (Normalized): Covid-mature-1 | Abundances (Normalized): Covid-mature-2 | Abundances (Normalized): Covid-mature-3 | Abundances (Normalized): Covid-mature-4 | Abundances (Normalized): Covid-mature-5 | Abundances (Normalized): Covid-mature-6 |
|---|---|---|---|---|---|---|
| Q4JQV9 | | 2874650.125 | 1802689.636 | | 433291.1656 | 119318019.5 |
| P22234 | | | | | | |
| O95394 | 2377856.34 | | | | | |
| P68402 | 734812.4425 | | | 1361525.129 | 1313069.221 | |
| P48163 | | | | | | |
| Q01581 | 622667.6696 | 526126.5383 | | 319850.4786 | 491541.163 | |
| O95164 | 674858.468 | | | | | |
| P10619 | | | | | | |
| P05997 | | | 1177494.736 | 685993.5203 | 495067.3442 | 2275440 |
| Q86V21 | | 831797.8577 | 1545404.619 | | 982989.9677 | |
| Q9NX62 | 520732.343 | | | | | 698065.71 |
| O94903 | | 1196595.85 | 3197136.042 | 1239309.251 | | |
| A7J390 | 1921793.813 | 3333731.461 | 1368235.105 | 3486577.153 | 2534597.799 | 5281160.529 |
| Q7L523 | | | 3954580.047 | 3473558.019 | 3703066.318 | |
| Q9Y6V0 | 2846073.061 | 203826.1669 | 2334369.37 | 2504239.709 | | 195367.2136 |
| O95445 | 523198.3874 | 753122.2441 | | 1010332.213 | | |
| Q9Y2A9 | | | | | | |
| Q9UBW5 | | | | | | |
| Q15293 | | | | | | |
| Q96HU8 | | | | | 1097551.517 | |
| Q13296 | | | | | | |
| Q92973 | | | | | | |
| Q9P0K7 | | | | | | |
| P62314 | | | | | | |
| P23246 | | | | | | |
| P22532 | 366832.8778 | | | | | |
| Q08554 | | | | | | |
| Q9UI42 | | | | | | |
| Q9HD42 | 1580023.359 | 4134225.11 | 2759227.507 | 1858588.304 | 1694117.418 | 35290159.67 |
| Q8IYJ3 | | | | | | |
| Q86WV6 | | | | | | |
| P61011 | 4405508.902 | 1944119.005 | 4487771.994 | | 2621648.979 | |
| P43686 | | | | | | |
| Q12840 | | | | | | |
| O75352 | | | | | 526558.7109 | 1617726.38 |
| Q8IY18 | | | | | | |
| Q00577 | | | | | | 872278645.9 |
| P00750 | | | | | | |
| Q96C24 | | | | | 901720.9982 | 790263.553 |
| Q16836 | 4828989.759 | 6268977.484 | 1509760.626 | 4386419.69 | | 2647192.721 |
| A0A075B6T7 | | | | | | 1375486.789 |
| P35998 | | | | | | 4026095.489 |
| Q86UL3 | | | | | | |
| O00182 | 449273.9468 | 757192.6219 | 792473.886 | | | 1173406.747 |
| Q86VP1 | | | | 37923608.07 | | |
| Q9UBM7 | | 736441.7177 | | 582197.2978 | | 3470687.299 |
| O75390 | | | | | | |
| P61601 | | | | 1342567.079 | | |
| Q96AC1 | | | | | | |
| P12956 | | | | | | |
| O75503 | | | | 544353.5081 | | |
| Q96HP4 | | | | | | |
| O94886 | 1188590.593 | 2403716.2 | 972672.9 | 773408.2828 | 741592.5232 | 2051218.667 |
| P15170 | | | | | | 876071.2433 |

Table S2. Enriched gene ontology (GO) terms associated with the differentially expressed proteins in COVID-19 colostrum and Ctrl colostrum sample groups

| Up/Down regulated | GO Term | % Associated Proteins | Associated Proteins Found |
|---|---|---|---|
| Up | humoral immune response mediated by circulating immunoglobulin | 34.25414276 | [C3, C4B, HLA-DRB1, HPX, IGHA1, IGHA2, IGHG3, IGHG4, IGHM, IGHV1-18, IGHV1-3, IGHV1-45, IGHV1-8, IGHV1OR15-1, IGHV2-26, IGHV2-5, IGHV3-11, IGHV3-13, IGHV3-15, IGHV3-16, IGHV3-20, IGHV3-21, IGHV3-23, IGHV3-30, IGHV3-35, IGHV3-38, IGHV3-43, IGHV3-49, IGHV3-64, IGHV3-64D, IGHV3-7, IGHV3-72, IGHV3-73, IGHV3-74, IGHV3-9, IGHV4-28, IGHV4-31, IGHV4-4, IGHV5-10-1, IGHV5-51, IGHV6-1, IGHV7-4-1, IGKC, IGKV1-12, IGKV1-16, IGKV1-17, IGKV1-5, IGKV1D-33, IGKV1D-39, IGKV2-30, IGKV3-20, IGKV3D-20, IGKV4-1, IGLC3, IGLL1, IGLV1-40, IGLV1-47, IGLV1-51, IGLV2-11, IGLV2-8, IGLV7-43, PSMA7] |
| Up | phagocytosis, recognition | 39.42307663 | [IGHA1, IGHA2, IGHG3, IGHG4, IGHM, IGHV1-18, IGHV1-3, IGHV1-45, IGHV1-8, IGHV1OR15-1, IGHV2-26, IGHV2-5, IGHV3-11, IGHV3-13, IGHV3-15, IGHV3-16, IGHV3-20, IGHV3-21, IGHV3-23, IGHV3-30, IGHV3-35, IGHV3-38, IGHV3-43, IGHV3-49, IGHV3-64, IGHV3-64D, IGHV3-7, IGHV3-72, IGHV3-73, IGHV3-74, IGHV3-9, IGHV4-28, IGHV4-31, IGHV4-4, IGHV5-10-1, IGHV5-51, IGHV6-1, IGHV7-4-1, IGKC, IGLC3, IGLL1] |
| Up | myeloid leukocyte mediated immunity | 10.29411793 | [ANPEP, ANXA3, ARPC5, ASAH1, AZU1, C3, CAMP, CHI3L1, CNN2, COTL1, CPNE3, CREG1, CTSB, CTSD, CTSG, CTSH, CTSZ, CYBB, DEFA4, ELANE, FTL, GM2A, GMFG, GSTP1, GUSB, GYG1, HK3, IGF2R, LAMP2, LCN2, LGALS3, LRG1, LTA4H, MNDA, NEU1, OLFM4, ORM1, ORM2, PA2G4, PGAM1, PIGR, PRCP, PRSS8, PRTN3, RAC2, RETN, RNASE3, RNASET2, S100A11, S100A12, S100A8, S100A9, S100P, SDCBP, SERPINA1, SERPINB1, SIRPA, SLPI, TINAGL1, TUBB4B, VCL, WDR1, XRCC5] |
| Up | regulation of humoral immune response | 23.44827652 | [C3, C4B, CFB, CFH, CXCL13, HPX, IGHG3, IGHG4, IGHV2-5, IGHV3-11, IGHV3-13, IGHV3-23, IGHV3-30, IGHV3-7, IGHV3-9, IGKC, IGKV1-12, IGKV1-16, IGKV1-17, IGKV1-5, IGKV1D-33, IGKV1D-39, IGKV2-30, IGKV3-20, IGKV3D-20, IGKV4-1, IGLC3, IGLV1-40, IGLV1-47, IGLV1-51, IGLV2-11, IGLV2-8, IGLV7-43, PSMA7] |
| Up | defense response to other organism | 7.24054718 | [ACTG1, ANXA3, APLP2, AZU1, C3, C4B, CAMP, CFB, CFH, CHID1, COTL1, CTSG, CXCL13, CYBB, DEFA4, DMBT1, ELANE, ERAP1, FLNA, FLNB, HLA-DRB1, HPX, IFI30, IFITM2, IGHA1, IGHA2, IGHG3, IGHG4, IGHM, IGHV1-18, IGHV1-3, IGHV1-45, IGHV1-8, IGHV1OR15-1, IGHV2-26, IGHV2-5, IGHV3-11, IGHV3-13, IGHV3-15, IGHV3-16, IGHV3-20, IGHV3-21, IGHV3-23, IGHV3-30, IGHV3-35, IGHV3-38, IGHV3-43, IGHV3-49, IGHV3-64, IGHV3-64D, IGHV3-7, IGHV3-72, IGHV3-73, IGHV3-74, IGHV3-9, IGHV4-28, IGHV4-31, IGHV4-4, IGHV5-10-1, IGHV5-51, IGHV6-1, IGHV7-4-1, IGKC, IGKV3-20, IGLC3, IGLL1, JCHAIN, LBP, LCN2, LGALS3, MNDA, MUC20, MUC4, PADI4, PSMA7, PSMB4, PSMB6, PSMB9, RNASE3, RNASET2, RPL13A, S100A12, S100A8, S100A9, SIRPA, SLPI, SMPDL3B, TINAGL1, TUBB4B, XRCC5] |
| Up | peptidase regulator activity | 5.737704754 | [APLP2, BIN1, C3, C4B, CTSH, KNG1, PI15, PRDX3, RARRES1, SERPINA1, SERPINA6, SERPINB1, SLPI, SPINT1] |
| Up | regulation of tumor necrosis factor production | 7.831325531 | [AZU1, CYBB, GPNMB, GSTP1, LBP, LCP1, ORM1, ORM2, S100A8, S100A9, SIRPA, THBS1, TSPO] |
| Up | tissue homeostasis | 5.78231287 | [ACTG1, AZGP1, CFH, CTSH, IGHA1, IGHA2, IGHG3, IGKC, JCHAIN, MUC4, PIGR, PIP, PROM1, RAC2, TNFRSF11B, VCL, ZG16B] |
| Up | myeloid leukocyte migration | 5.833333492 | [AZU1, CD9, CXCL13, LBP, LGALS3, PRTN3, RAC2, S100A12, S100A8, S100A9, SIRPA, SRP54, THBS1, WDR1] |
| Up | cell migration | 4.503393173 | [ACTG1, ADGRG1, ANXA3, APLP2, APOD, ARPC5, ATP5F1A, AZU1, CD9, CNN2, CPNE3, CTSH, CXCL13, ELANE, FERMT3, FLNA, GPNMB, GRB2, GSTP1, IGHA1, IGHA2, IGHM, IGHV2-5, IGHV3-11, IGHV3-13, IGHV3-23, IGHV3-30, IGHV3-7, IGHV3-9, IGKC, IGKV1-12, IGKV1-16, IGKV1-17, IGKV1-5, IGKV1D-33, IGKV1D-39, IGKV2-30, IGKV3-20, IGKV3D-20, IGKV4-1, IGLC3, IGLL1, IGLV1-40, IGLV1-47, IGLV1-51, IGLV2-11, IGLV2-8, IGLV7-43, JCHAIN, LBP, LCP1, LGALS3, MATN2, NARS1, NRP1, PRCP, PRSS8, PRTN3, PTPRF, RAC2, S100A11, S100A12, S100A8, S100A9, S100P, SDCBP, SIRPA, SOD2, SRP54, THBS1, TSPO, VCL, WDR1] |
| Up | defense response to bacterium | 15.34391499 | [ANXA3, AZU1, CAMP, CTSG, CXCL13, DEFA4, DMBT1, ELANE, IGHA1, IGHA2, IGHG3, IGHG4, IGHM, IGHV1-18, IGHV1-3, IGHV1-45, IGHV1-8, IGHV1OR15-1, IGHV2-26, IGHV2-5, IGHV3-11, IGHV3-13, IGHV3-15, IGHV3-16, IGHV3-20, IGHV3-21, IGHV3-23, IGHV3-30, IGHV3-35, IGHV3-38, IGHV3-43, IGHV3-49, IGHV3-64, IGHV3-64D, IGHV3-7, IGHV3-72, IGHV3-73, IGHV3-74, IGHV3-9, IGHV4-28, IGHV4-31, IGHV4-4, IGHV5-10-1, IGHV5-51, IGHV6-1, IGHV7-4-1, IGKC, IGKV3-20, IGLC3, IGLL1, JCHAIN, LBP, LCN2, RNASE3, S100A12, S100A8, S100A9, SLPI] |
| Up | regulation of chemokine production | 9.782608986 | [APOD, AZU1, ELANE, GSTP1, LBP, LCP1, S100A8, S100A9, SIRPA] |
| Up | peptidase activity | 4.118173599 | [ANPEP, APLP2, AZU1, BIN1, C3, C4B, CASP14, CFB, CHMP1A, CLC, CLCA4, CPM, CTSB, CTSD, CTSG, CTSH, CTSZ, ELANE, ERAP1, GGT5, KLK11, KNG1, LTA4H, OTUB1, PI15, PIP, PRCP, PRDX3, PRSS8, PRTN3, PSMA7, PSMB4, PSMB6, PSMB9, RARRES1, RNASE3, RNPEP, S100A8, S100A9, SERPINA1, SERPINA6, SERPINB1, SLPI, SPINT1, THBS1, TINAGL1] |
| Up | receptor-mediated endocytosis | 10.79545498 | [C3, CD9, DMBT1, EZR, GRB2, HNRNPK, HPX, IGF2R, IGHA1, IGHA2, IGHV2-5, IGHV3-11, IGHV3-13, IGHV3-23, IGHV3-30, IGHV3-7, IGHV3-9, IGKC, IGKV1-12, IGKV1-16, IGKV1-17, IGKV1-5, IGKV1D-33, IGKV1D-39, IGKV2-30, IGKV3-20, IGKV3D-20, IGKV4-1, IGLC3, IGLV1-40, IGLV1-47, IGLV1-51, IGLV2-11, IGLV2-8, IGLV7-43, JCHAIN, PRSS8, SDCBP] |
| Up | enzyme inhibitor activity | 4.567307472 | [ANXA3, ANXA4, APLP2, BIN1, C3, C4B, DMBT1, GMFG, KNG1, LGALS3, PI15, PRDX3, RARRES1, RNH1, SERPINA1, SERPINA6, SERPINB1, SLPI, SPINT1] |
| Up | retina homeostasis | 12.94117641 | [ACTG1, AZGP1, IGHA1, IGHA2, IGHG3, IGKC, JCHAIN, PIGR, PIP, PROM1, ZG16B] |
| Up | positive regulation of response to biotic stimulus | 5.223880768 | [HPX, LBP, MNDA, MUC20, MUC4, PSMA7, PSMB4, PSMB6, PSMB9, RNASE3, S100A8, S100A9, TINAGL1, XRCC5] |
| Up | positive regulation of vasculature development | 5.74162674 | [ANXA3, C3, CAMP, CHI3L1, CTSH, CYBB, ERAP1, LRG1, NRP1, PSMA7, SOD2, THBS1] |
| Up | cellular response to interleukin-12 | 15.51724148 | [CNN2, HNRNPA2B1, LCP1, LMNB1, S100A8, S100A9, SIRPA, SOD2, TALDO1] |
| Up | platelet degranulation | 10 | [APLP2, CD9, CHID1, FERMT3, FLNA, KNG1, LAMP2, LGALS3BP, ORM1, ORM2, SERPINA1, THBS1, VCL, WDR1] |
| Up | regulation of respiratory burst | 23.80952454 | [IGHA1, IGHA2, JCHAIN, LBP, RAC2] |
| Up | negative regulation of chemotaxis | 10.44776154 | [CXCL13, ELANE, GSTP1, NRP1, S100A8, S100A9, THBS1] |

Table S2. Cont.

| Up/Down regulated | GO Term | % Associated Proteins | Associated Proteins Found |
| --- | --- | --- | --- |
| Up | leukocyte mediated immunity | 12.79187775 | [ANPEP, ANXA3, ARPC5, ASAH1, AZGP1, AZU1, C3, C4B, CAMP, CHI3L1, CLC, CNN2, COTL1, CPNE3, CREG1, CTSB, CTSD, CTSG, CTSH, CTSZ, CYBB, DEFA4, ELANE, FTL, GM2A, GMFG, GSTP1, GUSB, GYG1, HK3, HLA-DRB1, HPX, IGF2R, IGHA1, IGHA2, IGHG3, IGHG4, IGHM, IGHV1-18, IGHV1-3, IGHV1-45, IGHV1-8, IGHV1OR15-1, IGHV2-26, IGHV2-5, IGHV3-11, IGHV3-13, IGHV3-15, IGHV3-16, IGHV3-20, IGHV3-21, IGHV3-23, IGHV3-30, IGHV3-35, IGHV3-38, IGHV3-43, IGHV3-49, IGHV3-64, IGHV3-64D, IGHV3-7, IGHV3-72, IGHV3-73, IGHV3-74, IGHV3-9, IGHV4-28, IGHV4-31, IGHV4-4, IGHV5-10-1, IGHV5-51, IGHV6-1, IGHV7-4-1, IGKC, IGKV1-12, IGKV1-16, IGKV1-17, IGKV1-5, IGKV1D-33, IGKV1D-39, IGKV2-30, IGKV3-20, IGKV3D-20, IGKV4-1, IGLC3, IGLL1, IGLV1-40, IGLV1-47, IGLV1-51, IGLV2-11, IGLV2-8, IGLV7-43, LAMP2, LCN2, LGALS3, LRG1, LTA4H, MNDA, NEU1, OLFM4, ORM1, ORM2, PA2G4, PGAM1, PIGR, PRCP, PRSS8, PRTN3, PSMA7, RAC2, RETN, RNASE3, RNASET2, S100A11, S100A12, S100A8, S100A9, S100P, SDCBP, SERPINA1, SERPINB1, SIRPA, SLPI, TINAGL1, TUBB4B, VCL, WDR1, XRCC5] |
| Up | leukocyte migration | 8.561019897 | [APOD, AZU1, CD9, CXCL13, ELANE, GRB2, IGHA1, IGHA2, IGHM, IGHV2-5, IGHV3-11, IGHV3-13, IGHV3-23, IGHV3-30, IGHV3-7, IGHV3-9, IGKC, IGKV1-12, IGKV1-16, IGKV1-17, IGKV1-5, IGKV1D-33, IGKV1D-39, IGKV2-30, IGKV3-20, IGKV3D-20, IGKV4-1, IGLC3, IGLL1, IGLV1-40, IGLV1-47, IGLV1-51, IGLV2-11, IGLV2-8, IGLV7-43, JCHAIN, LBP, LGALS3, PRTN3, RAC2, S100A12, S100A8, S100A9, SIRPA, SRP54, THBS1, WDR1] |
| Up | defense response to fungus | 18.18181801 | [COTL1, CTSG, DEFA4, ELANE, S100A12, S100A8, S100A9, TINAGL1] |
| Up | defense response to Gram-negative bacterium | 9.677419662 | [AZU1, CAMP, CTSG, DEFA4, DMBT1, ELANE, IGHM, LBP, RNASE3] |
| Up | leukocyte activation | 7.855626106 | [ANPEP, ANXA3, ARPC5, ASAH1, AZU1, C3, CAMP, CHI3L1, CLC, CNN2, COTL1, CPNE3, CREG1, CTSB, CTSD, CTSG, CTSH, CTSZ, CYBB, DEFA4, ELANE, FTL, GM2A, GMFG, GPNMB, GRB2, GSTP1, GUSB, GYG1, HK3, HLA-DRB1, IGF2R, IGHA1, IGHA2, IGHG3, IGHG4, IGHM, IGHV1-18, IGHV1-3, IGHV1-45, IGHV1-8, IGHV1OR15-1, IGHV2-26, IGHV2-5, IGHV3-11, IGHV3-13, IGHV3-15, IGHV3-16, IGHV3-20, IGHV3-21, IGHV3-23, IGHV3-30, IGHV3-35, IGHV3-38, IGHV3-43, IGHV3-49, IGHV3-64, IGHV3-64D, IGHV3-7, IGHV3-72, IGHV3-73, IGHV3-74, IGHV3-9, IGHV4-28, IGHV4-31, IGHV4-4, IGHV5-10-1, IGHV5-51, IGHV6-1, IGHV7-4-1, IGKC, IGLC3, IGLL1, LAMP2, LBP, LCN2, LCP1, LGALS1, LGALS3, LRG1, LTA4H, MNDA, NEU1, OLFM4, ORM1, ORM2, PA2G4, PGAM1, PIGR, PRCP, PRSS8, PRTN3, RAC2, RETN, RNASE3, RNASET2, S100A11, S100A12, S100A8, S100A9, S100P, SDCBP, SERPINA1, SERPINB1, SIRPA, SLPI, THBS1, TINAGL1, TUBB4B, VCL, XRCC5] |
| Up | regulation of interleukin-1 beta production | 8.695652008 | [AZU1, GSTP1, LCP1, MNDA, ORM1, ORM2, SERPINB1, SIRPA] |
| Up | positive regulation of response to external stimulus | 4.036697388 | [AZU1, C3, CXCL13, HPX, LBP, LCP1, MNDA, MUC20, MUC4, NRP1, PSMA7, PSMB4, PSMB6, PSMB9, RAC2, RNASE3, S100A12, S100A8, S100A9, THBS1, TINAGL1, XRCC5] |
| Up | antibacterial humoral response | 19.67213058 | [CAMP, CTSG, DEFA4, DMBT1, ELANE, IGHA1, IGHA2, IGHM, IGKV3-20, JCHAIN, RNASE3, SLPI] |
| Up | antimicrobial humoral response | 13.5714283 | [AZU1, CAMP, CTSG, CXCL13, DEFA4, DMBT1, ELANE, IGHA1, IGHA2, IGHM, IGKV3-20, JCHAIN, LCN2, PRTN3, RNASE3, S100A12, S100A8, S100A9, SLPI] |
| Up | vesicle-mediated transport | 6.173379898 | [ACTG1, ANPEP, ANXA3, APLP2, ARPC5, ASAH1, AZU1, BIN1, C3, C4B, CAMP, CD9, CHI3L1, CHID1, CHMP1A, CNN2, COTL1, CPNE3, CREG1, CTSB, CTSD, CTSG, CTSH, CTSZ, CYBB, DEFA4, DMBT1, ELANE, EZR, FERMT3, FLNA, FTL, GM2A, GMFG, GRB2, GSTP1, GUSB, GYG1, HK3, HNRNPK, HPX, HSPA13, IGF2R, IGHA1, IGHA2, IGHG3, IGHG4, IGHM, IGHV1-18, IGHV1-3, IGHV1-45, IGHV1-8, IGHV1OR15-1, IGHV2-26, IGHV2-5, IGHV3-11, IGHV3-13, IGHV3-15, IGHV3-16, IGHV3-20, IGHV3-21, IGHV3-23, IGHV3-30, IGHV3-35, IGHV3-38, IGHV3-43, IGHV3-49, IGHV3-64, IGHV3-64D, IGHV3-7, IGHV3-72, IGHV3-73, IGHV3-74, IGHV3-9, IGHV4-28, IGHV4-31, IGHV4-4, IGHV5-10-1, IGHV5-51, IGHV6-1, IGHV7-4-1, IGKC, IGKV1-12, IGKV1-16, IGKV1-17, IGKV1-5, IGKV1D-33, IGKV1D-39, IGKV2-30, IGKV3-20, IGKV3D-20, IGKV4-1, IGLC3, IGLL1, IGLV1-40, IGLV1-47, IGLV1-51, IGLV2-11, IGLV2-8, IGLV7-43, JCHAIN, KNG1, LAMP2, LBP, LCN2, LGALS3, LGALS3BP, LRG1, LTA4H, MNDA, NEU1, NRP1, OLFM4, ORM1, ORM2, PA2G4, PGAM1, PIGR, PRCP, PRSS8, PRTN3, RAC2, RETN, RNASE3, RNASET2, S100A11, S100A12, S100A8, S100A9, S100P, SDCBP, SERPINA1, SERPINB1, SIRPA, SLPI, THBS1, TINAGL1, TUBB4B, VCL, WDR1, XRCC5] |
| Up | serine-type peptidase activity | 4.838709831 | [APLP2, AZU1, CFB, CTSG, CTSH, ELANE, KLK11, PRCP, PRSS8, PRTN3, SERPINA1, SERPINA6, SERPINB1, SLPI, SPINT1] |
| Up | negative regulation of cytokine production | 5.86206913 | [ANXA4, APLP2, APOD, CHID1, ELANE, EZR, GPNMB, GSTP1, HLA-DRB1, LBP, NPTN, ORM1, SERPINB1, SIRPA, THBS1, TINAGL1, TSPO] |
| Down | ribosomal large subunit assembly | 9.677419662 | [RPL10, RPL5, RPL6] |
| Down | body fluid secretion | 4.424778938 | [CEL, CSN2, CSN3, SERPINC1, XDH] |
| Down | positive regulation of lipid storage | 11.11111069 | [APOB, PLIN2, PLIN3] |
| Down | ribosomal large subunit biogenesis | 6.578947544 | [RPL10, RPL10A, RPL14, RPL5, RPL6] |
| Down | nucleoside phosphate catabolic process | 5.194805145 | [CNP, HPRT1, NUDT5, XDH] |
| Down | positive regulation of lipid localization | 4.90196085 | [APOB, FABP3, PLIN2, PLIN3, SPP1] |
| Down | establishment of protein localization to endoplasmic reticulum | 8.064516068 | [RAB10, RPL10, RPL10A, RPL14, RPL27, RPL36AL, RPL5, RPL6, RPLP2, RPS24] |

Table S3. Enriched GO terms associated with the differentially expressed proteins in COVID-19 colostrum and COVID-19 mature sample groups

| Up/Down regulated | GOTerm | % Associated Proteins | Associated Proteins Found |
|---|---|---|---|
| Up | complement activation | 21.46 | [CFB, CRP, IGHA1, IGHG4, IGHV1-18, IGHV1-3, IGHV1-45, IGHV1-8, IGHV2-26, IGHV2-5, IGHV3-11, IGHV3-13, IGHV3-16, IGHV3-20, IGHV3-23, IGHV3-30, IGHV3-35, IGHV3-38, IGHV3-43, IGHV3-64, IGHV3-72, IGHV3-73, IGHV3-9, IGHV4-4, IGHV5-10-1, IGHV5-51, IGHV6-1, IGKV1-12, IGKV1-16, IGKV1-17, IGKV1-5, IGKV1D-33, IGKV1D-39, IGKV2-30, IGKV3-20, IGKV3D-20, IGKV4-1, IGLL1, IGLV1-40, IGLV1-47, IGLV1-51, IGLV2-8, IGLV7-43, PSMA7] |
| Up | phagocytosis, recognition | 25.00 | [IGHA1, IGHG4, IGHV1-18, IGHV1-3, IGHV1-45, IGHV1-8, IGHV2-26, IGHV2-5, IGHV3-11, IGHV3-13, IGHV3-16, IGHV3-20, IGHV3-23, IGHV3-30, IGHV3-35, IGHV3-38, IGHV3-43, IGHV3-64, IGHV3-72, IGHV3-73, IGHV3-9, IGHV4-4, IGHV5-10-1, IGHV5-51, IGHV6-1, IGLL1] |
| Up | regulation of complement activation | 19.05 | [CFB, IGHG4, IGHV2-5, IGHV3-11, IGHV3-13, IGHV3-23, IGHV3-30, IGHV3-9, IGKV1-12, IGKV1-16, IGKV1-17, IGKV1-5, IGKV1D-33, IGKV1D-39, IGKV2-30, IGKV3-20, IGKV3D-20, IGKV4-1, IGLV1-40, IGLV1-47, IGLV1-51, IGLV2-8, IGLV7-43, PSMA7] |
| Up | regulation of tumor necrosis factor production | 4.22 | [GPNMB, LBP, LCP1, ORM1, ORM2, S100A8, S100A9] |
| Up | leukocyte mediated immunity | 6.90 | [ASAH1, AZGP1, CHI3L1, CLC, CRP, CTSH, CTSZ, DEFA4, DPP7, FTL, GGH, GM2A, HK3, IGF2R, IGHA1, IGHG4, IGHV1-18, IGHV1-3, IGHV1-45, IGHV1-8, IGHV2-26, IGHV2-5, IGHV3-11, IGHV3-13, IGHV3-16, IGHV3-20, IGHV3-23, IGHV3-30, IGHV3-35, IGHV3-38, IGHV3-43, IGHV3-64, IGHV3-72, IGHV3-73, IGHV3-9, IGHV4-4, IGHV5-10-1, IGHV5-51, IGHV6-1, IGKV1-12, IGKV1-16, IGKV1-17, IGKV1-5, IGKV1D-33, IGKV1D-39, IGKV2-30, IGKV3-20, IGKV3D-20, IGKV4-1, IGLL1, IGLV1-40, IGLV1-47, IGLV1-51, IGLV2-8, IGLV7-43, LCN2, LTA4H, NEU1, ORM1, ORM2, PGAM1, PRSS8, PSMA7, RETN, S100A8, S100A9, TINAGL1, VCL] |
| Up | interleukin-12-mediated signaling pathway | 8.93 | [LCP1, LMNB1, S100A8, S100A9, SOD2] |
| Up | sphingolipid catabolic process | 10.53 | [ASAH1, GM2A, NEU1, SMPDL3B] |
| Up | antimicrobial humoral response | 5.00 | [DEFA4, DMBT1, IGHA1, IGKV3-20, LCN2, S100A8, S100A9] |
| Up | exopeptidase activity | 6.35 | [CPM, CTSH, CTSZ, DPP7, GGH, GGT5, LTA4H, RNPEP] |
| Up | leukocyte migration involved in inflammatory response | 15.79 | [LBP, S100A8, S100A9] |
| Up | leukocyte migration | 5.28 | [APOD, GP2, IGHA1, IGHV2-5, IGHV3-11, IGHV3-13, IGHV3-23, IGHV3-30, IGHV3-9, IGKV1-12, IGKV1-16, IGKV1-17, IGKV1-5, IGKV1D-33, IGKV1D-39, IGKV2-30, IGKV3-20, IGKV3D-20, IGKV4-1, IGLL1, IGLV1-40, IGLV1-47, IGLV1-51, IGLV2-8, IGLV7-43, LBP, S100A8, S100A9, SRP54] |
| Up | defense response to fungus | 9.09 | [DEFA4, S100A8, S100A9, TINAGL1] |
| Up | neutrophil degranulation | 4.27 | [ASAH1, CHI3L1, CTSH, CTSZ, DEFA4, DPP7, FTL, GGH, GM2A, HK3, IGF2R, LCN2, LTA4H, NEU1, ORM1, ORM2, PGAM1, PRSS8, RETN, S100A8, S100A9, TINAGL1, VCL] |
| Up | adaptive immune response | 7.99 | [AZGP1, CLC, CRP, CTSH, IGHA1, IGHG4, IGHV1-18, IGHV1-3, IGHV1-45, IGHV1-8, IGHV2-26, IGHV2-5, IGHV3-11, IGHV3-13, IGHV3-16, IGHV3-20, IGHV3-23, IGHV3-30, IGHV3-35, IGHV3-38, IGHV3-43, IGHV3-64, IGHV3-72, IGHV3-73, IGHV3-9, IGHV4-4, IGHV5-10-1, IGHV5-51, IGHV6-1, IGKV1-12, IGKV1-16, IGKV1-17, IGKV1-27, IGKV1-5, IGKV1-8, IGKV1D-13, IGKV1D-33, IGKV1D-39, IGKV2-30, IGKV2-40, IGKV2D-24, IGKV3-20, IGKV3D-11, IGKV3D-15, IGKV3D-20, IGKV4-1, IGKV6D-21, IGLL1, IGLV1-40, IGLV1-47, IGLV1-51, IGLV2-18, IGLV2-8, IGLV3-16, IGLV4-69, IGLV7-43, IGLV7-46, IGLV8-61, PSMA7, TINAGL1] |
| Down | SRP-dependent cotranslational protein targeting to membrane | 21.69811249 | [RPL10, RPL10A, RPL11, RPL13, RPL14, RPL17, RPL18A, RPL21, RPL23, RPL24, RPL27, RPL3, RPL35, RPL36, RPL36AL, RPL5, RPL6, RPLP2, RPS14, RPS19, RPS2, RPS20, RPS3A] |
| Down | ubiquitin ligase inhibitor activity | 50 | [RPL11, RPL17, RPL23, RPL5] |
| Down | ribosome assembly | 11.26760578 | [RPL10, RPL11, RPL24, RPL3, RPL5, RPL6, RPS14, RPS19] |
| Down | ribosome biogenesis | 4.088050365 | [RPL10, RPL10A, RPL11, RPL14, RPL24, RPL27, RPL3, RPL35, RPL5, RPL6, RPS14, RPS19, RPS2] |
| Down | sterol transport | 4.672897339 | [APOB, CEL, EGF, OSBP2, SCP2] |
| Down | positive regulation of lipid storage | 11.11111069 | [APOB, PLIN2, PLIN3] |
| Down | organic cyclic compound catabolic process | 4.087591171 | [CEL, CNP, HSPB1, NUDT5, RPL10, RPL10A, RPL11, RPL13, RPL14, RPL17, RPL18A, RPL21, RPL23, RPL24, RPL27, RPL3, RPL35, RPL36, RPL36AL, RPL5, RPL6, RPLP2, RPS14, RPS19, |
| Down | ribosomal large subunit biogenesis | 11.84210491 | [RPL10, RPL10A, RPL11, RPL14, RPL24, RPL3, RPL35, RPL5, RPL6] |

Table S4. Peptide to spectrum matches (PSMs) of identified glycopeptides

<table>
<tr><th>Query #:z</th><th>Protein Rank</th><th>Glycopeptide</th><th>Glycan</th><th>Modification Type(s)</th><th>Observed m/z</th><th>z</th><th>Observed (M+H)</th><th>Calc. mass (M+H)</th><th>Off-by-x error</th><th>Mass error (ppm)</th><th>StartPosition</th><th>Cleavage</th><th>Score</th><th>Delta</th><th>Delta Mod</th><th>|Log Prob|</th><th># of unique peptides</th><th>Protein Name</th><th>Protein DB number</th><th>Comment</th><th>Scan #</th><th>Scan Time</th></tr>
<tr><td>14439:4</td><td>1</td><td>R.RTAGWNVPIGTLRPFLN[+2570.925]WTGPPEPIEAAVAR.F</td><td>HexNAc(5)Hex(6)Fuc(2)NeuAc(1)</td><td>N[+2571]</td><td>1490.1959</td><td>4</td><td>5957.7619</td><td>5957.7371</td><td>0</td><td>4.2</td><td>140</td><td>Specific</td><td>311.26</td><td>311.26</td><td>311.26</td><td>5.97</td><td>51</td><td>>sp|P02788|TRFL_HUMAN Lactotransferrin OS=Homo sapiens OX=9606 GN=LTF PE=1 SV=6</td><td>1270</td><td>20201115_07413_1ng_glyco_hcd30_etd30_compound15.18041.18041.4</td><td>controllerType=0 controllerNumber=1 scan=18041</td><td>43.3873</td></tr>
<tr><td>13811:4</td><td>1</td><td>R.RTAGWNVPIGTLRPFLN[+2278.809]WTGPPEPIEAAVAR.F</td><td>HexNAc(5)Hex(6)NeuAc(1)</td><td>N[+2279]</td><td>1417.1694</td><td>4</td><td>5665.6559</td><td>5665.6213</td><td>0</td><td>6.1</td><td>140</td><td>Specific</td><td>332.81</td><td>332.81</td><td>332.81</td><td>4.77</td><td>51</td><td>>sp|P02788|TRFL_HUMAN Lactotransferrin OS=Homo sapiens OX=9606 GN=LTF PE=1 SV=6</td><td>1270</td><td>20201115_07413_1ng_glyco_hcd30_etd30_compound15.17372.17372.4</td><td>controllerType=0 controllerNumber=1 scan=17372</td><td>41.2616</td></tr>
<tr><td>14494:3</td><td>1</td><td>R.RTAGWNVPIGTLRPFLN[+2059.735]WTGPPEPIEAAVAR.F</td><td>HexNAc(4)Hex(5)Fuc(1)NeuAc(1)</td><td>N[+2060]</td><td>1816.2052</td><td>3</td><td>5446.6009</td><td>5446.547</td><td>0</td><td>9.9</td><td>140</td><td>Specific</td><td>397.69</td><td>387.38</td><td>387.38</td><td>5.57</td><td>51</td><td>>sp|P02788|TRFL_HUMAN Lactotransferrin OS=Homo sapiens OX=9606 GN=LTF PE=1 SV=6</td><td>1270</td><td>20201115_07413_1ng_glyco_hcd30_etd30_compound15.18099.18099.3</td><td>controllerType=0 controllerNumber=1 scan=18099</td><td>43.5649</td></tr>
<tr><td>14487:4</td><td>1</td><td>R.RTAGWNVPIGTLRPFLN[+2205.793]WTGPPEPIEAAVAR.F</td><td>HexNAc(4)Hex(5)Fuc(2)NeuAc(1)</td><td>N[+2206]</td><td>1398.9177</td><td>4</td><td>5592.6491</td><td>5592.6049</td><td>0</td><td>7.9</td><td>140</td><td>Specific</td><td>1047.44</td><td>976.15</td><td>976.15</td><td>17.24</td><td>51</td><td>>sp|P02788|TRFL_HUMAN Lactotransferrin OS=Homo sapiens OX=9606 GN=LTF PE=1 SV=6</td><td>1270</td><td>20201115_07413_1ng_glyco_hcd30_etd30_compound15.18092.18092.4</td><td>controllerType=0 controllerNumber=1 scan=18092</td><td>43.5496</td></tr>
<tr><td>14486:4</td><td>1</td><td>R.RTAGWNVPIGTLRPFLN[+2059.735]WTGPPEPIEAAVAR.F</td><td>HexNAc(4)Hex(5)Fuc(1)NeuAc(1)</td><td>N[+2060]</td><td>1362.4045</td><td>4</td><td>5446.5963</td><td>5446.547</td><td>0</td><td>9.1</td><td>140</td><td>Specific</td><td>1239.35</td><td>1113.13</td><td>1113.13</td><td>20.86</td><td>51</td><td>>sp|P02788|TRFL_HUMAN Lactotransferrin OS=Homo sapiens OX=9606 GN=LTF PE=1 SV=6</td><td>1270</td><td>20201115_07413_1ng_glyco_hcd30_etd30_compound15.18091.18091.4</td><td>controllerType=0 controllerNumber=1 scan=18091</td><td>43.5459</td></tr>
<tr><td>15355:3</td><td>1</td><td>R.TAGWNVPIGTLRPFLN[+2059.735]WTGPPEPIEAAVAR.F</td><td>HexNAc(4)Hex(5)Fuc(1)NeuAc(1)</td><td>N[+2060]</td><td>1764.1616</td><td>3</td><td>5290.4702</td><td>5290.4459</td><td>0</td><td>4.6</td><td>141</td><td>Specific</td><td>212.37</td><td>137.35</td><td>137.35</td><td>4.33</td><td>51</td><td>>sp|P02788|TRFL_HUMAN Lactotransferrin OS=Homo sapiens OX=9606 GN=LTF PE=1 SV=6</td><td>1270</td><td>20201115_07413_1ng_glyco_hcd30_etd30_compound15.19013.19013.2</td><td>controllerType=0 controllerNumber=1 scan=19013</td><td>46.3338</td></tr>
<tr><td>14624:4</td><td>1</td><td>R.TAGWNVPIGTLRPFLN[+2279.830]WTGPPEPIEAAVAR.F</td><td>HexNAc(5)Hex(6)Fuc(2)</td><td>N[+2280]</td><td>1378.3999</td><td>4</td><td>5510.5778</td><td>5510.5406</td><td>0</td><td>6.8</td><td>141</td><td>Specific</td><td>284.04</td><td>284.04</td><td>284.04</td><td>4.63</td><td>51</td><td>>sp|P02788|TRFL_HUMAN Lactotransferrin OS=Homo sapiens OX=9606 GN=LTF PE=1 SV=6</td><td>1270</td><td>20201115_07413_1ng_glyco_hcd30_etd30_compound15.18236.18236.4</td><td>controllerType=0 controllerNumber=1 scan=18236</td><td>43.948</td></tr>
<tr><td>14612:3</td><td>1</td><td>R.TAGWNVPIGTLRPFLN[+2279.830]WTGPPEPIEAAVAR.F</td><td>HexNAc(5)Hex(6)Fuc(2)</td><td>N[+2280]</td><td>1837.5318</td><td>3</td><td>5510.5809</td><td>5510.5406</td><td>0</td><td>7.3</td><td>141</td><td>Specific</td><td>311.85</td><td>264.91</td><td>264.91</td><td>4.16</td><td>51</td><td>>sp|P02788|TRFL_HUMAN Lactotransferrin OS=Homo sapiens OX=9606 GN=LTF PE=1 SV=6</td><td>1270</td><td>20201115_07413_1ng_glyco_hcd30_etd30_compound15.18224.18224.3</td><td>controllerType=0 controllerNumber=1 scan=18224</td><td>43.9078</td></tr>
<tr><td>15290:3</td><td>1</td><td>R.TAGWNVPIGTLRPFLN[+2424.867]WTGPPEPIEAAVAR.F</td><td>HexNAc(5)Hex(6)Fuc(1)NeuAc(1)</td><td>N[+2425]</td><td>1885.8693</td><td>3</td><td>5655.5935</td><td>5655.5781</td><td>0</td><td>2.7</td><td>141</td><td>Specific</td><td>317.42</td><td>216.91</td><td>216.91</td><td>4.86</td><td>51</td><td>>sp|P02788|TRFL_HUMAN Lactotransferrin OS=Homo sapiens OX=9606 GN=LTF PE=1 SV=6</td><td>1270</td><td>20201115_07413_1ng_glyco_hcd30_etd30_compound15.18943.18943.3</td><td>controllerType=0 controllerNumber=1 scan=18943</td><td>46.1116</td></tr>
<tr><td>15299:3</td><td>1</td><td>R.TAGWNVPIGTLRPFLN[+2570.925]WTGPPEPIEAAVAR.F</td><td>HexNAc(5)Hex(6)Fuc(2)NeuAc(1)</td><td>N[+2571]</td><td>1934.5553</td><td>3</td><td>5801.6513</td><td>5801.636</td><td>0</td><td>2.7</td><td>141</td><td>Specific</td><td>329.32</td><td>269.65</td><td>269.65</td><td>5.81</td><td>51</td><td>>sp|P02788|TRFL_HUMAN Lactotransferrin OS=Homo sapiens OX=9606 GN=LTF PE=1 SV=6</td><td>1270</td><td>20201115_07413_1ng_glyco_hcd30_etd30_compound15.18953.18953.3</td><td>controllerType=0 controllerNumber=1 scan=18953</td><td>46.1387</td></tr>
<tr><td>14661:3</td><td>1</td><td>R.TAGWNVPIGTLRPFLN[+1768.640]WTGPPEPIEAAVAR.F</td><td>HexNAc(4)Hex(5)Fuc(1)</td><td>N[+1769]</td><td>1667.12</td><td>3</td><td>4999.3453</td><td>4999.3504</td><td>0</td><td>-1</td><td>141</td><td>Specific</td><td>348.33</td><td>297.17</td><td>297.17</td><td>4.42</td><td>51</td><td>>sp|P02788|TRFL_HUMAN Lactotransferrin OS=Homo sapiens OX=9606 GN=LTF PE=1 SV=6</td><td>1270</td><td>20201115_07413_1ng_glyco_hcd30_etd30_compound15.18276.18276.3</td><td>controllerType=0 controllerNumber=1 scan=18276</td><td>44.0758</td></tr>
<tr><td>15245:3</td><td>1</td><td>R.TAGWNVPIGTLRPFLN[+2716.983]WTGPPEPIEAAVAR.F</td><td>HexNAc(5)Hex(6)Fuc(3)NeuAc(1)</td><td>N[+2717]</td><td>1983.2364</td><td>3</td><td>5947.6947</td><td>5947.6939</td><td>0</td><td>0.1</td><td>141</td><td>Specific</td><td>386.41</td><td>312.8</td><td>312.8</td><td>5.05</td><td>51</td><td>>sp|P02788|TRFL_HUMAN Lactotransferrin OS=Homo sapiens OX=9606 GN=LTF PE=1 SV=6</td><td>1270</td><td>20201115_07413_1ng_glyco_hcd30_etd30_compound15.18895.18895.3</td><td>controllerType=0 controllerNumber=1 scan=18895</td><td>45.9411</td></tr>
<tr><td>15353:3</td><td>1</td><td>R.TAGWNVPIGTLRPFLN[+2059.735]WTGPPEPIEAAVAR.F</td><td>HexNAc(4)Hex(5)Fuc(1)NeuAc(1)</td><td>N[+2060]</td><td>1764.1616</td><td>3</td><td>5290.4702</td><td>5290.4459</td><td>0</td><td>4.6</td><td>141</td><td>Specific</td><td>388.7</td><td>280.84</td><td>280.84</td><td>5.45</td><td>51</td><td>>sp|P02788|TRFL_HUMAN Lactotransferrin OS=Homo sapiens OX=9606 GN=LTF PE=1 SV=6</td><td>1270</td><td>20201115_07413_1ng_glyco_hcd30_etd30_compound15.19011.19011.2</td><td>controllerType=0 controllerNumber=1 scan=19011</td><td>46.3255</td></tr>
<tr><td>15338:3</td><td>1</td><td>R.TAGWNVPIGTLRPFLN[+2059.735]WTGPPEPIEAAVAR.F</td><td>HexNAc(4)Hex(5)Fuc(1)NeuAc(1)</td><td>N[+2060]</td><td>1764.1512</td><td>3</td><td>5290.4391</td><td>5290.4459</td><td>0</td><td>-1.3</td><td>141</td><td>Specific</td><td>449.13</td><td>327.24</td><td>327.24</td><td>4.68</td><td>51</td><td>>sp|P02788|TRFL_HUMAN Lactotransferrin OS=Homo sapiens OX=9606 GN=LTF PE=1 SV=6</td><td>1270</td><td>20201115_07413_1ng_glyco_hcd30_etd30_compound15.18995.18995.3</td><td>controllerType=0 controllerNumber=1 scan=18995</td><td>46.2633</td></tr>
<tr><td>15325:4</td><td>1</td><td>R.TAGWNVPIGTLRPFLN[+2351.851]WTGPPEPIEAAVAR.F</td><td>HexNAc(4)Hex(5)Fuc(3)NeuAc(1)</td><td>N[+2352]</td><td>1396.3976</td><td>4</td><td>5582.5685</td><td>5582.5617</td><td>0</td><td>1.2</td><td>141</td><td>Specific</td><td>512.92</td><td>508.71</td><td>508.71</td><td>6.96</td><td>51</td><td>>sp|P02788|TRFL_HUMAN Lactotransferrin OS=Homo sapiens OX=9606 GN=LTF PE=1 SV=6</td><td>1270</td><td>20201115_07413_1ng_glyco_hcd30_etd30_compound15.18981.18981.4</td><td>controllerType=0 controllerNumber=1 scan=18981</td><td>46.2288</td></tr>
<tr><td>14666:4</td><td>1</td><td>R.TAGWNVPIGTLRPFLN[+1768.640]WTGPPEPIEAAVAR.F</td><td>HexNAc(4)Hex(5)Fuc(1)</td><td>N[+1769]</td><td>1250.5927</td><td>4</td><td>4999.3488</td><td>4999.3504</td><td>0</td><td>-0.3</td><td>141</td><td>Specific</td><td>996.23</td><td>973.46</td><td>973.46</td><td>15.91</td><td>51</td><td>>sp|P02788|TRFL_HUMAN Lactotransferrin OS=Homo sapiens OX=9606 GN=LTF PE=1 SV=6</td><td>1270</td><td>20201115_07413_1ng_glyco_hcd30_etd30_compound15.18281.18281.4</td><td>controllerType=0 controllerNumber=1 scan=18281</td><td>44.0914</td></tr>
<tr><td>13874:3</td><td>1</td><td>R.TAGWNIPMGLLFN[+1460.529]QTGSC[+57.021]K.F</td><td>HexNAc(4)Hex(4)</td><td>C[+57], N[+1461]</td><td>1185.8566</td><td>3</td><td>3555.5553</td><td>3555.5387</td><td>0</td><td>4.7</td><td>485</td><td>Specific</td><td>209.19</td><td>188.72</td><td>188.72</td><td>3.22</td><td>51</td><td>>sp|P02788|TRFL_HUMAN Lactotransferrin OS=Homo sapiens OX=9606 GN=LTF PE=1 SV=6</td><td>1270</td><td>20201115_07413_1ng_glyco_hcd30_etd30_compound15.17439.17439.3</td><td>controllerType=0 controllerNumber=1 scan=17439</td><td>41.4695</td></tr>
<tr><td>15085:3</td><td>1</td><td>R.TAGWNIPMGLLFN[+1751.624]QTGSC[+57.021]K.F</td><td>HexNAc(4)Hex(4)NeuAc(1)</td><td>C[+57], N[+1752]</td><td>1282.8893</td><td>3</td><td>3846.6535</td><td>3846.6341</td><td>0</td><td>5</td><td>485</td><td>Specific</td><td>221.79</td><td>147.88</td><td>147.88</td><td>4.02</td><td>51</td><td>>sp|P02788|TRFL_HUMAN Lactotransferrin OS=Homo sapiens OX=9606 GN=LTF PE=1 SV=6</td><td>1270</td><td>20201115_07413_1ng_glyco_hcd30_etd30_compound15.18726.18726.3</td><td>controllerType=0 controllerNumber=1 scan=18726</td><td>45.4668</td></tr>
<tr><td>13848:3</td><td>1</td><td>R.TAGWNIPMGLLFN[+1419.502]QTGSC[+57.021]K.F</td><td>HexNAc(3)Hex(5)</td><td>C[+57], N[+1420]</td><td>1172.1796</td><td>3</td><td>3514.5242</td><td>3514.5122</td><td>0</td><td>3.4</td><td>485</td><td>Specific</td><td>272.94</td><td>266.45</td><td>146</td><td>4.56</td><td>51</td><td>>sp|P02788|TRFL_HUMAN Lactotransferrin OS=Homo sapiens OX=9606 GN=LTF PE=1 SV=6</td><td>1270</td><td>20201115_07413_1ng_glyco_hcd30_etd30_compound15.17411.17411.3</td><td>controllerType=0 controllerNumber=1 scan=17411</td><td>41.364</td></tr>
<tr><td>13500:3</td><td>1</td><td>R.TAGWNIPMGLLFN[+2133.772]QTGSC[+57.021]K.F</td><td>HexNAc(5)Hex(6)Fuc(1)</td><td>C[+57], N[+2134]</td><td>1410.2722</td><td>3</td><td>4228.8019</td><td>4228.7817</td><td>0</td><td>4.8</td><td>485</td><td>Specific</td><td>297.01</td><td>244.82</td><td>228.92</td><td>4.83</td><td>51</td><td>>sp|P02788|TRFL_HUMAN Lactotransferrin OS=Homo sapiens OX=9606 GN=LTF PE=1 SV=6</td><td>1270</td><td>20201115_07413_1ng_glyco_hcd30_etd30_compound15.17041.17041.3</td><td>controllerType=0 controllerNumber=1 scan=17041</td><td>40.2985</td></tr>
<tr><td>13771:3</td><td>1</td><td>R.TAGWNIPMGLLFN[+1581.555]QTGSC[+57.021]K.F</td><td>HexNAc(3)Hex(6)</td><td>C[+57], N[+1582]</td><td>1226.1966</td><td>3</td><td>3676.5752</td><td>3676.565</td><td>0</td><td>2.8</td><td>485</td><td>Specific</td><td>329.11</td><td>278.53</td><td>210.27</td><td>4.77</td><td>51</td><td>>sp|P02788|TRFL_HUMAN Lactotransferrin OS=Homo sapiens OX=9606 GN=LTF PE=1 SV=6</td><td>1270</td><td>20201115_07413_1ng_glyco_hcd30_etd30_compound15.17329.17329.3</td><td>controllerType=0 controllerNumber=1 scan=17329</td><td>41.1415</td></tr>
<tr><td>13703:3</td><td>1</td><td>R.TAGWNIPMGLLFN[+1606.587]QTGSC[+57.021]K.F</td><td>HexNAc(4)Hex(4)Fuc(1)</td><td>C[+57], N[+1607]</td><td>1234.5418</td><td>3</td><td>3701.6109</td><td>3701.5966</td><td>0</td><td>3.8</td><td>485</td><td>Specific</td><td>346.26</td><td>277.39</td><td>277.39</td><td>4.79</td><td>51</td><td>>sp|P02788|TRFL_HUMAN Lactotransferrin OS=Homo sapiens OX=9606 GN=LTF PE=1 SV=6</td><td>1270</td><td>20201115_07413_1ng_glyco_hcd30_etd30_compound15.17256.17256.3</td><td>controllerType=0 controllerNumber=1 scan=17256</td><td>40.8923</td></tr>
<tr><td>13589:3</td><td>1</td><td>R.TAGWNIPMGLLFN[+1768.640]QTGSC[+57.021]K.F</td><td>HexNAc(4)Hex(5)Fuc(1)</td><td>C[+57], N[+1769]</td><td>1288.5585</td><td>3</td><td>3863.661</td><td>3863.6495</td><td>0</td><td>3</td><td>485</td><td>Specific</td><td>375.13</td><td>264.91</td><td>264.91</td><td>5.48</td><td>51</td><td>>sp|P02788|TRFL_HUMAN Lactotransferrin OS=Homo sapiens OX=9606 GN=LTF PE=1 SV=6</td><td>1270</td><td>20201115_07413_1ng_glyco_hcd30_etd30_compound15.17135.17135.3</td><td>controllerType=0 controllerNumber=1 scan=17135</td><td>40.5485</td></tr>
<tr><td>13739:3</td><td>1</td><td>R.TAGWNIPMGLLFN[+1768.640]QTGSC[+57.021]K.F</td><td>HexNAc(4)Hex(5)Fuc(1)</td><td>C[+57], N[+1769]</td><td>1288.5578</td><td>3</td><td>3863.6588</td><td>3863.6495</td><td>0</td><td>2.4</td><td>485</td><td>Specific</td><td>376.82</td><td>244.72</td><td>244.72</td><td>5.25</td><td>51</td><td>>sp|P02788|TRFL_HUMAN Lactotransferrin OS=Homo sapiens OX=9606 GN=LTF PE=1 SV=6</td><td>1270</td><td>20201115_07413_1ng_glyco_hcd30_etd30_compound15.17295.17295.3</td><td>controllerType=0 controllerNumber=1 scan=17295</td><td>41.0069</td></tr>
<tr><td>13752:3</td><td>1</td><td>R.TAGWNIPMGLLFN[+1622.582]QTGSC[+57.021]K.F</td><td>HexNAc(4)Hex(5)</td><td>C[+57], N[+1623]</td><td>1239.8713</td><td>3</td><td>3717.5993</td><td>3717.5916</td><td>0</td><td>2.1</td><td>485</td><td>Specific</td><td>405.94</td><td>320.51</td><td>253.33</td><td>5.91</td><td>51</td><td>>sp|P02788|TRFL_HUMAN Lactotransferrin OS=Homo sapiens OX=9606 GN=LTF PE=1 SV=6</td><td>1270</td><td>20201115_07413_1ng_glyco_hcd30_etd30_compound15.17309.17309.3</td><td>controllerType=0 controllerNumber=1 scan=17309</td><td>41.0587</td></tr>
<tr><td>14979:3</td><td>1</td><td>R.TAGWNIPMGLLFN[+2059.735]QTGSC[+57.021]K.F</td><td>HexNAc(4)Hex(5)Fuc(1)NeuAc(1)</td><td>C[+57], N[+2060]</td><td>1385.5915</td><td>3</td><td>4154.7599</td><td>4154.7449</td><td>0</td><td>3.6</td><td>485</td><td>Specific</td><td>412.05</td><td>303.73</td><td>303.73</td><td>5.84</td><td>51</td><td>>sp|P02788|TRFL_HUMAN Lactotransferrin OS=Homo sapiens OX=9606 GN=LTF PE=1 SV=6</td><td>1270</td><td>20201115_07413_1ng_glyco_hcd30_etd30_compound15.18614.18614.3</td><td>controllerType=0 controllerNumber=1 scan=18614</td><td>45.0968</td></tr>
<tr><td>14891:3</td><td>1</td><td>R.TAGWNIPMGLLFN[+2059.735]QTGSC[+57.021]K.F</td><td>HexNAc(4)Hex(5)Fuc(1)NeuAc(1)</td><td>C[+57], N[+2060]</td><td>1385.5881</td><td>3</td><td>4154.7497</td><td>4154.7449</td><td>0</td><td>1.2</td><td>485</td><td>Specific</td><td>421.89</td><td>309.82</td><td>309.82</td><td>5.62</td><td>51</td><td>>sp|P02788|TRFL_HUMAN Lactotransferrin OS=Homo sapiens OX=9606 GN=LTF PE=1 SV=6</td><td>1270</td><td>20201115_07413_1ng_glyco_hcd30_etd30_compound15.18521.18521.3</td><td>controllerType=0 controllerNumber=1 scan=18521</td><td>44.8092</td></tr>
<tr><td>15068:3</td><td>1</td><td>R.TAGWNIPMGLLFN[+1751.624]QTGSC[+57.021]K.F</td><td>HexNAc(4)Hex(4)NeuAc(1)</td><td>C[+57], N[+1752]</td><td>1282.892</td><td>3</td><td>3846.6615</td><td>3846.6341</td><td>0</td><td>7.1</td><td>485</td><td>Specific</td><td>449.62</td><td>299.88</td><td>299.88</td><td>5.58</td><td>51</td><td>>sp|P02788|TRFL_HUMAN Lactotransferrin OS=Homo sapiens OX=9606 GN=LTF PE=1 SV=6</td><td>1270</td><td>20201115_07413_1ng_glyco_hcd30_etd30_compound15.18708.18708.3</td><td>controllerType=0 controllerNumber=1 scan=18708</td><td>45.4059</td></tr>
<tr><td>14988:3</td><td>1</td><td>R.TAGWNIPMGLLFN[+1913.677]QTGSC[+57.021]K.F</td><td>HexNAc(4)Hex(5)NeuAc(1)</td><td>C[+57], N[+1914]</td><td>1336.9017</td><td>3</td><td>4008.6905</td><td>4008.687</td><td>0</td><td>0.9</td><td>485</td><td>Specific</td><td>454.78</td><td>326.13</td><td>326.13</td><td>5.73</td><td>51</td><td>>sp|P02788|TRFL_HUMAN Lactotransferrin OS=Homo sapiens OX=9606 GN=LTF PE=1 SV=6</td><td>1270</td><td>20201115_07413_1ng_glyco_hcd30_etd30_compound15.18624.18624.3</td><td>controllerType=0 controllerNumber=1 scan=18624</td><td>45.1632</td></tr>
<tr><td>13865:3</td><td>1</td><td>R.TAGWNIPMGLLFN[+1216.423]QTGSC[+57.021]K.F</td><td>HexNAc(2)Hex(5)</td><td>C[+57], N[+1216]</td><td>1104.4877</td><td>3</td><td>3311.4485</td><td>3311.4328</td><td>0</td><td>4.8</td><td>485</td><td>Specific</td><td>462.52</td><td>440.88</td><td>219.19</td><td>6.74</td><td>51</td><td>>sp|P02788|TRFL_HUMAN Lactotransferrin OS=Homo sapiens OX=9606 GN=LTF PE=1 SV=6</td><td>1270</td><td>20201115_07413_1ng_glyco_hcd30_etd30_compound15.17430.17430.3</td><td>controllerType=0 controllerNumber=1 scan=17430</td><td>41.4231</td></tr>
<tr><td>16291:3</td><td>1</td><td>R.TAGWNIPMGLLFN[+2350.830]QTGSC[+57.021]K.F</td><td>HexNAc(4)Hex(5)Fuc(1)NeuAc(2)</td><td>C[+57], N[+2351]</td><td>1482.6236</td><td>3</td><td>4445.8562</td><td>4445.8403</td><td>0</td><td>3.6</td><td>485</td><td>Specific</td><td>469.62</td><td>312.57</td><td>312.57</td><td>5.96</td><td>51</td><td>>sp|P02788|TRFL_HUMAN Lactotransferrin OS=Homo sapiens OX=9606 GN=LTF PE=1 SV=6</td><td>1270</td><td>20201115_07413_1ng_glyco_hcd30_etd30_compound15.20013.20013.3</td><td>controllerType=0 controllerNumber=1 scan=20013</td><td>49.3595</td></tr>
<tr><td>13897:3</td><td>1</td><td>R.TAGWNIPMGLLFN[+1216.423]QTGSC[+57.021]K.F</td><td>HexNAc(2)Hex(5)</td><td>C[+57], N[+1216]</td><td>1104.4843</td><td>3</td><td>3311.4383</td><td>3311.4328</td><td>0</td><td>1.7</td><td>485</td><td>Specific</td><td>498.74</td><td>476.94</td><td>213.87</td><td>7.21</td><td>51</td><td>>sp|P02788|TRFL_HUMAN Lactotransferrin OS=Homo sapiens OX=9606 GN=LTF PE=1 SV=6</td><td>1270</td><td>20201115_07413_1ng_glyco_hcd30_etd30_compound15.17464.17464.3</td><td>controllerType=0 controllerNumber=1 scan=17464</td><td>41.5353</td></tr>
<tr><td>13873:2</td><td>1</td><td>R.TAGWNIPMGLLFN[+1216.423]QTGSC[+57.021]K.F</td><td>HexNAc(2)Hex(5)</td><td>C[+57], N[+1216]</td><td>1656.2265</td><td>2</td><td>3311.4457</td><td>3311.4328</td><td>0</td><td>3.9</td><td>485</td><td>Specific</td><td>545.99</td><td>529.08</td><td>177.18</td><td>7.3</td><td>51</td><td>>sp|P02788|TRFL_HUMAN Lactotransferrin OS=Homo sapiens OX=9606 GN=LTF PE=1 SV=6</td><td>1270</td><td>20201115_07413_1ng_glyco_hcd30_etd30_compound15.17438.17438.2</td><td>controllerType=0 controllerNumber=1 scan=17438</td><td>41.4625</td></tr>
<tr><td>14002:3</td><td>1</td><td>R.TAGWNIPM[+15.995]GLLFN[+1913.677]QTGSC[+57.021]K.F</td><td>HexNAc(4)Hex(5)NeuAc(1)</td><td>C[+57], M[+16], N[+1914]</td><td>1342.2361</td><td>3</td><td>4024.6939</td><td>4024.6819</td><td>0</td><td>3</td><td>485</td><td>Specific</td><td>210.04</td><td>99.84</td><td>99.84</td><td>3.29</td><td>51</td><td>>sp|P02788|TRFL_HUMAN Lactotransferrin OS=Homo sapiens OX=9606 GN=LTF PE=1 SV=6</td><td>1270</td><td>20201115_07413_1ng_glyco_hcd30_etd30_compound15.17575.17575.3</td><td>controllerType=0 controllerNumber=1 scan=17575</td><td>41.8586</td></tr>
<tr><td>13963:3</td><td>1</td><td>R.TAGWNIPM[+15.995]GLLFN[+2059.735]QTGSC[+57.021]K.F</td><td>HexNAc(4)Hex(5)Fuc(1)NeuAc(1)</td><td>C[+57], M[+16], N[+2060]</td><td>1390.9197</td><td>3</td><td>4170.7447</td><td>4170.7398</td><td>0</td><td>1.2</td><td>485</td><td>Specific</td><td>315.11</td><td>264.74</td><td>264.74</td><td>4.51</td><td>51</td><td>>sp|P02788|TRFL_HUMAN Lactotransferrin OS=Homo sapiens OX=9606 GN=LTF PE=1 SV=6</td><td>1270</td><td>20201115_07413_1ng_glyco_hcd30_etd30_compound15.17534.17534.3</td><td>controllerType=0 controllerNumber=1 scan=17534</td><td>41.7383</td></tr>
<tr><td>15137:3</td><td>1</td><td>R.TAGWNIPMGLLFN[+1913.677]QTGSC[+57.021]KFDEYFSQSC[+57.021]APGSDPR.S</td><td>HexNAc(4)Hex(5)NeuAc(1)</td><td>C[+57]*2, N[+1914]</td><td>1951.4845</td><td>3</td><td>5852.4388</td><td>5852.4341</td><td>0</td><td>0.8</td><td>485</td><td>Specific</td><td>298.88</td><td>215.35</td><td>215.35</td><td>4.23</td><td>51</td><td>>sp|P02788|TRFL_HUMAN Lactotransferrin OS=Homo sapiens OX=9606 GN=LTF PE=1 SV=6</td><td>1270</td><td>20201115_07413_1ng_glyco_hcd30_etd30_compound15.18781.18781.3</td><td>controllerType=0 controllerNumber=1 scan=18781</td><td>45.6216</td></tr>
<tr><td>16034:4</td><td>1</td><td>R.TAGWNIPMGLLFN[+2204.772]QTGSC[+57.021]KFDEYFSQSC[+57.021]APGSDPR.S</td><td>HexNAc(4)Hex(5)NeuAc(2)</td><td>C[+57]*2, N[+2205]</td><td>1536.6409</td><td>4</td><td>6143.5417</td><td>6143.5295</td><td>0</td><td>2</td><td>485</td><td>Specific</td><td>378.92</td><td>330.92</td><td>330.92</td><td>5.09</td><td>51</td><td>>sp|P02788|TRFL_HUMAN Lactotransferrin OS=Homo sapiens OX=9606 GN=LTF PE=1 SV=6</td><td>1270</td><td>20201115_07413_1ng_glyco_hcd30_etd30_compound15.19741.19741.4</td><td>controllerType=0 controllerNumber=1 scan=19741</td><td>48.5047</td></tr>
<tr><td>14355:3</td><td>1</td><td>R.TAGWNIPMGLLFN[+1768.640]QTGSC[+57.021]KFDEYFSQSC[+57.021]APGSDPR.S</td><td>HexNAc(4)Hex(5)Fuc(1)</td><td>C[+57]*2, N[+1769]</td><td>1903.1458</td><td>3</td><td>5707.423</td><td>5707.3966</td><td>0</td><td>4.6</td><td>485</td><td>Specific</td><td>536.45</td><td>419.11</td><td>419.11</td><td>6.37</td><td>51</td><td>>sp|P02788|TRFL_HUMAN Lactotransferrin OS=Homo sapiens OX=9606 GN=LTF PE=1 SV=6</td><td>1270</td><td>20201115_07413_1ng_glyco_hcd30_etd30_compound15.17952.17952.3</td><td>controllerType=0 controllerNumber=1 scan=17952</td><td>43.1264</td></tr>
<tr><td>15078:4</td><td>1</td><td>R.TAGWNIPMGLLFN[+2059.735]QTGSC[+57.021]KFDEYFSQSC[+57.021]APGSDPR.S</td><td>HexNAc(4)Hex(5)Fuc(1)NeuAc(1)</td><td>C[+57]*2, N[+2060]</td><td>1500.3879</td><td>4</td><td>5998.5299</td><td>5998.492</td><td>0</td><td>6.3</td><td>485</td><td>Specific</td><td>782.15</td><td>699.76</td><td>699.76</td><td>11.64</td><td>51</td><td>>sp|P02788|TRFL_HUMAN Lactotransferrin OS=Homo sapiens OX=9606 GN=LTF PE=1 SV=6</td><td>1270</td><td>20201115_07413_1ng_glyco_hcd30_etd30_compound15.18719.18719.4</td><td>controllerType=0 controllerNumber=1 scan=18719</td><td>45.4377</td></tr>
<tr><td>07625:3</td><td>1</td><td>R.N[+2059.735]GSDC[+57.021]PDKFC[+57.021]LFQSETK.N</td><td>HexNAc(4)Hex(5)Fuc(1)NeuAc(1)</td><td>C[+57]*2, N[+2060]</td><td>1364.8771</td><td>3</td><td>4092.6168</td><td>4092.6088</td><td>0</td><td>2</td><td>642</td><td>Specific</td><td>366.73</td><td>275.49</td><td>275.49</td><td>4.62</td><td>51</td><td>>sp|P02788|TRFL_HUMAN Lactotransferrin OS=Homo sapiens OX=9606 GN=LTF PE=1 SV=6</td><td>1270</td><td>20201115_07413_1ng_glyco_hcd30_etd30_compound15.10828.10828.3</td><td>controllerType=0 controllerNumber=1 scan=10828</td><td>26.6985</td></tr>
<tr><td>14197:3</td><td>2</td><td>R.LSLHRPALEDLLLGSEAN[+1501.555]LTC[+57.021]TLTGLR.D</td><td>HexNAc(5)Hex(3)</td><td>C[+57], N[+1502]</td><td>1489.0519</td><td>3</td><td>4465.1413</td><td>4465.1536</td><td>0</td><td>-2.8</td><td>127</td><td>Specific</td><td>441.97</td><td>372.89</td><td>372.89</td><td>5.25</td><td>22</td><td>>sp|P01876|IGHA1_HUMAN Immunoglobulin heavy constant alpha 1 OS=Homo sapiens OX=9606 GN=IGHA1 PE=1 SV=2</td><td>255</td><td>20201115_07413_1ng_glyco_hcd30_etd30_compound15.17782.17782.3</td><td>controllerType=0 controllerNumber=1 scan=17782</td><td>42.5926</td></tr>
<tr><td>14852:3</td><td>2</td><td>R.LSLHRPALEDLLLGSEAN[+1913.677]LTC[+57.021]TLTGLR.D</td><td>HexNAc(4)Hex(5)NeuAc(1)</td><td>C[+57], N[+1914]</td><td>1626.4366</td><td>3</td><td>4877.2952</td><td>4877.2753</td><td>0</td><td>4.1</td><td>127</td><td>Specific</td><td>442.97</td><td>365.66</td><td>365.66</td><td>6.6</td><td>22</td><td>>sp|P01876|IGHA1_HUMAN Immunoglobulin heavy constant alpha 1 OS=Homo sapiens OX=9606 GN=IGHA1 PE=1 SV=2</td><td>255</td><td>20201115_07413_1ng_glyco_hcd30_etd30_compound15.18479.18479.3</td><td>controllerType=0 controllerNumber=1 scan=18479</td><td>44.6885</td></tr>
<tr><td>14132:3</td><td>2</td><td>R.LSLHRPALEDLLLGSEAN[+1702.581]LTC[+57.021]TLTGLR.D</td><td>HexNAc(2)Hex(8)</td><td>C[+57], N[+1703]</td><td>1556.0772</td><td>3</td><td>4666.2171</td><td>4666.1796</td><td>0</td><td>8</td><td>127</td><td>Specific</td><td>478.92</td><td>468.37</td><td>468.37</td><td>6.23</td><td>22</td><td>>sp|P01876|IGHA1_HUMAN Immunoglobulin heavy constant alpha 1 OS=Homo sapiens OX=9606 GN=IGHA1 PE=1 SV=2</td><td>255</td><td>20201115_07413_1ng_glyco_hcd30_etd30_compound15.17712.17712.3</td><td>controllerType=0 controllerNumber=1 scan=17712</td><td>42.3082</td></tr>
<tr><td>14198:4</td><td>2</td><td>R.LSLHRPALEDLLLGSEAN[+1216.423]LTC[+57.021]TLTGLR.D</td><td>HexNAc(2)Hex(5)</td><td>C[+57], N[+1216]</td><td>1045.7599</td><td>4</td><td>4180.0179</td><td>4180.0211</td><td>0</td><td>-0.8</td><td>127</td><td>Specific</td><td>696.91</td><td>696.91</td><td>696.91</td><td>9.73</td><td>22</td><td>>sp|P01876|IGHA1_HUMAN Immunoglobulin heavy constant alpha 1 OS=Homo sapiens OX=9606 GN=IGHA1 PE=1 SV=2</td><td>255</td><td>20201115_07413_1ng_glyco_hcd30_etd30_compound15.17783.17783.4</td><td>controllerType=0 controllerNumber=1 scan=17783</td><td>42.5965</td></tr>
<tr><td>14846:4</td><td>2</td><td>R.LSLHRPALEDLLLGSEAN[+1913.677]LTC[+57.021]TLTGLR.D</td><td>HexNAc(4)Hex(5)NeuAc(1)</td><td>C[+57], N[+1914]</td><td>1220.0801</td><td>4</td><td>4877.2987</td><td>4877.2753</td><td>0</td><td>4.8</td><td>127</td><td>Specific</td><td>766.27</td><td>766.27</td><td>766.27</td><td>12.27</td><td>22</td><td>>sp|P01876|IGHA1_HUMAN Immunoglobulin heavy constant alpha 1 OS=Homo sapiens OX=9606 GN=IGHA1 PE=1 SV=2</td><td>255</td><td>20201115_07413_1ng_glyco_hcd30_etd30_compound15.18473.18473.4</td><td>controllerType=0 controllerNumber=1 scan=18473</td><td>44.6634</td></tr>
<tr><td>14118:4</td><td>2</td><td>R.LSLHRPALEDLLLGSEAN[+1702.581]LTC[+57.021]TLTGLR.D</td><td>HexNAc(2)Hex(8)</td><td>C[+57], N[+1703]</td><td>1167.3033</td><td>4</td><td>4666.1912</td><td>4666.1796</td><td>0</td><td>2.5</td><td>127</td><td>Specific</td><td>782.55</td><td>715.9</td><td>715.9</td><td>12.68</td><td>22</td><td>>sp|P01876|IGHA1_HUMAN Immunoglobulin heavy constant alpha 1 OS=Homo sapiens OX=9606 GN=IGHA1 PE=1 SV=2</td><td>255</td><td>20201115_07413_1ng_glyco_hcd30_etd30_compound15.17697.17697.4</td><td>controllerType=0 controllerNumber=1 scan=17697</td><td>42.2392</td></tr>
<tr><td>14124:4</td><td>2</td><td>R.LSLHRPALEDLLLGSEAN[+1702.581]LTC[+57.021]TLTGLR.D</td><td>HexNAc(2)Hex(8)</td><td>C[+57], N[+1703]</td><td>1167.309</td><td>4</td><td>4666.2142</td><td>4666.1796</td><td>0</td><td>7.4</td><td>127</td><td>Specific</td><td>844.58</td><td>844.58</td><td>844.58</td><td>13.24</td><td>22</td><td>>sp|P01876|IGHA1_HUMAN Immunoglobulin heavy constant alpha 1 OS=Homo sapiens OX=9606 GN=IGHA1 PE=1 SV=2</td><td>255</td><td>20201115_07413_1ng_glyco_hcd30_etd30_compound15.17704.17704.4</td><td>controllerType=0 controllerNumber=1 scan=17704</td><td>42.2686</td></tr>
<tr><td>14202:4</td><td>2</td><td>R.LSLHRPALEDLLLGSEAN[+1501.555]LTC[+57.021]TLTGLR.D</td><td>HexNAc(5)Hex(3)</td><td>C[+57], N[+1502]</td><td>1117.0479</td><td>4</td><td>4465.1697</td><td>4465.1536</td><td>0</td><td>3.6</td><td>127</td><td>Specific</td><td>925</td><td>920.21</td><td>920.21</td><td>15.61</td><td>22</td><td>>sp|P01876|IGHA1_HUMAN Immunoglobulin heavy constant alpha 1 OS=Homo sapiens OX=9606 GN=IGHA1 PE=1 SV=2</td><td>255</td><td>20201115_07413_1ng_glyco_hcd30_etd30_compound15.17788.17788.4</td><td>controllerType=0 controllerNumber=1 scan=17788</td><td>42.6106</td></tr>
<tr><td>14191:4</td><td>2</td><td>R.LSLHRPALEDLLLGSEAN[+1501.555]LTC[+57.021]TLTGLR.D</td><td>HexNAc(5)Hex(3)</td><td>C[+57], N[+1502]</td><td>1117.0436</td><td>4</td><td>4465.1527</td><td>4465.1536</td><td>0</td><td>-0.2</td><td>127</td><td>Specific</td><td>994.72</td><td>977.07</td><td>977.07</td><td>16.39</td><td>22</td><td>>sp|P01876|IGHA1_HUMAN Immunoglobulin heavy constant alpha 1 OS=Homo sapiens OX=9606 GN=IGHA1 PE=1 SV=2</td><td>255</td><td>20201115_07413_1ng_glyco_hcd30_etd30_compound15.17776.17776.4</td><td>controllerType=0 controllerNumber=1 scan=17776</td><td>42.5556</td></tr>
<tr><td>14142:4</td><td>2</td><td>R.LSLHRPALEDLLLGSEAN[+1864.634]LTC[+57.021]TLTGLR.D</td><td>HexNAc(2)Hex(9)</td><td>C[+57], N[+1865]</td><td>1207.8163</td><td>4</td><td>4828.2435</td><td>4828.2324</td><td>0</td><td>2.3</td><td>127</td><td>Specific</td><td>1041.25</td><td>1002.72</td><td>1002.72</td><td>17.68</td><td>22</td><td>>sp|P01876|IGHA1_HUMAN Immunoglobulin heavy constant alpha 1 OS=Homo sapiens OX=9606 GN=IGHA1 PE=1 SV=2</td><td>255</td><td>20201115_07413_1ng_glyco_hcd30_etd30_compound15.17723.17723.4</td><td>controllerType=0 controllerNumber=1 scan=17723</td><td>42.3488</td></tr>
<tr><td>14149:4</td><td>2</td><td>R.LSLHRPALEDLLLGSEAN[+1540.529]LTC[+57.021]TLTGLR.D</td><td>HexNAc(2)Hex(7)</td><td>C[+57], N[+1541]</td><td>1126.79</td><td>4</td><td>4504.138</td><td>4504.1268</td><td>0</td><td>2.5</td><td>127</td><td>Specific</td><td>1073.37</td><td>1005.55</td><td>1005.55</td><td>18.68</td><td>22</td><td>>sp|P01876|IGHA1_HUMAN Immunoglobulin heavy constant alpha 1 OS=Homo sapiens OX=9606 GN=IGHA1 PE=1 SV=2</td><td>255</td><td>20201115_07413_1ng_glyco_hcd30_etd30_compound15.17731.17731.4</td><td>controllerType=0 controllerNumber=1 scan=17731</td><td>42.3871</td></tr>
<tr><td>09536:3</td><td>2</td><td>R.LAGKPTHVN[+1501.555]VSVVMAEVDGTC[+57.021]Y.-</td><td>HexNAc(5)Hex(3)</td><td>C[+57], N[+1502]</td><td>1283.5762</td><td>3</td><td>3848.714</td><td>3848.6974</td><td>0</td><td>4.3</td><td>332</td><td>CRagged</td><td>215.57</td><td>138.42</td><td>138.42</td><td>2.78</td><td>22</td><td>>sp|P01876|IGHA1_HUMAN Immunoglobulin heavy constant alpha 1 OS=Homo sapiens OX=9606 GN=IGHA1 PE=1 SV=2</td><td>255</td><td>20201115_07413_1ng_glyco_hcd30_etd30_compound15.12826.12826.3</td><td>controllerType=0 controllerNumber=1 scan=12826</td><td>30.6109</td></tr>
<tr><td>09487:3</td><td>2</td><td>R.LAGKPTHVN[+1809.666]VSVVMAEVDGTC[+57.021]Y.-</td><td>HexNAc(5)Hex(4)Fuc(1)</td><td>C[+57], N[+1810]</td><td>1386.2776</td><td>3</td><td>4156.8182</td><td>4156.8081</td><td>0</td><td>2.4</td><td>332</td><td>CRagged</td><td>229.06</td><td>131.95</td><td>131.95</td><td>2.7</td><td>22</td><td>>sp|P01876|IGHA1_HUMAN Immunoglobulin heavy constant alpha 1 OS=Homo sapiens OX=9606 GN=IGHA1 PE=1 SV=2</td><td>255</td><td>20201115_07413_1ng_glyco_hcd30_etd30_compound15.12772.12772.3</td><td>controllerType=0 controllerNumber=1 scan=12772</td><td>30.4441</td></tr>
<tr><td>09484:3</td><td>2</td><td>R.LAGKPTHVN[+1663.608]VSVVMAEVDGTC[+57.021]Y.-</td><td>HexNAc(5)Hex(4)</td><td>C[+57], N[+1664]</td><td>1337.5919</td><td>3</td><td>4010.7611</td><td>4010.7502</td><td>0</td><td>2.7</td><td>332</td><td>CRagged</td><td>244.26</td><td>169.42</td><td>169.42</td><td>3.53</td><td>22</td><td>>sp|P01876|IGHA1_HUMAN Immunoglobulin heavy constant alpha 1 OS=Homo sapiens OX=9606 GN=IGHA1 PE=1 SV=2</td><td>255</td><td>20201115_07413_1ng_glyco_hcd30_etd30_compound15.12769.12769.3</td><td>controllerType=0 controllerNumber=1 scan=12769</td><td>30.4312</td></tr>
<tr><td>09541:3</td><td>2</td><td>R.LAGKPTHVN[+1647.613]VSVVMAEVDGTC[+57.021]Y.-</td><td>HexNAc(5)Hex(3)Fuc(1)</td><td>C[+57], N[+1648]</td><td>1332.2631</td><td>3</td><td>3994.7747</td><td>3994.7553</td><td>0</td><td>4.9</td><td>332</td><td>CRagged</td><td>259.84</td><td>151.81</td><td>151.81</td><td>3.24</td><td>22</td><td>>sp|P01876|IGHA1_HUMAN Immunoglobulin heavy constant alpha 1 OS=Homo sapiens OX=9606 GN=IGHA1 PE=1 SV=2</td><td>255</td><td>20201115_07413_1ng_glyco_hcd30_etd30_compound15.12831.12831.3</td><td>controllerType=0 controllerNumber=1 scan=12831</td><td>30.6333</td></tr>
<tr><td>09481:3</td><td>2</td><td>R.LAGKPTHVN[+1378.476]VSVVMAEVDGTC[+57.021]Y.-</td><td>HexNAc(2)Hex(6)</td><td>C[+57], N[+1378]</td><td>1242.5475</td><td>3</td><td>3725.6279</td><td>3725.6178</td><td>0</td><td>2.7</td><td>332</td><td>CRagged</td><td>264.17</td><td>203.63</td><td>190.83</td><td>3.8</td><td>22</td><td>>sp|P01876|IGHA1_HUMAN Immunoglobulin heavy constant alpha 1 OS=Homo sapiens OX=9606 GN=IGHA1 PE=1 SV=2</td><td>255</td><td>20201115_07413_1ng_glyco_hcd30_etd30_compound15.12766.12766.3</td><td>controllerType=0 controllerNumber=1 scan=12766</td><td>30.4224</td></tr>
<tr><td>10910:3</td><td>2</td><td>R.LAGKPTHVN[+1913.677]VSVVMAEVDGTC[+57.021]Y.-</td><td>HexNAc(4)Hex(5)NeuAc(1)</td><td>C[+57], N[+1914]</td><td>1420.9485</td><td>3</td><td>4260.8311</td><td>4260.8191</td><td>0</td><td>2.8</td><td>332</td><td>CRagged</td><td>268.76</td><td>161.02</td><td>161.02</td><td>3.53</td><td>22</td><td>>sp|P01876|IGHA1_HUMAN Immunoglobulin heavy constant alpha 1 OS=Homo sapiens OX=9606 GN=IGHA1 PE=1 SV=2</td><td>255</td><td>20201115_07413_1ng_glyco_hcd30_etd30_compound15.14284.14284.3</td><td>controllerType=0 controllerNumber=1 scan=14284</td><td>33.6471</td></tr>
<tr><td>10866:3</td><td>2</td><td>R.LAGKPTHVN[+2262.814]VSVVMAEVDGTC[+57.021]Y.-</td><td>HexNAc(5)Hex(5)Fuc(1)NeuAc(1)</td><td>C[+57], N[+2263]</td><td>1537.3272</td><td>3</td><td>4609.9671</td><td>4609.9564</td><td>0</td><td>2.3</td><td>332</td><td>CRagged</td><td>312.39</td><td>182.53</td><td>182.53</td><td>3.77</td><td>22</td><td>>sp|P01876|IGHA1_HUMAN Immunoglobulin heavy constant alpha 1 OS=Homo sapiens OX=9606 GN=IGHA1 PE=1 SV=2</td><td>255</td><td>20201115_07413_1ng_glyco_hcd30_etd30_compound15.14237.14237.3</td><td>controllerType=0 controllerNumber=1 scan=14237</td><td>33.5378</td></tr>
<tr><td>15490:3</td><td>3</td><td>R.GLSFDVSLEVSQGPGLLN[+2350.830]DTK.V</td><td>HexNAc(4)Hex(5)Fuc(1)NeuAc(2)</td><td>N[+2351]</td><td>1509.6573</td><td>3</td><td>4526.9573</td><td>4526.9436</td><td>0</td><td>3</td><td>118</td><td>Specific</td><td>471.15</td><td>327.67</td><td>327.67</td><td>5.99</td><td>22</td><td>>sp|P01833|PIGR_HUMAN Polymeric immunoglobulin receptor OS=Homo sapiens OX=9606 GN=PIGR PE=1 SV=4</td><td>902</td><td>20201115_07413_1ng_glyco_hcd30_etd30_compound15.19157.19157.3</td><td>controllerType=0 controllerNumber=1 scan=19157</td><td>46.5927</td></tr>
<tr><td>15013:3</td><td>3</td><td>K.QIGLYPVLVIDSSGYVNPN[+2205.793]YTGR.I</td><td>HexNAc(4)Hex(5)Fuc(2)NeuAc(1)</td><td>N[+2206]</td><td>1577.7053</td><td>3</td><td>4731.1013</td><td>4731.0963</td><td>0</td><td>1.1</td><td>168</td><td>Specific</td><td>267.42</td><td>198.42</td><td>198.42</td><td>4.83</td><td>22</td><td>>sp|P01833|PIGR_HUMAN Polymeric immunoglobulin receptor OS=Homo sapiens OX=9606 GN=PIGR PE=1 SV=4</td><td>902</td><td>20201115_07413_1ng_glyco_hcd30_etd30_compound15.18650.18650.3</td><td>controllerType=0 controllerNumber=1 scan=18650</td><td>45.2142</td></tr>
<tr><td>15036:3</td><td>3</td><td>K.QIGLYPVLVIDSSGYVNPN[+2059.735]YTGR.I</td><td>HexNAc(4)Hex(5)Fuc(1)NeuAc(1)</td><td>N[+2060]</td><td>1529.0245</td><td>3</td><td>4585.0589</td><td>4585.0384</td><td>0</td><td>4.5</td><td>168</td><td>Specific</td><td>581.09</td><td>489.94</td><td>489.94</td><td>7.99</td><td>22</td><td>>sp|P01833|PIGR_HUMAN Polymeric immunoglobulin receptor OS=Homo sapiens OX=9606 GN=PIGR PE=1 SV=4</td><td>902</td><td>20201115_07413_1ng_glyco_hcd30_etd30_compound15.18674.18674.3</td><td>controllerType=0 controllerNumber=1 scan=18674</td><td>45.2982</td></tr>
<tr><td>16098:3</td><td>3</td><td>R.LSLLEEPGN[+2350.830]GTFTVILNQLTSR.D</td><td>HexNAc(4)Hex(5)Fuc(1)NeuAc(2)</td><td>N[+2351]</td><td>1585.055</td><td>3</td><td>4753.1504</td><td>4753.1229</td><td>0</td><td>5.8</td><td>413</td><td>Specific</td><td>252.11</td><td>151.17</td><td>151.17</td><td>4.84</td><td>22</td><td>>sp|P01833|PIGR_HUMAN Polymeric immunoglobulin receptor OS=Homo sapiens OX=9606 GN=PIGR PE=1 SV=4</td><td>902</td><td>20201115_07413_1ng_glyco_hcd30_etd30_compound15.19809.19809.3</td><td>controllerType=0 controllerNumber=1 scan=19809</td><td>48.737</td></tr>
<tr><td>15984:3</td><td>3</td><td>R.LSLLEEPGN[+2569.905]GTFTVILNQLTSR.D</td><td>HexNAc(5)Hex(6)NeuAc(2)</td><td>N[+2570]</td><td>1658.0761</td><td>3</td><td>4972.2139</td><td>4972.1972</td><td>0</td><td>3.3</td><td>413</td><td>Specific</td><td>341.2</td><td>233.83</td><td>233.83</td><td>5.23</td><td>22</td><td>>sp|P01833|PIGR_HUMAN Polymeric immunoglobulin receptor OS=Homo sapiens OX=9606 GN=PIGR PE=1 SV=4</td><td>902</td><td>20201115_07413_1ng_glyco_hcd30_etd30_compound15.19687.19687.3</td><td>controllerType=0 controllerNumber=1 scan=19687</td><td>48.3603</td></tr>
<tr><td>16163:3</td><td>3</td><td>R.LSLLEEPGN[+2059.735]GTFTVILNQLTSR.D</td><td>HexNAc(4)Hex(5)Fuc(1)NeuAc(1)</td><td>N[+2060]</td><td>1488.0203</td><td>3</td><td>4462.0464</td><td>4462.0275</td><td>0</td><td>4.2</td><td>413</td><td>Specific</td><td>652.79</td><td>561.98</td><td>561.98</td><td>9.99</td><td>22</td><td>>sp|P01833|PIGR_HUMAN Polymeric immunoglobulin receptor OS=Homo sapiens OX=9606 GN=PIGR PE=1 SV=4</td><td>902</td><td>20201115_07413_1ng_glyco_hcd30_etd30_compound15.19878.19878.3</td><td>controllerType=0 controllerNumber=1 scan=19878</td><td>[illegible]</td></tr>
<tr><td>17663:3</td><td>3</td><td>R.LSLLEEPGN[+2350.830]GTFTVILNQLTSR.D</td><td>HexNAc(4)Hex(5)Fuc(1)NeuAc(2)</td><td>N[+2351]</td><td>1585.0493</td><td>3</td><td>4753.1332</td><td>4753.1229</td><td>0</td><td>2.2</td><td>413</td><td>Specific</td><td>690.12</td><td>489.81</td><td>489.81</td><td>10.4</td><td>22</td><td>>sp|P01833|PIGR_HUMAN Polymeric immunoglobulin receptor OS=Homo sapiens OX=9606 GN=PIGR PE=1 SV=4</td><td>902</td><td>20201115_07413_1ng_glyco_hcd30_etd30_compound15.21471.21471.3</td><td>controllerType=0 controllerNumber=1 scan=21471</td><td>[illegible]</td></tr>
<tr><td>13093:3</td><td>3</td><td>K.IIEGEPNLKVPGN[+2059.735]VTAVLGETLK.V</td><td>HexNAc(4)Hex(5)Fuc(1)NeuAc(1)</td><td>N[+2060]</td><td>1484.3687</td><td>3</td><td>4451.0916</td><td>4451.0843</td><td>0</td><td>1.6</td><td>457</td><td>Specific</td><td>349.08</td><td>318.07</td><td>318.07</td><td>4.16</td><td>22</td><td>>sp|P01833|PIGR_HUMAN Polymeric immunoglobulin receptor OS=Homo sapiens OX=9606 GN=PIGR PE=1 SV=4</td><td>902</td><td>20201115_07413_1ng_glyco_hcd30_etd30_compound15.16610.16610.3</td><td>controllerType=0 controllerNumber=1 scan=16610</td><td>[illegible]</td></tr>
<tr><td>13052:3</td><td>3</td><td>K.IIEGEPNLKVPGN[+2205.793]VTAVLGETLK.V</td><td>HexNAc(4)Hex(5)Fuc(2)NeuAc(1)</td><td>N[+2206]</td><td>1533.0562</td><td>3</td><td>4597.1541</td><td>4597.1422</td><td>0</td><td>2.6</td><td>457</td><td>Specific</td><td>407.53</td><td>328.17</td><td>328.17</td><td>4.23</td><td>22</td><td>>sp|P01833|PIGR_HUMAN Polymeric immunoglobulin receptor OS=Homo sapiens OX=9606 GN=PIGR PE=1 SV=4</td><td>902</td><td>20201115_07413_1ng_glyco_hcd30_etd30_compound15.16567.16567.3</td><td>controllerType=0 controllerNumber=1 scan=16567</td><td>39.1489</td></tr>
<tr><td>13057:3</td><td>3</td><td>K.IIEGEPNLKVPGN[+2205.793]VTAVLGETLK.V</td><td>HexNAc(4)Hex(5)Fuc(2)NeuAc(1)</td><td>N[+2206]</td><td>1533.0558</td><td>3</td><td>4597.153</td><td>4597.1422</td><td>0</td><td>2.3</td><td>457</td><td>Specific</td><td>413.11</td><td>343.56</td><td>343.56</td><td>3.86</td><td>22</td><td>>sp|P01833|PIGR_HUMAN Polymeric immunoglobulin receptor OS=Homo sapiens OX=9606 GN=PIGR PE=1 SV=4</td><td>902</td><td>20201115_07413_1ng_glyco_hcd30_etd30_compound15.16573.16573.3</td><td>controllerType=0 controllerNumber=1 scan=16573</td><td>39.1627</td></tr>
<tr><td>13101:3</td><td>3</td><td>K.IIEGEPNLKVPGN[+2059.735]VTAVLGETLK.V</td><td>HexNAc(4)Hex(5)Fuc(1)NeuAc(1)</td><td>N[+2060]</td><td>1484.3707</td><td>3</td><td>4451.0974</td><td>4451.0843</td><td>0</td><td>3</td><td>457</td><td>Specific</td><td>426.27</td><td>295.87</td><td>295.87</td><td>4.23</td><td>22</td><td>>sp|P01833|PIGR_HUMAN Polymeric immunoglobulin receptor OS=Homo sapiens OX=9606 GN=PIGR PE=1 SV=4</td><td>902</td><td>20201115_07413_1ng_glyco_hcd30_etd30_compound15.16620.16620.3</td><td>controllerType=0 controllerNumber=1 scan=16620</td><td>39.2912</td></tr>
<tr><td>12720:5</td><td>3</td><td>K.[+42.011]IIEGEPNLKVPGN[+1872.651]VTAVLGETLKVPC[+57.021]HFPC[+57.021]</td><td>HexNAc(3)Hex(6)NeuAc(1)</td><td>.[42], C[+57]*2, N[+1873]</td><td>1067.1028</td><td>5</td><td>5331.4848</td><td>5331.4679</td><td>0</td><td>3.2</td><td>457</td><td>Specific</td><td>225.14</td><td>217.93</td><td>217.93</td><td>2.64</td><td>22</td><td>>sp|P01833|PIGR_HUMAN Polymeric immunoglobulin receptor OS=Homo sapiens OX=9606 GN=PIGR PE=1 SV=4</td><td>902</td><td>20201115_07413_1ng_glyco_hcd30_etd30_compound15.16193.16193.5</td><td>controllerType=0 controllerNumber=1 scan=16193</td><td>38.2955</td></tr>
<tr><td>13756:4</td><td>3</td><td>K.IIEGEPNLKVPGN[+2350.830]VTAVLGETLKVPC[+57.021]HFPC[+57.021]K.F</td><td>HexNAc(4)Hex(5)Fuc(1)NeuAc(2)</td><td>C[+57]*2, N[+2351]</td><td>1442.6697</td><td>4</td><td>5767.6569</td><td>5767.6372</td><td>0</td><td>3.4</td><td>457</td><td>Specific</td><td>451.48</td><td>451.48</td><td>451.48</td><td>5.01</td><td>22</td><td>>sp|P01833|PIGR_HUMAN Polymeric immunoglobulin receptor OS=Homo sapiens OX=9606 GN=PIGR PE=1 SV=4</td><td>902</td><td>20201115_07413_1ng_glyco_hcd30_etd30_compound15.17313.17313.4</td><td>controllerType=0 controllerNumber=1 scan=17313</td><td>41.0671</td></tr>
<tr><td>12699:4</td><td>3</td><td>K.IIEGEPNLKVPGN[+2205.793]VTAVLGETLKVPC[+57.021]HFPC[+57.021]K.F</td><td>HexNAc(4)Hex(5)Fuc(2)NeuAc(1)</td><td>C[+57]*2, N[+2206]</td><td>1406.4188</td><td>4</td><td>5622.6535</td><td>5622.5997</td><td>0</td><td>9.6</td><td>457</td><td>Specific</td><td>488.88</td><td>488.88</td><td>488.88</td><td>4.33</td><td>22</td><td>>sp|P01833|PIGR_HUMAN Polymeric immunoglobulin receptor OS=Homo sapiens OX=9606 GN=PIGR PE=1 SV=4</td><td>902</td><td>20201115_07413_1ng_glyco_hcd30_etd30_compound15.16171.16171.4</td><td>controllerType=0 controllerNumber=1 scan=16171</td><td>38.2262</td></tr>
<tr><td>12710:4</td><td>3</td><td>K.IIEGEPNLKVPGN[+2059.735]VTAVLGETLKVPC[+57.021]HFPC[+57.021]K.F</td><td>HexNAc(4)Hex(5)Fuc(1)NeuAc(1)</td><td>C[+57]*2, N[+2060]</td><td>1369.8915</td><td>4</td><td>5476.5441</td><td>5476.5418</td><td>0</td><td>0.4</td><td>457</td><td>Specific</td><td>546.81</td><td>518.88</td><td>518.88</td><td>5.57</td><td>22</td><td>>sp|P01833|PIGR_HUMAN Polymeric immunoglobulin receptor OS=Homo sapiens OX=9606 GN=PIGR PE=1 SV=4</td><td>902</td><td>20201115_07413_1ng_glyco_hcd30_etd30_compound15.16183.16183.4</td><td>controllerType=0 controllerNumber=1 scan=16183</td><td>38.2499</td></tr>
<tr><td>13740:4</td><td>3</td><td>K.IIEGEPNLKVPGN[+2350.830]VTAVLGETLKVPC[+57.021]HFPC[+57.021]K.F</td><td>HexNAc(4)Hex(5)Fuc(1)NeuAc(2)</td><td>C[+57]*2, N[+2351]</td><td>1442.6721</td><td>4</td><td>5767.6667</td><td>5767.6372</td><td>0</td><td>5.1</td><td>457</td><td>Specific</td><td>558.9</td><td>558.9</td><td>558.9</td><td>5.45</td><td>22</td><td>>sp|P01833|PIGR_HUMAN Polymeric immunoglobulin receptor OS=Homo sapiens OX=9606 GN=PIGR PE=1 SV=4</td><td>902</td><td>20201115_07413_1ng_glyco_hcd30_etd30_compound15.17296.17296.2</td><td>controllerType=0 controllerNumber=1 scan=17296</td><td>41.0141</td></tr>
<tr><td>12662:5</td><td>3</td><td>K.IIEGEPNLKVPGN[+2059.735]VTAVLGETLKVPC[+57.021]HFPC[+57.021]K.F</td><td>HexNAc(4)Hex(5)Fuc(1)NeuAc(1)</td><td>C[+57]*2, N[+2060]</td><td>1096.1156</td><td>5</td><td>5476.5489</td><td>5476.5418</td><td>0</td><td>1.3</td><td>457</td><td>Specific</td><td>608.36</td><td>606.24</td><td>606.24</td><td>8.09</td><td>22</td><td>>sp|P01833|PIGR_HUMAN Polymeric immunoglobulin receptor OS=Homo sapiens OX=9606 GN=PIGR PE=1 SV=4</td><td>902</td><td>20201115_07413_1ng_glyco_hcd30_etd30_compound15.16131.16131.5</td><td>controllerType=0 controllerNumber=1 scan=16131</td><td>38.1165</td></tr>
<tr><td>13760:5</td><td>3</td><td>K.IIEGEPNLKVPGN[+2350.830]VTAVLGETLKVPC[+57.021]HFPC[+57.021]K.F</td><td>HexNAc(4)Hex(5)Fuc(1)NeuAc(2)</td><td>C[+57]*2, N[+2351]</td><td>1154.3339</td><td>5</td><td>5767.6402</td><td>5767.6372</td><td>0</td><td>0.5</td><td>457</td><td>Specific</td><td>651.83</td><td>651.83</td><td>651.83</td><td>8.8</td><td>22</td><td>>sp|P01833|PIGR_HUMAN Polymeric immunoglobulin receptor OS=Homo sapiens OX=9606 GN=PIGR PE=1 SV=4</td><td>902</td><td>20201115_07413_1ng_glyco_hcd30_etd30_compound15.17317.17317.5</td><td>controllerType=0 controllerNumber=1 scan=17317</td><td>41.0894</td></tr>
<tr><td>12691:5</td><td>3</td><td>K.IIEGEPNLKVPGN[+2205.793]VTAVLGETLKVPC[+57.021]HFPC[+57.021]K.F</td><td>HexNAc(4)Hex(5)Fuc(2)NeuAc(1)</td><td>C[+57]*2, N[+2206]</td><td>1125.3353</td><td>5</td><td>5622.6475</td><td>5622.5997</td><td>0</td><td>8.5</td><td>457</td><td>Specific</td><td>701.44</td><td>701.44</td><td>701.44</td><td>10.03</td><td>22</td><td>>sp|P01833|PIGR_HUMAN Polymeric immunoglobulin receptor OS=Homo sapiens OX=9606 GN=PIGR PE=1 SV=4</td><td>902</td><td>20201115_07413_1ng_glyco_hcd30_etd30_compound15.16163.16163.5</td><td>controllerType=0 controllerNumber=1 scan=16163</td><td>38.2059</td></tr>
<tr><td>12723:5</td><td>3</td><td>K.IIEGEPNLKVPGN[+2059.735]VTAVLGETLKVPC[+57.021]HFPC[+57.021]K.F</td><td>HexNAc(4)Hex(5)Fuc(1)NeuAc(1)</td><td>C[+57]*2, N[+2060]</td><td>1096.1169</td><td>5</td><td>5476.5556</td><td>5476.5418</td><td>0</td><td>2.5</td><td>457</td><td>Specific</td><td>741.6</td><td>617.92</td><td>617.92</td><td>10.48</td><td>22</td><td>>sp|P01833|PIGR_HUMAN Polymeric immunoglobulin receptor OS=Homo sapiens OX=9606 GN=PIGR PE=1 SV=4</td><td>902</td><td>20201115_07413_1ng_glyco_hcd30_etd30_compound15.16197.16197.5</td><td>controllerType=0 controllerNumber=1 scan=16197</td><td>38.3215</td></tr>
<tr><td>11347:3</td><td>3</td><td>K.VPGN[+2059.735]VTAVLGETLK.V</td><td>HexNAc(4)Hex(5)Fuc(1)NeuAc(1)</td><td>N[+2060]</td><td>1153.1874</td><td>3</td><td>3457.5478</td><td>3457.5348</td><td>0</td><td>3.7</td><td>466</td><td>Specific</td><td>313.82</td><td>196.87</td><td>196.87</td><td>4.75</td><td>22</td><td>>sp|P01833|PIGR_HUMAN Polymeric immunoglobulin receptor OS=Homo sapiens OX=9606 GN=PIGR PE=1 SV=4</td><td>902</td><td>20201115_07413_1ng_glyco_hcd30_etd30_compound15.14742.14742.3</td><td>controllerType=0 controllerNumber=1 scan=14742</td><td>34.6762</td></tr>
<tr><td>11283:4</td><td>3</td><td>K.VPGN[+2205.793]VTAVLGETLKVPC[+57.021]HFPC[+57.021]K.F</td><td>HexNAc(4)Hex(5)Fuc(2)NeuAc(1)</td><td>C[+57]*2, N[+2206]</td><td>1158.0212</td><td>4</td><td>4629.0628</td><td>4629.0502</td><td>0</td><td>2.7</td><td>466</td><td>Specific</td><td>357.51</td><td>311.38</td><td>311.38</td><td>3.58</td><td>22</td><td>>sp|P01833|PIGR_HUMAN Polymeric immunoglobulin receptor OS=Homo sapiens OX=9606 GN=PIGR PE=1 SV=4</td><td>902</td><td>20201115_07413_1ng_glyco_hcd30_etd30_compound15.14674.14674.4</td><td>controllerType=0 controllerNumber=1 scan=14674</td><td>34.5284</td></tr>
<tr><td>11306:4</td><td>3</td><td>K.VPGN[+2059.735]VTAVLGETLKVPC[+57.021]HFPC[+57.021]K.F</td><td>HexNAc(4)Hex(5)Fuc(1)NeuAc(1)</td><td>C[+57]*2, N[+2060]</td><td>1121.5071</td><td>4</td><td>4483.0067</td><td>4482.9923</td><td>0</td><td>3.2</td><td>466</td><td>Specific</td><td>527.78</td><td>468.96</td><td>468.96</td><td>5.85</td><td>22</td><td>>sp|P01833|PIGR_HUMAN Polymeric immunoglobulin receptor OS=Homo sapiens OX=9606 GN=PIGR PE=1 SV=4</td><td>902</td><td>20201115_07413_1ng_glyco_hcd30_etd30_compound15.14698.14698.4</td><td>controllerType=0 controllerNumber=1 scan=14698</td><td>34.5939</td></tr>
<tr><td>04825:3</td><td>3</td><td>K.WN[+1548.545]NTGC[+57.021]QALPSQDEGPSK.A</td><td>HexNAc(3)Hex(4)NeuAc(1)</td><td>C[+57], N[+1549]</td><td>1179.8234</td><td>3</td><td>3537.4557</td><td>3537.4215</td><td>0</td><td>9.7</td><td>498</td><td>Specific</td><td>242.05</td><td>97.31</td><td>97.31</td><td>3.68</td><td>22</td><td>>sp|P01833|PIGR_HUMAN Polymeric immunoglobulin receptor OS=Homo sapiens OX=9606 GN=PIGR PE=1 SV=4</td><td>902</td><td>20201115_07413_1ng_glyco_hcd30_etd30_compound15.7858.7858.3</td><td>controllerType=0 controllerNumber=1 scan=7858</td><td>21.1078</td></tr>
<tr><td>06479:3</td><td>3</td><td>K.WN[+2205.793]NTGC[+57.021]QALPSQDEGPSK.A</td><td>HexNAc(4)Hex(5)Fuc(2)NeuAc(1)</td><td>C[+57], N[+2206]</td><td>1398.8882</td><td>3</td><td>4194.65</td><td>4194.6695</td><td>0</td><td>-4.6</td><td>498</td><td>Specific</td><td>279.88</td><td>204.63</td><td>204.63</td><td>3.7</td><td>22</td><td>>sp|P01833|PIGR_HUMAN Polymeric immunoglobulin receptor OS=Homo sapiens OX=9606 GN=PIGR PE=1 SV=4</td><td>902</td><td>20201115_07413_1ng_glyco_hcd30_etd30_compound15.9608.9608.3</td><td>controllerType=0 controllerNumber=1 scan=9608</td><td>24.315</td></tr>
<tr><td>04724:3</td><td>3</td><td>K.WN[+1710.598]NTGC[+57.021]QALPSQDEGPSK.A</td><td>HexNAc(3)Hex(5)NeuAc(1)</td><td>C[+57], N[+1711]</td><td>1233.8336</td><td>3</td><td>3699.4861</td><td>3699.4743</td><td>0</td><td>3.2</td><td>498</td><td>Specific</td><td>425.46</td><td>322.01</td><td>322.01</td><td>5.84</td><td>22</td><td>>sp|P01833|PIGR_HUMAN Polymeric immunoglobulin receptor OS=Homo sapiens OX=9606 GN=PIGR PE=1 SV=4</td><td>902</td><td>20201115_07413_1ng_glyco_hcd30_etd30_compound15.7754.7754.3</td><td>controllerType=0 controllerNumber=1 scan=7754</td><td>20.9256</td></tr>
<tr><td>13870:3</td><td>5</td><td>K.RNIAAFGGDPNN[+1913.677]ITLFGESAGGASVSLQTLSPYNK.G</td><td>HexNAc(4)Hex(5)NeuAc(1)</td><td>N[+1914]</td><td>1827.5013</td><td>3</td><td>5480.4894</td><td>5480.4644</td><td>0</td><td>4.6</td><td>196</td><td>Specific</td><td>616.45</td><td>491.69</td><td>491.69</td><td>8.96</td><td>9</td><td>>sp|P19835|CEL_HUMAN Bile salt-activated lipase OS=Homo sapiens OX=9606 GN=CEL PE=1 SV=3</td><td>476</td><td>20201115_07413_1ng_glyco_hcd30_etd30_compound15.17435.17435.3</td><td>controllerType=0 controllerNumber=1 scan=17435</td><td>41.4485</td></tr>
<tr><td>13869:4</td><td>5</td><td>K.RNIAAFGGDPNN[+1913.677]ITLFGESAGGASVSLQTLSPYNK.G</td><td>HexNAc(4)Hex(5)NeuAc(1)</td><td>N[+1914]</td><td>1370.8774</td><td>4</td><td>5480.4879</td><td>5480.4644</td><td>0</td><td>4.3</td><td>196</td><td>Specific</td><td>726.04</td><td>675.29</td><td>675.29</td><td>11.14</td><td>9</td><td>>sp|P19835|CEL_HUMAN Bile salt-activated lipase OS=Homo sapiens OX=9606 GN=CEL PE=1 SV=3</td><td>476</td><td>20201115_07413_1ng_glyco_hcd30_etd30_compound15.17434.17434.4</td><td>controllerType=0 controllerNumber=1 scan=17434</td><td>41.4416</td></tr>
<tr><td>08381:3</td><td>6</td><td>K.TVLTPATNHMGN[+1540.529]VTFTIPANR.E</td><td>HexNAc(2)Hex(7)</td><td>N[+1541]</td><td>1265.9041</td><td>3</td><td>3795.6977</td><td>3795.6886</td><td>0</td><td>2.4</td><td>74</td><td>Specific</td><td>207.52</td><td>184.02</td><td>120.69</td><td>3.19</td><td>8</td><td>>sp|P01024|CO3_HUMAN Complement C3 OS=Homo sapiens OX=9606 GN=C3 PE=1 SV=2</td><td>492</td><td>20201115_07413_1ng_glyco_hcd30_etd30_compound15.11619.11619.3</td><td>controllerType=0 controllerNumber=1 scan=11619</td><td>28.2523</td></tr>
<tr><td>08369:3</td><td>6</td><td>K.TVLTPATNHMGN[+1540.529]VTFTIPANR.E</td><td>HexNAc(2)Hex(7)</td><td>N[+1541]</td><td>1265.9039</td><td>3</td><td>3795.697</td><td>3795.6886</td><td>0</td><td>2.2</td><td>74</td><td>Specific</td><td>247.7</td><td>186.85</td><td>135.44</td><td>3.72</td><td>8</td><td>>sp|P01024|CO3_HUMAN Complement C3 OS=Homo sapiens OX=9606 GN=C3 PE=1 SV=2</td><td>492</td><td>20201115_07413_1ng_glyco_hcd30_etd30_compound15.11606.11606.3</td><td>controllerType=0 controllerNumber=1 scan=11606</td><td>28.2133</td></tr>
<tr><td>08395:3</td><td>6</td><td>K.TVLTPATNHMGN[+1378.476]VTFTIPANR.E</td><td>HexNAc(2)Hex(6)</td><td>N[+1378]</td><td>1211.8851</td><td>3</td><td>3633.6406</td><td>3633.6358</td><td>0</td><td>1.3</td><td>74</td><td>Specific</td><td>364.11</td><td>280.18</td><td>154.48</td><td>5.41</td><td>8</td><td>>sp|P01024|CO3_HUMAN Complement C3 OS=Homo sapiens OX=9606 GN=C3 PE=1 SV=2</td><td>492</td><td>20201115_07413_1ng_glyco_hcd30_etd30_compound15.11635.11635.3</td><td>controllerType=0 controllerNumber=1 scan=11635</td><td>28.2885</td></tr>
<tr><td>07027:3</td><td>7</td><td>K.TAHC[+57.021]N[+2059.735]ESFYFLC[+57.021]K.R</td><td>HexNAc(4)Hex(5)Fuc(1)NeuAc(1)</td><td>C[+57]*2, N[+2060]</td><td>1246.1591</td><td>3</td><td>3736.4628</td><td>3736.4545</td><td>0</td><td>2.2</td><td>1201</td><td>Specific</td><td>207.62</td><td>123.26</td><td>123.26</td><td>3.81</td><td>10</td><td>>sp|P22897|MRC1_HUMAN Macrophage mannose receptor 1 OS=Homo sapiens OX=9606 GN=MRC1 PE=1 SV=1</td><td>1049</td><td>20201115_07413_1ng_glyco_hcd30_etd30_compound15.10192.10192.3</td><td>controllerType=0 controllerNumber=1 scan=10192</td><td>25.435</td></tr>
<tr><td>07590:4</td><td>8</td><td>R.QTGLAPGQEYEISLHIVKN[+2059.735]NTR.G</td><td>HexNAc(4)Hex(5)Fuc(1)NeuAc(1)</td><td>N[+2060]</td><td>1132.7727</td><td>4</td><td>4528.0692</td><td>4528.0241</td><td>0</td><td>9.9</td><td>770</td><td>Specific</td><td>397.65</td><td>356.97</td><td>356.97</td><td>3.31</td><td>11</td><td>>sp|P24821|TENA_HUMAN Tenascin OS=Homo sapiens OX=9606 GN=TNC PE=1 SV=3</td><td>486</td><td>20201115_07413_1ng_glyco_hcd30_etd30_compound15.10792.10792.4</td><td>controllerType=0 controllerNumber=1 scan=10792</td><td>26.6211</td></tr>
<tr><td>09573:3</td><td>14</td><td>R.IIVPLNNREN[+1694.603]ISDPTSPLR.T</td><td>HexNAc(3)Hex(4)Fuc(1)NeuAc(1)</td><td>N[+1695]</td><td>1281.6023</td><td>3</td><td>3842.7923</td><td>3842.7799</td><td>0</td><td>3.2</td><td>62</td><td>Specific</td><td>216.03</td><td>181.35</td><td>181.35</td><td>2.75</td><td>5</td><td>>sp|P01591|IGJ_HUMAN Immunoglobulin J chain OS=Homo sapiens OX=9606 GN=JCHAIN PE=1 SV=4</td><td>1392</td><td>20201115_07413_1ng_glyco_hcd30_etd30_compound15.12866.12866.3</td><td>controllerType=0 controllerNumber=1 scan=12866</td><td>30.7153</td></tr>
<tr><td>11566:4</td><td>15</td><td>K.LKELPGVC[+57.021]N[+2205.793]ETMMALWEEC[+57.021]KPC[+57.021]LK.Q</td><td>HexNAc(4)Hex(5)Fuc(2)NeuAc(1)</td><td>C[+57]*3, N[+2206]</td><td>1286.3113</td><td>4</td><td>5142.2234</td><td>5142.1776</td><td>0</td><td>8.9</td><td>95</td><td>Specific</td><td>424.82</td><td>271.3</td><td>271.3</td><td>2.85</td><td>9</td><td>>sp|P10909|CLUS_HUMAN Clusterin OS=Homo sapiens OX=9606 GN=CLU PE=1 SV=1</td><td>1467</td><td>20201115_07413_1ng_glyco_hcd30_etd30_compound15.14971.14971.4</td><td>controllerType=0 controllerNumber=1 scan=14971</td><td>35.2436</td></tr>
<tr><td>11490:4</td><td>15</td><td>K.LKELPGVC[+57.021]N[+2424.867]ETMMALWEEC[+57.021]KPC[+57.021]LK.Q</td><td>HexNAc(5)Hex(6)Fuc(1)NeuAc(1)</td><td>C[+57]*3, N[+2425]</td><td>1341.0758</td><td>4</td><td>5361.2814</td><td>5361.2518</td><td>0</td><td>5.5</td><td>95</td><td>Specific</td><td>426.95</td><td>306.59</td><td>33.12</td><td>3.4</td><td>9</td><td>>sp|P10909|CLUS_HUMAN Clusterin OS=Homo sapiens OX=9606 GN=CLU PE=1 SV=1</td><td>1467</td><td>20201115_07413_1ng_glyco_hcd30_etd30_compound15.14892.14892.4</td><td>controllerType=0 controllerNumber=1 scan=14892</td><td>35.0824</td></tr>
<tr><td>12780:4</td><td>15</td><td>K.LKELPGVC[+57.021]N[+2350.830]ETMMALWEEC[+57.021]KPC[+57.021]LK.Q</td><td>HexNAc(4)Hex(5)Fuc(1)NeuAc(2)</td><td>C[+57]*3, N[+2351]</td><td>1322.5608</td><td>4</td><td>5287.2213</td><td>5287.2151</td><td>0</td><td>1.2</td><td>95</td><td>Specific</td><td>453.8</td><td>325.44</td><td>325.44</td><td>3.76</td><td>9</td><td>>sp|P10909|CLUS_HUMAN Clusterin OS=Homo sapiens OX=9606 GN=CLU PE=1 SV=1</td><td>1467</td><td>20201115_07413_1ng_glyco_hcd30_etd30_compound15.16256.16256.4</td><td>controllerType=0 controllerNumber=1 scan=16256</td><td>38.4546</td></tr>
<tr><td>11610:3</td><td>15</td><td>K.LKELPGVC[+57.021]N[+2059.735]ETMMALWEEC[+57.021]KPC[+57.021]LK.Q</td><td>HexNAc(4)Hex(5)Fuc(1)NeuAc(1)</td><td>C[+57]*3, N[+2060]</td><td>1666.0582</td><td>3</td><td>4996.16</td><td>4996.1196</td><td>0</td><td>8.1</td><td>95</td><td>Specific</td><td>490.95</td><td>399.81</td><td>399.81</td><td>4.37</td><td>9</td><td>>sp|P10909|CLUS_HUMAN Clusterin OS=Homo sapiens OX=9606 GN=CLU PE=1 SV=1</td><td>1467</td><td>20201115_07413_1ng_glyco_hcd30_etd30_compound15.15019.15019.3</td><td>controllerType=0 controllerNumber=1 scan=15019</td><td>35.3578</td></tr>
<tr><td>11585:4</td><td>15</td><td>K.LKELPGVC[+57.021]N[+2059.735]ETMMALWEEC[+57.021]KPC[+57.021]LK.Q</td><td>HexNAc(4)Hex(5)Fuc(1)NeuAc(1)</td><td>C[+57]*3, N[+2060]</td><td>1249.7931</td><td>4</td><td>4996.1505</td><td>4996.1196</td><td>0</td><td>6.2</td><td>95</td><td>Specific</td><td>817.69</td><td>623.42</td><td>623.42</td><td>12.32</td><td>9</td><td>>sp|P10909|CLUS_HUMAN Clusterin OS=Homo sapiens OX=9606 GN=CLU PE=1 SV=1</td><td>1467</td><td>20201115_07413_1ng_glyco_hcd30_etd30_compound15.14993.14993.4</td><td>controllerType=0 controllerNumber=1 scan=14993</td><td>35.2982</td></tr>
<tr><td>18051:3</td><td>15</td><td>K.MLN[+2350.830]TSSLLEQLNEQFNWVSR.L</td><td>HexNAc(4)Hex(5)Fuc(1)NeuAc(2)</td><td>N[+2351]</td><td>1587.3473</td><td>3</td><td>4760.0275</td><td>4760.0171</td><td>0</td><td>2.2</td><td>352</td><td>Specific</td><td>468.35</td><td>301.4</td><td>301.4</td><td>6.03</td><td>9</td><td>>sp|P10909|CLUS_HUMAN Clusterin OS=Homo sapiens OX=9606 GN=CLU PE=1 SV=1</td><td>1467</td><td>20201115_07413_1ng_glyco_hcd30_etd30_compound15.21884.21884.3</td><td>controllerType=0 controllerNumber=1 scan=21884</td><td>54.3897</td></tr>
<tr><td>16397:3</td><td>15</td><td>K.MLN[+2059.735]TSSLLEQLNEQFNWVSR.L</td><td>HexNAc(4)Hex(5)Fuc(1)NeuAc(1)</td><td>N[+2060]</td><td>1490.3266</td><td>3</td><td>4468.9652</td><td>4468.9217</td><td>0</td><td>9.8</td><td>352</td><td>Specific</td><td>612.11</td><td>516.92</td><td>516.92</td><td>8.55</td><td>9</td><td>>sp|P10909|CLUS_HUMAN Clusterin OS=Homo sapiens OX=9606 GN=CLU PE=1 SV=1</td><td>1467</td><td>20201115_07413_1ng_glyco_hcd30_etd30_compound15.20126.20126.3</td><td>controllerType=0 controllerNumber=1 scan=20126</td><td>49.7684</td></tr>
<tr><td>08434:3</td><td>15</td><td>R.LAN[+2570.925]LTQGEDQYYLR.V</td><td>HexNAc(5)Hex(6)Fuc(2)NeuAc(1)</td><td>N[+2571]</td><td>1418.9368</td><td>3</td><td>4254.7959</td><td>4254.7587</td><td>0</td><td>8.7</td><td>372</td><td>Specific</td><td>206.44</td><td>138.37</td><td>138.37</td><td>3.23</td><td>9</td><td>>sp|P10909|CLUS_HUMAN Clusterin OS=Homo sapiens OX=9606 GN=CLU PE=1 SV=1</td><td>1467</td><td>20201115_07413_1ng_glyco_hcd30_etd30_compound15.11675.11675.3</td><td>controllerType=0 controllerNumber=1 scan=11675</td><td>28.3914</td></tr>
<tr><td>03761:3</td><td>17</td><td>K.TPLTAN[+2262.814]ITK.S</td><td>HexNAc(5)Hex(5)Fuc(1)NeuAc(1)</td><td>N[+2263]</td><td>1074.4672</td><td>3</td><td>3221.387</td><td>3221.3711</td><td>0</td><td>4.9</td><td>200</td><td>Specific</td><td>215.47</td><td>101</td><td>101</td><td>2.77</td><td>10</td><td>>sp|P01877|IGHA2_HUMAN Immunoglobulin heavy constant alpha 2 OS=Homo sapiens OX=9606 GN=IGHA2 PE=1 SV=4</td><td>576</td><td>20201115_07413_1ng_glyco_hcd30_etd30_compound15.6739.6739.3</td><td>controllerType=0 controllerNumber=1 scan=6739</td><td>19.0548</td></tr>
<tr><td>02269:2</td><td>17</td><td>K.TPLTAN[+1606.587]ITK.S</td><td>HexNAc(4)Hex(4)Fuc(1)</td><td>N[+1607]</td><td>1283.0783</td><td>2</td><td>2565.1493</td><td>2565.1435</td><td>0</td><td>2.3</td><td>200</td><td>Specific</td><td>229.41</td><td>58.94</td><td>58.94</td><td>2.63</td><td>10</td><td>>sp|P01877|IGHA2_HUMAN Immunoglobulin heavy constant alpha 2 OS=Homo sapiens OX=9606 GN=IGHA2 PE=1 SV=4</td><td>576</td><td>20201115_07413_1ng_glyco_hcd30_etd30_compound15.5138.5138.2</td><td>controllerType=0 controllerNumber=1 scan=5138</td><td>15.9397</td></tr>
<tr><td>02241:2</td><td>17</td><td>K.TPLTAN[+1971.719]ITK.S</td><td>HexNAc(5)Hex(5)Fuc(1)</td><td>N[+1972]</td><td>1465.6479</td><td>2</td><td>2930.2885</td><td>2930.2757</td><td>0</td><td>4.4</td><td>200</td><td>Specific</td><td>233.92</td><td>65.02</td><td>65.02</td><td>2.52</td><td>10</td><td>>sp|P01877|IGHA2_HUMAN Immunoglobulin heavy constant alpha 2 OS=Homo sapiens OX=9606 GN=IGHA2 PE=1 SV=4</td><td>576</td><td>20201115_07413_1ng_glyco_hcd30_etd30_compound15.5109.5109.2</td><td>controllerType=0 controllerNumber=1 scan=5109</td><td>15.8727</td></tr>
<tr><td>02236:3</td><td>17</td><td>K.TPLTAN[+1971.719]ITK.S</td><td>HexNAc(5)Hex(5)Fuc(1)</td><td>N[+1972]</td><td>977.4339</td><td>3</td><td>2930.287</td><td>2930.2757</td><td>0</td><td>3.9</td><td>200</td><td>Specific</td><td>293.36</td><td>133.22</td><td>133.22</td><td>3.21</td><td>10</td><td>>sp|P01877|IGHA2_HUMAN Immunoglobulin heavy constant alpha 2 OS=Homo sapiens OX=9606 GN=IGHA2 PE=1 SV=4</td><td>576</td><td>20201115_07413_1ng_glyco_hcd30_etd30_compound15.5104.5104.3</td><td>controllerType=0 controllerNumber=1 scan=5104</td><td>15.8589</td></tr>
<tr><td>02334:2</td><td>17</td><td>K.TPLTAN[+1501.555]ITK.S</td><td>HexNAc(5)Hex(3)</td><td>N[+1502]</td><td>1230.5633</td><td>2</td><td>2460.1193</td><td>2460.1121</td><td>0</td><td>2.9</td><td>200</td><td>Specific</td><td>295.43</td><td>85.38</td><td>85.38</td><td>2.91</td><td>10</td><td>>sp|P01877|IGHA2_HUMAN Immunoglobulin heavy constant alpha 2 OS=Homo sapiens OX=9606 GN=IGHA2 PE=1 SV=4</td><td>576</td><td>20201115_07413_1ng_glyco_hcd30_etd30_compound15.5207.5207.2</td><td>controllerType=0 controllerNumber=1 scan=5207</td><td>16.1044</td></tr>
<tr><td>02327:2</td><td>17</td><td>K.TPLTAN[+1647.613]ITK.S</td><td>HexNAc(5)Hex(3)Fuc(1)</td><td>N[+1648]</td><td>1303.5922</td><td>2</td><td>2606.1771</td><td>2606.17</td><td>0</td><td>2.7</td><td>200</td><td>Specific</td><td>299.77</td><td>92.23</td><td>92.23</td><td>2.83</td><td>10</td><td>>sp|P01877|IGHA2_HUMAN Immunoglobulin heavy constant alpha 2 OS=Homo sapiens OX=9606 GN=IGHA2 PE=1 SV=4</td><td>576</td><td>20201115_07413_1ng_glyco_hcd30_etd30_compound15.5200.5200.2</td><td>controllerType=0 controllerNumber=1 scan=5200</td><td>16.0879</td></tr>
<tr><td>02277:2</td><td>17</td><td>K.TPLTAN[+1809.666]ITK.S</td><td>HexNAc(5)Hex(4)Fuc(1)</td><td>N[+1810]</td><td>1384.6185</td><td>2</td><td>2768.2296</td><td>2768.2228</td><td>0</td><td>2.5</td><td>200</td><td>Specific</td><td>314.93</td><td>112.81</td><td>112.81</td><td>3.26</td><td>10</td><td>>sp|P01877|IGHA2_HUMAN Immunoglobulin heavy constant alpha 2 OS=Homo sapiens OX=9606 GN=IGHA2 PE=1 SV=4</td><td>576</td><td>20201115_07413_1ng_glyco_hcd30_etd30_compound15.5148.5148.2</td><td>controllerType=0 controllerNumber=1 scan=5148</td><td>15.9669</td></tr>
<tr><td>02278:3</td><td>17</td><td>K.TPLTAN[+1809.666]ITK.S</td><td>HexNAc(5)Hex(4)Fuc(1)</td><td>N[+1810]</td><td>923.4154</td><td>3</td><td>2768.2316</td><td>2768.2228</td><td>0</td><td>3.2</td><td>200</td><td>Specific</td><td>324.76</td><td>140.08</td><td>140.08</td><td>3.24</td><td>10</td><td>>sp|P01877|IGHA2_HUMAN Immunoglobulin heavy constant alpha 2 OS=Homo sapiens OX=9606 GN=IGHA2 PE=1 SV=4</td><td>576</td><td>20201115_07413_1ng_glyco_hcd30_etd30_compound15.5149.5149.3</td><td>controllerType=0 controllerNumber=1 scan=5149</td><td>15.9738</td></tr>
<tr><td>18644:4</td><td>19</td><td>R.QLAHQSN[+2350.830]STNIFFSPVSIATAFAMLSLGTK.A</td><td>HexNAc(4)Hex(5)Fuc(1)NeuAc(2)</td><td>N[+2351]</td><td>1383.8727</td><td>4</td><td>5532.4689</td><td>5532.4654</td><td>0</td><td>0.6</td><td>64</td><td>Specific</td><td>223.15</td><td>169.1</td><td>169.1</td><td>3.49</td><td>7</td><td>>sp|P01009|A1AT_HUMAN Alpha-1-antitrypsin OS=Homo sapiens OX=9606 GN=SERPINA1 PE=1 SV=3</td><td>642</td><td>20201115_07413_1ng_glyco_hcd30_etd30_compound15.22508.22508.2</td><td>controllerType=0 controllerNumber=1 scan=22508</td><td>55.972</td></tr>
<tr><td>18569:4</td><td>19</td><td>R.QLAHQSN[+2350.830]STNIFFSPVSIATAFAMLSLGTK.A</td><td>HexNAc(4)Hex(5)Fuc(1)NeuAc(2)</td><td>N[+2351]</td><td>1383.8843</td><td>4</td><td>5532.5153</td><td>5532.4654</td><td>0</td><td>9</td><td>64</td><td>Specific</td><td>233.33</td><td>197.42</td><td>197.42</td><td>3.28</td><td>7</td><td>>sp|P01009|A1AT_HUMAN Alpha-1-antitrypsin OS=Homo sapiens OX=9606 GN=SERPINA1 PE=1 SV=3</td><td>642</td><td>20201115_07413_1ng_glyco_hcd30_etd30_compound15.22428.22428.4</td><td>controllerType=0 controllerNumber=1 scan=22428</td><td>55.756</td></tr>
<tr><td>18574:3</td><td>19</td><td>R.QLAHQSN[+2204.772]STNIFFSPVSIATAFAMLSLGTK.A</td><td>HexNAc(4)Hex(5)NeuAc(2)</td><td>N[+2205]</td><td>1796.1435</td><td>3</td><td>5386.416</td><td>5386.4075</td><td>0</td><td>1.6</td><td>64</td><td>Specific</td><td>422.76</td><td>279</td><td>279</td><td>5.32</td><td>7</td><td>>sp|P01009|A1AT_HUMAN Alpha-1-antitrypsin OS=Homo sapiens OX=9606 GN=SERPINA1 PE=1 SV=3</td><td>642</td><td>20201115_07413_1ng_glyco_hcd30_etd30_compound15.22434.22434.3</td><td>controllerType=0 controllerNumber=1 scan=22434</td><td>55.7808</td></tr>
<tr><td>17632:4</td><td>19</td><td>K.ADTHDEILEGLNFN[+2861.000]LTEIPEAQIHEGFQELLR.T</td><td>HexNAc(5)Hex(6)NeuAc(3)</td><td>N[+2861]</td><td>1638.962</td><td>4</td><td>6552.8263</td><td>6552.8239</td><td>0</td><td>0.4</td><td>94</td><td>Specific</td><td>264.17</td><td>253.32</td><td>253.32</td><td>4.76</td><td>7</td><td>>sp|P01009|A1AT_HUMAN Alpha-1-antitrypsin OS=Homo sapiens OX=9606 GN=SERPINA1 PE=1 SV=3</td><td>642</td><td>20201115_07413_1ng_glyco_hcd30_etd30_compound15.21438.21438.4</td><td>controllerType=0 controllerNumber=1 scan=21438</td><td>53.0519</td></tr>
<tr><td>17626:4</td><td>19</td><td>K.ADTHDEILEGLNFN[+3007.058]LTEIPEAQIHEGFQELLR.T</td><td>HexNAc(5)Hex(6)Fuc(1)NeuAc(3)</td><td>N[+3007]</td><td>1675.4713</td><td>4</td><td>6698.8633</td><td>6698.8818</td><td>0</td><td>-2.8</td><td>94</td><td>Specific</td><td>435.29</td><td>435.29</td><td>435.29</td><td>5.61</td><td>7</td><td>>sp|P01009|A1AT_HUMAN Alpha-1-antitrypsin OS=Homo sapiens OX=9606 GN=SERPINA1 PE=1 SV=3</td><td>642</td><td>20201115_07413_1ng_glyco_hcd30_etd30_compound15.21431.21431.3</td><td>controllerType=0 controllerNumber=1 scan=21431</td><td>53.0274</td></tr>
<tr><td>15916:3</td><td>19</td><td>K.YLGN[+2204.772]ATAIFFLPDEGK.L</td><td>HexNAc(4)Hex(5)NeuAc(2)</td><td>N[+2205]</td><td>1320.9051</td><td>3</td><td>3960.7007</td><td>3960.6677</td><td>0</td><td>8.3</td><td>268</td><td>Specific</td><td>323.84</td><td>244.79</td><td>244.79</td><td>4.83</td><td>7</td><td>>sp|P01009|A1AT_HUMAN Alpha-1-antitrypsin OS=Homo sapiens OX=9606 GN=SERPINA1 PE=1 SV=3</td><td>642</td><td>20201115_07413_1ng_glyco_hcd30_etd30_compound15.19615.19615.3</td><td>controllerType=0 controllerNumber=1 scan=19615</td><td>48.1094</td></tr>
<tr><td>01503:3</td><td>24</td><td>R.LRN[+1216.423]VSWATGR.S</td><td>HexNAc(2)Hex(5)</td><td>N[+1216]</td><td>792.6911</td><td>3</td><td>2376.0588</td><td>2376.0559</td><td>0</td><td>1.2</td><td>149</td><td>Specific</td><td>281.24</td><td>149.28</td><td>149.28</td><td>2.55</td><td>3</td><td>>sp|P08571|CD14_HUMAN Monocyte differentiation antigen CD14 OS=Homo sapiens OX=9606 GN=CD14 PE=1 SV=2</td><td>1232</td><td>20201115_07413_1ng_glyco_hcd30_etd30_compound15.4319.4319.3</td><td>controllerType=0 controllerNumber=1 scan=4319</td><td>14.335</td></tr>
<tr><td>04574:3</td><td>38</td><td>R.LSPN[+2205.793]ASAEHLELR.W</td><td>HexNAc(4)Hex(5)Fuc(2)NeuAc(1)</td><td>N[+2206]</td><td>1214.8549</td><td>3</td><td>3642.5502</td><td>3642.5421</td><td>0</td><td>2.2</td><td>52</td><td>Specific</td><td>231.58</td><td>139.38</td><td>139.38</td><td>3.16</td><td>3</td><td>>sp|Q13410|BT1A1_HUMAN Butyrophilin subfamily 1 member A1 OS=Homo sapiens OX=9606 GN=BTN1A1 PE=1 SV=3</td><td>179</td><td>20201115_07413_1ng_glyco_hcd30_etd30_compound15.7597.7597.3</td><td>controllerType=0 controllerNumber=1 scan=7597</td><td>20.6321</td></tr>
<tr><td>00668:4</td><td>57</td><td>K.TKPREEQFN[+1444.534]STFR.V</td><td>HexNAc(4)Hex(3)Fuc(1)</td><td>N[+1445]</td><td>771.8447</td><td>4</td><td>3084.357</td><td>3084.3526</td><td>0</td><td>1.4</td><td>168</td><td>Specific</td><td>233.11</td><td>233.11</td><td>233.11</td><td>2.79</td><td>3</td><td>>sp|P01859|IGHG2_HUMAN Immunoglobulin heavy constant gamma 2 OS=Homo sapiens OX=9606 GN=IGHG2 PE=1 SV=2</td><td>387</td><td>20201115_07413_1ng_glyco_hcd30_etd30_compound15.3406.3406.4</td><td>controllerType=0 controllerNumber=1 scan=3406</td><td>12.5524</td></tr>
<tr><td>00615:4</td><td>57</td><td>K.TKPREEQFN[+1606.587]STFR.V</td><td>HexNAc(4)Hex(4)Fuc(1)</td><td>N[+1607]</td><td>812.3582</td><td>4</td><td>3246.4111</td><td>3246.4054</td><td>0</td><td>1.8</td><td>168</td><td>Specific</td><td>258.62</td><td>258.62</td><td>258.62</td><td>3.31</td><td>3</td><td>>sp|P01859|IGHG2_HUMAN Immunoglobulin heavy constant gamma 2 OS=Homo sapiens OX=9606 GN=IGHG2 PE=1 SV=2</td><td>387</td><td>20201115_07413_1ng_glyco_hcd30_etd30_compound15.3350.3350.4</td><td>controllerType=0 controllerNumber=1 scan=3350</td><td>12.4368</td></tr>
<tr><td>15784:5</td><td>1</td><td>R.RTAGWNVPIGTLRPFLN[+2059.735]WTGPPEPIEAAVAR.F</td><td>HexNAc(4)Hex(5)Fuc(1)NeuAc(1)</td><td>N[+2060]</td><td>1090.1202</td><td>5</td><td>5446.5721</td><td>5446.547</td><td>0</td><td>4.6</td><td>140</td><td>Specific</td><td>602.02</td><td>589.29</td><td>589.29</td><td>8.32</td><td>33</td><td>>sp|P02788|TRFL_HUMAN Lactotransferrin OS=Homo sapiens OX=9606 GN=LTF PE=1 SV=6</td><td>1270</td><td>20201115_07433_1ng_glyco_hcd30_etd30_compound15.19306.19306.5</td><td>controllerType=0 controllerNumber=1 scan=19306</td><td>43.6348</td></tr>
<tr><td>15778:4</td><td>1</td><td>R.RTAGWNVPIGTLRPFLN[+2059.735]WTGPPEPIEAAVAR.F</td><td>HexNAc(4)Hex(5)Fuc(1)NeuAc(1)</td><td>N[+2060]</td><td>1362.3969</td><td>4</td><td>5446.5656</td><td>5446.547</td><td>0</td><td>3.4</td><td>140</td><td>Specific</td><td>949.12</td><td>949.12</td><td>949.12</td><td>15.85</td><td>33</td><td>>sp|P02788|TRFL_HUMAN Lactotransferrin OS=Homo sapiens OX=9606 GN=LTF PE=1 SV=6</td><td>1270</td><td>20201115_07433_1ng_glyco_hcd30_etd30_compound15.19300.19300.4</td><td>controllerType=0 controllerNumber=1 scan=19300</td><td>43.6251</td></tr>
<tr><td>16578:3</td><td>1</td><td>R.TAGWNVPIGTLRPFLN[+2570.925]WTGPPEPIEAAVAR.F</td><td>HexNAc(5)Hex(6)Fuc(2)NeuAc(1)</td><td>N[+2571]</td><td>1934.5526</td><td>3</td><td>5801.6432</td><td>5801.636</td><td>0</td><td>1.2</td><td>141</td><td>Specific</td><td>284.5</td><td>178.51</td><td>178.51</td><td>4.69</td><td>33</td><td>>sp|P02788|TRFL_HUMAN Lactotransferrin OS=Homo sapiens OX=9606 GN=LTF PE=1 SV=6</td><td>1270</td><td>20201115_07433_1ng_glyco_hcd30_etd30_compound15.20143.20143.3</td><td>controllerType=0 controllerNumber=1 scan=20143</td><td>46.0015</td></tr>
<tr><td>16858:4</td><td>1</td><td>R.TAGWNVPIGTLRPFLN[+2059.735]WTGPPEPIEAAVAR.F</td><td>HexNAc(4)Hex(5)Fuc(1)NeuAc(1)</td><td>N[+2060]</td><td>1323.3748</td><td>4</td><td>5290.4772</td><td>5290.4459</td><td>0</td><td>5.9</td><td>141</td><td>Specific</td><td>287.51</td><td>287.51</td><td>287.51</td><td>3.96</td><td>33</td><td>>sp|P02788|TRFL_HUMAN Lactotransferrin OS=Homo sapiens OX=9606 GN=LTF PE=1 SV=6</td><td>1270</td><td>20201115_07433_1ng_glyco_hcd30_etd30_compound15.20443.20443.2</td><td>controllerType=0 controllerNumber=1 scan=20443</td><td>46.6566</td></tr>
<tr><td>16679:4</td><td>1</td><td>R.TAGWNVPIGTLRPFLN[+2059.735]WTGPPEPIEAAVAR.F</td><td>HexNAc(4)Hex(5)Fuc(1)NeuAc(1)</td><td>N[+2060]</td><td>1323.3709</td><td>4</td><td>5290.4616</td><td>5290.4459</td><td>0</td><td>3</td><td>141</td><td>Specific</td><td>340.19</td><td>340.19</td><td>340.19</td><td>4.65</td><td>33</td><td>>sp|P02788|TRFL_HUMAN Lactotransferrin OS=Homo sapiens OX=9606 GN=LTF PE=1 SV=6</td><td>1270</td><td>20201115_07433_1ng_glyco_hcd30_etd30_compound15.20251.20251.4</td><td>controllerType=0 controllerNumber=1 scan=20251</td><td>46.3187</td></tr>
<tr><td>16667:3</td><td>1</td><td>R.TAGWNVPIGTLRPFLN[+2059.735]WTGPPEPIEAAVAR.F</td><td>HexNAc(4)Hex(5)Fuc(1)NeuAc(1)</td><td>N[+2060]</td><td>1764.1594</td><td>3</td><td>5290.4636</td><td>5290.4459</td><td>0</td><td>3.4</td><td>141</td><td>Specific</td><td>418.07</td><td>310.88</td><td>310.88</td><td>5.08</td><td>33</td><td>>sp|P02788|TRFL_HUMAN Lactotransferrin OS=Homo sapiens OX=9606 GN=LTF PE=1 SV=6</td><td>1270</td><td>20201115_07433_1ng_glyco_hcd30_etd30_compound15.20238.20238.3</td><td>controllerType=0 controllerNumber=1 scan=20238</td><td>46.2712</td></tr>
<tr><td>15948:3</td><td>1</td><td>R.TAGWNVPIGTLRPFLN[+1768.640]WTGPPEPIEAAVAR.F</td><td>HexNAc(4)Hex(5)Fuc(1)</td><td>N[+1769]</td><td>1667.1309</td><td>3</td><td>4999.3783</td><td>4999.3504</td><td>0</td><td>5.6</td><td>141</td><td>Specific</td><td>419.75</td><td>373.21</td><td>373.21</td><td>5.96</td><td>33</td><td>>sp|P02788|TRFL_HUMAN Lactotransferrin OS=Homo sapiens OX=9606 GN=LTF PE=1 SV=6</td><td>1270</td><td>20201115_07433_1ng_glyco_hcd30_etd30_compound15.19478.19478.3</td><td>controllerType=0 controllerNumber=1 scan=19478</td><td>44.1104</td></tr>
<tr><td>16668:4</td><td>1</td><td>R.TAGWNVPIGTLRPFLN[+2351.851]WTGPPEPIEAAVAR.F</td><td>HexNAc(4)Hex(5)Fuc(3)NeuAc(1)</td><td>N[+2352]</td><td>1396.4001</td><td>4</td><td>5582.5788</td><td>5582.5617</td><td>0</td><td>3.1</td><td>141</td><td>Specific</td><td>509.57</td><td>509.57</td><td>509.57</td><td>7.7</td><td>33</td><td>>sp|P02788|TRFL_HUMAN Lactotransferrin OS=Homo sapiens OX=9606 GN=LTF PE=1 SV=6</td><td>1270</td><td>20201115_07433_1ng_glyco_hcd30_etd30_compound15.20239.20239.4</td><td>controllerType=0 controllerNumber=1 scan=20239</td><td>46.274</td></tr>
<tr><td>16658:3</td><td>1</td><td>R.TAGWNVPIGTLRPFLN[+2205.793]WTGPPEPIEAAVAR.F</td><td>HexNAc(4)Hex(5)Fuc(2)NeuAc(1)</td><td>N[+2206]</td><td>1812.8505</td><td>3</td><td>5436.5371</td><td>5436.5038</td><td>0</td><td>6.1</td><td>141</td><td>Specific</td><td>560.94</td><td>440.35</td><td>440.35</td><td>7.19</td><td>33</td><td>>sp|P02788|TRFL_HUMAN Lactotransferrin OS=Homo sapiens OX=9606 GN=LTF PE=1 SV=6</td><td>1270</td><td>20201115_07433_1ng_glyco_hcd30_etd30_compound15.20228.20228.3</td><td>controllerType=0 controllerNumber=1 scan=20228</td><td>46.213</td></tr>
<tr><td>16655:4</td><td>1</td><td>R.TAGWNVPIGTLRPFLN[+2205.793]WTGPPEPIEAAVAR.F</td><td>HexNAc(4)Hex(5)Fuc(2)NeuAc(1)</td><td>N[+2206]</td><td>1359.89</td><td>4</td><td>5436.5382</td><td>5436.5038</td><td>0</td><td>6.3</td><td>141</td><td>Specific</td><td>940.5</td><td>856.66</td><td>856.66</td><td>15.38</td><td>33</td><td>>sp|P02788|TRFL_HUMAN Lactotransferrin OS=Homo sapiens OX=9606 GN=LTF PE=1 SV=6</td><td>1270</td><td>20201115_07433_1ng_glyco_hcd30_etd30_compound15.20225.20225.4</td><td>controllerType=0 controllerNumber=1 scan=20225</td><td>46.2016</td></tr>
<tr><td>15947:4</td><td>1</td><td>R.TAGWNVPIGTLRPFLN[+1768.640]WTGPPEPIEAAVAR.F</td><td>HexNAc(4)Hex(5)Fuc(1)</td><td>N[+1769]</td><td>1250.5999</td><td>4</td><td>4999.3776</td><td>4999.3504</td><td>0</td><td>5.4</td><td>141</td><td>Specific</td><td>1061.19</td><td>1061.19</td><td>1061.19</td><td>18.56</td><td>33</td><td>>sp|P02788|TRFL_HUMAN Lactotransferrin OS=Homo sapiens OX=9606 GN=LTF PE=1 SV=6</td><td>1270</td><td>20201115_07433_1ng_glyco_hcd30_etd30_compound15.19477.19477.4</td><td>controllerType=0 controllerNumber=1 scan=19477</td><td>44.1053</td></tr>
<tr><td>14981:3</td><td>1</td><td>R.TAGWNIPMGLLFN[+1622.582]QTGSC[+57.021]K.F</td><td>HexNAc(4)Hex(5)</td><td>C[+57], N[+1623]</td><td>1239.8737</td><td>3</td><td>3717.6066</td><td>3717.5916</td><td>0</td><td>4</td><td>485</td><td>Specific</td><td>204.02</td><td>118.92</td><td>118.92</td><td>3.8</td><td>33</td><td>>sp|P02788|TRFL_HUMAN Lactotransferrin OS=Homo sapiens OX=9606 GN=LTF PE=1 SV=6</td><td>1270</td><td>20201115_07433_1ng_glyco_hcd30_etd30_compound15.18456.18456.3</td><td>controllerType=0 controllerNumber=1 scan=18456</td><td>41.1116</td></tr>
<tr><td>15133:3</td><td>1</td><td>R.TAGWNIPMGLLFN[+1216.423]QTGSC[+57.021]K.F</td><td>HexNAc(2)Hex(5)</td><td>C[+57], N[+1216]</td><td>1104.4852</td><td>3</td><td>3311.441</td><td>3311.4328</td><td>0</td><td>2.5</td><td>485</td><td>Specific</td><td>214.46</td><td>214.46</td><td>131.94</td><td>2.82</td><td>33</td><td>>sp|P02788|TRFL_HUMAN Lactotransferrin OS=Homo sapiens OX=9606 GN=LTF PE=1 SV=6</td><td>1270</td><td>20201115_07433_1ng_glyco_hcd30_etd30_compound15.18617.18617.3</td><td>controllerType=0 controllerNumber=1 scan=18617</td><td>41.5829</td></tr>
<tr><td>14919:3</td><td>1</td><td>R.TAGWNIPMGLLFN[+1606.587]QTGSC[+57.021]K.F</td><td>HexNAc(4)Hex(4)Fuc(1)</td><td>C[+57], N[+1607]</td><td>1234.5416</td><td>3</td><td>3701.6101</td><td>3701.5966</td><td>0</td><td>3.6</td><td>485</td><td>Specific</td><td>334.82</td><td>257.23</td><td>257.23</td><td>3.7</td><td>33</td><td>>sp|P02788|TRFL_HUMAN Lactotransferrin OS=Homo sapiens OX=9606 GN=LTF PE=1 SV=6</td><td>1270</td><td>20201115_07433_1ng_glyco_hcd30_etd30_compound15.18389.18389.3</td><td>controllerType=0 controllerNumber=1 scan=18389</td><td>40.9055</td></tr>
<tr><td>16181:3</td><td>1</td><td>R.TAGWNIPMGLLFN[+2205.793]QTGSC[+57.021]K.F</td><td>HexNAc(4)Hex(5)Fuc(2)NeuAc(1)</td><td>C[+57], N[+2206]</td><td>1434.2772</td><td>3</td><td>4300.8169</td><td>4300.8028</td><td>0</td><td>3.3</td><td>485</td><td>Specific</td><td>392.99</td><td>284.98</td><td>284.98</td><td>5.22</td><td>33</td><td>>sp|P02788|TRFL_HUMAN Lactotransferrin OS=Homo sapiens OX=9606 GN=LTF PE=1 SV=6</td><td>1270</td><td>20201115_07433_1ng_glyco_hcd30_etd30_compound15.19723.19723.3</td><td>controllerType=0 controllerNumber=1 scan=19723</td><td>44.8159</td></tr>
<tr><td>14835:3</td><td>1</td><td>R.TAGWNIPMGLLFN[+1768.640]QTGSC[+57.021]K.F</td><td>HexNAc(4)Hex(5)Fuc(1)</td><td>C[+57], N[+1769]</td><td>1288.5587</td><td>3</td><td>3863.6614</td><td>3863.6495</td><td>0</td><td>3.1</td><td>485</td><td>Specific</td><td>404.58</td><td>289.36</td><td>289.36</td><td>5.01</td><td>33</td><td>>sp|P02788|TRFL_HUMAN Lactotransferrin OS=Homo sapiens OX=9606 GN=LTF PE=1 SV=6</td><td>1270</td><td>20201115_07433_1ng_glyco_hcd30_etd30_compound15.18301.18301.3</td><td>controllerType=0 controllerNumber=1 scan=18301</td><td>40.6664</td></tr>
<tr><td>16186:3</td><td>1</td><td>R.TAGWNIPMGLLFN[+2059.735]QTGSC[+57.021]K.F</td><td>HexNAc(4)Hex(5)Fuc(1)NeuAc(1)</td><td>C[+57], N[+2060]</td><td>1385.5912</td><td>3</td><td>4154.7592</td><td>4154.7449</td><td>0</td><td>3.4</td><td>485</td><td>Specific</td><td>419.77</td><td>277.65</td><td>277.65</td><td>5.33</td><td>33</td><td>>sp|P02788|TRFL_HUMAN Lactotransferrin OS=Homo sapiens OX=9606 GN=LTF PE=1 SV=6</td><td>1270</td><td>20201115_07433_1ng_glyco_hcd30_etd30_compound15.19728.19728.3</td><td>controllerType=0 controllerNumber=1 scan=19728</td><td>44.83</td></tr>
</table>

Table S4. Cont.

| Query #:z | Protein Rank | Glycopeptide | Glycan | Modification Type(s) | Observed m/z | z | Observed (M+H) | Calc. mass (M+H) | Off-by-x error | Mass error (ppm) | StartPosition | Cleavage | Score | Delta | Delta Mod | \|Log Prob\| | # of unique peptides | Protein Name | Protein DB number | Comment | Scan # | Scan Time |
|---|---|---|---|---|---|---|---|---|---|---|---|---|---|---|---|---|---|---|---|---|---|---|
| 16318:4 | 1 | R.TAGWNIPMGLLFN[+1913.677]QTGSC[+57.021]K.F | HexNAc(4)Hex(5)NeuAc(1) | C[+57], N[+1914] | 1002.9306 | 4 | 4008.7008 | 4008.687 | 0 | 3.4 | 485 | Specific | 425.12 | 348.4 | 348.4 | 4.06 | 33 | >sp\|P02788\|TRFL_HUMAN Lactotransferrin OS=Homo sapiens OX=9606 GN=LTF PE=1 SV=6 | 1270 | 20201115_07433_1ng_glyco_hcd30_etd30_compound15.19868.19868.4 | controllerType=0 controllerNumber=1 scan=19868 | 45.2559 |
| 14852:4 | 1 | R.TAGWNIPMGLLFN[+1768.640]QTGSC[+57.021]K.F | HexNAc(4)Hex(5)Fuc(1) | C[+57], N[+1769] | 966.6708 | 4 | 3863.6614 | 3863.6495 | 0 | 3.1 | 485 | Specific | 447.13 | 433.88 | 433.88 | 4.72 | 33 | >sp\|P02788\|TRFL_HUMAN Lactotransferrin OS=Homo sapiens OX=9606 GN=LTF PE=1 SV=6 | 1270 | 20201115_07433_1ng_glyco_hcd30_etd30_compound15.18318.18318.4 | controllerType=0 controllerNumber=1 scan=18318 | 40.7004 |
| 16405:4 | 1 | R.TAGWNIPMGLLFN[+2205.793]QTGSC[+57.021]KFDEYFSQSC[+57.021]APGSDPR.S | HexNAc(4)Hex(5)Fuc(2)NeuAc(1) | C[+57]*2, N[+2206] | 1536.8971 | 4 | 6144.5666 | 6144.5499 | 0 | 2.7 | 485 | Specific | 396.64 | 359.12 | 359.12 | 5.6 | 33 | >sp\|P02788\|TRFL_HUMAN Lactotransferrin OS=Homo sapiens OX=9606 GN=LTF PE=1 SV=6 | 1270 | 20201115_07433_1ng_glyco_hcd30_etd30_compound15.19960.19960.4 | controllerType=0 controllerNumber=1 scan=19960 | 45.5201 |
| 15627:4 | 1 | R.TAGWNIPMGLLFN[+1768.640]QTGSC[+57.021]KFDEYFSQSC[+57.021]APGSDPR.S | HexNAc(4)Hex(5)Fuc(1) | C[+57]*2, N[+1769] | 1427.6117 | 4 | 5707.425 | 5707.3966 | 0 | 5 | 485 | Specific | 494.16 | 443.12 | 443.12 | 5.88 | 33 | >sp\|P02788\|TRFL_HUMAN Lactotransferrin OS=Homo sapiens OX=9606 GN=LTF PE=1 SV=6 | 1270 | 20201115_07433_1ng_glyco_hcd30_etd30_compound15.19141.19141.4 | controllerType=0 controllerNumber=1 scan=19141 | 43.1826 |
| 16394:4 | 1 | R.TAGWNIPMGLLFN[+2059.735]QTGSC[+57.021]KFDEYFSQSC[+57.021]APGSDPR.S | HexNAc(4)Hex(5)Fuc(1)NeuAc(1) | C[+57]*2, N[+2060] | 1500.3851 | 4 | 5998.5187 | 5998.492 | 0 | 4.4 | 485 | Specific | 579.27 | 503.51 | 503.51 | 8.64 | 33 | >sp\|P02788\|TRFL_HUMAN Lactotransferrin OS=Homo sapiens OX=9606 GN=LTF PE=1 SV=6 | 1270 | 20201115_07433_1ng_glyco_hcd30_etd30_compound15.19949.19949.4 | controllerType=0 controllerNumber=1 scan=19949 | 45.499 |
| 08273:3 | 1 | R.N[+2205.793]GSDC[+57.021]PDKFC[+57.021]LFQSETK.N | HexNAc(4)Hex(5)Fuc(2)NeuAc(1) | C[+57]*2, N[+2206] | 1413.5646 | 3 | 4238.6793 | 4238.6667 | 0 | 3 | 642 | Specific | 289.87 | 191.23 | 191.23 | 3.94 | 33 | >sp\|P02788\|TRFL_HUMAN Lactotransferrin OS=Homo sapiens OX=9606 GN=LTF PE=1 SV=6 | 1270 | 20201115_07433_1ng_glyco_hcd30_etd30_compound15.11403.11403.3 | controllerType=0 controllerNumber=1 scan=11403 | 26.597 |
| 08333:3 | 1 | R.N[+2059.735]GSDC[+57.021]PDKFC[+57.021]LFQSETK.N | HexNAc(4)Hex(5)Fuc(1)NeuAc(1) | C[+57]*2, N[+2060] | 1364.8837 | 3 | 4092.6366 | 4092.6088 | 0 | 6.8 | 642 | Specific | 338.63 | 250.2 | 250.2 | 3.87 | 33 | >sp\|P02788\|TRFL_HUMAN Lactotransferrin OS=Homo sapiens OX=9606 GN=LTF PE=1 SV=6 | 1270 | 20201115_07433_1ng_glyco_hcd30_etd30_compound15.11465.11465.3 | controllerType=0 controllerNumber=1 scan=11465 | 26.7072 |
| 16358:3 | 2 | R.NIAAFGGDPNN[+1913.677]ITLFGESAGGASVSLQTLSPYNK.G | HexNAc(4)Hex(5)NeuAc(1) | N[+1914] | 1775.467 | 3 | 5324.3865 | 5324.3633 | 0 | 4.3 | 197 | Specific | 644.44 | 487.82 | 487.82 | 9.92 | 18 | >sp\|P19835\|CEL_HUMAN Bile salt-activated lipase OS=Homo sapiens OX=9606 GN=CEL PE=1 SV=3 | 476 | 20201115_07433_1ng_glyco_hcd30_etd30_compound15.19911.19911.3 | controllerType=0 controllerNumber=1 scan=19911 | 45.3895 |
| 15481:4 | 3 | R.LSLHRPALEDLLLGSEAN[+1419.502]LTC[+57.021]TLTGLR.D | HexNAc(3)Hex(5) | C[+57], N[+1420] | 1096.5351 | 4 | 4383.1185 | 4383.1005 | 0 | 4.1 | 127 | Specific | 207.75 | 207.75 | 207.75 | 3.68 | 23 | >sp\|P01876\|IGHA1_HUMAN Immunoglobulin heavy constant alpha 1 OS=Homo sapiens OX=9606 GN=IGHA1 PE=1 SV=2 | 255 | 20201115_07433_1ng_glyco_hcd30_etd30_compound15.18985.18985.4 | controllerType=0 controllerNumber=1 scan=18985 | 42.6546 |
| 15477:3 | 3 | R.LSLHRPALEDLLLGSEAN[+1378.476]LTC[+57.021]TLTGLR.D | HexNAc(2)Hex(6) | C[+57], N[+1378] | 1448.0344 | 3 | 4342.0885 | 4342.074 | 0 | 3.4 | 127 | Specific | 324.22 | 317.67 | 317.67 | 5.96 | 23 | >sp\|P01876\|IGHA1_HUMAN Immunoglobulin heavy constant alpha 1 OS=Homo sapiens OX=9606 GN=IGHA1 PE=1 SV=2 | 255 | 20201115_07433_1ng_glyco_hcd30_etd30_compound15.18981.18981.3 | controllerType=0 controllerNumber=1 scan=18981 | 42.6452 |
| 15476:3 | 3 | R.LSLHRPALEDLLLGSEAN[+1501.555]LTC[+57.021]TLTGLR.D | HexNAc(5)Hex(3) | C[+57], N[+1502] | 1489.0617 | 3 | 4465.1706 | 4465.1536 | 0 | 3.8 | 127 | Specific | 398.93 | 371.52 | 371.52 | 7.5 | 23 | >sp\|P01876\|IGHA1_HUMAN Immunoglobulin heavy constant alpha 1 OS=Homo sapiens OX=9606 GN=IGHA1 PE=1 SV=2 | 255 | 20201115_07433_1ng_glyco_hcd30_etd30_compound15.18980.18980.3 | controllerType=0 controllerNumber=1 scan=18980 | 42.6422 |
| 15416:3 | 3 | R.LSLHRPALEDLLLGSEAN[+1864.634]LTC[+57.021]TLTGLR.D | HexNAc(2)Hex(9) | C[+57], N[+1865] | 1610.0868 | 3 | 4828.246 | 4828.2324 | 0 | 2.8 | 127 | Specific | 400.01 | 355.17 | 355.17 | 6.13 | 23 | >sp\|P01876\|IGHA1_HUMAN Immunoglobulin heavy constant alpha 1 OS=Homo sapiens OX=9606 GN=IGHA1 PE=1 SV=2 | 255 | 20201115_07433_1ng_glyco_hcd30_etd30_compound15.18916.18916.3 | controllerType=0 controllerNumber=1 scan=18916 | 42.4056 |
| 15430:3 | 3 | R.LSLHRPALEDLLLGSEAN[+1540.529]LTC[+57.021]TLTGLR.D | HexNAc(2)Hex(7) | C[+57], N[+1541] | 1502.0529 | 3 | 4504.1442 | 4504.1268 | 0 | 3.9 | 127 | Specific | 422.67 | 392.63 | 392.63 | 7.6 | 23 | >sp\|P01876\|IGHA1_HUMAN Immunoglobulin heavy constant alpha 1 OS=Homo sapiens OX=9606 GN=IGHA1 PE=1 SV=2 | 255 | 20201115_07433_1ng_glyco_hcd30_etd30_compound15.18931.18931.3 | controllerType=0 controllerNumber=1 scan=18931 | 42.4492 |
| 15417:3 | 3 | R.LSLHRPALEDLLLGSEAN[+1702.581]LTC[+57.021]TLTGLR.D | HexNAc(2)Hex(8) | C[+57], N[+1703] | 1556.0698 | 3 | 4666.1947 | 4666.1796 | 0 | 3.2 | 127 | Specific | 442.93 | 442.93 | 442.93 | 7.75 | 23 | >sp\|P01876\|IGHA1_HUMAN Immunoglobulin heavy constant alpha 1 OS=Homo sapiens OX=9606 GN=IGHA1 PE=1 SV=2 | 255 | 20201115_07433_1ng_glyco_hcd30_etd30_compound15.18917.18917.3 | controllerType=0 controllerNumber=1 scan=18917 | 42.4125 |
| 15493:3 | 3 | R.LSLHRPALEDLLLGSEAN[+1216.423]LTC[+57.021]TLTGLR.D | HexNAc(2)Hex(5) | C[+57], N[+1216] | 1394.0166 | 3 | 4180.0352 | 4180.0211 | 0 | 3.4 | 127 | Specific | 469.62 | 458.54 | 458.54 | 7.75 | 23 | >sp\|P01876\|IGHA1_HUMAN Immunoglobulin heavy constant alpha 1 OS=Homo sapiens OX=9606 GN=IGHA1 PE=1 SV=2 | 255 | 20201115_07433_1ng_glyco_hcd30_etd30_compound15.18998.18998.3 | controllerType=0 controllerNumber=1 scan=18998 | 42.7045 |
| 15475:4 | 3 | R.LSLHRPALEDLLLGSEAN[+1622.582]LTC[+57.021]TLTGLR.D | HexNAc(4)Hex(5) | C[+57], N[+1623] | 1147.3039 | 4 | 4586.1937 | 4586.1799 | 0 | 3 | 127 | Specific | 640.98 | 640.98 | 640.98 | 9.72 | 23 | >sp\|P01876\|IGHA1_HUMAN Immunoglobulin heavy constant alpha 1 OS=Homo sapiens OX=9606 GN=IGHA1 PE=1 SV=2 | 255 | 20201115_07433_1ng_glyco_hcd30_etd30_compound15.18979.18979.4 | controllerType=0 controllerNumber=1 scan=18979 | 42.6361 |
| 15411:4 | 3 | R.LSLHRPALEDLLLGSEAN[+1702.581]LTC[+57.021]TLTGLR.D | HexNAc(2)Hex(8) | C[+57], N[+1703] | 1167.3034 | 4 | 4666.1917 | 4666.1796 | 0 | 2.6 | 127 | Specific | 924.91 | 906.64 | 906.64 | 15.72 | 23 | >sp\|P01876\|IGHA1_HUMAN Immunoglobulin heavy constant alpha 1 OS=Homo sapiens OX=9606 GN=IGHA1 PE=1 SV=2 | 255 | 20201115_07433_1ng_glyco_hcd30_etd30_compound15.18911.18911.4 | controllerType=0 controllerNumber=1 scan=18911 | 42.3873 |
| 15470:4 | 3 | R.LSLHRPALEDLLLGSEAN[+1501.555]LTC[+57.021]TLTGLR.D | HexNAc(5)Hex(3) | C[+57], N[+1502] | 1117.0505 | 4 | 4465.18 | 4465.1536 | 0 | 5.9 | 127 | Specific | 933 | 933 | 933 | 15.3 | 23 | >sp\|P01876\|IGHA1_HUMAN Immunoglobulin heavy constant alpha 1 OS=Homo sapiens OX=9606 GN=IGHA1 PE=1 SV=2 | 255 | 20201115_07433_1ng_glyco_hcd30_etd30_compound15.18974.18974.4 | controllerType=0 controllerNumber=1 scan=18974 | 42.6256 |
| 15471:4 | 3 | R.LSLHRPALEDLLLGSEAN[+1378.476]LTC[+57.021]TLTGLR.D | HexNAc(2)Hex(6) | C[+57], N[+1378] | 1086.2787 | 4 | 4342.0931 | 4342.074 | 0 | 4.4 | 127 | Specific | 955.13 | 877.91 | 877.91 | 17.04 | 23 | >sp\|P01876\|IGHA1_HUMAN Immunoglobulin heavy constant alpha 1 OS=Homo sapiens OX=9606 GN=IGHA1 PE=1 SV=2 | 255 | 20201115_07433_1ng_glyco_hcd30_etd30_compound15.18975.18975.4 | controllerType=0 controllerNumber=1 scan=18975 | 42.6283 |
| 15410:4 | 3 | R.LSLHRPALEDLLLGSEAN[+1864.634]LTC[+57.021]TLTGLR.D | HexNAc(2)Hex(9) | C[+57], N[+1865] | 1207.8168 | 4 | 4828.2454 | 4828.2324 | 0 | 2.7 | 127 | Specific | 991.76 | 931.05 | 931.05 | 16.72 | 23 | >sp\|P01876\|IGHA1_HUMAN Immunoglobulin heavy constant alpha 1 OS=Homo sapiens OX=9606 GN=IGHA1 PE=1 SV=2 | 255 | 20201115_07433_1ng_glyco_hcd30_etd30_compound15.18910.18910.4 | controllerType=0 controllerNumber=1 scan=18910 | 42.3839 |
| 15482:4 | 3 | R.LSLHRPALEDLLLGSEAN[+1216.423]LTC[+57.021]TLTGLR.D | HexNAc(2)Hex(5) | C[+57], N[+1216] | 1045.7644 | 4 | 4180.036 | 4180.0211 | 0 | 3.5 | 127 | Specific | 1002.15 | 949.47 | 949.47 | 18.04 | 23 | >sp\|P01876\|IGHA1_HUMAN Immunoglobulin heavy constant alpha 1 OS=Homo sapiens OX=9606 GN=IGHA1 PE=1 SV=2 | 255 | 20201115_07433_1ng_glyco_hcd30_etd30_compound15.18986.18986.4 | controllerType=0 controllerNumber=1 scan=18986 | 42.6594 |
| 15424:4 | 3 | R.LSLHRPALEDLLLGSEAN[+1540.529]LTC[+57.021]TLTGLR.D | HexNAc(2)Hex(7) | C[+57], N[+1541] | 1126.7915 | 4 | 4504.1444 | 4504.1268 | 0 | 3.9 | 127 | Specific | 1080.1 | 994.05 | 994.05 | 19.04 | 23 | >sp\|P01876\|IGHA1_HUMAN Immunoglobulin heavy constant alpha 1 OS=Homo sapiens OX=9606 GN=IGHA1 PE=1 SV=2 | 255 | 20201115_07433_1ng_glyco_hcd30_etd30_compound15.18925.18925.4 | controllerType=0 controllerNumber=1 scan=18925 | 42.438 |
| 10308:3 | 3 | R.LAGKPTHVN[+1663.608]VSVVMAEVDGTC[+57.021]Y.- | HexNAc(5)Hex(4) | C[+57], N[+1664] | 1337.5936 | 3 | 4010.7661 | 4010.7502 | 0 | 4 | 332 | CRagged | 203.9 | 128.78 | 128.78 | 2.51 | 23 | >sp\|P01876\|IGHA1_HUMAN Immunoglobulin heavy constant alpha 1 OS=Homo sapiens OX=9606 GN=IGHA1 PE=1 SV=2 | 255 | 20201115_07433_1ng_glyco_hcd30_etd30_compound15.13514.13514.3 | controllerType=0 controllerNumber=1 scan=13514 | 30.5005 |
| 10346:3 | 3 | R.LAGKPTHVN[+1501.555]VSVVMAEVDGTC[+57.021]Y.- | HexNAc(5)Hex(3) | C[+57], N[+1502] | 1283.5745 | 3 | 3848.709 | 3848.6974 | 0 | 3 | 332 | CRagged | 209.24 | 140.38 | 140.38 | 2.57 | 23 | >sp\|P01876\|IGHA1_HUMAN Immunoglobulin heavy constant alpha 1 OS=Homo sapiens OX=9606 GN=IGHA1 PE=1 SV=2 | 255 | 20201115_07433_1ng_glyco_hcd30_etd30_compound15.13554.13554.3 | controllerType=0 controllerNumber=1 scan=13554 | 30.6447 |
| 10383:3 | 3 | R.LAGKPTHVN[+1647.613]VSVVMAEVDGTC[+57.021]Y.- | HexNAc(5)Hex(3)Fuc(1) | C[+57], N[+1648] | 1332.2613 | 3 | 3994.7693 | 3994.7553 | 0 | 3.5 | 332 | CRagged | 209.58 | 98.89 | 98.89 | 2.46 | 23 | >sp\|P01876\|IGHA1_HUMAN Immunoglobulin heavy constant alpha 1 OS=Homo sapiens OX=9606 GN=IGHA1 PE=1 SV=2 | 255 | 20201115_07433_1ng_glyco_hcd30_etd30_compound15.13593.13593.3 | controllerType=0 controllerNumber=1 scan=13593 | 30.726 |
| 10356:3 | 3 | R.LAGKPTHVN[+1216.423]VSVVMAEVDGTC[+57.021]Y.- | HexNAc(2)Hex(5) | C[+57], N[+1216] | 1188.5311 | 3 | 3563.5786 | 3563.5649 | 0 | 3.8 | 332 | CRagged | 224.6 | 211.94 | 181.85 | 3.07 | 23 | >sp\|P01876\|IGHA1_HUMAN Immunoglobulin heavy constant alpha 1 OS=Homo sapiens OX=9606 GN=IGHA1 PE=1 SV=2 | 255 | 20201115_07433_1ng_glyco_hcd30_etd30_compound15.13565.13565.3 | controllerType=0 controllerNumber=1 scan=13565 | 30.669 |
| 10293:3 | 3 | R.LAGKPTHVN[+1540.529]VSVVMAEVDGTC[+57.021]Y.- | HexNAc(2)Hex(7) | C[+57], N[+1541] | 1296.5679 | 3 | 3887.6892 | 3887.6706 | 0 | 4.8 | 332 | CRagged | 241.47 | 212.66 | 182.31 | 2.85 | 23 | >sp\|P01876\|IGHA1_HUMAN Immunoglobulin heavy constant alpha 1 OS=Homo sapiens OX=9606 GN=IGHA1 PE=1 SV=2 | 255 | 20201115_07433_1ng_glyco_hcd30_etd30_compound15.13498.13498.3 | controllerType=0 controllerNumber=1 scan=13498 | 30.4591 |
| 10288:3 | 3 | R.LAGKPTHVN[+1378.476]VSVVMAEVDGTC[+57.021]Y.- | HexNAc(2)Hex(6) | C[+57], N[+1378] | 1242.5497 | 3 | 3725.6347 | 3725.6178 | 0 | 4.5 | 332 | CRagged | 249.73 | 214.1 | 202.46 | 2.85 | 23 | >sp\|P01876\|IGHA1_HUMAN Immunoglobulin heavy constant alpha 1 OS=Homo sapiens OX=9606 GN=IGHA1 PE=1 SV=2 | 255 | 20201115_07433_1ng_glyco_hcd30_etd30_compound15.13493.13493.3 | controllerType=0 controllerNumber=1 scan=13493 | 30.45 |
| 10250:3 | 3 | R.LAGKPTHVN[+1702.581]VSVVMAEVDGTC[+57.021]Y.- | HexNAc(2)Hex(8) | C[+57], N[+1703] | 1350.5823 | 3 | 4049.7324 | 4049.7234 | 0 | 2.2 | 332 | CRagged | 261.28 | 205.06 | 189.41 | 3.11 | 23 | >sp\|P01876\|IGHA1_HUMAN Immunoglobulin heavy constant alpha 1 OS=Homo sapiens OX=9606 GN=IGHA1 PE=1 SV=2 | 255 | 20201115_07433_1ng_glyco_hcd30_etd30_compound15.13453.13453.3 | controllerType=0 controllerNumber=1 scan=13453 | 30.338 |
| 16853:3 | 5 | R.GLSFDVSLEVSQGPGLLN[+2350.830]DTK.V | HexNAc(4)Hex(5)Fuc(1)NeuAc(2) | N[+2351] | 1509.659 | 3 | 4526.9624 | 4526.9436 | 0 | 4.2 | 118 | Specific | 504.5 | 342.26 | 342.26 | 6.2 | 22 | >sp\|P01833\|PIGR_HUMAN Polymeric immunoglobulin receptor OS=Homo sapiens OX=9606 GN=PIGR PE=1 SV=4 | 902 | 20201115_07433_1ng_glyco_hcd30_etd30_compound15.20438.20438.3 | controllerType=0 controllerNumber=1 scan=20438 | 46.646 |
| 16385:3 | 5 | K.QIGLYPVLVIDSSGYVNPN[+1913.677]YTGR.I | HexNAc(4)Hex(5)NeuAc(1) | N[+1914] | 1480.3401 | 3 | 4439.0059 | 4438.9805 | 0 | 5.7 | 168 | Specific | 497.76 | 393.08 | 393.08 | 6.79 | 22 | >sp\|P01833\|PIGR_HUMAN Polymeric immunoglobulin receptor OS=Homo sapiens OX=9606 GN=PIGR PE=1 SV=4 | 902 | 20201115_07433_1ng_glyco_hcd30_etd30_compound15.19939.19939.3 | controllerType=0 controllerNumber=1 scan=19939 | 45.4646 |
| 16427:3 | 5 | R.LSLLEEPGN[+1913.677]GTFTVILNQLTSR.D | HexNAc(4)Hex(5)NeuAc(1) | N[+1914] | 1439.3383 | 3 | 4316.0004 | 4315.9696 | 0 | 7.1 | 413 | Specific | 345.49 | 307.49 | 307.49 | 4.6 | 22 | >sp\|P01833\|PIGR_HUMAN Polymeric immunoglobulin receptor OS=Homo sapiens OX=9606 GN=PIGR PE=1 SV=4 | 902 | 20201115_07433_1ng_glyco_hcd30_etd30_compound15.19983.19983.3 | controllerType=0 controllerNumber=1 scan=19983 | 45.5838 |
| 19520:4 | 5 | R.LSLLEEPGN[+2350.830]GTFTVILNQLTSR.D | HexNAc(4)Hex(5)Fuc(1)NeuAc(2) | N[+2351] | 1189.0408 | 4 | 4753.1414 | 4753.1229 | 0 | 3.9 | 413 | Specific | 361.49 | 302.28 | 302.28 | 3.75 | 22 | >sp\|P01833\|PIGR_HUMAN Polymeric immunoglobulin receptor OS=Homo sapiens OX=9606 GN=PIGR PE=1 SV=4 | 902 | 20201115_07433_1ng_glyco_hcd30_etd30_compound15.23246.23246.4 | controllerType=0 controllerNumber=1 scan=23246 | 53.3825 |
| 17657:3 | 5 | R.LSLLEEPGN[+2059.735]GTFTVILNQLTSR.D | HexNAc(4)Hex(5)Fuc(1)NeuAc(1) | N[+2060] | 1488.0223 | 3 | 4462.0523 | 4462.0275 | 0 | 5.6 | 413 | Specific | 510.05 | 407.87 | 407.87 | 6.79 | 22 | >sp\|P01833\|PIGR_HUMAN Polymeric immunoglobulin receptor OS=Homo sapiens OX=9606 GN=PIGR PE=1 SV=4 | 902 | 20201115_07433_1ng_glyco_hcd30_etd30_compound15.21287.21287.3 | controllerType=0 controllerNumber=1 scan=21287 | 48.9758 |
| 14242:3 | 5 | K.IIEGEPNLKVPGN[+2205.793]VTAVLGETLK.V | HexNAc(4)Hex(5)Fuc(2)NeuAc(1) | N[+2206] | 1533.0562 | 3 | 4597.1541 | 4597.1422 | 0 | 2.6 | 457 | Specific | 295.13 | 222.79 | 222.79 | 2.72 | 22 | >sp\|P01833\|PIGR_HUMAN Polymeric immunoglobulin receptor OS=Homo sapiens OX=9606 GN=PIGR PE=1 SV=4 | 902 | 20201115_07433_1ng_glyco_hcd30_etd30_compound15.17678.17678.3 | controllerType=0 controllerNumber=1 scan=17678 | 39.2613 |
| 14258:3 | 5 | K.IIEGEPNLKVPGN[+2059.735]VTAVLGETLK.V | HexNAc(4)Hex(5)Fuc(1)NeuAc(1) | N[+2060] | 1484.3725 | 3 | 4451.1029 | 4451.0843 | 0 | 4.2 | 457 | Specific | 363.6 | 318.33 | 318.33 | 3.89 | 22 | >sp\|P01833\|PIGR_HUMAN Polymeric immunoglobulin receptor OS=Homo sapiens OX=9606 GN=PIGR PE=1 SV=4 | 902 | 20201115_07433_1ng_glyco_hcd30_etd30_compound15.17695.17695.3 | controllerType=0 controllerNumber=1 scan=17695 | 39.3213 |
| 14235:4 | 5 | K.IIEGEPNLKVPGN[+2205.793]VTAVLGETLK.V | HexNAc(4)Hex(5)Fuc(2)NeuAc(1) | N[+2206] | 1150.0512 | 4 | 4597.1829 | 4597.1422 | 0 | 8.9 | 457 | Specific | 565.31 | 555.63 | 555.63 | 5.51 | 22 | >sp\|P01833\|PIGR_HUMAN Polymeric immunoglobulin receptor OS=Homo sapiens OX=9606 GN=PIGR PE=1 SV=4 | 902 | 20201115_07433_1ng_glyco_hcd30_etd30_compound15.17670.17670.2 | controllerType=0 controllerNumber=1 scan=17670 | 39.2319 |
| 13859:4 | 5 | K.IIEGEPNLKVPGN[+2059.735]VTAVLGETLKVPC[+57.021]HFPC[+57.021]K.F | HexNAc(4)Hex(5)Fuc(1)NeuAc(1) | C[+57]*2, N[+2060] | 1369.8864 | 4 | 5476.5236 | 5476.5418 | 0 | -3.3 | 457 | Specific | 319.82 | 319.82 | 319.82 | 2.63 | 22 | >sp\|P01833\|PIGR_HUMAN Polymeric immunoglobulin receptor OS=Homo sapiens OX=9606 GN=PIGR PE=1 SV=4 | 902 | 20201115_07433_1ng_glyco_hcd30_etd30_compound15.17238.17238.4 | controllerType=0 controllerNumber=1 scan=17238 | 38.3154 |
| 15010:4 | 5 | K.IIEGEPNLKVPGN[+2350.830]VTAVLGETLKVPC[+57.021]HFPC[+57.021]K.F | HexNAc(4)Hex(5)Fuc(1)NeuAc(2) | C[+57]*2, N[+2351] | 1442.673 | 4 | 5767.6701 | 5767.6372 | 0 | 5.7 | 457 | Specific | 336.85 | 336.85 | 336.85 | 3.83 | 22 | >sp\|P01833\|PIGR_HUMAN Polymeric immunoglobulin receptor OS=Homo sapiens OX=9606 GN=PIGR PE=1 SV=4 | 902 | 20201115_07433_1ng_glyco_hcd30_etd30_compound15.18486.18486.4 | controllerType=0 controllerNumber=1 scan=18486 | 41.1764 |
| 13845:4 | 5 | K.IIEGEPNLKVPGN[+2059.735]VTAVLGETLKVPC[+57.021]HFPC[+57.021]K.F | HexNAc(4)Hex(5)Fuc(1)NeuAc(1) | C[+57]*2, N[+2060] | 1369.8915 | 4 | 5476.5441 | 5476.5418 | 0 | 0.4 | 457 | Specific | 536.07 | 536.07 | 536.07 | 6.01 | 22 | >sp\|P01833\|PIGR_HUMAN Polymeric immunoglobulin receptor OS=Homo sapiens OX=9606 GN=PIGR PE=1 SV=4 | 902 | 20201115_07433_1ng_glyco_hcd30_etd30_compound15.17223.17223.4 | controllerType=0 controllerNumber=1 scan=17223 | 38.2595 |
| 13844:4 | 5 | K.IIEGEPNLKVPGN[+2205.793]VTAVLGETLKVPC[+57.021]HFPC[+57.021]K.F | HexNAc(4)Hex(5)Fuc(2)NeuAc(1) | C[+57]*2, N[+2206] | 1406.4117 | 4 | 5622.6252 | 5622.5997 | 0 | 4.5 | 457 | Specific | 549.85 | 549.85 | 549.85 | 6.53 | 22 | >sp\|P01833\|PIGR_HUMAN Polymeric immunoglobulin receptor OS=Homo sapiens OX=9606 GN=PIGR PE=1 SV=4 | 902 | 20201115_07433_1ng_glyco_hcd30_etd30_compound15.17222.17222.4 | controllerType=0 controllerNumber=1 scan=17222 | 38.2548 |
| 13839:5 | 5 | K.IIEGEPNLKVPGN[+2205.793]VTAVLGETLKVPC[+57.021]HFPC[+57.021]K.F | HexNAc(4)Hex(5)Fuc(2)NeuAc(1) | C[+57]*2, N[+2206] | 1125.3303 | 5 | 5622.6225 | 5622.5997 | 0 | 4.1 | 457 | Specific | 589.04 | 589.04 | 589.04 | 7.55 | 22 | >sp\|P01833\|PIGR_HUMAN Polymeric immunoglobulin receptor OS=Homo sapiens OX=9606 GN=PIGR PE=1 SV=4 | 902 | 20201115_07433_1ng_glyco_hcd30_etd30_compound15.17217.17217.5 | controllerType=0 controllerNumber=1 scan=17217 | 38.2433 |
| 13855:5 | 5 | K.IIEGEPNLKVPGN[+2059.735]VTAVLGETLKVPC[+57.021]HFPC[+57.021]K.F | HexNAc(4)Hex(5)Fuc(1)NeuAc(1) | C[+57]*2, N[+2060] | 1096.1169 | 5 | 5476.5556 | 5476.5418 | 0 | 2.5 | 457 | Specific | 664.39 | 664.39 | 664.39 | 9.02 | 22 | >sp\|P01833\|PIGR_HUMAN Polymeric immunoglobulin receptor OS=Homo sapiens OX=9606 GN=PIGR PE=1 SV=4 | 902 | 20201115_07433_1ng_glyco_hcd30_etd30_compound15.17234.17234.5 | controllerType=0 controllerNumber=1 scan=17234 | 38.3055 |
| 13828:4 | 5 | K.VPGN[+2351.851]VTAVLGETLKVPC[+57.021]HFPC[+57.021]K.F | HexNAc(4)Hex(5)Fuc(3)NeuAc(1) | C[+57]*2, N[+2352] | 1194.5334 | 4 | 4775.1116 | 4775.1081 | 0 | 0.7 | 466 | Specific | 234.56 | 234.56 | 234.56 | 2.43 | 22 | >sp\|P01833\|PIGR_HUMAN Polymeric immunoglobulin receptor OS=Homo sapiens OX=9606 GN=PIGR PE=1 SV=4 | 902 | 20201115_07433_1ng_glyco_hcd30_etd30_compound15.17205.17205.4 | controllerType=0 controllerNumber=1 scan=17205 | 38.1989 |
| 12340:4 | 5 | K.VPGN[+2205.793]VTAVLGETLKVPC[+57.021]HFPC[+57.021]K.F | HexNAc(4)Hex(5)Fuc(2)NeuAc(1) | C[+57]*2, N[+2206] | 1158.023 | 4 | 4629.0701 | 4629.0502 | 0 | 4.3 | 466 | Specific | 268.31 | 260.35 | 260.35 | 2.62 | 22 | >sp\|P01833\|PIGR_HUMAN Polymeric immunoglobulin receptor OS=Homo sapiens OX=9606 GN=PIGR PE=1 SV=4 | 902 | 20201115_07433_1ng_glyco_hcd30_etd30_compound15.15652.15652.4 | controllerType=0 controllerNumber=1 scan=15652 | 34.665 |
| 05281:3 | 5 | K.WN[+2059.735]NTGC[+57.021]QALPSQDEGPSK.A | HexNAc(4)Hex(5)Fuc(1)NeuAc(1) | C[+57], N[+2060] | 1350.2108 | 3 | 4048.6177 | 4048.6116 | 0 | 1.5 | 498 | Specific | 256.76 | 123.81 | 123.81 | 3.7 | 22 | >sp\|P01833\|PIGR_HUMAN Polymeric immunoglobulin receptor OS=Homo sapiens OX=9606 GN=PIGR PE=1 SV=4 | 902 | 20201115_07433_1ng_glyco_hcd30_etd30_compound15.8282.8282.3 | controllerType=0 controllerNumber=1 scan=8282 | 20.7514 |
| 05288:3 | 5 | K.WN[+1913.677]NTGC[+57.021]QALPSQDEGPSK.A | HexNAc(4)Hex(5)NeuAc(1) | C[+57], N[+1914] | 1301.525 | 3 | 3902.5603 | 3902.5537 | 0 | 1.7 | 498 | Specific | 564.56 | 369.69 | 369.69 | 7.97 | 22 | >sp\|P01833\|PIGR_HUMAN Polymeric immunoglobulin receptor OS=Homo sapiens OX=9606 GN=PIGR PE=1 SV=4 | 902 | 20201115_07433_1ng_glyco_hcd30_etd30_compound15.8289.8289.3 | controllerType=0 controllerNumber=1 scan=8289 | 20.7754 |
| 14546:3 | 8 | R.VWIALNSN[+2205.793]LTDNQYTWTDKWR.V | HexNAc(4)Hex(5)Fuc(2)NeuAc(1) | N[+2206] | 1610.7065 | 3 | 4830.1049 | 4830.0821 | 0 | 4.7 | 1153 | Specific | 304.82 | 204.79 | 204.79 | 2.88 | 11 | >sp\|P22897\|MRC1_HUMAN Macrophage mannose receptor 1 OS=Homo sapiens OX=9606 GN=MRC1 PE=1 SV=1 | 1049 | 20201115_07433_1ng_glyco_hcd30_etd30_compound15.17998.17998.3 | controllerType=0 controllerNumber=1 scan=17998 | 39.9387 |
| 07641:3 | 8 | K.TAHC[+57.021]N[+2205.793]ESFYFLC[+57.021]K.R | HexNAc(4)Hex(5)Fuc(2)NeuAc(1) | C[+57]*2, N[+2206] | 1294.8475 | 3 | 3882.5279 | 3882.5124 | 0 | 4 | 1201 | Specific | 252.1 | 120.34 | 120.34 | 4.1 | 11 | >sp\|P22897\|MRC1_HUMAN Macrophage mannose receptor 1 OS=Homo sapiens OX=9606 GN=MRC1 PE=1 SV=1 | 1049 | 20201115_07433_1ng_glyco_hcd30_etd30_compound15.10733.10733.3 | controllerType=0 controllerNumber=1 scan=10733 | 25.2564 |
| 07728:3 | 8 | K.TAHC[+57.021]N[+2059.735]ESFYFLC[+57.021]K.R | HexNAc(4)Hex(5)Fuc(1)NeuAc(1) | C[+57]*2, N[+2060] | 1246.158 | 3 | 3736.4595 | 3736.4545 | 0 | 1.3 | 1201 | Specific | 318.5 | 203.53 | 203.53 | 4.45 | 11 | >sp\|P22897\|MRC1_HUMAN Macrophage mannose receptor 1 OS=Homo sapiens OX=9606 GN=MRC1 PE=1 SV=1 | 1049 | 20201115_07433_1ng_glyco_hcd30_etd30_compound15.10824.10824.3 | controllerType=0 controllerNumber=1 scan=10824 | 25.438 |
| 07081:3 | 10 | R.LSPN[+2350.830]ASAEHLELR.W | HexNAc(4)Hex(5)Fuc(1)NeuAc(2) | N[+2351] | 1263.202 | 3 | 3787.5913 | 3787.5796 | 0 | 3.1 | 52 | Specific | 334.96 | 201.93 | 201.93 | 4.63 | 7 | >sp\|Q13410\|BT1A1_HUMAN Butyrophilin subfamily 1 member A1 OS=Homo sapiens OX=9606 GN=BTN1A1 PE=1 SV=3 | 179 | 20201115_07433_1ng_glyco_hcd30_etd30_compound15.10147.10147.3 | controllerType=0 controllerNumber=1 scan=10147 | 24.1193 |
| 05233:3 | 10 | R.LSPN[+2205.793]ASAEHLELR.W | HexNAc(4)Hex(5)Fuc(2)NeuAc(1) | N[+2206] | 1214.8563 | 3 | 3642.5542 | 3642.5421 | 0 | 3.3 | 52 | Specific | 393.8 | 250.03 | 250.03 | 5.04 | 7 | >sp\|Q13410\|BT1A1_HUMAN Butyrophilin subfamily 1 member A1 OS=Homo sapiens OX=9606 GN=BTN1A1 PE=1 SV=3 | 179 | 20201115_07433_1ng_glyco_hcd30_etd30_compound15.8232.8232.3 | controllerType=0 controllerNumber=1 scan=8232 | 20.6383 |
| 01953:4 | 12 | R.EKQTDEIKDTRN[+2059.735]ESTQNC[+57.021]VVAEPEK.M | HexNAc(4)Hex(5)Fuc(1)NeuAc(1) | C[+57], N[+2060] | 1252.7878 | 4 | 5008.1296 | 5008.1251 | 0 | 0.9 | 58 | Specific | 473.45 | 438.53 | 438.53 | 5.06 | 6 | >sp\|P47710\|CASA1_HUMAN Alpha-S1-casein OS=Homo sapiens OX=9606 GN=CSN1S1 PE=1 SV=1 | 989 | 20201115_07433_1ng_glyco_hcd30_etd30_compound15.4830.4830.4 | controllerType=0 controllerNumber=1 scan=4830 | 14.488 |
| 08600:4 | 12 | K.QTDEIKDTRN[+2059.735]ESTQNC[+57.021]VVAEPEKMESSISSSSEEMSLSK.C | HexNAc(4)Hex(5)Fuc(1)NeuAc(1) | C[+57], N[+2060] | 1613.4419 | 4 | 6450.7458 | 6450.7156 | 0 | 4.7 | 60 | Specific | 404.99 | 404.99 | 404.99 | 5.52 | 6 | >sp\|P47710\|CASA1_HUMAN Alpha-S1-casein OS=Homo sapiens OX=9606 GN=CSN1S1 PE=1 SV=1 | 989 | 20201115_07433_1ng_glyco_hcd30_etd30_compound15.11742.11742.4 | controllerType=0 controllerNumber=1 scan=11742 | 27.2144 |
| 00895:3 | 17 | K.TKPREEQFN[+1444.534]STFR.V | HexNAc(4)Hex(3)Fuc(1) | N[+1445] | 1028.7926 | 3 | 3084.3633 | 3084.3526 | 0 | 3.5 | 168 | Specific | 298.5 | 197.98 | 197.98 | 3.68 | 5 | >sp\|P01859\|IGHG2_HUMAN Immunoglobulin heavy constant gamma 2 OS=Homo sapiens OX=9606 GN=IGHG2 PE=1 SV=2 | 387 | 20201115_07433_1ng_glyco_hcd30_etd30_compound15.3730.3730.3 | controllerType=0 controllerNumber=1 scan=3730 | 12.5232 |
| 00858:4 | 17 | K.TKPREEQFN[+1606.587]STFR.V | HexNAc(4)Hex(4)Fuc(1) | N[+1607] | 812.3606 | 4 | 3246.4206 | 3246.4054 | 0 | 4.7 | 168 | Specific | 640.56 | 569.16 | 569.16 | 8.54 | 5 | >sp\|P01859\|IGHG2_HUMAN Immunoglobulin heavy constant gamma 2 OS=Homo sapiens OX=9606 GN=IGHG2 PE=1 SV=2 | 387 | 20201115_07433_1ng_glyco_hcd30_etd30_compound15.3691.3691.2 | controllerType=0 controllerNumber=1 scan=3691 | 12.4203 |
| 00890:4 | 17 | K.TKPREEQFN[+1444.534]STFR.V | HexNAc(4)Hex(3)Fuc(1) | N[+1445] | 771.8462 | 4 | 3084.363 | 3084.3526 | 0 | 3.4 | 168 | Specific | 680.92 | 565.28 | 565.28 | 8.99 | 5 | >sp\|P01859\|IGHG2_HUMAN Immunoglobulin heavy constant gamma 2 OS=Homo sapiens OX=9606 GN=IGHG2 PE=1 SV=2 | 387 | 20201115_07433_1ng_glyco_hcd30_etd30_compound15.3725.3725.4 | controllerType=0 controllerNumber=1 scan=3725 | 12.5079 |
| 20441:3 | 19 | R.QLAHQSN[+2204.772]STNIFFSPVSIATAFAMLSLGTK.A | HexNAc(4)Hex(5)NeuAc(2) | N[+2205] | 1796.1508 | 3 | 5386.438 | 5386.4075 | 0 | 5.7 | 64 | Specific | 413.99 | 289.42 | 289.42 | 5.37 | 4 | >sp\|P01009\|A1AT_HUMAN Alpha-1-antitrypsin OS=Homo sapiens OX=9606 GN=SERPINA1 PE=1 SV=3 | 642 | 20201115_07433_1ng_glyco_hcd30_etd30_compound15.24213.24213.3 | controllerType=0 controllerNumber=1 scan=24213 | 55.8069 |
| 20434:4 | 19 | R.QLAHQSN[+2204.772]STNIFFSPVSIATAFAMLSLGTK.A | HexNAc(4)Hex(5)NeuAc(2) | N[+2205] | 1347.364 | 4 | 5386.4342 | 5386.4075 | 0 | 5 | 64 | Specific | 619.29 | 555.68 | 555.68 | 8.45 | 4 | >sp\|P01009\|A1AT_HUMAN Alpha-1-antitrypsin OS=Homo sapiens OX=9606 GN=SERPINA1 PE=1 SV=3 | 642 | 20201115_07433_1ng_glyco_hcd30_etd30_compound15.24206.24206.4 | controllerType=0 controllerNumber=1 scan=24206 | 55.7786 |
| 17217:5 | 19 | K.YLGN[+2204.772]ATAIFFLPDEGKLQHLENELTHDIITK.F | HexNAc(4)Hex(5)NeuAc(2) | N[+2205] | 1149.9314 | 5 | 5745.6279 | 5745.6098 | 0 | 3.2 | 268 | Specific | 241.35 | 239.38 | 239.38 | 2.7 | 4 | >sp\|P01009\|A1AT_HUMAN Alpha-1-antitrypsin OS=Homo sapiens OX=9606 GN=SERPINA1 PE=1 SV=3 | 642 | 20201115_07433_1ng_glyco_hcd30_etd30_compound15.20822.20822.5 | controllerType=0 controllerNumber=1 scan=20822 | 47.6486 |
| 17211:4 | 19 | K.YLGN[+2204.772]ATAIFFLPDEGKLQHLENELTHDIITK.F | HexNAc(4)Hex(5)NeuAc(2) | N[+2205] | 1437.1642 | 4 | 5745.6349 | 5745.6098 | 0 | 4.4 | 268 | Specific | 396.91 | 396.91 | 396.91 | 5.98 | 4 | >sp\|P01009\|A1AT_HUMAN Alpha-1-antitrypsin OS=Homo sapiens OX=9606 GN=SERPINA1 PE=1 SV=3 | 642 | 20201115_07433_1ng_glyco_hcd30_etd30_compound15.20816.20816.4 | controllerType=0 controllerNumber=1 scan=20816 | 47.6267 |
| 02682:3 | 37 | K.TPLTAN[+1971.719]ITK.S | HexNAc(5)Hex(5)Fuc(1) | N[+1972] | 977.4326 | 3 | 2930.2832 | 2930.2757 | 0 | 2.6 | 200 | Specific | 221.8 | 116.97 | 116.97 | 2.42 | 7 | >sp\|P01877\|IGHA2_HUMAN Immunoglobulin heavy constant alpha 2 OS=Homo sapiens OX=9606 GN=IGHA2 PE=1 SV=4 | 576 | 20201115_07433_1ng_glyco_hcd30_etd30_compound15.5587.5587.3 | controllerType=0 controllerNumber=1 scan=5587 | 15.8993 |
| 02712:3 | 37 | K.TPLTAN[+1809.666]ITK.S | HexNAc(5)Hex(4)Fuc(1) | N[+1810] | 923.4148 | 3 | 2768.2299 | 2768.2228 | 0 | 2.6 | 200 | Specific | 293.04 | 129.53 | 129.53 | 2.46 | 7 | >sp\|P01877\|IGHA2_HUMAN Immunoglobulin heavy constant alpha 2 OS=Homo sapiens OX=9606 GN=IGHA2 PE=1 SV=4 | 576 | 20201115_07433_1ng_glyco_hcd30_etd30_compound15.5619.5619.3 | controllerType=0 controllerNumber=1 scan=5619 | 15.992 |
| 02751:2 | 37 | K.TPLTAN[+1647.613]ITK.S | HexNAc(5)Hex(3)Fuc(1) | N[+1648] | 1303.5918 | 2 | 2606.1764 | 2606.17 | 0 | 2.5 | 200 | Specific | 293.82 | 110.75 | 110.75 | 2.53 | 7 | >sp\|P01877\|IGHA2_HUMAN Immunoglobulin heavy constant alpha 2 OS=Homo sapiens OX=9606 GN=IGHA2 PE=1 SV=4 | 576 | 20201115_07433_1ng_glyco_hcd30_etd30_compound15.5660.5660.2 | controllerType=0 controllerNumber=1 scan=5660 | 16.1036 |
| 02717:2 | 37 | K.TPLTAN[+1809.666]ITK.S | HexNAc(5)Hex(4)Fuc(1) | N[+1810] | 1384.6178 | 2 | 2768.2284 | 2768.2228 | 0 | 2 | 200 | Specific | 302.18 | 108.74 | 108.74 | 2.43 | 7 | >sp\|P01877\|IGHA2_HUMAN Immunoglobulin heavy constant alpha 2 OS=Homo sapiens OX=9606 GN=IGHA2 PE=1 SV=4 | 576 | 20201115_07433_1ng_glyco_hcd30_etd30_compound15.5624.5624.2 | controllerType=0 controllerNumber=1 scan=5624 | 16.0157 |
| 02752:3 | 37 | K.TPLTAN[+1647.613]ITK.S | HexNAc(5)Hex(3)Fuc(1) | N[+1648] | 869.3972 | 3 | 2606.1772 | 2606.17 | 0 | 2.8 | 200 | Specific | 309.81 | 139.09 | 139.09 | 2.46 | 7 | >sp\|P01877\|IGHA2_HUMAN Immunoglobulin heavy constant alpha 2 OS=Homo sapiens OX=9606 GN=IGHA2 PE=1 SV=4 | 576 | 20201115_07433_1ng_glyco_hcd30_etd30_compound15.5661.5661.3 | controllerType=0 controllerNumber=1 scan=5661 | [illegible] |
| 02792:3 | 37 | K.TPLTAN[+1444.534]ITK.S | HexNAc(4)Hex(3)Fuc(1) | N[+1445] | 1202.0519 | 2 | 2403.0966 | 2403.0906 | 0 | 2.5 | 200 | Specific | 326.63 | 96.38 | 96.38 | 2.42 | 7 | >sp\|P01877\|IGHA2_HUMAN Immunoglobulin heavy constant alpha 2 OS=Homo sapiens OX=9606 GN=IGHA2 PE=1 SV=4 | 576 | 20201115_07433_1ng_glyco_hcd30_etd30_compound15.5703.5703.2 | controllerType=0 controllerNumber=1 scan=5703 | 16.2296 |
| 12910:3 | 41 | K.VVLHPN[+2205.793]YSQVDIGLIK.L | HexNAc(4)Hex(5)Fuc(2)NeuAc(1) | N[+2206] | 1334.2729 | 3 | 4000.8041 | 4000.8041 | 0 | 0 | 236 | Specific | 213.86 | 137.94 | 137.94 | 2.63 | 3 | >sp\|P00738\|HPT_HUMAN Haptoglobin OS=Homo sapiens OX=9606 GN=HP PE=1 SV=1 | 407 | 20201115_07433_1ng_glyco_hcd30_etd30_compound15.16245.16245.3 | controllerType=0 controllerNumber=1 scan=16245 | 35.9168 |
| 10274:3 | 60 | R.IIVPLNNREN[+1913.677]ISDPTSPLR.T | HexNAc(4)Hex(5)NeuAc(1) | N[+1914] | 1354.6279 | 3 | 4061.869 | 4061.8542 | 0 | 3.7 | 62 | Specific | 274.7 | 223.37 | 223.37 | 3.14 | 4 | >sp\|P01591\|IGJ_HUMAN Immunoglobulin J chain OS=Homo sapiens OX=9606 GN=JCHAIN PE=1 SV=4 | 1392 | 20201115_07433_1ng_glyco_hcd30_etd30_compound15.13478.13478.3 | controllerType=0 controllerNumber=1 scan=13478 | 30.4083 |
| 10243:3 | 60 | R.IIVPLNNREN[+2059.735]ISDPTSPLR.T | HexNAc(4)Hex(5)Fuc(1)NeuAc(1) | N[+2060] | 1403.3132 | 3 | 4207.9249 | 4207.9121 | 0 | 3.1 | 62 | Specific | 316.58 | 244.37 | 244.37 | 3.08 | 4 | >sp\|P01591\|IGJ_HUMAN Immunoglobulin J chain OS=Homo sapiens OX=9606 GN=JCHAIN PE=1 SV=4 | 1392 | 20201115_07433_1ng_glyco_hcd30_etd30_compound15.13445.13445.3 | controllerType=0 controllerNumber=1 scan=13445 | 30.3154 |
| 12655:4 | 61 | K.LKELPGVC[+57.021]N[+2059.735]ETMMALWEEC[+57.021]KPC[+57.021]LK.Q | HexNAc(4)Hex(5)Fuc(1)NeuAc(1) | C[+57]*3, N[+2060] | 1249.7904 | 4 | 4996.1398 | 4996.1196 | 0 | 4 | 95 | Specific | 432.12 | 331.13 | 331.13 | 3.88 | 3 | >sp\|P10909\|CLUS_HUMAN Clusterin OS=Homo sapiens OX=9606 GN=CLU PE=1 SV=1 | 1467 | 20201115_07433_1ng_glyco_hcd30_etd30_compound15.15980.15980.4 | controllerType=0 controllerNumber=1 scan=15980 | 35.3497 |
| 00256:4 | 61 | R.EIRHN[+2059.735]STGC[+57.021]LR.M | HexNAc(4)Hex(5)Fuc(1)NeuAc(1) | C[+57], N[+2060] | 851.359 | 4 | 3402.4143 | 3402.3994 | 0 | 4.4 | 287 | Specific | 342.19 | 241.88 | 241.88 | 3.86 | 3 | >sp\|P10909\|CLUS_HUMAN Clusterin OS=Homo sapiens OX=9606 GN=CLU PE=1 SV=1 | 1467 | 20201115_07433_1ng_glyco_hcd30_etd30_compound15.3062.3062.4 | controllerType=0 controllerNumber=1 scan=3062 | 11.2498 |
| 00284:4 | 72 | K.TKPREEQYN[+1606.587]STYR.V | HexNAc(4)Hex(4)Fuc(1) | N[+1607] | 820.3561 | 4 | 3278.4026 | 3278.3952 | 0 | 2.2 | 291 | Specific | 352.72 | 56.34 | 56.34 | 2.84 | 2 | >sp\|P0DOX5\|IGG1_HUMAN Immunoglobulin gamma-1 heavy chain OS=Homo sapiens OX=9606 PE=1 SV=2 | 914 | 20201115_07433_1ng_glyco_hcd30_etd30_compound15.3093.3093.4 | controllerType=0 controllerNumber=1 scan=3093 | 11.3405 |
| 14462:4 | 1 | R.RTAGWNVPIGTLRPFLN[+2350.830]WTGPPEPIEAAVAR.F | HexNAc(4)Hex(5)Fuc(1)NeuAc(2) | N[+2351] | 1435.1697 | 4 | 5737.6569 | 5737.6424 | 0 | 2.5 | 140 | Specific | 538.24 | 538.24 | 538.24 | 8.04 | 57 | >sp\|P02788\|TRFL_HUMAN Lactotransferrin OS=Homo sapiens OX=9606 GN=LTF PE=1 SV=6 | 1270 | 20201117_08512_1ng_glyco_hcd30_etd30_compound15.18041.18041.4 | controllerType=0 controllerNumber=1 scan=18041 | 46.0029 |
| 13642:4 | 1 | R.RTAGWNVPIGTLRPFLN[+2570.925]WTGPPEPIEAAVAR.F | HexNAc(5)Hex(6)Fuc(2)NeuAc(1) | N[+2571] | 1490.1947 | 4 | 5957.757 | 5957.7371 | 0 | 3.3 | 140 | Specific | 706.6 | 675.02 | 675.02 | 12.09 | 57 | >sp\|P02788\|TRFL_HUMAN Lactotransferrin OS=Homo sapiens OX=9606 GN=LTF PE=1 SV=6 | 1270 | 20201117_08512_1ng_glyco_hcd30_etd30_compound15.17173.17173.4 | controllerType=0 controllerNumber=1 scan=17173 | 43.3253 |
| 14405:5 | 1 | R.RTAGWNVPIGTLRPFLN[+2350.830]WTGPPEPIEAAVAR.F | HexNAc(4)Hex(5)Fuc(1)NeuAc(2) | N[+2351] | 1148.3342 | 5 | 5737.642 | 5737.6424 | 0 | -0.1 | 140 | Specific | 780.39 | 780.39 | 780.39 | 12.6 | 57 | >sp\|P02788\|TRFL_HUMAN Lactotransferrin OS=Homo sapiens OX=9606 GN=LTF PE=1 SV=6 | 1270 | 20201117_08512_1ng_glyco_hcd30_etd30_compound15.17981.17981.5 | controllerType=0 controllerNumber=1 scan=17981 | 45.8338 |
| 13723:4 | 1 | R.RTAGWNVPIGTLRPFLN[+2205.793]WTGPPEPIEAAVAR.F | HexNAc(4)Hex(5)Fuc(2)NeuAc(1) | N[+2206] | 1398.9076 | 4 | 5592.6086 | 5592.6049 | 0 | 0.7 | 140 | Specific | 882.98 | 853.26 | 853.26 | 15.08 | 57 | >sp\|P02788\|TRFL_HUMAN Lactotransferrin OS=Homo sapiens OX=9606 GN=LTF PE=1 SV=6 | 1270 | 20201117_08512_1ng_glyco_hcd30_etd30_compound15.17258.17258.4 | controllerType=0 controllerNumber=1 scan=17258 | 43.5648 |
| 14549:3 | 1 | R.TAGWNVPIGTLRPFLN[+2205.793]WTGPPEPIEAAVAR.F | HexNAc(4)Hex(5)Fuc(2)NeuAc(1) | N[+2206] | 1812.8438 | 3 | 5436.517 | 5436.5038 | 0 | 2.4 | 141 | Specific | 271.16 | 200.34 | 200.34 | 4.31 | 57 | >sp\|P02788\|TRFL_HUMAN Lactotransferrin OS=Homo sapiens OX=9606 GN=LTF PE=1 SV=6 | 1270 | 20201117_08512_1ng_glyco_hcd30_etd30_compound15.18133.18133.3 | controllerType=0 controllerNumber=1 scan=18133 | 46.3026 |
| 14495:3 | 1 | R.TAGWNVPIGTLRPFLN[+2570.925]WTGPPEPIEAAVAR.F | HexNAc(5)Hex(6)Fuc(2)NeuAc(1) | N[+2571] | 1934.5562 | 3 | 5801.6542 | 5801.636 | 0 | 3.1 | 141 | Specific | 328.18 | 246.94 | 246.94 | 4.88 | 57 | >sp\|P02788\|TRFL_HUMAN Lactotransferrin OS=Homo sapiens OX=9606 GN=LTF PE=1 SV=6 | 1270 | 20201117_08512_1ng_glyco_hcd30_etd30_compound15.18076.18076.3 | controllerType=0 controllerNumber=1 scan=18076 | 46.114 |
| 15446:3 | 1 | R.TAGWNVPIGTLRPFLN[+2350.830]WTGPPEPIEAAVAR.F | HexNAc(4)Hex(5)Fuc(1)NeuAc(2) | N[+2351] | 1861.1921 | 3 | 5581.5618 | 5581.5413 | 0 | 3.7 | 141 | Specific | 346.17 | 224.07 | 224.07 | 5.8 | 57 | >sp\|P02788\|TRFL_HUMAN Lactotransferrin OS=Homo sapiens OX=9606 GN=LTF PE=1 SV=6 | 1270 | 20201117_08512_1ng_glyco_hcd30_etd30_compound15.19086.19086.3 | controllerType=0 controllerNumber=1 scan=19086 | 49.3438 |
| 14604:3 | 1 | R.TAGWNVPIGTLRPFLN[+2059.735]WTGPPEPIEAAVAR.F | HexNAc(4)Hex(5)Fuc(1)NeuAc(1) | N[+2060] | 1764.1574 | 3 | 5290.4577 | 5290.4459 | 0 | 2.2 | 141 | Specific | 348.61 | 268.96 | 268.96 | 5.99 | 57 | >sp\|P02788\|TRFL_HUMAN Lactotransferrin OS=Homo sapiens OX=9606 GN=LTF PE=1 SV=6 | 1270 | 20201117_08512_1ng_glyco_hcd30_etd30_compound15.18191.18191.2 | controllerType=0 controllerNumber=1 scan=18191 | 46.4908 |
| 13875:3 | 1 | R.TAGWNVPIGTLRPFLN[+1768.640]WTGPPEPIEAAVAR.F | HexNAc(4)Hex(5)Fuc(1) | N[+1769] | 1667.1269 | 3 | 4999.3662 | 4999.3504 | 0 | 3.2 | 141 | Specific | 381.38 | 294.19 | 294.19 | 4.9 | 57 | >sp\|P02788\|TRFL_HUMAN Lactotransferrin OS=Homo sapiens OX=9606 GN=LTF PE=1 SV=6 | 1270 | 20201117_08512_1ng_glyco_hcd30_etd30_compound15.17421.17421.3 | controllerType=0 controllerNumber=1 scan=17421 | 44.0841 |
| 14550:3 | 1 | R.TAGWNVPIGTLRPFLN[+2059.735]WTGPPEPIEAAVAR.F | HexNAc(4)Hex(5)Fuc(1)NeuAc(1) | N[+2060] | 1764.1587 | 3 | 5290.4614 | 5290.4459 | 0 | 2.9 | 141 | Specific | 452.3 | 357.82 | 357.82 | 5.51 | 57 | >sp\|P02788\|TRFL_HUMAN Lactotransferrin OS=Homo sapiens OX=9606 GN=LTF PE=1 SV=6 | 1270 | 20201117_08512_1ng_glyco_hcd30_etd30_compound15.18134.18134.3 | controllerType=0 controllerNumber=1 scan=18134 | 46.3093 |
| 13880:4 | 1 | R.TAGWNVPIGTLRPFLN[+1913.677]WTGPPEPIEAAVAR.F | HexNAc(4)Hex(5)NeuAc(1) | N[+1914] | 1286.8615 | 4 | 5144.424 | 5144.388 | 0 | 7 | 141 | Specific | 471.87 | 471.87 | 471.87 | 5.51 | 57 | >sp\|P02788\|TRFL_HUMAN Lactotransferrin OS=Homo sapiens OX=9606 GN=LTF PE=1 SV=6 | 1270 | 20201117_08512_1ng_glyco_hcd30_etd30_compound15.17426.17426.4 | controllerType=0 controllerNumber=1 scan=17426 | 44.0947 |
| 14473:4 | 1 | R.TAGWNVPIGTLRPFLN[+2570.925]WTGPPEPIEAAVAR.F | HexNAc(5)Hex(6)Fuc(2)NeuAc(1) | N[+2571] | 1451.1692 | 4 | 5801.6549 | 5801.636 | 0 | 3.3 | 141 | Specific | 602.01 | 602.01 | 602.01 | 10.47 | 57 | >sp\|P02788\|TRFL_HUMAN Lactotransferrin OS=Homo sapiens OX=9606 GN=LTF PE=1 SV=6 | 1270 | 20201117_08512_1ng_glyco_hcd30_etd30_compound15.18053.18053.4 | controllerType=0 controllerNumber=1 scan=18053 | 46.0432 |
| 14489:4 | 1 | R.TAGWNVPIGTLRPFLN[+2570.925]WTGPPEPIEAAVAR.F | HexNAc(5)Hex(6)Fuc(2)NeuAc(1) | N[+2571] | 1451.1686 | 4 | 5801.6525 | 5801.636 | 0 | 2.8 | 141 | Specific | 660.51 | 653.52 | 653.52 | 11.47 | 57 | >sp\|P02788\|TRFL_HUMAN Lactotransferrin OS=Homo sapiens OX=9606 GN=LTF PE=1 SV=6 | 1270 | 20201117_08512_1ng_glyco_hcd30_etd30_compound15.18070.18070.4 | controllerType=0 controllerNumber=1 scan=18070 | 46.0976 |
| 14509:4 | 1 | R.TAGWNVPIGTLRPFLN[+2424.867]WTGPPEPIEAAVAR.F | HexNAc(5)Hex(6)Fuc(1)NeuAc(1) | N[+2425] | 1414.6542 | 4 | 5655.5949 | 5655.5781 | 0 | 3 | 141 | Specific | 784.66 | 678.1 | 678.1 | 13.47 | 57 | >sp\|P02788\|TRFL_HUMAN Lactotransferrin OS=Homo sapiens OX=9606 GN=LTF PE=1 SV=6 | 1270 | 20201117_08512_1ng_glyco_hcd30_etd30_compound15.18091.18091.4 | controllerType=0 controllerNumber=1 scan=18091 | 46.158 |
| 14544:4 | 1 | R.TAGWNVPIGTLRPFLN[+2205.793]WTGPPEPIEAAVAR.F | HexNAc(4)Hex(5)Fuc(2)NeuAc(1) | N[+2206] | 1359.884 | 4 | 5436.5143 | 5436.5038 | 0 | 1.9 | 141 | Specific | 830.86 | 779.17 | 779.17 | 14.46 | 57 | >sp\|P02788\|TRFL_HUMAN Lactotransferrin OS=Homo sapiens OX=9606 GN=LTF PE=1 SV=6 | 1270 | 20201117_08512_1ng_glyco_hcd30_etd30_compound15.18128.18128.4 | controllerType=0 controllerNumber=1 scan=18128 | 46.2863 |
| 15388:4 | 1 | R.TAGWNVPIGTLRPFLN[+2350.830]WTGPPEPIEAAVAR.F | HexNAc(4)Hex(5)Fuc(1)NeuAc(2) | N[+2351] | 1396.1438 | 4 | 5581.5534 | 5581.5413 | 0 | 2.2 | 141 | Specific | 882.29 | 865.74 | 865.74 | 15.46 | 57 | >sp\|P02788\|TRFL_HUMAN Lactotransferrin OS=Homo sapiens OX=9606 GN=LTF PE=1 SV=6 | 1270 | 20201117_08512_1ng_glyco_hcd30_etd30_compound15.19024.19024.4 | controllerType=0 controllerNumber=1 scan=19024 | 49.1202 |
| 14560:5 | 1 | R.TAGWNVPIGTLRPFLN[+2059.735]WTGPPEPIEAAVAR.F | HexNAc(4)Hex(5)Fuc(1)NeuAc(1) | N[+2060] | 1058.8976 | 5 | 5290.4588 | 5290.4459 | 0 | 2.4 | 141 | Specific | 942.03 | 874.89 | 874.89 | 14.95 | 57 | >sp\|P02788\|TRFL_HUMAN Lactotransferrin OS=Homo sapiens OX=9606 GN=LTF PE=1 SV=6 | 1270 | 20201117_08512_1ng_glyco_hcd30_etd30_compound15.18145.18145.5 | controllerType=0 controllerNumber=1 scan=18145 | 46.3485 |
| 14545:4 | 1 | R.TAGWNVPIGTLRPFLN[+2059.735]WTGPPEPIEAAVAR.F | HexNAc(4)Hex(5)Fuc(1)NeuAc(1) | N[+2060] | 1323.3723 | 4 | 5290.4674 | 5290.4459 | 0 | 4.1 | 141 | Specific | 975.06 | 949.22 | 949.22 | 17.47 | 57 | >sp\|P02788\|TRFL_HUMAN Lactotransferrin OS=Homo sapiens OX=9606 GN=LTF PE=1 SV=6 | 1270 | 20201117_08512_1ng_glyco_hcd30_etd30_compound15.18129.18129.4 | controllerType=0 controllerNumber=1 scan=18129 | 46.295 |
| 13876:4 | 1 | R.TAGWNVPIGTLRPFLN[+1768.640]WTGPPEPIEAAVAR.F | HexNAc(4)Hex(5)Fuc(1) | N[+1769] | 1250.597 | 4 | 4999.3664 | 4999.3504 | 0 | 3.2 | 141 | Specific | 1018.52 | 982 | 982 | 18.47 | 57 | >sp\|P02788\|TRFL_HUMAN Lactotransferrin OS=Homo sapiens OX=9606 GN=LTF PE=1 SV=6 | 1270 | 20201117_08512_1ng_glyco_hcd30_etd30_compound15.17422.17422.4 | controllerType=0 controllerNumber=1 scan=17422 | 44.0875 |
| 12828:3 | 1 | R.TAGWNIPMGLLFN[+1768.640]QTGSC[+57.021]K.F | HexNAc(4)Hex(5)Fuc(1) | C[+57], N[+1769] | 1288.5566 | 3 | 3863.6552 | 3863.6495 | 0 | 1.5 | 485 | Specific | 356.18 | 279.53 | 279.53 | 4.61 | 57 | >sp\|P02788\|TRFL_HUMAN Lactotransferrin OS=Homo sapiens OX=9606 GN=LTF PE=1 SV=6 | 1270 | 20201117_08512_1ng_glyco_hcd30_etd30_compound15.16312.16312.3 | controllerType=0 controllerNumber=1 scan=16312 | 40.7027 |
| 14211:3 | 1 | R.TAGWNIPMGLLFN[+1913.677]QTGSC[+57.021]K.F | HexNAc(4)Hex(5)NeuAc(1) | C[+57], N[+1914] | 1336.903 | 3 | 4008.6945 | 4008.687 | 0 | 1.9 | 485 | Specific | 409.24 | 275.88 | 275.88 | 5.69 | 57 | >sp\|P02788\|TRFL_HUMAN Lactotransferrin OS=Homo sapiens OX=9606 GN=LTF PE=1 SV=6 | 1270 | 20201117_08512_1ng_glyco_hcd30_etd30_compound15.17776.17776.3 | controllerType=0 controllerNumber=1 scan=17776 | 45.194 |
| 14108:3 | 1 | R.TAGWNIPMGLLFN[+2205.793]QTGSC[+57.021]K.F | HexNAc(4)Hex(5)Fuc(2)NeuAc(1) | C[+57], N[+2206] | 1434.2761 | 3 | 4300.8136 | 4300.8028 | 0 | 2.5 | 485 | Specific | 412.51 | 315.61 | 315.61 | 5.81 | 57 | >sp\|P02788\|TRFL_HUMAN Lactotransferrin OS=Homo sapiens OX=9606 GN=LTF PE=1 SV=6 | 1270 | 20201117_08512_1ng_glyco_hcd30_etd30_compound15.17666.17666.3 | controllerType=0 controllerNumber=1 scan=17666 | 44.806 |
| 15576:3 | 1 | R.TAGWNIPMGLLFN[+2204.772]QTGSC[+57.021]K.F | HexNAc(4)Hex(5)NeuAc(2) | C[+57], N[+2205] | 1433.9351 | 3 | 4299.7908 | 4299.7824 | 0 | 2 | 485 | Specific | 420.55 | 283.23 | 283.23 | 5.69 | 57 | >sp\|P02788\|TRFL_HUMAN Lactotransferrin OS=Homo sapiens OX=9606 GN=LTF PE=1 SV=6 | 1270 | 20201117_08512_1ng_glyco_hcd30_etd30_compound15.19224.19224.3 | controllerType=0 controllerNumber=1 scan=19224 | 49.8442 |
| 14122:3 | 1 | R.TAGWNIPMGLLFN[+2059.735]QTGSC[+57.021]K.F | HexNAc(4)Hex(5)Fuc(1)NeuAc(1) | C[+57], N[+2060] | 1385.591 | 3 | 4154.7584 | 4154.7449 | 0 | 3.3 | 485 | Specific | 425.22 | 304.22 | 304.22 | 5.81 | 57 | >sp\|P02788\|TRFL_HUMAN Lactotransferrin OS=Homo sapiens OX=9606 GN=LTF PE=1 SV=6 | 1270 | 20201117_08512_1ng_glyco_hcd30_etd30_compound15.17681.17681.3 | controllerType=0 controllerNumber=1 scan=17681 | 44.8566 |
| 13079:3 | 1 | R.TAGWNIPMGLLFN[+1216.423]QTGSC[+57.021]K.F | HexNAc(2)Hex(5) | C[+57], N[+1216] | 1104.4855 | 3 | 3311.442 | 3311.4328 | 0 | 2.8 | 485 | Specific | 475.18 | 444.89 | 226.28 | 7.2 | 57 | >sp\|P02788\|TRFL_HUMAN Lactotransferrin OS=Homo sapiens OX=9606 GN=LTF PE=1 SV=6 | 1270 | 20201117_08512_1ng_glyco_hcd30_etd30_compound15.16578.16578.3 | controllerType=0 controllerNumber=1 scan=16578 | 41.5168 |
| 15466:4 | 1 | R.TAGWNIPMGLLFN[+2350.830]QTGSC[+57.021]K.F | HexNAc(4)Hex(5)Fuc(1)NeuAc(2) | C[+57], N[+2351] | 1112.2177 | 4 | 4445.8489 | 4445.8403 | 0 | 1.9 | 485 | Specific | 503.28 | 443.12 | 443.12 | 6.75 | 57 | >sp\|P02788\|TRFL_HUMAN Lactotransferrin OS=Homo sapiens OX=9606 GN=LTF PE=1 SV=6 | 1270 | 20201117_08512_1ng_glyco_hcd30_etd30_compound15.19107.19107.4 | controllerType=0 controllerNumber=1 scan=19107 | 49.4191 |
| 13095:2 | 1 | R.TAGWNIPMGLLFN[+1216.423]QTGSC[+57.021]K.F | HexNAc(2)Hex(5) | C[+57], N[+1216] | 1656.2244 | 2 | 3311.4415 | 3311.4328 | 0 | 2.6 | 485 | Specific | 519.59 | 457.91 | 175.52 | 7.01 | 57 | >sp\|P02788\|TRFL_HUMAN Lactotransferrin OS=Homo sapiens OX=9606 GN=LTF PE=1 SV=6 | 1270 | 20201117_08512_1ng_glyco_hcd30_etd30_compound15.16595.16595.2 | controllerType=0 controllerNumber=1 scan=16595 | 41.5788 |
| 13025:3 | 1 | R.TAGWNIPM[+15.995]GLLFN[+2059.735]QTGSC[+57.021]K.F | HexNAc(4)Hex(5)Fuc(1)NeuAc(1) | C[+57], M[+16], N[+2060] | 1390.9229 | 3 | 4170.7542 | 4170.7398 | 0 | 3.4 | 485 | Specific | 353.88 | 295.76 | 295.76 | 4.73 | 57 | >sp\|P02788\|TRFL_HUMAN Lactotransferrin OS=Homo sapiens OX=9606 GN=LTF PE=1 SV=6 | 1270 | 20201117_08512_1ng_glyco_hcd30_etd30_compound15.16521.16521.3 | controllerType=0 controllerNumber=1 scan=16521 | 41.3312 |
| 13618:3 | 1 | R.TAGWNIPMGLLFN[+1768.640]QTGSC[+57.021]KFDEYFSQSC[+57.021]APGSDPR.S | HexNAc(4)Hex(5)Fuc(1) | C[+57]*2, N[+1769] | 1903.1413 | 3 | 5707.4094 | 5707.3966 | 0 | 2.2 | 485 | Specific | 271.01 | 235.75 | 235.75 | 5.47 | 57 | >sp\|P02788\|TRFL_HUMAN Lactotransferrin OS=Homo sapiens OX=9606 GN=LTF PE=1 SV=6 | 1270 | 20201117_08512_1ng_glyco_hcd30_etd30_compound15.17148.17148.3 | controllerType=0 controllerNumber=1 scan=17148 | 43.2474 |
| 14292:5 | 1 | R.TAGWNIPMGLLFN[+2059.735]QTGSC[+57.021]KFDEYFSQSC[+57.021]APGSDPR.S | HexNAc(4)Hex(5)Fuc(1)NeuAc(1) | C[+57]*2, N[+2060] | 1200.5066 | 5 | 5998.5039 | 5998.492 | 0 | 2 | 485 | Specific | 311.71 | 280.27 | 280.27 | 4.33 | 57 | >sp\|P02788\|TRFL_HUMAN Lactotransferrin OS=Homo sapiens OX=9606 GN=LTF PE=1 SV=6 | 1270 | 20201117_08512_1ng_glyco_hcd30_etd30_compound15.17863.17863.5 | controllerType=0 controllerNumber=1 scan=17863 | 45.5306 |
| 13727:3 | 1 | R.TAGWNIPMGLLFN[+1216.423]QTGSC[+57.021]KFDEYFSQSC[+57.021]APGSDPR.S | HexNAc(2)Hex(5) | C[+57]*2, N[+1216] | 1719.0702 | 3 | 5155.1959 | 5155.18 | 0 | 3.1 | 485 | Specific | 345.72 | 314.69 | 82.99 | 6.12 | 57 | >sp\|P02788\|TRFL_HUMAN Lactotransferrin OS=Homo sapiens OX=9606 GN=LTF PE=1 SV=6 | 1270 | 20201117_08512_1ng_glyco_hcd30_etd30_compound15.17263.17263.3 | controllerType=0 controllerNumber=1 scan=17263 | 43.5795 |
| 14275:4 | 1 | R.TAGWNIPMGLLFN[+2205.793]QTGSC[+57.021]KFDEYFSQSC[+57.021]APGSDPR.S | HexNAc(4)Hex(5)Fuc(2)NeuAc(1) | C[+57]*2, N[+2206] | 1536.8977 | 4 | 6144.569 | 6144.5499 | 0 | 3.1 | 485 | Specific | 390.76 | 311.79 | 311.79 | 6.43 | 57 | >sp\|P02788\|TRFL_HUMAN Lactotransferrin OS=Homo sapiens OX=9606 GN=LTF PE=1 SV=6 | 1270 | 20201117_08512_1ng_glyco_hcd30_etd30_compound15.17845.17845.4 | controllerType=0 controllerNumber=1 scan=17845 | 45.4672 |
| 15205:4 | 1 | R.TAGWNIPMGLLFN[+2350.830]QTGSC[+57.021]KFDEYFSQSC[+57.021]APGSDPR.S | HexNAc(4)Hex(5)Fuc(1)NeuAc(2) | C[+57]*2, N[+2351] | 1573.1565 | 4 | 6289.6042 | 6289.5875 | 0 | 2.7 | 485 | Specific | 405.96 | 368.79 | 368.79 | 8.08 | 57 | >sp\|P02788\|TRFL_HUMAN Lactotransferrin OS=Homo sapiens OX=9606 GN=LTF PE=1 SV=6 | 1270 | 20201117_08512_1ng_glyco_hcd30_etd30_compound15.18830.18830.4 | controllerType=0 controllerNumber=1 scan=18830 | 48.4816 |
| 13612:4 | 1 | R.TAGWNIPMGLLFN[+1768.640]QTGSC[+57.021]KFDEYFSQSC[+57.021]APGSDPR.S | HexNAc(4)Hex(5)Fuc(1) | C[+57]*2, N[+1769] | 1427.6084 | 4 | 5707.4118 | 5707.3966 | 0 | 2.7 | 485 | Specific | 424.06 | 359.55 | 359.55 | 5.65 | 57 | >sp\|P02788\|TRFL_HUMAN Lactotransferrin OS=Homo sapiens OX=9606 GN=LTF PE=1 SV=6 | 1270 | 20201117_08512_1ng_glyco_hcd30_etd30_compound15.17142.17142.4 | controllerType=0 controllerNumber=1 scan=17142 | 43.2316 |
| 13594:4 | 1 | R.TAGWNIPMGLLFN[+1768.640]QTGSC[+57.021]KFDEYFSQSC[+57.021]APGSDPR.S | HexNAc(4)Hex(5)Fuc(1) | C[+57]*2, N[+1769] | 1427.6083 | 4 | 5707.4113 | 5707.3966 | 0 | 2.6 | 485 | Specific | 517.18 | 458.49 | 458.49 | 7.01 | 57 | >sp\|P02788\|TRFL_HUMAN Lactotransferrin OS=Homo sapiens OX=9606 GN=LTF PE=1 SV=6 | 1270 | 20201117_08512_1ng_glyco_hcd30_etd30_compound15.17123.17123.4 | controllerType=0 controllerNumber=1 scan=17123 | 43.1761 |
| 13699:3 | 1 | R.TAGWNIPMGLLFN[+1216.423]QTGSC[+57.021]KFDEYFSQSC[+57.021]APGSDPR.S | HexNAc(2)Hex(5) | C[+57]*2, N[+1216] | 1719.068 | 3 | 5155.1893 | 5155.18 | 0 | 1.8 | 485 | Specific | 534.94 | 432.07 | 116.19 | 8.08 | 57 | >sp\|P02788\|TRFL_HUMAN Lactotransferrin OS=Homo sapiens OX=9606 GN=LTF PE=1 SV=6 | 1270 | 20201117_08512_1ng_glyco_hcd30_etd30_compound15.17233.17233.3 | controllerType=0 controllerNumber=1 scan=17233 | 43.4783 |
| 14274:4 | 1 | R.TAGWNIPMGLLFN[+2059.735]QTGSC[+57.021]KFDEYFSQSC[+57.021]APGSDPR.S | HexNAc(4)Hex(5)Fuc(1)NeuAc(1) | C[+57]*2, N[+2060] | 1500.3865 | 4 | 5998.5241 | 5998.492 | 0 | 5.3 | 485 | Specific | 575.68 | 535.1 | 535.1 | 9 | 57 | >sp\|P02788\|TRFL_HUMAN Lactotransferrin OS=Homo sapiens OX=9606 GN=LTF PE=1 SV=6 | 1270 | 20201117_08512_1ng_glyco_hcd30_etd30_compound15.17844.17844.4 | controllerType=0 controllerNumber=1 scan=17844 | 45.4644 |
| 14316:4 | 1 | R.TAGWNIPMGLLFN[+1913.677]QTGSC[+57.021]KFDEYFSQSC[+57.021]APGSDPR.S | HexNAc(4)Hex(5)NeuAc(1) | C[+57]*2, N[+1914] | 1463.8658 | 4 | 5852.4416 | 5852.4341 | 0 | 1.3 | 485 | Specific | 577.96 | 500.62 | 500.62 | 8.29 | 57 | >sp\|P02788\|TRFL_HUMAN Lactotransferrin OS=Homo sapiens OX=9606 GN=LTF PE=1 SV=6 | 1270 | 20201117_08512_1ng_glyco_hcd30_etd30_compound15.17888.17888.4 | controllerType=0 controllerNumber=1 scan=17888 | 45.6006 |
| 14440:4 | 1 | R.TAGWNIPMGLLFN[+1548.545]QTGSC[+57.021]KFDEYFSQSC[+57.021]APGSDPR.S | HexNAc(3)Hex(4)NeuAc(1) | C[+57]*2, N[+1549] | 1372.5848 | 4 | 5487.3175 | 5487.3019 | 0 | 2.8 | 485 | Specific | 740.48 | 629.99 | 629.99 | 11.56 | 57 | >sp\|P02788\|TRFL_HUMAN Lactotransferrin OS=Homo sapiens OX=9606 GN=LTF PE=1 SV=6 | 1270 | 20201117_08512_1ng_glyco_hcd30_etd30_compound15.18018.18018.4 | controllerType=0 controllerNumber=1 scan=18018 | 45.9281 |
| 13494:4 | 1 | R.TAGWNIPM[+15.995]GLLFN[+2059.735]QTGSC[+57.021]KFDEYFSQSC[+57.021]A | HexNAc(4)Hex(5)Fuc(1)NeuAc(1) | C[+57]*2, M[+16], N[+2060] | 1504.3824 | 4 | 6014.508 | 6014.487 | 0 | 3.5 | 485 | Specific | 249.7 | 190.77 | 190.77 | 4.64 | 57 | >sp\|P02788\|TRFL_HUMAN Lactotransferrin OS=Homo sapiens OX=9606 GN=LTF PE=1 SV=6 | 1270 | 20201117_08512_1ng_glyco_hcd30_etd30_compound15.17018.17018.4 | controllerType=0 controllerNumber=1 scan=17018 | 42.8602 |
| 08089:3 | 1 | K.FGRN[+2350.830]GSDC[+57.021]PDKFC[+57.021]LFQSETK.N | HexNAc(4)Hex(5)Fuc(1)NeuAc(2) | C[+57]*2, N[+2351] | 1581.9748 | 3 | 4743.9098 | 4743.8952 | 0 | 3.1 | 639 | Specific | 222.26 | 101.86 | 101.86 | 3.14 | 57 | >sp\|P02788\|TRFL_HUMAN Lactotransferrin OS=Homo sapiens OX=9606 GN=LTF PE=1 SV=6 | 1270 | 20201117_08512_1ng_glyco_hcd30_etd30_compound15.11326.11326.3 | controllerType=0 controllerNumber=1 scan=11326 | 28.0905 |
| 06863:3 | 1 | K.FGRN[+2059.735]GSDC[+57.021]PDKFC[+57.021]LFQSETK.N | HexNAc(4)Hex(5)Fuc(1)NeuAc(1) | C[+57]*2, N[+2060] | 1484.9439 | 3 | 4452.8172 | 4452.7998 | 0 | 3.9 | 639 | Specific | 231.12 | 144.42 | 144.42 | 2.83 | 57 | >sp\|P02788\|TRFL_HUMAN Lactotransferrin OS=Homo sapiens OX=9606 GN=LTF PE=1 SV=6 | 1270 | 20201117_08512_1ng_glyco_hcd30_etd30_compound15.10040.10040.3 | controllerType=0 controllerNumber=1 scan=10040 | 25.0331 |
| 06862:4 | 1 | K.FGRN[+2059.735]GSDC[+57.021]PDKFC[+57.021]LFQSETK.N | HexNAc(4)Hex(5)Fuc(1)NeuAc(1) | C[+57]*2, N[+2060] | 1113.9591 | 4 | 4452.8148 | 4452.7998 | 0 | 3.4 | 639 | Specific | 605.38 | 501.46 | 501.46 | 9.2 | 57 | >sp\|P02788\|TRFL_HUMAN Lactotransferrin OS=Homo sapiens OX=9606 GN=LTF PE=1 SV=6 | 1270 | 20201117_08512_1ng_glyco_hcd30_etd30_compound15.10039.10039.4 | controllerType=0 controllerNumber=1 scan=10039 | 25.0272 |
| 07517:3 | 2 | R.QTGLAPGQEYEISLHIVKN[+2059.735]NTR.G | HexNAc(4)Hex(5)Fuc(1)NeuAc(1) | N[+2060] | 1510.0173 | 3 | 4528.0373 | 4528.0241 | 0 | 2.9 | 770 | Specific | 265.89 | 171.53 | 171.53 | 3.25 | 43 | >sp\|P24821\|TENA_HUMAN Tenascin OS=Homo sapiens OX=9606 GN=TNC PE=1 SV=3 | 486 | 20201117_08512_1ng_glyco_hcd30_etd30_compound15.10726.10726.3 | controllerType=0 controllerNumber=1 scan=10726 | 26.6609 |
| 07484:3 | 2 | R.QTGLAPGQEYEISLHIVKN[+2059.735]NTR.G | HexNAc(4)Hex(5)Fuc(1)NeuAc(1) | N[+2060] | 1510.0174 | 3 | 4528.0376 | 4528.0241 | 0 | 3 | 770 | Specific | 319.99 | 205.23 | 205.23 | 3.8 | 43 | >sp\|P24821\|TENA_HUMAN Tenascin OS=Homo sapiens OX=9606 GN=TNC PE=1 SV=3 | 486 | 20201117_08512_1ng_glyco_hcd30_etd30_compound15.10691.10691.3 | controllerType=0 controllerNumber=1 scan=10691 | 26.568 |
| 08587:4 | 2 | R.QTGLAPGQEYEISLHIVKN[+2350.830]NTR.G | HexNAc(4)Hex(5)Fuc(1)NeuAc(2) | N[+2351] | 1205.5373 | 4 | 4819.1273 | 4819.1196 | 0 | 1.6 | 770 | Specific | 329.95 | 309.58 | 309.58 | 4.09 | 43 | >sp\|P24821\|TENA_HUMAN Tenascin OS=Homo sapiens OX=9606 GN=TNC PE=1 SV=3 | 486 | 20201117_08512_1ng_glyco_hcd30_etd30_compound15.11849.11849.4 | controllerType=0 controllerNumber=1 scan=11849 | 29.4549 |
| 07513:4 | 2 | R.QTGLAPGQEYEISLHIVKN[+2059.735]NTR.G | HexNAc(4)Hex(5)Fuc(1)NeuAc(1) | N[+2060] | 1132.7646 | 4 | 4528.0364 | 4528.0241 | 0 | 2.7 | 770 | Specific | 648.97 | 556.99 | 556.99 | 8.57 | 43 | >sp\|P24821\|TENA_HUMAN Tenascin OS=Homo sapiens OX=9606 GN=TNC PE=1 SV=3 | 486 | 20201117_08512_1ng_glyco_hcd30_etd30_compound15.10722.10722.4 | controllerType=0 controllerNumber=1 scan=10722 | 26.6516 |
| 11850:2 | 2 | R.LN[+1216.423]YSLPTGQWVGVQLPR.N | HexNAc(2)Hex(5) | N[+1216] | 1572.7395 | 2 | 3144.4718 | 3144.4617 | 0 | 3.2 | 1017 | Specific | 453.01 | 390.18 | 390.18 | 6.85 | 43 | >sp\|P24821\|TENA_HUMAN Tenascin OS=Homo sapiens OX=9606 GN=TNC PE=1 SV=3 | 486 | 20201117_08512_1ng_glyco_hcd30_etd30_compound15.15276.15276.2 | controllerType=0 controllerNumber=1 scan=15276 | 38.1693 |
| 11718:3 | 2 | R.LN[+1702.581]YSLPTGQWVGVQLPR.N | HexNAc(2)Hex(8) | N[+1703] | 1210.8802 | 3 | 3630.626 | 3630.6202 | 0 | 1.6 | 1017 | Specific | 539.11 | 448.44 | 448.44 | 7.6 | 43 | >sp\|P24821\|TENA_HUMAN Tenascin OS=Homo sapiens OX=9606 GN=TNC PE=1 SV=3 | 486 | 20201117_08512_1ng_glyco_hcd30_etd30_compound15.15136.15136.2 | controllerType=0 controllerNumber=1 scan=15136 | 37.7328 |
| 11726:3 | 2 | R.LN[+1702.581]YSLPTGQWVGVQLPR.N | HexNAc(2)Hex(8) | N[+1703] | 1210.8819 | 3 | 3630.6311 | 3630.6202 | 0 | 3 | 1017 | Specific | 557.26 | 471.94 | 471.94 | 7.6 | 43 | >sp\|P24821\|TENA_HUMAN Tenascin OS=Homo sapiens OX=9606 GN=TNC PE=1 SV=3 | 486 | 20201117_08512_1ng_glyco_hcd30_etd30_compound15.15145.15145.3 | controllerType=0 controllerNumber=1 scan=15145 | 37.754 |
| 11791:3 | 2 | R.LN[+1378.476]YSLPTGQWVGVQLPR.N | HexNAc(2)Hex(6) | N[+1378] | 1102.8461 | 3 | 3306.5237 | 3306.5145 | 0 | 2.8 | 1017 | Specific | 616.78 | 548.83 | 548.83 | 9.65 | 43 | >sp\|P24821\|TENA_HUMAN Tenascin OS=Homo sapiens OX=9606 GN=TNC PE=1 SV=3 | 486 | 20201117_08512_1ng_glyco_hcd30_etd30_compound15.15213.15213.3 | controllerType=0 controllerNumber=1 scan=15213 | 37.9577 |
| 11832:3 | 2 | R.LN[+1216.423]YSLPTGQWVGVQLPR.N | HexNAc(2)Hex(5) | N[+1216] | 1048.8296 | 3 | 3144.4743 | 3144.4617 | 0 | 4 | 1017 | Specific | 672.19 | 621.69 | 621.69 | 10.65 | 43 | >sp\|P24821\|TENA_HUMAN Tenascin OS=Homo sapiens OX=9606 GN=TNC PE=1 SV=3 | 486 | 20201117_08512_1ng_glyco_hcd30_etd30_compound15.15257.15257.3 | controllerType=0 controllerNumber=1 scan=15257 | 38.1059 |
| 03262:3 | 2 | R.N[+2059.735]TTSYVLR.G | HexNAc(4)Hex(5)Fuc(1)NeuAc(1) | N[+2060] | 1005.0866 | 3 | 3013.2452 | 3013.24 | 0 | 1.7 | 1034 | Specific | 264.85 | 128.36 | 128.36 | 2.94 | 43 | >sp\|P24821\|TENA_HUMAN Tenascin OS=Homo sapiens OX=9606 GN=TNC PE=1 SV=3 | 486 | 20201117_08512_1ng_glyco_hcd30_etd30_compound15.6301.6301.3 | controllerType=0 controllerNumber=1 scan=6301 | 17.8041 |
| 17083:5 | 3 | K.VFPLSLDSTPQDGNVVVAC[+57.021]LVQGFFPQEPLSVTWSESGQN[+2262.814]VTAI | HexNAc(5)Hex(5)Fuc(1)NeuAc(1) | C[+57], N[+2263] | 1408.6485 | 5 | 7039.2133 | 7039.1757 | 0 | 5.3 | 8 | Specific | 212.38 | 212.38 | 212.38 | 2.61 | 57 | >sp\|P01877\|IGHA2_HUMAN Immunoglobulin heavy constant alpha 2 OS=Homo sapiens OX=9606 GN=IGHA2 PE=1 SV=4 | 576 | 20201117_08512_1ng_glyco_hcd30_etd30_compound15.20829.20829.5 | controllerType=0 controllerNumber=1 scan=20829 | 55.0993 |
| 17089:4 | 3 | K.VFPLSLDSTPQDGNVVVAC[+57.021]LVQGFFPQEPLSVTWSESGQN[+2262.814]VTAI | HexNAc(5)Hex(5)Fuc(1)NeuAc(1) | C[+57], N[+2263] | 1760.5571 | 4 | 7039.2065 | 7039.1757 | 0 | 4.4 | 8 | Specific | 320.69 | 320.69 | 320.69 | 4.83 | 57 | >sp\|P01877\|IGHA2_HUMAN Immunoglobulin heavy constant alpha 2 OS=Homo sapiens OX=9606 GN=IGHA2 PE=1 SV=4 | 576 | 20201117_08512_1ng_glyco_hcd30_etd30_compound15.20835.20835.4 | controllerType=0 controllerNumber=1 scan=20835 | 55.1222 |
| 00862:3 | 3 | K.HYTN[+1971.719]SSQDVTVPC[+57.021]R.V | HexNAc(5)Hex(5)Fuc(1) | C[+57], N[+1972] | 1212.4982 | 3 | 3635.4801 | 3635.4682 | 0 | 3.3 | 89 | Specific | 275.2 | 220.32 | 220.32 | 4.66 | 57 | >sp\|P01877\|IGHA2_HUMAN Immunoglobulin heavy constant alpha 2 OS=Homo sapiens OX=9606 GN=IGHA2 PE=1 SV=4 | 576 | 20201117_08512_1ng_glyco_hcd30_etd30_compound15.3803.3803.3 | controllerType=0 controllerNumber=1 scan=3803 | 12.8647 |
| 01712:3 | 3 | K.HYTN[+2262.814]SSQDVTVPC[+57.021]R.V | HexNAc(5)Hex(5)Fuc(1)NeuAc(1) | C[+57], N[+2263] | 1309.5306 | 3 | 3926.5772 | 3926.5636 | 0 | 3.5 | 89 | Specific | 380.82 | 263.81 | 263.81 | 5.14 | 57 | >sp\|P01877\|IGHA2_HUMAN Immunoglobulin heavy constant alpha 2 OS=Homo sapiens OX=9606 GN=IGHA2 PE=1 SV=4 | 576 | 20201117_08512_1ng_glyco_hcd30_etd30_compound15.4687.4687.3 | controllerType=0 controllerNumber=1 scan=4687 | 14.4942 |
| 00903:3 | 3 | K.HYTN[+1444.534]SSQDVTVPC[+57.021]R.V | HexNAc(4)Hex(3)Fuc(1) | C[+57], N[+1445] | 1036.768 | 3 | 3108.2894 | 3108.2832 | 0 | 2 | 89 | Specific | 413.2 | 358 | 358 | 5.65 | 57 | >sp\|P01877\|IGHA2_HUMAN Immunoglobulin heavy constant alpha 2 OS=Homo sapiens OX=9606 GN=IGHA2 PE=1 SV=4 | 576 | 20201117_08512_1ng_glyco_hcd30_etd30_compound15.3845.3845.3 | controllerType=0 controllerNumber=1 scan=3845 | 12.9461 |
| 00907:3 | 3 | K.HYTN[+1809.666]SSQDVTVPC[+57.021]R.V | HexNAc(5)Hex(4)Fuc(1) | C[+57], N[+1810] | 1158.4792 | 3 | 3473.423 | 3473.4154 | 0 | 2.2 | 89 | Specific | 437.49 | 364.96 | 364.96 | 6.9 | 57 | >sp\|P01877\|IGHA2_HUMAN Immunoglobulin heavy constant alpha 2 OS=Homo sapiens OX=9606 GN=IGHA2 PE=1 SV=4 | 576 | 20201117_08512_1ng_glyco_hcd30_etd30_compound15.3850.3850.3 | controllerType=0 controllerNumber=1 scan=3850 | 12.9636 |
| 00913:3 | 3 | K.HYTN[+1647.613]SSQDVTVPC[+57.021]R.V | HexNAc(5)Hex(3)Fuc(1) | C[+57], N[+1648] | 1104.4611 | 3 | 3311.3687 | 3311.3625 | 0 | 1.9 | 89 | Specific | 519.54 | 461.82 | 461.82 | 7.01 | 57 | >sp\|P01877\|IGHA2_HUMAN Immunoglobulin heavy constant alpha 2 OS=Homo sapiens OX=9606 GN=IGHA2 PE=1 SV=4 | 576 | 20201117_08512_1ng_glyco_hcd30_etd30_compound15.3856.3856.3 | controllerType=0 controllerNumber=1 scan=3856 | 12.9778 |
| 02924:5 | 3 | K.HYTN[+1647.613]SSQDVTVPC[+57.021]RVPPPPPC[+57.021]C[+57.021]HPR.L | HexNAc(5)Hex(3)Fuc(1) | C[+57]*3, N[+1648] | 922.0023 | 5 | 4605.9823 | 4605.9689 | 0 | 2.9 | 89 | Specific | 222.37 | 222.37 | 222.37 | 2.62 | 57 | >sp\|P01877\|IGHA2_HUMAN Immunoglobulin heavy constant alpha 2 OS=Homo sapiens OX=9606 GN=IGHA2 PE=1 SV=4 | 576 | 20201117_08512_1ng_glyco_hcd30_etd30_compound15.5949.5949.5 | controllerType=0 controllerNumber=1 scan=5949 | 17.0809 |
| 02720:4 | 3 | K.HYTN[+1444.534]SSQDVTVPC[+57.021]RVPPPPPC[+57.021]C[+57.021]HPR.L | HexNAc(4)Hex(3)Fuc(1) | C[+57]*3, N[+1445] | 1101.4861 | 4 | 4402.9227 | 4402.8895 | 0 | 7.5 | 89 | Specific | 289.83 | 289.83 | 289.83 | 2.66 | 57 | >sp\|P01877\|IGHA2_HUMAN Immunoglobulin heavy constant alpha 2 OS=Homo sapiens OX=9606 GN=IGHA2 PE=1 SV=4 | 576 | 20201117_08512_1ng_glyco_hcd30_etd30_compound15.5735.5735.4 | controllerType=0 controllerNumber=1 scan=5735 | 16.6215 |

Table S4. Cont.

<table>
<tr><th>Query #:z</th><th>Protein Rank</th><th>Glycopeptide</th><th>Glycan</th><th>Modification Type(s)</th><th>Observed m/z</th><th>z</th><th>Observed (M+H)</th><th>Calc. mass (M+H)</th><th>Off-by-x error</th><th>Mass error (ppm)</th><th>StartPosition</th><th>Cleavage</th><th>Score</th><th>Delta</th><th>Delta Mod</th><th>|Log Prob|</th><th># of unique peptides</th><th>Protein Name</th><th>Protein DB number</th><th>Comment</th><th>Scan #</th><th>Scan Time</th></tr>
<tr><td>03239:4</td><td>3</td><td>K.HYTN[+2262.814]SSQDVTVPC[+57.021]RVPPPPPC[+57.021]C[+57.021]HPR.L</td><td>HexNAc(5)Hex(5)Fuc(1)NeuAc(1)</td><td>C[+57]*3, N[+2263]</td><td>1306.0502</td><td>4</td><td>5221.1789</td><td>5221.1699</td><td>0</td><td>1.7</td><td>89</td><td>Specific</td><td>359.65</td><td>356.15</td><td>356.15</td><td>4.52</td><td>57</td><td>>sp|P01877|IGHA2_HUMAN Immunoglobulin heavy constant alpha 2 OS=Homo sapiens OX=9606 GN=IGHA2 PE=1 SV=4</td><td>576</td><td>20201117_08512_1ng_glyco_hcd30_etd30_compound15.6277.6277.4</td><td>controllerType=0 controllerNumber=1 scan=6277</td><td>17.7472</td></tr>
<tr><td>03223:5</td><td>3</td><td>K.HYTN[+2262.814]SSQDVTVPC[+57.021]RVPPPPPC[+57.021]C[+57.021]HPR.L</td><td>HexNAc(5)Hex(5)Fuc(1)NeuAc(1)</td><td>C[+57]*3, N[+2263]</td><td>1045.0418</td><td>5</td><td>5221.1796</td><td>5221.1699</td><td>0</td><td>1.9</td><td>89</td><td>Specific</td><td>415.92</td><td>356.89</td><td>356.89</td><td>4.44</td><td>57</td><td>>sp|P01877|IGHA2_HUMAN Immunoglobulin heavy constant alpha 2 OS=Homo sapiens OX=9606 GN=IGHA2 PE=1 SV=4</td><td>576</td><td>20201117_08512_1ng_glyco_hcd30_etd30_compound15.6260.6260.5</td><td>controllerType=0 controllerNumber=1 scan=6260</td><td>17.7005</td></tr>
<tr><td>02648:4</td><td>3</td><td>K.HYTN[+1809.666]SSQDVTVPC[+57.021]RVPPPPPC[+57.021]C[+57.021]HPR.L</td><td>HexNAc(5)Hex(4)Fuc(1)</td><td>C[+57]*3, N[+1810]</td><td>1192.7636</td><td>4</td><td>4768.0325</td><td>4768.0217</td><td>0</td><td>2.3</td><td>89</td><td>Specific</td><td>497.5</td><td>401.6</td><td>401.6</td><td>6.04</td><td>57</td><td>>sp|P01877|IGHA2_HUMAN Immunoglobulin heavy constant alpha 2 OS=Homo sapiens OX=9606 GN=IGHA2 PE=1 SV=4</td><td>576</td><td>20201117_08512_1ng_glyco_hcd30_etd30_compound15.5660.5660.4</td><td>controllerType=0 controllerNumber=1 scan=5660</td><td>16.4408</td></tr>
<tr><td>02681:5</td><td>3</td><td>K.HYTN[+1647.613]SSQDVTVPC[+57.021]RVPPPPPC[+57.021]C[+57.021]HPR.L</td><td>HexNAc(5)Hex(3)Fuc(1)</td><td>C[+57]*3, N[+1648]</td><td>922.0023</td><td>5</td><td>4605.9823</td><td>4605.9689</td><td>0</td><td>2.9</td><td>89</td><td>Specific</td><td>511.98</td><td>476.94</td><td>476.94</td><td>6</td><td>57</td><td>>sp|P01877|IGHA2_HUMAN Immunoglobulin heavy constant alpha 2 OS=Homo sapiens OX=9606 GN=IGHA2 PE=1 SV=4</td><td>576</td><td>20201117_08512_1ng_glyco_hcd30_etd30_compound15.5695.5695.5</td><td>controllerType=0 controllerNumber=1 scan=5695</td><td>16.5305</td></tr>
<tr><td>02642:5</td><td>3</td><td>K.HYTN[+1809.666]SSQDVTVPC[+57.021]RVPPPPPC[+57.021]C[+57.021]HPR.L</td><td>HexNAc(5)Hex(4)Fuc(1)</td><td>C[+57]*3, N[+1810]</td><td>954.4126</td><td>5</td><td>4768.0337</td><td>4768.0217</td><td>0</td><td>2.5</td><td>89</td><td>Specific</td><td>535.23</td><td>535.23</td><td>535.23</td><td>6.04</td><td>57</td><td>>sp|P01877|IGHA2_HUMAN Immunoglobulin heavy constant alpha 2 OS=Homo sapiens OX=9606 GN=IGHA2 PE=1 SV=4</td><td>576</td><td>20201117_08512_1ng_glyco_hcd30_etd30_compound15.5654.5654.5</td><td>controllerType=0 controllerNumber=1 scan=5654</td><td>16.4263</td></tr>
<tr><td>02702:4</td><td>3</td><td>K.HYTN[+1647.613]SSQDVTVPC[+57.021]RVPPPPPC[+57.021]C[+57.021]HPR.L</td><td>HexNAc(5)Hex(3)Fuc(1)</td><td>C[+57]*3, N[+1648]</td><td>1152.2519</td><td>4</td><td>4605.9857</td><td>4605.9689</td><td>0</td><td>3.7</td><td>89</td><td>Specific</td><td>638.03</td><td>569.55</td><td>569.55</td><td>9.5</td><td>57</td><td>>sp|P01877|IGHA2_HUMAN Immunoglobulin heavy constant alpha 2 OS=Homo sapiens OX=9606 GN=IGHA2 PE=1 SV=4</td><td>576</td><td>20201117_08512_1ng_glyco_hcd30_etd30_compound15.5717.5717.4</td><td>controllerType=0 controllerNumber=1 scan=5717</td><td>16.5779</td></tr>
<tr><td>14166:3</td><td>3</td><td>R.LSLHRPALEDLLLGSEAN[+1710.598]LTC[+57.021]TLTGLR.D</td><td>HexNAc(3)Hex(5)NeuAc(1)</td><td>C[+57], N[+1711]</td><td>1558.7452</td><td>3</td><td>4674.221</td><td>4674.1959</td><td>0</td><td>5.4</td><td>114</td><td>Specific</td><td>284.61</td><td>226.5</td><td>226.5</td><td>4.21</td><td>57</td><td>>sp|P01877|IGHA2_HUMAN Immunoglobulin heavy constant alpha 2 OS=Homo sapiens OX=9606 GN=IGHA2 PE=1 SV=4</td><td>576</td><td>20201117_08512_1ng_glyco_hcd30_etd30_compound15.17727.17727.3</td><td>controllerType=0 controllerNumber=1 scan=17727</td><td>44.984</td></tr>
<tr><td>13440:3</td><td>3</td><td>R.LSLHRPALEDLLLGSEAN[+1622.582]LTC[+57.021]TLTGLR.D</td><td>HexNAc(4)Hex(5)</td><td>C[+57], N[+1623]</td><td>1529.4026</td><td>3</td><td>4586.1934</td><td>4586.1799</td><td>0</td><td>2.9</td><td>114</td><td>Specific</td><td>382.18</td><td>372.24</td><td>372.24</td><td>7.54</td><td>57</td><td>>sp|P01877|IGHA2_HUMAN Immunoglobulin heavy constant alpha 2 OS=Homo sapiens OX=9606 GN=IGHA2 PE=1 SV=4</td><td>576</td><td>20201117_08512_1ng_glyco_hcd30_etd30_compound15.16960.16960.3</td><td>controllerType=0 controllerNumber=1 scan=16960</td><td>42.6642</td></tr>
<tr><td>13477:3</td><td>3</td><td>R.LSLHRPALEDLLLGSEAN[+1501.555]LTC[+57.021]TLTGLR.D</td><td>HexNAc(5)Hex(3)</td><td>C[+57], N[+1502]</td><td>1489.0602</td><td>3</td><td>4465.1662</td><td>4465.1536</td><td>0</td><td>2.8</td><td>114</td><td>Specific</td><td>383.86</td><td>383.86</td><td>383.86</td><td>7.54</td><td>57</td><td>>sp|P01877|IGHA2_HUMAN Immunoglobulin heavy constant alpha 2 OS=Homo sapiens OX=9606 GN=IGHA2 PE=1 SV=4</td><td>576</td><td>20201117_08512_1ng_glyco_hcd30_etd30_compound15.17000.17000.6</td><td>controllerType=0 controllerNumber=1 scan=17000</td><td>42.813</td></tr>
<tr><td>13439:3</td><td>3</td><td>R.LSLHRPALEDLLLGSEAN[+1702.581]LTC[+57.021]TLTGLR.D</td><td>HexNAc(2)Hex(8)</td><td>C[+57], N[+1703]</td><td>1556.0734</td><td>3</td><td>4666.2057</td><td>4666.1796</td><td>0</td><td>5.6</td><td>114</td><td>Specific</td><td>384.09</td><td>345.88</td><td>345.88</td><td>5.82</td><td>57</td><td>>sp|P01877|IGHA2_HUMAN Immunoglobulin heavy constant alpha 2 OS=Homo sapiens OX=9606 GN=IGHA2 PE=1 SV=4</td><td>576</td><td>20201117_08512_1ng_glyco_hcd30_etd30_compound15.16959.16959.3</td><td>controllerType=0 controllerNumber=1 scan=16959</td><td>42.6559</td></tr>
<tr><td>14121:3</td><td>3</td><td>R.LSLHRPALEDLLLGSEAN[+1872.651]LTC[+57.021]TLTGLR.D</td><td>HexNAc(3)Hex(6)NeuAc(1)</td><td>C[+57], N[+1873]</td><td>1612.7601</td><td>3</td><td>4836.2656</td><td>4836.2487</td><td>0</td><td>3.5</td><td>114</td><td>Specific</td><td>390.44</td><td>319.63</td><td>319.63</td><td>6.09</td><td>57</td><td>>sp|P01877|IGHA2_HUMAN Immunoglobulin heavy constant alpha 2 OS=Homo sapiens OX=9606 GN=IGHA2 PE=1 SV=4</td><td>576</td><td>20201117_08512_1ng_glyco_hcd30_etd30_compound15.17680.17680.3</td><td>controllerType=0 controllerNumber=1 scan=17680</td><td>44.852</td></tr>
<tr><td>14138:4</td><td>3</td><td>R.LSLHRPALEDLLLGSEAN[+1913.677]LTC[+57.021]TLTGLR.D</td><td>HexNAc(4)Hex(5)NeuAc(1)</td><td>C[+57], N[+1914]</td><td>1220.0821</td><td>4</td><td>4877.3065</td><td>4877.2753</td><td>0</td><td>6.4</td><td>114</td><td>Specific</td><td>413.97</td><td>413.97</td><td>413.97</td><td>4.93</td><td>57</td><td>>sp|P01877|IGHA2_HUMAN Immunoglobulin heavy constant alpha 2 OS=Homo sapiens OX=9606 GN=IGHA2 PE=1 SV=4</td><td>576</td><td>20201117_08512_1ng_glyco_hcd30_etd30_compound15.17698.17698.4</td><td>controllerType=0 controllerNumber=1 scan=17698</td><td>44.9085</td></tr>
<tr><td>13386:3</td><td>3</td><td>R.LSLHRPALEDLLLGSEAN[+1864.634]LTC[+57.021]TLTGLR.D</td><td>HexNAc(2)Hex(9)</td><td>C[+57], N[+1865]</td><td>1610.0866</td><td>3</td><td>4828.2453</td><td>4828.2324</td><td>0</td><td>2.7</td><td>114</td><td>Specific</td><td>419.26</td><td>388.37</td><td>388.37</td><td>7.74</td><td>57</td><td>>sp|P01877|IGHA2_HUMAN Immunoglobulin heavy constant alpha 2 OS=Homo sapiens OX=9606 GN=IGHA2 PE=1 SV=4</td><td>576</td><td>20201117_08512_1ng_glyco_hcd30_etd30_compound15.16903.16903.3</td><td>controllerType=0 controllerNumber=1 scan=16903</td><td>42.4816</td></tr>
<tr><td>13465:3</td><td>3</td><td>R.LSLHRPALEDLLLGSEAN[+1460.529]LTC[+57.021]TLTGLR.D</td><td>HexNAc(4)Hex(4)</td><td>C[+57], N[+1461]</td><td>1475.3852</td><td>3</td><td>4424.141</td><td>4424.1271</td><td>0</td><td>3.2</td><td>114</td><td>Specific</td><td>422.47</td><td>372.22</td><td>372.22</td><td>7.74</td><td>57</td><td>>sp|P01877|IGHA2_HUMAN Immunoglobulin heavy constant alpha 2 OS=Homo sapiens OX=9606 GN=IGHA2 PE=1 SV=4</td><td>576</td><td>20201117_08512_1ng_glyco_hcd30_etd30_compound15.16987.16987.3</td><td>controllerType=0 controllerNumber=1 scan=16987</td><td>42.7662</td></tr>
<tr><td>14161:3</td><td>3</td><td>R.LSLHRPALEDLLLGSEAN[+1751.624]LTC[+57.021]TLTGLR.D</td><td>HexNAc(4)Hex(4)NeuAc(1)</td><td>C[+57], N[+1752]</td><td>1572.4191</td><td>3</td><td>4715.2428</td><td>4715.2225</td><td>0</td><td>4.3</td><td>114</td><td>Specific</td><td>425.62</td><td>390.72</td><td>390.72</td><td>7.27</td><td>57</td><td>>sp|P01877|IGHA2_HUMAN Immunoglobulin heavy constant alpha 2 OS=Homo sapiens OX=9606 GN=IGHA2 PE=1 SV=4</td><td>576</td><td>20201117_08512_1ng_glyco_hcd30_etd30_compound15.17722.17722.3</td><td>controllerType=0 controllerNumber=1 scan=17722</td><td>44.9658</td></tr>
<tr><td>13455:3</td><td>3</td><td>R.LSLHRPALEDLLLGSEAN[+1501.555]LTC[+57.021]TLTGLR.D</td><td>HexNAc(5)Hex(3)</td><td>C[+57], N[+1502]</td><td>1489.0605</td><td>3</td><td>4465.1669</td><td>4465.1536</td><td>0</td><td>3</td><td>114</td><td>Specific</td><td>433.72</td><td>404.15</td><td>404.15</td><td>7.74</td><td>57</td><td>>sp|P01877|IGHA2_HUMAN Immunoglobulin heavy constant alpha 2 OS=Homo sapiens OX=9606 GN=IGHA2 PE=1 SV=4</td><td>576</td><td>20201117_08512_1ng_glyco_hcd30_etd30_compound15.16976.16976.3</td><td>controllerType=0 controllerNumber=1 scan=16976</td><td>42.7315</td></tr>
<tr><td>13414:3</td><td>3</td><td>R.LSLHRPALEDLLLGSEAN[+1540.529]LTC[+57.021]TLTGLR.D</td><td>HexNAc(2)Hex(7)</td><td>C[+57], N[+1541]</td><td>1502.051</td><td>3</td><td>4504.1383</td><td>4504.1268</td><td>0</td><td>2.6</td><td>114</td><td>Specific</td><td>437.37</td><td>419.04</td><td>419.04</td><td>7.74</td><td>57</td><td>>sp|P01877|IGHA2_HUMAN Immunoglobulin heavy constant alpha 2 OS=Homo sapiens OX=9606 GN=IGHA2 PE=1 SV=4</td><td>576</td><td>20201117_08512_1ng_glyco_hcd30_etd30_compound15.16932.16932.3</td><td>controllerType=0 controllerNumber=1 scan=16932</td><td>42.5685</td></tr>
<tr><td>13424:3</td><td>3</td><td>R.LSLHRPALEDLLLGSEAN[+1378.476]LTC[+57.021]TLTGLR.D</td><td>HexNAc(2)Hex(6)</td><td>C[+57], N[+1378]</td><td>1448.0339</td><td>3</td><td>4342.0871</td><td>4342.074</td><td>0</td><td>3</td><td>114</td><td>Specific</td><td>472.13</td><td>472.13</td><td>472.13</td><td>7.69</td><td>57</td><td>>sp|P01877|IGHA2_HUMAN Immunoglobulin heavy constant alpha 2 OS=Homo sapiens OX=9606 GN=IGHA2 PE=1 SV=4</td><td>576</td><td>20201117_08512_1ng_glyco_hcd30_etd30_compound15.16943.16943.3</td><td>controllerType=0 controllerNumber=1 scan=16943</td><td>42.6081</td></tr>
<tr><td>13451:3</td><td>3</td><td>R.LSLHRPALEDLLLGSEAN[+1216.423]LTC[+57.021]TLTGLR.D</td><td>HexNAc(2)Hex(5)</td><td>C[+57], N[+1216]</td><td>1394.0151</td><td>3</td><td>4180.0307</td><td>4180.0211</td><td>0</td><td>2.3</td><td>114</td><td>Specific</td><td>543.54</td><td>533.26</td><td>533.26</td><td>8.09</td><td>57</td><td>>sp|P01877|IGHA2_HUMAN Immunoglobulin heavy constant alpha 2 OS=Homo sapiens OX=9606 GN=IGHA2 PE=1 SV=4</td><td>576</td><td>20201117_08512_1ng_glyco_hcd30_etd30_compound15.16972.16972.3</td><td>controllerType=0 controllerNumber=1 scan=16972</td><td>42.7199</td></tr>
<tr><td>13429:4</td><td>3</td><td>R.LSLHRPALEDLLLGSEAN[+1581.555]LTC[+57.021]TLTGLR.D</td><td>HexNAc(3)Hex(6)</td><td>C[+57], N[+1582]</td><td>1137.0466</td><td>4</td><td>4545.1644</td><td>4545.1533</td><td>0</td><td>2.4</td><td>114</td><td>Specific</td><td>602.8</td><td>583.64</td><td>583.64</td><td>10.12</td><td>57</td><td>>sp|P01877|IGHA2_HUMAN Immunoglobulin heavy constant alpha 2 OS=Homo sapiens OX=9606 GN=IGHA2 PE=1 SV=4</td><td>576</td><td>20201117_08512_1ng_glyco_hcd30_etd30_compound15.16948.16948.4</td><td>controllerType=0 controllerNumber=1 scan=16948</td><td>42.6226</td></tr>
<tr><td>13088:4</td><td>3</td><td>R.LSLHRPALEDLLLGSEAN[+1622.582]LTC[+57.021]TLTGLR.D</td><td>HexNAc(4)Hex(5)</td><td>C[+57], N[+1623]</td><td>1147.3035</td><td>4</td><td>4586.1922</td><td>4586.1799</td><td>0</td><td>2.7</td><td>114</td><td>Specific</td><td>607.39</td><td>607.39</td><td>607.39</td><td>10.12</td><td>57</td><td>>sp|P01877|IGHA2_HUMAN Immunoglobulin heavy constant alpha 2 OS=Homo sapiens OX=9606 GN=IGHA2 PE=1 SV=4</td><td>576</td><td>20201117_08512_1ng_glyco_hcd30_etd30_compound15.16588.16588.4</td><td>controllerType=0 controllerNumber=1 scan=16588</td><td>41.5417</td></tr>
<tr><td>14834:4</td><td>3</td><td>R.LSLHRPALEDLLLGSEAN[+2407.852]LTC[+57.021]TLTGLR.D</td><td>HexNAc(5)Hex(5)NeuAc(2)</td><td>C[+57], N[+2408]</td><td>1343.6179</td><td>4</td><td>5371.4499</td><td>5371.4501</td><td>0</td><td>0</td><td>114</td><td>Specific</td><td>657.59</td><td>609.93</td><td>609.93</td><td>9.36</td><td>57</td><td>>sp|P01877|IGHA2_HUMAN Immunoglobulin heavy constant alpha 2 OS=Homo sapiens OX=9606 GN=IGHA2 PE=1 SV=4</td><td>576</td><td>20201117_08512_1ng_glyco_hcd30_etd30_compound15.18435.18435.4</td><td>controllerType=0 controllerNumber=1 scan=18435</td><td>47.2124</td></tr>
<tr><td>13466:5</td><td>3</td><td>R.LSLHRPALEDLLLGSEAN[+1501.555]LTC[+57.021]TLTGLR.D</td><td>HexNAc(5)Hex(3)</td><td>C[+57], N[+1502]</td><td>893.8382</td><td>5</td><td>4465.162</td><td>4465.1536</td><td>0</td><td>1.9</td><td>114</td><td>Specific</td><td>758.26</td><td>753.97</td><td>753.97</td><td>12.54</td><td>57</td><td>>sp|P01877|IGHA2_HUMAN Immunoglobulin heavy constant alpha 2 OS=Homo sapiens OX=9606 GN=IGHA2 PE=1 SV=4</td><td>576</td><td>20201117_08512_1ng_glyco_hcd30_etd30_compound15.16988.16988.5</td><td>controllerType=0 controllerNumber=1 scan=16988</td><td>42.771</td></tr>
<tr><td>14156:4</td><td>3</td><td>R.LSLHRPALEDLLLGSEAN[+1751.624]LTC[+57.021]TLTGLR.D</td><td>HexNAc(4)Hex(4)NeuAc(1)</td><td>C[+57], N[+1752]</td><td>1179.5657</td><td>4</td><td>4715.241</td><td>4715.2225</td><td>0</td><td>3.9</td><td>114</td><td>Specific</td><td>825.82</td><td>799.92</td><td>799.92</td><td>14.12</td><td>57</td><td>>sp|P01877|IGHA2_HUMAN Immunoglobulin heavy constant alpha 2 OS=Homo sapiens OX=9606 GN=IGHA2 PE=1 SV=4</td><td>576</td><td>20201117_08512_1ng_glyco_hcd30_etd30_compound15.17717.17717.4</td><td>controllerType=0 controllerNumber=1 scan=17717</td><td>44.9546</td></tr>
<tr><td>13435:4</td><td>3</td><td>R.LSLHRPALEDLLLGSEAN[+1702.581]LTC[+57.021]TLTGLR.D</td><td>HexNAc(2)Hex(8)</td><td>C[+57], N[+1703]</td><td>1167.3024</td><td>4</td><td>4666.1878</td><td>4666.1796</td><td>0</td><td>1.8</td><td>114</td><td>Specific</td><td>847.89</td><td>804.95</td><td>804.95</td><td>14.11</td><td>57</td><td>>sp|P01877|IGHA2_HUMAN Immunoglobulin heavy constant alpha 2 OS=Homo sapiens OX=9606 GN=IGHA2 PE=1 SV=4</td><td>576</td><td>20201117_08512_1ng_glyco_hcd30_etd30_compound15.16955.16955.4</td><td>controllerType=0 controllerNumber=1 scan=16955</td><td>42.6482</td></tr>
<tr><td>13450:4</td><td>3</td><td>R.LSLHRPALEDLLLGSEAN[+1501.555]LTC[+57.021]TLTGLR.D</td><td>HexNAc(5)Hex(3)</td><td>C[+57], N[+1502]</td><td>1117.05</td><td>4</td><td>4465.178</td><td>4465.1536</td><td>0</td><td>5.5</td><td>114</td><td>Specific</td><td>854.73</td><td>854.73</td><td>854.73</td><td>14.65</td><td>57</td><td>>sp|P01877|IGHA2_HUMAN Immunoglobulin heavy constant alpha 2 OS=Homo sapiens OX=9606 GN=IGHA2 PE=1 SV=4</td><td>576</td><td>20201117_08512_1ng_glyco_hcd30_etd30_compound15.16971.16971.4</td><td>controllerType=0 controllerNumber=1 scan=16971</td><td>42.7095</td></tr>
<tr><td>13482:4</td><td>3</td><td>R.LSLHRPALEDLLLGSEAN[+1298.476]LTC[+57.021]TLTGLR.D</td><td>HexNAc(4)Hex(3)</td><td>C[+57], N[+1298]</td><td>1066.2754</td><td>4</td><td>4262.0799</td><td>4262.0742</td><td>0</td><td>1.3</td><td>114</td><td>Specific</td><td>859.73</td><td>859.73</td><td>859.73</td><td>15.11</td><td>57</td><td>>sp|P01877|IGHA2_HUMAN Immunoglobulin heavy constant alpha 2 OS=Homo sapiens OX=9606 GN=IGHA2 PE=1 SV=4</td><td>576</td><td>20201117_08512_1ng_glyco_hcd30_etd30_compound15.17005.17005.4</td><td>controllerType=0 controllerNumber=1 scan=17005</td><td>42.8265</td></tr>
<tr><td>13400:4</td><td>3</td><td>R.LSLHRPALEDLLLGSEAN[+1825.661]LTC[+57.021]TLTGLR.D</td><td>HexNAc(5)Hex(5)</td><td>C[+57], N[+1826]</td><td>1198.0733</td><td>4</td><td>4789.2713</td><td>4789.2592</td><td>0</td><td>2.5</td><td>114</td><td>Specific</td><td>887.14</td><td>855.59</td><td>855.59</td><td>15.12</td><td>57</td><td>>sp|P01877|IGHA2_HUMAN Immunoglobulin heavy constant alpha 2 OS=Homo sapiens OX=9606 GN=IGHA2 PE=1 SV=4</td><td>576</td><td>20201117_08512_1ng_glyco_hcd30_etd30_compound15.16917.16917.4</td><td>controllerType=0 controllerNumber=1 scan=16917</td><td>42.5185</td></tr>
<tr><td>13434:4</td><td>3</td><td>R.LSLHRPALEDLLLGSEAN[+1622.582]LTC[+57.021]TLTGLR.D</td><td>HexNAc(4)Hex(5)</td><td>C[+57], N[+1623]</td><td>1147.3044</td><td>4</td><td>4586.1956</td><td>4586.1799</td><td>0</td><td>3.4</td><td>114</td><td>Specific</td><td>947.21</td><td>846.22</td><td>846.22</td><td>16.12</td><td>57</td><td>>sp|P01877|IGHA2_HUMAN Immunoglobulin heavy constant alpha 2 OS=Homo sapiens OX=9606 GN=IGHA2 PE=1 SV=4</td><td>576</td><td>20201117_08512_1ng_glyco_hcd30_etd30_compound15.16954.16954.4</td><td>controllerType=0 controllerNumber=1 scan=16954</td><td>42.6453</td></tr>
<tr><td>13460:4</td><td>3</td><td>R.LSLHRPALEDLLLGSEAN[+1460.529]LTC[+57.021]TLTGLR.D</td><td>HexNAc(4)Hex(4)</td><td>C[+57], N[+1461]</td><td>1106.7922</td><td>4</td><td>4424.1468</td><td>4424.1271</td><td>0</td><td>4.5</td><td>114</td><td>Specific</td><td>954.91</td><td>878.61</td><td>878.61</td><td>16.65</td><td>57</td><td>>sp|P01877|IGHA2_HUMAN Immunoglobulin heavy constant alpha 2 OS=Homo sapiens OX=9606 GN=IGHA2 PE=1 SV=4</td><td>576</td><td>20201117_08512_1ng_glyco_hcd30_etd30_compound15.16982.16982.4</td><td>controllerType=0 controllerNumber=1 scan=16982</td><td>42.7523</td></tr>
<tr><td>13385:4</td><td>3</td><td>R.LSLHRPALEDLLLGSEAN[+1864.634]LTC[+57.021]TLTGLR.D</td><td>HexNAc(2)Hex(9)</td><td>C[+57], N[+1865]</td><td>1207.8172</td><td>4</td><td>4828.2469</td><td>4828.2324</td><td>0</td><td>3</td><td>114</td><td>Specific</td><td>1020.73</td><td>941.69</td><td>941.69</td><td>18.12</td><td>57</td><td>>sp|P01877|IGHA2_HUMAN Immunoglobulin heavy constant alpha 2 OS=Homo sapiens OX=9606 GN=IGHA2 PE=1 SV=4</td><td>576</td><td>20201117_08512_1ng_glyco_hcd30_etd30_compound15.16902.16902.4</td><td>controllerType=0 controllerNumber=1 scan=16902</td><td>42.4786</td></tr>
<tr><td>13447:4</td><td>3</td><td>R.LSLHRPALEDLLLGSEAN[+1216.423]LTC[+57.021]TLTGLR.D</td><td>HexNAc(2)Hex(5)</td><td>C[+57], N[+1216]</td><td>1045.7635</td><td>4</td><td>4180.0321</td><td>4180.0211</td><td>0</td><td>2.6</td><td>114</td><td>Specific</td><td>1022.86</td><td>993.55</td><td>993.55</td><td>18.12</td><td>57</td><td>>sp|P01877|IGHA2_HUMAN Immunoglobulin heavy constant alpha 2 OS=Homo sapiens OX=9606 GN=IGHA2 PE=1 SV=4</td><td>576</td><td>20201117_08512_1ng_glyco_hcd30_etd30_compound15.16967.16967.4</td><td>controllerType=0 controllerNumber=1 scan=16967</td><td>42.6926</td></tr>
<tr><td>13419:4</td><td>3</td><td>R.LSLHRPALEDLLLGSEAN[+1378.476]LTC[+57.021]TLTGLR.D</td><td>HexNAc(2)Hex(6)</td><td>C[+57], N[+1378]</td><td>1086.2767</td><td>4</td><td>4342.0848</td><td>4342.074</td><td>0</td><td>2.5</td><td>114</td><td>Specific</td><td>1087.38</td><td>1027.38</td><td>1027.38</td><td>19.12</td><td>57</td><td>>sp|P01877|IGHA2_HUMAN Immunoglobulin heavy constant alpha 2 OS=Homo sapiens OX=9606 GN=IGHA2 PE=1 SV=4</td><td>576</td><td>20201117_08512_1ng_glyco_hcd30_etd30_compound15.16938.16938.4</td><td>controllerType=0 controllerNumber=1 scan=16938</td><td>42.5922</td></tr>
<tr><td>13403:4</td><td>3</td><td>R.LSLHRPALEDLLLGSEAN[+1540.529]LTC[+57.021]TLTGLR.D</td><td>HexNAc(2)Hex(7)</td><td>C[+57], N[+1541]</td><td>1126.7901</td><td>4</td><td>4504.1385</td><td>4504.1268</td><td>0</td><td>2.6</td><td>114</td><td>Specific</td><td>1130.33</td><td>1045.56</td><td>1045.56</td><td>20.12</td><td>57</td><td>>sp|P01877|IGHA2_HUMAN Immunoglobulin heavy constant alpha 2 OS=Homo sapiens OX=9606 GN=IGHA2 PE=1 SV=4</td><td>576</td><td>20201117_08512_1ng_glyco_hcd30_etd30_compound15.16921.16921.4</td><td>controllerType=0 controllerNumber=1 scan=16921</td><td>42.532</td></tr>
<tr><td>02435:2</td><td>3</td><td>K.TPLTAN[+1971.719]ITK.S</td><td>HexNAc(5)Hex(5)Fuc(1)</td><td>N[+1972]</td><td>1465.6469</td><td>2</td><td>2930.2865</td><td>2930.2757</td><td>0</td><td>3.7</td><td>200</td><td>Specific</td><td>233.93</td><td>64.25</td><td>64.25</td><td>2.11</td><td>57</td><td>>sp|P01877|IGHA2_HUMAN Immunoglobulin heavy constant alpha 2 OS=Homo sapiens OX=9606 GN=IGHA2 PE=1 SV=4</td><td>576</td><td>20201117_08512_1ng_glyco_hcd30_etd30_compound15.5437.5437.2</td><td>controllerType=0 controllerNumber=1 scan=5437</td><td>15.8778</td></tr>
<tr><td>03943:3</td><td>3</td><td>K.TPLTAN[+2262.814]ITK.S</td><td>HexNAc(5)Hex(5)Fuc(1)NeuAc(1)</td><td>N[+2263]</td><td>1074.4629</td><td>3</td><td>3221.3742</td><td>3221.3711</td><td>0</td><td>1</td><td>200</td><td>Specific</td><td>235.23</td><td>116.65</td><td>116.65</td><td>2.73</td><td>57</td><td>>sp|P01877|IGHA2_HUMAN Immunoglobulin heavy constant alpha 2 OS=Homo sapiens OX=9606 GN=IGHA2 PE=1 SV=4</td><td>576</td><td>20201117_08512_1ng_glyco_hcd30_etd30_compound15.7008.7008.3</td><td>controllerType=0 controllerNumber=1 scan=7008</td><td>19.1229</td></tr>
<tr><td>02487:2</td><td>3</td><td>K.TPLTAN[+1606.587]ITK.S</td><td>HexNAc(4)Hex(4)Fuc(1)</td><td>N[+1607]</td><td>1283.0794</td><td>2</td><td>2565.1515</td><td>2565.1435</td><td>0</td><td>3.1</td><td>200</td><td>Specific</td><td>251.72</td><td>87.13</td><td>87.13</td><td>2.65</td><td>57</td><td>>sp|P01877|IGHA2_HUMAN Immunoglobulin heavy constant alpha 2 OS=Homo sapiens OX=9606 GN=IGHA2 PE=1 SV=4</td><td>576</td><td>20201117_08512_1ng_glyco_hcd30_etd30_compound15.5491.5491.2</td><td>controllerType=0 controllerNumber=1 scan=5491</td><td>16.0073</td></tr>
<tr><td>02460:3</td><td>3</td><td>K.TPLTAN[+1809.666]ITK.S</td><td>HexNAc(5)Hex(4)Fuc(1)</td><td>N[+1810]</td><td>923.4144</td><td>3</td><td>2768.2286</td><td>2768.2228</td><td>0</td><td>2.1</td><td>200</td><td>Specific</td><td>277.8</td><td>118.72</td><td>118.72</td><td>2.72</td><td>57</td><td>>sp|P01877|IGHA2_HUMAN Immunoglobulin heavy constant alpha 2 OS=Homo sapiens OX=9606 GN=IGHA2 PE=1 SV=4</td><td>576</td><td>20201117_08512_1ng_glyco_hcd30_etd30_compound15.5463.5463.3</td><td>controllerType=0 controllerNumber=1 scan=5463</td><td>15.9416</td></tr>
<tr><td>02422:3</td><td>3</td><td>K.TPLTAN[+1971.719]ITK.S</td><td>HexNAc(5)Hex(5)Fuc(1)</td><td>N[+1972]</td><td>977.4325</td><td>3</td><td>2930.2828</td><td>2930.2757</td><td>0</td><td>2.4</td><td>200</td><td>Specific</td><td>289.78</td><td>133.19</td><td>133.19</td><td>2.75</td><td>57</td><td>>sp|P01877|IGHA2_HUMAN Immunoglobulin heavy constant alpha 2 OS=Homo sapiens OX=9606 GN=IGHA2 PE=1 SV=4</td><td>576</td><td>20201117_08512_1ng_glyco_hcd30_etd30_compound15.5423.5423.3</td><td>controllerType=0 controllerNumber=1 scan=5423</td><td>15.8438</td></tr>
<tr><td>02521:3</td><td>3</td><td>K.TPLTAN[+1647.613]ITK.S</td><td>HexNAc(5)Hex(3)Fuc(1)</td><td>N[+1648]</td><td>869.3977</td><td>3</td><td>2606.1786</td><td>2606.17</td><td>0</td><td>3.3</td><td>200</td><td>Specific</td><td>293.27</td><td>120.62</td><td>120.62</td><td>2.69</td><td>57</td><td>>sp|P01877|IGHA2_HUMAN Immunoglobulin heavy constant alpha 2 OS=Homo sapiens OX=9606 GN=IGHA2 PE=1 SV=4</td><td>576</td><td>20201117_08512_1ng_glyco_hcd30_etd30_compound15.5527.5527.3</td><td>controllerType=0 controllerNumber=1 scan=5527</td><td>16.1195</td></tr>
<tr><td>02526:2</td><td>3</td><td>K.TPLTAN[+1501.555]ITK.S</td><td>HexNAc(5)Hex(3)</td><td>N[+1502]</td><td>1230.5645</td><td>2</td><td>2460.1217</td><td>2460.1121</td><td>0</td><td>3.9</td><td>200</td><td>Specific</td><td>296.4</td><td>87.75</td><td>87.75</td><td>2.63</td><td>57</td><td>>sp|P01877|IGHA2_HUMAN Immunoglobulin heavy constant alpha 2 OS=Homo sapiens OX=9606 GN=IGHA2 PE=1 SV=4</td><td>576</td><td>20201117_08512_1ng_glyco_hcd30_etd30_compound15.5532.5532.2</td><td>controllerType=0 controllerNumber=1 scan=5532</td><td>16.1322</td></tr>
<tr><td>02502:2</td><td>3</td><td>K.TPLTAN[+1647.613]ITK.S</td><td>HexNAc(5)Hex(3)Fuc(1)</td><td>N[+1648]</td><td>1303.5929</td><td>2</td><td>2606.1786</td><td>2606.17</td><td>0</td><td>3.3</td><td>200</td><td>Specific</td><td>305.52</td><td>114.37</td><td>114.37</td><td>2.75</td><td>57</td><td>>sp|P01877|IGHA2_HUMAN Immunoglobulin heavy constant alpha 2 OS=Homo sapiens OX=9606 GN=IGHA2 PE=1 SV=4</td><td>576</td><td>20201117_08512_1ng_glyco_hcd30_etd30_compound15.5507.5507.2</td><td>controllerType=0 controllerNumber=1 scan=5507</td><td>16.0543</td></tr>
<tr><td>02463:2</td><td>3</td><td>K.TPLTAN[+1809.666]ITK.S</td><td>HexNAc(5)Hex(4)Fuc(1)</td><td>N[+1810]</td><td>1384.6188</td><td>2</td><td>2768.2304</td><td>2768.2228</td><td>0</td><td>2.7</td><td>200</td><td>Specific</td><td>334.59</td><td>139.13</td><td>139.13</td><td>2.71</td><td>57</td><td>>sp|P01877|IGHA2_HUMAN Immunoglobulin heavy constant alpha 2 OS=Homo sapiens OX=9606 GN=IGHA2 PE=1 SV=4</td><td>576</td><td>20201117_08512_1ng_glyco_hcd30_etd30_compound15.5466.5466.2</td><td>controllerType=0 controllerNumber=1 scan=5466</td><td>15.951</td></tr>
<tr><td>02490:2</td><td>3</td><td>K.TPLTAN[+1444.534]ITK.S</td><td>HexNAc(4)Hex(3)Fuc(1)</td><td>N[+1445]</td><td>1202.0528</td><td>2</td><td>2403.0983</td><td>2403.0906</td><td>0</td><td>3.2</td><td>200</td><td>Specific</td><td>355.78</td><td>110.01</td><td>110.01</td><td>2.63</td><td>57</td><td>>sp|P01877|IGHA2_HUMAN Immunoglobulin heavy constant alpha 2 OS=Homo sapiens OX=9606 GN=IGHA2 PE=1 SV=4</td><td>576</td><td>20201117_08512_1ng_glyco_hcd30_etd30_compound15.5494.5494.2</td><td>controllerType=0 controllerNumber=1 scan=5494</td><td>16.0184</td></tr>
<tr><td>08351:3</td><td>3</td><td>R.MAGKPTHIN[+1809.666]VSVVMAEADGTC[+57.021]Y.-</td><td>HexNAc(5)Hex(4)Fuc(1)</td><td>C[+57], N[+1810]</td><td>1387.5899</td><td>3</td><td>4160.7551</td><td>4160.7489</td><td>0</td><td>1.5</td><td>319</td><td>CRagged</td><td>203.54</td><td>113.49</td><td>113.49</td><td>2.74</td><td>57</td><td>>sp|P01877|IGHA2_HUMAN Immunoglobulin heavy constant alpha 2 OS=Homo sapiens OX=9606 GN=IGHA2 PE=1 SV=4</td><td>576</td><td>20201117_08512_1ng_glyco_hcd30_etd30_compound15.11602.11602.3</td><td>controllerType=0 controllerNumber=1 scan=11602</td><td>28.8399</td></tr>
<tr><td>09598:3</td><td>3</td><td>R.MAGKPTHIN[+1710.598]VSVVMAEADGTC[+57.021]Y.-</td><td>HexNAc(3)Hex(5)NeuAc(1)</td><td>C[+57], N[+1711]</td><td>1354.5677</td><td>3</td><td>4061.6885</td><td>4061.6805</td><td>0</td><td>2</td><td>319</td><td>CRagged</td><td>212.44</td><td>118.28</td><td>118.28</td><td>2.81</td><td>57</td><td>>sp|P01877|IGHA2_HUMAN Immunoglobulin heavy constant alpha 2 OS=Homo sapiens OX=9606 GN=IGHA2 PE=1 SV=4</td><td>576</td><td>20201117_08512_1ng_glyco_hcd30_etd30_compound15.12907.12907.3</td><td>controllerType=0 controllerNumber=1 scan=12907</td><td>32.0493</td></tr>
<tr><td>09641:3</td><td>3</td><td>R.MAGKPTHIN[+1548.545]VSVVMAEADGTC[+57.021]Y.-</td><td>HexNAc(3)Hex(4)NeuAc(1)</td><td>C[+57], N[+1549]</td><td>1300.5505</td><td>3</td><td>3899.6369</td><td>3899.6277</td><td>0</td><td>2.4</td><td>319</td><td>CRagged</td><td>222.41</td><td>135.57</td><td>135.57</td><td>2.74</td><td>57</td><td>>sp|P01877|IGHA2_HUMAN Immunoglobulin heavy constant alpha 2 OS=Homo sapiens OX=9606 GN=IGHA2 PE=1 SV=4</td><td>576</td><td>20201117_08512_1ng_glyco_hcd30_etd30_compound15.12952.12952.3</td><td>controllerType=0 controllerNumber=1 scan=12952</td><td>32.1498</td></tr>
<tr><td>08276:3</td><td>3</td><td>R.MAGKPTHIN[+1702.581]VSVVMAEADGTC[+57.021]Y.-</td><td>HexNAc(2)Hex(8)</td><td>C[+57], N[+1703]</td><td>1351.8963</td><td>3</td><td>4053.6743</td><td>4053.6642</td><td>0</td><td>2.5</td><td>319</td><td>CRagged</td><td>246.17</td><td>246.17</td><td>196.29</td><td>3.65</td><td>57</td><td>>sp|P01877|IGHA2_HUMAN Immunoglobulin heavy constant alpha 2 OS=Homo sapiens OX=9606 GN=IGHA2 PE=1 SV=4</td><td>576</td><td>20201117_08512_1ng_glyco_hcd30_etd30_compound15.11523.11523.3</td><td>controllerType=0 controllerNumber=1 scan=11523</td><td>28.6226</td></tr>
<tr><td>08402:3</td><td>3</td><td>R.MAGKPTHIN[+1501.555]VSVVMAEADGTC[+57.021]Y.-</td><td>HexNAc(5)Hex(3)</td><td>C[+57], N[+1502]</td><td>1284.8876</td><td>3</td><td>3852.6483</td><td>3852.6382</td><td>0</td><td>2.6</td><td>319</td><td>CRagged</td><td>248.95</td><td>217.79</td><td>217.79</td><td>3.65</td><td>57</td><td>>sp|P01877|IGHA2_HUMAN Immunoglobulin heavy constant alpha 2 OS=Homo sapiens OX=9606 GN=IGHA2 PE=1 SV=4</td><td>576</td><td>20201117_08512_1ng_glyco_hcd30_etd30_compound15.11656.11656.3</td><td>controllerType=0 controllerNumber=1 scan=11656</td><td>28.9824</td></tr>
<tr><td>08316:3</td><td>3</td><td>R.MAGKPTHIN[+1378.476]VSVVMAEADGTC[+57.021]Y.-</td><td>HexNAc(2)Hex(6)</td><td>C[+57], N[+1378]</td><td>1243.8619</td><td>3</td><td>3729.5711</td><td>3729.5585</td><td>0</td><td>3.4</td><td>319</td><td>CRagged</td><td>274.28</td><td>229.02</td><td>197.78</td><td>3.65</td><td>57</td><td>>sp|P01877|IGHA2_HUMAN Immunoglobulin heavy constant alpha 2 OS=Homo sapiens OX=9606 GN=IGHA2 PE=1 SV=4</td><td>576</td><td>20201117_08512_1ng_glyco_hcd30_etd30_compound15.11565.11565.3</td><td>controllerType=0 controllerNumber=1 scan=11565</td><td>28.7343</td></tr>
<tr><td>08311:3</td><td>3</td><td>R.MAGKPTHIN[+1540.529]VSVVMAEADGTC[+57.021]Y.-</td><td>HexNAc(2)Hex(7)</td><td>C[+57], N[+1541]</td><td>1297.8802</td><td>3</td><td>3891.626</td><td>3891.6114</td><td>0</td><td>3.8</td><td>319</td><td>CRagged</td><td>277.03</td><td>277.03</td><td>193.35</td><td>4.09</td><td>57</td><td>>sp|P01877|IGHA2_HUMAN Immunoglobulin heavy constant alpha 2 OS=Homo sapiens OX=9606 GN=IGHA2 PE=1 SV=4</td><td>576</td><td>20201117_08512_1ng_glyco_hcd30_etd30_compound15.11560.11560.3</td><td>controllerType=0 controllerNumber=1 scan=11560</td><td>28.7206</td></tr>
<tr><td>08364:3</td><td>3</td><td>R.MAGKPTHIN[+1216.423]VSVVMAEADGTC[+57.021]Y.-</td><td>HexNAc(2)Hex(5)</td><td>C[+57], N[+1216]</td><td>1189.8423</td><td>3</td><td>3567.5125</td><td>3567.5057</td><td>0</td><td>1.9</td><td>319</td><td>CRagged</td><td>302.47</td><td>285.12</td><td>227.3</td><td>3.92</td><td>57</td><td>>sp|P01877|IGHA2_HUMAN Immunoglobulin heavy constant alpha 2 OS=Homo sapiens OX=9606 GN=IGHA2 PE=1 SV=4</td><td>576</td><td>20201117_08512_1ng_glyco_hcd30_etd30_compound15.11616.11616.3</td><td>controllerType=0 controllerNumber=1 scan=11616</td><td>28.8792</td></tr>
<tr><td>13430:3</td><td>4</td><td>R.GLSFDVSLEVSQGPGLLN[+2205.793]DTK.V</td><td>HexNAc(4)Hex(5)Fuc(2)NeuAc(1)</td><td>N[+2206]</td><td>1461.3118</td><td>3</td><td>4381.9209</td><td>4381.906</td><td>0</td><td>3.4</td><td>118</td><td>Specific</td><td>351.13</td><td>243.03</td><td>243.03</td><td>5.48</td><td>40</td><td>>sp|P01833|PIGR_HUMAN Polymeric immunoglobulin receptor OS=Homo sapiens OX=9606 GN=PIGR PE=1 SV=4</td><td>902</td><td>20201117_08512_1ng_glyco_hcd30_etd30_compound15.16949.16949.3</td><td>controllerType=0 controllerNumber=1 scan=16949</td><td>42.6267</td></tr>
<tr><td>13472:3</td><td>4</td><td>R.GLSFDVSLEVSQGPGLLN[+2059.735]DTK.V</td><td>HexNAc(4)Hex(5)Fuc(1)NeuAc(1)</td><td>N[+2060]</td><td>1412.6252</td><td>3</td><td>4235.861</td><td>4235.8481</td><td>0</td><td>3</td><td>118</td><td>Specific</td><td>376.37</td><td>264.43</td><td>264.43</td><td>5.64</td><td>40</td><td>>sp|P01833|PIGR_HUMAN Polymeric immunoglobulin receptor OS=Homo sapiens OX=9606 GN=PIGR PE=1 SV=4</td><td>902</td><td>20201117_08512_1ng_glyco_hcd30_etd30_compound15.16994.16994.3</td><td>controllerType=0 controllerNumber=1 scan=16994</td><td>42.7928</td></tr>
<tr><td>14680:4</td><td>4</td><td>R.GLSFDVSLEVSQGPGLLN[+2350.830]DTK.V</td><td>HexNAc(4)Hex(5)Fuc(1)NeuAc(2)</td><td>N[+2351]</td><td>1132.4943</td><td>4</td><td>4526.9554</td><td>4526.9436</td><td>0</td><td>2.6</td><td>118</td><td>Specific</td><td>467.73</td><td>430.27</td><td>430.27</td><td>5.57</td><td>40</td><td>>sp|P01833|PIGR_HUMAN Polymeric immunoglobulin receptor OS=Homo sapiens OX=9606 GN=PIGR PE=1 SV=4</td><td>902</td><td>20201117_08512_1ng_glyco_hcd30_etd30_compound15.18271.18271.4</td><td>controllerType=0 controllerNumber=1 scan=18271</td><td>46.69</td></tr>
<tr><td>14670:3</td><td>4</td><td>R.GLSFDVSLEVSQGPGLLN[+2350.830]DTK.V</td><td>HexNAc(4)Hex(5)Fuc(1)NeuAc(2)</td><td>N[+2351]</td><td>1509.6579</td><td>3</td><td>4526.9591</td><td>4526.9436</td><td>0</td><td>3.4</td><td>118</td><td>Specific</td><td>502.09</td><td>297.64</td><td>297.64</td><td>6.07</td><td>40</td><td>>sp|P01833|PIGR_HUMAN Polymeric immunoglobulin receptor OS=Homo sapiens OX=9606 GN=PIGR PE=1 SV=4</td><td>902</td><td>20201117_08512_1ng_glyco_hcd30_etd30_compound15.18261.18261.3</td><td>controllerType=0 controllerNumber=1 scan=18261</td><td>46.656</td></tr>
<tr><td>15605:4</td><td>4</td><td>K.QIGLYPVLVIDSSGYVNPN[+2204.772]YTGR.I</td><td>HexNAc(4)Hex(5)NeuAc(2)</td><td>N[+2205]</td><td>1183.2776</td><td>4</td><td>4730.0887</td><td>4730.0759</td><td>0</td><td>2.7</td><td>168</td><td>Specific</td><td>213.24</td><td>213.24</td><td>213.24</td><td>3.03</td><td>40</td><td>>sp|P01833|PIGR_HUMAN Polymeric immunoglobulin receptor OS=Homo sapiens OX=9606 GN=PIGR PE=1 SV=4</td><td>902</td><td>20201117_08512_1ng_glyco_hcd30_etd30_compound15.19255.19255.4</td><td>controllerType=0 controllerNumber=1 scan=19255</td><td>49.9551</td></tr>
<tr><td>15544:3</td><td>4</td><td>K.QIGLYPVLVIDSSGYVNPN[+2350.830]YTGR.I</td><td>HexNAc(4)Hex(5)Fuc(1)NeuAc(2)</td><td>N[+2351]</td><td>1626.056</td><td>3</td><td>4876.1533</td><td>4876.1338</td><td>0</td><td>4</td><td>168</td><td>Specific</td><td>390.11</td><td>279.3</td><td>279.3</td><td>5.64</td><td>40</td><td>>sp|P01833|PIGR_HUMAN Polymeric immunoglobulin receptor OS=Homo sapiens OX=9606 GN=PIGR PE=1 SV=4</td><td>902</td><td>20201117_08512_1ng_glyco_hcd30_etd30_compound15.19190.19190.3</td><td>controllerType=0 controllerNumber=1 scan=19190</td><td>49.7437</td></tr>
<tr><td>21132:3</td><td>4</td><td>R.[+42.011]LSLLEEPGN[+2351.851]GTFTVILNQLTSR.D</td><td>HexNAc(4)Hex(5)Fuc(3)NeuAc(1)</td><td>.[42], N[+2352]</td><td>1599.3898</td><td>3</td><td>4796.1549</td><td>4796.1539</td><td>0</td><td>0.2</td><td>413</td><td>Specific</td><td>260.26</td><td>182.79</td><td>182.79</td><td>4.02</td><td>40</td><td>>sp|P01833|PIGR_HUMAN Polymeric immunoglobulin receptor OS=Homo sapiens OX=9606 GN=PIGR PE=1 SV=4</td><td>902</td><td>20201117_08512_1ng_glyco_hcd30_etd30_compound15.25090.25090.2</td><td>controllerType=0 controllerNumber=1 scan=25090</td><td>65.3785</td></tr>
<tr><td>16886:3</td><td>4</td><td>R.LSLLEEPGN[+2350.830]GTFTVILNQLTSR.D</td><td>HexNAc(4)Hex(5)Fuc(1)NeuAc(2)</td><td>N[+2351]</td><td>1585.0449</td><td>3</td><td>4753.12</td><td>4753.1229</td><td>0</td><td>-0.6</td><td>413</td><td>Specific</td><td>331.14</td><td>181.63</td><td>181.63</td><td>3.47</td><td>40</td><td>>sp|P01833|PIGR_HUMAN Polymeric immunoglobulin receptor OS=Homo sapiens OX=9606 GN=PIGR PE=1 SV=4</td><td>902</td><td>20201117_08512_1ng_glyco_hcd30_etd30_compound15.20618.20618.3</td><td>controllerType=0 controllerNumber=1 scan=20618</td><td>54.3907</td></tr>
<tr><td>16590:4</td><td>4</td><td>R.LSLLEEPGN[+2350.830]GTFTVILNQLTSR.D</td><td>HexNAc(4)Hex(5)Fuc(1)NeuAc(2)</td><td>N[+2351]</td><td>1189.0406</td><td>4</td><td>4753.1405</td><td>4753.1229</td><td>0</td><td>3.7</td><td>413</td><td>Specific</td><td>443.42</td><td>389.83</td><td>389.83</td><td>5.57</td><td>40</td><td>>sp|P01833|PIGR_HUMAN Polymeric immunoglobulin receptor OS=Homo sapiens OX=9606 GN=PIGR PE=1 SV=4</td><td>902</td><td>20201117_08512_1ng_glyco_hcd30_etd30_compound15.20301.20301.4</td><td>controllerType=0 controllerNumber=1 scan=20301</td><td>53.2621</td></tr>
<tr><td>15339:3</td><td>4</td><td>R.LSLLEEPGN[+2076.750]GTFTVILNQLTSR.D</td><td>HexNAc(4)Hex(6)Fuc(2)</td><td>N[+2077]</td><td>1493.6944</td><td>3</td><td>4479.0686</td><td>4479.0428</td><td>0</td><td>5.8</td><td>413</td><td>Specific</td><td>451.89</td><td>373.31</td><td>373.31</td><td>5.96</td><td>40</td><td>>sp|P01833|PIGR_HUMAN Polymeric immunoglobulin receptor OS=Homo sapiens OX=9606 GN=PIGR PE=1 SV=4</td><td>902</td><td>20201117_08512_1ng_glyco_hcd30_etd30_compound15.18972.18972.2</td><td>controllerType=0 controllerNumber=1 scan=18972</td><td>48.9586</td></tr>
<tr><td>15304:4</td><td>4</td><td>R.LSLLEEPGN[+2205.793]GTFTVILNQLTSR.D</td><td>HexNAc(4)Hex(5)Fuc(2)NeuAc(1)</td><td>N[+2206]</td><td>1152.7797</td><td>4</td><td>4608.097</td><td>4608.0854</td><td>0</td><td>2.5</td><td>413</td><td>Specific</td><td>514.41</td><td>470.87</td><td>470.87</td><td>5.73</td><td>40</td><td>>sp|P01833|PIGR_HUMAN Polymeric immunoglobulin receptor OS=Homo sapiens OX=9606 GN=PIGR PE=1 SV=4</td><td>902</td><td>20201117_08512_1ng_glyco_hcd30_etd30_compound15.18934.18934.4</td><td>controllerType=0 controllerNumber=1 scan=18934</td><td>48.8042</td></tr>
<tr><td>15303:4</td><td>4</td><td>R.LSLLEEPGN[+2351.851]GTFTVILNQLTSR.D</td><td>HexNAc(4)Hex(5)Fuc(3)NeuAc(1)</td><td>N[+2352]</td><td>1189.2944</td><td>4</td><td>4754.1556</td><td>4754.1433</td><td>0</td><td>2.6</td><td>413</td><td>Specific</td><td>545.18</td><td>488.53</td><td>488.53</td><td>5.95</td><td>40</td><td>>sp|P01833|PIGR_HUMAN Polymeric immunoglobulin receptor OS=Homo sapiens OX=9606 GN=PIGR PE=1 SV=4</td><td>902</td><td>20201117_08512_1ng_glyco_hcd30_etd30_compound15.18933.18933.4</td><td>controllerType=0 controllerNumber=1 scan=18933</td><td>48.7969</td></tr>
<tr><td>15299:3</td><td>4</td><td>R.LSLLEEPGN[+2351.851]GTFTVILNQLTSR.D</td><td>HexNAc(4)Hex(5)Fuc(3)NeuAc(1)</td><td>N[+2352]</td><td>1585.3908</td><td>3</td><td>4754.1579</td><td>4754.1433</td><td>0</td><td>3.1</td><td>413</td><td>Specific</td><td>567.95</td><td>433.65</td><td>433.65</td><td>8.8</td><td>40</td><td>>sp|P01833|PIGR_HUMAN Polymeric immunoglobulin receptor OS=Homo sapiens OX=9606 GN=PIGR PE=1 SV=4</td><td>902</td><td>20201117_08512_1ng_glyco_hcd30_etd30_compound15.18929.18929.3</td><td>controllerType=0 controllerNumber=1 scan=18929</td><td>48.7873</td></tr>
<tr><td>16589:3</td><td>4</td><td>R.LSLLEEPGN[+2350.830]GTFTVILNQLTSR.D</td><td>HexNAc(4)Hex(5)Fuc(1)NeuAc(2)</td><td>N[+2351]</td><td>1585.0507</td><td>3</td><td>4753.1376</td><td>4753.1229</td><td>0</td><td>3.1</td><td>413</td><td>Specific</td><td>577.46</td><td>376.78</td><td>376.78</td><td>8.8</td><td>40</td><td>>sp|P01833|PIGR_HUMAN Polymeric immunoglobulin receptor OS=Homo sapiens OX=9606 GN=PIGR PE=1 SV=4</td><td>902</td><td>20201117_08512_1ng_glyco_hcd30_etd30_compound15.20300.20300.3</td><td>controllerType=0 controllerNumber=1 scan=20300</td><td>53.2579</td></tr>
<tr><td>15298:3</td><td>4</td><td>R.LSLLEEPGN[+2205.793]GTFTVILNQLTSR.D</td><td>HexNAc(4)Hex(5)Fuc(2)NeuAc(1)</td><td>N[+2206]</td><td>1536.7055</td><td>3</td><td>4608.102</td><td>4608.0854</td><td>0</td><td>3.6</td><td>413</td><td>Specific</td><td>584.41</td><td>426.8</td><td>426.8</td><td>8.8</td><td>40</td><td>>sp|P01833|PIGR_HUMAN Polymeric immunoglobulin receptor OS=Homo sapiens OX=9606 GN=PIGR PE=1 SV=4</td><td>902</td><td>20201117_08512_1ng_glyco_hcd30_etd30_compound15.18928.18928.3</td><td>controllerType=0 controllerNumber=1 scan=18928</td><td>48.7842</td></tr>
<tr><td>13636:3</td><td>4</td><td>K.IIEGEPNLKVPGN[+2350.830]VTAVLGETLK.V</td><td>HexNAc(4)Hex(5)Fuc(1)NeuAc(2)</td><td>N[+2351]</td><td>1581.4039</td><td>3</td><td>4742.197</td><td>4742.1797</td><td>0</td><td>3.7</td><td>457</td><td>Specific</td><td>308.16</td><td>243.94</td><td>243.94</td><td>3.77</td><td>40</td><td>>sp|P01833|PIGR_HUMAN Polymeric immunoglobulin receptor OS=Homo sapiens OX=9606 GN=PIGR PE=1 SV=4</td><td>902</td><td>20201117_08512_1ng_glyco_hcd30_etd30_compound15.17167.17167.3</td><td>controllerType=0 controllerNumber=1 scan=17167</td><td>43.3026</td></tr>
<tr><td>12230:3</td><td>4</td><td>K.IIEGEPNLKVPGN[+2205.793]VTAVLGETLK.V</td><td>HexNAc(4)Hex(5)Fuc(2)NeuAc(1)</td><td>N[+2206]</td><td>1533.0547</td><td>3</td><td>4597.1497</td><td>4597.1422</td><td>0</td><td>1.6</td><td>457</td><td>Specific</td><td>343.69</td><td>244.98</td><td>244.98</td><td>3.98</td><td>40</td><td>>sp|P01833|PIGR_HUMAN Polymeric immunoglobulin receptor OS=Homo sapiens OX=9606 GN=PIGR PE=1 SV=4</td><td>902</td><td>20201117_08512_1ng_glyco_hcd30_etd30_compound15.15683.15683.3</td><td>controllerType=0 controllerNumber=1 scan=15683</td><td>39.1422</td></tr>
<tr><td>12323:3</td><td>4</td><td>K.IIEGEPNLKVPGN[+2205.793]VTAVLGETLK.V</td><td>HexNAc(4)Hex(5)Fuc(2)NeuAc(1)</td><td>N[+2206]</td><td>1533.0563</td><td>3</td><td>4597.1544</td><td>4597.1422</td><td>0</td><td>2.7</td><td>457</td><td>Specific</td><td>349.55</td><td>281.65</td><td>281.65</td><td>4.18</td><td>40</td><td>>sp|P01833|PIGR_HUMAN Polymeric immunoglobulin receptor OS=Homo sapiens OX=9606 GN=PIGR PE=1 SV=4</td><td>902</td><td>20201117_08512_1ng_glyco_hcd30_etd30_compound15.15781.15781.3</td><td>controllerType=0 controllerNumber=1 scan=15781</td><td>39.38</td></tr>
<tr><td>12317:3</td><td>4</td><td>K.IIEGEPNLKVPGN[+2059.735]VTAVLGETLK.V</td><td>HexNAc(4)Hex(5)Fuc(1)NeuAc(1)</td><td>N[+2060]</td><td>1484.3705</td><td>3</td><td>4451.0971</td><td>4451.0843</td><td>0</td><td>2.9</td><td>457</td><td>Specific</td><td>358.97</td><td>270.51</td><td>270.51</td><td>4.18</td><td>40</td><td>>sp|P01833|PIGR_HUMAN Polymeric immunoglobulin receptor OS=Homo sapiens OX=9606 GN=PIGR PE=1 SV=4</td><td>902</td><td>20201117_08512_1ng_glyco_hcd30_etd30_compound15.15775.15775.3</td><td>controllerType=0 controllerNumber=1 scan=15775</td><td>39.3633</td></tr>
<tr><td>13593:4</td><td>4</td><td>K.IIEGEPNLKVPGN[+2350.830]VTAVLGETLK.V</td><td>HexNAc(4)Hex(5)Fuc(1)NeuAc(2)</td><td>N[+2351]</td><td>1186.3029</td><td>4</td><td>4742.1898</td><td>4742.1797</td><td>0</td><td>2.1</td><td>457</td><td>Specific</td><td>396.6</td><td>396.6</td><td>396.6</td><td>6.05</td><td>40</td><td>>sp|P01833|PIGR_HUMAN Polymeric immunoglobulin receptor OS=Homo sapiens OX=9606 GN=PIGR PE=1 SV=4</td><td>902</td><td>20201117_08512_1ng_glyco_hcd30_etd30_compound15.17122.17122.4</td><td>controllerType=0 controllerNumber=1 scan=17122</td><td>43.1707</td></tr>
<tr><td>13617:4</td><td>4</td><td>K.IIEGEPNLKVPGN[+2350.830]VTAVLGETLK.V</td><td>HexNAc(4)Hex(5)Fuc(1)NeuAc(2)</td><td>N[+2351]</td><td>1186.3033</td><td>4</td><td>4742.1912</td><td>4742.1797</td><td>0</td><td>2.4</td><td>457</td><td>Specific</td><td>510.96</td><td>506.33</td><td>506.33</td><td>6.37</td><td>40</td><td>>sp|P01833|PIGR_HUMAN Polymeric immunoglobulin receptor OS=Homo sapiens OX=9606 GN=PIGR PE=1 SV=4</td><td>902</td><td>20201117_08512_1ng_glyco_hcd30_etd30_compound15.17147.17147.4</td><td>controllerType=0 controllerNumber=1 scan=17147</td><td>43.243</td></tr>
<tr><td>12322:4</td><td>4</td><td>K.IIEGEPNLKVPGN[+2059.735]VTAVLGETLK.V</td><td>HexNAc(4)Hex(5)Fuc(1)NeuAc(1)</td><td>N[+2060]</td><td>1113.5289</td><td>4</td><td>4451.0936</td><td>4451.0843</td><td>0</td><td>2.1</td><td>457</td><td>Specific</td><td>550.97</td><td>504.14</td><td>504.14</td><td>6.59</td><td>40</td><td>>sp|P01833|PIGR_HUMAN Polymeric immunoglobulin receptor OS=Homo sapiens OX=9606 GN=PIGR PE=1 SV=4</td><td>902</td><td>20201117_08512_1ng_glyco_hcd30_etd30_compound15.15780.15780.4</td><td>controllerType=0 controllerNumber=1 scan=15780</td><td>39.3743</td></tr>
<tr><td>11911:3</td><td>4</td><td>K.IIEGEPNLKVPGN[+2059.735]VTAVLGETLKVPC[+57.021]HFPC[+57.021]K.F</td><td>HexNAc(4)Hex(5)Fuc(1)NeuAc(1)</td><td>C[+57]*2, N[+2060]</td><td>1826.1893</td><td>3</td><td>5476.5533</td><td>5476.5418</td><td>0</td><td>2.1</td><td>457</td><td>Specific</td><td>216.67</td><td>214.29</td><td>214.29</td><td>2.77</td><td>40</td><td>>sp|P01833|PIGR_HUMAN Polymeric immunoglobulin receptor OS=Homo sapiens OX=9606 GN=PIGR PE=1 SV=4</td><td>902</td><td>20201117_08512_1ng_glyco_hcd30_etd30_compound15.15340.15340.3</td><td>controllerType=0 controllerNumber=1 scan=15340</td><td>38.3465</td></tr>
<tr><td>12005:4</td><td>4</td><td>K.IIEGEPNLKVPGN[+2205.793]VTAVLGETLKVPC[+57.021]HFPC[+57.021]K.F</td><td>HexNAc(4)Hex(5)Fuc(2)NeuAc(1)</td><td>C[+57]*2, N[+2206]</td><td>1406.4081</td><td>4</td><td>5622.6105</td><td>5622.5997</td><td>0</td><td>1.9</td><td>457</td><td>Specific</td><td>217.54</td><td>217.54</td><td>217.54</td><td>2.92</td><td>40</td><td>>sp|P01833|PIGR_HUMAN Polymeric immunoglobulin receptor OS=Homo sapiens OX=9606 GN=PIGR PE=1 SV=4</td><td>902</td><td>20201117_08512_1ng_glyco_hcd30_etd30_compound15.15438.15438.4</td><td>controllerType=0 controllerNumber=1 scan=15438</td><td>38.5743</td></tr>
<tr><td>11967:4</td><td>4</td><td>K.IIEGEPNLKVPGN[+2205.793]VTAVLGETLKVPC[+57.021]HFPC[+57.021]K.F</td><td>HexNAc(4)Hex(5)Fuc(2)NeuAc(1)</td><td>C[+57]*2, N[+2206]</td><td>1406.41</td><td>4</td><td>5622.6183</td><td>5622.5997</td><td>0</td><td>3.3</td><td>457</td><td>Specific</td><td>317.16</td><td>317.16</td><td>317.16</td><td>4.44</td><td>40</td><td>>sp|P01833|PIGR_HUMAN Polymeric immunoglobulin receptor OS=Homo sapiens OX=9606 GN=PIGR PE=1 SV=4</td><td>902</td><td>20201117_08512_1ng_glyco_hcd30_etd30_compound15.15398.15398.2</td><td>controllerType=0 controllerNumber=1 scan=15398</td><td>38.4648</td></tr>
<tr><td>12987:6</td><td>4</td><td>K.IIEGEPNLKVPGN[+2350.830]VTAVLGETLKVPC[+57.021]HFPC[+57.021]K.F</td><td>HexNAc(4)Hex(5)Fuc(1)NeuAc(2)</td><td>C[+57]*2, N[+2351]</td><td>962.1135</td><td>6</td><td>5767.6447</td><td>5767.6372</td><td>0</td><td>1.3</td><td>457</td><td>Specific</td><td>329.94</td><td>329.94</td><td>329.94</td><td>4.98</td><td>40</td><td>>sp|P01833|PIGR_HUMAN Polymeric immunoglobulin receptor OS=Homo sapiens OX=9606 GN=PIGR PE=1 SV=4</td><td>902</td><td>20201117_08512_1ng_glyco_hcd30_etd30_compound15.16481.16481.6</td><td>controllerType=0 controllerNumber=1 scan=16481</td><td>41.1999</td></tr>
<tr><td>12928:4</td><td>4</td><td>K.IIEGEPNLKVPGN[+2350.830]VTAVLGETLKVPC[+57.021]HFPC[+57.021]K.F</td><td>HexNAc(4)Hex(5)Fuc(1)NeuAc(2)</td><td>C[+57]*2, N[+2351]</td><td>1442.6711</td><td>4</td><td>5767.6628</td><td>5767.6372</td><td>0</td><td>4.4</td><td>457</td><td>Specific</td><td>367.27</td><td>367.27</td><td>367.27</td><td>5.39</td><td>40</td><td>>sp|P01833|PIGR_HUMAN Polymeric immunoglobulin receptor OS=Homo sapiens OX=9606 GN=PIGR PE=1 SV=4</td><td>902</td><td>20201117_08512_1ng_glyco_hcd30_etd30_compound15.16419.16419.4</td><td>controllerType=0 controllerNumber=1 scan=16419</td><td>41.0231</td></tr>
<tr><td>12974:4</td><td>4</td><td>K.IIEGEPNLKVPGN[+2350.830]VTAVLGETLKVPC[+57.021]HFPC[+57.021]K.F</td><td>HexNAc(4)Hex(5)Fuc(1)NeuAc(2)</td><td>C[+57]*2, N[+2351]</td><td>1442.6687</td><td>4</td><td>5767.653</td><td>5767.6372</td><td>0</td><td>2.7</td><td>457</td><td>Specific</td><td>405.3</td><td>405.3</td><td>405.3</td><td>6.26</td><td>40</td><td>>sp|P01833|PIGR_HUMAN Polymeric immunoglobulin receptor OS=Homo sapiens OX=9606 GN=PIGR PE=1 SV=4</td><td>902</td><td>20201117_08512_1ng_glyco_hcd30_etd30_compound15.16468.16468.4</td><td>controllerType=0 controllerNumber=1 scan=16468</td><td>41.1628</td></tr>
<tr><td>11866:5</td><td>4</td><td>K.IIEGEPNLKVPGN[+2351.851]VTAVLGETLKVPC[+57.021]HFPC[+57.021]K.F</td><td>HexNAc(4)Hex(5)Fuc(3)NeuAc(1)</td><td>C[+57]*2, N[+2352]</td><td>1154.5409</td><td>5</td><td>5768.6754</td><td>5768.6576</td><td>0</td><td>3.1</td><td>457</td><td>Specific</td><td>422.27</td><td>422.27</td><td>422.27</td><td>6.05</td><td>40</td><td>>sp|P01833|PIGR_HUMAN Polymeric immunoglobulin receptor OS=Homo sapiens OX=9606 GN=PIGR PE=1 SV=4</td><td>902</td><td>20201117_08512_1ng_glyco_hcd30_etd30_compound15.15293.15293.5</td><td>controllerType=0 controllerNumber=1 scan=15293</td><td>38.228</td></tr>
<tr><td>12037:4</td><td>4</td><td>K.IIEGEPNLKVPGN[+2059.735]VTAVLGETLKVPC[+57.021]HFPC[+57.021]K.F</td><td>HexNAc(4)Hex(5)Fuc(1)NeuAc(1)</td><td>C[+57]*2, N[+2060]</td><td>1369.8951</td><td>4</td><td>5476.5587</td><td>5476.5418</td><td>0</td><td>3.1</td><td>457</td><td>Specific</td><td>445.52</td><td>430.67</td><td>430.67</td><td>6.01</td><td>40</td><td>>sp|P01833|PIGR_HUMAN Polymeric immunoglobulin receptor OS=Homo sapiens OX=9606 GN=PIGR PE=1 SV=4</td><td>902</td><td>20201117_08512_1ng_glyco_hcd30_etd30_compound15.15472.15472.4</td><td>controllerType=0 controllerNumber=1 scan=15472</td><td>38.6493</td></tr>
<tr><td>11874:4</td><td>4</td><td>K.IIEGEPNLKVPGN[+2351.851]VTAVLGETLKVPC[+57.021]HFPC[+57.021]K.F</td><td>HexNAc(4)Hex(5)Fuc(3)NeuAc(1)</td><td>C[+57]*2, N[+2352]</td><td>1442.9236</td><td>4</td><td>5768.6725</td><td>5768.6576</td><td>0</td><td>2.6</td><td>457</td><td>Specific</td><td>493.72</td><td>493.72</td><td>493.72</td><td>6.38</td><td>40</td><td>>sp|P01833|PIGR_HUMAN Polymeric immunoglobulin receptor OS=Homo sapiens OX=9606 GN=PIGR PE=1 SV=4</td><td>902</td><td>20201117_08512_1ng_glyco_hcd30_etd30_compound15.15301.15301.4</td><td>controllerType=0 controllerNumber=1 scan=15301</td><td>38.2545</td></tr>
<tr><td>11849:4</td><td>4</td><td>K.IIEGEPNLKVPGN[+2205.793]VTAVLGETLKVPC[+57.021]HFPC[+57.021]K.F</td><td>HexNAc(4)Hex(5)Fuc(2)NeuAc(1)</td><td>C[+57]*2, N[+2206]</td><td>1406.4099</td><td>4</td><td>5622.6178</td><td>5622.5997</td><td>0</td><td>3.2</td><td>457</td><td>Specific</td><td>549.71</td><td>549.71</td><td>549.71</td><td>6.61</td><td>40</td><td>>sp|P01833|PIGR_HUMAN Polymeric immunoglobulin receptor OS=Homo sapiens OX=9606 GN=PIGR PE=1 SV=4</td><td>902</td><td>20201117_08512_1ng_glyco_hcd30_etd30_compound15.15275.15275.4</td><td>controllerType=0 controllerNumber=1 scan=15275</td><td>38.1658</td></tr>
<tr><td>11867:4</td><td>4</td><td>K.IIEGEPNLKVPGN[+2059.735]VTAVLGETLKVPC[+57.021]HFPC[+57.021]K.F</td><td>HexNAc(4)Hex(5)Fuc(1)NeuAc(1)</td><td>C[+57]*2, N[+2060]</td><td>1369.8939</td><td>4</td><td>5476.5539</td><td>5476.5418</td><td>0</td><td>2.2</td><td>457</td><td>Specific</td><td>610.96</td><td>571.51</td><td>571.51</td><td>8.67</td><td>40</td><td>>sp|P01833|PIGR_HUMAN Polymeric immunoglobulin receptor OS=Homo sapiens OX=9606 GN=PIGR PE=1 SV=4</td><td>902</td><td>20201117_08512_1ng_glyco_hcd30_etd30_compound15.15294.15294.4</td><td>controllerType=0 controllerNumber=1 scan=15294</td><td>38.2325</td></tr>
<tr><td>12973:5</td><td>4</td><td>K.IIEGEPNLKVPGN[+2350.830]VTAVLGETLKVPC[+57.021]HFPC[+57.021]K.F</td><td>HexNAc(4)Hex(5)Fuc(1)NeuAc(2)</td><td>C[+57]*2, N[+2351]</td><td>1154.3359</td><td>5</td><td>5767.6506</td><td>5767.6372</td><td>0</td><td>2.3</td><td>457</td><td>Specific</td><td>640.88</td><td>640.88</td><td>640.88</td><td>8.66</td><td>40</td><td>>sp|P01833|PIGR_HUMAN Polymeric immunoglobulin receptor OS=Homo sapiens OX=9606 GN=PIGR PE=1 SV=4</td><td>902</td><td>20201117_08512_1ng_glyco_hcd30_etd30_compound15.16467.16467.5</td><td>controllerType=0 controllerNumber=1 scan=16467</td><td>41.1599</td></tr>
<tr><td>11865:5</td><td>4</td><td>K.IIEGEPNLKVPGN[+2059.735]VTAVLGETLKVPC[+57.021]HFPC[+57.021]K.F</td><td>HexNAc(4)Hex(5)Fuc(1)NeuAc(1)</td><td>C[+57]*2, N[+2060]</td><td>1096.1167</td><td>5</td><td>5476.5544</td><td>5476.5418</td><td>0</td><td>2.3</td><td>457</td><td>Specific</td><td>686.05</td><td>686.05</td><td>686.05</td><td>9.66</td><td>40</td><td>>sp|P01833|PIGR_HUMAN Polymeric immunoglobulin receptor OS=Homo sapiens OX=9606 GN=PIGR PE=1 SV=4</td><td>902</td><td>20201117_08512_1ng_glyco_hcd30_etd30_compound15.15292.15292.5</td><td>controllerType=0 controllerNumber=1 scan=15292</td><td>38.221</td></tr>
<tr><td>12926:5</td><td>4</td><td>K.IIEGEPNLKVPGN[+2350.830]VTAVLGETLKVPC[+57.021]HFPC[+57.021]K.F</td><td>HexNAc(4)Hex(5)Fuc(1)NeuAc(2)</td><td>C[+57]*2, N[+2351]</td><td>1154.3353</td><td>5</td><td>5767.6475</td><td>5767.6372</td><td>0</td><td>1.8</td><td>457</td><td>Specific</td><td>697.87</td><td>653.72</td><td>653.72</td><td>9.64</td><td>40</td><td>>sp|P01833|PIGR_HUMAN Polymeric immunoglobulin receptor OS=Homo sapiens OX=9606 GN=PIGR PE=1 SV=4</td><td>902</td><td>20201117_08512_1ng_glyco_hcd30_etd30_compound15.16417.16417.5</td><td>controllerType=0 controllerNumber=1 scan=16417</td><td>41.012</td></tr>
<tr><td>11846:5</td><td>4</td><td>K.IIEGEPNLKVPGN[+2205.793]VTAVLGETLKVPC[+57.021]HFPC[+57.021]K.F</td><td>HexNAc(4)Hex(5)Fuc(2)NeuAc(1)</td><td>C[+57]*2, N[+2206]</td><td>1125.3285</td><td>5</td><td>5622.6134</td><td>5622.5997</td><td>0</td><td>2.4</td><td>457</td><td>Specific</td><td>743.69</td><td>733.47</td><td>733.47</td><td>10.66</td><td>40</td><td>>sp|P01833|PIGR_HUMAN Polymeric immunoglobulin receptor OS=Homo sapiens OX=9606 GN=PIGR PE=1 SV=4</td><td>902</td><td>20201117_08512_1ng_glyco_hcd30_etd30_compound15.15272.15272.5</td><td>controllerType=0 controllerNumber=1 scan=15272</td><td>38.1562</td></tr>
<tr><td>10681:4</td><td>4</td><td>K.VPGN[+2059.735]VTAVLGETLKVPC[+57.021]HFPC[+57.021]K.F</td><td>HexNAc(4)Hex(5)Fuc(1)NeuAc(1)</td><td>C[+57]*2, N[+2060]</td><td>1121.506</td><td>4</td><td>4483.0023</td><td>4482.9923</td><td>0</td><td>2.2</td><td>466</td><td>Specific</td><td>380.23</td><td>380.23</td><td>380.23</td><td>5.68</td><td>40</td><td>>sp|P01833|PIGR_HUMAN Polymeric immunoglobulin receptor OS=Homo sapiens OX=9606 GN=PIGR PE=1 SV=4</td><td>902</td><td>20201117_08512_1ng_glyco_hcd30_etd30_compound15.14046.14046.4</td><td>controllerType=0 controllerNumber=1 scan=14046</td><td>34.7661</td></tr>
<tr><td>05041:3</td><td>4</td><td>K.WN[+1548.545]NTGC[+57.021]QALPSQDEGPSK.A</td><td>HexNAc(3)Hex(4)NeuAc(1)</td><td>C[+57], N[+1549]</td><td>1179.813</td><td>3</td><td>3537.4246</td><td>3537.4215</td><td>0</td><td>0.9</td><td>498</td><td>Specific</td><td>272.72</td><td>148.83</td><td>148.83</td><td>4.6</td><td>40</td><td>>sp|P01833|PIGR_HUMAN Polymeric immunoglobulin receptor OS=Homo sapiens OX=9606 GN=PIGR PE=1 SV=4</td><td>902</td><td>20201117_08512_1ng_glyco_hcd30_etd30_compound15.8146.8146.3</td><td>controllerType=0 controllerNumber=1 scan=8146</td><td>21.2187</td></tr>
<tr><td>04908:3</td><td>4</td><td>K.WN[+1872.651]NTGC[+57.021]QALPSQDEGPSK.A</td><td>HexNAc(3)Hex(6)NeuAc(1)</td><td>C[+57], N[+1873]</td><td>1287.8489</td><td>3</td><td>3861.5322</td><td>3861.5272</td><td>0</td><td>1.3</td><td>498</td><td>Specific</td><td>331.98</td><td>198.45</td><td>198.45</td><td>5.34</td><td>40</td><td>>sp|P01833|PIGR_HUMAN Polymeric immunoglobulin receptor OS=Homo sapiens OX=9606 GN=PIGR PE=1 SV=4</td><td>902</td><td>20201117_08512_1ng_glyco_hcd30_etd30_compound15.8008.8008.3</td><td>controllerType=0 controllerNumber=1 scan=8008</td><td>20.9456</td></tr>
<tr><td>03454:3</td><td>4</td><td>K.WN[+1216.423]NTGC[+57.021]QALPSQDEGPSK.A</td><td>HexNAc(2)Hex(5)</td><td>C[+57], N[+1216]</td><td>1069.1062</td><td>3</td><td>3205.3041</td><td>3205.2995</td><td>0</td><td>1.4</td><td>498</td><td>Specific</td><td>471.37</td><td>366.79</td><td>366.79</td><td>7.2</td><td>40</td><td>>sp|P01833|PIGR_HUMAN Polymeric immunoglobulin receptor OS=Homo sapiens OX=9606 GN=PIGR PE=1 SV=4</td><td>902</td><td>20201117_08512_1ng_glyco_hcd30_etd30_compound15.6501.6501.3</td><td>controllerType=0 controllerNumber=1 scan=6501</td><td>18.2303</td></tr>
<tr><td>04921:3</td><td>4</td><td>K.WN[+1710.598]NTGC[+57.021]QALPSQDEGPSK.A</td><td>HexNAc(3)Hex(5)NeuAc(1)</td><td>C[+57], N[+1711]</td><td>1233.8303</td><td>3</td><td>3699.4762</td><td>3699.4743</td><td>0</td><td>0.5</td><td>498</td><td>Specific</td><td>516.36</td><td>332.42</td><td>332.42</td><td>6.11</td><td>40</td><td>>sp|P01833|PIGR_HUMAN Polymeric immunoglobulin receptor OS=Homo sapiens OX=9606 GN=PIGR PE=1 SV=4</td><td>902</td><td>20201117_08512_1ng_glyco_hcd30_etd30_compound15.8021.8021.3</td><td>controllerType=0 controllerNumber=1 scan=8021</td><td>20.9717</td></tr>
<tr><td>04857:3</td><td>4</td><td>K.WN[+1913.677]NTGC[+57.021]QALPSQDEGPSK.A</td><td>HexNAc(4)Hex(5)NeuAc(1)</td><td>C[+57], N[+1914]</td><td>1301.5258</td><td>3</td><td>3902.5629</td><td>3902.5537</td><td>0</td><td>2.4</td><td>498</td><td>Specific</td><td>536.09</td><td>381.32</td><td>381.32</td><td>7.6</td><td>40</td><td>>sp|P01833|PIGR_HUMAN Polymeric immunoglobulin receptor OS=Homo sapiens OX=9606 GN=PIGR PE=1 SV=4</td><td>902</td><td>20201117_08512_1ng_glyco_hcd30_etd30_compound15.7955.7955.3</td><td>controllerType=0 controllerNumber=1 scan=7955</td><td>20.8398</td></tr>
<tr><td>08150:3</td><td>6</td><td>K.TVLTPATNHMGN[+1540.529]VTFTIPANR.E</td><td>HexNAc(2)Hex(7)</td><td>N[+1541]</td><td>1265.9025</td><td>3</td><td>3795.693</td><td>3795.6886</td><td>0</td><td>1.2</td><td>74</td><td>Specific</td><td>434.25</td><td>364.83</td><td>158.88</td><td>5.85</td><td>21</td><td>>sp|P01024|CO3_HUMAN Complement C3 OS=Homo sapiens OX=9606 GN=C3 PE=1 SV=2</td><td>492</td><td>20201117_08512_1ng_glyco_hcd30_etd30_compound15.11390.11390.3</td><td>controllerType=0 controllerNumber=1 scan=11390</td><td>28.2414</td></tr>
<tr><td>08185:3</td><td>6</td><td>K.TVLTPATNHMGN[+1378.476]VTFTIPANR.E</td><td>HexNAc(2)Hex(6)</td><td>N[+1378]</td><td>1211.8881</td><td>3</td><td>3633.6498</td><td>3633.6358</td><td>0</td><td>3.9</td><td>74</td><td>Specific</td><td>473.53</td><td>376.31</td><td>173.08</td><td>7</td><td>21</td><td>>sp|P01024|CO3_HUMAN Complement C3 OS=Homo sapiens OX=9606 GN=C3 PE=1 SV=2</td><td>492</td><td>20201117_08512_1ng_glyco_hcd30_etd30_compound15.11427.11427.3</td><td>controllerType=0 controllerNumber=1 scan=11427</td><td>28.3556</td></tr>
<tr><td>17289:4</td><td>12</td><td>R.QLAHQSN[+2350.830]STNIFFSPVSIATAFAMLSLGTK.A</td><td>HexNAc(4)Hex(5)Fuc(1)NeuAc(2)</td><td>N[+2351]</td><td>1383.8789</td><td>4</td><td>5532.4938</td><td>5532.4654</td><td>0</td><td>5.1</td><td>64</td><td>Specific</td><td>338.18</td><td>287.68</td><td>287.68</td><td>4.68</td><td>10</td><td>>sp|P01009|A1AT_HUMAN Alpha-1-antitrypsin OS=Homo sapiens OX=9606 GN=SERPINA1 PE=1 SV=3</td><td>642</td><td>20201117_08512_1ng_glyco_hcd30_etd30_compound15.21049.21049.4</td><td>controllerType=0 controllerNumber=1 scan=21049</td><td>55.8218</td></tr>
<tr><td>17337:4</td><td>12</td><td>R.QLAHQSN[+2204.772]STNIFFSPVSIATAFAMLSLGTK.A</td><td>HexNAc(4)Hex(5)NeuAc(2)</td><td>N[+2205]</td><td>1347.3606</td><td>4</td><td>5386.4206</td><td>5386.4075</td><td>0</td><td>2.4</td><td>64</td><td>Specific</td><td>347.56</td><td>285.76</td><td>269.05</td><td>5.15</td><td>10</td><td>>sp|P01009|A1AT_HUMAN Alpha-1-antitrypsin OS=Homo sapiens OX=9606 GN=SERPINA1 PE=1 SV=3</td><td>642</td><td>20201117_08512_1ng_glyco_hcd30_etd30_compound15.21100.21100.3</td><td>controllerType=0 controllerNumber=1 scan=21100</td><td>56.0165</td></tr>
<tr><td>17315:3</td><td>12</td><td>R.QLAHQSN[+2204.772]STNIFFSPVSIATAFAMLSLGTK.A</td><td>HexNAc(4)Hex(5)NeuAc(2)</td><td>N[+2205]</td><td>1796.1477</td><td>3</td><td>5386.4285</td><td>5386.4075</td><td>0</td><td>3.9</td><td>64</td><td>Specific</td><td>356.98</td><td>227.23</td><td>227.23</td><td>4.88</td><td>10</td><td>>sp|P01009|A1AT_HUMAN Alpha-1-antitrypsin OS=Homo sapiens OX=9606 GN=SERPINA1 PE=1 SV=3</td><td>642</td><td>20201117_08512_1ng_glyco_hcd30_etd30_compound15.21077.21077.3</td><td>controllerType=0 controllerNumber=1 scan=21077</td><td>55.9284</td></tr>
<tr><td>17314:4</td><td>12</td><td>R.QLAHQSN[+2204.772]STNIFFSPVSIATAFAMLSLGTK.A</td><td>HexNAc(4)Hex(5)NeuAc(2)</td><td>N[+2205]</td><td>1347.3626</td><td>4</td><td>5386.4284</td><td>5386.4075</td><td>0</td><td>3.9</td><td>64</td><td>Specific</td><td>732.79</td><td>618.15</td><td>618.15</td><td>12.47</td><td>10</td><td>>sp|P01009|A1AT_HUMAN Alpha-1-antitrypsin OS=Homo sapiens OX=9606 GN=SERPINA1 PE=1 SV=3</td><td>642</td><td>20201117_08512_1ng_glyco_hcd30_etd30_compound15.21076.21076.4</td><td>controllerType=0 controllerNumber=1 scan=21076</td><td>55.9238</td></tr>
<tr><td>16558:4</td><td>12</td><td>K.ADTHDEILEGLNFN[+2861.000]LTEIPEAQIHEGFQELLR.T</td><td>HexNAc(5)Hex(6)NeuAc(3)</td><td>N[+2861]</td><td>1638.97</td><td>4</td><td>6552.8581</td><td>6552.8239</td><td>0</td><td>5.2</td><td>94</td><td>Specific</td><td>204.22</td><td>204.22</td><td>204.22</td><td>4.18</td><td>10</td><td>>sp|P01009|A1AT_HUMAN Alpha-1-antitrypsin OS=Homo sapiens OX=9606 GN=SERPINA1 PE=1 SV=3</td><td>642</td><td>20201117_08512_1ng_glyco_hcd30_etd30_compound15.20266.20266.4</td><td>controllerType=0 controllerNumber=1 scan=20266</td><td>53.1273</td></tr>
<tr><td>16580:4</td><td>12</td><td>K.ADTHDEILEGLNFN[+2861.000]LTEIPEAQIHEGFQELLR.T</td><td>HexNAc(5)Hex(6)NeuAc(3)</td><td>N[+2861]</td><td>1638.9668</td><td>4</td><td>6552.8454</td><td>6552.8239</td><td>0</td><td>3.3</td><td>94</td><td>Specific</td><td>350.15</td><td>340.41</td><td>340.41</td><td>6.28</td><td>10</td><td>>sp|P01009|A1AT_HUMAN Alpha-1-antitrypsin OS=Homo sapiens OX=9606 GN=SERPINA1 PE=1 SV=3</td><td>642</td><td>20201117_08512_1ng_glyco_hcd30_etd30_compound15.20290.20290.4</td><td>controllerType=0 controllerNumber=1 scan=20290</td><td>53.2133</td></tr>
<tr><td>15005:4</td><td>12</td><td>K.YLGN[+2204.772]ATAIFFLPDEGKLQHLENELTHDIITK.F</td><td>HexNAc(4)Hex(5)NeuAc(2)</td><td>N[+2205]</td><td>1437.1608</td><td>4</td><td>5745.6212</td><td>5745.6098</td><td>0</td><td>2</td><td>268</td><td>Specific</td><td>534.96</td><td>528.32</td><td>528.32</td><td>6.91</td><td>10</td><td>>sp|P01009|A1AT_HUMAN Alpha-1-antitrypsin OS=Homo sapiens OX=9606 GN=SERPINA1 PE=1 SV=3</td><td>642</td><td>20201117_08512_1ng_glyco_hcd30_etd30_compound15.18616.18616.4</td><td>controllerType=0 controllerNumber=1 scan=18616</td><td>47.7467</td></tr>
<tr><td>15845:5</td><td>13</td><td>K.APDKNVIFSPLSISTALAFLSLGAHN[+2059.735]TTLTEILK.G</td><td>HexNAc(4)Hex(5)Fuc(1)NeuAc(1)</td><td>N[+2060]</td><td>1129.3496</td><td>5</td><td>5642.719</td><td>5642.7131</td><td>0</td><td>1</td><td>68</td><td>Specific</td><td>448.17</td><td>448.17</td><td>448.17</td><td>8.03</td><td>12</td><td>>sp|P01011|AACT_HUMAN Alpha-1-antichymotrypsin OS=Homo sapiens OX=9606 GN=SERPINA3 PE=1 SV=2</td><td>1314</td><td>20201117_08512_1ng_glyco_hcd30_etd30_compound15.19509.19509.5</td><td>controllerType=0 controllerNumber=1 scan=19509</td><td>50.7919</td></tr>
<tr><td>16668:4</td><td>13</td><td>K.APDKNVIFSPLSISTALAFLSLGAHN[+2350.830]TTLTEILK.G</td><td>HexNAc(4)Hex(5)Fuc(1)NeuAc(2)</td><td>N[+2351]</td><td>1484.2141</td><td>4</td><td>5933.8346</td><td>5933.8085</td><td>0</td><td>4.4</td><td>68</td><td>Specific</td><td>509.83</td><td>509.83</td><td>509.83</td><td>6.45</td><td>12</td><td>>sp|P01011|AACT_HUMAN Alpha-1-antichymotrypsin OS=Homo sapiens OX=9606 GN=SERPINA3 PE=1 SV=2</td><td>1314</td><td>20201117_08512_1ng_glyco_hcd30_etd30_compound15.20386.20386.4</td><td>controllerType=0 controllerNumber=1 scan=20386</td><td>53.5863</td></tr>
<tr><td>15844:4</td><td>13</td><td>K.APDKNVIFSPLSISTALAFLSLGAHN[+2059.735]TTLTEILK.G</td><td>HexNAc(4)Hex(5)Fuc(1)NeuAc(1)</td><td>N[+2060]</td><td>1411.4377</td><td>4</td><td>5642.7292</td><td>5642.7131</td><td>0</td><td>2.8</td><td>68</td><td>Specific</td><td>517.31</td><td>517.31</td><td>517.31</td><td>7.84</td><td>12</td><td>>sp|P01011|AACT_HUMAN Alpha-1-antichymotrypsin OS=Homo sapiens OX=9606 GN=SERPINA3 PE=1 SV=2</td><td>1314</td><td>20201117_08512_1ng_glyco_hcd30_etd30_compound15.19508.19508.4</td><td>controllerType=0 controllerNumber=1 scan=19508</td><td>50.7849</td></tr>
<tr><td>16655:5</td><td>13</td><td>K.APDKNVIFSPLSISTALAFLSLGAHN[+2350.830]TTLTEILK.G</td><td>HexNAc(4)Hex(5)Fuc(1)NeuAc(2)</td><td>N[+2351]</td><td>1187.573</td><td>5</td><td>5933.8359</td><td>5933.8085</td><td>0</td><td>4.6</td><td>68</td><td>Specific</td><td>775.92</td><td>775.92</td><td>775.92</td><td>13</td><td>12</td><td>>sp|P01011|AACT_HUMAN Alpha-1-antichymotrypsin OS=Homo sapiens OX=9606 GN=SERPINA3 PE=1 SV=2</td><td>1314</td><td>20201117_08512_1ng_glyco_hcd30_etd30_compound15.20372.20372.5</td><td>controllerType=0 controllerNumber=1 scan=20372</td><td>53.5294</td></tr>
<tr><td>10917:3</td><td>14</td><td>K.LKELPGVC[+57.021]N[+2059.735]ETMMALWEEC[+57.021]KPC[+57.021]LK.Q</td><td>HexNAc(4)Hex(5)Fuc(1)NeuAc(1)</td><td>C[+57]*3, N[+2060]</td><td>1666.0514</td><td>3</td><td>4996.1395</td><td>4996.1196</td><td>0</td><td>4</td><td>95</td><td>Specific</td><td>396.17</td><td>285.32</td><td>285.32</td><td>4.16</td><td>17</td><td>>sp|P10909|CLUS_HUMAN Clusterin OS=Homo sapiens OX=9606 GN=CLU PE=1 SV=1</td><td>1467</td><td>20201117_08512_1ng_glyco_hcd30_etd30_compound15.14294.14294.3</td><td>controllerType=0 controllerNumber=1 scan=14294</td><td>35.4638</td></tr>
<tr><td>10866:4</td><td>14</td><td>K.LKELPGVC[+57.021]N[+2351.851]ETMMALWEEC[+57.021]KPC[+57.021]LK.Q</td><td>HexNAc(4)Hex(5)Fuc(3)NeuAc(1)</td><td>C[+57]*3, N[+2352]</td><td>1322.8198</td><td>4</td><td>5288.2575</td><td>5288.2355</td><td>0</td><td>4.2</td><td>95</td><td>Specific</td><td>426.46</td><td>324.96</td><td>324.96</td><td>4.47</td><td>17</td><td>>sp|P10909|CLUS_HUMAN Clusterin OS=Homo sapiens OX=9606 GN=CLU PE=1 SV=1</td><td>1467</td><td>20201117_08512_1ng_glyco_hcd30_etd30_compound15.14241.14241.4</td><td>controllerType=0 controllerNumber=1 scan=14241</td><td>35.3209</td></tr>
<tr><td>12004:4</td><td>14</td><td>K.LKELPGVC[+57.021]N[+2350.830]ETMMALWEEC[+57.021]KPC[+57.021]LK.Q</td><td>HexNAc(4)Hex(5)Fuc(1)NeuAc(2)</td><td>C[+57]*3, N[+2351]</td><td>1322.5607</td><td>4</td><td>5287.2209</td><td>5287.2151</td><td>0</td><td>1.1</td><td>95</td><td>Specific</td><td>522.54</td><td>383.66</td><td>383.66</td><td>7.44</td><td>17</td><td>>sp|P10909|CLUS_HUMAN Clusterin OS=Homo sapiens OX=9606 GN=CLU PE=1 SV=1</td><td>1467</td><td>20201117_08512_1ng_glyco_hcd30_etd30_compound15.15437.15437.2</td><td>controllerType=0 controllerNumber=1 scan=15437</td><td>38.5697</td></tr>
<tr><td>10909:4</td><td>14</td><td>K.LKELPGVC[+57.021]N[+2059.735]ETMMALWEEC[+57.021]KPC[+57.021]LK.Q</td><td>HexNAc(4)Hex(5)Fuc(1)NeuAc(1)</td><td>C[+57]*3, N[+2060]</td><td>1249.7894</td><td>4</td><td>4996.1359</td><td>4996.1196</td><td>0</td><td>3.3</td><td>95</td><td>Specific</td><td>657.68</td><td>492.07</td><td>492.07</td><td>9.31</td><td>17</td><td>>sp|P10909|CLUS_HUMAN Clusterin OS=Homo sapiens OX=9606 GN=CLU PE=1 SV=1</td><td>1467</td><td>20201117_08512_1ng_glyco_hcd30_etd30_compound15.14286.14286.4</td><td>controllerType=0 controllerNumber=1 scan=14286</td><td>35.4372</td></tr>
<tr><td>00063:4</td><td>14</td><td>R.EIRHN[+2424.867]STGC[+57.021]LR.M</td><td>HexNAc(5)Hex(6)Fuc(1)NeuAc(1)</td><td>C[+57], N[+2425]</td><td>942.641</td><td>4</td><td>3767.5423</td><td>3767.5316</td><td>0</td><td>2.8</td><td>287</td><td>Specific</td><td>462.3</td><td>365.85</td><td>365.85</td><td>6.16</td><td>17</td><td>>sp|P10909|CLUS_HUMAN Clusterin OS=Homo sapiens OX=9606 GN=CLU PE=1 SV=1</td><td>1467</td><td>20201117_08512_1ng_glyco_hcd30_etd30_compound15.2972.2972.4</td><td>controllerType=0 controllerNumber=1 scan=2972</td><td>11.3018</td></tr>
<tr><td>00049:4</td><td>14</td><td>R.EIRHN[+2059.735]STGC[+57.021]LR.M</td><td>HexNAc(4)Hex(5)Fuc(1)NeuAc(1)</td><td>C[+57], N[+2060]</td><td>851.3582</td><td>4</td><td>3402.4111</td><td>3402.3994</td><td>0</td><td>3.5</td><td>287</td><td>Specific</td><td>483.47</td><td>380.28</td><td>380.28</td><td>6.32</td><td>17</td><td>>sp|P10909|CLUS_HUMAN Clusterin OS=Homo sapiens OX=9606 GN=CLU PE=1 SV=1</td><td>1467</td><td>20201117_08512_1ng_glyco_hcd30_etd30_compound15.2957.2957.4</td><td>controllerType=0 controllerNumber=1 scan=2957</td><td>11.2463</td></tr>
<tr><td>17097:3</td><td>14</td><td>K.MLN[+2204.772]TSSLLEQLNEQFNWVSR.L</td><td>HexNAc(4)Hex(5)NeuAc(2)</td><td>N[+2205]</td><td>1538.6642</td><td>3</td><td>4613.9782</td><td>4613.9592</td><td>0</td><td>4.1</td><td>352</td><td>Specific</td><td>255.97</td><td>148.99</td><td>148.99</td><td>4.76</td><td>17</td><td>>sp|P10909|CLUS_HUMAN Clusterin OS=Homo sapiens OX=9606 GN=CLU PE=1 SV=1</td><td>1467</td><td>20201117_08512_1ng_glyco_hcd30_etd30_compound15.20844.20844.3</td><td>controllerType=0 controllerNumber=1 scan=20844</td><td>55.1552</td></tr>
<tr><td>15726:3</td><td>14</td><td>K.MLN[+1913.677]TSSLLEQLNEQFNWVSR.L</td><td>HexNAc(4)Hex(5)NeuAc(1)</td><td>N[+1914]</td><td>1441.6326</td><td>3</td><td>4322.8833</td><td>4322.8637</td><td>0</td><td>4.5</td><td>352</td><td>Specific</td><td>559.87</td><td>400.48</td><td>400.48</td><td>7.28</td><td>17</td><td>>sp|P10909|CLUS_HUMAN Clusterin OS=Homo sapiens OX=9606 GN=CLU PE=1 SV=1</td><td>1467</td><td>20201117_08512_1ng_glyco_hcd30_etd30_compound15.19383.19383.3</td><td>controllerType=0 controllerNumber=1 scan=19383</td><td>50.3639</td></tr>
<tr><td>15559:3</td><td>14</td><td>K.MLN[+2059.735]TSSLLEQLNEQFNWVSR.L</td><td>HexNAc(4)Hex(5)Fuc(1)NeuAc(1)</td><td>N[+2060]</td><td>1490.3168</td><td>3</td><td>4468.9359</td><td>4468.9217</td><td>0</td><td>3.2</td><td>352</td><td>Specific</td><td>583.54</td><td>394.18</td><td>394.18</td><td>8.8</td><td>17</td><td>>sp|P10909|CLUS_HUMAN Clusterin OS=Homo sapiens OX=9606 GN=CLU PE=1 SV=1</td><td>1467</td><td>20201117_08512_1ng_glyco_hcd30_etd30_compound15.19206.19206.3</td><td>controllerType=0 controllerNumber=1 scan=19206</td><td>49.7965</td></tr>
<tr><td>09021:3</td><td>15</td><td>R.LAGKPTHVN[+1702.581]VSVVMAEVDGTC[+57.021]Y.-</td><td>HexNAc(2)Hex(8)</td><td>C[+57], N[+1703]</td><td>1350.5822</td><td>3</td><td>4049.7321</td><td>4049.7234</td><td>0</td><td>2.1</td><td>332</td><td>CRagged</td><td>206.66</td><td>166.31</td><td>160.84</td><td>2.68</td><td>20</td><td>>sp|P01876|IGHA1_HUMAN Immunoglobulin heavy constant alpha 1 OS=Homo sapiens OX=9606 GN=IGHA1 PE=1 SV=2</td><td>255</td><td>20201117_08512_1ng_glyco_hcd30_etd30_compound15.12303.12303.3</td><td>controllerType=0 controllerNumber=1 scan=12303</td><td>30.5108</td></tr>
<tr><td>10345:3</td><td>15</td><td>R.LAGKPTHVN[+2116.756]VSVVMAEVDGTC[+57.021]Y.-</td><td>HexNAc(5)Hex(5)NeuAc(1)</td><td>C[+57], N[+2117]</td><td>1488.639</td><td>3</td><td>4463.9024</td><td>4463.8985</td><td>0</td><td>0.9</td><td>332</td><td>CRagged</td><td>216.79</td><td>78.22</td><td>78.22</td><td>2.63</td><td>20</td><td>>sp|P01876|IGHA1_HUMAN Immunoglobulin heavy constant alpha 1 OS=Homo sapiens OX=9606 GN=IGHA1 PE=1 SV=2</td><td>255</td><td>20201117_08512_1ng_glyco_hcd30_etd30_compound15.13692.13692.3</td><td>controllerType=0 controllerNumber=1 scan=13692</td><td>33.7749</td></tr>
<tr><td>10332:3</td><td>15</td><td>R.LAGKPTHVN[+2262.814]VSVVMAEVDGTC[+57.021]Y.-</td><td>HexNAc(5)Hex(5)Fuc(1)NeuAc(1)</td><td>C[+57], N[+2263]</td><td>1537.3265</td><td>3</td><td>4609.9649</td><td>4609.9564</td><td>0</td><td>1.8</td><td>332</td><td>CRagged</td><td>216.85</td><td>41.81</td><td>41.81</td><td>2.38</td><td>20</td><td>>sp|P01876|IGHA1_HUMAN Immunoglobulin heavy constant alpha 1 OS=Homo sapiens OX=9606 GN=IGHA1 PE=1 SV=2</td><td>255</td><td>20201117_08512_1ng_glyco_hcd30_etd30_compound15.13679.13679.3</td><td>controllerType=0 controllerNumber=1 scan=13679</td><td>33.7451</td></tr>
<tr><td>09138:3</td><td>15</td><td>R.LAGKPTHVN[+1647.613]VSVVMAEVDGTC[+57.021]Y.-</td><td>HexNAc(5)Hex(3)Fuc(1)</td><td>C[+57], N[+1648]</td><td>1332.2599</td><td>3</td><td>3994.7653</td><td>3994.7553</td><td>0</td><td>2.5</td><td>332</td><td>CRagged</td><td>222.82</td><td>111.1</td><td>111.1</td><td>2.63</td><td>20</td><td>>sp|P01876|IGHA1_HUMAN Immunoglobulin heavy constant alpha 1 OS=Homo sapiens OX=9606 GN=IGHA1 PE=1 SV=2</td><td>255</td><td>20201117_08512_1ng_glyco_hcd30_etd30_compound15.12426.12426.3</td><td>controllerType=0 controllerNumber=1 scan=12426</td><td>30.857</td></tr>
<tr><td>09096:3</td><td>15</td><td>R.LAGKPTHVN[+1216.423]VSVVMAEVDGTC[+57.021]Y.-</td><td>HexNAc(2)Hex(5)</td><td>C[+57], N[+1216]</td><td>1188.529</td><td>3</td><td>3563.5724</td><td>3563.5649</td><td>0</td><td>2.1</td><td>332</td><td>CRagged</td><td>234.15</td><td>213.96</td><td>159.23</td><td>2.9</td><td>20</td><td>>sp|P01876|IGHA1_HUMAN Immunoglobulin heavy constant alpha 1 OS=Homo sapiens OX=9606 GN=IGHA1 PE=1 SV=2</td><td>255</td><td>20201117_08512_1ng_glyco_hcd30_etd30_compound15.12382.12382.3</td><td>controllerType=0 controllerNumber=1 scan=12382</td><td>30.7381</td></tr>
<tr><td>09044:3</td><td>15</td><td>R.LAGKPTHVN[+1378.476]VSVVMAEVDGTC[+57.021]Y.-</td><td>HexNAc(2)Hex(6)</td><td>C[+57], N[+1378]</td><td>1242.5474</td><td>3</td><td>3725.6277</td><td>3725.6178</td><td>0</td><td>2.7</td><td>332</td><td>CRagged</td><td>239.13</td><td>239.13</td><td>184.42</td><td>2.89</td><td>20</td><td>>sp|P01876|IGHA1_HUMAN Immunoglobulin heavy constant alpha 1 OS=Homo sapiens OX=9606 GN=IGHA1 PE=1 SV=2</td><td>255</td><td>20201117_08512_1ng_glyco_hcd30_etd30_compound15.12327.12327.3</td><td>controllerType=0 controllerNumber=1 scan=12327</td><td>30.5713</td></tr>
<tr><td>09133:3</td><td>15</td><td>R.LAGKPTHVN[+1501.555]VSVVMAEVDGTC[+57.021]Y.-</td><td>HexNAc(5)Hex(3)</td><td>C[+57], N[+1502]</td><td>1283.574</td><td>3</td><td>3848.7075</td><td>3848.6974</td><td>0</td><td>2.6</td><td>332</td><td>CRagged</td><td>241.89</td><td>145.62</td><td>145.62</td><td>3.15</td><td>20</td><td>>sp|P01876|IGHA1_HUMAN Immunoglobulin heavy constant alpha 1 OS=Homo sapiens OX=9606 GN=IGHA1 PE=1 SV=2</td><td>255</td><td>20201117_08512_1ng_glyco_hcd30_etd30_compound15.12421.12421.3</td><td>controllerType=0 controllerNumber=1 scan=12421</td><td>30.8384</td></tr>
<tr><td>08266:3</td><td>15</td><td>R.LAGKPTHVN[+1216.423]VSVVMAEVDGTC[+57.021]Y.-</td><td>HexNAc(2)Hex(5)</td><td>C[+57], N[+1216]</td><td>1188.5298</td><td>3</td><td>3563.575</td><td>3563.5649</td><td>0</td><td>2.8</td><td>332</td><td>CRagged</td><td>246.5</td><td>231.73</td><td>165.08</td><td>3.46</td><td>20</td><td>>sp|P01876|IGHA1_HUMAN Immunoglobulin heavy constant alpha 1 OS=Homo sapiens OX=9606 GN=IGHA1 PE=1 SV=2</td><td>255</td><td>20201117_08512_1ng_glyco_hcd30_etd30_compound15.11512.11512.3</td><td>controllerType=0 controllerNumber=1 scan=11512</td><td>28.5888</td></tr>
<tr><td>09050:3</td><td>15</td><td>R.LAGKPTHVN[+1971.719]VSVVMAEVDGTC[+57.021]Y.-</td><td>HexNAc(5)Hex(5)Fuc(1)</td><td>C[+57], N[+1972]</td><td>1440.2945</td><td>3</td><td>4318.8689</td><td>4318.861</td><td>0</td><td>1.8</td><td>332</td><td>CRagged</td><td>250.17</td><td>145.32</td><td>145.32</td><td>3.14</td><td>20</td><td>>sp|P01876|IGHA1_HUMAN Immunoglobulin heavy constant alpha 1 OS=Homo sapiens OX=9606 GN=IGHA1 PE=1 SV=2</td><td>255</td><td>20201117_08512_1ng_glyco_hcd30_etd30_compound15.12333.12333.3</td><td>controllerType=0 controllerNumber=1 scan=12333</td><td>30.5937</td></tr>
<tr><td>09060:3</td><td>15</td><td>R.LAGKPTHVN[+1809.666]VSVVMAEVDGTC[+57.021]Y.-</td><td>HexNAc(5)Hex(4)Fuc(1)</td><td>C[+57], N[+1810]</td><td>1386.2769</td><td>3</td><td>4156.8162</td><td>4156.8081</td><td>0</td><td>1.9</td><td>332</td><td>CRagged</td><td>254.96</td><td>201.43</td><td>201.43</td><td>3.46</td><td>20</td><td>>sp|P01876|IGHA1_HUMAN Immunoglobulin heavy constant alpha 1 OS=Homo sapiens OX=9606 GN=IGHA1 PE=1 SV=2</td><td>255</td><td>20201117_08512_1ng_glyco_hcd30_etd30_compound15.12344.12344.3</td><td>controllerType=0 controllerNumber=1 scan=12344</td><td>30.6322</td></tr>
<tr><td>09056:3</td><td>15</td><td>R.LAGKPTHVN[+1540.529]VSVVMAEVDGTC[+57.021]Y.-</td><td>HexNAc(2)Hex(7)</td><td>C[+57], N[+1541]</td><td>1296.5649</td><td>3</td><td>3887.6801</td><td>3887.6706</td><td>0</td><td>2.4</td><td>332</td><td>CRagged</td><td>270.52</td><td>205.8</td><td>205.16</td><td>3.47</td><td>20</td><td>>sp|P01876|IGHA1_HUMAN Immunoglobulin heavy constant alpha 1 OS=Homo sapiens OX=9606 GN=IGHA1 PE=1 SV=2</td><td>255</td><td>20201117_08512_1ng_glyco_hcd30_etd30_compound15.12340.12340.3</td><td>controllerType=0 controllerNumber=1 scan=12340</td><td>30.6199</td></tr>
<tr><td>11606:3</td><td>15</td><td>R.LAGKPTHVN[+2553.910]VSVVMAEVDGTC[+57.021]Y.-</td><td>HexNAc(5)Hex(5)Fuc(1)NeuAc(2)</td><td>C[+57], N[+2554]</td><td>1634.3605</td><td>3</td><td>4901.067</td><td>4901.0518</td><td>0</td><td>3.1</td><td>332</td><td>CRagged</td><td>290.97</td><td>100.58</td><td>100.58</td><td>2.97</td><td>20</td><td>>sp|P01876|IGHA1_HUMAN Immunoglobulin heavy constant alpha 1 OS=Homo sapiens OX=9606 GN=IGHA1 PE=1 SV=2</td><td>255</td><td>20201117_08512_1ng_glyco_hcd30_etd30_compound15.15019.15019.3</td><td>controllerType=0 controllerNumber=1 scan=15019</td><td>37.4127</td></tr>
<tr><td>09103:4</td><td>15</td><td>R.LAGKPTHVN[+1809.666]VSVVMAEVDGTC[+57.021]Y.-</td><td>HexNAc(5)Hex(4)Fuc(1)</td><td>C[+57], N[+1810]</td><td>1039.9591</td><td>4</td><td>4156.8148</td><td>4156.8081</td><td>0</td><td>1.6</td><td>332</td><td>CRagged</td><td>363.36</td><td>363.36</td><td>94.67</td><td>4.55</td><td>20</td><td>>sp|P01876|IGHA1_HUMAN Immunoglobulin heavy constant alpha 1 OS=Homo sapiens OX=9606 GN=IGHA1 PE=1 SV=2</td><td>255</td><td>20201117_08512_1ng_glyco_hcd30_etd30_compound15.12389.12389.4</td><td>controllerType=0 controllerNumber=1 scan=12389</td><td>30.7587</td></tr>
<tr><td>10341:4</td><td>15</td><td>R.LAGKPTHVN[+2262.814]VSVVMAEVDGTC[+57.021]Y.-</td><td>HexNAc(5)Hex(5)Fuc(1)NeuAc(1)</td><td>C[+57], N[+2263]</td><td>1153.2476</td><td>4</td><td>4609.9686</td><td>4609.9564</td><td>0</td><td>2.6</td><td>332</td><td>CRagged</td><td>428.27</td><td>344.13</td><td>344.13</td><td>4.76</td><td>20</td><td>>sp|P01876|IGHA1_HUMAN Immunoglobulin heavy constant alpha 1 OS=Homo sapiens OX=9606 GN=IGHA1 PE=1 SV=2</td><td>255</td><td>20201117_08512_1ng_glyco_hcd30_etd30_compound15.13688.13688.4</td><td>controllerType=0 controllerNumber=1 scan=13688</td><td>33.7666</td></tr>
<tr><td>11590:4</td><td>15</td><td>R.LAGKPTHVN[+2553.910]VSVVMAEVDGTC[+57.021]Y.-</td><td>HexNAc(5)Hex(5)Fuc(1)NeuAc(2)</td><td>C[+57], N[+2554]</td><td>1226.0215</td><td>4</td><td>4901.0643</td><td>4901.0518</td><td>0</td><td>2.5</td><td>332</td><td>CRagged</td><td>540.72</td><td>305.18</td><td>305.18</td><td>5.13</td><td>20</td><td>>sp|P01876|IGHA1_HUMAN Immunoglobulin heavy constant alpha 1 OS=Homo sapiens OX=9606 GN=IGHA1 PE=1 SV=2</td><td>255</td><td>20201117_08512_1ng_glyco_hcd30_etd30_compound15.15002.15002.4</td><td>controllerType=0 controllerNumber=1 scan=15002</td><td>37.3632</td></tr>
<tr><td>10306:4</td><td>20</td><td>R.FELLGC[+57.021]ELNGC[+57.021]ANPLGLKN[+1216.423]NSIPDKQITASSSYK.T</td><td>HexNAc(2)Hex(5)</td><td>C[+57]*2, N[+1216]</td><td>1239.5757</td><td>4</td><td>4955.2809</td><td>4955.2694</td><td>0</td><td>2.3</td><td>220</td><td>Specific</td><td>715.19</td><td>715.19</td><td>715.19</td><td>10.66</td><td>7</td><td>>sp|Q08431|MFGM_HUMAN Lactadherin OS=Homo sapiens OX=9606 GN=MFGE8 PE=1 SV=3</td><td>1064</td><td>20201117_08512_1ng_glyco_hcd30_etd30_compound15.13651.13651.4</td><td>controllerType=0 controllerNumber=1 scan=13651</td><td>33.6635</td></tr>
<tr><td>07180:3</td><td>20</td><td>K.VAYSN[+2059.735]DSANWTEYQDPR.T</td><td>HexNAc(4)Hex(5)Fuc(1)NeuAc(1)</td><td>N[+2060]</td><td>1359.2097</td><td>3</td><td>4075.6144</td><td>4075.6079</td><td>0</td><td>1.6</td><td>321</td><td>Specific</td><td>227.76</td><td>136.98</td><td>43.85</td><td>3.82</td><td>7</td><td>>sp|Q08431|MFGM_HUMAN Lactadherin OS=Homo sapiens OX=9606 GN=MFGE8 PE=1 SV=3</td><td>1064</td><td>20201117_08512_1ng_glyco_hcd30_etd30_compound15.10371.10371.3</td><td>controllerType=0 controllerNumber=1 scan=10371</td><td>25.7501</td></tr>
<tr><td>07793:4</td><td>21</td><td>K.QTDEIKDTRN[+2059.735]ESTQNC[+57.021]VVAEPEKMESSISSSSEEMSLSK.C</td><td>HexNAc(4)Hex(5)Fuc(1)NeuAc(1)</td><td>C[+57], N[+2060]</td><td>1613.4349</td><td>4</td><td>6450.7179</td><td>6450.7156</td><td>0</td><td>0.4</td><td>60</td><td>Specific</td><td>217.12</td><td>217.12</td><td>217.12</td><td>2.68</td><td>9</td><td>>sp|P47710|CASA1_HUMAN Alpha-S1-casein OS=Homo sapiens OX=9606 GN=CSN1S1 PE=1 SV=1</td><td>989</td><td>20201117_08512_1ng_glyco_hcd30_etd30_compound15.11016.11016.2</td><td>controllerType=0 controllerNumber=1 scan=11016</td><td>27.3399</td></tr>
</table>

Table S4. Cont.

| Query #:z | Protein Rank | Glycopeptide | Glycan | Modification Type(s) | Observed m/z | z | Observed (M+H) | Calc. mass (M+H) | Off-by-x error | Mass error (ppm) | StartPosition | Cleavage | Score | Delta | Delta Mod | \|Log Prob\| | # of unique peptides | Protein Name | Protein DB number | Comment | Scan # | Scan Time |
|---|---|---|---|---|---|---|---|---|---|---|---|---|---|---|---|---|---|---|---|---|---|---|
| 08984:3 | 26 | R.IIVPLNNREN[+2059.735]ISDPTSPLR.T | HexNAc(4)Hex(5)Fuc(1)NeuAc(1) | N[+2060] | 1403.3134 | 3 | 4207.9257 | 4207.9121 | 0 | 3.2 | 62 | Specific | 253.33 | 200.65 | 200.65 | 3.38 | 9 | >sp\|P01591\|IGJ_HUMAN Immunoglobulin J chain OS=Homo sapiens OX=9606 GN=JCHAIN PE=1 SV=4 | 1392 | 20201117_08512_1ng_glyco_hcd30_etd30_compound15.12264.12264.3 | controllerType=0 controllerNumber=1 scan=12264 | 30.3852 |
| 07913:3 | 26 | R.IIVPLNNREN[+1768.640]ISDPTSPLR.T | HexNAc(4)Hex(5)Fuc(1) | N[+1769] | 1306.2795 | 3 | 3916.8239 | 3916.8167 | 0 | 1.8 | 62 | Specific | 255.66 | 194.29 | 194.29 | 3.37 | 9 | >sp\|P01591\|IGJ_HUMAN Immunoglobulin J chain OS=Homo sapiens OX=9606 GN=JCHAIN PE=1 SV=4 | 1392 | 20201117_08512_1ng_glyco_hcd30_etd30_compound15.11142.11142.3 | controllerType=0 controllerNumber=1 scan=11142 | 27.6412 |
| 08985:3 | 26 | R.IIVPLNNREN[+1913.677]ISDPTSPLR.T | HexNAc(4)Hex(5)NeuAc(1) | N[+1914] | 1354.6273 | 3 | 4061.8672 | 4061.8542 | 0 | 3.2 | 62 | Specific | 286.25 | 227.83 | 227.83 | 3.46 | 9 | >sp\|P01591\|IGJ_HUMAN Immunoglobulin J chain OS=Homo sapiens OX=9606 GN=JCHAIN PE=1 SV=4 | 1392 | 20201117_08512_1ng_glyco_hcd30_etd30_compound15.12265.12265.3 | controllerType=0 controllerNumber=1 scan=12265 | 30.3891 |
| 07868:3 | 26 | R.IIVPLNNREN[+1663.608]ISDPTSPLR.T | HexNAc(5)Hex(4) | N[+1664] | 1271.2701 | 3 | 3811.7957 | 3811.7853 | 0 | 2.7 | 62 | Specific | 299.35 | 207.1 | 207.1 | 3.66 | 9 | >sp\|P01591\|IGJ_HUMAN Immunoglobulin J chain OS=Homo sapiens OX=9606 GN=JCHAIN PE=1 SV=4 | 1392 | 20201117_08512_1ng_glyco_hcd30_etd30_compound15.11095.11095.3 | controllerType=0 controllerNumber=1 scan=11095 | 27.5312 |
| 09027:3 | 26 | R.IIVPLNNREN[+1856.656]ISDPTSPLR.T | HexNAc(3)Hex(5)Fuc(1)NeuAc(1) | N[+1857] | 1335.6193 | 3 | 4004.8434 | 4004.8327 | 0 | 2.7 | 62 | Specific | 359.19 | 295.47 | 295.47 | 4.14 | 9 | >sp\|P01591\|IGJ_HUMAN Immunoglobulin J chain OS=Homo sapiens OX=9606 GN=JCHAIN PE=1 SV=4 | 1392 | 20201117_08512_1ng_glyco_hcd30_etd30_compound15.12309.12309.3 | controllerType=0 controllerNumber=1 scan=12309 | 30.5339 |
| 10389:3 | 26 | R.IIVPLNNREN[+2350.830]ISDPTSPLR.T | HexNAc(4)Hex(5)Fuc(1)NeuAc(2) | N[+2351] | 1500.3462 | 3 | 4499.0242 | 4499.0075 | 0 | 3.7 | 62 | Specific | 379.97 | 244.18 | 244.18 | 3.81 | 9 | >sp\|P01591\|IGJ_HUMAN Immunoglobulin J chain OS=Homo sapiens OX=9606 GN=JCHAIN PE=1 SV=4 | 1392 | 20201117_08512_1ng_glyco_hcd30_etd30_compound15.13739.13739.3 | controllerType=0 controllerNumber=1 scan=13739 | 33.9153 |
| 09007:4 | 26 | R.IIVPLNNREN[+1913.677]ISDPTSPLR.T | HexNAc(4)Hex(5)NeuAc(1) | N[+1914] | 1016.2229 | 4 | 4061.8697 | 4061.8542 | 0 | 3.8 | 62 | Specific | 525.4 | 525.4 | 525.4 | 6.46 | 9 | >sp\|P01591\|IGJ_HUMAN Immunoglobulin J chain OS=Homo sapiens OX=9606 GN=JCHAIN PE=1 SV=4 | 1392 | 20201117_08512_1ng_glyco_hcd30_etd30_compound15.12288.12288.4 | controllerType=0 controllerNumber=1 scan=12288 | 30.4498 |
| 00060:3 | 27 | K.YKN[+1647.613]NSDISSTR.G | HexNAc(5)Hex(3)Fuc(1) | N[+1648] | 978.0851 | 3 | 2932.2406 | 2932.2311 | 0 | 3.2 | 44 | Specific | 411.11 | 258.82 | 258.82 | 3.53 | 6 | >sp\|P01871\|IGHM_HUMAN Immunoglobulin heavy constant mu OS=Homo sapiens OX=9606 GN=IGHM PE=1 SV=4 | 328 | 20201117_08512_1ng_glyco_hcd30_etd30_compound15.2968.2968.3 | controllerType=0 controllerNumber=1 scan=2968 | 11.2877 |
| 00064:3 | 27 | K.YKN[+1647.613]NSDISSTR.G | HexNAc(5)Hex(3)Fuc(1) | N[+1648] | 978.0842 | 3 | 2932.238 | 2932.2311 | 0 | 2.4 | 44 | Specific | 411.76 | 274.41 | 274.41 | 3.51 | 6 | >sp\|P01871\|IGHM_HUMAN Immunoglobulin heavy constant mu OS=Homo sapiens OX=9606 GN=IGHM PE=1 SV=4 | 328 | 20201117_08512_1ng_glyco_hcd30_etd30_compound15.2973.2973.3 | controllerType=0 controllerNumber=1 scan=2973 | 11.3089 |
| 01713:3 | 28 | R.LRN[+1216.423]VSWATGR.S | HexNAc(2)Hex(5) | N[+1216] | 792.6927 | 3 | 2376.0634 | 2376.0559 | 0 | 3.1 | 149 | Specific | 328.52 | 143.61 | 143.61 | 2.61 | 5 | >sp\|P08571\|CD14_HUMAN Monocyte differentiation antigen CD14 OS=Homo sapiens OX=9606 GN=CD14 PE=1 SV=2 | 1232 | 20201117_08512_1ng_glyco_hcd30_etd30_compound15.4688.4688.3 | controllerType=0 controllerNumber=1 scan=4688 | 14.5016 |
| 04753:3 | 32 | R.LSPN[+2205.793]ASAEHLELR.W | HexNAc(4)Hex(5)Fuc(2)NeuAc(1) | N[+2206] | 1214.8552 | 3 | 3642.5509 | 3642.5421 | 0 | 2.4 | 52 | Specific | 276.65 | 181.44 | 181.44 | 2.94 | 7 | >sp\|Q13410\|BT1A1_HUMAN Butyrophilin subfamily 1 member A1 OS=Homo sapiens OX=9606 GN=BTN1A1 PE=1 SV=3 | 179 | 20201117_08512_1ng_glyco_hcd30_etd30_compound15.7847.7847.3 | controllerType=0 controllerNumber=1 scan=7847 | 20.6357 |
| 06513:3 | 32 | R.LSPN[+2350.830]ASAEHLELR.W | HexNAc(4)Hex(5)Fuc(1)NeuAc(2) | N[+2351] | 1263.2016 | 3 | 3787.5902 | 3787.5796 | 0 | 2.8 | 52 | Specific | 364.87 | 226.01 | 226.01 | 4.22 | 7 | >sp\|Q13410\|BT1A1_HUMAN Butyrophilin subfamily 1 member A1 OS=Homo sapiens OX=9606 GN=BTN1A1 PE=1 SV=3 | 179 | 20201117_08512_1ng_glyco_hcd30_etd30_compound15.9675.9675.3 | controllerType=0 controllerNumber=1 scan=9675 | 24.2071 |
| 07220:5 | 35 | K.SC[+57.021]QHN[+2059.735]GTMYQHGEIFSAHELFPSR.L | HexNAc(4)Hex(5)Fuc(1)NeuAc(1) | C[+57], N[+2060] | 979.4057 | 5 | 4892.9992 | 4892.9919 | 0 | 1.5 | 110 | Specific | 595.01 | 595.01 | 595.01 | 8.44 | 4 | >sp\|Q6WN34\|CRDL2_HUMAN Chordin-like protein 2 OS=Homo sapiens OX=9606 GN=CHRDL2 PE=1 SV=1 | 1189 | 20201117_08512_1ng_glyco_hcd30_etd30_compound15.10413.10413.5 | controllerType=0 controllerNumber=1 scan=10413 | 25.863 |
| 07200:4 | 35 | K.SC[+57.021]QHN[+2059.735]GTMYQHGEIFSAHELFPSR.L | HexNAc(4)Hex(5)Fuc(1)NeuAc(1) | C[+57], N[+2060] | 1224.0076 | 4 | 4893.0086 | 4892.9919 | 0 | 3.4 | 110 | Specific | 647.04 | 489.82 | 489.82 | 9.47 | 4 | >sp\|Q6WN34\|CRDL2_HUMAN Chordin-like protein 2 OS=Homo sapiens OX=9606 GN=CHRDL2 PE=1 SV=1 | 1189 | 20201117_08512_1ng_glyco_hcd30_etd30_compound15.10392.10392.4 | controllerType=0 controllerNumber=1 scan=10392 | 25.8109 |
| 15882:3 | 48 | R.QFWIFDVQNPQEVMMN[+2350.830]SSNIQVK.Q | HexNAc(4)Hex(5)Fuc(1)NeuAc(2) | N[+2351] | 1711.7353 | 3 | 5133.1914 | 5133.1631 | 0 | 5.5 | 64 | Specific | 364.88 | 268.25 | 268.25 | 5.17 | 3 | >sp\|P16671\|CD36_HUMAN Platelet glycoprotein 4 OS=Homo sapiens OX=9606 GN=CD36 PE=1 SV=2 | 812 | 20201117_08512_1ng_glyco_hcd30_etd30_compound15.19548.19548.3 | controllerType=0 controllerNumber=1 scan=19548 | 50.9185 |
| 13372:3 | 48 | R.NYIVPILWLN[+1864.634]ETGTIGDEKANMFR.S | HexNAc(2)Hex(9) | N[+1865] | 1553.697 | 3 | 4659.0763 | 4659.0574 | 0 | 4.1 | 408 | Specific | 420.57 | 377.38 | 377.38 | 6.17 | 3 | >sp\|P16671\|CD36_HUMAN Platelet glycoprotein 4 OS=Homo sapiens OX=9606 GN=CD36 PE=1 SV=2 | 812 | 20201117_08512_1ng_glyco_hcd30_etd30_compound15.16888.16888.3 | controllerType=0 controllerNumber=1 scan=16888 | 42.4234 |
| 12026:3 | 53 | R.KLPPGLLAN[+2059.735]FTLLR.T | HexNAc(4)Hex(5)Fuc(1)NeuAc(1) | N[+2060] | 1204.9061 | 3 | 3612.7036 | 3612.6923 | 0 | 3.1 | 178 | Specific | 355.17 | 309.8 | 309.8 | 4.12 | 2 | >sp\|P02750\|A2GL_HUMAN Leucine-rich alpha-2-glycoprotein OS=Homo sapiens OX=9606 GN=LRG1 PE=1 SV=2 | 518 | 20201117_08512_1ng_glyco_hcd30_etd30_compound15.15460.15460.3 | controllerType=0 controllerNumber=1 scan=15460 | 38.6242 |
| 00732:4 | 99 | K.TKPREEQFN[+1606.587]STFR.V | HexNAc(4)Hex(4)Fuc(1) | N[+1607] | 812.3591 | 4 | 3246.4145 | 3246.4054 | 0 | 2.8 | 168 | Specific | 325.3 | 325.3 | 325.3 | 3.48 | 1 | >sp\|P01859\|IGHG2_HUMAN Immunoglobulin heavy constant gamma 2 OS=Homo sapiens OX=9606 GN=IGHG2 PE=1 SV=2 | 387 | 20201117_08512_1ng_glyco_hcd30_etd30_compound15.3668.3668.4 | controllerType=0 controllerNumber=1 scan=3668 | 12.5948 |
| 15628:5 | 1 | R.RTAGWNVPIGTLRPFLN[+2204.772]WTGPPEPIEAAVAR.F | HexNAc(4)Hex(5)NeuAc(2) | N[+2205] | 1119.1271 | 5 | 5591.6063 | 5591.5845 | 0 | 3.9 | 140 | Specific | 366.7 | 366.7 | 366.7 | 5.66 | 33 | >sp\|P02788\|TRFL_HUMAN Lactotransferrin OS=Homo sapiens OX=9606 GN=LTF PE=1 SV=6 | 1270 | 20201119_08533_1ng_glyco_hcd30_etd30_compound15.19282.19282.3 | controllerType=0 controllerNumber=1 scan=19282 | 43.5372 |
| 15565:4 | 1 | R.RTAGWNVPIGTLRPFLN[+2716.983]WTGPPEPIEAAVAR.F | HexNAc(5)Hex(6)Fuc(3)NeuAc(1) | N[+2717] | 1526.7136 | 4 | 6103.8327 | 6103.795 | 0 | 6.2 | 140 | Specific | 734.19 | 734.19 | 734.19 | 10.44 | 33 | >sp\|P02788\|TRFL_HUMAN Lactotransferrin OS=Homo sapiens OX=9606 GN=LTF PE=1 SV=6 | 1270 | 20201119_08533_1ng_glyco_hcd30_etd30_compound15.19216.19216.4 | controllerType=0 controllerNumber=1 scan=19216 | 43.356 |
| 15661:5 | 1 | R.RTAGWNVPIGTLRPFLN[+2059.735]WTGPPEPIEAAVAR.F | HexNAc(4)Hex(5)Fuc(1)NeuAc(1) | N[+2060] | 1090.1185 | 5 | 5446.5636 | 5446.547 | 0 | 3 | 140 | Specific | 795.87 | 752.01 | 752.01 | 11.29 | 33 | >sp\|P02788\|TRFL_HUMAN Lactotransferrin OS=Homo sapiens OX=9606 GN=LTF PE=1 SV=6 | 1270 | 20201119_08533_1ng_glyco_hcd30_etd30_compound15.19317.19317.5 | controllerType=0 controllerNumber=1 scan=19317 | 43.6543 |
| 16511:3 | 1 | R.TAGWNVPIGTLRPFLN[+2351.851]WTGPPEPIEAAVAR.F | HexNAc(4)Hex(5)Fuc(3)NeuAc(1) | N[+2352] | 1861.5293 | 3 | 5582.5732 | 5582.5617 | 0 | 2.1 | 141 | Specific | 295.43 | 215.39 | 215.39 | 5.75 | 33 | >sp\|P02788\|TRFL_HUMAN Lactotransferrin OS=Homo sapiens OX=9606 GN=LTF PE=1 SV=6 | 1270 | 20201119_08533_1ng_glyco_hcd30_etd30_compound15.20214.20214.3 | controllerType=0 controllerNumber=1 scan=20214 | 46.2663 |
| 15825:4 | 1 | R.TAGWNVPIGTLRPFLN[+1768.640]WTGPPEPIEAAVAR.F | HexNAc(4)Hex(5)Fuc(1) | N[+1769] | 1250.5974 | 4 | 4999.3678 | 4999.3504 | 0 | 3.5 | 141 | Specific | 348.39 | 348.39 | 348.39 | 4.68 | 33 | >sp\|P02788\|TRFL_HUMAN Lactotransferrin OS=Homo sapiens OX=9606 GN=LTF PE=1 SV=6 | 1270 | 20201119_08533_1ng_glyco_hcd30_etd30_compound15.19489.19489.4 | controllerType=0 controllerNumber=1 scan=19489 | 44.1007 |
| 16544:3 | 1 | R.TAGWNVPIGTLRPFLN[+2059.735]WTGPPEPIEAAVAR.F | HexNAc(4)Hex(5)Fuc(1)NeuAc(1) | N[+2060] | 1764.1568 | 3 | 5290.4559 | 5290.4459 | 0 | 1.9 | 141 | Specific | 376.31 | 259.1 | 259.1 | 4.58 | 33 | >sp\|P02788\|TRFL_HUMAN Lactotransferrin OS=Homo sapiens OX=9606 GN=LTF PE=1 SV=6 | 1270 | 20201119_08533_1ng_glyco_hcd30_etd30_compound15.20249.20249.2 | controllerType=0 controllerNumber=1 scan=20249 | 46.3747 |
| 16479:4 | 1 | R.TAGWNVPIGTLRPFLN[+2716.983]WTGPPEPIEAAVAR.F | HexNAc(5)Hex(6)Fuc(3)NeuAc(1) | N[+2717] | 1487.6863 | 4 | 5947.7233 | 5947.6939 | 0 | 4.9 | 141 | Specific | 427.42 | 427.42 | 427.42 | 5.42 | 33 | >sp\|P02788\|TRFL_HUMAN Lactotransferrin OS=Homo sapiens OX=9606 GN=LTF PE=1 SV=6 | 1270 | 20201119_08533_1ng_glyco_hcd30_etd30_compound15.20180.20180.4 | controllerType=0 controllerNumber=1 scan=20180 | 46.1644 |
| 16487:4 | 1 | R.TAGWNVPIGTLRPFLN[+2570.925]WTGPPEPIEAAVAR.F | HexNAc(5)Hex(6)Fuc(2)NeuAc(1) | N[+2571] | 1451.1704 | 4 | 5801.6598 | 5801.636 | 0 | 4.1 | 141 | Specific | 494.02 | 494.02 | 494.02 | 6.6 | 33 | >sp\|P02788\|TRFL_HUMAN Lactotransferrin OS=Homo sapiens OX=9606 GN=LTF PE=1 SV=6 | 1270 | 20201119_08533_1ng_glyco_hcd30_etd30_compound15.20189.20189.4 | controllerType=0 controllerNumber=1 scan=20189 | 46.1935 |
| 16517:3 | 1 | R.TAGWNVPIGTLRPFLN[+2059.735]WTGPPEPIEAAVAR.F | HexNAc(4)Hex(5)Fuc(1)NeuAc(1) | N[+2060] | 1764.1583 | 3 | 5290.4603 | 5290.4459 | 0 | 2.7 | 141 | Specific | 551.39 | 436.31 | 436.31 | 7 | 33 | >sp\|P02788\|TRFL_HUMAN Lactotransferrin OS=Homo sapiens OX=9606 GN=LTF PE=1 SV=6 | 1270 | 20201119_08533_1ng_glyco_hcd30_etd30_compound15.20220.20220.3 | controllerType=0 controllerNumber=1 scan=20220 | 46.2883 |
| 16502:4 | 1 | R.TAGWNVPIGTLRPFLN[+2351.851]WTGPPEPIEAAVAR.F | HexNAc(4)Hex(5)Fuc(3)NeuAc(1) | N[+2352] | 1396.3984 | 4 | 5582.5719 | 5582.5617 | 0 | 1.8 | 141 | Specific | 747.66 | 747.66 | 747.66 | 11.65 | 33 | >sp\|P02788\|TRFL_HUMAN Lactotransferrin OS=Homo sapiens OX=9606 GN=LTF PE=1 SV=6 | 1270 | 20201119_08533_1ng_glyco_hcd30_etd30_compound15.20205.20205.4 | controllerType=0 controllerNumber=1 scan=20205 | 46.2469 |
| 16416:4 | 1 | R.TAGWNVPIGTLRPFLN[+2863.041]WTGPPEPIEAAVAR.F | HexNAc(5)Hex(6)Fuc(4)NeuAc(1) | N[+2863] | 1524.1974 | 4 | 6093.7677 | 6093.7518 | 0 | 2.6 | 141 | Specific | 758.33 | 735.23 | 735.23 | 12.66 | 33 | >sp\|P02788\|TRFL_HUMAN Lactotransferrin OS=Homo sapiens OX=9606 GN=LTF PE=1 SV=6 | 1270 | 20201119_08533_1ng_glyco_hcd30_etd30_compound15.20113.20113.4 | controllerType=0 controllerNumber=1 scan=20113 | 45.9646 |
| 16512:4 | 1 | R.TAGWNVPIGTLRPFLN[+2059.735]WTGPPEPIEAAVAR.F | HexNAc(4)Hex(5)Fuc(1)NeuAc(1) | N[+2060] | 1323.3705 | 4 | 5290.4601 | 5290.4459 | 0 | 2.7 | 141 | Specific | 846.28 | 784.45 | 784.45 | 13.66 | 33 | >sp\|P02788\|TRFL_HUMAN Lactotransferrin OS=Homo sapiens OX=9606 GN=LTF PE=1 SV=6 | 1270 | 20201119_08533_1ng_glyco_hcd30_etd30_compound15.20215.20215.4 | controllerType=0 controllerNumber=1 scan=20215 | 46.2756 |
| 14816:3 | 1 | R.TAGWNIPMGLLFN[+1606.587]QTGSC[+57.021]K.F | HexNAc(4)Hex(4)Fuc(1) | C[+57], N[+1607] | 1234.541 | 3 | 3701.6083 | 3701.5966 | 0 | 3.2 | 485 | Specific | 264.98 | 239.13 | 239.13 | 4.88 | 33 | >sp\|P02788\|TRFL_HUMAN Lactotransferrin OS=Homo sapiens OX=9606 GN=LTF PE=1 SV=6 | 1270 | 20201119_08533_1ng_glyco_hcd30_etd30_compound15.18423.18423.3 | controllerType=0 controllerNumber=1 scan=18423 | 40.9602 |
| 14694:3 | 1 | R.TAGWNIPMGLLFN[+1768.640]QTGSC[+57.021]K.F | HexNAc(4)Hex(5)Fuc(1) | C[+57], N[+1769] | 1288.5593 | 3 | 3863.6632 | 3863.6495 | 0 | 3.6 | 485 | Specific | 348.75 | 277.4 | 277.4 | 4.28 | 33 | >sp\|P02788\|TRFL_HUMAN Lactotransferrin OS=Homo sapiens OX=9606 GN=LTF PE=1 SV=6 | 1270 | 20201119_08533_1ng_glyco_hcd30_etd30_compound15.18295.18295.3 | controllerType=0 controllerNumber=1 scan=18295 | 40.6554 |
| 14705:3 | 1 | R.TAGWNIPMGLLFN[+1768.640]QTGSC[+57.021]K.F | HexNAc(4)Hex(5)Fuc(1) | C[+57], N[+1769] | 1288.5608 | 3 | 3863.668 | 3863.6495 | 0 | 4.8 | 485 | Specific | 386.28 | 288.65 | 288.65 | 4.68 | 33 | >sp\|P02788\|TRFL_HUMAN Lactotransferrin OS=Homo sapiens OX=9606 GN=LTF PE=1 SV=6 | 1270 | 20201119_08533_1ng_glyco_hcd30_etd30_compound15.18307.18307.3 | controllerType=0 controllerNumber=1 scan=18307 | 40.6899 |
| 16070:3 | 1 | R.TAGWNIPMGLLFN[+2059.735]QTGSC[+57.021]K.F | HexNAc(4)Hex(5)Fuc(1)NeuAc(1) | C[+57], N[+2060] | 1385.5912 | 3 | 4154.7592 | 4154.7449 | 0 | 3.4 | 485 | Specific | 395.51 | 283.75 | 283.75 | 5.3 | 33 | >sp\|P02788\|TRFL_HUMAN Lactotransferrin OS=Homo sapiens OX=9606 GN=LTF PE=1 SV=6 | 1270 | 20201119_08533_1ng_glyco_hcd30_etd30_compound15.19747.19747.3 | controllerType=0 controllerNumber=1 scan=19747 | 44.8276 |
| 16182:3 | 1 | R.TAGWNIPMGLLFN[+1913.677]QTGSC[+57.021]K.F | HexNAc(4)Hex(5)NeuAc(1) | C[+57], N[+1914] | 1336.912 | 3 | 4008.7216 | 4008.687 | 0 | 8.6 | 485 | Specific | 405.72 | 273.8 | 273.8 | 3.76 | 33 | >sp\|P02788\|TRFL_HUMAN Lactotransferrin OS=Homo sapiens OX=9606 GN=LTF PE=1 SV=6 | 1270 | 20201119_08533_1ng_glyco_hcd30_etd30_compound15.19865.19865.3 | controllerType=0 controllerNumber=1 scan=19865 | 45.237 |
| 16054:3 | 1 | R.TAGWNIPMGLLFN[+2205.793]QTGSC[+57.021]K.F | HexNAc(4)Hex(5)Fuc(2)NeuAc(1) | C[+57], N[+2206] | 1434.2764 | 3 | 4300.8147 | 4300.8028 | 0 | 2.8 | 485 | Specific | 440.7 | 276.51 | 276.51 | 5.45 | 33 | >sp\|P02788\|TRFL_HUMAN Lactotransferrin OS=Homo sapiens OX=9606 GN=LTF PE=1 SV=6 | 1270 | 20201119_08533_1ng_glyco_hcd30_etd30_compound15.19730.19730.3 | controllerType=0 controllerNumber=1 scan=19730 | 44.7855 |
| 17275:4 | 1 | R.TAGWNIPMGLLFN[+2350.830]QTGSC[+57.021]KFDEYFSQSC[+57.021]APGSDPR.S | HexNAc(4)Hex(5)Fuc(1)NeuAc(2) | C[+57]*2, N[+2351] | 1573.1591 | 4 | 6289.6144 | 6289.5875 | 0 | 4.3 | 485 | Specific | 273.38 | 249.04 | 249.04 | 4.91 | 33 | >sp\|P02788\|TRFL_HUMAN Lactotransferrin OS=Homo sapiens OX=9606 GN=LTF PE=1 SV=6 | 1270 | 20201119_08533_1ng_glyco_hcd30_etd30_compound15.21024.21024.4 | controllerType=0 controllerNumber=1 scan=21024 | 48.4785 |
| 17328:4 | 1 | R.TAGWNIPMGLLFN[+2204.772]QTGSC[+57.021]KFDEYFSQSC[+57.021]APGSDPR.S | HexNAc(4)Hex(5)NeuAc(2) | C[+57]*2, N[+2205] | 1536.6406 | 4 | 6143.5407 | 6143.5295 | 0 | 1.8 | 485 | Specific | 295.05 | 259.51 | 259.51 | 4.73 | 33 | >sp\|P02788\|TRFL_HUMAN Lactotransferrin OS=Homo sapiens OX=9606 GN=LTF PE=1 SV=6 | 1270 | 20201119_08533_1ng_glyco_hcd30_etd30_compound15.21081.21081.3 | controllerType=0 controllerNumber=1 scan=21081 | 48.6517 |
| 15505:4 | 1 | R.TAGWNIPMGLLFN[+1768.640]QTGSC[+57.021]KFDEYFSQSC[+57.021]APGSDPR.S | HexNAc(4)Hex(5)Fuc(1) | C[+57]*2, N[+1769] | 1427.6095 | 4 | 5707.4162 | 5707.3966 | 0 | 3.4 | 485 | Specific | 362.06 | 325.17 | 325.17 | 3.87 | 33 | >sp\|P02788\|TRFL_HUMAN Lactotransferrin OS=Homo sapiens OX=9606 GN=LTF PE=1 SV=6 | 1270 | 20201119_08533_1ng_glyco_hcd30_etd30_compound15.19153.19153.4 | controllerType=0 controllerNumber=1 scan=19153 | 43.1666 |
| 16262:4 | 1 | R.TAGWNIPMGLLFN[+2351.851]QTGSC[+57.021]KFDEYFSQSC[+57.021]APGSDPR.S | HexNAc(4)Hex(5)Fuc(3)NeuAc(1) | C[+57]*2, N[+2352] | 1573.4146 | 4 | 6290.6364 | 6290.6079 | 0 | 4.5 | 485 | Specific | 411.68 | 404.56 | 404.56 | 6.82 | 33 | >sp\|P02788\|TRFL_HUMAN Lactotransferrin OS=Homo sapiens OX=9606 GN=LTF PE=1 SV=6 | 1270 | 20201119_08533_1ng_glyco_hcd30_etd30_compound15.19950.19950.4 | controllerType=0 controllerNumber=1 scan=19950 | 45.5052 |
| 16244:4 | 1 | R.TAGWNIPMGLLFN[+2351.851]QTGSC[+57.021]KFDEYFSQSC[+57.021]APGSDPR.S | HexNAc(4)Hex(5)Fuc(3)NeuAc(1) | C[+57]*2, N[+2352] | 1573.4147 | 4 | 6290.6369 | 6290.6079 | 0 | 4.6 | 485 | Specific | 503.51 | 475.77 | 475.77 | 7.14 | 33 | >sp\|P02788\|TRFL_HUMAN Lactotransferrin OS=Homo sapiens OX=9606 GN=LTF PE=1 SV=6 | 1270 | 20201119_08533_1ng_glyco_hcd30_etd30_compound15.19931.19931.4 | controllerType=0 controllerNumber=1 scan=19931 | 45.4451 |
| 16272:4 | 1 | R.TAGWNIPMGLLFN[+2059.735]QTGSC[+57.021]KFDEYFSQSC[+57.021]APGSDPR.S | HexNAc(4)Hex(5)Fuc(1)NeuAc(1) | C[+57]*2, N[+2060] | 1500.3842 | 4 | 5998.5148 | 5998.492 | 0 | 3.8 | 485 | Specific | 638.83 | 555.96 | 555.96 | 9.31 | 33 | >sp\|P02788\|TRFL_HUMAN Lactotransferrin OS=Homo sapiens OX=9606 GN=LTF PE=1 SV=6 | 1270 | 20201119_08533_1ng_glyco_hcd30_etd30_compound15.19961.19961.4 | controllerType=0 controllerNumber=1 scan=19961 | 45.5444 |
| 16255:4 | 1 | R.TAGWNIPMGLLFN[+2205.793]QTGSC[+57.021]KFDEYFSQSC[+57.021]APGSDPR.S | HexNAc(4)Hex(5)Fuc(2)NeuAc(1) | C[+57]*2, N[+2206] | 1536.8981 | 4 | 6144.5705 | 6144.5499 | 0 | 3.3 | 485 | Specific | 654.77 | 615.33 | 615.33 | 10.06 | 33 | >sp\|P02788\|TRFL_HUMAN Lactotransferrin OS=Homo sapiens OX=9606 GN=LTF PE=1 SV=6 | 1270 | 20201119_08533_1ng_glyco_hcd30_etd30_compound15.19943.19943.4 | controllerType=0 controllerNumber=1 scan=19943 | 45.4911 |
| 08089:3 | 1 | R.N[+2205.793]GSDC[+57.021]PDKFC[+57.021]LFQSETK.N | HexNAc(4)Hex(5)Fuc(2)NeuAc(1) | C[+57]*2, N[+2206] | 1413.5668 | 3 | 4238.6859 | 4238.6667 | 0 | 4.5 | 642 | Specific | 204.72 | 97.56 | 97.56 | 4.01 | 33 | >sp\|P02788\|TRFL_HUMAN Lactotransferrin OS=Homo sapiens OX=9606 GN=LTF PE=1 SV=6 | 1270 | 20201119_08533_1ng_glyco_hcd30_etd30_compound15.11349.11349.3 | controllerType=0 controllerNumber=1 scan=11349 | 26.6131 |
| 15389:3 | 2 | R.LSLHRPALEDLLLGSEAN[+1419.502]LTC[+57.021]TLTGLR.D | HexNAc(3)Hex(5) | C[+57], N[+1420] | 1461.7075 | 3 | 4383.1078 | 4383.1005 | 0 | 1.7 | 127 | Specific | 232.67 | 179.51 | 179.51 | 3.52 | 28 | >sp\|P01876\|IGHA1_HUMAN Immunoglobulin heavy constant alpha 1 OS=Homo sapiens OX=9606 GN=IGHA1 PE=1 SV=2 | 255 | 20201119_08533_1ng_glyco_hcd30_etd30_compound15.19030.19030.3 | controllerType=0 controllerNumber=1 scan=19030 | 42.7855 |
| 15458:3 | 2 | R.LSLHRPALEDLLLGSEAN[+1095.397]LTC[+57.021]TLTGLR.D | HexNAc(3)Hex(3) | C[+57], N[+1095] | 1353.6694 | 3 | 4058.9936 | 4058.9949 | 0 | -0.3 | 127 | Specific | 257 | 224.13 | 224.13 | 3.93 | 28 | >sp\|P01876\|IGHA1_HUMAN Immunoglobulin heavy constant alpha 1 OS=Homo sapiens OX=9606 GN=IGHA1 PE=1 SV=2 | 255 | 20201119_08533_1ng_glyco_hcd30_etd30_compound15.19103.19103.3 | controllerType=0 controllerNumber=1 scan=19103 | 43.0008 |
| 15348:3 | 2 | R.LSLHRPALEDLLLGSEAN[+1540.529]LTC[+57.021]TLTGLR.D | HexNAc(2)Hex(7) | C[+57], N[+1541] | 1502.0538 | 3 | 4504.1468 | 4504.1268 | 0 | 4.4 | 127 | Specific | 425.4 | 410.98 | 410.98 | 7.36 | 28 | >sp\|P01876\|IGHA1_HUMAN Immunoglobulin heavy constant alpha 1 OS=Homo sapiens OX=9606 GN=IGHA1 PE=1 SV=2 | 255 | 20201119_08533_1ng_glyco_hcd30_etd30_compound15.18986.18986.3 | controllerType=0 controllerNumber=1 scan=18986 | 42.6449 |
| 15351:3 | 2 | R.LSLHRPALEDLLLGSEAN[+1663.608]LTC[+57.021]TLTGLR.D | HexNAc(5)Hex(4) | C[+57], N[+1664] | 1543.0789 | 3 | 4627.2222 | 4627.2064 | 0 | 3.4 | 127 | Specific | 429.87 | 394.35 | 394.35 | 7.12 | 28 | >sp\|P01876\|IGHA1_HUMAN Immunoglobulin heavy constant alpha 1 OS=Homo sapiens OX=9606 GN=IGHA1 PE=1 SV=2 | 255 | 20201119_08533_1ng_glyco_hcd30_etd30_compound15.18990.18990.3 | controllerType=0 controllerNumber=1 scan=18990 | 42.6564 |
| 15379:3 | 2 | R.LSLHRPALEDLLLGSEAN[+1501.555]LTC[+57.021]TLTGLR.D | HexNAc(5)Hex(3) | C[+57], N[+1502] | 1489.0589 | 3 | 4465.1621 | 4465.1536 | 0 | 1.9 | 127 | Specific | 440.32 | 388.8 | 388.8 | 7.16 | 28 | >sp\|P01876\|IGHA1_HUMAN Immunoglobulin heavy constant alpha 1 OS=Homo sapiens OX=9606 GN=IGHA1 PE=1 SV=2 | 255 | 20201119_08533_1ng_glyco_hcd30_etd30_compound15.19020.19020.3 | controllerType=0 controllerNumber=1 scan=19020 | 42.7655 |
| 15359:4 | 2 | R.LSLHRPALEDLLLGSEAN[+1581.555]LTC[+57.021]TLTGLR.D | HexNAc(3)Hex(6) | C[+57], N[+1582] | 1137.0464 | 4 | 4545.1639 | 4545.1533 | 0 | 2.3 | 127 | Specific | 454.04 | 454.04 | 454.04 | 6.51 | 28 | >sp\|P01876\|IGHA1_HUMAN Immunoglobulin heavy constant alpha 1 OS=Homo sapiens OX=9606 GN=IGHA1 PE=1 SV=2 | 255 | 20201119_08533_1ng_glyco_hcd30_etd30_compound15.18998.18998.4 | controllerType=0 controllerNumber=1 scan=18998 | 42.6879 |
| 15315:3 | 2 | R.LSLHRPALEDLLLGSEAN[+1702.581]LTC[+57.021]TLTGLR.D | HexNAc(2)Hex(8) | C[+57], N[+1703] | 1556.0735 | 3 | 4666.2061 | 4666.1796 | 0 | 5.7 | 127 | Specific | 462.92 | 438.81 | 438.81 | 6.55 | 28 | >sp\|P01876\|IGHA1_HUMAN Immunoglobulin heavy constant alpha 1 OS=Homo sapiens OX=9606 GN=IGHA1 PE=1 SV=2 | 255 | 20201119_08533_1ng_glyco_hcd30_etd30_compound15.18951.18951.3 | controllerType=0 controllerNumber=1 scan=18951 | 42.486 |
| 15368:3 | 2 | R.LSLHRPALEDLLLGSEAN[+1378.476]LTC[+57.021]TLTGLR.D | HexNAc(2)Hex(6) | C[+57], N[+1378] | 1448.0367 | 3 | 4342.0955 | 4342.074 | 0 | 5 | 127 | Specific | 505.7 | 476.53 | 476.53 | 6.82 | 28 | >sp\|P01876\|IGHA1_HUMAN Immunoglobulin heavy constant alpha 1 OS=Homo sapiens OX=9606 GN=IGHA1 PE=1 SV=2 | 255 | 20201119_08533_1ng_glyco_hcd30_etd30_compound15.19008.19008.3 | controllerType=0 controllerNumber=1 scan=19008 | 42.72 |
| 16076:4 | 2 | R.LSLHRPALEDLLLGSEAN[+1872.651]LTC[+57.021]TLTGLR.D | HexNAc(3)Hex(6)NeuAc(1) | C[+57], N[+1873] | 1209.8213 | 4 | 4836.2635 | 4836.2487 | 0 | 3.1 | 127 | Specific | 554.72 | 531.68 | 531.68 | 8.2 | 28 | >sp\|P01876\|IGHA1_HUMAN Immunoglobulin heavy constant alpha 1 OS=Homo sapiens OX=9606 GN=IGHA1 PE=1 SV=2 | 255 | 20201119_08533_1ng_glyco_hcd30_etd30_compound15.19753.19753.4 | controllerType=0 controllerNumber=1 scan=19753 | 44.8465 |
| 16105:4 | 2 | R.LSLHRPALEDLLLGSEAN[+1710.598]LTC[+57.021]TLTGLR.D | HexNAc(3)Hex(5)NeuAc(1) | C[+57], N[+1711] | 1169.3085 | 4 | 4674.2122 | 4674.1959 | 0 | 3.5 | 127 | Specific | 566.06 | 566.06 | 566.06 | 9.33 | 28 | >sp\|P01876\|IGHA1_HUMAN Immunoglobulin heavy constant alpha 1 OS=Homo sapiens OX=9606 GN=IGHA1 PE=1 SV=2 | 255 | 20201119_08533_1ng_glyco_hcd30_etd30_compound15.19784.19784.4 | controllerType=0 controllerNumber=1 scan=19784 | 44.952 |
| 15431:4 | 2 | R.LSLHRPALEDLLLGSEAN[+1216.423]LTC[+57.021]TLTGLR.D | HexNAc(2)Hex(5) | C[+57], N[+1216] | 1045.7619 | 4 | 4180.0257 | 4180.0211 | 0 | 1.1 | 127 | Specific | 701.1 | 701.1 | 701.1 | 12.07 | 28 | >sp\|P01876\|IGHA1_HUMAN Immunoglobulin heavy constant alpha 1 OS=Homo sapiens OX=9606 GN=IGHA1 PE=1 SV=2 | 255 | 20201119_08533_1ng_glyco_hcd30_etd30_compound15.19075.19075.2 | controllerType=0 controllerNumber=1 scan=19075 | 42.9425 |
| 15388:4 | 2 | R.LSLHRPALEDLLLGSEAN[+1257.449]LTC[+57.021]TLTGLR.D | HexNAc(3)Hex(4) | C[+57], N[+1257] | 1056.0196 | 4 | 4221.0565 | 4221.0477 | 0 | 2.1 | 127 | Specific | 849.79 | 849.79 | 849.79 | 14.07 | 28 | >sp\|P01876\|IGHA1_HUMAN Immunoglobulin heavy constant alpha 1 OS=Homo sapiens OX=9606 GN=IGHA1 PE=1 SV=2 | 255 | 20201119_08533_1ng_glyco_hcd30_etd30_compound15.19029.19029.4 | controllerType=0 controllerNumber=1 scan=19029 | 42.7814 |
| 15310:4 | 2 | R.LSLHRPALEDLLLGSEAN[+1702.581]LTC[+57.021]TLTGLR.D | HexNAc(2)Hex(8) | C[+57], N[+1703] | 1167.3069 | 4 | 4666.2059 | 4666.1796 | 0 | 5.6 | 127 | Specific | 865.18 | 853.17 | 853.17 | 14.47 | 28 | >sp\|P01876\|IGHA1_HUMAN Immunoglobulin heavy constant alpha 1 OS=Homo sapiens OX=9606 GN=IGHA1 PE=1 SV=2 | 255 | 20201119_08533_1ng_glyco_hcd30_etd30_compound15.18945.18945.4 | controllerType=0 controllerNumber=1 scan=18945 | 42.4521 |
| 15364:4 | 2 | R.LSLHRPALEDLLLGSEAN[+1501.555]LTC[+57.021]TLTGLR.D | HexNAc(5)Hex(3) | C[+57], N[+1502] | 1117.0497 | 4 | 4465.1771 | 4465.1536 | 0 | 5.3 | 127 | Specific | 887.18 | 887.18 | 887.18 | 14.47 | 28 | >sp\|P01876\|IGHA1_HUMAN Immunoglobulin heavy constant alpha 1 OS=Homo sapiens OX=9606 GN=IGHA1 PE=1 SV=2 | 255 | 20201119_08533_1ng_glyco_hcd30_etd30_compound15.19004.19004.4 | controllerType=0 controllerNumber=1 scan=19004 | 42.7114 |
| 15370:4 | 2 | R.LSLHRPALEDLLLGSEAN[+1419.502]LTC[+57.021]TLTGLR.D | HexNAc(3)Hex(5) | C[+57], N[+1420] | 1096.5358 | 4 | 4383.1214 | 4383.1005 | 0 | 4.8 | 127 | Specific | 930.2 | 869.95 | 869.95 | 15.47 | 28 | >sp\|P01876\|IGHA1_HUMAN Immunoglobulin heavy constant alpha 1 OS=Homo sapiens OX=9606 GN=IGHA1 PE=1 SV=2 | 255 | 20201119_08533_1ng_glyco_hcd30_etd30_compound15.19010.19010.4 | controllerType=0 controllerNumber=1 scan=19010 | 42.7297 |
| 15347:4 | 2 | R.LSLHRPALEDLLLGSEAN[+1663.608]LTC[+57.021]TLTGLR.D | HexNAc(5)Hex(4) | C[+57], N[+1664] | 1157.5619 | 4 | 4627.2259 | 4627.2064 | 0 | 4.2 | 127 | Specific | 987.83 | 985.99 | 985.99 | 17.33 | 28 | >sp\|P01876\|IGHA1_HUMAN Immunoglobulin heavy constant alpha 1 OS=Homo sapiens OX=9606 GN=IGHA1 PE=1 SV=2 | 255 | 20201119_08533_1ng_glyco_hcd30_etd30_compound15.18985.18985.4 | controllerType=0 controllerNumber=1 scan=18985 | 42.6418 |
| 15363:4 | 2 | R.LSLHRPALEDLLLGSEAN[+1378.476]LTC[+57.021]TLTGLR.D | HexNAc(2)Hex(6) | C[+57], N[+1378] | 1086.279 | 4 | 4342.0941 | 4342.074 | 0 | 4.6 | 127 | Specific | 1038.56 | 1000.39 | 1000.39 | 17.47 | 28 | >sp\|P01876\|IGHA1_HUMAN Immunoglobulin heavy constant alpha 1 OS=Homo sapiens OX=9606 GN=IGHA1 PE=1 SV=2 | 255 | 20201119_08533_1ng_glyco_hcd30_etd30_compound15.19003.19003.4 | controllerType=0 controllerNumber=1 scan=19003 | 42.7068 |
| 15376:4 | 2 | R.LSLHRPALEDLLLGSEAN[+1216.423]LTC[+57.021]TLTGLR.D | HexNAc(2)Hex(5) | C[+57], N[+1216] | 1045.7648 | 4 | 4180.0374 | 4180.0211 | 0 | 3.9 | 127 | Specific | 1075.23 | 1023.61 | 1023.61 | 19.33 | 28 | >sp\|P01876\|IGHA1_HUMAN Immunoglobulin heavy constant alpha 1 OS=Homo sapiens OX=9606 GN=IGHA1 PE=1 SV=2 | 255 | 20201119_08533_1ng_glyco_hcd30_etd30_compound15.19016.19016.4 | controllerType=0 controllerNumber=1 scan=19016 | 42.7532 |
| 15369:4 | 2 | R.LSLHRPALEDLLLGSEAN[+1460.529]LTC[+57.021]TLTGLR.D | HexNAc(4)Hex(4) | C[+57], N[+1461] | 1106.7941 | 4 | 4424.1546 | 4424.1271 | 0 | 6.2 | 127 | Specific | 1090.77 | 996.41 | 996.41 | 17.61 | 28 | >sp\|P01876\|IGHA1_HUMAN Immunoglobulin heavy constant alpha 1 OS=Homo sapiens OX=9606 GN=IGHA1 PE=1 SV=2 | 255 | 20201119_08533_1ng_glyco_hcd30_etd30_compound15.19009.19009.4 | controllerType=0 controllerNumber=1 scan=19009 | 42.7264 |
| 10138:3 | 2 | R.LAGKPTHVN[+1501.555]VSVVMAEVDGTC[+57.021]Y.- | HexNAc(5)Hex(3) | C[+57], N[+1502] | 1283.5754 | 3 | 3848.7116 | 3848.6974 | 0 | 3.7 | 332 | CRagged | 236.07 | 169.51 | 169.51 | 3.6 | 28 | >sp\|P01876\|IGHA1_HUMAN Immunoglobulin heavy constant alpha 1 OS=Homo sapiens OX=9606 GN=IGHA1 PE=1 SV=2 | 255 | 20201119_08533_1ng_glyco_hcd30_etd30_compound15.13480.13480.3 | controllerType=0 controllerNumber=1 scan=13480 | 30.8152 |
| 10096:3 | 2 | R.LAGKPTHVN[+1378.476]VSVVMAEVDGTC[+57.021]Y.- | HexNAc(2)Hex(6) | C[+57], N[+1378] | 1242.5479 | 3 | 3725.6292 | 3725.6178 | 0 | 3.1 | 332 | CRagged | 236.86 | 188.19 | 187.4 | 3.37 | 28 | >sp\|P01876\|IGHA1_HUMAN Immunoglobulin heavy constant alpha 1 OS=Homo sapiens OX=9606 GN=IGHA1 PE=1 SV=2 | 255 | 20201119_08533_1ng_glyco_hcd30_etd30_compound15.13435.13435.3 | controllerType=0 controllerNumber=1 scan=13435 | [illegible] |
| 10137:3 | 2 | R.LAGKPTHVN[+1647.613]VSVVMAEVDGTC[+57.021]Y.- | HexNAc(5)Hex(3)Fuc(1) | C[+57], N[+1648] | 1332.2606 | 3 | 3994.7671 | 3994.7553 | 0 | 3 | 332 | CRagged | 248.44 | 156.12 | 156.12 | 3.25 | 28 | >sp\|P01876\|IGHA1_HUMAN Immunoglobulin heavy constant alpha 1 OS=Homo sapiens OX=9606 GN=IGHA1 PE=1 SV=2 | 255 | 20201119_08533_1ng_glyco_hcd30_etd30_compound15.13479.13479.3 | controllerType=0 controllerNumber=1 scan=13479 | [illegible] |
| 10033:3 | 2 | R.LAGKPTHVN[+1702.581]VSVVMAEVDGTC[+57.021]Y.- | HexNAc(2)Hex(8) | C[+57], N[+1703] | 1350.5829 | 3 | 4049.7343 | 4049.7234 | 0 | 2.7 | 332 | CRagged | 251.42 | 191.69 | 191.69 | 3.76 | 28 | >sp\|P01876\|IGHA1_HUMAN Immunoglobulin heavy constant alpha 1 OS=Homo sapiens OX=9606 GN=IGHA1 PE=1 SV=2 | 255 | 20201119_08533_1ng_glyco_hcd30_etd30_compound15.13368.13368.3 | controllerType=0 controllerNumber=1 scan=13368 | 30.4721 |
| 10101:3 | 2 | R.LAGKPTHVN[+1663.608]VSVVMAEVDGTC[+57.021]Y.- | HexNAc(5)Hex(4) | C[+57], N[+1664] | 1337.5908 | 3 | 4010.7577 | 4010.7502 | 0 | 1.9 | 332 | CRagged | 257.69 | 222.34 | 175.61 | 3.62 | 28 | >sp\|P01876\|IGHA1_HUMAN Immunoglobulin heavy constant alpha 1 OS=Homo sapiens OX=9606 GN=IGHA1 PE=1 SV=2 | 255 | 20201119_08533_1ng_glyco_hcd30_etd30_compound15.13440.13440.3 | controllerType=0 controllerNumber=1 scan=13440 | 30.68 |
| 10097:3 | 2 | R.LAGKPTHVN[+1809.666]VSVVMAEVDGTC[+57.021]Y.- | HexNAc(5)Hex(4)Fuc(1) | C[+57], N[+1810] | 1386.278 | 3 | 4156.8195 | 4156.8081 | 0 | 2.7 | 332 | CRagged | 266.2 | 155.09 | 155.09 | 3.5 | 28 | >sp\|P01876\|IGHA1_HUMAN Immunoglobulin heavy constant alpha 1 OS=Homo sapiens OX=9606 GN=IGHA1 PE=1 SV=2 | 255 | 20201119_08533_1ng_glyco_hcd30_etd30_compound15.13436.13436.3 | controllerType=0 controllerNumber=1 scan=13436 | 30.6711 |
| 11657:4 | 2 | R.LAGKPTHVN[+2262.814]VSVVMAEVDGTC[+57.021]Y.- | HexNAc(5)Hex(5)Fuc(1)NeuAc(1) | C[+57], N[+2263] | 1153.2494 | 4 | 4609.9759 | 4609.9564 | 0 | 4.2 | 332 | CRagged | 279.02 | 187.83 | 187.83 | 3.59 | 28 | >sp\|P01876\|IGHA1_HUMAN Immunoglobulin heavy constant alpha 1 OS=Homo sapiens OX=9606 GN=IGHA1 PE=1 SV=2 | 255 | 20201119_08533_1ng_glyco_hcd30_etd30_compound15.15082.15082.4 | controllerType=0 controllerNumber=1 scan=15082 | 33.8441 |
| 10102:3 | 2 | R.LAGKPTHVN[+1216.423]VSVVMAEVDGTC[+57.021]Y.- | HexNAc(2)Hex(5) | C[+57], N[+1216] | 1188.5301 | 3 | 3563.5757 | 3563.5649 | 0 | 3 | 332 | CRagged | 282.96 | 282.96 | 219.45 | 4.22 | 28 | >sp\|P01876\|IGHA1_HUMAN Immunoglobulin heavy constant alpha 1 OS=Homo sapiens OX=9606 GN=IGHA1 PE=1 SV=2 | 255 | 20201119_08533_1ng_glyco_hcd30_etd30_compound15.13441.13441.3 | controllerType=0 controllerNumber=1 scan=13441 | 30.6868 |
| 04173:3 | 5 | R.FSDGLESN[+1864.634]SSTQFEVKK.Y | HexNAc(2)Hex(9) | N[+1865] | 1256.5215 | 3 | 3767.5501 | 3767.5421 | 0 | 2.1 | 219 | Specific | 203.22 | 129.6 | 129.6 | 3.7 | 10 | >sp\|P0C0L4\|CO4A_HUMAN Complement C4-A OS=Homo sapiens OX=9606 GN=C4A PE=1 SV=2 | 1235 | 20201119_08533_1ng_glyco_hcd30_etd30_compound15.7250.7250.3 | controllerType=0 controllerNumber=1 scan=7250 | 19.249 |
| 19106:3 | 6 | R.[+42.011]LSLLEEPGN[+2336.851]GTFTVILNQLTSR.D | HexNAc(6)Hex(6)Fuc(1) | .[42], N[+2337] | 1594.3808 | 3 | 4781.1278 | 4781.1542 | 0 | -5.5 | 413 | Specific | 305.36 | 207.16 | 207.16 | 2.69 | 18 | >sp\|P01833\|PIGR_HUMAN Polymeric immunoglobulin receptor OS=Homo sapiens OX=9606 GN=PIGR PE=1 SV=4 | 902 | 20201119_08533_1ng_glyco_hcd30_etd30_compound15.22963.22963.3 | controllerType=0 controllerNumber=1 scan=22963 | 53.5474 |
| 17373:3 | 6 | R.LSLLEEPGN[+2205.793]GTFTVILNQLTSR.D | HexNAc(4)Hex(5)Fuc(2)NeuAc(1) | N[+2206] | 1536.7075 | 3 | 4608.1078 | 4608.0854 | 0 | 4.9 | 413 | Specific | 490.89 | 323.59 | 323.59 | 5.01 | 18 | >sp\|P01833\|PIGR_HUMAN Polymeric immunoglobulin receptor OS=Homo sapiens OX=9606 GN=PIGR PE=1 SV=4 | 902 | 20201119_08533_1ng_glyco_hcd30_etd30_compound15.21128.21128.4 | controllerType=0 controllerNumber=1 scan=21128 | 48.8082 |
| 15541:3 | 6 | K.IIEGEPNLKVPGN[+2350.830]VTAVLGETLK.V | HexNAc(4)Hex(5)Fuc(1)NeuAc(2) | N[+2351] | 1581.4066 | 3 | 4742.2051 | 4742.1797 | 0 | 5.4 | 457 | Specific | 294.01 | 219.05 | 219.05 | 2.97 | 18 | >sp\|P01833\|PIGR_HUMAN Polymeric immunoglobulin receptor OS=Homo sapiens OX=9606 GN=PIGR PE=1 SV=4 | 902 | 20201119_08533_1ng_glyco_hcd30_etd30_compound15.19191.19191.3 | controllerType=0 controllerNumber=1 scan=19191 | 43.277 |
| 14101:3 | 6 | K.IIEGEPNLKVPGN[+2059.735]VTAVLGETLK.V | HexNAc(4)Hex(5)Fuc(1)NeuAc(1) | N[+2060] | 1484.3716 | 3 | 4451.1004 | 4451.0843 | 0 | 3.6 | 457 | Specific | 309.52 | 265.36 | 265.36 | 4.11 | 18 | >sp\|P01833\|PIGR_HUMAN Polymeric immunoglobulin receptor OS=Homo sapiens OX=9606 GN=PIGR PE=1 SV=4 | 902 | 20201119_08533_1ng_glyco_hcd30_etd30_compound15.17668.17668.3 | controllerType=0 controllerNumber=1 scan=17668 | 39.3943 |
| 15549:4 | 6 | K.IIEGEPNLKVPGN[+2350.830]VTAVLGETLK.V | HexNAc(4)Hex(5)Fuc(1)NeuAc(2) | N[+2351] | 1186.3072 | 4 | 4742.2069 | 4742.1797 | 0 | 5.7 | 457 | Specific | 352.67 | 340.24 | 340.24 | 3.07 | 18 | >sp\|P01833\|PIGR_HUMAN Polymeric immunoglobulin receptor OS=Homo sapiens OX=9606 GN=PIGR PE=1 SV=4 | 902 | 20201119_08533_1ng_glyco_hcd30_etd30_compound15.19199.19199.4 | controllerType=0 controllerNumber=1 scan=19199 | 43.2919 |
| 14043:3 | 6 | K.IIEGEPNLKVPGN[+2205.793]VTAVLGETLK.V | HexNAc(4)Hex(5)Fuc(2)NeuAc(1) | N[+2206] | 1533.0562 | 3 | 4597.1541 | 4597.1422 | 0 | 2.6 | 457 | Specific | 387.53 | 284.35 | 284.35 | 3.82 | 18 | >sp\|P01833\|PIGR_HUMAN Polymeric immunoglobulin receptor OS=Homo sapiens OX=9606 GN=PIGR PE=1 SV=4 | 902 | 20201119_08533_1ng_glyco_hcd30_etd30_compound15.17607.17607.3 | controllerType=0 controllerNumber=1 scan=17607 | 39.2311 |
| 14924:4 | 6 | K.IIEGEPNLKVPGN[+2350.830]VTAVLGETLKVPC[+57.021]HFPC[+57.021]K.F | HexNAc(4)Hex(5)Fuc(1)NeuAc(2) | C[+57]*2, N[+2351] | 1442.6718 | 4 | 5767.6652 | 5767.6372 | 0 | 4.8 | 457 | Specific | 401.95 | 401.95 | 401.95 | 4.02 | 18 | >sp\|P01833\|PIGR_HUMAN Polymeric immunoglobulin receptor OS=Homo sapiens OX=9606 GN=PIGR PE=1 SV=4 | 902 | 20201119_08533_1ng_glyco_hcd30_etd30_compound15.18539.18539.4 | controllerType=0 controllerNumber=1 scan=18539 | 41.3124 |
| 13653:4 | 6 | K.IIEGEPNLKVPGN[+2204.772]VTAVLGETLKVPC[+57.021]HFPC[+57.021]K.F | HexNAc(4)Hex(5)NeuAc(2) | C[+57]*2, N[+2205] | 1406.1595 | 4 | 5621.6164 | 5621.5793 | 0 | 6.6 | 457 | Specific | 445.46 | 445.46 | 445.46 | 3.58 | 18 | >sp\|P01833\|PIGR_HUMAN Polymeric immunoglobulin receptor OS=Homo sapiens OX=9606 GN=PIGR PE=1 SV=4 | 902 | 20201119_08533_1ng_glyco_hcd30_etd30_compound15.17166.17166.2 | controllerType=0 controllerNumber=1 scan=17166 | 38.3214 |
| 14919:5 | 6 | K.IIEGEPNLKVPGN[+2350.830]VTAVLGETLKVPC[+57.021]HFPC[+57.021]K.F | HexNAc(4)Hex(5)Fuc(1)NeuAc(2) | C[+57]*2, N[+2351] | 1154.3434 | 5 | 5767.6878 | 5767.6372 | 0 | 8.8 | 457 | Specific | 597.44 | 597.44 | 597.44 | 5.02 | 18 | >sp\|P01833\|PIGR_HUMAN Polymeric immunoglobulin receptor OS=Homo sapiens OX=9606 GN=PIGR PE=1 SV=4 | 902 | 20201119_08533_1ng_glyco_hcd30_etd30_compound15.18533.18533.5 | controllerType=0 controllerNumber=1 scan=18533 | 41.2893 |
| 13571:5 | 6 | K.IIEGEPNLKVPGN[+2205.793]VTAVLGETLKVPC[+57.021]HFPC[+57.021]K.F | HexNAc(4)Hex(5)Fuc(2)NeuAc(1) | C[+57]*2, N[+2206] | 1125.3317 | 5 | 5622.6292 | 5622.5997 | 0 | 5.2 | 457 | Specific | 759.18 | 741.52 | 741.52 | 10.27 | 18 | >sp\|P01833\|PIGR_HUMAN Polymeric immunoglobulin receptor OS=Homo sapiens OX=9606 GN=PIGR PE=1 SV=4 | 902 | 20201119_08533_1ng_glyco_hcd30_etd30_compound15.17080.17080.5 | controllerType=0 controllerNumber=1 scan=17080 | 38.1363 |
| 05073:3 | 6 | K.WN[+1872.651]NTGC[+57.021]QALPSQDEGPSK.A | HexNAc(3)Hex(6)NeuAc(1) | C[+57], N[+1873] | 1287.8525 | 3 | 3861.5429 | 3861.5272 | 0 | 4.1 | 498 | Specific | 224.01 | 103.96 | 103.96 | 4.01 | 18 | >sp\|P01833\|PIGR_HUMAN Polymeric immunoglobulin receptor OS=Homo sapiens OX=9606 GN=PIGR PE=1 SV=4 | 902 | 20201119_08533_1ng_glyco_hcd30_etd30_compound15.8181.8181.3 | controllerType=0 controllerNumber=1 scan=8181 | 20.872 |
| 05009:3 | 6 | K.WN[+2059.735]NTGC[+57.021]QALPSQDEGPSK.A | HexNAc(4)Hex(5)Fuc(1)NeuAc(1) | C[+57], N[+2060] | 1350.214 | 3 | 4048.6276 | 4048.6116 | 0 | 3.9 | 498 | Specific | 272.35 | 164.05 | 164.05 | 4.64 | 18 | >sp\|P01833\|PIGR_HUMAN Polymeric immunoglobulin receptor OS=Homo sapiens OX=9606 GN=PIGR PE=1 SV=4 | 902 | 20201119_08533_1ng_glyco_hcd30_etd30_compound15.8115.8115.3 | controllerType=0 controllerNumber=1 scan=8115 | 20.7482 |
| 05123:3 | 6 | K.WN[+1710.598]NTGC[+57.021]QALPSQDEGPSK.A | HexNAc(3)Hex(5)NeuAc(1) | C[+57], N[+1711] | 1233.8327 | 3 | 3699.4835 | 3699.4743 | 0 | 2.5 | 498 | Specific | 431.46 | 255.56 | 255.56 | 4.74 | 18 | >sp\|P01833\|PIGR_HUMAN Polymeric immunoglobulin receptor OS=Homo sapiens OX=9606 GN=PIGR PE=1 SV=4 | 902 | 20201119_08533_1ng_glyco_hcd30_etd30_compound15.8233.8233.3 | controllerType=0 controllerNumber=1 scan=8233 | 20.9746 |
| 05065:3 | 6 | K.WN[+1913.677]NTGC[+57.021]QALPSQDEGPSK.A | HexNAc(4)Hex(5)NeuAc(1) | C[+57], N[+1914] | 1301.5286 | 3 | 3902.5713 | 3902.5537 | 0 | 4.5 | 498 | Specific | 528.38 | 357.66 | 357.66 | 6.11 | 18 | >sp\|P01833\|PIGR_HUMAN Polymeric immunoglobulin receptor OS=Homo sapiens OX=9606 GN=PIGR PE=1 SV=4 | 902 | 20201119_08533_1ng_glyco_hcd30_etd30_compound15.8173.8173.3 | controllerType=0 controllerNumber=1 scan=8173 | 20.8548 |
| 00892:3 | 7 | K.HYTN[+1444.534]SSQDVTVPC[+57.021]R.V | HexNAc(4)Hex(3)Fuc(1) | C[+57], N[+1445] | 1036.7684 | 3 | 3108.2906 | 3108.2832 | 0 | 2.4 | 89 | Specific | 303.58 | 262.51 | 262.51 | 5.37 | 22 | >sp\|P01877\|IGHA2_HUMAN Immunoglobulin heavy constant alpha 2 OS=Homo sapiens OX=9606 GN=IGHA2 PE=1 SV=4 | 576 | 20201119_08533_1ng_glyco_hcd30_etd30_compound15.3838.3838.3 | controllerType=0 controllerNumber=1 scan=3838 | 12.9746 |
| 00935:3 | 7 | K.HYTN[+1647.613]SSQDVTVPC[+57.021]R.V | HexNAc(5)Hex(3)Fuc(1) | C[+57], N[+1648] | 1104.4616 | 3 | 3311.3702 | 3311.3625 | 0 | 2.3 | 89 | Specific | 394.06 | 335.03 | 335.03 | 5.59 | 22 | >sp\|P01877\|IGHA2_HUMAN Immunoglobulin heavy constant alpha 2 OS=Homo sapiens OX=9606 GN=IGHA2 PE=1 SV=4 | 576 | 20201119_08533_1ng_glyco_hcd30_etd30_compound15.3883.3883.2 | controllerType=0 controllerNumber=1 scan=3883 | 13.0676 |
| 00854:3 | 7 | K.HYTN[+1809.666]SSQDVTVPC[+57.021]R.V | HexNAc(5)Hex(4)Fuc(1) | C[+57], N[+1810] | 1158.4805 | 3 | 3473.427 | 3473.4154 | 0 | 3.4 | 89 | Specific | 397.32 | 329.65 | 329.65 | 5.3 | 22 | >sp\|P01877\|IGHA2_HUMAN Immunoglobulin heavy constant alpha 2 OS=Homo sapiens OX=9606 GN=IGHA2 PE=1 SV=4 | 576 | 20201119_08533_1ng_glyco_hcd30_etd30_compound15.3799.3799.3 | controllerType=0 controllerNumber=1 scan=3799 | 12.8885 |
| 00870:3 | 7 | K.HYTN[+1647.613]SSQDVTVPC[+57.021]R.V | HexNAc(5)Hex(3)Fuc(1) | C[+57], N[+1648] | 1104.4604 | 3 | 3311.3665 | 3311.3625 | 0 | 1.2 | 89 | Specific | 433.97 | 368.4 | 368.4 | 6.86 | 22 | >sp\|P01877\|IGHA2_HUMAN Immunoglobulin heavy constant alpha 2 OS=Homo sapiens OX=9606 GN=IGHA2 PE=1 SV=4 | 576 | 20201119_08533_1ng_glyco_hcd30_etd30_compound15.3816.3816.3 | controllerType=0 controllerNumber=1 scan=3816 | 12.9383 |
| 00867:3 | 7 | K.HYTN[+1647.613]SSQDVTVPC[+57.021]R.V | HexNAc(5)Hex(3)Fuc(1) | C[+57], N[+1648] | 1104.4633 | 3 | 3311.3753 | 3311.3625 | 0 | 3.9 | 89 | Specific | 490.17 | 419.75 | 419.75 | 7.44 | 22 | >sp\|P01877\|IGHA2_HUMAN Immunoglobulin heavy constant alpha 2 OS=Homo sapiens OX=9606 GN=IGHA2 PE=1 SV=4 | 576 | 20201119_08533_1ng_glyco_hcd30_etd30_compound15.3812.3812.3 | controllerType=0 controllerNumber=1 scan=3812 | 12.9225 |
| 02714:4 | 7 | K.HYTN[+1647.613]SSQDVTVPC[+57.021]RVPPPPPC[+57.021]C[+57.021]HPR.L | HexNAc(5)Hex(3)Fuc(1) | C[+57]*3, N[+1648] | 1152.252 | 4 | 4605.9862 | 4605.9689 | 0 | 3.8 | 89 | Specific | 462.78 | 402.91 | 402.91 | 5.57 | 22 | >sp\|P01877\|IGHA2_HUMAN Immunoglobulin heavy constant alpha 2 OS=Homo sapiens OX=9606 GN=IGHA2 PE=1 SV=4 | 576 | 20201119_08533_1ng_glyco_hcd30_etd30_compound15.5736.5736.4 | controllerType=0 controllerNumber=1 scan=5736 | 16.5772 |
| 02715:5 | 7 | K.HYTN[+1647.613]SSQDVTVPC[+57.021]RVPPPPPC[+57.021]C[+57.021]HPR.L | HexNAc(5)Hex(3)Fuc(1) | C[+57]*3, N[+1648] | 922.0027 | 5 | 4605.9844 | 4605.9689 | 0 | 3.4 | 89 | Specific | 484.64 | 458.13 | 458.13 | 5.8 | 22 | >sp\|P01877\|IGHA2_HUMAN Immunoglobulin heavy constant alpha 2 OS=Homo sapiens OX=9606 GN=IGHA2 PE=1 SV=4 | 576 | 20201119_08533_1ng_glyco_hcd30_etd30_compound15.5737.5737.5 | controllerType=0 controllerNumber=1 scan=5737 | 16.5835 |
| 02418:3 | 7 | K.TPLTAN[+1971.719]ITK.S | HexNAc(5)Hex(5)Fuc(1) | N[+1972] | 977.4337 | 3 | 2930.2867 | 2930.2757 | 0 | 3.8 | 200 | Specific | 243.68 | 119.08 | 119.08 | 2.66 | 22 | >sp\|P01877\|IGHA2_HUMAN Immunoglobulin heavy constant alpha 2 OS=Homo sapiens OX=9606 GN=IGHA2 PE=1 SV=4 | 576 | 20201119_08533_1ng_glyco_hcd30_etd30_compound15.5426.5426.3 | controllerType=0 controllerNumber=1 scan=5426 | 15.8634 |
| 02435:2 | 7 | K.TPLTAN[+1606.587]ITK.S | HexNAc(4)Hex(4)Fuc(1) | N[+1607] | 1283.0774 | 2 | 2565.1476 | 2565.1435 | 0 | 1.6 | 200 | Specific | 253.4 | 78.9 | 78.9 | 2.68 | 22 | >sp\|P01877\|IGHA2_HUMAN Immunoglobulin heavy constant alpha 2 OS=Homo sapiens OX=9606 GN=IGHA2 PE=1 SV=4 | 576 | 20201119_08533_1ng_glyco_hcd30_etd30_compound15.5444.5444.2 | controllerType=0 controllerNumber=1 scan=5444 | 15.8998 |
| 02510:3 | 7 | K.TPLTAN[+1647.613]ITK.S | HexNAc(5)Hex(3)Fuc(1) | N[+1648] | 869.3975 | 3 | 2606.1779 | 2606.17 | 0 | 3 | 200 | Specific | 270.67 | 117.22 | 117.22 | 2.68 | 22 | >sp\|P01877\|IGHA2_HUMAN Immunoglobulin heavy constant alpha 2 OS=Homo sapiens OX=9606 GN=IGHA2 PE=1 SV=4 | 576 | 20201119_08533_1ng_glyco_hcd30_etd30_compound15.5523.5523.3 | controllerType=0 controllerNumber=1 scan=5523 | 16.1094 |
| 02450:2 | 7 | K.TPLTAN[+1809.666]ITK.S | HexNAc(5)Hex(4)Fuc(1) | N[+1810] | 1384.6203 | 2 | 2768.2333 | 2768.2228 | 0 | 3.8 | 200 | Specific | 292.4 | 101.75 | 101.75 | 2.96 | 22 | >sp\|P01877\|IGHA2_HUMAN Immunoglobulin heavy constant alpha 2 OS=Homo sapiens OX=9606 GN=IGHA2 PE=1 SV=4 | 576 | 20201119_08533_1ng_glyco_hcd30_etd30_compound15.5460.5460.2 | controllerType=0 controllerNumber=1 scan=5460 | 15.9473 |
| 02447:3 | 7 | K.TPLTAN[+1809.666]ITK.S | HexNAc(5)Hex(4)Fuc(1) | N[+1810] | 923.416 | 3 | 2768.2336 | 2768.2228 | 0 | 3.9 | 200 | Specific | 298.15 | 129.07 | 129.07 | 2.82 | 22 | >sp\|P01877\|IGHA2_HUMAN Immunoglobulin heavy constant alpha 2 OS=Homo sapiens OX=9606 GN=IGHA2 PE=1 SV=4 | 576 | 20201119_08533_1ng_glyco_hcd30_etd30_compound15.5457.5457.3 | controllerType=0 controllerNumber=1 scan=5457 | 15.9362 |
| 02504:2 | 7 | K.TPLTAN[+1501.555]ITK.S | HexNAc(5)Hex(3) | N[+1502] | 1230.5654 | 2 | 2460.1234 | 2460.1121 | 0 | 4.6 | 200 | Specific | 308.66 | 97.08 | 97.08 | 2.96 | 22 | >sp\|P01877\|IGHA2_HUMAN Immunoglobulin heavy constant alpha 2 OS=Homo sapiens OX=9606 GN=IGHA2 PE=1 SV=4 | 576 | 20201119_08533_1ng_glyco_hcd30_etd30_compound15.5516.5516.2 | controllerType=0 controllerNumber=1 scan=5516 | 16.0842 |
| 02459:2 | 7 | K.TPLTAN[+1444.534]ITK.S | HexNAc(4)Hex(3)Fuc(1) | N[+1445] | 1202.0545 | 2 | 2403.1017 | 2403.0906 | 0 | 4.6 | 200 | Specific | 329.95 | 111.42 | 111.42 | 2.62 | 22 | >sp\|P01877\|IGHA2_HUMAN Immunoglobulin heavy constant alpha 2 OS=Homo sapiens OX=9606 GN=IGHA2 PE=1 SV=4 | 576 | 20201119_08533_1ng_glyco_hcd30_etd30_compound15.5469.5469.2 | controllerType=0 controllerNumber=1 scan=5469 | 15.9737 |
| 09102:3 | 7 | R.MAGKPTHIN[+1702.581]VSVVMAEADGTC[+57.021]Y.- | HexNAc(2)Hex(8) | C[+57], N[+1703] | 1351.8963 | 3 | 4053.6743 | 4053.6642 | 0 | 2.5 | 319 | CRagged | 212.85 | 212.85 | 176.89 | 3.43 | 22 | >sp\|P01877\|IGHA2_HUMAN Immunoglobulin heavy constant alpha 2 OS=Homo sapiens OX=9606 GN=IGHA2 PE=1 SV=4 | 576 | 20201119_08533_1ng_glyco_hcd30_etd30_compound15.12401.12401.3 | controllerType=0 controllerNumber=1 scan=12401 | 28.6356 |
| 09212:3 | 7 | R.MAGKPTHIN[+1501.555]VSVVMAEADGTC[+57.021]Y.- | HexNAc(5)Hex(3) | C[+57], N[+1502] | 1284.8886 | 3 | 3852.6513 | 3852.6382 | 0 | 3.4 | 319 | CRagged | 228.16 | 160.38 | 160.38 | 3.37 | 22 | >sp\|P01877\|IGHA2_HUMAN Immunoglobulin heavy constant alpha 2 OS=Homo sapiens OX=9606 GN=IGHA2 PE=1 SV=4 | 576 | 20201119_08533_1ng_glyco_hcd30_etd30_compound15.12516.12516.3 | controllerType=0 controllerNumber=1 scan=12516 | 28.916 |
| 09230:3 | 7 | R.MAGKPTHIN[+1647.613]VSVVMAEADGTC[+57.021]Y.- | HexNAc(5)Hex(3)Fuc(1) | C[+57], N[+1648] | 1333.5757 | 3 | 3998.7127 | 3998.6961 | 0 | 4.1 | 319 | CRagged | 236.79 | 149.43 | 149.43 | 3.34 | 22 | >sp\|P01877\|IGHA2_HUMAN Immunoglobulin heavy constant alpha 2 OS=Homo sapiens OX=9606 GN=IGHA2 PE=1 SV=4 | 576 | 20201119_08533_1ng_glyco_hcd30_etd30_compound15.12535.12535.3 | controllerType=0 controllerNumber=1 scan=12535 | 28.9533 |
| 09108:3 | 7 | R.MAGKPTHIN[+1540.529]VSVVMAEADGTC[+57.021]Y.- | HexNAc(2)Hex(7) | C[+57], N[+1541] | 1297.879 | 3 | 3891.6223 | 3891.6114 | 0 | 2.8 | 319 | CRagged | 267.11 | 267.11 | 198.69 | 4.14 | 22 | >sp\|P01877\|IGHA2_HUMAN Immunoglobulin heavy constant alpha 2 OS=Homo sapiens OX=9606 GN=IGHA2 PE=1 SV=4 | 576 | 20201119_08533_1ng_glyco_hcd30_etd30_compound15.12407.12407.3 | controllerType=0 controllerNumber=1 scan=12407 | 28.65 |
| 09130:3 | 7 | R.MAGKPTHIN[+1378.476]VSVVMAEADGTC[+57.021]Y.- | HexNAc(2)Hex(6) | C[+57], N[+1378] | 1243.8609 | 3 | 3729.5681 | 3729.5585 | 0 | 2.6 | 319 | CRagged | 270.72 | 259.63 | 203.82 | 4.11 | 22 | >sp\|P01877\|IGHA2_HUMAN Immunoglobulin heavy constant alpha 2 OS=Homo sapiens OX=9606 GN=IGHA2 PE=1 SV=4 | 576 | 20201119_08533_1ng_glyco_hcd30_etd30_compound15.12430.12430.3 | controllerType=0 controllerNumber=1 scan=12430 | 28.7102 |
| 09171:3 | 7 | R.MAGKPTHIN[+1216.423]VSVVMAEADGTC[+57.021]Y.- | HexNAc(2)Hex(5) | C[+57], N[+1216] | 1189.8426 | 3 | 3567.5132 | 3567.5057 | 0 | 2.1 | 319 | CRagged | 312.99 | 304.89 | 225.57 | 4.46 | 22 | >sp\|P01877\|IGHA2_HUMAN Immunoglobulin heavy constant alpha 2 OS=Homo sapiens OX=9606 GN=IGHA2 PE=1 SV=4 | 576 | 20201119_08533_1ng_glyco_hcd30_etd30_compound15.12473.12473.3 | controllerType=0 controllerNumber=1 scan=12473 | 28.8231 |
| 08939:3 | 9 | K.TVLTPATNHMGN[+1378.476]VTFTIPANR.E | HexNAc(2)Hex(6) | N[+1378] | 1211.8884 | 3 | 3633.6505 | 3633.6358 | 0 | 4.1 | 74 | Specific | 329.91 | 276.54 | 121.51 | 5.91 | 8 | >sp\|P01024\|CO3_HUMAN Complement C3 OS=Homo sapiens OX=9606 GN=C3 PE=1 SV=2 | 492 | 20201119_08533_1ng_glyco_hcd30_etd30_compound15.12231.12231.3 | controllerType=0 controllerNumber=1 scan=12231 | 28.3082 |
| 19102:5 | 12 | K.APDKNVIFSPLSISTALAFLSLGAHN[+2350.830]TTLTEILK.G | HexNAc(4)Hex(5)Fuc(1)NeuAc(2) | N[+2351] | 1187.571 | 5 | 5933.8261 | 5933.8085 | 0 | 3 | 68 | Specific | 225.01 | 225.01 | 225.01 | 2.91 | 6 | >sp\|P01011\|AACT_HUMAN Alpha-1-antichymotrypsin OS=Homo sapiens OX=9606 GN=SERPINA3 PE=1 SV=2 | 1314 | 20201119_08533_1ng_glyco_hcd30_etd30_compound15.22958.22958.5 | controllerType=0 controllerNumber=1 scan=22958 | 53.5318 |
| 19090:4 | 12 | K.APDKNVIFSPLSISTALAFLSLGAHN[+2350.830]TTLTEILK.G | HexNAc(4)Hex(5)Fuc(1)NeuAc(2) | N[+2351] | 1484.2116 | 4 | 5933.8244 | 5933.8085 | 0 | 2.7 | 68 | Specific | 594.5 | 581.95 | 581.95 | 8.49 | 6 | >sp\|P01011\|AACT_HUMAN Alpha-1-antichymotrypsin OS=Homo sapiens OX=9606 GN=SERPINA3 PE=1 SV=2 | 1314 | 20201119_08533_1ng_glyco_hcd30_etd30_compound15.22946.22946.4 | controllerType=0 controllerNumber=1 scan=22946 | 53.4908 |
| 06856:3 | 15 | R.LSPN[+2350.830]ASAEHLELR.W | HexNAc(4)Hex(5)Fuc(1)NeuAc(2) | N[+2351] | 1263.2001 | 3 | 3787.5858 | 3787.5796 | 0 | 1.7 | 52 | Specific | 390.42 | 256.94 | 256.94 | 5.05 | 5 | >sp\|Q13410\|BT1A1_HUMAN Butyrophilin subfamily 1 member A1 OS=Homo sapiens OX=9606 GN=BTN1A1 PE=1 SV=3 | 179 | 20201119_08533_1ng_glyco_hcd30_etd30_compound15.10030.10030.3 | controllerType=0 controllerNumber=1 scan=10030 | 24.106 |
| 07535:3 | 17 | K.TAHC[+57.021]N[+2059.735]ESFYFLC[+57.021]K.R | HexNAc(4)Hex(5)Fuc(1)NeuAc(1) | C[+57]*2, N[+2060] | 1246.1606 | 3 | 3736.4672 | 3736.4545 | 0 | 3.4 | 1201 | Specific | 272.45 | 180 | 180 | 4.82 | 4 | >sp\|P22897\|MRC1_HUMAN Macrophage mannose receptor 1 OS=Homo sapiens OX=9606 GN=MRC1 PE=1 SV=1 | 1049 | 20201119_08533_1ng_glyco_hcd30_etd30_compound15.10741.10741.3 | controllerType=0 controllerNumber=1 scan=10741 | 25.4445 |
| 07552:3 | 17 | K.TAHC[+57.021]N[+2059.735]ESFYFLC[+57.021]K.R | HexNAc(4)Hex(5)Fuc(1)NeuAc(1) | C[+57]*2, N[+2060] | 1246.1596 | 3 | 3736.4643 | 3736.4545 | 0 | 2.6 | 1201 | Specific | 307.16 | 204.24 | 204.24 | 5.04 | 4 | >sp\|P22897\|MRC1_HUMAN Macrophage mannose receptor 1 OS=Homo sapiens OX=9606 GN=MRC1 PE=1 SV=1 | 1049 | 20201119_08533_1ng_glyco_hcd30_etd30_compound15.10759.10759.3 | controllerType=0 controllerNumber=1 scan=10759 | 25.4776 |
| 19904:3 | 21 | R.QLAHQSN[+2204.772]STNIFFSPVSIATAFAMLSLGTK.A | HexNAc(4)Hex(5)NeuAc(2) | N[+2205] | 1796.1471 | 3 | 5386.4266 | 5386.4075 | 0 | 3.6 | 64 | Specific | 354.3 | 232.84 | 232.84 | 4.77 | 3 | >sp\|P01009\|A1AT_HUMAN Alpha-1-antitrypsin OS=Homo sapiens OX=9606 GN=SERPINA1 PE=1 SV=3 | 642 | 20201119_08533_1ng_glyco_hcd30_etd30_compound15.23806.23806.3 | controllerType=0 controllerNumber=1 scan=23806 | 55.8654 |
| 19883:4 | 21 | R.QLAHQSN[+2204.772]STNIFFSPVSIATAFAMLSLGTK.A | HexNAc(4)Hex(5)NeuAc(2) | N[+2205] | 1347.3573 | 4 | 5386.4074 | 5386.4075 | 0 | 0 | 64 | Specific | 564.91 | 485.06 | 372.68 | 6.6 | 3 | >sp\|P01009\|A1AT_HUMAN Alpha-1-antitrypsin OS=Homo sapiens OX=9606 GN=SERPINA1 PE=1 SV=3 | 642 | 20201119_08533_1ng_glyco_hcd30_etd30_compound15.23784.23784.4 | controllerType=0 controllerNumber=1 scan=23784 | 55.8002 |
| 19900:4 | 21 | R.QLAHQSN[+2204.772]STNIFFSPVSIATAFAMLSLGTK.A | HexNAc(4)Hex(5)NeuAc(2) | N[+2205] | 1347.3601 | 4 | 5386.4186 | 5386.4075 | 0 | 2.1 | 64 | Specific | 603.17 | 532.35 | 393.47 | 8.51 | 3 | >sp\|P01009\|A1AT_HUMAN Alpha-1-antitrypsin OS=Homo sapiens OX=9606 GN=SERPINA1 PE=1 SV=3 | 642 | 20201119_08533_1ng_glyco_hcd30_etd30_compound15.23802.23802.4 | controllerType=0 controllerNumber=1 scan=23802 | 55.855 |
| 19030:4 | 21 | K.ADTHDEILEGLNFN[+2861.000]LTEIPEAQIHEGFQELLR.T | HexNAc(5)Hex(6)NeuAc(3) | N[+2861] | 1638.9706 | 4 | 6552.8605 | 6552.8239 | 0 | 5.6 | 94 | Specific | 452.84 | 451.65 | 451.65 | 6.87 | 3 | >sp\|P01009\|A1AT_HUMAN Alpha-1-antitrypsin OS=Homo sapiens OX=9606 GN=SERPINA1 PE=1 SV=3 | 642 | 20201119_08533_1ng_glyco_hcd30_etd30_compound15.22882.22882.4 | controllerType=0 controllerNumber=1 scan=22882 | 53.2547 |
| 00062:4 | 25 | K.KKEDALN[+2059.735]ETRESETK.L | HexNAc(4)Hex(5)Fuc(1)NeuAc(1) | N[+2060] | 960.1668 | 4 | 3837.6453 | 3837.6276 | 0 | 4.6 | 80 | Specific | 477.66 | 402.67 | 402.67 | 4.66 | 6 | >sp\|P10909\|CLUS_HUMAN Clusterin OS=Homo sapiens OX=9606 GN=CLU PE=1 SV=1 | 1467 | 20201119_08533_1ng_glyco_hcd30_etd30_compound15.2973.2973.4 | controllerType=0 controllerNumber=1 scan=2973 | 11.301 |
| 00058:4 | 25 | K.KKEDALN[+2350.830]ETRESETK.L | HexNAc(4)Hex(5)Fuc(1)NeuAc(2) | N[+2351] | 1032.9412 | 4 | 4128.743 | 4128.723 | 0 | 4.8 | 80 | Specific | 528.15 | 407.28 | 407.28 | 5.55 | 6 | >sp\|P10909\|CLUS_HUMAN Clusterin OS=Homo sapiens OX=9606 GN=CLU PE=1 SV=1 | 1467 | 20201119_08533_1ng_glyco_hcd30_etd30_compound15.2969.2969.4 | controllerType=0 controllerNumber=1 scan=2969 | 11.2886 |
| 12285:4 | 25 | K.LKELPGVC[+57.021]N[+2205.793]ETMMALWEEC[+57.021]KPC[+57.021]LK.Q | HexNAc(4)Hex(5)Fuc(2)NeuAc(1) | C[+57]*3, N[+2206] | 1286.3053 | 4 | 5142.1994 | 5142.1776 | 0 | 4.2 | 95 | Specific | 444.79 | 359.94 | 359.94 | 4.75 | 6 | >sp\|P10909\|CLUS_HUMAN Clusterin OS=Homo sapiens OX=9606 GN=CLU PE=1 SV=1 | 1467 | 20201119_08533_1ng_glyco_hcd30_etd30_compound15.15738.15738.4 | controllerType=0 controllerNumber=1 scan=15738 | 35.3126 |
| 00065:3 | 25 | R.EIRHN[+2059.735]STGC[+57.021]LR.M | HexNAc(4)Hex(5)Fuc(1)NeuAc(1) | C[+57], N[+2060] | 1134.8085 | 3 | 3402.4109 | 3402.3994 | 0 | 3.4 | 287 | Specific | 271.65 | 141.59 | 141.59 | 2.57 | 6 | >sp\|P10909\|CLUS_HUMAN Clusterin OS=Homo sapiens OX=9606 GN=CLU PE=1 SV=1 | 1467 | 20201119_08533_1ng_glyco_hcd30_etd30_compound15.2977.2977.3 | controllerType=0 controllerNumber=1 scan=2977 | 11.3155 |
| 19422:3 | 25 | K.MLN[+2350.830]TSSLLEQLNEQFNWVSR.L | HexNAc(4)Hex(5)Fuc(1)NeuAc(2) | N[+2351] | 1587.3502 | 3 | 4760.0359 | 4760.0171 | 0 | 4 | 352 | Specific | 575.32 | 381.68 | 381.68 | 8.52 | 6 | >sp\|P10909\|CLUS_HUMAN Clusterin OS=Homo sapiens OX=9606 GN=CLU PE=1 SV=1 | 1467 | 20201119_08533_1ng_glyco_hcd30_etd30_compound15.23297.23297.3 | controllerType=0 controllerNumber=1 scan=23297 | 54.4981 |
| 11570:3 | 25 | R.LAN[+2350.830]LTQGEDQYYLR.V | HexNAc(4)Hex(5)Fuc(1)NeuAc(2) | N[+2351] | 1345.5641 | 3 | 4034.6779 | 4034.664 | 0 | 3.4 | 372 | Specific | 234.7 | 162.69 | 162.69 | 4.22 | 6 | >sp\|P10909\|CLUS_HUMAN Clusterin OS=Homo sapiens OX=9606 GN=CLU PE=1 SV=1 | 1467 | 20201119_08533_1ng_glyco_hcd30_etd30_compound15.14992.14992.3 | controllerType=0 controllerNumber=1 scan=14992 | 33.6378 |
| 08576:3 | 38 | R.IIVPLNNREN[+1460.529]ISDPTSPLR.T | HexNAc(4)Hex(4) | N[+1461] | 1203.5779 | 3 | 3608.7193 | 3608.7059 | 0 | 3.7 | 62 | Specific | 226.46 | 183.13 | 183.13 | 3.39 | 8 | >sp\|P01591\|IGJ_HUMAN Immunoglobulin J chain OS=Homo sapiens OX=9606 GN=JCHAIN PE=1 SV=4 | 1392 | 20201119_08533_1ng_glyco_hcd30_etd30_compound15.11854.11854.3 | controllerType=0 controllerNumber=1 scan=11854 | 27.5277 |
| 11710:3 | 38 | R.IIVPLNNREN[+2350.830]ISDPTSPLR.T | HexNAc(4)Hex(5)Fuc(1)NeuAc(2) | N[+2351] | 1500.3448 | 3 | 4499.0198 | 4499.0075 | 0 | 2.7 | 62 | Specific | 256.95 | 178.86 | 178.86 | 3.36 | 8 | >sp\|P01591\|IGJ_HUMAN Immunoglobulin J chain OS=Homo sapiens OX=9606 GN=JCHAIN PE=1 SV=4 | 1392 | 20201119_08533_1ng_glyco_hcd30_etd30_compound15.15137.15137.3 | controllerType=0 controllerNumber=1 scan=15137 | 33.9663 |
| 10030:3 | 38 | R.IIVPLNNREN[+1913.677]ISDPTSPLR.T | HexNAc(4)Hex(5)NeuAc(1) | N[+1914] | 1354.6274 | 3 | 4061.8676 | 4061.8542 | 0 | 3.3 | 62 | Specific | 261.87 | 221.04 | 221.04 | 3.49 | 8 | >sp\|P01591\|IGJ_HUMAN Immunoglobulin J chain OS=Homo sapiens OX=9606 GN=JCHAIN PE=1 SV=4 | 1392 | 20201119_08533_1ng_glyco_hcd30_etd30_compound15.13365.13365.3 | controllerType=0 controllerNumber=1 scan=13365 | 30.4584 |
| 08670:3 | 38 | R.IIVPLNNREN[+1501.555]ISDPTSPLR.T | HexNAc(5)Hex(3) | N[+1502] | 1217.2562 | 3 | 3649.7539 | 3649.7325 | 0 | 5.9 | 62 | Specific | 262.64 | 236.13 | 236.13 | 3.19 | 8 | >sp\|P01591\|IGJ_HUMAN Immunoglobulin J chain OS=Homo sapiens OX=9606 GN=JCHAIN PE=1 SV=4 | 1392 | 20201119_08533_1ng_glyco_hcd30_etd30_compound15.11952.11952.3 | controllerType=0 controllerNumber=1 scan=11952 | 27.721 |
| 08590:3 | 38 | R.IIVPLNNREN[+1663.608]ISDPTSPLR.T | HexNAc(5)Hex(4) | N[+1664] | 1271.2674 | 3 | 3811.7876 | 3811.7853 | 0 | 0.6 | 62 | Specific | 262.79 | 172.75 | 172.75 | 2.9 | 8 | >sp\|P01591\|IGJ_HUMAN Immunoglobulin J chain OS=Homo sapiens OX=9606 GN=JCHAIN PE=1 SV=4 | 1392 | 20201119_08533_1ng_glyco_hcd30_etd30_compound15.11869.11869.3 | controllerType=0 controllerNumber=1 scan=11869 | 27.56 |
| 08565:3 | 38 | R.IIVPLNNREN[+1663.608]ISDPTSPLR.T | HexNAc(5)Hex(4) | N[+1664] | 1271.2706 | 3 | 3811.7971 | 3811.7853 | 0 | 3.1 | 62 | Specific | 275.69 | 199.13 | 199.13 | 3.42 | 8 | >sp\|P01591\|IGJ_HUMAN Immunoglobulin J chain OS=Homo sapiens OX=9606 GN=JCHAIN PE=1 SV=4 | 1392 | 20201119_08533_1ng_glyco_hcd30_etd30_compound15.11843.11843.3 | controllerType=0 controllerNumber=1 scan=11843 | 27.5059 |
| 09949:3 | 38 | R.IIVPLNNREN[+2059.735]ISDPTSPLR.T | HexNAc(4)Hex(5)Fuc(1)NeuAc(1) | N[+2060] | 1403.3201 | 3 | 4207.9458 | 4207.9121 | 0 | 8 | 62 | Specific | 304.65 | 244.52 | 244.52 | 2.68 | 8 | >sp\|P01591\|IGJ_HUMAN Immunoglobulin J chain OS=Homo sapiens OX=9606 GN=JCHAIN PE=1 SV=4 | 1392 | 20201119_08533_1ng_glyco_hcd30_etd30_compound15.13280.13280.3 | controllerType=0 controllerNumber=1 scan=13280 | 30.2876 |
| 09960:3 | 38 | R.IIVPLNNREN[+2059.735]ISDPTSPLR.T | HexNAc(4)Hex(5)Fuc(1)NeuAc(1) | N[+2060] | 1403.3121 | 3 | 4207.9217 | 4207.9121 | 0 | 2.3 | 62 | Specific | 313.58 | 216.93 | 216.93 | 3.57 | 8 | >sp\|P01591\|IGJ_HUMAN Immunoglobulin J chain OS=Homo sapiens OX=9606 GN=JCHAIN PE=1 SV=4 | 1392 | 20201119_08533_1ng_glyco_hcd30_etd30_compound15.13292.13292.3 | controllerType=0 controllerNumber=1 scan=13292 | 30.3128 |
| 10065:3 | 38 | R.IIVPLNNREN[+1856.656]ISDPTSPLR.T | HexNAc(3)Hex(5)Fuc(1)NeuAc(1) | N[+1857] | 1335.619 | 3 | 4004.8423 | 4004.8327 | 0 | 2.4 | 62 | Specific | 323.08 | 239.6 | 239.6 | 3.45 | 8 | >sp\|P01591\|IGJ_HUMAN Immunoglobulin J chain OS=Homo sapiens OX=9606 GN=JCHAIN PE=1 SV=4 | 1392 | 20201119_08533_1ng_glyco_hcd30_etd30_compound15.13402.13402.3 | controllerType=0 controllerNumber=1 scan=13402 | 30.5808 |
| 00066:4 | 62 | K.TKPREEQYN[+1768.640]STYR.V | HexNAc(4)Hex(5)Fuc(1) | N[+1769] | 860.8701 | 4 | 3440.4585 | 3440.448 | 0 | 3 | 291 | Specific | 439.57 | 400.23 | 400.23 | 5.24 | 2 | >sp\|P0DOX5\|IGG1_HUMAN Immunoglobulin gamma-1 heavy chain OS=Homo sapiens OX=9606 PE=1 SV=2 | 914 | 20201119_08533_1ng_glyco_hcd30_etd30_compound15.2978.2978.4 | controllerType=0 controllerNumber=1 scan=2978 | 11.3227 |
| 00732:4 | 72 | K.TKPREEQFN[+1606.587]STFR.V | HexNAc(4)Hex(4)Fuc(1) | N[+1607] | 812.3602 | 4 | 3246.4189 | 3246.4054 | 0 | 4.2 | 168 | Specific | 314.29 | 314.29 | 314.29 | 3.9 | 2 | >sp\|P01859\|IGHG2_HUMAN Immunoglobulin heavy constant gamma 2 OS=Homo sapiens OX=9606 GN=IGHG2 PE=1 SV=2 | 387 | 20201119_08533_1ng_glyco_hcd30_etd30_compound15.3672.3672.4 | controllerType=0 controllerNumber=1 scan=3672 | 12.6093 |
| 00755:4 | 72 | K.TKPREEQFN[+1444.534]STFR.V | HexNAc(4)Hex(3)Fuc(1) | N[+1445] | 771.8436 | 4 | 3084.3528 | 3084.3526 | 0 | 0.1 | 168 | Specific | 347.35 | 344.46 | 344.46 | 3.17 | 2 | >sp\|P01859\|IGHG2_HUMAN Immunoglobulin heavy constant gamma 2 OS=Homo sapiens OX=9606 GN=IGHG2 PE=1 SV=2 | 387 | 20201119_08533_1ng_glyco_hcd30_etd30_compound15.3696.3696.4 | controllerType=0 controllerNumber=1 scan=3696 | 12.6669 |
| 00713:4 | 72 | K.TKPREEQFN[+1444.534]STFR.V | HexNAc(4)Hex(3)Fuc(1) | N[+1445] | 771.8457 | 4 | 3084.3611 | 3084.3526 | 0 | 2.8 | 168 | Specific | 435.52 | 417.58 | 417.58 | 5.24 | 2 | >sp\|P01859\|IGHG2_HUMAN Immunoglobulin heavy constant gamma 2 OS=Homo sapiens OX=9606 GN=IGHG2 PE=1 SV=2 | 387 | 20201119_08533_1ng_glyco_hcd30_etd30_compound15.3652.3652.2 | controllerType=0 controllerNumber=1 scan=3652 | 12.5592 |

Table S4. Cont.

| Query #:z | Protein Rank | Glycopeptide | Glycan | Modification Type(s) | Observed m/z | z | Observed (M+H) | Calc. mass (M+H) | Off-by-x error | Mass error (ppm) | StartPosition | Cleavage | Score | Delta | Delta Mod | \|Log Prob\| | # of unique peptides | Protein Name | Protein DB number | Comment | Scan # | Scan Time |
|---|---|---|---|---|---|---|---|---|---|---|---|---|---|---|---|---|---|---|---|---|---|---|
| 15665:4 | 1 | R.RTAGWNVPIGTLRPFLN[+2424.867]WTGPPEPIEAAVAR.F | HexNAc(5)Hex(6)Fuc(1)NeuAc(1) | N[+2425] | 1453.6804 | 4 | 5811.6999 | 5811.6792 | 0 | 3.6 | 140 | Specific | 356.8 | 356.8 | 356.8 | 5.14 | 56 | >sp\|P02788\|TRFL_HUMAN Lactotransferrin OS=Homo sapiens OX=9606 GN=LTF PE=1 SV=6 | 1270 | 20201115_10113_1ng_glyco_hcd30_etd30_compound15.18811.18811.4 | controllerType=0 controllerNumber=1 scan=18811 | 43.382 |
| 15701:4 | 1 | R.RTAGWNVPIGTLRPFLN[+2059.735]WTGPPEPIEAAVAR.F | HexNAc(4)Hex(5)Fuc(1)NeuAc(1) | N[+2060] | 1362.396 | 4 | 5446.5622 | 5446.547 | 0 | 2.8 | 140 | Specific | 371.87 | 371.87 | 371.87 | 7.83 | 56 | >sp\|P02788\|TRFL_HUMAN Lactotransferrin OS=Homo sapiens OX=9606 GN=LTF PE=1 SV=6 | 1270 | 20201115_10113_1ng_glyco_hcd30_etd30_compound15.18849.18849.2 | controllerType=0 controllerNumber=1 scan=18849 | 43.4744 |
| 15656:4 | 1 | R.RTAGWNVPIGTLRPFLN[+2570.925]WTGPPEPIEAAVAR.F | HexNAc(5)Hex(6)Fuc(2)NeuAc(1) | N[+2571] | 1490.1969 | 4 | 5957.7658 | 5957.7371 | 0 | 4.8 | 140 | Specific | 526.49 | 526.49 | 526.49 | 8.2 | 56 | >sp\|P02788\|TRFL_HUMAN Lactotransferrin OS=Homo sapiens OX=9606 GN=LTF PE=1 SV=6 | 1270 | 20201115_10113_1ng_glyco_hcd30_etd30_compound15.18802.18802.4 | controllerType=0 controllerNumber=1 scan=18802 | 43.3596 |
| 14913:4 | 1 | R.RTAGWNVPIGTLRPFLN[+2279.830]WTGPPEPIEAAVAR.F | HexNAc(5)Hex(6)Fuc(2) | N[+2280] | 1417.421 | 4 | 5666.6623 | 5666.6417 | 0 | 3.6 | 140 | Specific | 779.49 | 779.49 | 779.49 | 13.16 | 56 | >sp\|P02788\|TRFL_HUMAN Lactotransferrin OS=Homo sapiens OX=9606 GN=LTF PE=1 SV=6 | 1270 | 20201115_10113_1ng_glyco_hcd30_etd30_compound15.18018.18018.4 | controllerType=0 controllerNumber=1 scan=18018 | 41.1596 |
| 14988:4 | 1 | R.RTAGWNVPIGTLRPFLN[+1768.640]WTGPPEPIEAAVAR.F | HexNAc(4)Hex(5)Fuc(1) | N[+1769] | 1289.6216 | 4 | 5155.4645 | 5155.4516 | 0 | 2.5 | 140 | Specific | 1119.64 | 1056.85 | 1056.85 | 20.45 | 56 | >sp\|P02788\|TRFL_HUMAN Lactotransferrin OS=Homo sapiens OX=9606 GN=LTF PE=1 SV=6 | 1270 | 20201115_10113_1ng_glyco_hcd30_etd30_compound15.18098.18098.4 | controllerType=0 controllerNumber=1 scan=18098 | 41.3964 |
| 16519:4 | 1 | R.TAGWNVPIGTLRPFLN[+2716.983]WTGPPEPIEAAVAR.F | HexNAc(5)Hex(6)Fuc(3)NeuAc(1) | N[+2717] | 1487.6819 | 4 | 5947.7057 | 5947.6939 | 0 | 2 | 141 | Specific | 349.39 | 349.39 | 349.39 | 5.68 | 56 | >sp\|P02788\|TRFL_HUMAN Lactotransferrin OS=Homo sapiens OX=9606 GN=LTF PE=1 SV=6 | 1270 | 20201115_10113_1ng_glyco_hcd30_etd30_compound15.19715.19715.4 | controllerType=0 controllerNumber=1 scan=19715 | 46.0206 |
| 16586:3 | 1 | R.TAGWNVPIGTLRPFLN[+2059.735]WTGPPEPIEAAVAR.F | HexNAc(4)Hex(5)Fuc(1)NeuAc(1) | N[+2060] | 1764.1604 | 3 | 5290.4665 | 5290.4459 | 0 | 3.9 | 141 | Specific | 388.48 | 293.34 | 293.34 | 5.22 | 56 | >sp\|P02788\|TRFL_HUMAN Lactotransferrin OS=Homo sapiens OX=9606 GN=LTF PE=1 SV=6 | 1270 | 20201115_10113_1ng_glyco_hcd30_etd30_compound15.19787.19787.3 | controllerType=0 controllerNumber=1 scan=19787 | 46.2896 |
| 16577:3 | 1 | R.TAGWNVPIGTLRPFLN[+2205.793]WTGPPEPIEAAVAR.F | HexNAc(4)Hex(5)Fuc(2)NeuAc(1) | N[+2206] | 1812.8465 | 3 | 5436.525 | 5436.5038 | 0 | 3.9 | 141 | Specific | 457.35 | 356.99 | 356.99 | 5.71 | 56 | >sp\|P02788\|TRFL_HUMAN Lactotransferrin OS=Homo sapiens OX=9606 GN=LTF PE=1 SV=6 | 1270 | 20201115_10113_1ng_glyco_hcd30_etd30_compound15.19777.19777.3 | controllerType=0 controllerNumber=1 scan=19777 | 46.2526 |
| 16599:4 | 1 | R.TAGWNVPIGTLRPFLN[+2076.750]WTGPPEPIEAAVAR.F | HexNAc(4)Hex(6)Fuc(2) | N[+2077] | 1327.6255 | 4 | 5307.4801 | 5307.4612 | 0 | 3.6 | 141 | Specific | 480.43 | 480.43 | 480.43 | 8.15 | 56 | >sp\|P02788\|TRFL_HUMAN Lactotransferrin OS=Homo sapiens OX=9606 GN=LTF PE=1 SV=6 | 1270 | 20201115_10113_1ng_glyco_hcd30_etd30_compound15.19801.19801.4 | controllerType=0 controllerNumber=1 scan=19801 | 46.341 |
| 17352:4 | 1 | R.TAGWNVPIGTLRPFLN[+2862.021]WTGPPEPIEAAVAR.F | HexNAc(5)Hex(6)Fuc(2)NeuAc(2) | N[+2862] | 1523.9485 | 4 | 6092.7721 | 6092.7314 | 0 | 6.7 | 141 | Specific | 549 | 549 | 549 | 7.61 | 56 | >sp\|P02788\|TRFL_HUMAN Lactotransferrin OS=Homo sapiens OX=9606 GN=LTF PE=1 SV=6 | 1270 | 20201115_10113_1ng_glyco_hcd30_etd30_compound15.20596.20596.4 | controllerType=0 controllerNumber=1 scan=20596 | 48.6723 |
| 16508:4 | 1 | R.TAGWNVPIGTLRPFLN[+2570.925]WTGPPEPIEAAVAR.F | HexNAc(5)Hex(6)Fuc(2)NeuAc(1) | N[+2571] | 1451.1708 | 4 | 5801.6613 | 5801.636 | 0 | 4.4 | 141 | Specific | 701.06 | 680.4 | 680.4 | 12.42 | 56 | >sp\|P02788\|TRFL_HUMAN Lactotransferrin OS=Homo sapiens OX=9606 GN=LTF PE=1 SV=6 | 1270 | 20201115_10113_1ng_glyco_hcd30_etd30_compound15.19703.19703.4 | controllerType=0 controllerNumber=1 scan=19703 | 45.9803 |
| 16533:4 | 1 | R.TAGWNVPIGTLRPFLN[+2424.867]WTGPPEPIEAAVAR.F | HexNAc(5)Hex(6)Fuc(1)NeuAc(1) | N[+2425] | 1414.6544 | 4 | 5655.5959 | 5655.5781 | 0 | 3.1 | 141 | Specific | 746.31 | 745.96 | 745.96 | 12.71 | 56 | >sp\|P02788\|TRFL_HUMAN Lactotransferrin OS=Homo sapiens OX=9606 GN=LTF PE=1 SV=6 | 1270 | 20201115_10113_1ng_glyco_hcd30_etd30_compound15.19729.19729.4 | controllerType=0 controllerNumber=1 scan=19729 | 46.0515 |
| 16591:5 | 1 | R.TAGWNVPIGTLRPFLN[+2205.793]WTGPPEPIEAAVAR.F | HexNAc(4)Hex(5)Fuc(2)NeuAc(1) | N[+2206] | 1088.1091 | 5 | 5436.5166 | 5436.5038 | 0 | 2.4 | 141 | Specific | 767.67 | 767.67 | 767.67 | 13.11 | 56 | >sp\|P02788\|TRFL_HUMAN Lactotransferrin OS=Homo sapiens OX=9606 GN=LTF PE=1 SV=6 | 1270 | 20201115_10113_1ng_glyco_hcd30_etd30_compound15.19792.19792.5 | controllerType=0 controllerNumber=1 scan=19792 | 46.3034 |
| 16580:4 | 1 | R.TAGWNVPIGTLRPFLN[+2059.735]WTGPPEPIEAAVAR.F | HexNAc(4)Hex(5)Fuc(1)NeuAc(1) | N[+2060] | 1323.3723 | 4 | 5290.4674 | 5290.4459 | 0 | 4.1 | 141 | Specific | 1020.88 | 971.46 | 971.46 | 18.42 | 56 | >sp\|P02788\|TRFL_HUMAN Lactotransferrin OS=Homo sapiens OX=9606 GN=LTF PE=1 SV=6 | 1270 | 20201115_10113_1ng_glyco_hcd30_etd30_compound15.19781.19781.4 | controllerType=0 controllerNumber=1 scan=19781 | 46.2687 |
| 16595:5 | 1 | R.TAGWNVPIGTLRPFLN[+2059.735]WTGPPEPIEAAVAR.F | HexNAc(4)Hex(5)Fuc(1)NeuAc(1) | N[+2060] | 1058.8975 | 5 | 5290.4582 | 5290.4459 | 0 | 2.3 | 141 | Specific | 1102.16 | 1061.96 | 1061.96 | 20.11 | 56 | >sp\|P02788\|TRFL_HUMAN Lactotransferrin OS=Homo sapiens OX=9606 GN=LTF PE=1 SV=6 | 1270 | 20201115_10113_1ng_glyco_hcd30_etd30_compound15.19797.19797.5 | controllerType=0 controllerNumber=1 scan=19797 | 46.3196 |
| 15877:4 | 1 | R.TAGWNVPIGTLRPFLN[+1768.640]WTGPPEPIEAAVAR.F | HexNAc(4)Hex(5)Fuc(1) | N[+1769] | 1250.5978 | 4 | 4999.3693 | 4999.3504 | 0 | 3.8 | 141 | Specific | 1113.2 | 1073.99 | 1073.99 | 20.42 | 56 | >sp\|P02788\|TRFL_HUMAN Lactotransferrin OS=Homo sapiens OX=9606 GN=LTF PE=1 SV=6 | 1270 | 20201115_10113_1ng_glyco_hcd30_etd30_compound15.19036.19036.4 | controllerType=0 controllerNumber=1 scan=19036 | 44.0569 |
| 14774:2 | 1 | R.TAGWNIPMGLLFN[+1768.640]QTGSC[+57.021]K.F | HexNAc(4)Hex(5)Fuc(1) | C[+57], N[+1769] | 1932.335 | 2 | 3863.6628 | 3863.6495 | 0 | 3.5 | 485 | Specific | 207.47 | 100.51 | 100.51 | 3.79 | 56 | >sp\|P02788\|TRFL_HUMAN Lactotransferrin OS=Homo sapiens OX=9606 GN=LTF PE=1 SV=6 | 1270 | 20201115_10113_1ng_glyco_hcd30_etd30_compound15.17872.17872.2 | controllerType=0 controllerNumber=1 scan=17872 | 40.7818 |
| 16272:4 | 1 | R.TAGWNIPMGLLFN[+1913.677]QTGSC[+57.021]K.F | HexNAc(4)Hex(5)NeuAc(1) | C[+57], N[+1914] | 1002.9299 | 4 | 4008.6976 | 4008.687 | 0 | 2.6 | 485 | Specific | 245.9 | 213.82 | 213.82 | 3.13 | 56 | >sp\|P02788\|TRFL_HUMAN Lactotransferrin OS=Homo sapiens OX=9606 GN=LTF PE=1 SV=6 | 1270 | 20201115_10113_1ng_glyco_hcd30_etd30_compound15.19454.19454.4 | controllerType=0 controllerNumber=1 scan=19454 | 45.279 |
| 14808:3 | 1 | R.TAGWNIPMGLLFN[+1606.587]QTGSC[+57.021]K.F | HexNAc(4)Hex(4)Fuc(1) | C[+57], N[+1607] | 1234.54 | 3 | 3701.6054 | 3701.5966 | 0 | 2.4 | 485 | Specific | 278.71 | 252.84 | 252.84 | 4.52 | 56 | >sp\|P02788\|TRFL_HUMAN Lactotransferrin OS=Homo sapiens OX=9606 GN=LTF PE=1 SV=6 | 1270 | 20201115_10113_1ng_glyco_hcd30_etd30_compound15.17908.17908.3 | controllerType=0 controllerNumber=1 scan=17908 | 40.8811 |
| 14893:3 | 1 | R.TAGWNIPMGLLFN[+1622.582]QTGSC[+57.021]K.F | HexNAc(4)Hex(5) | C[+57], N[+1623] | 1239.873 | 3 | 3717.6044 | 3717.5916 | 0 | 3.5 | 485 | Specific | 334.15 | 266.33 | 228.84 | 5.72 | 56 | >sp\|P02788\|TRFL_HUMAN Lactotransferrin OS=Homo sapiens OX=9606 GN=LTF PE=1 SV=6 | 1270 | 20201115_10113_1ng_glyco_hcd30_etd30_compound15.17997.17997.3 | controllerType=0 controllerNumber=1 scan=17997 | 41.0895 |
| 14715:3 | 1 | R.TAGWNIPMGLLFN[+1768.640]QTGSC[+57.021]K.F | HexNAc(4)Hex(5)Fuc(1) | C[+57], N[+1769] | 1288.5597 | 3 | 3863.6647 | 3863.6495 | 0 | 3.9 | 485 | Specific | 372.87 | 286.79 | 286.79 | 5.45 | 56 | >sp\|P02788\|TRFL_HUMAN Lactotransferrin OS=Homo sapiens OX=9606 GN=LTF PE=1 SV=6 | 1270 | 20201115_10113_1ng_glyco_hcd30_etd30_compound15.17811.17811.3 | controllerType=0 controllerNumber=1 scan=17811 | 40.6347 |
| 16127:3 | 1 | R.TAGWNIPMGLLFN[+2351.851]QTGSC[+57.021]K.F | HexNAc(4)Hex(5)Fuc(3)NeuAc(1) | C[+57], N[+2352] | 1482.9715 | 3 | 4446.8999 | 4446.8607 | 0 | 8.8 | 485 | Specific | 381.89 | 274.93 | 274.93 | 4.42 | 56 | >sp\|P02788\|TRFL_HUMAN Lactotransferrin OS=Homo sapiens OX=9606 GN=LTF PE=1 SV=6 | 1270 | 20201115_10113_1ng_glyco_hcd30_etd30_compound15.19299.19299.3 | controllerType=0 controllerNumber=1 scan=19299 | 44.7437 |
| 14753:4 | 1 | R.TAGWNIPMGLLFN[+1768.640]QTGSC[+57.021]K.F | HexNAc(4)Hex(5)Fuc(1) | C[+57], N[+1769] | 966.6714 | 4 | 3863.6639 | 3863.6495 | 0 | 3.7 | 485 | Specific | 408.98 | 391.1 | 391.1 | 4.97 | 56 | >sp\|P02788\|TRFL_HUMAN Lactotransferrin OS=Homo sapiens OX=9606 GN=LTF PE=1 SV=6 | 1270 | 20201115_10113_1ng_glyco_hcd30_etd30_compound15.17850.17850.4 | controllerType=0 controllerNumber=1 scan=17850 | 40.72 |
| 16139:3 | 1 | R.TAGWNIPMGLLFN[+2059.735]QTGSC[+57.021]K.F | HexNAc(4)Hex(5)Fuc(1)NeuAc(1) | C[+57], N[+2060] | 1385.5895 | 3 | 4154.7541 | 4154.7449 | 0 | 2.2 | 485 | Specific | 423.43 | 283.85 | 283.85 | 5.84 | 56 | >sp\|P02788\|TRFL_HUMAN Lactotransferrin OS=Homo sapiens OX=9606 GN=LTF PE=1 SV=6 | 1270 | 20201115_10113_1ng_glyco_hcd30_etd30_compound15.19312.19312.3 | controllerType=0 controllerNumber=1 scan=19312 | 44.8071 |
| 16259:3 | 1 | R.TAGWNIPMGLLFN[+1913.677]QTGSC[+57.021]K.F | HexNAc(4)Hex(5)NeuAc(1) | C[+57], N[+1914] | 1336.9042 | 3 | 4008.6981 | 4008.687 | 0 | 2.8 | 485 | Specific | 424.62 | 281.15 | 281.15 | 5.84 | 56 | >sp\|P02788\|TRFL_HUMAN Lactotransferrin OS=Homo sapiens OX=9606 GN=LTF PE=1 SV=6 | 1270 | 20201115_10113_1ng_glyco_hcd30_etd30_compound15.19441.19441.3 | controllerType=0 controllerNumber=1 scan=19441 | 45.2457 |
| 17519:3 | 1 | R.TAGWNIPMGLLFN[+2350.830]QTGSC[+57.021]K.F | HexNAc(4)Hex(5)Fuc(1)NeuAc(2) | C[+57], N[+2351] | 1482.6257 | 3 | 4445.8624 | 4445.8403 | 0 | 5 | 485 | Specific | 472.52 | 322.28 | 322.28 | 5.99 | 56 | >sp\|P02788\|TRFL_HUMAN Lactotransferrin OS=Homo sapiens OX=9606 GN=LTF PE=1 SV=6 | 1270 | 20201115_10113_1ng_glyco_hcd30_etd30_compound15.20777.20777.3 | controllerType=0 controllerNumber=1 scan=20777 | 49.3213 |
| 15010:3 | 1 | R.TAGWNIPMGLLFN[+1216.423]QTGSC[+57.021]K.F | HexNAc(2)Hex(5) | C[+57], N[+1216] | 1104.4861 | 3 | 3311.4438 | 3311.4328 | 0 | 3.3 | 485 | Specific | 480.85 | 421.93 | 197.72 | 7.75 | 56 | >sp\|P02788\|TRFL_HUMAN Lactotransferrin OS=Homo sapiens OX=9606 GN=LTF PE=1 SV=6 | 1270 | 20201115_10113_1ng_glyco_hcd30_etd30_compound15.18121.18121.3 | controllerType=0 controllerNumber=1 scan=18121 | 41.451 |
| 15045:2 | 1 | R.TAGWNIPMGLLFN[+1216.423]QTGSC[+57.021]K.F | HexNAc(2)Hex(5) | C[+57], N[+1216] | 1656.2261 | 2 | 3311.445 | 3311.4328 | 0 | 3.7 | 485 | Specific | 522.65 | 521.17 | 177.2 | 7.69 | 56 | >sp\|P02788\|TRFL_HUMAN Lactotransferrin OS=Homo sapiens OX=9606 GN=LTF PE=1 SV=6 | 1270 | 20201115_10113_1ng_glyco_hcd30_etd30_compound15.18157.18157.2 | controllerType=0 controllerNumber=1 scan=18157 | 41.5458 |
| 15188:3 | 1 | R.TAGWNIPM[+15.995]GLLFN[+2059.735]QTGSC[+57.021]K.F | HexNAc(4)Hex(5)Fuc(1)NeuAc(1) | C[+57], M[+16], N[+2060] | 1390.9257 | 3 | 4170.7626 | 4170.7398 | 0 | 5.5 | 485 | Specific | 266.85 | 202.82 | 202.82 | 4.18 | 56 | >sp\|P02788\|TRFL_HUMAN Lactotransferrin OS=Homo sapiens OX=9606 GN=LTF PE=1 SV=6 | 1270 | 20201115_10113_1ng_glyco_hcd30_etd30_compound15.18307.18307.3 | controllerType=0 controllerNumber=1 scan=18307 | 41.9234 |
| 14935:3 | 1 | R.TAGWNIPM[+15.995]GLLFN[+2059.735]QTGSC[+57.021]K.F | HexNAc(4)Hex(5)Fuc(1)NeuAc(1) | C[+57], M[+16], N[+2060] | 1390.925 | 3 | 4170.7604 | 4170.7398 | 0 | 4.9 | 485 | Specific | 301.48 | 227.07 | 227.07 | 5.22 | 56 | >sp\|P02788\|TRFL_HUMAN Lactotransferrin OS=Homo sapiens OX=9606 GN=LTF PE=1 SV=6 | 1270 | 20201115_10113_1ng_glyco_hcd30_etd30_compound15.18041.18041.3 | controllerType=0 controllerNumber=1 scan=18041 | 41.2219 |
| 15599:3 | 1 | R.TAGWNIPMGLLFN[+1768.640]QTGSC[+57.021]KFDEYFSQSC[+57.021]APGSDPR.S | HexNAc(4)Hex(5)Fuc(1) | C[+57]*2, N[+1769] | 1903.1445 | 3 | 5707.4189 | 5707.3966 | 0 | 3.9 | 485 | Specific | 270.36 | 214.76 | 214.76 | 5.28 | 56 | >sp\|P02788\|TRFL_HUMAN Lactotransferrin OS=Homo sapiens OX=9606 GN=LTF PE=1 SV=6 | 1270 | 20201115_10113_1ng_glyco_hcd30_etd30_compound15.18742.18742.3 | controllerType=0 controllerNumber=1 scan=18742 | 43.2083 |
| 15707:3 | 1 | R.TAGWNIPMGLLFN[+1216.423]QTGSC[+57.021]KFDEYFSQSC[+57.021]APGSDPR.S | HexNAc(2)Hex(5) | C[+57]*2, N[+1216] | 1719.072 | 3 | 5155.2014 | 5155.18 | 0 | 4.2 | 485 | Specific | 282.35 | 231.91 | 113.49 | 4.36 | 56 | >sp\|P02788\|TRFL_HUMAN Lactotransferrin OS=Homo sapiens OX=9606 GN=LTF PE=1 SV=6 | 1270 | 20201115_10113_1ng_glyco_hcd30_etd30_compound15.18855.18855.3 | controllerType=0 controllerNumber=1 scan=18855 | 43.4915 |
| 15598:4 | 1 | R.TAGWNIPMGLLFN[+1768.640]QTGSC[+57.021]KFDEYFSQSC[+57.021]APGSDPR.S | HexNAc(4)Hex(5)Fuc(1) | C[+57]*2, N[+1769] | 1427.61 | 4 | 5707.4181 | 5707.3966 | 0 | 3.8 | 485 | Specific | 427.67 | 369.9 | 369.9 | 5.89 | 56 | >sp\|P02788\|TRFL_HUMAN Lactotransferrin OS=Homo sapiens OX=9606 GN=LTF PE=1 SV=6 | 1270 | 20201115_10113_1ng_glyco_hcd30_etd30_compound15.18741.18741.4 | controllerType=0 controllerNumber=1 scan=18741 | 43.205 |
| 16352:4 | 1 | R.TAGWNIPMGLLFN[+2059.735]QTGSC[+57.021]KFDEYFSQSC[+57.021]APGSDPR.S | HexNAc(4)Hex(5)Fuc(1)NeuAc(1) | C[+57]*2, N[+2060] | 1500.3817 | 4 | 5998.505 | 5998.492 | 0 | 2.2 | 485 | Specific | 551.29 | 498.98 | 498.98 | 8.38 | 56 | >sp\|P02788\|TRFL_HUMAN Lactotransferrin OS=Homo sapiens OX=9606 GN=LTF PE=1 SV=6 | 1270 | 20201115_10113_1ng_glyco_hcd30_etd30_compound15.19539.19539.4 | controllerType=0 controllerNumber=1 scan=19539 | 45.5233 |
| 08826:3 | 1 | R.N[+2205.793]GSDC[+57.021]PDKFC[+57.021]LFQSETK.N | HexNAc(4)Hex(5)Fuc(2)NeuAc(1) | C[+57]*2, N[+2206] | 1413.5666 | 3 | 4238.6852 | 4238.6667 | 0 | 4.4 | 642 | Specific | 235.52 | 167.4 | 167.4 | 3.59 | 56 | >sp\|P02788\|TRFL_HUMAN Lactotransferrin OS=Homo sapiens OX=9606 GN=LTF PE=1 SV=6 | 1270 | 20201115_10113_1ng_glyco_hcd30_etd30_compound15.11646.11646.3 | controllerType=0 controllerNumber=1 scan=11646 | 26.6549 |
| 08873:3 | 1 | R.N[+2059.735]GSDC[+57.021]PDKFC[+57.021]LFQSETK.N | HexNAc(4)Hex(5)Fuc(1)NeuAc(1) | C[+57]*2, N[+2060] | 1364.8784 | 3 | 4092.6205 | 4092.6088 | 0 | 2.9 | 642 | Specific | 337.56 | 212.16 | 212.16 | 4.99 | 56 | >sp\|P02788\|TRFL_HUMAN Lactotransferrin OS=Homo sapiens OX=9606 GN=LTF PE=1 SV=6 | 1270 | 20201115_10113_1ng_glyco_hcd30_etd30_compound15.11695.11695.3 | controllerType=0 controllerNumber=1 scan=11695 | 26.7609 |
| 15370:4 | 3 | R.LSLHRPALEDLLLGSEAN[+1864.634]LTC[+57.021]TLTGLR.D | HexNAc(2)Hex(9) | C[+57], N[+1865] | 1207.8174 | 4 | 4828.2479 | 4828.2324 | 0 | 3.2 | 127 | Specific | 259.52 | 259.52 | 259.52 | 5.06 | 34 | >sp\|P01876\|IGHA1_HUMAN Immunoglobulin heavy constant alpha 1 OS=Homo sapiens OX=9606 GN=IGHA1 PE=1 SV=2 | 255 | 20201115_10113_1ng_glyco_hcd30_etd30_compound15.18499.18499.4 | controllerType=0 controllerNumber=1 scan=18499 | 42.4517 |
| 16159:3 | 3 | R.LSLHRPALEDLLLGSEAN[+1751.624]LTC[+57.021]TLTGLR.D | HexNAc(4)Hex(4)NeuAc(1) | C[+57], N[+1752] | 1572.4172 | 3 | 4715.237 | 4715.2225 | 0 | 3.1 | 127 | Specific | 296.55 | 187.38 | 187.38 | 4.63 | 34 | >sp\|P01876\|IGHA1_HUMAN Immunoglobulin heavy constant alpha 1 OS=Homo sapiens OX=9606 GN=IGHA1 PE=1 SV=2 | 255 | 20201115_10113_1ng_glyco_hcd30_etd30_compound15.19333.19333.3 | controllerType=0 controllerNumber=1 scan=19333 | 44.8696 |
| 15434:3 | 3 | R.LSLHRPALEDLLLGSEAN[+1298.476]LTC[+57.021]TLTGLR.D | HexNAc(4)Hex(3) | C[+57], N[+1298] | 1421.3672 | 3 | 4262.0872 | 4262.0742 | 0 | 3 | 127 | Specific | 306.5 | 306.5 | 306.5 | 5.05 | 34 | >sp\|P01876\|IGHA1_HUMAN Immunoglobulin heavy constant alpha 1 OS=Homo sapiens OX=9606 GN=IGHA1 PE=1 SV=2 | 255 | 20201115_10113_1ng_glyco_hcd30_etd30_compound15.18566.18566.3 | controllerType=0 controllerNumber=1 scan=18566 | 42.6558 |
| 15359:5 | 3 | R.LSLHRPALEDLLLGSEAN[+1702.581]LTC[+57.021]TLTGLR.D | HexNAc(2)Hex(8) | C[+57], N[+1703] | 934.0419 | 5 | 4666.1804 | 4666.1796 | 0 | 0.2 | 127 | Specific | 370.9 | 368.16 | 368.16 | 5.24 | 34 | >sp\|P01876\|IGHA1_HUMAN Immunoglobulin heavy constant alpha 1 OS=Homo sapiens OX=9606 GN=IGHA1 PE=1 SV=2 | 255 | 20201115_10113_1ng_glyco_hcd30_etd30_compound15.18487.18487.5 | controllerType=0 controllerNumber=1 scan=18487 | 42.4128 |
| 15389:3 | 3 | R.LSLHRPALEDLLLGSEAN[+1378.476]LTC[+57.021]TLTGLR.D | HexNAc(2)Hex(6) | C[+57], N[+1378] | 1448.0324 | 3 | 4342.0827 | 4342.074 | 0 | 2 | 127 | Specific | 417.32 | 355.08 | 355.08 | 6.57 | 34 | >sp\|P01876\|IGHA1_HUMAN Immunoglobulin heavy constant alpha 1 OS=Homo sapiens OX=9606 GN=IGHA1 PE=1 SV=2 | 255 | 20201115_10113_1ng_glyco_hcd30_etd30_compound15.18519.18519.3 | controllerType=0 controllerNumber=1 scan=18519 | 42.5119 |
| 15412:3 | 3 | R.LSLHRPALEDLLLGSEAN[+1460.529]LTC[+57.021]TLTGLR.D | HexNAc(4)Hex(4) | C[+57], N[+1461] | 1475.3858 | 3 | 4424.1428 | 4424.1271 | 0 | 3.6 | 127 | Specific | 423.39 | 414.03 | 414.03 | 7.62 | 34 | >sp\|P01876\|IGHA1_HUMAN Immunoglobulin heavy constant alpha 1 OS=Homo sapiens OX=9606 GN=IGHA1 PE=1 SV=2 | 255 | 20201115_10113_1ng_glyco_hcd30_etd30_compound15.18543.18543.3 | controllerType=0 controllerNumber=1 scan=18543 | 42.5763 |
| 15376:3 | 3 | R.LSLHRPALEDLLLGSEAN[+1663.608]LTC[+57.021]TLTGLR.D | HexNAc(5)Hex(4) | C[+57], N[+1664] | 1543.0787 | 3 | 4627.2215 | 4627.2064 | 0 | 3.2 | 127 | Specific | 427.65 | 389.65 | 389.65 | 7.91 | 34 | >sp\|P01876\|IGHA1_HUMAN Immunoglobulin heavy constant alpha 1 OS=Homo sapiens OX=9606 GN=IGHA1 PE=1 SV=2 | 255 | 20201115_10113_1ng_glyco_hcd30_etd30_compound15.18505.18505.3 | controllerType=0 controllerNumber=1 scan=18505 | 42.4698 |
| 15371:3 | 3 | R.LSLHRPALEDLLLGSEAN[+1540.529]LTC[+57.021]TLTGLR.D | HexNAc(2)Hex(7) | C[+57], N[+1541] | 1502.053 | 3 | 4504.1446 | 4504.1268 | 0 | 3.9 | 127 | Specific | 434.06 | 421.01 | 421.01 | 7.62 | 34 | >sp\|P01876\|IGHA1_HUMAN Immunoglobulin heavy constant alpha 1 OS=Homo sapiens OX=9606 GN=IGHA1 PE=1 SV=2 | 255 | 20201115_10113_1ng_glyco_hcd30_etd30_compound15.18500.18500.3 | controllerType=0 controllerNumber=1 scan=18500 | 42.4557 |
| 15458:3 | 3 | R.LSLHRPALEDLLLGSEAN[+1216.423]LTC[+57.021]TLTGLR.D | HexNAc(2)Hex(5) | C[+57], N[+1216] | 1394.0153 | 3 | 4180.0314 | 4180.0211 | 0 | 2.5 | 127 | Specific | 443.63 | 419.91 | 419.91 | 8.18 | 34 | >sp\|P01876\|IGHA1_HUMAN Immunoglobulin heavy constant alpha 1 OS=Homo sapiens OX=9606 GN=IGHA1 PE=1 SV=2 | 255 | 20201115_10113_1ng_glyco_hcd30_etd30_compound15.18592.18592.3 | controllerType=0 controllerNumber=1 scan=18592 | 42.7351 |
| 15402:3 | 3 | R.LSLHRPALEDLLLGSEAN[+1501.555]LTC[+57.021]TLTGLR.D | HexNAc(5)Hex(3) | C[+57], N[+1502] | 1489.0613 | 3 | 4465.1695 | 4465.1536 | 0 | 3.6 | 127 | Specific | 445.22 | 413.7 | 413.7 | 8.18 | 34 | >sp\|P01876\|IGHA1_HUMAN Immunoglobulin heavy constant alpha 1 OS=Homo sapiens OX=9606 GN=IGHA1 PE=1 SV=2 | 255 | 20201115_10113_1ng_glyco_hcd30_etd30_compound15.18533.18533.3 | controllerType=0 controllerNumber=1 scan=18533 | 42.5504 |
| 16099:4 | 3 | R.LSLHRPALEDLLLGSEAN[+1872.651]LTC[+57.021]TLTGLR.D | HexNAc(3)Hex(6)NeuAc(1) | C[+57], N[+1873] | 1209.8213 | 4 | 4836.2635 | 4836.2487 | 0 | 3.1 | 127 | Specific | 460.43 | 405.36 | 405.36 | 7.34 | 34 | >sp\|P01876\|IGHA1_HUMAN Immunoglobulin heavy constant alpha 1 OS=Homo sapiens OX=9606 GN=IGHA1 PE=1 SV=2 | 255 | 20201115_10113_1ng_glyco_hcd30_etd30_compound15.19269.19269.3 | controllerType=0 controllerNumber=1 scan=19269 | 44.6236 |
| 15421:4 | 3 | R.LSLHRPALEDLLLGSEAN[+1419.502]LTC[+57.021]TLTGLR.D | HexNAc(3)Hex(5) | C[+57], N[+1420] | 1096.532 | 4 | 4383.1063 | 4383.1005 | 0 | 1.3 | 127 | Specific | 493.65 | 493.65 | 493.65 | 7.16 | 34 | >sp\|P01876\|IGHA1_HUMAN Immunoglobulin heavy constant alpha 1 OS=Homo sapiens OX=9606 GN=IGHA1 PE=1 SV=2 | 255 | 20201115_10113_1ng_glyco_hcd30_etd30_compound15.18553.18553.4 | controllerType=0 controllerNumber=1 scan=18553 | 42.6067 |
| 15346:3 | 3 | R.LSLHRPALEDLLLGSEAN[+1702.581]LTC[+57.021]TLTGLR.D | HexNAc(2)Hex(8) | C[+57], N[+1703] | 1556.0702 | 3 | 4666.1962 | 4666.1796 | 0 | 3.6 | 127 | Specific | 493.73 | 463.68 | 463.68 | 7.88 | 34 | >sp\|P01876\|IGHA1_HUMAN Immunoglobulin heavy constant alpha 1 OS=Homo sapiens OX=9606 GN=IGHA1 PE=1 SV=2 | 255 | 20201115_10113_1ng_glyco_hcd30_etd30_compound15.18474.18474.3 | controllerType=0 controllerNumber=1 scan=18474 | 42.3788 |
| 15352:3 | 3 | R.LSLHRPALEDLLLGSEAN[+1540.529]LTC[+57.021]TLTGLR.D | HexNAc(2)Hex(7) | C[+57], N[+1541] | 1502.0511 | 3 | 4504.1387 | 4504.1268 | 0 | 2.6 | 127 | Specific | 510.8 | 472.64 | 472.64 | 8.17 | 34 | >sp\|P01876\|IGHA1_HUMAN Immunoglobulin heavy constant alpha 1 OS=Homo sapiens OX=9606 GN=IGHA1 PE=1 SV=2 | 255 | 20201115_10113_1ng_glyco_hcd30_etd30_compound15.18480.18480.3 | controllerType=0 controllerNumber=1 scan=18480 | 42.3978 |
| 15397:4 | 3 | R.LSLHRPALEDLLLGSEAN[+1419.502]LTC[+57.021]TLTGLR.D | HexNAc(3)Hex(5) | C[+57], N[+1420] | 1096.5325 | 4 | 4383.1082 | 4383.1005 | 0 | 1.8 | 127 | Specific | 898.64 | 871.27 | 871.27 | 15.54 | 34 | >sp\|P01876\|IGHA1_HUMAN Immunoglobulin heavy constant alpha 1 OS=Homo sapiens OX=9606 GN=IGHA1 PE=1 SV=2 | 255 | 20201115_10113_1ng_glyco_hcd30_etd30_compound15.18527.18527.4 | controllerType=0 controllerNumber=1 scan=18527 | 42.5334 |
| 16151:4 | 3 | R.LSLHRPALEDLLLGSEAN[+1751.624]LTC[+57.021]TLTGLR.D | HexNAc(4)Hex(4)NeuAc(1) | C[+57], N[+1752] | 1179.5636 | 4 | 4715.2327 | 4715.2225 | 0 | 2.2 | 127 | Specific | 914.63 | 848.26 | 848.26 | 16.64 | 34 | >sp\|P01876\|IGHA1_HUMAN Immunoglobulin heavy constant alpha 1 OS=Homo sapiens OX=9606 GN=IGHA1 PE=1 SV=2 | 255 | 20201115_10113_1ng_glyco_hcd30_etd30_compound15.19325.19325.4 | controllerType=0 controllerNumber=1 scan=19325 | 44.8493 |
| 15345:4 | 3 | R.LSLHRPALEDLLLGSEAN[+1702.581]LTC[+57.021]TLTGLR.D | HexNAc(2)Hex(8) | C[+57], N[+1703] | 1167.3041 | 4 | 4666.1947 | 4666.1796 | 0 | 3.2 | 127 | Specific | 938.96 | 893.22 | 893.22 | 16.64 | 34 | >sp\|P01876\|IGHA1_HUMAN Immunoglobulin heavy constant alpha 1 OS=Homo sapiens OX=9606 GN=IGHA1 PE=1 SV=2 | 255 | 20201115_10113_1ng_glyco_hcd30_etd30_compound15.18473.18473.4 | controllerType=0 controllerNumber=1 scan=18473 | 42.3759 |
| 15453:4 | 3 | R.LSLHRPALEDLLLGSEAN[+1216.423]LTC[+57.021]TLTGLR.D | HexNAc(2)Hex(5) | C[+57], N[+1216] | 1045.7633 | 4 | 4180.0316 | 4180.0211 | 0 | 2.5 | 127 | Specific | 946.78 | 890.63 | 890.63 | 16.64 | 34 | >sp\|P01876\|IGHA1_HUMAN Immunoglobulin heavy constant alpha 1 OS=Homo sapiens OX=9606 GN=IGHA1 PE=1 SV=2 | 255 | 20201115_10113_1ng_glyco_hcd30_etd30_compound15.18587.18587.4 | controllerType=0 controllerNumber=1 scan=18587 | 42.7251 |
| 15401:4 | 3 | R.LSLHRPALEDLLLGSEAN[+1501.555]LTC[+57.021]TLTGLR.D | HexNAc(5)Hex(3) | C[+57], N[+1502] | 1117.0495 | 4 | 4465.1761 | 4465.1536 | 0 | 5 | 127 | Specific | 961.26 | 944.57 | 944.57 | 17.35 | 34 | >sp\|P01876\|IGHA1_HUMAN Immunoglobulin heavy constant alpha 1 OS=Homo sapiens OX=9606 GN=IGHA1 PE=1 SV=2 | 255 | 20201115_10113_1ng_glyco_hcd30_etd30_compound15.18532.18532.4 | controllerType=0 controllerNumber=1 scan=18532 | 42.5473 |
| 15426:4 | 3 | R.LSLHRPALEDLLLGSEAN[+1298.476]LTC[+57.021]TLTGLR.D | HexNAc(4)Hex(3) | C[+57], N[+1298] | 1066.2759 | 4 | 4262.0819 | 4262.0742 | 0 | 1.8 | 127 | Specific | 987.23 | 975.15 | 975.15 | 17.54 | 34 | >sp\|P01876\|IGHA1_HUMAN Immunoglobulin heavy constant alpha 1 OS=Homo sapiens OX=9606 GN=IGHA1 PE=1 SV=2 | 255 | 20201115_10113_1ng_glyco_hcd30_etd30_compound15.18558.18558.4 | controllerType=0 controllerNumber=1 scan=18558 | 42.6207 |
| 15396:4 | 3 | R.LSLHRPALEDLLLGSEAN[+1460.529]LTC[+57.021]TLTGLR.D | HexNAc(4)Hex(4) | C[+57], N[+1461] | 1106.7886 | 4 | 4424.1326 | 4424.1271 | 0 | 1.3 | 127 | Specific | 1004.86 | 909.23 | 909.23 | 18.54 | 34 | >sp\|P01876\|IGHA1_HUMAN Immunoglobulin heavy constant alpha 1 OS=Homo sapiens OX=9606 GN=IGHA1 PE=1 SV=2 | 255 | 20201115_10113_1ng_glyco_hcd30_etd30_compound15.18526.18526.4 | controllerType=0 controllerNumber=1 scan=18526 | 42.5304 |
| 15385:4 | 3 | R.LSLHRPALEDLLLGSEAN[+1378.476]LTC[+57.021]TLTGLR.D | HexNAc(2)Hex(6) | C[+57], N[+1378] | 1086.2764 | 4 | 4342.0838 | 4342.074 | 0 | 2.3 | 127 | Specific | 1011.07 | 970.73 | 970.73 | 18.64 | 34 | >sp\|P01876\|IGHA1_HUMAN Immunoglobulin heavy constant alpha 1 OS=Homo sapiens OX=9606 GN=IGHA1 PE=1 SV=2 | 255 | 20201115_10113_1ng_glyco_hcd30_etd30_compound15.18515.18515.4 | controllerType=0 controllerNumber=1 scan=18515 | 42.5028 |
| 15364:4 | 3 | R.LSLHRPALEDLLLGSEAN[+1540.529]LTC[+57.021]TLTGLR.D | HexNAc(2)Hex(7) | C[+57], N[+1541] | 1126.7908 | 4 | 4504.1414 | 4504.1268 | 0 | 3.3 | 127 | Specific | 1106.77 | 1002.04 | 1002.04 | 20.64 | 34 | >sp\|P01876\|IGHA1_HUMAN Immunoglobulin heavy constant alpha 1 OS=Homo sapiens OX=9606 GN=IGHA1 PE=1 SV=2 | 255 | 20201115_10113_1ng_glyco_hcd30_etd30_compound15.18493.18493.4 | controllerType=0 controllerNumber=1 scan=18493 | 42.4353 |
| 13429:3 | 3 | R.LAGKPTHVN[+2553.910]VSVVMAEVDGTC[+57.021]Y.- | HexNAc(5)Hex(5)Fuc(1)NeuAc(2) | C[+57], N[+2554] | 1634.3614 | 3 | 4901.0696 | 4901.0518 | 0 | 3.6 | 332 | CRagged | 202.37 | 42.42 | 42.42 | 2.56 | 34 | >sp\|P01876\|IGHA1_HUMAN Immunoglobulin heavy constant alpha 1 OS=Homo sapiens OX=9606 GN=IGHA1 PE=1 SV=2 | 255 | 20201115_10113_1ng_glyco_hcd30_etd30_compound15.16458.16458.3 | controllerType=0 controllerNumber=1 scan=16458 | 37.2836 |
| 10703:3 | 3 | R.LAGKPTHVN[+1647.613]VSVVMAEVDGTC[+57.021]Y.- | HexNAc(5)Hex(3)Fuc(1) | C[+57], N[+1648] | 1332.2596 | 3 | 3994.7642 | 3994.7553 | 0 | 2.2 | 332 | CRagged | 209.2 | 124.11 | 124.11 | 2.67 | 34 | >sp\|P01876\|IGHA1_HUMAN Immunoglobulin heavy constant alpha 1 OS=Homo sapiens OX=9606 GN=IGHA1 PE=1 SV=2 | 255 | 20201115_10113_1ng_glyco_hcd30_etd30_compound15.13604.13604.3 | controllerType=0 controllerNumber=1 scan=13604 | 30.7601 |
| 12051:3 | 3 | R.LAGKPTHVN[+2262.814]VSVVMAEVDGTC[+57.021]Y.- | HexNAc(5)Hex(5)Fuc(1)NeuAc(1) | C[+57], N[+2263] | 1537.3287 | 3 | 4609.9715 | 4609.9564 | 0 | 3.3 | 332 | CRagged | 215.02 | 52.93 | 52.93 | 2.56 | 34 | >sp\|P01876\|IGHA1_HUMAN Immunoglobulin heavy constant alpha 1 OS=Homo sapiens OX=9606 GN=IGHA1 PE=1 SV=2 | 255 | 20201115_10113_1ng_glyco_hcd30_etd30_compound15.15013.15013.3 | controllerType=0 controllerNumber=1 scan=15013 | 33.6199 |
| 10582:4 | 3 | R.LAGKPTHVN[+1702.581]VSVVMAEVDGTC[+57.021]Y.- | HexNAc(2)Hex(8) | C[+57], N[+1703] | 1013.1895 | 4 | 4049.7362 | 4049.7234 | 0 | 3.1 | 332 | CRagged | 216.29 | 216.29 | 69.71 | 3.14 | 34 | >sp\|P01876\|IGHA1_HUMAN Immunoglobulin heavy constant alpha 1 OS=Homo sapiens OX=9606 GN=IGHA1 PE=1 SV=2 | 255 | 20201115_10113_1ng_glyco_hcd30_etd30_compound15.13475.13475.4 | controllerType=0 controllerNumber=1 scan=13475 | 30.341 |
| 10609:3 | 3 | R.LAGKPTHVN[+1540.529]VSVVMAEVDGTC[+57.021]Y.- | HexNAc(2)Hex(7) | C[+57], N[+1541] | 1296.5655 | 3 | 3887.6819 | 3887.6706 | 0 | 2.9 | 332 | CRagged | 229.01 | 165.95 | 165.95 | 2.85 | 34 | >sp\|P01876\|IGHA1_HUMAN Immunoglobulin heavy constant alpha 1 OS=Homo sapiens OX=9606 GN=IGHA1 PE=1 SV=2 | 255 | 20201115_10113_1ng_glyco_hcd30_etd30_compound15.13504.13504.3 | controllerType=0 controllerNumber=1 scan=13504 | 30.4233 |
| 10623:3 | 3 | R.LAGKPTHVN[+1809.666]VSVVMAEVDGTC[+57.021]Y.- | HexNAc(5)Hex(4)Fuc(1) | C[+57], N[+1810] | 1386.2762 | 3 | 4156.814 | 4156.8081 | 0 | 1.4 | 332 | CRagged | 249.21 | 123.5 | 123.5 | 3.24 | 34 | >sp\|P01876\|IGHA1_HUMAN Immunoglobulin heavy constant alpha 1 OS=Homo sapiens OX=9606 GN=IGHA1 PE=1 SV=2 | 255 | 20201115_10113_1ng_glyco_hcd30_etd30_compound15.13519.13519.3 | controllerType=0 controllerNumber=1 scan=13519 | [illegible] |
| 10605:3 | 3 | R.LAGKPTHVN[+1378.476]VSVVMAEVDGTC[+57.021]Y.- | HexNAc(2)Hex(6) | C[+57], N[+1378] | 1242.5475 | 3 | 3725.6281 | 3725.6178 | 0 | 2.8 | 332 | CRagged | 261.88 | 202.82 | 196.95 | 3.92 | 34 | >sp\|P01876\|IGHA1_HUMAN Immunoglobulin heavy constant alpha 1 OS=Homo sapiens OX=9606 GN=IGHA1 PE=1 SV=2 | 255 | 20201115_10113_1ng_glyco_hcd30_etd30_compound15.13500.13500.3 | controllerType=0 controllerNumber=1 scan=13500 | [illegible] |
| 10573:3 | 3 | R.LAGKPTHVN[+1702.581]VSVVMAEVDGTC[+57.021]Y.- | HexNAc(2)Hex(8) | C[+57], N[+1703] | 1350.5826 | 3 | 4049.7332 | 4049.7234 | 0 | 2.4 | 332 | CRagged | 262.17 | 199.66 | 197.64 | 4.21 | 34 | >sp\|P01876\|IGHA1_HUMAN Immunoglobulin heavy constant alpha 1 OS=Homo sapiens OX=9606 GN=IGHA1 PE=1 SV=2 | 255 | 20201115_10113_1ng_glyco_hcd30_etd30_compound15.13466.13466.3 | controllerType=0 controllerNumber=1 scan=13466 | 30.3088 |
| 10624:3 | 3 | R.LAGKPTHVN[+1216.423]VSVVMAEVDGTC[+57.021]Y.- | HexNAc(2)Hex(5) | C[+57], N[+1216] | 1188.5289 | 3 | 3563.572 | 3563.5649 | 0 | 2 | 332 | CRagged | 277.27 | 264.63 | 204.65 | 4.15 | 34 | >sp\|P01876\|IGHA1_HUMAN Immunoglobulin heavy constant alpha 1 OS=Homo sapiens OX=9606 GN=IGHA1 PE=1 SV=2 | 255 | 20201115_10113_1ng_glyco_hcd30_etd30_compound15.13520.13520.3 | controllerType=0 controllerNumber=1 scan=13520 | 30.4813 |
| 12055:4 | 3 | R.LAGKPTHVN[+2262.814]VSVVMAEVDGTC[+57.021]Y.- | HexNAc(5)Hex(5)Fuc(1)NeuAc(1) | C[+57], N[+2263] | 1153.2486 | 4 | 4609.9725 | 4609.9564 | 0 | 3.5 | 332 | CRagged | 461.87 | 310.42 | 310.42 | 4.94 | 34 | >sp\|P01876\|IGHA1_HUMAN Immunoglobulin heavy constant alpha 1 OS=Homo sapiens OX=9606 GN=IGHA1 PE=1 SV=2 | 255 | 20201115_10113_1ng_glyco_hcd30_etd30_compound15.15017.15017.4 | controllerType=0 controllerNumber=1 scan=15017 | 33.6278 |
| 13409:4 | 3 | R.LAGKPTHVN[+2553.910]VSVVMAEVDGTC[+57.021]Y.- | HexNAc(5)Hex(5)Fuc(1)NeuAc(2) | C[+57], N[+2554] | 1226.0213 | 4 | 4901.0633 | 4901.0518 | 0 | 2.3 | 332 | CRagged | 477.1 | 251.56 | 251.56 | 4.7 | 34 | >sp\|P01876\|IGHA1_HUMAN Immunoglobulin heavy constant alpha 1 OS=Homo sapiens OX=9606 GN=IGHA1 PE=1 SV=2 | 255 | 20201115_10113_1ng_glyco_hcd30_etd30_compound15.16437.16437.4 | controllerType=0 controllerNumber=1 scan=16437 | 37.2353 |
| 13600:3 | 4 | R.LN[+1702.581]YSLPTGQWVGVQLPR.N | HexNAc(2)Hex(8) | N[+1703] | 1210.8812 | 3 | 3630.6289 | 3630.6202 | 0 | 2.4 | 1017 | Specific | 300.88 | 292.41 | 292.41 | 5.51 | 26 | >sp\|P24821\|TENA_HUMAN Tenascin OS=Homo sapiens OX=9606 GN=TNC PE=1 SV=3 | 486 | 20201115_10113_1ng_glyco_hcd30_etd30_compound15.16638.16638.3 | controllerType=0 controllerNumber=1 scan=16638 | 37.7527 |
| 13689:3 | 4 | R.LN[+1378.476]YSLPTGQWVGVQLPR.N | HexNAc(2)Hex(6) | N[+1378] | 1102.8462 | 3 | 3306.5241 | 3306.5145 | 0 | 2.9 | 1017 | Specific | 413.9 | 393.31 | 393.31 | 7.25 | 26 | >sp\|P24821\|TENA_HUMAN Tenascin OS=Homo sapiens OX=9606 GN=TNC PE=1 SV=3 | 486 | 20201115_10113_1ng_glyco_hcd30_etd30_compound15.16732.16732.3 | controllerType=0 controllerNumber=1 scan=16732 | 38.0177 |
| 13715:3 | 4 | R.LN[+1216.423]YSLPTGQWVGVQLPR.N | HexNAc(2)Hex(5) | N[+1216] | 1048.8283 | 3 | 3144.4703 | 3144.4617 | 0 | 2.7 | 1017 | Specific | 621.11 | 575.11 | 575.11 | 9.77 | 26 | >sp\|P24821\|TENA_HUMAN Tenascin OS=Homo sapiens OX=9606 GN=TNC PE=1 SV=3 | 486 | 20201115_10113_1ng_glyco_hcd30_etd30_compound15.16759.16759.3 | controllerType=0 controllerNumber=1 scan=16759 | 38.0797 |
| 09606:3 | 6 | K.TVLTPATNHMGN[+1540.529]VTFTIPANR.E | HexNAc(2)Hex(7) | N[+1541] | 1265.9029 | 3 | 3795.6941 | 3795.6886 | 0 | 1.4 | 74 | Specific | 388.53 | 308.67 | 138.03 | 5.76 | 21 | >sp\|P01024\|CO3_HUMAN Complement C3 OS=Homo sapiens OX=9606 GN=C3 PE=1 SV=2 | 492 | 20201115_10113_1ng_glyco_hcd30_etd30_compound15.12456.12456.3 | controllerType=0 controllerNumber=1 scan=12456 | 28.2103 |
| 09622:3 | 6 | K.TVLTPATNHMGN[+1540.529]VTFTIPANR.E | HexNAc(2)Hex(7) | N[+1541] | 1265.9037 | 3 | 3795.6967 | 3795.6886 | 0 | 2.1 | 74 | Specific | 399.83 | 312.13 | 159.5 | 4.73 | 21 | >sp\|P01024\|CO3_HUMAN Complement C3 OS=Homo sapiens OX=9606 GN=C3 PE=1 SV=2 | 492 | 20201115_10113_1ng_glyco_hcd30_etd30_compound15.12473.12473.3 | controllerType=0 controllerNumber=1 scan=12473 | 28.2486 |
| 09650:3 | 6 | K.TVLTPATNHMGN[+1378.476]VTFTIPANR.E | HexNAc(2)Hex(6) | N[+1378] | 1211.8863 | 3 | 3633.6443 | 3633.6358 | 0 | 2.3 | 74 | Specific | 412.54 | 291.3 | 175.03 | 5.94 | 21 | >sp\|P01024\|CO3_HUMAN Complement C3 OS=Homo sapiens OX=9606 GN=C3 PE=1 SV=2 | 492 | 20201115_10113_1ng_glyco_hcd30_etd30_compound15.12502.12502.3 | controllerType=0 controllerNumber=1 scan=12502 | 28.3081 |
| 15262:3 | 7 | R.GLSFDVSLEVSQGPGLLN[+2570.925]DTK.V | HexNAc(5)Hex(6)Fuc(2)NeuAc(1) | N[+2571] | 1583.0248 | 3 | 4747.06 | 4747.0382 | 0 | 4.6 | 118 | Specific | 371.05 | 219.74 | 219.74 | 5.39 | 31 | >sp\|P01833\|PIGR_HUMAN Polymeric immunoglobulin receptor OS=Homo sapiens OX=9606 GN=PIGR PE=1 SV=4 | 902 | 20201115_10113_1ng_glyco_hcd30_etd30_compound15.18385.18385.3 | controllerType=0 controllerNumber=1 scan=18385 | 42.1388 |
| 16291:4 | 7 | K.QIGLYPVLVIDSSGYVNPN[+2059.735]YTGR.I | HexNAc(4)Hex(5)Fuc(1)NeuAc(1) | N[+2060] | 1147.0175 | 4 | 4585.0482 | 4585.0384 | 0 | 2.1 | 168 | Specific | 210.52 | 210.52 | 210.52 | 2.72 | 31 | >sp\|P01833\|PIGR_HUMAN Polymeric immunoglobulin receptor OS=Homo sapiens OX=9606 GN=PIGR PE=1 SV=4 | 902 | 20201115_10113_1ng_glyco_hcd30_etd30_compound15.19474.19474.4 | controllerType=0 controllerNumber=1 scan=19474 | 45.3367 |
| 17426:3 | 7 | R.LSLLEEPGN[+2205.793]GTFTVILNQLTSR.D | HexNAc(4)Hex(5)Fuc(2)NeuAc(1) | N[+2206] | 1536.7086 | 3 | 4608.1111 | 4608.0854 | 0 | 5.6 | 413 | Specific | 258.62 | 206.59 | 206.59 | 4.18 | 31 | >sp\|P01833\|PIGR_HUMAN Polymeric immunoglobulin receptor OS=Homo sapiens OX=9606 GN=PIGR PE=1 SV=4 | 902 | 20201115_10113_1ng_glyco_hcd30_etd30_compound15.20676.20676.3 | controllerType=0 controllerNumber=1 scan=20676 | 48.9706 |
| 18708:4 | 7 | R.LSLLEEPGN[+2350.830]GTFTVILNQLTSR.D | HexNAc(4)Hex(5)Fuc(1)NeuAc(2) | N[+2351] | 1189.0418 | 4 | 4753.1453 | 4753.1229 | 0 | 4.7 | 413 | Specific | 602.6 | 522.24 | 522.24 | 8.09 | 31 | >sp\|P01833\|PIGR_HUMAN Polymeric immunoglobulin receptor OS=Homo sapiens OX=9606 GN=PIGR PE=1 SV=4 | 902 | 20201115_10113_1ng_glyco_hcd30_etd30_compound15.22041.22041.4 | controllerType=0 controllerNumber=1 scan=22041 | 53.2281 |
| 18707:3 | 7 | R.LSLLEEPGN[+2350.830]GTFTVILNQLTSR.D | HexNAc(4)Hex(5)Fuc(1)NeuAc(2) | N[+2351] | 1585.0539 | 3 | 4753.1471 | 4753.1229 | 0 | 5.1 | 413 | Specific | 639.7 | 444.16 | 444.16 | 9.72 | 31 | >sp\|P01833\|PIGR_HUMAN Polymeric immunoglobulin receptor OS=Homo sapiens OX=9606 GN=PIGR PE=1 SV=4 | 902 | 20201115_10113_1ng_glyco_hcd30_etd30_compound15.22040.22040.3 | controllerType=0 controllerNumber=1 scan=22040 | 53.2251 |
| 14159:3 | 7 | K.IIEGEPNLKVPGN[+2204.772]VTAVLGETLK.V | HexNAc(4)Hex(5)NeuAc(2) | N[+2205] | 1532.7216 | 3 | 4596.1503 | 4596.1218 | 0 | 6.2 | 457 | Specific | 357.55 | 282.91 | 282.91 | 3.12 | 31 | >sp\|P01833\|PIGR_HUMAN Polymeric immunoglobulin receptor OS=Homo sapiens OX=9606 GN=PIGR PE=1 SV=4 | 902 | 20201115_10113_1ng_glyco_hcd30_etd30_compound15.17226.17226.3 | controllerType=0 controllerNumber=1 scan=17226 | 39.2283 |
| 14887:5 | 7 | K.IIEGEPNLKVPGN[+2350.830]VTAVLGETLKVPC[+57.021]HFPC[+57.021]K.F | HexNAc(4)Hex(5)Fuc(1)NeuAc(2) | C[+57]*2, N[+2351] | 1154.3367 | 5 | 5767.6543 | 5767.6372 | 0 | 3 | 457 | Specific | 247.66 | 247.66 | 247.66 | 3.3 | 31 | >sp\|P01833\|PIGR_HUMAN Polymeric immunoglobulin receptor OS=Homo sapiens OX=9606 GN=PIGR PE=1 SV=4 | 902 | 20201115_10113_1ng_glyco_hcd30_etd30_compound15.17991.17991.5 | controllerType=0 controllerNumber=1 scan=17991 | 41.0719 |
| 14894:4 | 7 | K.IIEGEPNLKVPGN[+2350.830]VTAVLGETLKVPC[+57.021]HFPC[+57.021]K.F | HexNAc(4)Hex(5)Fuc(1)NeuAc(2) | C[+57]*2, N[+2351] | 1442.6708 | 4 | 5767.6613 | 5767.6372 | 0 | 4.2 | 457 | Specific | 379.85 | 377.58 | 377.58 | 4.7 | 31 | >sp\|P01833\|PIGR_HUMAN Polymeric immunoglobulin receptor OS=Homo sapiens OX=9606 GN=PIGR PE=1 SV=4 | 902 | 20201115_10113_1ng_glyco_hcd30_etd30_compound15.17998.17998.4 | controllerType=0 controllerNumber=1 scan=17998 | 41.0933 |
| 13784:5 | 7 | K.IIEGEPNLKVPGN[+2059.735]VTAVLGETLKVPC[+57.021]HFPC[+57.021]K.F | HexNAc(4)Hex(5)Fuc(1)NeuAc(1) | C[+57]*2, N[+2060] | 1096.1171 | 5 | 5476.5562 | 5476.5418 | 0 | 2.6 | 457 | Specific | 473.89 | 457.38 | 457.38 | 5.48 | 31 | >sp\|P01833\|PIGR_HUMAN Polymeric immunoglobulin receptor OS=Homo sapiens OX=9606 GN=PIGR PE=1 SV=4 | 902 | 20201115_10113_1ng_glyco_hcd30_etd30_compound15.16831.16831.3 | controllerType=0 controllerNumber=1 scan=16831 | 38.2686 |
| 13774:4 | 7 | K.IIEGEPNLKVPGN[+2205.793]VTAVLGETLKVPC[+57.021]HFPC[+57.021]K.F | HexNAc(4)Hex(5)Fuc(2)NeuAc(1) | C[+57]*2, N[+2206] | 1406.4103 | 4 | 5622.6193 | 5622.5997 | 0 | 3.5 | 457 | Specific | 485.05 | 462.19 | 462.19 | 5.35 | 31 | >sp\|P01833\|PIGR_HUMAN Polymeric immunoglobulin receptor OS=Homo sapiens OX=9606 GN=PIGR PE=1 SV=4 | 902 | 20201115_10113_1ng_glyco_hcd30_etd30_compound15.16821.16821.4 | controllerType=0 controllerNumber=1 scan=16821 | 38.2395 |
| 13858:5 | 7 | K.IIEGEPNLKVPGN[+2059.735]VTAVLGETLKVPC[+57.021]HFPC[+57.021]K.F | HexNAc(4)Hex(5)Fuc(1)NeuAc(1) | C[+57]*2, N[+2060] | 1096.1167 | 5 | 5476.5544 | 5476.5418 | 0 | 2.3 | 457 | Specific | 498.05 | 498.05 | 498.05 | 5.47 | 31 | >sp\|P01833\|PIGR_HUMAN Polymeric immunoglobulin receptor OS=Homo sapiens OX=9606 GN=PIGR PE=1 SV=4 | 902 | 20201115_10113_1ng_glyco_hcd30_etd30_compound15.16909.16909.5 | controllerType=0 controllerNumber=1 scan=16909 | 38.4449 |
| 13859:4 | 7 | K.IIEGEPNLKVPGN[+2059.735]VTAVLGETLKVPC[+57.021]HFPC[+57.021]K.F | HexNAc(4)Hex(5)Fuc(1)NeuAc(1) | C[+57]*2, N[+2060] | 1369.8943 | 4 | 5476.5563 | 5476.5418 | 0 | 2.5 | 457 | Specific | 511.88 | 511.88 | 511.88 | 5.64 | 31 | >sp\|P01833\|PIGR_HUMAN Polymeric immunoglobulin receptor OS=Homo sapiens OX=9606 GN=PIGR PE=1 SV=4 | 902 | 20201115_10113_1ng_glyco_hcd30_etd30_compound15.16910.16910.4 | controllerType=0 controllerNumber=1 scan=16910 | 38.4484 |
| 12389:4 | 7 | K.VPGN[+2205.793]VTAVLGETLKVPC[+57.021]HFPC[+57.021]K.F | HexNAc(4)Hex(5)Fuc(2)NeuAc(1) | C[+57]*2, N[+2206] | 1158.0202 | 4 | 4629.0589 | 4629.0502 | 0 | 1.9 | 466 | Specific | 275.76 | 275.76 | 275.76 | 3.62 | 31 | >sp\|P01833\|PIGR_HUMAN Polymeric immunoglobulin receptor OS=Homo sapiens OX=9606 GN=PIGR PE=1 SV=4 | 902 | 20201115_10113_1ng_glyco_hcd30_etd30_compound15.15367.15367.4 | controllerType=0 controllerNumber=1 scan=15367 | 34.5022 |
| 12479:4 | 7 | K.VPGN[+2059.735]VTAVLGETLKVPC[+57.021]HFPC[+57.021]K.F | HexNAc(4)Hex(5)Fuc(1)NeuAc(1) | C[+57]*2, N[+2060] | 1121.5046 | 4 | 4482.9964 | 4482.9923 | 0 | 0.9 | 466 | Specific | 311.22 | 260.45 | 260.45 | 3.63 | 31 | >sp\|P01833\|PIGR_HUMAN Polymeric immunoglobulin receptor OS=Homo sapiens OX=9606 GN=PIGR PE=1 SV=4 | 902 | 20201115_10113_1ng_glyco_hcd30_etd30_compound15.15462.15462.4 | controllerType=0 controllerNumber=1 scan=15462 | 34.7657 |
| 05832:3 | 7 | K.WN[+2059.735]NTGC[+57.021]QALPSQDEGPSK.A | HexNAc(4)Hex(5)Fuc(1)NeuAc(1) | C[+57], N[+2060] | 1350.2129 | 3 | 4048.6243 | 4048.6116 | 0 | 3.1 | 498 | Specific | 242 | 84.43 | 84.43 | 4.37 | 31 | >sp\|P01833\|PIGR_HUMAN Polymeric immunoglobulin receptor OS=Homo sapiens OX=9606 GN=PIGR PE=1 SV=4 | 902 | 20201115_10113_1ng_glyco_hcd30_etd30_compound15.8529.8529.3 | controllerType=0 controllerNumber=1 scan=8529 | 20.7578 |
| 05880:3 | 7 | K.WN[+1872.651]NTGC[+57.021]QALPSQDEGPSK.A | HexNAc(3)Hex(6)NeuAc(1) | C[+57], N[+1873] | 1287.8488 | 3 | 3861.5319 | 3861.5272 | 0 | 1.2 | 498 | Specific | 272.78 | 115.12 | 115.12 | 4.7 | 31 | >sp\|P01833\|PIGR_HUMAN Polymeric immunoglobulin receptor OS=Homo sapiens OX=9606 GN=PIGR PE=1 SV=4 | 902 | 20201115_10113_1ng_glyco_hcd30_etd30_compound15.8579.8579.3 | controllerType=0 controllerNumber=1 scan=8579 | 20.8578 |
| 05950:3 | 7 | K.WN[+1710.598]NTGC[+57.021]QALPSQDEGPSK.A | HexNAc(3)Hex(5)NeuAc(1) | C[+57], N[+1711] | 1233.8328 | 3 | 3699.4839 | 3699.4743 | 0 | 2.6 | 498 | Specific | 395.92 | 212.53 | 212.53 | 5.27 | 31 | >sp\|P01833\|PIGR_HUMAN Polymeric immunoglobulin receptor OS=Homo sapiens OX=9606 GN=PIGR PE=1 SV=4 | 902 | 20201115_10113_1ng_glyco_hcd30_etd30_compound15.8651.8651.3 | controllerType=0 controllerNumber=1 scan=8651 | 20.9814 |
| 05853:3 | 7 | K.WN[+1913.677]NTGC[+57.021]QALPSQDEGPSK.A | HexNAc(4)Hex(5)NeuAc(1) | C[+57], N[+1914] | 1301.5236 | 3 | 3902.5563 | 3902.5537 | 0 | 0.7 | 498 | Specific | 522.95 | 417.89 | 417.89 | 7.72 | 31 | >sp\|P01833\|PIGR_HUMAN Polymeric immunoglobulin receptor OS=Homo sapiens OX=9606 GN=PIGR PE=1 SV=4 | 902 | 20201115_10113_1ng_glyco_hcd30_etd30_compound15.8551.8551.3 | controllerType=0 controllerNumber=1 scan=8551 | 20.8045 |
| 15003:4 | 8 | K.RNIAAFGGDPNN[+1913.677]ITLFGESAGGASVSLQTLSPYNK.G | HexNAc(4)Hex(5)NeuAc(1) | N[+1914] | 1370.8749 | 4 | 5480.4777 | 5480.4644 | 0 | 2.4 | 196 | Specific | 250.08 | 250.08 | 250.08 | 4.29 | 11 | >sp\|P19835\|CEL_HUMAN Bile salt-activated lipase OS=Homo sapiens OX=9606 GN=CEL PE=1 SV=3 | 476 | 20201115_10113_1ng_glyco_hcd30_etd30_compound15.18113.18113.4 | controllerType=0 controllerNumber=1 scan=18113 | 41.4334 |
| 18239:4 | 14 | K.VFPLSLDSTPQDGNVVVAC[+57.021]LVQGFFPQEPLSVTWSESGQN[+1809.666]VTAI | HexNAc(5)Hex(4)Fuc(1) | C[+57], N[+1810] | 1647.2708 | 4 | 6586.0612 | 6586.0275 | 0 | 5.1 | 8 | Specific | 473.32 | 473.32 | 473.32 | 6.1 | 24 | >sp\|P01877\|IGHA2_HUMAN Immunoglobulin heavy constant alpha 2 OS=Homo sapiens OX=9606 GN=IGHA2 PE=1 SV=4 | 576 | 20201115_10113_1ng_glyco_hcd30_etd30_compound15.21544.21544.4 | controllerType=0 controllerNumber=1 scan=21544 | 51.7728 |
| 02531:3 | 14 | K.HYTN[+2262.814]SSQDVTVPC[+57.021]R.V | HexNAc(5)Hex(5)Fuc(1)NeuAc(1) | C[+57], N[+2263] | 1309.53 | 3 | 3926.5753 | 3926.5636 | 0 | 3 | 89 | Specific | 264.61 | 170.87 | 170.87 | 4.79 | 24 | >sp\|P01877\|IGHA2_HUMAN Immunoglobulin heavy constant alpha 2 OS=Homo sapiens OX=9606 GN=IGHA2 PE=1 SV=4 | 576 | 20201115_10113_1ng_glyco_hcd30_etd30_compound15.5105.5105.3 | controllerType=0 controllerNumber=1 scan=5105 | 14.4676 |
| 01726:3 | 14 | K.HYTN[+1647.613]SSQDVTVPC[+57.021]R.V | HexNAc(5)Hex(3)Fuc(1) | C[+57], N[+1648] | 1104.4624 | 3 | 3311.3727 | 3311.3625 | 0 | 3.1 | 89 | Specific | 460.1 | 384.82 | 384.82 | 7.45 | 24 | >sp\|P01877\|IGHA2_HUMAN Immunoglobulin heavy constant alpha 2 OS=Homo sapiens OX=9606 GN=IGHA2 PE=1 SV=4 | 576 | 20201115_10113_1ng_glyco_hcd30_etd30_compound15.4271.4271.3 | controllerType=0 controllerNumber=1 scan=4271 | 12.9892 |
| 03543:4 | 14 | K.HYTN[+1647.613]SSQDVTVPC[+57.021]RVPPPPPC[+57.021]C[+57.021]HPR.L | HexNAc(5)Hex(3)Fuc(1) | C[+57]*3, N[+1648] | 1152.2502 | 4 | 4605.9788 | 4605.9689 | 0 | 2.2 | 89 | Specific | 577.44 | 533.68 | 533.68 | 8.22 | 24 | >sp\|P01877\|IGHA2_HUMAN Immunoglobulin heavy constant alpha 2 OS=Homo sapiens OX=9606 GN=IGHA2 PE=1 SV=4 | 576 | 20201115_10113_1ng_glyco_hcd30_etd30_compound15.6157.6157.4 | controllerType=0 controllerNumber=1 scan=6157 | 16.5368 |
| 03521:5 | 14 | K.HYTN[+1647.613]SSQDVTVPC[+57.021]RVPPPPPC[+57.021]C[+57.021]HPR.L | HexNAc(5)Hex(3)Fuc(1) | C[+57]*3, N[+1648] | 922.0015 | 5 | 4605.9786 | 4605.9689 | 0 | 2.1 | 89 | Specific | 689.79 | 633.57 | 633.57 | 10.33 | 24 | >sp\|P01877\|IGHA2_HUMAN Immunoglobulin heavy constant alpha 2 OS=Homo sapiens OX=9606 GN=IGHA2 PE=1 SV=4 | 576 | 20201115_10113_1ng_glyco_hcd30_etd30_compound15.6134.6134.5 | controllerType=0 controllerNumber=1 scan=6134 | 16.4806 |
| 03312:3 | 14 | K.TPLTAN[+1971.719]ITK.S | HexNAc(5)Hex(5)Fuc(1) | N[+1972] | 977.4323 | 3 | 2930.2823 | 2930.2757 | 0 | 2.3 | 200 | Specific | 275.28 | 130.46 | 130.46 | 2.81 | 24 | >sp\|P01877\|IGHA2_HUMAN Immunoglobulin heavy constant alpha 2 OS=Homo sapiens OX=9606 GN=IGHA2 PE=1 SV=4 | 576 | 20201115_10113_1ng_glyco_hcd30_etd30_compound15.5914.5914.3 | controllerType=0 controllerNumber=1 scan=5914 | 15.8852 |
| 03392:3 | 14 | K.TPLTAN[+1647.613]ITK.S | HexNAc(5)Hex(3)Fuc(1) | N[+1648] | 869.3966 | 3 | 2606.1754 | 2606.17 | 0 | 2 | 200 | Specific | 291 | 139.55 | 139.55 | 2.78 | 24 | >sp\|P01877\|IGHA2_HUMAN Immunoglobulin heavy constant alpha 2 OS=Homo sapiens OX=9606 GN=IGHA2 PE=1 SV=4 | 576 | 20201115_10113_1ng_glyco_hcd30_etd30_compound15.5999.5999.3 | controllerType=0 controllerNumber=1 scan=5999 | 16.1438 |
| 03380:2 | 14 | K.TPLTAN[+1647.613]ITK.S | HexNAc(5)Hex(3)Fuc(1) | N[+1648] | 1303.5918 | 2 | 2606.1764 | 2606.17 | 0 | 2.5 | 200 | Specific | 299.28 | 118.88 | 118.88 | 2.76 | 24 | >sp\|P01877\|IGHA2_HUMAN Immunoglobulin heavy constant alpha 2 OS=Homo sapiens OX=9606 GN=IGHA2 PE=1 SV=4 | 576 | 20201115_10113_1ng_glyco_hcd30_etd30_compound15.5986.5986.2 | controllerType=0 controllerNumber=1 scan=5986 | 16.1009 |
| 03351:3 | 14 | K.TPLTAN[+1809.666]ITK.S | HexNAc(5)Hex(4)Fuc(1) | N[+1810] | 923.4152 | 3 | 2768.231 | 2768.2228 | 0 | 3 | 200 | Specific | 308.22 | 135.01 | 135.01 | 2.78 | 24 | >sp\|P01877\|IGHA2_HUMAN Immunoglobulin heavy constant alpha 2 OS=Homo sapiens OX=9606 GN=IGHA2 PE=1 SV=4 | 576 | 20201115_10113_1ng_glyco_hcd30_etd30_compound15.5955.5955.3 | controllerType=0 controllerNumber=1 scan=5955 | 16.0048 |
| 03348:2 | 14 | K.TPLTAN[+1809.666]ITK.S | HexNAc(5)Hex(4)Fuc(1) | N[+1810] | 1384.6189 | 2 | 2768.2306 | 2768.2228 | 0 | 2.8 | 200 | Specific | 314.55 | 111.16 | 111.16 | 2.76 | 24 | >sp\|P01877\|IGHA2_HUMAN Immunoglobulin heavy constant alpha 2 OS=Homo sapiens OX=9606 GN=IGHA2 PE=1 SV=4 | 576 | 20201115_10113_1ng_glyco_hcd30_etd30_compound15.5952.5952.2 | controllerType=0 controllerNumber=1 scan=5952 | 15.9948 |
| 03418:2 | 14 | K.TPLTAN[+1444.534]ITK.S | HexNAc(4)Hex(3)Fuc(1) | N[+1445] | 1202.0524 | 2 | 2403.0975 | 2403.0906 | 0 | 2.9 | 200 | Specific | 367.79 | 135.06 | 135.06 | 2.68 | 24 | >sp\|P01877\|IGHA2_HUMAN Immunoglobulin heavy constant alpha 2 OS=Homo sapiens OX=9606 GN=IGHA2 PE=1 SV=4 | 576 | 20201115_10113_1ng_glyco_hcd30_etd30_compound15.6026.6026.2 | controllerType=0 controllerNumber=1 scan=6026 | 16.2202 |
| 09722:3 | 14 | R.MAGKPTHIN[+1702.581]VSVVMAEADGTC[+57.021]Y.- | HexNAc(2)Hex(8) | C[+57], N[+1703] | 1351.8973 | 3 | 4053.6773 | 4053.6642 | 0 | 3.2 | 319 | CRagged | 246.27 | 246.27 | 193.46 | 4.39 | 24 | >sp\|P01877\|IGHA2_HUMAN Immunoglobulin heavy constant alpha 2 OS=Homo sapiens OX=9606 GN=IGHA2 PE=1 SV=4 | 576 | 20201115_10113_1ng_glyco_hcd30_etd30_compound15.12577.12577.3 | controllerType=0 controllerNumber=1 scan=12577 | 28.485 |
| 09792:3 | 14 | R.MAGKPTHIN[+1216.423]VSVVMAEADGTC[+57.021]Y.- | HexNAc(2)Hex(5) | C[+57], N[+1216] | 1189.8417 | 3 | 3567.5106 | 3567.5057 | 0 | 1.4 | 319 | CRagged | 268.88 | 262.7 | 196.89 | 4.06 | 24 | >sp\|P01877\|IGHA2_HUMAN Immunoglobulin heavy constant alpha 2 OS=Homo sapiens OX=9606 GN=IGHA2 PE=1 SV=4 | 576 | 20201115_10113_1ng_glyco_hcd30_etd30_compound15.12652.12652.3 | controllerType=0 controllerNumber=1 scan=12652 | 28.7224 |
| 09768:3 | 14 | R.MAGKPTHIN[+1378.476]VSVVMAEADGTC[+57.021]Y.- | HexNAc(2)Hex(6) | C[+57], N[+1378] | 1243.8692 | 3 | 3729.593 | 3729.5585 | 0 | 9.3 | 319 | CRagged | 283.35 | 260.08 | 216.1 | 3.24 | 24 | >sp\|P01877\|IGHA2_HUMAN Immunoglobulin heavy constant alpha 2 OS=Homo sapiens OX=9606 GN=IGHA2 PE=1 SV=4 | 576 | 20201115_10113_1ng_glyco_hcd30_etd30_compound15.12626.12626.3 | controllerType=0 controllerNumber=1 scan=12626 | 28.6212 |
| 19514:4 | 17 | R.QLAHQSN[+2205.793]STNIFFSPVSIATAFAMLSLGTK.A | HexNAc(4)Hex(5)Fuc(2)NeuAc(1) | N[+2206] | 1347.6141 | 4 | 5387.4347 | 5387.4279 | 0 | 1.3 | 64 | Specific | 425.38 | 385.69 | 385.69 | 7.05 | 8 | >sp\|P01009\|A1AT_HUMAN Alpha-1-antitrypsin OS=Homo sapiens OX=9606 GN=SERPINA1 PE=1 SV=3 | 642 | 20201115_10113_1ng_glyco_hcd30_etd30_compound15.22902.22902.4 | controllerType=0 controllerNumber=1 scan=22902 | 56.1364 |
| 19473:4 | 17 | R.QLAHQSN[+2204.772]STNIFFSPVSIATAFAMLSLGTK.A | HexNAc(4)Hex(5)NeuAc(2) | N[+2205] | 1347.3676 | 4 | 5386.4484 | 5386.4075 | 0 | 7.6 | 64 | Specific | 542.3 | 447.48 | 447.48 | 6.88 | 8 | >sp\|P01009\|A1AT_HUMAN Alpha-1-antitrypsin OS=Homo sapiens OX=9606 GN=SERPINA1 PE=1 SV=3 | 642 | 20201115_10113_1ng_glyco_hcd30_etd30_compound15.22859.22859.4 | controllerType=0 controllerNumber=1 scan=22859 | 55.9862 |
| 17900:4 | 18 | K.APDKNVIFSPLSISTALAFLSLGAHN[+2205.793]TTLTEILK.G | HexNAc(4)Hex(5)Fuc(2)NeuAc(1) | N[+2206] | 1447.9557 | 4 | 5788.8009 | 5788.771 | 0 | 5.2 | 68 | Specific | 280.27 | 280.27 | 280.27 | 3.4 | 11 | >sp\|P01011\|AACT_HUMAN Alpha-1-antichymotrypsin OS=Homo sapiens OX=9606 GN=SERPINA3 PE=1 SV=2 | 1314 | 20201115_10113_1ng_glyco_hcd30_etd30_compound15.21183.21183.4 | controllerType=0 controllerNumber=1 scan=21183 | 50.6681 |
| 17899:4 | 18 | K.APDKNVIFSPLSISTALAFLSLGAHN[+2059.735]TTLTEILK.G | HexNAc(4)Hex(5)Fuc(1)NeuAc(1) | N[+2060] | 1411.4396 | 4 | 5642.7365 | 5642.7131 | 0 | 4.1 | 68 | Specific | 593.79 | 593.79 | 593.79 | 8.98 | 11 | >sp\|P01011\|AACT_HUMAN Alpha-1-antichymotrypsin OS=Homo sapiens OX=9606 GN=SERPINA3 PE=1 SV=2 | 1314 | 20201115_10113_1ng_glyco_hcd30_etd30_compound15.21182.21182.4 | controllerType=0 controllerNumber=1 scan=21182 | 50.6645 |
| 17889:5 | 18 | K.APDKNVIFSPLSISTALAFLSLGAHN[+2059.735]TTLTEILK.G | HexNAc(4)Hex(5)Fuc(1)NeuAc(1) | N[+2060] | 1129.3519 | 5 | 5642.7306 | 5642.7131 | 0 | 3.1 | 68 | Specific | 616.37 | 616.37 | 616.37 | 10.34 | 11 | >sp\|P01011\|AACT_HUMAN Alpha-1-antichymotrypsin OS=Homo sapiens OX=9606 GN=SERPINA3 PE=1 SV=2 | 1314 | 20201115_10113_1ng_glyco_hcd30_etd30_compound15.21171.21171.5 | controllerType=0 controllerNumber=1 scan=21171 | 50.6286 |
| 13628:5 | 18 | K.GLKFN[+2350.830]LTETSEAEIHQSFQHLLR.T | HexNAc(4)Hex(5)Fuc(1)NeuAc(2) | N[+2351] | 1010.6538 | 5 | 5049.2396 | 5049.2251 | 0 | 2.9 | 102 | Specific | 437.54 | 437.54 | 437.54 | 5.6 | 11 | >sp\|P01011\|AACT_HUMAN Alpha-1-antichymotrypsin OS=Homo sapiens OX=9606 GN=SERPINA3 PE=1 SV=2 | 1314 | 20201115_10113_1ng_glyco_hcd30_etd30_compound15.16668.16668.5 | controllerType=0 controllerNumber=1 scan=16668 | 37.8284 |
| 05794:3 | 20 | R.LSPN[+2205.793]ASAEHLELR.W | HexNAc(4)Hex(5)Fuc(2)NeuAc(1) | N[+2206] | 1214.8552 | 3 | 3642.5509 | 3642.5421 | 0 | 2.4 | 52 | Specific | 283.5 | 196.91 | 196.91 | 3.92 | 8 | >sp\|Q13410\|BT1A1_HUMAN Butyrophilin subfamily 1 member A1 OS=Homo sapiens OX=9606 GN=BTN1A1 PE=1 SV=3 | 179 | 20201115_10113_1ng_glyco_hcd30_etd30_compound15.8490.8490.3 | controllerType=0 controllerNumber=1 scan=8490 | 20.6909 |
| 00858:4 | 22 | R.EIRHN[+2205.793]STGC[+57.021]LR.M | HexNAc(4)Hex(5)Fuc(2)NeuAc(1) | C[+57], N[+2206] | 887.8732 | 4 | 3548.471 | 3548.4573 | 0 | 3.9 | 287 | Specific | 434.08 | 348.89 | 348.89 | 3.4 | 13 | >sp\|P10909\|CLUS_HUMAN Clusterin OS=Homo sapiens OX=9606 GN=CLU PE=1 SV=1 | 1467 | 20201115_10113_1ng_glyco_hcd30_etd30_compound15.3370.3370.4 | controllerType=0 controllerNumber=1 scan=3370 | 11.2872 |
| 00847:4 | 22 | R.EIRHN[+2059.735]STGC[+57.021]LR.M | HexNAc(4)Hex(5)Fuc(1)NeuAc(1) | C[+57], N[+2060] | 851.3579 | 4 | 3402.4099 | 3402.3994 | 0 | 3.1 | 287 | Specific | 615.61 | 499.06 | 499.06 | 8.84 | 13 | >sp\|P10909\|CLUS_HUMAN Clusterin OS=Homo sapiens OX=9606 GN=CLU PE=1 SV=1 | 1467 | 20201115_10113_1ng_glyco_hcd30_etd30_compound15.3359.3359.4 | controllerType=0 controllerNumber=1 scan=3359 | 11.2519 |
| 17626:3 | 22 | K.MLN[+2205.793]TSSLLEQLNEQFNWVSR.L | HexNAc(4)Hex(5)Fuc(2)NeuAc(1) | N[+2206] | 1539.0045 | 3 | 4614.9988 | 4614.9796 | 0 | 4.2 | 352 | Specific | 317.54 | 204.05 | 204.05 | 3.72 | 13 | >sp\|P10909\|CLUS_HUMAN Clusterin OS=Homo sapiens OX=9606 GN=CLU PE=1 SV=1 | 1467 | 20201115_10113_1ng_glyco_hcd30_etd30_compound15.20891.20891.3 | controllerType=0 controllerNumber=1 scan=20891 | 49.6959 |
| 17642:4 | 22 | K.MLN[+2059.735]TSSLLEQLNEQFNWVSR.L | HexNAc(4)Hex(5)Fuc(1)NeuAc(1) | N[+2060] | 1117.9896 | 4 | 4468.9366 | 4468.9217 | 0 | 3.3 | 352 | Specific | 430.85 | 419.6 | 419.6 | 5.85 | 13 | >sp\|P10909\|CLUS_HUMAN Clusterin OS=Homo sapiens OX=9606 GN=CLU PE=1 SV=1 | 1467 | 20201115_10113_1ng_glyco_hcd30_etd30_compound15.20908.20908.4 | controllerType=0 controllerNumber=1 scan=20908 | 49.7566 |
| 08373:4 | 32 | K.SC[+57.021]QHN[+2059.735]GTMYQHGEIFSAHELFPSR.L | HexNAc(4)Hex(5)Fuc(1)NeuAc(1) | C[+57], N[+2060] | 1224.0094 | 4 | 4893.0159 | 4892.9919 | 0 | 4.9 | 110 | Specific | 588.34 | 511.48 | 511.48 | 8.45 | 5 | >sp\|Q6WN34\|CRDL2_HUMAN Chordin-like protein 2 OS=Homo sapiens OX=9606 GN=CHRDL2 PE=1 SV=1 | 1189 | 20201115_10113_1ng_glyco_hcd30_etd30_compound15.11170.11170.4 | controllerType=0 controllerNumber=1 scan=11170 | 25.6784 |
| 08364:5 | 32 | K.SC[+57.021]QHN[+2059.735]GTMYQHGEIFSAHELFPSR.L | HexNAc(4)Hex(5)Fuc(1)NeuAc(1) | C[+57], N[+2060] | 979.4078 | 5 | 4893.0099 | 4892.9919 | 0 | 3.7 | 110 | Specific | 678.03 | 657.4 | 657.4 | 10.65 | 5 | >sp\|Q6WN34\|CRDL2_HUMAN Chordin-like protein 2 OS=Homo sapiens OX=9606 GN=CHRDL2 PE=1 SV=1 | 1189 | 20201115_10113_1ng_glyco_hcd30_etd30_compound15.11161.11161.5 | controllerType=0 controllerNumber=1 scan=11161 | 25.6606 |
| 01971:4 | 36 | K.LENSLLDHRN[+2059.735]K.T | HexNAc(4)Hex(5)Fuc(1)NeuAc(1) | N[+2060] | 850.3687 | 4 | 3398.4529 | 3398.4474 | 0 | 1.6 | 247 | Specific | 308.23 | 232.56 | 232.56 | 3.29 | 3 | >sp\|P23280\|CAH6_HUMAN Carbonic anhydrase 6 OS=Homo sapiens OX=9606 GN=CA6 PE=1 SV=3 | 1012 | 20201115_10113_1ng_glyco_hcd30_etd30_compound15.4525.4525.4 | controllerType=0 controllerNumber=1 scan=4525 | 13.4732 |
| 12209:3 | 48 | R.IIVPLNNREN[+2350.830]ISDPTSPLR.T | HexNAc(4)Hex(5)Fuc(1)NeuAc(2) | N[+2351] | 1500.348 | 3 | 4499.0294 | 4499.0075 | 0 | 4.9 | 62 | Specific | 211.62 | 144.13 | 144.13 | 2.91 | 6 | >sp\|P01591\|IGJ_HUMAN Immunoglobulin J chain OS=Homo sapiens OX=9606 GN=JCHAIN PE=1 SV=4 | 1392 | 20201115_10113_1ng_glyco_hcd30_etd30_compound15.15178.15178.3 | controllerType=0 controllerNumber=1 scan=15178 | 34.0086 |
| 09263:3 | 48 | R.IIVPLNNREN[+1663.608]ISDPTSPLR.T | HexNAc(5)Hex(4) | N[+1664] | 1271.27 | 3 | 3811.7953 | 3811.7853 | 0 | 2.6 | 62 | Specific | 265.59 | 172.47 | 172.47 | 3.54 | 6 | >sp\|P01591\|IGJ_HUMAN Immunoglobulin J chain OS=Homo sapiens OX=9606 GN=JCHAIN PE=1 SV=4 | 1392 | 20201115_10113_1ng_glyco_hcd30_etd30_compound15.12101.12101.3 | controllerType=0 controllerNumber=1 scan=12101 | 27.5567 |
| 09251:3 | 48 | R.IIVPLNNREN[+1809.666]ISDPTSPLR.T | HexNAc(5)Hex(4)Fuc(1) | N[+1810] | 1319.9566 | 3 | 3957.8552 | 3957.8432 | 0 | 3 | 62 | Specific | 273.54 | 179.41 | 179.41 | 3.48 | 6 | >sp\|P01591\|IGJ_HUMAN Immunoglobulin J chain OS=Homo sapiens OX=9606 GN=JCHAIN PE=1 SV=4 | 1392 | 20201115_10113_1ng_glyco_hcd30_etd30_compound15.12089.12089.3 | controllerType=0 controllerNumber=1 scan=12089 | 27.5126 |
| 10590:3 | 48 | R.IIVPLNNREN[+2059.735]ISDPTSPLR.T | HexNAc(4)Hex(5)Fuc(1)NeuAc(1) | N[+2060] | 1403.3126 | 3 | 4207.9231 | 4207.9121 | 0 | 2.6 | 62 | Specific | 274.8 | 226.44 | 226.44 | 3.91 | 6 | >sp\|P01591\|IGJ_HUMAN Immunoglobulin J chain OS=Homo sapiens OX=9606 GN=JCHAIN PE=1 SV=4 | 1392 | 20201115_10113_1ng_glyco_hcd30_etd30_compound15.13484.13484.3 | controllerType=0 controllerNumber=1 scan=13484 | 30.3584 |
| 12228:3 | 48 | R.IIVPLNNREN[+2204.772]ISDPTSPLR.T | HexNAc(4)Hex(5)NeuAc(2) | N[+2205] | 1451.6594 | 3 | 4352.9635 | 4352.9496 | 0 | 3.2 | 62 | Specific | 339.88 | 221.61 | 221.61 | 4.38 | 6 | >sp\|P01591\|IGJ_HUMAN Immunoglobulin J chain OS=Homo sapiens OX=9606 GN=JCHAIN PE=1 SV=4 | 1392 | 20201115_10113_1ng_glyco_hcd30_etd30_compound15.15198.15198.3 | controllerType=0 controllerNumber=1 scan=15198 | 34.055 |
| 15346:4 | 1 | R.RTAGWNVPIGTLRPFLN[+2204.772]WTGPPEPIEAAVAR.F | HexNAc(4)Hex(5)NeuAc(2) | N[+2205] | 1398.6603 | 4 | 5591.6193 | 5591.5845 | 0 | 6.2 | 140 | Specific | 740.57 | 740.57 | 740.57 | 11.15 | 30 | >sp\|P02788\|TRFL_HUMAN Lactotransferrin OS=Homo sapiens OX=9606 GN=LTF PE=1 SV=6 | 1270 | 20201119_10132_1ng_glyco_hcd30_etd30_compound15.18991.18991.4 | controllerType=0 controllerNumber=1 scan=18991 | 43.659 |
| 15342:4 | 1 | R.RTAGWNVPIGTLRPFLN[+2059.735]WTGPPEPIEAAVAR.F | HexNAc(4)Hex(5)Fuc(1)NeuAc(1) | N[+2060] | 1362.4 | 4 | 5446.5783 | 5446.547 | 0 | 5.7 | 140 | Specific | 1023.81 | 988.83 | 988.83 | 16.57 | 30 | >sp\|P02788\|TRFL_HUMAN Lactotransferrin OS=Homo sapiens OX=9606 GN=LTF PE=1 SV=6 | 1270 | 20201119_10132_1ng_glyco_hcd30_etd30_compound15.18987.18987.4 | controllerType=0 controllerNumber=1 scan=18987 | 43.6508 |
| 15470:3 | 1 | R.TAGWNVPIGTLRPFLN[+2279.830]WTGPPEPIEAAVAR.F | HexNAc(5)Hex(6)Fuc(2) | N[+2280] | 1837.5268 | 3 | 5510.5659 | 5510.5406 | 0 | 4.6 | 141 | Specific | 248.46 | 195.15 | 195.15 | 4.14 | 30 | >sp\|P02788\|TRFL_HUMAN Lactotransferrin OS=Homo sapiens OX=9606 GN=LTF PE=1 SV=6 | 1270 | 20201119_10132_1ng_glyco_hcd30_etd30_compound15.19121.19121.3 | controllerType=0 controllerNumber=1 scan=19121 | 44.0008 |
| 16149:3 | 1 | R.TAGWNVPIGTLRPFLN[+2205.793]WTGPPEPIEAAVAR.F | HexNAc(4)Hex(5)Fuc(2)NeuAc(1) | N[+2206] | 1812.844 | 3 | 5436.5173 | 5436.5038 | 0 | 2.5 | 141 | Specific | 280.62 | 210.6 | 210.6 | 5.53 | 30 | >sp\|P02788\|TRFL_HUMAN Lactotransferrin OS=Homo sapiens OX=9606 GN=LTF PE=1 SV=6 | 1270 | 20201119_10132_1ng_glyco_hcd30_etd30_compound15.19842.19842.3 | controllerType=0 controllerNumber=1 scan=19842 | 46.2423 |
| 15493:4 | 1 | R.[+42.011]TAGWNVPIGTLRPFLN[+1872.651]WTGPPEPIEAAVAR.F | HexNAc(3)Hex(6)NeuAc(1) | .[42], N[+1873] | 1287.1108 | 4 | 5145.4215 | 5145.372 | 0 | 9.6 | 141 | Specific | 296.11 | 296.11 | 296.11 | 3.57 | 30 | >sp\|P02788\|TRFL_HUMAN Lactotransferrin OS=Homo sapiens OX=9606 GN=LTF PE=1 SV=6 | 1270 | 20201119_10132_1ng_glyco_hcd30_etd30_compound15.19146.19146.4 | controllerType=0 controllerNumber=1 scan=19146 | 44.0766 |
| 16076:3 | 1 | R.TAGWNVPIGTLRPFLN[+2570.925]WTGPPEPIEAAVAR.F | HexNAc(5)Hex(6)Fuc(2)NeuAc(1) | N[+2571] | 1934.551 | 3 | 5801.6384 | 5801.636 | 0 | 0.4 | 141 | Specific | 332.36 | 243.64 | 243.64 | 4.63 | 30 | >sp\|P02788\|TRFL_HUMAN Lactotransferrin OS=Homo sapiens OX=9606 GN=LTF PE=1 SV=6 | 1270 | 20201119_10132_1ng_glyco_hcd30_etd30_compound15.19764.19764.3 | controllerType=0 controllerNumber=1 scan=19764 | 46.0055 |
| 15498:3 | 1 | R.TAGWNVPIGTLRPFLN[+1768.640]WTGPPEPIEAAVAR.F | HexNAc(4)Hex(5)Fuc(1) | N[+1769] | 1667.1259 | 3 | 4999.3633 | 4999.3504 | 0 | 2.6 | 141 | Specific | 403.8 | 323.05 | 323.05 | 4.95 | 30 | >sp\|P02788\|TRFL_HUMAN Lactotransferrin OS=Homo sapiens OX=9606 GN=LTF PE=1 SV=6 | 1270 | 20201119_10132_1ng_glyco_hcd30_etd30_compound15.19151.19151.3 | controllerType=0 controllerNumber=1 scan=19151 | 44.0913 |

Table S4. Cont.

| Query #:z | Protein Rank | Glycopeptide | Glycan | Modification Type(s) | Observed m/z | z | Observed (M+H) | Calc. mass (M+H) | Off-by-x error | Mass error (ppm) | StartPosition | Cleavage | Score | Delta | Delta Mod | \|Log Prob\| | # of unique peptides | Protein Name | Protein DB number | Comment | Scan # | Scan Time |
|---|---|---|---|---|---|---|---|---|---|---|---|---|---|---|---|---|---|---|---|---|---|---|
| 16085:3 | 1 | R.TAGWNVPIGTLRPFLN[+2570.925]WTGPPEPIEAAVAR.F | HexNAc(5)Hex(6)Fuc(2)NeuAc(1) | N[+2571] | 1934.565 | 3 | 5801.6805 | 5801.636 | 0 | 7.7 | 141 | Specific | 418.77 | 347.29 | 347.29 | 4.39 | 30 | >sp\|P02788\|TRFL_HUMAN Lactotransferrin OS=Homo sapiens OX=9606 GN=LTF PE=1 SV=6 | 1270 | 20201119_10132_1ng_glyco_hcd30_etd30_compound15.19774.19774.3 | controllerType=0 controllerNumber=1 scan=19774 | 46.0266 |
| 15469:4 | 1 | R.TAGWNVPIGTLRPFLN[+2279.830]WTGPPEPIEAAVAR.F | HexNAc(5)Hex(6)Fuc(2) | N[+2280] | 1378.3969 | 4 | 5510.5656 | 5510.5406 | 0 | 4.5 | 141 | Specific | 576.83 | 569.43 | 569.43 | 7.49 | 30 | >sp\|P02788\|TRFL_HUMAN Lactotransferrin OS=Homo sapiens OX=9606 GN=LTF PE=1 SV=6 | 1270 | 20201119_10132_1ng_glyco_hcd30_etd30_compound15.19120.19120.4 | controllerType=0 controllerNumber=1 scan=19120 | 43.9959 |
| 16039:4 | 1 | R.TAGWNVPIGTLRPFLN[+2716.983]WTGPPEPIEAAVAR.F | HexNAc(5)Hex(6)Fuc(3)NeuAc(1) | N[+2717] | 1487.6805 | 4 | 5947.7004 | 5947.6939 | 0 | 1.1 | 141 | Specific | 620.71 | 620.71 | 620.71 | 9.2 | 30 | >sp\|P02788\|TRFL_HUMAN Lactotransferrin OS=Homo sapiens OX=9606 GN=LTF PE=1 SV=6 | 1270 | 20201119_10132_1ng_glyco_hcd30_etd30_compound15.19724.19724.2 | controllerType=0 controllerNumber=1 scan=19724 | 45.8908 |
| 16162:3 | 1 | R.TAGWNVPIGTLRPFLN[+2205.793]WTGPPEPIEAAVAR.F | HexNAc(4)Hex(5)Fuc(2)NeuAc(1) | N[+2206] | 1812.8479 | 3 | 5436.529 | 5436.5038 | 0 | 4.6 | 141 | Specific | 634.62 | 498.36 | 498.36 | 9.36 | 30 | >sp\|P02788\|TRFL_HUMAN Lactotransferrin OS=Homo sapiens OX=9606 GN=LTF PE=1 SV=6 | 1270 | 20201119_10132_1ng_glyco_hcd30_etd30_compound15.19856.19856.3 | controllerType=0 controllerNumber=1 scan=19856 | 46.2857 |
| 17119:4 | 1 | R.TAGWNVPIGTLRPFLN[+2350.830]WTGPPEPIEAAVAR.F | HexNAc(4)Hex(5)Fuc(1)NeuAc(2) | N[+2351] | 1396.1451 | 4 | 5581.5587 | 5581.5413 | 0 | 3.1 | 141 | Specific | 812.65 | 812.65 | 812.65 | 13.34 | 30 | >sp\|P02788\|TRFL_HUMAN Lactotransferrin OS=Homo sapiens OX=9606 GN=LTF PE=1 SV=6 | 1270 | 20201119_10132_1ng_glyco_hcd30_etd30_compound15.20869.20869.4 | controllerType=0 controllerNumber=1 scan=20869 | 49.1881 |
| 16048:4 | 1 | R.TAGWNVPIGTLRPFLN[+2716.983]WTGPPEPIEAAVAR.F | HexNAc(5)Hex(6)Fuc(3)NeuAc(1) | N[+2717] | 1487.6807 | 4 | 5947.701 | 5947.6939 | 0 | 1.2 | 141 | Specific | 814.46 | 814.46 | 814.46 | 13.2 | 30 | >sp\|P02788\|TRFL_HUMAN Lactotransferrin OS=Homo sapiens OX=9606 GN=LTF PE=1 SV=6 | 1270 | 20201119_10132_1ng_glyco_hcd30_etd30_compound15.19734.19734.4 | controllerType=0 controllerNumber=1 scan=19734 | 45.9161 |
| 16066:4 | 1 | R.TAGWNVPIGTLRPFLN[+2570.925]WTGPPEPIEAAVAR.F | HexNAc(5)Hex(6)Fuc(2)NeuAc(1) | N[+2571] | 1451.1677 | 4 | 5801.6491 | 5801.636 | 0 | 2.3 | 141 | Specific | 832.72 | 830.03 | 830.03 | 13.34 | 30 | >sp\|P02788\|TRFL_HUMAN Lactotransferrin OS=Homo sapiens OX=9606 GN=LTF PE=1 SV=6 | 1270 | 20201119_10132_1ng_glyco_hcd30_etd30_compound15.19754.19754.4 | controllerType=0 controllerNumber=1 scan=19754 | 45.9783 |
| 16148:4 | 1 | R.TAGWNVPIGTLRPFLN[+2205.793]WTGPPEPIEAAVAR.F | HexNAc(4)Hex(5)Fuc(2)NeuAc(1) | N[+2206] | 1359.8851 | 4 | 5436.5187 | 5436.5038 | 0 | 2.7 | 141 | Specific | 923.96 | 912.56 | 912.56 | 15.34 | 30 | >sp\|P02788\|TRFL_HUMAN Lactotransferrin OS=Homo sapiens OX=9606 GN=LTF PE=1 SV=6 | 1270 | 20201119_10132_1ng_glyco_hcd30_etd30_compound15.19841.19841.4 | controllerType=0 controllerNumber=1 scan=19841 | 46.2354 |
| 15499:4 | 1 | R.TAGWNVPIGTLRPFLN[+1768.640]WTGPPEPIEAAVAR.F | HexNAc(4)Hex(5)Fuc(1) | N[+1769] | 1250.5957 | 4 | 4999.361 | 4999.3504 | 0 | 2.1 | 141 | Specific | 930.88 | 927.29 | 927.29 | 15.34 | 30 | >sp\|P02788\|TRFL_HUMAN Lactotransferrin OS=Homo sapiens OX=9606 GN=LTF PE=1 SV=6 | 1270 | 20201119_10132_1ng_glyco_hcd30_etd30_compound15.19152.19152.4 | controllerType=0 controllerNumber=1 scan=19152 | 44.0945 |
| 14638:3 | 1 | R.TAGWNIPMGLLFN[+1622.582]QTGSC[+57.021]K.F | HexNAc(4)Hex(5) | C[+57], N[+1623] | 1239.879 | 3 | 3717.6223 | 3717.5916 | 0 | 8.3 | 485 | Specific | 230.39 | 178.07 | 178.07 | 3.03 | 30 | >sp\|P02788\|TRFL_HUMAN Lactotransferrin OS=Homo sapiens OX=9606 GN=LTF PE=1 SV=6 | 1270 | 20201119_10132_1ng_glyco_hcd30_etd30_compound15.18239.18239.6 | controllerType=0 controllerNumber=1 scan=18239 | 41.2927 |
| 14278:3 | 1 | R.TAGWNIPMGLLFN[+2279.830]QTGSC[+57.021]K.F | HexNAc(5)Hex(6)Fuc(2) | C[+57], N[+2280] | 1458.9589 | 3 | 4374.8622 | 4374.8396 | 0 | 5.2 | 485 | Specific | 263.7 | 195.4 | 195.4 | 4.39 | 30 | >sp\|P02788\|TRFL_HUMAN Lactotransferrin OS=Homo sapiens OX=9606 GN=LTF PE=1 SV=6 | 1270 | 20201119_10132_1ng_glyco_hcd30_etd30_compound15.17860.17860.3 | controllerType=0 controllerNumber=1 scan=17860 | 40.2315 |
| 14457:4 | 1 | R.TAGWNIPMGLLFN[+1768.640]QTGSC[+57.021]K.F | HexNAc(4)Hex(5)Fuc(1) | C[+57], N[+1769] | 966.6753 | 4 | 3863.6795 | 3863.6495 | 0 | 7.8 | 485 | Specific | 269.7 | 252.57 | 252.57 | 3.04 | 30 | >sp\|P02788\|TRFL_HUMAN Lactotransferrin OS=Homo sapiens OX=9606 GN=LTF PE=1 SV=6 | 1270 | 20201119_10132_1ng_glyco_hcd30_etd30_compound15.18048.18048.4 | controllerType=0 controllerNumber=1 scan=18048 | 40.727 |
| 14263:3 | 1 | R.TAGWNIPMGLLFN[+2279.830]QTGSC[+57.021]K.F | HexNAc(5)Hex(6)Fuc(2) | C[+57], N[+2280] | 1458.9577 | 3 | 4374.8585 | 4374.8396 | 0 | 4.3 | 485 | Specific | 272.83 | 192.49 | 192.49 | 5.18 | 30 | >sp\|P02788\|TRFL_HUMAN Lactotransferrin OS=Homo sapiens OX=9606 GN=LTF PE=1 SV=6 | 1270 | 20201119_10132_1ng_glyco_hcd30_etd30_compound15.17844.17844.3 | controllerType=0 controllerNumber=1 scan=17844 | 40.1932 |
| 14423:3 | 1 | R.TAGWNIPMGLLFN[+1768.640]QTGSC[+57.021]K.F | HexNAc(4)Hex(5)Fuc(1) | C[+57], N[+1769] | 1288.5612 | 3 | 3863.6691 | 3863.6495 | 0 | 5.1 | 485 | Specific | 369.88 | 268.94 | 268.94 | 3.98 | 30 | >sp\|P02788\|TRFL_HUMAN Lactotransferrin OS=Homo sapiens OX=9606 GN=LTF PE=1 SV=6 | 1270 | 20201119_10132_1ng_glyco_hcd30_etd30_compound15.18013.18013.3 | controllerType=0 controllerNumber=1 scan=18013 | 40.6344 |
| 14542:3 | 1 | R.TAGWNIPMGLLFN[+1622.582]QTGSC[+57.021]K.F | HexNAc(4)Hex(5) | C[+57], N[+1623] | 1239.8726 | 3 | 3717.6033 | 3717.5916 | 0 | 3.2 | 485 | Specific | 370.91 | 214.24 | 214.24 | 5.65 | 30 | >sp\|P02788\|TRFL_HUMAN Lactotransferrin OS=Homo sapiens OX=9606 GN=LTF PE=1 SV=6 | 1270 | 20201119_10132_1ng_glyco_hcd30_etd30_compound15.18138.18138.3 | controllerType=0 controllerNumber=1 scan=18138 | 41.0003 |
| 14556:3 | 1 | R.TAGWNIPMGLLFN[+1606.587]QTGSC[+57.021]K.F | HexNAc(4)Hex(4)Fuc(1) | C[+57], N[+1607] | 1234.5391 | 3 | 3701.6028 | 3701.5966 | 0 | 1.7 | 485 | Specific | 371.45 | 289.08 | 289.08 | 4.79 | 30 | >sp\|P02788\|TRFL_HUMAN Lactotransferrin OS=Homo sapiens OX=9606 GN=LTF PE=1 SV=6 | 1270 | 20201119_10132_1ng_glyco_hcd30_etd30_compound15.18153.18153.3 | controllerType=0 controllerNumber=1 scan=18153 | 41.0407 |
| 14444:2 | 1 | R.TAGWNIPMGLLFN[+1768.640]QTGSC[+57.021]K.F | HexNAc(4)Hex(5)Fuc(1) | C[+57], N[+1769] | 1932.3401 | 2 | 3863.6728 | 3863.6495 | 0 | 6 | 485 | Specific | 376.89 | 268.5 | 268.5 | 5.43 | 30 | >sp\|P02788\|TRFL_HUMAN Lactotransferrin OS=Homo sapiens OX=9606 GN=LTF PE=1 SV=6 | 1270 | 20201119_10132_1ng_glyco_hcd30_etd30_compound15.18035.18035.2 | controllerType=0 controllerNumber=1 scan=18035 | 40.6914 |
| 15717:3 | 1 | R.TAGWNIPMGLLFN[+2059.735]QTGSC[+57.021]K.F | HexNAc(4)Hex(5)Fuc(1)NeuAc(1) | C[+57], N[+2060] | 1385.5906 | 3 | 4154.7573 | 4154.7449 | 0 | 3 | 485 | Specific | 393.44 | 285.13 | 285.13 | 5.71 | 30 | >sp\|P02788\|TRFL_HUMAN Lactotransferrin OS=Homo sapiens OX=9606 GN=LTF PE=1 SV=6 | 1270 | 20201119_10132_1ng_glyco_hcd30_etd30_compound15.19383.19383.3 | controllerType=0 controllerNumber=1 scan=19383 | 44.8418 |
| 15694:3 | 1 | R.TAGWNIPMGLLFN[+2205.793]QTGSC[+57.021]K.F | HexNAc(4)Hex(5)Fuc(2)NeuAc(1) | C[+57], N[+2206] | 1434.2794 | 3 | 4300.8235 | 4300.8028 | 0 | 4.8 | 485 | Specific | 406.58 | 310.68 | 310.68 | 5.14 | 30 | >sp\|P02788\|TRFL_HUMAN Lactotransferrin OS=Homo sapiens OX=9606 GN=LTF PE=1 SV=6 | 1270 | 20201119_10132_1ng_glyco_hcd30_etd30_compound15.19359.19359.3 | controllerType=0 controllerNumber=1 scan=19359 | 44.7799 |
| 14392:3 | 1 | R.TAGWNIPMGLLFN[+1768.640]QTGSC[+57.021]K.F | HexNAc(4)Hex(5)Fuc(1) | C[+57], N[+1769] | 1288.5591 | 3 | 3863.6629 | 3863.6495 | 0 | 3.5 | 485 | Specific | 415.59 | 294.06 | 294.06 | 5.77 | 30 | >sp\|P02788\|TRFL_HUMAN Lactotransferrin OS=Homo sapiens OX=9606 GN=LTF PE=1 SV=6 | 1270 | 20201119_10132_1ng_glyco_hcd30_etd30_compound15.17980.17980.3 | controllerType=0 controllerNumber=1 scan=17980 | 40.5398 |
| 14699:3 | 1 | R.TAGWNIPMGLLFN[+1216.423]QTGSC[+57.021]K.F | HexNAc(2)Hex(5) | C[+57], N[+1216] | 1104.4853 | 3 | 3311.4412 | 3311.4328 | 0 | 2.5 | 485 | Specific | 482.24 | 464.17 | 223.77 | 7.56 | 30 | >sp\|P02788\|TRFL_HUMAN Lactotransferrin OS=Homo sapiens OX=9606 GN=LTF PE=1 SV=6 | 1270 | 20201119_10132_1ng_glyco_hcd30_etd30_compound15.18304.18304.3 | controllerType=0 controllerNumber=1 scan=18304 | 41.4828 |
| 12603:3 | 1 | R.TAGWNIPM[+15.995]GLLFN[+1768.640]QTGSC[+57.021]K.F | HexNAc(4)Hex(5)Fuc(1) | C[+57], M[+16], N[+1769] | 1293.8992 | 3 | 3879.6831 | 3879.6444 | 0 | 10 | 485 | Specific | 210.45 | 210.45 | 210.45 | 3.75 | 30 | >sp\|P02788\|TRFL_HUMAN Lactotransferrin OS=Homo sapiens OX=9606 GN=LTF PE=1 SV=6 | 1270 | 20201119_10132_1ng_glyco_hcd30_etd30_compound15.16065.16065.3 | controllerType=0 controllerNumber=1 scan=16065 | 36.3847 |
| 15198:4 | 1 | R.TAGWNIPMGLLFN[+1768.640]QTGSC[+57.021]KFDEYFSQSC[+57.021]APGSDPR.S | HexNAc(4)Hex(5)Fuc(1) | C[+57]*2, N[+1769] | 1427.6124 | 4 | 5707.4279 | 5707.3966 | 0 | 5.5 | 485 | Specific | 400.29 | 359.83 | 359.83 | 3.93 | 30 | >sp\|P02788\|TRFL_HUMAN Lactotransferrin OS=Homo sapiens OX=9606 GN=LTF PE=1 SV=6 | 1270 | 20201119_10132_1ng_glyco_hcd30_etd30_compound15.18834.18834.4 | controllerType=0 controllerNumber=1 scan=18834 | 43.1788 |
| 15237:4 | 1 | R.TAGWNIPMGLLFN[+1622.582]QTGSC[+57.021]KFDEYFSQSC[+57.021]APGSDPR.S | HexNAc(4)Hex(5) | C[+57]*2, N[+1623] | 1391.0986 | 4 | 5561.3727 | 5561.3387 | 0 | 6.1 | 485 | Specific | 415.1 | 409.61 | 409.61 | 5.23 | 30 | >sp\|P02788\|TRFL_HUMAN Lactotransferrin OS=Homo sapiens OX=9606 GN=LTF PE=1 SV=6 | 1270 | 20201119_10132_1ng_glyco_hcd30_etd30_compound15.18875.18875.4 | controllerType=0 controllerNumber=1 scan=18875 | 43.3027 |
| 15205:3 | 1 | R.TAGWNIPMGLLFN[+1768.640]QTGSC[+57.021]KFDEYFSQSC[+57.021]APGSDPR.S | HexNAc(4)Hex(5)Fuc(1) | C[+57]*2, N[+1769] | 1903.1451 | 3 | 5707.4208 | 5707.3966 | 0 | 4.2 | 485 | Specific | 468.42 | 400.46 | 400.46 | 6.68 | 30 | >sp\|P02788\|TRFL_HUMAN Lactotransferrin OS=Homo sapiens OX=9606 GN=LTF PE=1 SV=6 | 1270 | 20201119_10132_1ng_glyco_hcd30_etd30_compound15.18841.18841.3 | controllerType=0 controllerNumber=1 scan=18841 | 43.2008 |
| 15926:4 | 1 | R.TAGWNIPMGLLFN[+1913.677]QTGSC[+57.021]KFDEYFSQSC[+57.021]APGSDPR.S | HexNAc(4)Hex(5)NeuAc(1) | C[+57]*2, N[+1914] | 1463.8739 | 4 | 5852.4738 | 5852.4341 | 0 | 6.8 | 485 | Specific | 618.37 | 515.72 | 515.72 | 7.73 | 30 | >sp\|P02788\|TRFL_HUMAN Lactotransferrin OS=Homo sapiens OX=9606 GN=LTF PE=1 SV=6 | 1270 | 20201119_10132_1ng_glyco_hcd30_etd30_compound15.19606.19606.4 | controllerType=0 controllerNumber=1 scan=19606 | 45.6283 |
| 07895:3 | 1 | R.N[+2205.793]GSDC[+57.021]PDKFC[+57.021]LFQSETK.N | HexNAc(4)Hex(5)Fuc(2)NeuAc(1) | C[+57]*2, N[+2206] | 1413.5657 | 3 | 4238.6826 | 4238.6667 | 0 | 3.7 | 642 | Specific | 389.14 | 257.51 | 257.51 | 5.1 | 30 | >sp\|P02788\|TRFL_HUMAN Lactotransferrin OS=Homo sapiens OX=9606 GN=LTF PE=1 SV=6 | 1270 | 20201119_10132_1ng_glyco_hcd30_etd30_compound15.11135.11135.3 | controllerType=0 controllerNumber=1 scan=11135 | 26.5765 |
| 15103:3 | 2 | R.LSLHRPALEDLLLGSEAN[+1540.529]LTC[+57.021]TLTGLR.D | HexNAc(2)Hex(7) | C[+57], N[+1541] | 1502.0544 | 3 | 4504.1486 | 4504.1268 | 0 | 4.8 | 127 | Specific | 381.05 | 360.72 | 360.72 | 3.88 | 28 | >sp\|P01876\|IGHA1_HUMAN Immunoglobulin heavy constant alpha 1 OS=Homo sapiens OX=9606 GN=IGHA1 PE=1 SV=2 | 255 | 20201119_10132_1ng_glyco_hcd30_etd30_compound15.18730.18730.3 | controllerType=0 controllerNumber=1 scan=18730 | 42.72 |
| 15088:3 | 2 | R.LSLHRPALEDLLLGSEAN[+1419.502]LTC[+57.021]TLTGLR.D | HexNAc(3)Hex(5) | C[+57], N[+1420] | 1461.7189 | 3 | 4383.1422 | 4383.1005 | 0 | 9.5 | 127 | Specific | 385.15 | 367.65 | 367.65 | 6.05 | 28 | >sp\|P01876\|IGHA1_HUMAN Immunoglobulin heavy constant alpha 1 OS=Homo sapiens OX=9606 GN=IGHA1 PE=1 SV=2 | 255 | 20201119_10132_1ng_glyco_hcd30_etd30_compound15.18714.18714.3 | controllerType=0 controllerNumber=1 scan=18714 | 42.6473 |
| 15053:3 | 2 | R.LSLHRPALEDLLLGSEAN[+1702.581]LTC[+57.021]TLTGLR.D | HexNAc(2)Hex(8) | C[+57], N[+1703] | 1556.0698 | 3 | 4666.1947 | 4666.1796 | 0 | 3.2 | 127 | Specific | 441.87 | 427.22 | 427.22 | 6.72 | 28 | >sp\|P01876\|IGHA1_HUMAN Immunoglobulin heavy constant alpha 1 OS=Homo sapiens OX=9606 GN=IGHA1 PE=1 SV=2 | 255 | 20201119_10132_1ng_glyco_hcd30_etd30_compound15.18677.18677.3 | controllerType=0 controllerNumber=1 scan=18677 | 42.5202 |
| 15092:3 | 2 | R.LSLHRPALEDLLLGSEAN[+1622.582]LTC[+57.021]TLTGLR.D | HexNAc(4)Hex(5) | C[+57], N[+1623] | 1529.413 | 3 | 4586.2245 | 4586.1799 | 0 | 9.7 | 127 | Specific | 444.97 | 401.45 | 401.45 | 6.26 | 28 | >sp\|P01876\|IGHA1_HUMAN Immunoglobulin heavy constant alpha 1 OS=Homo sapiens OX=9606 GN=IGHA1 PE=1 SV=2 | 255 | 20201119_10132_1ng_glyco_hcd30_etd30_compound15.18718.18718.3 | controllerType=0 controllerNumber=1 scan=18718 | 42.6667 |
| 15025:3 | 2 | R.LSLHRPALEDLLLGSEAN[+1864.634]LTC[+57.021]TLTGLR.D | HexNAc(2)Hex(9) | C[+57], N[+1865] | 1610.0912 | 3 | 4828.2592 | 4828.2324 | 0 | 5.5 | 127 | Specific | 459.34 | 446.97 | 446.97 | 5.87 | 28 | >sp\|P01876\|IGHA1_HUMAN Immunoglobulin heavy constant alpha 1 OS=Homo sapiens OX=9606 GN=IGHA1 PE=1 SV=2 | 255 | 20201119_10132_1ng_glyco_hcd30_etd30_compound15.18647.18647.3 | controllerType=0 controllerNumber=1 scan=18647 | 42.438 |
| 15122:3 | 2 | R.LSLHRPALEDLLLGSEAN[+1216.423]LTC[+57.021]TLTGLR.D | HexNAc(2)Hex(5) | C[+57], N[+1216] | 1394.0216 | 3 | 4180.0502 | 4180.0211 | 0 | 7 | 127 | Specific | 485.3 | 482.53 | 482.53 | 6.73 | 28 | >sp\|P01876\|IGHA1_HUMAN Immunoglobulin heavy constant alpha 1 OS=Homo sapiens OX=9606 GN=IGHA1 PE=1 SV=2 | 255 | 20201119_10132_1ng_glyco_hcd30_etd30_compound15.18751.18751.3 | controllerType=0 controllerNumber=1 scan=18751 | 42.8399 |
| 15127:4 | 2 | R.LSLHRPALEDLLLGSEAN[+1257.449]LTC[+57.021]TLTGLR.D | HexNAc(3)Hex(4) | C[+57], N[+1257] | 1056.0248 | 4 | 4221.0774 | 4221.0477 | 0 | 7 | 127 | Specific | 688.28 | 688.28 | 688.28 | 10.15 | 28 | >sp\|P01876\|IGHA1_HUMAN Immunoglobulin heavy constant alpha 1 OS=Homo sapiens OX=9606 GN=IGHA1 PE=1 SV=2 | 255 | 20201119_10132_1ng_glyco_hcd30_etd30_compound15.18756.18756.4 | controllerType=0 controllerNumber=1 scan=18756 | 42.8592 |
| 15052:4 | 2 | R.LSLHRPALEDLLLGSEAN[+1702.581]LTC[+57.021]TLTGLR.D | HexNAc(2)Hex(8) | C[+57], N[+1703] | 1167.3036 | 4 | 4666.1927 | 4666.1796 | 0 | 2.8 | 127 | Specific | 740.71 | 740.71 | 740.71 | 11.42 | 28 | >sp\|P01876\|IGHA1_HUMAN Immunoglobulin heavy constant alpha 1 OS=Homo sapiens OX=9606 GN=IGHA1 PE=1 SV=2 | 255 | 20201119_10132_1ng_glyco_hcd30_etd30_compound15.18676.18676.4 | controllerType=0 controllerNumber=1 scan=18676 | 42.5171 |
| 15104:4 | 2 | R.LSLHRPALEDLLLGSEAN[+1419.502]LTC[+57.021]TLTGLR.D | HexNAc(3)Hex(5) | C[+57], N[+1420] | 1096.5365 | 4 | 4383.1243 | 4383.1005 | 0 | 5.4 | 127 | Specific | 864.24 | 864.24 | 864.24 | 13.57 | 28 | >sp\|P01876\|IGHA1_HUMAN Immunoglobulin heavy constant alpha 1 OS=Homo sapiens OX=9606 GN=IGHA1 PE=1 SV=2 | 255 | 20201119_10132_1ng_glyco_hcd30_etd30_compound15.18731.18731.4 | controllerType=0 controllerNumber=1 scan=18731 | 42.7319 |
| 15100:4 | 2 | R.LSLHRPALEDLLLGSEAN[+1622.582]LTC[+57.021]TLTGLR.D | HexNAc(4)Hex(5) | C[+57], N[+1623] | 1147.3069 | 4 | 4586.2059 | 4586.1799 | 0 | 5.7 | 127 | Specific | 903.22 | 819.96 | 819.96 | 14.57 | 28 | >sp\|P01876\|IGHA1_HUMAN Immunoglobulin heavy constant alpha 1 OS=Homo sapiens OX=9606 GN=IGHA1 PE=1 SV=2 | 255 | 20201119_10132_1ng_glyco_hcd30_etd30_compound15.18727.18727.4 | controllerType=0 controllerNumber=1 scan=18727 | 42.7095 |
| 15126:4 | 2 | R.LSLHRPALEDLLLGSEAN[+1298.476]LTC[+57.021]TLTGLR.D | HexNAc(4)Hex(3) | C[+57], N[+1298] | 1066.2811 | 4 | 4262.1026 | 4262.0742 | 0 | 6.6 | 127 | Specific | 909.99 | 909.99 | 909.99 | 15.15 | 28 | >sp\|P01876\|IGHA1_HUMAN Immunoglobulin heavy constant alpha 1 OS=Homo sapiens OX=9606 GN=IGHA1 PE=1 SV=2 | 255 | 20201119_10132_1ng_glyco_hcd30_etd30_compound15.18755.18755.4 | controllerType=0 controllerNumber=1 scan=18755 | 42.8517 |
| 15121:4 | 2 | R.LSLHRPALEDLLLGSEAN[+1216.423]LTC[+57.021]TLTGLR.D | HexNAc(2)Hex(5) | C[+57], N[+1216] | 1045.7679 | 4 | 4180.0498 | 4180.0211 | 0 | 6.9 | 127 | Specific | 940.28 | 922.87 | 922.87 | 15.15 | 28 | >sp\|P01876\|IGHA1_HUMAN Immunoglobulin heavy constant alpha 1 OS=Homo sapiens OX=9606 GN=IGHA1 PE=1 SV=2 | 255 | 20201119_10132_1ng_glyco_hcd30_etd30_compound15.18750.18750.4 | controllerType=0 controllerNumber=1 scan=18750 | 42.8329 |
| 15020:4 | 2 | R.LSLHRPALEDLLLGSEAN[+1864.634]LTC[+57.021]TLTGLR.D | HexNAc(2)Hex(9) | C[+57], N[+1865] | 1207.8199 | 4 | 4828.2576 | 4828.2324 | 0 | 5.2 | 127 | Specific | 1018.36 | 962.58 | 962.58 | 16.57 | 28 | >sp\|P01876\|IGHA1_HUMAN Immunoglobulin heavy constant alpha 1 OS=Homo sapiens OX=9606 GN=IGHA1 PE=1 SV=2 | 255 | 20201119_10132_1ng_glyco_hcd30_etd30_compound15.18642.18642.4 | controllerType=0 controllerNumber=1 scan=18642 | 42.4129 |
| 15099:4 | 2 | R.LSLHRPALEDLLLGSEAN[+1540.529]LTC[+57.021]TLTGLR.D | HexNAc(2)Hex(7) | C[+57], N[+1541] | 1126.7931 | 4 | 4504.1507 | 4504.1268 | 0 | 5.3 | 127 | Specific | 1019.5 | 950.03 | 950.03 | 16.57 | 28 | >sp\|P01876\|IGHA1_HUMAN Immunoglobulin heavy constant alpha 1 OS=Homo sapiens OX=9606 GN=IGHA1 PE=1 SV=2 | 255 | 20201119_10132_1ng_glyco_hcd30_etd30_compound15.18726.18726.4 | controllerType=0 controllerNumber=1 scan=18726 | 42.7025 |
| 15112:4 | 2 | R.LSLHRPALEDLLLGSEAN[+1378.476]LTC[+57.021]TLTGLR.D | HexNAc(2)Hex(6) | C[+57], N[+1378] | 1086.2823 | 4 | 4342.1072 | 4342.074 | 0 | 7.7 | 127 | Specific | 1028.39 | 1005.88 | 1005.88 | 16.66 | 28 | >sp\|P01876\|IGHA1_HUMAN Immunoglobulin heavy constant alpha 1 OS=Homo sapiens OX=9606 GN=IGHA1 PE=1 SV=2 | 255 | 20201119_10132_1ng_glyco_hcd30_etd30_compound15.18740.18740.4 | controllerType=0 controllerNumber=1 scan=18740 | 42.7779 |
| 15108:4 | 2 | R.LSLHRPALEDLLLGSEAN[+1460.529]LTC[+57.021]TLTGLR.D | HexNAc(4)Hex(4) | C[+57], N[+1461] | 1106.7937 | 4 | 4424.1531 | 4424.1271 | 0 | 5.9 | 127 | Specific | 1032.79 | 946.17 | 946.17 | 17.15 | 28 | >sp\|P01876\|IGHA1_HUMAN Immunoglobulin heavy constant alpha 1 OS=Homo sapiens OX=9606 GN=IGHA1 PE=1 SV=2 | 255 | 20201119_10132_1ng_glyco_hcd30_etd30_compound15.18735.18735.4 | controllerType=0 controllerNumber=1 scan=18735 | 42.7557 |
| 15107:4 | 2 | R.LSLHRPALEDLLLGSEAN[+1501.555]LTC[+57.021]TLTGLR.D | HexNAc(5)Hex(3) | C[+57], N[+1502] | 1117.0501 | 4 | 4465.1785 | 4465.1536 | 0 | 5.6 | 127 | Specific | 1265.65 | 1190.2 | 1190.2 | 21.57 | 28 | >sp\|P01876\|IGHA1_HUMAN Immunoglobulin heavy constant alpha 1 OS=Homo sapiens OX=9606 GN=IGHA1 PE=1 SV=2 | 255 | 20201119_10132_1ng_glyco_hcd30_etd30_compound15.18734.18734.4 | controllerType=0 controllerNumber=1 scan=18734 | 42.7492 |
| 09912:3 | 2 | R.LAGKPTHVN[+1663.608]VSVVMAEVDGTC[+57.021]Y.- | HexNAc(5)Hex(4) | C[+57], N[+1664] | 1337.5908 | 3 | 4010.7577 | 4010.7502 | 0 | 1.9 | 332 | CRagged | 211.88 | 173.73 | 173.73 | 2.88 | 28 | >sp\|P01876\|IGHA1_HUMAN Immunoglobulin heavy constant alpha 1 OS=Homo sapiens OX=9606 GN=IGHA1 PE=1 SV=2 | 255 | 20201119_10132_1ng_glyco_hcd30_etd30_compound15.13231.13231.3 | controllerType=0 controllerNumber=1 scan=13231 | 30.6243 |
| 10021:3 | 2 | R.LAGKPTHVN[+1095.397]VSVVMAEVDGTC[+57.021]Y.- | HexNAc(3)Hex(3) | C[+57], N[+1095] | 1148.1872 | 3 | 3442.5469 | 3442.5387 | 0 | 2.4 | 332 | CRagged | 227.55 | 166.93 | 166.93 | 3.03 | 28 | >sp\|P01876\|IGHA1_HUMAN Immunoglobulin heavy constant alpha 1 OS=Homo sapiens OX=9606 GN=IGHA1 PE=1 SV=2 | 255 | 20201119_10132_1ng_glyco_hcd30_etd30_compound15.13346.13346.3 | controllerType=0 controllerNumber=1 scan=13346 | 30.9428 |
| 09857:3 | 2 | R.LAGKPTHVN[+1702.581]VSVVMAEVDGTC[+57.021]Y.- | HexNAc(2)Hex(8) | C[+57], N[+1703] | 1350.5838 | 3 | 4049.7368 | 4049.7234 | 0 | 3.3 | 332 | CRagged | 246.45 | 205.53 | 175.8 | 4.11 | 28 | >sp\|P01876\|IGHA1_HUMAN Immunoglobulin heavy constant alpha 1 OS=Homo sapiens OX=9606 GN=IGHA1 PE=1 SV=2 | 255 | 20201119_10132_1ng_glyco_hcd30_etd30_compound15.13173.13173.3 | controllerType=0 controllerNumber=1 scan=13173 | 30.4448 |
| 09887:3 | 2 | R.LAGKPTHVN[+1378.476]VSVVMAEVDGTC[+57.021]Y.- | HexNAc(2)Hex(6) | C[+57], N[+1378] | 1242.5458 | 3 | 3725.6229 | 3725.6178 | 0 | 1.4 | 332 | CRagged | 258.9 | 215.39 | 197.62 | 4.03 | 28 | >sp\|P01876\|IGHA1_HUMAN Immunoglobulin heavy constant alpha 1 OS=Homo sapiens OX=9606 GN=IGHA1 PE=1 SV=2 | 255 | 20201119_10132_1ng_glyco_hcd30_etd30_compound15.13205.13205.3 | controllerType=0 controllerNumber=1 scan=13205 | 30.5485 |
| 09871:3 | 2 | R.LAGKPTHVN[+1540.529]VSVVMAEVDGTC[+57.021]Y.- | HexNAc(2)Hex(7) | C[+57], N[+1541] | 1296.5682 | 3 | 3887.69 | 3887.6706 | 0 | 5 | 332 | CRagged | 269.25 | 211.84 | 199.92 | 3.36 | 28 | >sp\|P01876\|IGHA1_HUMAN Immunoglobulin heavy constant alpha 1 OS=Homo sapiens OX=9606 GN=IGHA1 PE=1 SV=2 | 255 | 20201119_10132_1ng_glyco_hcd30_etd30_compound15.13188.13188.3 | controllerType=0 controllerNumber=1 scan=13188 | 30.4954 |
| 09945:3 | 2 | R.LAGKPTHVN[+1809.666]VSVVMAEVDGTC[+57.021]Y.- | HexNAc(5)Hex(4)Fuc(1) | C[+57], N[+1810] | 1386.28 | 3 | 4156.8253 | 4156.8081 | 0 | 4.1 | 332 | CRagged | 272.49 | 168.78 | 168.78 | 3.17 | 28 | >sp\|P01876\|IGHA1_HUMAN Immunoglobulin heavy constant alpha 1 OS=Homo sapiens OX=9606 GN=IGHA1 PE=1 SV=2 | 255 | 20201119_10132_1ng_glyco_hcd30_etd30_compound15.13266.13266.3 | controllerType=0 controllerNumber=1 scan=13266 | 30.7242 |
| 09962:3 | 2 | R.LAGKPTHVN[+1501.555]VSVVMAEVDGTC[+57.021]Y.- | HexNAc(5)Hex(3) | C[+57], N[+1502] | 1283.5748 | 3 | 3848.7097 | 3848.6974 | 0 | 3.2 | 332 | CRagged | 281 | 193.26 | 193.26 | 3.95 | 28 | >sp\|P01876\|IGHA1_HUMAN Immunoglobulin heavy constant alpha 1 OS=Homo sapiens OX=9606 GN=IGHA1 PE=1 SV=2 | 255 | 20201119_10132_1ng_glyco_hcd30_etd30_compound15.13284.13284.3 | controllerType=0 controllerNumber=1 scan=13284 | 30.78 |
| 09970:3 | 2 | R.LAGKPTHVN[+1647.613]VSVVMAEVDGTC[+57.021]Y.- | HexNAc(5)Hex(3)Fuc(1) | C[+57], N[+1648] | 1332.2604 | 3 | 3994.7667 | 3994.7553 | 0 | 2.9 | 332 | CRagged | 282.27 | 166.13 | 166.13 | 3.21 | 28 | >sp\|P01876\|IGHA1_HUMAN Immunoglobulin heavy constant alpha 1 OS=Homo sapiens OX=9606 GN=IGHA1 PE=1 SV=2 | 255 | 20201119_10132_1ng_glyco_hcd30_etd30_compound15.13292.13292.3 | controllerType=0 controllerNumber=1 scan=13292 | 30.798 |
| 09946:3 | 2 | R.LAGKPTHVN[+1216.423]VSVVMAEVDGTC[+57.021]Y.- | HexNAc(2)Hex(5) | C[+57], N[+1216] | 1188.5307 | 3 | 3563.5775 | 3563.5649 | 0 | 3.5 | 332 | CRagged | 289.59 | 273.9 | 211.52 | 4.2 | 28 | >sp\|P01876\|IGHA1_HUMAN Immunoglobulin heavy constant alpha 1 OS=Homo sapiens OX=9606 GN=IGHA1 PE=1 SV=2 | 255 | 20201119_10132_1ng_glyco_hcd30_etd30_compound15.13267.13267.3 | controllerType=0 controllerNumber=1 scan=13267 | 30.7279 |
| 15827:3 | 6 | K.QIGLYPVLVIDSSGYVNPN[+2059.735]YTGR.I | HexNAc(4)Hex(5)Fuc(1)NeuAc(1) | N[+2060] | 1529.024 | 3 | 4585.0574 | 4585.0384 | 0 | 4.1 | 168 | Specific | 607.76 | 543.48 | 543.48 | 9.16 | 17 | >sp\|P01833\|PIGR_HUMAN Polymeric immunoglobulin receptor OS=Homo sapiens OX=9606 GN=PIGR PE=1 SV=4 | 902 | 20201119_10132_1ng_glyco_hcd30_etd30_compound15.19501.19501.3 | controllerType=0 controllerNumber=1 scan=19501 | 45.2731 |
| 16984:4 | 6 | R.LSLLEEPGN[+2205.793]GTFTVILNQLTSR.D | HexNAc(4)Hex(5)Fuc(2)NeuAc(1) | N[+2206] | 1152.78 | 4 | 4608.098 | 4608.0854 | 0 | 2.7 | 413 | Specific | 380.58 | 328.48 | 328.48 | 5.68 | 17 | >sp\|P01833\|PIGR_HUMAN Polymeric immunoglobulin receptor OS=Homo sapiens OX=9606 GN=PIGR PE=1 SV=4 | 902 | 20201119_10132_1ng_glyco_hcd30_etd30_compound15.20726.20726.3 | controllerType=0 controllerNumber=1 scan=20726 | 48.7464 |
| 17001:4 | 6 | R.LSLLEEPGN[+2205.793]GTFTVILNQLTSR.D | HexNAc(4)Hex(5)Fuc(2)NeuAc(1) | N[+2206] | 1152.78 | 4 | 4608.098 | 4608.0854 | 0 | 2.7 | 413 | Specific | 408.17 | 338.98 | 338.98 | 5.21 | 17 | >sp\|P01833\|PIGR_HUMAN Polymeric immunoglobulin receptor OS=Homo sapiens OX=9606 GN=PIGR PE=1 SV=4 | 902 | 20201119_10132_1ng_glyco_hcd30_etd30_compound15.20744.20744.4 | controllerType=0 controllerNumber=1 scan=20744 | 48.8105 |
| 18860:4 | 6 | R.LSLLEEPGN[+2350.830]GTFTVILNQLTSR.D | HexNAc(4)Hex(5)Fuc(1)NeuAc(2) | N[+2351] | 1189.0447 | 4 | 4753.1571 | 4753.1229 | 0 | 7.2 | 413 | Specific | 427.12 | 337.09 | 337.09 | 4.94 | 17 | >sp\|P01833\|PIGR_HUMAN Polymeric immunoglobulin receptor OS=Homo sapiens OX=9606 GN=PIGR PE=1 SV=4 | 902 | 20201119_10132_1ng_glyco_hcd30_etd30_compound15.22696.22696.4 | controllerType=0 controllerNumber=1 scan=22696 | 53.3986 |
| 16997:3 | 6 | R.LSLLEEPGN[+2205.793]GTFTVILNQLTSR.D | HexNAc(4)Hex(5)Fuc(2)NeuAc(1) | N[+2206] | 1536.7057 | 3 | 4608.1027 | 4608.0854 | 0 | 3.7 | 413 | Specific | 605.12 | 484.68 | 484.68 | 9.16 | 17 | >sp\|P01833\|PIGR_HUMAN Polymeric immunoglobulin receptor OS=Homo sapiens OX=9606 GN=PIGR PE=1 SV=4 | 902 | 20201119_10132_1ng_glyco_hcd30_etd30_compound15.20740.20740.3 | controllerType=0 controllerNumber=1 scan=20740 | 48.7995 |
| 13811:3 | 6 | K.IIEGEPNLKVPGN[+2205.793]VTAVLGETLK.V | HexNAc(4)Hex(5)Fuc(2)NeuAc(1) | N[+2206] | 1533.0605 | 3 | 4597.1669 | 4597.1422 | 0 | 5.4 | 457 | Specific | 395.9 | 294.71 | 294.71 | 3.23 | 17 | >sp\|P01833\|PIGR_HUMAN Polymeric immunoglobulin receptor OS=Homo sapiens OX=9606 GN=PIGR PE=1 SV=4 | 902 | 20201119_10132_1ng_glyco_hcd30_etd30_compound15.17367.17367.3 | controllerType=0 controllerNumber=1 scan=17367 | 39.1525 |
| 13894:3 | 6 | K.IIEGEPNLKVPGN[+2059.735]VTAVLGETLK.V | HexNAc(4)Hex(5)Fuc(1)NeuAc(1) | N[+2060] | 1484.3714 | 3 | 4451.0996 | 4451.0843 | 0 | 3.4 | 457 | Specific | 403.94 | 307.33 | 307.33 | 4.08 | 17 | >sp\|P01833\|PIGR_HUMAN Polymeric immunoglobulin receptor OS=Homo sapiens OX=9606 GN=PIGR PE=1 SV=4 | 902 | 20201119_10132_1ng_glyco_hcd30_etd30_compound15.17455.17455.3 | controllerType=0 controllerNumber=1 scan=17455 | 39.3286 |
| 13453:4 | 6 | K.IIEGEPNLKVPGN[+2059.735]VTAVLGETLKVPC[+57.021]HFPC[+57.021]K.F | HexNAc(4)Hex(5)Fuc(1)NeuAc(1) | C[+57]*2, N[+2060] | 1369.9034 | 4 | 5476.592 | 5476.5418 | 0 | 9.2 | 457 | Specific | 330.84 | 330.84 | 330.84 | 2.76 | 17 | >sp\|P01833\|PIGR_HUMAN Polymeric immunoglobulin receptor OS=Homo sapiens OX=9606 GN=PIGR PE=1 SV=4 | 902 | 20201119_10132_1ng_glyco_hcd30_etd30_compound15.16955.16955.4 | controllerType=0 controllerNumber=1 scan=16955 | 38.3246 |
| 13427:4 | 6 | K.IIEGEPNLKVPGN[+2205.793]VTAVLGETLKVPC[+57.021]HFPC[+57.021]K.F | HexNAc(4)Hex(5)Fuc(2)NeuAc(1) | C[+57]*2, N[+2206] | 1406.4138 | 4 | 5622.6335 | 5622.5997 | 0 | 6 | 457 | Specific | 492.72 | 492.72 | 492.72 | 5.4 | 17 | >sp\|P01833\|PIGR_HUMAN Polymeric immunoglobulin receptor OS=Homo sapiens OX=9606 GN=PIGR PE=1 SV=4 | 902 | 20201119_10132_1ng_glyco_hcd30_etd30_compound15.16928.16928.4 | controllerType=0 controllerNumber=1 scan=16928 | 38.2517 |
| 13449:5 | 6 | K.IIEGEPNLKVPGN[+2059.735]VTAVLGETLKVPC[+57.021]HFPC[+57.021]K.F | HexNAc(4)Hex(5)Fuc(1)NeuAc(1) | C[+57]*2, N[+2060] | 1096.1227 | 5 | 5476.5843 | 5476.5418 | 0 | 7.8 | 457 | Specific | 510.59 | 467.37 | 467.37 | 4.72 | 17 | >sp\|P01833\|PIGR_HUMAN Polymeric immunoglobulin receptor OS=Homo sapiens OX=9606 GN=PIGR PE=1 SV=4 | 902 | 20201119_10132_1ng_glyco_hcd30_etd30_compound15.16951.16951.5 | controllerType=0 controllerNumber=1 scan=16951 | 38.3076 |
| 11743:3 | 6 | K.VPGN[+2205.793]VTAVLGETLK.V | HexNAc(4)Hex(5)Fuc(2)NeuAc(1) | N[+2206] | 1201.8724 | 3 | 3603.6026 | 3603.5927 | 0 | 2.7 | 466 | Specific | 325.33 | 232.22 | 232.22 | 5.34 | 17 | >sp\|P01833\|PIGR_HUMAN Polymeric immunoglobulin receptor OS=Homo sapiens OX=9606 GN=PIGR PE=1 SV=4 | 902 | 20201119_10132_1ng_glyco_hcd30_etd30_compound15.15166.15166.3 | controllerType=0 controllerNumber=1 scan=15166 | 34.4125 |
| 11832:4 | 6 | K.VPGN[+2059.735]VTAVLGETLKVPC[+57.021]HFPC[+57.021]K.F | HexNAc(4)Hex(5)Fuc(1)NeuAc(1) | C[+57]*2, N[+2060] | 1121.5115 | 4 | 4483.0242 | 4482.9923 | 0 | 7.1 | 466 | Specific | 343.75 | 316.45 | 316.45 | 3.19 | 17 | >sp\|P01833\|PIGR_HUMAN Polymeric immunoglobulin receptor OS=Homo sapiens OX=9606 GN=PIGR PE=1 SV=4 | 902 | 20201119_10132_1ng_glyco_hcd30_etd30_compound15.15259.15259.4 | controllerType=0 controllerNumber=1 scan=15259 | 34.6317 |
| 04919:3 | 6 | K.WN[+1913.677]NTGC[+57.021]QALPSQDEGPSK.A | HexNAc(4)Hex(5)NeuAc(1) | C[+57], N[+1914] | 1301.5259 | 3 | 3902.5633 | 3902.5537 | 0 | 2.4 | 498 | Specific | 360.16 | 230.1 | 230.1 | 5.65 | 17 | >sp\|P01833\|PIGR_HUMAN Polymeric immunoglobulin receptor OS=Homo sapiens OX=9606 GN=PIGR PE=1 SV=4 | 902 | 20201119_10132_1ng_glyco_hcd30_etd30_compound15.8021.8021.3 | controllerType=0 controllerNumber=1 scan=8021 | 20.7883 |
| 04844:3 | 6 | K.WN[+2059.735]NTGC[+57.021]QALPSQDEGPSK.A | HexNAc(4)Hex(5)Fuc(1)NeuAc(1) | C[+57], N[+2060] | 1350.2148 | 3 | 4048.6298 | 4048.6116 | 0 | 4.5 | 498 | Specific | 395.89 | 266.94 | 266.94 | 5.55 | 17 | >sp\|P01833\|PIGR_HUMAN Polymeric immunoglobulin receptor OS=Homo sapiens OX=9606 GN=PIGR PE=1 SV=4 | 902 | 20201119_10132_1ng_glyco_hcd30_etd30_compound15.7943.7943.3 | controllerType=0 controllerNumber=1 scan=7943 | 20.6409 |
| 18230:4 | 7 | K.VFPLSLDSTPQDGNVVVAC[+57.021]LVQGFFPQEPLSVTWSESGQN[+1647.613]VTAI | HexNAc(5)Hex(3)Fuc(1) | C[+57], N[+1648] | 1606.7529 | 4 | 6423.9899 | 6423.9746 | 0 | 2.4 | 8 | Specific | 267.62 | 267.62 | 267.62 | 4.48 | 17 | >sp\|P01877\|IGHA2_HUMAN Immunoglobulin heavy constant alpha 2 OS=Homo sapiens OX=9606 GN=IGHA2 PE=1 SV=4 | 576 | 20201119_10132_1ng_glyco_hcd30_etd30_compound15.22032.22032.4 | controllerType=0 controllerNumber=1 scan=22032 | 51.9301 |
| 00843:3 | 7 | K.HYTN[+1809.666]SSQDVTVPC[+57.021]R.V | HexNAc(5)Hex(4)Fuc(1) | C[+57], N[+1810] | 1158.4798 | 3 | 3473.4248 | 3473.4154 | 0 | 2.7 | 89 | Specific | 402.71 | 321.19 | 321.19 | 6.08 | 17 | >sp\|P01877\|IGHA2_HUMAN Immunoglobulin heavy constant alpha 2 OS=Homo sapiens OX=9606 GN=IGHA2 PE=1 SV=4 | 576 | 20201119_10132_1ng_glyco_hcd30_etd30_compound15.3782.3782.3 | controllerType=0 controllerNumber=1 scan=3782 | 12.8762 |
| 00863:3 | 7 | K.HYTN[+1647.613]SSQDVTVPC[+57.021]R.V | HexNAc(5)Hex(3)Fuc(1) | C[+57], N[+1648] | 1104.4613 | 3 | 3311.3694 | 3311.3625 | 0 | 2.1 | 89 | Specific | 493.85 | 426.6 | 426.6 | 7.65 | 17 | >sp\|P01877\|IGHA2_HUMAN Immunoglobulin heavy constant alpha 2 OS=Homo sapiens OX=9606 GN=IGHA2 PE=1 SV=4 | 576 | 20201119_10132_1ng_glyco_hcd30_etd30_compound15.3803.3803.3 | controllerType=0 controllerNumber=1 scan=3803 | 12.9381 |
| 02351:3 | 7 | K.TPLTAN[+1971.719]ITK.S | HexNAc(5)Hex(5)Fuc(1) | N[+1972] | 977.4333 | 3 | 2930.2854 | 2930.2757 | 0 | 3.3 | 200 | Specific | 221.05 | 126.04 | 126.04 | 2.65 | 17 | >sp\|P01877\|IGHA2_HUMAN Immunoglobulin heavy constant alpha 2 OS=Homo sapiens OX=9606 GN=IGHA2 PE=1 SV=4 | 576 | 20201119_10132_1ng_glyco_hcd30_etd30_compound15.5353.5353.3 | controllerType=0 controllerNumber=1 scan=5353 | 15.8353 |
| 02438:3 | 7 | K.TPLTAN[+1647.613]ITK.S | HexNAc(5)Hex(3)Fuc(1) | N[+1648] | 869.3979 | 3 | 2606.179 | 2606.17 | 0 | 3.5 | 200 | Specific | 253.69 | 127.16 | 127.16 | 2.63 | 17 | >sp\|P01877\|IGHA2_HUMAN Immunoglobulin heavy constant alpha 2 OS=Homo sapiens OX=9606 GN=IGHA2 PE=1 SV=4 | 576 | 20201119_10132_1ng_glyco_hcd30_etd30_compound15.5444.5444.3 | controllerType=0 controllerNumber=1 scan=5444 | 16.0495 |
| 02390:3 | 7 | K.TPLTAN[+1809.666]ITK.S | HexNAc(5)Hex(4)Fuc(1) | N[+1810] | 923.416 | 3 | 2768.2336 | 2768.2228 | 0 | 3.9 | 200 | Specific | 281.84 | 139.26 | 139.26 | 2.78 | 17 | >sp\|P01877\|IGHA2_HUMAN Immunoglobulin heavy constant alpha 2 OS=Homo sapiens OX=9606 GN=IGHA2 PE=1 SV=4 | 576 | 20201119_10132_1ng_glyco_hcd30_etd30_compound15.5394.5394.3 | controllerType=0 controllerNumber=1 scan=5394 | 15.9266 |
| 02382:2 | 7 | K.TPLTAN[+1809.666]ITK.S | HexNAc(5)Hex(4)Fuc(1) | N[+1810] | 1384.6198 | 2 | 2768.2323 | 2768.2228 | 0 | 3.4 | 200 | Specific | 299.92 | 114.69 | 114.69 | 2.83 | 17 | >sp\|P01877\|IGHA2_HUMAN Immunoglobulin heavy constant alpha 2 OS=Homo sapiens OX=9606 GN=IGHA2 PE=1 SV=4 | 576 | 20201119_10132_1ng_glyco_hcd30_etd30_compound15.5385.5385.2 | controllerType=0 controllerNumber=1 scan=5385 | 15.9034 |
| 02435:2 | 7 | K.TPLTAN[+1647.613]ITK.S | HexNAc(5)Hex(3)Fuc(1) | N[+1648] | 1303.5927 | 2 | 2606.1781 | 2606.17 | 0 | 3.1 | 200 | Specific | 313.5 | 111.38 | 111.38 | 2.84 | 17 | >sp\|P01877\|IGHA2_HUMAN Immunoglobulin heavy constant alpha 2 OS=Homo sapiens OX=9606 GN=IGHA2 PE=1 SV=4 | 576 | 20201119_10132_1ng_glyco_hcd30_etd30_compound15.5441.5441.2 | controllerType=0 controllerNumber=1 scan=5441 | 16.0403 |
| 02443:2 | 7 | K.TPLTAN[+1501.555]ITK.S | HexNAc(5)Hex(3) | N[+1502] | 1230.564 | 2 | 2460.1207 | 2460.1121 | 0 | 3.5 | 200 | Specific | 316.09 | 106.2 | 106.2 | 2.59 | 17 | >sp\|P01877\|IGHA2_HUMAN Immunoglobulin heavy constant alpha 2 OS=Homo sapiens OX=9606 GN=IGHA2 PE=1 SV=4 | 576 | 20201119_10132_1ng_glyco_hcd30_etd30_compound15.5449.5449.2 | controllerType=0 controllerNumber=1 scan=5449 | 16.0629 |
| 02398:2 | 7 | K.TPLTAN[+1444.534]ITK.S | HexNAc(4)Hex(3)Fuc(1) | N[+1445] | 1202.0539 | 2 | 2403.1005 | 2403.0906 | 0 | 4.1 | 200 | Specific | 330.68 | 102.54 | 102.54 | 2.49 | 17 | >sp\|P01877\|IGHA2_HUMAN Immunoglobulin heavy constant alpha 2 OS=Homo sapiens OX=9606 GN=IGHA2 PE=1 SV=4 | 576 | 20201119_10132_1ng_glyco_hcd30_etd30_compound15.5402.5402.2 | controllerType=0 controllerNumber=1 scan=5402 | 15.9446 |
| 08933:3 | 7 | R.MAGKPTHIN[+1702.581]VSVVMAEADGTC[+57.021]Y.- | HexNAc(2)Hex(8) | C[+57], N[+1703] | 1351.8947 | 3 | 4053.6696 | 4053.6642 | 0 | 1.3 | 319 | CRagged | 212.7 | 212.7 | 152.54 | 3.7 | 17 | >sp\|P01877\|IGHA2_HUMAN Immunoglobulin heavy constant alpha 2 OS=Homo sapiens OX=9606 GN=IGHA2 PE=1 SV=4 | 576 | 20201119_10132_1ng_glyco_hcd30_etd30_compound15.12213.12213.3 | controllerType=0 controllerNumber=1 scan=12213 | 28.5767 |
| 09092:3 | 7 | R.MAGKPTHIN[+1216.423]VSVVMAEADGTC[+57.021]Y.- | HexNAc(2)Hex(5) | C[+57], N[+1216] | 1189.842 | 3 | 3567.5114 | 3567.5057 | 0 | 1.6 | 319 | CRagged | 261.94 | 261.94 | 185.03 | 4.17 | 17 | >sp\|P01877\|IGHA2_HUMAN Immunoglobulin heavy constant alpha 2 OS=Homo sapiens OX=9606 GN=IGHA2 PE=1 SV=4 | 576 | 20201119_10132_1ng_glyco_hcd30_etd30_compound15.12379.12379.3 | controllerType=0 controllerNumber=1 scan=12379 | 28.9327 |
| 08999:3 | 7 | R.MAGKPTHIN[+1378.476]VSVVMAEADGTC[+57.021]Y.- | HexNAc(2)Hex(6) | C[+57], N[+1378] | 1243.8614 | 3 | 3729.5696 | 3729.5585 | 0 | 3 | 319 | CRagged | 275.62 | 241.73 | 181.86 | 4.13 | 17 | >sp\|P01877\|IGHA2_HUMAN Immunoglobulin heavy constant alpha 2 OS=Homo sapiens OX=9606 GN=IGHA2 PE=1 SV=4 | 576 | 20201119_10132_1ng_glyco_hcd30_etd30_compound15.12282.12282.3 | controllerType=0 controllerNumber=1 scan=12282 | 28.7306 |
| 07285:3 | 9 | K.TAHC[+57.021]N[+2205.793]ESFYFLC[+57.021]K.R | HexNAc(4)Hex(5)Fuc(2)NeuAc(1) | C[+57]*2, N[+2206] | 1294.8459 | 3 | 3882.5231 | 3882.5124 | 0 | 2.7 | 1201 | Specific | 208.86 | 141.16 | 141.16 | 3.52 | 7 | >sp\|P22897\|MRC1_HUMAN Macrophage mannose receptor 1 OS=Homo sapiens OX=9606 GN=MRC1 PE=1 SV=1 | 1049 | 20201119_10132_1ng_glyco_hcd30_etd30_compound15.10481.10481.3 | controllerType=0 controllerNumber=1 scan=10481 | 25.2596 |
| 07400:3 | 9 | K.TAHC[+57.021]N[+2059.735]ESFYFLC[+57.021]K.R | HexNAc(4)Hex(5)Fuc(1)NeuAc(1) | C[+57]*2, N[+2060] | 1246.16 | 3 | 3736.4654 | 3736.4545 | 0 | 2.9 | 1201 | Specific | 213.01 | 111.52 | 111.52 | 3.65 | 7 | >sp\|P22897\|MRC1_HUMAN Macrophage mannose receptor 1 OS=Homo sapiens OX=9606 GN=MRC1 PE=1 SV=1 | 1049 | 20201119_10132_1ng_glyco_hcd30_etd30_compound15.10600.10600.3 | controllerType=0 controllerNumber=1 scan=10600 | 25.4735 |
| 07383:3 | 9 | K.TAHC[+57.021]N[+2059.735]ESFYFLC[+57.021]K.R | HexNAc(4)Hex(5)Fuc(1)NeuAc(1) | C[+57]*2, N[+2060] | 1246.1597 | 3 | 3736.4646 | 3736.4545 | 0 | 2.7 | 1201 | Specific | 290.84 | 147.4 | 147.4 | 4.05 | 7 | >sp\|P22897\|MRC1_HUMAN Macrophage mannose receptor 1 OS=Homo sapiens OX=9606 GN=MRC1 PE=1 SV=1 | 1049 | 20201119_10132_1ng_glyco_hcd30_etd30_compound15.10582.10582.3 | controllerType=0 controllerNumber=1 scan=10582 | 25.4371 |
| 06706:3 | 13 | R.LSPN[+2350.830]ASAEHLELR.W | HexNAc(4)Hex(5)Fuc(1)NeuAc(2) | N[+2351] | 1263.2006 | 3 | 3787.5873 | 3787.5796 | 0 | 2 | 52 | Specific | 358.4 | 230.89 | 230.89 | 4.59 | 9 | >sp\|Q13410\|BT1A1_HUMAN Butyrophilin subfamily 1 member A1 OS=Homo sapiens OX=9606 GN=BTN1A1 PE=1 SV=3 | 179 | 20201119_10132_1ng_glyco_hcd30_etd30_compound15.9878.9878.3 | controllerType=0 controllerNumber=1 scan=9878 | 24.1447 |
| 04815:3 | 13 | R.LSPN[+2205.793]ASAEHLELR.W | HexNAc(4)Hex(5)Fuc(2)NeuAc(1) | N[+2206] | 1214.856 | 3 | 3642.5535 | 3642.5421 | 0 | 3.1 | 52 | Specific | 360.33 | 246.43 | 246.43 | 3.82 | 9 | >sp\|Q13410\|BT1A1_HUMAN Butyrophilin subfamily 1 member A1 OS=Homo sapiens OX=9606 GN=BTN1A1 PE=1 SV=3 | 179 | 20201119_10132_1ng_glyco_hcd30_etd30_compound15.7913.7913.3 | controllerType=0 controllerNumber=1 scan=7913 | 20.5804 |
| 09954:3 | 27 | R.IIVPLNNREN[+1694.603]ISDPTSPLR.T | HexNAc(3)Hex(4)Fuc(1)NeuAc(1) | N[+1695] | 1281.603 | 3 | 3842.7943 | 3842.7799 | 0 | 3.8 | 62 | Specific | 222.17 | 180.88 | 180.88 | 3.23 | 8 | >sp\|P01591\|IGJ_HUMAN Immunoglobulin J chain OS=Homo sapiens OX=9606 GN=JCHAIN PE=1 SV=4 | 1392 | 20201119_10132_1ng_glyco_hcd30_etd30_compound15.13275.13275.3 | controllerType=0 controllerNumber=1 scan=13275 | 30.742 |
| 08392:3 | 27 | R.IIVPLNNREN[+1663.608]ISDPTSPLR.T | HexNAc(5)Hex(4) | N[+1664] | 1271.2706 | 3 | 3811.7971 | 3811.7853 | 0 | 3.1 | 62 | Specific | 269.08 | 185.26 | 185.26 | 3.75 | 8 | >sp\|P01591\|IGJ_HUMAN Immunoglobulin J chain OS=Homo sapiens OX=9606 GN=JCHAIN PE=1 SV=4 | 1392 | 20201119_10132_1ng_glyco_hcd30_etd30_compound15.11652.11652.3 | controllerType=0 controllerNumber=1 scan=11652 | 27.5593 |
| 09810:3 | 27 | R.IIVPLNNREN[+2059.735]ISDPTSPLR.T | HexNAc(4)Hex(5)Fuc(1)NeuAc(1) | N[+2060] | 1403.3096 | 3 | 4207.9143 | 4207.9121 | 0 | 0.5 | 62 | Specific | 279.69 | 220.87 | 220.87 | 3.86 | 8 | >sp\|P01591\|IGJ_HUMAN Immunoglobulin J chain OS=Homo sapiens OX=9606 GN=JCHAIN PE=1 SV=4 | 1392 | 20201119_10132_1ng_glyco_hcd30_etd30_compound15.13124.13124.3 | controllerType=0 controllerNumber=1 scan=13124 | 30.3288 |
| 08351:3 | 27 | R.IIVPLNNREN[+1809.666]ISDPTSPLR.T | HexNAc(5)Hex(4)Fuc(1) | N[+1810] | 1319.9566 | 3 | 3957.8552 | 3957.8432 | 0 | 3 | 62 | Specific | 286.33 | 209.59 | 209.59 | 3.55 | 8 | >sp\|P01591\|IGJ_HUMAN Immunoglobulin J chain OS=Homo sapiens OX=9606 GN=JCHAIN PE=1 SV=4 | 1392 | 20201119_10132_1ng_glyco_hcd30_etd30_compound15.11609.11609.3 | controllerType=0 controllerNumber=1 scan=11609 | 27.4589 |
| 09852:3 | 27 | R.IIVPLNNREN[+1913.677]ISDPTSPLR.T | HexNAc(4)Hex(5)NeuAc(1) | N[+1914] | 1354.6269 | 3 | 4061.8661 | 4061.8542 | 0 | 2.9 | 62 | Specific | 308.45 | 232.79 | 232.79 | 3.86 | 8 | >sp\|P01591\|IGJ_HUMAN Immunoglobulin J chain OS=Homo sapiens OX=9606 GN=JCHAIN PE=1 SV=4 | 1392 | 20201119_10132_1ng_glyco_hcd30_etd30_compound15.13168.13168.3 | controllerType=0 controllerNumber=1 scan=13168 | 30.4283 |
| 08368:3 | 27 | R.IIVPLNNREN[+1809.666]ISDPTSPLR.T | HexNAc(5)Hex(4)Fuc(1) | N[+1810] | 1319.9571 | 3 | 3957.8567 | 3957.8432 | 0 | 3.4 | 62 | Specific | 318.07 | 231.77 | 231.77 | 3.5 | 8 | >sp\|P01591\|IGJ_HUMAN Immunoglobulin J chain OS=Homo sapiens OX=9606 GN=JCHAIN PE=1 SV=4 | 1392 | 20201119_10132_1ng_glyco_hcd30_etd30_compound15.11627.11627.3 | controllerType=0 controllerNumber=1 scan=11627 | 27.5016 |
| 09820:3 | 27 | R.IIVPLNNREN[+1913.677]ISDPTSPLR.T | HexNAc(4)Hex(5)NeuAc(1) | N[+1914] | 1354.6253 | 3 | 4061.8614 | 4061.8542 | 0 | 1.8 | 62 | Specific | 322.91 | 213.93 | 213.93 | 3.72 | 8 | >sp\|P01591\|IGJ_HUMAN Immunoglobulin J chain OS=Homo sapiens OX=9606 GN=JCHAIN PE=1 SV=4 | 1392 | 20201119_10132_1ng_glyco_hcd30_etd30_compound15.13134.13134.3 | controllerType=0 controllerNumber=1 scan=13134 | 30.3542 |
| 12134:4 | 29 | K.LKELPGVC[+57.021]N[+2204.772]ETMMALWEEC[+57.021]KPC[+57.021]LK.Q | HexNAc(4)Hex(5)NeuAc(2) | C[+57]*3, N[+2205] | 1286.0518 | 4 | 5141.1852 | 5141.1572 | 0 | 5.5 | 95 | Specific | 303.31 | 204.56 | 204.56 | 2.65 | 8 | >sp\|P10909\|CLUS_HUMAN Clusterin OS=Homo sapiens OX=9606 GN=CLU PE=1 SV=1 | 1467 | 20201119_10132_1ng_glyco_hcd30_etd30_compound15.15573.15573.4 | controllerType=0 controllerNumber=1 scan=15573 | 35.2666 |
| 12166:4 | 29 | K.LKELPGVC[+57.021]N[+2205.793]ETMMALWEEC[+57.021]KPC[+57.021]LK.Q | HexNAc(4)Hex(5)Fuc(2)NeuAc(1) | C[+57]*3, N[+2206] | 1286.3049 | 4 | 5142.1979 | 5142.1776 | 0 | 4 | 95 | Specific | 452.55 | 306.62 | 306.62 | 3.86 | 8 | >sp\|P10909\|CLUS_HUMAN Clusterin OS=Homo sapiens OX=9606 GN=CLU PE=1 SV=1 | 1467 | 20201119_10132_1ng_glyco_hcd30_etd30_compound15.15607.15607.4 | controllerType=0 controllerNumber=1 scan=15607 | 35.345 |
| 00076:4 | 29 | R.EIRHN[+2424.867]STGC[+57.021]LR.M | HexNAc(5)Hex(6)Fuc(1)NeuAc(1) | C[+57], N[+2425] | 942.6435 | 4 | 3767.5521 | 3767.5316 | 0 | 5.4 | 287 | Specific | 284.98 | 182.65 | 182.65 | 3.21 | 8 | >sp\|P10909\|CLUS_HUMAN Clusterin OS=Homo sapiens OX=9606 GN=CLU PE=1 SV=1 | 1467 | 20201119_10132_1ng_glyco_hcd30_etd30_compound15.2982.2982.2 | controllerType=0 controllerNumber=1 scan=2982 | 11.3023 |
| 00070:4 | 29 | R.EIRHN[+2059.735]STGC[+57.021]LR.M | HexNAc(4)Hex(5)Fuc(1)NeuAc(1) | C[+57], N[+2060] | 851.3589 | 4 | 3402.4138 | 3402.3994 | 0 | 4.2 | 287 | Specific | 449.79 | 336.18 | 336.18 | 4.94 | 8 | >sp\|P10909\|CLUS_HUMAN Clusterin OS=Homo sapiens OX=9606 GN=CLU PE=1 SV=1 | 1467 | 20201119_10132_1ng_glyco_hcd30_etd30_compound15.2976.2976.4 | controllerType=0 controllerNumber=1 scan=2976 | 11.281 |
| 19283:3 | 29 | K.MLN[+2350.830]TSSLLEQLNEQFNWVSR.L | HexNAc(4)Hex(5)Fuc(1)NeuAc(2) | N[+2351] | 1587.3532 | 3 | 4760.0451 | 4760.0171 | 0 | 5.9 | 352 | Specific | 332.04 | 213.13 | 213.13 | 4.29 | 8 | >sp\|P10909\|CLUS_HUMAN Clusterin OS=Homo sapiens OX=9606 GN=CLU PE=1 SV=1 | 1467 | 20201119_10132_1ng_glyco_hcd30_etd30_compound15.23140.23140.3 | controllerType=0 controllerNumber=1 scan=23140 | 54.4458 |
| 17325:3 | 29 | K.MLN[+2059.735]TSSLLEQLNEQFNWVSR.L | HexNAc(4)Hex(5)Fuc(1)NeuAc(1) | N[+2060] | 1490.3168 | 3 | 4468.9359 | 4468.9217 | 0 | 3.2 | 352 | Specific | 430.83 | 277.73 | 277.73 | 5.63 | 8 | >sp\|P10909\|CLUS_HUMAN Clusterin OS=Homo sapiens OX=9606 GN=CLU PE=1 SV=1 | 1467 | 20201119_10132_1ng_glyco_hcd30_etd30_compound15.21087.21087.3 | controllerType=0 controllerNumber=1 scan=21087 | 49.8427 |
| 00704:4 | 56 | K.TKPREEQFN[+1444.534]STFR.V | HexNAc(4)Hex(3)Fuc(1) | N[+1445] | 771.8461 | 4 | 3084.3625 | 3084.3526 | 0 | 3.2 | 168 | Specific | 462.7 | 412.23 | 412.23 | 5.21 | 3 | >sp\|P01859\|IGHG2_HUMAN Immunoglobulin heavy constant gamma 2 OS=Homo sapiens OX=9606 GN=IGHG2 PE=1 SV=2 | 387 | 20201119_10132_1ng_glyco_hcd30_etd30_compound15.3637.3637.4 | controllerType=0 controllerNumber=1 scan=3637 | 12.5787 |
| 00684:4 | 56 | K.TKPREEQFN[+1606.587]STFR.V | HexNAc(4)Hex(4)Fuc(1) | N[+1607] | 812.3606 | 4 | 3246.4204 | 3246.4054 | 0 | 4.6 | 168 | Specific | 465.47 | 428.64 | 428.64 | 4.36 | 3 | >sp\|P01859\|IGHG2_HUMAN Immunoglobulin heavy constant gamma 2 OS=Homo sapiens OX=9606 GN=IGHG2 PE=1 SV=2 | 387 | 20201119_10132_1ng_glyco_hcd30_etd30_compound15.3616.3616.4 | controllerType=0 controllerNumber=1 scan=3616 | 12.5258 |
| 12449:3 | 79 | K.VVLHPN[+2204.772]YSQVDIGLIK.L | HexNAc(4)Hex(5)NeuAc(2) | N[+2205] | 1333.9428 | 3 | 3999.8139 | 3999.7837 | 0 | 7.5 | 236 | Specific | 220.14 | 160.49 | 160.49 | 2.77 | 1 | >sp\|P00738\|HPT_HUMAN Haptoglobin OS=Homo sapiens OX=9606 GN=HP PE=1 SV=1 | 407 | 20201119_10132_1ng_glyco_hcd30_etd30_compound15.15904.15904.3 | controllerType=0 controllerNumber=1 scan=15904 | 36.0028 |
| 00073:4 | 80 | K.TKPREEQYN[+1606.587]STYR.V | HexNAc(4)Hex(4)Fuc(1) | N[+1607] | 820.3574 | 4 | 3278.4079 | 3278.3952 | 0 | 3.9 | 291 | Specific | 417.94 | 56.82 | 56.82 | 2.59 | 1 | >sp\|P0DOX5\|IGG1_HUMAN Immunoglobulin gamma-1 heavy chain OS=Homo sapiens OX=9606 PE=1 SV=2 | 914 | 20201119_10132_1ng_glyco_hcd30_etd30_compound15.2979.2979.4 | controllerType=0 controllerNumber=1 scan=2979 | 11.2917 |
| 16850:4 | 1 | R.[+42.011]RTAGWNVPIGTLRPFLN[+2336.851]WTGPPEPIEAAVAR.F | HexNAc(6)Hex(6)Fuc(1) | .[42], N[+2337] | 1442.1692 | 4 | 5765.6549 | 5765.6737 | 0 | -3.3 | 140 | Specific | 254.04 | 172.84 | 172.84 | 2.56 | 55 | >sp\|P02788\|TRFL_HUMAN Lactotransferrin OS=Homo sapiens OX=9606 GN=LTF PE=1 SV=6 | 1270 | 20201112_10613_1ng_glyco_hcd30_etd30_compound15.20226.20226.4 | controllerType=0 controllerNumber=1 scan=20226 | 45.9958 |
| 16142:4 | 1 | R.RTAGWNVPIGTLRPFLN[+2205.793]WTGPPEPIEAAVAR.F | HexNAc(4)Hex(5)Fuc(2)NeuAc(1) | N[+2206] | 1398.9075 | 4 | 5592.6081 | 5592.6049 | 0 | 0.6 | 140 | Specific | 657.75 | 642.79 | 642.79 | 10.37 | 55 | >sp\|P02788\|TRFL_HUMAN Lactotransferrin OS=Homo sapiens OX=9606 GN=LTF PE=1 SV=6 | 1270 | 20201112_10613_1ng_glyco_hcd30_etd30_compound15.19477.19477.4 | controllerType=0 controllerNumber=1 scan=19477 | 43.6956 |
| 16168:5 | 1 | R.RTAGWNVPIGTLRPFLN[+2059.735]WTGPPEPIEAAVAR.F | HexNAc(4)Hex(5)Fuc(1)NeuAc(1) | N[+2060] | 1090.1172 | 5 | 5446.5568 | 5446.547 | 0 | 1.8 | 140 | Specific | 781.56 | 781.56 | 781.56 | 12.47 | 55 | >sp\|P02788\|TRFL_HUMAN Lactotransferrin OS=Homo sapiens OX=9606 GN=LTF PE=1 SV=6 | 1270 | 20201112_10613_1ng_glyco_hcd30_etd30_compound15.19504.19504.5 | controllerType=0 controllerNumber=1 scan=19504 | 43.7733 |
| 16051:4 | 1 | R.RTAGWNVPIGTLRPFLN[+2570.925]WTGPPEPIEAAVAR.F | HexNAc(5)Hex(6)Fuc(2)NeuAc(1) | N[+2571] | 1490.1963 | 4 | 5957.7633 | 5957.7371 | 0 | 4.4 | 140 | Specific | 816.92 | 814.28 | 814.28 | 12.53 | 55 | >sp\|P02788\|TRFL_HUMAN Lactotransferrin OS=Homo sapiens OX=9606 GN=LTF PE=1 SV=6 | 1270 | 20201112_10613_1ng_glyco_hcd30_etd30_compound15.19381.19381.4 | controllerType=0 controllerNumber=1 scan=19381 | 43.4459 |
| 16157:4 | 1 | R.RTAGWNVPIGTLRPFLN[+2059.735]WTGPPEPIEAAVAR.F | HexNAc(4)Hex(5)Fuc(1)NeuAc(1) | N[+2060] | 1362.3948 | 4 | 5446.5573 | 5446.547 | 0 | 1.9 | 140 | Specific | 862.1 | 831.22 | 831.22 | 14.52 | 55 | >sp\|P02788\|TRFL_HUMAN Lactotransferrin OS=Homo sapiens OX=9606 GN=LTF PE=1 SV=6 | 1270 | 20201112_10613_1ng_glyco_hcd30_etd30_compound15.19493.19493.4 | controllerType=0 controllerNumber=1 scan=19493 | 43.7525 |
| 15528:4 | 1 | R.RTAGWNVPIGTLRPFLN[+1768.640]WTGPPEPIEAAVAR.F | HexNAc(4)Hex(5)Fuc(1) | N[+1769] | 1289.62 | 4 | 5155.4582 | 5155.4516 | 0 | 1.3 | 140 | Specific | 1154.62 | 1083.42 | 1083.42 | 20.52 | 55 | >sp\|P02788\|TRFL_HUMAN Lactotransferrin OS=Homo sapiens OX=9606 GN=LTF PE=1 SV=6 | 1270 | 20201112_10613_1ng_glyco_hcd30_etd30_compound15.18827.18827.4 | controllerType=0 controllerNumber=1 scan=18827 | 41.6541 |
| 16300:4 | 1 | R.TAGWNVPIGTLRPFLN[+2498.904]WTGPPEPIEAAVAR.F | HexNAc(6)Hex(7)Fuc(1) | N[+2499] | 1433.1639 | 4 | 5729.6339 | 5729.6148 | 0 | 3.3 | 141 | Specific | 247.04 | 247.04 | 247.04 | 3.38 | 55 | >sp\|P02788\|TRFL_HUMAN Lactotransferrin OS=Homo sapiens OX=9606 GN=LTF PE=1 SV=6 | 1270 | 20201112_10613_1ng_glyco_hcd30_etd30_compound15.19642.19642.4 | controllerType=0 controllerNumber=1 scan=19642 | 44.0966 |
| 16957:3 | 1 | R.TAGWNVPIGTLRPFLN[+2424.867]WTGPPEPIEAAVAR.F | HexNAc(5)Hex(6)Fuc(1)NeuAc(1) | N[+2425] | 1885.8673 | 3 | 5655.5873 | 5655.5781 | 0 | 1.6 | 141 | Specific | 289.76 | 225.18 | 225.18 | 4.48 | 55 | >sp\|P02788\|TRFL_HUMAN Lactotransferrin OS=Homo sapiens OX=9606 GN=LTF PE=1 SV=6 | 1270 | 20201112_10613_1ng_glyco_hcd30_etd30_compound15.20338.20338.3 | controllerType=0 controllerNumber=1 scan=20338 | 46.2845 |
| 17927:5 | 1 | R.TAGWNVPIGTLRPFLN[+2350.830]WTGPPEPIEAAVAR.F | HexNAc(4)Hex(5)Fuc(1)NeuAc(2) | N[+2351] | 1117.1169 | 5 | 5581.5556 | 5581.5413 | 0 | 2.6 | 141 | Specific | 308.59 | 308.59 | 308.59 | 3.74 | 55 | >sp\|P02788\|TRFL_HUMAN Lactotransferrin OS=Homo sapiens OX=9606 GN=LTF PE=1 SV=6 | 1270 | 20201112_10613_1ng_glyco_hcd30_etd30_compound15.21367.21367.5 | controllerType=0 controllerNumber=1 scan=21367 | 49.3294 |
| 17150:3 | 1 | R.TAGWNVPIGTLRPFLN[+2059.735]WTGPPEPIEAAVAR.F | HexNAc(4)Hex(5)Fuc(1)NeuAc(1) | N[+2060] | 1764.1521 | 3 | 5290.4416 | 5290.4459 | 0 | -0.8 | 141 | Specific | 358.12 | 227.88 | 227.88 | 3.62 | 55 | >sp\|P02788\|TRFL_HUMAN Lactotransferrin OS=Homo sapiens OX=9606 GN=LTF PE=1 SV=6 | 1270 | 20201112_10613_1ng_glyco_hcd30_etd30_compound15.20545.20545.3 | controllerType=0 controllerNumber=1 scan=20545 | 46.8125 |
| 16343:4 | 1 | R.[+42.011]TAGWNVPIGTLRPFLN[+1872.651]WTGPPEPIEAAVAR.F | HexNAc(3)Hex(6)NeuAc(1) | .[42], N[+1873] | 1287.1107 | 4 | 5145.4211 | 5145.372 | 0 | 9.5 | 141 | Specific | 409.25 | 409.25 | 409.25 | 4.6 | 55 | >sp\|P02788\|TRFL_HUMAN Lactotransferrin OS=Homo sapiens OX=9606 GN=LTF PE=1 SV=6 | 1270 | 20201112_10613_1ng_glyco_hcd30_etd30_compound15.19688.19688.4 | controllerType=0 controllerNumber=1 scan=19688 | 44.2425 |
| 17015:3 | 1 | R.TAGWNVPIGTLRPFLN[+2059.735]WTGPPEPIEAAVAR.F | HexNAc(4)Hex(5)Fuc(1)NeuAc(1) | N[+2060] | 1764.1578 | 3 | 5290.4588 | 5290.4459 | 0 | 2.5 | 141 | Specific | 423.3 | 316.2 | 316.2 | 5.48 | 55 | >sp\|P02788\|TRFL_HUMAN Lactotransferrin OS=Homo sapiens OX=9606 GN=LTF PE=1 SV=6 | 1270 | 20201112_10613_1ng_glyco_hcd30_etd30_compound15.20400.20400.3 | controllerType=0 controllerNumber=1 scan=20400 | 46.5009 |
| 17156:4 | 1 | R.TAGWNVPIGTLRPFLN[+2059.735]WTGPPEPIEAAVAR.F | HexNAc(4)Hex(5)Fuc(1)NeuAc(1) | N[+2060] | 1323.368 | 4 | 5290.4504 | 5290.4459 | 0 | 0.8 | 141 | Specific | 467.56 | 467.56 | 467.56 | 6.2 | 55 | >sp\|P02788\|TRFL_HUMAN Lactotransferrin OS=Homo sapiens OX=9606 GN=LTF PE=1 SV=6 | 1270 | 20201112_10613_1ng_glyco_hcd30_etd30_compound15.20551.20551.4 | controllerType=0 controllerNumber=1 scan=20551 | 46.8311 |
| 16861:4 | 1 | R.TAGWNVPIGTLRPFLN[+2936.057]WTGPPEPIEAAVAR.F | HexNAc(6)Hex(7)Fuc(2)NeuAc(1) | N[+2936] | 1542.4514 | 4 | 6166.7838 | 6166.7682 | 0 | 2.5 | 141 | Specific | 626.69 | 626.69 | 626.69 | 8.08 | 55 | >sp\|P02788\|TRFL_HUMAN Lactotransferrin OS=Homo sapiens OX=9606 GN=LTF PE=1 SV=6 | 1270 | 20201112_10613_1ng_glyco_hcd30_etd30_compound15.20237.20237.4 | controllerType=0 controllerNumber=1 scan=20237 | 46.0239 |
| 16956:4 | 1 | R.TAGWNVPIGTLRPFLN[+2424.867]WTGPPEPIEAAVAR.F | HexNAc(5)Hex(6)Fuc(1)NeuAc(1) | N[+2425] | 1414.6516 | 4 | 5655.5846 | 5655.5781 | 0 | 1.2 | 141 | Specific | 639.34 | 623.12 | 623.12 | 9.58 | 55 | >sp\|P02788\|TRFL_HUMAN Lactotransferrin OS=Homo sapiens OX=9606 GN=LTF PE=1 SV=6 | 1270 | 20201112_10613_1ng_glyco_hcd30_etd30_compound15.20337.20337.4 | controllerType=0 controllerNumber=1 scan=20337 | 46.2798 |
| 17014:4 | 1 | R.TAGWNVPIGTLRPFLN[+2059.735]WTGPPEPIEAAVAR.F | HexNAc(4)Hex(5)Fuc(1)NeuAc(1) | N[+2060] | 1323.3717 | 4 | 5290.465 | 5290.4459 | 0 | 3.6 | 141 | Specific | 693.2 | 664.66 | 664.66 | 9.6 | 55 | >sp\|P02788\|TRFL_HUMAN Lactotransferrin OS=Homo sapiens OX=9606 GN=LTF PE=1 SV=6 | 1270 | 20201112_10613_1ng_glyco_hcd30_etd30_compound15.20399.20399.4 | controllerType=0 controllerNumber=1 scan=20399 | 46.4982 |
| 16981:4 | 1 | R.TAGWNVPIGTLRPFLN[+2351.851]WTGPPEPIEAAVAR.F | HexNAc(4)Hex(5)Fuc(3)NeuAc(1) | N[+2352] | 1396.3972 | 4 | 5582.5671 | 5582.5617 | 0 | 1 | 141 | Specific | 735.42 | 702.19 | 702.19 | 11.58 | 55 | >sp\|P02788\|TRFL_HUMAN Lactotransferrin OS=Homo sapiens OX=9606 GN=LTF PE=1 SV=6 | 1270 | 20201112_10613_1ng_glyco_hcd30_etd30_compound15.20363.20363.4 | controllerType=0 controllerNumber=1 scan=20363 | 46.3685 |
| 16927:4 | 1 | R.TAGWNVPIGTLRPFLN[+2570.925]WTGPPEPIEAAVAR.F | HexNAc(5)Hex(6)Fuc(2)NeuAc(1) | N[+2571] | 1451.1678 | 4 | 5801.6496 | 5801.636 | 0 | 2.3 | 141 | Specific | 737.44 | 737.44 | 737.44 | 11.12 | 55 | >sp\|P02788\|TRFL_HUMAN Lactotransferrin OS=Homo sapiens OX=9606 GN=LTF PE=1 SV=6 | 1270 | 20201112_10613_1ng_glyco_hcd30_etd30_compound15.20307.20307.4 | controllerType=0 controllerNumber=1 scan=20307 | 46.2186 |
| 16369:4 | 1 | R.TAGWNVPIGTLRPFLN[+1768.640]WTGPPEPIEAAVAR.F | HexNAc(4)Hex(5)Fuc(1) | N[+1769] | 1250.5892 | 4 | 4999.3351 | 4999.3504 | 0 | -3.1 | 141 | Specific | 924.67 | 924.67 | 924.67 | 14.54 | 55 | >sp\|P02788\|TRFL_HUMAN Lactotransferrin OS=Homo sapiens OX=9606 GN=LTF PE=1 SV=6 | 1270 | 20201112_10613_1ng_glyco_hcd30_etd30_compound15.19716.19716.4 | controllerType=0 controllerNumber=1 scan=19716 | 44.3617 |
| 15565:3 | 1 | R.TAGWNIPMGLLFN[+1216.423]QTGSC[+57.021]K.F | HexNAc(2)Hex(5) | C[+57], N[+1216] | 1104.4833 | 3 | 3311.4354 | 3311.4328 | 0 | 0.8 | 485 | Specific | 257.63 | 242.4 | 119.89 | 3.93 | 55 | >sp\|P02788\|TRFL_HUMAN Lactotransferrin OS=Homo sapiens OX=9606 GN=LTF PE=1 SV=6 | 1270 | 20201112_10613_1ng_glyco_hcd30_etd30_compound15.18866.18866.3 | controllerType=0 controllerNumber=1 scan=18866 | 41.7572 |
| 17935:4 | 1 | R.TAGWNIPMGLLFN[+2350.830]QTGSC[+57.021]K.F | HexNAc(4)Hex(5)Fuc(1)NeuAc(2) | C[+57], N[+2351] | 1112.2193 | 4 | 4445.8553 | 4445.8403 | 0 | 3.4 | 485 | Specific | 275.67 | 267.96 | 267.96 | 3.25 | 55 | >sp\|P02788\|TRFL_HUMAN Lactotransferrin OS=Homo sapiens OX=9606 GN=LTF PE=1 SV=6 | 1270 | 20201112_10613_1ng_glyco_hcd30_etd30_compound15.21375.21375.4 | controllerType=0 controllerNumber=1 scan=21375 | 49.3488 |
| 16744:3 | 1 | R.TAGWNIPMGLLFN[+1913.677]QTGSC[+57.021]K.F | HexNAc(4)Hex(5)NeuAc(1) | C[+57], N[+1914] | 1336.9039 | 3 | 4008.697 | 4008.687 | 0 | 2.5 | 485 | Specific | 318.64 | 187.9 | 187.9 | 4.73 | 55 | >sp\|P02788\|TRFL_HUMAN Lactotransferrin OS=Homo sapiens OX=9606 GN=LTF PE=1 SV=6 | 1270 | 20201112_10613_1ng_glyco_hcd30_etd30_compound15.20113.20113.3 | controllerType=0 controllerNumber=1 scan=20113 | 45.6062 |
| 18085:3 | 1 | R.TAGWNIPMGLLFN[+2204.772]QTGSC[+57.021]K.F | HexNAc(4)Hex(5)NeuAc(2) | C[+57], N[+2205] | 1433.933 | 3 | 4299.7846 | 4299.7824 | 0 | 0.5 | 485 | Specific | 332.02 | 226.31 | 226.31 | 5.14 | 55 | >sp\|P02788\|TRFL_HUMAN Lactotransferrin OS=Homo sapiens OX=9606 GN=LTF PE=1 SV=6 | 1270 | 20201112_10613_1ng_glyco_hcd30_etd30_compound15.21534.21534.3 | controllerType=0 controllerNumber=1 scan=21534 | 49.8233 |
| 15429:3 | 1 | R.TAGWNIPMGLLFN[+1622.582]QTGSC[+57.021]K.F | HexNAc(4)Hex(5) | C[+57], N[+1623] | 1239.8775 | 3 | 3717.6179 | 3717.5916 | 0 | 7.1 | 485 | Specific | 360.73 | 263.12 | 243.54 | 2.97 | 55 | >sp\|P02788\|TRFL_HUMAN Lactotransferrin OS=Homo sapiens OX=9606 GN=LTF PE=1 SV=6 | 1270 | 20201112_10613_1ng_glyco_hcd30_etd30_compound15.18722.18722.3 | controllerType=0 controllerNumber=1 scan=18722 | 41.3211 |
| 15444:3 | 1 | R.TAGWNIPMGLLFN[+1622.582]QTGSC[+57.021]K.F | HexNAc(4)Hex(5) | C[+57], N[+1623] | 1239.8697 | 3 | 3717.5945 | 3717.5916 | 0 | 0.8 | 485 | Specific | 370.72 | 295.76 | 236.71 | 5.48 | 55 | >sp\|P02788\|TRFL_HUMAN Lactotransferrin OS=Homo sapiens OX=9606 GN=LTF PE=1 SV=6 | 1270 | 20201112_10613_1ng_glyco_hcd30_etd30_compound15.18738.18738.3 | controllerType=0 controllerNumber=1 scan=18738 | 41.3771 |
| 15281:3 | 1 | R.TAGWNIPMGLLFN[+1768.640]QTGSC[+57.021]K.F | HexNAc(4)Hex(5)Fuc(1) | C[+57], N[+1769] | 1288.5574 | 3 | 3863.6577 | 3863.6495 | 0 | 2.1 | 485 | Specific | 381.27 | 288.15 | 288.15 | 5.4 | 55 | >sp\|P02788\|TRFL_HUMAN Lactotransferrin OS=Homo sapiens OX=9606 GN=LTF PE=1 SV=6 | 1270 | 20201112_10613_1ng_glyco_hcd30_etd30_compound15.18566.18566.3 | controllerType=0 controllerNumber=1 scan=18566 | 40.8724 |
| 16547:3 | 1 | R.TAGWNIPMGLLFN[+2205.793]QTGSC[+57.021]K.F | HexNAc(4)Hex(5)Fuc(2)NeuAc(1) | C[+57], N[+2206] | 1434.2758 | 3 | 4300.8129 | 4300.8028 | 0 | 2.3 | 485 | Specific | 384.48 | 285.01 | 285.01 | 4.95 | 55 | >sp\|P02788\|TRFL_HUMAN Lactotransferrin OS=Homo sapiens OX=9606 GN=LTF PE=1 SV=6 | 1270 | 20201112_10613_1ng_glyco_hcd30_etd30_compound15.19903.19903.3 | controllerType=0 controllerNumber=1 scan=19903 | 44.8743 |
| 16556:3 | 1 | R.TAGWNIPMGLLFN[+2059.735]QTGSC[+57.021]K.F | HexNAc(4)Hex(5)Fuc(1)NeuAc(1) | C[+57], N[+2060] | 1385.5881 | 3 | 4154.7497 | 4154.7449 | 0 | 1.2 | 485 | Specific | 406.72 | 271.55 | 271.55 | 5.5 | 55 | >sp\|P02788\|TRFL_HUMAN Lactotransferrin OS=Homo sapiens OX=9606 GN=LTF PE=1 SV=6 | 1270 | 20201112_10613_1ng_glyco_hcd30_etd30_compound15.19913.19913.3 | controllerType=0 controllerNumber=1 scan=19913 | 44.9183 |
| 16663:3 | 1 | R.TAGWNIPMGLLFN[+1913.677]QTGSC[+57.021]K.F | HexNAc(4)Hex(5)NeuAc(1) | C[+57], N[+1914] | 1336.9015 | 3 | 4008.6901 | 4008.687 | 0 | 0.8 | 485 | Specific | 407.12 | 306.21 | 306.21 | 5.57 | 55 | >sp\|P02788\|TRFL_HUMAN Lactotransferrin OS=Homo sapiens OX=9606 GN=LTF PE=1 SV=6 | 1270 | 20201112_10613_1ng_glyco_hcd30_etd30_compound15.20026.20026.3 | controllerType=0 controllerNumber=1 scan=20026 | 45.274 |
| 17926:3 | 1 | R.TAGWNIPMGLLFN[+2350.830]QTGSC[+57.021]K.F | HexNAc(4)Hex(5)Fuc(1)NeuAc(2) | C[+57], N[+2351] | 1482.6231 | 3 | 4445.8548 | 4445.8403 | 0 | 3.3 | 485 | Specific | 417.48 | 197.51 | 197.51 | 4.95 | 55 | >sp\|P02788\|TRFL_HUMAN Lactotransferrin OS=Homo sapiens OX=9606 GN=LTF PE=1 SV=6 | 1270 | 20201112_10613_1ng_glyco_hcd30_etd30_compound15.21366.21366.3 | controllerType=0 controllerNumber=1 scan=21366 | 49.3263 |
| 15594:3 | 1 | R.TAGWNIPMGLLFN[+1216.423]QTGSC[+57.021]K.F | HexNAc(2)Hex(5) | C[+57], N[+1216] | 1104.4823 | 3 | 3311.4324 | 3311.4328 | 0 | -0.1 | 485 | Specific | 433.51 | 429.42 | 206.76 | 5.84 | 55 | >sp\|P02788\|TRFL_HUMAN Lactotransferrin OS=Homo sapiens OX=9606 GN=LTF PE=1 SV=6 | 1270 | 20201112_10613_1ng_glyco_hcd30_etd30_compound15.18897.18897.3 | controllerType=0 controllerNumber=1 scan=18897 | 41.8425 |
| 15129:3 | 1 | R.TAGWNIPM[+15.995]GLLFN[+1913.677]QTGSC[+57.021]K.F | HexNAc(4)Hex(5)NeuAc(1) | C[+57], M[+16], N[+1914] | 1342.233 | 3 | 4024.6843 | 4024.6819 | 0 | 0.6 | 485 | Specific | 229.27 | 147.33 | 147.33 | 4.52 | 55 | >sp\|P02788\|TRFL_HUMAN Lactotransferrin OS=Homo sapiens OX=9606 GN=LTF PE=1 SV=6 | 1270 | 20201112_10613_1ng_glyco_hcd30_etd30_compound15.18407.18407.3 | controllerType=0 controllerNumber=1 scan=18407 | 40.5086 |
| 14917:3 | 1 | R.TAGWNIPM[+15.995]GLLFN[+2059.735]QTGSC[+57.021]K.F | HexNAc(4)Hex(5)Fuc(1)NeuAc(1) | C[+57], M[+16], N[+2060] | 1390.9224 | 3 | 4170.7527 | 4170.7398 | 0 | 3.1 | 485 | Specific | 258.93 | 175.64 | 175.64 | 3.06 | 55 | >sp\|P02788\|TRFL_HUMAN Lactotransferrin OS=Homo sapiens OX=9606 GN=LTF PE=1 SV=6 | 1270 | 20201112_10613_1ng_glyco_hcd30_etd30_compound15.18185.18185.3 | controllerType=0 controllerNumber=1 scan=18185 | 39.9681 |

Table S4. Cont.

| Query #:z | Protein Rank | Glycopeptide | Glycan | Modification Type(s) | Observed m/z | z | Observed (M+H) | Calc. mass (M+H) | Off-by-x error | Mass error (ppm) | StartPosition | Cleavage | Score | Delta | Delta Mod | \|Log Prob\| | # of unique peptides | Protein Name | Protein DB number | Comment | Scan # | Scan Time |
|---|---|---|---|---|---|---|---|---|---|---|---|---|---|---|---|---|---|---|---|---|---|---|
| 15709:3 | 1 | R.TAGWNIPM[+15.995]GLLFN[+2059.735]QTGSC[+57.021]K.F | HexNAc(4)Hex(5)Fuc(1)NeuAc(1) | C[+57], M[+16], N[+2060] | 1390.9246 | 3 | 4170.7593 | 4170.7398 | 0 | 4.7 | 485 | Specific | 278.75 | 211.54 | 211.54 | 4.01 | 55 | >sp\|P02788\|TRFL_HUMAN Lactotransferrin OS=Homo sapiens OX=9606 GN=LTF PE=1 SV=6 | 1270 | 20201112_10613_1ng_glyco_hcd30_etd30_compound15.19019.19019.3 | controllerType=0 controllerNumber=1 scan=19019 | 42.2534 |
| 16025:3 | 1 | R.TAGWNIPMGLLFN[+1768.640]QTGSC[+57.021]KFDEYFSQSC[+57.021]APGSDPR.S | HexNAc(4)Hex(5)Fuc(1) | C[+57]*2, N[+1769] | 1903.1385 | 3 | 5707.401 | 5707.3966 | 0 | 0.8 | 485 | Specific | 286.78 | 258.57 | 258.57 | 5.04 | 55 | >sp\|P02788\|TRFL_HUMAN Lactotransferrin OS=Homo sapiens OX=9606 GN=LTF PE=1 SV=6 | 1270 | 20201112_10613_1ng_glyco_hcd30_etd30_compound15.19354.19354.3 | controllerType=0 controllerNumber=1 scan=19354 | 43.3638 |
| 16767:4 | 1 | R.TAGWNIPMGLLFN[+1913.677]QTGSC[+57.021]KFDEYFSQSC[+57.021]APGSDPR.S | HexNAc(4)Hex(5)NeuAc(1) | C[+57]*2, N[+1914] | 1463.8679 | 4 | 5852.4499 | 5852.4341 | 0 | 2.7 | 485 | Specific | 353.27 | 308.52 | 308.52 | 3.68 | 55 | >sp\|P02788\|TRFL_HUMAN Lactotransferrin OS=Homo sapiens OX=9606 GN=LTF PE=1 SV=6 | 1270 | 20201112_10613_1ng_glyco_hcd30_etd30_compound15.20138.20138.4 | controllerType=0 controllerNumber=1 scan=20138 | 45.7102 |
| 16057:4 | 1 | R.TAGWNIPMGLLFN[+1622.582]QTGSC[+57.021]KFDEYFSQSC[+57.021]APGSDPR.S | HexNAc(4)Hex(5) | C[+57]*2, N[+1623] | 1391.0949 | 4 | 5561.3576 | 5561.3387 | 0 | 3.4 | 485 | Specific | 361.5 | 326.54 | 70.89 | 3.62 | 55 | >sp\|P02788\|TRFL_HUMAN Lactotransferrin OS=Homo sapiens OX=9606 GN=LTF PE=1 SV=6 | 1270 | 20201112_10613_1ng_glyco_hcd30_etd30_compound15.19388.19388.4 | controllerType=0 controllerNumber=1 scan=19388 | 43.4604 |
| 16021:4 | 1 | R.TAGWNIPMGLLFN[+1768.640]QTGSC[+57.021]KFDEYFSQSC[+57.021]APGSDPR.S | HexNAc(4)Hex(5)Fuc(1) | C[+57]*2, N[+1769] | 1427.6072 | 4 | 5707.4069 | 5707.3966 | 0 | 1.8 | 485 | Specific | 461.25 | 406.39 | 406.39 | 7.09 | 55 | >sp\|P02788\|TRFL_HUMAN Lactotransferrin OS=Homo sapiens OX=9606 GN=LTF PE=1 SV=6 | 1270 | 20201112_10613_1ng_glyco_hcd30_etd30_compound15.19350.19350.4 | controllerType=0 controllerNumber=1 scan=19350 | 43.3561 |
| 16062:4 | 1 | R.TAGWNIPMGLLFN[+1606.587]QTGSC[+57.021]KFDEYFSQSC[+57.021]APGSDPR.S | HexNAc(4)Hex(4)Fuc(1) | C[+57]*2, N[+1607] | 1387.0935 | 4 | 5545.3522 | 5545.3438 | 0 | 1.5 | 485 | Specific | 484.23 | 432.85 | 432.85 | 5.84 | 55 | >sp\|P02788\|TRFL_HUMAN Lactotransferrin OS=Homo sapiens OX=9606 GN=LTF PE=1 SV=6 | 1270 | 20201112_10613_1ng_glyco_hcd30_etd30_compound15.19393.19393.4 | controllerType=0 controllerNumber=1 scan=19393 | 43.4707 |
| 16136:3 | 1 | R.TAGWNIPMGLLFN[+1216.423]QTGSC[+57.021]KFDEYFSQSC[+57.021]APGSDPR.S | HexNAc(2)Hex(5) | C[+57]*2, N[+1216] | 1719.0702 | 3 | 5155.1959 | 5155.18 | 0 | 3.1 | 485 | Specific | 503.23 | 419.92 | 135.72 | 6.68 | 55 | >sp\|P02788\|TRFL_HUMAN Lactotransferrin OS=Homo sapiens OX=9606 GN=LTF PE=1 SV=6 | 1270 | 20201112_10613_1ng_glyco_hcd30_etd30_compound15.19470.19470.3 | controllerType=0 controllerNumber=1 scan=19470 | 43.6681 |
| 17726:4 | 1 | R.TAGWNIPMGLLFN[+2204.772]QTGSC[+57.021]KFDEYFSQSC[+57.021]APGSDPR.S | HexNAc(4)Hex(5)NeuAc(2) | C[+57]*2, N[+2205] | 1536.6428 | 4 | 6143.5495 | 6143.5295 | 0 | 3.2 | 485 | Specific | 505.95 | 452.28 | 333.45 | 6.64 | 55 | >sp\|P02788\|TRFL_HUMAN Lactotransferrin OS=Homo sapiens OX=9606 GN=LTF PE=1 SV=6 | 1270 | 20201112_10613_1ng_glyco_hcd30_etd30_compound15.21153.21153.4 | controllerType=0 controllerNumber=1 scan=21153 | 48.5962 |
| 16736:4 | 1 | R.TAGWNIPMGLLFN[+2059.735]QTGSC[+57.021]KFDEYFSQSC[+57.021]APGSDPR.S | HexNAc(4)Hex(5)Fuc(1)NeuAc(1) | C[+57]*2, N[+2060] | 1500.3864 | 4 | 5998.5236 | 5998.492 | 0 | 5.3 | 485 | Specific | 569.29 | 505.61 | 505.61 | 6.48 | 55 | >sp\|P02788\|TRFL_HUMAN Lactotransferrin OS=Homo sapiens OX=9606 GN=LTF PE=1 SV=6 | 1270 | 20201112_10613_1ng_glyco_hcd30_etd30_compound15.20105.20105.4 | controllerType=0 controllerNumber=1 scan=20105 | 45.5784 |
| 08879:3 | 1 | R.N[+2205.793]GSDC[+57.021]PDKFC[+57.021]LFQSETK.N | HexNAc(4)Hex(5)Fuc(2)NeuAc(1) | C[+57]*2, N[+2206] | 1413.5611 | 3 | 4238.6687 | 4238.6667 | 0 | 0.5 | 642 | Specific | 255.04 | 184.73 | 184.73 | 4.59 | 55 | >sp\|P02788\|TRFL_HUMAN Lactotransferrin OS=Homo sapiens OX=9606 GN=LTF PE=1 SV=6 | 1270 | 20201112_10613_1ng_glyco_hcd30_etd30_compound15.11843.11843.3 | controllerType=0 controllerNumber=1 scan=11843 | 26.7307 |
| 10921:3 | 1 | R.N[+2350.830]GSDC[+57.021]PDKFC[+57.021]LFQSETK.N | HexNAc(4)Hex(5)Fuc(1)NeuAc(2) | C[+57]*2, N[+2351] | 1461.9085 | 3 | 4383.711 | 4383.7043 | 0 | 1.5 | 642 | Specific | 313.32 | 219.07 | 219.07 | 5.02 | 55 | >sp\|P02788\|TRFL_HUMAN Lactotransferrin OS=Homo sapiens OX=9606 GN=LTF PE=1 SV=6 | 1270 | 20201112_10613_1ng_glyco_hcd30_etd30_compound15.13966.13966.3 | controllerType=0 controllerNumber=1 scan=13966 | 30.9634 |
| 08962:4 | 2 | R.QTGLAPGQEYEISLHIVKN[+1913.677]NTR.G | HexNAc(4)Hex(5)NeuAc(1) | N[+1914] | 1096.2494 | 4 | 4381.9759 | 4381.9662 | 0 | 2.2 | 770 | Specific | 237.14 | 237.14 | 237.14 | 2.69 | 26 | >sp\|P24821\|TENA_HUMAN Tenascin OS=Homo sapiens OX=9606 GN=TNC PE=1 SV=3 | 486 | 20201112_10613_1ng_glyco_hcd30_etd30_compound15.11929.11929.4 | controllerType=0 controllerNumber=1 scan=11929 | 26.9173 |
| 08867:3 | 2 | R.QTGLAPGQEYEISLHIVKN[+2059.735]NTR.G | HexNAc(4)Hex(5)Fuc(1)NeuAc(1) | N[+2060] | 1510.0131 | 3 | 4528.0248 | 4528.0241 | 0 | 0.1 | 770 | Specific | 298.38 | 181.58 | 181.58 | 2.71 | 26 | >sp\|P24821\|TENA_HUMAN Tenascin OS=Homo sapiens OX=9606 GN=TNC PE=1 SV=3 | 486 | 20201112_10613_1ng_glyco_hcd30_etd30_compound15.11830.11830.3 | controllerType=0 controllerNumber=1 scan=11830 | 26.702 |
| 14296:3 | 2 | R.LN[+1216.423]YSLPTGQWVGVQLPR.N | HexNAc(2)Hex(5) | N[+1216] | 1048.8301 | 3 | 3144.4758 | 3144.4617 | 0 | 4.5 | 1017 | Specific | 286.13 | 248.02 | 248.02 | 4.33 | 26 | >sp\|P24821\|TENA_HUMAN Tenascin OS=Homo sapiens OX=9606 GN=TNC PE=1 SV=3 | 486 | 20201112_10613_1ng_glyco_hcd30_etd30_compound15.17503.17503.3 | controllerType=0 controllerNumber=1 scan=17503 | 38.4136 |
| 14246:3 | 2 | R.LN[+1378.476]YSLPTGQWVGVQLPR.N | HexNAc(2)Hex(6) | N[+1378] | 1102.8471 | 3 | 3306.5267 | 3306.5145 | 0 | 3.7 | 1017 | Specific | 515.06 | 469.84 | 469.84 | 6.24 | 26 | >sp\|P24821\|TENA_HUMAN Tenascin OS=Homo sapiens OX=9606 GN=TNC PE=1 SV=3 | 486 | 20201112_10613_1ng_glyco_hcd30_etd30_compound15.17451.17451.3 | controllerType=0 controllerNumber=1 scan=17451 | 38.2689 |
| 14138:3 | 2 | R.LN[+1702.581]YSLPTGQWVGVQLPR.N | HexNAc(2)Hex(8) | N[+1703] | 1210.8736 | 3 | 3630.6062 | 3630.6202 | 0 | -3.8 | 1017 | Specific | 530.56 | 429.07 | 429.07 | 6.5 | 26 | >sp\|P24821\|TENA_HUMAN Tenascin OS=Homo sapiens OX=9606 GN=TNC PE=1 SV=3 | 486 | 20201112_10613_1ng_glyco_hcd30_etd30_compound15.17336.17336.3 | controllerType=0 controllerNumber=1 scan=17336 | 37.9313 |
| 14186:3 | 2 | R.LN[+1540.529]YSLPTGQWVGVQLPR.N | HexNAc(2)Hex(7) | N[+1541] | 1156.8615 | 3 | 3468.57 | 3468.5674 | 0 | 0.7 | 1017 | Specific | 665.49 | 560.47 | 560.47 | 10.26 | 26 | >sp\|P24821\|TENA_HUMAN Tenascin OS=Homo sapiens OX=9606 GN=TNC PE=1 SV=3 | 486 | 20201112_10613_1ng_glyco_hcd30_etd30_compound15.17387.17387.3 | controllerType=0 controllerNumber=1 scan=17387 | 38.0743 |
| 04139:3 | 2 | R.N[+2059.735]TTSYVLR.G | HexNAc(4)Hex(5)Fuc(1)NeuAc(1) | N[+2060] | 1005.0861 | 3 | 3013.2437 | 3013.24 | 0 | 1.2 | 1034 | Specific | 312.47 | 183.54 | 183.54 | 4.41 | 26 | >sp\|P24821\|TENA_HUMAN Tenascin OS=Homo sapiens OX=9606 GN=TNC PE=1 SV=3 | 486 | 20201112_10613_1ng_glyco_hcd30_etd30_compound15.6917.6917.3 | controllerType=0 controllerNumber=1 scan=6917 | 17.8249 |
| 16521:3 | 3 | R.LSLHRPALEDLLLGSEAN[+1913.677]LTC[+57.021]TLTGLR.D | HexNAc(4)Hex(5)NeuAc(1) | C[+57], N[+1914] | 1626.4327 | 3 | 4877.2835 | 4877.2753 | 0 | 1.7 | 127 | Specific | 218.63 | 214.11 | 214.11 | 3.71 | 23 | >sp\|P01876\|IGHA1_HUMAN Immunoglobulin heavy constant alpha 1 OS=Homo sapiens OX=9606 GN=IGHA1 PE=1 SV=2 | 255 | 20201112_10613_1ng_glyco_hcd30_etd30_compound15.19875.19875.3 | controllerType=0 controllerNumber=1 scan=19875 | 44.7843 |
| 16791:3 | 3 | R.[+42.011]LSLHRPALEDLLLGSEAN[+1501.555]LTC[+57.021]TLTGLR.D | HexNAc(5)Hex(3) | .[42], C[+57], N[+1502] | 1503.0587 | 3 | 4507.1614 | 4507.1642 | 0 | -0.6 | 127 | Specific | 267.77 | 222.4 | 222.4 | 2.86 | 23 | >sp\|P01876\|IGHA1_HUMAN Immunoglobulin heavy constant alpha 1 OS=Homo sapiens OX=9606 GN=IGHA1 PE=1 SV=2 | 255 | 20201112_10613_1ng_glyco_hcd30_etd30_compound15.20163.20163.3 | controllerType=0 controllerNumber=1 scan=20163 | 45.779 |
| 15790:3 | 3 | R.LSLHRPALEDLLLGSEAN[+1864.634]LTC[+57.021]TLTGLR.D | HexNAc(2)Hex(9) | C[+57], N[+1865] | 1610.0836 | 3 | 4828.2361 | 4828.2324 | 0 | 0.8 | 127 | Specific | 338.86 | 338.86 | 338.86 | 5.38 | 23 | >sp\|P01876\|IGHA1_HUMAN Immunoglobulin heavy constant alpha 1 OS=Homo sapiens OX=9606 GN=IGHA1 PE=1 SV=2 | 255 | 20201112_10613_1ng_glyco_hcd30_etd30_compound15.19104.19104.3 | controllerType=0 controllerNumber=1 scan=19104 | 42.5098 |
| 15833:3 | 3 | R.LSLHRPALEDLLLGSEAN[+1540.529]LTC[+57.021]TLTGLR.D | HexNAc(2)Hex(7) | C[+57], N[+1541] | 1502.0491 | 3 | 4504.1328 | 4504.1268 | 0 | 1.3 | 127 | Specific | 377.95 | 337.25 | 337.25 | 5.67 | 23 | >sp\|P01876\|IGHA1_HUMAN Immunoglobulin heavy constant alpha 1 OS=Homo sapiens OX=9606 GN=IGHA1 PE=1 SV=2 | 255 | 20201112_10613_1ng_glyco_hcd30_etd30_compound15.19150.19150.3 | controllerType=0 controllerNumber=1 scan=19150 | 42.6543 |
| 16505:4 | 3 | R.LSLHRPALEDLLLGSEAN[+2116.756]LTC[+57.021]TLTGLR.D | HexNAc(5)Hex(5)NeuAc(1) | C[+57], N[+2117] | 1270.8444 | 4 | 5080.3556 | 5080.3547 | 0 | 0.2 | 127 | Specific | 382.13 | 382.13 | 382.13 | 5.55 | 23 | >sp\|P01876\|IGHA1_HUMAN Immunoglobulin heavy constant alpha 1 OS=Homo sapiens OX=9606 GN=IGHA1 PE=1 SV=2 | 255 | 20201112_10613_1ng_glyco_hcd30_etd30_compound15.19858.19858.4 | controllerType=0 controllerNumber=1 scan=19858 | 44.734 |
| 15861:3 | 3 | R.LSLHRPALEDLLLGSEAN[+1501.555]LTC[+57.021]TLTGLR.D | HexNAc(5)Hex(3) | C[+57], N[+1502] | 1489.0541 | 3 | 4465.1478 | 4465.1536 | 0 | -1.3 | 127 | Specific | 387.09 | 320.18 | 320.18 | 4.62 | 23 | >sp\|P01876\|IGHA1_HUMAN Immunoglobulin heavy constant alpha 1 OS=Homo sapiens OX=9606 GN=IGHA1 PE=1 SV=2 | 255 | 20201112_10613_1ng_glyco_hcd30_etd30_compound15.19180.19180.3 | controllerType=0 controllerNumber=1 scan=19180 | 42.7733 |
| 16563:4 | 3 | R.LSLHRPALEDLLLGSEAN[+1751.624]LTC[+57.021]TLTGLR.D | HexNAc(4)Hex(4)NeuAc(1) | C[+57], N[+1752] | 1179.5619 | 4 | 4715.2259 | 4715.2225 | 0 | 0.7 | 127 | Specific | 416.25 | 416.25 | 416.25 | 6.05 | 23 | >sp\|P01876\|IGHA1_HUMAN Immunoglobulin heavy constant alpha 1 OS=Homo sapiens OX=9606 GN=IGHA1 PE=1 SV=2 | 255 | 20201112_10613_1ng_glyco_hcd30_etd30_compound15.19920.19920.4 | controllerType=0 controllerNumber=1 scan=19920 | 44.933 |
| 15801:3 | 3 | R.LSLHRPALEDLLLGSEAN[+1702.581]LTC[+57.021]TLTGLR.D | HexNAc(2)Hex(8) | C[+57], N[+1703] | 1556.0689 | 3 | 4666.1922 | 4666.1796 | 0 | 2.7 | 127 | Specific | 419.24 | 367.02 | 367.02 | 6.44 | 23 | >sp\|P01876\|IGHA1_HUMAN Immunoglobulin heavy constant alpha 1 OS=Homo sapiens OX=9606 GN=IGHA1 PE=1 SV=2 | 255 | 20201112_10613_1ng_glyco_hcd30_etd30_compound15.19116.19116.3 | controllerType=0 controllerNumber=1 scan=19116 | 42.5418 |
| 16514:4 | 3 | R.LSLHRPALEDLLLGSEAN[+2116.756]LTC[+57.021]TLTGLR.D | HexNAc(5)Hex(5)NeuAc(1) | C[+57], N[+2117] | 1270.8461 | 4 | 5080.3625 | 5080.3547 | 0 | 1.5 | 127 | Specific | 523.84 | 523.84 | 523.84 | 6.5 | 23 | >sp\|P01876\|IGHA1_HUMAN Immunoglobulin heavy constant alpha 1 OS=Homo sapiens OX=9606 GN=IGHA1 PE=1 SV=2 | 255 | 20201112_10613_1ng_glyco_hcd30_etd30_compound15.19868.19868.4 | controllerType=0 controllerNumber=1 scan=19868 | 44.7635 |
| 15850:4 | 3 | R.LSLHRPALEDLLLGSEAN[+1419.502]LTC[+57.021]TLTGLR.D | HexNAc(3)Hex(5) | C[+57], N[+1420] | 1096.5323 | 4 | 4383.1072 | 4383.1005 | 0 | 1.5 | 127 | Specific | 670.09 | 670.09 | 670.09 | 10.62 | 23 | >sp\|P01876\|IGHA1_HUMAN Immunoglobulin heavy constant alpha 1 OS=Homo sapiens OX=9606 GN=IGHA1 PE=1 SV=2 | 255 | 20201112_10613_1ng_glyco_hcd30_etd30_compound15.19168.19168.4 | controllerType=0 controllerNumber=1 scan=19168 | 42.7252 |
| 15780:4 | 3 | R.LSLHRPALEDLLLGSEAN[+1864.634]LTC[+57.021]TLTGLR.D | HexNAc(2)Hex(9) | C[+57], N[+1865] | 1207.8146 | 4 | 4828.2366 | 4828.2324 | 0 | 0.9 | 127 | Specific | 781.46 | 752.5 | 752.5 | 12.62 | 23 | >sp\|P01876\|IGHA1_HUMAN Immunoglobulin heavy constant alpha 1 OS=Homo sapiens OX=9606 GN=IGHA1 PE=1 SV=2 | 255 | 20201112_10613_1ng_glyco_hcd30_etd30_compound15.19094.19094.4 | controllerType=0 controllerNumber=1 scan=19094 | 42.4742 |
| 16520:4 | 3 | R.LSLHRPALEDLLLGSEAN[+1913.677]LTC[+57.021]TLTGLR.D | HexNAc(4)Hex(5)NeuAc(1) | C[+57], N[+1914] | 1220.0747 | 4 | 4877.2772 | 4877.2753 | 0 | 0.4 | 127 | Specific | 816.24 | 783.19 | 783.19 | 13.47 | 23 | >sp\|P01876\|IGHA1_HUMAN Immunoglobulin heavy constant alpha 1 OS=Homo sapiens OX=9606 GN=IGHA1 PE=1 SV=2 | 255 | 20201112_10613_1ng_glyco_hcd30_etd30_compound15.19874.19874.4 | controllerType=0 controllerNumber=1 scan=19874 | 44.7804 |
| 15836:4 | 3 | R.LSLHRPALEDLLLGSEAN[+1663.608]LTC[+57.021]TLTGLR.D | HexNAc(5)Hex(4) | C[+57], N[+1664] | 1157.5586 | 4 | 4627.2127 | 4627.2064 | 0 | 1.4 | 127 | Specific | 823.61 | 823.61 | 823.61 | 13.62 | 23 | >sp\|P01876\|IGHA1_HUMAN Immunoglobulin heavy constant alpha 1 OS=Homo sapiens OX=9606 GN=IGHA1 PE=1 SV=2 | 255 | 20201112_10613_1ng_glyco_hcd30_etd30_compound15.19153.19153.4 | controllerType=0 controllerNumber=1 scan=19153 | 42.6712 |
| 15794:4 | 3 | R.LSLHRPALEDLLLGSEAN[+1702.581]LTC[+57.021]TLTGLR.D | HexNAc(2)Hex(8) | C[+57], N[+1703] | 1167.3027 | 4 | 4666.1888 | 4666.1796 | 0 | 2 | 127 | Specific | 844.6 | 838.49 | 838.49 | 13.62 | 23 | >sp\|P01876\|IGHA1_HUMAN Immunoglobulin heavy constant alpha 1 OS=Homo sapiens OX=9606 GN=IGHA1 PE=1 SV=2 | 255 | 20201112_10613_1ng_glyco_hcd30_etd30_compound15.19109.19109.4 | controllerType=0 controllerNumber=1 scan=19109 | 42.5232 |
| 15832:4 | 3 | R.LSLHRPALEDLLLGSEAN[+1540.529]LTC[+57.021]TLTGLR.D | HexNAc(2)Hex(7) | C[+57], N[+1541] | 1126.7884 | 4 | 4504.1317 | 4504.1268 | 0 | 1.1 | 127 | Specific | 917.53 | 867.37 | 867.37 | 15.62 | 23 | >sp\|P01876\|IGHA1_HUMAN Immunoglobulin heavy constant alpha 1 OS=Homo sapiens OX=9606 GN=IGHA1 PE=1 SV=2 | 255 | 20201112_10613_1ng_glyco_hcd30_etd30_compound15.19149.19149.4 | controllerType=0 controllerNumber=1 scan=19149 | 42.6504 |
| 15855:4 | 3 | R.LSLHRPALEDLLLGSEAN[+1216.423]LTC[+57.021]TLTGLR.D | HexNAc(2)Hex(5) | C[+57], N[+1216] | 1045.7622 | 4 | 4180.0272 | 4180.0211 | 0 | 1.4 | 127 | Specific | 996.22 | 969.69 | 969.69 | 16.62 | 23 | >sp\|P01876\|IGHA1_HUMAN Immunoglobulin heavy constant alpha 1 OS=Homo sapiens OX=9606 GN=IGHA1 PE=1 SV=2 | 255 | 20201112_10613_1ng_glyco_hcd30_etd30_compound15.19173.19173.4 | controllerType=0 controllerNumber=1 scan=19173 | 42.7468 |
| 15860:4 | 3 | R.LSLHRPALEDLLLGSEAN[+1501.555]LTC[+57.021]TLTGLR.D | HexNAc(5)Hex(3) | C[+57], N[+1502] | 1117.0422 | 4 | 4465.1468 | 4465.1536 | 0 | -1.5 | 127 | Specific | 1005.43 | 993.39 | 993.39 | 16.75 | 23 | >sp\|P01876\|IGHA1_HUMAN Immunoglobulin heavy constant alpha 1 OS=Homo sapiens OX=9606 GN=IGHA1 PE=1 SV=2 | 255 | 20201112_10613_1ng_glyco_hcd30_etd30_compound15.19179.19179.4 | controllerType=0 controllerNumber=1 scan=19179 | 42.7705 |
| 10901:3 | 3 | R.LAGKPTHVN[+1501.555]VSVVMAEVDGTC[+57.021]Y.- | HexNAc(5)Hex(3) | C[+57], N[+1502] | 1283.5735 | 3 | 3848.7061 | 3848.6974 | 0 | 2.3 | 332 | CRagged | 211.69 | 143 | 143 | 2.56 | 23 | >sp\|P01876\|IGHA1_HUMAN Immunoglobulin heavy constant alpha 1 OS=Homo sapiens OX=9606 GN=IGHA1 PE=1 SV=2 | 255 | 20201112_10613_1ng_glyco_hcd30_etd30_compound15.13945.13945.3 | controllerType=0 controllerNumber=1 scan=13945 | 30.9134 |
| 10780:3 | 3 | R.LAGKPTHVN[+1702.581]VSVVMAEVDGTC[+57.021]Y.- | HexNAc(2)Hex(8) | C[+57], N[+1703] | 1350.58 | 3 | 4049.7255 | 4049.7234 | 0 | 0.5 | 332 | CRagged | 220.2 | 165.93 | 165.93 | 2.87 | 23 | >sp\|P01876\|IGHA1_HUMAN Immunoglobulin heavy constant alpha 1 OS=Homo sapiens OX=9606 GN=IGHA1 PE=1 SV=2 | 255 | 20201112_10613_1ng_glyco_hcd30_etd30_compound15.13817.13817.3 | controllerType=0 controllerNumber=1 scan=13817 | 30.5379 |
| 10842:3 | 3 | R.LAGKPTHVN[+1216.423]VSVVMAEVDGTC[+57.021]Y.- | HexNAc(2)Hex(5) | C[+57], N[+1216] | 1188.5258 | 3 | 3563.5629 | 3563.5649 | 0 | -0.6 | 332 | CRagged | 232.68 | 210.64 | 172.55 | 2.58 | 23 | >sp\|P01876\|IGHA1_HUMAN Immunoglobulin heavy constant alpha 1 OS=Homo sapiens OX=9606 GN=IGHA1 PE=1 SV=2 | 255 | 20201112_10613_1ng_glyco_hcd30_etd30_compound15.13883.13883.3 | controllerType=0 controllerNumber=1 scan=13883 | 30.7425 |
| 10813:3 | 3 | R.LAGKPTHVN[+1809.666]VSVVMAEVDGTC[+57.021]Y.- | HexNAc(5)Hex(4)Fuc(1) | C[+57], N[+1810] | 1386.2753 | 3 | 4156.8114 | 4156.8081 | 0 | 0.8 | 332 | CRagged | 234.62 | 192.16 | 189.78 | 2.77 | 23 | >sp\|P01876\|IGHA1_HUMAN Immunoglobulin heavy constant alpha 1 OS=Homo sapiens OX=9606 GN=IGHA1 PE=1 SV=2 | 255 | 20201112_10613_1ng_glyco_hcd30_etd30_compound15.13853.13853.3 | controllerType=0 controllerNumber=1 scan=13853 | 30.6723 |
| 10896:3 | 3 | R.LAGKPTHVN[+1647.613]VSVVMAEVDGTC[+57.021]Y.- | HexNAc(5)Hex(3)Fuc(1) | C[+57], N[+1648] | 1332.2592 | 3 | 3994.7631 | 3994.7553 | 0 | 1.9 | 332 | CRagged | 237.16 | 119.11 | 119.11 | 2.63 | 23 | >sp\|P01876\|IGHA1_HUMAN Immunoglobulin heavy constant alpha 1 OS=Homo sapiens OX=9606 GN=IGHA1 PE=1 SV=2 | 255 | 20201112_10613_1ng_glyco_hcd30_etd30_compound15.13940.13940.3 | controllerType=0 controllerNumber=1 scan=13940 | 30.8957 |
| 10809:3 | 3 | R.LAGKPTHVN[+1378.476]VSVVMAEVDGTC[+57.021]Y.- | HexNAc(2)Hex(6) | C[+57], N[+1378] | 1242.5458 | 3 | 3725.6229 | 3725.6178 | 0 | 1.4 | 332 | CRagged | 263.26 | 263.26 | 192.05 | 3.2 | 23 | >sp\|P01876\|IGHA1_HUMAN Immunoglobulin heavy constant alpha 1 OS=Homo sapiens OX=9606 GN=IGHA1 PE=1 SV=2 | 255 | 20201112_10613_1ng_glyco_hcd30_etd30_compound15.13848.13848.3 | controllerType=0 controllerNumber=1 scan=13848 | 30.658 |
| 10807:3 | 3 | R.LAGKPTHVN[+1540.529]VSVVMAEVDGTC[+57.021]Y.- | HexNAc(2)Hex(7) | C[+57], N[+1541] | 1296.5626 | 3 | 3887.6731 | 3887.6706 | 0 | 0.7 | 332 | CRagged | 278.34 | 218.58 | 198.36 | 2.97 | 23 | >sp\|P01876\|IGHA1_HUMAN Immunoglobulin heavy constant alpha 1 OS=Homo sapiens OX=9606 GN=IGHA1 PE=1 SV=2 | 255 | 20201112_10613_1ng_glyco_hcd30_etd30_compound15.13846.13846.3 | controllerType=0 controllerNumber=1 scan=13846 | 30.6388 |
| 18055:4 | 4 | K.QIGLYPVLVIDSSGYVNPN[+2204.772]YTGR.I | HexNAc(4)Hex(5)NeuAc(2) | N[+2205] | 1183.2775 | 4 | 4730.0882 | 4730.0759 | 0 | 2.6 | 168 | Specific | 343.74 | 330.71 | 330.71 | 3.28 | 24 | >sp\|P01833\|PIGR_HUMAN Polymeric immunoglobulin receptor OS=Homo sapiens OX=9606 GN=PIGR PE=1 SV=4 | 902 | 20201112_10613_1ng_glyco_hcd30_etd30_compound15.21502.21502.4 | controllerType=0 controllerNumber=1 scan=21502 | 49.7238 |
| 18027:3 | 4 | K.QIGLYPVLVIDSSGYVNPN[+2204.772]YTGR.I | HexNAc(4)Hex(5)NeuAc(2) | N[+2205] | 1577.3582 | 3 | 4730.0601 | 4730.0759 | 0 | -3.3 | 168 | Specific | 375.1 | 281.52 | 281.52 | 3.83 | 24 | >sp\|P01833\|PIGR_HUMAN Polymeric immunoglobulin receptor OS=Homo sapiens OX=9606 GN=PIGR PE=1 SV=4 | 902 | 20201112_10613_1ng_glyco_hcd30_etd30_compound15.21473.21473.4 | controllerType=0 controllerNumber=1 scan=21473 | 49.6428 |
| 18045:3 | 4 | K.QIGLYPVLVIDSSGYVNPN[+2204.772]YTGR.I | HexNAc(4)Hex(5)NeuAc(2) | N[+2205] | 1577.363 | 3 | 4730.0744 | 4730.0759 | 0 | -0.3 | 168 | Specific | 533.16 | 411.25 | 411.25 | 6.84 | 24 | >sp\|P01833\|PIGR_HUMAN Polymeric immunoglobulin receptor OS=Homo sapiens OX=9606 GN=PIGR PE=1 SV=4 | 902 | 20201112_10613_1ng_glyco_hcd30_etd30_compound15.21492.21492.3 | controllerType=0 controllerNumber=1 scan=21492 | 49.6933 |
| 19423:3 | 4 | R.LSLLEEPGN[+2350.830]GTFTVILNQLTSR.D | HexNAc(4)Hex(5)Fuc(1)NeuAc(2) | N[+2351] | 1585.0519 | 3 | 4753.1413 | 4753.1229 | 0 | 3.9 | 413 | Specific | 383.18 | 243.8 | 243.8 | 3.92 | 24 | >sp\|P01833\|PIGR_HUMAN Polymeric immunoglobulin receptor OS=Homo sapiens OX=9606 GN=PIGR PE=1 SV=4 | 902 | 20201112_10613_1ng_glyco_hcd30_etd30_compound15.22947.22947.3 | controllerType=0 controllerNumber=1 scan=22947 | 53.557 |
| 16008:3 | 4 | K.IIEGEPNLKVPGN[+2350.830]VTAVLGETLK.V | HexNAc(4)Hex(5)Fuc(1)NeuAc(2) | N[+2351] | 1581.4033 | 3 | 4742.1952 | 4742.1797 | 0 | 3.3 | 457 | Specific | 253.8 | 174.51 | 174.51 | 2.64 | 24 | >sp\|P01833\|PIGR_HUMAN Polymeric immunoglobulin receptor OS=Homo sapiens OX=9606 GN=PIGR PE=1 SV=4 | 902 | 20201112_10613_1ng_glyco_hcd30_etd30_compound15.19336.19336.3 | controllerType=0 controllerNumber=1 scan=19336 | 43.3058 |
| 14707:3 | 4 | K.IIEGEPNLKVPGN[+2205.793]VTAVLGETLK.V | HexNAc(4)Hex(5)Fuc(2)NeuAc(1) | N[+2206] | 1533.051 | 3 | 4597.1383 | 4597.1422 | 0 | -0.8 | 457 | Specific | 346.51 | 281.28 | 281.28 | 2.74 | 24 | >sp\|P01833\|PIGR_HUMAN Polymeric immunoglobulin receptor OS=Homo sapiens OX=9606 GN=PIGR PE=1 SV=4 | 902 | 20201112_10613_1ng_glyco_hcd30_etd30_compound15.17963.17963.3 | controllerType=0 controllerNumber=1 scan=17963 | 39.4517 |
| 14715:3 | 4 | K.IIEGEPNLKVPGN[+2059.735]VTAVLGETLK.V | HexNAc(4)Hex(5)Fuc(1)NeuAc(1) | N[+2060] | 1484.3671 | 3 | 4451.0868 | 4451.0843 | 0 | 0.6 | 457 | Specific | 367.32 | 329.79 | 329.79 | 3.74 | 24 | >sp\|P01833\|PIGR_HUMAN Polymeric immunoglobulin receptor OS=Homo sapiens OX=9606 GN=PIGR PE=1 SV=4 | 902 | 20201112_10613_1ng_glyco_hcd30_etd30_compound15.17971.17971.3 | controllerType=0 controllerNumber=1 scan=17971 | 39.4739 |
| 14694:3 | 4 | K.IIEGEPNLKVPGN[+2205.793]VTAVLGETLK.V | HexNAc(4)Hex(5)Fuc(2)NeuAc(1) | N[+2206] | 1533.0519 | 3 | 4597.1413 | 4597.1422 | 0 | -0.2 | 457 | Specific | 401.02 | 304.18 | 304.18 | 3.65 | 24 | >sp\|P01833\|PIGR_HUMAN Polymeric immunoglobulin receptor OS=Homo sapiens OX=9606 GN=PIGR PE=1 SV=4 | 902 | 20201112_10613_1ng_glyco_hcd30_etd30_compound15.17947.17947.3 | controllerType=0 controllerNumber=1 scan=17947 | 39.4074 |
| 14727:3 | 4 | K.IIEGEPNLKVPGN[+2059.735]VTAVLGETLK.V | HexNAc(4)Hex(5)Fuc(1)NeuAc(1) | N[+2060] | 1484.3669 | 3 | 4451.0861 | 4451.0843 | 0 | 0.4 | 457 | Specific | 426.26 | 308.36 | 308.36 | 3.84 | 24 | >sp\|P01833\|PIGR_HUMAN Polymeric immunoglobulin receptor OS=Homo sapiens OX=9606 GN=PIGR PE=1 SV=4 | 902 | 20201112_10613_1ng_glyco_hcd30_etd30_compound15.17984.17984.3 | controllerType=0 controllerNumber=1 scan=17984 | 39.5047 |
| 14452:5 | 4 | K.IIEGEPNLKVPGN[+2059.735]VTAVLGETLKVPC[+57.021]HFPC[+57.021]K.F | HexNAc(4)Hex(5)Fuc(1)NeuAc(1) | C[+57]*2, N[+2060] | 1096.1147 | 5 | 5476.5446 | 5476.5418 | 0 | 0.5 | 457 | Specific | 249.77 | 170.96 | 170.96 | 2.78 | 24 | >sp\|P01833\|PIGR_HUMAN Polymeric immunoglobulin receptor OS=Homo sapiens OX=9606 GN=PIGR PE=1 SV=4 | 902 | 20201112_10613_1ng_glyco_hcd30_etd30_compound15.17666.17666.3 | controllerType=0 controllerNumber=1 scan=17666 | 38.7668 |
| 15449:4 | 4 | K.IIEGEPNLKVPGN[+2350.830]VTAVLGETLKVPC[+57.021]HFPC[+57.021]K.F | HexNAc(4)Hex(5)Fuc(1)NeuAc(2) | C[+57]*2, N[+2351] | 1442.6672 | 4 | 5767.6471 | 5767.6372 | 0 | 1.7 | 457 | Specific | 410.92 | 410.92 | 410.92 | 4.98 | 24 | >sp\|P01833\|PIGR_HUMAN Polymeric immunoglobulin receptor OS=Homo sapiens OX=9606 GN=PIGR PE=1 SV=4 | 902 | 20201112_10613_1ng_glyco_hcd30_etd30_compound15.18744.18744.4 | controllerType=0 controllerNumber=1 scan=18744 | 41.4008 |
| 15453:5 | 4 | K.IIEGEPNLKVPGN[+2350.830]VTAVLGETLKVPC[+57.021]HFPC[+57.021]K.F | HexNAc(4)Hex(5)Fuc(1)NeuAc(2) | C[+57]*2, N[+2351] | 1154.3344 | 5 | 5767.6427 | 5767.6372 | 0 | 0.9 | 457 | Specific | 579.28 | 579.28 | 579.28 | 6.66 | 24 | >sp\|P01833\|PIGR_HUMAN Polymeric immunoglobulin receptor OS=Homo sapiens OX=9606 GN=PIGR PE=1 SV=4 | 902 | 20201112_10613_1ng_glyco_hcd30_etd30_compound15.18748.18748.5 | controllerType=0 controllerNumber=1 scan=18748 | 41.4092 |
| 14403:4 | 4 | K.IIEGEPNLKVPGN[+2059.735]VTAVLGETLKVPC[+57.021]HFPC[+57.021]K.F | HexNAc(4)Hex(5)Fuc(1)NeuAc(1) | C[+57]*2, N[+2060] | 1369.8962 | 4 | 5476.5631 | 5476.5418 | 0 | 3.9 | 457 | Specific | 798.7 | 734.29 | 734.29 | 10.83 | 24 | >sp\|P01833\|PIGR_HUMAN Polymeric immunoglobulin receptor OS=Homo sapiens OX=9606 GN=PIGR PE=1 SV=4 | 902 | 20201112_10613_1ng_glyco_hcd30_etd30_compound15.17615.17615.2 | controllerType=0 controllerNumber=1 scan=17615 | 38.6523 |
| 12737:3 | 4 | K.VPGN[+2205.793]VTAVLGETLK.V | HexNAc(4)Hex(5)Fuc(2)NeuAc(1) | N[+2206] | 1201.8726 | 3 | 3603.6033 | 3603.5927 | 0 | 2.9 | 466 | Specific | 298.49 | 261.84 | 261.84 | 4.82 | 24 | >sp\|P01833\|PIGR_HUMAN Polymeric immunoglobulin receptor OS=Homo sapiens OX=9606 GN=PIGR PE=1 SV=4 | 902 | 20201112_10613_1ng_glyco_hcd30_etd30_compound15.15873.15873.3 | controllerType=0 controllerNumber=1 scan=15873 | 34.6038 |
| 15091:3 | 4 | K.VPGN[+2350.830]VTAVLGETLK.V | HexNAc(4)Hex(5)Fuc(1)NeuAc(2) | N[+2351] | 1250.2154 | 3 | 3748.6316 | 3748.6302 | 0 | 0.4 | 466 | Specific | 302.35 | 193.47 | 193.47 | 5.03 | 24 | >sp\|P01833\|PIGR_HUMAN Polymeric immunoglobulin receptor OS=Homo sapiens OX=9606 GN=PIGR PE=1 SV=4 | 902 | 20201112_10613_1ng_glyco_hcd30_etd30_compound15.18367.18367.3 | controllerType=0 controllerNumber=1 scan=18367 | 40.4066 |
| 12851:4 | 4 | K.[+42.011]VPGN[+2018.708]VTAVLGETLKVPC[+57.021]HFPC[+57.021]K.F | HexNAc(3)Hex(6)Fuc(1)NeuAc(1) | .[42], C[+57]*2, N[+2019] | 1121.756 | 4 | 4484.0023 | 4483.9763 | 0 | 5.8 | 466 | Specific | 207.24 | 207.24 | 207.24 | 2.5 | 24 | >sp\|P01833\|PIGR_HUMAN Polymeric immunoglobulin receptor OS=Homo sapiens OX=9606 GN=PIGR PE=1 SV=4 | 902 | 20201112_10613_1ng_glyco_hcd30_etd30_compound15.15993.15993.4 | controllerType=0 controllerNumber=1 scan=15993 | 34.9292 |
| 06004:3 | 4 | K.WN[+1548.545]NTGC[+57.021]QALPSQDEGPSK.A | HexNAc(3)Hex(4)NeuAc(1) | C[+57], N[+1549] | 1179.8137 | 3 | 3537.4264 | 3537.4215 | 0 | 1.4 | 498 | Specific | 218.92 | 123.6 | 123.6 | 4.67 | 24 | >sp\|P01833\|PIGR_HUMAN Polymeric immunoglobulin receptor OS=Homo sapiens OX=9606 GN=PIGR PE=1 SV=4 | 902 | 20201112_10613_1ng_glyco_hcd30_etd30_compound15.8849.8849.3 | controllerType=0 controllerNumber=1 scan=8849 | 21.1801 |
| 05829:3 | 4 | K.WN[+1872.651]NTGC[+57.021]QALPSQDEGPSK.A | HexNAc(3)Hex(6)NeuAc(1) | C[+57], N[+1873] | 1287.8472 | 3 | 3861.5271 | 3861.5272 | 0 | 0 | 498 | Specific | 345.35 | 209.43 | 209.43 | 5.01 | 24 | >sp\|P01833\|PIGR_HUMAN Polymeric immunoglobulin receptor OS=Homo sapiens OX=9606 GN=PIGR PE=1 SV=4 | 902 | 20201112_10613_1ng_glyco_hcd30_etd30_compound15.8667.8667.3 | controllerType=0 controllerNumber=1 scan=8667 | 20.8341 |
| 05912:3 | 4 | K.WN[+1710.598]NTGC[+57.021]QALPSQDEGPSK.A | HexNAc(3)Hex(5)NeuAc(1) | C[+57], N[+1711] | 1233.8317 | 3 | 3699.4806 | 3699.4743 | 0 | 1.7 | 498 | Specific | 359.17 | 230.68 | 230.68 | 5.29 | 24 | >sp\|P01833\|PIGR_HUMAN Polymeric immunoglobulin receptor OS=Homo sapiens OX=9606 GN=PIGR PE=1 SV=4 | 902 | 20201112_10613_1ng_glyco_hcd30_etd30_compound15.8754.8754.3 | controllerType=0 controllerNumber=1 scan=8754 | 21.0097 |
| 05897:3 | 4 | K.WN[+1710.598]NTGC[+57.021]QALPSQDEGPSK.A | HexNAc(3)Hex(5)NeuAc(1) | C[+57], N[+1711] | 1233.8322 | 3 | 3699.4821 | 3699.4743 | 0 | 2.1 | 498 | Specific | 411.96 | 242.56 | 242.56 | 5.53 | 24 | >sp\|P01833\|PIGR_HUMAN Polymeric immunoglobulin receptor OS=Homo sapiens OX=9606 GN=PIGR PE=1 SV=4 | 902 | 20201112_10613_1ng_glyco_hcd30_etd30_compound15.8738.8738.3 | controllerType=0 controllerNumber=1 scan=8738 | 20.9769 |
| 05861:3 | 4 | K.WN[+1913.677]NTGC[+57.021]QALPSQDEGPSK.A | HexNAc(4)Hex(5)NeuAc(1) | C[+57], N[+1914] | 1301.5236 | 3 | 3902.5563 | 3902.5537 | 0 | 0.7 | 498 | Specific | 463.25 | 300.14 | 300.14 | 5.76 | 24 | >sp\|P01833\|PIGR_HUMAN Polymeric immunoglobulin receptor OS=Homo sapiens OX=9606 GN=PIGR PE=1 SV=4 | 902 | 20201112_10613_1ng_glyco_hcd30_etd30_compound15.8701.8701.3 | controllerType=0 controllerNumber=1 scan=8701 | 20.9075 |
| 15508:4 | 5 | K.RNIAAFGGDPNN[+1913.677]ITLFGESAGGASVSLQTLSPYNK.G | HexNAc(4)Hex(5)NeuAc(1) | N[+1914] | 1370.874 | 4 | 5480.4743 | 5480.4644 | 0 | 1.8 | 196 | Specific | 651.29 | 631.53 | 631.53 | 9.43 | 9 | >sp\|P19835\|CEL_HUMAN Bile salt-activated lipase OS=Homo sapiens OX=9606 GN=CEL PE=1 SV=3 | 476 | 20201112_10613_1ng_glyco_hcd30_etd30_compound15.18806.18806.4 | controllerType=0 controllerNumber=1 scan=18806 | 41.5872 |
| 16716:3 | 5 | R.NIAAFGGDPNN[+1913.677]ITLFGESAGGASVSLQTLSPYNK.G | HexNAc(4)Hex(5)NeuAc(1) | N[+1914] | 1775.459 | 3 | 5324.3624 | 5324.3633 | 0 | -0.2 | 197 | Specific | 602.53 | 460.92 | 460.92 | 9.33 | 9 | >sp\|P19835\|CEL_HUMAN Bile salt-activated lipase OS=Homo sapiens OX=9606 GN=CEL PE=1 SV=3 | 476 | 20201112_10613_1ng_glyco_hcd30_etd30_compound15.20082.20082.3 | controllerType=0 controllerNumber=1 scan=20082 | 45.4663 |
| 09746:3 | 8 | K.TVLTPATNHMGN[+1540.529]VTFTIPANR.E | HexNAc(2)Hex(7) | N[+1541] | 1265.9026 | 3 | 3795.6934 | 3795.6886 | 0 | 1.3 | 74 | Specific | 235.72 | 202.15 | 123.59 | 3.04 | 8 | >sp\|P01024\|CO3_HUMAN Complement C3 OS=Homo sapiens OX=9606 GN=C3 PE=1 SV=2 | 492 | 20201112_10613_1ng_glyco_hcd30_etd30_compound15.12744.12744.2 | controllerType=0 controllerNumber=1 scan=12744 | 28.4823 |
| 08351:3 | 10 | K.TAHC[+57.021]N[+2059.735]ESFYFLC[+57.021]K.R | HexNAc(4)Hex(5)Fuc(1)NeuAc(1) | C[+57]*2, N[+2060] | 1246.1575 | 3 | 3736.458 | 3736.4545 | 0 | 0.9 | 1201 | Specific | 207.1 | 122.93 | 122.93 | 4.04 | 5 | >sp\|P22897\|MRC1_HUMAN Macrophage mannose receptor 1 OS=Homo sapiens OX=9606 GN=MRC1 PE=1 SV=1 | 1049 | 20201112_10613_1ng_glyco_hcd30_etd30_compound15.11288.11288.3 | controllerType=0 controllerNumber=1 scan=11288 | 25.6119 |
| 18159:4 | 15 | K.ADTHDEILEGLNFN[+2205.793]LTEIPEAQIHEGFQELLR.T | HexNAc(4)Hex(5)Fuc(2)NeuAc(1) | N[+2206] | 1475.1587 | 4 | 5897.6129 | 5897.6167 | 0 | -0.6 | 94 | Specific | 629.94 | 629.94 | 629.94 | 8.65 | 7 | >sp\|P01009\|A1AT_HUMAN Alpha-1-antitrypsin OS=Homo sapiens OX=9606 GN=SERPINA1 PE=1 SV=3 | 642 | 20201112_10613_1ng_glyco_hcd30_etd30_compound15.21612.21612.4 | controllerType=0 controllerNumber=1 scan=21612 | 50.0571 |
| 17447:5 | 15 | K.YLGN[+2204.772]ATAIFFLPDEGKLQHLENELTHDIITK.F | HexNAc(4)Hex(5)NeuAc(2) | N[+2205] | 1149.9294 | 5 | 5745.6181 | 5745.6098 | 0 | 1.5 | 268 | Specific | 326.92 | 301.37 | 301.37 | 3.46 | 7 | >sp\|P01009\|A1AT_HUMAN Alpha-1-antitrypsin OS=Homo sapiens OX=9606 GN=SERPINA1 PE=1 SV=3 | 642 | 20201112_10613_1ng_glyco_hcd30_etd30_compound15.20858.20858.5 | controllerType=0 controllerNumber=1 scan=20858 | 47.7016 |
| 17448:4 | 15 | K.YLGN[+2204.772]ATAIFFLPDEGKLQHLENELTHDIITK.F | HexNAc(4)Hex(5)NeuAc(2) | N[+2205] | 1437.1606 | 4 | 5745.6208 | 5745.6098 | 0 | 1.9 | 268 | Specific | 408.42 | 408.42 | 408.42 | 5.35 | 7 | >sp\|P01009\|A1AT_HUMAN Alpha-1-antitrypsin OS=Homo sapiens OX=9606 GN=SERPINA1 PE=1 SV=3 | 642 | 20201112_10613_1ng_glyco_hcd30_etd30_compound15.20859.20859.4 | controllerType=0 controllerNumber=1 scan=20859 | 47.7057 |
| 07616:3 | 18 | R.LSPN[+2350.830]ASAEHLELR.W | HexNAc(4)Hex(5)Fuc(1)NeuAc(2) | N[+2351] | 1263.2014 | 3 | 3787.5895 | 3787.5796 | 0 | 2.6 | 52 | Specific | 352.55 | 235.04 | 235.04 | 4.2 | 6 | >sp\|Q13410\|BT1A1_HUMAN Butyrophilin subfamily 1 member A1 OS=Homo sapiens OX=9606 GN=BTN1A1 PE=1 SV=3 | 179 | 20201112_10613_1ng_glyco_hcd30_etd30_compound15.10521.10521.3 | controllerType=0 controllerNumber=1 scan=10521 | 24.1362 |
| 05787:3 | 18 | R.LSPN[+2205.793]ASAEHLELR.W | HexNAc(4)Hex(5)Fuc(2)NeuAc(1) | N[+2206] | 1214.8537 | 3 | 3642.5465 | 3642.5421 | 0 | 1.2 | 52 | Specific | 375.62 | 259.06 | 259.06 | 3.74 | 6 | >sp\|Q13410\|BT1A1_HUMAN Butyrophilin subfamily 1 member A1 OS=Homo sapiens OX=9606 GN=BTN1A1 PE=1 SV=3 | 179 | 20201112_10613_1ng_glyco_hcd30_etd30_compound15.8624.8624.3 | controllerType=0 controllerNumber=1 scan=8624 | 20.7546 |
| 15817:3 | 18 | K.N[+1913.677]VSC[+57.021]YIQNLLLGQEK.K | HexNAc(4)Hex(5)NeuAc(1) | C[+57], N[+1914] | 1231.537 | 3 | 3692.5966 | 3692.5876 | 0 | 2.4 | 215 | Specific | 319.07 | 172.91 | 172.91 | 4.64 | 6 | >sp\|Q13410\|BT1A1_HUMAN Butyrophilin subfamily 1 member A1 OS=Homo sapiens OX=9606 GN=BTN1A1 PE=1 SV=3 | 179 | 20201112_10613_1ng_glyco_hcd30_etd30_compound15.19133.19133.3 | controllerType=0 controllerNumber=1 scan=19133 | 42.5962 |
| 04923:3 | 29 | K.TPLTAN[+2262.814]ITK.S | HexNAc(5)Hex(5)Fuc(1)NeuAc(1) | N[+2263] | 1074.4639 | 3 | 3221.3771 | 3221.3711 | 0 | 1.9 | 200 | Specific | 202.99 | 107.79 | 107.79 | 2.65 | 5 | >sp\|P01877\|IGHA2_HUMAN Immunoglobulin heavy constant alpha 2 OS=Homo sapiens OX=9606 GN=IGHA2 PE=1 SV=4 | 576 | 20201112_10613_1ng_glyco_hcd30_etd30_compound15.7730.7730.3 | controllerType=0 controllerNumber=1 scan=7730 | 19.2235 |
| 03261:3 | 29 | K.TPLTAN[+1971.719]ITK.S | HexNAc(5)Hex(5)Fuc(1) | N[+1972] | 977.4319 | 3 | 2930.281 | 2930.2757 | 0 | 1.8 | 200 | Specific | 235.38 | 120.13 | 120.13 | 2.66 | 5 | >sp\|P01877\|IGHA2_HUMAN Immunoglobulin heavy constant alpha 2 OS=Homo sapiens OX=9606 GN=IGHA2 PE=1 SV=4 | 576 | 20201112_10613_1ng_glyco_hcd30_etd30_compound15.6001.6001.3 | controllerType=0 controllerNumber=1 scan=6001 | 16.0685 |
| 03355:2 | 29 | K.TPLTAN[+1647.613]ITK.S | HexNAc(5)Hex(3)Fuc(1) | N[+1648] | 1303.59 | 2 | 2606.1727 | 2606.17 | 0 | 1 | 200 | Specific | 298.2 | 121.1 | 121.1 | 2.98 | 5 | >sp\|P01877\|IGHA2_HUMAN Immunoglobulin heavy constant alpha 2 OS=Homo sapiens OX=9606 GN=IGHA2 PE=1 SV=4 | 576 | 20201112_10613_1ng_glyco_hcd30_etd30_compound15.6099.6099.2 | controllerType=0 controllerNumber=1 scan=6099 | 16.2865 |
| 03362:3 | 29 | K.TPLTAN[+1647.613]ITK.S | HexNAc(5)Hex(3)Fuc(1) | N[+1648] | 869.3961 | 3 | 2606.1739 | 2606.17 | 0 | 1.5 | 200 | Specific | 306.34 | 127.06 | 127.06 | 2.81 | 5 | >sp\|P01877\|IGHA2_HUMAN Immunoglobulin heavy constant alpha 2 OS=Homo sapiens OX=9606 GN=IGHA2 PE=1 SV=4 | 576 | 20201112_10613_1ng_glyco_hcd30_etd30_compound15.6107.6107.3 | controllerType=0 controllerNumber=1 scan=6107 | 16.309 |
| 03310:2 | 29 | K.TPLTAN[+1809.666]ITK.S | HexNAc(5)Hex(4)Fuc(1) | N[+1810] | 1384.6167 | 2 | 2768.2262 | 2768.2228 | 0 | 1.2 | 200 | Specific | 311.15 | 126.96 | 126.96 | 2.98 | 5 | >sp\|P01877\|IGHA2_HUMAN Immunoglobulin heavy constant alpha 2 OS=Homo sapiens OX=9606 GN=IGHA2 PE=1 SV=4 | 576 | 20201112_10613_1ng_glyco_hcd30_etd30_compound15.6052.6052.2 | controllerType=0 controllerNumber=1 scan=6052 | 16.1836 |
| 03311:3 | 29 | K.TPLTAN[+1809.666]ITK.S | HexNAc(5)Hex(4)Fuc(1) | N[+1810] | 923.4135 | 3 | 2768.2261 | 2768.2228 | 0 | 1.2 | 200 | Specific | 318.09 | 134.68 | 134.68 | 2.81 | 5 | >sp\|P01877\|IGHA2_HUMAN Immunoglobulin heavy constant alpha 2 OS=Homo sapiens OX=9606 GN=IGHA2 PE=1 SV=4 | 576 | 20201112_10613_1ng_glyco_hcd30_etd30_compound15.6053.6053.3 | controllerType=0 controllerNumber=1 scan=6053 | 16.1905 |
| 15099:3 | 34 | R.NYIVPILWLN[+1864.634]ETGTIGDEKANM[+15.995]FR.S | HexNAc(2)Hex(9) | M[+16], N[+1865] | 1559.0229 | 3 | 4675.0541 | 4675.0523 | 0 | 0.4 | 408 | Specific | 581.48 | 557.51 | 557.51 | 7.55 | 2 | >sp\|P16671\|CD36_HUMAN Platelet glycoprotein 4 OS=Homo sapiens OX=9606 GN=CD36 PE=1 SV=2 | 812 | 20201112_10613_1ng_glyco_hcd30_etd30_compound15.18376.18376.3 | controllerType=0 controllerNumber=1 scan=18376 | 40.4344 |
| 08469:5 | 41 | K.[+42.011]SC[+57.021]QHN[+2018.708]GTMYQHGEIFSAHELFPSR.L | HexNAc(3)Hex(6)Fuc(1)NeuAc(1) | .[42], C[+57], N[+2019] | 979.6045 | 5 | 4893.9932 | 4893.9759 | 0 | 3.5 | 110 | Specific | 252.75 | 213.3 | 213.3 | 2.65 | 2 | >sp\|Q6WN34\|CRDL2_HUMAN Chordin-like protein 2 OS=Homo sapiens OX=9606 GN=CHRDL2 PE=1 SV=1 | 1189 | 20201112_10613_1ng_glyco_hcd30_etd30_compound15.11411.11411.5 | controllerType=0 controllerNumber=1 scan=11411 | 25.8399 |
| 13135:4 | 44 | K.LKELPGVC[+57.021]N[+2059.735]ETMMALWEEC[+57.021]KPC[+57.021]LK.Q | HexNAc(4)Hex(5)Fuc(1)NeuAc(1) | C[+57]*3, N[+2060] | 1249.7859 | 4 | 4996.1217 | 4996.1196 | 0 | 0.4 | 95 | Specific | 244.17 | 190.24 | 190.24 | 2.84 | 3 | >sp\|P10909\|CLUS_HUMAN Clusterin OS=Homo sapiens OX=9606 GN=CLU PE=1 SV=1 | 1467 | 20201112_10613_1ng_glyco_hcd30_etd30_compound15.16290.16290.4 | controllerType=0 controllerNumber=1 scan=16290 | 35.5915 |
| 13144:3 | 44 | K.LKELPGVC[+57.021]N[+2059.735]ETMMALWEEC[+57.021]KPC[+57.021]LK.Q | HexNAc(4)Hex(5)Fuc(1)NeuAc(1) | C[+57]*3, N[+2060] | 1666.0451 | 3 | 4996.1208 | 4996.1196 | 0 | 0.2 | 95 | Specific | 253.32 | 158.64 | 158.64 | 2.74 | 3 | >sp\|P10909\|CLUS_HUMAN Clusterin OS=Homo sapiens OX=9606 GN=CLU PE=1 SV=1 | 1467 | 20201112_10613_1ng_glyco_hcd30_etd30_compound15.16299.16299.3 | controllerType=0 controllerNumber=1 scan=16299 | 35.6126 |
| 00814:3 | 44 | R.EIRHN[+2059.735]STGC[+57.021]LR.M | HexNAc(4)Hex(5)Fuc(1)NeuAc(1) | C[+57], N[+2060] | 1134.8072 | 3 | 3402.4069 | 3402.3994 | 0 | 2.2 | 287 | Specific | 267.81 | 145.37 | 145.37 | 2.59 | 3 | >sp\|P10909\|CLUS_HUMAN Clusterin OS=Homo sapiens OX=9606 GN=CLU PE=1 SV=1 | 1467 | 20201112_10613_1ng_glyco_hcd30_etd30_compound15.3452.3452.3 | controllerType=0 controllerNumber=1 scan=3452 | 11.2524 |
| 00809:4 | 44 | R.EIRHN[+2059.735]STGC[+57.021]LR.M | HexNAc(4)Hex(5)Fuc(1)NeuAc(1) | C[+57], N[+2060] | 851.3551 | 4 | 3402.3987 | 3402.3994 | 0 | -0.2 | 287 | Specific | 294.12 | 225.54 | 225.54 | 4.33 | 3 | >sp\|P10909\|CLUS_HUMAN Clusterin OS=Homo sapiens OX=9606 GN=CLU PE=1 SV=1 | 1467 | 20201112_10613_1ng_glyco_hcd30_etd30_compound15.3445.3445.2 | controllerType=0 controllerNumber=1 scan=3445 | 11.2318 |
| 00815:4 | 44 | R.EIRHN[+2059.735]STGC[+57.021]LR.M | HexNAc(4)Hex(5)Fuc(1)NeuAc(1) | C[+57], N[+2060] | 851.3565 | 4 | 3402.4043 | 3402.3994 | 0 | 1.4 | 287 | Specific | 564.6 | 442.45 | 442.45 | 7.57 | 3 | >sp\|P10909\|CLUS_HUMAN Clusterin OS=Homo sapiens OX=9606 GN=CLU PE=1 SV=1 | 1467 | 20201112_10613_1ng_glyco_hcd30_etd30_compound15.3453.3453.4 | controllerType=0 controllerNumber=1 scan=3453 | 11.2601 |
| 18095:4 | 44 | K.MLN[+2059.735]TSSLLEQLNEQFNWVSR.L | HexNAc(4)Hex(5)Fuc(1)NeuAc(1) | N[+2060] | 1117.9876 | 4 | 4468.9285 | 4468.9217 | 0 | 1.5 | 352 | Specific | 347.69 | 326.41 | 326.41 | 3.47 | 3 | >sp\|P10909\|CLUS_HUMAN Clusterin OS=Homo sapiens OX=9606 GN=CLU PE=1 SV=1 | 1467 | 20201112_10613_1ng_glyco_hcd30_etd30_compound15.21545.21545.4 | controllerType=0 controllerNumber=1 scan=21545 | 49.8597 |
| 18084:3 | 44 | K.MLN[+2059.735]TSSLLEQLNEQFNWVSR.L | HexNAc(4)Hex(5)Fuc(1)NeuAc(1) | N[+2060] | 1490.3137 | 3 | 4468.9264 | 4468.9217 | 0 | 1.1 | 352 | Specific | 549.33 | 335.45 | 335.45 | 5.87 | 3 | >sp\|P10909\|CLUS_HUMAN Clusterin OS=Homo sapiens OX=9606 GN=CLU PE=1 SV=1 | 1467 | 20201112_10613_1ng_glyco_hcd30_etd30_compound15.21533.21533.3 | controllerType=0 controllerNumber=1 scan=21533 | 49.8202 |
| 12501:3 | 49 | R.IIVPLNNREN[+2204.772]ISDPTSPLR.T | HexNAc(4)Hex(5)NeuAc(2) | N[+2205] | 1451.6562 | 3 | 4352.954 | 4352.9496 | 0 | 1 | 62 | Specific | 269.52 | 199.87 | 199.87 | 3.42 | 4 | >sp\|P01591\|IGJ_HUMAN Immunoglobulin J chain OS=Homo sapiens OX=9606 GN=JCHAIN PE=1 SV=4 | 1392 | 20201112_10613_1ng_glyco_hcd30_etd30_compound15.15628.15628.3 | controllerType=0 controllerNumber=1 scan=15628 | 34.1266 |
| 10799:3 | 49 | R.IIVPLNNREN[+1913.677]ISDPTSPLR.T | HexNAc(4)Hex(5)NeuAc(1) | N[+1914] | 1354.6249 | 3 | 4061.8603 | 4061.8542 | 0 | 1.5 | 62 | Specific | 283.2 | 226.45 | 226.45 | 3.68 | 4 | >sp\|P01591\|IGJ_HUMAN Immunoglobulin J chain OS=Homo sapiens OX=9606 GN=JCHAIN PE=1 SV=4 | 1392 | 20201112_10613_1ng_glyco_hcd30_etd30_compound15.13837.13837.3 | controllerType=0 controllerNumber=1 scan=13837 | 30.5992 |
| 09200:4 | 51 | K.QTDEIKDTRN[+2059.735]ESTQNC[+57.021]VVAEPEKMESSISSSSEEMSLSK.C | HexNAc(4)Hex(5)Fuc(1)NeuAc(1) | C[+57], N[+2060] | 1613.4404 | 4 | 6450.7398 | 6450.7156 | 0 | 3.7 | 60 | Specific | 280.09 | 280.09 | 280.09 | 3.06 | 3 | >sp\|P47710\|CASA1_HUMAN Alpha-S1-casein OS=Homo sapiens OX=9606 GN=CSN1S1 PE=1 SV=1 | 989 | 20201112_10613_1ng_glyco_hcd30_etd30_compound15.12177.12177.4 | controllerType=0 controllerNumber=1 scan=12177 | 27.4191 |
| 16198:4 | 1 | R.RTAGWNVPIGTLRPFLN[+2205.793]WTGPPEPIEAAVAR.F | HexNAc(4)Hex(5)Fuc(2)NeuAc(1) | N[+2206] | 1398.9078 | 4 | 5592.6095 | 5592.6049 | 0 | 0.8 | 140 | Specific | 763.47 | 763.47 | 763.47 | 12.89 | 39 | >sp\|P02788\|TRFL_HUMAN Lactotransferrin OS=Homo sapiens OX=9606 GN=LTF PE=1 SV=6 | 1270 | 20201113_10633_1ng_glyco_hcd30_etd30_compound15.19816.19816.4 | controllerType=0 controllerNumber=1 scan=19816 | 43.6787 |
| 16252:4 | 1 | R.RTAGWNVPIGTLRPFLN[+2059.735]WTGPPEPIEAAVAR.F | HexNAc(4)Hex(5)Fuc(1)NeuAc(1) | N[+2060] | 1362.4021 | 4 | 5446.5866 | 5446.547 | 0 | 7.3 | 140 | Specific | 893.93 | 893.93 | 893.93 | 13.89 | 39 | >sp\|P02788\|TRFL_HUMAN Lactotransferrin OS=Homo sapiens OX=9606 GN=LTF PE=1 SV=6 | 1270 | 20201113_10633_1ng_glyco_hcd30_etd30_compound15.19873.19873.4 | controllerType=0 controllerNumber=1 scan=19873 | 43.8484 |
| 17012:3 | 1 | R.TAGWNVPIGTLRPFLN[+2570.925]WTGPPEPIEAAVAR.F | HexNAc(5)Hex(6)Fuc(2)NeuAc(1) | N[+2571] | 1934.5492 | 3 | 5801.6329 | 5801.636 | 0 | -0.5 | 141 | Specific | 200.16 | 154.94 | 154.94 | 4.98 | 39 | >sp\|P02788\|TRFL_HUMAN Lactotransferrin OS=Homo sapiens OX=9606 GN=LTF PE=1 SV=6 | 1270 | 20201113_10633_1ng_glyco_hcd30_etd30_compound15.20674.20674.3 | controllerType=0 controllerNumber=1 scan=20674 | 46.1558 |
| 16361:4 | 1 | R.[+42.011]TAGWNVPIGTLRPFLN[+2018.708]WTGPPEPIEAAVAR.F | HexNAc(3)Hex(6)Fuc(1)NeuAc(1) | .[42], N[+2019] | 1323.623 | 4 | 5291.4704 | 5291.4299 | 0 | 7.7 | 141 | Specific | 260.11 | 260.11 | 260.11 | 3.01 | 39 | >sp\|P02788\|TRFL_HUMAN Lactotransferrin OS=Homo sapiens OX=9606 GN=LTF PE=1 SV=6 | 1270 | 20201113_10633_1ng_glyco_hcd30_etd30_compound15.19987.19987.4 | controllerType=0 controllerNumber=1 scan=19987 | 44.1555 |
| 17130:3 | 1 | R.TAGWNVPIGTLRPFLN[+2059.735]WTGPPEPIEAAVAR.F | HexNAc(4)Hex(5)Fuc(1)NeuAc(1) | N[+2060] | 1764.155 | 3 | 5290.4504 | 5290.4459 | 0 | 0.9 | 141 | Specific | 299.8 | 207.99 | 207.99 | 5.55 | 39 | >sp\|P02788\|TRFL_HUMAN Lactotransferrin OS=Homo sapiens OX=9606 GN=LTF PE=1 SV=6 | 1270 | 20201113_10633_1ng_glyco_hcd30_etd30_compound15.20798.20798.3 | controllerType=0 controllerNumber=1 scan=20798 | 46.4948 |
| 17119:3 | 1 | R.TAGWNVPIGTLRPFLN[+2205.793]WTGPPEPIEAAVAR.F | HexNAc(4)Hex(5)Fuc(2)NeuAc(1) | N[+2206] | 1812.8427 | 3 | 5436.5137 | 5436.5038 | 0 | 1.8 | 141 | Specific | 356.99 | 290.87 | 290.87 | 6.01 | 39 | >sp\|P02788\|TRFL_HUMAN Lactotransferrin OS=Homo sapiens OX=9606 GN=LTF PE=1 SV=6 | 1270 | 20201113_10633_1ng_glyco_hcd30_etd30_compound15.20787.20787.3 | controllerType=0 controllerNumber=1 scan=20787 | 46.4672 |
| 16391:3 | 1 | R.TAGWNVPIGTLRPFLN[+1768.640]WTGPPEPIEAAVAR.F | HexNAc(4)Hex(5)Fuc(1) | N[+1769] | 1667.1253 | 3 | 4999.3614 | 4999.3504 | 0 | 2.2 | 141 | Specific | 413.37 | 346.68 | 346.68 | 5.28 | 39 | >sp\|P02788\|TRFL_HUMAN Lactotransferrin OS=Homo sapiens OX=9606 GN=LTF PE=1 SV=6 | 1270 | 20201113_10633_1ng_glyco_hcd30_etd30_compound15.20019.20019.3 | controllerType=0 controllerNumber=1 scan=20019 | 44.2522 |
| 17091:3 | 1 | R.TAGWNVPIGTLRPFLN[+2059.735]WTGPPEPIEAAVAR.F | HexNAc(4)Hex(5)Fuc(1)NeuAc(1) | N[+2060] | 1764.1576 | 3 | 5290.4581 | 5290.4459 | 0 | 2.3 | 141 | Specific | 458.3 | 339.11 | 339.11 | 5.9 | 39 | >sp\|P02788\|TRFL_HUMAN Lactotransferrin OS=Homo sapiens OX=9606 GN=LTF PE=1 SV=6 | 1270 | 20201113_10633_1ng_glyco_hcd30_etd30_compound15.20757.20757.3 | controllerType=0 controllerNumber=1 scan=20757 | 46.3778 |
| 17085:4 | 1 | R.TAGWNVPIGTLRPFLN[+2351.851]WTGPPEPIEAAVAR.F | HexNAc(4)Hex(5)Fuc(3)NeuAc(1) | N[+2352] | 1396.398 | 4 | 5582.57 | 5582.5617 | 0 | 1.5 | 141 | Specific | 657.34 | 657.34 | 657.34 | 10.89 | 39 | >sp\|P02788\|TRFL_HUMAN Lactotransferrin OS=Homo sapiens OX=9606 GN=LTF PE=1 SV=6 | 1270 | 20201113_10633_1ng_glyco_hcd30_etd30_compound15.20751.20751.4 | controllerType=0 controllerNumber=1 scan=20751 | 46.361 |
| 17007:4 | 1 | R.TAGWNVPIGTLRPFLN[+2570.925]WTGPPEPIEAAVAR.F | HexNAc(5)Hex(6)Fuc(2)NeuAc(1) | N[+2571] | 1451.167 | 4 | 5801.6462 | 5801.636 | 0 | 1.8 | 141 | Specific | 712.02 | 690.08 | 690.08 | 11.89 | 39 | >sp\|P02788\|TRFL_HUMAN Lactotransferrin OS=Homo sapiens OX=9606 GN=LTF PE=1 SV=6 | 1270 | 20201113_10633_1ng_glyco_hcd30_etd30_compound15.20669.20669.4 | controllerType=0 controllerNumber=1 scan=20669 | 46.1421 |
| 17075:4 | 1 | R.TAGWNVPIGTLRPFLN[+2205.793]WTGPPEPIEAAVAR.F | HexNAc(4)Hex(5)Fuc(2)NeuAc(1) | N[+2206] | 1359.8846 | 4 | 5436.5168 | 5436.5038 | 0 | 2.4 | 141 | Specific | 930.64 | 896.1 | 896.1 | 15.89 | 39 | >sp\|P02788\|TRFL_HUMAN Lactotransferrin OS=Homo sapiens OX=9606 GN=LTF PE=1 SV=6 | 1270 | 20201113_10633_1ng_glyco_hcd30_etd30_compound15.20740.20740.4 | controllerType=0 controllerNumber=1 scan=20740 | 46.3261 |
| 16382:4 | 1 | R.TAGWNVPIGTLRPFLN[+1768.640]WTGPPEPIEAAVAR.F | HexNAc(4)Hex(5)Fuc(1) | N[+1769] | 1250.5923 | 4 | 4999.3473 | 4999.3504 | 0 | -0.6 | 141 | Specific | 1099.6 | 1097.16 | 1097.16 | 18.66 | 39 | >sp\|P02788\|TRFL_HUMAN Lactotransferrin OS=Homo sapiens OX=9606 GN=LTF PE=1 SV=6 | 1270 | 20201113_10633_1ng_glyco_hcd30_etd30_compound15.20009.20009.4 | controllerType=0 controllerNumber=1 scan=20009 | 44.2226 |
| 15555:3 | 1 | R.TAGWNIPMGLLFN[+1460.529]QTGSC[+57.021]K.F | HexNAc(4)Hex(4) | C[+57], N[+1461] | 1185.8527 | 3 | 3555.5436 | 3555.5387 | 0 | 1.4 | 485 | Specific | 200.5 | 134.84 | 134.84 | 3.17 | 39 | >sp\|P02788\|TRFL_HUMAN Lactotransferrin OS=Homo sapiens OX=9606 GN=LTF PE=1 SV=6 | 1270 | 20201113_10633_1ng_glyco_hcd30_etd30_compound15.19136.19136.3 | controllerType=0 controllerNumber=1 scan=19136 | 41.6599 |
| 15649:3 | 1 | R.TAGWNIPMGLLFN[+1216.423]QTGSC[+57.021]K.F | HexNAc(2)Hex(5) | C[+57], N[+1216] | 1104.4846 | 3 | 3311.4394 | 3311.4328 | 0 | 2 | 485 | Specific | 275.8 | 257.62 | 178.76 | 3.97 | 39 | >sp\|P02788\|TRFL_HUMAN Lactotransferrin OS=Homo sapiens OX=9606 GN=LTF PE=1 SV=6 | 1270 | 20201113_10633_1ng_glyco_hcd30_etd30_compound15.19235.19235.3 | controllerType=0 controllerNumber=1 scan=19235 | 41.9544 |
| 16833:3 | 1 | R.TAGWNIPMGLLFN[+1751.624]QTGSC[+57.021]K.F | HexNAc(4)Hex(4)NeuAc(1) | C[+57], N[+1752] | 1282.8867 | 3 | 3846.6454 | 3846.6341 | 0 | 2.9 | 485 | Specific | 296.21 | 254.66 | 254.66 | 4.84 | 39 | >sp\|P02788\|TRFL_HUMAN Lactotransferrin OS=Homo sapiens OX=9606 GN=LTF PE=1 SV=6 | 1270 | 20201113_10633_1ng_glyco_hcd30_etd30_compound15.20484.20484.3 | controllerType=0 controllerNumber=1 scan=20484 | 45.6148 |
| 16671:3 | 1 | R.TAGWNIPMGLLFN[+2205.793]QTGSC[+57.021]K.F | HexNAc(4)Hex(5)Fuc(2)NeuAc(1) | C[+57], N[+2206] | 1434.2796 | 3 | 4300.8242 | 4300.8028 | 0 | 5 | 485 | Specific | 323.3 | 168.41 | 168.41 | 4.27 | 39 | >sp\|P02788\|TRFL_HUMAN Lactotransferrin OS=Homo sapiens OX=9606 GN=LTF PE=1 SV=6 | 1270 | 20201113_10633_1ng_glyco_hcd30_etd30_compound15.20312.20312.3 | controllerType=0 controllerNumber=1 scan=20312 | 45.027 |
| 15469:3 | 1 | R.TAGWNIPMGLLFN[+1622.582]QTGSC[+57.021]K.F | HexNAc(4)Hex(5) | C[+57], N[+1623] | 1239.8773 | 3 | 3717.6172 | 3717.5916 | 0 | 6.9 | 485 | Specific | 366.13 | 288.66 | 237.02 | 4.69 | 39 | >sp\|P02788\|TRFL_HUMAN Lactotransferrin OS=Homo sapiens OX=9606 GN=LTF PE=1 SV=6 | 1270 | 20201113_10633_1ng_glyco_hcd30_etd30_compound15.19046.19046.3 | controllerType=0 controllerNumber=1 scan=19046 | 41.3973 |
| 18170:3 | 1 | R.TAGWNIPMGLLFN[+2350.830]QTGSC[+57.021]K.F | HexNAc(4)Hex(5)Fuc(1)NeuAc(2) | C[+57], N[+2351] | 1482.6209 | 3 | 4445.8482 | 4445.8403 | 0 | 1.8 | 485 | Specific | 375.86 | 240.78 | 240.78 | 5.61 | 39 | >sp\|P02788\|TRFL_HUMAN Lactotransferrin OS=Homo sapiens OX=9606 GN=LTF PE=1 SV=6 | 1270 | 20201113_10633_1ng_glyco_hcd30_etd30_compound15.21895.21895.3 | controllerType=0 controllerNumber=1 scan=21895 | 49.3228 |
| 16655:3 | 1 | R.TAGWNIPMGLLFN[+2059.735]QTGSC[+57.021]K.F | HexNAc(4)Hex(5)Fuc(1)NeuAc(1) | C[+57], N[+2060] | 1385.5936 | 3 | 4154.7661 | 4154.7449 | 0 | 5.1 | 485 | Specific | 376.25 | 268.34 | 268.34 | 4.8 | 39 | >sp\|P02788\|TRFL_HUMAN Lactotransferrin OS=Homo sapiens OX=9606 GN=LTF PE=1 SV=6 | 1270 | 20201113_10633_1ng_glyco_hcd30_etd30_compound15.20296.20296.3 | controllerType=0 controllerNumber=1 scan=20296 | 44.9787 |
| 15394:3 | 1 | R.TAGWNIPMGLLFN[+1606.587]QTGSC[+57.021]K.F | HexNAc(4)Hex(4)Fuc(1) | C[+57], N[+1607] | 1234.5368 | 3 | 3701.5958 | 3701.5966 | 0 | -0.2 | 485 | Specific | 385.58 | 305.83 | 305.83 | 5.6 | 39 | >sp\|P02788\|TRFL_HUMAN Lactotransferrin OS=Homo sapiens OX=9606 GN=LTF PE=1 SV=6 | 1270 | 20201113_10633_1ng_glyco_hcd30_etd30_compound15.18966.18966.3 | controllerType=0 controllerNumber=1 scan=18966 | 41.1485 |
| 15300:3 | 1 | R.TAGWNIPMGLLFN[+1768.640]QTGSC[+57.021]K.F | HexNAc(4)Hex(5)Fuc(1) | C[+57], N[+1769] | 1288.5568 | 3 | 3863.6559 | 3863.6495 | 0 | 1.7 | 485 | Specific | 400.1 | 307.24 | 307.24 | 4.93 | 39 | >sp\|P02788\|TRFL_HUMAN Lactotransferrin OS=Homo sapiens OX=9606 GN=LTF PE=1 SV=6 | 1270 | 20201113_10633_1ng_glyco_hcd30_etd30_compound15.18868.18868.3 | controllerType=0 controllerNumber=1 scan=18868 | 40.8891 |
| 16741:3 | 1 | R.TAGWNIPMGLLFN[+1913.677]QTGSC[+57.021]K.F | HexNAc(4)Hex(5)NeuAc(1) | C[+57], N[+1914] | 1336.9029 | 3 | 4008.6941 | 4008.687 | 0 | 1.8 | 485 | Specific | 431.46 | 296.24 | 296.24 | 5.69 | 39 | >sp\|P02788\|TRFL_HUMAN Lactotransferrin OS=Homo sapiens OX=9606 GN=LTF PE=1 SV=6 | 1270 | 20201113_10633_1ng_glyco_hcd30_etd30_compound15.20387.20387.3 | controllerType=0 controllerNumber=1 scan=20387 | 45.2864 |
| 16871:4 | 1 | R.TAGWNIPMGLLFN[+1913.677]QTGSC[+57.021]KFDEYFSQSC[+57.021]APGSDPR.S | HexNAc(4)Hex(5)NeuAc(1) | C[+57]*2, N[+1914] | 1463.8696 | 4 | 5852.4567 | 5852.4341 | 0 | 3.9 | 485 | Specific | 311.69 | 271.59 | 271.59 | 4 | 39 | >sp\|P02788\|TRFL_HUMAN Lactotransferrin OS=Homo sapiens OX=9606 GN=LTF PE=1 SV=6 | 1270 | 20201113_10633_1ng_glyco_hcd30_etd30_compound15.20525.20525.4 | controllerType=0 controllerNumber=1 scan=20525 | 45.733 |
| 16112:4 | 1 | R.TAGWNIPMGLLFN[+1606.587]QTGSC[+57.021]KFDEYFSQSC[+57.021]APGSDPR.S | HexNAc(4)Hex(4)Fuc(1) | C[+57]*2, N[+1607] | 1387.0923 | 4 | 5545.3473 | 5545.3438 | 0 | 0.6 | 485 | Specific | 376.87 | 287.26 | 287.26 | 4.47 | 39 | >sp\|P02788\|TRFL_HUMAN Lactotransferrin OS=Homo sapiens OX=9606 GN=LTF PE=1 SV=6 | 1270 | 20201113_10633_1ng_glyco_hcd30_etd30_compound15.19726.19726.4 | controllerType=0 controllerNumber=1 scan=19726 | 43.4623 |
| 16095:4 | 1 | R.TAGWNIPMGLLFN[+1768.640]QTGSC[+57.021]KFDEYFSQSC[+57.021]APGSDPR.S | HexNAc(4)Hex(5)Fuc(1) | C[+57]*2, N[+1769] | 1427.6083 | 4 | 5707.4113 | 5707.3966 | 0 | 2.6 | 485 | Specific | 492.81 | 450.81 | 450.81 | 7.08 | 39 | >sp\|P02788\|TRFL_HUMAN Lactotransferrin OS=Homo sapiens OX=9606 GN=LTF PE=1 SV=6 | 1270 | 20201113_10633_1ng_glyco_hcd30_etd30_compound15.19708.19708.4 | controllerType=0 controllerNumber=1 scan=19708 | 43.4014 |
| 08412:3 | 1 | R.N[+2205.793]GSDC[+57.021]PDKFC[+57.021]LFQSETK.N | HexNAc(4)Hex(5)Fuc(2)NeuAc(1) | C[+57]*2, N[+2206] | 1413.5594 | 3 | 4238.6636 | 4238.6667 | 0 | -0.7 | 642 | Specific | 200.48 | 86.39 | 86.39 | 3.97 | 39 | >sp\|P02788\|TRFL_HUMAN Lactotransferrin OS=Homo sapiens OX=9606 GN=LTF PE=1 SV=6 | 1270 | 20201113_10633_1ng_glyco_hcd30_etd30_compound15.11627.11627.2 | controllerType=0 controllerNumber=1 scan=11627 | 26.7876 |
| 08502:3 | 1 | R.N[+2059.735]GSDC[+57.021]PDKFC[+57.021]LFQSETK.N | HexNAc(4)Hex(5)Fuc(1)NeuAc(1) | C[+57]*2, N[+2060] | 1364.8764 | 3 | 4092.6146 | 4092.6088 | 0 | 1.4 | 642 | Specific | 213.64 | 102.84 | 102.84 | 4.62 | 39 | >sp\|P02788\|TRFL_HUMAN Lactotransferrin OS=Homo sapiens OX=9606 GN=LTF PE=1 SV=6 | 1270 | 20201113_10633_1ng_glyco_hcd30_etd30_compound15.11721.11721.3 | controllerType=0 controllerNumber=1 scan=11721 | 26.9628 |
| 08381:3 | 1 | R.N[+2205.793]GSDC[+57.021]PDKFC[+57.021]LFQSETK.N | HexNAc(4)Hex(5)Fuc(2)NeuAc(1) | C[+57]*2, N[+2206] | 1413.5535 | 3 | 4238.646 | 4238.6667 | 0 | -4.9 | 642 | Specific | 287.82 | 187.02 | 187.02 | 3.41 | 39 | >sp\|P02788\|TRFL_HUMAN Lactotransferrin OS=Homo sapiens OX=9606 GN=LTF PE=1 SV=6 | 1270 | 20201113_10633_1ng_glyco_hcd30_etd30_compound15.11594.11594.3 | controllerType=0 controllerNumber=1 scan=11594 | 26.7291 |
| 15534:4 | 2 | K.RNIAAFGGDPNN[+1913.677]ITLFGESAGGASVSLQTLSPYNK.G | HexNAc(4)Hex(5)NeuAc(1) | N[+1914] | 1370.8721 | 4 | 5480.4665 | 5480.4644 | 0 | 0.4 | 196 | Specific | 387.92 | 383.76 | 383.76 | 6.14 | 13 | >sp\|P19835\|CEL_HUMAN Bile salt-activated lipase OS=Homo sapiens OX=9606 GN=CEL PE=1 SV=3 | 476 | 20201113_10633_1ng_glyco_hcd30_etd30_compound15.19114.19114.4 | controllerType=0 controllerNumber=1 scan=19114 | 41.6052 |
| 16789:3 | 2 | R.NIAAFGGDPNN[+1913.677]ITLFGESAGGASVSLQTLSPYNK.G | HexNAc(4)Hex(5)NeuAc(1) | N[+1914] | 1775.4611 | 3 | 5324.3689 | 5324.3633 | 0 | 1 | 197 | Specific | 469.54 | 324.2 | 324.2 | 6.37 | 13 | >sp\|P19835\|CEL_HUMAN Bile salt-activated lipase OS=Homo sapiens OX=9606 GN=CEL PE=1 SV=3 | 476 | 20201113_10633_1ng_glyco_hcd30_etd30_compound15.20438.20438.3 | controllerType=0 controllerNumber=1 scan=20438 | 45.4717 |
| 15931:3 | 4 | R.LSLHRPALEDLLLGSEAN[+1216.423]LTC[+57.021]TLTGLR.D | HexNAc(2)Hex(5) | C[+57], N[+1216] | 1394.0228 | 3 | 4180.0537 | 4180.0211 | 0 | 7.8 | 127 | Specific | 302.87 | 302.87 | 302.87 | 4.89 | 13 | >sp\|P01876\|IGHA1_HUMAN Immunoglobulin heavy constant alpha 1 OS=Homo sapiens OX=9606 GN=IGHA1 PE=1 SV=2 | 255 | 20201113_10633_1ng_glyco_hcd30_etd30_compound15.19534.19534.3 | controllerType=0 controllerNumber=1 scan=19534 | 42.8571 |
| 15914:5 | 4 | R.LSLHRPALEDLLLGSEAN[+1501.555]LTC[+57.021]TLTGLR.D | HexNAc(5)Hex(3) | C[+57], N[+1502] | 893.8385 | 5 | 4465.1635 | 4465.1536 | 0 | 2.2 | 127 | Specific | 346.47 | 346.47 | 346.47 | 4.85 | 13 | >sp\|P01876\|IGHA1_HUMAN Immunoglobulin heavy constant alpha 1 OS=Homo sapiens OX=9606 GN=IGHA1 PE=1 SV=2 | 255 | 20201113_10633_1ng_glyco_hcd30_etd30_compound15.19516.19516.3 | controllerType=0 controllerNumber=1 scan=19516 | 42.7916 |
| 15868:3 | 4 | R.LSLHRPALEDLLLGSEAN[+1540.529]LTC[+57.021]TLTGLR.D | HexNAc(2)Hex(7) | C[+57], N[+1541] | 1502.0478 | 3 | 4504.1288 | 4504.1268 | 0 | 0.5 | 127 | Specific | 348.61 | 289.64 | 289.64 | 5.62 | 13 | >sp\|P01876\|IGHA1_HUMAN Immunoglobulin heavy constant alpha 1 OS=Homo sapiens OX=9606 GN=IGHA1 PE=1 SV=2 | 255 | 20201113_10633_1ng_glyco_hcd30_etd30_compound15.19466.19466.3 | controllerType=0 controllerNumber=1 scan=19466 | 42.6026 |
| 15893:3 | 4 | R.LSLHRPALEDLLLGSEAN[+1663.608]LTC[+57.021]TLTGLR.D | HexNAc(5)Hex(4) | C[+57], N[+1664] | 1543.0883 | 3 | 4627.2503 | 4627.2064 | 0 | 9.5 | 127 | Specific | 381.36 | 366.49 | 366.49 | 5.52 | 13 | >sp\|P01876\|IGHA1_HUMAN Immunoglobulin heavy constant alpha 1 OS=Homo sapiens OX=9606 GN=IGHA1 PE=1 SV=2 | 255 | 20201113_10633_1ng_glyco_hcd30_etd30_compound15.19493.19493.3 | controllerType=0 controllerNumber=1 scan=19493 | 42.702 |
| 15925:3 | 4 | R.LSLHRPALEDLLLGSEAN[+1501.555]LTC[+57.021]TLTGLR.D | HexNAc(5)Hex(3) | C[+57], N[+1502] | 1489.0688 | 3 | 4465.1918 | 4465.1536 | 0 | 8.6 | 127 | Specific | 461.22 | 456.03 | 456.03 | 6.79 | 13 | >sp\|P01876\|IGHA1_HUMAN Immunoglobulin heavy constant alpha 1 OS=Homo sapiens OX=9606 GN=IGHA1 PE=1 SV=2 | 255 | 20201113_10633_1ng_glyco_hcd30_etd30_compound15.19528.19528.3 | controllerType=0 controllerNumber=1 scan=19528 | 42.8323 |
| 15859:4 | 4 | R.LSLHRPALEDLLLGSEAN[+1702.581]LTC[+57.021]TLTGLR.D | HexNAc(2)Hex(8) | C[+57], N[+1703] | 1167.3002 | 4 | 4666.179 | 4666.1796 | 0 | -0.1 | 127 | Specific | 809.85 | 764.45 | 764.45 | 13.66 | 13 | >sp\|P01876\|IGHA1_HUMAN Immunoglobulin heavy constant alpha 1 OS=Homo sapiens OX=9606 GN=IGHA1 PE=1 SV=2 | 255 | 20201113_10633_1ng_glyco_hcd30_etd30_compound15.19457.19457.4 | controllerType=0 controllerNumber=1 scan=19457 | 42.5764 |
| 15926:4 | 4 | R.LSLHRPALEDLLLGSEAN[+1216.423]LTC[+57.021]TLTGLR.D | HexNAc(2)Hex(5) | C[+57], N[+1216] | 1045.7675 | 4 | 4180.0482 | 4180.0211 | 0 | 6.5 | 127 | Specific | 820.85 | 780.84 | 780.84 | 12.89 | 13 | >sp\|P01876\|IGHA1_HUMAN Immunoglobulin heavy constant alpha 1 OS=Homo sapiens OX=9606 GN=IGHA1 PE=1 SV=2 | 255 | 20201113_10633_1ng_glyco_hcd30_etd30_compound15.19529.19529.4 | controllerType=0 controllerNumber=1 scan=19529 | 42.8392 |
| 15879:4 | 4 | R.LSLHRPALEDLLLGSEAN[+1501.555]LTC[+57.021]TLTGLR.D | HexNAc(5)Hex(3) | C[+57], N[+1502] | 1117.0546 | 4 | 4465.1966 | 4465.1536 | 0 | 9.6 | 127 | Specific | 888.01 | 884.76 | 884.76 | 14.04 | 13 | >sp\|P01876\|IGHA1_HUMAN Immunoglobulin heavy constant alpha 1 OS=Homo sapiens OX=9606 GN=IGHA1 PE=1 SV=2 | 255 | 20201113_10633_1ng_glyco_hcd30_etd30_compound15.19478.19478.4 | controllerType=0 controllerNumber=1 scan=19478 | 42.6295 |
| 15916:4 | 4 | R.LSLHRPALEDLLLGSEAN[+1298.476]LTC[+57.021]TLTGLR.D | HexNAc(4)Hex(3) | C[+57], N[+1298] | 1066.2758 | 4 | 4262.0814 | 4262.0742 | 0 | 1.7 | 127 | Specific | 932.53 | 932.53 | 932.53 | 15.89 | 13 | >sp\|P01876\|IGHA1_HUMAN Immunoglobulin heavy constant alpha 1 OS=Homo sapiens OX=9606 GN=IGHA1 PE=1 SV=2 | 255 | 20201113_10633_1ng_glyco_hcd30_etd30_compound15.19518.19518.4 | controllerType=0 controllerNumber=1 scan=19518 | 42.8031 |
| 15860:4 | 4 | R.LSLHRPALEDLLLGSEAN[+1540.529]LTC[+57.021]TLTGLR.D | HexNAc(2)Hex(7) | C[+57], N[+1541] | 1126.788 | 4 | 4504.1302 | 4504.1268 | 0 | 0.8 | 127 | Specific | 1006.84 | 921.64 | 921.64 | 17.89 | 13 | >sp\|P01876\|IGHA1_HUMAN Immunoglobulin heavy constant alpha 1 OS=Homo sapiens OX=9606 GN=IGHA1 PE=1 SV=2 | 255 | 20201113_10633_1ng_glyco_hcd30_etd30_compound15.19458.19458.4 | controllerType=0 controllerNumber=1 scan=19458 | 42.5817 |
| 10546:3 | 4 | R.LAGKPTHVN[+1501.555]VSVVMAEVDGTC[+57.021]Y.- | HexNAc(5)Hex(3) | C[+57], N[+1502] | 1283.5732 | 3 | 3848.705 | 3848.6974 | 0 | 2 | 332 | CRagged | 202.89 | 136.12 | 136.12 | 2.89 | 13 | >sp\|P01876\|IGHA1_HUMAN Immunoglobulin heavy constant alpha 1 OS=Homo sapiens OX=9606 GN=IGHA1 PE=1 SV=2 | 255 | 20201113_10633_1ng_glyco_hcd30_etd30_compound15.13841.13841.3 | controllerType=0 controllerNumber=1 scan=13841 | 30.9481 |
| 10458:3 | 4 | R.LAGKPTHVN[+1378.476]VSVVMAEVDGTC[+57.021]Y.- | HexNAc(2)Hex(6) | C[+57], N[+1378] | 1242.545 | 3 | 3725.6204 | 3725.6178 | 0 | 0.7 | 332 | CRagged | 237.87 | 187.6 | 187.6 | 2.95 | 13 | >sp\|P01876\|IGHA1_HUMAN Immunoglobulin heavy constant alpha 1 OS=Homo sapiens OX=9606 GN=IGHA1 PE=1 SV=2 | 255 | 20201113_10633_1ng_glyco_hcd30_etd30_compound15.13749.13749.3 | controllerType=0 controllerNumber=1 scan=13749 | 30.7292 |
| 10483:3 | 4 | R.LAGKPTHVN[+1216.423]VSVVMAEVDGTC[+57.021]Y.- | HexNAc(2)Hex(5) | C[+57], N[+1216] | 1188.5274 | 3 | 3563.5676 | 3563.5649 | 0 | 0.8 | 332 | CRagged | 247.53 | 235.29 | 178.42 | 3.7 | 13 | >sp\|P01876\|IGHA1_HUMAN Immunoglobulin heavy constant alpha 1 OS=Homo sapiens OX=9606 GN=IGHA1 PE=1 SV=2 | 255 | 20201113_10633_1ng_glyco_hcd30_etd30_compound15.13775.13775.3 | controllerType=0 controllerNumber=1 scan=13775 | 30.7829 |
| 15904:3 | 5 | R.GLSFDVSLEVSQGPGLLN[+2059.735]DTK.V | HexNAc(4)Hex(5)Fuc(1)NeuAc(1) | N[+2060] | 1412.6259 | 3 | 4235.8632 | 4235.8481 | 0 | 3.6 | 118 | Specific | 470.33 | 327.76 | 327.76 | 5.33 | 16 | >sp\|P01833\|PIGR_HUMAN Polymeric immunoglobulin receptor OS=Homo sapiens OX=9606 GN=PIGR PE=1 SV=4 | 902 | 20201113_10633_1ng_glyco_hcd30_etd30_compound15.19505.19505.3 | controllerType=0 controllerNumber=1 scan=19505 | 42.7547 |

Table S4. Cont.

| Query #:z | Protein Rank | Glycopeptide | Glycan | Modification Type(s) | Observed m/z | z | Observed (M+H) | Calc. mass (M+H) | Off-by-x error | Mass error (ppm) | StartPosition | Cleavage | Score | Delta | Delta Mod | \|Log Prob\| | # of unique peptides | Protein Name | Protein DB number | Comment | Scan # | Scan Time |
|---|---|---|---|---|---|---|---|---|---|---|---|---|---|---|---|---|---|---|---|---|---|---|
| 16759:3 | 5 | K.QIGLYPVLVIDSSGYVNPN[+2059.735]YTGR.I | HexNAc(4)Hex(5)Fuc(1)NeuAc(1) | N[+2060] | 1529.0241 | 3 | 4585.0578 | 4585.0384 | 0 | 4.2 | 168 | Specific | 472.15 | 385.66 | 385.66 | 6.54 | 16 | >sp\|P01833\|PIGR_HUMAN Polymeric immunoglobulin receptor OS=Homo sapiens OX=9606 GN=PIGR PE=1 SV=4 | 902 | 20201113_10633_1ng_glyco_hcd30_etd30_compound15.20406.20406.3 | controllerType=0 controllerNumber=1 scan=20406 | 45.3485 |
| 19931:4 | 5 | R.LSLLEEPGN[+2350.830]GTFTVILNQLTSR.D | HexNAc(4)Hex(5)Fuc(1)NeuAc(2) | N[+2351] | 1189.0414 | 4 | 4753.1439 | 4753.1229 | 0 | 4.4 | 413 | Specific | 546.91 | 479.22 | 479.22 | 5.84 | 16 | >sp\|P01833\|PIGR_HUMAN Polymeric immunoglobulin receptor OS=Homo sapiens OX=9606 GN=PIGR PE=1 SV=4 | 902 | 20201113_10633_1ng_glyco_hcd30_etd30_compound15.23750.23750.4 | controllerType=0 controllerNumber=1 scan=23750 | 53.2216 |
| 16086:3 | 5 | K.IIEGEPNLKVPGN[+2350.830]VTAVLGETLK.V | HexNAc(4)Hex(5)Fuc(1)NeuAc(2) | N[+2351] | 1581.4003 | 3 | 4742.1864 | 4742.1797 | 0 | 1.4 | 457 | Specific | 205.82 | 153.57 | 153.57 | 3.21 | 16 | >sp\|P01833\|PIGR_HUMAN Polymeric immunoglobulin receptor OS=Homo sapiens OX=9606 GN=PIGR PE=1 SV=4 | 902 | 20201113_10633_1ng_glyco_hcd30_etd30_compound15.19698.19698.3 | controllerType=0 controllerNumber=1 scan=19698 | 43.362 |
| 14279:4 | 5 | K.IIEGEPNLKVPGN[+2204.772]VTAVLGETLKVPC[+57.021]HFPC[+57.021]K.F | HexNAc(4)Hex(5)NeuAc(2) | C[+57]*2, N[+2205] | 1406.1525 | 4 | 5621.588 | 5621.5793 | 0 | 1.6 | 457 | Specific | 336.7 | 336.7 | 336.7 | 3.66 | 16 | >sp\|P01833\|PIGR_HUMAN Polymeric immunoglobulin receptor OS=Homo sapiens OX=9606 GN=PIGR PE=1 SV=4 | 902 | 20201113_10633_1ng_glyco_hcd30_etd30_compound15.17747.17747.4 | controllerType=0 controllerNumber=1 scan=17747 | 38.4827 |
| 15475:4 | 5 | K.IIEGEPNLKVPGN[+2350.830]VTAVLGETLKVPC[+57.021]HFPC[+57.021]K.F | HexNAc(4)Hex(5)Fuc(1)NeuAc(2) | C[+57]*2, N[+2351] | 1442.6766 | 4 | 5767.6847 | 5767.6372 | 0 | 8.2 | 457 | Specific | 389.77 | 389.77 | 389.77 | 3.9 | 16 | >sp\|P01833\|PIGR_HUMAN Polymeric immunoglobulin receptor OS=Homo sapiens OX=9606 GN=PIGR PE=1 SV=4 | 902 | 20201113_10633_1ng_glyco_hcd30_etd30_compound15.19052.19052.4 | controllerType=0 controllerNumber=1 scan=19052 | 41.4124 |
| 12509:3 | 5 | K.VPGN[+2205.793]VTAVLGETLK.V | HexNAc(4)Hex(5)Fuc(2)NeuAc(1) | N[+2206] | 1201.8714 | 3 | 3603.5996 | 3603.5927 | 0 | 1.9 | 466 | Specific | 202.05 | 153.14 | 153.14 | 4.3 | 16 | >sp\|P01833\|PIGR_HUMAN Polymeric immunoglobulin receptor OS=Homo sapiens OX=9606 GN=PIGR PE=1 SV=4 | 902 | 20201113_10633_1ng_glyco_hcd30_etd30_compound15.15905.15905.3 | controllerType=0 controllerNumber=1 scan=15905 | 34.6606 |
| 12607:4 | 5 | K.VPGN[+2059.735]VTAVLGETLKVPC[+57.021]HFPC[+57.021]K.F | HexNAc(4)Hex(5)Fuc(1)NeuAc(1) | C[+57]*2, N[+2060] | 1121.5044 | 4 | 4482.9959 | 4482.9923 | 0 | 0.8 | 466 | Specific | 361.91 | 306.67 | 306.67 | 4.24 | 16 | >sp\|P01833\|PIGR_HUMAN Polymeric immunoglobulin receptor OS=Homo sapiens OX=9606 GN=PIGR PE=1 SV=4 | 902 | 20201113_10633_1ng_glyco_hcd30_etd30_compound15.16007.16007.4 | controllerType=0 controllerNumber=1 scan=16007 | 34.8991 |
| 07186:3 | 5 | K.WN[+2204.772]NTGC[+57.021]QALPSQDEGPSK.A | HexNAc(4)Hex(5)NeuAc(2) | C[+57], N[+2205] | 1398.5549 | 3 | 4193.65 | 4193.6491 | 0 | 0.2 | 498 | Specific | 301.5 | 159.41 | 159.41 | 4.8 | 16 | >sp\|P01833\|PIGR_HUMAN Polymeric immunoglobulin receptor OS=Homo sapiens OX=9606 GN=PIGR PE=1 SV=4 | 902 | 20201113_10633_1ng_glyco_hcd30_etd30_compound15.10337.10337.3 | controllerType=0 controllerNumber=1 scan=10337 | 24.3869 |
| 05252:3 | 5 | K.WN[+2059.735]NTGC[+57.021]QALPSQDEGPSK.A | HexNAc(4)Hex(5)Fuc(1)NeuAc(1) | C[+57], N[+2060] | 1350.2115 | 3 | 4048.6199 | 4048.6116 | 0 | 2 | 498 | Specific | 459.18 | 305.68 | 305.68 | 6.12 | 16 | >sp\|P01833\|PIGR_HUMAN Polymeric immunoglobulin receptor OS=Homo sapiens OX=9606 GN=PIGR PE=1 SV=4 | 902 | 20201113_10633_1ng_glyco_hcd30_etd30_compound15.8329.8329.3 | controllerType=0 controllerNumber=1 scan=8329 | 20.7983 |
| 05305:3 | 5 | K.WN[+1913.677]NTGC[+57.021]QALPSQDEGPSK.A | HexNAc(4)Hex(5)NeuAc(1) | C[+57], N[+1914] | 1301.5248 | 3 | 3902.56 | 3902.5537 | 0 | 1.6 | 498 | Specific | 482.41 | 373.19 | 373.19 | 7.28 | 16 | >sp\|P01833\|PIGR_HUMAN Polymeric immunoglobulin receptor OS=Homo sapiens OX=9606 GN=PIGR PE=1 SV=4 | 902 | 20201113_10633_1ng_glyco_hcd30_etd30_compound15.8384.8384.3 | controllerType=0 controllerNumber=1 scan=8384 | 20.8994 |
| 07103:3 | 11 | R.LSPN[+2350.830]ASAEHLELR.W | HexNAc(4)Hex(5)Fuc(1)NeuAc(2) | N[+2351] | 1263.199 | 3 | 3787.5825 | 3787.5796 | 0 | 0.8 | 52 | Specific | 362.62 | 207.2 | 207.2 | 4.88 | 6 | >sp\|Q13410\|BT1A1_HUMAN Butyrophilin subfamily 1 member A1 OS=Homo sapiens OX=9606 GN=BTN1A1 PE=1 SV=3 | 179 | 20201113_10633_1ng_glyco_hcd30_etd30_compound15.10251.10251.3 | controllerType=0 controllerNumber=1 scan=10251 | 24.2222 |
| 05236:3 | 11 | R.LSPN[+2205.793]ASAEHLELR.W | HexNAc(4)Hex(5)Fuc(2)NeuAc(1) | N[+2206] | 1214.8548 | 3 | 3642.5498 | 3642.5421 | 0 | 2.1 | 52 | Specific | 375.8 | 269.94 | 269.94 | 5.22 | 6 | >sp\|Q13410\|BT1A1_HUMAN Butyrophilin subfamily 1 member A1 OS=Homo sapiens OX=9606 GN=BTN1A1 PE=1 SV=3 | 179 | 20201113_10633_1ng_glyco_hcd30_etd30_compound15.8312.8312.3 | controllerType=0 controllerNumber=1 scan=8312 | 20.7598 |
| 07824:3 | 12 | K.TAHC[+57.021]N[+2059.735]ESFYFLC[+57.021]K.R | HexNAc(4)Hex(5)Fuc(1)NeuAc(1) | C[+57]*2, N[+2060] | 1246.1584 | 3 | 3736.4606 | 3736.4545 | 0 | 1.6 | 1201 | Specific | 247.19 | 152.75 | 152.75 | 4.42 | 7 | >sp\|P22897\|MRC1_HUMAN Macrophage mannose receptor 1 OS=Homo sapiens OX=9606 GN=MRC1 PE=1 SV=1 | 1049 | 20201113_10633_1ng_glyco_hcd30_etd30_compound15.11010.11010.3 | controllerType=0 controllerNumber=1 scan=11010 | 25.6764 |
| 07811:3 | 12 | K.TAHC[+57.021]N[+2059.735]ESFYFLC[+57.021]K.R | HexNAc(4)Hex(5)Fuc(1)NeuAc(1) | C[+57]*2, N[+2060] | 1246.1576 | 3 | 3736.4584 | 3736.4545 | 0 | 1 | 1201 | Specific | 285.15 | 191.66 | 191.66 | 4.35 | 7 | >sp\|P22897\|MRC1_HUMAN Macrophage mannose receptor 1 OS=Homo sapiens OX=9606 GN=MRC1 PE=1 SV=1 | 1049 | 20201113_10633_1ng_glyco_hcd30_etd30_compound15.10996.10996.3 | controllerType=0 controllerNumber=1 scan=10996 | 25.6365 |
| 21103:3 | 17 | R.QLAHQSN[+2204.772]STNIFFSPVSIATAFAMLSLGTK.A | HexNAc(4)Hex(5)NeuAc(2) | N[+2205] | 1796.1432 | 3 | 5386.4149 | 5386.4075 | 0 | 1.4 | 64 | Specific | 342.24 | 215.8 | 215.8 | 5.66 | 3 | >sp\|P01009\|A1AT_HUMAN Alpha-1-antitrypsin OS=Homo sapiens OX=9606 GN=SERPINA1 PE=1 SV=3 | 642 | 20201113_10633_1ng_glyco_hcd30_etd30_compound15.24975.24975.3 | controllerType=0 controllerNumber=1 scan=24975 | 55.9549 |
| 21092:3 | 17 | R.QLAHQSN[+2204.772]STNIFFSPVSIATAFAMLSLGTK.A | HexNAc(4)Hex(5)NeuAc(2) | N[+2205] | 1796.1496 | 3 | 5386.4343 | 5386.4075 | 0 | 5 | 64 | Specific | 379.95 | 275.88 | 275.88 | 5.26 | 3 | >sp\|P01009\|A1AT_HUMAN Alpha-1-antitrypsin OS=Homo sapiens OX=9606 GN=SERPINA1 PE=1 SV=3 | 642 | 20201113_10633_1ng_glyco_hcd30_etd30_compound15.24963.24963.3 | controllerType=0 controllerNumber=1 scan=24963 | 55.9219 |
| 21099:4 | 17 | R.QLAHQSN[+2204.772]STNIFFSPVSIATAFAMLSLGTK.A | HexNAc(4)Hex(5)NeuAc(2) | N[+2205] | 1347.3557 | 4 | 5386.401 | 5386.4075 | 0 | -1.2 | 64 | Specific | 613.67 | 511.39 | 418.93 | 9.66 | 3 | >sp\|P01009\|A1AT_HUMAN Alpha-1-antitrypsin OS=Homo sapiens OX=9606 GN=SERPINA1 PE=1 SV=3 | 642 | 20201113_10633_1ng_glyco_hcd30_etd30_compound15.24971.24971.4 | controllerType=0 controllerNumber=1 scan=24971 | 55.9432 |
| 15069:3 | 36 | R.NYIVPILWLN[+1864.634]ETGTIGDEKANM[+15.995]FR.S | HexNAc(2)Hex(9) | M[+16], N[+1865] | 1559.0278 | 3 | 4675.0687 | 4675.0523 | 0 | 3.5 | 408 | Specific | 342.94 | 342.94 | 342.94 | 5.22 | 2 | >sp\|P16671\|CD36_HUMAN Platelet glycoprotein 4 OS=Homo sapiens OX=9606 GN=CD36 PE=1 SV=2 | 812 | 20201113_10633_1ng_glyco_hcd30_etd30_compound15.18626.18626.3 | controllerType=0 controllerNumber=1 scan=18626 | 40.3659 |
| 12899:4 | 38 | K.LKELPGVC[+57.021]N[+2205.793]ETMMALWEEC[+57.021]KPC[+57.021]LK.Q | HexNAc(4)Hex(5)Fuc(2)NeuAc(1) | C[+57]*3, N[+2206] | 1286.3014 | 4 | 5142.1837 | 5142.1776 | 0 | 1.2 | 95 | Specific | 284.77 | 214.29 | 214.29 | 3.21 | 4 | >sp\|P10909\|CLUS_HUMAN Clusterin OS=Homo sapiens OX=9606 GN=CLU PE=1 SV=1 | 1467 | 20201113_10633_1ng_glyco_hcd30_etd30_compound15.16311.16311.4 | controllerType=0 controllerNumber=1 scan=16311 | 35.4968 |
| 12955:4 | 38 | K.LKELPGVC[+57.021]N[+2059.735]ETMMALWEEC[+57.021]KPC[+57.021]LK.Q | HexNAc(4)Hex(5)Fuc(1)NeuAc(1) | C[+57]*3, N[+2060] | 1249.7882 | 4 | 4996.131 | 4996.1196 | 0 | 2.3 | 95 | Specific | 380.41 | 281.79 | 281.79 | 3.96 | 4 | >sp\|P10909\|CLUS_HUMAN Clusterin OS=Homo sapiens OX=9606 GN=CLU PE=1 SV=1 | 1467 | 20201113_10633_1ng_glyco_hcd30_etd30_compound15.16369.16369.4 | controllerType=0 controllerNumber=1 scan=16369 | 35.618 |
| 12941:4 | 38 | K.LKELPGVC[+57.021]N[+2059.735]ETMMALWEEC[+57.021]KPC[+57.021]LK.Q | HexNAc(4)Hex(5)Fuc(1)NeuAc(1) | C[+57]*3, N[+2060] | 1249.786 | 4 | 4996.1222 | 4996.1196 | 0 | 0.5 | 95 | Specific | 460.84 | 376.03 | 376.03 | 6 | 4 | >sp\|P10909\|CLUS_HUMAN Clusterin OS=Homo sapiens OX=9606 GN=CLU PE=1 SV=1 | 1467 | 20201113_10633_1ng_glyco_hcd30_etd30_compound15.16354.16354.4 | controllerType=0 controllerNumber=1 scan=16354 | 35.5875 |
| 00144:4 | 38 | R.EIRHN[+2059.735]STGC[+57.021]LR.M | HexNAc(4)Hex(5)Fuc(1)NeuAc(1) | C[+57], N[+2060] | 851.3563 | 4 | 3402.4035 | 3402.3994 | 0 | 1.2 | 287 | Specific | 347.62 | 269.3 | 269.3 | 5.16 | 4 | >sp\|P10909\|CLUS_HUMAN Clusterin OS=Homo sapiens OX=9606 GN=CLU PE=1 SV=1 | 1467 | 20201113_10633_1ng_glyco_hcd30_etd30_compound15.3027.3027.4 | controllerType=0 controllerNumber=1 scan=3027 | 11.3428 |
| 09515:3 | 38 | R.LAN[+2205.793]LTQGEDQYYLR.V | HexNAc(4)Hex(5)Fuc(2)NeuAc(1) | N[+2206] | 1297.2175 | 3 | 3889.6378 | 3889.6265 | 0 | 2.9 | 372 | Specific | 252.95 | 162.76 | 162.76 | 3.9 | 4 | >sp\|P10909\|CLUS_HUMAN Clusterin OS=Homo sapiens OX=9606 GN=CLU PE=1 SV=1 | 1467 | 20201113_10633_1ng_glyco_hcd30_etd30_compound15.12770.12770.3 | controllerType=0 controllerNumber=1 scan=12770 | 28.8966 |
| 10359:3 | 54 | R.IIVPLNNREN[+2059.735]ISDPTSPLR.T | HexNAc(4)Hex(5)Fuc(1)NeuAc(1) | N[+2060] | 1403.3113 | 3 | 4207.9195 | 4207.9121 | 0 | 1.8 | 62 | Specific | 235.15 | 206.38 | 206.38 | 3.39 | 3 | >sp\|P01591\|IGJ_HUMAN Immunoglobulin J chain OS=Homo sapiens OX=9606 GN=JCHAIN PE=1 SV=4 | 1392 | 20201113_10633_1ng_glyco_hcd30_etd30_compound15.13645.13645.3 | controllerType=0 controllerNumber=1 scan=13645 | 30.4704 |
| 02672:3 | 60 | K.TPLTAN[+1971.719]ITK.S | HexNAc(5)Hex(5)Fuc(1) | N[+1972] | 977.4323 | 3 | 2930.2825 | 2930.2757 | 0 | 2.3 | 200 | Specific | 223.64 | 114 | 114 | 2.7 | 3 | >sp\|P01877\|IGHA2_HUMAN Immunoglobulin heavy constant alpha 2 OS=Homo sapiens OX=9606 GN=IGHA2 PE=1 SV=4 | 576 | 20201113_10633_1ng_glyco_hcd30_etd30_compound15.5652.5652.3 | controllerType=0 controllerNumber=1 scan=5652 | 16.1222 |
| 02776:3 | 60 | K.TPLTAN[+1647.613]ITK.S | HexNAc(5)Hex(3)Fuc(1) | N[+1648] | 869.3968 | 3 | 2606.1757 | 2606.17 | 0 | 2.2 | 200 | Specific | 302.65 | 127.34 | 127.34 | 2.75 | 3 | >sp\|P01877\|IGHA2_HUMAN Immunoglobulin heavy constant alpha 2 OS=Homo sapiens OX=9606 GN=IGHA2 PE=1 SV=4 | 576 | 20201113_10633_1ng_glyco_hcd30_etd30_compound15.5761.5761.3 | controllerType=0 controllerNumber=1 scan=5761 | 16.3649 |
| 02752:2 | 60 | K.TPLTAN[+1647.613]ITK.S | HexNAc(5)Hex(3)Fuc(1) | N[+1648] | 1303.5904 | 2 | 2606.1735 | 2606.17 | 0 | 1.3 | 200 | Specific | 302.86 | 114.07 | 114.07 | 2.77 | 3 | >sp\|P01877\|IGHA2_HUMAN Immunoglobulin heavy constant alpha 2 OS=Homo sapiens OX=9606 GN=IGHA2 PE=1 SV=4 | 576 | 20201113_10633_1ng_glyco_hcd30_etd30_compound15.5736.5736.2 | controllerType=0 controllerNumber=1 scan=5736 | 16.3122 |
| 02716:3 | 60 | K.TPLTAN[+1809.666]ITK.S | HexNAc(5)Hex(4)Fuc(1) | N[+1810] | 923.4137 | 3 | 2768.2266 | 2768.2228 | 0 | 1.4 | 200 | Specific | 313.1 | 132.37 | 132.37 | 2.77 | 3 | >sp\|P01877\|IGHA2_HUMAN Immunoglobulin heavy constant alpha 2 OS=Homo sapiens OX=9606 GN=IGHA2 PE=1 SV=4 | 576 | 20201113_10633_1ng_glyco_hcd30_etd30_compound15.5698.5698.3 | controllerType=0 controllerNumber=1 scan=5698 | 16.2239 |
| 15322:4 | 1 | R.RTAGWNVPIGTLRPFLN[+2716.983]WTGPPEPIEAAVAR.F | HexNAc(5)Hex(6)Fuc(3)NeuAc(1) | N[+2717] | 1526.7117 | 4 | 6103.8249 | 6103.795 | 0 | 4.9 | 140 | Specific | 749.01 | 749.01 | 749.01 | 10.87 | 50 | >sp\|P02788\|TRFL_HUMAN Lactotransferrin OS=Homo sapiens OX=9606 GN=LTF PE=1 SV=6 | 1270 | 20201113_12413_1ng_glyco_hcd30_etd30_compound15.18898.18898.4 | controllerType=0 controllerNumber=1 scan=18898 | 43.3083 |
| 14738:4 | 1 | R.RTAGWNVPIGTLRPFLN[+1768.640]WTGPPEPIEAAVAR.F | HexNAc(4)Hex(5)Fuc(1) | N[+1769] | 1289.6223 | 4 | 5155.4674 | 5155.4516 | 0 | 3.1 | 140 | Specific | 752.25 | 752.25 | 752.25 | 12.45 | 50 | >sp\|P02788\|TRFL_HUMAN Lactotransferrin OS=Homo sapiens OX=9606 GN=LTF PE=1 SV=6 | 1270 | 20201113_12413_1ng_glyco_hcd30_etd30_compound15.18281.18281.4 | controllerType=0 controllerNumber=1 scan=18281 | 41.4834 |
| 15331:4 | 1 | R.RTAGWNVPIGTLRPFLN[+2570.925]WTGPPEPIEAAVAR.F | HexNAc(5)Hex(6)Fuc(2)NeuAc(1) | N[+2571] | 1490.1947 | 4 | 5957.757 | 5957.7371 | 0 | 3.3 | 140 | Specific | 848.71 | 801.51 | 801.51 | 13.29 | 50 | >sp\|P02788\|TRFL_HUMAN Lactotransferrin OS=Homo sapiens OX=9606 GN=LTF PE=1 SV=6 | 1270 | 20201113_12413_1ng_glyco_hcd30_etd30_compound15.18908.18908.4 | controllerType=0 controllerNumber=1 scan=18908 | 43.3424 |
| 15404:4 | 1 | R.RTAGWNVPIGTLRPFLN[+2059.735]WTGPPEPIEAAVAR.F | HexNAc(4)Hex(5)Fuc(1)NeuAc(1) | N[+2060] | 1362.3921 | 4 | 5446.5465 | 5446.547 | 0 | -0.1 | 140 | Specific | 1039.18 | 988.62 | 988.62 | 16.71 | 50 | >sp\|P02788\|TRFL_HUMAN Lactotransferrin OS=Homo sapiens OX=9606 GN=LTF PE=1 SV=6 | 1270 | 20201113_12413_1ng_glyco_hcd30_etd30_compound15.18985.18985.4 | controllerType=0 controllerNumber=1 scan=18985 | 43.5851 |
| 17203:3 | 1 | R.TAGWNVPIGTLRPFLN[+2350.830]WTGPPEPIEAAVAR.F | HexNAc(4)Hex(5)Fuc(1)NeuAc(2) | N[+2351] | 1861.187 | 3 | 5581.5464 | 5581.5413 | 0 | 0.9 | 141 | Specific | 205.77 | 114.93 | 114.93 | 4.04 | 50 | >sp\|P02788\|TRFL_HUMAN Lactotransferrin OS=Homo sapiens OX=9606 GN=LTF PE=1 SV=6 | 1270 | 20201113_12413_1ng_glyco_hcd30_etd30_compound15.20890.20890.3 | controllerType=0 controllerNumber=1 scan=20890 | 49.3233 |
| 15609:4 | 1 | R.TAGWNVPIGTLRPFLN[+1913.677]WTGPPEPIEAAVAR.F | HexNAc(4)Hex(5)NeuAc(1) | N[+1914] | 1286.8591 | 4 | 5144.4147 | 5144.388 | 0 | 5.2 | 141 | Specific | 215.97 | 215.97 | 215.97 | 3.05 | 50 | >sp\|P02788\|TRFL_HUMAN Lactotransferrin OS=Homo sapiens OX=9606 GN=LTF PE=1 SV=6 | 1270 | 20201113_12413_1ng_glyco_hcd30_etd30_compound15.19201.19201.4 | controllerType=0 controllerNumber=1 scan=19201 | 44.2312 |
| 16287:3 | 1 | R.TAGWNVPIGTLRPFLN[+2205.793]WTGPPEPIEAAVAR.F | HexNAc(4)Hex(5)Fuc(2)NeuAc(1) | N[+2206] | 1812.849 | 3 | 5436.5323 | 5436.5038 | 0 | 5.3 | 141 | Specific | 281.81 | 208.98 | 208.98 | 4.03 | 50 | >sp\|P02788\|TRFL_HUMAN Lactotransferrin OS=Homo sapiens OX=9606 GN=LTF PE=1 SV=6 | 1270 | 20201113_12413_1ng_glyco_hcd30_etd30_compound15.19920.19920.3 | controllerType=0 controllerNumber=1 scan=19920 | 46.4529 |
| 17209:4 | 1 | R.TAGWNVPIGTLRPFLN[+2350.830]WTGPPEPIEAAVAR.F | HexNAc(4)Hex(5)Fuc(1)NeuAc(2) | N[+2351] | 1396.1475 | 4 | 5581.568 | 5581.5413 | 0 | 4.8 | 141 | Specific | 362.15 | 362.15 | 362.15 | 4.6 | 50 | >sp\|P02788\|TRFL_HUMAN Lactotransferrin OS=Homo sapiens OX=9606 GN=LTF PE=1 SV=6 | 1270 | 20201113_12413_1ng_glyco_hcd30_etd30_compound15.20897.20897.4 | controllerType=0 controllerNumber=1 scan=20897 | 49.3444 |
| 16241:3 | 1 | R.TAGWNVPIGTLRPFLN[+2351.851]WTGPPEPIEAAVAR.F | HexNAc(4)Hex(5)Fuc(3)NeuAc(1) | N[+2352] | 1861.5256 | 3 | 5582.5622 | 5582.5617 | 0 | 0.1 | 141 | Specific | 400.74 | 340.28 | 340.28 | 4.64 | 50 | >sp\|P02788\|TRFL_HUMAN Lactotransferrin OS=Homo sapiens OX=9606 GN=LTF PE=1 SV=6 | 1270 | 20201113_12413_1ng_glyco_hcd30_etd30_compound15.19871.19871.3 | controllerType=0 controllerNumber=1 scan=19871 | 46.2878 |
| 16256:3 | 1 | R.TAGWNVPIGTLRPFLN[+2059.735]WTGPPEPIEAAVAR.F | HexNAc(4)Hex(5)Fuc(1)NeuAc(1) | N[+2060] | 1764.1588 | 3 | 5290.4618 | 5290.4459 | 0 | 3 | 141 | Specific | 513.48 | 378.64 | 378.64 | 6.69 | 50 | >sp\|P02788\|TRFL_HUMAN Lactotransferrin OS=Homo sapiens OX=9606 GN=LTF PE=1 SV=6 | 1270 | 20201113_12413_1ng_glyco_hcd30_etd30_compound15.19887.19887.3 | controllerType=0 controllerNumber=1 scan=19887 | 46.3595 |
| 16171:4 | 1 | R.TAGWNVPIGTLRPFLN[+2570.925]WTGPPEPIEAAVAR.F | HexNAc(5)Hex(6)Fuc(2)NeuAc(1) | N[+2571] | 1451.1663 | 4 | 5801.6432 | 5801.636 | 0 | 1.3 | 141 | Specific | 515.21 | 515.21 | 515.21 | 6.64 | 50 | >sp\|P02788\|TRFL_HUMAN Lactotransferrin OS=Homo sapiens OX=9606 GN=LTF PE=1 SV=6 | 1270 | 20201113_12413_1ng_glyco_hcd30_etd30_compound15.19797.19797.4 | controllerType=0 controllerNumber=1 scan=19797 | 46.0584 |
| 16242:3 | 1 | R.TAGWNVPIGTLRPFLN[+2059.735]WTGPPEPIEAAVAR.F | HexNAc(4)Hex(5)Fuc(1)NeuAc(1) | N[+2060] | 1764.1569 | 3 | 5290.4563 | 5290.4459 | 0 | 2 | 141 | Specific | 541 | 423.81 | 423.81 | 6.9 | 50 | >sp\|P02788\|TRFL_HUMAN Lactotransferrin OS=Homo sapiens OX=9606 GN=LTF PE=1 SV=6 | 1270 | 20201113_12413_1ng_glyco_hcd30_etd30_compound15.19872.19872.3 | controllerType=0 controllerNumber=1 scan=19872 | 46.299 |
| 16164:4 | 1 | R.TAGWNVPIGTLRPFLN[+2570.925]WTGPPEPIEAAVAR.F | HexNAc(5)Hex(6)Fuc(2)NeuAc(1) | N[+2571] | 1451.1676 | 4 | 5801.6486 | 5801.636 | 0 | 2.2 | 141 | Specific | 839.13 | 817.55 | 817.55 | 13.6 | 50 | >sp\|P02788\|TRFL_HUMAN Lactotransferrin OS=Homo sapiens OX=9606 GN=LTF PE=1 SV=6 | 1270 | 20201113_12413_1ng_glyco_hcd30_etd30_compound15.19789.19789.2 | controllerType=0 controllerNumber=1 scan=19789 | 46.0409 |
| 14776:2 | 1 | R.TAGWNIPMGLLFN[+1216.423]QTGSC[+57.021]K.F | HexNAc(2)Hex(5) | C[+57], N[+1216] | 1656.2229 | 2 | 3311.4385 | 3311.4328 | 0 | 1.7 | 485 | Specific | 236.22 | 227.83 | 114.64 | 4.82 | 50 | >sp\|P02788\|TRFL_HUMAN Lactotransferrin OS=Homo sapiens OX=9606 GN=LTF PE=1 SV=6 | 1270 | 20201113_12413_1ng_glyco_hcd30_etd30_compound15.18322.18322.2 | controllerType=0 controllerNumber=1 scan=18322 | 41.6409 |
| 16045:3 | 1 | R.TAGWNIPMGLLFN[+1548.545]QTGSC[+57.021]K.F | HexNAc(3)Hex(4)NeuAc(1) | C[+57], N[+1549] | 1215.1926 | 3 | 3643.5631 | 3643.5548 | 0 | 2.3 | 485 | Specific | 272.23 | 188.06 | 188.06 | 5.12 | 50 | >sp\|P02788\|TRFL_HUMAN Lactotransferrin OS=Homo sapiens OX=9606 GN=LTF PE=1 SV=6 | 1270 | 20201113_12413_1ng_glyco_hcd30_etd30_compound15.19664.19664.3 | controllerType=0 controllerNumber=1 scan=19664 | 45.6578 |
| 16008:3 | 1 | R.TAGWNIPMGLLFN[+1710.598]QTGSC[+57.021]K.F | HexNAc(3)Hex(5)NeuAc(1) | C[+57], N[+1711] | 1269.2075 | 3 | 3805.6078 | 3805.6076 | 0 | 0.1 | 485 | Specific | 277.03 | 190.79 | 190.79 | 4.38 | 50 | >sp\|P02788\|TRFL_HUMAN Lactotransferrin OS=Homo sapiens OX=9606 GN=LTF PE=1 SV=6 | 1270 | 20201113_12413_1ng_glyco_hcd30_etd30_compound15.19625.19625.3 | controllerType=0 controllerNumber=1 scan=19625 | 45.5534 |
| 14698:3 | 1 | R.TAGWNIPMGLLFN[+1622.582]QTGSC[+57.021]K.F | HexNAc(4)Hex(5) | C[+57], N[+1623] | 1239.8721 | 3 | 3717.6018 | 3717.5916 | 0 | 2.8 | 485 | Specific | 278.02 | 212.99 | 211.61 | 4.07 | 50 | >sp\|P02788\|TRFL_HUMAN Lactotransferrin OS=Homo sapiens OX=9606 GN=LTF PE=1 SV=6 | 1270 | 20201113_12413_1ng_glyco_hcd30_etd30_compound15.18239.18239.7 | controllerType=0 controllerNumber=1 scan=18239 | 41.3553 |
| 14746:3 | 1 | R.TAGWNIPMGLLFN[+1419.502]QTGSC[+57.021]K.F | HexNAc(3)Hex(5) | C[+57], N[+1420] | 1172.1799 | 3 | 3514.525 | 3514.5122 | 0 | 3.7 | 485 | Specific | 282.48 | 224.98 | 161.32 | 5.13 | 50 | >sp\|P02788\|TRFL_HUMAN Lactotransferrin OS=Homo sapiens OX=9606 GN=LTF PE=1 SV=6 | 1270 | 20201113_12413_1ng_glyco_hcd30_etd30_compound15.18289.18289.3 | controllerType=0 controllerNumber=1 scan=18289 | 41.5035 |
| 14383:3 | 1 | R.TAGWNIPMGLLFN[+2133.772]QTGSC[+57.021]K.F | HexNAc(5)Hex(6)Fuc(1) | C[+57], N[+2134] | 1410.2725 | 3 | 4228.803 | 4228.7817 | 0 | 5 | 485 | Specific | 302.27 | 165 | 165 | 4.61 | 50 | >sp\|P02788\|TRFL_HUMAN Lactotransferrin OS=Homo sapiens OX=9606 GN=LTF PE=1 SV=6 | 1270 | 20201113_12413_1ng_glyco_hcd30_etd30_compound15.17907.17907.3 | controllerType=0 controllerNumber=1 scan=17907 | 40.4466 |
| 14499:3 | 1 | R.TAGWNIPMGLLFN[+1768.640]QTGSC[+57.021]K.F | HexNAc(4)Hex(5)Fuc(1) | C[+57], N[+1769] | 1288.5605 | 3 | 3863.6669 | 3863.6495 | 0 | 4.5 | 485 | Specific | 350.13 | 301.11 | 301.11 | 5.42 | 50 | >sp\|P02788\|TRFL_HUMAN Lactotransferrin OS=Homo sapiens OX=9606 GN=LTF PE=1 SV=6 | 1270 | 20201113_12413_1ng_glyco_hcd30_etd30_compound15.18029.18029.3 | controllerType=0 controllerNumber=1 scan=18029 | 40.7469 |
| 14678:3 | 1 | R.TAGWNIPMGLLFN[+1581.555]QTGSC[+57.021]K.F | HexNAc(3)Hex(6) | C[+57], N[+1582] | 1226.1959 | 3 | 3676.573 | 3676.565 | 0 | 2.2 | 485 | Specific | 359.33 | 273.62 | 222.33 | 5.93 | 50 | >sp\|P02788\|TRFL_HUMAN Lactotransferrin OS=Homo sapiens OX=9606 GN=LTF PE=1 SV=6 | 1270 | 20201113_12413_1ng_glyco_hcd30_etd30_compound15.18218.18218.3 | controllerType=0 controllerNumber=1 scan=18218 | 41.2943 |
| 14575:3 | 1 | R.TAGWNIPMGLLFN[+1606.587]QTGSC[+57.021]K.F | HexNAc(4)Hex(4)Fuc(1) | C[+57], N[+1607] | 1234.5396 | 3 | 3701.6043 | 3701.5966 | 0 | 2.1 | 485 | Specific | 365.65 | 290.62 | 290.62 | 4.91 | 50 | >sp\|P02788\|TRFL_HUMAN Lactotransferrin OS=Homo sapiens OX=9606 GN=LTF PE=1 SV=6 | 1270 | 20201113_12413_1ng_glyco_hcd30_etd30_compound15.18109.18109.3 | controllerType=0 controllerNumber=1 scan=18109 | 40.9634 |
| 17214:3 | 1 | R.TAGWNIPMGLLFN[+2350.830]QTGSC[+57.021]K.F | HexNAc(4)Hex(5)Fuc(1)NeuAc(2) | C[+57], N[+2351] | 1482.6246 | 3 | 4445.8592 | 4445.8403 | 0 | 4.2 | 485 | Specific | 367.68 | 266.59 | 266.59 | 5.66 | 50 | >sp\|P02788\|TRFL_HUMAN Lactotransferrin OS=Homo sapiens OX=9606 GN=LTF PE=1 SV=6 | 1270 | 20201113_12413_1ng_glyco_hcd30_etd30_compound15.20902.20902.3 | controllerType=0 controllerNumber=1 scan=20902 | 49.3576 |
| 14667:3 | 1 | R.TAGWNIPMGLLFN[+1622.582]QTGSC[+57.021]K.F | HexNAc(4)Hex(5) | C[+57], N[+1623] | 1239.8716 | 3 | 3717.6004 | 3717.5916 | 0 | 2.4 | 485 | Specific | 371.47 | 294.65 | 236.29 | 5.89 | 50 | >sp\|P02788\|TRFL_HUMAN Lactotransferrin OS=Homo sapiens OX=9606 GN=LTF PE=1 SV=6 | 1270 | 20201113_12413_1ng_glyco_hcd30_etd30_compound15.18206.18206.3 | controllerType=0 controllerNumber=1 scan=18206 | 41.2475 |
| 15940:4 | 1 | R.TAGWNIPMGLLFN[+1913.677]QTGSC[+57.021]K.F | HexNAc(4)Hex(5)NeuAc(1) | C[+57], N[+1914] | 1002.9265 | 4 | 4008.6842 | 4008.687 | 0 | -0.7 | 485 | Specific | 372.33 | 296.91 | 296.91 | 3.94 | 50 | >sp\|P02788\|TRFL_HUMAN Lactotransferrin OS=Homo sapiens OX=9606 GN=LTF PE=1 SV=6 | 1270 | 20201113_12413_1ng_glyco_hcd30_etd30_compound15.19552.19552.4 | controllerType=0 controllerNumber=1 scan=19552 | 45.3186 |
| 15941:3 | 1 | R.TAGWNIPMGLLFN[+1694.603]QTGSC[+57.021]K.F | HexNAc(3)Hex(4)Fuc(1)NeuAc(1) | C[+57], N[+1695] | 1263.8755 | 3 | 3789.6121 | 3789.6127 | 0 | -0.2 | 485 | Specific | 383.91 | 273.58 | 273.58 | 4.17 | 50 | >sp\|P02788\|TRFL_HUMAN Lactotransferrin OS=Homo sapiens OX=9606 GN=LTF PE=1 SV=6 | 1270 | 20201113_12413_1ng_glyco_hcd30_etd30_compound15.19553.19553.3 | controllerType=0 controllerNumber=1 scan=19553 | 45.3303 |
| 14877:3 | 1 | R.TAGWNIPMGLLFN[+1095.397]QTGSC[+57.021]K.F | HexNAc(3)Hex(3) | C[+57], N[+1095] | 1064.1442 | 3 | 3190.418 | 3190.4065 | 0 | 3.6 | 485 | Specific | 407.68 | 354.44 | 354.44 | 5.81 | 50 | >sp\|P02788\|TRFL_HUMAN Lactotransferrin OS=Homo sapiens OX=9606 GN=LTF PE=1 SV=6 | 1270 | 20201113_12413_1ng_glyco_hcd30_etd30_compound15.18429.18429.3 | controllerType=0 controllerNumber=1 scan=18429 | 41.9417 |
| 15909:3 | 1 | R.TAGWNIPMGLLFN[+1913.677]QTGSC[+57.021]K.F | HexNAc(4)Hex(5)NeuAc(1) | C[+57], N[+1914] | 1336.8986 | 3 | 4008.6813 | 4008.687 | 0 | -1.4 | 485 | Specific | 413.19 | 291.28 | 291.28 | 4.65 | 50 | >sp\|P02788\|TRFL_HUMAN Lactotransferrin OS=Homo sapiens OX=9606 GN=LTF PE=1 SV=6 | 1270 | 20201113_12413_1ng_glyco_hcd30_etd30_compound15.19520.19520.3 | controllerType=0 controllerNumber=1 scan=19520 | 45.2343 |
| 15980:3 | 1 | R.TAGWNIPMGLLFN[+1751.624]QTGSC[+57.021]K.F | HexNAc(4)Hex(4)NeuAc(1) | C[+57], N[+1752] | 1282.8846 | 3 | 3846.6392 | 3846.6341 | 0 | 1.3 | 485 | Specific | 417.47 | 284.59 | 284.59 | 5.84 | 50 | >sp\|P02788\|TRFL_HUMAN Lactotransferrin OS=Homo sapiens OX=9606 GN=LTF PE=1 SV=6 | 1270 | 20201113_12413_1ng_glyco_hcd30_etd30_compound15.19595.19595.3 | controllerType=0 controllerNumber=1 scan=19595 | 45.4548 |
| 15816:3 | 1 | R.TAGWNIPMGLLFN[+2059.735]QTGSC[+57.021]K.F | HexNAc(4)Hex(5)Fuc(1)NeuAc(1) | C[+57], N[+2060] | 1385.5883 | 3 | 4154.7504 | 4154.7449 | 0 | 1.3 | 485 | Specific | 418.1 | 305.46 | 305.46 | 5.91 | 50 | >sp\|P02788\|TRFL_HUMAN Lactotransferrin OS=Homo sapiens OX=9606 GN=LTF PE=1 SV=6 | 1270 | 20201113_12413_1ng_glyco_hcd30_etd30_compound15.19421.19421.3 | controllerType=0 controllerNumber=1 scan=19421 | 44.8833 |
| 15944:3 | 1 | R.TAGWNIPMGLLFN[+1872.651]QTGSC[+57.021]K.F | HexNAc(3)Hex(6)NeuAc(1) | C[+57], N[+1873] | 1323.2276 | 3 | 3967.6682 | 3967.6604 | 0 | 2 | 485 | Specific | 430.71 | 296.14 | 250.38 | 5.92 | 50 | >sp\|P02788\|TRFL_HUMAN Lactotransferrin OS=Homo sapiens OX=9606 GN=LTF PE=1 SV=6 | 1270 | 20201113_12413_1ng_glyco_hcd30_etd30_compound15.19557.19557.3 | controllerType=0 controllerNumber=1 scan=19557 | 45.3459 |
| 14771:2 | 1 | R.TAGWNIPMGLLFN[+1216.423]QTGSC[+57.021]K.F | HexNAc(2)Hex(5) | C[+57], N[+1216] | 1656.2228 | 2 | 3311.4384 | 3311.4328 | 0 | 1.7 | 485 | Specific | 454.12 | 416.51 | 136.32 | 7.22 | 50 | >sp\|P02788\|TRFL_HUMAN Lactotransferrin OS=Homo sapiens OX=9606 GN=LTF PE=1 SV=6 | 1270 | 20201113_12413_1ng_glyco_hcd30_etd30_compound15.18316.18316.2 | controllerType=0 controllerNumber=1 scan=18316 | 41.6081 |
| 14762:3 | 1 | R.TAGWNIPMGLLFN[+1216.423]QTGSC[+57.021]K.F | HexNAc(2)Hex(5) | C[+57], N[+1216] | 1104.485 | 3 | 3311.4405 | 3311.4328 | 0 | 2.3 | 485 | Specific | 456.61 | 414.74 | 239.7 | 7.18 | 50 | >sp\|P02788\|TRFL_HUMAN Lactotransferrin OS=Homo sapiens OX=9606 GN=LTF PE=1 SV=6 | 1270 | 20201113_12413_1ng_glyco_hcd30_etd30_compound15.18307.18307.3 | controllerType=0 controllerNumber=1 scan=18307 | 41.5662 |
| 17326:3 | 1 | R.TAGWNIPMGLLFN[+2204.772]QTGSC[+57.021]K.F | HexNAc(4)Hex(5)NeuAc(2) | C[+57], N[+2205] | 1433.9341 | 3 | 4299.7879 | 4299.7824 | 0 | 1.3 | 485 | Specific | 469.53 | 289.75 | 289.75 | 5.99 | 50 | >sp\|P02788\|TRFL_HUMAN Lactotransferrin OS=Homo sapiens OX=9606 GN=LTF PE=1 SV=6 | 1270 | 20201113_12413_1ng_glyco_hcd30_etd30_compound15.21020.21020.3 | controllerType=0 controllerNumber=1 scan=21020 | 49.7355 |
| 15836:3 | 1 | R.TAGWNIPMGLLFN[+2205.793]QTGSC[+57.021]K.F | HexNAc(4)Hex(5)Fuc(2)NeuAc(1) | C[+57], N[+2206] | 1434.2741 | 3 | 4300.8078 | 4300.8028 | 0 | 1.2 | 485 | Specific | 480.04 | 327.94 | 327.94 | 6.15 | 50 | >sp\|P02788\|TRFL_HUMAN Lactotransferrin OS=Homo sapiens OX=9606 GN=LTF PE=1 SV=6 | 1270 | 20201113_12413_1ng_glyco_hcd30_etd30_compound15.19442.19442.3 | controllerType=0 controllerNumber=1 scan=19442 | 44.9458 |
| 14794:3 | 1 | R.TAGWNIPMGLLFN[+1216.423]QTGSC[+57.021]K.F | HexNAc(2)Hex(5) | C[+57], N[+1216] | 1104.4849 | 3 | 3311.4401 | 3311.4328 | 0 | 2.2 | 485 | Specific | 496.79 | 445.09 | 251.43 | 7.27 | 50 | >sp\|P02788\|TRFL_HUMAN Lactotransferrin OS=Homo sapiens OX=9606 GN=LTF PE=1 SV=6 | 1270 | 20201113_12413_1ng_glyco_hcd30_etd30_compound15.18341.18341.3 | controllerType=0 controllerNumber=1 scan=18341 | 41.6961 |
| 14714:3 | 1 | R.TAGWNIPM[+15.995]GLLFN[+2059.735]QTGSC[+57.021]K.F | HexNAc(4)Hex(5)Fuc(1)NeuAc(1) | C[+57], M[+16], N[+2060] | 1390.9243 | 3 | 4170.7582 | 4170.7398 | 0 | 4.4 | 485 | Specific | 274.83 | 244.29 | 244.29 | 3.72 | 50 | >sp\|P02788\|TRFL_HUMAN Lactotransferrin OS=Homo sapiens OX=9606 GN=LTF PE=1 SV=6 | 1270 | 20201113_12413_1ng_glyco_hcd30_etd30_compound15.18256.18256.3 | controllerType=0 controllerNumber=1 scan=18256 | 41.4152 |
| 16933:4 | 1 | R.TAGWNIPMGLLFN[+2350.830]QTGSC[+57.021]KFDEYFSQSC[+57.021]APGSDPR.S | HexNAc(4)Hex(5)Fuc(1)NeuAc(2) | C[+57]*2, N[+2351] | 1573.1598 | 4 | 6289.6173 | 6289.5875 | 0 | 4.8 | 485 | Specific | 327.43 | 292.88 | 292.88 | 4.9 | 50 | >sp\|P02788\|TRFL_HUMAN Lactotransferrin OS=Homo sapiens OX=9606 GN=LTF PE=1 SV=6 | 1270 | 20201113_12413_1ng_glyco_hcd30_etd30_compound15.20604.20604.4 | controllerType=0 controllerNumber=1 scan=20604 | 48.3242 |
| 16060:3 | 1 | R.TAGWNIPMGLLFN[+1913.677]QTGSC[+57.021]KFDEYFSQSC[+57.021]APGSDPR.S | HexNAc(4)Hex(5)NeuAc(1) | C[+57]*2, N[+1914] | 1951.5023 | 3 | 5852.4923 | 5852.4341 | 0 | 9.9 | 485 | Specific | 501.13 | 419.82 | 419.82 | 5.89 | 50 | >sp\|P02788\|TRFL_HUMAN Lactotransferrin OS=Homo sapiens OX=9606 GN=LTF PE=1 SV=6 | 1270 | 20201113_12413_1ng_glyco_hcd30_etd30_compound15.19680.19680.3 | controllerType=0 controllerNumber=1 scan=19680 | 45.7106 |
| 15317:4 | 1 | R.TAGWNIPMGLLFN[+1768.640]QTGSC[+57.021]KFDEYFSQSC[+57.021]APGSDPR.S | HexNAc(4)Hex(5)Fuc(1) | C[+57]*2, N[+1769] | 1427.6124 | 4 | 5707.4279 | 5707.3966 | 0 | 5.5 | 485 | Specific | 639.46 | 614.81 | 614.81 | 7.37 | 50 | >sp\|P02788\|TRFL_HUMAN Lactotransferrin OS=Homo sapiens OX=9606 GN=LTF PE=1 SV=6 | 1270 | 20201113_12413_1ng_glyco_hcd30_etd30_compound15.18893.18893.4 | controllerType=0 controllerNumber=1 scan=18893 | 43.2939 |
| 16059:4 | 1 | R.TAGWNIPMGLLFN[+1913.677]QTGSC[+57.021]KFDEYFSQSC[+57.021]APGSDPR.S | HexNAc(4)Hex(5)NeuAc(1) | C[+57]*2, N[+1914] | 1463.8785 | 4 | 5852.4923 | 5852.4341 | 0 | 9.9 | 485 | Specific | 792.12 | 719.96 | 719.96 | 10.89 | 50 | >sp\|P02788\|TRFL_HUMAN Lactotransferrin OS=Homo sapiens OX=9606 GN=LTF PE=1 SV=6 | 1270 | 20201113_12413_1ng_glyco_hcd30_etd30_compound15.19679.19679.4 | controllerType=0 controllerNumber=1 scan=19679 | 45.7051 |
| 15995:4 | 1 | R.TAGWNIPMGLLFN[+2059.735]QTGSC[+57.021]KFDEYFSQSC[+57.021]APGSDPR.S | HexNAc(4)Hex(5)Fuc(1)NeuAc(1) | C[+57]*2, N[+2060] | 1500.3801 | 4 | 5998.4987 | 5998.492 | 0 | 1.1 | 485 | Specific | 837.06 | 752.11 | 752.11 | 12.86 | 50 | >sp\|P02788\|TRFL_HUMAN Lactotransferrin OS=Homo sapiens OX=9606 GN=LTF PE=1 SV=6 | 1270 | 20201113_12413_1ng_glyco_hcd30_etd30_compound15.19611.19611.4 | controllerType=0 controllerNumber=1 scan=19611 | 45.5022 |
| 08151:3 | 1 | R.N[+2059.735]GSDC[+57.021]PDKFC[+57.021]LFQSETK.N | HexNAc(4)Hex(5)Fuc(1)NeuAc(1) | C[+57]*2, N[+2060] | 1364.879 | 3 | 4092.6223 | 4092.6088 | 0 | 3.3 | 642 | Specific | 296.29 | 196.03 | 196.03 | 4.67 | 50 | >sp\|P02788\|TRFL_HUMAN Lactotransferrin OS=Homo sapiens OX=9606 GN=LTF PE=1 SV=6 | 1270 | 20201113_12413_1ng_glyco_hcd30_etd30_compound15.11371.11371.3 | controllerType=0 controllerNumber=1 scan=11371 | 26.7667 |
| 08109:3 | 1 | R.N[+2205.793]GSDC[+57.021]PDKFC[+57.021]LFQSETK.N | HexNAc(4)Hex(5)Fuc(2)NeuAc(1) | C[+57]*2, N[+2206] | 1413.566 | 3 | 4238.6834 | 4238.6667 | 0 | 3.9 | 642 | Specific | 324.4 | 200.47 | 200.47 | 4.96 | 50 | >sp\|P02788\|TRFL_HUMAN Lactotransferrin OS=Homo sapiens OX=9606 GN=LTF PE=1 SV=6 | 1270 | 20201113_12413_1ng_glyco_hcd30_etd30_compound15.11328.11328.3 | controllerType=0 controllerNumber=1 scan=11328 | 26.6809 |
| 08094:3 | 1 | R.N[+2205.793]GSDC[+57.021]PDKFC[+57.021]LFQSETK.N | HexNAc(4)Hex(5)Fuc(2)NeuAc(1) | C[+57]*2, N[+2206] | 1413.5643 | 3 | 4238.6782 | 4238.6667 | 0 | 2.7 | 642 | Specific | 332.31 | 198.81 | 198.81 | 5.12 | 50 | >sp\|P02788\|TRFL_HUMAN Lactotransferrin OS=Homo sapiens OX=9606 GN=LTF PE=1 SV=6 | 1270 | 20201113_12413_1ng_glyco_hcd30_etd30_compound15.11312.11312.3 | controllerType=0 controllerNumber=1 scan=11312 | 26.6466 |
| 18773:3 | 3 | R.[+42.011]LSLLEEPGN[+2336.851]GTFTVILNQLTSR.D | HexNAc(6)Hex(6)Fuc(1) | .[42], N[+2337] | 1594.3928 | 3 | 4781.1637 | 4781.1542 | 0 | 2 | 413 | Specific | 301.07 | 244.61 | 244.61 | 3.97 | 25 | >sp\|P01833\|PIGR_HUMAN Polymeric immunoglobulin receptor OS=Homo sapiens OX=9606 GN=PIGR PE=1 SV=4 | 902 | 20201113_12413_1ng_glyco_hcd30_etd30_compound15.22548.22548.3 | controllerType=0 controllerNumber=1 scan=22548 | 53.4305 |
| 17045:3 | 3 | R.LSLLEEPGN[+2205.793]GTFTVILNQLTSR.D | HexNAc(4)Hex(5)Fuc(2)NeuAc(1) | N[+2206] | 1536.7042 | 3 | 4608.0979 | 4608.0854 | 0 | 2.7 | 413 | Specific | 330.6 | 240.65 | 240.65 | 4.09 | 25 | >sp\|P01833\|PIGR_HUMAN Polymeric immunoglobulin receptor OS=Homo sapiens OX=9606 GN=PIGR PE=1 SV=4 | 902 | 20201113_12413_1ng_glyco_hcd30_etd30_compound15.20722.20722.7 | controllerType=0 controllerNumber=1 scan=20722 | 48.7504 |
| 17139:3 | 3 | R.LSLLEEPGN[+2059.735]GTFTVILNQLTSR.D | HexNAc(4)Hex(5)Fuc(1)NeuAc(1) | N[+2060] | 1488.0224 | 3 | 4462.0526 | 4462.0275 | 0 | 5.6 | 413 | Specific | 499.08 | 396.03 | 396.03 | 6.42 | 25 | >sp\|P01833\|PIGR_HUMAN Polymeric immunoglobulin receptor OS=Homo sapiens OX=9606 GN=PIGR PE=1 SV=4 | 902 | 20201113_12413_1ng_glyco_hcd30_etd30_compound15.20822.20822.3 | controllerType=0 controllerNumber=1 scan=20822 | 49.0663 |
| 18723:4 | 3 | R.LSLLEEPGN[+2350.830]GTFTVILNQLTSR.D | HexNAc(4)Hex(5)Fuc(1)NeuAc(2) | N[+2351] | 1189.0418 | 4 | 4753.1453 | 4753.1229 | 0 | 4.7 | 413 | Specific | 546.45 | 457.05 | 457.05 | 5.72 | 25 | >sp\|P01833\|PIGR_HUMAN Polymeric immunoglobulin receptor OS=Homo sapiens OX=9606 GN=PIGR PE=1 SV=4 | 902 | 20201113_12413_1ng_glyco_hcd30_etd30_compound15.22494.22494.4 | controllerType=0 controllerNumber=1 scan=22494 | 53.2783 |
| 15311:3 | 3 | K.IIEGEPNLKVPGN[+2350.830]VTAVLGETLK.V | HexNAc(4)Hex(5)Fuc(1)NeuAc(2) | N[+2351] | 1581.4064 | 3 | 4742.2047 | 4742.1797 | 0 | 5.3 | 457 | Specific | 315.91 | 244 | 244 | 2.88 | 25 | >sp\|P01833\|PIGR_HUMAN Polymeric immunoglobulin receptor OS=Homo sapiens OX=9606 GN=PIGR PE=1 SV=4 | 902 | 20201113_12413_1ng_glyco_hcd30_etd30_compound15.18887.18887.3 | controllerType=0 controllerNumber=1 scan=18887 | 43.2755 |
| 15316:4 | 3 | K.IIEGEPNLKVPGN[+2350.830]VTAVLGETLK.V | HexNAc(4)Hex(5)Fuc(1)NeuAc(2) | N[+2351] | 1186.306 | 4 | 4742.202 | 4742.1797 | 0 | 4.7 | 457 | Specific | 328.8 | 287.22 | 287.22 | 3.02 | 25 | >sp\|P01833\|PIGR_HUMAN Polymeric immunoglobulin receptor OS=Homo sapiens OX=9606 GN=PIGR PE=1 SV=4 | 902 | 20201113_12413_1ng_glyco_hcd30_etd30_compound15.18892.18892.4 | controllerType=0 controllerNumber=1 scan=18892 | 43.2872 |
| 15301:3 | 3 | K.IIEGEPNLKVPGN[+2350.830]VTAVLGETLK.V | HexNAc(4)Hex(5)Fuc(1)NeuAc(2) | N[+2351] | 1581.4026 | 3 | 4742.1934 | 4742.1797 | 0 | 2.9 | 457 | Specific | 357.4 | 264.29 | 264.29 | 4.11 | 25 | >sp\|P01833\|PIGR_HUMAN Polymeric immunoglobulin receptor OS=Homo sapiens OX=9606 GN=PIGR PE=1 SV=4 | 902 | 20201113_12413_1ng_glyco_hcd30_etd30_compound15.18876.18876.2 | controllerType=0 controllerNumber=1 scan=18876 | 43.2462 |
| 13553:5 | 3 | K.IIEGEPNLKVPGN[+2205.793]VTAVLGETLKVPC[+57.021]HFPC[+57.021]K.F | HexNAc(4)Hex(5)Fuc(2)NeuAc(1) | C[+57]*2, N[+2206] | 1125.3289 | 5 | 5622.6152 | 5622.5997 | 0 | 2.8 | 457 | Specific | 312.75 | 312.75 | 312.75 | 3.58 | 25 | >sp\|P01833\|PIGR_HUMAN Polymeric immunoglobulin receptor OS=Homo sapiens OX=9606 GN=PIGR PE=1 SV=4 | 902 | 20201113_12413_1ng_glyco_hcd30_etd30_compound15.17020.17020.5 | controllerType=0 controllerNumber=1 scan=17020 | 38.4137 |
| 14662:4 | 3 | K.IIEGEPNLKVPGN[+2350.830]VTAVLGETLKVPC[+57.021]HFPC[+57.021]K.F | HexNAc(4)Hex(5)Fuc(1)NeuAc(2) | C[+57]*2, N[+2351] | 1442.6758 | 4 | 5767.6813 | 5767.6372 | 0 | 7.6 | 457 | Specific | 527.74 | 527.74 | 527.74 | 4.1 | 25 | >sp\|P01833\|PIGR_HUMAN Polymeric immunoglobulin receptor OS=Homo sapiens OX=9606 GN=PIGR PE=1 SV=4 | 902 | 20201113_12413_1ng_glyco_hcd30_etd30_compound15.18200.18200.4 | controllerType=0 controllerNumber=1 scan=18200 | 41.2161 |
| 14675:4 | 3 | K.IIEGEPNLKVPGN[+2350.830]VTAVLGETLKVPC[+57.021]HFPC[+57.021]K.F | HexNAc(4)Hex(5)Fuc(1)NeuAc(2) | C[+57]*2, N[+2351] | 1442.667 | 4 | 5767.6462 | 5767.6372 | 0 | 1.5 | 457 | Specific | 528.44 | 528.44 | 528.44 | 5.13 | 25 | >sp\|P01833\|PIGR_HUMAN Polymeric immunoglobulin receptor OS=Homo sapiens OX=9606 GN=PIGR PE=1 SV=4 | 902 | 20201113_12413_1ng_glyco_hcd30_etd30_compound15.18214.18214.4 | controllerType=0 controllerNumber=1 scan=18214 | 41.2799 |
| 14671:5 | 3 | K.IIEGEPNLKVPGN[+2350.830]VTAVLGETLKVPC[+57.021]HFPC[+57.021]K.F | HexNAc(4)Hex(5)Fuc(1)NeuAc(2) | C[+57]*2, N[+2351] | 1154.3366 | 5 | 5767.6536 | 5767.6372 | 0 | 2.8 | 457 | Specific | 686.53 | 686.53 | 686.53 | 8.52 | 25 | >sp\|P01833\|PIGR_HUMAN Polymeric immunoglobulin receptor OS=Homo sapiens OX=9606 GN=PIGR PE=1 SV=4 | 902 | 20201113_12413_1ng_glyco_hcd30_etd30_compound15.18210.18210.5 | controllerType=0 controllerNumber=1 scan=18210 | 41.2597 |
| 11952:3 | 3 | K.VPGN[+2205.793]VTAVLGETLK.V | HexNAc(4)Hex(5)Fuc(2)NeuAc(1) | N[+2206] | 1201.8693 | 3 | 3603.5934 | 3603.5927 | 0 | 0.2 | 466 | Specific | 258.59 | 159.39 | 159.39 | 4.23 | 25 | >sp\|P01833\|PIGR_HUMAN Polymeric immunoglobulin receptor OS=Homo sapiens OX=9606 GN=PIGR PE=1 SV=4 | 902 | 20201113_12413_1ng_glyco_hcd30_etd30_compound15.15345.15345.3 | controllerType=0 controllerNumber=1 scan=15345 | 34.5781 |
| 12008:3 | 3 | K.VPGN[+2059.735]VTAVLGETLK.V | HexNAc(4)Hex(5)Fuc(1)NeuAc(1) | N[+2060] | 1153.186 | 3 | 3457.5434 | 3457.5348 | 0 | 2.5 | 466 | Specific | 360.31 | 250.67 | 250.67 | 5.46 | 25 | >sp\|P01833\|PIGR_HUMAN Polymeric immunoglobulin receptor OS=Homo sapiens OX=9606 GN=PIGR PE=1 SV=4 | 902 | 20201113_12413_1ng_glyco_hcd30_etd30_compound15.15404.15404.3 | controllerType=0 controllerNumber=1 scan=15404 | 34.7471 |
| 13501:3 | 3 | K.VPGN[+2350.830]VTAVLGETLKVPC[+57.021]HFPC[+57.021]K.F | HexNAc(4)Hex(5)Fuc(1)NeuAc(2) | C[+57]*2, N[+2351] | 1592.0386 | 3 | 4774.1013 | 4774.0877 | 0 | 2.9 | 466 | Specific | 276.56 | 158.28 | 158.28 | 2.55 | 25 | >sp\|P01833\|PIGR_HUMAN Polymeric immunoglobulin receptor OS=Homo sapiens OX=9606 GN=PIGR PE=1 SV=4 | 902 | 20201113_12413_1ng_glyco_hcd30_etd30_compound15.16965.16965.3 | controllerType=0 controllerNumber=1 scan=16965 | 38.2675 |
| 11995:4 | 3 | K.VPGN[+2205.793]VTAVLGETLKVPC[+57.021]HFPC[+57.021]K.F | HexNAc(4)Hex(5)Fuc(2)NeuAc(1) | C[+57]*2, N[+2206] | 1158.0209 | 4 | 4629.0618 | 4629.0502 | 0 | 2.5 | 466 | Specific | 406.9 | 406.9 | 406.9 | 4.64 | 25 | >sp\|P01833\|PIGR_HUMAN Polymeric immunoglobulin receptor OS=Homo sapiens OX=9606 GN=PIGR PE=1 SV=4 | 902 | 20201113_12413_1ng_glyco_hcd30_etd30_compound15.15390.15390.4 | controllerType=0 controllerNumber=1 scan=15390 | 34.712 |
| 12020:4 | 3 | K.VPGN[+2059.735]VTAVLGETLKVPC[+57.021]HFPC[+57.021]K.F | HexNAc(4)Hex(5)Fuc(1)NeuAc(1) | C[+57]*2, N[+2060] | 1121.5059 | 4 | 4483.0018 | 4482.9923 | 0 | 2.1 | 466 | Specific | 484.47 | 457.32 | 457.32 | 5.28 | 25 | >sp\|P01833\|PIGR_HUMAN Polymeric immunoglobulin receptor OS=Homo sapiens OX=9606 GN=PIGR PE=1 SV=4 | 902 | 20201113_12413_1ng_glyco_hcd30_etd30_compound15.15416.15416.4 | controllerType=0 controllerNumber=1 scan=15416 | 34.7712 |
| 05114:3 | 3 | K.WN[+1872.651]NTGC[+57.021]QALPSQDEGPSK.A | HexNAc(3)Hex(6)NeuAc(1) | C[+57], N[+1873] | 1287.8506 | 3 | 3861.5374 | 3861.5272 | 0 | 2.6 | 498 | Specific | 201.58 | 108.44 | 108.44 | 4.14 | 25 | >sp\|P01833\|PIGR_HUMAN Polymeric immunoglobulin receptor OS=Homo sapiens OX=9606 GN=PIGR PE=1 SV=4 | 902 | 20201113_12413_1ng_glyco_hcd30_etd30_compound15.8207.8207.3 | controllerType=0 controllerNumber=1 scan=8207 | 20.9084 |
| 06993:3 | 3 | K.WN[+2204.772]NTGC[+57.021]QALPSQDEGPSK.A | HexNAc(4)Hex(5)NeuAc(2) | C[+57], N[+2205] | 1398.5607 | 3 | 4193.6676 | 4193.6491 | 0 | 4.4 | 498 | Specific | 240.15 | 129.99 | 129.99 | 4.76 | 25 | >sp\|P01833\|PIGR_HUMAN Polymeric immunoglobulin receptor OS=Homo sapiens OX=9606 GN=PIGR PE=1 SV=4 | 902 | 20201113_12413_1ng_glyco_hcd30_etd30_compound15.10161.10161.3 | controllerType=0 controllerNumber=1 scan=10161 | 24.4074 |
| 05035:3 | 3 | K.WN[+2059.735]NTGC[+57.021]QALPSQDEGPSK.A | HexNAc(4)Hex(5)Fuc(1)NeuAc(1) | C[+57], N[+2060] | 1350.2118 | 3 | 4048.621 | 4048.6116 | 0 | 2.3 | 498 | Specific | 249.77 | 139.51 | 139.51 | 4.92 | 25 | >sp\|P01833\|PIGR_HUMAN Polymeric immunoglobulin receptor OS=Homo sapiens OX=9606 GN=PIGR PE=1 SV=4 | 902 | 20201113_12413_1ng_glyco_hcd30_etd30_compound15.8125.8125.3 | controllerType=0 controllerNumber=1 scan=8125 | 20.7526 |
| 05063:3 | 3 | K.WN[+1913.677]NTGC[+57.021]QALPSQDEGPSK.A | HexNAc(4)Hex(5)NeuAc(1) | C[+57], N[+1914] | 1301.5257 | 3 | 3902.5625 | 3902.5537 | 0 | 2.3 | 498 | Specific | 502.64 | 317.64 | 317.64 | 6.16 | 25 | >sp\|P01833\|PIGR_HUMAN Polymeric immunoglobulin receptor OS=Homo sapiens OX=9606 GN=PIGR PE=1 SV=4 | 902 | 20201113_12413_1ng_glyco_hcd30_etd30_compound15.8154.8154.3 | controllerType=0 controllerNumber=1 scan=8154 | 20.8073 |
| 05144:3 | 3 | K.WN[+1710.598]NTGC[+57.021]QALPSQDEGPSK.A | HexNAc(3)Hex(5)NeuAc(1) | C[+57], N[+1711] | 1233.8321 | 3 | 3699.4817 | 3699.4743 | 0 | 2 | 498 | Specific | 504.31 | 308.81 | 308.81 | 6.16 | 25 | >sp\|P01833\|PIGR_HUMAN Polymeric immunoglobulin receptor OS=Homo sapiens OX=9606 GN=PIGR PE=1 SV=4 | 902 | 20201113_12413_1ng_glyco_hcd30_etd30_compound15.8238.8238.3 | controllerType=0 controllerNumber=1 scan=8238 | 20.9648 |
| 15764:3 | 4 | R.LSLHRPALEDLLLGSEAN[+2116.756]LTC[+57.021]TLTGLR.D | HexNAc(5)Hex(5)NeuAc(1) | C[+57], N[+2117] | 1694.1241 | 3 | 5080.3578 | 5080.3547 | 0 | 0.6 | 127 | Specific | 213.32 | 183.51 | 183.51 | 3.43 | 19 | >sp\|P01876\|IGHA1_HUMAN Immunoglobulin heavy constant alpha 1 OS=Homo sapiens OX=9606 GN=IGHA1 PE=1 SV=2 | 255 | 20201113_12413_1ng_glyco_hcd30_etd30_compound15.19365.19365.3 | controllerType=0 controllerNumber=1 scan=19365 | 44.6893 |
| 15121:3 | 4 | R.LSLHRPALEDLLLGSEAN[+1501.555]LTC[+57.021]TLTGLR.D | HexNAc(5)Hex(3) | C[+57], N[+1502] | 1489.0584 | 3 | 4465.1607 | 4465.1536 | 0 | 1.6 | 127 | Specific | 289.39 | 287.98 | 287.98 | 5.42 | 19 | >sp\|P01876\|IGHA1_HUMAN Immunoglobulin heavy constant alpha 1 OS=Homo sapiens OX=9606 GN=IGHA1 PE=1 SV=2 | 255 | 20201113_12413_1ng_glyco_hcd30_etd30_compound15.18686.18686.3 | controllerType=0 controllerNumber=1 scan=18686 | 42.6886 |
| 15101:4 | 4 | R.LSLHRPALEDLLLGSEAN[+1702.581]LTC[+57.021]TLTGLR.D | HexNAc(2)Hex(8) | C[+57], N[+1703] | 1167.3103 | 4 | 4666.2196 | 4666.1796 | 0 | 8.6 | 127 | Specific | 685.34 | 685.34 | 685.34 | 9.67 | 19 | >sp\|P01876\|IGHA1_HUMAN Immunoglobulin heavy constant alpha 1 OS=Homo sapiens OX=9606 GN=IGHA1 PE=1 SV=2 | 255 | 20201113_12413_1ng_glyco_hcd30_etd30_compound15.18664.18664.4 | controllerType=0 controllerNumber=1 scan=18664 | 42.5818 |
| 15081:4 | 4 | R.LSLHRPALEDLLLGSEAN[+1663.608]LTC[+57.021]TLTGLR.D | HexNAc(5)Hex(4) | C[+57], N[+1664] | 1157.5623 | 4 | 4627.2274 | 4627.2064 | 0 | 4.5 | 127 | Specific | 727.47 | 727.47 | 727.47 | 11.06 | 19 | >sp\|P01876\|IGHA1_HUMAN Immunoglobulin heavy constant alpha 1 OS=Homo sapiens OX=9606 GN=IGHA1 PE=1 SV=2 | 255 | 20201113_12413_1ng_glyco_hcd30_etd30_compound15.18643.18643.2 | controllerType=0 controllerNumber=1 scan=18643 | 42.4988 |
| 15075:4 | 4 | R.LSLHRPALEDLLLGSEAN[+1540.529]LTC[+57.021]TLTGLR.D | HexNAc(2)Hex(7) | C[+57], N[+1541] | 1126.7917 | 4 | 4504.1448 | 4504.1268 | 0 | 4 | 127 | Specific | 1120.23 | 1020.41 | 1020.41 | 19.49 | 19 | >sp\|P01876\|IGHA1_HUMAN Immunoglobulin heavy constant alpha 1 OS=Homo sapiens OX=9606 GN=IGHA1 PE=1 SV=2 | 255 | 20201113_12413_1ng_glyco_hcd30_etd30_compound15.18637.18637.4 | controllerType=0 controllerNumber=1 scan=18637 | 42.4854 |
| 10019:3 | 4 | R.LAGKPTHVN[+1540.529]VSVVMAEVDGTC[+57.021]Y.- | HexNAc(2)Hex(7) | C[+57], N[+1541] | 1296.5641 | 3 | 3887.6779 | 3887.6706 | 0 | 1.9 | 332 | CRagged | 234.31 | 192.29 | 166.44 | 3.07 | 19 | >sp\|P01876\|IGHA1_HUMAN Immunoglobulin heavy constant alpha 1 OS=Homo sapiens OX=9606 GN=IGHA1 PE=1 SV=2 | 255 | 20201113_12413_1ng_glyco_hcd30_etd30_compound15.13312.13312.3 | controllerType=0 controllerNumber=1 scan=13312 | 30.4982 |
| 10091:3 | 4 | R.LAGKPTHVN[+1501.555]VSVVMAEVDGTC[+57.021]Y.- | HexNAc(5)Hex(3) | C[+57], N[+1502] | 1283.571 | 3 | 3848.6984 | 3848.6974 | 0 | 0.3 | 332 | CRagged | 242.45 | 134.19 | 134.19 | 2.67 | 19 | >sp\|P01876\|IGHA1_HUMAN Immunoglobulin heavy constant alpha 1 OS=Homo sapiens OX=9606 GN=IGHA1 PE=1 SV=2 | 255 | 20201113_12413_1ng_glyco_hcd30_etd30_compound15.13388.13388.3 | controllerType=0 controllerNumber=1 scan=13388 | 30.7175 |
| 11570:3 | 4 | R.LAGKPTHVN[+2262.814]VSVVMAEVDGTC[+57.021]Y.- | HexNAc(5)Hex(5)Fuc(1)NeuAc(1) | C[+57], N[+2263] | 1537.3277 | 3 | 4609.9685 | 4609.9564 | 0 | 2.6 | 332 | CRagged | 250.1 | 65.89 | 65.89 | 2.63 | 19 | >sp\|P01876\|IGHA1_HUMAN Immunoglobulin heavy constant alpha 1 OS=Homo sapiens OX=9606 GN=IGHA1 PE=1 SV=2 | 255 | 20201113_12413_1ng_glyco_hcd30_etd30_compound15.14946.14946.3 | controllerType=0 controllerNumber=1 scan=14946 | 33.6814 |
| 10024:3 | 4 | R.LAGKPTHVN[+1378.476]VSVVMAEVDGTC[+57.021]Y.- | HexNAc(2)Hex(6) | C[+57], N[+1378] | 1242.5473 | 3 | 3725.6273 | 3725.6178 | 0 | 2.6 | 332 | CRagged | 256.88 | 189.02 | 189.02 | 3.69 | 19 | >sp\|P01876\|IGHA1_HUMAN Immunoglobulin heavy constant alpha 1 OS=Homo sapiens OX=9606 GN=IGHA1 PE=1 SV=2 | 255 | 20201113_12413_1ng_glyco_hcd30_etd30_compound15.13318.13318.3 | controllerType=0 controllerNumber=1 scan=13318 | 30.5263 |
| 11583:3 | 4 | R.LAGKPTHVN[+2059.735]VSVVMAEVDGTC[+57.021]Y.- | HexNAc(4)Hex(5)Fuc(1)NeuAc(1) | C[+57], N[+2060] | 1469.6343 | 3 | 4406.8885 | 4406.877 | 0 | 2.6 | 332 | CRagged | 260.65 | 128.2 | 128.2 | 3.18 | 19 | >sp\|P01876\|IGHA1_HUMAN Immunoglobulin heavy constant alpha 1 OS=Homo sapiens OX=9606 GN=IGHA1 PE=1 SV=2 | 255 | 20201113_12413_1ng_glyco_hcd30_etd30_compound15.14959.14959.3 | controllerType=0 controllerNumber=1 scan=14959 | 33.7215 |
| 10030:3 | 4 | R.LAGKPTHVN[+1809.666]VSVVMAEVDGTC[+57.021]Y.- | HexNAc(5)Hex(4)Fuc(1) | C[+57], N[+1810] | 1386.2773 | 3 | 4156.8173 | 4156.8081 | 0 | 2.2 | 332 | CRagged | 271.9 | 162.61 | 162.61 | 3.19 | 19 | >sp\|P01876\|IGHA1_HUMAN Immunoglobulin heavy constant alpha 1 OS=Homo sapiens OX=9606 GN=IGHA1 PE=1 SV=2 | 255 | 20201113_12413_1ng_glyco_hcd30_etd30_compound15.13324.13324.3 | controllerType=0 controllerNumber=1 scan=13324 | 30.5414 |
| 09993:3 | 4 | R.LAGKPTHVN[+1702.581]VSVVMAEVDGTC[+57.021]Y.- | HexNAc(2)Hex(8) | C[+57], N[+1703] | 1350.5833 | 3 | 4049.7354 | 4049.7234 | 0 | 3 | 332 | CRagged | 281.69 | 220.15 | 201.23 | 3.51 | 19 | >sp\|P01876\|IGHA1_HUMAN Immunoglobulin heavy constant alpha 1 OS=Homo sapiens OX=9606 GN=IGHA1 PE=1 SV=2 | 255 | 20201113_12413_1ng_glyco_hcd30_etd30_compound15.13285.13285.3 | controllerType=0 controllerNumber=1 scan=13285 | 30.4137 |
| 11538:3 | 4 | R.LAGKPTHVN[+2262.814]VSVVMAEVDGTC[+57.021]Y.- | HexNAc(5)Hex(5)Fuc(1)NeuAc(1) | C[+57], N[+2263] | 1537.332 | 3 | 4609.9813 | 4609.9564 | 0 | 5.4 | 332 | CRagged | 299.61 | 212.42 | 212.42 | 3.01 | 19 | >sp\|P01876\|IGHA1_HUMAN Immunoglobulin heavy constant alpha 1 OS=Homo sapiens OX=9606 GN=IGHA1 PE=1 SV=2 | 255 | 20201113_12413_1ng_glyco_hcd30_etd30_compound15.14912.14912.3 | controllerType=0 controllerNumber=1 scan=14912 | 33.5822 |
| 10052:3 | 4 | R.LAGKPTHVN[+1216.423]VSVVMAEVDGTC[+57.021]Y.- | HexNAc(2)Hex(5) | C[+57], N[+1216] | 1188.528 | 3 | 3563.5695 | 3563.5649 | 0 | 1.3 | 332 | CRagged | 301.51 | 301.51 | 203.56 | 4.43 | 19 | >sp\|P01876\|IGHA1_HUMAN Immunoglobulin heavy constant alpha 1 OS=Homo sapiens OX=9606 GN=IGHA1 PE=1 SV=2 | 255 | 20201113_12413_1ng_glyco_hcd30_etd30_compound15.13347.13347.3 | controllerType=0 controllerNumber=1 scan=13347 | 30.6029 |
| 11578:4 | 4 | R.LAGKPTHVN[+2262.814]VSVVMAEVDGTC[+57.021]Y.- | HexNAc(5)Hex(5)Fuc(1)NeuAc(1) | C[+57], N[+2263] | 1153.2482 | 4 | 4609.971 | 4609.9564 | 0 | 3.2 | 332 | CRagged | 387.18 | 344.52 | 344.52 | 3.94 | 19 | >sp\|P01876\|IGHA1_HUMAN Immunoglobulin heavy constant alpha 1 OS=Homo sapiens OX=9606 GN=IGHA1 PE=1 SV=2 | 255 | 20201113_12413_1ng_glyco_hcd30_etd30_compound15.14954.14954.4 | controllerType=0 controllerNumber=1 scan=14954 | 33.7076 |
| 08093:4 | 5 | R.QTGLAPGQEYEISLHIVKN[+2059.735]NTR.G | HexNAc(4)Hex(5)Fuc(1)NeuAc(1) | N[+2060] | 1132.764 | 4 | 4528.034 | 4528.0241 | 0 | 2.2 | 770 | Specific | 571.03 | 476.25 | 476.25 | 7.54 | 9 | >sp\|P24821\|TENA_HUMAN Tenascin OS=Homo sapiens OX=9606 GN=TNC PE=1 SV=3 | 486 | 20201113_12413_1ng_glyco_hcd30_etd30_compound15.11311.11311.4 | controllerType=0 controllerNumber=1 scan=11311 | 26.64 |
| 13376:3 | 5 | R.LN[+1702.581]YSLPTGQWVGVQLPR.N | HexNAc(2)Hex(8) | N[+1703] | 1210.8803 | 3 | 3630.6264 | 3630.6202 | 0 | 1.7 | 1017 | Specific | 275.79 | 258.02 | 258.02 | 4.52 | 9 | >sp\|P24821\|TENA_HUMAN Tenascin OS=Homo sapiens OX=9606 GN=TNC PE=1 SV=3 | 486 | 20201113_12413_1ng_glyco_hcd30_etd30_compound15.16833.16833.3 | controllerType=0 controllerNumber=1 scan=16833 | 37.9039 |
| 15986:3 | 6 | R.NIAAFGGDPNN[+1913.677]ITLFGESAGGASVSLQTLSPYNK.G | HexNAc(4)Hex(5)NeuAc(1) | N[+1914] | 1775.4652 | 3 | 5324.381 | 5324.3633 | 0 | 3.3 | 197 | Specific | 428.64 | 310.01 | 310.01 | 5.85 | 8 | >sp\|P19835\|CEL_HUMAN Bile salt-activated lipase OS=Homo sapiens OX=9606 GN=CEL PE=1 SV=3 | 476 | 20201113_12413_1ng_glyco_hcd30_etd30_compound15.19601.19601.3 | controllerType=0 controllerNumber=1 scan=19601 | 45.468 |
| 08952:3 | 8 | K.TVLTPATNHMGN[+1540.529]VTFTIPANR.E | HexNAc(2)Hex(7) | N[+1541] | 1265.9095 | 3 | 3795.7139 | 3795.6886 | 0 | 6.7 | 74 | Specific | 277.56 | 188 | 173.86 | 3.96 | 8 | >sp\|P01024\|CO3_HUMAN Complement C3 OS=Homo sapiens OX=9606 GN=C3 PE=1 SV=2 | 492 | 20201113_12413_1ng_glyco_hcd30_etd30_compound15.12204.12204.3 | controllerType=0 controllerNumber=1 scan=12204 | 28.3665 |
| 08971:3 | 8 | K.TVLTPATNHMGN[+1378.476]VTFTIPANR.E | HexNAc(2)Hex(6) | N[+1378] | 1211.8904 | 3 | 3633.6568 | 3633.6358 | 0 | 5.8 | 74 | Specific | 333.38 | 286.95 | 103.41 | 5.12 | 8 | >sp\|P01024\|CO3_HUMAN Complement C3 OS=Homo sapiens OX=9606 GN=C3 PE=1 SV=2 | 492 | 20201113_12413_1ng_glyco_hcd30_etd30_compound15.12223.12223.3 | controllerType=0 controllerNumber=1 scan=12223 | 28.401 |
| 18130:4 | 10 | K.VFPLSLDSTPQDGNVVVAC[+57.021]LVQGFFPQEPLSVTWSESGQN[+1809.666]VTAI | HexNAc(5)Hex(4)Fuc(1) | C[+57], N[+1810] | 1647.2723 | 4 | 6586.0675 | 6586.0275 | 0 | 6.1 | 8 | Specific | 396.93 | 396.93 | 396.93 | 5.48 | 18 | >sp\|P01877\|IGHA2_HUMAN Immunoglobulin heavy constant alpha 2 OS=Homo sapiens OX=9606 GN=IGHA2 PE=1 SV=4 | 576 | 20201113_12413_1ng_glyco_hcd30_etd30_compound15.21865.21865.4 | controllerType=0 controllerNumber=1 scan=21865 | 51.8719 |
| 01732:3 | 10 | K.HYTN[+2262.814]SSQDVTVPC[+57.021]R.V | HexNAc(5)Hex(5)Fuc(1)NeuAc(1) | C[+57], N[+2263] | 1309.529 | 3 | 3926.5724 | 3926.5636 | 0 | 2.2 | 89 | Specific | 340.35 | 226.57 | 226.57 | 5.61 | 18 | >sp\|P01877\|IGHA2_HUMAN Immunoglobulin heavy constant alpha 2 OS=Homo sapiens OX=9606 GN=IGHA2 PE=1 SV=4 | 576 | 20201113_12413_1ng_glyco_hcd30_etd30_compound15.4691.4691.3 | controllerType=0 controllerNumber=1 scan=4691 | 14.4996 |
| 00905:3 | 10 | K.HYTN[+1809.666]SSQDVTVPC[+57.021]R.V | HexNAc(5)Hex(4)Fuc(1) | C[+57], N[+1810] | 1158.4783 | 3 | 3473.4204 | 3473.4154 | 0 | 1.5 | 89 | Specific | 377.57 | 291.26 | 291.26 | 5.86 | 18 | >sp\|P01877\|IGHA2_HUMAN Immunoglobulin heavy constant alpha 2 OS=Homo sapiens OX=9606 GN=IGHA2 PE=1 SV=4 | 576 | 20201113_12413_1ng_glyco_hcd30_etd30_compound15.3832.3832.3 | controllerType=0 controllerNumber=1 scan=3832 | 12.9126 |
| 00924:3 | 10 | K.HYTN[+1647.613]SSQDVTVPC[+57.021]R.V | HexNAc(5)Hex(3)Fuc(1) | C[+57], N[+1648] | 1104.4621 | 3 | 3311.3716 | 3311.3625 | 0 | 2.7 | 89 | Specific | 424.68 | 350.18 | 350.18 | 6.02 | 18 | >sp\|P01877\|IGHA2_HUMAN Immunoglobulin heavy constant alpha 2 OS=Homo sapiens OX=9606 GN=IGHA2 PE=1 SV=4 | 576 | 20201113_12413_1ng_glyco_hcd30_etd30_compound15.3852.3852.3 | controllerType=0 controllerNumber=1 scan=3852 | 12.961 |
| 00920:3 | 10 | K.HYTN[+1647.613]SSQDVTVPC[+57.021]R.V | HexNAc(5)Hex(3)Fuc(1) | C[+57], N[+1648] | 1104.4606 | 3 | 3311.3672 | 3311.3625 | 0 | 1.4 | 89 | Specific | 486.44 | 404.57 | 404.57 | 7.31 | 18 | >sp\|P01877\|IGHA2_HUMAN Immunoglobulin heavy constant alpha 2 OS=Homo sapiens OX=9606 GN=IGHA2 PE=1 SV=4 | 576 | 20201113_12413_1ng_glyco_hcd30_etd30_compound15.3847.3847.3 | controllerType=0 controllerNumber=1 scan=3847 | 12.944 |
| 04151:2 | 10 | K.TPLTAN[+2059.735]ITK.S | HexNAc(4)Hex(5)Fuc(1)NeuAc(1) | N[+2060] | 1509.6519 | 2 | 3018.2965 | 3018.2917 | 0 | 1.6 | 200 | Specific | 210 | 78.6 | 78.6 | 3.23 | 18 | >sp\|P01877\|IGHA2_HUMAN Immunoglobulin heavy constant alpha 2 OS=Homo sapiens OX=9606 GN=IGHA2 PE=1 SV=4 | 576 | 20201113_12413_1ng_glyco_hcd30_etd30_compound15.7209.7209.2 | controllerType=0 controllerNumber=1 scan=7209 | 19.1454 |
| 02493:3 | 10 | K.TPLTAN[+1971.719]ITK.S | HexNAc(5)Hex(5)Fuc(1) | N[+1972] | 977.4328 | 3 | 2930.2839 | 2930.2757 | 0 | 2.8 | 200 | Specific | 237.62 | 127.71 | 127.71 | 2.67 | 18 | >sp\|P01877\|IGHA2_HUMAN Immunoglobulin heavy constant alpha 2 OS=Homo sapiens OX=9606 GN=IGHA2 PE=1 SV=4 | 576 | 20201113_12413_1ng_glyco_hcd30_etd30_compound15.5482.5482.3 | controllerType=0 controllerNumber=1 scan=5482 | 15.9865 |
| 02538:3 | 10 | K.TPLTAN[+1809.666]ITK.S | HexNAc(5)Hex(4)Fuc(1) | N[+1810] | 923.4147 | 3 | 2768.2296 | 2768.2228 | 0 | 2.4 | 200 | Specific | 299.33 | 137.08 | 137.08 | 3.01 | 18 | >sp\|P01877\|IGHA2_HUMAN Immunoglobulin heavy constant alpha 2 OS=Homo sapiens OX=9606 GN=IGHA2 PE=1 SV=4 | 576 | 20201113_12413_1ng_glyco_hcd30_etd30_compound15.5529.5529.3 | controllerType=0 controllerNumber=1 scan=5529 | 16.097 |
| 02584:2 | 10 | K.TPLTAN[+1647.613]ITK.S | HexNAc(5)Hex(3)Fuc(1) | N[+1648] | 1303.5934 | 2 | 2606.1796 | 2606.17 | 0 | 3.7 | 200 | Specific | 310.7 | 121.59 | 121.59 | 2.93 | 18 | >sp\|P01877\|IGHA2_HUMAN Immunoglobulin heavy constant alpha 2 OS=Homo sapiens OX=9606 GN=IGHA2 PE=1 SV=4 | 576 | 20201113_12413_1ng_glyco_hcd30_etd30_compound15.5577.5577.2 | controllerType=0 controllerNumber=1 scan=5577 | 16.1979 |
| 02609:3 | 10 | K.TPLTAN[+1647.613]ITK.S | HexNAc(5)Hex(3)Fuc(1) | N[+1648] | 869.3968 | 3 | 2606.1757 | 2606.17 | 0 | 2.2 | 200 | Specific | 326.6 | 151.68 | 151.68 | 3.3 | 18 | >sp\|P01877\|IGHA2_HUMAN Immunoglobulin heavy constant alpha 2 OS=Homo sapiens OX=9606 GN=IGHA2 PE=1 SV=4 | 576 | 20201113_12413_1ng_glyco_hcd30_etd30_compound15.5603.5603.3 | controllerType=0 controllerNumber=1 scan=5603 | 16.2453 |
| 02522:2 | 10 | K.TPLTAN[+1809.666]ITK.S | HexNAc(5)Hex(4)Fuc(1) | N[+1810] | 1384.6172 | 2 | 2768.2272 | 2768.2228 | 0 | 1.6 | 200 | Specific | 329.47 | 132.88 | 132.88 | 3.28 | 18 | >sp\|P01877\|IGHA2_HUMAN Immunoglobulin heavy constant alpha 2 OS=Homo sapiens OX=9606 GN=IGHA2 PE=1 SV=4 | 576 | 20201113_12413_1ng_glyco_hcd30_etd30_compound15.5512.5512.2 | controllerType=0 controllerNumber=1 scan=5512 | 16.0625 |
| 02573:2 | 10 | K.TPLTAN[+1444.534]ITK.S | HexNAc(4)Hex(3)Fuc(1) | N[+1445] | 1202.0513 | 2 | 2403.0953 | 2403.0906 | 0 | 2 | 200 | Specific | 334.96 | 111.29 | 111.29 | 2.68 | 18 | >sp\|P01877\|IGHA2_HUMAN Immunoglobulin heavy constant alpha 2 OS=Homo sapiens OX=9606 GN=IGHA2 PE=1 SV=4 | 576 | 20201113_12413_1ng_glyco_hcd30_etd30_compound15.5565.5565.2 | controllerType=0 controllerNumber=1 scan=5565 | 16.1678 |
| 09052:3 | 10 | R.MAGKPTHIN[+1702.581]VSVVMAEADGTC[+57.021]Y.- | HexNAc(2)Hex(8) | C[+57], N[+1703] | 1351.8978 | 3 | 4053.6787 | 4053.6642 | 0 | 3.6 | 319 | CRagged | 221.34 | 172.36 | 172.36 | 3.29 | 18 | >sp\|P01877\|IGHA2_HUMAN Immunoglobulin heavy constant alpha 2 OS=Homo sapiens OX=9606 GN=IGHA2 PE=1 SV=4 | 576 | 20201113_12413_1ng_glyco_hcd30_etd30_compound15.12308.12308.3 | controllerType=0 controllerNumber=1 scan=12308 | 28.5975 |
| 09058:3 | 10 | R.MAGKPTHIN[+1540.529]VSVVMAEADGTC[+57.021]Y.- | HexNAc(2)Hex(7) | C[+57], N[+1541] | 1297.8795 | 3 | 3891.6238 | 3891.6114 | 0 | 3.2 | 319 | CRagged | 250.34 | 185.77 | 160.1 | 3.56 | 18 | >sp\|P01877\|IGHA2_HUMAN Immunoglobulin heavy constant alpha 2 OS=Homo sapiens OX=9606 GN=IGHA2 PE=1 SV=4 | 576 | 20201113_12413_1ng_glyco_hcd30_etd30_compound15.12314.12314.3 | controllerType=0 controllerNumber=1 scan=12314 | 28.6131 |
| 09078:3 | 10 | R.MAGKPTHIN[+1378.476]VSVVMAEADGTC[+57.021]Y.- | HexNAc(2)Hex(6) | C[+57], N[+1378] | 1243.8616 | 3 | 3729.5703 | 3729.5585 | 0 | 3.2 | 319 | CRagged | 284.41 | 259.52 | 202.45 | 4.2 | 18 | >sp\|P01877\|IGHA2_HUMAN Immunoglobulin heavy constant alpha 2 OS=Homo sapiens OX=9606 GN=IGHA2 PE=1 SV=4 | 576 | 20201113_12413_1ng_glyco_hcd30_etd30_compound15.12335.12335.3 | controllerType=0 controllerNumber=1 scan=12335 | 28.6618 |
| 09130:3 | 10 | R.MAGKPTHIN[+1216.423]VSVVMAEADGTC[+57.021]Y.- | HexNAc(2)Hex(5) | C[+57], N[+1216] | 1189.8437 | 3 | 3567.5165 | 3567.5057 | 0 | 3 | 319 | CRagged | 323.3 | 294.92 | 212.4 | 4.72 | 18 | >sp\|P01877\|IGHA2_HUMAN Immunoglobulin heavy constant alpha 2 OS=Homo sapiens OX=9606 GN=IGHA2 PE=1 SV=4 | 576 | 20201113_12413_1ng_glyco_hcd30_etd30_compound15.12389.12389.3 | controllerType=0 controllerNumber=1 scan=12389 | 28.7745 |
| 10012:3 | 12 | R.IIVPLNNREN[+2059.735]ISDPTSPLR.T | HexNAc(4)Hex(5)Fuc(1)NeuAc(1) | N[+2060] | 1403.3102 | 3 | 4207.9162 | 4207.9121 | 0 | 1 | 62 | Specific | 211.81 | 193.84 | 193.84 | 3.32 | 6 | >sp\|P01591\|IGJ_HUMAN Immunoglobulin J chain OS=Homo sapiens OX=9606 GN=JCHAIN PE=1 SV=4 | 1392 | 20201113_12413_1ng_glyco_hcd30_etd30_compound15.13305.13305.3 | controllerType=0 controllerNumber=1 scan=13305 | 30.4729 |
| 11706:3 | 12 | R.IIVPLNNREN[+2350.830]ISDPTSPLR.T | HexNAc(4)Hex(5)Fuc(1)NeuAc(2) | N[+2351] | 1500.3462 | 3 | 4499.0242 | 4499.0075 | 0 | 3.7 | 62 | Specific | 245.43 | 173.1 | 173.1 | 3.18 | 6 | >sp\|P01591\|IGJ_HUMAN Immunoglobulin J chain OS=Homo sapiens OX=9606 GN=JCHAIN PE=1 SV=4 | 1392 | 20201113_12413_1ng_glyco_hcd30_etd30_compound15.15088.15088.3 | controllerType=0 controllerNumber=1 scan=15088 | 34.0063 |
| 10086:3 | 12 | R.IIVPLNNREN[+1856.656]ISDPTSPLR.T | HexNAc(3)Hex(5)Fuc(1)NeuAc(1) | N[+1857] | 1335.6192 | 3 | 4004.843 | 4004.8327 | 0 | 2.6 | 62 | Specific | 325.7 | 276.2 | 276.2 | 4.38 | 6 | >sp\|P01591\|IGJ_HUMAN Immunoglobulin J chain OS=Homo sapiens OX=9606 GN=JCHAIN PE=1 SV=4 | 1392 | 20201113_12413_1ng_glyco_hcd30_etd30_compound15.13383.13383.3 | controllerType=0 controllerNumber=1 scan=13383 | 30.6976 |
| 11668:3 | 12 | R.IIVPLNNREN[+2350.830]ISDPTSPLR.T | HexNAc(4)Hex(5)Fuc(1)NeuAc(2) | N[+2351] | 1500.3449 | 3 | 4499.0202 | 4499.0075 | 0 | 2.8 | 62 | Specific | 327.81 | 230.24 | 230.24 | 3.48 | 6 | >sp\|P01591\|IGJ_HUMAN Immunoglobulin J chain OS=Homo sapiens OX=9606 GN=JCHAIN PE=1 SV=4 | 1392 | 20201113_12413_1ng_glyco_hcd30_etd30_compound15.15048.15048.3 | controllerType=0 controllerNumber=1 scan=15048 | 33.9137 |
| 11730:3 | 12 | R.IIVPLNNREN[+2204.772]ISDPTSPLR.T | HexNAc(4)Hex(5)NeuAc(2) | N[+2205] | 1451.659 | 3 | 4352.9624 | 4352.9496 | 0 | 2.9 | 62 | Specific | 354.58 | 251.24 | 251.24 | 3.87 | 6 | >sp\|P01591\|IGJ_HUMAN Immunoglobulin J chain OS=Homo sapiens OX=9606 GN=JCHAIN PE=1 SV=4 | 1392 | 20201113_12413_1ng_glyco_hcd30_etd30_compound15.15113.15113.3 | controllerType=0 controllerNumber=1 scan=15113 | 34.0588 |

Table S4. Cont.

| Query #:z | Protein Rank | Glycopeptide | Glycan | Modification Type(s) | Observed m/z | z | Observed (M+H) | Calc. mass (M+H) | Off-by-x error | Mass error (ppm) | StartPosition | Cleavage | Score | Delta | Delta Mod | \|Log Prob\| | # of unique peptides | Protein Name | Protein DB number | Comment | Scan # | Scan Time |
|---|---|---|---|---|---|---|---|---|---|---|---|---|---|---|---|---|---|---|---|---|---|---|
| 18710:4 | 14 | K.ADTHDEILEGLNFN[+2861.000]LTEIPEAQIHEGFQELLR.T | HexNAc(5)Hex(6)NeuAc(3) | N[+2861] | 1638.9717 | 4 | 6552.8649 | 6552.8239 | 0 | 6.3 | 94 | Specific | 456.41 | 456.41 | 456.41 | 6.39 | 5 | >sp\|P01009\|A1AT_HUMAN Alpha-1-antitrypsin OS=Homo sapiens OX=9606 GN=SERPINA1 PE=1 SV=3 | 642 | 20201113_12413_1ng_glyco_hcd30_etd30_compound15.22481.22481.4 | controllerType=0 controllerNumber=1 scan=22481 | 53.2335 |
| 16716:5 | 14 | K.YLGN[+2204.772]ATAIFFLPDEGKLQHLENELTHDIITK.F | HexNAc(4)Hex(5)NeuAc(2) | N[+2205] | 1149.9304 | 5 | 5745.623 | 5745.6098 | 0 | 2.3 | 268 | Specific | 265.17 | 259.57 | 259.57 | 3.16 | 5 | >sp\|P01009\|A1AT_HUMAN Alpha-1-antitrypsin OS=Homo sapiens OX=9606 GN=SERPINA1 PE=1 SV=3 | 642 | 20201113_12413_1ng_glyco_hcd30_etd30_compound15.20375.20375.5 | controllerType=0 controllerNumber=1 scan=20375 | 47.7064 |
| 16715:4 | 14 | K.YLGN[+2204.772]ATAIFFLPDEGKLQHLENELTHDIITK.F | HexNAc(4)Hex(5)NeuAc(2) | N[+2205] | 1437.1616 | 4 | 5745.6247 | 5745.6098 | 0 | 2.6 | 268 | Specific | 705.86 | 700.98 | 700.98 | 10.36 | 5 | >sp\|P01009\|A1AT_HUMAN Alpha-1-antitrypsin OS=Homo sapiens OX=9606 GN=SERPINA1 PE=1 SV=3 | 642 | 20201113_12413_1ng_glyco_hcd30_etd30_compound15.20374.20374.4 | controllerType=0 controllerNumber=1 scan=20374 | 47.6994 |
| 12330:4 | 20 | K.LKELPGVC[+57.021]N[+2059.735]ETMMALWEEC[+57.021]KPC[+57.021]LK.Q | HexNAc(4)Hex(5)Fuc(1)NeuAc(1) | C[+57]*3, N[+2060] | 1249.7885 | 4 | 4996.132 | 4996.1196 | 0 | 2.5 | 95 | Specific | 641.22 | 495.36 | 495.36 | 7.62 | 8 | >sp\|P10909\|CLUS_HUMAN Clusterin OS=Homo sapiens OX=9606 GN=CLU PE=1 SV=1 | 1467 | 20201113_12413_1ng_glyco_hcd30_etd30_compound15.15740.15740.4 | controllerType=0 controllerNumber=1 scan=15740 | 35.4793 |
| 00065:4 | 20 | R.EIRHN[+2059.735]STGC[+57.021]LR.M | HexNAc(4)Hex(5)Fuc(1)NeuAc(1) | C[+57], N[+2060] | 851.3581 | 4 | 3402.4106 | 3402.3994 | 0 | 3.3 | 287 | Specific | 583.32 | 476.32 | 476.32 | 7.01 | 8 | >sp\|P10909\|CLUS_HUMAN Clusterin OS=Homo sapiens OX=9606 GN=CLU PE=1 SV=1 | 1467 | 20201113_12413_1ng_glyco_hcd30_etd30_compound15.2959.2959.4 | controllerType=0 controllerNumber=1 scan=2959 | 11.2578 |
| 17340:3 | 20 | K.MLN[+2059.735]TSSLLEQLNEQFNWVSR.L | HexNAc(4)Hex(5)Fuc(1)NeuAc(1) | N[+2060] | 1490.3134 | 3 | 4468.9257 | 4468.9217 | 0 | 0.9 | 352 | Specific | 311.7 | 181.43 | 181.43 | 4.9 | 8 | >sp\|P10909\|CLUS_HUMAN Clusterin OS=Homo sapiens OX=9606 GN=CLU PE=1 SV=1 | 1467 | 20201113_12413_1ng_glyco_hcd30_etd30_compound15.21035.21035.3 | controllerType=0 controllerNumber=1 scan=21035 | 49.7794 |
| 18818:5 | 21 | K.APDKNVIFSPLSISTALAFLSLGAHN[+2351.851]TTLTEILK.G | HexNAc(4)Hex(5)Fuc(3)NeuAc(1) | N[+2352] | 1187.7705 | 5 | 5934.8234 | 5934.8289 | 0 | -0.9 | 68 | Specific | 212.18 | 212.18 | 212.18 | 3.24 | 6 | >sp\|P01011\|AACT_HUMAN Alpha-1-antichymotrypsin OS=Homo sapiens OX=9606 GN=SERPINA3 PE=1 SV=2 | 1314 | 20201113_12413_1ng_glyco_hcd30_etd30_compound15.22596.22596.5 | controllerType=0 controllerNumber=1 scan=22596 | 53.5968 |
| 17764:4 | 21 | K.APDKNVIFSPLSISTALAFLSLGAHN[+2059.735]TTLTEILK.G | HexNAc(4)Hex(5)Fuc(1)NeuAc(1) | N[+2060] | 1411.4358 | 4 | 5642.7213 | 5642.7131 | 0 | 1.5 | 68 | Specific | 420.2 | 420.2 | 420.2 | 6.62 | 6 | >sp\|P01011\|AACT_HUMAN Alpha-1-antichymotrypsin OS=Homo sapiens OX=9606 GN=SERPINA3 PE=1 SV=2 | 1314 | 20201113_12413_1ng_glyco_hcd30_etd30_compound15.21480.21480.4 | controllerType=0 controllerNumber=1 scan=21480 | 50.8212 |
| 13391:4 | 21 | K.GLKFN[+2350.830]LTETSEAEIHQSFQHLLR.T | HexNAc(4)Hex(5)Fuc(1)NeuAc(2) | N[+2351] | 1263.0632 | 4 | 5049.231 | 5049.2251 | 0 | 1.2 | 102 | Specific | 248.53 | 183.64 | 183.64 | 2.56 | 6 | >sp\|P01011\|AACT_HUMAN Alpha-1-antichymotrypsin OS=Homo sapiens OX=9606 GN=SERPINA3 PE=1 SV=2 | 1314 | 20201113_12413_1ng_glyco_hcd30_etd30_compound15.16849.16849.4 | controllerType=0 controllerNumber=1 scan=16849 | 37.9491 |
| 13384:4 | 21 | K.GLKFN[+2350.830]LTETSEAEIHQSFQHLLR.T | HexNAc(4)Hex(5)Fuc(1)NeuAc(2) | N[+2351] | 1263.064 | 4 | 5049.234 | 5049.2251 | 0 | 1.8 | 102 | Specific | 365.24 | 324.59 | 324.59 | 3.39 | 6 | >sp\|P01011\|AACT_HUMAN Alpha-1-antichymotrypsin OS=Homo sapiens OX=9606 GN=SERPINA3 PE=1 SV=2 | 1314 | 20201113_12413_1ng_glyco_hcd30_etd30_compound15.16841.16841.4 | controllerType=0 controllerNumber=1 scan=16841 | 37.9221 |
| 07639:4 | 22 | K.SC[+57.021]QHN[+2059.735]GTMYQHGEIFSAHELFPSR.L | HexNAc(4)Hex(5)Fuc(1)NeuAc(1) | C[+57], N[+2060] | 1224.006 | 4 | 4893.0023 | 4892.9919 | 0 | 2.1 | 110 | Specific | 248.23 | 143.28 | 143.28 | 4.14 | 2 | >sp\|Q6WN34\|CRDL2_HUMAN Chordin-like protein 2 OS=Homo sapiens OX=9606 GN=CHRDL2 PE=1 SV=1 | 1189 | 20201113_12413_1ng_glyco_hcd30_etd30_compound15.10834.10834.2 | controllerType=0 controllerNumber=1 scan=10834 | 25.6288 |
| 07658:5 | 22 | K.SC[+57.021]QHN[+2059.735]GTMYQHGEIFSAHELFPSR.L | HexNAc(4)Hex(5)Fuc(1)NeuAc(1) | C[+57], N[+2060] | 979.4063 | 5 | 4893.0023 | 4892.9919 | 0 | 2.1 | 110 | Specific | 941.51 | 880.47 | 880.47 | 15.08 | 2 | >sp\|Q6WN34\|CRDL2_HUMAN Chordin-like protein 2 OS=Homo sapiens OX=9606 GN=CHRDL2 PE=1 SV=1 | 1189 | 20201113_12413_1ng_glyco_hcd30_etd30_compound15.10854.10854.5 | controllerType=0 controllerNumber=1 scan=10854 | 25.691 |
| 06848:3 | 24 | R.LSPN[+2350.830]ASAEHLELR.W | HexNAc(4)Hex(5)Fuc(1)NeuAc(2) | N[+2351] | 1263.2033 | 3 | 3787.5954 | 3787.5796 | 0 | 4.2 | 52 | Specific | 407.87 | 279.19 | 279.19 | 5.09 | 5 | >sp\|Q13410\|BT1A1_HUMAN Butyrophilin subfamily 1 member A1 OS=Homo sapiens OX=9606 GN=BTN1A1 PE=1 SV=3 | 179 | 20201113_12413_1ng_glyco_hcd30_etd30_compound15.10010.10010.3 | controllerType=0 controllerNumber=1 scan=10010 | 24.1046 |
| 08421:5 | 33 | K.QTDEIKDTRN[+2059.735]ESTQNC[+57.021]VVAEPEKMESSISSSSEEMSLSK.C | HexNAc(4)Hex(5)Fuc(1)NeuAc(1) | C[+57], N[+2060] | 1290.9535 | 5 | 6450.7384 | 6450.7156 | 0 | 3.5 | 60 | Specific | 215.64 | 215.64 | 215.64 | 2.65 | 5 | >sp\|P47710\|CASA1_HUMAN Alpha-S1-casein OS=Homo sapiens OX=9606 GN=CSN1S1 PE=1 SV=1 | 989 | 20201113_12413_1ng_glyco_hcd30_etd30_compound15.11652.11652.3 | controllerType=0 controllerNumber=1 scan=11652 | 27.3122 |
| 15812:4 | 1 | R.RTAGWNVPIGTLRPFLN[+2569.905]WTGPPEPIEAAVAR.F | HexNAc(5)Hex(6)NeuAc(2) | N[+2570] | 1489.942 | 4 | 5956.7462 | 5956.7167 | 0 | 5 | 140 | Specific | 216.27 | 216.27 | 216.27 | 2.86 | 42 | >sp\|P02788\|TRFL_HUMAN Lactotransferrin OS=Homo sapiens OX=9606 GN=LTF PE=1 SV=6 | 1270 | 20201113_12433_1ng_glyco_hcd30_etd30_compound15.19405.19405.4 | controllerType=0 controllerNumber=1 scan=19405 | 43.5799 |
| 15830:5 | 1 | R.RTAGWNVPIGTLRPFLN[+2205.793]WTGPPEPIEAAVAR.F | HexNAc(4)Hex(5)Fuc(2)NeuAc(1) | N[+2206] | 1119.3306 | 5 | 5592.6237 | 5592.6049 | 0 | 3.4 | 140 | Specific | 359.69 | 359.69 | 359.69 | 4.85 | 42 | >sp\|P02788\|TRFL_HUMAN Lactotransferrin OS=Homo sapiens OX=9606 GN=LTF PE=1 SV=6 | 1270 | 20201113_12433_1ng_glyco_hcd30_etd30_compound15.19424.19424.5 | controllerType=0 controllerNumber=1 scan=19424 | 43.6511 |
| 15742:4 | 1 | R.RTAGWNVPIGTLRPFLN[+2716.983]WTGPPEPIEAAVAR.F | HexNAc(5)Hex(6)Fuc(3)NeuAc(1) | N[+2717] | 1526.7089 | 4 | 6103.8136 | 6103.795 | 0 | 3.1 | 140 | Specific | 550.82 | 550.82 | 550.82 | 7.93 | 42 | >sp\|P02788\|TRFL_HUMAN Lactotransferrin OS=Homo sapiens OX=9606 GN=LTF PE=1 SV=6 | 1270 | 20201113_12433_1ng_glyco_hcd30_etd30_compound15.19332.19332.4 | controllerType=0 controllerNumber=1 scan=19332 | 43.3999 |
| 15748:4 | 1 | R.RTAGWNVPIGTLRPFLN[+2570.925]WTGPPEPIEAAVAR.F | HexNAc(5)Hex(6)Fuc(2)NeuAc(1) | N[+2571] | 1490.1953 | 4 | 5957.7594 | 5957.7371 | 0 | 3.8 | 140 | Specific | 881.23 | 872.71 | 872.71 | 14.78 | 42 | >sp\|P02788\|TRFL_HUMAN Lactotransferrin OS=Homo sapiens OX=9606 GN=LTF PE=1 SV=6 | 1270 | 20201113_12433_1ng_glyco_hcd30_etd30_compound15.19338.19338.4 | controllerType=0 controllerNumber=1 scan=19338 | 43.4124 |
| 15869:4 | 1 | R.RTAGWNVPIGTLRPFLN[+2059.735]WTGPPEPIEAAVAR.F | HexNAc(4)Hex(5)Fuc(1)NeuAc(1) | N[+2060] | 1362.3967 | 4 | 5446.5651 | 5446.547 | 0 | 3.3 | 140 | Specific | 977.36 | 971.28 | 971.28 | 16.78 | 42 | >sp\|P02788\|TRFL_HUMAN Lactotransferrin OS=Homo sapiens OX=9606 GN=LTF PE=1 SV=6 | 1270 | 20201113_12433_1ng_glyco_hcd30_etd30_compound15.19466.19466.4 | controllerType=0 controllerNumber=1 scan=19466 | 43.7784 |
| 16733:3 | 1 | R.TAGWNVPIGTLRPFLN[+2350.830]WTGPPEPIEAAVAR.F | HexNAc(4)Hex(5)Fuc(1)NeuAc(2) | N[+2351] | 1861.2004 | 3 | 5581.5867 | 5581.5413 | 0 | 8.1 | 141 | Specific | 262.09 | 179.38 | 179.38 | 4.33 | 42 | >sp\|P02788\|TRFL_HUMAN Lactotransferrin OS=Homo sapiens OX=9606 GN=LTF PE=1 SV=6 | 1270 | 20201113_12433_1ng_glyco_hcd30_etd30_compound15.20375.20375.3 | controllerType=0 controllerNumber=1 scan=20375 | 46.3918 |
| 16913:3 | 1 | R.TAGWNVPIGTLRPFLN[+2059.735]WTGPPEPIEAAVAR.F | HexNAc(4)Hex(5)Fuc(1)NeuAc(1) | N[+2060] | 1764.1689 | 3 | 5290.4922 | 5290.4459 | 0 | 8.8 | 141 | Specific | 329.45 | 246.95 | 246.95 | 4.36 | 42 | >sp\|P02788\|TRFL_HUMAN Lactotransferrin OS=Homo sapiens OX=9606 GN=LTF PE=1 SV=6 | 1270 | 20201113_12433_1ng_glyco_hcd30_etd30_compound15.20569.20569.3 | controllerType=0 controllerNumber=1 scan=20569 | 46.8131 |
| 16729:4 | 1 | R.TAGWNVPIGTLRPFLN[+2351.851]WTGPPEPIEAAVAR.F | HexNAc(4)Hex(5)Fuc(3)NeuAc(1) | N[+2352] | 1396.4009 | 4 | 5582.5817 | 5582.5617 | 0 | 3.6 | 141 | Specific | 340.9 | 340.9 | 340.9 | 4.64 | 42 | >sp\|P02788\|TRFL_HUMAN Lactotransferrin OS=Homo sapiens OX=9606 GN=LTF PE=1 SV=6 | 1270 | 20201113_12433_1ng_glyco_hcd30_etd30_compound15.20371.20371.4 | controllerType=0 controllerNumber=1 scan=20371 | 46.3834 |
| 16761:3 | 1 | R.TAGWNVPIGTLRPFLN[+2205.793]WTGPPEPIEAAVAR.F | HexNAc(4)Hex(5)Fuc(2)NeuAc(1) | N[+2206] | 1812.8447 | 3 | 5436.5195 | 5436.5038 | 0 | 2.9 | 141 | Specific | 413.34 | 309.14 | 309.14 | 5.5 | 42 | >sp\|P02788\|TRFL_HUMAN Lactotransferrin OS=Homo sapiens OX=9606 GN=LTF PE=1 SV=6 | 1270 | 20201113_12433_1ng_glyco_hcd30_etd30_compound15.20405.20405.3 | controllerType=0 controllerNumber=1 scan=20405 | 46.4867 |
| 16743:3 | 1 | R.TAGWNVPIGTLRPFLN[+2059.735]WTGPPEPIEAAVAR.F | HexNAc(4)Hex(5)Fuc(1)NeuAc(1) | N[+2060] | 1764.1602 | 3 | 5290.4662 | 5290.4459 | 0 | 3.8 | 141 | Specific | 442.32 | 354.5 | 354.5 | 6.45 | 42 | >sp\|P02788\|TRFL_HUMAN Lactotransferrin OS=Homo sapiens OX=9606 GN=LTF PE=1 SV=6 | 1270 | 20201113_12433_1ng_glyco_hcd30_etd30_compound15.20386.20386.2 | controllerType=0 controllerNumber=1 scan=20386 | 46.4366 |
| 17772:4 | 1 | R.TAGWNVPIGTLRPFLN[+2350.830]WTGPPEPIEAAVAR.F | HexNAc(4)Hex(5)Fuc(1)NeuAc(2) | N[+2351] | 1396.1506 | 4 | 5581.5807 | 5581.5413 | 0 | 7.1 | 141 | Specific | 664.49 | 626.29 | 626.29 | 9.83 | 42 | >sp\|P02788\|TRFL_HUMAN Lactotransferrin OS=Homo sapiens OX=9606 GN=LTF PE=1 SV=6 | 1270 | 20201113_12433_1ng_glyco_hcd30_etd30_compound15.21474.21474.3 | controllerType=0 controllerNumber=1 scan=21474 | 49.2999 |
| 16052:4 | 1 | R.TAGWNVPIGTLRPFLN[+1768.640]WTGPPEPIEAAVAR.F | HexNAc(4)Hex(5)Fuc(1) | N[+1769] | 1250.5991 | 4 | 4999.3747 | 4999.3504 | 0 | 4.8 | 141 | Specific | 806.87 | 794.37 | 794.37 | 12.74 | 42 | >sp\|P02788\|TRFL_HUMAN Lactotransferrin OS=Homo sapiens OX=9606 GN=LTF PE=1 SV=6 | 1270 | 20201113_12433_1ng_glyco_hcd30_etd30_compound15.19658.19658.4 | controllerType=0 controllerNumber=1 scan=19658 | 44.3078 |
| 16721:5 | 1 | R.TAGWNVPIGTLRPFLN[+2205.793]WTGPPEPIEAAVAR.F | HexNAc(4)Hex(5)Fuc(2)NeuAc(1) | N[+2206] | 1088.1113 | 5 | 5436.5275 | 5436.5038 | 0 | 4.4 | 141 | Specific | 823.69 | 723.08 | 723.08 | 11.89 | 42 | >sp\|P02788\|TRFL_HUMAN Lactotransferrin OS=Homo sapiens OX=9606 GN=LTF PE=1 SV=6 | 1270 | 20201113_12433_1ng_glyco_hcd30_etd30_compound15.20362.20362.4 | controllerType=0 controllerNumber=1 scan=20362 | 46.3468 |
| 16447:3 | 1 | R.TAGWNIPMGLLFN[+1751.624]QTGSC[+57.021]K.F | HexNAc(4)Hex(4)NeuAc(1) | C[+57], N[+1752] | 1282.8897 | 3 | 3846.6546 | 3846.6341 | 0 | 5.3 | 485 | Specific | 232.4 | 170.46 | 170.46 | 3.82 | 42 | >sp\|P02788\|TRFL_HUMAN Lactotransferrin OS=Homo sapiens OX=9606 GN=LTF PE=1 SV=6 | 1270 | 20201113_12433_1ng_glyco_hcd30_etd30_compound15.20076.20076.3 | controllerType=0 controllerNumber=1 scan=20076 | 45.6532 |
| 14853:3 | 1 | R.TAGWNIPMGLLFN[+1768.640]QTGSC[+57.021]K.F | HexNAc(4)Hex(5)Fuc(1) | C[+57], N[+1769] | 1288.5529 | 3 | 3863.6442 | 3863.6495 | 0 | -1.4 | 485 | Specific | 295.38 | 216.16 | 216.16 | 4.41 | 42 | >sp\|P02788\|TRFL_HUMAN Lactotransferrin OS=Homo sapiens OX=9606 GN=LTF PE=1 SV=6 | 1270 | 20201113_12433_1ng_glyco_hcd30_etd30_compound15.18394.18394.3 | controllerType=0 controllerNumber=1 scan=18394 | 40.6948 |
| 17930:3 | 1 | R.TAGWNIPMGLLFN[+2204.772]QTGSC[+57.021]K.F | HexNAc(4)Hex(5)NeuAc(2) | C[+57], N[+2205] | 1433.9402 | 3 | 4299.8062 | 4299.7824 | 0 | 5.5 | 485 | Specific | 327.33 | 195.26 | 195.26 | 4.27 | 42 | >sp\|P02788\|TRFL_HUMAN Lactotransferrin OS=Homo sapiens OX=9606 GN=LTF PE=1 SV=6 | 1270 | 20201113_12433_1ng_glyco_hcd30_etd30_compound15.21641.21641.3 | controllerType=0 controllerNumber=1 scan=21641 | 49.8053 |
| 16247:3 | 1 | R.TAGWNIPMGLLFN[+2205.793]QTGSC[+57.021]K.F | HexNAc(4)Hex(5)Fuc(2)NeuAc(1) | C[+57], N[+2206] | 1434.2747 | 3 | 4300.8096 | 4300.8028 | 0 | 1.6 | 485 | Specific | 376.76 | 267.79 | 267.79 | 5.71 | 42 | >sp\|P02788\|TRFL_HUMAN Lactotransferrin OS=Homo sapiens OX=9606 GN=LTF PE=1 SV=6 | 1270 | 20201113_12433_1ng_glyco_hcd30_etd30_compound15.19863.19863.3 | controllerType=0 controllerNumber=1 scan=19863 | 44.8936 |
| 15079:3 | 1 | R.TAGWNIPMGLLFN[+1622.582]QTGSC[+57.021]K.F | HexNAc(4)Hex(5) | C[+57], N[+1623] | 1239.8721 | 3 | 3717.6018 | 3717.5916 | 0 | 2.8 | 485 | Specific | 391 | 286.35 | 252.17 | 5.71 | 42 | >sp\|P02788\|TRFL_HUMAN Lactotransferrin OS=Homo sapiens OX=9606 GN=LTF PE=1 SV=6 | 1270 | 20201113_12433_1ng_glyco_hcd30_etd30_compound15.18631.18631.3 | controllerType=0 controllerNumber=1 scan=18631 | 41.3265 |
| 14907:3 | 1 | R.TAGWNIPMGLLFN[+1768.640]QTGSC[+57.021]K.F | HexNAc(4)Hex(5)Fuc(1) | C[+57], N[+1769] | 1288.561 | 3 | 3863.6684 | 3863.6495 | 0 | 4.9 | 485 | Specific | 392.44 | 271.18 | 271.18 | 4.63 | 42 | >sp\|P02788\|TRFL_HUMAN Lactotransferrin OS=Homo sapiens OX=9606 GN=LTF PE=1 SV=6 | 1270 | 20201113_12433_1ng_glyco_hcd30_etd30_compound15.18451.18451.3 | controllerType=0 controllerNumber=1 scan=18451 | 40.8517 |
| 14888:3 | 1 | R.TAGWNIPMGLLFN[+1768.640]QTGSC[+57.021]K.F | HexNAc(4)Hex(5)Fuc(1) | C[+57], N[+1769] | 1288.5572 | 3 | 3863.657 | 3863.6495 | 0 | 2 | 485 | Specific | 409 | 300.76 | 300.76 | 5.26 | 42 | >sp\|P02788\|TRFL_HUMAN Lactotransferrin OS=Homo sapiens OX=9606 GN=LTF PE=1 SV=6 | 1270 | 20201113_12433_1ng_glyco_hcd30_etd30_compound15.18430.18430.3 | controllerType=0 controllerNumber=1 scan=18430 | 40.789 |
| 16253:3 | 1 | R.TAGWNIPMGLLFN[+2059.735]QTGSC[+57.021]K.F | HexNAc(4)Hex(5)Fuc(1)NeuAc(1) | C[+57], N[+2060] | 1385.5879 | 3 | 4154.7493 | 4154.7449 | 0 | 1.1 | 485 | Specific | 427.56 | 301.35 | 301.35 | 5.23 | 42 | >sp\|P02788\|TRFL_HUMAN Lactotransferrin OS=Homo sapiens OX=9606 GN=LTF PE=1 SV=6 | 1270 | 20201113_12433_1ng_glyco_hcd30_etd30_compound15.19869.19869.3 | controllerType=0 controllerNumber=1 scan=19869 | 44.9069 |
| 16206:3 | 1 | R.TAGWNIPMGLLFN[+2205.793]QTGSC[+57.021]K.F | HexNAc(4)Hex(5)Fuc(2)NeuAc(1) | C[+57], N[+2206] | 1434.278 | 3 | 4300.8195 | 4300.8028 | 0 | 3.9 | 485 | Specific | 446.11 | 301.6 | 301.6 | 6.18 | 42 | >sp\|P02788\|TRFL_HUMAN Lactotransferrin OS=Homo sapiens OX=9606 GN=LTF PE=1 SV=6 | 1270 | 20201113_12433_1ng_glyco_hcd30_etd30_compound15.19819.19819.3 | controllerType=0 controllerNumber=1 scan=19819 | 44.7691 |
| 15761:3 | 1 | R.TAGWNIPMGLLFN[+1622.582]QTGSC[+57.021]KFDEYFSQSC[+57.021]APGSDPR.S | HexNAc(4)Hex(5) | C[+57]*2, N[+1623] | 1854.4581 | 3 | 5561.3597 | 5561.3387 | 0 | 3.8 | 485 | Specific | 255.45 | 182.39 | 89.44 | 4.97 | 42 | >sp\|P02788\|TRFL_HUMAN Lactotransferrin OS=Homo sapiens OX=9606 GN=LTF PE=1 SV=6 | 1270 | 20201113_12433_1ng_glyco_hcd30_etd30_compound15.19352.19352.3 | controllerType=0 controllerNumber=1 scan=19352 | 43.4528 |
| 15709:3 | 1 | R.TAGWNIPMGLLFN[+1768.640]QTGSC[+57.021]KFDEYFSQSC[+57.021]APGSDPR.S | HexNAc(4)Hex(5)Fuc(1) | C[+57]*2, N[+1769] | 1903.145 | 3 | 5707.4204 | 5707.3966 | 0 | 4.2 | 485 | Specific | 276.95 | 229.13 | 229.13 | 5.97 | 42 | >sp\|P02788\|TRFL_HUMAN Lactotransferrin OS=Homo sapiens OX=9606 GN=LTF PE=1 SV=6 | 1270 | 20201113_12433_1ng_glyco_hcd30_etd30_compound15.19297.19297.3 | controllerType=0 controllerNumber=1 scan=19297 | 43.2965 |
| 16453:4 | 1 | R.TAGWNIPMGLLFN[+1913.677]QTGSC[+57.021]KFDEYFSQSC[+57.021]APGSDPR.S | HexNAc(4)Hex(5)NeuAc(1) | C[+57]*2, N[+1914] | 1463.8712 | 4 | 5852.463 | 5852.4341 | 0 | 4.9 | 485 | Specific | 339.77 | 269.09 | 269.09 | 3.52 | 42 | >sp\|P02788\|TRFL_HUMAN Lactotransferrin OS=Homo sapiens OX=9606 GN=LTF PE=1 SV=6 | 1270 | 20201113_12433_1ng_glyco_hcd30_etd30_compound15.20083.20083.4 | controllerType=0 controllerNumber=1 scan=20083 | 45.6794 |
| 16431:4 | 1 | R.TAGWNIPMGLLFN[+2059.735]QTGSC[+57.021]KFDEYFSQSC[+57.021]APGSDPR.S | HexNAc(4)Hex(5)Fuc(1)NeuAc(1) | C[+57]*2, N[+2060] | 1500.3849 | 4 | 5998.5177 | 5998.492 | 0 | 4.3 | 485 | Specific | 411.77 | 380.97 | 380.97 | 7.88 | 42 | >sp\|P02788\|TRFL_HUMAN Lactotransferrin OS=Homo sapiens OX=9606 GN=LTF PE=1 SV=6 | 1270 | 20201113_12433_1ng_glyco_hcd30_etd30_compound15.20060.20060.4 | controllerType=0 controllerNumber=1 scan=20060 | 45.6228 |
| 15701:4 | 1 | R.TAGWNIPMGLLFN[+1768.640]QTGSC[+57.021]KFDEYFSQSC[+57.021]APGSDPR.S | HexNAc(4)Hex(5)Fuc(1) | C[+57]*2, N[+1769] | 1427.6106 | 4 | 5707.4206 | 5707.3966 | 0 | 4.2 | 485 | Specific | 759.7 | 697.39 | 697.39 | 13.18 | 42 | >sp\|P02788\|TRFL_HUMAN Lactotransferrin OS=Homo sapiens OX=9606 GN=LTF PE=1 SV=6 | 1270 | 20201113_12433_1ng_glyco_hcd30_etd30_compound15.19289.19289.4 | controllerType=0 controllerNumber=1 scan=19289 | 43.2764 |
| 10376:3 | 1 | R.N[+2350.830]GSDC[+57.021]PDKFC[+57.021]LFQSETK.N | HexNAc(4)Hex(5)Fuc(1)NeuAc(2) | C[+57]*2, N[+2351] | 1461.9117 | 3 | 4383.7205 | 4383.7043 | 0 | 3.7 | 642 | Specific | 200.05 | 103.78 | 103.78 | 4.88 | 42 | >sp\|P02788\|TRFL_HUMAN Lactotransferrin OS=Homo sapiens OX=9606 GN=LTF PE=1 SV=6 | 1270 | 20201113_12433_1ng_glyco_hcd30_etd30_compound15.13670.13670.3 | controllerType=0 controllerNumber=1 scan=13670 | 30.9959 |
| 08332:3 | 1 | R.N[+2059.735]GSDC[+57.021]PDKFC[+57.021]LFQSETK.N | HexNAc(4)Hex(5)Fuc(1)NeuAc(1) | C[+57]*2, N[+2060] | 1364.879 | 3 | 4092.6223 | 4092.6088 | 0 | 3.3 | 642 | Specific | 274.09 | 147.79 | 147.79 | 5.42 | 42 | >sp\|P02788\|TRFL_HUMAN Lactotransferrin OS=Homo sapiens OX=9606 GN=LTF PE=1 SV=6 | 1270 | 20201113_12433_1ng_glyco_hcd30_etd30_compound15.11547.11547.3 | controllerType=0 controllerNumber=1 scan=11547 | 26.8811 |
| 08223:3 | 1 | R.N[+2205.793]GSDC[+57.021]PDKFC[+57.021]LFQSETK.N | HexNAc(4)Hex(5)Fuc(2)NeuAc(1) | C[+57]*2, N[+2206] | 1413.5604 | 3 | 4238.6665 | 4238.6667 | 0 | -0.1 | 642 | Specific | 349.41 | 264.86 | 264.86 | 4.89 | 42 | >sp\|P02788\|TRFL_HUMAN Lactotransferrin OS=Homo sapiens OX=9606 GN=LTF PE=1 SV=6 | 1270 | 20201113_12433_1ng_glyco_hcd30_etd30_compound15.11434.11434.3 | controllerType=0 controllerNumber=1 scan=11434 | 26.671 |
| 15545:3 | 2 | R.LSLHRPALEDLLLGSEAN[+1378.476]LTC[+57.021]TLTGLR.D | HexNAc(2)Hex(6) | C[+57], N[+1378] | 1448.033 | 3 | 4342.0845 | 4342.074 | 0 | 2.4 | 127 | Specific | 205.79 | 205.79 | 205.79 | 5.17 | 23 | >sp\|P01876\|IGHA1_HUMAN Immunoglobulin heavy constant alpha 1 OS=Homo sapiens OX=9606 GN=IGHA1 PE=1 SV=2 | 255 | 20201113_12433_1ng_glyco_hcd30_etd30_compound15.19124.19124.3 | controllerType=0 controllerNumber=1 scan=19124 | 42.816 |
| 15486:4 | 2 | R.LSLHRPALEDLLLGSEAN[+1825.661]LTC[+57.021]TLTGLR.D | HexNAc(5)Hex(5) | C[+57], N[+1826] | 1198.0745 | 4 | 4789.2762 | 4789.2592 | 0 | 3.5 | 127 | Specific | 238.86 | 238.86 | 238.86 | 4.02 | 23 | >sp\|P01876\|IGHA1_HUMAN Immunoglobulin heavy constant alpha 1 OS=Homo sapiens OX=9606 GN=IGHA1 PE=1 SV=2 | 255 | 20201113_12433_1ng_glyco_hcd30_etd30_compound15.19061.19061.4 | controllerType=0 controllerNumber=1 scan=19061 | 42.5794 |
| 15527:3 | 2 | R.LSLHRPALEDLLLGSEAN[+1460.529]LTC[+57.021]TLTGLR.D | HexNAc(4)Hex(4) | C[+57], N[+1461] | 1475.3853 | 3 | 4424.1414 | 4424.1271 | 0 | 3.2 | 127 | Specific | 351.22 | 343.18 | 343.18 | 6.02 | 23 | >sp\|P01876\|IGHA1_HUMAN Immunoglobulin heavy constant alpha 1 OS=Homo sapiens OX=9606 GN=IGHA1 PE=1 SV=2 | 255 | 20201113_12433_1ng_glyco_hcd30_etd30_compound15.19105.19105.3 | controllerType=0 controllerNumber=1 scan=19105 | 42.7512 |
| 15517:3 | 2 | R.LSLHRPALEDLLLGSEAN[+1663.608]LTC[+57.021]TLTGLR.D | HexNAc(5)Hex(4) | C[+57], N[+1664] | 1543.0886 | 3 | 4627.2511 | 4627.2064 | 0 | 9.7 | 127 | Specific | 356.74 | 321.14 | 321.14 | 4.67 | 23 | >sp\|P01876\|IGHA1_HUMAN Immunoglobulin heavy constant alpha 1 OS=Homo sapiens OX=9606 GN=IGHA1 PE=1 SV=2 | 255 | 20201113_12433_1ng_glyco_hcd30_etd30_compound15.19094.19094.3 | controllerType=0 controllerNumber=1 scan=19094 | 42.7005 |
| 15570:3 | 2 | R.LSLHRPALEDLLLGSEAN[+1095.397]LTC[+57.021]TLTGLR.D | HexNAc(3)Hex(3) | C[+57], N[+1095] | 1353.6729 | 3 | 4059.0042 | 4058.9949 | 0 | 2.3 | 127 | Specific | 419.45 | 419.45 | 419.45 | 7.45 | 23 | >sp\|P01876\|IGHA1_HUMAN Immunoglobulin heavy constant alpha 1 OS=Homo sapiens OX=9606 GN=IGHA1 PE=1 SV=2 | 255 | 20201113_12433_1ng_glyco_hcd30_etd30_compound15.19151.19151.3 | controllerType=0 controllerNumber=1 scan=19151 | 42.8927 |
| 15454:3 | 2 | R.LSLHRPALEDLLLGSEAN[+1702.581]LTC[+57.021]TLTGLR.D | HexNAc(2)Hex(8) | C[+57], N[+1703] | 1556.0799 | 3 | 4666.2251 | 4666.1796 | 0 | 9.8 | 127 | Specific | 446.08 | 413.53 | 413.53 | 6.19 | 23 | >sp\|P01876\|IGHA1_HUMAN Immunoglobulin heavy constant alpha 1 OS=Homo sapiens OX=9606 GN=IGHA1 PE=1 SV=2 | 255 | 20201113_12433_1ng_glyco_hcd30_etd30_compound15.19027.19027.3 | controllerType=0 controllerNumber=1 scan=19027 | 42.4601 |
| 15503:3 | 2 | R.LSLHRPALEDLLLGSEAN[+1501.555]LTC[+57.021]TLTGLR.D | HexNAc(5)Hex(3) | C[+57], N[+1502] | 1489.0594 | 3 | 4465.1636 | 4465.1536 | 0 | 2.2 | 127 | Specific | 468.29 | 423.83 | 423.83 | 7.42 | 23 | >sp\|P01876\|IGHA1_HUMAN Immunoglobulin heavy constant alpha 1 OS=Homo sapiens OX=9606 GN=IGHA1 PE=1 SV=2 | 255 | 20201113_12433_1ng_glyco_hcd30_etd30_compound15.19079.19079.3 | controllerType=0 controllerNumber=1 scan=19079 | 42.6452 |
| 15491:3 | 2 | R.LSLHRPALEDLLLGSEAN[+1540.529]LTC[+57.021]TLTGLR.D | HexNAc(2)Hex(7) | C[+57], N[+1541] | 1502.0497 | 3 | 4504.1347 | 4504.1268 | 0 | 1.8 | 127 | Specific | 486.58 | 443.49 | 443.49 | 7.61 | 23 | >sp\|P01876\|IGHA1_HUMAN Immunoglobulin heavy constant alpha 1 OS=Homo sapiens OX=9606 GN=IGHA1 PE=1 SV=2 | 255 | 20201113_12433_1ng_glyco_hcd30_etd30_compound15.19067.19067.3 | controllerType=0 controllerNumber=1 scan=19067 | 42.597 |
| 15508:4 | 2 | R.LSLHRPALEDLLLGSEAN[+1663.608]LTC[+57.021]TLTGLR.D | HexNAc(5)Hex(4) | C[+57], N[+1664] | 1157.5671 | 4 | 4627.2464 | 4627.2064 | 0 | 8.6 | 127 | Specific | 856.59 | 856.59 | 856.59 | 13.59 | 23 | >sp\|P01876\|IGHA1_HUMAN Immunoglobulin heavy constant alpha 1 OS=Homo sapiens OX=9606 GN=IGHA1 PE=1 SV=2 | 255 | 20201113_12433_1ng_glyco_hcd30_etd30_compound15.19085.19085.4 | controllerType=0 controllerNumber=1 scan=19085 | 42.6709 |
| 15492:4 | 2 | R.LSLHRPALEDLLLGSEAN[+1501.555]LTC[+57.021]TLTGLR.D | HexNAc(5)Hex(3) | C[+57], N[+1502] | 1117.0462 | 4 | 4465.1629 | 4465.1536 | 0 | 2.1 | 127 | Specific | 928.15 | 901.61 | 901.61 | 15.82 | 23 | >sp\|P01876\|IGHA1_HUMAN Immunoglobulin heavy constant alpha 1 OS=Homo sapiens OX=9606 GN=IGHA1 PE=1 SV=2 | 255 | 20201113_12433_1ng_glyco_hcd30_etd30_compound15.19068.19068.4 | controllerType=0 controllerNumber=1 scan=19068 | 42.604 |
| 15539:4 | 2 | R.LSLHRPALEDLLLGSEAN[+1298.476]LTC[+57.021]TLTGLR.D | HexNAc(4)Hex(3) | C[+57], N[+1298] | 1066.2796 | 4 | 4262.0965 | 4262.0742 | 0 | 5.2 | 127 | Specific | 934.64 | 934.64 | 934.64 | 14.74 | 23 | >sp\|P01876\|IGHA1_HUMAN Immunoglobulin heavy constant alpha 1 OS=Homo sapiens OX=9606 GN=IGHA1 PE=1 SV=2 | 255 | 20201113_12433_1ng_glyco_hcd30_etd30_compound15.19118.19118.4 | controllerType=0 controllerNumber=1 scan=19118 | 42.7919 |
| 15522:4 | 2 | R.LSLHRPALEDLLLGSEAN[+1460.529]LTC[+57.021]TLTGLR.D | HexNAc(4)Hex(4) | C[+57], N[+1461] | 1106.7911 | 4 | 4424.1424 | 4424.1271 | 0 | 3.5 | 127 | Specific | 947.61 | 947.61 | 947.61 | 15.91 | 23 | >sp\|P01876\|IGHA1_HUMAN Immunoglobulin heavy constant alpha 1 OS=Homo sapiens OX=9606 GN=IGHA1 PE=1 SV=2 | 255 | 20201113_12433_1ng_glyco_hcd30_etd30_compound15.19100.19100.4 | controllerType=0 controllerNumber=1 scan=19100 | 42.7349 |
| 15447:4 | 2 | R.LSLHRPALEDLLLGSEAN[+1702.581]LTC[+57.021]TLTGLR.D | HexNAc(2)Hex(8) | C[+57], N[+1703] | 1167.3108 | 4 | 4666.2215 | 4666.1796 | 0 | 9 | 127 | Specific | 978.7 | 972.88 | 972.88 | 15.59 | 23 | >sp\|P01876\|IGHA1_HUMAN Immunoglobulin heavy constant alpha 1 OS=Homo sapiens OX=9606 GN=IGHA1 PE=1 SV=2 | 255 | 20201113_12433_1ng_glyco_hcd30_etd30_compound15.19020.19020.4 | controllerType=0 controllerNumber=1 scan=19020 | 42.4449 |
| 15458:4 | 2 | R.LSLHRPALEDLLLGSEAN[+1540.529]LTC[+57.021]TLTGLR.D | HexNAc(2)Hex(7) | C[+57], N[+1541] | 1126.797 | 4 | 4504.1663 | 4504.1268 | 0 | 8.8 | 127 | Specific | 984.26 | 925.5 | 925.5 | 15.59 | 23 | >sp\|P01876\|IGHA1_HUMAN Immunoglobulin heavy constant alpha 1 OS=Homo sapiens OX=9606 GN=IGHA1 PE=1 SV=2 | 255 | 20201113_12433_1ng_glyco_hcd30_etd30_compound15.19031.19031.4 | controllerType=0 controllerNumber=1 scan=19031 | 42.4746 |
| 15523:4 | 2 | R.LSLHRPALEDLLLGSEAN[+1216.423]LTC[+57.021]TLTGLR.D | HexNAc(2)Hex(5) | C[+57], N[+1216] | 1045.7633 | 4 | 4180.0316 | 4180.0211 | 0 | 2.5 | 127 | Specific | 1006.15 | 988.95 | 988.95 | 17.82 | 23 | >sp\|P01876\|IGHA1_HUMAN Immunoglobulin heavy constant alpha 1 OS=Homo sapiens OX=9606 GN=IGHA1 PE=1 SV=2 | 255 | 20201113_12433_1ng_glyco_hcd30_etd30_compound15.19101.19101.4 | controllerType=0 controllerNumber=1 scan=19101 | 42.738 |
| 15568:4 | 2 | R.LSLHRPALEDLLLGSEAN[+1095.397]LTC[+57.021]TLTGLR.D | HexNAc(3)Hex(3) | C[+57], N[+1095] | 1015.5076 | 4 | 4059.0086 | 4058.9949 | 0 | 3.4 | 127 | Specific | 1033.62 | 1033.62 | 1033.62 | 17.91 | 23 | >sp\|P01876\|IGHA1_HUMAN Immunoglobulin heavy constant alpha 1 OS=Homo sapiens OX=9606 GN=IGHA1 PE=1 SV=2 | 255 | 20201113_12433_1ng_glyco_hcd30_etd30_compound15.19149.19149.4 | controllerType=0 controllerNumber=1 scan=19149 | 42.8841 |
| 15440:4 | 2 | R.LSLHRPALEDLLLGSEAN[+1540.529]LTC[+57.021]TLTGLR.D | HexNAc(2)Hex(7) | C[+57], N[+1541] | 1126.7919 | 4 | 4504.1458 | 4504.1268 | 0 | 4.2 | 127 | Specific | 1079.02 | 1024.47 | 1024.47 | 18.91 | 23 | >sp\|P01876\|IGHA1_HUMAN Immunoglobulin heavy constant alpha 1 OS=Homo sapiens OX=9606 GN=IGHA1 PE=1 SV=2 | 255 | 20201113_12433_1ng_glyco_hcd30_etd30_compound15.19012.19012.2 | controllerType=0 controllerNumber=1 scan=19012 | 42.4154 |
| 10326:3 | 2 | R.LAGKPTHVN[+1501.555]VSVVMAEVDGTC[+57.021]Y.- | HexNAc(5)Hex(3) | C[+57], N[+1502] | 1283.5729 | 3 | 3848.7042 | 3848.6974 | 0 | 1.8 | 332 | CRagged | 201.01 | 113.49 | 113.49 | 2.57 | 23 | >sp\|P01876\|IGHA1_HUMAN Immunoglobulin heavy constant alpha 1 OS=Homo sapiens OX=9606 GN=IGHA1 PE=1 SV=2 | 255 | 20201113_12433_1ng_glyco_hcd30_etd30_compound15.13618.13618.3 | controllerType=0 controllerNumber=1 scan=13618 | 30.8814 |
| 10222:3 | 2 | R.LAGKPTHVN[+1809.666]VSVVMAEVDGTC[+57.021]Y.- | HexNAc(5)Hex(4)Fuc(1) | C[+57], N[+1810] | 1386.279 | 3 | 4156.8224 | 4156.8081 | 0 | 3.4 | 332 | CRagged | 201.36 | 136.31 | 136.31 | 2.69 | 23 | >sp\|P01876\|IGHA1_HUMAN Immunoglobulin heavy constant alpha 1 OS=Homo sapiens OX=9606 GN=IGHA1 PE=1 SV=2 | 255 | 20201113_12433_1ng_glyco_hcd30_etd30_compound15.13509.13509.3 | controllerType=0 controllerNumber=1 scan=13509 | 30.634 |
| 13469:4 | 2 | R.LAGKPTHVN[+2553.910]VSVVMAEVDGTC[+57.021]Y.- | HexNAc(5)Hex(5)Fuc(1)NeuAc(2) | C[+57], N[+2554] | 1226.022 | 4 | 4901.0662 | 4901.0518 | 0 | 2.9 | 332 | CRagged | 224.05 | 197.79 | 197.79 | 3.47 | 23 | >sp\|P01876\|IGHA1_HUMAN Immunoglobulin heavy constant alpha 1 OS=Homo sapiens OX=9606 GN=IGHA1 PE=1 SV=2 | 255 | 20201113_12433_1ng_glyco_hcd30_etd30_compound15.16910.16910.4 | controllerType=0 controllerNumber=1 scan=16910 | [illegible] |
| 10204:3 | 2 | R.LAGKPTHVN[+1378.476]VSVVMAEVDGTC[+57.021]Y.- | HexNAc(2)Hex(6) | C[+57], N[+1378] | 1242.5485 | 3 | 3725.631 | 3725.6178 | 0 | 3.6 | 332 | CRagged | 233.3 | 204.8 | 188.64 | 3.47 | 23 | >sp\|P01876\|IGHA1_HUMAN Immunoglobulin heavy constant alpha 1 OS=Homo sapiens OX=9606 GN=IGHA1 PE=1 SV=2 | 255 | 20201113_12433_1ng_glyco_hcd30_etd30_compound15.13491.13491.3 | controllerType=0 controllerNumber=1 scan=13491 | [illegible] |
| 10233:3 | 2 | R.LAGKPTHVN[+1809.666]VSVVMAEVDGTC[+57.021]Y.- | HexNAc(5)Hex(4)Fuc(1) | C[+57], N[+1810] | 1386.2763 | 3 | 4156.8144 | 4156.8081 | 0 | 1.5 | 332 | CRagged | 250.05 | 156.7 | 156.7 | 2.69 | 23 | >sp\|P01876\|IGHA1_HUMAN Immunoglobulin heavy constant alpha 1 OS=Homo sapiens OX=9606 GN=IGHA1 PE=1 SV=2 | 255 | 20201113_12433_1ng_glyco_hcd30_etd30_compound15.13521.13521.3 | controllerType=0 controllerNumber=1 scan=13521 | [illegible] |
| 10287:3 | 2 | R.LAGKPTHVN[+1501.555]VSVVMAEVDGTC[+57.021]Y.- | HexNAc(5)Hex(3) | C[+57], N[+1502] | 1283.5727 | 3 | 3848.7035 | 3848.6974 | 0 | 1.6 | 332 | CRagged | 252.44 | 162.37 | 162.37 | 3.35 | 23 | >sp\|P01876\|IGHA1_HUMAN Immunoglobulin heavy constant alpha 1 OS=Homo sapiens OX=9606 GN=IGHA1 PE=1 SV=2 | 255 | 20201113_12433_1ng_glyco_hcd30_etd30_compound15.13577.13577.3 | controllerType=0 controllerNumber=1 scan=13577 | 30.7813 |
| 10187:3 | 2 | R.LAGKPTHVN[+1540.529]VSVVMAEVDGTC[+57.021]Y.- | HexNAc(2)Hex(7) | C[+57], N[+1541] | 1296.5656 | 3 | 3887.6823 | 3887.6706 | 0 | 3 | 332 | CRagged | 253.35 | 210.39 | 204.39 | 3.79 | 23 | >sp\|P01876\|IGHA1_HUMAN Immunoglobulin heavy constant alpha 1 OS=Homo sapiens OX=9606 GN=IGHA1 PE=1 SV=2 | 255 | 20201113_12433_1ng_glyco_hcd30_etd30_compound15.13473.13473.3 | controllerType=0 controllerNumber=1 scan=13473 | 30.5453 |
| 10305:3 | 2 | R.LAGKPTHVN[+1647.613]VSVVMAEVDGTC[+57.021]Y.- | HexNAc(5)Hex(3)Fuc(1) | C[+57], N[+1648] | 1332.2586 | 3 | 3994.7613 | 3994.7553 | 0 | 1.5 | 332 | CRagged | 258.77 | 180.83 | 180.83 | 2.69 | 23 | >sp\|P01876\|IGHA1_HUMAN Immunoglobulin heavy constant alpha 1 OS=Homo sapiens OX=9606 GN=IGHA1 PE=1 SV=2 | 255 | 20201113_12433_1ng_glyco_hcd30_etd30_compound15.13596.13596.3 | controllerType=0 controllerNumber=1 scan=13596 | 30.8429 |
| 10247:3 | 2 | R.LAGKPTHVN[+1216.423]VSVVMAEVDGTC[+57.021]Y.- | HexNAc(2)Hex(5) | C[+57], N[+1216] | 1188.5289 | 3 | 3563.572 | 3563.5649 | 0 | 2 | 332 | CRagged | 276.51 | 255.09 | 192.78 | 3.94 | 23 | >sp\|P01876\|IGHA1_HUMAN Immunoglobulin heavy constant alpha 1 OS=Homo sapiens OX=9606 GN=IGHA1 PE=1 SV=2 | 255 | 20201113_12433_1ng_glyco_hcd30_etd30_compound15.13535.13535.3 | controllerType=0 controllerNumber=1 scan=13535 | 30.6935 |
| 12048:3 | 5 | R.TYNVLDMKN[+2059.735]TTC[+57.021]QDLQIEVTVK.G | HexNAc(4)Hex(5)Fuc(1)NeuAc(1) | C[+57], N[+2060] | 1558.3511 | 3 | 4673.0388 | 4673.0248 | 0 | 3 | 1383 | Specific | 241.07 | 105.68 | 105.68 | 3.59 | 11 | >sp\|P0C0L4\|CO4A_HUMAN Complement C4-A OS=Homo sapiens OX=9606 GN=C4A PE=1 SV=2 | 1235 | 20201113_12433_1ng_glyco_hcd30_etd30_compound15.15428.15428.3 | controllerType=0 controllerNumber=1 scan=15428 | 34.2458 |
| 09107:3 | 6 | K.TVLTPATNHMGN[+1540.529]VTFTIPANR.E | HexNAc(2)Hex(7) | N[+1541] | 1265.91 | 3 | 3795.7154 | 3795.6886 | 0 | 7 | 74 | Specific | 231.69 | 197.22 | 139.68 | 2.55 | 9 | >sp\|P01024\|CO3_HUMAN Complement C3 OS=Homo sapiens OX=9606 GN=C3 PE=1 SV=2 | 492 | 20201113_12433_1ng_glyco_hcd30_etd30_compound15.12351.12351.3 | controllerType=0 controllerNumber=1 scan=12351 | 28.3797 |
| 16907:3 | 7 | R.GLSFDVSLEVSQGPGLLN[+2351.851]DTK.V | HexNAc(4)Hex(5)Fuc(3)NeuAc(1) | N[+2352] | 1510.005 | 3 | 4528.0004 | 4527.964 | 0 | 8.1 | 118 | Specific | 290.86 | 225.32 | 225.32 | 4.03 | 19 | >sp\|P01833\|PIGR_HUMAN Polymeric immunoglobulin receptor OS=Homo sapiens OX=9606 GN=PIGR PE=1 SV=4 | 902 | 20201113_12433_1ng_glyco_hcd30_etd30_compound15.20562.20562.3 | controllerType=0 controllerNumber=1 scan=20562 | 46.7907 |
| 17910:3 | 7 | K.QIGLYPVLVIDSSGYVNPN[+2204.772]YTGR.I | HexNAc(4)Hex(5)NeuAc(2) | N[+2205] | 1577.3592 | 3 | 4730.063 | 4730.0759 | 0 | -2.7 | 168 | Specific | 567.33 | 458.9 | 458.9 | 6.6 | 19 | >sp\|P01833\|PIGR_HUMAN Polymeric immunoglobulin receptor OS=Homo sapiens OX=9606 GN=PIGR PE=1 SV=4 | 902 | 20201113_12433_1ng_glyco_hcd30_etd30_compound15.21620.21620.3 | controllerType=0 controllerNumber=1 scan=21620 | 49.7346 |
| 17615:4 | 7 | R.LSLLEEPGN[+2205.793]GTFTVILNQLTSR.D | HexNAc(4)Hex(5)Fuc(2)NeuAc(1) | N[+2206] | 1152.7801 | 4 | 4608.0985 | 4608.0854 | 0 | 2.8 | 413 | Specific | 353.88 | 328.71 | 328.71 | 4.57 | 19 | >sp\|P01833\|PIGR_HUMAN Polymeric immunoglobulin receptor OS=Homo sapiens OX=9606 GN=PIGR PE=1 SV=4 | 902 | 20201113_12433_1ng_glyco_hcd30_etd30_compound15.21308.21308.4 | controllerType=0 controllerNumber=1 scan=21308 | 48.8222 |
| 17604:3 | 7 | R.LSLLEEPGN[+2205.793]GTFTVILNQLTSR.D | HexNAc(4)Hex(5)Fuc(2)NeuAc(1) | N[+2206] | 1536.7032 | 3 | 4608.095 | 4608.0854 | 0 | 2.1 | 413 | Specific | 662.42 | 534.36 | 534.36 | 10.26 | 19 | >sp\|P01833\|PIGR_HUMAN Polymeric immunoglobulin receptor OS=Homo sapiens OX=9606 GN=PIGR PE=1 SV=4 | 902 | 20201113_12433_1ng_glyco_hcd30_etd30_compound15.21296.21296.3 | controllerType=0 controllerNumber=1 scan=21296 | 48.7872 |
| 15696:3 | 7 | K.IIEGEPNLKVPGN[+2350.830]VTAVLGETLK.V | HexNAc(4)Hex(5)Fuc(1)NeuAc(2) | N[+2351] | 1581.4003 | 3 | 4742.1864 | 4742.1797 | 0 | 1.4 | 457 | Specific | 255.9 | 212.9 | 212.9 | 3.26 | 19 | >sp\|P01833\|PIGR_HUMAN Polymeric immunoglobulin receptor OS=Homo sapiens OX=9606 GN=PIGR PE=1 SV=4 | 902 | 20201113_12433_1ng_glyco_hcd30_etd30_compound15.19283.19283.3 | controllerType=0 controllerNumber=1 scan=19283 | 43.2503 |
| 14306:3 | 7 | K.IIEGEPNLKVPGN[+2204.772]VTAVLGETLK.V | HexNAc(4)Hex(5)NeuAc(2) | N[+2205] | 1532.7205 | 3 | 4596.147 | 4596.1218 | 0 | 5.5 | 457 | Specific | 315.14 | 221.94 | 221.94 | 3.1 | 19 | >sp\|P01833\|PIGR_HUMAN Polymeric immunoglobulin receptor OS=Homo sapiens OX=9606 GN=PIGR PE=1 SV=4 | 902 | 20201113_12433_1ng_glyco_hcd30_etd30_compound15.17818.17818.3 | controllerType=0 controllerNumber=1 scan=17818 | 39.4278 |
| 15129:5 | 7 | K.IIEGEPNLKVPGN[+2350.830]VTAVLGETLKVPC[+57.021]HFPC[+57.021]K.F | HexNAc(4)Hex(5)Fuc(1)NeuAc(2) | C[+57]*2, N[+2351] | 1154.3362 | 5 | 5767.6518 | 5767.6372 | 0 | 2.5 | 457 | Specific | 374.17 | 374.17 | 374.17 | 4.77 | 19 | >sp\|P01833\|PIGR_HUMAN Polymeric immunoglobulin receptor OS=Homo sapiens OX=9606 GN=PIGR PE=1 SV=4 | 902 | 20201113_12433_1ng_glyco_hcd30_etd30_compound15.18684.18684.5 | controllerType=0 controllerNumber=1 scan=18684 | 41.4903 |
| 13872:4 | 7 | K.IIEGEPNLKVPGN[+2205.793]VTAVLGETLKVPC[+57.021]HFPC[+57.021]K.F | HexNAc(4)Hex(5)Fuc(2)NeuAc(1) | C[+57]*2, N[+2206] | 1406.4091 | 4 | 5622.6144 | 5622.5997 | 0 | 2.6 | 457 | Specific | 401.49 | 401.49 | 401.49 | 5.34 | 19 | >sp\|P01833\|PIGR_HUMAN Polymeric immunoglobulin receptor OS=Homo sapiens OX=9606 GN=PIGR PE=1 SV=4 | 902 | 20201113_12433_1ng_glyco_hcd30_etd30_compound15.17331.17331.4 | controllerType=0 controllerNumber=1 scan=17331 | 38.4023 |
| 13950:5 | 7 | K.IIEGEPNLKVPGN[+2059.735]VTAVLGETLKVPC[+57.021]HFPC[+57.021]K.F | HexNAc(4)Hex(5)Fuc(1)NeuAc(1) | C[+57]*2, N[+2060] | 1096.1161 | 5 | 5476.5514 | 5476.5418 | 0 | 1.7 | 457 | Specific | 445.31 | 445.31 | 445.31 | 4.92 | 19 | >sp\|P01833\|PIGR_HUMAN Polymeric immunoglobulin receptor OS=Homo sapiens OX=9606 GN=PIGR PE=1 SV=4 | 902 | 20201113_12433_1ng_glyco_hcd30_etd30_compound15.17412.17412.5 | controllerType=0 controllerNumber=1 scan=17412 | 38.5857 |
| 13873:5 | 7 | K.IIEGEPNLKVPGN[+2205.793]VTAVLGETLKVPC[+57.021]HFPC[+57.021]K.F | HexNAc(4)Hex(5)Fuc(2)NeuAc(1) | C[+57]*2, N[+2206] | 1125.3286 | 5 | 5622.614 | 5622.5997 | 0 | 2.5 | 457 | Specific | 523.99 | 523.99 | 523.99 | 5.64 | 19 | >sp\|P01833\|PIGR_HUMAN Polymeric immunoglobulin receptor OS=Homo sapiens OX=9606 GN=PIGR PE=1 SV=4 | 902 | 20201113_12433_1ng_glyco_hcd30_etd30_compound15.17332.17332.5 | controllerType=0 controllerNumber=1 scan=17332 | 38.4072 |
| 12223:3 | 7 | K.VPGN[+2205.793]VTAVLGETLK.V | HexNAc(4)Hex(5)Fuc(2)NeuAc(1) | N[+2206] | 1201.8727 | 3 | 3603.6037 | 3603.5927 | 0 | 3 | 466 | Specific | 254.5 | 170.74 | 170.74 | 4.46 | 19 | >sp\|P01833\|PIGR_HUMAN Polymeric immunoglobulin receptor OS=Homo sapiens OX=9606 GN=PIGR PE=1 SV=4 | 902 | 20201113_12433_1ng_glyco_hcd30_etd30_compound15.15610.15610.3 | controllerType=0 controllerNumber=1 scan=15610 | 34.5895 |
| 05179:3 | 7 | K.WN[+1913.677]NTGC[+57.021]QALPSQDEGPSK.A | HexNAc(4)Hex(5)NeuAc(1) | C[+57], N[+1914] | 1301.5278 | 3 | 3902.5687 | 3902.5537 | 0 | 3.9 | 498 | Specific | 226.23 | 133.62 | 133.62 | 4.67 | 19 | >sp\|P01833\|PIGR_HUMAN Polymeric immunoglobulin receptor OS=Homo sapiens OX=9606 GN=PIGR PE=1 SV=4 | 902 | 20201113_12433_1ng_glyco_hcd30_etd30_compound15.8273.8273.3 | controllerType=0 controllerNumber=1 scan=8273 | 21.0146 |
| 05125:3 | 7 | K.WN[+1872.651]NTGC[+57.021]QALPSQDEGPSK.A | HexNAc(3)Hex(6)NeuAc(1) | C[+57], N[+1873] | 1287.8503 | 3 | 3861.5363 | 3861.5272 | 0 | 2.4 | 498 | Specific | 351.34 | 284.26 | 284.26 | 5.52 | 19 | >sp\|P01833\|PIGR_HUMAN Polymeric immunoglobulin receptor OS=Homo sapiens OX=9606 GN=PIGR PE=1 SV=4 | 902 | 20201113_12433_1ng_glyco_hcd30_etd30_compound15.8217.8217.3 | controllerType=0 controllerNumber=1 scan=8217 | 20.9132 |
| 05044:3 | 7 | K.WN[+2059.735]NTGC[+57.021]QALPSQDEGPSK.A | HexNAc(4)Hex(5)Fuc(1)NeuAc(1) | C[+57], N[+2060] | 1350.2117 | 3 | 4048.6206 | 4048.6116 | 0 | 2.2 | 498 | Specific | 478.82 | 321.7 | 321.7 | 6.09 | 19 | >sp\|P01833\|PIGR_HUMAN Polymeric immunoglobulin receptor OS=Homo sapiens OX=9606 GN=PIGR PE=1 SV=4 | 902 | 20201113_12433_1ng_glyco_hcd30_etd30_compound15.8133.8133.3 | controllerType=0 controllerNumber=1 scan=8133 | 20.7608 |
| 05185:3 | 7 | K.WN[+1710.598]NTGC[+57.021]QALPSQDEGPSK.A | HexNAc(3)Hex(5)NeuAc(1) | C[+57], N[+1711] | 1233.8338 | 3 | 3699.4868 | 3699.4743 | 0 | 3.4 | 498 | Specific | 479.67 | 304.84 | 304.84 | 6.18 | 19 | >sp\|P01833\|PIGR_HUMAN Polymeric immunoglobulin receptor OS=Homo sapiens OX=9606 GN=PIGR PE=1 SV=4 | 902 | 20201113_12433_1ng_glyco_hcd30_etd30_compound15.8279.8279.3 | controllerType=0 controllerNumber=1 scan=8279 | 21.0299 |
| 18842:4 | 8 | K.VFPLSLDSTPQDGNVVVAC[+57.021]LVQGFFPQEPLSVTWSESGQN[+1647.613]VTAI | HexNAc(5)Hex(3)Fuc(1) | C[+57], N[+1648] | 1606.7564 | 4 | 6424.0036 | 6423.9746 | 0 | 4.5 | 8 | Specific | 482.63 | 482.63 | 482.63 | 6.02 | 18 | >sp\|P01877\|IGHA2_HUMAN Immunoglobulin heavy constant alpha 2 OS=Homo sapiens OX=9606 GN=IGHA2 PE=1 SV=4 | 576 | 20201113_12433_1ng_glyco_hcd30_etd30_compound15.22598.22598.2 | controllerType=0 controllerNumber=1 scan=22598 | 51.9584 |
| 00929:3 | 8 | K.HYTN[+1809.666]SSQDVTVPC[+57.021]R.V | HexNAc(5)Hex(4)Fuc(1) | C[+57], N[+1810] | 1158.4788 | 3 | 3473.4219 | 3473.4154 | 0 | 1.9 | 89 | Specific | 442.36 | 375.62 | 375.62 | 7.26 | 18 | >sp\|P01877\|IGHA2_HUMAN Immunoglobulin heavy constant alpha 2 OS=Homo sapiens OX=9606 GN=IGHA2 PE=1 SV=4 | 576 | 20201113_12433_1ng_glyco_hcd30_etd30_compound15.3864.3864.3 | controllerType=0 controllerNumber=1 scan=3864 | 12.9873 |
| 00935:3 | 8 | K.HYTN[+1647.613]SSQDVTVPC[+57.021]R.V | HexNAc(5)Hex(3)Fuc(1) | C[+57], N[+1648] | 1104.4615 | 3 | 3311.3698 | 3311.3625 | 0 | 2.2 | 89 | Specific | 502.35 | 435.13 | 435.13 | 7.45 | 18 | >sp\|P01877\|IGHA2_HUMAN Immunoglobulin heavy constant alpha 2 OS=Homo sapiens OX=9606 GN=IGHA2 PE=1 SV=4 | 576 | 20201113_12433_1ng_glyco_hcd30_etd30_compound15.3870.3870.3 | controllerType=0 controllerNumber=1 scan=3870 | 13.0028 |
| 02808:4 | 8 | K.HYTN[+1647.613]SSQDVTVPC[+57.021]RVPPPPPC[+57.021]C[+57.021]HPR.L | HexNAc(5)Hex(3)Fuc(1) | C[+57]*3, N[+1648] | 1152.2511 | 4 | 4605.9827 | 4605.9689 | 0 | 3 | 89 | Specific | 278.79 | 247.93 | 247.93 | 4.12 | 18 | >sp\|P01877\|IGHA2_HUMAN Immunoglobulin heavy constant alpha 2 OS=Homo sapiens OX=9606 GN=IGHA2 PE=1 SV=4 | 576 | 20201113_12433_1ng_glyco_hcd30_etd30_compound15.5816.5816.4 | controllerType=0 controllerNumber=1 scan=5816 | 16.65 |
| 04240:3 | 8 | K.TPLTAN[+2262.814]ITK.S | HexNAc(5)Hex(5)Fuc(1)NeuAc(1) | N[+2263] | 1074.4639 | 3 | 3221.3771 | 3221.3711 | 0 | 1.9 | 200 | Specific | 218.85 | 101.08 | 101.08 | 2.93 | 18 | >sp\|P01877\|IGHA2_HUMAN Immunoglobulin heavy constant alpha 2 OS=Homo sapiens OX=9606 GN=IGHA2 PE=1 SV=4 | 576 | 20201113_12433_1ng_glyco_hcd30_etd30_compound15.7300.7300.3 | controllerType=0 controllerNumber=1 scan=7300 | 19.2505 |
| 02617:2 | 8 | K.TPLTAN[+1647.613]ITK.S | HexNAc(5)Hex(3)Fuc(1) | N[+1648] | 1303.5907 | 2 | 2606.1742 | 2606.17 | 0 | 1.6 | 200 | Specific | 230.28 | 77.87 | 77.87 | 2.61 | 18 | >sp\|P01877\|IGHA2_HUMAN Immunoglobulin heavy constant alpha 2 OS=Homo sapiens OX=9606 GN=IGHA2 PE=1 SV=4 | 576 | 20201113_12433_1ng_glyco_hcd30_etd30_compound15.5617.5617.2 | controllerType=0 controllerNumber=1 scan=5617 | 16.232 |
| 02551:2 | 8 | K.TPLTAN[+1971.719]ITK.S | HexNAc(5)Hex(5)Fuc(1) | N[+1972] | 1465.6458 | 2 | 2930.2843 | 2930.2757 | 0 | 3 | 200 | Specific | 245.93 | 64.71 | 64.71 | 2.85 | 18 | >sp\|P01877\|IGHA2_HUMAN Immunoglobulin heavy constant alpha 2 OS=Homo sapiens OX=9606 GN=IGHA2 PE=1 SV=4 | 576 | 20201113_12433_1ng_glyco_hcd30_etd30_compound15.5548.5548.2 | controllerType=0 controllerNumber=1 scan=5548 | 16.0747 |
| 02534:3 | 8 | K.TPLTAN[+1971.719]ITK.S | HexNAc(5)Hex(5)Fuc(1) | N[+1972] | 977.4316 | 3 | 2930.2803 | 2930.2757 | 0 | 1.6 | 200 | Specific | 254.03 | 128.5 | 128.5 | 2.97 | 18 | >sp\|P01877\|IGHA2_HUMAN Immunoglobulin heavy constant alpha 2 OS=Homo sapiens OX=9606 GN=IGHA2 PE=1 SV=4 | 576 | 20201113_12433_1ng_glyco_hcd30_etd30_compound15.5530.5530.3 | controllerType=0 controllerNumber=1 scan=5530 | 16.0386 |
| 02565:3 | 8 | K.TPLTAN[+1809.666]ITK.S | HexNAc(5)Hex(4)Fuc(1) | N[+1810] | 923.4153 | 3 | 2768.2312 | 2768.2228 | 0 | 3 | 200 | Specific | 286.01 | 138.96 | 138.96 | 2.72 | 18 | >sp\|P01877\|IGHA2_HUMAN Immunoglobulin heavy constant alpha 2 OS=Homo sapiens OX=9606 GN=IGHA2 PE=1 SV=4 | 576 | 20201113_12433_1ng_glyco_hcd30_etd30_compound15.5563.5563.3 | controllerType=0 controllerNumber=1 scan=5563 | 16.1196 |
| 02572:2 | 8 | K.TPLTAN[+1444.534]ITK.S | HexNAc(4)Hex(3)Fuc(1) | N[+1445] | 1202.0529 | 2 | 2403.0985 | 2403.0906 | 0 | 3.3 | 200 | Specific | 295.26 | 63.28 | 63.28 | 2.39 | 18 | >sp\|P01877\|IGHA2_HUMAN Immunoglobulin heavy constant alpha 2 OS=Homo sapiens OX=9606 GN=IGHA2 PE=1 SV=4 | 576 | 20201113_12433_1ng_glyco_hcd30_etd30_compound15.5570.5570.2 | controllerType=0 controllerNumber=1 scan=5570 | 16.1482 |
| 02630:3 | 8 | K.TPLTAN[+1647.613]ITK.S | HexNAc(5)Hex(3)Fuc(1) | N[+1648] | 869.3979 | 3 | 2606.1792 | 2606.17 | 0 | 3.5 | 200 | Specific | 296.68 | 134.74 | 134.74 | 2.93 | 18 | >sp\|P01877\|IGHA2_HUMAN Immunoglobulin heavy constant alpha 2 OS=Homo sapiens OX=9606 GN=IGHA2 PE=1 SV=4 | 576 | 20201113_12433_1ng_glyco_hcd30_etd30_compound15.5631.5631.3 | controllerType=0 controllerNumber=1 scan=5631 | 16.2675 |
| 02561:2 | 8 | K.TPLTAN[+1809.666]ITK.S | HexNAc(5)Hex(4)Fuc(1) | N[+1810] | 1384.6189 | 2 | 2768.2306 | 2768.2228 | 0 | 2.8 | 200 | Specific | 305.84 | 114.61 | 114.61 | 2.75 | 18 | >sp\|P01877\|IGHA2_HUMAN Immunoglobulin heavy constant alpha 2 OS=Homo sapiens OX=9606 GN=IGHA2 PE=1 SV=4 | 576 | 20201113_12433_1ng_glyco_hcd30_etd30_compound15.5559.5559.2 | controllerType=0 controllerNumber=1 scan=5559 | 16.1069 |
| 09221:3 | 8 | R.MAGKPTHIN[+1702.581]VSVVMAEADGTC[+57.021]Y.- | HexNAc(2)Hex(8) | C[+57], N[+1703] | 1351.8956 | 3 | 4053.6721 | 4053.6642 | 0 | 2 | 319 | CRagged | 237.23 | 237.23 | 196.68 | 3.3 | 18 | >sp\|P01877\|IGHA2_HUMAN Immunoglobulin heavy constant alpha 2 OS=Homo sapiens OX=9606 GN=IGHA2 PE=1 SV=4 | 576 | 20201113_12433_1ng_glyco_hcd30_etd30_compound15.12470.12470.3 | controllerType=0 controllerNumber=1 scan=12470 | 28.6198 |
| 09247:3 | 8 | R.MAGKPTHIN[+1540.529]VSVVMAEADGTC[+57.021]Y.- | HexNAc(2)Hex(7) | C[+57], N[+1541] | 1297.8798 | 3 | 3891.6249 | 3891.6114 | 0 | 3.5 | 319 | CRagged | 285.35 | 279.89 | 211.01 | 4.29 | 18 | >sp\|P01877\|IGHA2_HUMAN Immunoglobulin heavy constant alpha 2 OS=Homo sapiens OX=9606 GN=IGHA2 PE=1 SV=4 | 576 | 20201113_12433_1ng_glyco_hcd30_etd30_compound15.12497.12497.3 | controllerType=0 controllerNumber=1 scan=12497 | 28.6757 |
| 09264:3 | 8 | R.MAGKPTHIN[+1378.476]VSVVMAEADGTC[+57.021]Y.- | HexNAc(2)Hex(6) | C[+57], N[+1378] | 1243.8609 | 3 | 3729.5681 | 3729.5585 | 0 | 2.6 | 319 | CRagged | 309.34 | 260.15 | 204.68 | 4.47 | 18 | >sp\|P01877\|IGHA2_HUMAN Immunoglobulin heavy constant alpha 2 OS=Homo sapiens OX=9606 GN=IGHA2 PE=1 SV=4 | 576 | 20201113_12433_1ng_glyco_hcd30_etd30_compound15.12515.12515.3 | controllerType=0 controllerNumber=1 scan=12515 | 28.7232 |
| 20500:3 | 11 | R.QLAHQSN[+2204.772]STNIFFSPVSIATAFAMLSLGTK.A | HexNAc(4)Hex(5)NeuAc(2) | N[+2205] | 1796.1499 | 3 | 5386.435 | 5386.4075 | 0 | 5.1 | 64 | Specific | 440.7 | 314.37 | 314.37 | 5.25 | 6 | >sp\|P01009\|A1AT_HUMAN Alpha-1-antitrypsin OS=Homo sapiens OX=9606 GN=SERPINA1 PE=1 SV=3 | 642 | 20201113_12433_1ng_glyco_hcd30_etd30_compound15.24345.24345.3 | controllerType=0 controllerNumber=1 scan=24345 | 55.944 |
| 20499:4 | 11 | R.QLAHQSN[+2204.772]STNIFFSPVSIATAFAMLSLGTK.A | HexNAc(4)Hex(5)NeuAc(2) | N[+2205] | 1347.3626 | 4 | 5386.4284 | 5386.4075 | 0 | 3.9 | 64 | Specific | 615.32 | 579.38 | 579.38 | 9.05 | 6 | >sp\|P01009\|A1AT_HUMAN Alpha-1-antitrypsin OS=Homo sapiens OX=9606 GN=SERPINA1 PE=1 SV=3 | 642 | 20201113_12433_1ng_glyco_hcd30_etd30_compound15.24344.24344.4 | controllerType=0 controllerNumber=1 scan=24344 | 55.9398 |
| 05029:3 | 12 | R.LSPN[+2205.793]ASAEHLELR.W | HexNAc(4)Hex(5)Fuc(2)NeuAc(1) | N[+2206] | 1214.8558 | 3 | 3642.5528 | 3642.5421 | 0 | 2.9 | 52 | Specific | 379.69 | 285.74 | 285.74 | 5.05 | 9 | >sp\|Q13410\|BT1A1_HUMAN Butyrophilin subfamily 1 member A1 OS=Homo sapiens OX=9606 GN=BTN1A1 PE=1 SV=3 | 179 | 20201113_12433_1ng_glyco_hcd30_etd30_compound15.8117.8117.3 | controllerType=0 controllerNumber=1 scan=8117 | 20.7055 |
| 06929:3 | 12 | R.LSPN[+2350.830]ASAEHLELR.W | HexNAc(4)Hex(5)Fuc(1)NeuAc(2) | N[+2351] | 1263.2006 | 3 | 3787.5873 | 3787.5796 | 0 | 2 | 52 | Specific | 396.27 | 228.09 | 228.09 | 4.81 | 9 | >sp\|Q13410\|BT1A1_HUMAN Butyrophilin subfamily 1 member A1 OS=Homo sapiens OX=9606 GN=BTN1A1 PE=1 SV=3 | 179 | 20201113_12433_1ng_glyco_hcd30_etd30_compound15.10085.10085.3 | controllerType=0 controllerNumber=1 scan=10085 | 24.2073 |
| 14578:3 | 14 | R.VWIALNSN[+2205.793]LTDNQYTWTDKWR.V | HexNAc(4)Hex(5)Fuc(2)NeuAc(1) | N[+2206] | 1610.7043 | 3 | 4830.0983 | 4830.0821 | 0 | 3.4 | 1153 | Specific | 380.98 | 254.94 | 254.94 | 4.37 | 9 | >sp\|P22897\|MRC1_HUMAN Macrophage mannose receptor 1 OS=Homo sapiens OX=9606 GN=MRC1 PE=1 SV=1 | 1049 | 20201113_12433_1ng_glyco_hcd30_etd30_compound15.18106.18106.3 | controllerType=0 controllerNumber=1 scan=18106 | 40.0249 |
| 07722:3 | 14 | K.TAHC[+57.021]N[+2059.735]ESFYFLC[+57.021]K.R | HexNAc(4)Hex(5)Fuc(1)NeuAc(1) | C[+57]*2, N[+2060] | 1246.1585 | 3 | 3736.461 | 3736.4545 | 0 | 1.7 | 1201 | Specific | 231.51 | 142.36 | 142.36 | 3.86 | 9 | >sp\|P22897\|MRC1_HUMAN Macrophage mannose receptor 1 OS=Homo sapiens OX=9606 GN=MRC1 PE=1 SV=1 | 1049 | 20201113_12433_1ng_glyco_hcd30_etd30_compound15.10910.10910.3 | controllerType=0 controllerNumber=1 scan=10910 | 25.653 |
| 00094:4 | 22 | K.KKEDALN[+2350.830]ETRESETK.L | HexNAc(4)Hex(5)Fuc(1)NeuAc(2) | N[+2351] | 1032.939 | 4 | 4128.7342 | 4128.723 | 0 | 2.7 | 80 | Specific | 265.77 | 223.76 | 223.76 | 3.35 | 7 | >sp\|P10909\|CLUS_HUMAN Clusterin OS=Homo sapiens OX=9606 GN=CLU PE=1 SV=1 | 1467 | 20201113_12433_1ng_glyco_hcd30_etd30_compound15.2995.2995.2 | controllerType=0 controllerNumber=1 scan=2995 | 11.3121 |
| 00097:4 | 22 | R.EIRHN[+2424.867]STGC[+57.021]LR.M | HexNAc(5)Hex(6)Fuc(1)NeuAc(1) | C[+57], N[+2425] | 942.6407 | 4 | 3767.5411 | 3767.5316 | 0 | 2.5 | 287 | Specific | 270.31 | 181.34 | 181.34 | 4.88 | 7 | >sp\|P10909\|CLUS_HUMAN Clusterin OS=Homo sapiens OX=9606 GN=CLU PE=1 SV=1 | 1467 | 20201113_12433_1ng_glyco_hcd30_etd30_compound15.2998.2998.4 | controllerType=0 controllerNumber=1 scan=2998 | 11.3228 |
| 00091:4 | 22 | R.EIRHN[+2205.793]STGC[+57.021]LR.M | HexNAc(4)Hex(5)Fuc(2)NeuAc(1) | C[+57], N[+2206] | 887.8738 | 4 | 3548.4735 | 3548.4573 | 0 | 4.6 | 287 | Specific | 366.91 | 276.66 | 276.66 | 2.96 | 7 | >sp\|P10909\|CLUS_HUMAN Clusterin OS=Homo sapiens OX=9606 GN=CLU PE=1 SV=1 | 1467 | 20201113_12433_1ng_glyco_hcd30_etd30_compound15.2992.2992.4 | controllerType=0 controllerNumber=1 scan=2992 | 11.301 |
| 00082:4 | 22 | R.EIRHN[+2059.735]STGC[+57.021]LR.M | HexNAc(4)Hex(5)Fuc(1)NeuAc(1) | C[+57], N[+2060] | 851.3577 | 4 | 3402.4089 | 3402.3994 | 0 | 2.8 | 287 | Specific | 429 | 326.58 | 326.58 | 5.77 | 7 | >sp\|P10909\|CLUS_HUMAN Clusterin OS=Homo sapiens OX=9606 GN=CLU PE=1 SV=1 | 1467 | 20201113_12433_1ng_glyco_hcd30_etd30_compound15.2982.2982.4 | controllerType=0 controllerNumber=1 scan=2982 | 11.2705 |
| 19851:3 | 22 | K.MLN[+2350.830]TSSLLEQLNEQFNWVSR.L | HexNAc(4)Hex(5)Fuc(1)NeuAc(2) | N[+2351] | 1587.3434 | 3 | 4760.0158 | 4760.0171 | 0 | -0.3 | 352 | Specific | 587.88 | 376.49 | 376.49 | 7.24 | 7 | >sp\|P10909\|CLUS_HUMAN Clusterin OS=Homo sapiens OX=9606 GN=CLU PE=1 SV=1 | 1467 | 20201113_12433_1ng_glyco_hcd30_etd30_compound15.23663.23663.3 | controllerType=0 controllerNumber=1 scan=23663 | 54.372 |
| 09330:3 | 22 | R.LAN[+2205.793]LTQGEDQYYLR.V | HexNAc(4)Hex(5)Fuc(2)NeuAc(1) | N[+2206] | 1297.2176 | 3 | 3889.6382 | 3889.6265 | 0 | 3 | 372 | Specific | 408.43 | 287.48 | 287.48 | 5.97 | 7 | >sp\|P10909\|CLUS_HUMAN Clusterin OS=Homo sapiens OX=9606 GN=CLU PE=1 SV=1 | 1467 | 20201113_12433_1ng_glyco_hcd30_etd30_compound15.12584.12584.3 | controllerType=0 controllerNumber=1 scan=12584 | 28.8936 |
| 00715:4 | 29 | K.TKPREEQFN[+1606.587]STFR.V | HexNAc(4)Hex(4)Fuc(1) | N[+1607] | 812.3598 | 4 | 3246.4172 | 3246.4054 | 0 | 3.6 | 168 | Specific | 292.67 | 292.67 | 292.67 | 5.39 | 5 | >sp\|P01859\|IGHG2_HUMAN Immunoglobulin heavy constant gamma 2 OS=Homo sapiens OX=9606 GN=IGHG2 PE=1 SV=2 | 387 | 20201113_12433_1ng_glyco_hcd30_etd30_compound15.3641.3641.2 | controllerType=0 controllerNumber=1 scan=3641 | 12.5561 |
| 10168:3 | 50 | R.IIVPLNNREN[+2059.735]ISDPTSPLR.T | HexNAc(4)Hex(5)Fuc(1)NeuAc(1) | N[+2060] | 1403.3121 | 3 | 4207.9217 | 4207.9121 | 0 | 2.3 | 62 | Specific | 207.71 | 185.59 | 185.59 | 3.95 | 4 | >sp\|P01591\|IGJ_HUMAN Immunoglobulin J chain OS=Homo sapiens OX=9606 GN=JCHAIN PE=1 SV=4 | 1392 | 20201113_12433_1ng_glyco_hcd30_etd30_compound15.13453.13453.3 | controllerType=0 controllerNumber=1 scan=13453 | 30.4947 |
| 08760:3 | 50 | R.IIVPLNNREN[+1663.608]ISDPTSPLR.T | HexNAc(5)Hex(4) | N[+1664] | 1271.2702 | 3 | 3811.796 | 3811.7853 | 0 | 2.8 | 62 | Specific | 231.6 | 160.53 | 160.53 | 3.84 | 4 | >sp\|P01591\|IGJ_HUMAN Immunoglobulin J chain OS=Homo sapiens OX=9606 GN=JCHAIN PE=1 SV=4 | 1392 | 20201113_12433_1ng_glyco_hcd30_etd30_compound15.11992.11992.3 | controllerType=0 controllerNumber=1 scan=11992 | 27.7389 |
| 11917:3 | 50 | R.IIVPLNNREN[+2350.830]ISDPTSPLR.T | HexNAc(4)Hex(5)Fuc(1)NeuAc(2) | N[+2351] | 1500.3451 | 3 | 4499.0209 | 4499.0075 | 0 | 3 | 62 | Specific | 247.39 | 167.37 | 167.37 | 4.26 | 4 | >sp\|P01591\|IGJ_HUMAN Immunoglobulin J chain OS=Homo sapiens OX=9606 GN=JCHAIN PE=1 SV=4 | 1392 | 20201113_12433_1ng_glyco_hcd30_etd30_compound15.15292.15292.3 | controllerType=0 controllerNumber=1 scan=15292 | 33.9617 |
| 10172:3 | 50 | R.IIVPLNNREN[+1913.677]ISDPTSPLR.T | HexNAc(4)Hex(5)NeuAc(1) | N[+1914] | 1354.6255 | 3 | 4061.8621 | 4061.8542 | 0 | 1.9 | 62 | Specific | 272.63 | 223.11 | 223.11 | 4.31 | 4 | >sp\|P01591\|IGJ_HUMAN Immunoglobulin J chain OS=Homo sapiens OX=9606 GN=JCHAIN PE=1 SV=4 | 1392 | 20201113_12433_1ng_glyco_hcd30_etd30_compound15.13457.13457.3 | controllerType=0 controllerNumber=1 scan=13457 | 30.5068 |
| 16337:3 | 1 | R.[+42.011]RTAGWNVPIGTLRPFLN[+1864.634]WTGPPEPIEAAVAR.F | HexNAc(2)Hex(9) | .[42], N[+1865] | 1765.1614 | 3 | 5293.4696 | 5293.4568 | 0 | 2.4 | 140 | Specific | 206.66 | 189.01 | 189.01 | 2.97 | 54 | >sp\|P02788\|TRFL_HUMAN Lactotransferrin OS=Homo sapiens OX=9606 GN=LTF PE=1 SV=6 | 1270 | 20201118_12812_1ng_glyco_hcd30_etd30_compound15.19976.19976.3 | controllerType=0 controllerNumber=1 scan=19976 | 46.5501 |
| 15381:4 | 1 | R.RTAGWNVPIGTLRPFLN[+2351.851]WTGPPEPIEAAVAR.F | HexNAc(4)Hex(5)Fuc(3)NeuAc(1) | N[+2352] | 1435.426 | 4 | 5738.6823 | 5738.6628 | 0 | 3.4 | 140 | Specific | 257.17 | 257.17 | 257.17 | 5.02 | 54 | >sp\|P02788\|TRFL_HUMAN Lactotransferrin OS=Homo sapiens OX=9606 GN=LTF PE=1 SV=6 | 1270 | 20201118_12812_1ng_glyco_hcd30_etd30_compound15.18968.18968.4 | controllerType=0 controllerNumber=1 scan=18968 | 43.5863 |
| 14698:4 | 1 | R.[+42.011]RTAGWNVPIGTLRPFLN[+1872.651]WTGPPEPIEAAVAR.F | HexNAc(3)Hex(6)NeuAc(1) | .[42], N[+1873] | 1326.1348 | 4 | 5301.5172 | 5301.4731 | 0 | 8.3 | 140 | Specific | 361.18 | 286.62 | 286.62 | 3.76 | 54 | >sp\|P02788\|TRFL_HUMAN Lactotransferrin OS=Homo sapiens OX=9606 GN=LTF PE=1 SV=6 | 1270 | 20201118_12812_1ng_glyco_hcd30_etd30_compound15.18245.18245.4 | controllerType=0 controllerNumber=1 scan=18245 | 41.3637 |
| 16094:4 | 1 | R.RTAGWNVPIGTLRPFLN[+2350.830]WTGPPEPIEAAVAR.F | HexNAc(4)Hex(5)Fuc(1)NeuAc(2) | N[+2351] | 1435.1694 | 4 | 5737.6559 | 5737.6424 | 0 | 2.4 | 140 | Specific | 662.4 | 662.4 | 662.4 | 10.43 | 54 | >sp\|P02788\|TRFL_HUMAN Lactotransferrin OS=Homo sapiens OX=9606 GN=LTF PE=1 SV=6 | 1270 | 20201118_12812_1ng_glyco_hcd30_etd30_compound15.19721.19721.4 | controllerType=0 controllerNumber=1 scan=19721 | 45.8599 |
| 14740:4 | 1 | R.RTAGWNVPIGTLRPFLN[+1768.640]WTGPPEPIEAAVAR.F | HexNAc(4)Hex(5)Fuc(1) | N[+1769] | 1289.6227 | 4 | 5155.4689 | 5155.4516 | 0 | 3.4 | 140 | Specific | 690.93 | 690.93 | 690.93 | 11.12 | 54 | >sp\|P02788\|TRFL_HUMAN Lactotransferrin OS=Homo sapiens OX=9606 GN=LTF PE=1 SV=6 | 1270 | 20201118_12812_1ng_glyco_hcd30_etd30_compound15.18289.18289.4 | controllerType=0 controllerNumber=1 scan=18289 | 41.5037 |
| 15296:4 | 1 | R.RTAGWNVPIGTLRPFLN[+2716.983]WTGPPEPIEAAVAR.F | HexNAc(5)Hex(6)Fuc(3)NeuAc(1) | N[+2717] | 1526.7122 | 4 | 6103.8268 | 6103.795 | 0 | 5.2 | 140 | Specific | 825.84 | 825.84 | 825.84 | 12.97 | 54 | >sp\|P02788\|TRFL_HUMAN Lactotransferrin OS=Homo sapiens OX=9606 GN=LTF PE=1 SV=6 | 1270 | 20201118_12812_1ng_glyco_hcd30_etd30_compound15.18879.18879.4 | controllerType=0 controllerNumber=1 scan=18879 | 43.3252 |
| 15423:4 | 1 | R.RTAGWNVPIGTLRPFLN[+2059.735]WTGPPEPIEAAVAR.F | HexNAc(4)Hex(5)Fuc(1)NeuAc(1) | N[+2060] | 1362.3967 | 4 | 5446.5651 | 5446.547 | 0 | 3.3 | 140 | Specific | 999.16 | 972.49 | 972.49 | 17.12 | 54 | >sp\|P02788\|TRFL_HUMAN Lactotransferrin OS=Homo sapiens OX=9606 GN=LTF PE=1 SV=6 | 1270 | 20201118_12812_1ng_glyco_hcd30_etd30_compound15.19013.19013.4 | controllerType=0 controllerNumber=1 scan=19013 | 43.721 |
| 16152:3 | 1 | R.TAGWNVPIGTLRPFLN[+2570.925]WTGPPEPIEAAVAR.F | HexNAc(5)Hex(6)Fuc(2)NeuAc(1) | N[+2571] | 1934.5549 | 3 | 5801.6501 | 5801.636 | 0 | 2.4 | 141 | Specific | 237.17 | 161.27 | 161.27 | 3.39 | 54 | >sp\|P02788\|TRFL_HUMAN Lactotransferrin OS=Homo sapiens OX=9606 GN=LTF PE=1 SV=6 | 1270 | 20201118_12812_1ng_glyco_hcd30_etd30_compound15.19782.19782.3 | controllerType=0 controllerNumber=1 scan=19782 | 46.0286 |
| 16233:3 | 1 | R.TAGWNVPIGTLRPFLN[+2351.851]WTGPPEPIEAAVAR.F | HexNAc(4)Hex(5)Fuc(3)NeuAc(1) | N[+2352] | 1861.5324 | 3 | 5582.5827 | 5582.5617 | 0 | 3.8 | 141 | Specific | 237.52 | 176.53 | 176.53 | 4.03 | 54 | >sp\|P02788\|TRFL_HUMAN Lactotransferrin OS=Homo sapiens OX=9606 GN=LTF PE=1 SV=6 | 1270 | 20201118_12812_1ng_glyco_hcd30_etd30_compound15.19867.19867.3 | controllerType=0 controllerNumber=1 scan=19867 | 46.2907 |
| 16255:3 | 1 | R.TAGWNVPIGTLRPFLN[+2205.793]WTGPPEPIEAAVAR.F | HexNAc(4)Hex(5)Fuc(2)NeuAc(1) | N[+2206] | 1812.848 | 3 | 5436.5294 | 5436.5038 | 0 | 4.7 | 141 | Specific | 355.86 | 273.75 | 273.75 | 3.94 | 54 | >sp\|P02788\|TRFL_HUMAN Lactotransferrin OS=Homo sapiens OX=9606 GN=LTF PE=1 SV=6 | 1270 | 20201118_12812_1ng_glyco_hcd30_etd30_compound15.19891.19891.3 | controllerType=0 controllerNumber=1 scan=19891 | 46.3749 |
| 16254:4 | 1 | R.TAGWNVPIGTLRPFLN[+2205.793]WTGPPEPIEAAVAR.F | HexNAc(4)Hex(5)Fuc(2)NeuAc(1) | N[+2206] | 1359.8838 | 4 | 5436.5133 | 5436.5038 | 0 | 1.8 | 141 | Specific | 359.48 | 359.48 | 359.48 | 4.13 | 54 | >sp\|P02788\|TRFL_HUMAN Lactotransferrin OS=Homo sapiens OX=9606 GN=LTF PE=1 SV=6 | 1270 | 20201118_12812_1ng_glyco_hcd30_etd30_compound15.19890.19890.4 | controllerType=0 controllerNumber=1 scan=19890 | 46.3712 |
| 17178:3 | 1 | R.TAGWNVPIGTLRPFLN[+2350.830]WTGPPEPIEAAVAR.F | HexNAc(4)Hex(5)Fuc(1)NeuAc(2) | N[+2351] | 1861.191 | 3 | 5581.5585 | 5581.5413 | 0 | 3.1 | 141 | Specific | 360.72 | 286.43 | 286.43 | 6.14 | 54 | >sp\|P02788\|TRFL_HUMAN Lactotransferrin OS=Homo sapiens OX=9606 GN=LTF PE=1 SV=6 | 1270 | 20201118_12812_1ng_glyco_hcd30_etd30_compound15.20867.20867.2 | controllerType=0 controllerNumber=1 scan=20867 | 49.2454 |
| 16239:3 | 1 | R.TAGWNVPIGTLRPFLN[+2059.735]WTGPPEPIEAAVAR.F | HexNAc(4)Hex(5)Fuc(1)NeuAc(1) | N[+2060] | 1764.1602 | 3 | 5290.4662 | 5290.4459 | 0 | 3.8 | 141 | Specific | 444.73 | 329.18 | 329.18 | 5.62 | 54 | >sp\|P02788\|TRFL_HUMAN Lactotransferrin OS=Homo sapiens OX=9606 GN=LTF PE=1 SV=6 | 1270 | 20201118_12812_1ng_glyco_hcd30_etd30_compound15.19874.19874.3 | controllerType=0 controllerNumber=1 scan=19874 | 46.3206 |
| 16151:4 | 1 | R.TAGWNVPIGTLRPFLN[+2570.925]WTGPPEPIEAAVAR.F | HexNAc(5)Hex(6)Fuc(2)NeuAc(1) | N[+2571] | 1451.1702 | 4 | 5801.6588 | 5801.636 | 0 | 3.9 | 141 | Specific | 716.47 | 716.47 | 716.47 | 10.99 | 54 | >sp\|P02788\|TRFL_HUMAN Lactotransferrin OS=Homo sapiens OX=9606 GN=LTF PE=1 SV=6 | 1270 | 20201118_12812_1ng_glyco_hcd30_etd30_compound15.19781.19781.4 | controllerType=0 controllerNumber=1 scan=19781 | 46.0256 |
| 16170:4 | 1 | R.TAGWNVPIGTLRPFLN[+2716.983]WTGPPEPIEAAVAR.F | HexNAc(5)Hex(6)Fuc(3)NeuAc(1) | N[+2717] | 1487.6843 | 4 | 5947.7155 | 5947.6939 | 0 | 3.6 | 141 | Specific | 785.77 | 785.77 | 785.77 | 12.96 | 54 | >sp\|P02788\|TRFL_HUMAN Lactotransferrin OS=Homo sapiens OX=9606 GN=LTF PE=1 SV=6 | 1270 | 20201118_12812_1ng_glyco_hcd30_etd30_compound15.19801.19801.4 | controllerType=0 controllerNumber=1 scan=19801 | 46.0882 |
| 16123:4 | 1 | R.TAGWNVPIGTLRPFLN[+2716.983]WTGPPEPIEAAVAR.F | HexNAc(5)Hex(6)Fuc(3)NeuAc(1) | N[+2717] | 1487.6845 | 4 | 5947.716 | 5947.6939 | 0 | 3.7 | 141 | Specific | 802.24 | 791.71 | 791.71 | 13.96 | 54 | >sp\|P02788\|TRFL_HUMAN Lactotransferrin OS=Homo sapiens OX=9606 GN=LTF PE=1 SV=6 | 1270 | 20201118_12812_1ng_glyco_hcd30_etd30_compound15.19751.19751.4 | controllerType=0 controllerNumber=1 scan=19751 | 45.9409 |

Table S4. Cont.

| Query #:z | Protein Rank | Glycopeptide | Glycan | Modification Type(s) | Observed m/z | z | Observed (M+H) | Calc. mass (M+H) | Off-by-x error | Mass error (ppm) | StartPosition | Cleavage | Score | Delta | Delta Mod | \|Log Prob\| | # of unique peptides | Protein Name | Protein DB number | Comment | Scan # | Scan Time |
|---|---|---|---|---|---|---|---|---|---|---|---|---|---|---|---|---|---|---|---|---|---|---|
| 16238:4 | 1 | R.TAGWNVPIGTLRPFLN[+2059.735]WTGPPEPIEAAVAR.F | HexNAc(4)Hex(5)Fuc(1)NeuAc(1) | N[+2060] | 1323.3728 | 4 | 5290.4694 | 5290.4459 | 0 | 4.4 | 141 | Specific | 867.21 | 845.84 | 845.84 | 14.96 | 54 | >sp\|P02788\|TRFL_HUMAN Lactotransferrin OS=Homo sapiens OX=9606 GN=LTF PE=1 SV=6 | 1270 | 20201118_12812_1ng_glyco_hcd30_etd30_compound15.19873.19873.4 | controllerType=0 controllerNumber=1 scan=19873 | 46.3178 |
| 17162:5 | 1 | R.TAGWNVPIGTLRPFLN[+2350.830]WTGPPEPIEAAVAR.F | HexNAc(4)Hex(5)Fuc(1)NeuAc(2) | N[+2351] | 1117.1168 | 5 | 5581.555 | 5581.5413 | 0 | 2.5 | 141 | Specific | 882.51 | 831.3 | 831.3 | 14.27 | 54 | >sp\|P02788\|TRFL_HUMAN Lactotransferrin OS=Homo sapiens OX=9606 GN=LTF PE=1 SV=6 | 1270 | 20201118_12812_1ng_glyco_hcd30_etd30_compound15.20850.20850.5 | controllerType=0 controllerNumber=1 scan=20850 | 49.1851 |
| 16223:4 | 1 | R.TAGWNVPIGTLRPFLN[+2351.851]WTGPPEPIEAAVAR.F | HexNAc(4)Hex(5)Fuc(3)NeuAc(1) | N[+2352] | 1396.3997 | 4 | 5582.5768 | 5582.5617 | 0 | 2.7 | 141 | Specific | 886.17 | 854.35 | 854.35 | 14.27 | 54 | >sp\|P02788\|TRFL_HUMAN Lactotransferrin OS=Homo sapiens OX=9606 GN=LTF PE=1 SV=6 | 1270 | 20201118_12812_1ng_glyco_hcd30_etd30_compound15.19857.19857.4 | controllerType=0 controllerNumber=1 scan=19857 | 46.2641 |
| 16008:3 | 1 | R.TAGWNIPMGLLFN[+1751.624]QTGSC[+57.021]K.F | HexNAc(4)Hex(4)NeuAc(1) | C[+57], N[+1752] | 1282.8887 | 3 | 3846.6516 | 3846.6341 | 0 | 4.5 | 485 | Specific | 210.79 | 127.22 | 127.22 | 4.26 | 54 | >sp\|P02788\|TRFL_HUMAN Lactotransferrin OS=Homo sapiens OX=9606 GN=LTF PE=1 SV=6 | 1270 | 20201118_12812_1ng_glyco_hcd30_etd30_compound15.19630.19630.3 | controllerType=0 controllerNumber=1 scan=19630 | 45.5859 |
| 14403:3 | 1 | R.TAGWNIPMGLLFN[+2133.772]QTGSC[+57.021]K.F | HexNAc(5)Hex(6)Fuc(1) | C[+57], N[+2134] | 1410.2689 | 3 | 4228.792 | 4228.7817 | 0 | 2.4 | 485 | Specific | 215.95 | 111.4 | 111.4 | 3.6 | 54 | >sp\|P02788\|TRFL_HUMAN Lactotransferrin OS=Homo sapiens OX=9606 GN=LTF PE=1 SV=6 | 1270 | 20201118_12812_1ng_glyco_hcd30_etd30_compound15.17932.17932.3 | controllerType=0 controllerNumber=1 scan=17932 | 40.4031 |
| 14371:3 | 1 | R.TAGWNIPMGLLFN[+2279.830]QTGSC[+57.021]K.F | HexNAc(5)Hex(6)Fuc(2) | C[+57], N[+2280] | 1458.9587 | 3 | 4374.8615 | 4374.8396 | 0 | 5 | 485 | Specific | 260.62 | 138.31 | 138.31 | 3.68 | 54 | >sp\|P02788\|TRFL_HUMAN Lactotransferrin OS=Homo sapiens OX=9606 GN=LTF PE=1 SV=6 | 1270 | 20201118_12812_1ng_glyco_hcd30_etd30_compound15.17898.17898.3 | controllerType=0 controllerNumber=1 scan=17898 | 40.3032 |
| 15802:3 | 1 | R.TAGWNIPMGLLFN[+2205.793]QTGSC[+57.021]K.F | HexNAc(4)Hex(5)Fuc(2)NeuAc(1) | C[+57], N[+2206] | 1434.2758 | 3 | 4300.8129 | 4300.8028 | 0 | 2.3 | 485 | Specific | 292.08 | 178.2 | 178.2 | 4.46 | 54 | >sp\|P02788\|TRFL_HUMAN Lactotransferrin OS=Homo sapiens OX=9606 GN=LTF PE=1 SV=6 | 1270 | 20201118_12812_1ng_glyco_hcd30_etd30_compound15.19413.19413.3 | controllerType=0 controllerNumber=1 scan=19413 | 44.8974 |
| 17245:4 | 1 | R.TAGWNIPMGLLFN[+2350.830]QTGSC[+57.021]K.F | HexNAc(4)Hex(5)Fuc(1)NeuAc(2) | C[+57], N[+2351] | 1112.2196 | 4 | 4445.8568 | 4445.8403 | 0 | 3.7 | 485 | Specific | 327.12 | 306.4 | 306.4 | 4.71 | 54 | >sp\|P02788\|TRFL_HUMAN Lactotransferrin OS=Homo sapiens OX=9606 GN=LTF PE=1 SV=6 | 1270 | 20201118_12812_1ng_glyco_hcd30_etd30_compound15.20938.20938.4 | controllerType=0 controllerNumber=1 scan=20938 | 49.4763 |
| 17374:3 | 1 | R.TAGWNIPMGLLFN[+2204.772]QTGSC[+57.021]K.F | HexNAc(4)Hex(5)NeuAc(2) | C[+57], N[+2205] | 1433.9401 | 3 | 4299.8058 | 4299.7824 | 0 | 5.4 | 485 | Specific | 349.65 | 256.54 | 256.54 | 4.41 | 54 | >sp\|P02788\|TRFL_HUMAN Lactotransferrin OS=Homo sapiens OX=9606 GN=LTF PE=1 SV=6 | 1270 | 20201118_12812_1ng_glyco_hcd30_etd30_compound15.21074.21074.3 | controllerType=0 controllerNumber=1 scan=21074 | 49.9056 |
| 14485:3 | 1 | R.TAGWNIPMGLLFN[+1768.640]QTGSC[+57.021]K.F | HexNAc(4)Hex(5)Fuc(1) | C[+57], N[+1769] | 1288.5585 | 3 | 3863.661 | 3863.6495 | 0 | 3 | 485 | Specific | 390.07 | 284.48 | 284.48 | 5.65 | 54 | >sp\|P02788\|TRFL_HUMAN Lactotransferrin OS=Homo sapiens OX=9606 GN=LTF PE=1 SV=6 | 1270 | 20201118_12812_1ng_glyco_hcd30_etd30_compound15.18019.18019.3 | controllerType=0 controllerNumber=1 scan=18019 | 40.6568 |
| 14646:3 | 1 | R.TAGWNIPMGLLFN[+1622.582]QTGSC[+57.021]K.F | HexNAc(4)Hex(5) | C[+57], N[+1623] | 1239.8734 | 3 | 3717.6056 | 3717.5916 | 0 | 3.8 | 485 | Specific | 390.53 | 305.55 | 260.59 | 4.82 | 54 | >sp\|P02788\|TRFL_HUMAN Lactotransferrin OS=Homo sapiens OX=9606 GN=LTF PE=1 SV=6 | 1270 | 20201118_12812_1ng_glyco_hcd30_etd30_compound15.18189.18189.3 | controllerType=0 controllerNumber=1 scan=18189 | 41.1744 |
| 14741:3 | 1 | R.TAGWNIPMGLLFN[+1216.423]QTGSC[+57.021]K.F | HexNAc(2)Hex(5) | C[+57], N[+1216] | 1104.4864 | 3 | 3311.4445 | 3311.4328 | 0 | 3.5 | 485 | Specific | 410.95 | 391.73 | 222.88 | 7.19 | 54 | >sp\|P02788\|TRFL_HUMAN Lactotransferrin OS=Homo sapiens OX=9606 GN=LTF PE=1 SV=6 | 1270 | 20201118_12812_1ng_glyco_hcd30_etd30_compound15.18290.18290.3 | controllerType=0 controllerNumber=1 scan=18290 | 41.5086 |
| 15901:3 | 1 | R.TAGWNIPMGLLFN[+1913.677]QTGSC[+57.021]K.F | HexNAc(4)Hex(5)NeuAc(1) | C[+57], N[+1914] | 1336.9044 | 3 | 4008.6985 | 4008.687 | 0 | 2.9 | 485 | Specific | 425.02 | 264.59 | 264.59 | 5.08 | 54 | >sp\|P02788\|TRFL_HUMAN Lactotransferrin OS=Homo sapiens OX=9606 GN=LTF PE=1 SV=6 | 1270 | 20201118_12812_1ng_glyco_hcd30_etd30_compound15.19518.19518.3 | controllerType=0 controllerNumber=1 scan=19518 | 45.2516 |
| 14468:3 | 1 | R.TAGWNIPMGLLFN[+1768.640]QTGSC[+57.021]K.F | HexNAc(4)Hex(5)Fuc(1) | C[+57], N[+1769] | 1288.5596 | 3 | 3863.6643 | 3863.6495 | 0 | 3.8 | 485 | Specific | 427.74 | 317.58 | 317.58 | 5.89 | 54 | >sp\|P02788\|TRFL_HUMAN Lactotransferrin OS=Homo sapiens OX=9606 GN=LTF PE=1 SV=6 | 1270 | 20201118_12812_1ng_glyco_hcd30_etd30_compound15.18001.18001.3 | controllerType=0 controllerNumber=1 scan=18001 | 40.6049 |
| 15787:3 | 1 | R.TAGWNIPMGLLFN[+2059.735]QTGSC[+57.021]K.F | HexNAc(4)Hex(5)Fuc(1)NeuAc(1) | C[+57], N[+2060] | 1385.5906 | 3 | 4154.7573 | 4154.7449 | 0 | 3 | 485 | Specific | 440.42 | 300.35 | 300.35 | 5.34 | 54 | >sp\|P02788\|TRFL_HUMAN Lactotransferrin OS=Homo sapiens OX=9606 GN=LTF PE=1 SV=6 | 1270 | 20201118_12812_1ng_glyco_hcd30_etd30_compound15.19397.19397.3 | controllerType=0 controllerNumber=1 scan=19397 | 44.8484 |
| 17223:3 | 1 | R.TAGWNIPMGLLFN[+2350.830]QTGSC[+57.021]K.F | HexNAc(4)Hex(5)Fuc(1)NeuAc(2) | C[+57], N[+2351] | 1482.6284 | 3 | 4445.8705 | 4445.8403 | 0 | 6.8 | 485 | Specific | 475.25 | 325.82 | 325.82 | 4.59 | 54 | >sp\|P02788\|TRFL_HUMAN Lactotransferrin OS=Homo sapiens OX=9606 GN=LTF PE=1 SV=6 | 1270 | 20201118_12812_1ng_glyco_hcd30_etd30_compound15.20915.20915.3 | controllerType=0 controllerNumber=1 scan=20915 | 49.4049 |
| 15912:4 | 1 | R.TAGWNIPMGLLFN[+1913.677]QTGSC[+57.021]K.F | HexNAc(4)Hex(5)NeuAc(1) | C[+57], N[+1914] | 1002.9294 | 4 | 4008.6956 | 4008.687 | 0 | 2.2 | 485 | Specific | 476.55 | 407.67 | 407.67 | 5.38 | 54 | >sp\|P02788\|TRFL_HUMAN Lactotransferrin OS=Homo sapiens OX=9606 GN=LTF PE=1 SV=6 | 1270 | 20201118_12812_1ng_glyco_hcd30_etd30_compound15.19529.19529.4 | controllerType=0 controllerNumber=1 scan=19529 | 45.282 |
| 14759:2 | 1 | R.TAGWNIPMGLLFN[+1216.423]QTGSC[+57.021]K.F | HexNAc(2)Hex(5) | C[+57], N[+1216] | 1656.2228 | 2 | 3311.4384 | 3311.4328 | 0 | 1.7 | 485 | Specific | 531.91 | 499.72 | 174.26 | 7.25 | 54 | >sp\|P02788\|TRFL_HUMAN Lactotransferrin OS=Homo sapiens OX=9606 GN=LTF PE=1 SV=6 | 1270 | 20201118_12812_1ng_glyco_hcd30_etd30_compound15.18309.18309.2 | controllerType=0 controllerNumber=1 scan=18309 | 41.5588 |
| 14591:3 | 1 | R.TAGWNIPM[+15.995]GLLFN[+2059.735]QTGSC[+57.021]K.F | HexNAc(4)Hex(5)Fuc(1)NeuAc(1) | C[+57], M[+16], N[+2060] | 1390.9233 | 3 | 4170.7553 | 4170.7398 | 0 | 3.7 | 485 | Specific | 287.35 | 240.36 | 240.36 | 4.6 | 54 | >sp\|P02788\|TRFL_HUMAN Lactotransferrin OS=Homo sapiens OX=9606 GN=LTF PE=1 SV=6 | 1270 | 20201118_12812_1ng_glyco_hcd30_etd30_compound15.18131.18131.3 | controllerType=0 controllerNumber=1 scan=18131 | 40.9843 |
| 14622:3 | 1 | R.TAGWNIPM[+15.995]GLLFN[+2059.735]QTGSC[+57.021]K.F | HexNAc(4)Hex(5)Fuc(1)NeuAc(1) | C[+57], M[+16], N[+2060] | 1390.9256 | 3 | 4170.7622 | 4170.7398 | 0 | 5.4 | 485 | Specific | 316.21 | 272.44 | 272.44 | 3.12 | 54 | >sp\|P02788\|TRFL_HUMAN Lactotransferrin OS=Homo sapiens OX=9606 GN=LTF PE=1 SV=6 | 1270 | 20201118_12812_1ng_glyco_hcd30_etd30_compound15.18163.18163.3 | controllerType=0 controllerNumber=1 scan=18163 | 41.0726 |
| 15998:4 | 1 | R.TAGWNIPMGLLFN[+2076.750]QTGSC[+57.021]KFDEYFSQSC[+57.021]APGSDPR.S | HexNAc(4)Hex(6)Fuc(2) | C[+57]*2, N[+2077] | 1504.6415 | 4 | 6015.5441 | 6015.5074 | 0 | 6.1 | 485 | Specific | 248.2 | 248.2 | 248.2 | 3.31 | 54 | >sp\|P02788\|TRFL_HUMAN Lactotransferrin OS=Homo sapiens OX=9606 GN=LTF PE=1 SV=6 | 1270 | 20201118_12812_1ng_glyco_hcd30_etd30_compound15.19620.19620.4 | controllerType=0 controllerNumber=1 scan=19620 | 45.5579 |
| 15264:3 | 1 | R.TAGWNIPMGLLFN[+1768.640]QTGSC[+57.021]KFDEYFSQSC[+57.021]APGSDPR.S | HexNAc(4)Hex(5)Fuc(1) | C[+57]*2, N[+1769] | 1903.1407 | 3 | 5707.4076 | 5707.3966 | 0 | 1.9 | 485 | Specific | 279.75 | 237.91 | 237.91 | 4.84 | 54 | >sp\|P02788\|TRFL_HUMAN Lactotransferrin OS=Homo sapiens OX=9606 GN=LTF PE=1 SV=6 | 1270 | 20201118_12812_1ng_glyco_hcd30_etd30_compound15.18845.18845.3 | controllerType=0 controllerNumber=1 scan=18845 | 43.2204 |
| 16057:4 | 1 | R.[+42.011]TAGWNIPMGLLFN[+2018.708]QTGSC[+57.021]KFDEYFSQSC[+57.021]A | HexNAc(3)Hex(6)Fuc(1)NeuAc(1) | .[42], C[+57]*2, N[+2019] | 1500.6379 | 4 | 5999.5299 | 5999.4761 | 0 | 9 | 485 | Specific | 320.24 | 294.06 | 294.06 | 3.29 | 54 | >sp\|P02788\|TRFL_HUMAN Lactotransferrin OS=Homo sapiens OX=9606 GN=LTF PE=1 SV=6 | 1270 | 20201118_12812_1ng_glyco_hcd30_etd30_compound15.19682.19682.4 | controllerType=0 controllerNumber=1 scan=19682 | 45.7309 |
| 16033:4 | 1 | R.TAGWNIPMGLLFN[+1913.677]QTGSC[+57.021]KFDEYFSQSC[+57.021]APGSDPR.S | HexNAc(4)Hex(5)NeuAc(1) | C[+57]*2, N[+1914] | 1463.866 | 4 | 5852.4421 | 5852.4341 | 0 | 1.4 | 485 | Specific | 384.29 | 298.15 | 298.15 | 3.76 | 54 | >sp\|P02788\|TRFL_HUMAN Lactotransferrin OS=Homo sapiens OX=9606 GN=LTF PE=1 SV=6 | 1270 | 20201118_12812_1ng_glyco_hcd30_etd30_compound15.19657.19657.4 | controllerType=0 controllerNumber=1 scan=19657 | 45.6758 |
| 15257:4 | 1 | R.TAGWNIPMGLLFN[+1768.640]QTGSC[+57.021]KFDEYFSQSC[+57.021]APGSDPR.S | HexNAc(4)Hex(5)Fuc(1) | C[+57]*2, N[+1769] | 1427.6099 | 4 | 5707.4176 | 5707.3966 | 0 | 3.7 | 485 | Specific | 603.33 | 568.12 | 568.12 | 9.53 | 54 | >sp\|P02788\|TRFL_HUMAN Lactotransferrin OS=Homo sapiens OX=9606 GN=LTF PE=1 SV=6 | 1270 | 20201118_12812_1ng_glyco_hcd30_etd30_compound15.18838.18838.4 | controllerType=0 controllerNumber=1 scan=18838 | 43.2045 |
| 15977:4 | 1 | R.TAGWNIPMGLLFN[+2059.735]QTGSC[+57.021]KFDEYFSQSC[+57.021]APGSDPR.S | HexNAc(4)Hex(5)Fuc(1)NeuAc(1) | C[+57]*2, N[+2060] | 1500.3855 | 4 | 5998.5202 | 5998.492 | 0 | 4.7 | 485 | Specific | 627.13 | 554.88 | 554.88 | 8.39 | 54 | >sp\|P02788\|TRFL_HUMAN Lactotransferrin OS=Homo sapiens OX=9606 GN=LTF PE=1 SV=6 | 1270 | 20201118_12812_1ng_glyco_hcd30_etd30_compound15.19598.19598.4 | controllerType=0 controllerNumber=1 scan=19598 | 45.484 |
| 08321:3 | 1 | R.N[+2059.735]GSDC[+57.021]PDKFC[+57.021]LFQSETK.N | HexNAc(4)Hex(5)Fuc(1)NeuAc(1) | C[+57]*2, N[+2060] | 1364.8787 | 3 | 4092.6216 | 4092.6088 | 0 | 3.1 | 642 | Specific | 226.51 | 169 | 169 | 4.22 | 54 | >sp\|P02788\|TRFL_HUMAN Lactotransferrin OS=Homo sapiens OX=9606 GN=LTF PE=1 SV=6 | 1270 | 20201118_12812_1ng_glyco_hcd30_etd30_compound15.11541.11541.3 | controllerType=0 controllerNumber=1 scan=11541 | 26.8114 |
| 08188:3 | 1 | R.N[+2204.772]GSDC[+57.021]PDKFC[+57.021]LFQSETK.N | HexNAc(4)Hex(5)NeuAc(2) | C[+57]*2, N[+2205] | 1413.2322 | 3 | 4237.6822 | 4237.6463 | 0 | 8.4 | 642 | Specific | 348.62 | 243.25 | 243.25 | 4.33 | 54 | >sp\|P02788\|TRFL_HUMAN Lactotransferrin OS=Homo sapiens OX=9606 GN=LTF PE=1 SV=6 | 1270 | 20201118_12812_1ng_glyco_hcd30_etd30_compound15.11402.11402.3 | controllerType=0 controllerNumber=1 scan=11402 | 26.5235 |
| 15835:3 | 2 | R.LSLHRPALEDLLLGSEAN[+1751.624]LTC[+57.021]TLTGLR.D | HexNAc(4)Hex(4)NeuAc(1) | C[+57], N[+1752] | 1572.4167 | 3 | 4715.2355 | 4715.2225 | 0 | 2.8 | 127 | Specific | 306.99 | 236.92 | 236.92 | 5.08 | 26 | >sp\|P01876\|IGHA1_HUMAN Immunoglobulin heavy constant alpha 1 OS=Homo sapiens OX=9606 GN=IGHA1 PE=1 SV=2 | 255 | 20201118_12812_1ng_glyco_hcd30_etd30_compound15.19447.19447.3 | controllerType=0 controllerNumber=1 scan=19447 | 45.0022 |
| 15071:3 | 2 | R.LSLHRPALEDLLLGSEAN[+1864.634]LTC[+57.021]TLTGLR.D | HexNAc(2)Hex(9) | C[+57], N[+1865] | 1610.0866 | 3 | 4828.2453 | 4828.2324 | 0 | 2.7 | 127 | Specific | 356.73 | 322.68 | 322.68 | 5.2 | 26 | >sp\|P01876\|IGHA1_HUMAN Immunoglobulin heavy constant alpha 1 OS=Homo sapiens OX=9606 GN=IGHA1 PE=1 SV=2 | 255 | 20201118_12812_1ng_glyco_hcd30_etd30_compound15.18639.18639.3 | controllerType=0 controllerNumber=1 scan=18639 | 42.536 |
| 15150:3 | 2 | R.LSLHRPALEDLLLGSEAN[+1460.529]LTC[+57.021]TLTGLR.D | HexNAc(4)Hex(4) | C[+57], N[+1461] | 1475.3897 | 3 | 4424.1546 | 4424.1271 | 0 | 6.2 | 127 | Specific | 405.06 | 371.97 | 371.97 | 6.26 | 26 | >sp\|P01876\|IGHA1_HUMAN Immunoglobulin heavy constant alpha 1 OS=Homo sapiens OX=9606 GN=IGHA1 PE=1 SV=2 | 255 | 20201118_12812_1ng_glyco_hcd30_etd30_compound15.18723.18723.3 | controllerType=0 controllerNumber=1 scan=18723 | 42.8143 |
| 15115:3 | 2 | R.LSLHRPALEDLLLGSEAN[+1378.476]LTC[+57.021]TLTGLR.D | HexNAc(2)Hex(6) | C[+57], N[+1378] | 1448.0355 | 3 | 4342.0918 | 4342.074 | 0 | 4.1 | 127 | Specific | 415.07 | 385.76 | 385.76 | 7.4 | 26 | >sp\|P01876\|IGHA1_HUMAN Immunoglobulin heavy constant alpha 1 OS=Homo sapiens OX=9606 GN=IGHA1 PE=1 SV=2 | 255 | 20201118_12812_1ng_glyco_hcd30_etd30_compound15.18686.18686.3 | controllerType=0 controllerNumber=1 scan=18686 | 42.6892 |
| 15137:3 | 2 | R.LSLHRPALEDLLLGSEAN[+1216.423]LTC[+57.021]TLTGLR.D | HexNAc(2)Hex(5) | C[+57], N[+1216] | 1394.0167 | 3 | 4180.0354 | 4180.0211 | 0 | 3.4 | 127 | Specific | 472.84 | 463.09 | 463.09 | 7.53 | 26 | >sp\|P01876\|IGHA1_HUMAN Immunoglobulin heavy constant alpha 1 OS=Homo sapiens OX=9606 GN=IGHA1 PE=1 SV=2 | 255 | 20201118_12812_1ng_glyco_hcd30_etd30_compound15.18709.18709.3 | controllerType=0 controllerNumber=1 scan=18709 | 42.7785 |
| 15084:3 | 2 | R.LSLHRPALEDLLLGSEAN[+1540.529]LTC[+57.021]TLTGLR.D | HexNAc(2)Hex(7) | C[+57], N[+1541] | 1502.0527 | 3 | 4504.1435 | 4504.1268 | 0 | 3.7 | 127 | Specific | 473.7 | 465.86 | 465.86 | 7.53 | 26 | >sp\|P01876\|IGHA1_HUMAN Immunoglobulin heavy constant alpha 1 OS=Homo sapiens OX=9606 GN=IGHA1 PE=1 SV=2 | 255 | 20201118_12812_1ng_glyco_hcd30_etd30_compound15.18653.18653.3 | controllerType=0 controllerNumber=1 scan=18653 | 42.5819 |
| 15130:3 | 2 | R.LSLHRPALEDLLLGSEAN[+1501.555]LTC[+57.021]TLTGLR.D | HexNAc(5)Hex(3) | C[+57], N[+1502] | 1489.0616 | 3 | 4465.1702 | 4465.1536 | 0 | 3.7 | 127 | Specific | 476.59 | 460.15 | 460.15 | 7.53 | 26 | >sp\|P01876\|IGHA1_HUMAN Immunoglobulin heavy constant alpha 1 OS=Homo sapiens OX=9606 GN=IGHA1 PE=1 SV=2 | 255 | 20201118_12812_1ng_glyco_hcd30_etd30_compound15.18702.18702.3 | controllerType=0 controllerNumber=1 scan=18702 | 42.7524 |
| 15075:3 | 2 | R.LSLHRPALEDLLLGSEAN[+1702.581]LTC[+57.021]TLTGLR.D | HexNAc(2)Hex(8) | C[+57], N[+1703] | 1556.0691 | 3 | 4666.1929 | 4666.1796 | 0 | 2.8 | 127 | Specific | 483.06 | 434.24 | 434.24 | 6.94 | 26 | >sp\|P01876\|IGHA1_HUMAN Immunoglobulin heavy constant alpha 1 OS=Homo sapiens OX=9606 GN=IGHA1 PE=1 SV=2 | 255 | 20201118_12812_1ng_glyco_hcd30_etd30_compound15.18643.18643.3 | controllerType=0 controllerNumber=1 scan=18643 | 42.5487 |
| 15120:3 | 2 | R.LSLHRPALEDLLLGSEAN[+1622.582]LTC[+57.021]TLTGLR.D | HexNAc(4)Hex(5) | C[+57], N[+1623] | 1529.4035 | 3 | 4586.1959 | 4586.1799 | 0 | 3.5 | 127 | Specific | 484.36 | 416.81 | 416.81 | 7.62 | 26 | >sp\|P01876\|IGHA1_HUMAN Immunoglobulin heavy constant alpha 1 OS=Homo sapiens OX=9606 GN=IGHA1 PE=1 SV=2 | 255 | 20201118_12812_1ng_glyco_hcd30_etd30_compound15.18691.18691.3 | controllerType=0 controllerNumber=1 scan=18691 | 42.7015 |
| 15186:4 | 2 | R.LSLHRPALEDLLLGSEAN[+1298.476]LTC[+57.021]TLTGLR.D | HexNAc(4)Hex(3) | C[+57], N[+1298] | 1066.2775 | 4 | 4262.0882 | 4262.0742 | 0 | 3.3 | 127 | Specific | 524.72 | 524.72 | 524.72 | 8.26 | 26 | >sp\|P01876\|IGHA1_HUMAN Immunoglobulin heavy constant alpha 1 OS=Homo sapiens OX=9606 GN=IGHA1 PE=1 SV=2 | 255 | 20201118_12812_1ng_glyco_hcd30_etd30_compound15.18762.18762.4 | controllerType=0 controllerNumber=1 scan=18762 | 42.9717 |
| 15825:4 | 2 | R.LSLHRPALEDLLLGSEAN[+1751.624]LTC[+57.021]TLTGLR.D | HexNAc(4)Hex(4)NeuAc(1) | C[+57], N[+1752] | 1179.5645 | 4 | 4715.2362 | 4715.2225 | 0 | 2.9 | 127 | Specific | 612.13 | 602.48 | 602.48 | 9.43 | 26 | >sp\|P01876\|IGHA1_HUMAN Immunoglobulin heavy constant alpha 1 OS=Homo sapiens OX=9606 GN=IGHA1 PE=1 SV=2 | 255 | 20201118_12812_1ng_glyco_hcd30_etd30_compound15.19437.19437.4 | controllerType=0 controllerNumber=1 scan=19437 | 44.9757 |
| 15141:4 | 2 | R.LSLHRPALEDLLLGSEAN[+1663.608]LTC[+57.021]TLTGLR.D | HexNAc(5)Hex(4) | C[+57], N[+1664] | 1157.5619 | 4 | 4627.2259 | 4627.2064 | 0 | 4.2 | 127 | Specific | 894.53 | 894.53 | 894.53 | 15.12 | 26 | >sp\|P01876\|IGHA1_HUMAN Immunoglobulin heavy constant alpha 1 OS=Homo sapiens OX=9606 GN=IGHA1 PE=1 SV=2 | 255 | 20201118_12812_1ng_glyco_hcd30_etd30_compound15.18714.18714.4 | controllerType=0 controllerNumber=1 scan=18714 | 42.7899 |
| 15109:4 | 2 | R.LSLHRPALEDLLLGSEAN[+1540.529]LTC[+57.021]TLTGLR.D | HexNAc(2)Hex(7) | C[+57], N[+1541] | 1126.7902 | 4 | 4504.139 | 4504.1268 | 0 | 2.7 | 127 | Specific | 947.11 | 872.52 | 872.52 | 15.43 | 26 | >sp\|P01876\|IGHA1_HUMAN Immunoglobulin heavy constant alpha 1 OS=Homo sapiens OX=9606 GN=IGHA1 PE=1 SV=2 | 255 | 20201118_12812_1ng_glyco_hcd30_etd30_compound15.18680.18680.4 | controllerType=0 controllerNumber=1 scan=18680 | 42.6661 |
| 15047:4 | 2 | R.LSLHRPALEDLLLGSEAN[+1864.634]LTC[+57.021]TLTGLR.D | HexNAc(2)Hex(9) | C[+57], N[+1865] | 1207.8179 | 4 | 4828.2498 | 4828.2324 | 0 | 3.6 | 127 | Specific | 993.42 | 933.61 | 933.61 | 17.12 | 26 | >sp\|P01876\|IGHA1_HUMAN Immunoglobulin heavy constant alpha 1 OS=Homo sapiens OX=9606 GN=IGHA1 PE=1 SV=2 | 255 | 20201118_12812_1ng_glyco_hcd30_etd30_compound15.18614.18614.4 | controllerType=0 controllerNumber=1 scan=18614 | 42.4426 |
| 15069:4 | 2 | R.LSLHRPALEDLLLGSEAN[+1702.581]LTC[+57.021]TLTGLR.D | HexNAc(2)Hex(8) | C[+57], N[+1703] | 1167.3039 | 4 | 4666.1937 | 4666.1796 | 0 | 3 | 127 | Specific | 1012.95 | 978.66 | 978.66 | 18.12 | 26 | >sp\|P01876\|IGHA1_HUMAN Immunoglobulin heavy constant alpha 1 OS=Homo sapiens OX=9606 GN=IGHA1 PE=1 SV=2 | 255 | 20201118_12812_1ng_glyco_hcd30_etd30_compound15.18637.18637.4 | controllerType=0 controllerNumber=1 scan=18637 | 42.5263 |
| 15110:4 | 2 | R.LSLHRPALEDLLLGSEAN[+1378.476]LTC[+57.021]TLTGLR.D | HexNAc(2)Hex(6) | C[+57], N[+1378] | 1086.2782 | 4 | 4342.0911 | 4342.074 | 0 | 4 | 127 | Specific | 1020.78 | 993.3 | 993.3 | 18.12 | 26 | >sp\|P01876\|IGHA1_HUMAN Immunoglobulin heavy constant alpha 1 OS=Homo sapiens OX=9606 GN=IGHA1 PE=1 SV=2 | 255 | 20201118_12812_1ng_glyco_hcd30_etd30_compound15.18681.18681.4 | controllerType=0 controllerNumber=1 scan=18681 | 42.6697 |
| 15095:4 | 2 | R.LSLHRPALEDLLLGSEAN[+1622.582]LTC[+57.021]TLTGLR.D | HexNAc(4)Hex(5) | C[+57], N[+1623] | 1147.3042 | 4 | 4586.1951 | 4586.1799 | 0 | 3.3 | 127 | Specific | 1022.26 | 891.31 | 891.31 | 18.12 | 26 | >sp\|P01876\|IGHA1_HUMAN Immunoglobulin heavy constant alpha 1 OS=Homo sapiens OX=9606 GN=IGHA1 PE=1 SV=2 | 255 | 20201118_12812_1ng_glyco_hcd30_etd30_compound15.18665.18665.4 | controllerType=0 controllerNumber=1 scan=18665 | 42.6181 |
| 15129:4 | 2 | R.LSLHRPALEDLLLGSEAN[+1216.423]LTC[+57.021]TLTGLR.D | HexNAc(2)Hex(5) | C[+57], N[+1216] | 1045.7638 | 4 | 4180.0335 | 4180.0211 | 0 | 3 | 127 | Specific | 1046.87 | 989.59 | 989.59 | 18.12 | 26 | >sp\|P01876\|IGHA1_HUMAN Immunoglobulin heavy constant alpha 1 OS=Homo sapiens OX=9606 GN=IGHA1 PE=1 SV=2 | 255 | 20201118_12812_1ng_glyco_hcd30_etd30_compound15.18701.18701.4 | controllerType=0 controllerNumber=1 scan=18701 | 42.7435 |
| 15124:4 | 2 | R.LSLHRPALEDLLLGSEAN[+1501.555]LTC[+57.021]TLTGLR.D | HexNAc(5)Hex(3) | C[+57], N[+1502] | 1117.0489 | 4 | 4465.1737 | 4465.1536 | 0 | 4.5 | 127 | Specific | 1048.19 | 1048.19 | 1048.19 | 18.12 | 26 | >sp\|P01876\|IGHA1_HUMAN Immunoglobulin heavy constant alpha 1 OS=Homo sapiens OX=9606 GN=IGHA1 PE=1 SV=2 | 255 | 20201118_12812_1ng_glyco_hcd30_etd30_compound15.18696.18696.4 | controllerType=0 controllerNumber=1 scan=18696 | 42.7263 |
| 10219:3 | 2 | R.LAGKPTHVN[+1809.666]VSVVMAEVDGTC[+57.021]Y.- | HexNAc(5)Hex(4)Fuc(1) | C[+57], N[+1810] | 1386.2776 | 3 | 4156.8184 | 4156.8081 | 0 | 2.5 | 332 | CRagged | 201.76 | 84.16 | 84.16 | 2.59 | 26 | >sp\|P01876\|IGHA1_HUMAN Immunoglobulin heavy constant alpha 1 OS=Homo sapiens OX=9606 GN=IGHA1 PE=1 SV=2 | 255 | 20201118_12812_1ng_glyco_hcd30_etd30_compound15.13515.13515.3 | controllerType=0 controllerNumber=1 scan=13515 | 30.6308 |
| 11798:3 | 2 | R.LAGKPTHVN[+2262.814]VSVVMAEVDGTC[+57.021]Y.- | HexNAc(5)Hex(5)Fuc(1)NeuAc(1) | C[+57], N[+2263] | 1537.3301 | 3 | 4609.9759 | 4609.9564 | 0 | 4.2 | 332 | CRagged | 203.21 | 71.28 | 71.28 | 2.58 | 26 | >sp\|P01876\|IGHA1_HUMAN Immunoglobulin heavy constant alpha 1 OS=Homo sapiens OX=9606 GN=IGHA1 PE=1 SV=2 | 255 | 20201118_12812_1ng_glyco_hcd30_etd30_compound15.15171.15171.3 | controllerType=0 controllerNumber=1 scan=15171 | 33.8139 |
| 10187:3 | 2 | R.LAGKPTHVN[+1378.476]VSVVMAEVDGTC[+57.021]Y.- | HexNAc(2)Hex(6) | C[+57], N[+1378] | 1242.5469 | 3 | 3725.6262 | 3725.6178 | 0 | 2.3 | 332 | CRagged | 218.22 | 218.22 | 169.26 | 3.37 | 26 | >sp\|P01876\|IGHA1_HUMAN Immunoglobulin heavy constant alpha 1 OS=Homo sapiens OX=9606 GN=IGHA1 PE=1 SV=2 | 255 | 20201118_12812_1ng_glyco_hcd30_etd30_compound15.13481.13481.3 | controllerType=0 controllerNumber=1 scan=13481 | 30.5399 |
| 11767:3 | 2 | R.LAGKPTHVN[+2262.814]VSVVMAEVDGTC[+57.021]Y.- | HexNAc(5)Hex(5)Fuc(1)NeuAc(1) | C[+57], N[+2263] | 1537.326 | 3 | 4609.9634 | 4609.9564 | 0 | 1.5 | 332 | CRagged | 234.94 | 91.17 | 91.17 | 2.59 | 26 | >sp\|P01876\|IGHA1_HUMAN Immunoglobulin heavy constant alpha 1 OS=Homo sapiens OX=9606 GN=IGHA1 PE=1 SV=2 | 255 | 20201118_12812_1ng_glyco_hcd30_etd30_compound15.15138.15138.3 | controllerType=0 controllerNumber=1 scan=15138 | 33.7354 |
| 10148:3 | 2 | R.LAGKPTHVN[+1702.581]VSVVMAEVDGTC[+57.021]Y.- | HexNAc(2)Hex(8) | C[+57], N[+1703] | 1350.5849 | 3 | 4049.7401 | 4049.7234 | 0 | 4.1 | 332 | CRagged | 237.46 | 191.56 | 181.8 | 3.93 | 26 | >sp\|P01876\|IGHA1_HUMAN Immunoglobulin heavy constant alpha 1 OS=Homo sapiens OX=9606 GN=IGHA1 PE=1 SV=2 | 255 | 20201118_12812_1ng_glyco_hcd30_etd30_compound15.13440.13440.3 | controllerType=0 controllerNumber=1 scan=13440 | 30.4157 |
| 10282:3 | 2 | R.LAGKPTHVN[+1501.555]VSVVMAEVDGTC[+57.021]Y.- | HexNAc(5)Hex(3) | C[+57], N[+1502] | 1283.5754 | 3 | 3848.7116 | 3848.6974 | 0 | 3.7 | 332 | CRagged | 246.01 | 173.24 | 173.24 | 3.71 | 26 | >sp\|P01876\|IGHA1_HUMAN Immunoglobulin heavy constant alpha 1 OS=Homo sapiens OX=9606 GN=IGHA1 PE=1 SV=2 | 255 | 20201118_12812_1ng_glyco_hcd30_etd30_compound15.13581.13581.3 | controllerType=0 controllerNumber=1 scan=13581 | 30.8259 |
| 10274:3 | 2 | R.LAGKPTHVN[+1647.613]VSVVMAEVDGTC[+57.021]Y.- | HexNAc(5)Hex(3)Fuc(1) | C[+57], N[+1648] | 1332.2608 | 3 | 3994.7678 | 3994.7553 | 0 | 3.1 | 332 | CRagged | 266.21 | 161.32 | 161.32 | 4.13 | 26 | >sp\|P01876\|IGHA1_HUMAN Immunoglobulin heavy constant alpha 1 OS=Homo sapiens OX=9606 GN=IGHA1 PE=1 SV=2 | 255 | 20201118_12812_1ng_glyco_hcd30_etd30_compound15.13572.13572.3 | controllerType=0 controllerNumber=1 scan=13572 | 30.8108 |
| 10234:3 | 2 | R.LAGKPTHVN[+1216.423]VSVVMAEVDGTC[+57.021]Y.- | HexNAc(2)Hex(5) | C[+57], N[+1216] | 1188.5296 | 3 | 3563.5742 | 3563.5649 | 0 | 2.6 | 332 | CRagged | 266.84 | 258.01 | 194.77 | 3.96 | 26 | >sp\|P01876\|IGHA1_HUMAN Immunoglobulin heavy constant alpha 1 OS=Homo sapiens OX=9606 GN=IGHA1 PE=1 SV=2 | 255 | 20201118_12812_1ng_glyco_hcd30_etd30_compound15.13531.13531.3 | controllerType=0 controllerNumber=1 scan=13531 | 30.7015 |
| 10182:3 | 2 | R.LAGKPTHVN[+1540.529]VSVVMAEVDGTC[+57.021]Y.- | HexNAc(2)Hex(7) | C[+57], N[+1541] | 1296.56 | 3 | 3887.6654 | 3887.6706 | 0 | -1.3 | 332 | CRagged | 305.18 | 229.81 | 201.18 | 3.37 | 26 | >sp\|P01876\|IGHA1_HUMAN Immunoglobulin heavy constant alpha 1 OS=Homo sapiens OX=9606 GN=IGHA1 PE=1 SV=2 | 255 | 20201118_12812_1ng_glyco_hcd30_etd30_compound15.13476.13476.3 | controllerType=0 controllerNumber=1 scan=13476 | 30.5227 |
| 14986:3 | 5 | R.GLSFDVSLEVSQGPGLLN[+2569.905]DTK.V | HexNAc(5)Hex(6)NeuAc(2) | N[+2570] | 1582.6907 | 3 | 4746.0576 | 4746.0178 | 0 | 8.4 | 118 | Specific | 332.11 | 201.98 | 201.98 | 4.88 | 33 | >sp\|P01833\|PIGR_HUMAN Polymeric immunoglobulin receptor OS=Homo sapiens OX=9606 GN=PIGR PE=1 SV=4 | 902 | 20201118_12812_1ng_glyco_hcd30_etd30_compound15.18549.18549.3 | controllerType=0 controllerNumber=1 scan=18549 | 42.2414 |
| 14916:3 | 5 | R.GLSFDVSLEVSQGPGLLN[+2716.983]DTK.V | HexNAc(5)Hex(6)Fuc(3)NeuAc(1) | N[+2717] | 1631.7103 | 3 | 4893.1162 | 4893.0962 | 0 | 4.1 | 118 | Specific | 362.02 | 229.91 | 229.91 | 6.03 | 33 | >sp\|P01833\|PIGR_HUMAN Polymeric immunoglobulin receptor OS=Homo sapiens OX=9606 GN=PIGR PE=1 SV=4 | 902 | 20201118_12812_1ng_glyco_hcd30_etd30_compound15.18475.18475.3 | controllerType=0 controllerNumber=1 scan=18475 | 42.0417 |
| 16404:3 | 5 | R.GLSFDVSLEVSQGPGLLN[+2350.830]DTK.V | HexNAc(4)Hex(5)Fuc(1)NeuAc(2) | N[+2351] | 1509.6561 | 3 | 4526.9536 | 4526.9436 | 0 | 2.2 | 118 | Specific | 497.83 | 296.61 | 296.61 | 5.85 | 33 | >sp\|P01833\|PIGR_HUMAN Polymeric immunoglobulin receptor OS=Homo sapiens OX=9606 GN=PIGR PE=1 SV=4 | 902 | 20201118_12812_1ng_glyco_hcd30_etd30_compound15.20047.20047.3 | controllerType=0 controllerNumber=1 scan=20047 | 46.7141 |
| 16396:3 | 5 | R.GLSFDVSLEVSQGPGLLN[+2350.830]DTK.V | HexNAc(4)Hex(5)Fuc(1)NeuAc(2) | N[+2351] | 1509.6567 | 3 | 4526.9555 | 4526.9436 | 0 | 2.6 | 118 | Specific | 500.99 | 316.14 | 316.14 | 5.85 | 33 | >sp\|P01833\|PIGR_HUMAN Polymeric immunoglobulin receptor OS=Homo sapiens OX=9606 GN=PIGR PE=1 SV=4 | 902 | 20201118_12812_1ng_glyco_hcd30_etd30_compound15.20038.20038.3 | controllerType=0 controllerNumber=1 scan=20038 | 46.6935 |
| 17322:3 | 5 | K.QIGLYPVLVIDSSGYVNPN[+2350.830]YTGR.I | HexNAc(4)Hex(5)Fuc(1)NeuAc(2) | N[+2351] | 1626.06 | 3 | 4876.1654 | 4876.1338 | 0 | 6.5 | 168 | Specific | 362.9 | 265.89 | 265.89 | 4.64 | 33 | >sp\|P01833\|PIGR_HUMAN Polymeric immunoglobulin receptor OS=Homo sapiens OX=9606 GN=PIGR PE=1 SV=4 | 902 | 20201118_12812_1ng_glyco_hcd30_etd30_compound15.21019.21019.3 | controllerType=0 controllerNumber=1 scan=21019 | 49.7429 |
| 15971:3 | 5 | K.QIGLYPVLVIDSSGYVNPN[+1913.677]YTGR.I | HexNAc(4)Hex(5)NeuAc(1) | N[+1914] | 1480.3377 | 3 | 4438.9986 | 4438.9805 | 0 | 4.1 | 168 | Specific | 522.83 | 432.7 | 432.7 | 8.19 | 33 | >sp\|P01833\|PIGR_HUMAN Polymeric immunoglobulin receptor OS=Homo sapiens OX=9606 GN=PIGR PE=1 SV=4 | 902 | 20201118_12812_1ng_glyco_hcd30_etd30_compound15.19591.19591.3 | controllerType=0 controllerNumber=1 scan=19591 | 45.4624 |
| 17089:3 | 5 | R.LSLLEEPGN[+2204.772]GTFTVILNQLTSR.D | HexNAc(4)Hex(5)NeuAc(2) | N[+2205] | 1536.3746 | 3 | 4607.1092 | 4607.065 | 0 | 9.6 | 413 | Specific | 395.74 | 234.66 | 234.66 | 5.13 | 33 | >sp\|P01833\|PIGR_HUMAN Polymeric immunoglobulin receptor OS=Homo sapiens OX=9606 GN=PIGR PE=1 SV=4 | 902 | 20201118_12812_1ng_glyco_hcd30_etd30_compound15.20773.20773.3 | controllerType=0 controllerNumber=1 scan=20773 | 48.9706 |
| 18728:3 | 5 | R.LSLLEEPGN[+2350.830]GTFTVILNQLTSR.D | HexNAc(4)Hex(5)Fuc(1)NeuAc(2) | N[+2351] | 1585.0527 | 3 | 4753.1435 | 4753.1229 | 0 | 4.3 | 413 | Specific | 425.06 | 281.56 | 281.56 | 6.19 | 33 | >sp\|P01833\|PIGR_HUMAN Polymeric immunoglobulin receptor OS=Homo sapiens OX=9606 GN=PIGR PE=1 SV=4 | 902 | 20201118_12812_1ng_glyco_hcd30_etd30_compound15.22505.22505.3 | controllerType=0 controllerNumber=1 scan=22505 | 53.8059 |
| 17091:3 | 5 | R.LSLLEEPGN[+2076.750]GTFTVILNQLTSR.D | HexNAc(4)Hex(6)Fuc(2) | N[+2077] | 1493.6968 | 3 | 4479.076 | 4479.0428 | 0 | 7.4 | 413 | Specific | 448.68 | 384.86 | 384.86 | 6.31 | 33 | >sp\|P01833\|PIGR_HUMAN Polymeric immunoglobulin receptor OS=Homo sapiens OX=9606 GN=PIGR PE=1 SV=4 | 902 | 20201118_12812_1ng_glyco_hcd30_etd30_compound15.20775.20775.3 | controllerType=0 controllerNumber=1 scan=20775 | [illegible] |
| 17041:3 | 5 | R.LSLLEEPGN[+2205.793]GTFTVILNQLTSR.D | HexNAc(4)Hex(5)Fuc(2)NeuAc(1) | N[+2206] | 1536.7056 | 3 | 4608.1023 | 4608.0854 | 0 | 3.7 | 413 | Specific | 582.02 | 475.92 | 475.92 | 9.05 | 33 | >sp\|P01833\|PIGR_HUMAN Polymeric immunoglobulin receptor OS=Homo sapiens OX=9606 GN=PIGR PE=1 SV=4 | 902 | 20201118_12812_1ng_glyco_hcd30_etd30_compound15.20722.20722.3 | controllerType=0 controllerNumber=1 scan=20722 | [illegible] |
| 17046:4 | 5 | R.LSLLEEPGN[+2205.793]GTFTVILNQLTSR.D | HexNAc(4)Hex(5)Fuc(2)NeuAc(1) | N[+2206] | 1152.7801 | 4 | 4608.0985 | 4608.0854 | 0 | 2.8 | 413 | Specific | 671.64 | 611.14 | 611.14 | 9.4 | 33 | >sp\|P01833\|PIGR_HUMAN Polymeric immunoglobulin receptor OS=Homo sapiens OX=9606 GN=PIGR PE=1 SV=4 | 902 | 20201118_12812_1ng_glyco_hcd30_etd30_compound15.20727.20727.4 | controllerType=0 controllerNumber=1 scan=20727 | 48.81 |
| 15310:3 | 5 | K.IIEGEPNLKVPGN[+2350.830]VTAVLGETLK.V | HexNAc(4)Hex(5)Fuc(1)NeuAc(2) | N[+2351] | 1581.4058 | 3 | 4742.2029 | 4742.1797 | 0 | 4.9 | 457 | Specific | 254.81 | 203.56 | 203.56 | 2.97 | 33 | >sp\|P01833\|PIGR_HUMAN Polymeric immunoglobulin receptor OS=Homo sapiens OX=9606 GN=PIGR PE=1 SV=4 | 902 | 20201118_12812_1ng_glyco_hcd30_etd30_compound15.18894.18894.3 | controllerType=0 controllerNumber=1 scan=18894 | 43.3688 |
| 13941:3 | 5 | K.IIEGEPNLKVPGN[+2059.735]VTAVLGETLK.V | HexNAc(4)Hex(5)Fuc(1)NeuAc(1) | N[+2060] | 1484.3703 | 3 | 4451.0963 | 4451.0843 | 0 | 2.7 | 457 | Specific | 351.94 | 276.34 | 276.34 | 3.59 | 33 | >sp\|P01833\|PIGR_HUMAN Polymeric immunoglobulin receptor OS=Homo sapiens OX=9606 GN=PIGR PE=1 SV=4 | 902 | 20201118_12812_1ng_glyco_hcd30_etd30_compound15.17444.17444.3 | controllerType=0 controllerNumber=1 scan=17444 | 39.2875 |
| 13888:3 | 5 | K.IIEGEPNLKVPGN[+2204.772]VTAVLGETLK.V | HexNAc(4)Hex(5)NeuAc(2) | N[+2205] | 1532.7214 | 3 | 4596.1496 | 4596.1218 | 0 | 6 | 457 | Specific | 365.68 | 287.63 | 287.63 | 3.23 | 33 | >sp\|P01833\|PIGR_HUMAN Polymeric immunoglobulin receptor OS=Homo sapiens OX=9606 GN=PIGR PE=1 SV=4 | 902 | 20201118_12812_1ng_glyco_hcd30_etd30_compound15.17387.17387.3 | controllerType=0 controllerNumber=1 scan=17387 | 39.1424 |
| 13931:3 | 5 | K.IIEGEPNLKVPGN[+2059.735]VTAVLGETLK.V | HexNAc(4)Hex(5)Fuc(1)NeuAc(1) | N[+2060] | 1484.3691 | 3 | 4451.0927 | 4451.0843 | 0 | 1.9 | 457 | Specific | 378.84 | 258.37 | 258.37 | 3.69 | 33 | >sp\|P01833\|PIGR_HUMAN Polymeric immunoglobulin receptor OS=Homo sapiens OX=9606 GN=PIGR PE=1 SV=4 | 902 | 20201118_12812_1ng_glyco_hcd30_etd30_compound15.17433.17433.3 | controllerType=0 controllerNumber=1 scan=17433 | 39.2551 |
| 13905:3 | 5 | K.IIEGEPNLKVPGN[+2205.793]VTAVLGETLK.V | HexNAc(4)Hex(5)Fuc(2)NeuAc(1) | N[+2206] | 1533.0554 | 3 | 4597.1515 | 4597.1422 | 0 | 2 | 457 | Specific | 422.38 | 338.11 | 338.11 | 4.17 | 33 | >sp\|P01833\|PIGR_HUMAN Polymeric immunoglobulin receptor OS=Homo sapiens OX=9606 GN=PIGR PE=1 SV=4 | 902 | 20201118_12812_1ng_glyco_hcd30_etd30_compound15.17406.17406.3 | controllerType=0 controllerNumber=1 scan=17406 | 39.1787 |
| 13947:4 | 5 | K.IIEGEPNLKVPGN[+2059.735]VTAVLGETLK.V | HexNAc(4)Hex(5)Fuc(1)NeuAc(1) | N[+2060] | 1113.5295 | 4 | 4451.096 | 4451.0843 | 0 | 2.6 | 457 | Specific | 551.55 | 496.2 | 496.2 | 5.8 | 33 | >sp\|P01833\|PIGR_HUMAN Polymeric immunoglobulin receptor OS=Homo sapiens OX=9606 GN=PIGR PE=1 SV=4 | 902 | 20201118_12812_1ng_glyco_hcd30_etd30_compound15.17450.17450.4 | controllerType=0 controllerNumber=1 scan=17450 | 39.2993 |
| 13524:4 | 5 | K.IIEGEPNLKVPGN[+2205.793]VTAVLGETLKVPC[+57.021]HFPC[+57.021]K.F | HexNAc(4)Hex(5)Fuc(2)NeuAc(1) | C[+57]*2, N[+2206] | 1406.41 | 4 | 5622.6183 | 5622.5997 | 0 | 3.3 | 457 | Specific | 215.72 | 215.72 | 215.72 | 2.74 | 33 | >sp\|P01833\|PIGR_HUMAN Polymeric immunoglobulin receptor OS=Homo sapiens OX=9606 GN=PIGR PE=1 SV=4 | 902 | 20201118_12812_1ng_glyco_hcd30_etd30_compound15.16979.16979.4 | controllerType=0 controllerNumber=1 scan=16979 | 38.299 |
| 14668:4 | 5 | K.IIEGEPNLKVPGN[+2350.830]VTAVLGETLKVPC[+57.021]HFPC[+57.021]K.F | HexNAc(4)Hex(5)Fuc(1)NeuAc(2) | C[+57]*2, N[+2351] | 1442.6709 | 4 | 5767.6618 | 5767.6372 | 0 | 4.3 | 457 | Specific | 405.47 | 405.47 | 405.47 | 5.36 | 33 | >sp\|P01833\|PIGR_HUMAN Polymeric immunoglobulin receptor OS=Homo sapiens OX=9606 GN=PIGR PE=1 SV=4 | 902 | 20201118_12812_1ng_glyco_hcd30_etd30_compound15.18213.18213.4 | controllerType=0 controllerNumber=1 scan=18213 | 41.2632 |
| 13527:5 | 5 | K.IIEGEPNLKVPGN[+2059.735]VTAVLGETLKVPC[+57.021]HFPC[+57.021]K.F | HexNAc(4)Hex(5)Fuc(1)NeuAc(1) | C[+57]*2, N[+2060] | 1096.1179 | 5 | 5476.5605 | 5476.5418 | 0 | 3.4 | 457 | Specific | 496.83 | 421.33 | 421.33 | 5.43 | 33 | >sp\|P01833\|PIGR_HUMAN Polymeric immunoglobulin receptor OS=Homo sapiens OX=9606 GN=PIGR PE=1 SV=4 | 902 | 20201118_12812_1ng_glyco_hcd30_etd30_compound15.16983.16983.5 | controllerType=0 controllerNumber=1 scan=16983 | 38.312 |
| 13531:4 | 5 | K.IIEGEPNLKVPGN[+2059.735]VTAVLGETLKVPC[+57.021]HFPC[+57.021]K.F | HexNAc(4)Hex(5)Fuc(1)NeuAc(1) | C[+57]*2, N[+2060] | 1369.8959 | 4 | 5476.5617 | 5476.5418 | 0 | 3.6 | 457 | Specific | 517.6 | 517.6 | 517.6 | 5.59 | 33 | >sp\|P01833\|PIGR_HUMAN Polymeric immunoglobulin receptor OS=Homo sapiens OX=9606 GN=PIGR PE=1 SV=4 | 902 | 20201118_12812_1ng_glyco_hcd30_etd30_compound15.16987.16987.4 | controllerType=0 controllerNumber=1 scan=16987 | 38.322 |
| 14660:5 | 5 | K.IIEGEPNLKVPGN[+2350.830]VTAVLGETLKVPC[+57.021]HFPC[+57.021]K.F | HexNAc(4)Hex(5)Fuc(1)NeuAc(2) | C[+57]*2, N[+2351] | 1154.3372 | 5 | 5767.6567 | 5767.6372 | 0 | 3.4 | 457 | Specific | 541.4 | 541.4 | 541.4 | 6.22 | 33 | >sp\|P01833\|PIGR_HUMAN Polymeric immunoglobulin receptor OS=Homo sapiens OX=9606 GN=PIGR PE=1 SV=4 | 902 | 20201118_12812_1ng_glyco_hcd30_etd30_compound15.18205.18205.5 | controllerType=0 controllerNumber=1 scan=18205 | 41.2357 |
| 13521:5 | 5 | K.IIEGEPNLKVPGN[+2059.735]VTAVLGETLKVPC[+57.021]HFPC[+57.021]K.F | HexNAc(4)Hex(5)Fuc(1)NeuAc(1) | C[+57]*2, N[+2060] | 1096.1173 | 5 | 5476.5575 | 5476.5418 | 0 | 2.9 | 457 | Specific | 597.42 | 597.42 | 597.42 | 6.45 | 33 | >sp\|P01833\|PIGR_HUMAN Polymeric immunoglobulin receptor OS=Homo sapiens OX=9606 GN=PIGR PE=1 SV=4 | 902 | 20201118_12812_1ng_glyco_hcd30_etd30_compound15.16976.16976.5 | controllerType=0 controllerNumber=1 scan=16976 | 38.2897 |
| 13519:5 | 5 | K.IIEGEPNLKVPGN[+2205.793]VTAVLGETLKVPC[+57.021]HFPC[+57.021]K.F | HexNAc(4)Hex(5)Fuc(2)NeuAc(1) | C[+57]*2, N[+2206] | 1125.3284 | 5 | 5622.6128 | 5622.5997 | 0 | 2.3 | 457 | Specific | 678.78 | 678.78 | 678.78 | 8.45 | 33 | >sp\|P01833\|PIGR_HUMAN Polymeric immunoglobulin receptor OS=Homo sapiens OX=9606 GN=PIGR PE=1 SV=4 | 902 | 20201118_12812_1ng_glyco_hcd30_etd30_compound15.16974.16974.5 | controllerType=0 controllerNumber=1 scan=16974 | 38.2807 |
| 12032:3 | 5 | K.VPGN[+2205.793]VTAVLGETLK.V | HexNAc(4)Hex(5)Fuc(2)NeuAc(1) | N[+2206] | 1201.8733 | 3 | 3603.6055 | 3603.5927 | 0 | 3.5 | 466 | Specific | 276.08 | 207.41 | 207.41 | 3.47 | 33 | >sp\|P01833\|PIGR_HUMAN Polymeric immunoglobulin receptor OS=Homo sapiens OX=9606 GN=PIGR PE=1 SV=4 | 902 | 20201118_12812_1ng_glyco_hcd30_etd30_compound15.15416.15416.3 | controllerType=0 controllerNumber=1 scan=15416 | 34.4469 |
| 12073:4 | 5 | K.VPGN[+2205.793]VTAVLGETLKVPC[+57.021]HFPC[+57.021]K.F | HexNAc(4)Hex(5)Fuc(2)NeuAc(1) | C[+57]*2, N[+2206] | 1158.0212 | 4 | 4629.0628 | 4629.0502 | 0 | 2.7 | 466 | Specific | 232.8 | 232.8 | 232.8 | 2.75 | 33 | >sp\|P01833\|PIGR_HUMAN Polymeric immunoglobulin receptor OS=Homo sapiens OX=9606 GN=PIGR PE=1 SV=4 | 902 | 20201118_12812_1ng_glyco_hcd30_etd30_compound15.15460.15460.4 | controllerType=0 controllerNumber=1 scan=15460 | 34.5718 |
| 12148:4 | 5 | K.VPGN[+2059.735]VTAVLGETLKVPC[+57.021]HFPC[+57.021]K.F | HexNAc(4)Hex(5)Fuc(1)NeuAc(1) | C[+57]*2, N[+2060] | 1121.5061 | 4 | 4483.0028 | 4482.9923 | 0 | 2.3 | 466 | Specific | 254.17 | 215.83 | 215.83 | 2.99 | 33 | >sp\|P01833\|PIGR_HUMAN Polymeric immunoglobulin receptor OS=Homo sapiens OX=9606 GN=PIGR PE=1 SV=4 | 902 | 20201118_12812_1ng_glyco_hcd30_etd30_compound15.15539.15539.4 | controllerType=0 controllerNumber=1 scan=15539 | 34.8121 |
| 13532:4 | 5 | K.VPGN[+2350.830]VTAVLGETLKVPC[+57.021]HFPC[+57.021]K.F | HexNAc(4)Hex(5)Fuc(1)NeuAc(2) | C[+57]*2, N[+2351] | 1194.2828 | 4 | 4774.1092 | 4774.0877 | 0 | 4.5 | 466 | Specific | 308.76 | 250.93 | 250.93 | 4.21 | 33 | >sp\|P01833\|PIGR_HUMAN Polymeric immunoglobulin receptor OS=Homo sapiens OX=9606 GN=PIGR PE=1 SV=4 | 902 | 20201118_12812_1ng_glyco_hcd30_etd30_compound15.16988.16988.4 | controllerType=0 controllerNumber=1 scan=16988 | 38.3286 |
| 05137:3 | 5 | K.WN[+2059.735]NTGC[+57.021]QALPSQDEGPSK.A | HexNAc(4)Hex(5)Fuc(1)NeuAc(1) | C[+57], N[+2060] | 1350.2118 | 3 | 4048.621 | 4048.6116 | 0 | 2.3 | 498 | Specific | 369.77 | 233.2 | 233.2 | 4.93 | 33 | >sp\|P01833\|PIGR_HUMAN Polymeric immunoglobulin receptor OS=Homo sapiens OX=9606 GN=PIGR PE=1 SV=4 | 902 | 20201118_12812_1ng_glyco_hcd30_etd30_compound15.8219.8219.3 | controllerType=0 controllerNumber=1 scan=8219 | 20.6837 |
| 05221:3 | 5 | K.WN[+1913.677]NTGC[+57.021]QALPSQDEGPSK.A | HexNAc(4)Hex(5)NeuAc(1) | C[+57], N[+1914] | 1301.5259 | 3 | 3902.5633 | 3902.5537 | 0 | 2.4 | 498 | Specific | 492.49 | 311.34 | 311.34 | 5.43 | 33 | >sp\|P01833\|PIGR_HUMAN Polymeric immunoglobulin receptor OS=Homo sapiens OX=9606 GN=PIGR PE=1 SV=4 | 902 | 20201118_12812_1ng_glyco_hcd30_etd30_compound15.8306.8306.3 | controllerType=0 controllerNumber=1 scan=8306 | 20.846 |
| 09091:3 | 7 | K.TVLTPATNHMGN[+1378.476]VTFTIPANR.E | HexNAc(2)Hex(6) | N[+1378] | 1211.8879 | 3 | 3633.6491 | 3633.6358 | 0 | 3.7 | 74 | Specific | 404.11 | 319.54 | 150.82 | 6.47 | 10 | >sp\|P01024\|CO3_HUMAN Complement C3 OS=Homo sapiens OX=9606 GN=C3 PE=1 SV=2 | 492 | 20201118_12812_1ng_glyco_hcd30_etd30_compound15.12341.12341.3 | controllerType=0 controllerNumber=1 scan=12341 | 28.3068 |
| 15961:4 | 8 | R.NIAAFGGDPNN[+1913.677]ITLFGESAGGASVSLQTLSPYNK.G | HexNAc(4)Hex(5)NeuAc(1) | N[+1914] | 1331.8535 | 4 | 5324.3922 | 5324.3633 | 0 | 5.4 | 197 | Specific | 264.4 | 243.76 | 243.76 | 2.85 | 9 | >sp\|P19835\|CEL_HUMAN Bile salt-activated lipase OS=Homo sapiens OX=9606 GN=CEL PE=1 SV=3 | 476 | 20201118_12812_1ng_glyco_hcd30_etd30_compound15.19581.19581.4 | controllerType=0 controllerNumber=1 scan=19581 | 45.4341 |
| 15954:4 | 8 | R.NIAAFGGDPNN[+1913.677]ITLFGESAGGASVSLQTLSPYNK.G | HexNAc(4)Hex(5)NeuAc(1) | N[+1914] | 1331.8505 | 4 | 5324.38 | 5324.3633 | 0 | 3.1 | 197 | Specific | 391.65 | 369.99 | 369.99 | 5.92 | 9 | >sp\|P19835\|CEL_HUMAN Bile salt-activated lipase OS=Homo sapiens OX=9606 GN=CEL PE=1 SV=3 | 476 | 20201118_12812_1ng_glyco_hcd30_etd30_compound15.19573.19573.4 | controllerType=0 controllerNumber=1 scan=19573 | 45.4198 |
| 15970:3 | 8 | R.NIAAFGGDPNN[+1913.677]ITLFGESAGGASVSLQTLSPYNK.G | HexNAc(4)Hex(5)NeuAc(1) | N[+1914] | 1775.4704 | 3 | 5324.3967 | 5324.3633 | 0 | 6.3 | 197 | Specific | 522.81 | 411.05 | 411.05 | 6.97 | 9 | >sp\|P19835\|CEL_HUMAN Bile salt-activated lipase OS=Homo sapiens OX=9606 GN=CEL PE=1 SV=3 | 476 | 20201118_12812_1ng_glyco_hcd30_etd30_compound15.19590.19590.3 | controllerType=0 controllerNumber=1 scan=19590 | 45.4585 |
| 12588:3 | 11 | R.ALGFEN[+2350.830]ATQALGR.A | HexNAc(4)Hex(5)Fuc(1)NeuAc(2) | N[+2351] | 1233.5193 | 3 | 3698.5435 | 3698.5319 | 0 | 3.1 | 64 | Specific | 260.9 | 153.5 | 153.5 | 4.41 | 9 | >sp\|Q08380\|LG3BP_HUMAN Galectin-3-binding protein OS=Homo sapiens OX=9606 GN=LGALS3BP PE=1 SV=1 | 1562 | 20201118_12812_1ng_glyco_hcd30_etd30_compound15.15998.15998.3 | controllerType=0 controllerNumber=1 scan=15998 | 35.8299 |
| 05281:3 | 11 | K.AAIPSALDTN[+2059.735]SSK.S | HexNAc(4)Hex(5)Fuc(1)NeuAc(1) | N[+2060] | 1112.1386 | 3 | 3334.4012 | 3334.3936 | 0 | 2.3 | 542 | Specific | 204.09 | 101.46 | 101.46 | 2.27 | 9 | >sp\|Q08380\|LG3BP_HUMAN Galectin-3-binding protein OS=Homo sapiens OX=9606 GN=LGALS3BP PE=1 SV=1 | 1562 | 20201118_12812_1ng_glyco_hcd30_etd30_compound15.8368.8368.3 | controllerType=0 controllerNumber=1 scan=8368 | 20.968 |
| 03482:2 | 11 | K.AAIPSALDTN[+1216.423]SSK.S | HexNAc(2)Hex(5) | N[+1216] | 1246.0469 | 2 | 2491.0866 | 2491.0815 | 0 | 2 | 542 | Specific | 281.38 | 113.72 | 113.72 | 2.54 | 9 | >sp\|Q08380\|LG3BP_HUMAN Galectin-3-binding protein OS=Homo sapiens OX=9606 GN=LGALS3BP PE=1 SV=1 | 1562 | 20201118_12812_1ng_glyco_hcd30_etd30_compound15.6502.6502.2 | controllerType=0 controllerNumber=1 scan=6502 | 17.6694 |
| 19471:4 | 14 | R.QLAHQSN[+2350.830]STNIFFSPVSIATAFAMLSLGTK.A | HexNAc(4)Hex(5)Fuc(1)NeuAc(2) | N[+2351] | 1383.876 | 4 | 5532.4821 | 5532.4654 | 0 | 3 | 64 | Specific | 266.49 | 229.2 | 229.2 | 4.34 | 10 | >sp\|P01009\|A1AT_HUMAN Alpha-1-antitrypsin OS=Homo sapiens OX=9606 GN=SERPINA1 PE=1 SV=3 | 642 | 20201118_12812_1ng_glyco_hcd30_etd30_compound15.23290.23290.4 | controllerType=0 controllerNumber=1 scan=23290 | 55.9023 |
| 19500:4 | 14 | R.[+42.011]QLAHQSN[+2190.793]STNIFFSPVSIATAFAMLSLGTK.A | HexNAc(6)Hex(6) | .[42], N[+2191] | 1354.359 | 4 | 5414.4142 | 5414.4388 | 0 | -4.5 | 64 | Specific | 349.68 | 276.06 | 276.06 | 2.61 | 10 | >sp\|P01009\|A1AT_HUMAN Alpha-1-antitrypsin OS=Homo sapiens OX=9606 GN=SERPINA1 PE=1 SV=3 | 642 | 20201118_12812_1ng_glyco_hcd30_etd30_compound15.23321.23321.4 | controllerType=0 controllerNumber=1 scan=23321 | 56.0127 |
| 19483:3 | 14 | R.QLAHQSN[+2204.772]STNIFFSPVSIATAFAMLSLGTK.A | HexNAc(4)Hex(5)NeuAc(2) | N[+2205] | 1796.1456 | 3 | 5386.4222 | 5386.4075 | 0 | 2.7 | 64 | Specific | 386.72 | 263.92 | 263.92 | 5.45 | 10 | >sp\|P01009\|A1AT_HUMAN Alpha-1-antitrypsin OS=Homo sapiens OX=9606 GN=SERPINA1 PE=1 SV=3 | 642 | 20201118_12812_1ng_glyco_hcd30_etd30_compound15.23303.23303.3 | controllerType=0 controllerNumber=1 scan=23303 | 55.9533 |
| 19482:4 | 14 | R.QLAHQSN[+2204.772]STNIFFSPVSIATAFAMLSLGTK.A | HexNAc(4)Hex(5)NeuAc(2) | N[+2205] | 1347.3604 | 4 | 5386.4196 | 5386.4075 | 0 | 2.2 | 64 | Specific | 618.52 | 494.2 | 388.74 | 8.31 | 10 | >sp\|P01009\|A1AT_HUMAN Alpha-1-antitrypsin OS=Homo sapiens OX=9606 GN=SERPINA1 PE=1 SV=3 | 642 | 20201118_12812_1ng_glyco_hcd30_etd30_compound15.23302.23302.4 | controllerType=0 controllerNumber=1 scan=23302 | 55.9498 |
| 19445:4 | 14 | R.QLAHQSN[+2204.772]STNIFFSPVSIATAFAMLSLGTK.A | HexNAc(4)Hex(5)NeuAc(2) | N[+2205] | 1347.3578 | 4 | 5386.4093 | 5386.4075 | 0 | 0.3 | 64 | Specific | 670.2 | 591.73 | 406 | 10.16 | 10 | >sp\|P01009\|A1AT_HUMAN Alpha-1-antitrypsin OS=Homo sapiens OX=9606 GN=SERPINA1 PE=1 SV=3 | 642 | 20201118_12812_1ng_glyco_hcd30_etd30_compound15.23262.23262.4 | controllerType=0 controllerNumber=1 scan=23262 | 55.8168 |
| 18803:4 | 14 | R.QLAHQSN[+2204.772]STNIFFSPVSIATAFAM[+15.995]LSLGTK.A | HexNAc(4)Hex(5)NeuAc(2) | M[+16], N[+2205] | 1351.3599 | 4 | 5402.4176 | 5402.4024 | 0 | 2.8 | 64 | Specific | 309.54 | 239.27 | 239.27 | 3.73 | 10 | >sp\|P01009\|A1AT_HUMAN Alpha-1-antitrypsin OS=Homo sapiens OX=9606 GN=SERPINA1 PE=1 SV=3 | 642 | 20201118_12812_1ng_glyco_hcd30_etd30_compound15.22584.22584.4 | controllerType=0 controllerNumber=1 scan=22584 | 54.0155 |
| 16889:3 | 14 | K.YLGN[+2204.772]ATAIFFLPDEGK.L | HexNAc(4)Hex(5)NeuAc(2) | N[+2205] | 1320.8989 | 3 | 3960.682 | 3960.6677 | 0 | 3.6 | 268 | Specific | 267.07 | 202.3 | 202.3 | 5.32 | 10 | >sp\|P01009\|A1AT_HUMAN Alpha-1-antitrypsin OS=Homo sapiens OX=9606 GN=SERPINA1 PE=1 SV=3 | 642 | 20201118_12812_1ng_glyco_hcd30_etd30_compound15.20561.20561.3 | controllerType=0 controllerNumber=1 scan=20561 | 48.3121 |
| 16708:5 | 14 | K.YLGN[+2204.772]ATAIFFLPDEGKLQHLENELTHDIITK.F | HexNAc(4)Hex(5)NeuAc(2) | N[+2205] | 1149.9312 | 5 | 5745.6267 | 5745.6098 | 0 | 2.9 | 268 | Specific | 312.3 | 305.21 | 305.21 | 3.16 | 10 | >sp\|P01009\|A1AT_HUMAN Alpha-1-antitrypsin OS=Homo sapiens OX=9606 GN=SERPINA1 PE=1 SV=3 | 642 | 20201118_12812_1ng_glyco_hcd30_etd30_compound15.20369.20369.5 | controllerType=0 controllerNumber=1 scan=20369 | 47.6952 |
| 16709:4 | 14 | K.YLGN[+2204.772]ATAIFFLPDEGKLQHLENELTHDIITK.F | HexNAc(4)Hex(5)NeuAc(2) | N[+2205] | 1437.1624 | 4 | 5745.6276 | 5745.6098 | 0 | 3.1 | 268 | Specific | 471.46 | 471.46 | 471.46 | 5.83 | 10 | >sp\|P01009\|A1AT_HUMAN Alpha-1-antitrypsin OS=Homo sapiens OX=9606 GN=SERPINA1 PE=1 SV=3 | 642 | 20201118_12812_1ng_glyco_hcd30_etd30_compound15.20370.20370.4 | controllerType=0 controllerNumber=1 scan=20370 | 47.6987 |
| 05091:3 | 15 | R.LSPN[+2205.793]ASAEHLELR.W | HexNAc(4)Hex(5)Fuc(2)NeuAc(1) | N[+2206] | 1214.8555 | 3 | 3642.552 | 3642.5421 | 0 | 2.7 | 52 | Specific | 331.1 | 210.04 | 210.04 | 3.31 | 10 | >sp\|Q13410\|BT1A1_HUMAN Butyrophilin subfamily 1 member A1 OS=Homo sapiens OX=9606 GN=BTN1A1 PE=1 SV=3 | 179 | 20201118_12812_1ng_glyco_hcd30_etd30_compound15.8171.8171.3 | controllerType=0 controllerNumber=1 scan=8171 | 20.5835 |
| 07012:3 | 15 | R.LSPN[+2350.830]ASAEHLELR.W | HexNAc(4)Hex(5)Fuc(1)NeuAc(2) | N[+2351] | 1263.2026 | 3 | 3787.5932 | 3787.5796 | 0 | 3.6 | 52 | Specific | 348.77 | 224.67 | 224.67 | 5.03 | 10 | >sp\|Q13410\|BT1A1_HUMAN Butyrophilin subfamily 1 member A1 OS=Homo sapiens OX=9606 GN=BTN1A1 PE=1 SV=3 | 179 | 20201118_12812_1ng_glyco_hcd30_etd30_compound15.10162.10162.3 | controllerType=0 controllerNumber=1 scan=10162 | 24.1392 |
| 15089:3 | 15 | K.N[+1913.677]VSC[+57.021]YIQNLLLGQEK.K | HexNAc(4)Hex(5)NeuAc(1) | C[+57], N[+1914] | 1231.5389 | 3 | 3692.6021 | 3692.5876 | 0 | 3.9 | 215 | Specific | 206.33 | 74.86 | 74.86 | 3.2 | 10 | >sp\|Q13410\|BT1A1_HUMAN Butyrophilin subfamily 1 member A1 OS=Homo sapiens OX=9606 GN=BTN1A1 PE=1 SV=3 | 179 | 20201118_12812_1ng_glyco_hcd30_etd30_compound15.18658.18658.3 | controllerType=0 controllerNumber=1 scan=18658 | 42.5913 |
| 02555:3 | 26 | K.TPLTAN[+1971.719]ITK.S | HexNAc(5)Hex(5)Fuc(1) | N[+1972] | 977.4334 | 3 | 2930.2856 | 2930.2757 | 0 | 3.4 | 315 | Specific | 235.42 | 118.8 | 118.8 | 2.51 | 8 | >sp\|P0DOX2\|IGA2_HUMAN Immunoglobulin alpha-2 heavy chain OS=Homo sapiens OX=9606 PE=1 SV=2 | 631 | 20201118_12812_1ng_glyco_hcd30_etd30_compound15.5539.5539.3 | controllerType=0 controllerNumber=1 scan=5539 | 15.8438 |
| 02658:3 | 26 | K.TPLTAN[+1647.613]ITK.S | HexNAc(5)Hex(3)Fuc(1) | N[+1648] | 869.3979 | 3 | 2606.179 | 2606.17 | 0 | 3.5 | 315 | Specific | 262.75 | 130.77 | 130.77 | 2.66 | 8 | >sp\|P0DOX2\|IGA2_HUMAN Immunoglobulin alpha-2 heavy chain OS=Homo sapiens OX=9606 PE=1 SV=2 | 631 | 20201118_12812_1ng_glyco_hcd30_etd30_compound15.5647.5647.3 | controllerType=0 controllerNumber=1 scan=5647 | 16.1007 |
| 02651:2 | 26 | K.TPLTAN[+1647.613]ITK.S | HexNAc(5)Hex(3)Fuc(1) | N[+1648] | 1303.5928 | 2 | 2606.1784 | 2606.17 | 0 | 3.2 | 315 | Specific | 290.36 | 118.26 | 118.26 | 3.05 | 8 | >sp\|P0DOX2\|IGA2_HUMAN Immunoglobulin alpha-2 heavy chain OS=Homo sapiens OX=9606 PE=1 SV=2 | 631 | 20201118_12812_1ng_glyco_hcd30_etd30_compound15.5639.5639.2 | controllerType=0 controllerNumber=1 scan=5639 | 16.0731 |
| 02603:3 | 26 | K.TPLTAN[+1809.666]ITK.S | HexNAc(5)Hex(4)Fuc(1) | N[+1810] | 923.4156 | 3 | 2768.2321 | 2768.2228 | 0 | 3.4 | 315 | Specific | 292.88 | 143.41 | 143.41 | 2.94 | 8 | >sp\|P0DOX2\|IGA2_HUMAN Immunoglobulin alpha-2 heavy chain OS=Homo sapiens OX=9606 PE=1 SV=2 | 631 | 20201118_12812_1ng_glyco_hcd30_etd30_compound15.5589.5589.3 | controllerType=0 controllerNumber=1 scan=5589 | 15.9394 |
| 02599:2 | 26 | K.TPLTAN[+1809.666]ITK.S | HexNAc(5)Hex(4)Fuc(1) | N[+1810] | 1384.6193 | 2 | 2768.2313 | 2768.2228 | 0 | 3.1 | 315 | Specific | 304.59 | 121.68 | 121.68 | 2.78 | 8 | >sp\|P0DOX2\|IGA2_HUMAN Immunoglobulin alpha-2 heavy chain OS=Homo sapiens OX=9606 PE=1 SV=2 | 631 | 20201118_12812_1ng_glyco_hcd30_etd30_compound15.5585.5585.2 | controllerType=0 controllerNumber=1 scan=5585 | 15.9266 |
| 02627:2 | 26 | K.TPLTAN[+1444.534]ITK.S | HexNAc(4)Hex(3)Fuc(1) | N[+1445] | 1202.0509 | 2 | 2403.0946 | 2403.0906 | 0 | 1.7 | 315 | Specific | 336.49 | 102.05 | 102.05 | 2.5 | 8 | >sp\|P0DOX2\|IGA2_HUMAN Immunoglobulin alpha-2 heavy chain OS=Homo sapiens OX=9606 PE=1 SV=2 | 631 | 20201118_12812_1ng_glyco_hcd30_etd30_compound15.5614.5614.2 | controllerType=0 controllerNumber=1 scan=5614 | 15.9996 |
| 10152:3 | 29 | R.IIVPLNNREN[+1913.677]ISDPTSPLR.T | HexNAc(4)Hex(5)NeuAc(1) | N[+1914] | 1354.628 | 3 | 4061.8694 | 4061.8542 | 0 | 3.8 | 62 | Specific | 267.85 | 225.13 | 225.13 | 3.76 | 6 | >sp\|P01591\|IGJ_HUMAN Immunoglobulin J chain OS=Homo sapiens OX=9606 GN=JCHAIN PE=1 SV=4 | 1392 | 20201118_12812_1ng_glyco_hcd30_etd30_compound15.13444.13444.3 | controllerType=0 controllerNumber=1 scan=13444 | 30.4253 |
| 10128:3 | 29 | R.IIVPLNNREN[+2059.735]ISDPTSPLR.T | HexNAc(4)Hex(5)Fuc(1)NeuAc(1) | N[+2060] | 1403.3135 | 3 | 4207.926 | 4207.9121 | 0 | 3.3 | 62 | Specific | 281.68 | 205.47 | 205.47 | 3.28 | 6 | >sp\|P01591\|IGJ_HUMAN Immunoglobulin J chain OS=Homo sapiens OX=9606 GN=JCHAIN PE=1 SV=4 | 1392 | 20201118_12812_1ng_glyco_hcd30_etd30_compound15.13419.13419.3 | controllerType=0 controllerNumber=1 scan=13419 | 30.3546 |
| 10208:3 | 29 | R.IIVPLNNREN[+1856.656]ISDPTSPLR.T | HexNAc(3)Hex(5)Fuc(1)NeuAc(1) | N[+1857] | 1335.6177 | 3 | 4004.8386 | 4004.8327 | 0 | 1.5 | 62 | Specific | 324.06 | 219.11 | 219.11 | 3.15 | 6 | >sp\|P01591\|IGJ_HUMAN Immunoglobulin J chain OS=Homo sapiens OX=9606 GN=JCHAIN PE=1 SV=4 | 1392 | 20201118_12812_1ng_glyco_hcd30_etd30_compound15.13503.13503.3 | controllerType=0 controllerNumber=1 scan=13503 | 30.585 |
| 09087:4 | 31 | K.SC[+57.021]QHN[+2350.830]GTMYQHGEIFSAHELFPSR.L | HexNAc(4)Hex(5)Fuc(1)NeuAc(2) | C[+57], N[+2351] | 1296.783 | 4 | 5184.11 | 5184.0873 | 0 | 4.4 | 110 | Specific | 294.03 | 184.79 | 184.79 | 4.76 | 5 | >sp\|Q6WN34\|CRDL2_HUMAN Chordin-like protein 2 OS=Homo sapiens OX=9606 GN=CHRDL2 PE=1 SV=1 | 1189 | 20201118_12812_1ng_glyco_hcd30_etd30_compound15.12337.12337.4 | controllerType=0 controllerNumber=1 scan=12337 | 28.2945 |
| 07831:4 | 31 | K.SC[+57.021]QHN[+2059.735]GTMYQHGEIFSAHELFPSR.L | HexNAc(4)Hex(5)Fuc(1)NeuAc(1) | C[+57], N[+2060] | 1224.0088 | 4 | 4893.0135 | 4892.9919 | 0 | 4.4 | 110 | Specific | 623.15 | 542.35 | 542.35 | 9.93 | 5 | >sp\|Q6WN34\|CRDL2_HUMAN Chordin-like protein 2 OS=Homo sapiens OX=9606 GN=CHRDL2 PE=1 SV=1 | 1189 | 20201118_12812_1ng_glyco_hcd30_etd30_compound15.11019.11019.4 | controllerType=0 controllerNumber=1 scan=11019 | 25.7237 |
| 00102:4 | 32 | R.EIRHN[+2059.735]STGC[+57.021]LR.M | HexNAc(4)Hex(5)Fuc(1)NeuAc(1) | C[+57], N[+2060] | 851.3585 | 4 | 3402.4121 | 3402.3994 | 0 | 3.7 | 287 | Specific | 477.1 | 358.55 | 358.55 | 5.18 | 5 | >sp\|P10909\|CLUS_HUMAN Clusterin OS=Homo sapiens OX=9606 GN=CLU PE=1 SV=1 | 1467 | 20201118_12812_1ng_glyco_hcd30_etd30_compound15.2988.2988.2 | controllerType=0 controllerNumber=1 scan=2988 | 11.2457 |
| 00105:4 | 32 | R.EIRHN[+2059.735]STGC[+57.021]LR.M | HexNAc(4)Hex(5)Fuc(1)NeuAc(1) | C[+57], N[+2060] | 851.3579 | 4 | 3402.4097 | 3402.3994 | 0 | 3 | 287 | Specific | 667.9 | 492.74 | 492.74 | 9.72 | 5 | >sp\|P10909\|CLUS_HUMAN Clusterin OS=Homo sapiens OX=9606 GN=CLU PE=1 SV=1 | 1467 | 20201118_12812_1ng_glyco_hcd30_etd30_compound15.2992.2992.4 | controllerType=0 controllerNumber=1 scan=2992 | 11.2605 |
| 18945:4 | 32 | K.MLN[+2350.830]TSSLLEQLNEQFNWVSR.L | HexNAc(4)Hex(5)Fuc(1)NeuAc(2) | N[+2351] | 1190.7662 | 4 | 4760.043 | 4760.0171 | 0 | 5.4 | 352 | Specific | 252.03 | 229.07 | 229.07 | 2.97 | 5 | >sp\|P10909\|CLUS_HUMAN Clusterin OS=Homo sapiens OX=9606 GN=CLU PE=1 SV=1 | 1467 | 20201118_12812_1ng_glyco_hcd30_etd30_compound15.22736.22736.4 | controllerType=0 controllerNumber=1 scan=22736 | 54.4931 |
| 01864:3 | 34 | K.N[+1216.423]NSIPDKQITASSSYK.T | HexNAc(2)Hex(5) | N[+1216] | 990.4404 | 3 | 2969.3066 | 2969.2991 | 0 | 2.5 | 238 | Specific | 227.94 | 158.84 | 158.84 | 2.51 | 4 | >sp\|Q08431\|MFGM_HUMAN Lactadherin OS=Homo sapiens OX=9606 GN=MFGE8 PE=1 SV=3 | 1064 | 20201118_12812_1ng_glyco_hcd30_etd30_compound15.4820.4820.3 | controllerType=0 controllerNumber=1 scan=4820 | 14.5698 |
| 08513:4 | 36 | K.QTDEIKDTRN[+2059.735]ESTQNC[+57.021]VVAEPEKMESSISSSSEEMSLSK.C | HexNAc(4)Hex(5)Fuc(1)NeuAc(1) | C[+57], N[+2060] | 1613.443 | 4 | 6450.7502 | 6450.7156 | 0 | 5.4 | 60 | Specific | 253.7 | 253.7 | 253.7 | 2.59 | 3 | >sp\|P47710\|CASA1_HUMAN Alpha-S1-casein OS=Homo sapiens OX=9606 GN=CSN1S1 PE=1 SV=1 | 989 | 20201118_12812_1ng_glyco_hcd30_etd30_compound15.11741.11741.2 | controllerType=0 controllerNumber=1 scan=11741 | 27.1913 |
| 12810:4 | 69 | K.MVSHHN[+2204.772]LTTGATLINEQWLLTTAK.N | HexNAc(4)Hex(5)NeuAc(2) | N[+2205] | 1221.8003 | 4 | 4884.1795 | 4884.1647 | 0 | 3 | 179 | Specific | 543.78 | 499.19 | 499.19 | 7.09 | 2 | >sp\|P00738\|HPT_HUMAN Haptoglobin OS=Homo sapiens OX=9606 GN=HP PE=1 SV=1 | 407 | 20201118_12812_1ng_glyco_hcd30_etd30_compound15.16232.16232.4 | controllerType=0 controllerNumber=1 scan=16232 | 36.4863 |
| 12633:3 | 69 | K.VVLHPN[+2204.772]YSQVDIGLIK.L | HexNAc(4)Hex(5)NeuAc(2) | N[+2205] | 1333.9385 | 3 | 3999.801 | 3999.7837 | 0 | 4.3 | 236 | Specific | 286.01 | 189.44 | 189.44 | 4.15 | 2 | >sp\|P00738\|HPT_HUMAN Haptoglobin OS=Homo sapiens OX=9606 GN=HP PE=1 SV=1 | 407 | 20201118_12812_1ng_glyco_hcd30_etd30_compound15.16046.16046.3 | controllerType=0 controllerNumber=1 scan=16046 | 35.968 |
| 01787:3 | 71 | R.LRN[+1216.423]VSWATGR.S | HexNAc(2)Hex(5) | N[+1216] | 792.6936 | 3 | 2376.0663 | 2376.0559 | 0 | 4.4 | 149 | Specific | 302.12 | 166.27 | 166.27 | 2.68 | 2 | >sp\|P08571\|CD14_HUMAN Monocyte differentiation antigen CD14 OS=Homo sapiens OX=9606 GN=CD14 PE=1 SV=2 | 1232 | 20201118_12812_1ng_glyco_hcd30_etd30_compound15.4740.4740.3 | controllerType=0 controllerNumber=1 scan=4740 | 14.4257 |
| 19964:3 | 84 | R.AVN[+2204.772]ITSENLIDDVVSLIR.G | HexNAc(4)Hex(5)NeuAc(2) | N[+2205] | 1392.6263 | 3 | 4175.8643 | 4175.8481 | 0 | 3.9 | 413 | Specific | 288.76 | 167.26 | 167.26 | 5.14 | 1 | >sp\|P02748\|CO9_HUMAN Complement component C9 OS=Homo sapiens OX=9606 GN=C9 PE=1 SV=2 | 1547 | 20201118_12812_1ng_glyco_hcd30_etd30_compound15.23809.23809.3 | controllerType=0 controllerNumber=1 scan=23809 | 57.228 |
| 15979:4 | 1 | R.RTAGWNVPIGTLRPFLN[+2350.830]WTGPPEPIEAAVAR.F | HexNAc(4)Hex(5)Fuc(1)NeuAc(2) | N[+2351] | 1435.1704 | 4 | 5737.6598 | 5737.6424 | 0 | 3 | 140 | Specific | 718.25 | 718.25 | 718.25 | 12.03 | 36 | >sp\|P02788\|TRFL_HUMAN Lactotransferrin OS=Homo sapiens OX=9606 GN=LTF PE=1 SV=6 | 1270 | 20201118_12833_1ng_glyco_hcd30_etd30_compound15.19612.19612.4 | controllerType=0 controllerNumber=1 scan=19612 | 45.8691 |
| 15285:4 | 1 | R.RTAGWNVPIGTLRPFLN[+2351.851]WTGPPEPIEAAVAR.F | HexNAc(4)Hex(5)Fuc(3)NeuAc(1) | N[+2352] | 1435.4262 | 4 | 5738.6828 | 5738.6628 | 0 | 3.5 | 140 | Specific | 793.82 | 556.75 | 556.75 | 13.03 | 36 | >sp\|P02788\|TRFL_HUMAN Lactotransferrin OS=Homo sapiens OX=9606 GN=LTF PE=1 SV=6 | 1270 | 20201118_12833_1ng_glyco_hcd30_etd30_compound15.18878.18878.4 | controllerType=0 controllerNumber=1 scan=18878 | 43.516 |
| 15334:4 | 1 | R.RTAGWNVPIGTLRPFLN[+2059.735]WTGPPEPIEAAVAR.F | HexNAc(4)Hex(5)Fuc(1)NeuAc(1) | N[+2060] | 1362.3964 | 4 | 5446.5636 | 5446.547 | 0 | 3.1 | 140 | Specific | 1016.79 | 947.24 | 947.24 | 18.03 | 36 | >sp\|P02788\|TRFL_HUMAN Lactotransferrin OS=Homo sapiens OX=9606 GN=LTF PE=1 SV=6 | 1270 | 20201118_12833_1ng_glyco_hcd30_etd30_compound15.18930.18930.4 | controllerType=0 controllerNumber=1 scan=18930 | 43.6999 |
| 17048:3 | 1 | R.TAGWNVPIGTLRPFLN[+2350.830]WTGPPEPIEAAVAR.F | HexNAc(4)Hex(5)Fuc(1)NeuAc(2) | N[+2351] | 1861.1995 | 3 | 5581.5841 | 5581.5413 | 0 | 7.7 | 141 | Specific | 211.58 | 127.62 | 127.62 | 2.99 | 36 | >sp\|P02788\|TRFL_HUMAN Lactotransferrin OS=Homo sapiens OX=9606 GN=LTF PE=1 SV=6 | 1270 | 20201118_12833_1ng_glyco_hcd30_etd30_compound15.20748.20748.3 | controllerType=0 controllerNumber=1 scan=20748 | 49.3003 |
| 16117:3 | 1 | R.TAGWNVPIGTLRPFLN[+2205.793]WTGPPEPIEAAVAR.F | HexNAc(4)Hex(5)Fuc(2)NeuAc(1) | N[+2206] | 1812.8449 | 3 | 5436.5202 | 5436.5038 | 0 | 3 | 141 | Specific | 441.05 | 337.29 | 337.29 | 5.43 | 36 | >sp\|P02788\|TRFL_HUMAN Lactotransferrin OS=Homo sapiens OX=9606 GN=LTF PE=1 SV=6 | 1270 | 20201118_12833_1ng_glyco_hcd30_etd30_compound15.19758.19758.3 | controllerType=0 controllerNumber=1 scan=19758 | 46.3271 |
| 15472:3 | 1 | R.TAGWNVPIGTLRPFLN[+1768.640]WTGPPEPIEAAVAR.F | HexNAc(4)Hex(5)Fuc(1) | N[+1769] | 1667.129 | 3 | 4999.3724 | 4999.3504 | 0 | 4.4 | 141 | Specific | 491.82 | 414.27 | 414.27 | 6.72 | 36 | >sp\|P02788\|TRFL_HUMAN Lactotransferrin OS=Homo sapiens OX=9606 GN=LTF PE=1 SV=6 | 1270 | 20201118_12833_1ng_glyco_hcd30_etd30_compound15.19075.19075.3 | controllerType=0 controllerNumber=1 scan=19075 | 44.1253 |
| 16097:3 | 1 | R.TAGWNVPIGTLRPFLN[+2205.793]WTGPPEPIEAAVAR.F | HexNAc(4)Hex(5)Fuc(2)NeuAc(1) | N[+2206] | 1812.8471 | 3 | 5436.5268 | 5436.5038 | 0 | 4.2 | 141 | Specific | 517.42 | 408.9 | 408.9 | 7.87 | 36 | >sp\|P02788\|TRFL_HUMAN Lactotransferrin OS=Homo sapiens OX=9606 GN=LTF PE=1 SV=6 | 1270 | 20201118_12833_1ng_glyco_hcd30_etd30_compound15.19736.19736.3 | controllerType=0 controllerNumber=1 scan=19736 | 46.2451 |
| 16109:3 | 1 | R.TAGWNVPIGTLRPFLN[+2059.735]WTGPPEPIEAAVAR.F | HexNAc(4)Hex(5)Fuc(1)NeuAc(1) | N[+2060] | 1764.1611 | 3 | 5290.4687 | 5290.4459 | 0 | 4.3 | 141 | Specific | 552.77 | 446.99 | 446.99 | 6.86 | 36 | >sp\|P02788\|TRFL_HUMAN Lactotransferrin OS=Homo sapiens OX=9606 GN=LTF PE=1 SV=6 | 1270 | 20201118_12833_1ng_glyco_hcd30_etd30_compound15.19749.19749.3 | controllerType=0 controllerNumber=1 scan=19749 | 46.3001 |
| 16118:5 | 1 | R.TAGWNVPIGTLRPFLN[+2059.735]WTGPPEPIEAAVAR.F | HexNAc(4)Hex(5)Fuc(1)NeuAc(1) | N[+2060] | 1058.8987 | 5 | 5290.4643 | 5290.4459 | 0 | 3.5 | 141 | Specific | 748.13 | 707.89 | 707.89 | 10.89 | 36 | >sp\|P02788\|TRFL_HUMAN Lactotransferrin OS=Homo sapiens OX=9606 GN=LTF PE=1 SV=6 | 1270 | 20201118_12833_1ng_glyco_hcd30_etd30_compound15.19759.19759.5 | controllerType=0 controllerNumber=1 scan=19759 | 46.3302 |
| 16091:4 | 1 | R.TAGWNVPIGTLRPFLN[+2205.793]WTGPPEPIEAAVAR.F | HexNAc(4)Hex(5)Fuc(2)NeuAc(1) | N[+2206] | 1359.8873 | 4 | 5436.5275 | 5436.5038 | 0 | 4.4 | 141 | Specific | 874.85 | 855.53 | 855.53 | 14.38 | 36 | >sp\|P02788\|TRFL_HUMAN Lactotransferrin OS=Homo sapiens OX=9606 GN=LTF PE=1 SV=6 | 1270 | 20201118_12833_1ng_glyco_hcd30_etd30_compound15.19730.19730.4 | controllerType=0 controllerNumber=1 scan=19730 | 46.2202 |

Table S4. Cont.

| Query #:z | Protein Rank | Glycopeptide | Glycan | Modification Type(s) | Observed m/z | z | Observed (M+H) | Calc. mass (M+H) | Off-by-x error | Mass error (ppm) | StartPosition | Cleavage | Score | Delta | Delta Mod | \|Log Prob\| | # of unique peptides | Protein Name | Protein DB number | Comment | Scan # | Scan Time |
|---|---|---|---|---|---|---|---|---|---|---|---|---|---|---|---|---|---|---|---|---|---|---|
| 16101:4 | 1 | R.TAGWNVPIGTLRPFLN[+2059.735]WTGPPEPIEAAVAR.F | HexNAc(4)Hex(5)Fuc(1)NeuAc(1) | N[+2060] | 1323.3726 | 4 | 5290.4684 | 5290.4459 | 0 | 4.3 | 141 | Specific | 942.77 | 937.38 | 937.38 | 15.38 | 36 | >sp\|P02788\|TRFL_HUMAN Lactotransferrin OS=Homo sapiens OX=9606 GN=LTF PE=1 SV=6 | 1270 | 20201118_12833_1ng_glyco_hcd30_etd30_compound15.19741.19741.4 | controllerType=0 controllerNumber=1 scan=19741 | 46.2674 |
| 15473:4 | 1 | R.TAGWNVPIGTLRPFLN[+1768.640]WTGPPEPIEAAVAR.F | HexNAc(4)Hex(5)Fuc(1) | N[+1769] | 1250.5984 | 4 | 4999.3717 | 4999.3504 | 0 | 4.3 | 141 | Specific | 1052.92 | 1033.34 | 1033.34 | 18.71 | 36 | >sp\|P02788\|TRFL_HUMAN Lactotransferrin OS=Homo sapiens OX=9606 GN=LTF PE=1 SV=6 | 1270 | 20201118_12833_1ng_glyco_hcd30_etd30_compound15.19076.19076.4 | controllerType=0 controllerNumber=1 scan=19076 | 44.1321 |
| 14422:3 | 1 | R.TAGWNIPMGLLFN[+1768.640]QTGSC[+57.021]K.F | HexNAc(4)Hex(5)Fuc(1) | C[+57], N[+1769] | 1288.5569 | 3 | 3863.6563 | 3863.6495 | 0 | 1.8 | 485 | Specific | 384.97 | 305.09 | 305.09 | 4.86 | 36 | >sp\|P02788\|TRFL_HUMAN Lactotransferrin OS=Homo sapiens OX=9606 GN=LTF PE=1 SV=6 | 1270 | 20201118_12833_1ng_glyco_hcd30_etd30_compound15.17965.17965.3 | controllerType=0 controllerNumber=1 scan=17965 | 40.6804 |
| 15683:3 | 1 | R.TAGWNIPMGLLFN[+2059.735]QTGSC[+57.021]K.F | HexNAc(4)Hex(5)Fuc(1)NeuAc(1) | C[+57], N[+2060] | 1385.5887 | 3 | 4154.7515 | 4154.7449 | 0 | 1.6 | 485 | Specific | 403.73 | 258.59 | 258.59 | 3.91 | 36 | >sp\|P02788\|TRFL_HUMAN Lactotransferrin OS=Homo sapiens OX=9606 GN=LTF PE=1 SV=6 | 1270 | 20201118_12833_1ng_glyco_hcd30_etd30_compound15.19299.19299.3 | controllerType=0 controllerNumber=1 scan=19299 | 44.8486 |
| 15674:3 | 1 | R.TAGWNIPMGLLFN[+2205.793]QTGSC[+57.021]K.F | HexNAc(4)Hex(5)Fuc(2)NeuAc(1) | C[+57], N[+2206] | 1434.2805 | 3 | 4300.8268 | 4300.8028 | 0 | 5.6 | 485 | Specific | 422.69 | 325.88 | 325.88 | 4.62 | 36 | >sp\|P02788\|TRFL_HUMAN Lactotransferrin OS=Homo sapiens OX=9606 GN=LTF PE=1 SV=6 | 1270 | 20201118_12833_1ng_glyco_hcd30_etd30_compound15.19289.19289.3 | controllerType=0 controllerNumber=1 scan=19289 | 44.8049 |
| 15803:3 | 1 | R.TAGWNIPMGLLFN[+1913.677]QTGSC[+57.021]K.F | HexNAc(4)Hex(5)NeuAc(1) | C[+57], N[+1914] | 1336.9048 | 3 | 4008.7 | 4008.687 | 0 | 3.2 | 485 | Specific | 458.22 | 309.28 | 309.28 | 5.2 | 36 | >sp\|P02788\|TRFL_HUMAN Lactotransferrin OS=Homo sapiens OX=9606 GN=LTF PE=1 SV=6 | 1270 | 20201118_12833_1ng_glyco_hcd30_etd30_compound15.19426.19426.3 | controllerType=0 controllerNumber=1 scan=19426 | 45.2719 |
| 15883:5 | 1 | R.TAGWNIPMGLLFN[+2059.735]QTGSC[+57.021]KFDEYFSQSC[+57.021]APGSDPR.S | HexNAc(4)Hex(5)Fuc(1)NeuAc(1) | C[+57]*2, N[+2060] | 1200.5079 | 5 | 5998.5106 | 5998.492 | 0 | 3.1 | 485 | Specific | 329.94 | 309.84 | 309.84 | 3.71 | 36 | >sp\|P02788\|TRFL_HUMAN Lactotransferrin OS=Homo sapiens OX=9606 GN=LTF PE=1 SV=6 | 1270 | 20201118_12833_1ng_glyco_hcd30_etd30_compound15.19511.19511.5 | controllerType=0 controllerNumber=1 scan=19511 | 45.5734 |
| 15912:3 | 1 | R.TAGWNIPMGLLFN[+1913.677]QTGSC[+57.021]KFDEYFSQSC[+57.021]APGSDPR.S | HexNAc(4)Hex(5)NeuAc(1) | C[+57]*2, N[+1914] | 1951.4893 | 3 | 5852.4535 | 5852.4341 | 0 | 3.3 | 485 | Specific | 384.63 | 310.8 | 310.8 | 5.46 | 36 | >sp\|P02788\|TRFL_HUMAN Lactotransferrin OS=Homo sapiens OX=9606 GN=LTF PE=1 SV=6 | 1270 | 20201118_12833_1ng_glyco_hcd30_etd30_compound15.19541.19541.3 | controllerType=0 controllerNumber=1 scan=19541 | 45.6582 |
| 15903:4 | 1 | R.TAGWNIPMGLLFN[+1913.677]QTGSC[+57.021]KFDEYFSQSC[+57.021]APGSDPR.S | HexNAc(4)Hex(5)NeuAc(1) | C[+57]*2, N[+1914] | 1463.8672 | 4 | 5852.4469 | 5852.4341 | 0 | 2.2 | 485 | Specific | 424.77 | 306.93 | 306.93 | 4.54 | 36 | >sp\|P02788\|TRFL_HUMAN Lactotransferrin OS=Homo sapiens OX=9606 GN=LTF PE=1 SV=6 | 1270 | 20201118_12833_1ng_glyco_hcd30_etd30_compound15.19532.19532.4 | controllerType=0 controllerNumber=1 scan=19532 | 45.6361 |
| 15845:4 | 1 | R.TAGWNIPMGLLFN[+2205.793]QTGSC[+57.021]KFDEYFSQSC[+57.021]APGSDPR.S | HexNAc(4)Hex(5)Fuc(2)NeuAc(1) | C[+57]*2, N[+2206] | 1536.9004 | 4 | 6144.5797 | 6144.5499 | 0 | 4.8 | 485 | Specific | 502.54 | 482.18 | 482.18 | 6.76 | 36 | >sp\|P02788\|TRFL_HUMAN Lactotransferrin OS=Homo sapiens OX=9606 GN=LTF PE=1 SV=6 | 1270 | 20201118_12833_1ng_glyco_hcd30_etd30_compound15.19471.19471.4 | controllerType=0 controllerNumber=1 scan=19471 | 45.439 |
| 15180:3 | 1 | R.TAGWNIPMGLLFN[+1768.640]QTGSC[+57.021]KFDEYFSQSC[+57.021]APGSDPR.S | HexNAc(4)Hex(5)Fuc(1) | C[+57]*2, N[+1769] | 1903.1485 | 3 | 5707.431 | 5707.3966 | 0 | 6 | 485 | Specific | 521.21 | 447.99 | 447.99 | 6.61 | 36 | >sp\|P02788\|TRFL_HUMAN Lactotransferrin OS=Homo sapiens OX=9606 GN=LTF PE=1 SV=6 | 1270 | 20201118_12833_1ng_glyco_hcd30_etd30_compound15.18767.18767.3 | controllerType=0 controllerNumber=1 scan=18767 | 43.1793 |
| 15882:4 | 1 | R.TAGWNIPMGLLFN[+2059.735]QTGSC[+57.021]KFDEYFSQSC[+57.021]APGSDPR.S | HexNAc(4)Hex(5)Fuc(1)NeuAc(1) | C[+57]*2, N[+2060] | 1500.3872 | 4 | 5998.527 | 5998.492 | 0 | 5.8 | 485 | Specific | 537.95 | 501.13 | 501.13 | 6.29 | 36 | >sp\|P02788\|TRFL_HUMAN Lactotransferrin OS=Homo sapiens OX=9606 GN=LTF PE=1 SV=6 | 1270 | 20201118_12833_1ng_glyco_hcd30_etd30_compound15.19510.19510.4 | controllerType=0 controllerNumber=1 scan=19510 | 45.5705 |
| 15223:4 | 1 | R.TAGWNIPMGLLFN[+1622.582]QTGSC[+57.021]KFDEYFSQSC[+57.021]APGSDPR.S | HexNAc(4)Hex(5) | C[+57]*2, N[+1623] | 1391.0955 | 4 | 5561.36 | 5561.3387 | 0 | 3.8 | 485 | Specific | 626.33 | 556.34 | 138.29 | 8.14 | 36 | >sp\|P02788\|TRFL_HUMAN Lactotransferrin OS=Homo sapiens OX=9606 GN=LTF PE=1 SV=6 | 1270 | 20201118_12833_1ng_glyco_hcd30_etd30_compound15.18813.18813.4 | controllerType=0 controllerNumber=1 scan=18813 | 43.3453 |
| 16773:4 | 1 | R.TAGWNIPMGLLFN[+2350.830]QTGSC[+57.021]KFDEYFSQSC[+57.021]APGSDPR.S | HexNAc(4)Hex(5)Fuc(1)NeuAc(2) | C[+57]*2, N[+2351] | 1573.1586 | 4 | 6289.6125 | 6289.5875 | 0 | 4 | 485 | Specific | 684.69 | 633.91 | 633.91 | 10.26 | 36 | >sp\|P02788\|TRFL_HUMAN Lactotransferrin OS=Homo sapiens OX=9606 GN=LTF PE=1 SV=6 | 1270 | 20201118_12833_1ng_glyco_hcd30_etd30_compound15.20456.20456.4 | controllerType=0 controllerNumber=1 scan=20456 | 48.3483 |
| 15173:4 | 1 | R.TAGWNIPMGLLFN[+1768.640]QTGSC[+57.021]KFDEYFSQSC[+57.021]APGSDPR.S | HexNAc(4)Hex(5)Fuc(1) | C[+57]*2, N[+1769] | 1427.6158 | 4 | 5707.4416 | 5707.3966 | 0 | 7.9 | 485 | Specific | 828.16 | 752.86 | 752.86 | 12.37 | 36 | >sp\|P02788\|TRFL_HUMAN Lactotransferrin OS=Homo sapiens OX=9606 GN=LTF PE=1 SV=6 | 1270 | 20201118_12833_1ng_glyco_hcd30_etd30_compound15.18760.18760.4 | controllerType=0 controllerNumber=1 scan=18760 | 43.1651 |
| 15220:4 | 1 | R.TAGWNIPM[+15.995]GLLFN[+2059.735]QTGSC[+57.021]KFDEYFSQSC[+57.021]A | HexNAc(4)Hex(5)Fuc(1)NeuAc(1) | C[+57]*2, M[+16], N[+2060] | 1504.3851 | 4 | 6014.5187 | 6014.487 | 0 | 5.3 | 485 | Specific | 417.54 | 328.98 | 328.98 | 4.96 | 36 | >sp\|P02788\|TRFL_HUMAN Lactotransferrin OS=Homo sapiens OX=9606 GN=LTF PE=1 SV=6 | 1270 | 20201118_12833_1ng_glyco_hcd30_etd30_compound15.18809.18809.4 | controllerType=0 controllerNumber=1 scan=18809 | 43.3345 |
| 07242:3 | 1 | K.FGRN[+2205.793]GSDC[+57.021]PDKFC[+57.021]LFQSETK.N | HexNAc(4)Hex(5)Fuc(2)NeuAc(1) | C[+57]*2, N[+2206] | 1533.6277 | 3 | 4598.8687 | 4598.8577 | 0 | 2.4 | 639 | Specific | 234.22 | 159.84 | 159.84 | 3.51 | 36 | >sp\|P02788\|TRFL_HUMAN Lactotransferrin OS=Homo sapiens OX=9606 GN=LTF PE=1 SV=6 | 1270 | 20201118_12833_1ng_glyco_hcd30_etd30_compound15.10419.10419.3 | controllerType=0 controllerNumber=1 scan=10419 | 24.8151 |
| 07312:4 | 1 | K.FGRN[+2059.735]GSDC[+57.021]PDKFC[+57.021]LFQSETK.N | HexNAc(4)Hex(5)Fuc(1)NeuAc(1) | C[+57]*2, N[+2060] | 1113.9572 | 4 | 4452.807 | 4452.7998 | 0 | 1.6 | 639 | Specific | 469.05 | 303.06 | 303.06 | 5.39 | 36 | >sp\|P02788\|TRFL_HUMAN Lactotransferrin OS=Homo sapiens OX=9606 GN=LTF PE=1 SV=6 | 1270 | 20201118_12833_1ng_glyco_hcd30_etd30_compound15.10492.10492.4 | controllerType=0 controllerNumber=1 scan=10492 | 24.9653 |
| 08066:3 | 1 | R.N[+2205.793]GSDC[+57.021]PDKFC[+57.021]LFQSETK.N | HexNAc(4)Hex(5)Fuc(2)NeuAc(1) | C[+57]*2, N[+2206] | 1413.5527 | 3 | 4238.6435 | 4238.6667 | 0 | -5.5 | 642 | Specific | 215.1 | 151.18 | 151.18 | 2.82 | 36 | >sp\|P02788\|TRFL_HUMAN Lactotransferrin OS=Homo sapiens OX=9606 GN=LTF PE=1 SV=6 | 1270 | 20201118_12833_1ng_glyco_hcd30_etd30_compound15.11282.11282.3 | controllerType=0 controllerNumber=1 scan=11282 | 26.5257 |
| 08093:3 | 1 | R.N[+2205.793]GSDC[+57.021]PDKFC[+57.021]LFQSETK.N | HexNAc(4)Hex(5)Fuc(2)NeuAc(1) | C[+57]*2, N[+2206] | 1413.5645 | 3 | 4238.679 | 4238.6667 | 0 | 2.9 | 642 | Specific | 255.1 | 189.31 | 189.31 | 4.4 | 36 | >sp\|P02788\|TRFL_HUMAN Lactotransferrin OS=Homo sapiens OX=9606 GN=LTF PE=1 SV=6 | 1270 | 20201118_12833_1ng_glyco_hcd30_etd30_compound15.11311.11311.3 | controllerType=0 controllerNumber=1 scan=11311 | 26.599 |
| 08079:3 | 1 | R.N[+2205.793]GSDC[+57.021]PDKFC[+57.021]LFQSETK.N | HexNAc(4)Hex(5)Fuc(2)NeuAc(1) | C[+57]*2, N[+2206] | 1413.5628 | 3 | 4238.6739 | 4238.6667 | 0 | 1.7 | 642 | Specific | 278.94 | 177.08 | 177.08 | 4.45 | 36 | >sp\|P02788\|TRFL_HUMAN Lactotransferrin OS=Homo sapiens OX=9606 GN=LTF PE=1 SV=6 | 1270 | 20201118_12833_1ng_glyco_hcd30_etd30_compound15.11296.11296.3 | controllerType=0 controllerNumber=1 scan=11296 | 26.561 |
| 15085:3 | 2 | R.LSLHRPALEDLLLGSEAN[+1257.449]LTC[+57.021]TLTGLR.D | HexNAc(3)Hex(4) | C[+57], N[+1257] | 1407.6921 | 3 | 4221.0617 | 4221.0477 | 0 | 3.3 | 127 | Specific | 287.37 | 280.41 | 280.41 | 5.37 | 30 | >sp\|P01876\|IGHA1_HUMAN Immunoglobulin heavy constant alpha 1 OS=Homo sapiens OX=9606 GN=IGHA1 PE=1 SV=2 | 255 | 20201118_12833_1ng_glyco_hcd30_etd30_compound15.18666.18666.3 | controllerType=0 controllerNumber=1 scan=18666 | 42.8559 |
| 15100:3 | 2 | R.LSLHRPALEDLLLGSEAN[+1298.476]LTC[+57.021]TLTGLR.D | HexNAc(4)Hex(3) | C[+57], N[+1298] | 1421.3693 | 3 | 4262.0934 | 4262.0742 | 0 | 4.5 | 127 | Specific | 363.94 | 363.94 | 363.94 | 5.46 | 30 | >sp\|P01876\|IGHA1_HUMAN Immunoglobulin heavy constant alpha 1 OS=Homo sapiens OX=9606 GN=IGHA1 PE=1 SV=2 | 255 | 20201118_12833_1ng_glyco_hcd30_etd30_compound15.18682.18682.3 | controllerType=0 controllerNumber=1 scan=18682 | 42.9173 |
| 15049:3 | 2 | R.LSLHRPALEDLLLGSEAN[+1378.476]LTC[+57.021]TLTGLR.D | HexNAc(2)Hex(6) | C[+57], N[+1378] | 1448.0344 | 3 | 4342.0885 | 4342.074 | 0 | 3.4 | 127 | Specific | 379.5 | 373.05 | 373.05 | 7 | 30 | >sp\|P01876\|IGHA1_HUMAN Immunoglobulin heavy constant alpha 1 OS=Homo sapiens OX=9606 GN=IGHA1 PE=1 SV=2 | 255 | 20201118_12833_1ng_glyco_hcd30_etd30_compound15.18628.18628.3 | controllerType=0 controllerNumber=1 scan=18628 | 42.7303 |
| 15055:3 | 2 | R.LSLHRPALEDLLLGSEAN[+1501.555]LTC[+57.021]TLTGLR.D | HexNAc(5)Hex(3) | C[+57], N[+1502] | 1489.0606 | 3 | 4465.1673 | 4465.1536 | 0 | 3.1 | 127 | Specific | 383.68 | 348.38 | 348.38 | 5.79 | 30 | >sp\|P01876\|IGHA1_HUMAN Immunoglobulin heavy constant alpha 1 OS=Homo sapiens OX=9606 GN=IGHA1 PE=1 SV=2 | 255 | 20201118_12833_1ng_glyco_hcd30_etd30_compound15.18634.18634.3 | controllerType=0 controllerNumber=1 scan=18634 | 42.7541 |
| 15008:3 | 2 | R.LSLHRPALEDLLLGSEAN[+1702.581]LTC[+57.021]TLTGLR.D | HexNAc(2)Hex(8) | C[+57], N[+1703] | 1556.0674 | 3 | 4666.1878 | 4666.1796 | 0 | 1.8 | 127 | Specific | 399.89 | 361.66 | 361.66 | 5.84 | 30 | >sp\|P01876\|IGHA1_HUMAN Immunoglobulin heavy constant alpha 1 OS=Homo sapiens OX=9606 GN=IGHA1 PE=1 SV=2 | 255 | 20201118_12833_1ng_glyco_hcd30_etd30_compound15.18584.18584.3 | controllerType=0 controllerNumber=1 scan=18584 | 42.5495 |
| 15074:5 | 2 | R.LSLHRPALEDLLLGSEAN[+1524.534]LTC[+57.021]TLTGLR.D | HexNAc(2)Hex(6)Fuc(1) | C[+57], N[+1525] | 898.4302 | 5 | 4488.1219 | 4488.1319 | 0 | -2.2 | 127 | Specific | 436.33 | 436.33 | 436.33 | 3.96 | 30 | >sp\|P01876\|IGHA1_HUMAN Immunoglobulin heavy constant alpha 1 OS=Homo sapiens OX=9606 GN=IGHA1 PE=1 SV=2 | 255 | 20201118_12833_1ng_glyco_hcd30_etd30_compound15.18655.18655.5 | controllerType=0 controllerNumber=1 scan=18655 | 42.8197 |
| 15033:3 | 2 | R.LSLHRPALEDLLLGSEAN[+1663.608]LTC[+57.021]TLTGLR.D | HexNAc(5)Hex(4) | C[+57], N[+1664] | 1543.0781 | 3 | 4627.2196 | 4627.2064 | 0 | 2.9 | 127 | Specific | 439.86 | 387.96 | 387.96 | 7.15 | 30 | >sp\|P01876\|IGHA1_HUMAN Immunoglobulin heavy constant alpha 1 OS=Homo sapiens OX=9606 GN=IGHA1 PE=1 SV=2 | 255 | 20201118_12833_1ng_glyco_hcd30_etd30_compound15.18611.18611.3 | controllerType=0 controllerNumber=1 scan=18611 | 42.6493 |
| 15723:4 | 2 | R.LSLHRPALEDLLLGSEAN[+1710.598]LTC[+57.021]TLTGLR.D | HexNAc(3)Hex(5)NeuAc(1) | C[+57], N[+1711] | 1169.3088 | 4 | 4674.2132 | 4674.1959 | 0 | 3.7 | 127 | Specific | 455.47 | 455.47 | 455.47 | 7.76 | 30 | >sp\|P01876\|IGHA1_HUMAN Immunoglobulin heavy constant alpha 1 OS=Homo sapiens OX=9606 GN=IGHA1 PE=1 SV=2 | 255 | 20201118_12833_1ng_glyco_hcd30_etd30_compound15.19341.19341.4 | controllerType=0 controllerNumber=1 scan=19341 | 44.9689 |
| 15060:3 | 2 | R.LSLHRPALEDLLLGSEAN[+1216.423]LTC[+57.021]TLTGLR.D | HexNAc(2)Hex(5) | C[+57], N[+1216] | 1394.0148 | 3 | 4180.0299 | 4180.0211 | 0 | 2.1 | 127 | Specific | 510.89 | 497.93 | 497.93 | 7.56 | 30 | >sp\|P01876\|IGHA1_HUMAN Immunoglobulin heavy constant alpha 1 OS=Homo sapiens OX=9606 GN=IGHA1 PE=1 SV=2 | 255 | 20201118_12833_1ng_glyco_hcd30_etd30_compound15.18640.18640.3 | controllerType=0 controllerNumber=1 scan=18640 | 42.7716 |
| 15018:4 | 2 | R.LSLHRPALEDLLLGSEAN[+1864.634]LTC[+57.021]TLTGLR.D | HexNAc(2)Hex(9) | C[+57], N[+1865] | 1207.8178 | 4 | 4828.2493 | 4828.2324 | 0 | 3.5 | 127 | Specific | 530.91 | 530.91 | 530.91 | 7.19 | 30 | >sp\|P01876\|IGHA1_HUMAN Immunoglobulin heavy constant alpha 1 OS=Homo sapiens OX=9606 GN=IGHA1 PE=1 SV=2 | 255 | 20201118_12833_1ng_glyco_hcd30_etd30_compound15.18595.18595.4 | controllerType=0 controllerNumber=1 scan=18595 | 42.5945 |
| 15120:4 | 2 | R.LSLHRPALEDLLLGSEAN[+1460.529]LTC[+57.021]TLTGLR.D | HexNAc(4)Hex(4) | C[+57], N[+1461] | 1106.7912 | 4 | 4424.1429 | 4424.1271 | 0 | 3.6 | 127 | Specific | 555.86 | 555.86 | 555.86 | 8.01 | 30 | >sp\|P01876\|IGHA1_HUMAN Immunoglobulin heavy constant alpha 1 OS=Homo sapiens OX=9606 GN=IGHA1 PE=1 SV=2 | 255 | 20201118_12833_1ng_glyco_hcd30_etd30_compound15.18703.18703.4 | controllerType=0 controllerNumber=1 scan=18703 | 42.9751 |
| 15066:4 | 2 | R.LSLHRPALEDLLLGSEAN[+1622.582]LTC[+57.021]TLTGLR.D | HexNAc(4)Hex(5) | C[+57], N[+1623] | 1147.3022 | 4 | 4586.1868 | 4586.1799 | 0 | 1.5 | 127 | Specific | 673.62 | 673.62 | 673.62 | 11.09 | 30 | >sp\|P01876\|IGHA1_HUMAN Immunoglobulin heavy constant alpha 1 OS=Homo sapiens OX=9606 GN=IGHA1 PE=1 SV=2 | 255 | 20201118_12833_1ng_glyco_hcd30_etd30_compound15.18646.18646.4 | controllerType=0 controllerNumber=1 scan=18646 | 42.7904 |
| 15075:4 | 2 | R.LSLHRPALEDLLLGSEAN[+1524.534]LTC[+57.021]TLTGLR.D | HexNAc(2)Hex(6)Fuc(1) | C[+57], N[+1525] | 1122.7843 | 4 | 4488.1155 | 4488.1319 | 0 | -3.6 | 127 | Specific | 721.27 | 721.27 | 721.27 | 10.84 | 30 | >sp\|P01876\|IGHA1_HUMAN Immunoglobulin heavy constant alpha 1 OS=Homo sapiens OX=9606 GN=IGHA1 PE=1 SV=2 | 255 | 20201118_12833_1ng_glyco_hcd30_etd30_compound15.18656.18656.4 | controllerType=0 controllerNumber=1 scan=18656 | 42.8243 |
| 15080:4 | 2 | R.LSLHRPALEDLLLGSEAN[+1257.449]LTC[+57.021]TLTGLR.D | HexNAc(3)Hex(4) | C[+57], N[+1257] | 1056.0187 | 4 | 4221.053 | 4221.0477 | 0 | 1.3 | 127 | Specific | 754.5 | 754.5 | 754.5 | 12.32 | 30 | >sp\|P01876\|IGHA1_HUMAN Immunoglobulin heavy constant alpha 1 OS=Homo sapiens OX=9606 GN=IGHA1 PE=1 SV=2 | 255 | 20201118_12833_1ng_glyco_hcd30_etd30_compound15.18661.18661.4 | controllerType=0 controllerNumber=1 scan=18661 | 42.8386 |
| 15045:4 | 2 | R.LSLHRPALEDLLLGSEAN[+1622.582]LTC[+57.021]TLTGLR.D | HexNAc(4)Hex(5) | C[+57], N[+1623] | 1147.3033 | 4 | 4586.1912 | 4586.1799 | 0 | 2.5 | 127 | Specific | 843 | 773.62 | 773.62 | 14.09 | 30 | >sp\|P01876\|IGHA1_HUMAN Immunoglobulin heavy constant alpha 1 OS=Homo sapiens OX=9606 GN=IGHA1 PE=1 SV=2 | 255 | 20201118_12833_1ng_glyco_hcd30_etd30_compound15.18624.18624.4 | controllerType=0 controllerNumber=1 scan=18624 | 42.7216 |
| 15003:4 | 2 | R.LSLHRPALEDLLLGSEAN[+1702.581]LTC[+57.021]TLTGLR.D | HexNAc(2)Hex(8) | C[+57], N[+1703] | 1167.3033 | 4 | 4666.1912 | 4666.1796 | 0 | 2.5 | 127 | Specific | 847.23 | 847.23 | 847.23 | 14.09 | 30 | >sp\|P01876\|IGHA1_HUMAN Immunoglobulin heavy constant alpha 1 OS=Homo sapiens OX=9606 GN=IGHA1 PE=1 SV=2 | 255 | 20201118_12833_1ng_glyco_hcd30_etd30_compound15.18579.18579.4 | controllerType=0 controllerNumber=1 scan=18579 | 42.5378 |
| 15032:4 | 2 | R.LSLHRPALEDLLLGSEAN[+1663.608]LTC[+57.021]TLTGLR.D | HexNAc(5)Hex(4) | C[+57], N[+1664] | 1157.5608 | 4 | 4627.2215 | 4627.2064 | 0 | 3.3 | 127 | Specific | 929.34 | 929.34 | 929.34 | 16.03 | 30 | >sp\|P01876\|IGHA1_HUMAN Immunoglobulin heavy constant alpha 1 OS=Homo sapiens OX=9606 GN=IGHA1 PE=1 SV=2 | 255 | 20201118_12833_1ng_glyco_hcd30_etd30_compound15.18610.18610.4 | controllerType=0 controllerNumber=1 scan=18610 | 42.6461 |
| 14970:4 | 2 | R.LSLHRPALEDLLLGSEAN[+1864.634]LTC[+57.021]TLTGLR.D | HexNAc(2)Hex(9) | C[+57], N[+1865] | 1207.8172 | 4 | 4828.2469 | 4828.2324 | 0 | 3 | 127 | Specific | 940.9 | 904.87 | 904.87 | 16.03 | 30 | >sp\|P01876\|IGHA1_HUMAN Immunoglobulin heavy constant alpha 1 OS=Homo sapiens OX=9606 GN=IGHA1 PE=1 SV=2 | 255 | 20201118_12833_1ng_glyco_hcd30_etd30_compound15.18543.18543.4 | controllerType=0 controllerNumber=1 scan=18543 | 42.4036 |
| 15069:4 | 2 | R.LSLHRPALEDLLLGSEAN[+1298.476]LTC[+57.021]TLTGLR.D | HexNAc(4)Hex(3) | C[+57], N[+1298] | 1066.2767 | 4 | 4262.0848 | 4262.0742 | 0 | 2.5 | 127 | Specific | 958.99 | 958.99 | 958.99 | 17.09 | 30 | >sp\|P01876\|IGHA1_HUMAN Immunoglobulin heavy constant alpha 1 OS=Homo sapiens OX=9606 GN=IGHA1 PE=1 SV=2 | 255 | 20201118_12833_1ng_glyco_hcd30_etd30_compound15.18649.18649.4 | controllerType=0 controllerNumber=1 scan=18649 | 42.7998 |
| 15044:4 | 2 | R.LSLHRPALEDLLLGSEAN[+1501.555]LTC[+57.021]TLTGLR.D | HexNAc(5)Hex(3) | C[+57], N[+1502] | 1117.049 | 4 | 4465.1741 | 4465.1536 | 0 | 4.6 | 127 | Specific | 963.34 | 955.34 | 955.34 | 16.7 | 30 | >sp\|P01876\|IGHA1_HUMAN Immunoglobulin heavy constant alpha 1 OS=Homo sapiens OX=9606 GN=IGHA1 PE=1 SV=2 | 255 | 20201118_12833_1ng_glyco_hcd30_etd30_compound15.18623.18623.4 | controllerType=0 controllerNumber=1 scan=18623 | 42.7188 |
| 15017:4 | 2 | R.LSLHRPALEDLLLGSEAN[+1540.529]LTC[+57.021]TLTGLR.D | HexNAc(2)Hex(7) | C[+57], N[+1541] | 1126.7917 | 4 | 4504.1448 | 4504.1268 | 0 | 4 | 127 | Specific | 982.43 | 927.37 | 927.37 | 17.03 | 30 | >sp\|P01876\|IGHA1_HUMAN Immunoglobulin heavy constant alpha 1 OS=Homo sapiens OX=9606 GN=IGHA1 PE=1 SV=2 | 255 | 20201118_12833_1ng_glyco_hcd30_etd30_compound15.18594.18594.4 | controllerType=0 controllerNumber=1 scan=18594 | 42.5911 |
| 15050:4 | 2 | R.LSLHRPALEDLLLGSEAN[+1216.423]LTC[+57.021]TLTGLR.D | HexNAc(2)Hex(5) | C[+57], N[+1216] | 1045.7636 | 4 | 4180.0325 | 4180.0211 | 0 | 2.7 | 127 | Specific | 1014.57 | 976.14 | 976.14 | 18.09 | 30 | >sp\|P01876\|IGHA1_HUMAN Immunoglobulin heavy constant alpha 1 OS=Homo sapiens OX=9606 GN=IGHA1 PE=1 SV=2 | 255 | 20201118_12833_1ng_glyco_hcd30_etd30_compound15.18629.18629.4 | controllerType=0 controllerNumber=1 scan=18629 | 42.737 |
| 15056:4 | 2 | R.LSLHRPALEDLLLGSEAN[+1460.529]LTC[+57.021]TLTGLR.D | HexNAc(4)Hex(4) | C[+57], N[+1461] | 1106.7934 | 4 | 4424.1517 | 4424.1271 | 0 | 5.6 | 127 | Specific | 1016.92 | 958.95 | 958.95 | 17.7 | 30 | >sp\|P01876\|IGHA1_HUMAN Immunoglobulin heavy constant alpha 1 OS=Homo sapiens OX=9606 GN=IGHA1 PE=1 SV=2 | 255 | 20201118_12833_1ng_glyco_hcd30_etd30_compound15.18635.18635.4 | controllerType=0 controllerNumber=1 scan=18635 | 42.7573 |
| 15022:4 | 2 | R.LSLHRPALEDLLLGSEAN[+1378.476]LTC[+57.021]TLTGLR.D | HexNAc(2)Hex(6) | C[+57], N[+1378] | 1086.2784 | 4 | 4342.0916 | 4342.074 | 0 | 4.1 | 127 | Specific | 1038.24 | 1000.46 | 1000.46 | 18.03 | 30 | >sp\|P01876\|IGHA1_HUMAN Immunoglobulin heavy constant alpha 1 OS=Homo sapiens OX=9606 GN=IGHA1 PE=1 SV=2 | 255 | 20201118_12833_1ng_glyco_hcd30_etd30_compound15.18599.18599.4 | controllerType=0 controllerNumber=1 scan=18599 | 42.6099 |
| 10142:3 | 2 | R.LAGKPTHVN[+1663.608]VSVVMAEVDGTC[+57.021]Y.- | HexNAc(5)Hex(4) | C[+57], N[+1664] | 1337.5911 | 3 | 4010.7588 | 4010.7502 | 0 | 2.1 | 332 | CRagged | 203.27 | 113.41 | 113.41 | 2.75 | 30 | >sp\|P01876\|IGHA1_HUMAN Immunoglobulin heavy constant alpha 1 OS=Homo sapiens OX=9606 GN=IGHA1 PE=1 SV=2 | 255 | 20201118_12833_1ng_glyco_hcd30_etd30_compound15.13440.13440.3 | controllerType=0 controllerNumber=1 scan=13440 | 30.7058 |
| 10059:3 | 2 | R.LAGKPTHVN[+1702.581]VSVVMAEVDGTC[+57.021]Y.- | HexNAc(2)Hex(8) | C[+57], N[+1703] | 1350.5822 | 3 | 4049.7321 | 4049.7234 | 0 | 2.1 | 332 | CRagged | 226.2 | 175.43 | 160.37 | 3.02 | 30 | >sp\|P01876\|IGHA1_HUMAN Immunoglobulin heavy constant alpha 1 OS=Homo sapiens OX=9606 GN=IGHA1 PE=1 SV=2 | 255 | 20201118_12833_1ng_glyco_hcd30_etd30_compound15.13352.13352.3 | controllerType=0 controllerNumber=1 scan=13352 | 30.4587 |
| 11681:3 | 2 | R.LAGKPTHVN[+1751.624]VSVVMAEVDGTC[+57.021]Y.- | HexNAc(4)Hex(4)NeuAc(1) | C[+57], N[+1752] | 1366.9312 | 3 | 4098.7791 | 4098.7663 | 0 | 3.1 | 332 | CRagged | 238.68 | 124.81 | 124.81 | 2.72 | 30 | >sp\|P01876\|IGHA1_HUMAN Immunoglobulin heavy constant alpha 1 OS=Homo sapiens OX=9606 GN=IGHA1 PE=1 SV=2 | 255 | 20201118_12833_1ng_glyco_hcd30_etd30_compound15.15056.15056.3 | controllerType=0 controllerNumber=1 scan=15056 | 33.9395 |
| 10176:3 | 2 | R.LAGKPTHVN[+1501.555]VSVVMAEVDGTC[+57.021]Y.- | HexNAc(5)Hex(3) | C[+57], N[+1502] | 1283.5722 | 3 | 3848.702 | 3848.6974 | 0 | 1.2 | 332 | CRagged | 240.1 | 157.22 | 157.22 | 2.81 | 30 | >sp\|P01876\|IGHA1_HUMAN Immunoglobulin heavy constant alpha 1 OS=Homo sapiens OX=9606 GN=IGHA1 PE=1 SV=2 | 255 | 20201118_12833_1ng_glyco_hcd30_etd30_compound15.13476.13476.3 | controllerType=0 controllerNumber=1 scan=13476 | 30.8019 |
| 10071:3 | 2 | R.LAGKPTHVN[+1540.529]VSVVMAEVDGTC[+57.021]Y.- | HexNAc(2)Hex(7) | C[+57], N[+1541] | 1296.5649 | 3 | 3887.6801 | 3887.6706 | 0 | 2.4 | 332 | CRagged | 244.32 | 180.56 | 180.56 | 3.46 | 30 | >sp\|P01876\|IGHA1_HUMAN Immunoglobulin heavy constant alpha 1 OS=Homo sapiens OX=9606 GN=IGHA1 PE=1 SV=2 | 255 | 20201118_12833_1ng_glyco_hcd30_etd30_compound15.13365.13365.3 | controllerType=0 controllerNumber=1 scan=13365 | 30.4986 |
| 10239:3 | 2 | R.LAGKPTHVN[+1095.397]VSVVMAEVDGTC[+57.021]Y.- | HexNAc(3)Hex(3) | C[+57], N[+1095] | 1148.189 | 3 | 3442.5524 | 3442.5387 | 0 | 4 | 332 | CRagged | 244.41 | 180.48 | 180.48 | 3.41 | 30 | >sp\|P01876\|IGHA1_HUMAN Immunoglobulin heavy constant alpha 1 OS=Homo sapiens OX=9606 GN=IGHA1 PE=1 SV=2 | 255 | 20201118_12833_1ng_glyco_hcd30_etd30_compound15.13542.13542.3 | controllerType=0 controllerNumber=1 scan=13542 | 30.9763 |
| 11613:3 | 2 | R.LAGKPTHVN[+1913.677]VSVVMAEVDGTC[+57.021]Y.- | HexNAc(4)Hex(5)NeuAc(1) | C[+57], N[+1914] | 1420.9506 | 3 | 4260.8373 | 4260.8191 | 0 | 4.3 | 332 | CRagged | 246.43 | 130.61 | 130.61 | 2.83 | 30 | >sp\|P01876\|IGHA1_HUMAN Immunoglobulin heavy constant alpha 1 OS=Homo sapiens OX=9606 GN=IGHA1 PE=1 SV=2 | 255 | 20201118_12833_1ng_glyco_hcd30_etd30_compound15.14985.14985.3 | controllerType=0 controllerNumber=1 scan=14985 | 33.7588 |
| 10105:3 | 2 | R.LAGKPTHVN[+1378.476]VSVVMAEVDGTC[+57.021]Y.- | HexNAc(2)Hex(6) | C[+57], N[+1378] | 1242.5466 | 3 | 3725.6251 | 3725.6178 | 0 | 2 | 332 | CRagged | 247.61 | 193.52 | 174.58 | 3.46 | 30 | >sp\|P01876\|IGHA1_HUMAN Immunoglobulin heavy constant alpha 1 OS=Homo sapiens OX=9606 GN=IGHA1 PE=1 SV=2 | 255 | 20201118_12833_1ng_glyco_hcd30_etd30_compound15.13401.13401.3 | controllerType=0 controllerNumber=1 scan=13401 | 30.598 |
| 10180:3 | 2 | R.LAGKPTHVN[+1647.613]VSVVMAEVDGTC[+57.021]Y.- | HexNAc(5)Hex(3)Fuc(1) | C[+57], N[+1648] | 1332.258 | 3 | 3994.7594 | 3994.7553 | 0 | 1 | 332 | CRagged | 284.45 | 151.62 | 151.62 | 2.84 | 30 | >sp\|P01876\|IGHA1_HUMAN Immunoglobulin heavy constant alpha 1 OS=Homo sapiens OX=9606 GN=IGHA1 PE=1 SV=2 | 255 | 20201118_12833_1ng_glyco_hcd30_etd30_compound15.13480.13480.3 | controllerType=0 controllerNumber=1 scan=13480 | 30.8142 |
| 10140:3 | 2 | R.LAGKPTHVN[+1216.423]VSVVMAEVDGTC[+57.021]Y.- | HexNAc(2)Hex(5) | C[+57], N[+1216] | 1188.5286 | 3 | 3563.5713 | 3563.5649 | 0 | 1.8 | 332 | CRagged | 285.51 | 257.21 | 200.93 | 4.01 | 30 | >sp\|P01876\|IGHA1_HUMAN Immunoglobulin heavy constant alpha 1 OS=Homo sapiens OX=9606 GN=IGHA1 PE=1 SV=2 | 255 | 20201118_12833_1ng_glyco_hcd30_etd30_compound15.13438.13438.3 | controllerType=0 controllerNumber=1 scan=13438 | 30.6931 |
| 13106:4 | 2 | R.LAGKPTHVN[+2553.910]VSVVMAEVDGTC[+57.021]Y.- | HexNAc(5)Hex(5)Fuc(1)NeuAc(2) | C[+57], N[+2554] | 1226.0215 | 4 | 4901.0643 | 4901.0518 | 0 | 2.5 | 332 | CRagged | 349.82 | 214.63 | 214.63 | 3.25 | 30 | >sp\|P01876\|IGHA1_HUMAN Immunoglobulin heavy constant alpha 1 OS=Homo sapiens OX=9606 GN=IGHA1 PE=1 SV=2 | 255 | 20201118_12833_1ng_glyco_hcd30_etd30_compound15.16547.16547.4 | controllerType=0 controllerNumber=1 scan=16547 | 37.3759 |
| 04267:3 | 5 | R.FSDGLESN[+1864.634]SSTQFEVKK.Y | HexNAc(2)Hex(9) | N[+1865] | 1256.5212 | 3 | 3767.549 | 3767.5421 | 0 | 1.8 | 219 | Specific | 252.06 | 174.74 | 174.74 | 3.66 | 12 | >sp\|P0C0L5\|CO4B_HUMAN Complement C4-B OS=Homo sapiens OX=9606 GN=C4B PE=1 SV=2 | 1424 | 20201118_12833_1ng_glyco_hcd30_etd30_compound15.7331.7331.3 | controllerType=0 controllerNumber=1 scan=7331 | 19.2181 |
| 16277:3 | 8 | R.GLSFDVSLEVSQGPGLLN[+2350.830]DTK.V | HexNAc(4)Hex(5)Fuc(1)NeuAc(2) | N[+2351] | 1509.6568 | 3 | 4526.9558 | 4526.9436 | 0 | 2.7 | 118 | Specific | 477.58 | 274.81 | 274.81 | 5.46 | 14 | >sp\|P01833\|PIGR_HUMAN Polymeric immunoglobulin receptor OS=Homo sapiens OX=9606 GN=PIGR PE=1 SV=4 | 902 | 20201118_12833_1ng_glyco_hcd30_etd30_compound15.19928.19928.3 | controllerType=0 controllerNumber=1 scan=19928 | 46.7138 |
| 15870:3 | 8 | K.QIGLYPVLVIDSSGYVNPN[+1913.677]YTGR.I | HexNAc(4)Hex(5)NeuAc(1) | N[+1914] | 1480.3382 | 3 | 4439 | 4438.9805 | 0 | 4.4 | 168 | Specific | 204.83 | 163.57 | 163.57 | 3.65 | 14 | >sp\|P01833\|PIGR_HUMAN Polymeric immunoglobulin receptor OS=Homo sapiens OX=9606 GN=PIGR PE=1 SV=4 | 902 | 20201118_12833_1ng_glyco_hcd30_etd30_compound15.19497.19497.3 | controllerType=0 controllerNumber=1 scan=19497 | 45.5178 |
| 15823:4 | 8 | K.QIGLYPVLVIDSSGYVNPN[+2059.735]YTGR.I | HexNAc(4)Hex(5)Fuc(1)NeuAc(1) | N[+2060] | 1147.0204 | 4 | 4585.0599 | 4585.0384 | 0 | 4.7 | 168 | Specific | 400.67 | 363.42 | 363.42 | 3.8 | 14 | >sp\|P01833\|PIGR_HUMAN Polymeric immunoglobulin receptor OS=Homo sapiens OX=9606 GN=PIGR PE=1 SV=4 | 902 | 20201118_12833_1ng_glyco_hcd30_etd30_compound15.19447.19447.4 | controllerType=0 controllerNumber=1 scan=19447 | 45.3451 |
| 18645:3 | 8 | R.LSLLEEPGN[+2350.830]GTFTVILNQLTSR.D | HexNAc(4)Hex(5)Fuc(1)NeuAc(2) | N[+2351] | 1585.0513 | 3 | 4753.1394 | 4753.1229 | 0 | 3.5 | 413 | Specific | 391.05 | 254.9 | 254.9 | 4.82 | 14 | >sp\|P01833\|PIGR_HUMAN Polymeric immunoglobulin receptor OS=Homo sapiens OX=9606 GN=PIGR PE=1 SV=4 | 902 | 20201118_12833_1ng_glyco_hcd30_etd30_compound15.22435.22435.3 | controllerType=0 controllerNumber=1 scan=22435 | 53.5282 |
| 18588:4 | 8 | R.LSLLEEPGN[+2350.830]GTFTVILNQLTSR.D | HexNAc(4)Hex(5)Fuc(1)NeuAc(2) | N[+2351] | 1189.0414 | 4 | 4753.1439 | 4753.1229 | 0 | 4.4 | 413 | Specific | 492.37 | 452.81 | 452.81 | 5.34 | 14 | >sp\|P01833\|PIGR_HUMAN Polymeric immunoglobulin receptor OS=Homo sapiens OX=9606 GN=PIGR PE=1 SV=4 | 902 | 20201118_12833_1ng_glyco_hcd30_etd30_compound15.22374.22374.4 | controllerType=0 controllerNumber=1 scan=22374 | 53.3103 |
| 16931:4 | 8 | R.LSLLEEPGN[+2059.735]GTFTVILNQLTSR.D | HexNAc(4)Hex(5)Fuc(1)NeuAc(1) | N[+2060] | 1116.2657 | 4 | 4462.0408 | 4462.0275 | 0 | 3 | 413 | Specific | 578.29 | 524.82 | 524.82 | 7.41 | 14 | >sp\|P01833\|PIGR_HUMAN Polymeric immunoglobulin receptor OS=Homo sapiens OX=9606 GN=PIGR PE=1 SV=4 | 902 | 20201118_12833_1ng_glyco_hcd30_etd30_compound15.20624.20624.4 | controllerType=0 controllerNumber=1 scan=20624 | 48.9059 |
| 16918:3 | 8 | R.LSLLEEPGN[+2059.735]GTFTVILNQLTSR.D | HexNAc(4)Hex(5)Fuc(1)NeuAc(1) | N[+2060] | 1488.0202 | 3 | 4462.046 | 4462.0275 | 0 | 4.2 | 413 | Specific | 697.7 | 572.68 | 572.68 | 10.23 | 14 | >sp\|P01833\|PIGR_HUMAN Polymeric immunoglobulin receptor OS=Homo sapiens OX=9606 GN=PIGR PE=1 SV=4 | 902 | 20201118_12833_1ng_glyco_hcd30_etd30_compound15.20610.20610.3 | controllerType=0 controllerNumber=1 scan=20610 | 48.8591 |
| 13829:3 | 8 | K.IIEGEPNLKVPGN[+2205.793]VTAVLGETLK.V | HexNAc(4)Hex(5)Fuc(2)NeuAc(1) | N[+2206] | 1533.0517 | 3 | 4597.1405 | 4597.1422 | 0 | -0.4 | 457 | Specific | 336.89 | 286.18 | 286.18 | 2.74 | 14 | >sp\|P01833\|PIGR_HUMAN Polymeric immunoglobulin receptor OS=Homo sapiens OX=9606 GN=PIGR PE=1 SV=4 | 902 | 20201118_12833_1ng_glyco_hcd30_etd30_compound15.17334.17334.3 | controllerType=0 controllerNumber=1 scan=17334 | [illegible] |
| 13837:3 | 8 | K.IIEGEPNLKVPGN[+2204.772]VTAVLGETLK.V | HexNAc(4)Hex(5)NeuAc(2) | N[+2205] | 1532.7209 | 3 | 4596.1481 | 4596.1218 | 0 | 5.7 | 457 | Specific | 387.86 | 297.3 | 297.3 | 2.96 | 14 | >sp\|P01833\|PIGR_HUMAN Polymeric immunoglobulin receptor OS=Homo sapiens OX=9606 GN=PIGR PE=1 SV=4 | 902 | 20201118_12833_1ng_glyco_hcd30_etd30_compound15.17343.17343.3 | controllerType=0 controllerNumber=1 scan=17343 | [illegible] |
| 14596:4 | 8 | K.IIEGEPNLKVPGN[+2350.830]VTAVLGETLKVPC[+57.021]HFPC[+57.021]K.F | HexNAc(4)Hex(5)Fuc(1)NeuAc(2) | C[+57]*2, N[+2351] | 1442.6687 | 4 | 5767.653 | 5767.6372 | 0 | 2.7 | 457 | Specific | 449.06 | 448.64 | 448.64 | 5.74 | 14 | >sp\|P01833\|PIGR_HUMAN Polymeric immunoglobulin receptor OS=Homo sapiens OX=9606 GN=PIGR PE=1 SV=4 | 902 | 20201118_12833_1ng_glyco_hcd30_etd30_compound15.18148.18148.4 | controllerType=0 controllerNumber=1 scan=18148 | 41.1951 |
| 14592:5 | 8 | K.IIEGEPNLKVPGN[+2350.830]VTAVLGETLKVPC[+57.021]HFPC[+57.021]K.F | HexNAc(4)Hex(5)Fuc(1)NeuAc(2) | C[+57]*2, N[+2351] | 1154.3357 | 5 | 5767.6494 | 5767.6372 | 0 | 2.1 | 457 | Specific | 487.11 | 487.11 | 487.11 | 6.07 | 14 | >sp\|P01833\|PIGR_HUMAN Polymeric immunoglobulin receptor OS=Homo sapiens OX=9606 GN=PIGR PE=1 SV=4 | 902 | 20201118_12833_1ng_glyco_hcd30_etd30_compound15.18144.18144.5 | controllerType=0 controllerNumber=1 scan=18144 | 41.1865 |
| 13506:4 | 8 | K.IIEGEPNLKVPGN[+2205.793]VTAVLGETLKVPC[+57.021]HFPC[+57.021]K.F | HexNAc(4)Hex(5)Fuc(2)NeuAc(1) | C[+57]*2, N[+2206] | 1406.407 | 4 | 5622.6061 | 5622.5997 | 0 | 1.1 | 457 | Specific | 522.79 | 522.79 | 522.79 | 5.36 | 14 | >sp\|P01833\|PIGR_HUMAN Polymeric immunoglobulin receptor OS=Homo sapiens OX=9606 GN=PIGR PE=1 SV=4 | 902 | 20201118_12833_1ng_glyco_hcd30_etd30_compound15.16969.16969.4 | controllerType=0 controllerNumber=1 scan=16969 | 38.3881 |
| 05087:3 | 8 | K.WN[+2059.735]NTGC[+57.021]QALPSQDEGPSK.A | HexNAc(4)Hex(5)Fuc(1)NeuAc(1) | C[+57], N[+2060] | 1350.2129 | 3 | 4048.6243 | 4048.6116 | 0 | 3.1 | 498 | Specific | 250.5 | 130.23 | 130.23 | 3.97 | 14 | >sp\|P01833\|PIGR_HUMAN Polymeric immunoglobulin receptor OS=Homo sapiens OX=9606 GN=PIGR PE=1 SV=4 | 902 | 20201118_12833_1ng_glyco_hcd30_etd30_compound15.8180.8180.3 | controllerType=0 controllerNumber=1 scan=8180 | 20.7249 |
| 11287:6 | 21 | K.SAVQGPPERDLC[+57.021]GC[+57.021]YSVSSVLPGC[+57.021]AQPWNHGETFTC[+ | HexNAc(5)Hex(3)Fuc(1) | C[+57]*4, N[+1648] | 1243.9009 | 6 | 7458.3692 | 7458.3591 | 0 | 1.4 | 156 | Specific | 678.92 | 678.92 | 678.92 | 9.13 | 7 | >sp\|P01877\|IGHA2_HUMAN Immunoglobulin heavy constant alpha 2 OS=Homo sapiens OX=9606 GN=IGHA2 PE=1 SV=4 | 576 | 20201118_12833_1ng_glyco_hcd30_etd30_compound15.14631.14631.6 | controllerType=0 controllerNumber=1 scan=14631 | 33.0639 |
| 02500:3 | 21 | K.TPLTAN[+1971.719]ITK.S | HexNAc(5)Hex(5)Fuc(1) | N[+1972] | 977.4334 | 3 | 2930.2858 | 2930.2757 | 0 | 3.4 | 200 | Specific | 245.41 | 126.49 | 126.49 | 2.54 | 7 | >sp\|P01877\|IGHA2_HUMAN Immunoglobulin heavy constant alpha 2 OS=Homo sapiens OX=9606 GN=IGHA2 PE=1 SV=4 | 576 | 20201118_12833_1ng_glyco_hcd30_etd30_compound15.5494.5494.3 | controllerType=0 controllerNumber=1 scan=5494 | 15.8593 |
| 02593:3 | 21 | K.TPLTAN[+1647.613]ITK.S | HexNAc(5)Hex(3)Fuc(1) | N[+1648] | 869.3963 | 3 | 2606.1744 | 2606.17 | 0 | 1.7 | 200 | Specific | 294.64 | 137.65 | 137.65 | 2.63 | 7 | >sp\|P01877\|IGHA2_HUMAN Immunoglobulin heavy constant alpha 2 OS=Homo sapiens OX=9606 GN=IGHA2 PE=1 SV=4 | 576 | 20201118_12833_1ng_glyco_hcd30_etd30_compound15.5591.5591.3 | controllerType=0 controllerNumber=1 scan=5591 | 16.0933 |
| 02581:2 | 21 | K.TPLTAN[+1647.613]ITK.S | HexNAc(5)Hex(3)Fuc(1) | N[+1648] | 1303.5911 | 2 | 2606.1749 | 2606.17 | 0 | 1.9 | 200 | Specific | 302.65 | 115.85 | 115.85 | 2.73 | 7 | >sp\|P01877\|IGHA2_HUMAN Immunoglobulin heavy constant alpha 2 OS=Homo sapiens OX=9606 GN=IGHA2 PE=1 SV=4 | 576 | 20201118_12833_1ng_glyco_hcd30_etd30_compound15.5579.5579.2 | controllerType=0 controllerNumber=1 scan=5579 | 16.062 |
| 02546:3 | 21 | K.TPLTAN[+1809.666]ITK.S | HexNAc(5)Hex(4)Fuc(1) | N[+1810] | 923.4153 | 3 | 2768.2312 | 2768.2228 | 0 | 3 | 200 | Specific | 306.73 | 130.13 | 130.13 | 2.59 | 7 | >sp\|P01877\|IGHA2_HUMAN Immunoglobulin heavy constant alpha 2 OS=Homo sapiens OX=9606 GN=IGHA2 PE=1 SV=4 | 576 | 20201118_12833_1ng_glyco_hcd30_etd30_compound15.5542.5542.3 | controllerType=0 controllerNumber=1 scan=5542 | 15.9633 |
| 19591:3 | 24 | R.QLAHQSN[+2204.772]STNIFFSPVSIATAFAMLSLGTK.A | HexNAc(4)Hex(5)NeuAc(2) | N[+2205] | 1796.1478 | 3 | 5386.4288 | 5386.4075 | 0 | 4 | 64 | Specific | 282.62 | 166.07 | 166.07 | 5.34 | 4 | >sp\|P01009\|A1AT_HUMAN Alpha-1-antitrypsin OS=Homo sapiens OX=9606 GN=SERPINA1 PE=1 SV=3 | 642 | 20201118_12833_1ng_glyco_hcd30_etd30_compound15.23428.23428.3 | controllerType=0 controllerNumber=1 scan=23428 | 55.9033 |
| 19590:4 | 24 | R.QLAHQSN[+2204.772]STNIFFSPVSIATAFAMLSLGTK.A | HexNAc(4)Hex(5)NeuAc(2) | N[+2205] | 1347.3606 | 4 | 5386.4206 | 5386.4075 | 0 | 2.4 | 64 | Specific | 614.28 | 498.37 | 426.84 | 8.95 | 4 | >sp\|P01009\|A1AT_HUMAN Alpha-1-antitrypsin OS=Homo sapiens OX=9606 GN=SERPINA1 PE=1 SV=3 | 642 | 20201118_12833_1ng_glyco_hcd30_etd30_compound15.23427.23427.4 | controllerType=0 controllerNumber=1 scan=23427 | 55.8986 |
| 05029:3 | 30 | R.LSPN[+2351.851]ASAEHLELR.W | HexNAc(4)Hex(5)Fuc(3)NeuAc(1) | N[+2352] | 1263.542 | 3 | 3788.6116 | 3788.6 | 0 | 3.1 | 52 | Specific | 216.98 | 156.29 | 156.29 | 2.88 | 7 | >sp\|Q13410\|BT1A1_HUMAN Butyrophilin subfamily 1 member A1 OS=Homo sapiens OX=9606 GN=BTN1A1 PE=1 SV=3 | 179 | 20201118_12833_1ng_glyco_hcd30_etd30_compound15.8119.8119.3 | controllerType=0 controllerNumber=1 scan=8119 | 20.5979 |
| 05058:3 | 30 | R.LSPN[+2205.793]ASAEHLELR.W | HexNAc(4)Hex(5)Fuc(2)NeuAc(1) | N[+2206] | 1214.8559 | 3 | 3642.5531 | 3642.5421 | 0 | 3 | 52 | Specific | 337.39 | 218.47 | 218.47 | 3.3 | 7 | >sp\|Q13410\|BT1A1_HUMAN Butyrophilin subfamily 1 member A1 OS=Homo sapiens OX=9606 GN=BTN1A1 PE=1 SV=3 | 179 | 20201118_12833_1ng_glyco_hcd30_etd30_compound15.8150.8150.3 | controllerType=0 controllerNumber=1 scan=8150 | 20.6628 |
| 06891:3 | 30 | R.LSPN[+2350.830]ASAEHLELR.W | HexNAc(4)Hex(5)Fuc(1)NeuAc(2) | N[+2351] | 1263.2001 | 3 | 3787.5858 | 3787.5796 | 0 | 1.7 | 52 | Specific | 349.75 | 187.07 | 187.07 | 4.24 | 7 | >sp\|Q13410\|BT1A1_HUMAN Butyrophilin subfamily 1 member A1 OS=Homo sapiens OX=9606 GN=BTN1A1 PE=1 SV=3 | 179 | 20201118_12833_1ng_glyco_hcd30_etd30_compound15.10051.10051.3 | controllerType=0 controllerNumber=1 scan=10051 | 24.0977 |
| 06882:3 | 30 | R.LSPN[+2350.830]ASAEHLELR.W | HexNAc(4)Hex(5)Fuc(1)NeuAc(2) | N[+2351] | 1263.2004 | 3 | 3787.5866 | 3787.5796 | 0 | 1.8 | 52 | Specific | 379.52 | 249.12 | 249.12 | 4.42 | 7 | >sp\|Q13410\|BT1A1_HUMAN Butyrophilin subfamily 1 member A1 OS=Homo sapiens OX=9606 GN=BTN1A1 PE=1 SV=3 | 179 | 20201118_12833_1ng_glyco_hcd30_etd30_compound15.10041.10041.3 | controllerType=0 controllerNumber=1 scan=10041 | 24.0747 |
| 08553:3 | 33 | R.IIVPLNNREN[+1809.666]ISDPTSPLR.T | HexNAc(5)Hex(4)Fuc(1) | N[+1810] | 1319.9556 | 3 | 3957.8523 | 3957.8432 | 0 | 2.3 | 62 | Specific | 214.38 | 155.57 | 155.57 | 2.75 | 4 | >sp\|P01591\|IGJ_HUMAN Immunoglobulin J chain OS=Homo sapiens OX=9606 GN=JCHAIN PE=1 SV=4 | 1392 | 20201118_12833_1ng_glyco_hcd30_etd30_compound15.11789.11789.3 | controllerType=0 controllerNumber=1 scan=11789 | 27.5262 |
| 08566:3 | 33 | R.IIVPLNNREN[+1663.608]ISDPTSPLR.T | HexNAc(5)Hex(4) | N[+1664] | 1271.2681 | 3 | 3811.7898 | 3811.7853 | 0 | 1.2 | 62 | Specific | 244.28 | 186.63 | 186.63 | 2.68 | 4 | >sp\|P01591\|IGJ_HUMAN Immunoglobulin J chain OS=Homo sapiens OX=9606 GN=JCHAIN PE=1 SV=4 | 1392 | 20201118_12833_1ng_glyco_hcd30_etd30_compound15.11803.11803.3 | controllerType=0 controllerNumber=1 scan=11803 | 27.5581 |
| 11740:3 | 33 | R.IIVPLNNREN[+2204.772]ISDPTSPLR.T | HexNAc(4)Hex(5)NeuAc(2) | N[+2205] | 1451.6589 | 3 | 4352.9621 | 4352.9496 | 0 | 2.9 | 62 | Specific | 258.27 | 178.48 | 178.48 | 3.39 | 4 | >sp\|P01591\|IGJ_HUMAN Immunoglobulin J chain OS=Homo sapiens OX=9606 GN=JCHAIN PE=1 SV=4 | 1392 | 20201118_12833_1ng_glyco_hcd30_etd30_compound15.15118.15118.3 | controllerType=0 controllerNumber=1 scan=15118 | 34.0999 |
| 12699:4 | 35 | K.MVSHHN[+2204.772]LTTGATLINEQWLLTTAK.N | HexNAc(4)Hex(5)NeuAc(2) | N[+2205] | 1221.8013 | 4 | 4884.1834 | 4884.1647 | 0 | 3.8 | 179 | Specific | 383.94 | 371.05 | 371.05 | 5.21 | 3 | >sp\|P00738\|HPT_HUMAN Haptoglobin OS=Homo sapiens OX=9606 GN=HP PE=1 SV=1 | 407 | 20201118_12833_1ng_glyco_hcd30_etd30_compound15.16122.16122.3 | controllerType=0 controllerNumber=1 scan=16122 | 36.3933 |
| 12523:3 | 35 | K.VVLHPN[+2204.772]YSQVDIGLIK.L | HexNAc(4)Hex(5)NeuAc(2) | N[+2205] | 1333.938 | 3 | 3999.7996 | 3999.7837 | 0 | 4 | 236 | Specific | 284.57 | 187.42 | 187.42 | 3.31 | 3 | >sp\|P00738\|HPT_HUMAN Haptoglobin OS=Homo sapiens OX=9606 GN=HP PE=1 SV=1 | 407 | 20201118_12833_1ng_glyco_hcd30_etd30_compound15.15937.15937.3 | controllerType=0 controllerNumber=1 scan=15937 | 35.9233 |
| 00093:4 | 47 | K.KKEDALN[+2350.830]ETRESETK.L | HexNAc(4)Hex(5)Fuc(1)NeuAc(2) | N[+2351] | 1032.9411 | 4 | 4128.7425 | 4128.723 | 0 | 4.7 | 80 | Specific | 210.35 | 177.87 | 177.87 | 2.71 | 4 | >sp\|P10909\|CLUS_HUMAN Clusterin OS=Homo sapiens OX=9606 GN=CLU PE=1 SV=1 | 1467 | 20201118_12833_1ng_glyco_hcd30_etd30_compound15.2992.2992.4 | controllerType=0 controllerNumber=1 scan=2992 | 11.307 |
| 12276:4 | 47 | K.LKELPGVC[+57.021]N[+2059.735]ETMMALWEEC[+57.021]KPC[+57.021]LK.Q | HexNAc(4)Hex(5)Fuc(1)NeuAc(1) | C[+57]*3, N[+2060] | 1249.7872 | 4 | 4996.1271 | 4996.1196 | 0 | 1.5 | 95 | Specific | 438.47 | 307.61 | 307.61 | 4.34 | 4 | >sp\|P10909\|CLUS_HUMAN Clusterin OS=Homo sapiens OX=9606 GN=CLU PE=1 SV=1 | 1467 | 20201118_12833_1ng_glyco_hcd30_etd30_compound15.15679.15679.4 | controllerType=0 controllerNumber=1 scan=15679 | 35.4198 |
| 00081:3 | 47 | R.EIRHN[+2059.735]STGC[+57.021]LR.M | HexNAc(4)Hex(5)Fuc(1)NeuAc(1) | C[+57], N[+2060] | 1134.8087 | 3 | 3402.4117 | 3402.3994 | 0 | 3.6 | 287 | Specific | 223.95 | 131.03 | 131.03 | 2.56 | 4 | >sp\|P10909\|CLUS_HUMAN Clusterin OS=Homo sapiens OX=9606 GN=CLU PE=1 SV=1 | 1467 | 20201118_12833_1ng_glyco_hcd30_etd30_compound15.2980.2980.3 | controllerType=0 controllerNumber=1 scan=2980 | 11.2634 |
| 00082:4 | 47 | R.EIRHN[+2059.735]STGC[+57.021]LR.M | HexNAc(4)Hex(5)Fuc(1)NeuAc(1) | C[+57], N[+2060] | 851.3577 | 4 | 3402.4089 | 3402.3994 | 0 | 2.8 | 287 | Specific | 652.57 | 491.95 | 491.95 | 9.64 | 4 | >sp\|P10909\|CLUS_HUMAN Clusterin OS=Homo sapiens OX=9606 GN=CLU PE=1 SV=1 | 1467 | 20201118_12833_1ng_glyco_hcd30_etd30_compound15.2981.2981.4 | controllerType=0 controllerNumber=1 scan=2981 | 11.2705 |
| 00788:4 | 71 | K.TKPREEQFN[+1444.534]STFR.V | HexNAc(4)Hex(3)Fuc(1) | N[+1445] | 771.8458 | 4 | 3084.3613 | 3084.3526 | 0 | 2.8 | 168 | Specific | 364.91 | 354.24 | 354.24 | 3.63 | 1 | >sp\|P01859\|IGHG2_HUMAN Immunoglobulin heavy constant gamma 2 OS=Homo sapiens OX=9606 GN=IGHG2 PE=1 SV=2 | 387 | 20201118_12833_1ng_glyco_hcd30_etd30_compound15.3716.3716.4 | controllerType=0 controllerNumber=1 scan=3716 | 12.6347 |
| 00078:4 | 79 | K.TKPREEQYN[+1606.587]STYR.V | HexNAc(4)Hex(4)Fuc(1) | N[+1607] | 820.3566 | 4 | 3278.4045 | 3278.3952 | 0 | 2.8 | 291 | Specific | 395.94 | 55.48 | 55.48 | 2.11 | 1 | >sp\|P0DOX5\|IGG1_HUMAN Immunoglobulin gamma-1 heavy chain OS=Homo sapiens OX=9606 PE=1 SV=2 | 914 | 20201118_12833_1ng_glyco_hcd30_etd30_compound15.2976.2976.4 | controllerType=0 controllerNumber=1 scan=2976 | 11.249 |
| 16356:4 | 1 | R.RTAGWNVPIGTLRPFLN[+2350.830]WTGPPEPIEAAVAR.F | HexNAc(4)Hex(5)Fuc(1)NeuAc(2) | N[+2351] | 1435.1672 | 4 | 5737.6471 | 5737.6424 | 0 | 0.8 | 140 | Specific | 800.81 | 800.81 | 800.81 | 12.54 | 39 | >sp\|P02788\|TRFL_HUMAN Lactotransferrin OS=Homo sapiens OX=9606 GN=LTF PE=1 SV=6 | 1270 | 20201114_13813_1ng_glyco_hcd30_etd30_compound15.19975.19975.4 | controllerType=0 controllerNumber=1 scan=19975 | 45.7548 |
| 15013:4 | 1 | R.RTAGWNVPIGTLRPFLN[+1768.640]WTGPPEPIEAAVAR.F | HexNAc(4)Hex(5)Fuc(1) | N[+1769] | 1289.6176 | 4 | 5155.4484 | 5155.4516 | 0 | -0.6 | 140 | Specific | 1036.72 | 1035.25 | 1035.25 | 16.54 | 39 | >sp\|P02788\|TRFL_HUMAN Lactotransferrin OS=Homo sapiens OX=9606 GN=LTF PE=1 SV=6 | 1270 | 20201114_13813_1ng_glyco_hcd30_etd30_compound15.18549.18549.4 | controllerType=0 controllerNumber=1 scan=18549 | 41.3808 |
| 15859:3 | 1 | R.TAGWNVPIGTLRPFLN[+1768.640]WTGPPEPIEAAVAR.F | HexNAc(4)Hex(5)Fuc(1) | N[+1769] | 1667.1308 | 3 | 4999.3779 | 4999.3504 | 0 | 5.5 | 141 | Specific | 234.83 | 179.59 | 179.59 | 3.66 | 39 | >sp\|P02788\|TRFL_HUMAN Lactotransferrin OS=Homo sapiens OX=9606 GN=LTF PE=1 SV=6 | 1270 | 20201114_13813_1ng_glyco_hcd30_etd30_compound15.19447.19447.3 | controllerType=0 controllerNumber=1 scan=19447 | 44.0983 |
| 16555:3 | 1 | R.TAGWNVPIGTLRPFLN[+2059.735]WTGPPEPIEAAVAR.F | HexNAc(4)Hex(5)Fuc(1)NeuAc(1) | N[+2060] | 1764.1555 | 3 | 5290.4519 | 5290.4459 | 0 | 1.1 | 141 | Specific | 320.75 | 228.27 | 228.27 | 6.32 | 39 | >sp\|P02788\|TRFL_HUMAN Lactotransferrin OS=Homo sapiens OX=9606 GN=LTF PE=1 SV=6 | 1270 | 20201114_13813_1ng_glyco_hcd30_etd30_compound15.20187.20187.3 | controllerType=0 controllerNumber=1 scan=20187 | 46.4092 |
| 16552:3 | 1 | R.TAGWNVPIGTLRPFLN[+2205.793]WTGPPEPIEAAVAR.F | HexNAc(4)Hex(5)Fuc(2)NeuAc(1) | N[+2206] | 1812.8498 | 3 | 5436.5349 | 5436.5038 | 0 | 5.7 | 141 | Specific | 364.08 | 284.5 | 284.5 | 4.75 | 39 | >sp\|P02788\|TRFL_HUMAN Lactotransferrin OS=Homo sapiens OX=9606 GN=LTF PE=1 SV=6 | 1270 | 20201114_13813_1ng_glyco_hcd30_etd30_compound15.20183.20183.2 | controllerType=0 controllerNumber=1 scan=20183 | 46.3998 |
| 16483:4 | 1 | R.TAGWNVPIGTLRPFLN[+2716.983]WTGPPEPIEAAVAR.F | HexNAc(5)Hex(6)Fuc(3)NeuAc(1) | N[+2717] | 1487.6899 | 4 | 5947.7379 | 5947.6939 | 0 | 7.4 | 141 | Specific | 364.55 | 364.55 | 364.55 | 6.21 | 39 | >sp\|P02788\|TRFL_HUMAN Lactotransferrin OS=Homo sapiens OX=9606 GN=LTF PE=1 SV=6 | 1270 | 20201114_13813_1ng_glyco_hcd30_etd30_compound15.20109.20109.4 | controllerType=0 controllerNumber=1 scan=20109 | 46.1363 |
| 16433:4 | 1 | R.TAGWNVPIGTLRPFLN[+2936.057]WTGPPEPIEAAVAR.F | HexNAc(6)Hex(7)Fuc(2)NeuAc(1) | N[+2936] | 1542.4475 | 4 | 6166.7682 | 6166.7682 | 0 | 0 | 141 | Specific | 531.11 | 531.11 | 531.11 | 7.77 | 39 | >sp\|P02788\|TRFL_HUMAN Lactotransferrin OS=Homo sapiens OX=9606 GN=LTF PE=1 SV=6 | 1270 | 20201114_13813_1ng_glyco_hcd30_etd30_compound15.20056.20056.2 | controllerType=0 controllerNumber=1 scan=20056 | 45.963 |
| 16708:3 | 1 | R.TAGWNVPIGTLRPFLN[+2059.735]WTGPPEPIEAAVAR.F | HexNAc(4)Hex(5)Fuc(1)NeuAc(1) | N[+2060] | 1764.1684 | 3 | 5290.4907 | 5290.4459 | 0 | 8.5 | 141 | Specific | 533.1 | 431.58 | 431.58 | 6.35 | 39 | >sp\|P02788\|TRFL_HUMAN Lactotransferrin OS=Homo sapiens OX=9606 GN=LTF PE=1 SV=6 | 1270 | 20201114_13813_1ng_glyco_hcd30_etd30_compound15.20350.20350.3 | controllerType=0 controllerNumber=1 scan=20350 | 46.7832 |
| 16545:3 | 1 | R.TAGWNVPIGTLRPFLN[+2059.735]WTGPPEPIEAAVAR.F | HexNAc(4)Hex(5)Fuc(1)NeuAc(1) | N[+2060] | 1764.1634 | 3 | 5290.4757 | 5290.4459 | 0 | 5.6 | 141 | Specific | 580.61 | 400.56 | 400.56 | 8.43 | 39 | >sp\|P02788\|TRFL_HUMAN Lactotransferrin OS=Homo sapiens OX=9606 GN=LTF PE=1 SV=6 | 1270 | 20201114_13813_1ng_glyco_hcd30_etd30_compound15.20176.20176.2 | controllerType=0 controllerNumber=1 scan=20176 | 46.3815 |
| 15800:4 | 1 | R.TAGWNVPIGTLRPFLN[+2425.887]WTGPPEPIEAAVAR.F | HexNAc(5)Hex(6)Fuc(3) | N[+2426] | 1414.9117 | 4 | 5656.6252 | 5656.5985 | 0 | 4.7 | 141 | Specific | 735.85 | 512.09 | 512.09 | 11.65 | 39 | >sp\|P02788\|TRFL_HUMAN Lactotransferrin OS=Homo sapiens OX=9606 GN=LTF PE=1 SV=6 | 1270 | 20201114_13813_1ng_glyco_hcd30_etd30_compound15.19384.19384.4 | controllerType=0 controllerNumber=1 scan=19384 | 43.9052 |
| 15865:4 | 1 | R.TAGWNVPIGTLRPFLN[+1768.640]WTGPPEPIEAAVAR.F | HexNAc(4)Hex(5)Fuc(1) | N[+1769] | 1250.5996 | 4 | 4999.3766 | 4999.3504 | 0 | 5.2 | 141 | Specific | 989.4 | 989.4 | 989.4 | 16.65 | 39 | >sp\|P02788\|TRFL_HUMAN Lactotransferrin OS=Homo sapiens OX=9606 GN=LTF PE=1 SV=6 | 1270 | 20201114_13813_1ng_glyco_hcd30_etd30_compound15.19453.19453.4 | controllerType=0 controllerNumber=1 scan=19453 | 44.1205 |
| 14818:2 | 1 | R.TAGWNIPMGLLFN[+1768.640]QTGSC[+57.021]K.F | HexNAc(4)Hex(5)Fuc(1) | C[+57], N[+1769] | 1932.3298 | 2 | 3863.6523 | 3863.6495 | 0 | 0.7 | 485 | Specific | 215.99 | 92.73 | 92.73 | 4.74 | 39 | >sp\|P02788\|TRFL_HUMAN Lactotransferrin OS=Homo sapiens OX=9606 GN=LTF PE=1 SV=6 | 1270 | 20201114_13813_1ng_glyco_hcd30_etd30_compound15.18342.18342.2 | controllerType=0 controllerNumber=1 scan=18342 | 40.7334 |
| 15091:2 | 1 | R.TAGWNIPMGLLFN[+1216.423]QTGSC[+57.021]K.F | HexNAc(2)Hex(5) | C[+57], N[+1216] | 1656.2308 | 2 | 3311.4543 | 3311.4328 | 0 | 6.5 | 485 | Specific | 216.49 | 216.49 | 111.1 | 5.65 | 39 | >sp\|P02788\|TRFL_HUMAN Lactotransferrin OS=Homo sapiens OX=9606 GN=LTF PE=1 SV=6 | 1270 | 20201114_13813_1ng_glyco_hcd30_etd30_compound15.18632.18632.2 | controllerType=0 controllerNumber=1 scan=18632 | 41.6333 |
| 16438:3 | 1 | R.TAGWNIPMGLLFN[+2059.735]QTGSC[+57.021]K.F | HexNAc(4)Hex(5)Fuc(1)NeuAc(1) | C[+57], N[+2060] | 1385.5755 | 3 | 4154.7119 | 4154.7449 | 0 | -7.9 | 485 | Specific | 236.71 | 113.78 | 113.78 | 4.1 | 39 | >sp\|P02788\|TRFL_HUMAN Lactotransferrin OS=Homo sapiens OX=9606 GN=LTF PE=1 SV=6 | 1270 | 20201114_13813_1ng_glyco_hcd30_etd30_compound15.20061.20061.3 | controllerType=0 controllerNumber=1 scan=20061 | 45.9765 |
| 14627:3 | 1 | R.TAGWNIPMGLLFN[+2279.830]QTGSC[+57.021]K.F | HexNAc(5)Hex(6)Fuc(2) | C[+57], N[+2280] | 1458.955 | 3 | 4374.8505 | 4374.8396 | 0 | 2.5 | 485 | Specific | 260.9 | 186.57 | 186.57 | 6.52 | 39 | >sp\|P02788\|TRFL_HUMAN Lactotransferrin OS=Homo sapiens OX=9606 GN=LTF PE=1 SV=6 | 1270 | 20201114_13813_1ng_glyco_hcd30_etd30_compound15.18139.18139.3 | controllerType=0 controllerNumber=1 scan=18139 | 40.2024 |
| 16278:3 | 1 | R.TAGWNIPMGLLFN[+1751.624]QTGSC[+57.021]K.F | HexNAc(4)Hex(4)NeuAc(1) | C[+57], N[+1752] | 1282.8954 | 3 | 3846.6718 | 3846.6341 | 0 | 9.8 | 485 | Specific | 265.89 | 211.47 | 211.47 | 5.71 | 39 | >sp\|P02788\|TRFL_HUMAN Lactotransferrin OS=Homo sapiens OX=9606 GN=LTF PE=1 SV=6 | 1270 | 20201114_13813_1ng_glyco_hcd30_etd30_compound15.19892.19892.3 | controllerType=0 controllerNumber=1 scan=19892 | 45.4952 |
| 17699:3 | 1 | R.TAGWNIPMGLLFN[+2204.772]QTGSC[+57.021]K.F | HexNAc(4)Hex(5)NeuAc(2) | C[+57], N[+2205] | 1433.9383 | 3 | 4299.8003 | 4299.7824 | 0 | 4.2 | 485 | Specific | 299.82 | 214.28 | 214.28 | 6.27 | 39 | >sp\|P02788\|TRFL_HUMAN Lactotransferrin OS=Homo sapiens OX=9606 GN=LTF PE=1 SV=6 | 1270 | 20201114_13813_1ng_glyco_hcd30_etd30_compound15.21402.21402.3 | controllerType=0 controllerNumber=1 scan=21402 | 49.8776 |
| 15987:3 | 1 | R.TAGWNIPMGLLFN[+2424.867]QTGSC[+57.021]K.F | HexNAc(5)Hex(6)Fuc(1)NeuAc(1) | C[+57], N[+2425] | 1507.2982 | 3 | 4519.8799 | 4519.8771 | 0 | 0.6 | 485 | Specific | 327.54 | 206.99 | 206.99 | 6.04 | 39 | >sp\|P02788\|TRFL_HUMAN Lactotransferrin OS=Homo sapiens OX=9606 GN=LTF PE=1 SV=6 | 1270 | 20201114_13813_1ng_glyco_hcd30_etd30_compound15.19582.19582.3 | controllerType=0 controllerNumber=1 scan=19582 | 44.4581 |
| 14885:3 | 1 | R.TAGWNIPMGLLFN[+1768.640]QTGSC[+57.021]K.F | HexNAc(4)Hex(5)Fuc(1) | C[+57], N[+1769] | 1288.5568 | 3 | 3863.6559 | 3863.6495 | 0 | 1.7 | 485 | Specific | 333.99 | 277.19 | 277.19 | 5.95 | 39 | >sp\|P02788\|TRFL_HUMAN Lactotransferrin OS=Homo sapiens OX=9606 GN=LTF PE=1 SV=6 | 1270 | 20201114_13813_1ng_glyco_hcd30_etd30_compound15.18413.18413.3 | controllerType=0 controllerNumber=1 scan=18413 | 40.9443 |
| 14688:3 | 1 | R.TAGWNIPMGLLFN[+2133.772]QTGSC[+57.021]K.F | HexNAc(5)Hex(6)Fuc(1) | C[+57], N[+2134] | 1410.2735 | 3 | 4228.8059 | 4228.7817 | 0 | 5.7 | 485 | Specific | 334.12 | 296.46 | 223.32 | 6.25 | 39 | >sp\|P02788\|TRFL_HUMAN Lactotransferrin OS=Homo sapiens OX=9606 GN=LTF PE=1 SV=6 | 1270 | 20201114_13813_1ng_glyco_hcd30_etd30_compound15.18203.18203.3 | controllerType=0 controllerNumber=1 scan=18203 | 40.3548 |
| 14869:3 | 1 | R.TAGWNIPMGLLFN[+1606.587]QTGSC[+57.021]K.F | HexNAc(4)Hex(4)Fuc(1) | C[+57], N[+1607] | 1234.5399 | 3 | 3701.6051 | 3701.5966 | 0 | 2.3 | 485 | Specific | 343.34 | 272.13 | 272.13 | 6.95 | 39 | >sp\|P02788\|TRFL_HUMAN Lactotransferrin OS=Homo sapiens OX=9606 GN=LTF PE=1 SV=6 | 1270 | 20201114_13813_1ng_glyco_hcd30_etd30_compound15.18396.18396.3 | controllerType=0 controllerNumber=1 scan=18396 | 40.8929 |
| 17632:3 | 1 | R.TAGWNIPMGLLFN[+2204.772]QTGSC[+57.021]K.F | HexNAc(4)Hex(5)NeuAc(2) | C[+57], N[+2205] | 1433.9374 | 3 | 4299.7978 | 4299.7824 | 0 | 3.6 | 485 | Specific | 348.67 | 239.2 | 239.2 | 6.33 | 39 | >sp\|P02788\|TRFL_HUMAN Lactotransferrin OS=Homo sapiens OX=9606 GN=LTF PE=1 SV=6 | 1270 | 20201114_13813_1ng_glyco_hcd30_etd30_compound15.21332.21332.3 | controllerType=0 controllerNumber=1 scan=21332 | 49.6515 |
| 14933:3 | 1 | R.TAGWNIPMGLLFN[+1622.582]QTGSC[+57.021]K.F | HexNAc(4)Hex(5) | C[+57], N[+1623] | 1239.8727 | 3 | 3717.6037 | 3717.5916 | 0 | 3.3 | 485 | Specific | 370.4 | 285.63 | 257.33 | 5.59 | 39 | >sp\|P02788\|TRFL_HUMAN Lactotransferrin OS=Homo sapiens OX=9606 GN=LTF PE=1 SV=6 | 1270 | 20201114_13813_1ng_glyco_hcd30_etd30_compound15.18464.18464.3 | controllerType=0 controllerNumber=1 scan=18464 | 41.1084 |
| 14789:3 | 1 | R.TAGWNIPMGLLFN[+1768.640]QTGSC[+57.021]K.F | HexNAc(4)Hex(5)Fuc(1) | C[+57], N[+1769] | 1288.5579 | 3 | 3863.6592 | 3863.6495 | 0 | 2.5 | 485 | Specific | 386.55 | 261.91 | 261.91 | 6.02 | 39 | >sp\|P02788\|TRFL_HUMAN Lactotransferrin OS=Homo sapiens OX=9606 GN=LTF PE=1 SV=6 | 1270 | 20201114_13813_1ng_glyco_hcd30_etd30_compound15.18312.18312.3 | controllerType=0 controllerNumber=1 scan=18312 | 40.6571 |
| 16104:3 | 1 | R.TAGWNIPMGLLFN[+2059.735]QTGSC[+57.021]K.F | HexNAc(4)Hex(5)Fuc(1)NeuAc(1) | C[+57], N[+2060] | 1385.589 | 3 | 4154.7526 | 4154.7449 | 0 | 1.9 | 485 | Specific | 390.79 | 289.99 | 289.99 | 7.02 | 39 | >sp\|P02788\|TRFL_HUMAN Lactotransferrin OS=Homo sapiens OX=9606 GN=LTF PE=1 SV=6 | 1270 | 20201114_13813_1ng_glyco_hcd30_etd30_compound15.19706.19706.3 | controllerType=0 controllerNumber=1 scan=19706 | 44.8352 |
| 16184:3 | 1 | R.TAGWNIPMGLLFN[+1913.677]QTGSC[+57.021]K.F | HexNAc(4)Hex(5)NeuAc(1) | C[+57], N[+1914] | 1336.91 | 3 | 4008.7154 | 4008.687 | 0 | 7.1 | 485 | Specific | 408.36 | 267.47 | 267.47 | 6.96 | 39 | >sp\|P02788\|TRFL_HUMAN Lactotransferrin OS=Homo sapiens OX=9606 GN=LTF PE=1 SV=6 | 1270 | 20201114_13813_1ng_glyco_hcd30_etd30_compound15.19792.19792.3 | controllerType=0 controllerNumber=1 scan=19792 | 45.1368 |
| 15061:3 | 1 | R.TAGWNIPMGLLFN[+1216.423]QTGSC[+57.021]K.F | HexNAc(2)Hex(5) | C[+57], N[+1216] | 1104.4911 | 3 | 3311.4588 | 3311.4328 | 0 | 7.8 | 485 | Specific | 408.74 | 386.69 | 211.43 | 8.1 | 39 | >sp\|P02788\|TRFL_HUMAN Lactotransferrin OS=Homo sapiens OX=9606 GN=LTF PE=1 SV=6 | 1270 | 20201114_13813_1ng_glyco_hcd30_etd30_compound15.18601.18601.3 | controllerType=0 controllerNumber=1 scan=18601 | 41.5456 |
| 16144:3 | 1 | R.TAGWNIPMGLLFN[+2059.735]QTGSC[+57.021]K.F | HexNAc(4)Hex(5)Fuc(1)NeuAc(1) | C[+57], N[+2060] | 1385.591 | 3 | 4154.7584 | 4154.7449 | 0 | 3.3 | 485 | Specific | 416.69 | 295.31 | 295.31 | 6.73 | 39 | >sp\|P02788\|TRFL_HUMAN Lactotransferrin OS=Homo sapiens OX=9606 GN=LTF PE=1 SV=6 | 1270 | 20201114_13813_1ng_glyco_hcd30_etd30_compound15.19749.19749.3 | controllerType=0 controllerNumber=1 scan=19749 | 44.969 |
| 16074:3 | 1 | R.TAGWNIPMGLLFN[+2205.793]QTGSC[+57.021]K.F | HexNAc(4)Hex(5)Fuc(2)NeuAc(1) | C[+57], N[+2206] | 1434.2756 | 3 | 4300.8122 | 4300.8028 | 0 | 2.2 | 485 | Specific | 419.16 | 277.88 | 277.88 | 7.19 | 39 | >sp\|P02788\|TRFL_HUMAN Lactotransferrin OS=Homo sapiens OX=9606 GN=LTF PE=1 SV=6 | 1270 | 20201114_13813_1ng_glyco_hcd30_etd30_compound15.19674.19674.3 | controllerType=0 controllerNumber=1 scan=19674 | 44.7377 |
| 15049:3 | 1 | R.TAGWNIPMGLLFN[+1216.423]QTGSC[+57.021]K.F | HexNAc(2)Hex(5) | C[+57], N[+1216] | 1104.487 | 3 | 3311.4463 | 3311.4328 | 0 | 4.1 | 485 | Specific | 425 | 406.44 | 211.78 | 7.9 | 39 | >sp\|P02788\|TRFL_HUMAN Lactotransferrin OS=Homo sapiens OX=9606 GN=LTF PE=1 SV=6 | 1270 | 20201114_13813_1ng_glyco_hcd30_etd30_compound15.18588.18588.3 | controllerType=0 controllerNumber=1 scan=18588 | 41.5038 |
| 17524:3 | 1 | R.TAGWNIPMGLLFN[+2350.830]QTGSC[+57.021]K.F | HexNAc(4)Hex(5)Fuc(1)NeuAc(2) | C[+57], N[+2351] | 1482.6242 | 3 | 4445.8581 | 4445.8403 | 0 | 4 | 485 | Specific | 460.03 | 319.88 | 319.88 | 6.9 | 39 | >sp\|P02788\|TRFL_HUMAN Lactotransferrin OS=Homo sapiens OX=9606 GN=LTF PE=1 SV=6 | 1270 | 20201114_13813_1ng_glyco_hcd30_etd30_compound15.21216.21216.3 | controllerType=0 controllerNumber=1 scan=21216 | 49.2466 |
| 14953:3 | 1 | R.TAGWNIPM[+15.995]GLLFN[+2059.735]QTGSC[+57.021]K.F | HexNAc(4)Hex(5)Fuc(1)NeuAc(1) | C[+57], M[+16], N[+2060] | 1390.9277 | 3 | 4170.7685 | 4170.7398 | 0 | 6.9 | 485 | Specific | 264.89 | 233.52 | 233.52 | 4.98 | 39 | >sp\|P02788\|TRFL_HUMAN Lactotransferrin OS=Homo sapiens OX=9606 GN=LTF PE=1 SV=6 | 1270 | 20201114_13813_1ng_glyco_hcd30_etd30_compound15.18486.18486.3 | controllerType=0 controllerNumber=1 scan=18486 | 41.1815 |
| 15166:3 | 1 | R.TAGWNIPM[+15.995]GLLFN[+2059.735]QTGSC[+57.021]K.F | HexNAc(4)Hex(5)Fuc(1)NeuAc(1) | C[+57], M[+16], N[+2060] | 1390.924 | 3 | 4170.7575 | 4170.7398 | 0 | 4.2 | 485 | Specific | 323.28 | 257.23 | 257.23 | 5.11 | 39 | >sp\|P02788\|TRFL_HUMAN Lactotransferrin OS=Homo sapiens OX=9606 GN=LTF PE=1 SV=6 | 1270 | 20201114_13813_1ng_glyco_hcd30_etd30_compound15.18712.18712.3 | controllerType=0 controllerNumber=1 scan=18712 | 41.8475 |
| 16316:4 | 1 | R.TAGWNIPMGLLFN[+1913.677]QTGSC[+57.021]KFDEYFSQSC[+57.021]APGSDPR.S | HexNAc(4)Hex(5)NeuAc(1) | C[+57]*2, N[+1914] | 1463.8762 | 4 | 5852.4831 | 5852.4341 | 0 | 8.4 | 485 | Specific | 707.72 | 637.52 | 637.52 | 9.87 | 39 | >sp\|P02788\|TRFL_HUMAN Lactotransferrin OS=Homo sapiens OX=9606 GN=LTF PE=1 SV=6 | 1270 | 20201114_13813_1ng_glyco_hcd30_etd30_compound15.19933.19933.4 | controllerType=0 controllerNumber=1 scan=19933 | 45.6268 |
| 08117:3 | 1 | R.N[+2059.735]GSDC[+57.021]PDKFC[+57.021]LFQSETK.N | HexNAc(4)Hex(5)Fuc(1)NeuAc(1) | C[+57]*2, N[+2060] | 1364.8763 | 3 | 4092.6143 | 4092.6088 | 0 | 1.3 | 642 | Specific | 259.1 | 161.91 | 161.91 | 5.41 | 39 | >sp\|P02788\|TRFL_HUMAN Lactotransferrin OS=Homo sapiens OX=9606 GN=LTF PE=1 SV=6 | 1270 | 20201114_13813_1ng_glyco_hcd30_etd30_compound15.11321.11321.3 | controllerType=0 controllerNumber=1 scan=11321 | 26.6884 |
| 08084:3 | 1 | R.N[+2205.793]GSDC[+57.021]PDKFC[+57.021]LFQSETK.N | HexNAc(4)Hex(5)Fuc(2)NeuAc(1) | C[+57]*2, N[+2206] | 1413.5649 | 3 | 4238.6801 | 4238.6667 | 0 | 3.1 | 642 | Specific | 313.39 | 215.95 | 215.95 | 5.35 | 39 | >sp\|P02788\|TRFL_HUMAN Lactotransferrin OS=Homo sapiens OX=9606 GN=LTF PE=1 SV=6 | 1270 | 20201114_13813_1ng_glyco_hcd30_etd30_compound15.11287.11287.3 | controllerType=0 controllerNumber=1 scan=11287 | 26.6226 |
| 16077:4 | 2 | R.LSLHRPALEDLLLGSEAN[+1872.651]LTC[+57.021]TLTGLR.D | HexNAc(3)Hex(6)NeuAc(1) | C[+57], N[+1873] | 1209.8244 | 4 | 4836.2757 | 4836.2487 | 0 | 5.6 | 127 | Specific | 297.91 | 297.91 | 297.91 | 4.45 | 21 | >sp\|P01876\|IGHA1_HUMAN Immunoglobulin heavy constant alpha 1 OS=Homo sapiens OX=9606 GN=IGHA1 PE=1 SV=2 | 255 | 20201114_13813_1ng_glyco_hcd30_etd30_compound15.19677.19677.4 | controllerType=0 controllerNumber=1 scan=19677 | 44.7451 |
| 15405:3 | 2 | R.LSLHRPALEDLLLGSEAN[+1501.555]LTC[+57.021]TLTGLR.D | HexNAc(5)Hex(3) | C[+57], N[+1502] | 1489.0628 | 3 | 4465.1739 | 4465.1536 | 0 | 4.5 | 127 | Specific | 298.77 | 298.77 | 298.77 | 5.37 | 21 | >sp\|P01876\|IGHA1_HUMAN Immunoglobulin heavy constant alpha 1 OS=Homo sapiens OX=9606 GN=IGHA1 PE=1 SV=2 | 255 | 20201114_13813_1ng_glyco_hcd30_etd30_compound15.18967.18967.3 | controllerType=0 controllerNumber=1 scan=18967 | 42.714 |
| 15367:3 | 2 | R.LSLHRPALEDLLLGSEAN[+1501.555]LTC[+57.021]TLTGLR.D | HexNAc(5)Hex(3) | C[+57], N[+1502] | 1489.0578 | 3 | 4465.1588 | 4465.1536 | 0 | 1.2 | 127 | Specific | 484.3 | 429.5 | 429.5 | 7.69 | 21 | >sp\|P01876\|IGHA1_HUMAN Immunoglobulin heavy constant alpha 1 OS=Homo sapiens OX=9606 GN=IGHA1 PE=1 SV=2 | 255 | 20201114_13813_1ng_glyco_hcd30_etd30_compound15.18926.18926.3 | controllerType=0 controllerNumber=1 scan=18926 | 42.5759 |
| 15307:4 | 2 | R.LSLHRPALEDLLLGSEAN[+1663.608]LTC[+57.021]TLTGLR.D | HexNAc(5)Hex(4) | C[+57], N[+1664] | 1157.5678 | 4 | 4627.2493 | 4627.2064 | 0 | 9.3 | 127 | Specific | 759.02 | 759.02 | 759.02 | 12.27 | 21 | >sp\|P01876\|IGHA1_HUMAN Immunoglobulin heavy constant alpha 1 OS=Homo sapiens OX=9606 GN=IGHA1 PE=1 SV=2 | 255 | 20201114_13813_1ng_glyco_hcd30_etd30_compound15.18861.18861.2 | controllerType=0 controllerNumber=1 scan=18861 | 42.3162 |
| 15362:4 | 2 | R.LSLHRPALEDLLLGSEAN[+1216.423]LTC[+57.021]TLTGLR.D | HexNAc(2)Hex(5) | C[+57], N[+1216] | 1045.7613 | 4 | 4180.0233 | 4180.0211 | 0 | 0.5 | 127 | Specific | 866.12 | 859.14 | 859.14 | 14.47 | 21 | >sp\|P01876\|IGHA1_HUMAN Immunoglobulin heavy constant alpha 1 OS=Homo sapiens OX=9606 GN=IGHA1 PE=1 SV=2 | 255 | 20201114_13813_1ng_glyco_hcd30_etd30_compound15.18921.18921.4 | controllerType=0 controllerNumber=1 scan=18921 | 42.5577 |
| 15312:4 | 2 | R.LSLHRPALEDLLLGSEAN[+1864.634]LTC[+57.021]TLTGLR.D | HexNAc(2)Hex(9) | C[+57], N[+1865] | 1207.8243 | 4 | 4828.2752 | 4828.2324 | 0 | 8.9 | 127 | Specific | 904.55 | 899.2 | 899.2 | 15.27 | 21 | >sp\|P01876\|IGHA1_HUMAN Immunoglobulin heavy constant alpha 1 OS=Homo sapiens OX=9606 GN=IGHA1 PE=1 SV=2 | 255 | 20201114_13813_1ng_glyco_hcd30_etd30_compound15.18867.18867.4 | controllerType=0 controllerNumber=1 scan=18867 | 42.3356 |
| 15372:4 | 2 | R.LSLHRPALEDLLLGSEAN[+1663.608]LTC[+57.021]TLTGLR.D | HexNAc(5)Hex(4) | C[+57], N[+1664] | 1157.568 | 4 | 4627.2503 | 4627.2064 | 0 | 9.5 | 127 | Specific | 917.23 | 909.51 | 909.51 | 15.27 | 21 | >sp\|P01876\|IGHA1_HUMAN Immunoglobulin heavy constant alpha 1 OS=Homo sapiens OX=9606 GN=IGHA1 PE=1 SV=2 | 255 | 20201114_13813_1ng_glyco_hcd30_etd30_compound15.18932.18932.4 | controllerType=0 controllerNumber=1 scan=18932 | 42.5979 |

Table S4. Cont.

| Query #:z | Protein Rank | Glycopeptide | Glycan | Modification Type(s) | Observed m/z | z | Observed (M+H) | Calc. mass (M+H) | Off-by-x error | Mass error (ppm) | StartPosition | Cleavage | Score | Delta | Delta Mod | \|Log Prob\| | # of unique peptides | Protein Name | Protein DB number | Comment | Scan # | Scan Time |
|---|---|---|---|---|---|---|---|---|---|---|---|---|---|---|---|---|---|---|---|---|---|---|
| 15357:4 | 2 | R.LSLHRPALEDLLLGSEAN[+1501.555]LTC[+57.021]TLTGLR.D | HexNAc(5)Hex(3) | C[+57], N[+1502] | 1117.0442 | 4 | 4465.1551 | 4465.1536 | 0 | 0.3 | 127 | Specific | 924.87 | 913.81 | 913.81 | 15.47 | 21 | >sp\|P01876\|IGHA1_HUMAN Immunoglobulin heavy constant alpha 1 OS=Homo sapiens OX=9606 GN=IGHA1 PE=1 SV=2 | 255 | 20201114_13813_1ng_glyco_hcd30_etd30_compound15.18916.18916.4 | controllerType=0 controllerNumber=1 scan=18916 | 42.5358 |
| 15401:4 | 2 | R.LSLHRPALEDLLLGSEAN[+1501.555]LTC[+57.021]TLTGLR.D | HexNAc(5)Hex(3) | C[+57], N[+1502] | 1117.0483 | 4 | 4465.1712 | 4465.1536 | 0 | 3.9 | 127 | Specific | 947.82 | 943.37 | 943.37 | 15.66 | 21 | >sp\|P01876\|IGHA1_HUMAN Immunoglobulin heavy constant alpha 1 OS=Homo sapiens OX=9606 GN=IGHA1 PE=1 SV=2 | 255 | 20201114_13813_1ng_glyco_hcd30_etd30_compound15.18963.18963.4 | controllerType=0 controllerNumber=1 scan=18963 | 42.7066 |
| 15369:4 | 2 | R.LSLHRPALEDLLLGSEAN[+1460.529]LTC[+57.021]TLTGLR.D | HexNAc(4)Hex(4) | C[+57], N[+1461] | 1106.787 | 4 | 4424.1263 | 4424.1271 | 0 | -0.2 | 127 | Specific | 1020.43 | 927.08 | 927.08 | 17.47 | 21 | >sp\|P01876\|IGHA1_HUMAN Immunoglobulin heavy constant alpha 1 OS=Homo sapiens OX=9606 GN=IGHA1 PE=1 SV=2 | 255 | 20201114_13813_1ng_glyco_hcd30_etd30_compound15.18928.18928.4 | controllerType=0 controllerNumber=1 scan=18928 | 42.5868 |
| 15324:4 | 2 | R.LSLHRPALEDLLLGSEAN[+1540.529]LTC[+57.021]TLTGLR.D | HexNAc(2)Hex(7) | C[+57], N[+1541] | 1126.7884 | 4 | 4504.1317 | 4504.1268 | 0 | 1.1 | 127 | Specific | 1259.19 | 1143.74 | 1143.74 | 23.09 | 21 | >sp\|P01876\|IGHA1_HUMAN Immunoglobulin heavy constant alpha 1 OS=Homo sapiens OX=9606 GN=IGHA1 PE=1 SV=2 | 255 | 20201114_13813_1ng_glyco_hcd30_etd30_compound15.18880.18880.4 | controllerType=0 controllerNumber=1 scan=18880 | 42.3987 |
| 15378:4 | 2 | R.LSLHRPALEDLLLGSEAN[+1216.423]LTC[+57.021]TLTGLR.D | HexNAc(2)Hex(5) | C[+57], N[+1216] | 1045.7701 | 4 | 4180.0584 | 4180.0211 | 0 | 8.9 | 127 | Specific | 1301.62 | 1222.44 | 1222.44 | 23.27 | 21 | >sp\|P01876\|IGHA1_HUMAN Immunoglobulin heavy constant alpha 1 OS=Homo sapiens OX=9606 GN=IGHA1 PE=1 SV=2 | 255 | 20201114_13813_1ng_glyco_hcd30_etd30_compound15.18938.18938.4 | controllerType=0 controllerNumber=1 scan=18938 | 42.6168 |
| 11675:3 | 2 | R.LAGKPTHVN[+2262.814]VSVVMAEVDGTC[+57.021]Y.- | HexNAc(5)Hex(5)Fuc(1)NeuAc(1) | C[+57], N[+2263] | 1537.325 | 3 | 4609.9605 | 4609.9564 | 0 | 0.9 | 332 | CRagged | 202.86 | 52.07 | 52.07 | 2.57 | 21 | >sp\|P01876\|IGHA1_HUMAN Immunoglobulin heavy constant alpha 1 OS=Homo sapiens OX=9606 GN=IGHA1 PE=1 SV=2 | 255 | 20201114_13813_1ng_glyco_hcd30_etd30_compound15.15037.15037.3 | controllerType=0 controllerNumber=1 scan=15037 | 33.6336 |
| 10054:3 | 2 | R.LAGKPTHVN[+1702.581]VSVVMAEVDGTC[+57.021]Y.- | HexNAc(2)Hex(8) | C[+57], N[+1703] | 1350.5816 | 3 | 4049.7302 | 4049.7234 | 0 | 1.7 | 332 | CRagged | 210.19 | 205.57 | 171.55 | 3.55 | 21 | >sp\|P01876\|IGHA1_HUMAN Immunoglobulin heavy constant alpha 1 OS=Homo sapiens OX=9606 GN=IGHA1 PE=1 SV=2 | 255 | 20201114_13813_1ng_glyco_hcd30_etd30_compound15.13334.13334.3 | controllerType=0 controllerNumber=1 scan=13334 | 30.3697 |
| 10069:3 | 2 | R.LAGKPTHVN[+1378.476]VSVVMAEVDGTC[+57.021]Y.- | HexNAc(2)Hex(6) | C[+57], N[+1378] | 1242.5486 | 3 | 3725.6314 | 3725.6178 | 0 | 3.7 | 332 | CRagged | 227.5 | 227.5 | 171.59 | 3.19 | 21 | >sp\|P01876\|IGHA1_HUMAN Immunoglobulin heavy constant alpha 1 OS=Homo sapiens OX=9606 GN=IGHA1 PE=1 SV=2 | 255 | 20201114_13813_1ng_glyco_hcd30_etd30_compound15.13350.13350.3 | controllerType=0 controllerNumber=1 scan=13350 | 30.422 |
| 10073:3 | 2 | R.LAGKPTHVN[+1540.529]VSVVMAEVDGTC[+57.021]Y.- | HexNAc(2)Hex(7) | C[+57], N[+1541] | 1296.5677 | 3 | 3887.6885 | 3887.6706 | 0 | 4.6 | 332 | CRagged | 235.81 | 235.81 | 179.93 | 3.01 | 21 | >sp\|P01876\|IGHA1_HUMAN Immunoglobulin heavy constant alpha 1 OS=Homo sapiens OX=9606 GN=IGHA1 PE=1 SV=2 | 255 | 20201114_13813_1ng_glyco_hcd30_etd30_compound15.13354.13354.3 | controllerType=0 controllerNumber=1 scan=13354 | 30.4325 |
| 10117:3 | 2 | R.LAGKPTHVN[+1809.666]VSVVMAEVDGTC[+57.021]Y.- | HexNAc(5)Hex(4)Fuc(1) | C[+57], N[+1810] | 1386.2761 | 3 | 4156.8136 | 4156.8081 | 0 | 1.3 | 332 | CRagged | 252.53 | 154.1 | 154.1 | 3.45 | 21 | >sp\|P01876\|IGHA1_HUMAN Immunoglobulin heavy constant alpha 1 OS=Homo sapiens OX=9606 GN=IGHA1 PE=1 SV=2 | 255 | 20201114_13813_1ng_glyco_hcd30_etd30_compound15.13401.13401.3 | controllerType=0 controllerNumber=1 scan=13401 | 30.5435 |
| 10163:3 | 2 | R.LAGKPTHVN[+1647.613]VSVVMAEVDGTC[+57.021]Y.- | HexNAc(5)Hex(3)Fuc(1) | C[+57], N[+1648] | 1332.2588 | 3 | 3994.762 | 3994.7553 | 0 | 1.7 | 332 | CRagged | 261.18 | 144.78 | 144.78 | 3.38 | 21 | >sp\|P01876\|IGHA1_HUMAN Immunoglobulin heavy constant alpha 1 OS=Homo sapiens OX=9606 GN=IGHA1 PE=1 SV=2 | 255 | 20201114_13813_1ng_glyco_hcd30_etd30_compound15.13451.13451.3 | controllerType=0 controllerNumber=1 scan=13451 | 30.6925 |
| 10160:3 | 2 | R.LAGKPTHVN[+1647.613]VSVVMAEVDGTC[+57.021]Y.- | HexNAc(5)Hex(3)Fuc(1) | C[+57], N[+1648] | 1332.2576 | 3 | 3994.7583 | 3994.7553 | 0 | 0.8 | 332 | CRagged | 265.13 | 166.32 | 166.32 | 3.14 | 21 | >sp\|P01876\|IGHA1_HUMAN Immunoglobulin heavy constant alpha 1 OS=Homo sapiens OX=9606 GN=IGHA1 PE=1 SV=2 | 255 | 20201114_13813_1ng_glyco_hcd30_etd30_compound15.13447.13447.3 | controllerType=0 controllerNumber=1 scan=13447 | 30.6776 |
| 16257:3 | 6 | K.QIGLYPVLVIDSSGYVNPN[+1913.677]YTGR.I | HexNAc(4)Hex(5)NeuAc(1) | N[+1914] | 1480.3358 | 3 | 4438.9927 | 4438.9805 | 0 | 2.8 | 168 | Specific | 476.9 | 391.13 | 391.13 | 7.13 | 21 | >sp\|P01833\|PIGR_HUMAN Polymeric immunoglobulin receptor OS=Homo sapiens OX=9606 GN=PIGR PE=1 SV=4 | 902 | 20201114_13813_1ng_glyco_hcd30_etd30_compound15.19870.19870.3 | controllerType=0 controllerNumber=1 scan=19870 | 45.4203 |
| 17304:3 | 6 | R.LSLLEEPGN[+2570.925]GTFTVILNQLTSR.D | HexNAc(5)Hex(6)Fuc(2)NeuAc(1) | N[+2571] | 1658.4204 | 3 | 4973.2466 | 4973.2176 | 0 | 5.8 | 413 | Specific | 271.02 | 171.15 | 171.15 | 3.58 | 21 | >sp\|P01833\|PIGR_HUMAN Polymeric immunoglobulin receptor OS=Homo sapiens OX=9606 GN=PIGR PE=1 SV=4 | 902 | 20201114_13813_1ng_glyco_hcd30_etd30_compound15.20981.20981.3 | controllerType=0 controllerNumber=1 scan=20981 | 48.4948 |
| 19048:3 | 6 | R.LSLLEEPGN[+2350.830]GTFTVILNQLTSR.D | HexNAc(4)Hex(5)Fuc(1)NeuAc(2) | N[+2351] | 1585.0506 | 3 | 4753.1372 | 4753.1229 | 0 | 3 | 413 | Specific | 596.28 | 434.23 | 434.23 | 8.15 | 21 | >sp\|P01833\|PIGR_HUMAN Polymeric immunoglobulin receptor OS=Homo sapiens OX=9606 GN=PIGR PE=1 SV=4 | 902 | 20201114_13813_1ng_glyco_hcd30_etd30_compound15.22829.22829.3 | controllerType=0 controllerNumber=1 scan=22829 | 53.1389 |
| 15534:3 | 6 | K.IIEGEPNLKVPGN[+2350.830]VTAVLGETLK.V | HexNAc(4)Hex(5)Fuc(1)NeuAc(2) | N[+2351] | 1581.398 | 3 | 4742.1795 | 4742.1797 | 0 | -0.1 | 457 | Specific | 356.67 | 284.86 | 284.86 | 3.35 | 21 | >sp\|P01833\|PIGR_HUMAN Polymeric immunoglobulin receptor OS=Homo sapiens OX=9606 GN=PIGR PE=1 SV=4 | 902 | 20201114_13813_1ng_glyco_hcd30_etd30_compound15.19103.19103.2 | controllerType=0 controllerNumber=1 scan=19103 | 43.1313 |
| 14220:3 | 6 | K.IIEGEPNLKVPGN[+2205.793]VTAVLGETLK.V | HexNAc(4)Hex(5)Fuc(2)NeuAc(1) | N[+2206] | 1533.0638 | 3 | 4597.1768 | 4597.1422 | 0 | 7.5 | 457 | Specific | 373.9 | 280.23 | 280.23 | 3.55 | 21 | >sp\|P01833\|PIGR_HUMAN Polymeric immunoglobulin receptor OS=Homo sapiens OX=9606 GN=PIGR PE=1 SV=4 | 902 | 20201114_13813_1ng_glyco_hcd30_etd30_compound15.17710.17710.3 | controllerType=0 controllerNumber=1 scan=17710 | 39.2166 |
| 14980:5 | 6 | K.IIEGEPNLKVPGN[+2350.830]VTAVLGETLKVPC[+57.021]HFPC[+57.021]K.F | HexNAc(4)Hex(5)Fuc(1)NeuAc(2) | C[+57]*2, N[+2351] | 1154.3447 | 5 | 5767.6945 | 5767.6372 | 0 | 9.9 | 457 | Specific | 459.83 | 459.83 | 459.83 | 3.89 | 21 | >sp\|P01833\|PIGR_HUMAN Polymeric immunoglobulin receptor OS=Homo sapiens OX=9606 GN=PIGR PE=1 SV=4 | 902 | 20201114_13813_1ng_glyco_hcd30_etd30_compound15.18514.18514.5 | controllerType=0 controllerNumber=1 scan=18514 | 41.2819 |
| 12198:3 | 6 | K.VPGN[+2059.735]VTAVLGETLK.V | HexNAc(4)Hex(5)Fuc(1)NeuAc(1) | N[+2060] | 1153.1844 | 3 | 3457.5386 | 3457.5348 | 0 | 1.1 | 466 | Specific | 273.93 | 187.96 | 187.96 | 6.52 | 21 | >sp\|P01833\|PIGR_HUMAN Polymeric immunoglobulin receptor OS=Homo sapiens OX=9606 GN=PIGR PE=1 SV=4 | 902 | 20201114_13813_1ng_glyco_hcd30_etd30_compound15.15582.15582.3 | controllerType=0 controllerNumber=1 scan=15582 | 34.7307 |
| 12093:3 | 6 | K.VPGN[+2205.793]VTAVLGETLK.V | HexNAc(4)Hex(5)Fuc(2)NeuAc(1) | N[+2206] | 1201.8722 | 3 | 3603.6022 | 3603.5927 | 0 | 2.6 | 466 | Specific | 346.74 | 248.14 | 248.14 | 6.76 | 21 | >sp\|P01833\|PIGR_HUMAN Polymeric immunoglobulin receptor OS=Homo sapiens OX=9606 GN=PIGR PE=1 SV=4 | 902 | 20201114_13813_1ng_glyco_hcd30_etd30_compound15.15472.15472.3 | controllerType=0 controllerNumber=1 scan=15472 | 34.4626 |
| 12141:4 | 6 | K.VPGN[+2205.793]VTAVLGETLKVPC[+57.021]HFPC[+57.021]K.F | HexNAc(4)Hex(5)Fuc(2)NeuAc(1) | C[+57]*2, N[+2206] | 1158.0241 | 4 | 4629.0745 | 4629.0502 | 0 | 5.3 | 466 | Specific | 388.92 | 386.34 | 386.34 | 4.14 | 21 | >sp\|P01833\|PIGR_HUMAN Polymeric immunoglobulin receptor OS=Homo sapiens OX=9606 GN=PIGR PE=1 SV=4 | 902 | 20201114_13813_1ng_glyco_hcd30_etd30_compound15.15522.15522.4 | controllerType=0 controllerNumber=1 scan=15522 | 34.5908 |
| 05032:3 | 6 | K.WN[+1872.651]NTGC[+57.021]QALPSQDEGPSK.A | HexNAc(3)Hex(6)NeuAc(1) | C[+57], N[+1873] | 1287.8499 | 3 | 3861.5352 | 3861.5272 | 0 | 2.1 | 498 | Specific | 302.36 | 164.5 | 164.5 | 6.51 | 21 | >sp\|P01833\|PIGR_HUMAN Polymeric immunoglobulin receptor OS=Homo sapiens OX=9606 GN=PIGR PE=1 SV=4 | 902 | 20201114_13813_1ng_glyco_hcd30_etd30_compound15.8101.8101.3 | controllerType=0 controllerNumber=1 scan=8101 | 20.7683 |
| 05208:3 | 6 | K.WN[+1548.545]NTGC[+57.021]QALPSQDEGPSK.A | HexNAc(3)Hex(4)NeuAc(1) | C[+57], N[+1549] | 1179.8132 | 3 | 3537.4249 | 3537.4215 | 0 | 1 | 498 | Specific | 312.9 | 176.01 | 176.01 | 6.51 | 21 | >sp\|P01833\|PIGR_HUMAN Polymeric immunoglobulin receptor OS=Homo sapiens OX=9606 GN=PIGR PE=1 SV=4 | 902 | 20201114_13813_1ng_glyco_hcd30_etd30_compound15.8283.8283.3 | controllerType=0 controllerNumber=1 scan=8283 | 21.107 |
| 05101:3 | 6 | K.WN[+1710.598]NTGC[+57.021]QALPSQDEGPSK.A | HexNAc(3)Hex(5)NeuAc(1) | C[+57], N[+1711] | 1233.8307 | 3 | 3699.4777 | 3699.4743 | 0 | 0.9 | 498 | Specific | 382.36 | 218.08 | 218.08 | 6.73 | 21 | >sp\|P01833\|PIGR_HUMAN Polymeric immunoglobulin receptor OS=Homo sapiens OX=9606 GN=PIGR PE=1 SV=4 | 902 | 20201114_13813_1ng_glyco_hcd30_etd30_compound15.8173.8173.3 | controllerType=0 controllerNumber=1 scan=8173 | 20.9105 |
| 06900:3 | 6 | K.WN[+2204.772]NTGC[+57.021]QALPSQDEGPSK.A | HexNAc(4)Hex(5)NeuAc(2) | C[+57], N[+2205] | 1398.5555 | 3 | 4193.6519 | 4193.6491 | 0 | 0.7 | 498 | Specific | 388.47 | 268.04 | 268.04 | 6.4 | 21 | >sp\|P01833\|PIGR_HUMAN Polymeric immunoglobulin receptor OS=Homo sapiens OX=9606 GN=PIGR PE=1 SV=4 | 902 | 20201114_13813_1ng_glyco_hcd30_etd30_compound15.10042.10042.3 | controllerType=0 controllerNumber=1 scan=10042 | 24.2719 |
| 04964:3 | 6 | K.WN[+2059.735]NTGC[+57.021]QALPSQDEGPSK.A | HexNAc(4)Hex(5)Fuc(1)NeuAc(1) | C[+57], N[+2060] | 1350.2094 | 3 | 4048.6137 | 4048.6116 | 0 | 0.5 | 498 | Specific | 428.24 | 311.56 | 311.56 | 6.54 | 21 | >sp\|P01833\|PIGR_HUMAN Polymeric immunoglobulin receptor OS=Homo sapiens OX=9606 GN=PIGR PE=1 SV=4 | 902 | 20201114_13813_1ng_glyco_hcd30_etd30_compound15.8031.8031.3 | controllerType=0 controllerNumber=1 scan=8031 | 20.6417 |
| 05022:3 | 6 | K.WN[+1913.677]NTGC[+57.021]QALPSQDEGPSK.A | HexNAc(4)Hex(5)NeuAc(1) | C[+57], N[+1914] | 1301.5246 | 3 | 3902.5592 | 3902.5537 | 0 | 1.4 | 498 | Specific | 446.35 | 248.56 | 248.56 | 7.18 | 21 | >sp\|P01833\|PIGR_HUMAN Polymeric immunoglobulin receptor OS=Homo sapiens OX=9606 GN=PIGR PE=1 SV=4 | 902 | 20201114_13813_1ng_glyco_hcd30_etd30_compound15.8091.8091.3 | controllerType=0 controllerNumber=1 scan=8091 | 20.746 |
| 00958:3 | 10 | K.HYTN[+1647.613]SSQDVTVPC[+57.021]R.V | HexNAc(5)Hex(3)Fuc(1) | C[+57], N[+1648] | 1104.4599 | 3 | 3311.365 | 3311.3625 | 0 | 0.8 | 89 | Specific | 201.35 | 134.33 | 134.33 | 3.37 | 13 | >sp\|P01877\|IGHA2_HUMAN Immunoglobulin heavy constant alpha 2 OS=Homo sapiens OX=9606 GN=IGHA2 PE=1 SV=4 | 576 | 20201114_13813_1ng_glyco_hcd30_etd30_compound15.3853.3853.2 | controllerType=0 controllerNumber=1 scan=3853 | 12.9414 |
| 00917:3 | 10 | K.HYTN[+1809.666]SSQDVTVPC[+57.021]R.V | HexNAc(5)Hex(4)Fuc(1) | C[+57], N[+1810] | 1158.4782 | 3 | 3473.4201 | 3473.4154 | 0 | 1.4 | 89 | Specific | 328.86 | 273.96 | 273.96 | 6.73 | 13 | >sp\|P01877\|IGHA2_HUMAN Immunoglobulin heavy constant alpha 2 OS=Homo sapiens OX=9606 GN=IGHA2 PE=1 SV=4 | 576 | 20201114_13813_1ng_glyco_hcd30_etd30_compound15.3810.3810.3 | controllerType=0 controllerNumber=1 scan=3810 | 12.8543 |
| 00949:3 | 10 | K.HYTN[+1809.666]SSQDVTVPC[+57.021]R.V | HexNAc(5)Hex(4)Fuc(1) | C[+57], N[+1810] | 1158.475 | 3 | 3473.4105 | 3473.4154 | 0 | -1.4 | 89 | Specific | 392.65 | 301.08 | 301.08 | 6.06 | 13 | >sp\|P01877\|IGHA2_HUMAN Immunoglobulin heavy constant alpha 2 OS=Homo sapiens OX=9606 GN=IGHA2 PE=1 SV=4 | 576 | 20201114_13813_1ng_glyco_hcd30_etd30_compound15.3844.3844.3 | controllerType=0 controllerNumber=1 scan=3844 | 12.9206 |
| 00964:3 | 10 | K.HYTN[+1647.613]SSQDVTVPC[+57.021]R.V | HexNAc(5)Hex(3)Fuc(1) | C[+57], N[+1648] | 1104.4599 | 3 | 3311.365 | 3311.3625 | 0 | 0.8 | 89 | Specific | 510.74 | 430.14 | 430.14 | 7.77 | 13 | >sp\|P01877\|IGHA2_HUMAN Immunoglobulin heavy constant alpha 2 OS=Homo sapiens OX=9606 GN=IGHA2 PE=1 SV=4 | 576 | 20201114_13813_1ng_glyco_hcd30_etd30_compound15.3859.3859.3 | controllerType=0 controllerNumber=1 scan=3859 | 12.9708 |
| 02523:3 | 10 | K.TPLTAN[+1809.666]ITK.S | HexNAc(5)Hex(4)Fuc(1) | N[+1810] | 923.4145 | 3 | 2768.229 | 2768.2228 | 0 | 2.2 | 200 | Specific | 273.05 | 128.72 | 128.72 | 2.73 | 13 | >sp\|P01877\|IGHA2_HUMAN Immunoglobulin heavy constant alpha 2 OS=Homo sapiens OX=9606 GN=IGHA2 PE=1 SV=4 | 576 | 20201114_13813_1ng_glyco_hcd30_etd30_compound15.5492.5492.3 | controllerType=0 controllerNumber=1 scan=5492 | 16.0666 |
| 02554:2 | 10 | K.TPLTAN[+1647.613]ITK.S | HexNAc(5)Hex(3)Fuc(1) | N[+1648] | 1303.5905 | 2 | 2606.1737 | 2606.17 | 0 | 1.4 | 200 | Specific | 292.44 | 108.69 | 108.69 | 2.91 | 13 | >sp\|P01877\|IGHA2_HUMAN Immunoglobulin heavy constant alpha 2 OS=Homo sapiens OX=9606 GN=IGHA2 PE=1 SV=4 | 576 | 20201114_13813_1ng_glyco_hcd30_etd30_compound15.5524.5524.2 | controllerType=0 controllerNumber=1 scan=5524 | 16.1257 |
| 02518:2 | 10 | K.TPLTAN[+1809.666]ITK.S | HexNAc(5)Hex(4)Fuc(1) | N[+1810] | 1384.6178 | 2 | 2768.2284 | 2768.2228 | 0 | 2 | 200 | Specific | 301.53 | 108.16 | 108.16 | 2.91 | 13 | >sp\|P01877\|IGHA2_HUMAN Immunoglobulin heavy constant alpha 2 OS=Homo sapiens OX=9606 GN=IGHA2 PE=1 SV=4 | 576 | 20201114_13813_1ng_glyco_hcd30_etd30_compound15.5487.5487.2 | controllerType=0 controllerNumber=1 scan=5487 | 16.0531 |
| 02589:3 | 10 | K.TPLTAN[+1647.613]ITK.S | HexNAc(5)Hex(3)Fuc(1) | N[+1648] | 869.3953 | 3 | 2606.1713 | 2606.17 | 0 | 0.5 | 200 | Specific | 309.71 | 122.28 | 122.28 | 2.57 | 13 | >sp\|P01877\|IGHA2_HUMAN Immunoglobulin heavy constant alpha 2 OS=Homo sapiens OX=9606 GN=IGHA2 PE=1 SV=4 | 576 | 20201114_13813_1ng_glyco_hcd30_etd30_compound15.5561.5561.3 | controllerType=0 controllerNumber=1 scan=5561 | 16.211 |
| 12496:4 | 13 | K.LKELPGVC[+57.021]N[+2190.793]ETM[+15.995]MALWEEC[+57.021]KPC[+57.021 | HexNAc(6)Hex(6) | C[+57]*3, M[+16], N[+2191] | 1286.554 | 4 | 5143.194 | 5143.1728 | 0 | 4.1 | 95 | Specific | 228.61 | 170.88 | 0 | 2.54 | 9 | >sp\|P10909\|CLUS_HUMAN Clusterin OS=Homo sapiens OX=9606 GN=CLU PE=1 SV=1 | 1467 | 20201114_13813_1ng_glyco_hcd30_etd30_compound15.15891.15891.4 | controllerType=0 controllerNumber=1 scan=15891 | 35.3308 |
| 00132:4 | 13 | R.EIRHN[+2424.867]STGC[+57.021]LR.M | HexNAc(5)Hex(6)Fuc(1)NeuAc(1) | C[+57], N[+2425] | 942.6413 | 4 | 3767.5433 | 3767.5316 | 0 | 3.1 | 287 | Specific | 543.94 | 373.76 | 373.76 | 5.78 | 9 | >sp\|P10909\|CLUS_HUMAN Clusterin OS=Homo sapiens OX=9606 GN=CLU PE=1 SV=1 | 1467 | 20201114_13813_1ng_glyco_hcd30_etd30_compound15.2992.2992.4 | controllerType=0 controllerNumber=1 scan=2992 | 11.2744 |
| 00129:4 | 13 | R.EIRHN[+2059.735]STGC[+57.021]LR.M | HexNAc(4)Hex(5)Fuc(1)NeuAc(1) | C[+57], N[+2060] | 851.358 | 4 | 3402.4101 | 3402.3994 | 0 | 3.2 | 287 | Specific | 724.34 | 559.53 | 559.53 | 9.71 | 9 | >sp\|P10909\|CLUS_HUMAN Clusterin OS=Homo sapiens OX=9606 GN=CLU PE=1 SV=1 | 1467 | 20201114_13813_1ng_glyco_hcd30_etd30_compound15.2989.2989.4 | controllerType=0 controllerNumber=1 scan=2989 | 11.2627 |
| 17631:3 | 13 | K.MLN[+2059.735]TSSLLEQLNEQFNWVSR.L | HexNAc(4)Hex(5)Fuc(1)NeuAc(1) | N[+2060] | 1490.317 | 3 | 4468.9363 | 4468.9217 | 0 | 3.3 | 352 | Specific | 655.99 | 422.78 | 422.78 | 10.15 | 9 | >sp\|P10909\|CLUS_HUMAN Clusterin OS=Homo sapiens OX=9606 GN=CLU PE=1 SV=1 | 1467 | 20201114_13813_1ng_glyco_hcd30_etd30_compound15.21331.21331.3 | controllerType=0 controllerNumber=1 scan=21331 | 49.6481 |
| 09181:3 | 13 | R.LAN[+2205.793]LTQGEDQYYLR.V | HexNAc(4)Hex(5)Fuc(2)NeuAc(1) | N[+2206] | 1297.2154 | 3 | 3889.6316 | 3889.6265 | 0 | 1.3 | 372 | Specific | 268.92 | 190.29 | 190.29 | 6.52 | 9 | >sp\|P10909\|CLUS_HUMAN Clusterin OS=Homo sapiens OX=9606 GN=CLU PE=1 SV=1 | 1467 | 20201114_13813_1ng_glyco_hcd30_etd30_compound15.12427.12427.3 | controllerType=0 controllerNumber=1 scan=12427 | 28.7348 |
| 20106:3 | 15 | R.QLAHQSN[+2204.772]STNIFFSPVSIATAFAMLSLGTK.A | HexNAc(4)Hex(5)NeuAc(2) | N[+2205] | 1796.1441 | 3 | 5386.4178 | 5386.4075 | 0 | 1.9 | 64 | Specific | 357.63 | 234.12 | 234.12 | 6.32 | 5 | >sp\|P01009\|A1AT_HUMAN Alpha-1-antitrypsin OS=Homo sapiens OX=9606 GN=SERPINA1 PE=1 SV=3 | 642 | 20201114_13813_1ng_glyco_hcd30_etd30_compound15.23946.23946.3 | controllerType=0 controllerNumber=1 scan=23946 | 55.8364 |
| 19024:4 | 15 | K.ADTHDEILEGLNFN[+2861.000]LTEIPEAQIHEGFQELLR.T | HexNAc(5)Hex(6)NeuAc(3) | N[+2861] | 1638.9622 | 4 | 6552.8268 | 6552.8239 | 0 | 0.4 | 94 | Specific | 302.53 | 302.53 | 302.53 | 4.97 | 5 | >sp\|P01009\|A1AT_HUMAN Alpha-1-antitrypsin OS=Homo sapiens OX=9606 GN=SERPINA1 PE=1 SV=3 | 642 | 20201114_13813_1ng_glyco_hcd30_etd30_compound15.22804.22804.4 | controllerType=0 controllerNumber=1 scan=22804 | 53.0707 |
| 06843:3 | 17 | K.WEC[+57.021]KN[+2059.735]DTLLGIK.G | HexNAc(4)Hex(5)Fuc(1)NeuAc(1) | C[+57], N[+2060] | 1179.5024 | 3 | 3536.4926 | 3536.4865 | 0 | 1.7 | 100 | Specific | 289.11 | 174.1 | 174.1 | 3.06 | 7 | >sp\|P22897\|MRC1_HUMAN Macrophage mannose receptor 1 OS=Homo sapiens OX=9606 GN=MRC1 PE=1 SV=1 | 1049 | 20201114_13813_1ng_glyco_hcd30_etd30_compound15.9983.9983.3 | controllerType=0 controllerNumber=1 scan=9983 | 24.1422 |
| 04949:3 | 19 | R.LSPN[+2205.793]ASAEHLELR.W | HexNAc(4)Hex(5)Fuc(2)NeuAc(1) | N[+2206] | 1214.8537 | 3 | 3642.5465 | 3642.5421 | 0 | 1.2 | 52 | Specific | 350.97 | 235.36 | 235.36 | 3.91 | 6 | >sp\|Q13410\|BT1A1_HUMAN Butyrophilin subfamily 1 member A1 OS=Homo sapiens OX=9606 GN=BTN1A1 PE=1 SV=3 | 179 | 20201114_13813_1ng_glyco_hcd30_etd30_compound15.8015.8015.3 | controllerType=0 controllerNumber=1 scan=8015 | 20.6073 |
| 06809:3 | 19 | R.LSPN[+2350.830]ASAEHLELR.W | HexNAc(4)Hex(5)Fuc(1)NeuAc(2) | N[+2351] | 1263.201 | 3 | 3787.5884 | 3787.5796 | 0 | 2.3 | 52 | Specific | 381.24 | 242.2 | 242.2 | 5.14 | 6 | >sp\|Q13410\|BT1A1_HUMAN Butyrophilin subfamily 1 member A1 OS=Homo sapiens OX=9606 GN=BTN1A1 PE=1 SV=3 | 179 | 20201114_13813_1ng_glyco_hcd30_etd30_compound15.9948.9948.3 | controllerType=0 controllerNumber=1 scan=9948 | 24.0812 |
| 02162:4 | 23 | K.QTDEIKDTRN[+2059.735]ESTQNC[+57.021]VVAEPEK.M | HexNAc(4)Hex(5)Fuc(1)NeuAc(1) | C[+57], N[+2060] | 1188.5069 | 4 | 4751.0057 | 4750.9876 | 0 | 3.8 | 60 | Specific | 614.16 | 524.91 | 524.91 | 8.23 | 3 | >sp\|P47710\|CASA1_HUMAN Alpha-S1-casein OS=Homo sapiens OX=9606 GN=CSN1S1 PE=1 SV=1 | 989 | 20201114_13813_1ng_glyco_hcd30_etd30_compound15.5113.5113.4 | controllerType=0 controllerNumber=1 scan=5113 | 15.2879 |
| 00739:4 | 52 | K.TKPREEQFN[+1444.534]STFR.V | HexNAc(4)Hex(3)Fuc(1) | N[+1445] | 771.847 | 4 | 3084.3662 | 3084.3526 | 0 | 4.4 | 168 | Specific | 270.06 | 270.06 | 270.06 | 3.1 | 4 | >sp\|P01859\|IGHG2_HUMAN Immunoglobulin heavy constant gamma 2 OS=Homo sapiens OX=9606 GN=IGHG2 PE=1 SV=2 | 387 | 20201114_13813_1ng_glyco_hcd30_etd30_compound15.3626.3626.4 | controllerType=0 controllerNumber=1 scan=3626 | 12.5193 |
| 00720:4 | 52 | K.TKPREEQFN[+1606.587]STFR.V | HexNAc(4)Hex(4)Fuc(1) | N[+1607] | 812.3578 | 4 | 3246.4094 | 3246.4054 | 0 | 1.2 | 168 | Specific | 311.07 | 311.07 | 311.07 | 3.6 | 4 | >sp\|P01859\|IGHG2_HUMAN Immunoglobulin heavy constant gamma 2 OS=Homo sapiens OX=9606 GN=IGHG2 PE=1 SV=2 | 387 | 20201114_13813_1ng_glyco_hcd30_etd30_compound15.3606.3606.2 | controllerType=0 controllerNumber=1 scan=3606 | 12.473 |
| 12722:3 | 60 | K.VVLHPN[+2204.772]YSQVDIGLIK.L | HexNAc(4)Hex(5)NeuAc(2) | N[+2205] | 1333.935 | 3 | 3999.7904 | 3999.7837 | 0 | 1.7 | 236 | Specific | 352.16 | 235.51 | 235.51 | 5.54 | 1 | >sp\|P00738\|HPT_HUMAN Haptoglobin OS=Homo sapiens OX=9606 GN=HP PE=1 SV=1 | 407 | 20201114_13813_1ng_glyco_hcd30_etd30_compound15.16129.16129.3 | controllerType=0 controllerNumber=1 scan=16129 | 35.8568 |
| 08846:4 | 64 | K.SC[+57.021]QHN[+2350.830]GTMYQHGEIFSAHELFPSR.L | HexNAc(4)Hex(5)Fuc(1)NeuAc(2) | C[+57], N[+2351] | 1296.7783 | 4 | 5184.0915 | 5184.0873 | 0 | 0.8 | 110 | Specific | 250.03 | 238.01 | 238.01 | 4.36 | 1 | >sp\|Q6WN34\|CRDL2_HUMAN Chordin-like protein 2 OS=Homo sapiens OX=9606 GN=CHRDL2 PE=1 SV=1 | 1189 | 20201114_13813_1ng_glyco_hcd30_etd30_compound15.12077.12077.4 | controllerType=0 controllerNumber=1 scan=12077 | 28.032 |
| 15931:4 | 1 | R.RTAGWNVPIGTLRPFLN[+1768.640]WTGPPEPIEAAVAR.F | HexNAc(4)Hex(5)Fuc(1) | N[+1769] | 1289.6238 | 4 | 5155.4733 | 5155.4516 | 0 | 4.2 | 140 | Specific | 858.03 | 858.03 | 858.03 | 14.83 | 37 | >sp\|P02788\|TRFL_HUMAN Lactotransferrin OS=Homo sapiens OX=9606 GN=LTF PE=1 SV=6 | 1270 | 20201116_13832_1ng_glyco_hcd30_etd30_compound15.19416.19416.4 | controllerType=0 controllerNumber=1 scan=19416 | 41.4518 |
| 17519:3 | 1 | R.TAGWNVPIGTLRPFLN[+2205.793]WTGPPEPIEAAVAR.F | HexNAc(4)Hex(5)Fuc(2)NeuAc(1) | N[+2206] | 1812.8437 | 3 | 5436.5166 | 5436.5038 | 0 | 2.4 | 141 | Specific | 219.03 | 141.89 | 141.89 | 3.63 | 37 | >sp\|P02788\|TRFL_HUMAN Lactotransferrin OS=Homo sapiens OX=9606 GN=LTF PE=1 SV=6 | 1270 | 20201116_13832_1ng_glyco_hcd30_etd30_compound15.21093.21093.2 | controllerType=0 controllerNumber=1 scan=21093 | 46.3565 |
| 17548:3 | 1 | R.TAGWNVPIGTLRPFLN[+2059.735]WTGPPEPIEAAVAR.F | HexNAc(4)Hex(5)Fuc(1)NeuAc(1) | N[+2060] | 1764.163 | 3 | 5290.4746 | 5290.4459 | 0 | 5.4 | 141 | Specific | 254.65 | 191.24 | 191.24 | 5.21 | 37 | >sp\|P02788\|TRFL_HUMAN Lactotransferrin OS=Homo sapiens OX=9606 GN=LTF PE=1 SV=6 | 1270 | 20201116_13832_1ng_glyco_hcd30_etd30_compound15.21123.21123.2 | controllerType=0 controllerNumber=1 scan=21123 | 46.4188 |
| 17502:3 | 1 | R.TAGWNVPIGTLRPFLN[+2205.793]WTGPPEPIEAAVAR.F | HexNAc(4)Hex(5)Fuc(2)NeuAc(1) | N[+2206] | 1812.8463 | 3 | 5436.5243 | 5436.5038 | 0 | 3.8 | 141 | Specific | 306.27 | 202.89 | 202.89 | 5.51 | 37 | >sp\|P02788\|TRFL_HUMAN Lactotransferrin OS=Homo sapiens OX=9606 GN=LTF PE=1 SV=6 | 1270 | 20201116_13832_1ng_glyco_hcd30_etd30_compound15.21075.21075.2 | controllerType=0 controllerNumber=1 scan=21075 | 46.315 |
| 16793:4 | 1 | R.[+42.011]TAGWNVPIGTLRPFLN[+2018.708]WTGPPEPIEAAVAR.F | HexNAc(3)Hex(6)Fuc(1)NeuAc(1) | .[42], N[+2019] | 1323.6257 | 4 | 5291.4811 | 5291.4299 | 0 | 9.7 | 141 | Specific | 307.47 | 307.47 | 307.47 | 2.79 | 37 | >sp\|P02788\|TRFL_HUMAN Lactotransferrin OS=Homo sapiens OX=9606 GN=LTF PE=1 SV=6 | 1270 | 20201116_13832_1ng_glyco_hcd30_etd30_compound15.20326.20326.4 | controllerType=0 controllerNumber=1 scan=20326 | 44.0651 |
| 18525:3 | 1 | R.TAGWNVPIGTLRPFLN[+2350.830]WTGPPEPIEAAVAR.F | HexNAc(4)Hex(5)Fuc(1)NeuAc(2) | N[+2351] | 1861.192 | 3 | 5581.5614 | 5581.5413 | 0 | 3.6 | 141 | Specific | 386.98 | 309.05 | 309.05 | 6.02 | 37 | >sp\|P02788\|TRFL_HUMAN Lactotransferrin OS=Homo sapiens OX=9606 GN=LTF PE=1 SV=6 | 1270 | 20201116_13832_1ng_glyco_hcd30_etd30_compound15.22157.22157.3 | controllerType=0 controllerNumber=1 scan=22157 | 49.1597 |
| 16830:4 | 1 | R.TAGWNVPIGTLRPFLN[+1768.640]WTGPPEPIEAAVAR.F | HexNAc(4)Hex(5)Fuc(1) | N[+1769] | 1250.5973 | 4 | 4999.3673 | 4999.3504 | 0 | 3.4 | 141 | Specific | 427.54 | 427.54 | 427.54 | 5.69 | 37 | >sp\|P02788\|TRFL_HUMAN Lactotransferrin OS=Homo sapiens OX=9606 GN=LTF PE=1 SV=6 | 1270 | 20201116_13832_1ng_glyco_hcd30_etd30_compound15.20365.20365.4 | controllerType=0 controllerNumber=1 scan=20365 | 44.1898 |
| 17484:3 | 1 | R.TAGWNVPIGTLRPFLN[+2059.735]WTGPPEPIEAAVAR.F | HexNAc(4)Hex(5)Fuc(1)NeuAc(1) | N[+2060] | 1764.159 | 3 | 5290.4625 | 5290.4459 | 0 | 3.1 | 141 | Specific | 459.32 | 348.18 | 348.18 | 4.74 | 37 | >sp\|P02788\|TRFL_HUMAN Lactotransferrin OS=Homo sapiens OX=9606 GN=LTF PE=1 SV=6 | 1270 | 20201116_13832_1ng_glyco_hcd30_etd30_compound15.21056.21056.3 | controllerType=0 controllerNumber=1 scan=21056 | 46.259 |
| 18560:4 | 1 | R.TAGWNVPIGTLRPFLN[+2350.830]WTGPPEPIEAAVAR.F | HexNAc(4)Hex(5)Fuc(1)NeuAc(2) | N[+2351] | 1396.1423 | 4 | 5581.5475 | 5581.5413 | 0 | 1.1 | 141 | Specific | 462.2 | 462.2 | 462.2 | 6.25 | 37 | >sp\|P02788\|TRFL_HUMAN Lactotransferrin OS=Homo sapiens OX=9606 GN=LTF PE=1 SV=6 | 1270 | 20201116_13832_1ng_glyco_hcd30_etd30_compound15.22194.22194.4 | controllerType=0 controllerNumber=1 scan=22194 | 49.2785 |
| 18519:4 | 1 | R.TAGWNVPIGTLRPFLN[+2350.830]WTGPPEPIEAAVAR.F | HexNAc(4)Hex(5)Fuc(1)NeuAc(2) | N[+2351] | 1396.145 | 4 | 5581.5583 | 5581.5413 | 0 | 3 | 141 | Specific | 665.16 | 651.46 | 651.46 | 10.41 | 37 | >sp\|P02788\|TRFL_HUMAN Lactotransferrin OS=Homo sapiens OX=9606 GN=LTF PE=1 SV=6 | 1270 | 20201116_13832_1ng_glyco_hcd30_etd30_compound15.22151.22151.4 | controllerType=0 controllerNumber=1 scan=22151 | 49.1442 |
| 17480:5 | 1 | R.TAGWNVPIGTLRPFLN[+2205.793]WTGPPEPIEAAVAR.F | HexNAc(4)Hex(5)Fuc(2)NeuAc(1) | N[+2206] | 1088.1088 | 5 | 5436.5147 | 5436.5038 | 0 | 2 | 141 | Specific | 862.71 | 829.45 | 829.45 | 14.11 | 37 | >sp\|P02788\|TRFL_HUMAN Lactotransferrin OS=Homo sapiens OX=9606 GN=LTF PE=1 SV=6 | 1270 | 20201116_13832_1ng_glyco_hcd30_etd30_compound15.21052.21052.5 | controllerType=0 controllerNumber=1 scan=21052 | 46.2436 |
| 17492:4 | 1 | R.TAGWNVPIGTLRPFLN[+2059.735]WTGPPEPIEAAVAR.F | HexNAc(4)Hex(5)Fuc(1)NeuAc(1) | N[+2060] | 1323.3737 | 4 | 5290.4728 | 5290.4459 | 0 | 5.1 | 141 | Specific | 900.3 | 880.25 | 880.25 | 15.47 | 37 | >sp\|P02788\|TRFL_HUMAN Lactotransferrin OS=Homo sapiens OX=9606 GN=LTF PE=1 SV=6 | 1270 | 20201116_13832_1ng_glyco_hcd30_etd30_compound15.21065.21065.4 | controllerType=0 controllerNumber=1 scan=21065 | 46.2871 |
| 17461:4 | 1 | R.TAGWNVPIGTLRPFLN[+2205.793]WTGPPEPIEAAVAR.F | HexNAc(4)Hex(5)Fuc(2)NeuAc(1) | N[+2206] | 1359.8861 | 4 | 5436.5226 | 5436.5038 | 0 | 3.5 | 141 | Specific | 929.2 | 909.75 | 909.75 | 15.41 | 37 | >sp\|P02788\|TRFL_HUMAN Lactotransferrin OS=Homo sapiens OX=9606 GN=LTF PE=1 SV=6 | 1270 | 20201116_13832_1ng_glyco_hcd30_etd30_compound15.21032.21032.4 | controllerType=0 controllerNumber=1 scan=21032 | 46.1855 |
| 15863:3 | 1 | R.TAGWNIPMGLLFN[+1622.582]QTGSC[+57.021]K.F | HexNAc(4)Hex(5) | C[+57], N[+1623] | 1239.8764 | 3 | 3717.6146 | 3717.5916 | 0 | 6.2 | 485 | Specific | 294.91 | 250.27 | 179.61 | 4.65 | 37 | >sp\|P02788\|TRFL_HUMAN Lactotransferrin OS=Homo sapiens OX=9606 GN=LTF PE=1 SV=6 | 1270 | 20201116_13832_1ng_glyco_hcd30_etd30_compound15.19344.19344.3 | controllerType=0 controllerNumber=1 scan=19344 | 41.2202 |
| 15665:3 | 1 | R.TAGWNIPMGLLFN[+1768.640]QTGSC[+57.021]K.F | HexNAc(4)Hex(5)Fuc(1) | C[+57], N[+1769] | 1288.559 | 3 | 3863.6625 | 3863.6495 | 0 | 3.4 | 485 | Specific | 358.36 | 269.53 | 269.53 | 5.4 | 37 | >sp\|P02788\|TRFL_HUMAN Lactotransferrin OS=Homo sapiens OX=9606 GN=LTF PE=1 SV=6 | 1270 | 20201116_13832_1ng_glyco_hcd30_etd30_compound15.19136.19136.3 | controllerType=0 controllerNumber=1 scan=19136 | 40.[illegible] |
| 15676:3 | 1 | R.TAGWNIPMGLLFN[+1768.640]QTGSC[+57.021]K.F | HexNAc(4)Hex(5)Fuc(1) | C[+57], N[+1769] | 1288.5605 | 3 | 3863.6669 | 3863.6495 | 0 | 4.5 | 485 | Specific | 393.17 | 283.48 | 283.48 | 4.71 | 37 | >sp\|P02788\|TRFL_HUMAN Lactotransferrin OS=Homo sapiens OX=9606 GN=LTF PE=1 SV=6 | 1270 | 20201116_13832_1ng_glyco_hcd30_etd30_compound15.19148.19148.3 | controllerType=0 controllerNumber=1 scan=19148 | 40.[illegible] |
| 17170:3 | 1 | R.TAGWNIPMGLLFN[+1913.677]QTGSC[+57.021]K.F | HexNAc(4)Hex(5)NeuAc(1) | C[+57], N[+1914] | 1336.9059 | 3 | 4008.7033 | 4008.687 | 0 | 4.1 | 485 | Specific | 410.7 | 279.03 | 279.03 | 5.85 | 37 | >sp\|P02788\|TRFL_HUMAN Lactotransferrin OS=Homo sapiens OX=9606 GN=LTF PE=1 SV=6 | 1270 | 20201116_13832_1ng_glyco_hcd30_etd30_compound15.20725.20725.3 | controllerType=0 controllerNumber=1 scan=20725 | 45.2804 |
| 17056:3 | 1 | R.TAGWNIPMGLLFN[+2059.735]QTGSC[+57.021]K.F | HexNAc(4)Hex(5)Fuc(1)NeuAc(1) | C[+57], N[+2060] | 1385.5936 | 3 | 4154.7661 | 4154.7449 | 0 | 5.1 | 485 | Specific | 443.57 | 322.08 | 322.08 | 5.87 | 37 | >sp\|P02788\|TRFL_HUMAN Lactotransferrin OS=Homo sapiens OX=9606 GN=LTF PE=1 SV=6 | 1270 | 20201116_13832_1ng_glyco_hcd30_etd30_compound15.20603.20603.3 | controllerType=0 controllerNumber=1 scan=20603 | 44.8659 |
| 16567:3 | 1 | R.TAGWNIPMGLLFN[+1622.582]QTGSC[+57.021]KFDEYFSQSC[+57.021]APGSDPR.S | HexNAc(4)Hex(5) | C[+57]*2, N[+1623] | 1854.4555 | 3 | 5561.352 | 5561.3387 | 0 | 2.4 | 485 | Specific | 223.67 | 189.89 | 118.57 | 3.56 | 37 | >sp\|P02788\|TRFL_HUMAN Lactotransferrin OS=Homo sapiens OX=9606 GN=LTF PE=1 SV=6 | 1270 | 20201116_13832_1ng_glyco_hcd30_etd30_compound15.20088.20088.3 | controllerType=0 controllerNumber=1 scan=20088 | 43.3732 |
| 16531:3 | 1 | R.TAGWNIPMGLLFN[+1768.640]QTGSC[+57.021]KFDEYFSQSC[+57.021]APGSDPR.S | HexNAc(4)Hex(5)Fuc(1) | C[+57]*2, N[+1769] | 1903.1457 | 3 | 5707.4226 | 5707.3966 | 0 | 4.6 | 485 | Specific | 258.09 | 217.25 | 217.25 | 5.23 | 37 | >sp\|P02788\|TRFL_HUMAN Lactotransferrin OS=Homo sapiens OX=9606 GN=LTF PE=1 SV=6 | 1270 | 20201116_13832_1ng_glyco_hcd30_etd30_compound15.20050.20050.3 | controllerType=0 controllerNumber=1 scan=20050 | 43.2571 |
| 17216:4 | 1 | R.TAGWNIPMGLLFN[+2205.793]QTGSC[+57.021]KFDEYFSQSC[+57.021]APGSDPR.S | HexNAc(4)Hex(5)Fuc(2)NeuAc(1) | C[+57]*2, N[+2206] | 1536.9008 | 4 | 6144.5812 | 6144.5499 | 0 | 5.1 | 485 | Specific | 354.71 | 329.53 | 329.53 | 5.43 | 37 | >sp\|P02788\|TRFL_HUMAN Lactotransferrin OS=Homo sapiens OX=9606 GN=LTF PE=1 SV=6 | 1270 | 20201116_13832_1ng_glyco_hcd30_etd30_compound15.20774.20774.4 | controllerType=0 controllerNumber=1 scan=20774 | 45.4429 |
| 16482:4 | 1 | R.[+42.011]TAGWNIPMGLLFN[+1872.651]QTGSC[+57.021]KFDEYFSQSC[+57.021]A | HexNAc(3)Hex(6)NeuAc(1) | .[42], C[+57]*2, N[+1873] | 1464.1228 | 4 | 5853.4694 | 5853.4181 | 0 | 8.8 | 485 | Specific | 360.31 | 299.12 | 299.12 | 2.95 | 37 | >sp\|P02788\|TRFL_HUMAN Lactotransferrin OS=Homo sapiens OX=9606 GN=LTF PE=1 SV=6 | 1270 | 20201116_13832_1ng_glyco_hcd30_etd30_compound15.19998.19998.4 | controllerType=0 controllerNumber=1 scan=19998 | 43.1216 |
| 16556:4 | 1 | R.TAGWNIPMGLLFN[+1606.587]QTGSC[+57.021]KFDEYFSQSC[+57.021]APGSDPR.S | HexNAc(4)Hex(4)Fuc(1) | C[+57]*2, N[+1607] | 1387.0942 | 4 | 5545.3551 | 5545.3438 | 0 | 2 | 485 | Specific | 364.56 | 320.91 | 320.91 | 3.87 | 37 | >sp\|P02788\|TRFL_HUMAN Lactotransferrin OS=Homo sapiens OX=9606 GN=LTF PE=1 SV=6 | 1270 | 20201116_13832_1ng_glyco_hcd30_etd30_compound15.20076.20076.4 | controllerType=0 controllerNumber=1 scan=20076 | 43.3319 |
| 17243:4 | 1 | R.TAGWNIPMGLLFN[+2059.735]QTGSC[+57.021]KFDEYFSQSC[+57.021]APGSDPR.S | HexNAc(4)Hex(5)Fuc(1)NeuAc(1) | C[+57]*2, N[+2060] | 1500.386 | 4 | 5998.5221 | 5998.492 | 0 | 5 | 485 | Specific | 455.54 | 422.08 | 422.08 | 6 | 37 | >sp\|P02788\|TRFL_HUMAN Lactotransferrin OS=Homo sapiens OX=9606 GN=LTF PE=1 SV=6 | 1270 | 20201116_13832_1ng_glyco_hcd30_etd30_compound15.20803.20803.4 | controllerType=0 controllerNumber=1 scan=20803 | 45.5479 |
| 17266:4 | 1 | R.TAGWNIPMGLLFN[+1913.677]QTGSC[+57.021]KFDEYFSQSC[+57.021]APGSDPR.S | HexNAc(4)Hex(5)NeuAc(1) | C[+57]*2, N[+1914] | 1463.8687 | 4 | 5852.4528 | 5852.4341 | 0 | 3.2 | 485 | Specific | 584.99 | 510.87 | 510.87 | 8.16 | 37 | >sp\|P02788\|TRFL_HUMAN Lactotransferrin OS=Homo sapiens OX=9606 GN=LTF PE=1 SV=6 | 1270 | 20201116_13832_1ng_glyco_hcd30_etd30_compound15.20827.20827.4 | controllerType=0 controllerNumber=1 scan=20827 | 45.6134 |
| 16547:4 | 1 | R.TAGWNIPMGLLFN[+1622.582]QTGSC[+57.021]KFDEYFSQSC[+57.021]APGSDPR.S | HexNAc(4)Hex(5) | C[+57]*2, N[+1623] | 1391.0962 | 4 | 5561.3629 | 5561.3387 | 0 | 4.4 | 485 | Specific | 595.66 | 594.29 | 166.45 | 8.53 | 37 | >sp\|P02788\|TRFL_HUMAN Lactotransferrin OS=Homo sapiens OX=9606 GN=LTF PE=1 SV=6 | 1270 | 20201116_13832_1ng_glyco_hcd30_etd30_compound15.20067.20067.4 | controllerType=0 controllerNumber=1 scan=20067 | 43.3097 |
| 16526:4 | 1 | R.TAGWNIPMGLLFN[+1768.640]QTGSC[+57.021]KFDEYFSQSC[+57.021]APGSDPR.S | HexNAc(4)Hex(5)Fuc(1) | C[+57]*2, N[+1769] | 1427.6134 | 4 | 5707.4318 | 5707.3966 | 0 | 6.2 | 485 | Specific | 645.19 | 611.59 | 611.59 | 8.52 | 37 | >sp\|P02788\|TRFL_HUMAN Lactotransferrin OS=Homo sapiens OX=9606 GN=LTF PE=1 SV=6 | 1270 | 20201116_13832_1ng_glyco_hcd30_etd30_compound15.20045.20045.4 | controllerType=0 controllerNumber=1 scan=20045 | 43.2488 |
| 07677:4 | 1 | K.FGRN[+2205.793]GSDC[+57.021]PDKFC[+57.021]LFQSETK.N | HexNAc(4)Hex(5)Fuc(2)NeuAc(1) | C[+57]*2, N[+2206] | 1150.4731 | 4 | 4598.8704 | 4598.8577 | 0 | 2.8 | 639 | Specific | 242.1 | 179.13 | 179.13 | 5.06 | 37 | >sp\|P02788\|TRFL_HUMAN Lactotransferrin OS=Homo sapiens OX=9606 GN=LTF PE=1 SV=6 | 1270 | 20201116_13832_1ng_glyco_hcd30_etd30_compound15.10764.10764.2 | controllerType=0 controllerNumber=1 scan=10764 | 24.8326 |
| 07682:4 | 1 | K.FGRN[+2205.793]GSDC[+57.021]PDKFC[+57.021]LFQSETK.N | HexNAc(4)Hex(5)Fuc(2)NeuAc(1) | C[+57]*2, N[+2206] | 1150.4739 | 4 | 4598.8738 | 4598.8577 | 0 | 3.5 | 639 | Specific | 389.72 | 304.57 | 304.57 | 5.93 | 37 | >sp\|P02788\|TRFL_HUMAN Lactotransferrin OS=Homo sapiens OX=9606 GN=LTF PE=1 SV=6 | 1270 | 20201116_13832_1ng_glyco_hcd30_etd30_compound15.10770.10770.4 | controllerType=0 controllerNumber=1 scan=10770 | 24.8494 |
| 16385:3 | 2 | R.LSLHRPALEDLLLGSEAN[+1216.423]LTC[+57.021]TLTGLR.D | HexNAc(2)Hex(5) | C[+57], N[+1216] | 1394.0195 | 3 | 4180.0438 | 4180.0211 | 0 | 5.4 | 127 | Specific | 297.21 | 294.52 | 294.52 | 5.48 | 26 | >sp\|P01876\|IGHA1_HUMAN Immunoglobulin heavy constant alpha 1 OS=Homo sapiens OX=9606 GN=IGHA1 PE=1 SV=2 | 255 | 20201116_13832_1ng_glyco_hcd30_etd30_compound15.19895.19895.3 | controllerType=0 controllerNumber=1 scan=19895 | 42.7957 |
| 17039:4 | 2 | R.LSLHRPALEDLLLGSEAN[+1872.651]LTC[+57.021]TLTGLR.D | HexNAc(3)Hex(6)NeuAc(1) | C[+57], N[+1873] | 1209.8232 | 4 | 4836.2708 | 4836.2487 | 0 | 4.6 | 127 | Specific | 401.21 | 401.21 | 401.21 | 5.8 | 26 | >sp\|P01876\|IGHA1_HUMAN Immunoglobulin heavy constant alpha 1 OS=Homo sapiens OX=9606 GN=IGHA1 PE=1 SV=2 | 255 | 20201116_13832_1ng_glyco_hcd30_etd30_compound15.20585.20585.4 | controllerType=0 controllerNumber=1 scan=20585 | 44.8182 |
| 16310:3 | 2 | R.LSLHRPALEDLLLGSEAN[+1702.581]LTC[+57.021]TLTGLR.D | HexNAc(2)Hex(8) | C[+57], N[+1703] | 1556.0718 | 3 | 4666.201 | 4666.1796 | 0 | 4.6 | 127 | Specific | 410.42 | 339.92 | 339.92 | 5.79 | 26 | >sp\|P01876\|IGHA1_HUMAN Immunoglobulin heavy constant alpha 1 OS=Homo sapiens OX=9606 GN=IGHA1 PE=1 SV=2 | 255 | 20201116_13832_1ng_glyco_hcd30_etd30_compound15.19815.19815.3 | controllerType=0 controllerNumber=1 scan=19815 | 42.4912 |
| 16371:3 | 2 | R.LSLHRPALEDLLLGSEAN[+1501.555]LTC[+57.021]TLTGLR.D | HexNAc(5)Hex(3) | C[+57], N[+1502] | 1489.0627 | 3 | 4465.1735 | 4465.1536 | 0 | 4.5 | 127 | Specific | 425.01 | 413.02 | 413.02 | 7.26 | 26 | >sp\|P01876\|IGHA1_HUMAN Immunoglobulin heavy constant alpha 1 OS=Homo sapiens OX=9606 GN=IGHA1 PE=1 SV=2 | 255 | 20201116_13832_1ng_glyco_hcd30_etd30_compound15.19880.19880.3 | controllerType=0 controllerNumber=1 scan=19880 | 42.7295 |
| 16298:3 | 2 | R.LSLHRPALEDLLLGSEAN[+1864.634]LTC[+57.021]TLTGLR.D | HexNAc(2)Hex(9) | C[+57], N[+1865] | 1610.0889 | 3 | 4828.2522 | 4828.2324 | 0 | 4.1 | 127 | Specific | 431.9 | 390.7 | 390.7 | 7.26 | 26 | >sp\|P01876\|IGHA1_HUMAN Immunoglobulin heavy constant alpha 1 OS=Homo sapiens OX=9606 GN=IGHA1 PE=1 SV=2 | 255 | 20201116_13832_1ng_glyco_hcd30_etd30_compound15.19802.19802.3 | controllerType=0 controllerNumber=1 scan=19802 | 42.4441 |
| 16336:3 | 2 | R.LSLHRPALEDLLLGSEAN[+1378.476]LTC[+57.021]TLTGLR.D | HexNAc(2)Hex(6) | C[+57], N[+1378] | 1448.038 | 3 | 4342.0995 | 4342.074 | 0 | 5.9 | 127 | Specific | 443.61 | 408.84 | 408.84 | 6.41 | 26 | >sp\|P01876\|IGHA1_HUMAN Immunoglobulin heavy constant alpha 1 OS=Homo sapiens OX=9606 GN=IGHA1 PE=1 SV=2 | 255 | 20201116_13832_1ng_glyco_hcd30_etd30_compound15.19843.19843.3 | controllerType=0 controllerNumber=1 scan=19843 | 42.5837 |
| 16324:3 | 2 | R.LSLHRPALEDLLLGSEAN[+1540.529]LTC[+57.021]TLTGLR.D | HexNAc(2)Hex(7) | C[+57], N[+1541] | 1502.0525 | 3 | 4504.1431 | 4504.1268 | 0 | 3.6 | 127 | Specific | 444.14 | 416.83 | 416.83 | 7.42 | 26 | >sp\|P01876\|IGHA1_HUMAN Immunoglobulin heavy constant alpha 1 OS=Homo sapiens OX=9606 GN=IGHA1 PE=1 SV=2 | 255 | 20201116_13832_1ng_glyco_hcd30_etd30_compound15.19830.19830.3 | controllerType=0 controllerNumber=1 scan=19830 | 42.5432 |
| 17079:4 | 2 | R.LSLHRPALEDLLLGSEAN[+1710.598]LTC[+57.021]TLTGLR.D | HexNAc(3)Hex(5)NeuAc(1) | C[+57], N[+1711] | 1169.3088 | 4 | 4674.2132 | 4674.1959 | 0 | 3.7 | 127 | Specific | 457 | 457 | 457 | 6.61 | 26 | >sp\|P01876\|IGHA1_HUMAN Immunoglobulin heavy constant alpha 1 OS=Homo sapiens OX=9606 GN=IGHA1 PE=1 SV=2 | 255 | 20201116_13832_1ng_glyco_hcd30_etd30_compound15.20627.20627.4 | controllerType=0 controllerNumber=1 scan=20627 | 44.9307 |
| 17031:4 | 2 | R.LSLHRPALEDLLLGSEAN[+1913.677]LTC[+57.021]TLTGLR.D | HexNAc(4)Hex(5)NeuAc(1) | C[+57], N[+1914] | 1220.0801 | 4 | 4877.2987 | 4877.2753 | 0 | 4.8 | 127 | Specific | 569.93 | 569.93 | 569.93 | 8.52 | 26 | >sp\|P01876\|IGHA1_HUMAN Immunoglobulin heavy constant alpha 1 OS=Homo sapiens OX=9606 GN=IGHA1 PE=1 SV=2 | 255 | 20201116_13832_1ng_glyco_hcd30_etd30_compound15.20577.20577.4 | controllerType=0 controllerNumber=1 scan=20577 | 44.7905 |
| 16383:4 | 2 | R.LSLHRPALEDLLLGSEAN[+1663.608]LTC[+57.021]TLTGLR.D | HexNAc(5)Hex(4) | C[+57], N[+1664] | 1157.5649 | 4 | 4627.2376 | 4627.2064 | 0 | 6.7 | 127 | Specific | 664.04 | 664.04 | 664.04 | 9.82 | 26 | >sp\|P01876\|IGHA1_HUMAN Immunoglobulin heavy constant alpha 1 OS=Homo sapiens OX=9606 GN=IGHA1 PE=1 SV=2 | 255 | 20201116_13832_1ng_glyco_hcd30_etd30_compound15.19893.19893.4 | controllerType=0 controllerNumber=1 scan=19893 | 42.7819 |
| 16993:4 | 2 | R.LSLHRPALEDLLLGSEAN[+1913.677]LTC[+57.021]TLTGLR.D | HexNAc(4)Hex(5)NeuAc(1) | C[+57], N[+1914] | 1220.079 | 4 | 4877.2943 | 4877.2753 | 0 | 3.9 | 127 | Specific | 715.59 | 715.59 | 715.59 | 11.83 | 26 | >sp\|P01876\|IGHA1_HUMAN Immunoglobulin heavy constant alpha 1 OS=Homo sapiens OX=9606 GN=IGHA1 PE=1 SV=2 | 255 | 20201116_13832_1ng_glyco_hcd30_etd30_compound15.20537.20537.4 | controllerType=0 controllerNumber=1 scan=20537 | 44.6701 |
| 17013:4 | 2 | R.LSLHRPALEDLLLGSEAN[+2116.756]LTC[+57.021]TLTGLR.D | HexNAc(5)Hex(5)NeuAc(1) | C[+57], N[+2117] | 1270.8496 | 4 | 5080.3766 | 5080.3547 | 0 | 4.3 | 127 | Specific | 717.26 | 683.84 | 683.84 | 10.99 | 26 | >sp\|P01876\|IGHA1_HUMAN Immunoglobulin heavy constant alpha 1 OS=Homo sapiens OX=9606 GN=IGHA1 PE=1 SV=2 | 255 | 20201116_13832_1ng_glyco_hcd30_etd30_compound15.20558.20558.4 | controllerType=0 controllerNumber=1 scan=20558 | 44.7314 |
| 16364:4 | 2 | R.LSLHRPALEDLLLGSEAN[+1622.582]LTC[+57.021]TLTGLR.D | HexNAc(4)Hex(5) | C[+57], N[+1623] | 1147.305 | 4 | 4586.1981 | 4586.1799 | 0 | 4 | 127 | Specific | 729.17 | 729.17 | 729.17 | 11.83 | 26 | >sp\|P01876\|IGHA1_HUMAN Immunoglobulin heavy constant alpha 1 OS=Homo sapiens OX=9606 GN=IGHA1 PE=1 SV=2 | 255 | 20201116_13832_1ng_glyco_hcd30_etd30_compound15.19873.19873.4 | controllerType=0 controllerNumber=1 scan=19873 | 42.6994 |
| 17012:4 | 2 | R.LSLHRPALEDLLLGSEAN[+1913.677]LTC[+57.021]TLTGLR.D | HexNAc(4)Hex(5)NeuAc(1) | C[+57], N[+1914] | 1220.079 | 4 | 4877.2943 | 4877.2753 | 0 | 3.9 | 127 | Specific | 776.39 | 769.94 | 769.94 | 12.83 | 26 | >sp\|P01876\|IGHA1_HUMAN Immunoglobulin heavy constant alpha 1 OS=Homo sapiens OX=9606 GN=IGHA1 PE=1 SV=2 | 255 | 20201116_13832_1ng_glyco_hcd30_etd30_compound15.20557.20557.4 | controllerType=0 controllerNumber=1 scan=20557 | 44.7266 |
| 16398:4 | 2 | R.LSLHRPALEDLLLGSEAN[+1298.476]LTC[+57.021]TLTGLR.D | HexNAc(4)Hex(3) | C[+57], N[+1298] | 1066.2781 | 4 | 4262.0906 | 4262.0742 | 0 | 3.9 | 127 | Specific | 812.11 | 812.11 | 812.11 | 13.83 | 26 | >sp\|P01876\|IGHA1_HUMAN Immunoglobulin heavy constant alpha 1 OS=Homo sapiens OX=9606 GN=IGHA1 PE=1 SV=2 | 255 | 20201116_13832_1ng_glyco_hcd30_etd30_compound15.19909.19909.4 | controllerType=0 controllerNumber=1 scan=19909 | 42.8443 |
| 16294:4 | 2 | R.LSLHRPALEDLLLGSEAN[+1702.581]LTC[+57.021]TLTGLR.D | HexNAc(2)Hex(8) | C[+57], N[+1703] | 1167.3051 | 4 | 4666.1986 | 4666.1796 | 0 | 4.1 | 127 | Specific | 893.73 | 854.07 | 854.07 | 14.83 | 26 | >sp\|P01876\|IGHA1_HUMAN Immunoglobulin heavy constant alpha 1 OS=Homo sapiens OX=9606 GN=IGHA1 PE=1 SV=2 | 255 | 20201116_13832_1ng_glyco_hcd30_etd30_compound15.19798.19798.4 | controllerType=0 controllerNumber=1 scan=19798 | 42.4293 |
| 16307:4 | 2 | R.LSLHRPALEDLLLGSEAN[+1540.529]LTC[+57.021]TLTGLR.D | HexNAc(2)Hex(7) | C[+57], N[+1541] | 1126.7925 | 4 | 4504.1483 | 4504.1268 | 0 | 4.8 | 127 | Specific | 910.45 | 875.07 | 875.07 | 15.52 | 26 | >sp\|P01876\|IGHA1_HUMAN Immunoglobulin heavy constant alpha 1 OS=Homo sapiens OX=9606 GN=IGHA1 PE=1 SV=2 | 255 | 20201116_13832_1ng_glyco_hcd30_etd30_compound15.19812.19812.4 | controllerType=0 controllerNumber=1 scan=19812 | 42.4829 |
| 16378:4 | 2 | R.LSLHRPALEDLLLGSEAN[+1216.423]LTC[+57.021]TLTGLR.D | HexNAc(2)Hex(5) | C[+57], N[+1216] | 1045.7659 | 4 | 4180.0418 | 4180.0211 | 0 | 4.9 | 127 | Specific | 927.35 | 894.45 | 894.45 | 15.52 | 26 | >sp\|P01876\|IGHA1_HUMAN Immunoglobulin heavy constant alpha 1 OS=Homo sapiens OX=9606 GN=IGHA1 PE=1 SV=2 | 255 | 20201116_13832_1ng_glyco_hcd30_etd30_compound15.19888.19888.4 | controllerType=0 controllerNumber=1 scan=19888 | 42.7658 |
| 16350:4 | 2 | R.LSLHRPALEDLLLGSEAN[+1501.555]LTC[+57.021]TLTGLR.D | HexNAc(5)Hex(3) | C[+57], N[+1502] | 1117.0498 | 4 | 4465.1776 | 4465.1536 | 0 | 5.4 | 127 | Specific | 964.52 | 940.76 | 940.76 | 16.52 | 26 | >sp\|P01876\|IGHA1_HUMAN Immunoglobulin heavy constant alpha 1 OS=Homo sapiens OX=9606 GN=IGHA1 PE=1 SV=2 | 255 | 20201116_13832_1ng_glyco_hcd30_etd30_compound15.19858.19858.4 | controllerType=0 controllerNumber=1 scan=19858 | 42.6334 |
| 16360:4 | 2 | R.LSLHRPALEDLLLGSEAN[+1460.529]LTC[+57.021]TLTGLR.D | HexNAc(4)Hex(4) | C[+57], N[+1461] | 1106.7926 | 4 | 4424.1488 | 4424.1271 | 0 | 4.9 | 127 | Specific | 1049.4 | 983.5 | 983.5 | 17.52 | 26 | >sp\|P01876\|IGHA1_HUMAN Immunoglobulin heavy constant alpha 1 OS=Homo sapiens OX=9606 GN=IGHA1 PE=1 SV=2 | 255 | 20201116_13832_1ng_glyco_hcd30_etd30_compound15.19868.19868.4 | controllerType=0 controllerNumber=1 scan=19868 | 42.6768 |
| 10853:3 | 2 | R.LAGKPTHVN[+1501.555]VSVVMAEVDGTC[+57.021]Y.- | HexNAc(5)Hex(3) | C[+57], N[+1502] | 1283.5756 | 3 | 3848.7123 | 3848.6974 | 0 | 3.9 | 332 | CRagged | 203.09 | 127.18 | 127.18 | 3 | 26 | >sp\|P01876\|IGHA1_HUMAN Immunoglobulin heavy constant alpha 1 OS=Homo sapiens OX=9606 GN=IGHA1 PE=1 SV=2 | 255 | 20201116_13832_1ng_glyco_hcd30_etd30_compound15.14064.14064.3 | controllerType=0 controllerNumber=1 scan=14064 | 30.8003 |
| 10804:3 | 2 | R.LAGKPTHVN[+1216.423]VSVVMAEVDGTC[+57.021]Y.- | HexNAc(2)Hex(5) | C[+57], N[+1216] | 1188.5292 | 3 | 3563.5731 | 3563.5649 | 0 | 2.3 | 332 | CRagged | 225.58 | 209.31 | 167.77 | 3.52 | 26 | >sp\|P01876\|IGHA1_HUMAN Immunoglobulin heavy constant alpha 1 OS=Homo sapiens OX=9606 GN=IGHA1 PE=1 SV=2 | 255 | 20201116_13832_1ng_glyco_hcd30_etd30_compound15.14013.14013.3 | controllerType=0 controllerNumber=1 scan=14013 | 30.6979 |
| 10772:3 | 2 | R.LAGKPTHVN[+1378.476]VSVVMAEVDGTC[+57.021]Y.- | HexNAc(2)Hex(6) | C[+57], N[+1378] | 1242.5474 | 3 | 3725.6277 | 3725.6178 | 0 | 2.7 | 332 | CRagged | 237.12 | 202.48 | 180.73 | 3.27 | 26 | >sp\|P01876\|IGHA1_HUMAN Immunoglobulin heavy constant alpha 1 OS=Homo sapiens OX=9606 GN=IGHA1 PE=1 SV=2 | 255 | 20201116_13832_1ng_glyco_hcd30_etd30_compound15.13980.13980.3 | controllerType=0 controllerNumber=1 scan=13980 | 30.6083 |
| 10812:3 | 2 | R.LAGKPTHVN[+1809.666]VSVVMAEVDGTC[+57.021]Y.- | HexNAc(5)Hex(4)Fuc(1) | C[+57], N[+1810] | 1386.2784 | 3 | 4156.8206 | 4156.8081 | 0 | 3 | 332 | CRagged | 240.08 | 147.49 | 147.49 | 3.28 | 26 | >sp\|P01876\|IGHA1_HUMAN Immunoglobulin heavy constant alpha 1 OS=Homo sapiens OX=9606 GN=IGHA1 PE=1 SV=2 | 255 | 20201116_13832_1ng_glyco_hcd30_etd30_compound15.14022.14022.3 | controllerType=0 controllerNumber=1 scan=14022 | 30.7229 |
| 10708:3 | 2 | R.LAGKPTHVN[+1702.581]VSVVMAEVDGTC[+57.021]Y.- | HexNAc(2)Hex(8) | C[+57], N[+1703] | 1350.5847 | 3 | 4049.7394 | 4049.7234 | 0 | 3.9 | 332 | CRagged | 260.79 | 214 | 171.77 | 4.07 | 26 | >sp\|P01876\|IGHA1_HUMAN Immunoglobulin heavy constant alpha 1 OS=Homo sapiens OX=9606 GN=IGHA1 PE=1 SV=2 | 255 | 20201116_13832_1ng_glyco_hcd30_etd30_compound15.13913.13913.3 | controllerType=0 controllerNumber=1 scan=13913 | 30.446 |
| 10813:3 | 2 | R.LAGKPTHVN[+1216.423]VSVVMAEVDGTC[+57.021]Y.- | HexNAc(2)Hex(5) | C[+57], N[+1216] | 1188.5307 | 3 | 3563.5775 | 3563.5649 | 0 | 3.5 | 332 | CRagged | 279.02 | 267.88 | 170.96 | 4.23 | 26 | >sp\|P01876\|IGHA1_HUMAN Immunoglobulin heavy constant alpha 1 OS=Homo sapiens OX=9606 GN=IGHA1 PE=1 SV=2 | 255 | 20201116_13832_1ng_glyco_hcd30_etd30_compound15.14023.14023.3 | controllerType=0 controllerNumber=1 scan=14023 | 30.7309 |
| 10861:3 | 2 | R.LAGKPTHVN[+1647.613]VSVVMAEVDGTC[+57.021]Y.- | HexNAc(5)Hex(3)Fuc(1) | C[+57], N[+1648] | 1332.2623 | 3 | 3994.7722 | 3994.7553 | 0 | 4.2 | 332 | CRagged | 282.59 | 153.17 | 153.17 | 3.63 | 26 | >sp\|P01876\|IGHA1_HUMAN Immunoglobulin heavy constant alpha 1 OS=Homo sapiens OX=9606 GN=IGHA1 PE=1 SV=2 | 255 | 20201116_13832_1ng_glyco_hcd30_etd30_compound15.14073.14073.3 | controllerType=0 controllerNumber=1 scan=14073 | 30.8234 |
| 10884:4 | 2 | R.LAGKPTHVN[+1647.613]VSVVMAEVDGTC[+57.021]Y.- | HexNAc(5)Hex(3)Fuc(1) | C[+57], N[+1648] | 999.449 | 4 | 3994.7742 | 3994.7553 | 0 | 4.7 | 332 | CRagged | 300.98 | 300.98 | 300.98 | 3.7 | 26 | >sp\|P01876\|IGHA1_HUMAN Immunoglobulin heavy constant alpha 1 OS=Homo sapiens OX=9606 GN=IGHA1 PE=1 SV=2 | 255 | 20201116_13832_1ng_glyco_hcd30_etd30_compound15.14097.14097.4 | controllerType=0 controllerNumber=1 scan=14097 | 30.8766 |
| 17685:4 | 5 | R.GLSFDVSLEVSQGPGLLN[+2350.830]DTK.V | HexNAc(4)Hex(5)Fuc(1)NeuAc(2) | N[+2351] | 1132.4947 | 4 | 4526.9569 | 4526.9436 | 0 | 2.9 | 118 | Specific | 251.84 | 243.14 | 243.14 | 4.83 | 18 | >sp\|P01833\|PIGR_HUMAN Polymeric immunoglobulin receptor OS=Homo sapiens OX=9606 GN=PIGR PE=1 SV=4 | 902 | 20201116_13832_1ng_glyco_hcd30_etd30_compound15.21269.21269.2 | controllerType=0 controllerNumber=1 scan=21269 | 46.6548 |
| 16332:4 | 5 | R.GLSFDVSLEVSQGPGLLN[+2205.793]DTK.V | HexNAc(4)Hex(5)Fuc(2)NeuAc(1) | N[+2206] | 1096.2353 | 4 | 4381.9193 | 4381.906 | 0 | 3 | 118 | Specific | 274.66 | 274.66 | 274.66 | 3.52 | 18 | >sp\|P01833\|PIGR_HUMAN Polymeric immunoglobulin receptor OS=Homo sapiens OX=9606 GN=PIGR PE=1 SV=4 | 902 | 20201116_13832_1ng_glyco_hcd30_etd30_compound15.19838.19838.4 | controllerType=0 controllerNumber=1 scan=19838 | 42.5694 |
| 16320:3 | 5 | R.GLSFDVSLEVSQGPGLLN[+2205.793]DTK.V | HexNAc(4)Hex(5)Fuc(2)NeuAc(1) | N[+2206] | 1461.3129 | 3 | 4381.9242 | 4381.906 | 0 | 4.1 | 118 | Specific | 456.89 | 294.73 | 294.73 | 5.89 | 18 | >sp\|P01833\|PIGR_HUMAN Polymeric immunoglobulin receptor OS=Homo sapiens OX=9606 GN=PIGR PE=1 SV=4 | 902 | 20201116_13832_1ng_glyco_hcd30_etd30_compound15.19826.19826.3 | controllerType=0 controllerNumber=1 scan=19826 | 42.5335 |
| 16952:3 | 5 | R.GLSFDVSLEVSQGPGLLN[+2205.793]DTKVYTVDLGR.T | HexNAc(4)Hex(5)Fuc(2)NeuAc(1) | N[+2206] | 1762.4764 | 3 | 5285.4147 | 5285.3875 | 0 | 5.1 | 118 | Specific | 211.46 | 139.53 | 139.53 | 3.9 | 18 | >sp\|P01833\|PIGR_HUMAN Polymeric immunoglobulin receptor OS=Homo sapiens OX=9606 GN=PIGR PE=1 SV=4 | 902 | 20201116_13832_1ng_glyco_hcd30_etd30_compound15.20493.20493.3 | controllerType=0 controllerNumber=1 scan=20493 | 44.5482 |
| 17907:4 | 5 | R.GLSFDVSLEVSQGPGLLN[+2351.851]DTKVYTVDLGR.T | HexNAc(4)Hex(5)Fuc(3)NeuAc(1) | N[+2352] | 1358.6163 | 4 | 5431.4434 | 5431.4454 | 0 | -0.4 | 118 | Specific | 348.32 | 348.32 | 348.32 | 2.82 | 18 | >sp\|P01833\|PIGR_HUMAN Polymeric immunoglobulin receptor OS=Homo sapiens OX=9606 GN=PIGR PE=1 SV=4 | 902 | 20201116_13832_1ng_glyco_hcd30_etd30_compound15.21502.21502.4 | controllerType=0 controllerNumber=1 scan=21502 | 47.2649 |
| 17167:4 | 5 | K.QIGLYPVLVIDSSGYVNPN[+2059.735]YTGR.I | HexNAc(4)Hex(5)Fuc(1)NeuAc(1) | N[+2060] | 1147.0181 | 4 | 4585.0506 | 4585.0384 | 0 | 2.7 | 168 | Specific | 449.02 | 439.11 | 439.11 | 5.63 | 18 | >sp\|P01833\|PIGR_HUMAN Polymeric immunoglobulin receptor OS=Homo sapiens OX=9606 GN=PIGR PE=1 SV=4 | 902 | 20201116_13832_1ng_glyco_hcd30_etd30_compound15.20721.20721.4 | controllerType=0 controllerNumber=1 scan=20721 | 45.2657 |
| 17176:4 | 5 | K.QIGLYPVLVIDSSGYVNPN[+2059.735]YTGR.I | HexNAc(4)Hex(5)Fuc(1)NeuAc(1) | N[+2060] | 1147.0187 | 4 | 4585.053 | 4585.0384 | 0 | 3.2 | 168 | Specific | 494.23 | 470.95 | 470.95 | 5.94 | 18 | >sp\|P01833\|PIGR_HUMAN Polymeric immunoglobulin receptor OS=Homo sapiens OX=9606 GN=PIGR PE=1 SV=4 | 902 | 20201116_13832_1ng_glyco_hcd30_etd30_compound15.20731.20731.4 | controllerType=0 controllerNumber=1 scan=20731 | 45.306 |
| 17185:3 | 5 | K.QIGLYPVLVIDSSGYVNPN[+2059.735]YTGR.I | HexNAc(4)Hex(5)Fuc(1)NeuAc(1) | N[+2060] | 1529.0247 | 3 | 4585.0596 | 4585.0384 | 0 | 4.6 | 168 | Specific | 579.66 | 490.3 | 490.3 | 8.24 | 18 | >sp\|P01833\|PIGR_HUMAN Polymeric immunoglobulin receptor OS=Homo sapiens OX=9606 GN=PIGR PE=1 SV=4 | 902 | 20201116_13832_1ng_glyco_hcd30_etd30_compound15.20741.20741.3 | controllerType=0 controllerNumber=1 scan=20741 | 45.333 |
| 20321:3 | 5 | R.LSLLEEPGN[+2350.830]GTFTVILNQLTSR.D | HexNAc(4)Hex(5)Fuc(1)NeuAc(2) | N[+2351] | 1585.0502 | 3 | 4753.1361 | 4753.1229 | 0 | 2.8 | 413 | Specific | 493.87 | 361.78 | 361.78 | 7.12 | 18 | >sp\|P01833\|PIGR_HUMAN Polymeric immunoglobulin receptor OS=Homo sapiens OX=9606 GN=PIGR PE=1 SV=4 | 902 | 20201116_13832_1ng_glyco_hcd30_etd30_compound15.24050.24050.3 | controllerType=0 controllerNumber=1 scan=24050 | 53.408 |
| 18408:4 | 5 | R.LSLLEEPGN[+2205.793]GTFTVILNQLTSR.D | HexNAc(4)Hex(5)Fuc(2)NeuAc(1) | N[+2206] | 1152.7825 | 4 | 4608.1082 | 4608.0854 | 0 | 5 | 413 | Specific | 561.21 | 498.21 | 498.21 | 7.06 | 18 | >sp\|P01833\|PIGR_HUMAN Polymeric immunoglobulin receptor OS=Homo sapiens OX=9606 GN=PIGR PE=1 SV=4 | 902 | 20201116_13832_1ng_glyco_hcd30_etd30_compound15.22032.22032.4 | controllerType=0 controllerNumber=1 scan=22032 | 48.7745 |
| 18391:3 | 5 | R.LSLLEEPGN[+2205.793]GTFTVILNQLTSR.D | HexNAc(4)Hex(5)Fuc(2)NeuAc(1) | N[+2206] | 1536.7077 | 3 | 4608.1085 | 4608.0854 | 0 | 5 | 413 | Specific | 633.39 | 507.08 | 507.08 | 9.24 | 18 | >sp\|P01833\|PIGR_HUMAN Polymeric immunoglobulin receptor OS=Homo sapiens OX=9606 GN=PIGR PE=1 SV=4 | 902 | 20201116_13832_1ng_glyco_hcd30_etd30_compound15.22014.22014.3 | controllerType=0 controllerNumber=1 scan=22014 | 48.7159 |
| 15061:3 | 5 | K.IIEGEPNLKVPGN[+2205.793]VTAVLGETLK.V | HexNAc(4)Hex(5)Fuc(2)NeuAc(1) | N[+2206] | 1533.0547 | 3 | 4597.1497 | 4597.1422 | 0 | 1.6 | 457 | Specific | 212.9 | 182.49 | 182.49 | 3.48 | 18 | >sp\|P01833\|PIGR_HUMAN Polymeric immunoglobulin receptor OS=Homo sapiens OX=9606 GN=PIGR PE=1 SV=4 | 902 | 20201116_13832_1ng_glyco_hcd30_etd30_compound15.18501.18501.3 | controllerType=0 controllerNumber=1 scan=18501 | 39.329 |
| 15027:3 | 5 | K.IIEGEPNLKVPGN[+2205.793]VTAVLGETLK.V | HexNAc(4)Hex(5)Fuc(2)NeuAc(1) | N[+2206] | 1533.0506 | 3 | 4597.1372 | 4597.1422 | 0 | -1.1 | 457 | Specific | 344.02 | 264.02 | 264.02 | 2.61 | 18 | >sp\|P01833\|PIGR_HUMAN Polymeric immunoglobulin receptor OS=Homo sapiens OX=9606 GN=PIGR PE=1 SV=4 | 902 | 20201116_13832_1ng_glyco_hcd30_etd30_compound15.18464.18464.3 | controllerType=0 controllerNumber=1 scan=18464 | 39.2375 |
| 14580:4 | 5 | K.IIEGEPNLKVPGN[+2205.793]VTAVLGETLKVPC[+57.021]HFPC[+57.021]K.F | HexNAc(4)Hex(5)Fuc(2)NeuAc(1) | C[+57]*2, N[+2206] | 1406.4103 | 4 | 5622.6193 | 5622.5997 | 0 | 3.5 | 457 | Specific | 348.55 | 348.55 | 348.55 | 4.35 | 18 | >sp\|P01833\|PIGR_HUMAN Polymeric immunoglobulin receptor OS=Homo sapiens OX=9606 GN=PIGR PE=1 SV=4 | 902 | 20201116_13832_1ng_glyco_hcd30_etd30_compound15.17959.17959.4 | controllerType=0 controllerNumber=1 scan=17959 | 38.2635 |
| 14579:5 | 5 | K.IIEGEPNLKVPGN[+2059.735]VTAVLGETLKVPC[+57.021]HFPC[+57.021]K.F | HexNAc(4)Hex(5)Fuc(1)NeuAc(1) | C[+57]*2, N[+2060] | 1096.1173 | 5 | 5476.5575 | 5476.5418 | 0 | 2.9 | 457 | Specific | 445.4 | 445.4 | 445.4 | 5.39 | 18 | >sp\|P01833\|PIGR_HUMAN Polymeric immunoglobulin receptor OS=Homo sapiens OX=9606 GN=PIGR PE=1 SV=4 | 902 | 20201116_13832_1ng_glyco_hcd30_etd30_compound15.17958.17958.5 | controllerType=0 controllerNumber=1 scan=17958 | 38.2591 |
| 14568:5 | 5 | K.IIEGEPNLKVPGN[+2059.735]VTAVLGETLKVPC[+57.021]HFPC[+57.021]K.F | HexNAc(4)Hex(5)Fuc(1)NeuAc(1) | C[+57]*2, N[+2060] | 1096.119 | 5 | 5476.566 | 5476.5418 | 0 | 4.4 | 457 | Specific | 489.82 | 489.82 | 489.82 | 6.1 | 18 | >sp\|P01833\|PIGR_HUMAN Polymeric immunoglobulin receptor OS=Homo sapiens OX=9606 GN=PIGR PE=1 SV=4 | 902 | 20201116_13832_1ng_glyco_hcd30_etd30_compound15.17946.17946.5 | controllerType=0 controllerNumber=1 scan=17946 | 38.2253 |
| 14554:5 | 5 | K.IIEGEPNLKVPGN[+2205.793]VTAVLGETLKVPC[+57.021]HFPC[+57.021]K.F | HexNAc(4)Hex(5)Fuc(2)NeuAc(1) | C[+57]*2, N[+2206] | 1125.3293 | 5 | 5622.6176 | 5622.5997 | 0 | 3.2 | 457 | Specific | 755.51 | 755.51 | 755.51 | 11.47 | 18 | >sp\|P01833\|PIGR_HUMAN Polymeric immunoglobulin receptor OS=Homo sapiens OX=9606 GN=PIGR PE=1 SV=4 | 902 | 20201116_13832_1ng_glyco_hcd30_etd30_compound15.17931.17931.5 | controllerType=0 controllerNumber=1 scan=17931 | 38.1879 |
| 05354:3 | 5 | K.WN[+2059.735]NTGC[+57.021]QALPSQDEGPSK.A | HexNAc(4)Hex(5)Fuc(1)NeuAc(1) | C[+57], N[+2060] | 1350.2129 | 3 | 4048.6243 | 4048.6116 | 0 | 3.1 | 498 | Specific | 308.56 | 198.39 | 198.39 | 5.15 | 18 | >sp\|P01833\|PIGR_HUMAN Polymeric immunoglobulin receptor OS=Homo sapiens OX=9606 GN=PIGR PE=1 SV=4 | 902 | 20201116_13832_1ng_glyco_hcd30_etd30_compound15.8358.8358.3 | controllerType=0 controllerNumber=1 scan=8358 | 20.7101 |
| 15940:4 | 6 | K.RNIAAFGGDPNN[+1913.677]ITLFGESAGGASVSLQTLSPYNK.G | HexNAc(4)Hex(5)NeuAc(1) | N[+1914] | 1370.8763 | 4 | 5480.4836 | 5480.4644 | 0 | 3.5 | 196 | Specific | 547.84 | 515.29 | 515.29 | 6.91 | 9 | >sp\|P19835\|CEL_HUMAN Bile salt-activated lipase OS=Homo sapiens OX=9606 GN=CEL PE=1 SV=3 | 476 | 20201116_13832_1ng_glyco_hcd30_etd30_compound15.19426.19426.4 | controllerType=0 controllerNumber=1 scan=19426 | 41.4831 |
| 17206:3 | 6 | R.NIAAFGGDPNN[+1913.677]ITLFGESAGGASVSLQTLSPYNK.G | HexNAc(4)Hex(5)NeuAc(1) | N[+1914] | 1775.4654 | 3 | 5324.3817 | 5324.3633 | 0 | 3.5 | 197 | Specific | 709.25 | 588.1 | 588.1 | 11.65 | 9 | >sp\|P19835\|CEL_HUMAN Bile salt-activated lipase OS=Homo sapiens OX=9606 GN=CEL PE=1 SV=3 | 476 | 20201116_13832_1ng_glyco_hcd30_etd30_compound15.20763.20763.3 | controllerType=0 controllerNumber=1 scan=20763 | 45.407 |
| 12357:4 | 8 | K.KGN[+2204.772]TTLNSFVIPSESDVPTHC[+57.021]PSQWWPYAGHC[+57.021]YK.I | HexNAc(4)Hex(5)NeuAc(2) | C[+57]*2, N[+2205] | 1542.9132 | 4 | 6168.631 | 6168.5942 | 0 | 6 | 342 | Specific | 231.13 | 217.21 | 217.21 | 3.33 | 7 | >sp\|P22897\|MRC1_HUMAN Macrophage mannose receptor 1 OS=Homo sapiens OX=9606 GN=MRC1 PE=1 SV=1 | 1049 | 20201116_13832_1ng_glyco_hcd30_etd30_compound15.15646.15646.4 | controllerType=0 controllerNumber=1 scan=15646 | 33.6269 |
| 05348:3 | 17 | R.LSPN[+2205.793]ASAEHLELR.W | HexNAc(4)Hex(5)Fuc(2)NeuAc(1) | N[+2206] | 1214.8558 | 3 | 3642.5528 | 3642.5421 | 0 | 2.9 | 52 | Specific | 265.3 | 170.26 | 170.26 | 2.97 | 3 | >sp\|Q13410\|BT1A1_HUMAN Butyrophilin subfamily 1 member A1 OS=Homo sapiens OX=9606 GN=BTN1A1 PE=1 SV=3 | 179 | 20201116_13832_1ng_glyco_hcd30_etd30_compound15.8352.8352.3 | controllerType=0 controllerNumber=1 scan=8352 | 20.6946 |
| 07319:3 | 17 | R.LSPN[+2350.830]ASAEHLELR.W | HexNAc(4)Hex(5)Fuc(1)NeuAc(2) | N[+2351] | 1263.2003 | 3 | 3787.5862 | 3787.5796 | 0 | 1.8 | 52 | Specific | 279.74 | 161.2 | 161.2 | 3.56 | 3 | >sp\|Q13410\|BT1A1_HUMAN Butyrophilin subfamily 1 member A1 OS=Homo sapiens OX=9606 GN=BTN1A1 PE=1 SV=3 | 179 | 20201116_13832_1ng_glyco_hcd30_etd30_compound15.10391.10391.3 | controllerType=0 controllerNumber=1 scan=10391 | 24.1764 |
| 21336:3 | 20 | R.QLAHQSN[+2204.772]STNIFFSPVSIATAFAMLSLGTK.A | HexNAc(4)Hex(5)NeuAc(2) | N[+2205] | 1796.1488 | 3 | 5386.4317 | 5386.4075 | 0 | 4.5 | 64 | Specific | 277.94 | 172.56 | 172.56 | 4.35 | 3 | >sp\|P01009\|A1AT_HUMAN Alpha-1-antitrypsin OS=Homo sapiens OX=9606 GN=SERPINA1 PE=1 SV=3 | 642 | 20201116_13832_1ng_glyco_hcd30_etd30_compound15.25116.25116.3 | controllerType=0 controllerNumber=1 scan=25116 | 55.9102 |
| 21320:4 | 20 | R.QLAHQSN[+2204.772]STNIFFSPVSIATAFAMLSLGTK.A | HexNAc(4)Hex(5)NeuAc(2) | N[+2205] | 1347.3616 | 4 | 5386.4245 | 5386.4075 | 0 | 3.2 | 64 | Specific | 426.54 | 385.45 | 385.45 | 6.88 | 3 | >sp\|P01009\|A1AT_HUMAN Alpha-1-antitrypsin OS=Homo sapiens OX=9606 GN=SERPINA1 PE=1 SV=3 | 642 | 20201116_13832_1ng_glyco_hcd30_etd30_compound15.25099.25099.2 | controllerType=0 controllerNumber=1 scan=25099 | 55.8483 |
| 21314:3 | 20 | R.QLAHQSN[+2204.772]STNIFFSPVSIATAFAMLSLGTK.A | HexNAc(4)Hex(5)NeuAc(2) | N[+2205] | 1796.1485 | 3 | 5386.431 | 5386.4075 | 0 | 4.4 | 64 | Specific | 474.45 | 344.1 | 344.1 | 6.28 | 3 | >sp\|P01009\|A1AT_HUMAN Alpha-1-antitrypsin OS=Homo sapiens OX=9606 GN=SERPINA1 PE=1 SV=3 | 642 | 20201116_13832_1ng_glyco_hcd30_etd30_compound15.25092.25092.3 | controllerType=0 controllerNumber=1 scan=25092 | 55.8259 |
| 21335:4 | 20 | R.QLAHQSN[+2204.772]STNIFFSPVSIATAFAMLSLGTK.A | HexNAc(4)Hex(5)NeuAc(2) | N[+2205] | 1347.3617 | 4 | 5386.425 | 5386.4075 | 0 | 3.2 | 64 | Specific | 514.43 | 432.27 | 432.27 | 6.18 | 3 | >sp\|P01009\|A1AT_HUMAN Alpha-1-antitrypsin OS=Homo sapiens OX=9606 GN=SERPINA1 PE=1 SV=3 | 642 | 20201116_13832_1ng_glyco_hcd30_etd30_compound15.25115.25115.4 | controllerType=0 controllerNumber=1 scan=25115 | 55.9056 |
| 18051:5 | 20 | K.YLGN[+2204.772]ATAIFFLPDEGKLQHLENELTHDIITK.F | HexNAc(4)Hex(5)NeuAc(2) | N[+2205] | 1149.9343 | 5 | 5745.6425 | 5745.6098 | 0 | 5.7 | 268 | Specific | 227.68 | 227.68 | 227.68 | 2.67 | 3 | >sp\|P01009\|A1AT_HUMAN Alpha-1-antitrypsin OS=Homo sapiens OX=9606 GN=SERPINA1 PE=1 SV=3 | 642 | 20201116_13832_1ng_glyco_hcd30_etd30_compound15.21655.21655.5 | controllerType=0 controllerNumber=1 scan=21655 | 47.6906 |

Table S4. Cont.

| Query #:z | Protein Rank | Glycopeptide | Glycan | Modification Type(s) | Observed m/z | z | Observed (M+H) | Calc. mass (M+H) | Off-by-x error | Mass error (ppm) | StartPosition | Cleavage | Score | Delta | Delta Mod | \|Log Prob\| | # of unique peptides | Protein Name | Protein DB number | Comment | Scan # | Scan Time |
|---|---|---|---|---|---|---|---|---|---|---|---|---|---|---|---|---|---|---|---|---|---|---|
| 18047:4 | 20 | K.YLGN[+2204.772]ATAIFFLPDEGKLQHLENELTHDIITK.F | HexNAc(4)Hex(5)NeuAc(2) | N[+2205] | 1437.1671 | 4 | 5745.6466 | 5745.6098 | 0 | 6.4 | 268 | Specific | 515.46 | 515.46 | 515.46 | 5.76 | 3 | >sp\|P01009\|A1AT_HUMAN Alpha-1-antitrypsin OS=Homo sapiens OX=9606 GN=SERPINA1 PE=1 SV=3 | 642 | 20201116_13832_1ng_glyco_hcd30_etd30_compound15.21651.21651.4 | controllerType=0 controllerNumber=1 scan=21651 | 47.6811 |
| 02769:2 | 23 | K.TPLTAN[+1809.666]ITK.S | HexNAc(5)Hex(4)Fuc(1) | N[+1810] | 1384.6203 | 2 | 2768.2333 | 2768.2228 | 0 | 3.8 | 200 | Specific | 220.95 | 62.04 | 62.04 | 2.58 | 8 | >sp\|P01877\|IGHA2_HUMAN Immunoglobulin heavy constant alpha 2 OS=Homo sapiens OX=9606 GN=IGHA2 PE=1 SV=4 | 576 | 20201116_13832_1ng_glyco_hcd30_etd30_compound15.5681.5681.2 | controllerType=0 controllerNumber=1 scan=5681 | 16.0228 |
| 02764:3 | 23 | K.TPLTAN[+1809.666]ITK.S | HexNAc(5)Hex(4)Fuc(1) | N[+1810] | 923.4164 | 3 | 2768.2347 | 2768.2228 | 0 | 4.3 | 200 | Specific | 294.67 | 132.4 | 132.4 | 2.67 | 8 | >sp\|P01877\|IGHA2_HUMAN Immunoglobulin heavy constant alpha 2 OS=Homo sapiens OX=9606 GN=IGHA2 PE=1 SV=4 | 576 | 20201116_13832_1ng_glyco_hcd30_etd30_compound15.5676.5676.3 | controllerType=0 controllerNumber=1 scan=5676 | 16.0089 |
| 02805:2 | 23 | K.TPLTAN[+1647.613]ITK.S | HexNAc(5)Hex(3)Fuc(1) | N[+1648] | 1303.5942 | 2 | 2606.181 | 2606.17 | 0 | 4.2 | 200 | Specific | 307.87 | 123.84 | 123.84 | 2.61 | 8 | >sp\|P01877\|IGHA2_HUMAN Immunoglobulin heavy constant alpha 2 OS=Homo sapiens OX=9606 GN=IGHA2 PE=1 SV=4 | 576 | 20201116_13832_1ng_glyco_hcd30_etd30_compound15.5719.5719.2 | controllerType=0 controllerNumber=1 scan=5719 | 16.1127 |
| 02813:3 | 23 | K.TPLTAN[+1647.613]ITK.S | HexNAc(5)Hex(3)Fuc(1) | N[+1648] | 869.3985 | 3 | 2606.1808 | 2606.17 | 0 | 4.2 | 200 | Specific | 316.12 | 146.94 | 146.94 | 2.67 | 8 | >sp\|P01877\|IGHA2_HUMAN Immunoglobulin heavy constant alpha 2 OS=Homo sapiens OX=9606 GN=IGHA2 PE=1 SV=4 | 576 | 20201116_13832_1ng_glyco_hcd30_etd30_compound15.5727.5727.3 | controllerType=0 controllerNumber=1 scan=5727 | 16.1313 |
| 09799:3 | 23 | R.MAGKPTHIN[+1378.476]VSVVMAEADGTC[+57.021]Y.- | HexNAc(2)Hex(6) | C[+57], N[+1378] | 1243.8611 | 3 | 3729.5689 | 3729.5585 | 0 | 2.8 | 319 | CRagged | 215.27 | 200.26 | 152.26 | 3.56 | 8 | >sp\|P01877\|IGHA2_HUMAN Immunoglobulin heavy constant alpha 2 OS=Homo sapiens OX=9606 GN=IGHA2 PE=1 SV=4 | 576 | 20201116_13832_1ng_glyco_hcd30_etd30_compound15.12971.12971.3 | controllerType=0 controllerNumber=1 scan=12971 | 28.7751 |
| 13213:4 | 24 | K.LKELPGVC[+57.021]N[+2205.793]ETMMALWEEC[+57.021]KPC[+57.021]LK.Q | HexNAc(4)Hex(5)Fuc(2)NeuAc(1) | C[+57]*3, N[+2206] | 1286.304 | 4 | 5142.194 | 5142.1776 | 0 | 3.2 | 95 | Specific | 547.49 | 462.89 | 462.89 | 6.03 | 5 | >sp\|P10909\|CLUS_HUMAN Clusterin OS=Homo sapiens OX=9606 GN=CLU PE=1 SV=1 | 1467 | 20201116_13832_1ng_glyco_hcd30_etd30_compound15.16535.16535.4 | controllerType=0 controllerNumber=1 scan=16535 | 35.3137 |
| 13268:4 | 24 | K.LKELPGVC[+57.021]N[+2059.735]ETMMALWEEC[+57.021]KPC[+57.021]LK.Q | HexNAc(4)Hex(5)Fuc(1)NeuAc(1) | C[+57]*3, N[+2060] | 1249.7908 | 4 | 4996.1413 | 4996.1196 | 0 | 4.3 | 95 | Specific | 571.62 | 409.38 | 409.38 | 7.3 | 5 | >sp\|P10909\|CLUS_HUMAN Clusterin OS=Homo sapiens OX=9606 GN=CLU PE=1 SV=1 | 1467 | 20201116_13832_1ng_glyco_hcd30_etd30_compound15.16592.16592.4 | controllerType=0 controllerNumber=1 scan=16592 | 35.4344 |
| 13190:4 | 24 | K.LKELPGVC[+57.021]N[+2205.793]ETMMALWEEC[+57.021]KPC[+57.021]LK.Q | HexNAc(4)Hex(5)Fuc(2)NeuAc(1) | C[+57]*3, N[+2206] | 1286.3041 | 4 | 5142.1945 | 5142.1776 | 0 | 3.3 | 95 | Specific | 601.52 | 455.07 | 455.07 | 7.91 | 5 | >sp\|P10909\|CLUS_HUMAN Clusterin OS=Homo sapiens OX=9606 GN=CLU PE=1 SV=1 | 1467 | 20201116_13832_1ng_glyco_hcd30_etd30_compound15.16511.16511.4 | controllerType=0 controllerNumber=1 scan=16511 | 35.2676 |
| 20706:3 | 24 | K.MLN[+2350.830]TSSLLEQLNEQFNWVSR.L | HexNAc(4)Hex(5)Fuc(1)NeuAc(2) | N[+2351] | 1587.3498 | 3 | 4760.0348 | 4760.0171 | 0 | 3.7 | 352 | Specific | 517.89 | 319.8 | 319.8 | 6.5 | 5 | >sp\|P10909\|CLUS_HUMAN Clusterin OS=Homo sapiens OX=9606 GN=CLU PE=1 SV=1 | 1467 | 20201116_13832_1ng_glyco_hcd30_etd30_compound15.24454.24454.3 | controllerType=0 controllerNumber=1 scan=24454 | 54.3846 |
| 00294:4 | 57 | K.TKPREEQYN[+1606.587]STYR.V | HexNAc(4)Hex(4)Fuc(1) | N[+1607] | 820.3572 | 4 | 3278.407 | 3278.3952 | 0 | 3.6 | 291 | Specific | 200.01 | 50.1 | 50.1 | 2.47 | 2 | >sp\|P0DOX5\|IGG1_HUMAN Immunoglobulin gamma-1 heavy chain OS=Homo sapiens OX=9606 PE=1 SV=2 | 914 | 20201116_13832_1ng_glyco_hcd30_etd30_compound15.3109.3109.4 | controllerType=0 controllerNumber=1 scan=3109 | 11.3075 |
| 00955:4 | 69 | K.TKPREEQFN[+1444.534]STFR.V | HexNAc(4)Hex(3)Fuc(1) | N[+1445] | 771.8474 | 4 | 3084.3677 | 3084.3526 | 0 | 4.9 | 168 | Specific | 386.81 | 338.08 | 338.08 | 3.4 | 2 | >sp\|P01859\|IGHG2_HUMAN Immunoglobulin heavy constant gamma 2 OS=Homo sapiens OX=9606 GN=IGHG2 PE=1 SV=2 | 387 | 20201116_13832_1ng_glyco_hcd30_etd30_compound15.3797.3797.4 | controllerType=0 controllerNumber=1 scan=3797 | 12.5687 |
| 00961:4 | 69 | K.TKPREEQFN[+1444.534]STFR.V | HexNAc(4)Hex(3)Fuc(1) | N[+1445] | 771.8465 | 4 | 3084.364 | 3084.3526 | 0 | 3.7 | 168 | Specific | 406.38 | 348.44 | 348.44 | 3.69 | 2 | >sp\|P01859\|IGHG2_HUMAN Immunoglobulin heavy constant gamma 2 OS=Homo sapiens OX=9606 GN=IGHG2 PE=1 SV=2 | 387 | 20201116_13832_1ng_glyco_hcd30_etd30_compound15.3804.3804.4 | controllerType=0 controllerNumber=1 scan=3804 | 12.5872 |
| 14957:4 | 1 | R.RTAGWNVPIGTLRPFLN[+2279.830]WTGPPEPIEAAVAR.F | HexNAc(5)Hex(6)Fuc(2) | N[+2280] | 1417.4196 | 4 | 5666.6564 | 5666.6417 | 0 | 2.6 | 140 | Specific | 373.26 | 373.26 | 373.26 | 6.9 | 49 | >sp\|P02788\|TRFL_HUMAN Lactotransferrin OS=Homo sapiens OX=9606 GN=LTF PE=1 SV=6 | 1270 | 20201115_15212_1ng_glyco_hcd30_etd30_compound15.18321.18321.4 | controllerType=0 controllerNumber=1 scan=18321 | 41.2962 |
| 15617:4 | 1 | R.RTAGWNVPIGTLRPFLN[+2570.925]WTGPPEPIEAAVAR.F | HexNAc(5)Hex(6)Fuc(2)NeuAc(1) | N[+2571] | 1490.1943 | 4 | 5957.7555 | 5957.7371 | 0 | 3.1 | 140 | Specific | 657.61 | 657.61 | 657.61 | 10.46 | 49 | >sp\|P02788\|TRFL_HUMAN Lactotransferrin OS=Homo sapiens OX=9606 GN=LTF PE=1 SV=6 | 1270 | 20201115_15212_1ng_glyco_hcd30_etd30_compound15.19018.19018.4 | controllerType=0 controllerNumber=1 scan=19018 | 43.3165 |
| 15752:4 | 1 | R.RTAGWNVPIGTLRPFLN[+2059.735]WTGPPEPIEAAVAR.F | HexNAc(4)Hex(5)Fuc(1)NeuAc(1) | N[+2060] | 1362.3966 | 4 | 5446.5646 | 5446.547 | 0 | 3.2 | 140 | Specific | 907.75 | 885.93 | 885.93 | 15.46 | 49 | >sp\|P02788\|TRFL_HUMAN Lactotransferrin OS=Homo sapiens OX=9606 GN=LTF PE=1 SV=6 | 1270 | 20201115_15212_1ng_glyco_hcd30_etd30_compound15.19160.19160.4 | controllerType=0 controllerNumber=1 scan=19160 | 43.6573 |
| 14980:4 | 1 | R.RTAGWNVPIGTLRPFLN[+1768.640]WTGPPEPIEAAVAR.F | HexNAc(4)Hex(5)Fuc(1) | N[+1769] | 1289.6216 | 4 | 5155.4645 | 5155.4516 | 0 | 2.5 | 140 | Specific | 1048.16 | 1048.16 | 1048.16 | 17.51 | 49 | >sp\|P02788\|TRFL_HUMAN Lactotransferrin OS=Homo sapiens OX=9606 GN=LTF PE=1 SV=6 | 1270 | 20201115_15212_1ng_glyco_hcd30_etd30_compound15.18346.18346.4 | controllerType=0 controllerNumber=1 scan=18346 | 41.3713 |
| 15854:4 | 1 | R.TAGWNVPIGTLRPFLN[+2498.904]WTGPPEPIEAAVAR.F | HexNAc(6)Hex(7)Fuc(1) | N[+2499] | 1433.1626 | 4 | 5729.6286 | 5729.6148 | 0 | 2.4 | 141 | Specific | 236.14 | 236.14 | 236.14 | 3.71 | 49 | >sp\|P02788\|TRFL_HUMAN Lactotransferrin OS=Homo sapiens OX=9606 GN=LTF PE=1 SV=6 | 1270 | 20201115_15212_1ng_glyco_hcd30_etd30_compound15.19266.19266.4 | controllerType=0 controllerNumber=1 scan=19266 | 43.8845 |
| 17667:3 | 1 | R.TAGWNVPIGTLRPFLN[+2350.830]WTGPPEPIEAAVAR.F | HexNAc(4)Hex(5)Fuc(1)NeuAc(2) | N[+2351] | 1861.1909 | 3 | 5581.5581 | 5581.5413 | 0 | 3 | 141 | Specific | 335.85 | 235.33 | 235.33 | 5.23 | 49 | >sp\|P02788\|TRFL_HUMAN Lactotransferrin OS=Homo sapiens OX=9606 GN=LTF PE=1 SV=6 | 1270 | 20201115_15212_1ng_glyco_hcd30_etd30_compound15.21177.21177.3 | controllerType=0 controllerNumber=1 scan=21177 | 49.1806 |
| 16624:3 | 1 | R.TAGWNVPIGTLRPFLN[+2205.793]WTGPPEPIEAAVAR.F | HexNAc(4)Hex(5)Fuc(2)NeuAc(1) | N[+2206] | 1812.8446 | 3 | 5436.5192 | 5436.5038 | 0 | 2.8 | 141 | Specific | 336.78 | 247.96 | 247.96 | 4.87 | 49 | >sp\|P02788\|TRFL_HUMAN Lactotransferrin OS=Homo sapiens OX=9606 GN=LTF PE=1 SV=6 | 1270 | 20201115_15212_1ng_glyco_hcd30_etd30_compound15.20078.20078.3 | controllerType=0 controllerNumber=1 scan=20078 | 46.2351 |
| 16486:4 | 1 | R.TAGWNVPIGTLRPFLN[+2716.983]WTGPPEPIEAAVAR.F | HexNAc(5)Hex(6)Fuc(3)NeuAc(1) | N[+2717] | 1487.6846 | 4 | 5947.7165 | 5947.6939 | 0 | 3.8 | 141 | Specific | 344 | 344 | 344 | 5.17 | 49 | >sp\|P02788\|TRFL_HUMAN Lactotransferrin OS=Homo sapiens OX=9606 GN=LTF PE=1 SV=6 | 1270 | 20201115_15212_1ng_glyco_hcd30_etd30_compound15.19933.19933.2 | controllerType=0 controllerNumber=1 scan=19933 | 45.8712 |
| 16557:3 | 1 | R.TAGWNVPIGTLRPFLN[+2424.867]WTGPPEPIEAAVAR.F | HexNAc(5)Hex(6)Fuc(1)NeuAc(1) | N[+2425] | 1885.8718 | 3 | 5655.6008 | 5655.5781 | 0 | 4 | 141 | Specific | 389.58 | 284.48 | 284.48 | 5.52 | 49 | >sp\|P02788\|TRFL_HUMAN Lactotransferrin OS=Homo sapiens OX=9606 GN=LTF PE=1 SV=6 | 1270 | 20201115_15212_1ng_glyco_hcd30_etd30_compound15.20007.20007.3 | controllerType=0 controllerNumber=1 scan=20007 | 46.045 |
| 16660:4 | 1 | R.TAGWNVPIGTLRPFLN[+2076.750]WTGPPEPIEAAVAR.F | HexNAc(4)Hex(6)Fuc(2) | N[+2077] | 1327.6277 | 4 | 5307.4889 | 5307.4612 | 0 | 5.2 | 141 | Specific | 407.81 | 407.81 | 407.81 | 6.64 | 49 | >sp\|P02788\|TRFL_HUMAN Lactotransferrin OS=Homo sapiens OX=9606 GN=LTF PE=1 SV=6 | 1270 | 20201115_15212_1ng_glyco_hcd30_etd30_compound15.20116.20116.4 | controllerType=0 controllerNumber=1 scan=20116 | 46.3517 |
| 16649:3 | 1 | R.TAGWNVPIGTLRPFLN[+2059.735]WTGPPEPIEAAVAR.F | HexNAc(4)Hex(5)Fuc(1)NeuAc(1) | N[+2060] | 1764.1595 | 3 | 5290.464 | 5290.4459 | 0 | 3.4 | 141 | Specific | 408.36 | 341.19 | 341.19 | 5.59 | 49 | >sp\|P02788\|TRFL_HUMAN Lactotransferrin OS=Homo sapiens OX=9606 GN=LTF PE=1 SV=6 | 1270 | 20201115_15212_1ng_glyco_hcd30_etd30_compound15.20104.20104.2 | controllerType=0 controllerNumber=1 scan=20104 | 46.3239 |
| 16677:3 | 1 | R.TAGWNVPIGTLRPFLN[+2059.735]WTGPPEPIEAAVAR.F | HexNAc(4)Hex(5)Fuc(1)NeuAc(1) | N[+2060] | 1764.1576 | 3 | 5290.4581 | 5290.4459 | 0 | 2.3 | 141 | Specific | 426.77 | 320.92 | 320.92 | 5.1 | 49 | >sp\|P02788\|TRFL_HUMAN Lactotransferrin OS=Homo sapiens OX=9606 GN=LTF PE=1 SV=6 | 1270 | 20201115_15212_1ng_glyco_hcd30_etd30_compound15.20134.20134.3 | controllerType=0 controllerNumber=1 scan=20134 | 46.3963 |
| 15910:3 | 1 | R.TAGWNVPIGTLRPFLN[+1768.640]WTGPPEPIEAAVAR.F | HexNAc(4)Hex(5)Fuc(1) | N[+1769] | 1667.1256 | 3 | 4999.3622 | 4999.3504 | 0 | 2.3 | 141 | Specific | 442.98 | 385.87 | 385.87 | 6.9 | 49 | >sp\|P02788\|TRFL_HUMAN Lactotransferrin OS=Homo sapiens OX=9606 GN=LTF PE=1 SV=6 | 1270 | 20201115_15212_1ng_glyco_hcd30_etd30_compound15.19326.19326.3 | controllerType=0 controllerNumber=1 scan=19326 | 44.0835 |
| 16654:3 | 1 | R.TAGWNVPIGTLRPFLN[+2059.735]WTGPPEPIEAAVAR.F | HexNAc(4)Hex(5)Fuc(1)NeuAc(1) | N[+2060] | 1764.1588 | 3 | 5290.4618 | 5290.4459 | 0 | 3 | 141 | Specific | 473.78 | 359.25 | 359.25 | 6.13 | 49 | >sp\|P02788\|TRFL_HUMAN Lactotransferrin OS=Homo sapiens OX=9606 GN=LTF PE=1 SV=6 | 1270 | 20201115_15212_1ng_glyco_hcd30_etd30_compound15.20110.20110.3 | controllerType=0 controllerNumber=1 scan=20110 | 46.3397 |
| 16631:5 | 1 | R.TAGWNVPIGTLRPFLN[+2205.793]WTGPPEPIEAAVAR.F | HexNAc(4)Hex(5)Fuc(2)NeuAc(1) | N[+2206] | 1088.1085 | 5 | 5436.5135 | 5436.5038 | 0 | 1.8 | 141 | Specific | 541.65 | 541.65 | 541.65 | 7.05 | 49 | >sp\|P02788\|TRFL_HUMAN Lactotransferrin OS=Homo sapiens OX=9606 GN=LTF PE=1 SV=6 | 1270 | 20201115_15212_1ng_glyco_hcd30_etd30_compound15.20085.20085.5 | controllerType=0 controllerNumber=1 scan=20085 | 46.2548 |
| 17666:4 | 1 | R.TAGWNVPIGTLRPFLN[+2350.830]WTGPPEPIEAAVAR.F | HexNAc(4)Hex(5)Fuc(1)NeuAc(2) | N[+2351] | 1396.1444 | 4 | 5581.5558 | 5581.5413 | 0 | 2.6 | 141 | Specific | 648.6 | 648.6 | 648.6 | 9.84 | 49 | >sp\|P02788\|TRFL_HUMAN Lactotransferrin OS=Homo sapiens OX=9606 GN=LTF PE=1 SV=6 | 1270 | 20201115_15212_1ng_glyco_hcd30_etd30_compound15.21176.21176.4 | controllerType=0 controllerNumber=1 scan=21176 | 49.1778 |
| 15909:4 | 1 | R.TAGWNVPIGTLRPFLN[+1768.640]WTGPPEPIEAAVAR.F | HexNAc(4)Hex(5)Fuc(1) | N[+1769] | 1250.5969 | 4 | 4999.3659 | 4999.3504 | 0 | 3.1 | 141 | Specific | 676.55 | 676.55 | 676.55 | 10.79 | 49 | >sp\|P02788\|TRFL_HUMAN Lactotransferrin OS=Homo sapiens OX=9606 GN=LTF PE=1 SV=6 | 1270 | 20201115_15212_1ng_glyco_hcd30_etd30_compound15.19325.19325.4 | controllerType=0 controllerNumber=1 scan=19325 | 44.0805 |
| 17671:5 | 1 | R.TAGWNVPIGTLRPFLN[+2350.830]WTGPPEPIEAAVAR.F | HexNAc(4)Hex(5)Fuc(1)NeuAc(2) | N[+2351] | 1117.1162 | 5 | 5581.552 | 5581.5413 | 0 | 1.9 | 141 | Specific | 679.6 | 638.17 | 638.17 | 10.43 | 49 | >sp\|P02788\|TRFL_HUMAN Lactotransferrin OS=Homo sapiens OX=9606 GN=LTF PE=1 SV=6 | 1270 | 20201115_15212_1ng_glyco_hcd30_etd30_compound15.21181.21181.5 | controllerType=0 controllerNumber=1 scan=21181 | 49.1895 |
| 16653:4 | 1 | R.TAGWNVPIGTLRPFLN[+2059.735]WTGPPEPIEAAVAR.F | HexNAc(4)Hex(5)Fuc(1)NeuAc(1) | N[+2060] | 1323.3717 | 4 | 5290.465 | 5290.4459 | 0 | 3.6 | 141 | Specific | 699.2 | 687.48 | 687.48 | 10.79 | 49 | >sp\|P02788\|TRFL_HUMAN Lactotransferrin OS=Homo sapiens OX=9606 GN=LTF PE=1 SV=6 | 1270 | 20201115_15212_1ng_glyco_hcd30_etd30_compound15.20109.20109.4 | controllerType=0 controllerNumber=1 scan=20109 | 46.3369 |
| 16595:4 | 1 | R.TAGWNVPIGTLRPFLN[+2424.867]WTGPPEPIEAAVAR.F | HexNAc(5)Hex(6)Fuc(1)NeuAc(1) | N[+2425] | 1414.6547 | 4 | 5655.5968 | 5655.5781 | 0 | 3.3 | 141 | Specific | 702.71 | 702.71 | 702.71 | 11.79 | 49 | >sp\|P02788\|TRFL_HUMAN Lactotransferrin OS=Homo sapiens OX=9606 GN=LTF PE=1 SV=6 | 1270 | 20201115_15212_1ng_glyco_hcd30_etd30_compound15.20047.20047.4 | controllerType=0 controllerNumber=1 scan=20047 | 46.1357 |
| 16623:4 | 1 | R.TAGWNVPIGTLRPFLN[+2205.793]WTGPPEPIEAAVAR.F | HexNAc(4)Hex(5)Fuc(2)NeuAc(1) | N[+2206] | 1359.8846 | 4 | 5436.5168 | 5436.5038 | 0 | 2.4 | 141 | Specific | 841.66 | 840.7 | 840.7 | 13.84 | 49 | >sp\|P02788\|TRFL_HUMAN Lactotransferrin OS=Homo sapiens OX=9606 GN=LTF PE=1 SV=6 | 1270 | 20201115_15212_1ng_glyco_hcd30_etd30_compound15.20077.20077.4 | controllerType=0 controllerNumber=1 scan=20077 | 46.2323 |
| 15027:3 | 1 | R.TAGWNIPMGLLFN[+1216.423]QTGSC[+57.021]K.F | HexNAc(2)Hex(5) | C[+57], N[+1216] | 1104.4848 | 3 | 3311.4398 | 3311.4328 | 0 | 2.1 | 485 | Specific | 214.32 | 166.11 | 129.98 | 3.23 | 49 | >sp\|P02788\|TRFL_HUMAN Lactotransferrin OS=Homo sapiens OX=9606 GN=LTF PE=1 SV=6 | 1270 | 20201115_15212_1ng_glyco_hcd30_etd30_compound15.18396.18396.3 | controllerType=0 controllerNumber=1 scan=18396 | 41.52 |
| 16209:4 | 1 | R.TAGWNIPMGLLFN[+2059.735]QTGSC[+57.021]K.F | HexNAc(4)Hex(5)Fuc(1)NeuAc(1) | C[+57], N[+2060] | 1039.4463 | 4 | 4154.7635 | 4154.7449 | 0 | 4.5 | 485 | Specific | 249.53 | 204.51 | 204.51 | 3.88 | 49 | >sp\|P02788\|TRFL_HUMAN Lactotransferrin OS=Homo sapiens OX=9606 GN=LTF PE=1 SV=6 | 1270 | 20201115_15212_1ng_glyco_hcd30_etd30_compound15.19640.19640.3 | controllerType=0 controllerNumber=1 scan=19640 | 44.9636 |
| 14590:3 | 1 | R.TAGWNIPMGLLFN[+2279.830]QTGSC[+57.021]K.F | HexNAc(5)Hex(6)Fuc(2) | C[+57], N[+2280] | 1458.9554 | 3 | 4374.8516 | 4374.8396 | 0 | 2.7 | 485 | Specific | 258.48 | 192.27 | 192.13 | 4.98 | 49 | >sp\|P02788\|TRFL_HUMAN Lactotransferrin OS=Homo sapiens OX=9606 GN=LTF PE=1 SV=6 | 1270 | 20201115_15212_1ng_glyco_hcd30_etd30_compound15.17936.17936.3 | controllerType=0 controllerNumber=1 scan=17936 | 40.2735 |
| 14817:3 | 1 | R.TAGWNIPMGLLFN[+1606.587]QTGSC[+57.021]K.F | HexNAc(4)Hex(4)Fuc(1) | C[+57], N[+1607] | 1234.5401 | 3 | 3701.6057 | 3701.5966 | 0 | 2.5 | 485 | Specific | 363.55 | 303.43 | 303.43 | 5.06 | 49 | >sp\|P02788\|TRFL_HUMAN Lactotransferrin OS=Homo sapiens OX=9606 GN=LTF PE=1 SV=6 | 1270 | 20201115_15212_1ng_glyco_hcd30_etd30_compound15.18174.18174.3 | controllerType=0 controllerNumber=1 scan=18174 | 40.8917 |
| 14893:3 | 1 | R.TAGWNIPMGLLFN[+1622.582]QTGSC[+57.021]K.F | HexNAc(4)Hex(5) | C[+57], N[+1623] | 1239.8726 | 3 | 3717.6033 | 3717.5916 | 0 | 3.2 | 485 | Specific | 381.44 | 279.22 | 265.36 | 5.33 | 49 | >sp\|P02788\|TRFL_HUMAN Lactotransferrin OS=Homo sapiens OX=9606 GN=LTF PE=1 SV=6 | 1270 | 20201115_15212_1ng_glyco_hcd30_etd30_compound15.18254.18254.3 | controllerType=0 controllerNumber=1 scan=18254 | 41.0974 |
| 16343:3 | 1 | R.TAGWNIPMGLLFN[+1751.624]QTGSC[+57.021]K.F | HexNAc(4)Hex(4)NeuAc(1) | C[+57], N[+1752] | 1282.8863 | 3 | 3846.6443 | 3846.6341 | 0 | 2.6 | 485 | Specific | 397.46 | 270.03 | 270.03 | 5.38 | 49 | >sp\|P02788\|TRFL_HUMAN Lactotransferrin OS=Homo sapiens OX=9606 GN=LTF PE=1 SV=6 | 1270 | 20201115_15212_1ng_glyco_hcd30_etd30_compound15.19784.19784.3 | controllerType=0 controllerNumber=1 scan=19784 | 45.4794 |
| 16318:3 | 1 | R.TAGWNIPMGLLFN[+1913.677]QTGSC[+57.021]K.F | HexNAc(4)Hex(5)NeuAc(1) | C[+57], N[+1914] | 1336.9078 | 3 | 4008.7088 | 4008.687 | 0 | 5.4 | 485 | Specific | 416.02 | 268.58 | 268.58 | 4.78 | 49 | >sp\|P02788\|TRFL_HUMAN Lactotransferrin OS=Homo sapiens OX=9606 GN=LTF PE=1 SV=6 | 1270 | 20201115_15212_1ng_glyco_hcd30_etd30_compound15.19757.19757.3 | controllerType=0 controllerNumber=1 scan=19757 | 45.3814 |
| 16148:3 | 1 | R.TAGWNIPMGLLFN[+2205.793]QTGSC[+57.021]K.F | HexNAc(4)Hex(5)Fuc(2)NeuAc(1) | C[+57], N[+2206] | 1434.2765 | 3 | 4300.8151 | 4300.8028 | 0 | 2.9 | 485 | Specific | 445.52 | 290.93 | 290.93 | 5.62 | 49 | >sp\|P02788\|TRFL_HUMAN Lactotransferrin OS=Homo sapiens OX=9606 GN=LTF PE=1 SV=6 | 1270 | 20201115_15212_1ng_glyco_hcd30_etd30_compound15.19576.19576.3 | controllerType=0 controllerNumber=1 scan=19576 | 44.7797 |
| 16291:4 | 1 | R.TAGWNIPMGLLFN[+1913.677]QTGSC[+57.021]K.F | HexNAc(4)Hex(5)NeuAc(1) | C[+57], N[+1914] | 1002.9292 | 4 | 4008.6949 | 4008.687 | 0 | 2 | 485 | Specific | 455.14 | 396.82 | 396.82 | 6.3 | 49 | >sp\|P02788\|TRFL_HUMAN Lactotransferrin OS=Homo sapiens OX=9606 GN=LTF PE=1 SV=6 | 1270 | 20201115_15212_1ng_glyco_hcd30_etd30_compound15.19727.19727.4 | controllerType=0 controllerNumber=1 scan=19727 | 45.2462 |
| 16269:3 | 1 | R.TAGWNIPMGLLFN[+1913.677]QTGSC[+57.021]K.F | HexNAc(4)Hex(5)NeuAc(1) | C[+57], N[+1914] | 1336.903 | 3 | 4008.6944 | 4008.687 | 0 | 1.9 | 485 | Specific | 512.2 | 331.21 | 331.21 | 6.15 | 49 | >sp\|P02788\|TRFL_HUMAN Lactotransferrin OS=Homo sapiens OX=9606 GN=LTF PE=1 SV=6 | 1270 | 20201115_15212_1ng_glyco_hcd30_etd30_compound15.19703.19703.3 | controllerType=0 controllerNumber=1 scan=19703 | 45.1522 |
| 14971:3 | 1 | R.TAGWNIPM[+15.995]GLLFN[+2059.735]QTGSC[+57.021]K.F | HexNAc(4)Hex(5)Fuc(1)NeuAc(1) | C[+57], M[+16], N[+2060] | 1390.9245 | 3 | 4170.7589 | 4170.7398 | 0 | 4.6 | 485 | Specific | 249.72 | 210.59 | 210.59 | 3.51 | 49 | >sp\|P02788\|TRFL_HUMAN Lactotransferrin OS=Homo sapiens OX=9606 GN=LTF PE=1 SV=6 | 1270 | 20201115_15212_1ng_glyco_hcd30_etd30_compound15.18336.18336.3 | controllerType=0 controllerNumber=1 scan=18336 | 41.3418 |
| 15586:4 | 1 | R.TAGWNIPMGLLFN[+1768.640]QTGSC[+57.021]KFDEYFSQSC[+57.021]APGSDPR.S | HexNAc(4)Hex(5)Fuc(1) | C[+57]*2, N[+1769] | 1427.6115 | 4 | 5707.424 | 5707.3966 | 0 | 4.8 | 485 | Specific | 532.75 | 479.07 | 479.07 | 5.75 | 49 | >sp\|P02788\|TRFL_HUMAN Lactotransferrin OS=Homo sapiens OX=9606 GN=LTF PE=1 SV=6 | 1270 | 20201115_15212_1ng_glyco_hcd30_etd30_compound15.18985.18985.4 | controllerType=0 controllerNumber=1 scan=18985 | 43.2034 |
| 16342:4 | 1 | R.TAGWNIPMGLLFN[+2059.735]QTGSC[+57.021]KFDEYFSQSC[+57.021]APGSDPR.S | HexNAc(4)Hex(5)Fuc(1)NeuAc(1) | C[+57]*2, N[+2060] | 1500.3831 | 4 | 5998.5104 | 5998.492 | 0 | 3.1 | 485 | Specific | 577.4 | 503.41 | 503.41 | 8.2 | 49 | >sp\|P02788\|TRFL_HUMAN Lactotransferrin OS=Homo sapiens OX=9606 GN=LTF PE=1 SV=6 | 1270 | 20201115_15212_1ng_glyco_hcd30_etd30_compound15.19783.19783.4 | controllerType=0 controllerNumber=1 scan=19783 | 45.4755 |
| 08366:3 | 1 | R.N[+2205.793]GSDC[+57.021]PDKFC[+57.021]LFQSETK.N | HexNAc(4)Hex(5)Fuc(2)NeuAc(1) | C[+57]*2, N[+2206] | 1413.564 | 3 | 4238.6775 | 4238.6667 | 0 | 2.5 | 642 | Specific | 263.36 | 140.28 | 140.28 | 3.67 | 49 | >sp\|P02788\|TRFL_HUMAN Lactotransferrin OS=Homo sapiens OX=9606 GN=LTF PE=1 SV=6 | 1270 | 20201115_15212_1ng_glyco_hcd30_etd30_compound15.11405.11405.3 | controllerType=0 controllerNumber=1 scan=11405 | 26.605 |
| 08461:3 | 1 | R.N[+2059.735]GSDC[+57.021]PDKFC[+57.021]LFQSETK.N | HexNAc(4)Hex(5)Fuc(1)NeuAc(1) | C[+57]*2, N[+2060] | 1364.8776 | 3 | 4092.6183 | 4092.6088 | 0 | 2.3 | 642 | Specific | 312.94 | 195.25 | 195.25 | 4.16 | 49 | >sp\|P02788\|TRFL_HUMAN Lactotransferrin OS=Homo sapiens OX=9606 GN=LTF PE=1 SV=6 | 1270 | 20201115_15212_1ng_glyco_hcd30_etd30_compound15.11504.11504.3 | controllerType=0 controllerNumber=1 scan=11504 | 26.7982 |
| 16105:4 | 2 | R.LSLHRPALEDLLLGSEAN[+1872.651]LTC[+57.021]TLTGLR.D | HexNAc(3)Hex(6)NeuAc(1) | C[+57], N[+1873] | 1209.8223 | 4 | 4836.2674 | 4836.2487 | 0 | 3.9 | 127 | Specific | 308.98 | 308.98 | 308.98 | 4.53 | 29 | >sp\|P01876\|IGHA1_HUMAN Immunoglobulin heavy constant alpha 1 OS=Homo sapiens OX=9606 GN=IGHA1 PE=1 SV=2 | 255 | 20201115_15212_1ng_glyco_hcd30_etd30_compound15.19530.19530.3 | controllerType=0 controllerNumber=1 scan=19530 | 44.6251 |
| 15338:3 | 2 | R.LSLHRPALEDLLLGSEAN[+1864.634]LTC[+57.021]TLTGLR.D | HexNAc(2)Hex(9) | C[+57], N[+1865] | 1610.0875 | 3 | 4828.2479 | 4828.2324 | 0 | 3.2 | 127 | Specific | 341.99 | 341.99 | 341.99 | 5.47 | 29 | >sp\|P01876\|IGHA1_HUMAN Immunoglobulin heavy constant alpha 1 OS=Homo sapiens OX=9606 GN=IGHA1 PE=1 SV=2 | 255 | 20201115_15212_1ng_glyco_hcd30_etd30_compound15.18723.18723.3 | controllerType=0 controllerNumber=1 scan=18723 | 42.3859 |
| 15407:3 | 2 | R.LSLHRPALEDLLLGSEAN[+1501.555]LTC[+57.021]TLTGLR.D | HexNAc(5)Hex(3) | C[+57], N[+1502] | 1489.0589 | 3 | 4465.1621 | 4465.1536 | 0 | 1.9 | 127 | Specific | 363.75 | 340.82 | 340.82 | 5.58 | 29 | >sp\|P01876\|IGHA1_HUMAN Immunoglobulin heavy constant alpha 1 OS=Homo sapiens OX=9606 GN=IGHA1 PE=1 SV=2 | 255 | 20201115_15212_1ng_glyco_hcd30_etd30_compound15.18796.18796.3 | controllerType=0 controllerNumber=1 scan=18796 | 42.6254 |
| 15413:3 | 2 | R.LSLHRPALEDLLLGSEAN[+1460.529]LTC[+57.021]TLTGLR.D | HexNAc(4)Hex(4) | C[+57], N[+1461] | 1475.3812 | 3 | 4424.1289 | 4424.1271 | 0 | 0.4 | 127 | Specific | 368.53 | 305.1 | 305.1 | 4.99 | 29 | >sp\|P01876\|IGHA1_HUMAN Immunoglobulin heavy constant alpha 1 OS=Homo sapiens OX=9606 GN=IGHA1 PE=1 SV=2 | 255 | 20201115_15212_1ng_glyco_hcd30_etd30_compound15.18802.18802.3 | controllerType=0 controllerNumber=1 scan=18802 | 42.6458 |
| 15385:3 | 2 | R.LSLHRPALEDLLLGSEAN[+1622.582]LTC[+57.021]TLTGLR.D | HexNAc(4)Hex(5) | C[+57], N[+1623] | 1529.4044 | 3 | 4586.1985 | 4586.1799 | 0 | 4.1 | 127 | Specific | 380.94 | 380.16 | 380.16 | 6.67 | 29 | >sp\|P01876\|IGHA1_HUMAN Immunoglobulin heavy constant alpha 1 OS=Homo sapiens OX=9606 GN=IGHA1 PE=1 SV=2 | 255 | 20201115_15212_1ng_glyco_hcd30_etd30_compound15.18772.18772.3 | controllerType=0 controllerNumber=1 scan=18772 | 42.5495 |
| 15323:3 | 2 | R.LSLHRPALEDLLLGSEAN[+1702.581]LTC[+57.021]TLTGLR.D | HexNAc(2)Hex(8) | C[+57], N[+1703] | 1556.0706 | 3 | 4666.1973 | 4666.1796 | 0 | 3.8 | 127 | Specific | 401.5 | 353.04 | 353.04 | 5.24 | 29 | >sp\|P01876\|IGHA1_HUMAN Immunoglobulin heavy constant alpha 1 OS=Homo sapiens OX=9606 GN=IGHA1 PE=1 SV=2 | 255 | 20201115_15212_1ng_glyco_hcd30_etd30_compound15.18707.18707.3 | controllerType=0 controllerNumber=1 scan=18707 | 42.3382 |
| 15379:3 | 2 | R.LSLHRPALEDLLLGSEAN[+1378.476]LTC[+57.021]TLTGLR.D | HexNAc(2)Hex(6) | C[+57], N[+1378] | 1448.0358 | 3 | 4342.0929 | 4342.074 | 0 | 4.4 | 127 | Specific | 518.66 | 518.66 | 518.66 | 6.42 | 29 | >sp\|P01876\|IGHA1_HUMAN Immunoglobulin heavy constant alpha 1 OS=Homo sapiens OX=9606 GN=IGHA1 PE=1 SV=2 | 255 | 20201115_15212_1ng_glyco_hcd30_etd30_compound15.18766.18766.3 | controllerType=0 controllerNumber=1 scan=18766 | 42.5273 |
| 15344:5 | 2 | R.LSLHRPALEDLLLGSEAN[+1864.634]LTC[+57.021]TLTGLR.D | HexNAc(2)Hex(9) | C[+57], N[+1865] | 966.4554 | 5 | 4828.2479 | 4828.2324 | 0 | 3.2 | 127 | Specific | 544.59 | 544.59 | 544.59 | 7.68 | 29 | >sp\|P01876\|IGHA1_HUMAN Immunoglobulin heavy constant alpha 1 OS=Homo sapiens OX=9606 GN=IGHA1 PE=1 SV=2 | 255 | 20201115_15212_1ng_glyco_hcd30_etd30_compound15.18729.18729.5 | controllerType=0 controllerNumber=1 scan=18729 | 42.4069 |
| 15367:4 | 2 | R.LSLHRPALEDLLLGSEAN[+1864.634]LTC[+57.021]TLTGLR.D | HexNAc(2)Hex(9) | C[+57], N[+1865] | 1207.8178 | 4 | 4828.2493 | 4828.2324 | 0 | 3.5 | 127 | Specific | 654.63 | 654.63 | 654.63 | 10.63 | 29 | >sp\|P01876\|IGHA1_HUMAN Immunoglobulin heavy constant alpha 1 OS=Homo sapiens OX=9606 GN=IGHA1 PE=1 SV=2 | 255 | 20201115_15212_1ng_glyco_hcd30_etd30_compound15.18753.18753.2 | controllerType=0 controllerNumber=1 scan=18753 | 42.4811 |
| 16187:4 | 2 | R.LSLHRPALEDLLLGSEAN[+1710.598]LTC[+57.021]TLTGLR.D | HexNAc(3)Hex(5)NeuAc(1) | C[+57], N[+1711] | 1169.3099 | 4 | 4674.2176 | 4674.1959 | 0 | 4.6 | 127 | Specific | 676.58 | 659.93 | 659.93 | 9.97 | 29 | >sp\|P01876\|IGHA1_HUMAN Immunoglobulin heavy constant alpha 1 OS=Homo sapiens OX=9606 GN=IGHA1 PE=1 SV=2 | 255 | 20201115_15212_1ng_glyco_hcd30_etd30_compound15.19617.19617.4 | controllerType=0 controllerNumber=1 scan=19617 | 44.895 |
| 16126:4 | 2 | R.LSLHRPALEDLLLGSEAN[+1872.651]LTC[+57.021]TLTGLR.D | HexNAc(3)Hex(6)NeuAc(1) | C[+57], N[+1873] | 1209.821 | 4 | 4836.262 | 4836.2487 | 0 | 2.7 | 127 | Specific | 712.22 | 649.17 | 649.17 | 11.68 | 29 | >sp\|P01876\|IGHA1_HUMAN Immunoglobulin heavy constant alpha 1 OS=Homo sapiens OX=9606 GN=IGHA1 PE=1 SV=2 | 255 | 20201115_15212_1ng_glyco_hcd30_etd30_compound15.19552.19552.4 | controllerType=0 controllerNumber=1 scan=19552 | 44.7084 |
| 15406:4 | 2 | R.LSLHRPALEDLLLGSEAN[+1501.555]LTC[+57.021]TLTGLR.D | HexNAc(5)Hex(3) | C[+57], N[+1502] | 1117.0473 | 4 | 4465.1673 | 4465.1536 | 0 | 3.1 | 127 | Specific | 899.81 | 876.24 | 876.24 | 14.63 | 29 | >sp\|P01876\|IGHA1_HUMAN Immunoglobulin heavy constant alpha 1 OS=Homo sapiens OX=9606 GN=IGHA1 PE=1 SV=2 | 255 | 20201115_15212_1ng_glyco_hcd30_etd30_compound15.18795.18795.4 | controllerType=0 controllerNumber=1 scan=18795 | 42.6188 |
| 15372:4 | 2 | R.LSLHRPALEDLLLGSEAN[+1663.608]LTC[+57.021]TLTGLR.D | HexNAc(5)Hex(4) | C[+57], N[+1664] | 1157.5613 | 4 | 4627.2235 | 4627.2064 | 0 | 3.7 | 127 | Specific | 917.8 | 873.08 | 873.08 | 15.63 | 29 | >sp\|P01876\|IGHA1_HUMAN Immunoglobulin heavy constant alpha 1 OS=Homo sapiens OX=9606 GN=IGHA1 PE=1 SV=2 | 255 | 20201115_15212_1ng_glyco_hcd30_etd30_compound15.18759.18759.4 | controllerType=0 controllerNumber=1 scan=18759 | 42.5024 |
| 15357:4 | 2 | R.LSLHRPALEDLLLGSEAN[+1702.581]LTC[+57.021]TLTGLR.D | HexNAc(2)Hex(8) | C[+57], N[+1703] | 1167.3034 | 4 | 4666.1917 | 4666.1796 | 0 | 2.6 | 127 | Specific | 932.29 | 892.35 | 892.35 | 15.68 | 29 | >sp\|P01876\|IGHA1_HUMAN Immunoglobulin heavy constant alpha 1 OS=Homo sapiens OX=9606 GN=IGHA1 PE=1 SV=2 | 255 | 20201115_15212_1ng_glyco_hcd30_etd30_compound15.18743.18743.2 | controllerType=0 controllerNumber=1 scan=18743 | 42.4549 |
| 15318:4 | 2 | R.LSLHRPALEDLLLGSEAN[+1702.581]LTC[+57.021]TLTGLR.D | HexNAc(2)Hex(8) | C[+57], N[+1703] | 1167.3041 | 4 | 4666.1947 | 4666.1796 | 0 | 3.2 | 127 | Specific | 986.99 | 956.67 | 956.67 | 16.63 | 29 | >sp\|P01876\|IGHA1_HUMAN Immunoglobulin heavy constant alpha 1 OS=Homo sapiens OX=9606 GN=IGHA1 PE=1 SV=2 | 255 | 20201115_15212_1ng_glyco_hcd30_etd30_compound15.18702.18702.4 | controllerType=0 controllerNumber=1 scan=18702 | 42.3279 |
| 15388:4 | 2 | R.LSLHRPALEDLLLGSEAN[+1460.529]LTC[+57.021]TLTGLR.D | HexNAc(4)Hex(4) | C[+57], N[+1461] | 1106.7911 | 4 | 4424.1424 | 4424.1271 | 0 | 3.5 | 127 | Specific | 992.34 | 992.34 | 992.34 | 16.63 | 29 | >sp\|P01876\|IGHA1_HUMAN Immunoglobulin heavy constant alpha 1 OS=Homo sapiens OX=9606 GN=IGHA1 PE=1 SV=2 | 255 | 20201115_15212_1ng_glyco_hcd30_etd30_compound15.18776.18776.4 | controllerType=0 controllerNumber=1 scan=18776 | 42.5623 |
| 15417:4 | 2 | R.LSLHRPALEDLLLGSEAN[+1298.476]LTC[+57.021]TLTGLR.D | HexNAc(4)Hex(3) | C[+57], N[+1298] | 1066.2753 | 4 | 4262.0794 | 4262.0742 | 0 | 1.2 | 127 | Specific | 1004.86 | 981.86 | 981.86 | 17.36 | 29 | >sp\|P01876\|IGHA1_HUMAN Immunoglobulin heavy constant alpha 1 OS=Homo sapiens OX=9606 GN=IGHA1 PE=1 SV=2 | 255 | 20201115_15212_1ng_glyco_hcd30_etd30_compound15.18806.18806.4 | controllerType=0 controllerNumber=1 scan=18806 | 42.6571 |
| 15309:4 | 2 | R.LSLHRPALEDLLLGSEAN[+1540.529]LTC[+57.021]TLTGLR.D | HexNAc(2)Hex(7) | C[+57], N[+1541] | 1126.7914 | 4 | 4504.1439 | 4504.1268 | 0 | 3.8 | 127 | Specific | 1031.8 | 993.71 | 993.71 | 17.63 | 29 | >sp\|P01876\|IGHA1_HUMAN Immunoglobulin heavy constant alpha 1 OS=Homo sapiens OX=9606 GN=IGHA1 PE=1 SV=2 | 255 | 20201115_15212_1ng_glyco_hcd30_etd30_compound15.18692.18692.2 | controllerType=0 controllerNumber=1 scan=18692 | 42.2906 |
| 15377:4 | 2 | R.LSLHRPALEDLLLGSEAN[+1378.476]LTC[+57.021]TLTGLR.D | HexNAc(2)Hex(6) | C[+57], N[+1378] | 1086.2786 | 4 | 4342.0926 | 4342.074 | 0 | 4.3 | 127 | Specific | 1102.17 | 1038.97 | 1038.97 | 18.97 | 29 | >sp\|P01876\|IGHA1_HUMAN Immunoglobulin heavy constant alpha 1 OS=Homo sapiens OX=9606 GN=IGHA1 PE=1 SV=2 | 255 | 20201115_15212_1ng_glyco_hcd30_etd30_compound15.18764.18764.4 | controllerType=0 controllerNumber=1 scan=18764 | 42.5167 |
| 15358:4 | 2 | R.LSLHRPALEDLLLGSEAN[+1540.529]LTC[+57.021]TLTGLR.D | HexNAc(2)Hex(7) | C[+57], N[+1541] | 1126.7912 | 4 | 4504.1429 | 4504.1268 | 0 | 3.6 | 127 | Specific | 1117.16 | 1067.27 | 1067.27 | 19.63 | 29 | >sp\|P01876\|IGHA1_HUMAN Immunoglobulin heavy constant alpha 1 OS=Homo sapiens OX=9606 GN=IGHA1 PE=1 SV=2 | 255 | 20201115_15212_1ng_glyco_hcd30_etd30_compound15.18744.18744.2 | controllerType=0 controllerNumber=1 scan=18744 | 42.4596 |
| 11938:3 | 2 | R.LAGKPTHVN[+1751.624]VSVVMAEVDGTC[+57.021]Y.- | HexNAc(4)Hex(4)NeuAc(1) | C[+57], N[+1752] | 1366.9277 | 3 | 4098.7685 | 4098.7663 | 0 | 0.5 | 332 | CRagged | 230.69 | 121.71 | 121.71 | 2.87 | 29 | >sp\|P01876\|IGHA1_HUMAN Immunoglobulin heavy constant alpha 1 OS=Homo sapiens OX=9606 GN=IGHA1 PE=1 SV=2 | 255 | 20201115_15212_1ng_glyco_hcd30_etd30_compound15.15141.15141.3 | controllerType=0 controllerNumber=1 scan=15141 | 33.915 |
| 10219:3 | 2 | R.LAGKPTHVN[+1378.476]VSVVMAEVDGTC[+57.021]Y.- | HexNAc(2)Hex(6) | C[+57], N[+1378] | 1242.5479 | 3 | 3725.6292 | 3725.6178 | 0 | 3.1 | 332 | CRagged | 232.89 | 190.02 | 172.92 | 3.23 | 29 | >sp\|P01876\|IGHA1_HUMAN Immunoglobulin heavy constant alpha 1 OS=Homo sapiens OX=9606 GN=IGHA1 PE=1 SV=2 | 255 | 20201115_15212_1ng_glyco_hcd30_etd30_compound15.13332.13332.3 | controllerType=0 controllerNumber=1 scan=13332 | 30.4279 |
| 11795:3 | 2 | R.LAGKPTHVN[+2262.814]VSVVMAEVDGTC[+57.021]Y.- | HexNAc(5)Hex(5)Fuc(1)NeuAc(1) | C[+57], N[+2263] | 1537.3278 | 3 | 4609.9689 | 4609.9564 | 0 | 2.7 | 332 | CRagged | 234.9 | 69.86 | 69.86 | 2.62 | 29 | >sp\|P01876\|IGHA1_HUMAN Immunoglobulin heavy constant alpha 1 OS=Homo sapiens OX=9606 GN=IGHA1 PE=1 SV=2 | 255 | 20201115_15212_1ng_glyco_hcd30_etd30_compound15.14992.14992.3 | controllerType=0 controllerNumber=1 scan=14992 | 33.6008 |
| 13367:3 | 2 | R.LAGKPTHVN[+2553.910]VSVVMAEVDGTC[+57.021]Y.- | HexNAc(5)Hex(5)Fuc(1)NeuAc(2) | C[+57], N[+2554] | 1634.3605 | 3 | 4901.067 | 4901.0518 | 0 | 3.1 | 332 | CRagged | 241.56 | 60.39 | 60.39 | 2.82 | 29 | >sp\|P01876\|IGHA1_HUMAN Immunoglobulin heavy constant alpha 1 OS=Homo sapiens OX=9606 GN=IGHA1 PE=1 SV=2 | 255 | 20201115_15212_1ng_glyco_hcd30_etd30_compound15.16632.16632.3 | controllerType=0 controllerNumber=1 scan=16632 | 37.3414 |
| 10189:3 | 2 | R.LAGKPTHVN[+1702.581]VSVVMAEVDGTC[+57.021]Y.- | HexNAc(2)Hex(8) | C[+57], N[+1703] | 1350.5821 | 3 | 4049.7317 | 4049.7234 | 0 | 2 | 332 | CRagged | 243.67 | 201.67 | 191.13 | 4.02 | 29 | >sp\|P01876\|IGHA1_HUMAN Immunoglobulin heavy constant alpha 1 OS=Homo sapiens OX=9606 GN=IGHA1 PE=1 SV=2 | 255 | 20201115_15212_1ng_glyco_hcd30_etd30_compound15.13301.13301.3 | controllerType=0 controllerNumber=1 scan=13301 | 30.3202 |
| 10252:3 | 2 | R.LAGKPTHVN[+1216.423]VSVVMAEVDGTC[+57.021]Y.- | HexNAc(2)Hex(5) | C[+57], N[+1216] | 1188.5292 | 3 | 3563.5731 | 3563.5649 | 0 | 2.3 | 332 | CRagged | 246.22 | 246.22 | 190.49 | 4 | 29 | >sp\|P01876\|IGHA1_HUMAN Immunoglobulin heavy constant alpha 1 OS=Homo sapiens OX=9606 GN=IGHA1 PE=1 SV=2 | 255 | 20201115_15212_1ng_glyco_hcd30_etd30_compound15.13367.13367.3 | controllerType=0 controllerNumber=1 scan=13367 | 30.5025 |
| 10230:3 | 2 | R.LAGKPTHVN[+1809.666]VSVVMAEVDGTC[+57.021]Y.- | HexNAc(5)Hex(4)Fuc(1) | C[+57], N[+1810] | 1386.2763 | 3 | 4156.8144 | 4156.8081 | 0 | 1.5 | 332 | CRagged | 247.96 | 121.79 | 121.79 | 2.96 | 29 | >sp\|P01876\|IGHA1_HUMAN Immunoglobulin heavy constant alpha 1 OS=Homo sapiens OX=9606 GN=IGHA1 PE=1 SV=2 | 255 | 20201115_15212_1ng_glyco_hcd30_etd30_compound15.13344.13344.3 | controllerType=0 controllerNumber=1 scan=13344 | 30.4479 |
| 10204:3 | 2 | R.LAGKPTHVN[+1540.529]VSVVMAEVDGTC[+57.021]Y.- | HexNAc(2)Hex(7) | C[+57], N[+1541] | 1296.5655 | 3 | 3887.6819 | 3887.6706 | 0 | 2.9 | 332 | CRagged | 252.11 | 200.84 | 200.84 | 4.02 | 29 | >sp\|P01876\|IGHA1_HUMAN Immunoglobulin heavy constant alpha 1 OS=Homo sapiens OX=9606 GN=IGHA1 PE=1 SV=2 | 255 | 20201115_15212_1ng_glyco_hcd30_etd30_compound15.13317.13317.3 | controllerType=0 controllerNumber=1 scan=13317 | 30.3739 |
| 10317:3 | 2 | R.LAGKPTHVN[+1647.613]VSVVMAEVDGTC[+57.021]Y.- | HexNAc(5)Hex(3)Fuc(1) | C[+57], N[+1648] | 1332.2598 | 3 | 3994.7649 | 3994.7553 | 0 | 2.4 | 332 | CRagged | 263.58 | 113.93 | 113.93 | 3.03 | 29 | >sp\|P01876\|IGHA1_HUMAN Immunoglobulin heavy constant alpha 1 OS=Homo sapiens OX=9606 GN=IGHA1 PE=1 SV=2 | 255 | 20201115_15212_1ng_glyco_hcd30_etd30_compound15.13435.13435.3 | controllerType=0 controllerNumber=1 scan=13435 | 30.6637 |
| 11848:3 | 2 | R.LAGKPTHVN[+2262.814]VSVVMAEVDGTC[+57.021]Y.- | HexNAc(5)Hex(5)Fuc(1)NeuAc(1) | C[+57], N[+2263] | 1537.326 | 3 | 4609.9634 | 4609.9564 | 0 | 1.5 | 332 | CRagged | 281.53 | 91.41 | 91.41 | 2.9 | 29 | >sp\|P01876\|IGHA1_HUMAN Immunoglobulin heavy constant alpha 1 OS=Homo sapiens OX=9606 GN=IGHA1 PE=1 SV=2 | 255 | 20201115_15212_1ng_glyco_hcd30_etd30_compound15.15047.15047.3 | controllerType=0 controllerNumber=1 scan=15047 | 33.6953 |
| 13363:4 | 2 | R.LAGKPTHVN[+2553.910]VSVVMAEVDGTC[+57.021]Y.- | HexNAc(5)Hex(5)Fuc(1)NeuAc(2) | C[+57], N[+2554] | 1226.0226 | 4 | 4901.0687 | 4901.0518 | 0 | 3.4 | 332 | CRagged | 308.25 | 205.76 | 205.76 | 4.21 | 29 | >sp\|P01876\|IGHA1_HUMAN Immunoglobulin heavy constant alpha 1 OS=Homo sapiens OX=9606 GN=IGHA1 PE=1 SV=2 | 255 | 20201115_15212_1ng_glyco_hcd30_etd30_compound15.16628.16628.4 | controllerType=0 controllerNumber=1 scan=16628 | 37.3302 |
| 15246:3 | 4 | R.GLSFDVSLEVSQGPGLLN[+2715.963]DTK.V | HexNAc(5)Hex(6)Fuc(1)NeuAc(2) | N[+2716] | 1631.3748 | 3 | 4892.1099 | 4892.0758 | 0 | 7 | 118 | Specific | 428.22 | 248.49 | 248.49 | 4.14 | 31 | >sp\|P01833\|PIGR_HUMAN Polymeric immunoglobulin receptor OS=Homo sapiens OX=9606 GN=PIGR PE=1 SV=4 | 902 | 20201115_15212_1ng_glyco_hcd30_etd30_compound15.18626.18626.3 | controllerType=0 controllerNumber=1 scan=18626 | 42.1359 |
| 16798:3 | 4 | R.GLSFDVSLEVSQGPGLLN[+2350.830]DTK.V | HexNAc(4)Hex(5)Fuc(1)NeuAc(2) | N[+2351] | 1509.6544 | 3 | 4526.9485 | 4526.9436 | 0 | 1.1 | 118 | Specific | 442.98 | 256.86 | 256.86 | 5.3 | 31 | >sp\|P01833\|PIGR_HUMAN Polymeric immunoglobulin receptor OS=Homo sapiens OX=9606 GN=PIGR PE=1 SV=4 | 902 | 20201115_15212_1ng_glyco_hcd30_etd30_compound15.20263.20263.3 | controllerType=0 controllerNumber=1 scan=20263 | 46.6022 |
| 16837:3 | 4 | R.GLSFDVSLEVSQGPGLLN[+2350.830]DTK.V | HexNAc(4)Hex(5)Fuc(1)NeuAc(2) | N[+2351] | 1509.6578 | 3 | 4526.9588 | 4526.9436 | 0 | 3.4 | 118 | Specific | 443.83 | 271.56 | 271.56 | 5.57 | 31 | >sp\|P01833\|PIGR_HUMAN Polymeric immunoglobulin receptor OS=Homo sapiens OX=9606 GN=PIGR PE=1 SV=4 | 902 | 20201115_15212_1ng_glyco_hcd30_etd30_compound15.20304.20304.3 | controllerType=0 controllerNumber=1 scan=20304 | 46.72 |
| 17818:4 | 4 | K.QIGLYPVLVIDSSGYVNPN[+2204.772]YTGR.I | HexNAc(4)Hex(5)NeuAc(2) | N[+2205] | 1183.2803 | 4 | 4730.0994 | 4730.0759 | 0 | 5 | 168 | Specific | 409.52 | 406.94 | 406.94 | 5.87 | 31 | >sp\|P01833\|PIGR_HUMAN Polymeric immunoglobulin receptor OS=Homo sapiens OX=9606 GN=PIGR PE=1 SV=4 | 902 | 20201115_15212_1ng_glyco_hcd30_etd30_compound15.21337.21337.4 | controllerType=0 controllerNumber=1 scan=21337 | 49.7538 |
| 16290:3 | 4 | K.QIGLYPVLVIDSSGYVNPN[+2059.735]YTGR.I | HexNAc(4)Hex(5)Fuc(1)NeuAc(1) | N[+2060] | 1529.0228 | 3 | 4585.0537 | 4585.0384 | 0 | 3.3 | 168 | Specific | 605.99 | 515.86 | 515.86 | 9.44 | 31 | >sp\|P01833\|PIGR_HUMAN Polymeric immunoglobulin receptor OS=Homo sapiens OX=9606 GN=PIGR PE=1 SV=4 | 902 | 20201115_15212_1ng_glyco_hcd30_etd30_compound15.19726.19726.3 | controllerType=0 controllerNumber=1 scan=19726 | 45.2427 |
| 19109:3 | 4 | R.[+42.011]LSLLEEPGN[+2336.851]GTFTVILNQLTSR.D | HexNAc(6)Hex(6)Fuc(1) | .[42], N[+2337] | 1594.3836 | 3 | 4781.1363 | 4781.1542 | 0 | -3.8 | 413 | Specific | 311.61 | 191.62 | 191.62 | 2.94 | 31 | >sp\|P01833\|PIGR_HUMAN Polymeric immunoglobulin receptor OS=Homo sapiens OX=9606 GN=PIGR PE=1 SV=4 | 902 | 20201115_15212_1ng_glyco_hcd30_etd30_compound15.22707.22707.3 | controllerType=0 controllerNumber=1 scan=22707 | 53.4087 |
| 17525:4 | 4 | R.LSLLEEPGN[+2205.793]GTFTVILNQLTSR.D | HexNAc(4)Hex(5)Fuc(2)NeuAc(1) | N[+2206] | 1152.7812 | 4 | 4608.1029 | 4608.0854 | 0 | 3.8 | 413 | Specific | 414.96 | 385.1 | 385.1 | 6.26 | 31 | >sp\|P01833\|PIGR_HUMAN Polymeric immunoglobulin receptor OS=Homo sapiens OX=9606 GN=PIGR PE=1 SV=4 | 902 | 20201115_15212_1ng_glyco_hcd30_etd30_compound15.21027.21027.4 | controllerType=0 controllerNumber=1 scan=21027 | 48.7524 |
| 17591:3 | 4 | R.LSLLEEPGN[+2059.735]GTFTVILNQLTSR.D | HexNAc(4)Hex(5)Fuc(1)NeuAc(1) | N[+2060] | 1488.0171 | 3 | 4462.0369 | 4462.0275 | 0 | 2.1 | 413 | Specific | 532.03 | 430.62 | 430.62 | 7.37 | 31 | >sp\|P01833\|PIGR_HUMAN Polymeric immunoglobulin receptor OS=Homo sapiens OX=9606 GN=PIGR PE=1 SV=4 | 902 | 20201115_15212_1ng_glyco_hcd30_etd30_compound15.21097.21097.3 | controllerType=0 controllerNumber=1 scan=21097 | 48.961 |
| 19073:4 | 4 | R.LSLLEEPGN[+2350.830]GTFTVILNQLTSR.D | HexNAc(4)Hex(5)Fuc(1)NeuAc(2) | N[+2351] | 1189.0429 | 4 | 4753.1497 | 4753.1229 | 0 | 5.6 | 413 | Specific | 605.41 | 541.29 | 541.29 | 8.41 | 31 | >sp\|P01833\|PIGR_HUMAN Polymeric immunoglobulin receptor OS=Homo sapiens OX=9606 GN=PIGR PE=1 SV=4 | 902 | 20201115_15212_1ng_glyco_hcd30_etd30_compound15.22669.22669.4 | controllerType=0 controllerNumber=1 scan=22669 | 53.293 |
| 19041:3 | 4 | R.LSLLEEPGN[+2350.830]GTFTVILNQLTSR.D | HexNAc(4)Hex(5)Fuc(1)NeuAc(2) | N[+2351] | 1585.0532 | 3 | 4753.1449 | 4753.1229 | 0 | 4.6 | 413 | Specific | 631.19 | 441.74 | 441.74 | 8.77 | 31 | >sp\|P01833\|PIGR_HUMAN Polymeric immunoglobulin receptor OS=Homo sapiens OX=9606 GN=PIGR PE=1 SV=4 | 902 | 20201115_15212_1ng_glyco_hcd30_etd30_compound15.22635.22635.3 | controllerType=0 controllerNumber=1 scan=22635 | 53.178 |
| 14194:3 | 4 | K.IIEGEPNLKVPGN[+2205.793]VTAVLGETLK.V | HexNAc(4)Hex(5)Fuc(2)NeuAc(1) | N[+2206] | 1533.0574 | 3 | 4597.1576 | 4597.1422 | 0 | 3.4 | 457 | Specific | 259.53 | 183.88 | 183.88 | 3.11 | 31 | >sp\|P01833\|PIGR_HUMAN Polymeric immunoglobulin receptor OS=Homo sapiens OX=9606 GN=PIGR PE=1 SV=4 | 902 | 20201115_15212_1ng_glyco_hcd30_etd30_compound15.17519.17519.3 | controllerType=0 controllerNumber=1 scan=17519 | 39.4092 |
| 14169:4 | 4 | K.IIEGEPNLKVPGN[+2059.735]VTAVLGETLK.V | HexNAc(4)Hex(5)Fuc(1)NeuAc(1) | N[+2060] | 1113.5275 | 4 | 4451.0882 | 4451.0843 | 0 | 0.9 | 457 | Specific | 370.37 | 370.37 | 370.37 | 5.45 | 31 | >sp\|P01833\|PIGR_HUMAN Polymeric immunoglobulin receptor OS=Homo sapiens OX=9606 GN=PIGR PE=1 SV=4 | 902 | 20201115_15212_1ng_glyco_hcd30_etd30_compound15.17492.17492.2 | controllerType=0 controllerNumber=1 scan=17492 | 39.3123 |
| 13743:4 | 4 | K.IIEGEPNLKVPGN[+2059.735]VTAVLGETLKVPC[+57.021]HFPC[+57.021]K.F | HexNAc(4)Hex(5)Fuc(1)NeuAc(1) | C[+57]*2, N[+2060] | 1369.8903 | 4 | 5476.5392 | 5476.5418 | 0 | -0.5 | 457 | Specific | 281.61 | 281.61 | 281.61 | 2.63 | 31 | >sp\|P01833\|PIGR_HUMAN Polymeric immunoglobulin receptor OS=Homo sapiens OX=9606 GN=PIGR PE=1 SV=4 | 902 | 20201115_15212_1ng_glyco_hcd30_etd30_compound15.17026.17026.4 | controllerType=0 controllerNumber=1 scan=17026 | 38.3057 |
| 14906:4 | 4 | K.IIEGEPNLKVPGN[+2350.830]VTAVLGETLKVPC[+57.021]HFPC[+57.021]K.F | HexNAc(4)Hex(5)Fuc(1)NeuAc(2) | C[+57]*2, N[+2351] | 1442.6681 | 4 | 5767.6505 | 5767.6372 | 0 | 2.3 | 457 | Specific | 310.47 | 302.61 | 302.61 | 3.45 | 31 | >sp\|P01833\|PIGR_HUMAN Polymeric immunoglobulin receptor OS=Homo sapiens OX=9606 GN=PIGR PE=1 SV=4 | 902 | 20201115_15212_1ng_glyco_hcd30_etd30_compound15.18267.18267.4 | controllerType=0 controllerNumber=1 scan=18267 | 41.1331 |
| 13802:5 | 4 | K.IIEGEPNLKVPGN[+2059.735]VTAVLGETLKVPC[+57.021]HFPC[+57.021]K.F | HexNAc(4)Hex(5)Fuc(1)NeuAc(1) | C[+57]*2, N[+2060] | 1096.1171 | 5 | 5476.5562 | 5476.5418 | 0 | 2.6 | 457 | Specific | 312.94 | 312.94 | 312.94 | 3.56 | 31 | >sp\|P01833\|PIGR_HUMAN Polymeric immunoglobulin receptor OS=Homo sapiens OX=9606 GN=PIGR PE=1 SV=4 | 902 | 20201115_15212_1ng_glyco_hcd30_etd30_compound15.17087.17087.5 | controllerType=0 controllerNumber=1 scan=17087 | 38.4257 |
| 13795:5 | 4 | K.IIEGEPNLKVPGN[+2204.772]VTAVLGETLKVPC[+57.021]HFPC[+57.021]K.F | HexNAc(4)Hex(5)NeuAc(2) | C[+57]*2, N[+2205] | 1125.1267 | 5 | 5621.6045 | 5621.5793 | 0 | 4.5 | 457 | Specific | 376.43 | 376.43 | 376.43 | 4.55 | 31 | >sp\|P01833\|PIGR_HUMAN Polymeric immunoglobulin receptor OS=Homo sapiens OX=9606 GN=PIGR PE=1 SV=4 | 902 | 20201115_15212_1ng_glyco_hcd30_etd30_compound15.17080.17080.5 | controllerType=0 controllerNumber=1 scan=17080 | 38.4122 |
| 13708:5 | 4 | K.IIEGEPNLKVPGN[+2205.793]VTAVLGETLKVPC[+57.021]HFPC[+57.021]K.F | HexNAc(4)Hex(5)Fuc(2)NeuAc(1) | C[+57]*2, N[+2206] | 1125.3285 | 5 | 5622.6134 | 5622.5997 | 0 | 2.4 | 457 | Specific | 486.91 | 486.91 | 486.91 | 5.98 | 31 | >sp\|P01833\|PIGR_HUMAN Polymeric immunoglobulin receptor OS=Homo sapiens OX=9606 GN=PIGR PE=1 SV=4 | 902 | 20201115_15212_1ng_glyco_hcd30_etd30_compound15.16989.16989.5 | controllerType=0 controllerNumber=1 scan=16989 | 38.2111 |
| 14934:5 | 4 | K.IIEGEPNLKVPGN[+2350.830]VTAVLGETLKVPC[+57.021]HFPC[+57.021]K.F | HexNAc(4)Hex(5)Fuc(1)NeuAc(2) | C[+57]*2, N[+2351] | 1154.3366 | 5 | 5767.6536 | 5767.6372 | 0 | 2.8 | 457 | Specific | 521.57 | 521.57 | 521.57 | 6.07 | 31 | >sp\|P01833\|PIGR_HUMAN Polymeric immunoglobulin receptor OS=Homo sapiens OX=9606 GN=PIGR PE=1 SV=4 | 902 | 20201115_15212_1ng_glyco_hcd30_etd30_compound15.18297.18297.5 | controllerType=0 controllerNumber=1 scan=18297 | 41.2169 |
| 13730:4 | 4 | K.IIEGEPNLKVPGN[+2204.772]VTAVLGETLKVPC[+57.021]HFPC[+57.021]K.F | HexNAc(4)Hex(5)NeuAc(2) | C[+57]*2, N[+2205] | 1406.1576 | 4 | 5621.6086 | 5621.5793 | 0 | 5.2 | 457 | Specific | 539.46 | 539.46 | 539.46 | 5.31 | 31 | >sp\|P01833\|PIGR_HUMAN Polymeric immunoglobulin receptor OS=Homo sapiens OX=9606 GN=PIGR PE=1 SV=4 | 902 | 20201115_15212_1ng_glyco_hcd30_etd30_compound15.17012.17012.4 | controllerType=0 controllerNumber=1 scan=17012 | 38.2749 |
| 05446:3 | 4 | K.WN[+1913.677]NTGC[+57.021]QALPSQDEGPSK.A | HexNAc(4)Hex(5)NeuAc(1) | C[+57], N[+1914] | 1301.5272 | 3 | 3902.5669 | 3902.5537 | 0 | 3.4 | 498 | Specific | 266.89 | 135.78 | 135.78 | 4.19 | 31 | >sp\|P01833\|PIGR_HUMAN Polymeric immunoglobulin receptor OS=Homo sapiens OX=9606 GN=PIGR PE=1 SV=4 | 902 | 20201115_15212_1ng_glyco_hcd30_etd30_compound15.8355.8355.3 | controllerType=0 controllerNumber=1 scan=8355 | 20.9129 |
| 05335:3 | 4 | K.WN[+2059.735]NTGC[+57.021]QALPSQDEGPSK.A | HexNAc(4)Hex(5)Fuc(1)NeuAc(1) | C[+57], N[+2060] | 1350.2144 | 3 | 4048.6287 | 4048.6116 | 0 | 4.2 | 498 | Specific | 447.94 | 307.02 | 307.02 | 5.34 | 31 | >sp\|P01833\|PIGR_HUMAN Polymeric immunoglobulin receptor OS=Homo sapiens OX=9606 GN=PIGR PE=1 SV=4 | 902 | 20201115_15212_1ng_glyco_hcd30_etd30_compound15.8240.8240.3 | controllerType=0 controllerNumber=1 scan=8240 | 20.6892 |
| 09257:3 | 6 | K.TVLTPATNHMGN[+1378.476]VTFTIPANR.E | HexNAc(2)Hex(6) | N[+1378] | 1211.887 | 3 | 3633.6465 | 3633.6358 | 0 | 2.9 | 74 | Specific | 215.31 | 175.02 | 109.36 | 3.39 | 15 | >sp\|P01024\|CO3_HUMAN Complement C3 OS=Homo sapiens OX=9606 GN=C3 PE=1 SV=2 | 492 | 20201115_15212_1ng_glyco_hcd30_etd30_compound15.12331.12331.3 | controllerType=0 controllerNumber=1 scan=12331 | 28.348 |
| 09209:3 | 6 | K.TVLTPATNHMGN[+1540.529]VTFTIPANR.E | HexNAc(2)Hex(7) | N[+1541] | 1265.9051 | 3 | 3795.7007 | 3795.6886 | 0 | 3.2 | 74 | Specific | 379.74 | 336.75 | 112.88 | 5.53 | 15 | >sp\|P01024\|CO3_HUMAN Complement C3 OS=Homo sapiens OX=9606 GN=C3 PE=1 SV=2 | 492 | 20201115_15212_1ng_glyco_hcd30_etd30_compound15.12280.12280.3 | controllerType=0 controllerNumber=1 scan=12280 | 28.2014 |
| 15010:4 | 7 | K.RNIAAFGGDPNN[+1913.677]ITLFGESAGGASVSLQTLSPYNK.G | HexNAc(4)Hex(5)NeuAc(1) | N[+1914] | 1370.8777 | 4 | 5480.4889 | 5480.4644 | 0 | 4.5 | 196 | Specific | 588.91 | 573.19 | 573.19 | 7.29 | 10 | >sp\|P19835\|CEL_HUMAN Bile salt-activated lipase OS=Homo sapiens OX=9606 GN=CEL PE=1 SV=3 | 476 | 20201115_15212_1ng_glyco_hcd30_etd30_compound15.18378.18378.4 | controllerType=0 controllerNumber=1 scan=18378 | 41.4659 |
| 19851:3 | 8 | R.QLAHQSN[+2204.772]STNIFFSPVSIATAFAMLSLGTK.A | HexNAc(4)Hex(5)NeuAc(2) | N[+2205] | 1796.1461 | 3 | 5386.4237 | 5386.4075 | 0 | 3 | 64 | Specific | 439.6 | 315.13 | 315.13 | 5.73 | 9 | >sp\|P01009\|A1AT_HUMAN Alpha-1-antitrypsin OS=Homo sapiens OX=9606 GN=SERPINA1 PE=1 SV=3 | 642 | 20201115_15212_1ng_glyco_hcd30_etd30_compound15.23496.23496.3 | controllerType=0 controllerNumber=1 scan=23496 | 55.7824 |
| 19845:4 | 8 | R.QLAHQSN[+2204.772]STNIFFSPVSIATAFAMLSLGTK.A | HexNAc(4)Hex(5)NeuAc(2) | N[+2205] | 1347.3611 | 4 | 5386.4225 | 5386.4075 | 0 | 2.8 | 64 | Specific | 619.75 | 503.5 | 434.26 | 9.84 | 9 | >sp\|P01009\|A1AT_HUMAN Alpha-1-antitrypsin OS=Homo sapiens OX=9606 GN=SERPINA1 PE=1 SV=3 | 642 | 20201115_15212_1ng_glyco_hcd30_etd30_compound15.23490.23490.4 | controllerType=0 controllerNumber=1 scan=23490 | 55.7692 |
| 19017:4 | 8 | K.ADTHDEILEGLNFN[+2861.000]LTEIPEAQIHEGFQELLR.T | HexNAc(5)Hex(6)NeuAc(3) | N[+2861] | 1638.963 | 4 | 6552.8302 | 6552.8239 | 0 | 1 | 94 | Specific | 269.1 | 269.1 | 269.1 | 4.89 | 9 | >sp\|P01009\|A1AT_HUMAN Alpha-1-antitrypsin OS=Homo sapiens OX=9606 GN=SERPINA1 PE=1 SV=3 | 642 | 20201115_15212_1ng_glyco_hcd30_etd30_compound15.22609.22609.4 | controllerType=0 controllerNumber=1 scan=22609 | 53.0959 |
| 19629:4 | 11 | K.VFPLSLDSTPQDGNVVVAC[+57.021]LVQGFFPQEPLSVTWSESGQN[+2263.835]VTAI | HexNAc(5)Hex(5)Fuc(3) | C[+57], N[+2264] | 1760.8114 | 4 | 7040.2238 | 7040.1961 | 0 | 3.9 | 8 | Specific | 253.71 | 253.71 | 253.71 | 4.86 | 22 | >sp\|P01877\|IGHA2_HUMAN Immunoglobulin heavy constant alpha 2 OS=Homo sapiens OX=9606 GN=IGHA2 PE=1 SV=4 | 576 | 20201115_15212_1ng_glyco_hcd30_etd30_compound15.23260.23260.4 | controllerType=0 controllerNumber=1 scan=23260 | 55.0526 |
| 01249:3 | 11 | K.HYTN[+1971.719]SSQDVTVPC[+57.021]R.V | HexNAc(5)Hex(5)Fuc(1) | C[+57], N[+1972] | 1212.4994 | 3 | 3635.4836 | 3635.4682 | 0 | 4.3 | 89 | Specific | 248.24 | 186.66 | 186.66 | 4.45 | 22 | >sp\|P01877\|IGHA2_HUMAN Immunoglobulin heavy constant alpha 2 OS=Homo sapiens OX=9606 GN=IGHA2 PE=1 SV=4 | 576 | 20201115_15212_1ng_glyco_hcd30_etd30_compound15.3997.3997.3 | controllerType=0 controllerNumber=1 scan=3997 | 12.8422 |
| 02066:3 | 11 | K.HYTN[+2262.814]SSQDVTVPC[+57.021]R.V | HexNAc(5)Hex(5)Fuc(1)NeuAc(1) | C[+57], N[+2263] | 1309.5318 | 3 | 3926.5808 | 3926.5636 | 0 | 4.4 | 89 | Specific | 358.2 | 249.22 | 249.22 | 3.9 | 22 | >sp\|P01877\|IGHA2_HUMAN Immunoglobulin heavy constant alpha 2 OS=Homo sapiens OX=9606 GN=IGHA2 PE=1 SV=4 | 576 | 20201115_15212_1ng_glyco_hcd30_etd30_compound15.4845.4845.3 | controllerType=0 controllerNumber=1 scan=4845 | 14.4575 |
| 01250:3 | 11 | K.HYTN[+1809.666]SSQDVTVPC[+57.021]R.V | HexNAc(5)Hex(4)Fuc(1) | C[+57], N[+1810] | 1158.4785 | 3 | 3473.4208 | 3473.4154 | 0 | 1.6 | 89 | Specific | 396.74 | 332.31 | 332.31 | 5.25 | 22 | >sp\|P01877\|IGHA2_HUMAN Immunoglobulin heavy constant alpha 2 OS=Homo sapiens OX=9606 GN=IGHA2 PE=1 SV=4 | 576 | 20201115_15212_1ng_glyco_hcd30_etd30_compound15.3998.3998.3 | controllerType=0 controllerNumber=1 scan=3998 | 12.8508 |
| 01272:3 | 11 | K.HYTN[+1647.613]SSQDVTVPC[+57.021]R.V | HexNAc(5)Hex(3)Fuc(1) | C[+57], N[+1648] | 1104.4607 | 3 | 3311.3676 | 3311.3625 | 0 | 1.5 | 89 | Specific | 443.39 | 394.66 | 394.66 | 6.31 | 22 | >sp\|P01877\|IGHA2_HUMAN Immunoglobulin heavy constant alpha 2 OS=Homo sapiens OX=9606 GN=IGHA2 PE=1 SV=4 | 576 | 20201115_15212_1ng_glyco_hcd30_etd30_compound15.4021.4021.3 | controllerType=0 controllerNumber=1 scan=4021 | 12.9016 |
| 03044:4 | 11 | K.HYTN[+1647.613]SSQDVTVPC[+57.021]RVPPPPPC[+57.021]C[+57.021]HPR.L | HexNAc(5)Hex(3)Fuc(1) | C[+57]*3, N[+1648] | 1152.252 | 4 | 4605.9862 | 4605.9689 | 0 | 3.8 | 89 | Specific | 315.98 | 287.99 | 287.99 | 3.41 | 22 | >sp\|P01877\|IGHA2_HUMAN Immunoglobulin heavy constant alpha 2 OS=Homo sapiens OX=9606 GN=IGHA2 PE=1 SV=4 | 576 | 20201115_15212_1ng_glyco_hcd30_etd30_compound15.5864.5864.4 | controllerType=0 controllerNumber=1 scan=5864 | 16.5235 |
| 03041:5 | 11 | K.HYTN[+1647.613]SSQDVTVPC[+57.021]RVPPPPPC[+57.021]C[+57.021]HPR.L | HexNAc(5)Hex(3)Fuc(1) | C[+57]*3, N[+1648] | 922.0031 | 5 | 4605.9866 | 4605.9689 | 0 | 3.8 | 89 | Specific | 534.18 | 495.31 | 495.31 | 6.11 | 22 | >sp\|P01877\|IGHA2_HUMAN Immunoglobulin heavy constant alpha 2 OS=Homo sapiens OX=9606 GN=IGHA2 PE=1 SV=4 | 576 | 20201115_15212_1ng_glyco_hcd30_etd30_compound15.5861.5861.5 | controllerType=0 controllerNumber=1 scan=5861 | 16.5124 |
| 02789:3 | 11 | K.TPLTAN[+1971.719]ITK.S | HexNAc(5)Hex(5)Fuc(1) | N[+1972] | 977.4339 | 3 | 2930.287 | 2930.2757 | 0 | 3.9 | 200 | Specific | 267.1 | 121.72 | 121.72 | 2.69 | 22 | >sp\|P01877\|IGHA2_HUMAN Immunoglobulin heavy constant alpha 2 OS=Homo sapiens OX=9606 GN=IGHA2 PE=1 SV=4 | 576 | 20201115_15212_1ng_glyco_hcd30_etd30_compound15.5597.5597.3 | controllerType=0 controllerNumber=1 scan=5597 | 15.862 |
| 02884:3 | 11 | K.TPLTAN[+1647.613]ITK.S | HexNAc(5)Hex(3)Fuc(1) | N[+1648] | 869.3982 | 3 | 2606.1801 | 2606.17 | 0 | 3.9 | 200 | Specific | 290.73 | 130.16 | 130.16 | 2.72 | 22 | >sp\|P01877\|IGHA2_HUMAN Immunoglobulin heavy constant alpha 2 OS=Homo sapiens OX=9606 GN=IGHA2 PE=1 SV=4 | 576 | 20201115_15212_1ng_glyco_hcd30_etd30_compound15.5697.5697.3 | controllerType=0 controllerNumber=1 scan=5697 | 16.136 |
| 02878:2 | 11 | K.TPLTAN[+1647.613]ITK.S | HexNAc(5)Hex(3)Fuc(1) | N[+1648] | 1303.5935 | 2 | 2606.1798 | 2606.17 | 0 | 3.8 | 200 | Specific | 300.22 | 117.59 | 117.59 | 2.89 | 22 | >sp\|P01877\|IGHA2_HUMAN Immunoglobulin heavy constant alpha 2 OS=Homo sapiens OX=9606 GN=IGHA2 PE=1 SV=4 | 576 | 20201115_15212_1ng_glyco_hcd30_etd30_compound15.5691.5691.2 | controllerType=0 controllerNumber=1 scan=5691 | 16.1169 |
| 02830:3 | 11 | K.TPLTAN[+1809.666]ITK.S | HexNAc(5)Hex(4)Fuc(1) | N[+1810] | 923.416 | 3 | 2768.2336 | 2768.2228 | 0 | 3.9 | 200 | Specific | 301.39 | 134.9 | 134.9 | 2.81 | 22 | >sp\|P01877\|IGHA2_HUMAN Immunoglobulin heavy constant alpha 2 OS=Homo sapiens OX=9606 GN=IGHA2 PE=1 SV=4 | 576 | 20201115_15212_1ng_glyco_hcd30_etd30_compound15.5640.5640.3 | controllerType=0 controllerNumber=1 scan=5640 | 15.9799 |
| 02836:2 | 11 | K.TPLTAN[+1809.666]ITK.S | HexNAc(5)Hex(4)Fuc(1) | N[+1810] | 1384.6208 | 2 | 2768.2343 | 2768.2228 | 0 | 4.1 | 200 | Specific | 309 | 111.49 | 111.49 | 2.68 | 22 | >sp\|P01877\|IGHA2_HUMAN Immunoglobulin heavy constant alpha 2 OS=Homo sapiens OX=9606 GN=IGHA2 PE=1 SV=4 | 576 | 20201115_15212_1ng_glyco_hcd30_etd30_compound15.5646.5646.2 | controllerType=0 controllerNumber=1 scan=5646 | 15.9938 |
| 02848:2 | 11 | K.TPLTAN[+1444.534]ITK.S | HexNAc(4)Hex(3)Fuc(1) | N[+1445] | 1202.0514 | 2 | 2403.0956 | 2403.0906 | 0 | 2.1 | 200 | Specific | 341.85 | 107.39 | 107.39 | 2.64 | 22 | >sp\|P01877\|IGHA2_HUMAN Immunoglobulin heavy constant alpha 2 OS=Homo sapiens OX=9606 GN=IGHA2 PE=1 SV=4 | 576 | 20201115_15212_1ng_glyco_hcd30_etd30_compound15.5659.5659.2 | controllerType=0 controllerNumber=1 scan=5659 | 16.0244 |
| 09337:3 | 11 | R.MAGKPTHIN[+1540.529]VSVVMAEADGTC[+57.021]Y.- | HexNAc(2)Hex(7) | C[+57], N[+1541] | 1297.879 | 3 | 3891.6223 | 3891.6114 | 0 | 2.8 | 319 | CRagged | 258.77 | 258.77 | 186.1 | 4.36 | 22 | >sp\|P01877\|IGHA2_HUMAN Immunoglobulin heavy constant alpha 2 OS=Homo sapiens OX=9606 GN=IGHA2 PE=1 SV=4 | 576 | 20201115_15212_1ng_glyco_hcd30_etd30_compound15.12415.12415.3 | controllerType=0 controllerNumber=1 scan=12415 | 28.5662 |
| 09314:3 | 11 | R.MAGKPTHIN[+1702.581]VSVVMAEADGTC[+57.021]Y.- | HexNAc(2)Hex(8) | C[+57], N[+1703] | 1351.8992 | 3 | 4053.6831 | 4053.6642 | 0 | 4.7 | 319 | CRagged | 272.43 | 271.44 | 151.35 | 3.55 | 22 | >sp\|P01877\|IGHA2_HUMAN Immunoglobulin heavy constant alpha 2 OS=Homo sapiens OX=9606 GN=IGHA2 PE=1 SV=4 | 576 | 20201115_15212_1ng_glyco_hcd30_etd30_compound15.12391.12391.3 | controllerType=0 controllerNumber=1 scan=12391 | 28.501 |
| 09349:3 | 11 | R.MAGKPTHIN[+1378.476]VSVVMAEADGTC[+57.021]Y.- | HexNAc(2)Hex(6) | C[+57], N[+1378] | 1243.8609 | 3 | 3729.5681 | 3729.5585 | 0 | 2.6 | 319 | CRagged | 300.16 | 258.38 | 220.06 | 4.7 | 22 | >sp\|P01877\|IGHA2_HUMAN Immunoglobulin heavy constant alpha 2 OS=Homo sapiens OX=9606 GN=IGHA2 PE=1 SV=4 | 576 | 20201115_15212_1ng_glyco_hcd30_etd30_compound15.12428.12428.3 | controllerType=0 controllerNumber=1 scan=12428 | 28.6082 |
| 09394:3 | 11 | R.MAGKPTHIN[+1216.423]VSVVMAEADGTC[+57.021]Y.- | HexNAc(2)Hex(5) | C[+57], N[+1216] | 1189.8442 | 3 | 3567.518 | 3567.5057 | 0 | 3.4 | 319 | CRagged | 300.39 | 283.86 | 217.7 | 4.39 | 22 | >sp\|P01877\|IGHA2_HUMAN Immunoglobulin heavy constant alpha 2 OS=Homo sapiens OX=9606 GN=IGHA2 PE=1 SV=4 | 576 | 20201115_15212_1ng_glyco_hcd30_etd30_compound15.12475.12475.3 | controllerType=0 controllerNumber=1 scan=12475 | 28.7233 |
| 12790:3 | 23 | K.VVLHPN[+2204.772]YSQVDIGLIK.L | HexNAc(4)Hex(5)NeuAc(2) | N[+2205] | 1333.9356 | 3 | 3999.7923 | 3999.7837 | 0 | 2.1 | 236 | Specific | 283.58 | 195.16 | 195.16 | 4.17 | 5 | >sp\|P00738\|HPT_HUMAN Haptoglobin OS=Homo sapiens OX=9606 GN=HP PE=1 SV=1 | 407 | 20201115_15212_1ng_glyco_hcd30_etd30_compound15.16030.16030.3 | controllerType=0 controllerNumber=1 scan=16030 | 35.9537 |
| 10295:3 | 25 | R.IIVPLNNREN[+1856.656]ISDPTSPLR.T | HexNAc(3)Hex(5)Fuc(1)NeuAc(1) | N[+1857] | 1335.622 | 3 | 4004.8515 | 4004.8327 | 0 | 4.7 | 62 | Specific | 249.38 | 200.71 | 200.71 | 3.03 | 8 | >sp\|P01591\|IGJ_HUMAN Immunoglobulin J chain OS=Homo sapiens OX=9606 GN=JCHAIN PE=1 SV=4 | 1392 | 20201115_15212_1ng_glyco_hcd30_etd30_compound15.13412.13412.3 | controllerType=0 controllerNumber=1 scan=13412 | 30.6007 |
| 11960:3 | 25 | R.IIVPLNNREN[+2350.830]ISDPTSPLR.T | HexNAc(4)Hex(5)Fuc(1)NeuAc(2) | N[+2351] | 1500.3425 | 3 | 4499.0128 | 4499.0075 | 0 | 1.2 | 62 | Specific | 318.28 | 234.22 | 234.22 | 3.39 | 8 | >sp\|P01591\|IGJ_HUMAN Immunoglobulin J chain OS=Homo sapiens OX=9606 GN=JCHAIN PE=1 SV=4 | 1392 | 20201115_15212_1ng_glyco_hcd30_etd30_compound15.15164.15164.3 | controllerType=0 controllerNumber=1 scan=15164 | 33.9619 |
| 07197:3 | 28 | R.LSPN[+2350.830]ASAEHLELR.W | HexNAc(4)Hex(5)Fuc(1)NeuAc(2) | N[+2351] | 1263.2012 | 3 | 3787.5891 | 3787.5796 | 0 | 2.5 | 52 | Specific | 288.48 | 148.68 | 148.68 | 3.76 | 5 | >sp\|Q13410\|BT1A1_HUMAN Butyrophilin subfamily 1 member A1 OS=Homo sapiens OX=9606 GN=BTN1A1 PE=1 SV=3 | 179 | 20201115_15212_1ng_glyco_hcd30_etd30_compound15.10172.10172.3 | controllerType=0 controllerNumber=1 scan=10172 | 24.1469 |
| 05315:3 | 28 | R.LSPN[+2205.793]ASAEHLELR.W | HexNAc(4)Hex(5)Fuc(2)NeuAc(1) | N[+2206] | 1214.8541 | 3 | 3642.5476 | 3642.5421 | 0 | 1.5 | 52 | Specific | 300.07 | 195.91 | 195.91 | 3.86 | 5 | >sp\|Q13410\|BT1A1_HUMAN Butyrophilin subfamily 1 member A1 OS=Homo sapiens OX=9606 GN=BTN1A1 PE=1 SV=3 | 179 | 20201115_15212_1ng_glyco_hcd30_etd30_compound15.8219.8219.3 | controllerType=0 controllerNumber=1 scan=8219 | 20.6496 |
| 07206:3 | 28 | R.LSPN[+2350.830]ASAEHLELR.W | HexNAc(4)Hex(5)Fuc(1)NeuAc(2) | N[+2351] | 1263.1993 | 3 | 3787.5833 | 3787.5796 | 0 | 1 | 52 | Specific | 304.1 | 185.64 | 185.64 | 3.61 | 5 | >sp\|Q13410\|BT1A1_HUMAN Butyrophilin subfamily 1 member A1 OS=Homo sapiens OX=9606 GN=BTN1A1 PE=1 SV=3 | 179 | 20201115_15212_1ng_glyco_hcd30_etd30_compound15.10182.10182.3 | controllerType=0 controllerNumber=1 scan=10182 | 24.1701 |
| 00457:4 | 30 | R.EIRHN[+2424.867]STGC[+57.021]LR.M | HexNAc(5)Hex(6)Fuc(1)NeuAc(1) | C[+57], N[+2425] | 942.6409 | 4 | 3767.5418 | 3767.5316 | 0 | 2.7 | 287 | Specific | 442.08 | 281.55 | 281.55 | 4.25 | 8 | >sp\|P10909\|CLUS_HUMAN Clusterin OS=Homo sapiens OX=9606 GN=CLU PE=1 SV=1 | 1467 | 20201115_15212_1ng_glyco_hcd30_etd30_compound15.3173.3173.4 | controllerType=0 controllerNumber=1 scan=3173 | 11.2546 |
| 00468:4 | 30 | R.EIRHN[+2059.735]STGC[+57.021]LR.M | HexNAc(4)Hex(5)Fuc(1)NeuAc(1) | C[+57], N[+2060] | 851.3573 | 4 | 3402.4075 | 3402.3994 | 0 | 2.4 | 287 | Specific | 663.2 | 500.99 | 500.99 | 9.11 | 8 | >sp\|P10909\|CLUS_HUMAN Clusterin OS=Homo sapiens OX=9606 GN=CLU PE=1 SV=1 | 1467 | 20201115_15212_1ng_glyco_hcd30_etd30_compound15.3185.3185.4 | controllerType=0 controllerNumber=1 scan=3185 | 11.3014 |
| 02084:4 | 33 | R.EKQTDEIKDTRN[+2059.735]ESTQNC[+57.021]VVAEPEK.M | HexNAc(4)Hex(5)Fuc(1)NeuAc(1) | C[+57], N[+2060] | 1252.7898 | 4 | 5008.1374 | 5008.1251 | 0 | 2.4 | 58 | Specific | 439.12 | 403.29 | 403.29 | 5.15 | 5 | >sp\|P47710\|CASA1_HUMAN Alpha-S1-casein OS=Homo sapiens OX=9606 GN=CSN1S1 PE=1 SV=1 | 989 | 20201115_15212_1ng_glyco_hcd30_etd30_compound15.4864.4864.4 | controllerType=0 controllerNumber=1 scan=4864 | 14.5065 |
| 02483:4 | 33 | K.QTDEIKDTRN[+2059.735]ESTQNC[+57.021]VVAEPEK.M | HexNAc(4)Hex(5)Fuc(1)NeuAc(1) | C[+57], N[+2060] | 1188.5063 | 4 | 4751.0032 | 4750.9876 | 0 | 3.3 | 60 | Specific | 620.27 | 511.91 | 511.91 | 8.93 | 5 | >sp\|P47710\|CASA1_HUMAN Alpha-S1-casein OS=Homo sapiens OX=9606 GN=CSN1S1 PE=1 SV=1 | 989 | 20201115_15212_1ng_glyco_hcd30_etd30_compound15.5278.5278.4 | controllerType=0 controllerNumber=1 scan=5278 | 15.282 |
| 07909:5 | 39 | K.SC[+57.021]QHN[+2059.735]GTMYQHGEIFSAHELFPSR.L | HexNAc(4)Hex(5)Fuc(1)NeuAc(1) | C[+57], N[+2060] | 979.4132 | 5 | 4893.037 | 4892.9919 | 0 | 9.2 | 110 | Specific | 662.82 | 601.98 | 601.98 | 9.08 | 2 | >sp\|Q6WN34\|CRDL2_HUMAN Chordin-like protein 2 OS=Homo sapiens OX=9606 GN=CHRDL2 PE=1 SV=1 | 1189 | 20201115_15212_1ng_glyco_hcd30_etd30_compound15.10918.10918.3 | controllerType=0 controllerNumber=1 scan=10918 | 25.6261 |

Table S4. Cont.

| Query #:z | Protein Rank | Glycopeptide | Glycan | Modification Type(s) | Observed m/z | z | Observed (M+H) | Calc. mass (M+H) | Off-by-x error | Mass error (ppm) | StartPosition | Cleavage | Score | Delta | Delta Mod | \|Log Prob\| | # of unique peptides | Protein Name | Protein DB number | Comment | Scan # | Scan Time |
|---|---|---|---|---|---|---|---|---|---|---|---|---|---|---|---|---|---|---|---|---|---|---|
| 01030:4 | 82 | K.TKPREEQFN[+1768.640]STFR.V | HexNAc(4)Hex(5)Fuc(1) | N[+1769] | 852.8727 | 4 | 3408.469 | 3408.4582 | 0 | 3.2 | 168 | Specific | 206.59 | 206.59 | 206.59 | 3.1 | 3 | >sp\|P01859\|IGHG2_HUMAN Immunoglobulin heavy constant gamma 2 OS=Homo sapiens OX=9606 GN=IGHG2 PE=1 SV=2 | 387 | 20201115_15212_1ng_glyco_hcd30_etd30_compound15.3769.3769.4 | controllerType=0 controllerNumber=1 scan=3769 | 12.3996 |
| 01080:4 | 82 | K.TKPREEQFN[+1606.587]STFR.V | HexNAc(4)Hex(4)Fuc(1) | N[+1607] | 812.3589 | 4 | 3246.4138 | 3246.4054 | 0 | 2.6 | 168 | Specific | 274.84 | 274.84 | 274.84 | 3.64 | 3 | >sp\|P01859\|IGHG2_HUMAN Immunoglobulin heavy constant gamma 2 OS=Homo sapiens OX=9606 GN=IGHG2 PE=1 SV=2 | 387 | 20201115_15212_1ng_glyco_hcd30_etd30_compound15.3821.3821.4 | controllerType=0 controllerNumber=1 scan=3821 | 12.5045 |
| 15729:5 | 1 | R.RTAGWNVPIGTLRPFLN[+2059.735]WTGPPEPIEAAVAR.F | HexNAc(4)Hex(5)Fuc(1)NeuAc(1) | N[+2060] | 1090.1168 | 5 | 5446.555 | 5446.547 | 0 | 1.5 | 140 | Specific | 330.13 | 330.13 | 330.13 | 4.15 | 53 | >sp\|P02788\|TRFL_HUMAN Lactotransferrin OS=Homo sapiens OX=9606 GN=LTF PE=1 SV=6 | 1270 | 20201119_15232_1ng_glyco_hcd30_etd30_compound15.19372.19372.5 | controllerType=0 controllerNumber=1 scan=19372 | 43.7086 |
| 15676:5 | 1 | R.RTAGWNVPIGTLRPFLN[+2204.772]WTGPPEPIEAAVAR.F | HexNAc(4)Hex(5)NeuAc(2) | N[+2205] | 1119.1292 | 5 | 5591.6167 | 5591.5845 | 0 | 5.8 | 140 | Specific | 468.1 | 447.31 | 447.31 | 6.1 | 53 | >sp\|P02788\|TRFL_HUMAN Lactotransferrin OS=Homo sapiens OX=9606 GN=LTF PE=1 SV=6 | 1270 | 20201119_15232_1ng_glyco_hcd30_etd30_compound15.19316.19316.6 | controllerType=0 controllerNumber=1 scan=19316 | 43.5329 |
| 15633:4 | 1 | R.RTAGWNVPIGTLRPFLN[+2570.925]WTGPPEPIEAAVAR.F | HexNAc(5)Hex(6)Fuc(2)NeuAc(1) | N[+2571] | 1490.1952 | 4 | 5957.7589 | 5957.7371 | 0 | 3.7 | 140 | Specific | 512.88 | 512.88 | 512.88 | 6.55 | 53 | >sp\|P02788\|TRFL_HUMAN Lactotransferrin OS=Homo sapiens OX=9606 GN=LTF PE=1 SV=6 | 1270 | 20201119_15232_1ng_glyco_hcd30_etd30_compound15.19271.19271.4 | controllerType=0 controllerNumber=1 scan=19271 | 43.4133 |
| 16417:4 | 1 | R.RTAGWNVPIGTLRPFLN[+2350.830]WTGPPEPIEAAVAR.F | HexNAc(4)Hex(5)Fuc(1)NeuAc(2) | N[+2351] | 1435.1694 | 4 | 5737.6558 | 5737.6424 | 0 | 2.3 | 140 | Specific | 926.06 | 926.06 | 926.06 | 15.25 | 53 | >sp\|P02788\|TRFL_HUMAN Lactotransferrin OS=Homo sapiens OX=9606 GN=LTF PE=1 SV=6 | 1270 | 20201119_15232_1ng_glyco_hcd30_etd30_compound15.20099.20099.4 | controllerType=0 controllerNumber=1 scan=20099 | 45.8451 |
| 15041:4 | 1 | R.RTAGWNVPIGTLRPFLN[+1768.640]WTGPPEPIEAAVAR.F | HexNAc(4)Hex(5)Fuc(1) | N[+1769] | 1289.6221 | 4 | 5155.4665 | 5155.4516 | 0 | 2.9 | 140 | Specific | 975.76 | 969.73 | 969.73 | 16.25 | 53 | >sp\|P02788\|TRFL_HUMAN Lactotransferrin OS=Homo sapiens OX=9606 GN=LTF PE=1 SV=6 | 1270 | 20201119_15232_1ng_glyco_hcd30_etd30_compound15.18644.18644.4 | controllerType=0 controllerNumber=1 scan=18644 | 41.4477 |
| 15720:4 | 1 | R.RTAGWNVPIGTLRPFLN[+2059.735]WTGPPEPIEAAVAR.F | HexNAc(4)Hex(5)Fuc(1)NeuAc(1) | N[+2060] | 1362.3973 | 4 | 5446.5675 | 5446.547 | 0 | 3.8 | 140 | Specific | 1014.96 | 984.04 | 984.04 | 17.82 | 53 | >sp\|P02788\|TRFL_HUMAN Lactotransferrin OS=Homo sapiens OX=9606 GN=LTF PE=1 SV=6 | 1270 | 20201119_15232_1ng_glyco_hcd30_etd30_compound15.19363.19363.4 | controllerType=0 controllerNumber=1 scan=19363 | 43.6854 |
| 16546:5 | 1 | R.TAGWNVPIGTLRPFLN[+2205.793]WTGPPEPIEAAVAR.F | HexNAc(4)Hex(5)Fuc(2)NeuAc(1) | N[+2206] | 1088.1115 | 5 | 5436.5282 | 5436.5038 | 0 | 4.5 | 141 | Specific | 262.23 | 249.89 | 249.89 | 3.74 | 53 | >sp\|P02788\|TRFL_HUMAN Lactotransferrin OS=Homo sapiens OX=9606 GN=LTF PE=1 SV=6 | 1270 | 20201119_15232_1ng_glyco_hcd30_etd30_compound15.20235.20235.4 | controllerType=0 controllerNumber=1 scan=20235 | 46.2501 |
| 16595:3 | 1 | R.TAGWNVPIGTLRPFLN[+2204.772]WTGPPEPIEAAVAR.F | HexNAc(4)Hex(5)NeuAc(2) | N[+2205] | 1812.5102 | 3 | 5435.5161 | 5435.4834 | 0 | 6 | 141 | Specific | 296.91 | 207.65 | 207.65 | 4.8 | 53 | >sp\|P02788\|TRFL_HUMAN Lactotransferrin OS=Homo sapiens OX=9606 GN=LTF PE=1 SV=6 | 1270 | 20201119_15232_1ng_glyco_hcd30_etd30_compound15.20287.20287.3 | controllerType=0 controllerNumber=1 scan=20287 | 46.3921 |
| 16597:3 | 1 | R.TAGWNVPIGTLRPFLN[+2059.735]WTGPPEPIEAAVAR.F | HexNAc(4)Hex(5)Fuc(1)NeuAc(1) | N[+2060] | 1764.1602 | 3 | 5290.4662 | 5290.4459 | 0 | 3.8 | 141 | Specific | 319.23 | 221.84 | 221.84 | 5.74 | 53 | >sp\|P02788\|TRFL_HUMAN Lactotransferrin OS=Homo sapiens OX=9606 GN=LTF PE=1 SV=6 | 1270 | 20201119_15232_1ng_glyco_hcd30_etd30_compound15.20289.20289.3 | controllerType=0 controllerNumber=1 scan=20289 | 46.3962 |
| 16503:4 | 1 | R.TAGWNVPIGTLRPFLN[+2716.983]WTGPPEPIEAAVAR.F | HexNAc(5)Hex(6)Fuc(3)NeuAc(1) | N[+2717] | 1487.6854 | 4 | 5947.7199 | 5947.6939 | 0 | 4.4 | 141 | Specific | 322.82 | 322.82 | 322.82 | 5.92 | 53 | >sp\|P02788\|TRFL_HUMAN Lactotransferrin OS=Homo sapiens OX=9606 GN=LTF PE=1 SV=6 | 1270 | 20201119_15232_1ng_glyco_hcd30_etd30_compound15.20190.20190.4 | controllerType=0 controllerNumber=1 scan=20190 | 46.1244 |
| 17585:3 | 1 | R.TAGWNVPIGTLRPFLN[+2350.830]WTGPPEPIEAAVAR.F | HexNAc(4)Hex(5)Fuc(1)NeuAc(2) | N[+2351] | 1861.1995 | 3 | 5581.5841 | 5581.5413 | 0 | 7.7 | 141 | Specific | 376.78 | 307.53 | 307.53 | 4.96 | 53 | >sp\|P02788\|TRFL_HUMAN Lactotransferrin OS=Homo sapiens OX=9606 GN=LTF PE=1 SV=6 | 1270 | 20201119_15232_1ng_glyco_hcd30_etd30_compound15.21333.21333.3 | controllerType=0 controllerNumber=1 scan=21333 | 49.2875 |
| 15903:4 | 1 | R.TAGWNVPIGTLRPFLN[+1913.677]WTGPPEPIEAAVAR.F | HexNAc(4)Hex(5)NeuAc(1) | N[+1914] | 1286.8613 | 4 | 5144.4235 | 5144.388 | 0 | 6.9 | 141 | Specific | 379.26 | 379.26 | 379.26 | 5.9 | 53 | >sp\|P02788\|TRFL_HUMAN Lactotransferrin OS=Homo sapiens OX=9606 GN=LTF PE=1 SV=6 | 1270 | 20201119_15232_1ng_glyco_hcd30_etd30_compound15.19555.19555.4 | controllerType=0 controllerNumber=1 scan=19555 | 44.212 |
| 16549:3 | 1 | R.TAGWNVPIGTLRPFLN[+2059.735]WTGPPEPIEAAVAR.F | HexNAc(4)Hex(5)Fuc(1)NeuAc(1) | N[+2060] | 1764.1594 | 3 | 5290.4636 | 5290.4459 | 0 | 3.4 | 141 | Specific | 382.11 | 281.24 | 281.24 | 5.77 | 53 | >sp\|P02788\|TRFL_HUMAN Lactotransferrin OS=Homo sapiens OX=9606 GN=LTF PE=1 SV=6 | 1270 | 20201119_15232_1ng_glyco_hcd30_etd30_compound15.20239.20239.3 | controllerType=0 controllerNumber=1 scan=20239 | 46.2627 |
| 17607:3 | 1 | R.TAGWNVPIGTLRPFLN[+2351.851]WTGPPEPIEAAVAR.F | HexNAc(4)Hex(5)Fuc(3)NeuAc(1) | N[+2352] | 1861.5223 | 3 | 5582.5523 | 5582.5617 | 0 | -1.7 | 141 | Specific | 396.91 | 332.84 | 332.84 | 4.54 | 53 | >sp\|P02788\|TRFL_HUMAN Lactotransferrin OS=Homo sapiens OX=9606 GN=LTF PE=1 SV=6 | 1270 | 20201119_15232_1ng_glyco_hcd30_etd30_compound15.21356.21356.3 | controllerType=0 controllerNumber=1 scan=21356 | 49.3479 |
| 16534:3 | 1 | R.TAGWNVPIGTLRPFLN[+2205.793]WTGPPEPIEAAVAR.F | HexNAc(4)Hex(5)Fuc(2)NeuAc(1) | N[+2206] | 1812.8475 | 3 | 5436.5279 | 5436.5038 | 0 | 4.4 | 141 | Specific | 445.96 | 362.45 | 362.45 | 6.81 | 53 | >sp\|P02788\|TRFL_HUMAN Lactotransferrin OS=Homo sapiens OX=9606 GN=LTF PE=1 SV=6 | 1270 | 20201119_15232_1ng_glyco_hcd30_etd30_compound15.20223.20223.3 | controllerType=0 controllerNumber=1 scan=20223 | 46.2093 |
| 15875:3 | 1 | R.TAGWNVPIGTLRPFLN[+1768.640]WTGPPEPIEAAVAR.F | HexNAc(4)Hex(5)Fuc(1) | N[+1769] | 1667.1279 | 3 | 4999.3691 | 4999.3504 | 0 | 3.7 | 141 | Specific | 450.13 | 363.94 | 363.94 | 7.27 | 53 | >sp\|P02788\|TRFL_HUMAN Lactotransferrin OS=Homo sapiens OX=9606 GN=LTF PE=1 SV=6 | 1270 | 20201119_15232_1ng_glyco_hcd30_etd30_compound15.19526.19526.3 | controllerType=0 controllerNumber=1 scan=19526 | 44.1239 |
| 16723:4 | 1 | R.TAGWNVPIGTLRPFLN[+2059.735]WTGPPEPIEAAVAR.F | HexNAc(4)Hex(5)Fuc(1)NeuAc(1) | N[+2060] | 1323.3712 | 4 | 5290.463 | 5290.4459 | 0 | 3.2 | 141 | Specific | 530.53 | 509.69 | 509.69 | 8.63 | 53 | >sp\|P02788\|TRFL_HUMAN Lactotransferrin OS=Homo sapiens OX=9606 GN=LTF PE=1 SV=6 | 1270 | 20201119_15232_1ng_glyco_hcd30_etd30_compound15.20422.20422.4 | controllerType=0 controllerNumber=1 scan=20422 | 46.6219 |
| 17584:4 | 1 | R.TAGWNVPIGTLRPFLN[+2350.830]WTGPPEPIEAAVAR.F | HexNAc(4)Hex(5)Fuc(1)NeuAc(2) | N[+2351] | 1396.1442 | 4 | 5581.5548 | 5581.5413 | 0 | 2.4 | 141 | Specific | 589.94 | 589.94 | 589.94 | 9.01 | 53 | >sp\|P02788\|TRFL_HUMAN Lactotransferrin OS=Homo sapiens OX=9606 GN=LTF PE=1 SV=6 | 1270 | 20201119_15232_1ng_glyco_hcd30_etd30_compound15.21332.21332.4 | controllerType=0 controllerNumber=1 scan=21332 | 49.2844 |
| 16471:4 | 1 | R.TAGWNVPIGTLRPFLN[+2570.925]WTGPPEPIEAAVAR.F | HexNAc(5)Hex(6)Fuc(2)NeuAc(1) | N[+2571] | 1451.1733 | 4 | 5801.6715 | 5801.636 | 0 | 6.1 | 141 | Specific | 831.84 | 827.04 | 827.04 | 13.63 | 53 | >sp\|P02788\|TRFL_HUMAN Lactotransferrin OS=Homo sapiens OX=9606 GN=LTF PE=1 SV=6 | 1270 | 20201119_15232_1ng_glyco_hcd30_etd30_compound15.20157.20157.4 | controllerType=0 controllerNumber=1 scan=20157 | 46.0224 |
| 15876:4 | 1 | R.TAGWNVPIGTLRPFLN[+1768.640]WTGPPEPIEAAVAR.F | HexNAc(4)Hex(5)Fuc(1) | N[+1769] | 1250.5985 | 4 | 4999.3722 | 4999.3504 | 0 | 4.4 | 141 | Specific | 858.51 | 858.51 | 858.51 | 15.58 | 53 | >sp\|P02788\|TRFL_HUMAN Lactotransferrin OS=Homo sapiens OX=9606 GN=LTF PE=1 SV=6 | 1270 | 20201119_15232_1ng_glyco_hcd30_etd30_compound15.19527.19527.4 | controllerType=0 controllerNumber=1 scan=19527 | 44.1278 |
| 16535:4 | 1 | R.TAGWNVPIGTLRPFLN[+2205.793]WTGPPEPIEAAVAR.F | HexNAc(4)Hex(5)Fuc(2)NeuAc(1) | N[+2206] | 1359.8877 | 4 | 5436.529 | 5436.5038 | 0 | 4.6 | 141 | Specific | 875.72 | 838.08 | 838.08 | 14.52 | 53 | >sp\|P02788\|TRFL_HUMAN Lactotransferrin OS=Homo sapiens OX=9606 GN=LTF PE=1 SV=6 | 1270 | 20201119_15232_1ng_glyco_hcd30_etd30_compound15.20224.20224.4 | controllerType=0 controllerNumber=1 scan=20224 | 46.2145 |
| 16242:4 | 1 | R.TAGWNIPMGLLFN[+1913.677]QTGSC[+57.021]K.F | HexNAc(4)Hex(5)NeuAc(1) | C[+57], N[+1914] | 1002.9311 | 4 | 4008.7025 | 4008.687 | 0 | 3.9 | 485 | Specific | 211.81 | 171.57 | 171.57 | 3.44 | 53 | >sp\|P02788\|TRFL_HUMAN Lactotransferrin OS=Homo sapiens OX=9606 GN=LTF PE=1 SV=6 | 1270 | 20201119_15232_1ng_glyco_hcd30_etd30_compound15.19914.19914.4 | controllerType=0 controllerNumber=1 scan=19914 | 45.2991 |
| 16306:3 | 1 | R.TAGWNIPMGLLFN[+1751.624]QTGSC[+57.021]K.F | HexNAc(4)Hex(4)NeuAc(1) | C[+57], N[+1752] | 1282.8871 | 3 | 3846.6469 | 3846.6341 | 0 | 3.3 | 485 | Specific | 363.73 | 243.31 | 243.31 | 5.85 | 53 | >sp\|P02788\|TRFL_HUMAN Lactotransferrin OS=Homo sapiens OX=9606 GN=LTF PE=1 SV=6 | 1270 | 20201119_15232_1ng_glyco_hcd30_etd30_compound15.19982.19982.3 | controllerType=0 controllerNumber=1 scan=19982 | 45.5167 |
| 14788:3 | 1 | R.TAGWNIPMGLLFN[+1768.640]QTGSC[+57.021]K.F | HexNAc(4)Hex(5)Fuc(1) | C[+57], N[+1769] | 1288.5591 | 3 | 3863.6629 | 3863.6495 | 0 | 3.5 | 485 | Specific | 376.56 | 280.53 | 280.53 | 5.85 | 53 | >sp\|P02788\|TRFL_HUMAN Lactotransferrin OS=Homo sapiens OX=9606 GN=LTF PE=1 SV=6 | 1270 | 20201119_15232_1ng_glyco_hcd30_etd30_compound15.18375.18375.3 | controllerType=0 controllerNumber=1 scan=18375 | 40.6395 |
| 14894:3 | 1 | R.TAGWNIPMGLLFN[+1606.587]QTGSC[+57.021]K.F | HexNAc(4)Hex(4)Fuc(1) | C[+57], N[+1607] | 1234.5406 | 3 | 3701.6072 | 3701.5966 | 0 | 2.9 | 485 | Specific | 382.65 | 294.89 | 294.89 | 5.04 | 53 | >sp\|P02788\|TRFL_HUMAN Lactotransferrin OS=Homo sapiens OX=9606 GN=LTF PE=1 SV=6 | 1270 | 20201119_15232_1ng_glyco_hcd30_etd30_compound15.18487.18487.3 | controllerType=0 controllerNumber=1 scan=18487 | 40.9483 |
| 14999:3 | 1 | R.TAGWNIPMGLLFN[+1460.529]QTGSC[+57.021]K.F | HexNAc(4)Hex(4) | C[+57], N[+1461] | 1185.8547 | 3 | 3555.5495 | 3555.5387 | 0 | 3 | 485 | Specific | 387.62 | 311.07 | 227.21 | 4.58 | 53 | >sp\|P02788\|TRFL_HUMAN Lactotransferrin OS=Homo sapiens OX=9606 GN=LTF PE=1 SV=6 | 1270 | 20201119_15232_1ng_glyco_hcd30_etd30_compound15.18600.18600.3 | controllerType=0 controllerNumber=1 scan=18600 | 41.3136 |
| 14931:3 | 1 | R.TAGWNIPMGLLFN[+1622.582]QTGSC[+57.021]K.F | HexNAc(4)Hex(5) | C[+57], N[+1623] | 1239.8731 | 3 | 3717.6048 | 3717.5916 | 0 | 3.6 | 485 | Specific | 389.42 | 302.64 | 259.51 | 6.21 | 53 | >sp\|P02788\|TRFL_HUMAN Lactotransferrin OS=Homo sapiens OX=9606 GN=LTF PE=1 SV=6 | 1270 | 20201119_15232_1ng_glyco_hcd30_etd30_compound15.18528.18528.3 | controllerType=0 controllerNumber=1 scan=18528 | 41.0775 |
| 16094:3 | 1 | R.TAGWNIPMGLLFN[+2205.793]QTGSC[+57.021]K.F | HexNAc(4)Hex(5)Fuc(2)NeuAc(1) | C[+57], N[+2206] | 1434.277 | 3 | 4300.8166 | 4300.8028 | 0 | 3.2 | 485 | Specific | 409.47 | 286.54 | 286.54 | 6.29 | 53 | >sp\|P02788\|TRFL_HUMAN Lactotransferrin OS=Homo sapiens OX=9606 GN=LTF PE=1 SV=6 | 1270 | 20201119_15232_1ng_glyco_hcd30_etd30_compound15.19757.19757.3 | controllerType=0 controllerNumber=1 scan=19757 | 44.8083 |
| 16114:3 | 1 | R.TAGWNIPMGLLFN[+2059.735]QTGSC[+57.021]K.F | HexNAc(4)Hex(5)Fuc(1)NeuAc(1) | C[+57], N[+2060] | 1385.5912 | 3 | 4154.7592 | 4154.7449 | 0 | 3.4 | 485 | Specific | 414.88 | 317.9 | 317.9 | 6.29 | 53 | >sp\|P02788\|TRFL_HUMAN Lactotransferrin OS=Homo sapiens OX=9606 GN=LTF PE=1 SV=6 | 1270 | 20201119_15232_1ng_glyco_hcd30_etd30_compound15.19779.19779.3 | controllerType=0 controllerNumber=1 scan=19779 | 44.8704 |
| 16217:3 | 1 | R.TAGWNIPMGLLFN[+1913.677]QTGSC[+57.021]K.F | HexNAc(4)Hex(5)NeuAc(1) | C[+57], N[+1914] | 1336.9052 | 3 | 4008.7011 | 4008.687 | 0 | 3.5 | 485 | Specific | 440.1 | 310.6 | 310.6 | 6.51 | 53 | >sp\|P02788\|TRFL_HUMAN Lactotransferrin OS=Homo sapiens OX=9606 GN=LTF PE=1 SV=6 | 1270 | 20201119_15232_1ng_glyco_hcd30_etd30_compound15.19888.19888.3 | controllerType=0 controllerNumber=1 scan=19888 | 45.2131 |
| 17644:3 | 1 | R.TAGWNIPMGLLFN[+2350.830]QTGSC[+57.021]K.F | HexNAc(4)Hex(5)Fuc(1)NeuAc(2) | C[+57], N[+2351] | 1482.6243 | 3 | 4445.8584 | 4445.8403 | 0 | 4.1 | 485 | Specific | 470.93 | 300.23 | 300.23 | 6.51 | 53 | >sp\|P02788\|TRFL_HUMAN Lactotransferrin OS=Homo sapiens OX=9606 GN=LTF PE=1 SV=6 | 1270 | 20201119_15232_1ng_glyco_hcd30_etd30_compound15.21395.21395.3 | controllerType=0 controllerNumber=1 scan=21395 | 49.4561 |
| 15050:3 | 1 | R.TAGWNIPMGLLFN[+1216.423]QTGSC[+57.021]K.F | HexNAc(2)Hex(5) | C[+57], N[+1216] | 1104.486 | 3 | 3311.4434 | 3311.4328 | 0 | 3.2 | 485 | Specific | 476.88 | 451.12 | 250.35 | 7.71 | 53 | >sp\|P02788\|TRFL_HUMAN Lactotransferrin OS=Homo sapiens OX=9606 GN=LTF PE=1 SV=6 | 1270 | 20201119_15232_1ng_glyco_hcd30_etd30_compound15.18654.18654.3 | controllerType=0 controllerNumber=1 scan=18654 | 41.482 |
| 15064:2 | 1 | R.TAGWNIPMGLLFN[+1216.423]QTGSC[+57.021]K.F | HexNAc(2)Hex(5) | C[+57], N[+1216] | 1656.2236 | 2 | 3311.4398 | 3311.4328 | 0 | 2.1 | 485 | Specific | 552.29 | 552.29 | 162.73 | 6.97 | 53 | >sp\|P02788\|TRFL_HUMAN Lactotransferrin OS=Homo sapiens OX=9606 GN=LTF PE=1 SV=6 | 1270 | 20201119_15232_1ng_glyco_hcd30_etd30_compound15.18669.18669.2 | controllerType=0 controllerNumber=1 scan=18669 | 41.5236 |
| 14962:3 | 1 | R.TAGWNIPM[+15.995]GLLFN[+2059.735]QTGSC[+57.021]K.F | HexNAc(4)Hex(5)Fuc(1)NeuAc(1) | C[+57], M[+16], N[+2060] | 1390.9238 | 3 | 4170.7567 | 4170.7398 | 0 | 4.1 | 485 | Specific | 312.08 | 268.64 | 268.64 | 5.59 | 53 | >sp\|P02788\|TRFL_HUMAN Lactotransferrin OS=Homo sapiens OX=9606 GN=LTF PE=1 SV=6 | 1270 | 20201119_15232_1ng_glyco_hcd30_etd30_compound15.18560.18560.3 | controllerType=0 controllerNumber=1 scan=18560 | 41.1827 |
| 15622:3 | 1 | R.TAGWNIPMGLLFN[+1622.582]QTGSC[+57.021]KFDEYFSQSC[+57.021]APGSDPR.S | HexNAc(4)Hex(5) | C[+57]*2, N[+1623] | 1854.4578 | 3 | 5561.359 | 5561.3387 | 0 | 3.6 | 485 | Specific | 257.57 | 201.2 | 201.2 | 5.28 | 53 | >sp\|P02788\|TRFL_HUMAN Lactotransferrin OS=Homo sapiens OX=9606 GN=LTF PE=1 SV=6 | 1270 | 20201119_15232_1ng_glyco_hcd30_etd30_compound15.19259.19259.3 | controllerType=0 controllerNumber=1 scan=19259 | 43.3852 |
| 16318:5 | 1 | R.TAGWNIPMGLLFN[+2059.735]QTGSC[+57.021]KFDEYFSQSC[+57.021]APGSDPR.S | HexNAc(4)Hex(5)Fuc(1)NeuAc(1) | C[+57]*2, N[+2060] | 1200.509 | 5 | 5998.5161 | 5998.492 | 0 | 4 | 485 | Specific | 304.49 | 304.49 | 304.49 | 5.33 | 53 | >sp\|P02788\|TRFL_HUMAN Lactotransferrin OS=Homo sapiens OX=9606 GN=LTF PE=1 SV=6 | 1270 | 20201119_15232_1ng_glyco_hcd30_etd30_compound15.19995.19995.5 | controllerType=0 controllerNumber=1 scan=19995 | 45.5495 |
| 16369:4 | 1 | R.TAGWNIPMGLLFN[+1913.677]QTGSC[+57.021]KFDEYFSQSC[+57.021]APGSDPR.S | HexNAc(4)Hex(5)NeuAc(1) | C[+57]*2, N[+1914] | 1463.8665 | 4 | 5852.444 | 5852.4341 | 0 | 1.7 | 485 | Specific | 386.02 | 326.13 | 326.13 | 5.01 | 53 | >sp\|P02788\|TRFL_HUMAN Lactotransferrin OS=Homo sapiens OX=9606 GN=LTF PE=1 SV=6 | 1270 | 20201119_15232_1ng_glyco_hcd30_etd30_compound15.20049.20049.4 | controllerType=0 controllerNumber=1 scan=20049 | 45.7152 |
| 15609:4 | 1 | R.TAGWNIPMGLLFN[+1622.582]QTGSC[+57.021]KFDEYFSQSC[+57.021]APGSDPR.S | HexNAc(4)Hex(5) | C[+57]*2, N[+1623] | 1391.0972 | 4 | 5561.3669 | 5561.3387 | 0 | 5.1 | 485 | Specific | 471.91 | 429.06 | 127.64 | 5.96 | 53 | >sp\|P02788\|TRFL_HUMAN Lactotransferrin OS=Homo sapiens OX=9606 GN=LTF PE=1 SV=6 | 1270 | 20201119_15232_1ng_glyco_hcd30_etd30_compound15.19246.19246.4 | controllerType=0 controllerNumber=1 scan=19246 | 43.3415 |
| 15596:4 | 1 | R.TAGWNIPMGLLFN[+1606.587]QTGSC[+57.021]KFDEYFSQSC[+57.021]APGSDPR.S | HexNAc(4)Hex(4)Fuc(1) | C[+57]*2, N[+1607] | 1387.0974 | 4 | 5545.3678 | 5545.3438 | 0 | 4.3 | 485 | Specific | 566.8 | 522.71 | 522.71 | 8.33 | 53 | >sp\|P02788\|TRFL_HUMAN Lactotransferrin OS=Homo sapiens OX=9606 GN=LTF PE=1 SV=6 | 1270 | 20201119_15232_1ng_glyco_hcd30_etd30_compound15.19232.19232.4 | controllerType=0 controllerNumber=1 scan=19232 | 43.295 |
| 16304:4 | 1 | R.TAGWNIPMGLLFN[+2205.793]QTGSC[+57.021]KFDEYFSQSC[+57.021]APGSDPR.S | HexNAc(4)Hex(5)Fuc(2)NeuAc(1) | C[+57]*2, N[+2206] | 1536.8981 | 4 | 6144.5705 | 6144.5499 | 0 | 3.3 | 485 | Specific | 575.19 | 534.32 | 534.32 | 9.32 | 53 | >sp\|P02788\|TRFL_HUMAN Lactotransferrin OS=Homo sapiens OX=9606 GN=LTF PE=1 SV=6 | 1270 | 20201119_15232_1ng_glyco_hcd30_etd30_compound15.19980.19980.4 | controllerType=0 controllerNumber=1 scan=19980 | 45.5078 |
| 16299:4 | 1 | R.TAGWNIPMGLLFN[+2059.735]QTGSC[+57.021]KFDEYFSQSC[+57.021]APGSDPR.S | HexNAc(4)Hex(5)Fuc(1)NeuAc(1) | C[+57]*2, N[+2060] | 1500.386 | 4 | 5998.5221 | 5998.492 | 0 | 5 | 485 | Specific | 733.95 | 658.46 | 658.46 | 11.27 | 53 | >sp\|P02788\|TRFL_HUMAN Lactotransferrin OS=Homo sapiens OX=9606 GN=LTF PE=1 SV=6 | 1270 | 20201119_15232_1ng_glyco_hcd30_etd30_compound15.19975.19975.4 | controllerType=0 controllerNumber=1 scan=19975 | 45.4975 |
| 15575:4 | 1 | R.TAGWNIPMGLLFN[+1768.640]QTGSC[+57.021]KFDEYFSQSC[+57.021]APGSDPR.S | HexNAc(4)Hex(5)Fuc(1) | C[+57]*2, N[+1769] | 1427.6127 | 4 | 5707.4289 | 5707.3966 | 0 | 5.6 | 485 | Specific | 812.68 | 739.37 | 739.37 | 13.27 | 53 | >sp\|P02788\|TRFL_HUMAN Lactotransferrin OS=Homo sapiens OX=9606 GN=LTF PE=1 SV=6 | 1270 | 20201119_15232_1ng_glyco_hcd30_etd30_compound15.19210.19210.4 | controllerType=0 controllerNumber=1 scan=19210 | 43.2244 |
| 10326:3 | 1 | R.N[+2351.851]GSDC[+57.021]PDKFC[+57.021]LFQSETK.N | HexNAc(4)Hex(5)Fuc(3)NeuAc(1) | C[+57]*2, N[+2352] | 1462.2418 | 3 | 4384.7107 | 4384.7247 | 0 | -3.2 | 642 | Specific | 258.11 | 211.51 | 211.51 | 3.11 | 53 | >sp\|P02788\|TRFL_HUMAN Lactotransferrin OS=Homo sapiens OX=9606 GN=LTF PE=1 SV=6 | 1270 | 20201119_15232_1ng_glyco_hcd30_etd30_compound15.13661.13661.3 | controllerType=0 controllerNumber=1 scan=13661 | 30.948 |
| 08207:3 | 1 | R.N[+2204.772]GSDC[+57.021]PDKFC[+57.021]LFQSETK.N | HexNAc(4)Hex(5)NeuAc(2) | C[+57]*2, N[+2205] | 1413.231 | 3 | 4237.6785 | 4237.6463 | 0 | 7.6 | 642 | Specific | 260.26 | 152.66 | 152.66 | 3.01 | 53 | >sp\|P02788\|TRFL_HUMAN Lactotransferrin OS=Homo sapiens OX=9606 GN=LTF PE=1 SV=6 | 1270 | 20201119_15232_1ng_glyco_hcd30_etd30_compound15.11459.11459.3 | controllerType=0 controllerNumber=1 scan=11459 | 26.6715 |
| 08227:3 | 1 | R.N[+2059.735]GSDC[+57.021]PDKFC[+57.021]LFQSETK.N | HexNAc(4)Hex(5)Fuc(1)NeuAc(1) | C[+57]*2, N[+2060] | 1364.8781 | 3 | 4092.6198 | 4092.6088 | 0 | 2.7 | 642 | Specific | 270.28 | 159.83 | 159.83 | 4.32 | 53 | >sp\|P02788\|TRFL_HUMAN Lactotransferrin OS=Homo sapiens OX=9606 GN=LTF PE=1 SV=6 | 1270 | 20201119_15232_1ng_glyco_hcd30_etd30_compound15.11479.11479.3 | controllerType=0 controllerNumber=1 scan=11479 | 26.7084 |
| 08141:3 | 1 | R.N[+2205.793]GSDC[+57.021]PDKFC[+57.021]LFQSETK.N | HexNAc(4)Hex(5)Fuc(2)NeuAc(1) | C[+57]*2, N[+2206] | 1413.5622 | 3 | 4238.672 | 4238.6667 | 0 | 1.2 | 642 | Specific | 309.31 | 190.81 | 190.81 | 3.94 | 53 | >sp\|P02788\|TRFL_HUMAN Lactotransferrin OS=Homo sapiens OX=9606 GN=LTF PE=1 SV=6 | 1270 | 20201119_15232_1ng_glyco_hcd30_etd30_compound15.11389.11389.3 | controllerType=0 controllerNumber=1 scan=11389 | 26.5188 |
| 08255:3 | 1 | R.N[+2059.735]GSDC[+57.021]PDKFC[+57.021]LFQSETK.N | HexNAc(4)Hex(5)Fuc(1)NeuAc(1) | C[+57]*2, N[+2060] | 1364.8791 | 3 | 4092.6227 | 4092.6088 | 0 | 3.4 | 642 | Specific | 336.22 | 242.05 | 242.05 | 5.46 | 53 | >sp\|P02788\|TRFL_HUMAN Lactotransferrin OS=Homo sapiens OX=9606 GN=LTF PE=1 SV=6 | 1270 | 20201119_15232_1ng_glyco_hcd30_etd30_compound15.11509.11509.3 | controllerType=0 controllerNumber=1 scan=11509 | 26.7769 |
| 15433:3 | 2 | R.LSLHRPALEDLLLGSEAN[+1501.555]LTC[+57.021]TLTGLR.D | HexNAc(5)Hex(3) | C[+57], N[+1502] | 1489.0613 | 3 | 4465.1695 | 4465.1536 | 0 | 3.6 | 127 | Specific | 282.36 | 206.05 | 206.05 | 5.74 | 33 | >sp\|P01876\|IGHA1_HUMAN Immunoglobulin heavy constant alpha 1 OS=Homo sapiens OX=9606 GN=IGHA1 PE=1 SV=2 | 255 | 20201119_15232_1ng_glyco_hcd30_etd30_compound15.19059.19059.3 | controllerType=0 controllerNumber=1 scan=19059 | 42.728 |
| 15466:3 | 2 | R.LSLHRPALEDLLLGSEAN[+1298.476]LTC[+57.021]TLTGLR.D | HexNAc(4)Hex(3) | C[+57], N[+1298] | 1421.3699 | 3 | 4262.0952 | 4262.0742 | 0 | 4.9 | 127 | Specific | 318.2 | 318.2 | 318.2 | 5.06 | 33 | >sp\|P01876\|IGHA1_HUMAN Immunoglobulin heavy constant alpha 1 OS=Homo sapiens OX=9606 GN=IGHA1 PE=1 SV=2 | 255 | 20201119_15232_1ng_glyco_hcd30_etd30_compound15.19094.19094.3 | controllerType=0 controllerNumber=1 scan=19094 | 42.8477 |
| 16162:3 | 2 | R.LSLHRPALEDLLLGSEAN[+1751.624]LTC[+57.021]TLTGLR.D | HexNAc(4)Hex(4)NeuAc(1) | C[+57], N[+1752] | 1572.4201 | 3 | 4715.2457 | 4715.2225 | 0 | 4.9 | 127 | Specific | 323.47 | 301.04 | 301.04 | 5.35 | 33 | >sp\|P01876\|IGHA1_HUMAN Immunoglobulin heavy constant alpha 1 OS=Homo sapiens OX=9606 GN=IGHA1 PE=1 SV=2 | 255 | 20201119_15232_1ng_glyco_hcd30_etd30_compound15.19829.19829.3 | controllerType=0 controllerNumber=1 scan=19829 | 44.9932 |
| 15467:5 | 2 | R.LSLHRPALEDLLLGSEAN[+1524.534]LTC[+57.021]TLTGLR.D | HexNAc(2)Hex(6)Fuc(1) | C[+57], N[+1525] | 898.4312 | 5 | 4488.1271 | 4488.1319 | 0 | -1.1 | 127 | Specific | 329 | 329 | 329 | 3.32 | 33 | >sp\|P01876\|IGHA1_HUMAN Immunoglobulin heavy constant alpha 1 OS=Homo sapiens OX=9606 GN=IGHA1 PE=1 SV=2 | 255 | 20201119_15232_1ng_glyco_hcd30_etd30_compound15.19095.19095.5 | controllerType=0 controllerNumber=1 scan=19095 | 42.8514 |
| 15417:3 | 2 | R.LSLHRPALEDLLLGSEAN[+1663.608]LTC[+57.021]TLTGLR.D | HexNAc(5)Hex(4) | C[+57], N[+1664] | 1543.0801 | 3 | 4627.2259 | 4627.2064 | 0 | 4.2 | 127 | Specific | 385.31 | 365.96 | 365.96 | 7.58 | 33 | >sp\|P01876\|IGHA1_HUMAN Immunoglobulin heavy constant alpha 1 OS=Homo sapiens OX=9606 GN=IGHA1 PE=1 SV=2 | 255 | 20201119_15232_1ng_glyco_hcd30_etd30_compound15.19042.19042.3 | controllerType=0 controllerNumber=1 scan=19042 | 42.6737 |
| 15465:5 | 2 | R.LSLHRPALEDLLLGSEAN[+1501.555]LTC[+57.021]TLTGLR.D | HexNAc(5)Hex(3) | C[+57], N[+1502] | 893.8404 | 5 | 4465.1727 | 4465.1536 | 0 | 4.3 | 127 | Specific | 389.46 | 389.46 | 389.46 | 7.05 | 33 | >sp\|P01876\|IGHA1_HUMAN Immunoglobulin heavy constant alpha 1 OS=Homo sapiens OX=9606 GN=IGHA1 PE=1 SV=2 | 255 | 20201119_15232_1ng_glyco_hcd30_etd30_compound15.19093.19093.5 | controllerType=0 controllerNumber=1 scan=19093 | 42.8418 |
| 15440:3 | 2 | R.LSLHRPALEDLLLGSEAN[+1622.582]LTC[+57.021]TLTGLR.D | HexNAc(4)Hex(5) | C[+57], N[+1623] | 1529.3998 | 3 | 4586.185 | 4586.1799 | 0 | 1.1 | 127 | Specific | 402.24 | 333.47 | 333.47 | 5.49 | 33 | >sp\|P01876\|IGHA1_HUMAN Immunoglobulin heavy constant alpha 1 OS=Homo sapiens OX=9606 GN=IGHA1 PE=1 SV=2 | 255 | 20201119_15232_1ng_glyco_hcd30_etd30_compound15.19066.19066.3 | controllerType=0 controllerNumber=1 scan=19066 | 42.7509 |
| 15434:4 | 2 | R.LSLHRPALEDLLLGSEAN[+1419.502]LTC[+57.021]TLTGLR.D | HexNAc(3)Hex(5) | C[+57], N[+1420] | 1096.5345 | 4 | 4383.116 | 4383.1005 | 0 | 3.5 | 127 | Specific | 441.39 | 441.39 | 441.39 | 7.88 | 33 | >sp\|P01876\|IGHA1_HUMAN Immunoglobulin heavy constant alpha 1 OS=Homo sapiens OX=9606 GN=IGHA1 PE=1 SV=2 | 255 | 20201119_15232_1ng_glyco_hcd30_etd30_compound15.19060.19060.4 | controllerType=0 controllerNumber=1 scan=19060 | 42.7312 |
| 15402:3 | 2 | R.LSLHRPALEDLLLGSEAN[+1540.529]LTC[+57.021]TLTGLR.D | HexNAc(2)Hex(7) | C[+57], N[+1541] | 1502.0517 | 3 | 4504.1405 | 4504.1268 | 0 | 3.1 | 127 | Specific | 444.69 | 417.2 | 417.2 | 7.32 | 33 | >sp\|P01876\|IGHA1_HUMAN Immunoglobulin heavy constant alpha 1 OS=Homo sapiens OX=9606 GN=IGHA1 PE=1 SV=2 | 255 | 20201119_15232_1ng_glyco_hcd30_etd30_compound15.19026.19026.3 | controllerType=0 controllerNumber=1 scan=19026 | 42.6073 |
| 15416:3 | 2 | R.LSLHRPALEDLLLGSEAN[+1378.476]LTC[+57.021]TLTGLR.D | HexNAc(2)Hex(6) | C[+57], N[+1378] | 1448.0362 | 3 | 4342.094 | 4342.074 | 0 | 4.6 | 127 | Specific | 460.09 | 453.31 | 453.31 | 6.82 | 33 | >sp\|P01876\|IGHA1_HUMAN Immunoglobulin heavy constant alpha 1 OS=Homo sapiens OX=9606 GN=IGHA1 PE=1 SV=2 | 255 | 20201119_15232_1ng_glyco_hcd30_etd30_compound15.19041.19041.3 | controllerType=0 controllerNumber=1 scan=19041 | 42.6649 |
| 15488:3 | 2 | R.LSLHRPALEDLLLGSEAN[+1095.397]LTC[+57.021]TLTGLR.D | HexNAc(3)Hex(3) | C[+57], N[+1095] | 1353.673 | 3 | 4059.0045 | 4058.9949 | 0 | 2.4 | 127 | Specific | 468.84 | 408 | 408 | 7.32 | 33 | >sp\|P01876\|IGHA1_HUMAN Immunoglobulin heavy constant alpha 1 OS=Homo sapiens OX=9606 GN=IGHA1 PE=1 SV=2 | 255 | 20201119_15232_1ng_glyco_hcd30_etd30_compound15.19117.19117.3 | controllerType=0 controllerNumber=1 scan=19117 | 42.9264 |
| 15444:3 | 2 | R.LSLHRPALEDLLLGSEAN[+1216.423]LTC[+57.021]TLTGLR.D | HexNAc(2)Hex(5) | C[+57], N[+1216] | 1394.0178 | 3 | 4180.0387 | 4180.0211 | 0 | 4.2 | 127 | Specific | 479.6 | 479.6 | 479.6 | 7.88 | 33 | >sp\|P01876\|IGHA1_HUMAN Immunoglobulin heavy constant alpha 1 OS=Homo sapiens OX=9606 GN=IGHA1 PE=1 SV=2 | 255 | 20201119_15232_1ng_glyco_hcd30_etd30_compound15.19071.19071.3 | controllerType=0 controllerNumber=1 scan=19071 | 42.7674 |
| 15377:3 | 2 | R.LSLHRPALEDLLLGSEAN[+1702.581]LTC[+57.021]TLTGLR.D | HexNAc(2)Hex(8) | C[+57], N[+1703] | 1556.0723 | 3 | 4666.2024 | 4666.1796 | 0 | 4.9 | 127 | Specific | 482.25 | 451.37 | 451.37 | 6.87 | 33 | >sp\|P01876\|IGHA1_HUMAN Immunoglobulin heavy constant alpha 1 OS=Homo sapiens OX=9606 GN=IGHA1 PE=1 SV=2 | 255 | 20201119_15232_1ng_glyco_hcd30_etd30_compound15.18999.18999.3 | controllerType=0 controllerNumber=1 scan=18999 | 42.5019 |
| 15363:3 | 2 | R.LSLHRPALEDLLLGSEAN[+1864.634]LTC[+57.021]TLTGLR.D | HexNAc(2)Hex(9) | C[+57], N[+1865] | 1610.0889 | 3 | 4828.2522 | 4828.2324 | 0 | 4.1 | 127 | Specific | 526.83 | 492.11 | 492.11 | 8.16 | 33 | >sp\|P01876\|IGHA1_HUMAN Immunoglobulin heavy constant alpha 1 OS=Homo sapiens OX=9606 GN=IGHA1 PE=1 SV=2 | 255 | 20201119_15232_1ng_glyco_hcd30_etd30_compound15.18984.18984.3 | controllerType=0 controllerNumber=1 scan=18984 | 42.4464 |
| 16157:4 | 2 | R.LSLHRPALEDLLLGSEAN[+1751.624]LTC[+57.021]TLTGLR.D | HexNAc(4)Hex(4)NeuAc(1) | C[+57], N[+1752] | 1179.5647 | 4 | 4715.2371 | 4715.2225 | 0 | 3.1 | 127 | Specific | 555.37 | 553.54 | 553.54 | 7.6 | 33 | >sp\|P01876\|IGHA1_HUMAN Immunoglobulin heavy constant alpha 1 OS=Homo sapiens OX=9606 GN=IGHA1 PE=1 SV=2 | 255 | 20201119_15232_1ng_glyco_hcd30_etd30_compound15.19824.19824.4 | controllerType=0 controllerNumber=1 scan=19824 | 44.982 |
| 16944:4 | 2 | R.LSLHRPALEDLLLGSEAN[+2204.772]LTC[+57.021]TLTGLR.D | HexNAc(4)Hex(5)NeuAc(2) | C[+57], N[+2205] | 1292.8506 | 4 | 5168.3805 | 5168.3707 | 0 | 1.9 | 127 | Specific | 557.52 | 527.83 | 527.83 | 6.99 | 33 | >sp\|P01876\|IGHA1_HUMAN Immunoglobulin heavy constant alpha 1 OS=Homo sapiens OX=9606 GN=IGHA1 PE=1 SV=2 | 255 | 20201119_15232_1ng_glyco_hcd30_etd30_compound15.20655.20655.4 | controllerType=0 controllerNumber=1 scan=20655 | 47.2945 |
| 16075:4 | 2 | R.LSLHRPALEDLLLGSEAN[+1913.677]LTC[+57.021]TLTGLR.D | HexNAc(4)Hex(5)NeuAc(1) | C[+57], N[+1914] | 1220.0778 | 4 | 4877.2894 | 4877.2753 | 0 | 2.9 | 127 | Specific | 795.64 | 770.64 | 770.64 | 12.55 | 33 | >sp\|P01876\|IGHA1_HUMAN Immunoglobulin heavy constant alpha 1 OS=Homo sapiens OX=9606 GN=IGHA1 PE=1 SV=2 | 255 | 20201119_15232_1ng_glyco_hcd30_etd30_compound15.19737.19737.4 | controllerType=0 controllerNumber=1 scan=19737 | 44.7314 |
| 16126:4 | 2 | R.LSLHRPALEDLLLGSEAN[+1710.598]LTC[+57.021]TLTGLR.D | HexNAc(3)Hex(5)NeuAc(1) | C[+57], N[+1711] | 1169.3083 | 4 | 4674.2113 | 4674.1959 | 0 | 3.3 | 127 | Specific | 812.89 | 780.44 | 780.44 | 14.12 | 33 | >sp\|P01876\|IGHA1_HUMAN Immunoglobulin heavy constant alpha 1 OS=Homo sapiens OX=9606 GN=IGHA1 PE=1 SV=2 | 255 | 20201119_15232_1ng_glyco_hcd30_etd30_compound15.19791.19791.4 | controllerType=0 controllerNumber=1 scan=19791 | 44.9026 |
| 15455:4 | 2 | R.LSLHRPALEDLLLGSEAN[+1257.449]LTC[+57.021]TLTGLR.D | HexNAc(3)Hex(4) | C[+57], N[+1257] | 1056.0213 | 4 | 4221.0633 | 4221.0477 | 0 | 3.7 | 127 | Specific | 874.72 | 874.72 | 874.72 | 15.12 | 33 | >sp\|P01876\|IGHA1_HUMAN Immunoglobulin heavy constant alpha 1 OS=Homo sapiens OX=9606 GN=IGHA1 PE=1 SV=2 | 255 | 20201119_15232_1ng_glyco_hcd30_etd30_compound15.19082.19082.4 | controllerType=0 controllerNumber=1 scan=19082 | 42.8083 |
| 15411:4 | 2 | R.LSLHRPALEDLLLGSEAN[+1663.608]LTC[+57.021]TLTGLR.D | HexNAc(5)Hex(4) | C[+57], N[+1664] | 1157.5617 | 4 | 4627.2249 | 4627.2064 | 0 | 4 | 127 | Specific | 939.55 | 926.81 | 926.81 | 16.12 | 33 | >sp\|P01876\|IGHA1_HUMAN Immunoglobulin heavy constant alpha 1 OS=Homo sapiens OX=9606 GN=IGHA1 PE=1 SV=2 | 255 | 20201119_15232_1ng_glyco_hcd30_etd30_compound15.19036.19036.4 | controllerType=0 controllerNumber=1 scan=19036 | 42.6513 |
| 15371:4 | 2 | R.LSLHRPALEDLLLGSEAN[+1702.581]LTC[+57.021]TLTGLR.D | HexNAc(2)Hex(8) | C[+57], N[+1703] | 1167.3061 | 4 | 4666.2025 | 4666.1796 | 0 | 4.9 | 127 | Specific | 967.91 | 944.23 | 944.23 | 16.06 | 33 | >sp\|P01876\|IGHA1_HUMAN Immunoglobulin heavy constant alpha 1 OS=Homo sapiens OX=9606 GN=IGHA1 PE=1 SV=2 | 255 | 20201119_15232_1ng_glyco_hcd30_etd30_compound15.18993.18993.4 | controllerType=0 controllerNumber=1 scan=18993 | 42.4796 |
| 15482:4 | 2 | R.LSLHRPALEDLLLGSEAN[+1095.397]LTC[+57.021]TLTGLR.D | HexNAc(3)Hex(3) | C[+57], N[+1095] | 1015.5069 | 4 | 4059.0059 | 4058.9949 | 0 | 2.7 | 127 | Specific | 973.56 | 973.56 | 973.56 | 16.55 | 33 | >sp\|P01876\|IGHA1_HUMAN Immunoglobulin heavy constant alpha 1 OS=Homo sapiens OX=9606 GN=IGHA1 PE=1 SV=2 | 255 | 20201119_15232_1ng_glyco_hcd30_etd30_compound15.19111.19111.4 | controllerType=0 controllerNumber=1 scan=19111 | 42.9061 |
| 15459:4 | 2 | R.LSLHRPALEDLLLGSEAN[+1298.476]LTC[+57.021]TLTGLR.D | HexNAc(4)Hex(3) | C[+57], N[+1298] | 1066.2781 | 4 | 4262.0906 | 4262.0742 | 0 | 3.9 | 127 | Specific | 975.75 | 975.75 | 975.75 | 17.12 | 33 | >sp\|P01876\|IGHA1_HUMAN Immunoglobulin heavy constant alpha 1 OS=Homo sapiens OX=9606 GN=IGHA1 PE=1 SV=2 | 255 | 20201119_15232_1ng_glyco_hcd30_etd30_compound15.19087.19087.4 | controllerType=0 controllerNumber=1 scan=19087 | 42.8247 |
| 15443:4 | 2 | R.LSLHRPALEDLLLGSEAN[+1216.423]LTC[+57.021]TLTGLR.D | HexNAc(2)Hex(5) | C[+57], N[+1216] | 1045.7653 | 4 | 4180.0394 | 4180.0211 | 0 | 4.4 | 127 | Specific | 1044.41 | 1026.02 | 1026.02 | 17.06 | 33 | >sp\|P01876\|IGHA1_HUMAN Immunoglobulin heavy constant alpha 1 OS=Homo sapiens OX=9606 GN=IGHA1 PE=1 SV=2 | 255 | 20201119_15232_1ng_glyco_hcd30_etd30_compound15.19070.19070.4 | controllerType=0 controllerNumber=1 scan=19070 | 42.7643 |
| 15412:4 | 2 | R.LSLHRPALEDLLLGSEAN[+1378.476]LTC[+57.021]TLTGLR.D | HexNAc(2)Hex(6) | C[+57], N[+1378] | 1086.2786 | 4 | 4342.0926 | 4342.074 | 0 | 4.3 | 127 | Specific | 1057.26 | 1013.83 | 1013.83 | 19.12 | 33 | >sp\|P01876\|IGHA1_HUMAN Immunoglobulin heavy constant alpha 1 OS=Homo sapiens OX=9606 GN=IGHA1 PE=1 SV=2 | 255 | 20201119_15232_1ng_glyco_hcd30_etd30_compound15.19037.19037.4 | controllerType=0 controllerNumber=1 scan=19037 | 42.6546 |
| 15398:4 | 2 | R.LSLHRPALEDLLLGSEAN[+1540.529]LTC[+57.021]TLTGLR.D | HexNAc(2)Hex(7) | C[+57], N[+1541] | 1126.7906 | 4 | 4504.1405 | 4504.1268 | 0 | 3 | 127 | Specific | 1062.93 | 981.22 | 981.22 | 18.55 | 33 | >sp\|P01876\|IGHA1_HUMAN Immunoglobulin heavy constant alpha 1 OS=Homo sapiens OX=9606 GN=IGHA1 PE=1 SV=2 | 255 | 20201119_15232_1ng_glyco_hcd30_etd30_compound15.19022.19022.4 | controllerType=0 controllerNumber=1 scan=19022 | 42.5891 |
| 15358:4 | 2 | R.LSLHRPALEDLLLGSEAN[+1864.634]LTC[+57.021]TLTGLR.D | HexNAc(2)Hex(9) | C[+57], N[+1865] | 1207.8185 | 4 | 4828.2523 | 4828.2324 | 0 | 4.1 | 127 | Specific | 1064.2 | 1019.44 | 1019.44 | 19.12 | 33 | >sp\|P01876\|IGHA1_HUMAN Immunoglobulin heavy constant alpha 1 OS=Homo sapiens OX=9606 GN=IGHA1 PE=1 SV=2 | 255 | 20201119_15232_1ng_glyco_hcd30_etd30_compound15.18979.18979.4 | controllerType=0 controllerNumber=1 scan=18979 | 42.4251 |
| 10255:3 | 2 | R.LAGKPTHVN[+1809.666]VSVVMAEVDGTC[+57.021]Y.- | HexNAc(5)Hex(4)Fuc(1) | C[+57], N[+1810] | 1386.2781 | 3 | 4156.8197 | 4156.8081 | 0 | 2.8 | 332 | CRagged | 218.16 | 136.06 | 136.06 | 2.63 | 33 | >sp\|P01876\|IGHA1_HUMAN Immunoglobulin heavy constant alpha 1 OS=Homo sapiens OX=9606 GN=IGHA1 PE=1 SV=2 | 255 | 20201119_15232_1ng_glyco_hcd30_etd30_compound15.13586.13586.3 | controllerType=0 controllerNumber=1 scan=13586 | 30.7568 |
| 10201:3 | 2 | R.LAGKPTHVN[+1378.476]VSVVMAEVDGTC[+57.021]Y.- | HexNAc(2)Hex(6) | C[+57], N[+1378] | 1242.5455 | 3 | 3725.6218 | 3725.6178 | 0 | 1.1 | 332 | CRagged | 246.7 | 194.76 | 194.3 | 2.66 | 33 | >sp\|P01876\|IGHA1_HUMAN Immunoglobulin heavy constant alpha 1 OS=Homo sapiens OX=9606 GN=IGHA1 PE=1 SV=2 | 255 | 20201119_15232_1ng_glyco_hcd30_etd30_compound15.13529.13529.3 | controllerType=0 controllerNumber=1 scan=13529 | 30.6016 |
| 10172:3 | 2 | R.LAGKPTHVN[+1540.529]VSVVMAEVDGTC[+57.021]Y.- | HexNAc(2)Hex(7) | C[+57], N[+1541] | 1296.5668 | 3 | 3887.6859 | 3887.6706 | 0 | 3.9 | 332 | CRagged | 251.89 | 188.49 | 188.49 | 3.46 | 33 | >sp\|P01876\|IGHA1_HUMAN Immunoglobulin heavy constant alpha 1 OS=Homo sapiens OX=9606 GN=IGHA1 PE=1 SV=2 | 255 | 20201119_15232_1ng_glyco_hcd30_etd30_compound15.13499.13499.3 | controllerType=0 controllerNumber=1 scan=13499 | 30.536 |
| 10294:3 | 2 | R.LAGKPTHVN[+1501.555]VSVVMAEVDGTC[+57.021]Y.- | HexNAc(5)Hex(3) | C[+57], N[+1502] | 1283.5738 | 3 | 3848.7068 | 3848.6974 | 0 | 2.4 | 332 | CRagged | 255.14 | 182.41 | 182.41 | 3.01 | 33 | >sp\|P01876\|IGHA1_HUMAN Immunoglobulin heavy constant alpha 1 OS=Homo sapiens OX=9606 GN=IGHA1 PE=1 SV=2 | 255 | 20201119_15232_1ng_glyco_hcd30_etd30_compound15.13628.13628.3 | controllerType=0 controllerNumber=1 scan=13628 | 30.8618 |
| 10272:3 | 2 | R.LAGKPTHVN[+1647.613]VSVVMAEVDGTC[+57.021]Y.- | HexNAc(5)Hex(3)Fuc(1) | C[+57], N[+1648] | 1332.2591 | 3 | 3994.7627 | 3994.7553 | 0 | 1.9 | 332 | CRagged | 256.13 | 133.65 | 133.65 | 2.73 | 33 | >sp\|P01876\|IGHA1_HUMAN Immunoglobulin heavy constant alpha 1 OS=Homo sapiens OX=9606 GN=IGHA1 PE=1 SV=2 | 255 | 20201119_15232_1ng_glyco_hcd30_etd30_compound15.13605.13605.3 | controllerType=0 controllerNumber=1 scan=13605 | 30.8066 |
| 10138:3 | 2 | R.LAGKPTHVN[+1702.581]VSVVMAEVDGTC[+57.021]Y.- | HexNAc(2)Hex(8) | C[+57], N[+1703] | 1350.5849 | 3 | 4049.7401 | 4049.7234 | 0 | 4.1 | 332 | CRagged | 268.62 | 254.89 | 200.71 | 3.67 | 33 | >sp\|P01876\|IGHA1_HUMAN Immunoglobulin heavy constant alpha 1 OS=Homo sapiens OX=9606 GN=IGHA1 PE=1 SV=2 | 255 | 20201119_15232_1ng_glyco_hcd30_etd30_compound15.13463.13463.3 | controllerType=0 controllerNumber=1 scan=13463 | 30.418 |
| 10253:3 | 2 | R.LAGKPTHVN[+1216.423]VSVVMAEVDGTC[+57.021]Y.- | HexNAc(2)Hex(5) | C[+57], N[+1216] | 1188.5302 | 3 | 3563.576 | 3563.5649 | 0 | 3.1 | 332 | CRagged | 270.44 | 270.44 | 193.71 | 3.18 | 33 | >sp\|P01876\|IGHA1_HUMAN Immunoglobulin heavy constant alpha 1 OS=Homo sapiens OX=9606 GN=IGHA1 PE=1 SV=2 | 255 | 20201119_15232_1ng_glyco_hcd30_etd30_compound15.13584.13584.3 | controllerType=0 controllerNumber=1 scan=13584 | 30.7488 |
| 10264:4 | 2 | R.LAGKPTHVN[+1809.666]VSVVMAEVDGTC[+57.021]Y.- | HexNAc(5)Hex(4)Fuc(1) | C[+57], N[+1810] | 1039.9604 | 4 | 4156.8198 | 4156.8081 | 0 | 2.8 | 332 | CRagged | 299.16 | 299.16 | 89.66 | 3.38 | 33 | >sp\|P01876\|IGHA1_HUMAN Immunoglobulin heavy constant alpha 1 OS=Homo sapiens OX=9606 GN=IGHA1 PE=1 SV=2 | 255 | 20201119_15232_1ng_glyco_hcd30_etd30_compound15.13595.13595.4 | controllerType=0 controllerNumber=1 scan=13595 | 30.7816 |
| 13409:4 | 2 | R.LAGKPTHVN[+2553.910]VSVVMAEVDGTC[+57.021]Y.- | HexNAc(5)Hex(5)Fuc(1)NeuAc(2) | C[+57], N[+2554] | 1226.0231 | 4 | 4901.0706 | 4901.0518 | 0 | 3.8 | 332 | CRagged | 350.31 | 175.32 | 175.32 | 4.16 | 33 | >sp\|P01876\|IGHA1_HUMAN Immunoglobulin heavy constant alpha 1 OS=Homo sapiens OX=9606 GN=IGHA1 PE=1 SV=2 | 255 | 20201119_15232_1ng_glyco_hcd30_etd30_compound15.16891.16891.4 | controllerType=0 controllerNumber=1 scan=16891 | 37.3794 |
| 16247:3 | 4 | K.QIGLYPVLVIDSSGYVNPN[+2059.735]YTGR.I | HexNAc(4)Hex(5)Fuc(1)NeuAc(1) | N[+2060] | 1529.0242 | 3 | 4585.0581 | 4585.0384 | 0 | 4.3 | 168 | Specific | 523.26 | 440.57 | 440.57 | 7.7 | 19 | >sp\|P01833\|PIGR_HUMAN Polymeric immunoglobulin receptor OS=Homo sapiens OX=9606 GN=PIGR PE=1 SV=4 | 902 | 20201119_15232_1ng_glyco_hcd30_etd30_compound15.19919.19919.3 | controllerType=0 controllerNumber=1 scan=19919 | 45.3163 |
| 19271:3 | 4 | R.LSLLEEPGN[+2350.830]GTFTVILNQLTSR.D | HexNAc(4)Hex(5)Fuc(1)NeuAc(2) | N[+2351] | 1585.0563 | 3 | 4753.1544 | 4753.1229 | 0 | 6.6 | 413 | Specific | 409.11 | 303.89 | 303.89 | 5.05 | 19 | >sp\|P01833\|PIGR_HUMAN Polymeric immunoglobulin receptor OS=Homo sapiens OX=9606 GN=PIGR PE=1 SV=4 | 902 | 20201119_15232_1ng_glyco_hcd30_etd30_compound15.23109.23109.3 | controllerType=0 controllerNumber=1 scan=23109 | 53.5651 |
| 17413:3 | 4 | R.LSLLEEPGN[+2205.793]GTFTVILNQLTSR.D | HexNAc(4)Hex(5)Fuc(2)NeuAc(1) | N[+2206] | 1536.7101 | 3 | 4608.1159 | 4608.0854 | 0 | 6.6 | 413 | Specific | 527.99 | 464.22 | 464.22 | 6.75 | 19 | >sp\|P01833\|PIGR_HUMAN Polymeric immunoglobulin receptor OS=Homo sapiens OX=9606 GN=PIGR PE=1 SV=4 | 902 | 20201119_15232_1ng_glyco_hcd30_etd30_compound15.21150.21150.4 | controllerType=0 controllerNumber=1 scan=21150 | 48.7178 |
| 17475:4 | 4 | R.LSLLEEPGN[+2059.735]GTFTVILNQLTSR.D | HexNAc(4)Hex(5)Fuc(1)NeuAc(1) | N[+2060] | 1116.2665 | 4 | 4462.0443 | 4462.0275 | 0 | 3.8 | 413 | Specific | 546.13 | 521.97 | 521.97 | 7.09 | 19 | >sp\|P01833\|PIGR_HUMAN Polymeric immunoglobulin receptor OS=Homo sapiens OX=9606 GN=PIGR PE=1 SV=4 | 902 | 20201119_15232_1ng_glyco_hcd30_etd30_compound15.21216.21216.4 | controllerType=0 controllerNumber=1 scan=21216 | 48.9472 |
| 19196:4 | 4 | R.LSLLEEPGN[+2350.830]GTFTVILNQLTSR.D | HexNAc(4)Hex(5)Fuc(1)NeuAc(2) | N[+2351] | 1189.0413 | 4 | 4753.1434 | 4753.1229 | 0 | 4.3 | 413 | Specific | 575.43 | 482.73 | 482.73 | 8.04 | 19 | >sp\|P01833\|PIGR_HUMAN Polymeric immunoglobulin receptor OS=Homo sapiens OX=9606 GN=PIGR PE=1 SV=4 | 902 | 20201119_15232_1ng_glyco_hcd30_etd30_compound15.23029.23029.4 | controllerType=0 controllerNumber=1 scan=23029 | 53.2829 |
| 17423:3 | 4 | R.LSLLEEPGN[+2205.793]GTFTVILNQLTSR.D | HexNAc(4)Hex(5)Fuc(2)NeuAc(1) | N[+2206] | 1536.7066 | 3 | 4608.1052 | 4608.0854 | 0 | 4.3 | 413 | Specific | 648.78 | 526.52 | 526.52 | 9.65 | 19 | >sp\|P01833\|PIGR_HUMAN Polymeric immunoglobulin receptor OS=Homo sapiens OX=9606 GN=PIGR PE=1 SV=4 | 902 | 20201119_15232_1ng_glyco_hcd30_etd30_compound15.21161.21161.3 | controllerType=0 controllerNumber=1 scan=21161 | 48.7601 |
| 14245:3 | 4 | K.IIEGEPNLKVPGN[+2205.793]VTAVLGETLK.V | HexNAc(4)Hex(5)Fuc(2)NeuAc(1) | N[+2206] | 1533.0565 | 3 | 4597.1548 | 4597.1422 | 0 | 2.7 | 457 | Specific | 261.42 | 205.33 | 205.33 | 3.2 | 19 | >sp\|P01833\|PIGR_HUMAN Polymeric immunoglobulin receptor OS=Homo sapiens OX=9606 GN=PIGR PE=1 SV=4 | 902 | 20201119_15232_1ng_glyco_hcd30_etd30_compound15.17800.17800.3 | controllerType=0 controllerNumber=1 scan=17800 | 39.3718 |
| 14242:3 | 4 | K.IIEGEPNLKVPGN[+2059.735]VTAVLGETLK.V | HexNAc(4)Hex(5)Fuc(1)NeuAc(1) | N[+2060] | 1484.3707 | 3 | 4451.0974 | 4451.0843 | 0 | 3 | 457 | Specific | 323.52 | 260.04 | 260.04 | 3.64 | 19 | >sp\|P01833\|PIGR_HUMAN Polymeric immunoglobulin receptor OS=Homo sapiens OX=9606 GN=PIGR PE=1 SV=4 | 902 | 20201119_15232_1ng_glyco_hcd30_etd30_compound15.17797.17797.3 | controllerType=0 controllerNumber=1 scan=17797 | 39.3623 |
| 14164:3 | 4 | K.IIEGEPNLKVPGN[+2205.793]VTAVLGETLK.V | HexNAc(4)Hex(5)Fuc(2)NeuAc(1) | N[+2206] | 1533.0552 | 3 | 4597.1511 | 4597.1422 | 0 | 1.9 | 457 | Specific | 407.37 | 349.17 | 349.17 | 4.09 | 19 | >sp\|P01833\|PIGR_HUMAN Polymeric immunoglobulin receptor OS=Homo sapiens OX=9606 GN=PIGR PE=1 SV=4 | 902 | 20201119_15232_1ng_glyco_hcd30_etd30_compound15.17715.17715.3 | controllerType=0 controllerNumber=1 scan=17715 | 39.1723 |
| 14967:5 | 4 | K.IIEGEPNLKVPGN[+2350.830]VTAVLGETLKVPC[+57.021]HFPC[+57.021]K.F | HexNAc(4)Hex(5)Fuc(1)NeuAc(2) | C[+57]*2, N[+2351] | 1154.3373 | 5 | 5767.6573 | 5767.6372 | 0 | 3.5 | 457 | Specific | 366.55 | 366.55 | 366.55 | 5.3 | 19 | >sp\|P01833\|PIGR_HUMAN Polymeric immunoglobulin receptor OS=Homo sapiens OX=9606 GN=PIGR PE=1 SV=4 | 902 | 20201119_15232_1ng_glyco_hcd30_etd30_compound15.18566.18566.5 | controllerType=0 controllerNumber=1 scan=18566 | 41.2043 |
| 14955:5 | 4 | K.IIEGEPNLKVPGN[+2350.830]VTAVLGETLKVPC[+57.021]HFPC[+57.021]K.F | HexNAc(4)Hex(5)Fuc(1)NeuAc(2) | C[+57]*2, N[+2351] | 1154.3363 | 5 | 5767.6524 | 5767.6372 | 0 | 2.6 | 457 | Specific | 387.08 | 387.08 | 387.08 | 5.06 | 19 | >sp\|P01833\|PIGR_HUMAN Polymeric immunoglobulin receptor OS=Homo sapiens OX=9606 GN=PIGR PE=1 SV=4 | 902 | 20201119_15232_1ng_glyco_hcd30_etd30_compound15.18553.18553.5 | controllerType=0 controllerNumber=1 scan=18553 | 41.1617 |
| 14973:4 | 4 | K.IIEGEPNLKVPGN[+2350.830]VTAVLGETLKVPC[+57.021]HFPC[+57.021]K.F | HexNAc(4)Hex(5)Fuc(1)NeuAc(2) | C[+57]*2, N[+2351] | 1442.6718 | 4 | 5767.6652 | 5767.6372 | 0 | 4.8 | 457 | Specific | 403.97 | 403.97 | 403.97 | 5.03 | 19 | >sp\|P01833\|PIGR_HUMAN Polymeric immunoglobulin receptor OS=Homo sapiens OX=9606 GN=PIGR PE=1 SV=4 | 902 | 20201119_15232_1ng_glyco_hcd30_etd30_compound15.18572.18572.4 | controllerType=0 controllerNumber=1 scan=18572 | 41.2215 |
| 13758:5 | 4 | K.IIEGEPNLKVPGN[+2059.735]VTAVLGETLKVPC[+57.021]HFPC[+57.021]K.F | HexNAc(4)Hex(5)Fuc(1)NeuAc(1) | C[+57]*2, N[+2060] | 1096.1178 | 5 | 5476.5599 | 5476.5418 | 0 | 3.3 | 457 | Specific | 523.35 | 514.05 | 514.05 | 6.66 | 19 | >sp\|P01833\|PIGR_HUMAN Polymeric immunoglobulin receptor OS=Homo sapiens OX=9606 GN=PIGR PE=1 SV=4 | 902 | 20201119_15232_1ng_glyco_hcd30_etd30_compound15.17255.17255.5 | controllerType=0 controllerNumber=1 scan=17255 | 38.1678 |
| 05153:3 | 4 | K.WN[+1913.677]NTGC[+57.021]QALPSQDEGPSK.A | HexNAc(4)Hex(5)NeuAc(1) | C[+57], N[+1914] | 1301.5287 | 3 | 3902.5717 | 3902.5537 | 0 | 4.6 | 498 | Specific | 370.45 | 211.04 | 211.04 | 4.78 | 19 | >sp\|P01833\|PIGR_HUMAN Polymeric immunoglobulin receptor OS=Homo sapiens OX=9606 GN=PIGR PE=1 SV=4 | 902 | 20201119_15232_1ng_glyco_hcd30_etd30_compound15.8261.8261.3 | controllerType=0 controllerNumber=1 scan=8261 | 20.9067 |
| 05044:3 | 4 | K.WN[+2059.735]NTGC[+57.021]QALPSQDEGPSK.A | HexNAc(4)Hex(5)Fuc(1)NeuAc(1) | C[+57], N[+2060] | 1350.2125 | 3 | 4048.6228 | 4048.6116 | 0 | 2.8 | 498 | Specific | 400.36 | 282.35 | 282.35 | 5.73 | 19 | >sp\|P01833\|PIGR_HUMAN Polymeric immunoglobulin receptor OS=Homo sapiens OX=9606 GN=PIGR PE=1 SV=4 | 902 | 20201119_15232_1ng_glyco_hcd30_etd30_compound15.8148.8148.3 | controllerType=0 controllerNumber=1 scan=8148 | 20.698 |
| 15068:3 | 6 | K.RNIAAFGGDPNN[+1913.677]ITLFGESAGGASVSLQTLSPYNK.G | HexNAc(4)Hex(5)NeuAc(1) | N[+1914] | 1827.5014 | 3 | 5480.4897 | 5480.4644 | 0 | 4.6 | 196 | Specific | 395.72 | 302.32 | 302.32 | 5.03 | 12 | >sp\|P19835\|CEL_HUMAN Bile salt-activated lipase OS=Homo sapiens OX=9606 GN=CEL PE=1 SV=3 | 476 | 20201119_15232_1ng_glyco_hcd30_etd30_compound15.18673.18673.3 | controllerType=0 controllerNumber=1 scan=18673 | 41.5355 |
| 16275:3 | 6 | R.NIAAFGGDPNN[+1913.677]ITLFGESAGGASVSLQTLSPYNK.G | HexNAc(4)Hex(5)NeuAc(1) | N[+1914] | 1775.4687 | 3 | 5324.3916 | 5324.3633 | 0 | 5.3 | 197 | Specific | 813.47 | 657.67 | 657.67 | 13.22 | 12 | >sp\|P19835\|CEL_HUMAN Bile salt-activated lipase OS=Homo sapiens OX=9606 GN=CEL PE=1 SV=3 | 476 | 20201119_15232_1ng_glyco_hcd30_etd30_compound15.19949.19949.3 | controllerType=0 controllerNumber=1 scan=19949 | 45.4118 |
| 09038:3 | 7 | K.TVLTPATNHMGN[+1378.476]VTFTIPANR.E | HexNAc(2)Hex(6) | N[+1378] | 1211.8887 | 3 | 3633.6516 | 3633.6358 | 0 | 4.4 | 74 | Specific | 276.26 | 253.15 | 138.35 | 4.84 | 8 | >sp\|P01024\|CO3_HUMAN Complement C3 OS=Homo sapiens OX=9606 GN=C3 PE=1 SV=2 | 492 | 20201119_15232_1ng_glyco_hcd30_etd30_compound15.12322.12322.3 | controllerType=0 controllerNumber=1 scan=12322 | 28.3081 |
| 05020:3 | 8 | R.LSPN[+2205.793]ASAEHLELR.W | HexNAc(4)Hex(5)Fuc(2)NeuAc(1) | N[+2206] | 1214.8565 | 3 | 3642.555 | 3642.5421 | 0 | 3.5 | 52 | Specific | 363.71 | 256.37 | 256.37 | 4.04 | 7 | >sp\|Q13410\|BT1A1_HUMAN Butyrophilin subfamily 1 member A1 OS=Homo sapiens OX=9606 GN=BTN1A1 PE=1 SV=3 | 179 | 20201119_15232_1ng_glyco_hcd30_etd30_compound15.8123.8123.3 | controllerType=0 controllerNumber=1 scan=8123 | 20.6456 |
| 06927:3 | 8 | R.LSPN[+2350.830]ASAEHLELR.W | HexNAc(4)Hex(5)Fuc(1)NeuAc(2) | N[+2351] | 1263.2015 | 3 | 3787.5899 | 3787.5796 | 0 | 2.7 | 52 | Specific | 386.8 | 252.84 | 252.84 | 4.08 | 7 | >sp\|Q13410\|BT1A1_HUMAN Butyrophilin subfamily 1 member A1 OS=Homo sapiens OX=9606 GN=BTN1A1 PE=1 SV=3 | 179 | 20201119_15232_1ng_glyco_hcd30_etd30_compound15.10100.10100.3 | controllerType=0 controllerNumber=1 scan=10100 | 24.1059 |
| 18635:4 | 9 | K.VFPLSLDSTPQDGNVVVAC[+57.021]LVQGFFPQEPLSVTWSESGQN[+1647.613]VTAI | HexNAc(5)Hex(3)Fuc(1) | C[+57], N[+1648] | 1606.7544 | 4 | 6423.9958 | 6423.9746 | 0 | 3.3 | 8 | Specific | 654.96 | 654.96 | 654.96 | 10.91 | 17 | >sp\|P01877\|IGHA2_HUMAN Immunoglobulin heavy constant alpha 2 OS=Homo sapiens OX=9606 GN=IGHA2 PE=1 SV=4 | 576 | 20201119_15232_1ng_glyco_hcd30_etd30_compound15.22434.22434.2 | controllerType=0 controllerNumber=1 scan=22434 | 51.9254 |
| 00858:3 | 9 | K.HYTN[+1809.666]SSQDVTVPC[+57.021]R.V | HexNAc(5)Hex(4)Fuc(1) | C[+57], N[+1810] | 1158.4802 | 3 | 3473.4259 | 3473.4154 | 0 | 3 | 89 | Specific | 354.63 | 290.72 | 290.72 | 5.21 | 17 | >sp\|P01877\|IGHA2_HUMAN Immunoglobulin heavy constant alpha 2 OS=Homo sapiens OX=9606 GN=IGHA2 PE=1 SV=4 | 576 | 20201119_15232_1ng_glyco_hcd30_etd30_compound15.3800.3800.3 | controllerType=0 controllerNumber=1 scan=3800 | 12.8806 |
| 00893:3 | 9 | K.HYTN[+1647.613]SSQDVTVPC[+57.021]R.V | HexNAc(5)Hex(3)Fuc(1) | C[+57], N[+1648] | 1104.4621 | 3 | 3311.3716 | 3311.3625 | 0 | 2.7 | 89 | Specific | 482.97 | 404.26 | 404.26 | 6.73 | 17 | >sp\|P01877\|IGHA2_HUMAN Immunoglobulin heavy constant alpha 2 OS=Homo sapiens OX=9606 GN=IGHA2 PE=1 SV=4 | 576 | 20201119_15232_1ng_glyco_hcd30_etd30_compound15.3837.3837.3 | controllerType=0 controllerNumber=1 scan=3837 | 12.9572 |
| 02744:4 | 9 | K.HYTN[+1647.613]SSQDVTVPC[+57.021]RVPPPPPC[+57.021]C[+57.021]HPR.L | HexNAc(5)Hex(3)Fuc(1) | C[+57]*3, N[+1648] | 1152.2502 | 4 | 4605.9788 | 4605.9689 | 0 | 2.2 | 89 | Specific | 272.96 | 271.53 | 271.53 | 3.02 | 17 | >sp\|P01877\|IGHA2_HUMAN Immunoglobulin heavy constant alpha 2 OS=Homo sapiens OX=9606 GN=IGHA2 PE=1 SV=4 | 576 | 20201119_15232_1ng_glyco_hcd30_etd30_compound15.5763.5763.4 | controllerType=0 controllerNumber=1 scan=5763 | 16.5383 |
| 02745:5 | 9 | K.HYTN[+1647.613]SSQDVTVPC[+57.021]RVPPPPPC[+57.021]C[+57.021]HPR.L | HexNAc(5)Hex(3)Fuc(1) | C[+57]*3, N[+1648] | 922.0018 | 5 | 4605.9799 | 4605.9689 | 0 | 2.4 | 89 | Specific | 524.17 | 497.51 | 497.51 | 5.27 | 17 | >sp\|P01877\|IGHA2_HUMAN Immunoglobulin heavy constant alpha 2 OS=Homo sapiens OX=9606 GN=IGHA2 PE=1 SV=4 | 576 | 20201119_15232_1ng_glyco_hcd30_etd30_compound15.5764.5764.5 | controllerType=0 controllerNumber=1 scan=5764 | 16.5471 |
| 02444:3 | 9 | K.TPLTAN[+1971.719]ITK.S | HexNAc(5)Hex(5)Fuc(1) | N[+1972] | 977.4329 | 3 | 2930.2841 | 2930.2757 | 0 | 2.9 | 200 | Specific | 228.56 | 126.19 | 126.19 | 2.71 | 17 | >sp\|P01877\|IGHA2_HUMAN Immunoglobulin heavy constant alpha 2 OS=Homo sapiens OX=9606 GN=IGHA2 PE=1 SV=4 | 576 | 20201119_15232_1ng_glyco_hcd30_etd30_compound15.5450.5450.3 | controllerType=0 controllerNumber=1 scan=5450 | 15.8554 |
| 02488:3 | 9 | K.TPLTAN[+1809.666]ITK.S | HexNAc(5)Hex(4)Fuc(1) | N[+1810] | 923.4156 | 3 | 2768.2321 | 2768.2228 | 0 | 3.4 | 200 | Specific | 250.41 | 126.28 | 126.28 | 2.96 | 17 | >sp\|P01877\|IGHA2_HUMAN Immunoglobulin heavy constant alpha 2 OS=Homo sapiens OX=9606 GN=IGHA2 PE=1 SV=4 | 576 | 20201119_15232_1ng_glyco_hcd30_etd30_compound15.5496.5496.3 | controllerType=0 controllerNumber=1 scan=5496 | 15.9591 |
| 02544:3 | 9 | K.TPLTAN[+1647.613]ITK.S | HexNAc(5)Hex(3)Fuc(1) | N[+1648] | 869.3975 | 3 | 2606.1779 | 2606.17 | 0 | 3 | 200 | Specific | 260.53 | 124.88 | 124.88 | 2.82 | 17 | >sp\|P01877\|IGHA2_HUMAN Immunoglobulin heavy constant alpha 2 OS=Homo sapiens OX=9606 GN=IGHA2 PE=1 SV=4 | 576 | 20201119_15232_1ng_glyco_hcd30_etd30_compound15.5554.5554.3 | controllerType=0 controllerNumber=1 scan=5554 | 16.1036 |
| 02487:2 | 9 | K.TPLTAN[+1809.666]ITK.S | HexNAc(5)Hex(4)Fuc(1) | N[+1810] | 1384.6194 | 2 | 2768.2316 | 2768.2228 | 0 | 3.2 | 200 | Specific | 295.69 | 109.93 | 109.93 | 2.76 | 17 | >sp\|P01877\|IGHA2_HUMAN Immunoglobulin heavy constant alpha 2 OS=Homo sapiens OX=9606 GN=IGHA2 PE=1 SV=4 | 576 | 20201119_15232_1ng_glyco_hcd30_etd30_compound15.5495.5495.2 | controllerType=0 controllerNumber=1 scan=5495 | 15.9529 |
| 02528:2 | 9 | K.TPLTAN[+1647.613]ITK.S | HexNAc(5)Hex(3)Fuc(1) | N[+1648] | 1303.5923 | 2 | 2606.1774 | 2606.17 | 0 | 2.8 | 200 | Specific | 315.33 | 122.37 | 122.37 | 2.65 | 17 | >sp\|P01877\|IGHA2_HUMAN Immunoglobulin heavy constant alpha 2 OS=Homo sapiens OX=9606 GN=IGHA2 PE=1 SV=4 | 576 | 20201119_15232_1ng_glyco_hcd30_etd30_compound15.5538.5538.2 | controllerType=0 controllerNumber=1 scan=5538 | 16.0679 |
| 02491:2 | 9 | K.TPLTAN[+1444.534]ITK.S | HexNAc(4)Hex(3)Fuc(1) | N[+1445] | 1202.0527 | 2 | 2403.098 | 2403.0906 | 0 | 3.1 | 200 | Specific | 339.55 | 111.77 | 111.77 | 2.6 | 17 | >sp\|P01877\|IGHA2_HUMAN Immunoglobulin heavy constant alpha 2 OS=Homo sapiens OX=9606 GN=IGHA2 PE=1 SV=4 | 576 | 20201119_15232_1ng_glyco_hcd30_etd30_compound15.5499.5499.2 | controllerType=0 controllerNumber=1 scan=5499 | 15.9682 |
| 09299:3 | 9 | R.MAGKPTHIN[+1216.423]VSVVMAEADGTC[+57.021]Y.- | HexNAc(2)Hex(5) | C[+57], N[+1216] | 1189.8433 | 3 | 3567.5154 | 3567.5057 | 0 | 2.7 | 319 | CRagged | 230.57 | 230.57 | 165.27 | 2.94 | 17 | >sp\|P01877\|IGHA2_HUMAN Immunoglobulin heavy constant alpha 2 OS=Homo sapiens OX=9606 GN=IGHA2 PE=1 SV=4 | 576 | 20201119_15232_1ng_glyco_hcd30_etd30_compound15.12594.12594.3 | controllerType=0 controllerNumber=1 scan=12594 | 28.8723 |
| 09214:3 | 9 | R.MAGKPTHIN[+1702.581]VSVVMAEADGTC[+57.021]Y.- | HexNAc(2)Hex(8) | C[+57], N[+1703] | 1351.8967 | 3 | 4053.6754 | 4053.6642 | 0 | 2.8 | 319 | CRagged | 234.34 | 177.03 | 177.03 | 2.79 | 17 | >sp\|P01877\|IGHA2_HUMAN Immunoglobulin heavy constant alpha 2 OS=Homo sapiens OX=9606 GN=IGHA2 PE=1 SV=4 | 576 | 20201119_15232_1ng_glyco_hcd30_etd30_compound15.12505.12505.3 | controllerType=0 controllerNumber=1 scan=12505 | 28.6637 |
| 09254:3 | 9 | R.MAGKPTHIN[+1378.476]VSVVMAEADGTC[+57.021]Y.- | HexNAc(2)Hex(6) | C[+57], N[+1378] | 1243.8614 | 3 | 3729.5696 | 3729.5585 | 0 | 3 | 319 | CRagged | 276.31 | 263.46 | 220.39 | 3.34 | 17 | >sp\|P01877\|IGHA2_HUMAN Immunoglobulin heavy constant alpha 2 OS=Homo sapiens OX=9606 GN=IGHA2 PE=1 SV=4 | 576 | 20201119_15232_1ng_glyco_hcd30_etd30_compound15.12547.12547.3 | controllerType=0 controllerNumber=1 scan=12547 | 28.767 |
| 00065:3 | 13 | K.YKN[+1647.613]NSDISSTR.G | HexNAc(5)Hex(3)Fuc(1) | N[+1648] | 978.085 | 3 | 2932.2404 | 2932.2311 | 0 | 3.2 | 44 | Specific | 310.78 | 195.9 | 195.9 | 3.12 | 5 | >sp\|P01871\|IGHM_HUMAN Immunoglobulin heavy constant mu OS=Homo sapiens OX=9606 GN=IGHM PE=1 SV=4 | 328 | 20201119_15232_1ng_glyco_hcd30_etd30_compound15.2974.2974.3 | controllerType=0 controllerNumber=1 scan=2974 | 11.2951 |
| 07626:3 | 16 | K.TAHC[+57.021]N[+2059.735]ESFYFLC[+57.021]K.R | HexNAc(4)Hex(5)Fuc(1)NeuAc(1) | C[+57]*2, N[+2060] | 1246.1576 | 3 | 3736.4584 | 3736.4545 | 0 | 1 | 1201 | Specific | 227.93 | 154.39 | 154.39 | 2.83 | 7 | >sp\|P22897\|MRC1_HUMAN Macrophage mannose receptor 1 OS=Homo sapiens OX=9606 GN=MRC1 PE=1 SV=1 | 1049 | 20201119_15232_1ng_glyco_hcd30_etd30_compound15.10832.10832.3 | controllerType=0 controllerNumber=1 scan=10832 | 25.4655 |
| 20173:3 | 23 | R.QLAHQSN[+2204.772]STNIFFSPVSIATAFAMLSLGTK.A | HexNAc(4)Hex(5)NeuAc(2) | N[+2205] | 1796.148 | 3 | 5386.4296 | 5386.4075 | 0 | 4.1 | 64 | Specific | 383.97 | 273.54 | 273.54 | 6.02 | 3 | >sp\|P01009\|A1AT_HUMAN Alpha-1-antitrypsin OS=Homo sapiens OX=9606 GN=SERPINA1 PE=1 SV=3 | 642 | 20201119_15232_1ng_glyco_hcd30_etd30_compound15.24057.24057.3 | controllerType=0 controllerNumber=1 scan=24057 | 55.9078 |
| 20153:4 | 23 | R.QLAHQSN[+2204.772]STNIFFSPVSIATAFAMLSLGTK.A | HexNAc(4)Hex(5)NeuAc(2) | N[+2205] | 1347.3585 | 4 | 5386.4123 | 5386.4075 | 0 | 0.9 | 64 | Specific | 581.94 | 502.14 | 368.27 | 7.94 | 3 | >sp\|P01009\|A1AT_HUMAN Alpha-1-antitrypsin OS=Homo sapiens OX=9606 GN=SERPINA1 PE=1 SV=3 | 642 | 20201119_15232_1ng_glyco_hcd30_etd30_compound15.24036.24036.4 | controllerType=0 controllerNumber=1 scan=24036 | 55.8511 |
| 20172:4 | 23 | R.QLAHQSN[+2204.772]STNIFFSPVSIATAFAMLSLGTK.A | HexNAc(4)Hex(5)NeuAc(2) | N[+2205] | 1347.3608 | 4 | 5386.4215 | 5386.4075 | 0 | 2.6 | 64 | Specific | 614.96 | 528.79 | 417.92 | 8.95 | 3 | >sp\|P01009\|A1AT_HUMAN Alpha-1-antitrypsin OS=Homo sapiens OX=9606 GN=SERPINA1 PE=1 SV=3 | 642 | 20201119_15232_1ng_glyco_hcd30_etd30_compound15.24056.24056.4 | controllerType=0 controllerNumber=1 scan=24056 | 55.9031 |
| 19257:5 | 28 | K.APDKNVIFSPLSISTALAFLSLGAHN[+2350.830]TTLTEILK.G | HexNAc(4)Hex(5)Fuc(1)NeuAc(2) | N[+2351] | 1187.5728 | 5 | 5933.8347 | 5933.8085 | 0 | 4.4 | 68 | Specific | 293.41 | 293.41 | 293.41 | 5.33 | 4 | >sp\|P01011\|AACT_HUMAN Alpha-1-antichymotrypsin OS=Homo sapiens OX=9606 GN=SERPINA3 PE=1 SV=2 | 1314 | 20201119_15232_1ng_glyco_hcd30_etd30_compound15.23094.23094.5 | controllerType=0 controllerNumber=1 scan=23094 | 53.5201 |
| 08703:3 | 30 | R.IIVPLNNREN[+1501.555]ISDPTSPLR.T | HexNAc(5)Hex(3) | N[+1502] | 1217.2576 | 3 | 3649.7583 | 3649.7325 | 0 | 7.1 | 62 | Specific | 217.92 | 217.92 | 217.92 | 2.69 | 5 | >sp\|P01591\|IGJ_HUMAN Immunoglobulin J chain OS=Homo sapiens OX=9606 GN=JCHAIN PE=1 SV=4 | 1392 | 20201119_15232_1ng_glyco_hcd30_etd30_compound15.11975.11975.2 | controllerType=0 controllerNumber=1 scan=11975 | 27.668 |
| 08706:3 | 30 | R.IIVPLNNREN[+1647.613]ISDPTSPLR.T | HexNAc(5)Hex(3)Fuc(1) | N[+1648] | 1265.9401 | 3 | 3795.8058 | 3795.7904 | 0 | 4.1 | 62 | Specific | 228.37 | 157.59 | 157.59 | 3.43 | 5 | >sp\|P01591\|IGJ_HUMAN Immunoglobulin J chain OS=Homo sapiens OX=9606 GN=JCHAIN PE=1 SV=4 | 1392 | 20201119_15232_1ng_glyco_hcd30_etd30_compound15.11978.11978.3 | controllerType=0 controllerNumber=1 scan=11978 | 27.6787 |
| 08667:3 | 30 | R.IIVPLNNREN[+1663.608]ISDPTSPLR.T | HexNAc(5)Hex(4) | N[+1664] | 1271.2681 | 3 | 3811.7898 | 3811.7853 | 0 | 1.2 | 62 | Specific | 267.91 | 174.76 | 174.76 | 3.07 | 5 | >sp\|P01591\|IGJ_HUMAN Immunoglobulin J chain OS=Homo sapiens OX=9606 GN=JCHAIN PE=1 SV=4 | 1392 | 20201119_15232_1ng_glyco_hcd30_etd30_compound15.11938.11938.3 | controllerType=0 controllerNumber=1 scan=11938 | 27.6042 |
| 10118:3 | 30 | R.IIVPLNNREN[+2059.735]ISDPTSPLR.T | HexNAc(4)Hex(5)Fuc(1)NeuAc(1) | N[+2060] | 1403.3135 | 3 | 4207.926 | 4207.9121 | 0 | 3.3 | 62 | Specific | 293.45 | 229.81 | 229.81 | 3.98 | 5 | >sp\|P01591\|IGJ_HUMAN Immunoglobulin J chain OS=Homo sapiens OX=9606 GN=JCHAIN PE=1 SV=4 | 1392 | 20201119_15232_1ng_glyco_hcd30_etd30_compound15.13442.13442.3 | controllerType=0 controllerNumber=1 scan=13442 | 30.3679 |
| 12481:4 | 32 | K.LKELPGVC[+57.021]N[+2205.793]ETMMALWEEC[+57.021]KPC[+57.021]LK.Q | HexNAc(4)Hex(5)Fuc(2)NeuAc(1) | C[+57]*3, N[+2206] | 1286.3036 | 4 | 5142.1925 | 5142.1776 | 0 | 2.9 | 95 | Specific | 502 | 411.72 | 411.72 | 5.42 | 6 | >sp\|P10909\|CLUS_HUMAN Clusterin OS=Homo sapiens OX=9606 GN=CLU PE=1 SV=1 | 1467 | 20201119_15232_1ng_glyco_hcd30_etd30_compound15.15923.15923.4 | controllerType=0 controllerNumber=1 scan=15923 | 35.2898 |
| 12549:4 | 32 | K.LKELPGVC[+57.021]N[+2059.735]ETMMALWEEC[+57.021]KPC[+57.021]LK.Q | HexNAc(4)Hex(5)Fuc(1)NeuAc(1) | C[+57]*3, N[+2060] | 1249.7903 | 4 | 4996.1393 | 4996.1196 | 0 | 3.9 | 95 | Specific | 525.24 | 375.04 | 375.04 | 6.22 | 6 | >sp\|P10909\|CLUS_HUMAN Clusterin OS=Homo sapiens OX=9606 GN=CLU PE=1 SV=1 | 1467 | 20201119_15232_1ng_glyco_hcd30_etd30_compound15.15994.15994.4 | controllerType=0 controllerNumber=1 scan=15994 | 35.4303 |
| 00062:4 | 32 | R.EIRHN[+2424.867]STGC[+57.021]LR.M | HexNAc(5)Hex(6)Fuc(1)NeuAc(1) | C[+57], N[+2425] | 942.6414 | 4 | 3767.5438 | 3767.5316 | 0 | 3.2 | 287 | Specific | 370.73 | 263.52 | 263.52 | 4.08 | 6 | >sp\|P10909\|CLUS_HUMAN Clusterin OS=Homo sapiens OX=9606 GN=CLU PE=1 SV=1 | 1467 | 20201119_15232_1ng_glyco_hcd30_etd30_compound15.2971.2971.4 | controllerType=0 controllerNumber=1 scan=2971 | 11.2839 |
| 00058:4 | 32 | R.EIRHN[+2059.735]STGC[+57.021]LR.M | HexNAc(4)Hex(5)Fuc(1)NeuAc(1) | C[+57], N[+2060] | 851.3591 | 4 | 3402.4145 | 3402.3994 | 0 | 4.5 | 287 | Specific | 589.29 | 440.9 | 440.9 | 6.72 | 6 | >sp\|P10909\|CLUS_HUMAN Clusterin OS=Homo sapiens OX=9606 GN=CLU PE=1 SV=1 | 1467 | 20201119_15232_1ng_glyco_hcd30_etd30_compound15.2967.2967.4 | controllerType=0 controllerNumber=1 scan=2967 | 11.2656 |
| 00709:4 | 71 | K.TKPREEQFN[+1606.587]STFR.V | HexNAc(4)Hex(4)Fuc(1) | N[+1607] | 812.3587 | 4 | 3246.4128 | 3246.4054 | 0 | 2.3 | 168 | Specific | 428.1 | 377.77 | 377.77 | 4.7 | 2 | >sp\|P01859\|IGHG2_HUMAN Immunoglobulin heavy constant gamma 2 OS=Homo sapiens OX=9606 GN=IGHG2 PE=1 SV=2 | 387 | 20201119_15232_1ng_glyco_hcd30_etd30_compound15.3645.3645.4 | controllerType=0 controllerNumber=1 scan=3645 | 12.5605 |
| 15548:4 | 1 | R.RTAGWNVPIGTLRPFLN[+2716.983]WTGPPEPIEAAVAR.F | HexNAc(5)Hex(6)Fuc(3)NeuAc(1) | N[+2717] | 1526.7075 | 4 | 6103.8083 | 6103.795 | 0 | 2.2 | 140 | Specific | 799.09 | 781.59 | 781.59 | 12.81 | 40 | >sp\|P02788\|TRFL_HUMAN Lactotransferrin OS=Homo sapiens OX=9606 GN=LTF PE=1 SV=6 | 1270 | 20201113_15413_1ng_glyco_hcd30_etd30_compound15.19138.19138.4 | controllerType=0 controllerNumber=1 scan=19138 | 43.3697 |

Table S4. Cont.

| Query #:z | Protein Rank | Glycopeptide | Glycan | Modification Type(s) | Observed m/z | z | Observed (M+H) | Calc. mass (M+H) | Off-by-x error | Mass error (ppm) | StartPosition | Cleavage | Score | Delta | Delta Mod | \|Log Prob\| | # of unique peptides | Protein Name | Protein DB number | Comment | Scan # | Scan Time |
|---|---|---|---|---|---|---|---|---|---|---|---|---|---|---|---|---|---|---|---|---|---|---|
| 15555:4 | 1 | R.RTAGWNVPIGTLRPFLN[+2570.925]WTGPPEPIEAAVAR.F | HexNAc(5)Hex(6)Fuc(2)NeuAc(1) | N[+2571] | 1490.1948 | 4 | 5957.7575 | 5957.7371 | 0 | 3.4 | 140 | Specific | 804.75 | 772.71 | 772.71 | 13.34 | 40 | >sp\|P02788\|TRFL_HUMAN Lactotransferrin OS=Homo sapiens OX=9606 GN=LTF PE=1 SV=6 | 1270 | 20201113_15413_1ng_glyco_hcd30_etd30_compound15.19145.19145.4 | controllerType=0 controllerNumber=1 scan=19145 | 43.3883 |
| 15640:4 | 1 | R.RTAGWNVPIGTLRPFLN[+2205.793]WTGPPEPIEAAVAR.F | HexNAc(4)Hex(5)Fuc(2)NeuAc(1) | N[+2206] | 1398.9117 | 4 | 5592.6252 | 5592.6049 | 0 | 3.6 | 140 | Specific | 893.13 | 893.13 | 893.13 | 14.34 | 40 | >sp\|P02788\|TRFL_HUMAN Lactotransferrin OS=Homo sapiens OX=9606 GN=LTF PE=1 SV=6 | 1270 | 20201113_15413_1ng_glyco_hcd30_etd30_compound15.19235.19235.4 | controllerType=0 controllerNumber=1 scan=19235 | 43.6457 |
| 15902:3 | 1 | R.TAGWNVPIGTLRPFLN[+1768.640]WTGPPEPIEAAVAR.F | HexNAc(4)Hex(5)Fuc(1) | N[+1769] | 1667.1265 | 3 | 4999.3649 | 4999.3504 | 0 | 2.9 | 141 | Specific | 222.91 | 194.68 | 194.68 | 3.24 | 40 | >sp\|P02788\|TRFL_HUMAN Lactotransferrin OS=Homo sapiens OX=9606 GN=LTF PE=1 SV=6 | 1270 | 20201113_15413_1ng_glyco_hcd30_etd30_compound15.19510.19510.3 | controllerType=0 controllerNumber=1 scan=19510 | 44.3934 |
| 16709:3 | 1 | R.TAGWNVPIGTLRPFLN[+2059.735]WTGPPEPIEAAVAR.F | HexNAc(4)Hex(5)Fuc(1)NeuAc(1) | N[+2060] | 1764.1699 | 3 | 5290.4951 | 5290.4459 | 0 | 9.3 | 141 | Specific | 291.61 | 231.19 | 231.19 | 5.33 | 40 | >sp\|P02788\|TRFL_HUMAN Lactotransferrin OS=Homo sapiens OX=9606 GN=LTF PE=1 SV=6 | 1270 | 20201113_15413_1ng_glyco_hcd30_etd30_compound15.20363.20363.3 | controllerType=0 controllerNumber=1 scan=20363 | 46.8594 |
| 17528:3 | 1 | R.TAGWNVPIGTLRPFLN[+2350.830]WTGPPEPIEAAVAR.F | HexNAc(4)Hex(5)Fuc(1)NeuAc(2) | N[+2351] | 1861.1993 | 3 | 5581.5834 | 5581.5413 | 0 | 7.5 | 141 | Specific | 330.6 | 274.67 | 274.67 | 4.92 | 40 | >sp\|P02788\|TRFL_HUMAN Lactotransferrin OS=Homo sapiens OX=9606 GN=LTF PE=1 SV=6 | 1270 | 20201113_15413_1ng_glyco_hcd30_etd30_compound15.21228.21228.3 | controllerType=0 controllerNumber=1 scan=21228 | 49.4422 |
| 16560:3 | 1 | R.TAGWNVPIGTLRPFLN[+2059.735]WTGPPEPIEAAVAR.F | HexNAc(4)Hex(5)Fuc(1)NeuAc(1) | N[+2060] | 1764.1604 | 3 | 5290.4665 | 5290.4459 | 0 | 3.9 | 141 | Specific | 361.92 | 263.16 | 263.16 | 5.73 | 40 | >sp\|P02788\|TRFL_HUMAN Lactotransferrin OS=Homo sapiens OX=9606 GN=LTF PE=1 SV=6 | 1270 | 20201113_15413_1ng_glyco_hcd30_etd30_compound15.20205.20205.2 | controllerType=0 controllerNumber=1 scan=20205 | 46.4579 |
| 16305:4 | 1 | R.TAGWNVPIGTLRPFLN[+3227.153]WTGPPEPIEAAVAR.F | HexNAc(6)Hex(7)Fuc(2)NeuAc(2) | N[+3227] | 1615.2313 | 4 | 6457.9035 | 6457.8636 | 0 | 6.2 | 141 | Specific | 492.9 | 492.9 | 492.9 | 7.17 | 40 | >sp\|P02788\|TRFL_HUMAN Lactotransferrin OS=Homo sapiens OX=9606 GN=LTF PE=1 SV=6 | 1270 | 20201113_15413_1ng_glyco_hcd30_etd30_compound15.19936.19936.2 | controllerType=0 controllerNumber=1 scan=19936 | 45.7009 |
| 16516:3 | 1 | R.TAGWNVPIGTLRPFLN[+2205.793]WTGPPEPIEAAVAR.F | HexNAc(4)Hex(5)Fuc(2)NeuAc(1) | N[+2206] | 1812.8562 | 3 | 5436.5539 | 5436.5038 | 0 | 9.2 | 141 | Specific | 519.29 | 418.76 | 418.76 | 7.1 | 40 | >sp\|P02788\|TRFL_HUMAN Lactotransferrin OS=Homo sapiens OX=9606 GN=LTF PE=1 SV=6 | 1270 | 20201113_15413_1ng_glyco_hcd30_etd30_compound15.20158.20158.3 | controllerType=0 controllerNumber=1 scan=20158 | 46.305 |
| 16428:4 | 1 | R.TAGWNVPIGTLRPFLN[+2863.041]WTGPPEPIEAAVAR.F | HexNAc(5)Hex(6)Fuc(4)NeuAc(1) | N[+2863] | 1524.2007 | 4 | 6093.7809 | 6093.7518 | 0 | 4.8 | 141 | Specific | 556.9 | 556.9 | 556.9 | 6.21 | 40 | >sp\|P02788\|TRFL_HUMAN Lactotransferrin OS=Homo sapiens OX=9606 GN=LTF PE=1 SV=6 | 1270 | 20201113_15413_1ng_glyco_hcd30_etd30_compound15.20065.20065.4 | controllerType=0 controllerNumber=1 scan=20065 | 46.0365 |
| 16472:4 | 1 | R.TAGWNVPIGTLRPFLN[+2570.925]WTGPPEPIEAAVAR.F | HexNAc(5)Hex(6)Fuc(2)NeuAc(1) | N[+2571] | 1451.1675 | 4 | 5801.6481 | 5801.636 | 0 | 2.1 | 141 | Specific | 626.1 | 626.1 | 626.1 | 9.89 | 40 | >sp\|P02788\|TRFL_HUMAN Lactotransferrin OS=Homo sapiens OX=9606 GN=LTF PE=1 SV=6 | 1270 | 20201113_15413_1ng_glyco_hcd30_etd30_compound15.20112.20112.4 | controllerType=0 controllerNumber=1 scan=20112 | 46.1649 |
| 17518:4 | 1 | R.TAGWNVPIGTLRPFLN[+2350.830]WTGPPEPIEAAVAR.F | HexNAc(4)Hex(5)Fuc(1)NeuAc(2) | N[+2351] | 1396.1534 | 4 | 5581.592 | 5581.5413 | 0 | 9.1 | 141 | Specific | 739.57 | 704.78 | 704.78 | 11.31 | 40 | >sp\|P02788\|TRFL_HUMAN Lactotransferrin OS=Homo sapiens OX=9606 GN=LTF PE=1 SV=6 | 1270 | 20201113_15413_1ng_glyco_hcd30_etd30_compound15.21218.21218.4 | controllerType=0 controllerNumber=1 scan=21218 | 49.411 |
| 16503:4 | 1 | R.TAGWNVPIGTLRPFLN[+2424.867]WTGPPEPIEAAVAR.F | HexNAc(5)Hex(6)Fuc(1)NeuAc(1) | N[+2425] | 1414.6531 | 4 | 5655.5905 | 5655.5781 | 0 | 2.2 | 141 | Specific | 829.27 | 825.4 | 825.4 | 13.89 | 40 | >sp\|P02788\|TRFL_HUMAN Lactotransferrin OS=Homo sapiens OX=9606 GN=LTF PE=1 SV=6 | 1270 | 20201113_15413_1ng_glyco_hcd30_etd30_compound15.20144.20144.4 | controllerType=0 controllerNumber=1 scan=20144 | 46.2481 |
| 16515:4 | 1 | R.TAGWNVPIGTLRPFLN[+2205.793]WTGPPEPIEAAVAR.F | HexNAc(4)Hex(5)Fuc(2)NeuAc(1) | N[+2206] | 1359.8949 | 4 | 5436.5578 | 5436.5038 | 0 | 9.9 | 141 | Specific | 924.73 | 878.16 | 878.16 | 15.31 | 40 | >sp\|P02788\|TRFL_HUMAN Lactotransferrin OS=Homo sapiens OX=9606 GN=LTF PE=1 SV=6 | 1270 | 20201113_15413_1ng_glyco_hcd30_etd30_compound15.20157.20157.4 | controllerType=0 controllerNumber=1 scan=20157 | 46.3011 |
| 14866:3 | 1 | R.TAGWNIPMGLLFN[+1768.640]QTGSC[+57.021]K.F | HexNAc(4)Hex(5)Fuc(1) | C[+57], N[+1769] | 1288.5569 | 3 | 3863.6563 | 3863.6495 | 0 | 1.8 | 485 | Specific | 284.14 | 240.89 | 240.89 | 5.3 | 40 | >sp\|P02788\|TRFL_HUMAN Lactotransferrin OS=Homo sapiens OX=9606 GN=LTF PE=1 SV=6 | 1270 | 20201113_15413_1ng_glyco_hcd30_etd30_compound15.18417.18417.3 | controllerType=0 controllerNumber=1 scan=18417 | 41.2375 |
| 14809:3 | 1 | R.TAGWNIPMGLLFN[+1606.587]QTGSC[+57.021]K.F | HexNAc(4)Hex(4)Fuc(1) | C[+57], N[+1607] | 1234.5413 | 3 | 3701.6094 | 3701.5966 | 0 | 3.4 | 485 | Specific | 337.49 | 250.57 | 250.57 | 3.75 | 40 | >sp\|P02788\|TRFL_HUMAN Lactotransferrin OS=Homo sapiens OX=9606 GN=LTF PE=1 SV=6 | 1270 | 20201113_15413_1ng_glyco_hcd30_etd30_compound15.18357.18357.3 | controllerType=0 controllerNumber=1 scan=18357 | 41.0577 |
| 14880:3 | 1 | R.TAGWNIPMGLLFN[+1622.582]QTGSC[+57.021]K.F | HexNAc(4)Hex(5) | C[+57], N[+1623] | 1239.8737 | 3 | 3717.6066 | 3717.5916 | 0 | 4 | 485 | Specific | 342.81 | 262.93 | 229.95 | 5.13 | 40 | >sp\|P02788\|TRFL_HUMAN Lactotransferrin OS=Homo sapiens OX=9606 GN=LTF PE=1 SV=6 | 1270 | 20201113_15413_1ng_glyco_hcd30_etd30_compound15.18432.18432.3 | controllerType=0 controllerNumber=1 scan=18432 | 41.2774 |
| 17523:4 | 1 | R.TAGWNIPMGLLFN[+2350.830]QTGSC[+57.021]K.F | HexNAc(4)Hex(5)Fuc(1)NeuAc(2) | C[+57], N[+2351] | 1112.2229 | 4 | 4445.8699 | 4445.8403 | 0 | 6.7 | 485 | Specific | 351.24 | 346.43 | 346.43 | 3.43 | 40 | >sp\|P02788\|TRFL_HUMAN Lactotransferrin OS=Homo sapiens OX=9606 GN=LTF PE=1 SV=6 | 1270 | 20201113_15413_1ng_glyco_hcd30_etd30_compound15.21223.21223.4 | controllerType=0 controllerNumber=1 scan=21223 | 49.4205 |
| 16220:3 | 1 | R.TAGWNIPMGLLFN[+1872.651]QTGSC[+57.021]K.F | HexNAc(3)Hex(6)NeuAc(1) | C[+57], N[+1873] | 1323.2313 | 3 | 3967.6792 | 3967.6604 | 0 | 4.7 | 485 | Specific | 360.3 | 259.75 | 226.39 | 4.78 | 40 | >sp\|P02788\|TRFL_HUMAN Lactotransferrin OS=Homo sapiens OX=9606 GN=LTF PE=1 SV=6 | 1270 | 20201113_15413_1ng_glyco_hcd30_etd30_compound15.19846.19846.3 | controllerType=0 controllerNumber=1 scan=19846 | 45.4043 |
| 14737:3 | 1 | R.TAGWNIPMGLLFN[+1768.640]QTGSC[+57.021]K.F | HexNAc(4)Hex(5)Fuc(1) | C[+57], N[+1769] | 1288.5589 | 3 | 3863.6621 | 3863.6495 | 0 | 3.3 | 485 | Specific | 365.92 | 284.81 | 284.81 | 4.1 | 40 | >sp\|P02788\|TRFL_HUMAN Lactotransferrin OS=Homo sapiens OX=9606 GN=LTF PE=1 SV=6 | 1270 | 20201113_15413_1ng_glyco_hcd30_etd30_compound15.18281.18281.3 | controllerType=0 controllerNumber=1 scan=18281 | 40.8261 |
| 16047:3 | 1 | R.TAGWNIPMGLLFN[+2205.793]QTGSC[+57.021]K.F | HexNAc(4)Hex(5)Fuc(2)NeuAc(1) | C[+57], N[+2206] | 1434.2786 | 3 | 4300.8213 | 4300.8028 | 0 | 4.3 | 485 | Specific | 374.4 | 268.85 | 268.85 | 5.11 | 40 | >sp\|P02788\|TRFL_HUMAN Lactotransferrin OS=Homo sapiens OX=9606 GN=LTF PE=1 SV=6 | 1270 | 20201113_15413_1ng_glyco_hcd30_etd30_compound15.19663.19663.3 | controllerType=0 controllerNumber=1 scan=19663 | 44.8228 |
| 16248:3 | 1 | R.TAGWNIPMGLLFN[+1751.624]QTGSC[+57.021]K.F | HexNAc(4)Hex(4)NeuAc(1) | C[+57], N[+1752] | 1282.8947 | 3 | 3846.6696 | 3846.6341 | 0 | 9.2 | 485 | Specific | 394.82 | 284.08 | 284.08 | 4.99 | 40 | >sp\|P02788\|TRFL_HUMAN Lactotransferrin OS=Homo sapiens OX=9606 GN=LTF PE=1 SV=6 | 1270 | 20201113_15413_1ng_glyco_hcd30_etd30_compound15.19876.19876.3 | controllerType=0 controllerNumber=1 scan=19876 | 45.4992 |
| 16170:3 | 1 | R.TAGWNIPMGLLFN[+1913.677]QTGSC[+57.021]K.F | HexNAc(4)Hex(5)NeuAc(1) | C[+57], N[+1914] | 1336.9074 | 3 | 4008.7077 | 4008.687 | 0 | 5.2 | 485 | Specific | 433.78 | 290.18 | 290.18 | 5.11 | 40 | >sp\|P02788\|TRFL_HUMAN Lactotransferrin OS=Homo sapiens OX=9606 GN=LTF PE=1 SV=6 | 1270 | 20201113_15413_1ng_glyco_hcd30_etd30_compound15.19793.19793.3 | controllerType=0 controllerNumber=1 scan=19793 | 45.2188 |
| 16065:3 | 1 | R.TAGWNIPMGLLFN[+2059.735]QTGSC[+57.021]K.F | HexNAc(4)Hex(5)Fuc(1)NeuAc(1) | C[+57], N[+2060] | 1385.5887 | 3 | 4154.7515 | 4154.7449 | 0 | 1.6 | 485 | Specific | 458.12 | 378.96 | 378.96 | 7 | 40 | >sp\|P02788\|TRFL_HUMAN Lactotransferrin OS=Homo sapiens OX=9606 GN=LTF PE=1 SV=6 | 1270 | 20201113_15413_1ng_glyco_hcd30_etd30_compound15.19682.19682.3 | controllerType=0 controllerNumber=1 scan=19682 | 44.8843 |
| 15179:3 | 1 | R.TAGWNIPM[+15.995]GLLFN[+2059.735]QTGSC[+57.021]K.F | HexNAc(4)Hex(5)Fuc(1)NeuAc(1) | C[+57], M[+16], N[+2060] | 1390.9251 | 3 | 4170.7608 | 4170.7398 | 0 | 5 | 485 | Specific | 258.89 | 224.97 | 224.97 | 3.2 | 40 | >sp\|P02788\|TRFL_HUMAN Lactotransferrin OS=Homo sapiens OX=9606 GN=LTF PE=1 SV=6 | 1270 | 20201113_15413_1ng_glyco_hcd30_etd30_compound15.18747.18747.3 | controllerType=0 controllerNumber=1 scan=18747 | 42.1458 |
| 15079:3 | 1 | R.TAGWNIPM[+15.995]GLLFN[+2059.735]QTGSC[+57.021]K.F | HexNAc(4)Hex(5)Fuc(1)NeuAc(1) | C[+57], M[+16], N[+2060] | 1390.9241 | 3 | 4170.7578 | 4170.7398 | 0 | 4.3 | 485 | Specific | 276.75 | 205.92 | 205.92 | 3.36 | 40 | >sp\|P02788\|TRFL_HUMAN Lactotransferrin OS=Homo sapiens OX=9606 GN=LTF PE=1 SV=6 | 1270 | 20201113_15413_1ng_glyco_hcd30_etd30_compound15.18641.18641.3 | controllerType=0 controllerNumber=1 scan=18641 | 41.84 |
| 14816:3 | 1 | R.TAGWNIPM[+15.995]GLLFN[+2059.735]QTGSC[+57.021]K.F | HexNAc(4)Hex(5)Fuc(1)NeuAc(1) | C[+57], M[+16], N[+2060] | 1390.9272 | 3 | 4170.767 | 4170.7398 | 0 | 6.5 | 485 | Specific | 319.19 | 274.6 | 274.6 | 3.09 | 40 | >sp\|P02788\|TRFL_HUMAN Lactotransferrin OS=Homo sapiens OX=9606 GN=LTF PE=1 SV=6 | 1270 | 20201113_15413_1ng_glyco_hcd30_etd30_compound15.18364.18364.3 | controllerType=0 controllerNumber=1 scan=18364 | 41.0859 |
| 17487:4 | 1 | R.[+42.011]TAGWNIPMGLLFN[+2059.735]QTGSC[+57.021]KFDEYFSQSC[+57.021]A | HexNAc(4)Hex(5)Fuc(1)NeuAc(1) | .[42], C[+57]*2, N[+2060] | 1510.8833 | 4 | 6040.5114 | 6040.5026 | 0 | 1.5 | 485 | Specific | 218.86 | 176.57 | 176.57 | 2.88 | 40 | >sp\|P02788\|TRFL_HUMAN Lactotransferrin OS=Homo sapiens OX=9606 GN=LTF PE=1 SV=6 | 1270 | 20201113_15413_1ng_glyco_hcd30_etd30_compound15.21185.21185.4 | controllerType=0 controllerNumber=1 scan=21185 | 49.2898 |
| 10271:3 | 1 | R.N[+2350.830]GSDC[+57.021]PDKFC[+57.021]LFQSETK.N | HexNAc(4)Hex(5)Fuc(1)NeuAc(2) | C[+57]*2, N[+2351] | 1461.9108 | 3 | 4383.7179 | 4383.7043 | 0 | 3.1 | 642 | Specific | 255.61 | 158.87 | 158.87 | 3.67 | 40 | >sp\|P02788\|TRFL_HUMAN Lactotransferrin OS=Homo sapiens OX=9606 GN=LTF PE=1 SV=6 | 1270 | 20201113_15413_1ng_glyco_hcd30_etd30_compound15.13580.13580.3 | controllerType=0 controllerNumber=1 scan=13580 | 30.9854 |
| 08172:3 | 1 | R.N[+2205.793]GSDC[+57.021]PDKFC[+57.021]LFQSETK.N | HexNAc(4)Hex(5)Fuc(2)NeuAc(1) | C[+57]*2, N[+2206] | 1413.5635 | 3 | 4238.676 | 4238.6667 | 0 | 2.2 | 642 | Specific | 330.72 | 223.03 | 223.03 | 4.48 | 40 | >sp\|P02788\|TRFL_HUMAN Lactotransferrin OS=Homo sapiens OX=9606 GN=LTF PE=1 SV=6 | 1270 | 20201113_15413_1ng_glyco_hcd30_etd30_compound15.11399.11399.3 | controllerType=0 controllerNumber=1 scan=11399 | 26.6858 |
| 15393:3 | 2 | R.LSLHRPALEDLLLGSEAN[+1095.397]LTC[+57.021]TLTGLR.D | HexNAc(3)Hex(3) | C[+57], N[+1095] | 1353.6734 | 3 | 4059.0056 | 4058.9949 | 0 | 2.7 | 127 | Specific | 227.46 | 227.46 | 227.46 | 4.85 | 28 | >sp\|P01876\|IGHA1_HUMAN Immunoglobulin heavy constant alpha 1 OS=Homo sapiens OX=9606 GN=IGHA1 PE=1 SV=2 | 255 | 20201113_15413_1ng_glyco_hcd30_etd30_compound15.18974.18974.3 | controllerType=0 controllerNumber=1 scan=18974 | 42.882 |
| 16095:3 | 2 | R.LSLHRPALEDLLLGSEAN[+1751.624]LTC[+57.021]TLTGLR.D | HexNAc(4)Hex(4)NeuAc(1) | C[+57], N[+1752] | 1572.4138 | 3 | 4715.2267 | 4715.2225 | 0 | 0.9 | 127 | Specific | 255.4 | 180.25 | 180.25 | 4.41 | 28 | >sp\|P01876\|IGHA1_HUMAN Immunoglobulin heavy constant alpha 1 OS=Homo sapiens OX=9606 GN=IGHA1 PE=1 SV=2 | 255 | 20201113_15413_1ng_glyco_hcd30_etd30_compound15.19713.19713.3 | controllerType=0 controllerNumber=1 scan=19713 | 44.9606 |
| 16060:3 | 2 | R.LSLHRPALEDLLLGSEAN[+1872.651]LTC[+57.021]TLTGLR.D | HexNAc(3)Hex(6)NeuAc(1) | C[+57], N[+1873] | 1612.7634 | 3 | 4836.2755 | 4836.2487 | 0 | 5.5 | 127 | Specific | 256.55 | 199.68 | 199.68 | 4.52 | 28 | >sp\|P01876\|IGHA1_HUMAN Immunoglobulin heavy constant alpha 1 OS=Homo sapiens OX=9606 GN=IGHA1 PE=1 SV=2 | 255 | 20201113_15413_1ng_glyco_hcd30_etd30_compound15.19676.19676.3 | controllerType=0 controllerNumber=1 scan=19676 | 44.8554 |
| 15379:4 | 2 | R.LSLHRPALEDLLLGSEAN[+1460.529]LTC[+57.021]TLTGLR.D | HexNAc(4)Hex(4) | C[+57], N[+1461] | 1106.7928 | 4 | 4424.1492 | 4424.1271 | 0 | 5 | 127 | Specific | 298.12 | 298.12 | 298.12 | 4.42 | 28 | >sp\|P01876\|IGHA1_HUMAN Immunoglobulin heavy constant alpha 1 OS=Homo sapiens OX=9606 GN=IGHA1 PE=1 SV=2 | 255 | 20201113_15413_1ng_glyco_hcd30_etd30_compound15.18959.18959.4 | controllerType=0 controllerNumber=1 scan=18959 | 42.8311 |
| 15381:4 | 2 | R.LSLHRPALEDLLLGSEAN[+1298.476]LTC[+57.021]TLTGLR.D | HexNAc(4)Hex(3) | C[+57], N[+1298] | 1066.2774 | 4 | 4262.0877 | 4262.0742 | 0 | 3.2 | 127 | Specific | 410.21 | 410.21 | 410.21 | 7.09 | 28 | >sp\|P01876\|IGHA1_HUMAN Immunoglobulin heavy constant alpha 1 OS=Homo sapiens OX=9606 GN=IGHA1 PE=1 SV=2 | 255 | 20201113_15413_1ng_glyco_hcd30_etd30_compound15.18961.18961.4 | controllerType=0 controllerNumber=1 scan=18961 | 42.8451 |
| 15324:3 | 2 | R.LSLHRPALEDLLLGSEAN[+1622.582]LTC[+57.021]TLTGLR.D | HexNAc(4)Hex(5) | C[+57], N[+1623] | 1529.4006 | 3 | 4586.1872 | 4586.1799 | 0 | 1.6 | 127 | Specific | 418.9 | 369.68 | 369.68 | 7.08 | 28 | >sp\|P01876\|IGHA1_HUMAN Immunoglobulin heavy constant alpha 1 OS=Homo sapiens OX=9606 GN=IGHA1 PE=1 SV=2 | 255 | 20201113_15413_1ng_glyco_hcd30_etd30_compound15.18900.18900.3 | controllerType=0 controllerNumber=1 scan=18900 | 42.6205 |
| 15305:3 | 2 | R.LSLHRPALEDLLLGSEAN[+1540.529]LTC[+57.021]TLTGLR.D | HexNAc(2)Hex(7) | C[+57], N[+1541] | 1502.0502 | 3 | 4504.1361 | 4504.1268 | 0 | 2.1 | 127 | Specific | 437.24 | 392.98 | 392.98 | 7.08 | 28 | >sp\|P01876\|IGHA1_HUMAN Immunoglobulin heavy constant alpha 1 OS=Homo sapiens OX=9606 GN=IGHA1 PE=1 SV=2 | 255 | 20201113_15413_1ng_glyco_hcd30_etd30_compound15.18880.18880.3 | controllerType=0 controllerNumber=1 scan=18880 | 42.5463 |
| 15357:3 | 2 | R.LSLHRPALEDLLLGSEAN[+1378.476]LTC[+57.021]TLTGLR.D | HexNAc(2)Hex(6) | C[+57], N[+1378] | 1448.0346 | 3 | 4342.0893 | 4342.074 | 0 | 3.5 | 127 | Specific | 440.68 | 440.68 | 440.68 | 6.67 | 28 | >sp\|P01876\|IGHA1_HUMAN Immunoglobulin heavy constant alpha 1 OS=Homo sapiens OX=9606 GN=IGHA1 PE=1 SV=2 | 255 | 20201113_15413_1ng_glyco_hcd30_etd30_compound15.18935.18935.3 | controllerType=0 controllerNumber=1 scan=18935 | 42.7397 |
| 15355:3 | 2 | R.LSLHRPALEDLLLGSEAN[+1216.423]LTC[+57.021]TLTGLR.D | HexNAc(2)Hex(5) | C[+57], N[+1216] | 1394.0156 | 3 | 4180.0321 | 4180.0211 | 0 | 2.6 | 127 | Specific | 448.02 | 435.03 | 435.03 | 7.14 | 28 | >sp\|P01876\|IGHA1_HUMAN Immunoglobulin heavy constant alpha 1 OS=Homo sapiens OX=9606 GN=IGHA1 PE=1 SV=2 | 255 | 20201113_15413_1ng_glyco_hcd30_etd30_compound15.18933.18933.3 | controllerType=0 controllerNumber=1 scan=18933 | 42.731 |
| 15353:4 | 2 | R.LSLHRPALEDLLLGSEAN[+1216.423]LTC[+57.021]TLTGLR.D | HexNAc(2)Hex(5) | C[+57], N[+1216] | 1045.7635 | 4 | 4180.0321 | 4180.0211 | 0 | 2.6 | 127 | Specific | 607.13 | 607.13 | 607.13 | 9.81 | 28 | >sp\|P01876\|IGHA1_HUMAN Immunoglobulin heavy constant alpha 1 OS=Homo sapiens OX=9606 GN=IGHA1 PE=1 SV=2 | 255 | 20201113_15413_1ng_glyco_hcd30_etd30_compound15.18931.18931.4 | controllerType=0 controllerNumber=1 scan=18931 | 42.721 |
| 15321:4 | 2 | R.LSLHRPALEDLLLGSEAN[+1622.582]LTC[+57.021]TLTGLR.D | HexNAc(4)Hex(5) | C[+57], N[+1623] | 1147.3027 | 4 | 4586.1888 | 4586.1799 | 0 | 1.9 | 127 | Specific | 617.94 | 561.88 | 561.88 | 9.81 | 28 | >sp\|P01876\|IGHA1_HUMAN Immunoglobulin heavy constant alpha 1 OS=Homo sapiens OX=9606 GN=IGHA1 PE=1 SV=2 | 255 | 20201113_15413_1ng_glyco_hcd30_etd30_compound15.18897.18897.4 | controllerType=0 controllerNumber=1 scan=18897 | 42.6061 |
| 15409:4 | 2 | R.LSLHRPALEDLLLGSEAN[+1095.397]LTC[+57.021]TLTGLR.D | HexNAc(3)Hex(3) | C[+57], N[+1095] | 1015.5094 | 4 | 4059.0159 | 4058.9949 | 0 | 5.2 | 127 | Specific | 685.34 | 685.34 | 685.34 | 10.21 | 28 | >sp\|P01876\|IGHA1_HUMAN Immunoglobulin heavy constant alpha 1 OS=Homo sapiens OX=9606 GN=IGHA1 PE=1 SV=2 | 255 | 20201113_15413_1ng_glyco_hcd30_etd30_compound15.18991.18991.4 | controllerType=0 controllerNumber=1 scan=18991 | 42.9258 |
| 15257:4 | 2 | R.LSLHRPALEDLLLGSEAN[+1540.529]LTC[+57.021]TLTGLR.D | HexNAc(2)Hex(7) | C[+57], N[+1541] | 1126.7903 | 4 | 4504.1395 | 4504.1268 | 0 | 2.8 | 127 | Specific | 832.44 | 800.99 | 800.99 | 13.34 | 28 | >sp\|P01876\|IGHA1_HUMAN Immunoglobulin heavy constant alpha 1 OS=Homo sapiens OX=9606 GN=IGHA1 PE=1 SV=2 | 255 | 20201113_15413_1ng_glyco_hcd30_etd30_compound15.18829.18829.2 | controllerType=0 controllerNumber=1 scan=18829 | 42.3757 |
| 15368:4 | 2 | R.LSLHRPALEDLLLGSEAN[+1378.476]LTC[+57.021]TLTGLR.D | HexNAc(2)Hex(6) | C[+57], N[+1378] | 1086.2781 | 4 | 4342.0906 | 4342.074 | 0 | 3.8 | 127 | Specific | 851.93 | 812.04 | 812.04 | 14.34 | 28 | >sp\|P01876\|IGHA1_HUMAN Immunoglobulin heavy constant alpha 1 OS=Homo sapiens OX=9606 GN=IGHA1 PE=1 SV=2 | 255 | 20201113_15413_1ng_glyco_hcd30_etd30_compound15.18947.18947.4 | controllerType=0 controllerNumber=1 scan=18947 | 42.7834 |
| 16018:4 | 2 | R.LSLHRPALEDLLLGSEAN[+1913.677]LTC[+57.021]TLTGLR.D | HexNAc(4)Hex(5)NeuAc(1) | C[+57], N[+1914] | 1220.073 | 4 | 4877.2703 | 4877.2753 | 0 | -1 | 127 | Specific | 899.2 | 860.91 | 860.91 | 13.85 | 28 | >sp\|P01876\|IGHA1_HUMAN Immunoglobulin heavy constant alpha 1 OS=Homo sapiens OX=9606 GN=IGHA1 PE=1 SV=2 | 255 | 20201113_15413_1ng_glyco_hcd30_etd30_compound15.19632.19632.4 | controllerType=0 controllerNumber=1 scan=19632 | 44.7207 |
| 16061:4 | 2 | R.LSLHRPALEDLLLGSEAN[+1751.624]LTC[+57.021]TLTGLR.D | HexNAc(4)Hex(4)NeuAc(1) | C[+57], N[+1752] | 1179.5643 | 4 | 4715.2352 | 4715.2225 | 0 | 2.7 | 127 | Specific | 912.36 | 826.89 | 826.89 | 15.81 | 28 | >sp\|P01876\|IGHA1_HUMAN Immunoglobulin heavy constant alpha 1 OS=Homo sapiens OX=9606 GN=IGHA1 PE=1 SV=2 | 255 | 20201113_15413_1ng_glyco_hcd30_etd30_compound15.19677.19677.4 | controllerType=0 controllerNumber=1 scan=19677 | 44.8631 |
| 15347:4 | 2 | R.LSLHRPALEDLLLGSEAN[+1501.555]LTC[+57.021]TLTGLR.D | HexNAc(5)Hex(3) | C[+57], N[+1502] | 1117.0472 | 4 | 4465.1668 | 4465.1536 | 0 | 3 | 127 | Specific | 912.4 | 912.4 | 912.4 | 15.34 | 28 | >sp\|P01876\|IGHA1_HUMAN Immunoglobulin heavy constant alpha 1 OS=Homo sapiens OX=9606 GN=IGHA1 PE=1 SV=2 | 255 | 20201113_15413_1ng_glyco_hcd30_etd30_compound15.18925.18925.4 | controllerType=0 controllerNumber=1 scan=18925 | 42.6966 |
| 15299:4 | 2 | R.LSLHRPALEDLLLGSEAN[+1540.529]LTC[+57.021]TLTGLR.D | HexNAc(2)Hex(7) | C[+57], N[+1541] | 1126.7893 | 4 | 4504.1356 | 4504.1268 | 0 | 2 | 127 | Specific | 1093.34 | 968.39 | 968.39 | 18.81 | 28 | >sp\|P01876\|IGHA1_HUMAN Immunoglobulin heavy constant alpha 1 OS=Homo sapiens OX=9606 GN=IGHA1 PE=1 SV=2 | 255 | 20201113_15413_1ng_glyco_hcd30_etd30_compound15.18874.18874.4 | controllerType=0 controllerNumber=1 scan=18874 | 42.5202 |
| 11703:4 | 2 | R.LAGKPTHVN[+2262.814]VSVVMAEVDGTC[+57.021]Y.- | HexNAc(5)Hex(5)Fuc(1)NeuAc(1) | C[+57], N[+2263] | 1153.2496 | 4 | 4609.9764 | 4609.9564 | 0 | 4.3 | 332 | CRagged | 200.32 | 151.7 | 151.7 | 2.65 | 28 | >sp\|P01876\|IGHA1_HUMAN Immunoglobulin heavy constant alpha 1 OS=Homo sapiens OX=9606 GN=IGHA1 PE=1 SV=2 | 255 | 20201113_15413_1ng_glyco_hcd30_etd30_compound15.15085.15085.4 | controllerType=0 controllerNumber=1 scan=15085 | 33.7906 |
| 09162:3 | 2 | R.LAGKPTHVN[+1216.423]VSVVMAEVDGTC[+57.021]Y.- | HexNAc(2)Hex(5) | C[+57], N[+1216] | 1188.5252 | 3 | 3563.5611 | 3563.5649 | 0 | -1.1 | 332 | CRagged | 205.71 | 196.18 | 169.7 | 2.77 | 28 | >sp\|P01876\|IGHA1_HUMAN Immunoglobulin heavy constant alpha 1 OS=Homo sapiens OX=9606 GN=IGHA1 PE=1 SV=2 | 255 | 20201113_15413_1ng_glyco_hcd30_etd30_compound15.12426.12426.3 | controllerType=0 controllerNumber=1 scan=12426 | 28.6329 |
| 10136:3 | 2 | R.LAGKPTHVN[+1809.666]VSVVMAEVDGTC[+57.021]Y.- | HexNAc(5)Hex(4)Fuc(1) | C[+57], N[+1810] | 1386.2769 | 3 | 4156.8162 | 4156.8081 | 0 | 1.9 | 332 | CRagged | 208.34 | 91 | 91 | 2.62 | 28 | >sp\|P01876\|IGHA1_HUMAN Immunoglobulin heavy constant alpha 1 OS=Homo sapiens OX=9606 GN=IGHA1 PE=1 SV=2 | 255 | 20201113_15413_1ng_glyco_hcd30_etd30_compound15.13439.13439.3 | controllerType=0 controllerNumber=1 scan=13439 | 30.6315 |
| 11694:3 | 2 | R.LAGKPTHVN[+2262.814]VSVVMAEVDGTC[+57.021]Y.- | HexNAc(5)Hex(5)Fuc(1)NeuAc(1) | C[+57], N[+2263] | 1537.3296 | 3 | 4609.9744 | 4609.9564 | 0 | 3.9 | 332 | CRagged | 212.61 | 54.65 | 54.65 | 2.16 | 28 | >sp\|P01876\|IGHA1_HUMAN Immunoglobulin heavy constant alpha 1 OS=Homo sapiens OX=9606 GN=IGHA1 PE=1 SV=2 | 255 | 20201113_15413_1ng_glyco_hcd30_etd30_compound15.15076.15076.3 | controllerType=0 controllerNumber=1 scan=15076 | 33.7652 |
| 10195:3 | 2 | R.LAGKPTHVN[+1501.555]VSVVMAEVDGTC[+57.021]Y.- | HexNAc(5)Hex(3) | C[+57], N[+1502] | 1283.5734 | 3 | 3848.7057 | 3848.6974 | 0 | 2.2 | 332 | CRagged | 232.72 | 146.49 | 146.49 | 2.87 | 28 | >sp\|P01876\|IGHA1_HUMAN Immunoglobulin heavy constant alpha 1 OS=Homo sapiens OX=9606 GN=IGHA1 PE=1 SV=2 | 255 | 20201113_15413_1ng_glyco_hcd30_etd30_compound15.13501.13501.3 | controllerType=0 controllerNumber=1 scan=13501 | 30.807 |
| 10161:3 | 2 | R.LAGKPTHVN[+1216.423]VSVVMAEVDGTC[+57.021]Y.- | HexNAc(2)Hex(5) | C[+57], N[+1216] | 1188.5285 | 3 | 3563.5709 | 3563.5649 | 0 | 1.7 | 332 | CRagged | 244.13 | 231.82 | 167.91 | 3.9 | 28 | >sp\|P01876\|IGHA1_HUMAN Immunoglobulin heavy constant alpha 1 OS=Homo sapiens OX=9606 GN=IGHA1 PE=1 SV=2 | 255 | 20201113_15413_1ng_glyco_hcd30_etd30_compound15.13465.13465.3 | controllerType=0 controllerNumber=1 scan=13465 | 30.7033 |
| 10118:3 | 2 | R.LAGKPTHVN[+1378.476]VSVVMAEVDGTC[+57.021]Y.- | HexNAc(2)Hex(6) | C[+57], N[+1378] | 1242.5467 | 3 | 3725.6255 | 3725.6178 | 0 | 2.1 | 332 | CRagged | 249.78 | 190.94 | 183.19 | 3.6 | 28 | >sp\|P01876\|IGHA1_HUMAN Immunoglobulin heavy constant alpha 1 OS=Homo sapiens OX=9606 GN=IGHA1 PE=1 SV=2 | 255 | 20201113_15413_1ng_glyco_hcd30_etd30_compound15.13420.13420.3 | controllerType=0 controllerNumber=1 scan=13420 | 30.5797 |
| 10078:3 | 2 | R.LAGKPTHVN[+1702.581]VSVVMAEVDGTC[+57.021]Y.- | HexNAc(2)Hex(8) | C[+57], N[+1703] | 1350.5826 | 3 | 4049.7332 | 4049.7234 | 0 | 2.4 | 332 | CRagged | 251.14 | 187.31 | 187.31 | 3.35 | 28 | >sp\|P01876\|IGHA1_HUMAN Immunoglobulin heavy constant alpha 1 OS=Homo sapiens OX=9606 GN=IGHA1 PE=1 SV=2 | 255 | 20201113_15413_1ng_glyco_hcd30_etd30_compound15.13378.13378.3 | controllerType=0 controllerNumber=1 scan=13378 | 30.4693 |
| 10100:3 | 2 | R.LAGKPTHVN[+1540.529]VSVVMAEVDGTC[+57.021]Y.- | HexNAc(2)Hex(7) | C[+57], N[+1541] | 1296.5652 | 3 | 3887.6812 | 3887.6706 | 0 | 2.7 | 332 | CRagged | 279.87 | 202.01 | 202.01 | 3.45 | 28 | >sp\|P01876\|IGHA1_HUMAN Immunoglobulin heavy constant alpha 1 OS=Homo sapiens OX=9606 GN=IGHA1 PE=1 SV=2 | 255 | 20201113_15413_1ng_glyco_hcd30_etd30_compound15.13401.13401.3 | controllerType=0 controllerNumber=1 scan=13401 | 30.5285 |
| 10210:3 | 2 | R.LAGKPTHVN[+1647.613]VSVVMAEVDGTC[+57.021]Y.- | HexNAc(5)Hex(3)Fuc(1) | C[+57], N[+1648] | 1332.2593 | 3 | 3994.7635 | 3994.7553 | 0 | 2 | 332 | CRagged | 287.89 | 155.16 | 155.16 | 3.33 | 28 | >sp\|P01876\|IGHA1_HUMAN Immunoglobulin heavy constant alpha 1 OS=Homo sapiens OX=9606 GN=IGHA1 PE=1 SV=2 | 255 | 20201113_15413_1ng_glyco_hcd30_etd30_compound15.13517.13517.3 | controllerType=0 controllerNumber=1 scan=13517 | 30.8493 |
| 13335:4 | 2 | R.LAGKPTHVN[+2553.910]VSVVMAEVDGTC[+57.021]Y.- | HexNAc(5)Hex(5)Fuc(1)NeuAc(2) | C[+57], N[+2554] | 1226.0223 | 4 | 4901.0672 | 4901.0518 | 0 | 3.1 | 332 | CRagged | 302.36 | 256.99 | 256.99 | 3.62 | 28 | >sp\|P01876\|IGHA1_HUMAN Immunoglobulin heavy constant alpha 1 OS=Homo sapiens OX=9606 GN=IGHA1 PE=1 SV=2 | 255 | 20201113_15413_1ng_glyco_hcd30_etd30_compound15.16785.16785.2 | controllerType=0 controllerNumber=1 scan=16785 | 37.3733 |
| 13314:4 | 2 | R.LAGKPTHVN[+2553.910]VSVVMAEVDGTC[+57.021]Y.- | HexNAc(5)Hex(5)Fuc(1)NeuAc(2) | C[+57], N[+2554] | 1226.0192 | 4 | 4901.055 | 4901.0518 | 0 | 0.7 | 332 | CRagged | 382.68 | 222.08 | 222.08 | 3.44 | 28 | >sp\|P01876\|IGHA1_HUMAN Immunoglobulin heavy constant alpha 1 OS=Homo sapiens OX=9606 GN=IGHA1 PE=1 SV=2 | 255 | 20201113_15413_1ng_glyco_hcd30_etd30_compound15.16763.16763.3 | controllerType=0 controllerNumber=1 scan=16763 | 37.3022 |
| 13607:3 | 4 | R.LN[+1540.529]YSLPTGQWVGVQLPR.N | HexNAc(2)Hex(7) | N[+1541] | 1156.8639 | 3 | 3468.5773 | 3468.5674 | 0 | 2.9 | 1017 | Specific | 415.31 | 394.34 | 394.34 | 6.47 | 10 | >sp\|P24821\|TENA_HUMAN Tenascin OS=Homo sapiens OX=9606 GN=TNC PE=1 SV=3 | 486 | 20201113_15413_1ng_glyco_hcd30_etd30_compound15.17069.17069.3 | controllerType=0 controllerNumber=1 scan=17069 | 38.0401 |
| 04291:3 | 5 | R.FSDGLESN[+1864.634]SSTQFEVKK.Y | HexNAc(2)Hex(9) | N[+1865] | 1256.5204 | 3 | 3767.5468 | 3767.5421 | 0 | 1.2 | 219 | Specific | 200.25 | 137.46 | 137.46 | 2.75 | 11 | >sp\|P0C0L4\|CO4A_HUMAN Complement C4-A OS=Homo sapiens OX=9606 GN=C4A PE=1 SV=2 | 1235 | 20201113_15413_1ng_glyco_hcd30_etd30_compound15.7366.7366.3 | controllerType=0 controllerNumber=1 scan=7366 | 19.3738 |
| 15127:3 | 6 | R.GLSFDVSLEVSQGPGLLN[+2863.041]DTK.V | HexNAc(5)Hex(6)Fuc(4)NeuAc(1) | N[+2863] | 1680.3997 | 3 | 5039.1847 | 5039.1541 | 0 | 6.1 | 118 | Specific | 204.95 | 125.63 | 125.63 | 3.72 | 20 | >sp\|P01833\|PIGR_HUMAN Polymeric immunoglobulin receptor OS=Homo sapiens OX=9606 GN=PIGR PE=1 SV=4 | 902 | 20201113_15413_1ng_glyco_hcd30_etd30_compound15.18691.18691.3 | controllerType=0 controllerNumber=1 scan=18691 | 41.967 |
| 15332:3 | 6 | R.GLSFDVSLEVSQGPGLLN[+2205.793]DTK.V | HexNAc(4)Hex(5)Fuc(2)NeuAc(1) | N[+2206] | 1461.3128 | 3 | 4381.9239 | 4381.906 | 0 | 4.1 | 118 | Specific | 238.54 | 154.26 | 154.26 | 2.8 | 20 | >sp\|P01833\|PIGR_HUMAN Polymeric immunoglobulin receptor OS=Homo sapiens OX=9606 GN=PIGR PE=1 SV=4 | 902 | 20201113_15413_1ng_glyco_hcd30_etd30_compound15.18909.18909.3 | controllerType=0 controllerNumber=1 scan=18909 | 42.6467 |
| 15160:3 | 6 | R.GLSFDVSLEVSQGPGLLN[+2715.963]DTK.V | HexNAc(5)Hex(6)Fuc(1)NeuAc(2) | N[+2716] | 1631.373 | 3 | 4892.1044 | 4892.0758 | 0 | 5.9 | 118 | Specific | 406.12 | 221.27 | 221.27 | 4.79 | 20 | >sp\|P01833\|PIGR_HUMAN Polymeric immunoglobulin receptor OS=Homo sapiens OX=9606 GN=PIGR PE=1 SV=4 | 902 | 20201113_15413_1ng_glyco_hcd30_etd30_compound15.18726.18726.3 | controllerType=0 controllerNumber=1 scan=18726 | 42.0771 |
| 17624:4 | 6 | K.QIGLYPVLVIDSSGYVNPN[+2204.772]YTGR.I | HexNAc(4)Hex(5)NeuAc(2) | N[+2205] | 1183.2775 | 4 | 4730.0882 | 4730.0759 | 0 | 2.6 | 168 | Specific | 429.8 | 378.78 | 378.78 | 5.77 | 20 | >sp\|P01833\|PIGR_HUMAN Polymeric immunoglobulin receptor OS=Homo sapiens OX=9606 GN=PIGR PE=1 SV=4 | 902 | 20201113_15413_1ng_glyco_hcd30_etd30_compound15.21331.21331.4 | controllerType=0 controllerNumber=1 scan=21331 | 49.7954 |
| 17403:4 | 6 | R.LSLLEEPGN[+2059.735]GTFTVILNQLTSR.D | HexNAc(4)Hex(5)Fuc(1)NeuAc(1) | N[+2060] | 1116.2692 | 4 | 4462.055 | 4462.0275 | 0 | 6.2 | 413 | Specific | 372.12 | 370.21 | 370.21 | 5.03 | 20 | >sp\|P01833\|PIGR_HUMAN Polymeric immunoglobulin receptor OS=Homo sapiens OX=9606 GN=PIGR PE=1 SV=4 | 902 | 20201113_15413_1ng_glyco_hcd30_etd30_compound15.21096.21096.4 | controllerType=0 controllerNumber=1 scan=21096 | 48.9966 |
| 19083:4 | 6 | R.LSLLEEPGN[+2350.830]GTFTVILNQLTSR.D | HexNAc(4)Hex(5)Fuc(1)NeuAc(2) | N[+2351] | 1189.0407 | 4 | 4753.1409 | 4753.1229 | 0 | 3.8 | 413 | Specific | 405.22 | 368.53 | 368.53 | 5.3 | 20 | >sp\|P01833\|PIGR_HUMAN Polymeric immunoglobulin receptor OS=Homo sapiens OX=9606 GN=PIGR PE=1 SV=4 | 902 | 20201113_15413_1ng_glyco_hcd30_etd30_compound15.22859.22859.4 | controllerType=0 controllerNumber=1 scan=22859 | 53.2696 |
| 19074:3 | 6 | R.LSLLEEPGN[+2350.830]GTFTVILNQLTSR.D | HexNAc(4)Hex(5)Fuc(1)NeuAc(2) | N[+2351] | 1585.0551 | 3 | 4753.1508 | 4753.1229 | 0 | 5.9 | 413 | Specific | 584.59 | 400.94 | 400.94 | 7.62 | 20 | >sp\|P01833\|PIGR_HUMAN Polymeric immunoglobulin receptor OS=Homo sapiens OX=9606 GN=PIGR PE=1 SV=4 | 902 | 20201113_15413_1ng_glyco_hcd30_etd30_compound15.22850.22850.3 | controllerType=0 controllerNumber=1 scan=22850 | 53.2363 |
| 15510:3 | 6 | K.IIEGEPNLKVPGN[+2350.830]VTAVLGETLK.V | HexNAc(4)Hex(5)Fuc(1)NeuAc(2) | N[+2351] | 1581.404 | 3 | 4742.1974 | 4742.1797 | 0 | 3.7 | 457 | Specific | 299.4 | 236.65 | 236.65 | 2.97 | 20 | >sp\|P01833\|PIGR_HUMAN Polymeric immunoglobulin receptor OS=Homo sapiens OX=9606 GN=PIGR PE=1 SV=4 | 902 | 20201113_15413_1ng_glyco_hcd30_etd30_compound15.19097.19097.3 | controllerType=0 controllerNumber=1 scan=19097 | 43.2369 |
| 14189:3 | 6 | K.IIEGEPNLKVPGN[+2059.735]VTAVLGETLK.V | HexNAc(4)Hex(5)Fuc(1)NeuAc(1) | N[+2060] | 1484.3699 | 3 | 4451.0952 | 4451.0843 | 0 | 2.5 | 457 | Specific | 338.4 | 223.53 | 223.53 | 3.53 | 20 | >sp\|P01833\|PIGR_HUMAN Polymeric immunoglobulin receptor OS=Homo sapiens OX=9606 GN=PIGR PE=1 SV=4 | 902 | 20201113_15413_1ng_glyco_hcd30_etd30_compound15.17702.17702.3 | controllerType=0 controllerNumber=1 scan=17702 | 39.4619 |
| 14132:3 | 6 | K.IIEGEPNLKVPGN[+2205.793]VTAVLGETLK.V | HexNAc(4)Hex(5)Fuc(2)NeuAc(1) | N[+2206] | 1533.0521 | 3 | 4597.1417 | 4597.1422 | 0 | -0.1 | 457 | Specific | 356.2 | 322.67 | 322.67 | 2.85 | 20 | >sp\|P01833\|PIGR_HUMAN Polymeric immunoglobulin receptor OS=Homo sapiens OX=9606 GN=PIGR PE=1 SV=4 | 902 | 20201113_15413_1ng_glyco_hcd30_etd30_compound15.17639.17639.3 | controllerType=0 controllerNumber=1 scan=17639 | 39.2846 |
| 14164:3 | 6 | K.IIEGEPNLKVPGN[+2205.793]VTAVLGETLK.V | HexNAc(4)Hex(5)Fuc(2)NeuAc(1) | N[+2206] | 1533.0549 | 3 | 4597.15 | 4597.1422 | 0 | 1.7 | 457 | Specific | 397.5 | 292.87 | 292.87 | 3.61 | 20 | >sp\|P01833\|PIGR_HUMAN Polymeric immunoglobulin receptor OS=Homo sapiens OX=9606 GN=PIGR PE=1 SV=4 | 902 | 20201113_15413_1ng_glyco_hcd30_etd30_compound15.17675.17675.3 | controllerType=0 controllerNumber=1 scan=17675 | 39.3759 |
| 12067:3 | 6 | K.VPGN[+2205.793]VTAVLGETLK.V | HexNAc(4)Hex(5)Fuc(2)NeuAc(1) | N[+2206] | 1201.873 | 3 | 3603.6044 | 3603.5927 | 0 | 3.2 | 466 | Specific | 279.16 | 189.15 | 189.15 | 3.36 | 20 | >sp\|P01833\|PIGR_HUMAN Polymeric immunoglobulin receptor OS=Homo sapiens OX=9606 GN=PIGR PE=1 SV=4 | 902 | 20201113_15413_1ng_glyco_hcd30_etd30_compound15.15464.15464.3 | controllerType=0 controllerNumber=1 scan=15464 | 34.568 |
| 14583:3 | 6 | K.VPGN[+2350.830]VTAVLGETLK.V | HexNAc(4)Hex(5)Fuc(1)NeuAc(2) | N[+2351] | 1250.2177 | 3 | 3748.6386 | 3748.6302 | 0 | 2.2 | 466 | Specific | 287.47 | 196.79 | 196.79 | 5.05 | 20 | >sp\|P01833\|PIGR_HUMAN Polymeric immunoglobulin receptor OS=Homo sapiens OX=9606 GN=PIGR PE=1 SV=4 | 902 | 20201113_15413_1ng_glyco_hcd30_etd30_compound15.18118.18118.3 | controllerType=0 controllerNumber=1 scan=18118 | 40.3886 |
| 05158:3 | 6 | K.WN[+1913.677]NTGC[+57.021]QALPSQDEGPSK.A | HexNAc(4)Hex(5)NeuAc(1) | C[+57], N[+1914] | 1301.5264 | 3 | 3902.5647 | 3902.5537 | 0 | 2.8 | 498 | Specific | 244.29 | 114.66 | 114.66 | 4.08 | 20 | >sp\|P01833\|PIGR_HUMAN Polymeric immunoglobulin receptor OS=Homo sapiens OX=9606 GN=PIGR PE=1 SV=4 | 902 | 20201113_15413_1ng_glyco_hcd30_etd30_compound15.8265.8265.3 | controllerType=0 controllerNumber=1 scan=8265 | 20.9973 |
| 05185:3 | 6 | K.WN[+1710.598]NTGC[+57.021]QALPSQDEGPSK.A | HexNAc(3)Hex(5)NeuAc(1) | C[+57], N[+1711] | 1233.8322 | 3 | 3699.4821 | 3699.4743 | 0 | 2.1 | 498 | Specific | 364.35 | 217.02 | 217.02 | 5.23 | 20 | >sp\|P01833\|PIGR_HUMAN Polymeric immunoglobulin receptor OS=Homo sapiens OX=9606 GN=PIGR PE=1 SV=4 | 902 | 20201113_15413_1ng_glyco_hcd30_etd30_compound15.8293.8293.3 | controllerType=0 controllerNumber=1 scan=8293 | 21.0556 |
| 05035:3 | 6 | K.WN[+2059.735]NTGC[+57.021]QALPSQDEGPSK.A | HexNAc(4)Hex(5)Fuc(1)NeuAc(1) | C[+57], N[+2060] | 1350.212 | 3 | 4048.6214 | 4048.6116 | 0 | 2.4 | 498 | Specific | 455.43 | 288.08 | 288.08 | 5.77 | 20 | >sp\|P01833\|PIGR_HUMAN Polymeric immunoglobulin receptor OS=Homo sapiens OX=9606 GN=PIGR PE=1 SV=4 | 902 | 20201113_15413_1ng_glyco_hcd30_etd30_compound15.8137.8137.3 | controllerType=0 controllerNumber=1 scan=8137 | 20.7505 |
| 06967:3 | 7 | K.WEC[+57.021]KN[+2059.735]DTLLGIK.G | HexNAc(4)Hex(5)Fuc(1)NeuAc(1) | C[+57], N[+2060] | 1179.502 | 3 | 3536.4915 | 3536.4865 | 0 | 1.4 | 100 | Specific | 212.12 | 111.89 | 111.89 | 2.56 | 11 | >sp\|P22897\|MRC1_HUMAN Macrophage mannose receptor 1 OS=Homo sapiens OX=9606 GN=MRC1 PE=1 SV=1 | 1049 | 20201113_15413_1ng_glyco_hcd30_etd30_compound15.10142.10142.3 | controllerType=0 controllerNumber=1 scan=10142 | 24.3429 |
| 06851:3 | 7 | K.WEC[+57.021]KN[+2205.793]DTLLGIK.G | HexNAc(4)Hex(5)Fuc(2)NeuAc(1) | C[+57], N[+2206] | 1228.1885 | 3 | 3682.551 | 3682.5444 | 0 | 1.8 | 100 | Specific | 230.27 | 123.86 | 123.86 | 2.62 | 11 | >sp\|P22897\|MRC1_HUMAN Macrophage mannose receptor 1 OS=Homo sapiens OX=9606 GN=MRC1 PE=1 SV=1 | 1049 | 20201113_15413_1ng_glyco_hcd30_etd30_compound15.10022.10022.3 | controllerType=0 controllerNumber=1 scan=10022 | 24.1152 |
| 11691:4 | 7 | K.KGN[+2205.793]TTLNSFVIPSESDVPTHC[+57.021]PSQWWPYAGHC[+57.021]YK.I | HexNAc(4)Hex(5)Fuc(2)NeuAc(1) | C[+57]*2, N[+2206] | 1543.1627 | 4 | 6169.6291 | 6169.6146 | 0 | 2.3 | 342 | Specific | 324.32 | 262.58 | 262.58 | 3.99 | 11 | >sp\|P22897\|MRC1_HUMAN Macrophage mannose receptor 1 OS=Homo sapiens OX=9606 GN=MRC1 PE=1 SV=1 | 1049 | 20201113_15413_1ng_glyco_hcd30_etd30_compound15.15072.15072.2 | controllerType=0 controllerNumber=1 scan=15072 | 33.7494 |
| 14989:4 | 8 | K.RNIAAFGGDPNN[+1913.677]ITLFGESAGGASVSLQTLSPYNK.G | HexNAc(4)Hex(5)NeuAc(1) | N[+1914] | 1370.8827 | 4 | 5480.5089 | 5480.4644 | 0 | 8.1 | 196 | Specific | 316.78 | 295.05 | 295.05 | 3.87 | 9 | >sp\|P19835\|CEL_HUMAN Bile salt-activated lipase OS=Homo sapiens OX=9606 GN=CEL PE=1 SV=3 | 476 | 20201113_15413_1ng_glyco_hcd30_etd30_compound15.18546.18546.4 | controllerType=0 controllerNumber=1 scan=18546 | 41.5663 |
| 09090:3 | 12 | K.TVLTPATNHMGN[+1378.476]VTFTIPANR.E | HexNAc(2)Hex(6) | N[+1378] | 1211.8853 | 3 | 3633.6414 | 3633.6358 | 0 | 1.5 | 74 | Specific | 311.17 | 256.48 | 126.26 | 4.05 | 6 | >sp\|P01024\|CO3_HUMAN Complement C3 OS=Homo sapiens OX=9606 GN=C3 PE=1 SV=2 | 492 | 20201113_15413_1ng_glyco_hcd30_etd30_compound15.12351.12351.3 | controllerType=0 controllerNumber=1 scan=12351 | 28.4701 |
| 18499:4 | 13 | K.VFPLSLDSTPQDGNVVVAC[+57.021]LVQGFFPQEPLSVTWSESGQN[+1647.613]VTAI | HexNAc(5)Hex(3)Fuc(1) | C[+57], N[+1648] | 1606.7596 | 4 | 6424.0168 | 6423.9746 | 0 | 6.6 | 8 | Specific | 423.16 | 423.16 | 423.16 | 5.69 | 16 | >sp\|P01877\|IGHA2_HUMAN Immunoglobulin heavy constant alpha 2 OS=Homo sapiens OX=9606 GN=IGHA2 PE=1 SV=4 | 576 | 20201113_15413_1ng_glyco_hcd30_etd30_compound15.22246.22246.2 | controllerType=0 controllerNumber=1 scan=22246 | 51.9079 |
| 01752:3 | 13 | K.HYTN[+2262.814]SSQDVTVPC[+57.021]R.V | HexNAc(5)Hex(5)Fuc(1)NeuAc(1) | C[+57], N[+2263] | 1309.5296 | 3 | 3926.5742 | 3926.5636 | 0 | 2.7 | 89 | Specific | 268.37 | 134.26 | 134.26 | 4.55 | 16 | >sp\|P01877\|IGHA2_HUMAN Immunoglobulin heavy constant alpha 2 OS=Homo sapiens OX=9606 GN=IGHA2 PE=1 SV=4 | 576 | 20201113_15413_1ng_glyco_hcd30_etd30_compound15.4725.4725.3 | controllerType=0 controllerNumber=1 scan=4725 | 14.5382 |
| 00980:3 | 13 | K.HYTN[+1647.613]SSQDVTVPC[+57.021]R.V | HexNAc(5)Hex(3)Fuc(1) | C[+57], N[+1648] | 1104.4602 | 3 | 3311.3661 | 3311.3625 | 0 | 1.1 | 89 | Specific | 345.89 | 305.08 | 305.08 | 4.35 | 16 | >sp\|P01877\|IGHA2_HUMAN Immunoglobulin heavy constant alpha 2 OS=Homo sapiens OX=9606 GN=IGHA2 PE=1 SV=4 | 576 | 20201113_15413_1ng_glyco_hcd30_etd30_compound15.3923.3923.2 | controllerType=0 controllerNumber=1 scan=3923 | 13.0872 |
| 00926:3 | 13 | K.HYTN[+1809.666]SSQDVTVPC[+57.021]R.V | HexNAc(5)Hex(4)Fuc(1) | C[+57], N[+1810] | 1158.4802 | 3 | 3473.4259 | 3473.4154 | 0 | 3 | 89 | Specific | 415.92 | 345.42 | 345.42 | 5.18 | 16 | >sp\|P01877\|IGHA2_HUMAN Immunoglobulin heavy constant alpha 2 OS=Homo sapiens OX=9606 GN=IGHA2 PE=1 SV=4 | 576 | 20201113_15413_1ng_glyco_hcd30_etd30_compound15.3867.3867.3 | controllerType=0 controllerNumber=1 scan=3867 | 12.9599 |
| 00944:3 | 13 | K.HYTN[+1647.613]SSQDVTVPC[+57.021]R.V | HexNAc(5)Hex(3)Fuc(1) | C[+57], N[+1648] | 1104.4611 | 3 | 3311.3687 | 3311.3625 | 0 | 1.9 | 89 | Specific | 447.98 | 391.79 | 391.79 | 6.84 | 16 | >sp\|P01877\|IGHA2_HUMAN Immunoglobulin heavy constant alpha 2 OS=Homo sapiens OX=9606 GN=IGHA2 PE=1 SV=4 | 576 | 20201113_15413_1ng_glyco_hcd30_etd30_compound15.3886.3886.3 | controllerType=0 controllerNumber=1 scan=3886 | 13.0072 |
| 02519:3 | 13 | K.TPLTAN[+1971.719]ITK.S | HexNAc(5)Hex(5)Fuc(1) | N[+1972] | 977.4301 | 3 | 2930.2759 | 2930.2757 | 0 | 0.1 | 200 | Specific | 243.46 | 113.43 | 113.43 | 2.66 | 16 | >sp\|P01877\|IGHA2_HUMAN Immunoglobulin heavy constant alpha 2 OS=Homo sapiens OX=9606 GN=IGHA2 PE=1 SV=4 | 576 | 20201113_15413_1ng_glyco_hcd30_etd30_compound15.5525.5525.3 | controllerType=0 controllerNumber=1 scan=5525 | 16.0229 |
| 04205:3 | 13 | K.TPLTAN[+2262.814]ITK.S | HexNAc(5)Hex(5)Fuc(1)NeuAc(1) | N[+2263] | 1074.4639 | 3 | 3221.3771 | 3221.3711 | 0 | 1.9 | 200 | Specific | 254.16 | 109.59 | 109.59 | 3.48 | 16 | >sp\|P01877\|IGHA2_HUMAN Immunoglobulin heavy constant alpha 2 OS=Homo sapiens OX=9606 GN=IGHA2 PE=1 SV=4 | 576 | 20201113_15413_1ng_glyco_hcd30_etd30_compound15.7277.7277.3 | controllerType=0 controllerNumber=1 scan=7277 | 19.215 |
| 02620:3 | 13 | K.TPLTAN[+1647.613]ITK.S | HexNAc(5)Hex(3)Fuc(1) | N[+1648] | 869.3979 | 3 | 2606.1792 | 2606.17 | 0 | 3.5 | 200 | Specific | 286.93 | 121.99 | 121.99 | 2.58 | 16 | >sp\|P01877\|IGHA2_HUMAN Immunoglobulin heavy constant alpha 2 OS=Homo sapiens OX=9606 GN=IGHA2 PE=1 SV=4 | 576 | 20201113_15413_1ng_glyco_hcd30_etd30_compound15.5630.5630.3 | controllerType=0 controllerNumber=1 scan=5630 | 16.2668 |
| 02560:3 | 13 | K.TPLTAN[+1809.666]ITK.S | HexNAc(5)Hex(4)Fuc(1) | N[+1810] | 923.4152 | 3 | 2768.231 | 2768.2228 | 0 | 3 | 200 | Specific | 295.49 | 139.12 | 139.12 | 2.74 | 16 | >sp\|P01877\|IGHA2_HUMAN Immunoglobulin heavy constant alpha 2 OS=Homo sapiens OX=9606 GN=IGHA2 PE=1 SV=4 | 576 | 20201113_15413_1ng_glyco_hcd30_etd30_compound15.5568.5568.3 | controllerType=0 controllerNumber=1 scan=5568 | 16.1374 |
| 02605:2 | 13 | K.TPLTAN[+1647.613]ITK.S | HexNAc(5)Hex(3)Fuc(1) | N[+1648] | 1303.594 | 2 | 2606.1808 | 2606.17 | 0 | 4.1 | 200 | Specific | 305.31 | 123.7 | 123.7 | 2.87 | 16 | >sp\|P01877\|IGHA2_HUMAN Immunoglobulin heavy constant alpha 2 OS=Homo sapiens OX=9606 GN=IGHA2 PE=1 SV=4 | 576 | 20201113_15413_1ng_glyco_hcd30_etd30_compound15.5615.5615.2 | controllerType=0 controllerNumber=1 scan=5615 | 16.2394 |
| 02559:2 | 13 | K.TPLTAN[+1809.666]ITK.S | HexNAc(5)Hex(4)Fuc(1) | N[+1810] | 1384.6189 | 2 | 2768.2306 | 2768.2228 | 0 | 2.8 | 200 | Specific | 315.03 | 118.84 | 118.84 | 2.62 | 16 | >sp\|P01877\|IGHA2_HUMAN Immunoglobulin heavy constant alpha 2 OS=Homo sapiens OX=9606 GN=IGHA2 PE=1 SV=4 | 576 | 20201113_15413_1ng_glyco_hcd30_etd30_compound15.5567.5567.2 | controllerType=0 controllerNumber=1 scan=5567 | 16.1306 |
| 09151:3 | 13 | R.MAGKPTHIN[+1702.581]VSVVMAEADGTC[+57.021]Y.- | HexNAc(2)Hex(8) | C[+57], N[+1703] | 1351.8913 | 3 | 4053.6593 | 4053.6642 | 0 | -1.2 | 319 | CRagged | 237.2 | 237.2 | 171.65 | 2.72 | 16 | >sp\|P01877\|IGHA2_HUMAN Immunoglobulin heavy constant alpha 2 OS=Homo sapiens OX=9606 GN=IGHA2 PE=1 SV=4 | 576 | 20201113_15413_1ng_glyco_hcd30_etd30_compound15.12415.12415.3 | controllerType=0 controllerNumber=1 scan=12415 | 28.6042 |
| 09221:3 | 13 | R.MAGKPTHIN[+1378.476]VSVVMAEADGTC[+57.021]Y.- | HexNAc(2)Hex(6) | C[+57], N[+1378] | 1243.8622 | 3 | 3729.5722 | 3729.5585 | 0 | 3.7 | 319 | CRagged | 238.21 | 233.39 | 151.21 | 2.85 | 16 | >sp\|P01877\|IGHA2_HUMAN Immunoglobulin heavy constant alpha 2 OS=Homo sapiens OX=9606 GN=IGHA2 PE=1 SV=4 | 576 | 20201113_15413_1ng_glyco_hcd30_etd30_compound15.12488.12488.3 | controllerType=0 controllerNumber=1 scan=12488 | 28.7803 |
| 09193:3 | 13 | R.MAGKPTHIN[+1540.529]VSVVMAEADGTC[+57.021]Y.- | HexNAc(2)Hex(7) | C[+57], N[+1541] | 1297.8795 | 3 | 3891.6238 | 3891.6114 | 0 | 3.2 | 319 | CRagged | 253.07 | 194.66 | 177.68 | 3.27 | 16 | >sp\|P01877\|IGHA2_HUMAN Immunoglobulin heavy constant alpha 2 OS=Homo sapiens OX=9606 GN=IGHA2 PE=1 SV=4 | 576 | 20201113_15413_1ng_glyco_hcd30_etd30_compound15.12459.12459.3 | controllerType=0 controllerNumber=1 scan=12459 | 28.7077 |
| 09242:3 | 13 | R.MAGKPTHIN[+1216.423]VSVVMAEADGTC[+57.021]Y.- | HexNAc(2)Hex(5) | C[+57], N[+1216] | 1189.8436 | 3 | 3567.5161 | 3567.5057 | 0 | 2.9 | 319 | CRagged | 278.19 | 255.5 | 193.98 | 3.41 | 16 | >sp\|P01877\|IGHA2_HUMAN Immunoglobulin heavy constant alpha 2 OS=Homo sapiens OX=9606 GN=IGHA2 PE=1 SV=4 | 576 | 20201113_15413_1ng_glyco_hcd30_etd30_compound15.12510.12510.3 | controllerType=0 controllerNumber=1 scan=12510 | 28.8505 |
| 08681:3 | 15 | R.IIVPLNNREN[+1809.666]ISDPTSPLR.T | HexNAc(5)Hex(4)Fuc(1) | N[+1810] | 1319.9557 | 3 | 3957.8527 | 3957.8432 | 0 | 2.4 | 62 | Specific | 210.21 | 170.75 | 170.75 | 2.59 | 9 | >sp\|P01591\|IGJ_HUMAN Immunoglobulin J chain OS=Homo sapiens OX=9606 GN=JCHAIN PE=1 SV=4 | 1392 | 20201113_15413_1ng_glyco_hcd30_etd30_compound15.11926.11926.3 | controllerType=0 controllerNumber=1 scan=11926 | 27.6649 |
| 08719:3 | 15 | R.IIVPLNNREN[+1663.608]ISDPTSPLR.T | HexNAc(5)Hex(4) | N[+1664] | 1271.2685 | 3 | 3811.7909 | 3811.7853 | 0 | 1.5 | 62 | Specific | 248.57 | 156.76 | 156.76 | 2.87 | 9 | >sp\|P01591\|IGJ_HUMAN Immunoglobulin J chain OS=Homo sapiens OX=9606 GN=JCHAIN PE=1 SV=4 | 1392 | 20201113_15413_1ng_glyco_hcd30_etd30_compound15.11966.11966.3 | controllerType=0 controllerNumber=1 scan=11966 | 27.7332 |
| 10152:3 | 15 | R.IIVPLNNREN[+1856.656]ISDPTSPLR.T | HexNAc(3)Hex(5)Fuc(1)NeuAc(1) | N[+1857] | 1335.6186 | 3 | 4004.8412 | 4004.8327 | 0 | 2.1 | 62 | Specific | 257.99 | 194.28 | 194.28 | 3.07 | 9 | >sp\|P01591\|IGJ_HUMAN Immunoglobulin J chain OS=Homo sapiens OX=9606 GN=JCHAIN PE=1 SV=4 | 1392 | 20201113_15413_1ng_glyco_hcd30_etd30_compound15.13455.13455.3 | controllerType=0 controllerNumber=1 scan=13455 | 30.6831 |
| 11778:3 | 15 | R.IIVPLNNREN[+2350.830]ISDPTSPLR.T | HexNAc(4)Hex(5)Fuc(1)NeuAc(2) | N[+2351] | 1500.342 | 3 | 4499.0114 | 4499.0075 | 0 | 0.9 | 62 | Specific | 285.37 | 219.37 | 219.37 | 2.76 | 9 | >sp\|P01591\|IGJ_HUMAN Immunoglobulin J chain OS=Homo sapiens OX=9606 GN=JCHAIN PE=1 SV=4 | 1392 | 20201113_15413_1ng_glyco_hcd30_etd30_compound15.15163.15163.3 | controllerType=0 controllerNumber=1 scan=15163 | 33.9678 |
| 11857:3 | 15 | R.IIVPLNNREN[+2204.772]ISDPTSPLR.T | HexNAc(4)Hex(5)NeuAc(2) | N[+2205] | 1451.6598 | 3 | 4352.965 | 4352.9496 | 0 | 3.5 | 62 | Specific | 296.96 | 229.14 | 229.14 | 3.14 | 9 | >sp\|P01591\|IGJ_HUMAN Immunoglobulin J chain OS=Homo sapiens OX=9606 GN=JCHAIN PE=1 SV=4 | 1392 | 20201113_15413_1ng_glyco_hcd30_etd30_compound15.15245.15245.3 | controllerType=0 controllerNumber=1 scan=15245 | 34.1266 |
| 10095:3 | 15 | R.IIVPLNNREN[+1913.677]ISDPTSPLR.T | HexNAc(4)Hex(5)NeuAc(1) | N[+1914] | 1354.6255 | 3 | 4061.8621 | 4061.8542 | 0 | 1.9 | 62 | Specific | 305.56 | 240.65 | 240.65 | 3.33 | 9 | >sp\|P01591\|IGJ_HUMAN Immunoglobulin J chain OS=Homo sapiens OX=9606 GN=JCHAIN PE=1 SV=4 | 1392 | 20201113_15413_1ng_glyco_hcd30_etd30_compound15.13395.13395.3 | controllerType=0 controllerNumber=1 scan=13395 | 30.5036 |
| 07606:3 | 16 | K.VAYSN[+2204.772]DSANWTEYQDPR.T | HexNAc(4)Hex(5)NeuAc(2) | N[+2205] | 1407.5634 | 3 | 4220.6757 | 4220.6454 | 0 | 7.2 | 321 | Specific | 430.36 | 283 | 66.66 | 4.42 | 4 | >sp\|Q08431\|MFGM_HUMAN Lactadherin OS=Homo sapiens OX=9606 GN=MFGE8 PE=1 SV=3 | 1064 | 20201113_15413_1ng_glyco_hcd30_etd30_compound15.10807.10807.3 | controllerType=0 controllerNumber=1 scan=10807 | 25.5134 |
| 05056:3 | 17 | R.LSPN[+2205.793]ASAEHLELR.W | HexNAc(4)Hex(5)Fuc(2)NeuAc(1) | N[+2206] | 1214.8543 | 3 | 3642.5484 | 3642.5421 | 0 | 1.7 | 52 | Specific | 290.72 | 176.47 | 176.47 | 3.86 | 7 | >sp\|Q13410\|BT1A1_HUMAN Butyrophilin subfamily 1 member A1 OS=Homo sapiens OX=9606 GN=BTN1A1 PE=1 SV=3 | 179 | 20201113_15413_1ng_glyco_hcd30_etd30_compound15.8159.8159.3 | controllerType=0 controllerNumber=1 scan=8159 | 20.7922 |
| 06875:3 | 17 | R.LSPN[+2350.830]ASAEHLELR.W | HexNAc(4)Hex(5)Fuc(1)NeuAc(2) | N[+2351] | 1263.2007 | 3 | 3787.5877 | 3787.5796 | 0 | 2.1 | 52 | Specific | 395.61 | 248.18 | 248.18 | 5.22 | 7 | >sp\|Q13410\|BT1A1_HUMAN Butyrophilin subfamily 1 member A1 OS=Homo sapiens OX=9606 GN=BTN1A1 PE=1 SV=3 | 179 | 20201113_15413_1ng_glyco_hcd30_etd30_compound15.10047.10047.3 | controllerType=0 controllerNumber=1 scan=10047 | 24.1696 |
| 01765:4 | 20 | R.EKQTDEIKDTRN[+2059.735]ESTQNC[+57.021]VVAEPEK.M | HexNAc(4)Hex(5)Fuc(1)NeuAc(1) | C[+57], N[+2060] | 1252.7902 | 4 | 5008.1388 | 5008.1251 | 0 | 2.7 | 58 | Specific | 580.23 | 500.33 | 500.33 | 6.05 | 6 | >sp\|P47710\|CASA1_HUMAN Alpha-S1-casein OS=Homo sapiens OX=9606 GN=CSN1S1 PE=1 SV=1 | 989 | 20201113_15413_1ng_glyco_hcd30_etd30_compound15.4739.4739.4 | controllerType=0 controllerNumber=1 scan=4739 | 14.5698 |
| 02205:4 | 20 | K.QTDEIKDTRN[+2059.735]ESTQNC[+57.021]VVAEPEK.M | HexNAc(4)Hex(5)Fuc(1)NeuAc(1) | C[+57], N[+2060] | 1188.5061 | 4 | 4751.0028 | 4750.9876 | 0 | 3.2 | 60 | Specific | 548.45 | 482.71 | 482.71 | 6.07 | 6 | >sp\|P47710\|CASA1_HUMAN Alpha-S1-casein OS=Homo sapiens OX=9606 GN=CSN1S1 PE=1 SV=1 | 989 | 20201113_15413_1ng_glyco_hcd30_etd30_compound15.5195.5195.4 | controllerType=0 controllerNumber=1 scan=5195 | 15.3982 |
| 13810:4 | 21 | K.LKELPGVC[+57.021]N[+2350.830]ETMMALWEEC[+57.021]KPC[+57.021]LK.Q | HexNAc(4)Hex(5)Fuc(1)NeuAc(2) | C[+57]*3, N[+2351] | 1322.5616 | 4 | 5287.2248 | 5287.2151 | 0 | 1.8 | 95 | Specific | 431.75 | 294 | 294 | 3.48 | 7 | >sp\|P10909\|CLUS_HUMAN Clusterin OS=Homo sapiens OX=9606 GN=CLU PE=1 SV=1 | 1467 | 20201113_15413_1ng_glyco_hcd30_etd30_compound15.17282.17282.4 | controllerType=0 controllerNumber=1 scan=17282 | 38.5745 |
| 00081:3 | 21 | R.EIRHN[+2059.735]STGC[+57.021]LR.M | HexNAc(4)Hex(5)Fuc(1)NeuAc(1) | C[+57], N[+2060] | 1134.8068 | 3 | 3402.4058 | 3402.3994 | 0 | 1.9 | 287 | Specific | 224.49 | 121.35 | 121.35 | 2.49 | 7 | >sp\|P10909\|CLUS_HUMAN Clusterin OS=Homo sapiens OX=9606 GN=CLU PE=1 SV=1 | 1467 | 20201113_15413_1ng_glyco_hcd30_etd30_compound15.2990.2990.3 | controllerType=0 controllerNumber=1 scan=2990 | 11.3431 |
| 00068:4 | 21 | R.EIRHN[+2059.735]STGC[+57.021]LR.M | HexNAc(4)Hex(5)Fuc(1)NeuAc(1) | C[+57], N[+2060] | 851.3575 | 4 | 3402.4082 | 3402.3994 | 0 | 2.6 | 287 | Specific | 530.81 | 403.81 | 403.81 | 6.28 | 7 | >sp\|P10909\|CLUS_HUMAN Clusterin OS=Homo sapiens OX=9606 GN=CLU PE=1 SV=1 | 1467 | 20201113_15413_1ng_glyco_hcd30_etd30_compound15.2976.2976.4 | controllerType=0 controllerNumber=1 scan=2976 | 11.2954 |
| 17625:3 | 21 | K.MLN[+2059.735]TSSLLEQLNEQFNWVSR.L | HexNAc(4)Hex(5)Fuc(1)NeuAc(1) | N[+2060] | 1490.3167 | 3 | 4468.9356 | 4468.9217 | 0 | 3.1 | 352 | Specific | 612.55 | 423.09 | 423.09 | 8.72 | 7 | >sp\|P10909\|CLUS_HUMAN Clusterin OS=Homo sapiens OX=9606 GN=CLU PE=1 SV=1 | 1467 | 20201113_15413_1ng_glyco_hcd30_etd30_compound15.21332.21332.3 | controllerType=0 controllerNumber=1 scan=21332 | 49.8013 |
| 15227:4 | 1 | R.LSLHRPALEDLLLGSEAN[+1419.502]LTC[+57.021]TLTGLR.D | HexNAc(3)Hex(5) | C[+57], N[+1420] | 1096.535 | 4 | 4383.118 | 4383.1005 | 0 | 4 | 127 | Specific | 220.32 | 220.32 | 220.32 | 4.32 | 36 | >sp\|P01876\|IGHA1_HUMAN Immunoglobulin heavy constant alpha 1 OS=Homo sapiens OX=9606 GN=IGHA1 PE=1 SV=2 | 255 | 20201119_15432_1ng_glyco_hcd30_etd30_compound15.18840.18840.4 | controllerType=0 controllerNumber=1 scan=18840 | 42.8509 |
| 15143:3 | 1 | R.LSLHRPALEDLLLGSEAN[+1702.581]LTC[+57.021]TLTGLR.D | HexNAc(2)Hex(8) | C[+57], N[+1703] | 1556.0704 | 3 | 4666.1966 | 4666.1796 | 0 | 3.6 | 127 | Specific | 369.18 | 329.87 | 329.87 | 5.12 | 36 | >sp\|P01876\|IGHA1_HUMAN Immunoglobulin heavy constant alpha 1 OS=Homo sapiens OX=9606 GN=IGHA1 PE=1 SV=2 | 255 | 20201119_15432_1ng_glyco_hcd30_etd30_compound15.18751.18751.3 | controllerType=0 controllerNumber=1 scan=18751 | 42.5506 |
| 15140:3 | 1 | R.LSLHRPALEDLLLGSEAN[+1864.634]LTC[+57.021]TLTGLR.D | HexNAc(2)Hex(9) | C[+57], N[+1865] | 1610.0873 | 3 | 4828.2475 | 4828.2324 | 0 | 3.1 | 127 | Specific | 430.38 | 382.65 | 382.65 | 6.41 | 36 | >sp\|P01876\|IGHA1_HUMAN Immunoglobulin heavy constant alpha 1 OS=Homo sapiens OX=9606 GN=IGHA1 PE=1 SV=2 | 255 | 20201119_15432_1ng_glyco_hcd30_etd30_compound15.18748.18748.3 | controllerType=0 controllerNumber=1 scan=18748 | 42.5399 |
| 15892:4 | 1 | R.LSLHRPALEDLLLGSEAN[+2116.756]LTC[+57.021]TLTGLR.D | HexNAc(5)Hex(5)NeuAc(1) | C[+57], N[+2117] | 1270.8502 | 4 | 5080.3791 | 5080.3547 | 0 | 4.8 | 127 | Specific | 437.1 | 437.1 | 437.1 | 6.21 | 36 | >sp\|P01876\|IGHA1_HUMAN Immunoglobulin heavy constant alpha 1 OS=Homo sapiens OX=9606 GN=IGHA1 PE=1 SV=2 | 255 | 20201119_15432_1ng_glyco_hcd30_etd30_compound15.19542.19542.4 | controllerType=0 controllerNumber=1 scan=19542 | 44.8819 |
| 15214:3 | 1 | R.LSLHRPALEDLLLGSEAN[+1501.555]LTC[+57.021]TLTGLR.D | HexNAc(5)Hex(3) | C[+57], N[+1502] | 1489.0621 | 3 | 4465.1717 | 4465.1536 | 0 | 4 | 127 | Specific | 444.91 | 444.91 | 444.91 | 7.02 | 36 | >sp\|P01876\|IGHA1_HUMAN Immunoglobulin heavy constant alpha 1 OS=Homo sapiens OX=9606 GN=IGHA1 PE=1 SV=2 | 255 | 20201119_15432_1ng_glyco_hcd30_etd30_compound15.18827.18827.3 | controllerType=0 controllerNumber=1 scan=18827 | 42.8076 |
| 15166:3 | 1 | R.LSLHRPALEDLLLGSEAN[+1540.529]LTC[+57.021]TLTGLR.D | HexNAc(2)Hex(7) | C[+57], N[+1541] | 1502.0524 | 3 | 4504.1427 | 4504.1268 | 0 | 3.5 | 127 | Specific | 454.47 | 410.69 | 410.69 | 7.02 | 36 | >sp\|P01876\|IGHA1_HUMAN Immunoglobulin heavy constant alpha 1 OS=Homo sapiens OX=9606 GN=IGHA1 PE=1 SV=2 | 255 | 20201119_15432_1ng_glyco_hcd30_etd30_compound15.18776.18776.3 | controllerType=0 controllerNumber=1 scan=18776 | 42.6328 |
| 15186:3 | 1 | R.LSLHRPALEDLLLGSEAN[+1378.476]LTC[+57.021]TLTGLR.D | HexNAc(2)Hex(6) | C[+57], N[+1378] | 1448.0366 | 3 | 4342.0951 | 4342.074 | 0 | 4.9 | 127 | Specific | 463.37 | 439.57 | 439.57 | 6.37 | 36 | >sp\|P01876\|IGHA1_HUMAN Immunoglobulin heavy constant alpha 1 OS=Homo sapiens OX=9606 GN=IGHA1 PE=1 SV=2 | 255 | 20201119_15432_1ng_glyco_hcd30_etd30_compound15.18797.18797.3 | controllerType=0 controllerNumber=1 scan=18797 | 42.7089 |
| 15220:3 | 1 | R.LSLHRPALEDLLLGSEAN[+1216.423]LTC[+57.021]TLTGLR.D | HexNAc(2)Hex(5) | C[+57], N[+1216] | 1394.0168 | 3 | 4180.0358 | 4180.0211 | 0 | 3.5 | 127 | Specific | 484.8 | 481.54 | 481.54 | 7.1 | 36 | >sp\|P01876\|IGHA1_HUMAN Immunoglobulin heavy constant alpha 1 OS=Homo sapiens OX=9606 GN=IGHA1 PE=1 SV=2 | 255 | 20201119_15432_1ng_glyco_hcd30_etd30_compound15.18833.18833.3 | controllerType=0 controllerNumber=1 scan=18833 | 42.8254 |
| 15919:4 | 1 | R.LSLHRPALEDLLLGSEAN[+1710.598]LTC[+57.021]TLTGLR.D | HexNAc(3)Hex(5)NeuAc(1) | C[+57], N[+1711] | 1169.3089 | 4 | 4674.2137 | 4674.1959 | 0 | 3.8 | 127 | Specific | 540.07 | 540.07 | 540.07 | 7.83 | 36 | >sp\|P01876\|IGHA1_HUMAN Immunoglobulin heavy constant alpha 1 OS=Homo sapiens OX=9606 GN=IGHA1 PE=1 SV=2 | 255 | 20201119_15432_1ng_glyco_hcd30_etd30_compound15.19570.19570.4 | controllerType=0 controllerNumber=1 scan=19570 | 44.9662 |
| 15243:4 | 1 | R.LSLHRPALEDLLLGSEAN[+1257.449]LTC[+57.021]TLTGLR.D | HexNAc(3)Hex(4) | C[+57], N[+1257] | 1056.0206 | 4 | 4221.0604 | 4221.0477 | 0 | 3 | 127 | Specific | 650.8 | 650.8 | 650.8 | 10.39 | 36 | >sp\|P01876\|IGHA1_HUMAN Immunoglobulin heavy constant alpha 1 OS=Homo sapiens OX=9606 GN=IGHA1 PE=1 SV=2 | 255 | 20201119_15432_1ng_glyco_hcd30_etd30_compound15.18857.18857.4 | controllerType=0 controllerNumber=1 scan=18857 | 42.9071 |
| 15914:4 | 1 | R.LSLHRPALEDLLLGSEAN[+1751.624]LTC[+57.021]TLTGLR.D | HexNAc(4)Hex(4)NeuAc(1) | C[+57], N[+1752] | 1179.566 | 4 | 4715.242 | 4715.2225 | 0 | 4.1 | 127 | Specific | 737.51 | 718.36 | 718.36 | 11.87 | 36 | >sp\|P01876\|IGHA1_HUMAN Immunoglobulin heavy constant alpha 1 OS=Homo sapiens OX=9606 GN=IGHA1 PE=1 SV=2 | 255 | 20201119_15432_1ng_glyco_hcd30_etd30_compound15.19565.19565.4 | controllerType=0 controllerNumber=1 scan=19565 | 44.9498 |
| 15241:4 | 1 | R.LSLHRPALEDLLLGSEAN[+1524.534]LTC[+57.021]TLTGLR.D | HexNAc(2)Hex(6)Fuc(1) | C[+57], N[+1525] | 1122.785 | 4 | 4488.118 | 4488.1319 | 0 | -3.1 | 127 | Specific | 755.81 | 755.81 | 755.81 | 11.18 | 36 | >sp\|P01876\|IGHA1_HUMAN Immunoglobulin heavy constant alpha 1 OS=Homo sapiens OX=9606 GN=IGHA1 PE=1 SV=2 | 255 | 20201119_15432_1ng_glyco_hcd30_etd30_compound15.18855.18855.4 | controllerType=0 controllerNumber=1 scan=18855 | 42.8964 |
| 15236:4 | 1 | R.LSLHRPALEDLLLGSEAN[+1298.476]LTC[+57.021]TLTGLR.D | HexNAc(4)Hex(3) | C[+57], N[+1298] | 1066.2774 | 4 | 4262.0877 | 4262.0742 | 0 | 3.2 | 127 | Specific | 872.59 | 872.59 | 872.59 | 14.87 | 36 | >sp\|P01876\|IGHA1_HUMAN Immunoglobulin heavy constant alpha 1 OS=Homo sapiens OX=9606 GN=IGHA1 PE=1 SV=2 | 255 | 20201119_15432_1ng_glyco_hcd30_etd30_compound15.18850.18850.4 | controllerType=0 controllerNumber=1 scan=18850 | 42.8846 |
| 15215:4 | 1 | R.LSLHRPALEDLLLGSEAN[+1622.582]LTC[+57.021]TLTGLR.D | HexNAc(4)Hex(5) | C[+57], N[+1623] | 1147.3036 | 4 | 4586.1927 | 4586.1799 | 0 | 2.8 | 127 | Specific | 882.32 | 800.94 | 800.94 | 14.39 | 36 | >sp\|P01876\|IGHA1_HUMAN Immunoglobulin heavy constant alpha 1 OS=Homo sapiens OX=9606 GN=IGHA1 PE=1 SV=2 | 255 | 20201119_15432_1ng_glyco_hcd30_etd30_compound15.18828.18828.4 | controllerType=0 controllerNumber=1 scan=18828 | 42.8105 |
| 15195:4 | 1 | R.LSLHRPALEDLLLGSEAN[+1663.608]LTC[+57.021]TLTGLR.D | HexNAc(5)Hex(4) | C[+57], N[+1664] | 1157.5611 | 4 | 4627.2225 | 4627.2064 | 0 | 3.5 | 127 | Specific | 888.17 | 888.17 | 888.17 | 14.87 | 36 | >sp\|P01876\|IGHA1_HUMAN Immunoglobulin heavy constant alpha 1 OS=Homo sapiens OX=9606 GN=IGHA1 PE=1 SV=2 | 255 | 20201119_15432_1ng_glyco_hcd30_etd30_compound15.18807.18807.4 | controllerType=0 controllerNumber=1 scan=18807 | 42.7459 |
| 15201:4 | 1 | R.LSLHRPALEDLLLGSEAN[+1501.555]LTC[+57.021]TLTGLR.D | HexNAc(5)Hex(3) | C[+57], N[+1502] | 1117.0478 | 4 | 4465.1693 | 4465.1536 | 0 | 3.5 | 127 | Specific | 906.89 | 906.89 | 906.89 | 15.87 | 36 | >sp\|P01876\|IGHA1_HUMAN Immunoglobulin heavy constant alpha 1 OS=Homo sapiens OX=9606 GN=IGHA1 PE=1 SV=2 | 255 | 20201119_15432_1ng_glyco_hcd30_etd30_compound15.18813.18813.4 | controllerType=0 controllerNumber=1 scan=18813 | 42.7678 |
| 15181:4 | 1 | R.LSLHRPALEDLLLGSEAN[+1378.476]LTC[+57.021]TLTGLR.D | HexNAc(2)Hex(6) | C[+57], N[+1378] | 1086.2787 | 4 | 4342.0931 | 4342.074 | 0 | 4.4 | 127 | Specific | 942.22 | 912.93 | 912.93 | 15.22 | 36 | >sp\|P01876\|IGHA1_HUMAN Immunoglobulin heavy constant alpha 1 OS=Homo sapiens OX=9606 GN=IGHA1 PE=1 SV=2 | 255 | 20201119_15432_1ng_glyco_hcd30_etd30_compound15.18792.18792.4 | controllerType=0 controllerNumber=1 scan=18792 | 42.6926 |
| 15135:4 | 1 | R.LSLHRPALEDLLLGSEAN[+1702.581]LTC[+57.021]TLTGLR.D | HexNAc(2)Hex(8) | C[+57], N[+1703] | 1167.3041 | 4 | 4666.1947 | 4666.1796 | 0 | 3.2 | 127 | Specific | 990.73 | 935.16 | 935.16 | 16.87 | 36 | >sp\|P01876\|IGHA1_HUMAN Immunoglobulin heavy constant alpha 1 OS=Homo sapiens OX=9606 GN=IGHA1 PE=1 SV=2 | 255 | 20201119_15432_1ng_glyco_hcd30_etd30_compound15.18743.18743.4 | controllerType=0 controllerNumber=1 scan=18743 | 42.5216 |
| 15134:4 | 1 | R.LSLHRPALEDLLLGSEAN[+1864.634]LTC[+57.021]TLTGLR.D | HexNAc(2)Hex(9) | C[+57], N[+1865] | 1207.8179 | 4 | 4828.2498 | 4828.2324 | 0 | 3.6 | 127 | Specific | 998.71 | 948.62 | 948.62 | 16.87 | 36 | >sp\|P01876\|IGHA1_HUMAN Immunoglobulin heavy constant alpha 1 OS=Homo sapiens OX=9606 GN=IGHA1 PE=1 SV=2 | 255 | 20201119_15432_1ng_glyco_hcd30_etd30_compound15.18742.18742.4 | controllerType=0 controllerNumber=1 scan=18742 | 42.5172 |
| 15219:4 | 1 | R.LSLHRPALEDLLLGSEAN[+1460.529]LTC[+57.021]TLTGLR.D | HexNAc(4)Hex(4) | C[+57], N[+1461] | 1106.7944 | 4 | 4424.1556 | 4424.1271 | 0 | 6.5 | 127 | Specific | 1054.15 | 953.8 | 953.8 | 17.36 | 36 | >sp\|P01876\|IGHA1_HUMAN Immunoglobulin heavy constant alpha 1 OS=Homo sapiens OX=9606 GN=IGHA1 PE=1 SV=2 | 255 | 20201119_15432_1ng_glyco_hcd30_etd30_compound15.18832.18832.4 | controllerType=0 controllerNumber=1 scan=18832 | 42.8195 |
| 15154:4 | 1 | R.LSLHRPALEDLLLGSEAN[+1540.529]LTC[+57.021]TLTGLR.D | HexNAc(2)Hex(7) | C[+57], N[+1541] | 1126.79 | 4 | 4504.138 | 4504.1268 | 0 | 2.5 | 127 | Specific | 1076.88 | 972.38 | 972.38 | 18.39 | 36 | >sp\|P01876\|IGHA1_HUMAN Immunoglobulin heavy constant alpha 1 OS=Homo sapiens OX=9606 GN=IGHA1 PE=1 SV=2 | 255 | 20201119_15432_1ng_glyco_hcd30_etd30_compound15.18763.18763.4 | controllerType=0 controllerNumber=1 scan=18763 | 42.5908 |
| 11649:3 | 1 | R.LAGKPTHVN[+1913.677]VSVVMAEVDGTC[+57.021]Y.- | HexNAc(4)Hex(5)NeuAc(1) | C[+57], N[+1914] | 1420.9506 | 3 | 4260.8373 | 4260.8191 | 0 | 4.3 | 332 | CRagged | 216.85 | 100.71 | 100.71 | 3.54 | 36 | >sp\|P01876\|IGHA1_HUMAN Immunoglobulin heavy constant alpha 1 OS=Homo sapiens OX=9606 GN=IGHA1 PE=1 SV=2 | 255 | 20201119_15432_1ng_glyco_hcd30_etd30_compound15.15051.15051.3 | controllerType=0 controllerNumber=1 scan=15051 | 33.8315 |
| 10234:3 | 1 | R.LAGKPTHVN[+1095.397]VSVVMAEVDGTC[+57.021]Y.- | HexNAc(3)Hex(3) | C[+57], N[+1095] | 1148.1881 | 3 | 3442.5499 | 3442.5387 | 0 | 3.3 | 332 | CRagged | 222.21 | 168.74 | 168.74 | 3.72 | 36 | >sp\|P01876\|IGHA1_HUMAN Immunoglobulin heavy constant alpha 1 OS=Homo sapiens OX=9606 GN=IGHA1 PE=1 SV=2 | 255 | 20201119_15432_1ng_glyco_hcd30_etd30_compound15.13553.13553.3 | controllerType=0 controllerNumber=1 scan=13553 | 31.002 |

Table S4. Cont.

| Query #:z | Protein Rank | Glycopeptide | Glycan | Modification Type(s) | Observed m/z | z | Observed (M+H) | Calc. mass (M+H) | Off-by-x error | Mass error (ppm) | StartPosition | Cleavage | Score | Delta | Delta Mod | \|Log Prob\| | # of unique peptides | Protein Name | Protein DB number | Comment | Scan # | Scan Time |
|---|---|---|---|---|---|---|---|---|---|---|---|---|---|---|---|---|---|---|---|---|---|---|
| 10117:3 | 1 | R.LAGKPTHVN[+1540.529]VSVVMAEVDGTC[+57.021]Y.- | HexNAc(2)Hex(7) | C[+57], N[+1541] | 1296.564 | 3 | 3887.6775 | 3887.6706 | 0 | 1.8 | 332 | CRagged | 238.17 | 174.25 | 174.25 | 3.28 | 36 | >sp\|P01876\|IGHA1_HUMAN Immunoglobulin heavy constant alpha 1 OS=Homo sapiens OX=9606 GN=IGHA1 PE=1 SV=2 | 255 | 20201119_15432_1ng_glyco_hcd30_etd30_compound15.13430.13430.3 | controllerType=0 controllerNumber=1 scan=13430 | 30.6641 |
| 10134:3 | 1 | R.LAGKPTHVN[+1378.476]VSVVMAEVDGTC[+57.021]Y.- | HexNAc(2)Hex(6) | C[+57], N[+1378] | 1242.5493 | 3 | 3725.6332 | 3725.6178 | 0 | 4.1 | 332 | CRagged | 239.19 | 239.19 | 195.88 | 3.97 | 36 | >sp\|P01876\|IGHA1_HUMAN Immunoglobulin heavy constant alpha 1 OS=Homo sapiens OX=9606 GN=IGHA1 PE=1 SV=2 | 255 | 20201119_15432_1ng_glyco_hcd30_etd30_compound15.13448.13448.3 | controllerType=0 controllerNumber=1 scan=13448 | 30.722 |
| 10209:3 | 1 | R.LAGKPTHVN[+1501.555]VSVVMAEVDGTC[+57.021]Y.- | HexNAc(5)Hex(3) | C[+57], N[+1502] | 1283.5734 | 3 | 3848.7057 | 3848.6974 | 0 | 2.2 | 332 | CRagged | 252.07 | 166.04 | 166.04 | 3.78 | 36 | >sp\|P01876\|IGHA1_HUMAN Immunoglobulin heavy constant alpha 1 OS=Homo sapiens OX=9606 GN=IGHA1 PE=1 SV=2 | 255 | 20201119_15432_1ng_glyco_hcd30_etd30_compound15.13527.13527.3 | controllerType=0 controllerNumber=1 scan=13527 | 30.9319 |
| 10076:3 | 1 | R.LAGKPTHVN[+1702.581]VSVVMAEVDGTC[+57.021]Y.- | HexNAc(2)Hex(8) | C[+57], N[+1703] | 1350.5829 | 3 | 4049.7343 | 4049.7234 | 0 | 2.7 | 332 | CRagged | 264.74 | 253.81 | 197.46 | 4.22 | 36 | >sp\|P01876\|IGHA1_HUMAN Immunoglobulin heavy constant alpha 1 OS=Homo sapiens OX=9606 GN=IGHA1 PE=1 SV=2 | 255 | 20201119_15432_1ng_glyco_hcd30_etd30_compound15.13387.13387.3 | controllerType=0 controllerNumber=1 scan=13387 | 30.5631 |
| 10150:3 | 1 | R.LAGKPTHVN[+1216.423]VSVVMAEVDGTC[+57.021]Y.- | HexNAc(2)Hex(5) | C[+57], N[+1216] | 1188.5308 | 3 | 3563.5779 | 3563.5649 | 0 | 3.6 | 332 | CRagged | 279.8 | 248.02 | 213.81 | 4.7 | 36 | >sp\|P01876\|IGHA1_HUMAN Immunoglobulin heavy constant alpha 1 OS=Homo sapiens OX=9606 GN=IGHA1 PE=1 SV=2 | 255 | 20201119_15432_1ng_glyco_hcd30_etd30_compound15.13465.13465.3 | controllerType=0 controllerNumber=1 scan=13465 | 30.7761 |
| 10190:3 | 1 | R.LAGKPTHVN[+1647.613]VSVVMAEVDGTC[+57.021]Y.- | HexNAc(5)Hex(3)Fuc(1) | C[+57], N[+1648] | 1332.2601 | 3 | 3994.7656 | 3994.7553 | 0 | 2.6 | 332 | CRagged | 299.69 | 158.15 | 158.15 | 3.89 | 36 | >sp\|P01876\|IGHA1_HUMAN Immunoglobulin heavy constant alpha 1 OS=Homo sapiens OX=9606 GN=IGHA1 PE=1 SV=2 | 255 | 20201119_15432_1ng_glyco_hcd30_etd30_compound15.13507.13507.3 | controllerType=0 controllerNumber=1 scan=13507 | 30.8794 |
| 13215:4 | 1 | R.LAGKPTHVN[+2553.910]VSVVMAEVDGTC[+57.021]Y.- | HexNAc(5)Hex(5)Fuc(1)NeuAc(2) | C[+57], N[+2554] | 1226.0226 | 4 | 4901.0687 | 4901.0518 | 0 | 3.4 | 332 | CRagged | 310.47 | 244.16 | 244.16 | 4.24 | 36 | >sp\|P01876\|IGHA1_HUMAN Immunoglobulin heavy constant alpha 1 OS=Homo sapiens OX=9606 GN=IGHA1 PE=1 SV=2 | 255 | 20201119_15432_1ng_glyco_hcd30_etd30_compound15.16690.16690.4 | controllerType=0 controllerNumber=1 scan=16690 | 37.4921 |
| 11668:4 | 1 | R.LAGKPTHVN[+2262.814]VSVVMAEVDGTC[+57.021]Y.- | HexNAc(5)Hex(5)Fuc(1)NeuAc(1) | C[+57], N[+2263] | 1153.2492 | 4 | 4609.9749 | 4609.9564 | 0 | 4 | 332 | CRagged | 455.63 | 299.5 | 299.5 | 4.7 | 36 | >sp\|P01876\|IGHA1_HUMAN Immunoglobulin heavy constant alpha 1 OS=Homo sapiens OX=9606 GN=IGHA1 PE=1 SV=2 | 255 | 20201119_15432_1ng_glyco_hcd30_etd30_compound15.15071.15071.4 | controllerType=0 controllerNumber=1 scan=15071 | 33.8942 |
| 14761:4 | 2 | R.[+42.011]RTAGWNVPIGTLRPFLN[+1872.651]WTGPPEPIEAAVAR.F | HexNAc(3)Hex(6)NeuAc(1) | .[42], N[+1873] | 1326.1354 | 4 | 5301.5197 | 5301.4731 | 0 | 8.8 | 140 | Specific | 203.88 | 184.03 | 184.03 | 2.68 | 32 | >sp\|P02788\|TRFL_HUMAN Lactotransferrin OS=Homo sapiens OX=9606 GN=LTF PE=1 SV=6 | 1270 | 20201119_15432_1ng_glyco_hcd30_etd30_compound15.18348.18348.4 | controllerType=0 controllerNumber=1 scan=18348 | 41.3999 |
| 15441:5 | 2 | R.RTAGWNVPIGTLRPFLN[+2205.793]WTGPPEPIEAAVAR.F | HexNAc(4)Hex(5)Fuc(2)NeuAc(1) | N[+2206] | 1119.3322 | 5 | 5592.6317 | 5592.6049 | 0 | 4.8 | 140 | Specific | 952.52 | 889.89 | 889.89 | 15.62 | 32 | >sp\|P02788\|TRFL_HUMAN Lactotransferrin OS=Homo sapiens OX=9606 GN=LTF PE=1 SV=6 | 1270 | 20201119_15432_1ng_glyco_hcd30_etd30_compound15.19066.19066.2 | controllerType=0 controllerNumber=1 scan=19066 | 43.5188 |
| 14769:4 | 2 | R.RTAGWNVPIGTLRPFLN[+1768.640]WTGPPEPIEAAVAR.F | HexNAc(4)Hex(5)Fuc(1) | N[+1769] | 1289.6211 | 4 | 5155.4626 | 5155.4516 | 0 | 2.1 | 140 | Specific | 973.87 | 973.87 | 973.87 | 16.39 | 32 | >sp\|P02788\|TRFL_HUMAN Lactotransferrin OS=Homo sapiens OX=9606 GN=LTF PE=1 SV=6 | 1270 | 20201119_15432_1ng_glyco_hcd30_etd30_compound15.18356.18356.4 | controllerType=0 controllerNumber=1 scan=18356 | 41.4266 |
| 16277:3 | 2 | R.TAGWNVPIGTLRPFLN[+2570.925]WTGPPEPIEAAVAR.F | HexNAc(5)Hex(6)Fuc(2)NeuAc(1) | N[+2571] | 1934.5579 | 3 | 5801.6593 | 5801.636 | 0 | 4 | 141 | Specific | 264.94 | 179.25 | 179.25 | 4.94 | 32 | >sp\|P02788\|TRFL_HUMAN Lactotransferrin OS=Homo sapiens OX=9606 GN=LTF PE=1 SV=6 | 1270 | 20201119_15432_1ng_glyco_hcd30_etd30_compound15.19950.19950.3 | controllerType=0 controllerNumber=1 scan=19950 | 46.1547 |
| 16323:3 | 2 | R.TAGWNVPIGTLRPFLN[+2205.793]WTGPPEPIEAAVAR.F | HexNAc(4)Hex(5)Fuc(2)NeuAc(1) | N[+2206] | 1812.8479 | 3 | 5436.529 | 5436.5038 | 0 | 4.6 | 141 | Specific | 332.93 | 218.99 | 218.99 | 4.75 | 32 | >sp\|P02788\|TRFL_HUMAN Lactotransferrin OS=Homo sapiens OX=9606 GN=LTF PE=1 SV=6 | 1270 | 20201119_15432_1ng_glyco_hcd30_etd30_compound15.19998.19998.2 | controllerType=0 controllerNumber=1 scan=19998 | 46.2942 |
| 17308:3 | 2 | R.TAGWNVPIGTLRPFLN[+2350.830]WTGPPEPIEAAVAR.F | HexNAc(4)Hex(5)Fuc(1)NeuAc(2) | N[+2351] | 1861.1915 | 3 | 5581.5599 | 5581.5413 | 0 | 3.3 | 141 | Specific | 366.08 | 290.17 | 290.17 | 5.34 | 32 | >sp\|P02788\|TRFL_HUMAN Lactotransferrin OS=Homo sapiens OX=9606 GN=LTF PE=1 SV=6 | 1270 | 20201119_15432_1ng_glyco_hcd30_etd30_compound15.21040.21040.3 | controllerType=0 controllerNumber=1 scan=21040 | 49.1674 |
| 15636:3 | 2 | R.TAGWNVPIGTLRPFLN[+1768.640]WTGPPEPIEAAVAR.F | HexNAc(4)Hex(5)Fuc(1) | N[+1769] | 1667.1296 | 3 | 4999.3743 | 4999.3504 | 0 | 4.8 | 141 | Specific | 385.04 | 333.36 | 333.36 | 4.92 | 32 | >sp\|P02788\|TRFL_HUMAN Lactotransferrin OS=Homo sapiens OX=9606 GN=LTF PE=1 SV=6 | 1270 | 20201119_15432_1ng_glyco_hcd30_etd30_compound15.19272.19272.3 | controllerType=0 controllerNumber=1 scan=19272 | 44.0994 |
| 16299:5 | 2 | R.TAGWNVPIGTLRPFLN[+2205.793]WTGPPEPIEAAVAR.F | HexNAc(4)Hex(5)Fuc(2)NeuAc(1) | N[+2206] | 1088.1096 | 5 | 5436.519 | 5436.5038 | 0 | 2.8 | 141 | Specific | 406.08 | 406.08 | 406.08 | 5.56 | 32 | >sp\|P02788\|TRFL_HUMAN Lactotransferrin OS=Homo sapiens OX=9606 GN=LTF PE=1 SV=6 | 1270 | 20201119_15432_1ng_glyco_hcd30_etd30_compound15.19973.19973.4 | controllerType=0 controllerNumber=1 scan=19973 | 46.2131 |
| 16371:3 | 2 | R.TAGWNVPIGTLRPFLN[+2059.735]WTGPPEPIEAAVAR.F | HexNAc(4)Hex(5)Fuc(1)NeuAc(1) | N[+2060] | 1764.1595 | 3 | 5290.464 | 5290.4459 | 0 | 3.4 | 141 | Specific | 478.22 | 359.14 | 359.14 | 6.61 | 32 | >sp\|P02788\|TRFL_HUMAN Lactotransferrin OS=Homo sapiens OX=9606 GN=LTF PE=1 SV=6 | 1270 | 20201119_15432_1ng_glyco_hcd30_etd30_compound15.20049.20049.2 | controllerType=0 controllerNumber=1 scan=20049 | 46.427 |
| 16272:4 | 2 | R.TAGWNVPIGTLRPFLN[+2570.925]WTGPPEPIEAAVAR.F | HexNAc(5)Hex(6)Fuc(2)NeuAc(1) | N[+2571] | 1451.1702 | 4 | 5801.6588 | 5801.636 | 0 | 3.9 | 141 | Specific | 551 | 551 | 551 | 7.23 | 32 | >sp\|P02788\|TRFL_HUMAN Lactotransferrin OS=Homo sapiens OX=9606 GN=LTF PE=1 SV=6 | 1270 | 20201119_15432_1ng_glyco_hcd30_etd30_compound15.19945.19945.4 | controllerType=0 controllerNumber=1 scan=19945 | 46.1407 |
| 17305:4 | 2 | R.TAGWNVPIGTLRPFLN[+2350.830]WTGPPEPIEAAVAR.F | HexNAc(4)Hex(5)Fuc(1)NeuAc(2) | N[+2351] | 1396.1443 | 4 | 5581.5553 | 5581.5413 | 0 | 2.5 | 141 | Specific | 720.64 | 720.64 | 720.64 | 10.79 | 32 | >sp\|P02788\|TRFL_HUMAN Lactotransferrin OS=Homo sapiens OX=9606 GN=LTF PE=1 SV=6 | 1270 | 20201119_15432_1ng_glyco_hcd30_etd30_compound15.21037.21037.4 | controllerType=0 controllerNumber=1 scan=21037 | 49.1603 |
| 15635:4 | 2 | R.TAGWNVPIGTLRPFLN[+1768.640]WTGPPEPIEAAVAR.F | HexNAc(4)Hex(5)Fuc(1) | N[+1769] | 1250.5999 | 4 | 4999.3776 | 4999.3504 | 0 | 5.4 | 141 | Specific | 971.54 | 894.99 | 894.99 | 15.62 | 32 | >sp\|P02788\|TRFL_HUMAN Lactotransferrin OS=Homo sapiens OX=9606 GN=LTF PE=1 SV=6 | 1270 | 20201119_15432_1ng_glyco_hcd30_etd30_compound15.19271.19271.4 | controllerType=0 controllerNumber=1 scan=19271 | 44.096 |
| 14765:3 | 2 | R.TAGWNIPMGLLFN[+1460.529]QTGSC[+57.021]K.F | HexNAc(4)Hex(4) | C[+57], N[+1461] | 1185.8535 | 3 | 3555.5458 | 3555.5387 | 0 | 2 | 485 | Specific | 207.23 | 147.24 | 142.32 | 2.57 | 32 | >sp\|P02788\|TRFL_HUMAN Lactotransferrin OS=Homo sapiens OX=9606 GN=LTF PE=1 SV=6 | 1270 | 20201119_15432_1ng_glyco_hcd30_etd30_compound15.18352.18352.3 | controllerType=0 controllerNumber=1 scan=18352 | 41.4089 |
| 14643:3 | 2 | R.TAGWNIPMGLLFN[+1768.640]QTGSC[+57.021]K.F | HexNAc(4)Hex(5)Fuc(1) | C[+57], N[+1769] | 1288.5604 | 3 | 3863.6665 | 3863.6495 | 0 | 4.4 | 485 | Specific | 324.49 | 234.18 | 234.18 | 4.51 | 32 | >sp\|P02788\|TRFL_HUMAN Lactotransferrin OS=Homo sapiens OX=9606 GN=LTF PE=1 SV=6 | 1270 | 20201119_15432_1ng_glyco_hcd30_etd30_compound15.18223.18223.3 | controllerType=0 controllerNumber=1 scan=18223 | 41.0285 |
| 14799:3 | 2 | R.TAGWNIPMGLLFN[+1216.423]QTGSC[+57.021]K.F | HexNAc(2)Hex(5) | C[+57], N[+1216] | 1104.4876 | 3 | 3311.4482 | 3311.4328 | 0 | 4.6 | 485 | Specific | 334.69 | 314.35 | 199.27 | 3.4 | 32 | >sp\|P02788\|TRFL_HUMAN Lactotransferrin OS=Homo sapiens OX=9606 GN=LTF PE=1 SV=6 | 1270 | 20201119_15432_1ng_glyco_hcd30_etd30_compound15.18388.18388.3 | controllerType=0 controllerNumber=1 scan=18388 | 41.5205 |
| 14677:3 | 2 | R.TAGWNIPMGLLFN[+1622.582]QTGSC[+57.021]K.F | HexNAc(4)Hex(5) | C[+57], N[+1623] | 1239.8727 | 3 | 3717.6037 | 3717.5916 | 0 | 3.3 | 485 | Specific | 343.2 | 252.7 | 243.43 | 4.51 | 32 | >sp\|P02788\|TRFL_HUMAN Lactotransferrin OS=Homo sapiens OX=9606 GN=LTF PE=1 SV=6 | 1270 | 20201119_15432_1ng_glyco_hcd30_etd30_compound15.18259.18259.3 | controllerType=0 controllerNumber=1 scan=18259 | 41.1373 |
| 15852:3 | 2 | R.TAGWNIPMGLLFN[+2205.793]QTGSC[+57.021]K.F | HexNAc(4)Hex(5)Fuc(2)NeuAc(1) | C[+57], N[+2206] | 1434.2772 | 3 | 4300.8169 | 4300.8028 | 0 | 3.3 | 485 | Specific | 377.92 | 271.86 | 271.86 | 4.32 | 32 | >sp\|P02788\|TRFL_HUMAN Lactotransferrin OS=Homo sapiens OX=9606 GN=LTF PE=1 SV=6 | 1270 | 20201119_15432_1ng_glyco_hcd30_etd30_compound15.19500.19500.3 | controllerType=0 controllerNumber=1 scan=19500 | 44.7559 |
| 16066:3 | 2 | R.TAGWNIPMGLLFN[+1751.624]QTGSC[+57.021]K.F | HexNAc(4)Hex(4)NeuAc(1) | C[+57], N[+1752] | 1282.8859 | 3 | 3846.6432 | 3846.6341 | 0 | 2.4 | 485 | Specific | 378.21 | 257.31 | 257.31 | 3.85 | 32 | >sp\|P02788\|TRFL_HUMAN Lactotransferrin OS=Homo sapiens OX=9606 GN=LTF PE=1 SV=6 | 1270 | 20201119_15432_1ng_glyco_hcd30_etd30_compound15.19727.19727.3 | controllerType=0 controllerNumber=1 scan=19727 | 45.5234 |
| 14651:3 | 2 | R.TAGWNIPMGLLFN[+1622.582]QTGSC[+57.021]K.F | HexNAc(4)Hex(5) | C[+57], N[+1623] | 1239.874 | 3 | 3717.6073 | 3717.5916 | 0 | 4.2 | 485 | Specific | 393.76 | 307.21 | 269.66 | 4.61 | 32 | >sp\|P02788\|TRFL_HUMAN Lactotransferrin OS=Homo sapiens OX=9606 GN=LTF PE=1 SV=6 | 1270 | 20201119_15432_1ng_glyco_hcd30_etd30_compound15.18231.18231.3 | controllerType=0 controllerNumber=1 scan=18231 | 41.0538 |
| 14601:3 | 2 | R.TAGWNIPMGLLFN[+1606.587]QTGSC[+57.021]K.F | HexNAc(4)Hex(4)Fuc(1) | C[+57], N[+1607] | 1234.5405 | 3 | 3701.6068 | 3701.5966 | 0 | 2.8 | 485 | Specific | 393.89 | 301.93 | 301.93 | 3.73 | 32 | >sp\|P02788\|TRFL_HUMAN Lactotransferrin OS=Homo sapiens OX=9606 GN=LTF PE=1 SV=6 | 1270 | 20201119_15432_1ng_glyco_hcd30_etd30_compound15.18179.18179.3 | controllerType=0 controllerNumber=1 scan=18179 | 40.9097 |
| 14509:3 | 2 | R.TAGWNIPMGLLFN[+1768.640]QTGSC[+57.021]K.F | HexNAc(4)Hex(5)Fuc(1) | C[+57], N[+1769] | 1288.5591 | 3 | 3863.6629 | 3863.6495 | 0 | 3.5 | 485 | Specific | 408.48 | 272.53 | 272.53 | 4.52 | 32 | >sp\|P02788\|TRFL_HUMAN Lactotransferrin OS=Homo sapiens OX=9606 GN=LTF PE=1 SV=6 | 1270 | 20201119_15432_1ng_glyco_hcd30_etd30_compound15.18082.18082.3 | controllerType=0 controllerNumber=1 scan=18082 | 40.6349 |
| 15885:3 | 2 | R.TAGWNIPMGLLFN[+2059.735]QTGSC[+57.021]K.F | HexNAc(4)Hex(5)Fuc(1)NeuAc(1) | C[+57], N[+2060] | 1385.5911 | 3 | 4154.7588 | 4154.7449 | 0 | 3.4 | 485 | Specific | 438.99 | 314.5 | 314.5 | 4.8 | 32 | >sp\|P02788\|TRFL_HUMAN Lactotransferrin OS=Homo sapiens OX=9606 GN=LTF PE=1 SV=6 | 1270 | 20201119_15432_1ng_glyco_hcd30_etd30_compound15.19535.19535.3 | controllerType=0 controllerNumber=1 scan=19535 | 44.8666 |
| 15991:3 | 2 | R.TAGWNIPMGLLFN[+1913.677]QTGSC[+57.021]K.F | HexNAc(4)Hex(5)NeuAc(1) | C[+57], N[+1914] | 1336.9045 | 3 | 4008.6989 | 4008.687 | 0 | 3 | 485 | Specific | 450.86 | 290.53 | 290.53 | 4.24 | 32 | >sp\|P02788\|TRFL_HUMAN Lactotransferrin OS=Homo sapiens OX=9606 GN=LTF PE=1 SV=6 | 1270 | 20201119_15432_1ng_glyco_hcd30_etd30_compound15.19647.19647.3 | controllerType=0 controllerNumber=1 scan=19647 | 45.2512 |
| 17398:3 | 2 | R.TAGWNIPMGLLFN[+2350.830]QTGSC[+57.021]K.F | HexNAc(4)Hex(5)Fuc(1)NeuAc(2) | C[+57], N[+2351] | 1482.6243 | 3 | 4445.8584 | 4445.8403 | 0 | 4.1 | 485 | Specific | 472.65 | 316.84 | 316.84 | 5.01 | 32 | >sp\|P02788\|TRFL_HUMAN Lactotransferrin OS=Homo sapiens OX=9606 GN=LTF PE=1 SV=6 | 1270 | 20201119_15432_1ng_glyco_hcd30_etd30_compound15.21135.21135.3 | controllerType=0 controllerNumber=1 scan=21135 | 49.4539 |
| 15357:3 | 2 | R.TAGWNIPMGLLFN[+1768.640]QTGSC[+57.021]KFDEYFSQSC[+57.021]APGSDPR.S | HexNAc(4)Hex(5)Fuc(1) | C[+57]*2, N[+1769] | 1903.1438 | 3 | 5707.4167 | 5707.3966 | 0 | 3.5 | 485 | Specific | 280.81 | 231.61 | 231.61 | 5.24 | 32 | >sp\|P02788\|TRFL_HUMAN Lactotransferrin OS=Homo sapiens OX=9606 GN=LTF PE=1 SV=6 | 1270 | 20201119_15432_1ng_glyco_hcd30_etd30_compound15.18978.18978.3 | controllerType=0 controllerNumber=1 scan=18978 | 43.2753 |
| 15352:4 | 2 | R.TAGWNIPMGLLFN[+1768.640]QTGSC[+57.021]KFDEYFSQSC[+57.021]APGSDPR.S | HexNAc(4)Hex(5)Fuc(1) | C[+57]*2, N[+1769] | 1427.6119 | 4 | 5707.4259 | 5707.3966 | 0 | 5.1 | 485 | Specific | 611.95 | 544.77 | 544.77 | 7.5 | 32 | >sp\|P02788\|TRFL_HUMAN Lactotransferrin OS=Homo sapiens OX=9606 GN=LTF PE=1 SV=6 | 1270 | 20201119_15432_1ng_glyco_hcd30_etd30_compound15.18973.18973.4 | controllerType=0 controllerNumber=1 scan=18973 | 43.2665 |
| 16065:5 | 2 | R.TAGWNIPMGLLFN[+2059.735]QTGSC[+57.021]KFDEYFSQSC[+57.021]APGSDPR.S | HexNAc(4)Hex(5)Fuc(1)NeuAc(1) | C[+57]*2, N[+2060] | 1200.5098 | 5 | 5998.5197 | 5998.492 | 0 | 4.6 | 485 | Specific | 638.8 | 548.16 | 548.16 | 7.5 | 32 | >sp\|P02788\|TRFL_HUMAN Lactotransferrin OS=Homo sapiens OX=9606 GN=LTF PE=1 SV=6 | 1270 | 20201119_15432_1ng_glyco_hcd30_etd30_compound15.19726.19726.5 | controllerType=0 controllerNumber=1 scan=19726 | 45.5186 |
| 16053:4 | 2 | R.TAGWNIPMGLLFN[+2059.735]QTGSC[+57.021]KFDEYFSQSC[+57.021]APGSDPR.S | HexNAc(4)Hex(5)Fuc(1)NeuAc(1) | C[+57]*2, N[+2060] | 1500.3848 | 4 | 5998.5172 | 5998.492 | 0 | 4.2 | 485 | Specific | 841.7 | 771.92 | 771.92 | 12.56 | 32 | >sp\|P02788\|TRFL_HUMAN Lactotransferrin OS=Homo sapiens OX=9606 GN=LTF PE=1 SV=6 | 1270 | 20201119_15432_1ng_glyco_hcd30_etd30_compound15.19714.19714.4 | controllerType=0 controllerNumber=1 scan=19714 | 45.4787 |
| 16048:4 | 2 | R.TAGWNIPMGLLFN[+2205.793]QTGSC[+57.021]KFDEYFSQSC[+57.021]APGSDPR.S | HexNAc(4)Hex(5)Fuc(2)NeuAc(1) | C[+57]*2, N[+2206] | 1536.8977 | 4 | 6144.569 | 6144.5499 | 0 | 3.1 | 485 | Specific | 866.35 | 791.17 | 791.17 | 13.56 | 32 | >sp\|P02788\|TRFL_HUMAN Lactotransferrin OS=Homo sapiens OX=9606 GN=LTF PE=1 SV=6 | 1270 | 20201119_15432_1ng_glyco_hcd30_etd30_compound15.19707.19707.4 | controllerType=0 controllerNumber=1 scan=19707 | 45.4559 |
| 08152:3 | 2 | R.N[+2205.793]GSDC[+57.021]PDKFC[+57.021]LFQSETK.N | HexNAc(4)Hex(5)Fuc(2)NeuAc(1) | C[+57]*2, N[+2206] | 1413.5624 | 3 | 4238.6728 | 4238.6667 | 0 | 1.4 | 642 | Specific | 294.44 | 207.12 | 207.12 | 4.14 | 32 | >sp\|P02788\|TRFL_HUMAN Lactotransferrin OS=Homo sapiens OX=9606 GN=LTF PE=1 SV=6 | 1270 | 20201119_15432_1ng_glyco_hcd30_etd30_compound15.11386.11386.3 | controllerType=0 controllerNumber=1 scan=11386 | 26.6043 |
| 08147:3 | 2 | R.N[+2205.793]GSDC[+57.021]PDKFC[+57.021]LFQSETK.N | HexNAc(4)Hex(5)Fuc(2)NeuAc(1) | C[+57]*2, N[+2206] | 1413.5656 | 3 | 4238.6823 | 4238.6667 | 0 | 3.7 | 642 | Specific | 328.1 | 221.63 | 221.63 | 5.06 | 32 | >sp\|P02788\|TRFL_HUMAN Lactotransferrin OS=Homo sapiens OX=9606 GN=LTF PE=1 SV=6 | 1270 | 20201119_15432_1ng_glyco_hcd30_etd30_compound15.11380.11380.3 | controllerType=0 controllerNumber=1 scan=11380 | 26.5861 |
| 06628:3 | 4 | R.FSDGLESN[+1864.634]SSTQFEVK.K | HexNAc(2)Hex(9) | N[+1865] | 1213.8199 | 3 | 3639.4451 | 3639.4472 | 0 | -0.6 | 219 | Specific | 213.72 | 213.72 | 213.72 | 2.58 | 14 | >sp\|P0C0L4\|CO4A_HUMAN Complement C4-A OS=Homo sapiens OX=9606 GN=C4A PE=1 SV=2 | 1235 | 20201119_15432_1ng_glyco_hcd30_etd30_compound15.9788.9788.3 | controllerType=0 controllerNumber=1 scan=9788 | 23.5713 |
| 04209:3 | 4 | R.FSDGLESN[+1864.634]SSTQFEVKK.Y | HexNAc(2)Hex(9) | N[+1865] | 1256.5231 | 3 | 3767.5548 | 3767.5421 | 0 | 3.4 | 219 | Specific | 212.15 | 153.64 | 153.64 | 3.55 | 14 | >sp\|P0C0L4\|CO4A_HUMAN Complement C4-A OS=Homo sapiens OX=9606 GN=C4A PE=1 SV=2 | 1235 | 20201119_15432_1ng_glyco_hcd30_etd30_compound15.7284.7284.3 | controllerType=0 controllerNumber=1 scan=7284 | 19.2343 |
| 07654:3 | 5 | K.TAHC[+57.021]N[+2059.735]ESFYFLC[+57.021]K.R | HexNAc(4)Hex(5)Fuc(1)NeuAc(1) | C[+57]*2, N[+2060] | 1246.16 | 3 | 3736.4654 | 3736.4545 | 0 | 2.9 | 1201 | Specific | 240.8 | 135.46 | 135.46 | 3.51 | 12 | >sp\|P22897\|MRC1_HUMAN Macrophage mannose receptor 1 OS=Homo sapiens OX=9606 GN=MRC1 PE=1 SV=1 | 1049 | 20201119_15432_1ng_glyco_hcd30_etd30_compound15.10859.10859.3 | controllerType=0 controllerNumber=1 scan=10859 | 25.557 |
| 07530:3 | 5 | K.TAHC[+57.021]N[+2205.793]ESFYFLC[+57.021]K.R | HexNAc(4)Hex(5)Fuc(2)NeuAc(1) | C[+57]*2, N[+2206] | 1294.8449 | 3 | 3882.5202 | 3882.5124 | 0 | 2 | 1201 | Specific | 245.61 | 166.48 | 166.48 | 3.74 | 12 | >sp\|P22897\|MRC1_HUMAN Macrophage mannose receptor 1 OS=Homo sapiens OX=9606 GN=MRC1 PE=1 SV=1 | 1049 | 20201119_15432_1ng_glyco_hcd30_etd30_compound15.10730.10730.3 | controllerType=0 controllerNumber=1 scan=10730 | 25.2958 |
| 07627:3 | 5 | K.TAHC[+57.021]N[+2059.735]ESFYFLC[+57.021]K.R | HexNAc(4)Hex(5)Fuc(1)NeuAc(1) | C[+57]*2, N[+2060] | 1246.1579 | 3 | 3736.4591 | 3736.4545 | 0 | 1.2 | 1201 | Specific | 289.65 | 197.09 | 197.09 | 3.66 | 12 | >sp\|P22897\|MRC1_HUMAN Macrophage mannose receptor 1 OS=Homo sapiens OX=9606 GN=MRC1 PE=1 SV=1 | 1049 | 20201119_15432_1ng_glyco_hcd30_etd30_compound15.10831.10831.3 | controllerType=0 controllerNumber=1 scan=10831 | 25.501 |
| 16015:4 | 6 | K.QIGLYPVLVIDSSGYVNPN[+2059.735]YTGR.I | HexNAc(4)Hex(5)Fuc(1)NeuAc(1) | N[+2060] | 1147.0208 | 4 | 4585.0613 | 4585.0384 | 0 | 5 | 168 | Specific | 436.46 | 426.86 | 426.86 | 5.1 | 17 | >sp\|P01833\|PIGR_HUMAN Polymeric immunoglobulin receptor OS=Homo sapiens OX=9606 GN=PIGR PE=1 SV=4 | 902 | 20201119_15432_1ng_glyco_hcd30_etd30_compound15.19672.19672.4 | controllerType=0 controllerNumber=1 scan=19672 | 45.3331 |
| 16022:3 | 6 | K.QIGLYPVLVIDSSGYVNPN[+2059.735]YTGR.I | HexNAc(4)Hex(5)Fuc(1)NeuAc(1) | N[+2060] | 1529.0237 | 3 | 4585.0567 | 4585.0384 | 0 | 4 | 168 | Specific | 563.21 | 490.03 | 490.03 | 7.79 | 17 | >sp\|P01833\|PIGR_HUMAN Polymeric immunoglobulin receptor OS=Homo sapiens OX=9606 GN=PIGR PE=1 SV=4 | 902 | 20201119_15432_1ng_glyco_hcd30_etd30_compound15.19680.19680.3 | controllerType=0 controllerNumber=1 scan=19680 | 45.3618 |
| 19071:3 | 6 | R.[+42.011]LSLLEEPGN[+2336.851]GTFTVILNQLTSR.D | HexNAc(6)Hex(6)Fuc(1) | .[42], N[+2337] | 1594.3868 | 3 | 4781.1458 | 4781.1542 | 0 | -1.8 | 413 | Specific | 230.61 | 136.63 | 136.63 | 2.46 | 17 | >sp\|P01833\|PIGR_HUMAN Polymeric immunoglobulin receptor OS=Homo sapiens OX=9606 GN=PIGR PE=1 SV=4 | 902 | 20201119_15432_1ng_glyco_hcd30_etd30_compound15.22890.22890.3 | controllerType=0 controllerNumber=1 scan=22890 | 53.4504 |
| 17225:3 | 6 | R.LSLLEEPGN[+2205.793]GTFTVILNQLTSR.D | HexNAc(4)Hex(5)Fuc(2)NeuAc(1) | N[+2206] | 1536.7084 | 3 | 4608.1107 | 4608.0854 | 0 | 5.5 | 413 | Specific | 619.4 | 524.64 | 524.64 | 8.14 | 17 | >sp\|P01833\|PIGR_HUMAN Polymeric immunoglobulin receptor OS=Homo sapiens OX=9606 GN=PIGR PE=1 SV=4 | 902 | 20201119_15432_1ng_glyco_hcd30_etd30_compound15.20953.20953.3 | controllerType=0 controllerNumber=1 scan=20953 | 48.9255 |
| 13922:3 | 6 | K.IIEGEPNLKVPGN[+2204.772]VTAVLGETLK.V | HexNAc(4)Hex(5)NeuAc(2) | N[+2205] | 1532.7211 | 3 | 4596.1488 | 4596.1218 | 0 | 5.9 | 457 | Specific | 332.35 | 252.71 | 252.71 | 3.3 | 17 | >sp\|P01833\|PIGR_HUMAN Polymeric immunoglobulin receptor OS=Homo sapiens OX=9606 GN=PIGR PE=1 SV=4 | 902 | 20201119_15432_1ng_glyco_hcd30_etd30_compound15.17458.17458.3 | controllerType=0 controllerNumber=1 scan=17458 | 39.1695 |
| 13991:4 | 6 | K.IIEGEPNLKVPGN[+2059.735]VTAVLGETLK.V | HexNAc(4)Hex(5)Fuc(1)NeuAc(1) | N[+2060] | 1113.5287 | 4 | 4451.0931 | 4451.0843 | 0 | 2 | 457 | Specific | 363.84 | 344.75 | 344.75 | 3.95 | 17 | >sp\|P01833\|PIGR_HUMAN Polymeric immunoglobulin receptor OS=Homo sapiens OX=9606 GN=PIGR PE=1 SV=4 | 902 | 20201119_15432_1ng_glyco_hcd30_etd30_compound15.17533.17533.4 | controllerType=0 controllerNumber=1 scan=17533 | 39.3546 |
| 13975:3 | 6 | K.IIEGEPNLKVPGN[+2059.735]VTAVLGETLK.V | HexNAc(4)Hex(5)Fuc(1)NeuAc(1) | N[+2060] | 1484.3705 | 3 | 4451.0971 | 4451.0843 | 0 | 2.9 | 457 | Specific | 366 | 304.29 | 304.29 | 4.39 | 17 | >sp\|P01833\|PIGR_HUMAN Polymeric immunoglobulin receptor OS=Homo sapiens OX=9606 GN=PIGR PE=1 SV=4 | 902 | 20201119_15432_1ng_glyco_hcd30_etd30_compound15.17516.17516.3 | controllerType=0 controllerNumber=1 scan=17516 | 39.3015 |
| 14713:4 | 6 | K.IIEGEPNLKVPGN[+2350.830]VTAVLGETLKVPC[+57.021]HFPC[+57.021]K.F | HexNAc(4)Hex(5)Fuc(1)NeuAc(2) | C[+57]*2, N[+2351] | 1442.6726 | 4 | 5767.6686 | 5767.6372 | 0 | 5.4 | 457 | Specific | 203.3 | 203.3 | 203.3 | 2.5 | 17 | >sp\|P01833\|PIGR_HUMAN Polymeric immunoglobulin receptor OS=Homo sapiens OX=9606 GN=PIGR PE=1 SV=4 | 902 | 20201119_15432_1ng_glyco_hcd30_etd30_compound15.18297.18297.4 | controllerType=0 controllerNumber=1 scan=18297 | 41.252 |
| 13527:4 | 6 | K.IIEGEPNLKVPGN[+2205.793]VTAVLGETLKVPC[+57.021]HFPC[+57.021]K.F | HexNAc(4)Hex(5)Fuc(2)NeuAc(1) | C[+57]*2, N[+2206] | 1406.4074 | 4 | 5622.6076 | 5622.5997 | 0 | 1.4 | 457 | Specific | 492.88 | 492.88 | 492.88 | 4.79 | 17 | >sp\|P01833\|PIGR_HUMAN Polymeric immunoglobulin receptor OS=Homo sapiens OX=9606 GN=PIGR PE=1 SV=4 | 902 | 20201119_15432_1ng_glyco_hcd30_etd30_compound15.17017.17017.4 | controllerType=0 controllerNumber=1 scan=17017 | 38.2455 |
| 11931:3 | 6 | K.VPGN[+2205.793]VTAVLGETLK.V | HexNAc(4)Hex(5)Fuc(2)NeuAc(1) | N[+2206] | 1201.8742 | 3 | 3603.6081 | 3603.5927 | 0 | 4.3 | 466 | Specific | 298.12 | 227.45 | 227.45 | 3.6 | 17 | >sp\|P01833\|PIGR_HUMAN Polymeric immunoglobulin receptor OS=Homo sapiens OX=9606 GN=PIGR PE=1 SV=4 | 902 | 20201119_15432_1ng_glyco_hcd30_etd30_compound15.15345.15345.3 | controllerType=0 controllerNumber=1 scan=15345 | 34.5033 |
| 05067:3 | 6 | K.WN[+2059.735]NTGC[+57.021]QALPSQDEGPSK.A | HexNAc(4)Hex(5)Fuc(1)NeuAc(1) | C[+57], N[+2060] | 1350.2121 | 3 | 4048.6217 | 4048.6116 | 0 | 2.5 | 498 | Specific | 323.65 | 189.94 | 189.94 | 3.9 | 17 | >sp\|P01833\|PIGR_HUMAN Polymeric immunoglobulin receptor OS=Homo sapiens OX=9606 GN=PIGR PE=1 SV=4 | 902 | 20201119_15432_1ng_glyco_hcd30_etd30_compound15.8172.8172.3 | controllerType=0 controllerNumber=1 scan=8172 | 20.7772 |
| 08980:3 | 7 | K.TVLTPATNHMGN[+1378.476]VTFTIPANR.E | HexNAc(2)Hex(6) | N[+1378] | 1211.8876 | 3 | 3633.6483 | 3633.6358 | 0 | 3.5 | 74 | Specific | 447.97 | 366.13 | 190.53 | 7.22 | 8 | >sp\|P01024\|CO3_HUMAN Complement C3 OS=Homo sapiens OX=9606 GN=C3 PE=1 SV=2 | 492 | 20201119_15432_1ng_glyco_hcd30_etd30_compound15.12247.12247.3 | controllerType=0 controllerNumber=1 scan=12247 | 28.3098 |
| 18404:4 | 9 | K.VFPLSLDSTPQDGNVVVAC[+57.021]LVQGFFPQEPLSVTWSESGQN[+1647.613]VTAI | HexNAc(5)Hex(3)Fuc(1) | C[+57], N[+1648] | 1606.7617 | 4 | 6424.0251 | 6423.9746 | 0 | 7.8 | 8 | Specific | 331.27 | 331.27 | 331.27 | 3.69 | 16 | >sp\|P01877\|IGHA2_HUMAN Immunoglobulin heavy constant alpha 2 OS=Homo sapiens OX=9606 GN=IGHA2 PE=1 SV=4 | 576 | 20201119_15432_1ng_glyco_hcd30_etd30_compound15.22186.22186.5 | controllerType=0 controllerNumber=1 scan=22186 | 51.864 |
| 00919:3 | 9 | K.HYTN[+1809.666]SSQDVTVPC[+57.021]R.V | HexNAc(5)Hex(4)Fuc(1) | C[+57], N[+1810] | 1158.48 | 3 | 3473.4256 | 3473.4154 | 0 | 2.9 | 89 | Specific | 340.63 | 276.32 | 276.32 | 3.95 | 16 | >sp\|P01877\|IGHA2_HUMAN Immunoglobulin heavy constant alpha 2 OS=Homo sapiens OX=9606 GN=IGHA2 PE=1 SV=4 | 576 | 20201119_15432_1ng_glyco_hcd30_etd30_compound15.3861.3861.3 | controllerType=0 controllerNumber=1 scan=3861 | 12.9545 |
| 00911:3 | 9 | K.HYTN[+1809.666]SSQDVTVPC[+57.021]R.V | HexNAc(5)Hex(4)Fuc(1) | C[+57], N[+1810] | 1158.4798 | 3 | 3473.4248 | 3473.4154 | 0 | 2.7 | 89 | Specific | 403.89 | 347.25 | 347.25 | 4.55 | 16 | >sp\|P01877\|IGHA2_HUMAN Immunoglobulin heavy constant alpha 2 OS=Homo sapiens OX=9606 GN=IGHA2 PE=1 SV=4 | 576 | 20201119_15432_1ng_glyco_hcd30_etd30_compound15.3852.3852.3 | controllerType=0 controllerNumber=1 scan=3852 | 12.9256 |
| 00927:3 | 9 | K.HYTN[+1647.613]SSQDVTVPC[+57.021]R.V | HexNAc(5)Hex(3)Fuc(1) | C[+57], N[+1648] | 1104.4623 | 3 | 3311.3724 | 3311.3625 | 0 | 3 | 89 | Specific | 476.14 | 415.27 | 415.27 | 5.84 | 16 | >sp\|P01877\|IGHA2_HUMAN Immunoglobulin heavy constant alpha 2 OS=Homo sapiens OX=9606 GN=IGHA2 PE=1 SV=4 | 576 | 20201119_15432_1ng_glyco_hcd30_etd30_compound15.3869.3869.3 | controllerType=0 controllerNumber=1 scan=3869 | 12.9847 |
| 02771:5 | 9 | K.HYTN[+1647.613]SSQDVTVPC[+57.021]RVPPPPPC[+57.021]C[+57.021]HPR.L | HexNAc(5)Hex(3)Fuc(1) | C[+57]*3, N[+1648] | 922.0025 | 5 | 4605.9835 | 4605.9689 | 0 | 3.2 | 89 | Specific | 421.99 | 397.77 | 397.77 | 5.7 | 16 | >sp\|P01877\|IGHA2_HUMAN Immunoglobulin heavy constant alpha 2 OS=Homo sapiens OX=9606 GN=IGHA2 PE=1 SV=4 | 576 | 20201119_15432_1ng_glyco_hcd30_etd30_compound15.5789.5789.5 | controllerType=0 controllerNumber=1 scan=5789 | 16.5847 |
| 02487:2 | 9 | K.TPLTAN[+1606.587]ITK.S | HexNAc(4)Hex(4)Fuc(1) | N[+1607] | 1283.0782 | 2 | 2565.1491 | 2565.1435 | 0 | 2.2 | 200 | Specific | 261.58 | 95.29 | 95.29 | 2.87 | 16 | >sp\|P01877\|IGHA2_HUMAN Immunoglobulin heavy constant alpha 2 OS=Homo sapiens OX=9606 GN=IGHA2 PE=1 SV=4 | 576 | 20201119_15432_1ng_glyco_hcd30_etd30_compound15.5493.5493.2 | controllerType=0 controllerNumber=1 scan=5493 | 15.9457 |
| 02491:3 | 9 | K.TPLTAN[+1809.666]ITK.S | HexNAc(5)Hex(4)Fuc(1) | N[+1810] | 923.4157 | 3 | 2768.2327 | 2768.2228 | 0 | 3.5 | 200 | Specific | 280.92 | 121.03 | 121.03 | 2.63 | 16 | >sp\|P01877\|IGHA2_HUMAN Immunoglobulin heavy constant alpha 2 OS=Homo sapiens OX=9606 GN=IGHA2 PE=1 SV=4 | 576 | 20201119_15432_1ng_glyco_hcd30_etd30_compound15.5498.5498.3 | controllerType=0 controllerNumber=1 scan=5498 | 15.9617 |
| 02531:2 | 9 | K.TPLTAN[+1647.613]ITK.S | HexNAc(5)Hex(3)Fuc(1) | N[+1648] | 1303.5926 | 2 | 2606.1779 | 2606.17 | 0 | 3 | 200 | Specific | 286 | 109.76 | 109.76 | 2.83 | 16 | >sp\|P01877\|IGHA2_HUMAN Immunoglobulin heavy constant alpha 2 OS=Homo sapiens OX=9606 GN=IGHA2 PE=1 SV=4 | 576 | 20201119_15432_1ng_glyco_hcd30_etd30_compound15.5540.5540.2 | controllerType=0 controllerNumber=1 scan=5540 | 16.0697 |
| 02554:2 | 9 | K.TPLTAN[+1501.555]ITK.S | HexNAc(5)Hex(3) | N[+1502] | 1230.5645 | 2 | 2460.1217 | 2460.1121 | 0 | 3.9 | 200 | Specific | 302.94 | 83.93 | 83.93 | 2.83 | 16 | >sp\|P01877\|IGHA2_HUMAN Immunoglobulin heavy constant alpha 2 OS=Homo sapiens OX=9606 GN=IGHA2 PE=1 SV=4 | 576 | 20201119_15432_1ng_glyco_hcd30_etd30_compound15.5564.5564.2 | controllerType=0 controllerNumber=1 scan=5564 | 16.[illegible]337 |
| 02492:2 | 9 | K.TPLTAN[+1809.666]ITK.S | HexNAc(5)Hex(4)Fuc(1) | N[+1810] | 1384.6198 | 2 | 2768.2323 | 2768.2228 | 0 | 3.4 | 200 | Specific | 306.3 | 122.23 | 122.23 | 2.92 | 16 | >sp\|P01877\|IGHA2_HUMAN Immunoglobulin heavy constant alpha 2 OS=Homo sapiens OX=9606 GN=IGHA2 PE=1 SV=4 | 576 | 20201119_15432_1ng_glyco_hcd30_etd30_compound15.5499.5499.2 | controllerType=0 controllerNumber=1 scan=5499 | 15.[illegible]84 |
| 02523:2 | 9 | K.TPLTAN[+1444.534]ITK.S | HexNAc(5)Hex(3)Fuc(1) | N[+1445] | 1202.0523 | 2 | 2403.0973 | 2403.0906 | 0 | 2.8 | 200 | Specific | 340.56 | 107.11 | 107.11 | 2.62 | 16 | >sp\|P01877\|IGHA2_HUMAN Immunoglobulin heavy constant alpha 2 OS=Homo sapiens OX=9606 GN=IGHA2 PE=1 SV=4 | 576 | 20201119_15432_1ng_glyco_hcd30_etd30_compound15.5531.5531.2 | controllerType=0 controllerNumber=1 scan=5531 | 16.0451 |
| 09275:3 | 9 | R.MAGKPTHIN[+1216.423]VSVVMAEADGTC[+57.021]Y.- | HexNAc(2)Hex(5) | C[+57], N[+1216] | 1189.8438 | 3 | 3567.5169 | 3567.5057 | 0 | 3.1 | 319 | CRagged | 283.72 | 271.14 | 189.9 | 4.69 | 16 | >sp\|P01877\|IGHA2_HUMAN Immunoglobulin heavy constant alpha 2 OS=Homo sapiens OX=9606 GN=IGHA2 PE=1 SV=4 | 576 | 20201119_15432_1ng_glyco_hcd30_etd30_compound15.12555.12555.3 | controllerType=0 controllerNumber=1 scan=12555 | 28.9638 |
| 09201:3 | 9 | R.MAGKPTHIN[+1378.476]VSVVMAEADGTC[+57.021]Y.- | HexNAc(2)Hex(6) | C[+57], N[+1378] | 1243.8615 | 3 | 3729.57 | 3729.5585 | 0 | 3.1 | 319 | CRagged | 288.29 | 244.46 | 210.04 | 4.35 | 16 | >sp\|P01877\|IGHA2_HUMAN Immunoglobulin heavy constant alpha 2 OS=Homo sapiens OX=9606 GN=IGHA2 PE=1 SV=4 | 576 | 20201119_15432_1ng_glyco_hcd30_etd30_compound15.12477.12477.3 | controllerType=0 controllerNumber=1 scan=12477 | 28.7756 |
| 14805:4 | 10 | K.RNIAAFGGDPNN[+1913.677]ITLFGESAGGASVSLQTLSPYNK.G | HexNAc(4)Hex(5)NeuAc(1) | N[+1914] | 1370.8812 | 4 | 5480.5031 | 5480.4644 | 0 | 7.1 | 196 | Specific | 380.3 | 379.32 | 379.32 | 3.69 | 5 | >sp\|P19835\|CEL_HUMAN Bile salt-activated lipase OS=Homo sapiens OX=9606 GN=CEL PE=1 SV=3 | 476 | 20201119_15432_1ng_glyco_hcd30_etd30_compound15.18394.18394.4 | controllerType=0 controllerNumber=1 scan=18394 | 41.5356 |
| 04975:3 | 15 | R.LSPN[+2205.793]ASAEHLELR.W | HexNAc(4)Hex(5)Fuc(2)NeuAc(1) | N[+2206] | 1214.8564 | 3 | 3642.5546 | 3642.5421 | 0 | 3.4 | 52 | Specific | 381.68 | 280.8 | 280.8 | 4.38 | 7 | >sp\|Q13410\|BT1A1_HUMAN Butyrophilin subfamily 1 member A1 OS=Homo sapiens OX=9606 GN=BTN1A1 PE=1 SV=3 | 179 | 20201119_15432_1ng_glyco_hcd30_etd30_compound15.8076.8076.3 | controllerType=0 controllerNumber=1 scan=8076 | 20.5912 |
| 06929:3 | 15 | R.LSPN[+2350.830]ASAEHLELR.W | HexNAc(4)Hex(5)Fuc(1)NeuAc(2) | N[+2351] | 1263.201 | 3 | 3787.5884 | 3787.5796 | 0 | 2.3 | 52 | Specific | 412.3 | 269.52 | 269.52 | 4.1 | 7 | >sp\|Q13410\|BT1A1_HUMAN Butyrophilin subfamily 1 member A1 OS=Homo sapiens OX=9606 GN=BTN1A1 PE=1 SV=3 | 179 | 20201119_15432_1ng_glyco_hcd30_etd30_compound15.10100.10100.3 | controllerType=0 controllerNumber=1 scan=10100 | 24.1402 |
| 12308:4 | 20 | K.LKELPGVC[+57.021]N[+2205.793]ETMMALWEEC[+57.021]KPC[+57.021]LK.Q | HexNAc(4)Hex(5)Fuc(2)NeuAc(1) | C[+57]*3, N[+2206] | 1286.3047 | 4 | 5142.1969 | 5142.1776 | 0 | 3.8 | 95 | Specific | 543.67 | 412.99 | 412.99 | 6.64 | 7 | >sp\|P10909\|CLUS_HUMAN Clusterin OS=Homo sapiens OX=9606 GN=CLU PE=1 SV=1 | 1467 | 20201119_15432_1ng_glyco_hcd30_etd30_compound15.15740.15740.4 | controllerType=0 controllerNumber=1 scan=15740 | 35.4004 |
| 12333:4 | 20 | K.LKELPGVC[+57.021]N[+2059.735]ETMMALWEEC[+57.021]KPC[+57.021]LK.Q | HexNAc(4)Hex(5)Fuc(1)NeuAc(1) | C[+57]*3, N[+2060] | 1249.7913 | 4 | 4996.1432 | 4996.1196 | 0 | 4.7 | 95 | Specific | 632.57 | 457.62 | 457.62 | 8.06 | 7 | >sp\|P10909\|CLUS_HUMAN Clusterin OS=Homo sapiens OX=9606 GN=CLU PE=1 SV=1 | 1467 | 20201119_15432_1ng_glyco_hcd30_etd30_compound15.15766.15766.4 | controllerType=0 controllerNumber=1 scan=15766 | 35.4808 |
| 00062:4 | 20 | R.EIRHN[+2059.735]STGC[+57.021]LR.M | HexNAc(4)Hex(5)Fuc(1)NeuAc(1) | C[+57], N[+2060] | 851.3576 | 4 | 3402.4084 | 3402.3994 | 0 | 2.7 | 287 | Specific | 415.68 | 324.14 | 324.14 | 4.33 | 7 | >sp\|P10909\|CLUS_HUMAN Clusterin OS=Homo sapiens OX=9606 GN=CLU PE=1 SV=1 | 1467 | 20201119_15432_1ng_glyco_hcd30_etd30_compound15.2968.2968.4 | controllerType=0 controllerNumber=1 scan=2968 | 11.253 |
| 00072:4 | 20 | R.EIRHN[+2205.793]STGC[+57.021]LR.M | HexNAc(4)Hex(5)Fuc(2)NeuAc(1) | C[+57], N[+2206] | 887.8749 | 4 | 3548.4776 | 3548.4573 | 0 | 5.7 | 287 | Specific | 583.27 | 467.25 | 467.25 | 6.81 | 7 | >sp\|P10909\|CLUS_HUMAN Clusterin OS=Homo sapiens OX=9606 GN=CLU PE=1 SV=1 | 1467 | 20201119_15432_1ng_glyco_hcd30_etd30_compound15.2979.2979.4 | controllerType=0 controllerNumber=1 scan=2979 | 11.2971 |
| 19440:4 | 20 | K.MLN[+2350.830]TSSLLEQLNEQFNWVSR.L | HexNAc(4)Hex(5)Fuc(1)NeuAc(2) | N[+2351] | 1190.7646 | 4 | 4760.0364 | 4760.0171 | 0 | 4.1 | 352 | Specific | 278.93 | 244.89 | 244.89 | 3.59 | 7 | >sp\|P10909\|CLUS_HUMAN Clusterin OS=Homo sapiens OX=9606 GN=CLU PE=1 SV=1 | 1467 | 20201119_15432_1ng_glyco_hcd30_etd30_compound15.23277.23277.5 | controllerType=0 controllerNumber=1 scan=23277 | 54.4173 |
| 17496:3 | 20 | K.MLN[+2205.793]TSSLLEQLNEQFNWVSR.L | HexNAc(4)Hex(5)Fuc(2)NeuAc(1) | N[+2206] | 1539.0045 | 3 | 4614.9988 | 4614.9796 | 0 | 4.2 | 352 | Specific | 285.84 | 167.25 | 167.25 | 3.65 | 7 | >sp\|P10909\|CLUS_HUMAN Clusterin OS=Homo sapiens OX=9606 GN=CLU PE=1 SV=1 | 1467 | 20201119_15432_1ng_glyco_hcd30_etd30_compound15.21238.21238.3 | controllerType=0 controllerNumber=1 scan=21238 | 49.7612 |
| 08658:3 | 22 | R.IIVPLNNREN[+1663.608]ISDPTSPLR.T | HexNAc(5)Hex(4) | N[+1664] | 1271.264 | 3 | 3811.7774 | 3811.7853 | 0 | -2.1 | 62 | Specific | 240.45 | 168.06 | 168.06 | 2.8 | 6 | >sp\|P01591\|IGJ_HUMAN Immunoglobulin J chain OS=Homo sapiens OX=9606 GN=JCHAIN PE=1 SV=4 | 1392 | 20201119_15432_1ng_glyco_hcd30_etd30_compound15.11913.11913.3 | controllerType=0 controllerNumber=1 scan=11913 | 27.6402 |
| 10099:3 | 22 | R.IIVPLNNREN[+1856.656]ISDPTSPLR.T | HexNAc(3)Hex(5)Fuc(1)NeuAc(1) | N[+1857] | 1335.6193 | 3 | 4004.8434 | 4004.8327 | 0 | 2.7 | 62 | Specific | 250.7 | 204.27 | 204.27 | 3.53 | 6 | >sp\|P01591\|IGJ_HUMAN Immunoglobulin J chain OS=Homo sapiens OX=9606 GN=JCHAIN PE=1 SV=4 | 1392 | 20201119_15432_1ng_glyco_hcd30_etd30_compound15.13411.13411.3 | controllerType=0 controllerNumber=1 scan=13411 | 30.6104 |
| 08617:3 | 22 | R.IIVPLNNREN[+1809.666]ISDPTSPLR.T | HexNAc(5)Hex(4)Fuc(1) | N[+1810] | 1319.9572 | 3 | 3957.8571 | 3957.8432 | 0 | 3.5 | 62 | Specific | 265.53 | 204.56 | 204.56 | 4.18 | 6 | >sp\|P01591\|IGJ_HUMAN Immunoglobulin J chain OS=Homo sapiens OX=9606 GN=JCHAIN PE=1 SV=4 | 1392 | 20201119_15432_1ng_glyco_hcd30_etd30_compound15.11870.11870.3 | controllerType=0 controllerNumber=1 scan=11870 | 27.5465 |
| 09994:3 | 22 | R.IIVPLNNREN[+2059.735]ISDPTSPLR.T | HexNAc(4)Hex(5)Fuc(1)NeuAc(1) | N[+2060] | 1403.3135 | 3 | 4207.926 | 4207.9121 | 0 | 3.3 | 62 | Specific | 279.37 | 223.68 | 223.68 | 4.13 | 6 | >sp\|P01591\|IGJ_HUMAN Immunoglobulin J chain OS=Homo sapiens OX=9606 GN=JCHAIN PE=1 SV=4 | 1392 | 20201119_15432_1ng_glyco_hcd30_etd30_compound15.13301.13301.3 | controllerType=0 controllerNumber=1 scan=13301 | 30.3531 |
| 08604:3 | 22 | R.IIVPLNNREN[+1809.666]ISDPTSPLR.T | HexNAc(5)Hex(4)Fuc(1) | N[+1810] | 1319.9576 | 3 | 3957.8582 | 3957.8432 | 0 | 3.8 | 62 | Specific | 293.58 | 208.78 | 208.78 | 4.11 | 6 | >sp\|P01591\|IGJ_HUMAN Immunoglobulin J chain OS=Homo sapiens OX=9606 GN=JCHAIN PE=1 SV=4 | 1392 | 20201119_15432_1ng_glyco_hcd30_etd30_compound15.11856.11856.3 | controllerType=0 controllerNumber=1 scan=11856 | 27.5112 |
| 10033:3 | 22 | R.IIVPLNNREN[+1913.677]ISDPTSPLR.T | HexNAc(4)Hex(5)NeuAc(1) | N[+1914] | 1354.6263 | 3 | 4061.8643 | 4061.8542 | 0 | 2.5 | 62 | Specific | 315.25 | 213.39 | 213.39 | 3.82 | 6 | >sp\|P01591\|IGJ_HUMAN Immunoglobulin J chain OS=Homo sapiens OX=9606 GN=JCHAIN PE=1 SV=4 | 1392 | 20201119_15432_1ng_glyco_hcd30_etd30_compound15.13342.13342.3 | controllerType=0 controllerNumber=1 scan=13342 | 30.4618 |
| 20084:3 | 25 | R.QLAHQSN[+2204.772]STNIFFSPVSIATAFAMLSLGTK.A | HexNAc(4)Hex(5)NeuAc(2) | N[+2205] | 1796.1478 | 3 | 5386.4288 | 5386.4075 | 0 | 4 | 64 | Specific | 264.99 | 154.17 | 154.17 | 5.17 | 3 | >sp\|P01009\|A1AT_HUMAN Alpha-1-antitrypsin OS=Homo sapiens OX=9606 GN=SERPINA1 PE=1 SV=3 | 642 | 20201119_15432_1ng_glyco_hcd30_etd30_compound15.23951.23951.3 | controllerType=0 controllerNumber=1 scan=23951 | 55.9712 |
| 20044:3 | 25 | R.QLAHQSN[+2204.772]STNIFFSPVSIATAFAMLSLGTK.A | HexNAc(4)Hex(5)NeuAc(2) | N[+2205] | 1796.1462 | 3 | 5386.4241 | 5386.4075 | 0 | 3.1 | 64 | Specific | 297.55 | 185.89 | 185.89 | 4.93 | 3 | >sp\|P01009\|A1AT_HUMAN Alpha-1-antitrypsin OS=Homo sapiens OX=9606 GN=SERPINA1 PE=1 SV=3 | 642 | 20201119_15432_1ng_glyco_hcd30_etd30_compound15.23908.23908.3 | controllerType=0 controllerNumber=1 scan=23908 | 55.8424 |
| 20048:3 | 25 | R.QLAHQSN[+2204.772]STNIFFSPVSIATAFAMLSLGTK.A | HexNAc(4)Hex(5)NeuAc(2) | N[+2205] | 1796.148 | 3 | 5386.4296 | 5386.4075 | 0 | 4.1 | 64 | Specific | 329.51 | 210.8 | 210.8 | 5.8 | 3 | >sp\|P01009\|A1AT_HUMAN Alpha-1-antitrypsin OS=Homo sapiens OX=9606 GN=SERPINA1 PE=1 SV=3 | 642 | 20201119_15432_1ng_glyco_hcd30_etd30_compound15.23913.23913.3 | controllerType=0 controllerNumber=1 scan=23913 | 55.8578 |
| 20081:4 | 25 | R.QLAHQSN[+2204.772]STNIFFSPVSIATAFAMLSLGTK.A | HexNAc(4)Hex(5)NeuAc(2) | N[+2205] | 1347.361 | 4 | 5386.422 | 5386.4075 | 0 | 2.7 | 64 | Specific | 559.06 | 481.87 | 338.99 | 6.1 | 3 | >sp\|P01009\|A1AT_HUMAN Alpha-1-antitrypsin OS=Homo sapiens OX=9606 GN=SERPINA1 PE=1 SV=3 | 642 | 20201119_15432_1ng_glyco_hcd30_etd30_compound15.23948.23948.4 | controllerType=0 controllerNumber=1 scan=23948 | 55.9609 |
| 12780:4 | 40 | K.MVSHHN[+2204.772]LTTGATLINEQWLLTTAK.N | HexNAc(4)Hex(5)NeuAc(2) | N[+2205] | 1221.8029 | 4 | 4884.1898 | 4884.1647 | 0 | 5.1 | 179 | Specific | 289.5 | 285.45 | 285.45 | 4.04 | 3 | >sp\|P00738\|HPT_HUMAN Haptoglobin OS=Homo sapiens OX=9606 GN=HP PE=1 SV=1 | 407 | 20201119_15432_1ng_glyco_hcd30_etd30_compound15.16234.16234.2 | controllerType=0 controllerNumber=1 scan=16234 | 36.5205 |
| 07741:4 | 77 | K.SC[+57.021]QHN[+2205.793]GTMYQHGEIFSAHELFPSR.L | HexNAc(4)Hex(5)Fuc(2)NeuAc(1) | C[+57], N[+2206] | 1260.5237 | 4 | 5039.0729 | 5039.0498 | 0 | 4.6 | 110 | Specific | 469.66 | 386.77 | 386.77 | 5.89 | 1 | >sp\|Q6WN34\|CRDL2_HUMAN Chordin-like protein 2 OS=Homo sapiens OX=9606 GN=CHRDL2 PE=1 SV=1 | 1189 | 20201119_15432_1ng_glyco_hcd30_etd30_compound15.10949.10949.4 | controllerType=0 controllerNumber=1 scan=10949 | 25.7306 |
| 15349:5 | 1 | R.RTAGWNVPIGTLRPFLN[+2350.830]WTGPPEPIEAAVAR.F | HexNAc(4)Hex(5)Fuc(1)NeuAc(2) | N[+2351] | 1148.3375 | 5 | 5737.6585 | 5737.6424 | 0 | 2.8 | 140 | Specific | 376.5 | 376.5 | 376.5 | 6.34 | 58 | >sp\|P02788\|TRFL_HUMAN Lactotransferrin OS=Homo sapiens OX=9606 GN=LTF PE=1 SV=6 | 1270 | 20201119_16913_1ng_glyco_hcd30_etd30_compound15.18974.18974.5 | controllerType=0 controllerNumber=1 scan=18974 | 45.8459 |
| 14507:4 | 1 | R.RTAGWNVPIGTLRPFLN[+2863.041]WTGPPEPIEAAVAR.F | HexNAc(5)Hex(6)Fuc(4)NeuAc(1) | N[+2863] | 1563.224 | 4 | 6249.8742 | 6249.8529 | 0 | 3.4 | 140 | Specific | 433.63 | 433.63 | 433.63 | 7.39 | 58 | >sp\|P02788\|TRFL_HUMAN Lactotransferrin OS=Homo sapiens OX=9606 GN=LTF PE=1 SV=6 | 1270 | 20201119_16913_1ng_glyco_hcd30_etd30_compound15.18086.18086.4 | controllerType=0 controllerNumber=1 scan=18086 | 43.1986 |
| 15408:4 | 1 | R.RTAGWNVPIGTLRPFLN[+2350.830]WTGPPEPIEAAVAR.F | HexNAc(4)Hex(5)Fuc(1)NeuAc(2) | N[+2351] | 1435.1686 | 4 | 5737.6525 | 5737.6424 | 0 | 1.8 | 140 | Specific | 593.74 | 593.74 | 593.74 | 8.08 | 58 | >sp\|P02788\|TRFL_HUMAN Lactotransferrin OS=Homo sapiens OX=9606 GN=LTF PE=1 SV=6 | 1270 | 20201119_16913_1ng_glyco_hcd30_etd30_compound15.19036.19036.4 | controllerType=0 controllerNumber=1 scan=19036 | 45.9998 |
| 14564:4 | 1 | R.RTAGWNVPIGTLRPFLN[+2716.983]WTGPPEPIEAAVAR.F | HexNAc(5)Hex(6)Fuc(3)NeuAc(1) | N[+2717] | 1526.7113 | 4 | 6103.8234 | 6103.795 | 0 | 4.7 | 140 | Specific | 639.76 | 639.76 | 639.76 | 9.38 | 58 | >sp\|P02788\|TRFL_HUMAN Lactotransferrin OS=Homo sapiens OX=9606 GN=LTF PE=1 SV=6 | 1270 | 20201119_16913_1ng_glyco_hcd30_etd30_compound15.18146.18146.4 | controllerType=0 controllerNumber=1 scan=18146 | 43.3609 |
| 14636:4 | 1 | R.RTAGWNVPIGTLRPFLN[+2351.851]WTGPPEPIEAAVAR.F | HexNAc(4)Hex(5)Fuc(3)NeuAc(1) | N[+2352] | 1435.4285 | 4 | 5738.6921 | 5738.6628 | 0 | 5.1 | 140 | Specific | 680.44 | 555.13 | 555.13 | 10.38 | 58 | >sp\|P02788\|TRFL_HUMAN Lactotransferrin OS=Homo sapiens OX=9606 GN=LTF PE=1 SV=6 | 1270 | 20201119_16913_1ng_glyco_hcd30_etd30_compound15.18221.18221.4 | controllerType=0 controllerNumber=1 scan=18221 | 43.5673 |
| 15385:4 | 1 | R.RTAGWNVPIGTLRPFLN[+2350.830]WTGPPEPIEAAVAR.F | HexNAc(4)Hex(5)Fuc(1)NeuAc(2) | N[+2351] | 1435.1702 | 4 | 5737.6588 | 5737.6424 | 0 | 2.9 | 140 | Specific | 712.55 | 712.55 | 712.55 | 11.94 | 58 | >sp\|P02788\|TRFL_HUMAN Lactotransferrin OS=Homo sapiens OX=9606 GN=LTF PE=1 SV=6 | 1270 | 20201119_16913_1ng_glyco_hcd30_etd30_compound15.19012.19012.4 | controllerType=0 controllerNumber=1 scan=19012 | 45.9468 |
| 14530:4 | 1 | R.RTAGWNVPIGTLRPFLN[+2863.041]WTGPPEPIEAAVAR.F | HexNAc(5)Hex(6)Fuc(4)NeuAc(1) | N[+2863] | 1563.2241 | 4 | 6249.8747 | 6249.8529 | 0 | 3.5 | 140 | Specific | 764.74 | 764.74 | 764.74 | 12.94 | 58 | >sp\|P02788\|TRFL_HUMAN Lactotransferrin OS=Homo sapiens OX=9606 GN=LTF PE=1 SV=6 | 1270 | 20201119_16913_1ng_glyco_hcd30_etd30_compound15.18110.18110.4 | controllerType=0 controllerNumber=1 scan=18110 | 43.2541 |
| 13954:4 | 1 | R.RTAGWNVPIGTLRPFLN[+1768.640]WTGPPEPIEAAVAR.F | HexNAc(4)Hex(5)Fuc(1) | N[+1769] | 1289.6215 | 4 | 5155.464 | 5155.4516 | 0 | 2.4 | 140 | Specific | 880.41 | 880.41 | 880.41 | 14.94 | 58 | >sp\|P02788\|TRFL_HUMAN Lactotransferrin OS=Homo sapiens OX=9606 GN=LTF PE=1 SV=6 | 1270 | 20201119_16913_1ng_glyco_hcd30_etd30_compound15.17500.17500.4 | controllerType=0 controllerNumber=1 scan=17500 | 41.4007 |
| 14539:4 | 1 | R.RTAGWNVPIGTLRPFLN[+2570.925]WTGPPEPIEAAVAR.F | HexNAc(5)Hex(6)Fuc(2)NeuAc(1) | N[+2571] | 1490.1952 | 4 | 5957.7589 | 5957.7371 | 0 | 3.7 | 140 | Specific | 896.4 | 869.39 | 869.39 | 14.94 | 58 | >sp\|P02788\|TRFL_HUMAN Lactotransferrin OS=Homo sapiens OX=9606 GN=LTF PE=1 SV=6 | 1270 | 20201119_16913_1ng_glyco_hcd30_etd30_compound15.18119.18119.4 | controllerType=0 controllerNumber=1 scan=18119 | 43.2798 |
| 14662:4 | 1 | R.RTAGWNVPIGTLRPFLN[+2059.735]WTGPPEPIEAAVAR.F | HexNAc(4)Hex(5)Fuc(1)NeuAc(1) | N[+2060] | 1362.3953 | 4 | 5446.5592 | 5446.547 | 0 | 2.3 | 140 | Specific | 1100.65 | 1037.96 | 1037.96 | 19.94 | 58 | >sp\|P02788\|TRFL_HUMAN Lactotransferrin OS=Homo sapiens OX=9606 GN=LTF PE=1 SV=6 | 1270 | 20201119_16913_1ng_glyco_hcd30_etd30_compound15.18249.18249.4 | controllerType=0 controllerNumber=1 scan=18249 | 43.6675 |
| 14819:4 | 1 | R.TAGWNVPIGTLRPFLN[+2059.735]WTGPPEPIEAAVAR.F | HexNAc(4)Hex(5)Fuc(1)NeuAc(1) | N[+2060] | 1323.373 | 4 | 5290.4704 | 5290.4459 | 0 | 4.6 | 141 | Specific | 220.17 | 220.17 | 220.17 | 3.24 | 58 | >sp\|P02788\|TRFL_HUMAN Lactotransferrin OS=Homo sapiens OX=9606 GN=LTF PE=1 SV=6 | 1270 | 20201119_16913_1ng_glyco_hcd30_etd30_compound15.18412.18412.4 | controllerType=0 controllerNumber=1 scan=18412 | 44.0493 |
| 15266:4 | 1 | R.TAGWNVPIGTLRPFLN[+3227.153]WTGPPEPIEAAVAR.F | HexNAc(6)Hex(7)Fuc(2)NeuAc(2) | N[+3227] | 1615.2303 | 4 | 6457.8996 | 6457.8636 | 0 | 5.6 | 141 | Specific | 222.89 | 222.89 | 222.89 | 4.06 | 58 | >sp\|P02788\|TRFL_HUMAN Lactotransferrin OS=Homo sapiens OX=9606 GN=LTF PE=1 SV=6 | 1270 | 20201119_16913_1ng_glyco_hcd30_etd30_compound15.18887.18887.4 | controllerType=0 controllerNumber=1 scan=18887 | 45.6104 |
| 15607:3 | 1 | R.[+42.011]TAGWNVPIGTLRPFLN[+2018.708]WTGPPEPIEAAVAR.F | HexNAc(3)Hex(6)Fuc(1)NeuAc(1) | .[42], N[+2019] | 1764.4989 | 3 | 5291.4821 | 5291.4299 | 0 | 9.9 | 141 | Specific | 284.26 | 256.29 | 256.29 | 3.4 | 58 | >sp\|P02788\|TRFL_HUMAN Lactotransferrin OS=Homo sapiens OX=9606 GN=LTF PE=1 SV=6 | 1270 | 20201119_16913_1ng_glyco_hcd30_etd30_compound15.19245.19245.3 | controllerType=0 controllerNumber=1 scan=19245 | 46.5512 |
| 15406:4 | 1 | R.TAGWNVPIGTLRPFLN[+2791.020]WTGPPEPIEAAVAR.F | HexNAc(6)Hex(7)Fuc(3) | N[+2791] | 1506.1835 | 4 | 6021.7121 | 6021.7307 | 0 | -3.1 | 141 | Specific | 292.38 | 292.38 | 292.38 | 4 | 58 | >sp\|P02788\|TRFL_HUMAN Lactotransferrin OS=Homo sapiens OX=9606 GN=LTF PE=1 SV=6 | 1270 | 20201119_16913_1ng_glyco_hcd30_etd30_compound15.19034.19034.4 | controllerType=0 controllerNumber=1 scan=19034 | 45.9938 |
| 15500:3 | 1 | R.TAGWNVPIGTLRPFLN[+2205.793]WTGPPEPIEAAVAR.F | HexNAc(4)Hex(5)Fuc(2)NeuAc(1) | N[+2206] | 1812.8458 | 3 | 5436.5228 | 5436.5038 | 0 | 3.5 | 141 | Specific | 309.04 | 221.87 | 221.87 | 5.58 | 58 | >sp\|P02788\|TRFL_HUMAN Lactotransferrin OS=Homo sapiens OX=9606 GN=LTF PE=1 SV=6 | 1270 | 20201119_16913_1ng_glyco_hcd30_etd30_compound15.19134.19134.2 | controllerType=0 controllerNumber=1 scan=19134 | 46.3237 |
| 15661:3 | 1 | R.TAGWNVPIGTLRPFLN[+2059.735]WTGPPEPIEAAVAR.F | HexNAc(4)Hex(5)Fuc(1)NeuAc(1) | N[+2060] | 1764.1602 | 3 | 5290.4662 | 5290.4459 | 0 | 3.8 | 141 | Specific | 325.7 | 216.92 | 216.92 | 6.02 | 58 | >sp\|P02788\|TRFL_HUMAN Lactotransferrin OS=Homo sapiens OX=9606 GN=LTF PE=1 SV=6 | 1270 | 20201119_16913_1ng_glyco_hcd30_etd30_compound15.19302.19302.2 | controllerType=0 controllerNumber=1 scan=19302 | 46.682 |
| 16462:3 | 1 | R.TAGWNVPIGTLRPFLN[+2350.830]WTGPPEPIEAAVAR.F | HexNAc(4)Hex(5)Fuc(1)NeuAc(2) | N[+2351] | 1861.1904 | 3 | 5581.5566 | 5581.5413 | 0 | 2.7 | 141 | Specific | 398.66 | 321.66 | 321.66 | 6.52 | 58 | >sp\|P02788\|TRFL_HUMAN Lactotransferrin OS=Homo sapiens OX=9606 GN=LTF PE=1 SV=6 | 1270 | 20201119_16913_1ng_glyco_hcd30_etd30_compound15.20148.20148.3 | controllerType=0 controllerNumber=1 scan=20148 | 49.1684 |
| 15483:3 | 1 | R.TAGWNVPIGTLRPFLN[+2205.793]WTGPPEPIEAAVAR.F | HexNAc(4)Hex(5)Fuc(2)NeuAc(1) | N[+2206] | 1812.8441 | 3 | 5436.5177 | 5436.5038 | 0 | 2.6 | 141 | Specific | 457.54 | 351.8 | 351.8 | 5.79 | 58 | >sp\|P02788\|TRFL_HUMAN Lactotransferrin OS=Homo sapiens OX=9606 GN=LTF PE=1 SV=6 | 1270 | 20201119_16913_1ng_glyco_hcd30_etd30_compound15.19116.19116.3 | controllerType=0 controllerNumber=1 scan=19116 | 46.2447 |
| 15488:3 | 1 | R.TAGWNVPIGTLRPFLN[+2059.735]WTGPPEPIEAAVAR.F | HexNAc(4)Hex(5)Fuc(1)NeuAc(1) | N[+2060] | 1764.1584 | 3 | 5290.4607 | 5290.4459 | 0 | 2.8 | 141 | Specific | 578.99 | 454.81 | 454.81 | 9.34 | 58 | >sp\|P02788\|TRFL_HUMAN Lactotransferrin OS=Homo sapiens OX=9606 GN=LTF PE=1 SV=6 | 1270 | 20201119_16913_1ng_glyco_hcd30_etd30_compound15.19121.19121.3 | controllerType=0 controllerNumber=1 scan=19121 | 46.2692 |
| 15387:4 | 1 | R.TAGWNVPIGTLRPFLN[+2863.041]WTGPPEPIEAAVAR.F | HexNAc(5)Hex(6)Fuc(4)NeuAc(1) | N[+2863] | 1524.1949 | 4 | 6093.758 | 6093.7518 | 0 | 1 | 141 | Specific | 607.32 | 607.32 | 607.32 | 9.45 | 58 | >sp\|P02788\|TRFL_HUMAN Lactotransferrin OS=Homo sapiens OX=9606 GN=LTF PE=1 SV=6 | 1270 | 20201119_16913_1ng_glyco_hcd30_etd30_compound15.19014.19014.4 | controllerType=0 controllerNumber=1 scan=19014 | 45.9548 |
| 16318:4 | 1 | R.TAGWNVPIGTLRPFLN[+2862.021]WTGPPEPIEAAVAR.F | HexNAc(5)Hex(6)Fuc(2)NeuAc(2) | N[+2862] | 1523.9456 | 4 | 6092.7604 | 6092.7314 | 0 | 4.8 | 141 | Specific | 674.59 | 674.59 | 674.59 | 10.75 | 58 | >sp\|P02788\|TRFL_HUMAN Lactotransferrin OS=Homo sapiens OX=9606 GN=LTF PE=1 SV=6 | 1270 | 20201119_16913_1ng_glyco_hcd30_etd30_compound15.19995.19995.4 | controllerType=0 controllerNumber=1 scan=19995 | 48.6884 |
| 15820:4 | 1 | R.TAGWNVPIGTLRPFLN[+2059.735]WTGPPEPIEAAVAR.F | HexNAc(4)Hex(5)Fuc(1)NeuAc(1) | N[+2060] | 1323.3794 | 4 | 5290.4958 | 5290.4459 | 0 | 9.4 | 141 | Specific | 721.65 | 721.65 | 721.65 | 11.12 | 58 | >sp\|P02788\|TRFL_HUMAN Lactotransferrin OS=Homo sapiens OX=9606 GN=LTF PE=1 SV=6 | 1270 | 20201119_16913_1ng_glyco_hcd30_etd30_compound15.19468.19468.4 | controllerType=0 controllerNumber=1 scan=19468 | 47.1158 |
| 16461:4 | 1 | R.TAGWNVPIGTLRPFLN[+2350.830]WTGPPEPIEAAVAR.F | HexNAc(4)Hex(5)Fuc(1)NeuAc(2) | N[+2351] | 1396.1448 | 4 | 5581.5573 | 5581.5413 | 0 | 2.9 | 141 | Specific | 790.25 | 789.05 | 789.05 | 13.31 | 58 | >sp\|P02788\|TRFL_HUMAN Lactotransferrin OS=Homo sapiens OX=9606 GN=LTF PE=1 SV=6 | 1270 | 20201119_16913_1ng_glyco_hcd30_etd30_compound15.20147.20147.4 | controllerType=0 controllerNumber=1 scan=20147 | 49.1647 |
| 15356:4 | 1 | R.TAGWNVPIGTLRPFLN[+2863.041]WTGPPEPIEAAVAR.F | HexNAc(5)Hex(6)Fuc(4)NeuAc(1) | N[+2863] | 1524.1973 | 4 | 6093.7672 | 6093.7518 | 0 | 2.5 | 141 | Specific | 790.31 | 790.31 | 790.31 | 13.31 | 58 | >sp\|P02788\|TRFL_HUMAN Lactotransferrin OS=Homo sapiens OX=9606 GN=LTF PE=1 SV=6 | 1270 | 20201119_16913_1ng_glyco_hcd30_etd30_compound15.18981.18981.4 | controllerType=0 controllerNumber=1 scan=18981 | 45.8705 |
| 16473:5 | 1 | R.TAGWNVPIGTLRPFLN[+2350.830]WTGPPEPIEAAVAR.F | HexNAc(4)Hex(5)Fuc(1)NeuAc(2) | N[+2351] | 1117.1176 | 5 | 5581.5587 | 5581.5413 | 0 | 3.1 | 141 | Specific | 944.4 | 902.23 | 902.23 | 16.31 | 58 | >sp\|P02788\|TRFL_HUMAN Lactotransferrin OS=Homo sapiens OX=9606 GN=LTF PE=1 SV=6 | 1270 | 20201119_16913_1ng_glyco_hcd30_etd30_compound15.20160.20160.5 | controllerType=0 controllerNumber=1 scan=20160 | 49.2175 |
| 15482:4 | 1 | R.TAGWNVPIGTLRPFLN[+2205.793]WTGPPEPIEAAVAR.F | HexNAc(4)Hex(5)Fuc(2)NeuAc(1) | N[+2206] | 1359.8849 | 4 | 5436.5177 | 5436.5038 | 0 | 2.6 | 141 | Specific | 962.81 | 962.81 | 962.81 | 17.31 | 58 | >sp\|P02788\|TRFL_HUMAN Lactotransferrin OS=Homo sapiens OX=9606 GN=LTF PE=1 SV=6 | 1270 | 20201119_16913_1ng_glyco_hcd30_etd30_compound15.19115.19115.4 | controllerType=0 controllerNumber=1 scan=19115 | 46.2417 |
| 15487:4 | 1 | R.TAGWNVPIGTLRPFLN[+2059.735]WTGPPEPIEAAVAR.F | HexNAc(4)Hex(5)Fuc(1)NeuAc(1) | N[+2060] | 1323.3691 | 4 | 5290.4547 | 5290.4459 | 0 | 1.7 | 141 | Specific | 1059.07 | 981.64 | 981.64 | 18.45 | 58 | >sp\|P02788\|TRFL_HUMAN Lactotransferrin OS=Homo sapiens OX=9606 GN=LTF PE=1 SV=6 | 1270 | 20201119_16913_1ng_glyco_hcd30_etd30_compound15.19120.19120.4 | controllerType=0 controllerNumber=1 scan=19120 | 46.2553 |
| 13721:3 | 1 | R.TAGWNIPMGLLFN[+1768.640]QTGSC[+57.021]K.F | HexNAc(4)Hex(5)Fuc(1) | C[+57], N[+1769] | 1288.5567 | 3 | 3863.6555 | 3863.6495 | 0 | 1.6 | 485 | Specific | 308.51 | 237.06 | 237.06 | 4.66 | 58 | >sp\|P02788\|TRFL_HUMAN Lactotransferrin OS=Homo sapiens OX=9606 GN=LTF PE=1 SV=6 | 1270 | 20201119_16913_1ng_glyco_hcd30_etd30_compound15.17254.17254.3 | controllerType=0 controllerNumber=1 scan=17254 | 40.6635 |
| 15043:3 | 1 | R.TAGWNIPMGLLFN[+2351.851]QTGSC[+57.021]K.F | HexNAc(4)Hex(5)Fuc(3)NeuAc(1) | C[+57], N[+2352] | 1482.9665 | 3 | 4446.8849 | 4446.8607 | 0 | 5.4 | 485 | Specific | 313.18 | 227.24 | 227.24 | 4.96 | 58 | >sp\|P02788\|TRFL_HUMAN Lactotransferrin OS=Homo sapiens OX=9606 GN=LTF PE=1 SV=6 | 1270 | 20201119_16913_1ng_glyco_hcd30_etd30_compound15.18650.18650.3 | controllerType=0 controllerNumber=1 scan=18650 | 44.7835 |
| 15082:4 | 1 | R.TAGWNIPMGLLFN[+2059.735]QTGSC[+57.021]K.F | HexNAc(4)Hex(5)Fuc(1)NeuAc(1) | C[+57], N[+2060] | 1039.4467 | 4 | 4154.765 | 4154.7449 | 0 | 4.8 | 485 | Specific | 321.8 | 302.66 | 302.66 | 4.44 | 58 | >sp\|P02788\|TRFL_HUMAN Lactotransferrin OS=Homo sapiens OX=9606 GN=LTF PE=1 SV=6 | 1270 | 20201119_16913_1ng_glyco_hcd30_etd30_compound15.18691.18691.4 | controllerType=0 controllerNumber=1 scan=18691 | 44.9154 |
| 16522:3 | 1 | R.TAGWNIPMGLLFN[+2350.830]QTGSC[+57.021]K.F | HexNAc(4)Hex(5)Fuc(1)NeuAc(2) | C[+57], N[+2351] | 1482.6247 | 3 | 4445.8595 | 4445.8403 | 0 | 4.3 | 485 | Specific | 353.91 | 253.22 | 253.22 | 5.02 | 58 | >sp\|P02788\|TRFL_HUMAN Lactotransferrin OS=Homo sapiens OX=9606 GN=LTF PE=1 SV=6 | 1270 | 20201119_16913_1ng_glyco_hcd30_etd30_compound15.20214.20214.3 | controllerType=0 controllerNumber=1 scan=20214 | 49.4611 |
| 13583:3 | 1 | R.TAGWNIPMGLLFN[+2279.830]QTGSC[+57.021]K.F | HexNAc(5)Hex(6)Fuc(2) | C[+57], N[+2280] | 1458.9529 | 3 | 4374.8443 | 4374.8396 | 0 | 1.1 | 485 | Specific | 368.07 | 273.17 | 242.42 | 4.76 | 58 | >sp\|P02788\|TRFL_HUMAN Lactotransferrin OS=Homo sapiens OX=9606 GN=LTF PE=1 SV=6 | 1270 | 20201119_16913_1ng_glyco_hcd30_etd30_compound15.17108.17108.3 | controllerType=0 controllerNumber=1 scan=17108 | 40.2704 |
| 13600:3 | 1 | R.TAGWNIPMGLLFN[+2133.772]QTGSC[+57.021]K.F | HexNAc(5)Hex(6)Fuc(1) | C[+57], N[+2134] | 1410.2702 | 3 | 4228.796 | 4228.7817 | 0 | 3.4 | 485 | Specific | 374.48 | 279.11 | 248.28 | 4.96 | 58 | >sp\|P02788\|TRFL_HUMAN Lactotransferrin OS=Homo sapiens OX=9606 GN=LTF PE=1 SV=6 | 1270 | 20201119_16913_1ng_glyco_hcd30_etd30_compound15.17126.17126.3 | controllerType=0 controllerNumber=1 scan=17126 | 40.3183 |
| 15060:3 | 1 | R.TAGWNIPMGLLFN[+2059.735]QTGSC[+57.021]K.F | HexNAc(4)Hex(5)Fuc(1)NeuAc(1) | C[+57], N[+2060] | 1385.593 | 3 | 4154.7643 | 4154.7449 | 0 | 4.7 | 485 | Specific | 382.61 | 270.14 | 270.14 | 5.05 | 58 | >sp\|P02788\|TRFL_HUMAN Lactotransferrin OS=Homo sapiens OX=9606 GN=LTF PE=1 SV=6 | 1270 | 20201119_16913_1ng_glyco_hcd30_etd30_compound15.18668.18668.3 | controllerType=0 controllerNumber=1 scan=18668 | 44.8416 |
| 15056:3 | 1 | R.TAGWNIPMGLLFN[+2205.793]QTGSC[+57.021]K.F | HexNAc(4)Hex(5)Fuc(2)NeuAc(1) | C[+57], N[+2206] | 1434.2791 | 3 | 4300.8228 | 4300.8028 | 0 | 4.6 | 485 | Specific | 397.42 | 311.33 | 311.33 | 5.25 | 58 | >sp\|P02788\|TRFL_HUMAN Lactotransferrin OS=Homo sapiens OX=9606 GN=LTF PE=1 SV=6 | 1270 | 20201119_16913_1ng_glyco_hcd30_etd30_compound15.18664.18664.3 | controllerType=0 controllerNumber=1 scan=18664 | 44.832 |
| 16633:3 | 1 | R.TAGWNIPMGLLFN[+2204.772]QTGSC[+57.021]K.F | HexNAc(4)Hex(5)NeuAc(2) | C[+57], N[+2205] | 1433.9377 | 3 | 4299.7985 | 4299.7824 | 0 | 3.7 | 485 | Specific | 429.18 | 338.1 | 338.1 | 5.84 | 58 | >sp\|P02788\|TRFL_HUMAN Lactotransferrin OS=Homo sapiens OX=9606 GN=LTF PE=1 SV=6 | 1270 | 20201119_16913_1ng_glyco_hcd30_etd30_compound15.20331.20331.3 | controllerType=0 controllerNumber=1 scan=20331 | 49.8133 |
| 15189:3 | 1 | R.TAGWNIPMGLLFN[+1872.651]QTGSC[+57.021]K.F | HexNAc(3)Hex(6)NeuAc(1) | C[+57], N[+1873] | 1323.2357 | 3 | 3967.6924 | 3967.6604 | 0 | 8.1 | 485 | Specific | 433.2 | 319.26 | 216.22 | 4.54 | 58 | >sp\|P02788\|TRFL_HUMAN Lactotransferrin OS=Homo sapiens OX=9606 GN=LTF PE=1 SV=6 | 1270 | 20201119_16913_1ng_glyco_hcd30_etd30_compound15.18806.18806.3 | controllerType=0 controllerNumber=1 scan=18806 | 45.3622 |
| 13994:2 | 1 | R.TAGWNIPMGLLFN[+1216.423]QTGSC[+57.021]K.F | HexNAc(2)Hex(5) | C[+57], N[+1216] | 1656.2198 | 2 | 3311.4323 | 3311.4328 | 0 | -0.2 | 485 | Specific | 442.68 | 423.61 | 145.58 | 6.24 | 58 | >sp\|P02788\|TRFL_HUMAN Lactotransferrin OS=Homo sapiens OX=9606 GN=LTF PE=1 SV=6 | 1270 | 20201119_16913_1ng_glyco_hcd30_etd30_compound15.17542.17542.2 | controllerType=0 controllerNumber=1 scan=17542 | 41.5328 |
| 13967:3 | 1 | R.TAGWNIPMGLLFN[+1216.423]QTGSC[+57.021]K.F | HexNAc(2)Hex(5) | C[+57], N[+1216] | 1104.4876 | 3 | 3311.4482 | 3311.4328 | 0 | 4.6 | 485 | Specific | 475.05 | 447.3 | 240.22 | 6.87 | 58 | >sp\|P02788\|TRFL_HUMAN Lactotransferrin OS=Homo sapiens OX=9606 GN=LTF PE=1 SV=6 | 1270 | 20201119_16913_1ng_glyco_hcd30_etd30_compound15.17514.17514.3 | controllerType=0 controllerNumber=1 scan=17514 | 41.4384 |
| 15138:3 | 1 | R.TAGWNIPMGLLFN[+1913.677]QTGSC[+57.021]K.F | HexNAc(4)Hex(5)NeuAc(1) | C[+57], N[+1914] | 1336.9068 | 3 | 4008.7058 | 4008.687 | 0 | 4.7 | 485 | Specific | 477.59 | 307.7 | 307.7 | 5.51 | 58 | >sp\|P02788\|TRFL_HUMAN Lactotransferrin OS=Homo sapiens OX=9606 GN=LTF PE=1 SV=6 | 1270 | 20201119_16913_1ng_glyco_hcd30_etd30_compound15.18753.18753.3 | controllerType=0 controllerNumber=1 scan=18753 | 45.2153 |
| 16506:3 | 1 | R.TAGWNIPMGLLFN[+2350.830]QTGSC[+57.021]K.F | HexNAc(4)Hex(5)Fuc(1)NeuAc(2) | C[+57], N[+2351] | 1482.6309 | 3 | 4445.8782 | 4445.8403 | 0 | 8.5 | 485 | Specific | 591.46 | 413.24 | 413.24 | 7.45 | 58 | >sp\|P02788\|TRFL_HUMAN Lactotransferrin OS=Homo sapiens OX=9606 GN=LTF PE=1 SV=6 | 1270 | 20201119_16913_1ng_glyco_hcd30_etd30_compound15.20196.20196.3 | controllerType=0 controllerNumber=1 scan=20196 | 49.3422 |
| 14323:3 | 1 | R.TAGWNIPM[+15.995]GLLFN[+2059.735]QTGSC[+57.021]K.F | HexNAc(4)Hex(5)Fuc(1)NeuAc(1) | C[+57], M[+16], N[+2060] | 1390.9277 | 3 | 4170.7685 | 4170.7398 | 0 | 6.9 | 485 | Specific | 287.13 | 225.83 | 225.83 | 3.6 | 58 | >sp\|P02788\|TRFL_HUMAN Lactotransferrin OS=Homo sapiens OX=9606 GN=LTF PE=1 SV=6 | 1270 | 20201119_16913_1ng_glyco_hcd30_etd30_compound15.17890.17890.3 | controllerType=0 controllerNumber=1 scan=17890 | 42.5297 |
| 16219:4 | 1 | R.TAGWNIPMGLLFN[+2350.830]QTGSC[+57.021]KFDEYFSQSC[+57.021]APGSDPR.S | HexNAc(4)Hex(5)Fuc(1)NeuAc(2) | C[+57]*2, N[+2351] | 1573.1589 | 4 | 6289.6139 | 6289.5875 | 0 | 4.2 | 485 | Specific | 543.8 | 503.21 | 503.21 | 6.82 | 58 | >sp\|P02788\|TRFL_HUMAN Lactotransferrin OS=Homo sapiens OX=9606 GN=LTF PE=1 SV=6 | 1270 | 20201119_16913_1ng_glyco_hcd30_etd30_compound15.19891.19891.4 | controllerType=0 controllerNumber=1 scan=19891 | 48.4086 |
| 15234:4 | 1 | R.TAGWNIPMGLLFN[+2059.735]QTGSC[+57.021]KFDEYFSQSC[+57.021]APGSDPR.S | HexNAc(4)Hex(5)Fuc(1)NeuAc(1) | C[+57]*2, N[+2060] | 1500.3862 | 4 | 5998.5231 | 5998.492 | 0 | 5.2 | 485 | Specific | 675.32 | 604.02 | 604.02 | 9.85 | 58 | >sp\|P02788\|TRFL_HUMAN Lactotransferrin OS=Homo sapiens OX=9606 GN=LTF PE=1 SV=6 | 1270 | 20201119_16913_1ng_glyco_hcd30_etd30_compound15.18854.18854.4 | controllerType=0 controllerNumber=1 scan=18854 | 45.5175 |
| 09927:3 | 1 | R.N[+2350.830]GSDC[+57.021]PDKFC[+57.021]LFQSETK.N | HexNAc(4)Hex(5)Fuc(1)NeuAc(2) | C[+57]*2, N[+2351] | 1461.9139 | 3 | 4383.7271 | 4383.7043 | 0 | 5.2 | 642 | Specific | 316.85 | 193.12 | 193.12 | 4.23 | 58 | >sp\|P02788\|TRFL_HUMAN Lactotransferrin OS=Homo sapiens OX=9606 GN=LTF PE=1 SV=6 | 1270 | 20201119_16913_1ng_glyco_hcd30_etd30_compound15.13253.13253.3 | controllerType=0 controllerNumber=1 scan=13253 | 31.002 |

Table S4. Cont.

| Query #:z | Protein Rank | Glycopeptide | Glycan | Modification Type(s) | Observed m/z | z | Observed (M+H) | Calc. mass (M+H) | Off-by-x error | Mass error (ppm) | StartPosition | Cleavage | Score | Delta | Delta Mod | \|Log Prob\| | # of unique peptides | Protein Name | Protein DB number | Comment | Scan # | Scan Time |
|---|---|---|---|---|---|---|---|---|---|---|---|---|---|---|---|---|---|---|---|---|---|---|
| 08044:3 | 1 | R.N[+2059.735]GSDC[+57.021]PDKFC[+57.021]LFQSETK.N | HexNAc(4)Hex(5)Fuc(1)NeuAc(1) | C[+57]*2, N[+2060] | 1364.879 | 3 | 4092.6223 | 4092.6088 | 0 | 3.3 | 642 | Specific | 371.69 | 266.1 | 266.1 | 5.15 | 58 | >sp\|P02788\|TRFL_HUMAN Lactotransferrin OS=Homo sapiens OX=9606 GN=LTF PE=1 SV=6 | 1270 | 20201119_16913_1ng_glyco_hcd30_etd30_compound15.11288.11288.3 | controllerType=0 controllerNumber=1 scan=11288 | 26.7786 |
| 07221:3 | 2 | R.GN[+2059.735]FSTEGC[+57.021]GC[+57.021]VC[+57.021]EPGWK.G | HexNAc(4)Hex(5)Fuc(1)NeuAc(1) | C[+57]*3, N[+2060] | 1335.5101 | 3 | 4004.5156 | 4004.5023 | 0 | 3.3 | 165 | Specific | 315.94 | 206.15 | 206.15 | 5.25 | 31 | >sp\|P24821\|TENA_HUMAN Tenascin OS=Homo sapiens OX=9606 GN=TNC PE=1 SV=3 | 486 | 20201119_16913_1ng_glyco_hcd30_etd30_compound15.10422.10422.3 | controllerType=0 controllerNumber=1 scan=10422 | 25.0104 |
| 01379:3 | 2 | K.GPN[+2059.735]C[+57.021]SEPEC[+57.021]PGNC[+57.021]HLR.G | HexNAc(4)Hex(5)Fuc(1)NeuAc(1) | C[+57]*3, N[+2060] | 1315.1735 | 3 | 3943.506 | 3943.4931 | 0 | 3.3 | 182 | Specific | 282.99 | 144.22 | 144.22 | 5.12 | 31 | >sp\|P24821\|TENA_HUMAN Tenascin OS=Homo sapiens OX=9606 GN=TNC PE=1 SV=3 | 486 | 20201119_16913_1ng_glyco_hcd30_etd30_compound15.4347.4347.3 | controllerType=0 controllerNumber=1 scan=4347 | 13.917 |
| 12679:3 | 2 | R.LN[+1702.581]YSLPTGQWVGVQLPR.N | HexNAc(2)Hex(8) | N[+1703] | 1210.8882 | 3 | 3630.6502 | 3630.6202 | 0 | 8.3 | 1017 | Specific | 467.45 | 438.59 | 438.59 | 6.24 | 31 | >sp\|P24821\|TENA_HUMAN Tenascin OS=Homo sapiens OX=9606 GN=TNC PE=1 SV=3 | 486 | 20201119_16913_1ng_glyco_hcd30_etd30_compound15.16145.16145.3 | controllerType=0 controllerNumber=1 scan=16145 | 37.783 |
| 12801:2 | 2 | R.LN[+1216.423]YSLPTGQWVGVQLPR.N | HexNAc(2)Hex(5) | N[+1216] | 1572.7382 | 2 | 3144.4691 | 3144.4617 | 0 | 2.4 | 1017 | Specific | 542.13 | 467.86 | 467.86 | 7.63 | 31 | >sp\|P24821\|TENA_HUMAN Tenascin OS=Homo sapiens OX=9606 GN=TNC PE=1 SV=3 | 486 | 20201119_16913_1ng_glyco_hcd30_etd30_compound15.16273.16273.2 | controllerType=0 controllerNumber=1 scan=16273 | 38.1598 |
| 12779:3 | 2 | R.LN[+1216.423]YSLPTGQWVGVQLPR.N | HexNAc(2)Hex(5) | N[+1216] | 1048.8291 | 3 | 3144.4728 | 3144.4617 | 0 | 3.5 | 1017 | Specific | 567.61 | 527.72 | 527.72 | 8.63 | 31 | >sp\|P24821\|TENA_HUMAN Tenascin OS=Homo sapiens OX=9606 GN=TNC PE=1 SV=3 | 486 | 20201119_16913_1ng_glyco_hcd30_etd30_compound15.16250.16250.3 | controllerType=0 controllerNumber=1 scan=16250 | 38.0881 |
| 12744:3 | 2 | R.LN[+1378.476]YSLPTGQWVGVQLPR.N | HexNAc(2)Hex(6) | N[+1378] | 1102.8462 | 3 | 3306.5241 | 3306.5145 | 0 | 2.9 | 1017 | Specific | 607.14 | 548.89 | 548.89 | 9.63 | 31 | >sp\|P24821\|TENA_HUMAN Tenascin OS=Homo sapiens OX=9606 GN=TNC PE=1 SV=3 | 486 | 20201119_16913_1ng_glyco_hcd30_etd30_compound15.16213.16213.3 | controllerType=0 controllerNumber=1 scan=16213 | 37.9775 |
| 06894:3 | 2 | R.N[+2059.735]LTVPGSLR.A | HexNAc(4)Hex(5)Fuc(1)NeuAc(1) | N[+2060] | 1006.1049 | 3 | 3016.3003 | 3016.2873 | 0 | 4.3 | 1119 | Specific | 271.93 | 131.81 | 131.81 | 3 | 31 | >sp\|P24821\|TENA_HUMAN Tenascin OS=Homo sapiens OX=9606 GN=TNC PE=1 SV=3 | 486 | 20201119_16913_1ng_glyco_hcd30_etd30_compound15.10080.10080.3 | controllerType=0 controllerNumber=1 scan=10080 | 24.3172 |
| 06854:3 | 2 | R.N[+2059.735]LTVPGSLR.A | HexNAc(4)Hex(5)Fuc(1)NeuAc(1) | N[+2060] | 1006.1026 | 3 | 3016.2931 | 3016.2873 | 0 | 1.9 | 1119 | Specific | 276.9 | 151.83 | 151.83 | 2.77 | 31 | >sp\|P24821\|TENA_HUMAN Tenascin OS=Homo sapiens OX=9606 GN=TNC PE=1 SV=3 | 486 | 20201119_16913_1ng_glyco_hcd30_etd30_compound15.10038.10038.2 | controllerType=0 controllerNumber=1 scan=10038 | 24.2329 |
| 14357:4 | 3 | R.LSLHRPALEDLLLGSEAN[+1663.608]LTC[+57.021]TLTGLR.D | HexNAc(5)Hex(4) | C[+57], N[+1664] | 1157.5625 | 4 | 4627.2283 | 4627.2064 | 0 | 4.7 | 127 | Specific | 271.06 | 271.06 | 271.06 | 4.99 | 35 | >sp\|P01876\|IGHA1_HUMAN Immunoglobulin heavy constant alpha 1 OS=Homo sapiens OX=9606 GN=IGHA1 PE=1 SV=2 | 255 | 20201119_16913_1ng_glyco_hcd30_etd30_compound15.17927.17927.4 | controllerType=0 controllerNumber=1 scan=17927 | 42.6657 |
| 14345:4 | 3 | R.LSLHRPALEDLLLGSEAN[+1378.476]LTC[+57.021]TLTGLR.D | HexNAc(2)Hex(6) | C[+57], N[+1378] | 1086.2765 | 4 | 4342.0843 | 4342.074 | 0 | 2.4 | 127 | Specific | 325.58 | 325.58 | 325.58 | 6.36 | 35 | >sp\|P01876\|IGHA1_HUMAN Immunoglobulin heavy constant alpha 1 OS=Homo sapiens OX=9606 GN=IGHA1 PE=1 SV=2 | 255 | 20201119_16913_1ng_glyco_hcd30_etd30_compound15.17914.17914.4 | controllerType=0 controllerNumber=1 scan=17914 | 42.6208 |
| 14354:3 | 3 | R.LSLHRPALEDLLLGSEAN[+1378.476]LTC[+57.021]TLTGLR.D | HexNAc(2)Hex(6) | C[+57], N[+1378] | 1448.0355 | 3 | 4342.0918 | 4342.074 | 0 | 4.1 | 127 | Specific | 416.47 | 407.14 | 407.14 | 6.61 | 35 | >sp\|P01876\|IGHA1_HUMAN Immunoglobulin heavy constant alpha 1 OS=Homo sapiens OX=9606 GN=IGHA1 PE=1 SV=2 | 255 | 20201119_16913_1ng_glyco_hcd30_etd30_compound15.17923.17923.3 | controllerType=0 controllerNumber=1 scan=17923 | 42.6561 |
| 14360:3 | 3 | R.LSLHRPALEDLLLGSEAN[+1663.608]LTC[+57.021]TLTGLR.D | HexNAc(5)Hex(4) | C[+57], N[+1664] | 1543.0825 | 3 | 4627.2328 | 4627.2064 | 0 | 5.7 | 127 | Specific | 424.86 | 375.77 | 375.77 | 5.96 | 35 | >sp\|P01876\|IGHA1_HUMAN Immunoglobulin heavy constant alpha 1 OS=Homo sapiens OX=9606 GN=IGHA1 PE=1 SV=2 | 255 | 20201119_16913_1ng_glyco_hcd30_etd30_compound15.17930.17930.3 | controllerType=0 controllerNumber=1 scan=17930 | 42.6716 |
| 14364:3 | 3 | R.LSLHRPALEDLLLGSEAN[+1378.476]LTC[+57.021]TLTGLR.D | HexNAc(2)Hex(6) | C[+57], N[+1378] | 1448.039 | 3 | 4342.1024 | 4342.074 | 0 | 6.6 | 127 | Specific | 457.24 | 451.8 | 451.8 | 6.4 | 35 | >sp\|P01876\|IGHA1_HUMAN Immunoglobulin heavy constant alpha 1 OS=Homo sapiens OX=9606 GN=IGHA1 PE=1 SV=2 | 255 | 20201119_16913_1ng_glyco_hcd30_etd30_compound15.17934.17934.3 | controllerType=0 controllerNumber=1 scan=17934 | 42.6802 |
| 14366:4 | 3 | R.LSLHRPALEDLLLGSEAN[+1702.581]LTC[+57.021]TLTGLR.D | HexNAc(2)Hex(8) | C[+57], N[+1703] | 1167.3046 | 4 | 4666.1966 | 4666.1796 | 0 | 3.6 | 127 | Specific | 484.14 | 468.32 | 468.32 | 7.18 | 35 | >sp\|P01876\|IGHA1_HUMAN Immunoglobulin heavy constant alpha 1 OS=Homo sapiens OX=9606 GN=IGHA1 PE=1 SV=2 | 255 | 20201119_16913_1ng_glyco_hcd30_etd30_compound15.17936.17936.4 | controllerType=0 controllerNumber=1 scan=17936 | 42.6882 |
| 14385:3 | 3 | R.LSLHRPALEDLLLGSEAN[+1216.423]LTC[+57.021]TLTGLR.D | HexNAc(2)Hex(5) | C[+57], N[+1216] | 1394.0175 | 3 | 4180.038 | 4180.0211 | 0 | 4 | 127 | Specific | 504.32 | 504.32 | 504.32 | 6.83 | 35 | >sp\|P01876\|IGHA1_HUMAN Immunoglobulin heavy constant alpha 1 OS=Homo sapiens OX=9606 GN=IGHA1 PE=1 SV=2 | 255 | 20201119_16913_1ng_glyco_hcd30_etd30_compound15.17956.17956.3 | controllerType=0 controllerNumber=1 scan=17956 | 42.7511 |
| 14375:4 | 3 | R.LSLHRPALEDLLLGSEAN[+1216.423]LTC[+57.021]TLTGLR.D | HexNAc(2)Hex(5) | C[+57], N[+1216] | 1045.7652 | 4 | 4180.0389 | 4180.0211 | 0 | 4.2 | 127 | Specific | 555.03 | 555.03 | 555.03 | 7.75 | 35 | >sp\|P01876\|IGHA1_HUMAN Immunoglobulin heavy constant alpha 1 OS=Homo sapiens OX=9606 GN=IGHA1 PE=1 SV=2 | 255 | 20201119_16913_1ng_glyco_hcd30_etd30_compound15.17946.17946.4 | controllerType=0 controllerNumber=1 scan=17946 | 42.7242 |
| 14386:4 | 3 | R.LSLHRPALEDLLLGSEAN[+1540.529]LTC[+57.021]TLTGLR.D | HexNAc(2)Hex(7) | C[+57], N[+1541] | 1126.7865 | 4 | 4504.1243 | 4504.1268 | 0 | -0.5 | 127 | Specific | 570.36 | 567.25 | 567.25 | 7.46 | 35 | >sp\|P01876\|IGHA1_HUMAN Immunoglobulin heavy constant alpha 1 OS=Homo sapiens OX=9606 GN=IGHA1 PE=1 SV=2 | 255 | 20201119_16913_1ng_glyco_hcd30_etd30_compound15.17957.17957.4 | controllerType=0 controllerNumber=1 scan=17957 | 42.7559 |
| 14430:4 | 3 | R.LSLHRPALEDLLLGSEAN[+1501.555]LTC[+57.021]TLTGLR.D | HexNAc(5)Hex(3) | C[+57], N[+1502] | 1117.0472 | 4 | 4465.1668 | 4465.1536 | 0 | 3 | 127 | Specific | 592.8 | 592.8 | 592.8 | 9.34 | 35 | >sp\|P01876\|IGHA1_HUMAN Immunoglobulin heavy constant alpha 1 OS=Homo sapiens OX=9606 GN=IGHA1 PE=1 SV=2 | 255 | 20201119_16913_1ng_glyco_hcd30_etd30_compound15.18004.18004.4 | controllerType=0 controllerNumber=1 scan=18004 | 42.9449 |
| 14452:4 | 3 | R.LSLHRPALEDLLLGSEAN[+1298.476]LTC[+57.021]TLTGLR.D | HexNAc(4)Hex(3) | C[+57], N[+1298] | 1066.2786 | 4 | 4262.0926 | 4262.0742 | 0 | 4.3 | 127 | Specific | 721.09 | 721.09 | 721.09 | 11.78 | 35 | >sp\|P01876\|IGHA1_HUMAN Immunoglobulin heavy constant alpha 1 OS=Homo sapiens OX=9606 GN=IGHA1 PE=1 SV=2 | 255 | 20201119_16913_1ng_glyco_hcd30_etd30_compound15.18027.18027.4 | controllerType=0 controllerNumber=1 scan=18027 | 43.0109 |
| 14396:4 | 3 | R.LSLHRPALEDLLLGSEAN[+1257.449]LTC[+57.021]TLTGLR.D | HexNAc(3)Hex(4) | C[+57], N[+1257] | 1056.0201 | 4 | 4221.0584 | 4221.0477 | 0 | 2.5 | 127 | Specific | 772.94 | 765.2 | 765.2 | 13.34 | 35 | >sp\|P01876\|IGHA1_HUMAN Immunoglobulin heavy constant alpha 1 OS=Homo sapiens OX=9606 GN=IGHA1 PE=1 SV=2 | 255 | 20201119_16913_1ng_glyco_hcd30_etd30_compound15.17968.17968.4 | controllerType=0 controllerNumber=1 scan=17968 | 42.8006 |
| 15076:4 | 3 | R.LSLHRPALEDLLLGSEAN[+1710.598]LTC[+57.021]TLTGLR.D | HexNAc(3)Hex(5)NeuAc(1) | C[+57], N[+1711] | 1169.3094 | 4 | 4674.2158 | 4674.1959 | 0 | 4.2 | 127 | Specific | 851.35 | 815.8 | 815.8 | 14.78 | 35 | >sp\|P01876\|IGHA1_HUMAN Immunoglobulin heavy constant alpha 1 OS=Homo sapiens OX=9606 GN=IGHA1 PE=1 SV=2 | 255 | 20201119_16913_1ng_glyco_hcd30_etd30_compound15.18685.18685.4 | controllerType=0 controllerNumber=1 scan=18685 | 44.8877 |
| 14390:4 | 3 | R.LSLHRPALEDLLLGSEAN[+1622.582]LTC[+57.021]TLTGLR.D | HexNAc(4)Hex(5) | C[+57], N[+1623] | 1147.3052 | 4 | 4586.199 | 4586.1799 | 0 | 4.2 | 127 | Specific | 867.7 | 776.57 | 776.57 | 14.78 | 35 | >sp\|P01876\|IGHA1_HUMAN Immunoglobulin heavy constant alpha 1 OS=Homo sapiens OX=9606 GN=IGHA1 PE=1 SV=2 | 255 | 20201119_16913_1ng_glyco_hcd30_etd30_compound15.17962.17962.4 | controllerType=0 controllerNumber=1 scan=17962 | 42.7699 |
| 15078:4 | 3 | R.LSLHRPALEDLLLGSEAN[+1751.624]LTC[+57.021]TLTGLR.D | HexNAc(4)Hex(4)NeuAc(1) | C[+57], N[+1752] | 1179.5663 | 4 | 4715.2434 | 4715.2225 | 0 | 4.4 | 127 | Specific | 878.15 | 878.15 | 878.15 | 14.78 | 35 | >sp\|P01876\|IGHA1_HUMAN Immunoglobulin heavy constant alpha 1 OS=Homo sapiens OX=9606 GN=IGHA1 PE=1 SV=2 | 255 | 20201119_16913_1ng_glyco_hcd30_etd30_compound15.18687.18687.4 | controllerType=0 controllerNumber=1 scan=18687 | 44.9031 |
| 15068:4 | 3 | R.LSLHRPALEDLLLGSEAN[+1872.651]LTC[+57.021]TLTGLR.D | HexNAc(3)Hex(6)NeuAc(1) | C[+57], N[+1873] | 1209.8228 | 4 | 4836.2694 | 4836.2487 | 0 | 4.3 | 127 | Specific | 878.57 | 728.37 | 728.37 | 14.78 | 35 | >sp\|P01876\|IGHA1_HUMAN Immunoglobulin heavy constant alpha 1 OS=Homo sapiens OX=9606 GN=IGHA1 PE=1 SV=2 | 255 | 20201119_16913_1ng_glyco_hcd30_etd30_compound15.18676.18676.4 | controllerType=0 controllerNumber=1 scan=18676 | 44.8686 |
| 14329:4 | 3 | R.LSLHRPALEDLLLGSEAN[+1702.581]LTC[+57.021]TLTGLR.D | HexNAc(2)Hex(8) | C[+57], N[+1703] | 1167.3095 | 4 | 4666.2161 | 4666.1796 | 0 | 7.8 | 127 | Specific | 948.53 | 947.65 | 947.65 | 15.22 | 35 | >sp\|P01876\|IGHA1_HUMAN Immunoglobulin heavy constant alpha 1 OS=Homo sapiens OX=9606 GN=IGHA1 PE=1 SV=2 | 255 | 20201119_16913_1ng_glyco_hcd30_etd30_compound15.17897.17897.4 | controllerType=0 controllerNumber=1 scan=17897 | 42.5654 |
| 14374:4 | 3 | R.LSLHRPALEDLLLGSEAN[+1501.555]LTC[+57.021]TLTGLR.D | HexNAc(5)Hex(3) | C[+57], N[+1502] | 1117.0514 | 4 | 4465.1839 | 4465.1536 | 0 | 6.8 | 127 | Specific | 972.52 | 972.52 | 972.52 | 16.22 | 35 | >sp\|P01876\|IGHA1_HUMAN Immunoglobulin heavy constant alpha 1 OS=Homo sapiens OX=9606 GN=IGHA1 PE=1 SV=2 | 255 | 20201119_16913_1ng_glyco_hcd30_etd30_compound15.17945.17945.4 | controllerType=0 controllerNumber=1 scan=17945 | 42.7192 |
| 14315:4 | 3 | R.LSLHRPALEDLLLGSEAN[+1864.634]LTC[+57.021]TLTGLR.D | HexNAc(2)Hex(9) | C[+57], N[+1865] | 1207.8177 | 4 | 4828.2488 | 4828.2324 | 0 | 3.4 | 127 | Specific | 992.64 | 952.06 | 952.06 | 17.34 | 35 | >sp\|P01876\|IGHA1_HUMAN Immunoglobulin heavy constant alpha 1 OS=Homo sapiens OX=9606 GN=IGHA1 PE=1 SV=2 | 255 | 20201119_16913_1ng_glyco_hcd30_etd30_compound15.17882.17882.4 | controllerType=0 controllerNumber=1 scan=17882 | 42.4928 |
| 14380:4 | 3 | R.LSLHRPALEDLLLGSEAN[+1460.529]LTC[+57.021]TLTGLR.D | HexNAc(4)Hex(4) | C[+57], N[+1461] | 1106.7956 | 4 | 4424.1605 | 4424.1271 | 0 | 7.6 | 127 | Specific | 1084.52 | 986.29 | 986.29 | 18.22 | 35 | >sp\|P01876\|IGHA1_HUMAN Immunoglobulin heavy constant alpha 1 OS=Homo sapiens OX=9606 GN=IGHA1 PE=1 SV=2 | 255 | 20201119_16913_1ng_glyco_hcd30_etd30_compound15.17951.17951.4 | controllerType=0 controllerNumber=1 scan=17951 | 42.7373 |
| 11227:4 | 3 | R.LAGKPTHVN[+2262.814]VSVVMAEVDGTC[+57.021]Y.- | HexNAc(5)Hex(5)Fuc(1)NeuAc(1) | C[+57], N[+2263] | 1153.2492 | 4 | 4609.9749 | 4609.9564 | 0 | 4 | 332 | CRagged | 218.19 | 131.94 | 131.94 | 2.95 | 35 | >sp\|P01876\|IGHA1_HUMAN Immunoglobulin heavy constant alpha 1 OS=Homo sapiens OX=9606 GN=IGHA1 PE=1 SV=2 | 255 | 20201119_16913_1ng_glyco_hcd30_etd30_compound15.14620.14620.4 | controllerType=0 controllerNumber=1 scan=14620 | 33.905 |
| 09861:3 | 3 | R.LAGKPTHVN[+1501.555]VSVVMAEVDGTC[+57.021]Y.- | HexNAc(5)Hex(3) | C[+57], N[+1502] | 1283.5712 | 3 | 3848.6991 | 3848.6974 | 0 | 0.4 | 332 | CRagged | 236.63 | 149.05 | 149.05 | 2.59 | 35 | >sp\|P01876\|IGHA1_HUMAN Immunoglobulin heavy constant alpha 1 OS=Homo sapiens OX=9606 GN=IGHA1 PE=1 SV=2 | 255 | 20201119_16913_1ng_glyco_hcd30_etd30_compound15.13183.13183.3 | controllerType=0 controllerNumber=1 scan=13183 | 30.8006 |
| 09821:3 | 3 | R.LAGKPTHVN[+1809.666]VSVVMAEVDGTC[+57.021]Y.- | HexNAc(5)Hex(4)Fuc(1) | C[+57], N[+1810] | 1386.2794 | 3 | 4156.8235 | 4156.8081 | 0 | 3.7 | 332 | CRagged | 236.74 | 157.82 | 157.82 | 2.91 | 35 | >sp\|P01876\|IGHA1_HUMAN Immunoglobulin heavy constant alpha 1 OS=Homo sapiens OX=9606 GN=IGHA1 PE=1 SV=2 | 255 | 20201119_16913_1ng_glyco_hcd30_etd30_compound15.13141.13141.3 | controllerType=0 controllerNumber=1 scan=13141 | 30.6683 |
| 09836:3 | 3 | R.LAGKPTHVN[+1809.666]VSVVMAEVDGTC[+57.021]Y.- | HexNAc(5)Hex(4)Fuc(1) | C[+57], N[+1810] | 1386.2781 | 3 | 4156.8198 | 4156.8081 | 0 | 2.8 | 332 | CRagged | 237.31 | 116.09 | 116.09 | 2.88 | 35 | >sp\|P01876\|IGHA1_HUMAN Immunoglobulin heavy constant alpha 1 OS=Homo sapiens OX=9606 GN=IGHA1 PE=1 SV=2 | 255 | 20201119_16913_1ng_glyco_hcd30_etd30_compound15.13157.13157.3 | controllerType=0 controllerNumber=1 scan=13157 | 30.7192 |
| 09787:3 | 3 | R.LAGKPTHVN[+1378.476]VSVVMAEVDGTC[+57.021]Y.- | HexNAc(2)Hex(6) | C[+57], N[+1378] | 1242.5471 | 3 | 3725.6266 | 3725.6178 | 0 | 2.4 | 332 | CRagged | 249.02 | 204.78 | 184.78 | 3.83 | 35 | >sp\|P01876\|IGHA1_HUMAN Immunoglobulin heavy constant alpha 1 OS=Homo sapiens OX=9606 GN=IGHA1 PE=1 SV=2 | 255 | 20201119_16913_1ng_glyco_hcd30_etd30_compound15.13105.13105.3 | controllerType=0 controllerNumber=1 scan=13105 | 30.5573 |
| 09763:3 | 3 | R.LAGKPTHVN[+1702.581]VSVVMAEVDGTC[+57.021]Y.- | HexNAc(2)Hex(8) | C[+57], N[+1703] | 1350.5806 | 3 | 4049.7273 | 4049.7234 | 0 | 1 | 332 | CRagged | 249.44 | 180.56 | 180.56 | 3.24 | 35 | >sp\|P01876\|IGHA1_HUMAN Immunoglobulin heavy constant alpha 1 OS=Homo sapiens OX=9606 GN=IGHA1 PE=1 SV=2 | 255 | 20201119_16913_1ng_glyco_hcd30_etd30_compound15.13080.13080.3 | controllerType=0 controllerNumber=1 scan=13080 | 30.4928 |
| 09875:3 | 3 | R.LAGKPTHVN[+1647.613]VSVVMAEVDGTC[+57.021]Y.- | HexNAc(5)Hex(3)Fuc(1) | C[+57], N[+1648] | 1332.2603 | 3 | 3994.7664 | 3994.7553 | 0 | 2.8 | 332 | CRagged | 281.43 | 145.01 | 145.01 | 3.59 | 35 | >sp\|P01876\|IGHA1_HUMAN Immunoglobulin heavy constant alpha 1 OS=Homo sapiens OX=9606 GN=IGHA1 PE=1 SV=2 | 255 | 20201119_16913_1ng_glyco_hcd30_etd30_compound15.13198.13198.3 | controllerType=0 controllerNumber=1 scan=13198 | 30.8445 |
| 09840:3 | 3 | R.LAGKPTHVN[+1216.423]VSVVMAEVDGTC[+57.021]Y.- | HexNAc(2)Hex(5) | C[+57], N[+1216] | 1188.5306 | 3 | 3563.5772 | 3563.5649 | 0 | 3.4 | 332 | CRagged | 292.97 | 274.5 | 213.4 | 3.88 | 35 | >sp\|P01876\|IGHA1_HUMAN Immunoglobulin heavy constant alpha 1 OS=Homo sapiens OX=9606 GN=IGHA1 PE=1 SV=2 | 255 | 20201119_16913_1ng_glyco_hcd30_etd30_compound15.13161.13161.3 | controllerType=0 controllerNumber=1 scan=13161 | 30.7308 |
| 09786:3 | 3 | R.LAGKPTHVN[+1540.529]VSVVMAEVDGTC[+57.021]Y.- | HexNAc(2)Hex(7) | C[+57], N[+1541] | 1296.5635 | 3 | 3887.676 | 3887.6706 | 0 | 1.4 | 332 | CRagged | 294.2 | 226.25 | 217.69 | 3.88 | 35 | >sp\|P01876\|IGHA1_HUMAN Immunoglobulin heavy constant alpha 1 OS=Homo sapiens OX=9606 GN=IGHA1 PE=1 SV=2 | 255 | 20201119_16913_1ng_glyco_hcd30_etd30_compound15.13104.13104.3 | controllerType=0 controllerNumber=1 scan=13104 | 30.5475 |
| 11204:4 | 3 | R.LAGKPTHVN[+2263.835]VSVVMAEVDGTC[+57.021]Y.- | HexNAc(5)Hex(5)Fuc(3) | C[+57], N[+2264] | 1153.4996 | 4 | 4610.9764 | 4610.9768 | 0 | -0.1 | 332 | CRagged | 410.25 | 360.75 | 360.75 | 3.15 | 35 | >sp\|P01876\|IGHA1_HUMAN Immunoglobulin heavy constant alpha 1 OS=Homo sapiens OX=9606 GN=IGHA1 PE=1 SV=2 | 255 | 20201119_16913_1ng_glyco_hcd30_etd30_compound15.14596.14596.4 | controllerType=0 controllerNumber=1 scan=14596 | 33.8414 |
| 16599:3 | 4 | K.QIGLYPVLVIDSSGYVNPN[+2350.830]YTGR.I | HexNAc(4)Hex(5)Fuc(1)NeuAc(2) | N[+2351] | 1626.0554 | 3 | 4876.1515 | 4876.1338 | 0 | 3.6 | 168 | Specific | 365.33 | 235.55 | 235.55 | 5.74 | 38 | >sp\|P01833\|PIGR_HUMAN Polymeric immunoglobulin receptor OS=Homo sapiens OX=9606 GN=PIGR PE=1 SV=4 | 902 | 20201119_16913_1ng_glyco_hcd30_etd30_compound15.20295.20295.3 | controllerType=0 controllerNumber=1 scan=20295 | 49.7034 |
| 16660:4 | 4 | K.QIGLYPVLVIDSSGYVNPN[+2204.772]YTGR.I | HexNAc(4)Hex(5)NeuAc(2) | N[+2205] | 1183.2787 | 4 | 4730.0931 | 4730.0759 | 0 | 3.6 | 168 | Specific | 387.16 | 360.58 | 360.58 | 5.27 | 38 | >sp\|P01833\|PIGR_HUMAN Polymeric immunoglobulin receptor OS=Homo sapiens OX=9606 GN=PIGR PE=1 SV=4 | 902 | 20201119_16913_1ng_glyco_hcd30_etd30_compound15.20359.20359.4 | controllerType=0 controllerNumber=1 scan=20359 | 49.8996 |
| 15188:4 | 4 | K.QIGLYPVLVIDSSGYVNPN[+2059.735]YTGR.I | HexNAc(4)Hex(5)Fuc(1)NeuAc(1) | N[+2060] | 1147.0254 | 4 | 4585.0799 | 4585.0384 | 0 | 9.1 | 168 | Specific | 390.36 | 390.36 | 390.36 | 4.88 | 38 | >sp\|P01833\|PIGR_HUMAN Polymeric immunoglobulin receptor OS=Homo sapiens OX=9606 GN=PIGR PE=1 SV=4 | 902 | 20201119_16913_1ng_glyco_hcd30_etd30_compound15.18805.18805.4 | controllerType=0 controllerNumber=1 scan=18805 | 45.3565 |
| 15223:3 | 4 | K.QIGLYPVLVIDSSGYVNPN[+1913.677]YTGR.I | HexNAc(4)Hex(5)NeuAc(1) | N[+1914] | 1480.3373 | 3 | 4438.9975 | 4438.9805 | 0 | 3.8 | 168 | Specific | 590.92 | 496.7 | 496.7 | 8.63 | 38 | >sp\|P01833\|PIGR_HUMAN Polymeric immunoglobulin receptor OS=Homo sapiens OX=9606 GN=PIGR PE=1 SV=4 | 902 | 20201119_16913_1ng_glyco_hcd30_etd30_compound15.18842.18842.3 | controllerType=0 controllerNumber=1 scan=18842 | 45.4766 |
| 16448:3 | 4 | R.[+42.011]LSLLEEPGN[+2018.708]GTFTVILNQLTSR.D | HexNAc(3)Hex(6)Fuc(1)NeuAc(1) | .[42], N[+2019] | 1488.3528 | 3 | 4463.044 | 4463.0115 | 0 | 7.3 | 413 | Specific | 247.94 | 199.23 | 199.23 | 3.44 | 38 | >sp\|P01833\|PIGR_HUMAN Polymeric immunoglobulin receptor OS=Homo sapiens OX=9606 GN=PIGR PE=1 SV=4 | 902 | 20201119_16913_1ng_glyco_hcd30_etd30_compound15.20133.20133.3 | controllerType=0 controllerNumber=1 scan=20133 | 49.1053 |
| 18799:3 | 4 | R.[+42.011]LSLLEEPGN[+2205.793]GTFTVILNQLTSR.D | HexNAc(4)Hex(5)Fuc(2)NeuAc(1) | .[42], N[+2206] | 1550.7044 | 3 | 4650.0987 | 4650.096 | 0 | 0.6 | 413 | Specific | 264.78 | 177.1 | 177.1 | 4.23 | 38 | >sp\|P01833\|PIGR_HUMAN Polymeric immunoglobulin receptor OS=Homo sapiens OX=9606 GN=PIGR PE=1 SV=4 | 902 | 20201119_16913_1ng_glyco_hcd30_etd30_compound15.22631.22631.3 | controllerType=0 controllerNumber=1 scan=22631 | 57.0217 |
| 21667:4 | 4 | R.[+42.011]LSLLEEPGN[+2351.851]GTFTVILNQLTSR.D | HexNAc(4)Hex(5)Fuc(3)NeuAc(1) | .[42], N[+2352] | 1199.7935 | 4 | 4796.1522 | 4796.1539 | 0 | -0.4 | 413 | Specific | 294.07 | 262.24 | 262.24 | 2.86 | 38 | >sp\|P01833\|PIGR_HUMAN Polymeric immunoglobulin receptor OS=Homo sapiens OX=9606 GN=PIGR PE=1 SV=4 | 902 | 20201119_16913_1ng_glyco_hcd30_etd30_compound15.25669.25669.4 | controllerType=0 controllerNumber=1 scan=25669 | 65.3712 |
| 17984:3 | 4 | R.LSLLEEPGN[+2350.830]GTFTVILNQLTSR.D | HexNAc(4)Hex(5)Fuc(1)NeuAc(2) | N[+2351] | 1585.0578 | 3 | 4753.1588 | 4753.1229 | 0 | 7.6 | 413 | Specific | 329.62 | 239.68 | 239.68 | 4.46 | 38 | >sp\|P01833\|PIGR_HUMAN Polymeric immunoglobulin receptor OS=Homo sapiens OX=9606 GN=PIGR PE=1 SV=4 | 902 | 20201119_16913_1ng_glyco_hcd30_etd30_compound15.21760.21760.3 | controllerType=0 controllerNumber=1 scan=21760 | 54.0509 |
| 16400:4 | 4 | R.LSLLEEPGN[+2059.735]GTFTVILNQLTSR.D | HexNAc(4)Hex(5)Fuc(1)NeuAc(1) | N[+2060] | 1116.2654 | 4 | 4462.0399 | 4462.0275 | 0 | 2.8 | 413 | Specific | 444.12 | 412.82 | 412.82 | 6.6 | 38 | >sp\|P01833\|PIGR_HUMAN Polymeric immunoglobulin receptor OS=Homo sapiens OX=9606 GN=PIGR PE=1 SV=4 | 902 | 20201119_16913_1ng_glyco_hcd30_etd30_compound15.20083.20083.4 | controllerType=0 controllerNumber=1 scan=20083 | 48.9808 |
| 16353:4 | 4 | R.LSLLEEPGN[+2205.793]GTFTVILNQLTSR.D | HexNAc(4)Hex(5)Fuc(2)NeuAc(1) | N[+2206] | 1152.7819 | 4 | 4608.1058 | 4608.0854 | 0 | 4.4 | 413 | Specific | 568.64 | 523.41 | 523.41 | 7.24 | 38 | >sp\|P01833\|PIGR_HUMAN Polymeric immunoglobulin receptor OS=Homo sapiens OX=9606 GN=PIGR PE=1 SV=4 | 902 | 20201119_16913_1ng_glyco_hcd30_etd30_compound15.20033.20033.4 | controllerType=0 controllerNumber=1 scan=20033 | 48.8101 |
| 16396:3 | 4 | R.LSLLEEPGN[+2059.735]GTFTVILNQLTSR.D | HexNAc(4)Hex(5)Fuc(1)NeuAc(1) | N[+2060] | 1488.0201 | 3 | 4462.0457 | 4462.0275 | 0 | 4.1 | 413 | Specific | 623.96 | 534.92 | 534.92 | 9.07 | 38 | >sp\|P01833\|PIGR_HUMAN Polymeric immunoglobulin receptor OS=Homo sapiens OX=9606 GN=PIGR PE=1 SV=4 | 902 | 20201119_16913_1ng_glyco_hcd30_etd30_compound15.20079.20079.3 | controllerType=0 controllerNumber=1 scan=20079 | 48.9734 |
| 17755:4 | 4 | R.LSLLEEPGN[+2350.830]GTFTVILNQLTSR.D | HexNAc(4)Hex(5)Fuc(1)NeuAc(2) | N[+2351] | 1189.0411 | 4 | 4753.1424 | 4753.1229 | 0 | 4.1 | 413 | Specific | 640.66 | 542.47 | 542.47 | 9.07 | 38 | >sp\|P01833\|PIGR_HUMAN Polymeric immunoglobulin receptor OS=Homo sapiens OX=9606 GN=PIGR PE=1 SV=4 | 902 | 20201119_16913_1ng_glyco_hcd30_etd30_compound15.21516.21516.3 | controllerType=0 controllerNumber=1 scan=21516 | 53.2297 |
| 17829:3 | 4 | R.LSLLEEPGN[+2350.830]GTFTVILNQLTSR.D | HexNAc(4)Hex(5)Fuc(1)NeuAc(2) | N[+2351] | 1585.05 | 3 | 4753.1354 | 4753.1229 | 0 | 2.6 | 413 | Specific | 642.54 | 462.03 | 462.03 | 9.63 | 38 | >sp\|P01833\|PIGR_HUMAN Polymeric immunoglobulin receptor OS=Homo sapiens OX=9606 GN=PIGR PE=1 SV=4 | 902 | 20201119_16913_1ng_glyco_hcd30_etd30_compound15.21595.21595.3 | controllerType=0 controllerNumber=1 scan=21595 | 53.4906 |
| 13267:4 | 4 | K.IIEGEPNLKVPGN[+2059.735]VTAVLGETLK.V | HexNAc(4)Hex(5)Fuc(1)NeuAc(1) | N[+2060] | 1113.5336 | 4 | 4451.1126 | 4451.0843 | 0 | 6.4 | 457 | Specific | 226.06 | 226.06 | 226.06 | 2.54 | 38 | >sp\|P01833\|PIGR_HUMAN Polymeric immunoglobulin receptor OS=Homo sapiens OX=9606 GN=PIGR PE=1 SV=4 | 902 | 20201119_16913_1ng_glyco_hcd30_etd30_compound15.16776.16776.2 | controllerType=0 controllerNumber=1 scan=16776 | 39.4005 |
| 13170:3 | 4 | K.IIEGEPNLKVPGN[+2205.793]VTAVLGETLK.V | HexNAc(4)Hex(5)Fuc(2)NeuAc(1) | N[+2206] | 1533.0566 | 3 | 4597.1552 | 4597.1422 | 0 | 2.8 | 457 | Specific | 293.91 | 235.01 | 235.01 | 4.41 | 38 | >sp\|P01833\|PIGR_HUMAN Polymeric immunoglobulin receptor OS=Homo sapiens OX=9606 GN=PIGR PE=1 SV=4 | 902 | 20201119_16913_1ng_glyco_hcd30_etd30_compound15.16674.16674.3 | controllerType=0 controllerNumber=1 scan=16674 | 39.1346 |
| 13212:3 | 4 | K.IIEGEPNLKVPGN[+2205.793]VTAVLGETLK.V | HexNAc(4)Hex(5)Fuc(2)NeuAc(1) | N[+2206] | 1533.0573 | 3 | 4597.1574 | 4597.1422 | 0 | 3.3 | 457 | Specific | 396.43 | 326.49 | 326.49 | 4.25 | 38 | >sp\|P01833\|PIGR_HUMAN Polymeric immunoglobulin receptor OS=Homo sapiens OX=9606 GN=PIGR PE=1 SV=4 | 902 | 20201119_16913_1ng_glyco_hcd30_etd30_compound15.16719.16719.3 | controllerType=0 controllerNumber=1 scan=16719 | 39.2491 |
| 13216:3 | 4 | K.IIEGEPNLKVPGN[+2059.735]VTAVLGETLK.V | HexNAc(4)Hex(5)Fuc(1)NeuAc(1) | N[+2060] | 1484.3709 | 3 | 4451.0982 | 4451.0843 | 0 | 3.1 | 457 | Specific | 457.5 | 382.09 | 382.09 | 6.97 | 38 | >sp\|P01833\|PIGR_HUMAN Polymeric immunoglobulin receptor OS=Homo sapiens OX=9606 GN=PIGR PE=1 SV=4 | 902 | 20201119_16913_1ng_glyco_hcd30_etd30_compound15.16723.16723.3 | controllerType=0 controllerNumber=1 scan=16723 | 39.2584 |
| 13220:4 | 4 | K.IIEGEPNLKVPGN[+2205.793]VTAVLGETLK.V | HexNAc(4)Hex(5)Fuc(2)NeuAc(1) | N[+2206] | 1150.0448 | 4 | 4597.1575 | 4597.1422 | 0 | 3.3 | 457 | Specific | 617.53 | 497.46 | 497.46 | 8.03 | 38 | >sp\|P01833\|PIGR_HUMAN Polymeric immunoglobulin receptor OS=Homo sapiens OX=9606 GN=PIGR PE=1 SV=4 | 902 | 20201119_16913_1ng_glyco_hcd30_etd30_compound15.16727.16727.4 | controllerType=0 controllerNumber=1 scan=16727 | 39.2666 |
| 13846:4 | 4 | K.IIEGEPNLKVPGN[+2350.830]VTAVLGETLKVPC[+57.021]HFPC[+57.021]K.F | HexNAc(4)Hex(5)Fuc(1)NeuAc(2) | C[+57]*2, N[+2351] | 1442.6713 | 4 | 5767.6632 | 5767.6372 | 0 | 4.5 | 457 | Specific | 274.71 | 274.71 | 274.71 | 2.86 | 38 | >sp\|P01833\|PIGR_HUMAN Polymeric immunoglobulin receptor OS=Homo sapiens OX=9606 GN=PIGR PE=1 SV=4 | 902 | 20201119_16913_1ng_glyco_hcd30_etd30_compound15.17386.17386.2 | controllerType=0 controllerNumber=1 scan=17386 | 41.0611 |
| 12895:5 | 4 | K.IIEGEPNLKVPGN[+2205.793]VTAVLGETLKVPC[+57.021]HFPC[+57.021]K.F | HexNAc(4)Hex(5)Fuc(2)NeuAc(1) | C[+57]*2, N[+2206] | 1125.3291 | 5 | 5622.6164 | 5622.5997 | 0 | 3 | 457 | Specific | 336.92 | 336.92 | 336.92 | 3.49 | 38 | >sp\|P01833\|PIGR_HUMAN Polymeric immunoglobulin receptor OS=Homo sapiens OX=9606 GN=PIGR PE=1 SV=4 | 902 | 20201119_16913_1ng_glyco_hcd30_etd30_compound15.16372.16372.5 | controllerType=0 controllerNumber=1 scan=16372 | 38.4507 |
| 12873:5 | 4 | K.IIEGEPNLKVPGN[+2205.793]VTAVLGETLKVPC[+57.021]HFPC[+57.021]K.F | HexNAc(4)Hex(5)Fuc(2)NeuAc(1) | C[+57]*2, N[+2206] | 1125.3306 | 5 | 5622.6237 | 5622.5997 | 0 | 4.3 | 457 | Specific | 595.58 | 595.58 | 595.58 | 6.36 | 38 | >sp\|P01833\|PIGR_HUMAN Polymeric immunoglobulin receptor OS=Homo sapiens OX=9606 GN=PIGR PE=1 SV=4 | 902 | 20201119_16913_1ng_glyco_hcd30_etd30_compound15.16349.16349.5 | controllerType=0 controllerNumber=1 scan=16349 | 38.3742 |
| 13638:3 | 4 | K.VPGN[+2350.830]VTAVLGETLK.V | HexNAc(4)Hex(5)Fuc(1)NeuAc(2) | N[+2351] | 1250.2205 | 3 | 3748.647 | 3748.6302 | 0 | 4.5 | 466 | Specific | 329.48 | 190.91 | 190.91 | 4.75 | 38 | >sp\|P01833\|PIGR_HUMAN Polymeric immunoglobulin receptor OS=Homo sapiens OX=9606 GN=PIGR PE=1 SV=4 | 902 | 20201119_16913_1ng_glyco_hcd30_etd30_compound15.17166.17166.3 | controllerType=0 controllerNumber=1 scan=17166 | 40.4347 |
| 11447:3 | 4 | K.VPGN[+2205.793]VTAVLGETLK.V | HexNAc(4)Hex(5)Fuc(2)NeuAc(1) | N[+2206] | 1201.8755 | 3 | 3603.6121 | 3603.5927 | 0 | 5.4 | 466 | Specific | 335.35 | 227.46 | 227.46 | 5.01 | 38 | >sp\|P01833\|PIGR_HUMAN Polymeric immunoglobulin receptor OS=Homo sapiens OX=9606 GN=PIGR PE=1 SV=4 | 902 | 20201119_16913_1ng_glyco_hcd30_etd30_compound15.14851.14851.3 | controllerType=0 controllerNumber=1 scan=14851 | 34.4839 |
| 11507:3 | 4 | K.VPGN[+2059.735]VTAVLGETLK.V | HexNAc(4)Hex(5)Fuc(1)NeuAc(1) | N[+2060] | 1153.1865 | 3 | 3457.5448 | 3457.5348 | 0 | 2.9 | 466 | Specific | 344.9 | 236.88 | 236.88 | 5.57 | 38 | >sp\|P01833\|PIGR_HUMAN Polymeric immunoglobulin receptor OS=Homo sapiens OX=9606 GN=PIGR PE=1 SV=4 | 902 | 20201119_16913_1ng_glyco_hcd30_etd30_compound15.14914.14914.3 | controllerType=0 controllerNumber=1 scan=14914 | 34.6757 |
| 12846:3 | 4 | K.VPGN[+2350.830]VTAVLGETLKVPC[+57.021]HFPC[+57.021]K.F | HexNAc(4)Hex(5)Fuc(1)NeuAc(2) | C[+57]*2, N[+2351] | 1592.0362 | 3 | 4774.094 | 4774.0877 | 0 | 1.3 | 466 | Specific | 334.75 | 244.58 | 244.58 | 2.99 | 38 | >sp\|P01833\|PIGR_HUMAN Polymeric immunoglobulin receptor OS=Homo sapiens OX=9606 GN=PIGR PE=1 SV=4 | 902 | 20201119_16913_1ng_glyco_hcd30_etd30_compound15.16320.16320.3 | controllerType=0 controllerNumber=1 scan=16320 | [illegible] |
| 06986:3 | 4 | K.WN[+2204.772]NTGC[+57.021]QALPSQDEGPSK.A | HexNAc(4)Hex(5)NeuAc(2) | C[+57], N[+2205] | 1398.5579 | 3 | 4193.6592 | 4193.6491 | 0 | 2.4 | 498 | Specific | 344.52 | 209.17 | 209.17 | 5.3 | 38 | >sp\|P01833\|PIGR_HUMAN Polymeric immunoglobulin receptor OS=Homo sapiens OX=9606 GN=PIGR PE=1 SV=4 | 902 | 20201119_16913_1ng_glyco_hcd30_etd30_compound15.10177.10177.3 | controllerType=0 controllerNumber=1 scan=10177 | [illegible] |
| 06980:3 | 4 | K.WN[+2204.772]NTGC[+57.021]QALPSQDEGPSK.A | HexNAc(4)Hex(5)NeuAc(2) | C[+57], N[+2205] | 1398.5551 | 3 | 4193.6508 | 4193.6491 | 0 | 0.4 | 498 | Specific | 361.52 | 252.34 | 252.34 | 4.87 | 38 | >sp\|P01833\|PIGR_HUMAN Polymeric immunoglobulin receptor OS=Homo sapiens OX=9606 GN=PIGR PE=1 SV=4 | 902 | 20201119_16913_1ng_glyco_hcd30_etd30_compound15.10170.10170.3 | controllerType=0 controllerNumber=1 scan=10170 | [illegible] |
| 04950:3 | 4 | K.WN[+2059.735]NTGC[+57.021]QALPSQDEGPSK.A | HexNAc(4)Hex(5)Fuc(1)NeuAc(1) | C[+57], N[+2060] | 1350.2139 | 3 | 4048.6272 | 4048.6116 | 0 | 3.9 | 498 | Specific | 417.06 | 306.29 | 306.29 | 5.71 | 38 | >sp\|P01833\|PIGR_HUMAN Polymeric immunoglobulin receptor OS=Homo sapiens OX=9606 GN=PIGR PE=1 SV=4 | 902 | 20201119_16913_1ng_glyco_hcd30_etd30_compound15.8060.8060.3 | controllerType=0 controllerNumber=1 scan=8060 | 20.6952 |
| 05110:3 | 4 | K.WN[+1710.598]NTGC[+57.021]QALPSQDEGPSK.A | HexNAc(3)Hex(5)NeuAc(1) | C[+57], N[+1711] | 1233.8321 | 3 | 3699.4817 | 3699.4743 | 0 | 2 | 498 | Specific | 446.08 | 276.59 | 276.59 | 5.01 | 38 | >sp\|P01833\|PIGR_HUMAN Polymeric immunoglobulin receptor OS=Homo sapiens OX=9606 GN=PIGR PE=1 SV=4 | 902 | 20201119_16913_1ng_glyco_hcd30_etd30_compound15.8227.8227.3 | controllerType=0 controllerNumber=1 scan=8227 | 21.0198 |
| 05017:3 | 4 | K.WN[+1913.677]NTGC[+57.021]QALPSQDEGPSK.A | HexNAc(4)Hex(5)NeuAc(1) | C[+57], N[+1914] | 1301.5268 | 3 | 3902.5658 | 3902.5537 | 0 | 3.1 | 498 | Specific | 530.59 | 349.97 | 349.97 | 6.35 | 38 | >sp\|P01833\|PIGR_HUMAN Polymeric immunoglobulin receptor OS=Homo sapiens OX=9606 GN=PIGR PE=1 SV=4 | 902 | 20201119_16913_1ng_glyco_hcd30_etd30_compound15.8130.8130.3 | controllerType=0 controllerNumber=1 scan=8130 | 20.8259 |
| 05105:3 | 4 | K.WN[+1710.598]NTGC[+57.021]QALPSQDEGPSK.A | HexNAc(3)Hex(5)NeuAc(1) | C[+57], N[+1711] | 1233.8331 | 3 | 3699.4846 | 3699.4743 | 0 | 2.8 | 498 | Specific | 556.42 | 399.91 | 399.91 | 7.59 | 38 | >sp\|P01833\|PIGR_HUMAN Polymeric immunoglobulin receptor OS=Homo sapiens OX=9606 GN=PIGR PE=1 SV=4 | 902 | 20201119_16913_1ng_glyco_hcd30_etd30_compound15.8221.8221.3 | controllerType=0 controllerNumber=1 scan=8221 | 21.0015 |
| 06524:3 | 5 | R.FSDGLESN[+1864.634]SSTQFEVK.K | HexNAc(2)Hex(9) | N[+1865] | 1213.8256 | 3 | 3639.4623 | 3639.4472 | 0 | 4.2 | 219 | Specific | 306.47 | 306.47 | 306.47 | 5.03 | 18 | >sp\|P0C0L4\|CO4A_HUMAN Complement C4-A OS=Homo sapiens OX=9606 GN=C4A PE=1 SV=2 | 1235 | 20201119_16913_1ng_glyco_hcd30_etd30_compound15.9696.9696.3 | controllerType=0 controllerNumber=1 scan=9696 | 23.5708 |
| 08701:3 | 7 | K.TVLTPATNHMGN[+1702.581]VTFTIPANR.E | HexNAc(2)Hex(8) | N[+1703] | 1319.9269 | 3 | 3957.7661 | 3957.7414 | 0 | 6.2 | 74 | Specific | 266.59 | 211.5 | 146.96 | 4.26 | 17 | >sp\|P01024\|CO3_HUMAN Complement C3 OS=Homo sapiens OX=9606 GN=C3 PE=1 SV=2 | 492 | 20201119_16913_1ng_glyco_hcd30_etd30_compound15.11972.11972.3 | controllerType=0 controllerNumber=1 scan=11972 | 28.1234 |
| 08736:3 | 7 | K.TVLTPATNHMGN[+1540.529]VTFTIPANR.E | HexNAc(2)Hex(7) | N[+1541] | 1265.9057 | 3 | 3795.7025 | 3795.6886 | 0 | 3.7 | 74 | Specific | 377.18 | 283.41 | 149.17 | 4.09 | 17 | >sp\|P01024\|CO3_HUMAN Complement C3 OS=Homo sapiens OX=9606 GN=C3 PE=1 SV=2 | 492 | 20201119_16913_1ng_glyco_hcd30_etd30_compound15.12009.12009.3 | controllerType=0 controllerNumber=1 scan=12009 | 28.2041 |
| 08754:3 | 7 | K.TVLTPATNHMGN[+1540.529]VTFTIPANR.E | HexNAc(2)Hex(7) | N[+1541] | 1265.9033 | 3 | 3795.6952 | 3795.6886 | 0 | 1.7 | 74 | Specific | 385.01 | 338.89 | 130.05 | 3.57 | 17 | >sp\|P01024\|CO3_HUMAN Complement C3 OS=Homo sapiens OX=9606 GN=C3 PE=1 SV=2 | 492 | 20201119_16913_1ng_glyco_hcd30_etd30_compound15.12028.12028.3 | controllerType=0 controllerNumber=1 scan=12028 | 28.2403 |
| 13987:4 | 8 | K.RNIAAFGGDPNN[+1913.677]ITLFGESAGGASVSLQTLSPYNK.G | HexNAc(4)Hex(5)NeuAc(1) | N[+1914] | 1370.8735 | 4 | 5480.4723 | 5480.4644 | 0 | 1.4 | 196 | Specific | 468.53 | 451.82 | 451.82 | 6.19 | 13 | >sp\|P19835\|CEL_HUMAN Bile salt-activated lipase OS=Homo sapiens OX=9606 GN=CEL PE=1 SV=3 | 476 | 20201119_16913_1ng_glyco_hcd30_etd30_compound15.17535.17535.4 | controllerType=0 controllerNumber=1 scan=17535 | 41.5066 |
| 17498:4 | 10 | K.VFPLSLDSTPQDGNVVVAC[+57.021]LVQGFFPQEPLSVTWSESGQN[+1647.613]VTAI | HexNAc(5)Hex(3)Fuc(1) | C[+57], N[+1648] | 1606.7562 | 4 | 6424.0031 | 6423.9746 | 0 | 4.4 | 8 | Specific | 395 | 395 | 395 | 6.93 | 23 | >sp\|P01877\|IGHA2_HUMAN Immunoglobulin heavy constant alpha 2 OS=Homo sapiens OX=9606 GN=IGHA2 PE=1 SV=4 | 576 | 20201119_16913_1ng_glyco_hcd30_etd30_compound15.21245.21245.4 | controllerType=0 controllerNumber=1 scan=21245 | 52.4076 |
| 17283:4 | 10 | K.VFPLSLDSTPQDGNVVVAC[+57.021]LVQGFFPQEPLSVTWSESGQN[+1809.666]VTAI | HexNAc(5)Hex(4)Fuc(1) | C[+57], N[+1810] | 1647.2665 | 4 | 6586.0441 | 6586.0275 | 0 | 2.5 | 8 | Specific | 509.03 | 509.03 | 509.03 | 7.62 | 23 | >sp\|P01877\|IGHA2_HUMAN Immunoglobulin heavy constant alpha 2 OS=Homo sapiens OX=9606 GN=IGHA2 PE=1 SV=4 | 576 | 20201119_16913_1ng_glyco_hcd30_etd30_compound15.21018.21018.4 | controllerType=0 controllerNumber=1 scan=21018 | 51.8505 |
| 01699:3 | 10 | K.HYTN[+2262.814]SSQDVTVPC[+57.021]R.V | HexNAc(5)Hex(5)Fuc(1)NeuAc(1) | C[+57], N[+2263] | 1309.5313 | 3 | 3926.5794 | 3926.5636 | 0 | 4 | 89 | Specific | 234.39 | 130.4 | 130.4 | 4.37 | 23 | >sp\|P01877\|IGHA2_HUMAN Immunoglobulin heavy constant alpha 2 OS=Homo sapiens OX=9606 GN=IGHA2 PE=1 SV=4 | 576 | 20201119_16913_1ng_glyco_hcd30_etd30_compound15.4682.4682.3 | controllerType=0 controllerNumber=1 scan=4682 | 14.5311 |
| 00953:3 | 10 | K.HYTN[+1647.613]SSQDVTVPC[+57.021]R.V | HexNAc(5)Hex(3)Fuc(1) | C[+57], N[+1648] | 1104.463 | 3 | 3311.3746 | 3311.3625 | 0 | 3.6 | 89 | Specific | 234.47 | 202.27 | 202.27 | 3.88 | 23 | >sp\|P01877\|IGHA2_HUMAN Immunoglobulin heavy constant alpha 2 OS=Homo sapiens OX=9606 GN=IGHA2 PE=1 SV=4 | 576 | 20201119_16913_1ng_glyco_hcd30_etd30_compound15.3906.3906.2 | controllerType=0 controllerNumber=1 scan=3906 | 13.1444 |
| 00834:3 | 10 | K.HYTN[+1971.719]SSQDVTVPC[+57.021]R.V | HexNAc(5)Hex(5)Fuc(1) | C[+57], N[+1972] | 1212.4974 | 3 | 3635.4776 | 3635.4682 | 0 | 2.6 | 89 | Specific | 308.67 | 235.28 | 235.28 | 5.55 | 23 | >sp\|P01877\|IGHA2_HUMAN Immunoglobulin heavy constant alpha 2 OS=Homo sapiens OX=9606 GN=IGHA2 PE=1 SV=4 | 576 | 20201119_16913_1ng_glyco_hcd30_etd30_compound15.3782.3782.3 | controllerType=0 controllerNumber=1 scan=3782 | 12.8952 |
| 00877:3 | 10 | K.HYTN[+1444.534]SSQDVTVPC[+57.021]R.V | HexNAc(4)Hex(3)Fuc(1) | C[+57], N[+1445] | 1036.7687 | 3 | 3108.2916 | 3108.2832 | 0 | 2.7 | 89 | Specific | 366.43 | 282.25 | 282.25 | 4.99 | 23 | >sp\|P01877\|IGHA2_HUMAN Immunoglobulin heavy constant alpha 2 OS=Homo sapiens OX=9606 GN=IGHA2 PE=1 SV=4 | 576 | 20201119_16913_1ng_glyco_hcd30_etd30_compound15.3827.3827.3 | controllerType=0 controllerNumber=1 scan=3827 | 12.9874 |
| 00848:3 | 10 | K.HYTN[+1809.666]SSQDVTVPC[+57.021]R.V | HexNAc(5)Hex(4)Fuc(1) | C[+57], N[+1810] | 1158.4803 | 3 | 3473.4263 | 3473.4154 | 0 | 3.1 | 89 | Specific | 455.78 | 388.87 | 388.87 | 7.46 | 23 | >sp\|P01877\|IGHA2_HUMAN Immunoglobulin heavy constant alpha 2 OS=Homo sapiens OX=9606 GN=IGHA2 PE=1 SV=4 | 576 | 20201119_16913_1ng_glyco_hcd30_etd30_compound15.3797.3797.3 | controllerType=0 controllerNumber=1 scan=3797 | 12.9279 |
| 00873:3 | 10 | K.HYTN[+1647.613]SSQDVTVPC[+57.021]R.V | HexNAc(5)Hex(3)Fuc(1) | C[+57], N[+1648] | 1104.4624 | 3 | 3311.3727 | 3311.3625 | 0 | 3.1 | 89 | Specific | 473.74 | 408.86 | 408.86 | 7.46 | 23 | >sp\|P01877\|IGHA2_HUMAN Immunoglobulin heavy constant alpha 2 OS=Homo sapiens OX=9606 GN=IGHA2 PE=1 SV=4 | 576 | 20201119_16913_1ng_glyco_hcd30_etd30_compound15.3823.3823.3 | controllerType=0 controllerNumber=1 scan=3823 | 12.9777 |
| 02431:2 | 10 | K.TPLTAN[+1971.719]ITK.S | HexNAc(5)Hex(5)Fuc(1) | N[+1972] | 1465.6451 | 2 | 2930.2828 | 2930.2757 | 0 | 2.5 | 200 | Specific | 231.25 | 67.13 | 67.13 | 2.62 | 23 | >sp\|P01877\|IGHA2_HUMAN Immunoglobulin heavy constant alpha 2 OS=Homo sapiens OX=9606 GN=IGHA2 PE=1 SV=4 | 576 | 20201119_16913_1ng_glyco_hcd30_etd30_compound15.5442.5442.2 | controllerType=0 controllerNumber=1 scan=5442 | 15.892 |
| 04109:3 | 10 | K.TPLTAN[+2262.814]ITK.S | HexNAc(5)Hex(5)Fuc(1)NeuAc(1) | N[+2263] | 1074.4644 | 3 | 3221.3786 | 3221.3711 | 0 | 2.3 | 200 | Specific | 256.37 | 114.88 | 114.88 | 4.51 | 23 | >sp\|P01877\|IGHA2_HUMAN Immunoglobulin heavy constant alpha 2 OS=Homo sapiens OX=9606 GN=IGHA2 PE=1 SV=4 | 576 | 20201119_16913_1ng_glyco_hcd30_etd30_compound15.7188.7188.3 | controllerType=0 controllerNumber=1 scan=7188 | 19.1056 |
| 02402:3 | 10 | K.TPLTAN[+1971.719]ITK.S | HexNAc(5)Hex(5)Fuc(1) | N[+1972] | 977.4325 | 3 | 2930.2828 | 2930.2757 | 0 | 2.4 | 200 | Specific | 258.93 | 130.3 | 130.3 | 3.06 | 23 | >sp\|P01877\|IGHA2_HUMAN Immunoglobulin heavy constant alpha 2 OS=Homo sapiens OX=9606 GN=IGHA2 PE=1 SV=4 | 576 | 20201119_16913_1ng_glyco_hcd30_etd30_compound15.5412.5412.3 | controllerType=0 controllerNumber=1 scan=5412 | 15.8319 |
| 02505:2 | 10 | K.TPLTAN[+1501.555]ITK.S | HexNAc(5)Hex(3) | N[+1502] | 1230.564 | 2 | 2460.1207 | 2460.1121 | 0 | 3.5 | 200 | Specific | 270.29 | 54.82 | 54.82 | 2.58 | 23 | >sp\|P01877\|IGHA2_HUMAN Immunoglobulin heavy constant alpha 2 OS=Homo sapiens OX=9606 GN=IGHA2 PE=1 SV=4 | 576 | 20201119_16913_1ng_glyco_hcd30_etd30_compound15.5519.5519.2 | controllerType=0 controllerNumber=1 scan=5519 | 16.0793 |
| 02511:3 | 10 | K.TPLTAN[+1647.613]ITK.S | HexNAc(5)Hex(3)Fuc(1) | N[+1648] | 869.3977 | 3 | 2606.1785 | 2606.17 | 0 | 3.2 | 200 | Specific | 275.32 | 127.65 | 127.65 | 3.06 | 23 | >sp\|P01877\|IGHA2_HUMAN Immunoglobulin heavy constant alpha 2 OS=Homo sapiens OX=9606 GN=IGHA2 PE=1 SV=4 | 576 | 20201119_16913_1ng_glyco_hcd30_etd30_compound15.5526.5526.3 | controllerType=0 controllerNumber=1 scan=5526 | 16.0996 |
| 04114:3 | 10 | K.TPLTAN[+2262.814]ITK.S | HexNAc(5)Hex(5)Fuc(1)NeuAc(1) | N[+2263] | 1074.4668 | 3 | 3221.3859 | 3221.3711 | 0 | 4.6 | 200 | Specific | 275.68 | 112.79 | 112.79 | 3.96 | 23 | >sp\|P01877\|IGHA2_HUMAN Immunoglobulin heavy constant alpha 2 OS=Homo sapiens OX=9606 GN=IGHA2 PE=1 SV=4 | 576 | 20201119_16913_1ng_glyco_hcd30_etd30_compound15.7194.7194.3 | controllerType=0 controllerNumber=1 scan=7194 | 19.1223 |
| 02454:3 | 10 | K.TPLTAN[+1809.666]ITK.S | HexNAc(5)Hex(4)Fuc(1) | N[+1810] | 923.4155 | 3 | 2768.2319 | 2768.2228 | 0 | 3.3 | 200 | Specific | 286.83 | 127.88 | 127.88 | 3 | 23 | >sp\|P01877\|IGHA2_HUMAN Immunoglobulin heavy constant alpha 2 OS=Homo sapiens OX=9606 GN=IGHA2 PE=1 SV=4 | 576 | 20201119_16913_1ng_glyco_hcd30_etd30_compound15.5466.5466.3 | controllerType=0 controllerNumber=1 scan=5466 | 15.9622 |
| 02483:2 | 10 | K.TPLTAN[+1647.613]ITK.S | HexNAc(5)Hex(3)Fuc(1) | N[+1648] | 1303.5929 | 2 | 2606.1786 | 2606.17 | 0 | 3.3 | 200 | Specific | 312.28 | 126.49 | 126.49 | 3.03 | 23 | >sp\|P01877\|IGHA2_HUMAN Immunoglobulin heavy constant alpha 2 OS=Homo sapiens OX=9606 GN=IGHA2 PE=1 SV=4 | 576 | 20201119_16913_1ng_glyco_hcd30_etd30_compound15.5497.5497.2 | controllerType=0 controllerNumber=1 scan=5497 | 16.0383 |
| 02467:2 | 10 | K.TPLTAN[+1444.534]ITK.S | HexNAc(4)Hex(3)Fuc(1) | N[+1445] | 1202.0538 | 2 | 2403.1002 | 2403.0906 | 0 | 4 | 200 | Specific | 320.2 | 84.21 | 84.21 | 2.54 | 23 | >sp\|P01877\|IGHA2_HUMAN Immunoglobulin heavy constant alpha 2 OS=Homo sapiens OX=9606 GN=IGHA2 PE=1 SV=4 | 576 | 20201119_16913_1ng_glyco_hcd30_etd30_compound15.5480.5480.2 | controllerType=0 controllerNumber=1 scan=5480 | 15.9905 |
| 02446:2 | 10 | K.TPLTAN[+1809.666]ITK.S | HexNAc(5)Hex(4)Fuc(1) | N[+1810] | 1384.6193 | 2 | 2768.2313 | 2768.2228 | 0 | 3.1 | 200 | Specific | 322.32 | 126.86 | 126.86 | 3.06 | 23 | >sp\|P01877\|IGHA2_HUMAN Immunoglobulin heavy constant alpha 2 OS=Homo sapiens OX=9606 GN=IGHA2 PE=1 SV=4 | 576 | 20201119_16913_1ng_glyco_hcd30_etd30_compound15.5458.5458.2 | controllerType=0 controllerNumber=1 scan=5458 | 15.9369 |
| 08922:3 | 10 | R.MAGKPTHIN[+1702.581]VSVVMAEADGTC[+57.021]Y.- | HexNAc(2)Hex(8) | C[+57], N[+1703] | 1351.8961 | 3 | 4053.6736 | 4053.6642 | 0 | 2.3 | 319 | CRagged | 248.44 | 248.44 | 167.84 | 4.82 | 23 | >sp\|P01877\|IGHA2_HUMAN Immunoglobulin heavy constant alpha 2 OS=Homo sapiens OX=9606 GN=IGHA2 PE=1 SV=4 | 576 | 20201119_16913_1ng_glyco_hcd30_etd30_compound15.12203.12203.3 | controllerType=0 controllerNumber=1 scan=12203 | 28.6435 |
| 09023:3 | 10 | R.MAGKPTHIN[+1216.423]VSVVMAEADGTC[+57.021]Y.- | HexNAc(2)Hex(5) | C[+57], N[+1216] | 1189.846 | 3 | 3567.5235 | 3567.5057 | 0 | 5 | 319 | CRagged | 271.01 | 265.58 | 189.96 | 3.88 | 23 | >sp\|P01877\|IGHA2_HUMAN Immunoglobulin heavy constant alpha 2 OS=Homo sapiens OX=9606 GN=IGHA2 PE=1 SV=4 | 576 | 20201119_16913_1ng_glyco_hcd30_etd30_compound15.12309.12309.3 | controllerType=0 controllerNumber=1 scan=12309 | 28.8888 |
| 08993:3 | 10 | R.MAGKPTHIN[+1378.476]VSVVMAEADGTC[+57.021]Y.- | HexNAc(2)Hex(6) | C[+57], N[+1378] | 1243.862 | 3 | 3729.5714 | 3729.5585 | 0 | 3.5 | 319 | CRagged | 328.07 | 277.89 | 215.25 | 4.66 | 23 | >sp\|P01877\|IGHA2_HUMAN Immunoglobulin heavy constant alpha 2 OS=Homo sapiens OX=9606 GN=IGHA2 PE=1 SV=4 | 576 | 20201119_16913_1ng_glyco_hcd30_etd30_compound15.12278.12278.3 | controllerType=0 controllerNumber=1 scan=12278 | 28.8113 |
| 16849:5 | 11 | K.APDKNVIFSPLSISTALAFLSLGAHN[+2424.867]TTLTEILK.G | HexNAc(5)Hex(6)Fuc(1)NeuAc(1) | N[+2425] | 1202.3824 | 5 | 6007.8831 | 6007.8453 | 0 | 6.3 | 68 | Specific | 296.34 | 296.34 | 296.34 | 3.23 | 11 | >sp\|P01011\|AACT_HUMAN Alpha-1-antichymotrypsin OS=Homo sapiens OX=9606 GN=SERPINA3 PE=1 SV=2 | 1314 | 20201119_16913_1ng_glyco_hcd30_etd30_compound15.20558.20558.2 | controllerType=0 controllerNumber=1 scan=20558 | 50.4569 |
| 07478:3 | 12 | K.TAHC[+57.021]N[+2059.735]ESFYFLC[+57.021]K.R | HexNAc(4)Hex(5)Fuc(1)NeuAc(1) | C[+57]*2, N[+2060] | 1246.1591 | 3 | 3736.4628 | 3736.4545 | 0 | 2.2 | 1201 | Specific | 221.06 | 130.06 | 130.06 | 4.37 | 10 | >sp\|P22897\|MRC1_HUMAN Macrophage mannose receptor 1 OS=Homo sapiens OX=9606 GN=MRC1 PE=1 SV=1 | 1049 | 20201119_16913_1ng_glyco_hcd30_etd30_compound15.10690.10690.3 | controllerType=0 controllerNumber=1 scan=10690 | 25.5387 |
| 11781:3 | 14 | K.LKELPGVC[+57.021]N[+2059.735]ETMMALWEEC[+57.021]KPC[+57.021]LK.Q | HexNAc(4)Hex(5)Fuc(1)NeuAc(1) | C[+57]*3, N[+2060] | 1666.0511 | 3 | 4996.1388 | 4996.1196 | 0 | 3.8 | 95 | Specific | 274.55 | 190 | 190 | 3.65 | 19 | >sp\|P10909\|CLUS_HUMAN Clusterin OS=Homo sapiens OX=9606 GN=CLU PE=1 SV=1 | 1467 | 20201119_16913_1ng_glyco_hcd30_etd30_compound15.15203.15203.3 | controllerType=0 controllerNumber=1 scan=15203 | 35.4595 |
| 12944:4 | 14 | K.LKELPGVC[+57.021]N[+2350.830]ETMMALWEEC[+57.021]KPC[+57.021]LK.Q | HexNAc(4)Hex(5)Fuc(1)NeuAc(2) | C[+57]*3, N[+2351] | 1322.5696 | 4 | 5287.2565 | 5287.2151 | 0 | 7.8 | 95 | Specific | 365.02 | 216.36 | 216.36 | 2.72 | 19 | >sp\|P10909\|CLUS_HUMAN Clusterin OS=Homo sapiens OX=9606 GN=CLU PE=1 SV=1 | 1467 | 20201119_16913_1ng_glyco_hcd30_etd30_compound15.16424.16424.4 | controllerType=0 controllerNumber=1 scan=16424 | 38.5972 |
| 11799:3 | 14 | K.LKELPGVC[+57.021]N[+2059.735]ETMMALWEEC[+57.021]KPC[+57.021]LK.Q | HexNAc(4)Hex(5)Fuc(1)NeuAc(1) | C[+57]*3, N[+2060] | 1666.051 | 3 | 4996.1384 | 4996.1196 | 0 | 3.8 | 95 | Specific | 367.4 | 311.23 | 311.23 | 4.09 | 19 | >sp\|P10909\|CLUS_HUMAN Clusterin OS=Homo sapiens OX=9606 GN=CLU PE=1 SV=1 | 1467 | 20201119_16913_1ng_glyco_hcd30_etd30_compound15.15222.15222.3 | controllerType=0 controllerNumber=1 scan=15222 | 35.509 |
| 11714:4 | 14 | K.LKELPGVC[+57.021]N[+2205.793]ETMMALWEEC[+57.021]KPC[+57.021]LK.Q | HexNAc(4)Hex(5)Fuc(2)NeuAc(1) | C[+57]*3, N[+2206] | 1286.3025 | 4 | 5142.1881 | 5142.1776 | 0 | 2.1 | 95 | Specific | 662.73 | 474.41 | 474.41 | 8.48 | 19 | >sp\|P10909\|CLUS_HUMAN Clusterin OS=Homo sapiens OX=9606 GN=CLU PE=1 SV=1 | 1467 | 20201119_16913_1ng_glyco_hcd30_etd30_compound15.15133.15133.4 | controllerType=0 controllerNumber=1 scan=15133 | 35.282 |
| 11792:4 | 14 | K.LKELPGVC[+57.021]N[+2059.735]ETMMALWEEC[+57.021]KPC[+57.021]LK.Q | HexNAc(4)Hex(5)Fuc(1)NeuAc(1) | C[+57]*3, N[+2060] | 1249.7897 | 4 | 4996.1369 | 4996.1196 | 0 | 3.4 | 95 | Specific | 705.17 | 547.98 | 547.98 | 10.52 | 19 | >sp\|P10909\|CLUS_HUMAN Clusterin OS=Homo sapiens OX=9606 GN=CLU PE=1 SV=1 | 1467 | 20201119_16913_1ng_glyco_hcd30_etd30_compound15.15215.15215.4 | controllerType=0 controllerNumber=1 scan=15215 | 35.491 |
| 00104:4 | 14 | R.EIRHN[+2350.830]STGC[+57.021]LR.M | HexNAc(4)Hex(5)Fuc(1)NeuAc(2) | C[+57], N[+2351] | 924.1335 | 4 | 3693.5123 | 3693.4948 | 0 | 4.7 | 287 | Specific | 328.37 | 218.8 | 218.8 | 3.08 | 19 | >sp\|P10909\|CLUS_HUMAN Clusterin OS=Homo sapiens OX=9606 GN=CLU PE=1 SV=1 | 1467 | 20201119_16913_1ng_glyco_hcd30_etd30_compound15.3022.3022.4 | controllerType=0 controllerNumber=1 scan=3022 | 11.4593 |
| 00048:4 | 14 | R.EIRHN[+2205.793]STGC[+57.021]LR.M | HexNAc(4)Hex(5)Fuc(2)NeuAc(1) | C[+57], N[+2206] | 887.8747 | 4 | 3548.4771 | 3548.4573 | 0 | 5.6 | 287 | Specific | 395.87 | 305.48 | 305.48 | 3.12 | 19 | >sp\|P10909\|CLUS_HUMAN Clusterin OS=Homo sapiens OX=9606 GN=CLU PE=1 SV=1 | 1467 | 20201119_16913_1ng_glyco_hcd30_etd30_compound15.2963.2963.4 | controllerType=0 controllerNumber=1 scan=2963 | 11.3187 |
| 00040:4 | 14 | R.EIRHN[+2424.867]STGC[+57.021]LR.M | HexNAc(5)Hex(6)Fuc(1)NeuAc(1) | C[+57], N[+2425] | 942.6422 | 4 | 3767.5469 | 3767.5316 | 0 | 4.1 | 287 | Specific | 511.02 | 326.13 | 326.13 | 4.25 | 19 | >sp\|P10909\|CLUS_HUMAN Clusterin OS=Homo sapiens OX=9606 GN=CLU PE=1 SV=1 | 1467 | 20201119_16913_1ng_glyco_hcd30_etd30_compound15.2955.2955.4 | controllerType=0 controllerNumber=1 scan=2955 | 11.2813 |
| 00034:4 | 14 | R.EIRHN[+2059.735]STGC[+57.021]LR.M | HexNAc(4)Hex(5)Fuc(1)NeuAc(1) | C[+57], N[+2060] | 851.359 | 4 | 3402.4143 | 3402.3994 | 0 | 4.4 | 287 | Specific | 619.46 | 461.06 | 461.06 | 7.93 | 19 | >sp\|P10909\|CLUS_HUMAN Clusterin OS=Homo sapiens OX=9606 GN=CLU PE=1 SV=1 | 1467 | 20201119_16913_1ng_glyco_hcd30_etd30_compound15.2948.2948.4 | controllerType=0 controllerNumber=1 scan=2948 | 11.2492 |
| 16659:4 | 14 | K.MLN[+2059.735]TSSLLEQLNEQFNWVSR.L | HexNAc(4)Hex(5)Fuc(1)NeuAc(1) | N[+2060] | 1117.9904 | 4 | 4468.9398 | 4468.9217 | 0 | 4.1 | 352 | Specific | 272.44 | 246.74 | 246.74 | 3.97 | 19 | >sp\|P10909\|CLUS_HUMAN Clusterin OS=Homo sapiens OX=9606 GN=CLU PE=1 SV=1 | 1467 | 20201119_16913_1ng_glyco_hcd30_etd30_compound15.20358.20358.4 | controllerType=0 controllerNumber=1 scan=20358 | 49.8914 |
| 16649:3 | 14 | K.MLN[+2059.735]TSSLLEQLNEQFNWVSR.L | HexNAc(4)Hex(5)Fuc(1)NeuAc(1) | N[+2060] | 1490.3174 | 3 | 4468.9378 | 4468.9217 | 0 | 3.6 | 352 | Specific | 615.69 | 443.09 | 443.09 | 9.63 | 19 | >sp\|P10909\|CLUS_HUMAN Clusterin OS=Homo sapiens OX=9606 GN=CLU PE=1 SV=1 | 1467 | 20201119_16913_1ng_glyco_hcd30_etd30_compound15.20348.20348.3 | controllerType=0 controllerNumber=1 scan=20348 | 49.8658 |
| 16364:3 | 14 | K.M[+15.995]LN[+2059.735]TSSLLEQLNEQFNWVSR.L | HexNAc(4)Hex(5)Fuc(1)NeuAc(1) | M[+16], N[+2060] | 1495.6553 | 3 | 4484.9514 | 4484.9166 | 0 | 7.8 | 352 | Specific | 437.72 | 297.16 | 297.16 | 4.6 | 19 | >sp\|P10909\|CLUS_HUMAN Clusterin OS=Homo sapiens OX=9606 GN=CLU PE=1 SV=1 | 1467 | 20201119_16913_1ng_glyco_hcd30_etd30_compound15.20044.20044.3 | controllerType=0 controllerNumber=1 scan=20044 | 48.8362 |
| 11129:3 | 14 | R.LAN[+2350.830]LTQGEDQYYLR.V | HexNAc(4)Hex(5)Fuc(1)NeuAc(2) | N[+2351] | 1345.5629 | 3 | 4034.6742 | 4034.664 | 0 | 2.5 | 372 | Specific | 350.52 | 247.31 | 247.31 | 5.57 | 19 | >sp\|P10909\|CLUS_HUMAN Clusterin OS=Homo sapiens OX=9606 GN=CLU PE=1 SV=1 | 1467 | 20201119_16913_1ng_glyco_hcd30_etd30_compound15.14518.14518.3 | controllerType=0 controllerNumber=1 scan=14518 | 33.6668 |
| 18541:4 | 17 | R.QLAHQSN[+2350.830]STNIFFSPVSIATAFAMLSLGTK.A | HexNAc(4)Hex(5)Fuc(1)NeuAc(2) | N[+2351] | 1383.8721 | 4 | 5532.4665 | 5532.4654 | 0 | 0.2 | 64 | Specific | 262.38 | 214.79 | 214.79 | 3.62 | 8 | >sp\|P01009\|A1AT_HUMAN Alpha-1-antitrypsin OS=Homo sapiens OX=9606 GN=SERPINA1 PE=1 SV=3 | 642 | 20201119_16913_1ng_glyco_hcd30_etd30_compound15.22356.22356.4 | controllerType=0 controllerNumber=1 scan=22356 | 56.1067 |
| 18487:3 | 17 | R.QLAHQSN[+2204.772]STNIFFSPVSIATAFAMLSLGTK.A | HexNAc(4)Hex(5)NeuAc(2) | N[+2205] | 1796.1494 | 3 | 5386.4336 | 5386.4075 | 0 | 4.8 | 64 | Specific | 440.73 | 304.94 | 304.94 | 6.22 | 8 | >sp\|P01009\|A1AT_HUMAN Alpha-1-antitrypsin OS=Homo sapiens OX=9606 GN=SERPINA1 PE=1 SV=3 | 642 | 20201119_16913_1ng_glyco_hcd30_etd30_compound15.22299.22299.3 | controllerType=0 controllerNumber=1 scan=22299 | 55.9169 |
| 18475:4 | 17 | R.QLAHQSN[+2350.830]STNIFFSPVSIATAFAMLSLGTK.A | HexNAc(4)Hex(5)Fuc(1)NeuAc(2) | N[+2351] | 1383.876 | 4 | 5532.4821 | 5532.4654 | 0 | 3 | 64 | Specific | 568.54 | 483.64 | 483.64 | 8.65 | 8 | >sp\|P01009\|A1AT_HUMAN Alpha-1-antitrypsin OS=Homo sapiens OX=9606 GN=SERPINA1 PE=1 SV=3 | 642 | 20201119_16913_1ng_glyco_hcd30_etd30_compound15.22286.22286.4 | controllerType=0 controllerNumber=1 scan=22286 | 55.8632 |
| 18486:4 | 17 | R.QLAHQSN[+2204.772]STNIFFSPVSIATAFAMLSLGTK.A | HexNAc(4)Hex(5)NeuAc(2) | N[+2205] | 1347.3634 | 4 | 5386.4318 | 5386.4075 | 0 | 4.5 | 64 | Specific | 678.36 | 569.56 | 569.56 | 10.09 | 8 | >sp\|P01009\|A1AT_HUMAN Alpha-1-antitrypsin OS=Homo sapiens OX=9606 GN=SERPINA1 PE=1 SV=3 | 642 | 20201119_16913_1ng_glyco_hcd30_etd30_compound15.22298.22298.4 | controllerType=0 controllerNumber=1 scan=22298 | 55.9135 |
| 17752:4 | 17 | K.ADTHDEILEGLNFN[+2861.000]LTEIPEAQIHEGFQELLR.T | HexNAc(5)Hex(6)NeuAc(3) | N[+2861] | 1638.9711 | 4 | 6552.8625 | 6552.8239 | 0 | 5.9 | 94 | Specific | 310.51 | 310.51 | 310.51 | 5.34 | 8 | >sp\|P01009\|A1AT_HUMAN Alpha-1-antitrypsin OS=Homo sapiens OX=9606 GN=SERPINA1 PE=1 SV=3 | 642 | 20201119_16913_1ng_glyco_hcd30_etd30_compound15.21513.21513.4 | controllerType=0 controllerNumber=1 scan=21513 | 53.2169 |
| 05376:3 | 18 | K.KLLN[+2059.735]LTVR.I | HexNAc(4)Hex(5)Fuc(1)NeuAc(1) | N[+2060] | 1006.1289 | 3 | 3016.3723 | 3016.3601 | 0 | 4 | 133 | Specific | 249.83 | 171.46 | 171.46 | 2.72 | 9 | >sp\|Q6UX06\|OLFM4_HUMAN Olfactomedin-4 OS=Homo sapiens OX=9606 GN=OLFM4 PE=1 SV=1 | 624 | 20201119_16913_1ng_glyco_hcd30_etd30_compound15.8502.8502.3 | controllerType=0 controllerNumber=1 scan=8502 | 21.4899 |
| 09924:2 | 18 | R.VN[+1216.423]LTTNTIAVTQTLPNAAYNNR.F | HexNAc(2)Hex(5) | N[+1216] | 1803.3442 | 2 | 3605.6811 | 3605.6699 | 0 | 3.1 | 351 | Specific | 571.25 | 513.88 | 513.88 | 8.67 | 9 | >sp\|Q6UX06\|OLFM4_HUMAN Olfactomedin-4 OS=Homo sapiens OX=9606 GN=OLFM4 PE=1 SV=1 | 624 | 20201119_16913_1ng_glyco_hcd30_etd30_compound15.13249.13249.2 | controllerType=0 controllerNumber=1 scan=13249 | 30.9872 |
| 09914:3 | 18 | R.VN[+1216.423]LTTNTIAVTQTLPNAAYNNR.F | HexNAc(2)Hex(5) | N[+1216] | 1202.5655 | 3 | 3605.6819 | 3605.6699 | 0 | 3.3 | 351 | Specific | 618.68 | 561.95 | 561.95 | 9.63 | 9 | >sp\|Q6UX06\|OLFM4_HUMAN Olfactomedin-4 OS=Homo sapiens OX=9606 GN=OLFM4 PE=1 SV=1 | 624 | 20201119_16913_1ng_glyco_hcd30_etd30_compound15.13239.13239.3 | controllerType=0 controllerNumber=1 scan=13239 | 30.9569 |
| 02989:3 | 21 | R.LRN[+1913.677]VSWATGR.S | HexNAc(4)Hex(5)NeuAc(1) | N[+1914] | 1025.111 | 3 | 3073.3184 | 3073.3101 | 0 | 2.7 | 149 | Specific | 211.24 | 140.41 | 140.41 | 2.64 | 7 | >sp\|P08571\|CD14_HUMAN Monocyte differentiation antigen CD14 OS=Homo sapiens OX=9606 GN=CD14 PE=1 SV=2 | 1232 | 20201119_16913_1ng_glyco_hcd30_etd30_compound15.6023.6023.3 | controllerType=0 controllerNumber=1 scan=6023 | 17.0243 |
| 01704:3 | 21 | R.LRN[+1216.423]VSWATGR.S | HexNAc(2)Hex(5) | N[+1216] | 792.6935 | 3 | 2376.066 | 2376.0559 | 0 | 4.2 | 149 | Specific | 243.97 | 126.41 | 126.41 | 2.58 | 7 | >sp\|P08571\|CD14_HUMAN Monocyte differentiation antigen CD14 OS=Homo sapiens OX=9606 GN=CD14 PE=1 SV=2 | 1232 | 20201119_16913_1ng_glyco_hcd30_etd30_compound15.4687.4687.3 | controllerType=0 controllerNumber=1 scan=4687 | 14.545 |
| 00047:3 | 22 | K.YKN[+2059.735]NSDISSTR.G | HexNAc(4)Hex(5)Fuc(1)NeuAc(1) | N[+2060] | 1115.4616 | 3 | 3344.3702 | 3344.3528 | 0 | 5.2 | 44 | Specific | 244.42 | 131.5 | 131.5 | 3.53 | 4 | >sp\|P01871\|IGHM_HUMAN Immunoglobulin heavy constant mu OS=Homo sapiens OX=9606 GN=IGHM PE=1 SV=4 | 328 | 20201119_16913_1ng_glyco_hcd30_etd30_compound15.2962.2962.3 | controllerType=0 controllerNumber=1 scan=2962 | 11.3105 |
| 19086:4 | 24 | R.QLDMLDLSN[+2204.772]NSLASVPEGLWASLGQPNWDMR.D | HexNAc(4)Hex(5)NeuAc(2) | N[+2205] | 1416.3674 | 4 | 5662.4479 | 5662.4239 | 0 | 4.2 | 261 | Specific | 289.33 | 289.33 | 289.33 | 3.57 | 4 | >sp\|P02750\|A2GL_HUMAN Leucine-rich alpha-2-glycoprotein OS=Homo sapiens OX=9606 GN=LRG1 PE=1 SV=2 | 518 | 20201119_16913_1ng_glyco_hcd30_etd30_compound15.22935.22935.4 | controllerType=0 controllerNumber=1 scan=22935 | 57.9816 |
| 12148:4 | 28 | K.MVSHHN[+2204.772]LTTGATLINEQWLLTTAK.N | HexNAc(4)Hex(5)NeuAc(2) | N[+2205] | 1221.8003 | 4 | 4884.1795 | 4884.1647 | 0 | 3 | 179 | Specific | 628.2 | 625.99 | 625.99 | 9.01 | 5 | >sp\|P00738\|HPT_HUMAN Haptoglobin OS=Homo sapiens OX=9606 GN=HP PE=1 SV=1 | 407 | 20201119_16913_1ng_glyco_hcd30_etd30_compound15.15588.15588.3 | controllerType=0 controllerNumber=1 scan=15588 | 36.4519 |
| 09735:3 | 29 | R.IIVPLNNREN[+1913.677]ISDPTSPLR.T | HexNAc(4)Hex(5)NeuAc(1) | N[+1914] | 1354.6253 | 3 | 4061.8614 | 4061.8542 | 0 | 1.8 | 62 | Specific | 251.26 | 219.07 | 219.07 | 3.06 | 5 | >sp\|P01591\|IGJ_HUMAN Immunoglobulin J chain OS=Homo sapiens OX=9606 GN=JCHAIN PE=1 SV=4 | 1392 | 20201119_16913_1ng_glyco_hcd30_etd30_compound15.13050.13050.3 | controllerType=0 controllerNumber=1 scan=13050 | 30.3958 |
| 12603:3 | 30 | K.RNYIVPILWLN[+1864.634]ETGTIGDEK.A | HexNAc(2)Hex(9) | N[+1865] | 1399.2967 | 3 | 4195.8755 | 4195.8685 | 0 | 1.7 | 407 | Specific | 235.2 | 197.34 | 197.34 | 3.7 | 4 | >sp\|P16671\|CD36_HUMAN Platelet glycoprotein 4 OS=Homo sapiens OX=9606 GN=CD36 PE=1 SV=2 | 812 | 20201119_16913_1ng_glyco_hcd30_etd30_compound15.16064.16064.3 | controllerType=0 controllerNumber=1 scan=16064 | 37.5486 |
| 04909:3 | 33 | R.LSPN[+2205.793]ASAEHLELR.W | HexNAc(4)Hex(5)Fuc(2)NeuAc(1) | N[+2206] | 1214.8555 | 3 | 3642.552 | 3642.5421 | 0 | 2.7 | 52 | Specific | 277.9 | 181.29 | 181.29 | 4.02 | 4 | >sp\|Q13410\|BT1A1_HUMAN Butyrophilin subfamily 1 member A1 OS=Homo sapiens OX=9606 GN=BTN1A1 PE=1 SV=3 | 179 | 20201119_16913_1ng_glyco_hcd30_etd30_compound15.8018.8018.3 | controllerType=0 controllerNumber=1 scan=8018 | 20.6179 |
| 06840:3 | 33 | R.LSPN[+2350.830]ASAEHLELR.W | HexNAc(4)Hex(5)Fuc(1)NeuAc(2) | N[+2351] | 1263.2012 | 3 | 3787.5891 | 3787.5796 | 0 | 2.5 | 52 | Specific | 390.85 | 225.18 | 225.18 | 5.77 | 4 | >sp\|Q13410\|BT1A1_HUMAN Butyrophilin subfamily 1 member A1 OS=Homo sapiens OX=9606 GN=BTN1A1 PE=1 SV=3 | 179 | 20201119_16913_1ng_glyco_hcd30_etd30_compound15.10024.10024.3 | controllerType=0 controllerNumber=1 scan=10024 | 24.1985 |
| 01727:3 | 34 | K.N[+1216.423]NSIPDKQITASSSYK.T | HexNAc(2)Hex(5) | N[+1216] | 990.4431 | 3 | 2969.3149 | 2969.2991 | 0 | 5.3 | 238 | Specific | 388.58 | 218.42 | 218.42 | 3.03 | 6 | >sp\|Q08431\|MFGM_HUMAN Lactadherin OS=Homo sapiens OX=9606 GN=MFGE8 PE=1 SV=3 | 1064 | 20201119_16913_1ng_glyco_hcd30_etd30_compound15.4711.4711.3 | controllerType=0 controllerNumber=1 scan=4711 | 14.5955 |
| 00724:4 | 93 | K.TKPREEQFN[+1606.587]STFR.V | HexNAc(4)Hex(4)Fuc(1) | N[+1607] | 812.3614 | 4 | 3246.4238 | 3246.4054 | 0 | 5.7 | 168 | Specific | 286.5 | 285.09 | 285.09 | 3.74 | 2 | >sp\|P01859\|IGHG2_HUMAN Immunoglobulin heavy constant gamma 2 OS=Homo sapiens OX=9606 GN=IGHG2 PE=1 SV=2 | 387 | 20201119_16913_1ng_glyco_hcd30_etd30_compound15.3667.3667.4 | controllerType=0 controllerNumber=1 scan=3667 | 12.6742 |
| 00700:4 | 93 | K.TKPREEQFN[+1606.587]STFR.V | HexNAc(4)Hex(4)Fuc(1) | N[+1607] | 812.3606 | 4 | 3246.4204 | 3246.4054 | 0 | 4.6 | 168 | Specific | 449.23 | 420.28 | 420.28 | 6.41 | 2 | >sp\|P01859\|IGHG2_HUMAN Immunoglobulin heavy constant gamma 2 OS=Homo sapiens OX=9606 GN=IGHG2 PE=1 SV=2 | 387 | 20201119_16913_1ng_glyco_hcd30_etd30_compound15.3642.3642.4 | controllerType=0 controllerNumber=1 scan=3642 | 12.616 |
| 15991:4 | 1 | R.RTAGWNVPIGTLRPFLN[+2863.041]WTGPPEPIEAAVAR.F | HexNAc(5)Hex(6)Fuc(4)NeuAc(1) | N[+2863] | 1563.2322 | 4 | 6249.9069 | 6249.8529 | 0 | 8.6 | 140 | Specific | 273.06 | 273.06 | 273.06 | 4.92 | 38 | >sp\|P02788\|TRFL_HUMAN Lactotransferrin OS=Homo sapiens OX=9606 GN=LTF PE=1 SV=6 | 1270 | 20201114_16933_1ng_glyco_hcd30_etd30_compound15.19564.19564.4 | controllerType=0 controllerNumber=1 scan=19564 | 43.3108 |
| 16026:4 | 1 | R.RTAGWNVPIGTLRPFLN[+2571.946]WTGPPEPIEAAVAR.F | HexNAc(5)Hex(6)Fuc(4) | N[+2572] | 1490.4462 | 4 | 5958.7629 | 5958.7575 | 0 | 0.9 | 140 | Specific | 352.05 | 352.05 | 352.05 | 5.92 | 38 | >sp\|P02788\|TRFL_HUMAN Lactotransferrin OS=Homo sapiens OX=9606 GN=LTF PE=1 SV=6 | 1270 | 20201114_16933_1ng_glyco_hcd30_etd30_compound15.19601.19601.4 | controllerType=0 controllerNumber=1 scan=19601 | 43.4227 |
| 16780:4 | 1 | R.RTAGWNVPIGTLRPFLN[+2351.851]WTGPPEPIEAAVAR.F | HexNAc(4)Hex(5)Fuc(3)NeuAc(1) | N[+2352] | 1435.4185 | 4 | 5738.6522 | 5738.6628 | 0 | -1.9 | 140 | Specific | 477.72 | 477.72 | 477.72 | 6.16 | 38 | >sp\|P02788\|TRFL_HUMAN Lactotransferrin OS=Homo sapiens OX=9606 GN=LTF PE=1 SV=6 | 1270 | 20201114_16933_1ng_glyco_hcd30_etd30_compound15.20403.20403.4 | controllerType=0 controllerNumber=1 scan=20403 | 45.8861 |
| 16011:4 | 1 | R.RTAGWNVPIGTLRPFLN[+2570.925]WTGPPEPIEAAVAR.F | HexNAc(5)Hex(6)Fuc(2)NeuAc(1) | N[+2571] | 1490.191 | 4 | 5957.7423 | 5957.7371 | 0 | 0.9 | 140 | Specific | 632.55 | 632.55 | 632.55 | 9.44 | 38 | >sp\|P02788\|TRFL_HUMAN Lactotransferrin OS=Homo sapiens OX=9606 GN=LTF PE=1 SV=6 | 1270 | 20201114_16933_1ng_glyco_hcd30_etd30_compound15.19585.19585.4 | controllerType=0 controllerNumber=1 scan=19585 | 43.3712 |

Table S4. Cont.

| Query #:z | Protein Rank | Glycopeptide | Glycan | Modification Type(s) | Observed m/z | z | Observed (M+H) | Calc. mass (M+H) | Off-by-x error | Mass error (ppm) | StartPosition | Cleavage | Score | Delta | Delta Mod | \|Log Prob\| | # of unique peptides | Protein Name | Protein DB number | Comment | Scan # | Scan Time |
|---|---|---|---|---|---|---|---|---|---|---|---|---|---|---|---|---|---|---|---|---|---|---|
| 16754:4 | 1 | R.RTAGWNVPIGTLRPFLN[+2350.830]WTGPPEPIEAAVAR.F | HexNAc(4)Hex(5)Fuc(1)NeuAc(2) | N[+2351] | 1435.1737 | 4 | 5737.673 | 5737.6424 | 0 | 5.3 | 140 | Specific | 875.06 | 875.06 | 875.06 | 13.98 | 38 | >sp\|P02788\|TRFL_HUMAN Lactotransferrin OS=Homo sapiens OX=9606 GN=LTF PE=1 SV=6 | 1270 | 20201114_16933_1ng_glyco_hcd30_etd30_compound15.20375.20375.4 | controllerType=0 controllerNumber=1 scan=20375 | 45.8056 |
| 16903:3 | 1 | R.TAGWNVPIGTLRPFLN[+2351.851]WTGPPEPIEAAVAR.F | HexNAc(4)Hex(5)Fuc(3)NeuAc(1) | N[+2352] | 1861.5286 | 3 | 5582.5714 | 5582.5617 | 0 | 1.7 | 141 | Specific | 218.97 | 155.1 | 155.1 | 4.96 | 38 | >sp\|P02788\|TRFL_HUMAN Lactotransferrin OS=Homo sapiens OX=9606 GN=LTF PE=1 SV=6 | 1270 | 20201114_16933_1ng_glyco_hcd30_etd30_compound15.20534.20534.3 | controllerType=0 controllerNumber=1 scan=20534 | 46.2804 |
| 16930:3 | 1 | R.TAGWNVPIGTLRPFLN[+2205.793]WTGPPEPIEAAVAR.F | HexNAc(4)Hex(5)Fuc(2)NeuAc(1) | N[+2206] | 1812.8454 | 3 | 5436.5217 | 5436.5038 | 0 | 3.3 | 141 | Specific | 364.03 | 281.69 | 281.69 | 6.25 | 38 | >sp\|P02788\|TRFL_HUMAN Lactotransferrin OS=Homo sapiens OX=9606 GN=LTF PE=1 SV=6 | 1270 | 20201114_16933_1ng_glyco_hcd30_etd30_compound15.20562.20562.2 | controllerType=0 controllerNumber=1 scan=20562 | 46.3627 |
| 16781:4 | 1 | R.TAGWNVPIGTLRPFLN[+2716.983]WTGPPEPIEAAVAR.F | HexNAc(5)Hex(6)Fuc(3)NeuAc(1) | N[+2717] | 1487.6815 | 4 | 5947.7043 | 5947.6939 | 0 | 1.7 | 141 | Specific | 398.25 | 398.25 | 398.25 | 6.36 | 38 | >sp\|P02788\|TRFL_HUMAN Lactotransferrin OS=Homo sapiens OX=9606 GN=LTF PE=1 SV=6 | 1270 | 20201114_16933_1ng_glyco_hcd30_etd30_compound15.20404.20404.4 | controllerType=0 controllerNumber=1 scan=20404 | 45.892 |
| 16898:3 | 1 | R.TAGWNVPIGTLRPFLN[+2059.735]WTGPPEPIEAAVAR.F | HexNAc(4)Hex(5)Fuc(1)NeuAc(1) | N[+2060] | 1764.1567 | 3 | 5290.4556 | 5290.4459 | 0 | 1.8 | 141 | Specific | 426.98 | 310.49 | 310.49 | 6.59 | 38 | >sp\|P02788\|TRFL_HUMAN Lactotransferrin OS=Homo sapiens OX=9606 GN=LTF PE=1 SV=6 | 1270 | 20201114_16933_1ng_glyco_hcd30_etd30_compound15.20529.20529.3 | controllerType=0 controllerNumber=1 scan=20529 | 46.2622 |
| 16878:3 | 1 | R.TAGWNVPIGTLRPFLN[+2205.793]WTGPPEPIEAAVAR.F | HexNAc(4)Hex(5)Fuc(2)NeuAc(1) | N[+2206] | 1812.8575 | 3 | 5436.558 | 5436.5038 | 0 | 10 | 141 | Specific | 450.19 | 389.23 | 389.23 | 5.78 | 38 | >sp\|P02788\|TRFL_HUMAN Lactotransferrin OS=Homo sapiens OX=9606 GN=LTF PE=1 SV=6 | 1270 | 20201114_16933_1ng_glyco_hcd30_etd30_compound15.20508.20508.3 | controllerType=0 controllerNumber=1 scan=20508 | 46.2056 |
| 17897:3 | 1 | R.TAGWNVPIGTLRPFLN[+2350.830]WTGPPEPIEAAVAR.F | HexNAc(4)Hex(5)Fuc(1)NeuAc(2) | N[+2351] | 1861.1904 | 3 | 5581.5566 | 5581.5413 | 0 | 2.7 | 141 | Specific | 469.16 | 391.03 | 391.03 | 7.97 | 38 | >sp\|P02788\|TRFL_HUMAN Lactotransferrin OS=Homo sapiens OX=9606 GN=LTF PE=1 SV=6 | 1270 | 20201114_16933_1ng_glyco_hcd30_etd30_compound15.21588.21588.3 | controllerType=0 controllerNumber=1 scan=21588 | 49.1688 |
| 16266:3 | 1 | R.TAGWNVPIGTLRPFLN[+1768.640]WTGPPEPIEAAVAR.F | HexNAc(4)Hex(5)Fuc(1) | N[+1769] | 1667.1302 | 3 | 4999.3761 | 4999.3504 | 0 | 5.1 | 141 | Specific | 473.19 | 390.01 | 390.01 | 7.22 | 38 | >sp\|P02788\|TRFL_HUMAN Lactotransferrin OS=Homo sapiens OX=9606 GN=LTF PE=1 SV=6 | 1270 | 20201114_16933_1ng_glyco_hcd30_etd30_compound15.19854.19854.3 | controllerType=0 controllerNumber=1 scan=19854 | 44.1077 |
| 16902:4 | 1 | R.TAGWNVPIGTLRPFLN[+2351.851]WTGPPEPIEAAVAR.F | HexNAc(4)Hex(5)Fuc(3)NeuAc(1) | N[+2352] | 1396.3971 | 4 | 5582.5666 | 5582.5617 | 0 | 0.9 | 141 | Specific | 551.02 | 541.21 | 541.21 | 7.08 | 38 | >sp\|P02788\|TRFL_HUMAN Lactotransferrin OS=Homo sapiens OX=9606 GN=LTF PE=1 SV=6 | 1270 | 20201114_16933_1ng_glyco_hcd30_etd30_compound15.20533.20533.4 | controllerType=0 controllerNumber=1 scan=20533 | 46.2748 |
| 17877:4 | 1 | R.TAGWNVPIGTLRPFLN[+2350.830]WTGPPEPIEAAVAR.F | HexNAc(4)Hex(5)Fuc(1)NeuAc(2) | N[+2351] | 1396.1456 | 4 | 5581.5607 | 5581.5413 | 0 | 3.5 | 141 | Specific | 873.66 | 853.9 | 853.9 | 14.83 | 38 | >sp\|P02788\|TRFL_HUMAN Lactotransferrin OS=Homo sapiens OX=9606 GN=LTF PE=1 SV=6 | 1270 | 20201114_16933_1ng_glyco_hcd30_etd30_compound15.21566.21566.4 | controllerType=0 controllerNumber=1 scan=21566 | 49.1051 |
| 16899:4 | 1 | R.TAGWNVPIGTLRPFLN[+2059.735]WTGPPEPIEAAVAR.F | HexNAc(4)Hex(5)Fuc(1)NeuAc(1) | N[+2060] | 1323.3698 | 4 | 5290.4572 | 5290.4459 | 0 | 2.1 | 141 | Specific | 1060.12 | 1048.25 | 1048.25 | 19.01 | 38 | >sp\|P02788\|TRFL_HUMAN Lactotransferrin OS=Homo sapiens OX=9606 GN=LTF PE=1 SV=6 | 1270 | 20201114_16933_1ng_glyco_hcd30_etd30_compound15.20530.20530.4 | controllerType=0 controllerNumber=1 scan=20530 | 46.2686 |
| 16658:3 | 1 | R.TAGWNIPMGLLFN[+1751.624]QTGSC[+57.021]K.F | HexNAc(4)Hex(4)NeuAc(1) | C[+57], N[+1752] | 1282.8823 | 3 | 3846.6322 | 3846.6341 | 0 | -0.5 | 485 | Specific | 252.37 | 188.27 | 188.27 | 4.97 | 38 | >sp\|P02788\|TRFL_HUMAN Lactotransferrin OS=Homo sapiens OX=9606 GN=LTF PE=1 SV=6 | 1270 | 20201114_16933_1ng_glyco_hcd30_etd30_compound15.20273.20273.3 | controllerType=0 controllerNumber=1 scan=20273 | 45.4886 |
| 15432:3 | 1 | R.TAGWNIPMGLLFN[+1216.423]QTGSC[+57.021]K.F | HexNAc(2)Hex(5) | C[+57], N[+1216] | 1104.4853 | 3 | 3311.4412 | 3311.4328 | 0 | 2.5 | 485 | Specific | 289.58 | 285.97 | 146.98 | 4.65 | 38 | >sp\|P02788\|TRFL_HUMAN Lactotransferrin OS=Homo sapiens OX=9606 GN=LTF PE=1 SV=6 | 1270 | 20201114_16933_1ng_glyco_hcd30_etd30_compound15.18973.18973.3 | controllerType=0 controllerNumber=1 scan=18973 | 41.581 |
| 16634:3 | 1 | R.TAGWNIPMGLLFN[+2059.735]QTGSC[+57.021]K.F | HexNAc(4)Hex(5)Fuc(1)NeuAc(1) | C[+57], N[+2060] | 1385.5911 | 3 | 4154.7588 | 4154.7449 | 0 | 3.4 | 485 | Specific | 298.46 | 228.02 | 228.02 | 5.67 | 38 | >sp\|P02788\|TRFL_HUMAN Lactotransferrin OS=Homo sapiens OX=9606 GN=LTF PE=1 SV=6 | 1270 | 20201114_16933_1ng_glyco_hcd30_etd30_compound15.20248.20248.3 | controllerType=0 controllerNumber=1 scan=20248 | 45.4156 |
| 15259:3 | 1 | R.TAGWNIPMGLLFN[+1622.582]QTGSC[+57.021]K.F | HexNAc(4)Hex(5) | C[+57], N[+1623] | 1239.8733 | 3 | 3717.6053 | 3717.5916 | 0 | 3.7 | 485 | Specific | 307.45 | 231.97 | 231.97 | 5.77 | 38 | >sp\|P02788\|TRFL_HUMAN Lactotransferrin OS=Homo sapiens OX=9606 GN=LTF PE=1 SV=6 | 1270 | 20201114_16933_1ng_glyco_hcd30_etd30_compound15.18790.18790.3 | controllerType=0 controllerNumber=1 scan=18790 | 41.0859 |
| 15094:3 | 1 | R.TAGWNIPMGLLFN[+1768.640]QTGSC[+57.021]K.F | HexNAc(4)Hex(5)Fuc(1) | C[+57], N[+1769] | 1288.564 | 3 | 3863.6775 | 3863.6495 | 0 | 7.3 | 485 | Specific | 330.56 | 256.83 | 256.83 | 4.76 | 38 | >sp\|P02788\|TRFL_HUMAN Lactotransferrin OS=Homo sapiens OX=9606 GN=LTF PE=1 SV=6 | 1270 | 20201114_16933_1ng_glyco_hcd30_etd30_compound15.18616.18616.3 | controllerType=0 controllerNumber=1 scan=18616 | 40.6438 |
| 15221:3 | 1 | R.TAGWNIPMGLLFN[+1768.640]QTGSC[+57.021]K.F | HexNAc(4)Hex(5)Fuc(1) | C[+57], N[+1769] | 1288.565 | 3 | 3863.6804 | 3863.6495 | 0 | 8 | 485 | Specific | 335.38 | 249.75 | 249.75 | 4.76 | 38 | >sp\|P02788\|TRFL_HUMAN Lactotransferrin OS=Homo sapiens OX=9606 GN=LTF PE=1 SV=6 | 1270 | 20201114_16933_1ng_glyco_hcd30_etd30_compound15.18750.18750.3 | controllerType=0 controllerNumber=1 scan=18750 | 40.9803 |
| 16445:3 | 1 | R.TAGWNIPMGLLFN[+2350.830]QTGSC[+57.021]K.F | HexNAc(4)Hex(5)Fuc(1)NeuAc(2) | C[+57], N[+2351] | 1482.6259 | 3 | 4445.8632 | 4445.8403 | 0 | 5.1 | 485 | Specific | 348.36 | 147.78 | 147.78 | 4.82 | 38 | >sp\|P02788\|TRFL_HUMAN Lactotransferrin OS=Homo sapiens OX=9606 GN=LTF PE=1 SV=6 | 1270 | 20201114_16933_1ng_glyco_hcd30_etd30_compound15.20045.20045.3 | controllerType=0 controllerNumber=1 scan=20045 | 44.7038 |
| 16530:3 | 1 | R.TAGWNIPMGLLFN[+2059.735]QTGSC[+57.021]K.F | HexNAc(4)Hex(5)Fuc(1)NeuAc(1) | C[+57], N[+2060] | 1385.5879 | 3 | 4154.7493 | 4154.7449 | 0 | 1.1 | 485 | Specific | 356.28 | 202.08 | 202.08 | 5.78 | 38 | >sp\|P02788\|TRFL_HUMAN Lactotransferrin OS=Homo sapiens OX=9606 GN=LTF PE=1 SV=6 | 1270 | 20201114_16933_1ng_glyco_hcd30_etd30_compound15.20137.20137.3 | controllerType=0 controllerNumber=1 scan=20137 | 45.0368 |
| 18075:3 | 1 | R.TAGWNIPMGLLFN[+2204.772]QTGSC[+57.021]K.F | HexNAc(4)Hex(5)NeuAc(2) | C[+57], N[+2205] | 1433.9327 | 3 | 4299.7835 | 4299.7824 | 0 | 0.3 | 485 | Specific | 358.46 | 236.13 | 236.13 | 5.88 | 38 | >sp\|P02788\|TRFL_HUMAN Lactotransferrin OS=Homo sapiens OX=9606 GN=LTF PE=1 SV=6 | 1270 | 20201114_16933_1ng_glyco_hcd30_etd30_compound15.21775.21775.3 | controllerType=0 controllerNumber=1 scan=21775 | 49.6825 |
| 16619:3 | 1 | R.TAGWNIPMGLLFN[+1913.677]QTGSC[+57.021]K.F | HexNAc(4)Hex(5)NeuAc(1) | C[+57], N[+1914] | 1336.8992 | 3 | 4008.6831 | 4008.687 | 0 | -1 | 485 | Specific | 375.68 | 184.68 | 184.68 | 5.19 | 38 | >sp\|P02788\|TRFL_HUMAN Lactotransferrin OS=Homo sapiens OX=9606 GN=LTF PE=1 SV=6 | 1270 | 20201114_16933_1ng_glyco_hcd30_etd30_compound15.20232.20232.3 | controllerType=0 controllerNumber=1 scan=20232 | 45.372 |
| 15399:3 | 1 | R.TAGWNIPMGLLFN[+1216.423]QTGSC[+57.021]K.F | HexNAc(2)Hex(5) | C[+57], N[+1216] | 1104.4848 | 3 | 3311.4398 | 3311.4328 | 0 | 2.1 | 485 | Specific | 385.35 | 357.05 | 202.24 | 6.63 | 38 | >sp\|P02788\|TRFL_HUMAN Lactotransferrin OS=Homo sapiens OX=9606 GN=LTF PE=1 SV=6 | 1270 | 20201114_16933_1ng_glyco_hcd30_etd30_compound15.18938.18938.3 | controllerType=0 controllerNumber=1 scan=18938 | 41.4962 |
| 16493:3 | 1 | R.TAGWNIPMGLLFN[+2059.735]QTGSC[+57.021]K.F | HexNAc(4)Hex(5)Fuc(1)NeuAc(1) | C[+57], N[+2060] | 1385.5904 | 3 | 4154.7566 | 4154.7449 | 0 | 2.8 | 485 | Specific | 406.91 | 287.74 | 287.74 | 6.37 | 38 | >sp\|P02788\|TRFL_HUMAN Lactotransferrin OS=Homo sapiens OX=9606 GN=LTF PE=1 SV=6 | 1270 | 20201114_16933_1ng_glyco_hcd30_etd30_compound15.20097.20097.3 | controllerType=0 controllerNumber=1 scan=20097 | 44.861 |
| 17971:3 | 1 | R.TAGWNIPMGLLFN[+2350.830]QTGSC[+57.021]K.F | HexNAc(4)Hex(5)Fuc(1)NeuAc(2) | C[+57], N[+2351] | 1482.6215 | 3 | 4445.85 | 4445.8403 | 0 | 2.2 | 485 | Specific | 415.5 | 269.06 | 269.06 | 6.55 | 38 | >sp\|P02788\|TRFL_HUMAN Lactotransferrin OS=Homo sapiens OX=9606 GN=LTF PE=1 SV=6 | 1270 | 20201114_16933_1ng_glyco_hcd30_etd30_compound15.21666.21666.3 | controllerType=0 controllerNumber=1 scan=21666 | 49.3826 |
| 17900:3 | 1 | R.TAGWNIPMGLLFN[+2350.830]QTGSC[+57.021]K.F | HexNAc(4)Hex(5)Fuc(1)NeuAc(2) | C[+57], N[+2351] | 1482.6231 | 3 | 4445.8548 | 4445.8403 | 0 | 3.3 | 485 | Specific | 422.15 | 285.95 | 285.95 | 6.37 | 38 | >sp\|P02788\|TRFL_HUMAN Lactotransferrin OS=Homo sapiens OX=9606 GN=LTF PE=1 SV=6 | 1270 | 20201114_16933_1ng_glyco_hcd30_etd30_compound15.21591.21591.3 | controllerType=0 controllerNumber=1 scan=21591 | 49.1762 |
| 16441:3 | 1 | R.TAGWNIPMGLLFN[+2205.793]QTGSC[+57.021]K.F | HexNAc(4)Hex(5)Fuc(2)NeuAc(1) | C[+57], N[+2206] | 1434.2725 | 3 | 4300.803 | 4300.8028 | 0 | 0.1 | 485 | Specific | 438.15 | 319.14 | 319.14 | 6.28 | 38 | >sp\|P02788\|TRFL_HUMAN Lactotransferrin OS=Homo sapiens OX=9606 GN=LTF PE=1 SV=6 | 1270 | 20201114_16933_1ng_glyco_hcd30_etd30_compound15.20041.20041.3 | controllerType=0 controllerNumber=1 scan=20041 | 44.6945 |
| 16566:3 | 1 | R.TAGWNIPMGLLFN[+1913.677]QTGSC[+57.021]K.F | HexNAc(4)Hex(5)NeuAc(1) | C[+57], N[+1914] | 1336.9114 | 3 | 4008.7197 | 4008.687 | 0 | 8.2 | 485 | Specific | 463.51 | 322.43 | 322.43 | 6.08 | 38 | >sp\|P02788\|TRFL_HUMAN Lactotransferrin OS=Homo sapiens OX=9606 GN=LTF PE=1 SV=6 | 1270 | 20201114_16933_1ng_glyco_hcd30_etd30_compound15.20176.20176.3 | controllerType=0 controllerNumber=1 scan=20176 | 45.1919 |
| 15980:2 | 1 | R.TAGWNIPMGLLFN[+203.079]QTGSC[+57.021]K.F | HexNAc(1) | C[+57], N[+203] | 1149.5567 | 2 | 2298.1061 | 2298.0893 | 0 | 7.3 | 485 | Specific | 655.56 | 636.69 | 636.69 | 9.58 | 38 | >sp\|P02788\|TRFL_HUMAN Lactotransferrin OS=Homo sapiens OX=9606 GN=LTF PE=1 SV=6 | 1270 | 20201114_16933_1ng_glyco_hcd30_etd30_compound15.19552.19552.2 | controllerType=0 controllerNumber=1 scan=19552 | 43.2809 |
| 15203:3 | 1 | R.TAGWNIPM[+15.995]GLLFN[+2059.735]QTGSC[+57.021]K.F | HexNAc(4)Hex(5)Fuc(1)NeuAc(1) | C[+57], M[+16], N[+2060] | 1390.9221 | 3 | 4170.7516 | 4170.7398 | 0 | 2.8 | 485 | Specific | 300.38 | 239.43 | 239.43 | 5.77 | 38 | >sp\|P02788\|TRFL_HUMAN Lactotransferrin OS=Homo sapiens OX=9606 GN=LTF PE=1 SV=6 | 1270 | 20201114_16933_1ng_glyco_hcd30_etd30_compound15.18731.18731.3 | controllerType=0 controllerNumber=1 scan=18731 | 40.9216 |
| 15185:3 | 1 | R.TAGWNIPM[+15.995]GLLFN[+2059.735]QTGSC[+57.021]K.F | HexNAc(4)Hex(5)Fuc(1)NeuAc(1) | C[+57], M[+16], N[+2060] | 1390.925 | 3 | 4170.7604 | 4170.7398 | 0 | 4.9 | 485 | Specific | 306.61 | 245.49 | 245.49 | 5.2 | 38 | >sp\|P02788\|TRFL_HUMAN Lactotransferrin OS=Homo sapiens OX=9606 GN=LTF PE=1 SV=6 | 1270 | 20201114_16933_1ng_glyco_hcd30_etd30_compound15.18712.18712.3 | controllerType=0 controllerNumber=1 scan=18712 | 40.8628 |
| 16717:4 | 1 | R.TAGWNIPMGLLFN[+1913.677]QTGSC[+57.021]KFDEYFSQSC[+57.021]APGSDPR.S | HexNAc(4)Hex(5)NeuAc(1) | C[+57]*2, N[+1914] | 1463.8665 | 4 | 5852.444 | 5852.4341 | 0 | 1.7 | 485 | Specific | 322.23 | 256.48 | 256.48 | 4.43 | 38 | >sp\|P02788\|TRFL_HUMAN Lactotransferrin OS=Homo sapiens OX=9606 GN=LTF PE=1 SV=6 | 1270 | 20201114_16933_1ng_glyco_hcd30_etd30_compound15.20336.20336.4 | controllerType=0 controllerNumber=1 scan=20336 | 45.6805 |
| 16688:4 | 1 | R.TAGWNIPMGLLFN[+1913.677]QTGSC[+57.021]KFDEYFSQSC[+57.021]APGSDPR.S | HexNAc(4)Hex(5)NeuAc(1) | C[+57]*2, N[+1914] | 1463.8655 | 4 | 5852.4401 | 5852.4341 | 0 | 1 | 485 | Specific | 571.63 | 498.67 | 498.67 | 7.79 | 38 | >sp\|P02788\|TRFL_HUMAN Lactotransferrin OS=Homo sapiens OX=9606 GN=LTF PE=1 SV=6 | 1270 | 20201114_16933_1ng_glyco_hcd30_etd30_compound15.20305.20305.4 | controllerType=0 controllerNumber=1 scan=20305 | 45.5883 |
| 15955:4 | 1 | R.TAGWNIPMGLLFN[+1768.640]QTGSC[+57.021]KFDEYFSQSC[+57.021]APGSDPR.S | HexNAc(4)Hex(5)Fuc(1) | C[+57]*2, N[+1769] | 1427.6067 | 4 | 5707.4049 | 5707.3966 | 0 | 1.5 | 485 | Specific | 637.57 | 597.75 | 597.75 | 9.07 | 38 | >sp\|P02788\|TRFL_HUMAN Lactotransferrin OS=Homo sapiens OX=9606 GN=LTF PE=1 SV=6 | 1270 | 20201114_16933_1ng_glyco_hcd30_etd30_compound15.19526.19526.4 | controllerType=0 controllerNumber=1 scan=19526 | 43.1942 |
| 16644:4 | 1 | R.TAGWNIPMGLLFN[+2059.735]QTGSC[+57.021]KFDEYFSQSC[+57.021]APGSDPR.S | HexNAc(4)Hex(5)Fuc(1)NeuAc(1) | C[+57]*2, N[+2060] | 1500.3789 | 4 | 5998.4938 | 5998.492 | 0 | 0.3 | 485 | Specific | 769.26 | 695.64 | 695.64 | 11.79 | 38 | >sp\|P02788\|TRFL_HUMAN Lactotransferrin OS=Homo sapiens OX=9606 GN=LTF PE=1 SV=6 | 1270 | 20201114_16933_1ng_glyco_hcd30_etd30_compound15.20259.20259.4 | controllerType=0 controllerNumber=1 scan=20259 | 45.4493 |
| 08115:3 | 1 | R.N[+2205.793]GSDC[+57.021]PDKFC[+57.021]LFQSETK.N | HexNAc(4)Hex(5)Fuc(2)NeuAc(1) | C[+57]*2, N[+2206] | 1413.5671 | 3 | 4238.6867 | 4238.6667 | 0 | 4.7 | 642 | Specific | 232.96 | 168.93 | 168.93 | 3.94 | 38 | >sp\|P02788\|TRFL_HUMAN Lactotransferrin OS=Homo sapiens OX=9606 GN=LTF PE=1 SV=6 | 1270 | 20201114_16933_1ng_glyco_hcd30_etd30_compound15.11303.11303.3 | controllerType=0 controllerNumber=1 scan=11303 | 26.5371 |
| 08125:3 | 1 | R.N[+2205.793]GSDC[+57.021]PDKFC[+57.021]LFQSETK.N | HexNAc(4)Hex(5)Fuc(2)NeuAc(1) | C[+57]*2, N[+2206] | 1413.5635 | 3 | 4238.676 | 4238.6667 | 0 | 2.2 | 642 | Specific | 294.66 | 223.76 | 223.76 | 5.2 | 38 | >sp\|P02788\|TRFL_HUMAN Lactotransferrin OS=Homo sapiens OX=9606 GN=LTF PE=1 SV=6 | 1270 | 20201114_16933_1ng_glyco_hcd30_etd30_compound15.11314.11314.3 | controllerType=0 controllerNumber=1 scan=11314 | 26.562 |
| 16482:3 | 2 | R.LSLHRPALEDLLLGSEAN[+1751.624]LTC[+57.021]TLTGLR.D | HexNAc(4)Hex(4)NeuAc(1) | C[+57], N[+1752] | 1572.4203 | 3 | 4715.2463 | 4715.2225 | 0 | 5.1 | 127 | Specific | 287.55 | 203.34 | 203.34 | 4.66 | 20 | >sp\|P01876\|IGHA1_HUMAN Immunoglobulin heavy constant alpha 1 OS=Homo sapiens OX=9606 GN=IGHA1 PE=1 SV=2 | 255 | 20201114_16933_1ng_glyco_hcd30_etd30_compound15.20084.20084.3 | controllerType=0 controllerNumber=1 scan=20084 | 44.8112 |
| 15681:3 | 2 | R.LSLHRPALEDLLLGSEAN[+1702.581]LTC[+57.021]TLTGLR.D | HexNAc(2)Hex(8) | C[+57], N[+1703] | 1556.0704 | 3 | 4666.1966 | 4666.1796 | 0 | 3.6 | 127 | Specific | 320.89 | 317.49 | 317.49 | 5.64 | 20 | >sp\|P01876\|IGHA1_HUMAN Immunoglobulin heavy constant alpha 1 OS=Homo sapiens OX=9606 GN=IGHA1 PE=1 SV=2 | 255 | 20201114_16933_1ng_glyco_hcd30_etd30_compound15.19236.19236.3 | controllerType=0 controllerNumber=1 scan=19236 | 42.3616 |
| 15804:4 | 2 | R.LSLHRPALEDLLLGSEAN[+1298.476]LTC[+57.021]TLTGLR.D | HexNAc(4)Hex(3) | C[+57], N[+1298] | 1066.2743 | 4 | 4262.0755 | 4262.0742 | 0 | 0.3 | 127 | Specific | 434.05 | 434.05 | 434.05 | 7.59 | 20 | >sp\|P01876\|IGHA1_HUMAN Immunoglobulin heavy constant alpha 1 OS=Homo sapiens OX=9606 GN=IGHA1 PE=1 SV=2 | 255 | 20201114_16933_1ng_glyco_hcd30_etd30_compound15.19367.19367.4 | controllerType=0 controllerNumber=1 scan=19367 | 42.7898 |
| 16488:4 | 2 | R.LSLHRPALEDLLLGSEAN[+1710.598]LTC[+57.021]TLTGLR.D | HexNAc(3)Hex(5)NeuAc(1) | C[+57], N[+1711] | 1169.3109 | 4 | 4674.2218 | 4674.1959 | 0 | 5.5 | 127 | Specific | 496.01 | 496.01 | 496.01 | 7.24 | 20 | >sp\|P01876\|IGHA1_HUMAN Immunoglobulin heavy constant alpha 1 OS=Homo sapiens OX=9606 GN=IGHA1 PE=1 SV=2 | 255 | 20201114_16933_1ng_glyco_hcd30_etd30_compound15.20090.20090.4 | controllerType=0 controllerNumber=1 scan=20090 | 44.8279 |
| 15746:4 | 2 | R.LSLHRPALEDLLLGSEAN[+1216.423]LTC[+57.021]TLTGLR.D | HexNAc(2)Hex(5) | C[+57], N[+1216] | 1045.7626 | 4 | 4180.0286 | 4180.0211 | 0 | 1.8 | 127 | Specific | 972.86 | 950.77 | 950.77 | 17.01 | 20 | >sp\|P01876\|IGHA1_HUMAN Immunoglobulin heavy constant alpha 1 OS=Homo sapiens OX=9606 GN=IGHA1 PE=1 SV=2 | 255 | 20201114_16933_1ng_glyco_hcd30_etd30_compound15.19304.19304.4 | controllerType=0 controllerNumber=1 scan=19304 | 42.5746 |
| 15740:4 | 2 | R.LSLHRPALEDLLLGSEAN[+1460.529]LTC[+57.021]TLTGLR.D | HexNAc(4)Hex(4) | C[+57], N[+1461] | 1106.7892 | 4 | 4424.1351 | 4424.1271 | 0 | 1.8 | 127 | Specific | 1017.61 | 934.1 | 934.1 | 18.01 | 20 | >sp\|P01876\|IGHA1_HUMAN Immunoglobulin heavy constant alpha 1 OS=Homo sapiens OX=9606 GN=IGHA1 PE=1 SV=2 | 255 | 20201114_16933_1ng_glyco_hcd30_etd30_compound15.19298.19298.4 | controllerType=0 controllerNumber=1 scan=19298 | 42.5417 |
| 15695:4 | 2 | R.LSLHRPALEDLLLGSEAN[+1702.581]LTC[+57.021]TLTGLR.D | HexNAc(2)Hex(8) | C[+57], N[+1703] | 1167.3014 | 4 | 4666.1839 | 4666.1796 | 0 | 0.9 | 127 | Specific | 1025.83 | 980.02 | 980.02 | 17.73 | 20 | >sp\|P01876\|IGHA1_HUMAN Immunoglobulin heavy constant alpha 1 OS=Homo sapiens OX=9606 GN=IGHA1 PE=1 SV=2 | 255 | 20201114_16933_1ng_glyco_hcd30_etd30_compound15.19251.19251.4 | controllerType=0 controllerNumber=1 scan=19251 | 42.4056 |
| 15735:4 | 2 | R.LSLHRPALEDLLLGSEAN[+1378.476]LTC[+57.021]TLTGLR.D | HexNAc(2)Hex(6) | C[+57], N[+1378] | 1086.2767 | 4 | 4342.0848 | 4342.074 | 0 | 2.5 | 127 | Specific | 1086.14 | 1040.57 | 1040.57 | 19.01 | 20 | >sp\|P01876\|IGHA1_HUMAN Immunoglobulin heavy constant alpha 1 OS=Homo sapiens OX=9606 GN=IGHA1 PE=1 SV=2 | 255 | 20201114_16933_1ng_glyco_hcd30_etd30_compound15.19293.19293.4 | controllerType=0 controllerNumber=1 scan=19293 | 42.5272 |
| 15736:4 | 2 | R.LSLHRPALEDLLLGSEAN[+1501.555]LTC[+57.021]TLTGLR.D | HexNAc(5)Hex(3) | C[+57], N[+1502] | 1117.0458 | 4 | 4465.1614 | 4465.1536 | 0 | 1.8 | 127 | Specific | 1145.52 | 1096.14 | 1096.14 | 20.01 | 20 | >sp\|P01876\|IGHA1_HUMAN Immunoglobulin heavy constant alpha 1 OS=Homo sapiens OX=9606 GN=IGHA1 PE=1 SV=2 | 255 | 20201114_16933_1ng_glyco_hcd30_etd30_compound15.19294.19294.4 | controllerType=0 controllerNumber=1 scan=19294 | 42.5311 |
| 15779:4 | 2 | R.LSLHRPALEDLLLGSEAN[+1501.555]LTC[+57.021]TLTGLR.D | HexNAc(5)Hex(3) | C[+57], N[+1502] | 1117.0544 | 4 | 4465.1956 | 4465.1536 | 0 | 9.4 | 127 | Specific | 1181.52 | 1146.52 | 1146.52 | 20.01 | 20 | >sp\|P01876\|IGHA1_HUMAN Immunoglobulin heavy constant alpha 1 OS=Homo sapiens OX=9606 GN=IGHA1 PE=1 SV=2 | 255 | 20201114_16933_1ng_glyco_hcd30_etd30_compound15.19340.19340.2 | controllerType=0 controllerNumber=1 scan=19340 | 42.6884 |
| 10102:3 | 2 | R.LAGKPTHVN[+1702.581]VSVVMAEVDGTC[+57.021]Y.- | HexNAc(2)Hex(8) | C[+57], N[+1703] | 1350.5833 | 3 | 4049.7354 | 4049.7234 | 0 | 3 | 332 | CRagged | 228.25 | 168.32 | 168.32 | 3.38 | 20 | >sp\|P01876\|IGHA1_HUMAN Immunoglobulin heavy constant alpha 1 OS=Homo sapiens OX=9606 GN=IGHA1 PE=1 SV=2 | 255 | 20201114_16933_1ng_glyco_hcd30_etd30_compound15.13366.13366.3 | controllerType=0 controllerNumber=1 scan=13366 | 30.3137 |
| 10134:3 | 2 | R.LAGKPTHVN[+1540.529]VSVVMAEVDGTC[+57.021]Y.- | HexNAc(2)Hex(7) | C[+57], N[+1541] | 1296.5627 | 3 | 3887.6735 | 3887.6706 | 0 | 0.7 | 332 | CRagged | 229.09 | 169.03 | 169.03 | 2.97 | 20 | >sp\|P01876\|IGHA1_HUMAN Immunoglobulin heavy constant alpha 1 OS=Homo sapiens OX=9606 GN=IGHA1 PE=1 SV=2 | 255 | 20201114_16933_1ng_glyco_hcd30_etd30_compound15.13400.13400.3 | controllerType=0 controllerNumber=1 scan=13400 | 30.3827 |
| 10223:3 | 2 | R.LAGKPTHVN[+1501.555]VSVVMAEVDGTC[+57.021]Y.- | HexNAc(5)Hex(3) | C[+57], N[+1502] | 1283.5735 | 3 | 3848.7061 | 3848.6974 | 0 | 2.3 | 332 | CRagged | 242.84 | 148.65 | 148.65 | 3.67 | 20 | >sp\|P01876\|IGHA1_HUMAN Immunoglobulin heavy constant alpha 1 OS=Homo sapiens OX=9606 GN=IGHA1 PE=1 SV=2 | 255 | 20201114_16933_1ng_glyco_hcd30_etd30_compound15.13493.13493.3 | controllerType=0 controllerNumber=1 scan=13493 | 30.6201 |
| 10106:3 | 2 | R.LAGKPTHVN[+1702.581]VSVVMAEVDGTC[+57.021]Y.- | HexNAc(2)Hex(8) | C[+57], N[+1703] | 1350.5855 | 3 | 4049.742 | 4049.7234 | 0 | 4.6 | 332 | CRagged | 251.49 | 203.82 | 199.92 | 3.67 | 20 | >sp\|P01876\|IGHA1_HUMAN Immunoglobulin heavy constant alpha 1 OS=Homo sapiens OX=9606 GN=IGHA1 PE=1 SV=2 | 255 | 20201114_16933_1ng_glyco_hcd30_etd30_compound15.13371.13371.3 | controllerType=0 controllerNumber=1 scan=13371 | 30.3298 |
| 10159:3 | 2 | R.LAGKPTHVN[+1378.476]VSVVMAEVDGTC[+57.021]Y.- | HexNAc(2)Hex(6) | C[+57], N[+1378] | 1242.5475 | 3 | 3725.6281 | 3725.6178 | 0 | 2.8 | 332 | CRagged | 267.12 | 219.08 | 196.48 | 4.04 | 20 | >sp\|P01876\|IGHA1_HUMAN Immunoglobulin heavy constant alpha 1 OS=Homo sapiens OX=9606 GN=IGHA1 PE=1 SV=2 | 255 | 20201114_16933_1ng_glyco_hcd30_etd30_compound15.13426.13426.3 | controllerType=0 controllerNumber=1 scan=13426 | 30.4426 |
| 10179:3 | 2 | R.LAGKPTHVN[+1809.666]VSVVMAEVDGTC[+57.021]Y.- | HexNAc(5)Hex(4)Fuc(1) | C[+57], N[+1810] | 1386.2773 | 3 | 4156.8173 | 4156.8081 | 0 | 2.2 | 332 | CRagged | 273.83 | 165.63 | 165.63 | 4.04 | 20 | >sp\|P01876\|IGHA1_HUMAN Immunoglobulin heavy constant alpha 1 OS=Homo sapiens OX=9606 GN=IGHA1 PE=1 SV=2 | 255 | 20201114_16933_1ng_glyco_hcd30_etd30_compound15.13447.13447.3 | controllerType=0 controllerNumber=1 scan=13447 | 30.4935 |
| 10201:3 | 2 | R.LAGKPTHVN[+1216.423]VSVVMAEVDGTC[+57.021]Y.- | HexNAc(2)Hex(5) | C[+57], N[+1216] | 1188.529 | 3 | 3563.5724 | 3563.5649 | 0 | 2.1 | 332 | CRagged | 274.62 | 246.84 | 202.85 | 4.32 | 20 | >sp\|P01876\|IGHA1_HUMAN Immunoglobulin heavy constant alpha 1 OS=Homo sapiens OX=9606 GN=IGHA1 PE=1 SV=2 | 255 | 20201114_16933_1ng_glyco_hcd30_etd30_compound15.13470.13470.3 | controllerType=0 controllerNumber=1 scan=13470 | 30.5536 |
| 10214:3 | 2 | R.LAGKPTHVN[+1216.423]VSVVMAEVDGTC[+57.021]Y.- | HexNAc(2)Hex(5) | C[+57], N[+1216] | 1188.5291 | 3 | 3563.5728 | 3563.5649 | 0 | 2.2 | 332 | CRagged | 277.96 | 277.96 | 202.2 | 4.39 | 20 | >sp\|P01876\|IGHA1_HUMAN Immunoglobulin heavy constant alpha 1 OS=Homo sapiens OX=9606 GN=IGHA1 PE=1 SV=2 | 255 | 20201114_16933_1ng_glyco_hcd30_etd30_compound15.13484.13484.3 | controllerType=0 controllerNumber=1 scan=13484 | 30.5979 |
| 06536:3 | 4 | R.FSDGLESN[+1864.634]SSTQFEVK.K | HexNAc(2)Hex(9) | N[+1865] | 1213.8311 | 3 | 3639.4788 | 3639.4472 | 0 | 8.7 | 219 | Specific | 255.06 | 255.06 | 255.06 | 5.11 | 9 | >sp\|P0C0L4\|CO4A_HUMAN Complement C4-A OS=Homo sapiens OX=9606 GN=C4A PE=1 SV=2 | 1235 | 20201114_16933_1ng_glyco_hcd30_etd30_compound15.9644.9644.3 | controllerType=0 controllerNumber=1 scan=9644 | 23.5116 |
| 19697:3 | 9 | R.LSLLEEPGN[+2350.830]GTFTVILNQLTSR.D | HexNAc(4)Hex(5)Fuc(1)NeuAc(2) | N[+2351] | 1585.0511 | 3 | 4753.1387 | 4753.1229 | 0 | 3.3 | 413 | Specific | 358.96 | 237.48 | 237.48 | 5.56 | 14 | >sp\|P01833\|PIGR_HUMAN Polymeric immunoglobulin receptor OS=Homo sapiens OX=9606 GN=PIGR PE=1 SV=4 | 902 | 20201114_16933_1ng_glyco_hcd30_etd30_compound15.23486.23486.3 | controllerType=0 controllerNumber=1 scan=23486 | 53.4283 |
| 17746:3 | 9 | R.LSLLEEPGN[+2205.793]GTFTVILNQLTSR.D | HexNAc(4)Hex(5)Fuc(2)NeuAc(1) | N[+2206] | 1536.6994 | 3 | 4608.0836 | 4608.0854 | 0 | -0.4 | 413 | Specific | 658.14 | 501.45 | 501.45 | 9.64 | 14 | >sp\|P01833\|PIGR_HUMAN Polymeric immunoglobulin receptor OS=Homo sapiens OX=9606 GN=PIGR PE=1 SV=4 | 902 | 20201114_16933_1ng_glyco_hcd30_etd30_compound15.21429.21429.3 | controllerType=0 controllerNumber=1 scan=21429 | 48.713 |
| 14465:4 | 9 | K.IIEGEPNLKVPGN[+2205.793]VTAVLGETLK.V | HexNAc(4)Hex(5)Fuc(2)NeuAc(1) | N[+2206] | 1150.0435 | 4 | 4597.1522 | 4597.1422 | 0 | 2.2 | 457 | Specific | 233.19 | 233.19 | 233.19 | 3.11 | 14 | >sp\|P01833\|PIGR_HUMAN Polymeric immunoglobulin receptor OS=Homo sapiens OX=9606 GN=PIGR PE=1 SV=4 | 902 | 20201114_16933_1ng_glyco_hcd30_etd30_compound15.17954.17954.4 | controllerType=0 controllerNumber=1 scan=17954 | 39.2651 |
| 14442:3 | 9 | K.IIEGEPNLKVPGN[+2059.735]VTAVLGETLK.V | HexNAc(4)Hex(5)Fuc(1)NeuAc(1) | N[+2060] | 1484.3702 | 3 | 4451.096 | 4451.0843 | 0 | 2.6 | 457 | Specific | 287.13 | 228.75 | 228.75 | 3.43 | 14 | >sp\|P01833\|PIGR_HUMAN Polymeric immunoglobulin receptor OS=Homo sapiens OX=9606 GN=PIGR PE=1 SV=4 | 902 | 20201114_16933_1ng_glyco_hcd30_etd30_compound15.17930.17930.3 | controllerType=0 controllerNumber=1 scan=17930 | 39.2073 |
| 14446:3 | 9 | K.IIEGEPNLKVPGN[+2204.772]VTAVLGETLK.V | HexNAc(4)Hex(5)NeuAc(2) | N[+2205] | 1532.7199 | 3 | 4596.1452 | 4596.1218 | 0 | 5.1 | 457 | Specific | 394.24 | 309.46 | 309.46 | 3.29 | 14 | >sp\|P01833\|PIGR_HUMAN Polymeric immunoglobulin receptor OS=Homo sapiens OX=9606 GN=PIGR PE=1 SV=4 | 902 | 20201114_16933_1ng_glyco_hcd30_etd30_compound15.17935.17935.3 | controllerType=0 controllerNumber=1 scan=17935 | 39.2249 |
| 14105:4 | 9 | K.IIEGEPNLKVPGN[+2205.793]VTAVLGETLKVPC[+57.021]HFPC[+57.021]K.F | HexNAc(4)Hex(5)Fuc(2)NeuAc(1) | C[+57]*2, N[+2206] | 1406.4069 | 4 | 5622.6058 | 5622.5997 | 0 | 1.1 | 457 | Specific | 278.53 | 278.53 | 278.53 | 2.66 | 14 | >sp\|P01833\|PIGR_HUMAN Polymeric immunoglobulin receptor OS=Homo sapiens OX=9606 GN=PIGR PE=1 SV=4 | 902 | 20201114_16933_1ng_glyco_hcd30_etd30_compound15.17554.17554.4 | controllerType=0 controllerNumber=1 scan=17554 | 38.442 |
| 15286:5 | 9 | K.IIEGEPNLKVPGN[+2350.830]VTAVLGETLKVPC[+57.021]HFPC[+57.021]K.F | HexNAc(4)Hex(5)Fuc(1)NeuAc(2) | C[+57]*2, N[+2351] | 1154.3435 | 5 | 5767.6884 | 5767.6372 | 0 | 8.9 | 457 | Specific | 499.65 | 499.65 | 499.65 | 4.22 | 14 | >sp\|P01833\|PIGR_HUMAN Polymeric immunoglobulin receptor OS=Homo sapiens OX=9606 GN=PIGR PE=1 SV=4 | 902 | 20201114_16933_1ng_glyco_hcd30_etd30_compound15.18819.18819.5 | controllerType=0 controllerNumber=1 scan=18819 | 41.1797 |
| 12225:3 | 9 | K.VPGN[+2205.793]VTAVLGETLK.V | HexNAc(4)Hex(5)Fuc(2)NeuAc(1) | N[+2206] | 1201.8705 | 3 | 3603.5971 | 3603.5927 | 0 | 1.2 | 466 | Specific | 286.81 | 183.92 | 183.92 | 3.75 | 14 | >sp\|P01833\|PIGR_HUMAN Polymeric immunoglobulin receptor OS=Homo sapiens OX=9606 GN=PIGR PE=1 SV=4 | 902 | 20201114_16933_1ng_glyco_hcd30_etd30_compound15.15593.15593.3 | controllerType=0 controllerNumber=1 scan=15593 | 34.4705 |
| 12271:4 | 9 | K.VPGN[+2205.793]VTAVLGETLKVPC[+57.021]HFPC[+57.021]K.F | HexNAc(4)Hex(5)Fuc(2)NeuAc(1) | C[+57]*2, N[+2206] | 1158.0213 | 4 | 4629.0633 | 4629.0502 | 0 | 2.8 | 466 | Specific | 330.07 | 322.58 | 322.58 | 3.6 | 14 | >sp\|P01833\|PIGR_HUMAN Polymeric immunoglobulin receptor OS=Homo sapiens OX=9606 GN=PIGR PE=1 SV=4 | 902 | 20201114_16933_1ng_glyco_hcd30_etd30_compound15.15641.15641.3 | controllerType=0 controllerNumber=1 scan=15641 | 34.5726 |
| 04972:3 | 9 | K.WN[+2059.735]NTGC[+57.021]QALPSQDEGPSK.A | HexNAc(4)Hex(5)Fuc(1)NeuAc(1) | C[+57], N[+2060] | 1350.2118 | 3 | 4048.621 | 4048.6116 | 0 | 2.3 | 498 | Specific | 351.22 | 213.3 | 213.3 | 6.07 | 14 | >sp\|P01833\|PIGR_HUMAN Polymeric immunoglobulin receptor OS=Homo sapiens OX=9606 GN=PIGR PE=1 SV=4 | 902 | 20201114_16933_1ng_glyco_hcd30_etd30_compound15.8021.8021.3 | controllerType=0 controllerNumber=1 scan=8021 | 20.6695 |
| 05053:3 | 9 | K.WN[+1913.677]NTGC[+57.021]QALPSQDEGPSK.A | HexNAc(4)Hex(5)NeuAc(1) | C[+57], N[+1914] | 1301.525 | 3 | 3902.5603 | 3902.5537 | 0 | 1.7 | 498 | Specific | 410.08 | 275.73 | 275.73 | 6.55 | 14 | >sp\|P01833\|PIGR_HUMAN Polymeric immunoglobulin receptor OS=Homo sapiens OX=9606 GN=PIGR PE=1 SV=4 | 902 | 20201114_16933_1ng_glyco_hcd30_etd30_compound15.8105.8105.3 | controllerType=0 controllerNumber=1 scan=8105 | 20.8[illegible]33 |
| 00870:3 | 15 | K.HYTN[+1809.666]SSQDVTVPC[+57.021]R.V | HexNAc(5)Hex(4)Fuc(1) | C[+57], N[+1810] | 1158.4776 | 3 | 3473.4182 | 3473.4154 | 0 | 0.8 | 89 | Specific | 221.37 | 164.95 | 164.95 | 4.15 | 10 | >sp\|P01877\|IGHA2_HUMAN Immunoglobulin heavy constant alpha 2 OS=Homo sapiens OX=9606 GN=IGHA2 PE=1 SV=4 | 576 | 20201114_16933_1ng_glyco_hcd30_etd30_compound15.3746.3746.3 | controllerType=0 controllerNumber=1 scan=3746 | 12.8643 |
| 00886:3 | 15 | K.HYTN[+1647.613]SSQDVTVPC[+57.021]R.V | HexNAc(5)Hex(3)Fuc(1) | C[+57], N[+1648] | 1104.4602 | 3 | 3311.3661 | 3311.3625 | 0 | 1.1 | 89 | Specific | 464.84 | 391 | 391 | 6.77 | 10 | >sp\|P01877\|IGHA2_HUMAN Immunoglobulin heavy constant alpha 2 OS=Homo sapiens OX=9606 GN=IGHA2 PE=1 SV=4 | 576 | 20201114_16933_1ng_glyco_hcd30_etd30_compound15.3763.3763.3 | controllerType=0 controllerNumber=1 scan=3763 | 12.9017 |
| 02480:2 | 15 | K.TPLTAN[+1809.666]ITK.S | HexNAc(5)Hex(4)Fuc(1) | N[+1810] | 1384.618 | 2 | 2768.2286 | 2768.2228 | 0 | 2.1 | 200 | Specific | 252.38 | 63.86 | 63.86 | 2.57 | 10 | >sp\|P01877\|IGHA2_HUMAN Immunoglobulin heavy constant alpha 2 OS=Homo sapiens OX=9606 GN=IGHA2 PE=1 SV=4 | 576 | 20201114_16933_1ng_glyco_hcd30_etd30_compound15.5429.5429.2 | controllerType=0 controllerNumber=1 scan=5429 | 16.0342 |
| 02536:3 | 15 | K.TPLTAN[+1647.613]ITK.S | HexNAc(5)Hex(3)Fuc(1) | N[+1648] | 869.3971 | 3 | 2606.1767 | 2606.17 | 0 | 2.6 | 200 | Specific | 269.59 | 112.19 | 112.19 | 2.72 | 10 | >sp\|P01877\|IGHA2_HUMAN Immunoglobulin heavy constant alpha 2 OS=Homo sapiens OX=9606 GN=IGHA2 PE=1 SV=4 | 576 | 20201114_16933_1ng_glyco_hcd30_etd30_compound15.5487.5487.3 | controllerType=0 controllerNumber=1 scan=5487 | 16.1602 |
| 02488:3 | 15 | K.TPLTAN[+1809.666]ITK.S | HexNAc(5)Hex(4)Fuc(1) | N[+1810] | 923.4148 | 3 | 2768.2297 | 2768.2228 | 0 | 2.5 | 200 | Specific | 283.16 | 132.95 | 132.95 | 2.86 | 10 | >sp\|P01877\|IGHA2_HUMAN Immunoglobulin heavy constant alpha 2 OS=Homo sapiens OX=9606 GN=IGHA2 PE=1 SV=4 | 576 | 20201114_16933_1ng_glyco_hcd30_etd30_compound15.5437.5437.3 | controllerType=0 controllerNumber=1 scan=5437 | 16.0523 |
| 02528:2 | 15 | K.TPLTAN[+1647.613]ITK.S | HexNAc(5)Hex(3)Fuc(1) | N[+1648] | 1303.592 | 2 | 2606.1767 | 2606.17 | 0 | 2.6 | 200 | Specific | 292.41 | 112.67 | 112.67 | 3.06 | 10 | >sp\|P01877\|IGHA2_HUMAN Immunoglobulin heavy constant alpha 2 OS=Homo sapiens OX=9606 GN=IGHA2 PE=1 SV=4 | 576 | 20201114_16933_1ng_glyco_hcd30_etd30_compound15.5479.5479.2 | controllerType=0 controllerNumber=1 scan=5479 | 16.1447 |
| 02542:2 | 15 | K.TPLTAN[+1501.555]ITK.S | HexNAc(5)Hex(3) | N[+1502] | 1230.5634 | 2 | 2460.1195 | 2460.1121 | 0 | 3 | 200 | Specific | 302.36 | 81.48 | 81.48 | 2.56 | 10 | >sp\|P01877\|IGHA2_HUMAN Immunoglobulin heavy constant alpha 2 OS=Homo sapiens OX=9606 GN=IGHA2 PE=1 SV=4 | 576 | 20201114_16933_1ng_glyco_hcd30_etd30_compound15.5493.5493.2 | controllerType=0 controllerNumber=1 scan=5493 | 16.1755 |
| 02474:2 | 15 | K.TPLTAN[+1444.534]ITK.S | HexNAc(4)Hex(3)Fuc(1) | N[+1445] | 1202.0513 | 2 | 2403.0953 | 2403.0906 | 0 | 2 | 200 | Specific | 323.02 | 110.93 | 110.93 | 3.25 | 10 | >sp\|P01877\|IGHA2_HUMAN Immunoglobulin heavy constant alpha 2 OS=Homo sapiens OX=9606 GN=IGHA2 PE=1 SV=4 | 576 | 20201114_16933_1ng_glyco_hcd30_etd30_compound15.5422.5422.2 | controllerType=0 controllerNumber=1 scan=5422 | 16.0163 |
| 09154:3 | 15 | R.MAGKPTHIN[+1540.529]VSVVMAEADGTC[+57.021]Y.- | HexNAc(2)Hex(7) | C[+57], N[+1541] | 1297.8806 | 3 | 3891.6271 | 3891.6114 | 0 | 4 | 319 | CRagged | 214.8 | 214.8 | 176.17 | 4.01 | 10 | >sp\|P01877\|IGHA2_HUMAN Immunoglobulin heavy constant alpha 2 OS=Homo sapiens OX=9606 GN=IGHA2 PE=1 SV=4 | 576 | 20201114_16933_1ng_glyco_hcd30_etd30_compound15.12384.12384.3 | controllerType=0 controllerNumber=1 scan=12384 | 28.5894 |
| 09184:3 | 15 | R.MAGKPTHIN[+1378.476]VSVVMAEADGTC[+57.021]Y.- | HexNAc(2)Hex(6) | C[+57], N[+1378] | 1243.8603 | 3 | 3729.5663 | 3729.5585 | 0 | 2.1 | 319 | CRagged | 252.75 | 210.26 | 192.57 | 4.15 | 10 | >sp\|P01877\|IGHA2_HUMAN Immunoglobulin heavy constant alpha 2 OS=Homo sapiens OX=9606 GN=IGHA2 PE=1 SV=4 | 576 | 20201114_16933_1ng_glyco_hcd30_etd30_compound15.12416.12416.3 | controllerType=0 controllerNumber=1 scan=12416 | 28.6829 |
| 04935:3 | 17 | R.LSPN[+2205.793]ASAEHLELR.W | HexNAc(4)Hex(5)Fuc(2)NeuAc(1) | N[+2206] | 1214.8543 | 3 | 3642.5484 | 3642.5421 | 0 | 1.7 | 52 | Specific | 295.59 | 170.6 | 170.6 | 4.03 | 6 | >sp\|Q13410\|BT1A1_HUMAN Butyrophilin subfamily 1 member A1 OS=Homo sapiens OX=9606 GN=BTN1A1 PE=1 SV=3 | 179 | 20201114_16933_1ng_glyco_hcd30_etd30_compound15.7982.7982.3 | controllerType=0 controllerNumber=1 scan=7982 | 20.5908 |
| 06834:3 | 17 | R.LSPN[+2350.830]ASAEHLELR.W | HexNAc(4)Hex(5)Fuc(1)NeuAc(2) | N[+2351] | 1263.1999 | 3 | 3787.5851 | 3787.5796 | 0 | 1.5 | 52 | Specific | 391 | 259.75 | 259.75 | 4.62 | 6 | >sp\|Q13410\|BT1A1_HUMAN Butyrophilin subfamily 1 member A1 OS=Homo sapiens OX=9606 GN=BTN1A1 PE=1 SV=3 | 179 | 20201114_16933_1ng_glyco_hcd30_etd30_compound15.9953.9953.3 | controllerType=0 controllerNumber=1 scan=9953 | 24.0874 |
| 15722:3 | 17 | K.N[+1913.677]VSC[+57.021]YIQNLLLGQEK.K | HexNAc(4)Hex(5)NeuAc(1) | C[+57], N[+1914] | 1231.5428 | 3 | 3692.6138 | 3692.5876 | 0 | 7.1 | 215 | Specific | 221.45 | 109.76 | 109.76 | 4.3 | 6 | >sp\|Q13410\|BT1A1_HUMAN Butyrophilin subfamily 1 member A1 OS=Homo sapiens OX=9606 GN=BTN1A1 PE=1 SV=3 | 179 | 20201114_16933_1ng_glyco_hcd30_etd30_compound15.19279.19279.3 | controllerType=0 controllerNumber=1 scan=19279 | 42.4716 |
| 14132:4 | 24 | K.LKELPGVC[+57.021]N[+2350.830]ETMMALWEEC[+57.021]KPC[+57.021]LK.Q | HexNAc(4)Hex(5)Fuc(1)NeuAc(2) | C[+57]*3, N[+2351] | 1322.5614 | 4 | 5287.2238 | 5287.2151 | 0 | 1.7 | 95 | Specific | 321.41 | 198.54 | 198.54 | 3.64 | 5 | >sp\|P10909\|CLUS_HUMAN Clusterin OS=Homo sapiens OX=9606 GN=CLU PE=1 SV=1 | 1467 | 20201114_16933_1ng_glyco_hcd30_etd30_compound15.17583.17583.4 | controllerType=0 controllerNumber=1 scan=17583 | 38.5201 |
| 15763:3 | 24 | K.ELPGVC[+57.021]N[+2350.830]ETMMALWEEC[+57.021]KPC[+57.021]LK.Q | HexNAc(4)Hex(5)Fuc(1)NeuAc(2) | C[+57]*3, N[+2351] | 1682.6853 | 3 | 5046.0413 | 5046.036 | 0 | 1.1 | 97 | Specific | 218.31 | 124.55 | 124.55 | 4.18 | 5 | >sp\|P10909\|CLUS_HUMAN Clusterin OS=Homo sapiens OX=9606 GN=CLU PE=1 SV=1 | 1467 | 20201114_16933_1ng_glyco_hcd30_etd30_compound15.19323.19323.3 | controllerType=0 controllerNumber=1 scan=19323 | 42.6411 |
| 00107:4 | 24 | R.EIRHN[+2059.735]STGC[+57.021]LR.M | HexNAc(4)Hex(5)Fuc(1)NeuAc(1) | C[+57], N[+2060] | 851.3582 | 4 | 3402.4111 | 3402.3994 | 0 | 3.5 | 287 | Specific | 353.86 | 280.66 | 280.66 | 4.52 | 5 | >sp\|P10909\|CLUS_HUMAN Clusterin OS=Homo sapiens OX=9606 GN=CLU PE=1 SV=1 | 1467 | 20201114_16933_1ng_glyco_hcd30_etd30_compound15.2948.2948.4 | controllerType=0 controllerNumber=1 scan=2948 | 11.3019 |
| 18105:3 | 24 | K.MLN[+2059.735]TSSLLEQLNEQFNWVSR.L | HexNAc(4)Hex(5)Fuc(1)NeuAc(1) | N[+2060] | 1490.3163 | 3 | 4468.9345 | 4468.9217 | 0 | 2.9 | 352 | Specific | 405.21 | 263.87 | 263.87 | 5.86 | 5 | >sp\|P10909\|CLUS_HUMAN Clusterin OS=Homo sapiens OX=9606 GN=CLU PE=1 SV=1 | 1467 | 20201114_16933_1ng_glyco_hcd30_etd30_compound15.21806.21806.5 | controllerType=0 controllerNumber=1 scan=21806 | 49.7779 |
| 20705:4 | 30 | R.QLAHQSN[+2350.830]STNIFFSPVSIATAFAMLSLGTK.A | HexNAc(4)Hex(5)Fuc(1)NeuAc(2) | N[+2351] | 1383.8723 | 4 | 5532.4674 | 5532.4654 | 0 | 0.4 | 64 | Specific | 232.99 | 180.82 | 180.82 | 3.71 | 2 | >sp\|P01009\|A1AT_HUMAN Alpha-1-antitrypsin OS=Homo sapiens OX=9606 GN=SERPINA1 PE=1 SV=3 | 642 | 20201114_16933_1ng_glyco_hcd30_etd30_compound15.24550.24550.4 | controllerType=0 controllerNumber=1 scan=24550 | 55.748 |
| 08641:3 | 50 | R.IIVPLNNREN[+1809.666]ISDPTSPLR.T | HexNAc(5)Hex(4)Fuc(1) | N[+1810] | 1319.9551 | 3 | 3957.8509 | 3957.8432 | 0 | 1.9 | 62 | Specific | 204.69 | 163.13 | 163.13 | 3.02 | 3 | >sp\|P01591\|IGJ_HUMAN Immunoglobulin J chain OS=Homo sapiens OX=9606 GN=JCHAIN PE=1 SV=4 | 1392 | 20201114_16933_1ng_glyco_hcd30_etd30_compound15.11850.11850.3 | controllerType=0 controllerNumber=1 scan=11850 | 27.5321 |
| 08645:3 | 50 | R.IIVPLNNREN[+1663.608]ISDPTSPLR.T | HexNAc(5)Hex(4) | N[+1664] | 1271.2623 | 3 | 3811.7722 | 3811.7853 | 0 | -3.4 | 62 | Specific | 268.59 | 190.74 | 190.74 | 2.75 | 3 | >sp\|P01591\|IGJ_HUMAN Immunoglobulin J chain OS=Homo sapiens OX=9606 GN=JCHAIN PE=1 SV=4 | 1392 | 20201114_16933_1ng_glyco_hcd30_etd30_compound15.11854.11854.3 | controllerType=0 controllerNumber=1 scan=11854 | 27.5442 |
| 15839:3 | 52 | R.NYIVPILWLN[+1864.634]ETGTIGDEK.A | HexNAc(2)Hex(9) | N[+1865] | 1347.2675 | 3 | 4039.788 | 4039.7674 | 0 | 5.1 | 408 | Specific | 296.99 | 296.99 | 296.99 | 5.28 | 1 | >sp\|P16671\|CD36_HUMAN Platelet glycoprotein 4 OS=Homo sapiens OX=9606 GN=CD36 PE=1 SV=2 | 812 | 20201114_16933_1ng_glyco_hcd30_etd30_compound15.19404.19404.3 | controllerType=0 controllerNumber=1 scan=19404 | 42.9007 |
| 01103:3 | 1 | K.GPN[+2059.735]C[+57.021]SEPEC[+57.021]PGNC[+57.021]HLR.G | HexNAc(4)Hex(5)Fuc(1)NeuAc(1) | C[+57]*3, N[+2060] | 1315.1757 | 3 | 3943.5126 | 3943.4931 | 0 | 4.9 | 182 | Specific | 384.24 | 266.1 | 266.1 | 4.91 | 59 | >sp\|P24821\|TENA_HUMAN Tenascin OS=Homo sapiens OX=9606 GN=TNC PE=1 SV=3 | 486 | 20201116_fqg_colostrum_1ng_glyco_hcd30_etd30_compound15.4037.4037.3 | controllerType=0 controllerNumber=1 scan=4037 | 13.8384 |
| 12847:2 | 1 | R.LN[+1702.581]YSLPTGQWVGVQLPR.N | HexNAc(2)Hex(8) | N[+1703] | 1815.8231 | 2 | 3630.6389 | 3630.6202 | 0 | 5.2 | 1017 | Specific | 301.13 | 270.54 | 270.54 | 4.65 | 59 | >sp\|P24821\|TENA_HUMAN Tenascin OS=Homo sapiens OX=9606 GN=TNC PE=1 SV=3 | 486 | 20201116_fqg_colostrum_1ng_glyco_hcd30_etd30_compound15.16253.16253.2 | controllerType=0 controllerNumber=1 scan=16253 | 37.7101 |
| 12809:3 | 1 | R.LN[+1702.581]YSLPTGQWVGVQLPR.N | HexNAc(2)Hex(8) | N[+1703] | 1210.8807 | 3 | 3630.6275 | 3630.6202 | 0 | 2 | 1017 | Specific | 374.86 | 303.31 | 303.31 | 4.46 | 59 | >sp\|P24821\|TENA_HUMAN Tenascin OS=Homo sapiens OX=9606 GN=TNC PE=1 SV=3 | 486 | 20201116_fqg_colostrum_1ng_glyco_hcd30_etd30_compound15.16214.16214.2 | controllerType=0 controllerNumber=1 scan=16214 | 37.6221 |
| 12834:3 | 1 | R.LN[+1702.581]YSLPTGQWVGVQLPR.N | HexNAc(2)Hex(8) | N[+1703] | 1210.8807 | 3 | 3630.6275 | 3630.6202 | 0 | 2 | 1017 | Specific | 378.74 | 362.57 | 362.57 | 4.67 | 59 | >sp\|P24821\|TENA_HUMAN Tenascin OS=Homo sapiens OX=9606 GN=TNC PE=1 SV=3 | 486 | 20201116_fqg_colostrum_1ng_glyco_hcd30_etd30_compound15.16240.16240.3 | controllerType=0 controllerNumber=1 scan=16240 | 37.6813 |
| 12975:2 | 1 | R.LN[+1378.476]YSLPTGQWVGVQLPR.N | HexNAc(2)Hex(6) | N[+1378] | 1653.7653 | 2 | 3306.5233 | 3306.5145 | 0 | 2.7 | 1017 | Specific | 476 | 416.85 | 416.85 | 7.47 | 59 | >sp\|P24821\|TENA_HUMAN Tenascin OS=Homo sapiens OX=9606 GN=TNC PE=1 SV=3 | 486 | 20201116_fqg_colostrum_1ng_glyco_hcd30_etd30_compound15.16386.16386.2 | controllerType=0 controllerNumber=1 scan=16386 | 37.993 |
| 12987:3 | 1 | R.LN[+1216.423]YSLPTGQWVGVQLPR.N | HexNAc(2)Hex(5) | N[+1216] | 1048.8336 | 3 | 3144.4864 | 3144.4617 | 0 | 7.8 | 1017 | Specific | 582.96 | 563.56 | 563.56 | 7.78 | 59 | >sp\|P24821\|TENA_HUMAN Tenascin OS=Homo sapiens OX=9606 GN=TNC PE=1 SV=3 | 486 | 20201116_fqg_colostrum_1ng_glyco_hcd30_etd30_compound15.16400.16400.3 | controllerType=0 controllerNumber=1 scan=16400 | 38.0582 |
| 12983:2 | 1 | R.LN[+1216.423]YSLPTGQWVGVQLPR.N | HexNAc(2)Hex(5) | N[+1216] | 1572.7453 | 2 | 3144.4833 | 3144.4617 | 0 | 6.9 | 1017 | Specific | 589.87 | 572.36 | 572.36 | 7.63 | 59 | >sp\|P24821\|TENA_HUMAN Tenascin OS=Homo sapiens OX=9606 GN=TNC PE=1 SV=3 | 486 | 20201116_fqg_colostrum_1ng_glyco_hcd30_etd30_compound15.16396.16396.2 | controllerType=0 controllerNumber=1 scan=16396 | 38.0358 |
| 01379:3 | 1 | R.N[+1768.640]TTSYVLR.G | HexNAc(4)Hex(5)Fuc(1) | N[+1769] | 908.0554 | 3 | 2722.1518 | 2722.1446 | 0 | 2.6 | 1034 | Specific | 261.02 | 128.73 | 128.73 | 3.38 | 59 | >sp\|P24821\|TENA_HUMAN Tenascin OS=Homo sapiens OX=9606 GN=TNC PE=1 SV=3 | 486 | 20201116_fqg_colostrum_1ng_glyco_hcd30_etd30_compound15.4324.4324.3 | controllerType=0 controllerNumber=1 scan=4324 | 14.4677 |
| 02773:3 | 1 | R.N[+2059.735]TTSYVLR.G | HexNAc(4)Hex(5)Fuc(1)NeuAc(1) | N[+2060] | 1005.0891 | 3 | 3013.2527 | 3013.24 | 0 | 4.2 | 1034 | Specific | 265.61 | 142.55 | 142.55 | 3.6 | 59 | >sp\|P24821\|TENA_HUMAN Tenascin OS=Homo sapiens OX=9606 GN=TNC PE=1 SV=3 | 486 | 20201116_fqg_colostrum_1ng_glyco_hcd30_etd30_compound15.5777.5777.3 | controllerType=0 controllerNumber=1 scan=5777 | 17.7095 |
| 06210:2 | 1 | R.N[+2059.735]LTVPGSLR.A | HexNAc(4)Hex(5)Fuc(1)NeuAc(1) | N[+2060] | 1508.6497 | 2 | 3016.2921 | 3016.2873 | 0 | 1.6 | 1119 | Specific | 212.07 | 107.62 | 107.62 | 2.67 | 59 | >sp\|P24821\|TENA_HUMAN Tenascin OS=Homo sapiens OX=9606 GN=TNC PE=1 SV=3 | 486 | 20201116_fqg_colostrum_1ng_glyco_hcd30_etd30_compound15.9359.9359.2 | controllerType=0 controllerNumber=1 scan=9359 | 24.2505 |
| 30605:4 | 2 | R.LSLHRPALEDLLLGSEAN[+1501.555]LTC[+57.021]TLTGLR.D | HexNAc(5)Hex(3) | C[+57], N[+1502] | 1117.0462 | 4 | 4465.1629 | 4465.1536 | 0 | 2.1 | 127 | Specific | 202 | 202 | 202 | 4.71 | 58 | >sp\|P01876\|IGHA1_HUMAN Immunoglobulin heavy constant alpha 1 OS=Homo sapiens OX=9606 GN=IGHA1 PE=1 SV=2 | 255 | 20201116_fqg_colostrum_1ng_glyco_hcd30_etd30_compound15.36995.36995.2 | controllerType=0 controllerNumber=1 scan=36995 | 91.1122 |
| 21712:4 | 2 | R.LSLHRPALEDLLLGSEAN[+1501.555]LTC[+57.021]TLTGLR.D | HexNAc(5)Hex(3) | C[+57], N[+1502] | 1117.0401 | 4 | 4465.1385 | 4465.1536 | 0 | -3.4 | 127 | Specific | 222.19 | 222.19 | 222.19 | 3.1 | 58 | >sp\|P01876\|IGHA1_HUMAN Immunoglobulin heavy constant alpha 1 OS=Homo sapiens OX=9606 GN=IGHA1 PE=1 SV=2 | 255 | 20201116_fqg_colostrum_1ng_glyco_hcd30_etd30_compound15.25634.25634.4 | controllerType=0 controllerNumber=1 scan=25634 | 65.0836 |
| 14976:3 | 2 | R.LSLHRPALEDLLLGSEAN[+1298.476]LTC[+57.021]TLTGLR.D | HexNAc(4)Hex(3) | C[+57], N[+1298] | 1421.3683 | 3 | 4262.0905 | 4262.0742 | 0 | 3.8 | 127 | Specific | 223.18 | 223.18 | 223.18 | 3.61 | 58 | >sp\|P01876\|IGHA1_HUMAN Immunoglobulin heavy constant alpha 1 OS=Homo sapiens OX=9606 GN=IGHA1 PE=1 SV=2 | 255 | 20201116_fqg_colostrum_1ng_glyco_hcd30_etd30_compound15.18477.18477.2 | controllerType=0 controllerNumber=1 scan=18477 | 42.7408 |
| 30284:4 | 2 | R.LSLHRPALEDLLLGSEAN[+1216.423]LTC[+57.021]TLTGLR.D | HexNAc(2)Hex(5) | C[+57], N[+1216] | 1045.764 | 4 | 4180.034 | 4180.0211 | 0 | 3.1 | 127 | Specific | 226.8 | 226.8 | 226.8 | 4.12 | 58 | >sp\|P01876\|IGHA1_HUMAN Immunoglobulin heavy constant alpha 1 OS=Homo sapiens OX=9606 GN=IGHA1 PE=1 SV=2 | 255 | 20201116_fqg_colostrum_1ng_glyco_hcd30_etd30_compound15.36656.36656.4 | controllerType=0 controllerNumber=1 scan=36656 | 90.2607 |
| 30212:4 | 2 | R.LSLHRPALEDLLLGSEAN[+1501.555]LTC[+57.021]TLTGLR.D | HexNAc(5)Hex(3) | C[+57], N[+1502] | 1117.0441 | 4 | 4465.1546 | 4465.1536 | 0 | 0.2 | 127 | Specific | 243.3 | 243.3 | 243.3 | 5.19 | 58 | >sp\|P01876\|IGHA1_HUMAN Immunoglobulin heavy constant alpha 1 OS=Homo sapiens OX=9606 GN=IGHA1 PE=1 SV=2 | 255 | 20201116_fqg_colostrum_1ng_glyco_hcd30_etd30_compound15.36580.36580.4 | controllerType=0 controllerNumber=1 scan=36580 | 90.0616 |
| 14949:5 | 2 | R.LSLHRPALEDLLLGSEAN[+1298.476]LTC[+57.021]TLTGLR.D | HexNAc(4)Hex(3) | C[+57], N[+1298] | 853.2212 | 5 | 4262.077 | 4262.0742 | 0 | 0.6 | 127 | Specific | 293.77 | 293.77 | 293.77 | 4.75 | 58 | >sp\|P01876\|IGHA1_HUMAN Immunoglobulin heavy constant alpha 1 OS=Homo sapiens OX=9606 GN=IGHA1 PE=1 SV=2 | 255 | 20201116_fqg_colostrum_1ng_glyco_hcd30_etd30_compound15.18448.18448.5 | controllerType=0 controllerNumber=1 scan=18448 | 42.6549 |
| 30120:4 | 2 | R.LSLHRPALEDLLLGSEAN[+1216.423]LTC[+57.021]TLTGLR.D | HexNAc(2)Hex(5) | C[+57], N[+1216] | 1045.7625 | 4 | 4180.0281 | 4180.0211 | 0 | 1.7 | 127 | Specific | 294.58 | 294.58 | 294.58 | 6.05 | 58 | >sp\|P01876\|IGHA1_HUMAN Immunoglobulin heavy constant alpha 1 OS=Homo sapiens OX=9606 GN=IGHA1 PE=1 SV=2 | 255 | 20201116_fqg_colostrum_1ng_glyco_hcd30_etd30_compound15.36483.36483.4 | controllerType=0 controllerNumber=1 scan=36483 | 89.8023 |
| 29963:4 | 2 | R.LSLHRPALEDLLLGSEAN[+1216.423]LTC[+57.021]TLTGLR.D | HexNAc(2)Hex(5) | C[+57], N[+1216] | 1045.763 | 4 | 4180.0301 | 4180.0211 | 0 | 2.1 | 127 | Specific | 297.22 | 297.22 | 297.22 | 6.05 | 58 | >sp\|P01876\|IGHA1_HUMAN Immunoglobulin heavy constant alpha 1 OS=Homo sapiens OX=9606 GN=IGHA1 PE=1 SV=2 | 255 | 20201116_fqg_colostrum_1ng_glyco_hcd30_etd30_compound15.36318.36318.4 | controllerType=0 controllerNumber=1 scan=36318 | 89.3916 |
| 14862:3 | 2 | R.LSLHRPALEDLLLGSEAN[+1419.502]LTC[+57.021]TLTGLR.D | HexNAc(3)Hex(5) | C[+57], N[+1420] | 1461.7087 | 3 | 4383.1115 | 4383.1005 | 0 | 2.5 | 127 | Specific | 366.29 | 314.55 | 314.55 | 6.37 | 58 | >sp\|P01876\|IGHA1_HUMAN Immunoglobulin heavy constant alpha 1 OS=Homo sapiens OX=9606 GN=IGHA1 PE=1 SV=2 | 255 | 20201116_fqg_colostrum_1ng_glyco_hcd30_etd30_compound15.18357.18357.3 | controllerType=0 controllerNumber=1 scan=18357 | 42.3887 |
| 14881:3 | 2 | R.LSLHRPALEDLLLGSEAN[+1622.582]LTC[+57.021]TLTGLR.D | HexNAc(4)Hex(5) | C[+57], N[+1623] | 1529.4036 | 3 | 4586.1963 | 4586.1799 | 0 | 3.6 | 127 | Specific | 369 | 332.98 | 332.98 | 5.38 | 58 | >sp\|P01876\|IGHA1_HUMAN Immunoglobulin heavy constant alpha 1 OS=Homo sapiens OX=9606 GN=IGHA1 PE=1 SV=2 | 255 | 20201116_fqg_colostrum_1ng_glyco_hcd30_etd30_compound15.18377.18377.3 | controllerType=0 controllerNumber=1 scan=18377 | 42.4408 |
| 14942:3 | 2 | R.LSLHRPALEDLLLGSEAN[+1257.449]LTC[+57.021]TLTGLR.D | HexNAc(3)Hex(4) | C[+57], N[+1257] | 1407.6916 | 3 | 4221.0602 | 4221.0477 | 0 | 3 | 127 | Specific | 379.71 | 352.23 | 352.23 | 6.57 | 58 | >sp\|P01876\|IGHA1_HUMAN Immunoglobulin heavy constant alpha 1 OS=Homo sapiens OX=9606 GN=IGHA1 PE=1 SV=2 | 255 | 20201116_fqg_colostrum_1ng_glyco_hcd30_etd30_compound15.18441.18441.3 | controllerType=0 controllerNumber=1 scan=18441 | 42.6345 |
| 14863:3 | 2 | R.LSLHRPALEDLLLGSEAN[+1460.529]LTC[+57.021]TLTGLR.D | HexNAc(4)Hex(4) | C[+57], N[+1461] | 1475.3848 | 3 | 4424.1399 | 4424.1271 | 0 | 2.9 | 127 | Specific | 396.53 | 369.36 | 369.36 | 7.78 | 58 | >sp\|P01876\|IGHA1_HUMAN Immunoglobulin heavy constant alpha 1 OS=Homo sapiens OX=9606 GN=IGHA1 PE=1 SV=2 | 255 | 20201116_fqg_colostrum_1ng_glyco_hcd30_etd30_compound15.18358.18358.6 | controllerType=0 controllerNumber=1 scan=18358 | 42.3922 |
| 15674:3 | 2 | R.LSLHRPALEDLLLGSEAN[+1751.624]LTC[+57.021]TLTGLR.D | HexNAc(4)Hex(4)NeuAc(1) | C[+57], N[+1752] | 1572.417 | 3 | 4715.2366 | 4715.2225 | 0 | 3 | 127 | Specific | 405.1 | 295.38 | 295.38 | 6.5 | 58 | >sp\|P01876\|IGHA1_HUMAN Immunoglobulin heavy constant alpha 1 OS=Homo sapiens OX=9606 GN=IGHA1 PE=1 SV=2 | 255 | 20201116_fqg_colostrum_1ng_glyco_hcd30_etd30_compound15.19213.19213.3 | controllerType=0 controllerNumber=1 scan=19213 | 44.7863 |
| 15649:3 | 2 | R.LSLHRPALEDLLLGSEAN[+1872.651]LTC[+57.021]TLTGLR.D | HexNAc(3)Hex(6)NeuAc(1) | C[+57], N[+1873] | 1612.7587 | 3 | 4836.2616 | 4836.2487 | 0 | 2.7 | 127 | Specific | 407.25 | 312.62 | 312.62 | 5.31 | 58 | >sp\|P01876\|IGHA1_HUMAN Immunoglobulin heavy constant alpha 1 OS=Homo sapiens OX=9606 GN=IGHA1 PE=1 SV=2 | 255 | 20201116_fqg_colostrum_1ng_glyco_hcd30_etd30_compound15.19187.19187.3 | controllerType=0 controllerNumber=1 scan=19187 | 44.7173 |
| 14820:3 | 2 | R.LSLHRPALEDLLLGSEAN[+1663.608]LTC[+57.021]TLTGLR.D | HexNAc(5)Hex(4) | C[+57], N[+1664] | 1543.0766 | 3 | 4627.2152 | 4627.2064 | 0 | 1.9 | 127 | Specific | 407.69 | 358.21 | 358.21 | 6.36 | 58 | >sp\|P01876\|IGHA1_HUMAN Immunoglobulin heavy constant alpha 1 OS=Homo sapiens OX=9606 GN=IGHA1 PE=1 SV=2 | 255 | 20201116_fqg_colostrum_1ng_glyco_hcd30_etd30_compound15.18313.18313.3 | controllerType=0 controllerNumber=1 scan=18313 | 42.2576 |
| 14846:3 | 2 | R.LSLHRPALEDLLLGSEAN[+1460.529]LTC[+57.021]TLTGLR.D | HexNAc(4)Hex(4) | C[+57], N[+1461] | 1475.3829 | 3 | 4424.1341 | 4424.1271 | 0 | 1.6 | 127 | Specific | 424.64 | 364.36 | 364.36 | 7.57 | 58 | >sp\|P01876\|IGHA1_HUMAN Immunoglobulin heavy constant alpha 1 OS=Homo sapiens OX=9606 GN=IGHA1 PE=1 SV=2 | 255 | 20201116_fqg_colostrum_1ng_glyco_hcd30_etd30_compound15.18340.18340.3 | controllerType=0 controllerNumber=1 scan=18340 | 42.339 |
| 14775:3 | 2 | R.LSLHRPALEDLLLGSEAN[+1864.634]LTC[+57.021]TLTGLR.D | HexNAc(2)Hex(9) | C[+57], N[+1865] | 1610.0842 | 3 | 4828.2379 | 4828.2324 | 0 | 1.1 | 127 | Specific | 427.53 | 382.61 | 382.61 | 6.37 | 58 | >sp\|P01876\|IGHA1_HUMAN Immunoglobulin heavy constant alpha 1 OS=Homo sapiens OX=9606 GN=IGHA1 PE=1 SV=2 | 255 | 20201116_fqg_colostrum_1ng_glyco_hcd30_etd30_compound15.18266.18266.3 | controllerType=0 controllerNumber=1 scan=18266 | 42.1463 |
| 15669:3 | 2 | R.LSLHRPALEDLLLGSEAN[+1710.598]LTC[+57.021]TLTGLR.D | HexNAc(3)Hex(5)NeuAc(1) | C[+57], N[+1711] | 1558.7408 | 3 | 4674.2078 | 4674.1959 | 0 | 2.5 | 127 | Specific | 429.06 | 328.12 | 328.12 | 6.5 | 58 | >sp\|P01876\|IGHA1_HUMAN Immunoglobulin heavy constant alpha 1 OS=Homo sapiens OX=9606 GN=IGHA1 PE=1 SV=2 | 255 | 20201116_fqg_colostrum_1ng_glyco_hcd30_etd30_compound15.19208.19208.3 | controllerType=0 controllerNumber=1 scan=19208 | 44.7762 |
| 14936:3 | 2 | R.LSLHRPALEDLLLGSEAN[+1095.397]LTC[+57.021]TLTGLR.D | HexNAc(3)Hex(3) | C[+57], N[+1095] | 1353.6713 | 3 | 4058.9994 | 4058.9949 | 0 | 1.1 | 127 | Specific | 440.38 | 440.38 | 440.38 | 7.64 | 58 | >sp\|P01876\|IGHA1_HUMAN Immunoglobulin heavy constant alpha 1 OS=Homo sapiens OX=9606 GN=IGHA1 PE=1 SV=2 | 255 | 20201116_fqg_colostrum_1ng_glyco_hcd30_etd30_compound15.18435.18435.3 | controllerType=0 controllerNumber=1 scan=18435 | 42.6175 |
| 14871:3 | 2 | R.LSLHRPALEDLLLGSEAN[+1501.555]LTC[+57.021]TLTGLR.D | HexNAc(5)Hex(3) | C[+57], N[+1502] | 1489.0616 | 3 | 4465.1702 | 4465.1536 | 0 | 3.7 | 127 | Specific | 442.87 | 408.48 | 408.48 | 7.99 | 58 | >sp\|P01876\|IGHA1_HUMAN Immunoglobulin heavy constant alpha 1 OS=Homo sapiens OX=9606 GN=IGHA1 PE=1 SV=2 | 255 | 20201116_fqg_colostrum_1ng_glyco_hcd30_etd30_compound15.18367.18367.3 | controllerType=0 controllerNumber=1 scan=18367 | 42.4196 |
| 14883:3 | 2 | R.LSLHRPALEDLLLGSEAN[+1298.476]LTC[+57.021]TLTGLR.D | HexNAc(4)Hex(3) | C[+57], N[+1298] | 1421.3686 | 3 | 4262.0912 | 4262.0742 | 0 | 4 | 127 | Specific | 447.51 | 447.51 | 447.51 | 7.99 | 58 | >sp\|P01876\|IGHA1_HUMAN Immunoglobulin heavy constant alpha 1 OS=Homo sapiens OX=9606 GN=IGHA1 PE=1 SV=2 | 255 | 20201116_fqg_colostrum_1ng_glyco_hcd30_etd30_compound15.18379.18379.6 | controllerType=0 controllerNumber=1 scan=18379 | 42.449 |
| 14795:3 | 2 | R.LSLHRPALEDLLLGSEAN[+1864.634]LTC[+57.021]TLTGLR.D | HexNAc(2)Hex(9) | C[+57], N[+1865] | 1610.086 | 3 | 4828.2434 | 4828.2324 | 0 | 2.3 | 127 | Specific | 448.33 | 396.04 | 396.04 | 6.45 | 58 | >sp\|P01876\|IGHA1_HUMAN Immunoglobulin heavy constant alpha 1 OS=Homo sapiens OX=9606 GN=IGHA1 PE=1 SV=2 | 255 | 20201116_fqg_colostrum_1ng_glyco_hcd30_etd30_compound15.18287.18287.3 | controllerType=0 controllerNumber=1 scan=18287 | 42.1898 |
| 16579:3 | 2 | R.LSLHRPALEDLLLGSEAN[+2204.772]LTC[+57.021]TLTGLR.D | HexNAc(4)Hex(5)NeuAc(2) | C[+57], N[+2205] | 1723.4692 | 3 | 5168.3931 | 5168.3707 | 0 | 4.3 | 127 | Specific | 453.27 | 325.72 | 325.72 | 4.65 | 58 | >sp\|P01876\|IGHA1_HUMAN Immunoglobulin heavy constant alpha 1 OS=Homo sapiens OX=9606 GN=IGHA1 PE=1 SV=2 | 255 | 20201116_fqg_colostrum_1ng_glyco_hcd30_etd30_compound15.20167.20167.3 | controllerType=0 controllerNumber=1 scan=20167 | 47.2059 |
| 15647:5 | 2 | R.LSLHRPALEDLLLGSEAN[+1913.677]LTC[+57.021]TLTGLR.D | HexNAc(4)Hex(5)NeuAc(1) | C[+57], N[+1914] | 976.2624 | 5 | 4877.283 | 4877.2753 | 0 | 1.6 | 127 | Specific | 456.14 | 456.14 | 456.14 | 6.64 | 58 | >sp\|P01876\|IGHA1_HUMAN Immunoglobulin heavy constant alpha 1 OS=Homo sapiens OX=9606 GN=IGHA1 PE=1 SV=2 | 255 | 20201116_fqg_colostrum_1ng_glyco_hcd30_etd30_compound15.19185.19185.5 | controllerType=0 controllerNumber=1 scan=19185 | 44.7096 |
| 15707:3 | 2 | R.LSLHRPALEDLLLGSEAN[+1548.545]LTC[+57.021]TLTGLR.D | HexNAc(3)Hex(4)NeuAc(1) | C[+57], N[+1549] | 1504.7231 | 3 | 4512.1547 | 4512.1431 | 0 | 2.6 | 127 | Specific | 462.49 | 378.73 | 378.73 | 7.99 | 58 | >sp\|P01876\|IGHA1_HUMAN Immunoglobulin heavy constant alpha 1 OS=Homo sapiens OX=9606 GN=IGHA1 PE=1 SV=2 | 255 | 20201116_fqg_colostrum_1ng_glyco_hcd30_etd30_compound15.19248.19248.3 | controllerType=0 controllerNumber=1 scan=19248 | 44.8693 |
| 14851:3 | 2 | R.LSLHRPALEDLLLGSEAN[+1216.423]LTC[+57.021]TLTGLR.D | HexNAc(2)Hex(5) | C[+57], N[+1216] | 1394.0156 | 3 | 4180.0321 | 4180.0211 | 0 | 2.6 | 127 | Specific | 487.76 | 476.76 | 476.76 | 8.05 | 58 | >sp\|P01876\|IGHA1_HUMAN Immunoglobulin heavy constant alpha 1 OS=Homo sapiens OX=9606 GN=IGHA1 PE=1 SV=2 | 255 | 20201116_fqg_colostrum_1ng_glyco_hcd30_etd30_compound15.18346.18346.3 | controllerType=0 controllerNumber=1 scan=18346 | 42.3624 |
| 14796:3 | 2 | R.LSLHRPALEDLLLGSEAN[+1702.581]LTC[+57.021]TLTGLR.D | HexNAc(2)Hex(8) | C[+57], N[+1703] | 1556.0677 | 3 | 4666.1885 | 4666.1796 | 0 | 1.9 | 127 | Specific | 497.67 | 451.9 | 451.9 | 7.7 | 58 | >sp\|P01876\|IGHA1_HUMAN Immunoglobulin heavy constant alpha 1 OS=Homo sapiens OX=9606 GN=IGHA1 PE=1 SV=2 | 255 | 20201116_fqg_colostrum_1ng_glyco_hcd30_etd30_compound15.18288.18288.3 | controllerType=0 controllerNumber=1 scan=18288 | 42.1938 |
| 14969:3 | 2 | R.LSLHRPALEDLLLGSEAN[+892.317]LTC[+57.021]TLTGLR.D | HexNAc(2)Hex(3) | C[+57], N[+892] | 1285.9803 | 3 | 3855.9263 | 3855.9155 | 0 | 2.8 | 127 | Specific | 513.29 | 513.29 | 513.29 | 8.05 | 58 | >sp\|P01876\|IGHA1_HUMAN Immunoglobulin heavy constant alpha 1 OS=Homo sapiens OX=9606 GN=IGHA1 PE=1 SV=2 | 255 | 20201116_fqg_colostrum_1ng_glyco_hcd30_etd30_compound15.18470.18470.3 | controllerType=0 controllerNumber=1 scan=18470 | 42.7223 |
| 14918:3 | 2 | R.LSLHRPALEDLLLGSEAN[+1054.370]LTC[+57.021]TLTGLR.D | HexNAc(2)Hex(4) | C[+57], N[+1054] | 1339.9975 | 3 | 4017.9779 | 4017.9683 | 0 | 2.4 | 127 | Specific | 513.8 | 503.84 | 503.84 | 8.05 | 58 | >sp\|P01876\|IGHA1_HUMAN Immunoglobulin heavy constant alpha 1 OS=Homo sapiens OX=9606 GN=IGHA1 PE=1 SV=2 | 255 | 20201116_fqg_colostrum_1ng_glyco_hcd30_etd30_compound15.18416.18416.3 | controllerType=0 controllerNumber=1 scan=18416 | 42.5603 |
| 14842:3 | 2 | R.LSLHRPALEDLLLGSEAN[+1378.476]LTC[+57.021]TLTGLR.D | HexNAc(2)Hex(6) | C[+57], N[+1378] | 1448.0322 | 3 | 4342.0819 | 4342.074 | 0 | 1.8 | 127 | Specific | 564.45 | 540.25 | 540.25 | 8.9 | 58 | >sp\|P01876\|IGHA1_HUMAN Immunoglobulin heavy constant alpha 1 OS=Homo sapiens OX=9606 GN=IGHA1 PE=1 SV=2 | 255 | 20201116_fqg_colostrum_1ng_glyco_hcd30_etd30_compound15.18336.18336.3 | controllerType=0 controllerNumber=1 scan=18336 | 42.3285 |
| 15408:4 | 2 | R.LSLHRPALEDLLLGSEAN[+1501.555]LTC[+57.021]TLTGLR.D | HexNAc(5)Hex(3) | C[+57], N[+1502] | 1117.0411 | 4 | 4465.1424 | 4465.1536 | 0 | -2.5 | 127 | Specific | 631.56 | 581.79 | 581.79 | 9.05 | 58 | >sp\|P01876\|IGHA1_HUMAN Immunoglobulin heavy constant alpha 1 OS=Homo sapiens OX=9606 GN=IGHA1 PE=1 SV=2 | 255 | 20201116_fqg_colostrum_1ng_glyco_hcd30_etd30_compound15.18929.18929.2 | controllerType=0 controllerNumber=1 scan=18929 | 43.8604 |
| 29722:4 | 2 | R.LSLHRPALEDLLLGSEAN[+1501.555]LTC[+57.021]TLTGLR.D | HexNAc(5)Hex(3) | C[+57], N[+1502] | 1117.0452 | 4 | 4465.159 | 4465.1536 | 0 | 1.2 | 127 | Specific | 667.42 | 667.42 | 667.42 | 11.19 | 58 | >sp\|P01876\|IGHA1_HUMAN Immunoglobulin heavy constant alpha 1 OS=Homo sapiens OX=9606 GN=IGHA1 PE=1 SV=2 | 255 | 20201116_fqg_colostrum_1ng_glyco_hcd30_etd30_compound15.36058.36058.4 | controllerType=0 controllerNumber=1 scan=36058 | 88.6822 |
| 15642:4 | 2 | R.LSLHRPALEDLLLGSEAN[+1872.651]LTC[+57.021]TLTGLR.D | HexNAc(3)Hex(6)NeuAc(1) | C[+57], N[+1873] | 1209.8215 | 4 | 4836.264 | 4836.2487 | 0 | 3.2 | 127 | Specific | 679.27 | 612.43 | 612.43 | 11.54 | 58 | >sp\|P01876\|IGHA1_HUMAN Immunoglobulin heavy constant alpha 1 OS=Homo sapiens OX=9606 GN=IGHA1 PE=1 SV=2 | 255 | 20201116_fqg_colostrum_1ng_glyco_hcd30_etd30_compound15.19180.19180.4 | controllerType=0 controllerNumber=1 scan=19180 | 44.6992 |
| 14943:4 | 2 | R.LSLHRPALEDLLLGSEAN[+1524.534]LTC[+57.021]TLTGLR.D | HexNAc(2)Hex(6)Fuc(1) | C[+57], N[+1525] | 1122.7868 | 4 | 4488.1253 | 4488.1319 | 0 | -1.5 | 127 | Specific | 710.74 | 710.74 | 710.74 | 11.05 | 58 | >sp\|P01876\|IGHA1_HUMAN Immunoglobulin heavy constant alpha 1 OS=Homo sapiens OX=9606 GN=IGHA1 PE=1 SV=2 | 255 | 20201116_fqg_colostrum_1ng_glyco_hcd30_etd30_compound15.18442.18442.4 | controllerType=0 controllerNumber=1 scan=18442 | 42.6378 |
| 14868:5 | 2 | R.LSLHRPALEDLLLGSEAN[+1216.423]LTC[+57.021]TLTGLR.D | HexNAc(2)Hex(5) | C[+57], N[+1216] | 836.8124 | 5 | 4180.0329 | 4180.0211 | 0 | 2.8 | 127 | Specific | 727.26 | 717.55 | 717.55 | 12.69 | 58 | >sp\|P01876\|IGHA1_HUMAN Immunoglobulin heavy constant alpha 1 OS=Homo sapiens OX=9606 GN=IGHA1 PE=1 SV=2 | 255 | 20201116_fqg_colostrum_1ng_glyco_hcd30_etd30_compound15.18363.18363.5 | controllerType=0 controllerNumber=1 scan=18363 | 42.4074 |
| 14899:4 | 2 | R.LSLHRPALEDLLLGSEAN[+1257.449]LTC[+57.021]TLTGLR.D | HexNAc(3)Hex(4) | C[+57], N[+1257] | 1056.0173 | 4 | 4221.0472 | 4221.0477 | 0 | -0.1 | 127 | Specific | 730.2 | 730.2 | 730.2 | 11.46 | 58 | >sp\|P01876\|IGHA1_HUMAN Immunoglobulin heavy constant alpha 1 OS=Homo sapiens OX=9606 GN=IGHA1 PE=1 SV=2 | 255 | 20201116_fqg_colostrum_1ng_glyco_hcd30_etd30_compound15.18396.18396.4 | controllerType=0 controllerNumber=1 scan=18396 | 42.5078 |
| 14932:4 | 2 | R.LSLHRPALEDLLLGSEAN[+1524.534]LTC[+57.021]TLTGLR.D | HexNAc(2)Hex(6)Fuc(1) | C[+57], N[+1525] | 1122.7848 | 4 | 4488.1175 | 4488.1319 | 0 | -3.2 | 127 | Specific | 758.82 | 758.82 | 758.82 | 12.05 | 58 | >sp\|P01876\|IGHA1_HUMAN Immunoglobulin heavy constant alpha 1 OS=Homo sapiens OX=9606 GN=IGHA1 PE=1 SV=2 | 255 | 20201116_fqg_colostrum_1ng_glyco_hcd30_etd30_compound15.18430.18430.4 | controllerType=0 controllerNumber=1 scan=18430 | 42.6011 |
| 15663:4 | 2 | R.LSLHRPALEDLLLGSEAN[+1710.598]LTC[+57.021]TLTGLR.D | HexNAc(3)Hex(5)NeuAc(1) | C[+57], N[+1711] | 1169.3081 | 4 | 4674.2108 | 4674.1959 | 0 | 3.2 | 127 | Specific | 766.7 | 736.34 | 736.34 | 13.54 | 58 | >sp\|P01876\|IGHA1_HUMAN Immunoglobulin heavy constant alpha 1 OS=Homo sapiens OX=9606 GN=IGHA1 PE=1 SV=2 | 255 | 20201116_fqg_colostrum_1ng_glyco_hcd30_etd30_compound15.19202.19202.4 | controllerType=0 controllerNumber=1 scan=19202 | 44.7607 |

Table S4. Cont.

| Query #:z | Protein Rank | Glycopeptide | Glycan | Modification Type(s) | Observed m/z | z | Observed (M+H) | Calc. mass (M+H) | Off-by-x error | Mass error (ppm) | StartPosition | Cleavage | Score | Delta | Delta Mod | \|Log Prob\| | # of unique peptides | Protein Name | Protein DB number | Comment | Scan # | Scan Time |
|---|---|---|---|---|---|---|---|---|---|---|---|---|---|---|---|---|---|---|---|---|---|---|
| 14882:5 | 2 | R.LSLHRPALEDLLLGSEAN[+1501.555]LTC[+57.021]TLTGLR.D | HexNAc(5)Hex(3) | C[+57], N[+1502] | 893.8394 | 5 | 4465.1678 | 4465.1536 | 0 | 3.2 | 127 | Specific | 777.04 | 768.82 | 768.82 | 13.69 | 58 | >sp\|P01876\|IGHA1_HUMAN Immunoglobulin heavy constant alpha 1 OS=Homo sapiens OX=9606 GN=IGHA1 PE=1 SV=2 | 255 | 20201116_fqg_colostrum_1ng_glyco_hcd30_etd30_compound15.18378.18378.5 | controllerType=0 controllerNumber=1 scan=18378 | 42.446 |
| 15694:4 | 2 | R.LSLHRPALEDLLLGSEAN[+1548.545]LTC[+57.021]TLTGLR.D | HexNAc(3)Hex(4)NeuAc(1) | C[+57], N[+1549] | 1128.7942 | 4 | 4512.1551 | 4512.1431 | 0 | 2.7 | 127 | Specific | 820.97 | 814.99 | 814.99 | 14.54 | 58 | >sp\|P01876\|IGHA1_HUMAN Immunoglobulin heavy constant alpha 1 OS=Homo sapiens OX=9606 GN=IGHA1 PE=1 SV=2 | 255 | 20201116_fqg_colostrum_1ng_glyco_hcd30_etd30_compound15.19234.19234.4 | controllerType=0 controllerNumber=1 scan=19234 | 44.8289 |
| 14770:4 | 2 | R.LSLHRPALEDLLLGSEAN[+1864.634]LTC[+57.021]TLTGLR.D | HexNAc(2)Hex(9) | C[+57], N[+1865] | 1207.8157 | 4 | 4828.241 | 4828.2324 | 0 | 1.8 | 127 | Specific | 858.92 | 841.95 | 841.95 | 15.19 | 58 | >sp\|P01876\|IGHA1_HUMAN Immunoglobulin heavy constant alpha 1 OS=Homo sapiens OX=9606 GN=IGHA1 PE=1 SV=2 | 255 | 20201116_fqg_colostrum_1ng_glyco_hcd30_etd30_compound15.18261.18261.4 | controllerType=0 controllerNumber=1 scan=18261 | 42.136 |
| 15667:4 | 2 | R.LSLHRPALEDLLLGSEAN[+1751.624]LTC[+57.021]TLTGLR.D | HexNAc(4)Hex(4)NeuAc(1) | C[+57], N[+1752] | 1179.5638 | 4 | 4715.2332 | 4715.2225 | 0 | 2.3 | 127 | Specific | 887.56 | 830.58 | 830.58 | 15.19 | 58 | >sp\|P01876\|IGHA1_HUMAN Immunoglobulin heavy constant alpha 1 OS=Homo sapiens OX=9606 GN=IGHA1 PE=1 SV=2 | 255 | 20201116_fqg_colostrum_1ng_glyco_hcd30_etd30_compound15.19206.19206.4 | controllerType=0 controllerNumber=1 scan=19206 | 44.7685 |
| 14913:4 | 2 | R.LSLHRPALEDLLLGSEAN[+1501.555]LTC[+57.021]TLTGLR.D | HexNAc(5)Hex(3) | C[+57], N[+1502] | 1117.0469 | 4 | 4465.1658 | 4465.1536 | 0 | 2.7 | 127 | Specific | 927.37 | 893.5 | 893.5 | 16.54 | 58 | >sp\|P01876\|IGHA1_HUMAN Immunoglobulin heavy constant alpha 1 OS=Homo sapiens OX=9606 GN=IGHA1 PE=1 SV=2 | 255 | 20201116_fqg_colostrum_1ng_glyco_hcd30_etd30_compound15.18411.18411.4 | controllerType=0 controllerNumber=1 scan=18411 | 42.5473 |
| 14831:4 | 2 | R.LSLHRPALEDLLLGSEAN[+1540.529]LTC[+57.021]TLTGLR.D | HexNAc(2)Hex(7) | C[+57], N[+1541] | 1126.7889 | 4 | 4504.1336 | 4504.1268 | 0 | 1.5 | 127 | Specific | 931.5 | 927.77 | 927.77 | 16.19 | 58 | >sp\|P01876\|IGHA1_HUMAN Immunoglobulin heavy constant alpha 1 OS=Homo sapiens OX=9606 GN=IGHA1 PE=1 SV=2 | 255 | 20201116_fqg_colostrum_1ng_glyco_hcd30_etd30_compound15.18324.18324.4 | controllerType=0 controllerNumber=1 scan=18324 | 42.2863 |
| 14852:4 | 2 | R.LSLHRPALEDLLLGSEAN[+1419.502]LTC[+57.021]TLTGLR.D | HexNAc(3)Hex(5) | C[+57], N[+1420] | 1096.5337 | 4 | 4383.1131 | 4383.1005 | 0 | 2.9 | 127 | Specific | 977.15 | 920.79 | 920.79 | 17.54 | 58 | >sp\|P01876\|IGHA1_HUMAN Immunoglobulin heavy constant alpha 1 OS=Homo sapiens OX=9606 GN=IGHA1 PE=1 SV=2 | 255 | 20201116_fqg_colostrum_1ng_glyco_hcd30_etd30_compound15.18347.18347.4 | controllerType=0 controllerNumber=1 scan=18347 | 42.3652 |
| 14819:4 | 2 | R.LSLHRPALEDLLLGSEAN[+1702.581]LTC[+57.021]TLTGLR.D | HexNAc(2)Hex(8) | C[+57], N[+1703] | 1167.3025 | 4 | 4666.1883 | 4666.1796 | 0 | 1.9 | 127 | Specific | 979.36 | 946.79 | 946.79 | 17.19 | 58 | >sp\|P01876\|IGHA1_HUMAN Immunoglobulin heavy constant alpha 1 OS=Homo sapiens OX=9606 GN=IGHA1 PE=1 SV=2 | 255 | 20201116_fqg_colostrum_1ng_glyco_hcd30_etd30_compound15.18312.18312.4 | controllerType=0 controllerNumber=1 scan=18312 | 42.252 |
| 14904:4 | 2 | R.LSLHRPALEDLLLGSEAN[+1501.555]LTC[+57.021]TLTGLR.D | HexNAc(5)Hex(3) | C[+57], N[+1502] | 1117.0461 | 4 | 4465.1624 | 4465.1536 | 0 | 2 | 127 | Specific | 988.31 | 988.31 | 988.31 | 17.19 | 58 | >sp\|P01876\|IGHA1_HUMAN Immunoglobulin heavy constant alpha 1 OS=Homo sapiens OX=9606 GN=IGHA1 PE=1 SV=2 | 255 | 20201116_fqg_colostrum_1ng_glyco_hcd30_etd30_compound15.18401.18401.2 | controllerType=0 controllerNumber=1 scan=18401 | 42.5195 |
| 14825:4 | 2 | R.LSLHRPALEDLLLGSEAN[+1663.608]LTC[+57.021]TLTGLR.D | HexNAc(5)Hex(4) | C[+57], N[+1664] | 1157.5594 | 4 | 4627.2156 | 4627.2064 | 0 | 2 | 127 | Specific | 991.55 | 983.99 | 983.99 | 17.19 | 58 | >sp\|P01876\|IGHA1_HUMAN Immunoglobulin heavy constant alpha 1 OS=Homo sapiens OX=9606 GN=IGHA1 PE=1 SV=2 | 255 | 20201116_fqg_colostrum_1ng_glyco_hcd30_etd30_compound15.18318.18318.4 | controllerType=0 controllerNumber=1 scan=18318 | 42.269 |
| 14835:4 | 2 | R.LSLHRPALEDLLLGSEAN[+1540.529]LTC[+57.021]TLTGLR.D | HexNAc(2)Hex(7) | C[+57], N[+1541] | 1126.7887 | 4 | 4504.1331 | 4504.1268 | 0 | 1.4 | 127 | Specific | 1009.65 | 919.22 | 919.22 | 18.19 | 58 | >sp\|P01876\|IGHA1_HUMAN Immunoglobulin heavy constant alpha 1 OS=Homo sapiens OX=9606 GN=IGHA1 PE=1 SV=2 | 255 | 20201116_fqg_colostrum_1ng_glyco_hcd30_etd30_compound15.18329.18329.4 | controllerType=0 controllerNumber=1 scan=18329 | 42.3005 |
| 14876:4 | 2 | R.LSLHRPALEDLLLGSEAN[+1460.529]LTC[+57.021]TLTGLR.D | HexNAc(4)Hex(4) | C[+57], N[+1461] | 1106.7915 | 4 | 4424.1444 | 4424.1271 | 0 | 3.9 | 127 | Specific | 1014.45 | 966.67 | 966.67 | 18.54 | 58 | >sp\|P01876\|IGHA1_HUMAN Immunoglobulin heavy constant alpha 1 OS=Homo sapiens OX=9606 GN=IGHA1 PE=1 SV=2 | 255 | 20201116_fqg_colostrum_1ng_glyco_hcd30_etd30_compound15.18372.18372.4 | controllerType=0 controllerNumber=1 scan=18372 | 42.431 |
| 14892:4 | 2 | R.LSLHRPALEDLLLGSEAN[+1298.476]LTC[+57.021]TLTGLR.D | HexNAc(4)Hex(3) | C[+57], N[+1298] | 1066.2753 | 4 | 4262.0794 | 4262.0742 | 0 | 1.2 | 127 | Specific | 1023.93 | 995.45 | 995.45 | 18.19 | 58 | >sp\|P01876\|IGHA1_HUMAN Immunoglobulin heavy constant alpha 1 OS=Homo sapiens OX=9606 GN=IGHA1 PE=1 SV=2 | 255 | 20201116_fqg_colostrum_1ng_glyco_hcd30_etd30_compound15.18389.18389.4 | controllerType=0 controllerNumber=1 scan=18389 | 42.4919 |
| 14840:4 | 2 | R.LSLHRPALEDLLLGSEAN[+1216.423]LTC[+57.021]TLTGLR.D | HexNAc(2)Hex(5) | C[+57], N[+1216] | 1045.7626 | 4 | 4180.0286 | 4180.0211 | 0 | 1.8 | 127 | Specific | 1043.72 | 1022.6 | 1022.6 | 18.19 | 58 | >sp\|P01876\|IGHA1_HUMAN Immunoglobulin heavy constant alpha 1 OS=Homo sapiens OX=9606 GN=IGHA1 PE=1 SV=2 | 255 | 20201116_fqg_colostrum_1ng_glyco_hcd30_etd30_compound15.18334.18334.4 | controllerType=0 controllerNumber=1 scan=18334 | 42.3172 |
| 14993:4 | 2 | R.LSLHRPALEDLLLGSEAN[+892.317]LTC[+57.021]TLTGLR.D | HexNAc(2)Hex(3) | C[+57], N[+892] | 964.7366 | 4 | 3855.9244 | 3855.9155 | 0 | 2.3 | 127 | Specific | 1065.03 | 1065.03 | 1065.03 | 19.19 | 58 | >sp\|P01876\|IGHA1_HUMAN Immunoglobulin heavy constant alpha 1 OS=Homo sapiens OX=9606 GN=IGHA1 PE=1 SV=2 | 255 | 20201116_fqg_colostrum_1ng_glyco_hcd30_etd30_compound15.18495.18495.4 | controllerType=0 controllerNumber=1 scan=18495 | 42.7947 |
| 14836:4 | 2 | R.LSLHRPALEDLLLGSEAN[+1378.476]LTC[+57.021]TLTGLR.D | HexNAc(2)Hex(6) | C[+57], N[+1378] | 1086.2756 | 4 | 4342.0804 | 4342.074 | 0 | 1.5 | 127 | Specific | 1074.69 | 1037.29 | 1037.29 | 19.19 | 58 | >sp\|P01876\|IGHA1_HUMAN Immunoglobulin heavy constant alpha 1 OS=Homo sapiens OX=9606 GN=IGHA1 PE=1 SV=2 | 255 | 20201116_fqg_colostrum_1ng_glyco_hcd30_etd30_compound15.18330.18330.4 | controllerType=0 controllerNumber=1 scan=18330 | 42.3052 |
| 14935:4 | 2 | R.LSLHRPALEDLLLGSEAN[+1054.370]LTC[+57.021]TLTGLR.D | HexNAc(2)Hex(4) | C[+57], N[+1054] | 1005.2492 | 4 | 4017.9752 | 4017.9683 | 0 | 1.7 | 127 | Specific | 1105.02 | 1091.16 | 1091.16 | 20.19 | 58 | >sp\|P01876\|IGHA1_HUMAN Immunoglobulin heavy constant alpha 1 OS=Homo sapiens OX=9606 GN=IGHA1 PE=1 SV=2 | 255 | 20201116_fqg_colostrum_1ng_glyco_hcd30_etd30_compound15.18434.18434.4 | controllerType=0 controllerNumber=1 scan=18434 | 42.614 |
| 08143:3 | 2 | R.LAGKPTHVN[+1663.608]VSVVMAEVDGTC[+57.021]Y.- | HexNAc(5)Hex(4) | C[+57], N[+1664] | 1337.5906 | 3 | 4010.7573 | 4010.7502 | 0 | 1.8 | 332 | CRagged | 218.78 | 142.31 | 140.48 | 2.73 | 58 | >sp\|P01876\|IGHA1_HUMAN Immunoglobulin heavy constant alpha 1 OS=Homo sapiens OX=9606 GN=IGHA1 PE=1 SV=2 | 255 | 20201116_fqg_colostrum_1ng_glyco_hcd30_etd30_compound15.11368.11368.3 | controllerType=0 controllerNumber=1 scan=11368 | 28.2805 |
| 09040:3 | 2 | R.LAGKPTHVN[+1378.476]VSVVMAEVDGTC[+57.021]Y.- | HexNAc(2)Hex(6) | C[+57], N[+1378] | 1242.5472 | 3 | 3725.627 | 3725.6178 | 0 | 2.5 | 332 | CRagged | 223.72 | 223.72 | 161.89 | 3.6 | 58 | >sp\|P01876\|IGHA1_HUMAN Immunoglobulin heavy constant alpha 1 OS=Homo sapiens OX=9606 GN=IGHA1 PE=1 SV=2 | 255 | 20201116_fqg_colostrum_1ng_glyco_hcd30_etd30_compound15.12301.12301.3 | controllerType=0 controllerNumber=1 scan=12301 | 30.2724 |
| 08172:3 | 2 | R.LAGKPTHVN[+1606.587]VSVVMAEVDGTC[+57.021]Y.- | HexNAc(4)Hex(4)Fuc(1) | C[+57], N[+1607] | 1318.5856 | 3 | 3953.7423 | 3953.7288 | 0 | 3.4 | 332 | CRagged | 232.19 | 157.55 | 157.55 | 3.13 | 58 | >sp\|P01876\|IGHA1_HUMAN Immunoglobulin heavy constant alpha 1 OS=Homo sapiens OX=9606 GN=IGHA1 PE=1 SV=2 | 255 | 20201116_fqg_colostrum_1ng_glyco_hcd30_etd30_compound15.11398.11398.3 | controllerType=0 controllerNumber=1 scan=11398 | 28.3562 |
| 08146:3 | 2 | R.LAGKPTHVN[+1378.476]VSVVMAEVDGTC[+57.021]Y.- | HexNAc(2)Hex(6) | C[+57], N[+1378] | 1242.5466 | 3 | 3725.6251 | 3725.6178 | 0 | 2 | 332 | CRagged | 241.47 | 194.91 | 194.91 | 3.94 | 58 | >sp\|P01876\|IGHA1_HUMAN Immunoglobulin heavy constant alpha 1 OS=Homo sapiens OX=9606 GN=IGHA1 PE=1 SV=2 | 255 | 20201116_fqg_colostrum_1ng_glyco_hcd30_etd30_compound15.11371.11371.3 | controllerType=0 controllerNumber=1 scan=11371 | 28.2888 |
| 09157:3 | 2 | R.LAGKPTHVN[+1647.613]VSVVMAEVDGTC[+57.021]Y.- | HexNAc(5)Hex(3)Fuc(1) | C[+57], N[+1648] | 1332.2586 | 3 | 3994.7613 | 3994.7553 | 0 | 1.5 | 332 | CRagged | 243.16 | 94.16 | 94.16 | 3.22 | 58 | >sp\|P01876\|IGHA1_HUMAN Immunoglobulin heavy constant alpha 1 OS=Homo sapiens OX=9606 GN=IGHA1 PE=1 SV=2 | 255 | 20201116_fqg_colostrum_1ng_glyco_hcd30_etd30_compound15.12423.12423.3 | controllerType=0 controllerNumber=1 scan=12423 | 30.5194 |
| 09025:3 | 2 | R.LAGKPTHVN[+1702.581]VSVVMAEVDGTC[+57.021]Y.- | HexNAc(2)Hex(8) | C[+57], N[+1703] | 1350.5829 | 3 | 4049.7343 | 4049.7234 | 0 | 2.7 | 332 | CRagged | 243.56 | 187.9 | 187.9 | 4.27 | 58 | >sp\|P01876\|IGHA1_HUMAN Immunoglobulin heavy constant alpha 1 OS=Homo sapiens OX=9606 GN=IGHA1 PE=1 SV=2 | 255 | 20201116_fqg_colostrum_1ng_glyco_hcd30_etd30_compound15.12285.12285.3 | controllerType=0 controllerNumber=1 scan=12285 | 30.2156 |
| 10788:3 | 2 | R.LAGKPTHVN[+2262.814]VSVVMAEVDGTC[+57.021]Y.- | HexNAc(5)Hex(5)Fuc(1)NeuAc(1) | C[+57], N[+2263] | 1537.3274 | 3 | 4609.9678 | 4609.9564 | 0 | 2.5 | 332 | CRagged | 247.55 | 117.51 | 117.51 | 4.03 | 58 | >sp\|P01876\|IGHA1_HUMAN Immunoglobulin heavy constant alpha 1 OS=Homo sapiens OX=9606 GN=IGHA1 PE=1 SV=2 | 255 | 20201116_fqg_colostrum_1ng_glyco_hcd30_etd30_compound15.14113.14113.3 | controllerType=0 controllerNumber=1 scan=14113 | 33.6022 |
| 09211:3 | 2 | R.LAGKPTHVN[+1444.534]VSVVMAEVDGTC[+57.021]Y.- | HexNAc(4)Hex(3)Fuc(1) | C[+57], N[+1445] | 1264.5672 | 3 | 3791.687 | 3791.6759 | 0 | 2.9 | 332 | CRagged | 248.18 | 144.77 | 144.77 | 4.03 | 58 | >sp\|P01876\|IGHA1_HUMAN Immunoglobulin heavy constant alpha 1 OS=Homo sapiens OX=9606 GN=IGHA1 PE=1 SV=2 | 255 | 20201116_fqg_colostrum_1ng_glyco_hcd30_etd30_compound15.12479.12479.3 | controllerType=0 controllerNumber=1 scan=12479 | 30.6344 |
| 10782:3 | 2 | R.LAGKPTHVN[+1913.677]VSVVMAEVDGTC[+57.021]Y.- | HexNAc(4)Hex(5)NeuAc(1) | C[+57], N[+1914] | 1420.9479 | 3 | 4260.8292 | 4260.8191 | 0 | 2.4 | 332 | CRagged | 249.67 | 180.61 | 180.61 | 3.62 | 58 | >sp\|P01876\|IGHA1_HUMAN Immunoglobulin heavy constant alpha 1 OS=Homo sapiens OX=9606 GN=IGHA1 PE=1 SV=2 | 255 | 20201116_fqg_colostrum_1ng_glyco_hcd30_etd30_compound15.14107.14107.3 | controllerType=0 controllerNumber=1 scan=14107 | 33.5915 |
| 08156:3 | 2 | R.LAGKPTHVN[+1216.423]VSVVMAEVDGTC[+57.021]Y.- | HexNAc(2)Hex(5) | C[+57], N[+1216] | 1188.5313 | 3 | 3563.5794 | 3563.5649 | 0 | 4 | 332 | CRagged | 271.03 | 248.43 | 183.58 | 4.52 | 58 | >sp\|P01876\|IGHA1_HUMAN Immunoglobulin heavy constant alpha 1 OS=Homo sapiens OX=9606 GN=IGHA1 PE=1 SV=2 | 255 | 20201116_fqg_colostrum_1ng_glyco_hcd30_etd30_compound15.11382.11382.3 | controllerType=0 controllerNumber=1 scan=11382 | 28.331 |
| 09142:3 | 2 | R.LAGKPTHVN[+1054.370]VSVVMAEVDGTC[+57.021]Y.- | HexNAc(2)Hex(4) | C[+57], N[+1054] | 1134.5114 | 3 | 3401.5196 | 3401.5121 | 0 | 2.2 | 332 | CRagged | 289.02 | 252.72 | 196.07 | 4.14 | 58 | >sp\|P01876\|IGHA1_HUMAN Immunoglobulin heavy constant alpha 1 OS=Homo sapiens OX=9606 GN=IGHA1 PE=1 SV=2 | 255 | 20201116_fqg_colostrum_1ng_glyco_hcd30_etd30_compound15.12407.12407.3 | controllerType=0 controllerNumber=1 scan=12407 | 30.4766 |
| 09093:2 | 2 | R.LAGKPTHVN[+1216.423]VSVVMAEVDGTC[+57.021]Y.- | HexNAc(2)Hex(5) | C[+57], N[+1216] | 1782.2887 | 2 | 3563.57 | 3563.5649 | 0 | 1.4 | 332 | CRagged | 337.46 | 337.46 | 136.22 | 5.71 | 58 | >sp\|P01876\|IGHA1_HUMAN Immunoglobulin heavy constant alpha 1 OS=Homo sapiens OX=9606 GN=IGHA1 PE=1 SV=2 | 255 | 20201116_fqg_colostrum_1ng_glyco_hcd30_etd30_compound15.12356.12356.2 | controllerType=0 controllerNumber=1 scan=12356 | 30.3832 |
| 18178:5 | 3 | K.VFPLSLDSTPQDGNVVVAC[+57.021]LVQGFFPQEPLSVTWSESGQN[+1444.534]VTAI | HexNAc(4)Hex(3)Fuc(1) | C[+57], N[+1445] | 1244.9893 | 5 | 6220.9172 | 6220.8953 | 0 | 3.5 | 8 | Specific | 202.25 | 202.25 | 202.25 | 3.68 | 65 | >sp\|P01877\|IGHA2_HUMAN Immunoglobulin heavy constant alpha 2 OS=Homo sapiens OX=9606 GN=IGHA2 PE=1 SV=4 | 576 | 20201116_fqg_colostrum_1ng_glyco_hcd30_etd30_compound15.21855.21855.5 | controllerType=0 controllerNumber=1 scan=21855 | 52.133 |
| 18547:5 | 3 | K.VFPLSLDSTPQDGNVVVAC[+57.021]LVQGFFPQEPLSVTWSESGQN[+1444.534]VTAI | HexNAc(4)Hex(3)Fuc(1) | C[+57], N[+1445] | 1244.9893 | 5 | 6220.9172 | 6220.8953 | 0 | 3.5 | 8 | Specific | 233.26 | 233.26 | 233.26 | 3.92 | 65 | >sp\|P01877\|IGHA2_HUMAN Immunoglobulin heavy constant alpha 2 OS=Homo sapiens OX=9606 GN=IGHA2 PE=1 SV=4 | 576 | 20201116_fqg_colostrum_1ng_glyco_hcd30_etd30_compound15.22246.22246.5 | controllerType=0 controllerNumber=1 scan=22246 | 53.3709 |
| 19423:4 | 3 | K.VFPLSLDSTPQDGNVVVAC[+57.021]LVQGFFPQEPLSVTWSESGQN[+2262.814]VTAI | HexNAc(5)Hex(5)Fuc(1)NeuAc(1) | C[+57], N[+2263] | 1760.5539 | 4 | 7039.1938 | 7039.1757 | 0 | 2.6 | 8 | Specific | 233.85 | 233.85 | 233.85 | 5.52 | 65 | >sp\|P01877\|IGHA2_HUMAN Immunoglobulin heavy constant alpha 2 OS=Homo sapiens OX=9606 GN=IGHA2 PE=1 SV=4 | 576 | 20201116_fqg_colostrum_1ng_glyco_hcd30_etd30_compound15.23177.23177.4 | controllerType=0 controllerNumber=1 scan=23177 | 56.5027 |
| 18062:4 | 3 | K.VFPLSLDSTPQDGNVVVAC[+57.021]LVQGFFPQEPLSVTWSESGQN[+1809.666]VTAI | HexNAc(5)Hex(4)Fuc(1) | C[+57], N[+1810] | 1647.2683 | 4 | 6586.0514 | 6586.0275 | 0 | 3.6 | 8 | Specific | 260.13 | 260.13 | 260.13 | 6.45 | 65 | >sp\|P01877\|IGHA2_HUMAN Immunoglobulin heavy constant alpha 2 OS=Homo sapiens OX=9606 GN=IGHA2 PE=1 SV=4 | 576 | 20201116_fqg_colostrum_1ng_glyco_hcd30_etd30_compound15.21733.21733.4 | controllerType=0 controllerNumber=1 scan=21733 | 51.7821 |
| 19182:4 | 3 | K.VFPLSLDSTPQDGNVVVAC[+57.021]LVQGFFPQEPLSVTWSESGQN[+2059.735]VTAI | HexNAc(4)Hex(5)Fuc(1)NeuAc(1) | C[+57], N[+2060] | 1709.787 | 4 | 6836.126 | 6836.0963 | 0 | 4.3 | 8 | Specific | 265.19 | 265.19 | 265.19 | 5.72 | 65 | >sp\|P01877\|IGHA2_HUMAN Immunoglobulin heavy constant alpha 2 OS=Homo sapiens OX=9606 GN=IGHA2 PE=1 SV=4 | 576 | 20201116_fqg_colostrum_1ng_glyco_hcd30_etd30_compound15.22920.22920.4 | controllerType=0 controllerNumber=1 scan=22920 | 55.5827 |
| 19033:5 | 3 | K.VFPLSLDSTPQDGNVVVAC[+57.021]LVQGFFPQEPLSVTWSESGQN[+2262.814]VTAI | HexNAc(5)Hex(5)Fuc(1)NeuAc(1) | C[+57], N[+2263] | 1408.6481 | 5 | 7039.2115 | 7039.1757 | 0 | 5.1 | 8 | Specific | 301.83 | 301.83 | 301.83 | 4.68 | 65 | >sp\|P01877\|IGHA2_HUMAN Immunoglobulin heavy constant alpha 2 OS=Homo sapiens OX=9606 GN=IGHA2 PE=1 SV=4 | 576 | 20201116_fqg_colostrum_1ng_glyco_hcd30_etd30_compound15.22762.22762.5 | controllerType=0 controllerNumber=1 scan=22762 | 55.0465 |
| 18096:4 | 3 | K.VFPLSLDSTPQDGNVVVAC[+57.021]LVQGFFPQEPLSVTWSESGQN[+1971.719]VTAI | HexNAc(5)Hex(5)Fuc(1) | C[+57], N[+1972] | 1687.7851 | 4 | 6748.1186 | 6748.0803 | 0 | 5.7 | 8 | Specific | 313.48 | 313.48 | 313.48 | 5.87 | 65 | >sp\|P01877\|IGHA2_HUMAN Immunoglobulin heavy constant alpha 2 OS=Homo sapiens OX=9606 GN=IGHA2 PE=1 SV=4 | 576 | 20201116_fqg_colostrum_1ng_glyco_hcd30_etd30_compound15.21769.21769.4 | controllerType=0 controllerNumber=1 scan=21769 | 51.8901 |
| 19525:4 | 3 | K.VFPLSLDSTPQDGNVVVAC[+57.021]LVQGFFPQEPLSVTWSESGQN[+2059.735]VTAI | HexNAc(4)Hex(5)Fuc(1)NeuAc(1) | C[+57], N[+2060] | 1709.7891 | 4 | 6836.1348 | 6836.0963 | 0 | 5.6 | 8 | Specific | 339.77 | 339.77 | 339.77 | 6.13 | 65 | >sp\|P01877\|IGHA2_HUMAN Immunoglobulin heavy constant alpha 2 OS=Homo sapiens OX=9606 GN=IGHA2 PE=1 SV=4 | 576 | 20201116_fqg_colostrum_1ng_glyco_hcd30_etd30_compound15.23286.23286.4 | controllerType=0 controllerNumber=1 scan=23286 | 56.921 |
| 18118:4 | 3 | K.VFPLSLDSTPQDGNVVVAC[+57.021]LVQGFFPQEPLSVTWSESGQN[+1606.587]VTAI | HexNAc(4)Hex(4)Fuc(1) | C[+57], N[+1607] | 1596.5016 | 4 | 6382.9845 | 6382.9481 | 0 | 5.7 | 8 | Specific | 368.32 | 368.32 | 368.32 | 7.56 | 65 | >sp\|P01877\|IGHA2_HUMAN Immunoglobulin heavy constant alpha 2 OS=Homo sapiens OX=9606 GN=IGHA2 PE=1 SV=4 | 576 | 20201116_fqg_colostrum_1ng_glyco_hcd30_etd30_compound15.21792.21792.4 | controllerType=0 controllerNumber=1 scan=21792 | 51.9653 |
| 20299:6 | 3 | K.VFPLSLDSTPQDGNVVVAC[+57.021]LVQGFFPQEPLSVTWSESGQN[+2351.851]VTAI | HexNAc(4)Hex(5)Fuc(3)NeuAc(1) | C[+57], N[+2352] | 1188.8741 | 6 | 7128.2081 | 7128.2121 | 0 | -0.6 | 8 | Specific | 375.76 | 375.76 | 375.76 | 7.22 | 65 | >sp\|P01877\|IGHA2_HUMAN Immunoglobulin heavy constant alpha 2 OS=Homo sapiens OX=9606 GN=IGHA2 PE=1 SV=4 | 576 | 20201116_fqg_colostrum_1ng_glyco_hcd30_etd30_compound15.24118.24118.6 | controllerType=0 controllerNumber=1 scan=24118 | 60.0013 |
| 19537:4 | 3 | K.VFPLSLDSTPQDGNVVVAC[+57.021]LVQGFFPQEPLSVTWSESGQN[+1694.603]VTAI | HexNAc(3)Hex(4)Fuc(1)NeuAc(1) | C[+57], N[+1695] | 1618.5022 | 4 | 6470.987 | 6470.9641 | 0 | 3.5 | 8 | Specific | 376.72 | 376.72 | 376.72 | 8.29 | 65 | >sp\|P01877\|IGHA2_HUMAN Immunoglobulin heavy constant alpha 2 OS=Homo sapiens OX=9606 GN=IGHA2 PE=1 SV=4 | 576 | 20201116_fqg_colostrum_1ng_glyco_hcd30_etd30_compound15.23299.23299.4 | controllerType=0 controllerNumber=1 scan=23299 | 56.9881 |
| 18630:4 | 3 | K.VFPLSLDSTPQDGNVVVAC[+57.021]LVQGFFPQEPLSVTWSESGQN[+1038.375]VTAI | HexNAc(2)Hex(3)Fuc(1) | C[+57], N[+1038] | 1454.4441 | 4 | 5814.7546 | 5814.7365 | 0 | 3.1 | 8 | Specific | 386.49 | 386.49 | 386.49 | 6.84 | 65 | >sp\|P01877\|IGHA2_HUMAN Immunoglobulin heavy constant alpha 2 OS=Homo sapiens OX=9606 GN=IGHA2 PE=1 SV=4 | 576 | 20201116_fqg_colostrum_1ng_glyco_hcd30_etd30_compound15.22333.22333.4 | controllerType=0 controllerNumber=1 scan=22333 | 53.6574 |
| 18035:4 | 3 | K.VFPLSLDSTPQDGNVVVAC[+57.021]LVQGFFPQEPLSVTWSESGQN[+1809.666]VTAI | HexNAc(5)Hex(4)Fuc(1) | C[+57], N[+1810] | 1647.2671 | 4 | 6586.0465 | 6586.0275 | 0 | 2.9 | 8 | Specific | 387.26 | 387.26 | 387.26 | 8.29 | 65 | >sp\|P01877\|IGHA2_HUMAN Immunoglobulin heavy constant alpha 2 OS=Homo sapiens OX=9606 GN=IGHA2 PE=1 SV=4 | 576 | 20201116_fqg_colostrum_1ng_glyco_hcd30_etd30_compound15.21704.21704.4 | controllerType=0 controllerNumber=1 scan=21704 | 51.6876 |
| 19037:4 | 3 | K.VFPLSLDSTPQDGNVVVAC[+57.021]LVQGFFPQEPLSVTWSESGQN[+2262.814]VTAI | HexNAc(5)Hex(5)Fuc(1)NeuAc(1) | C[+57], N[+2263] | 1760.5577 | 4 | 7039.209 | 7039.1757 | 0 | 4.7 | 8 | Specific | 392.97 | 392.97 | 392.97 | 7.56 | 65 | >sp\|P01877\|IGHA2_HUMAN Immunoglobulin heavy constant alpha 2 OS=Homo sapiens OX=9606 GN=IGHA2 PE=1 SV=4 | 576 | 20201116_fqg_colostrum_1ng_glyco_hcd30_etd30_compound15.22766.22766.4 | controllerType=0 controllerNumber=1 scan=22766 | 55.0561 |
| 18100:4 | 3 | K.VFPLSLDSTPQDGNVVVAC[+57.021]LVQGFFPQEPLSVTWSESGQN[+1647.613]VTAI | HexNAc(5)Hex(3)Fuc(1) | C[+57], N[+1648] | 1606.7557 | 4 | 6424.0011 | 6423.9746 | 0 | 4.1 | 8 | Specific | 405.3 | 405.3 | 405.3 | 8.43 | 65 | >sp\|P01877\|IGHA2_HUMAN Immunoglobulin heavy constant alpha 2 OS=Homo sapiens OX=9606 GN=IGHA2 PE=1 SV=4 | 576 | 20201116_fqg_colostrum_1ng_glyco_hcd30_etd30_compound15.21773.21773.4 | controllerType=0 controllerNumber=1 scan=21773 | 51.9072 |
| 18480:4 | 3 | K.VFPLSLDSTPQDGNVVVAC[+57.021]LVQGFFPQEPLSVTWSESGQN[+1444.534]VTAI | HexNAc(4)Hex(3)Fuc(1) | C[+57], N[+1445] | 1555.9882 | 4 | 6220.9308 | 6220.8953 | 0 | 5.7 | 8 | Specific | 468.83 | 468.83 | 468.83 | 7.76 | 65 | >sp\|P01877\|IGHA2_HUMAN Immunoglobulin heavy constant alpha 2 OS=Homo sapiens OX=9606 GN=IGHA2 PE=1 SV=4 | 576 | 20201116_fqg_colostrum_1ng_glyco_hcd30_etd30_compound15.22176.22176.4 | controllerType=0 controllerNumber=1 scan=22176 | 53.1895 |
| 18138:4 | 3 | K.VFPLSLDSTPQDGNVVVAC[+57.021]LVQGFFPQEPLSVTWSESGQN[+1444.534]VTAI | HexNAc(4)Hex(3)Fuc(1) | C[+57], N[+1445] | 1555.9805 | 4 | 6220.9001 | 6220.8953 | 0 | 0.8 | 8 | Specific | 602.57 | 602.57 | 602.57 | 10.41 | 65 | >sp\|P01877\|IGHA2_HUMAN Immunoglobulin heavy constant alpha 2 OS=Homo sapiens OX=9606 GN=IGHA2 PE=1 SV=4 | 576 | 20201116_fqg_colostrum_1ng_glyco_hcd30_etd30_compound15.21813.21813.4 | controllerType=0 controllerNumber=1 scan=21813 | 52.0256 |
| 00756:2 | 3 | K.HYTN[+1809.666]SSQDVTVPC[+57.021]R.V | HexNAc(5)Hex(4)Fuc(1) | C[+57], N[+1810] | 1737.2171 | 2 | 3473.427 | 3473.4154 | 0 | 3.3 | 89 | Specific | 200.14 | 125.26 | 125.26 | 3.73 | 65 | >sp\|P01877\|IGHA2_HUMAN Immunoglobulin heavy constant alpha 2 OS=Homo sapiens OX=9606 GN=IGHA2 PE=1 SV=4 | 576 | 20201116_fqg_colostrum_1ng_glyco_hcd30_etd30_compound15.3673.3673.2 | controllerType=0 controllerNumber=1 scan=3673 | 12.9303 |
| 00729:2 | 3 | K.HYTN[+1444.534]SSQDVTVPC[+57.021]R.V | HexNAc(4)Hex(3)Fuc(1) | C[+57], N[+1445] | 1554.6497 | 2 | 3108.2921 | 3108.2832 | 0 | 2.9 | 89 | Specific | 263.08 | 194.14 | 194.14 | 3.75 | 65 | >sp\|P01877\|IGHA2_HUMAN Immunoglobulin heavy constant alpha 2 OS=Homo sapiens OX=9606 GN=IGHA2 PE=1 SV=4 | 576 | 20201116_fqg_colostrum_1ng_glyco_hcd30_etd30_compound15.3645.3645.2 | controllerType=0 controllerNumber=1 scan=3645 | 12.8585 |
| 00718:2 | 3 | K.HYTN[+1606.587]SSQDVTVPC[+57.021]R.V | HexNAc(4)Hex(4)Fuc(1) | C[+57], N[+1607] | 1635.6752 | 2 | 3270.3432 | 3270.336 | 0 | 2.2 | 89 | Specific | 275.14 | 227.63 | 227.63 | 3.66 | 65 | >sp\|P01877\|IGHA2_HUMAN Immunoglobulin heavy constant alpha 2 OS=Homo sapiens OX=9606 GN=IGHA2 PE=1 SV=4 | 576 | 20201116_fqg_colostrum_1ng_glyco_hcd30_etd30_compound15.3633.3633.2 | controllerType=0 controllerNumber=1 scan=3633 | 12.8332 |
| 00767:2 | 3 | K.HYTN[+1647.613]SSQDVTVPC[+57.021]R.V | HexNAc(5)Hex(3)Fuc(1) | C[+57], N[+1648] | 1656.1921 | 2 | 3311.3768 | 3311.3625 | 0 | 4.3 | 89 | Specific | 291.51 | 221.21 | 221.21 | 3.11 | 65 | >sp\|P01877\|IGHA2_HUMAN Immunoglobulin heavy constant alpha 2 OS=Homo sapiens OX=9606 GN=IGHA2 PE=1 SV=4 | 576 | 20201116_fqg_colostrum_1ng_glyco_hcd30_etd30_compound15.3685.3685.2 | controllerType=0 controllerNumber=1 scan=3685 | 12.9659 |
| 00794:2 | 3 | K.HYTN[+876.322]SSQDVTVPC[+57.021]R.V | HexNAc(2)Hex(2)Fuc(1) | C[+57], N[+876] | 1270.5441 | 2 | 2540.0809 | 2540.0716 | 0 | 3.7 | 89 | Specific | 340.04 | 276.57 | 276.57 | 3.72 | 65 | >sp\|P01877\|IGHA2_HUMAN Immunoglobulin heavy constant alpha 2 OS=Homo sapiens OX=9606 GN=IGHA2 PE=1 SV=4 | 576 | 20201116_fqg_colostrum_1ng_glyco_hcd30_etd30_compound15.3713.3713.2 | controllerType=0 controllerNumber=1 scan=3713 | 13.0574 |
| 00702:3 | 3 | K.HYTN[+1971.719]SSQDVTVPC[+57.021]R.V | HexNAc(5)Hex(5)Fuc(1) | C[+57], N[+1972] | 1212.4987 | 3 | 3635.4816 | 3635.4682 | 0 | 3.7 | 89 | Specific | 371.82 | 296.68 | 296.68 | 5.84 | 65 | >sp\|P01877\|IGHA2_HUMAN Immunoglobulin heavy constant alpha 2 OS=Homo sapiens OX=9606 GN=IGHA2 PE=1 SV=4 | 576 | 20201116_fqg_colostrum_1ng_glyco_hcd30_etd30_compound15.3616.3616.3 | controllerType=0 controllerNumber=1 scan=3616 | 12.7901 |
| 00670:3 | 3 | K.HYTN[+1768.640]SSQDVTVPC[+57.021]R.V | HexNAc(4)Hex(5)Fuc(1) | C[+57], N[+1769] | 1144.8048 | 3 | 3432.4 | 3432.3888 | 0 | 3.2 | 89 | Specific | 374.73 | 312.94 | 312.94 | 5.84 | 65 | >sp\|P01877\|IGHA2_HUMAN Immunoglobulin heavy constant alpha 2 OS=Homo sapiens OX=9606 GN=IGHA2 PE=1 SV=4 | 576 | 20201116_fqg_colostrum_1ng_glyco_hcd30_etd30_compound15.3582.3582.3 | controllerType=0 controllerNumber=1 scan=3582 | 12.667 |
| 00743:2 | 3 | K.HYTN[+1038.375]SSQDVTVPC[+57.021]R.V | HexNAc(2)Hex(3)Fuc(1) | C[+57], N[+1038] | 1351.5693 | 2 | 2702.1312 | 2702.1244 | 0 | 2.5 | 89 | Specific | 381.76 | 310.69 | 310.69 | 4.37 | 65 | >sp\|P01877\|IGHA2_HUMAN Immunoglobulin heavy constant alpha 2 OS=Homo sapiens OX=9606 GN=IGHA2 PE=1 SV=4 | 576 | 20201116_fqg_colostrum_1ng_glyco_hcd30_etd30_compound15.3659.3659.2 | controllerType=0 controllerNumber=1 scan=3659 | 12.894 |
| 00710:3 | 3 | K.HYTN[+1809.666]SSQDVTVPC[+57.021]R.V | HexNAc(5)Hex(4)Fuc(1) | C[+57], N[+1810] | 1158.4787 | 3 | 3473.4215 | 3473.4154 | 0 | 1.8 | 89 | Specific | 384.5 | 308.18 | 308.18 | 5.5 | 65 | >sp\|P01877\|IGHA2_HUMAN Immunoglobulin heavy constant alpha 2 OS=Homo sapiens OX=9606 GN=IGHA2 PE=1 SV=4 | 576 | 20201116_fqg_colostrum_1ng_glyco_hcd30_etd30_compound15.3625.3625.3 | controllerType=0 controllerNumber=1 scan=3625 | 12.8101 |
| 01386:3 | 3 | K.HYTN[+2262.814]SSQDVTVPC[+57.021]R.V | HexNAc(5)Hex(5)Fuc(1)NeuAc(1) | C[+57], N[+2263] | 1309.5297 | 3 | 3926.5746 | 3926.5636 | 0 | 2.8 | 89 | Specific | 397.35 | 279.29 | 279.29 | 5.65 | 65 | >sp\|P01877\|IGHA2_HUMAN Immunoglobulin heavy constant alpha 2 OS=Homo sapiens OX=9606 GN=IGHA2 PE=1 SV=4 | 576 | 20201116_fqg_colostrum_1ng_glyco_hcd30_etd30_compound15.4332.4332.3 | controllerType=0 controllerNumber=1 scan=4332 | 14.4942 |
| 01329:3 | 3 | K.HYTN[+2059.735]SSQDVTVPC[+57.021]R.V | HexNAc(4)Hex(5)Fuc(1)NeuAc(1) | C[+57], N[+2060] | 1241.8383 | 3 | 3723.5004 | 3723.4842 | 0 | 4.3 | 89 | Specific | 457.21 | 335.65 | 335.65 | 5.52 | 65 | >sp\|P01877\|IGHA2_HUMAN Immunoglobulin heavy constant alpha 2 OS=Homo sapiens OX=9606 GN=IGHA2 PE=1 SV=4 | 576 | 20201116_fqg_colostrum_1ng_glyco_hcd30_etd30_compound15.4272.4272.3 | controllerType=0 controllerNumber=1 scan=4272 | 14.3348 |
| 00730:3 | 3 | K.HYTN[+1647.613]SSQDVTVPC[+57.021]R.V | HexNAc(5)Hex(3)Fuc(1) | C[+57], N[+1648] | 1104.4619 | 3 | 3311.3713 | 3311.3625 | 0 | 2.6 | 89 | Specific | 465.69 | 394.64 | 394.64 | 7.47 | 65 | >sp\|P01877\|IGHA2_HUMAN Immunoglobulin heavy constant alpha 2 OS=Homo sapiens OX=9606 GN=IGHA2 PE=1 SV=4 | 576 | 20201116_fqg_colostrum_1ng_glyco_hcd30_etd30_compound15.3646.3646.3 | controllerType=0 controllerNumber=1 scan=3646 | 12.8656 |
| 01114:3 | 3 | K.HYTN[+1444.534]SSQDVTVPC[+57.021]R.V | HexNAc(4)Hex(3)Fuc(1) | C[+57], N[+1445] | 1036.7681 | 3 | 3108.2897 | 3108.2832 | 0 | 2.1 | 89 | Specific | 506.78 | 425.71 | 425.71 | 7.18 | 65 | >sp\|P01877\|IGHA2_HUMAN Immunoglobulin heavy constant alpha 2 OS=Homo sapiens OX=9606 GN=IGHA2 PE=1 SV=4 | 576 | 20201116_fqg_colostrum_1ng_glyco_hcd30_etd30_compound15.4048.4048.3 | controllerType=0 controllerNumber=1 scan=4048 | 13.8717 |
| 00676:3 | 3 | K.HYTN[+1606.587]SSQDVTVPC[+57.021]R.V | HexNAc(4)Hex(4)Fuc(1) | C[+57], N[+1607] | 1090.7864 | 3 | 3270.3447 | 3270.336 | 0 | 2.7 | 89 | Specific | 530.44 | 457.87 | 457.87 | 7.74 | 65 | >sp\|P01877\|IGHA2_HUMAN Immunoglobulin heavy constant alpha 2 OS=Homo sapiens OX=9606 GN=IGHA2 PE=1 SV=4 | 576 | 20201116_fqg_colostrum_1ng_glyco_hcd30_etd30_compound15.3589.3589.3 | controllerType=0 controllerNumber=1 scan=3589 | 12.6879 |
| 01400:3 | 3 | K.HYTN[+1694.603]SSQDVTVPC[+57.021]R.V | HexNAc(3)Hex(4)Fuc(1)NeuAc(1) | C[+57], N[+1695] | 1120.1258 | 3 | 3358.3627 | 3358.352 | 0 | 3.2 | 89 | Specific | 532.43 | 413.41 | 413.41 | 7.74 | 65 | >sp\|P01877\|IGHA2_HUMAN Immunoglobulin heavy constant alpha 2 OS=Homo sapiens OX=9606 GN=IGHA2 PE=1 SV=4 | 576 | 20201116_fqg_colostrum_1ng_glyco_hcd30_etd30_compound15.4346.4346.3 | controllerType=0 controllerNumber=1 scan=4346 | 14.5324 |
| 00683:3 | 3 | K.HYTN[+1403.507]SSQDVTVPC[+57.021]R.V | HexNAc(3)Hex(4)Fuc(1) | C[+57], N[+1404] | 1023.0931 | 3 | 3067.2648 | 3067.2566 | 0 | 2.7 | 89 | Specific | 534.68 | 445.76 | 445.76 | 7.74 | 65 | >sp\|P01877\|IGHA2_HUMAN Immunoglobulin heavy constant alpha 2 OS=Homo sapiens OX=9606 GN=IGHA2 PE=1 SV=4 | 576 | 20201116_fqg_colostrum_1ng_glyco_hcd30_etd30_compound15.3596.3596.3 | controllerType=0 controllerNumber=1 scan=3596 | 12.7102 |
| 00692:3 | 3 | K.HYTN[+1444.534]SSQDVTVPC[+57.021]R.V | HexNAc(4)Hex(3)Fuc(1) | C[+57], N[+1445] | 1036.7698 | 3 | 3108.2949 | 3108.2832 | 0 | 3.8 | 89 | Specific | 552.15 | 481.66 | 481.66 | 7.74 | 65 | >sp\|P01877\|IGHA2_HUMAN Immunoglobulin heavy constant alpha 2 OS=Homo sapiens OX=9606 GN=IGHA2 PE=1 SV=4 | 576 | 20201116_fqg_colostrum_1ng_glyco_hcd30_etd30_compound15.3606.3606.3 | controllerType=0 controllerNumber=1 scan=3606 | 12.7465 |
| 00775:3 | 3 | K.HYTN[+876.322]SSQDVTVPC[+57.021]R.V | HexNAc(2)Hex(2)Fuc(1) | C[+57], N[+876] | 847.365 | 3 | 2540.0803 | 2540.0716 | 0 | 3.4 | 89 | Specific | 579.82 | 419.09 | 419.09 | 7.53 | 65 | >sp\|P01877\|IGHA2_HUMAN Immunoglobulin heavy constant alpha 2 OS=Homo sapiens OX=9606 GN=IGHA2 PE=1 SV=4 | 576 | 20201116_fqg_colostrum_1ng_glyco_hcd30_etd30_compound15.3693.3693.3 | controllerType=0 controllerNumber=1 scan=3693 | 13.0018 |
| 01373:4 | 3 | K.HYTN[+2262.814]SSQDVTVPC[+57.021]R.V | HexNAc(5)Hex(5)Fuc(1)NeuAc(1) | C[+57], N[+2263] | 982.3988 | 4 | 3926.5733 | 3926.5636 | 0 | 2.5 | 89 | Specific | 601.56 | 520.51 | 520.51 | 9.87 | 65 | >sp\|P01877\|IGHA2_HUMAN Immunoglobulin heavy constant alpha 2 OS=Homo sapiens OX=9606 GN=IGHA2 PE=1 SV=4 | 576 | 20201116_fqg_colostrum_1ng_glyco_hcd30_etd30_compound15.4318.4318.4 | controllerType=0 controllerNumber=1 scan=4318 | [illegible] |
| 00711:3 | 3 | K.HYTN[+1241.454]SSQDVTVPC[+57.021]R.V | HexNAc(3)Hex(3)Fuc(1) | C[+57], N[+1241] | 969.0751 | 3 | 2905.2108 | 2905.2038 | 0 | 2.4 | 89 | Specific | 613.97 | 479 | 479 | 9.37 | 65 | >sp\|P01877\|IGHA2_HUMAN Immunoglobulin heavy constant alpha 2 OS=Homo sapiens OX=9606 GN=IGHA2 PE=1 SV=4 | 576 | 20201116_fqg_colostrum_1ng_glyco_hcd30_etd30_compound15.3626.3626.3 | controllerType=0 controllerNumber=1 scan=3626 | [illegible] |
| 00717:3 | 3 | K.HYTN[+1038.375]SSQDVTVPC[+57.021]R.V | HexNAc(2)Hex(3)Fuc(1) | C[+57], N[+1038] | 901.381 | 3 | 2702.1285 | 2702.1244 | 0 | 1.5 | 89 | Specific | 625.54 | 507.82 | 507.82 | 9.37 | 65 | >sp\|P01877\|IGHA2_HUMAN Immunoglobulin heavy constant alpha 2 OS=Homo sapiens OX=9606 GN=IGHA2 PE=1 SV=4 | 576 | 20201116_fqg_colostrum_1ng_glyco_hcd30_etd30_compound15.3632.3632.3 | controllerType=0 controllerNumber=1 scan=3632 | 12.8299 |
| 02234:5 | 3 | K.HYTN[+1647.613]SSQDVTVPC[+57.021]RVPPPPPC[+57.021]C[+57.021]HPR.L | HexNAc(5)Hex(3)Fuc(1) | C[+57]*3, N[+1648] | 922.0018 | 5 | 4605.9799 | 4605.9689 | 0 | 2.4 | 89 | Specific | 253.29 | 234.3 | 234.3 | 3.75 | 65 | >sp\|P01877\|IGHA2_HUMAN Immunoglobulin heavy constant alpha 2 OS=Homo sapiens OX=9606 GN=IGHA2 PE=1 SV=4 | 576 | 20201116_fqg_colostrum_1ng_glyco_hcd30_etd30_compound15.5216.5216.5 | controllerType=0 controllerNumber=1 scan=5216 | 16.4439 |
| 02239:4 | 3 | K.HYTN[+1647.613]SSQDVTVPC[+57.021]RVPPPPPC[+57.021]C[+57.021]HPR.L | HexNAc(5)Hex(3)Fuc(1) | C[+57]*3, N[+1648] | 1152.2509 | 4 | 4605.9818 | 4605.9689 | 0 | 2.8 | 89 | Specific | 575.83 | 570.6 | 570.6 | 7.68 | 65 | >sp\|P01877\|IGHA2_HUMAN Immunoglobulin heavy constant alpha 2 OS=Homo sapiens OX=9606 GN=IGHA2 PE=1 SV=4 | 576 | 20201116_fqg_colostrum_1ng_glyco_hcd30_etd30_compound15.5221.5221.4 | controllerType=0 controllerNumber=1 scan=5221 | 16.4597 |
| 03516:3 | 3 | K.TPLTAN[+2262.814]ITK.S | HexNAc(5)Hex(5)Fuc(1)NeuAc(1) | N[+2263] | 1074.4649 | 3 | 3221.3801 | 3221.3711 | 0 | 2.8 | 200 | Specific | 225.92 | 106.43 | 106.43 | 2.77 | 65 | >sp\|P01877\|IGHA2_HUMAN Immunoglobulin heavy constant alpha 2 OS=Homo sapiens OX=9606 GN=IGHA2 PE=1 SV=4 | 576 | 20201116_fqg_colostrum_1ng_glyco_hcd30_etd30_compound15.6556.6556.3 | controllerType=0 controllerNumber=1 scan=6556 | 19.0432 |
| 01912:2 | 3 | K.TPLTAN[+1768.640]ITK.S | HexNAc(4)Hex(5)Fuc(1) | N[+1769] | 1364.1051 | 2 | 2727.203 | 2727.1963 | 0 | 2.5 | 200 | Specific | 233.7 | 96.49 | 96.49 | 2.7 | 65 | >sp\|P01877\|IGHA2_HUMAN Immunoglobulin heavy constant alpha 2 OS=Homo sapiens OX=9606 GN=IGHA2 PE=1 SV=4 | 576 | 20201116_fqg_colostrum_1ng_glyco_hcd30_etd30_compound15.4878.4878.2 | controllerType=0 controllerNumber=1 scan=4878 | 15.5771 |
| 03510:2 | 3 | K.TPLTAN[+2059.735]ITK.S | HexNAc(4)Hex(5)Fuc(1)NeuAc(1) | N[+2060] | 1509.6541 | 2 | 3018.3009 | 3018.2917 | 0 | 3.1 | 200 | Specific | 253.75 | 80.45 | 80.45 | 3.15 | 65 | >sp\|P01877\|IGHA2_HUMAN Immunoglobulin heavy constant alpha 2 OS=Homo sapiens OX=9606 GN=IGHA2 PE=1 SV=4 | 576 | 20201116_fqg_colostrum_1ng_glyco_hcd30_etd30_compound15.6550.6550.2 | controllerType=0 controllerNumber=1 scan=6550 | 19.0246 |
| 03540:2 | 3 | K.TPLTAN[+2262.814]ITK.S | HexNAc(5)Hex(5)Fuc(1)NeuAc(1) | N[+2263] | 1611.195 | 2 | 3221.3827 | 3221.3711 | 0 | 3.6 | 200 | Specific | 262.65 | 85.05 | 85.05 | 4.2 | 65 | >sp\|P01877\|IGHA2_HUMAN Immunoglobulin heavy constant alpha 2 OS=Homo sapiens OX=9606 GN=IGHA2 PE=1 SV=4 | 576 | 20201116_fqg_colostrum_1ng_glyco_hcd30_etd30_compound15.6581.6581.2 | controllerType=0 controllerNumber=1 scan=6581 | 19.1104 |
| 01978:3 | 3 | K.TPLTAN[+1971.719]ITK.S | HexNAc(5)Hex(5)Fuc(1) | N[+1972] | 977.4342 | 3 | 2930.2881 | 2930.2757 | 0 | 4.3 | 200 | Specific | 275.64 | 124.27 | 124.27 | 3.24 | 65 | >sp\|P01877\|IGHA2_HUMAN Immunoglobulin heavy constant alpha 2 OS=Homo sapiens OX=9606 GN=IGHA2 PE=1 SV=4 | 576 | 20201116_fqg_colostrum_1ng_glyco_hcd30_etd30_compound15.4947.4947.3 | controllerType=0 controllerNumber=1 scan=4947 | 15.7316 |
| 03502:3 | 3 | K.TPLTAN[+2059.735]ITK.S | HexNAc(4)Hex(5)Fuc(1)NeuAc(1) | N[+2060] | 1006.7721 | 3 | 3018.3018 | 3018.2917 | 0 | 3.3 | 200 | Specific | 281.14 | 122.87 | 122.87 | 3.57 | 65 | >sp\|P01877\|IGHA2_HUMAN Immunoglobulin heavy constant alpha 2 OS=Homo sapiens OX=9606 GN=IGHA2 PE=1 SV=4 | 576 | 20201116_fqg_colostrum_1ng_glyco_hcd30_etd30_compound15.6541.6541.3 | controllerType=0 controllerNumber=1 scan=6541 | 18.9986 |
| 02034:2 | 3 | K.TPLTAN[+1663.608]ITK.S | HexNAc(5)Hex(4) | N[+1664] | 1311.5898 | 2 | 2622.1723 | 2622.1649 | 0 | 2.8 | 200 | Specific | 281.54 | 98.96 | 98.96 | 2.89 | 65 | >sp\|P01877\|IGHA2_HUMAN Immunoglobulin heavy constant alpha 2 OS=Homo sapiens OX=9606 GN=IGHA2 PE=1 SV=4 | 576 | 20201116_fqg_colostrum_1ng_glyco_hcd30_etd30_compound15.5006.5006.2 | controllerType=0 controllerNumber=1 scan=5006 | 15.9095 |
| 02070:3 | 3 | K.TPLTAN[+1647.613]ITK.S | HexNAc(5)Hex(3)Fuc(1) | N[+1648] | 869.3963 | 3 | 2606.1744 | 2606.17 | 0 | 1.7 | 200 | Specific | 284.71 | 123.23 | 123.23 | 2.69 | 65 | >sp\|P01877\|IGHA2_HUMAN Immunoglobulin heavy constant alpha 2 OS=Homo sapiens OX=9606 GN=IGHA2 PE=1 SV=4 | 576 | 20201116_fqg_colostrum_1ng_glyco_hcd30_etd30_compound15.5044.5044.3 | controllerType=0 controllerNumber=1 scan=5044 | 16.0317 |
| 02060:2 | 3 | K.TPLTAN[+1647.613]ITK.S | HexNAc(5)Hex(3)Fuc(1) | N[+1648] | 1303.5907 | 2 | 2606.1742 | 2606.17 | 0 | 1.6 | 200 | Specific | 286.55 | 105.66 | 105.66 | 2.77 | 65 | >sp\|P01877\|IGHA2_HUMAN Immunoglobulin heavy constant alpha 2 OS=Homo sapiens OX=9606 GN=IGHA2 PE=1 SV=4 | 576 | 20201116_fqg_colostrum_1ng_glyco_hcd30_etd30_compound15.5034.5034.2 | controllerType=0 controllerNumber=1 scan=5034 | 16.0083 |
| 01991:2 | 3 | K.TPLTAN[+1971.719]ITK.S | HexNAc(5)Hex(5)Fuc(1) | N[+1972] | 1465.6466 | 2 | 2930.286 | 2930.2757 | 0 | 3.5 | 200 | Specific | 297.93 | 95.78 | 95.78 | 2.73 | 65 | >sp\|P01877\|IGHA2_HUMAN Immunoglobulin heavy constant alpha 2 OS=Homo sapiens OX=9606 GN=IGHA2 PE=1 SV=4 | 576 | 20201116_fqg_colostrum_1ng_glyco_hcd30_etd30_compound15.4961.4961.2 | controllerType=0 controllerNumber=1 scan=4961 | 15.7711 |
| 02024:3 | 3 | K.TPLTAN[+1809.666]ITK.S | HexNAc(5)Hex(4)Fuc(1) | N[+1810] | 923.4151 | 3 | 2768.2308 | 2768.2228 | 0 | 2.9 | 200 | Specific | 302.58 | 147.51 | 147.51 | 2.89 | 65 | >sp\|P01877\|IGHA2_HUMAN Immunoglobulin heavy constant alpha 2 OS=Homo sapiens OX=9606 GN=IGHA2 PE=1 SV=4 | 576 | 20201116_fqg_colostrum_1ng_glyco_hcd30_etd30_compound15.4996.4996.3 | controllerType=0 controllerNumber=1 scan=4996 | 15.8876 |
| 02023:2 | 3 | K.TPLTAN[+1809.666]ITK.S | HexNAc(5)Hex(4)Fuc(1) | N[+1810] | 1384.6194 | 2 | 2768.2316 | 2768.2228 | 0 | 3.2 | 200 | Specific | 303.67 | 107.08 | 107.08 | 2.89 | 65 | >sp\|P01877\|IGHA2_HUMAN Immunoglobulin heavy constant alpha 2 OS=Homo sapiens OX=9606 GN=IGHA2 PE=1 SV=4 | 576 | 20201116_fqg_colostrum_1ng_glyco_hcd30_etd30_compound15.4995.4995.2 | controllerType=0 controllerNumber=1 scan=4995 | 15.8847 |
| 01974:2 | 3 | K.TPLTAN[+1606.587]ITK.S | HexNAc(4)Hex(4)Fuc(1) | N[+1607] | 1283.0807 | 2 | 2565.1542 | 2565.1435 | 0 | 4.2 | 200 | Specific | 303.69 | 113.53 | 113.53 | 2.97 | 65 | >sp\|P01877\|IGHA2_HUMAN Immunoglobulin heavy constant alpha 2 OS=Homo sapiens OX=9606 GN=IGHA2 PE=1 SV=4 | 576 | 20201116_fqg_colostrum_1ng_glyco_hcd30_etd30_compound15.4943.4943.2 | controllerType=0 controllerNumber=1 scan=4943 | 15.7203 |
| 01951:2 | 3 | K.TPLTAN[+1216.423]ITK.S | HexNAc(2)Hex(5) | N[+1216] | 1087.9967 | 2 | 2174.9862 | 2174.9796 | 0 | 3 | 200 | Specific | 305.1 | 109.89 | 109.89 | 2.89 | 65 | >sp\|P01877\|IGHA2_HUMAN Immunoglobulin heavy constant alpha 2 OS=Homo sapiens OX=9606 GN=IGHA2 PE=1 SV=4 | 576 | 20201116_fqg_colostrum_1ng_glyco_hcd30_etd30_compound15.4919.4919.2 | controllerType=0 controllerNumber=1 scan=4919 | 15.6675 |
| 02097:2 | 3 | K.TPLTAN[+1647.613]ITK.S | HexNAc(5)Hex(3)Fuc(1) | N[+1648] | 1303.5911 | 2 | 2606.1749 | 2606.17 | 0 | 1.9 | 200 | Specific | 320.4 | 99.87 | 99.87 | 2.79 | 65 | >sp\|P01877\|IGHA2_HUMAN Immunoglobulin heavy constant alpha 2 OS=Homo sapiens OX=9606 GN=IGHA2 PE=1 SV=4 | 576 | 20201116_fqg_colostrum_1ng_glyco_hcd30_etd30_compound15.5072.5072.2 | controllerType=0 controllerNumber=1 scan=5072 | 16.0957 |
| 03725:2 | 3 | K.TPLTAN[+1694.603]ITK.S | HexNAc(3)Hex(4)Fuc(1)NeuAc(1) | N[+1695] | 1327.0892 | 2 | 2653.171 | 2653.1595 | 0 | 4.3 | 200 | Specific | 321.17 | 117.17 | 117.17 | 2.96 | 65 | >sp\|P01877\|IGHA2_HUMAN Immunoglobulin heavy constant alpha 2 OS=Homo sapiens OX=9606 GN=IGHA2 PE=1 SV=4 | 576 | 20201116_fqg_colostrum_1ng_glyco_hcd30_etd30_compound15.6773.6773.2 | controllerType=0 controllerNumber=1 scan=6773 | 19.4951 |
| 02065:2 | 3 | K.TPLTAN[+1241.454]ITK.S | HexNAc(3)Hex(3)Fuc(1) | N[+1241] | 1100.5114 | 2 | 2200.0155 | 2200.0113 | 0 | 1.9 | 200 | Specific | 321.5 | 103.18 | 103.18 | 2.79 | 65 | >sp\|P01877\|IGHA2_HUMAN Immunoglobulin heavy constant alpha 2 OS=Homo sapiens OX=9606 GN=IGHA2 PE=1 SV=4 | 576 | 20201116_fqg_colostrum_1ng_glyco_hcd30_etd30_compound15.5039.5039.2 | controllerType=0 controllerNumber=1 scan=5039 | 16.0189 |
| 02020:3 | 3 | K.TPLTAN[+1606.587]ITK.S | HexNAc(4)Hex(4)Fuc(1) | N[+1607] | 855.7213 | 3 | 2565.1495 | 2565.1435 | 0 | 2.3 | 200 | Specific | 324.49 | 131.24 | 131.24 | 3.02 | 65 | >sp\|P01877\|IGHA2_HUMAN Immunoglobulin heavy constant alpha 2 OS=Homo sapiens OX=9606 GN=IGHA2 PE=1 SV=4 | 576 | 20201116_fqg_colostrum_1ng_glyco_hcd30_etd30_compound15.4991.4991.3 | controllerType=0 controllerNumber=1 scan=4991 | 15.8713 |
| 02030:2 | 3 | K.TPLTAN[+1444.534]ITK.S | HexNAc(4)Hex(3)Fuc(1) | N[+1445] | 1202.0533 | 2 | 2403.0993 | 2403.0906 | 0 | 3.6 | 200 | Specific | 346.05 | 128 | 128 | 2.85 | 65 | >sp\|P01877\|IGHA2_HUMAN Immunoglobulin heavy constant alpha 2 OS=Homo sapiens OX=9606 GN=IGHA2 PE=1 SV=4 | 576 | 20201116_fqg_colostrum_1ng_glyco_hcd30_etd30_compound15.5002.5002.2 | controllerType=0 controllerNumber=1 scan=5002 | 15.9008 |
| 02083:2 | 3 | K.TPLTAN[+1038.375]ITK.S | HexNAc(2)Hex(3)Fuc(1) | N[+1038] | 998.9716 | 2 | 1996.9359 | 1996.9319 | 0 | 2 | 200 | Specific | 361.58 | 162.63 | 162.63 | 3.23 | 65 | >sp\|P01877\|IGHA2_HUMAN Immunoglobulin heavy constant alpha 2 OS=Homo sapiens OX=9606 GN=IGHA2 PE=1 SV=4 | 576 | 20201116_fqg_colostrum_1ng_glyco_hcd30_etd30_compound15.5058.5058.2 | controllerType=0 controllerNumber=1 scan=5058 | 16.0625 |
| 01995:2 | 3 | K.TPLTAN[+1403.507]ITK.S | HexNAc(3)Hex(4)Fuc(1) | N[+1404] | 1181.5395 | 2 | 2362.0717 | 2362.0641 | 0 | 3.2 | 200 | Specific | 361.66 | 133.37 | 133.37 | 2.94 | 65 | >sp\|P01877\|IGHA2_HUMAN Immunoglobulin heavy constant alpha 2 OS=Homo sapiens OX=9606 GN=IGHA2 PE=1 SV=4 | 576 | 20201116_fqg_colostrum_1ng_glyco_hcd30_etd30_compound15.4965.4965.2 | controllerType=0 controllerNumber=1 scan=4965 | 15.7883 |
| 02057:2 | 3 | K.TPLTAN[+1298.476]ITK.S | HexNAc(4)Hex(3) | N[+1298] | 1129.0236 | 2 | 2257.0399 | 2257.0327 | 0 | 3.2 | 200 | Specific | 364.73 | 134.41 | 134.41 | 3.3 | 65 | >sp\|P01877\|IGHA2_HUMAN Immunoglobulin heavy constant alpha 2 OS=Homo sapiens OX=9606 GN=IGHA2 PE=1 SV=4 | 576 | 20201116_fqg_colostrum_1ng_glyco_hcd30_etd30_compound15.5030.5030.2 | controllerType=0 controllerNumber=1 scan=5030 | 15.9946 |
| 02055:2 | 3 | K.TPLTAN[+1501.555]ITK.S | HexNAc(5)Hex(3) | N[+1502] | 1230.5635 | 2 | 2460.1198 | 2460.1121 | 0 | 3.1 | 200 | Specific | 369.81 | 127.33 | 127.33 | 3.44 | 65 | >sp\|P01877\|IGHA2_HUMAN Immunoglobulin heavy constant alpha 2 OS=Homo sapiens OX=9606 GN=IGHA2 PE=1 SV=4 | 576 | 20201116_fqg_colostrum_1ng_glyco_hcd30_etd30_compound15.5028.5028.2 | controllerType=0 controllerNumber=1 scan=5028 | 15.9755 |
| 02082:3 | 3 | K.TPLTAN[+1444.534]ITK.S | HexNAc(4)Hex(3)Fuc(1) | N[+1445] | 801.7038 | 3 | 2403.0969 | 2403.0906 | 0 | 2.6 | 200 | Specific | 487.42 | 291.49 | 291.49 | 4.26 | 65 | >sp\|P01877\|IGHA2_HUMAN Immunoglobulin heavy constant alpha 2 OS=Homo sapiens OX=9606 GN=IGHA2 PE=1 SV=4 | 576 | 20201116_fqg_colostrum_1ng_glyco_hcd30_etd30_compound15.5057.5057.3 | controllerType=0 controllerNumber=1 scan=5057 | 16.0585 |
| 07306:3 | 3 | R.MAGKPTHIN[+1216.423]VSVVMAEADGTC[+57.021].Y | HexNAc(2)Hex(5) | C[+57], N[+1216] | 1135.491 | 3 | 3404.4584 | 3404.4424 | 0 | 4.7 | 319 | CRagged | 229.36 | 191.23 | 165 | 2.91 | 65 | >sp\|P01877\|IGHA2_HUMAN Immunoglobulin heavy constant alpha 2 OS=Homo sapiens OX=9606 GN=IGHA2 PE=1 SV=4 | 576 | 20201116_fqg_colostrum_1ng_glyco_hcd30_etd30_compound15.10498.10498.3 | controllerType=0 controllerNumber=1 scan=10498 | 26.5371 |
| 06447:3 | 3 | R.MAGKPTHIN[+1216.423]VSVVMAEADGTC[+57.021].Y | HexNAc(2)Hex(5) | C[+57], N[+1216] | 1135.489 | 3 | 3404.4526 | 3404.4424 | 0 | 3 | 319 | CRagged | 279.4 | 247.2 | 193.69 | 4.52 | 65 | >sp\|P01877\|IGHA2_HUMAN Immunoglobulin heavy constant alpha 2 OS=Homo sapiens OX=9606 GN=IGHA2 PE=1 SV=4 | 576 | 20201116_fqg_colostrum_1ng_glyco_hcd30_etd30_compound15.9607.9607.3 | controllerType=0 controllerNumber=1 scan=9607 | 24.7936 |
| 06398:3 | 3 | R.MAGKPTHIN[+1378.476]VSVVMAEADGTC[+57.021].Y | HexNAc(2)Hex(6) | C[+57], N[+1378] | 1189.5084 | 3 | 3566.5105 | 3566.4952 | 0 | 4.3 | 319 | CRagged | 281.8 | 258.34 | 177.3 | 3.74 | 65 | >sp\|P01877\|IGHA2_HUMAN Immunoglobulin heavy constant alpha 2 OS=Homo sapiens OX=9606 GN=IGHA2 PE=1 SV=4 | 576 | 20201116_fqg_colostrum_1ng_glyco_hcd30_etd30_compound15.9555.9555.3 | controllerType=0 controllerNumber=1 scan=9555 | 24.6791 |
| 05832:3 | 3 | R.M[+15.995]AGKPTHIN[+1216.423]VSVVMAEADGTC[+57.021].Y | HexNAc(2)Hex(5) | C[+57], M[+16], N[+1216] | 1140.8202 | 3 | 3420.4461 | 3420.4373 | 0 | 2.6 | 319 | CRagged | 214.76 | 184.68 | 119.88 | 3.25 | 65 | >sp\|P01877\|IGHA2_HUMAN Immunoglobulin heavy constant alpha 2 OS=Homo sapiens OX=9606 GN=IGHA2 PE=1 SV=4 | 576 | 20201116_fqg_colostrum_1ng_glyco_hcd30_etd30_compound15.8968.8968.3 | controllerType=0 controllerNumber=1 scan=8968 | 23.5209 |
| 07627:3 | 3 | R.MAGKPTHIN[+1378.476]VSVVMAEADGTC[+57.021]Y.- | HexNAc(2)Hex(6) | C[+57], N[+1378] | 1243.8616 | 3 | 3729.5703 | 3729.5585 | 0 | 3.2 | 319 | CRagged | 206.99 | 185.1 | 121.32 | 3.25 | 65 | >sp\|P01877\|IGHA2_HUMAN Immunoglobulin heavy constant alpha 2 OS=Homo sapiens OX=9606 GN=IGHA2 PE=1 SV=4 | 576 | 20201116_fqg_colostrum_1ng_glyco_hcd30_etd30_compound15.10831.10831.3 | controllerType=0 controllerNumber=1 scan=10831 | 27.1583 |
| 08183:3 | 3 | R.MAGKPTHIN[+1702.581]VSVVMAEADGTC[+57.021]Y.- | HexNAc(2)Hex(8) | C[+57], N[+1703] | 1351.8984 | 3 | 4053.6806 | 4053.6642 | 0 | 4 | 319 | CRagged | 246.18 | 246.18 | 193.56 | 4.78 | 65 | >sp\|P01877\|IGHA2_HUMAN Immunoglobulin heavy constant alpha 2 OS=Homo sapiens OX=9606 GN=IGHA2 PE=1 SV=4 | 576 | 20201116_fqg_colostrum_1ng_glyco_hcd30_etd30_compound15.11410.11410.3 | controllerType=0 controllerNumber=1 scan=11410 | 28.3968 |
| 08241:3 | 3 | R.MAGKPTHIN[+1378.476]VSVVMAEADGTC[+57.021]Y.- | HexNAc(2)Hex(6) | C[+57], N[+1378] | 1243.8615 | 3 | 3729.57 | 3729.5585 | 0 | 3.1 | 319 | CRagged | 288.13 | 234.48 | 205.09 | 4.48 | 65 | >sp\|P01877\|IGHA2_HUMAN Immunoglobulin heavy constant alpha 2 OS=Homo sapiens OX=9606 GN=IGHA2 PE=1 SV=4 | 576 | 20201116_fqg_colostrum_1ng_glyco_hcd30_etd30_compound15.11471.11471.3 | controllerType=0 controllerNumber=1 scan=11471 | 28.5703 |
| 08268:3 | 3 | R.MAGKPTHIN[+1216.423]VSVVMAEADGTC[+57.021]Y.- | HexNAc(2)Hex(5) | C[+57], N[+1216] | 1189.8426 | 3 | 3567.5132 | 3567.5057 | 0 | 2.1 | 319 | CRagged | 327.33 | 272.86 | 214.37 | 4.16 | 65 | >sp\|P01877\|IGHA2_HUMAN Immunoglobulin heavy constant alpha 2 OS=Homo sapiens OX=9606 GN=IGHA2 PE=1 SV=4 | 576 | 20201116_fqg_colostrum_1ng_glyco_hcd30_etd30_compound15.11499.11499.3 | controllerType=0 controllerNumber=1 scan=11499 | 28.629 |
| 08313:3 | 3 | R.MAGKPTHIN[+1054.370]VSVVMAEADGTC[+57.021]Y.- | HexNAc(2)Hex(4) | C[+57], N[+1054] | 1135.8317 | 3 | 3405.4805 | 3405.4529 | 0 | 8.1 | 319 | CRagged | 369.69 | 320.09 | 236.73 | 3.75 | 65 | >sp\|P01877\|IGHA2_HUMAN Immunoglobulin heavy constant alpha 2 OS=Homo sapiens OX=9606 GN=IGHA2 PE=1 SV=4 | 576 | 20201116_fqg_colostrum_1ng_glyco_hcd30_etd30_compound15.11546.11546.3 | controllerType=0 controllerNumber=1 scan=11546 | 28.7519 |
| 07794:3 | 3 | R.M[+15.995]AGKPTHIN[+1378.476]VSVVMAEADGTC[+57.021]Y.- | HexNAc(2)Hex(6) | C[+57], M[+16], N[+1378] | 1249.1972 | 3 | 3745.577 | 3745.5535 | 0 | 6.3 | 319 | CRagged | 205.22 | 173.75 | 136.57 | 2.74 | 65 | >sp\|P01877\|IGHA2_HUMAN Immunoglobulin heavy constant alpha 2 OS=Homo sapiens OX=9606 GN=IGHA2 PE=1 SV=4 | 576 | 20201116_fqg_colostrum_1ng_glyco_hcd30_etd30_compound15.11005.11005.3 | controllerType=0 controllerNumber=1 scan=11005 | 27.5474 |
| 15145:4 | 4 | R.RTAGWNVPIGTLRPFLN[+2716.983]WTGPPEPIEAAVAR.F | HexNAc(5)Hex(6)Fuc(3)NeuAc(1) | N[+2717] | 1526.7106 | 4 | 6103.8205 | 6103.795 | 0 | 4.2 | 140 | Specific | 622.82 | 601.56 | 601.56 | 10.21 | 32 | >sp\|P02788\|TRFL_HUMAN Lactotransferrin OS=Homo sapiens OX=9606 GN=LTF PE=1 SV=6 | 1270 | 20201116_fqg_colostrum_1ng_glyco_hcd30_etd30_compound15.18655.18655.4 | controllerType=0 controllerNumber=1 scan=18655 | 43.2258 |
| 15268:4 | 4 | R.RTAGWNVPIGTLRPFLN[+2059.735]WTGPPEPIEAAVAR.F | HexNAc(4)Hex(5)Fuc(1)NeuAc(1) | N[+2060] | 1362.398 | 4 | 5446.57 | 5446.547 | 0 | 4.2 | 140 | Specific | 1010.25 | 966.29 | 966.29 | 18.21 | 32 | >sp\|P02788\|TRFL_HUMAN Lactotransferrin OS=Homo sapiens OX=9606 GN=LTF PE=1 SV=6 | 1270 | 20201116_fqg_colostrum_1ng_glyco_hcd30_etd30_compound15.18783.18783.4 | controllerType=0 controllerNumber=1 scan=18783 | 43.5299 |
| 16275:4 | 4 | R.[+42.011]TAGWNVPIGTLRPFLN[+2190.793]WTGPPEPIEAAVAR.F | HexNAc(6)Hex(6) | .[42], N[+2191] | 1366.6333 | 4 | 5463.5114 | 5463.5147 | 0 | -0.6 | 141 | Specific | 210.61 | 130.82 | 130.82 | 2.79 | 32 | >sp\|P02788\|TRFL_HUMAN Lactotransferrin OS=Homo sapiens OX=9606 GN=LTF PE=1 SV=6 | 1270 | 20201116_fqg_colostrum_1ng_glyco_hcd30_etd30_compound15.19848.19848.4 | controllerType=0 controllerNumber=1 scan=19848 | 46.3495 |
| 16313:4 | 4 | R.TAGWNVPIGTLRPFLN[+2205.793]WTGPPEPIEAAVAR.F | HexNAc(4)Hex(5)Fuc(2)NeuAc(1) | N[+2206] | 1359.8838 | 4 | 5436.5133 | 5436.5038 | 0 | 1.8 | 141 | Specific | 216.65 | 216.65 | 216.65 | 3.33 | 32 | >sp\|P02788\|TRFL_HUMAN Lactotransferrin OS=Homo sapiens OX=9606 GN=LTF PE=1 SV=6 | 1270 | 20201116_fqg_colostrum_1ng_glyco_hcd30_etd30_compound15.19888.19888.4 | controllerType=0 controllerNumber=1 scan=19888 | 46.467 |
| 16256:3 | 4 | R.TAGWNVPIGTLRPFLN[+2205.793]WTGPPEPIEAAVAR.F | HexNAc(4)Hex(5)Fuc(2)NeuAc(1) | N[+2206] | 1812.8451 | 3 | 5436.5206 | 5436.5038 | 0 | 3.1 | 141 | Specific | 241.62 | 165.19 | 165.19 | 4.4 | 32 | >sp\|P02788\|TRFL_HUMAN Lactotransferrin OS=Homo sapiens OX=9606 GN=LTF PE=1 SV=6 | 1270 | 20201116_fqg_colostrum_1ng_glyco_hcd30_etd30_compound15.19828.19828.3 | controllerType=0 controllerNumber=1 scan=19828 | 46.2923 |
| 17258:5 | 4 | R.TAGWNVPIGTLRPFLN[+2350.830]WTGPPEPIEAAVAR.F | HexNAc(4)Hex(5)Fuc(1)NeuAc(2) | N[+2351] | 1117.1179 | 5 | 5581.5605 | 5581.5413 | 0 | 3.4 | 141 | Specific | 242.36 | 242.36 | 242.36 | 4.8 | 32 | >sp\|P02788\|TRFL_HUMAN Lactotransferrin OS=Homo sapiens OX=9606 GN=LTF PE=1 SV=6 | 1270 | 20201116_fqg_colostrum_1ng_glyco_hcd30_etd30_compound15.20883.20883.5 | controllerType=0 controllerNumber=1 scan=20883 | 49.2094 |
| 17242:3 | 4 | R.TAGWNVPIGTLRPFLN[+2350.830]WTGPPEPIEAAVAR.F | HexNAc(4)Hex(5)Fuc(1)NeuAc(2) | N[+2351] | 1861.1922 | 3 | 5581.5621 | 5581.5413 | 0 | 3.7 | 141 | Specific | 332.22 | 264.99 | 264.99 | 6.25 | 32 | >sp\|P02788\|TRFL_HUMAN Lactotransferrin OS=Homo sapiens OX=9606 GN=LTF PE=1 SV=6 | 1270 | 20201116_fqg_colostrum_1ng_glyco_hcd30_etd30_compound15.20866.20866.3 | controllerType=0 controllerNumber=1 scan=20866 | 49.1587 |
| 17358:4 | 4 | R.TAGWNVPIGTLRPFLN[+2350.830]WTGPPEPIEAAVAR.F | HexNAc(4)Hex(5)Fuc(1)NeuAc(2) | N[+2351] | 1396.1454 | 4 | 5581.5597 | 5581.5413 | 0 | 3.3 | 141 | Specific | 354.26 | 354.26 | 354.26 | 5.36 | 32 | >sp\|P02788\|TRFL_HUMAN Lactotransferrin OS=Homo sapiens OX=9606 GN=LTF PE=1 SV=6 | 1270 | 20201116_fqg_colostrum_1ng_glyco_hcd30_etd30_compound15.20988.20988.4 | controllerType=0 controllerNumber=1 scan=20988 | 49.509 |
| 16251:4 | 4 | R.TAGWNVPIGTLRPFLN[+2205.793]WTGPPEPIEAAVAR.F | HexNAc(4)Hex(5)Fuc(2)NeuAc(1) | N[+2206] | 1359.8849 | 4 | 5436.5177 | 5436.5038 | 0 | 2.6 | 141 | Specific | 694.52 | 694.52 | 694.52 | 11.69 | 32 | >sp\|P02788\|TRFL_HUMAN Lactotransferrin OS=Homo sapiens OX=9606 GN=LTF PE=1 SV=6 | 1270 | 20201116_fqg_colostrum_1ng_glyco_hcd30_etd30_compound15.19823.19823.4 | controllerType=0 controllerNumber=1 scan=19823 | 46.2742 |
| 16267:5 | 4 | R.TAGWNVPIGTLRPFLN[+2059.735]WTGPPEPIEAAVAR.F | HexNAc(4)Hex(5)Fuc(1)NeuAc(1) | N[+2060] | 1058.8989 | 5 | 5290.4655 | 5290.4459 | 0 | 3.7 | 141 | Specific | 887.35 | 877.1 | 877.1 | 15.69 | 32 | >sp\|P02788\|TRFL_HUMAN Lactotransferrin OS=Homo sapiens OX=9606 GN=LTF PE=1 SV=6 | 1270 | 20201116_fqg_colostrum_1ng_glyco_hcd30_etd30_compound15.19840.19840.5 | controllerType=0 controllerNumber=1 scan=19840 | 46.333 |
| 16252:4 | 4 | R.TAGWNVPIGTLRPFLN[+2059.735]WTGPPEPIEAAVAR.F | HexNAc(4)Hex(5)Fuc(1)NeuAc(1) | N[+2060] | 1323.3709 | 4 | 5290.4616 | 5290.4459 | 0 | 3 | 141 | Specific | 909.68 | 870.2 | 870.2 | 16.69 | 32 | >sp\|P02788\|TRFL_HUMAN Lactotransferrin OS=Homo sapiens OX=9606 GN=LTF PE=1 SV=6 | 1270 | 20201116_fqg_colostrum_1ng_glyco_hcd30_etd30_compound15.19824.19824.4 | controllerType=0 controllerNumber=1 scan=19824 | 46.2839 |
| 15829:3 | 4 | R.TAGWNIPMGLLFN[+1913.677]QTGSC[+57.021]K.F | HexNAc(4)Hex(5)NeuAc(1) | C[+57], N[+1914] | 1336.9042 | 3 | 4008.6981 | 4008.687 | 0 | 2.8 | 485 | Specific | 311.99 | 191.82 | 191.82 | 5.83 | 32 | >sp\|P02788\|TRFL_HUMAN Lactotransferrin OS=Homo sapiens OX=9606 GN=LTF PE=1 SV=6 | 1270 | 20201116_fqg_colostrum_1ng_glyco_hcd30_etd30_compound15.19376.19376.3 | controllerType=0 controllerNumber=1 scan=19376 | 45.2141 |
| 15722:3 | 4 | R.TAGWNIPMGLLFN[+2059.735]QTGSC[+57.021]K.F | HexNAc(4)Hex(5)Fuc(1)NeuAc(1) | C[+57], N[+2060] | 1385.5897 | 3 | 4154.7544 | 4154.7449 | 0 | 2.3 | 485 | Specific | 342.21 | 256.29 | 256.29 | 5.79 | 32 | >sp\|P02788\|TRFL_HUMAN Lactotransferrin OS=Homo sapiens OX=9606 GN=LTF PE=1 SV=6 | 1270 | 20201116_fqg_colostrum_1ng_glyco_hcd30_etd30_compound15.19264.19264.3 | controllerType=0 controllerNumber=1 scan=19264 | 44.9167 |
| 17303:3 | 4 | R.TAGWNIPMGLLFN[+2350.830]QTGSC[+57.021]K.F | HexNAc(4)Hex(5)Fuc(1)NeuAc(2) | C[+57], N[+2351] | 1482.6244 | 3 | 4445.8588 | 4445.8403 | 0 | 4.2 | 485 | Specific | 401.62 | 281.05 | 281.05 | 6.45 | 32 | >sp\|P02788\|TRFL_HUMAN Lactotransferrin OS=Homo sapiens OX=9606 GN=LTF PE=1 SV=6 | 1270 | 20201116_fqg_colostrum_1ng_glyco_hcd30_etd30_compound15.20930.20930.3 | controllerType=0 controllerNumber=1 scan=20930 | 49.3494 |
| 08058:3 | 5 | K.TVLTPATNHMGN[+1540.529]VTFTIPANR.E | HexNAc(2)Hex(7) | N[+1541] | 1265.9045 | 3 | 3795.6989 | 3795.6886 | 0 | 2.7 | 74 | Specific | 401.88 | 348.4 | 138.42 | 5.5 | 30 | >sp\|P01024\|CO3_HUMAN Complement C3 OS=Homo sapiens OX=9606 GN=C3 PE=1 SV=2 | 492 | 20201116_fqg_colostrum_1ng_glyco_hcd30_etd30_compound15.11279.11279.3 | controllerType=0 controllerNumber=1 scan=11279 | 28.0913 |
| 15442:3 | 6 | K.QIGLYPVLVIDSSGYVNPN[+1257.449]YTGR.I | HexNAc(3)Hex(4) | N[+1257] | 1261.583 | 3 | 3782.7344 | 3782.7529 | 0 | -4.9 | 168 | Specific | 262.02 | 262.02 | 262.02 | 3.74 | 29 | >sp\|P01833\|PIGR_HUMAN Polymeric immunoglobulin receptor OS=Homo sapiens OX=9606 GN=PIGR PE=1 SV=4 | 902 | 20201116_fqg_colostrum_1ng_glyco_hcd30_etd30_compound15.18965.18965.3 | controllerType=0 controllerNumber=1 scan=18965 | 43.9454 |
| 15828:3 | 6 | K.QIGLYPVLVIDSSGYVNPN[+2205.793]YTGR.I | HexNAc(4)Hex(5)Fuc(2)NeuAc(1) | N[+2206] | 1577.7093 | 3 | 4731.1133 | 4731.0963 | 0 | 3.6 | 168 | Specific | 377.24 | 340.52 | 340.52 | 6.72 | 29 | >sp\|P01833\|PIGR_HUMAN Polymeric immunoglobulin receptor OS=Homo sapiens OX=9606 GN=PIGR PE=1 SV=4 | 902 | 20201116_fqg_colostrum_1ng_glyco_hcd30_etd30_compound15.19375.19375.3 | controllerType=0 controllerNumber=1 scan=19375 | 45.2111 |
| 21740:3 | 6 | R.[+42.011]LSLLEEPGN[+2351.851]GTFTVILNQLTSR.D | HexNAc(4)Hex(5)Fuc(3)NeuAc(1) | .[42], N[+2352] | 1599.394 | 3 | 4796.1674 | 4796.1539 | 0 | 2.8 | 413 | Specific | 301.92 | 190.73 | 190.73 | 5.83 | 29 | >sp\|P01833\|PIGR_HUMAN Polymeric immunoglobulin receptor OS=Homo sapiens OX=9606 GN=PIGR PE=1 SV=4 | 902 | 20201116_fqg_colostrum_1ng_glyco_hcd30_etd30_compound15.25665.25665.3 | controllerType=0 controllerNumber=1 scan=25665 | 65.1869 |
| 18567:4 | 6 | R.LSLLEEPGN[+2350.830]GTFTVILNQLTSR.D | HexNAc(4)Hex(5)Fuc(1)NeuAc(2) | N[+2351] | 1189.0424 | 4 | 4753.1478 | 4753.1229 | 0 | 5.2 | 413 | Specific | 305.85 | 273.28 | 273.28 | 3.88 | 29 | >sp\|P01833\|PIGR_HUMAN Polymeric immunoglobulin receptor OS=Homo sapiens OX=9606 GN=PIGR PE=1 SV=4 | 902 | 20201116_fqg_colostrum_1ng_glyco_hcd30_etd30_compound15.22267.22267.4 | controllerType=0 controllerNumber=1 scan=22267 | 53.4312 |
| 21755:3 | 6 | R.[+42.011]LSLLEEPGN[+2351.851]GTFTVILNQLTSR.D | HexNAc(4)Hex(5)Fuc(3)NeuAc(1) | .[42], N[+2352] | 1599.3824 | 3 | 4796.1326 | 4796.1539 | 0 | -4.4 | 413 | Specific | 353.9 | 223.27 | 223.27 | 4.94 | 29 | >sp\|P01833\|PIGR_HUMAN Polymeric immunoglobulin receptor OS=Homo sapiens OX=9606 GN=PIGR PE=1 SV=4 | 902 | 20201116_fqg_colostrum_1ng_glyco_hcd30_etd30_compound15.25681.25681.3 | controllerType=0 controllerNumber=1 scan=25681 | 65.2441 |
| 18595:3 | 6 | R.LSLLEEPGN[+2496.888]GTFTVILNQLTSR.D | HexNAc(4)Hex(5)Fuc(2)NeuAc(2) | N[+2497] | 1633.7411 | 3 | 4899.2089 | 4899.1808 | 0 | 5.7 | 413 | Specific | 400.57 | 292.66 | 292.66 | 5.71 | 29 | >sp\|P01833\|PIGR_HUMAN Polymeric immunoglobulin receptor OS=Homo sapiens OX=9606 GN=PIGR PE=1 SV=4 | 902 | 20201116_fqg_colostrum_1ng_glyco_hcd30_etd30_compound15.22296.22296.3 | controllerType=0 controllerNumber=1 scan=22296 | 53.5325 |
| 17112:4 | 6 | R.LSLLEEPGN[+2205.793]GTFTVILNQLTSR.D | HexNAc(4)Hex(5)Fuc(2)NeuAc(1) | N[+2206] | 1152.781 | 4 | 4608.1024 | 4608.0854 | 0 | 3.7 | 413 | Specific | 456.09 | 391.48 | 391.48 | 6.94 | 29 | >sp\|P01833\|PIGR_HUMAN Polymeric immunoglobulin receptor OS=Homo sapiens OX=9606 GN=PIGR PE=1 SV=4 | 902 | 20201116_fqg_colostrum_1ng_glyco_hcd30_etd30_compound15.20729.20729.4 | controllerType=0 controllerNumber=1 scan=20729 | 48.7697 |
| 18583:3 | 6 | R.LSLLEEPGN[+2350.830]GTFTVILNQLTSR.D | HexNAc(4)Hex(5)Fuc(1)NeuAc(2) | N[+2351] | 1585.0589 | 3 | 4753.1621 | 4753.1229 | 0 | 8.3 | 413 | Specific | 533.72 | 394.56 | 394.56 | 6.96 | 29 | >sp\|P01833\|PIGR_HUMAN Polymeric immunoglobulin receptor OS=Homo sapiens OX=9606 GN=PIGR PE=1 SV=4 | 902 | 20201116_fqg_colostrum_1ng_glyco_hcd30_etd30_compound15.22284.22284.3 | controllerType=0 controllerNumber=1 scan=22284 | 53.4838 |
| 17101:3 | 6 | R.LSLLEEPGN[+2205.793]GTFTVILNQLTSR.D | HexNAc(4)Hex(5)Fuc(2)NeuAc(1) | N[+2206] | 1536.7067 | 3 | 4608.1056 | 4608.0854 | 0 | 4.4 | 413 | Specific | 561.27 | 470.35 | 470.35 | 8.65 | 29 | >sp\|P01833\|PIGR_HUMAN Polymeric immunoglobulin receptor OS=Homo sapiens OX=9606 GN=PIGR PE=1 SV=4 | 902 | 20201116_fqg_colostrum_1ng_glyco_hcd30_etd30_compound15.20717.20717.3 | controllerType=0 controllerNumber=1 scan=20717 | 48.7354 |
| 17090:3 | 6 | R.LSLLEEPGN[+2351.851]GTFTVILNQLTSR.D | HexNAc(4)Hex(5)Fuc(3)NeuAc(1) | N[+2352] | 1585.3925 | 3 | 4754.163 | 4754.1433 | 0 | 4.1 | 413 | Specific | 582.65 | 467.08 | 467.08 | 9.39 | 29 | >sp\|P01833\|PIGR_HUMAN Polymeric immunoglobulin receptor OS=Homo sapiens OX=9606 GN=PIGR PE=1 SV=4 | 902 | 20201116_fqg_colostrum_1ng_glyco_hcd30_etd30_compound15.20706.20706.3 | controllerType=0 controllerNumber=1 scan=20706 | 48.7055 |
| 17096:4 | 6 | R.LSLLEEPGN[+2351.851]GTFTVILNQLTSR.D | HexNAc(4)Hex(5)Fuc(3)NeuAc(1) | N[+2352] | 1189.2962 | 4 | 4754.1629 | 4754.1433 | 0 | 4.1 | 413 | Specific | 622.64 | 575.49 | 575.49 | 9.2 | 29 | >sp\|P01833\|PIGR_HUMAN Polymeric immunoglobulin receptor OS=Homo sapiens OX=9606 GN=PIGR PE=1 SV=4 | 902 | 20201116_fqg_colostrum_1ng_glyco_hcd30_etd30_compound15.20712.20712.4 | controllerType=0 controllerNumber=1 scan=20712 | 48.7213 |
| 13544:3 | 6 | K.IIEGEPNLKVPGN[+2059.735]VTAVLGETLK.V | HexNAc(4)Hex(5)Fuc(1)NeuAc(1) | N[+2060] | 1484.3698 | 3 | 4451.0949 | 4451.0843 | 0 | 2.4 | 457 | Specific | 321.11 | 254.95 | 254.95 | 4.09 | 29 | >sp\|P01833\|PIGR_HUMAN Polymeric immunoglobulin receptor OS=Homo sapiens OX=9606 GN=PIGR PE=1 SV=4 | 902 | 20201116_fqg_colostrum_1ng_glyco_hcd30_etd30_compound15.16981.16981.3 | controllerType=0 controllerNumber=1 scan=16981 | 39.3208 |
| 13445:3 | 6 | K.IIEGEPNLKVPGN[+2351.851]VTAVLGETLK.V | HexNAc(4)Hex(5)Fuc(3)NeuAc(1) | N[+2352] | 1581.743 | 3 | 4743.2144 | 4743.2001 | 0 | 3 | 457 | Specific | 351.3 | 270.62 | 270.62 | 4.15 | 29 | >sp\|P01833\|PIGR_HUMAN Polymeric immunoglobulin receptor OS=Homo sapiens OX=9606 GN=PIGR PE=1 SV=4 | 902 | 20201116_fqg_colostrum_1ng_glyco_hcd30_etd30_compound15.16877.16877.3 | controllerType=0 controllerNumber=1 scan=16877 | 39.0489 |
| 13471:3 | 6 | K.IIEGEPNLKVPGN[+2205.793]VTAVLGETLK.V | HexNAc(4)Hex(5)Fuc(2)NeuAc(1) | N[+2206] | 1533.0555 | 3 | 4597.1519 | 4597.1422 | 0 | 2.1 | 457 | Specific | 418.72 | 335.14 | 335.14 | 4.79 | 29 | >sp\|P01833\|PIGR_HUMAN Polymeric immunoglobulin receptor OS=Homo sapiens OX=9606 GN=PIGR PE=1 SV=4 | 902 | 20201116_fqg_colostrum_1ng_glyco_hcd30_etd30_compound15.16904.16904.3 | controllerType=0 controllerNumber=1 scan=16904 | 39.1016 |
| 13040:4 | 6 | K.IIEGEPNLKVPGN[+2205.793]VTAVLGETLKVPC[+57.021]HFPC[+57.021]K.F | HexNAc(4)Hex(5)Fuc(2)NeuAc(1) | C[+57]*2, N[+2206] | 1406.4124 | 4 | 5622.6276 | 5622.5997 | 0 | 5 | 457 | Specific | 262 | 262 | 262 | 2.87 | 29 | >sp\|P01833\|PIGR_HUMAN Polymeric immunoglobulin receptor OS=Homo sapiens OX=9606 GN=PIGR PE=1 SV=4 | 902 | 20201116_fqg_colostrum_1ng_glyco_hcd30_etd30_compound15.16456.16456.4 | controllerType=0 controllerNumber=1 scan=16456 | 38.2162 |
| 14359:5 | 6 | K.IIEGEPNLKVPGN[+2350.830]VTAVLGETLKVPC[+57.021]HFPC[+57.021]K.F | HexNAc(4)Hex(5)Fuc(1)NeuAc(2) | C[+57]*2, N[+2351] | 1154.3358 | 5 | 5767.65 | 5767.6372 | 0 | 2.2 | 457 | Specific | 419.59 | 419.59 | 419.59 | 4.97 | 29 | >sp\|P01833\|PIGR_HUMAN Polymeric immunoglobulin receptor OS=Homo sapiens OX=9606 GN=PIGR PE=1 SV=4 | 902 | 20201116_fqg_colostrum_1ng_glyco_hcd30_etd30_compound15.17832.17832.5 | controllerType=0 controllerNumber=1 scan=17832 | 41.1182 |
| 04472:2 | 6 | K.WN[+1710.598]NTGC[+57.021]QALPSQDEGPSK.A | HexNAc(3)Hex(5)NeuAc(1) | C[+57], N[+1711] | 1850.247 | 2 | 3699.4868 | 3699.4743 | 0 | 3.4 | 498 | Specific | 268.98 | 149.18 | 149.18 | 4.95 | 29 | >sp\|P01833\|PIGR_HUMAN Polymeric immunoglobulin receptor OS=Homo sapiens OX=9606 GN=PIGR PE=1 SV=4 | 902 | 20201116_fqg_colostrum_1ng_glyco_hcd30_etd30_compound15.7547.7547.2 | controllerType=0 controllerNumber=1 scan=7547 | 20.9561 |

Table S4. Cont.

| Query #:z | Protein Rank | Glycopeptide | Glycan | Modification Type(s) | Observed m/z | z | Observed (M+H) | Calc. mass (M+H) | Off-by-x error | Mass error (ppm) | StartPosition | Cleavage | Score | Delta | Delta Mod | \|Log Prob\| | # of unique peptides | Protein Name | Protein DB number | Comment | Scan # | Scan Time |
|---|---|---|---|---|---|---|---|---|---|---|---|---|---|---|---|---|---|---|---|---|---|---|
| 06270:3 | 6 | K.WN[+2204.772]NTGC[+57.021]QALPSQDEGPSK.A | HexNAc(4)Hex(5)NeuAc(2) | C[+57], N[+2205] | 1398.5577 | 3 | 4193.6585 | 4193.6491 | 0 | 2.2 | 498 | Specific | 297.23 | 154.72 | 154.72 | 4.8 | 29 | >sp\|P01833\|PIGR_HUMAN Polymeric immunoglobulin receptor OS=Homo sapiens OX=9606 GN=PIGR PE=1 SV=4 | 902 | 20201116_fqg_colostrum_1ng_glyco_hcd30_etd30_compound15.9422.9422.3 | controllerType=0 controllerNumber=1 scan=9422 | 24.3888 |
| 04468:3 | 6 | K.WN[+1710.598]NTGC[+57.021]QALPSQDEGPSK.A | HexNAc(3)Hex(5)NeuAc(1) | C[+57], N[+1711] | 1233.8337 | 3 | 3699.4865 | 3699.4743 | 0 | 3.3 | 498 | Specific | 333.73 | 196.23 | 196.23 | 5.37 | 29 | >sp\|P01833\|PIGR_HUMAN Polymeric immunoglobulin receptor OS=Homo sapiens OX=9606 GN=PIGR PE=1 SV=4 | 902 | 20201116_fqg_colostrum_1ng_glyco_hcd30_etd30_compound15.7543.7543.3 | controllerType=0 controllerNumber=1 scan=7543 | 20.9474 |
| 04267:3 | 6 | K.WN[+2205.793]NTGC[+57.021]QALPSQDEGPSK.A | HexNAc(4)Hex(5)Fuc(2)NeuAc(1) | C[+57], N[+2206] | 1398.8997 | 3 | 4194.6846 | 4194.6695 | 0 | 3.6 | 498 | Specific | 363.55 | 246.13 | 246.13 | 5.83 | 29 | >sp\|P01833\|PIGR_HUMAN Polymeric immunoglobulin receptor OS=Homo sapiens OX=9606 GN=PIGR PE=1 SV=4 | 902 | 20201116_fqg_colostrum_1ng_glyco_hcd30_etd30_compound15.7335.7335.3 | controllerType=0 controllerNumber=1 scan=7335 | 20.5803 |
| 04369:3 | 6 | K.WN[+1872.651]NTGC[+57.021]QALPSQDEGPSK.A | HexNAc(3)Hex(6)NeuAc(1) | C[+57], N[+1873] | 1287.8505 | 3 | 3861.537 | 3861.5272 | 0 | 2.6 | 498 | Specific | 405.54 | 328.81 | 328.81 | 6.33 | 29 | >sp\|P01833\|PIGR_HUMAN Polymeric immunoglobulin receptor OS=Homo sapiens OX=9606 GN=PIGR PE=1 SV=4 | 902 | 20201116_fqg_colostrum_1ng_glyco_hcd30_etd30_compound15.7441.7441.3 | controllerType=0 controllerNumber=1 scan=7441 | 20.7804 |
| 04354:3 | 6 | K.WN[+1913.677]NTGC[+57.021]QALPSQDEGPSK.A | HexNAc(4)Hex(5)NeuAc(1) | C[+57], N[+1914] | 1301.5254 | 3 | 3902.5618 | 3902.5537 | 0 | 2.1 | 498 | Specific | 522.87 | 337.81 | 337.81 | 6.34 | 29 | >sp\|P01833\|PIGR_HUMAN Polymeric immunoglobulin receptor OS=Homo sapiens OX=9606 GN=PIGR PE=1 SV=4 | 902 | 20201116_fqg_colostrum_1ng_glyco_hcd30_etd30_compound15.7425.7425.3 | controllerType=0 controllerNumber=1 scan=7425 | 20.749 |
| 19304:4 | 9 | R.[+42.011]QLAHQSN[+2190.793]STNIFFSPVSIATAFAMLSLGTK.A | HexNAc(6)Hex(6) | .[42], N[+2191] | 1354.3613 | 4 | 5414.4235 | 5414.4388 | 0 | -2.8 | 64 | Specific | 308.04 | 250.97 | 250.97 | 2.98 | 11 | >sp\|P01009\|A1AT_HUMAN Alpha-1-antitrypsin OS=Homo sapiens OX=9606 GN=SERPINA1 PE=1 SV=3 | 642 | 20201116_fqg_colostrum_1ng_glyco_hcd30_etd30_compound15.23050.23050.4 | controllerType=0 controllerNumber=1 scan=23050 | 56.0357 |
| 19247:3 | 9 | R.QLAHQSN[+2204.772]STNIFFSPVSIATAFAMLSLGTK.A | HexNAc(4)Hex(5)NeuAc(2) | N[+2205] | 1796.1464 | 3 | 5386.4248 | 5386.4075 | 0 | 3.2 | 64 | Specific | 502.12 | 373.65 | 373.65 | 8.34 | 11 | >sp\|P01009\|A1AT_HUMAN Alpha-1-antitrypsin OS=Homo sapiens OX=9606 GN=SERPINA1 PE=1 SV=3 | 642 | 20201116_fqg_colostrum_1ng_glyco_hcd30_etd30_compound15.22989.22989.3 | controllerType=0 controllerNumber=1 scan=22989 | 55.8207 |
| 19241:4 | 9 | R.QLAHQSN[+2204.772]STNIFFSPVSIATAFAMLSLGTK.A | HexNAc(4)Hex(5)NeuAc(2) | N[+2205] | 1347.3606 | 4 | 5386.4206 | 5386.4075 | 0 | 2.4 | 64 | Specific | 733.96 | 631 | 470.72 | 12.34 | 11 | >sp\|P01009\|A1AT_HUMAN Alpha-1-antitrypsin OS=Homo sapiens OX=9606 GN=SERPINA1 PE=1 SV=3 | 642 | 20201116_fqg_colostrum_1ng_glyco_hcd30_etd30_compound15.22983.22983.4 | controllerType=0 controllerNumber=1 scan=22983 | 55.8011 |
| 18490:4 | 9 | K.ADTHDEILEGLNFN[+2861.000]LTEIPEAQIHEGFQELLR.T | HexNAc(5)Hex(6)NeuAc(3) | N[+2861] | 1638.9735 | 4 | 6552.8722 | 6552.8239 | 0 | 7.4 | 94 | Specific | 202.06 | 202.06 | 202.06 | 3.67 | 11 | >sp\|P01009\|A1AT_HUMAN Alpha-1-antitrypsin OS=Homo sapiens OX=9606 GN=SERPINA1 PE=1 SV=3 | 642 | 20201116_fqg_colostrum_1ng_glyco_hcd30_etd30_compound15.22186.22186.4 | controllerType=0 controllerNumber=1 scan=22186 | 53.2184 |
| 16727:4 | 9 | K.YLGN[+2204.772]ATAIFFLPDEGKLQHLENELTHDIITK.F | HexNAc(4)Hex(5)NeuAc(2) | N[+2205] | 1437.1631 | 4 | 5745.6305 | 5745.6098 | 0 | 3.6 | 268 | Specific | 396.29 | 396.29 | 396.29 | 5.64 | 11 | >sp\|P01009\|A1AT_HUMAN Alpha-1-antitrypsin OS=Homo sapiens OX=9606 GN=SERPINA1 PE=1 SV=3 | 642 | 20201116_fqg_colostrum_1ng_glyco_hcd30_etd30_compound15.20324.20324.4 | controllerType=0 controllerNumber=1 scan=20324 | 47.6793 |
| 02742:2 | 11 | K.AAIPSALDTN[+1216.423]SSK.S | HexNAc(2)Hex(5) | N[+1216] | 1246.0477 | 2 | 2491.088 | 2491.0815 | 0 | 2.6 | 542 | Specific | 384.52 | 201.12 | 201.12 | 3.99 | 7 | >sp\|Q08380\|LG3BP_HUMAN Galectin-3-binding protein OS=Homo sapiens OX=9606 GN=LGALS3BP PE=1 SV=1 | 1562 | 20201116_fqg_colostrum_1ng_glyco_hcd30_etd30_compound15.5744.5744.2 | controllerType=0 controllerNumber=1 scan=5744 | 17.6293 |
| 07801:3 | 14 | R.IIVPLNNREN[+1647.613]ISDPTSPLR.T | HexNAc(5)Hex(3)Fuc(1) | N[+1648] | 1265.9416 | 3 | 3795.8102 | 3795.7904 | 0 | 5.2 | 62 | Specific | 264.54 | 188.48 | 188.48 | 3.27 | 16 | >sp\|P01591\|IGJ_HUMAN Immunoglobulin J chain OS=Homo sapiens OX=9606 GN=JCHAIN PE=1 SV=4 | 1392 | 20201116_fqg_colostrum_1ng_glyco_hcd30_etd30_compound15.11012.11012.3 | controllerType=0 controllerNumber=1 scan=11012 | 27.5614 |
| 10948:3 | 14 | R.IIVPLNNREN[+2350.830]ISDPTSPLR.T | HexNAc(4)Hex(5)Fuc(1)NeuAc(2) | N[+2351] | 1500.3428 | 3 | 4499.0139 | 4499.0075 | 0 | 1.4 | 62 | Specific | 274.32 | 194.34 | 194.34 | 3.6 | 16 | >sp\|P01591\|IGJ_HUMAN Immunoglobulin J chain OS=Homo sapiens OX=9606 GN=JCHAIN PE=1 SV=4 | 1392 | 20201116_fqg_colostrum_1ng_glyco_hcd30_etd30_compound15.14281.14281.3 | controllerType=0 controllerNumber=1 scan=14281 | 33.9088 |
| 10971:3 | 14 | R.IIVPLNNREN[+2350.830]ISDPTSPLR.T | HexNAc(4)Hex(5)Fuc(1)NeuAc(2) | N[+2351] | 1500.3422 | 3 | 4499.0121 | 4499.0075 | 0 | 1 | 62 | Specific | 280.24 | 180.73 | 180.73 | 3.28 | 16 | >sp\|P01591\|IGJ_HUMAN Immunoglobulin J chain OS=Homo sapiens OX=9606 GN=JCHAIN PE=1 SV=4 | 1392 | 20201116_fqg_colostrum_1ng_glyco_hcd30_etd30_compound15.14305.14305.3 | controllerType=0 controllerNumber=1 scan=14305 | 33.9576 |
| 07810:3 | 14 | R.IIVPLNNREN[+1501.555]ISDPTSPLR.T | HexNAc(5)Hex(3) | N[+1502] | 1217.256 | 3 | 3649.7536 | 3649.7325 | 0 | 5.8 | 62 | Specific | 284.36 | 244.01 | 244.01 | 3.45 | 16 | >sp\|P01591\|IGJ_HUMAN Immunoglobulin J chain OS=Homo sapiens OX=9606 GN=JCHAIN PE=1 SV=4 | 1392 | 20201116_fqg_colostrum_1ng_glyco_hcd30_etd30_compound15.11021.11021.3 | controllerType=0 controllerNumber=1 scan=11021 | 27.5871 |
| 09204:3 | 14 | R.IIVPLNNREN[+1694.603]ISDPTSPLR.T | HexNAc(3)Hex(4)Fuc(1)NeuAc(1) | N[+1695] | 1281.6009 | 3 | 3842.7881 | 3842.7799 | 0 | 2.1 | 62 | Specific | 291.97 | 215.05 | 215.05 | 3.57 | 16 | >sp\|P01591\|IGJ_HUMAN Immunoglobulin J chain OS=Homo sapiens OX=9606 GN=JCHAIN PE=1 SV=4 | 1392 | 20201116_fqg_colostrum_1ng_glyco_hcd30_etd30_compound15.12471.12471.3 | controllerType=0 controllerNumber=1 scan=12471 | 30.6101 |
| 07740:3 | 14 | R.IIVPLNNREN[+1663.608]ISDPTSPLR.T | HexNAc(5)Hex(4) | N[+1664] | 1271.2708 | 3 | 3811.7979 | 3811.7853 | 0 | 3.3 | 62 | Specific | 298.43 | 227.71 | 227.71 | 4.13 | 16 | >sp\|P01591\|IGJ_HUMAN Immunoglobulin J chain OS=Homo sapiens OX=9606 GN=JCHAIN PE=1 SV=4 | 1392 | 20201116_fqg_colostrum_1ng_glyco_hcd30_etd30_compound15.10949.10949.3 | controllerType=0 controllerNumber=1 scan=10949 | 27.434 |
| 07726:3 | 14 | R.IIVPLNNREN[+1606.587]ISDPTSPLR.T | HexNAc(4)Hex(4)Fuc(1) | N[+1607] | 1252.2614 | 3 | 3754.7697 | 3754.7638 | 0 | 1.6 | 62 | Specific | 303.28 | 226.61 | 226.61 | 4.04 | 16 | >sp\|P01591\|IGJ_HUMAN Immunoglobulin J chain OS=Homo sapiens OX=9606 GN=JCHAIN PE=1 SV=4 | 1392 | 20201116_fqg_colostrum_1ng_glyco_hcd30_etd30_compound15.10934.10934.3 | controllerType=0 controllerNumber=1 scan=10934 | 27.3959 |
| 07944:3 | 14 | R.IIVPLNNREN[+1038.375]ISDPTSPLR.T | HexNAc(2)Hex(3)Fuc(1) | N[+1038] | 1062.8582 | 3 | 3186.56 | 3186.5523 | 0 | 2.4 | 62 | Specific | 314.06 | 257.75 | 257.75 | 4.35 | 16 | >sp\|P01591\|IGJ_HUMAN Immunoglobulin J chain OS=Homo sapiens OX=9606 GN=JCHAIN PE=1 SV=4 | 1392 | 20201116_fqg_colostrum_1ng_glyco_hcd30_etd30_compound15.11160.11160.3 | controllerType=0 controllerNumber=1 scan=11160 | 27.8557 |
| 07774:3 | 14 | R.IIVPLNNREN[+1768.640]ISDPTSPLR.T | HexNAc(4)Hex(5)Fuc(1) | N[+1769] | 1306.2805 | 3 | 3916.8268 | 3916.8167 | 0 | 2.6 | 62 | Specific | 330.32 | 242.49 | 242.49 | 4.43 | 16 | >sp\|P01591\|IGJ_HUMAN Immunoglobulin J chain OS=Homo sapiens OX=9606 GN=JCHAIN PE=1 SV=4 | 1392 | 20201116_fqg_colostrum_1ng_glyco_hcd30_etd30_compound15.10984.10984.3 | controllerType=0 controllerNumber=1 scan=10984 | 27.5068 |
| 07751:3 | 14 | R.IIVPLNNREN[+1460.529]ISDPTSPLR.T | HexNAc(4)Hex(4) | N[+1461] | 1203.5768 | 3 | 3608.716 | 3608.7059 | 0 | 2.8 | 62 | Specific | 332.53 | 251.88 | 251.88 | 4.43 | 16 | >sp\|P01591\|IGJ_HUMAN Immunoglobulin J chain OS=Homo sapiens OX=9606 GN=JCHAIN PE=1 SV=4 | 1392 | 20201116_fqg_colostrum_1ng_glyco_hcd30_etd30_compound15.10960.10960.3 | controllerType=0 controllerNumber=1 scan=10960 | 27.4546 |
| 09020:3 | 14 | R.IIVPLNNREN[+2059.735]ISDPTSPLR.T | HexNAc(4)Hex(5)Fuc(1)NeuAc(1) | N[+2060] | 1403.3132 | 3 | 4207.9249 | 4207.9121 | 0 | 3.1 | 62 | Specific | 374.08 | 299.25 | 299.25 | 4.8 | 16 | >sp\|P01591\|IGJ_HUMAN Immunoglobulin J chain OS=Homo sapiens OX=9606 GN=JCHAIN PE=1 SV=4 | 1392 | 20201116_fqg_colostrum_1ng_glyco_hcd30_etd30_compound15.12280.12280.3 | controllerType=0 controllerNumber=1 scan=12280 | 30.1988 |
| 07769:3 | 14 | R.IIVPLNNREN[+1257.449]ISDPTSPLR.T | HexNAc(3)Hex(4) | N[+1257] | 1135.8842 | 3 | 3405.638 | 3405.6266 | 0 | 3.4 | 62 | Specific | 393.62 | 327.2 | 327.2 | 6 | 16 | >sp\|P01591\|IGJ_HUMAN Immunoglobulin J chain OS=Homo sapiens OX=9606 GN=JCHAIN PE=1 SV=4 | 1392 | 20201116_fqg_colostrum_1ng_glyco_hcd30_etd30_compound15.10979.10979.3 | controllerType=0 controllerNumber=1 scan=10979 | 27.4907 |
| 17623:4 | 15 | K.APDKNVIFSPLSISTALAFLSLGAHN[+2569.905]TTLTEILK.G | HexNAc(5)Hex(6)NeuAc(2) | N[+2570] | 1538.9861 | 4 | 6152.9225 | 6152.8828 | 0 | 6.5 | 68 | Specific | 425.05 | 425.05 | 425.05 | 6.42 | 11 | >sp\|P01011\|AACT_HUMAN Alpha-1-antichymotrypsin OS=Homo sapiens OX=9606 GN=SERPINA3 PE=1 SV=2 | 1314 | 20201116_fqg_colostrum_1ng_glyco_hcd30_etd30_compound15.21267.21267.3 | controllerType=0 controllerNumber=1 scan=21267 | 50.3722 |
| 00087:4 | 23 | R.EIRHN[+2059.735]STGC[+57.021]LR.M | HexNAc(4)Hex(5)Fuc(1)NeuAc(1) | C[+57], N[+2060] | 851.3583 | 4 | 3402.4114 | 3402.3994 | 0 | 3.5 | 287 | Specific | 607.65 | 487.69 | 487.69 | 8.99 | 7 | >sp\|P10909\|CLUS_HUMAN Clusterin OS=Homo sapiens OX=9606 GN=CLU PE=1 SV=1 | 1467 | 20201116_fqg_colostrum_1ng_glyco_hcd30_etd30_compound15.2968.2968.4 | controllerType=0 controllerNumber=1 scan=2968 | 11.2431 |
| 17446:4 | 23 | K.MLN[+2059.735]TSSLLEQLNEQFNWVSR.L | HexNAc(4)Hex(5)Fuc(1)NeuAc(1) | N[+2060] | 1117.9887 | 4 | 4468.9329 | 4468.9217 | 0 | 2.5 | 352 | Specific | 319.04 | 311.74 | 311.74 | 4.8 | 7 | >sp\|P10909\|CLUS_HUMAN Clusterin OS=Homo sapiens OX=9606 GN=CLU PE=1 SV=1 | 1467 | 20201116_fqg_colostrum_1ng_glyco_hcd30_etd30_compound15.21081.21081.4 | controllerType=0 controllerNumber=1 scan=21081 | 49.8115 |
| 17436:3 | 23 | K.MLN[+2059.735]TSSLLEQLNEQFNWVSR.L | HexNAc(4)Hex(5)Fuc(1)NeuAc(1) | N[+2060] | 1490.3196 | 3 | 4468.9444 | 4468.9217 | 0 | 5.1 | 352 | Specific | 598.16 | 402.77 | 402.77 | 8.65 | 7 | >sp\|P10909\|CLUS_HUMAN Clusterin OS=Homo sapiens OX=9606 GN=CLU PE=1 SV=1 | 1467 | 20201116_fqg_colostrum_1ng_glyco_hcd30_etd30_compound15.21070.21070.3 | controllerType=0 controllerNumber=1 scan=21070 | 49.7721 |
| 18857:3 | 23 | K.MLN[+2350.830]TSSLLEQLNEQFNWVSR.L | HexNAc(4)Hex(5)Fuc(1)NeuAc(2) | N[+2351] | 1587.3506 | 3 | 4760.0374 | 4760.0171 | 0 | 4.3 | 352 | Specific | 608.16 | 425.2 | 425.2 | 9.65 | 7 | >sp\|P10909\|CLUS_HUMAN Clusterin OS=Homo sapiens OX=9606 GN=CLU PE=1 SV=1 | 1467 | 20201116_fqg_colostrum_1ng_glyco_hcd30_etd30_compound15.22573.22573.3 | controllerType=0 controllerNumber=1 scan=22573 | 54.3705 |
| 01396:3 | 29 | K.N[+1216.423]NSIPDKQITASSSYK.T | HexNAc(2)Hex(5) | N[+1216] | 990.4408 | 3 | 2969.3077 | 2969.2991 | 0 | 2.9 | 238 | Specific | 459.28 | 290.93 | 290.93 | 4.05 | 5 | >sp\|Q08431\|MFGM_HUMAN Lactadherin OS=Homo sapiens OX=9606 GN=MFGE8 PE=1 SV=3 | 1064 | 20201116_fqg_colostrum_1ng_glyco_hcd30_etd30_compound15.4342.4342.3 | controllerType=0 controllerNumber=1 scan=4342 | 14.5135 |
| 01301:3 | 33 | R.VVLGAHN[+730.264]LSR.R | HexNAc(2)Hex(2) | N[+730] | 599.2992 | 3 | 1795.883 | 1795.8807 | 0 | 1.3 | 82 | Specific | 227.26 | 205.06 | 205.06 | 2.72 | 4 | >sp\|P08246\|ELNE_HUMAN Neutrophil elastase OS=Homo sapiens OX=9606 GN=ELANE PE=1 SV=1 | 1269 | 20201116_fqg_colostrum_1ng_glyco_hcd30_etd30_compound15.4242.4242.3 | controllerType=0 controllerNumber=1 scan=4242 | 14.2491 |
| 16219:2 | 33 | R.GIASVLQELN[+876.322]VTVVTSLC[+57.021]R.R | HexNAc(2)Hex(2)Fuc(1) | C[+57], N[+876] | 1468.231 | 2 | 2935.4547 | 2935.4439 | 0 | 3.7 | 164 | Specific | 668.27 | 616.28 | 616.28 | 11.24 | 4 | >sp\|P08246\|ELNE_HUMAN Neutrophil elastase OS=Homo sapiens OX=9606 GN=ELANE PE=1 SV=1 | 1269 | 20201116_fqg_colostrum_1ng_glyco_hcd30_etd30_compound15.19789.19789.2 | controllerType=0 controllerNumber=1 scan=19789 | 46.1762 |
| 16197:3 | 33 | R.GIASVLQELN[+876.322]VTVVTSLC[+57.021]R.R | HexNAc(2)Hex(2)Fuc(1) | C[+57], N[+876] | 979.1555 | 3 | 2935.4519 | 2935.4439 | 0 | 2.7 | 164 | Specific | 870.79 | 761.09 | 761.09 | 15.39 | 4 | >sp\|P08246\|ELNE_HUMAN Neutrophil elastase OS=Homo sapiens OX=9606 GN=ELANE PE=1 SV=1 | 1269 | 20201116_fqg_colostrum_1ng_glyco_hcd30_etd30_compound15.19766.19766.3 | controllerType=0 controllerNumber=1 scan=19766 | 46.1022 |
| 00612:3 | 41 | K.TKPREEQFN[+1606.587]STFR.V | HexNAc(4)Hex(4)Fuc(1) | N[+1607] | 1082.8101 | 3 | 3246.4157 | 3246.4054 | 0 | 3.2 | 168 | Specific | 210.18 | 147.48 | 147.48 | 2.71 | 5 | >sp\|P01859\|IGHG2_HUMAN Immunoglobulin heavy constant gamma 2 OS=Homo sapiens OX=9606 GN=IGHG2 PE=1 SV=2 | 387 | 20201116_fqg_colostrum_1ng_glyco_hcd30_etd30_compound15.3522.3522.3 | controllerType=0 controllerNumber=1 scan=3522 | 12.5376 |
| 00578:4 | 41 | K.TKPREEQFN[+1768.640]STFR.V | HexNAc(4)Hex(5)Fuc(1) | N[+1769] | 852.8732 | 4 | 3408.4709 | 3408.4582 | 0 | 3.7 | 168 | Specific | 311.12 | 256.47 | 256.47 | 3.64 | 5 | >sp\|P01859\|IGHG2_HUMAN Immunoglobulin heavy constant gamma 2 OS=Homo sapiens OX=9606 GN=IGHG2 PE=1 SV=2 | 387 | 20201116_fqg_colostrum_1ng_glyco_hcd30_etd30_compound15.3486.3486.4 | controllerType=0 controllerNumber=1 scan=3486 | 12.4423 |
| 00602:4 | 41 | K.TKPREEQFN[+1606.587]STFR.V | HexNAc(4)Hex(4)Fuc(1) | N[+1607] | 812.3587 | 4 | 3246.4131 | 3246.4054 | 0 | 2.4 | 168 | Specific | 605.66 | 503.84 | 503.84 | 7.81 | 5 | >sp\|P01859\|IGHG2_HUMAN Immunoglobulin heavy constant gamma 2 OS=Homo sapiens OX=9606 GN=IGHG2 PE=1 SV=2 | 387 | 20201116_fqg_colostrum_1ng_glyco_hcd30_etd30_compound15.3511.3511.4 | controllerType=0 controllerNumber=1 scan=3511 | 12.5007 |
| 02848:2 | 44 | R.SC[+57.021]PAC[+57.021]PGSN[+876.322]ITIR.N | HexNAc(2)Hex(2)Fuc(1) | C[+57]*2, N[+876] | 1155.0015 | 2 | 2308.9957 | 2308.9894 | 0 | 2.7 | 315 | Specific | 366.02 | 247.31 | 247.31 | 4.24 | 4 | >sp\|P05164\|PERM_HUMAN Myeloperoxidase OS=Homo sapiens OX=9606 GN=MPO PE=1 SV=1 | 12 | 20201116_fqg_colostrum_1ng_glyco_hcd30_etd30_compound15.5854.5854.2 | controllerType=0 controllerNumber=1 scan=5854 | 17.8384 |
| 00593:2 | 44 | R.SYN[+1038.375]DSVDPR.I | HexNAc(2)Hex(3)Fuc(1) | N[+1038] | 1045.9259 | 2 | 2090.8446 | 2090.8395 | 0 | 2.5 | 481 | Specific | 346.72 | 244.51 | 244.51 | 3.93 | 4 | >sp\|P05164\|PERM_HUMAN Myeloperoxidase OS=Homo sapiens OX=9606 GN=MPO PE=1 SV=1 | 12 | 20201116_fqg_colostrum_1ng_glyco_hcd30_etd30_compound15.3502.3502.2 | controllerType=0 controllerNumber=1 scan=3502 | 12.483 |
| 01342:3 | 46 | R.LRN[+1216.423]VSWATGR.S | HexNAc(2)Hex(5) | N[+1216] | 792.6936 | 3 | 2376.0663 | 2376.0559 | 0 | 4.4 | 149 | Specific | 329.88 | 163.76 | 163.76 | 2.65 | 4 | >sp\|P08571\|CD14_HUMAN Monocyte differentiation antigen CD14 OS=Homo sapiens OX=9606 GN=CD14 PE=1 SV=2 | 1232 | 20201116_fqg_colostrum_1ng_glyco_hcd30_etd30_compound15.4285.4285.3 | controllerType=0 controllerNumber=1 scan=4285 | 14.3607 |
| 15616:5 | 115 | R.VGAHDPPAQVLQELN[+876.322]VTVVTFFC[+57.021]RPHNIC[+57.021]TFVPR.R | HexNAc(2)Hex(2)Fuc(1) | C[+57]*2, N[+876] | 959.8775 | 5 | 4795.3585 | 4795.3117 | 0 | 9.8 | 160 | Specific | 462.19 | 462.19 | 462.19 | 5.31 | 2 | >sp\|P24158\|PRTN3_HUMAN Myeloblastin OS=Homo sapiens OX=9606 GN=PRTN3 PE=1 SV=3 | 931 | 20201116_fqg_colostrum_1ng_glyco_hcd30_etd30_compound15.19151.19151.5 | controllerType=0 controllerNumber=1 scan=19151 | 44.5389 |
| 07170:3 | 134 | R.FVN[+876.322]VTVTPEDQC[+57.021]RPNNVC[+57.021]TGVLTR.R | HexNAc(2)Hex(2)Fuc(1) | C[+57]*2, N[+876] | 1218.2341 | 3 | 3652.6876 | 3652.6728 | 0 | 4.1 | 169 | Specific | 376.74 | 328.28 | 328.28 | 4.91 | 2 | >sp\|P20160\|CAP7_HUMAN Azurocidin OS=Homo sapiens OX=9606 GN=AZU1 PE=1 SV=3 | 850 | 20201116_fqg_colostrum_1ng_glyco_hcd30_etd30_compound15.10356.10356.3 | controllerType=0 controllerNumber=1 scan=10356 | 26.2262 |
| 15160:5 | 1 | R.LSLHRPALEDLLLGSEAN[+1501.555]LTC[+57.021]TLTGLR.D | HexNAc(5)Hex(3) | C[+57], N[+1502] | 893.839 | 5 | 4465.166 | 4465.1536 | 0 | 2.8 | 127 | Specific | 216.84 | 216.84 | 216.84 | 3.17 | 31 | >sp\|P01876\|IGHA1_HUMAN Immunoglobulin heavy constant alpha 1 OS=Homo sapiens OX=9606 GN=IGHA1 PE=1 SV=2 | 255 | 20201117_fqg_milk_1ng_glyco_hcd30_etd30_compound15.18751.18751.5 | controllerType=0 controllerNumber=1 scan=18751 | 42.7951 |
| 15142:3 | 1 | R.LSLHRPALEDLLLGSEAN[+1460.529]LTC[+57.021]TLTGLR.D | HexNAc(4)Hex(4) | C[+57], N[+1461] | 1475.3827 | 3 | 4424.1337 | 4424.1271 | 0 | 1.5 | 127 | Specific | 372.48 | 312 | 312 | 5.32 | 31 | >sp\|P01876\|IGHA1_HUMAN Immunoglobulin heavy constant alpha 1 OS=Homo sapiens OX=9606 GN=IGHA1 PE=1 SV=2 | 255 | 20201117_fqg_milk_1ng_glyco_hcd30_etd30_compound15.18732.18732.3 | controllerType=0 controllerNumber=1 scan=18732 | 42.7339 |
| 15122:3 | 1 | R.LSLHRPALEDLLLGSEAN[+1378.476]LTC[+57.021]TLTGLR.D | HexNAc(2)Hex(6) | C[+57], N[+1378] | 1448.0319 | 3 | 4342.0812 | 4342.074 | 0 | 1.7 | 127 | Specific | 397.35 | 368.04 | 368.04 | 6.48 | 31 | >sp\|P01876\|IGHA1_HUMAN Immunoglobulin heavy constant alpha 1 OS=Homo sapiens OX=9606 GN=IGHA1 PE=1 SV=2 | 255 | 20201117_fqg_milk_1ng_glyco_hcd30_etd30_compound15.18711.18711.3 | controllerType=0 controllerNumber=1 scan=18711 | 42.668 |
| 15155:3 | 1 | R.LSLHRPALEDLLLGSEAN[+1501.555]LTC[+57.021]TLTGLR.D | HexNAc(5)Hex(3) | C[+57], N[+1502] | 1489.0605 | 3 | 4465.1669 | 4465.1536 | 0 | 3 | 127 | Specific | 411.51 | 392.3 | 392.3 | 7.08 | 31 | >sp\|P01876\|IGHA1_HUMAN Immunoglobulin heavy constant alpha 1 OS=Homo sapiens OX=9606 GN=IGHA1 PE=1 SV=2 | 255 | 20201117_fqg_milk_1ng_glyco_hcd30_etd30_compound15.18746.18746.3 | controllerType=0 controllerNumber=1 scan=18746 | 42.7809 |
| 15069:3 | 1 | R.LSLHRPALEDLLLGSEAN[+1864.634]LTC[+57.021]TLTGLR.D | HexNAc(2)Hex(9) | C[+57], N[+1865] | 1610.086 | 3 | 4828.2434 | 4828.2324 | 0 | 2.3 | 127 | Specific | 430.62 | 374.38 | 374.38 | 5.98 | 31 | >sp\|P01876\|IGHA1_HUMAN Immunoglobulin heavy constant alpha 1 OS=Homo sapiens OX=9606 GN=IGHA1 PE=1 SV=2 | 255 | 20201117_fqg_milk_1ng_glyco_hcd30_etd30_compound15.18654.18654.3 | controllerType=0 controllerNumber=1 scan=18654 | 42.4336 |
| 15127:4 | 1 | R.LSLHRPALEDLLLGSEAN[+1419.502]LTC[+57.021]TLTGLR.D | HexNAc(3)Hex(5) | C[+57], N[+1420] | 1096.5329 | 4 | 4383.1097 | 4383.1005 | 0 | 2.1 | 127 | Specific | 444.81 | 444.81 | 444.81 | 6.48 | 31 | >sp\|P01876\|IGHA1_HUMAN Immunoglobulin heavy constant alpha 1 OS=Homo sapiens OX=9606 GN=IGHA1 PE=1 SV=2 | 255 | 20201117_fqg_milk_1ng_glyco_hcd30_etd30_compound15.18716.18716.4 | controllerType=0 controllerNumber=1 scan=18716 | 42.6845 |
| 15802:4 | 1 | R.LSLHRPALEDLLLGSEAN[+2116.756]LTC[+57.021]TLTGLR.D | HexNAc(5)Hex(5)NeuAc(1) | C[+57], N[+2117] | 1270.8496 | 4 | 5080.3766 | 5080.3547 | 0 | 4.3 | 127 | Specific | 457.12 | 457.12 | 457.12 | 5.93 | 31 | >sp\|P01876\|IGHA1_HUMAN Immunoglobulin heavy constant alpha 1 OS=Homo sapiens OX=9606 GN=IGHA1 PE=1 SV=2 | 255 | 20201117_fqg_milk_1ng_glyco_hcd30_etd30_compound15.19429.19429.4 | controllerType=0 controllerNumber=1 scan=19429 | 44.7447 |
| 15128:3 | 1 | R.LSLHRPALEDLLLGSEAN[+1216.423]LTC[+57.021]TLTGLR.D | HexNAc(2)Hex(5) | C[+57], N[+1216] | 1394.014 | 3 | 4180.0274 | 4180.0211 | 0 | 1.5 | 127 | Specific | 481.41 | 481.41 | 481.41 | 7.34 | 31 | >sp\|P01876\|IGHA1_HUMAN Immunoglobulin heavy constant alpha 1 OS=Homo sapiens OX=9606 GN=IGHA1 PE=1 SV=2 | 255 | 20201117_fqg_milk_1ng_glyco_hcd30_etd30_compound15.18717.18717.3 | controllerType=0 controllerNumber=1 scan=18717 | 42.6882 |
| 15091:3 | 1 | R.LSLHRPALEDLLLGSEAN[+1540.529]LTC[+57.021]TLTGLR.D | HexNAc(2)Hex(7) | C[+57], N[+1541] | 1502.0503 | 3 | 4504.1365 | 4504.1268 | 0 | 2.2 | 127 | Specific | 518.69 | 434.08 | 434.08 | 7.08 | 31 | >sp\|P01876\|IGHA1_HUMAN Immunoglobulin heavy constant alpha 1 OS=Homo sapiens OX=9606 GN=IGHA1 PE=1 SV=2 | 255 | 20201117_fqg_milk_1ng_glyco_hcd30_etd30_compound15.18677.18677.3 | controllerType=0 controllerNumber=1 scan=18677 | 42.5084 |
| 15803:4 | 1 | R.LSLHRPALEDLLLGSEAN[+1872.651]LTC[+57.021]TLTGLR.D | HexNAc(3)Hex(6)NeuAc(1) | C[+57], N[+1873] | 1209.8234 | 4 | 4836.2718 | 4836.2487 | 0 | 4.8 | 127 | Specific | 674.64 | 586.45 | 586.45 | 10.2 | 31 | >sp\|P01876\|IGHA1_HUMAN Immunoglobulin heavy constant alpha 1 OS=Homo sapiens OX=9606 GN=IGHA1 PE=1 SV=2 | 255 | 20201117_fqg_milk_1ng_glyco_hcd30_etd30_compound15.19430.19430.4 | controllerType=0 controllerNumber=1 scan=19430 | 44.7538 |
| 15185:4 | 1 | R.LSLHRPALEDLLLGSEAN[+1054.370]LTC[+57.021]TLTGLR.D | HexNAc(2)Hex(4) | C[+57], N[+1054] | 1005.2489 | 4 | 4017.9737 | 4017.9683 | 0 | 1.3 | 127 | Specific | 696.07 | 696.07 | 696.07 | 11.01 | 31 | >sp\|P01876\|IGHA1_HUMAN Immunoglobulin heavy constant alpha 1 OS=Homo sapiens OX=9606 GN=IGHA1 PE=1 SV=2 | 255 | 20201117_fqg_milk_1ng_glyco_hcd30_etd30_compound15.18777.18777.4 | controllerType=0 controllerNumber=1 scan=18777 | 42.8841 |
| 15144:4 | 1 | R.LSLHRPALEDLLLGSEAN[+1257.449]LTC[+57.021]TLTGLR.D | HexNAc(3)Hex(4) | C[+57], N[+1257] | 1056.0186 | 4 | 4221.0526 | 4221.0477 | 0 | 1.2 | 127 | Specific | 729.83 | 729.83 | 729.83 | 12.01 | 31 | >sp\|P01876\|IGHA1_HUMAN Immunoglobulin heavy constant alpha 1 OS=Homo sapiens OX=9606 GN=IGHA1 PE=1 SV=2 | 255 | 20201117_fqg_milk_1ng_glyco_hcd30_etd30_compound15.18734.18734.4 | controllerType=0 controllerNumber=1 scan=18734 | 42.7456 |
| 15840:4 | 1 | R.LSLHRPALEDLLLGSEAN[+1710.598]LTC[+57.021]TLTGLR.D | HexNAc(3)Hex(5)NeuAc(1) | C[+57], N[+1711] | 1169.3079 | 4 | 4674.2098 | 4674.1959 | 0 | 3 | 127 | Specific | 787.67 | 773.75 | 773.75 | 12.95 | 31 | >sp\|P01876\|IGHA1_HUMAN Immunoglobulin heavy constant alpha 1 OS=Homo sapiens OX=9606 GN=IGHA1 PE=1 SV=2 | 255 | 20201117_fqg_milk_1ng_glyco_hcd30_etd30_compound15.19469.19469.4 | controllerType=0 controllerNumber=1 scan=19469 | 44.8673 |
| 15143:4 | 1 | R.LSLHRPALEDLLLGSEAN[+1524.534]LTC[+57.021]TLTGLR.D | HexNAc(2)Hex(6)Fuc(1) | C[+57], N[+1525] | 1122.7828 | 4 | 4488.1092 | 4488.1319 | 0 | -5 | 127 | Specific | 802.62 | 791.98 | 791.98 | 12.89 | 31 | >sp\|P01876\|IGHA1_HUMAN Immunoglobulin heavy constant alpha 1 OS=Homo sapiens OX=9606 GN=IGHA1 PE=1 SV=2 | 255 | 20201117_fqg_milk_1ng_glyco_hcd30_etd30_compound15.18733.18733.4 | controllerType=0 controllerNumber=1 scan=18733 | 42.7418 |
| 15175:4 | 1 | R.LSLHRPALEDLLLGSEAN[+1095.397]LTC[+57.021]TLTGLR.D | HexNAc(3)Hex(3) | C[+57], N[+1095] | 1015.5058 | 4 | 4059.0015 | 4058.9949 | 0 | 1.6 | 127 | Specific | 807.03 | 807.03 | 807.03 | 13.74 | 31 | >sp\|P01876\|IGHA1_HUMAN Immunoglobulin heavy constant alpha 1 OS=Homo sapiens OX=9606 GN=IGHA1 PE=1 SV=2 | 255 | 20201117_fqg_milk_1ng_glyco_hcd30_etd30_compound15.18767.18767.4 | controllerType=0 controllerNumber=1 scan=18767 | 42.8491 |
| 15154:4 | 1 | R.LSLHRPALEDLLLGSEAN[+1501.555]LTC[+57.021]TLTGLR.D | HexNAc(5)Hex(3) | C[+57], N[+1502] | 1117.049 | 4 | 4465.1741 | 4465.1536 | 0 | 4.6 | 127 | Specific | 946.47 | 946.47 | 946.47 | 15.2 | 31 | >sp\|P01876\|IGHA1_HUMAN Immunoglobulin heavy constant alpha 1 OS=Homo sapiens OX=9606 GN=IGHA1 PE=1 SV=2 | 255 | 20201117_fqg_milk_1ng_glyco_hcd30_etd30_compound15.18745.18745.4 | controllerType=0 controllerNumber=1 scan=18745 | 42.7781 |
| 15121:4 | 1 | R.LSLHRPALEDLLLGSEAN[+1216.423]LTC[+57.021]TLTGLR.D | HexNAc(2)Hex(5) | C[+57], N[+1216] | 1045.7624 | 4 | 4180.0277 | 4180.0211 | 0 | 1.6 | 127 | Specific | 968.65 | 942.41 | 942.41 | 16.74 | 31 | >sp\|P01876\|IGHA1_HUMAN Immunoglobulin heavy constant alpha 1 OS=Homo sapiens OX=9606 GN=IGHA1 PE=1 SV=2 | 255 | 20201117_fqg_milk_1ng_glyco_hcd30_etd30_compound15.18710.18710.4 | controllerType=0 controllerNumber=1 scan=18710 | 42.665 |
| 15095:4 | 1 | R.LSLHRPALEDLLLGSEAN[+1663.608]LTC[+57.021]TLTGLR.D | HexNAc(5)Hex(4) | C[+57], N[+1664] | 1157.5595 | 4 | 4627.2161 | 4627.2064 | 0 | 2.1 | 127 | Specific | 975.8 | 969.91 | 969.91 | 16.74 | 31 | >sp\|P01876\|IGHA1_HUMAN Immunoglobulin heavy constant alpha 1 OS=Homo sapiens OX=9606 GN=IGHA1 PE=1 SV=2 | 255 | 20201117_fqg_milk_1ng_glyco_hcd30_etd30_compound15.18681.18681.4 | controllerType=0 controllerNumber=1 scan=18681 | 42.5223 |
| 15138:4 | 1 | R.LSLHRPALEDLLLGSEAN[+1298.476]LTC[+57.021]TLTGLR.D | HexNAc(4)Hex(3) | C[+57], N[+1298] | 1066.2758 | 4 | 4262.0814 | 4262.0742 | 0 | 1.7 | 127 | Specific | 982.75 | 982.75 | 982.75 | 16.74 | 31 | >sp\|P01876\|IGHA1_HUMAN Immunoglobulin heavy constant alpha 1 OS=Homo sapiens OX=9606 GN=IGHA1 PE=1 SV=2 | 255 | 20201117_fqg_milk_1ng_glyco_hcd30_etd30_compound15.18728.18728.4 | controllerType=0 controllerNumber=1 scan=18728 | 42.7264 |
| 15126:4 | 1 | R.LSLHRPALEDLLLGSEAN[+1460.529]LTC[+57.021]TLTGLR.D | HexNAc(4)Hex(4) | C[+57], N[+1461] | 1106.7906 | 4 | 4424.1405 | 4424.1271 | 0 | 3 | 127 | Specific | 986.97 | 923.57 | 923.57 | 16.95 | 31 | >sp\|P01876\|IGHA1_HUMAN Immunoglobulin heavy constant alpha 1 OS=Homo sapiens OX=9606 GN=IGHA1 PE=1 SV=2 | 255 | 20201117_fqg_milk_1ng_glyco_hcd30_etd30_compound15.18715.18715.4 | controllerType=0 controllerNumber=1 scan=18715 | 42.6781 |
| 15065:4 | 1 | R.LSLHRPALEDLLLGSEAN[+1864.634]LTC[+57.021]TLTGLR.D | HexNAc(2)Hex(9) | C[+57], N[+1865] | 1207.8161 | 4 | 4828.2425 | 4828.2324 | 0 | 2.1 | 127 | Specific | 1001.16 | 927.13 | 927.13 | 17.74 | 31 | >sp\|P01876\|IGHA1_HUMAN Immunoglobulin heavy constant alpha 1 OS=Homo sapiens OX=9606 GN=IGHA1 PE=1 SV=2 | 255 | 20201117_fqg_milk_1ng_glyco_hcd30_etd30_compound15.18650.18650.4 | controllerType=0 controllerNumber=1 scan=18650 | 42.4262 |
| 15113:4 | 1 | R.LSLHRPALEDLLLGSEAN[+1378.476]LTC[+57.021]TLTGLR.D | HexNAc(2)Hex(6) | C[+57], N[+1378] | 1086.2768 | 4 | 4342.0853 | 4342.074 | 0 | 2.6 | 127 | Specific | 1009.55 | 946.37 | 946.37 | 17.95 | 31 | >sp\|P01876\|IGHA1_HUMAN Immunoglobulin heavy constant alpha 1 OS=Homo sapiens OX=9606 GN=IGHA1 PE=1 SV=2 | 255 | 20201117_fqg_milk_1ng_glyco_hcd30_etd30_compound15.18701.18701.4 | controllerType=0 controllerNumber=1 scan=18701 | 42.6101 |
| 15083:4 | 1 | R.LSLHRPALEDLLLGSEAN[+1540.529]LTC[+57.021]TLTGLR.D | HexNAc(2)Hex(7) | C[+57], N[+1541] | 1126.79 | 4 | 4504.138 | 4504.1268 | 0 | 2.5 | 127 | Specific | 1021.43 | 942.75 | 942.75 | 17.74 | 31 | >sp\|P01876\|IGHA1_HUMAN Immunoglobulin heavy constant alpha 1 OS=Homo sapiens OX=9606 GN=IGHA1 PE=1 SV=2 | 255 | 20201117_fqg_milk_1ng_glyco_hcd30_etd30_compound15.18669.18669.4 | controllerType=0 controllerNumber=1 scan=18669 | 42.491 |
| 10018:3 | 1 | R.LAGKPTHVN[+1663.608]VSVVMAEVDGTC[+57.021]Y.- | HexNAc(5)Hex(4) | C[+57], N[+1664] | 1337.5914 | 3 | 4010.7595 | 4010.7502 | 0 | 2.3 | 332 | CRagged | 209.68 | 124.36 | 124.36 | 2.66 | 31 | >sp\|P01876\|IGHA1_HUMAN Immunoglobulin heavy constant alpha 1 OS=Homo sapiens OX=9606 GN=IGHA1 PE=1 SV=2 | 255 | 20201117_fqg_milk_1ng_glyco_hcd30_etd30_compound15.13320.13320.3 | controllerType=0 controllerNumber=1 scan=13320 | 30.547 |
| 11691:3 | 1 | R.LAGKPTHVN[+1751.624]VSVVMAEVDGTC[+57.021]Y.- | HexNAc(4)Hex(4)NeuAc(1) | C[+57], N[+1752] | 1366.931 | 3 | 4098.7783 | 4098.7663 | 0 | 2.9 | 332 | CRagged | 233.78 | 139.81 | 139.81 | 3.14 | 31 | >sp\|P01876\|IGHA1_HUMAN Immunoglobulin heavy constant alpha 1 OS=Homo sapiens OX=9606 GN=IGHA1 PE=1 SV=2 | 255 | 20201117_fqg_milk_1ng_glyco_hcd30_etd30_compound15.15075.15075.3 | controllerType=0 controllerNumber=1 scan=15075 | 33.9224 |
| 10081:3 | 1 | R.LAGKPTHVN[+1501.555]VSVVMAEVDGTC[+57.021]Y.- | HexNAc(5)Hex(3) | C[+57], N[+1502] | 1283.5742 | 3 | 3848.7079 | 3848.6974 | 0 | 2.7 | 332 | CRagged | 235.33 | 173.28 | 173.28 | 3.08 | 31 | >sp\|P01876\|IGHA1_HUMAN Immunoglobulin heavy constant alpha 1 OS=Homo sapiens OX=9606 GN=IGHA1 PE=1 SV=2 | 255 | 20201117_fqg_milk_1ng_glyco_hcd30_etd30_compound15.13387.13387.3 | controllerType=0 controllerNumber=1 scan=13387 | 30.7272 |
| 10062:3 | 1 | R.LAGKPTHVN[+1216.423]VSVVMAEVDGTC[+57.021]Y.- | HexNAc(2)Hex(5) | C[+57], N[+1216] | 1188.5287 | 3 | 3563.5717 | 3563.5649 | 0 | 1.9 | 332 | CRagged | 243.32 | 222.43 | 171.31 | 3.53 | 31 | >sp\|P01876\|IGHA1_HUMAN Immunoglobulin heavy constant alpha 1 OS=Homo sapiens OX=9606 GN=IGHA1 PE=1 SV=2 | 255 | 20201117_fqg_milk_1ng_glyco_hcd30_etd30_compound15.13367.13367.3 | controllerType=0 controllerNumber=1 scan=13367 | 30.6779 |
| 10009:3 | 1 | R.LAGKPTHVN[+1378.476]VSVVMAEVDGTC[+57.021]Y.- | HexNAc(2)Hex(6) | C[+57], N[+1378] | 1242.5477 | 3 | 3725.6284 | 3725.6178 | 0 | 2.9 | 332 | CRagged | 245.17 | 211.91 | 187.53 | 3.98 | 31 | >sp\|P01876\|IGHA1_HUMAN Immunoglobulin heavy constant alpha 1 OS=Homo sapiens OX=9606 GN=IGHA1 PE=1 SV=2 | 255 | 20201117_fqg_milk_1ng_glyco_hcd30_etd30_compound15.13311.13311.3 | controllerType=0 controllerNumber=1 scan=13311 | 30.5238 |
| 10061:3 | 1 | R.LAGKPTHVN[+1809.666]VSVVMAEVDGTC[+57.021]Y.- | HexNAc(5)Hex(4)Fuc(1) | C[+57], N[+1810] | 1386.2769 | 3 | 4156.8162 | 4156.8081 | 0 | 1.9 | 332 | CRagged | 254.39 | 147.32 | 147.32 | 3.1 | 31 | >sp\|P01876\|IGHA1_HUMAN Immunoglobulin heavy constant alpha 1 OS=Homo sapiens OX=9606 GN=IGHA1 PE=1 SV=2 | 255 | 20201117_fqg_milk_1ng_glyco_hcd30_etd30_compound15.13366.13366.3 | controllerType=0 controllerNumber=1 scan=13366 | 30.6712 |
| 13162:4 | 1 | R.LAGKPTHVN[+2553.910]VSVVMAEVDGTC[+57.021]Y.- | HexNAc(5)Hex(5)Fuc(1)NeuAc(2) | C[+57], N[+2554] | 1226.0219 | 4 | 4901.0657 | 4901.0518 | 0 | 2.8 | 332 | CRagged | 318.88 | 252.89 | 252.89 | 4.56 | 31 | >sp\|P01876\|IGHA1_HUMAN Immunoglobulin heavy constant alpha 1 OS=Homo sapiens OX=9606 GN=IGHA1 PE=1 SV=2 | 255 | 20201117_fqg_milk_1ng_glyco_hcd30_etd30_compound15.16610.16610.4 | controllerType=0 controllerNumber=1 scan=16610 | 37.3312 |
| 15409:5 | 2 | R.RTAGWNVPIGTLRPFLN[+2059.735]WTGPPEPIEAAVAR.F | HexNAc(4)Hex(5)Fuc(1)NeuAc(1) | N[+2060] | 1090.118 | 5 | 5446.5611 | 5446.547 | 0 | 2.6 | 140 | Specific | 500.62 | 485.91 | 485.91 | 6.04 | 34 | >sp\|P02788\|TRFL_HUMAN Lactotransferrin OS=Homo sapiens OX=9606 GN=LTF PE=1 SV=6 | 1270 | 20201117_fqg_milk_1ng_glyco_hcd30_etd30_compound15.19015.19015.5 | controllerType=0 controllerNumber=1 scan=19015 | 43.615 |
| 15402:4 | 2 | R.RTAGWNVPIGTLRPFLN[+2059.735]WTGPPEPIEAAVAR.F | HexNAc(4)Hex(5)Fuc(1)NeuAc(1) | N[+2060] | 1362.3955 | 4 | 5446.5602 | 5446.547 | 0 | 2.4 | 140 | Specific | 925.82 | 893.31 | 893.31 | 15.74 | 34 | >sp\|P02788\|TRFL_HUMAN Lactotransferrin OS=Homo sapiens OX=9606 GN=LTF PE=1 SV=6 | 1270 | 20201117_fqg_milk_1ng_glyco_hcd30_etd30_compound15.19008.19008.4 | controllerType=0 controllerNumber=1 scan=19008 | 43.5829 |
| 16366:3 | 2 | R.[+42.011]TAGWNVPIGTLRPFLN[+2018.708]WTGPPEPIEAAVAR.F | HexNAc(3)Hex(6)Fuc(1)NeuAc(1) | .[42], N[+2019] | 1764.4882 | 3 | 5291.45 | 5291.4299 | 0 | 3.8 | 141 | Specific | 224.59 | 181.98 | 181.98 | 2.84 | 34 | >sp\|P02788\|TRFL_HUMAN Lactotransferrin OS=Homo sapiens OX=9606 GN=LTF PE=1 SV=6 | 1270 | 20201117_fqg_milk_1ng_glyco_hcd30_etd30_compound15.20025.20025.3 | controllerType=0 controllerNumber=1 scan=20025 | 46.4517 |
| 17321:3 | 2 | R.TAGWNVPIGTLRPFLN[+2350.830]WTGPPEPIEAAVAR.F | HexNAc(4)Hex(5)Fuc(1)NeuAc(2) | N[+2351] | 1861.1877 | 3 | 5581.5486 | 5581.5413 | 0 | 1.3 | 141 | Specific | 342.86 | 260.23 | 260.23 | 4.98 | 34 | >sp\|P02788\|TRFL_HUMAN Lactotransferrin OS=Homo sapiens OX=9606 GN=LTF PE=1 SV=6 | 1270 | 20201117_fqg_milk_1ng_glyco_hcd30_etd30_compound15.21037.21037.3 | controllerType=0 controllerNumber=1 scan=21037 | 49.183 |
| 17336:5 | 2 | R.TAGWNVPIGTLRPFLN[+2350.830]WTGPPEPIEAAVAR.F | HexNAc(4)Hex(5)Fuc(1)NeuAc(2) | N[+2351] | 1117.1168 | 5 | 5581.555 | 5581.5413 | 0 | 2.5 | 141 | Specific | 352.49 | 349.98 | 349.98 | 4.21 | 34 | >sp\|P02788\|TRFL_HUMAN Lactotransferrin OS=Homo sapiens OX=9606 GN=LTF PE=1 SV=6 | 1270 | 20201117_fqg_milk_1ng_glyco_hcd30_etd30_compound15.21053.21053.5 | controllerType=0 controllerNumber=1 scan=21053 | 49.2171 |
| 15588:3 | 2 | R.TAGWNVPIGTLRPFLN[+1768.640]WTGPPEPIEAAVAR.F | HexNAc(4)Hex(5)Fuc(1) | N[+1769] | 1667.1294 | 3 | 4999.3735 | 4999.3504 | 0 | 4.6 | 141 | Specific | 456.06 | 370.77 | 370.77 | 5.93 | 34 | >sp\|P02788\|TRFL_HUMAN Lactotransferrin OS=Homo sapiens OX=9606 GN=LTF PE=1 SV=6 | 1270 | 20201117_fqg_milk_1ng_glyco_hcd30_etd30_compound15.19203.19203.3 | controllerType=0 controllerNumber=1 scan=19203 | 44.0921 |
| 16296:5 | 2 | R.TAGWNVPIGTLRPFLN[+2059.735]WTGPPEPIEAAVAR.F | HexNAc(4)Hex(5)Fuc(1)NeuAc(1) | N[+2060] | 1058.8999 | 5 | 5290.4704 | 5290.4459 | 0 | 4.6 | 141 | Specific | 517.32 | 502.64 | 502.64 | 5.97 | 34 | >sp\|P02788\|TRFL_HUMAN Lactotransferrin OS=Homo sapiens OX=9606 GN=LTF PE=1 SV=6 | 1270 | 20201117_fqg_milk_1ng_glyco_hcd30_etd30_compound15.19951.19951.5 | controllerType=0 controllerNumber=1 scan=19951 | 46.3015 |
| 17315:4 | 2 | R.TAGWNVPIGTLRPFLN[+2350.830]WTGPPEPIEAAVAR.F | HexNAc(4)Hex(5)Fuc(1)NeuAc(2) | N[+2351] | 1396.1433 | 4 | 5581.5514 | 5581.5413 | 0 | 1.8 | 141 | Specific | 682.86 | 662.54 | 662.54 | 10.66 | 34 | >sp\|P02788\|TRFL_HUMAN Lactotransferrin OS=Homo sapiens OX=9606 GN=LTF PE=1 SV=6 | 1270 | 20201117_fqg_milk_1ng_glyco_hcd30_etd30_compound15.21031.21031.4 | controllerType=0 controllerNumber=1 scan=21031 | 49.1598 |
| 15587:4 | 2 | R.TAGWNVPIGTLRPFLN[+1768.640]WTGPPEPIEAAVAR.F | HexNAc(4)Hex(5)Fuc(1) | N[+1769] | 1250.5986 | 4 | 4999.3727 | 4999.3504 | 0 | 4.5 | 141 | Specific | 870.24 | 862.03 | 862.03 | 14.11 | 34 | >sp\|P02788\|TRFL_HUMAN Lactotransferrin OS=Homo sapiens OX=9606 GN=LTF PE=1 SV=6 | 1270 | 20201117_fqg_milk_1ng_glyco_hcd30_etd30_compound15.19202.19202.4 | controllerType=0 controllerNumber=1 scan=19202 | 44.0885 |
| 14429:3 | 2 | R.TAGWNIPMGLLFN[+1913.677]QTGSC[+57.021]K.F | HexNAc(4)Hex(5)NeuAc(1) | C[+57], N[+1914] | 1336.9059 | 3 | 4008.7033 | 4008.687 | 0 | 4.1 | 485 | Specific | 225.05 | 158.67 | 158.67 | 2.89 | 34 | >sp\|P02788\|TRFL_HUMAN Lactotransferrin OS=Homo sapiens OX=9606 GN=LTF PE=1 SV=6 | 1270 | 20201117_fqg_milk_1ng_glyco_hcd30_etd30_compound15.17979.17979.3 | controllerType=0 controllerNumber=1 scan=17979 | 40.4888 |
| 14733:3 | 2 | R.TAGWNIPMGLLFN[+1460.529]QTGSC[+57.021]K.F | HexNAc(4)Hex(4) | C[+57], N[+1461] | 1185.8542 | 3 | 3555.548 | 3555.5387 | 0 | 2.6 | 485 | Specific | 244.93 | 168.67 | 168.67 | 3.49 | 34 | >sp\|P02788\|TRFL_HUMAN Lactotransferrin OS=Homo sapiens OX=9606 GN=LTF PE=1 SV=6 | 1270 | 20201117_fqg_milk_1ng_glyco_hcd30_etd30_compound15.18299.18299.3 | controllerType=0 controllerNumber=1 scan=18299 | 41.3862 |
| 15835:3 | 2 | R.TAGWNIPMGLLFN[+2205.793]QTGSC[+57.021]K.F | HexNAc(4)Hex(5)Fuc(2)NeuAc(1) | C[+57], N[+2206] | 1434.2763 | 3 | 4300.8144 | 4300.8028 | 0 | 2.7 | 485 | Specific | 304.98 | 224.12 | 224.12 | 5.22 | 34 | >sp\|P02788\|TRFL_HUMAN Lactotransferrin OS=Homo sapiens OX=9606 GN=LTF PE=1 SV=6 | 1270 | 20201117_fqg_milk_1ng_glyco_hcd30_etd30_compound15.19464.19464.3 | controllerType=0 controllerNumber=1 scan=19464 | 44.8547 |
| 17404:3 | 2 | R.TAGWNIPMGLLFN[+2350.830]QTGSC[+57.021]K.F | HexNAc(4)Hex(5)Fuc(1)NeuAc(2) | C[+57], N[+2351] | 1482.6241 | 3 | 4445.8577 | 4445.8403 | 0 | 3.9 | 485 | Specific | 330.09 | 201.87 | 201.87 | 4.44 | 34 | >sp\|P02788\|TRFL_HUMAN Lactotransferrin OS=Homo sapiens OX=9606 GN=LTF PE=1 SV=6 | 1270 | 20201117_fqg_milk_1ng_glyco_hcd30_etd30_compound15.21124.21124.3 | controllerType=0 controllerNumber=1 scan=21124 | 49.3943 |
| 14646:3 | 2 | R.TAGWNIPMGLLFN[+1622.582]QTGSC[+57.021]K.F | HexNAc(4)Hex(5) | C[+57], N[+1623] | 1239.871 | 3 | 3717.5985 | 3717.5916 | 0 | 1.9 | 485 | Specific | 341.63 | 275.51 | 223.83 | 5.24 | 34 | >sp\|P02788\|TRFL_HUMAN Lactotransferrin OS=Homo sapiens OX=9606 GN=LTF PE=1 SV=6 | 1270 | 20201117_fqg_milk_1ng_glyco_hcd30_etd30_compound15.18207.18207.3 | controllerType=0 controllerNumber=1 scan=18207 | 41.1104 |
| 15830:3 | 2 | R.TAGWNIPMGLLFN[+2059.735]QTGSC[+57.021]K.F | HexNAc(4)Hex(5)Fuc(1)NeuAc(1) | C[+57], N[+2060] | 1385.5905 | 3 | 4154.757 | 4154.7449 | 0 | 2.9 | 485 | Specific | 354.02 | 249.03 | 249.03 | 5.42 | 34 | >sp\|P02788\|TRFL_HUMAN Lactotransferrin OS=Homo sapiens OX=9606 GN=LTF PE=1 SV=6 | 1270 | 20201117_fqg_milk_1ng_glyco_hcd30_etd30_compound15.19459.19459.3 | controllerType=0 controllerNumber=1 scan=19459 | 44.8401 |
| 14486:3 | 2 | R.TAGWNIPMGLLFN[+1768.640]QTGSC[+57.021]K.F | HexNAc(4)Hex(5)Fuc(1) | C[+57], N[+1769] | 1288.558 | 3 | 3863.6596 | 3863.6495 | 0 | 2.6 | 485 | Specific | 356.06 | 272.33 | 272.33 | 5.24 | 34 | >sp\|P02788\|TRFL_HUMAN Lactotransferrin OS=Homo sapiens OX=9606 GN=LTF PE=1 SV=6 | 1270 | 20201117_fqg_milk_1ng_glyco_hcd30_etd30_compound15.18039.18039.3 | controllerType=0 controllerNumber=1 scan=18039 | 40.6254 |
| 14481:3 | 2 | R.TAGWNIPMGLLFN[+1768.640]QTGSC[+57.021]K.F | HexNAc(4)Hex(5)Fuc(1) | C[+57], N[+1769] | 1288.5569 | 3 | 3863.6563 | 3863.6495 | 0 | 1.8 | 485 | Specific | 382.34 | 297.59 | 297.59 | 4.27 | 34 | >sp\|P02788\|TRFL_HUMAN Lactotransferrin OS=Homo sapiens OX=9606 GN=LTF PE=1 SV=6 | 1270 | 20201117_fqg_milk_1ng_glyco_hcd30_etd30_compound15.18033.18033.3 | controllerType=0 controllerNumber=1 scan=18033 | 40.6105 |
| 14568:3 | 2 | R.TAGWNIPMGLLFN[+1606.587]QTGSC[+57.021]K.F | HexNAc(4)Hex(4)Fuc(1) | C[+57], N[+1607] | 1234.5408 | 3 | 3701.6079 | 3701.5966 | 0 | 3.1 | 485 | Specific | 382.37 | 309.87 | 309.87 | 5.46 | 34 | >sp\|P02788\|TRFL_HUMAN Lactotransferrin OS=Homo sapiens OX=9606 GN=LTF PE=1 SV=6 | 1270 | 20201117_fqg_milk_1ng_glyco_hcd30_etd30_compound15.18125.18125.3 | controllerType=0 controllerNumber=1 scan=18125 | 40.8827 |
| 14767:3 | 2 | R.TAGWNIPMGLLFN[+1216.423]QTGSC[+57.021]K.F | HexNAc(2)Hex(5) | C[+57], N[+1216] | 1104.4854 | 3 | 3311.4416 | 3311.4328 | 0 | 2.7 | 485 | Specific | 383.36 | 375.4 | 183.73 | 5.62 | 34 | >sp\|P02788\|TRFL_HUMAN Lactotransferrin OS=Homo sapiens OX=9606 GN=LTF PE=1 SV=6 | 1270 | 20201117_fqg_milk_1ng_glyco_hcd30_etd30_compound15.18335.18335.3 | controllerType=0 controllerNumber=1 scan=18335 | 41.4917 |
| 16003:3 | 2 | R.TAGWNIPMGLLFN[+1751.624]QTGSC[+57.021]K.F | HexNAc(4)Hex(4)NeuAc(1) | C[+57], N[+1752] | 1282.8864 | 3 | 3846.6447 | 3846.6341 | 0 | 2.7 | 485 | Specific | 410.68 | 246.94 | 246.94 | 5.73 | 34 | >sp\|P02788\|TRFL_HUMAN Lactotransferrin OS=Homo sapiens OX=9606 GN=LTF PE=1 SV=6 | 1270 | 20201117_fqg_milk_1ng_glyco_hcd30_etd30_compound15.19643.19643.3 | controllerType=0 controllerNumber=1 scan=19643 | 45.4464 |
| 15927:3 | 2 | R.TAGWNIPMGLLFN[+1913.677]QTGSC[+57.021]K.F | HexNAc(4)Hex(5)NeuAc(1) | C[+57], N[+1914] | 1336.9056 | 3 | 4008.7022 | 4008.687 | 0 | 3.8 | 485 | Specific | 420.44 | 271.01 | 271.01 | 5.75 | 34 | >sp\|P02788\|TRFL_HUMAN Lactotransferrin OS=Homo sapiens OX=9606 GN=LTF PE=1 SV=6 | 1270 | 20201117_fqg_milk_1ng_glyco_hcd30_etd30_compound15.19562.19562.3 | controllerType=0 controllerNumber=1 scan=19562 | 45.1778 |
| 15941:4 | 2 | R.TAGWNIPMGLLFN[+1913.677]QTGSC[+57.021]K.F | HexNAc(4)Hex(5)NeuAc(1) | C[+57], N[+1914] | 1002.9295 | 4 | 4008.6964 | 4008.687 | 0 | 2.3 | 485 | Specific | 455.36 | 400.11 | 400.11 | 5.88 | 34 | >sp\|P02788\|TRFL_HUMAN Lactotransferrin OS=Homo sapiens OX=9606 GN=LTF PE=1 SV=6 | 1270 | 20201117_fqg_milk_1ng_glyco_hcd30_etd30_compound15.19577.19577.4 | controllerType=0 controllerNumber=1 scan=19577 | 45.2263 |
| 14712:3 | 2 | R.TAGWNIPM[+15.995]GLLFN[+2059.735]QTGSC[+57.021]K.F | HexNAc(4)Hex(5)Fuc(1)NeuAc(1) | C[+57], M[+16], N[+2060] | 1390.9225 | 3 | 4170.7531 | 4170.7398 | 0 | 3.2 | 485 | Specific | 294.68 | 246.62 | 246.62 | 3.92 | 34 | >sp\|P02788\|TRFL_HUMAN Lactotransferrin OS=Homo sapiens OX=9606 GN=LTF PE=1 SV=6 | 1270 | 20201117_fqg_milk_1ng_glyco_hcd30_etd30_compound15.18277.18277.3 | controllerType=0 controllerNumber=1 scan=18277 | 41.3173 |
| 14676:3 | 2 | R.TAGWNIPM[+15.995]GLLFN[+2059.735]QTGSC[+57.021]K.F | HexNAc(4)Hex(5)Fuc(1)NeuAc(1) | C[+57], M[+16], N[+2060] | 1390.9211 | 3 | 4170.7487 | 4170.7398 | 0 | 2.1 | 485 | Specific | 314.66 | 233.58 | 233.58 | 5.24 | 34 | >sp\|P02788\|TRFL_HUMAN Lactotransferrin OS=Homo sapiens OX=9606 GN=LTF PE=1 SV=6 | 1270 | 20201117_fqg_milk_1ng_glyco_hcd30_etd30_compound15.18239.18239.3 | controllerType=0 controllerNumber=1 scan=18239 | 41.1999 |
| 14822:3 | 2 | R.TAGWNIPM[+15.995]GLLFN[+2059.735]QTGSC[+57.021]K.F | HexNAc(4)Hex(5)Fuc(1)NeuAc(1) | C[+57], M[+16], N[+2060] | 1390.9224 | 3 | 4170.7527 | 4170.7398 | 0 | 3.1 | 485 | Specific | 314.97 | 242.72 | 242.72 | 4.19 | 34 | >sp\|P02788\|TRFL_HUMAN Lactotransferrin OS=Homo sapiens OX=9606 GN=LTF PE=1 SV=6 | 1270 | 20201117_fqg_milk_1ng_glyco_hcd30_etd30_compound15.18393.18393.2 | controllerType=0 controllerNumber=1 scan=18393 | 41.6501 |
| 15276:4 | 2 | R.TAGWNIPMGLLFN[+1768.640]QTGSC[+57.021]KFDEYFSQSC[+57.021]APGSDPR.S | HexNAc(4)Hex(5)Fuc(1) | C[+57]*2, N[+1769] | 1427.6067 | 4 | 5707.4049 | 5707.3966 | 0 | 1.5 | 485 | Specific | 377.2 | 336.55 | 336.55 | 3.54 | 34 | >sp\|P02788\|TRFL_HUMAN Lactotransferrin OS=Homo sapiens OX=9606 GN=LTF PE=1 SV=6 | 1270 | 20201117_fqg_milk_1ng_glyco_hcd30_etd30_compound15.18874.18874.4 | controllerType=0 controllerNumber=1 scan=18874 | 43.1768 |
| 16007:4 | 2 | R.TAGWNIPMGLLFN[+2059.735]QTGSC[+57.021]KFDEYFSQSC[+57.021]APGSDPR.S | HexNAc(4)Hex(5)Fuc(1)NeuAc(1) | C[+57]*2, N[+2060] | 1500.3835 | 4 | 5998.5124 | 5998.492 | 0 | 3.4 | 485 | Specific | 706.13 | 649.59 | 649.59 | 10.72 | 34 | >sp\|P02788\|TRFL_HUMAN Lactotransferrin OS=Homo sapiens OX=9606 GN=LTF PE=1 SV=6 | 1270 | 20201117_fqg_milk_1ng_glyco_hcd30_etd30_compound15.19647.19647.4 | controllerType=0 controllerNumber=1 scan=19647 | 45.4633 |
| 16018:4 | 2 | R.TAGWNIPMGLLFN[+2059.735]QTGSC[+57.021]KFDEYFSQSC[+57.021]APGSDPR.S | HexNAc(4)Hex(5)Fuc(1)NeuAc(1) | C[+57]*2, N[+2060] | 1500.3835 | 4 | 5998.5124 | 5998.492 | 0 | 3.4 | 485 | Specific | 724.88 | 634.69 | 634.69 | 10.72 | 34 | >sp\|P02788\|TRFL_HUMAN Lactotransferrin OS=Homo sapiens OX=9606 GN=LTF PE=1 SV=6 | 1270 | 20201117_fqg_milk_1ng_glyco_hcd30_etd30_compound15.19659.19659.4 | controllerType=0 controllerNumber=1 scan=19659 | 45.5007 |
| 10154:3 | 2 | R.N[+2350.830]GSDC[+57.021]PDKFC[+57.021]LFQSETK.N | HexNAc(4)Hex(5)Fuc(1)NeuAc(2) | C[+57]*2, N[+2351] | 1461.9101 | 3 | 4383.7157 | 4383.7043 | 0 | 2.6 | 642 | Specific | 226.97 | 161.67 | 161.67 | 3.51 | 34 | >sp\|P02788\|TRFL_HUMAN Lactotransferrin OS=Homo sapiens OX=9606 GN=LTF PE=1 SV=6 | 1270 | 20201117_fqg_milk_1ng_glyco_hcd30_etd30_compound15.13463.13463.3 | controllerType=0 controllerNumber=1 scan=13463 | 30.9074 |
| 08063:3 | 2 | R.N[+2205.793]GSDC[+57.021]PDKFC[+57.021]LFQSETK.N | HexNAc(4)Hex(5)Fuc(2)NeuAc(1) | C[+57]*2, N[+2206] | 1413.561 | 3 | 4238.6684 | 4238.6667 | 0 | 0.4 | 642 | Specific | 234.24 | 118.7 | 118.7 | 3.64 | 34 | >sp\|P02788\|TRFL_HUMAN Lactotransferrin OS=Homo sapiens OX=9606 GN=LTF PE=1 SV=6 | 1270 | 20201117_fqg_milk_1ng_glyco_hcd30_etd30_compound15.11289.11289.3 | controllerType=0 controllerNumber=1 scan=11289 | 26.5205 |
| 08165:3 | 2 | R.N[+2059.735]GSDC[+57.021]PDKFC[+57.021]LFQSETK.N | HexNAc(4)Hex(5)Fuc(1)NeuAc(1) | C[+57]*2, N[+2060] | 1364.8764 | 3 | 4092.6146 | 4092.6088 | 0 | 1.4 | 642 | Specific | 317.92 | 222.55 | 222.55 | 4.62 | 34 | >sp\|P02788\|TRFL_HUMAN Lactotransferrin OS=Homo sapiens OX=9606 GN=LTF PE=1 SV=6 | 1270 | 20201117_fqg_milk_1ng_glyco_hcd30_etd30_compound15.11396.11396.3 | controllerType=0 controllerNumber=1 scan=11396 | 26.7313 |
| 08086:3 | 2 | R.N[+2205.793]GSDC[+57.021]PDKFC[+57.021]LFQSETK.N | HexNAc(4)Hex(5)Fuc(2)NeuAc(1) | C[+57]*2, N[+2206] | 1413.5641 | 3 | 4238.6779 | 4238.6667 | 0 | 2.6 | 642 | Specific | 335.47 | 226.59 | 226.59 | 4.34 | 34 | >sp\|P02788\|TRFL_HUMAN Lactotransferrin OS=Homo sapiens OX=9606 GN=LTF PE=1 SV=6 | 1270 | 20201117_fqg_milk_1ng_glyco_hcd30_etd30_compound15.11314.11314.3 | controllerType=0 controllerNumber=1 scan=11314 | 26.5687 |
| 16459:3 | 5 | R.GLSFDVSLEVSQGPGLLN[+2350.830]DTK.V | HexNAc(4)Hex(5)Fuc(1)NeuAc(2) | N[+2351] | 1509.6566 | 3 | 4526.9551 | 4526.9436 | 0 | 2.6 | 118 | Specific | 485.36 | 279.37 | 279.37 | 5.91 | 16 | >sp\|P01833\|PIGR_HUMAN Polymeric immunoglobulin receptor OS=Homo sapiens OX=9606 GN=PIGR PE=1 SV=4 | 902 | 20201117_fqg_milk_1ng_glyco_hcd30_etd30_compound15.20128.20128.3 | controllerType=0 controllerNumber=1 scan=20128 | 46.6113 |
| 16496:3 | 5 | R.GLSFDVSLEVSQGPGLLN[+2350.830]DTK.V | HexNAc(4)Hex(5)Fuc(1)NeuAc(2) | N[+2351] | 1509.6541 | 3 | 4526.9478 | 4526.9436 | 0 | 0.9 | 118 | Specific | 499.42 | 294.24 | 294.24 | 6.18 | 16 | >sp\|P01833\|PIGR_HUMAN Polymeric immunoglobulin receptor OS=Homo sapiens OX=9606 GN=PIGR PE=1 SV=4 | 902 | 20201117_fqg_milk_1ng_glyco_hcd30_etd30_compound15.20168.20168.3 | controllerType=0 controllerNumber=1 scan=20168 | 46.7272 |
| 19117:4 | 5 | R.LSLLEEPGN[+2350.830]GTFTVILNQLTSR.D | HexNAc(4)Hex(5)Fuc(1)NeuAc(2) | N[+2351] | 1189.0431 | 4 | 4753.1507 | 4753.1229 | 0 | 5.8 | 413 | Specific | 489.18 | 422.68 | 422.68 | 5.96 | 16 | >sp\|P01833\|PIGR_HUMAN Polymeric immunoglobulin receptor OS=Homo sapiens OX=9606 GN=PIGR PE=1 SV=4 | 902 | 20201117_fqg_milk_1ng_glyco_hcd30_etd30_compound15.22923.22923.4 | controllerType=0 controllerNumber=1 scan=22923 | 53.3324 |
| 17203:4 | 5 | R.LSLLEEPGN[+2205.793]GTFTVILNQLTSR.D | HexNAc(4)Hex(5)Fuc(2)NeuAc(1) | N[+2206] | 1152.7796 | 4 | 4608.0965 | 4608.0854 | 0 | 2.4 | 413 | Specific | 556.76 | 478.75 | 478.75 | 6.53 | 16 | >sp\|P01833\|PIGR_HUMAN Polymeric immunoglobulin receptor OS=Homo sapiens OX=9606 GN=PIGR PE=1 SV=4 | 902 | 20201117_fqg_milk_1ng_glyco_hcd30_etd30_compound15.20913.20913.4 | controllerType=0 controllerNumber=1 scan=20913 | 48.8382 |
| 19116:3 | 5 | R.LSLLEEPGN[+2350.830]GTFTVILNQLTSR.D | HexNAc(4)Hex(5)Fuc(1)NeuAc(2) | N[+2351] | 1585.0532 | 3 | 4753.1449 | 4753.1229 | 0 | 4.6 | 413 | Specific | 610.17 | 422.56 | 422.56 | 8.97 | 16 | >sp\|P01833\|PIGR_HUMAN Polymeric immunoglobulin receptor OS=Homo sapiens OX=9606 GN=PIGR PE=1 SV=4 | 902 | 20201117_fqg_milk_1ng_glyco_hcd30_etd30_compound15.22922.22922.3 | controllerType=0 controllerNumber=1 scan=22922 | 53.329 |
| 17175:3 | 5 | R.LSLLEEPGN[+2205.793]GTFTVILNQLTSR.D | HexNAc(4)Hex(5)Fuc(2)NeuAc(1) | N[+2206] | 1536.7045 | 3 | 4608.099 | 4608.0854 | 0 | 3 | 413 | Specific | 621.7 | 497.56 | 497.56 | 9.73 | 16 | >sp\|P01833\|PIGR_HUMAN Polymeric immunoglobulin receptor OS=Homo sapiens OX=9606 GN=PIGR PE=1 SV=4 | 902 | 20201117_fqg_milk_1ng_glyco_hcd30_etd30_compound15.20883.20883.3 | controllerType=0 controllerNumber=1 scan=20883 | 48.7276 |
| 17217:3 | 5 | R.LSLLEEPGN[+2059.735]GTFTVILNQLTSR.D | HexNAc(4)Hex(5)Fuc(1)NeuAc(1) | N[+2060] | 1488.0186 | 3 | 4462.0413 | 4462.0275 | 0 | 3.1 | 413 | Specific | 648.9 | 509.87 | 509.87 | 9.73 | 16 | >sp\|P01833\|PIGR_HUMAN Polymeric immunoglobulin receptor OS=Homo sapiens OX=9606 GN=PIGR PE=1 SV=4 | 902 | 20201117_fqg_milk_1ng_glyco_hcd30_etd30_compound15.20928.20928.3 | controllerType=0 controllerNumber=1 scan=20928 | 48.8966 |
| 13899:4 | 5 | K.IIEGEPNLKVPGN[+2205.793]VTAVLGETLK.V | HexNAc(4)Hex(5)Fuc(2)NeuAc(1) | N[+2206] | 1150.0419 | 4 | 4597.1458 | 4597.1422 | 0 | 0.8 | 457 | Specific | 256.77 | 247.51 | 247.51 | 3.48 | 16 | >sp\|P01833\|PIGR_HUMAN Polymeric immunoglobulin receptor OS=Homo sapiens OX=9606 GN=PIGR PE=1 SV=4 | 902 | 20201117_fqg_milk_1ng_glyco_hcd30_etd30_compound15.17419.17419.2 | controllerType=0 controllerNumber=1 scan=17419 | 39.2253 |
| 13907:4 | 5 | K.IIEGEPNLKVPGN[+2204.772]VTAVLGETLK.V | HexNAc(4)Hex(5)NeuAc(2) | N[+2205] | 1149.7926 | 4 | 4596.1488 | 4596.1218 | 0 | 5.9 | 457 | Specific | 395.13 | 392.95 | 392.95 | 4.41 | 16 | >sp\|P01833\|PIGR_HUMAN Polymeric immunoglobulin receptor OS=Homo sapiens OX=9606 GN=PIGR PE=1 SV=4 | 902 | 20201117_fqg_milk_1ng_glyco_hcd30_etd30_compound15.17428.17428.4 | controllerType=0 controllerNumber=1 scan=17428 | 39.2519 |
| 13926:3 | 5 | K.IIEGEPNLKVPGN[+2205.793]VTAVLGETLK.V | HexNAc(4)Hex(5)Fuc(2)NeuAc(1) | N[+2206] | 1533.0562 | 3 | 4597.1541 | 4597.1422 | 0 | 2.6 | 457 | Specific | 396.88 | 301.48 | 301.48 | 3.73 | 16 | >sp\|P01833\|PIGR_HUMAN Polymeric immunoglobulin receptor OS=Homo sapiens OX=9606 GN=PIGR PE=1 SV=4 | 902 | 20201117_fqg_milk_1ng_glyco_hcd30_etd30_compound15.17448.17448.3 | controllerType=0 controllerNumber=1 scan=17448 | 39.2993 |
| 13892:3 | 5 | K.IIEGEPNLKVPGN[+2205.793]VTAVLGETLK.V | HexNAc(4)Hex(5)Fuc(2)NeuAc(1) | N[+2206] | 1533.0561 | 3 | 4597.1537 | 4597.1422 | 0 | 2.5 | 457 | Specific | 412.28 | 327.63 | 327.63 | 3.9 | 16 | >sp\|P01833\|PIGR_HUMAN Polymeric immunoglobulin receptor OS=Homo sapiens OX=9606 GN=PIGR PE=1 SV=4 | 902 | 20201117_fqg_milk_1ng_glyco_hcd30_etd30_compound15.17411.17411.3 | controllerType=0 controllerNumber=1 scan=17411 | 39.2099 |
| 13531:4 | 5 | K.IIEGEPNLKVPGN[+2205.793]VTAVLGETLKVPC[+57.021]HFPC[+57.021]K.F | HexNAc(4)Hex(5)Fuc(2)NeuAc(1) | C[+57]*2, N[+2206] | 1406.4077 | 4 | 5622.609 | 5622.5997 | 0 | 1.7 | 457 | Specific | 313.27 | 313.27 | 313.27 | 3.07 | 16 | >sp\|P01833\|PIGR_HUMAN Polymeric immunoglobulin receptor OS=Homo sapiens OX=9606 GN=PIGR PE=1 SV=4 | 902 | 20201117_fqg_milk_1ng_glyco_hcd30_etd30_compound15.16998.16998.4 | controllerType=0 controllerNumber=1 scan=16998 | 38.3051 |
| 13510:5 | 5 | K.IIEGEPNLKVPGN[+2205.793]VTAVLGETLKVPC[+57.021]HFPC[+57.021]K.F | HexNAc(4)Hex(5)Fuc(2)NeuAc(1) | C[+57]*2, N[+2206] | 1125.3273 | 5 | 5622.6073 | 5622.5997 | 0 | 1.3 | 457 | Specific | 518.71 | 518.71 | 518.71 | 5.71 | 16 | >sp\|P01833\|PIGR_HUMAN Polymeric immunoglobulin receptor OS=Homo sapiens OX=9606 GN=PIGR PE=1 SV=4 | 902 | 20201117_fqg_milk_1ng_glyco_hcd30_etd30_compound15.16976.16976.5 | controllerType=0 controllerNumber=1 scan=16976 | 38.2342 |
| 13524:5 | 5 | K.IIEGEPNLKVPGN[+2205.793]VTAVLGETLKVPC[+57.021]HFPC[+57.021]K.F | HexNAc(4)Hex(5)Fuc(2)NeuAc(1) | C[+57]*2, N[+2206] | 1125.3282 | 5 | 5622.6121 | 5622.5997 | 0 | 2.2 | 457 | Specific | 633.47 | 633.47 | 633.47 | 7.53 | 16 | >sp\|P01833\|PIGR_HUMAN Polymeric immunoglobulin receptor OS=Homo sapiens OX=9606 GN=PIGR PE=1 SV=4 | 902 | 20201117_fqg_milk_1ng_glyco_hcd30_etd30_compound15.16991.16991.5 | controllerType=0 controllerNumber=1 scan=16991 | 38.2819 |
| 11930:3 | 5 | K.VPGN[+2205.793]VTAVLGETLK.V | HexNAc(4)Hex(5)Fuc(2)NeuAc(1) | N[+2206] | 1201.8725 | 3 | 3603.6029 | 3603.5927 | 0 | 2.8 | 466 | Specific | 294.83 | 204.42 | 204.42 | 4.52 | 16 | >sp\|P01833\|PIGR_HUMAN Polymeric immunoglobulin receptor OS=Homo sapiens OX=9606 GN=PIGR PE=1 SV=4 | 902 | 20201117_fqg_milk_1ng_glyco_hcd30_etd30_compound15.15325.15325.3 | controllerType=0 controllerNumber=1 scan=15325 | 34.5055 |
| 04975:3 | 5 | K.WN[+2059.735]NTGC[+57.021]QALPSQDEGPSK.A | HexNAc(4)Hex(5)Fuc(1)NeuAc(1) | C[+57], N[+2060] | 1350.2114 | 3 | 4048.6195 | 4048.6116 | 0 | 2 | 498 | Specific | 452.95 | 322.37 | 322.37 | 5.73 | 16 | >sp\|P01833\|PIGR_HUMAN Polymeric immunoglobulin receptor OS=Homo sapiens OX=9606 GN=PIGR PE=1 SV=4 | 902 | 20201117_fqg_milk_1ng_glyco_hcd30_etd30_compound15.8068.8068.3 | controllerType=0 controllerNumber=1 scan=8068 | 20.6119 |
| 05060:3 | 5 | K.WN[+1913.677]NTGC[+57.021]QALPSQDEGPSK.A | HexNAc(4)Hex(5)NeuAc(1) | C[+57], N[+1914] | 1301.5242 | 3 | 3902.5581 | 3902.5537 | 0 | 1.1 | 498 | Specific | 518.46 | 402.68 | 402.68 | 7.54 | 16 | >sp\|P01833\|PIGR_HUMAN Polymeric immunoglobulin receptor OS=Homo sapiens OX=9606 GN=PIGR PE=1 SV=4 | 902 | 20201117_fqg_milk_1ng_glyco_hcd30_etd30_compound15.8156.8156.3 | controllerType=0 controllerNumber=1 scan=8156 | 20.7729 |
| 20086:3 | 7 | R.QLAHQSN[+2204.772]STNIFFSPVSIATAFAMLSLGTK.A | HexNAc(4)Hex(5)NeuAc(2) | N[+2205] | 1796.1421 | 3 | 5386.4116 | 5386.4075 | 0 | 0.8 | 64 | Specific | 341.51 | 237.07 | 237.07 | 5.87 | 7 | >sp\|P01009\|A1AT_HUMAN Alpha-1-antitrypsin OS=Homo sapiens OX=9606 GN=SERPINA1 PE=1 SV=3 | 642 | 20201117_fqg_milk_1ng_glyco_hcd30_etd30_compound15.23941.23941.3 | controllerType=0 controllerNumber=1 scan=23941 | 55.9156 |
| 20083:4 | 7 | R.QLAHQSN[+2204.772]STNIFFSPVSIATAFAMLSLGTK.A | HexNAc(4)Hex(5)NeuAc(2) | N[+2205] | 1347.3571 | 4 | 5386.4064 | 5386.4075 | 0 | -0.2 | 64 | Specific | 462.5 | 377.51 | 345.11 | 5.28 | 7 | >sp\|P01009\|A1AT_HUMAN Alpha-1-antitrypsin OS=Homo sapiens OX=9606 GN=SERPINA1 PE=1 SV=3 | 642 | 20201117_fqg_milk_1ng_glyco_hcd30_etd30_compound15.23938.23938.4 | controllerType=0 controllerNumber=1 scan=23938 | 55.9077 |
| 20054:4 | 7 | R.QLAHQSN[+2204.772]STNIFFSPVSIATAFAMLSLGTK.A | HexNAc(4)Hex(5)NeuAc(2) | N[+2205] | 1347.3582 | 4 | 5386.4108 | 5386.4075 | 0 | 0.6 | 64 | Specific | 607.83 | 488.07 | 357.87 | 9.92 | 7 | >sp\|P01009\|A1AT_HUMAN Alpha-1-antitrypsin OS=Homo sapiens OX=9606 GN=SERPINA1 PE=1 SV=3 | 642 | 20201117_fqg_milk_1ng_glyco_hcd30_etd30_compound15.23907.23907.4 | controllerType=0 controllerNumber=1 scan=23907 | 55.8129 |
| 17005:3 | 7 | K.YLGN[+2204.772]ATAIFFLPDEGK.L | HexNAc(4)Hex(5)NeuAc(2) | N[+2205] | 1320.8963 | 3 | 3960.6743 | 3960.6677 | 0 | 1.7 | 268 | Specific | 288.62 | 198.66 | 198.66 | 5.01 | 7 | >sp\|P01009\|A1AT_HUMAN Alpha-1-antitrypsin OS=Homo sapiens OX=9606 GN=SERPINA1 PE=1 SV=3 | 642 | 20201117_fqg_milk_1ng_glyco_hcd30_etd30_compound15.20704.20704.3 | controllerType=0 controllerNumber=1 scan=20704 | 48.2217 |
| 18442:4 | 8 | K.VFPLSLDSTPQDGNVVVAC[+57.021]LVQGFFPQEPLSVTWSESGQN[+1647.613]VTAI | HexNAc(5)Hex(3)Fuc(1) | C[+57], N[+1648] | 1606.7489 | 4 | 6423.9738 | 6423.9746 | 0 | -0.1 | 8 | Specific | 269.66 | 269.66 | 269.66 | 3.9 | 24 | >sp\|P01877\|IGHA2_HUMAN Immunoglobulin heavy constant alpha 2 OS=Homo sapiens OX=9606 GN=IGHA2 PE=1 SV=4 | 576 | 20201117_fqg_milk_1ng_glyco_hcd30_etd30_compound15.22209.22209.4 | controllerType=0 controllerNumber=1 scan=22209 | 51.8353 |
| 01706:3 | 8 | K.HYTN[+2262.814]SSQDVTVPC[+57.021]R.V | HexNAc(5)Hex(5)Fuc(1)NeuAc(1) | C[+57], N[+2263] | 1309.5294 | 3 | 3926.5735 | 3926.5636 | 0 | 2.5 | 89 | Specific | 252.77 | 140.28 | 140.28 | 3.94 | 24 | >sp\|P01877\|IGHA2_HUMAN Immunoglobulin heavy constant alpha 2 OS=Homo sapiens OX=9606 GN=IGHA2 PE=1 SV=4 | 576 | 20201117_fqg_milk_1ng_glyco_hcd30_etd30_compound15.4670.4670.3 | controllerType=0 controllerNumber=1 scan=4670 | 14.4633 |
| 00933:3 | 8 | K.HYTN[+1647.613]SSQDVTVPC[+57.021]R.V | HexNAc(5)Hex(3)Fuc(1) | C[+57], N[+1648] | 1104.4611 | 3 | 3311.3687 | 3311.3625 | 0 | 1.9 | 89 | Specific | 269.12 | 252.11 | 252.11 | 3.4 | 24 | >sp\|P01877\|IGHA2_HUMAN Immunoglobulin heavy constant alpha 2 OS=Homo sapiens OX=9606 GN=IGHA2 PE=1 SV=4 | 576 | 20201117_fqg_milk_1ng_glyco_hcd30_etd30_compound15.3867.3867.2 | controllerType=0 controllerNumber=1 scan=3867 | 12.9779 |
| 00896:3 | 8 | K.HYTN[+1647.613]SSQDVTVPC[+57.021]R.V | HexNAc(5)Hex(3)Fuc(1) | C[+57], N[+1648] | 1104.4622 | 3 | 3311.372 | 3311.3625 | 0 | 2.9 | 89 | Specific | 490.38 | 422.23 | 422.23 | 6.86 | 24 | >sp\|P01877\|IGHA2_HUMAN Immunoglobulin heavy constant alpha 2 OS=Homo sapiens OX=9606 GN=IGHA2 PE=1 SV=4 | 576 | 20201117_fqg_milk_1ng_glyco_hcd30_etd30_compound15.3828.3828.3 | controllerType=0 controllerNumber=1 scan=3828 | 12.8798 |
| 00886:3 | 8 | K.HYTN[+1809.666]SSQDVTVPC[+57.021]R.V | HexNAc(5)Hex(4)Fuc(1) | C[+57], N[+1810] | 1158.4794 | 3 | 3473.4237 | 3473.4154 | 0 | 2.4 | 89 | Specific | 507.78 | 430.45 | 430.45 | 6.66 | 24 | >sp\|P01877\|IGHA2_HUMAN Immunoglobulin heavy constant alpha 2 OS=Homo sapiens OX=9606 GN=IGHA2 PE=1 SV=4 | 576 | 20201117_fqg_milk_1ng_glyco_hcd30_etd30_compound15.3818.3818.3 | controllerType=0 controllerNumber=1 scan=3818 | 12.8567 |
| 02415:2 | 8 | K.TPLTAN[+1606.587]ITK.S | HexNAc(4)Hex(4)Fuc(1) | N[+1607] | 1283.0789 | 2 | 2565.1505 | 2565.1435 | 0 | 2.8 | 200 | Specific | 206.39 | 63.86 | 63.86 | 2.56 | 24 | >sp\|P01877\|IGHA2_HUMAN Immunoglobulin heavy constant alpha 2 OS=Homo sapiens OX=9606 GN=IGHA2 PE=1 SV=4 | 576 | 20201117_fqg_milk_1ng_glyco_hcd30_etd30_compound15.5412.5412.2 | controllerType=0 controllerNumber=1 scan=5412 | 15.8143 |
| 02530:3 | 8 | K.TPLTAN[+1647.613]ITK.S | HexNAc(5)Hex(3)Fuc(1) | N[+1648] | 869.3961 | 3 | 2606.1739 | 2606.17 | 0 | 1.5 | 200 | Specific | 283.65 | 121.68 | 121.68 | 2.65 | 24 | >sp\|P01877\|IGHA2_HUMAN Immunoglobulin heavy constant alpha 2 OS=Homo sapiens OX=9606 GN=IGHA2 PE=1 SV=4 | 576 | 20201117_fqg_milk_1ng_glyco_hcd30_etd30_compound15.5532.5532.3 | controllerType=0 controllerNumber=1 scan=5532 | 16.0963 |
| 02518:2 | 8 | K.TPLTAN[+1647.613]ITK.S | HexNAc(5)Hex(3)Fuc(1) | N[+1648] | 1303.5906 | 2 | 2606.174 | 2606.17 | 0 | 1.5 | 200 | Specific | 298.54 | 116.77 | 116.77 | 2.66 | 24 | >sp\|P01877\|IGHA2_HUMAN Immunoglobulin heavy constant alpha 2 OS=Homo sapiens OX=9606 GN=IGHA2 PE=1 SV=4 | 576 | 20201117_fqg_milk_1ng_glyco_hcd30_etd30_compound15.5520.5520.2 | controllerType=0 controllerNumber=1 scan=5520 | 16.061 |
| 02475:2 | 8 | K.TPLTAN[+1809.666]ITK.S | HexNAc(5)Hex(4)Fuc(1) | N[+1810] | 1384.6197 | 2 | 2768.2321 | 2768.2228 | 0 | 3.3 | 200 | Specific | 304.66 | 122.26 | 122.26 | 2.74 | 24 | >sp\|P01877\|IGHA2_HUMAN Immunoglobulin heavy constant alpha 2 OS=Homo sapiens OX=9606 GN=IGHA2 PE=1 SV=4 | 576 | 20201117_fqg_milk_1ng_glyco_hcd30_etd30_compound15.5475.5475.2 | controllerType=0 controllerNumber=1 scan=5475 | 15.9534 |
| 02476:3 | 8 | K.TPLTAN[+1809.666]ITK.S | HexNAc(5)Hex(4)Fuc(1) | N[+1810] | 923.4155 | 3 | 2768.2319 | 2768.2228 | 0 | 3.3 | 200 | Specific | 317.51 | 135.95 | 135.95 | 2.76 | 24 | >sp\|P01877\|IGHA2_HUMAN Immunoglobulin heavy constant alpha 2 OS=Homo sapiens OX=9606 GN=IGHA2 PE=1 SV=4 | 576 | 20201117_fqg_milk_1ng_glyco_hcd30_etd30_compound15.5476.5476.3 | controllerType=0 controllerNumber=1 scan=5476 | 15.9616 |
| 02527:2 | 8 | K.TPLTAN[+1501.555]ITK.S | HexNAc(5)Hex(3) | N[+1502] | 1230.5621 | 2 | 2460.1168 | 2460.1121 | 0 | 1.9 | 200 | Specific | 318.13 | 88.34 | 88.34 | 2.49 | 24 | >sp\|P01877\|IGHA2_HUMAN Immunoglobulin heavy constant alpha 2 OS=Homo sapiens OX=9606 GN=IGHA2 PE=1 SV=4 | 576 | 20201117_fqg_milk_1ng_glyco_hcd30_etd30_compound15.5529.5529.2 | controllerType=0 controllerNumber=1 scan=5529 | 16.084 |
| 02479:2 | 8 | K.TPLTAN[+1444.534]ITK.S | HexNAc(4)Hex(3)Fuc(1) | N[+1445] | 1202.0524 | 2 | 2403.0975 | 2403.0906 | 0 | 2.9 | 200 | Specific | 318.79 | 91.79 | 91.79 | 2.49 | 24 | >sp\|P01877\|IGHA2_HUMAN Immunoglobulin heavy constant alpha 2 OS=Homo sapiens OX=9606 GN=IGHA2 PE=1 SV=4 | 576 | 20201117_fqg_milk_1ng_glyco_hcd30_etd30_compound15.5479.5479.2 | controllerType=0 controllerNumber=1 scan=5479 | 15.9723 |

Table S4. Cont.

| Query #:z | Protein Rank | Glycopeptide | Glycan | Modification Type(s) | Observed m/z | z | Observed (M+H) | Calc. mass (M+H) | Off-by-x error | Mass error (ppm) | StartPosition | Cleavage | Score | Delta | Delta Mod | \|Log Prob\| | # of unique peptides | Protein Name | Protein DB number | Comment | Scan # | Scan Time |
|---|---|---|---|---|---|---|---|---|---|---|---|---|---|---|---|---|---|---|---|---|---|---|
| 09205:3 | 8 | R.MAGKPTHIN[+1501.555]VSVVMAEADGTC[+57.021]Y.- | HexNAc(5)Hex(3) | C[+57], N[+1502] | 1284.8864 | 3 | 3852.6447 | 3852.6382 | 0 | 1.7 | 319 | CRagged | 209.88 | 152.77 | 152.77 | 2.82 | 24 | >sp\|P01877\|IGHA2_HUMAN Immunoglobulin heavy constant alpha 2 OS=Homo sapiens OX=9606 GN=IGHA2 PE=1 SV=4 | 576 | 20201117_fqg_milk_1ng_glyco_hcd30_etd30_compound15.12476.12476.3 | controllerType=0 controllerNumber=1 scan=12476 | 28.8735 |
| 09078:3 | 8 | R.MAGKPTHIN[+1702.581]VSVVMAEADGTC[+57.021]Y.- | HexNAc(2)Hex(8) | C[+57], N[+1703] | 1351.8962 | 3 | 4053.674 | 4053.6642 | 0 | 2.4 | 319 | CRagged | 237.75 | 237.75 | 181.01 | 3.81 | 24 | >sp\|P01877\|IGHA2_HUMAN Immunoglobulin heavy constant alpha 2 OS=Homo sapiens OX=9606 GN=IGHA2 PE=1 SV=4 | 576 | 20201117_fqg_milk_1ng_glyco_hcd30_etd30_compound15.12343.12343.3 | controllerType=0 controllerNumber=1 scan=12343 | 28.5263 |
| 09127:3 | 8 | R.MAGKPTHIN[+1378.476]VSVVMAEADGTC[+57.021]Y.- | HexNAc(2)Hex(6) | C[+57], N[+1378] | 1243.8609 | 3 | 3729.5681 | 3729.5585 | 0 | 2.6 | 319 | CRagged | 286.38 | 219.22 | 211.31 | 4.21 | 24 | >sp\|P01877\|IGHA2_HUMAN Immunoglobulin heavy constant alpha 2 OS=Homo sapiens OX=9606 GN=IGHA2 PE=1 SV=4 | 576 | 20201117_fqg_milk_1ng_glyco_hcd30_etd30_compound15.12394.12394.3 | controllerType=0 controllerNumber=1 scan=12394 | 28.6592 |
| 09098:3 | 8 | R.MAGKPTHIN[+1540.529]VSVVMAEADGTC[+57.021]Y.- | HexNAc(2)Hex(7) | C[+57], N[+1541] | 1297.8771 | 3 | 3891.6168 | 3891.6114 | 0 | 1.4 | 319 | CRagged | 302.9 | 299.13 | 177.68 | 4.72 | 24 | >sp\|P01877\|IGHA2_HUMAN Immunoglobulin heavy constant alpha 2 OS=Homo sapiens OX=9606 GN=IGHA2 PE=1 SV=4 | 576 | 20201117_fqg_milk_1ng_glyco_hcd30_etd30_compound15.12364.12364.3 | controllerType=0 controllerNumber=1 scan=12364 | 28.5789 |
| 09167:3 | 8 | R.MAGKPTHIN[+1216.423]VSVVMAEADGTC[+57.021]Y.- | HexNAc(2)Hex(5) | C[+57], N[+1216] | 1189.8433 | 3 | 3567.5154 | 3567.5057 | 0 | 2.7 | 319 | CRagged | 328.57 | 277.64 | 223.97 | 4.8 | 24 | >sp\|P01877\|IGHA2_HUMAN Immunoglobulin heavy constant alpha 2 OS=Homo sapiens OX=9606 GN=IGHA2 PE=1 SV=4 | 576 | 20201117_fqg_milk_1ng_glyco_hcd30_etd30_compound15.12436.12436.3 | controllerType=0 controllerNumber=1 scan=12436 | 28.786 |
| 04974:3 | 15 | R.LSPN[+2205.793]ASAEHLELR.W | HexNAc(4)Hex(5)Fuc(2)NeuAc(1) | N[+2206] | 1214.8549 | 3 | 3642.5502 | 3642.5421 | 0 | 2.2 | 52 | Specific | 397.2 | 285.97 | 285.97 | 4.54 | 9 | >sp\|Q13410\|BT1A1_HUMAN Butyrophilin subfamily 1 member A1 OS=Homo sapiens OX=9606 GN=BTN1A1 PE=1 SV=3 | 179 | 20201117_fqg_milk_1ng_glyco_hcd30_etd30_compound15.8067.8067.3 | controllerType=0 controllerNumber=1 scan=8067 | 20.6065 |
| 06863:3 | 15 | R.LSPN[+2350.830]ASAEHLELR.W | HexNAc(4)Hex(5)Fuc(1)NeuAc(2) | N[+2351] | 1263.202 | 3 | 3787.5913 | 3787.5796 | 0 | 3.1 | 52 | Specific | 414.04 | 261.86 | 261.86 | 5.12 | 9 | >sp\|Q13410\|BT1A1_HUMAN Butyrophilin subfamily 1 member A1 OS=Homo sapiens OX=9606 GN=BTN1A1 PE=1 SV=3 | 179 | 20201117_fqg_milk_1ng_glyco_hcd30_etd30_compound15.10027.10027.3 | controllerType=0 controllerNumber=1 scan=10027 | 24.0711 |
| 15092:3 | 15 | K.N[+1913.677]VSC[+57.021]YIQNLLLGQEK.K | HexNAc(4)Hex(5)NeuAc(1) | C[+57], N[+1914] | 1231.5362 | 3 | 3692.594 | 3692.5876 | 0 | 1.7 | 215 | Specific | 312.58 | 170.45 | 170.45 | 4.94 | 9 | >sp\|Q13410\|BT1A1_HUMAN Butyrophilin subfamily 1 member A1 OS=Homo sapiens OX=9606 GN=BTN1A1 PE=1 SV=3 | 179 | 20201117_fqg_milk_1ng_glyco_hcd30_etd30_compound15.18678.18678.3 | controllerType=0 controllerNumber=1 scan=18678 | 42.5122 |
| 02158:4 | 19 | K.QTDEIKDTRN[+2059.735]ESTQNC[+57.021]VVAEPEK.M | HexNAc(4)Hex(5)Fuc(1)NeuAc(1) | C[+57], N[+2060] | 1188.5048 | 4 | 4750.9974 | 4750.9876 | 0 | 2.1 | 60 | Specific | 431.91 | 340.37 | 340.37 | 5.04 | 5 | >sp\|P47710\|CASA1_HUMAN Alpha-S1-casein OS=Homo sapiens OX=9606 GN=CSN1S1 PE=1 SV=1 | 989 | 20201117_fqg_milk_1ng_glyco_hcd30_etd30_compound15.5139.5139.4 | controllerType=0 controllerNumber=1 scan=5139 | 15.3124 |
| 02134:4 | 19 | K.QTDEIKDTRN[+2059.735]ESTQNC[+57.021]VVAEPEK.M | HexNAc(4)Hex(5)Fuc(1)NeuAc(1) | C[+57], N[+2060] | 1188.5059 | 4 | 4751.0018 | 4750.9876 | 0 | 3 | 60 | Specific | 690.33 | 587.7 | 587.7 | 9.86 | 5 | >sp\|P47710\|CASA1_HUMAN Alpha-S1-casein OS=Homo sapiens OX=9606 GN=CSN1S1 PE=1 SV=1 | 989 | 20201117_fqg_milk_1ng_glyco_hcd30_etd30_compound15.5114.5114.4 | controllerType=0 controllerNumber=1 scan=5114 | 15.2626 |
| 11754:3 | 21 | R.IIVPLNNREN[+2204.772]ISDPTSPLR.T | HexNAc(4)Hex(5)NeuAc(2) | N[+2205] | 1451.6576 | 3 | 4352.9584 | 4352.9496 | 0 | 2 | 62 | Specific | 218.71 | 121.74 | 121.74 | 2.68 | 8 | >sp\|P01591\|IGJ_HUMAN Immunoglobulin J chain OS=Homo sapiens OX=9606 GN=JCHAIN PE=1 SV=4 | 1392 | 20201117_fqg_milk_1ng_glyco_hcd30_etd30_compound15.15141.15141.3 | controllerType=0 controllerNumber=1 scan=15141 | 34.1146 |
| 10055:3 | 21 | R.IIVPLNNREN[+1856.656]ISDPTSPLR.T | HexNAc(3)Hex(5)Fuc(1)NeuAc(1) | N[+1857] | 1335.617 | 3 | 4004.8364 | 4004.8327 | 0 | 0.9 | 62 | Specific | 231.98 | 192.06 | 192.06 | 2.86 | 8 | >sp\|P01591\|IGJ_HUMAN Immunoglobulin J chain OS=Homo sapiens OX=9606 GN=JCHAIN PE=1 SV=4 | 1392 | 20201117_fqg_milk_1ng_glyco_hcd30_etd30_compound15.13359.13359.3 | controllerType=0 controllerNumber=1 scan=13359 | 30.6457 |
| 08556:3 | 21 | R.IIVPLNNREN[+1460.529]ISDPTSPLR.T | HexNAc(4)Hex(4) | N[+1461] | 1203.5761 | 3 | 3608.7138 | 3608.7059 | 0 | 2.2 | 62 | Specific | 254.63 | 181.65 | 181.65 | 2.96 | 8 | >sp\|P01591\|IGJ_HUMAN Immunoglobulin J chain OS=Homo sapiens OX=9606 GN=JCHAIN PE=1 SV=4 | 1392 | 20201117_fqg_milk_1ng_glyco_hcd30_etd30_compound15.11801.11801.3 | controllerType=0 controllerNumber=1 scan=11801 | 27.4972 |
| 08590:3 | 21 | R.IIVPLNNREN[+1663.608]ISDPTSPLR.T | HexNAc(5)Hex(4) | N[+1664] | 1271.2647 | 3 | 3811.7796 | 3811.7853 | 0 | -1.5 | 62 | Specific | 263.68 | 180.98 | 180.98 | 2.52 | 8 | >sp\|P01591\|IGJ_HUMAN Immunoglobulin J chain OS=Homo sapiens OX=9606 GN=JCHAIN PE=1 SV=4 | 1392 | 20201117_fqg_milk_1ng_glyco_hcd30_etd30_compound15.11837.11837.3 | controllerType=0 controllerNumber=1 scan=11837 | 27.5807 |
| 09952:3 | 21 | R.IIVPLNNREN[+1913.677]ISDPTSPLR.T | HexNAc(4)Hex(5)NeuAc(1) | N[+1914] | 1354.626 | 3 | 4061.8635 | 4061.8542 | 0 | 2.3 | 62 | Specific | 283.25 | 219.76 | 219.76 | 2.97 | 8 | >sp\|P01591\|IGJ_HUMAN Immunoglobulin J chain OS=Homo sapiens OX=9606 GN=JCHAIN PE=1 SV=4 | 1392 | 20201117_fqg_milk_1ng_glyco_hcd30_etd30_compound15.13252.13252.3 | controllerType=0 controllerNumber=1 scan=13252 | 30.3834 |
| 11735:3 | 21 | R.IIVPLNNREN[+2204.772]ISDPTSPLR.T | HexNAc(4)Hex(5)NeuAc(2) | N[+2205] | 1451.6561 | 3 | 4352.9536 | 4352.9496 | 0 | 0.9 | 62 | Specific | 284.89 | 211.6 | 211.6 | 3.75 | 8 | >sp\|P01591\|IGJ_HUMAN Immunoglobulin J chain OS=Homo sapiens OX=9606 GN=JCHAIN PE=1 SV=4 | 1392 | 20201117_fqg_milk_1ng_glyco_hcd30_etd30_compound15.15121.15121.3 | controllerType=0 controllerNumber=1 scan=15121 | 34.072 |
| 07638:4 | 34 | K.SC[+57.021]QHN[+2205.793]GTMYQHGEIFSAHELFPSR.L | HexNAc(4)Hex(5)Fuc(2)NeuAc(1) | C[+57], N[+2206] | 1260.522 | 4 | 5039.0661 | 5039.0498 | 0 | 3.2 | 110 | Specific | 412.91 | 375.17 | 375.17 | 6.94 | 3 | >sp\|Q6WN34\|CRDL2_HUMAN Chordin-like protein 2 OS=Homo sapiens OX=9606 GN=CHRDL2 PE=1 SV=1 | 1189 | 20201117_fqg_milk_1ng_glyco_hcd30_etd30_compound15.10839.10839.4 | controllerType=0 controllerNumber=1 scan=10839 | 25.6346 |
| 19495:3 | 76 | K.MLN[+2351.851]TSSLLEQLNEQFNWVSR.L | HexNAc(4)Hex(5)Fuc(3)NeuAc(1) | N[+2352] | 1587.6848 | 3 | 4761.0398 | 4761.0375 | 0 | 0.5 | 352 | Specific | 306.11 | 184.48 | 184.48 | 5.21 | 2 | >sp\|P10909\|CLUS_HUMAN Clusterin OS=Homo sapiens OX=9606 GN=CLU PE=1 SV=1 | 1467 | 20201117_fqg_milk_1ng_glyco_hcd30_etd30_compound15.23321.23321.3 | controllerType=0 controllerNumber=1 scan=23321 | 54.4462 |
| 01357:3 | 1 | K.GPN[+2059.735]C[+57.021]SEPEC[+57.021]PGNC[+57.021]HLR.G | HexNAc(4)Hex(5)Fuc(1)NeuAc(1) | C[+57]*3, N[+2060] | 1315.1728 | 3 | 3943.5038 | 3943.4931 | 0 | 2.7 | 182 | Specific | 226.86 | 117.9 | 117.9 | 3.28 | 52 | >sp\|P24821\|TENA_HUMAN Tenascin OS=Homo sapiens OX=9606 GN=TNC PE=1 SV=3 | 486 | 20201118_lss_colostrum_1ng_glyco_hcd30_etd30_compound15.4306.4306.3 | controllerType=0 controllerNumber=1 scan=4306 | 13.7999 |
| 12380:3 | 1 | R.LN[+1540.529]YSLPTGQWVGVQLPR.N | HexNAc(2)Hex(7) | N[+1541] | 1156.8682 | 3 | 3468.5901 | 3468.5674 | 0 | 6.6 | 1017 | Specific | 494.95 | 460.33 | 460.33 | 5.88 | 52 | >sp\|P24821\|TENA_HUMAN Tenascin OS=Homo sapiens OX=9606 GN=TNC PE=1 SV=3 | 486 | 20201118_lss_colostrum_1ng_glyco_hcd30_etd30_compound15.15793.15793.3 | controllerType=0 controllerNumber=1 scan=15793 | 37.8846 |
| 12434:2 | 1 | R.LN[+1216.423]YSLPTGQWVGVQLPR.N | HexNAc(2)Hex(5) | N[+1216] | 1572.7394 | 2 | 3144.4716 | 3144.4617 | 0 | 3.1 | 1017 | Specific | 527.43 | 456.71 | 456.71 | 7.1 | 52 | >sp\|P24821\|TENA_HUMAN Tenascin OS=Homo sapiens OX=9606 GN=TNC PE=1 SV=3 | 486 | 20201118_lss_colostrum_1ng_glyco_hcd30_etd30_compound15.15850.15850.2 | controllerType=0 controllerNumber=1 scan=15850 | 38.0567 |
| 12420:3 | 1 | R.LN[+1216.423]YSLPTGQWVGVQLPR.N | HexNAc(2)Hex(5) | N[+1216] | 1048.8308 | 3 | 3144.478 | 3144.4617 | 0 | 5.2 | 1017 | Specific | 637.25 | 585.69 | 585.69 | 8.82 | 52 | >sp\|P24821\|TENA_HUMAN Tenascin OS=Homo sapiens OX=9606 GN=TNC PE=1 SV=3 | 486 | 20201118_lss_colostrum_1ng_glyco_hcd30_etd30_compound15.15835.15835.3 | controllerType=0 controllerNumber=1 scan=15835 | 38.0164 |
| 01716:2 | 1 | R.N[+1216.423]TTSYVLR.G | HexNAc(2)Hex(5) | N[+1216] | 1085.4698 | 2 | 2169.9323 | 2169.9279 | 0 | 2 | 1034 | Specific | 233.97 | 81.7 | 81.7 | 2.59 | 52 | >sp\|P24821\|TENA_HUMAN Tenascin OS=Homo sapiens OX=9606 GN=TNC PE=1 SV=3 | 486 | 20201118_lss_colostrum_1ng_glyco_hcd30_etd30_compound15.4679.4679.2 | controllerType=0 controllerNumber=1 scan=4679 | 14.5032 |
| 03318:3 | 1 | R.N[+2059.735]TTSYVLR.G | HexNAc(4)Hex(5)Fuc(1)NeuAc(1) | N[+2060] | 1005.0876 | 3 | 3013.2481 | 3013.24 | 0 | 2.7 | 1034 | Specific | 334.62 | 172.31 | 172.31 | 4.2 | 52 | >sp\|P24821\|TENA_HUMAN Tenascin OS=Homo sapiens OX=9606 GN=TNC PE=1 SV=3 | 486 | 20201118_lss_colostrum_1ng_glyco_hcd30_etd30_compound15.6345.6345.3 | controllerType=0 controllerNumber=1 scan=6345 | 17.7229 |
| 15201:5 | 2 | R.RTAGWNVPIGTLRPFLN[+2350.830]WTGPPEPIEAAVAR.F | HexNAc(4)Hex(5)Fuc(1)NeuAc(2) | N[+2351] | 1148.337 | 5 | 5737.6561 | 5737.6424 | 0 | 2.4 | 140 | Specific | 351.58 | 351.58 | 351.58 | 4.99 | 57 | >sp\|P02788\|TRFL_HUMAN Lactotransferrin OS=Homo sapiens OX=9606 GN=LTF PE=1 SV=6 | 1270 | 20201118_lss_colostrum_1ng_glyco_hcd30_etd30_compound15.18759.18759.5 | controllerType=0 controllerNumber=1 scan=18759 | 45.8384 |
| 14437:4 | 2 | R.RTAGWNVPIGTLRPFLN[+2059.735]WTGPPEPIEAAVAR.F | HexNAc(4)Hex(5)Fuc(1)NeuAc(1) | N[+2060] | 1362.3969 | 4 | 5446.5656 | 5446.547 | 0 | 3.4 | 140 | Specific | 775.29 | 775.29 | 775.29 | 12.86 | 57 | >sp\|P02788\|TRFL_HUMAN Lactotransferrin OS=Homo sapiens OX=9606 GN=LTF PE=1 SV=6 | 1270 | 20201118_lss_colostrum_1ng_glyco_hcd30_etd30_compound15.17957.17957.4 | controllerType=0 controllerNumber=1 scan=17957 | 43.6777 |
| 14298:4 | 2 | R.RTAGWNVPIGTLRPFLN[+2716.983]WTGPPEPIEAAVAR.F | HexNAc(5)Hex(6)Fuc(3)NeuAc(1) | N[+2717] | 1526.71 | 4 | 6103.818 | 6103.795 | 0 | 3.8 | 140 | Specific | 805.93 | 805.93 | 805.93 | 13.86 | 57 | >sp\|P02788\|TRFL_HUMAN Lactotransferrin OS=Homo sapiens OX=9606 GN=LTF PE=1 SV=6 | 1270 | 20201118_lss_colostrum_1ng_glyco_hcd30_etd30_compound15.17811.17811.4 | controllerType=0 controllerNumber=1 scan=17811 | 43.2851 |
| 14384:4 | 2 | R.RTAGWNVPIGTLRPFLN[+2351.851]WTGPPEPIEAAVAR.F | HexNAc(4)Hex(5)Fuc(3)NeuAc(1) | N[+2352] | 1435.4265 | 4 | 5738.6842 | 5738.6628 | 0 | 3.7 | 140 | Specific | 868.2 | 616.11 | 616.11 | 14.86 | 57 | >sp\|P02788\|TRFL_HUMAN Lactotransferrin OS=Homo sapiens OX=9606 GN=LTF PE=1 SV=6 | 1270 | 20201118_lss_colostrum_1ng_glyco_hcd30_etd30_compound15.17901.17901.4 | controllerType=0 controllerNumber=1 scan=17901 | 43.5176 |
| 14389:4 | 2 | R.RTAGWNVPIGTLRPFLN[+2205.793]WTGPPEPIEAAVAR.F | HexNAc(4)Hex(5)Fuc(2)NeuAc(1) | N[+2206] | 1398.9114 | 4 | 5592.6237 | 5592.6049 | 0 | 3.4 | 140 | Specific | 950.21 | 905.87 | 905.87 | 16.86 | 57 | >sp\|P02788\|TRFL_HUMAN Lactotransferrin OS=Homo sapiens OX=9606 GN=LTF PE=1 SV=6 | 1270 | 20201118_lss_colostrum_1ng_glyco_hcd30_etd30_compound15.17906.17906.4 | controllerType=0 controllerNumber=1 scan=17906 | 43.5402 |
| 13689:4 | 2 | R.RTAGWNVPIGTLRPFLN[+1768.640]WTGPPEPIEAAVAR.F | HexNAc(4)Hex(5)Fuc(1) | N[+1769] | 1289.6209 | 4 | 5155.4616 | 5155.4516 | 0 | 1.9 | 140 | Specific | 978.75 | 978.75 | 978.75 | 16.75 | 57 | >sp\|P02788\|TRFL_HUMAN Lactotransferrin OS=Homo sapiens OX=9606 GN=LTF PE=1 SV=6 | 1270 | 20201118_lss_colostrum_1ng_glyco_hcd30_etd30_compound15.17167.17167.4 | controllerType=0 controllerNumber=1 scan=17167 | 41.3827 |
| 15342:3 | 2 | R.TAGWNVPIGTLRPFLN[+2350.830]WTGPPEPIEAAVAR.F | HexNAc(4)Hex(5)Fuc(1)NeuAc(2) | N[+2351] | 1861.1989 | 3 | 5581.5823 | 5581.5413 | 0 | 7.3 | 141 | Specific | 239.59 | 141.12 | 141.12 | 3.39 | 57 | >sp\|P02788\|TRFL_HUMAN Lactotransferrin OS=Homo sapiens OX=9606 GN=LTF PE=1 SV=6 | 1270 | 20201118_lss_colostrum_1ng_glyco_hcd30_etd30_compound15.18909.18909.3 | controllerType=0 controllerNumber=1 scan=18909 | 46.3443 |
| 16227:3 | 2 | R.TAGWNVPIGTLRPFLN[+2350.830]WTGPPEPIEAAVAR.F | HexNAc(4)Hex(5)Fuc(1)NeuAc(2) | N[+2351] | 1861.1922 | 3 | 5581.5621 | 5581.5413 | 0 | 3.7 | 141 | Specific | 268.67 | 196.16 | 196.16 | 5.57 | 57 | >sp\|P02788\|TRFL_HUMAN Lactotransferrin OS=Homo sapiens OX=9606 GN=LTF PE=1 SV=6 | 1270 | 20201118_lss_colostrum_1ng_glyco_hcd30_etd30_compound15.19844.19844.3 | controllerType=0 controllerNumber=1 scan=19844 | 49.2527 |
| 15348:3 | 2 | R.TAGWNVPIGTLRPFLN[+2205.793]WTGPPEPIEAAVAR.F | HexNAc(4)Hex(5)Fuc(2)NeuAc(1) | N[+2206] | 1812.8462 | 3 | 5436.5239 | 5436.5038 | 0 | 3.7 | 141 | Specific | 279.77 | 181.01 | 181.01 | 4.79 | 57 | >sp\|P02788\|TRFL_HUMAN Lactotransferrin OS=Homo sapiens OX=9606 GN=LTF PE=1 SV=6 | 1270 | 20201118_lss_colostrum_1ng_glyco_hcd30_etd30_compound15.18915.18915.3 | controllerType=0 controllerNumber=1 scan=18915 | 46.3538 |
| 14586:4 | 2 | R.[+42.011]TAGWNVPIGTLRPFLN[+1872.651]WTGPPEPIEAAVAR.F | HexNAc(3)Hex(6)NeuAc(1) | .[42], N[+1873] | 1287.1108 | 4 | 5145.4215 | 5145.372 | 0 | 9.6 | 141 | Specific | 293.3 | 293.3 | 293.3 | 3.43 | 57 | >sp\|P02788\|TRFL_HUMAN Lactotransferrin OS=Homo sapiens OX=9606 GN=LTF PE=1 SV=6 | 1270 | 20201118_lss_colostrum_1ng_glyco_hcd30_etd30_compound15.18113.18113.4 | controllerType=0 controllerNumber=1 scan=18113 | 44.0562 |
| 16282:4 | 2 | R.[+42.011]TAGWNVPIGTLRPFLN[+2336.851]WTGPPEPIEAAVAR.F | HexNAc(6)Hex(6)Fuc(1) | .[42], N[+2337] | 1403.1444 | 4 | 5609.5558 | 5609.5726 | 0 | -3 | 141 | Specific | 297.16 | 297.16 | 297.16 | 3.74 | 57 | >sp\|P02788\|TRFL_HUMAN Lactotransferrin OS=Homo sapiens OX=9606 GN=LTF PE=1 SV=6 | 1270 | 20201118_lss_colostrum_1ng_glyco_hcd30_etd30_compound15.19902.19902.4 | controllerType=0 controllerNumber=1 scan=19902 | 49.4304 |
| 14603:3 | 2 | R.TAGWNVPIGTLRPFLN[+1768.640]WTGPPEPIEAAVAR.F | HexNAc(4)Hex(5)Fuc(1) | N[+1769] | 1667.1273 | 3 | 4999.3673 | 4999.3504 | 0 | 3.4 | 141 | Specific | 357.09 | 308.01 | 308.01 | 5.35 | 57 | >sp\|P02788\|TRFL_HUMAN Lactotransferrin OS=Homo sapiens OX=9606 GN=LTF PE=1 SV=6 | 1270 | 20201118_lss_colostrum_1ng_glyco_hcd30_etd30_compound15.18130.18130.3 | controllerType=0 controllerNumber=1 scan=18130 | 44.0969 |
| 15358:3 | 2 | R.TAGWNVPIGTLRPFLN[+2205.793]WTGPPEPIEAAVAR.F | HexNAc(4)Hex(5)Fuc(2)NeuAc(1) | N[+2206] | 1812.8427 | 3 | 5436.5137 | 5436.5038 | 0 | 1.8 | 141 | Specific | 371.47 | 297.1 | 297.1 | 5.88 | 57 | >sp\|P02788\|TRFL_HUMAN Lactotransferrin OS=Homo sapiens OX=9606 GN=LTF PE=1 SV=6 | 1270 | 20201118_lss_colostrum_1ng_glyco_hcd30_etd30_compound15.18926.18926.3 | controllerType=0 controllerNumber=1 scan=18926 | 46.3883 |
| 15463:3 | 2 | R.TAGWNVPIGTLRPFLN[+2059.735]WTGPPEPIEAAVAR.F | HexNAc(4)Hex(5)Fuc(1)NeuAc(1) | N[+2060] | 1764.1588 | 3 | 5290.4618 | 5290.4459 | 0 | 3 | 141 | Specific | 397.55 | 318.09 | 318.09 | 5.99 | 57 | >sp\|P02788\|TRFL_HUMAN Lactotransferrin OS=Homo sapiens OX=9606 GN=LTF PE=1 SV=6 | 1270 | 20201118_lss_colostrum_1ng_glyco_hcd30_etd30_compound15.19036.19036.3 | controllerType=0 controllerNumber=1 scan=19036 | 46.7019 |
| 14555:4 | 2 | R.TAGWNVPIGTLRPFLN[+2425.887]WTGPPEPIEAAVAR.F | HexNAc(5)Hex(6)Fuc(3) | N[+2426] | 1414.9092 | 4 | 5656.6149 | 5656.5985 | 0 | 2.9 | 141 | Specific | 500.25 | 321.11 | 321.11 | 6.33 | 57 | >sp\|P02788\|TRFL_HUMAN Lactotransferrin OS=Homo sapiens OX=9606 GN=LTF PE=1 SV=6 | 1270 | 20201118_lss_colostrum_1ng_glyco_hcd30_etd30_compound15.18080.18080.4 | controllerType=0 controllerNumber=1 scan=18080 | 43.9584 |
| 15335:4 | 2 | R.TAGWNVPIGTLRPFLN[+2351.851]WTGPPEPIEAAVAR.F | HexNAc(4)Hex(5)Fuc(3)NeuAc(1) | N[+2352] | 1396.4015 | 4 | 5582.5841 | 5582.5617 | 0 | 4 | 141 | Specific | 560.21 | 560.21 | 560.21 | 8.94 | 57 | >sp\|P02788\|TRFL_HUMAN Lactotransferrin OS=Homo sapiens OX=9606 GN=LTF PE=1 SV=6 | 1270 | 20201118_lss_colostrum_1ng_glyco_hcd30_etd30_compound15.18902.18902.4 | controllerType=0 controllerNumber=1 scan=18902 | 46.3249 |
| 16231:5 | 2 | R.TAGWNVPIGTLRPFLN[+2350.830]WTGPPEPIEAAVAR.F | HexNAc(4)Hex(5)Fuc(1)NeuAc(2) | N[+2351] | 1117.1179 | 5 | 5581.5605 | 5581.5413 | 0 | 3.4 | 141 | Specific | 583.38 | 581.15 | 581.15 | 8.16 | 57 | >sp\|P02788\|TRFL_HUMAN Lactotransferrin OS=Homo sapiens OX=9606 GN=LTF PE=1 SV=6 | 1270 | 20201118_lss_colostrum_1ng_glyco_hcd30_etd30_compound15.19848.19848.5 | controllerType=0 controllerNumber=1 scan=19848 | 49.2605 |
| 15357:4 | 2 | R.TAGWNVPIGTLRPFLN[+2205.793]WTGPPEPIEAAVAR.F | HexNAc(4)Hex(5)Fuc(2)NeuAc(1) | N[+2206] | 1359.8827 | 4 | 5436.5089 | 5436.5038 | 0 | 1 | 141 | Specific | 662.28 | 662.28 | 662.28 | 10.83 | 57 | >sp\|P02788\|TRFL_HUMAN Lactotransferrin OS=Homo sapiens OX=9606 GN=LTF PE=1 SV=6 | 1270 | 20201118_lss_colostrum_1ng_glyco_hcd30_etd30_compound15.18925.18925.4 | controllerType=0 controllerNumber=1 scan=18925 | 46.3853 |
| 16226:4 | 2 | R.TAGWNVPIGTLRPFLN[+2350.830]WTGPPEPIEAAVAR.F | HexNAc(4)Hex(5)Fuc(1)NeuAc(2) | N[+2351] | 1396.147 | 4 | 5581.5661 | 5581.5413 | 0 | 4.4 | 141 | Specific | 666.41 | 666.41 | 666.41 | 11 | 57 | >sp\|P02788\|TRFL_HUMAN Lactotransferrin OS=Homo sapiens OX=9606 GN=LTF PE=1 SV=6 | 1270 | 20201118_lss_colostrum_1ng_glyco_hcd30_etd30_compound15.19843.19843.4 | controllerType=0 controllerNumber=1 scan=19843 | 49.2498 |
| 15218:4 | 2 | R.TAGWNVPIGTLRPFLN[+2716.983]WTGPPEPIEAAVAR.F | HexNAc(5)Hex(6)Fuc(3)NeuAc(1) | N[+2717] | 1487.6846 | 4 | 5947.7165 | 5947.6939 | 0 | 3.8 | 141 | Specific | 683.04 | 671.51 | 671.51 | 10.94 | 57 | >sp\|P02788\|TRFL_HUMAN Lactotransferrin OS=Homo sapiens OX=9606 GN=LTF PE=1 SV=6 | 1270 | 20201118_lss_colostrum_1ng_glyco_hcd30_etd30_compound15.18777.18777.4 | controllerType=0 controllerNumber=1 scan=18777 | 45.8983 |
| 15298:4 | 2 | R.TAGWNVPIGTLRPFLN[+2570.925]WTGPPEPIEAAVAR.F | HexNAc(5)Hex(6)Fuc(2)NeuAc(1) | N[+2571] | 1451.1755 | 4 | 5801.6803 | 5801.636 | 0 | 7.6 | 141 | Specific | 807.66 | 807.66 | 807.66 | 13.26 | 57 | >sp\|P02788\|TRFL_HUMAN Lactotransferrin OS=Homo sapiens OX=9606 GN=LTF PE=1 SV=6 | 1270 | 20201118_lss_colostrum_1ng_glyco_hcd30_etd30_compound15.18862.18862.4 | controllerType=0 controllerNumber=1 scan=18862 | 46.191 |
| 14598:4 | 2 | R.TAGWNVPIGTLRPFLN[+1768.640]WTGPPEPIEAAVAR.F | HexNAc(4)Hex(5)Fuc(1) | N[+1769] | 1250.5974 | 4 | 4999.3678 | 4999.3504 | 0 | 3.5 | 141 | Specific | 893.17 | 872.14 | 872.14 | 15.48 | 57 | >sp\|P02788\|TRFL_HUMAN Lactotransferrin OS=Homo sapiens OX=9606 GN=LTF PE=1 SV=6 | 1270 | 20201118_lss_colostrum_1ng_glyco_hcd30_etd30_compound15.18125.18125.4 | controllerType=0 controllerNumber=1 scan=18125 | 44.0877 |
| 13462:3 | 2 | R.TAGWNIPMGLLFN[+1768.640]QTGSC[+57.021]K.F | HexNAc(4)Hex(5)Fuc(1) | C[+57], N[+1769] | 1288.5604 | 3 | 3863.6665 | 3863.6495 | 0 | 4.4 | 485 | Specific | 272.93 | 199.5 | 199.5 | 3.7 | 57 | >sp\|P02788\|TRFL_HUMAN Lactotransferrin OS=Homo sapiens OX=9606 GN=LTF PE=1 SV=6 | 1270 | 20201118_lss_colostrum_1ng_glyco_hcd30_etd30_compound15.16928.16928.3 | controllerType=0 controllerNumber=1 scan=16928 | 40.699 |
| 14929:3 | 2 | R.TAGWNIPMGLLFN[+2205.793]QTGSC[+57.021]K.F | HexNAc(4)Hex(5)Fuc(2)NeuAc(1) | C[+57], N[+2206] | 1434.2769 | 3 | 4300.8162 | 4300.8028 | 0 | 3.1 | 485 | Specific | 275.13 | 165.95 | 165.95 | 4.88 | 57 | >sp\|P02788\|TRFL_HUMAN Lactotransferrin OS=Homo sapiens OX=9606 GN=LTF PE=1 SV=6 | 1270 | 20201118_lss_colostrum_1ng_glyco_hcd30_etd30_compound15.18473.18473.3 | controllerType=0 controllerNumber=1 scan=18473 | 45.0506 |
| 16289:4 | 2 | R.TAGWNIPMGLLFN[+2350.830]QTGSC[+57.021]K.F | HexNAc(4)Hex(5)Fuc(1)NeuAc(2) | C[+57], N[+2351] | 1112.2187 | 4 | 4445.8529 | 4445.8403 | 0 | 2.8 | 485 | Specific | 306.44 | 250.6 | 250.6 | 4.77 | 57 | >sp\|P02788\|TRFL_HUMAN Lactotransferrin OS=Homo sapiens OX=9606 GN=LTF PE=1 SV=6 | 1270 | 20201118_lss_colostrum_1ng_glyco_hcd30_etd30_compound15.19909.19909.4 | controllerType=0 controllerNumber=1 scan=19909 | 49.4467 |
| 16295:3 | 2 | R.TAGWNIPMGLLFN[+2350.830]QTGSC[+57.021]K.F | HexNAc(4)Hex(5)Fuc(1)NeuAc(2) | C[+57], N[+2351] | 1482.6244 | 3 | 4445.8588 | 4445.8403 | 0 | 4.2 | 485 | Specific | 329.02 | 226.99 | 226.99 | 5.44 | 57 | >sp\|P02788\|TRFL_HUMAN Lactotransferrin OS=Homo sapiens OX=9606 GN=LTF PE=1 SV=6 | 1270 | 20201118_lss_colostrum_1ng_glyco_hcd30_etd30_compound15.19916.19916.3 | controllerType=0 controllerNumber=1 scan=19916 | 49.4804 |
| 13434:3 | 2 | R.TAGWNIPMGLLFN[+1768.640]QTGSC[+57.021]K.F | HexNAc(4)Hex(5)Fuc(1) | C[+57], N[+1769] | 1288.5587 | 3 | 3863.6614 | 3863.6495 | 0 | 3.1 | 485 | Specific | 380.92 | 271.39 | 271.39 | 5.36 | 57 | >sp\|P02788\|TRFL_HUMAN Lactotransferrin OS=Homo sapiens OX=9606 GN=LTF PE=1 SV=6 | 1270 | 20201118_lss_colostrum_1ng_glyco_hcd30_etd30_compound15.16898.16898.3 | controllerType=0 controllerNumber=1 scan=16898 | 40.6354 |
| 14833:3 | 2 | R.TAGWNIPMGLLFN[+2351.851]QTGSC[+57.021]K.F | HexNAc(4)Hex(5)Fuc(3)NeuAc(1) | C[+57], N[+2352] | 1482.9645 | 3 | 4446.8791 | 4446.8607 | 0 | 4.1 | 485 | Specific | 382.88 | 273.49 | 273.49 | 5.9 | 57 | >sp\|P02788\|TRFL_HUMAN Lactotransferrin OS=Homo sapiens OX=9606 GN=LTF PE=1 SV=6 | 1270 | 20201118_lss_colostrum_1ng_glyco_hcd30_etd30_compound15.18372.18372.3 | controllerType=0 controllerNumber=1 scan=18372 | 44.7705 |
| 14852:3 | 2 | R.TAGWNIPMGLLFN[+2059.735]QTGSC[+57.021]K.F | HexNAc(4)Hex(5)Fuc(1)NeuAc(1) | C[+57], N[+2060] | 1385.592 | 3 | 4154.7614 | 4154.7449 | 0 | 4 | 485 | Specific | 395.55 | 304.48 | 304.48 | 5.89 | 57 | >sp\|P02788\|TRFL_HUMAN Lactotransferrin OS=Homo sapiens OX=9606 GN=LTF PE=1 SV=6 | 1270 | 20201118_lss_colostrum_1ng_glyco_hcd30_etd30_compound15.18392.18392.3 | controllerType=0 controllerNumber=1 scan=18392 | 44.8276 |
| 14991:3 | 2 | R.TAGWNIPMGLLFN[+1913.677]QTGSC[+57.021]K.F | HexNAc(4)Hex(5)NeuAc(1) | C[+57], N[+1914] | 1336.905 | 3 | 4008.7003 | 4008.687 | 0 | 3.3 | 485 | Specific | 455.56 | 303.84 | 303.84 | 5.92 | 57 | >sp\|P02788\|TRFL_HUMAN Lactotransferrin OS=Homo sapiens OX=9606 GN=LTF PE=1 SV=6 | 1270 | 20201118_lss_colostrum_1ng_glyco_hcd30_etd30_compound15.18539.18539.3 | controllerType=0 controllerNumber=1 scan=18539 | 45.2606 |
| 13716:3 | 2 | R.TAGWNIPMGLLFN[+1216.423]QTGSC[+57.021]K.F | HexNAc(2)Hex(5) | C[+57], N[+1216] | 1104.485 | 3 | 3311.4405 | 3311.4328 | 0 | 2.3 | 485 | Specific | 493.63 | 437.84 | 230.83 | 7.44 | 57 | >sp\|P02788\|TRFL_HUMAN Lactotransferrin OS=Homo sapiens OX=9606 GN=LTF PE=1 SV=6 | 1270 | 20201118_lss_colostrum_1ng_glyco_hcd30_etd30_compound15.17195.17195.3 | controllerType=0 controllerNumber=1 scan=17195 | 41.4539 |
| 13737:2 | 2 | R.TAGWNIPMGLLFN[+1216.423]QTGSC[+57.021]K.F | HexNAc(2)Hex(5) | C[+57], N[+1216] | 1656.226 | 2 | 3311.4447 | 3311.4328 | 0 | 3.6 | 485 | Specific | 565.87 | 565.87 | 200.31 | 8.65 | 57 | >sp\|P02788\|TRFL_HUMAN Lactotransferrin OS=Homo sapiens OX=9606 GN=LTF PE=1 SV=6 | 1270 | 20201118_lss_colostrum_1ng_glyco_hcd30_etd30_compound15.17217.17217.2 | controllerType=0 controllerNumber=1 scan=17217 | 41.5234 |
| 13201:3 | 2 | R.TAGWNIPM[+15.995]GLLFN[+2059.735]QTGSC[+57.021]K.F | HexNAc(4)Hex(5)Fuc(1)NeuAc(1) | C[+57], M[+16], N[+2060] | 1390.925 | 3 | 4170.7604 | 4170.7398 | 0 | 4.9 | 485 | Specific | 217.5 | 187.08 | 187.08 | 2.79 | 57 | >sp\|P02788\|TRFL_HUMAN Lactotransferrin OS=Homo sapiens OX=9606 GN=LTF PE=1 SV=6 | 1270 | 20201118_lss_colostrum_1ng_glyco_hcd30_etd30_compound15.16654.16654.3 | controllerType=0 controllerNumber=1 scan=16654 | 39.9791 |
| 14264:4 | 2 | R.TAGWNIPMGLLFN[+1768.640]QTGSC[+57.021]KFDEYFSQSC[+57.021]APGSDPR.S | HexNAc(4)Hex(5)Fuc(1) | C[+57]*2, N[+1769] | 1427.61 | 4 | 5707.4181 | 5707.3966 | 0 | 3.8 | 485 | Specific | 319.31 | 274.15 | 274.15 | 4.6 | 57 | >sp\|P02788\|TRFL_HUMAN Lactotransferrin OS=Homo sapiens OX=9606 GN=LTF PE=1 SV=6 | 1270 | 20201118_lss_colostrum_1ng_glyco_hcd30_etd30_compound15.17775.17775.4 | controllerType=0 controllerNumber=1 scan=17775 | 43.1697 |
| 14388:4 | 2 | R.TAGWNIPMGLLFN[+1216.423]QTGSC[+57.021]KFDEYFSQSC[+57.021]APGSDPR.S | HexNAc(2)Hex(5) | C[+57]*2, N[+1216] | 1289.5555 | 4 | 5155.2004 | 5155.18 | 0 | 4 | 485 | Specific | 326.38 | 291.51 | 99.98 | 4.65 | 57 | >sp\|P02788\|TRFL_HUMAN Lactotransferrin OS=Homo sapiens OX=9606 GN=LTF PE=1 SV=6 | 1270 | 20201118_lss_colostrum_1ng_glyco_hcd30_etd30_compound15.17905.17905.4 | controllerType=0 controllerNumber=1 scan=17905 | 43.5313 |
| 15127:4 | 2 | R.TAGWNIPMGLLFN[+1913.677]QTGSC[+57.021]KFDEYFSQSC[+57.021]APGSDPR.S | HexNAc(4)Hex(5)NeuAc(1) | C[+57]*2, N[+1914] | 1463.8676 | 4 | 5852.4484 | 5852.4341 | 0 | 2.4 | 485 | Specific | 335.24 | 252.57 | 252.57 | 5.09 | 57 | >sp\|P02788\|TRFL_HUMAN Lactotransferrin OS=Homo sapiens OX=9606 GN=LTF PE=1 SV=6 | 1270 | 20201118_lss_colostrum_1ng_glyco_hcd30_etd30_compound15.18682.18682.4 | controllerType=0 controllerNumber=1 scan=18682 | 45.6598 |
| 15131:3 | 2 | R.TAGWNIPMGLLFN[+1913.677]QTGSC[+57.021]KFDEYFSQSC[+57.021]APGSDPR.S | HexNAc(4)Hex(5)NeuAc(1) | C[+57]*2, N[+1914] | 1951.4896 | 3 | 5852.4542 | 5852.4341 | 0 | 3.4 | 485 | Specific | 348.64 | 278.42 | 278.42 | 5.54 | 57 | >sp\|P02788\|TRFL_HUMAN Lactotransferrin OS=Homo sapiens OX=9606 GN=LTF PE=1 SV=6 | 1270 | 20201118_lss_colostrum_1ng_glyco_hcd30_etd30_compound15.18686.18686.3 | controllerType=0 controllerNumber=1 scan=18686 | 45.6685 |
| 14390:3 | 2 | R.TAGWNIPMGLLFN[+1216.423]QTGSC[+57.021]KFDEYFSQSC[+57.021]APGSDPR.S | HexNAc(2)Hex(5) | C[+57]*2, N[+1216] | 1719.0705 | 3 | 5155.197 | 5155.18 | 0 | 3.3 | 485 | Specific | 438.45 | 384.87 | 119.34 | 7.21 | 57 | >sp\|P02788\|TRFL_HUMAN Lactotransferrin OS=Homo sapiens OX=9606 GN=LTF PE=1 SV=6 | 1270 | 20201118_lss_colostrum_1ng_glyco_hcd30_etd30_compound15.17907.17907.3 | controllerType=0 controllerNumber=1 scan=17907 | 43.5432 |
| 15039:4 | 2 | R.TAGWNIPMGLLFN[+2351.851]QTGSC[+57.021]KFDEYFSQSC[+57.021]APGSDPR.S | HexNAc(4)Hex(5)Fuc(3)NeuAc(1) | C[+57]*2, N[+2352] | 1573.4136 | 4 | 6290.6325 | 6290.6079 | 0 | 3.9 | 485 | Specific | 470.62 | 443.3 | 443.3 | 7.26 | 57 | >sp\|P02788\|TRFL_HUMAN Lactotransferrin OS=Homo sapiens OX=9606 GN=LTF PE=1 SV=6 | 1270 | 20201118_lss_colostrum_1ng_glyco_hcd30_etd30_compound15.18590.18590.4 | controllerType=0 controllerNumber=1 scan=18590 | 45.4369 |
| 16011:4 | 2 | R.TAGWNIPMGLLFN[+2350.830]QTGSC[+57.021]KFDEYFSQSC[+57.021]APGSDPR.S | HexNAc(4)Hex(5)Fuc(1)NeuAc(2) | C[+57]*2, N[+2351] | 1573.1636 | 4 | 6289.6325 | 6289.5875 | 0 | 7.2 | 485 | Specific | 737.43 | 713.23 | 713.23 | 10.45 | 57 | >sp\|P02788\|TRFL_HUMAN Lactotransferrin OS=Homo sapiens OX=9606 GN=LTF PE=1 SV=6 | 1270 | 20201118_lss_colostrum_1ng_glyco_hcd30_etd30_compound15.19615.19615.4 | controllerType=0 controllerNumber=1 scan=19615 | 48.4986 |
| 14107:4 | 3 | R.LSLHRPALEDLLLGSEAN[+1825.661]LTC[+57.021]TLTGLR.D | HexNAc(5)Hex(5) | C[+57], N[+1826] | 1198.0762 | 4 | 4789.283 | 4789.2592 | 0 | 5 | 127 | Specific | 334.12 | 334.12 | 334.12 | 4.93 | 45 | >sp\|P01876\|IGHA1_HUMAN Immunoglobulin heavy constant alpha 1 OS=Homo sapiens OX=9606 GN=IGHA1 PE=1 SV=2 | 255 | 20201118_lss_colostrum_1ng_glyco_hcd30_etd30_compound15.17607.17607.4 | controllerType=0 controllerNumber=1 scan=17607 | [illegible] |
| 14837:3 | 3 | R.LSLHRPALEDLLLGSEAN[+2116.756]LTC[+57.021]TLTGLR.D | HexNAc(5)Hex(5)NeuAc(1) | C[+57], N[+2117] | 1694.1264 | 3 | 5080.3647 | 5080.3547 | 0 | 2 | 127 | Specific | 334.33 | 253.47 | 253.47 | 5.89 | 45 | >sp\|P01876\|IGHA1_HUMAN Immunoglobulin heavy constant alpha 1 OS=Homo sapiens OX=9606 GN=IGHA1 PE=1 SV=2 | 255 | 20201118_lss_colostrum_1ng_glyco_hcd30_etd30_compound15.18377.18377.3 | controllerType=0 controllerNumber=1 scan=18377 | [illegible] |
| 14938:4 | 3 | R.LSLHRPALEDLLLGSEAN[+1548.545]LTC[+57.021]TLTGLR.D | HexNAc(3)Hex(4)NeuAc(1) | C[+57], N[+1549] | 1128.7952 | 4 | 4512.159 | 4512.1431 | 0 | 3.5 | 127 | Specific | 367.24 | 367.24 | 367.24 | 7.16 | 45 | >sp\|P01876\|IGHA1_HUMAN Immunoglobulin heavy constant alpha 1 OS=Homo sapiens OX=9606 GN=IGHA1 PE=1 SV=2 | 255 | 20201118_lss_colostrum_1ng_glyco_hcd30_etd30_compound15.18483.18483.4 | controllerType=0 controllerNumber=1 scan=18483 | 45.081 |
| 14106:3 | 3 | R.LSLHRPALEDLLLGSEAN[+1540.529]LTC[+57.021]TLTGLR.D | HexNAc(2)Hex(7) | C[+57], N[+1541] | 1502.0523 | 3 | 4504.1424 | 4504.1268 | 0 | 3.5 | 127 | Specific | 367.31 | 367.31 | 367.31 | 7.59 | 45 | >sp\|P01876\|IGHA1_HUMAN Immunoglobulin heavy constant alpha 1 OS=Homo sapiens OX=9606 GN=IGHA1 PE=1 SV=2 | 255 | 20201118_lss_colostrum_1ng_glyco_hcd30_etd30_compound15.17606.17606.3 | controllerType=0 controllerNumber=1 scan=17606 | 42.6238 |
| 14067:3 | 3 | R.LSLHRPALEDLLLGSEAN[+1702.581]LTC[+57.021]TLTGLR.D | HexNAc(2)Hex(8) | C[+57], N[+1703] | 1556.0696 | 3 | 4666.1944 | 4666.1796 | 0 | 3.2 | 127 | Specific | 393.72 | 361.72 | 361.72 | 7.59 | 45 | >sp\|P01876\|IGHA1_HUMAN Immunoglobulin heavy constant alpha 1 OS=Homo sapiens OX=9606 GN=IGHA1 PE=1 SV=2 | 255 | 20201118_lss_colostrum_1ng_glyco_hcd30_etd30_compound15.17565.17565.3 | controllerType=0 controllerNumber=1 scan=17565 | 42.5021 |
| 14827:3 | 3 | R.LSLHRPALEDLLLGSEAN[+1913.677]LTC[+57.021]TLTGLR.D | HexNAc(4)Hex(5)NeuAc(1) | C[+57], N[+1914] | 1626.4361 | 3 | 4877.2937 | 4877.2753 | 0 | 3.8 | 127 | Specific | 397.52 | 329.67 | 329.67 | 6.36 | 45 | >sp\|P01876\|IGHA1_HUMAN Immunoglobulin heavy constant alpha 1 OS=Homo sapiens OX=9606 GN=IGHA1 PE=1 SV=2 | 255 | 20201118_lss_colostrum_1ng_glyco_hcd30_etd30_compound15.18366.18366.3 | controllerType=0 controllerNumber=1 scan=18366 | 44.732 |
| 14101:3 | 3 | R.LSLHRPALEDLLLGSEAN[+1622.582]LTC[+57.021]TLTGLR.D | HexNAc(4)Hex(5) | C[+57], N[+1623] | 1529.4046 | 3 | 4586.1992 | 4586.1799 | 0 | 4.2 | 127 | Specific | 403.1 | 342.33 | 342.33 | 5.95 | 45 | >sp\|P01876\|IGHA1_HUMAN Immunoglobulin heavy constant alpha 1 OS=Homo sapiens OX=9606 GN=IGHA1 PE=1 SV=2 | 255 | 20201118_lss_colostrum_1ng_glyco_hcd30_etd30_compound15.17601.17601.3 | controllerType=0 controllerNumber=1 scan=17601 | 42.6107 |
| 14125:3 | 3 | R.LSLHRPALEDLLLGSEAN[+1663.608]LTC[+57.021]TLTGLR.D | HexNAc(5)Hex(4) | C[+57], N[+1664] | 1543.0784 | 3 | 4627.2207 | 4627.2064 | 0 | 3.1 | 127 | Specific | 407.92 | 374.23 | 374.23 | 7.67 | 45 | >sp\|P01876\|IGHA1_HUMAN Immunoglobulin heavy constant alpha 1 OS=Homo sapiens OX=9606 GN=IGHA1 PE=1 SV=2 | 255 | 20201118_lss_colostrum_1ng_glyco_hcd30_etd30_compound15.17626.17626.3 | controllerType=0 controllerNumber=1 scan=17626 | 42.6898 |
| 14047:3 | 3 | R.LSLHRPALEDLLLGSEAN[+1864.634]LTC[+57.021]TLTGLR.D | HexNAc(2)Hex(9) | C[+57], N[+1865] | 1610.0884 | 3 | 4828.2508 | 4828.2324 | 0 | 3.8 | 127 | Specific | 419.72 | 364.9 | 364.9 | 7.67 | 45 | >sp\|P01876\|IGHA1_HUMAN Immunoglobulin heavy constant alpha 1 OS=Homo sapiens OX=9606 GN=IGHA1 PE=1 SV=2 | 255 | 20201118_lss_colostrum_1ng_glyco_hcd30_etd30_compound15.17544.17544.3 | controllerType=0 controllerNumber=1 scan=17544 | 42.435 |
| 14124:3 | 3 | R.LSLHRPALEDLLLGSEAN[+1216.423]LTC[+57.021]TLTGLR.D | HexNAc(2)Hex(5) | C[+57], N[+1216] | 1394.0154 | 3 | 4180.0318 | 4180.0211 | 0 | 2.5 | 127 | Specific | 460.01 | 446.68 | 446.68 | 7.61 | 45 | >sp\|P01876\|IGHA1_HUMAN Immunoglobulin heavy constant alpha 1 OS=Homo sapiens OX=9606 GN=IGHA1 PE=1 SV=2 | 255 | 20201118_lss_colostrum_1ng_glyco_hcd30_etd30_compound15.17625.17625.3 | controllerType=0 controllerNumber=1 scan=17625 | 42.6853 |
| 14863:4 | 3 | R.LSLHRPALEDLLLGSEAN[+2116.756]LTC[+57.021]TLTGLR.D | HexNAc(5)Hex(5)NeuAc(1) | C[+57], N[+2117] | 1270.8472 | 4 | 5080.3669 | 5080.3547 | 0 | 2.4 | 127 | Specific | 472.77 | 462.26 | 462.26 | 7.18 | 45 | >sp\|P01876\|IGHA1_HUMAN Immunoglobulin heavy constant alpha 1 OS=Homo sapiens OX=9606 GN=IGHA1 PE=1 SV=2 | 255 | 20201118_lss_colostrum_1ng_glyco_hcd30_etd30_compound15.18404.18404.4 | controllerType=0 controllerNumber=1 scan=18404 | 44.8619 |
| 14095:3 | 3 | R.LSLHRPALEDLLLGSEAN[+1378.476]LTC[+57.021]TLTGLR.D | HexNAc(2)Hex(6) | C[+57], N[+1378] | 1448.0344 | 3 | 4342.0885 | 4342.074 | 0 | 3.4 | 127 | Specific | 532 | 526.72 | 526.72 | 7.93 | 45 | >sp\|P01876\|IGHA1_HUMAN Immunoglobulin heavy constant alpha 1 OS=Homo sapiens OX=9606 GN=IGHA1 PE=1 SV=2 | 255 | 20201118_lss_colostrum_1ng_glyco_hcd30_etd30_compound15.17594.17594.3 | controllerType=0 controllerNumber=1 scan=17594 | 42.5853 |
| 14862:4 | 3 | R.LSLHRPALEDLLLGSEAN[+1913.677]LTC[+57.021]TLTGLR.D | HexNAc(4)Hex(5)NeuAc(1) | C[+57], N[+1914] | 1220.0794 | 4 | 4877.2957 | 4877.2753 | 0 | 4.2 | 127 | Specific | 591.19 | 591.19 | 591.19 | 9.39 | 45 | >sp\|P01876\|IGHA1_HUMAN Immunoglobulin heavy constant alpha 1 OS=Homo sapiens OX=9606 GN=IGHA1 PE=1 SV=2 | 255 | 20201118_lss_colostrum_1ng_glyco_hcd30_etd30_compound15.18403.18403.4 | controllerType=0 controllerNumber=1 scan=18403 | 44.8564 |
| 14822:4 | 3 | R.LSLHRPALEDLLLGSEAN[+1913.677]LTC[+57.021]TLTGLR.D | HexNAc(4)Hex(5)NeuAc(1) | C[+57], N[+1914] | 1220.0787 | 4 | 4877.2928 | 4877.2753 | 0 | 3.6 | 127 | Specific | 778.45 | 763.59 | 763.59 | 13.39 | 45 | >sp\|P01876\|IGHA1_HUMAN Immunoglobulin heavy constant alpha 1 OS=Homo sapiens OX=9606 GN=IGHA1 PE=1 SV=2 | 255 | 20201118_lss_colostrum_1ng_glyco_hcd30_etd30_compound15.18361.18361.4 | controllerType=0 controllerNumber=1 scan=18361 | 44.7232 |
| 14884:3 | 3 | R.LSLHRPALEDLLLGSEAN[+1710.598]LTC[+57.021]TLTGLR.D | HexNAc(3)Hex(5)NeuAc(1) | C[+57], N[+1711] | 1169.3094 | 4 | 4674.2156 | 4674.1959 | 0 | 4.2 | 127 | Specific | 813.71 | 813.71 | 813.71 | 13.9 | 45 | >sp\|P01876\|IGHA1_HUMAN Immunoglobulin heavy constant alpha 1 OS=Homo sapiens OX=9606 GN=IGHA1 PE=1 SV=2 | 255 | 20201118_lss_colostrum_1ng_glyco_hcd30_etd30_compound15.18426.18426.4 | controllerType=0 controllerNumber=1 scan=18426 | 44.9168 |
| 14120:4 | 3 | R.LSLHRPALEDLLLGSEAN[+1622.582]LTC[+57.021]TLTGLR.D | HexNAc(4)Hex(5) | C[+57], N[+1623] | 1147.3025 | 4 | 4586.1883 | 4586.1799 | 0 | 1.8 | 127 | Specific | 829.84 | 827.77 | 827.77 | 14.28 | 45 | >sp\|P01876\|IGHA1_HUMAN Immunoglobulin heavy constant alpha 1 OS=Homo sapiens OX=9606 GN=IGHA1 PE=1 SV=2 | 255 | 20201118_lss_colostrum_1ng_glyco_hcd30_etd30_compound15.17621.17621.4 | controllerType=0 controllerNumber=1 scan=17621 | 42.6771 |
| 14883:4 | 3 | R.LSLHRPALEDLLLGSEAN[+1751.624]LTC[+57.021]TLTGLR.D | HexNAc(4)Hex(4)NeuAc(1) | C[+57], N[+1752] | 1179.5649 | 4 | 4715.2376 | 4715.2225 | 0 | 3.2 | 127 | Specific | 915.08 | 906.32 | 906.32 | 16.39 | 45 | >sp\|P01876\|IGHA1_HUMAN Immunoglobulin heavy constant alpha 1 OS=Homo sapiens OX=9606 GN=IGHA1 PE=1 SV=2 | 255 | 20201118_lss_colostrum_1ng_glyco_hcd30_etd30_compound15.18425.18425.4 | controllerType=0 controllerNumber=1 scan=18425 | 44.9122 |
| 14100:4 | 3 | R.LSLHRPALEDLLLGSEAN[+1540.529]LTC[+57.021]TLTGLR.D | HexNAc(2)Hex(7) | C[+57], N[+1541] | 1126.7891 | 4 | 4504.1346 | 4504.1268 | 0 | 1.7 | 127 | Specific | 929.28 | 877.49 | 877.49 | 16.28 | 45 | >sp\|P01876\|IGHA1_HUMAN Immunoglobulin heavy constant alpha 1 OS=Homo sapiens OX=9606 GN=IGHA1 PE=1 SV=2 | 255 | 20201118_lss_colostrum_1ng_glyco_hcd30_etd30_compound15.17600.17600.4 | controllerType=0 controllerNumber=1 scan=17600 | 42.6076 |
| 14043:4 | 3 | R.LSLHRPALEDLLLGSEAN[+1864.634]LTC[+57.021]TLTGLR.D | HexNAc(2)Hex(9) | C[+57], N[+1865] | 1207.8171 | 4 | 4828.2464 | 4828.2324 | 0 | 2.9 | 127 | Specific | 984.01 | 956.46 | 956.46 | 17.39 | 45 | >sp\|P01876\|IGHA1_HUMAN Immunoglobulin heavy constant alpha 1 OS=Homo sapiens OX=9606 GN=IGHA1 PE=1 SV=2 | 255 | 20201118_lss_colostrum_1ng_glyco_hcd30_etd30_compound15.17540.17540.4 | controllerType=0 controllerNumber=1 scan=17540 | 42.4253 |
| 14061:4 | 3 | R.LSLHRPALEDLLLGSEAN[+1702.581]LTC[+57.021]TLTGLR.D | HexNAc(2)Hex(8) | C[+57], N[+1703] | 1167.3039 | 4 | 4666.1937 | 4666.1796 | 0 | 3 | 127 | Specific | 1017.09 | 947.18 | 947.18 | 18.39 | 45 | >sp\|P01876\|IGHA1_HUMAN Immunoglobulin heavy constant alpha 1 OS=Homo sapiens OX=9606 GN=IGHA1 PE=1 SV=2 | 255 | 20201118_lss_colostrum_1ng_glyco_hcd30_etd30_compound15.17559.17559.4 | controllerType=0 controllerNumber=1 scan=17559 | 42.4817 |
| 14074:4 | 3 | R.LSLHRPALEDLLLGSEAN[+1663.608]LTC[+57.021]TLTGLR.D | HexNAc(5)Hex(4) | C[+57], N[+1664] | 1157.5617 | 4 | 4627.2249 | 4627.2064 | 0 | 4 | 127 | Specific | 1017.14 | 995.12 | 995.12 | 18.39 | 45 | >sp\|P01876\|IGHA1_HUMAN Immunoglobulin heavy constant alpha 1 OS=Homo sapiens OX=9606 GN=IGHA1 PE=1 SV=2 | 255 | 20201118_lss_colostrum_1ng_glyco_hcd30_etd30_compound15.17572.17572.4 | controllerType=0 controllerNumber=1 scan=17572 | 42.524 |
| 14154:4 | 3 | R.LSLHRPALEDLLLGSEAN[+1095.397]LTC[+57.021]TLTGLR.D | HexNAc(3)Hex(3) | C[+57], N[+1095] | 1015.5063 | 4 | 4059.0032 | 4058.9949 | 0 | 2.1 | 127 | Specific | 1054.69 | 1054.69 | 1054.69 | 19.28 | 45 | >sp\|P01876\|IGHA1_HUMAN Immunoglobulin heavy constant alpha 1 OS=Homo sapiens OX=9606 GN=IGHA1 PE=1 SV=2 | 255 | 20201118_lss_colostrum_1ng_glyco_hcd30_etd30_compound15.17658.17658.4 | controllerType=0 controllerNumber=1 scan=17658 | 42.8039 |
| 14119:4 | 3 | R.LSLHRPALEDLLLGSEAN[+1216.423]LTC[+57.021]TLTGLR.D | HexNAc(2)Hex(5) | C[+57], N[+1216] | 1045.7637 | 4 | 4180.033 | 4180.0211 | 0 | 2.8 | 127 | Specific | 1098.17 | 1063.28 | 1063.28 | 19.39 | 45 | >sp\|P01876\|IGHA1_HUMAN Immunoglobulin heavy constant alpha 1 OS=Homo sapiens OX=9606 GN=IGHA1 PE=1 SV=2 | 255 | 20201118_lss_colostrum_1ng_glyco_hcd30_etd30_compound15.17620.17620.4 | controllerType=0 controllerNumber=1 scan=17620 | 42.6742 |
| 14085:4 | 3 | R.LSLHRPALEDLLLGSEAN[+1378.476]LTC[+57.021]TLTGLR.D | HexNAc(2)Hex(6) | C[+57], N[+1378] | 1086.2768 | 4 | 4342.0853 | 4342.074 | 0 | 2.6 | 127 | Specific | 1194.73 | 1108.61 | 1108.61 | 21.39 | 45 | >sp\|P01876\|IGHA1_HUMAN Immunoglobulin heavy constant alpha 1 OS=Homo sapiens OX=9606 GN=IGHA1 PE=1 SV=2 | 255 | 20201118_lss_colostrum_1ng_glyco_hcd30_etd30_compound15.17584.17584.4 | controllerType=0 controllerNumber=1 scan=17584 | 42.5623 |
| 10920:3 | 3 | R.LAGKPTHVN[+2059.735]VSVVMAEVDGTC[+57.021]Y.- | HexNAc(4)Hex(5)Fuc(1)NeuAc(1) | C[+57], N[+2060] | 1469.6342 | 3 | 4406.8881 | 4406.877 | 0 | 2.5 | 332 | CRagged | 208.96 | 115.57 | 115.57 | 2.75 | 45 | >sp\|P01876\|IGHA1_HUMAN Immunoglobulin heavy constant alpha 1 OS=Homo sapiens OX=9606 GN=IGHA1 PE=1 SV=2 | 255 | 20201118_lss_colostrum_1ng_glyco_hcd30_etd30_compound15.14258.14258.3 | controllerType=0 controllerNumber=1 scan=14258 | 33.834 |
| 09557:3 | 3 | R.LAGKPTHVN[+1663.608]VSVVMAEVDGTC[+57.021]Y.- | HexNAc(5)Hex(4) | C[+57], N[+1664] | 1337.5922 | 3 | 4010.7621 | 4010.7502 | 0 | 3 | 332 | CRagged | 215.98 | 114.28 | 114.28 | 2.75 | 45 | >sp\|P01876\|IGHA1_HUMAN Immunoglobulin heavy constant alpha 1 OS=Homo sapiens OX=9606 GN=IGHA1 PE=1 SV=2 | 255 | 20201118_lss_colostrum_1ng_glyco_hcd30_etd30_compound15.12836.12836.3 | controllerType=0 controllerNumber=1 scan=12836 | 30.5742 |
| 08733:3 | 3 | R.LAGKPTHVN[+1378.476]VSVVMAEVDGTC[+57.021]Y.- | HexNAc(2)Hex(6) | C[+57], N[+1378] | 1242.5489 | 3 | 3725.6321 | 3725.6178 | 0 | 3.8 | 332 | CRagged | 220.04 | 220.04 | 160.92 | 3.21 | 45 | >sp\|P01876\|IGHA1_HUMAN Immunoglobulin heavy constant alpha 1 OS=Homo sapiens OX=9606 GN=IGHA1 PE=1 SV=2 | 255 | 20201118_lss_colostrum_1ng_glyco_hcd30_etd30_compound15.11975.11975.3 | controllerType=0 controllerNumber=1 scan=11975 | 28.4822 |
| 09638:3 | 3 | R.LAGKPTHVN[+1501.555]VSVVMAEVDGTC[+57.021]Y.- | HexNAc(5)Hex(3) | C[+57], N[+1502] | 1283.5754 | 3 | 3848.7116 | 3848.6974 | 0 | 3.7 | 332 | CRagged | 231.95 | 143.38 | 143.38 | 2.82 | 45 | >sp\|P01876\|IGHA1_HUMAN Immunoglobulin heavy constant alpha 1 OS=Homo sapiens OX=9606 GN=IGHA1 PE=1 SV=2 | 255 | 20201118_lss_colostrum_1ng_glyco_hcd30_etd30_compound15.12921.12921.3 | controllerType=0 controllerNumber=1 scan=12921 | 30.819 |
| 09506:3 | 3 | R.LAGKPTHVN[+1702.581]VSVVMAEVDGTC[+57.021]Y.- | HexNAc(2)Hex(8) | C[+57], N[+1703] | 1350.5851 | 3 | 4049.7409 | 4049.7234 | 0 | 4.3 | 332 | CRagged | 233.84 | 166.78 | 166.78 | 2.68 | 45 | >sp\|P01876\|IGHA1_HUMAN Immunoglobulin heavy constant alpha 1 OS=Homo sapiens OX=9606 GN=IGHA1 PE=1 SV=2 | 255 | 20201118_lss_colostrum_1ng_glyco_hcd30_etd30_compound15.12782.12782.3 | controllerType=0 controllerNumber=1 scan=12782 | 30.4297 |
| 09755:3 | 3 | R.LAGKPTHVN[+1216.423]VSVVMAEVDGTC[+57.021]Y.- | HexNAc(2)Hex(5) | C[+57], N[+1216] | 1188.5314 | 3 | 3563.5797 | 3563.5649 | 0 | 4.2 | 332 | CRagged | 236.21 | 236.21 | 164.21 | 3.47 | 45 | >sp\|P01876\|IGHA1_HUMAN Immunoglobulin heavy constant alpha 1 OS=Homo sapiens OX=9606 GN=IGHA1 PE=1 SV=2 | 255 | 20201118_lss_colostrum_1ng_glyco_hcd30_etd30_compound15.13043.13043.3 | controllerType=0 controllerNumber=1 scan=13043 | 31.1464 |
| 08757:3 | 3 | R.LAGKPTHVN[+1216.423]VSVVMAEVDGTC[+57.021]Y.- | HexNAc(2)Hex(5) | C[+57], N[+1216] | 1188.5303 | 3 | 3563.5764 | 3563.5649 | 0 | 3.2 | 332 | CRagged | 243.65 | 218.71 | 186.6 | 3.82 | 45 | >sp\|P01876\|IGHA1_HUMAN Immunoglobulin heavy constant alpha 1 OS=Homo sapiens OX=9606 GN=IGHA1 PE=1 SV=2 | 255 | 20201118_lss_colostrum_1ng_glyco_hcd30_etd30_compound15.12000.12000.3 | controllerType=0 controllerNumber=1 scan=12000 | 28.5667 |
| 09556:3 | 3 | R.LAGKPTHVN[+1378.476]VSVVMAEVDGTC[+57.021]Y.- | HexNAc(2)Hex(6) | C[+57], N[+1378] | 1242.5475 | 3 | 3725.6281 | 3725.6178 | 0 | 2.8 | 332 | CRagged | 246.01 | 246.01 | 197.4 | 4.34 | 45 | >sp\|P01876\|IGHA1_HUMAN Immunoglobulin heavy constant alpha 1 OS=Homo sapiens OX=9606 GN=IGHA1 PE=1 SV=2 | 255 | 20201118_lss_colostrum_1ng_glyco_hcd30_etd30_compound15.12835.12835.3 | controllerType=0 controllerNumber=1 scan=12835 | 30.5712 |
| 09568:3 | 3 | R.LAGKPTHVN[+1809.666]VSVVMAEVDGTC[+57.021]Y.- | HexNAc(5)Hex(4)Fuc(1) | C[+57], N[+1810] | 1386.2775 | 3 | 4156.818 | 4156.8081 | 0 | 2.4 | 332 | CRagged | 257.7 | 165.92 | 165.92 | 3.38 | 45 | >sp\|P01876\|IGHA1_HUMAN Immunoglobulin heavy constant alpha 1 OS=Homo sapiens OX=9606 GN=IGHA1 PE=1 SV=2 | 255 | 20201118_lss_colostrum_1ng_glyco_hcd30_etd30_compound15.12847.12847.3 | controllerType=0 controllerNumber=1 scan=12847 | 30.5933 |
| 09578:3 | 3 | R.LAGKPTHVN[+1216.423]VSVVMAEVDGTC[+57.021]Y.- | HexNAc(2)Hex(5) | C[+57], N[+1216] | 1188.5301 | 3 | 3563.5757 | 3563.5649 | 0 | 3 | 332 | CRagged | 262.93 | 249.52 | 187.79 | 4.45 | 45 | >sp\|P01876\|IGHA1_HUMAN Immunoglobulin heavy constant alpha 1 OS=Homo sapiens OX=9606 GN=IGHA1 PE=1 SV=2 | 255 | 20201118_lss_colostrum_1ng_glyco_hcd30_etd30_compound15.12858.12858.3 | controllerType=0 controllerNumber=1 scan=12858 | 30.6321 |
| 09519:3 | 3 | R.LAGKPTHVN[+1540.529]VSVVMAEVDGTC[+57.021]Y.- | HexNAc(2)Hex(7) | C[+57], N[+1541] | 1296.5652 | 3 | 3887.6812 | 3887.6706 | 0 | 2.7 | 332 | CRagged | 269.45 | 204.64 | 204.64 | 3.87 | 45 | >sp\|P01876\|IGHA1_HUMAN Immunoglobulin heavy constant alpha 1 OS=Homo sapiens OX=9606 GN=IGHA1 PE=1 SV=2 | 255 | 20201118_lss_colostrum_1ng_glyco_hcd30_etd30_compound15.12796.12796.3 | controllerType=0 controllerNumber=1 scan=12796 | 30.4699 |
| 09661:3 | 3 | R.LAGKPTHVN[+1647.613]VSVVMAEVDGTC[+57.021]Y.- | HexNAc(5)Hex(3)Fuc(1) | C[+57], N[+1648] | 1332.259 | 3 | 3994.7624 | 3994.7553 | 0 | 1.8 | 332 | CRagged | 295.28 | 157.82 | 157.82 | 3.73 | 45 | >sp\|P01876\|IGHA1_HUMAN Immunoglobulin heavy constant alpha 1 OS=Homo sapiens OX=9606 GN=IGHA1 PE=1 SV=2 | 255 | 20201118_lss_colostrum_1ng_glyco_hcd30_etd30_compound15.12945.12945.3 | controllerType=0 controllerNumber=1 scan=12945 | 30.8823 |
| 10880:3 | 3 | R.LAGKPTHVN[+2262.814]VSVVMAEVDGTC[+57.021]Y.- | HexNAc(5)Hex(5)Fuc(1)NeuAc(1) | C[+57], N[+2263] | 1537.3276 | 3 | 4609.9682 | 4609.9564 | 0 | 2.6 | 332 | CRagged | 317.66 | 130.69 | 130.69 | 3.83 | 45 | >sp\|P01876\|IGHA1_HUMAN Immunoglobulin heavy constant alpha 1 OS=Homo sapiens OX=9606 GN=IGHA1 PE=1 SV=2 | 255 | 20201118_lss_colostrum_1ng_glyco_hcd30_etd30_compound15.14217.14217.3 | controllerType=0 controllerNumber=1 scan=14217 | 33.7303 |
| 10879:4 | 3 | R.LAGKPTHVN[+2262.814]VSVVMAEVDGTC[+57.021]Y.- | HexNAc(5)Hex(5)Fuc(1)NeuAc(1) | C[+57], N[+2263] | 1153.2471 | 4 | 4609.9666 | 4609.9564 | 0 | 2.2 | 332 | CRagged | 339.65 | 262.88 | 262.88 | 4.51 | 45 | >sp\|P01876\|IGHA1_HUMAN Immunoglobulin heavy constant alpha 1 OS=Homo sapiens OX=9606 GN=IGHA1 PE=1 SV=2 | 255 | 20201118_lss_colostrum_1ng_glyco_hcd30_etd30_compound15.14216.14216.4 | controllerType=0 controllerNumber=1 scan=14216 | 33.7269 |
| 12202:4 | 3 | R.LAGKPTHVN[+2553.910]VSVVMAEVDGTC[+57.021]Y.- | HexNAc(5)Hex(5)Fuc(1)NeuAc(2) | C[+57], N[+2554] | 1226.0226 | 4 | 4901.0687 | 4901.0518 | 0 | 3.4 | 332 | CRagged | 431.6 | 235.81 | 235.81 | 4.62 | 45 | >sp\|P01876\|IGHA1_HUMAN Immunoglobulin heavy constant alpha 1 OS=Homo sapiens OX=9606 GN=IGHA1 PE=1 SV=2 | 255 | 20201118_lss_colostrum_1ng_glyco_hcd30_etd30_compound15.15605.15605.4 | controllerType=0 controllerNumber=1 scan=15605 | 37.3512 |
| 07613:3 | 3 | R.LAGKPTHVN[+1216.423]VSVVM[+15.995]AEVDGTC[+57.021]Y.- | HexNAc(2)Hex(5) | C[+57], M[+16], N[+1216] | 1193.8633 | 3 | 3579.5755 | 3579.5599 | 0 | 4.4 | 332 | CRagged | 223.65 | 180.96 | 180.96 | 2.76 | 45 | >sp\|P01876\|IGHA1_HUMAN Immunoglobulin heavy constant alpha 1 OS=Homo sapiens OX=9606 GN=IGHA1 PE=1 SV=2 | 255 | 20201118_lss_colostrum_1ng_glyco_hcd30_etd30_compound15.10805.10805.3 | controllerType=0 controllerNumber=1 scan=10805 | 25.9695 |
| 08636:3 | 4 | K.TVLTPATNHMGN[+1378.476]VTFTIPANR.E | HexNAc(2)Hex(6) | N[+1378] | 1211.8868 | 3 | 3633.6458 | 3633.6358 | 0 | 2.7 | 74 | Specific | 493.96 | 389.15 | 162.57 | 7.44 | 26 | >sp\|P01024\|CO3_HUMAN Complement C3 OS=Homo sapiens OX=9606 GN=C3 PE=1 SV=2 | 492 | 20201118_lss_colostrum_1ng_glyco_hcd30_etd30_compound15.11874.11874.3 | controllerType=0 controllerNumber=1 scan=11874 | 28.2624 |
| 13965:3 | 5 | R.GLSFDVSLEVSQGPGLLN[+2715.963]DTK.V | HexNAc(5)Hex(6)Fuc(1)NeuAc(2) | N[+2716] | 1631.3793 | 3 | 4892.1234 | 4892.0758 | 0 | 9.7 | 118 | Specific | 362.28 | 253.07 | 253.07 | 4.27 | 45 | >sp\|P01833\|PIGR_HUMAN Polymeric immunoglobulin receptor OS=Homo sapiens OX=9606 GN=PIGR PE=1 SV=4 | 902 | 20201118_lss_colostrum_1ng_glyco_hcd30_etd30_compound15.17456.17456.3 | controllerType=0 controllerNumber=1 scan=17456 | 42.1501 |
| 14997:4 | 5 | K.QIGLYPVLVIDSSGYVNPN[+2205.793]YTGR.I | HexNAc(4)Hex(5)Fuc(2)NeuAc(1) | N[+2206] | 1183.5339 | 4 | 4731.1136 | 4731.0963 | 0 | 3.7 | 168 | Specific | 409.79 | 409.79 | 409.79 | 6.06 | 45 | >sp\|P01833\|PIGR_HUMAN Polymeric immunoglobulin receptor OS=Homo sapiens OX=9606 GN=PIGR PE=1 SV=4 | 902 | 20201118_lss_colostrum_1ng_glyco_hcd30_etd30_compound15.18545.18545.4 | controllerType=0 controllerNumber=1 scan=18545 | 45.2777 |
| 16361:3 | 5 | K.QIGLYPVLVIDSSGYVNPN[+2350.830]YTGR.I | HexNAc(4)Hex(5)Fuc(1)NeuAc(2) | N[+2351] | 1626.0527 | 3 | 4876.1435 | 4876.1338 | 0 | 2 | 168 | Specific | 470.17 | 314.65 | 314.65 | 5.81 | 45 | >sp\|P01833\|PIGR_HUMAN Polymeric immunoglobulin receptor OS=Homo sapiens OX=9606 GN=PIGR PE=1 SV=4 | 902 | 20201118_lss_colostrum_1ng_glyco_hcd30_etd30_compound15.19986.19986.3 | controllerType=0 controllerNumber=1 scan=19986 | 49.7263 |
| 15008:4 | 5 | K.QIGLYPVLVIDSSGYVNPN[+2059.735]YTGR.I | HexNAc(4)Hex(5)Fuc(1)NeuAc(1) | N[+2060] | 1147.0187 | 4 | 4585.053 | 4585.0384 | 0 | 3.2 | 168 | Specific | 488.21 | 428.56 | 428.56 | 6.42 | 45 | >sp\|P01833\|PIGR_HUMAN Polymeric immunoglobulin receptor OS=Homo sapiens OX=9606 GN=PIGR PE=1 SV=4 | 902 | 20201118_lss_colostrum_1ng_glyco_hcd30_etd30_compound15.18557.18557.4 | controllerType=0 controllerNumber=1 scan=18557 | 45.3164 |
| 15007:3 | 5 | K.QIGLYPVLVIDSSGYVNPN[+2059.735]YTGR.I | HexNAc(4)Hex(5)Fuc(1)NeuAc(1) | N[+2060] | 1529.024 | 3 | 4585.0574 | 4585.0384 | 0 | 4.1 | 168 | Specific | 522.29 | 460.57 | 460.57 | 7.64 | 45 | >sp\|P01833\|PIGR_HUMAN Polymeric immunoglobulin receptor OS=Homo sapiens OX=9606 GN=PIGR PE=1 SV=4 | 902 | 20201118_lss_colostrum_1ng_glyco_hcd30_etd30_compound15.18556.18556.3 | controllerType=0 controllerNumber=1 scan=18556 | 45.3133 |
| 15040:3 | 5 | K.QIGLYPVLVIDSSGYVNPN[+1913.677]YTGR.I | HexNAc(4)Hex(5)NeuAc(1) | N[+1914] | 1480.3392 | 3 | 4439.003 | 4438.9805 | 0 | 5.1 | 168 | Specific | 574.86 | 518.69 | 518.69 | 8.26 | 45 | >sp\|P01833\|PIGR_HUMAN Polymeric immunoglobulin receptor OS=Homo sapiens OX=9606 GN=PIGR PE=1 SV=4 | 902 | 20201118_lss_colostrum_1ng_glyco_hcd30_etd30_compound15.18591.18591.3 | controllerType=0 controllerNumber=1 scan=18591 | 45.445 |
| 21210:3 | 5 | R.[+42.011]LSLLEEPGN[+2351.851]GTFTVILNQLTSR.D | HexNAc(4)Hex(5)Fuc(3)NeuAc(1) | .[42], N[+2352] | 1599.3859 | 3 | 4796.1432 | 4796.1539 | 0 | -2.2 | 413 | Specific | 250.04 | 131.82 | 131.82 | 3.38 | 45 | >sp\|P01833\|PIGR_HUMAN Polymeric immunoglobulin receptor OS=Homo sapiens OX=9606 GN=PIGR PE=1 SV=4 | 902 | 20201118_lss_colostrum_1ng_glyco_hcd30_etd30_compound15.25132.25132.3 | controllerType=0 controllerNumber=1 scan=25132 | 65.4402 |
| 16024:3 | 5 | R.LSLLEEPGN[+2570.925]GTFTVILNQLTSR.D | HexNAc(5)Hex(6)Fuc(2)NeuAc(1) | N[+2571] | 1658.4237 | 3 | 4973.2565 | 4973.2176 | 0 | 7.8 | 413 | Specific | 438.27 | 283.59 | 283.59 | 3.98 | 45 | >sp\|P01833\|PIGR_HUMAN Polymeric immunoglobulin receptor OS=Homo sapiens OX=9606 GN=PIGR PE=1 SV=4 | 902 | 20201118_lss_colostrum_1ng_glyco_hcd30_etd30_compound15.19628.19628.3 | controllerType=0 controllerNumber=1 scan=19628 | 48.5435 |
| 17354:4 | 5 | R.LSLLEEPGN[+2350.830]GTFTVILNQLTSR.D | HexNAc(4)Hex(5)Fuc(1)NeuAc(2) | N[+2351] | 1189.0426 | 4 | 4753.1488 | 4753.1229 | 0 | 5.4 | 413 | Specific | 522.4 | 442.95 | 442.95 | 6.03 | 45 | >sp\|P01833\|PIGR_HUMAN Polymeric immunoglobulin receptor OS=Homo sapiens OX=9606 GN=PIGR PE=1 SV=4 | 902 | 20201118_lss_colostrum_1ng_glyco_hcd30_etd30_compound15.21042.21042.4 | controllerType=0 controllerNumber=1 scan=21042 | 53.2471 |
| 16087:3 | 5 | R.LSLLEEPGN[+2205.793]GTFTVILNQLTSR.D | HexNAc(4)Hex(5)Fuc(2)NeuAc(1) | N[+2206] | 1536.7084 | 3 | 4608.1107 | 4608.0854 | 0 | 5.5 | 413 | Specific | 556.67 | 463.21 | 463.21 | 7.15 | 45 | >sp\|P01833\|PIGR_HUMAN Polymeric immunoglobulin receptor OS=Homo sapiens OX=9606 GN=PIGR PE=1 SV=4 | 902 | 20201118_lss_colostrum_1ng_glyco_hcd30_etd30_compound15.19695.19695.3 | controllerType=0 controllerNumber=1 scan=19695 | 48.7781 |
| 16078:4 | 5 | R.LSLLEEPGN[+2351.851]GTFTVILNQLTSR.D | HexNAc(4)Hex(5)Fuc(3)NeuAc(1) | N[+2352] | 1189.2967 | 4 | 4754.1649 | 4754.1433 | 0 | 4.5 | 413 | Specific | 629.9 | 546.37 | 546.37 | 8.38 | 45 | >sp\|P01833\|PIGR_HUMAN Polymeric immunoglobulin receptor OS=Homo sapiens OX=9606 GN=PIGR PE=1 SV=4 | 902 | 20201118_lss_colostrum_1ng_glyco_hcd30_etd30_compound15.19686.19686.4 | controllerType=0 controllerNumber=1 scan=19686 | 48.7427 |
| 16109:4 | 5 | R.LSLLEEPGN[+2059.735]GTFTVILNQLTSR.D | HexNAc(4)Hex(5)Fuc(1)NeuAc(1) | N[+2060] | 1116.2663 | 4 | 4462.0433 | 4462.0275 | 0 | 3.5 | 413 | Specific | 633.71 | 567.08 | 567.08 | 8.86 | 45 | >sp\|P01833\|PIGR_HUMAN Polymeric immunoglobulin receptor OS=Homo sapiens OX=9606 GN=PIGR PE=1 SV=4 | 902 | 20201118_lss_colostrum_1ng_glyco_hcd30_etd30_compound15.19719.19719.4 | controllerType=0 controllerNumber=1 scan=19719 | 48.8619 |
| 17347:3 | 5 | R.LSLLEEPGN[+2350.830]GTFTVILNQLTSR.D | HexNAc(4)Hex(5)Fuc(1)NeuAc(2) | N[+2351] | 1585.0534 | 3 | 4753.1457 | 4753.1229 | 0 | 4.8 | 413 | Specific | 664.64 | 467.37 | 467.37 | 10.26 | 45 | >sp\|P01833\|PIGR_HUMAN Polymeric immunoglobulin receptor OS=Homo sapiens OX=9606 GN=PIGR PE=1 SV=4 | 902 | 20201118_lss_colostrum_1ng_glyco_hcd30_etd30_compound15.21035.21035.3 | controllerType=0 controllerNumber=1 scan=21035 | 53.2271 |
| 16103:3 | 5 | R.LSLLEEPGN[+2059.735]GTFTVILNQLTSR.D | HexNAc(4)Hex(5)Fuc(1)NeuAc(1) | N[+2060] | 1488.0204 | 3 | 4462.0468 | 4462.0275 | 0 | 4.3 | 413 | Specific | 669.34 | 529.54 | 529.54 | 10.26 | 45 | >sp\|P01833\|PIGR_HUMAN Polymeric immunoglobulin receptor OS=Homo sapiens OX=9606 GN=PIGR PE=1 SV=4 | 902 | 20201118_lss_colostrum_1ng_glyco_hcd30_etd30_compound15.19713.19713.3 | controllerType=0 controllerNumber=1 scan=19713 | 48.8405 |
| 12951:3 | 5 | K.IIEGEPNLKVPGN[+2205.793]VTAVLGETLK.V | HexNAc(4)Hex(5)Fuc(2)NeuAc(1) | N[+2206] | 1533.0566 | 3 | 4597.1552 | 4597.1422 | 0 | 2.8 | 457 | Specific | 210.72 | 179.45 | 179.45 | 3.63 | 45 | >sp\|P01833\|PIGR_HUMAN Polymeric immunoglobulin receptor OS=Homo sapiens OX=9606 GN=PIGR PE=1 SV=4 | 902 | 20201118_lss_colostrum_1ng_glyco_hcd30_etd30_compound15.16393.16393.3 | controllerType=0 controllerNumber=1 scan=16393 | 39.4383 |
| 12933:3 | 5 | K.IIEGEPNLKVPGN[+2205.793]VTAVLGETLK.V | HexNAc(4)Hex(5)Fuc(2)NeuAc(1) | N[+2206] | 1533.0578 | 3 | 4597.1588 | 4597.1422 | 0 | 3.6 | 457 | Specific | 295.68 | 227.68 | 227.68 | 4.04 | 45 | >sp\|P01833\|PIGR_HUMAN Polymeric immunoglobulin receptor OS=Homo sapiens OX=9606 GN=PIGR PE=1 SV=4 | 902 | 20201118_lss_colostrum_1ng_glyco_hcd30_etd30_compound15.16374.16374.3 | controllerType=0 controllerNumber=1 scan=16374 | 39.3596 |
| 12848:3 | 5 | K.IIEGEPNLKVPGN[+2351.851]VTAVLGETLK.V | HexNAc(4)Hex(5)Fuc(3)NeuAc(1) | N[+2352] | 1581.7424 | 3 | 4743.2125 | 4743.2001 | 0 | 2.6 | 457 | Specific | 304.18 | 263.57 | 263.57 | 4.3 | 45 | >sp\|P01833\|PIGR_HUMAN Polymeric immunoglobulin receptor OS=Homo sapiens OX=9606 GN=PIGR PE=1 SV=4 | 902 | 20201118_lss_colostrum_1ng_glyco_hcd30_etd30_compound15.16285.16285.3 | controllerType=0 controllerNumber=1 scan=16285 | 39.1481 |
| 14270:3 | 5 | K.IIEGEPNLKVPGN[+2350.830]VTAVLGETLK.V | HexNAc(4)Hex(5)Fuc(1)NeuAc(2) | N[+2351] | 1581.4036 | 3 | 4742.1963 | 4742.1797 | 0 | 3.5 | 457 | Specific | 326.61 | 245.79 | 245.79 | 4.36 | 45 | >sp\|P01833\|PIGR_HUMAN Polymeric immunoglobulin receptor OS=Homo sapiens OX=9606 GN=PIGR PE=1 SV=4 | 902 | 20201118_lss_colostrum_1ng_glyco_hcd30_etd30_compound15.17781.17781.3 | controllerType=0 controllerNumber=1 scan=17781 | 43.1861 |
| 12927:3 | 5 | K.IIEGEPNLKVPGN[+2059.735]VTAVLGETLK.V | HexNAc(4)Hex(5)Fuc(1)NeuAc(1) | N[+2060] | 1484.3698 | 3 | 4451.0949 | 4451.0843 | 0 | 2.4 | 457 | Specific | 363.35 | 291.13 | 291.13 | 4.44 | 45 | >sp\|P01833\|PIGR_HUMAN Polymeric immunoglobulin receptor OS=Homo sapiens OX=9606 GN=PIGR PE=1 SV=4 | 902 | 20201118_lss_colostrum_1ng_glyco_hcd30_etd30_compound15.16368.16368.3 | controllerType=0 controllerNumber=1 scan=16368 | 39.3477 |
| 14304:4 | 5 | K.IIEGEPNLKVPGN[+2350.830]VTAVLGETLK.V | HexNAc(4)Hex(5)Fuc(1)NeuAc(2) | N[+2351] | 1186.305 | 4 | 4742.1981 | 4742.1797 | 0 | 3.9 | 457 | Specific | 463.33 | 448.71 | 448.71 | 5.95 | 45 | >sp\|P01833\|PIGR_HUMAN Polymeric immunoglobulin receptor OS=Homo sapiens OX=9606 GN=PIGR PE=1 SV=4 | 902 | 20201118_lss_colostrum_1ng_glyco_hcd30_etd30_compound15.17817.17817.4 | controllerType=0 controllerNumber=1 scan=17817 | 43.2953 |
| 12913:4 | 5 | K.IIEGEPNLKVPGN[+2059.735]VTAVLGETLK.V | HexNAc(4)Hex(5)Fuc(1)NeuAc(1) | N[+2060] | 1113.5303 | 4 | 4451.0994 | 4451.0843 | 0 | 3.4 | 457 | Specific | 489.89 | 485.39 | 485.39 | 6.48 | 45 | >sp\|P01833\|PIGR_HUMAN Polymeric immunoglobulin receptor OS=Homo sapiens OX=9606 GN=PIGR PE=1 SV=4 | 902 | 20201118_lss_colostrum_1ng_glyco_hcd30_etd30_compound15.16353.16353.4 | controllerType=0 controllerNumber=1 scan=16353 | 39.3094 |
| 11678:5 | 5 | K.IIEGEPNLKVPGN[+2424.867]VTAVLGETLKVPC[+57.021]HFPC[+57.021]K.F | HexNAc(5)Hex(6)Fuc(1)NeuAc(1) | C[+57]*2, N[+2425] | 1169.1466 | 5 | 5841.7039 | 5841.674 | 0 | 5.1 | 457 | Specific | 213.25 | 213.25 | 213.25 | 2.63 | 45 | >sp\|P01833\|PIGR_HUMAN Polymeric immunoglobulin receptor OS=Homo sapiens OX=9606 GN=PIGR PE=1 SV=4 | 902 | 20201118_lss_colostrum_1ng_glyco_hcd30_etd30_compound15.15054.15054.5 | controllerType=0 controllerNumber=1 scan=15054 | 35.9291 |
| 12584:4 | 5 | K.[+42.011]IIEGEPNLKVPGN[+2018.708]VTAVLGETLKVPC[+57.021]HFPC[+57.021]K | HexNAc(3)Hex(6)Fuc(1)NeuAc(1) | .[42], C[+57]*2, N[+2019] | 1370.1432 | 4 | 5477.5509 | 5477.5258 | 0 | 4.6 | 457 | Specific | 257.99 | 257.99 | 257.99 | 3.08 | 45 | >sp\|P01833\|PIGR_HUMAN Polymeric immunoglobulin receptor OS=Homo sapiens OX=9606 GN=PIGR PE=1 SV=4 | 902 | 20201118_lss_colostrum_1ng_glyco_hcd30_etd30_compound15.16007.16007.4 | controllerType=0 controllerNumber=1 scan=16007 | 38.4508 |
| 11736:4 | 5 | K.IIEGEPNLKVPGN[+2059.735]VTAVLGETLKVPC[+57.021]HFPC[+57.021]K.F | HexNAc(4)Hex(5)Fuc(1)NeuAc(1) | C[+57]*2, N[+2060] | 1369.8986 | 4 | 5476.5724 | 5476.5418 | 0 | 5.6 | 457 | Specific | 320.94 | 320.94 | 320.94 | 3.2 | 45 | >sp\|P01833\|PIGR_HUMAN Polymeric immunoglobulin receptor OS=Homo sapiens OX=9606 GN=PIGR PE=1 SV=4 | 902 | 20201118_lss_colostrum_1ng_glyco_hcd30_etd30_compound15.15115.15115.4 | controllerType=0 controllerNumber=1 scan=15115 | 36.0991 |
| 12613:5 | 5 | K.IIEGEPNLKVPGN[+2205.793]VTAVLGETLKVPC[+57.021]HFPC[+57.021]K.F | HexNAc(4)Hex(5)Fuc(2)NeuAc(1) | C[+57]*2, N[+2206] | 1125.3292 | 5 | 5622.617 | 5622.5997 | 0 | 3.1 | 457 | Specific | 381.85 | 381.85 | 381.85 | 5.52 | 45 | >sp\|P01833\|PIGR_HUMAN Polymeric immunoglobulin receptor OS=Homo sapiens OX=9606 GN=PIGR PE=1 SV=4 | 902 | 20201118_lss_colostrum_1ng_glyco_hcd30_etd30_compound15.16038.16038.5 | controllerType=0 controllerNumber=1 scan=16038 | 38.5283 |
| 12453:4 | 5 | K.IIEGEPNLKVPGN[+2205.793]VTAVLGETLKVPC[+57.021]HFPC[+57.021]K.F | HexNAc(4)Hex(5)Fuc(2)NeuAc(1) | C[+57]*2, N[+2206] | 1406.4097 | 4 | 5622.6169 | 5622.5997 | 0 | 3 | 457 | Specific | 391.05 | 391.05 | 391.05 | 5.44 | 45 | >sp\|P01833\|PIGR_HUMAN Polymeric immunoglobulin receptor OS=Homo sapiens OX=9606 GN=PIGR PE=1 SV=4 | 902 | 20201118_lss_colostrum_1ng_glyco_hcd30_etd30_compound15.15870.15870.4 | controllerType=0 controllerNumber=1 scan=15870 | 38.1165 |
| 13615:4 | 5 | K.IIEGEPNLKVPGN[+2350.830]VTAVLGETLKVPC[+57.021]HFPC[+57.021]K.F | HexNAc(4)Hex(5)Fuc(1)NeuAc(2) | C[+57]*2, N[+2351] | 1442.6688 | 4 | 5767.6535 | 5767.6372 | 0 | 2.8 | 457 | Specific | 417.72 | 417.72 | 417.72 | 5.63 | 45 | >sp\|P01833\|PIGR_HUMAN Polymeric immunoglobulin receptor OS=Homo sapiens OX=9606 GN=PIGR PE=1 SV=4 | 902 | 20201118_lss_colostrum_1ng_glyco_hcd30_etd30_compound15.17089.17089.4 | controllerType=0 controllerNumber=1 scan=17089 | 41.1474 |
| 13614:5 | 5 | K.IIEGEPNLKVPGN[+2350.830]VTAVLGETLKVPC[+57.021]HFPC[+57.021]K.F | HexNAc(4)Hex(5)Fuc(1)NeuAc(2) | C[+57]*2, N[+2351] | 1154.3361 | 5 | 5767.6512 | 5767.6372 | 0 | 2.4 | 457 | Specific | 509.39 | 509.39 | 509.39 | 5.76 | 45 | >sp\|P01833\|PIGR_HUMAN Polymeric immunoglobulin receptor OS=Homo sapiens OX=9606 GN=PIGR PE=1 SV=4 | 902 | 20201118_lss_colostrum_1ng_glyco_hcd30_etd30_compound15.17088.17088.5 | controllerType=0 controllerNumber=1 scan=17088 | 41.1438 |
| 12449:5 | 5 | K.IIEGEPNLKVPGN[+2205.793]VTAVLGETLKVPC[+57.021]HFPC[+57.021]K.F | HexNAc(4)Hex(5)Fuc(2)NeuAc(1) | C[+57]*2, N[+2206] | 1125.3289 | 5 | 5622.6152 | 5622.5997 | 0 | 2.8 | 457 | Specific | 524.31 | 524.31 | 524.31 | 6.13 | 45 | >sp\|P01833\|PIGR_HUMAN Polymeric immunoglobulin receptor OS=Homo sapiens OX=9606 GN=PIGR PE=1 SV=4 | 902 | 20201118_lss_colostrum_1ng_glyco_hcd30_etd30_compound15.15866.15866.5 | controllerType=0 controllerNumber=1 scan=15866 | 38.1072 |
| 12473:5 | 5 | K.IIEGEPNLKVPGN[+2351.851]VTAVLGETLKVPC[+57.021]HFPC[+57.021]K.F | HexNAc(4)Hex(5)Fuc(3)NeuAc(1) | C[+57]*2, N[+2352] | 1154.5388 | 5 | 5768.665 | 5768.6576 | 0 | 1.3 | 457 | Specific | 561.53 | 561.53 | 561.53 | 7.24 | 45 | >sp\|P01833\|PIGR_HUMAN Polymeric immunoglobulin receptor OS=Homo sapiens OX=9606 GN=PIGR PE=1 SV=4 | 902 | 20201118_lss_colostrum_1ng_glyco_hcd30_etd30_compound15.15891.15891.5 | controllerType=0 controllerNumber=1 scan=15891 | 38.1743 |

Table S4. Cont.

| Query #:z | Protein Rank | Glycopeptide | Glycan | Modification Type(s) | Observed m/z | z | Observed (M+H) | Calc. mass (M+H) | Off-by-x error | Mass error (ppm) | StartPosition | Cleavage | Score | Delta | Delta Mod | \|Log Prob\| | # of unique peptides | Protein Name | Protein DB number | Comment | Scan # | Scan Time |
|---|---|---|---|---|---|---|---|---|---|---|---|---|---|---|---|---|---|---|---|---|---|---|
| 12465:5 | 5 | K.IIEGEPNLKVPGN[+2059.735]VTAVLGETLKVPC[+57.021]HFPC[+57.021]K.F | HexNAc(4)Hex(5)Fuc(1)NeuAc(1) | C[+57]*2, N[+2060] | 1096.1169 | 5 | 5476.5556 | 5476.5418 | 0 | 2.5 | 457 | Specific | 586.22 | 583.05 | 583.05 | 7.35 | 45 | >sp\|P01833\|PIGR_HUMAN Polymeric immunoglobulin receptor OS=Homo sapiens OX=9606 GN=PIGR PE=1 SV=4 | 902 | 20201118_lss_colostrum_1ng_glyco_hcd30_etd30_compound15.15882.15882.5 | controllerType=0 controllerNumber=1 scan=15882 | 38.1506 |
| 12483:4 | 5 | K.IIEGEPNLKVPGN[+2059.735]VTAVLGETLKVPC[+57.021]HFPC[+57.021]K.F | HexNAc(4)Hex(5)Fuc(1)NeuAc(1) | C[+57]*2, N[+2060] | 1369.8931 | 4 | 5476.5504 | 5476.5418 | 0 | 1.6 | 457 | Specific | 586.6 | 570.18 | 570.18 | 7.1 | 45 | >sp\|P01833\|PIGR_HUMAN Polymeric immunoglobulin receptor OS=Homo sapiens OX=9606 GN=PIGR PE=1 SV=4 | 902 | 20201118_lss_colostrum_1ng_glyco_hcd30_etd30_compound15.15901.15901.4 | controllerType=0 controllerNumber=1 scan=15901 | 38.2062 |
| 11179:4 | 5 | K.VPGN[+2205.793]VTAVLGETLKVPC[+57.021]HFPC[+57.021]K.F | HexNAc(4)Hex(5)Fuc(2)NeuAc(1) | C[+57]*2, N[+2206] | 1158.0212 | 4 | 4629.0628 | 4629.0502 | 0 | 2.7 | 466 | Specific | 250.33 | 247.67 | 247.67 | 3.5 | 45 | >sp\|P01833\|PIGR_HUMAN Polymeric immunoglobulin receptor OS=Homo sapiens OX=9606 GN=PIGR PE=1 SV=4 | 902 | 20201118_lss_colostrum_1ng_glyco_hcd30_etd30_compound15.14531.14531.4 | controllerType=0 controllerNumber=1 scan=14531 | 34.571 |
| 04858:3 | 5 | K.WN[+2204.772]NTGC[+57.021]QALPSQDEGPSK.A | HexNAc(4)Hex(5)NeuAc(2) | C[+57], N[+2205] | 1398.5627 | 3 | 4193.6735 | 4193.6491 | 0 | 5.8 | 498 | Specific | 288.32 | 160.27 | 160.27 | 4.09 | 45 | >sp\|P01833\|PIGR_HUMAN Polymeric immunoglobulin receptor OS=Homo sapiens OX=9606 GN=PIGR PE=1 SV=4 | 902 | 20201118_lss_colostrum_1ng_glyco_hcd30_etd30_compound15.7944.7944.3 | controllerType=0 controllerNumber=1 scan=7944 | 20.6511 |
| 03517:3 | 5 | K.WN[+1216.423]NTGC[+57.021]QALPSQDEGPSK.A | HexNAc(2)Hex(5) | C[+57], N[+1216] | 1069.0988 | 3 | 3205.2818 | 3205.2995 | 0 | -5.5 | 498 | Specific | 321.09 | 239.97 | 239.97 | 3.52 | 45 | >sp\|P01833\|PIGR_HUMAN Polymeric immunoglobulin receptor OS=Homo sapiens OX=9606 GN=PIGR PE=1 SV=4 | 902 | 20201118_lss_colostrum_1ng_glyco_hcd30_etd30_compound15.6553.6553.3 | controllerType=0 controllerNumber=1 scan=6553 | 18.1673 |
| 03489:2 | 5 | K.WN[+1216.423]NTGC[+57.021]QALPSQDEGPSK.A | HexNAc(2)Hex(5) | C[+57], N[+1216] | 1603.1552 | 2 | 3205.3031 | 3205.2995 | 0 | 1.1 | 498 | Specific | 344.24 | 278.91 | 278.91 | 5.07 | 45 | >sp\|P01833\|PIGR_HUMAN Polymeric immunoglobulin receptor OS=Homo sapiens OX=9606 GN=PIGR PE=1 SV=4 | 902 | 20201118_lss_colostrum_1ng_glyco_hcd30_etd30_compound15.6524.6524.2 | controllerType=0 controllerNumber=1 scan=6524 | 18.1116 |
| 04875:3 | 5 | K.WN[+2059.735]NTGC[+57.021]QALPSQDEGPSK.A | HexNAc(4)Hex(5)Fuc(1)NeuAc(1) | C[+57], N[+2060] | 1350.2118 | 3 | 4048.621 | 4048.6116 | 0 | 2.3 | 498 | Specific | 354.09 | 254.72 | 254.72 | 5.17 | 45 | >sp\|P01833\|PIGR_HUMAN Polymeric immunoglobulin receptor OS=Homo sapiens OX=9606 GN=PIGR PE=1 SV=4 | 902 | 20201118_lss_colostrum_1ng_glyco_hcd30_etd30_compound15.7962.7962.3 | controllerType=0 controllerNumber=1 scan=7962 | 20.6924 |
| 05050:3 | 5 | K.WN[+1710.598]NTGC[+57.021]QALPSQDEGPSK.A | HexNAc(3)Hex(5)NeuAc(1) | C[+57], N[+1711] | 1233.8323 | 3 | 3699.4824 | 3699.4743 | 0 | 2.2 | 498 | Specific | 377.32 | 298.12 | 298.12 | 5.36 | 45 | >sp\|P01833\|PIGR_HUMAN Polymeric immunoglobulin receptor OS=Homo sapiens OX=9606 GN=PIGR PE=1 SV=4 | 902 | 20201118_lss_colostrum_1ng_glyco_hcd30_etd30_compound15.8144.8144.3 | controllerType=0 controllerNumber=1 scan=8144 | 21.0504 |
| 05082:3 | 5 | K.WN[+1548.545]NTGC[+57.021]QALPSQDEGPSK.A | HexNAc(3)Hex(4)NeuAc(1) | C[+57], N[+1549] | 1179.814 | 3 | 3537.4275 | 3537.4215 | 0 | 1.7 | 498 | Specific | 443.22 | 363.25 | 363.25 | 6.79 | 45 | >sp\|P01833\|PIGR_HUMAN Polymeric immunoglobulin receptor OS=Homo sapiens OX=9606 GN=PIGR PE=1 SV=4 | 902 | 20201118_lss_colostrum_1ng_glyco_hcd30_etd30_compound15.8177.8177.3 | controllerType=0 controllerNumber=1 scan=8177 | 21.115 |
| 04920:3 | 5 | K.WN[+1872.651]NTGC[+57.021]QALPSQDEGPSK.A | HexNAc(3)Hex(6)NeuAc(1) | C[+57], N[+1873] | 1287.8494 | 3 | 3861.5337 | 3861.5272 | 0 | 1.7 | 498 | Specific | 490.59 | 333.2 | 333.2 | 5.77 | 45 | >sp\|P01833\|PIGR_HUMAN Polymeric immunoglobulin receptor OS=Homo sapiens OX=9606 GN=PIGR PE=1 SV=4 | 902 | 20201118_lss_colostrum_1ng_glyco_hcd30_etd30_compound15.8009.8009.3 | controllerType=0 controllerNumber=1 scan=8009 | 20.7917 |
| 04921:3 | 5 | K.WN[+1913.677]NTGC[+57.021]QALPSQDEGPSK.A | HexNAc(4)Hex(5)NeuAc(1) | C[+57], N[+1914] | 1301.5253 | 3 | 3902.5614 | 3902.5537 | 0 | 2 | 498 | Specific | 517.24 | 343.99 | 343.99 | 5.77 | 45 | >sp\|P01833\|PIGR_HUMAN Polymeric immunoglobulin receptor OS=Homo sapiens OX=9606 GN=PIGR PE=1 SV=4 | 902 | 20201118_lss_colostrum_1ng_glyco_hcd30_etd30_compound15.8010.8010.3 | controllerType=0 controllerNumber=1 scan=8010 | 20.7971 |
| 04085:3 | 6 | R.FSDGLESN[+1864.634]SSTQFEVKK.Y | HexNAc(2)Hex(9) | N[+1865] | 1256.5239 | 3 | 3767.557 | 3767.5421 | 0 | 4 | 219 | Specific | 267.07 | 211.62 | 211.62 | 3.93 | 22 | >sp\|P0C0L4\|CO4A_HUMAN Complement C4-A OS=Homo sapiens OX=9606 GN=C4A PE=1 SV=2 | 1235 | 20201118_lss_colostrum_1ng_glyco_hcd30_etd30_compound15.7143.7143.3 | controllerType=0 controllerNumber=1 scan=7143 | 19.1897 |
| 00086:4 | 10 | K.KKEDALN[+2059.735]ETRESETK.L | HexNAc(4)Hex(5)Fuc(1)NeuAc(1) | N[+2060] | 960.1651 | 4 | 3837.6387 | 3837.6276 | 0 | 2.9 | 80 | Specific | 706 | 590.04 | 590.04 | 10.34 | 28 | >sp\|P10909\|CLUS_HUMAN Clusterin OS=Homo sapiens OX=9606 GN=CLU PE=1 SV=1 | 1467 | 20201118_lss_colostrum_1ng_glyco_hcd30_etd30_compound15.2985.2985.4 | controllerType=0 controllerNumber=1 scan=2985 | 11.3154 |
| 11397:4 | 10 | K.LKELPGVC[+57.021]N[+2570.925]ETMMALWEEC[+57.021]KPC[+57.021]LK.Q | HexNAc(5)Hex(6)Fuc(2)NeuAc(1) | C[+57]*3, N[+2571] | 1377.5873 | 4 | 5507.3273 | 5507.3097 | 0 | 3.2 | 95 | Specific | 256.57 | 194.13 | 194.13 | 3.52 | 28 | >sp\|P10909\|CLUS_HUMAN Clusterin OS=Homo sapiens OX=9606 GN=CLU PE=1 SV=1 | 1467 | 20201118_lss_colostrum_1ng_glyco_hcd30_etd30_compound15.14759.14759.4 | controllerType=0 controllerNumber=1 scan=14759 | 35.1515 |
| 10401:4 | 10 | K.LKELPGVC[+57.021]N[+2279.830]ETMMALWEEC[+57.021]KPC[+57.021]LK.Q | HexNAc(5)Hex(6)Fuc(2) | C[+57]*3, N[+2280] | 1304.8108 | 4 | 5216.2213 | 5216.2143 | 0 | 1.3 | 95 | Specific | 287.51 | 241.34 | 27.55 | 3.83 | 28 | >sp\|P10909\|CLUS_HUMAN Clusterin OS=Homo sapiens OX=9606 GN=CLU PE=1 SV=1 | 1467 | 20201118_lss_colostrum_1ng_glyco_hcd30_etd30_compound15.13717.13717.2 | controllerType=0 controllerNumber=1 scan=13717 | 32.6242 |
| 11517:3 | 10 | K.LKELPGVC[+57.021]N[+2059.735]ETMMALWEEC[+57.021]KPC[+57.021]LK.Q | HexNAc(4)Hex(5)Fuc(1)NeuAc(1) | C[+57]*3, N[+2060] | 1666.0466 | 3 | 4996.1252 | 4996.1196 | 0 | 1.1 | 95 | Specific | 305.21 | 229 | 229 | 4.03 | 28 | >sp\|P10909\|CLUS_HUMAN Clusterin OS=Homo sapiens OX=9606 GN=CLU PE=1 SV=1 | 1467 | 20201118_lss_colostrum_1ng_glyco_hcd30_etd30_compound15.14885.14885.3 | controllerType=0 controllerNumber=1 scan=14885 | 35.4782 |
| 11452:4 | 10 | K.LKELPGVC[+57.021]N[+2205.793]ETMMALWEEC[+57.021]KPC[+57.021]LK.Q | HexNAc(4)Hex(5)Fuc(2)NeuAc(1) | C[+57]*3, N[+2206] | 1286.3031 | 4 | 5142.1906 | 5142.1776 | 0 | 2.5 | 95 | Specific | 402.17 | 386.81 | 386.81 | 5.91 | 28 | >sp\|P10909\|CLUS_HUMAN Clusterin OS=Homo sapiens OX=9606 GN=CLU PE=1 SV=1 | 1467 | 20201118_lss_colostrum_1ng_glyco_hcd30_etd30_compound15.14817.14817.4 | controllerType=0 controllerNumber=1 scan=14817 | 35.3043 |
| 11497:3 | 10 | K.LKELPGVC[+57.021]N[+2059.735]ETMMALWEEC[+57.021]KPC[+57.021]LK.Q | HexNAc(4)Hex(5)Fuc(1)NeuAc(1) | C[+57]*3, N[+2060] | 1666.0488 | 3 | 4996.1318 | 4996.1196 | 0 | 2.4 | 95 | Specific | 434.18 | 339.35 | 339.35 | 4.65 | 28 | >sp\|P10909\|CLUS_HUMAN Clusterin OS=Homo sapiens OX=9606 GN=CLU PE=1 SV=1 | 1467 | 20201118_lss_colostrum_1ng_glyco_hcd30_etd30_compound15.14864.14864.3 | controllerType=0 controllerNumber=1 scan=14864 | 35.4339 |
| 11328:4 | 10 | K.LKELPGVC[+57.021]N[+2716.983]ETMMALWEEC[+57.021]KPC[+57.021]LK.Q | HexNAc(5)Hex(6)Fuc(3)NeuAc(1) | C[+57]*3, N[+2717] | 1414.1019 | 4 | 5653.3859 | 5653.3677 | 0 | 3.2 | 95 | Specific | 473.75 | 396.79 | 396.79 | 5.97 | 28 | >sp\|P10909\|CLUS_HUMAN Clusterin OS=Homo sapiens OX=9606 GN=CLU PE=1 SV=1 | 1467 | 20201118_lss_colostrum_1ng_glyco_hcd30_etd30_compound15.14687.14687.4 | controllerType=0 controllerNumber=1 scan=14687 | 34.9969 |
| 11470:4 | 10 | K.LKELPGVC[+57.021]N[+2059.735]ETMMALWEEC[+57.021]KPC[+57.021]LK.Q | HexNAc(4)Hex(5)Fuc(1)NeuAc(1) | C[+57]*3, N[+2060] | 1249.7883 | 4 | 4996.1315 | 4996.1196 | 0 | 2.4 | 95 | Specific | 567.72 | 426.44 | 426.44 | 7.49 | 28 | >sp\|P10909\|CLUS_HUMAN Clusterin OS=Homo sapiens OX=9606 GN=CLU PE=1 SV=1 | 1467 | 20201118_lss_colostrum_1ng_glyco_hcd30_etd30_compound15.14836.14836.4 | controllerType=0 controllerNumber=1 scan=14836 | 35.3629 |
| 00079:3 | 10 | R.EIRHN[+2424.867]STGC[+57.021]LR.M | HexNAc(5)Hex(6)Fuc(1)NeuAc(1) | C[+57], N[+2425] | 1256.5197 | 3 | 3767.5446 | 3767.5316 | 0 | 3.5 | 287 | Specific | 223.78 | 103.95 | 103.95 | 2.6 | 28 | >sp\|P10909\|CLUS_HUMAN Clusterin OS=Homo sapiens OX=9606 GN=CLU PE=1 SV=1 | 1467 | 20201118_lss_colostrum_1ng_glyco_hcd30_etd30_compound15.2977.2977.3 | controllerType=0 controllerNumber=1 scan=2977 | 11.2834 |
| 00076:3 | 10 | R.EIRHN[+2351.851]STGC[+57.021]LR.M | HexNAc(4)Hex(5)Fuc(3)NeuAc(1) | C[+57], N[+2352] | 1232.18 | 3 | 3694.5254 | 3694.5152 | 0 | 2.8 | 287 | Specific | 232.33 | 139.77 | 139.77 | 2.6 | 28 | >sp\|P10909\|CLUS_HUMAN Clusterin OS=Homo sapiens OX=9606 GN=CLU PE=1 SV=1 | 1467 | 20201118_lss_colostrum_1ng_glyco_hcd30_etd30_compound15.2974.2974.3 | controllerType=0 controllerNumber=1 scan=2974 | 11.272 |
| 00065:4 | 10 | R.EIRHN[+2059.735]STGC[+57.021]LR.M | HexNAc(4)Hex(5)Fuc(1)NeuAc(1) | C[+57], N[+2060] | 851.3552 | 4 | 3402.3992 | 3402.3994 | 0 | -0.1 | 287 | Specific | 312.54 | 241.16 | 241.16 | 3.01 | 28 | >sp\|P10909\|CLUS_HUMAN Clusterin OS=Homo sapiens OX=9606 GN=CLU PE=1 SV=1 | 1467 | 20201118_lss_colostrum_1ng_glyco_hcd30_etd30_compound15.2961.2961.4 | controllerType=0 controllerNumber=1 scan=2961 | 11.2176 |
| 00091:4 | 10 | R.EIRHN[+2570.925]STGC[+57.021]LR.M | HexNAc(5)Hex(6)Fuc(2)NeuAc(1) | C[+57], N[+2571] | 979.1569 | 4 | 3913.6058 | 3913.5895 | 0 | 4.2 | 287 | Specific | 377.56 | 304.94 | 304.94 | 4.6 | 28 | >sp\|P10909\|CLUS_HUMAN Clusterin OS=Homo sapiens OX=9606 GN=CLU PE=1 SV=1 | 1467 | 20201118_lss_colostrum_1ng_glyco_hcd30_etd30_compound15.2990.2990.4 | controllerType=0 controllerNumber=1 scan=2990 | 11.3359 |
| 00070:4 | 10 | R.EIRHN[+2424.867]STGC[+57.021]LR.M | HexNAc(5)Hex(6)Fuc(1)NeuAc(1) | C[+57], N[+2425] | 942.643 | 4 | 3767.5504 | 3767.5316 | 0 | 5 | 287 | Specific | 510.49 | 390.18 | 390.18 | 5.76 | 28 | >sp\|P10909\|CLUS_HUMAN Clusterin OS=Homo sapiens OX=9606 GN=CLU PE=1 SV=1 | 1467 | 20201118_lss_colostrum_1ng_glyco_hcd30_etd30_compound15.2968.2968.4 | controllerType=0 controllerNumber=1 scan=2968 | 11.2452 |
| 00080:4 | 10 | R.EIRHN[+2351.851]STGC[+57.021]LR.M | HexNAc(4)Hex(5)Fuc(3)NeuAc(1) | C[+57], N[+2352] | 924.3869 | 4 | 3694.5259 | 3694.5152 | 0 | 2.9 | 287 | Specific | 577.02 | 420.68 | 420.68 | 7.45 | 28 | >sp\|P10909\|CLUS_HUMAN Clusterin OS=Homo sapiens OX=9606 GN=CLU PE=1 SV=1 | 1467 | 20201118_lss_colostrum_1ng_glyco_hcd30_etd30_compound15.2978.2978.4 | controllerType=0 controllerNumber=1 scan=2978 | 11.2905 |
| 00075:4 | 10 | R.EIRHN[+2279.830]STGC[+57.021]LR.M | HexNAc(5)Hex(6)Fuc(2) | C[+57], N[+2280] | 906.3821 | 4 | 3622.5064 | 3622.4941 | 0 | 3.4 | 287 | Specific | 586.55 | 486.39 | 486.39 | 7.45 | 28 | >sp\|P10909\|CLUS_HUMAN Clusterin OS=Homo sapiens OX=9606 GN=CLU PE=1 SV=1 | 1467 | 20201118_lss_colostrum_1ng_glyco_hcd30_etd30_compound15.2973.2973.4 | controllerType=0 controllerNumber=1 scan=2973 | 11.2649 |
| 16349:4 | 10 | K.MLN[+2351.851]TSSLLEQLNEQFNWVSR.L | HexNAc(4)Hex(5)Fuc(3)NeuAc(1) | N[+2352] | 1191.0202 | 4 | 4761.0589 | 4761.0375 | 0 | 4.5 | 352 | Specific | 221.63 | 210.05 | 210.05 | 2.7 | 28 | >sp\|P10909\|CLUS_HUMAN Clusterin OS=Homo sapiens OX=9606 GN=CLU PE=1 SV=1 | 1467 | 20201118_lss_colostrum_1ng_glyco_hcd30_etd30_compound15.19973.19973.4 | controllerType=0 controllerNumber=1 scan=19973 | 49.6715 |
| 16365:3 | 10 | K.MLN[+2424.867]TSSLLEQLNEQFNWVSR.L | HexNAc(5)Hex(6)Fuc(1)NeuAc(1) | N[+2425] | 1612.0227 | 3 | 4834.0535 | 4834.0539 | 0 | -0.1 | 352 | Specific | 229.4 | 99.97 | 71.38 | 2.66 | 28 | >sp\|P10909\|CLUS_HUMAN Clusterin OS=Homo sapiens OX=9606 GN=CLU PE=1 SV=1 | 1467 | 20201118_lss_colostrum_1ng_glyco_hcd30_etd30_compound15.19990.19990.3 | controllerType=0 controllerNumber=1 scan=19990 | 49.7377 |
| 17639:4 | 10 | K.MLN[+2350.830]TSSLLEQLNEQFNWVSR.L | HexNAc(4)Hex(5)Fuc(1)NeuAc(2) | N[+2351] | 1190.7649 | 4 | 4760.0379 | 4760.0171 | 0 | 4.4 | 352 | Specific | 370.44 | 320.37 | 320.37 | 3.99 | 28 | >sp\|P10909\|CLUS_HUMAN Clusterin OS=Homo sapiens OX=9606 GN=CLU PE=1 SV=1 | 1467 | 20201118_lss_colostrum_1ng_glyco_hcd30_etd30_compound15.21347.21347.4 | controllerType=0 controllerNumber=1 scan=21347 | 54.3748 |
| 16401:4 | 10 | K.MLN[+2059.735]TSSLLEQLNEQFNWVSR.L | HexNAc(4)Hex(5)Fuc(1)NeuAc(1) | N[+2060] | 1117.99 | 4 | 4468.9383 | 4468.9217 | 0 | 3.7 | 352 | Specific | 451.75 | 419.48 | 419.48 | 6.21 | 28 | >sp\|P10909\|CLUS_HUMAN Clusterin OS=Homo sapiens OX=9606 GN=CLU PE=1 SV=1 | 1467 | 20201118_lss_colostrum_1ng_glyco_hcd30_etd30_compound15.20029.20029.4 | controllerType=0 controllerNumber=1 scan=20029 | 49.8794 |
| 16777:3 | 10 | K.MLN[+1458.442]TSSLLEQLNEQFNWVSR.L | HexNAc(2)Hex(6)Phospho(1) | N[+1458] | 1289.8868 | 3 | 3867.6458 | 3867.6287 | 0 | 4.4 | 352 | Specific | 532.68 | 417.83 | 417.83 | 7.15 | 28 | >sp\|P10909\|CLUS_HUMAN Clusterin OS=Homo sapiens OX=9606 GN=CLU PE=1 SV=1 | 1467 | 20201118_lss_colostrum_1ng_glyco_hcd30_etd30_compound15.20429.20429.3 | controllerType=0 controllerNumber=1 scan=20429 | 51.2376 |
| 16565:3 | 10 | K.MLN[+1872.651]TSSLLEQLNEQFNWVSR.L | HexNAc(3)Hex(6)NeuAc(1) | N[+1873] | 1427.9583 | 3 | 4281.8604 | 4281.8372 | 0 | 5.4 | 352 | Specific | 536.52 | 366.52 | 136.31 | 7.15 | 28 | >sp\|P10909\|CLUS_HUMAN Clusterin OS=Homo sapiens OX=9606 GN=CLU PE=1 SV=1 | 1467 | 20201118_lss_colostrum_1ng_glyco_hcd30_etd30_compound15.20203.20203.3 | controllerType=0 controllerNumber=1 scan=20203 | 50.4607 |
| 16550:3 | 10 | K.MLN[+1913.677]TSSLLEQLNEQFNWVSR.L | HexNAc(4)Hex(5)NeuAc(1) | N[+1914] | 1441.6324 | 3 | 4322.8826 | 4322.8637 | 0 | 4.4 | 352 | Specific | 550.72 | 400.69 | 400.69 | 7.15 | 28 | >sp\|P10909\|CLUS_HUMAN Clusterin OS=Homo sapiens OX=9606 GN=CLU PE=1 SV=1 | 1467 | 20201118_lss_colostrum_1ng_glyco_hcd30_etd30_compound15.20187.20187.3 | controllerType=0 controllerNumber=1 scan=20187 | 50.3984 |
| 16336:3 | 10 | K.MLN[+2351.851]TSSLLEQLNEQFNWVSR.L | HexNAc(4)Hex(5)Fuc(3)NeuAc(1) | N[+2352] | 1587.692 | 3 | 4761.0613 | 4761.0375 | 0 | 5 | 352 | Specific | 595.46 | 400.29 | 400.29 | 8.26 | 28 | >sp\|P10909\|CLUS_HUMAN Clusterin OS=Homo sapiens OX=9606 GN=CLU PE=1 SV=1 | 1467 | 20201118_lss_colostrum_1ng_glyco_hcd30_etd30_compound15.19959.19959.3 | controllerType=0 controllerNumber=1 scan=19959 | 49.622 |
| 16400:3 | 10 | K.MLN[+2059.735]TSSLLEQLNEQFNWVSR.L | HexNAc(4)Hex(5)Fuc(1)NeuAc(1) | N[+2060] | 1490.3187 | 3 | 4468.9414 | 4468.9217 | 0 | 4.4 | 352 | Specific | 608.41 | 430.64 | 430.64 | 9.26 | 28 | >sp\|P10909\|CLUS_HUMAN Clusterin OS=Homo sapiens OX=9606 GN=CLU PE=1 SV=1 | 1467 | 20201118_lss_colostrum_1ng_glyco_hcd30_etd30_compound15.20028.20028.3 | controllerType=0 controllerNumber=1 scan=20028 | 49.8751 |
| 17665:3 | 10 | K.MLN[+2350.830]TSSLLEQLNEQFNWVSR.L | HexNAc(4)Hex(5)Fuc(1)NeuAc(2) | N[+2351] | 1587.3528 | 3 | 4760.044 | 4760.0171 | 0 | 5.7 | 352 | Specific | 657.76 | 460.78 | 460.78 | 10.26 | 28 | >sp\|P10909\|CLUS_HUMAN Clusterin OS=Homo sapiens OX=9606 GN=CLU PE=1 SV=1 | 1467 | 20201118_lss_colostrum_1ng_glyco_hcd30_etd30_compound15.21375.21375.3 | controllerType=0 controllerNumber=1 scan=21375 | 54.4864 |
| 16086:4 | 10 | K.M[+15.995]LN[+2059.735]TSSLLEQLNEQFNWVSR.L | HexNAc(4)Hex(5)Fuc(1)NeuAc(1) | M[+16], N[+2060] | 1121.99 | 4 | 4484.9383 | 4484.9166 | 0 | 4.8 | 352 | Specific | 200.74 | 179.84 | 179.84 | 2.61 | 28 | >sp\|P10909\|CLUS_HUMAN Clusterin OS=Homo sapiens OX=9606 GN=CLU PE=1 SV=1 | 1467 | 20201118_lss_colostrum_1ng_glyco_hcd30_etd30_compound15.19694.19694.4 | controllerType=0 controllerNumber=1 scan=19694 | 48.7689 |
| 16080:3 | 10 | K.M[+15.995]LN[+2059.735]TSSLLEQLNEQFNWVSR.L | HexNAc(4)Hex(5)Fuc(1)NeuAc(1) | M[+16], N[+2060] | 1495.6515 | 3 | 4484.9401 | 4484.9166 | 0 | 5.2 | 352 | Specific | 589.66 | 435.29 | 435.29 | 8.26 | 28 | >sp\|P10909\|CLUS_HUMAN Clusterin OS=Homo sapiens OX=9606 GN=CLU PE=1 SV=1 | 1467 | 20201118_lss_colostrum_1ng_glyco_hcd30_etd30_compound15.19688.19688.3 | controllerType=0 controllerNumber=1 scan=19688 | 48.7498 |
| 10820:3 | 10 | R.LAN[+2350.830]LTQGEDQYYLR.V | HexNAc(4)Hex(5)Fuc(1)NeuAc(2) | N[+2351] | 1345.564 | 3 | 4034.6775 | 4034.664 | 0 | 3.3 | 372 | Specific | 386.98 | 264.34 | 264.34 | 5.47 | 28 | >sp\|P10909\|CLUS_HUMAN Clusterin OS=Homo sapiens OX=9606 GN=CLU PE=1 SV=1 | 1467 | 20201118_lss_colostrum_1ng_glyco_hcd30_etd30_compound15.14154.14154.3 | controllerType=0 controllerNumber=1 scan=14154 | 33.5901 |
| 17817:5 | 11 | K.VFPLSLDSTPQDGNVVVAC[+57.021]LVQGFFPQEPLSVTWSESGQN[+2262.814]VTAI | HexNAc(5)Hex(5)Fuc(1)NeuAc(1) | C[+57], N[+2263] | 1408.6467 | 5 | 7039.2041 | 7039.1757 | 0 | 4 | 8 | Specific | 246.38 | 246.38 | 246.38 | 4.36 | 26 | >sp\|P01877\|IGHA2_HUMAN Immunoglobulin heavy constant alpha 2 OS=Homo sapiens OX=9606 GN=IGHA2 PE=1 SV=4 | 576 | 20201118_lss_colostrum_1ng_glyco_hcd30_etd30_compound15.21538.21538.5 | controllerType=0 controllerNumber=1 scan=21538 | 55.119 |
| 16941:4 | 11 | K.VFPLSLDSTPQDGNVVVAC[+57.021]LVQGFFPQEPLSVTWSESGQN[+1809.666]VTAI | HexNAc(5)Hex(4)Fuc(1) | C[+57], N[+1810] | 1647.2698 | 4 | 6586.0573 | 6586.0275 | 0 | 4.5 | 8 | Specific | 318.11 | 318.11 | 318.11 | 4.96 | 26 | >sp\|P01877\|IGHA2_HUMAN Immunoglobulin heavy constant alpha 2 OS=Homo sapiens OX=9606 GN=IGHA2 PE=1 SV=4 | 576 | 20201118_lss_colostrum_1ng_glyco_hcd30_etd30_compound15.20603.20603.4 | controllerType=0 controllerNumber=1 scan=20603 | 51.8216 |
| 17072:4 | 11 | K.VFPLSLDSTPQDGNVVVAC[+57.021]LVQGFFPQEPLSVTWSESGQN[+1809.666]VTAI | HexNAc(5)Hex(4)Fuc(1) | C[+57], N[+1810] | 1647.2692 | 4 | 6586.0548 | 6586.0275 | 0 | 4.2 | 8 | Specific | 451.51 | 451.51 | 451.51 | 7.14 | 26 | >sp\|P01877\|IGHA2_HUMAN Immunoglobulin heavy constant alpha 2 OS=Homo sapiens OX=9606 GN=IGHA2 PE=1 SV=4 | 576 | 20201118_lss_colostrum_1ng_glyco_hcd30_etd30_compound15.20743.20743.4 | controllerType=0 controllerNumber=1 scan=20743 | 52.265 |
| 17816:4 | 11 | K.VFPLSLDSTPQDGNVVVAC[+57.021]LVQGFFPQEPLSVTWSESGQN[+2262.814]VTAI | HexNAc(5)Hex(5)Fuc(1)NeuAc(1) | C[+57], N[+2263] | 1760.5586 | 4 | 7039.2124 | 7039.1757 | 0 | 5.2 | 8 | Specific | 472.26 | 472.26 | 472.26 | 6.65 | 26 | >sp\|P01877\|IGHA2_HUMAN Immunoglobulin heavy constant alpha 2 OS=Homo sapiens OX=9606 GN=IGHA2 PE=1 SV=4 | 576 | 20201118_lss_colostrum_1ng_glyco_hcd30_etd30_compound15.21537.21537.4 | controllerType=0 controllerNumber=1 scan=21537 | 55.113 |
| 00811:3 | 11 | K.HYTN[+1971.719]SSQDVTVPC[+57.021]R.V | HexNAc(5)Hex(5)Fuc(1) | C[+57], N[+1972] | 1212.4966 | 3 | 3635.4754 | 3635.4682 | 0 | 2 | 89 | Specific | 375.45 | 305.31 | 305.31 | 5.26 | 26 | >sp\|P01877\|IGHA2_HUMAN Immunoglobulin heavy constant alpha 2 OS=Homo sapiens OX=9606 GN=IGHA2 PE=1 SV=4 | 576 | 20201118_lss_colostrum_1ng_glyco_hcd30_etd30_compound15.3740.3740.3 | controllerType=0 controllerNumber=1 scan=3740 | 12.7794 |
| 00862:3 | 11 | K.HYTN[+1444.534]SSQDVTVPC[+57.021]R.V | HexNAc(4)Hex(3)Fuc(1) | C[+57], N[+1445] | 1036.7687 | 3 | 3108.2916 | 3108.2832 | 0 | 2.7 | 89 | Specific | 420.52 | 325.73 | 325.73 | 5.62 | 26 | >sp\|P01877\|IGHA2_HUMAN Immunoglobulin heavy constant alpha 2 OS=Homo sapiens OX=9606 GN=IGHA2 PE=1 SV=4 | 576 | 20201118_lss_colostrum_1ng_glyco_hcd30_etd30_compound15.3793.3793.3 | controllerType=0 controllerNumber=1 scan=3793 | 12.8819 |
| 00836:3 | 11 | K.HYTN[+1809.666]SSQDVTVPC[+57.021]R.V | HexNAc(5)Hex(4)Fuc(1) | C[+57], N[+1810] | 1158.4786 | 3 | 3473.4212 | 3473.4154 | 0 | 1.7 | 89 | Specific | 423.75 | 347.51 | 347.51 | 5.51 | 26 | >sp\|P01877\|IGHA2_HUMAN Immunoglobulin heavy constant alpha 2 OS=Homo sapiens OX=9606 GN=IGHA2 PE=1 SV=4 | 576 | 20201118_lss_colostrum_1ng_glyco_hcd30_etd30_compound15.3766.3766.3 | controllerType=0 controllerNumber=1 scan=3766 | 12.8337 |
| 00875:3 | 11 | K.HYTN[+1647.613]SSQDVTVPC[+57.021]R.V | HexNAc(5)Hex(3)Fuc(1) | C[+57], N[+1648] | 1104.4637 | 3 | 3311.3764 | 3311.3625 | 0 | 4.2 | 89 | Specific | 527.04 | 455.29 | 455.29 | 7.1 | 26 | >sp\|P01877\|IGHA2_HUMAN Immunoglobulin heavy constant alpha 2 OS=Homo sapiens OX=9606 GN=IGHA2 PE=1 SV=4 | 576 | 20201118_lss_colostrum_1ng_glyco_hcd30_etd30_compound15.3807.3807.3 | controllerType=0 controllerNumber=1 scan=3807 | 12.9196 |
| 02725:4 | 11 | K.HYTN[+1647.613]SSQDVTVPC[+57.021]RVPPPPPC[+57.021]C[+57.021]HPR.L | HexNAc(5)Hex(3)Fuc(1) | C[+57]*3, N[+1648] | 1152.2513 | 4 | 4605.9832 | 4605.9689 | 0 | 3.1 | 89 | Specific | 378.74 | 378.74 | 378.74 | 6.22 | 26 | >sp\|P01877\|IGHA2_HUMAN Immunoglobulin heavy constant alpha 2 OS=Homo sapiens OX=9606 GN=IGHA2 PE=1 SV=4 | 576 | 20201118_lss_colostrum_1ng_glyco_hcd30_etd30_compound15.5728.5728.4 | controllerType=0 controllerNumber=1 scan=5728 | 16.5532 |
| 02716:5 | 11 | K.HYTN[+1647.613]SSQDVTVPC[+57.021]RVPPPPPC[+57.021]C[+57.021]HPR.L | HexNAc(5)Hex(3)Fuc(1) | C[+57]*3, N[+1648] | 922.0019 | 5 | 4605.9802 | 4605.9689 | 0 | 2.5 | 89 | Specific | 487.97 | 459.02 | 459.02 | 6.53 | 26 | >sp\|P01877\|IGHA2_HUMAN Immunoglobulin heavy constant alpha 2 OS=Homo sapiens OX=9606 GN=IGHA2 PE=1 SV=4 | 576 | 20201118_lss_colostrum_1ng_glyco_hcd30_etd30_compound15.5719.5719.5 | controllerType=0 controllerNumber=1 scan=5719 | 16.5285 |
| 04041:2 | 11 | K.TPLTAN[+2059.735]ITK.S | HexNAc(4)Hex(5)Fuc(1)NeuAc(1) | N[+2060] | 1509.6529 | 2 | 3018.2985 | 3018.2917 | 0 | 2.2 | 200 | Specific | 205.3 | 57.28 | 57.28 | 2.71 | 26 | >sp\|P01877\|IGHA2_HUMAN Immunoglobulin heavy constant alpha 2 OS=Homo sapiens OX=9606 GN=IGHA2 PE=1 SV=4 | 576 | 20201118_lss_colostrum_1ng_glyco_hcd30_etd30_compound15.7097.7097.2 | controllerType=0 controllerNumber=1 scan=7097 | 19.0978 |
| 04014:3 | 11 | K.TPLTAN[+2262.814]ITK.S | HexNAc(5)Hex(5)Fuc(1)NeuAc(1) | N[+2263] | 1074.4656 | 3 | 3221.3823 | 3221.3711 | 0 | 3.5 | 200 | Specific | 258.97 | 117.31 | 117.31 | 3.31 | 26 | >sp\|P01877\|IGHA2_HUMAN Immunoglobulin heavy constant alpha 2 OS=Homo sapiens OX=9606 GN=IGHA2 PE=1 SV=4 | 576 | 20201118_lss_colostrum_1ng_glyco_hcd30_etd30_compound15.7069.7069.3 | controllerType=0 controllerNumber=1 scan=7069 | 19.0371 |
| 02452:2 | 11 | K.TPLTAN[+1971.719]ITK.S | HexNAc(5)Hex(5)Fuc(1) | N[+1972] | 1465.6463 | 2 | 2930.2853 | 2930.2757 | 0 | 3.3 | 200 | Specific | 262.05 | 62.48 | 62.48 | 2.65 | 26 | >sp\|P01877\|IGHA2_HUMAN Immunoglobulin heavy constant alpha 2 OS=Homo sapiens OX=9606 GN=IGHA2 PE=1 SV=4 | 576 | 20201118_lss_colostrum_1ng_glyco_hcd30_etd30_compound15.5442.5442.2 | controllerType=0 controllerNumber=1 scan=5442 | 15.8661 |
| 02426:3 | 11 | K.TPLTAN[+1971.719]ITK.S | HexNAc(5)Hex(5)Fuc(1) | N[+1972] | 977.4331 | 3 | 2930.2847 | 2930.2757 | 0 | 3.1 | 200 | Specific | 290.86 | 133.33 | 133.33 | 2.95 | 26 | >sp\|P01877\|IGHA2_HUMAN Immunoglobulin heavy constant alpha 2 OS=Homo sapiens OX=9606 GN=IGHA2 PE=1 SV=4 | 576 | 20201118_lss_colostrum_1ng_glyco_hcd30_etd30_compound15.5415.5415.3 | controllerType=0 controllerNumber=1 scan=5415 | 15.7877 |
| 02517:2 | 11 | K.TPLTAN[+1647.613]ITK.S | HexNAc(5)Hex(3)Fuc(1) | N[+1648] | 1303.5917 | 2 | 2606.1762 | 2606.17 | 0 | 2.4 | 200 | Specific | 303.6 | 115.93 | 115.93 | 2.81 | 26 | >sp\|P01877\|IGHA2_HUMAN Immunoglobulin heavy constant alpha 2 OS=Homo sapiens OX=9606 GN=IGHA2 PE=1 SV=4 | 576 | 20201118_lss_colostrum_1ng_glyco_hcd30_etd30_compound15.5511.5511.2 | controllerType=0 controllerNumber=1 scan=5511 | 16.0393 |
| 02481:2 | 11 | K.TPLTAN[+1809.666]ITK.S | HexNAc(5)Hex(4)Fuc(1) | N[+1810] | 1384.6181 | 2 | 2768.2289 | 2768.2228 | 0 | 2.2 | 200 | Specific | 304.58 | 115.95 | 115.95 | 2.77 | 26 | >sp\|P01877\|IGHA2_HUMAN Immunoglobulin heavy constant alpha 2 OS=Homo sapiens OX=9606 GN=IGHA2 PE=1 SV=4 | 576 | 20201118_lss_colostrum_1ng_glyco_hcd30_etd30_compound15.5473.5473.2 | controllerType=0 controllerNumber=1 scan=5473 | 15.9372 |
| 02535:3 | 11 | K.TPLTAN[+1647.613]ITK.S | HexNAc(5)Hex(3)Fuc(1) | N[+1648] | 869.3973 | 3 | 2606.1774 | 2606.17 | 0 | 2.8 | 200 | Specific | 306.61 | 134.72 | 134.72 | 2.79 | 26 | >sp\|P01877\|IGHA2_HUMAN Immunoglobulin heavy constant alpha 2 OS=Homo sapiens OX=9606 GN=IGHA2 PE=1 SV=4 | 576 | 20201118_lss_colostrum_1ng_glyco_hcd30_etd30_compound15.5530.5530.3 | controllerType=0 controllerNumber=1 scan=5530 | 16.0949 |
| 02548:2 | 11 | K.TPLTAN[+1501.555]ITK.S | HexNAc(5)Hex(3) | N[+1502] | 1230.5641 | 2 | 2460.121 | 2460.1121 | 0 | 3.6 | 200 | Specific | 308.19 | 102.24 | 102.24 | 2.81 | 26 | >sp\|P01877\|IGHA2_HUMAN Immunoglobulin heavy constant alpha 2 OS=Homo sapiens OX=9606 GN=IGHA2 PE=1 SV=4 | 576 | 20201118_lss_colostrum_1ng_glyco_hcd30_etd30_compound15.5543.5543.2 | controllerType=0 controllerNumber=1 scan=5543 | 16.1247 |
| 02485:3 | 11 | K.TPLTAN[+1809.666]ITK.S | HexNAc(5)Hex(4)Fuc(1) | N[+1810] | 923.414 | 3 | 2768.2275 | 2768.2228 | 0 | 1.7 | 200 | Specific | 310.43 | 138.96 | 138.96 | 2.75 | 26 | >sp\|P01877\|IGHA2_HUMAN Immunoglobulin heavy constant alpha 2 OS=Homo sapiens OX=9606 GN=IGHA2 PE=1 SV=4 | 576 | 20201118_lss_colostrum_1ng_glyco_hcd30_etd30_compound15.5477.5477.3 | controllerType=0 controllerNumber=1 scan=5477 | 15.9483 |
| 02538:2 | 11 | K.TPLTAN[+1444.534]ITK.S | HexNAc(4)Hex(3)Fuc(1) | N[+1445] | 1202.053 | 2 | 2403.0988 | 2403.0906 | 0 | 3.4 | 200 | Specific | 351.63 | 126.16 | 126.16 | 2.72 | 26 | >sp\|P01877\|IGHA2_HUMAN Immunoglobulin heavy constant alpha 2 OS=Homo sapiens OX=9606 GN=IGHA2 PE=1 SV=4 | 576 | 20201118_lss_colostrum_1ng_glyco_hcd30_etd30_compound15.5533.5533.2 | controllerType=0 controllerNumber=1 scan=5533 | 16.1046 |
| 08772:3 | 11 | R.MAGKPTHIN[+1702.581]VSVVMAEADGTC[+57.021]Y.- | HexNAc(2)Hex(8) | C[+57], N[+1703] | 1351.8974 | 3 | 4053.6776 | 4053.6642 | 0 | 3.3 | 319 | CRagged | 253.5 | 253.5 | 185.7 | 4.65 | 26 | >sp\|P01877\|IGHA2_HUMAN Immunoglobulin heavy constant alpha 2 OS=Homo sapiens OX=9606 GN=IGHA2 PE=1 SV=4 | 576 | 20201118_lss_colostrum_1ng_glyco_hcd30_etd30_compound15.12016.12016.3 | controllerType=0 controllerNumber=1 scan=12016 | 28.6164 |
| 08816:3 | 11 | R.MAGKPTHIN[+1378.476]VSVVMAEADGTC[+57.021]Y.- | HexNAc(2)Hex(6) | C[+57], N[+1378] | 1243.8635 | 3 | 3729.5758 | 3729.5585 | 0 | 4.6 | 319 | CRagged | 280.6 | 268.77 | 217.71 | 4.09 | 26 | >sp\|P01877\|IGHA2_HUMAN Immunoglobulin heavy constant alpha 2 OS=Homo sapiens OX=9606 GN=IGHA2 PE=1 SV=4 | 576 | 20201118_lss_colostrum_1ng_glyco_hcd30_etd30_compound15.12062.12062.3 | controllerType=0 controllerNumber=1 scan=12062 | 28.729 |
| 08846:3 | 11 | R.MAGKPTHIN[+1216.423]VSVVMAEADGTC[+57.021]Y.- | HexNAc(2)Hex(5) | C[+57], N[+1216] | 1189.8448 | 3 | 3567.5198 | 3567.5057 | 0 | 3.9 | 319 | CRagged | 353.28 | 305.43 | 223.09 | 4.82 | 26 | >sp\|P01877\|IGHA2_HUMAN Immunoglobulin heavy constant alpha 2 OS=Homo sapiens OX=9606 GN=IGHA2 PE=1 SV=4 | 576 | 20201118_lss_colostrum_1ng_glyco_hcd30_etd30_compound15.12093.12093.3 | controllerType=0 controllerNumber=1 scan=12093 | 28.8185 |
| 18370:4 | 12 | K.APDKNVIFSPLSISTALAFLSLGAHN[+2861.000]TTLTEILK.G | HexNAc(5)Hex(6)NeuAc(3) | N[+2861] | 1611.7576 | 4 | 6444.0085 | 6443.9782 | 0 | 4.7 | 68 | Specific | 206.35 | 206.35 | 206.35 | 3.67 | 14 | >sp\|P01011\|AACT_HUMAN Alpha-1-antichymotrypsin OS=Homo sapiens OX=9606 GN=SERPINA3 PE=1 SV=2 | 1314 | 20201118_lss_colostrum_1ng_glyco_hcd30_etd30_compound15.22127.22127.4 | controllerType=0 controllerNumber=1 scan=22127 | 57.1479 |
| 17448:4 | 12 | K.APDKNVIFSPLSISTALAFLSLGAHN[+2350.830]TTLTEILK.G | HexNAc(4)Hex(5)Fuc(1)NeuAc(2) | N[+2351] | 1484.2123 | 4 | 5933.8273 | 5933.8085 | 0 | 3.2 | 68 | Specific | 278.79 | 278.79 | 278.79 | 4.82 | 14 | >sp\|P01011\|AACT_HUMAN Alpha-1-antichymotrypsin OS=Homo sapiens OX=9606 GN=SERPINA3 PE=1 SV=2 | 1314 | 20201118_lss_colostrum_1ng_glyco_hcd30_etd30_compound15.21143.21143.4 | controllerType=0 controllerNumber=1 scan=21143 | 53.6044 |
| 16643:4 | 12 | K.APDKNVIFSPLSISTALAFLSLGAHN[+2059.735]TTLTEILK.G | HexNAc(4)Hex(5)Fuc(1)NeuAc(1) | N[+2060] | 1411.4382 | 4 | 5642.7311 | 5642.7131 | 0 | 3.2 | 68 | Specific | 412.27 | 412.27 | 412.27 | 6.67 | 14 | >sp\|P01011\|AACT_HUMAN Alpha-1-antichymotrypsin OS=Homo sapiens OX=9606 GN=SERPINA3 PE=1 SV=2 | 1314 | 20201118_lss_colostrum_1ng_glyco_hcd30_etd30_compound15.20286.20286.4 | controllerType=0 controllerNumber=1 scan=20286 | [illegible] |
| 18059:4 | 14 | R.QLAHQSN[+2350.830]STNIFFSPVSIATAFAMLSLGTK.A | HexNAc(4)Hex(5)Fuc(1)NeuAc(2) | N[+2351] | 1383.8737 | 4 | 5532.4728 | 5532.4654 | 0 | 1.3 | 64 | Specific | 321.6 | 291.5 | 291.5 | 5.55 | 13 | >sp\|P01009\|A1AT_HUMAN Alpha-1-antitrypsin OS=Homo sapiens OX=9606 GN=SERPINA1 PE=1 SV=3 | 642 | 20201118_lss_colostrum_1ng_glyco_hcd30_etd30_compound15.21796.21796.4 | controllerType=0 controllerNumber=1 scan=21796 | 56.0431 |
| 18025:3 | 14 | R.QLAHQSN[+2204.772]STNIFFSPVSIATAFAMLSLGTK.A | HexNAc(4)Hex(5)NeuAc(2) | N[+2205] | 1796.1462 | 3 | 5386.4241 | 5386.4075 | 0 | 3.1 | 64 | Specific | 492.82 | 381.75 | 381.75 | 8.18 | 13 | >sp\|P01009\|A1AT_HUMAN Alpha-1-antitrypsin OS=Homo sapiens OX=9606 GN=SERPINA1 PE=1 SV=3 | 642 | 20201118_lss_colostrum_1ng_glyco_hcd30_etd30_compound15.21760.21760.3 | controllerType=0 controllerNumber=1 scan=21760 | 55.9138 |
| 18026:5 | 14 | R.QLAHQSN[+2204.772]STNIFFSPVSIATAFAMLSLGTK.A | HexNAc(4)Hex(5)NeuAc(2) | N[+2205] | 1078.0906 | 5 | 5386.4238 | 5386.4075 | 0 | 3 | 64 | Specific | 644.79 | 543.98 | 543.98 | 9.6 | 13 | >sp\|P01009\|A1AT_HUMAN Alpha-1-antitrypsin OS=Homo sapiens OX=9606 GN=SERPINA1 PE=1 SV=3 | 642 | 20201118_lss_colostrum_1ng_glyco_hcd30_etd30_compound15.21761.21761.5 | controllerType=0 controllerNumber=1 scan=21761 | 55.9178 |
| 18021:4 | 14 | R.QLAHQSN[+2204.772]STNIFFSPVSIATAFAMLSLGTK.A | HexNAc(4)Hex(5)NeuAc(2) | N[+2205] | 1347.3617 | 4 | 5386.425 | 5386.4075 | 0 | 3.2 | 64 | Specific | 690.04 | 553.44 | 553.44 | 11.48 | 13 | >sp\|P01009\|A1AT_HUMAN Alpha-1-antitrypsin OS=Homo sapiens OX=9606 GN=SERPINA1 PE=1 SV=3 | 642 | 20201118_lss_colostrum_1ng_glyco_hcd30_etd30_compound15.21756.21756.4 | controllerType=0 controllerNumber=1 scan=21756 | 55.9042 |
| 17560:4 | 14 | R.QLAHQSN[+2204.772]STNIFFSPVSIATAFAM[+15.995]LSLGTK.A | HexNAc(4)Hex(5)NeuAc(2) | M[+16], N[+2205] | 1351.3572 | 4 | 5402.4069 | 5402.4024 | 0 | 0.8 | 64 | Specific | 224.23 | 216.93 | 216.93 | 3.85 | 13 | >sp\|P01009\|A1AT_HUMAN Alpha-1-antitrypsin OS=Homo sapiens OX=9606 GN=SERPINA1 PE=1 SV=3 | 642 | 20201118_lss_colostrum_1ng_glyco_hcd30_etd30_compound15.21262.21262.4 | controllerType=0 controllerNumber=1 scan=21262 | 54.0458 |
| 17537:4 | 14 | R.QLAHQSN[+2204.772]STNIFFSPVSIATAFAM[+15.995]LSLGTK.A | HexNAc(4)Hex(5)NeuAc(2) | M[+16], N[+2205] | 1351.3618 | 4 | 5402.4254 | 5402.4024 | 0 | 4.3 | 64 | Specific | 509.04 | 395.85 | 395.85 | 6.81 | 13 | >sp\|P01009\|A1AT_HUMAN Alpha-1-antitrypsin OS=Homo sapiens OX=9606 GN=SERPINA1 PE=1 SV=3 | 642 | 20201118_lss_colostrum_1ng_glyco_hcd30_etd30_compound15.21238.21238.4 | controllerType=0 controllerNumber=1 scan=21238 | 53.9487 |
| 17332:4 | 14 | K.ADTHDEILEGLNFN[+2861.000]LTEIPEAQIHEGFQELLR.T | HexNAc(5)Hex(6)NeuAc(3) | N[+2861] | 1638.9686 | 4 | 6552.8527 | 6552.8239 | 0 | 4.4 | 94 | Specific | 319.26 | 319.26 | 319.26 | 5.26 | 13 | >sp\|P01009\|A1AT_HUMAN Alpha-1-antitrypsin OS=Homo sapiens OX=9606 GN=SERPINA1 PE=1 SV=3 | 642 | 20201118_lss_colostrum_1ng_glyco_hcd30_etd30_compound15.21019.21019.4 | controllerType=0 controllerNumber=1 scan=21019 | 53.169 |
| 16461:4 | 14 | K.ADTHDEILEGLNFN[+2204.772]LTEIPEAQIHEGFQELLR.T | HexNAc(4)Hex(5)NeuAc(2) | N[+2205] | 1474.9108 | 4 | 5896.6212 | 5896.5963 | 0 | 4.2 | 94 | Specific | 504.98 | 477.32 | 477.32 | 6.87 | 13 | >sp\|P01009\|A1AT_HUMAN Alpha-1-antitrypsin OS=Homo sapiens OX=9606 GN=SERPINA1 PE=1 SV=3 | 642 | 20201118_lss_colostrum_1ng_glyco_hcd30_etd30_compound15.20093.20093.4 | controllerType=0 controllerNumber=1 scan=20093 | 50.0931 |
| 10833:4 | 16 | R.FELLGC[+57.021]ELNGC[+57.021]ANPLGLKN[+1216.423]NSIPDKQITASSSYK.T | HexNAc(2)Hex(5) | C[+57]*2, N[+1216] | 1239.5758 | 4 | 4955.2814 | 4955.2694 | 0 | 2.4 | 220 | Specific | 855.9 | 855.9 | 855.9 | 14.23 | 10 | >sp\|Q08431\|MFGM_HUMAN Lactadherin OS=Homo sapiens OX=9606 GN=MFGE8 PE=1 SV=3 | 1064 | 20201118_lss_colostrum_1ng_glyco_hcd30_etd30_compound15.14168.14168.4 | controllerType=0 controllerNumber=1 scan=14168 | 33.6199 |
| 01610:3 | 16 | K.N[+1540.529]NSIPDKQITASSSYK.T | HexNAc(2)Hex(7) | N[+1541] | 1098.4761 | 3 | 3293.4138 | 3293.4048 | 0 | 2.7 | 238 | Specific | 269.2 | 208.89 | 208.89 | 3.93 | 10 | >sp\|Q08431\|MFGM_HUMAN Lactadherin OS=Homo sapiens OX=9606 GN=MFGE8 PE=1 SV=3 | 1064 | 20201118_lss_colostrum_1ng_glyco_hcd30_etd30_compound15.4569.4569.3 | controllerType=0 controllerNumber=1 scan=4569 | 14.2664 |
| 01703:3 | 16 | K.N[+1216.423]NSIPDKQITASSSYK.T | HexNAc(2)Hex(5) | N[+1216] | 990.44 | 3 | 2969.3055 | 2969.2991 | 0 | 2.2 | 238 | Specific | 405.88 | 242.67 | 242.67 | 3.74 | 10 | >sp\|Q08431\|MFGM_HUMAN Lactadherin OS=Homo sapiens OX=9606 GN=MFGE8 PE=1 SV=3 | 1064 | 20201118_lss_colostrum_1ng_glyco_hcd30_etd30_compound15.4666.4666.3 | controllerType=0 controllerNumber=1 scan=4666 | 14.4772 |
| 03277:2 | 17 | K.AAIPSALDTN[+1216.423]SSK.S | HexNAc(2)Hex(5) | N[+1216] | 1246.0451 | 2 | 2491.0829 | 2491.0815 | 0 | 0.5 | 542 | Specific | 431.55 | 250.47 | 250.47 | 4.37 | 7 | >sp\|Q08380\|LG3BP_HUMAN Galectin-3-binding protein OS=Homo sapiens OX=9606 GN=LGALS3BP PE=1 SV=1 | 1562 | 20201118_lss_colostrum_1ng_glyco_hcd30_etd30_compound15.6302.6302.2 | controllerType=0 controllerNumber=1 scan=6302 | 17.6339 |
| 01660:3 | 22 | R.LRN[+1216.423]VSWATGR.S | HexNAc(2)Hex(5) | N[+1216] | 792.6923 | 3 | 2376.0623 | 2376.0559 | 0 | 2.7 | 149 | Specific | 384.93 | 173.31 | 173.31 | 2.77 | 6 | >sp\|P08571\|CD14_HUMAN Monocyte differentiation antigen CD14 OS=Homo sapiens OX=9606 GN=CD14 PE=1 SV=2 | 1232 | 20201118_lss_colostrum_1ng_glyco_hcd30_etd30_compound15.4621.4621.3 | controllerType=0 controllerNumber=1 scan=4621 | 14.3629 |
| 09609:3 | 24 | R.IIVPLNNREN[+1694.603]ISDPTSPLR.T | HexNAc(3)Hex(4)Fuc(1)NeuAc(1) | N[+1695] | 1281.6038 | 3 | 3842.7969 | 3842.7799 | 0 | 4.4 | 62 | Specific | 255.3 | 205.65 | 205.65 | 3.38 | 11 | >sp\|P01591\|IGJ_HUMAN Immunoglobulin J chain OS=Homo sapiens OX=9606 GN=JCHAIN PE=1 SV=4 | 1392 | 20201118_lss_colostrum_1ng_glyco_hcd30_etd30_compound15.12890.12890.3 | controllerType=0 controllerNumber=1 scan=12890 | 30.7365 |
| 09500:3 | 24 | R.IIVPLNNREN[+1913.677]ISDPTSPLR.T | HexNAc(4)Hex(5)NeuAc(1) | N[+1914] | 1354.6286 | 3 | 4061.8712 | 4061.8542 | 0 | 4.2 | 62 | Specific | 284.85 | 229.21 | 229.21 | 4.04 | 11 | >sp\|P01591\|IGJ_HUMAN Immunoglobulin J chain OS=Homo sapiens OX=9606 GN=JCHAIN PE=1 SV=4 | 1392 | 20201118_lss_colostrum_1ng_glyco_hcd30_etd30_compound15.12776.12776.3 | controllerType=0 controllerNumber=1 scan=12776 | 30.4132 |
| 08316:3 | 24 | R.IIVPLNNREN[+1809.666]ISDPTSPLR.T | HexNAc(5)Hex(4)Fuc(1) | N[+1810] | 1319.9557 | 3 | 3957.8527 | 3957.8432 | 0 | 2.4 | 62 | Specific | 287.16 | 201.18 | 201.18 | 3.63 | 11 | >sp\|P01591\|IGJ_HUMAN Immunoglobulin J chain OS=Homo sapiens OX=9606 GN=JCHAIN PE=1 SV=4 | 1392 | 20201118_lss_colostrum_1ng_glyco_hcd30_etd30_compound15.11540.11540.3 | controllerType=0 controllerNumber=1 scan=11540 | 27.5178 |
| 09543:3 | 24 | R.IIVPLNNREN[+1856.656]ISDPTSPLR.T | HexNAc(3)Hex(5)Fuc(1)NeuAc(1) | N[+1857] | 1335.6196 | 3 | 4004.8441 | 4004.8327 | 0 | 2.9 | 62 | Specific | 291.78 | 224.11 | 224.11 | 4.04 | 11 | >sp\|P01591\|IGJ_HUMAN Immunoglobulin J chain OS=Homo sapiens OX=9606 GN=JCHAIN PE=1 SV=4 | 1392 | 20201118_lss_colostrum_1ng_glyco_hcd30_etd30_compound15.12821.12821.3 | controllerType=0 controllerNumber=1 scan=12821 | 30.5298 |
| 11009:3 | 24 | R.IIVPLNNREN[+2204.772]ISDPTSPLR.T | HexNAc(4)Hex(5)NeuAc(2) | N[+2205] | 1451.6585 | 3 | 4352.961 | 4352.9496 | 0 | 2.6 | 62 | Specific | 310.14 | 222.22 | 222.22 | 4.04 | 11 | >sp\|P01591\|IGJ_HUMAN Immunoglobulin J chain OS=Homo sapiens OX=9606 GN=JCHAIN PE=1 SV=4 | 1392 | 20201118_lss_colostrum_1ng_glyco_hcd30_etd30_compound15.14352.14352.3 | controllerType=0 controllerNumber=1 scan=14352 | 34.0887 |
| 08317:3 | 24 | R.IIVPLNNREN[+1663.608]ISDPTSPLR.T | HexNAc(5)Hex(4) | N[+1664] | 1271.2693 | 3 | 3811.7935 | 3811.7853 | 0 | 2.1 | 62 | Specific | 354.4 | 246.9 | 246.9 | 4.25 | 11 | >sp\|P01591\|IGJ_HUMAN Immunoglobulin J chain OS=Homo sapiens OX=9606 GN=JCHAIN PE=1 SV=4 | 1392 | 20201118_lss_colostrum_1ng_glyco_hcd30_etd30_compound15.11541.11541.3 | controllerType=0 controllerNumber=1 scan=11541 | 27.5228 |
| 10587:4 | 31 | K.MVSHHN[+2059.735]LTTGATLINEQWLLTTAK.N | HexNAc(4)Hex(5)Fuc(1)NeuAc(1) | N[+2060] | 1185.5424 | 4 | 4739.1478 | 4739.1272 | 0 | 4.3 | 179 | Specific | 384.54 | 359.21 | 359.21 | 4.71 | 7 | >sp\|P00738\|HPT_HUMAN Haptoglobin OS=Homo sapiens OX=9606 GN=HP PE=1 SV=1 | 407 | 20201118_lss_colostrum_1ng_glyco_hcd30_etd30_compound15.13911.13911.4 | controllerType=0 controllerNumber=1 scan=13911 | 33.0438 |
| 04857:3 | 38 | R.LSPN[+2205.793]ASAEHLELR.W | HexNAc(4)Hex(5)Fuc(2)NeuAc(1) | N[+2206] | 1214.855 | 3 | 3642.5506 | 3642.5421 | 0 | 2.3 | 52 | Specific | 231.85 | 148.98 | 148.98 | 2.96 | 5 | >sp\|Q13410\|BT1A1_HUMAN Butyrophilin subfamily 1 member A1 OS=Homo sapiens OX=9606 GN=BTN1A1 PE=1 SV=3 | 179 | 20201118_lss_colostrum_1ng_glyco_hcd30_etd30_compound15.7943.7943.3 | controllerType=0 controllerNumber=1 scan=7943 | 20.6436 |
| 06744:3 | 38 | R.LSPN[+2350.830]ASAEHLELR.W | HexNAc(4)Hex(5)Fuc(1)NeuAc(2) | N[+2351] | 1263.2017 | 3 | 3787.5906 | 3787.5796 | 0 | 2.9 | 52 | Specific | 270.05 | 146.55 | 146.55 | 3.82 | 5 | >sp\|Q13410\|BT1A1_HUMAN Butyrophilin subfamily 1 member A1 OS=Homo sapiens OX=9606 GN=BTN1A1 PE=1 SV=3 | 179 | 20201118_lss_colostrum_1ng_glyco_hcd30_etd30_compound15.9900.9900.3 | controllerType=0 controllerNumber=1 scan=9900 | 24.171 |
| 12726:3 | 45 | R.KLPPGLLAN[+2059.735]FTLLR.T | HexNAc(4)Hex(5)Fuc(1)NeuAc(1) | N[+2060] | 1204.905 | 3 | 3612.7003 | 3612.6923 | 0 | 2.2 | 178 | Specific | 243.73 | 204.74 | 204.74 | 3.29 | 3 | >sp\|P02750\|A2GL_HUMAN Leucine-rich alpha-2-glycoprotein OS=Homo sapiens OX=9606 GN=LRG1 PE=1 SV=2 | 518 | 20201118_lss_colostrum_1ng_glyco_hcd30_etd30_compound15.16156.16156.3 | controllerType=0 controllerNumber=1 scan=16156 | 38.8518 |
| 12700:3 | 45 | R.KLPPGLLAN[+2059.735]FTLLR.T | HexNAc(4)Hex(5)Fuc(1)NeuAc(1) | N[+2060] | 1204.9045 | 3 | 3612.6989 | 3612.6923 | 0 | 1.8 | 178 | Specific | 281.61 | 245.17 | 245.17 | 4.04 | 3 | >sp\|P02750\|A2GL_HUMAN Leucine-rich alpha-2-glycoprotein OS=Homo sapiens OX=9606 GN=LRG1 PE=1 SV=2 | 518 | 20201118_lss_colostrum_1ng_glyco_hcd30_etd30_compound15.16129.16129.3 | controllerType=0 controllerNumber=1 scan=16129 | 38.7788 |
| 18369:3 | 51 | R.AVN[+2204.772]ITSENLIDDVVSLIR.G | HexNAc(4)Hex(5)NeuAc(2) | N[+2205] | 1392.6274 | 3 | 4175.8676 | 4175.8481 | 0 | 4.7 | 413 | Specific | 351.4 | 246.31 | 246.31 | 4.79 | 2 | >sp\|P02748\|CO9_HUMAN Complement component C9 OS=Homo sapiens OX=9606 GN=C9 PE=1 SV=2 | 1547 | 20201118_lss_colostrum_1ng_glyco_hcd30_etd30_compound15.22126.22126.3 | controllerType=0 controllerNumber=1 scan=22126 | 57.1402 |
| 14027:3 | 53 | R.NYIVPILWLN[+1864.634]ETGTIGDEKANMFR.S | HexNAc(2)Hex(9) | N[+1865] | 1553.6959 | 3 | 4659.073 | 4659.0574 | 0 | 3.3 | 408 | Specific | 411.89 | 411.89 | 411.89 | 7.01 | 3 | >sp\|P16671\|CD36_HUMAN Platelet glycoprotein 4 OS=Homo sapiens OX=9606 GN=CD36 PE=1 SV=2 | 812 | 20201118_lss_colostrum_1ng_glyco_hcd30_etd30_compound15.17523.17523.3 | controllerType=0 controllerNumber=1 scan=17523 | 42.3616 |
| 00696:4 | 89 | K.TKPREEQFN[+1606.587]STFR.V | HexNAc(4)Hex(4)Fuc(1) | N[+1607] | 812.359 | 4 | 3246.4143 | 3246.4054 | 0 | 2.7 | 168 | Specific | 250.94 | 245.77 | 245.77 | 3.96 | 3 | >sp\|P01859\|IGHG2_HUMAN Immunoglobulin heavy constant gamma 2 OS=Homo sapiens OX=9606 GN=IGHG2 PE=1 SV=2 | 387 | 20201118_lss_colostrum_1ng_glyco_hcd30_etd30_compound15.3621.3621.4 | controllerType=0 controllerNumber=1 scan=3621 | 12.5572 |
| 00654:4 | 89 | K.TKPREEQFN[+1768.640]STFR.V | HexNAc(4)Hex(5)Fuc(1) | N[+1769] | 852.8724 | 4 | 3408.4678 | 3408.4582 | 0 | 2.8 | 168 | Specific | 252.17 | 252.17 | 252.17 | 4.19 | 3 | >sp\|P01859\|IGHG2_HUMAN Immunoglobulin heavy constant gamma 2 OS=Homo sapiens OX=9606 GN=IGHG2 PE=1 SV=2 | 387 | 20201118_lss_colostrum_1ng_glyco_hcd30_etd30_compound15.3577.3577.4 | controllerType=0 controllerNumber=1 scan=3577 | 12.4613 |
| 00630:4 | 89 | K.TKPREEQFN[+1768.640]STFR.V | HexNAc(4)Hex(5)Fuc(1) | N[+1769] | 852.8712 | 4 | 3408.4631 | 3408.4582 | 0 | 1.4 | 168 | Specific | 459.56 | 400.93 | 400.93 | 6.41 | 3 | >sp\|P01859\|IGHG2_HUMAN Immunoglobulin heavy constant gamma 2 OS=Homo sapiens OX=9606 GN=IGHG2 PE=1 SV=2 | 387 | 20201118_lss_colostrum_1ng_glyco_hcd30_etd30_compound15.3552.3552.4 | controllerType=0 controllerNumber=1 scan=3552 | 12.4071 |
| 15171:4 | 1 | R.RTAGWNVPIGTLRPFLN[+2351.851]WTGPPEPIEAAVAR.F | HexNAc(4)Hex(5)Fuc(3)NeuAc(1) | N[+2352] | 1435.4248 | 4 | 5738.6774 | 5738.6628 | 0 | 2.5 | 140 | Specific | 620.19 | 620.19 | 620.19 | 9.15 | 48 | >sp\|P02788\|TRFL_HUMAN Lactotransferrin OS=Homo sapiens OX=9606 GN=LTF PE=1 SV=6 | 1270 | 20201118_lss_milk_1ng_glyco_hcd30_etd30_compound15.18750.18750.4 | controllerType=0 controllerNumber=1 scan=18750 | 43.5506 |
| 15202:5 | 1 | R.RTAGWNVPIGTLRPFLN[+2059.735]WTGPPEPIEAAVAR.F | HexNAc(4)Hex(5)Fuc(1)NeuAc(1) | N[+2060] | 1090.1173 | 5 | 5446.5575 | 5446.547 | 0 | 1.9 | 140 | Specific | 735.09 | 735.09 | 735.09 | 10.32 | 48 | >sp\|P02788\|TRFL_HUMAN Lactotransferrin OS=Homo sapiens OX=9606 GN=LTF PE=1 SV=6 | 1270 | 20201118_lss_milk_1ng_glyco_hcd30_etd30_compound15.18783.18783.5 | controllerType=0 controllerNumber=1 scan=18783 | 43.6563 |
| 15163:5 | 1 | R.RTAGWNVPIGTLRPFLN[+2205.793]WTGPPEPIEAAVAR.F | HexNAc(4)Hex(5)Fuc(2)NeuAc(1) | N[+2206] | 1119.3296 | 5 | 5592.6189 | 5592.6049 | 0 | 2.5 | 140 | Specific | 856.92 | 803.9 | 803.9 | 13.6 | 48 | >sp\|P02788\|TRFL_HUMAN Lactotransferrin OS=Homo sapiens OX=9606 GN=LTF PE=1 SV=6 | 1270 | 20201118_lss_milk_1ng_glyco_hcd30_etd30_compound15.18741.18741.3 | controllerType=0 controllerNumber=1 scan=18741 | 43.5245 |
| 14460:4 | 1 | R.RTAGWNVPIGTLRPFLN[+1768.640]WTGPPEPIEAAVAR.F | HexNAc(4)Hex(5)Fuc(1) | N[+1769] | 1289.6221 | 4 | 5155.4665 | 5155.4516 | 0 | 2.9 | 140 | Specific | 889.98 | 889.98 | 889.98 | 14.15 | 48 | >sp\|P02788\|TRFL_HUMAN Lactotransferrin OS=Homo sapiens OX=9606 GN=LTF PE=1 SV=6 | 1270 | 20201118_lss_milk_1ng_glyco_hcd30_etd30_compound15.18001.18001.4 | controllerType=0 controllerNumber=1 scan=18001 | 41.3957 |
| 15198:4 | 1 | R.RTAGWNVPIGTLRPFLN[+2059.735]WTGPPEPIEAAVAR.F | HexNAc(4)Hex(5)Fuc(1)NeuAc(1) | N[+2060] | 1362.3951 | 4 | 5446.5587 | 5446.547 | 0 | 2.2 | 140 | Specific | 1082.66 | 1026.18 | 1026.18 | 18.15 | 48 | >sp\|P02788\|TRFL_HUMAN Lactotransferrin OS=Homo sapiens OX=9606 GN=LTF PE=1 SV=6 | 1270 | 20201118_lss_milk_1ng_glyco_hcd30_etd30_compound15.18779.18779.4 | controllerType=0 controllerNumber=1 scan=18779 | 43.6491 |
| 16048:3 | 1 | R.TAGWNVPIGTLRPFLN[+2715.963]WTGPPEPIEAAVAR.F | HexNAc(5)Hex(6)Fuc(1)NeuAc(2) | N[+2716] | 1982.9094 | 3 | 5946.7136 | 5946.6735 | 0 | 6.7 | 141 | Specific | 229.65 | 109.74 | 109.74 | 2.7 | 48 | >sp\|P02788\|TRFL_HUMAN Lactotransferrin OS=Homo sapiens OX=9606 GN=LTF PE=1 SV=6 | 1270 | 20201118_lss_milk_1ng_glyco_hcd30_etd30_compound15.19671.19671.3 | controllerType=0 controllerNumber=1 scan=19671 | 46.1057 |
| 16104:3 | 1 | R.TAGWNVPIGTLRPFLN[+2205.793]WTGPPEPIEAAVAR.F | HexNAc(4)Hex(5)Fuc(2)NeuAc(1) | N[+2206] | 1812.8455 | 3 | 5436.5221 | 5436.5038 | 0 | 3.4 | 141 | Specific | 408.08 | 295.92 | 295.92 | 5.85 | 48 | >sp\|P02788\|TRFL_HUMAN Lactotransferrin OS=Homo sapiens OX=9606 GN=LTF PE=1 SV=6 | 1270 | 20201118_lss_milk_1ng_glyco_hcd30_etd30_compound15.19731.19731.3 | controllerType=0 controllerNumber=1 scan=19731 | 46.2889 |
| 15366:3 | 1 | R.TAGWNVPIGTLRPFLN[+1768.640]WTGPPEPIEAAVAR.F | HexNAc(4)Hex(5)Fuc(1) | N[+1769] | 1667.1253 | 3 | 4999.3614 | 4999.3504 | 0 | 2.2 | 141 | Specific | 409.53 | 340.28 | 340.28 | 5.09 | 48 | >sp\|P02788\|TRFL_HUMAN Lactotransferrin OS=Homo sapiens OX=9606 GN=LTF PE=1 SV=6 | 1270 | 20201118_lss_milk_1ng_glyco_hcd30_etd30_compound15.18954.18954.3 | controllerType=0 controllerNumber=1 scan=18954 | 44.0927 |
| 16110:3 | 1 | R.TAGWNVPIGTLRPFLN[+2059.735]WTGPPEPIEAAVAR.F | HexNAc(4)Hex(5)Fuc(1)NeuAc(1) | N[+2060] | 1764.161 | 3 | 5290.4684 | 5290.4459 | 0 | 4.3 | 141 | Specific | 443.02 | 329.77 | 329.77 | 5.67 | 48 | >sp\|P02788\|TRFL_HUMAN Lactotransferrin OS=Homo sapiens OX=9606 GN=LTF PE=1 SV=6 | 1270 | 20201118_lss_milk_1ng_glyco_hcd30_etd30_compound15.19737.19737.3 | controllerType=0 controllerNumber=1 scan=19737 | 46.3051 |
| 17050:3 | 1 | R.TAGWNVPIGTLRPFLN[+2350.830]WTGPPEPIEAAVAR.F | HexNAc(4)Hex(5)Fuc(1)NeuAc(2) | N[+2351] | 1861.1899 | 3 | 5581.5552 | 5581.5413 | 0 | 2.5 | 141 | Specific | 459.77 | 356.02 | 356.02 | 6.27 | 48 | >sp\|P02788\|TRFL_HUMAN Lactotransferrin OS=Homo sapiens OX=9606 GN=LTF PE=1 SV=6 | 1270 | 20201118_lss_milk_1ng_glyco_hcd30_etd30_compound15.20729.20729.3 | controllerType=0 controllerNumber=1 scan=20729 | 49.1727 |
| 16116:5 | 1 | R.TAGWNVPIGTLRPFLN[+2205.793]WTGPPEPIEAAVAR.F | HexNAc(4)Hex(5)Fuc(2)NeuAc(1) | N[+2206] | 1088.109 | 5 | 5436.516 | 5436.5038 | 0 | 2.2 | 141 | Specific | 592.17 | 592.17 | 592.17 | 8.8 | 48 | >sp\|P02788\|TRFL_HUMAN Lactotransferrin OS=Homo sapiens OX=9606 GN=LTF PE=1 SV=6 | 1270 | 20201118_lss_milk_1ng_glyco_hcd30_etd30_compound15.19743.19743.5 | controllerType=0 controllerNumber=1 scan=19743 | 46.319 |
| 16060:4 | 1 | R.TAGWNVPIGTLRPFLN[+2716.983]WTGPPEPIEAAVAR.F | HexNAc(5)Hex(6)Fuc(3)NeuAc(1) | N[+2717] | 1487.684 | 4 | 5947.714 | 5947.6939 | 0 | 3.4 | 141 | Specific | 606.52 | 583.46 | 583.46 | 9.2 | 48 | >sp\|P02788\|TRFL_HUMAN Lactotransferrin OS=Homo sapiens OX=9606 GN=LTF PE=1 SV=6 | 1270 | 20201118_lss_milk_1ng_glyco_hcd30_etd30_compound15.19684.19684.4 | controllerType=0 controllerNumber=1 scan=19684 | 46.1354 |
| 15329:4 | 1 | R.TAGWNVPIGTLRPFLN[+2279.830]WTGPPEPIEAAVAR.F | HexNAc(5)Hex(6)Fuc(2) | N[+2280] | 1378.3944 | 4 | 5510.5558 | 5510.5406 | 0 | 2.8 | 141 | Specific | 660.68 | 617.4 | 617.4 | 10.8 | 48 | >sp\|P02788\|TRFL_HUMAN Lactotransferrin OS=Homo sapiens OX=9606 GN=LTF PE=1 SV=6 | 1270 | 20201118_lss_milk_1ng_glyco_hcd30_etd30_compound15.18915.18915.4 | controllerType=0 controllerNumber=1 scan=18915 | 43.9622 |
| 17049:4 | 1 | R.TAGWNVPIGTLRPFLN[+2350.830]WTGPPEPIEAAVAR.F | HexNAc(4)Hex(5)Fuc(1)NeuAc(2) | N[+2351] | 1396.1438 | 4 | 5581.5534 | 5581.5413 | 0 | 2.2 | 141 | Specific | 679.89 | 674.16 | 674.16 | 10.8 | 48 | >sp\|P02788\|TRFL_HUMAN Lactotransferrin OS=Homo sapiens OX=9606 GN=LTF PE=1 SV=6 | 1270 | 20201118_lss_milk_1ng_glyco_hcd30_etd30_compound15.20728.20728.4 | controllerType=0 controllerNumber=1 scan=20728 | 49.1698 |
| 16065:4 | 1 | R.TAGWNVPIGTLRPFLN[+2424.867]WTGPPEPIEAAVAR.F | HexNAc(5)Hex(6)Fuc(1)NeuAc(1) | N[+2425] | 1414.6541 | 4 | 5655.5944 | 5655.5781 | 0 | 2.9 | 141 | Specific | 696.16 | 673.1 | 673.1 | 10.8 | 48 | >sp\|P02788\|TRFL_HUMAN Lactotransferrin OS=Homo sapiens OX=9606 GN=LTF PE=1 SV=6 | 1270 | 20201118_lss_milk_1ng_glyco_hcd30_etd30_compound15.19689.19689.4 | controllerType=0 controllerNumber=1 scan=19689 | 46.147 |
| 16103:4 | 1 | R.TAGWNVPIGTLRPFLN[+2205.793]WTGPPEPIEAAVAR.F | HexNAc(4)Hex(5)Fuc(2)NeuAc(1) | N[+2206] | 1359.8853 | 4 | 5436.5192 | 5436.5038 | 0 | 2.8 | 141 | Specific | 779.52 | 779.52 | 779.52 | 12.8 | 48 | >sp\|P02788\|TRFL_HUMAN Lactotransferrin OS=Homo sapiens OX=9606 GN=LTF PE=1 SV=6 | 1270 | 20201118_lss_milk_1ng_glyco_hcd30_etd30_compound15.19730.19730.4 | controllerType=0 controllerNumber=1 scan=19730 | 46.286 |
| 17046:4 | 1 | R.TAGWNVPIGTLRPFLN[+2350.830]WTGPPEPIEAAVAR.F | HexNAc(4)Hex(5)Fuc(1)NeuAc(2) | N[+2351] | 1396.1431 | 4 | 5581.5504 | 5581.5413 | 0 | 1.6 | 141 | Specific | 840.97 | 784.03 | 784.03 | 13.8 | 48 | >sp\|P02788\|TRFL_HUMAN Lactotransferrin OS=Homo sapiens OX=9606 GN=LTF PE=1 SV=6 | 1270 | 20201118_lss_milk_1ng_glyco_hcd30_etd30_compound15.20724.20724.4 | controllerType=0 controllerNumber=1 scan=20724 | 49.1577 |
| 16109:4 | 1 | R.TAGWNVPIGTLRPFLN[+2059.735]WTGPPEPIEAAVAR.F | HexNAc(4)Hex(5)Fuc(1)NeuAc(1) | N[+2060] | 1323.3729 | 4 | 5290.4699 | 5290.4459 | 0 | 4.5 | 141 | Specific | 938.28 | 923.32 | 923.32 | 15.7 | 48 | >sp\|P02788\|TRFL_HUMAN Lactotransferrin OS=Homo sapiens OX=9606 GN=LTF PE=1 SV=6 | 1270 | 20201118_lss_milk_1ng_glyco_hcd30_etd30_compound15.19736.19736.4 | controllerType=0 controllerNumber=1 scan=19736 | 46.3017 |
| 15361:4 | 1 | R.TAGWNVPIGTLRPFLN[+1768.640]WTGPPEPIEAAVAR.F | HexNAc(4)Hex(5)Fuc(1) | N[+1769] | 1250.5957 | 4 | 4999.361 | 4999.3504 | 0 | 2.1 | 141 | Specific | 1053.44 | 1046.84 | 1046.84 | 18.8 | 48 | >sp\|P02788\|TRFL_HUMAN Lactotransferrin OS=Homo sapiens OX=9606 GN=LTF PE=1 SV=6 | 1270 | 20201118_lss_milk_1ng_glyco_hcd30_etd30_compound15.18949.18949.4 | controllerType=0 controllerNumber=1 scan=18949 | 44.0794 |
| 15647:4 | 1 | R.TAGWNIPMGLLFN[+2205.793]QTGSC[+57.021]K.F | HexNAc(4)Hex(5)Fuc(2)NeuAc(1) | C[+57], N[+2206] | 1075.9595 | 4 | 4300.8162 | 4300.8028 | 0 | 3.1 | 485 | Specific | 236.11 | 213.6 | 213.6 | 2.93 | 48 | >sp\|P02788\|TRFL_HUMAN Lactotransferrin OS=Homo sapiens OX=9606 GN=LTF PE=1 SV=6 | 1270 | 20201118_lss_milk_1ng_glyco_hcd30_etd30_compound15.19248.19248.4 | controllerType=0 controllerNumber=1 scan=19248 | 44.8693 |
| 14196:3 | 1 | R.TAGWNIPMGLLFN[+1913.677]QTGSC[+57.021]K.F | HexNAc(4)Hex(5)NeuAc(1) | C[+57], N[+1914] | 1336.9068 | 3 | 4008.7058 | 4008.687 | 0 | 4.7 | 485 | Specific | 246.29 | 220.21 | 220.21 | 3.94 | 48 | >sp\|P02788\|TRFL_HUMAN Lactotransferrin OS=Homo sapiens OX=9606 GN=LTF PE=1 SV=6 | 1270 | 20201118_lss_milk_1ng_glyco_hcd30_etd30_compound15.17722.17722.3 | controllerType=0 controllerNumber=1 scan=17722 | 40.5786 |
| 17252:3 | 1 | R.TAGWNIPMGLLFN[+2204.772]QTGSC[+57.021]K.F | HexNAc(4)Hex(5)NeuAc(2) | C[+57], N[+2205] | 1433.9362 | 3 | 4299.7941 | 4299.7824 | 0 | 2.7 | 485 | Specific | 320.15 | 233.53 | 233.53 | 4.81 | 48 | >sp\|P02788\|TRFL_HUMAN Lactotransferrin OS=Homo sapiens OX=9606 GN=LTF PE=1 SV=6 | 1270 | 20201118_lss_milk_1ng_glyco_hcd30_etd30_compound15.20944.20944.3 | controllerType=0 controllerNumber=1 scan=20944 | 49.8814 |
| 14501:3 | 1 | R.TAGWNIPMGLLFN[+1216.423]QTGSC[+57.021]K.F | HexNAc(2)Hex(5) | C[+57], N[+1216] | 1104.4855 | 3 | 3311.442 | 3311.4328 | 0 | 2.8 | 485 | Specific | 321.53 | 312.37 | 171.23 | 4.22 | 48 | >sp\|P02788\|TRFL_HUMAN Lactotransferrin OS=Homo sapiens OX=9606 GN=LTF PE=1 SV=6 | 1270 | 20201118_lss_milk_1ng_glyco_hcd30_etd30_compound15.18044.18044.3 | controllerType=0 controllerNumber=1 scan=18044 | 41.5255 |
| 17144:4 | 1 | R.TAGWNIPMGLLFN[+2350.830]QTGSC[+57.021]K.F | HexNAc(4)Hex(5)Fuc(1)NeuAc(2) | C[+57], N[+2351] | 1112.2179 | 4 | 4445.8499 | 4445.8403 | 0 | 2.2 | 485 | Specific | 335.26 | 283.13 | 283.13 | 4.14 | 48 | >sp\|P02788\|TRFL_HUMAN Lactotransferrin OS=Homo sapiens OX=9606 GN=LTF PE=1 SV=6 | 1270 | 20201118_lss_milk_1ng_glyco_hcd30_etd30_compound15.20828.20828.4 | controllerType=0 controllerNumber=1 scan=20828 | 49.474 |
| 14240:3 | 1 | R.TAGWNIPMGLLFN[+1768.640]QTGSC[+57.021]K.F | HexNAc(4)Hex(5)Fuc(1) | C[+57], N[+1769] | 1288.5641 | 3 | 3863.6779 | 3863.6495 | 0 | 7.4 | 485 | Specific | 346.71 | 259.71 | 259.71 | 3.59 | 48 | >sp\|P02788\|TRFL_HUMAN Lactotransferrin OS=Homo sapiens OX=9606 GN=LTF PE=1 SV=6 | 1270 | 20201118_lss_milk_1ng_glyco_hcd30_etd30_compound15.17769.17769.3 | controllerType=0 controllerNumber=1 scan=17769 | 40.7067 |
| 15617:3 | 1 | R.TAGWNIPMGLLFN[+2205.793]QTGSC[+57.021]K.F | HexNAc(4)Hex(5)Fuc(2)NeuAc(1) | C[+57], N[+2206] | 1434.2753 | 3 | 4300.8114 | 4300.8028 | 0 | 2 | 485 | Specific | 362.77 | 264.52 | 264.52 | 4.98 | 48 | >sp\|P02788\|TRFL_HUMAN Lactotransferrin OS=Homo sapiens OX=9606 GN=LTF PE=1 SV=6 | 1270 | 20201118_lss_milk_1ng_glyco_hcd30_etd30_compound15.19217.19217.3 | controllerType=0 controllerNumber=1 scan=19217 | 44.7822 |
| 15743:3 | 1 | R.TAGWNIPMGLLFN[+1913.677]QTGSC[+57.021]K.F | HexNAc(4)Hex(5)NeuAc(1) | C[+57], N[+1914] | 1336.9037 | 3 | 4008.6967 | 4008.687 | 0 | 2.4 | 485 | Specific | 421.79 | 285.54 | 285.54 | 5.06 | 48 | >sp\|P02788\|TRFL_HUMAN Lactotransferrin OS=Homo sapiens OX=9606 GN=LTF PE=1 SV=6 | 1270 | 20201118_lss_milk_1ng_glyco_hcd30_etd30_compound15.19350.19350.3 | controllerType=0 controllerNumber=1 scan=19350 | 45.2084 |
| 15770:4 | 1 | R.TAGWNIPMGLLFN[+1913.677]QTGSC[+57.021]K.F | HexNAc(4)Hex(5)NeuAc(1) | C[+57], N[+1914] | 1002.9294 | 4 | 4008.6956 | 4008.687 | 0 | 2.2 | 485 | Specific | 440.65 | 372.32 | 372.32 | 5.65 | 48 | >sp\|P02788\|TRFL_HUMAN Lactotransferrin OS=Homo sapiens OX=9606 GN=LTF PE=1 SV=6 | 1270 | 20201118_lss_milk_1ng_glyco_hcd30_etd30_compound15.19379.19379.4 | controllerType=0 controllerNumber=1 scan=19379 | 45.2954 |
| 17125:3 | 1 | R.TAGWNIPMGLLFN[+2350.830]QTGSC[+57.021]K.F | HexNAc(4)Hex(5)Fuc(1)NeuAc(2) | C[+57], N[+2351] | 1482.6243 | 3 | 4445.8584 | 4445.8403 | 0 | 4.1 | 485 | Specific | 467.77 | 321.12 | 321.12 | 5.24 | 48 | >sp\|P02788\|TRFL_HUMAN Lactotransferrin OS=Homo sapiens OX=9606 GN=LTF PE=1 SV=6 | 1270 | 20201118_lss_milk_1ng_glyco_hcd30_etd30_compound15.20808.20808.3 | controllerType=0 controllerNumber=1 scan=20808 | 49.3971 |
| 15044:3 | 1 | R.TAGWNIPMGLLFN[+1768.640]QTGSC[+57.021]KFDEYFSQSC[+57.021]APGSDPR. | HexNAc(4)Hex(5)Fuc(1) | C[+57]*2, N[+1769] | 1903.1428 | 3 | 5707.4138 | 5707.3966 | 0 | 3 | 485 | Specific | 299.79 | 243.4 | 243.4 | 4.68 | 48 | >sp\|P02788\|TRFL_HUMAN Lactotransferrin OS=Homo sapiens OX=9606 GN=LTF PE=1 SV=6 | 1270 | 20201118_lss_milk_1ng_glyco_hcd30_etd30_compound15.18616.18616.3 | controllerType=0 controllerNumber=1 scan=18616 | 43.1633 |
| 16810:4 | 1 | R.TAGWNIPMGLLFN[+2350.830]QTGSC[+57.021]KFDEYFSQSC[+57.021]APGSDPR. | HexNAc(4)Hex(5)Fuc(1)NeuAc(2) | C[+57]*2, N[+2351] | 1573.1582 | 4 | 6289.611 | 6289.5875 | 0 | 3.7 | 485 | Specific | 438.14 | 415.68 | 415.68 | 6.24 | 48 | >sp\|P02788\|TRFL_HUMAN Lactotransferrin OS=Homo sapiens OX=9606 GN=LTF PE=1 SV=6 | 1270 | 20201118_lss_milk_1ng_glyco_hcd30_etd30_compound15.20476.20476.4 | controllerType=0 controllerNumber=1 scan=20476 | 48.4125 |
| 16871:4 | 1 | R.TAGWNIPMGLLFN[+2204.772]QTGSC[+57.021]KFDEYFSQSC[+57.021]APGSDPR. | HexNAc(4)Hex(5)NeuAc(2) | C[+57]*2, N[+2205] | 1536.6439 | 4 | 6143.5539 | 6143.5295 | 0 | 4 | 485 | Specific | 493.19 | 438.74 | 438.74 | 6.55 | 48 | >sp\|P02788\|TRFL_HUMAN Lactotransferrin OS=Homo sapiens OX=9606 GN=LTF PE=1 SV=6 | 1270 | 20201118_lss_milk_1ng_glyco_hcd30_etd30_compound15.20540.20540.4 | controllerType=0 controllerNumber=1 scan=20540 | 48.6 |
| 15842:4 | 1 | R.TAGWNIPMGLLFN[+2205.793]QTGSC[+57.021]KFDEYFSQSC[+57.021]APGSDPR. | HexNAc(4)Hex(5)Fuc(2)NeuAc(1) | C[+57]*2, N[+2206] | 1536.8986 | 4 | 6144.5724 | 6144.5499 | 0 | 3.7 | 485 | Specific | 495.52 | 461.5 | 461.5 | 6.55 | 48 | >sp\|P02788\|TRFL_HUMAN Lactotransferrin OS=Homo sapiens OX=9606 GN=LTF PE=1 SV=6 | 1270 | 20201118_lss_milk_1ng_glyco_hcd30_etd30_compound15.19455.19455.4 | controllerType=0 controllerNumber=1 scan=19455 | 45.5156 |
| 15158:3 | 1 | R.TAGWNIPMGLLFN[+1216.423]QTGSC[+57.021]KFDEYFSQSC[+57.021]APGSDPR. | HexNAc(2)Hex(5) | C[+57]*2, N[+1216] | 1719.0711 | 3 | 5155.1988 | 5155.18 | 0 | 3.7 | 485 | Specific | 556.72 | 462.38 | 149.43 | 6.69 | 48 | >sp\|P02788\|TRFL_HUMAN Lactotransferrin OS=Homo sapiens OX=9606 GN=LTF PE=1 SV=6 | 1270 | 20201118_lss_milk_1ng_glyco_hcd30_etd30_compound15.18736.18736.3 | controllerType=0 controllerNumber=1 scan=18736 | 43.5036 |
| 15038:4 | 1 | R.TAGWNIPMGLLFN[+1768.640]QTGSC[+57.021]KFDEYFSQSC[+57.021]APGSDPR. | HexNAc(4)Hex(5)Fuc(1) | C[+57]*2, N[+1769] | 1427.6104 | 4 | 5707.4196 | 5707.3966 | 0 | 4 | 485 | Specific | 619.84 | 573.07 | 573.07 | 8.75 | 48 | >sp\|P02788\|TRFL_HUMAN Lactotransferrin OS=Homo sapiens OX=9606 GN=LTF PE=1 SV=6 | 1270 | 20201118_lss_milk_1ng_glyco_hcd30_etd30_compound15.18610.18610.4 | controllerType=0 controllerNumber=1 scan=18610 | 43.1477 |
| 15837:4 | 1 | R.TAGWNIPMGLLFN[+2059.735]QTGSC[+57.021]KFDEYFSQSC[+57.021]APGSDPR. | HexNAc(4)Hex(5)Fuc(1)NeuAc(1) | C[+57]*2, N[+2060] | 1500.3872 | 4 | 5998.527 | 5998.492 | 0 | 5.8 | 485 | Specific | 844.75 | 785.68 | 785.68 | 12.28 | 48 | >sp\|P02788\|TRFL_HUMAN Lactotransferrin OS=Homo sapiens OX=9606 GN=LTF PE=1 SV=6 | 1270 | 20201118_lss_milk_1ng_glyco_hcd30_etd30_compound15.19450.19450.4 | controllerType=0 controllerNumber=1 scan=19450 | 45.5053 |
| 08046:3 | 1 | R.N[+2205.793]GSDC[+57.021]PDKFC[+57.021]LFQSETK.N | HexNAc(4)Hex(5)Fuc(2)NeuAc(1) | C[+57]*2, N[+2206] | 1413.5629 | 3 | 4238.6742 | 4238.6667 | 0 | 1.8 | 642 | Specific | 271.73 | 160.64 | 160.64 | 3.95 | 48 | >sp\|P02788\|TRFL_HUMAN Lactotransferrin OS=Homo sapiens OX=9606 GN=LTF PE=1 SV=6 | 1270 | 20201118_lss_milk_1ng_glyco_hcd30_etd30_compound15.11278.11278.3 | controllerType=0 controllerNumber=1 scan=11278 | 26.5291 |
| 08117:3 | 1 | R.N[+2059.735]GSDC[+57.021]PDKFC[+57.021]LFQSETK.N | HexNAc(4)Hex(5)Fuc(1)NeuAc(1) | C[+57]*2, N[+2060] | 1364.8788 | 3 | 4092.622 | 4092.6088 | 0 | 3.2 | 642 | Specific | 343.15 | 255.04 | 255.04 | 4.38 | 48 | >sp\|P02788\|TRFL_HUMAN Lactotransferrin OS=Homo sapiens OX=9606 GN=LTF PE=1 SV=6 | 1270 | 20201118_lss_milk_1ng_glyco_hcd30_etd30_compound15.11353.11353.3 | controllerType=0 controllerNumber=1 scan=11353 | 26.6912 |
| 14941:3 | 2 | R.LSLHRPALEDLLLGSEAN[+1298.476]LTC[+57.021]TLTGLR.D | HexNAc(4)Hex(3) | C[+57], N[+1298] | 1421.3636 | 3 | 4262.0762 | 4262.0742 | 0 | 0.5 | 127 | Specific | 316.33 | 249.99 | 249.99 | 4.87 | 31 | >sp\|P01876\|IGHA1_HUMAN Immunoglobulin heavy constant alpha 1 OS=Homo sapiens OX=9606 GN=IGHA1 PE=1 SV=2 | 255 | 20201118_lss_milk_1ng_glyco_hcd30_etd30_compound15.18507.18507.2 | controllerType=0 controllerNumber=1 scan=18507 | 42.8413 |
| 15685:4 | 2 | R.LSLHRPALEDLLLGSEAN[+1751.624]LTC[+57.021]TLTGLR.D | HexNAc(4)Hex(4)NeuAc(1) | C[+57], N[+1752] | 1179.5649 | 4 | 4715.2376 | 4715.2225 | 0 | 3.2 | 127 | Specific | 341.93 | 341.93 | 341.93 | 4.72 | 31 | >sp\|P01876\|IGHA1_HUMAN Immunoglobulin heavy constant alpha 1 OS=Homo sapiens OX=9606 GN=IGHA1 PE=1 SV=2 | 255 | 20201118_lss_milk_1ng_glyco_hcd30_etd30_compound15.19289.19289.4 | controllerType=0 controllerNumber=1 scan=19289 | 44.995 |

Table S4. Cont.

| Query #:z | Protein Rank | Glycopeptide | Glycan | Modification Type(s) | Observed m/z | z | Observed (M+H) | Calc. mass (M+H) | Off-by-x error | Mass error (ppm) | StartPosition | Cleavage | Score | Delta | Delta Mod | \|Log Prob\| | # of unique peptides | Protein Name | Protein DB number | Comment | Scan # | Scan Time |
|---|---|---|---|---|---|---|---|---|---|---|---|---|---|---|---|---|---|---|---|---|---|---|
| 14899:3 | 2 | R.LSLHRPALEDLLLGSEAN[+1501.555]LTC[+57.021]TLTGLR.D | HexNAc(5)Hex(3) | C[+57], N[+1502] | 1489.0588 | 3 | 4465.1618 | 4465.1536 | 0 | 1.8 | 127 | Specific | 391.69 | 346.82 | 346.82 | 5.72 | 31 | >sp\|P01876\|IGHA1_HUMAN Immunoglobulin heavy constant alpha 1 OS=Homo sapiens OX=9606 GN=IGHA1 PE=1 SV=2 | 255 | 20201118_lss_milk_1ng_glyco_hcd30_etd30_compound15.18463.18463.3 | controllerType=0 controllerNumber=1 scan=18463 | 42.7062 |
| 14886:3 | 2 | R.LSLHRPALEDLLLGSEAN[+1622.582]LTC[+57.021]TLTGLR.D | HexNAc(4)Hex(5) | C[+57], N[+1623] | 1529.4024 | 3 | 4586.1926 | 4586.1799 | 0 | 2.8 | 127 | Specific | 398.51 | 345.69 | 345.69 | 5.72 | 31 | >sp\|P01876\|IGHA1_HUMAN Immunoglobulin heavy constant alpha 1 OS=Homo sapiens OX=9606 GN=IGHA1 PE=1 SV=2 | 255 | 20201118_lss_milk_1ng_glyco_hcd30_etd30_compound15.18449.18449.3 | controllerType=0 controllerNumber=1 scan=18449 | 42.6536 |
| 14954:3 | 2 | R.LSLHRPALEDLLLGSEAN[+1216.423]LTC[+57.021]TLTGLR.D | HexNAc(2)Hex(5) | C[+57], N[+1216] | 1394.018 | 3 | 4180.0395 | 4180.0211 | 0 | 4.4 | 127 | Specific | 403.84 | 395.91 | 395.91 | 6.97 | 31 | >sp\|P01876\|IGHA1_HUMAN Immunoglobulin heavy constant alpha 1 OS=Homo sapiens OX=9606 GN=IGHA1 PE=1 SV=2 | 255 | 20201118_lss_milk_1ng_glyco_hcd30_etd30_compound15.18521.18521.3 | controllerType=0 controllerNumber=1 scan=18521 | 42.8791 |
| 14875:3 | 2 | R.LSLHRPALEDLLLGSEAN[+1663.608]LTC[+57.021]TLTGLR.D | HexNAc(5)Hex(4) | C[+57], N[+1664] | 1543.0778 | 3 | 4627.2189 | 4627.2064 | 0 | 2.7 | 127 | Specific | 430.41 | 371.09 | 371.09 | 6.58 | 31 | >sp\|P01876\|IGHA1_HUMAN Immunoglobulin heavy constant alpha 1 OS=Homo sapiens OX=9606 GN=IGHA1 PE=1 SV=2 | 255 | 20201118_lss_milk_1ng_glyco_hcd30_etd30_compound15.18437.18437.3 | controllerType=0 controllerNumber=1 scan=18437 | 42.6062 |
| 14832:3 | 2 | R.LSLHRPALEDLLLGSEAN[+1864.634]LTC[+57.021]TLTGLR.D | HexNAc(2)Hex(9) | C[+57], N[+1865] | 1610.0878 | 3 | 4828.2489 | 4828.2324 | 0 | 3.4 | 127 | Specific | 441.97 | 416.79 | 416.79 | 6.68 | 31 | >sp\|P01876\|IGHA1_HUMAN Immunoglobulin heavy constant alpha 1 OS=Homo sapiens OX=9606 GN=IGHA1 PE=1 SV=2 | 255 | 20201118_lss_milk_1ng_glyco_hcd30_etd30_compound15.18391.18391.3 | controllerType=0 controllerNumber=1 scan=18391 | 42.4446 |
| 14884:3 | 2 | R.LSLHRPALEDLLLGSEAN[+1378.476]LTC[+57.021]TLTGLR.D | HexNAc(2)Hex(6) | C[+57], N[+1378] | 1448.0342 | 3 | 4342.0882 | 4342.074 | 0 | 3.3 | 127 | Specific | 454.88 | 431.51 | 431.51 | 7.17 | 31 | >sp\|P01876\|IGHA1_HUMAN Immunoglobulin heavy constant alpha 1 OS=Homo sapiens OX=9606 GN=IGHA1 PE=1 SV=2 | 255 | 20201118_lss_milk_1ng_glyco_hcd30_etd30_compound15.18447.18447.3 | controllerType=0 controllerNumber=1 scan=18447 | 42.6382 |
| 14905:3 | 2 | R.LSLHRPALEDLLLGSEAN[+1216.423]LTC[+57.021]TLTGLR.D | HexNAc(2)Hex(5) | C[+57], N[+1216] | 1394.0142 | 3 | 4180.0281 | 4180.0211 | 0 | 1.7 | 127 | Specific | 484.89 | 476.16 | 476.16 | 7.38 | 31 | >sp\|P01876\|IGHA1_HUMAN Immunoglobulin heavy constant alpha 1 OS=Homo sapiens OX=9606 GN=IGHA1 PE=1 SV=2 | 255 | 20201118_lss_milk_1ng_glyco_hcd30_etd30_compound15.18469.18469.3 | controllerType=0 controllerNumber=1 scan=18469 | 42.7248 |
| 14907:4 | 2 | R.LSLHRPALEDLLLGSEAN[+1540.529]LTC[+57.021]TLTGLR.D | HexNAc(2)Hex(7) | C[+57], N[+1541] | 1126.784 | 4 | 4504.1141 | 4504.1268 | 0 | -2.8 | 127 | Specific | 580.07 | 554.1 | 554.1 | 6.71 | 31 | >sp\|P01876\|IGHA1_HUMAN Immunoglobulin heavy constant alpha 1 OS=Homo sapiens OX=9606 GN=IGHA1 PE=1 SV=2 | 255 | 20201118_lss_milk_1ng_glyco_hcd30_etd30_compound15.18471.18471.4 | controllerType=0 controllerNumber=1 scan=18471 | 42.734 |
| 15628:4 | 2 | R.LSLHRPALEDLLLGSEAN[+1913.677]LTC[+57.021]TLTGLR.D | HexNAc(4)Hex(5)NeuAc(1) | C[+57], N[+1914] | 1220.0766 | 4 | 4877.2845 | 4877.2753 | 0 | 1.9 | 127 | Specific | 720.25 | 720.25 | 720.25 | 11.85 | 31 | >sp\|P01876\|IGHA1_HUMAN Immunoglobulin heavy constant alpha 1 OS=Homo sapiens OX=9606 GN=IGHA1 PE=1 SV=2 | 255 | 20201118_lss_milk_1ng_glyco_hcd30_etd30_compound15.19229.19229.4 | controllerType=0 controllerNumber=1 scan=19229 | 44.8193 |
| 14964:4 | 2 | R.LSLHRPALEDLLLGSEAN[+1095.397]LTC[+57.021]TLTGLR.D | HexNAc(3)Hex(3) | C[+57], N[+1095] | 1015.506 | 4 | 4059.002 | 4058.9949 | 0 | 1.8 | 127 | Specific | 802.72 | 802.72 | 802.72 | 13.85 | 31 | >sp\|P01876\|IGHA1_HUMAN Immunoglobulin heavy constant alpha 1 OS=Homo sapiens OX=9606 GN=IGHA1 PE=1 SV=2 | 255 | 20201118_lss_milk_1ng_glyco_hcd30_etd30_compound15.18532.18532.4 | controllerType=0 controllerNumber=1 scan=18532 | 42.9219 |
| 14943:4 | 2 | R.LSLHRPALEDLLLGSEAN[+1298.476]LTC[+57.021]TLTGLR.D | HexNAc(4)Hex(3) | C[+57], N[+1298] | 1066.2759 | 4 | 4262.0819 | 4262.0742 | 0 | 1.8 | 127 | Specific | 834.94 | 834.94 | 834.94 | 13.85 | 31 | >sp\|P01876\|IGHA1_HUMAN Immunoglobulin heavy constant alpha 1 OS=Homo sapiens OX=9606 GN=IGHA1 PE=1 SV=2 | 255 | 20201118_lss_milk_1ng_glyco_hcd30_etd30_compound15.18509.18509.4 | controllerType=0 controllerNumber=1 scan=18509 | 42.8484 |
| 14880:4 | 2 | R.LSLHRPALEDLLLGSEAN[+1622.582]LTC[+57.021]TLTGLR.D | HexNAc(4)Hex(5) | C[+57], N[+1623] | 1147.3031 | 4 | 4586.1907 | 4586.1799 | 0 | 2.4 | 127 | Specific | 866.15 | 764.67 | 764.67 | 14.85 | 31 | >sp\|P01876\|IGHA1_HUMAN Immunoglobulin heavy constant alpha 1 OS=Homo sapiens OX=9606 GN=IGHA1 PE=1 SV=2 | 255 | 20201118_lss_milk_1ng_glyco_hcd30_etd30_compound15.18443.18443.4 | controllerType=0 controllerNumber=1 scan=18443 | 42.6286 |
| 14894:4 | 2 | R.LSLHRPALEDLLLGSEAN[+1501.555]LTC[+57.021]TLTGLR.D | HexNAc(5)Hex(3) | C[+57], N[+1502] | 1117.0469 | 4 | 4465.1658 | 4465.1536 | 0 | 2.7 | 127 | Specific | 882.18 | 882.18 | 882.18 | 14.85 | 31 | >sp\|P01876\|IGHA1_HUMAN Immunoglobulin heavy constant alpha 1 OS=Homo sapiens OX=9606 GN=IGHA1 PE=1 SV=2 | 255 | 20201118_lss_milk_1ng_glyco_hcd30_etd30_compound15.18458.18458.4 | controllerType=0 controllerNumber=1 scan=18458 | 42.6913 |
| 14860:4 | 2 | R.LSLHRPALEDLLLGSEAN[+1663.608]LTC[+57.021]TLTGLR.D | HexNAc(5)Hex(4) | C[+57], N[+1664] | 1157.5599 | 4 | 4627.2176 | 4627.2064 | 0 | 2.4 | 127 | Specific | 924.58 | 920.93 | 920.93 | 15.85 | 31 | >sp\|P01876\|IGHA1_HUMAN Immunoglobulin heavy constant alpha 1 OS=Homo sapiens OX=9606 GN=IGHA1 PE=1 SV=2 | 255 | 20201118_lss_milk_1ng_glyco_hcd30_etd30_compound15.18421.18421.4 | controllerType=0 controllerNumber=1 scan=18421 | 42.5581 |
| 14844:4 | 2 | R.LSLHRPALEDLLLGSEAN[+1702.581]LTC[+57.021]TLTGLR.D | HexNAc(2)Hex(8) | C[+57], N[+1703] | 1167.3025 | 4 | 4666.1883 | 4666.1796 | 0 | 1.9 | 127 | Specific | 938.39 | 896.9 | 896.9 | 15.85 | 31 | >sp\|P01876\|IGHA1_HUMAN Immunoglobulin heavy constant alpha 1 OS=Homo sapiens OX=9606 GN=IGHA1 PE=1 SV=2 | 255 | 20201118_lss_milk_1ng_glyco_hcd30_etd30_compound15.18404.18404.4 | controllerType=0 controllerNumber=1 scan=18404 | 42.4908 |
| 14901:4 | 2 | R.LSLHRPALEDLLLGSEAN[+1460.529]LTC[+57.021]TLTGLR.D | HexNAc(4)Hex(4) | C[+57], N[+1461] | 1106.7903 | 4 | 4424.1395 | 4424.1271 | 0 | 2.8 | 127 | Specific | 944.7 | 883.28 | 883.28 | 15.85 | 31 | >sp\|P01876\|IGHA1_HUMAN Immunoglobulin heavy constant alpha 1 OS=Homo sapiens OX=9606 GN=IGHA1 PE=1 SV=2 | 255 | 20201118_lss_milk_1ng_glyco_hcd30_etd30_compound15.18465.18465.4 | controllerType=0 controllerNumber=1 scan=18465 | 42.7149 |
| 14895:4 | 2 | R.LSLHRPALEDLLLGSEAN[+1216.423]LTC[+57.021]TLTGLR.D | HexNAc(2)Hex(5) | C[+57], N[+1216] | 1045.7625 | 4 | 4180.0281 | 4180.0211 | 0 | 1.7 | 127 | Specific | 1022.88 | 974.59 | 974.59 | 17.85 | 31 | >sp\|P01876\|IGHA1_HUMAN Immunoglobulin heavy constant alpha 1 OS=Homo sapiens OX=9606 GN=IGHA1 PE=1 SV=2 | 255 | 20201118_lss_milk_1ng_glyco_hcd30_etd30_compound15.18459.18459.4 | controllerType=0 controllerNumber=1 scan=18459 | 42.6959 |
| 14822:4 | 2 | R.LSLHRPALEDLLLGSEAN[+1864.634]LTC[+57.021]TLTGLR.D | HexNAc(2)Hex(9) | C[+57], N[+1865] | 1207.8175 | 4 | 4828.2484 | 4828.2324 | 0 | 3.3 | 127 | Specific | 1054.82 | 1006.86 | 1006.86 | 18.75 | 31 | >sp\|P01876\|IGHA1_HUMAN Immunoglobulin heavy constant alpha 1 OS=Homo sapiens OX=9606 GN=IGHA1 PE=1 SV=2 | 255 | 20201118_lss_milk_1ng_glyco_hcd30_etd30_compound15.18381.18381.4 | controllerType=0 controllerNumber=1 scan=18381 | 42.409 |
| 14870:4 | 2 | R.LSLHRPALEDLLLGSEAN[+1378.476]LTC[+57.021]TLTGLR.D | HexNAc(2)Hex(6) | C[+57], N[+1378] | 1086.2764 | 4 | 4342.0838 | 4342.074 | 0 | 2.3 | 127 | Specific | 1116.18 | 1032.59 | 1032.59 | 19.85 | 31 | >sp\|P01876\|IGHA1_HUMAN Immunoglobulin heavy constant alpha 1 OS=Homo sapiens OX=9606 GN=IGHA1 PE=1 SV=2 | 255 | 20201118_lss_milk_1ng_glyco_hcd30_etd30_compound15.18432.18432.4 | controllerType=0 controllerNumber=1 scan=18432 | 42.5949 |
| 09961:3 | 2 | R.LAGKPTHVN[+1663.608]VSVVMAEVDGTC[+57.021]Y.- | HexNAc(5)Hex(4) | C[+57], N[+1664] | 1337.5904 | 3 | 4010.7566 | 4010.7502 | 0 | 1.6 | 332 | CRagged | 234.71 | 154.45 | 154.45 | 2.71 | 31 | >sp\|P01876\|IGHA1_HUMAN Immunoglobulin heavy constant alpha 1 OS=Homo sapiens OX=9606 GN=IGHA1 PE=1 SV=2 | 255 | 20201118_lss_milk_1ng_glyco_hcd30_etd30_compound15.13271.13271.3 | controllerType=0 controllerNumber=1 scan=13271 | 30.5082 |
| 09949:3 | 2 | R.LAGKPTHVN[+1540.529]VSVVMAEVDGTC[+57.021]Y.- | HexNAc(2)Hex(7) | C[+57], N[+1541] | 1296.5655 | 3 | 3887.6819 | 3887.6706 | 0 | 2.9 | 332 | CRagged | 235.61 | 177.73 | 177.73 | 2.91 | 31 | >sp\|P01876\|IGHA1_HUMAN Immunoglobulin heavy constant alpha 1 OS=Homo sapiens OX=9606 GN=IGHA1 PE=1 SV=2 | 255 | 20201118_lss_milk_1ng_glyco_hcd30_etd30_compound15.13258.13258.3 | controllerType=0 controllerNumber=1 scan=13258 | 30.4771 |
| 10002:3 | 2 | R.LAGKPTHVN[+1378.476]VSVVMAEVDGTC[+57.021]Y.- | HexNAc(2)Hex(6) | C[+57], N[+1378] | 1242.5552 | 3 | 3725.6511 | 3725.6178 | 0 | 9 | 332 | CRagged | 244.38 | 244.38 | 193.82 | 2.73 | 31 | >sp\|P01876\|IGHA1_HUMAN Immunoglobulin heavy constant alpha 1 OS=Homo sapiens OX=9606 GN=IGHA1 PE=1 SV=2 | 255 | 20201118_lss_milk_1ng_glyco_hcd30_etd30_compound15.13314.13314.3 | controllerType=0 controllerNumber=1 scan=13314 | 30.6156 |
| 11537:3 | 2 | R.LAGKPTHVN[+1872.651]VSVVMAEVDGTC[+57.021]Y.- | HexNAc(3)Hex(6)NeuAc(1) | C[+57], N[+1873] | 1407.274 | 3 | 4219.8074 | 4219.7926 | 0 | 3.5 | 332 | CRagged | 244.68 | 157.24 | 143.53 | 3.31 | 31 | >sp\|P01876\|IGHA1_HUMAN Immunoglobulin heavy constant alpha 1 OS=Homo sapiens OX=9606 GN=IGHA1 PE=1 SV=2 | 255 | 20201118_lss_milk_1ng_glyco_hcd30_etd30_compound15.14920.14920.3 | controllerType=0 controllerNumber=1 scan=14920 | 33.8525 |
| 09975:3 | 2 | R.LAGKPTHVN[+1809.666]VSVVMAEVDGTC[+57.021]Y.- | HexNAc(5)Hex(4)Fuc(1) | C[+57], N[+1810] | 1386.275 | 3 | 4156.8103 | 4156.8081 | 0 | 0.5 | 332 | CRagged | 245.38 | 139.47 | 139.47 | 2.83 | 31 | >sp\|P01876\|IGHA1_HUMAN Immunoglobulin heavy constant alpha 1 OS=Homo sapiens OX=9606 GN=IGHA1 PE=1 SV=2 | 255 | 20201118_lss_milk_1ng_glyco_hcd30_etd30_compound15.13285.13285.3 | controllerType=0 controllerNumber=1 scan=13285 | 30.5477 |
| 10036:3 | 2 | R.LAGKPTHVN[+1647.613]VSVVMAEVDGTC[+57.021]Y.- | HexNAc(5)Hex(3)Fuc(1) | C[+57], N[+1648] | 1332.2603 | 3 | 3994.7664 | 3994.7553 | 0 | 2.8 | 332 | CRagged | 251.5 | 143.71 | 143.71 | 3.19 | 31 | >sp\|P01876\|IGHA1_HUMAN Immunoglobulin heavy constant alpha 1 OS=Homo sapiens OX=9606 GN=IGHA1 PE=1 SV=2 | 255 | 20201118_lss_milk_1ng_glyco_hcd30_etd30_compound15.13350.13350.3 | controllerType=0 controllerNumber=1 scan=13350 | 30.757 |
| 11579:3 | 2 | R.LAGKPTHVN[+1751.624]VSVVMAEVDGTC[+57.021]Y.- | HexNAc(4)Hex(4)NeuAc(1) | C[+57], N[+1752] | 1366.9263 | 3 | 4098.7644 | 4098.7663 | 0 | -0.5 | 332 | CRagged | 260.93 | 131.61 | 131.61 | 2.6 | 31 | >sp\|P01876\|IGHA1_HUMAN Immunoglobulin heavy constant alpha 1 OS=Homo sapiens OX=9606 GN=IGHA1 PE=1 SV=2 | 255 | 20201118_lss_milk_1ng_glyco_hcd30_etd30_compound15.14964.14964.3 | controllerType=0 controllerNumber=1 scan=14964 | 33.968 |
| 10020:3 | 2 | R.LAGKPTHVN[+1216.423]VSVVMAEVDGTC[+57.021]Y.- | HexNAc(2)Hex(5) | C[+57], N[+1216] | 1188.5283 | 3 | 3563.5702 | 3563.5649 | 0 | 1.5 | 332 | CRagged | 292.67 | 255.17 | 205.05 | 4.19 | 31 | >sp\|P01876\|IGHA1_HUMAN Immunoglobulin heavy constant alpha 1 OS=Homo sapiens OX=9606 GN=IGHA1 PE=1 SV=2 | 255 | 20201118_lss_milk_1ng_glyco_hcd30_etd30_compound15.13333.13333.3 | controllerType=0 controllerNumber=1 scan=13333 | 30.6957 |
| 04141:3 | 4 | R.FSDGLESN[+1864.634]SSTQFEVKK.Y | HexNAc(2)Hex(9) | N[+1865] | 1256.5204 | 3 | 3767.5468 | 3767.5421 | 0 | 1.2 | 219 | Specific | 257.32 | 205.05 | 205.05 | 3.22 | 17 | >sp\|P0C0L4\|CO4A_HUMAN Complement C4-A OS=Homo sapiens OX=9606 GN=C4A PE=1 SV=2 | 1235 | 20201118_lss_milk_1ng_glyco_hcd30_etd30_compound15.7213.7213.3 | controllerType=0 controllerNumber=1 scan=7213 | 19.1317 |
| 16318:3 | 6 | R.GLSFDVSLEVSQGPGLLN[+2350.830]DTK.V | HexNAc(4)Hex(5)Fuc(1)NeuAc(2) | N[+2351] | 1509.6585 | 3 | 4526.961 | 4526.9436 | 0 | 3.8 | 118 | Specific | 340.48 | 240.47 | 240.47 | 4.7 | 25 | >sp\|P01833\|PIGR_HUMAN Polymeric immunoglobulin receptor OS=Homo sapiens OX=9606 GN=PIGR PE=1 SV=4 | 902 | 20201118_lss_milk_1ng_glyco_hcd30_etd30_compound15.19956.19956.3 | controllerType=0 controllerNumber=1 scan=19956 | 46.8652 |
| 15769:4 | 6 | K.QIGLYPVLVIDSSGYVNPN[+2059.735]YTGR.I | HexNAc(4)Hex(5)Fuc(1)NeuAc(1) | N[+2060] | 1147.0185 | 4 | 4585.0521 | 4585.0384 | 0 | 3 | 168 | Specific | 479 | 454.11 | 454.11 | 5.96 | 25 | >sp\|P01833\|PIGR_HUMAN Polymeric immunoglobulin receptor OS=Homo sapiens OX=9606 GN=PIGR PE=1 SV=4 | 902 | 20201118_lss_milk_1ng_glyco_hcd30_etd30_compound15.19378.19378.4 | controllerType=0 controllerNumber=1 scan=19378 | 45.2916 |
| 18405:4 | 6 | R.LSLLEEPGN[+2350.830]GTFTVILNQLTSR.D | HexNAc(4)Hex(5)Fuc(1)NeuAc(2) | N[+2351] | 1189.04 | 4 | 4753.138 | 4753.1229 | 0 | 3.2 | 413 | Specific | 514.45 | 423.9 | 423.9 | 6.24 | 25 | >sp\|P01833\|PIGR_HUMAN Polymeric immunoglobulin receptor OS=Homo sapiens OX=9606 GN=PIGR PE=1 SV=4 | 902 | 20201118_lss_milk_1ng_glyco_hcd30_etd30_compound15.22165.22165.4 | controllerType=0 controllerNumber=1 scan=22165 | 53.392 |
| 18452:3 | 6 | R.LSLLEEPGN[+2350.830]GTFTVILNQLTSR.D | HexNAc(4)Hex(5)Fuc(1)NeuAc(2) | N[+2351] | 1585.0517 | 3 | 4753.1405 | 4753.1229 | 0 | 3.7 | 413 | Specific | 557.89 | 394.88 | 394.88 | 6.82 | 25 | >sp\|P01833\|PIGR_HUMAN Polymeric immunoglobulin receptor OS=Homo sapiens OX=9606 GN=PIGR PE=1 SV=4 | 902 | 20201118_lss_milk_1ng_glyco_hcd30_etd30_compound15.22215.22215.3 | controllerType=0 controllerNumber=1 scan=22215 | 53.5319 |
| 15055:3 | 6 | K.IIEGEPNLKVPGN[+2350.830]VTAVLGETLK.V | HexNAc(4)Hex(5)Fuc(1)NeuAc(2) | N[+2351] | 1581.4033 | 3 | 4742.1952 | 4742.1797 | 0 | 3.3 | 457 | Specific | 254.53 | 211.1 | 211.1 | 3.4 | 25 | >sp\|P01833\|PIGR_HUMAN Polymeric immunoglobulin receptor OS=Homo sapiens OX=9606 GN=PIGR PE=1 SV=4 | 902 | 20201118_lss_milk_1ng_glyco_hcd30_etd30_compound15.18628.18628.3 | controllerType=0 controllerNumber=1 scan=18628 | 43.2081 |
| 13703:4 | 6 | K.IIEGEPNLKVPGN[+2205.793]VTAVLGETLK.V | HexNAc(4)Hex(5)Fuc(2)NeuAc(1) | N[+2206] | 1150.0442 | 4 | 4597.1551 | 4597.1422 | 0 | 2.8 | 457 | Specific | 346.74 | 341.02 | 341.02 | 3.89 | 25 | >sp\|P01833\|PIGR_HUMAN Polymeric immunoglobulin receptor OS=Homo sapiens OX=9606 GN=PIGR PE=1 SV=4 | 902 | 20201118_lss_milk_1ng_glyco_hcd30_etd30_compound15.17204.17204.4 | controllerType=0 controllerNumber=1 scan=17204 | 39.3494 |
| 13669:3 | 6 | K.IIEGEPNLKVPGN[+2205.793]VTAVLGETLK.V | HexNAc(4)Hex(5)Fuc(2)NeuAc(1) | N[+2206] | 1533.0572 | 3 | 4597.157 | 4597.1422 | 0 | 3.2 | 457 | Specific | 377.38 | 291.89 | 291.89 | 3.82 | 25 | >sp\|P01833\|PIGR_HUMAN Polymeric immunoglobulin receptor OS=Homo sapiens OX=9606 GN=PIGR PE=1 SV=4 | 902 | 20201118_lss_milk_1ng_glyco_hcd30_etd30_compound15.17169.17169.3 | controllerType=0 controllerNumber=1 scan=17169 | 39.2662 |
| 13633:3 | 6 | K.IIEGEPNLKVPGN[+2205.793]VTAVLGETLK.V | HexNAc(4)Hex(5)Fuc(2)NeuAc(1) | N[+2206] | 1533.0545 | 3 | 4597.1489 | 4597.1422 | 0 | 1.5 | 457 | Specific | 377.75 | 275.62 | 275.62 | 3.83 | 25 | >sp\|P01833\|PIGR_HUMAN Polymeric immunoglobulin receptor OS=Homo sapiens OX=9606 GN=PIGR PE=1 SV=4 | 902 | 20201118_lss_milk_1ng_glyco_hcd30_etd30_compound15.17130.17130.3 | controllerType=0 controllerNumber=1 scan=17130 | 39.1404 |
| 13270:4 | 6 | K.IIEGEPNLKVPGN[+2059.735]VTAVLGETLKVPC[+57.021]HFPC[+57.021]K.F | HexNAc(4)Hex(5)Fuc(1)NeuAc(1) | C[+57]*2, N[+2060] | 1369.8911 | 4 | 5476.5426 | 5476.5418 | 0 | 0.2 | 457 | Specific | 311.54 | 311.54 | 311.54 | 2.87 | 25 | >sp\|P01833\|PIGR_HUMAN Polymeric immunoglobulin receptor OS=Homo sapiens OX=9606 GN=PIGR PE=1 SV=4 | 902 | 20201118_lss_milk_1ng_glyco_hcd30_etd30_compound15.16737.16737.4 | controllerType=0 controllerNumber=1 scan=16737 | 38.2652 |
| 14379:5 | 6 | K.IIEGEPNLKVPGN[+2350.830]VTAVLGETLKVPC[+57.021]HFPC[+57.021]K.F | HexNAc(4)Hex(5)Fuc(1)NeuAc(2) | C[+57]*2, N[+2351] | 1154.3364 | 5 | 5767.653 | 5767.6372 | 0 | 2.7 | 457 | Specific | 369.88 | 369.88 | 369.88 | 4.3 | 25 | >sp\|P01833\|PIGR_HUMAN Polymeric immunoglobulin receptor OS=Homo sapiens OX=9606 GN=PIGR PE=1 SV=4 | 902 | 20201118_lss_milk_1ng_glyco_hcd30_etd30_compound15.17915.17915.5 | controllerType=0 controllerNumber=1 scan=17915 | 41.118 |
| 03598:3 | 6 | K.WN[+1216.423]NTGC[+57.021]QALPSQDEGPSK.A | HexNAc(2)Hex(5) | C[+57], N[+1216] | 1069.1064 | 3 | 3205.3046 | 3205.2995 | 0 | 1.6 | 498 | Specific | 203.26 | 128.88 | 128.88 | 3.08 | 25 | >sp\|P01833\|PIGR_HUMAN Polymeric immunoglobulin receptor OS=Homo sapiens OX=9606 GN=PIGR PE=1 SV=4 | 902 | 20201118_lss_milk_1ng_glyco_hcd30_etd30_compound15.6648.6648.3 | controllerType=0 controllerNumber=1 scan=6648 | 18.1442 |
| 05045:3 | 6 | K.WN[+1872.651]NTGC[+57.021]QALPSQDEGPSK.A | HexNAc(3)Hex(6)NeuAc(1) | C[+57], N[+1873] | 1287.8498 | 3 | 3861.5348 | 3861.5272 | 0 | 2 | 498 | Specific | 278.59 | 143.27 | 143.27 | 4.13 | 25 | >sp\|P01833\|PIGR_HUMAN Polymeric immunoglobulin receptor OS=Homo sapiens OX=9606 GN=PIGR PE=1 SV=4 | 902 | 20201118_lss_milk_1ng_glyco_hcd30_etd30_compound15.8149.8149.3 | controllerType=0 controllerNumber=1 scan=8149 | 20.8192 |
| 04982:3 | 6 | K.WN[+2059.735]NTGC[+57.021]QALPSQDEGPSK.A | HexNAc(4)Hex(5)Fuc(1)NeuAc(1) | C[+57], N[+2060] | 1350.2117 | 3 | 4048.6206 | 4048.6116 | 0 | 2.2 | 498 | Specific | 355.07 | 191.5 | 191.5 | 4.39 | 25 | >sp\|P01833\|PIGR_HUMAN Polymeric immunoglobulin receptor OS=Homo sapiens OX=9606 GN=PIGR PE=1 SV=4 | 902 | 20201118_lss_milk_1ng_glyco_hcd30_etd30_compound15.8084.8084.3 | controllerType=0 controllerNumber=1 scan=8084 | 20.6904 |
| 05111:3 | 6 | K.WN[+1710.598]NTGC[+57.021]QALPSQDEGPSK.A | HexNAc(3)Hex(5)NeuAc(1) | C[+57], N[+1711] | 1233.8325 | 3 | 3699.4828 | 3699.4743 | 0 | 2.3 | 498 | Specific | 388.24 | 305.16 | 305.16 | 4.84 | 25 | >sp\|P01833\|PIGR_HUMAN Polymeric immunoglobulin receptor OS=Homo sapiens OX=9606 GN=PIGR PE=1 SV=4 | 902 | 20201118_lss_milk_1ng_glyco_hcd30_etd30_compound15.8218.8218.3 | controllerType=0 controllerNumber=1 scan=8218 | 20.9567 |
| 05049:3 | 6 | K.WN[+1913.677]NTGC[+57.021]QALPSQDEGPSK.A | HexNAc(4)Hex(5)NeuAc(1) | C[+57], N[+1914] | 1301.5252 | 3 | 3902.5611 | 3902.5537 | 0 | 1.9 | 498 | Specific | 461.82 | 299.17 | 299.17 | 5.11 | 25 | >sp\|P01833\|PIGR_HUMAN Polymeric immunoglobulin receptor OS=Homo sapiens OX=9606 GN=PIGR PE=1 SV=4 | 902 | 20201118_lss_milk_1ng_glyco_hcd30_etd30_compound15.8154.8154.3 | controllerType=0 controllerNumber=1 scan=8154 | 20.8356 |
| 08914:3 | 7 | K.TVLTPATNHMGN[+1378.476]VTFTIPANR.E | HexNAc(2)Hex(6) | N[+1378] | 1211.8848 | 3 | 3633.6399 | 3633.6358 | 0 | 1.1 | 74 | Specific | 405.6 | 329.56 | 154.53 | 4.73 | 14 | >sp\|P01024\|CO3_HUMAN Complement C3 OS=Homo sapiens OX=9606 GN=C3 PE=1 SV=2 | 492 | 20201118_lss_milk_1ng_glyco_hcd30_etd30_compound15.12183.12183.3 | controllerType=0 controllerNumber=1 scan=12183 | 28.3224 |
| 18117:4 | 9 | K.VFPLSLDSTPQDGNVVVAC[+57.021]LVQGFFPQEPLSVTWSESGQN[+1647.613]VTAI | HexNAc(5)Hex(3)Fuc(1) | C[+57], N[+1648] | 1606.7527 | 4 | 6423.9889 | 6423.9746 | 0 | 2.2 | 8 | Specific | 277.46 | 277.46 | 277.46 | 5.09 | 24 | >sp\|P01877\|IGHA2_HUMAN Immunoglobulin heavy constant alpha 2 OS=Homo sapiens OX=9606 GN=IGHA2 PE=1 SV=4 | 576 | 20201118_lss_milk_1ng_glyco_hcd30_etd30_compound15.21860.21860.4 | controllerType=0 controllerNumber=1 scan=21860 | 52.4683 |
| 18934:4 | 9 | K.VFPLSLDSTPQDGNVVVAC[+57.021]LVQGFFPQEPLSVTWSESGQN[+2262.814]VTAI | HexNAc(5)Hex(5)Fuc(1)NeuAc(1) | C[+57], N[+2263] | 1760.5549 | 4 | 7039.1978 | 7039.1757 | 0 | 3.1 | 8 | Specific | 314.47 | 314.47 | 314.47 | 5.09 | 24 | >sp\|P01877\|IGHA2_HUMAN Immunoglobulin heavy constant alpha 2 OS=Homo sapiens OX=9606 GN=IGHA2 PE=1 SV=4 | 576 | 20201118_lss_milk_1ng_glyco_hcd30_etd30_compound15.22726.22726.4 | controllerType=0 controllerNumber=1 scan=22726 | 55.1346 |
| 17903:4 | 9 | K.VFPLSLDSTPQDGNVVVAC[+57.021]LVQGFFPQEPLSVTWSESGQN[+1809.666]VTAI | HexNAc(5)Hex(4)Fuc(1) | C[+57], N[+1810] | 1647.2697 | 4 | 6586.0568 | 6586.0275 | 0 | 4.5 | 8 | Specific | 412.88 | 412.88 | 412.88 | 6.63 | 24 | >sp\|P01877\|IGHA2_HUMAN Immunoglobulin heavy constant alpha 2 OS=Homo sapiens OX=9606 GN=IGHA2 PE=1 SV=4 | 576 | 20201118_lss_milk_1ng_glyco_hcd30_etd30_compound15.21632.21632.4 | controllerType=0 controllerNumber=1 scan=21632 | 51.8895 |
| 17889:4 | 9 | K.VFPLSLDSTPQDGNVVVAC[+57.021]LVQGFFPQEPLSVTWSESGQN[+1647.613]VTAI | HexNAc(5)Hex(3)Fuc(1) | C[+57], N[+1648] | 1606.7518 | 4 | 6423.9855 | 6423.9746 | 0 | 1.7 | 8 | Specific | 483.51 | 483.51 | 483.51 | 7.05 | 24 | >sp\|P01877\|IGHA2_HUMAN Immunoglobulin heavy constant alpha 2 OS=Homo sapiens OX=9606 GN=IGHA2 PE=1 SV=4 | 576 | 20201118_lss_milk_1ng_glyco_hcd30_etd30_compound15.21617.21617.5 | controllerType=0 controllerNumber=1 scan=21617 | 51.8325 |
| 01720:3 | 9 | K.HYTN[+2262.814]SSQDVTVPC[+57.021]R.V | HexNAc(5)Hex(5)Fuc(1)NeuAc(1) | C[+57], N[+2263] | 1309.5284 | 3 | 3926.5706 | 3926.5636 | 0 | 1.8 | 89 | Specific | 235.33 | 124.02 | 124.02 | 3.12 | 24 | >sp\|P01877\|IGHA2_HUMAN Immunoglobulin heavy constant alpha 2 OS=Homo sapiens OX=9606 GN=IGHA2 PE=1 SV=4 | 576 | 20201118_lss_milk_1ng_glyco_hcd30_etd30_compound15.4696.4696.3 | controllerType=0 controllerNumber=1 scan=4696 | 14.4893 |
| 00869:3 | 9 | K.HYTN[+1809.666]SSQDVTVPC[+57.021]R.V | HexNAc(5)Hex(4)Fuc(1) | C[+57], N[+1810] | 1158.4789 | 3 | 3473.4223 | 3473.4154 | 0 | 2 | 89 | Specific | 405.45 | 332.06 | 332.06 | 5.07 | 24 | >sp\|P01877\|IGHA2_HUMAN Immunoglobulin heavy constant alpha 2 OS=Homo sapiens OX=9606 GN=IGHA2 PE=1 SV=4 | 576 | 20201118_lss_milk_1ng_glyco_hcd30_etd30_compound15.3810.3810.3 | controllerType=0 controllerNumber=1 scan=3810 | 12.8765 |
| 00889:3 | 9 | K.HYTN[+1647.613]SSQDVTVPC[+57.021]R.V | HexNAc(5)Hex(3)Fuc(1) | C[+57], N[+1648] | 1104.4597 | 3 | 3311.3647 | 3311.3625 | 0 | 0.6 | 89 | Specific | 490.04 | 429.39 | 429.39 | 6.08 | 24 | >sp\|P01877\|IGHA2_HUMAN Immunoglobulin heavy constant alpha 2 OS=Homo sapiens OX=9606 GN=IGHA2 PE=1 SV=4 | 576 | 20201118_lss_milk_1ng_glyco_hcd30_etd30_compound15.3831.3831.3 | controllerType=0 controllerNumber=1 scan=3831 | 12.9247 |
| 02774:5 | 9 | K.HYTN[+1647.613]SSQDVTVPC[+57.021]RVPPPPPC[+57.021]C[+57.021]HPR.L | HexNAc(5)Hex(3)Fuc(1) | C[+57]*3, N[+1648] | 922.0017 | 5 | 4605.9795 | 4605.9689 | 0 | 2.3 | 89 | Specific | 529.66 | 516.78 | 516.78 | 6.52 | 24 | >sp\|P01877\|IGHA2_HUMAN Immunoglobulin heavy constant alpha 2 OS=Homo sapiens OX=9606 GN=IGHA2 PE=1 SV=4 | 576 | 20201118_lss_milk_1ng_glyco_hcd30_etd30_compound15.5793.5793.5 | controllerType=0 controllerNumber=1 scan=5793 | 16.617 |
| 04118:3 | 9 | K.TPLTAN[+2262.814]ITK.S | HexNAc(5)Hex(5)Fuc(1)NeuAc(1) | N[+2263] | 1074.4635 | 3 | 3221.376 | 3221.3711 | 0 | 1.5 | 200 | Specific | 227.71 | 105.24 | 105.24 | 2.58 | 24 | >sp\|P01877\|IGHA2_HUMAN Immunoglobulin heavy constant alpha 2 OS=Homo sapiens OX=9606 GN=IGHA2 PE=1 SV=4 | 576 | 20201118_lss_milk_1ng_glyco_hcd30_etd30_compound15.7189.7189.3 | controllerType=0 controllerNumber=1 scan=7189 | 19.0759 |
| 04119:3 | 9 | K.TPLTAN[+2059.735]ITK.S | HexNAc(4)Hex(5)Fuc(1)NeuAc(1) | N[+2060] | 1006.7705 | 3 | 3018.2971 | 3018.2917 | 0 | 1.8 | 200 | Specific | 230.55 | 115.14 | 115.14 | 2.75 | 24 | >sp\|P01877\|IGHA2_HUMAN Immunoglobulin heavy constant alpha 2 OS=Homo sapiens OX=9606 GN=IGHA2 PE=1 SV=4 | 576 | 20201118_lss_milk_1ng_glyco_hcd30_etd30_compound15.7190.7190.3 | controllerType=0 controllerNumber=1 scan=7190 | 19.0834 |
| 02462:2 | 9 | K.TPLTAN[+1971.719]ITK.S | HexNAc(5)Hex(5)Fuc(1) | N[+1972] | 1465.6446 | 2 | 2930.2819 | 2930.2757 | 0 | 2.1 | 200 | Specific | 239.03 | 30.17 | 30.17 | 2.57 | 24 | >sp\|P01877\|IGHA2_HUMAN Immunoglobulin heavy constant alpha 2 OS=Homo sapiens OX=9606 GN=IGHA2 PE=1 SV=4 | 576 | 20201118_lss_milk_1ng_glyco_hcd30_etd30_compound15.5467.5467.2 | controllerType=0 controllerNumber=1 scan=5467 | 15.8838 |
| 02542:3 | 9 | K.TPLTAN[+1647.613]ITK.S | HexNAc(5)Hex(3)Fuc(1) | N[+1648] | 869.3962 | 3 | 2606.1741 | 2606.17 | 0 | 1.6 | 200 | Specific | 290.5 | 136.78 | 136.78 | 2.65 | 24 | >sp\|P01877\|IGHA2_HUMAN Immunoglobulin heavy constant alpha 2 OS=Homo sapiens OX=9606 GN=IGHA2 PE=1 SV=4 | 576 | 20201118_lss_milk_1ng_glyco_hcd30_etd30_compound15.5551.5551.3 | controllerType=0 controllerNumber=1 scan=5551 | 16.0838 |
| 02506:2 | 9 | K.TPLTAN[+1444.534]ITK.S | HexNAc(4)Hex(3)Fuc(1) | N[+1445] | 1202.0517 | 2 | 2403.0961 | 2403.0906 | 0 | 2.3 | 200 | Specific | 299.31 | 72.29 | 72.29 | 2.6 | 24 | >sp\|P01877\|IGHA2_HUMAN Immunoglobulin heavy constant alpha 2 OS=Homo sapiens OX=9606 GN=IGHA2 PE=1 SV=4 | 576 | 20201118_lss_milk_1ng_glyco_hcd30_etd30_compound15.5513.5513.2 | controllerType=0 controllerNumber=1 scan=5513 | 15.991 |
| 02440:3 | 9 | K.TPLTAN[+1971.719]ITK.S | HexNAc(5)Hex(5)Fuc(1) | N[+1972] | 977.4315 | 3 | 2930.2801 | 2930.2757 | 0 | 1.5 | 200 | Specific | 303.9 | 149.86 | 149.86 | 2.8 | 24 | >sp\|P01877\|IGHA2_HUMAN Immunoglobulin heavy constant alpha 2 OS=Homo sapiens OX=9606 GN=IGHA2 PE=1 SV=4 | 576 | 20201118_lss_milk_1ng_glyco_hcd30_etd30_compound15.5444.5444.3 | controllerType=0 controllerNumber=1 scan=5444 | 15.8403 |
| 02481:2 | 9 | K.TPLTAN[+1809.666]ITK.S | HexNAc(5)Hex(4)Fuc(1) | N[+1810] | 1384.6175 | 2 | 2768.2277 | 2768.2228 | 0 | 1.7 | 200 | Specific | 307.39 | 110.13 | 110.13 | 2.65 | 24 | >sp\|P01877\|IGHA2_HUMAN Immunoglobulin heavy constant alpha 2 OS=Homo sapiens OX=9606 GN=IGHA2 PE=1 SV=4 | 576 | 20201118_lss_milk_1ng_glyco_hcd30_etd30_compound15.5487.5487.2 | controllerType=0 controllerNumber=1 scan=5487 | 15.9358 |
| 02551:2 | 9 | K.TPLTAN[+1501.555]ITK.S | HexNAc(5)Hex(3) | N[+1502] | 1230.5617 | 2 | 2460.1161 | 2460.1121 | 0 | 1.6 | 200 | Specific | 312.89 | 88.39 | 88.39 | 2.66 | 24 | >sp\|P01877\|IGHA2_HUMAN Immunoglobulin heavy constant alpha 2 OS=Homo sapiens OX=9606 GN=IGHA2 PE=1 SV=4 | 576 | 20201118_lss_milk_1ng_glyco_hcd30_etd30_compound15.5560.5560.2 | controllerType=0 controllerNumber=1 scan=5560 | 16.1095 |
| 02480:3 | 9 | K.TPLTAN[+1809.666]ITK.S | HexNAc(5)Hex(4)Fuc(1) | N[+1810] | 923.4146 | 3 | 2768.2292 | 2768.2228 | 0 | 2.3 | 200 | Specific | 314.96 | 150.34 | 150.34 | 2.8 | 24 | >sp\|P01877\|IGHA2_HUMAN Immunoglobulin heavy constant alpha 2 OS=Homo sapiens OX=9606 GN=IGHA2 PE=1 SV=4 | 576 | 20201118_lss_milk_1ng_glyco_hcd30_etd30_compound15.5486.5486.3 | controllerType=0 controllerNumber=1 scan=5486 | 15.9288 |
| 02535:2 | 9 | K.TPLTAN[+1647.613]ITK.S | HexNAc(5)Hex(3)Fuc(1) | N[+1648] | 1303.5907 | 2 | 2606.1742 | 2606.17 | 0 | 1.6 | 200 | Specific | 350.06 | 93.55 | 93.55 | 2.62 | 24 | >sp\|P01877\|IGHA2_HUMAN Immunoglobulin heavy constant alpha 2 OS=Homo sapiens OX=9606 GN=IGHA2 PE=1 SV=4 | 576 | 20201118_lss_milk_1ng_glyco_hcd30_etd30_compound15.5543.5543.2 | controllerType=0 controllerNumber=1 scan=5543 | 16.0623 |
| 09020:3 | 9 | R.MAGKPTHIN[+1702.581]VSVVMAEADGTC[+57.021]Y.- | HexNAc(2)Hex(8) | C[+57], N[+1703] | 1351.8964 | 3 | 4053.6747 | 4053.6642 | 0 | 2.6 | 319 | CRagged | 254.12 | 206.41 | 206.41 | 3.91 | 24 | >sp\|P01877\|IGHA2_HUMAN Immunoglobulin heavy constant alpha 2 OS=Homo sapiens OX=9606 GN=IGHA2 PE=1 SV=4 | 576 | 20201118_lss_milk_1ng_glyco_hcd30_etd30_compound15.12293.12293.3 | controllerType=0 controllerNumber=1 scan=12293 | 28.5411 |
| 09061:3 | 9 | R.MAGKPTHIN[+1540.529]VSVVMAEADGTC[+57.021]Y.- | HexNAc(2)Hex(7) | C[+57], N[+1541] | 1297.8787 | 3 | 3891.6216 | 3891.6114 | 0 | 2.6 | 319 | CRagged | 269.46 | 269.46 | 199.67 | 4.18 | 24 | >sp\|P01877\|IGHA2_HUMAN Immunoglobulin heavy constant alpha 2 OS=Homo sapiens OX=9606 GN=IGHA2 PE=1 SV=4 | 576 | 20201118_lss_milk_1ng_glyco_hcd30_etd30_compound15.12336.12336.3 | controllerType=0 controllerNumber=1 scan=12336 | 28.[illegible]361 |
| 09074:3 | 9 | R.MAGKPTHIN[+1378.476]VSVVMAEADGTC[+57.021]Y.- | HexNAc(2)Hex(6) | C[+57], N[+1378] | 1243.8591 | 3 | 3729.5626 | 3729.5585 | 0 | 1.1 | 319 | CRagged | 271.59 | 260.05 | 185.29 | 3.63 | 24 | >sp\|P01877\|IGHA2_HUMAN Immunoglobulin heavy constant alpha 2 OS=Homo sapiens OX=9606 GN=IGHA2 PE=1 SV=4 | 576 | 20201118_lss_milk_1ng_glyco_hcd30_etd30_compound15.12350.12350.3 | controllerType=0 controllerNumber=1 scan=12350 | 28.[illegible]63 |
| 09120:3 | 9 | R.MAGKPTHIN[+1216.423]VSVVMAEADGTC[+57.021]Y.- | HexNAc(2)Hex(5) | C[+57], N[+1216] | 1189.8454 | 3 | 3567.5216 | 3567.5057 | 0 | 4.5 | 319 | CRagged | 298.35 | 273.31 | 224.54 | 3.89 | 24 | >sp\|P01877\|IGHA2_HUMAN Immunoglobulin heavy constant alpha 2 OS=Homo sapiens OX=9606 GN=IGHA2 PE=1 SV=4 | 576 | 20201118_lss_milk_1ng_glyco_hcd30_etd30_compound15.12398.12398.3 | controllerType=0 controllerNumber=1 scan=12398 | 28.[illegible]394 |
| 04926:3 | 11 | R.LSPN[+2205.793]ASAEHLELR.W | HexNAc(4)Hex(5)Fuc(2)NeuAc(1) | N[+2206] | 1214.8533 | 3 | 3642.5454 | 3642.5421 | 0 | 0.9 | 52 | Specific | 352.67 | 253.18 | 253.18 | 3.67 | 11 | >sp\|Q13410\|BT1A1_HUMAN Butyrophilin subfamily 1 member A1 OS=Homo sapiens OX=9606 GN=BTN1A1 PE=1 SV=3 | 179 | 20201118_lss_milk_1ng_glyco_hcd30_etd30_compound15.8025.8025.3 | controllerType=0 controllerNumber=1 scan=8025 | 20.5623 |
| 06877:3 | 11 | R.LSPN[+2350.830]ASAEHLELR.W | HexNAc(4)Hex(5)Fuc(1)NeuAc(2) | N[+2351] | 1263.2 | 3 | 3787.5855 | 3787.5796 | 0 | 1.6 | 52 | Specific | 364.84 | 224.28 | 224.28 | 4.71 | 11 | >sp\|Q13410\|BT1A1_HUMAN Butyrophilin subfamily 1 member A1 OS=Homo sapiens OX=9606 GN=BTN1A1 PE=1 SV=3 | 179 | 20201118_lss_milk_1ng_glyco_hcd30_etd30_compound15.10047.10047.3 | controllerType=0 controllerNumber=1 scan=10047 | 24.1402 |
| 04929:3 | 11 | R.LSPN[+2205.793]ASAEHLELR.W | HexNAc(4)Hex(5)Fuc(2)NeuAc(1) | N[+2206] | 1214.8542 | 3 | 3642.548 | 3642.5421 | 0 | 1.6 | 52 | Specific | 403.48 | 279.8 | 279.8 | 4.88 | 11 | >sp\|Q13410\|BT1A1_HUMAN Butyrophilin subfamily 1 member A1 OS=Homo sapiens OX=9606 GN=BTN1A1 PE=1 SV=3 | 179 | 20201118_lss_milk_1ng_glyco_hcd30_etd30_compound15.8029.8029.3 | controllerType=0 controllerNumber=1 scan=8029 | 20.577 |
| 09921:3 | 17 | R.IIVPLNNREN[+2059.735]ISDPTSPLR.T | HexNAc(4)Hex(5)Fuc(1)NeuAc(1) | N[+2060] | 1403.3115 | 3 | 4207.9198 | 4207.9121 | 0 | 1.8 | 62 | Specific | 214.06 | 194.18 | 194.18 | 2.96 | 9 | >sp\|P01591\|IGJ_HUMAN Immunoglobulin J chain OS=Homo sapiens OX=9606 GN=JCHAIN PE=1 SV=4 | 1392 | 20201118_lss_milk_1ng_glyco_hcd30_etd30_compound15.13229.13229.3 | controllerType=0 controllerNumber=1 scan=13229 | 30.4024 |
| 11622:3 | 17 | R.IIVPLNNREN[+2204.772]ISDPTSPLR.T | HexNAc(4)Hex(5)NeuAc(2) | N[+2205] | 1451.6579 | 3 | 4352.9591 | 4352.9496 | 0 | 2.2 | 62 | Specific | 274.62 | 192.29 | 192.29 | 3.4 | 9 | >sp\|P01591\|IGJ_HUMAN Immunoglobulin J chain OS=Homo sapiens OX=9606 GN=JCHAIN PE=1 SV=4 | 1392 | 20201118_lss_milk_1ng_glyco_hcd30_etd30_compound15.15009.15009.3 | controllerType=0 controllerNumber=1 scan=15009 | 34.0805 |
| 09901:3 | 17 | R.IIVPLNNREN[+1913.677]ISDPTSPLR.T | HexNAc(4)Hex(5)NeuAc(1) | N[+1914] | 1354.6237 | 3 | 4061.8566 | 4061.8542 | 0 | 0.6 | 62 | Specific | 288.94 | 212.91 | 212.91 | 3.18 | 9 | >sp\|P01591\|IGJ_HUMAN Immunoglobulin J chain OS=Homo sapiens OX=9606 GN=JCHAIN PE=1 SV=4 | 1392 | 20201118_lss_milk_1ng_glyco_hcd30_etd30_compound15.13208.13208.3 | controllerType=0 controllerNumber=1 scan=13208 | 30.3623 |
| 08514:3 | 17 | R.IIVPLNNREN[+1663.608]ISDPTSPLR.T | HexNAc(5)Hex(4) | N[+1664] | 1271.2648 | 3 | 3811.7799 | 3811.7853 | 0 | -1.4 | 62 | Specific | 309.57 | 236.48 | 236.48 | 2.89 | 9 | >sp\|P01591\|IGJ_HUMAN Immunoglobulin J chain OS=Homo sapiens OX=9606 GN=JCHAIN PE=1 SV=4 | 1392 | 20201118_lss_milk_1ng_glyco_hcd30_etd30_compound15.11767.11767.3 | controllerType=0 controllerNumber=1 scan=11767 | 27.4795 |
| 08469:3 | 17 | R.IIVPLNNREN[+1809.666]ISDPTSPLR.T | HexNAc(5)Hex(4)Fuc(1) | N[+1810] | 1319.954 | 3 | 3957.8476 | 3957.8432 | 0 | 1.1 | 62 | Specific | 314.78 | 225.55 | 225.55 | 3.18 | 9 | >sp\|P01591\|IGJ_HUMAN Immunoglobulin J chain OS=Homo sapiens OX=9606 GN=JCHAIN PE=1 SV=4 | 1392 | 20201118_lss_milk_1ng_glyco_hcd30_etd30_compound15.11720.11720.3 | controllerType=0 controllerNumber=1 scan=11720 | 27.3772 |
| 08591:3 | 17 | R.IIVPLNNREN[+1647.613]ISDPTSPLR.T | HexNAc(5)Hex(3)Fuc(1) | N[+1648] | 1265.9372 | 3 | 3795.797 | 3795.7904 | 0 | 1.7 | 62 | Specific | 331.48 | 254.58 | 254.58 | 3.95 | 9 | >sp\|P01591\|IGJ_HUMAN Immunoglobulin J chain OS=Homo sapiens OX=9606 GN=JCHAIN PE=1 SV=4 | 1392 | 20201118_lss_milk_1ng_glyco_hcd30_etd30_compound15.11847.11847.3 | controllerType=0 controllerNumber=1 scan=11847 | 27.6458 |
| 01692:3 | 22 | R.LRN[+1216.423]VSWATGR.S | HexNAc(2)Hex(5) | N[+1216] | 792.6933 | 3 | 2376.0652 | 2376.0559 | 0 | 3.9 | 149 | Specific | 334.66 | 171.48 | 171.48 | 2.58 | 5 | >sp\|P08571\|CD14_HUMAN Monocyte differentiation antigen CD14 OS=Homo sapiens OX=9606 GN=CD14 PE=1 SV=2 | 1232 | 20201118_lss_milk_1ng_glyco_hcd30_etd30_compound15.4666.4666.3 | controllerType=0 controllerNumber=1 scan=4666 | 14.4205 |
| 12082:4 | 24 | K.LKELPGVC[+57.021]N[+2351.851]ETMMALWEEC[+57.021]KPC[+57.021]LK.Q | HexNAc(4)Hex(5)Fuc(3)NeuAc(1) | C[+57]*3, N[+2352] | 1322.8182 | 4 | 5288.2511 | 5288.2355 | 0 | 3 | 95 | Specific | 208.19 | 166.01 | 166.01 | 2.65 | 9 | >sp\|P10909\|CLUS_HUMAN Clusterin OS=Homo sapiens OX=9606 GN=CLU PE=1 SV=1 | 1467 | 20201118_lss_milk_1ng_glyco_hcd30_etd30_compound15.15491.15491.4 | controllerType=0 controllerNumber=1 scan=15491 | 35.2573 |
| 12005:4 | 24 | K.LKELPGVC[+57.021]N[+2570.925]ETMMALWEEC[+57.021]KPC[+57.021]LK.Q | HexNAc(5)Hex(6)Fuc(2)NeuAc(1) | C[+57]*3, N[+2571] | 1377.5878 | 4 | 5507.3293 | 5507.3097 | 0 | 3.5 | 95 | Specific | 251.08 | 221.94 | 221.94 | 3.57 | 9 | >sp\|P10909\|CLUS_HUMAN Clusterin OS=Homo sapiens OX=9606 GN=CLU PE=1 SV=1 | 1467 | 20201118_lss_milk_1ng_glyco_hcd30_etd30_compound15.15410.15410.4 | controllerType=0 controllerNumber=1 scan=15410 | 35.0768 |
| 12100:4 | 24 | K.LKELPGVC[+57.021]N[+2205.793]ETMMALWEEC[+57.021]KPC[+57.021]LK.Q | HexNAc(4)Hex(5)Fuc(2)NeuAc(1) | C[+57]*3, N[+2206] | 1286.3043 | 4 | 5142.1955 | 5142.1776 | 0 | 3.5 | 95 | Specific | 555.01 | 403.56 | 403.56 | 5.87 | 9 | >sp\|P10909\|CLUS_HUMAN Clusterin OS=Homo sapiens OX=9606 GN=CLU PE=1 SV=1 | 1467 | 20201118_lss_milk_1ng_glyco_hcd30_etd30_compound15.15510.15510.4 | controllerType=0 controllerNumber=1 scan=15510 | 35.3132 |
| 12126:4 | 24 | K.LKELPGVC[+57.021]N[+2059.735]ETMMALWEEC[+57.021]KPC[+57.021]LK.Q | HexNAc(4)Hex(5)Fuc(1)NeuAc(1) | C[+57]*3, N[+2060] | 1249.7885 | 4 | 4996.132 | 4996.1196 | 0 | 2.5 | 95 | Specific | 720.28 | 584.41 | 584.41 | 10.36 | 9 | >sp\|P10909\|CLUS_HUMAN Clusterin OS=Homo sapiens OX=9606 GN=CLU PE=1 SV=1 | 1467 | 20201118_lss_milk_1ng_glyco_hcd30_etd30_compound15.15537.15537.4 | controllerType=0 controllerNumber=1 scan=15537 | 35.375 |
| 00057:4 | 24 | R.EIRHN[+2424.867]STGC[+57.021]LR.M | HexNAc(5)Hex(6)Fuc(1)NeuAc(1) | C[+57], N[+2425] | 942.6396 | 4 | 3767.5367 | 3767.5316 | 0 | 1.4 | 287 | Specific | 346.05 | 257.78 | 257.78 | 3.33 | 9 | >sp\|P10909\|CLUS_HUMAN Clusterin OS=Homo sapiens OX=9606 GN=CLU PE=1 SV=1 | 1467 | 20201118_lss_milk_1ng_glyco_hcd30_etd30_compound15.2965.2965.4 | controllerType=0 controllerNumber=1 scan=2965 | 11.2706 |
| 00052:4 | 24 | R.EIRHN[+2059.735]STGC[+57.021]LR.M | HexNAc(4)Hex(5)Fuc(1)NeuAc(1) | C[+57], N[+2060] | 851.357 | 4 | 3402.4062 | 3402.3994 | 0 | 2 | 287 | Specific | 694.68 | 541.06 | 541.06 | 8.98 | 9 | >sp\|P10909\|CLUS_HUMAN Clusterin OS=Homo sapiens OX=9606 GN=CLU PE=1 SV=1 | 1467 | 20201118_lss_milk_1ng_glyco_hcd30_etd30_compound15.2960.2960.4 | controllerType=0 controllerNumber=1 scan=2960 | 11.2569 |
| 18742:3 | 24 | K.MLN[+2350.830]TSSLLEQLNEQFNWVSR.L | HexNAc(4)Hex(5)Fuc(1)NeuAc(2) | N[+2351] | 1587.3504 | 3 | 4760.0366 | 4760.0171 | 0 | 4.1 | 352 | Specific | 488.06 | 292.05 | 292.05 | 5.36 | 9 | >sp\|P10909\|CLUS_HUMAN Clusterin OS=Homo sapiens OX=9606 GN=CLU PE=1 SV=1 | 1467 | 20201118_lss_milk_1ng_glyco_hcd30_etd30_compound15.22523.22523.3 | controllerType=0 controllerNumber=1 scan=22523 | 54.5584 |
| 11419:3 | 24 | R.LAN[+2350.830]LTQGEDQYYLR.V | HexNAc(4)Hex(5)Fuc(1)NeuAc(2) | N[+2351] | 1345.5617 | 3 | 4034.6706 | 4034.664 | 0 | 1.6 | 372 | Specific | 364.55 | 251.13 | 251.13 | 4.75 | 9 | >sp\|P10909\|CLUS_HUMAN Clusterin OS=Homo sapiens OX=9606 GN=CLU PE=1 SV=1 | 1467 | 20201118_lss_milk_1ng_glyco_hcd30_etd30_compound15.14796.14796.3 | controllerType=0 controllerNumber=1 scan=14796 | 33.5498 |
| 19144:3 | 29 | R.QLAHQSN[+2204.772]STNIFFSPVSIATAFAMLSLGTK.A | HexNAc(4)Hex(5)NeuAc(2) | N[+2205] | 1796.1411 | 3 | 5386.4087 | 5386.4075 | 0 | 0.2 | 64 | Specific | 440.92 | 315.91 | 315.91 | 5.58 | 4 | >sp\|P01009\|A1AT_HUMAN Alpha-1-antitrypsin OS=Homo sapiens OX=9606 GN=SERPINA1 PE=1 SV=3 | 642 | 20201118_lss_milk_1ng_glyco_hcd30_etd30_compound15.22950.22950.3 | controllerType=0 controllerNumber=1 scan=22950 | 55.8583 |
| 19168:4 | 29 | R.QLAHQSN[+2204.772]STNIFFSPVSIATAFAMLSLGTK.A | HexNAc(4)Hex(5)NeuAc(2) | N[+2205] | 1347.3613 | 4 | 5386.4235 | 5386.4075 | 0 | 3 | 64 | Specific | 685.1 | 624.6 | 457.22 | 10.8 | 4 | >sp\|P01009\|A1AT_HUMAN Alpha-1-antitrypsin OS=Homo sapiens OX=9606 GN=SERPINA1 PE=1 SV=3 | 642 | 20201118_lss_milk_1ng_glyco_hcd30_etd30_compound15.22977.22977.3 | controllerType=0 controllerNumber=1 scan=22977 | 55.9594 |
| 19140:4 | 29 | R.QLAHQSN[+2204.772]STNIFFSPVSIATAFAMLSLGTK.A | HexNAc(4)Hex(5)NeuAc(2) | N[+2205] | 1347.3577 | 4 | 5386.409 | 5386.4075 | 0 | 0.3 | 64 | Specific | 735 | 610.59 | 508.71 | 11.22 | 4 | >sp\|P01009\|A1AT_HUMAN Alpha-1-antitrypsin OS=Homo sapiens OX=9606 GN=SERPINA1 PE=1 SV=3 | 642 | 20201118_lss_milk_1ng_glyco_hcd30_etd30_compound15.22946.22946.4 | controllerType=0 controllerNumber=1 scan=22946 | 55.8397 |
| 18357:4 | 29 | K.ADTHDEILEGLNFN[+2861.000]LTEIPEAQIHEGFQELLR.T | HexNAc(5)Hex(6)NeuAc(3) | N[+2861] | 1638.9701 | 4 | 6552.8586 | 6552.8239 | 0 | 5.3 | 94 | Specific | 263.41 | 263.41 | 263.41 | 4.42 | 4 | >sp\|P01009\|A1AT_HUMAN Alpha-1-antitrypsin OS=Homo sapiens OX=9606 GN=SERPINA1 PE=1 SV=3 | 642 | 20201118_lss_milk_1ng_glyco_hcd30_etd30_compound15.22114.22114.4 | controllerType=0 controllerNumber=1 scan=22114 | 53.2286 |
| 14959:3 | 50 | R.NYIVPILWLN[+1864.634]ETGTIGDEK.A | HexNAc(2)Hex(9) | N[+1865] | 1347.2698 | 3 | 4039.7949 | 4039.7674 | 0 | 6.8 | 408 | Specific | 282.37 | 282.37 | 282.37 | 4.06 | 2 | >sp\|P16671\|CD36_HUMAN Platelet glycoprotein 4 OS=Homo sapiens OX=9606 GN=CD36 PE=1 SV=2 | 812 | 20201118_lss_milk_1ng_glyco_hcd30_etd30_compound15.18526.18526.3 | controllerType=0 controllerNumber=1 scan=18526 | 42.9 |
| 00749:4 | 65 | K.TKPREEQFN[+1444.534]STFR.V | HexNAc(4)Hex(3)Fuc(1) | N[+1445] | 771.8452 | 4 | 3084.3589 | 3084.3526 | 0 | 2 | 168 | Specific | 405.08 | 339.87 | 339.87 | 4.08 | 3 | >sp\|P01859\|IGHG2_HUMAN Immunoglobulin heavy constant gamma 2 OS=Homo sapiens OX=9606 GN=IGHG2 PE=1 SV=2 | 387 | 20201118_lss_milk_1ng_glyco_hcd30_etd30_compound15.3685.3685.4 | controllerType=0 controllerNumber=1 scan=3685 | 12.62 |
| 00684:4 | 65 | K.TKPREEQFN[+1606.587]STFR.V | HexNAc(4)Hex(4)Fuc(1) | N[+1607] | 812.359 | 4 | 3246.4143 | 3246.4054 | 0 | 2.7 | 168 | Specific | 507.22 | 460.26 | 460.26 | 5.93 | 3 | >sp\|P01859\|IGHG2_HUMAN Immunoglobulin heavy constant gamma 2 OS=Homo sapiens OX=9606 GN=IGHG2 PE=1 SV=2 | 387 | 20201118_lss_milk_1ng_glyco_hcd30_etd30_compound15.3617.3617.4 | controllerType=0 controllerNumber=1 scan=3617 | 12.4902 |
| 00075:4 | 87 | K.TKPREEQYN[+1444.534]STYR.V | HexNAc(4)Hex(3)Fuc(1) | N[+1445] | 779.8442 | 4 | 3116.355 | 3116.3424 | 0 | 4 | 291 | Specific | 327.43 | 40.58 | 40.58 | 2.74 | 2 | >sp\|P0DOX5\|IGG1_HUMAN Immunoglobulin gamma-1 heavy chain OS=Homo sapiens OX=9606 PE=1 SV=2 | 914 | 20201118_lss_milk_1ng_glyco_hcd30_etd30_compound15.2984.2984.4 | controllerType=0 controllerNumber=1 scan=2984 | 11.3331 |
| 00062:4 | 87 | K.TKPREEQYN[+1606.587]STYR.V | HexNAc(4)Hex(4)Fuc(1) | N[+1607] | 820.3556 | 4 | 3278.4004 | 3278.3952 | 0 | 1.6 | 291 | Specific | 673.32 | 63.99 | 63.99 | 6.38 | 2 | >sp\|P0DOX5\|IGG1_HUMAN Immunoglobulin gamma-1 heavy chain OS=Homo sapiens OX=9606 PE=1 SV=2 | 914 | 20201118_lss_milk_1ng_glyco_hcd30_etd30_compound15.2970.2970.4 | controllerType=0 controllerNumber=1 scan=2970 | 11.2972 |
| 14023:3 | 1 | R.LSLHRPALEDLLLGSEAN[+2116.756]LTC[+57.021]TLTGLR.D | HexNAc(5)Hex(5)NeuAc(1) | C[+57], N[+2117] | 1694.1276 | 3 | 5080.3684 | 5080.3547 | 0 | 2.7 | 127 | Specific | 314.19 | 187.15 | 187.15 | 5.73 | 72 | >sp\|P01876\|IGHA1_HUMAN Immunoglobulin heavy constant alpha 1 OS=Homo sapiens OX=9606 GN=IGHA1 PE=1 SV=2 | 255 | 20201116_xd_colostrum_1ng_glyco_hcd30_etd30_compound15.17335.17335.2 | controllerType=0 controllerNumber=1 scan=17335 | 44.7257 |
| 13424:3 | 1 | R.LSLHRPALEDLLLGSEAN[+1095.397]LTC[+57.021]TLTGLR.D | HexNAc(3)Hex(3) | C[+57], N[+1095] | 1353.671 | 3 | 4058.9983 | 4058.9949 | 0 | 0.9 | 127 | Specific | 343.27 | 321.64 | 321.64 | 5.48 | 72 | >sp\|P01876\|IGHA1_HUMAN Immunoglobulin heavy constant alpha 1 OS=Homo sapiens OX=9606 GN=IGHA1 PE=1 SV=2 | 255 | 20201116_xd_colostrum_1ng_glyco_hcd30_etd30_compound15.16703.16703.3 | controllerType=0 controllerNumber=1 scan=16703 | 42.8734 |
| 13986:3 | 1 | R.LSLHRPALEDLLLGSEAN[+2116.756]LTC[+57.021]TLTGLR.D | HexNAc(5)Hex(5)NeuAc(1) | C[+57], N[+2117] | 1694.1281 | 3 | 5080.3699 | 5080.3547 | 0 | 3 | 127 | Specific | 365.26 | 287.43 | 287.43 | 5.26 | 72 | >sp\|P01876\|IGHA1_HUMAN Immunoglobulin heavy constant alpha 1 OS=Homo sapiens OX=9606 GN=IGHA1 PE=1 SV=2 | 255 | 20201116_xd_colostrum_1ng_glyco_hcd30_etd30_compound15.17296.17296.3 | controllerType=0 controllerNumber=1 scan=17296 | 44.5998 |
| 13387:3 | 1 | R.LSLHRPALEDLLLGSEAN[+1501.555]LTC[+57.021]TLTGLR.D | HexNAc(5)Hex(3) | C[+57], N[+1502] | 1489.0599 | 3 | 4465.1651 | 4465.1536 | 0 | 2.6 | 127 | Specific | 371.29 | 302.11 | 302.11 | 5.38 | 72 | >sp\|P01876\|IGHA1_HUMAN Immunoglobulin heavy constant alpha 1 OS=Homo sapiens OX=9606 GN=IGHA1 PE=1 SV=2 | 255 | 20201116_xd_colostrum_1ng_glyco_hcd30_etd30_compound15.16664.16664.6 | controllerType=0 controllerNumber=1 scan=16664 | 42.7463 |
| 13297:3 | 1 | R.LSLHRPALEDLLLGSEAN[+1663.608]LTC[+57.021]TLTGLR.D | HexNAc(5)Hex(4) | C[+57], N[+1664] | 1543.0793 | 3 | 4627.2233 | 4627.2064 | 0 | 3.6 | 127 | Specific | 382.67 | 325.63 | 325.63 | 5.68 | 72 | >sp\|P01876\|IGHA1_HUMAN Immunoglobulin heavy constant alpha 1 OS=Homo sapiens OX=9606 GN=IGHA1 PE=1 SV=2 | 255 | 20201116_xd_colostrum_1ng_glyco_hcd30_etd30_compound15.16569.16569.3 | controllerType=0 controllerNumber=1 scan=16569 | 42.4396 |
| 14047:3 | 1 | R.LSLHRPALEDLLLGSEAN[+1872.651]LTC[+57.021]TLTGLR.D | HexNAc(3)Hex(6)NeuAc(1) | C[+57], N[+1873] | 1612.7614 | 3 | 4836.2697 | 4836.2487 | 0 | 4.3 | 127 | Specific | 387.63 | 301.36 | 301.36 | 5.68 | 72 | >sp\|P01876\|IGHA1_HUMAN Immunoglobulin heavy constant alpha 1 OS=Homo sapiens OX=9606 GN=IGHA1 PE=1 SV=2 | 255 | 20201116_xd_colostrum_1ng_glyco_hcd30_etd30_compound15.17361.17361.3 | controllerType=0 controllerNumber=1 scan=17361 | 44.8099 |
| 14797:5 | 1 | R.LSLHRPALEDLLLGSEAN[+2407.852]LTC[+57.021]TLTGLR.D | HexNAc(5)Hex(5)NeuAc(2) | C[+57], N[+2408] | 1075.0975 | 5 | 5371.4586 | 5371.4501 | 0 | 1.6 | 127 | Specific | 389 | 384.69 | 384.69 | 7.08 | 72 | >sp\|P01876\|IGHA1_HUMAN Immunoglobulin heavy constant alpha 1 OS=Homo sapiens OX=9606 GN=IGHA1 PE=1 SV=2 | 255 | 20201116_xd_colostrum_1ng_glyco_hcd30_etd30_compound15.18155.18155.5 | controllerType=0 controllerNumber=1 scan=18155 | 47.1669 |
| 13359:3 | 1 | R.LSLHRPALEDLLLGSEAN[+1298.476]LTC[+57.021]TLTGLR.D | HexNAc(4)Hex(3) | C[+57], N[+1298] | 1421.3655 | 3 | 4262.0821 | 4262.0742 | 0 | 1.8 | 127 | Specific | 393.01 | 354.07 | 354.07 | 6.47 | 72 | >sp\|P01876\|IGHA1_HUMAN Immunoglobulin heavy constant alpha 1 OS=Homo sapiens OX=9606 GN=IGHA1 PE=1 SV=2 | 255 | 20201116_xd_colostrum_1ng_glyco_hcd30_etd30_compound15.16634.16634.3 | controllerType=0 controllerNumber=1 scan=16634 | 42.644 |
| 13355:4 | 1 | R.LSLHRPALEDLLLGSEAN[+1419.502]LTC[+57.021]TLTGLR.D | HexNAc(3)Hex(5) | C[+57], N[+1420] | 1096.5312 | 4 | 4383.1029 | 4383.1005 | 0 | 0.5 | 127 | Specific | 396.06 | 396.06 | 396.06 | 7.34 | 72 | >sp\|P01876\|IGHA1_HUMAN Immunoglobulin heavy constant alpha 1 OS=Homo sapiens OX=9606 GN=IGHA1 PE=1 SV=2 | 255 | 20201116_xd_colostrum_1ng_glyco_hcd30_etd30_compound15.16630.16630.4 | controllerType=0 controllerNumber=1 scan=16630 | 42.6331 |
| 13338:3 | 1 | R.LSLHRPALEDLLLGSEAN[+1460.529]LTC[+57.021]TLTGLR.D | HexNAc(4)Hex(4) | C[+57], N[+1461] | 1475.3847 | 3 | 4424.1395 | 4424.1271 | 0 | 2.8 | 127 | Specific | 398.11 | 362.14 | 362.14 | 6.47 | 72 | >sp\|P01876\|IGHA1_HUMAN Immunoglobulin heavy constant alpha 1 OS=Homo sapiens OX=9606 GN=IGHA1 PE=1 SV=2 | 255 | 20201116_xd_colostrum_1ng_glyco_hcd30_etd30_compound15.16612.16612.3 | controllerType=0 controllerNumber=1 scan=16612 | 42.5801 |
| 13307:3 | 1 | R.LSLHRPALEDLLLGSEAN[+1702.581]LTC[+57.021]TLTGLR.D | HexNAc(2)Hex(8) | C[+57], N[+1703] | 1556.0726 | 3 | 4666.2031 | 4666.1796 | 0 | 5 | 127 | Specific | 398.73 | 355.44 | 355.44 | 4.96 | 72 | >sp\|P01876\|IGHA1_HUMAN Immunoglobulin heavy constant alpha 1 OS=Homo sapiens OX=9606 GN=IGHA1 PE=1 SV=2 | 255 | 20201116_xd_colostrum_1ng_glyco_hcd30_etd30_compound15.16579.16579.3 | controllerType=0 controllerNumber=1 scan=16579 | 42.4629 |
| 14011:3 | 1 | R.LSLHRPALEDLLLGSEAN[+1913.677]LTC[+57.021]TLTGLR.D | HexNAc(4)Hex(5)NeuAc(1) | C[+57], N[+1914] | 1626.4343 | 3 | 4877.2882 | 4877.2753 | 0 | 2.7 | 127 | Specific | 434.52 | 317.27 | 317.27 | 6.35 | 72 | >sp\|P01876\|IGHA1_HUMAN Immunoglobulin heavy constant alpha 1 OS=Homo sapiens OX=9606 GN=IGHA1 PE=1 SV=2 | 255 | 20201116_xd_colostrum_1ng_glyco_hcd30_etd30_compound15.17323.17323.3 | controllerType=0 controllerNumber=1 scan=17323 | 44.6871 |
| 13344:3 | 1 | R.LSLHRPALEDLLLGSEAN[+1501.555]LTC[+57.021]TLTGLR.D | HexNAc(5)Hex(3) | C[+57], N[+1502] | 1489.0589 | 3 | 4465.1621 | 4465.1536 | 0 | 1.9 | 127 | Specific | 468.12 | 406.11 | 406.11 | 7.86 | 72 | >sp\|P01876\|IGHA1_HUMAN Immunoglobulin heavy constant alpha 1 OS=Homo sapiens OX=9606 GN=IGHA1 PE=1 SV=2 | 255 | 20201116_xd_colostrum_1ng_glyco_hcd30_etd30_compound15.16619.16619.3 | controllerType=0 controllerNumber=1 scan=16619 | 42.6016 |
| 14048:3 | 1 | R.LSLHRPALEDLLLGSEAN[+1710.598]LTC[+57.021]TLTGLR.D | HexNAc(3)Hex(5)NeuAc(1) | C[+57], N[+1711] | 1558.7437 | 3 | 4674.2166 | 4674.1959 | 0 | 4.4 | 127 | Specific | 468.52 | 403.1 | 403.1 | 7.27 | 72 | >sp\|P01876\|IGHA1_HUMAN Immunoglobulin heavy constant alpha 1 OS=Homo sapiens OX=9606 GN=IGHA1 PE=1 SV=2 | 255 | 20201116_xd_colostrum_1ng_glyco_hcd30_etd30_compound15.17362.17362.3 | controllerType=0 controllerNumber=1 scan=17362 | 44.8139 |
| 13607:4 | 1 | R.LSLHRPALEDLLLGSEAN[+1501.555]LTC[+57.021]TLTGLR.D | HexNAc(5)Hex(3) | C[+57], N[+1502] | 1117.0368 | 4 | 4465.1253 | 4465.1536 | 0 | -6.3 | 127 | Specific | 468.79 | 468.79 | 468.79 | 5.63 | 72 | >sp\|P01876\|IGHA1_HUMAN Immunoglobulin heavy constant alpha 1 OS=Homo sapiens OX=9606 GN=IGHA1 PE=1 SV=2 | 255 | 20201116_xd_colostrum_1ng_glyco_hcd30_etd30_compound15.16897.16897.4 | controllerType=0 controllerNumber=1 scan=16897 | 43.4538 |
| 13381:3 | 1 | R.LSLHRPALEDLLLGSEAN[+1054.370]LTC[+57.021]TLTGLR.D | HexNAc(2)Hex(4) | C[+57], N[+1054] | 1339.9966 | 3 | 4017.9754 | 4017.9683 | 0 | 1.8 | 127 | Specific | 470.61 | 448.16 | 448.16 | 7.86 | 72 | >sp\|P01876\|IGHA1_HUMAN Immunoglobulin heavy constant alpha 1 OS=Homo sapiens OX=9606 GN=IGHA1 PE=1 SV=2 | 255 | 20201116_xd_colostrum_1ng_glyco_hcd30_etd30_compound15.16658.16658.3 | controllerType=0 controllerNumber=1 scan=16658 | 42.7256 |
| 13289:3 | 1 | R.LSLHRPALEDLLLGSEAN[+1540.529]LTC[+57.021]TLTGLR.D | HexNAc(2)Hex(7) | C[+57], N[+1541] | 1502.0505 | 3 | 4504.1369 | 4504.1268 | 0 | 2.2 | 127 | Specific | 476.05 | 434.68 | 434.68 | 7.86 | 72 | >sp\|P01876\|IGHA1_HUMAN Immunoglobulin heavy constant alpha 1 OS=Homo sapiens OX=9606 GN=IGHA1 PE=1 SV=2 | 255 | 20201116_xd_colostrum_1ng_glyco_hcd30_etd30_compound15.16560.16560.3 | controllerType=0 controllerNumber=1 scan=16560 | 42.4145 |
| 13423:3 | 1 | R.LSLHRPALEDLLLGSEAN[+892.317]LTC[+57.021]TLTGLR.D | HexNAc(2)Hex(3) | C[+57], N[+892] | 1285.9803 | 3 | 3855.9263 | 3855.9155 | 0 | 2.8 | 127 | Specific | 493.62 | 484.89 | 484.89 | 8 | 72 | >sp\|P01876\|IGHA1_HUMAN Immunoglobulin heavy constant alpha 1 OS=Homo sapiens OX=9606 GN=IGHA1 PE=1 SV=2 | 255 | 20201116_xd_colostrum_1ng_glyco_hcd30_etd30_compound15.16702.16702.3 | controllerType=0 controllerNumber=1 scan=16702 | 42.868 |
| 13630:4 | 1 | R.LSLHRPALEDLLLGSEAN[+1501.555]LTC[+57.021]TLTGLR.D | HexNAc(5)Hex(3) | C[+57], N[+1502] | 1117.0378 | 4 | 4465.1292 | 4465.1536 | 0 | -5.5 | 127 | Specific | 495.01 | 495.01 | 495.01 | 6.49 | 72 | >sp\|P01876\|IGHA1_HUMAN Immunoglobulin heavy constant alpha 1 OS=Homo sapiens OX=9606 GN=IGHA1 PE=1 SV=2 | 255 | 20201116_xd_colostrum_1ng_glyco_hcd30_etd30_compound15.16921.16921.4 | controllerType=0 controllerNumber=1 scan=16921 | 43.5277 |
| 13322:3 | 1 | R.LSLHRPALEDLLLGSEAN[+1216.423]LTC[+57.021]TLTGLR.D | HexNAc(2)Hex(5) | C[+57], N[+1216] | 1394.0154 | 3 | 4180.0318 | 4180.0211 | 0 | 2.5 | 127 | Specific | 536.28 | 536.28 | 536.28 | 8.11 | 72 | >sp\|P01876\|IGHA1_HUMAN Immunoglobulin heavy constant alpha 1 OS=Homo sapiens OX=9606 GN=IGHA1 PE=1 SV=2 | 255 | 20201116_xd_colostrum_1ng_glyco_hcd30_etd30_compound15.16596.16596.3 | controllerType=0 controllerNumber=1 scan=16596 | 42.5419 |
| 13425:4 | 1 | R.LSLHRPALEDLLLGSEAN[+1622.582]LTC[+57.021]TLTGLR.D | HexNAc(4)Hex(5) | C[+57], N[+1623] | 1147.3035 | 4 | 4586.1922 | 4586.1799 | 0 | 2.7 | 127 | Specific | 543.14 | 543.14 | 543.14 | 7.62 | 72 | >sp\|P01876\|IGHA1_HUMAN Immunoglobulin heavy constant alpha 1 OS=Homo sapiens OX=9606 GN=IGHA1 PE=1 SV=2 | 255 | 20201116_xd_colostrum_1ng_glyco_hcd30_etd30_compound15.16704.16704.4 | controllerType=0 controllerNumber=1 scan=16704 | 42.8787 |
| 13303:5 | 1 | R.LSLHRPALEDLLLGSEAN[+1663.608]LTC[+57.021]TLTGLR.D | HexNAc(5)Hex(4) | C[+57], N[+1664] | 926.249 | 5 | 4627.2161 | 4627.2064 | 0 | 2.1 | 127 | Specific | 587.27 | 587.27 | 587.27 | 8.88 | 72 | >sp\|P01876\|IGHA1_HUMAN Immunoglobulin heavy constant alpha 1 OS=Homo sapiens OX=9606 GN=IGHA1 PE=1 SV=2 | 255 | 20201116_xd_colostrum_1ng_glyco_hcd30_etd30_compound15.16575.16575.5 | controllerType=0 controllerNumber=1 scan=16575 | 42.4549 |
| 13364:5 | 1 | R.LSLHRPALEDLLLGSEAN[+1216.423]LTC[+57.021]TLTGLR.D | HexNAc(2)Hex(5) | C[+57], N[+1216] | 836.8109 | 5 | 4180.0256 | 4180.0211 | 0 | 1.1 | 127 | Specific | 645.74 | 624.31 | 624.31 | 10.12 | 72 | >sp\|P01876\|IGHA1_HUMAN Immunoglobulin heavy constant alpha 1 OS=Homo sapiens OX=9606 GN=IGHA1 PE=1 SV=2 | 255 | 20201116_xd_colostrum_1ng_glyco_hcd30_etd30_compound15.16640.16640.5 | controllerType=0 controllerNumber=1 scan=16640 | 42.6595 |
| 14110:4 | 1 | R.LSLHRPALEDLLLGSEAN[+1751.624]LTC[+57.021]TLTGLR.D | HexNAc(4)Hex(4)NeuAc(1) | C[+57], N[+1752] | 1179.5628 | 4 | 4715.2293 | 4715.2225 | 0 | 1.5 | 127 | Specific | 662.94 | 652.75 | 652.75 | 9.93 | 72 | >sp\|P01876\|IGHA1_HUMAN Immunoglobulin heavy constant alpha 1 OS=Homo sapiens OX=9606 GN=IGHA1 PE=1 SV=2 | 255 | 20201116_xd_colostrum_1ng_glyco_hcd30_etd30_compound15.17427.17427.4 | controllerType=0 controllerNumber=1 scan=17427 | 45.0136 |
| 13403:4 | 1 | R.LSLHRPALEDLLLGSEAN[+1257.449]LTC[+57.021]TLTGLR.D | HexNAc(3)Hex(4) | C[+57], N[+1257] | 1056.0207 | 4 | 4221.0609 | 4221.0477 | 0 | 3.1 | 127 | Specific | 668.19 | 668.19 | 668.19 | 11.59 | 72 | >sp\|P01876\|IGHA1_HUMAN Immunoglobulin heavy constant alpha 1 OS=Homo sapiens OX=9606 GN=IGHA1 PE=1 SV=2 | 255 | 20201116_xd_colostrum_1ng_glyco_hcd30_etd30_compound15.16681.16681.4 | controllerType=0 controllerNumber=1 scan=16681 | 42.8046 |
| 13277:4 | 1 | R.LSLHRPALEDLLLGSEAN[+1864.634]LTC[+57.021]TLTGLR.D | HexNAc(2)Hex(9) | C[+57], N[+1865] | 1207.814 | 4 | 4828.2342 | 4828.2324 | 0 | 0.4 | 127 | Specific | 784.19 | 784.19 | 784.19 | 12.83 | 72 | >sp\|P01876\|IGHA1_HUMAN Immunoglobulin heavy constant alpha 1 OS=Homo sapiens OX=9606 GN=IGHA1 PE=1 SV=2 | 255 | 20201116_xd_colostrum_1ng_glyco_hcd30_etd30_compound15.16548.16548.4 | controllerType=0 controllerNumber=1 scan=16548 | 42.3757 |
| 13987:5 | 1 | R.LSLHRPALEDLLLGSEAN[+2116.756]LTC[+57.021]TLTGLR.D | HexNAc(5)Hex(5)NeuAc(1) | C[+57], N[+2117] | 1016.8798 | 5 | 5080.3697 | 5080.3547 | 0 | 3 | 127 | Specific | 800.41 | 719.61 | 719.61 | 14.17 | 72 | >sp\|P01876\|IGHA1_HUMAN Immunoglobulin heavy constant alpha 1 OS=Homo sapiens OX=9606 GN=IGHA1 PE=1 SV=2 | 255 | 20201116_xd_colostrum_1ng_glyco_hcd30_etd30_compound15.17297.17297.5 | controllerType=0 controllerNumber=1 scan=17297 | 44.6057 |
| 13349:5 | 1 | R.LSLHRPALEDLLLGSEAN[+1501.555]LTC[+57.021]TLTGLR.D | HexNAc(5)Hex(3) | C[+57], N[+1502] | 893.8379 | 5 | 4465.1605 | 4465.1536 | 0 | 1.5 | 127 | Specific | 823.13 | 820.83 | 820.83 | 13.23 | 72 | >sp\|P01876\|IGHA1_HUMAN Immunoglobulin heavy constant alpha 1 OS=Homo sapiens OX=9606 GN=IGHA1 PE=1 SV=2 | 255 | 20201116_xd_colostrum_1ng_glyco_hcd30_etd30_compound15.16624.16624.5 | controllerType=0 controllerNumber=1 scan=16624 | 42.6134 |
| 14015:5 | 1 | R.LSLHRPALEDLLLGSEAN[+1913.677]LTC[+57.021]TLTGLR.D | HexNAc(4)Hex(5)NeuAc(1) | C[+57], N[+1914] | 976.262 | 5 | 4877.2811 | 4877.2753 | 0 | 1.2 | 127 | Specific | 828.22 | 755.35 | 755.35 | 13.23 | 72 | >sp\|P01876\|IGHA1_HUMAN Immunoglobulin heavy constant alpha 1 OS=Homo sapiens OX=9606 GN=IGHA1 PE=1 SV=2 | 255 | 20201116_xd_colostrum_1ng_glyco_hcd30_etd30_compound15.17327.17327.5 | controllerType=0 controllerNumber=1 scan=17327 | 44.6953 |
| 14010:4 | 1 | R.LSLHRPALEDLLLGSEAN[+1913.677]LTC[+57.021]TLTGLR.D | HexNAc(4)Hex(5)NeuAc(1) | C[+57], N[+1914] | 1220.0778 | 4 | 4877.2894 | 4877.2753 | 0 | 2.9 | 127 | Specific | 831.02 | 814.41 | 814.41 | 14.59 | 72 | >sp\|P01876\|IGHA1_HUMAN Immunoglobulin heavy constant alpha 1 OS=Homo sapiens OX=9606 GN=IGHA1 PE=1 SV=2 | 255 | 20201116_xd_colostrum_1ng_glyco_hcd30_etd30_compound15.17322.17322.4 | controllerType=0 controllerNumber=1 scan=17322 | 44.6842 |
| 14042:4 | 1 | R.LSLHRPALEDLLLGSEAN[+1872.651]LTC[+57.021]TLTGLR.D | HexNAc(3)Hex(6)NeuAc(1) | C[+57], N[+1873] | 1209.8245 | 4 | 4836.2762 | 4836.2487 | 0 | 5.7 | 127 | Specific | 832.86 | 752.79 | 752.79 | 13.8 | 72 | >sp\|P01876\|IGHA1_HUMAN Immunoglobulin heavy constant alpha 1 OS=Homo sapiens OX=9606 GN=IGHA1 PE=1 SV=2 | 255 | 20201116_xd_colostrum_1ng_glyco_hcd30_etd30_compound15.17356.17356.4 | controllerType=0 controllerNumber=1 scan=17356 | 44.7963 |
| 13302:4 | 1 | R.LSLHRPALEDLLLGSEAN[+1581.555]LTC[+57.021]TLTGLR.D | HexNAc(3)Hex(6) | C[+57], N[+1582] | 1137.0486 | 4 | 4545.1727 | 4545.1533 | 0 | 4.3 | 127 | Specific | 834.74 | 834.74 | 834.74 | 14 | 72 | >sp\|P01876\|IGHA1_HUMAN Immunoglobulin heavy constant alpha 1 OS=Homo sapiens OX=9606 GN=IGHA1 PE=1 SV=2 | 255 | 20201116_xd_colostrum_1ng_glyco_hcd30_etd30_compound15.16574.16574.4 | controllerType=0 controllerNumber=1 scan=16574 | 42.4518 |
| 13980:4 | 1 | R.LSLHRPALEDLLLGSEAN[+2116.756]LTC[+57.021]TLTGLR.D | HexNAc(5)Hex(5)NeuAc(1) | C[+57], N[+2117] | 1270.8466 | 4 | 5080.3644 | 5080.3547 | 0 | 1.9 | 127 | Specific | 839.99 | 782.28 | 782.28 | 14.59 | 72 | >sp\|P01876\|IGHA1_HUMAN Immunoglobulin heavy constant alpha 1 OS=Homo sapiens OX=9606 GN=IGHA1 PE=1 SV=2 | 255 | 20201116_xd_colostrum_1ng_glyco_hcd30_etd30_compound15.17290.17290.4 | controllerType=0 controllerNumber=1 scan=17290 | 44.5881 |
| 13343:4 | 1 | R.LSLHRPALEDLLLGSEAN[+1501.555]LTC[+57.021]TLTGLR.D | HexNAc(5)Hex(3) | C[+57], N[+1502] | 1117.0466 | 4 | 4465.1644 | 4465.1536 | 0 | 2.4 | 127 | Specific | 855.04 | 855.04 | 855.04 | 15.59 | 72 | >sp\|P01876\|IGHA1_HUMAN Immunoglobulin heavy constant alpha 1 OS=Homo sapiens OX=9606 GN=IGHA1 PE=1 SV=2 | 255 | 20201116_xd_colostrum_1ng_glyco_hcd30_etd30_compound15.16618.16618.4 | controllerType=0 controllerNumber=1 scan=16618 | 42.5988 |
| 14046:4 | 1 | R.LSLHRPALEDLLLGSEAN[+1710.598]LTC[+57.021]TLTGLR.D | HexNAc(3)Hex(5)NeuAc(1) | C[+57], N[+1711] | 1169.3096 | 4 | 4674.2166 | 4674.1959 | 0 | 4.4 | 127 | Specific | 868.69 | 868.69 | 868.69 | 15 | 72 | >sp\|P01876\|IGHA1_HUMAN Immunoglobulin heavy constant alpha 1 OS=Homo sapiens OX=9606 GN=IGHA1 PE=1 SV=2 | 255 | 20201116_xd_colostrum_1ng_glyco_hcd30_etd30_compound15.17360.17360.4 | controllerType=0 controllerNumber=1 scan=17360 | 44.805 |
| 13282:4 | 1 | R.LSLHRPALEDLLLGSEAN[+1581.555]LTC[+57.021]TLTGLR.D | HexNAc(3)Hex(6) | C[+57], N[+1582] | 1137.0462 | 4 | 4545.1629 | 4545.1533 | 0 | 2.1 | 127 | Specific | 936.33 | 878.94 | 878.94 | 16.59 | 72 | >sp\|P01876\|IGHA1_HUMAN Immunoglobulin heavy constant alpha 1 OS=Homo sapiens OX=9606 GN=IGHA1 PE=1 SV=2 | 255 | 20201116_xd_colostrum_1ng_glyco_hcd30_etd30_compound15.16553.16553.4 | controllerType=0 controllerNumber=1 scan=16553 | 42.3904 |
| 13328:4 | 1 | R.LSLHRPALEDLLLGSEAN[+1378.476]LTC[+57.021]TLTGLR.D | HexNAc(2)Hex(6) | C[+57], N[+1378] | 1086.277 | 4 | 4342.0863 | 4342.074 | 0 | 2.8 | 127 | Specific | 969.92 | 942.08 | 942.08 | 17.59 | 72 | >sp\|P01876\|IGHA1_HUMAN Immunoglobulin heavy constant alpha 1 OS=Homo sapiens OX=9606 GN=IGHA1 PE=1 SV=2 | 255 | 20201116_xd_colostrum_1ng_glyco_hcd30_etd30_compound15.16602.16602.4 | controllerType=0 controllerNumber=1 scan=16602 | 42.5569 |
| 13313:4 | 1 | R.LSLHRPALEDLLLGSEAN[+1216.423]LTC[+57.021]TLTGLR.D | HexNAc(2)Hex(5) | C[+57], N[+1216] | 1045.7637 | 4 | 4180.033 | 4180.0211 | 0 | 2.8 | 127 | Specific | 1021.89 | 955.03 | 955.03 | 18.59 | 72 | >sp\|P01876\|IGHA1_HUMAN Immunoglobulin heavy constant alpha 1 OS=Homo sapiens OX=9606 GN=IGHA1 PE=1 SV=2 | 255 | 20201116_xd_colostrum_1ng_glyco_hcd30_etd30_compound15.16586.16586.4 | controllerType=0 controllerNumber=1 scan=16586 | 42.5087 |
| 13356:4 | 1 | R.LSLHRPALEDLLLGSEAN[+1298.476]LTC[+57.021]TLTGLR.D | HexNAc(4)Hex(3) | C[+57], N[+1298] | 1066.2753 | 4 | 4262.0794 | 4262.0742 | 0 | 1.2 | 127 | Specific | 1045.54 | 1045.54 | 1045.54 | 17.83 | 72 | >sp\|P01876\|IGHA1_HUMAN Immunoglobulin heavy constant alpha 1 OS=Homo sapiens OX=9606 GN=IGHA1 PE=1 SV=2 | 255 | 20201116_xd_colostrum_1ng_glyco_hcd30_etd30_compound15.16631.16631.4 | controllerType=0 controllerNumber=1 scan=16631 | 42.6367 |
| 13416:4 | 1 | R.LSLHRPALEDLLLGSEAN[+892.317]LTC[+57.021]TLTGLR.D | HexNAc(2)Hex(3) | C[+57], N[+892] | 964.7352 | 4 | 3855.9188 | 3855.9155 | 0 | 0.9 | 127 | Specific | 1046.3 | 1043.34 | 1043.34 | 17.83 | 72 | >sp\|P01876\|IGHA1_HUMAN Immunoglobulin heavy constant alpha 1 OS=Homo sapiens OX=9606 GN=IGHA1 PE=1 SV=2 | 255 | 20201116_xd_colostrum_1ng_glyco_hcd30_etd30_compound15.16695.16695.4 | controllerType=0 controllerNumber=1 scan=16695 | 42.8471 |
| 13319:4 | 1 | R.LSLHRPALEDLLLGSEAN[+1460.529]LTC[+57.021]TLTGLR.D | HexNAc(4)Hex(4) | C[+57], N[+1461] | 1106.7903 | 4 | 4424.1395 | 4424.1271 | 0 | 2.8 | 127 | Specific | 1057.31 | 933.49 | 933.49 | 19.59 | 72 | >sp\|P01876\|IGHA1_HUMAN Immunoglobulin heavy constant alpha 1 OS=Homo sapiens OX=9606 GN=IGHA1 PE=1 SV=2 | 255 | 20201116_xd_colostrum_1ng_glyco_hcd30_etd30_compound15.16592.16592.4 | controllerType=0 controllerNumber=1 scan=16592 | 42.5295 |
| 13278:4 | 1 | R.LSLHRPALEDLLLGSEAN[+1540.529]LTC[+57.021]TLTGLR.D | HexNAc(2)Hex(7) | C[+57], N[+1541] | 1126.789 | 4 | 4504.1341 | 4504.1268 | 0 | 1.6 | 127 | Specific | 1079.97 | 1003.18 | 1003.18 | 19.59 | 72 | >sp\|P01876\|IGHA1_HUMAN Immunoglobulin heavy constant alpha 1 OS=Homo sapiens OX=9606 GN=IGHA1 PE=1 SV=2 | 255 | 20201116_xd_colostrum_1ng_glyco_hcd30_etd30_compound15.16549.16549.4 | controllerType=0 controllerNumber=1 scan=16549 | 42.3807 |

Table S4. Cont.

<table>
<tr><th>Query #:z</th><th>Protein Rank</th><th>Glycopeptide</th><th>Glycan</th><th>Modification Type(s)</th><th>Observed m/z</th><th>z</th><th>Observed (M+H)</th><th>Calc. mass (M+H)</th><th>Off-by-x error</th><th>Mass error (ppm)</th><th>StartPosition</th><th>Cleavage</th><th>Score</th><th>Delta</th><th>Delta Mod</th><th>|Log Prob|</th><th># of unique peptides</th><th>Protein Name</th><th>Protein DB number</th><th>Comment</th><th>Scan #</th><th>Scan Time</th></tr>
<tr><td>13380:4</td><td>1</td><td>R.LSLHRPALEDLLLGSEAN[+1054.370]LTC[+57.021]TLTGLR.D</td><td>HexNAc(2)Hex(4)</td><td>C[+57], N[+1054]</td><td>1005.2499</td><td>4</td><td>4017.9779</td><td>4017.9683</td><td>0</td><td>2.4</td><td>127</td><td>Specific</td><td>1180.39</td><td>1141.43</td><td>1141.43</td><td>21.59</td><td>72</td><td>>sp|P01876|IGHA1_HUMAN Immunoglobulin heavy constant alpha 1 OS=Homo sapiens OX=9606 GN=IGHA1 PE=1 SV=2</td><td>255</td><td>20201116_xd_colostrum_1ng_glyco_hcd30_etd30_compound15.16657.16657.4</td><td>controllerType=0 controllerNumber=1 scan=16657</td><td>42.7225</td></tr>
<tr><td>09266:4</td><td>1</td><td>K.TIDRLAGKPTHVN[+1216.423]VSVVMAEVDGTC[+57.021]Y.-</td><td>HexNAc(2)Hex(5)</td><td>C[+57], N[+1216]</td><td>1012.9666</td><td>4</td><td>4048.8446</td><td>4048.8247</td><td>0</td><td>4.9</td><td>328</td><td>CRagged</td><td>277.15</td><td>277.15</td><td>88.83</td><td>4</td><td>72</td><td>>sp|P01876|IGHA1_HUMAN Immunoglobulin heavy constant alpha 1 OS=Homo sapiens OX=9606 GN=IGHA1 PE=1 SV=2</td><td>255</td><td>20201116_xd_colostrum_1ng_glyco_hcd30_etd30_compound15.12335.12335.4</td><td>controllerType=0 controllerNumber=1 scan=12335</td><td>31.1945</td></tr>
<tr><td>07409:3</td><td>1</td><td>R.LAGKPTHVN[+1216.423]VSVVMAEVDGTC[+57.021].Y</td><td>HexNAc(2)Hex(5)</td><td>C[+57], N[+1216]</td><td>1134.1759</td><td>3</td><td>3400.5132</td><td>3400.5016</td><td>0</td><td>3.4</td><td>332</td><td>CRagged</td><td>303.58</td><td>237.34</td><td>219.42</td><td>4.26</td><td>72</td><td>>sp|P01876|IGHA1_HUMAN Immunoglobulin heavy constant alpha 1 OS=Homo sapiens OX=9606 GN=IGHA1 PE=1 SV=2</td><td>255</td><td>20201116_xd_colostrum_1ng_glyco_hcd30_etd30_compound15.10385.10385.3</td><td>controllerType=0 controllerNumber=1 scan=10385</td><td>26.1008</td></tr>
<tr><td>08244:3</td><td>1</td><td>R.LAGKPTHVN[+1702.581]VSVVMAEVDGTC[+57.021]Y.-</td><td>HexNAc(2)Hex(8)</td><td>C[+57], N[+1703]</td><td>1350.5815</td><td>3</td><td>4049.7299</td><td>4049.7234</td><td>0</td><td>1.6</td><td>332</td><td>CRagged</td><td>204</td><td>163.59</td><td>118.03</td><td>3.5</td><td>72</td><td>>sp|P01876|IGHA1_HUMAN Immunoglobulin heavy constant alpha 1 OS=Homo sapiens OX=9606 GN=IGHA1 PE=1 SV=2</td><td>255</td><td>20201116_xd_colostrum_1ng_glyco_hcd30_etd30_compound15.11260.11260.3</td><td>controllerType=0 controllerNumber=1 scan=11260</td><td>28.2312</td></tr>
<tr><td>09040:4</td><td>1</td><td>R.LAGKPTHVN[+1971.719]VSVVMAEVDGTC[+57.021]Y.-</td><td>HexNAc(5)Hex(5)Fuc(1)</td><td>C[+57], N[+1972]</td><td>1080.4731</td><td>4</td><td>4318.8704</td><td>4318.861</td><td>0</td><td>2.2</td><td>332</td><td>CRagged</td><td>205.83</td><td>205.83</td><td>99.73</td><td>3.66</td><td>72</td><td>>sp|P01876|IGHA1_HUMAN Immunoglobulin heavy constant alpha 1 OS=Homo sapiens OX=9606 GN=IGHA1 PE=1 SV=2</td><td>255</td><td>20201116_xd_colostrum_1ng_glyco_hcd30_etd30_compound15.12097.12097.4</td><td>controllerType=0 controllerNumber=1 scan=12097</td><td>30.5141</td></tr>
<tr><td>11579:3</td><td>1</td><td>R.LAGKPTHVN[+2350.830]VSVVMAEVDGTC[+57.021]Y.-</td><td>HexNAc(4)Hex(5)Fuc(1)NeuAc(2)</td><td>C[+57], N[+2351]</td><td>1566.6667</td><td>3</td><td>4697.9855</td><td>4697.9724</td><td>0</td><td>2.8</td><td>332</td><td>CRagged</td><td>219.53</td><td>60.9</td><td>60.9</td><td>2.84</td><td>72</td><td>>sp|P01876|IGHA1_HUMAN Immunoglobulin heavy constant alpha 1 OS=Homo sapiens OX=9606 GN=IGHA1 PE=1 SV=2</td><td>255</td><td>20201116_xd_colostrum_1ng_glyco_hcd30_etd30_compound15.14758.14758.3</td><td>controllerType=0 controllerNumber=1 scan=14758</td><td>37.2917</td></tr>
<tr><td>08284:3</td><td>1</td><td>R.LAGKPTHVN[+1663.608]VSVVMAEVDGTC[+57.021]Y.-</td><td>HexNAc(5)Hex(4)</td><td>C[+57], N[+1664]</td><td>1337.5914</td><td>3</td><td>4010.7595</td><td>4010.7502</td><td>0</td><td>2.3</td><td>332</td><td>CRagged</td><td>220.91</td><td>136.39</td><td>136.39</td><td>3.22</td><td>72</td><td>>sp|P01876|IGHA1_HUMAN Immunoglobulin heavy constant alpha 1 OS=Homo sapiens OX=9606 GN=IGHA1 PE=1 SV=2</td><td>255</td><td>20201116_xd_colostrum_1ng_glyco_hcd30_etd30_compound15.11303.11303.3</td><td>controllerType=0 controllerNumber=1 scan=11303</td><td>28.3604</td></tr>
<tr><td>08252:3</td><td>1</td><td>R.LAGKPTHVN[+1702.581]VSVVMAEVDGTC[+57.021]Y.-</td><td>HexNAc(2)Hex(8)</td><td>C[+57], N[+1703]</td><td>1350.5836</td><td>3</td><td>4049.7361</td><td>4049.7234</td><td>0</td><td>3.1</td><td>332</td><td>CRagged</td><td>224.79</td><td>172.61</td><td>168.44</td><td>3.56</td><td>72</td><td>>sp|P01876|IGHA1_HUMAN Immunoglobulin heavy constant alpha 1 OS=Homo sapiens OX=9606 GN=IGHA1 PE=1 SV=2</td><td>255</td><td>20201116_xd_colostrum_1ng_glyco_hcd30_etd30_compound15.11269.11269.3</td><td>controllerType=0 controllerNumber=1 scan=11269</td><td>28.2554</td></tr>
<tr><td>10286:3</td><td>1</td><td>R.LAGKPTHVN[+1872.651]VSVVMAEVDGTC[+57.021]Y.-</td><td>HexNAc(3)Hex(6)NeuAc(1)</td><td>C[+57], N[+1873]</td><td>1407.2729</td><td>3</td><td>4219.8041</td><td>4219.7926</td><td>0</td><td>2.7</td><td>332</td><td>CRagged</td><td>227.89</td><td>103.35</td><td>103.35</td><td>3.13</td><td>72</td><td>>sp|P01876|IGHA1_HUMAN Immunoglobulin heavy constant alpha 1 OS=Homo sapiens OX=9606 GN=IGHA1 PE=1 SV=2</td><td>255</td><td>20201116_xd_colostrum_1ng_glyco_hcd30_etd30_compound15.13402.13402.3</td><td>controllerType=0 controllerNumber=1 scan=13402</td><td>33.7881</td></tr>
<tr><td>08756:3</td><td>1</td><td>R.LAGKPTHVN[+1216.423]VSVVMAEVDGTC[+57.021]Y.-</td><td>HexNAc(2)Hex(5)</td><td>C[+57], N[+1216]</td><td>1188.5297</td><td>3</td><td>3563.5746</td><td>3563.5649</td><td>0</td><td>2.7</td><td>332</td><td>CRagged</td><td>232.57</td><td>221.41</td><td>175.1</td><td>3.67</td><td>72</td><td>>sp|P01876|IGHA1_HUMAN Immunoglobulin heavy constant alpha 1 OS=Homo sapiens OX=9606 GN=IGHA1 PE=1 SV=2</td><td>255</td><td>20201116_xd_colostrum_1ng_glyco_hcd30_etd30_compound15.11800.11800.3</td><td>controllerType=0 controllerNumber=1 scan=11800</td><td>29.7732</td></tr>
<tr><td>09020:4</td><td>1</td><td>R.LAGKPTHVN[+1825.661]VSVVMAEVDGTC[+57.021]Y.-</td><td>HexNAc(5)Hex(5)</td><td>C[+57], N[+1826]</td><td>1043.9615</td><td>4</td><td>4172.824</td><td>4172.8031</td><td>0</td><td>5</td><td>332</td><td>CRagged</td><td>238.82</td><td>238.82</td><td>43.59</td><td>3.25</td><td>72</td><td>>sp|P01876|IGHA1_HUMAN Immunoglobulin heavy constant alpha 1 OS=Homo sapiens OX=9606 GN=IGHA1 PE=1 SV=2</td><td>255</td><td>20201116_xd_colostrum_1ng_glyco_hcd30_etd30_compound15.12076.12076.4</td><td>controllerType=0 controllerNumber=1 scan=12076</td><td>30.4548</td></tr>
<tr><td>09006:3</td><td>1</td><td>R.LAGKPTHVN[+1971.719]VSVVMAEVDGTC[+57.021]Y.-</td><td>HexNAc(5)Hex(5)Fuc(1)</td><td>C[+57], N[+1972]</td><td>1440.2951</td><td>3</td><td>4318.8707</td><td>4318.861</td><td>0</td><td>2.3</td><td>332</td><td>CRagged</td><td>239.61</td><td>113.67</td><td>113.67</td><td>3.13</td><td>72</td><td>>sp|P01876|IGHA1_HUMAN Immunoglobulin heavy constant alpha 1 OS=Homo sapiens OX=9606 GN=IGHA1 PE=1 SV=2</td><td>255</td><td>20201116_xd_colostrum_1ng_glyco_hcd30_etd30_compound15.12061.12061.3</td><td>controllerType=0 controllerNumber=1 scan=12061</td><td>30.4156</td></tr>
<tr><td>10266:3</td><td>1</td><td>R.LAGKPTHVN[+2262.814]VSVVMAEVDGTC[+57.021]Y.-</td><td>HexNAc(5)Hex(5)Fuc(1)NeuAc(1)</td><td>C[+57], N[+2263]</td><td>1537.3281</td><td>3</td><td>4609.9696</td><td>4609.9564</td><td>0</td><td>2.9</td><td>332</td><td>CRagged</td><td>241.37</td><td>69.57</td><td>69.57</td><td>3.12</td><td>72</td><td>>sp|P01876|IGHA1_HUMAN Immunoglobulin heavy constant alpha 1 OS=Homo sapiens OX=9606 GN=IGHA1 PE=1 SV=2</td><td>255</td><td>20201116_xd_colostrum_1ng_glyco_hcd30_etd30_compound15.13381.13381.3</td><td>controllerType=0 controllerNumber=1 scan=13381</td><td>33.7363</td></tr>
<tr><td>08971:3</td><td>1</td><td>R.LAGKPTHVN[+1702.581]VSVVMAEVDGTC[+57.021]Y.-</td><td>HexNAc(2)Hex(8)</td><td>C[+57], N[+1703]</td><td>1350.5854</td><td>3</td><td>4049.7416</td><td>4049.7234</td><td>0</td><td>4.5</td><td>332</td><td>CRagged</td><td>241.77</td><td>187.29</td><td>187.29</td><td>4.11</td><td>72</td><td>>sp|P01876|IGHA1_HUMAN Immunoglobulin heavy constant alpha 1 OS=Homo sapiens OX=9606 GN=IGHA1 PE=1 SV=2</td><td>255</td><td>20201116_xd_colostrum_1ng_glyco_hcd30_etd30_compound15.12025.12025.3</td><td>controllerType=0 controllerNumber=1 scan=12025</td><td>30.3059</td></tr>
<tr><td>08273:3</td><td>1</td><td>R.LAGKPTHVN[+1971.719]VSVVMAEVDGTC[+57.021]Y.-</td><td>HexNAc(5)Hex(5)Fuc(1)</td><td>C[+57], N[+1972]</td><td>1440.2949</td><td>3</td><td>4318.87</td><td>4318.861</td><td>0</td><td>2.1</td><td>332</td><td>CRagged</td><td>244.75</td><td>190.55</td><td>157.7</td><td>4.38</td><td>72</td><td>>sp|P01876|IGHA1_HUMAN Immunoglobulin heavy constant alpha 1 OS=Homo sapiens OX=9606 GN=IGHA1 PE=1 SV=2</td><td>255</td><td>20201116_xd_colostrum_1ng_glyco_hcd30_etd30_compound15.11291.11291.3</td><td>controllerType=0 controllerNumber=1 scan=11291</td><td>28.3218</td></tr>
<tr><td>08300:3</td><td>1</td><td>R.LAGKPTHVN[+1809.666]VSVVMAEVDGTC[+57.021]Y.-</td><td>HexNAc(5)Hex(4)Fuc(1)</td><td>C[+57], N[+1810]</td><td>1386.2824</td><td>3</td><td>4156.8327</td><td>4156.8081</td><td>0</td><td>5.9</td><td>332</td><td>CRagged</td><td>246.1</td><td>160.13</td><td>160.13</td><td>3.64</td><td>72</td><td>>sp|P01876|IGHA1_HUMAN Immunoglobulin heavy constant alpha 1 OS=Homo sapiens OX=9606 GN=IGHA1 PE=1 SV=2</td><td>255</td><td>20201116_xd_colostrum_1ng_glyco_hcd30_etd30_compound15.11320.11320.3</td><td>controllerType=0 controllerNumber=1 scan=11320</td><td>28.412</td></tr>
<tr><td>09046:3</td><td>1</td><td>R.LAGKPTHVN[+1460.529]VSVVMAEVDGTC[+57.021]Y.-</td><td>HexNAc(4)Hex(4)</td><td>C[+57], N[+1461]</td><td>1269.8968</td><td>3</td><td>3807.6758</td><td>3807.6709</td><td>0</td><td>1.3</td><td>332</td><td>CRagged</td><td>247.88</td><td>163.84</td><td>163.84</td><td>3.73</td><td>72</td><td>>sp|P01876|IGHA1_HUMAN Immunoglobulin heavy constant alpha 1 OS=Homo sapiens OX=9606 GN=IGHA1 PE=1 SV=2</td><td>255</td><td>20201116_xd_colostrum_1ng_glyco_hcd30_etd30_compound15.12103.12103.3</td><td>controllerType=0 controllerNumber=1 scan=12103</td><td>30.53</td></tr>
<tr><td>09076:2</td><td>1</td><td>R.LAGKPTHVN[+1216.423]VSVVMAEVDGTC[+57.021]Y.-</td><td>HexNAc(2)Hex(5)</td><td>C[+57], N[+1216]</td><td>1782.2912</td><td>2</td><td>3563.5752</td><td>3563.5649</td><td>0</td><td>2.9</td><td>332</td><td>CRagged</td><td>249.17</td><td>241.29</td><td>138.17</td><td>5.45</td><td>72</td><td>>sp|P01876|IGHA1_HUMAN Immunoglobulin heavy constant alpha 1 OS=Homo sapiens OX=9606 GN=IGHA1 PE=1 SV=2</td><td>255</td><td>20201116_xd_colostrum_1ng_glyco_hcd30_etd30_compound15.12135.12135.2</td><td>controllerType=0 controllerNumber=1 scan=12135</td><td>30.6268</td></tr>
<tr><td>10309:3</td><td>1</td><td>R.LAGKPTHVN[+1710.598]VSVVMAEVDGTC[+57.021]Y.-</td><td>HexNAc(3)Hex(5)NeuAc(1)</td><td>C[+57], N[+1711]</td><td>1353.2546</td><td>3</td><td>4057.7492</td><td>4057.7397</td><td>0</td><td>2.3</td><td>332</td><td>CRagged</td><td>249.74</td><td>103.58</td><td>103.58</td><td>3.94</td><td>72</td><td>>sp|P01876|IGHA1_HUMAN Immunoglobulin heavy constant alpha 1 OS=Homo sapiens OX=9606 GN=IGHA1 PE=1 SV=2</td><td>255</td><td>20201116_xd_colostrum_1ng_glyco_hcd30_etd30_compound15.13426.13426.3</td><td>controllerType=0 controllerNumber=1 scan=13426</td><td>33.8719</td></tr>
<tr><td>09036:3</td><td>1</td><td>R.LAGKPTHVN[+1606.587]VSVVMAEVDGTC[+57.021]Y.-</td><td>HexNAc(4)Hex(4)Fuc(1)</td><td>C[+57], N[+1607]</td><td>1318.5834</td><td>3</td><td>3953.7357</td><td>3953.7288</td><td>0</td><td>1.8</td><td>332</td><td>CRagged</td><td>249.83</td><td>155.05</td><td>155.05</td><td>4.01</td><td>72</td><td>>sp|P01876|IGHA1_HUMAN Immunoglobulin heavy constant alpha 1 OS=Homo sapiens OX=9606 GN=IGHA1 PE=1 SV=2</td><td>255</td><td>20201116_xd_colostrum_1ng_glyco_hcd30_etd30_compound15.12093.12093.3</td><td>controllerType=0 controllerNumber=1 scan=12093</td><td>30.5041</td></tr>
<tr><td>09107:3</td><td>1</td><td>R.LAGKPTHVN[+1647.613]VSVVMAEVDGTC[+57.021]Y.-</td><td>HexNAc(5)Hex(3)Fuc(1)</td><td>C[+57], N[+1648]</td><td>1332.2621</td><td>3</td><td>3994.7719</td><td>3994.7553</td><td>0</td><td>4.1</td><td>332</td><td>CRagged</td><td>253.09</td><td>159.92</td><td>159.92</td><td>3.89</td><td>72</td><td>>sp|P01876|IGHA1_HUMAN Immunoglobulin heavy constant alpha 1 OS=Homo sapiens OX=9606 GN=IGHA1 PE=1 SV=2</td><td>255</td><td>20201116_xd_colostrum_1ng_glyco_hcd30_etd30_compound15.12168.12168.3</td><td>controllerType=0 controllerNumber=1 scan=12168</td><td>30.7224</td></tr>
<tr><td>08276:3</td><td>1</td><td>R.LAGKPTHVN[+1378.476]VSVVMAEVDGTC[+57.021]Y.-</td><td>HexNAc(2)Hex(6)</td><td>C[+57], N[+1378]</td><td>1242.5468</td><td>3</td><td>3725.6259</td><td>3725.6178</td><td>0</td><td>2.2</td><td>332</td><td>CRagged</td><td>253.17</td><td>197.86</td><td>197.86</td><td>4.35</td><td>72</td><td>>sp|P01876|IGHA1_HUMAN Immunoglobulin heavy constant alpha 1 OS=Homo sapiens OX=9606 GN=IGHA1 PE=1 SV=2</td><td>255</td><td>20201116_xd_colostrum_1ng_glyco_hcd30_etd30_compound15.11294.11294.3</td><td>controllerType=0 controllerNumber=1 scan=11294</td><td>28.3355</td></tr>
<tr><td>08993:3</td><td>1</td><td>R.LAGKPTHVN[+1540.529]VSVVMAEVDGTC[+57.021]Y.-</td><td>HexNAc(2)Hex(7)</td><td>C[+57], N[+1541]</td><td>1296.5644</td><td>3</td><td>3887.6786</td><td>3887.6706</td><td>0</td><td>2.1</td><td>332</td><td>CRagged</td><td>256.67</td><td>198.63</td><td>183.46</td><td>4.58</td><td>72</td><td>>sp|P01876|IGHA1_HUMAN Immunoglobulin heavy constant alpha 1 OS=Homo sapiens OX=9606 GN=IGHA1 PE=1 SV=2</td><td>255</td><td>20201116_xd_colostrum_1ng_glyco_hcd30_etd30_compound15.12048.12048.3</td><td>controllerType=0 controllerNumber=1 scan=12048</td><td>30.3712</td></tr>
<tr><td>08301:3</td><td>1</td><td>R.LAGKPTHVN[+1606.587]VSVVMAEVDGTC[+57.021]Y.-</td><td>HexNAc(4)Hex(4)Fuc(1)</td><td>C[+57], N[+1607]</td><td>1318.5893</td><td>3</td><td>3953.7533</td><td>3953.7288</td><td>0</td><td>6.2</td><td>332</td><td>CRagged</td><td>257.14</td><td>156.89</td><td>156.89</td><td>3.38</td><td>72</td><td>>sp|P01876|IGHA1_HUMAN Immunoglobulin heavy constant alpha 1 OS=Homo sapiens OX=9606 GN=IGHA1 PE=1 SV=2</td><td>255</td><td>20201116_xd_colostrum_1ng_glyco_hcd30_etd30_compound15.11321.11321.3</td><td>controllerType=0 controllerNumber=1 scan=11321</td><td>28.4189</td></tr>
<tr><td>08272:3</td><td>1</td><td>R.LAGKPTHVN[+1540.529]VSVVMAEVDGTC[+57.021]Y.-</td><td>HexNAc(2)Hex(7)</td><td>C[+57], N[+1541]</td><td>1296.5648</td><td>3</td><td>3887.6797</td><td>3887.6706</td><td>0</td><td>2.3</td><td>332</td><td>CRagged</td><td>258.11</td><td>201.41</td><td>200.13</td><td>4.35</td><td>72</td><td>>sp|P01876|IGHA1_HUMAN Immunoglobulin heavy constant alpha 1 OS=Homo sapiens OX=9606 GN=IGHA1 PE=1 SV=2</td><td>255</td><td>20201116_xd_colostrum_1ng_glyco_hcd30_etd30_compound15.11290.11290.3</td><td>controllerType=0 controllerNumber=1 scan=11290</td><td>28.3164</td></tr>
<tr><td>09094:3</td><td>1</td><td>R.LAGKPTHVN[+1501.555]VSVVMAEVDGTC[+57.021]Y.-</td><td>HexNAc(5)Hex(3)</td><td>C[+57], N[+1502]</td><td>1283.5744</td><td>3</td><td>3848.7086</td><td>3848.6974</td><td>0</td><td>2.9</td><td>332</td><td>CRagged</td><td>258.35</td><td>176.61</td><td>176.61</td><td>4.45</td><td>72</td><td>>sp|P01876|IGHA1_HUMAN Immunoglobulin heavy constant alpha 1 OS=Homo sapiens OX=9606 GN=IGHA1 PE=1 SV=2</td><td>255</td><td>20201116_xd_colostrum_1ng_glyco_hcd30_etd30_compound15.12154.12154.3</td><td>controllerType=0 controllerNumber=1 scan=12154</td><td>30.6758</td></tr>
<tr><td>10210:3</td><td>1</td><td>R.LAGKPTHVN[+2116.756]VSVVMAEVDGTC[+57.021]Y.-</td><td>HexNAc(5)Hex(5)NeuAc(1)</td><td>C[+57], N[+2117]</td><td>1488.6429</td><td>3</td><td>4463.9141</td><td>4463.8985</td><td>0</td><td>3.5</td><td>332</td><td>CRagged</td><td>258.39</td><td>108.04</td><td>108.04</td><td>3.43</td><td>72</td><td>>sp|P01876|IGHA1_HUMAN Immunoglobulin heavy constant alpha 1 OS=Homo sapiens OX=9606 GN=IGHA1 PE=1 SV=2</td><td>255</td><td>20201116_xd_colostrum_1ng_glyco_hcd30_etd30_compound15.13322.13322.3</td><td>controllerType=0 controllerNumber=1 scan=13322</td><td>33.5703</td></tr>
<tr><td>10235:4</td><td>1</td><td>R.LAGKPTHVN[+1913.677]VSVVMAEVDGTC[+57.021]Y.-</td><td>HexNAc(4)Hex(5)NeuAc(1)</td><td>C[+57], N[+1914]</td><td>1065.9628</td><td>4</td><td>4260.8294</td><td>4260.8191</td><td>0</td><td>2.4</td><td>332</td><td>CRagged</td><td>277.56</td><td>251.04</td><td>251.04</td><td>4.9</td><td>72</td><td>>sp|P01876|IGHA1_HUMAN Immunoglobulin heavy constant alpha 1 OS=Homo sapiens OX=9606 GN=IGHA1 PE=1 SV=2</td><td>255</td><td>20201116_xd_colostrum_1ng_glyco_hcd30_etd30_compound15.13348.13348.4</td><td>controllerType=0 controllerNumber=1 scan=13348</td><td>33.6361</td></tr>
<tr><td>09030:3</td><td>1</td><td>R.LAGKPTHVN[+1809.666]VSVVMAEVDGTC[+57.021]Y.-</td><td>HexNAc(5)Hex(4)Fuc(1)</td><td>C[+57], N[+1810]</td><td>1386.2767</td><td>3</td><td>4156.8155</td><td>4156.8081</td><td>0</td><td>1.8</td><td>332</td><td>CRagged</td><td>282.01</td><td>164.34</td><td>164.34</td><td>4.38</td><td>72</td><td>>sp|P01876|IGHA1_HUMAN Immunoglobulin heavy constant alpha 1 OS=Homo sapiens OX=9606 GN=IGHA1 PE=1 SV=2</td><td>255</td><td>20201116_xd_colostrum_1ng_glyco_hcd30_etd30_compound15.12087.12087.3</td><td>controllerType=0 controllerNumber=1 scan=12087</td><td>30.4883</td></tr>
<tr><td>09009:3</td><td>1</td><td>R.LAGKPTHVN[+1378.476]VSVVMAEVDGTC[+57.021]Y.-</td><td>HexNAc(2)Hex(6)</td><td>C[+57], N[+1378]</td><td>1242.5505</td><td>3</td><td>3725.6369</td><td>3725.6178</td><td>0</td><td>5.1</td><td>332</td><td>CRagged</td><td>285.37</td><td>285.37</td><td>216.69</td><td>4.07</td><td>72</td><td>>sp|P01876|IGHA1_HUMAN Immunoglobulin heavy constant alpha 1 OS=Homo sapiens OX=9606 GN=IGHA1 PE=1 SV=2</td><td>255</td><td>20201116_xd_colostrum_1ng_glyco_hcd30_etd30_compound15.12065.12065.3</td><td>controllerType=0 controllerNumber=1 scan=12065</td><td>30.4309</td></tr>
<tr><td>08344:3</td><td>1</td><td>R.LAGKPTHVN[+1054.370]VSVVMAEVDGTC[+57.021]Y.-</td><td>HexNAc(2)Hex(4)</td><td>C[+57], N[+1054]</td><td>1134.5132</td><td>3</td><td>3401.5251</td><td>3401.5121</td><td>0</td><td>3.8</td><td>332</td><td>CRagged</td><td>285.43</td><td>276.49</td><td>167.32</td><td>4.27</td><td>72</td><td>>sp|P01876|IGHA1_HUMAN Immunoglobulin heavy constant alpha 1 OS=Homo sapiens OX=9606 GN=IGHA1 PE=1 SV=2</td><td>255</td><td>20201116_xd_colostrum_1ng_glyco_hcd30_etd30_compound15.11367.11367.3</td><td>controllerType=0 controllerNumber=1 scan=11367</td><td>28.5944</td></tr>
<tr><td>11566:3</td><td>1</td><td>R.LAGKPTHVN[+2553.910]VSVVMAEVDGTC[+57.021]Y.-</td><td>HexNAc(5)Hex(5)Fuc(1)NeuAc(2)</td><td>C[+57], N[+2554]</td><td>1634.3622</td><td>3</td><td>4901.0722</td><td>4901.0518</td><td>0</td><td>4.2</td><td>332</td><td>CRagged</td><td>288.88</td><td>162.9</td><td>162.9</td><td>3.82</td><td>72</td><td>>sp|P01876|IGHA1_HUMAN Immunoglobulin heavy constant alpha 1 OS=Homo sapiens OX=9606 GN=IGHA1 PE=1 SV=2</td><td>255</td><td>20201116_xd_colostrum_1ng_glyco_hcd30_etd30_compound15.14744.14744.3</td><td>controllerType=0 controllerNumber=1 scan=14744</td><td>37.2374</td></tr>
<tr><td>09144:3</td><td>1</td><td>R.LAGKPTHVN[+1444.534]VSVVMAEVDGTC[+57.021]Y.-</td><td>HexNAc(4)Hex(3)Fuc(1)</td><td>C[+57], N[+1445]</td><td>1264.5673</td><td>3</td><td>3791.6874</td><td>3791.6759</td><td>0</td><td>3</td><td>332</td><td>CRagged</td><td>294.07</td><td>191.65</td><td>191.65</td><td>4.68</td><td>72</td><td>>sp|P01876|IGHA1_HUMAN Immunoglobulin heavy constant alpha 1 OS=Homo sapiens OX=9606 GN=IGHA1 PE=1 SV=2</td><td>255</td><td>20201116_xd_colostrum_1ng_glyco_hcd30_etd30_compound15.12207.12207.3</td><td>controllerType=0 controllerNumber=1 scan=12207</td><td>30.8423</td></tr>
<tr><td>10255:3</td><td>1</td><td>R.LAGKPTHVN[+2059.735]VSVVMAEVDGTC[+57.021]Y.-</td><td>HexNAc(4)Hex(5)Fuc(1)NeuAc(1)</td><td>C[+57], N[+2060]</td><td>1469.6357</td><td>3</td><td>4406.8925</td><td>4406.877</td><td>0</td><td>3.5</td><td>332</td><td>CRagged</td><td>306.96</td><td>141.6</td><td>141.6</td><td>3.54</td><td>72</td><td>>sp|P01876|IGHA1_HUMAN Immunoglobulin heavy constant alpha 1 OS=Homo sapiens OX=9606 GN=IGHA1 PE=1 SV=2</td><td>255</td><td>20201116_xd_colostrum_1ng_glyco_hcd30_etd30_compound15.13369.13369.3</td><td>controllerType=0 controllerNumber=1 scan=13369</td><td>33.693</td></tr>
<tr><td>09106:3</td><td>1</td><td>R.LAGKPTHVN[+1054.370]VSVVMAEVDGTC[+57.021]Y.-</td><td>HexNAc(2)Hex(4)</td><td>C[+57], N[+1054]</td><td>1134.5137</td><td>3</td><td>3401.5265</td><td>3401.5121</td><td>0</td><td>4.2</td><td>332</td><td>CRagged</td><td>330.62</td><td>311.88</td><td>203.93</td><td>4.72</td><td>72</td><td>>sp|P01876|IGHA1_HUMAN Immunoglobulin heavy constant alpha 1 OS=Homo sapiens OX=9606 GN=IGHA1 PE=1 SV=2</td><td>255</td><td>20201116_xd_colostrum_1ng_glyco_hcd30_etd30_compound15.12167.12167.3</td><td>controllerType=0 controllerNumber=1 scan=12167</td><td>30.7195</td></tr>
<tr><td>10291:4</td><td>1</td><td>R.LAGKPTHVN[+1872.651]VSVVMAEVDGTC[+57.021]Y.-</td><td>HexNAc(3)Hex(6)NeuAc(1)</td><td>C[+57], N[+1873]</td><td>1055.7068</td><td>4</td><td>4219.8055</td><td>4219.7926</td><td>0</td><td>3.1</td><td>332</td><td>CRagged</td><td>341.15</td><td>334.36</td><td>39.41</td><td>5.37</td><td>72</td><td>>sp|P01876|IGHA1_HUMAN Immunoglobulin heavy constant alpha 1 OS=Homo sapiens OX=9606 GN=IGHA1 PE=1 SV=2</td><td>255</td><td>20201116_xd_colostrum_1ng_glyco_hcd30_etd30_compound15.13407.13407.4</td><td>controllerType=0 controllerNumber=1 scan=13407</td><td>33.7971</td></tr>
<tr><td>10265:4</td><td>1</td><td>R.LAGKPTHVN[+2262.814]VSVVMAEVDGTC[+57.021]Y.-</td><td>HexNAc(5)Hex(5)Fuc(1)NeuAc(1)</td><td>C[+57], N[+2263]</td><td>1153.2475</td><td>4</td><td>4609.9681</td><td>4609.9564</td><td>0</td><td>2.5</td><td>332</td><td>CRagged</td><td>346.43</td><td>259.81</td><td>259.81</td><td>5.36</td><td>72</td><td>>sp|P01876|IGHA1_HUMAN Immunoglobulin heavy constant alpha 1 OS=Homo sapiens OX=9606 GN=IGHA1 PE=1 SV=2</td><td>255</td><td>20201116_xd_colostrum_1ng_glyco_hcd30_etd30_compound15.13380.13380.4</td><td>controllerType=0 controllerNumber=1 scan=13380</td><td>33.7332</td></tr>
<tr><td>08305:3</td><td>1</td><td>R.LAGKPTHVN[+1216.423]VSVVMAEVDGTC[+57.021]Y.-</td><td>HexNAc(2)Hex(5)</td><td>C[+57], N[+1216]</td><td>1188.5341</td><td>3</td><td>3563.5878</td><td>3563.5649</td><td>0</td><td>6.4</td><td>332</td><td>CRagged</td><td>387.06</td><td>373.6</td><td>238.89</td><td>5.65</td><td>72</td><td>>sp|P01876|IGHA1_HUMAN Immunoglobulin heavy constant alpha 1 OS=Homo sapiens OX=9606 GN=IGHA1 PE=1 SV=2</td><td>255</td><td>20201116_xd_colostrum_1ng_glyco_hcd30_etd30_compound15.11325.11325.3</td><td>controllerType=0 controllerNumber=1 scan=11325</td><td>28.4347</td></tr>
<tr><td>11639:4</td><td>1</td><td>R.LAGKPTHVN[+2204.772]VSVVMAEVDGTC[+57.021]Y.-</td><td>HexNAc(4)Hex(5)NeuAc(2)</td><td>C[+57], N[+2205]</td><td>1138.7362</td><td>4</td><td>4551.9232</td><td>4551.9145</td><td>0</td><td>1.9</td><td>332</td><td>CRagged</td><td>388.52</td><td>222.61</td><td>222.61</td><td>5.11</td><td>72</td><td>>sp|P01876|IGHA1_HUMAN Immunoglobulin heavy constant alpha 1 OS=Homo sapiens OX=9606 GN=IGHA1 PE=1 SV=2</td><td>255</td><td>20201116_xd_colostrum_1ng_glyco_hcd30_etd30_compound15.14821.14821.4</td><td>controllerType=0 controllerNumber=1 scan=14821</td><td>37.4561</td></tr>
<tr><td>10230:4</td><td>1</td><td>R.LAGKPTHVN[+2116.756]VSVVMAEVDGTC[+57.021]Y.-</td><td>HexNAc(5)Hex(5)NeuAc(1)</td><td>C[+57], N[+2117]</td><td>1116.7322</td><td>4</td><td>4463.9071</td><td>4463.8985</td><td>0</td><td>1.9</td><td>332</td><td>CRagged</td><td>439.05</td><td>271.4</td><td>271.4</td><td>5.75</td><td>72</td><td>>sp|P01876|IGHA1_HUMAN Immunoglobulin heavy constant alpha 1 OS=Homo sapiens OX=9606 GN=IGHA1 PE=1 SV=2</td><td>255</td><td>20201116_xd_colostrum_1ng_glyco_hcd30_etd30_compound15.13343.13343.4</td><td>controllerType=0 controllerNumber=1 scan=13343</td><td>33.6238</td></tr>
<tr><td>08333:4</td><td>1</td><td>R.LAGKPTHVN[+1216.423]VSVVMAEVDGTC[+57.021]Y.-</td><td>HexNAc(2)Hex(5)</td><td>C[+57], N[+1216]</td><td>891.651</td><td>4</td><td>3563.5821</td><td>3563.5649</td><td>0</td><td>4.8</td><td>332</td><td>CRagged</td><td>454.08</td><td>454.08</td><td>102.72</td><td>6.06</td><td>72</td><td>>sp|P01876|IGHA1_HUMAN Immunoglobulin heavy constant alpha 1 OS=Homo sapiens OX=9606 GN=IGHA1 PE=1 SV=2</td><td>255</td><td>20201116_xd_colostrum_1ng_glyco_hcd30_etd30_compound15.11355.11355.4</td><td>controllerType=0 controllerNumber=1 scan=11355</td><td>28.5429</td></tr>
<tr><td>11575:4</td><td>1</td><td>R.LAGKPTHVN[+2350.830]VSVVMAEVDGTC[+57.021]Y.-</td><td>HexNAc(4)Hex(5)Fuc(1)NeuAc(2)</td><td>C[+57], N[+2351]</td><td>1175.2514</td><td>4</td><td>4697.9837</td><td>4697.9724</td><td>0</td><td>2.4</td><td>332</td><td>CRagged</td><td>480.55</td><td>407.59</td><td>407.59</td><td>7.24</td><td>72</td><td>>sp|P01876|IGHA1_HUMAN Immunoglobulin heavy constant alpha 1 OS=Homo sapiens OX=9606 GN=IGHA1 PE=1 SV=2</td><td>255</td><td>20201116_xd_colostrum_1ng_glyco_hcd30_etd30_compound15.14754.14754.4</td><td>controllerType=0 controllerNumber=1 scan=14754</td><td>37.2823</td></tr>
<tr><td>11560:4</td><td>1</td><td>R.LAGKPTHVN[+2553.910]VSVVMAEVDGTC[+57.021]Y.-</td><td>HexNAc(5)Hex(5)Fuc(1)NeuAc(2)</td><td>C[+57], N[+2554]</td><td>1226.0231</td><td>4</td><td>4901.0706</td><td>4901.0518</td><td>0</td><td>3.8</td><td>332</td><td>CRagged</td><td>488.71</td><td>267.72</td><td>267.72</td><td>4.86</td><td>72</td><td>>sp|P01876|IGHA1_HUMAN Immunoglobulin heavy constant alpha 1 OS=Homo sapiens OX=9606 GN=IGHA1 PE=1 SV=2</td><td>255</td><td>20201116_xd_colostrum_1ng_glyco_hcd30_etd30_compound15.14738.14738.4</td><td>controllerType=0 controllerNumber=1 scan=14738</td><td>37.2268</td></tr>
<tr><td>07323:3</td><td>1</td><td>R.LAGKPTHVN[+1378.476]VSVVM[+15.995]AEVDGTC[+57.021]Y.-</td><td>HexNAc(2)Hex(6)</td><td>C[+57], M[+16], N[+1378]</td><td>1247.8804</td><td>3</td><td>3741.6267</td><td>3741.6127</td><td>0</td><td>3.8</td><td>332</td><td>CRagged</td><td>207.1</td><td>138.67</td><td>138.67</td><td>2.96</td><td>72</td><td>>sp|P01876|IGHA1_HUMAN Immunoglobulin heavy constant alpha 1 OS=Homo sapiens OX=9606 GN=IGHA1 PE=1 SV=2</td><td>255</td><td>20201116_xd_colostrum_1ng_glyco_hcd30_etd30_compound15.10295.10295.3</td><td>controllerType=0 controllerNumber=1 scan=10295</td><td>25.876</td></tr>
<tr><td>07620:3</td><td>1</td><td>R.LAGKPTHVN[+1216.423]VSVVM[+15.995]AEVDGTC[+57.021]Y.-</td><td>HexNAc(2)Hex(5)</td><td>C[+57], M[+16], N[+1216]</td><td>1193.8644</td><td>3</td><td>3579.5788</td><td>3579.5599</td><td>0</td><td>5.3</td><td>332</td><td>CRagged</td><td>212.35</td><td>161.76</td><td>161.76</td><td>3.03</td><td>72</td><td>>sp|P01876|IGHA1_HUMAN Immunoglobulin heavy constant alpha 1 OS=Homo sapiens OX=9606 GN=IGHA1 PE=1 SV=2</td><td>255</td><td>20201116_xd_colostrum_1ng_glyco_hcd30_etd30_compound15.10608.10608.3</td><td>controllerType=0 controllerNumber=1 scan=10608</td><td>26.656</td></tr>
<tr><td>07096:3</td><td>1</td><td>R.LAGKPTHVN[+1216.423]VSVVM[+15.995]AEVDGTC[+57.021]Y.-</td><td>HexNAc(2)Hex(5)</td><td>C[+57], M[+16], N[+1216]</td><td>1193.8631</td><td>3</td><td>3579.5747</td><td>3579.5599</td><td>0</td><td>4.2</td><td>332</td><td>CRagged</td><td>225.95</td><td>200.39</td><td>200.39</td><td>3.08</td><td>72</td><td>>sp|P01876|IGHA1_HUMAN Immunoglobulin heavy constant alpha 1 OS=Homo sapiens OX=9606 GN=IGHA1 PE=1 SV=2</td><td>255</td><td>20201116_xd_colostrum_1ng_glyco_hcd30_etd30_compound15.10058.10058.3</td><td>controllerType=0 controllerNumber=1 scan=10058</td><td>25.3285</td></tr>
<tr><td>12897:4</td><td>2</td><td>R.RTAGWNVPIGTLRPFLN[+2424.867]WTGPPEPIEAAVAR.F</td><td>HexNAc(5)Hex(6)Fuc(1)NeuAc(1)</td><td>N[+2425]</td><td>1453.6797</td><td>4</td><td>5811.6969</td><td>5811.6792</td><td>0</td><td>3.1</td><td>140</td><td>Specific</td><td>407.01</td><td>380.63</td><td>380.63</td><td>7.03</td><td>64</td><td>>sp|P02788|TRFL_HUMAN Lactotransferrin OS=Homo sapiens OX=9606 GN=LTF PE=1 SV=6</td><td>1270</td><td>20201116_xd_colostrum_1ng_glyco_hcd30_etd30_compound15.16146.16146.4</td><td>controllerType=0 controllerNumber=1 scan=16146</td><td>41.1816</td></tr>
<tr><td>13624:5</td><td>2</td><td>R.RTAGWNVPIGTLRPFLN[+2351.851]WTGPPEPIEAAVAR.F</td><td>HexNAc(4)Hex(5)Fuc(3)NeuAc(1)</td><td>N[+2352]</td><td>1148.542</td><td>5</td><td>5738.6809</td><td>5738.6628</td><td>0</td><td>3.1</td><td>140</td><td>Specific</td><td>494.35</td><td>494.35</td><td>494.35</td><td>8.08</td><td>64</td><td>>sp|P02788|TRFL_HUMAN Lactotransferrin OS=Homo sapiens OX=9606 GN=LTF PE=1 SV=6</td><td>1270</td><td>20201116_xd_colostrum_1ng_glyco_hcd30_etd30_compound15.16915.16915.5</td><td>controllerType=0 controllerNumber=1 scan=16915</td><td>43.5008</td></tr>
<tr><td>13559:4</td><td>2</td><td>R.RTAGWNVPIGTLRPFLN[+2716.983]WTGPPEPIEAAVAR.F</td><td>HexNAc(5)Hex(6)Fuc(3)NeuAc(1)</td><td>N[+2717]</td><td>1526.7062</td><td>4</td><td>6103.8029</td><td>6103.795</td><td>0</td><td>1.3</td><td>140</td><td>Specific</td><td>647.92</td><td>647.92</td><td>647.92</td><td>9.93</td><td>64</td><td>>sp|P02788|TRFL_HUMAN Lactotransferrin OS=Homo sapiens OX=9606 GN=LTF PE=1 SV=6</td><td>1270</td><td>20201116_xd_colostrum_1ng_glyco_hcd30_etd30_compound15.16847.16847.4</td><td>controllerType=0 controllerNumber=1 scan=16847</td><td>43.3313</td></tr>
<tr><td>13620:4</td><td>2</td><td>R.RTAGWNVPIGTLRPFLN[+2351.851]WTGPPEPIEAAVAR.F</td><td>HexNAc(4)Hex(5)Fuc(3)NeuAc(1)</td><td>N[+2352]</td><td>1435.4265</td><td>4</td><td>5738.6842</td><td>5738.6628</td><td>0</td><td>3.7</td><td>140</td><td>Specific</td><td>816.57</td><td>576.03</td><td>576.03</td><td>14.11</td><td>64</td><td>>sp|P02788|TRFL_HUMAN Lactotransferrin OS=Homo sapiens OX=9606 GN=LTF PE=1 SV=6</td><td>1270</td><td>20201116_xd_colostrum_1ng_glyco_hcd30_etd30_compound15.16911.16911.4</td><td>controllerType=0 controllerNumber=1 scan=16911</td><td>43.4932</td></tr>
<tr><td>13652:4</td><td>2</td><td>R.RTAGWNVPIGTLRPFLN[+2059.735]WTGPPEPIEAAVAR.F</td><td>HexNAc(4)Hex(5)Fuc(1)NeuAc(1)</td><td>N[+2060]</td><td>1362.3943</td><td>4</td><td>5446.5553</td><td>5446.547</td><td>0</td><td>1.5</td><td>140</td><td>Specific</td><td>925.62</td><td>910.62</td><td>910.62</td><td>15.93</td><td>64</td><td>>sp|P02788|TRFL_HUMAN Lactotransferrin OS=Homo sapiens OX=9606 GN=LTF PE=1 SV=6</td><td>1270</td><td>20201116_xd_colostrum_1ng_glyco_hcd30_etd30_compound15.16945.16945.4</td><td>controllerType=0 controllerNumber=1 scan=16945</td><td>43.6122</td></tr>
<tr><td>13821:4</td><td>2</td><td>R.[+42.011]TAGWNVPIGTLRPFLN[+1872.651]WTGPPEPIEAAVAR.F</td><td>HexNAc(3)Hex(6)NeuAc(1)</td><td>.[42], N[+1873]</td><td>1287.1108</td><td>4</td><td>5145.4215</td><td>5145.372</td><td>0</td><td>9.6</td><td>141</td><td>Specific</td><td>241.04</td><td>241.04</td><td>241.04</td><td>4.04</td><td>64</td><td>>sp|P02788|TRFL_HUMAN Lactotransferrin OS=Homo sapiens OX=9606 GN=LTF PE=1 SV=6</td><td>1270</td><td>20201116_xd_colostrum_1ng_glyco_hcd30_etd30_compound15.17122.17122.4</td><td>controllerType=0 controllerNumber=1 scan=17122</td><td>44.0764</td></tr>
<tr><td>14529:3</td><td>2</td><td>R.TAGWNVPIGTLRPFLN[+2205.793]WTGPPEPIEAAVAR.F</td><td>HexNAc(4)Hex(5)Fuc(2)NeuAc(1)</td><td>N[+2206]</td><td>1812.8451</td><td>3</td><td>5436.5206</td><td>5436.5038</td><td>0</td><td>3.1</td><td>141</td><td>Specific</td><td>252.36</td><td>191.37</td><td>191.37</td><td>5.25</td><td>64</td><td>>sp|P02788|TRFL_HUMAN Lactotransferrin OS=Homo sapiens OX=9606 GN=LTF PE=1 SV=6</td><td>1270</td><td>20201116_xd_colostrum_1ng_glyco_hcd30_etd30_compound15.17872.17872.2</td><td>controllerType=0 controllerNumber=1 scan=17872</td><td>46.3114</td></tr>
<tr><td>14516:3</td><td>2</td><td>R.TAGWNVPIGTLRPFLN[+2205.793]WTGPPEPIEAAVAR.F</td><td>HexNAc(4)Hex(5)Fuc(2)NeuAc(1)</td><td>N[+2206]</td><td>1812.8452</td><td>3</td><td>5436.521</td><td>5436.5038</td><td>0</td><td>3.2</td><td>141</td><td>Specific</td><td>348.03</td><td>262.15</td><td>262.15</td><td>5.82</td><td>64</td><td>>sp|P02788|TRFL_HUMAN Lactotransferrin OS=Homo sapiens OX=9606 GN=LTF PE=1 SV=6</td><td>1270</td><td>20201116_xd_colostrum_1ng_glyco_hcd30_etd30_compound15.17858.17858.3</td><td>controllerType=0 controllerNumber=1 scan=17858</td><td>46.2494</td></tr>
<tr><td>14501:3</td><td>2</td><td>R.TAGWNVPIGTLRPFLN[+2351.851]WTGPPEPIEAAVAR.F</td><td>HexNAc(4)Hex(5)Fuc(3)NeuAc(1)</td><td>N[+2352]</td><td>1861.5323</td><td>3</td><td>5582.5824</td><td>5582.5617</td><td>0</td><td>3.7</td><td>141</td><td>Specific</td><td>398.64</td><td>331.81</td><td>331.81</td><td>6.29</td><td>64</td><td>>sp|P02788|TRFL_HUMAN Lactotransferrin OS=Homo sapiens OX=9606 GN=LTF PE=1 SV=6</td><td>1270</td><td>20201116_xd_colostrum_1ng_glyco_hcd30_etd30_compound15.17842.17842.3</td><td>controllerType=0 controllerNumber=1 scan=17842</td><td>46.1979</td></tr>
<tr><td>14540:3</td><td>2</td><td>R.TAGWNVPIGTLRPFLN[+2059.735]WTGPPEPIEAAVAR.F</td><td>HexNAc(4)Hex(5)Fuc(1)NeuAc(1)</td><td>N[+2060]</td><td>1764.1602</td><td>3</td><td>5290.4662</td><td>5290.4459</td><td>0</td><td>3.8</td><td>141</td><td>Specific</td><td>412.36</td><td>306.09</td><td>306.09</td><td>5.45</td><td>64</td><td>>sp|P02788|TRFL_HUMAN Lactotransferrin OS=Homo sapiens OX=9606 GN=LTF PE=1 SV=6</td><td>1270</td><td>20201116_xd_colostrum_1ng_glyco_hcd30_etd30_compound15.17884.17884.3</td><td>controllerType=0 controllerNumber=1 scan=17884</td><td>46.3537</td></tr>
<tr><td>15368:3</td><td>2</td><td>R.TAGWNVPIGTLRPFLN[+2496.888]WTGPPEPIEAAVAR.F</td><td>HexNAc(4)Hex(5)Fuc(2)NeuAc(2)</td><td>N[+2497]</td><td>1909.8776</td><td>3</td><td>5727.6184</td><td>5727.5992</td><td>0</td><td>3.4</td><td>141</td><td>Specific</td><td>444.27</td><td>356.47</td><td>356.47</td><td>6.44</td><td>64</td><td>>sp|P02788|TRFL_HUMAN Lactotransferrin OS=Homo sapiens OX=9606 GN=LTF PE=1 SV=6</td><td>1270</td><td>20201116_xd_colostrum_1ng_glyco_hcd30_etd30_compound15.18764.18764.3</td><td>controllerType=0 controllerNumber=1 scan=18764</td><td>49.2781</td></tr>
<tr><td>14536:5</td><td>2</td><td>R.TAGWNVPIGTLRPFLN[+2498.904]WTGPPEPIEAAVAR.F</td><td>HexNAc(6)Hex(7)Fuc(1)</td><td>N[+2499]</td><td>1146.7327</td><td>5</td><td>5729.6342</td><td>5729.6148</td><td>0</td><td>3.4</td><td>141</td><td>Specific</td><td>534.66</td><td>531.32</td><td>531.32</td><td>8.22</td><td>64</td><td>>sp|P02788|TRFL_HUMAN Lactotransferrin OS=Homo sapiens OX=9606 GN=LTF PE=1 SV=6</td><td>1270</td><td>20201116_xd_colostrum_1ng_glyco_hcd30_etd30_compound15.17879.17879.5</td><td>controllerType=0 controllerNumber=1 scan=17879</td><td>46.3362</td></tr>
<tr><td>14376:4</td><td>2</td><td>R.TAGWNVPIGTLRPFLN[+2863.041]WTGPPEPIEAAVAR.F</td><td>HexNAc(5)Hex(6)Fuc(4)NeuAc(1)</td><td>N[+2863]</td><td>1524.197</td><td>4</td><td>6093.7663</td><td>6093.7518</td><td>0</td><td>2.4</td><td>141</td><td>Specific</td><td>648.7</td><td>648.7</td><td>648.7</td><td>10.89</td><td>64</td><td>>sp|P02788|TRFL_HUMAN Lactotransferrin OS=Homo sapiens OX=9606 GN=LTF PE=1 SV=6</td><td>1270</td><td>20201116_xd_colostrum_1ng_glyco_hcd30_etd30_compound15.17710.17710.4</td><td>controllerType=0 controllerNumber=1 scan=17710</td><td>45.8244</td></tr>
<tr><td>15337:4</td><td>2</td><td>R.TAGWNVPIGTLRPFLN[+2496.888]WTGPPEPIEAAVAR.F</td><td>HexNAc(4)Hex(5)Fuc(2)NeuAc(2)</td><td>N[+2497]</td><td>1432.6587</td><td>4</td><td>5727.6129</td><td>5727.5992</td><td>0</td><td>2.4</td><td>141</td><td>Specific</td><td>706.7</td><td>691.62</td><td>691.62</td><td>12.89</td><td>64</td><td>>sp|P02788|TRFL_HUMAN Lactotransferrin OS=Homo sapiens OX=9606 GN=LTF PE=1 SV=6</td><td>1270</td><td>20201116_xd_colostrum_1ng_glyco_hcd30_etd30_compound15.18730.18730.4</td><td>controllerType=0 controllerNumber=1 scan=18730</td><td>49.1372</td></tr>
<tr><td>14431:4</td><td>2</td><td>R.TAGWNVPIGTLRPFLN[+2570.925]WTGPPEPIEAAVAR.F</td><td>HexNAc(5)Hex(6)Fuc(2)NeuAc(1)</td><td>N[+2571]</td><td>1451.1704</td><td>4</td><td>5801.6598</td><td>5801.636</td><td>0</td><td>4.1</td><td>141</td><td>Specific</td><td>739.94</td><td>723.39</td><td>723.39</td><td>12.3</td><td>64</td><td>>sp|P02788|TRFL_HUMAN Lactotransferrin OS=Homo sapiens OX=9606 GN=LTF PE=1 SV=6</td><td>1270</td><td>20201116_xd_colostrum_1ng_glyco_hcd30_etd30_compound15.17768.17768.4</td><td>controllerType=0 controllerNumber=1 scan=17768</td><td>45.9738</td></tr>
<tr><td>13840:4</td><td>2</td><td>R.TAGWNVPIGTLRPFLN[+1768.640]WTGPPEPIEAAVAR.F</td><td>HexNAc(4)Hex(5)Fuc(1)</td><td>N[+1769]</td><td>1250.5966</td><td>4</td><td>4999.3644</td><td>4999.3504</td><td>0</td><td>2.8</td><td>141</td><td>Specific</td><td>749.67</td><td>749.67</td><td>749.67</td><td>12.89</td><td>64</td><td>>sp|P02788|TRFL_HUMAN Lactotransferrin OS=Homo sapiens OX=9606 GN=LTF PE=1 SV=6</td><td>1270</td><td>20201116_xd_colostrum_1ng_glyco_hcd30_etd30_compound15.17142.17142.4</td><td>controllerType=0 controllerNumber=1 scan=17142</td><td>44.1314</td></tr>
<tr><td>14649:4</td><td>2</td><td>R.TAGWNVPIGTLRPFLN[+2351.851]WTGPPEPIEAAVAR.F</td><td>HexNAc(4)Hex(5)Fuc(3)NeuAc(1)</td><td>N[+2352]</td><td>1396.3993</td><td>4</td><td>5582.5754</td><td>5582.5617</td><td>0</td><td>2.4</td><td>141</td><td>Specific</td><td>753.03</td><td>744.3</td><td>744.3</td><td>13.89</td><td>64</td><td>>sp|P02788|TRFL_HUMAN Lactotransferrin OS=Homo sapiens OX=9606 GN=LTF PE=1 SV=6</td><td>1270</td><td>20201116_xd_colostrum_1ng_glyco_hcd30_etd30_compound15.17998.17998.4</td><td>controllerType=0 controllerNumber=1 scan=17998</td><td>46.6682</td></tr>
<tr><td>14523:5</td><td>2</td><td>R.TAGWNVPIGTLRPFLN[+2205.793]WTGPPEPIEAAVAR.F</td><td>HexNAc(4)Hex(5)Fuc(2)NeuAc(1)</td><td>N[+2206]</td><td>1088.1101</td><td>5</td><td>5436.5214</td><td>5436.5038</td><td>0</td><td>3.3</td><td>141</td><td>Specific</td><td>806.52</td><td>804.48</td><td>804.48</td><td>14.3</td><td>64</td><td>>sp|P02788|TRFL_HUMAN Lactotransferrin OS=Homo sapiens OX=9606 GN=LTF PE=1 SV=6</td><td>1270</td><td>20201116_xd_colostrum_1ng_glyco_hcd30_etd30_compound15.17865.17865.5</td><td>controllerType=0 controllerNumber=1 scan=17865</td><td>46.2784</td></tr>
<tr><td>14512:4</td><td>2</td><td>R.TAGWNVPIGTLRPFLN[+2205.793]WTGPPEPIEAAVAR.F</td><td>HexNAc(4)Hex(5)Fuc(2)NeuAc(1)</td><td>N[+2206]</td><td>1359.8861</td><td>4</td><td>5436.5226</td><td>5436.5038</td><td>0</td><td>3.5</td><td>141</td><td>Specific</td><td>861.49</td><td>851.1</td><td>851.1</td><td>15.3</td><td>64</td><td>>sp|P02788|TRFL_HUMAN Lactotransferrin OS=Homo sapiens OX=9606 GN=LTF PE=1 SV=6</td><td>1270</td><td>20201116_xd_colostrum_1ng_glyco_hcd30_etd30_compound15.17854.17854.4</td><td>controllerType=0 controllerNumber=1 scan=17854</td><td>[illegible]</td></tr>
<tr><td>14508:4</td><td>2</td><td>R.TAGWNVPIGTLRPFLN[+2205.793]WTGPPEPIEAAVAR.F</td><td>HexNAc(4)Hex(5)Fuc(2)NeuAc(1)</td><td>N[+2206]</td><td>1359.8888</td><td>4</td><td>5436.5334</td><td>5436.5038</td><td>0</td><td>5.4</td><td>141</td><td>Specific</td><td>868.95</td><td>831.82</td><td>831.82</td><td>15.09</td><td>64</td><td>>sp|P02788|TRFL_HUMAN Lactotransferrin OS=Homo sapiens OX=9606 GN=LTF PE=1 SV=6</td><td>1270</td><td>20201116_xd_colostrum_1ng_glyco_hcd30_etd30_compound15.17849.17849.2</td><td>controllerType=0 controllerNumber=1 scan=17849</td><td>[illegible]</td></tr>
<tr><td>15323:4</td><td>2</td><td>R.TAGWNVPIGTLRPFLN[+2350.830]WTGPPEPIEAAVAR.F</td><td>HexNAc(4)Hex(5)Fuc(1)NeuAc(2)</td><td>N[+2351]</td><td>1396.1464</td><td>4</td><td>5581.5636</td><td>5581.5413</td><td>0</td><td>4</td><td>141</td><td>Specific</td><td>874.43</td><td>857.11</td><td>857.11</td><td>15.3</td><td>64</td><td>>sp|P02788|TRFL_HUMAN Lactotransferrin OS=Homo sapiens OX=9606 GN=LTF PE=1 SV=6</td><td>1270</td><td>20201116_xd_colostrum_1ng_glyco_hcd30_etd30_compound15.18715.18715.4</td><td>controllerType=0 controllerNumber=1 scan=18715</td><td>49.0867</td></tr>
<tr><td>14539:4</td><td>2</td><td>R.TAGWNVPIGTLRPFLN[+2059.735]WTGPPEPIEAAVAR.F</td><td>HexNAc(4)Hex(5)Fuc(1)NeuAc(1)</td><td>N[+2060]</td><td>1323.3722</td><td>4</td><td>5290.467</td><td>5290.4459</td><td>0</td><td>4</td><td>141</td><td>Specific</td><td>903.41</td><td>859.95</td><td>859.95</td><td>16.3</td><td>64</td><td>>sp|P02788|TRFL_HUMAN Lactotransferrin OS=Homo sapiens OX=9606 GN=LTF PE=1 SV=6</td><td>1270</td><td>20201116_xd_colostrum_1ng_glyco_hcd30_etd30_compound15.17883.17883.4</td><td>controllerType=0 controllerNumber=1 scan=17883</td><td>46.3509</td></tr>
<tr><td>14545:5</td><td>2</td><td>R.TAGWNVPIGTLRPFLN[+2059.735]WTGPPEPIEAAVAR.F</td><td>HexNAc(4)Hex(5)Fuc(1)NeuAc(1)</td><td>N[+2060]</td><td>1058.8988</td><td>5</td><td>5290.4649</td><td>5290.4459</td><td>0</td><td>3.6</td><td>141</td><td>Specific</td><td>921.64</td><td>921.64</td><td>921.64</td><td>16.3</td><td>64</td><td>>sp|P02788|TRFL_HUMAN Lactotransferrin OS=Homo sapiens OX=9606 GN=LTF PE=1 SV=6</td><td>1270</td><td>20201116_xd_colostrum_1ng_glyco_hcd30_etd30_compound15.17889.17889.5</td><td>controllerType=0 controllerNumber=1 scan=17889</td><td>46.3697</td></tr>
<tr><td>14496:4</td><td>2</td><td>R.TAGWNVPIGTLRPFLN[+2351.851]WTGPPEPIEAAVAR.F</td><td>HexNAc(4)Hex(5)Fuc(3)NeuAc(1)</td><td>N[+2352]</td><td>1396.4037</td><td>4</td><td>5582.5929</td><td>5582.5617</td><td>0</td><td>5.6</td><td>141</td><td>Specific</td><td>965.21</td><td>965.21</td><td>965.21</td><td>17.09</td><td>64</td><td>>sp|P02788|TRFL_HUMAN Lactotransferrin OS=Homo sapiens OX=9606 GN=LTF PE=1 SV=6</td><td>1270</td><td>20201116_xd_colostrum_1ng_glyco_hcd30_etd30_compound15.17837.17837.4</td><td>controllerType=0 controllerNumber=1 scan=17837</td><td>46.1868</td></tr>
<tr><td>14503:5</td><td>2</td><td>R.TAGWNVPIGTLRPFLN[+2351.851]WTGPPEPIEAAVAR.F</td><td>HexNAc(4)Hex(5)Fuc(3)NeuAc(1)</td><td>N[+2352]</td><td>1117.3229</td><td>5</td><td>5582.5853</td><td>5582.5617</td><td>0</td><td>4.2</td><td>141</td><td>Specific</td><td>1041.68</td><td>964.93</td><td>964.93</td><td>18.3</td><td>64</td><td>>sp|P02788|TRFL_HUMAN Lactotransferrin OS=Homo sapiens OX=9606 GN=LTF PE=1 SV=6</td><td>1270</td><td>20201116_xd_colostrum_1ng_glyco_hcd30_etd30_compound15.17844.17844.5</td><td>controllerType=0 controllerNumber=1 scan=17844</td><td>46.2087</td></tr>
<tr><td>12615:3</td><td>2</td><td>R.TAGWNIPMGLLFN[+2351.851]QTGSC[+57.021]K.F</td><td>HexNAc(4)Hex(5)Fuc(3)NeuAc(1)</td><td>C[+57], N[+2352]</td><td>1482.9704</td><td>3</td><td>4446.8966</td><td>4446.8607</td><td>0</td><td>8.1</td><td>485</td><td>Specific</td><td>202.4</td><td>180.55</td><td>180.55</td><td>3.12</td><td>64</td><td>>sp|P02788|TRFL_HUMAN Lactotransferrin OS=Homo sapiens OX=9606 GN=LTF PE=1 SV=6</td><td>1270</td><td>20201116_xd_colostrum_1ng_glyco_hcd30_etd30_compound15.15848.15848.3</td><td>controllerType=0 controllerNumber=1 scan=15848</td><td>40.2637</td></tr>
<tr><td>14158:3</td><td>2</td><td>R.TAGWNIPMGLLFN[+1856.656]QTGSC[+57.021]K.F</td><td>HexNAc(3)Hex(5)Fuc(1)NeuAc(1)</td><td>C[+57], N[+1857]</td><td>1317.8985</td><td>3</td><td>3951.6809</td><td>3951.6655</td><td>0</td><td>3.9</td><td>485</td><td>Specific</td><td>265.67</td><td>172.36</td><td>172.36</td><td>5.39</td><td>64</td><td>>sp|P02788|TRFL_HUMAN Lactotransferrin OS=Homo sapiens OX=9606 GN=LTF PE=1 SV=6</td><td>1270</td><td>20201116_xd_colostrum_1ng_glyco_hcd30_etd30_compound15.17479.17479.3</td><td>controllerType=0 controllerNumber=1 scan=17479</td><td>45.1903</td></tr>
<tr><td>14026:3</td><td>2</td><td>R.TAGWNIPMGLLFN[+2351.851]QTGSC[+57.021]K.F</td><td>HexNAc(4)Hex(5)Fuc(3)NeuAc(1)</td><td>C[+57], N[+2352]</td><td>1482.9636</td><td>3</td><td>4446.8761</td><td>4446.8607</td><td>0</td><td>3.5</td><td>485</td><td>Specific</td><td>338</td><td>236.04</td><td>236.04</td><td>4.88</td><td>64</td><td>>sp|P02788|TRFL_HUMAN Lactotransferrin OS=Homo sapiens OX=9606 GN=LTF PE=1 SV=6</td><td>1270</td><td>20201116_xd_colostrum_1ng_glyco_hcd30_etd30_compound15.17339.17339.3</td><td>controllerType=0 controllerNumber=1 scan=17339</td><td>44.7358</td></tr>
<tr><td>15499:3</td><td>2</td><td>R.TAGWNIPMGLLFN[+2204.772]QTGSC[+57.021]K.F</td><td>HexNAc(4)Hex(5)NeuAc(2)</td><td>C[+57], N[+2205]</td><td>1433.9361</td><td>3</td><td>4299.7937</td><td>4299.7824</td><td>0</td><td>2.6</td><td>485</td><td>Specific</td><td>349.92</td><td>231.14</td><td>231.14</td><td>6.36</td><td>64</td><td>>sp|P02788|TRFL_HUMAN Lactotransferrin OS=Homo sapiens OX=9606 GN=LTF PE=1 SV=6</td><td>1270</td><td>20201116_xd_colostrum_1ng_glyco_hcd30_etd30_compound15.18906.18906.3</td><td>controllerType=0 controllerNumber=1 scan=18906</td><td>49.8191</td></tr>
<tr><td>15374:3</td><td>2</td><td>R.TAGWNIPMGLLFN[+2350.830]QTGSC[+57.021]K.F</td><td>HexNAc(4)Hex(5)Fuc(1)NeuAc(2)</td><td>C[+57], N[+2351]</td><td>1482.624</td><td>3</td><td>4445.8573</td><td>4445.8403</td><td>0</td><td>3.8</td><td>485</td><td>Specific</td><td>371.98</td><td>233.55</td><td>233.55</td><td>5.8</td><td>64</td><td>>sp|P02788|TRFL_HUMAN Lactotransferrin OS=Homo sapiens OX=9606 GN=LTF PE=1 SV=6</td><td>1270</td><td>20201116_xd_colostrum_1ng_glyco_hcd30_etd30_compound15.18771.18771.3</td><td>controllerType=0 controllerNumber=1 scan=18771</td><td>49.3057</td></tr>
<tr><td>15384:4</td><td>2</td><td>R.TAGWNIPMGLLFN[+2350.830]QTGSC[+57.021]K.F</td><td>HexNAc(4)Hex(5)Fuc(1)NeuAc(2)</td><td>C[+57], N[+2351]</td><td>1112.2189</td><td>4</td><td>4445.8538</td><td>4445.8403</td><td>0</td><td>3</td><td>485</td><td>Specific</td><td>421.18</td><td>365.71</td><td>365.71</td><td>8.01</td><td>64</td><td>>sp|P02788|TRFL_HUMAN Lactotransferrin OS=Homo sapiens OX=9606 GN=LTF PE=1 SV=6</td><td>1270</td><td>20201116_xd_colostrum_1ng_glyco_hcd30_etd30_compound15.18781.18781.4</td><td>controllerType=0 controllerNumber=1 scan=18781</td><td>49.341</td></tr>
<tr><td>14198:3</td><td>2</td><td>R.TAGWNIPMGLLFN[+1872.651]QTGSC[+57.021]K.F</td><td>HexNAc(3)Hex(6)NeuAc(1)</td><td>C[+57], N[+1873]</td><td>1323.2287</td><td>3</td><td>3967.6715</td><td>3967.6604</td><td>0</td><td>2.8</td><td>485</td><td>Specific</td><td>434.56</td><td>309.93</td><td>223.27</td><td>6.59</td><td>64</td><td>>sp|P02788|TRFL_HUMAN Lactotransferrin OS=Homo sapiens OX=9606 GN=LTF PE=1 SV=6</td><td>1270</td><td>20201116_xd_colostrum_1ng_glyco_hcd30_etd30_compound15.17521.17521.3</td><td>controllerType=0 controllerNumber=1 scan=17521</td><td>45.3342</td></tr>
<tr><td>14066:3</td><td>2</td><td>R.TAGWNIPMGLLFN[+2059.735]QTGSC[+57.021]K.F</td><td>HexNAc(4)Hex(5)Fuc(1)NeuAc(1)</td><td>C[+57], N[+2060]</td><td>1385.5884</td><td>3</td><td>4154.7508</td><td>4154.7449</td><td>0</td><td>1.4</td><td>485</td><td>Specific</td><td>456.27</td><td>303.04</td><td>303.04</td><td>5.91</td><td>64</td><td>>sp|P02788|TRFL_HUMAN Lactotransferrin OS=Homo sapiens OX=9606 GN=LTF PE=1 SV=6</td><td>1270</td><td>20201116_xd_colostrum_1ng_glyco_hcd30_etd30_compound15.17381.17381.3</td><td>controllerType=0 controllerNumber=1 scan=17381</td><td>44.8681</td></tr>
<tr><td>14027:3</td><td>2</td><td>R.TAGWNIPMGLLFN[+2205.793]QTGSC[+57.021]K.F</td><td>HexNAc(4)Hex(5)Fuc(2)NeuAc(1)</td><td>C[+57], N[+2206]</td><td>1434.2772</td><td>3</td><td>4300.8169</td><td>4300.8028</td><td>0</td><td>3.3</td><td>485</td><td>Specific</td><td>465.02</td><td>310.47</td><td>310.47</td><td>6.09</td><td>64</td><td>>sp|P02788|TRFL_HUMAN Lactotransferrin OS=Homo sapiens OX=9606 GN=LTF PE=1 SV=6</td><td>1270</td><td>20201116_xd_colostrum_1ng_glyco_hcd30_etd30_compound15.17340.17340.3</td><td>controllerType=0 controllerNumber=1 scan=17340</td><td>44.7391</td></tr>
<tr><td>15175:4</td><td>2</td><td>R.TAGWNIPMGLLFN[+2204.772]QTGSC[+57.021]KFDEYFSQSC[+57.021]APGSDPR.S</td><td>HexNAc(4)Hex(5)NeuAc(2)</td><td>C[+57]*2, N[+2205]</td><td>1536.6393</td><td>4</td><td>6143.5353</td><td>6143.5295</td><td>0</td><td>0.9</td><td>485</td><td>Specific</td><td>226.99</td><td>183.44</td><td>183.44</td><td>3.84</td><td>64</td><td>>sp|P02788|TRFL_HUMAN Lactotransferrin OS=Homo sapiens OX=9606 GN=LTF PE=1 SV=6</td><td>1270</td><td>20201116_xd_colostrum_1ng_glyco_hcd30_etd30_compound15.18558.18558.4</td><td>controllerType=0 controllerNumber=1 scan=18558</td><td>48.5692</td></tr>
<tr><td>14203:4</td><td>2</td><td>R.TAGWNIPMGLLFN[+2351.851]QTGSC[+57.021]KFDEYFSQSC[+57.021]APGSDPR.S</td><td>HexNAc(4)Hex(5)Fuc(3)NeuAc(1)</td><td>C[+57]*2, N[+2352]</td><td>1573.412</td><td>4</td><td>6290.6261</td><td>6290.6079</td><td>0</td><td>2.9</td><td>485</td><td>Specific</td><td>334.42</td><td>307.78</td><td>307.78</td><td>6.76</td><td>64</td><td>>sp|P02788|TRFL_HUMAN Lactotransferrin OS=Homo sapiens OX=9606 GN=LTF PE=1 SV=6</td><td>1270</td><td>20201116_xd_colostrum_1ng_glyco_hcd30_etd30_compound15.17527.17527.4</td><td>controllerType=0 controllerNumber=1 scan=17527</td><td>45.3537</td></tr>
<tr><td>14375:4</td><td>2</td><td>R.TAGWNIPMGLLFN[+1710.598]QTGSC[+57.021]KFDEYFSQSC[+57.021]APGSDPR.S</td><td>HexNAc(3)Hex(5)NeuAc(1)</td><td>C[+57]*2, N[+1711]</td><td>1413.094</td><td>4</td><td>5649.3542</td><td>5649.3548</td><td>0</td><td>-0.1</td><td>485</td><td>Specific</td><td>373.18</td><td>280.29</td><td>280.29</td><td>4.26</td><td>64</td><td>>sp|P02788|TRFL_HUMAN Lactotransferrin OS=Homo sapiens OX=9606 GN=LTF PE=1 SV=6</td><td>1270</td><td>20201116_xd_colostrum_1ng_glyco_hcd30_etd30_compound15.17709.17709.4</td><td>controllerType=0 controllerNumber=1 scan=17709</td><td>45.818</td></tr>
<tr><td>15154:4</td><td>2</td><td>R.TAGWNIPMGLLFN[+2204.772]QTGSC[+57.021]KFDEYFSQSC[+57.021]APGSDPR.S</td><td>HexNAc(4)Hex(5)NeuAc(2)</td><td>C[+57]*2, N[+2205]</td><td>1536.6401</td><td>4</td><td>6143.5387</td><td>6143.5295</td><td>0</td><td>1.5</td><td>485</td><td>Specific</td><td>448.76</td><td>404.67</td><td>336.23</td><td>7.6</td><td>64</td><td>>sp|P02788|TRFL_HUMAN Lactotransferrin OS=Homo sapiens OX=9606 GN=LTF PE=1 SV=6</td><td>1270</td><td>20201116_xd_colostrum_1ng_glyco_hcd30_etd30_compound15.18536.18536.4</td><td>controllerType=0 controllerNumber=1 scan=18536</td><td>48.509</td></tr>
<tr><td>15101:4</td><td>2</td><td>R.TAGWNIPMGLLFN[+2350.830]QTGSC[+57.021]KFDEYFSQSC[+57.021]APGSDPR.S</td><td>HexNAc(4)Hex(5)Fuc(1)NeuAc(2)</td><td>C[+57]*2, N[+2351]</td><td>1573.1589</td><td>4</td><td>6289.6139</td><td>6289.5875</td><td>0</td><td>4.2</td><td>485</td><td>Specific</td><td>519.24</td><td>489.19</td><td>489.19</td><td>7.92</td><td>64</td><td>>sp|P02788|TRFL_HUMAN Lactotransferrin OS=Homo sapiens OX=9606 GN=LTF PE=1 SV=6</td><td>1270</td><td>20201116_xd_colostrum_1ng_glyco_hcd30_etd30_compound15.18480.18480.4</td><td>controllerType=0 controllerNumber=1 scan=18480</td><td>48.3344</td></tr>
<tr><td>13522:4</td><td>2</td><td>R.TAGWNIPMGLLFN[+1768.640]QTGSC[+57.021]KFDEYFSQSC[+57.021]APGSDPR.S</td><td>HexNAc(4)Hex(5)Fuc(1)</td><td>C[+57]*2, N[+1769]</td><td>1427.6115</td><td>4</td><td>5707.424</td><td>5707.3966</td><td>0</td><td>4.8</td><td>485</td><td>Specific</td><td>726.2</td><td>639.33</td><td>639.33</td><td>11.9</td><td>64</td><td>>sp|P02788|TRFL_HUMAN Lactotransferrin OS=Homo sapiens OX=9606 GN=LTF PE=1 SV=6</td><td>1270</td><td>20201116_xd_colostrum_1ng_glyco_hcd30_etd30_compound15.16808.16808.4</td><td>controllerType=0 controllerNumber=1 scan=16808</td><td>43.2134</td></tr>
<tr><td>07577:3</td><td>2</td><td>R.N[+2351.851]GSDC[+57.021]PDKFC[+57.021]LFQSETK.N</td><td>HexNAc(4)Hex(5)Fuc(3)NeuAc(1)</td><td>C[+57]*2, N[+2352]</td><td>1462.2526</td><td>3</td><td>4384.7433</td><td>4384.7247</td><td>0</td><td>4.3</td><td>642</td><td>Specific</td><td>308.28</td><td>221.15</td><td>221.15</td><td>4.1</td><td>64</td><td>>sp|P02788|TRFL_HUMAN Lactotransferrin OS=Homo sapiens OX=9606 GN=LTF PE=1 SV=6</td><td>1270</td><td>20201116_xd_colostrum_1ng_glyco_hcd30_etd30_compound15.10563.10563.3</td><td>controllerType=0 controllerNumber=1 scan=10563</td><td>26.5417</td></tr>
<tr><td>08267:3</td><td>3</td><td>K.TVLTPATNHMGN[+1378.476]VTFTIPANR.E</td><td>HexNAc(2)Hex(6)</td><td>N[+1378]</td><td>1211.8864</td><td>3</td><td>3633.6447</td><td>3633.6358</td><td>0</td><td>2.4</td><td>74</td><td>Specific</td><td>496.82</td><td>386.25</td><td>157.17</td><td>7.95</td><td>47</td><td>>sp|P01024|CO3_HUMAN Complement C3 OS=Homo sapiens OX=9606 GN=C3 PE=1 SV=2</td><td>492</td><td>20201116_xd_colostrum_1ng_glyco_hcd30_etd30_compound15.11285.11285.3</td><td>controllerType=0 controllerNumber=1 scan=11285</td><td>28.3045</td></tr>
<tr><td>08240:3</td><td>3</td><td>K.TVLTPATNHMGN[+1540.529]VTFTIPANR.E</td><td>HexNAc(2)Hex(7)</td><td>N[+1541]</td><td>1265.9035</td><td>3</td><td>3795.6959</td><td>3795.6886</td><td>0</td><td>1.9</td><td>74</td><td>Specific</td><td>507.76</td><td>390.32</td><td>189.46</td><td>7.95</td><td>47</td><td>>sp|P01024|CO3_HUMAN Complement C3 OS=Homo sapiens OX=9606 GN=C3 PE=1 SV=2</td><td>492</td><td>20201116_xd_colostrum_1ng_glyco_hcd30_etd30_compound15.11256.11256.3</td><td>controllerType=0 controllerNumber=1 scan=11256</td><td>28.2188</td></tr>
<tr><td>11819:3</td><td>4</td><td>R.LN[+1216.423]YSLPTGQWVGVQLPR.N</td><td>HexNAc(2)Hex(5)</td><td>N[+1216]</td><td>1048.8314</td><td>3</td><td>3144.4798</td><td>3144.4617</td><td>0</td><td>5.7</td><td>1017</td><td>Specific</td><td>382.78</td><td>342.36</td><td>342.36</td><td>5.3</td><td>37</td><td>>sp|P24821|TENA_HUMAN Tenascin OS=Homo sapiens OX=9606 GN=TNC PE=1 SV=3</td><td>486</td><td>20201116_xd_colostrum_1ng_glyco_hcd30_etd30_compound15.15011.15011.3</td><td>controllerType=0 controllerNumber=1 scan=15011</td><td>38.0303</td></tr>
<tr><td>11846:2</td><td>4</td><td>R.LN[+1216.423]YSLPTGQWVGVQLPR.N</td><td>HexNAc(2)Hex(5)</td><td>N[+1216]</td><td>1572.7397</td><td>2</td><td>3144.4721</td><td>3144.4617</td><td>0</td><td>3.3</td><td>1017</td><td>Specific</td><td>531.63</td><td>473.41</td><td>473.41</td><td>6.85</td><td>37</td><td>>sp|P24821|TENA_HUMAN Tenascin OS=Homo sapiens OX=9606 GN=TNC PE=1 SV=3</td><td>486</td><td>20201116_xd_colostrum_1ng_glyco_hcd30_etd30_compound15.15040.15040.2</td><td>controllerType=0 controllerNumber=1 scan=15040</td><td>38.1312</td></tr>
<tr><td>11831:3</td><td>4</td><td>R.LN[+1216.423]YSLPTGQWVGVQLPR.N</td><td>HexNAc(2)Hex(5)</td><td>N[+1216]</td><td>1048.8327</td><td>3</td><td>3144.4835</td><td>3144.4617</td><td>0</td><td>6.9</td><td>1017</td><td>Specific</td><td>626.81</td><td>577.38</td><td>577.38</td><td>8.71</td><td>37</td><td>>sp|P24821|TENA_HUMAN Tenascin OS=Homo sapiens OX=9606 GN=TNC PE=1 SV=3</td><td>486</td><td>20201116_xd_colostrum_1ng_glyco_hcd30_etd30_compound15.15024.15024.3</td><td>controllerType=0 controllerNumber=1 scan=15024</td><td>38.084</td></tr>
<tr><td>03583:3</td><td>4</td><td>R.N[+2351.851]TTSYVLR.G</td><td>HexNAc(4)Hex(5)Fuc(3)NeuAc(1)</td><td>N[+2352]</td><td>1102.4606</td><td>3</td><td>3305.3672</td><td>3305.3558</td><td>0</td><td>3.5</td><td>1034</td><td>Specific</td><td>205.09</td><td>103.7</td><td>103.7</td><td>2.85</td><td>37</td><td>>sp|P24821|TENA_HUMAN Tenascin OS=Homo sapiens OX=9606 GN=TNC PE=1 SV=3</td><td>486</td><td>20201116_xd_colostrum_1ng_glyco_hcd30_etd30_compound15.6391.6391.3</td><td>controllerType=0 controllerNumber=1 scan=6391</td><td>17.5091</td></tr>
<tr><td>03730:3</td><td>4</td><td>R.N[+2059.735]TTSYVLR.G</td><td>HexNAc(4)Hex(5)Fuc(1)NeuAc(1)</td><td>N[+2060]</td><td>1005.0865</td><td>3</td><td>3013.245</td><td>3013.24</td><td>0</td><td>1.7</td><td>1034</td><td>Specific</td><td>275.69</td><td>171.64</td><td>171.64</td><td>5</td><td>37</td><td>>sp|P24821|TENA_HUMAN Tenascin OS=Homo sapiens OX=9606 GN=TNC PE=1 SV=3</td><td>486</td><td>20201116_xd_colostrum_1ng_glyco_hcd30_etd30_compound15.6544.6544.3</td><td>controllerType=0 controllerNumber=1 scan=6544</td><td>17.8347</td></tr>
<tr><td>13103:3</td><td>5</td><td>R.GLSFDVSLEVSQGPGLLN[+2862.021]DTK.V</td><td>HexNAc(5)Hex(6)Fuc(2)NeuAc(2)</td><td>N[+2862]</td><td>1680.0603</td><td>3</td><td>5038.1662</td><td>5038.1337</td><td>0</td><td>6.5</td><td>118</td><td>Specific</td><td>346.36</td><td>173.26</td><td>173.26</td><td>5.1</td><td>60</td><td>>sp|P01833|PIGR_HUMAN Polymeric immunoglobulin receptor OS=Homo sapiens OX=9606 GN=PIGR PE=1 SV=4</td><td>902</td><td>20201116_xd_colostrum_1ng_glyco_hcd30_etd30_compound15.16364.16364.3</td><td>controllerType=0 controllerNumber=1 scan=16364</td><td>41.8505</td></tr>
<tr><td>13308:3</td><td>5</td><td>R.GLSFDVSLEVSQGPGLLN[+2351.851]DTK.V</td><td>HexNAc(4)Hex(5)Fuc(3)NeuAc(1)</td><td>N[+2352]</td><td>1509.9985</td><td>3</td><td>4527.9809</td><td>4527.964</td><td>0</td><td>3.7</td><td>118</td><td>Specific</td><td>446.5</td><td>245.77</td><td>245.77</td><td>5.96</td><td>60</td><td>>sp|P01833|PIGR_HUMAN Polymeric immunoglobulin receptor OS=Homo sapiens OX=9606 GN=PIGR PE=1 SV=4</td><td>902</td><td>20201116_xd_colostrum_1ng_glyco_hcd30_etd30_compound15.16580.16580.3</td><td>controllerType=0 controllerNumber=1 scan=16580</td><td>42.4692</td></tr>
<tr><td>14634:3</td><td>5</td><td>R.GLSFDVSLEVSQGPGLLN[+2350.830]DTK.V</td><td>HexNAc(4)Hex(5)Fuc(1)NeuAc(2)</td><td>N[+2351]</td><td>1509.6585</td><td>3</td><td>4526.961</td><td>4526.9436</td><td>0</td><td>3.8</td><td>118</td><td>Specific</td><td>510.69</td><td>357.12</td><td>357.12</td><td>6.43</td><td>60</td><td>>sp|P01833|PIGR_HUMAN Polymeric immunoglobulin receptor OS=Homo sapiens OX=9606 GN=PIGR PE=1 SV=4</td><td>902</td><td>20201116_xd_colostrum_1ng_glyco_hcd30_etd30_compound15.17983.17983.3</td><td>controllerType=0 controllerNumber=1 scan=17983</td><td>46.625</td></tr>
<tr><td>14884:4</td><td>5</td><td>R.GLSFDVSLEVSQGPGLLN[+2350.830]DTKVYTVDLGR.T</td><td>HexNAc(4)Hex(5)Fuc(1)NeuAc(2)</td><td>N[+2351]</td><td>1358.3658</td><td>4</td><td>5430.4416</td><td>5430.425</td><td>0</td><td>3.1</td><td>118</td><td>Specific</td><td>257.66</td><td>245.74</td><td>245.74</td><td>4.99</td><td>60</td><td>>sp|P01833|PIGR_HUMAN Polymeric immunoglobulin receptor OS=Homo sapiens OX=9606 GN=PIGR PE=1 SV=4</td><td>902</td><td>20201116_xd_colostrum_1ng_glyco_hcd30_etd30_compound15.18247.18247.4</td><td>controllerType=0 controllerNumber=1 scan=18247</td><td>47.4503</td></tr>
<tr><td>13975:3</td><td>5</td><td>R.GLSFDVSLEVSQGPGLLN[+2351.851]DTKVYTVDLGR.T</td><td>HexNAc(4)Hex(5)Fuc(3)NeuAc(1)</td><td>N[+2352]</td><td>1811.1577</td><td>3</td><td>5431.4585</td><td>5431.4454</td><td>0</td><td>2.4</td><td>118</td><td>Specific</td><td>395.31</td><td>295.49</td><td>295.49</td><td>5.55</td><td>60</td><td>>sp|P01833|PIGR_HUMAN Polymeric immunoglobulin receptor OS=Homo sapiens OX=9606 GN=PIGR PE=1 SV=4</td><td>902</td><td>20201116_xd_colostrum_1ng_glyco_hcd30_etd30_compound15.17284.17284.3</td><td>controllerType=0 controllerNumber=1 scan=17284</td><td>44.5667</td></tr>
<tr><td>14038:3</td><td>5</td><td>K.QIGLYPVLVIDSSGYVNPN[+2716.983]YTGR.I</td><td>HexNAc(5)Hex(6)Fuc(3)NeuAc(1)</td><td>N[+2717]</td><td>1748.1062</td><td>3</td><td>5242.3039</td><td>5242.2864</td><td>0</td><td>3.3</td><td>168</td><td>Specific</td><td>298.2</td><td>233.05</td><td>233.05</td><td>5.55</td><td>60</td><td>>sp|P01833|PIGR_HUMAN Polymeric immunoglobulin receptor OS=Homo sapiens OX=9606 GN=PIGR PE=1 SV=4</td><td>902</td><td>20201116_xd_colostrum_1ng_glyco_hcd30_etd30_compound15.17351.17351.3</td><td>controllerType=0 controllerNumber=1 scan=17351</td><td>44.7808</td></tr>
<tr><td>16566:3</td><td>5</td><td>K.QIGLYPVLVIDSSGYVNPN[+2188.740]YTGR.I</td><td>HexNAc(2)Hex(11)</td><td>N[+2189]</td><td>1572.0341</td><td>3</td><td>4714.0878</td><td>4714.0433</td><td>0</td><td>9.4</td><td>168</td><td>Specific</td><td>368.01</td><td>280.3</td><td>280.3</td><td>5.13</td><td>60</td><td>>sp|P01833|PIGR_HUMAN Polymeric immunoglobulin receptor OS=Homo sapiens OX=9606 GN=PIGR PE=1 SV=4</td><td>902</td><td>20201116_xd_colostrum_1ng_glyco_hcd30_etd30_compound15.20050.20050.3</td><td>controllerType=0 controllerNumber=1 scan=20050</td><td>53.9145</td></tr>
<tr><td>14239:3</td><td>5</td><td>K.QIGLYPVLVIDSSGYVNPN[+1872.651]YTGR.I</td><td>HexNAc(3)Hex(6)NeuAc(1)</td><td>N[+1873]</td><td>1466.663</td><td>3</td><td>4397.9745</td><td>4397.9539</td><td>0</td><td>4.7</td><td>168</td><td>Specific</td><td>381.53</td><td>284.94</td><td>284.94</td><td>5.8</td><td>60</td><td>>sp|P01833|PIGR_HUMAN Polymeric immunoglobulin receptor OS=Homo sapiens OX=9606 GN=PIGR PE=1 SV=4</td><td>902</td><td>20201116_xd_colostrum_1ng_glyco_hcd30_etd30_compound15.17565.17565.3</td><td>controllerType=0 controllerNumber=1 scan=17565</td><td>45.4621</td></tr>
<tr><td>15470:3</td><td>5</td><td>K.QIGLYPVLVIDSSGYVNPN[+2350.830]YTGR.I</td><td>HexNAc(4)Hex(5)Fuc(1)NeuAc(2)</td><td>N[+2351]</td><td>1626.0554</td><td>3</td><td>4876.1515</td><td>4876.1338</td><td>0</td><td>3.6</td><td>168</td><td>Specific</td><td>448.68</td><td>351.34</td><td>351.34</td><td>6.28</td><td>60</td><td>>sp|P01833|PIGR_HUMAN Polymeric immunoglobulin receptor OS=Homo sapiens OX=9606 GN=PIGR PE=1 SV=4</td><td>902</td><td>20201116_xd_colostrum_1ng_glyco_hcd30_etd30_compound15.18874.18874.3</td><td>controllerType=0 controllerNumber=1 scan=18874</td><td>49.6969</td></tr>
<tr><td>15471:4</td><td>5</td><td>K.QIGLYPVLVIDSSGYVNPN[+2350.830]YTGR.I</td><td>HexNAc(4)Hex(5)Fuc(1)NeuAc(2)</td><td>N[+2351]</td><td>1219.7934</td><td>4</td><td>4876.1517</td><td>4876.1338</td><td>0</td><td>3.7</td><td>168</td><td>Specific</td><td>455.09</td><td>400.63</td><td>400.63</td><td>6.8</td><td>60</td><td>>sp|P01833|PIGR_HUMAN Polymeric immunoglobulin receptor OS=Homo sapiens OX=9606 GN=PIGR PE=1 SV=4</td><td>902</td><td>20201116_xd_colostrum_1ng_glyco_hcd30_etd30_compound15.18875.18875.4</td><td>controllerType=0 controllerNumber=1 scan=18875</td><td>49.7067</td></tr>
<tr><td>14157:4</td><td>5</td><td>K.QIGLYPVLVIDSSGYVNPN[+2205.793]YTGR.I</td><td>HexNAc(4)Hex(5)Fuc(2)NeuAc(1)</td><td>N[+2206]</td><td>1183.5334</td><td>4</td><td>4731.1116</td><td>4731.0963</td><td>0</td><td>3.2</td><td>168</td><td>Specific</td><td>516.6</td><td>492.15</td><td>492.15</td><td>6.94</td><td>60</td><td>>sp|P01833|PIGR_HUMAN Polymeric immunoglobulin receptor OS=Homo sapiens OX=9606 GN=PIGR PE=1 SV=4</td><td>902</td><td>20201116_xd_colostrum_1ng_glyco_hcd30_etd30_compound15.17478.17478.4</td><td>controllerType=0 controllerNumber=1 scan=17478</td><td>45.1869</td></tr>
<tr><td>14148:3</td><td>5</td><td>K.QIGLYPVLVIDSSGYVNPN[+2351.851]YTGR.I</td><td>HexNAc(4)Hex(5)Fuc(3)NeuAc(1)</td><td>N[+2352]</td><td>1626.3997</td><td>3</td><td>4877.1846</td><td>4877.1542</td><td>0</td><td>6.2</td><td>168</td><td>Specific</td><td>548.8</td><td>445.65</td><td>445.65</td><td>7.56</td><td>60</td><td>>sp|P01833|PIGR_HUMAN Polymeric immunoglobulin receptor OS=Homo sapiens OX=9606 GN=PIGR PE=1 SV=4</td><td>902</td><td>20201116_xd_colostrum_1ng_glyco_hcd30_etd30_compound15.17468.17468.3</td><td>controllerType=0 controllerNumber=1 scan=17468</td><td>45.1465</td></tr>
<tr><td>14169:3</td><td>5</td><td>K.QIGLYPVLVIDSSGYVNPN[+2059.735]YTGR.I</td><td>HexNAc(4)Hex(5)Fuc(1)NeuAc(1)</td><td>N[+2060]</td><td>1529.0234</td><td>3</td><td>4585.0556</td><td>4585.0384</td><td>0</td><td>3.7</td><td>168</td><td>Specific</td><td>582.19</td><td>508.3</td><td>508.3</td><td>8.84</td><td>60</td><td>>sp|P01833|PIGR_HUMAN Polymeric immunoglobulin receptor OS=Homo sapiens OX=9606 GN=PIGR PE=1 SV=4</td><td>902</td><td>20201116_xd_colostrum_1ng_glyco_hcd30_etd30_compound15.17490.17490.3</td><td>controllerType=0 controllerNumber=1 scan=17490</td><td>45.23</td></tr>
<tr><td>14170:4</td><td>5</td><td>K.QIGLYPVLVIDSSGYVNPN[+2059.735]YTGR.I</td><td>HexNAc(4)Hex(5)Fuc(1)NeuAc(1)</td><td>N[+2060]</td><td>1147.0177</td><td>4</td><td>4585.0491</td><td>4585.0384</td><td>0</td><td>2.3</td><td>168</td><td>Specific</td><td>585.49</td><td>525.08</td><td>525.08</td><td>8.54</td><td>60</td><td>>sp|P01833|PIGR_HUMAN Polymeric immunoglobulin receptor OS=Homo sapiens OX=9606 GN=PIGR PE=1 SV=4</td><td>902</td><td>20201116_xd_colostrum_1ng_glyco_hcd30_etd30_compound15.17491.17491.4</td><td>controllerType=0 controllerNumber=1 scan=17491</td><td>45.2333</td></tr>
<tr><td>14227:3</td><td>5</td><td>K.QIGLYPVLVIDSSGYVNPN[+1913.677]YTGR.I</td><td>HexNAc(4)Hex(5)NeuAc(1)</td><td>N[+1914]</td><td>1480.3362</td><td>3</td><td>4438.9942</td><td>4438.9805</td><td>0</td><td>3.1</td><td>168</td><td>Specific</td><td>592.01</td><td>478.83</td><td>478.83</td><td>9.43</td><td>60</td><td>>sp|P01833|PIGR_HUMAN Polymeric immunoglobulin receptor OS=Homo sapiens OX=9606 GN=PIGR PE=1 SV=4</td><td>902</td><td>20201116_xd_colostrum_1ng_glyco_hcd30_etd30_compound15.17552.17552.3</td><td>controllerType=0 controllerNumber=1 scan=17552</td><td>45.4235</td></tr>
<tr><td>16402:3</td><td>5</td><td>K.AQYEGRLSLLEEPGN[+1663.608]GTFTVILNQLTSR.D</td><td>HexNAc(5)Hex(4)</td><td>N[+1664]</td><td>1590.7343</td><td>3</td><td>4770.1884</td><td>4770.2249</td><td>0</td><td>-7.7</td><td>407</td><td>Specific</td><td>215.11</td><td>193.41</td><td>193.41</td><td>2.32</td><td>60</td><td>>sp|P01833|PIGR_HUMAN Polymeric immunoglobulin receptor OS=Homo sapiens OX=9606 GN=PIGR PE=1 SV=4</td><td>902</td><td>20201116_xd_colostrum_1ng_glyco_hcd30_etd30_compound15.19873.19873.3</td><td>controllerType=0 controllerNumber=1 scan=19873</td><td>53.2459</td></tr>
<tr><td>16678:3</td><td>5</td><td>R.LSLLEEPGN[+2351.851]GTFTVILNQLTSR.D</td><td>HexNAc(4)Hex(5)Fuc(3)NeuAc(1)</td><td>N[+2352]</td><td>1585.3819</td><td>3</td><td>4754.1311</td><td>4754.1433</td><td>0</td><td>-2.6</td><td>413</td><td>Specific</td><td>249.95</td><td>113.15</td><td>113.15</td><td>3.26</td><td>60</td><td>>sp|P01833|PIGR_HUMAN Polymeric immunoglobulin receptor OS=Homo sapiens OX=9606 GN=PIGR PE=1 SV=4</td><td>902</td><td>20201116_xd_colostrum_1ng_glyco_hcd30_etd30_compound15.20170.20170.3</td><td>controllerType=0 controllerNumber=1 scan=20170</td><td>54.3534</td></tr>
<tr><td>15091:4</td><td>5</td><td>R.LSLLEEPGN[+2863.041]GTFTVILNQLTSR.D</td><td>HexNAc(5)Hex(6)Fuc(4)NeuAc(1)</td><td>N[+2863]</td><td>1317.0916</td><td>4</td><td>5265.3444</td><td>5265.3334</td><td>0</td><td>2.1</td><td>413</td><td>Specific</td><td>309.99</td><td>216.76</td><td>216.76</td><td>4.56</td><td>60</td><td>>sp|P01833|PIGR_HUMAN Polymeric immunoglobulin receptor OS=Homo sapiens OX=9606 GN=PIGR PE=1 SV=4</td><td>902</td><td>20201116_xd_colostrum_1ng_glyco_hcd30_etd30_compound15.18469.18469.4</td><td>controllerType=0 controllerNumber=1 scan=18469</td><td>48.2822</td></tr>
<tr><td>15088:3</td><td>5</td><td>R.LSLLEEPGN[+2862.021]GTFTVILNQLTSR.D</td><td>HexNAc(5)Hex(6)Fuc(2)NeuAc(2)</td><td>N[+2862]</td><td>1755.4587</td><td>3</td><td>5264.3616</td><td>5264.313</td><td>0</td><td>9.2</td><td>413</td><td>Specific</td><td>385.36</td><td>246.41</td><td>246.41</td><td>5.12</td><td>60</td><td>>sp|P01833|PIGR_HUMAN Polymeric immunoglobulin receptor OS=Homo sapiens OX=9606 GN=PIGR PE=1 SV=4</td><td>902</td><td>20201116_xd_colostrum_1ng_glyco_hcd30_etd30_compound15.18466.18466.3</td><td>controllerType=0 controllerNumber=1 scan=18466</td><td>48.2697</td></tr>
<tr><td>15052:3</td><td>5</td><td>R.LSLLEEPGN[+2863.041]GTFTVILNQLTSR.D</td><td>HexNAc(5)Hex(6)Fuc(4)NeuAc(1)</td><td>N[+2863]</td><td>1755.7887</td><td>3</td><td>5265.3514</td><td>5265.3334</td><td>0</td><td>3.4</td><td>413</td><td>Specific</td><td>545.09</td><td>258.36</td><td>258.36</td><td>6.21</td><td>60</td><td>>sp|P01833|PIGR_HUMAN Polymeric immunoglobulin receptor OS=Homo sapiens OX=9606 GN=PIGR PE=1 SV=4</td><td>902</td><td>20201116_xd_colostrum_1ng_glyco_hcd30_etd30_compound15.18428.18428.3</td><td>controllerType=0 controllerNumber=1 scan=18428</td><td>48.1348</td></tr>
<tr><td>16385:4</td><td>5</td><td>R.LSLLEEPGN[+2350.830]GTFTVILNQLTSR.D</td><td>HexNAc(4)Hex(5)Fuc(1)NeuAc(2)</td><td>N[+2351]</td><td>1189.0422</td><td>4</td><td>4753.1468</td><td>4753.1229</td><td>0</td><td>5</td><td>413</td><td>Specific</td><td>559.06</td><td>450.99</td><td>450.99</td><td>6.84</td><td>60</td><td>>sp|P01833|PIGR_HUMAN Polymeric immunoglobulin receptor OS=Homo sapiens OX=9606 GN=PIGR PE=1 SV=4</td><td>902</td><td>20201116_xd_colostrum_1ng_glyco_hcd30_etd30_compound15.19855.19855.4</td><td>controllerType=0 controllerNumber=1 scan=19855</td><td>53.1895</td></tr>
<tr><td>15297:3</td><td>5</td><td>R.LSLLEEPGN[+2350.830]GTFTVILNQLTSR.D</td><td>HexNAc(4)Hex(5)Fuc(1)NeuAc(2)</td><td>N[+2351]</td><td>1585.0568</td><td>3</td><td>4753.1559</td><td>4753.1229</td><td>0</td><td>6.9</td><td>413</td><td>Specific</td><td>560.81</td><td>406.59</td><td>406.59</td><td>8.33</td><td>60</td><td>>sp|P01833|PIGR_HUMAN Polymeric immunoglobulin receptor OS=Homo sapiens OX=9606 GN=PIGR PE=1 SV=4</td><td>902</td><td>20201116_xd_colostrum_1ng_glyco_hcd30_etd30_compound15.18687.18687.3</td><td>controllerType=0 controllerNumber=1 scan=18687</td><td>48.9725</td></tr>
<tr><td>15238:4</td><td>5</td><td>R.LSLLEEPGN[+2205.793]GTFTVILNQLTSR.D</td><td>HexNAc(4)Hex(5)Fuc(2)NeuAc(1)</td><td>N[+2206]</td><td>1152.7824</td><td>4</td><td>4608.1077</td><td>4608.0854</td><td>0</td><td>4.8</td><td>413</td><td>Specific</td><td>589.55</td><td>502.91</td><td>502.91</td><td>7.63</td><td>60</td><td>>sp|P01833|PIGR_HUMAN Polymeric immunoglobulin receptor OS=Homo sapiens OX=9606 GN=PIGR PE=1 SV=4</td><td>902</td><td>20201116_xd_colostrum_1ng_glyco_hcd30_etd30_compound15.18624.18624.4</td><td>controllerType=0 controllerNumber=1 scan=18624</td><td>48.7667</td></tr>
<tr><td>16382:3</td><td>5</td><td>R.LSLLEEPGN[+2350.830]GTFTVILNQLTSR.D</td><td>HexNAc(4)Hex(5)Fuc(1)NeuAc(2)</td><td>N[+2351]</td><td>1585.0529</td><td>3</td><td>4753.1442</td><td>4753.1229</td><td>0</td><td>4.5</td><td>413</td><td>Specific</td><td>617.89</td><td>424.02</td><td>424.02</td><td>9.84</td><td>60</td><td>>sp|P01833|PIGR_HUMAN Polymeric immunoglobulin receptor OS=Homo sapiens OX=9606 GN=PIGR PE=1 SV=4</td><td>902</td><td>20201116_xd_colostrum_1ng_glyco_hcd30_etd30_compound15.19852.19852.3</td><td>controllerType=0 controllerNumber=1 scan=19852</td><td>53.1776</td></tr>
<tr><td>16431:3</td><td>5</td><td>R.LSLLEEPGN[+2496.888]GTFTVILNQLTSR.D</td><td>HexNAc(4)Hex(5)Fuc(2)NeuAc(2)</td><td>N[+2497]</td><td>1633.7378</td><td>3</td><td>4899.199</td><td>4899.1808</td><td>0</td><td>3.7</td><td>413</td><td>Specific</td><td>645.71</td><td>511.58</td><td>511.58</td><td>9.84</td><td>60</td><td>>sp|P01833|PIGR_HUMAN Polymeric immunoglobulin receptor OS=Homo sapiens OX=9606 GN=PIGR PE=1 SV=4</td><td>902</td><td>20201116_xd_colostrum_1ng_glyco_hcd30_etd30_compound15.19904.19904.3</td><td>controllerType=0 controllerNumber=1 scan=19904</td><td>53.3725</td></tr>
<tr><td>15227:3</td><td>5</td><td>R.LSLLEEPGN[+2205.793]GTFTVILNQLTSR.D</td><td>HexNAc(4)Hex(5)Fuc(2)NeuAc(1)</td><td>N[+2206]</td><td>1536.7088</td><td>3</td><td>4608.1118</td><td>4608.0854</td><td>0</td><td>5.7</td><td>413</td><td>Specific</td><td>701.29</td><td>584.64</td><td>584.64</td><td>11.64</td><td>60</td><td>>sp|P01833|PIGR_HUMAN Polymeric immunoglobulin receptor OS=Homo sapiens OX=9606 GN=PIGR PE=1 SV=4</td><td>902</td><td>20201116_xd_colostrum_1ng_glyco_hcd30_etd30_compound15.18613.18613.3</td><td>controllerType=0 controllerNumber=1 scan=18613</td><td>48.74</td></tr>
<tr><td>12250:4</td><td>5</td><td>K.IIEGEPNLKVPGN[+2205.793]VTAVLGETLK.V</td><td>HexNAc(4)Hex(5)Fuc(2)NeuAc(1)</td><td>N[+2206]</td><td>1150.0467</td><td>4</td><td>4597.1649</td><td>4597.1422</td><td>0</td><td>4.9</td><td>457</td><td>Specific</td><td>248.37</td><td>248.37</td><td>248.37</td><td>3.78</td><td>60</td><td>>sp|P01833|PIGR_HUMAN Polymeric immunoglobulin receptor OS=Homo sapiens OX=9606 GN=PIGR PE=1 SV=4</td><td>902</td><td>20201116_xd_colostrum_1ng_glyco_hcd30_etd30_compound15.15465.15465.4</td><td>controllerType=0 controllerNumber=1 scan=15465</td><td>39.2715</td></tr>
<tr><td>13517:4</td><td>5</td><td>K.IIEGEPNLKVPGN[+2350.830]VTAVLGETLK.V</td><td>HexNAc(4)Hex(5)Fuc(1)NeuAc(2)</td><td>N[+2351]</td><td>1186.3047</td><td>4</td><td>4742.1971</td><td>4742.1797</td><td>0</td><td>3.7</td><td>457</td><td>Specific</td><td>339.94</td><td>333.45</td><td>333.45</td><td>4.57</td><td>60</td><td>>sp|P01833|PIGR_HUMAN Polymeric immunoglobulin receptor OS=Homo sapiens OX=9606 GN=PIGR PE=1 SV=4</td><td>902</td><td>20201116_xd_colostrum_1ng_glyco_hcd30_etd30_compound15.16802.16802.4</td><td>controllerType=0 controllerNumber=1 scan=16802</td><td>43.1945</td></tr>
<tr><td>12244:3</td><td>5</td><td>K.IIEGEPNLKVPGN[+2205.793]VTAVLGETLK.V</td><td>HexNAc(4)Hex(5)Fuc(2)NeuAc(1)</td><td>N[+2206]</td><td>1533.056</td><td>3</td><td>4597.1533</td><td>4597.1422</td><td>0</td><td>2.4</td><td>457</td><td>Specific</td><td>377.84</td><td>296.21</td><td>296.21</td><td>5.22</td><td>60</td><td>>sp|P01833|PIGR_HUMAN Polymeric immunoglobulin receptor OS=Homo sapiens OX=9606 GN=PIGR PE=1 SV=4</td><td>902</td><td>20201116_xd_colostrum_1ng_glyco_hcd30_etd30_compound15.15459.15459.3</td><td>controllerType=0 controllerNumber=1 scan=15459</td><td>39.2601</td></tr>
<tr><td>11734:5</td><td>5</td><td>K.IIEGEPNLKVPGN[+2863.041]VTAVLGETLKVPC[+57.021]HFPC[+57.021]K.F</td><td>HexNAc(5)Hex(6)Fuc(4)NeuAc(1)</td><td>C[+57]*2, N[+2863]</td><td>1256.7775</td><td>5</td><td>6279.8582</td><td>6279.8477</td><td>0</td><td>1.7</td><td>457</td><td>Specific</td><td>305.73</td><td>289.35</td><td>289.35</td><td>4.5</td><td>60</td><td>>sp|P01833|PIGR_HUMAN Polymeric immunoglobulin receptor OS=Homo sapiens OX=9606 GN=PIGR PE=1 SV=4</td><td>902</td><td>20201116_xd_colostrum_1ng_glyco_hcd30_etd30_compound15.14921.14921.5</td><td>controllerType=0 controllerNumber=1 scan=14921</td><td>37.7387</td></tr>
<tr><td>12888:4</td><td>5</td><td>K.IIEGEPNLKVPGN[+2350.830]VTAVLGETLKVPC[+57.021]HFPC[+57.021]K.F</td><td>HexNAc(4)Hex(5)Fuc(1)NeuAc(2)</td><td>C[+57]*2, N[+2351]</td><td>1442.6704</td><td>4</td><td>5767.6598</td><td>5767.6372</td><td>0</td><td>3.9</td><td>457</td><td>Specific</td><td>323.91</td><td>323.91</td><td>323.91</td><td>3.95</td><td>60</td><td>>sp|P01833|PIGR_HUMAN Polymeric immunoglobulin receptor OS=Homo sapiens OX=9606 GN=PIGR PE=1 SV=4</td><td>902</td><td>20201116_xd_colostrum_1ng_glyco_hcd30_etd30_compound15.16137.16137.4</td><td>controllerType=0 controllerNumber=1 scan=16137</td><td>41.1574</td></tr>
<tr><td>11878:4</td><td>5</td><td>K.IIEGEPNLKVPGN[+2205.793]VTAVLGETLKVPC[+57.021]HFPC[+57.021]K.F</td><td>HexNAc(4)Hex(5)Fuc(2)NeuAc(1)</td><td>C[+57]*2, N[+2206]</td><td>1406.4106</td><td>4</td><td>5622.6208</td><td>5622.5997</td><td>0</td><td>3.7</td><td>457</td><td>Specific</td><td>401.61</td><td>401.61</td><td>401.61</td><td>5.68</td><td>60</td><td>>sp|P01833|PIGR_HUMAN Polymeric immunoglobulin receptor OS=Homo sapiens OX=9606 GN=PIGR PE=1 SV=4</td><td>902</td><td>20201116_xd_colostrum_1ng_glyco_hcd30_etd30_compound15.15074.15074.4</td><td>controllerType=0 controllerNumber=1 scan=15074</td><td>38.2391</td></tr>
<tr><td>11887:4</td><td>5</td><td>K.IIEGEPNLKVPGN[+2059.735]VTAVLGETLKVPC[+57.021]HFPC[+57.021]K.F</td><td>HexNAc(4)Hex(5)Fuc(1)NeuAc(1)</td><td>C[+57]*2, N[+2060]</td><td>1369.8949</td><td>4</td><td>5476.5578</td><td>5476.5418</td><td>0</td><td>2.9</td><td>457</td><td>Specific</td><td>410.53</td><td>410.53</td><td>410.53</td><td>6.03</td><td>60</td><td>>sp|P01833|PIGR_HUMAN Polymeric immunoglobulin receptor OS=Homo sapiens OX=9606 GN=PIGR PE=1 SV=4</td><td>902</td><td>20201116_xd_colostrum_1ng_glyco_hcd30_etd30_compound15.15083.15083.4</td><td>controllerType=0 controllerNumber=1 scan=15083</td><td>38.2652</td></tr>
<tr><td>11861:4</td><td>5</td><td>K.IIEGEPNLKVPGN[+2351.851]VTAVLGETLKVPC[+57.021]HFPC[+57.021]K.F</td><td>HexNAc(4)Hex(5)Fuc(3)NeuAc(1)</td><td>C[+57]*2, N[+2352]</td><td>1442.9247</td><td>4</td><td>5768.6769</td><td>5768.6576</td><td>0</td><td>3.3</td><td>457</td><td>Specific</td><td>471.85</td><td>471.85</td><td>471.85</td><td>5.77</td><td>60</td><td>>sp|P01833|PIGR_HUMAN Polymeric immunoglobulin receptor OS=Homo sapiens OX=9606 GN=PIGR PE=1 SV=4</td><td>902</td><td>20201116_xd_colostrum_1ng_glyco_hcd30_etd30_compound15.15056.15056.4</td><td>controllerType=0 controllerNumber=1 scan=15056</td><td>38.1859</td></tr>
<tr><td>12887:5</td><td>5</td><td>K.IIEGEPNLKVPGN[+2350.830]VTAVLGETLKVPC[+57.021]HFPC[+57.021]K.F</td><td>HexNAc(4)Hex(5)Fuc(1)NeuAc(2)</td><td>C[+57]*2, N[+2351]</td><td>1154.3376</td><td>5</td><td>5767.6591</td><td>5767.6372</td><td>0</td><td>3.8</td><td>457</td><td>Specific</td><td>645.41</td><td>645.41</td><td>645.41</td><td>8.5</td><td>60</td><td>>sp|P01833|PIGR_HUMAN Polymeric immunoglobulin receptor OS=Homo sapiens OX=9606 GN=PIGR PE=1 SV=4</td><td>902</td><td>20201116_xd_colostrum_1ng_glyco_hcd30_etd30_compound15.16136.16136.5</td><td>controllerType=0 controllerNumber=1 scan=16136</td><td>41.1543</td></tr>
<tr><td>11877:5</td><td>5</td><td>K.IIEGEPNLKVPGN[+2205.793]VTAVLGETLKVPC[+57.021]HFPC[+57.021]K.F</td><td>HexNAc(4)Hex(5)Fuc(2)NeuAc(1)</td><td>C[+57]*2, N[+2206]</td><td>1125.3293</td><td>5</td><td>5622.6176</td><td>5622.5997</td><td>0</td><td>3.2</td><td>457</td><td>Specific</td><td>647.34</td><td>647.34</td><td>647.34</td><td>8.8</td><td>60</td><td>>sp|P01833|PIGR_HUMAN Polymeric immunoglobulin receptor OS=Homo sapiens OX=9606 GN=PIGR PE=1 SV=4</td><td>902</td><td>20201116_xd_colostrum_1ng_glyco_hcd30_etd30_compound15.15073.15073.5</td><td>controllerType=0 controllerNumber=1 scan=15073</td><td>38.2361</td></tr>
<tr><td>11882:5</td><td>5</td><td>K.IIEGEPNLKVPGN[+2059.735]VTAVLGETLKVPC[+57.021]HFPC[+57.021]K.F</td><td>HexNAc(4)Hex(5)Fuc(1)NeuAc(1)</td><td>C[+57]*2, N[+2060]</td><td>1096.1174</td><td>5</td><td>5476.5581</td><td>5476.5418</td><td>0</td><td>3</td><td>457</td><td>Specific</td><td>697.38</td><td>633.17</td><td>633.17</td><td>10.09</td><td>60</td><td>>sp|P01833|PIGR_HUMAN Polymeric immunoglobulin receptor OS=Homo sapiens OX=9606 GN=PIGR PE=1 SV=4</td><td>902</td><td>20201116_xd_colostrum_1ng_glyco_hcd30_etd30_compound15.15078.15078.5</td><td>controllerType=0 controllerNumber=1 scan=15078</td><td>38.2493</td></tr>
<tr><td>12858:5</td><td>5</td><td>K.IIEGEPNLKVPGN[+2350.830]VTAVLGETLKVPC[+57.021]HFPC[+57.021]K.F</td><td>HexNAc(4)Hex(5)Fuc(1)NeuAc(2)</td><td>C[+57]*2, N[+2351]</td><td>1154.3359</td><td>5</td><td>5767.6506</td><td>5767.6372</td><td>0</td><td>2.3</td><td>457</td><td>Specific</td><td>708.12</td><td>702.87</td><td>702.87</td><td>11.21</td><td>60</td><td>>sp|P01833|PIGR_HUMAN Polymeric immunoglobulin receptor OS=Homo sapiens OX=9606 GN=PIGR PE=1 SV=4</td><td>902</td><td>20201116_xd_colostrum_1ng_glyco_hcd30_etd30_compound15.16105.16105.5</td><td>controllerType=0 controllerNumber=1 scan=16105</td><td>41.0343</td></tr>
<tr><td>11860:5</td><td>5</td><td>K.IIEGEPNLKVPGN[+2351.851]VTAVLGETLKVPC[+57.021]HFPC[+57.021]K.F</td><td>HexNAc(4)Hex(5)Fuc(3)NeuAc(1)</td><td>C[+57]*2, N[+2352]</td><td>1154.5404</td><td>5</td><td>5768.6729</td><td>5768.6576</td><td>0</td><td>2.7</td><td>457</td><td>Specific</td><td>710.2</td><td>710.2</td><td>710.2</td><td>11.39</td><td>60</td><td>>sp|P01833|PIGR_HUMAN Polymeric immunoglobulin receptor OS=Homo sapiens OX=9606 GN=PIGR PE=1 SV=4</td><td>902</td><td>20201116_xd_colostrum_1ng_glyco_hcd30_etd30_compound15.15055.15055.5</td><td>controllerType=0 controllerNumber=1 scan=15055</td><td>38.183</td></tr>
<tr><td>10549:4</td><td>5</td><td>K.VPGN[+2351.851]VTAVLGETLKVPC[+57.021]HFPC[+57.021]K.F</td><td>HexNAc(4)Hex(5)Fuc(3)NeuAc(1)</td><td>C[+57]*2, N[+2352]</td><td>1194.5352</td><td>4</td><td>4775.119</td><td>4775.1081</td><td>0</td><td>2.3</td><td>466</td><td>Specific</td><td>217.71</td><td>217.71</td><td>217.71</td><td>3.01</td><td>60</td><td>>sp|P01833|PIGR_HUMAN Polymeric immunoglobulin receptor OS=Homo sapiens OX=9606 GN=PIGR PE=1 SV=4</td><td>902</td><td>20201116_xd_colostrum_1ng_glyco_hcd30_etd30_compound15.13678.13678.4</td><td>controllerType=0 controllerNumber=1 scan=13678</td><td>34.5222</td></tr>
<tr><td>10555:4</td><td>5</td><td>K.VPGN[+2205.793]VTAVLGETLKVPC[+57.021]HFPC[+57.021]K.F</td><td>HexNAc(4)Hex(5)Fuc(2)NeuAc(1)</td><td>C[+57]*2, N[+2206]</td><td>1158.021</td><td>4</td><td>4629.0623</td><td>4629.0502</td><td>0</td><td>2.6</td><td>466</td><td>Specific</td><td>234.69</td><td>234.69</td><td>234.69</td><td>3.45</td><td>60</td><td>>sp|P01833|PIGR_HUMAN Polymeric immunoglobulin receptor OS=Homo sapiens OX=9606 GN=PIGR PE=1 SV=4</td><td>902</td><td>20201116_xd_colostrum_1ng_glyco_hcd30_etd30_compound15.13684.13684.4</td><td>controllerType=0 controllerNumber=1 scan=13684</td><td>34.5432</td></tr>
</table>

Table S4. Cont.

| Query #:z | Protein Rank | Glycopeptide | Glycan | Modification Type(s) | Observed m/z | z | Observed (M+H) | Calc. mass (M+H) | Off-by-x error | Mass error (ppm) | StartPosition | Cleavage | Score | Delta | Delta Mod | \|Log Prob\| | # of unique peptides | Protein Name | Protein DB number | Comment | Scan # | Scan Time |
|---|---|---|---|---|---|---|---|---|---|---|---|---|---|---|---|---|---|---|---|---|---|---|
| 11892:4 | 5 | K.VPGN[+2350.830]VTAVLGETLKVPC[+57.021]HFPC[+57.021]K.F | HexNAc(4)Hex(5)Fuc(1)NeuAc(2) | C[+57]*2, N[+2351] | 1194.2808 | 4 | 4774.1014 | 4774.0877 | 0 | 2.9 | 466 | Specific | 335.91 | 284.33 | 284.33 | 4.48 | 60 | >sp\|P01833\|PIGR_HUMAN Polymeric immunoglobulin receptor OS=Homo sapiens OX=9606 GN=PIGR PE=1 SV=4 | 902 | 20201116_xd_colostrum_1ng_glyco_hcd30_etd30_compound15.15088.15088.4 | controllerType=0 controllerNumber=1 scan=15088 | 38.2747 |
| 06749:3 | 5 | K.WN[+2204.772]NTGC[+57.021]QALPSQDEGPSK.A | HexNAc(4)Hex(5)NeuAc(2) | C[+57], N[+2205] | 1398.5577 | 3 | 4193.6585 | 4193.6491 | 0 | 2.2 | 498 | Specific | 219.58 | 81.27 | 81.27 | 3.13 | 60 | >sp\|P01833\|PIGR_HUMAN Polymeric immunoglobulin receptor OS=Homo sapiens OX=9606 GN=PIGR PE=1 SV=4 | 902 | 20201116_xd_colostrum_1ng_glyco_hcd30_etd30_compound15.9694.9694.3 | controllerType=0 controllerNumber=1 scan=9694 | 24.4275 |
| 05060:3 | 5 | K.WN[+2205.793]NTGC[+57.021]QALPSQDEGPSK.A | HexNAc(4)Hex(5)Fuc(2)NeuAc(1) | C[+57], N[+2206] | 1398.8978 | 3 | 4194.6787 | 4194.6695 | 0 | 2.2 | 498 | Specific | 443.54 | 294.03 | 294.03 | 5.94 | 60 | >sp\|P01833\|PIGR_HUMAN Polymeric immunoglobulin receptor OS=Homo sapiens OX=9606 GN=PIGR PE=1 SV=4 | 902 | 20201116_xd_colostrum_1ng_glyco_hcd30_etd30_compound15.7931.7931.3 | controllerType=0 controllerNumber=1 scan=7931 | 20.5887 |
| 05102:3 | 5 | K.WN[+2059.735]NTGC[+57.021]QALPSQDEGPSK.A | HexNAc(4)Hex(5)Fuc(1)NeuAc(1) | C[+57], N[+2060] | 1350.2122 | 3 | 4048.6221 | 4048.6116 | 0 | 2.6 | 498 | Specific | 444.74 | 309.95 | 309.95 | 6.06 | 60 | >sp\|P01833\|PIGR_HUMAN Polymeric immunoglobulin receptor OS=Homo sapiens OX=9606 GN=PIGR PE=1 SV=4 | 902 | 20201116_xd_colostrum_1ng_glyco_hcd30_etd30_compound15.7975.7975.3 | controllerType=0 controllerNumber=1 scan=7975 | 20.6885 |
| 05193:3 | 5 | K.WN[+1710.598]NTGC[+57.021]QALPSQDEGPSK.A | HexNAc(3)Hex(5)NeuAc(1) | C[+57], N[+1711] | 1233.8336 | 3 | 3699.4861 | 3699.4743 | 0 | 3.2 | 498 | Specific | 499.77 | 408.48 | 408.48 | 7.04 | 60 | >sp\|P01833\|PIGR_HUMAN Polymeric immunoglobulin receptor OS=Homo sapiens OX=9606 GN=PIGR PE=1 SV=4 | 902 | 20201116_xd_colostrum_1ng_glyco_hcd30_etd30_compound15.8070.8070.3 | controllerType=0 controllerNumber=1 scan=8070 | 20.9089 |
| 05123:3 | 5 | K.WN[+1872.651]NTGC[+57.021]QALPSQDEGPSK.A | HexNAc(3)Hex(6)NeuAc(1) | C[+57], N[+1873] | 1287.8516 | 3 | 3861.5403 | 3861.5272 | 0 | 3.4 | 498 | Specific | 525.1 | 332.22 | 332.22 | 5.92 | 60 | >sp\|P01833\|PIGR_HUMAN Polymeric immunoglobulin receptor OS=Homo sapiens OX=9606 GN=PIGR PE=1 SV=4 | 902 | 20201116_xd_colostrum_1ng_glyco_hcd30_etd30_compound15.7997.7997.3 | controllerType=0 controllerNumber=1 scan=7997 | 20.741 |
| 17100:3 | 8 | R.QLAHQSN[+2204.772]STNIFFSPVSIATAFAMLSLGTK.A | HexNAc(4)Hex(5)NeuAc(2) | N[+2205] | 1796.1458 | 3 | 5386.423 | 5386.4075 | 0 | 2.9 | 64 | Specific | 279.92 | 173.45 | 173.45 | 6.14 | 16 | >sp\|P01009\|A1AT_HUMAN Alpha-1-antitrypsin OS=Homo sapiens OX=9606 GN=SERPINA1 PE=1 SV=3 | 642 | 20201116_xd_colostrum_1ng_glyco_hcd30_etd30_compound15.20624.20624.3 | controllerType=0 controllerNumber=1 scan=20624 | 56.0306 |
| 17030:3 | 8 | R.QLAHQSN[+2350.830]STNIFFSPVSIATAFAMLSLGTK.A | HexNAc(4)Hex(5)Fuc(1)NeuAc(2) | N[+2351] | 1844.8291 | 3 | 5532.4726 | 5532.4654 | 0 | 1.3 | 64 | Specific | 298.74 | 213.94 | 213.94 | 5.27 | 16 | >sp\|P01009\|A1AT_HUMAN Alpha-1-antitrypsin OS=Homo sapiens OX=9606 GN=SERPINA1 PE=1 SV=3 | 642 | 20201116_xd_colostrum_1ng_glyco_hcd30_etd30_compound15.20549.20549.3 | controllerType=0 controllerNumber=1 scan=20549 | 55.7532 |
| 17029:4 | 8 | R.QLAHQSN[+2350.830]STNIFFSPVSIATAFAMLSLGTK.A | HexNAc(4)Hex(5)Fuc(1)NeuAc(2) | N[+2351] | 1383.8738 | 4 | 5532.4733 | 5532.4654 | 0 | 1.4 | 64 | Specific | 576.64 | 516 | 516 | 9.12 | 16 | >sp\|P01009\|A1AT_HUMAN Alpha-1-antitrypsin OS=Homo sapiens OX=9606 GN=SERPINA1 PE=1 SV=3 | 642 | 20201116_xd_colostrum_1ng_glyco_hcd30_etd30_compound15.20548.20548.4 | controllerType=0 controllerNumber=1 scan=20548 | 55.7491 |
| 17099:4 | 8 | R.QLAHQSN[+2204.772]STNIFFSPVSIATAFAMLSLGTK.A | HexNAc(4)Hex(5)NeuAc(2) | N[+2205] | 1347.3617 | 4 | 5386.425 | 5386.4075 | 0 | 3.2 | 64 | Specific | 670.98 | 570.79 | 570.79 | 11.3 | 16 | >sp\|P01009\|A1AT_HUMAN Alpha-1-antitrypsin OS=Homo sapiens OX=9606 GN=SERPINA1 PE=1 SV=3 | 642 | 20201116_xd_colostrum_1ng_glyco_hcd30_etd30_compound15.20623.20623.4 | controllerType=0 controllerNumber=1 scan=20623 | 56.0236 |
| 16579:4 | 8 | R.QLAHQSN[+2204.772]STNIFFSPVSIATAFAM[+15.995]LSLGTK.A | HexNAc(4)Hex(5)NeuAc(2) | M[+16], N[+2205] | 1351.3605 | 4 | 5402.4201 | 5402.4024 | 0 | 3.3 | 64 | Specific | 476.01 | 342.52 | 342.52 | 5.73 | 16 | >sp\|P01009\|A1AT_HUMAN Alpha-1-antitrypsin OS=Homo sapiens OX=9606 GN=SERPINA1 PE=1 SV=3 | 642 | 20201116_xd_colostrum_1ng_glyco_hcd30_etd30_compound15.20064.20064.4 | controllerType=0 controllerNumber=1 scan=20064 | 53.9706 |
| 14927:5 | 8 | K.YLGN[+2204.772]ATAIFFLPDEGKLQHLENELTHDIITK.F | HexNAc(4)Hex(5)NeuAc(2) | N[+2205] | 1149.9373 | 5 | 5745.6572 | 5745.6098 | 0 | 8.3 | 268 | Specific | 598.84 | 548.33 | 548.33 | 7.43 | 16 | >sp\|P01009\|A1AT_HUMAN Alpha-1-antitrypsin OS=Homo sapiens OX=9606 GN=SERPINA1 PE=1 SV=3 | 642 | 20201116_xd_colostrum_1ng_glyco_hcd30_etd30_compound15.18293.18293.5 | controllerType=0 controllerNumber=1 scan=18293 | 47.6476 |
| 14922:4 | 8 | K.YLGN[+2204.772]ATAIFFLPDEGKLQHLENELTHDIITK.F | HexNAc(4)Hex(5)NeuAc(2) | N[+2205] | 1437.1705 | 4 | 5745.6603 | 5745.6098 | 0 | 8.8 | 268 | Specific | 717.24 | 717.24 | 717.24 | 10.74 | 16 | >sp\|P01009\|A1AT_HUMAN Alpha-1-antitrypsin OS=Homo sapiens OX=9606 GN=SERPINA1 PE=1 SV=3 | 642 | 20201116_xd_colostrum_1ng_glyco_hcd30_etd30_compound15.18288.18288.4 | controllerType=0 controllerNumber=1 scan=18288 | 47.6083 |
| 16168:4 | 10 | K.VFPLSLDSTPQDGNVVVAC[+57.021]LVQGFFPQEPLSVTWSESGQN[+1444.534]VTAI | HexNAc(4)Hex(3)Fuc(1) | C[+57], N[+1445] | 1555.9873 | 4 | 6220.9274 | 6220.8953 | 0 | 5.2 | 8 | Specific | 242.66 | 242.66 | 242.66 | 4.67 | 24 | >sp\|P01877\|IGHA2_HUMAN Immunoglobulin heavy constant alpha 2 OS=Homo sapiens OX=9606 GN=IGHA2 PE=1 SV=4 | 576 | 20201116_xd_colostrum_1ng_glyco_hcd30_etd30_compound15.19623.19623.4 | controllerType=0 controllerNumber=1 scan=19623 | 52.3823 |
| 16872:5 | 10 | K.VFPLSLDSTPQDGNVVVAC[+57.021]LVQGFFPQEPLSVTWSESGQN[+2262.814]VTAI | HexNAc(5)Hex(5)Fuc(1)NeuAc(1) | C[+57], N[+2263] | 1408.6487 | 5 | 7039.2145 | 7039.1757 | 0 | 5.5 | 8 | Specific | 321.61 | 321.61 | 321.61 | 5.02 | 24 | >sp\|P01877\|IGHA2_HUMAN Immunoglobulin heavy constant alpha 2 OS=Homo sapiens OX=9606 GN=IGHA2 PE=1 SV=4 | 576 | 20201116_xd_colostrum_1ng_glyco_hcd30_etd30_compound15.20380.20380.5 | controllerType=0 controllerNumber=1 scan=20380 | 55.114 |
| 16315:4 | 10 | K.VFPLSLDSTPQDGNVVVAC[+57.021]LVQGFFPQEPLSVTWSESGQN[+1809.666]VTAI | HexNAc(5)Hex(4)Fuc(1) | C[+57], N[+1810] | 1647.2684 | 4 | 6586.0519 | 6586.0275 | 0 | 3.7 | 8 | Specific | 327.94 | 327.94 | 327.94 | 6.23 | 24 | >sp\|P01877\|IGHA2_HUMAN Immunoglobulin heavy constant alpha 2 OS=Homo sapiens OX=9606 GN=IGHA2 PE=1 SV=4 | 576 | 20201116_xd_colostrum_1ng_glyco_hcd30_etd30_compound15.19780.19780.4 | controllerType=0 controllerNumber=1 scan=19780 | 52.9105 |
| 16050:4 | 10 | K.VFPLSLDSTPQDGNVVVAC[+57.021]LVQGFFPQEPLSVTWSESGQN[+1606.587]VTAI | HexNAc(4)Hex(4)Fuc(1) | C[+57], N[+1607] | 1596.4968 | 4 | 6382.9655 | 6382.9481 | 0 | 2.7 | 8 | Specific | 339.07 | 339.07 | 339.07 | 6.31 | 24 | >sp\|P01877\|IGHA2_HUMAN Immunoglobulin heavy constant alpha 2 OS=Homo sapiens OX=9606 GN=IGHA2 PE=1 SV=4 | 576 | 20201116_xd_colostrum_1ng_glyco_hcd30_etd30_compound15.19496.19496.4 | controllerType=0 controllerNumber=1 scan=19496 | 51.9155 |
| 15982:4 | 10 | K.VFPLSLDSTPQDGNVVVAC[+57.021]LVQGFFPQEPLSVTWSESGQN[+1971.719]VTAI | HexNAc(5)Hex(5)Fuc(1) | C[+57], N[+1972] | 1687.7807 | 4 | 6748.1011 | 6748.0803 | 0 | 3.1 | 8 | Specific | 370.21 | 370.21 | 370.21 | 8.27 | 24 | >sp\|P01877\|IGHA2_HUMAN Immunoglobulin heavy constant alpha 2 OS=Homo sapiens OX=9606 GN=IGHA2 PE=1 SV=4 | 576 | 20201116_xd_colostrum_1ng_glyco_hcd30_etd30_compound15.19424.19424.4 | controllerType=0 controllerNumber=1 scan=19424 | 51.686 |
| 16876:6 | 10 | K.VFPLSLDSTPQDGNVVVAC[+57.021]LVQGFFPQEPLSVTWSESGQN[+2262.814]VTAI | HexNAc(5)Hex(5)Fuc(1)NeuAc(1) | C[+57], N[+2263] | 1174.0403 | 6 | 7039.2057 | 7039.1757 | 0 | 4.3 | 8 | Specific | 390.4 | 390.4 | 390.4 | 7.68 | 24 | >sp\|P01877\|IGHA2_HUMAN Immunoglobulin heavy constant alpha 2 OS=Homo sapiens OX=9606 GN=IGHA2 PE=1 SV=4 | 576 | 20201116_xd_colostrum_1ng_glyco_hcd30_etd30_compound15.20384.20384.6 | controllerType=0 controllerNumber=1 scan=20384 | 55.1325 |
| 16079:4 | 10 | K.VFPLSLDSTPQDGNVVVAC[+57.021]LVQGFFPQEPLSVTWSESGQN[+1809.666]VTAI | HexNAc(5)Hex(4)Fuc(1) | C[+57], N[+1810] | 1647.2699 | 4 | 6586.0578 | 6586.0275 | 0 | 4.6 | 8 | Specific | 399.44 | 399.44 | 399.44 | 7.68 | 24 | >sp\|P01877\|IGHA2_HUMAN Immunoglobulin heavy constant alpha 2 OS=Homo sapiens OX=9606 GN=IGHA2 PE=1 SV=4 | 576 | 20201116_xd_colostrum_1ng_glyco_hcd30_etd30_compound15.19527.19527.4 | controllerType=0 controllerNumber=1 scan=19527 | 52.0167 |
| 16089:4 | 10 | K.VFPLSLDSTPQDGNVVVAC[+57.021]LVQGFFPQEPLSVTWSESGQN[+1606.587]VTAI | HexNAc(4)Hex(4)Fuc(1) | C[+57], N[+1607] | 1596.4977 | 4 | 6382.9689 | 6382.9481 | 0 | 3.3 | 8 | Specific | 509.55 | 509.55 | 509.55 | 7.98 | 24 | >sp\|P01877\|IGHA2_HUMAN Immunoglobulin heavy constant alpha 2 OS=Homo sapiens OX=9606 GN=IGHA2 PE=1 SV=4 | 576 | 20201116_xd_colostrum_1ng_glyco_hcd30_etd30_compound15.19537.19537.4 | controllerType=0 controllerNumber=1 scan=19537 | 52.0466 |
| 16135:4 | 10 | K.VFPLSLDSTPQDGNVVVAC[+57.021]LVQGFFPQEPLSVTWSESGQN[+1647.613]VTAI | HexNAc(5)Hex(3)Fuc(1) | C[+57], N[+1648] | 1606.7567 | 4 | 6424.005 | 6423.9746 | 0 | 4.7 | 8 | Specific | 546.46 | 546.46 | 546.46 | 8.09 | 24 | >sp\|P01877\|IGHA2_HUMAN Immunoglobulin heavy constant alpha 2 OS=Homo sapiens OX=9606 GN=IGHA2 PE=1 SV=4 | 576 | 20201116_xd_colostrum_1ng_glyco_hcd30_etd30_compound15.19588.19588.4 | controllerType=0 controllerNumber=1 scan=19588 | 52.2361 |
| 16873:4 | 10 | K.VFPLSLDSTPQDGNVVVAC[+57.021]LVQGFFPQEPLSVTWSESGQN[+2262.814]VTAI | HexNAc(5)Hex(5)Fuc(1)NeuAc(1) | C[+57], N[+2263] | 1760.561 | 4 | 7039.2222 | 7039.1757 | 0 | 6.6 | 8 | Specific | 552.75 | 552.75 | 552.75 | 7.58 | 24 | >sp\|P01877\|IGHA2_HUMAN Immunoglobulin heavy constant alpha 2 OS=Homo sapiens OX=9606 GN=IGHA2 PE=1 SV=4 | 576 | 20201116_xd_colostrum_1ng_glyco_hcd30_etd30_compound15.20381.20381.4 | controllerType=0 controllerNumber=1 scan=20381 | 55.1205 |
| 09984:6 | 10 | K.SAVQGPPERDLC[+57.021]GC[+57.021]YSVSSVLPGC[+57.021]AQPWNHGETFTC[+! | HexNAc(5)Hex(4)Fuc(1) | C[+57]*4, N[+1810] | 1270.9107 | 6 | 7620.4278 | 7620.4119 | 0 | 2.1 | 156 | Specific | 566.46 | 502.52 | 502.52 | 7.55 | 24 | >sp\|P01877\|IGHA2_HUMAN Immunoglobulin heavy constant alpha 2 OS=Homo sapiens OX=9606 GN=IGHA2 PE=1 SV=4 | 576 | 20201116_xd_colostrum_1ng_glyco_hcd30_etd30_compound15.13086.13086.6 | controllerType=0 controllerNumber=1 scan=13086 | 32.9951 |
| 04404:2 | 10 | K.TPLTAN[+2262.814]ITK.S | HexNAc(5)Hex(5)Fuc(1)NeuAc(1) | N[+2263] | 1611.1933 | 2 | 3221.3793 | 3221.3711 | 0 | 2.5 | 200 | Specific | 232.15 | 74.32 | 74.32 | 2.88 | 24 | >sp\|P01877\|IGHA2_HUMAN Immunoglobulin heavy constant alpha 2 OS=Homo sapiens OX=9606 GN=IGHA2 PE=1 SV=4 | 576 | 20201116_xd_colostrum_1ng_glyco_hcd30_etd30_compound15.7248.7248.2 | controllerType=0 controllerNumber=1 scan=7248 | 19.1188 |
| 02781:3 | 10 | K.TPLTAN[+2117.777]ITK.S | HexNAc(5)Hex(5)Fuc(2) | N[+2118] | 1026.1183 | 3 | 3076.3404 | 3076.3336 | 0 | 2.2 | 200 | Specific | 242.38 | 111.8 | 111.8 | 3.79 | 24 | >sp\|P01877\|IGHA2_HUMAN Immunoglobulin heavy constant alpha 2 OS=Homo sapiens OX=9606 GN=IGHA2 PE=1 SV=4 | 576 | 20201116_xd_colostrum_1ng_glyco_hcd30_etd30_compound15.5553.5553.3 | controllerType=0 controllerNumber=1 scan=5553 | 15.5738 |
| 02762:3 | 10 | K.TPLTAN[+2117.777]ITK.S | HexNAc(5)Hex(5)Fuc(2) | N[+2118] | 1026.1184 | 3 | 3076.3408 | 3076.3336 | 0 | 2.3 | 200 | Specific | 246.43 | 118.86 | 118.86 | 3.35 | 24 | >sp\|P01877\|IGHA2_HUMAN Immunoglobulin heavy constant alpha 2 OS=Homo sapiens OX=9606 GN=IGHA2 PE=1 SV=4 | 576 | 20201116_xd_colostrum_1ng_glyco_hcd30_etd30_compound15.5533.5533.3 | controllerType=0 controllerNumber=1 scan=5533 | 15.5354 |
| 04387:3 | 10 | K.TPLTAN[+2262.814]ITK.S | HexNAc(5)Hex(5)Fuc(1)NeuAc(1) | N[+2263] | 1074.4649 | 3 | 3221.3801 | 3221.3711 | 0 | 2.8 | 200 | Specific | 269.82 | 111.13 | 111.13 | 3.79 | 24 | >sp\|P01877\|IGHA2_HUMAN Immunoglobulin heavy constant alpha 2 OS=Homo sapiens OX=9606 GN=IGHA2 PE=1 SV=4 | 576 | 20201116_xd_colostrum_1ng_glyco_hcd30_etd30_compound15.7230.7230.3 | controllerType=0 controllerNumber=1 scan=7230 | 19.0655 |
| 02900:3 | 10 | K.TPLTAN[+1971.719]ITK.S | HexNAc(5)Hex(5)Fuc(1) | N[+1972] | 977.4337 | 3 | 2930.2867 | 2930.2757 | 0 | 3.8 | 200 | Specific | 276.74 | 134.69 | 134.69 | 2.91 | 24 | >sp\|P01877\|IGHA2_HUMAN Immunoglobulin heavy constant alpha 2 OS=Homo sapiens OX=9606 GN=IGHA2 PE=1 SV=4 | 576 | 20201116_xd_colostrum_1ng_glyco_hcd30_etd30_compound15.5676.5676.3 | controllerType=0 controllerNumber=1 scan=5676 | 15.7994 |
| 04393:3 | 10 | K.TPLTAN[+2059.735]ITK.S | HexNAc(4)Hex(5)Fuc(1)NeuAc(1) | N[+2060] | 1006.7713 | 3 | 3018.2994 | 3018.2917 | 0 | 2.6 | 200 | Specific | 283.17 | 128.43 | 128.43 | 3.66 | 24 | >sp\|P01877\|IGHA2_HUMAN Immunoglobulin heavy constant alpha 2 OS=Homo sapiens OX=9606 GN=IGHA2 PE=1 SV=4 | 576 | 20201116_xd_colostrum_1ng_glyco_hcd30_etd30_compound15.7236.7236.3 | controllerType=0 controllerNumber=1 scan=7236 | 19.0818 |
| 03031:2 | 10 | K.TPLTAN[+1038.375]ITK.S | HexNAc(2)Hex(3)Fuc(1) | N[+1038] | 998.973 | 2 | 1996.9387 | 1996.9319 | 0 | 3.4 | 200 | Specific | 284.95 | 88.92 | 88.92 | 2.89 | 24 | >sp\|P01877\|IGHA2_HUMAN Immunoglobulin heavy constant alpha 2 OS=Homo sapiens OX=9606 GN=IGHA2 PE=1 SV=4 | 576 | 20201116_xd_colostrum_1ng_glyco_hcd30_etd30_compound15.5815.5815.2 | controllerType=0 controllerNumber=1 scan=5815 | 16.2613 |
| 02919:2 | 10 | K.TPLTAN[+1971.719]ITK.S | HexNAc(5)Hex(5)Fuc(1) | N[+1972] | 1465.6464 | 2 | 2930.2855 | 2930.2757 | 0 | 3.4 | 200 | Specific | 294.22 | 89.97 | 89.97 | 2.89 | 24 | >sp\|P01877\|IGHA2_HUMAN Immunoglobulin heavy constant alpha 2 OS=Homo sapiens OX=9606 GN=IGHA2 PE=1 SV=4 | 576 | 20201116_xd_colostrum_1ng_glyco_hcd30_etd30_compound15.5696.5696.2 | controllerType=0 controllerNumber=1 scan=5696 | 15.8627 |
| 02996:3 | 10 | K.TPLTAN[+1647.613]ITK.S | HexNAc(5)Hex(3)Fuc(1) | N[+1648] | 869.398 | 3 | 2606.1796 | 2606.17 | 0 | 3.7 | 200 | Specific | 295.76 | 137.01 | 137.01 | 3.04 | 24 | >sp\|P01877\|IGHA2_HUMAN Immunoglobulin heavy constant alpha 2 OS=Homo sapiens OX=9606 GN=IGHA2 PE=1 SV=4 | 576 | 20201116_xd_colostrum_1ng_glyco_hcd30_etd30_compound15.5778.5778.3 | controllerType=0 controllerNumber=1 scan=5778 | 16.1355 |
| 02988:2 | 10 | K.TPLTAN[+1647.613]ITK.S | HexNAc(5)Hex(3)Fuc(1) | N[+1648] | 1303.5937 | 2 | 2606.1801 | 2606.17 | 0 | 3.9 | 200 | Specific | 299.46 | 113.88 | 113.88 | 2.89 | 24 | >sp\|P01877\|IGHA2_HUMAN Immunoglobulin heavy constant alpha 2 OS=Homo sapiens OX=9606 GN=IGHA2 PE=1 SV=4 | 576 | 20201116_xd_colostrum_1ng_glyco_hcd30_etd30_compound15.5770.5770.2 | controllerType=0 controllerNumber=1 scan=5770 | 16.108 |
| 02928:2 | 10 | K.TPLTAN[+1606.587]ITK.S | HexNAc(4)Hex(4)Fuc(1) | N[+1607] | 1283.0784 | 2 | 2565.1495 | 2565.1435 | 0 | 2.4 | 200 | Specific | 305.37 | 107.21 | 107.21 | 3.16 | 24 | >sp\|P01877\|IGHA2_HUMAN Immunoglobulin heavy constant alpha 2 OS=Homo sapiens OX=9606 GN=IGHA2 PE=1 SV=4 | 576 | 20201116_xd_colostrum_1ng_glyco_hcd30_etd30_compound15.5706.5706.2 | controllerType=0 controllerNumber=1 scan=5706 | 15.8984 |
| 02945:3 | 10 | K.TPLTAN[+1809.666]ITK.S | HexNAc(5)Hex(4)Fuc(1) | N[+1810] | 923.4159 | 3 | 2768.233 | 2768.2228 | 0 | 3.7 | 200 | Specific | 306.16 | 130.85 | 130.85 | 2.91 | 24 | >sp\|P01877\|IGHA2_HUMAN Immunoglobulin heavy constant alpha 2 OS=Homo sapiens OX=9606 GN=IGHA2 PE=1 SV=4 | 576 | 20201116_xd_colostrum_1ng_glyco_hcd30_etd30_compound15.5724.5724.3 | controllerType=0 controllerNumber=1 scan=5724 | 15.9598 |
| 02955:2 | 10 | K.TPLTAN[+1809.666]ITK.S | HexNAc(5)Hex(4)Fuc(1) | N[+1810] | 1384.6203 | 2 | 2768.2333 | 2768.2228 | 0 | 3.8 | 200 | Specific | 313.44 | 115.43 | 115.43 | 2.89 | 24 | >sp\|P01877\|IGHA2_HUMAN Immunoglobulin heavy constant alpha 2 OS=Homo sapiens OX=9606 GN=IGHA2 PE=1 SV=4 | 576 | 20201116_xd_colostrum_1ng_glyco_hcd30_etd30_compound15.5735.5735.2 | controllerType=0 controllerNumber=1 scan=5735 | 15.999 |
| 02951:2 | 10 | K.TPLTAN[+1403.507]ITK.S | HexNAc(3)Hex(4)Fuc(1) | N[+1404] | 1181.5397 | 2 | 2362.0722 | 2362.0641 | 0 | 3.4 | 200 | Specific | 317.5 | 92.64 | 92.64 | 2.89 | 24 | >sp\|P01877\|IGHA2_HUMAN Immunoglobulin heavy constant alpha 2 OS=Homo sapiens OX=9606 GN=IGHA2 PE=1 SV=4 | 576 | 20201116_xd_colostrum_1ng_glyco_hcd30_etd30_compound15.5730.5730.2 | controllerType=0 controllerNumber=1 scan=5730 | 15.9822 |
| 02941:3 | 10 | K.TPLTAN[+1606.587]ITK.S | HexNAc(4)Hex(4)Fuc(1) | N[+1607] | 855.7224 | 3 | 2565.1526 | 2565.1435 | 0 | 3.6 | 200 | Specific | 341.19 | 140.27 | 140.27 | 3.16 | 24 | >sp\|P01877\|IGHA2_HUMAN Immunoglobulin heavy constant alpha 2 OS=Homo sapiens OX=9606 GN=IGHA2 PE=1 SV=4 | 576 | 20201116_xd_colostrum_1ng_glyco_hcd30_etd30_compound15.5720.5720.3 | controllerType=0 controllerNumber=1 scan=5720 | 15.9492 |
| 02987:2 | 10 | K.TPLTAN[+1444.534]ITK.S | HexNAc(4)Hex(3)Fuc(1) | N[+1445] | 1202.0538 | 2 | 2403.1002 | 2403.0906 | 0 | 4 | 200 | Specific | 350.48 | 109.53 | 109.53 | 2.95 | 24 | >sp\|P01877\|IGHA2_HUMAN Immunoglobulin heavy constant alpha 2 OS=Homo sapiens OX=9606 GN=IGHA2 PE=1 SV=4 | 576 | 20201116_xd_colostrum_1ng_glyco_hcd30_etd30_compound15.5769.5769.2 | controllerType=0 controllerNumber=1 scan=5769 | 16.103 |
| 03015:2 | 10 | K.TPLTAN[+1241.454]ITK.S | HexNAc(3)Hex(3)Fuc(1) | N[+1241] | 1100.5133 | 2 | 2200.0194 | 2200.0113 | 0 | 3.7 | 200 | Specific | 356.33 | 133.79 | 133.79 | 3.01 | 24 | >sp\|P01877\|IGHA2_HUMAN Immunoglobulin heavy constant alpha 2 OS=Homo sapiens OX=9606 GN=IGHA2 PE=1 SV=4 | 576 | 20201116_xd_colostrum_1ng_glyco_hcd30_etd30_compound15.5798.5798.2 | controllerType=0 controllerNumber=1 scan=5798 | 16.2028 |
| 03005:3 | 10 | K.TPLTAN[+1444.534]ITK.S | HexNAc(4)Hex(3)Fuc(1) | N[+1445] | 801.7044 | 3 | 2403.0986 | 2403.0906 | 0 | 3.3 | 200 | Specific | 401.69 | 214.58 | 214.58 | 3.47 | 24 | >sp\|P01877\|IGHA2_HUMAN Immunoglobulin heavy constant alpha 2 OS=Homo sapiens OX=9606 GN=IGHA2 PE=1 SV=4 | 576 | 20201116_xd_colostrum_1ng_glyco_hcd30_etd30_compound15.5788.5788.3 | controllerType=0 controllerNumber=1 scan=5788 | 16.161 |
| 03666:2 | 12 | K.AAIPSALDTN[+1216.423]SSK.S | HexNAc(2)Hex(5) | N[+1216] | 1246.0475 | 2 | 2491.0878 | 2491.0815 | 0 | 2.5 | 542 | Specific | 458.95 | 254.66 | 254.66 | 4.92 | 11 | >sp\|Q08380\|LG3BP_HUMAN Galectin-3-binding protein OS=Homo sapiens OX=9606 GN=LGALS3BP PE=1 SV=1 | 1562 | 20201116_xd_colostrum_1ng_glyco_hcd30_etd30_compound15.6477.6477.2 | controllerType=0 controllerNumber=1 scan=6477 | 17.7 |
| 08047:3 | 16 | R.IIVPLNNREN[+1501.555]ISDPTSPLR.T | HexNAc(5)Hex(3) | N[+1502] | 1217.2477 | 3 | 3649.7287 | 3649.7325 | 0 | -1 | 62 | Specific | 228.18 | 169.27 | 169.27 | 2.81 | 12 | >sp\|P01591\|IGJ_HUMAN Immunoglobulin J chain OS=Homo sapiens OX=9606 GN=JCHAIN PE=1 SV=4 | 1392 | 20201116_xd_colostrum_1ng_glyco_hcd30_etd30_compound15.11055.11055.3 | controllerType=0 controllerNumber=1 scan=11055 | 27.7585 |
| 07973:3 | 16 | R.IIVPLNNREN[+1663.608]ISDPTSPLR.T | HexNAc(5)Hex(4) | N[+1664] | 1271.2706 | 3 | 3811.7971 | 3811.7853 | 0 | 3.1 | 62 | Specific | 267.57 | 172.61 | 172.61 | 4.17 | 12 | >sp\|P01591\|IGJ_HUMAN Immunoglobulin J chain OS=Homo sapiens OX=9606 GN=JCHAIN PE=1 SV=4 | 1392 | 20201116_xd_colostrum_1ng_glyco_hcd30_etd30_compound15.10978.10978.3 | controllerType=0 controllerNumber=1 scan=10978 | 27.5615 |
| 08957:3 | 16 | R.IIVPLNNREN[+2059.735]ISDPTSPLR.T | HexNAc(4)Hex(5)Fuc(1)NeuAc(1) | N[+2060] | 1403.314 | 3 | 4207.9275 | 4207.9121 | 0 | 3.7 | 62 | Specific | 314.54 | 227.53 | 227.53 | 3.77 | 12 | >sp\|P01591\|IGJ_HUMAN Immunoglobulin J chain OS=Homo sapiens OX=9606 GN=JCHAIN PE=1 SV=4 | 1392 | 20201116_xd_colostrum_1ng_glyco_hcd30_etd30_compound15.12010.12010.3 | controllerType=0 controllerNumber=1 scan=12010 | 30.2493 |
| 09057:3 | 16 | R.IIVPLNNREN[+1856.656]ISDPTSPLR.T | HexNAc(3)Hex(5)Fuc(1)NeuAc(1) | N[+1857] | 1335.6177 | 3 | 4004.8386 | 4004.8327 | 0 | 1.5 | 62 | Specific | 354.26 | 274.19 | 274.19 | 4.03 | 12 | >sp\|P01591\|IGJ_HUMAN Immunoglobulin J chain OS=Homo sapiens OX=9606 GN=JCHAIN PE=1 SV=4 | 1392 | 20201116_xd_colostrum_1ng_glyco_hcd30_etd30_compound15.12115.12115.3 | controllerType=0 controllerNumber=1 scan=12115 | 30.5703 |
| 07940:3 | 16 | R.IIVPLNNREN[+1809.666]ISDPTSPLR.T | HexNAc(5)Hex(4)Fuc(1) | N[+1810] | 1319.9575 | 3 | 3957.8578 | 3957.8432 | 0 | 3.7 | 62 | Specific | 357.11 | 243.55 | 243.55 | 4.48 | 12 | >sp\|P01591\|IGJ_HUMAN Immunoglobulin J chain OS=Homo sapiens OX=9606 GN=JCHAIN PE=1 SV=4 | 1392 | 20201116_xd_colostrum_1ng_glyco_hcd30_etd30_compound15.10943.10943.3 | controllerType=0 controllerNumber=1 scan=10943 | 27.4524 |
| 09015:4 | 16 | R.IIVPLNNREN[+1913.677]ISDPTSPLR.T | HexNAc(4)Hex(5)NeuAc(1) | N[+1914] | 1016.2247 | 4 | 4061.8768 | 4061.8542 | 0 | 5.6 | 62 | Specific | 362.47 | 362.47 | 362.47 | 4.63 | 12 | >sp\|P01591\|IGJ_HUMAN Immunoglobulin J chain OS=Homo sapiens OX=9606 GN=JCHAIN PE=1 SV=4 | 1392 | 20201116_xd_colostrum_1ng_glyco_hcd30_etd30_compound15.12071.12071.4 | controllerType=0 controllerNumber=1 scan=12071 | 30.4456 |
| 07981:4 | 16 | R.IIVPLNNREN[+1663.608]ISDPTSPLR.T | HexNAc(5)Hex(4) | N[+1664] | 953.7038 | 4 | 3811.7935 | 3811.7853 | 0 | 2.2 | 62 | Specific | 468.9 | 447.62 | 447.62 | 7.09 | 12 | >sp\|P01591\|IGJ_HUMAN Immunoglobulin J chain OS=Homo sapiens OX=9606 GN=JCHAIN PE=1 SV=4 | 1392 | 20201116_xd_colostrum_1ng_glyco_hcd30_etd30_compound15.10986.10986.4 | controllerType=0 controllerNumber=1 scan=10986 | 27.5735 |
| 00540:3 | 18 | K.YKN[+2262.814]NSDISSTR.G | HexNAc(5)Hex(5)Fuc(1)NeuAc(1) | N[+2263] | 1183.1551 | 3 | 3547.4507 | 3547.4322 | 0 | 5.2 | 44 | Specific | 265.46 | 92.24 | 92.24 | 2.85 | 9 | >sp\|P01871\|IGHM_HUMAN Immunoglobulin heavy constant mu OS=Homo sapiens OX=9606 GN=IGHM PE=1 SV=4 | 328 | 20201116_xd_colostrum_1ng_glyco_hcd30_etd30_compound15.3224.3224.3 | controllerType=0 controllerNumber=1 scan=3224 | 11.2793 |
| 09232:3 | 18 | R.TVDKSTGKPTLYN[+1378.476]VSLVMSDTAGTC[+57.021]Y.- | HexNAc(2)Hex(6) | C[+57], N[+1378] | 1396.2846 | 3 | 4186.8393 | 4186.8187 | 0 | 4.9 | 428 | CRagged | 256.41 | 236.11 | 147.12 | 3.93 | 9 | >sp\|P01871\|IGHM_HUMAN Immunoglobulin heavy constant mu OS=Homo sapiens OX=9606 GN=IGHM PE=1 SV=4 | 328 | 20201116_xd_colostrum_1ng_glyco_hcd30_etd30_compound15.12299.12299.3 | controllerType=0 controllerNumber=1 scan=12299 | 31.0872 |
| 02253:3 | 20 | K.N[+1216.423]NSIPDKQITASSSYK.T | HexNAc(2)Hex(5) | N[+1216] | 990.4427 | 3 | 2969.3134 | 2969.2991 | 0 | 4.8 | 238 | Specific | 384.51 | 211.4 | 211.4 | 3.13 | 7 | >sp\|Q08431\|MFGM_HUMAN Lactadherin OS=Homo sapiens OX=9606 GN=MFGE8 PE=1 SV=3 | 1064 | 20201116_xd_colostrum_1ng_glyco_hcd30_etd30_compound15.5005.5005.3 | controllerType=0 controllerNumber=1 scan=5005 | 14.568 |
| 09213:3 | 24 | R.VN[+1216.423]LTTNTIAVTQTLPNAAYNNR.F | HexNAc(2)Hex(5) | N[+1216] | 1202.5663 | 3 | 3605.6845 | 3605.6699 | 0 | 4 | 351 | Specific | 527.84 | 498.34 | 498.34 | 7.76 | 5 | >sp\|Q6UX06\|OLFM4_HUMAN Olfactomedin-4 OS=Homo sapiens OX=9606 GN=OLFM4 PE=1 SV=1 | 624 | 20201116_xd_colostrum_1ng_glyco_hcd30_etd30_compound15.12279.12279.3 | controllerType=0 controllerNumber=1 scan=12279 | 31.0334 |
| 10828:4 | 27 | K.LKELPGVC[+57.021]N[+2351.851]ETMMALWEEC[+57.021]KPC[+57.021]LK.Q | HexNAc(4)Hex(5)Fuc(3)NeuAc(1) | C[+57]*3, N[+2352] | 1322.8173 | 4 | 5288.2472 | 5288.2355 | 0 | 2.2 | 95 | Specific | 406.43 | 246.31 | 246.31 | 4.63 | 9 | >sp\|P10909\|CLUS_HUMAN Clusterin OS=Homo sapiens OX=9606 GN=CLU PE=1 SV=1 | 1467 | 20201116_xd_colostrum_1ng_glyco_hcd30_etd30_compound15.13970.13970.4 | controllerType=0 controllerNumber=1 scan=13970 | 35.2537 |
| 00560:4 | 27 | R.EIRHN[+2059.735]STGC[+57.021]LR.M | HexNAc(4)Hex(5)Fuc(1)NeuAc(1) | C[+57], N[+2060] | 851.3572 | 4 | 3402.407 | 3402.3994 | 0 | 2.2 | 287 | Specific | 319.11 | 236.11 | 236.11 | 4.58 | 9 | >sp\|P10909\|CLUS_HUMAN Clusterin OS=Homo sapiens OX=9606 GN=CLU PE=1 SV=1 | 1467 | 20201116_xd_colostrum_1ng_glyco_hcd30_etd30_compound15.3245.3245.4 | controllerType=0 controllerNumber=1 scan=3245 | 11.355 |
| 00536:4 | 27 | R.EIRHN[+2351.851]STGC[+57.021]LR.M | HexNAc(4)Hex(5)Fuc(3)NeuAc(1) | C[+57], N[+2352] | 924.3863 | 4 | 3694.5233 | 3694.5152 | 0 | 2.2 | 287 | Specific | 560.5 | 403.38 | 403.38 | 7.87 | 9 | >sp\|P10909\|CLUS_HUMAN Clusterin OS=Homo sapiens OX=9606 GN=CLU PE=1 SV=1 | 1467 | 20201116_xd_colostrum_1ng_glyco_hcd30_etd30_compound15.3220.3220.4 | controllerType=0 controllerNumber=1 scan=3220 | 11.2667 |
| 15480:3 | 27 | K.MLN[+2059.735]TSSLLEQLNEQFNWVSR.L | HexNAc(4)Hex(5)Fuc(1)NeuAc(1) | N[+2060] | 1490.3173 | 3 | 4468.9373 | 4468.9217 | 0 | 3.5 | 352 | Specific | 647.33 | 424.98 | 424.98 | 9.84 | 9 | >sp\|P10909\|CLUS_HUMAN Clusterin OS=Homo sapiens OX=9606 GN=CLU PE=1 SV=1 | 1467 | 20201116_xd_colostrum_1ng_glyco_hcd30_etd30_compound15.18885.18885.3 | controllerType=0 controllerNumber=1 scan=18885 | 49.7363 |
| 11698:3 | 33 | K.TFYWDFYTN[+2351.851]R.T | HexNAc(4)Hex(5)Fuc(3)NeuAc(1) | N[+2352] | 1255.5015 | 3 | 3764.49 | 3764.4777 | 0 | 3.3 | 97 | Specific | 211.62 | 148.25 | 148.25 | 3.24 | 4 | >sp\|P12273\|PIP_HUMAN Prolactin-inducible protein OS=Homo sapiens OX=9606 GN=PIP PE=1 SV=1 | 647 | 20201116_xd_colostrum_1ng_glyco_hcd30_etd30_compound15.14883.14883.3 | controllerType=0 controllerNumber=1 scan=14883 | 37.6279 |
| 15518:5 | 34 | R.SDHGSSISC[+57.021]QPPAEIPGYLPADTVHLAVEFFN[+2350.830]LTHLPANLLQGASI | HexNAc(4)Hex(5)Fuc(1)NeuAc(2) | C[+57], N[+2351] | 1450.8682 | 5 | 7250.3119 | 7250.2714 | 0 | 5.6 | 48 | Specific | 326.01 | 326.01 | 326.01 | 4.73 | 5 | >sp\|P02750\|A2GL_HUMAN Leucine-rich alpha-2-glycoprotein OS=Homo sapiens OX=9606 GN=LRG1 PE=1 SV=2 | 518 | 20201116_xd_colostrum_1ng_glyco_hcd30_etd30_compound15.18927.18927.5 | controllerType=0 controllerNumber=1 scan=18927 | 49.9143 |
| 08725:3 | 49 | R.ADGTVNQIEGEATPVN[+1054.370]LTEPAKLEVK.F | HexNAc(2)Hex(4) | N[+1054] | 1259.9352 | 3 | 3777.7912 | 3777.7798 | 0 | 3 | 83 | Specific | 241.62 | 210.88 | 210.88 | 4.74 | 3 | >sp\|P05090\|APOD_HUMAN Apolipoprotein D OS=Homo sapiens OX=9606 GN=APOD PE=1 SV=1 | 594 | 20201116_xd_colostrum_1ng_glyco_hcd30_etd30_compound15.11768.11768.3 | controllerType=0 controllerNumber=1 scan=11768 | 29.6918 |
| 14747:4 | 1 | R.RTAGWNVPIGTLRPFLN[+2278.809]WTGPPEPIEAAVAR.F | HexNAc(5)Hex(6)NeuAc(1) | N[+2279] | 1417.1693 | 4 | 5665.6554 | 5665.6213 | 0 | 6 | 140 | Specific | 279.46 | 279.46 | 279.46 | 3.44 | 46 | >sp\|P02788\|TRFL_HUMAN Lactotransferrin OS=Homo sapiens OX=9606 GN=LTF PE=1 SV=6 | 1270 | 20201113_xd_milk_1ng_glyco_hcd30_etd30_compound15.18248.18248.4 | controllerType=0 controllerNumber=1 scan=18248 | 41.4604 |
| 15390:4 | 1 | R.RTAGWNVPIGTLRPFLN[+2716.983]WTGPPEPIEAAVAR.F | HexNAc(5)Hex(6)Fuc(3)NeuAc(1) | N[+2717] | 1526.7089 | 4 | 6103.8136 | 6103.795 | 0 | 3.1 | 140 | Specific | 664.55 | 654.35 | 654.35 | 10.57 | 46 | >sp\|P02788\|TRFL_HUMAN Lactotransferrin OS=Homo sapiens OX=9606 GN=LTF PE=1 SV=6 | 1270 | 20201113_xd_milk_1ng_glyco_hcd30_etd30_compound15.18927.18927.4 | controllerType=0 controllerNumber=1 scan=18927 | 43.4254 |
| 15454:4 | 1 | R.RTAGWNVPIGTLRPFLN[+2351.851]WTGPPEPIEAAVAR.F | HexNAc(4)Hex(5)Fuc(3)NeuAc(1) | N[+2352] | 1435.4257 | 4 | 5738.6808 | 5738.6628 | 0 | 3.1 | 140 | Specific | 871.13 | 644.89 | 644.89 | 14.57 | 46 | >sp\|P02788\|TRFL_HUMAN Lactotransferrin OS=Homo sapiens OX=9606 GN=LTF PE=1 SV=6 | 1270 | 20201113_xd_milk_1ng_glyco_hcd30_etd30_compound15.18994.18994.4 | controllerType=0 controllerNumber=1 scan=18994 | 43.5994 |
| 15466:4 | 1 | R.RTAGWNVPIGTLRPFLN[+2059.735]WTGPPEPIEAAVAR.F | HexNAc(4)Hex(5)Fuc(1)NeuAc(1) | N[+2060] | 1362.3944 | 4 | 5446.5558 | 5446.547 | 0 | 1.6 | 140 | Specific | 1112.55 | 1084.34 | 1084.34 | 19.54 | 46 | >sp\|P02788\|TRFL_HUMAN Lactotransferrin OS=Homo sapiens OX=9606 GN=LTF PE=1 SV=6 | 1270 | 20201113_xd_milk_1ng_glyco_hcd30_etd30_compound15.19006.19006.4 | controllerType=0 controllerNumber=1 scan=19006 | 43.6431 |
| 15693:3 | 1 | R.TAGWNVPIGTLRPFLN[+1768.640]WTGPPEPIEAAVAR.F | HexNAc(4)Hex(5)Fuc(1) | N[+1769] | 1667.1268 | 3 | 4999.3658 | 4999.3504 | 0 | 3.1 | 141 | Specific | 286.02 | 282.91 | 282.91 | 4.57 | 46 | >sp\|P02788\|TRFL_HUMAN Lactotransferrin OS=Homo sapiens OX=9606 GN=LTF PE=1 SV=6 | 1270 | 20201113_xd_milk_1ng_glyco_hcd30_etd30_compound15.19244.19244.3 | controllerType=0 controllerNumber=1 scan=19244 | [illegible] |
| 16358:3 | 1 | R.TAGWNVPIGTLRPFLN[+2059.735]WTGPPEPIEAAVAR.F | HexNAc(4)Hex(5)Fuc(1)NeuAc(1) | N[+2060] | 1764.1549 | 3 | 5290.4501 | 5290.4459 | 0 | 0.8 | 141 | Specific | 413.32 | 317.89 | 317.89 | 6.09 | 46 | >sp\|P02788\|TRFL_HUMAN Lactotransferrin OS=Homo sapiens OX=9606 GN=LTF PE=1 SV=6 | 1270 | 20201113_xd_milk_1ng_glyco_hcd30_etd30_compound15.19948.19948.3 | controllerType=0 controllerNumber=1 scan=19948 | 46.4348 |
| 16333:3 | 1 | R.TAGWNVPIGTLRPFLN[+2205.793]WTGPPEPIEAAVAR.F | HexNAc(4)Hex(5)Fuc(2)NeuAc(1) | N[+2206] | 1812.8481 | 3 | 5436.5298 | 5436.5038 | 0 | 4.8 | 141 | Specific | 569.32 | 447.55 | 447.55 | 7.73 | 46 | >sp\|P02788\|TRFL_HUMAN Lactotransferrin OS=Homo sapiens OX=9606 GN=LTF PE=1 SV=6 | 1270 | 20201113_xd_milk_1ng_glyco_hcd30_etd30_compound15.19921.19921.3 | controllerType=0 controllerNumber=1 scan=19921 | 46.3285 |
| 17299:4 | 1 | R.TAGWNVPIGTLRPFLN[+2350.830]WTGPPEPIEAAVAR.F | HexNAc(4)Hex(5)Fuc(1)NeuAc(2) | N[+2351] | 1396.1416 | 4 | 5581.5446 | 5581.5413 | 0 | 0.6 | 141 | Specific | 596.98 | 596.98 | 596.98 | 7.97 | 46 | >sp\|P02788\|TRFL_HUMAN Lactotransferrin OS=Homo sapiens OX=9606 GN=LTF PE=1 SV=6 | 1270 | 20201113_xd_milk_1ng_glyco_hcd30_etd30_compound15.20943.20943.4 | controllerType=0 controllerNumber=1 scan=20943 | 49.3423 |
| 16357:4 | 1 | R.TAGWNVPIGTLRPFLN[+2059.735]WTGPPEPIEAAVAR.F | HexNAc(4)Hex(5)Fuc(1)NeuAc(1) | N[+2060] | 1323.3693 | 4 | 5290.4552 | 5290.4459 | 0 | 1.8 | 141 | Specific | 841.07 | 823.55 | 823.55 | 14.29 | 46 | >sp\|P02788\|TRFL_HUMAN Lactotransferrin OS=Homo sapiens OX=9606 GN=LTF PE=1 SV=6 | 1270 | 20201113_xd_milk_1ng_glyco_hcd30_etd30_compound15.19947.19947.4 | controllerType=0 controllerNumber=1 scan=19947 | 46.4321 |
| 16321:4 | 1 | R.TAGWNVPIGTLRPFLN[+2570.925]WTGPPEPIEAAVAR.F | HexNAc(5)Hex(6)Fuc(2)NeuAc(1) | N[+2571] | 1451.1683 | 4 | 5801.6514 | 5801.636 | 0 | 2.7 | 141 | Specific | 876.01 | 871.38 | 871.38 | 15.31 | 46 | >sp\|P02788\|TRFL_HUMAN Lactotransferrin OS=Homo sapiens OX=9606 GN=LTF PE=1 SV=6 | 1270 | 20201113_xd_milk_1ng_glyco_hcd30_etd30_compound15.19907.19907.4 | controllerType=0 controllerNumber=1 scan=19907 | 46.2754 |
| 16332:4 | 1 | R.TAGWNVPIGTLRPFLN[+2205.793]WTGPPEPIEAAVAR.F | HexNAc(4)Hex(5)Fuc(2)NeuAc(1) | N[+2206] | 1359.8881 | 4 | 5436.5304 | 5436.5038 | 0 | 4.9 | 141 | Specific | 971.55 | 965.77 | 965.77 | 17.1 | 46 | >sp\|P02788\|TRFL_HUMAN Lactotransferrin OS=Homo sapiens OX=9606 GN=LTF PE=1 SV=6 | 1270 | 20201113_xd_milk_1ng_glyco_hcd30_etd30_compound15.19920.19920.4 | controllerType=0 controllerNumber=1 scan=19920 | 46.3233 |
| 14409:3 | 1 | R.TAGWNIPMGLLFN[+2133.772]QTGSC[+57.021]K.F | HexNAc(5)Hex(6)Fuc(1) | C[+57], N[+2134] | 1410.2691 | 3 | 4228.7927 | 4228.7817 | 0 | 2.6 | 485 | Specific | 310.39 | 251.15 | 204.19 | 5.19 | 46 | >sp\|P02788\|TRFL_HUMAN Lactotransferrin OS=Homo sapiens OX=9606 GN=LTF PE=1 SV=6 | 1270 | 20201113_xd_milk_1ng_glyco_hcd30_etd30_compound15.17893.17893.3 | controllerType=0 controllerNumber=1 scan=17893 | 40.5107 |
| 14549:2 | 1 | R.TAGWNIPMGLLFN[+1768.640]QTGSC[+57.021]K.F | HexNAc(4)Hex(5)Fuc(1) | C[+57], N[+1769] | 1932.3321 | 2 | 3863.657 | 3863.6495 | 0 | 1.9 | 485 | Specific | 345.24 | 210.45 | 210.45 | 5.04 | 46 | >sp\|P02788\|TRFL_HUMAN Lactotransferrin OS=Homo sapiens OX=9606 GN=LTF PE=1 SV=6 | 1270 | 20201113_xd_milk_1ng_glyco_hcd30_etd30_compound15.18039.18039.2 | controllerType=0 controllerNumber=1 scan=18039 | 40.8715 |
| 15860:3 | 1 | R.TAGWNIPMGLLFN[+2205.793]QTGSC[+57.021]K.F | HexNAc(4)Hex(5)Fuc(2)NeuAc(1) | C[+57], N[+2206] | 1434.2866 | 3 | 4300.8451 | 4300.8028 | 0 | 9.8 | 485 | Specific | 373.73 | 283.59 | 283.59 | 4.33 | 46 | >sp\|P02788\|TRFL_HUMAN Lactotransferrin OS=Homo sapiens OX=9606 GN=LTF PE=1 SV=6 | 1270 | 20201113_xd_milk_1ng_glyco_hcd30_etd30_compound15.19421.19421.3 | controllerType=0 controllerNumber=1 scan=19421 | 44.8142 |
| 14680:3 | 1 | R.TAGWNIPMGLLFN[+1622.582]QTGSC[+57.021]K.F | HexNAc(4)Hex(5) | C[+57], N[+1623] | 1239.8715 | 3 | 3717.6 | 3717.5916 | 0 | 2.3 | 485 | Specific | 375.87 | 291.3 | 230.8 | 5.5 | 46 | >sp\|P02788\|TRFL_HUMAN Lactotransferrin OS=Homo sapiens OX=9606 GN=LTF PE=1 SV=6 | 1270 | 20201113_xd_milk_1ng_glyco_hcd30_etd30_compound15.18178.18178.3 | controllerType=0 controllerNumber=1 scan=18178 | 41.2794 |
| 15878:3 | 1 | R.TAGWNIPMGLLFN[+2059.735]QTGSC[+57.021]K.F | HexNAc(4)Hex(5)Fuc(1)NeuAc(1) | C[+57], N[+2060] | 1385.5914 | 3 | 4154.7595 | 4154.7449 | 0 | 3.5 | 485 | Specific | 382.95 | 260.16 | 260.16 | 5.33 | 46 | >sp\|P02788\|TRFL_HUMAN Lactotransferrin OS=Homo sapiens OX=9606 GN=LTF PE=1 SV=6 | 1270 | 20201113_xd_milk_1ng_glyco_hcd30_etd30_compound15.19440.19440.3 | controllerType=0 controllerNumber=1 scan=19440 | 44.8759 |
| 14388:3 | 1 | R.TAGWNIPMGLLFN[+2279.830]QTGSC[+57.021]K.F | HexNAc(5)Hex(6)Fuc(2) | C[+57], N[+2280] | 1458.9557 | 3 | 4374.8527 | 4374.8396 | 0 | 3 | 485 | Specific | 386.1 | 326.34 | 238.96 | 5.57 | 46 | >sp\|P02788\|TRFL_HUMAN Lactotransferrin OS=Homo sapiens OX=9606 GN=LTF PE=1 SV=6 | 1270 | 20201113_xd_milk_1ng_glyco_hcd30_etd30_compound15.17871.17871.3 | controllerType=0 controllerNumber=1 scan=17871 | 40.4547 |
| 16028:3 | 1 | R.TAGWNIPMGLLFN[+1872.651]QTGSC[+57.021]K.F | HexNAc(3)Hex(6)NeuAc(1) | C[+57], N[+1873] | 1323.2287 | 3 | 3967.6715 | 3967.6604 | 0 | 2.8 | 485 | Specific | 393.9 | 260.94 | 244.24 | 5.5 | 46 | >sp\|P02788\|TRFL_HUMAN Lactotransferrin OS=Homo sapiens OX=9606 GN=LTF PE=1 SV=6 | 1270 | 20201113_xd_milk_1ng_glyco_hcd30_etd30_compound15.19599.19599.3 | controllerType=0 controllerNumber=1 scan=19599 | 45.3976 |
| 14531:3 | 1 | R.TAGWNIPMGLLFN[+1768.640]QTGSC[+57.021]K.F | HexNAc(4)Hex(5)Fuc(1) | C[+57], N[+1769] | 1288.5584 | 3 | 3863.6607 | 3863.6495 | 0 | 2.9 | 485 | Specific | 403.96 | 288.37 | 288.37 | 5.47 | 46 | >sp\|P02788\|TRFL_HUMAN Lactotransferrin OS=Homo sapiens OX=9606 GN=LTF PE=1 SV=6 | 1270 | 20201113_xd_milk_1ng_glyco_hcd30_etd30_compound15.18021.18021.3 | controllerType=0 controllerNumber=1 scan=18021 | 40.8292 |
| 14562:4 | 1 | R.TAGWNIPMGLLFN[+1768.640]QTGSC[+57.021]K.F | HexNAc(4)Hex(5)Fuc(1) | C[+57], N[+1769] | 966.6687 | 4 | 3863.6529 | 3863.6495 | 0 | 0.9 | 485 | Specific | 411.19 | 363.26 | 363.26 | 4.16 | 46 | >sp\|P02788\|TRFL_HUMAN Lactotransferrin OS=Homo sapiens OX=9606 GN=LTF PE=1 SV=6 | 1270 | 20201113_xd_milk_1ng_glyco_hcd30_etd30_compound15.18053.18053.4 | controllerType=0 controllerNumber=1 scan=18053 | 40.9106 |
| 16018:4 | 1 | R.TAGWNIPMGLLFN[+1913.677]QTGSC[+57.021]K.F | HexNAc(4)Hex(5)NeuAc(1) | C[+57], N[+1914] | 1002.9292 | 4 | 4008.6951 | 4008.687 | 0 | 2 | 485 | Specific | 427.26 | 341.62 | 341.62 | 4.9 | 46 | >sp\|P02788\|TRFL_HUMAN Lactotransferrin OS=Homo sapiens OX=9606 GN=LTF PE=1 SV=6 | 1270 | 20201113_xd_milk_1ng_glyco_hcd30_etd30_compound15.19588.19588.4 | controllerType=0 controllerNumber=1 scan=19588 | 45.3646 |
| 17415:3 | 1 | R.TAGWNIPMGLLFN[+2204.772]QTGSC[+57.021]K.F | HexNAc(4)Hex(5)NeuAc(2) | C[+57], N[+2205] | 1433.9354 | 3 | 4299.7915 | 4299.7824 | 0 | 2.1 | 485 | Specific | 428.11 | 292.7 | 292.7 | 5.45 | 46 | >sp\|P02788\|TRFL_HUMAN Lactotransferrin OS=Homo sapiens OX=9606 GN=LTF PE=1 SV=6 | 1270 | 20201113_xd_milk_1ng_glyco_hcd30_etd30_compound15.21065.21065.3 | controllerType=0 controllerNumber=1 scan=21065 | 49.7053 |
| 17300:3 | 1 | R.TAGWNIPMGLLFN[+2350.830]QTGSC[+57.021]K.F | HexNAc(4)Hex(5)Fuc(1)NeuAc(2) | C[+57], N[+2351] | 1482.6216 | 3 | 4445.8504 | 4445.8403 | 0 | 2.3 | 485 | Specific | 433.97 | 257.72 | 257.72 | 5.3 | 46 | >sp\|P02788\|TRFL_HUMAN Lactotransferrin OS=Homo sapiens OX=9606 GN=LTF PE=1 SV=6 | 1270 | 20201113_xd_milk_1ng_glyco_hcd30_etd30_compound15.20944.20944.3 | controllerType=0 controllerNumber=1 scan=20944 | 49.3459 |
| 15989:3 | 1 | R.TAGWNIPMGLLFN[+1913.677]QTGSC[+57.021]K.F | HexNAc(4)Hex(5)NeuAc(1) | C[+57], N[+1914] | 1336.9056 | 3 | 4008.7022 | 4008.687 | 0 | 3.8 | 485 | Specific | 465.63 | 336.07 | 336.07 | 5.97 | 46 | >sp\|P02788\|TRFL_HUMAN Lactotransferrin OS=Homo sapiens OX=9606 GN=LTF PE=1 SV=6 | 1270 | 20201113_xd_milk_1ng_glyco_hcd30_etd30_compound15.19558.19558.3 | controllerType=0 controllerNumber=1 scan=19558 | 45.27 |
| 14387:3 | 1 | R.TAGWNIPM[+15.995]GLLFN[+1913.677]QTGSC[+57.021]K.F | HexNAc(4)Hex(5)NeuAc(1) | C[+57], M[+16], N[+1914] | 1342.2357 | 3 | 4024.6924 | 4024.6819 | 0 | 2.6 | 485 | Specific | 300.53 | 222.58 | 222.58 | 5.09 | 46 | >sp\|P02788\|TRFL_HUMAN Lactotransferrin OS=Homo sapiens OX=9606 GN=LTF PE=1 SV=6 | 1270 | 20201113_xd_milk_1ng_glyco_hcd30_etd30_compound15.17870.17870.3 | controllerType=0 controllerNumber=1 scan=17870 | 40.4504 |
| 16134:4 | 1 | R.TAGWNIPMGLLFN[+2059.735]QTGSC[+57.021]KFDEYFSQSC[+57.021]APGSDPR.S | HexNAc(4)Hex(5)Fuc(1)NeuAc(1) | C[+57]*2, N[+2060] | 1500.3804 | 4 | 5998.4997 | 5998.492 | 0 | 1.3 | 485 | Specific | 307.57 | 238.76 | 238.76 | 3.21 | 46 | >sp\|P02788\|TRFL_HUMAN Lactotransferrin OS=Homo sapiens OX=9606 GN=LTF PE=1 SV=6 | 1270 | 20201113_xd_milk_1ng_glyco_hcd30_etd30_compound15.19711.19711.4 | controllerType=0 controllerNumber=1 scan=19711 | 45.75 |
| 17014:4 | 1 | R.TAGWNIPMGLLFN[+2350.830]QTGSC[+57.021]KFDEYFSQSC[+57.021]APGSDPR.S | HexNAc(4)Hex(5)Fuc(1)NeuAc(2) | C[+57]*2, N[+2351] | 1573.1578 | 4 | 6289.6095 | 6289.5875 | 0 | 3.5 | 485 | Specific | 336.76 | 276.27 | 276.27 | 4.72 | 46 | >sp\|P02788\|TRFL_HUMAN Lactotransferrin OS=Homo sapiens OX=9606 GN=LTF PE=1 SV=6 | 1270 | 20201113_xd_milk_1ng_glyco_hcd30_etd30_compound15.20642.20642.4 | controllerType=0 controllerNumber=1 scan=20642 | 48.454 |
| 15328:3 | 1 | R.TAGWNIPMGLLFN[+1768.640]QTGSC[+57.021]KFDEYFSQSC[+57.021]APGSDPR.S | HexNAc(4)Hex(5)Fuc(1) | C[+57]*2, N[+1769] | 1903.1443 | 3 | 5707.4182 | 5707.3966 | 0 | 3.8 | 485 | Specific | 366.21 | 299.73 | 299.73 | 4.85 | 46 | >sp\|P02788\|TRFL_HUMAN Lactotransferrin OS=Homo sapiens OX=9606 GN=LTF PE=1 SV=6 | 1270 | 20201113_xd_milk_1ng_glyco_hcd30_etd30_compound15.18862.18862.3 | controllerType=0 controllerNumber=1 scan=18862 | 43.2681 |
| 17058:4 | 1 | R.TAGWNIPMGLLFN[+2204.772]QTGSC[+57.021]KFDEYFSQSC[+57.021]APGSDPR.S | HexNAc(4)Hex(5)NeuAc(2) | C[+57]*2, N[+2205] | 1536.6427 | 4 | 6143.549 | 6143.5295 | 0 | 3.2 | 485 | Specific | 485.24 | 450.54 | 301.06 | 6.76 | 46 | >sp\|P02788\|TRFL_HUMAN Lactotransferrin OS=Homo sapiens OX=9606 GN=LTF PE=1 SV=6 | 1270 | 20201113_xd_milk_1ng_glyco_hcd30_etd30_compound15.20688.20688.4 | controllerType=0 controllerNumber=1 scan=20688 | 48.5863 |
| 16061:4 | 1 | R.TAGWNIPMGLLFN[+2205.793]QTGSC[+57.021]KFDEYFSQSC[+57.021]APGSDPR.S | HexNAc(4)Hex(5)Fuc(2)NeuAc(1) | C[+57]*2, N[+2206] | 1536.8998 | 4 | 6144.5773 | 6144.5499 | 0 | 4.5 | 485 | Specific | 519.52 | 466.53 | 466.53 | 6.55 | 46 | >sp\|P02788\|TRFL_HUMAN Lactotransferrin OS=Homo sapiens OX=9606 GN=LTF PE=1 SV=6 | 1270 | 20201113_xd_milk_1ng_glyco_hcd30_etd30_compound15.19634.19634.4 | controllerType=0 controllerNumber=1 scan=19634 | 45.5138 |
| 16097:4 | 1 | R.TAGWNIPMGLLFN[+2059.735]QTGSC[+57.021]KFDEYFSQSC[+57.021]APGSDPR.S | HexNAc(4)Hex(5)Fuc(1)NeuAc(1) | C[+57]*2, N[+2060] | 1500.3906 | 4 | 5998.5407 | 5998.492 | 0 | 8.1 | 485 | Specific | 763.85 | 712.11 | 712.11 | 11.04 | 46 | >sp\|P02788\|TRFL_HUMAN Lactotransferrin OS=Homo sapiens OX=9606 GN=LTF PE=1 SV=6 | 1270 | 20201113_xd_milk_1ng_glyco_hcd30_etd30_compound15.19672.19672.4 | controllerType=0 controllerNumber=1 scan=19672 | 45.6248 |
| 15322:4 | 1 | R.TAGWNIPMGLLFN[+1768.640]QTGSC[+57.021]KFDEYFSQSC[+57.021]APGSDPR.S | HexNAc(4)Hex(5)Fuc(1) | C[+57]*2, N[+1769] | 1427.6111 | 4 | 5707.4225 | 5707.3966 | 0 | 4.5 | 485 | Specific | 769.97 | 682.23 | 682.23 | 11.8 | 46 | >sp\|P02788\|TRFL_HUMAN Lactotransferrin OS=Homo sapiens OX=9606 GN=LTF PE=1 SV=6 | 1270 | 20201113_xd_milk_1ng_glyco_hcd30_etd30_compound15.18856.18856.4 | controllerType=0 controllerNumber=1 scan=18856 | 43.2521 |
| 08292:3 | 1 | R.N[+2205.793]GSDC[+57.021]PDKFC[+57.021]LFQSETK.N | HexNAc(4)Hex(5)Fuc(2)NeuAc(1) | C[+57]*2, N[+2206] | 1413.5652 | 3 | 4238.6812 | 4238.6667 | 0 | 3.4 | 642 | Specific | 274.63 | 174.7 | 174.7 | 4.18 | 46 | >sp\|P02788\|TRFL_HUMAN Lactotransferrin OS=Homo sapiens OX=9606 GN=LTF PE=1 SV=6 | 1270 | 20201113_xd_milk_1ng_glyco_hcd30_etd30_compound15.11485.11485.3 | controllerType=0 controllerNumber=1 scan=11485 | 26.7532 |
| 15839:3 | 2 | R.LSLHRPALEDLLLGSEAN[+2116.756]LTC[+57.021]TLTGLR.D | HexNAc(5)Hex(5)NeuAc(1) | C[+57], N[+2117] | 1694.1328 | 3 | 5080.3838 | 5080.3547 | 0 | 5.7 | 127 | Specific | 280.09 | 231.4 | 231.4 | 4.81 | 28 | >sp\|P01876\|IGHA1_HUMAN Immunoglobulin heavy constant alpha 1 OS=Homo sapiens OX=9606 GN=IGHA1 PE=1 SV=2 | 255 | 20201113_xd_milk_1ng_glyco_hcd30_etd30_compound15.19398.19398.3 | controllerType=0 controllerNumber=1 scan=19398 | 44.7386 |
| 15129:3 | 2 | R.LSLHRPALEDLLLGSEAN[+1501.555]LTC[+57.021]TLTGLR.D | HexNAc(5)Hex(3) | C[+57], N[+1502] | 1489.0623 | 3 | 4465.1724 | 4465.1536 | 0 | 4.2 | 127 | Specific | 304.75 | 304.75 | 304.75 | 5.16 | 28 | >sp\|P01876\|IGHA1_HUMAN Immunoglobulin heavy constant alpha 1 OS=Homo sapiens OX=9606 GN=IGHA1 PE=1 SV=2 | 255 | 20201113_xd_milk_1ng_glyco_hcd30_etd30_compound15.18653.18653.3 | controllerType=0 controllerNumber=1 scan=18653 | 42.6575 |
| 15153:3 | 2 | R.LSLHRPALEDLLLGSEAN[+1460.529]LTC[+57.021]TLTGLR.D | HexNAc(4)Hex(4) | C[+57], N[+1461] | 1475.386 | 3 | 4424.1436 | 4424.1271 | 0 | 3.7 | 127 | Specific | 313.33 | 256.55 | 256.55 | 5.13 | 28 | >sp\|P01876\|IGHA1_HUMAN Immunoglobulin heavy constant alpha 1 OS=Homo sapiens OX=9606 GN=IGHA1 PE=1 SV=2 | 255 | 20201113_xd_milk_1ng_glyco_hcd30_etd30_compound15.18678.18678.3 | controllerType=0 controllerNumber=1 scan=18678 | 42.723 |
| 15083:3 | 2 | R.LSLHRPALEDLLLGSEAN[+1702.581]LTC[+57.021]TLTGLR.D | HexNAc(2)Hex(8) | C[+57], N[+1703] | 1556.0683 | 3 | 4666.1903 | 4666.1796 | 0 | 2.3 | 127 | Specific | 372.26 | 350.71 | 350.71 | 5.82 | 28 | >sp\|P01876\|IGHA1_HUMAN Immunoglobulin heavy constant alpha 1 OS=Homo sapiens OX=9606 GN=IGHA1 PE=1 SV=2 | 255 | 20201113_xd_milk_1ng_glyco_hcd30_etd30_compound15.18604.18604.3 | controllerType=0 controllerNumber=1 scan=18604 | 42.5158 |
| 15170:5 | 2 | R.LSLHRPALEDLLLGSEAN[+1501.555]LTC[+57.021]TLTGLR.D | HexNAc(5)Hex(3) | C[+57], N[+1502] | 893.843 | 5 | 4465.1861 | 4465.1536 | 0 | 7.3 | 127 | Specific | 400.53 | 400.53 | 400.53 | 5.03 | 28 | >sp\|P01876\|IGHA1_HUMAN Immunoglobulin heavy constant alpha 1 OS=Homo sapiens OX=9606 GN=IGHA1 PE=1 SV=2 | 255 | 20201113_xd_milk_1ng_glyco_hcd30_etd30_compound15.18696.18696.5 | controllerType=0 controllerNumber=1 scan=18696 | 42.7824 |
| 15105:3 | 2 | R.LSLHRPALEDLLLGSEAN[+1622.582]LTC[+57.021]TLTGLR.D | HexNAc(4)Hex(5) | C[+57], N[+1623] | 1529.4015 | 3 | 4586.1901 | 4586.1799 | 0 | 2.2 | 127 | Specific | 437.07 | 380.11 | 380.11 | 6.96 | 28 | >sp\|P01876\|IGHA1_HUMAN Immunoglobulin heavy constant alpha 1 OS=Homo sapiens OX=9606 GN=IGHA1 PE=1 SV=2 | 255 | 20201113_xd_milk_1ng_glyco_hcd30_etd30_compound15.18627.18627.3 | controllerType=0 controllerNumber=1 scan=18627 | 42.5773 |
| 15084:3 | 2 | R.LSLHRPALEDLLLGSEAN[+1540.529]LTC[+57.021]TLTGLR.D | HexNAc(2)Hex(7) | C[+57], N[+1541] | 1502.0522 | 3 | 4504.142 | 4504.1268 | 0 | 3.4 | 127 | Specific | 444.08 | 444.08 | 444.08 | 7.32 | 28 | >sp\|P01876\|IGHA1_HUMAN Immunoglobulin heavy constant alpha 1 OS=Homo sapiens OX=9606 GN=IGHA1 PE=1 SV=2 | 255 | 20201113_xd_milk_1ng_glyco_hcd30_etd30_compound15.18605.18605.3 | controllerType=0 controllerNumber=1 scan=18605 | 42.5204 |
| 15049:3 | 2 | R.LSLHRPALEDLLLGSEAN[+1864.634]LTC[+57.021]TLTGLR.D | HexNAc(2)Hex(9) | C[+57], N[+1865] | 1610.0854 | 3 | 4828.2416 | 4828.2324 | 0 | 1.9 | 127 | Specific | 447.26 | 414.83 | 414.83 | 7.3 | 28 | >sp\|P01876\|IGHA1_HUMAN Immunoglobulin heavy constant alpha 1 OS=Homo sapiens OX=9606 GN=IGHA1 PE=1 SV=2 | 255 | 20201113_xd_milk_1ng_glyco_hcd30_etd30_compound15.18568.18568.3 | controllerType=0 controllerNumber=1 scan=18568 | 42.4034 |
| 15090:3 | 2 | R.LSLHRPALEDLLLGSEAN[+1663.608]LTC[+57.021]TLTGLR.D | HexNAc(5)Hex(4) | C[+57], N[+1664] | 1543.0788 | 3 | 4627.2218 | 4627.2064 | 0 | 3.3 | 127 | Specific | 458.21 | 427.22 | 427.22 | 7.32 | 28 | >sp\|P01876\|IGHA1_HUMAN Immunoglobulin heavy constant alpha 1 OS=Homo sapiens OX=9606 GN=IGHA1 PE=1 SV=2 | 255 | 20201113_xd_milk_1ng_glyco_hcd30_etd30_compound15.18611.18611.3 | controllerType=0 controllerNumber=1 scan=18611 | 42.5357 |
| 15148:4 | 2 | R.LSLHRPALEDLLLGSEAN[+1460.529]LTC[+57.021]TLTGLR.D | HexNAc(4)Hex(4) | C[+57], N[+1461] | 1106.7914 | 4 | 4424.1439 | 4424.1271 | 0 | 3.8 | 127 | Specific | 473.21 | 473.21 | 473.21 | 7.74 | 28 | >sp\|P01876\|IGHA1_HUMAN Immunoglobulin heavy constant alpha 1 OS=Homo sapiens OX=9606 GN=IGHA1 PE=1 SV=2 | 255 | 20201113_xd_milk_1ng_glyco_hcd30_etd30_compound15.18673.18673.4 | controllerType=0 controllerNumber=1 scan=18673 | 42.7098 |
| 15104:3 | 2 | R.LSLHRPALEDLLLGSEAN[+1378.476]LTC[+57.021]TLTGLR.D | HexNAc(2)Hex(6) | C[+57], N[+1378] | 1448.0323 | 3 | 4342.0823 | 4342.074 | 0 | 1.9 | 127 | Specific | 474.1 | 466.26 | 466.26 | 7.3 | 28 | >sp\|P01876\|IGHA1_HUMAN Immunoglobulin heavy constant alpha 1 OS=Homo sapiens OX=9606 GN=IGHA1 PE=1 SV=2 | 255 | 20201113_xd_milk_1ng_glyco_hcd30_etd30_compound15.18626.18626.3 | controllerType=0 controllerNumber=1 scan=18626 | 42.574 |
| 15134:3 | 2 | R.LSLHRPALEDLLLGSEAN[+1216.423]LTC[+57.021]TLTGLR.D | HexNAc(2)Hex(5) | C[+57], N[+1216] | 1394.0164 | 3 | 4180.0347 | 4180.0211 | 0 | 3.2 | 127 | Specific | 510.91 | 510.91 | 510.91 | 7.36 | 28 | >sp\|P01876\|IGHA1_HUMAN Immunoglobulin heavy constant alpha 1 OS=Homo sapiens OX=9606 GN=IGHA1 PE=1 SV=2 | 255 | 20201113_xd_milk_1ng_glyco_hcd30_etd30_compound15.18658.18658.3 | controllerType=0 controllerNumber=1 scan=18658 | 42.671 |
| 15819:4 | 2 | R.LSLHRPALEDLLLGSEAN[+1913.677]LTC[+57.021]TLTGLR.D | HexNAc(4)Hex(5)NeuAc(1) | C[+57], N[+1914] | 1220.0791 | 4 | 4877.2947 | 4877.2753 | 0 | 4 | 127 | Specific | 738.65 | 691.18 | 691.18 | 12.04 | 28 | >sp\|P01876\|IGHA1_HUMAN Immunoglobulin heavy constant alpha 1 OS=Homo sapiens OX=9606 GN=IGHA1 PE=1 SV=2 | 255 | 20201113_xd_milk_1ng_glyco_hcd30_etd30_compound15.19377.19377.4 | controllerType=0 controllerNumber=1 scan=19377 | 44.6715 |
| 15077:4 | 2 | R.LSLHRPALEDLLLGSEAN[+1702.581]LTC[+57.021]TLTGLR.D | HexNAc(2)Hex(8) | C[+57], N[+1703] | 1167.3028 | 4 | 4666.1893 | 4666.1796 | 0 | 2.1 | 127 | Specific | 761.78 | 761.78 | 761.78 | 13.23 | 28 | >sp\|P01876\|IGHA1_HUMAN Immunoglobulin heavy constant alpha 1 OS=Homo sapiens OX=9606 GN=IGHA1 PE=1 SV=2 | 255 | 20201113_xd_milk_1ng_glyco_hcd30_etd30_compound15.18598.18598.4 | controllerType=0 controllerNumber=1 scan=18598 | 42.5008 |
| 15099:4 | 2 | R.LSLHRPALEDLLLGSEAN[+1622.582]LTC[+57.021]TLTGLR.D | HexNAc(4)Hex(5) | C[+57], N[+1623] | 1147.303 | 4 | 4586.1903 | 4586.1799 | 0 | 2.3 | 127 | Specific | 789.03 | 680.75 | 680.75 | 13.25 | 28 | >sp\|P01876\|IGHA1_HUMAN Immunoglobulin heavy constant alpha 1 OS=Homo sapiens OX=9606 GN=IGHA1 PE=1 SV=2 | 255 | 20201113_xd_milk_1ng_glyco_hcd30_etd30_compound15.18621.18621.4 | controllerType=0 controllerNumber=1 scan=18621 | 42.5616 |
| 15078:4 | 2 | R.LSLHRPALEDLLLGSEAN[+1663.608]LTC[+57.021]TLTGLR.D | HexNAc(5)Hex(4) | C[+57], N[+1664] | 1157.56 | 4 | 4627.2181 | 4627.2064 | 0 | 2.5 | 127 | Specific | 795.75 | 795.75 | 795.75 | 13.25 | 28 | >sp\|P01876\|IGHA1_HUMAN Immunoglobulin heavy constant alpha 1 OS=Homo sapiens OX=9606 GN=IGHA1 PE=1 SV=2 | 255 | 20201113_xd_milk_1ng_glyco_hcd30_etd30_compound15.18599.18599.4 | controllerType=0 controllerNumber=1 scan=18599 | 42.504 |
| 15828:4 | 2 | R.LSLHRPALEDLLLGSEAN[+2116.756]LTC[+57.021]TLTGLR.D | HexNAc(5)Hex(5)NeuAc(1) | C[+57], N[+2117] | 1270.8533 | 4 | 5080.3913 | 5080.3547 | 0 | 7.2 | 127 | Specific | 795.88 | 763.89 | 763.89 | 12.28 | 28 | >sp\|P01876\|IGHA1_HUMAN Immunoglobulin heavy constant alpha 1 OS=Homo sapiens OX=9606 GN=IGHA1 PE=1 SV=2 | 255 | 20201113_xd_milk_1ng_glyco_hcd30_etd30_compound15.19387.19387.4 | controllerType=0 controllerNumber=1 scan=19387 | 44.7039 |
| 15161:4 | 2 | R.LSLHRPALEDLLLGSEAN[+1298.476]LTC[+57.021]TLTGLR.D | HexNAc(4)Hex(3) | C[+57], N[+1298] | 1066.2776 | 4 | 4262.0887 | 4262.0742 | 0 | 3.4 | 127 | Specific | 806.28 | 806.28 | 806.28 | 14.25 | 28 | >sp\|P01876\|IGHA1_HUMAN Immunoglobulin heavy constant alpha 1 OS=Homo sapiens OX=9606 GN=IGHA1 PE=1 SV=2 | 255 | 20201113_xd_milk_1ng_glyco_hcd30_etd30_compound15.18686.18686.4 | controllerType=0 controllerNumber=1 scan=18686 | 42.7586 |
| 15128:4 | 2 | R.LSLHRPALEDLLLGSEAN[+1501.555]LTC[+57.021]TLTGLR.D | HexNAc(5)Hex(3) | C[+57], N[+1502] | 1117.0506 | 4 | 4465.1805 | 4465.1536 | 0 | 6 | 127 | Specific | 943.74 | 931.41 | 931.41 | 15.94 | 28 | >sp\|P01876\|IGHA1_HUMAN Immunoglobulin heavy constant alpha 1 OS=Homo sapiens OX=9606 GN=IGHA1 PE=1 SV=2 | 255 | 20201113_xd_milk_1ng_glyco_hcd30_etd30_compound15.18652.18652.4 | controllerType=0 controllerNumber=1 scan=18652 | 42.6544 |
| 15053:4 | 2 | R.LSLHRPALEDLLLGSEAN[+1864.634]LTC[+57.021]TLTGLR.D | HexNAc(2)Hex(9) | C[+57], N[+1865] | 1207.8157 | 4 | 4828.241 | 4828.2324 | 0 | 1.8 | 127 | Specific | 953.2 | 905.75 | 905.75 | 17.23 | 28 | >sp\|P01876\|IGHA1_HUMAN Immunoglobulin heavy constant alpha 1 OS=Homo sapiens OX=9606 GN=IGHA1 PE=1 SV=2 | 255 | 20201113_xd_milk_1ng_glyco_hcd30_etd30_compound15.18572.18572.4 | controllerType=0 controllerNumber=1 scan=18572 | 42.4164 |
| 15094:4 | 2 | R.LSLHRPALEDLLLGSEAN[+1540.529]LTC[+57.021]TLTGLR.D | HexNAc(2)Hex(7) | C[+57], N[+1541] | 1126.7891 | 4 | 4504.1346 | 4504.1268 | 0 | 1.7 | 127 | Specific | 955.27 | 912.77 | 912.77 | 17.23 | 28 | >sp\|P01876\|IGHA1_HUMAN Immunoglobulin heavy constant alpha 1 OS=Homo sapiens OX=9606 GN=IGHA1 PE=1 SV=2 | 255 | 20201113_xd_milk_1ng_glyco_hcd30_etd30_compound15.18616.18616.4 | controllerType=0 controllerNumber=1 scan=18616 | 42.5497 |
| 15133:4 | 2 | R.LSLHRPALEDLLLGSEAN[+1216.423]LTC[+57.021]TLTGLR.D | HexNAc(2)Hex(5) | C[+57], N[+1216] | 1045.7643 | 4 | 4180.0355 | 4180.0211 | 0 | 3.4 | 127 | Specific | 1003.84 | 939.11 | 939.11 | 18.25 | 28 | >sp\|P01876\|IGHA1_HUMAN Immunoglobulin heavy constant alpha 1 OS=Homo sapiens OX=9606 GN=IGHA1 PE=1 SV=2 | 255 | 20201113_xd_milk_1ng_glyco_hcd30_etd30_compound15.18657.18657.4 | controllerType=0 controllerNumber=1 scan=18657 | 42.6652 |
| 15095:4 | 2 | R.LSLHRPALEDLLLGSEAN[+1378.476]LTC[+57.021]TLTGLR.D | HexNAc(2)Hex(6) | C[+57], N[+1378] | 1086.2759 | 4 | 4342.0819 | 4342.074 | 0 | 1.8 | 127 | Specific | 1037.78 | 974.54 | 974.54 | 18.23 | 28 | >sp\|P01876\|IGHA1_HUMAN Immunoglobulin heavy constant alpha 1 OS=Homo sapiens OX=9606 GN=IGHA1 PE=1 SV=2 | 255 | 20201113_xd_milk_1ng_glyco_hcd30_etd30_compound15.18617.18617.4 | controllerType=0 controllerNumber=1 scan=18617 | 42.5532 |
| 10098:3 | 2 | R.LAGKPTHVN[+1702.581]VSVVMAEVDGTC[+57.021]Y.- | HexNAc(2)Hex(8) | C[+57], N[+1703] | 1350.5827 | 3 | 4049.7335 | 4049.7234 | 0 | 2.5 | 332 | CRagged | 206.62 | 166.53 | 159.06 | 2.81 | 28 | >sp\|P01876\|IGHA1_HUMAN Immunoglobulin heavy constant alpha 1 OS=Homo sapiens OX=9606 GN=IGHA1 PE=1 SV=2 | 255 | 20201113_xd_milk_1ng_glyco_hcd30_etd30_compound15.13362.13362.3 | controllerType=0 controllerNumber=1 scan=13362 | 30.4413 |
| 10156:3 | 2 | R.LAGKPTHVN[+1809.666]VSVVMAEVDGTC[+57.021]Y.- | HexNAc(5)Hex(4)Fuc(1) | C[+57], N[+1810] | 1386.2783 | 3 | 4156.8202 | 4156.8081 | 0 | 2.9 | 332 | CRagged | 215.03 | 114.18 | 114.18 | 2.46 | 28 | >sp\|P01876\|IGHA1_HUMAN Immunoglobulin heavy constant alpha 1 OS=Homo sapiens OX=9606 GN=IGHA1 PE=1 SV=2 | 255 | 20201113_xd_milk_1ng_glyco_hcd30_etd30_compound15.13423.13423.3 | controllerType=0 controllerNumber=1 scan=13423 | 30.6222 |
| 10251:3 | 2 | R.LAGKPTHVN[+1501.555]VSVVMAEVDGTC[+57.021]Y.- | HexNAc(5)Hex(3) | C[+57], N[+1502] | 1283.5729 | 3 | 3848.7042 | 3848.6974 | 0 | 1.8 | 332 | CRagged | 221.61 | 147.03 | 147.03 | 2.73 | 28 | >sp\|P01876\|IGHA1_HUMAN Immunoglobulin heavy constant alpha 1 OS=Homo sapiens OX=9606 GN=IGHA1 PE=1 SV=2 | 255 | 20201113_xd_milk_1ng_glyco_hcd30_etd30_compound15.13522.13522.3 | controllerType=0 controllerNumber=1 scan=13522 | 30.8395 |
| 11587:3 | 2 | R.LAGKPTHVN[+2262.814]VSVVMAEVDGTC[+57.021]Y.- | HexNAc(5)Hex(5)Fuc(1)NeuAc(1) | C[+57], N[+2263] | 1537.3278 | 3 | 4609.9689 | 4609.9564 | 0 | 2.7 | 332 | CRagged | 232.04 | 97.94 | 97.94 | 2.51 | 28 | >sp\|P01876\|IGHA1_HUMAN Immunoglobulin heavy constant alpha 1 OS=Homo sapiens OX=9606 GN=IGHA1 PE=1 SV=2 | 255 | 20201113_xd_milk_1ng_glyco_hcd30_etd30_compound15.14922.14922.3 | controllerType=0 controllerNumber=1 scan=14922 | 33.6264 |
| 10124:3 | 2 | R.LAGKPTHVN[+1540.529]VSVVMAEVDGTC[+57.021]Y.- | HexNAc(2)Hex(7) | C[+57], N[+1541] | 1296.5635 | 3 | 3887.676 | 3887.6706 | 0 | 1.4 | 332 | CRagged | 236.62 | 186.47 | 163.93 | 3 | 28 | >sp\|P01876\|IGHA1_HUMAN Immunoglobulin heavy constant alpha 1 OS=Homo sapiens OX=9606 GN=IGHA1 PE=1 SV=2 | 255 | 20201113_xd_milk_1ng_glyco_hcd30_etd30_compound15.13389.13389.3 | controllerType=0 controllerNumber=1 scan=13389 | 30.527 |
| 10134:3 | 2 | R.LAGKPTHVN[+1378.476]VSVVMAEVDGTC[+57.021]Y.- | HexNAc(2)Hex(6) | C[+57], N[+1378] | 1242.5458 | 3 | 3725.6229 | 3725.6178 | 0 | 1.4 | 332 | CRagged | 253.02 | 253.02 | 198.37 | 3.55 | 28 | >sp\|P01876\|IGHA1_HUMAN Immunoglobulin heavy constant alpha 1 OS=Homo sapiens OX=9606 GN=IGHA1 PE=1 SV=2 | 255 | 20201113_xd_milk_1ng_glyco_hcd30_etd30_compound15.13400.13400.3 | controllerType=0 controllerNumber=1 scan=13400 | 30.5596 |
| 10208:3 | 2 | R.LAGKPTHVN[+1647.613]VSVVMAEVDGTC[+57.021]Y.- | HexNAc(5)Hex(3)Fuc(1) | C[+57], N[+1648] | 1332.2608 | 3 | 3994.7678 | 3994.7553 | 0 | 3.1 | 332 | CRagged | 256.97 | 134.61 | 134.61 | 3.05 | 28 | >sp\|P01876\|IGHA1_HUMAN Immunoglobulin heavy constant alpha 1 OS=Homo sapiens OX=9606 GN=IGHA1 PE=1 SV=2 | 255 | 20201113_xd_milk_1ng_glyco_hcd30_etd30_compound15.13477.13477.3 | controllerType=0 controllerNumber=1 scan=13477 | 30.7311 |
| 10140:3 | 2 | R.LAGKPTHVN[+1216.423]VSVVMAEVDGTC[+57.021]Y.- | HexNAc(2)Hex(5) | C[+57], N[+1216] | 1188.5284 | 3 | 3563.5706 | 3563.5649 | 0 | 1.6 | 332 | CRagged | 272.78 | 243.65 | 194.22 | 3.55 | 28 | >sp\|P01876\|IGHA1_HUMAN Immunoglobulin heavy constant alpha 1 OS=Homo sapiens OX=9606 GN=IGHA1 PE=1 SV=2 | 255 | 20201113_xd_milk_1ng_glyco_hcd30_etd30_compound15.13406.13406.3 | controllerType=0 controllerNumber=1 scan=13406 | 30.5741 |
| 11619:4 | 2 | R.LAGKPTHVN[+2262.814]VSVVMAEVDGTC[+57.021]Y.- | HexNAc(5)Hex(5)Fuc(1)NeuAc(1) | C[+57], N[+2263] | 1153.2454 | 4 | 4609.9598 | 4609.9564 | 0 | 0.7 | 332 | CRagged | 447.88 | 367.04 | 367.04 | 5.62 | 28 | >sp\|P01876\|IGHA1_HUMAN Immunoglobulin heavy constant alpha 1 OS=Homo sapiens OX=9606 GN=IGHA1 PE=1 SV=2 | 255 | 20201113_xd_milk_1ng_glyco_hcd30_etd30_compound15.14955.14955.4 | controllerType=0 controllerNumber=1 scan=14955 | 33.6888 |
| 06783:3 | 3 | R.FSDGLESN[+1864.634]SSTQFEVK.K | HexNAc(2)Hex(9) | N[+1865] | 1213.822 | 3 | 3639.4513 | 3639.4472 | 0 | 1.1 | 219 | Specific | 370.74 | 370.74 | 370.74 | 7.06 | 24 | >sp\|P0C0L5\|CO4B_HUMAN Complement C4-B OS=Homo sapiens OX=9606 GN=C4B PE=1 SV=2 | 1424 | 20201113_xd_milk_1ng_glyco_hcd30_etd30_compound15.9908.9908.3 | controllerType=0 controllerNumber=1 scan=9908 | 23.6719 |
| 04462:3 | 3 | R.FSDGLESN[+1864.634]SSTQFEVKK.Y | HexNAc(2)Hex(9) | N[+1865] | 1256.5222 | 3 | 3767.5519 | 3767.5421 | 0 | 2.6 | 219 | Specific | 242.7 | 184.48 | 184.48 | 3.31 | 24 | >sp\|P0C0L5\|CO4B_HUMAN Complement C4-B OS=Homo sapiens OX=9606 GN=C4B PE=1 SV=2 | 1424 | 20201113_xd_milk_1ng_glyco_hcd30_etd30_compound15.7503.7503.3 | controllerType=0 controllerNumber=1 scan=7503 | 19.4107 |
| 15986:4 | 5 | K.QIGLYPVLVIDSSGYVNPN[+2059.735]YTGR.I | HexNAc(4)Hex(5)Fuc(1)NeuAc(1) | N[+2060] | 1147.0186 | 4 | 4585.0526 | 4585.0384 | 0 | 3.1 | 168 | Specific | 344.19 | 344.19 | 344.19 | 4.39 | 29 | >sp\|P01833\|PIGR_HUMAN Polymeric immunoglobulin receptor OS=Homo sapiens OX=9606 GN=PIGR PE=1 SV=4 | 902 | 20201113_xd_milk_1ng_glyco_hcd30_etd30_compound15.19554.19554.3 | controllerType=0 controllerNumber=1 scan=19554 | 45.2588 |
| 16007:3 | 5 | K.QIGLYPVLVIDSSGYVNPN[+2059.735]YTGR.I | HexNAc(4)Hex(5)Fuc(1)NeuAc(1) | N[+2060] | 1529.0228 | 3 | 4585.0537 | 4585.0384 | 0 | 3.3 | 168 | Specific | 599.98 | 476.51 | 476.51 | 8.62 | 29 | >sp\|P01833\|PIGR_HUMAN Polymeric immunoglobulin receptor OS=Homo sapiens OX=9606 GN=PIGR PE=1 SV=4 | 902 | 20201113_xd_milk_1ng_glyco_hcd30_etd30_compound15.19577.19577.3 | controllerType=0 controllerNumber=1 scan=19577 | 45.3359 |
| 17167:3 | 5 | R.LSLLEEPGN[+2076.750]GTFTVILNQLTSR.D | HexNAc(4)Hex(6)Fuc(2) | N[+2077] | 1493.692 | 3 | 4479.0613 | 4479.0428 | 0 | 4.1 | 413 | Specific | 406.82 | 358.02 | 358.02 | 4.66 | 29 | >sp\|P01833\|PIGR_HUMAN Polymeric immunoglobulin receptor OS=Homo sapiens OX=9606 GN=PIGR PE=1 SV=4 | 902 | 20201113_xd_milk_1ng_glyco_hcd30_etd30_compound15.20803.20803.3 | controllerType=0 controllerNumber=1 scan=20803 | 48.9444 |
| 17120:3 | 5 | R.LSLLEEPGN[+2205.793]GTFTVILNQLTSR.D | HexNAc(4)Hex(5)Fuc(2)NeuAc(1) | N[+2206] | 1536.7066 | 3 | 4608.1052 | 4608.0854 | 0 | 4.3 | 413 | Specific | 526.4 | 428.51 | 428.51 | 7.41 | 29 | >sp\|P01833\|PIGR_HUMAN Polymeric immunoglobulin receptor OS=Homo sapiens OX=9606 GN=PIGR PE=1 SV=4 | 902 | 20201113_xd_milk_1ng_glyco_hcd30_etd30_compound15.20754.20754.3 | controllerType=0 controllerNumber=1 scan=20754 | 48.7829 |
| 17121:4 | 5 | R.LSLLEEPGN[+2205.793]GTFTVILNQLTSR.D | HexNAc(4)Hex(5)Fuc(2)NeuAc(1) | N[+2206] | 1152.7804 | 4 | 4608.0999 | 4608.0854 | 0 | 3.1 | 413 | Specific | 550.41 | 507.74 | 507.74 | 6.7 | 29 | >sp\|P01833\|PIGR_HUMAN Polymeric immunoglobulin receptor OS=Homo sapiens OX=9606 GN=PIGR PE=1 SV=4 | 902 | 20201113_xd_milk_1ng_glyco_hcd30_etd30_compound15.20755.20755.4 | controllerType=0 controllerNumber=1 scan=20755 | 48.7871 |
| 17188:3 | 5 | R.LSLLEEPGN[+2059.735]GTFTVILNQLTSR.D | HexNAc(4)Hex(5)Fuc(1)NeuAc(1) | N[+2060] | 1488.017 | 3 | 4462.0365 | 4462.0275 | 0 | 2 | 413 | Specific | 634.98 | 499.94 | 499.94 | 9.6 | 29 | >sp\|P01833\|PIGR_HUMAN Polymeric immunoglobulin receptor OS=Homo sapiens OX=9606 GN=PIGR PE=1 SV=4 | 902 | 20201113_xd_milk_1ng_glyco_hcd30_etd30_compound15.20826.20826.3 | controllerType=0 controllerNumber=1 scan=20826 | 49.0133 |
| 13966:4 | 5 | K.IIEGEPNLKVPGN[+2205.793]VTAVLGETLK.V | HexNAc(4)Hex(5)Fuc(2)NeuAc(1) | N[+2206] | 1150.0441 | 4 | 4597.1546 | 4597.1422 | 0 | 2.7 | 457 | Specific | 275.74 | 258.63 | 258.63 | 2.78 | 29 | >sp\|P01833\|PIGR_HUMAN Polymeric immunoglobulin receptor OS=Homo sapiens OX=9606 GN=PIGR PE=1 SV=4 | 902 | 20201113_xd_milk_1ng_glyco_hcd30_etd30_compound15.17429.17429.4 | controllerType=0 controllerNumber=1 scan=17429 | 39.4972 |

Table S4. Cont.

| Query #:z | Protein Rank | Glycopeptide | Glycan | Modification Type(s) | Observed m/z | z | Observed (M+H) | Calc. mass (M+H) | Off-by-x error | Mass error (ppm) | StartPosition | Cleavage | Score | Delta | Delta Mod | \|Log Prob\| | # of unique peptides | Protein Name | Protein DB number | Comment | Scan # | Scan Time |
|---|---|---|---|---|---|---|---|---|---|---|---|---|---|---|---|---|---|---|---|---|---|---|
| 13961:3 | 5 | K.IIEGEPNLKVPGN[+2205.793]VTAVLGETLK.V | HexNAc(4)Hex(5)Fuc(2)NeuAc(1) | N[+2206] | 1533.0556 | 3 | 4597.1522 | 4597.1422 | 0 | 2.2 | 457 | Specific | 293.38 | 242.98 | 242.98 | 3.16 | 29 | >sp\|P01833\|PIGR_HUMAN Polymeric immunoglobulin receptor OS=Homo sapiens OX=9606 GN=PIGR PE=1 SV=4 | 902 | 20201113_xd_milk_1ng_glyco_hcd30_etd30_compound15.17424.17424.3 | controllerType=0 controllerNumber=1 scan=17424 | 39.4832 |
| 13944:3 | 5 | K.IIEGEPNLKVPGN[+1913.677]VTAVLGETLK.V | HexNAc(4)Hex(5)NeuAc(1) | N[+1914] | 1435.6808 | 3 | 4305.028 | 4305.0264 | 0 | 0.4 | 457 | Specific | 298.65 | 234.37 | 234.37 | 2.87 | 29 | >sp\|P01833\|PIGR_HUMAN Polymeric immunoglobulin receptor OS=Homo sapiens OX=9606 GN=PIGR PE=1 SV=4 | 902 | 20201113_xd_milk_1ng_glyco_hcd30_etd30_compound15.17406.17406.3 | controllerType=0 controllerNumber=1 scan=17406 | 39.4281 |
| 13884:3 | 5 | K.IIEGEPNLKVPGN[+2205.793]VTAVLGETLK.V | HexNAc(4)Hex(5)Fuc(2)NeuAc(1) | N[+2206] | 1533.051 | 3 | 4597.1383 | 4597.1422 | 0 | -0.8 | 457 | Specific | 345.85 | 262.78 | 262.78 | 2.84 | 29 | >sp\|P01833\|PIGR_HUMAN Polymeric immunoglobulin receptor OS=Homo sapiens OX=9606 GN=PIGR PE=1 SV=4 | 902 | 20201113_xd_milk_1ng_glyco_hcd30_etd30_compound15.17341.17341.3 | controllerType=0 controllerNumber=1 scan=17341 | 39.2481 |
| 13928:4 | 5 | K.IIEGEPNLKVPGN[+2205.793]VTAVLGETLK.V | HexNAc(4)Hex(5)Fuc(2)NeuAc(1) | N[+2206] | 1150.0442 | 4 | 4597.1551 | 4597.1422 | 0 | 2.8 | 457 | Specific | 586.53 | 535.32 | 535.32 | 7.09 | 29 | >sp\|P01833\|PIGR_HUMAN Polymeric immunoglobulin receptor OS=Homo sapiens OX=9606 GN=PIGR PE=1 SV=4 | 902 | 20201113_xd_milk_1ng_glyco_hcd30_etd30_compound15.17389.17389.4 | controllerType=0 controllerNumber=1 scan=17389 | 39.382 |
| 13607:5 | 5 | K.IIEGEPNLKVPGN[+2059.735]VTAVLGETLKVPC[+57.021]HFPC[+57.021]K.F | HexNAc(4)Hex(5)Fuc(1)NeuAc(1) | C[+57]*2, N[+2060] | 1096.1136 | 5 | 5476.5391 | 5476.5418 | 0 | -0.5 | 457 | Specific | 333.14 | 320.87 | 320.87 | 2.53 | 29 | >sp\|P01833\|PIGR_HUMAN Polymeric immunoglobulin receptor OS=Homo sapiens OX=9606 GN=PIGR PE=1 SV=4 | 902 | 20201113_xd_milk_1ng_glyco_hcd30_etd30_compound15.17033.17033.5 | controllerType=0 controllerNumber=1 scan=17033 | 38.5923 |
| 13556:5 | 5 | K.IIEGEPNLKVPGN[+2204.772]VTAVLGETLKVPC[+57.021]HFPC[+57.021]K.F | HexNAc(4)Hex(5)NeuAc(2) | C[+57]*2, N[+2205] | 1125.1273 | 5 | 5621.6075 | 5621.5793 | 0 | 5 | 457 | Specific | 399.75 | 399.75 | 399.75 | 4.09 | 29 | >sp\|P01833\|PIGR_HUMAN Polymeric immunoglobulin receptor OS=Homo sapiens OX=9606 GN=PIGR PE=1 SV=4 | 902 | 20201113_xd_milk_1ng_glyco_hcd30_etd30_compound15.16980.16980.5 | controllerType=0 controllerNumber=1 scan=16980 | 38.484 |
| 14386:3 | 5 | K.VPGN[+2350.830]VTAVLGETLK.V | HexNAc(4)Hex(5)Fuc(1)NeuAc(2) | N[+2351] | 1250.2172 | 3 | 3748.6371 | 3748.6302 | 0 | 1.8 | 466 | Specific | 217.06 | 144.39 | 144.39 | 3.37 | 29 | >sp\|P01833\|PIGR_HUMAN Polymeric immunoglobulin receptor OS=Homo sapiens OX=9606 GN=PIGR PE=1 SV=4 | 902 | 20201113_xd_milk_1ng_glyco_hcd30_etd30_compound15.17869.17869.3 | controllerType=0 controllerNumber=1 scan=17869 | 40.4442 |
| 12057:3 | 5 | K.VPGN[+2059.735]VTAVLGETLK.V | HexNAc(4)Hex(5)Fuc(1)NeuAc(1) | N[+2060] | 1153.1862 | 3 | 3457.5441 | 3457.5348 | 0 | 2.7 | 466 | Specific | 240.96 | 161.53 | 161.53 | 3.51 | 29 | >sp\|P01833\|PIGR_HUMAN Polymeric immunoglobulin receptor OS=Homo sapiens OX=9606 GN=PIGR PE=1 SV=4 | 902 | 20201113_xd_milk_1ng_glyco_hcd30_etd30_compound15.15414.15414.3 | controllerType=0 controllerNumber=1 scan=15414 | 34.826 |
| 11965:3 | 5 | K.VPGN[+2205.793]VTAVLGETLK.V | HexNAc(4)Hex(5)Fuc(2)NeuAc(1) | N[+2206] | 1201.8719 | 3 | 3603.6011 | 3603.5927 | 0 | 2.3 | 466 | Specific | 309.37 | 212.64 | 212.64 | 3.8 | 29 | >sp\|P01833\|PIGR_HUMAN Polymeric immunoglobulin receptor OS=Homo sapiens OX=9606 GN=PIGR PE=1 SV=4 | 902 | 20201113_xd_milk_1ng_glyco_hcd30_etd30_compound15.15317.15317.3 | controllerType=0 controllerNumber=1 scan=15317 | 34.5543 |
| 12064:4 | 5 | K.VPGN[+2059.735]VTAVLGETLKVPC[+57.021]HFPC[+57.021]K.F | HexNAc(4)Hex(5)Fuc(1)NeuAc(1) | C[+57]*2, N[+2060] | 1121.5085 | 4 | 4483.012 | 4482.9923 | 0 | 4.4 | 466 | Specific | 239.1 | 226.02 | 226.02 | 2.5 | 29 | >sp\|P01833\|PIGR_HUMAN Polymeric immunoglobulin receptor OS=Homo sapiens OX=9606 GN=PIGR PE=1 SV=4 | 902 | 20201113_xd_milk_1ng_glyco_hcd30_etd30_compound15.15421.15421.4 | controllerType=0 controllerNumber=1 scan=15421 | 34.848 |
| 05282:3 | 5 | K.WN[+1913.677]NTGC[+57.021]QALPSQDEGPSK.A | HexNAc(4)Hex(5)NeuAc(1) | C[+57], N[+1914] | 1301.5254 | 3 | 3902.5618 | 3902.5537 | 0 | 2.1 | 498 | Specific | 425.89 | 265.31 | 265.31 | 5.45 | 29 | >sp\|P01833\|PIGR_HUMAN Polymeric immunoglobulin receptor OS=Homo sapiens OX=9606 GN=PIGR PE=1 SV=4 | 902 | 20201113_xd_milk_1ng_glyco_hcd30_etd30_compound15.8352.8352.3 | controllerType=0 controllerNumber=1 scan=8352 | 20.8969 |
| 05207:3 | 5 | K.WN[+2059.735]NTGC[+57.021]QALPSQDEGPSK.A | HexNAc(4)Hex(5)Fuc(1)NeuAc(1) | C[+57], N[+2060] | 1350.2099 | 3 | 4048.6151 | 4048.6116 | 0 | 0.9 | 498 | Specific | 435.71 | 312.05 | 312.05 | 5.52 | 29 | >sp\|P01833\|PIGR_HUMAN Polymeric immunoglobulin receptor OS=Homo sapiens OX=9606 GN=PIGR PE=1 SV=4 | 902 | 20201113_xd_milk_1ng_glyco_hcd30_etd30_compound15.8274.8274.3 | controllerType=0 controllerNumber=1 scan=8274 | 20.7436 |
| 05359:3 | 5 | K.WN[+1710.598]NTGC[+57.021]QALPSQDEGPSK.A | HexNAc(3)Hex(5)NeuAc(1) | C[+57], N[+1711] | 1233.8321 | 3 | 3699.4817 | 3699.4743 | 0 | 2 | 498 | Specific | 540.73 | 445.18 | 445.18 | 7.66 | 29 | >sp\|P01833\|PIGR_HUMAN Polymeric immunoglobulin receptor OS=Homo sapiens OX=9606 GN=PIGR PE=1 SV=4 | 902 | 20201113_xd_milk_1ng_glyco_hcd30_etd30_compound15.8432.8432.3 | controllerType=0 controllerNumber=1 scan=8432 | 21.054 |
| 09209:3 | 7 | K.TVLTPATNHMGN[+1378.476]VTFTIPANR.E | HexNAc(2)Hex(6) | N[+1378] | 1211.8867 | 3 | 3633.6454 | 3633.6358 | 0 | 2.6 | 74 | Specific | 454.51 | 361.97 | 152.53 | 6.06 | 14 | >sp\|P01024\|CO3_HUMAN Complement C3 OS=Homo sapiens OX=9606 GN=C3 PE=1 SV=2 | 492 | 20201113_xd_milk_1ng_glyco_hcd30_etd30_compound15.12437.12437.3 | controllerType=0 controllerNumber=1 scan=12437 | 28.5281 |
| 09165:3 | 7 | K.TVLTPATNHMGN[+1540.529]VTFTIPANR.E | HexNAc(2)Hex(7) | N[+1541] | 1265.9046 | 3 | 3795.6992 | 3795.6886 | 0 | 2.8 | 74 | Specific | 466.85 | 363.22 | 168.03 | 6.06 | 14 | >sp\|P01024\|CO3_HUMAN Complement C3 OS=Homo sapiens OX=9606 GN=C3 PE=1 SV=2 | 492 | 20201113_xd_milk_1ng_glyco_hcd30_etd30_compound15.12390.12390.3 | controllerType=0 controllerNumber=1 scan=12390 | 28.3893 |
| 16029:3 | 8 | R.NIAAFGGDPNN[+1913.677]ITLFGESAGGASVSLQTLSPYNK.G | HexNAc(4)Hex(5)NeuAc(1) | N[+1914] | 1775.467 | 3 | 5324.3865 | 5324.3633 | 0 | 4.3 | 197 | Specific | 563.96 | 436.74 | 436.74 | 8.41 | 12 | >sp\|P19835\|CEL_HUMAN Bile salt-activated lipase OS=Homo sapiens OX=9606 GN=CEL PE=1 SV=3 | 476 | 20201113_xd_milk_1ng_glyco_hcd30_etd30_compound15.19600.19600.3 | controllerType=0 controllerNumber=1 scan=19600 | 45.4021 |
| 16916:3 | 11 | K.YLGN[+2204.772]ATAIFFLPDEGK.L | HexNAc(4)Hex(5)NeuAc(2) | N[+2205] | 1320.8965 | 3 | 3960.6751 | 3960.6677 | 0 | 1.9 | 268 | Specific | 205.22 | 92.66 | 92.66 | 3.78 | 6 | >sp\|P01009\|A1AT_HUMAN Alpha-1-antitrypsin OS=Homo sapiens OX=9606 GN=SERPINA1 PE=1 SV=3 | 642 | 20201113_xd_milk_1ng_glyco_hcd30_etd30_compound15.20538.20538.2 | controllerType=0 controllerNumber=1 scan=20538 | 48.1182 |
| 07079:3 | 12 | R.LSPN[+2350.830]ASAEHLELR.W | HexNAc(4)Hex(5)Fuc(1)NeuAc(2) | N[+2351] | 1263.201 | 3 | 3787.5884 | 3787.5796 | 0 | 2.3 | 52 | Specific | 303.48 | 156.07 | 156.07 | 3.24 | 8 | >sp\|Q13410\|BT1A1_HUMAN Butyrophilin subfamily 1 member A1 OS=Homo sapiens OX=9606 GN=BTN1A1 PE=1 SV=3 | 179 | 20201113_xd_milk_1ng_glyco_hcd30_etd30_compound15.10215.10215.3 | controllerType=0 controllerNumber=1 scan=10215 | 24.258 |
| 05201:3 | 12 | R.LSPN[+2205.793]ASAEHLELR.W | HexNAc(4)Hex(5)Fuc(2)NeuAc(1) | N[+2206] | 1214.8532 | 3 | 3642.5451 | 3642.5421 | 0 | 0.8 | 52 | Specific | 392.43 | 262.56 | 262.56 | 3.41 | 8 | >sp\|Q13410\|BT1A1_HUMAN Butyrophilin subfamily 1 member A1 OS=Homo sapiens OX=9606 GN=BTN1A1 PE=1 SV=3 | 179 | 20201113_xd_milk_1ng_glyco_hcd30_etd30_compound15.8268.8268.3 | controllerType=0 controllerNumber=1 scan=8268 | 20.7312 |
| 11739:3 | 14 | R.IIVPLNNREN[+2350.830]ISDPTSPLR.T | HexNAc(4)Hex(5)Fuc(1)NeuAc(2) | N[+2351] | 1500.3443 | 3 | 4499.0183 | 4499.0075 | 0 | 2.4 | 62 | Specific | 201.34 | 162.18 | 162.18 | 2.68 | 7 | >sp\|P01591\|IGJ_HUMAN Immunoglobulin J chain OS=Homo sapiens OX=9606 GN=JCHAIN PE=1 SV=4 | 1392 | 20201113_xd_milk_1ng_glyco_hcd30_etd30_compound15.15080.15080.3 | controllerType=0 controllerNumber=1 scan=15080 | 33.9921 |
| 10123:3 | 14 | R.IIVPLNNREN[+1913.677]ISDPTSPLR.T | HexNAc(4)Hex(5)NeuAc(1) | N[+1914] | 1354.6254 | 3 | 4061.8617 | 4061.8542 | 0 | 1.9 | 62 | Specific | 265.17 | 198.03 | 198.03 | 3.04 | 7 | >sp\|P01591\|IGJ_HUMAN Immunoglobulin J chain OS=Homo sapiens OX=9606 GN=JCHAIN PE=1 SV=4 | 1392 | 20201113_xd_milk_1ng_glyco_hcd30_etd30_compound15.13388.13388.3 | controllerType=0 controllerNumber=1 scan=13388 | 30.5234 |
| 10097:3 | 14 | R.IIVPLNNREN[+2059.735]ISDPTSPLR.T | HexNAc(4)Hex(5)Fuc(1)NeuAc(1) | N[+2060] | 1403.3121 | 3 | 4207.9217 | 4207.9121 | 0 | 2.3 | 62 | Specific | 269.16 | 219.03 | 219.03 | 2.81 | 7 | >sp\|P01591\|IGJ_HUMAN Immunoglobulin J chain OS=Homo sapiens OX=9606 GN=JCHAIN PE=1 SV=4 | 1392 | 20201113_xd_milk_1ng_glyco_hcd30_etd30_compound15.13361.13361.3 | controllerType=0 controllerNumber=1 scan=13361 | 30.4376 |
| 08778:3 | 14 | R.IIVPLNNREN[+1809.666]ISDPTSPLR.T | HexNAc(5)Hex(4)Fuc(1) | N[+1810] | 1319.9539 | 3 | 3957.8472 | 3957.8432 | 0 | 1 | 62 | Specific | 271.94 | 177.85 | 177.85 | 2.74 | 7 | >sp\|P01591\|IGJ_HUMAN Immunoglobulin J chain OS=Homo sapiens OX=9606 GN=JCHAIN PE=1 SV=4 | 1392 | 20201113_xd_milk_1ng_glyco_hcd30_etd30_compound15.11989.11989.3 | controllerType=0 controllerNumber=1 scan=11989 | 27.6407 |
| 08824:3 | 14 | R.IIVPLNNREN[+1663.608]ISDPTSPLR.T | HexNAc(5)Hex(4) | N[+1664] | 1271.2685 | 3 | 3811.7909 | 3811.7853 | 0 | 1.5 | 62 | Specific | 274.89 | 201.83 | 201.83 | 3 | 7 | >sp\|P01591\|IGJ_HUMAN Immunoglobulin J chain OS=Homo sapiens OX=9606 GN=JCHAIN PE=1 SV=4 | 1392 | 20201113_xd_milk_1ng_glyco_hcd30_etd30_compound15.12037.12037.3 | controllerType=0 controllerNumber=1 scan=12037 | 27.7448 |
| 18726:4 | 20 | K.APDKNVIFSPLSISTALAFLSLGAHN[+2350.830]TTLTEILK.G | HexNAc(4)Hex(5)Fuc(1)NeuAc(2) | N[+2351] | 1484.2135 | 4 | 5933.8322 | 5933.8085 | 0 | 4 | 68 | Specific | 275.93 | 275.93 | 275.93 | 3.38 | 7 | >sp\|P01011\|AACT_HUMAN Alpha-1-antichymotrypsin OS=Homo sapiens OX=9606 GN=SERPINA3 PE=1 SV=2 | 1314 | 20201113_xd_milk_1ng_glyco_hcd30_etd30_compound15.22447.22447.2 | controllerType=0 controllerNumber=1 scan=22447 | 53.566 |
| 17694:4 | 20 | K.APDKNVIFSPLSISTALAFLSLGAHN[+2716.983]TTLTEILK.G | HexNAc(5)Hex(6)Fuc(3)NeuAc(1) | N[+2717] | 1575.7505 | 4 | 6299.9801 | 6299.9611 | 0 | 3 | 68 | Specific | 304.75 | 304.75 | 304.75 | 4.71 | 7 | >sp\|P01011\|AACT_HUMAN Alpha-1-antichymotrypsin OS=Homo sapiens OX=9606 GN=SERPINA3 PE=1 SV=2 | 1314 | 20201113_xd_milk_1ng_glyco_hcd30_etd30_compound15.21358.21358.4 | controllerType=0 controllerNumber=1 scan=21358 | 50.5214 |
| 14345:3 | 22 | K.AALAAFNAQNN[+2204.772]GSNFQLEEISR.A | HexNAc(4)Hex(5)NeuAc(2) | N[+2205] | 1523.9849 | 3 | 4569.9402 | 4569.9256 | 0 | 3.2 | 166 | Specific | 541.21 | 397.42 | 397.42 | 7.62 | 4 | >sp\|P02765\|FETUA_HUMAN Alpha-2-HS-glycoprotein OS=Homo sapiens OX=9606 GN=AHSG PE=1 SV=2 | 135 | 20201113_xd_milk_1ng_glyco_hcd30_etd30_compound15.17826.17826.3 | controllerType=0 controllerNumber=1 scan=17826 | 40.3491 |
| 19226:4 | 23 | K.VFPLSLDSTPQDGNVVVAC[+57.021]LVQGFFPQEPLSVTWSESGQN[+2262.814]VTAI | HexNAc(5)Hex(5)Fuc(1)NeuAc(1) | C[+57], N[+2263] | 1760.5522 | 4 | 7039.187 | 7039.1757 | 0 | 1.6 | 8 | Specific | 285.78 | 285.78 | 285.78 | 5.4 | 11 | >sp\|P01877\|IGHA2_HUMAN Immunoglobulin heavy constant alpha 2 OS=Homo sapiens OX=9606 GN=IGHA2 PE=1 SV=4 | 576 | 20201113_xd_milk_1ng_glyco_hcd30_etd30_compound15.22976.22976.4 | controllerType=0 controllerNumber=1 scan=22976 | 55.117 |
| 04364:3 | 23 | K.TPLTAN[+2262.814]ITK.S | HexNAc(5)Hex(5)Fuc(1)NeuAc(1) | N[+2263] | 1074.4635 | 3 | 3221.376 | 3221.3711 | 0 | 1.5 | 200 | Specific | 221.01 | 81.59 | 81.59 | 2.43 | 11 | >sp\|P01877\|IGHA2_HUMAN Immunoglobulin heavy constant alpha 2 OS=Homo sapiens OX=9606 GN=IGHA2 PE=1 SV=4 | 576 | 20201113_xd_milk_1ng_glyco_hcd30_etd30_compound15.7402.7402.3 | controllerType=0 controllerNumber=1 scan=7402 | 19.2289 |
| 02689:2 | 23 | K.TPLTAN[+1971.719]ITK.S | HexNAc(5)Hex(5)Fuc(1) | N[+1972] | 1465.6473 | 2 | 2930.2872 | 2930.2757 | 0 | 4 | 200 | Specific | 260.05 | 84.17 | 84.17 | 2.45 | 11 | >sp\|P01877\|IGHA2_HUMAN Immunoglobulin heavy constant alpha 2 OS=Homo sapiens OX=9606 GN=IGHA2 PE=1 SV=4 | 576 | 20201113_xd_milk_1ng_glyco_hcd30_etd30_compound15.5661.5661.2 | controllerType=0 controllerNumber=1 scan=5661 | 16.0661 |
| 02681:3 | 23 | K.TPLTAN[+1971.719]ITK.S | HexNAc(5)Hex(5)Fuc(1) | N[+1972] | 977.4335 | 3 | 2930.2859 | 2930.2757 | 0 | 3.5 | 200 | Specific | 268.33 | 126.62 | 126.62 | 2.51 | 11 | >sp\|P01877\|IGHA2_HUMAN Immunoglobulin heavy constant alpha 2 OS=Homo sapiens OX=9606 GN=IGHA2 PE=1 SV=4 | 576 | 20201113_xd_milk_1ng_glyco_hcd30_etd30_compound15.5653.5653.3 | controllerType=0 controllerNumber=1 scan=5653 | 16.0499 |
| 02795:2 | 23 | K.TPLTAN[+1501.555]ITK.S | HexNAc(5)Hex(3) | N[+1502] | 1230.5632 | 2 | 2460.119 | 2460.1121 | 0 | 2.8 | 200 | Specific | 288.62 | 90.23 | 90.23 | 2.63 | 11 | >sp\|P01877\|IGHA2_HUMAN Immunoglobulin heavy constant alpha 2 OS=Homo sapiens OX=9606 GN=IGHA2 PE=1 SV=4 | 576 | 20201113_xd_milk_1ng_glyco_hcd30_etd30_compound15.5772.5772.2 | controllerType=0 controllerNumber=1 scan=5772 | 16.3444 |
| 02782:3 | 23 | K.TPLTAN[+1647.613]ITK.S | HexNAc(5)Hex(3)Fuc(1) | N[+1648] | 869.3968 | 3 | 2606.1757 | 2606.17 | 0 | 2.2 | 200 | Specific | 297.09 | 138.9 | 138.9 | 2.67 | 11 | >sp\|P01877\|IGHA2_HUMAN Immunoglobulin heavy constant alpha 2 OS=Homo sapiens OX=9606 GN=IGHA2 PE=1 SV=4 | 576 | 20201113_xd_milk_1ng_glyco_hcd30_etd30_compound15.5759.5759.3 | controllerType=0 controllerNumber=1 scan=5759 | 16.3106 |
| 02716:2 | 23 | K.TPLTAN[+1809.666]ITK.S | HexNAc(5)Hex(4)Fuc(1) | N[+1810] | 1384.6183 | 2 | 2768.2294 | 2768.2228 | 0 | 2.4 | 200 | Specific | 301.58 | 118.22 | 118.22 | 2.5 | 11 | >sp\|P01877\|IGHA2_HUMAN Immunoglobulin heavy constant alpha 2 OS=Homo sapiens OX=9606 GN=IGHA2 PE=1 SV=4 | 576 | 20201113_xd_milk_1ng_glyco_hcd30_etd30_compound15.5689.5689.2 | controllerType=0 controllerNumber=1 scan=5689 | 16.1308 |
| 02767:2 | 23 | K.TPLTAN[+1647.613]ITK.S | HexNAc(5)Hex(3)Fuc(1) | N[+1648] | 1303.5909 | 2 | 2606.1744 | 2606.17 | 0 | 1.7 | 200 | Specific | 303.05 | 107.75 | 107.75 | 2.49 | 11 | >sp\|P01877\|IGHA2_HUMAN Immunoglobulin heavy constant alpha 2 OS=Homo sapiens OX=9606 GN=IGHA2 PE=1 SV=4 | 576 | 20201113_xd_milk_1ng_glyco_hcd30_etd30_compound15.5743.5743.2 | controllerType=0 controllerNumber=1 scan=5743 | 16.267 |
| 02724:3 | 23 | K.TPLTAN[+1809.666]ITK.S | HexNAc(5)Hex(4)Fuc(1) | N[+1810] | 923.4155 | 3 | 2768.2319 | 2768.2228 | 0 | 3.3 | 200 | Specific | 303.28 | 128.25 | 128.25 | 2.55 | 11 | >sp\|P01877\|IGHA2_HUMAN Immunoglobulin heavy constant alpha 2 OS=Homo sapiens OX=9606 GN=IGHA2 PE=1 SV=4 | 576 | 20201113_xd_milk_1ng_glyco_hcd30_etd30_compound15.5698.5698.3 | controllerType=0 controllerNumber=1 scan=5698 | 16.153 |
| 02673:3 | 23 | K.TPLTAN[+1971.719]ITK.S | HexNAc(5)Hex(5)Fuc(1) | N[+1972] | 977.4333 | 3 | 2930.2854 | 2930.2757 | 0 | 3.3 | 200 | Specific | 304.19 | 131.44 | 131.44 | 2.67 | 11 | >sp\|P01877\|IGHA2_HUMAN Immunoglobulin heavy constant alpha 2 OS=Homo sapiens OX=9606 GN=IGHA2 PE=1 SV=4 | 576 | 20201113_xd_milk_1ng_glyco_hcd30_etd30_compound15.5644.5644.3 | controllerType=0 controllerNumber=1 scan=5644 | 16.0262 |
| 01928:4 | 24 | R.EKQTDEIKDTRN[+2059.735]ESTQNC[+57.021]VVAEPEK.M | HexNAc(4)Hex(5)Fuc(1)NeuAc(1) | C[+57], N[+2060] | 1252.7896 | 4 | 5008.1364 | 5008.1251 | 0 | 2.2 | 58 | Specific | 679.41 | 588.12 | 588.12 | 8.91 | 6 | >sp\|P47710\|CASA1_HUMAN Alpha-S1-casein OS=Homo sapiens OX=9606 GN=CSN1S1 PE=1 SV=1 | 989 | 20201113_xd_milk_1ng_glyco_hcd30_etd30_compound15.4871.4871.4 | controllerType=0 controllerNumber=1 scan=4871 | 14.6515 |
| 01854:3 | 25 | K.N[+1540.529]NSIPDKQITASSSYK.T | HexNAc(2)Hex(7) | N[+1541] | 1098.475 | 3 | 3293.4105 | 3293.4048 | 0 | 1.7 | 238 | Specific | 215.24 | 141.42 | 141.42 | 3.61 | 3 | >sp\|Q08431\|MFGM_HUMAN Lactadherin OS=Homo sapiens OX=9606 GN=MFGE8 PE=1 SV=3 | 1064 | 20201113_xd_milk_1ng_glyco_hcd30_etd30_compound15.4794.4794.3 | controllerType=0 controllerNumber=1 scan=4794 | 14.5008 |
| 01850:3 | 27 | R.LRN[+1216.423]VSWATGR.S | HexNAc(2)Hex(5) | N[+1216] | 792.6914 | 3 | 2376.0598 | 2376.0559 | 0 | 1.6 | 149 | Specific | 363.66 | 182.65 | 182.65 | 2.46 | 5 | >sp\|P08571\|CD14_HUMAN Monocyte differentiation antigen CD14 OS=Homo sapiens OX=9606 GN=CD14 PE=1 SV=2 | 1232 | 20201113_xd_milk_1ng_glyco_hcd30_etd30_compound15.4790.4790.3 | controllerType=0 controllerNumber=1 scan=4790 | 14.489 |
| 00150:4 | 36 | K.KKEDALN[+2059.735]ETRESETK.L | HexNAc(4)Hex(5)Fuc(1)NeuAc(1) | N[+2060] | 960.1647 | 4 | 3837.6368 | 3837.6276 | 0 | 2.4 | 80 | Specific | 343.09 | 328.31 | 328.31 | 3.09 | 7 | >sp\|P10909\|CLUS_HUMAN Clusterin OS=Homo sapiens OX=9606 GN=CLU PE=1 SV=1 | 1467 | 20201113_xd_milk_1ng_glyco_hcd30_etd30_compound15.3022.3022.2 | controllerType=0 controllerNumber=1 scan=3022 | 11.3169 |
| 00143:4 | 36 | R.EIRHN[+2059.735]STGC[+57.021]LR.M | HexNAc(4)Hex(5)Fuc(1)NeuAc(1) | C[+57], N[+2060] | 851.3577 | 4 | 3402.4092 | 3402.3994 | 0 | 2.9 | 287 | Specific | 648.83 | 506.99 | 506.99 | 8.27 | 7 | >sp\|P10909\|CLUS_HUMAN Clusterin OS=Homo sapiens OX=9606 GN=CLU PE=1 SV=1 | 1467 | 20201113_xd_milk_1ng_glyco_hcd30_etd30_compound15.3015.3015.4 | controllerType=0 controllerNumber=1 scan=3015 | 11.2936 |
| 18980:3 | 36 | K.MLN[+2350.830]TSSLLEQLNEQFNWVSR.L | HexNAc(4)Hex(5)Fuc(1)NeuAc(2) | N[+2351] | 1587.3491 | 3 | 4760.0326 | 4760.0171 | 0 | 3.3 | 352 | Specific | 379.84 | 224.36 | 224.36 | 5.17 | 7 | >sp\|P10909\|CLUS_HUMAN Clusterin OS=Homo sapiens OX=9606 GN=CLU PE=1 SV=1 | 1467 | 20201113_xd_milk_1ng_glyco_hcd30_etd30_compound15.22717.22717.3 | controllerType=0 controllerNumber=1 scan=22717 | 54.3969 |
| 17431:3 | 36 | K.MLN[+2205.793]TSSLLEQLNEQFNWVSR.L | HexNAc(4)Hex(5)Fuc(2)NeuAc(1) | N[+2206] | 1539.0092 | 3 | 4615.0131 | 4614.9796 | 0 | 7.3 | 352 | Specific | 441.75 | 296.22 | 296.22 | 4.84 | 7 | >sp\|P10909\|CLUS_HUMAN Clusterin OS=Homo sapiens OX=9606 GN=CLU PE=1 SV=1 | 1467 | 20201113_xd_milk_1ng_glyco_hcd30_etd30_compound15.21082.21082.3 | controllerType=0 controllerNumber=1 scan=21082 | 49.7787 |
| 17445:3 | 36 | K.MLN[+2059.735]TSSLLEQLNEQFNWVSR.L | HexNAc(4)Hex(5)Fuc(1)NeuAc(1) | N[+2060] | 1490.3231 | 3 | 4468.9546 | 4468.9217 | 0 | 7.4 | 352 | Specific | 545.83 | 370.64 | 370.64 | 6.65 | 7 | >sp\|P10909\|CLUS_HUMAN Clusterin OS=Homo sapiens OX=9606 GN=CLU PE=1 SV=1 | 1467 | 20201113_xd_milk_1ng_glyco_hcd30_etd30_compound15.21097.21097.3 | controllerType=0 controllerNumber=1 scan=21097 | 49.8544 |
| 06845:4 | 1 | R.QTGLAPGQEYEISLHIVKN[+2351.851]NTR.G | HexNAc(4)Hex(5)Fuc(3)NeuAc(1) | N[+2352] | 1205.7929 | 4 | 4820.1497 | 4820.14 | 0 | 2 | 770 | Specific | 461.9 | 379.61 | 379.61 | 6.11 | 68 | >sp\|P24821\|TENA_HUMAN Tenascin OS=Homo sapiens OX=9606 GN=TNC PE=1 SV=3 | 486 | 20201118_ly_colostrum_1ng_glyco_hcd30_etd30_compound15.10033.10033.4 | controllerType=0 controllerNumber=1 scan=10033 | 26.4898 |
| 07862:4 | 1 | R.QTGLAPGQEYEISLHIVKN[+2350.830]NTR.G | HexNAc(4)Hex(5)Fuc(1)NeuAc(2) | N[+2351] | 1205.5387 | 4 | 4819.1331 | 4819.1196 | 0 | 2.8 | 770 | Specific | 522.71 | 459.3 | 459.3 | 6.68 | 68 | >sp\|P24821\|TENA_HUMAN Tenascin OS=Homo sapiens OX=9606 GN=TNC PE=1 SV=3 | 486 | 20201118_ly_colostrum_1ng_glyco_hcd30_etd30_compound15.11102.11102.4 | controllerType=0 controllerNumber=1 scan=11102 | 29.397 |
| 06883:4 | 1 | R.QTGLAPGQEYEISLHIVKN[+2059.735]NTR.G | HexNAc(4)Hex(5)Fuc(1)NeuAc(1) | N[+2060] | 1132.7659 | 4 | 4528.0418 | 4528.0241 | 0 | 3.9 | 770 | Specific | 560.57 | 476.23 | 476.23 | 7.54 | 68 | >sp\|P24821\|TENA_HUMAN Tenascin OS=Homo sapiens OX=9606 GN=TNC PE=1 SV=3 | 486 | 20201118_ly_colostrum_1ng_glyco_hcd30_etd30_compound15.10073.10073.4 | controllerType=0 controllerNumber=1 scan=10073 | 26.5982 |
| 06004:5 | 1 | R.QTGLAPGQEYEISLHIVKN[+2351.851]NTRGPGLKR.V | HexNAc(4)Hex(5)Fuc(3)NeuAc(1) | N[+2352] | 1086.5122 | 5 | 5428.5319 | 5428.5158 | 0 | 3 | 770 | Specific | 586.93 | 586.93 | 586.93 | 7.77 | 68 | >sp\|P24821\|TENA_HUMAN Tenascin OS=Homo sapiens OX=9606 GN=TNC PE=1 SV=3 | 486 | 20201118_ly_colostrum_1ng_glyco_hcd30_etd30_compound15.9150.9150.5 | controllerType=0 controllerNumber=1 scan=9150 | 24.211 |
| 09848:4 | 1 | K.FDRYRLN[+1216.423]YSLPTGQWVGVQLPR.N | HexNAc(2)Hex(5) | N[+1216] | 971.2144 | 4 | 3881.8358 | 3881.8226 | 0 | 3.4 | 1012 | Specific | 510.96 | 510.96 | 510.96 | 6.15 | 68 | >sp\|P24821\|TENA_HUMAN Tenascin OS=Homo sapiens OX=9606 GN=TNC PE=1 SV=3 | 486 | 20201118_ly_colostrum_1ng_glyco_hcd30_etd30_compound15.13184.13184.4 | controllerType=0 controllerNumber=1 scan=13184 | 34.8226 |
| 10985:2 | 1 | R.LN[+1216.423]YSLPTGQWVGVQLPR.N | HexNAc(2)Hex(5) | N[+1216] | 1572.7375 | 2 | 3144.4677 | 3144.4617 | 0 | 1.9 | 1017 | Specific | 500.07 | 444.59 | 444.59 | 7.3 | 68 | >sp\|P24821\|TENA_HUMAN Tenascin OS=Homo sapiens OX=9606 GN=TNC PE=1 SV=3 | 486 | 20201118_ly_colostrum_1ng_glyco_hcd30_etd30_compound15.14381.14381.2 | controllerType=0 controllerNumber=1 scan=14381 | 38.0818 |
| 10978:3 | 1 | R.LN[+1216.423]YSLPTGQWVGVQLPR.N | HexNAc(2)Hex(5) | N[+1216] | 1048.8289 | 3 | 3144.4721 | 3144.4617 | 0 | 3.3 | 1017 | Specific | 632.81 | 560.57 | 560.57 | 9.41 | 68 | >sp\|P24821\|TENA_HUMAN Tenascin OS=Homo sapiens OX=9606 GN=TNC PE=1 SV=3 | 486 | 20201118_ly_colostrum_1ng_glyco_hcd30_etd30_compound15.14374.14374.3 | controllerType=0 controllerNumber=1 scan=14374 | 38.0567 |
| 03096:3 | 1 | R.N[+2351.851]TTSYVLR.G | HexNAc(4)Hex(5)Fuc(3)NeuAc(1) | N[+2352] | 1102.4593 | 3 | 3305.3632 | 3305.3558 | 0 | 2.2 | 1034 | Specific | 210.09 | 103.17 | 103.17 | 2.83 | 68 | >sp\|P24821\|TENA_HUMAN Tenascin OS=Homo sapiens OX=9606 GN=TNC PE=1 SV=3 | 486 | 20201118_ly_colostrum_1ng_glyco_hcd30_etd30_compound15.6116.6116.3 | controllerType=0 controllerNumber=1 scan=6116 | 17.5131 |
| 03206:3 | 1 | R.N[+2059.735]TTSYVLR.G | HexNAc(4)Hex(5)Fuc(1)NeuAc(1) | N[+2060] | 1005.0892 | 3 | 3013.253 | 3013.24 | 0 | 4.3 | 1034 | Specific | 240.34 | 116.39 | 116.39 | 3.07 | 68 | >sp\|P24821\|TENA_HUMAN Tenascin OS=Homo sapiens OX=9606 GN=TNC PE=1 SV=3 | 486 | 20201118_ly_colostrum_1ng_glyco_hcd30_etd30_compound15.6232.6232.3 | controllerType=0 controllerNumber=1 scan=6232 | 17.8298 |
| 01734:3 | 1 | R.N[+1768.640]TTSYVLR.G | HexNAc(4)Hex(5)Fuc(1) | N[+1769] | 908.0564 | 3 | 2722.1547 | 2722.1446 | 0 | 3.7 | 1034 | Specific | 262.89 | 126.09 | 126.09 | 2.99 | 68 | >sp\|P24821\|TENA_HUMAN Tenascin OS=Homo sapiens OX=9606 GN=TNC PE=1 SV=3 | 486 | 20201118_ly_colostrum_1ng_glyco_hcd30_etd30_compound15.4696.4696.3 | controllerType=0 controllerNumber=1 scan=4696 | 14.5039 |
| 13247:3 | 2 | R.LSLHRPALEDLLLGSEAN[+1751.624]LTC[+57.021]TLTGLR.D | HexNAc(4)Hex(4)NeuAc(1) | C[+57], N[+1752] | 1572.416 | 3 | 4715.2333 | 4715.2225 | 0 | 2.3 | 127 | Specific | 268.55 | 204.53 | 204.53 | 5.1 | 65 | >sp\|P01876\|IGHA1_HUMAN Immunoglobulin heavy constant alpha 1 OS=Homo sapiens OX=9606 GN=IGHA1 PE=1 SV=2 | 255 | 20201118_ly_colostrum_1ng_glyco_hcd30_etd30_compound15.16765.16765.3 | controllerType=0 controllerNumber=1 scan=16765 | 45.0184 |
| 13195:5 | 2 | R.LSLHRPALEDLLLGSEAN[+1913.677]LTC[+57.021]TLTGLR.D | HexNAc(4)Hex(5)NeuAc(1) | C[+57], N[+1914] | 976.2627 | 5 | 4877.2842 | 4877.2753 | 0 | 1.8 | 127 | Specific | 289.89 | 289.89 | 289.89 | 4.95 | 65 | >sp\|P01876\|IGHA1_HUMAN Immunoglobulin heavy constant alpha 1 OS=Homo sapiens OX=9606 GN=IGHA1 PE=1 SV=2 | 255 | 20201118_ly_colostrum_1ng_glyco_hcd30_etd30_compound15.16710.16710.5 | controllerType=0 controllerNumber=1 scan=16710 | 44.8396 |
| 12556:3 | 2 | R.LSLHRPALEDLLLGSEAN[+1501.555]LTC[+57.021]TLTGLR.D | HexNAc(5)Hex(3) | C[+57], N[+1502] | 1489.0607 | 3 | 4465.1676 | 4465.1536 | 0 | 3.1 | 127 | Specific | 322.64 | 281.85 | 281.85 | 6.38 | 65 | >sp\|P01876\|IGHA1_HUMAN Immunoglobulin heavy constant alpha 1 OS=Homo sapiens OX=9606 GN=IGHA1 PE=1 SV=2 | 255 | 20201118_ly_colostrum_1ng_glyco_hcd30_etd30_compound15.16037.16037.6 | controllerType=0 controllerNumber=1 scan=16037 | 42.8573 |
| 25808:4 | 2 | R.LSLHRPALEDLLLGSEAN[+1501.555]LTC[+57.021]TLTGLR.D | HexNAc(5)Hex(3) | C[+57], N[+1502] | 1117.0459 | 4 | 4465.1619 | 4465.1536 | 0 | 1.9 | 127 | Specific | 335.16 | 335.16 | 335.16 | 6.75 | 65 | >sp\|P01876\|IGHA1_HUMAN Immunoglobulin heavy constant alpha 1 OS=Homo sapiens OX=9606 GN=IGHA1 PE=1 SV=2 | 255 | 20201118_ly_colostrum_1ng_glyco_hcd30_etd30_compound15.32909.32909.4 | controllerType=0 controllerNumber=1 scan=32909 | 88.7636 |
| 13953:3 | 2 | R.LSLHRPALEDLLLGSEAN[+2204.772]LTC[+57.021]TLTGLR.D | HexNAc(4)Hex(5)NeuAc(2) | C[+57], N[+2205] | 1723.4654 | 3 | 5168.3817 | 5168.3707 | 0 | 2.1 | 127 | Specific | 337.76 | 228.34 | 228.34 | 6.33 | 65 | >sp\|P01876\|IGHA1_HUMAN Immunoglobulin heavy constant alpha 1 OS=Homo sapiens OX=9606 GN=IGHA1 PE=1 SV=2 | 255 | 20201118_ly_colostrum_1ng_glyco_hcd30_etd30_compound15.17514.17514.3 | controllerType=0 controllerNumber=1 scan=17514 | 47.3379 |
| 13243:3 | 2 | R.LSLHRPALEDLLLGSEAN[+1548.545]LTC[+57.021]TLTGLR.D | HexNAc(3)Hex(4)NeuAc(1) | C[+57], N[+1549] | 1504.7245 | 3 | 4512.1591 | 4512.1431 | 0 | 3.5 | 127 | Specific | 340.51 | 258.57 | 258.57 | 6.38 | 65 | >sp\|P01876\|IGHA1_HUMAN Immunoglobulin heavy constant alpha 1 OS=Homo sapiens OX=9606 GN=IGHA1 PE=1 SV=2 | 255 | 20201118_ly_colostrum_1ng_glyco_hcd30_etd30_compound15.16761.16761.3 | controllerType=0 controllerNumber=1 scan=16761 | 45.0115 |
| 13189:3 | 2 | R.LSLHRPALEDLLLGSEAN[+2116.756]LTC[+57.021]TLTGLR.D | HexNAc(5)Hex(5)NeuAc(1) | C[+57], N[+2117] | 1694.1301 | 3 | 5080.3757 | 5080.3547 | 0 | 4.1 | 127 | Specific | 352.08 | 300.27 | 300.27 | 5.33 | 65 | >sp\|P01876\|IGHA1_HUMAN Immunoglobulin heavy constant alpha 1 OS=Homo sapiens OX=9606 GN=IGHA1 PE=1 SV=2 | 255 | 20201118_ly_colostrum_1ng_glyco_hcd30_etd30_compound15.16704.16704.3 | controllerType=0 controllerNumber=1 scan=16704 | 44.8215 |
| 25991:4 | 2 | R.LSLHRPALEDLLLGSEAN[+1501.555]LTC[+57.021]TLTGLR.D | HexNAc(5)Hex(3) | C[+57], N[+1502] | 1117.047 | 4 | 4465.1663 | 4465.1536 | 0 | 2.9 | 127 | Specific | 353.17 | 353.17 | 353.17 | 6.75 | 65 | >sp\|P01876\|IGHA1_HUMAN Immunoglobulin heavy constant alpha 1 OS=Homo sapiens OX=9606 GN=IGHA1 PE=1 SV=2 | 255 | 20201118_ly_colostrum_1ng_glyco_hcd30_etd30_compound15.33112.33112.4 | controllerType=0 controllerNumber=1 scan=33112 | 89.2132 |
| 12558:3 | 2 | R.LSLHRPALEDLLLGSEAN[+1054.370]LTC[+57.021]TLTGLR.D | HexNAc(2)Hex(4) | C[+57], N[+1054] | 1339.9974 | 3 | 4017.9776 | 4017.9683 | 0 | 2.3 | 127 | Specific | 376.02 | 366.58 | 366.58 | 8.16 | 65 | >sp\|P01876\|IGHA1_HUMAN Immunoglobulin heavy constant alpha 1 OS=Homo sapiens OX=9606 GN=IGHA1 PE=1 SV=2 | 255 | 20201118_ly_colostrum_1ng_glyco_hcd30_etd30_compound15.16039.16039.3 | controllerType=0 controllerNumber=1 scan=16039 | 42.8661 |
| 12497:3 | 2 | R.LSLHRPALEDLLLGSEAN[+1216.423]LTC[+57.021]TLTGLR.D | HexNAc(2)Hex(5) | C[+57], N[+1216] | 1394.0147 | 3 | 4180.0296 | 4180.0211 | 0 | 2 | 127 | Specific | 381.39 | 381.39 | 381.39 | 8.16 | 65 | >sp\|P01876\|IGHA1_HUMAN Immunoglobulin heavy constant alpha 1 OS=Homo sapiens OX=9606 GN=IGHA1 PE=1 SV=2 | 255 | 20201118_ly_colostrum_1ng_glyco_hcd30_etd30_compound15.15975.15975.6 | controllerType=0 controllerNumber=1 scan=15975 | 42.6626 |
| 15867:4 | 2 | R.LSLHRPALEDLLLGSEAN[+1501.555]LTC[+57.021]TLTGLR.D | HexNAc(5)Hex(3) | C[+57], N[+1502] | 1117.0474 | 4 | 4465.1678 | 4465.1536 | 0 | 3.2 | 127 | Specific | 385.84 | 385.84 | 385.84 | 7.79 | 65 | >sp\|P01876\|IGHA1_HUMAN Immunoglobulin heavy constant alpha 1 OS=Homo sapiens OX=9606 GN=IGHA1 PE=1 SV=2 | 255 | 20201118_ly_colostrum_1ng_glyco_hcd30_etd30_compound15.19560.19560.3 | controllerType=0 controllerNumber=1 scan=19560 | 54.[illegible]589 |
| 13219:3 | 2 | R.LSLHRPALEDLLLGSEAN[+1710.598]LTC[+57.021]TLTGLR.D | HexNAc(3)Hex(5)NeuAc(1) | C[+57], N[+1711] | 1558.7422 | 3 | 4674.2122 | 4674.1959 | 0 | 3.5 | 127 | Specific | 393.04 | 270.46 | 270.46 | 6.39 | 65 | >sp\|P01876\|IGHA1_HUMAN Immunoglobulin heavy constant alpha 1 OS=Homo sapiens OX=9606 GN=IGHA1 PE=1 SV=2 | 255 | 20201118_ly_colostrum_1ng_glyco_hcd30_etd30_compound15.16736.16736.3 | controllerType=0 controllerNumber=1 scan=16736 | 44.9195 |
| 12442:3 | 2 | R.LSLHRPALEDLLLGSEAN[+1663.608]LTC[+57.021]TLTGLR.D | HexNAc(5)Hex(4) | C[+57], N[+1664] | 1543.0772 | 3 | 4627.2171 | 4627.2064 | 0 | 2.3 | 127 | Specific | 394.57 | 368.23 | 368.23 | 8.16 | 65 | >sp\|P01876\|IGHA1_HUMAN Immunoglobulin heavy constant alpha 1 OS=Homo sapiens OX=9606 GN=IGHA1 PE=1 SV=2 | 255 | 20201118_ly_colostrum_1ng_glyco_hcd30_etd30_compound15.15916.15916.3 | controllerType=0 controllerNumber=1 scan=15916 | 42.4814 |
| 13205:3 | 2 | R.LSLHRPALEDLLLGSEAN[+1872.651]LTC[+57.021]TLTGLR.D | HexNAc(3)Hex(6)NeuAc(1) | C[+57], N[+1873] | 1612.7607 | 3 | 4836.2675 | 4836.2487 | 0 | 3.9 | 127 | Specific | 397.84 | 341.04 | 341.04 | 6.68 | 65 | >sp\|P01876\|IGHA1_HUMAN Immunoglobulin heavy constant alpha 1 OS=Homo sapiens OX=9606 GN=IGHA1 PE=1 SV=2 | 255 | 20201118_ly_colostrum_1ng_glyco_hcd30_etd30_compound15.16721.16721.3 | controllerType=0 controllerNumber=1 scan=16721 | 44.8744 |
| 12483:3 | 2 | R.LSLHRPALEDLLLGSEAN[+1460.529]LTC[+57.021]TLTGLR.D | HexNAc(4)Hex(4) | C[+57], N[+1461] | 1475.3864 | 3 | 4424.1447 | 4424.1271 | 0 | 4 | 127 | Specific | 403.26 | 394.61 | 394.61 | 8.16 | 65 | >sp\|P01876\|IGHA1_HUMAN Immunoglobulin heavy constant alpha 1 OS=Homo sapiens OX=9606 GN=IGHA1 PE=1 SV=2 | 255 | 20201118_ly_colostrum_1ng_glyco_hcd30_etd30_compound15.15960.15960.3 | controllerType=0 controllerNumber=1 scan=15960 | 42.6252 |
| 12545:3 | 2 | R.LSLHRPALEDLLLGSEAN[+1298.476]LTC[+57.021]TLTGLR.D | HexNAc(4)Hex(3) | C[+57], N[+1298] | 1421.3682 | 3 | 4262.0901 | 4262.0742 | 0 | 3.7 | 127 | Specific | 404.38 | 398.49 | 398.49 | 8.16 | 65 | >sp\|P01876\|IGHA1_HUMAN Immunoglobulin heavy constant alpha 1 OS=Homo sapiens OX=9606 GN=IGHA1 PE=1 SV=2 | 255 | 20201118_ly_colostrum_1ng_glyco_hcd30_etd30_compound15.16026.16026.3 | controllerType=0 controllerNumber=1 scan=16026 | 42.8316 |
| 13188:3 | 2 | R.LSLHRPALEDLLLGSEAN[+1913.677]LTC[+57.021]TLTGLR.D | HexNAc(4)Hex(5)NeuAc(1) | C[+57], N[+1914] | 1626.4368 | 3 | 4877.2959 | 4877.2753 | 0 | 4.2 | 127 | Specific | 405.06 | 345.91 | 345.91 | 5.65 | 65 | >sp\|P01876\|IGHA1_HUMAN Immunoglobulin heavy constant alpha 1 OS=Homo sapiens OX=9606 GN=IGHA1 PE=1 SV=2 | 255 | 20201118_ly_colostrum_1ng_glyco_hcd30_etd30_compound15.16703.16703.3 | controllerType=0 controllerNumber=1 scan=16703 | 44.8137 |
| 14803:4 | 2 | R.LSLHRPALEDLLLGSEAN[+1501.555]LTC[+57.021]TLTGLR.D | HexNAc(5)Hex(3) | C[+57], N[+1502] | 1117.0467 | 4 | 4465.1649 | 4465.1536 | 0 | 2.5 | 127 | Specific | 426.14 | 426.14 | 426.14 | 8.24 | 65 | >sp\|P01876\|IGHA1_HUMAN Immunoglobulin heavy constant alpha 1 OS=Homo sapiens OX=9606 GN=IGHA1 PE=1 SV=2 | 255 | 20201118_ly_colostrum_1ng_glyco_hcd30_etd30_compound15.18421.18421.4 | controllerType=0 controllerNumber=1 scan=18421 | 50.5826 |
| 12466:3 | 2 | R.LSLHRPALEDLLLGSEAN[+1663.608]LTC[+57.021]TLTGLR.D | HexNAc(5)Hex(4) | C[+57], N[+1664] | 1543.0794 | 3 | 4627.2237 | 4627.2064 | 0 | 3.7 | 127 | Specific | 430.8 | 395.18 | 395.18 | 8.16 | 65 | >sp\|P01876\|IGHA1_HUMAN Immunoglobulin heavy constant alpha 1 OS=Homo sapiens OX=9606 GN=IGHA1 PE=1 SV=2 | 255 | 20201118_ly_colostrum_1ng_glyco_hcd30_etd30_compound15.15942.15942.3 | controllerType=0 controllerNumber=1 scan=15942 | 42.5679 |
| 12595:3 | 2 | R.LSLHRPALEDLLLGSEAN[+892.317]LTC[+57.021]TLTGLR.D | HexNAc(2)Hex(3) | C[+57], N[+892] | 1285.9782 | 3 | 3855.9201 | 3855.9155 | 0 | 1.2 | 127 | Specific | 455.22 | 453.62 | 453.62 | 8.34 | 65 | >sp\|P01876\|IGHA1_HUMAN Immunoglobulin heavy constant alpha 1 OS=Homo sapiens OX=9606 GN=IGHA1 PE=1 SV=2 | 255 | 20201118_ly_colostrum_1ng_glyco_hcd30_etd30_compound15.16078.16078.3 | controllerType=0 controllerNumber=1 scan=16078 | 42.9743 |
| 12400:3 | 2 | R.LSLHRPALEDLLLGSEAN[+1864.634]LTC[+57.021]TLTGLR.D | HexNAc(2)Hex(9) | C[+57], N[+1865] | 1610.0864 | 3 | 4828.2445 | 4828.2324 | 0 | 2.5 | 127 | Specific | 463.81 | 419.55 | 419.55 | 8.34 | 65 | >sp\|P01876\|IGHA1_HUMAN Immunoglobulin heavy constant alpha 1 OS=Homo sapiens OX=9606 GN=IGHA1 PE=1 SV=2 | 255 | 20201118_ly_colostrum_1ng_glyco_hcd30_etd30_compound15.15872.15872.3 | controllerType=0 controllerNumber=1 scan=15872 | 42.3454 |
| 12477:3 | 2 | R.LSLHRPALEDLLLGSEAN[+1501.555]LTC[+57.021]TLTGLR.D | HexNAc(5)Hex(3) | C[+57], N[+1502] | 1489.0607 | 3 | 4465.1676 | 4465.1536 | 0 | 3.1 | 127 | Specific | 466.45 | 440.53 | 440.53 | 8.11 | 65 | >sp\|P01876\|IGHA1_HUMAN Immunoglobulin heavy constant alpha 1 OS=Homo sapiens OX=9606 GN=IGHA1 PE=1 SV=2 | 255 | 20201118_ly_colostrum_1ng_glyco_hcd30_etd30_compound15.15954.15954.3 | controllerType=0 controllerNumber=1 scan=15954 | 42.6079 |
| 12418:3 | 2 | R.LSLHRPALEDLLLGSEAN[+1702.581]LTC[+57.021]TLTGLR.D | HexNAc(2)Hex(8) | C[+57], N[+1703] | 1556.0693 | 3 | 4666.1933 | 4666.1796 | 0 | 2.9 | 127 | Specific | 467.04 | 422.33 | 422.33 | 8.11 | 65 | >sp\|P01876\|IGHA1_HUMAN Immunoglobulin heavy constant alpha 1 OS=Homo sapiens OX=9606 GN=IGHA1 PE=1 SV=2 | 255 | 20201118_ly_colostrum_1ng_glyco_hcd30_etd30_compound15.15891.15891.3 | controllerType=0 controllerNumber=1 scan=15891 | 42.3952 |
| 12473:5 | 2 | R.LSLHRPALEDLLLGSEAN[+1663.608]LTC[+57.021]TLTGLR.D | HexNAc(5)Hex(4) | C[+57], N[+1664] | 926.2487 | 5 | 4627.2146 | 4627.2064 | 0 | 1.8 | 127 | Specific | 468.57 | 468.57 | 468.57 | 6.79 | 65 | >sp\|P01876\|IGHA1_HUMAN Immunoglobulin heavy constant alpha 1 OS=Homo sapiens OX=9606 GN=IGHA1 PE=1 SV=2 | 255 | 20201118_ly_colostrum_1ng_glyco_hcd30_etd30_compound15.15949.15949.5 | controllerType=0 controllerNumber=1 scan=15949 | 42.588 |
| 12467:3 | 2 | R.LSLHRPALEDLLLGSEAN[+1378.476]LTC[+57.021]TLTGLR.D | HexNAc(2)Hex(6) | C[+57], N[+1378] | 1448.0352 | 3 | 4342.0911 | 4342.074 | 0 | 3.9 | 127 | Specific | 486.14 | 486.14 | 486.14 | 8.29 | 65 | >sp\|P01876\|IGHA1_HUMAN Immunoglobulin heavy constant alpha 1 OS=Homo sapiens OX=9606 GN=IGHA1 PE=1 SV=2 | 255 | 20201118_ly_colostrum_1ng_glyco_hcd30_etd30_compound15.15943.15943.3 | controllerType=0 controllerNumber=1 scan=15943 | 42.571 |
| 12435:3 | 2 | R.LSLHRPALEDLLLGSEAN[+1540.529]LTC[+57.021]TLTGLR.D | HexNAc(2)Hex(7) | C[+57], N[+1541] | 1502.0507 | 3 | 4504.1376 | 4504.1268 | 0 | 2.4 | 127 | Specific | 492.26 | 462.34 | 462.34 | 8.51 | 65 | >sp\|P01876\|IGHA1_HUMAN Immunoglobulin heavy constant alpha 1 OS=Homo sapiens OX=9606 GN=IGHA1 PE=1 SV=2 | 255 | 20201118_ly_colostrum_1ng_glyco_hcd30_etd30_compound15.15909.15909.3 | controllerType=0 controllerNumber=1 scan=15909 | 42.4544 |
| 12546:3 | 2 | R.LSLHRPALEDLLLGSEAN[+1095.397]LTC[+57.021]TLTGLR.D | HexNAc(3)Hex(3) | C[+57], N[+1095] | 1353.6729 | 3 | 4059.0042 | 4058.9949 | 0 | 2.3 | 127 | Specific | 500.65 | 491.1 | 491.1 | 8.51 | 65 | >sp\|P01876\|IGHA1_HUMAN Immunoglobulin heavy constant alpha 1 OS=Homo sapiens OX=9606 GN=IGHA1 PE=1 SV=2 | 255 | 20201118_ly_colostrum_1ng_glyco_hcd30_etd30_compound15.16027.16027.3 | controllerType=0 controllerNumber=1 scan=16027 | 42.8348 |
| 12476:3 | 2 | R.LSLHRPALEDLLLGSEAN[+1216.423]LTC[+57.021]TLTGLR.D | HexNAc(2)Hex(5) | C[+57], N[+1216] | 1394.0154 | 3 | 4180.0318 | 4180.0211 | 0 | 2.5 | 127 | Specific | 505.71 | 505.71 | 505.71 | 8.51 | 65 | >sp\|P01876\|IGHA1_HUMAN Immunoglobulin heavy constant alpha 1 OS=Homo sapiens OX=9606 GN=IGHA1 PE=1 SV=2 | 255 | 20201118_ly_colostrum_1ng_glyco_hcd30_etd30_compound15.15953.15953.3 | controllerType=0 controllerNumber=1 scan=15953 | 42.602 |
| 13952:4 | 2 | R.LSLHRPALEDLLLGSEAN[+2204.772]LTC[+57.021]TLTGLR.D | HexNAc(4)Hex(5)NeuAc(2) | C[+57], N[+2205] | 1292.8513 | 4 | 5168.3835 | 5168.3707 | 0 | 2.5 | 127 | Specific | 519.35 | 519.35 | 519.35 | 7.12 | 65 | >sp\|P01876\|IGHA1_HUMAN Immunoglobulin heavy constant alpha 1 OS=Homo sapiens OX=9606 GN=IGHA1 PE=1 SV=2 | 255 | 20201118_ly_colostrum_1ng_glyco_hcd30_etd30_compound15.17513.17513.4 | controllerType=0 controllerNumber=1 scan=17513 | 47.3345 |
| 12551:4 | 2 | R.LSLHRPALEDLLLGSEAN[+1647.613]LTC[+57.021]TLTGLR.D | HexNAc(5)Hex(3)Fuc(1) | C[+57], N[+1648] | 1153.5611 | 4 | 4611.2225 | 4611.2115 | 0 | 2.4 | 127 | Specific | 525.37 | 524.15 | 524.15 | 8.6 | 65 | >sp\|P01876\|IGHA1_HUMAN Immunoglobulin heavy constant alpha 1 OS=Homo sapiens OX=9606 GN=IGHA1 PE=1 SV=2 | 255 | 20201118_ly_colostrum_1ng_glyco_hcd30_etd30_compound15.16032.16032.4 | controllerType=0 controllerNumber=1 scan=16032 | 42.8466 |
| 14764:4 | 2 | R.LSLHRPALEDLLLGSEAN[+1501.555]LTC[+57.021]TLTGLR.D | HexNAc(5)Hex(3) | C[+57], N[+1502] | 1117.0478 | 4 | 4465.1693 | 4465.1536 | 0 | 3.5 | 127 | Specific | 536.57 | 536.57 | 536.57 | 7.13 | 65 | >sp\|P01876\|IGHA1_HUMAN Immunoglobulin heavy constant alpha 1 OS=Homo sapiens OX=9606 GN=IGHA1 PE=1 SV=2 | 255 | 20201118_ly_colostrum_1ng_glyco_hcd30_etd30_compound15.18379.18379.4 | controllerType=0 controllerNumber=1 scan=18379 | 50.4104 |
| 13908:4 | 2 | R.LSLHRPALEDLLLGSEAN[+2407.852]LTC[+57.021]TLTGLR.D | HexNAc(5)Hex(5)NeuAc(2) | C[+57], N[+2408] | 1343.6201 | 4 | 5371.4587 | 5371.4501 | 0 | 1.6 | 127 | Specific | 587.81 | 511.71 | 511.71 | 8.41 | 65 | >sp\|P01876\|IGHA1_HUMAN Immunoglobulin heavy constant alpha 1 OS=Homo sapiens OX=9606 GN=IGHA1 PE=1 SV=2 | 255 | 20201118_ly_colostrum_1ng_glyco_hcd30_etd30_compound15.17467.17467.4 | controllerType=0 controllerNumber=1 scan=17467 | 47.2109 |
| 12215:4 | 2 | R.LSLHRPALEDLLLGSEAN[+1501.555]LTC[+57.021]TLTGLR.D | HexNAc(5)Hex(3) | C[+57], N[+1502] | 1117.0447 | 4 | 4465.1571 | 4465.1536 | 0 | 0.8 | 127 | Specific | 643.97 | 643.72 | 643.72 | 9.54 | 65 | >sp\|P01876\|IGHA1_HUMAN Immunoglobulin heavy constant alpha 1 OS=Homo sapiens OX=9606 GN=IGHA1 PE=1 SV=2 | 255 | 20201118_ly_colostrum_1ng_glyco_hcd30_etd30_compound15.15676.15676.4 | controllerType=0 controllerNumber=1 scan=15676 | 41.7657 |
| 12533:4 | 2 | R.LSLHRPALEDLLLGSEAN[+1257.449]LTC[+57.021]TLTGLR.D | HexNAc(3)Hex(4) | C[+57], N[+1257] | 1056.0175 | 4 | 4221.0482 | 4221.0477 | 0 | 0.1 | 127 | Specific | 644.7 | 644.7 | 644.7 | 9.54 | 65 | >sp\|P01876\|IGHA1_HUMAN Immunoglobulin heavy constant alpha 1 OS=Homo sapiens OX=9606 GN=IGHA1 PE=1 SV=2 | 255 | 20201118_ly_colostrum_1ng_glyco_hcd30_etd30_compound15.16013.16013.4 | controllerType=0 controllerNumber=1 scan=16013 | 42.7931 |
| 12481:4 | 2 | R.LSLHRPALEDLLLGSEAN[+1419.502]LTC[+57.021]TLTGLR.D | HexNAc(3)Hex(5) | C[+57], N[+1420] | 1096.5336 | 4 | 4383.1126 | 4383.1005 | 0 | 2.8 | 127 | Specific | 708.93 | 694.45 | 694.45 | 12.66 | 65 | >sp\|P01876\|IGHA1_HUMAN Immunoglobulin heavy constant alpha 1 OS=Homo sapiens OX=9606 GN=IGHA1 PE=1 SV=2 | 255 | 20201118_ly_colostrum_1ng_glyco_hcd30_etd30_compound15.15958.15958.4 | controllerType=0 controllerNumber=1 scan=15958 | 42.6155 |
| 13200:4 | 2 | R.LSLHRPALEDLLLGSEAN[+1872.651]LTC[+57.021]TLTGLR.D | HexNAc(3)Hex(6)NeuAc(1) | C[+57], N[+1873] | 1209.8228 | 4 | 4836.2694 | 4836.2487 | 0 | 4.3 | 127 | Specific | 715.64 | 685.83 | 685.83 | 12.44 | 65 | >sp\|P01876\|IGHA1_HUMAN Immunoglobulin heavy constant alpha 1 OS=Homo sapiens OX=9606 GN=IGHA1 PE=1 SV=2 | 255 | 20201118_ly_colostrum_1ng_glyco_hcd30_etd30_compound15.16716.16716.4 | controllerType=0 controllerNumber=1 scan=16716 | 44.8604 |
| 13184:4 | 2 | R.LSLHRPALEDLLLGSEAN[+2116.756]LTC[+57.021]TLTGLR.D | HexNAc(5)Hex(5)NeuAc(1) | C[+57], N[+2117] | 1270.8481 | 4 | 5080.3708 | 5080.3547 | 0 | 3.2 | 127 | Specific | 750.05 | 660.12 | 660.12 | 13.44 | 65 | >sp\|P01876\|IGHA1_HUMAN Immunoglobulin heavy constant alpha 1 OS=Homo sapiens OX=9606 GN=IGHA1 PE=1 SV=2 | 255 | 20201118_ly_colostrum_1ng_glyco_hcd30_etd30_compound15.16699.16699.4 | controllerType=0 controllerNumber=1 scan=16699 | 44.8068 |
| 13150:5 | 2 | R.LSLHRPALEDLLLGSEAN[+2116.756]LTC[+57.021]TLTGLR.D | HexNAc(5)Hex(5)NeuAc(1) | C[+57], N[+2117] | 1016.8794 | 5 | 5080.3679 | 5080.3547 | 0 | 2.6 | 127 | Specific | 766.84 | 741.27 | 741.27 | 12.59 | 65 | >sp\|P01876\|IGHA1_HUMAN Immunoglobulin heavy constant alpha 1 OS=Homo sapiens OX=9606 GN=IGHA1 PE=1 SV=2 | 255 | 20201118_ly_colostrum_1ng_glyco_hcd30_etd30_compound15.16663.16663.5 | controllerType=0 controllerNumber=1 scan=16663 | 44.6944 |
| 13235:4 | 2 | R.LSLHRPALEDLLLGSEAN[+1548.545]LTC[+57.021]TLTGLR.D | HexNAc(3)Hex(4)NeuAc(1) | C[+57], N[+1549] | 1128.7948 | 4 | 4512.1575 | 4512.1431 | 0 | 3.2 | 127 | Specific | 778.33 | 748.74 | 748.74 | 13.44 | 65 | >sp\|P01876\|IGHA1_HUMAN Immunoglobulin heavy constant alpha 1 OS=Homo sapiens OX=9606 GN=IGHA1 PE=1 SV=2 | 255 | 20201118_ly_colostrum_1ng_glyco_hcd30_etd30_compound15.16753.16753.4 | controllerType=0 controllerNumber=1 scan=16753 | 44.9807 |
| 12498:5 | 2 | R.LSLHRPALEDLLLGSEAN[+1216.423]LTC[+57.021]TLTGLR.D | HexNAc(2)Hex(5) | C[+57], N[+1216] | 836.8114 | 5 | 4180.0277 | 4180.0211 | 0 | 1.6 | 127 | Specific | 782.21 | 740.25 | 740.25 | 13.71 | 65 | >sp\|P01876\|IGHA1_HUMAN Immunoglobulin heavy constant alpha 1 OS=Homo sapiens OX=9606 GN=IGHA1 PE=1 SV=2 | 255 | 20201118_ly_colostrum_1ng_glyco_hcd30_etd30_compound15.15976.15976.5 | controllerType=0 controllerNumber=1 scan=15976 | 42.6696 |
| 12471:4 | 2 | R.LSLHRPALEDLLLGSEAN[+1581.555]LTC[+57.021]TLTGLR.D | HexNAc(3)Hex(6) | C[+57], N[+1582] | 1137.0475 | 4 | 4545.1683 | 4545.1533 | 0 | 3.3 | 127 | Specific | 796.65 | 768.7 | 768.7 | 13.44 | 65 | >sp\|P01876\|IGHA1_HUMAN Immunoglobulin heavy constant alpha 1 OS=Homo sapiens OX=9606 GN=IGHA1 PE=1 SV=2 | 255 | 20201118_ly_colostrum_1ng_glyco_hcd30_etd30_compound15.15947.15947.4 | controllerType=0 controllerNumber=1 scan=15947 | 42.5779 |
| 12511:5 | 2 | R.LSLHRPALEDLLLGSEAN[+1501.555]LTC[+57.021]TLTGLR.D | HexNAc(5)Hex(3) | C[+57], N[+1502] | 893.838 | 5 | 4465.1611 | 4465.1536 | 0 | 1.7 | 127 | Specific | 832.03 | 810.85 | 810.85 | 13.45 | 65 | >sp\|P01876\|IGHA1_HUMAN Immunoglobulin heavy constant alpha 1 OS=Homo sapiens OX=9606 GN=IGHA1 PE=1 SV=2 | 255 | 20201118_ly_colostrum_1ng_glyco_hcd30_etd30_compound15.15990.15990.5 | controllerType=0 controllerNumber=1 scan=15990 | 42.7131 |
| 13183:4 | 2 | R.LSLHRPALEDLLLGSEAN[+1913.677]LTC[+57.021]TLTGLR.D | HexNAc(4)Hex(5)NeuAc(1) | C[+57], N[+1914] | 1220.0796 | 4 | 4877.2967 | 4877.2753 | 0 | 4.4 | 127 | Specific | 884.02 | 821.61 | 821.61 | 15.44 | 65 | >sp\|P01876\|IGHA1_HUMAN Immunoglobulin heavy constant alpha 1 OS=Homo sapiens OX=9606 GN=IGHA1 PE=1 SV=2 | 255 | 20201118_ly_colostrum_1ng_glyco_hcd30_etd30_compound15.16698.16698.4 | controllerType=0 controllerNumber=1 scan=16698 | 44.8036 |
| 12505:4 | 2 | R.LSLHRPALEDLLLGSEAN[+1298.476]LTC[+57.021]TLTGLR.D | HexNAc(4)Hex(3) | C[+57], N[+1298] | 1066.2773 | 4 | 4262.0872 | 4262.0742 | 0 | 3.1 | 127 | Specific | 894.85 | 894.85 | 894.85 | 15.44 | 65 | >sp\|P01876\|IGHA1_HUMAN Immunoglobulin heavy constant alpha 1 OS=Homo sapiens OX=9606 GN=IGHA1 PE=1 SV=2 | 255 | 20201118_ly_colostrum_1ng_glyco_hcd30_etd30_compound15.15983.15983.4 | controllerType=0 controllerNumber=1 scan=15983 | 42.6897 |
| 12540:4 | 2 | R.LSLHRPALEDLLLGSEAN[+1501.555]LTC[+57.021]TLTGLR.D | HexNAc(5)Hex(3) | C[+57], N[+1502] | 1117.0492 | 4 | 4465.1751 | 4465.1536 | 0 | 4.8 | 127 | Specific | 907 | 907 | 907 | 15.85 | 65 | >sp\|P01876\|IGHA1_HUMAN Immunoglobulin heavy constant alpha 1 OS=Homo sapiens OX=9606 GN=IGHA1 PE=1 SV=2 | 255 | 20201118_ly_colostrum_1ng_glyco_hcd30_etd30_compound15.16021.16021.4 | controllerType=0 controllerNumber=1 scan=16021 | 42.8215 |
| 12751:4 | 2 | R.LSLHRPALEDLLLGSEAN[+1501.555]LTC[+57.021]TLTGLR.D | HexNAc(5)Hex(3) | C[+57], N[+1502] | 1117.0475 | 4 | 4465.1683 | 4465.1536 | 0 | 3.3 | 127 | Specific | 931.11 | 922.08 | 922.08 | 16.44 | 65 | >sp\|P01876\|IGHA1_HUMAN Immunoglobulin heavy constant alpha 1 OS=Homo sapiens OX=9606 GN=IGHA1 PE=1 SV=2 | 255 | 20201118_ly_colostrum_1ng_glyco_hcd30_etd30_compound15.16243.16243.3 | controllerType=0 controllerNumber=1 scan=16243 | 43.4525 |
| 13190:4 | 2 | R.LSLHRPALEDLLLGSEAN[+1710.598]LTC[+57.021]TLTGLR.D | HexNAc(3)Hex(5)NeuAc(1) | C[+57], N[+1711] | 1169.3089 | 4 | 4674.2137 | 4674.1959 | 0 | 3.8 | 127 | Specific | 956.87 | 945.1 | 945.1 | 17.44 | 65 | >sp\|P01876\|IGHA1_HUMAN Immunoglobulin heavy constant alpha 1 OS=Homo sapiens OX=9606 GN=IGHA1 PE=1 SV=2 | 255 | 20201118_ly_colostrum_1ng_glyco_hcd30_etd30_compound15.16705.16705.4 | controllerType=0 controllerNumber=1 scan=16705 | 44.8268 |
| 12460:4 | 2 | R.LSLHRPALEDLLLGSEAN[+1663.608]LTC[+57.021]TLTGLR.D | HexNAc(5)Hex(4) | C[+57], N[+1664] | 1157.5606 | 4 | 4627.2205 | 4627.2064 | 0 | 3 | 127 | Specific | 966.62 | 946.04 | 946.04 | 17.44 | 65 | >sp\|P01876\|IGHA1_HUMAN Immunoglobulin heavy constant alpha 1 OS=Homo sapiens OX=9606 GN=IGHA1 PE=1 SV=2 | 255 | 20201118_ly_colostrum_1ng_glyco_hcd30_etd30_compound15.15936.15936.4 | controllerType=0 controllerNumber=1 scan=15936 | 42.545 |
| 12413:4 | 2 | R.LSLHRPALEDLLLGSEAN[+1702.581]LTC[+57.021]TLTGLR.D | HexNAc(2)Hex(8) | C[+57], N[+1703] | 1167.3029 | 4 | 4666.1898 | 4666.1796 | 0 | 2.2 | 127 | Specific | 992.92 | 937.34 | 937.34 | 17.66 | 65 | >sp\|P01876\|IGHA1_HUMAN Immunoglobulin heavy constant alpha 1 OS=Homo sapiens OX=9606 GN=IGHA1 PE=1 SV=2 | 255 | 20201118_ly_colostrum_1ng_glyco_hcd30_etd30_compound15.15886.15886.4 | controllerType=0 controllerNumber=1 scan=15886 | 42.3823 |
| 12465:4 | 2 | R.LSLHRPALEDLLLGSEAN[+1216.423]LTC[+57.021]TLTGLR.D | HexNAc(2)Hex(5) | C[+57], N[+1216] | 1045.7642 | 4 | 4180.035 | 4180.0211 | 0 | 3.3 | 127 | Specific | 1019.69 | 992.17 | 992.17 | 18.44 | 65 | >sp\|P01876\|IGHA1_HUMAN Immunoglobulin heavy constant alpha 1 OS=Homo sapiens OX=9606 GN=IGHA1 PE=1 SV=2 | 255 | 20201118_ly_colostrum_1ng_glyco_hcd30_etd30_compound15.15941.15941.4 | controllerType=0 controllerNumber=1 scan=15941 | 42.5612 |
| 12492:4 | 2 | R.LSLHRPALEDLLLGSEAN[+1460.529]LTC[+57.021]TLTGLR.D | HexNAc(4)Hex(4) | C[+57], N[+1461] | 1106.7914 | 4 | 4424.1439 | 4424.1271 | 0 | 3.8 | 127 | Specific | 1039.13 | 930.51 | 930.51 | 18.44 | 65 | >sp\|P01876\|IGHA1_HUMAN Immunoglobulin heavy constant alpha 1 OS=Homo sapiens OX=9606 GN=IGHA1 PE=1 SV=2 | 255 | 20201118_ly_colostrum_1ng_glyco_hcd30_etd30_compound15.15970.15970.4 | controllerType=0 controllerNumber=1 scan=15970 | 42.6513 |
| 12428:4 | 2 | R.LSLHRPALEDLLLGSEAN[+1540.529]LTC[+57.021]TLTGLR.D | HexNAc(2)Hex(7) | C[+57], N[+1541] | 1126.7902 | 4 | 4504.139 | 4504.1268 | 0 | 2.7 | 127 | Specific | 1042.77 | 960.29 | 960.29 | 18.66 | 65 | >sp\|P01876\|IGHA1_HUMAN Immunoglobulin heavy constant alpha 1 OS=Homo sapiens OX=9606 GN=IGHA1 PE=1 SV=2 | 255 | 20201118_ly_colostrum_1ng_glyco_hcd30_etd30_compound15.15902.15902.4 | controllerType=0 controllerNumber=1 scan=15902 | 42.4362 |
| 12461:4 | 2 | R.LSLHRPALEDLLLGSEAN[+1378.476]LTC[+57.021]TLTGLR.D | HexNAc(2)Hex(6) | C[+57], N[+1378] | 1086.2778 | 4 | 4342.0892 | 4342.074 | 0 | 3.5 | 127 | Specific | 1062.07 | 997.34 | 997.34 | 19.44 | 65 | >sp\|P01876\|IGHA1_HUMAN Immunoglobulin heavy constant alpha 1 OS=Homo sapiens OX=9606 GN=IGHA1 PE=1 SV=2 | 255 | 20201118_ly_colostrum_1ng_glyco_hcd30_etd30_compound15.15937.15937.4 | controllerType=0 controllerNumber=1 scan=15937 | 42.5487 |
| 12531:4 | 2 | R.LSLHRPALEDLLLGSEAN[+1054.370]LTC[+57.021]TLTGLR.D | HexNAc(2)Hex(4) | C[+57], N[+1054] | 1005.2495 | 4 | 4017.9761 | 4017.9683 | 0 | 2 | 127 | Specific | 1104.15 | 1060.12 | 1060.12 | 20.66 | 65 | >sp\|P01876\|IGHA1_HUMAN Immunoglobulin heavy constant alpha 1 OS=Homo sapiens OX=9606 GN=IGHA1 PE=1 SV=2 | 255 | 20201118_ly_colostrum_1ng_glyco_hcd30_etd30_compound15.16011.16011.4 | controllerType=0 controllerNumber=1 scan=16011 | 42.7829 |
| 13329:4 | 2 | R.LSLHRPALEDLLLGSEAN[+1216.423]LTC[+57.021]TLTGLRDASGVTFTWTPSSGK.S | HexNAc(2)Hex(5) | C[+57], N[+1216] | 1426.1948 | 4 | 5701.7575 | 5701.7311 | 0 | 4.6 | 127 | Specific | 736.46 | 736.46 | 736.46 | 11.47 | 65 | >sp\|P01876\|IGHA1_HUMAN Immunoglobulin heavy constant alpha 1 OS=Homo sapiens OX=9606 GN=IGHA1 PE=1 SV=2 | 255 | 20201118_ly_colostrum_1ng_glyco_hcd30_etd30_compound15.16852.16852.4 | controllerType=0 controllerNumber=1 scan=16852 | 45.3104 |
| 13313:5 | 2 | R.LSLHRPALEDLLLGSEAN[+1216.423]LTC[+57.021]TLTGLRDASGVTFTWTPSSGK.S | HexNAc(2)Hex(5) | C[+57], N[+1216] | 1141.156 | 5 | 5701.7509 | 5701.7311 | 0 | 3.5 | 127 | Specific | 876.46 | 876.46 | 876.46 | 14.92 | 65 | >sp\|P01876\|IGHA1_HUMAN Immunoglobulin heavy constant alpha 1 OS=Homo sapiens OX=9606 GN=IGHA1 PE=1 SV=2 | 255 | 20201118_ly_colostrum_1ng_glyco_hcd30_etd30_compound15.16835.16835.5 | controllerType=0 controllerNumber=1 scan=16835 | 45.2501 |
| 13316:5 | 2 | R.LSLHRPALEDLLLGSEAN[+1216.423]LTC[+57.021]TLTGLRDASGVTFTWTPSSGK.S | HexNAc(2)Hex(5) | C[+57], N[+1216] | 1141.1571 | 5 | 5701.7564 | 5701.7311 | 0 | 4.4 | 127 | Specific | 974.08 | 974.08 | 974.08 | 16.92 | 65 | >sp\|P01876\|IGHA1_HUMAN Immunoglobulin heavy constant alpha 1 OS=Homo sapiens OX=9606 GN=IGHA1 PE=1 SV=2 | 255 | 20201118_ly_colostrum_1ng_glyco_hcd30_etd30_compound15.16839.16839.5 | controllerType=0 controllerNumber=1 scan=16839 | 45.2661 |
| 08509:3 | 2 | K.TIDRLAGKPTHVN[+1216.423]VSVVMAEVDGTC[+57.021]Y.- | HexNAc(2)Hex(5) | C[+57], N[+1216] | 1350.282 | 3 | 4048.8316 | 4048.8247 | 0 | 1.7 | 328 | CRagged | 212.81 | 190.71 | 98 | 3.78 | 65 | >sp\|P01876\|IGHA1_HUMAN Immunoglobulin heavy constant alpha 1 OS=Homo sapiens OX=9606 GN=IGHA1 PE=1 SV=2 | 255 | 20201118_ly_colostrum_1ng_glyco_hcd30_etd30_compound15.11781.11781.3 | controllerType=0 controllerNumber=1 scan=11781 | 31.2138 |
| 08488:3 | 2 | K.TIDRLAGKPTHVN[+1378.476]VSVVMAEVDGTC[+57.021]Y.- | HexNAc(2)Hex(6) | C[+57], N[+1378] | 1404.3022 | 3 | 4210.892 | 4210.8776 | 0 | 3.4 | 328 | CRagged | 230.42 | 190.59 | 121.78 | 3.59 | 65 | >sp\|P01876\|IGHA1_HUMAN Immunoglobulin heavy constant alpha 1 OS=Homo sapiens OX=9606 GN=IGHA1 PE=1 SV=2 | 255 | 20201118_ly_colostrum_1ng_glyco_hcd30_etd30_compound15.11758.11758.3 | controllerType=0 controllerNumber=1 scan=11758 | 31.1317 |
| 08497:4 | 2 | K.TIDRLAGKPTHVN[+1663.608]VSVVMAEVDGTC[+57.021]Y.- | HexNAc(5)Hex(4) | C[+57], N[+1664] | 1124.7608 | 4 | 4496.0213 | 4496.01 | 0 | 2.5 | 328 | CRagged | 484.73 | 484.73 | 98.1 | 7.12 | 65 | >sp\|P01876\|IGHA1_HUMAN Immunoglobulin heavy constant alpha 1 OS=Homo sapiens OX=9606 GN=IGHA1 PE=1 SV=2 | 255 | 20201118_ly_colostrum_1ng_glyco_hcd30_etd30_compound15.11768.11768.4 | controllerType=0 controllerNumber=1 scan=11768 | 31.1665 |
| 09455:4 | 2 | K.TIDRLAGKPTHVN[+2262.814]VSVVMAEVDGTC[+57.021]Y.- | HexNAc(5)Hex(5)Fuc(1)NeuAc(1) | C[+57], N[+2263] | 1274.5627 | 4 | 5095.2292 | 5095.2162 | 0 | 2.5 | 328 | CRagged | 538.85 | 528.94 | 528.94 | 7.19 | 65 | >sp\|P01876\|IGHA1_HUMAN Immunoglobulin heavy constant alpha 1 OS=Homo sapiens OX=9606 GN=IGHA1 PE=1 SV=2 | 255 | 20201118_ly_colostrum_1ng_glyco_hcd30_etd30_compound15.12771.12771.4 | controllerType=0 controllerNumber=1 scan=12771 | 33.6736 |
| 08498:4 | 2 | K.TIDRLAGKPTHVN[+1216.423]VSVVMAEVDGTC[+57.021]Y.- | HexNAc(2)Hex(5) | C[+57], N[+1216] | 1012.9648 | 4 | 4048.8375 | 4048.8247 | 0 | 3.1 | 328 | CRagged | 641.14 | 641.14 | 147.62 | 10.31 | 65 | >sp\|P01876\|IGHA1_HUMAN Immunoglobulin heavy constant alpha 1 OS=Homo sapiens OX=9606 GN=IGHA1 PE=1 SV=2 | 255 | 20201118_ly_colostrum_1ng_glyco_hcd30_etd30_compound15.11769.11769.4 | controllerType=0 controllerNumber=1 scan=11769 | 31.1742 |
| 08478:4 | 2 | K.TIDRLAGKPTHVN[+1378.476]VSVVMAEVDGTC[+57.021]Y.- | HexNAc(2)Hex(6) | C[+57], N[+1378] | 1053.4775 | 4 | 4210.888 | 4210.8776 | 0 | 2.5 | 328 | CRagged | 668.11 | 653.94 | 172.96 | 10.28 | 65 | >sp\|P01876\|IGHA1_HUMAN Immunoglobulin heavy constant alpha 1 OS=Homo sapiens OX=9606 GN=IGHA1 PE=1 SV=2 | 255 | 20201118_ly_colostrum_1ng_glyco_hcd30_etd30_compound15.11748.11748.4 | controllerType=0 controllerNumber=1 scan=11748 | 31.0988 |
| 08346:2 | 2 | R.LAGKPTHVN[+1216.423]VSVVMAEVDGTC[+57.021]Y.- | HexNAc(2)Hex(5) | C[+57], N[+1216] | 1782.2883 | 2 | 3563.5693 | 3563.5649 | 0 | 1.2 | 332 | CRagged | 201.41 | 201.41 | 121.31 | 4.95 | 65 | >sp\|P01876\|IGHA1_HUMAN Immunoglobulin heavy constant alpha 1 OS=Homo sapiens OX=9606 GN=IGHA1 PE=1 SV=2 | 255 | 20201118_ly_colostrum_1ng_glyco_hcd30_etd30_compound15.11610.11610.2 | controllerType=0 controllerNumber=1 scan=11610 | 30.7278 |
| 07609:3 | 2 | R.LAGKPTHVN[+1647.613]VSVVMAEVDGTC[+57.021]Y.- | HexNAc(5)Hex(3)Fuc(1) | C[+57], N[+1648] | 1332.2626 | 3 | 3994.7733 | 3994.7553 | 0 | 4.5 | 332 | CRagged | 206.38 | 132.14 | 132.14 | 3.05 | 65 | >sp\|P01876\|IGHA1_HUMAN Immunoglobulin heavy constant alpha 1 OS=Homo sapiens OX=9606 GN=IGHA1 PE=1 SV=2 | 255 | 20201118_ly_colostrum_1ng_glyco_hcd30_etd30_compound15.10836.10836.3 | controllerType=0 controllerNumber=1 scan=10836 | 28.6438 |
| 08406:3 | 2 | R.LAGKPTHVN[+1095.397]VSVVMAEVDGTC[+57.021]Y.- | HexNAc(3)Hex(3) | C[+57], N[+1095] | 1148.1894 | 3 | 3442.5535 | 3442.5387 | 0 | 4.3 | 332 | CRagged | 206.43 | 119.87 | 119.87 | 2.99 | 65 | >sp\|P01876\|IGHA1_HUMAN Immunoglobulin heavy constant alpha 1 OS=Homo sapiens OX=9606 GN=IGHA1 PE=1 SV=2 | 255 | 20201118_ly_colostrum_1ng_glyco_hcd30_etd30_compound15.11673.11673.3 | controllerType=0 controllerNumber=1 scan=11673 | 30.9071 |
| 07545:3 | 2 | R.LAGKPTHVN[+1378.476]VSVVMAEVDGTC[+57.021]Y.- | HexNAc(2)Hex(6) | C[+57], N[+1378] | 1242.5479 | 3 | 3725.6292 | 3725.6178 | 0 | 3.1 | 332 | CRagged | 207.84 | 164.93 | 158.8 | 3.13 | 65 | >sp\|P01876\|IGHA1_HUMAN Immunoglobulin heavy constant alpha 1 OS=Homo sapiens OX=9606 GN=IGHA1 PE=1 SV=2 | 255 | 20201118_ly_colostrum_1ng_glyco_hcd30_etd30_compound15.10768.10768.3 | controllerType=0 controllerNumber=1 scan=10768 | 28.4214 |
| 08282:3 | 2 | R.LAGKPTHVN[+1540.529]VSVVMAEVDGTC[+57.021]Y.- | HexNAc(2)Hex(7) | C[+57], N[+1541] | 1296.5676 | 3 | 3887.6881 | 3887.6706 | 0 | 4.5 | 332 | CRagged | 215.65 | 168.17 | 168.17 | 3.13 | 65 | >sp\|P01876\|IGHA1_HUMAN Immunoglobulin heavy constant alpha 1 OS=Homo sapiens OX=9606 GN=IGHA1 PE=1 SV=2 | 255 | 20201118_ly_colostrum_1ng_glyco_hcd30_etd30_compound15.11543.11543.3 | controllerType=0 controllerNumber=1 scan=11543 | 30.5502 |
| 09492:3 | 2 | R.LAGKPTHVN[+1913.677]VSVVMAEVDGTC[+57.021]Y.- | HexNAc(4)Hex(5)NeuAc(1) | C[+57], N[+1914] | 1420.9474 | 3 | 4260.8278 | 4260.8191 | 0 | 2 | 332 | CRagged | 229.42 | 106.25 | 106.25 | 3.02 | 65 | >sp\|P01876\|IGHA1_HUMAN Immunoglobulin heavy constant alpha 1 OS=Homo sapiens OX=9606 GN=IGHA1 PE=1 SV=2 | 255 | 20201118_ly_colostrum_1ng_glyco_hcd30_etd30_compound15.12810.12810.3 | controllerType=0 controllerNumber=1 scan=12810 | 33.7755 |
| 08243:3 | 2 | R.LAGKPTHVN[+1702.581]VSVVMAEVDGTC[+57.021]Y.- | HexNAc(2)Hex(8) | C[+57], N[+1703] | 1350.5843 | 3 | 4049.7383 | 4049.7234 | 0 | 3.7 | 332 | CRagged | 235.01 | 174.55 | 174.13 | 3.42 | 65 | >sp\|P01876\|IGHA1_HUMAN Immunoglobulin heavy constant alpha 1 OS=Homo sapiens OX=9606 GN=IGHA1 PE=1 SV=2 | 255 | 20201118_ly_colostrum_1ng_glyco_hcd30_etd30_compound15.11502.11502.3 | controllerType=0 controllerNumber=1 scan=11502 | 30.4114 |
| 08277:3 | 2 | R.LAGKPTHVN[+1378.476]VSVVMAEVDGTC[+57.021]Y.- | HexNAc(2)Hex(6) | C[+57], N[+1378] | 1242.5467 | 3 | 3725.6255 | 3725.6178 | 0 | 2.1 | 332 | CRagged | 242.81 | 190.58 | 179.41 | 4.26 | 65 | >sp\|P01876\|IGHA1_HUMAN Immunoglobulin heavy constant alpha 1 OS=Homo sapiens OX=9606 GN=IGHA1 PE=1 SV=2 | 255 | 20201118_ly_colostrum_1ng_glyco_hcd30_etd30_compound15.11538.11538.3 | controllerType=0 controllerNumber=1 scan=11538 | 30.5419 |
| 08341:3 | 2 | R.LAGKPTHVN[+1501.555]VSVVMAEVDGTC[+57.021]Y.- | HexNAc(5)Hex(3) | C[+57], N[+1502] | 1283.5742 | 3 | 3848.7079 | 3848.6974 | 0 | 2.7 | 332 | CRagged | 244.25 | 161.13 | 161.13 | 3.87 | 65 | >sp\|P01876\|IGHA1_HUMAN Immunoglobulin heavy constant alpha 1 OS=Homo sapiens OX=9606 GN=IGHA1 PE=1 SV=2 | 255 | 20201118_ly_colostrum_1ng_glyco_hcd30_etd30_compound15.11605.11605.3 | controllerType=0 controllerNumber=1 scan=11605 | 30.7167 |
| 08400:3 | 2 | R.LAGKPTHVN[+1444.534]VSVVMAEVDGTC[+57.021]Y.- | HexNAc(4)Hex(3)Fuc(1) | C[+57], N[+1445] | 1264.5663 | 3 | 3791.6845 | 3791.6759 | 0 | 2.2 | 332 | CRagged | 245.86 | 162.75 | 162.75 | 3.87 | 65 | >sp\|P01876\|IGHA1_HUMAN Immunoglobulin heavy constant alpha 1 OS=Homo sapiens OX=9606 GN=IGHA1 PE=1 SV=2 | 255 | 20201118_ly_colostrum_1ng_glyco_hcd30_etd30_compound15.11666.11666.3 | controllerType=0 controllerNumber=1 scan=11666 | 30.8812 |
| 08360:3 | 2 | R.LAGKPTHVN[+1647.613]VSVVMAEVDGTC[+57.021]Y.- | HexNAc(5)Hex(3)Fuc(1) | C[+57], N[+1648] | 1332.2596 | 3 | 3994.7642 | 3994.7553 | 0 | 2.2 | 332 | CRagged | 252.71 | 158.42 | 158.42 | 4.01 | 65 | >sp\|P01876\|IGHA1_HUMAN Immunoglobulin heavy constant alpha 1 OS=Homo sapiens OX=9606 GN=IGHA1 PE=1 SV=2 | 255 | 20201118_ly_colostrum_1ng_glyco_hcd30_etd30_compound15.11625.11625.3 | controllerType=0 controllerNumber=1 scan=11625 | 30.7746 |
| 09472:3 | 2 | R.LAGKPTHVN[+2262.814]VSVVMAEVDGTC[+57.021]Y.- | HexNAc(5)Hex(5)Fuc(1)NeuAc(1) | C[+57], N[+2263] | 1537.3283 | 3 | 4609.9704 | 4609.9564 | 0 | 3 | 332 | CRagged | 253.54 | 139.97 | 139.97 | 3.67 | 65 | >sp\|P01876\|IGHA1_HUMAN Immunoglobulin heavy constant alpha 1 OS=Homo sapiens OX=9606 GN=IGHA1 PE=1 SV=2 | 255 | 20201118_ly_colostrum_1ng_glyco_hcd30_etd30_compound15.12789.12789.3 | controllerType=0 controllerNumber=1 scan=12789 | 33.724 |
| 10740:3 | 2 | R.LAGKPTHVN[+2350.830]VSVVMAEVDGTC[+57.021]Y.- | HexNAc(4)Hex(5)Fuc(1)NeuAc(2) | C[+57], N[+2351] | 1566.6681 | 3 | 4697.9899 | 4697.9724 | 0 | 3.7 | 332 | CRagged | 265.82 | 71.4 | 71.4 | 3.32 | 65 | >sp\|P01876\|IGHA1_HUMAN Immunoglobulin heavy constant alpha 1 OS=Homo sapiens OX=9606 GN=IGHA1 PE=1 SV=2 | 255 | 20201118_ly_colostrum_1ng_glyco_hcd30_etd30_compound15.14123.14123.3 | controllerType=0 controllerNumber=1 scan=14123 | 37.3385 |
| 08395:3 | 2 | R.LAGKPTHVN[+1095.397]VSVVMAEVDGTC[+57.021]Y.- | HexNAc(3)Hex(3) | C[+57], N[+1095] | 1148.1864 | 3 | 3442.5447 | 3442.5387 | 0 | 1.8 | 332 | CRagged | 276.38 | 192.86 | 192.86 | 4.42 | 65 | >sp\|P01876\|IGHA1_HUMAN Immunoglobulin heavy constant alpha 1 OS=Homo sapiens OX=9606 GN=IGHA1 PE=1 SV=2 | 255 | 20201118_ly_colostrum_1ng_glyco_hcd30_etd30_compound15.11661.11661.3 | controllerType=0 controllerNumber=1 scan=11661 | 30.8645 |
| 08386:3 | 2 | R.LAGKPTHVN[+1054.370]VSVVMAEVDGTC[+57.021]Y.- | HexNAc(2)Hex(4) | C[+57], N[+1054] | 1134.5109 | 3 | 3401.5181 | 3401.5121 | 0 | 1.8 | 332 | CRagged | 280.78 | 255.07 | 194.73 | 4.82 | 65 | >sp\|P01876\|IGHA1_HUMAN Immunoglobulin heavy constant alpha 1 OS=Homo sapiens OX=9606 GN=IGHA1 PE=1 SV=2 | 255 | 20201118_ly_colostrum_1ng_glyco_hcd30_etd30_compound15.11652.11652.3 | controllerType=0 controllerNumber=1 scan=11652 | 30.8473 |
| 08319:3 | 2 | R.LAGKPTHVN[+1216.423]VSVVMAEVDGTC[+57.021]Y.- | HexNAc(2)Hex(5) | C[+57], N[+1216] | 1188.5298 | 3 | 3563.575 | 3563.5649 | 0 | 2.8 | 332 | CRagged | 284.26 | 253.68 | 215.74 | 4.65 | 65 | >sp\|P01876\|IGHA1_HUMAN Immunoglobulin heavy constant alpha 1 OS=Homo sapiens OX=9606 GN=IGHA1 PE=1 SV=2 | 255 | 20201118_ly_colostrum_1ng_glyco_hcd30_etd30_compound15.11582.11582.3 | controllerType=0 controllerNumber=1 scan=11582 | 30.6589 |
| 07571:3 | 2 | R.LAGKPTHVN[+1216.423]VSVVMAEVDGTC[+57.021]Y.- | HexNAc(2)Hex(5) | C[+57], N[+1216] | 1188.5303 | 3 | 3563.5764 | 3563.5649 | 0 | 3.2 | 332 | CRagged | 288.73 | 259.81 | 203.64 | 4.44 | 65 | >sp\|P01876\|IGHA1_HUMAN Immunoglobulin heavy constant alpha 1 OS=Homo sapiens OX=9606 GN=IGHA1 PE=1 SV=2 | 255 | 20201118_ly_colostrum_1ng_glyco_hcd30_etd30_compound15.10795.10795.3 | controllerType=0 controllerNumber=1 scan=10795 | 28.507 |

Table S4. Cont.

| Query #:z | Protein Rank | Glycopeptide | Glycan | Modification Type(s) | Observed m/z | z | Observed (M+H) | Calc. mass (M+H) | Off-by-x error | Mass error (ppm) | StartPosition | Cleavage | Score | Delta | Delta Mod | \|Log Prob\| | # of unique peptides | Protein Name | Protein DB number | Comment | Scan # | Scan Time |
|---|---|---|---|---|---|---|---|---|---|---|---|---|---|---|---|---|---|---|---|---|---|---|
| 10816:3 | 2 | R.LAGKPTHVN[+2204.772]VSVVMAEVDGTC[+57.021]Y.- | HexNAc(4)Hex(5)NeuAc(2) | C[+57], N[+2205] | 1517.9803 | 3 | 4551.9263 | 4551.9145 | 0 | 2.6 | 332 | CRagged | 291.99 | 146.24 | 146.24 | 4.09 | 65 | >sp\|P01876\|IGHA1_HUMAN Immunoglobulin heavy constant alpha 1 OS=Homo sapiens OX=9606 GN=IGHA1 PE=1 SV=2 | 255 | 20201118_ly_colostrum_1ng_glyco_hcd30_etd30_compound15.14203.14203.3 | controllerType=0 controllerNumber=1 scan=14203 | 37.5528 |
| 09476:4 | 2 | R.LAGKPTHVN[+2262.814]VSVVMAEVDGTC[+57.021]Y.- | HexNAc(5)Hex(5)Fuc(1)NeuAc(1) | C[+57], N[+2263] | 1153.2481 | 4 | 4609.9705 | 4609.9564 | 0 | 3.1 | 332 | CRagged | 516.79 | 349.51 | 349.51 | 5.14 | 65 | >sp\|P01876\|IGHA1_HUMAN Immunoglobulin heavy constant alpha 1 OS=Homo sapiens OX=9606 GN=IGHA1 PE=1 SV=2 | 255 | 20201118_ly_colostrum_1ng_glyco_hcd30_etd30_compound15.12793.12793.4 | controllerType=0 controllerNumber=1 scan=12793 | 33.7319 |
| 06682:3 | 2 | R.LAGKPTHVN[+1216.423]VSVVM[+15.995]AEVDGTC[+57.021]Y.- | HexNAc(2)Hex(5) | C[+57], M[+16], N[+1216] | 1193.8611 | 3 | 3579.5689 | 3579.5599 | 0 | 2.5 | 332 | CRagged | 224.75 | 181.77 | 181.77 | 3.46 | 65 | >sp\|P01876\|IGHA1_HUMAN Immunoglobulin heavy constant alpha 1 OS=Homo sapiens OX=9606 GN=IGHA1 PE=1 SV=2 | 255 | 20201118_ly_colostrum_1ng_glyco_hcd30_etd30_compound15.9861.9861.3 | controllerType=0 controllerNumber=1 scan=9861 | 26.0155 |
| 12783:5 | 3 | R.RTAGWNVPIGTLRPFLN[+2351.851]WTGPPEPIEAAVAR.F | HexNAc(4)Hex(5)Fuc(3)NeuAc(1) | N[+2352] | 1148.5414 | 5 | 5738.6778 | 5738.6628 | 0 | 2.6 | 140 | Specific | 582.85 | 563.97 | 563.97 | 8.46 | 49 | >sp\|P02788\|TRFL_HUMAN Lactotransferrin OS=Homo sapiens OX=9606 GN=LTF PE=1 SV=6 | 1270 | 20201118_ly_colostrum_1ng_glyco_hcd30_etd30_compound15.16276.16276.5 | controllerType=0 controllerNumber=1 scan=16276 | 43.5279 |
| 12767:4 | 3 | R.RTAGWNVPIGTLRPFLN[+2351.851]WTGPPEPIEAAVAR.F | HexNAc(4)Hex(5)Fuc(3)NeuAc(1) | N[+2352] | 1435.4266 | 4 | 5738.6847 | 5738.6628 | 0 | 3.8 | 140 | Specific | 787.77 | 589.38 | 589.38 | 13.31 | 49 | >sp\|P02788\|TRFL_HUMAN Lactotransferrin OS=Homo sapiens OX=9606 GN=LTF PE=1 SV=6 | 1270 | 20201118_ly_colostrum_1ng_glyco_hcd30_etd30_compound15.16260.16260.4 | controllerType=0 controllerNumber=1 scan=16260 | 43.4941 |
| 13669:3 | 3 | R.TAGWNVPIGTLRPFLN[+2205.793]WTGPPEPIEAAVAR.F | HexNAc(4)Hex(5)Fuc(2)NeuAc(1) | N[+2206] | 1812.8455 | 3 | 5436.5221 | 5436.5038 | 0 | 3.4 | 141 | Specific | 208.03 | 133.53 | 133.53 | 3.49 | 49 | >sp\|P02788\|TRFL_HUMAN Lactotransferrin OS=Homo sapiens OX=9606 GN=LTF PE=1 SV=6 | 1270 | 20201118_ly_colostrum_1ng_glyco_hcd30_etd30_compound15.17212.17212.3 | controllerType=0 controllerNumber=1 scan=17212 | 46.3619 |
| 14448:3 | 3 | R.TAGWNVPIGTLRPFLN[+2350.830]WTGPPEPIEAAVAR.F | HexNAc(4)Hex(5)Fuc(1)NeuAc(2) | N[+2351] | 1861.1925 | 3 | 5581.5629 | 5581.5413 | 0 | 3.9 | 141 | Specific | 368.58 | 308.93 | 308.93 | 5.05 | 49 | >sp\|P02788\|TRFL_HUMAN Lactotransferrin OS=Homo sapiens OX=9606 GN=LTF PE=1 SV=6 | 1270 | 20201118_ly_colostrum_1ng_glyco_hcd30_etd30_compound15.18042.18042.3 | controllerType=0 controllerNumber=1 scan=18042 | 49.1632 |
| 14486:3 | 3 | R.TAGWNVPIGTLRPFLN[+2496.888]WTGPPEPIEAAVAR.F | HexNAc(4)Hex(5)Fuc(2)NeuAc(2) | N[+2497] | 1909.8756 | 3 | 5727.6122 | 5727.5992 | 0 | 2.3 | 141 | Specific | 417.27 | 294.57 | 294.57 | 6.75 | 49 | >sp\|P02788\|TRFL_HUMAN Lactotransferrin OS=Homo sapiens OX=9606 GN=LTF PE=1 SV=6 | 1270 | 20201118_ly_colostrum_1ng_glyco_hcd30_etd30_compound15.18082.18082.3 | controllerType=0 controllerNumber=1 scan=18082 | 49.3008 |
| 13611:5 | 3 | R.TAGWNVPIGTLRPFLN[+2350.830]WTGPPEPIEAAVAR.F | HexNAc(4)Hex(5)Fuc(1)NeuAc(2) | N[+2351] | 1117.1217 | 5 | 5581.5794 | 5581.5413 | 0 | 6.8 | 141 | Specific | 452.76 | 444.7 | 444.7 | 5.59 | 49 | >sp\|P02788\|TRFL_HUMAN Lactotransferrin OS=Homo sapiens OX=9606 GN=LTF PE=1 SV=6 | 1270 | 20201118_ly_colostrum_1ng_glyco_hcd30_etd30_compound15.17150.17150.3 | controllerType=0 controllerNumber=1 scan=17150 | 46.1447 |
| 13552:4 | 3 | R.TAGWNVPIGTLRPFLN[+2716.983]WTGPPEPIEAAVAR.F | HexNAc(5)Hex(6)Fuc(3)NeuAc(1) | N[+2717] | 1487.6829 | 4 | 5947.7096 | 5947.6939 | 0 | 2.6 | 141 | Specific | 507.87 | 500.31 | 500.31 | 7.3 | 49 | >sp\|P02788\|TRFL_HUMAN Lactotransferrin OS=Homo sapiens OX=9606 GN=LTF PE=1 SV=6 | 1270 | 20201118_ly_colostrum_1ng_glyco_hcd30_etd30_compound15.17088.17088.4 | controllerType=0 controllerNumber=1 scan=17088 | 45.959 |
| 13716:4 | 3 | R.TAGWNVPIGTLRPFLN[+2059.735]WTGPPEPIEAAVAR.F | HexNAc(4)Hex(5)Fuc(1)NeuAc(1) | N[+2060] | 1323.3704 | 4 | 5290.4596 | 5290.4459 | 0 | 2.6 | 141 | Specific | 673.32 | 673.32 | 673.32 | 11.71 | 49 | >sp\|P02788\|TRFL_HUMAN Lactotransferrin OS=Homo sapiens OX=9606 GN=LTF PE=1 SV=6 | 1270 | 20201118_ly_colostrum_1ng_glyco_hcd30_etd30_compound15.17262.17262.4 | controllerType=0 controllerNumber=1 scan=17262 | 46.5271 |
| 13668:4 | 3 | R.TAGWNVPIGTLRPFLN[+2205.793]WTGPPEPIEAAVAR.F | HexNAc(4)Hex(5)Fuc(2)NeuAc(1) | N[+2206] | 1359.8856 | 4 | 5436.5207 | 5436.5038 | 0 | 3.1 | 141 | Specific | 772.19 | 752.55 | 752.55 | 13.48 | 49 | >sp\|P02788\|TRFL_HUMAN Lactotransferrin OS=Homo sapiens OX=9606 GN=LTF PE=1 SV=6 | 1270 | 20201118_ly_colostrum_1ng_glyco_hcd30_etd30_compound15.17211.17211.4 | controllerType=0 controllerNumber=1 scan=17211 | 46.3587 |
| 14460:5 | 3 | R.TAGWNVPIGTLRPFLN[+2350.830]WTGPPEPIEAAVAR.F | HexNAc(4)Hex(5)Fuc(1)NeuAc(2) | N[+2351] | 1117.1184 | 5 | 5581.5629 | 5581.5413 | 0 | 3.9 | 141 | Specific | 786.1 | 754.17 | 754.17 | 12.24 | 49 | >sp\|P02788\|TRFL_HUMAN Lactotransferrin OS=Homo sapiens OX=9606 GN=LTF PE=1 SV=6 | 1270 | 20201118_ly_colostrum_1ng_glyco_hcd30_etd30_compound15.18054.18054.5 | controllerType=0 controllerNumber=1 scan=18054 | 49.2027 |
| 14469:4 | 3 | R.TAGWNVPIGTLRPFLN[+2496.888]WTGPPEPIEAAVAR.F | HexNAc(4)Hex(5)Fuc(2)NeuAc(2) | N[+2497] | 1432.6591 | 4 | 5727.6144 | 5727.5992 | 0 | 2.7 | 141 | Specific | 809.37 | 773.69 | 773.69 | 14.71 | 49 | >sp\|P02788\|TRFL_HUMAN Lactotransferrin OS=Homo sapiens OX=9606 GN=LTF PE=1 SV=6 | 1270 | 20201118_ly_colostrum_1ng_glyco_hcd30_etd30_compound15.18064.18064.4 | controllerType=0 controllerNumber=1 scan=18064 | 49.238 |
| 14447:4 | 3 | R.TAGWNVPIGTLRPFLN[+2350.830]WTGPPEPIEAAVAR.F | HexNAc(4)Hex(5)Fuc(1)NeuAc(2) | N[+2351] | 1396.1467 | 4 | 5581.5651 | 5581.5413 | 0 | 4.3 | 141 | Specific | 906.88 | 874.88 | 874.88 | 16.48 | 49 | >sp\|P02788\|TRFL_HUMAN Lactotransferrin OS=Homo sapiens OX=9606 GN=LTF PE=1 SV=6 | 1270 | 20201118_ly_colostrum_1ng_glyco_hcd30_etd30_compound15.18041.18041.4 | controllerType=0 controllerNumber=1 scan=18041 | 49.1603 |
| 13163:3 | 3 | R.TAGWNIPMGLLFN[+2351.851]QTGSC[+57.021]K.F | HexNAc(4)Hex(5)Fuc(3)NeuAc(1) | C[+57], N[+2352] | 1482.9628 | 3 | 4446.8739 | 4446.8607 | 0 | 3 | 485 | Specific | 390.35 | 267.72 | 267.72 | 5.22 | 49 | >sp\|P02788\|TRFL_HUMAN Lactotransferrin OS=Homo sapiens OX=9606 GN=LTF PE=1 SV=6 | 1270 | 20201118_ly_colostrum_1ng_glyco_hcd30_etd30_compound15.16676.16676.3 | controllerType=0 controllerNumber=1 scan=16676 | 44.7391 |
| 14523:3 | 3 | R.TAGWNIPMGLLFN[+2350.830]QTGSC[+57.021]K.F | HexNAc(4)Hex(5)Fuc(1)NeuAc(2) | C[+57], N[+2351] | 1482.6248 | 3 | 4445.8599 | 4445.8403 | 0 | 4.4 | 485 | Specific | 428.93 | 279.47 | 279.47 | 5.44 | 49 | >sp\|P02788\|TRFL_HUMAN Lactotransferrin OS=Homo sapiens OX=9606 GN=LTF PE=1 SV=6 | 1270 | 20201118_ly_colostrum_1ng_glyco_hcd30_etd30_compound15.18122.18122.3 | controllerType=0 controllerNumber=1 scan=18122 | 49.4397 |
| 13371:4 | 3 | R.TAGWNIPMGLLFN[+2205.793]QTGSC[+57.021]KFDEYFSQSC[+57.021]APGSDPR.S | HexNAc(4)Hex(5)Fuc(2)NeuAc(1) | C[+57]*2, N[+2206] | 1536.903 | 4 | 6144.59 | 6144.5499 | 0 | 6.5 | 485 | Specific | 210.98 | 210.98 | 210.98 | 3.01 | 49 | >sp\|P02788\|TRFL_HUMAN Lactotransferrin OS=Homo sapiens OX=9606 GN=LTF PE=1 SV=6 | 1270 | 20201118_ly_colostrum_1ng_glyco_hcd30_etd30_compound15.16897.16897.3 | controllerType=0 controllerNumber=1 scan=16897 | 45.4581 |
| 12683:4 | 3 | R.TAGWNIPMGLLFN[+1768.640]QTGSC[+57.021]KFDEYFSQSC[+57.021]APGSDPR.S | HexNAc(4)Hex(5)Fuc(1) | C[+57]*2, N[+1769] | 1427.6106 | 4 | 5707.4206 | 5707.3966 | 0 | 4.2 | 485 | Specific | 239.26 | 234.02 | 234.02 | 3.56 | 49 | >sp\|P02788\|TRFL_HUMAN Lactotransferrin OS=Homo sapiens OX=9606 GN=LTF PE=1 SV=6 | 1270 | 20201118_ly_colostrum_1ng_glyco_hcd30_etd30_compound15.16172.16172.4 | controllerType=0 controllerNumber=1 scan=16172 | 43.2622 |
| 13384:4 | 3 | R.TAGWNIPMGLLFN[+2059.735]QTGSC[+57.021]KFDEYFSQSC[+57.021]APGSDPR.S | HexNAc(4)Hex(5)Fuc(1)NeuAc(1) | C[+57]*2, N[+2060] | 1500.3839 | 4 | 5998.5138 | 5998.492 | 0 | 3.6 | 485 | Specific | 290.35 | 275.33 | 275.33 | 4.89 | 49 | >sp\|P02788\|TRFL_HUMAN Lactotransferrin OS=Homo sapiens OX=9606 GN=LTF PE=1 SV=6 | 1270 | 20201118_ly_colostrum_1ng_glyco_hcd30_etd30_compound15.16911.16911.4 | controllerType=0 controllerNumber=1 scan=16911 | 45.4994 |
| 12670:3 | 3 | R.TAGWNIPMGLLFN[+1768.640]QTGSC[+57.021]KFDEYFSQSC[+57.021]APGSDPR.S | HexNAc(4)Hex(5)Fuc(1) | C[+57]*2, N[+1769] | 1903.1419 | 3 | 5707.4112 | 5707.3966 | 0 | 2.6 | 485 | Specific | 363.83 | 304.92 | 304.92 | 6.26 | 49 | >sp\|P02788\|TRFL_HUMAN Lactotransferrin OS=Homo sapiens OX=9606 GN=LTF PE=1 SV=6 | 1270 | 20201118_ly_colostrum_1ng_glyco_hcd30_etd30_compound15.16158.16158.3 | controllerType=0 controllerNumber=1 scan=16158 | 43.2186 |
| 14248:4 | 3 | R.TAGWNIPMGLLFN[+2350.830]QTGSC[+57.021]KFDEYFSQSC[+57.021]APGSDPR.S | HexNAc(4)Hex(5)Fuc(1)NeuAc(2) | C[+57]*2, N[+2351] | 1573.1578 | 4 | 6289.6095 | 6289.5875 | 0 | 3.5 | 485 | Specific | 375.15 | 343.34 | 343.34 | 6.31 | 49 | >sp\|P02788\|TRFL_HUMAN Lactotransferrin OS=Homo sapiens OX=9606 GN=LTF PE=1 SV=6 | 1270 | 20201118_ly_colostrum_1ng_glyco_hcd30_etd30_compound15.17829.17829.4 | controllerType=0 controllerNumber=1 scan=17829 | 48.4314 |
| 13263:4 | 3 | R.TAGWNIPMGLLFN[+2863.041]QTGSC[+57.021]KFDEYFSQSC[+57.021]APGSDPR.S | HexNAc(5)Hex(6)Fuc(4)NeuAc(1) | C[+57]*2, N[+2863] | 1701.2136 | 4 | 6801.8325 | 6801.798 | 0 | 5.1 | 485 | Specific | 382.02 | 348.27 | 348.27 | 5.72 | 49 | >sp\|P02788\|TRFL_HUMAN Lactotransferrin OS=Homo sapiens OX=9606 GN=LTF PE=1 SV=6 | 1270 | 20201118_ly_colostrum_1ng_glyco_hcd30_etd30_compound15.16782.16782.4 | controllerType=0 controllerNumber=1 scan=16782 | 45.0686 |
| 13353:4 | 3 | R.TAGWNIPMGLLFN[+2351.851]QTGSC[+57.021]KFDEYFSQSC[+57.021]APGSDPR.S | HexNAc(4)Hex(5)Fuc(3)NeuAc(1) | C[+57]*2, N[+2352] | 1573.4132 | 4 | 6290.631 | 6290.6079 | 0 | 3.7 | 485 | Specific | 619.24 | 586.75 | 586.75 | 10.22 | 49 | >sp\|P02788\|TRFL_HUMAN Lactotransferrin OS=Homo sapiens OX=9606 GN=LTF PE=1 SV=6 | 1270 | 20201118_ly_colostrum_1ng_glyco_hcd30_etd30_compound15.16878.16878.4 | controllerType=0 controllerNumber=1 scan=16878 | 45.392 |
| 13788:4 | 4 | R.GLSFDVSLEVSQGPGLLN[+2350.830]DTK.V | HexNAc(4)Hex(5)Fuc(1)NeuAc(2) | N[+2351] | 1132.4942 | 4 | 4526.9549 | 4526.9436 | 0 | 2.5 | 118 | Specific | 349.33 | 315.3 | 315.3 | 4.63 | 63 | >sp\|P01833\|PIGR_HUMAN Polymeric immunoglobulin receptor OS=Homo sapiens OX=9606 GN=PIGR PE=1 SV=4 | 902 | 20201118_ly_colostrum_1ng_glyco_hcd30_etd30_compound15.17339.17339.2 | controllerType=0 controllerNumber=1 scan=17339 | 46.7719 |
| 13774:4 | 4 | R.GLSFDVSLEVSQGPGLLN[+2351.851]DTK.V | HexNAc(4)Hex(5)Fuc(3)NeuAc(1) | N[+2352] | 1132.7425 | 4 | 4527.9481 | 4527.964 | 0 | -3.5 | 118 | Specific | 362.77 | 332.19 | 332.19 | 3.84 | 63 | >sp\|P01833\|PIGR_HUMAN Polymeric immunoglobulin receptor OS=Homo sapiens OX=9606 GN=PIGR PE=1 SV=4 | 902 | 20201118_ly_colostrum_1ng_glyco_hcd30_etd30_compound15.17324.17324.4 | controllerType=0 controllerNumber=1 scan=17324 | 46.7203 |
| 13787:3 | 4 | R.GLSFDVSLEVSQGPGLLN[+2350.830]DTK.V | HexNAc(4)Hex(5)Fuc(1)NeuAc(2) | N[+2351] | 1509.6572 | 3 | 4526.9569 | 4526.9436 | 0 | 3 | 118 | Specific | 525.19 | 333.32 | 333.32 | 6.65 | 63 | >sp\|P01833\|PIGR_HUMAN Polymeric immunoglobulin receptor OS=Homo sapiens OX=9606 GN=PIGR PE=1 SV=4 | 902 | 20201118_ly_colostrum_1ng_glyco_hcd30_etd30_compound15.17338.17338.3 | controllerType=0 controllerNumber=1 scan=17338 | 46.766 |
| 14004:4 | 4 | R.GLSFDVSLEVSQGPGLLN[+2350.830]DTKVYTVDLGR.T | HexNAc(4)Hex(5)Fuc(1)NeuAc(2) | N[+2351] | 1358.366 | 4 | 5430.4421 | 5430.425 | 0 | 3.1 | 118 | Specific | 411.18 | 387.34 | 387.34 | 6.49 | 63 | >sp\|P01833\|PIGR_HUMAN Polymeric immunoglobulin receptor OS=Homo sapiens OX=9606 GN=PIGR PE=1 SV=4 | 902 | 20201118_ly_colostrum_1ng_glyco_hcd30_etd30_compound15.17568.17568.4 | controllerType=0 controllerNumber=1 scan=17568 | 47.5044 |
| 14776:3 | 4 | K.QIGLYPVLVIDSSGYVNPN[+2204.772]YTGR.I | HexNAc(4)Hex(5)NeuAc(2) | N[+2205] | 1577.3683 | 3 | 4730.0905 | 4730.0759 | 0 | 3.1 | 168 | Specific | 206.86 | 105.61 | 105.61 | 4.1 | 63 | >sp\|P01833\|PIGR_HUMAN Polymeric immunoglobulin receptor OS=Homo sapiens OX=9606 GN=PIGR PE=1 SV=4 | 902 | 20201118_ly_colostrum_1ng_glyco_hcd30_etd30_compound15.18392.18392.3 | controllerType=0 controllerNumber=1 scan=18392 | 50.4625 |
| 14717:3 | 4 | K.QIGLYPVLVIDSSGYVNPN[+2350.830]YTGR.I | HexNAc(4)Hex(5)Fuc(1)NeuAc(2) | N[+2351] | 1626.0535 | 3 | 4876.146 | 4876.1338 | 0 | 2.5 | 168 | Specific | 237.2 | 162.2 | 162.2 | 4.59 | 63 | >sp\|P01833\|PIGR_HUMAN Polymeric immunoglobulin receptor OS=Homo sapiens OX=9606 GN=PIGR PE=1 SV=4 | 902 | 20201118_ly_colostrum_1ng_glyco_hcd30_etd30_compound15.18329.18329.3 | controllerType=0 controllerNumber=1 scan=18329 | 50.2169 |
| 14742:3 | 4 | K.QIGLYPVLVIDSSGYVNPN[+2204.772]YTGR.I | HexNAc(4)Hex(5)NeuAc(2) | N[+2205] | 1577.3671 | 3 | 4730.0868 | 4730.0759 | 0 | 2.3 | 168 | Specific | 501.96 | 390.51 | 390.51 | 7.9 | 63 | >sp\|P01833\|PIGR_HUMAN Polymeric immunoglobulin receptor OS=Homo sapiens OX=9606 GN=PIGR PE=1 SV=4 | 902 | 20201118_ly_colostrum_1ng_glyco_hcd30_etd30_compound15.18356.18356.3 | controllerType=0 controllerNumber=1 scan=18356 | 50.3132 |
| 14593:4 | 4 | K.QIGLYPVLVIDSSGYVNPN[+2350.830]YTGR.I | HexNAc(4)Hex(5)Fuc(1)NeuAc(2) | N[+2351] | 1219.7929 | 4 | 4876.1497 | 4876.1338 | 0 | 3.3 | 168 | Specific | 535.16 | 506.46 | 506.46 | 6.81 | 63 | >sp\|P01833\|PIGR_HUMAN Polymeric immunoglobulin receptor OS=Homo sapiens OX=9606 GN=PIGR PE=1 SV=4 | 902 | 20201118_ly_colostrum_1ng_glyco_hcd30_etd30_compound15.18196.18196.4 | controllerType=0 controllerNumber=1 scan=18196 | 49.7042 |
| 14589:3 | 4 | K.QIGLYPVLVIDSSGYVNPN[+2350.830]YTGR.I | HexNAc(4)Hex(5)Fuc(1)NeuAc(2) | N[+2351] | 1626.054 | 3 | 4876.1475 | 4876.1338 | 0 | 2.8 | 168 | Specific | 560.5 | 470.71 | 470.71 | 9.19 | 63 | >sp\|P01833\|PIGR_HUMAN Polymeric immunoglobulin receptor OS=Homo sapiens OX=9606 GN=PIGR PE=1 SV=4 | 902 | 20201118_ly_colostrum_1ng_glyco_hcd30_etd30_compound15.18192.18192.3 | controllerType=0 controllerNumber=1 scan=18192 | 49.6891 |
| 15505:4 | 4 | R.LSLLEEPGN[+2496.888]GTFTVILNQLTSR.D | HexNAc(4)Hex(5)Fuc(2)NeuAc(2) | N[+2497] | 1225.5523 | 4 | 4899.1873 | 4899.1808 | 0 | 1.3 | 413 | Specific | 280.18 | 264.31 | 264.31 | 4.55 | 63 | >sp\|P01833\|PIGR_HUMAN Polymeric immunoglobulin receptor OS=Homo sapiens OX=9606 GN=PIGR PE=1 SV=4 | 902 | 20201118_ly_colostrum_1ng_glyco_hcd30_etd30_compound15.19171.19171.4 | controllerType=0 controllerNumber=1 scan=19171 | 53.272 |
| 15535:4 | 4 | R.LSLLEEPGN[+2553.910]GTFTVILNQLTSR.D | HexNAc(5)Hex(5)Fuc(1)NeuAc(2) | N[+2554] | 1239.8126 | 4 | 4956.2287 | 4956.2023 | 0 | 5.3 | 413 | Specific | 329.3 | 321.15 | 321.15 | 3.84 | 63 | >sp\|P01833\|PIGR_HUMAN Polymeric immunoglobulin receptor OS=Homo sapiens OX=9606 GN=PIGR PE=1 SV=4 | 902 | 20201118_ly_colostrum_1ng_glyco_hcd30_etd30_compound15.19203.19203.4 | controllerType=0 controllerNumber=1 scan=19203 | 53.4089 |
| 14376:4 | 4 | R.LSLLEEPGN[+2076.750]GTFTVILNQLTSR.D | HexNAc(4)Hex(6)Fuc(2) | N[+2077] | 1120.5238 | 4 | 4479.0736 | 4479.0428 | 0 | 6.9 | 413 | Specific | 330.08 | 330.08 | 330.08 | 4.14 | 63 | >sp\|P01833\|PIGR_HUMAN Polymeric immunoglobulin receptor OS=Homo sapiens OX=9606 GN=PIGR PE=1 SV=4 | 902 | 20201118_ly_colostrum_1ng_glyco_hcd30_etd30_compound15.17965.17965.4 | controllerType=0 controllerNumber=1 scan=17965 | 48.8835 |
| 14657:3 | 4 | R.LSLLEEPGN[+1200.428]GTFTVILNQLTSR.D | HexNAc(2)Hex(4)Fuc(1) | N[+1200] | 1201.5844 | 3 | 3602.7387 | 3602.7205 | 0 | 5 | 413 | Specific | 475.62 | 475.62 | 475.62 | 6.91 | 63 | >sp\|P01833\|PIGR_HUMAN Polymeric immunoglobulin receptor OS=Homo sapiens OX=9606 GN=PIGR PE=1 SV=4 | 902 | 20201118_ly_colostrum_1ng_glyco_hcd30_etd30_compound15.18265.18265.2 | controllerType=0 controllerNumber=1 scan=18265 | 49.964 |
| 15493:4 | 4 | R.LSLLEEPGN[+2350.830]GTFTVILNQLTSR.D | HexNAc(4)Hex(5)Fuc(1)NeuAc(2) | N[+2351] | 1189.0436 | 4 | 4753.1527 | 4753.1229 | 0 | 6.3 | 413 | Specific | 529.24 | 436.36 | 436.36 | 6.05 | 63 | >sp\|P01833\|PIGR_HUMAN Polymeric immunoglobulin receptor OS=Homo sapiens OX=9606 GN=PIGR PE=1 SV=4 | 902 | 20201118_ly_colostrum_1ng_glyco_hcd30_etd30_compound15.19158.19158.4 | controllerType=0 controllerNumber=1 scan=19158 | 53.221 |
| 14348:4 | 4 | R.LSLLEEPGN[+2351.851]GTFTVILNQLTSR.D | HexNAc(4)Hex(5)Fuc(3)NeuAc(1) | N[+2352] | 1189.2969 | 4 | 4754.1658 | 4754.1433 | 0 | 4.7 | 413 | Specific | 564.98 | 488.81 | 488.81 | 7.27 | 63 | >sp\|P01833\|PIGR_HUMAN Polymeric immunoglobulin receptor OS=Homo sapiens OX=9606 GN=PIGR PE=1 SV=4 | 902 | 20201118_ly_colostrum_1ng_glyco_hcd30_etd30_compound15.17934.17934.4 | controllerType=0 controllerNumber=1 scan=17934 | 48.7622 |
| 15603:3 | 4 | R.LSLLEEPGN[+2350.830]GTFTVILNQLTSR.D | HexNAc(4)Hex(5)Fuc(1)NeuAc(2) | N[+2351] | 1585.0569 | 3 | 4753.1563 | 4753.1229 | 0 | 7 | 413 | Specific | 585.68 | 448.99 | 448.99 | 8.13 | 63 | >sp\|P01833\|PIGR_HUMAN Polymeric immunoglobulin receptor OS=Homo sapiens OX=9606 GN=PIGR PE=1 SV=4 | 902 | 20201118_ly_colostrum_1ng_glyco_hcd30_etd30_compound15.19276.19276.3 | controllerType=0 controllerNumber=1 scan=19276 | 53.6773 |
| 15567:4 | 4 | R.LSLLEEPGN[+2496.888]GTFTVILNQLTSR.D | HexNAc(4)Hex(5)Fuc(2)NeuAc(2) | N[+2497] | 1225.5523 | 4 | 4899.1873 | 4899.1808 | 0 | 1.3 | 413 | Specific | 623.47 | 529.24 | 529.24 | 9.09 | 63 | >sp\|P01833\|PIGR_HUMAN Polymeric immunoglobulin receptor OS=Homo sapiens OX=9606 GN=PIGR PE=1 SV=4 | 902 | 20201118_ly_colostrum_1ng_glyco_hcd30_etd30_compound15.19237.19237.4 | controllerType=0 controllerNumber=1 scan=19237 | 53.5241 |
| 15561:3 | 4 | R.LSLLEEPGN[+2496.888]GTFTVILNQLTSR.D | HexNAc(4)Hex(5)Fuc(2)NeuAc(2) | N[+2497] | 1633.7358 | 3 | 4899.1928 | 4899.1808 | 0 | 2.4 | 413 | Specific | 632.27 | 429.12 | 429.12 | 10.19 | 63 | >sp\|P01833\|PIGR_HUMAN Polymeric immunoglobulin receptor OS=Homo sapiens OX=9606 GN=PIGR PE=1 SV=4 | 902 | 20201118_ly_colostrum_1ng_glyco_hcd30_etd30_compound15.19231.19231.3 | controllerType=0 controllerNumber=1 scan=19231 | 53.5067 |
| 14342:3 | 4 | R.LSLLEEPGN[+2351.851]GTFTVILNQLTSR.D | HexNAc(4)Hex(5)Fuc(3)NeuAc(1) | N[+2352] | 1585.3928 | 3 | 4754.1637 | 4754.1433 | 0 | 4.3 | 413 | Specific | 638.1 | 525.7 | 525.7 | 9.97 | 63 | >sp\|P01833\|PIGR_HUMAN Polymeric immunoglobulin receptor OS=Homo sapiens OX=9606 GN=PIGR PE=1 SV=4 | 902 | 20201118_ly_colostrum_1ng_glyco_hcd30_etd30_compound15.17928.17928.3 | controllerType=0 controllerNumber=1 scan=17928 | 48.7436 |
| 14349:3 | 4 | R.LSLLEEPGN[+2205.793]GTFTVILNQLTSR.D | HexNAc(4)Hex(5)Fuc(2)NeuAc(1) | N[+2206] | 1536.7071 | 3 | 4608.1067 | 4608.0854 | 0 | 4.6 | 413 | Specific | 689.65 | 561.49 | 561.49 | 10.38 | 63 | >sp\|P01833\|PIGR_HUMAN Polymeric immunoglobulin receptor OS=Homo sapiens OX=9606 GN=PIGR PE=1 SV=4 | 902 | 20201118_ly_colostrum_1ng_glyco_hcd30_etd30_compound15.17935.17935.3 | controllerType=0 controllerNumber=1 scan=17935 | 48.7652 |
| 11310:3 | 4 | K.IIEGEPNLKVPGN[+2351.851]VTAVLGETLK.V | HexNAc(4)Hex(5)Fuc(3)NeuAc(1) | N[+2352] | 1581.7426 | 3 | 4743.2133 | 4743.2001 | 0 | 2.8 | 457 | Specific | 331.07 | 283.14 | 283.14 | 4.55 | 63 | >sp\|P01833\|PIGR_HUMAN Polymeric immunoglobulin receptor OS=Homo sapiens OX=9606 GN=PIGR PE=1 SV=4 | 902 | 20201118_ly_colostrum_1ng_glyco_hcd30_etd30_compound15.14724.14724.3 | controllerType=0 controllerNumber=1 scan=14724 | 39.1353 |
| 10913:4 | 4 | K.IIEGEPNLKVPGN[+2863.041]VTAVLGETLKVPC[+57.021]HFPC[+57.021]K.F | HexNAc(5)Hex(6)Fuc(4)NeuAc(1) | C[+57]*2, N[+2863] | 1570.7217 | 4 | 6279.8649 | 6279.8477 | 0 | 2.7 | 457 | Specific | 230.39 | 230.39 | 230.39 | 3.89 | 63 | >sp\|P01833\|PIGR_HUMAN Polymeric immunoglobulin receptor OS=Homo sapiens OX=9606 GN=PIGR PE=1 SV=4 | 902 | 20201118_ly_colostrum_1ng_glyco_hcd30_etd30_compound15.14305.14305.4 | controllerType=0 controllerNumber=1 scan=14305 | 37.838 |
| 10292:4 | 4 | K.[+42.011]IIEGEPNLKVPGN[+2018.708]VTAVLGETLKVPC[+57.021]HFPC[+57.021]K | HexNAc(3)Hex(6)Fuc(1)NeuAc(1) | .[42], C[+57]*2, N[+2019] | 1370.1505 | 4 | 5477.5802 | 5477.5258 | 0 | 9.9 | 457 | Specific | 231.28 | 201.24 | 201.24 | 2.29 | 63 | >sp\|P01833\|PIGR_HUMAN Polymeric immunoglobulin receptor OS=Homo sapiens OX=9606 GN=PIGR PE=1 SV=4 | 902 | 20201118_ly_colostrum_1ng_glyco_hcd30_etd30_compound15.13650.13650.4 | controllerType=0 controllerNumber=1 scan=13650 | 36.0239 |
| 11039:6 | 4 | K.IIEGEPNLKVPGN[+2205.793]VTAVLGETLKVPC[+57.021]HFPC[+57.021]K.F | HexNAc(4)Hex(5)Fuc(2)NeuAc(1) | C[+57]*2, N[+2206] | 937.9405 | 6 | 5622.6065 | 5622.5997 | 0 | 1.2 | 457 | Specific | 236.44 | 236.44 | 236.44 | 2.99 | 63 | >sp\|P01833\|PIGR_HUMAN Polymeric immunoglobulin receptor OS=Homo sapiens OX=9606 GN=PIGR PE=1 SV=4 | 902 | 20201118_ly_colostrum_1ng_glyco_hcd30_etd30_compound15.14438.14438.6 | controllerType=0 controllerNumber=1 scan=14438 | 38.2794 |
| 11115:5 | 4 | K.IIEGEPNLKVPGN[+2350.830]VTAVLGETLKVPC[+57.021]HFPC[+57.021]K.F | HexNAc(4)Hex(5)Fuc(1)NeuAc(2) | C[+57]*2, N[+2351] | 1154.339 | 5 | 5767.6659 | 5767.6372 | 0 | 5 | 457 | Specific | 264.23 | 264.23 | 264.23 | 3.01 | 63 | >sp\|P01833\|PIGR_HUMAN Polymeric immunoglobulin receptor OS=Homo sapiens OX=9606 GN=PIGR PE=1 SV=4 | 902 | 20201118_ly_colostrum_1ng_glyco_hcd30_etd30_compound15.14519.14519.5 | controllerType=0 controllerNumber=1 scan=14519 | 38.5299 |
| 11135:4 | 4 | K.IIEGEPNLKVPGN[+2059.735]VTAVLGETLKVPC[+57.021]HFPC[+57.021]K.F | HexNAc(4)Hex(5)Fuc(1)NeuAc(1) | C[+57]*2, N[+2060] | 1369.8978 | 4 | 5476.5695 | 5476.5418 | 0 | 5.1 | 457 | Specific | 272.89 | 272.89 | 272.89 | 4.07 | 63 | >sp\|P01833\|PIGR_HUMAN Polymeric immunoglobulin receptor OS=Homo sapiens OX=9606 GN=PIGR PE=1 SV=4 | 902 | 20201118_ly_colostrum_1ng_glyco_hcd30_etd30_compound15.14540.14540.4 | controllerType=0 controllerNumber=1 scan=14540 | 38.5784 |
| 10963:5 | 4 | K.IIEGEPNLKVPGN[+2863.041]VTAVLGETLKVPC[+57.021]HFPC[+57.021]K.F | HexNAc(5)Hex(6)Fuc(4)NeuAc(1) | C[+57]*2, N[+2863] | 1256.7784 | 5 | 6279.8631 | 6279.8477 | 0 | 2.4 | 457 | Specific | 338.51 | 338.51 | 338.51 | 4.94 | 63 | >sp\|P01833\|PIGR_HUMAN Polymeric immunoglobulin receptor OS=Homo sapiens OX=9606 GN=PIGR PE=1 SV=4 | 902 | 20201118_ly_colostrum_1ng_glyco_hcd30_etd30_compound15.14358.14358.5 | controllerType=0 controllerNumber=1 scan=14358 | 38.0022 |
| 10282:5 | 4 | K.IIEGEPNLKVPGN[+2263.835]VTAVLGETLKVPC[+57.021]HFPC[+57.021]K.F | HexNAc(5)Hex(5)Fuc(3) | C[+57]*2, N[+2264] | 1136.9381 | 5 | 5680.6615 | 5680.6416 | 0 | 3.5 | 457 | Specific | 393.82 | 393.82 | 393.82 | 5.83 | 63 | >sp\|P01833\|PIGR_HUMAN Polymeric immunoglobulin receptor OS=Homo sapiens OX=9606 GN=PIGR PE=1 SV=4 | 902 | 20201118_ly_colostrum_1ng_glyco_hcd30_etd30_compound15.13639.13639.5 | controllerType=0 controllerNumber=1 scan=13639 | 35.9782 |
| 11989:4 | 4 | K.IIEGEPNLKVPGN[+2350.830]VTAVLGETLKVPC[+57.021]HFPC[+57.021]K.F | HexNAc(4)Hex(5)Fuc(1)NeuAc(2) | C[+57]*2, N[+2351] | 1442.6696 | 4 | 5767.6564 | 5767.6372 | 0 | 3.3 | 457 | Specific | 429.47 | 429.47 | 429.47 | 6.35 | 63 | >sp\|P01833\|PIGR_HUMAN Polymeric immunoglobulin receptor OS=Homo sapiens OX=9606 GN=PIGR PE=1 SV=4 | 902 | 20201118_ly_colostrum_1ng_glyco_hcd30_etd30_compound15.15438.15438.4 | controllerType=0 controllerNumber=1 scan=15438 | 41.054 |
| 11031:4 | 4 | K.IIEGEPNLKVPGN[+2351.851]VTAVLGETLKVPC[+57.021]HFPC[+57.021]K.F | HexNAc(4)Hex(5)Fuc(3)NeuAc(1) | C[+57]*2, N[+2352] | 1442.9241 | 4 | 5768.6745 | 5768.6576 | 0 | 2.9 | 457 | Specific | 494.21 | 494.21 | 494.21 | 6.65 | 63 | >sp\|P01833\|PIGR_HUMAN Polymeric immunoglobulin receptor OS=Homo sapiens OX=9606 GN=PIGR PE=1 SV=4 | 902 | 20201118_ly_colostrum_1ng_glyco_hcd30_etd30_compound15.14430.14430.4 | controllerType=0 controllerNumber=1 scan=14430 | 38.2437 |
| 11977:4 | 4 | K.IIEGEPNLKVPGN[+2350.830]VTAVLGETLKVPC[+57.021]HFPC[+57.021]K.F | HexNAc(4)Hex(5)Fuc(1)NeuAc(2) | C[+57]*2, N[+2351] | 1442.6681 | 4 | 5767.6505 | 5767.6372 | 0 | 2.3 | 457 | Specific | 506.73 | 506.73 | 506.73 | 6.87 | 63 | >sp\|P01833\|PIGR_HUMAN Polymeric immunoglobulin receptor OS=Homo sapiens OX=9606 GN=PIGR PE=1 SV=4 | 902 | 20201118_ly_colostrum_1ng_glyco_hcd30_etd30_compound15.15425.15425.4 | controllerType=0 controllerNumber=1 scan=15425 | 40.9945 |
| 11027:5 | 4 | K.IIEGEPNLKVPGN[+2351.851]VTAVLGETLKVPC[+57.021]HFPC[+57.021]K.F | HexNAc(4)Hex(5)Fuc(3)NeuAc(1) | C[+57]*2, N[+2352] | 1154.5402 | 5 | 5768.6717 | 5768.6576 | 0 | 2.4 | 457 | Specific | 525.06 | 525.06 | 525.06 | 6.79 | 63 | >sp\|P01833\|PIGR_HUMAN Polymeric immunoglobulin receptor OS=Homo sapiens OX=9606 GN=PIGR PE=1 SV=4 | 902 | 20201118_ly_colostrum_1ng_glyco_hcd30_etd30_compound15.14426.14426.5 | controllerType=0 controllerNumber=1 scan=14426 | 38.2326 |
| 11985:5 | 4 | K.IIEGEPNLKVPGN[+2350.830]VTAVLGETLKVPC[+57.021]HFPC[+57.021]K.F | HexNAc(4)Hex(5)Fuc(1)NeuAc(2) | C[+57]*2, N[+2351] | 1154.3364 | 5 | 5767.653 | 5767.6372 | 0 | 2.7 | 457 | Specific | 582.94 | 582.94 | 582.94 | 7.85 | 63 | >sp\|P01833\|PIGR_HUMAN Polymeric immunoglobulin receptor OS=Homo sapiens OX=9606 GN=PIGR PE=1 SV=4 | 902 | 20201118_ly_colostrum_1ng_glyco_hcd30_etd30_compound15.15434.15434.5 | controllerType=0 controllerNumber=1 scan=15434 | 41.0371 |
| 11010:4 | 4 | K.IIEGEPNLKVPGN[+2205.793]VTAVLGETLKVPC[+57.021]HFPC[+57.021]K.F | HexNAc(4)Hex(5)Fuc(2)NeuAc(1) | C[+57]*2, N[+2206] | 1406.4094 | 4 | 5622.6159 | 5622.5997 | 0 | 2.9 | 457 | Specific | 598.28 | 598.28 | 598.28 | 8.1 | 63 | >sp\|P01833\|PIGR_HUMAN Polymeric immunoglobulin receptor OS=Homo sapiens OX=9606 GN=PIGR PE=1 SV=4 | 902 | 20201118_ly_colostrum_1ng_glyco_hcd30_etd30_compound15.14408.14408.4 | controllerType=0 controllerNumber=1 scan=14408 | 38.171 |
| 11042:4 | 4 | K.IIEGEPNLKVPGN[+2059.735]VTAVLGETLKVPC[+57.021]HFPC[+57.021]K.F | HexNAc(4)Hex(5)Fuc(1)NeuAc(1) | C[+57]*2, N[+2060] | 1369.8936 | 4 | 5476.5524 | 5476.5418 | 0 | 1.9 | 457 | Specific | 658.65 | 603.62 | 603.62 | 10.32 | 63 | >sp\|P01833\|PIGR_HUMAN Polymeric immunoglobulin receptor OS=Homo sapiens OX=9606 GN=PIGR PE=1 SV=4 | 902 | 20201118_ly_colostrum_1ng_glyco_hcd30_etd30_compound15.14442.14442.4 | controllerType=0 controllerNumber=1 scan=14442 | 38.2949 |
| 11015:5 | 4 | K.IIEGEPNLKVPGN[+2059.735]VTAVLGETLKVPC[+57.021]HFPC[+57.021]K.F | HexNAc(4)Hex(5)Fuc(1)NeuAc(1) | C[+57]*2, N[+2060] | 1096.1158 | 5 | 5476.5501 | 5476.5418 | 0 | 1.5 | 457 | Specific | 667.3 | 667.3 | 667.3 | 10.18 | 63 | >sp\|P01833\|PIGR_HUMAN Polymeric immunoglobulin receptor OS=Homo sapiens OX=9606 GN=PIGR PE=1 SV=4 | 902 | 20201118_ly_colostrum_1ng_glyco_hcd30_etd30_compound15.14413.14413.5 | controllerType=0 controllerNumber=1 scan=14413 | 38.1836 |
| 11009:5 | 4 | K.IIEGEPNLKVPGN[+2205.793]VTAVLGETLKVPC[+57.021]HFPC[+57.021]K.F | HexNAc(4)Hex(5)Fuc(2)NeuAc(1) | C[+57]*2, N[+2206] | 1125.3281 | 5 | 5622.6115 | 5622.5997 | 0 | 2.1 | 457 | Specific | 690.29 | 690.29 | 690.29 | 10.04 | 63 | >sp\|P01833\|PIGR_HUMAN Polymeric immunoglobulin receptor OS=Homo sapiens OX=9606 GN=PIGR PE=1 SV=4 | 902 | 20201118_ly_colostrum_1ng_glyco_hcd30_etd30_compound15.14407.14407.5 | controllerType=0 controllerNumber=1 scan=14407 | 38.1678 |
| 09755:4 | 4 | K.VPGN[+2205.793]VTAVLGETLKVPC[+57.021]HFPC[+57.021]K.F | HexNAc(4)Hex(5)Fuc(2)NeuAc(1) | C[+57]*2, N[+2206] | 1158.0218 | 4 | 4629.0653 | 4629.0502 | 0 | 3.2 | 466 | Specific | 201.93 | 201.93 | 201.93 | 2.87 | 63 | >sp\|P01833\|PIGR_HUMAN Polymeric immunoglobulin receptor OS=Homo sapiens OX=9606 GN=PIGR PE=1 SV=4 | 902 | 20201118_ly_colostrum_1ng_glyco_hcd30_etd30_compound15.13086.13086.4 | controllerType=0 controllerNumber=1 scan=13086 | 34.5533 |
| 07485:4 | 4 | K.FSSYEKYWC[+57.021]KWN[+1710.598]NTGC[+57.021]QALPSQDEGPSK.A | HexNAc(3)Hex(5)NeuAc(1) | C[+57]*2, N[+1711] | 1270.2765 | 4 | 5078.0841 | 5078.0759 | 0 | 1.6 | 488 | Specific | 497.51 | 415.82 | 415.82 | 7.02 | 63 | >sp\|P01833\|PIGR_HUMAN Polymeric immunoglobulin receptor OS=Homo sapiens OX=9606 GN=PIGR PE=1 SV=4 | 902 | 20201118_ly_colostrum_1ng_glyco_hcd30_etd30_compound15.10705.10705.4 | controllerType=0 controllerNumber=1 scan=10705 | 28.267 |
| 07390:4 | 4 | K.FSSYEKYWC[+57.021]KWN[+2205.793]NTGC[+57.021]QALPSQDEGPSK.A | HexNAc(4)Hex(5)Fuc(2)NeuAc(1) | C[+57]*2, N[+2206] | 1394.077 | 4 | 5573.2863 | 5573.2711 | 0 | 2.7 | 488 | Specific | 506.81 | 395.29 | 395.29 | 8.14 | 63 | >sp\|P01833\|PIGR_HUMAN Polymeric immunoglobulin receptor OS=Homo sapiens OX=9606 GN=PIGR PE=1 SV=4 | 902 | 20201118_ly_colostrum_1ng_glyco_hcd30_etd30_compound15.10605.10605.4 | controllerType=0 controllerNumber=1 scan=10605 | 27.9918 |
| 07478:4 | 4 | K.FSSYEKYWC[+57.021]KWN[+1710.598]NTGC[+57.021]QALPSQDEGPSK.A | HexNAc(3)Hex(5)NeuAc(1) | C[+57]*2, N[+1711] | 1270.2772 | 4 | 5078.0871 | 5078.0759 | 0 | 2.2 | 488 | Specific | 539.92 | 412.73 | 412.73 | 8.37 | 63 | >sp\|P01833\|PIGR_HUMAN Polymeric immunoglobulin receptor OS=Homo sapiens OX=9606 GN=PIGR PE=1 SV=4 | 902 | 20201118_ly_colostrum_1ng_glyco_hcd30_etd30_compound15.10697.10697.4 | controllerType=0 controllerNumber=1 scan=10697 | 28.2449 |
| 05954:3 | 4 | K.YWC[+57.021]KWN[+2205.793]NTGC[+57.021]QALPSQDEGPSK.A | HexNAc(4)Hex(5)Fuc(2)NeuAc(1) | C[+57]*2, N[+2206] | 1611.3227 | 3 | 4831.9535 | 4831.9378 | 0 | 3.3 | 494 | Specific | 318.28 | 187.26 | 187.26 | 5.42 | 63 | >sp\|P01833\|PIGR_HUMAN Polymeric immunoglobulin receptor OS=Homo sapiens OX=9606 GN=PIGR PE=1 SV=4 | 902 | 20201118_ly_colostrum_1ng_glyco_hcd30_etd30_compound15.9097.9097.3 | controllerType=0 controllerNumber=1 scan=9097 | 24.0661 |
| 06080:3 | 4 | K.YWC[+57.021]KWN[+1710.598]NTGC[+57.021]QALPSQDEGPSK.A | HexNAc(3)Hex(5)NeuAc(1) | C[+57]*2, N[+1711] | 1446.261 | 3 | 4336.7686 | 4336.7426 | 0 | 6 | 494 | Specific | 344.93 | 258.47 | 258.47 | 5.01 | 63 | >sp\|P01833\|PIGR_HUMAN Polymeric immunoglobulin receptor OS=Homo sapiens OX=9606 GN=PIGR PE=1 SV=4 | 902 | 20201118_ly_colostrum_1ng_glyco_hcd30_etd30_compound15.9230.9230.3 | controllerType=0 controllerNumber=1 scan=9230 | 24.4382 |
| 04575:3 | 4 | K.WN[+1913.677]NTGC[+57.021]QALPSQDEGPSK.A | HexNAc(4)Hex(5)NeuAc(1) | C[+57], N[+1914] | 1301.5276 | 3 | 3902.5684 | 3902.5537 | 0 | 3.8 | 498 | Specific | 367 | 196.6 | 196.6 | 5.06 | 63 | >sp\|P01833\|PIGR_HUMAN Polymeric immunoglobulin receptor OS=Homo sapiens OX=9606 GN=PIGR PE=1 SV=4 | 902 | 20201118_ly_colostrum_1ng_glyco_hcd30_etd30_compound15.7659.7659.3 | controllerType=0 controllerNumber=1 scan=7659 | 20.8846 |
| 04617:3 | 4 | K.WN[+1710.598]NTGC[+57.021]QALPSQDEGPSK.A | HexNAc(3)Hex(5)NeuAc(1) | C[+57], N[+1711] | 1233.8339 | 3 | 3699.4872 | 3699.4743 | 0 | 3.5 | 498 | Specific | 472.09 | 286.67 | 286.67 | 5.45 | 63 | >sp\|P01833\|PIGR_HUMAN Polymeric immunoglobulin receptor OS=Homo sapiens OX=9606 GN=PIGR PE=1 SV=4 | 902 | 20201118_ly_colostrum_1ng_glyco_hcd30_etd30_compound15.7703.7703.3 | controllerType=0 controllerNumber=1 scan=7703 | 20.981 |
| 06438:4 | 4 | K.WN[+1872.651]NTGC[+57.021]QALPSQDEGPSKAFVNC[+57.021]DENSR.L | HexNAc(3)Hex(6)NeuAc(1) | C[+57]*2, N[+1873] | 1264.2649 | 4 | 5054.0378 | 5054.0203 | 0 | 3.5 | 498 | Specific | 319.5 | 276.29 | 276.29 | 4.88 | 63 | >sp\|P01833\|PIGR_HUMAN Polymeric immunoglobulin receptor OS=Homo sapiens OX=9606 GN=PIGR PE=1 SV=4 | 902 | 20201118_ly_colostrum_1ng_glyco_hcd30_etd30_compound15.9605.9605.4 | controllerType=0 controllerNumber=1 scan=9605 | 25.3866 |
| 17425:5 | 6 | K.VFPLSLDSTPQDGNVVVAC[+57.021]LVQGFFPQEPLSVTWSESGQN[+2553.910]VTAI | HexNAc(5)Hex(5)Fuc(1)NeuAc(2) | C[+57], N[+2554] | 1466.8669 | 5 | 7330.3052 | 7330.2711 | 0 | 4.7 | 8 | Specific | 225.28 | 225.28 | 225.28 | 3.3 | 61 | >sp\|P01877\|IGHA2_HUMAN Immunoglobulin heavy constant alpha 2 OS=Homo sapiens OX=9606 GN=IGHA2 PE=1 SV=4 | 576 | 20201118_ly_colostrum_1ng_glyco_hcd30_etd30_compound15.21215.21215.5 | controllerType=0 controllerNumber=1 scan=21215 | 59.9793 |
| 15474:4 | 6 | K.VFPLSLDSTPQDGNVVVAC[+57.021]LVQGFFPQEPLSVTWSESGQN[+1809.666]VTAI | HexNAc(5)Hex(4)Fuc(1) | C[+57], N[+1810] | 1647.2695 | 4 | 6586.0563 | 6586.0275 | 0 | 4.4 | 8 | Specific | 270.07 | 270.07 | 270.07 | 6.01 | 61 | >sp\|P01877\|IGHA2_HUMAN Immunoglobulin heavy constant alpha 2 OS=Homo sapiens OX=9606 GN=IGHA2 PE=1 SV=4 | 576 | 20201118_ly_colostrum_1ng_glyco_hcd30_etd30_compound15.19138.19138.4 | controllerType=0 controllerNumber=1 scan=19138 | 53.1568 |
| 15146:4 | 6 | K.VFPLSLDSTPQDGNVVVAC[+57.021]LVQGFFPQEPLSVTWSESGQN[+1809.666]VTAI | HexNAc(5)Hex(4)Fuc(1) | C[+57], N[+1810] | 1647.2697 | 4 | 6586.0568 | 6586.0275 | 0 | 4.5 | 8 | Specific | 322.71 | 322.71 | 322.71 | 6.17 | 61 | >sp\|P01877\|IGHA2_HUMAN Immunoglobulin heavy constant alpha 2 OS=Homo sapiens OX=9606 GN=IGHA2 PE=1 SV=4 | 576 | 20201118_ly_colostrum_1ng_glyco_hcd30_etd30_compound15.18787.18787.4 | controllerType=0 controllerNumber=1 scan=18787 | 51.8427 |
| 15967:5 | 6 | K.VFPLSLDSTPQDGNVVVAC[+57.021]LVQGFFPQEPLSVTWSESGQN[+2262.814]VTAI | HexNAc(5)Hex(5)Fuc(1)NeuAc(1) | C[+57], N[+2263] | 1408.6526 | 5 | 7039.2341 | 7039.1757 | 0 | 8.3 | 8 | Specific | 326.4 | 326.4 | 326.4 | 4.12 | 61 | >sp\|P01877\|IGHA2_HUMAN Immunoglobulin heavy constant alpha 2 OS=Homo sapiens OX=9606 GN=IGHA2 PE=1 SV=4 | 576 | 20201118_ly_colostrum_1ng_glyco_hcd30_etd30_compound15.19667.19667.5 | controllerType=0 controllerNumber=1 scan=19667 | 55.1564 |
| 15497:4 | 6 | K.VFPLSLDSTPQDGNVVVAC[+57.021]LVQGFFPQEPLSVTWSESGQN[+1647.613]VTAI | HexNAc(5)Hex(3)Fuc(1) | C[+57], N[+1648] | 1606.7498 | 4 | 6423.9772 | 6423.9746 | 0 | 0.4 | 8 | Specific | 399.04 | 399.04 | 399.04 | 6.76 | 61 | >sp\|P01877\|IGHA2_HUMAN Immunoglobulin heavy constant alpha 2 OS=Homo sapiens OX=9606 GN=IGHA2 PE=1 SV=4 | 576 | 20201118_ly_colostrum_1ng_glyco_hcd30_etd30_compound15.19162.19162.4 | controllerType=0 controllerNumber=1 scan=19162 | 53.24 |
| 15970:6 | 6 | K.VFPLSLDSTPQDGNVVVAC[+57.021]LVQGFFPQEPLSVTWSESGQN[+2262.814]VTAI | HexNAc(5)Hex(5)Fuc(1)NeuAc(1) | C[+57], N[+2263] | 1174.0434 | 6 | 7039.224 | 7039.1757 | 0 | 6.9 | 8 | Specific | 431.2 | 431.2 | 431.2 | 7.05 | 61 | >sp\|P01877\|IGHA2_HUMAN Immunoglobulin heavy constant alpha 2 OS=Homo sapiens OX=9606 GN=IGHA2 PE=1 SV=4 | 576 | 20201118_ly_colostrum_1ng_glyco_hcd30_etd30_compound15.19670.19670.6 | controllerType=0 controllerNumber=1 scan=19670 | 55.1676 |
| 15966:4 | 6 | K.VFPLSLDSTPQDGNVVVAC[+57.021]LVQGFFPQEPLSVTWSESGQN[+2262.814]VTAI | HexNAc(5)Hex(5)Fuc(1)NeuAc(1) | C[+57], N[+2263] | 1760.5637 | 4 | 7039.2329 | 7039.1757 | 0 | 8.1 | 8 | Specific | 467.29 | 467.29 | 467.29 | 7 | 61 | >sp\|P01877\|IGHA2_HUMAN Immunoglobulin heavy constant alpha 2 OS=Homo sapiens OX=9606 GN=IGHA2 PE=1 SV=4 | 576 | 20201118_ly_colostrum_1ng_glyco_hcd30_etd30_compound15.19666.19666.4 | controllerType=0 controllerNumber=1 scan=19666 | 55.1488 |
| 15177:4 | 6 | K.VFPLSLDSTPQDGNVVVAC[+57.021]LVQGFFPQEPLSVTWSESGQN[+1647.613]VTAI | HexNAc(5)Hex(3)Fuc(1) | C[+57], N[+1648] | 1606.7582 | 4 | 6424.0109 | 6423.9746 | 0 | 5.6 | 8 | Specific | 624.36 | 624.36 | 624.36 | 9.72 | 61 | >sp\|P01877\|IGHA2_HUMAN Immunoglobulin heavy constant alpha 2 OS=Homo sapiens OX=9606 GN=IGHA2 PE=1 SV=4 | 576 | 20201118_ly_colostrum_1ng_glyco_hcd30_etd30_compound15.18820.18820.4 | controllerType=0 controllerNumber=1 scan=18820 | 51.9651 |
| 01688:3 | 6 | K.HYTN[+2059.735]SSQDVTVPC[+57.021]R.V | HexNAc(4)Hex(5)Fuc(1)NeuAc(1) | C[+57], N[+2060] | 1241.835 | 3 | 3723.4905 | 3723.4842 | 0 | 1.7 | 89 | Specific | 211.87 | 119.1 | 119.1 | 3.75 | 61 | >sp\|P01877\|IGHA2_HUMAN Immunoglobulin heavy constant alpha 2 OS=Homo sapiens OX=9606 GN=IGHA2 PE=1 SV=4 | 576 | 20201118_ly_colostrum_1ng_glyco_hcd30_etd30_compound15.4648.4648.3 | controllerType=0 controllerNumber=1 scan=4648 | 14.3963 |
| 00851:3 | 6 | K.HYTN[+1971.719]SSQDVTVPC[+57.021]R.V | HexNAc(5)Hex(5)Fuc(1) | C[+57], N[+1972] | 1212.4971 | 3 | 3635.4768 | 3635.4682 | 0 | 2.4 | 89 | Specific | 439.79 | 350.78 | 350.78 | 5.91 | 61 | >sp\|P01877\|IGHA2_HUMAN Immunoglobulin heavy constant alpha 2 OS=Homo sapiens OX=9606 GN=IGHA2 PE=1 SV=4 | 576 | 20201118_ly_colostrum_1ng_glyco_hcd30_etd30_compound15.3780.3780.3 | controllerType=0 controllerNumber=1 scan=3780 | 12.8269 |
| 01709:3 | 6 | K.HYTN[+2262.814]SSQDVTVPC[+57.021]R.V | HexNAc(5)Hex(5)Fuc(1)NeuAc(1) | C[+57], N[+2263] | 1309.5308 | 3 | 3926.5779 | 3926.5636 | 0 | 3.6 | 89 | Specific | 440.31 | 283.52 | 283.52 | 5.35 | 61 | >sp\|P01877\|IGHA2_HUMAN Immunoglobulin heavy constant alpha 2 OS=Homo sapiens OX=9606 GN=IGHA2 PE=1 SV=4 | 576 | 20201118_ly_colostrum_1ng_glyco_hcd30_etd30_compound15.4670.4670.3 | controllerType=0 controllerNumber=1 scan=4670 | 14.4499 |
| 00858:3 | 6 | K.HYTN[+1809.666]SSQDVTVPC[+57.021]R.V | HexNAc(5)Hex(4)Fuc(1) | C[+57], N[+1810] | 1158.4793 | 3 | 3473.4234 | 3473.4154 | 0 | 2.3 | 89 | Specific | 459.45 | 390.79 | 390.79 | 7.12 | 61 | >sp\|P01877\|IGHA2_HUMAN Immunoglobulin heavy constant alpha 2 OS=Homo sapiens OX=9606 GN=IGHA2 PE=1 SV=4 | 576 | 20201118_ly_colostrum_1ng_glyco_hcd30_etd30_compound15.3787.3787.3 | controllerType=0 controllerNumber=1 scan=3787 | 12.845 |
| 00834:3 | 6 | K.HYTN[+1606.587]SSQDVTVPC[+57.021]R.V | HexNAc(4)Hex(4)Fuc(1) | C[+57], N[+1607] | 1090.7864 | 3 | 3270.3447 | 3270.336 | 0 | 2.7 | 89 | Specific | 510.42 | 428.85 | 428.85 | 7.3 | 61 | >sp\|P01877\|IGHA2_HUMAN Immunoglobulin heavy constant alpha 2 OS=Homo sapiens OX=9606 GN=IGHA2 PE=1 SV=4 | 576 | 20201118_ly_colostrum_1ng_glyco_hcd30_etd30_compound15.3762.3762.3 | controllerType=0 controllerNumber=1 scan=3762 | 12.7848 |
| 00868:3 | 6 | K.HYTN[+1444.534]SSQDVTVPC[+57.021]R.V | HexNAc(4)Hex(3)Fuc(1) | C[+57], N[+1445] | 1036.7681 | 3 | 3108.2897 | 3108.2832 | 0 | 2.1 | 89 | Specific | 515.48 | 429.08 | 429.08 | 7.3 | 61 | >sp\|P01877\|IGHA2_HUMAN Immunoglobulin heavy constant alpha 2 OS=Homo sapiens OX=9606 GN=IGHA2 PE=1 SV=4 | 576 | 20201118_ly_colostrum_1ng_glyco_hcd30_etd30_compound15.3798.3798.3 | controllerType=0 controllerNumber=1 scan=3798 | 12.8839 |
| 00879:3 | 6 | K.HYTN[+1647.613]SSQDVTVPC[+57.021]R.V | HexNAc(5)Hex(3)Fuc(1) | C[+57], N[+1648] | 1104.4611 | 3 | 3311.3687 | 3311.3625 | 0 | 1.9 | 89 | Specific | 526.04 | 460.48 | 460.48 | 7.53 | 61 | >sp\|P01877\|IGHA2_HUMAN Immunoglobulin heavy constant alpha 2 OS=Homo sapiens OX=9606 GN=IGHA2 PE=1 SV=4 | 576 | 20201118_ly_colostrum_1ng_glyco_hcd30_etd30_compound15.3809.3809.3 | controllerType=0 controllerNumber=1 scan=3809 | 12.9087 |
| 03456:4 | 6 | K.HYTN[+2262.814]SSQDVTVPC[+57.021]RVPPPPPC[+57.021]C[+57.021]HPR.L | HexNAc(5)Hex(5)Fuc(1)NeuAc(1) | C[+57]*3, N[+2263] | 1306.0505 | 4 | 5221.1803 | 5221.1699 | 0 | 2 | 89 | Specific | 203.22 | 193.5 | 193.5 | 3.47 | 61 | >sp\|P01877\|IGHA2_HUMAN Immunoglobulin heavy constant alpha 2 OS=Homo sapiens OX=9606 GN=IGHA2 PE=1 SV=4 | 576 | 20201118_ly_colostrum_1ng_glyco_hcd30_etd30_compound15.6492.6492.4 | controllerType=0 controllerNumber=1 scan=6492 | 18.3946 |
| 02706:3 | 6 | K.HYTN[+1647.613]SSQDVTVPC[+57.021]RVPPPPPC[+57.021]C[+57.021]HPR.L | HexNAc(5)Hex(3)Fuc(1) | C[+57]*3, N[+1648] | 1535.9999 | 3 | 4605.9853 | 4605.9689 | 0 | 3.6 | 89 | Specific | 227.32 | 153.62 | 153.62 | 2.89 | 61 | >sp\|P01877\|IGHA2_HUMAN Immunoglobulin heavy constant alpha 2 OS=Homo sapiens OX=9606 GN=IGHA2 PE=1 SV=4 | 576 | 20201118_ly_colostrum_1ng_glyco_hcd30_etd30_compound15.5707.5707.3 | controllerType=0 controllerNumber=1 scan=5707 | 16.5673 |
| 02690:4 | 6 | K.HYTN[+1971.719]SSQDVTVPC[+57.021]RVPPPPPC[+57.021]C[+57.021]HPR.L | HexNAc(5)Hex(5)Fuc(1) | C[+57]*3, N[+1972] | 1233.278 | 4 | 4930.0902 | 4930.0745 | 0 | 3.2 | 89 | Specific | 375.07 | 317.88 | 317.88 | 4.53 | 61 | >sp\|P01877\|IGHA2_HUMAN Immunoglobulin heavy constant alpha 2 OS=Homo sapiens OX=9606 GN=IGHA2 PE=1 SV=4 | 576 | 20201118_ly_colostrum_1ng_glyco_hcd30_etd30_compound15.5690.5690.4 | controllerType=0 controllerNumber=1 scan=5690 | 16.519 |
| 03935:4 | 6 | K.HYTN[+2553.910]SSQDVTVPC[+57.021]RVPPPPPC[+57.021]C[+57.021]HPR.L | HexNAc(5)Hex(5)Fuc(1)NeuAc(2) | C[+57]*3, N[+2554] | 1378.8261 | 4 | 5512.2824 | 5512.2653 | 0 | 3.1 | 89 | Specific | 396.9 | 323.9 | 323.9 | 6.03 | 61 | >sp\|P01877\|IGHA2_HUMAN Immunoglobulin heavy constant alpha 2 OS=Homo sapiens OX=9606 GN=IGHA2 PE=1 SV=4 | 576 | 20201118_ly_colostrum_1ng_glyco_hcd30_etd30_compound15.6993.6993.4 | controllerType=0 controllerNumber=1 scan=6993 | 19.5029 |
| 02688:5 | 6 | K.HYTN[+1971.719]SSQDVTVPC[+57.021]RVPPPPPC[+57.021]C[+57.021]HPR.L | HexNAc(5)Hex(5)Fuc(1) | C[+57]*3, N[+1972] | 986.8239 | 5 | 4930.0906 | 4930.0745 | 0 | 3.3 | 89 | Specific | 446.07 | 445.27 | 445.27 | 6.05 | 61 | >sp\|P01877\|IGHA2_HUMAN Immunoglobulin heavy constant alpha 2 OS=Homo sapiens OX=9606 GN=IGHA2 PE=1 SV=4 | 576 | 20201118_ly_colostrum_1ng_glyco_hcd30_etd30_compound15.5688.5688.5 | controllerType=0 controllerNumber=1 scan=5688 | 16.5099 |
| 02682:5 | 6 | K.HYTN[+1647.613]SSQDVTVPC[+57.021]RVPPPPPC[+57.021]C[+57.021]HPR.L | HexNAc(5)Hex(3)Fuc(1) | C[+57]*3, N[+1648] | 922.0029 | 5 | 4605.9853 | 4605.9689 | 0 | 3.6 | 89 | Specific | 462.21 | 457.13 | 457.13 | 5.89 | 61 | >sp\|P01877\|IGHA2_HUMAN Immunoglobulin heavy constant alpha 2 OS=Homo sapiens OX=9606 GN=IGHA2 PE=1 SV=4 | 576 | 20201118_ly_colostrum_1ng_glyco_hcd30_etd30_compound15.5682.5682.5 | controllerType=0 controllerNumber=1 scan=5682 | 16.4973 |
| 02639:5 | 6 | K.HYTN[+1809.666]SSQDVTVPC[+57.021]RVPPPPPC[+57.021]C[+57.021]HPR.L | HexNAc(5)Hex(4)Fuc(1) | C[+57]*3, N[+1810] | 954.4119 | 5 | 4768.0303 | 4768.0217 | 0 | 1.8 | 89 | Specific | 469.08 | 410.23 | 410.23 | 6.43 | 61 | >sp\|P01877\|IGHA2_HUMAN Immunoglobulin heavy constant alpha 2 OS=Homo sapiens OX=9606 GN=IGHA2 PE=1 SV=4 | 576 | 20201118_ly_colostrum_1ng_glyco_hcd30_etd30_compound15.5636.5636.5 | controllerType=0 controllerNumber=1 scan=5636 | 16.3593 |
| 02695:5 | 6 | K.HYTN[+1606.587]SSQDVTVPC[+57.021]RVPPPPPC[+57.021]C[+57.021]HPR.L | HexNAc(4)Hex(4)Fuc(1) | C[+57]*3, N[+1607] | 913.7978 | 5 | 4564.9597 | 4564.9423 | 0 | 3.8 | 89 | Specific | 487.49 | 454.66 | 454.66 | 6.06 | 61 | >sp\|P01877\|IGHA2_HUMAN Immunoglobulin heavy constant alpha 2 OS=Homo sapiens OX=9606 GN=IGHA2 PE=1 SV=4 | 576 | 20201118_ly_colostrum_1ng_glyco_hcd30_etd30_compound15.5695.5695.5 | controllerType=0 controllerNumber=1 scan=5695 | 16.5358 |
| 02659:5 | 6 | K.HYTN[+1647.613]SSQDVTVPC[+57.021]RVPPPPPC[+57.021]C[+57.021]HPR.L | HexNAc(5)Hex(3)Fuc(1) | C[+57]*3, N[+1648] | 922.0006 | 5 | 4605.9741 | 4605.9689 | 0 | 1.1 | 89 | Specific | 488.93 | 440.77 | 440.77 | 6.45 | 61 | >sp\|P01877\|IGHA2_HUMAN Immunoglobulin heavy constant alpha 2 OS=Homo sapiens OX=9606 GN=IGHA2 PE=1 SV=4 | 576 | 20201118_ly_colostrum_1ng_glyco_hcd30_etd30_compound15.5657.5657.5 | controllerType=0 controllerNumber=1 scan=5657 | 16.4227 |
| 02683:4 | 6 | K.HYTN[+1647.613]SSQDVTVPC[+57.021]RVPPPPPC[+57.021]C[+57.021]HPR.L | HexNAc(5)Hex(3)Fuc(1) | C[+57]*3, N[+1648] | 1152.2515 | 4 | 4605.9842 | 4605.9689 | 0 | 3.3 | 89 | Specific | 513.73 | 469.84 | 469.84 | 6.53 | 61 | >sp\|P01877\|IGHA2_HUMAN Immunoglobulin heavy constant alpha 2 OS=Homo sapiens OX=9606 GN=IGHA2 PE=1 SV=4 | 576 | 20201118_ly_colostrum_1ng_glyco_hcd30_etd30_compound15.5683.5683.4 | controllerType=0 controllerNumber=1 scan=5683 | 16.5007 |
| 03149:4 | 6 | K.HYTN[+2262.814]SSQDVTVPC[+57.021]RVPPPPPC[+57.021]C[+57.021]HPR.L | HexNAc(5)Hex(5)Fuc(1)NeuAc(1) | C[+57]*3, N[+2263] | 1306.0521 | 4 | 5221.1867 | 5221.1699 | 0 | 3.2 | 89 | Specific | 564.99 | 449.95 | 449.95 | 8.94 | 61 | >sp\|P01877\|IGHA2_HUMAN Immunoglobulin heavy constant alpha 2 OS=Homo sapiens OX=9606 GN=IGHA2 PE=1 SV=4 | 576 | 20201118_ly_colostrum_1ng_glyco_hcd30_etd30_compound15.6171.6171.4 | controllerType=0 controllerNumber=1 scan=6171 | 17.6613 |
| 02696:4 | 6 | K.HYTN[+1606.587]SSQDVTVPC[+57.021]RVPPPPPC[+57.021]C[+57.021]HPR.L | HexNAc(4)Hex(4)Fuc(1) | C[+57]*3, N[+1607] | 1141.9938 | 4 | 4564.9534 | 4564.9423 | 0 | 2.4 | 89 | Specific | 570.5 | 537.66 | 537.66 | 9.16 | 61 | >sp\|P01877\|IGHA2_HUMAN Immunoglobulin heavy constant alpha 2 OS=Homo sapiens OX=9606 GN=IGHA2 PE=1 SV=4 | 576 | 20201118_ly_colostrum_1ng_glyco_hcd30_etd30_compound15.5696.5696.4 | controllerType=0 controllerNumber=1 scan=5696 | 16.5423 |
| 03145:5 | 6 | K.HYTN[+2262.814]SSQDVTVPC[+57.021]RVPPPPPC[+57.021]C[+57.021]HPR.L | HexNAc(5)Hex(5)Fuc(1)NeuAc(1) | C[+57]*3, N[+2263] | 1045.0428 | 5 | 5221.1851 | 5221.1699 | 0 | 2.9 | 89 | Specific | 578.38 | 483.54 | 483.54 | 8.8 | 61 | >sp\|P01877\|IGHA2_HUMAN Immunoglobulin heavy constant alpha 2 OS=Homo sapiens OX=9606 GN=IGHA2 PE=1 SV=4 | 576 | 20201118_ly_colostrum_1ng_glyco_hcd30_etd30_compound15.6167.6167.5 | controllerType=0 controllerNumber=1 scan=6167 | 17.6441 |
| 02689:5 | 6 | K.HYTN[+1444.534]SSQDVTVPC[+57.021]RVPPPPPC[+57.021]C[+57.021]HPR.L | HexNAc(4)Hex(3)Fuc(1) | C[+57]*3, N[+1445] | 881.3866 | 5 | 4402.904 | 4402.8895 | 0 | 3.3 | 89 | Specific | 614.18 | 563.11 | 563.11 | 8.52 | 61 | >sp\|P01877\|IGHA2_HUMAN Immunoglobulin heavy constant alpha 2 OS=Homo sapiens OX=9606 GN=IGHA2 PE=1 SV=4 | 576 | 20201118_ly_colostrum_1ng_glyco_hcd30_etd30_compound15.5689.5689.5 | controllerType=0 controllerNumber=1 scan=5689 | 16.5153 |
| 02640:4 | 6 | K.HYTN[+1809.666]SSQDVTVPC[+57.021]RVPPPPPC[+57.021]C[+57.021]HPR.L | HexNAc(5)Hex(4)Fuc(1) | C[+57]*3, N[+1810] | 1192.764 | 4 | 4768.034 | 4768.0217 | 0 | 2.6 | 89 | Specific | 711.23 | 630.76 | 630.76 | 12.16 | 61 | >sp\|P01877\|IGHA2_HUMAN Immunoglobulin heavy constant alpha 2 OS=Homo sapiens OX=9606 GN=IGHA2 PE=1 SV=4 | 576 | 20201118_ly_colostrum_1ng_glyco_hcd30_etd30_compound15.5637.5637.4 | controllerType=0 controllerNumber=1 scan=5637 | 16.3632 |
| 02694:4 | 6 | K.HYTN[+1444.534]SSQDVTVPC[+57.021]RVPPPPPC[+57.021]C[+57.021]HPR.L | HexNAc(4)Hex(3)Fuc(1) | C[+57]*3, N[+1445] | 1101.481 | 4 | 4402.9022 | 4402.8895 | 0 | 2.9 | 89 | Specific | 726.85 | 651.27 | 651.27 | 11.94 | 61 | >sp\|P01877\|IGHA2_HUMAN Immunoglobulin heavy constant alpha 2 OS=Homo sapiens OX=9606 GN=IGHA2 PE=1 SV=4 | 576 | 20201118_ly_colostrum_1ng_glyco_hcd30_etd30_compound15.5694.5694.4 | controllerType=0 controllerNumber=1 scan=5694 | 16.53 |
| 13151:5 | 6 | R.LSLHRPALEDLLLGSEAN[+1378.476]LTC[+57.021]TLTGLRDASGATFTWTPSSGK.S | HexNAc(2)Hex(6) | C[+57], N[+1378] | 1167.9581 | 5 | 5835.7616 | 5835.7526 | 0 | 1.5 | 114 | Specific | 500.47 | 500.47 | 500.47 | 6.57 | 61 | >sp\|P01877\|IGHA2_HUMAN Immunoglobulin heavy constant alpha 2 OS=Homo sapiens OX=9606 GN=IGHA2 PE=1 SV=4 | 576 | 20201118_ly_colostrum_1ng_glyco_hcd30_etd30_compound15.16664.16664.5 | controllerType=0 controllerNumber=1 scan=16664 | 44.6975 |
| 13134:5 | 6 | R.LSLHRPALEDLLLGSEAN[+1540.529]LTC[+57.021]TLTGLRDASGATFTWTPSSGK.S | HexNAc(2)Hex(7) | C[+57], N[+1541] | 1200.3694 | 5 | 5997.8178 | 5997.8054 | 0 | 2.1 | 114 | Specific | 768.91 | 768.91 | 768.91 | 11.88 | 61 | >sp\|P01877\|IGHA2_HUMAN Immunoglobulin heavy constant alpha 2 OS=Homo sapiens OX=9606 GN=IGHA2 PE=1 SV=4 | 576 | 20201118_ly_colostrum_1ng_glyco_hcd30_etd30_compound15.16646.16646.5 | controllerType=0 controllerNumber=1 scan=16646 | 44.6378 |
| 13171:4 | 6 | R.LSLHRPALEDLLLGSEAN[+1216.423]LTC[+57.021]TLTGLRDASGATFTWTPSSGK.S | HexNAc(2)Hex(5) | C[+57], N[+1216] | 1419.1865 | 4 | 5673.7243 | 5673.6998 | 0 | 4.3 | 114 | Specific | 829.51 | 829.51 | 829.51 | 14.06 | 61 | >sp\|P01877\|IGHA2_HUMAN Immunoglobulin heavy constant alpha 2 OS=Homo sapiens OX=9606 GN=IGHA2 PE=1 SV=4 | 576 | 20201118_ly_colostrum_1ng_glyco_hcd30_etd30_compound15.16685.16685.4 | controllerType=0 controllerNumber=1 scan=16685 | 44.7613 |
| 13144:4 | 6 | R.LSLHRPALEDLLLGSEAN[+1378.476]LTC[+57.021]TLTGLRDASGATFTWTPSSGK.S | HexNAc(2)Hex(6) | C[+57], N[+1378] | 1459.6985 | 4 | 5835.7721 | 5835.7526 | 0 | 3.3 | 114 | Specific | 855.06 | 855.06 | 855.06 | 15.06 | 61 | >sp\|P01877\|IGHA2_HUMAN Immunoglobulin heavy constant alpha 2 OS=Homo sapiens OX=9606 GN=IGHA2 PE=1 SV=4 | 576 | 20201118_ly_colostrum_1ng_glyco_hcd30_etd30_compound15.16656.16656.4 | controllerType=0 controllerNumber=1 scan=16656 | 44.6646 |
| 13145:5 | 6 | R.LSLHRPALEDLLLGSEAN[+1501.555]LTC[+57.021]TLTGLRDASGATFTWTPSSGK.S | HexNAc(5)Hex(3) | C[+57], N[+1502] | 1192.5759 | 5 | 5958.8505 | 5958.8323 | 0 | 3.1 | 114 | Specific | 921.94 | 911.98 | 911.98 | 15.92 | 61 | >sp\|P01877\|IGHA2_HUMAN Immunoglobulin heavy constant alpha 2 OS=Homo sapiens OX=9606 GN=IGHA2 PE=1 SV=4 | 576 | 20201118_ly_colostrum_1ng_glyco_hcd30_etd30_compound15.16657.16657.5 | controllerType=0 controllerNumber=1 scan=16657 | 44.6704 |
| 13166:5 | 6 | R.LSLHRPALEDLLLGSEAN[+1216.423]LTC[+57.021]TLTGLRDASGATFTWTPSSGK.S | HexNAc(2)Hex(5) | C[+57], N[+1216] | 1135.549 | 5 | 5673.7157 | 5673.6998 | 0 | 2.8 | 114 | Specific | 947.37 | 947.37 | 947.37 | 16.14 | 61 | >sp\|P01877\|IGHA2_HUMAN Immunoglobulin heavy constant alpha 2 OS=Homo sapiens OX=9606 GN=IGHA2 PE=1 SV=4 | 576 | 20201118_ly_colostrum_1ng_glyco_hcd30_etd30_compound15.16680.16680.5 | controllerType=0 controllerNumber=1 scan=16680 | 44.7508 |
| 13156:4 | 6 | R.LSLHRPALEDLLLGSEAN[+1501.555]LTC[+57.021]TLTGLRDASGATFTWTPSSGK.S | HexNAc(5)Hex(3) | C[+57], N[+1502] | 1490.4677 | 4 | 5958.8488 | 5958.8323 | 0 | 2.8 | 114 | Specific | 954.97 | 954.97 | 954.97 | 17.28 | 61 | >sp\|P01877\|IGHA2_HUMAN Immunoglobulin heavy constant alpha 2 OS=Homo sapiens OX=9606 GN=IGHA2 PE=1 SV=4 | 576 | 20201118_ly_colostrum_1ng_glyco_hcd30_etd30_compound15.16669.16669.4 | controllerType=0 controllerNumber=1 scan=16669 | 44.7153 |
| 03799:2 | 6 | K.TPLTAN[+2262.814]ITK.S | HexNAc(5)Hex(5)Fuc(1)NeuAc(1) | N[+2263] | 1611.1927 | 2 | 3221.3781 | 3221.3711 | 0 | 2.2 | 200 | Specific | 211.58 | 34.14 | 34.14 | 2.86 | 61 | >sp\|P01877\|IGHA2_HUMAN Immunoglobulin heavy constant alpha 2 OS=Homo sapiens OX=9606 GN=IGHA2 PE=1 SV=4 | 576 | 20201118_ly_colostrum_1ng_glyco_hcd30_etd30_compound15.6850.6850.2 | controllerType=0 controllerNumber=1 scan=6850 | 19.1376 |
| 03785:2 | 6 | K.TPLTAN[+2059.735]ITK.S | HexNAc(4)Hex(5)Fuc(1)NeuAc(1) | N[+2060] | 1509.6546 | 2 | 3018.3019 | 3018.2917 | 0 | 3.4 | 200 | Specific | 228.75 | 74.22 | 74.22 | 3.01 | 61 | >sp\|P01877\|IGHA2_HUMAN Immunoglobulin heavy constant alpha 2 OS=Homo sapiens OX=9606 GN=IGHA2 PE=1 SV=4 | 576 | 20201118_ly_colostrum_1ng_glyco_hcd30_etd30_compound15.6836.6836.2 | controllerType=0 controllerNumber=1 scan=6836 | 19.1055 |
| 03770:3 | 6 | K.TPLTAN[+2262.814]ITK.S | HexNAc(5)Hex(5)Fuc(1)NeuAc(1) | N[+2263] | 1074.4648 | 3 | 3221.3797 | 3221.3711 | 0 | 2.7 | 200 | Specific | 257.81 | 115.46 | 115.46 | 3.19 | 61 | >sp\|P01877\|IGHA2_HUMAN Immunoglobulin heavy constant alpha 2 OS=Homo sapiens OX=9606 GN=IGHA2 PE=1 SV=4 | 576 | 20201118_ly_colostrum_1ng_glyco_hcd30_etd30_compound15.6820.6820.3 | controllerType=0 controllerNumber=1 scan=6820 | 19.071 |
| 03773:3 | 6 | K.TPLTAN[+2059.735]ITK.S | HexNAc(4)Hex(5)Fuc(1)NeuAc(1) | N[+2060] | 1006.7715 | 3 | 3018.2998 | 3018.2917 | 0 | 2.7 | 200 | Specific | 261.98 | 133.14 | 133.14 | 3.16 | 61 | >sp\|P01877\|IGHA2_HUMAN Immunoglobulin heavy constant alpha 2 OS=Homo sapiens OX=9606 GN=IGHA2 PE=1 SV=4 | 576 | 20201118_ly_colostrum_1ng_glyco_hcd30_etd30_compound15.6823.6823.3 | controllerType=0 controllerNumber=1 scan=6823 | 19.0795 |
| 02546:3 | 6 | K.TPLTAN[+1647.613]ITK.S | HexNAc(5)Hex(3)Fuc(1) | N[+1648] | 869.3974 | 3 | 2606.1777 | 2606.17 | 0 | 3 | 200 | Specific | 269.86 | 131.69 | 131.69 | 2.89 | 61 | >sp\|P01877\|IGHA2_HUMAN Immunoglobulin heavy constant alpha 2 OS=Homo sapiens OX=9606 GN=IGHA2 PE=1 SV=4 | 576 | 20201118_ly_colostrum_1ng_glyco_hcd30_etd30_compound15.5538.5538.3 | controllerType=0 controllerNumber=1 scan=5538 | 16.0798 |
| 02458:3 | 6 | K.TPLTAN[+1971.719]ITK.S | HexNAc(5)Hex(5)Fuc(1) | N[+1972] | 977.4335 | 3 | 2930.2859 | 2930.2757 | 0 | 3.5 | 200 | Specific | 287.87 | 131.67 | 131.67 | 2.8 | 61 | >sp\|P01877\|IGHA2_HUMAN Immunoglobulin heavy constant alpha 2 OS=Homo sapiens OX=9606 GN=IGHA2 PE=1 SV=4 | 576 | 20201118_ly_colostrum_1ng_glyco_hcd30_etd30_compound15.5445.5445.3 | controllerType=0 controllerNumber=1 scan=5445 | 15.789 |
| 02578:2 | 6 | K.TPLTAN[+1647.613]ITK.S | HexNAc(5)Hex(3)Fuc(1) | N[+1648] | 1303.5942 | 2 | 2606.181 | 2606.17 | 0 | 4.2 | 200 | Specific | 298.68 | 93.53 | 93.53 | 2.84 | 61 | >sp\|P01877\|IGHA2_HUMAN Immunoglobulin heavy constant alpha 2 OS=Homo sapiens OX=9606 GN=IGHA2 PE=1 SV=4 | 576 | 20201118_ly_colostrum_1ng_glyco_hcd30_etd30_compound15.5572.5572.2 | controllerType=0 controllerNumber=1 scan=5572 | 16.1862 |
| 02497:2 | 6 | K.TPLTAN[+1809.666]ITK.S | HexNAc(5)Hex(4)Fuc(1) | N[+1810] | 1384.618 | 2 | 2768.2286 | 2768.2228 | 0 | 2.1 | 200 | Specific | 300.06 | 117.73 | 117.73 | 2.97 | 61 | >sp\|P01877\|IGHA2_HUMAN Immunoglobulin heavy constant alpha 2 OS=Homo sapiens OX=9606 GN=IGHA2 PE=1 SV=4 | 576 | 20201118_ly_colostrum_1ng_glyco_hcd30_etd30_compound15.5486.5486.2 | controllerType=0 controllerNumber=1 scan=5486 | 15.9027 |
| 02539:2 | 6 | K.TPLTAN[+1647.613]ITK.S | HexNAc(5)Hex(3)Fuc(1) | N[+1648] | 1303.5923 | 2 | 2606.1774 | 2606.17 | 0 | 2.8 | 200 | Specific | 308.59 | 126.31 | 126.31 | 2.95 | 61 | >sp\|P01877\|IGHA2_HUMAN Immunoglobulin heavy constant alpha 2 OS=Homo sapiens OX=9606 GN=IGHA2 PE=1 SV=4 | 576 | 20201118_ly_colostrum_1ng_glyco_hcd30_etd30_compound15.5531.5531.2 | controllerType=0 controllerNumber=1 scan=5531 | 16.0572 |
| 02505:3 | 6 | K.TPLTAN[+1809.666]ITK.S | HexNAc(5)Hex(4)Fuc(1) | N[+1810] | 923.4148 | 3 | 2768.2299 | 2768.2228 | 0 | 2.6 | 200 | Specific | 308.85 | 136.7 | 136.7 | 2.87 | 61 | >sp\|P01877\|IGHA2_HUMAN Immunoglobulin heavy constant alpha 2 OS=Homo sapiens OX=9606 GN=IGHA2 PE=1 SV=4 | 576 | 20201118_ly_colostrum_1ng_glyco_hcd30_etd30_compound15.5494.5494.3 | controllerType=0 controllerNumber=1 scan=5494 | 15.9407 |
| 02493:2 | 6 | K.TPLTAN[+1606.587]ITK.S | HexNAc(4)Hex(4)Fuc(1) | N[+1607] | 1283.0785 | 2 | 2565.1498 | 2565.1435 | 0 | 2.5 | 200 | Specific | 315.24 | 132.75 | 132.75 | 3.19 | 61 | >sp\|P01877\|IGHA2_HUMAN Immunoglobulin heavy constant alpha 2 OS=Homo sapiens OX=9606 GN=IGHA2 PE=1 SV=4 | 576 | 20201118_ly_colostrum_1ng_glyco_hcd30_etd30_compound15.5482.5482.2 | controllerType=0 controllerNumber=1 scan=5482 | 15.8847 |
| 02555:2 | 6 | K.TPLTAN[+1444.534]ITK.S | HexNAc(4)Hex(3)Fuc(1) | N[+1445] | 1202.0529 | 2 | 2403.0985 | 2403.0906 | 0 | 3.3 | 200 | Specific | 331.78 | 83.48 | 83.48 | 2.79 | 61 | >sp\|P01877\|IGHA2_HUMAN Immunoglobulin heavy constant alpha 2 OS=Homo sapiens OX=9606 GN=IGHA2 PE=1 SV=4 | 576 | 20201118_ly_colostrum_1ng_glyco_hcd30_etd30_compound15.5548.5548.2 | controllerType=0 controllerNumber=1 scan=5548 | 16.1142 |
| 07584:3 | 6 | R.MAGKPTHIN[+1864.634]VSVVMAEADGTC[+57.021]Y.- | HexNAc(2)Hex(9) | C[+57], N[+1865] | 1405.9166 | 3 | 4215.7351 | 4215.717 | 0 | 4.3 | 319 | CRagged | 204.52 | 204.52 | 204.52 | 3.42 | 61 | >sp\|P01877\|IGHA2_HUMAN Immunoglobulin heavy constant alpha 2 OS=Homo sapiens OX=9606 GN=IGHA2 PE=1 SV=4 | 576 | 20201118_ly_colostrum_1ng_glyco_hcd30_etd30_compound15.10809.10809.3 | controllerType=0 controllerNumber=1 scan=10809 | 28.5527 |
| 07135:3 | 6 | R.MAGKPTHIN[+1378.476]VSVVMAEADGTC[+57.021]Y.- | HexNAc(2)Hex(6) | C[+57], N[+1378] | 1243.8614 | 3 | 3729.5696 | 3729.5585 | 0 | 3 | 319 | CRagged | 228.86 | 173.94 | 157.41 | 3.42 | 61 | >sp\|P01877\|IGHA2_HUMAN Immunoglobulin heavy constant alpha 2 OS=Homo sapiens OX=9606 GN=IGHA2 PE=1 SV=4 | 576 | 20201118_ly_colostrum_1ng_glyco_hcd30_etd30_compound15.10337.10337.3 | controllerType=0 controllerNumber=1 scan=10337 | 27.2866 |
| 07628:3 | 6 | R.MAGKPTHIN[+1663.608]VSVVMAEADGTC[+57.021]Y.- | HexNAc(5)Hex(4) | C[+57], N[+1664] | 1338.9066 | 3 | 4014.7051 | 4014.691 | 0 | 3.5 | 319 | CRagged | 234.68 | 115.59 | 115.59 | 2.94 | 61 | >sp\|P01877\|IGHA2_HUMAN Immunoglobulin heavy constant alpha 2 OS=Homo sapiens OX=9606 GN=IGHA2 PE=1 SV=4 | 576 | 20201118_ly_colostrum_1ng_glyco_hcd30_etd30_compound15.10856.10856.3 | controllerType=0 controllerNumber=1 scan=10856 | 28.6985 |
| 07659:3 | 6 | R.MAGKPTHIN[+1216.423]VSVVMAEADGTC[+57.021]Y.- | HexNAc(2)Hex(5) | C[+57], N[+1216] | 1189.8428 | 3 | 3567.5139 | 3567.5057 | 0 | 2.3 | 319 | CRagged | 250.32 | 217.78 | 191.94 | 4.46 | 61 | >sp\|P01877\|IGHA2_HUMAN Immunoglobulin heavy constant alpha 2 OS=Homo sapiens OX=9606 GN=IGHA2 PE=1 SV=4 | 576 | 20201118_ly_colostrum_1ng_glyco_hcd30_etd30_compound15.10889.10889.3 | controllerType=0 controllerNumber=1 scan=10889 | 28.7926 |
| 07576:3 | 6 | R.MAGKPTHIN[+1702.581]VSVVMAEADGTC[+57.021]Y.- | HexNAc(2)Hex(8) | C[+57], N[+1703] | 1351.8984 | 3 | 4053.6806 | 4053.6642 | 0 | 4 | 319 | CRagged | 252.95 | 252.95 | 195.07 | 4.36 | 61 | >sp\|P01877\|IGHA2_HUMAN Immunoglobulin heavy constant alpha 2 OS=Homo sapiens OX=9606 GN=IGHA2 PE=1 SV=4 | 576 | 20201118_ly_colostrum_1ng_glyco_hcd30_etd30_compound15.10801.10801.3 | controllerType=0 controllerNumber=1 scan=10801 | 28.5326 |
| 08799:3 | 6 | R.MAGKPTHIN[+1710.598]VSVVMAEADGTC[+57.021]Y.- | HexNAc(3)Hex(5)NeuAc(1) | C[+57], N[+1711] | 1354.5687 | 3 | 4061.6914 | 4061.6805 | 0 | 2.7 | 319 | CRagged | 253.92 | 181.26 | 156.75 | 4.42 | 61 | >sp\|P01877\|IGHA2_HUMAN Immunoglobulin heavy constant alpha 2 OS=Homo sapiens OX=9606 GN=IGHA2 PE=1 SV=4 | 576 | 20201118_ly_colostrum_1ng_glyco_hcd30_etd30_compound15.12084.12084.3 | controllerType=0 controllerNumber=1 scan=12084 | 31.9595 |
| 07198:3 | 6 | R.MAGKPTHIN[+1216.423]VSVVMAEADGTC[+57.021]Y.- | HexNAc(2)Hex(5) | C[+57], N[+1216] | 1189.8437 | 3 | 3567.5165 | 3567.5057 | 0 | 3 | 319 | CRagged | 265.04 | 252.23 | 194.39 | 4.36 | 61 | >sp\|P01877\|IGHA2_HUMAN Immunoglobulin heavy constant alpha 2 OS=Homo sapiens OX=9606 GN=IGHA2 PE=1 SV=4 | 576 | 20201118_ly_colostrum_1ng_glyco_hcd30_etd30_compound15.10403.10403.3 | controllerType=0 controllerNumber=1 scan=10403 | 27.4562 |
| 08838:3 | 6 | R.MAGKPTHIN[+1548.545]VSVVMAEADGTC[+57.021]Y.- | HexNAc(3)Hex(4)NeuAc(1) | C[+57], N[+1549] | 1300.5514 | 3 | 3899.6398 | 3899.6277 | 0 | 3.1 | 319 | CRagged | 266.83 | 172.6 | 172.6 | 4.21 | 61 | >sp\|P01877\|IGHA2_HUMAN Immunoglobulin heavy constant alpha 2 OS=Homo sapiens OX=9606 GN=IGHA2 PE=1 SV=4 | 576 | 20201118_ly_colostrum_1ng_glyco_hcd30_etd30_compound15.12125.12125.3 | controllerType=0 controllerNumber=1 scan=12125 | 32.0678 |
| 07702:3 | 6 | R.MAGKPTHIN[+1647.613]VSVVMAEADGTC[+57.021]Y.- | HexNAc(5)Hex(3)Fuc(1) | C[+57], N[+1648] | 1333.5744 | 3 | 3998.7086 | 3998.6961 | 0 | 3.1 | 319 | CRagged | 269.75 | 122.52 | 122.52 | 3.43 | 61 | >sp\|P01877\|IGHA2_HUMAN Immunoglobulin heavy constant alpha 2 OS=Homo sapiens OX=9606 GN=IGHA2 PE=1 SV=4 | 576 | 20201118_ly_colostrum_1ng_glyco_hcd30_etd30_compound15.10934.10934.3 | controllerType=0 controllerNumber=1 scan=10934 | 28.9105 |
| 07622:3 | 6 | R.MAGKPTHIN[+1378.476]VSVVMAEADGTC[+57.021]Y.- | HexNAc(2)Hex(6) | C[+57], N[+1378] | 1243.8624 | 3 | 3729.5725 | 3729.5585 | 0 | 3.8 | 319 | CRagged | 280.2 | 219.51 | 172.25 | 4.19 | 61 | >sp\|P01877\|IGHA2_HUMAN Immunoglobulin heavy constant alpha 2 OS=Homo sapiens OX=9606 GN=IGHA2 PE=1 SV=4 | 576 | 20201118_ly_colostrum_1ng_glyco_hcd30_etd30_compound15.10850.10850.3 | controllerType=0 controllerNumber=1 scan=10850 | 28.6878 |
| 07690:3 | 6 | R.MAGKPTHIN[+1501.555]VSVVMAEADGTC[+57.021]Y.- | HexNAc(5)Hex(3) | C[+57], N[+1502] | 1284.8874 | 3 | 3852.6476 | 3852.6382 | 0 | 2.4 | 319 | CRagged | 303.12 | 252.81 | 252.81 | 4.65 | 61 | >sp\|P01877\|IGHA2_HUMAN Immunoglobulin heavy constant alpha 2 OS=Homo sapiens OX=9606 GN=IGHA2 PE=1 SV=4 | 576 | 20201118_ly_colostrum_1ng_glyco_hcd30_etd30_compound15.10921.10921.3 | controllerType=0 controllerNumber=1 scan=10921 | 28.8717 |
| 07592:3 | 6 | R.MAGKPTHIN[+1540.529]VSVVMAEADGTC[+57.021]Y.- | HexNAc(2)Hex(7) | C[+57], N[+1541] | 1297.8796 | 3 | 3891.6242 | 3891.6114 | 0 | 3.3 | 319 | CRagged | 303.41 | 303.41 | 185.28 | 5.77 | 61 | >sp\|P01877\|IGHA2_HUMAN Immunoglobulin heavy constant alpha 2 OS=Homo sapiens OX=9606 GN=IGHA2 PE=1 SV=4 | 576 | 20201118_ly_colostrum_1ng_glyco_hcd30_etd30_compound15.10818.10818.3 | controllerType=0 controllerNumber=1 scan=10818 | 28.5895 |
| 07302:3 | 6 | R.M[+15.995]AGKPTHIN[+1540.529]VSVVMAEADGTC[+57.021]Y.- | HexNAc(2)Hex(7) | C[+57], M[+16], N[+1541] | 1303.2075 | 3 | 3907.6078 | 3907.6063 | 0 | 0.4 | 319 | CRagged | 201.42 | 180.23 | 109.87 | 2.93 | 61 | >sp\|P01877\|IGHA2_HUMAN Immunoglobulin heavy constant alpha 2 OS=Homo sapiens OX=9606 GN=IGHA2 PE=1 SV=4 | 576 | 20201118_ly_colostrum_1ng_glyco_hcd30_etd30_compound15.10512.10512.3 | controllerType=0 controllerNumber=1 scan=10512 | 27.7569 |
| 07379:3 | 6 | R.M[+15.995]AGKPTHIN[+1216.423]VSVVMAEADGTC[+57.021]Y.- | HexNAc(2)Hex(5) | C[+57], M[+16], N[+1216] | 1195.1749 | 3 | 3583.51 | 3583.5006 | 0 | 2.6 | 319 | CRagged | 217.5 | 168.95 | 130.65 | 3.27 | 61 | >sp\|P01877\|IGHA2_HUMAN Immunoglobulin heavy constant alpha 2 OS=Homo sapiens OX=9606 GN=IGHA2 PE=1 SV=4 | 576 | 20201118_ly_colostrum_1ng_glyco_hcd30_etd30_compound15.10593.10593.3 | controllerType=0 controllerNumber=1 scan=10593 | 27.9569 |
| 07344:3 | 6 | R.M[+15.995]AGKPTHIN[+1216.423]VSVVMAEADGTC[+57.021]Y.- | HexNAc(2)Hex(5) | C[+57], M[+16], N[+1216] | 1195.1733 | 3 | 3583.5053 | 3583.5006 | 0 | 1.3 | 319 | CRagged | 274.5 | 234.01 | 171.33 | 4.46 | 61 | >sp\|P01877\|IGHA2_HUMAN Immunoglobulin heavy constant alpha 2 OS=Homo sapiens OX=9606 GN=IGHA2 PE=1 SV=4 | 576 | 20201118_ly_colostrum_1ng_glyco_hcd30_etd30_compound15.10556.10556.3 | controllerType=0 controllerNumber=1 scan=10556 | 27.8503 |

Table S4. Cont.

| Query #:z | Protein Rank | Glycopeptide | Glycan | Modification Type(s) | Observed m/z | z | Observed (M+H) | Calc. mass (M+H) | Off-by-x error | Mass error (ppm) | StartPosition | Cleavage | Score | Delta | Delta Mod | \|Log Prob\| | # of unique peptides | Protein Name | Protein DB number | Comment | Scan # | Scan Time |
|---|---|---|---|---|---|---|---|---|---|---|---|---|---|---|---|---|---|---|---|---|---|---|
| 16127:3 | 9 | R.QLAHQSN[+2350.830]STNIFFSPVSIATAFAMLSLGTK.A | HexNAc(4)Hex(5)Fuc(1)NeuAc(2) | N[+2351] | 1844.8343 | 3 | 5532.4884 | 5532.4654 | 0 | 4.2 | 64 | Specific | 322.97 | 216.81 | 216.81 | 5.96 | 15 | >sp\|P01009\|A1AT_HUMAN Alpha-1-antitrypsin OS=Homo sapiens OX=9606 GN=SERPINA1 PE=1 SV=3 | 642 | 20201118_ly_colostrum_1ng_glyco_hcd30_etd30_compound15.19839.19839.3 | controllerType=0 controllerNumber=1 scan=19839 | 55.8006 |
| 16151:3 | 9 | R.QLAHQSN[+2204.772]STNIFFSPVSIATAFAMLSLGTK.A | HexNAc(4)Hex(5)NeuAc(2) | N[+2205] | 1796.1464 | 3 | 5386.4246 | 5386.4075 | 0 | 3.2 | 64 | Specific | 466.95 | 336.81 | 336.81 | 6.71 | 15 | >sp\|P01009\|A1AT_HUMAN Alpha-1-antitrypsin OS=Homo sapiens OX=9606 GN=SERPINA1 PE=1 SV=3 | 642 | 20201118_ly_colostrum_1ng_glyco_hcd30_etd30_compound15.19865.19865.3 | controllerType=0 controllerNumber=1 scan=19865 | 55.8919 |
| 16133:4 | 9 | R.QLAHQSN[+2350.830]STNIFFSPVSIATAFAMLSLGTK.A | HexNAc(4)Hex(5)Fuc(1)NeuAc(2) | N[+2351] | 1383.8774 | 4 | 5532.4879 | 5532.4654 | 0 | 4.1 | 64 | Specific | 824.91 | 732.05 | 732.05 | 14.48 | 15 | >sp\|P01009\|A1AT_HUMAN Alpha-1-antitrypsin OS=Homo sapiens OX=9606 GN=SERPINA1 PE=1 SV=3 | 642 | 20201118_ly_colostrum_1ng_glyco_hcd30_etd30_compound15.19846.19846.4 | controllerType=0 controllerNumber=1 scan=19846 | 55.8289 |
| 11127:4 | 13 | K.LKELPGVC[+57.021]N[+2350.830]ETMMALWEEC[+57.021]KPC[+57.021]LK.Q | HexNAc(4)Hex(5)Fuc(1)NeuAc(2) | C[+57]*3, N[+2351] | 1322.562 | 4 | 5287.2262 | 5287.2151 | 0 | 2.1 | 95 | Specific | 323.34 | 196.14 | 196.14 | 4.23 | 17 | >sp\|P10909\|CLUS_HUMAN Clusterin OS=Homo sapiens OX=9606 GN=CLU PE=1 SV=1 | 1467 | 20201118_ly_colostrum_1ng_glyco_hcd30_etd30_compound15.14531.14531.4 | controllerType=0 controllerNumber=1 scan=14531 | 38.5617 |
| 10027:4 | 13 | K.LKELPGVC[+57.021]N[+2205.793]ETMMALWEEC[+57.021]KPC[+57.021]LK.Q | HexNAc(4)Hex(5)Fuc(2)NeuAc(1) | C[+57]*3, N[+2206] | 1286.3029 | 4 | 5142.1896 | 5142.1776 | 0 | 2.3 | 95 | Specific | 423.55 | 289.21 | 289.21 | 4.83 | 17 | >sp\|P10909\|CLUS_HUMAN Clusterin OS=Homo sapiens OX=9606 GN=CLU PE=1 SV=1 | 1467 | 20201118_ly_colostrum_1ng_glyco_hcd30_etd30_compound15.13371.13371.4 | controllerType=0 controllerNumber=1 scan=13371 | 35.2797 |
| 10013:3 | 13 | K.LKELPGVC[+57.021]N[+2351.851]ETMMALWEEC[+57.021]KPC[+57.021]LK.Q | HexNAc(4)Hex(5)Fuc(3)NeuAc(1) | C[+57]*3, N[+2352] | 1763.4209 | 3 | 5288.248 | 5288.2355 | 0 | 2.4 | 95 | Specific | 437.12 | 300.56 | 300.56 | 5 | 17 | >sp\|P10909\|CLUS_HUMAN Clusterin OS=Homo sapiens OX=9606 GN=CLU PE=1 SV=1 | 1467 | 20201118_ly_colostrum_1ng_glyco_hcd30_etd30_compound15.13356.13356.3 | controllerType=0 controllerNumber=1 scan=13356 | 35.2411 |
| 10016:4 | 13 | K.LKELPGVC[+57.021]N[+2351.851]ETMMALWEEC[+57.021]KPC[+57.021]LK.Q | HexNAc(4)Hex(5)Fuc(3)NeuAc(1) | C[+57]*3, N[+2352] | 1322.8175 | 4 | 5288.2482 | 5288.2355 | 0 | 2.4 | 95 | Specific | 612.42 | 481.73 | 481.73 | 10.02 | 17 | >sp\|P10909\|CLUS_HUMAN Clusterin OS=Homo sapiens OX=9606 GN=CLU PE=1 SV=1 | 1467 | 20201118_ly_colostrum_1ng_glyco_hcd30_etd30_compound15.13360.13360.4 | controllerType=0 controllerNumber=1 scan=13360 | 35.2556 |
| 00080:3 | 13 | R.EIRHN[+2059.735]STGC[+57.021]LR.M | HexNAc(4)Hex(5)Fuc(1)NeuAc(1) | C[+57], N[+2060] | 1134.8072 | 3 | 3402.4069 | 3402.3994 | 0 | 2.2 | 287 | Specific | 209.43 | 99.63 | 99.63 | 2.85 | 17 | >sp\|P10909\|CLUS_HUMAN Clusterin OS=Homo sapiens OX=9606 GN=CLU PE=1 SV=1 | 1467 | 20201118_ly_colostrum_1ng_glyco_hcd30_etd30_compound15.2975.2975.3 | controllerType=0 controllerNumber=1 scan=2975 | 11.2416 |
| 00084:4 | 13 | R.EIRHN[+2351.851]STGC[+57.021]LR.M | HexNAc(4)Hex(5)Fuc(3)NeuAc(1) | C[+57], N[+2352] | 924.3871 | 4 | 3694.5264 | 3694.5152 | 0 | 3 | 287 | Specific | 634.22 | 480.9 | 480.9 | 8.71 | 17 | >sp\|P10909\|CLUS_HUMAN Clusterin OS=Homo sapiens OX=9606 GN=CLU PE=1 SV=1 | 1467 | 20201118_ly_colostrum_1ng_glyco_hcd30_etd30_compound15.2980.2980.4 | controllerType=0 controllerNumber=1 scan=2980 | 11.259 |
| 00092:4 | 13 | R.EIRHN[+2205.793]STGC[+57.021]LR.M | HexNAc(4)Hex(5)Fuc(2)NeuAc(1) | C[+57], N[+2206] | 887.8733 | 4 | 3548.4713 | 3548.4573 | 0 | 3.9 | 287 | Specific | 654.4 | 499.17 | 499.17 | 9.71 | 17 | >sp\|P10909\|CLUS_HUMAN Clusterin OS=Homo sapiens OX=9606 GN=CLU PE=1 SV=1 | 1467 | 20201118_ly_colostrum_1ng_glyco_hcd30_etd30_compound15.2988.2988.4 | controllerType=0 controllerNumber=1 scan=2988 | 11.2908 |
| 00104:4 | 13 | R.EIRHN[+2059.735]STGC[+57.021]LR.M | HexNAc(4)Hex(5)Fuc(1)NeuAc(1) | C[+57], N[+2060] | 851.3568 | 4 | 3402.4055 | 3402.3994 | 0 | 1.8 | 287 | Specific | 684.58 | 514.91 | 514.91 | 9.93 | 17 | >sp\|P10909\|CLUS_HUMAN Clusterin OS=Homo sapiens OX=9606 GN=CLU PE=1 SV=1 | 1467 | 20201118_ly_colostrum_1ng_glyco_hcd30_etd30_compound15.3001.3001.4 | controllerType=0 controllerNumber=1 scan=3001 | 11.3377 |
| 15774:4 | 13 | K.MLN[+2350.830]TSSLLEQLNEQFNWVSR.L | HexNAc(4)Hex(5)Fuc(1)NeuAc(2) | N[+2351] | 1190.7669 | 4 | 4760.0457 | 4760.0171 | 0 | 6 | 352 | Specific | 315.87 | 279.75 | 279.75 | 3.77 | 17 | >sp\|P10909\|CLUS_HUMAN Clusterin OS=Homo sapiens OX=9606 GN=CLU PE=1 SV=1 | 1467 | 20201118_ly_colostrum_1ng_glyco_hcd30_etd30_compound15.19460.19460.4 | controllerType=0 controllerNumber=1 scan=19460 | 54.3782 |
| 14769:3 | 13 | K.MLN[+1872.651]TSSLLEQLNEQFNWVSR.L | HexNAc(3)Hex(6)NeuAc(1) | N[+1873] | 1427.9586 | 3 | 4281.8611 | 4281.8372 | 0 | 5.6 | 352 | Specific | 477 | 335.54 | 133.89 | 5.65 | 17 | >sp\|P10909\|CLUS_HUMAN Clusterin OS=Homo sapiens OX=9606 GN=CLU PE=1 SV=1 | 1467 | 20201118_ly_colostrum_1ng_glyco_hcd30_etd30_compound15.18385.18385.3 | controllerType=0 controllerNumber=1 scan=18385 | 50.435 |
| 14601:3 | 13 | K.MLN[+2205.793]TSSLLEQLNEQFNWVSR.L | HexNAc(4)Hex(5)Fuc(2)NeuAc(1) | N[+2206] | 1539.0036 | 3 | 4614.9962 | 4614.9796 | 0 | 3.6 | 352 | Specific | 569.8 | 387.44 | 387.44 | 8.97 | 17 | >sp\|P10909\|CLUS_HUMAN Clusterin OS=Homo sapiens OX=9606 GN=CLU PE=1 SV=1 | 1467 | 20201118_ly_colostrum_1ng_glyco_hcd30_etd30_compound15.18205.18205.3 | controllerType=0 controllerNumber=1 scan=18205 | 49.7421 |
| 15765:3 | 13 | K.MLN[+2350.830]TSSLLEQLNEQFNWVSR.L | HexNAc(4)Hex(5)Fuc(1)NeuAc(2) | N[+2351] | 1587.3575 | 3 | 4760.0579 | 4760.0171 | 0 | 8.6 | 352 | Specific | 592.49 | 399.71 | 399.71 | 7.78 | 17 | >sp\|P10909\|CLUS_HUMAN Clusterin OS=Homo sapiens OX=9606 GN=CLU PE=1 SV=1 | 1467 | 20201118_ly_colostrum_1ng_glyco_hcd30_etd30_compound15.19451.19451.3 | controllerType=0 controllerNumber=1 scan=19451 | 54.3434 |
| 14628:3 | 13 | K.MLN[+2059.735]TSSLLEQLNEQFNWVSR.L | HexNAc(4)Hex(5)Fuc(1)NeuAc(1) | N[+2060] | 1490.3156 | 3 | 4468.9323 | 4468.9217 | 0 | 2.4 | 352 | Specific | 644.86 | 412.15 | 412.15 | 10.19 | 17 | >sp\|P10909\|CLUS_HUMAN Clusterin OS=Homo sapiens OX=9606 GN=CLU PE=1 SV=1 | 1467 | 20201118_ly_colostrum_1ng_glyco_hcd30_etd30_compound15.18234.18234.3 | controllerType=0 controllerNumber=1 scan=18234 | 49.8485 |
| 03858:3 | 14 | R.FSDGLESN[+1702.581]SSTQFEVKK.Y | HexNAc(2)Hex(8) | N[+1703] | 1202.5046 | 3 | 3605.4992 | 3605.4893 | 0 | 2.7 | 219 | Specific | 221.14 | 168.19 | 168.19 | 3.89 | 12 | >sp\|P0C0L4\|CO4A_HUMAN Complement C4-A OS=Homo sapiens OX=9606 GN=C4A PE=1 SV=2 | 1235 | 20201118_ly_colostrum_1ng_glyco_hcd30_etd30_compound15.6912.6912.3 | controllerType=0 controllerNumber=1 scan=6912 | 19.2939 |
| 09610:4 | 15 | R.IIVPLNNREN[+2204.772]ISDPTSPLR.T | HexNAc(4)Hex(5)NeuAc(2) | N[+2205] | 1088.9952 | 4 | 4352.9588 | 4352.9496 | 0 | 2.1 | 62 | Specific | 227.69 | 227.69 | 227.69 | 3.07 | 13 | >sp\|P01591\|IGJ_HUMAN Immunoglobulin J chain OS=Homo sapiens OX=9606 GN=JCHAIN PE=1 SV=4 | 1392 | 20201118_ly_colostrum_1ng_glyco_hcd30_etd30_compound15.12933.12933.4 | controllerType=0 controllerNumber=1 scan=12933 | 34.1248 |
| 07289:3 | 15 | R.IIVPLNNREN[+1663.608]ISDPTSPLR.T | HexNAc(5)Hex(4) | N[+1664] | 1271.2696 | 3 | 3811.7942 | 3811.7853 | 0 | 2.3 | 62 | Specific | 234.94 | 162.81 | 162.81 | 2.99 | 13 | >sp\|P01591\|IGJ_HUMAN Immunoglobulin J chain OS=Homo sapiens OX=9606 GN=JCHAIN PE=1 SV=4 | 1392 | 20201118_ly_colostrum_1ng_glyco_hcd30_etd30_compound15.10499.10499.3 | controllerType=0 controllerNumber=1 scan=10499 | 27.7132 |
| 07249:3 | 15 | R.IIVPLNNREN[+1809.666]ISDPTSPLR.T | HexNAc(5)Hex(4)Fuc(1) | N[+1810] | 1319.9577 | 3 | 3957.8585 | 3957.8432 | 0 | 3.9 | 62 | Specific | 241.89 | 145.97 | 145.97 | 3.25 | 13 | >sp\|P01591\|IGJ_HUMAN Immunoglobulin J chain OS=Homo sapiens OX=9606 GN=JCHAIN PE=1 SV=4 | 1392 | 20201118_ly_colostrum_1ng_glyco_hcd30_etd30_compound15.10457.10457.3 | controllerType=0 controllerNumber=1 scan=10457 | 27.6002 |
| 08242:3 | 15 | R.IIVPLNNREN[+1913.677]ISDPTSPLR.T | HexNAc(4)Hex(5)NeuAc(1) | N[+1914] | 1354.6277 | 3 | 4061.8687 | 4061.8542 | 0 | 3.6 | 62 | Specific | 274.05 | 224.98 | 224.98 | 3.35 | 13 | >sp\|P01591\|IGJ_HUMAN Immunoglobulin J chain OS=Homo sapiens OX=9606 GN=JCHAIN PE=1 SV=4 | 1392 | 20201118_ly_colostrum_1ng_glyco_hcd30_etd30_compound15.11501.11501.3 | controllerType=0 controllerNumber=1 scan=11501 | 30.4077 |
| 07181:3 | 15 | R.IIVPLNNREN[+1809.666]ISDPTSPLR.T | HexNAc(5)Hex(4)Fuc(1) | N[+1810] | 1319.9562 | 3 | 3957.8541 | 3957.8432 | 0 | 2.8 | 62 | Specific | 282.63 | 184.55 | 184.55 | 3.7 | 13 | >sp\|P01591\|IGJ_HUMAN Immunoglobulin J chain OS=Homo sapiens OX=9606 GN=JCHAIN PE=1 SV=4 | 1392 | 20201118_ly_colostrum_1ng_glyco_hcd30_etd30_compound15.10385.10385.3 | controllerType=0 controllerNumber=1 scan=10385 | 27.4064 |
| 09600:3 | 15 | R.IIVPLNNREN[+2204.772]ISDPTSPLR.T | HexNAc(4)Hex(5)NeuAc(2) | N[+2205] | 1451.6581 | 3 | 4352.9599 | 4352.9496 | 0 | 2.4 | 62 | Specific | 287.55 | 222.57 | 222.57 | 3.59 | 13 | >sp\|P01591\|IGJ_HUMAN Immunoglobulin J chain OS=Homo sapiens OX=9606 GN=JCHAIN PE=1 SV=4 | 1392 | 20201118_ly_colostrum_1ng_glyco_hcd30_etd30_compound15.12923.12923.3 | controllerType=0 controllerNumber=1 scan=12923 | 34.0867 |
| 08184:3 | 15 | R.IIVPLNNREN[+2059.735]ISDPTSPLR.T | HexNAc(4)Hex(5)Fuc(1)NeuAc(1) | N[+2060] | 1403.3127 | 3 | 4207.9235 | 4207.9121 | 0 | 2.7 | 62 | Specific | 302.54 | 261.97 | 261.97 | 3.99 | 13 | >sp\|P01591\|IGJ_HUMAN Immunoglobulin J chain OS=Homo sapiens OX=9606 GN=JCHAIN PE=1 SV=4 | 1392 | 20201118_ly_colostrum_1ng_glyco_hcd30_etd30_compound15.11440.11440.3 | controllerType=0 controllerNumber=1 scan=11440 | 30.2418 |
| 08298:3 | 15 | R.IIVPLNNREN[+1856.656]ISDPTSPLR.T | HexNAc(3)Hex(5)Fuc(1)NeuAc(1) | N[+1857] | 1335.6191 | 3 | 4004.8427 | 4004.8327 | 0 | 2.5 | 62 | Specific | 304.25 | 216.15 | 216.15 | 3.73 | 13 | >sp\|P01591\|IGJ_HUMAN Immunoglobulin J chain OS=Homo sapiens OX=9606 GN=JCHAIN PE=1 SV=4 | 1392 | 20201118_ly_colostrum_1ng_glyco_hcd30_etd30_compound15.11560.11560.3 | controllerType=0 controllerNumber=1 scan=11560 | 30.595 |
| 08247:4 | 15 | R.IIVPLNNREN[+1913.677]ISDPTSPLR.T | HexNAc(4)Hex(5)NeuAc(1) | N[+1914] | 1016.222 | 4 | 4061.866 | 4061.8542 | 0 | 2.9 | 62 | Specific | 506.07 | 506.07 | 506.07 | 5.88 | 13 | >sp\|P01591\|IGJ_HUMAN Immunoglobulin J chain OS=Homo sapiens OX=9606 GN=JCHAIN PE=1 SV=4 | 1392 | 20201118_ly_colostrum_1ng_glyco_hcd30_etd30_compound15.11506.11506.4 | controllerType=0 controllerNumber=1 scan=11506 | 30.4198 |
| 08222:4 | 15 | R.IIVPLNNREN[+2059.735]ISDPTSPLR.T | HexNAc(4)Hex(5)Fuc(1)NeuAc(1) | N[+2060] | 1052.7364 | 4 | 4207.9237 | 4207.9121 | 0 | 2.8 | 62 | Specific | 524.47 | 506.08 | 506.08 | 6.33 | 13 | >sp\|P01591\|IGJ_HUMAN Immunoglobulin J chain OS=Homo sapiens OX=9606 GN=JCHAIN PE=1 SV=4 | 1392 | 20201118_ly_colostrum_1ng_glyco_hcd30_etd30_compound15.11480.11480.4 | controllerType=0 controllerNumber=1 scan=11480 | 30.3483 |
| 03166:2 | 17 | K.AAIPSALDTN[+1216.423]SSK.S | HexNAc(2)Hex(5) | N[+1216] | 1246.0487 | 2 | 2491.0902 | 2491.0815 | 0 | 3.5 | 542 | Specific | 345.22 | 155.63 | 155.63 | 3.27 | 8 | >sp\|Q08380\|LG3BP_HUMAN Galectin-3-binding protein OS=Homo sapiens OX=9606 GN=LGALS3BP PE=1 SV=1 | 1562 | 20201118_ly_colostrum_1ng_glyco_hcd30_etd30_compound15.6189.6189.2 | controllerType=0 controllerNumber=1 scan=6189 | 17.7081 |
| 08477:3 | 20 | R.TVDKSTGKPTLYN[+1378.476]VSLVMSDTAGTC[+57.021]Y.- | HexNAc(2)Hex(6) | C[+57], N[+1378] | 1396.2816 | 3 | 4186.8301 | 4186.8187 | 0 | 2.7 | 428 | CRagged | 323.32 | 271.33 | 159.82 | 4.89 | 9 | >sp\|P01871\|IGHM_HUMAN Immunoglobulin heavy constant mu OS=Homo sapiens OX=9606 GN=IGHM PE=1 SV=4 | 328 | 20201118_ly_colostrum_1ng_glyco_hcd30_etd30_compound15.11747.11747.3 | controllerType=0 controllerNumber=1 scan=11747 | 31.0942 |
| 00729:4 | 64 | K.TKPREEQFN[+1606.587]STFR.V | HexNAc(4)Hex(4)Fuc(1) | N[+1607] | 812.3587 | 4 | 3246.4131 | 3246.4054 | 0 | 2.4 | 168 | Specific | 503.59 | 436.81 | 436.81 | 5.38 | 3 | >sp\|P01859\|IGHG2_HUMAN Immunoglobulin heavy constant gamma 2 OS=Homo sapiens OX=9606 GN=IGHG2 PE=1 SV=2 | 387 | 20201118_ly_colostrum_1ng_glyco_hcd30_etd30_compound15.3653.3653.4 | controllerType=0 controllerNumber=1 scan=3653 | 12.5613 |
| 00719:4 | 64 | K.TKPREEQFN[+1606.587]STFR.V | HexNAc(4)Hex(4)Fuc(1) | N[+1607] | 812.3593 | 4 | 3246.4155 | 3246.4054 | 0 | 3.1 | 168 | Specific | 640.66 | 536.08 | 536.08 | 7.59 | 3 | >sp\|P01859\|IGHG2_HUMAN Immunoglobulin heavy constant gamma 2 OS=Homo sapiens OX=9606 GN=IGHG2 PE=1 SV=2 | 387 | 20201118_ly_colostrum_1ng_glyco_hcd30_etd30_compound15.3642.3642.4 | controllerType=0 controllerNumber=1 scan=3642 | 12.5307 |
| 06805:3 | 100 | R.FVN[+876.322]VTVTPEDQC[+57.021]RPNNVC[+57.021]TGVLTR.R | HexNAc(2)Hex(2)Fuc(1) | C[+57]*2, N[+876] | 1218.2336 | 3 | 3652.6862 | 3652.6728 | 0 | 3.7 | 169 | Specific | 256.94 | 214.62 | 214.62 | 2.63 | 1 | >sp\|P20160\|CAP7_HUMAN Azurocidin OS=Homo sapiens OX=9606 GN=AZU1 PE=1 SV=3 | 850 | 20201118_ly_colostrum_1ng_glyco_hcd30_etd30_compound15.9990.9990.3 | controllerType=0 controllerNumber=1 scan=9990 | 26.3575 |
| 15072:4 | 1 | R.[+42.011]RTAGWNVPIGTLRPFLN[+1872.651]WTGPPEPIEAAVAR.F | HexNAc(3)Hex(6)NeuAc(1) | .[42], N[+1873] | 1326.1359 | 4 | 5301.5216 | 5301.4731 | 0 | 9.2 | 140 | Specific | 391.05 | 323.19 | 323.19 | 3.1 | 38 | >sp\|P02788\|TRFL_HUMAN Lactotransferrin OS=Homo sapiens OX=9606 GN=LTF PE=1 SV=6 | 1270 | 20201117_ly_milk_1ng_glyco_hcd30_etd30_compound15.18642.18642.4 | controllerType=0 controllerNumber=1 scan=18642 | 41.3105 |
| 16472:4 | 1 | R.RTAGWNVPIGTLRPFLN[+2350.830]WTGPPEPIEAAVAR.F | HexNAc(4)Hex(5)Fuc(1)NeuAc(2) | N[+2351] | 1435.1752 | 4 | 5737.6789 | 5737.6424 | 0 | 6.4 | 140 | Specific | 715.62 | 715.62 | 715.62 | 10.43 | 38 | >sp\|P02788\|TRFL_HUMAN Lactotransferrin OS=Homo sapiens OX=9606 GN=LTF PE=1 SV=6 | 1270 | 20201117_ly_milk_1ng_glyco_hcd30_etd30_compound15.20121.20121.4 | controllerType=0 controllerNumber=1 scan=20121 | 45.8316 |
| 15113:4 | 1 | R.RTAGWNVPIGTLRPFLN[+1768.640]WTGPPEPIEAAVAR.F | HexNAc(4)Hex(5)Fuc(1) | N[+1769] | 1289.6243 | 4 | 5155.4753 | 5155.4516 | 0 | 4.6 | 140 | Specific | 927.42 | 910.22 | 910.22 | 15.42 | 38 | >sp\|P02788\|TRFL_HUMAN Lactotransferrin OS=Homo sapiens OX=9606 GN=LTF PE=1 SV=6 | 1270 | 20201117_ly_milk_1ng_glyco_hcd30_etd30_compound15.18685.18685.4 | controllerType=0 controllerNumber=1 scan=18685 | 41.4469 |
| 15963:3 | 1 | R.TAGWNVPIGTLRPFLN[+1768.640]WTGPPEPIEAAVAR.F | HexNAc(4)Hex(5)Fuc(1) | N[+1769] | 1667.1276 | 3 | 4999.3684 | 4999.3504 | 0 | 3.6 | 141 | Specific | 386.57 | 328.79 | 328.79 | 4.92 | 38 | >sp\|P02788\|TRFL_HUMAN Lactotransferrin OS=Homo sapiens OX=9606 GN=LTF PE=1 SV=6 | 1270 | 20201117_ly_milk_1ng_glyco_hcd30_etd30_compound15.19583.19583.3 | controllerType=0 controllerNumber=1 scan=19583 | 44.1303 |
| 16629:3 | 1 | R.TAGWNVPIGTLRPFLN[+2059.735]WTGPPEPIEAAVAR.F | HexNAc(4)Hex(5)Fuc(1)NeuAc(1) | N[+2060] | 1764.1628 | 3 | 5290.4739 | 5290.4459 | 0 | 5.3 | 141 | Specific | 399.15 | 267.98 | 267.98 | 4.54 | 38 | >sp\|P02788\|TRFL_HUMAN Lactotransferrin OS=Homo sapiens OX=9606 GN=LTF PE=1 SV=6 | 1270 | 20201117_ly_milk_1ng_glyco_hcd30_etd30_compound15.20286.20286.3 | controllerType=0 controllerNumber=1 scan=20286 | 46.3172 |
| 16660:3 | 1 | R.TAGWNVPIGTLRPFLN[+2204.772]WTGPPEPIEAAVAR.F | HexNAc(4)Hex(5)NeuAc(2) | N[+2205] | 1812.5135 | 3 | 5435.5259 | 5435.4834 | 0 | 7.8 | 141 | Specific | 472.76 | 352.44 | 352.44 | 4.63 | 38 | >sp\|P02788\|TRFL_HUMAN Lactotransferrin OS=Homo sapiens OX=9606 GN=LTF PE=1 SV=6 | 1270 | 20201117_ly_milk_1ng_glyco_hcd30_etd30_compound15.20319.20319.3 | controllerType=0 controllerNumber=1 scan=20319 | 46.4074 |
| 16608:3 | 1 | R.TAGWNVPIGTLRPFLN[+2205.793]WTGPPEPIEAAVAR.F | HexNAc(4)Hex(5)Fuc(2)NeuAc(1) | N[+2206] | 1812.8466 | 3 | 5436.5254 | 5436.5038 | 0 | 4 | 141 | Specific | 472.81 | 362.8 | 362.8 | 5.47 | 38 | >sp\|P02788\|TRFL_HUMAN Lactotransferrin OS=Homo sapiens OX=9606 GN=LTF PE=1 SV=6 | 1270 | 20201117_ly_milk_1ng_glyco_hcd30_etd30_compound15.20264.20264.3 | controllerType=0 controllerNumber=1 scan=20264 | 46.243 |
| 16630:5 | 1 | R.TAGWNVPIGTLRPFLN[+2205.793]WTGPPEPIEAAVAR.F | HexNAc(4)Hex(5)Fuc(2)NeuAc(1) | N[+2206] | 1088.1111 | 5 | 5436.5263 | 5436.5038 | 0 | 4.1 | 141 | Specific | 542.91 | 505.81 | 505.81 | 6.9 | 38 | >sp\|P02788\|TRFL_HUMAN Lactotransferrin OS=Homo sapiens OX=9606 GN=LTF PE=1 SV=6 | 1270 | 20201117_ly_milk_1ng_glyco_hcd30_etd30_compound15.20287.20287.5 | controllerType=0 controllerNumber=1 scan=20287 | 46.3237 |
| 16631:5 | 1 | R.TAGWNVPIGTLRPFLN[+2059.735]WTGPPEPIEAAVAR.F | HexNAc(4)Hex(5)Fuc(1)NeuAc(1) | N[+2060] | 1058.8998 | 5 | 5290.4698 | 5290.4459 | 0 | 4.5 | 141 | Specific | 608.96 | 590.13 | 590.13 | 8.87 | 38 | >sp\|P02788\|TRFL_HUMAN Lactotransferrin OS=Homo sapiens OX=9606 GN=LTF PE=1 SV=6 | 1270 | 20201117_ly_milk_1ng_glyco_hcd30_etd30_compound15.20288.20288.5 | controllerType=0 controllerNumber=1 scan=20288 | 46.3276 |
| 16513:4 | 1 | R.TAGWNVPIGTLRPFLN[+2716.983]WTGPPEPIEAAVAR.F | HexNAc(5)Hex(6)Fuc(3)NeuAc(1) | N[+2717] | 1487.6864 | 4 | 5947.7238 | 5947.6939 | 0 | 5 | 141 | Specific | 771.79 | 771.79 | 771.79 | 12.28 | 38 | >sp\|P02788\|TRFL_HUMAN Lactotransferrin OS=Homo sapiens OX=9606 GN=LTF PE=1 SV=6 | 1270 | 20201117_ly_milk_1ng_glyco_hcd30_etd30_compound15.20164.20164.4 | controllerType=0 controllerNumber=1 scan=20164 | 45.9472 |
| 16604:4 | 1 | R.TAGWNVPIGTLRPFLN[+2205.793]WTGPPEPIEAAVAR.F | HexNAc(4)Hex(5)Fuc(2)NeuAc(1) | N[+2206] | 1359.8877 | 4 | 5436.529 | 5436.5038 | 0 | 4.6 | 141 | Specific | 833.27 | 813.16 | 813.16 | 13.68 | 38 | >sp\|P02788\|TRFL_HUMAN Lactotransferrin OS=Homo sapiens OX=9606 GN=LTF PE=1 SV=6 | 1270 | 20201117_ly_milk_1ng_glyco_hcd30_etd30_compound15.20260.20260.4 | controllerType=0 controllerNumber=1 scan=20260 | 46.2338 |
| 16623:4 | 1 | R.TAGWNVPIGTLRPFLN[+2059.735]WTGPPEPIEAAVAR.F | HexNAc(4)Hex(5)Fuc(1)NeuAc(1) | N[+2060] | 1323.3751 | 4 | 5290.4787 | 5290.4459 | 0 | 6.2 | 141 | Specific | 844.63 | 766.87 | 766.87 | 12.68 | 38 | >sp\|P02788\|TRFL_HUMAN Lactotransferrin OS=Homo sapiens OX=9606 GN=LTF PE=1 SV=6 | 1270 | 20201117_ly_milk_1ng_glyco_hcd30_etd30_compound15.20280.20280.4 | controllerType=0 controllerNumber=1 scan=20280 | 46.305 |
| 15962:4 | 1 | R.TAGWNVPIGTLRPFLN[+1768.640]WTGPPEPIEAAVAR.F | HexNAc(4)Hex(5)Fuc(1) | N[+1769] | 1250.5978 | 4 | 4999.3693 | 4999.3504 | 0 | 3.8 | 141 | Specific | 1086.41 | 1086.41 | 1086.41 | 18.68 | 38 | >sp\|P02788\|TRFL_HUMAN Lactotransferrin OS=Homo sapiens OX=9606 GN=LTF PE=1 SV=6 | 1270 | 20201117_ly_milk_1ng_glyco_hcd30_etd30_compound15.19582.19582.4 | controllerType=0 controllerNumber=1 scan=19582 | 44.1257 |
| 14992:3 | 1 | R.TAGWNIPMGLLFN[+1768.640]QTGSC[+57.021]K.F | HexNAc(4)Hex(5)Fuc(1) | C[+57], N[+1769] | 1288.5576 | 3 | 3863.6581 | 3863.6495 | 0 | 2.2 | 485 | Specific | 249.19 | 229.93 | 229.93 | 5.22 | 38 | >sp\|P02788\|TRFL_HUMAN Lactotransferrin OS=Homo sapiens OX=9606 GN=LTF PE=1 SV=6 | 1270 | 20201117_ly_milk_1ng_glyco_hcd30_etd30_compound15.18557.18557.3 | controllerType=0 controllerNumber=1 scan=18557 | 41.0643 |
| 15005:3 | 1 | R.TAGWNIPMGLLFN[+1622.582]QTGSC[+57.021]K.F | HexNAc(4)Hex(5) | C[+57], N[+1623] | 1239.871 | 3 | 3717.5985 | 3717.5916 | 0 | 1.9 | 485 | Specific | 342.93 | 273.75 | 231.9 | 5.56 | 38 | >sp\|P02788\|TRFL_HUMAN Lactotransferrin OS=Homo sapiens OX=9606 GN=LTF PE=1 SV=6 | 1270 | 20201117_ly_milk_1ng_glyco_hcd30_etd30_compound15.18571.18571.3 | controllerType=0 controllerNumber=1 scan=18571 | 41.0973 |
| 17813:3 | 1 | R.TAGWNIPMGLLFN[+2204.772]QTGSC[+57.021]K.F | HexNAc(4)Hex(5)NeuAc(2) | C[+57], N[+2205] | 1433.9369 | 3 | 4299.7963 | 4299.7824 | 0 | 3.2 | 485 | Specific | 350.85 | 273.46 | 273.46 | 5.62 | 38 | >sp\|P02788\|TRFL_HUMAN Lactotransferrin OS=Homo sapiens OX=9606 GN=LTF PE=1 SV=6 | 1270 | 20201117_ly_milk_1ng_glyco_hcd30_etd30_compound15.21539.21539.3 | controllerType=0 controllerNumber=1 scan=21539 | 49.814 |
| 14831:3 | 1 | R.TAGWNIPMGLLFN[+1768.640]QTGSC[+57.021]K.F | HexNAc(4)Hex(5)Fuc(1) | C[+57], N[+1769] | 1288.5595 | 3 | 3863.664 | 3863.6495 | 0 | 3.8 | 485 | Specific | 389.75 | 296.23 | 296.23 | 5.67 | 38 | >sp\|P02788\|TRFL_HUMAN Lactotransferrin OS=Homo sapiens OX=9606 GN=LTF PE=1 SV=6 | 1270 | 20201117_ly_milk_1ng_glyco_hcd30_etd30_compound15.18388.18388.3 | controllerType=0 controllerNumber=1 scan=18388 | 40.6093 |
| 15124:3 | 1 | R.TAGWNIPMGLLFN[+1216.423]QTGSC[+57.021]K.F | HexNAc(2)Hex(5) | C[+57], N[+1216] | 1104.4868 | 3 | 3311.446 | 3311.4328 | 0 | 4 | 485 | Specific | 402.41 | 376.83 | 204.46 | 5.86 | 38 | >sp\|P02788\|TRFL_HUMAN Lactotransferrin OS=Homo sapiens OX=9606 GN=LTF PE=1 SV=6 | 1270 | 20201117_ly_milk_1ng_glyco_hcd30_etd30_compound15.18697.18697.3 | controllerType=0 controllerNumber=1 scan=18697 | 41.4873 |
| 14925:3 | 1 | R.TAGWNIPMGLLFN[+1606.587]QTGSC[+57.021]K.F | HexNAc(4)Hex(4)Fuc(1) | C[+57], N[+1607] | 1234.5416 | 3 | 3701.6101 | 3701.5966 | 0 | 3.6 | 485 | Specific | 405.28 | 296.42 | 296.42 | 4.96 | 38 | >sp\|P02788\|TRFL_HUMAN Lactotransferrin OS=Homo sapiens OX=9606 GN=LTF PE=1 SV=6 | 1270 | 20201117_ly_milk_1ng_glyco_hcd30_etd30_compound15.18487.18487.3 | controllerType=0 controllerNumber=1 scan=18487 | 40.891 |
| 16182:3 | 1 | R.TAGWNIPMGLLFN[+2059.735]QTGSC[+57.021]K.F | HexNAc(4)Hex(5)Fuc(1)NeuAc(1) | C[+57], N[+2060] | 1385.5906 | 3 | 4154.7573 | 4154.7449 | 0 | 3 | 485 | Specific | 423.25 | 286.81 | 286.81 | 5.78 | 38 | >sp\|P02788\|TRFL_HUMAN Lactotransferrin OS=Homo sapiens OX=9606 GN=LTF PE=1 SV=6 | 1270 | 20201117_ly_milk_1ng_glyco_hcd30_etd30_compound15.19813.19813.3 | controllerType=0 controllerNumber=1 scan=19813 | 44.8145 |
| 16168:3 | 1 | R.TAGWNIPMGLLFN[+2205.793]QTGSC[+57.021]K.F | HexNAc(4)Hex(5)Fuc(2)NeuAc(1) | C[+57], N[+2206] | 1434.2764 | 3 | 4300.8147 | 4300.8028 | 0 | 2.8 | 485 | Specific | 441.19 | 322.12 | 322.12 | 6.16 | 38 | >sp\|P02788\|TRFL_HUMAN Lactotransferrin OS=Homo sapiens OX=9606 GN=LTF PE=1 SV=6 | 1270 | 20201117_ly_milk_1ng_glyco_hcd30_etd30_compound15.19799.19799.3 | controllerType=0 controllerNumber=1 scan=19799 | 44.7687 |
| 16283:3 | 1 | R.TAGWNIPMGLLFN[+1913.677]QTGSC[+57.021]K.F | HexNAc(4)Hex(5)NeuAc(1) | C[+57], N[+1914] | 1336.9044 | 3 | 4008.6985 | 4008.687 | 0 | 2.9 | 485 | Specific | 449.96 | 291.39 | 291.39 | 6.16 | 38 | >sp\|P02788\|TRFL_HUMAN Lactotransferrin OS=Homo sapiens OX=9606 GN=LTF PE=1 SV=6 | 1270 | 20201117_ly_milk_1ng_glyco_hcd30_etd30_compound15.19921.19921.3 | controllerType=0 controllerNumber=1 scan=19921 | 45.1932 |
| 16308:4 | 1 | R.TAGWNIPMGLLFN[+1913.677]QTGSC[+57.021]K.F | HexNAc(4)Hex(5)NeuAc(1) | C[+57], N[+1914] | 1002.9311 | 4 | 4008.7027 | 4008.687 | 0 | 3.9 | 485 | Specific | 455.83 | 390.04 | 390.04 | 5.59 | 38 | >sp\|P02788\|TRFL_HUMAN Lactotransferrin OS=Homo sapiens OX=9606 GN=LTF PE=1 SV=6 | 1270 | 20201117_ly_milk_1ng_glyco_hcd30_etd30_compound15.19947.19947.4 | controllerType=0 controllerNumber=1 scan=19947 | 45.273 |
| 14542:3 | 1 | R.TAGWNIPM[+15.995]GLLFN[+2059.735]QTGSC[+57.021]K.F | HexNAc(4)Hex(5)Fuc(1)NeuAc(1) | C[+57], M[+16], N[+2060] | 1390.9296 | 3 | 4170.7743 | 4170.7398 | 0 | 8.3 | 485 | Specific | 232.6 | 163.46 | 163.46 | 2.59 | 38 | >sp\|P02788\|TRFL_HUMAN Lactotransferrin OS=Homo sapiens OX=9606 GN=LTF PE=1 SV=6 | 1270 | 20201117_ly_milk_1ng_glyco_hcd30_etd30_compound15.18083.18083.3 | controllerType=0 controllerNumber=1 scan=18083 | 39.8489 |
| 15055:3 | 1 | R.TAGWNIPM[+15.995]GLLFN[+2059.735]QTGSC[+57.021]K.F | HexNAc(4)Hex(5)Fuc(1)NeuAc(1) | C[+57], M[+16], N[+2060] | 1390.9244 | 3 | 4170.7586 | 4170.7398 | 0 | 4.5 | 485 | Specific | 394.42 | 294.9 | 294.9 | 5.67 | 38 | >sp\|P02788\|TRFL_HUMAN Lactotransferrin OS=Homo sapiens OX=9606 GN=LTF PE=1 SV=6 | 1270 | 20201117_ly_milk_1ng_glyco_hcd30_etd30_compound15.18624.18624.3 | controllerType=0 controllerNumber=1 scan=18624 | 41.2571 |
| 16418:4 | 1 | R.TAGWNIPMGLLFN[+1913.677]QTGSC[+57.021]KFDEYFSQSC[+57.021]APGSDPR.S | HexNAc(4)Hex(5)NeuAc(1) | C[+57]*2, N[+1914] | 1463.8667 | 4 | 5852.445 | 5852.4341 | 0 | 1.9 | 485 | Specific | 357.85 | 282.75 | 282.75 | 5.21 | 38 | >sp\|P02788\|TRFL_HUMAN Lactotransferrin OS=Homo sapiens OX=9606 GN=LTF PE=1 SV=6 | 1270 | 20201117_ly_milk_1ng_glyco_hcd30_etd30_compound15.20064.20064.4 | controllerType=0 controllerNumber=1 scan=20064 | 45.6767 |
| 15662:4 | 1 | R.TAGWNIPMGLLFN[+1768.640]QTGSC[+57.021]KFDEYFSQSC[+57.021]APGSDPR.S | HexNAc(4)Hex(5)Fuc(1) | C[+57]*2, N[+1769] | 1427.6078 | 4 | 5707.4093 | 5707.3966 | 0 | 2.2 | 485 | Specific | 419.53 | 308.19 | 308.19 | 4.33 | 38 | >sp\|P02788\|TRFL_HUMAN Lactotransferrin OS=Homo sapiens OX=9606 GN=LTF PE=1 SV=6 | 1270 | 20201117_ly_milk_1ng_glyco_hcd30_etd30_compound15.19266.19266.4 | controllerType=0 controllerNumber=1 scan=19266 | 43.2302 |
| 16364:4 | 1 | R.TAGWNIPMGLLFN[+2059.735]QTGSC[+57.021]KFDEYFSQSC[+57.021]APGSDPR.S | HexNAc(4)Hex(5)Fuc(1)NeuAc(1) | C[+57]*2, N[+2060] | 1500.3866 | 4 | 5998.5246 | 5998.492 | 0 | 5.4 | 485 | Specific | 721.8 | 668.25 | 668.25 | 10.77 | 38 | >sp\|P02788\|TRFL_HUMAN Lactotransferrin OS=Homo sapiens OX=9606 GN=LTF PE=1 SV=6 | 1270 | 20201117_ly_milk_1ng_glyco_hcd30_etd30_compound15.20007.20007.4 | controllerType=0 controllerNumber=1 scan=20007 | 45.4891 |
| 10442:3 | 1 | R.N[+2350.830]GSDC[+57.021]PDKFC[+57.021]LFQSETK.N | HexNAc(4)Hex(5)Fuc(1)NeuAc(2) | C[+57]*2, N[+2351] | 1461.9119 | 3 | 4383.7212 | 4383.7043 | 0 | 3.9 | 642 | Specific | 222.24 | 121.43 | 121.43 | 4.23 | 38 | >sp\|P02788\|TRFL_HUMAN Lactotransferrin OS=Homo sapiens OX=9606 GN=LTF PE=1 SV=6 | 1270 | 20201117_ly_milk_1ng_glyco_hcd30_etd30_compound15.13765.13765.3 | controllerType=0 controllerNumber=1 scan=13765 | 30.9195 |
| 08215:3 | 1 | R.N[+2205.793]GSDC[+57.021]PDKFC[+57.021]LFQSETK.N | HexNAc(4)Hex(5)Fuc(2)NeuAc(1) | C[+57]*2, N[+2206] | 1413.5627 | 3 | 4238.6735 | 4238.6667 | 0 | 1.6 | 642 | Specific | 232.78 | 156.36 | 156.36 | 4.45 | 38 | >sp\|P02788\|TRFL_HUMAN Lactotransferrin OS=Homo sapiens OX=9606 GN=LTF PE=1 SV=6 | 1270 | 20201117_ly_milk_1ng_glyco_hcd30_etd30_compound15.11453.11453.3 | controllerType=0 controllerNumber=1 scan=11453 | 26.5251 |
| 08238:3 | 1 | R.N[+2205.793]GSDC[+57.021]PDKFC[+57.021]LFQSETK.N | HexNAc(4)Hex(5)Fuc(2)NeuAc(1) | C[+57]*2, N[+2206] | 1413.5641 | 3 | 4238.6779 | 4238.6667 | 0 | 2.6 | 642 | Specific | 294.68 | 188.52 | 188.52 | 5.17 | 38 | >sp\|P02788\|TRFL_HUMAN Lactotransferrin OS=Homo sapiens OX=9606 GN=LTF PE=1 SV=6 | 1270 | 20201117_ly_milk_1ng_glyco_hcd30_etd30_compound15.11478.11478.3 | controllerType=0 controllerNumber=1 scan=11478 | 26.5859 |
| 08319:3 | 1 | R.N[+2059.735]GSDC[+57.021]PDKFC[+57.021]LFQSETK.N | HexNAc(4)Hex(5)Fuc(1)NeuAc(1) | C[+57]*2, N[+2060] | 1364.8786 | 3 | 4092.6212 | 4092.6088 | 0 | 3 | 642 | Specific | 316.39 | 190.53 | 190.53 | 5.17 | 38 | >sp\|P02788\|TRFL_HUMAN Lactotransferrin OS=Homo sapiens OX=9606 GN=LTF PE=1 SV=6 | 1270 | 20201117_ly_milk_1ng_glyco_hcd30_etd30_compound15.11562.11562.3 | controllerType=0 controllerNumber=1 scan=11562 | 26.7531 |
| 15503:3 | 2 | R.LSLHRPALEDLLLGSEAN[+1622.582]LTC[+57.021]TLTGLR.D | HexNAc(4)Hex(5) | C[+57], N[+1623] | 1529.4045 | 3 | 4586.1989 | 4586.1799 | 0 | 4.1 | 127 | Specific | 321.1 | 321.1 | 321.1 | 5.79 | 29 | >sp\|P01876\|IGHA1_HUMAN Immunoglobulin heavy constant alpha 1 OS=Homo sapiens OX=9606 GN=IGHA1 PE=1 SV=2 | 255 | 20201117_ly_milk_1ng_glyco_hcd30_etd30_compound15.19097.19097.3 | controllerType=0 controllerNumber=1 scan=19097 | 42.6928 |
| 15513:3 | 2 | R.LSLHRPALEDLLLGSEAN[+1460.529]LTC[+57.021]TLTGLR.D | HexNAc(4)Hex(4) | C[+57], N[+1461] | 1475.3891 | 3 | 4424.1527 | 4424.1271 | 0 | 5.8 | 127 | Specific | 361.93 | 309.93 | 309.93 | 5.46 | 29 | >sp\|P01876\|IGHA1_HUMAN Immunoglobulin heavy constant alpha 1 OS=Homo sapiens OX=9606 GN=IGHA1 PE=1 SV=2 | 255 | 20201117_ly_milk_1ng_glyco_hcd30_etd30_compound15.19108.19108.3 | controllerType=0 controllerNumber=1 scan=19108 | 42.[illegible]322 |
| 16181:4 | 2 | R.LSLHRPALEDLLLGSEAN[+1872.651]LTC[+57.021]TLTGLR.D | HexNAc(3)Hex(6)NeuAc(1) | C[+57], N[+1873] | 1209.823 | 4 | 4836.2703 | 4836.2487 | 0 | 4.5 | 127 | Specific | 382.2 | 382.2 | 382.2 | 7.15 | 29 | >sp\|P01876\|IGHA1_HUMAN Immunoglobulin heavy constant alpha 1 OS=Homo sapiens OX=9606 GN=IGHA1 PE=1 SV=2 | 255 | 20201117_ly_milk_1ng_glyco_hcd30_etd30_compound15.19812.19812.4 | controllerType=0 controllerNumber=1 scan=19812 | 44.8103 |
| 15442:3 | 2 | R.LSLHRPALEDLLLGSEAN[+1702.581]LTC[+57.021]TLTGLR.D | HexNAc(2)Hex(8) | C[+57], N[+1703] | 1556.07 | 3 | 4666.1955 | 4666.1796 | 0 | 3.4 | 127 | Specific | 390.27 | 384.44 | 384.44 | 7.28 | 29 | >sp\|P01876\|IGHA1_HUMAN Immunoglobulin heavy constant alpha 1 OS=Homo sapiens OX=9606 GN=IGHA1 PE=1 SV=2 | 255 | 20201117_ly_milk_1ng_glyco_hcd30_etd30_compound15.19033.19033.3 | controllerType=0 controllerNumber=1 scan=19033 | 42.4946 |
| 15480:3 | 2 | R.LSLHRPALEDLLLGSEAN[+1663.608]LTC[+57.021]TLTGLR.D | HexNAc(5)Hex(4) | C[+57], N[+1664] | 1543.0816 | 3 | 4627.2302 | 4627.2064 | 0 | 5.1 | 127 | Specific | 399.29 | 374.38 | 374.38 | 6.74 | 29 | >sp\|P01876\|IGHA1_HUMAN Immunoglobulin heavy constant alpha 1 OS=Homo sapiens OX=9606 GN=IGHA1 PE=1 SV=2 | 255 | 20201117_ly_milk_1ng_glyco_hcd30_etd30_compound15.19073.19073.3 | controllerType=0 controllerNumber=1 scan=19073 | 42.6063 |
| 15443:3 | 2 | R.LSLHRPALEDLLLGSEAN[+1864.634]LTC[+57.021]TLTGLR.D | HexNAc(2)Hex(9) | C[+57], N[+1865] | 1610.0876 | 3 | 4828.2482 | 4828.2324 | 0 | 3.3 | 127 | Specific | 467.14 | 423.92 | 423.92 | 7.61 | 29 | >sp\|P01876\|IGHA1_HUMAN Immunoglobulin heavy constant alpha 1 OS=Homo sapiens OX=9606 GN=IGHA1 PE=1 SV=2 | 255 | 20201117_ly_milk_1ng_glyco_hcd30_etd30_compound15.19034.19034.3 | controllerType=0 controllerNumber=1 scan=19034 | 42.4979 |
| 15447:3 | 2 | R.LSLHRPALEDLLLGSEAN[+1540.529]LTC[+57.021]TLTGLR.D | HexNAc(2)Hex(7) | C[+57], N[+1541] | 1502.0525 | 3 | 4504.1431 | 4504.1268 | 0 | 3.6 | 127 | Specific | 468.93 | 429.08 | 429.08 | 7.61 | 29 | >sp\|P01876\|IGHA1_HUMAN Immunoglobulin heavy constant alpha 1 OS=Homo sapiens OX=9606 GN=IGHA1 PE=1 SV=2 | 255 | 20201117_ly_milk_1ng_glyco_hcd30_etd30_compound15.19038.19038.3 | controllerType=0 controllerNumber=1 scan=19038 | 42.5102 |
| 15491:3 | 2 | R.LSLHRPALEDLLLGSEAN[+1501.555]LTC[+57.021]TLTGLR.D | HexNAc(5)Hex(3) | C[+57], N[+1502] | 1489.0627 | 3 | 4465.1735 | 4465.1536 | 0 | 4.5 | 127 | Specific | 473.05 | 430.1 | 430.1 | 7.46 | 29 | >sp\|P01876\|IGHA1_HUMAN Immunoglobulin heavy constant alpha 1 OS=Homo sapiens OX=9606 GN=IGHA1 PE=1 SV=2 | 255 | 20201117_ly_milk_1ng_glyco_hcd30_etd30_compound15.19085.19085.3 | controllerType=0 controllerNumber=1 scan=19085 | 42.6517 |
| 15476:3 | 2 | R.LSLHRPALEDLLLGSEAN[+1378.476]LTC[+57.021]TLTGLR.D | HexNAc(2)Hex(6) | C[+57], N[+1378] | 1448.0378 | 3 | 4342.0988 | 4342.074 | 0 | 5.7 | 127 | Specific | 496.35 | 469.22 | 469.22 | 7.06 | 29 | >sp\|P01876\|IGHA1_HUMAN Immunoglobulin heavy constant alpha 1 OS=Homo sapiens OX=9606 GN=IGHA1 PE=1 SV=2 | 255 | 20201117_ly_milk_1ng_glyco_hcd30_etd30_compound15.19069.19069.3 | controllerType=0 controllerNumber=1 scan=19069 | 42.5977 |
| 15507:3 | 2 | R.LSLHRPALEDLLLGSEAN[+1216.423]LTC[+57.021]TLTGLR.D | HexNAc(2)Hex(5) | C[+57], N[+1216] | 1394.0189 | 3 | 4180.042 | 4180.0211 | 0 | 5 | 127 | Specific | 512.74 | 503.31 | 503.31 | 7.06 | 29 | >sp\|P01876\|IGHA1_HUMAN Immunoglobulin heavy constant alpha 1 OS=Homo sapiens OX=9606 GN=IGHA1 PE=1 SV=2 | 255 | 20201117_ly_milk_1ng_glyco_hcd30_etd30_compound15.19102.19102.3 | controllerType=0 controllerNumber=1 scan=19102 | 42.7091 |
| 16151:4 | 2 | R.LSLHRPALEDLLLGSEAN[+2116.756]LTC[+57.021]TLTGLR.D | HexNAc(5)Hex(5)NeuAc(1) | C[+57], N[+2117] | 1270.8484 | 4 | 5080.3717 | 5080.3547 | 0 | 3.4 | 127 | Specific | 724 | 606.4 | 606.4 | 11.87 | 29 | >sp\|P01876\|IGHA1_HUMAN Immunoglobulin heavy constant alpha 1 OS=Homo sapiens OX=9606 GN=IGHA1 PE=1 SV=2 | 255 | 20201117_ly_milk_1ng_glyco_hcd30_etd30_compound15.19781.19781.4 | controllerType=0 controllerNumber=1 scan=19781 | 44.7163 |
| 15487:4 | 2 | R.LSLHRPALEDLLLGSEAN[+1419.502]LTC[+57.021]TLTGLR.D | HexNAc(3)Hex(5) | C[+57], N[+1420] | 1096.5362 | 4 | 4383.1229 | 4383.1005 | 0 | 5.1 | 127 | Specific | 758.8 | 758.8 | 758.8 | 12.32 | 29 | >sp\|P01876\|IGHA1_HUMAN Immunoglobulin heavy constant alpha 1 OS=Homo sapiens OX=9606 GN=IGHA1 PE=1 SV=2 | 255 | 20201117_ly_milk_1ng_glyco_hcd30_etd30_compound15.19080.19080.4 | controllerType=0 controllerNumber=1 scan=19080 | 42.6346 |
| 15495:4 | 2 | R.LSLHRPALEDLLLGSEAN[+1622.582]LTC[+57.021]TLTGLR.D | HexNAc(4)Hex(5) | C[+57], N[+1623] | 1147.3051 | 4 | 4586.1986 | 4586.1799 | 0 | 4.1 | 127 | Specific | 854.04 | 854.04 | 854.04 | 14.72 | 29 | >sp\|P01876\|IGHA1_HUMAN Immunoglobulin heavy constant alpha 1 OS=Homo sapiens OX=9606 GN=IGHA1 PE=1 SV=2 | 255 | 20201117_ly_milk_1ng_glyco_hcd30_etd30_compound15.19089.19089.4 | controllerType=0 controllerNumber=1 scan=19089 | 42.6596 |
| 15490:4 | 2 | R.LSLHRPALEDLLLGSEAN[+1501.555]LTC[+57.021]TLTGLR.D | HexNAc(5)Hex(3) | C[+57], N[+1502] | 1117.05 | 4 | 4465.178 | 4465.1536 | 0 | 5.5 | 127 | Specific | 864.41 | 864.41 | 864.41 | 14.32 | 29 | >sp\|P01876\|IGHA1_HUMAN Immunoglobulin heavy constant alpha 1 OS=Homo sapiens OX=9606 GN=IGHA1 PE=1 SV=2 | 255 | 20201117_ly_milk_1ng_glyco_hcd30_etd30_compound15.19084.19084.4 | controllerType=0 controllerNumber=1 scan=19084 | 42.6481 |
| 15475:4 | 2 | R.LSLHRPALEDLLLGSEAN[+1663.608]LTC[+57.021]TLTGLR.D | HexNAc(5)Hex(4) | C[+57], N[+1664] | 1157.5628 | 4 | 4627.2293 | 4627.2064 | 0 | 4.9 | 127 | Specific | 909.62 | 894.71 | 894.71 | 15.32 | 29 | >sp\|P01876\|IGHA1_HUMAN Immunoglobulin heavy constant alpha 1 OS=Homo sapiens OX=9606 GN=IGHA1 PE=1 SV=2 | 255 | 20201117_ly_milk_1ng_glyco_hcd30_etd30_compound15.19068.19068.4 | controllerType=0 controllerNumber=1 scan=19068 | 42.5945 |
| 16141:4 | 2 | R.LSLHRPALEDLLLGSEAN[+1913.677]LTC[+57.021]TLTGLR.D | HexNAc(4)Hex(5)NeuAc(1) | C[+57], N[+1914] | 1220.0776 | 4 | 4877.2884 | 4877.2753 | 0 | 2.7 | 127 | Specific | 930.82 | 896.38 | 896.38 | 15.87 | 29 | >sp\|P01876\|IGHA1_HUMAN Immunoglobulin heavy constant alpha 1 OS=Homo sapiens OX=9606 GN=IGHA1 PE=1 SV=2 | 255 | 20201117_ly_milk_1ng_glyco_hcd30_etd30_compound15.19770.19770.4 | controllerType=0 controllerNumber=1 scan=19770 | 44.6792 |
| 15437:4 | 2 | R.LSLHRPALEDLLLGSEAN[+1864.634]LTC[+57.021]TLTGLR.D | HexNAc(2)Hex(9) | C[+57], N[+1865] | 1207.8179 | 4 | 4828.2498 | 4828.2324 | 0 | 3.6 | 127 | Specific | 937.87 | 887.73 | 887.73 | 15.72 | 29 | >sp\|P01876\|IGHA1_HUMAN Immunoglobulin heavy constant alpha 1 OS=Homo sapiens OX=9606 GN=IGHA1 PE=1 SV=2 | 255 | 20201117_ly_milk_1ng_glyco_hcd30_etd30_compound15.19028.19028.4 | controllerType=0 controllerNumber=1 scan=19028 | 42.483 |
| 15438:4 | 2 | R.LSLHRPALEDLLLGSEAN[+1702.581]LTC[+57.021]TLTGLR.D | HexNAc(2)Hex(8) | C[+57], N[+1703] | 1167.3039 | 4 | 4666.1937 | 4666.1796 | 0 | 3 | 127 | Specific | 972.59 | 930.19 | 930.19 | 16.87 | 29 | >sp\|P01876\|IGHA1_HUMAN Immunoglobulin heavy constant alpha 1 OS=Homo sapiens OX=9606 GN=IGHA1 PE=1 SV=2 | 255 | 20201117_ly_milk_1ng_glyco_hcd30_etd30_compound15.19029.19029.4 | controllerType=0 controllerNumber=1 scan=19029 | 42.4874 |
| 15506:4 | 2 | R.LSLHRPALEDLLLGSEAN[+1460.529]LTC[+57.021]TLTGLR.D | HexNAc(4)Hex(4) | C[+57], N[+1461] | 1106.7962 | 4 | 4424.1629 | 4424.1271 | 0 | 8.1 | 127 | Specific | 1037.26 | 937.14 | 937.14 | 16.87 | 29 | >sp\|P01876\|IGHA1_HUMAN Immunoglobulin heavy constant alpha 1 OS=Homo sapiens OX=9606 GN=IGHA1 PE=1 SV=2 | 255 | 20201117_ly_milk_1ng_glyco_hcd30_etd30_compound15.19101.19101.4 | controllerType=0 controllerNumber=1 scan=19101 | 42.7056 |
| 15466:4 | 2 | R.LSLHRPALEDLLLGSEAN[+1378.476]LTC[+57.021]TLTGLR.D | HexNAc(2)Hex(6) | C[+57], N[+1378] | 1086.2774 | 4 | 4342.0877 | 4342.074 | 0 | 3.2 | 127 | Specific | 1042.53 | 982.82 | 982.82 | 17.87 | 29 | >sp\|P01876\|IGHA1_HUMAN Immunoglobulin heavy constant alpha 1 OS=Homo sapiens OX=9606 GN=IGHA1 PE=1 SV=2 | 255 | 20201117_ly_milk_1ng_glyco_hcd30_etd30_compound15.19058.19058.4 | controllerType=0 controllerNumber=1 scan=19058 | 42.5705 |
| 15502:4 | 2 | R.LSLHRPALEDLLLGSEAN[+1216.423]LTC[+57.021]TLTGLR.D | HexNAc(2)Hex(5) | C[+57], N[+1216] | 1045.7653 | 4 | 4180.0394 | 4180.0211 | 0 | 4.4 | 127 | Specific | 1072.87 | 1052.78 | 1052.78 | 18.72 | 29 | >sp\|P01876\|IGHA1_HUMAN Immunoglobulin heavy constant alpha 1 OS=Homo sapiens OX=9606 GN=IGHA1 PE=1 SV=2 | 255 | 20201117_ly_milk_1ng_glyco_hcd30_etd30_compound15.19096.19096.4 | controllerType=0 controllerNumber=1 scan=19096 | 42.6894 |
| 15433:4 | 2 | R.LSLHRPALEDLLLGSEAN[+1540.529]LTC[+57.021]TLTGLR.D | HexNAc(2)Hex(7) | C[+57], N[+1541] | 1126.7892 | 4 | 4504.1351 | 4504.1268 | 0 | 1.8 | 127 | Specific | 1133.66 | 996 | 996 | 19.81 | 29 | >sp\|P01876\|IGHA1_HUMAN Immunoglobulin heavy constant alpha 1 OS=Homo sapiens OX=9606 GN=IGHA1 PE=1 SV=2 | 255 | 20201117_ly_milk_1ng_glyco_hcd30_etd30_compound15.19023.19023.4 | controllerType=0 controllerNumber=1 scan=19023 | 42.4668 |
| 10323:4 | 2 | R.LAGKPTHVN[+1809.666]VSVVMAEVDGTC[+57.021]Y.- | HexNAc(5)Hex(4)Fuc(1) | C[+57], N[+1810] | 1039.9596 | 4 | 4156.8167 | 4156.8081 | 0 | 2.1 | 332 | CRagged | 206.55 | 206.55 | 46.93 | 2.91 | 29 | >sp\|P01876\|IGHA1_HUMAN Immunoglobulin heavy constant alpha 1 OS=Homo sapiens OX=9606 GN=IGHA1 PE=1 SV=2 | 255 | 20201117_ly_milk_1ng_glyco_hcd30_etd30_compound15.13640.13640.2 | controllerType=0 controllerNumber=1 scan=13640 | 30.6156 |
| 10331:3 | 2 | R.LAGKPTHVN[+1809.666]VSVVMAEVDGTC[+57.021]Y.- | HexNAc(5)Hex(4)Fuc(1) | C[+57], N[+1810] | 1386.2795 | 3 | 4156.8239 | 4156.8081 | 0 | 3.8 | 332 | CRagged | 206.71 | 89.52 | 89.52 | 2.57 | 29 | >sp\|P01876\|IGHA1_HUMAN Immunoglobulin heavy constant alpha 1 OS=Homo sapiens OX=9606 GN=IGHA1 PE=1 SV=2 | 255 | 20201117_ly_milk_1ng_glyco_hcd30_etd30_compound15.13649.13649.3 | controllerType=0 controllerNumber=1 scan=13649 | 30.6509 |
| 10237:3 | 2 | R.LAGKPTHVN[+1702.581]VSVVMAEVDGTC[+57.021]Y.- | HexNAc(2)Hex(8) | C[+57], N[+1703] | 1350.5832 | 3 | 4049.735 | 4049.7234 | 0 | 2.9 | 332 | CRagged | 233.18 | 186.62 | 174.92 | 3.13 | 29 | >sp\|P01876\|IGHA1_HUMAN Immunoglobulin heavy constant alpha 1 OS=Homo sapiens OX=9606 GN=IGHA1 PE=1 SV=2 | 255 | 20201117_ly_milk_1ng_glyco_hcd30_etd30_compound15.13550.13550.3 | controllerType=0 controllerNumber=1 scan=13550 | 30.4075 |
| 10299:3 | 2 | R.LAGKPTHVN[+1663.608]VSVVMAEVDGTC[+57.021]Y.- | HexNAc(5)Hex(4) | C[+57], N[+1664] | 1337.5895 | 3 | 4010.7541 | 4010.7502 | 0 | 1 | 332 | CRagged | 236.39 | 174.4 | 144.81 | 2.5 | 29 | >sp\|P01876\|IGHA1_HUMAN Immunoglobulin heavy constant alpha 1 OS=Homo sapiens OX=9606 GN=IGHA1 PE=1 SV=2 | 255 | 20201117_ly_milk_1ng_glyco_hcd30_etd30_compound15.13615.13615.3 | controllerType=0 controllerNumber=1 scan=13615 | 30.5524 |
| 10312:3 | 2 | R.LAGKPTHVN[+1809.666]VSVVMAEVDGTC[+57.021]Y.- | HexNAc(5)Hex(4)Fuc(1) | C[+57], N[+1810] | 1386.2765 | 3 | 4156.8151 | 4156.8081 | 0 | 1.7 | 332 | CRagged | 237.14 | 181 | 180.11 | 2.79 | 29 | >sp\|P01876\|IGHA1_HUMAN Immunoglobulin heavy constant alpha 1 OS=Homo sapiens OX=9606 GN=IGHA1 PE=1 SV=2 | 255 | 20201117_ly_milk_1ng_glyco_hcd30_etd30_compound15.13629.13629.3 | controllerType=0 controllerNumber=1 scan=13629 | 30.5971 |
| 10351:3 | 2 | R.LAGKPTHVN[+1501.555]VSVVMAEVDGTC[+57.021]Y.- | HexNAc(5)Hex(3) | C[+57], N[+1502] | 1283.5744 | 3 | 3848.7086 | 3848.6974 | 0 | 2.9 | 332 | CRagged | 254.74 | 172.33 | 172.33 | 3.22 | 29 | >sp\|P01876\|IGHA1_HUMAN Immunoglobulin heavy constant alpha 1 OS=Homo sapiens OX=9606 GN=IGHA1 PE=1 SV=2 | 255 | 20201117_ly_milk_1ng_glyco_hcd30_etd30_compound15.13670.13670.3 | controllerType=0 controllerNumber=1 scan=13670 | 30.7166 |
| 10272:3 | 2 | R.LAGKPTHVN[+1540.529]VSVVMAEVDGTC[+57.021]Y.- | HexNAc(2)Hex(7) | C[+57], N[+1541] | 1296.5646 | 3 | 3887.6793 | 3887.6706 | 0 | 2.3 | 332 | CRagged | 257.78 | 195.12 | 195.12 | 3.61 | 29 | >sp\|P01876\|IGHA1_HUMAN Immunoglobulin heavy constant alpha 1 OS=Homo sapiens OX=9606 GN=IGHA1 PE=1 SV=2 | 255 | 20201117_ly_milk_1ng_glyco_hcd30_etd30_compound15.13587.13587.3 | controllerType=0 controllerNumber=1 scan=13587 | 30.4852 |
| 10369:3 | 2 | R.LAGKPTHVN[+1647.613]VSVVMAEVDGTC[+57.021]Y.- | HexNAc(5)Hex(3)Fuc(1) | C[+57], N[+1648] | 1332.2603 | 3 | 3994.7664 | 3994.7553 | 0 | 2.8 | 332 | CRagged | 260.84 | 166.87 | 166.87 | 3.22 | 29 | >sp\|P01876\|IGHA1_HUMAN Immunoglobulin heavy constant alpha 1 OS=Homo sapiens OX=9606 GN=IGHA1 PE=1 SV=2 | 255 | 20201117_ly_milk_1ng_glyco_hcd30_etd30_compound15.13689.13689.3 | controllerType=0 controllerNumber=1 scan=13689 | 30.7579 |
| 10291:3 | 2 | R.LAGKPTHVN[+1378.476]VSVVMAEVDGTC[+57.021]Y.- | HexNAc(2)Hex(6) | C[+57], N[+1378] | 1242.5466 | 3 | 3725.6251 | 3725.6178 | 0 | 2 | 332 | CRagged | 262.68 | 262.68 | 183.82 | 3.82 | 29 | >sp\|P01876\|IGHA1_HUMAN Immunoglobulin heavy constant alpha 1 OS=Homo sapiens OX=9606 GN=IGHA1 PE=1 SV=2 | 255 | 20201117_ly_milk_1ng_glyco_hcd30_etd30_compound15.13607.13607.3 | controllerType=0 controllerNumber=1 scan=13607 | 30.5359 |
| 10335:3 | 2 | R.LAGKPTHVN[+1216.423]VSVVMAEVDGTC[+57.021]Y.- | HexNAc(2)Hex(5) | C[+57], N[+1216] | 1188.5305 | 3 | 3563.5768 | 3563.5649 | 0 | 3.3 | 332 | CRagged | 268.52 | 241.07 | 208.64 | 3.43 | 29 | >sp\|P01876\|IGHA1_HUMAN Immunoglobulin heavy constant alpha 1 OS=Homo sapiens OX=9606 GN=IGHA1 PE=1 SV=2 | 255 | 20201117_ly_milk_1ng_glyco_hcd30_etd30_compound15.13653.13653.3 | controllerType=0 controllerNumber=1 scan=13653 | 30.6616 |
| 10391:4 | 2 | R.LAGKPTHVN[+1647.613]VSVVMAEVDGTC[+57.021]Y.- | HexNAc(5)Hex(3)Fuc(1) | C[+57], N[+1648] | 999.4471 | 4 | 3994.7667 | 3994.7553 | 0 | 2.8 | 332 | CRagged | 325.75 | 325.75 | 325.75 | 3.64 | 29 | >sp\|P01876\|IGHA1_HUMAN Immunoglobulin heavy constant alpha 1 OS=Homo sapiens OX=9606 GN=IGHA1 PE=1 SV=2 | 255 | 20201117_ly_milk_1ng_glyco_hcd30_etd30_compound15.13712.13712.4 | controllerType=0 controllerNumber=1 scan=13712 | 30.8135 |
| 15481:3 | 6 | R.GLSFDVSLEVSQGPGLLN[+2205.793]DTK.V | HexNAc(4)Hex(5)Fuc(2)NeuAc(1) | N[+2206] | 1461.3151 | 3 | 4381.9308 | 4381.906 | 0 | 5.7 | 118 | Specific | 285.87 | 190.06 | 190.06 | 3.41 | 23 | >sp\|P01833\|PIGR_HUMAN Polymeric immunoglobulin receptor OS=Homo sapiens OX=9606 GN=PIGR PE=1 SV=4 | 902 | 20201117_ly_milk_1ng_glyco_hcd30_etd30_compound15.19074.19074.3 | controllerType=0 controllerNumber=1 scan=19074 | 42.6149 |
| 16806:4 | 6 | R.GLSFDVSLEVSQGPGLLN[+2350.830]DTK.V | HexNAc(4)Hex(5)Fuc(1)NeuAc(2) | N[+2351] | 1132.4952 | 4 | 4526.9588 | 4526.9436 | 0 | 3.4 | 118 | Specific | 441.8 | 402.89 | 402.89 | 5.98 | 23 | >sp\|P01833\|PIGR_HUMAN Polymeric immunoglobulin receptor OS=Homo sapiens OX=9606 GN=PIGR PE=1 SV=4 | 902 | 20201117_ly_milk_1ng_glyco_hcd30_etd30_compound15.20473.20473.4 | controllerType=0 controllerNumber=1 scan=20473 | 46.7002 |
| 16826:3 | 6 | R.GLSFDVSLEVSQGPGLLN[+2350.830]DTK.V | HexNAc(4)Hex(5)Fuc(1)NeuAc(2) | N[+2351] | 1509.6583 | 3 | 4526.9602 | 4526.9436 | 0 | 3.7 | 118 | Specific | 521.4 | 338.42 | 338.42 | 6.09 | 23 | >sp\|P01833\|PIGR_HUMAN Polymeric immunoglobulin receptor OS=Homo sapiens OX=9606 GN=PIGR PE=1 SV=4 | 902 | 20201117_ly_milk_1ng_glyco_hcd30_etd30_compound15.20495.20495.3 | controllerType=0 controllerNumber=1 scan=20495 | 46.7648 |
| 16796:3 | 6 | R.GLSFDVSLEVSQGPGLLN[+2350.830]DTK.V | HexNAc(4)Hex(5)Fuc(1)NeuAc(2) | N[+2351] | 1509.6581 | 3 | 4526.9599 | 4526.9436 | 0 | 3.6 | 118 | Specific | 548.86 | 357.35 | 357.35 | 6.09 | 23 | >sp\|P01833\|PIGR_HUMAN Polymeric immunoglobulin receptor OS=Homo sapiens OX=9606 GN=PIGR PE=1 SV=4 | 902 | 20201117_ly_milk_1ng_glyco_hcd30_etd30_compound15.20463.20463.3 | controllerType=0 controllerNumber=1 scan=20463 | 46.6682 |
| 16361:3 | 6 | K.QIGLYPVLVIDSSGYVNPN[+1913.677]YTGR.I | HexNAc(4)Hex(5)NeuAc(1) | N[+1914] | 1480.3387 | 3 | 4439.0015 | 4438.9805 | 0 | 4.7 | 168 | Specific | 410.07 | 323.01 | 323.01 | 5.51 | 23 | >sp\|P01833\|PIGR_HUMAN Polymeric immunoglobulin receptor OS=Homo sapiens OX=9606 GN=PIGR PE=1 SV=4 | 902 | 20201117_ly_milk_1ng_glyco_hcd30_etd30_compound15.20003.20003.3 | controllerType=0 controllerNumber=1 scan=20003 | 45.4742 |
| 16333:3 | 6 | K.QIGLYPVLVIDSSGYVNPN[+2059.735]YTGR.I | HexNAc(4)Hex(5)Fuc(1)NeuAc(1) | N[+2060] | 1529.0233 | 3 | 4585.0552 | 4585.0384 | 0 | 3.7 | 168 | Specific | 525.11 | 432.71 | 432.71 | 7.29 | 23 | >sp\|P01833\|PIGR_HUMAN Polymeric immunoglobulin receptor OS=Homo sapiens OX=9606 GN=PIGR PE=1 SV=4 | 902 | 20201117_ly_milk_1ng_glyco_hcd30_etd30_compound15.19974.19974.3 | controllerType=0 controllerNumber=1 scan=19974 | 45.3722 |
| 17513:3 | 6 | R.LSLLEEPGN[+2205.793]GTFTVILNQLTSR.D | HexNAc(4)Hex(5)Fuc(2)NeuAc(1) | N[+2206] | 1536.7059 | 3 | 4608.103 | 4608.0854 | 0 | 3.8 | 413 | Specific | 612.86 | 506.72 | 506.72 | 9.25 | 23 | >sp\|P01833\|PIGR_HUMAN Polymeric immunoglobulin receptor OS=Homo sapiens OX=9606 GN=PIGR PE=1 SV=4 | 902 | 20201117_ly_milk_1ng_glyco_hcd30_etd30_compound15.21223.21223.3 | controllerType=0 controllerNumber=1 scan=21223 | 48.8717 |
| 19231:3 | 6 | R.LSLLEEPGN[+2350.830]GTFTVILNQLTSR.D | HexNAc(4)Hex(5)Fuc(1)NeuAc(2) | N[+2351] | 1585.0525 | 3 | 4753.1431 | 4753.1229 | 0 | 4.2 | 413 | Specific | 618.39 | 450.66 | 450.66 | 9.25 | 23 | >sp\|P01833\|PIGR_HUMAN Polymeric immunoglobulin receptor OS=Homo sapiens OX=9606 GN=PIGR PE=1 SV=4 | 902 | 20201117_ly_milk_1ng_glyco_hcd30_etd30_compound15.23029.23029.3 | controllerType=0 controllerNumber=1 scan=23029 | 53.2326 |
| 14265:3 | 6 | K.IIEGEPNLKVPGN[+2205.793]VTAVLGETLK.V | HexNAc(4)Hex(5)Fuc(2)NeuAc(1) | N[+2206] | 1533.0568 | 3 | 4597.1559 | 4597.1422 | 0 | 3 | 457 | Specific | 286.87 | 234 | 234 | 3.52 | 23 | >sp\|P01833\|PIGR_HUMAN Polymeric immunoglobulin receptor OS=Homo sapiens OX=9606 GN=PIGR PE=1 SV=4 | 902 | 20201117_ly_milk_1ng_glyco_hcd30_etd30_compound15.17792.17792.3 | controllerType=0 controllerNumber=1 scan=17792 | 39.2405 |
| 15644:3 | 6 | K.IIEGEPNLKVPGN[+2350.830]VTAVLGETLK.V | HexNAc(4)Hex(5)Fuc(1)NeuAc(2) | N[+2351] | 1581.4046 | 3 | 4742.1992 | 4742.1797 | 0 | 4.1 | 457 | Specific | 306.48 | 269.76 | 269.76 | 3.59 | 23 | >sp\|P01833\|PIGR_HUMAN Polymeric immunoglobulin receptor OS=Homo sapiens OX=9606 GN=PIGR PE=1 SV=4 | 902 | 20201117_ly_milk_1ng_glyco_hcd30_etd30_compound15.19247.19247.3 | controllerType=0 controllerNumber=1 scan=19247 | 43.1796 |
| 14218:3 | 6 | K.IIEGEPNLKVPGN[+2205.793]VTAVLGETLK.V | HexNAc(4)Hex(5)Fuc(2)NeuAc(1) | N[+2206] | 1533.0557 | 3 | 4597.1526 | 4597.1422 | 0 | 2.3 | 457 | Specific | 365.27 | 276.45 | 276.45 | 3.78 | 23 | >sp\|P01833\|PIGR_HUMAN Polymeric immunoglobulin receptor OS=Homo sapiens OX=9606 GN=PIGR PE=1 SV=4 | 902 | 20201117_ly_milk_1ng_glyco_hcd30_etd30_compound15.17741.17741.3 | controllerType=0 controllerNumber=1 scan=17741 | 39.1234 |
| 13854:4 | 6 | K.IIEGEPNLKVPGN[+2205.793]VTAVLGETLKVPC[+57.021]HFPC[+57.021]K.F | HexNAc(4)Hex(5)Fuc(2)NeuAc(1) | C[+57]*2, N[+2206] | 1406.4095 | 4 | 5622.6164 | 5622.5997 | 0 | 3 | 457 | Specific | 307.22 | 307.22 | 307.22 | 3.09 | 23 | >sp\|P01833\|PIGR_HUMAN Polymeric immunoglobulin receptor OS=Homo sapiens OX=9606 GN=PIGR PE=1 SV=4 | 902 | 20201117_ly_milk_1ng_glyco_hcd30_etd30_compound15.17332.17332.4 | controllerType=0 controllerNumber=1 scan=17332 | 38.2671 |
| 13874:4 | 6 | K.IIEGEPNLKVPGN[+2205.793]VTAVLGETLKVPC[+57.021]HFPC[+57.021]K.F | HexNAc(4)Hex(5)Fuc(2)NeuAc(1) | C[+57]*2, N[+2206] | 1406.4111 | 4 | 5622.6227 | 5622.5997 | 0 | 4.1 | 457 | Specific | 330.77 | 330.77 | 330.77 | 3.22 | 23 | >sp\|P01833\|PIGR_HUMAN Polymeric immunoglobulin receptor OS=Homo sapiens OX=9606 GN=PIGR PE=1 SV=4 | 902 | 20201117_ly_milk_1ng_glyco_hcd30_etd30_compound15.17353.17353.4 | controllerType=0 controllerNumber=1 scan=17353 | 38.3187 |
| 13870:5 | 6 | K.IIEGEPNLKVPGN[+2205.793]VTAVLGETLKVPC[+57.021]HFPC[+57.021]K.F | HexNAc(4)Hex(5)Fuc(2)NeuAc(1) | C[+57]*2, N[+2206] | 1125.3304 | 5 | 5622.6231 | 5622.5997 | 0 | 4.2 | 457 | Specific | 677.58 | 677.58 | 677.58 | 8.48 | 23 | >sp\|P01833\|PIGR_HUMAN Polymeric immunoglobulin receptor OS=Homo sapiens OX=9606 GN=PIGR PE=1 SV=4 | 902 | 20201117_ly_milk_1ng_glyco_hcd30_etd30_compound15.17349.17349.5 | controllerType=0 controllerNumber=1 scan=17349 | 38.3063 |
| 14772:3 | 6 | K.VPGN[+2350.830]VTAVLGETLK.V | HexNAc(4)Hex(5)Fuc(1)NeuAc(2) | N[+2351] | 1250.2202 | 3 | 3748.646 | 3748.6302 | 0 | 4.2 | 466 | Specific | 322.62 | 187.36 | 187.36 | 4.43 | 23 | >sp\|P01833\|PIGR_HUMAN Polymeric immunoglobulin receptor OS=Homo sapiens OX=9606 GN=PIGR PE=1 SV=4 | 902 | 20201117_ly_milk_1ng_glyco_hcd30_etd30_compound15.18324.18324.3 | controllerType=0 controllerNumber=1 scan=18324 | 40.4081 |
| 12285:3 | 6 | K.VPGN[+2059.735]VTAVLGETLK.V | HexNAc(4)Hex(5)Fuc(1)NeuAc(1) | N[+2060] | 1153.1857 | 3 | 3457.5426 | 3457.5348 | 0 | 2.3 | 466 | Specific | 344.24 | 234.67 | 234.67 | 5.33 | 23 | >sp\|P01833\|PIGR_HUMAN Polymeric immunoglobulin receptor OS=Homo sapiens OX=9606 GN=PIGR PE=1 SV=4 | 902 | 20201117_ly_milk_1ng_glyco_hcd30_etd30_compound15.15696.15696.3 | controllerType=0 controllerNumber=1 scan=15696 | 34.6555 |
| 05109:3 | 6 | K.WN[+2059.735]NTGC[+57.021]QALPSQDEGPSK.A | HexNAc(4)Hex(5)Fuc(1)NeuAc(1) | C[+57], N[+2060] | 1350.2131 | 3 | 4048.6247 | 4048.6116 | 0 | 3.2 | 498 | Specific | 219.99 | 81.3 | 81.3 | 4.33 | 23 | >sp\|P01833\|PIGR_HUMAN Polymeric immunoglobulin receptor OS=Homo sapiens OX=9606 GN=PIGR PE=1 SV=4 | 902 | 20201117_ly_milk_1ng_glyco_hcd30_etd30_compound15.8215.8215.3 | controllerType=0 controllerNumber=1 scan=8215 | 20.7238 |
| 05242:3 | 6 | K.WN[+1710.598]NTGC[+57.021]QALPSQDEGPSK.A | HexNAc(3)Hex(5)NeuAc(1) | C[+57], N[+1711] | 1233.8323 | 3 | 3699.4824 | 3699.4743 | 0 | 2.2 | 498 | Specific | 266.7 | 135.44 | 135.44 | 4.84 | 23 | >sp\|P01833\|PIGR_HUMAN Polymeric immunoglobulin receptor OS=Homo sapiens OX=9606 GN=PIGR PE=1 SV=4 | 902 | 20201117_ly_milk_1ng_glyco_hcd30_etd30_compound15.8352.8352.3 | controllerType=0 controllerNumber=1 scan=8352 | 20.9662 |
| 05197:3 | 6 | K.WN[+1913.677]NTGC[+57.021]QALPSQDEGPSK.A | HexNAc(4)Hex(5)NeuAc(1) | C[+57], N[+1914] | 1301.5234 | 3 | 3902.5556 | 3902.5537 | 0 | 0.5 | 498 | Specific | 283.49 | 133.73 | 133.73 | 3.92 | 23 | >sp\|P01833\|PIGR_HUMAN Polymeric immunoglobulin receptor OS=Homo sapiens OX=9606 GN=PIGR PE=1 SV=4 | 902 | 20201117_ly_milk_1ng_glyco_hcd30_etd30_compound15.8306.8306.3 | controllerType=0 controllerNumber=1 scan=8306 | 20.887 |
| 05101:3 | 6 | K.WN[+2059.735]NTGC[+57.021]QALPSQDEGPSK.A | HexNAc(4)Hex(5)Fuc(1)NeuAc(1) | C[+57], N[+2060] | 1350.2136 | 3 | 4048.6261 | 4048.6116 | 0 | 3.6 | 498 | Specific | 339.37 | 189.67 | 189.67 | 5.24 | 23 | >sp\|P01833\|PIGR_HUMAN Polymeric immunoglobulin receptor OS=Homo sapiens OX=9606 GN=PIGR PE=1 SV=4 | 902 | 20201117_ly_milk_1ng_glyco_hcd30_etd30_compound15.8206.8206.3 | controllerType=0 controllerNumber=1 scan=8206 | 20.7016 |
| 05134:3 | 6 | K.WN[+1913.677]NTGC[+57.021]QALPSQDEGPSK.A | HexNAc(4)Hex(5)NeuAc(1) | C[+57], N[+1914] | 1301.5262 | 3 | 3902.564 | 3902.5537 | 0 | 2.6 | 498 | Specific | 488.57 | 298.64 | 298.64 | 6.15 | 23 | >sp\|P01833\|PIGR_HUMAN Polymeric immunoglobulin receptor OS=Homo sapiens OX=9606 GN=PIGR PE=1 SV=4 | 902 | 20201117_ly_milk_1ng_glyco_hcd30_etd30_compound15.8240.8240.3 | controllerType=0 controllerNumber=1 scan=8240 | 20.7692 |
| 07602:3 | 8 | K.TAHC[+57.021]N[+2205.793]ESFYFLC[+57.021]K.R | HexNAc(4)Hex(5)Fuc(2)NeuAc(1) | C[+57]*2, N[+2206] | 1294.8492 | 3 | 3882.533 | 3882.5124 | 0 | 5.3 | 1201 | Specific | 201.63 | 122.1 | 122.1 | 2.97 | 11 | >sp\|P22897\|MRC1_HUMAN Macrophage mannose receptor 1 OS=Homo sapiens OX=9606 GN=MRC1 PE=1 SV=1 | 1049 | 20201117_ly_milk_1ng_glyco_hcd30_etd30_compound15.10802.10802.3 | controllerType=0 controllerNumber=1 scan=10802 | 25.2386 |
| 07703:3 | 8 | K.TAHC[+57.021]N[+2059.735]ESFYFLC[+57.021]K.R | HexNAc(4)Hex(5)Fuc(1)NeuAc(1) | C[+57]*2, N[+2060] | 1246.159 | 3 | 3736.4624 | 3736.4545 | 0 | 2.1 | 1201 | Specific | 289.88 | 180.64 | 180.64 | 3.87 | 11 | >sp\|P22897\|MRC1_HUMAN Macrophage mannose receptor 1 OS=Homo sapiens OX=9606 GN=MRC1 PE=1 SV=1 | 1049 | 20201117_ly_milk_1ng_glyco_hcd30_etd30_compound15.10907.10907.3 | controllerType=0 controllerNumber=1 scan=10907 | 25.4256 |
| 05036:3 | 12 | R.LSPN[+2205.793]ASAEHLELR.W | HexNAc(4)Hex(5)Fuc(2)NeuAc(1) | N[+2206] | 1214.8559 | 3 | 3642.5531 | 3642.5421 | 0 | 3 | 52 | Specific | 362.04 | 257.38 | 257.38 | 3.7 | 9 | >sp\|Q13410\|BT1A1_HUMAN Butyrophilin subfamily 1 member A1 OS=Homo sapiens OX=9606 GN=BTN1A1 PE=1 SV=3 | 179 | 20201117_ly_milk_1ng_glyco_hcd30_etd30_compound15.8138.8138.3 | controllerType=0 controllerNumber=1 scan=8138 | 20.5686 |
| 06959:3 | 12 | R.LSPN[+2350.830]ASAEHLELR.W | HexNAc(4)Hex(5)Fuc(1)NeuAc(2) | N[+2351] | 1263.2015 | 3 | 3787.5899 | 3787.5796 | 0 | 2.7 | 52 | Specific | 372.95 | 237.06 | 237.06 | 4.27 | 9 | >sp\|Q13410\|BT1A1_HUMAN Butyrophilin subfamily 1 member A1 OS=Homo sapiens OX=9606 GN=BTN1A1 PE=1 SV=3 | 179 | 20201117_ly_milk_1ng_glyco_hcd30_etd30_compound15.10132.10132.3 | controllerType=0 controllerNumber=1 scan=10132 | 24.0618 |
| 15465:3 | 12 | K.N[+1913.677]VSC[+57.021]YIQNLLLGQEK.K | HexNAc(4)Hex(5)NeuAc(1) | C[+57], N[+1914] | 1231.5379 | 3 | 3692.5991 | 3692.5876 | 0 | 3.1 | 215 | Specific | 320.2 | 209.46 | 209.46 | 5.39 | 9 | >sp\|Q13410\|BT1A1_HUMAN Butyrophilin subfamily 1 member A1 OS=Homo sapiens OX=9606 GN=BTN1A1 PE=1 SV=3 | 179 | 20201117_ly_milk_1ng_glyco_hcd30_etd30_compound15.19057.19057.3 | controllerType=0 controllerNumber=1 scan=19057 | 42.5648 |
| 00861:3 | 13 | K.HYTN[+1809.666]SSQDVTVPC[+57.021]R.V | HexNAc(5)Hex(4)Fuc(1) | C[+57], N[+1810] | 1158.4791 | 3 | 3473.4226 | 3473.4154 | 0 | 2.1 | 89 | Specific | 395.1 | 321.37 | 321.37 | 5.81 | 16 | >sp\|P01877\|IGHA2_HUMAN Immunoglobulin heavy constant alpha 2 OS=Homo sapiens OX=9606 GN=IGHA2 PE=1 SV=4 | 576 | 20201117_ly_milk_1ng_glyco_hcd30_etd30_compound15.3802.3802.3 | controllerType=0 controllerNumber=1 scan=3802 | 12.8323 |
| 00876:3 | 13 | K.HYTN[+1647.613]SSQDVTVPC[+57.021]R.V | HexNAc(5)Hex(3)Fuc(1) | C[+57], N[+1648] | 1104.4613 | 3 | 3311.3694 | 3311.3625 | 0 | 2.1 | 89 | Specific | 489.75 | 408.86 | 408.86 | 7.4 | 16 | >sp\|P01877\|IGHA2_HUMAN Immunoglobulin heavy constant alpha 2 OS=Homo sapiens OX=9606 GN=IGHA2 PE=1 SV=4 | 576 | 20201117_ly_milk_1ng_glyco_hcd30_etd30_compound15.3817.3817.3 | controllerType=0 controllerNumber=1 scan=3817 | 12.8641 |
| 04229:3 | 13 | K.TPLTAN[+2262.814]ITK.S | HexNAc(5)Hex(5)Fuc(1)NeuAc(1) | N[+2263] | 1074.4638 | 3 | 3221.3768 | 3221.3711 | 0 | 1.8 | 200 | Specific | 203.83 | 103.94 | 103.94 | 3.13 | 16 | >sp\|P01877\|IGHA2_HUMAN Immunoglobulin heavy constant alpha 2 OS=Homo sapiens OX=9606 GN=IGHA2 PE=1 SV=4 | 576 | 20201117_ly_milk_1ng_glyco_hcd30_etd30_compound15.7303.7303.3 | controllerType=0 controllerNumber=1 scan=7303 | 19.1439 |
| 02518:3 | 13 | K.TPLTAN[+1809.666]ITK.S | HexNAc(5)Hex(4)Fuc(1) | N[+1810] | 923.4145 | 3 | 2768.2288 | 2768.2228 | 0 | 2.2 | 200 | Specific | 297.74 | 145.69 | 145.69 | 2.54 | 16 | >sp\|P01877\|IGHA2_HUMAN Immunoglobulin heavy constant alpha 2 OS=Homo sapiens OX=9606 GN=IGHA2 PE=1 SV=4 | 576 | 20201117_ly_milk_1ng_glyco_hcd30_etd30_compound15.5526.5526.3 | controllerType=0 controllerNumber=1 scan=5526 | 15.9728 |
| 02521:2 | 13 | K.TPLTAN[+1809.666]ITK.S | HexNAc(5)Hex(4)Fuc(1) | N[+1810] | 1384.6191 | 2 | 2768.2308 | 2768.2228 | 0 | 2.9 | 200 | Specific | 299.73 | 110.98 | 110.98 | 2.56 | 16 | >sp\|P01877\|IGHA2_HUMAN Immunoglobulin heavy constant alpha 2 OS=Homo sapiens OX=9606 GN=IGHA2 PE=1 SV=4 | 576 | 20201117_ly_milk_1ng_glyco_hcd30_etd30_compound15.5530.5530.2 | controllerType=0 controllerNumber=1 scan=5530 | 15.987 |
| 02571:3 | 13 | K.TPLTAN[+1647.613]ITK.S | HexNAc(5)Hex(3)Fuc(1) | N[+1648] | 869.3976 | 3 | 2606.1783 | 2606.17 | 0 | 3.2 | 200 | Specific | 303.33 | 131.13 | 131.13 | 2.53 | 16 | >sp\|P01877\|IGHA2_HUMAN Immunoglobulin heavy constant alpha 2 OS=Homo sapiens OX=9606 GN=IGHA2 PE=1 SV=4 | 576 | 20201117_ly_milk_1ng_glyco_hcd30_etd30_compound15.5582.5582.3 | controllerType=0 controllerNumber=1 scan=5582 | 16.1168 |
| 02565:2 | 13 | K.TPLTAN[+1647.613]ITK.S | HexNAc(5)Hex(3)Fuc(1) | N[+1648] | 1303.5932 | 2 | 2606.1791 | 2606.17 | 0 | 3.5 | 200 | Specific | 312.12 | 116.99 | 116.99 | 2.61 | 16 | >sp\|P01877\|IGHA2_HUMAN Immunoglobulin heavy constant alpha 2 OS=Homo sapiens OX=9606 GN=IGHA2 PE=1 SV=4 | 576 | 20201117_ly_milk_1ng_glyco_hcd30_etd30_compound15.5576.5576.2 | controllerType=0 controllerNumber=1 scan=5576 | 16.1021 |
| 02620:2 | 13 | K.TPLTAN[+1444.534]ITK.S | HexNAc(4)Hex(3)Fuc(1) | N[+1445] | 1202.0536 | 2 | 2403.1 | 2403.0906 | 0 | 3.9 | 200 | Specific | 330.2 | 99.65 | 99.65 | 2.5 | 16 | >sp\|P01877\|IGHA2_HUMAN Immunoglobulin heavy constant alpha 2 OS=Homo sapiens OX=9606 GN=IGHA2 PE=1 SV=4 | 576 | 20201117_ly_milk_1ng_glyco_hcd30_etd30_compound15.5633.5633.2 | controllerType=0 controllerNumber=1 scan=5633 | 16.2309 |
| 09450:3 | 13 | R.MAGKPTHIN[+1647.613]VSVVMAEADGTC[+57.021]Y.- | HexNAc(5)Hex(3)Fuc(1) | C[+57], N[+1648] | 1333.5723 | 3 | 3998.7024 | 3998.6961 | 0 | 1.6 | 319 | CRagged | 222.41 | 148.98 | 148.98 | 2.91 | 16 | >sp\|P01877\|IGHA2_HUMAN Immunoglobulin heavy constant alpha 2 OS=Homo sapiens OX=9606 GN=IGHA2 PE=1 SV=4 | 576 | 20201117_ly_milk_1ng_glyco_hcd30_etd30_compound15.12735.12735.3 | controllerType=0 controllerNumber=1 scan=12735 | 28.9373 |
| 09428:3 | 13 | R.MAGKPTHIN[+1501.555]VSVVMAEADGTC[+57.021]Y.- | HexNAc(5)Hex(3) | C[+57], N[+1502] | 1284.8865 | 3 | 3852.645 | 3852.6382 | 0 | 1.8 | 319 | CRagged | 231.89 | 160.51 | 160.51 | 2.91 | 16 | >sp\|P01877\|IGHA2_HUMAN Immunoglobulin heavy constant alpha 2 OS=Homo sapiens OX=9606 GN=IGHA2 PE=1 SV=4 | 576 | 20201117_ly_milk_1ng_glyco_hcd30_etd30_compound15.12712.12712.3 | controllerType=0 controllerNumber=1 scan=12712 | 28.8933 |
| 09315:3 | 13 | R.MAGKPTHIN[+1540.529]VSVVMAEADGTC[+57.021]Y.- | HexNAc(2)Hex(7) | C[+57], N[+1541] | 1297.8788 | 3 | 3891.622 | 3891.6114 | 0 | 2.7 | 319 | CRagged | 240 | 238.18 | 154.8 | 2.98 | 16 | >sp\|P01877\|IGHA2_HUMAN Immunoglobulin heavy constant alpha 2 OS=Homo sapiens OX=9606 GN=IGHA2 PE=1 SV=4 | 576 | 20201117_ly_milk_1ng_glyco_hcd30_etd30_compound15.12595.12595.3 | controllerType=0 controllerNumber=1 scan=12595 | 28.6205 |
| 09414:3 | 13 | R.MAGKPTHIN[+1216.423]VSVVMAEADGTC[+57.021]Y.- | HexNAc(2)Hex(5) | C[+57], N[+1216] | 1189.8425 | 3 | 3567.5128 | 3567.5057 | 0 | 2 | 319 | CRagged | 241.72 | 235.62 | 191.55 | 3.61 | 16 | >sp\|P01877\|IGHA2_HUMAN Immunoglobulin heavy constant alpha 2 OS=Homo sapiens OX=9606 GN=IGHA2 PE=1 SV=4 | 576 | 20201117_ly_milk_1ng_glyco_hcd30_etd30_compound15.12698.12698.3 | controllerType=0 controllerNumber=1 scan=12698 | 28.8485 |

Table S4. Cont.

| Query #:z | Protein Rank | Glycopeptide | Glycan | Modification Type(s) | Observed m/z | z | Observed (M+H) | Calc. mass (M+H) | Off-by-x error | Mass error (ppm) | StartPosition | Cleavage | Score | Delta | Delta Mod | \|Log Prob\| | # of unique peptides | Protein Name | Protein DB number | Comment | Scan # | Scan Time |
|---|---|---|---|---|---|---|---|---|---|---|---|---|---|---|---|---|---|---|---|---|---|---|
| 09332:3 | 13 | R.MAGKPTHIN[+1378.476]VSVVMAEADGTC[+57.021]Y.- | HexNAc(2)Hex(6) | C[+57], N[+1378] | 1243.8622 | 3 | 3729.5722 | 3729.5585 | 0 | 3.7 | 319 | CRagged | 260.45 | 232.53 | 200.05 | 3.52 | 16 | >sp\|P01877\|IGHA2_HUMAN Immunoglobulin heavy constant alpha 2 OS=Homo sapiens OX=9606 GN=IGHA2 PE=1 SV=4 | 576 | 20201117_ly_milk_1ng_glyco_hcd30_etd30_compound15.12613.12613.3 | controllerType=0 controllerNumber=1 scan=12613 | 28.6697 |
| 09291:3 | 13 | R.MAGKPTHIN[+1702.581]VSVVMAEADGTC[+57.021]Y.- | HexNAc(2)Hex(8) | C[+57], N[+1703] | 1351.8978 | 3 | 4053.6787 | 4053.6642 | 0 | 3.6 | 319 | CRagged | 280.08 | 280.08 | 181.99 | 4.4 | 16 | >sp\|P01877\|IGHA2_HUMAN Immunoglobulin heavy constant alpha 2 OS=Homo sapiens OX=9606 GN=IGHA2 PE=1 SV=4 | 576 | 20201117_ly_milk_1ng_glyco_hcd30_etd30_compound15.12569.12569.3 | controllerType=0 controllerNumber=1 scan=12569 | 28.5468 |
| 20199:3 | 25 | R.QLAHQSN[+2204.772]STNIFFSPVSIATAFAMLSLGTK.A | HexNAc(4)Hex(5)NeuAc(2) | N[+2205] | 1796.1482 | 3 | 5386.4299 | 5386.4075 | 0 | 4.2 | 64 | Specific | 359.61 | 252.97 | 252.97 | 4.64 | 3 | >sp\|P01009\|A1AT_HUMAN Alpha-1-antitrypsin OS=Homo sapiens OX=9606 GN=SERPINA1 PE=1 SV=3 | 642 | 20201117_ly_milk_1ng_glyco_hcd30_etd30_compound15.24049.24049.3 | controllerType=0 controllerNumber=1 scan=24049 | 55.8665 |
| 20208:5 | 25 | R.QLAHQSN[+2204.772]STNIFFSPVSIATAFAMLSLGTK.A | HexNAc(4)Hex(5)NeuAc(2) | N[+2205] | 1078.0966 | 5 | 5386.4537 | 5386.4075 | 0 | 8.6 | 64 | Specific | 381.29 | 381.29 | 381.29 | 4.84 | 3 | >sp\|P01009\|A1AT_HUMAN Alpha-1-antitrypsin OS=Homo sapiens OX=9606 GN=SERPINA1 PE=1 SV=3 | 642 | 20201117_ly_milk_1ng_glyco_hcd30_etd30_compound15.24059.24059.5 | controllerType=0 controllerNumber=1 scan=24059 | 55.9067 |
| 20194:4 | 25 | R.QLAHQSN[+2204.772]STNIFFSPVSIATAFAMLSLGTK.A | HexNAc(4)Hex(5)NeuAc(2) | N[+2205] | 1347.3606 | 4 | 5386.4206 | 5386.4075 | 0 | 2.4 | 64 | Specific | 681.7 | 579.21 | 430.98 | 9.96 | 3 | >sp\|P01009\|A1AT_HUMAN Alpha-1-antitrypsin OS=Homo sapiens OX=9606 GN=SERPINA1 PE=1 SV=3 | 642 | 20201117_ly_milk_1ng_glyco_hcd30_etd30_compound15.24044.24044.4 | controllerType=0 controllerNumber=1 scan=24044 | 55.8549 |
| 13085:4 | 28 | K.MVSHHN[+2204.772]LTTGATLINEQWLLTTAK.N | HexNAc(4)Hex(5)NeuAc(2) | N[+2205] | 1221.8012 | 4 | 4884.1829 | 4884.1647 | 0 | 3.7 | 179 | Specific | 264.3 | 264.3 | 264.3 | 3.96 | 4 | >sp\|P00738\|HPT_HUMAN Haptoglobin OS=Homo sapiens OX=9606 GN=HP PE=1 SV=1 | 407 | 20201117_ly_milk_1ng_glyco_hcd30_etd30_compound15.16529.16529.3 | controllerType=0 controllerNumber=1 scan=16529 | 36.4604 |
| 08734:3 | 29 | R.IIVPLNNREN[+1809.666]ISDPTSPLR.T | HexNAc(5)Hex(4)Fuc(1) | N[+1810] | 1319.9545 | 3 | 3957.849 | 3957.8432 | 0 | 1.5 | 62 | Specific | 234.4 | 133.55 | 133.55 | 2.74 | 4 | >sp\|P01591\|IGJ_HUMAN Immunoglobulin J chain OS=Homo sapiens OX=9606 GN=JCHAIN PE=1 SV=4 | 1392 | 20201117_ly_milk_1ng_glyco_hcd30_etd30_compound15.11992.11992.3 | controllerType=0 controllerNumber=1 scan=11992 | 27.5097 |
| 11983:3 | 29 | R.IIVPLNNREN[+2350.830]ISDPTSPLR.T | HexNAc(4)Hex(5)Fuc(1)NeuAc(2) | N[+2351] | 1500.3438 | 3 | 4499.0169 | 4499.0075 | 0 | 2.1 | 62 | Specific | 258.22 | 200.83 | 200.83 | 4.24 | 4 | >sp\|P01591\|IGJ_HUMAN Immunoglobulin J chain OS=Homo sapiens OX=9606 GN=JCHAIN PE=1 SV=4 | 1392 | 20201117_ly_milk_1ng_glyco_hcd30_etd30_compound15.15381.15381.3 | controllerType=0 controllerNumber=1 scan=15381 | 33.9565 |
| 10216:3 | 29 | R.IIVPLNNREN[+2059.735]ISDPTSPLR.T | HexNAc(4)Hex(5)Fuc(1)NeuAc(1) | N[+2060] | 1403.3124 | 3 | 4207.9228 | 4207.9121 | 0 | 2.5 | 62 | Specific | 281.76 | 212.21 | 212.21 | 3.99 | 4 | >sp\|P01591\|IGJ_HUMAN Immunoglobulin J chain OS=Homo sapiens OX=9606 GN=JCHAIN PE=1 SV=4 | 1392 | 20201117_ly_milk_1ng_glyco_hcd30_etd30_compound15.13529.13529.3 | controllerType=0 controllerNumber=1 scan=13529 | 30.3624 |
| 12628:4 | 42 | K.LKELPGVC[+57.021]N[+2059.735]ETMMALWEEC[+57.021]KPC[+57.021]LK.Q | HexNAc(4)Hex(5)Fuc(1)NeuAc(1) | C[+57]*3, N[+2060] | 1249.7893 | 4 | 4996.1354 | 4996.1196 | 0 | 3.2 | 95 | Specific | 468.69 | 348.99 | 348.99 | 4.37 | 7 | >sp\|P10909\|CLUS_HUMAN Clusterin OS=Homo sapiens OX=9606 GN=CLU PE=1 SV=1 | 1467 | 20201117_ly_milk_1ng_glyco_hcd30_etd30_compound15.16053.16053.4 | controllerType=0 controllerNumber=1 scan=16053 | 35.4221 |
| 00074:4 | 42 | R.EIRHN[+2059.735]STGC[+57.021]LR.M | HexNAc(4)Hex(5)Fuc(1)NeuAc(1) | C[+57], N[+2060] | 851.3571 | 4 | 3402.4067 | 3402.3994 | 0 | 2.2 | 287 | Specific | 360.57 | 274.14 | 274.14 | 4.22 | 7 | >sp\|P10909\|CLUS_HUMAN Clusterin OS=Homo sapiens OX=9606 GN=CLU PE=1 SV=1 | 1467 | 20201117_ly_milk_1ng_glyco_hcd30_etd30_compound15.2984.2984.4 | controllerType=0 controllerNumber=1 scan=2984 | 11.3055 |
| 19638:3 | 42 | K.MLN[+2350.830]TSSLLEQLNEQFNWVSR.L | HexNAc(4)Hex(5)Fuc(1)NeuAc(2) | N[+2351] | 1587.35 | 3 | 4760.0355 | 4760.0171 | 0 | 3.9 | 352 | Specific | 489.94 | 344.84 | 344.84 | 5.8 | 7 | >sp\|P10909\|CLUS_HUMAN Clusterin OS=Homo sapiens OX=9606 GN=CLU PE=1 SV=1 | 1467 | 20201117_ly_milk_1ng_glyco_hcd30_etd30_compound15.23461.23461.3 | controllerType=0 controllerNumber=1 scan=23461 | 54.4874 |
| 11817:3 | 42 | R.LAN[+2350.830]LTQGEDQYYLR.V | HexNAc(4)Hex(5)Fuc(1)NeuAc(2) | N[+2351] | 1345.5613 | 3 | 4034.6695 | 4034.664 | 0 | 1.3 | 372 | Specific | 224.26 | 131.42 | 131.42 | 3.7 | 7 | >sp\|P10909\|CLUS_HUMAN Clusterin OS=Homo sapiens OX=9606 GN=CLU PE=1 SV=1 | 1467 | 20201117_ly_milk_1ng_glyco_hcd30_etd30_compound15.15207.15207.3 | controllerType=0 controllerNumber=1 scan=15207 | 33.5429 |
| 08619:4 | 47 | K.QTDEIKDTRN[+2059.735]ESTQNC[+57.021]VVAEPEKMESSISSSSEEMSLSK.C | HexNAc(4)Hex(5)Fuc(1)NeuAc(1) | C[+57], N[+2060] | 1613.4377 | 4 | 6450.7292 | 6450.7156 | 0 | 2.1 | 60 | Specific | 216.19 | 216.19 | 216.19 | 2.74 | 5 | >sp\|P47710\|CASA1_HUMAN Alpha-S1-casein OS=Homo sapiens OX=9606 GN=CSN1S1 PE=1 SV=1 | 989 | 20201117_ly_milk_1ng_glyco_hcd30_etd30_compound15.11873.11873.4 | controllerType=0 controllerNumber=1 scan=11873 | 27.2918 |
| 13519:3 | 1 | R.LSLHRPALEDLLLGSEAN[+1702.581]LTC[+57.021]TLTGLR.D | HexNAc(2)Hex(8) | C[+57], N[+1703] | 1556.07 | 3 | 4666.1955 | 4666.1796 | 0 | 3.4 | 127 | Specific | 298.68 | 287.86 | 287.86 | 5.14 | 64 | >sp\|P01876\|IGHA1_HUMAN Immunoglobulin heavy constant alpha 1 OS=Homo sapiens OX=9606 GN=IGHA1 PE=1 SV=2 | 255 | 20201113_gy_colo_milk_1ng_glyco_hcd30_etd30_compound15.16966.16966.3 | controllerType=0 controllerNumber=1 scan=16966 | 42.5365 |
| 13607:3 | 1 | R.LSLHRPALEDLLLGSEAN[+1460.529]LTC[+57.021]TLTGLR.D | HexNAc(4)Hex(4) | C[+57], N[+1461] | 1475.3849 | 3 | 4424.1401 | 4424.1271 | 0 | 3 | 127 | Specific | 313.99 | 291.49 | 291.49 | 6.28 | 64 | >sp\|P01876\|IGHA1_HUMAN Immunoglobulin heavy constant alpha 1 OS=Homo sapiens OX=9606 GN=IGHA1 PE=1 SV=2 | 255 | 20201113_gy_colo_milk_1ng_glyco_hcd30_etd30_compound15.17059.17059.3 | controllerType=0 controllerNumber=1 scan=17059 | 42.8422 |
| 14977:4 | 1 | R.LSLHRPALEDLLLGSEAN[+2204.772]LTC[+57.021]TLTGLR.D | HexNAc(4)Hex(5)NeuAc(2) | C[+57], N[+2205] | 1292.8519 | 4 | 5168.3859 | 5168.3707 | 0 | 2.9 | 127 | Specific | 334.83 | 334.83 | 334.83 | 5.52 | 64 | >sp\|P01876\|IGHA1_HUMAN Immunoglobulin heavy constant alpha 1 OS=Homo sapiens OX=9606 GN=IGHA1 PE=1 SV=2 | 255 | 20201113_gy_colo_milk_1ng_glyco_hcd30_etd30_compound15.18509.18509.4 | controllerType=0 controllerNumber=1 scan=18509 | 47.2925 |
| 14168:3 | 1 | R.LSLHRPALEDLLLGSEAN[+2116.756]LTC[+57.021]TLTGLR.D | HexNAc(5)Hex(5)NeuAc(1) | C[+57], N[+2117] | 1694.1257 | 3 | 5080.3625 | 5080.3547 | 0 | 1.5 | 127 | Specific | 382.81 | 270.74 | 270.74 | 5.29 | 64 | >sp\|P01876\|IGHA1_HUMAN Immunoglobulin heavy constant alpha 1 OS=Homo sapiens OX=9606 GN=IGHA1 PE=1 SV=2 | 255 | 20201113_gy_colo_milk_1ng_glyco_hcd30_etd30_compound15.17654.17654.3 | controllerType=0 controllerNumber=1 scan=17654 | 44.7146 |
| 13500:3 | 1 | R.LSLHRPALEDLLLGSEAN[+1663.608]LTC[+57.021]TLTGLR.D | HexNAc(5)Hex(4) | C[+57], N[+1664] | 1543.0766 | 3 | 4627.2152 | 4627.2064 | 0 | 1.9 | 127 | Specific | 398.36 | 345.33 | 345.33 | 6.66 | 64 | >sp\|P01876\|IGHA1_HUMAN Immunoglobulin heavy constant alpha 1 OS=Homo sapiens OX=9606 GN=IGHA1 PE=1 SV=2 | 255 | 20201113_gy_colo_milk_1ng_glyco_hcd30_etd30_compound15.16946.16946.3 | controllerType=0 controllerNumber=1 scan=16946 | 42.4724 |
| 13562:3 | 1 | R.LSLHRPALEDLLLGSEAN[+1298.476]LTC[+57.021]TLTGLR.D | HexNAc(4)Hex(3) | C[+57], N[+1298] | 1421.367 | 3 | 4262.0864 | 4262.0742 | 0 | 2.9 | 127 | Specific | 408.17 | 408.17 | 408.17 | 7.86 | 64 | >sp\|P01876\|IGHA1_HUMAN Immunoglobulin heavy constant alpha 1 OS=Homo sapiens OX=9606 GN=IGHA1 PE=1 SV=2 | 255 | 20201113_gy_colo_milk_1ng_glyco_hcd30_etd30_compound15.17011.17011.3 | controllerType=0 controllerNumber=1 scan=17011 | 42.681 |
| 13538:3 | 1 | R.LSLHRPALEDLLLGSEAN[+1501.555]LTC[+57.021]TLTGLR.D | HexNAc(5)Hex(3) | C[+57], N[+1502] | 1489.0654 | 3 | 4465.1815 | 4465.1536 | 0 | 6.3 | 127 | Specific | 410.9 | 409.93 | 409.93 | 6.6 | 64 | >sp\|P01876\|IGHA1_HUMAN Immunoglobulin heavy constant alpha 1 OS=Homo sapiens OX=9606 GN=IGHA1 PE=1 SV=2 | 255 | 20201113_gy_colo_milk_1ng_glyco_hcd30_etd30_compound15.16986.16986.3 | controllerType=0 controllerNumber=1 scan=16986 | 42.6056 |
| 13528:5 | 1 | R.LSLHRPALEDLLLGSEAN[+1663.608]LTC[+57.021]TLTGLR.D | HexNAc(5)Hex(4) | C[+57], N[+1664] | 926.2478 | 5 | 4627.21 | 4627.2064 | 0 | 0.8 | 127 | Specific | 414.71 | 400.88 | 400.88 | 6.32 | 64 | >sp\|P01876\|IGHA1_HUMAN Immunoglobulin heavy constant alpha 1 OS=Homo sapiens OX=9606 GN=IGHA1 PE=1 SV=2 | 255 | 20201113_gy_colo_milk_1ng_glyco_hcd30_etd30_compound15.16975.16975.3 | controllerType=0 controllerNumber=1 scan=16975 | 42.5675 |
| 14964:3 | 1 | R.LSLHRPALEDLLLGSEAN[+2204.772]LTC[+57.021]TLTGLR.D | HexNAc(4)Hex(5)NeuAc(2) | C[+57], N[+2205] | 1723.4715 | 3 | 5168.4 | 5168.3707 | 0 | 5.7 | 127 | Specific | 427.25 | 309.52 | 309.52 | 4.21 | 64 | >sp\|P01876\|IGHA1_HUMAN Immunoglobulin heavy constant alpha 1 OS=Homo sapiens OX=9606 GN=IGHA1 PE=1 SV=2 | 255 | 20201113_gy_colo_milk_1ng_glyco_hcd30_etd30_compound15.18495.18495.3 | controllerType=0 controllerNumber=1 scan=18495 | 47.2483 |
| 13566:3 | 1 | R.LSLHRPALEDLLLGSEAN[+1378.476]LTC[+57.021]TLTGLR.D | HexNAc(2)Hex(6) | C[+57], N[+1378] | 1448.0366 | 3 | 4342.0951 | 4342.074 | 0 | 4.9 | 127 | Specific | 429.44 | 377.76 | 377.76 | 6.6 | 64 | >sp\|P01876\|IGHA1_HUMAN Immunoglobulin heavy constant alpha 1 OS=Homo sapiens OX=9606 GN=IGHA1 PE=1 SV=2 | 255 | 20201113_gy_colo_milk_1ng_glyco_hcd30_etd30_compound15.17016.17016.3 | controllerType=0 controllerNumber=1 scan=17016 | 42.6944 |
| 13467:3 | 1 | R.LSLHRPALEDLLLGSEAN[+1864.634]LTC[+57.021]TLTGLR.D | HexNAc(2)Hex(9) | C[+57], N[+1865] | 1610.0859 | 3 | 4828.2431 | 4828.2324 | 0 | 2.2 | 127 | Specific | 430.02 | 381.03 | 381.03 | 7.86 | 64 | >sp\|P01876\|IGHA1_HUMAN Immunoglobulin heavy constant alpha 1 OS=Homo sapiens OX=9606 GN=IGHA1 PE=1 SV=2 | 255 | 20201113_gy_colo_milk_1ng_glyco_hcd30_etd30_compound15.16911.16911.3 | controllerType=0 controllerNumber=1 scan=16911 | 42.3772 |
| 14204:3 | 1 | R.LSLHRPALEDLLLGSEAN[+1710.598]LTC[+57.021]TLTGLR.D | HexNAc(3)Hex(5)NeuAc(1) | C[+57], N[+1711] | 1558.7494 | 3 | 4674.2338 | 4674.1959 | 0 | 8.1 | 127 | Specific | 433.41 | 321.98 | 321.98 | 4.87 | 64 | >sp\|P01876\|IGHA1_HUMAN Immunoglobulin heavy constant alpha 1 OS=Homo sapiens OX=9606 GN=IGHA1 PE=1 SV=2 | 255 | 20201113_gy_colo_milk_1ng_glyco_hcd30_etd30_compound15.17692.17692.3 | controllerType=0 controllerNumber=1 scan=17692 | 44.828 |
| 13496:3 | 1 | R.LSLHRPALEDLLLGSEAN[+1540.529]LTC[+57.021]TLTGLR.D | HexNAc(2)Hex(7) | C[+57], N[+1541] | 1502.0497 | 3 | 4504.1347 | 4504.1268 | 0 | 1.8 | 127 | Specific | 436.14 | 396.77 | 396.77 | 7.86 | 64 | >sp\|P01876\|IGHA1_HUMAN Immunoglobulin heavy constant alpha 1 OS=Homo sapiens OX=9606 GN=IGHA1 PE=1 SV=2 | 255 | 20201113_gy_colo_milk_1ng_glyco_hcd30_etd30_compound15.16942.16942.3 | controllerType=0 controllerNumber=1 scan=16942 | 42.4651 |
| 14140:3 | 1 | R.LSLHRPALEDLLLGSEAN[+1913.677]LTC[+57.021]TLTGLR.D | HexNAc(4)Hex(5)NeuAc(1) | C[+57], N[+1914] | 1626.4413 | 3 | 4877.3095 | 4877.2753 | 0 | 7 | 127 | Specific | 441.12 | 344.21 | 344.21 | 4.25 | 64 | >sp\|P01876\|IGHA1_HUMAN Immunoglobulin heavy constant alpha 1 OS=Homo sapiens OX=9606 GN=IGHA1 PE=1 SV=2 | 255 | 20201113_gy_colo_milk_1ng_glyco_hcd30_etd30_compound15.17624.17624.4 | controllerType=0 controllerNumber=1 scan=17624 | 44.6145 |
| 13597:3 | 1 | R.LSLHRPALEDLLLGSEAN[+1095.397]LTC[+57.021]TLTGLR.D | HexNAc(3)Hex(3) | C[+57], N[+1095] | 1353.6723 | 3 | 4059.0023 | 4058.9949 | 0 | 1.8 | 127 | Specific | 452.15 | 451.83 | 451.83 | 8.03 | 64 | >sp\|P01876\|IGHA1_HUMAN Immunoglobulin heavy constant alpha 1 OS=Homo sapiens OX=9606 GN=IGHA1 PE=1 SV=2 | 255 | 20201113_gy_colo_milk_1ng_glyco_hcd30_etd30_compound15.17048.17048.3 | controllerType=0 controllerNumber=1 scan=17048 | 42.805 |
| 13544:3 | 1 | R.LSLHRPALEDLLLGSEAN[+1216.423]LTC[+57.021]TLTGLR.D | HexNAc(2)Hex(5) | C[+57], N[+1216] | 1394.0196 | 3 | 4180.0442 | 4180.0211 | 0 | 5.5 | 127 | Specific | 463.16 | 463.16 | 463.16 | 6.78 | 64 | >sp\|P01876\|IGHA1_HUMAN Immunoglobulin heavy constant alpha 1 OS=Homo sapiens OX=9606 GN=IGHA1 PE=1 SV=2 | 255 | 20201113_gy_colo_milk_1ng_glyco_hcd30_etd30_compound15.16992.16992.3 | controllerType=0 controllerNumber=1 scan=16992 | 42.6228 |
| 13571:3 | 1 | R.LSLHRPALEDLLLGSEAN[+1216.423]LTC[+57.021]TLTGLR.D | HexNAc(2)Hex(5) | C[+57], N[+1216] | 1394.0141 | 3 | 4180.0277 | 4180.0211 | 0 | 1.6 | 127 | Specific | 477.13 | 432.07 | 432.07 | 8.03 | 64 | >sp\|P01876\|IGHA1_HUMAN Immunoglobulin heavy constant alpha 1 OS=Homo sapiens OX=9606 GN=IGHA1 PE=1 SV=2 | 255 | 20201113_gy_colo_milk_1ng_glyco_hcd30_etd30_compound15.17021.17021.2 | controllerType=0 controllerNumber=1 scan=17021 | 42.7059 |
| 14241:3 | 1 | R.LSLHRPALEDLLLGSEAN[+1548.545]LTC[+57.021]TLTGLR.D | HexNAc(3)Hex(4)NeuAc(1) | C[+57], N[+1549] | 1504.7219 | 3 | 4512.151 | 4512.1431 | 0 | 1.8 | 127 | Specific | 482.94 | 381.6 | 381.6 | 8.22 | 64 | >sp\|P01876\|IGHA1_HUMAN Immunoglobulin heavy constant alpha 1 OS=Homo sapiens OX=9606 GN=IGHA1 PE=1 SV=2 | 255 | 20201113_gy_colo_milk_1ng_glyco_hcd30_etd30_compound15.17731.17731.3 | controllerType=0 controllerNumber=1 scan=17731 | 44.9534 |
| 14199:3 | 1 | R.LSLHRPALEDLLLGSEAN[+1751.624]LTC[+57.021]TLTGLR.D | HexNAc(4)Hex(4)NeuAc(1) | C[+57], N[+1752] | 1572.4241 | 3 | 4715.2578 | 4715.2225 | 0 | 7.5 | 127 | Specific | 492.13 | 455.48 | 455.48 | 6.61 | 64 | >sp\|P01876\|IGHA1_HUMAN Immunoglobulin heavy constant alpha 1 OS=Homo sapiens OX=9606 GN=IGHA1 PE=1 SV=2 | 255 | 20201113_gy_colo_milk_1ng_glyco_hcd30_etd30_compound15.17687.17687.3 | controllerType=0 controllerNumber=1 scan=17687 | 44.8202 |
| 13636:3 | 1 | R.LSLHRPALEDLLLGSEAN[+892.317]LTC[+57.021]TLTGLR.D | HexNAc(2)Hex(3) | C[+57], N[+892] | 1285.9805 | 3 | 3855.927 | 3855.9155 | 0 | 3 | 127 | Specific | 539.66 | 535.44 | 535.44 | 8.27 | 64 | >sp\|P01876\|IGHA1_HUMAN Immunoglobulin heavy constant alpha 1 OS=Homo sapiens OX=9606 GN=IGHA1 PE=1 SV=2 | 255 | 20201113_gy_colo_milk_1ng_glyco_hcd30_etd30_compound15.17090.17090.3 | controllerType=0 controllerNumber=1 scan=17090 | 42.9504 |
| 13590:4 | 1 | R.LSLHRPALEDLLLGSEAN[+1647.613]LTC[+57.021]TLTGLR.D | HexNAc(5)Hex(3)Fuc(1) | C[+57], N[+1648] | 1153.56 | 4 | 4611.2181 | 4611.2115 | 0 | 1.4 | 127 | Specific | 598.4 | 598.4 | 598.4 | 9.63 | 64 | >sp\|P01876\|IGHA1_HUMAN Immunoglobulin heavy constant alpha 1 OS=Homo sapiens OX=9606 GN=IGHA1 PE=1 SV=2 | 255 | 20201113_gy_colo_milk_1ng_glyco_hcd30_etd30_compound15.17041.17041.4 | controllerType=0 controllerNumber=1 scan=17041 | 42.7779 |
| 13553:4 | 1 | R.LSLHRPALEDLLLGSEAN[+1419.502]LTC[+57.021]TLTGLR.D | HexNAc(3)Hex(5) | C[+57], N[+1420] | 1096.5335 | 4 | 4383.1121 | 4383.1005 | 0 | 2.7 | 127 | Specific | 648.3 | 648.3 | 648.3 | 10.63 | 64 | >sp\|P01876\|IGHA1_HUMAN Immunoglobulin heavy constant alpha 1 OS=Homo sapiens OX=9606 GN=IGHA1 PE=1 SV=2 | 255 | 20201113_gy_colo_milk_1ng_glyco_hcd30_etd30_compound15.17002.17002.4 | controllerType=0 controllerNumber=1 scan=17002 | 42.6498 |
| 13539:4 | 1 | R.LSLHRPALEDLLLGSEAN[+1622.582]LTC[+57.021]TLTGLR.D | HexNAc(4)Hex(5) | C[+57], N[+1623] | 1147.3067 | 4 | 4586.2049 | 4586.1799 | 0 | 5.5 | 127 | Specific | 688.7 | 677.93 | 677.93 | 10.38 | 64 | >sp\|P01876\|IGHA1_HUMAN Immunoglobulin heavy constant alpha 1 OS=Homo sapiens OX=9606 GN=IGHA1 PE=1 SV=2 | 255 | 20201113_gy_colo_milk_1ng_glyco_hcd30_etd30_compound15.16987.16987.4 | controllerType=0 controllerNumber=1 scan=16987 | 42.6084 |
| 13585:4 | 1 | R.LSLHRPALEDLLLGSEAN[+1054.370]LTC[+57.021]TLTGLR.D | HexNAc(2)Hex(4) | C[+57], N[+1054] | 1005.2496 | 4 | 4017.9764 | 4017.9683 | 0 | 2 | 127 | Specific | 778.45 | 778.45 | 778.45 | 13.63 | 64 | >sp\|P01876\|IGHA1_HUMAN Immunoglobulin heavy constant alpha 1 OS=Homo sapiens OX=9606 GN=IGHA1 PE=1 SV=2 | 255 | 20201113_gy_colo_milk_1ng_glyco_hcd30_etd30_compound15.17036.17036.4 | controllerType=0 controllerNumber=1 scan=17036 | 42.7582 |
| 13560:5 | 1 | R.LSLHRPALEDLLLGSEAN[+1501.555]LTC[+57.021]TLTGLR.D | HexNAc(5)Hex(3) | C[+57], N[+1502] | 893.8385 | 5 | 4465.1635 | 4465.1536 | 0 | 2.2 | 127 | Specific | 795.77 | 771.7 | 771.7 | 14.02 | 64 | >sp\|P01876\|IGHA1_HUMAN Immunoglobulin heavy constant alpha 1 OS=Homo sapiens OX=9606 GN=IGHA1 PE=1 SV=2 | 255 | 20201113_gy_colo_milk_1ng_glyco_hcd30_etd30_compound15.17009.17009.5 | controllerType=0 controllerNumber=1 scan=17009 | 42.67 |
| 14173:4 | 1 | R.LSLHRPALEDLLLGSEAN[+1872.651]LTC[+57.021]TLTGLR.D | HexNAc(3)Hex(6)NeuAc(1) | C[+57], N[+1873] | 1209.8211 | 4 | 4836.2625 | 4836.2487 | 0 | 2.8 | 127 | Specific | 804.69 | 702.56 | 702.56 | 14.63 | 64 | >sp\|P01876\|IGHA1_HUMAN Immunoglobulin heavy constant alpha 1 OS=Homo sapiens OX=9606 GN=IGHA1 PE=1 SV=2 | 255 | 20201113_gy_colo_milk_1ng_glyco_hcd30_etd30_compound15.17659.17659.4 | controllerType=0 controllerNumber=1 scan=17659 | 44.7327 |
| 14198:4 | 1 | R.LSLHRPALEDLLLGSEAN[+1751.624]LTC[+57.021]TLTGLR.D | HexNAc(4)Hex(4)NeuAc(1) | C[+57], N[+1752] | 1179.571 | 4 | 4715.262 | 4715.2225 | 0 | 8.4 | 127 | Specific | 841.46 | 841.46 | 841.46 | 12.76 | 64 | >sp\|P01876\|IGHA1_HUMAN Immunoglobulin heavy constant alpha 1 OS=Homo sapiens OX=9606 GN=IGHA1 PE=1 SV=2 | 255 | 20201113_gy_colo_milk_1ng_glyco_hcd30_etd30_compound15.17686.17686.4 | controllerType=0 controllerNumber=1 scan=17686 | 44.8173 |
| 13514:4 | 1 | R.LSLHRPALEDLLLGSEAN[+1702.581]LTC[+57.021]TLTGLR.D | HexNAc(2)Hex(8) | C[+57], N[+1703] | 1167.3028 | 4 | 4666.1893 | 4666.1796 | 0 | 2.1 | 127 | Specific | 856.29 | 848.87 | 848.87 | 15.63 | 64 | >sp\|P01876\|IGHA1_HUMAN Immunoglobulin heavy constant alpha 1 OS=Homo sapiens OX=9606 GN=IGHA1 PE=1 SV=2 | 255 | 20201113_gy_colo_milk_1ng_glyco_hcd30_etd30_compound15.16961.16961.4 | controllerType=0 controllerNumber=1 scan=16961 | 42.5263 |
| 14234:4 | 1 | R.LSLHRPALEDLLLGSEAN[+1548.545]LTC[+57.021]TLTGLR.D | HexNAc(3)Hex(4)NeuAc(1) | C[+57], N[+1549] | 1128.7931 | 4 | 4512.1507 | 4512.1431 | 0 | 1.7 | 127 | Specific | 862.62 | 809.18 | 809.18 | 15.63 | 64 | >sp\|P01876\|IGHA1_HUMAN Immunoglobulin heavy constant alpha 1 OS=Homo sapiens OX=9606 GN=IGHA1 PE=1 SV=2 | 255 | 20201113_gy_colo_milk_1ng_glyco_hcd30_etd30_compound15.17724.17724.4 | controllerType=0 controllerNumber=1 scan=17724 | 44.9379 |
| 14205:4 | 1 | R.LSLHRPALEDLLLGSEAN[+1710.598]LTC[+57.021]TLTGLR.D | HexNAc(3)Hex(5)NeuAc(1) | C[+57], N[+1711] | 1169.3146 | 4 | 4674.2366 | 4674.1959 | 0 | 8.7 | 127 | Specific | 895.06 | 879.96 | 879.96 | 13.76 | 64 | >sp\|P01876\|IGHA1_HUMAN Immunoglobulin heavy constant alpha 1 OS=Homo sapiens OX=9606 GN=IGHA1 PE=1 SV=2 | 255 | 20201113_gy_colo_milk_1ng_glyco_hcd30_etd30_compound15.17693.17693.4 | controllerType=0 controllerNumber=1 scan=17693 | 44.8355 |
| 13545:4 | 1 | R.LSLHRPALEDLLLGSEAN[+1460.529]LTC[+57.021]TLTGLR.D | HexNAc(4)Hex(4) | C[+57], N[+1461] | 1106.7948 | 4 | 4424.1575 | 4424.1271 | 0 | 6.9 | 127 | Specific | 905.55 | 854.24 | 854.24 | 15.02 | 64 | >sp\|P01876\|IGHA1_HUMAN Immunoglobulin heavy constant alpha 1 OS=Homo sapiens OX=9606 GN=IGHA1 PE=1 SV=2 | 255 | 20201113_gy_colo_milk_1ng_glyco_hcd30_etd30_compound15.16993.16993.4 | controllerType=0 controllerNumber=1 scan=16993 | 42.6257 |
| 13495:4 | 1 | R.LSLHRPALEDLLLGSEAN[+1540.529]LTC[+57.021]TLTGLR.D | HexNAc(2)Hex(7) | C[+57], N[+1541] | 1126.7892 | 4 | 4504.1351 | 4504.1268 | 0 | 1.8 | 127 | Specific | 909.33 | 825.94 | 825.94 | 16.63 | 64 | >sp\|P01876\|IGHA1_HUMAN Immunoglobulin heavy constant alpha 1 OS=Homo sapiens OX=9606 GN=IGHA1 PE=1 SV=2 | 255 | 20201113_gy_colo_milk_1ng_glyco_hcd30_etd30_compound15.16941.16941.4 | controllerType=0 controllerNumber=1 scan=16941 | 42.462 |
| 13521:4 | 1 | R.LSLHRPALEDLLLGSEAN[+1501.555]LTC[+57.021]TLTGLR.D | HexNAc(5)Hex(3) | C[+57], N[+1502] | 1117.0467 | 4 | 4465.1649 | 4465.1536 | 0 | 2.5 | 127 | Specific | 921.77 | 921.77 | 921.77 | 16.63 | 64 | >sp\|P01876\|IGHA1_HUMAN Immunoglobulin heavy constant alpha 1 OS=Homo sapiens OX=9606 GN=IGHA1 PE=1 SV=2 | 255 | 20201113_gy_colo_milk_1ng_glyco_hcd30_etd30_compound15.16968.16968.4 | controllerType=0 controllerNumber=1 scan=16968 | 42.5472 |
| 13515:4 | 1 | R.LSLHRPALEDLLLGSEAN[+1663.608]LTC[+57.021]TLTGLR.D | HexNAc(5)Hex(4) | C[+57], N[+1664] | 1157.5595 | 4 | 4627.2161 | 4627.2064 | 0 | 2.1 | 127 | Specific | 927.74 | 911.59 | 911.59 | 16.63 | 64 | >sp\|P01876\|IGHA1_HUMAN Immunoglobulin heavy constant alpha 1 OS=Homo sapiens OX=9606 GN=IGHA1 PE=1 SV=2 | 255 | 20201113_gy_colo_milk_1ng_glyco_hcd30_etd30_compound15.16962.16962.4 | controllerType=0 controllerNumber=1 scan=16962 | 42.5296 |
| 13565:4 | 1 | R.LSLHRPALEDLLLGSEAN[+1378.476]LTC[+57.021]TLTGLR.D | HexNAc(2)Hex(6) | C[+57], N[+1378] | 1086.2751 | 4 | 4342.0784 | 4342.074 | 0 | 1 | 127 | Specific | 951.76 | 926.32 | 926.32 | 16.75 | 64 | >sp\|P01876\|IGHA1_HUMAN Immunoglobulin heavy constant alpha 1 OS=Homo sapiens OX=9606 GN=IGHA1 PE=1 SV=2 | 255 | 20201113_gy_colo_milk_1ng_glyco_hcd30_etd30_compound15.17015.17015.4 | controllerType=0 controllerNumber=1 scan=17015 | 42.6903 |
| 13554:4 | 1 | R.LSLHRPALEDLLLGSEAN[+1298.476]LTC[+57.021]TLTGLR.D | HexNAc(4)Hex(3) | C[+57], N[+1298] | 1066.2756 | 4 | 4262.0804 | 4262.0742 | 0 | 1.4 | 127 | Specific | 1015.26 | 1001.03 | 1001.03 | 18.63 | 64 | >sp\|P01876\|IGHA1_HUMAN Immunoglobulin heavy constant alpha 1 OS=Homo sapiens OX=9606 GN=IGHA1 PE=1 SV=2 | 255 | 20201113_gy_colo_milk_1ng_glyco_hcd30_etd30_compound15.17003.17003.4 | controllerType=0 controllerNumber=1 scan=17003 | 42.6539 |
| 13537:4 | 1 | R.LSLHRPALEDLLLGSEAN[+1216.423]LTC[+57.021]TLTGLR.D | HexNAc(2)Hex(5) | C[+57], N[+1216] | 1045.7674 | 4 | 4180.0477 | 4180.0211 | 0 | 6.4 | 127 | Specific | 1035.52 | 994.83 | 994.83 | 17.38 | 64 | >sp\|P01876\|IGHA1_HUMAN Immunoglobulin heavy constant alpha 1 OS=Homo sapiens OX=9606 GN=IGHA1 PE=1 SV=2 | 255 | 20201113_gy_colo_milk_1ng_glyco_hcd30_etd30_compound15.16985.16985.4 | controllerType=0 controllerNumber=1 scan=16985 | 42.5991 |
| 09050:3 | 1 | R.LAGKPTHVN[+1095.397]VSVVMAEVDGTC[+57.021]Y.- | HexNAc(3)Hex(3) | C[+57], N[+1095] | 1148.1873 | 3 | 3442.5473 | 3442.5387 | 0 | 2.5 | 332 | CRagged | 200.06 | 154.79 | 154.79 | 3.37 | 64 | >sp\|P01876\|IGHA1_HUMAN Immunoglobulin heavy constant alpha 1 OS=Homo sapiens OX=9606 GN=IGHA1 PE=1 SV=2 | 255 | 20201113_gy_colo_milk_1ng_glyco_hcd30_etd30_compound15.12280.12280.3 | controllerType=0 controllerNumber=1 scan=12280 | 31.0459 |
| 08083:3 | 1 | R.LAGKPTHVN[+1378.476]VSVVMAEVDGTC[+57.021]Y.- | HexNAc(2)Hex(6) | C[+57], N[+1378] | 1242.5479 | 3 | 3725.6292 | 3725.6178 | 0 | 3.1 | 332 | CRagged | 206.26 | 159.41 | 159.41 | 3.31 | 64 | >sp\|P01876\|IGHA1_HUMAN Immunoglobulin heavy constant alpha 1 OS=Homo sapiens OX=9606 GN=IGHA1 PE=1 SV=2 | 255 | 20201113_gy_colo_milk_1ng_glyco_hcd30_etd30_compound15.11266.11266.3 | controllerType=0 controllerNumber=1 scan=11266 | 28.5073 |
| 09049:3 | 1 | R.LAGKPTHVN[+1216.423]VSVVMAEVDGTC[+57.021]Y.- | HexNAc(2)Hex(5) | C[+57], N[+1216] | 1188.5287 | 3 | 3563.5717 | 3563.5649 | 0 | 1.9 | 332 | CRagged | 206.96 | 194.89 | 157.29 | 3.42 | 64 | >sp\|P01876\|IGHA1_HUMAN Immunoglobulin heavy constant alpha 1 OS=Homo sapiens OX=9606 GN=IGHA1 PE=1 SV=2 | 255 | 20201113_gy_colo_milk_1ng_glyco_hcd30_etd30_compound15.12279.12279.3 | controllerType=0 controllerNumber=1 scan=12279 | 31.0417 |
| 08906:3 | 1 | R.LAGKPTHVN[+1663.608]VSVVMAEVDGTC[+57.021]Y.- | HexNAc(5)Hex(4) | C[+57], N[+1664] | 1337.5922 | 3 | 4010.7621 | 4010.7502 | 0 | 3 | 332 | CRagged | 207.65 | 152.83 | 152.83 | 3.05 | 64 | >sp\|P01876\|IGHA1_HUMAN Immunoglobulin heavy constant alpha 1 OS=Homo sapiens OX=9606 GN=IGHA1 PE=1 SV=2 | 255 | 20201113_gy_colo_milk_1ng_glyco_hcd30_etd30_compound15.12128.12128.3 | controllerType=0 controllerNumber=1 scan=12128 | 30.6162 |
| 08933:4 | 1 | R.LAGKPTHVN[+1971.719]VSVVMAEVDGTC[+57.021]Y.- | HexNAc(5)Hex(5)Fuc(1) | C[+57], N[+1972] | 1080.4733 | 4 | 4318.8714 | 4318.861 | 0 | 2.4 | 332 | CRagged | 220.02 | 220.02 | 122.22 | 3.8 | 64 | >sp\|P01876\|IGHA1_HUMAN Immunoglobulin heavy constant alpha 1 OS=Homo sapiens OX=9606 GN=IGHA1 PE=1 SV=2 | 255 | 20201113_gy_colo_milk_1ng_glyco_hcd30_etd30_compound15.12156.12156.4 | controllerType=0 controllerNumber=1 scan=12156 | 30.7069 |
| 10367:3 | 1 | R.LAGKPTHVN[+1751.624]VSVVMAEVDGTC[+57.021]Y.- | HexNAc(4)Hex(4)NeuAc(1) | C[+57], N[+1752] | 1366.9312 | 3 | 4098.7791 | 4098.7663 | 0 | 3.1 | 332 | CRagged | 220.37 | 85.86 | 85.86 | 2.62 | 64 | >sp\|P01876\|IGHA1_HUMAN Immunoglobulin heavy constant alpha 1 OS=Homo sapiens OX=9606 GN=IGHA1 PE=1 SV=2 | 255 | 20201113_gy_colo_milk_1ng_glyco_hcd30_etd30_compound15.13654.13654.3 | controllerType=0 controllerNumber=1 scan=13654 | 34.0672 |
| 10249:3 | 1 | R.LAGKPTHVN[+2262.814]VSVVMAEVDGTC[+57.021]Y.- | HexNAc(5)Hex(5)Fuc(1)NeuAc(1) | C[+57], N[+2263] | 1537.3292 | 3 | 4609.9729 | 4609.9564 | 0 | 3.6 | 332 | CRagged | 231.89 | 73.68 | 73.68 | 2.62 | 64 | >sp\|P01876\|IGHA1_HUMAN Immunoglobulin heavy constant alpha 1 OS=Homo sapiens OX=9606 GN=IGHA1 PE=1 SV=2 | 255 | 20201113_gy_colo_milk_1ng_glyco_hcd30_etd30_compound15.13529.13529.3 | controllerType=0 controllerNumber=1 scan=13529 | 33.7275 |
| 08982:2 | 1 | R.LAGKPTHVN[+1216.423]VSVVMAEVDGTC[+57.021]Y.- | HexNAc(2)Hex(5) | C[+57], N[+1216] | 1782.2896 | 2 | 3563.572 | 3563.5649 | 0 | 2 | 332 | CRagged | 236.77 | 236.77 | 93.71 | 4.67 | 64 | >sp\|P01876\|IGHA1_HUMAN Immunoglobulin heavy constant alpha 1 OS=Homo sapiens OX=9606 GN=IGHA1 PE=1 SV=2 | 255 | 20201113_gy_colo_milk_1ng_glyco_hcd30_etd30_compound15.12208.12208.2 | controllerType=0 controllerNumber=1 scan=12208 | 30.8563 |
| 08916:3 | 1 | R.LAGKPTHVN[+1809.666]VSVVMAEVDGTC[+57.021]Y.- | HexNAc(5)Hex(4)Fuc(1) | C[+57], N[+1810] | 1386.2778 | 3 | 4156.8187 | 4156.8081 | 0 | 2.6 | 332 | CRagged | 252.08 | 135.94 | 135.94 | 3.84 | 64 | >sp\|P01876\|IGHA1_HUMAN Immunoglobulin heavy constant alpha 1 OS=Homo sapiens OX=9606 GN=IGHA1 PE=1 SV=2 | 255 | 20201113_gy_colo_milk_1ng_glyco_hcd30_etd30_compound15.12138.12138.3 | controllerType=0 controllerNumber=1 scan=12138 | 30.6463 |
| 08862:3 | 1 | R.LAGKPTHVN[+1702.581]VSVVMAEVDGTC[+57.021]Y.- | HexNAc(2)Hex(8) | C[+57], N[+1703] | 1350.5821 | 3 | 4049.7317 | 4049.7234 | 0 | 2 | 332 | CRagged | 253.54 | 201.32 | 182.09 | 4.32 | 64 | >sp\|P01876\|IGHA1_HUMAN Immunoglobulin heavy constant alpha 1 OS=Homo sapiens OX=9606 GN=IGHA1 PE=1 SV=2 | 255 | 20201113_gy_colo_milk_1ng_glyco_hcd30_etd30_compound15.12082.12082.3 | controllerType=0 controllerNumber=1 scan=12082 | 30.4773 |
| 08971:3 | 1 | R.LAGKPTHVN[+1501.555]VSVVMAEVDGTC[+57.021]Y.- | HexNAc(5)Hex(3) | C[+57], N[+1502] | 1283.5727 | 3 | 3848.7035 | 3848.6974 | 0 | 1.6 | 332 | CRagged | 255.12 | 158.07 | 158.07 | 4.27 | 64 | >sp\|P01876\|IGHA1_HUMAN Immunoglobulin heavy constant alpha 1 OS=Homo sapiens OX=9606 GN=IGHA1 PE=1 SV=2 | 255 | 20201113_gy_colo_milk_1ng_glyco_hcd30_etd30_compound15.12197.12197.3 | controllerType=0 controllerNumber=1 scan=12197 | 30.8278 |
| 08904:3 | 1 | R.LAGKPTHVN[+1378.476]VSVVMAEVDGTC[+57.021]Y.- | HexNAc(2)Hex(6) | C[+57], N[+1378] | 1242.5478 | 3 | 3725.6288 | 3725.6178 | 0 | 3 | 332 | CRagged | 262.1 | 199.71 | 187.05 | 4.54 | 64 | >sp\|P01876\|IGHA1_HUMAN Immunoglobulin heavy constant alpha 1 OS=Homo sapiens OX=9606 GN=IGHA1 PE=1 SV=2 | 255 | 20201113_gy_colo_milk_1ng_glyco_hcd30_etd30_compound15.12126.12126.3 | controllerType=0 controllerNumber=1 scan=12126 | 30.6028 |
| 10353:3 | 1 | R.LAGKPTHVN[+1710.598]VSVVMAEVDGTC[+57.021]Y.- | HexNAc(3)Hex(5)NeuAc(1) | C[+57], N[+1711] | 1353.2536 | 3 | 4057.7462 | 4057.7397 | 0 | 1.6 | 332 | CRagged | 263.4 | 209.12 | 174.21 | 4.32 | 64 | >sp\|P01876\|IGHA1_HUMAN Immunoglobulin heavy constant alpha 1 OS=Homo sapiens OX=9606 GN=IGHA1 PE=1 SV=2 | 255 | 20201113_gy_colo_milk_1ng_glyco_hcd30_etd30_compound15.13639.13639.3 | controllerType=0 controllerNumber=1 scan=13639 | 34.0229 |
| 08972:3 | 1 | R.LAGKPTHVN[+1647.613]VSVVMAEVDGTC[+57.021]Y.- | HexNAc(5)Hex(3)Fuc(1) | C[+57], N[+1648] | 1332.259 | 3 | 3994.7624 | 3994.7553 | 0 | 1.8 | 332 | CRagged | 264.66 | 133.47 | 133.47 | 3.84 | 64 | >sp\|P01876\|IGHA1_HUMAN Immunoglobulin heavy constant alpha 1 OS=Homo sapiens OX=9606 GN=IGHA1 PE=1 SV=2 | 255 | 20201113_gy_colo_milk_1ng_glyco_hcd30_etd30_compound15.12198.12198.3 | controllerType=0 controllerNumber=1 scan=12198 | [illegible] |
| 08881:3 | 1 | R.LAGKPTHVN[+1540.529]VSVVMAEVDGTC[+57.021]Y.- | HexNAc(2)Hex(7) | C[+57], N[+1541] | 1296.5643 | 3 | 3887.6782 | 3887.6706 | 0 | 2 | 332 | CRagged | 269.83 | 205.84 | 205.84 | 4.54 | 64 | >sp\|P01876\|IGHA1_HUMAN Immunoglobulin heavy constant alpha 1 OS=Homo sapiens OX=9606 GN=IGHA1 PE=1 SV=2 | 255 | 20201113_gy_colo_milk_1ng_glyco_hcd30_etd30_compound15.12101.12101.3 | controllerType=0 controllerNumber=1 scan=12101 | [illegible] |
| 11692:3 | 1 | R.LAGKPTHVN[+2204.772]VSVVMAEVDGTC[+57.021]Y.- | HexNAc(4)Hex(5)NeuAc(2) | C[+57], N[+2205] | 1517.9804 | 3 | 4551.9267 | 4551.9145 | 0 | 2.7 | 332 | CRagged | 270.73 | 167.44 | 167.44 | 4.49 | 64 | >sp\|P01876\|IGHA1_HUMAN Immunoglobulin heavy constant alpha 1 OS=Homo sapiens OX=9606 GN=IGHA1 PE=1 SV=2 | 255 | 20201113_gy_colo_milk_1ng_glyco_hcd30_etd30_compound15.15043.15043.3 | controllerType=0 controllerNumber=1 scan=15043 | 37.4987 |
| 08169:3 | 1 | R.LAGKPTHVN[+1054.370]VSVVMAEVDGTC[+57.021]Y.- | HexNAc(2)Hex(4) | C[+57], N[+1054] | 1134.5093 | 3 | 3401.5134 | 3401.5121 | 0 | 0.4 | 332 | CRagged | 271.2 | 258.13 | 161.63 | 3.92 | 64 | >sp\|P01876\|IGHA1_HUMAN Immunoglobulin heavy constant alpha 1 OS=Homo sapiens OX=9606 GN=IGHA1 PE=1 SV=2 | 255 | 20201113_gy_colo_milk_1ng_glyco_hcd30_etd30_compound15.11356.11356.3 | controllerType=0 controllerNumber=1 scan=11356 | 28.7702 |
| 08105:3 | 1 | R.LAGKPTHVN[+1216.423]VSVVMAEVDGTC[+57.021]Y.- | HexNAc(2)Hex(5) | C[+57], N[+1216] | 1188.5286 | 3 | 3563.5713 | 3563.5649 | 0 | 1.8 | 332 | CRagged | 275.74 | 240.71 | 180.84 | 4.57 | 64 | >sp\|P01876\|IGHA1_HUMAN Immunoglobulin heavy constant alpha 1 OS=Homo sapiens OX=9606 GN=IGHA1 PE=1 SV=2 | 255 | 20201113_gy_colo_milk_1ng_glyco_hcd30_etd30_compound15.11289.11289.3 | controllerType=0 controllerNumber=1 scan=11289 | 28.582 |
| 08993:3 | 1 | R.LAGKPTHVN[+1444.534]VSVVMAEVDGTC[+57.021]Y.- | HexNAc(4)Hex(3)Fuc(1) | C[+57], N[+1445] | 1264.5645 | 3 | 3791.679 | 3791.6759 | 0 | 0.8 | 332 | CRagged | 276.42 | 151.93 | 151.93 | 3.38 | 64 | >sp\|P01876\|IGHA1_HUMAN Immunoglobulin heavy constant alpha 1 OS=Homo sapiens OX=9606 GN=IGHA1 PE=1 SV=2 | 255 | 20201113_gy_colo_milk_1ng_glyco_hcd30_etd30_compound15.12220.12220.3 | controllerType=0 controllerNumber=1 scan=12220 | 30.8935 |
| 10250:3 | 1 | R.LAGKPTHVN[+1913.677]VSVVMAEVDGTC[+57.021]Y.- | HexNAc(4)Hex(5)NeuAc(1) | C[+57], N[+1914] | 1420.9496 | 3 | 4260.8344 | 4260.8191 | 0 | 3.6 | 332 | CRagged | 282.56 | 187.48 | 187.48 | 3.31 | 64 | >sp\|P01876\|IGHA1_HUMAN Immunoglobulin heavy constant alpha 1 OS=Homo sapiens OX=9606 GN=IGHA1 PE=1 SV=2 | 255 | 20201113_gy_colo_milk_1ng_glyco_hcd30_etd30_compound15.13530.13530.3 | controllerType=0 controllerNumber=1 scan=13530 | 33.7307 |
| 08087:3 | 1 | R.LAGKPTHVN[+1809.666]VSVVMAEVDGTC[+57.021]Y.- | HexNAc(5)Hex(4)Fuc(1) | C[+57], N[+1810] | 1386.2781 | 3 | 4156.8198 | 4156.8081 | 0 | 2.8 | 332 | CRagged | 286.21 | 157.95 | 157.95 | 4.32 | 64 | >sp\|P01876\|IGHA1_HUMAN Immunoglobulin heavy constant alpha 1 OS=Homo sapiens OX=9606 GN=IGHA1 PE=1 SV=2 | 255 | 20201113_gy_colo_milk_1ng_glyco_hcd30_etd30_compound15.11270.11270.3 | controllerType=0 controllerNumber=1 scan=11270 | 28.5172 |
| 10269:3 | 1 | R.LAGKPTHVN[+2059.735]VSVVMAEVDGTC[+57.021]Y.- | HexNAc(4)Hex(5)Fuc(1)NeuAc(1) | C[+57], N[+2060] | 1469.6359 | 3 | 4406.8932 | 4406.877 | 0 | 3.7 | 332 | CRagged | 289.3 | 132.34 | 132.34 | 3.01 | 64 | >sp\|P01876\|IGHA1_HUMAN Immunoglobulin heavy constant alpha 1 OS=Homo sapiens OX=9606 GN=IGHA1 PE=1 SV=2 | 255 | 20201113_gy_colo_milk_1ng_glyco_hcd30_etd30_compound15.13550.13550.3 | controllerType=0 controllerNumber=1 scan=13550 | 33.8019 |
| 08962:2 | 1 | R.LAGKPTHVN[+1216.423]VSVVMAEVDGTC[+57.021]Y.- | HexNAc(2)Hex(5) | C[+57], N[+1216] | 1782.2893 | 2 | 3563.5713 | 3563.5649 | 0 | 1.8 | 332 | CRagged | 291.84 | 291.84 | 148.11 | 5.99 | 64 | >sp\|P01876\|IGHA1_HUMAN Immunoglobulin heavy constant alpha 1 OS=Homo sapiens OX=9606 GN=IGHA1 PE=1 SV=2 | 255 | 20201113_gy_colo_milk_1ng_glyco_hcd30_etd30_compound15.12187.12187.2 | controllerType=0 controllerNumber=1 scan=12187 | 30.7922 |
| 11635:4 | 1 | R.LAGKPTHVN[+2350.830]VSVVMAEVDGTC[+57.021]Y.- | HexNAc(4)Hex(5)Fuc(1)NeuAc(2) | C[+57], N[+2351] | 1175.2509 | 4 | 4697.9818 | 4697.9724 | 0 | 2 | 332 | CRagged | 303.03 | 247.09 | 247.09 | 4.79 | 64 | >sp\|P01876\|IGHA1_HUMAN Immunoglobulin heavy constant alpha 1 OS=Homo sapiens OX=9606 GN=IGHA1 PE=1 SV=2 | 255 | 20201113_gy_colo_milk_1ng_glyco_hcd30_etd30_compound15.14984.14984.4 | controllerType=0 controllerNumber=1 scan=14984 | 37.3604 |
| 08994:3 | 1 | R.LAGKPTHVN[+1054.370]VSVVMAEVDGTC[+57.021]Y.- | HexNAc(2)Hex(4) | C[+57], N[+1054] | 1134.5103 | 3 | 3401.5163 | 3401.5121 | 0 | 1.2 | 332 | CRagged | 308.5 | 292.92 | 215.05 | 4.24 | 64 | >sp\|P01876\|IGHA1_HUMAN Immunoglobulin heavy constant alpha 1 OS=Homo sapiens OX=9606 GN=IGHA1 PE=1 SV=2 | 255 | 20201113_gy_colo_milk_1ng_glyco_hcd30_etd30_compound15.12221.12221.3 | controllerType=0 controllerNumber=1 scan=12221 | 30.9 |
| 09015:4 | 1 | R.LAGKPTHVN[+1647.613]VSVVMAEVDGTC[+57.021]Y.- | HexNAc(5)Hex(3)Fuc(1) | C[+57], N[+1648] | 999.4461 | 4 | 3994.7628 | 3994.7553 | 0 | 1.9 | 332 | CRagged | 323.97 | 323.97 | 323.97 | 5.09 | 64 | >sp\|P01876\|IGHA1_HUMAN Immunoglobulin heavy constant alpha 1 OS=Homo sapiens OX=9606 GN=IGHA1 PE=1 SV=2 | 255 | 20201113_gy_colo_milk_1ng_glyco_hcd30_etd30_compound15.12243.12243.4 | controllerType=0 controllerNumber=1 scan=12243 | 30.9465 |
| 11639:4 | 1 | R.LAGKPTHVN[+2553.910]VSVVMAEVDGTC[+57.021]Y.- | HexNAc(5)Hex(5)Fuc(1)NeuAc(2) | C[+57], N[+2554] | 1226.0216 | 4 | 4901.0648 | 4901.0518 | 0 | 2.6 | 332 | CRagged | 347.66 | 207.5 | 207.5 | 5.06 | 64 | >sp\|P01876\|IGHA1_HUMAN Immunoglobulin heavy constant alpha 1 OS=Homo sapiens OX=9606 GN=IGHA1 PE=1 SV=2 | 255 | 20201113_gy_colo_milk_1ng_glyco_hcd30_etd30_compound15.14988.14988.4 | controllerType=0 controllerNumber=1 scan=14988 | 37.3685 |
| 09046:3 | 1 | R.LAGKPTHVN[+892.317]VSVVMAEVDGTC[+57.021]Y.- | HexNAc(2)Hex(3) | C[+57], N[+892] | 1080.4958 | 3 | 3239.4727 | 3239.4593 | 0 | 4.1 | 332 | CRagged | 378.44 | 373.38 | 164.54 | 5.7 | 64 | >sp\|P01876\|IGHA1_HUMAN Immunoglobulin heavy constant alpha 1 OS=Homo sapiens OX=9606 GN=IGHA1 PE=1 SV=2 | 255 | 20201113_gy_colo_milk_1ng_glyco_hcd30_etd30_compound15.12275.12275.3 | controllerType=0 controllerNumber=1 scan=12275 | 31.0288 |
| 10279:4 | 1 | R.LAGKPTHVN[+2059.735]VSVVMAEVDGTC[+57.021]Y.- | HexNAc(4)Hex(5)Fuc(1)NeuAc(1) | C[+57], N[+2060] | 1102.4775 | 4 | 4406.888 | 4406.877 | 0 | 2.5 | 332 | CRagged | 417.6 | 372.22 | 372.22 | 7.07 | 64 | >sp\|P01876\|IGHA1_HUMAN Immunoglobulin heavy constant alpha 1 OS=Homo sapiens OX=9606 GN=IGHA1 PE=1 SV=2 | 255 | 20201113_gy_colo_milk_1ng_glyco_hcd30_etd30_compound15.13560.13560.4 | controllerType=0 controllerNumber=1 scan=13560 | 33.8313 |
| 10259:4 | 1 | R.LAGKPTHVN[+2262.814]VSVVMAEVDGTC[+57.021]Y.- | HexNAc(5)Hex(5)Fuc(1)NeuAc(1) | C[+57], N[+2263] | 1153.2489 | 4 | 4609.9739 | 4609.9564 | 0 | 3.8 | 332 | CRagged | 548.24 | 399.63 | 399.63 | 6.18 | 64 | >sp\|P01876\|IGHA1_HUMAN Immunoglobulin heavy constant alpha 1 OS=Homo sapiens OX=9606 GN=IGHA1 PE=1 SV=2 | 255 | 20201113_gy_colo_milk_1ng_glyco_hcd30_etd30_compound15.13539.13539.4 | controllerType=0 controllerNumber=1 scan=13539 | 33.767 |
| 07079:3 | 1 | R.LAGKPTHVN[+1647.613]VSVVM[+15.995]AEVDGTC[+57.021]Y.- | HexNAc(5)Hex(3)Fuc(1) | C[+57], M[+16], N[+1648] | 1337.5911 | 3 | 4010.7588 | 4010.7502 | 0 | 2.1 | 332 | CRagged | 205.87 | 111.32 | 111.32 | 3.04 | 64 | >sp\|P01876\|IGHA1_HUMAN Immunoglobulin heavy constant alpha 1 OS=Homo sapiens OX=9606 GN=IGHA1 PE=1 SV=2 | 255 | 20201113_gy_colo_milk_1ng_glyco_hcd30_etd30_compound15.10217.10217.3 | controllerType=0 controllerNumber=1 scan=10217 | 26.1456 |
| 06616:3 | 2 | R.GN[+2059.735]FSTEGC[+57.021]GC[+57.021]VC[+57.021]EPGWK.G | HexNAc(4)Hex(5)Fuc(1)NeuAc(1) | C[+57]*3, N[+2060] | 1335.5097 | 3 | 4004.5145 | 4004.5023 | 0 | 3.1 | 165 | Specific | 364.62 | 259.49 | 259.49 | 6.32 | 44 | >sp\|P24821\|TENA_HUMAN Tenascin OS=Homo sapiens OX=9606 GN=TNC PE=1 SV=3 | 486 | 20201113_gy_colo_milk_1ng_glyco_hcd30_etd30_compound15.9734.9734.3 | controllerType=0 controllerNumber=1 scan=9734 | 25.0946 |
| 08508:4 | 2 | R.QTGLAPGQEYEISLHIVKN[+2350.830]NTR.G | HexNAc(4)Hex(5)Fuc(1)NeuAc(2) | N[+2351] | 1205.5389 | 4 | 4819.1336 | 4819.1196 | 0 | 2.9 | 770 | Specific | 351.33 | 306.47 | 306.47 | 3.88 | 44 | >sp\|P24821\|TENA_HUMAN Tenascin OS=Homo sapiens OX=9606 GN=TNC PE=1 SV=3 | 486 | 20201113_gy_colo_milk_1ng_glyco_hcd30_etd30_compound15.11710.11710.4 | controllerType=0 controllerNumber=1 scan=11710 | 29.5551 |
| 07316:4 | 2 | R.QTGLAPGQEYEISLHIVKN[+2059.735]NTR.G | HexNAc(4)Hex(5)Fuc(1)NeuAc(1) | N[+2060] | 1132.763 | 4 | 4528.0301 | 4528.0241 | 0 | 1.3 | 770 | Specific | 375.05 | 374.93 | 374.93 | 4.92 | 44 | >sp\|P24821\|TENA_HUMAN Tenascin OS=Homo sapiens OX=9606 GN=TNC PE=1 SV=3 | 486 | 20201113_gy_colo_milk_1ng_glyco_hcd30_etd30_compound15.10465.10465.4 | controllerType=0 controllerNumber=1 scan=10465 | 26.7141 |
| 11839:3 | 2 | R.LN[+1702.581]YSLPTGQWVGVQLPR.N | HexNAc(2)Hex(8) | N[+1703] | 1210.8784 | 3 | 3630.6205 | 3630.6202 | 0 | 0.1 | 1017 | Specific | 383.22 | 327.23 | 327.23 | 5.78 | 44 | >sp\|P24821\|TENA_HUMAN Tenascin OS=Homo sapiens OX=9606 GN=TNC PE=1 SV=3 | 486 | 20201113_gy_colo_milk_1ng_glyco_hcd30_etd30_compound15.15197.15197.2 | controllerType=0 controllerNumber=1 scan=15197 | 37.8424 |
| 11973:2 | 2 | R.LN[+1378.476]YSLPTGQWVGVQLPR.N | HexNAc(2)Hex(6) | N[+1378] | 1653.767 | 2 | 3306.5267 | 3306.5145 | 0 | 3.7 | 1017 | Specific | 420.91 | 330.51 | 330.51 | 5.17 | 44 | >sp\|P24821\|TENA_HUMAN Tenascin OS=Homo sapiens OX=9606 GN=TNC PE=1 SV=3 | 486 | 20201113_gy_colo_milk_1ng_glyco_hcd30_etd30_compound15.15338.15338.2 | controllerType=0 controllerNumber=1 scan=15338 | 38.2313 |
| 11993:2 | 2 | R.LN[+1216.423]YSLPTGQWVGVQLPR.N | HexNAc(2)Hex(5) | N[+1216] | 1572.7364 | 2 | 3144.4655 | 3144.4617 | 0 | 1.2 | 1017 | Specific | 508.91 | 446.07 | 446.07 | 6.95 | 44 | >sp\|P24821\|TENA_HUMAN Tenascin OS=Homo sapiens OX=9606 GN=TNC PE=1 SV=3 | 486 | 20201113_gy_colo_milk_1ng_glyco_hcd30_etd30_compound15.15359.15359.2 | controllerType=0 controllerNumber=1 scan=15359 | 38.2917 |
| 11921:3 | 2 | R.LN[+1540.529]YSLPTGQWVGVQLPR.N | HexNAc(2)Hex(7) | N[+1541] | 1156.8669 | 3 | 3468.5861 | 3468.5674 | 0 | 5.4 | 1017 | Specific | 559.57 | 507.59 | 507.59 | 7.01 | 44 | >sp\|P24821\|TENA_HUMAN Tenascin OS=Homo sapiens OX=9606 GN=TNC PE=1 SV=3 | 486 | 20201113_gy_colo_milk_1ng_glyco_hcd30_etd30_compound15.15283.15283.3 | controllerType=0 controllerNumber=1 scan=15283 | 38.0674 |
| 11861:3 | 2 | R.LN[+1702.581]YSLPTGQWVGVQLPR.N | HexNAc(2)Hex(8) | N[+1703] | 1210.8792 | 3 | 3630.6231 | 3630.6202 | 0 | 0.8 | 1017 | Specific | 574.49 | 482.05 | 482.05 | 8.39 | 44 | >sp\|P24821\|TENA_HUMAN Tenascin OS=Homo sapiens OX=9606 GN=TNC PE=1 SV=3 | 486 | 20201113_gy_colo_milk_1ng_glyco_hcd30_etd30_compound15.15220.15220.3 | controllerType=0 controllerNumber=1 scan=15220 | 37.9045 |
| 11979:3 | 2 | R.LN[+1216.423]YSLPTGQWVGVQLPR.N | HexNAc(2)Hex(5) | N[+1216] | 1048.831 | 3 | 3144.4783 | 3144.4617 | 0 | 5.3 | 1017 | Specific | 640.16 | 604.65 | 604.65 | 9.02 | 44 | >sp\|P24821\|TENA_HUMAN Tenascin OS=Homo sapiens OX=9606 GN=TNC PE=1 SV=3 | 486 | 20201113_gy_colo_milk_1ng_glyco_hcd30_etd30_compound15.15344.15344.3 | controllerType=0 controllerNumber=1 scan=15344 | 38.2534 |
| 02907:3 | 2 | R.N[+2059.735]TTSYVLR.G | HexNAc(4)Hex(5)Fuc(1)NeuAc(1) | N[+2060] | 1005.088 | 3 | 3013.2496 | 3013.24 | 0 | 3.2 | 1034 | Specific | 319.35 | 177.13 | 177.13 | 4.32 | 44 | >sp\|P24821\|TENA_HUMAN Tenascin OS=Homo sapiens OX=9606 GN=TNC PE=1 SV=3 | 486 | 20201113_gy_colo_milk_1ng_glyco_hcd30_etd30_compound15.5876.5876.3 | controllerType=0 controllerNumber=1 scan=5876 | 17.8597 |
| 13510:4 | 2 | R.LN[+2059.735]WTAADQAYEHFIIQVQEANKVEAAR.N | HexNAc(4)Hex(5)Fuc(1)NeuAc(1) | N[+2060] | 1294.5818 | 4 | 5175.3053 | 5175.2945 | 0 | 2.1 | 1092 | Specific | 208.5 | 208.5 | 208.5 | 2.65 | 44 | >sp\|P24821\|TENA_HUMAN Tenascin OS=Homo sapiens OX=9606 GN=TNC PE=1 SV=3 | 486 | 20201113_gy_colo_milk_1ng_glyco_hcd30_etd30_compound15.16956.16956.4 | controllerType=0 controllerNumber=1 scan=16956 | 42.5085 |
| 06320:3 | 2 | R.N[+2059.735]LTVPGSLR.A | HexNAc(4)Hex(5)Fuc(1)NeuAc(1) | N[+2060] | 1006.1026 | 3 | 3016.2931 | 3016.2873 | 0 | 1.9 | 1119 | Specific | 285.83 | 151.3 | 151.3 | 4.16 | 44 | >sp\|P24821\|TENA_HUMAN Tenascin OS=Homo sapiens OX=9606 GN=TNC PE=1 SV=3 | 486 | 20201113_gy_colo_milk_1ng_glyco_hcd30_etd30_compound15.9425.9425.3 | controllerType=0 controllerNumber=1 scan=9425 | 24.4156 |
| 13839:4 | 3 | R.RTAGWNVPIGTLRPFLN[+2351.851]WTGPPEPIEAAVAR.F | HexNAc(4)Hex(5)Fuc(3)NeuAc(1) | N[+2352] | 1435.4247 | 4 | 5738.6769 | 5738.6628 | 0 | 2.5 | 140 | Specific | 275.07 | 275.07 | 275.07 | 5.29 | 45 | >sp\|P02788\|TRFL_HUMAN Lactotransferrin OS=Homo sapiens OX=9606 GN=LTF PE=1 SV=6 | 1270 | 20201113_gy_colo_milk_1ng_glyco_hcd30_etd30_compound15.17305.17305.4 | controllerType=0 controllerNumber=1 scan=17305 | 43.6384 |
| 13171:4 | 3 | R.RTAGWNVPIGTLRPFLN[+2278.809]WTGPPEPIEAAVAR.F | HexNAc(5)Hex(6)NeuAc(1) | N[+2279] | 1417.1704 | 4 | 5665.6598 | 5665.6213 | 0 | 6.8 | 140 | Specific | 476.15 | 476.15 | 476.15 | 5.81 | 45 | >sp\|P02788\|TRFL_HUMAN Lactotransferrin OS=Homo sapiens OX=9606 GN=LTF PE=1 SV=6 | 1270 | 20201113_gy_colo_milk_1ng_glyco_hcd30_etd30_compound15.16599.16599.4 | controllerType=0 controllerNumber=1 scan=16599 | 41.5096 |
| 13794:4 | 3 | R.RTAGWNVPIGTLRPFLN[+2570.925]WTGPPEPIEAAVAR.F | HexNAc(5)Hex(6)Fuc(2)NeuAc(1) | N[+2571] | 1490.194 | 4 | 5957.7541 | 5957.7371 | 0 | 2.9 | 140 | Specific | 630.25 | 630.25 | 630.25 | 10.69 | 45 | >sp\|P02788\|TRFL_HUMAN Lactotransferrin OS=Homo sapiens OX=9606 GN=LTF PE=1 SV=6 | 1270 | 20201113_gy_colo_milk_1ng_glyco_hcd30_etd30_compound15.17258.17258.4 | controllerType=0 controllerNumber=1 scan=17258 | 43.522 |
| 13747:4 | 3 | R.RTAGWNVPIGTLRPFLN[+2863.041]WTGPPEPIEAAVAR.F | HexNAc(5)Hex(6)Fuc(4)NeuAc(1) | N[+2863] | 1563.2235 | 4 | 6249.8722 | 6249.8529 | 0 | 3.1 | 140 | Specific | 687.15 | 687.15 | 687.15 | 11.69 | 45 | >sp\|P02788\|TRFL_HUMAN Lactotransferrin OS=Homo sapiens OX=9606 GN=LTF PE=1 SV=6 | 1270 | 20201113_gy_colo_milk_1ng_glyco_hcd30_etd30_compound15.17208.17208.4 | controllerType=0 controllerNumber=1 scan=17208 | 43.3509 |
| 14525:4 | 3 | R.RTAGWNVPIGTLRPFLN[+2350.830]WTGPPEPIEAAVAR.F | HexNAc(4)Hex(5)Fuc(1)NeuAc(2) | N[+2351] | 1435.1702 | 4 | 5737.6588 | 5737.6424 | 0 | 2.9 | 140 | Specific | 827.47 | 827.47 | 827.47 | 14.69 | 45 | >sp\|P02788\|TRFL_HUMAN Lactotransferrin OS=Homo sapiens OX=9606 GN=LTF PE=1 SV=6 | 1270 | 20201113_gy_colo_milk_1ng_glyco_hcd30_etd30_compound15.18031.18031.3 | controllerType=0 controllerNumber=1 scan=18031 | 45.8618 |
| 15616:4 | 3 | R.[+42.011]TAGWNVPIGTLRPFLN[+2336.851]WTGPPEPIEAAVAR.F | HexNAc(6)Hex(6)Fuc(1) | .[42], N[+2337] | 1403.1437 | 4 | 5609.5529 | 5609.5726 | 0 | -3.5 | 141 | Specific | 218.59 | 218.59 | 218.59 | 3.03 | 45 | >sp\|P02788\|TRFL_HUMAN Lactotransferrin OS=Homo sapiens OX=9606 GN=LTF PE=1 SV=6 | 1270 | 20201113_gy_colo_milk_1ng_glyco_hcd30_etd30_compound15.19188.19188.4 | controllerType=0 controllerNumber=1 scan=19188 | 49.4106 |
| 15608:3 | 3 | R.TAGWNVPIGTLRPFLN[+2350.830]WTGPPEPIEAAVAR.F | HexNAc(4)Hex(5)Fuc(1)NeuAc(2) | N[+2351] | 1861.1858 | 3 | 5581.5427 | 5581.5413 | 0 | 0.3 | 141 | Specific | 244.94 | 164.07 | 164.07 | 5.31 | 45 | >sp\|P02788\|TRFL_HUMAN Lactotransferrin OS=Homo sapiens OX=9606 GN=LTF PE=1 SV=6 | 1270 | 20201113_gy_colo_milk_1ng_glyco_hcd30_etd30_compound15.19179.19179.2 | controllerType=0 controllerNumber=1 scan=19179 | 49.3834 |
| 14625:3 | 3 | R.TAGWNVPIGTLRPFLN[+2570.925]WTGPPEPIEAAVAR.F | HexNAc(5)Hex(6)Fuc(2)NeuAc(1) | N[+2571] | 1934.5547 | 3 | 5801.6494 | 5801.636 | 0 | 2.3 | 141 | Specific | 248.14 | 160.98 | 160.98 | 6.18 | 45 | >sp\|P02788\|TRFL_HUMAN Lactotransferrin OS=Homo sapiens OX=9606 GN=LTF PE=1 SV=6 | 1270 | 20201113_gy_colo_milk_1ng_glyco_hcd30_etd30_compound15.18137.18137.3 | controllerType=0 controllerNumber=1 scan=18137 | 46.1806 |
| 14715:3 | 3 | R.TAGWNVPIGTLRPFLN[+2205.793]WTGPPEPIEAAVAR.F | HexNAc(4)Hex(5)Fuc(2)NeuAc(1) | N[+2206] | 1812.8449 | 3 | 5436.5202 | 5436.5038 | 0 | 3 | 141 | Specific | 303.9 | 232.58 | 232.58 | 5.29 | 45 | >sp\|P02788\|TRFL_HUMAN Lactotransferrin OS=Homo sapiens OX=9606 GN=LTF PE=1 SV=6 | 1270 | 20201113_gy_colo_milk_1ng_glyco_hcd30_etd30_compound15.18232.18232.3 | controllerType=0 controllerNumber=1 scan=18232 | 46.4616 |
| 14701:3 | 3 | R.TAGWNVPIGTLRPFLN[+2350.830]WTGPPEPIEAAVAR.F | HexNAc(4)Hex(5)Fuc(1)NeuAc(2) | N[+2351] | 1861.202 | 3 | 5581.5914 | 5581.5413 | 0 | 9 | 141 | Specific | 337.75 | 243.01 | 243.01 | 3.35 | 45 | >sp\|P02788\|TRFL_HUMAN Lactotransferrin OS=Homo sapiens OX=9606 GN=LTF PE=1 SV=6 | 1270 | 20201113_gy_colo_milk_1ng_glyco_hcd30_etd30_compound15.18217.18217.3 | controllerType=0 controllerNumber=1 scan=18217 | 46.4095 |
| 14055:4 | 3 | R.TAGWNVPIGTLRPFLN[+1768.640]WTGPPEPIEAAVAR.F | HexNAc(4)Hex(5)Fuc(1) | N[+1769] | 1250.5974 | 4 | 4999.3678 | 4999.3504 | 0 | 3.5 | 141 | Specific | 361.03 | 361.03 | 361.03 | 5.02 | 45 | >sp\|P02788\|TRFL_HUMAN Lactotransferrin OS=Homo sapiens OX=9606 GN=LTF PE=1 SV=6 | 1270 | 20201113_gy_colo_milk_1ng_glyco_hcd30_etd30_compound15.17532.17532.4 | controllerType=0 controllerNumber=1 scan=17532 | 44.3181 |
| 14041:3 | 3 | R.TAGWNVPIGTLRPFLN[+1768.640]WTGPPEPIEAAVAR.F | HexNAc(4)Hex(5)Fuc(1) | N[+1769] | 1667.1325 | 3 | 4999.383 | 4999.3504 | 0 | 6.5 | 141 | Specific | 453.63 | 401.56 | 401.56 | 6.11 | 45 | >sp\|P02788\|TRFL_HUMAN Lactotransferrin OS=Homo sapiens OX=9606 GN=LTF PE=1 SV=6 | 1270 | 20201113_gy_colo_milk_1ng_glyco_hcd30_etd30_compound15.17517.17517.3 | controllerType=0 controllerNumber=1 scan=17517 | 44.2441 |
| 15565:3 | 3 | R.TAGWNVPIGTLRPFLN[+2350.830]WTGPPEPIEAAVAR.F | HexNAc(4)Hex(5)Fuc(1)NeuAc(2) | N[+2351] | 1861.2038 | 3 | 5581.5969 | 5581.5413 | 0 | 10 | 141 | Specific | 511.14 | 415.34 | 415.34 | 5.68 | 45 | >sp\|P02788\|TRFL_HUMAN Lactotransferrin OS=Homo sapiens OX=9606 GN=LTF PE=1 SV=6 | 1270 | 20201113_gy_colo_milk_1ng_glyco_hcd30_etd30_compound15.19133.19133.3 | controllerType=0 controllerNumber=1 scan=19133 | 49.2156 |
| 14736:4 | 3 | R.TAGWNVPIGTLRPFLN[+2076.750]WTGPPEPIEAAVAR.F | HexNAc(4)Hex(6)Fuc(2) | N[+2077] | 1327.6266 | 4 | 5307.4845 | 5307.4612 | 0 | 4.4 | 141 | Specific | 530.57 | 530.57 | 530.57 | 7.88 | 45 | >sp\|P02788\|TRFL_HUMAN Lactotransferrin OS=Homo sapiens OX=9606 GN=LTF PE=1 SV=6 | 1270 | 20201113_gy_colo_milk_1ng_glyco_hcd30_etd30_compound15.18254.18254.4 | controllerType=0 controllerNumber=1 scan=18254 | 46.5183 |
| 14697:4 | 3 | R.TAGWNVPIGTLRPFLN[+2350.830]WTGPPEPIEAAVAR.F | HexNAc(4)Hex(5)Fuc(1)NeuAc(2) | N[+2351] | 1396.1484 | 4 | 5581.5719 | 5581.5413 | 0 | 5.5 | 141 | Specific | 552.54 | 552.54 | 552.54 | 6.74 | 45 | >sp\|P02788\|TRFL_HUMAN Lactotransferrin OS=Homo sapiens OX=9606 GN=LTF PE=1 SV=6 | 1270 | 20201113_gy_colo_milk_1ng_glyco_hcd30_etd30_compound15.18213.18213.4 | controllerType=0 controllerNumber=1 scan=18213 | 46.4024 |
| 14621:4 | 3 | R.TAGWNVPIGTLRPFLN[+2570.925]WTGPPEPIEAAVAR.F | HexNAc(5)Hex(6)Fuc(2)NeuAc(1) | N[+2571] | 1451.1689 | 4 | 5801.654 | 5801.636 | 0 | 3.1 | 141 | Specific | 764.14 | 746.69 | 746.69 | 14.02 | 45 | >sp\|P02788\|TRFL_HUMAN Lactotransferrin OS=Homo sapiens OX=9606 GN=LTF PE=1 SV=6 | 1270 | 20201113_gy_colo_milk_1ng_glyco_hcd30_etd30_compound15.18133.18133.4 | controllerType=0 controllerNumber=1 scan=18133 | 46.1733 |
| 14376:3 | 3 | R.TAGWNIPMGLLFN[+1694.603]QTGSC[+57.021]K.F | HexNAc(3)Hex(4)Fuc(1)NeuAc(1) | C[+57], N[+1695] | 1263.8781 | 3 | 3789.6198 | 3789.6127 | 0 | 1.9 | 485 | Specific | 285.47 | 217.56 | 217.56 | 6.43 | 45 | >sp\|P02788\|TRFL_HUMAN Lactotransferrin OS=Homo sapiens OX=9606 GN=LTF PE=1 SV=6 | 1270 | 20201113_gy_colo_milk_1ng_glyco_hcd30_etd30_compound15.17874.17874.3 | controllerType=0 controllerNumber=1 scan=17874 | 45.4442 |
| 14412:3 | 3 | R.TAGWNIPMGLLFN[+1710.598]QTGSC[+57.021]K.F | HexNAc(3)Hex(5)NeuAc(1) | C[+57], N[+1711] | 1269.2081 | 3 | 3805.6096 | 3805.6076 | 0 | 0.5 | 485 | Specific | 299.09 | 200.81 | 200.81 | 5.53 | 45 | >sp\|P02788\|TRFL_HUMAN Lactotransferrin OS=Homo sapiens OX=9606 GN=LTF PE=1 SV=6 | 1270 | 20201113_gy_colo_milk_1ng_glyco_hcd30_etd30_compound15.17912.17912.3 | controllerType=0 controllerNumber=1 scan=17912 | 45.5554 |
| 13266:2 | 3 | R.TAGWNIPMGLLFN[+1216.423]QTGSC[+57.021]K.F | HexNAc(2)Hex(5) | C[+57], N[+1216] | 1656.2282 | 2 | 3311.4491 | 3311.4328 | 0 | 4.9 | 485 | Specific | 345.57 | 331.65 | 140.92 | 5.4 | 45 | >sp\|P02788\|TRFL_HUMAN Lactotransferrin OS=Homo sapiens OX=9606 GN=LTF PE=1 SV=6 | 1270 | 20201113_gy_colo_milk_1ng_glyco_hcd30_etd30_compound15.16699.16699.2 | controllerType=0 controllerNumber=1 scan=16699 | 41.8158 |
| 13110:3 | 3 | R.TAGWNIPMGLLFN[+1622.582]QTGSC[+57.021]K.F | HexNAc(4)Hex(5) | C[+57], N[+1623] | 1239.8736 | 3 | 3717.6062 | 3717.5916 | 0 | 3.9 | 485 | Specific | 349.36 | 263.29 | 253.75 | 5.57 | 45 | >sp\|P02788\|TRFL_HUMAN Lactotransferrin OS=Homo sapiens OX=9606 GN=LTF PE=1 SV=6 | 1270 | 20201113_gy_colo_milk_1ng_glyco_hcd30_etd30_compound15.16535.16535.3 | controllerType=0 controllerNumber=1 scan=16535 | 41.3308 |
| 12948:3 | 3 | R.TAGWNIPMGLLFN[+1768.640]QTGSC[+57.021]K.F | HexNAc(4)Hex(5)Fuc(1) | C[+57], N[+1769] | 1288.5593 | 3 | 3863.6632 | 3863.6495 | 0 | 3.6 | 485 | Specific | 360.14 | 256.56 | 256.56 | 5.65 | 45 | >sp\|P02788\|TRFL_HUMAN Lactotransferrin OS=Homo sapiens OX=9606 GN=LTF PE=1 SV=6 | 1270 | 20201113_gy_colo_milk_1ng_glyco_hcd30_etd30_compound15.16364.16364.3 | controllerType=0 controllerNumber=1 scan=16364 | 40.8293 |
| 13107:3 | 3 | R.TAGWNIPMGLLFN[+1622.582]QTGSC[+57.021]K.F | HexNAc(4)Hex(5) | C[+57], N[+1623] | 1239.8749 | 3 | 3717.6103 | 3717.5916 | 0 | 5 | 485 | Specific | 360.48 | 302.29 | 233.06 | 5.65 | 45 | >sp\|P02788\|TRFL_HUMAN Lactotransferrin OS=Homo sapiens OX=9606 GN=LTF PE=1 SV=6 | 1270 | 20201113_gy_colo_milk_1ng_glyco_hcd30_etd30_compound15.16531.16531.3 | controllerType=0 controllerNumber=1 scan=16531 | 41.3158 |
| 15593:4 | 3 | R.TAGWNIPMGLLFN[+2350.830]QTGSC[+57.021]K.F | HexNAc(4)Hex(5)Fuc(1)NeuAc(2) | C[+57], N[+2351] | 1112.2228 | 4 | 4445.8695 | 4445.8403 | 0 | 6.6 | 485 | Specific | 415.9 | 347.86 | 347.86 | 4.82 | 45 | >sp\|P02788\|TRFL_HUMAN Lactotransferrin OS=Homo sapiens OX=9606 GN=LTF PE=1 SV=6 | 1270 | 20201113_gy_colo_milk_1ng_glyco_hcd30_etd30_compound15.19163.19163.4 | controllerType=0 controllerNumber=1 scan=19163 | 49.3322 |
| 14226:3 | 3 | R.TAGWNIPMGLLFN[+2059.735]QTGSC[+57.021]K.F | HexNAc(4)Hex(5)Fuc(1)NeuAc(1) | C[+57], N[+2060] | 1385.5921 | 3 | 4154.7617 | 4154.7449 | 0 | 4.1 | 485 | Specific | 420.71 | 391.03 | 391.03 | 6.15 | 45 | >sp\|P02788\|TRFL_HUMAN Lactotransferrin OS=Homo sapiens OX=9606 GN=LTF PE=1 SV=6 | 1270 | 20201113_gy_colo_milk_1ng_glyco_hcd30_etd30_compound15.17715.17715.3 | controllerType=0 controllerNumber=1 scan=17715 | 44.9066 |
| 14190:3 | 3 | R.TAGWNIPMGLLFN[+2351.851]QTGSC[+57.021]K.F | HexNAc(4)Hex(5)Fuc(3)NeuAc(1) | C[+57], N[+2352] | 1482.9622 | 3 | 4446.8721 | 4446.8607 | 0 | 2.6 | 485 | Specific | 422.69 | 323.14 | 323.14 | 6.95 | 45 | >sp\|P02788\|TRFL_HUMAN Lactotransferrin OS=Homo sapiens OX=9606 GN=LTF PE=1 SV=6 | 1270 | 20201113_gy_colo_milk_1ng_glyco_hcd30_etd30_compound15.17677.17677.3 | controllerType=0 controllerNumber=1 scan=17677 | 44.7807 |
| 15613:3 | 3 | R.TAGWNIPMGLLFN[+2350.830]QTGSC[+57.021]K.F | HexNAc(4)Hex(5)Fuc(1)NeuAc(2) | C[+57], N[+2351] | 1482.6238 | 3 | 4445.857 | 4445.8403 | 0 | 3.7 | 485 | Specific | 438.56 | 282.97 | 282.97 | 5.7 | 45 | >sp\|P02788\|TRFL_HUMAN Lactotransferrin OS=Homo sapiens OX=9606 GN=LTF PE=1 SV=6 | 1270 | 20201113_gy_colo_milk_1ng_glyco_hcd30_etd30_compound15.19185.19185.3 | controllerType=0 controllerNumber=1 scan=19185 | 49.4041 |
| 15702:3 | 3 | R.TAGWNIPMGLLFN[+2204.772]QTGSC[+57.021]K.F | HexNAc(4)Hex(5)NeuAc(2) | C[+57], N[+2205] | 1433.9371 | 3 | 4299.7967 | 4299.7824 | 0 | 3.3 | 485 | Specific | 439 | 334.03 | 334.03 | 6.04 | 45 | >sp\|P02788\|TRFL_HUMAN Lactotransferrin OS=Homo sapiens OX=9606 GN=LTF PE=1 SV=6 | 1270 | 20201113_gy_colo_milk_1ng_glyco_hcd30_etd30_compound15.19280.19280.3 | controllerType=0 controllerNumber=1 scan=19280 | 49.7334 |
| 14316:3 | 3 | R.TAGWNIPMGLLFN[+1913.677]QTGSC[+57.021]K.F | HexNAc(4)Hex(5)NeuAc(1) | C[+57], N[+1914] | 1336.9037 | 3 | 4008.6967 | 4008.687 | 0 | 2.4 | 485 | Specific | 445.2 | 306.12 | 306.12 | 7.13 | 45 | >sp\|P02788\|TRFL_HUMAN Lactotransferrin OS=Homo sapiens OX=9606 GN=LTF PE=1 SV=6 | 1270 | 20201113_gy_colo_milk_1ng_glyco_hcd30_etd30_compound15.17811.17811.3 | controllerType=0 controllerNumber=1 scan=17811 | 45.2439 |
| 13010:2 | 3 | R.TAGWNIPMGLLFN[+203.079]QTGSC[+57.021]K.F | HexNAc(1) | C[+57], N[+203] | 1149.5402 | 2 | 2298.0731 | 2298.0893 | 0 | -7 | 485 | Specific | 470.87 | 334.27 | 334.27 | 3.08 | 45 | >sp\|P02788\|TRFL_HUMAN Lactotransferrin OS=Homo sapiens OX=9606 GN=LTF PE=1 SV=6 | 1270 | 20201113_gy_colo_milk_1ng_glyco_hcd30_etd30_compound15.16429.16429.2 | controllerType=0 controllerNumber=1 scan=16429 | 41.0226 |
| 14195:3 | 3 | R.TAGWNIPMGLLFN[+2205.793]QTGSC[+57.021]K.F | HexNAc(4)Hex(5)Fuc(2)NeuAc(1) | C[+57], N[+2206] | 1434.2767 | 3 | 4300.8155 | 4300.8028 | 0 | 2.9 | 485 | Specific | 474.83 | 330.4 | 330.4 | 7.36 | 45 | >sp\|P02788\|TRFL_HUMAN Lactotransferrin OS=Homo sapiens OX=9606 GN=LTF PE=1 SV=6 | 1270 | 20201113_gy_colo_milk_1ng_glyco_hcd30_etd30_compound15.17682.17682.3 | controllerType=0 controllerNumber=1 scan=17682 | 44.8059 |
| 13220:3 | 3 | R.TAGWNIPMGLLFN[+1216.423]QTGSC[+57.021]K.F | HexNAc(2)Hex(5) | C[+57], N[+1216] | 1104.4859 | 3 | 3311.4431 | 3311.4328 | 0 | 3.1 | 485 | Specific | 478.29 | 395.33 | 217.88 | 8.5 | 45 | >sp\|P02788\|TRFL_HUMAN Lactotransferrin OS=Homo sapiens OX=9606 GN=LTF PE=1 SV=6 | 1270 | 20201113_gy_colo_milk_1ng_glyco_hcd30_etd30_compound15.16651.16651.3 | controllerType=0 controllerNumber=1 scan=16651 | 41.6596 |
| 14356:3 | 3 | R.TAGWNIPMGLLFN[+1872.651]QTGSC[+57.021]K.F | HexNAc(3)Hex(6)NeuAc(1) | C[+57], N[+1873] | 1323.2285 | 3 | 3967.6708 | 3967.6604 | 0 | 2.6 | 485 | Specific | 504.67 | 348.96 | 277.03 | 7.55 | 45 | >sp\|P02788\|TRFL_HUMAN Lactotransferrin OS=Homo sapiens OX=9606 GN=LTF PE=1 SV=6 | 1270 | 20201113_gy_colo_milk_1ng_glyco_hcd30_etd30_compound15.17853.17853.3 | controllerType=0 controllerNumber=1 scan=17853 | 45.3628 |
| 14338:4 | 3 | R.TAGWNIPMGLLFN[+1913.677]QTGSC[+57.021]K.F | HexNAc(4)Hex(5)NeuAc(1) | C[+57], N[+1914] | 1002.9311 | 4 | 4008.7027 | 4008.687 | 0 | 3.9 | 485 | Specific | 513.8 | 446.59 | 446.59 | 5.54 | 45 | >sp\|P02788\|TRFL_HUMAN Lactotransferrin OS=Homo sapiens OX=9606 GN=LTF PE=1 SV=6 | 1270 | 20201113_gy_colo_milk_1ng_glyco_hcd30_etd30_compound15.17834.17834.4 | controllerType=0 controllerNumber=1 scan=17834 | 45.3011 |
| 12671:3 | 3 | R.TAGWNIPM[+15.995]GLLFN[+2059.735]QTGSC[+57.021]K.F | HexNAc(4)Hex(5)Fuc(1)NeuAc(1) | C[+57], M[+16], N[+2060] | 1390.925 | 3 | 4170.7604 | 4170.7398 | 0 | 4.9 | 485 | Specific | 222.06 | 190.34 | 190.34 | 2.81 | 45 | >sp\|P02788\|TRFL_HUMAN Lactotransferrin OS=Homo sapiens OX=9606 GN=LTF PE=1 SV=6 | 1270 | 20201113_gy_colo_milk_1ng_glyco_hcd30_etd30_compound15.16073.16073.3 | controllerType=0 controllerNumber=1 scan=16073 | 40.0437 |
| 14420:4 | 3 | R.TAGWNIPMGLLFN[+2059.735]QTGSC[+57.021]KFDEYFSQSC[+57.021]APGSDPR.S | HexNAc(4)Hex(5)Fuc(1)NeuAc(1) | C[+57]*2, N[+2060] | 1500.3826 | 4 | 5998.5085 | 5998.492 | 0 | 2.7 | 485 | Specific | 372.35 | 329.54 | 329.54 | 6.65 | 45 | >sp\|P02788\|TRFL_HUMAN Lactotransferrin OS=Homo sapiens OX=9606 GN=LTF PE=1 SV=6 | 1270 | 20201113_gy_colo_milk_1ng_glyco_hcd30_etd30_compound15.17921.17921.4 | controllerType=0 controllerNumber=1 scan=17921 | 45.5818 |
| 07206:3 | 3 | R.N[+2569.905]GSDC[+57.021]PDKFC[+57.021]LFQSETK.N | HexNAc(5)Hex(6)NeuAc(2) | C[+57]*2, N[+2570] | 1534.9399 | 3 | 4602.8051 | 4602.7785 | 0 | 5.8 | 642 | Specific | 258.97 | 157.19 | 157.19 | 3.41 | 45 | >sp\|P02788\|TRFL_HUMAN Lactotransferrin OS=Homo sapiens OX=9606 GN=LTF PE=1 SV=6 | 1270 | 20201113_gy_colo_milk_1ng_glyco_hcd30_etd30_compound15.10350.10350.3 | controllerType=0 controllerNumber=1 scan=10350 | 26.4593 |
| 06010:3 | 4 | R.FSDGLESN[+1864.634]SSTQFEVK.K | HexNAc(2)Hex(9) | N[+1865] | 1213.824 | 3 | 3639.4575 | 3639.4472 | 0 | 2.9 | 219 | Specific | 444.31 | 420.65 | 420.65 | 8.03 | 22 | >sp\|P0C0L4\|CO4A_HUMAN Complement C4-A OS=Homo sapiens OX=9606 GN=C4A PE=1 SV=2 | 1235 | 20201113_gy_colo_milk_1ng_glyco_hcd30_etd30_compound15.9100.9100.3 | controllerType=0 controllerNumber=1 scan=9100 | 23.709 |
| 03711:3 | 4 | R.FSDGLESN[+1864.634]SSTQFEVKK.Y | HexNAc(2)Hex(9) | N[+1865] | 1256.5214 | 3 | 3767.5497 | 3767.5421 | 0 | 2 | 219 | Specific | 232.01 | 168.71 | 168.71 | 2.9 | 22 | >sp\|P0C0L4\|CO4A_HUMAN Complement C4-A OS=Homo sapiens OX=9606 GN=C4A PE=1 SV=2 | 1235 | 20201113_gy_colo_milk_1ng_glyco_hcd30_etd30_compound15.6714.6714.3 | controllerType=0 controllerNumber=1 scan=6714 | 19.3827 |
| 15688:4 | 5 | K.QIGLYPVLVIDSSGYVNPN[+2204.772]YTGR.I | HexNAc(4)Hex(5)NeuAc(2) | N[+2205] | 1183.2748 | 4 | 4730.0775 | 4730.0759 | 0 | 0.3 | 168 | Specific | 519.61 | 464.61 | 464.61 | 5.81 | 35 | >sp\|P01833\|PIGR_HUMAN Polymeric immunoglobulin receptor OS=Homo sapiens OX=9606 GN=PIGR PE=1 SV=4 | 902 | 20201113_gy_colo_milk_1ng_glyco_hcd30_etd30_compound15.19265.19265.4 | controllerType=0 controllerNumber=1 scan=19265 | 49.6828 |
| 15655:3 | 5 | K.QIGLYPVLVIDSSGYVNPN[+2350.830]YTGR.I | HexNAc(4)Hex(5)Fuc(1)NeuAc(2) | N[+2351] | 1626.0554 | 3 | 4876.1515 | 4876.1338 | 0 | 3.6 | 168 | Specific | 551.44 | 456.12 | 456.12 | 7.6 | 35 | >sp\|P01833\|PIGR_HUMAN Polymeric immunoglobulin receptor OS=Homo sapiens OX=9606 GN=PIGR PE=1 SV=4 | 902 | 20201113_gy_colo_milk_1ng_glyco_hcd30_etd30_compound15.19229.19229.3 | controllerType=0 controllerNumber=1 scan=19229 | 49.5513 |
| 15687:3 | 5 | K.QIGLYPVLVIDSSGYVNPN[+2204.772]YTGR.I | HexNAc(4)Hex(5)NeuAc(2) | N[+2205] | 1577.3643 | 3 | 4730.0784 | 4730.0759 | 0 | 0.5 | 168 | Specific | 569.24 | 457.88 | 457.88 | 8.87 | 35 | >sp\|P01833\|PIGR_HUMAN Polymeric immunoglobulin receptor OS=Homo sapiens OX=9606 GN=PIGR PE=1 SV=4 | 902 | 20201113_gy_colo_milk_1ng_glyco_hcd30_etd30_compound15.19264.19264.3 | controllerType=0 controllerNumber=1 scan=19264 | 49.6783 |

Table S4. Cont.

| Query #:z | Protein Rank | Glycopeptide | Glycan | Modification Type(s) | Observed m/z | z | Observed (M+H) | Calc. mass (M+H) | Off-by-x error | Mass error (ppm) | StartPosition | Cleavage | Score | Delta | Delta Mod | \|Log Prob\| | # of unique peptides | Protein Name | Protein DB number | Comment | Scan # | Scan Time |
|---|---|---|---|---|---|---|---|---|---|---|---|---|---|---|---|---|---|---|---|---|---|---|
| 14317:3 | 5 | K.QIGLYPVLVIDSSGYVNPN[+2205.793]YTGR.I | HexNAc(4)Hex(5)Fuc(2)NeuAc(1) | N[+2206] | 1577.7075 | 3 | 4731.1078 | 4731.0963 | 0 | 2.4 | 168 | Specific | 572.54 | 465.61 | 465.61 | 9.74 | 35 | >sp\|P01833\|PIGR_HUMAN Polymeric immunoglobulin receptor OS=Homo sapiens OX=9606 GN=PIGR PE=1 SV=4 | 902 | 20201113_gy_colo_milk_1ng_glyco_hcd30_etd30_compound15.17812.17812.3 | controllerType=0 controllerNumber=1 scan=17812 | 45.2467 |
| 17029:3 | 5 | R.LSLLEEPGN[+2351.851]GTFTVILNQLTSR.D | HexNAc(4)Hex(5)Fuc(3)NeuAc(1) | N[+2352] | 1585.3798 | 3 | 4754.1249 | 4754.1433 | 0 | -3.9 | 413 | Specific | 206.97 | 64.49 | 64.49 | 2.96 | 35 | >sp\|P01833\|PIGR_HUMAN Polymeric immunoglobulin receptor OS=Homo sapiens OX=9606 GN=PIGR PE=1 SV=4 | 902 | 20201113_gy_colo_milk_1ng_glyco_hcd30_etd30_compound15.20690.20690.3 | controllerType=0 controllerNumber=1 scan=20690 | 54.3935 |
| 20852:3 | 5 | R.[+42.011]LSLLEEPGN[+2351.851]GTFTVILNQLTSR.D | HexNAc(4)Hex(5)Fuc(3)NeuAc(1) | .[42], N[+2352] | 1599.393 | 3 | 4796.1645 | 4796.1539 | 0 | 2.2 | 413 | Specific | 255.27 | 165.68 | 165.68 | 6.3 | 35 | >sp\|P01833\|PIGR_HUMAN Polymeric immunoglobulin receptor OS=Homo sapiens OX=9606 GN=PIGR PE=1 SV=4 | 902 | 20201113_gy_colo_milk_1ng_glyco_hcd30_etd30_compound15.24739.24739.3 | controllerType=0 controllerNumber=1 scan=24739 | 64.8912 |
| 15509:3 | 5 | R.LSLLEEPGN[+2205.793]GTFTVILNQLTSR.D | HexNAc(4)Hex(5)Fuc(2)NeuAc(1) | N[+2206] | 1536.7051 | 3 | 4608.1009 | 4608.0854 | 0 | 3.4 | 413 | Specific | 517.7 | 381.83 | 381.83 | 7.55 | 35 | >sp\|P01833\|PIGR_HUMAN Polymeric immunoglobulin receptor OS=Homo sapiens OX=9606 GN=PIGR PE=1 SV=4 | 902 | 20201113_gy_colo_milk_1ng_glyco_hcd30_etd30_compound15.19074.19074.3 | controllerType=0 controllerNumber=1 scan=19074 | 49.026 |
| 16763:3 | 5 | R.LSLLEEPGN[+2496.888]GTFTVILNQLTSR.D | HexNAc(4)Hex(5)Fuc(2)NeuAc(2) | N[+2497] | 1633.7354 | 3 | 4899.1917 | 4899.1808 | 0 | 2.2 | 413 | Specific | 588.12 | 419.62 | 419.62 | 9.74 | 35 | >sp\|P01833\|PIGR_HUMAN Polymeric immunoglobulin receptor OS=Homo sapiens OX=9606 GN=PIGR PE=1 SV=4 | 902 | 20201113_gy_colo_milk_1ng_glyco_hcd30_etd30_compound15.20407.20407.3 | controllerType=0 controllerNumber=1 scan=20407 | 53.4421 |
| 15443:3 | 5 | R.LSLLEEPGN[+2351.851]GTFTVILNQLTSR.D | HexNAc(4)Hex(5)Fuc(3)NeuAc(1) | N[+2352] | 1585.3899 | 3 | 4754.1553 | 4754.1433 | 0 | 2.5 | 413 | Specific | 590.08 | 455.99 | 455.99 | 9.74 | 35 | >sp\|P01833\|PIGR_HUMAN Polymeric immunoglobulin receptor OS=Homo sapiens OX=9606 GN=PIGR PE=1 SV=4 | 902 | 20201113_gy_colo_milk_1ng_glyco_hcd30_etd30_compound15.19004.19004.3 | controllerType=0 controllerNumber=1 scan=19004 | 48.7934 |
| 15449:4 | 5 | R.LSLLEEPGN[+2351.851]GTFTVILNQLTSR.D | HexNAc(4)Hex(5)Fuc(3)NeuAc(1) | N[+2352] | 1189.2951 | 4 | 4754.1585 | 4754.1433 | 0 | 3.2 | 413 | Specific | 600.19 | 481.14 | 481.14 | 7.59 | 35 | >sp\|P01833\|PIGR_HUMAN Polymeric immunoglobulin receptor OS=Homo sapiens OX=9606 GN=PIGR PE=1 SV=4 | 902 | 20201113_gy_colo_milk_1ng_glyco_hcd30_etd30_compound15.19010.19010.4 | controllerType=0 controllerNumber=1 scan=19010 | 48.8052 |
| 12397:3 | 5 | K.IIEGEPNLKVPGN[+2351.851]VTAVLGETLK.V | HexNAc(4)Hex(5)Fuc(3)NeuAc(1) | N[+2352] | 1581.7426 | 3 | 4743.2133 | 4743.2001 | 0 | 2.8 | 457 | Specific | 227.29 | 182.66 | 182.66 | 2.85 | 35 | >sp\|P01833\|PIGR_HUMAN Polymeric immunoglobulin receptor OS=Homo sapiens OX=9606 GN=PIGR PE=1 SV=4 | 902 | 20201113_gy_colo_milk_1ng_glyco_hcd30_etd30_compound15.15785.15785.3 | controllerType=0 controllerNumber=1 scan=15785 | 39.355 |
| 12427:3 | 5 | K.IIEGEPNLKVPGN[+1913.677]VTAVLGETLK.V | HexNAc(4)Hex(5)NeuAc(1) | N[+1914] | 1435.6856 | 3 | 4305.0423 | 4305.0264 | 0 | 3.7 | 457 | Specific | 234.41 | 189.12 | 189.12 | 2.65 | 35 | >sp\|P01833\|PIGR_HUMAN Polymeric immunoglobulin receptor OS=Homo sapiens OX=9606 GN=PIGR PE=1 SV=4 | 902 | 20201113_gy_colo_milk_1ng_glyco_hcd30_etd30_compound15.15816.15816.3 | controllerType=0 controllerNumber=1 scan=15816 | 39.4172 |
| 12379:3 | 5 | K.IIEGEPNLKVPGN[+2351.851]VTAVLGETLK.V | HexNAc(4)Hex(5)Fuc(3)NeuAc(1) | N[+2352] | 1581.7411 | 3 | 4743.2089 | 4743.2001 | 0 | 1.9 | 457 | Specific | 276.55 | 267.75 | 267.75 | 3.96 | 35 | >sp\|P01833\|PIGR_HUMAN Polymeric immunoglobulin receptor OS=Homo sapiens OX=9606 GN=PIGR PE=1 SV=4 | 902 | 20201113_gy_colo_milk_1ng_glyco_hcd30_etd30_compound15.15766.15766.3 | controllerType=0 controllerNumber=1 scan=15766 | 39.3115 |
| 12359:3 | 5 | K.IIEGEPNLKVPGN[+2205.793]VTAVLGETLK.V | HexNAc(4)Hex(5)Fuc(2)NeuAc(1) | N[+2206] | 1533.0608 | 3 | 4597.168 | 4597.1422 | 0 | 5.6 | 457 | Specific | 353.88 | 314.72 | 314.72 | 2.95 | 35 | >sp\|P01833\|PIGR_HUMAN Polymeric immunoglobulin receptor OS=Homo sapiens OX=9606 GN=PIGR PE=1 SV=4 | 902 | 20201113_gy_colo_milk_1ng_glyco_hcd30_etd30_compound15.15745.15745.3 | controllerType=0 controllerNumber=1 scan=15745 | 39.2569 |
| 12348:3 | 5 | K.IIEGEPNLKVPGN[+2351.851]VTAVLGETLK.V | HexNAc(4)Hex(5)Fuc(3)NeuAc(1) | N[+2352] | 1581.7446 | 3 | 4743.2191 | 4743.2001 | 0 | 4 | 457 | Specific | 406.7 | 330.03 | 330.03 | 3.81 | 35 | >sp\|P01833\|PIGR_HUMAN Polymeric immunoglobulin receptor OS=Homo sapiens OX=9606 GN=PIGR PE=1 SV=4 | 902 | 20201113_gy_colo_milk_1ng_glyco_hcd30_etd30_compound15.15733.15733.3 | controllerType=0 controllerNumber=1 scan=15733 | 39.2291 |
| 12070:5 | 5 | K.IIEGEPNLKVPGN[+2204.772]VTAVLGETLKVPC[+57.021]HFPC[+57.021]K.F | HexNAc(4)Hex(5)NeuAc(2) | C[+57]*2, N[+2205] | 1125.131 | 5 | 5621.6258 | 5621.5793 | 0 | 8.3 | 457 | Specific | 296.02 | 296.02 | 296.02 | 2.6 | 35 | >sp\|P01833\|PIGR_HUMAN Polymeric immunoglobulin receptor OS=Homo sapiens OX=9606 GN=PIGR PE=1 SV=4 | 902 | 20201113_gy_colo_milk_1ng_glyco_hcd30_etd30_compound15.15440.15440.5 | controllerType=0 controllerNumber=1 scan=15440 | 38.5222 |
| 12840:3 | 5 | K.VPGN[+2350.830]VTAVLGETLK.V | HexNAc(4)Hex(5)Fuc(1)NeuAc(2) | N[+2351] | 1250.2176 | 3 | 3748.6382 | 3748.6302 | 0 | 2.1 | 466 | Specific | 232.48 | 136.27 | 136.27 | 3.33 | 35 | >sp\|P01833\|PIGR_HUMAN Polymeric immunoglobulin receptor OS=Homo sapiens OX=9606 GN=PIGR PE=1 SV=4 | 902 | 20201113_gy_colo_milk_1ng_glyco_hcd30_etd30_compound15.16251.16251.3 | controllerType=0 controllerNumber=1 scan=16251 | 40.5284 |
| 10555:3 | 5 | K.VPGN[+2205.793]VTAVLGETLK.V | HexNAc(4)Hex(5)Fuc(2)NeuAc(1) | N[+2206] | 1201.872 | 3 | 3603.6015 | 3603.5927 | 0 | 2.4 | 466 | Specific | 282.13 | 174.34 | 174.34 | 5.93 | 35 | >sp\|P01833\|PIGR_HUMAN Polymeric immunoglobulin receptor OS=Homo sapiens OX=9606 GN=PIGR PE=1 SV=4 | 902 | 20201113_gy_colo_milk_1ng_glyco_hcd30_etd30_compound15.13854.13854.3 | controllerType=0 controllerNumber=1 scan=13854 | 34.6251 |
| 12024:4 | 5 | K.VPGN[+2350.830]VTAVLGETLKVPC[+57.021]HFPC[+57.021]K.F | HexNAc(4)Hex(5)Fuc(1)NeuAc(2) | C[+57]*2, N[+2351] | 1194.2785 | 4 | 4774.0921 | 4774.0877 | 0 | 0.9 | 466 | Specific | 317.87 | 273.36 | 273.36 | 3.07 | 35 | >sp\|P01833\|PIGR_HUMAN Polymeric immunoglobulin receptor OS=Homo sapiens OX=9606 GN=PIGR PE=1 SV=4 | 902 | 20201113_gy_colo_milk_1ng_glyco_hcd30_etd30_compound15.15391.15391.4 | controllerType=0 controllerNumber=1 scan=15391 | 38.3804 |
| 04678:3 | 5 | K.WN[+1548.545]NTGC[+57.021]QALPSQDEGPSK.A | HexNAc(3)Hex(4)NeuAc(1) | C[+57], N[+1549] | 1179.8145 | 3 | 3537.429 | 3537.4215 | 0 | 2.1 | 498 | Specific | 302.07 | 153.13 | 153.13 | 5.88 | 35 | >sp\|P01833\|PIGR_HUMAN Polymeric immunoglobulin receptor OS=Homo sapiens OX=9606 GN=PIGR PE=1 SV=4 | 902 | 20201113_gy_colo_milk_1ng_glyco_hcd30_etd30_compound15.7717.7717.3 | controllerType=0 controllerNumber=1 scan=7717 | 21.2417 |
| 06326:3 | 5 | K.WN[+2204.772]NTGC[+57.021]QALPSQDEGPSK.A | HexNAc(4)Hex(5)NeuAc(2) | C[+57], N[+2205] | 1398.5573 | 3 | 4193.6574 | 4193.6491 | 0 | 2 | 498 | Specific | 331.38 | 199.07 | 199.07 | 6 | 35 | >sp\|P01833\|PIGR_HUMAN Polymeric immunoglobulin receptor OS=Homo sapiens OX=9606 GN=PIGR PE=1 SV=4 | 902 | 20201113_gy_colo_milk_1ng_glyco_hcd30_etd30_compound15.9431.9431.3 | controllerType=0 controllerNumber=1 scan=9431 | 24.4276 |
| 04418:3 | 5 | K.WN[+2205.793]NTGC[+57.021]QALPSQDEGPSK.A | HexNAc(4)Hex(5)Fuc(2)NeuAc(1) | C[+57], N[+2206] | 1398.8981 | 3 | 4194.6798 | 4194.6695 | 0 | 2.5 | 498 | Specific | 354.21 | 189.51 | 189.51 | 6 | 35 | >sp\|P01833\|PIGR_HUMAN Polymeric immunoglobulin receptor OS=Homo sapiens OX=9606 GN=PIGR PE=1 SV=4 | 902 | 20201113_gy_colo_milk_1ng_glyco_hcd30_etd30_compound15.7447.7447.3 | controllerType=0 controllerNumber=1 scan=7447 | 20.7384 |
| 04430:3 | 5 | K.WN[+2059.735]NTGC[+57.021]QALPSQDEGPSK.A | HexNAc(4)Hex(5)Fuc(1)NeuAc(1) | C[+57], N[+2060] | 1350.2129 | 3 | 4048.6243 | 4048.6116 | 0 | 3.1 | 498 | Specific | 401.91 | 267.73 | 267.73 | 6.37 | 35 | >sp\|P01833\|PIGR_HUMAN Polymeric immunoglobulin receptor OS=Homo sapiens OX=9606 GN=PIGR PE=1 SV=4 | 902 | 20201113_gy_colo_milk_1ng_glyco_hcd30_etd30_compound15.7460.7460.3 | controllerType=0 controllerNumber=1 scan=7460 | 20.7668 |
| 06294:3 | 5 | K.WN[+2204.772]NTGC[+57.021]QALPSQDEGPSK.A | HexNAc(4)Hex(5)NeuAc(2) | C[+57], N[+2205] | 1398.5566 | 3 | 4193.6552 | 4193.6491 | 0 | 1.4 | 498 | Specific | 430.46 | 333.45 | 333.45 | 6.71 | 35 | >sp\|P01833\|PIGR_HUMAN Polymeric immunoglobulin receptor OS=Homo sapiens OX=9606 GN=PIGR PE=1 SV=4 | 902 | 20201113_gy_colo_milk_1ng_glyco_hcd30_etd30_compound15.9397.9397.3 | controllerType=0 controllerNumber=1 scan=9397 | 24.3485 |
| 04527:3 | 5 | K.WN[+1913.677]NTGC[+57.021]QALPSQDEGPSK.A | HexNAc(4)Hex(5)NeuAc(1) | C[+57], N[+1914] | 1301.5267 | 3 | 3902.5655 | 3902.5537 | 0 | 3 | 498 | Specific | 439.86 | 285.48 | 285.48 | 6.37 | 35 | >sp\|P01833\|PIGR_HUMAN Polymeric immunoglobulin receptor OS=Homo sapiens OX=9606 GN=PIGR PE=1 SV=4 | 902 | 20201113_gy_colo_milk_1ng_glyco_hcd30_etd30_compound15.7561.7561.3 | controllerType=0 controllerNumber=1 scan=7561 | 20.9599 |
| 03128:3 | 5 | K.WN[+1216.423]NTGC[+57.021]QALPSQDEGPSK.A | HexNAc(2)Hex(5) | C[+57], N[+1216] | 1069.1052 | 3 | 3205.301 | 3205.2995 | 0 | 0.5 | 498 | Specific | 477.55 | 374.21 | 374.21 | 7.15 | 35 | >sp\|P01833\|PIGR_HUMAN Polymeric immunoglobulin receptor OS=Homo sapiens OX=9606 GN=PIGR PE=1 SV=4 | 902 | 20201113_gy_colo_milk_1ng_glyco_hcd30_etd30_compound15.6106.6106.3 | controllerType=0 controllerNumber=1 scan=6106 | 18.3243 |
| 04503:3 | 5 | K.WN[+1872.651]NTGC[+57.021]QALPSQDEGPSK.A | HexNAc(3)Hex(6)NeuAc(1) | C[+57], N[+1873] | 1287.8519 | 3 | 3861.541 | 3861.5272 | 0 | 3.6 | 498 | Specific | 479.73 | 302.06 | 302.06 | 5.51 | 35 | >sp\|P01833\|PIGR_HUMAN Polymeric immunoglobulin receptor OS=Homo sapiens OX=9606 GN=PIGR PE=1 SV=4 | 902 | 20201113_gy_colo_milk_1ng_glyco_hcd30_etd30_compound15.7536.7536.3 | controllerType=0 controllerNumber=1 scan=7536 | 20.9086 |
| 03105:3 | 5 | K.WN[+1216.423]NTGC[+57.021]QALPSQDEGPSK.A | HexNAc(2)Hex(5) | C[+57], N[+1216] | 1069.1067 | 3 | 3205.3056 | 3205.2995 | 0 | 1.9 | 498 | Specific | 493.15 | 406.38 | 406.38 | 8.22 | 35 | >sp\|P01833\|PIGR_HUMAN Polymeric immunoglobulin receptor OS=Homo sapiens OX=9606 GN=PIGR PE=1 SV=4 | 902 | 20201113_gy_colo_milk_1ng_glyco_hcd30_etd30_compound15.6082.6082.3 | controllerType=0 controllerNumber=1 scan=6082 | 18.2799 |
| 04581:3 | 5 | K.WN[+1710.598]NTGC[+57.021]QALPSQDEGPSK.A | HexNAc(3)Hex(5)NeuAc(1) | C[+57], N[+1711] | 1233.8326 | 3 | 3699.4832 | 3699.4743 | 0 | 2.4 | 498 | Specific | 496.07 | 325.47 | 325.47 | 7.07 | 35 | >sp\|P01833\|PIGR_HUMAN Polymeric immunoglobulin receptor OS=Homo sapiens OX=9606 GN=PIGR PE=1 SV=4 | 902 | 20201113_gy_colo_milk_1ng_glyco_hcd30_etd30_compound15.7617.7617.3 | controllerType=0 controllerNumber=1 scan=7617 | 21.0668 |
| 08078:3 | 7 | K.TVLTPATNHMGN[+1540.529]VTFTIPANR.E | HexNAc(2)Hex(7) | N[+1541] | 1265.903 | 3 | 3795.6945 | 3795.6886 | 0 | 1.5 | 74 | Specific | 323.61 | 262.03 | 135.54 | 6.31 | 12 | >sp\|P01024\|CO3_HUMAN Complement C3 OS=Homo sapiens OX=9606 GN=C3 PE=1 SV=2 | 492 | 20201113_gy_colo_milk_1ng_glyco_hcd30_etd30_compound15.11260.11260.3 | controllerType=0 controllerNumber=1 scan=11260 | 28.4766 |
| 08088:3 | 7 | K.TVLTPATNHMGN[+1378.476]VTFTIPANR.E | HexNAc(2)Hex(6) | N[+1378] | 1211.887 | 3 | 3633.6465 | 3633.6358 | 0 | 2.9 | 74 | Specific | 327.51 | 295.63 | 124.13 | 5.39 | 12 | >sp\|P01024\|CO3_HUMAN Complement C3 OS=Homo sapiens OX=9606 GN=C3 PE=1 SV=2 | 492 | 20201113_gy_colo_milk_1ng_glyco_hcd30_etd30_compound15.11271.11271.3 | controllerType=0 controllerNumber=1 scan=11271 | 28.5229 |
| 17460:3 | 9 | R.QLAHQSN[+2350.830]STNIFFSPVSIATAFAMLSLGTK.A | HexNAc(4)Hex(5)Fuc(1)NeuAc(2) | N[+2351] | 1844.8294 | 3 | 5532.4737 | 5532.4654 | 0 | 1.5 | 64 | Specific | 218.87 | 131.01 | 131.01 | 4.86 | 10 | >sp\|P01009\|A1AT_HUMAN Alpha-1-antitrypsin OS=Homo sapiens OX=9606 GN=SERPINA1 PE=1 SV=3 | 642 | 20201113_gy_colo_milk_1ng_glyco_hcd30_etd30_compound15.21147.21147.3 | controllerType=0 controllerNumber=1 scan=21147 | 55.8644 |
| 19382:4 | 9 | R.QLAHQSN[+2861.000]STNIFFSPVSIATAFAMLSLGTK.A | HexNAc(5)Hex(6)NeuAc(3) | N[+2861] | 1511.4202 | 4 | 6042.6588 | 6042.6351 | 0 | 3.9 | 64 | Specific | 235.99 | 233.78 | 233.78 | 3.38 | 10 | >sp\|P01009\|A1AT_HUMAN Alpha-1-antitrypsin OS=Homo sapiens OX=9606 GN=SERPINA1 PE=1 SV=3 | 642 | 20201113_gy_colo_milk_1ng_glyco_hcd30_etd30_compound15.23177.23177.4 | controllerType=0 controllerNumber=1 scan=23177 | 61.2257 |
| 17517:4 | 9 | R.QLAHQSN[+2350.830]STNIFFSPVSIATAFAMLSLGTK.A | HexNAc(4)Hex(5)Fuc(1)NeuAc(2) | N[+2351] | 1383.876 | 4 | 5532.4821 | 5532.4654 | 0 | 3 | 64 | Specific | 323.07 | 269.79 | 269.79 | 5.96 | 10 | >sp\|P01009\|A1AT_HUMAN Alpha-1-antitrypsin OS=Homo sapiens OX=9606 GN=SERPINA1 PE=1 SV=3 | 642 | 20201113_gy_colo_milk_1ng_glyco_hcd30_etd30_compound15.21208.21208.4 | controllerType=0 controllerNumber=1 scan=21208 | 56.0808 |
| 17470:3 | 9 | R.QLAHQSN[+2204.772]STNIFFSPVSIATAFAMLSLGTK.A | HexNAc(4)Hex(5)NeuAc(2) | N[+2205] | 1796.1523 | 3 | 5386.4424 | 5386.4075 | 0 | 6.5 | 64 | Specific | 439.71 | 330.52 | 330.52 | 4.79 | 10 | >sp\|P01009\|A1AT_HUMAN Alpha-1-antitrypsin OS=Homo sapiens OX=9606 GN=SERPINA1 PE=1 SV=3 | 642 | 20201113_gy_colo_milk_1ng_glyco_hcd30_etd30_compound15.21158.21158.3 | controllerType=0 controllerNumber=1 scan=21158 | 55.9061 |
| 17453:4 | 9 | R.QLAHQSN[+2350.830]STNIFFSPVSIATAFAMLSLGTK.A | HexNAc(4)Hex(5)Fuc(1)NeuAc(2) | N[+2351] | 1383.8737 | 4 | 5532.4728 | 5532.4654 | 0 | 1.3 | 64 | Specific | 498.64 | 419.63 | 419.63 | 6.85 | 10 | >sp\|P01009\|A1AT_HUMAN Alpha-1-antitrypsin OS=Homo sapiens OX=9606 GN=SERPINA1 PE=1 SV=3 | 642 | 20201113_gy_colo_milk_1ng_glyco_hcd30_etd30_compound15.21140.21140.4 | controllerType=0 controllerNumber=1 scan=21140 | 55.8433 |
| 17471:5 | 9 | R.QLAHQSN[+2204.772]STNIFFSPVSIATAFAMLSLGTK.A | HexNAc(4)Hex(5)NeuAc(2) | N[+2205] | 1078.0901 | 5 | 5386.4213 | 5386.4075 | 0 | 2.6 | 64 | Specific | 707.74 | 623.92 | 505.18 | 12.27 | 10 | >sp\|P01009\|A1AT_HUMAN Alpha-1-antitrypsin OS=Homo sapiens OX=9606 GN=SERPINA1 PE=1 SV=3 | 642 | 20201113_gy_colo_milk_1ng_glyco_hcd30_etd30_compound15.21159.21159.5 | controllerType=0 controllerNumber=1 scan=21159 | 55.9129 |
| 17466:4 | 9 | R.QLAHQSN[+2204.772]STNIFFSPVSIATAFAMLSLGTK.A | HexNAc(4)Hex(5)NeuAc(2) | N[+2205] | 1347.3661 | 4 | 5386.4425 | 5386.4075 | 0 | 6.5 | 64 | Specific | 751.88 | 627.48 | 627.48 | 12.77 | 10 | >sp\|P01009\|A1AT_HUMAN Alpha-1-antitrypsin OS=Homo sapiens OX=9606 GN=SERPINA1 PE=1 SV=3 | 642 | 20201113_gy_colo_milk_1ng_glyco_hcd30_etd30_compound15.21154.21154.4 | controllerType=0 controllerNumber=1 scan=21154 | 55.8869 |
| 15104:5 | 9 | K.YLGN[+2204.772]ATAIFFLPDEGKLQHLENELTHDIITK.F | HexNAc(4)Hex(5)NeuAc(2) | N[+2205] | 1149.9325 | 5 | 5745.6334 | 5745.6098 | 0 | 4.1 | 268 | Specific | 280.85 | 243.08 | 243.08 | 2.65 | 10 | >sp\|P01009\|A1AT_HUMAN Alpha-1-antitrypsin OS=Homo sapiens OX=9606 GN=SERPINA1 PE=1 SV=3 | 642 | 20201113_gy_colo_milk_1ng_glyco_hcd30_etd30_compound15.18644.18644.5 | controllerType=0 controllerNumber=1 scan=18644 | 47.7306 |
| 15103:4 | 9 | K.YLGN[+2204.772]ATAIFFLPDEGKLQHLENELTHDIITK.F | HexNAc(4)Hex(5)NeuAc(2) | N[+2205] | 1437.1642 | 4 | 5745.6349 | 5745.6098 | 0 | 4.4 | 268 | Specific | 573.54 | 573.54 | 573.54 | 7.36 | 10 | >sp\|P01009\|A1AT_HUMAN Alpha-1-antitrypsin OS=Homo sapiens OX=9606 GN=SERPINA1 PE=1 SV=3 | 642 | 20201113_gy_colo_milk_1ng_glyco_hcd30_etd30_compound15.18643.18643.4 | controllerType=0 controllerNumber=1 scan=18643 | 47.7273 |
| 16447:4 | 17 | K.VFPLSLDSTPQDGNVVVAC[+57.021]LVQGFFPQEPLSVTWSESGQN[+1444.534]VTAI | HexNAc(4)Hex(3)Fuc(1) | C[+57], N[+1445] | 1555.9849 | 4 | 6220.9176 | 6220.8953 | 0 | 3.6 | 8 | Specific | 255.2 | 255.2 | 255.2 | 4.88 | 13 | >sp\|P01877\|IGHA2_HUMAN Immunoglobulin heavy constant alpha 2 OS=Homo sapiens OX=9606 GN=IGHA2 PE=1 SV=4 | 576 | 20201113_gy_colo_milk_1ng_glyco_hcd30_etd30_compound15.20071.20071.2 | controllerType=0 controllerNumber=1 scan=20071 | 52.3128 |
| 16330:4 | 17 | K.VFPLSLDSTPQDGNVVVAC[+57.021]LVQGFFPQEPLSVTWSESGQN[+1809.666]VTAI | HexNAc(5)Hex(4)Fuc(1) | C[+57], N[+1810] | 1647.2715 | 4 | 6586.0641 | 6586.0275 | 0 | 5.6 | 8 | Specific | 424.05 | 424.05 | 424.05 | 6.51 | 13 | >sp\|P01877\|IGHA2_HUMAN Immunoglobulin heavy constant alpha 2 OS=Homo sapiens OX=9606 GN=IGHA2 PE=1 SV=4 | 576 | 20201113_gy_colo_milk_1ng_glyco_hcd30_etd30_compound15.19947.19947.4 | controllerType=0 controllerNumber=1 scan=19947 | 51.9219 |
| 16400:4 | 17 | K.VFPLSLDSTPQDGNVVVAC[+57.021]LVQGFFPQEPLSVTWSESGQN[+1647.613]VTAI | HexNAc(5)Hex(3)Fuc(1) | C[+57], N[+1648] | 1606.7527 | 4 | 6423.9889 | 6423.9746 | 0 | 2.2 | 8 | Specific | 444.29 | 444.29 | 444.29 | 7.94 | 13 | >sp\|P01877\|IGHA2_HUMAN Immunoglobulin heavy constant alpha 2 OS=Homo sapiens OX=9606 GN=IGHA2 PE=1 SV=4 | 576 | 20201113_gy_colo_milk_1ng_glyco_hcd30_etd30_compound15.20021.20021.4 | controllerType=0 controllerNumber=1 scan=20021 | 52.1589 |
| 02065:3 | 17 | K.TPLTAN[+1971.719]ITK.S | HexNAc(5)Hex(5)Fuc(1) | N[+1972] | 977.4306 | 3 | 2930.2773 | 2930.2757 | 0 | 0.6 | 200 | Specific | 216.81 | 101.14 | 101.14 | 2.64 | 13 | >sp\|P01877\|IGHA2_HUMAN Immunoglobulin heavy constant alpha 2 OS=Homo sapiens OX=9606 GN=IGHA2 PE=1 SV=4 | 576 | 20201113_gy_colo_milk_1ng_glyco_hcd30_etd30_compound15.4959.4959.3 | controllerType=0 controllerNumber=1 scan=4959 | 15.9992 |
| 03633:3 | 17 | K.TPLTAN[+2262.814]ITK.S | HexNAc(5)Hex(5)Fuc(1)NeuAc(1) | N[+2263] | 1074.4644 | 3 | 3221.3786 | 3221.3711 | 0 | 2.3 | 200 | Specific | 218.22 | 97.63 | 97.63 | 2.78 | 13 | >sp\|P01877\|IGHA2_HUMAN Immunoglobulin heavy constant alpha 2 OS=Homo sapiens OX=9606 GN=IGHA2 PE=1 SV=4 | 576 | 20201113_gy_colo_milk_1ng_glyco_hcd30_etd30_compound15.6633.6633.3 | controllerType=0 controllerNumber=1 scan=6633 | 19.2244 |
| 02114:2 | 17 | K.TPLTAN[+1444.534]ITK.S | HexNAc(4)Hex(3)Fuc(1) | N[+1445] | 1202.0464 | 2 | 2403.0856 | 2403.0906 | 0 | -2.1 | 200 | Specific | 245.85 | 64.99 | 64.99 | 2.6 | 13 | >sp\|P01877\|IGHA2_HUMAN Immunoglobulin heavy constant alpha 2 OS=Homo sapiens OX=9606 GN=IGHA2 PE=1 SV=4 | 576 | 20201113_gy_colo_milk_1ng_glyco_hcd30_etd30_compound15.5013.5013.2 | controllerType=0 controllerNumber=1 scan=5013 | 16.1259 |
| 03629:2 | 17 | K.TPLTAN[+2059.735]ITK.S | HexNAc(4)Hex(5)Fuc(1)NeuAc(1) | N[+2060] | 1509.6526 | 2 | 3018.298 | 3018.2917 | 0 | 2.1 | 200 | Specific | 253.1 | 81.29 | 81.29 | 4.33 | 13 | >sp\|P01877\|IGHA2_HUMAN Immunoglobulin heavy constant alpha 2 OS=Homo sapiens OX=9606 GN=IGHA2 PE=1 SV=4 | 576 | 20201113_gy_colo_milk_1ng_glyco_hcd30_etd30_compound15.6629.6629.2 | controllerType=0 controllerNumber=1 scan=6629 | 19.2118 |
| 03618:3 | 17 | K.TPLTAN[+2059.735]ITK.S | HexNAc(4)Hex(5)Fuc(1)NeuAc(1) | N[+2060] | 1006.772 | 3 | 3018.3015 | 3018.2917 | 0 | 3.2 | 200 | Specific | 253.79 | 127.74 | 127.74 | 2.89 | 13 | >sp\|P01877\|IGHA2_HUMAN Immunoglobulin heavy constant alpha 2 OS=Homo sapiens OX=9606 GN=IGHA2 PE=1 SV=4 | 576 | 20201113_gy_colo_milk_1ng_glyco_hcd30_etd30_compound15.6617.6617.3 | controllerType=0 controllerNumber=1 scan=6617 | 19.186 |
| 02195:2 | 17 | K.TPLTAN[+1241.454]ITK.S | HexNAc(3)Hex(3)Fuc(1) | N[+1241] | 1100.5098 | 2 | 2200.0123 | 2200.0113 | 0 | 0.5 | 200 | Specific | 323.01 | 81.38 | 81.38 | 2.66 | 13 | >sp\|P01877\|IGHA2_HUMAN Immunoglobulin heavy constant alpha 2 OS=Homo sapiens OX=9606 GN=IGHA2 PE=1 SV=4 | 576 | 20201113_gy_colo_milk_1ng_glyco_hcd30_etd30_compound15.5102.5102.2 | controllerType=0 controllerNumber=1 scan=5102 | 16.3302 |
| 02158:2 | 17 | K.TPLTAN[+1444.534]ITK.S | HexNAc(4)Hex(3)Fuc(1) | N[+1445] | 1202.0512 | 2 | 2403.0951 | 2403.0906 | 0 | 1.9 | 200 | Specific | 344.87 | 112.76 | 112.76 | 2.92 | 13 | >sp\|P01877\|IGHA2_HUMAN Immunoglobulin heavy constant alpha 2 OS=Homo sapiens OX=9606 GN=IGHA2 PE=1 SV=4 | 576 | 20201113_gy_colo_milk_1ng_glyco_hcd30_etd30_compound15.5062.5062.2 | controllerType=0 controllerNumber=1 scan=5062 | 16.2446 |
| 07702:3 | 25 | R.IIVPLNNREN[+1809.666]ISDPTSPLR.T | HexNAc(5)Hex(4)Fuc(1) | N[+1810] | 1319.9546 | 3 | 3957.8494 | 3957.8432 | 0 | 1.6 | 62 | Specific | 223.91 | 171.71 | 171.71 | 2.8 | 12 | >sp\|P01591\|IGJ_HUMAN Immunoglobulin J chain OS=Homo sapiens OX=9606 GN=JCHAIN PE=1 SV=4 | 1392 | 20201113_gy_colo_milk_1ng_glyco_hcd30_etd30_compound15.10868.10868.3 | controllerType=0 controllerNumber=1 scan=10868 | 27.6153 |
| 07833:3 | 25 | R.IIVPLNNREN[+1501.555]ISDPTSPLR.T | HexNAc(5)Hex(3) | N[+1502] | 1217.2524 | 3 | 3649.7426 | 3649.7325 | 0 | 2.8 | 62 | Specific | 236.04 | 153.78 | 153.78 | 2.83 | 12 | >sp\|P01591\|IGJ_HUMAN Immunoglobulin J chain OS=Homo sapiens OX=9606 GN=JCHAIN PE=1 SV=4 | 1392 | 20201113_gy_colo_milk_1ng_glyco_hcd30_etd30_compound15.11004.11004.3 | controllerType=0 controllerNumber=1 scan=11004 | 27.9058 |
| 08866:3 | 25 | R.IIVPLNNREN[+1913.677]ISDPTSPLR.T | HexNAc(4)Hex(5)NeuAc(1) | N[+1914] | 1354.6246 | 3 | 4061.8592 | 4061.8542 | 0 | 1.2 | 62 | Specific | 280.69 | 229.86 | 229.86 | 2.89 | 12 | >sp\|P01591\|IGJ_HUMAN Immunoglobulin J chain OS=Homo sapiens OX=9606 GN=JCHAIN PE=1 SV=4 | 1392 | 20201113_gy_colo_milk_1ng_glyco_hcd30_etd30_compound15.12086.12086.3 | controllerType=0 controllerNumber=1 scan=12086 | 30.4851 |
| 07748:3 | 25 | R.IIVPLNNREN[+1663.608]ISDPTSPLR.T | HexNAc(5)Hex(4) | N[+1664] | 1271.2692 | 3 | 3811.7931 | 3811.7853 | 0 | 2 | 62 | Specific | 286.11 | 197.98 | 197.98 | 3.57 | 12 | >sp\|P01591\|IGJ_HUMAN Immunoglobulin J chain OS=Homo sapiens OX=9606 GN=JCHAIN PE=1 SV=4 | 1392 | 20201113_gy_colo_milk_1ng_glyco_hcd30_etd30_compound15.10916.10916.3 | controllerType=0 controllerNumber=1 scan=10916 | 27.7167 |
| 08981:3 | 25 | R.IIVPLNNREN[+1694.603]ISDPTSPLR.T | HexNAc(3)Hex(4)Fuc(1)NeuAc(1) | N[+1695] | 1281.6006 | 3 | 3842.7874 | 3842.7799 | 0 | 2 | 62 | Specific | 305.52 | 231.56 | 231.56 | 3.81 | 12 | >sp\|P01591\|IGJ_HUMAN Immunoglobulin J chain OS=Homo sapiens OX=9606 GN=JCHAIN PE=1 SV=4 | 1392 | 20201113_gy_colo_milk_1ng_glyco_hcd30_etd30_compound15.12207.12207.3 | controllerType=0 controllerNumber=1 scan=12207 | 30.8527 |
| 05598:3 | 25 | R.EN[+1913.677]ISDPTSPLR.T | HexNAc(4)Hex(5)NeuAc(1) | N[+1914] | 1048.105 | 3 | 3142.3005 | 3142.2938 | 0 | 2.1 | 70 | Specific | 300.68 | 171.31 | 171.31 | 4.38 | 12 | >sp\|P01591\|IGJ_HUMAN Immunoglobulin J chain OS=Homo sapiens OX=9606 GN=JCHAIN PE=1 SV=4 | 1392 | 20201113_gy_colo_milk_1ng_glyco_hcd30_etd30_compound15.8671.8671.3 | controllerType=0 controllerNumber=1 scan=8671 | 22.9035 |
| 04436:3 | 39 | R.LSPN[+2205.793]ASAEHLELR.W | HexNAc(4)Hex(5)Fuc(2)NeuAc(1) | N[+2206] | 1214.8565 | 3 | 3642.555 | 3642.5421 | 0 | 3.5 | 52 | Specific | 238.17 | 167.64 | 167.64 | 2.7 | 4 | >sp\|Q13410\|BT1A1_HUMAN Butyrophilin subfamily 1 member A1 OS=Homo sapiens OX=9606 GN=BTN1A1 PE=1 SV=3 | 179 | 20201113_gy_colo_milk_1ng_glyco_hcd30_etd30_compound15.7466.7466.3 | controllerType=0 controllerNumber=1 scan=7466 | 20.782 |
| 14647:3 | 41 | R.GIASVLQELN[+876.322]VTVVTSLC[+57.021]R.R | HexNAc(2)Hex(2)Fuc(1) | C[+57], N[+876] | 979.1519 | 3 | 2935.4411 | 2935.4439 | 0 | -1 | 164 | Specific | 253.19 | 251.13 | 251.13 | 3.18 | 3 | >sp\|P08246\|ELNE_HUMAN Neutrophil elastase OS=Homo sapiens OX=9606 GN=ELANE PE=1 SV=1 | 1269 | 20201113_gy_colo_milk_1ng_glyco_hcd30_etd30_compound15.18160.18160.3 | controllerType=0 controllerNumber=1 scan=18160 | 46.2463 |
| 15715:3 | 48 | K.MLN[+2059.735]TSSLLEQLNEQFNWVSR.L | HexNAc(4)Hex(5)Fuc(1)NeuAc(1) | N[+2060] | 1490.3174 | 3 | 4468.9378 | 4468.9217 | 0 | 3.6 | 352 | Specific | 519.2 | 300.87 | 300.87 | 6.17 | 6 | >sp\|P10909\|CLUS_HUMAN Clusterin OS=Homo sapiens OX=9606 GN=CLU PE=1 SV=1 | 1467 | 20201113_gy_colo_milk_1ng_glyco_hcd30_etd30_compound15.19293.19293.3 | controllerType=0 controllerNumber=1 scan=19293 | 49.7822 |
| 11085:3 | 110 | R.EDFVLTAAHC[+57.021]WGSNIN[+203.079]VTLGAHNIQR.R | HexNAc(1) | C[+57], N[+203] | 1042.846 | 3 | 3126.5234 | 3126.5062 | 0 | 5.5 | 56 | Specific | 295.92 | 258.55 | 258.55 | 3.35 | 1 | >sp\|P08311\|CATG_HUMAN Cathepsin G OS=Homo sapiens OX=9606 GN=CTSG PE=1 SV=2 | 1574 | 20201113_gy_colo_milk_1ng_glyco_hcd30_etd30_compound15.14408.14408.3 | controllerType=0 controllerNumber=1 scan=14408 | 35.9671 |
| 08573:3 | 112 | R.ADGTVNQIEGEATPVN[+1216.423]LTEPAKLEVK.F | HexNAc(2)Hex(5) | N[+1216] | 1313.9501 | 3 | 3939.8358 | 3939.8326 | 0 | 0.8 | 83 | Specific | 269.09 | 269.09 | 269.09 | 3.23 | 1 | >sp\|P05090\|APOD_HUMAN Apolipoprotein D OS=Homo sapiens OX=9606 GN=APOD PE=1 SV=1 | 594 | 20201113_gy_colo_milk_1ng_glyco_hcd30_etd30_compound15.11777.11777.3 | controllerType=0 controllerNumber=1 scan=11777 | 29.7126 |
| 15330:4 | 1 | R.[+42.011]RTAGWNVPIGTLRPFLN[+2018.708]WTGPPEPIEAAVAR.F | HexNAc(3)Hex(6)Fuc(1)NeuAc(1) | .[42], N[+2019] | 1362.6503 | 4 | 5447.5793 | 5447.531 | 0 | 8.9 | 140 | Specific | 224.19 | 218.47 | 218.47 | 2.75 | 50 | >sp\|P02788\|TRFL_HUMAN Lactotransferrin OS=Homo sapiens OX=9606 GN=LTF PE=1 SV=6 | 1270 | 20201113_gy_milk_1ng_glyco_hcd30_etd30_compound15.18864.18864.4 | controllerType=0 controllerNumber=1 scan=18864 | 41.3288 |
| 15974:4 | 1 | R.RTAGWNVPIGTLRPFLN[+2569.905]WTGPPEPIEAAVAR.F | HexNAc(5)Hex(6)NeuAc(2) | N[+2570] | 1489.9441 | 4 | 5956.7546 | 5956.7167 | 0 | 6.4 | 140 | Specific | 653.44 | 653.44 | 653.44 | 10.41 | 50 | >sp\|P02788\|TRFL_HUMAN Lactotransferrin OS=Homo sapiens OX=9606 GN=LTF PE=1 SV=6 | 1270 | 20201113_gy_milk_1ng_glyco_hcd30_etd30_compound15.19547.19547.4 | controllerType=0 controllerNumber=1 scan=19547 | 43.4227 |
| 15376:4 | 1 | R.RTAGWNVPIGTLRPFLN[+1768.640]WTGPPEPIEAAVAR.F | HexNAc(4)Hex(5)Fuc(1) | N[+1769] | 1289.6209 | 4 | 5155.4616 | 5155.4516 | 0 | 1.9 | 140 | Specific | 709.91 | 709.91 | 709.91 | 12.2 | 50 | >sp\|P02788\|TRFL_HUMAN Lactotransferrin OS=Homo sapiens OX=9606 GN=LTF PE=1 SV=6 | 1270 | 20201113_gy_milk_1ng_glyco_hcd30_etd30_compound15.18912.18912.4 | controllerType=0 controllerNumber=1 scan=18912 | 41.4667 |
| 15364:4 | 1 | R.RTAGWNVPIGTLRPFLN[+1913.677]WTGPPEPIEAAVAR.F | HexNAc(4)Hex(5)NeuAc(1) | N[+1914] | 1325.8844 | 4 | 5300.5158 | 5300.4891 | 0 | 5 | 140 | Specific | 922.91 | 849.7 | 849.7 | 15.41 | 50 | >sp\|P02788\|TRFL_HUMAN Lactotransferrin OS=Homo sapiens OX=9606 GN=LTF PE=1 SV=6 | 1270 | 20201113_gy_milk_1ng_glyco_hcd30_etd30_compound15.18900.18900.4 | controllerType=0 controllerNumber=1 scan=18900 | 41.4373 |
| 16234:4 | 1 | R.TAGWNVPIGTLRPFLN[+1768.640]WTGPPEPIEAAVAR.F | HexNAc(4)Hex(5)Fuc(1) | N[+1769] | 1250.5966 | 4 | 4999.3644 | 4999.3504 | 0 | 2.8 | 141 | Specific | 204.97 | 204.97 | 204.97 | 3.92 | 50 | >sp\|P02788\|TRFL_HUMAN Lactotransferrin OS=Homo sapiens OX=9606 GN=LTF PE=1 SV=6 | 1270 | 20201113_gy_milk_1ng_glyco_hcd30_etd30_compound15.19820.19820.4 | controllerType=0 controllerNumber=1 scan=19820 | 44.1865 |
| 16200:3 | 1 | R.[+42.011]TAGWNVPIGTLRPFLN[+2018.708]WTGPPEPIEAAVAR.F | HexNAc(3)Hex(6)Fuc(1)NeuAc(1) | .[42], N[+2019] | 1764.4987 | 3 | 5291.4817 | 5291.4299 | 0 | 9.8 | 141 | Specific | 214.57 | 149.18 | 149.18 | 2.66 | 50 | >sp\|P02788\|TRFL_HUMAN Lactotransferrin OS=Homo sapiens OX=9606 GN=LTF PE=1 SV=6 | 1270 | 20201113_gy_milk_1ng_glyco_hcd30_etd30_compound15.19784.19784.3 | controllerType=0 controllerNumber=1 scan=19784 | 44.0334 |
| 16917:3 | 1 | R.TAGWNVPIGTLRPFLN[+2059.735]WTGPPEPIEAAVAR.F | HexNAc(4)Hex(5)Fuc(1)NeuAc(1) | N[+2060] | 1764.1578 | 3 | 5290.4588 | 5290.4459 | 0 | 2.5 | 141 | Specific | 240.83 | 139.62 | 139.62 | 4.94 | 50 | >sp\|P02788\|TRFL_HUMAN Lactotransferrin OS=Homo sapiens OX=9606 GN=LTF PE=1 SV=6 | 1270 | 20201113_gy_milk_1ng_glyco_hcd30_etd30_compound15.20544.20544.3 | controllerType=0 controllerNumber=1 scan=20544 | 46.3813 |
| 17974:4 | 1 | R.[+42.011]TAGWNVPIGTLRPFLN[+2336.851]WTGPPEPIEAAVAR.F | HexNAc(6)Hex(6)Fuc(1) | .[42], N[+2337] | 1403.142 | 4 | 5609.5461 | 5609.5726 | 0 | -4.7 | 141 | Specific | 251.35 | 251.35 | 251.35 | 3.3 | 50 | >sp\|P02788\|TRFL_HUMAN Lactotransferrin OS=Homo sapiens OX=9606 GN=LTF PE=1 SV=6 | 1270 | 20201113_gy_milk_1ng_glyco_hcd30_etd30_compound15.21661.21661.4 | controllerType=0 controllerNumber=1 scan=21661 | 49.2147 |
| 16889:3 | 1 | R.TAGWNVPIGTLRPFLN[+2351.851]WTGPPEPIEAAVAR.F | HexNAc(4)Hex(5)Fuc(3)NeuAc(1) | N[+2352] | 1861.5305 | 3 | 5582.5769 | 5582.5617 | 0 | 2.7 | 141 | Specific | 292.76 | 244.04 | 244.04 | 4.89 | 50 | >sp\|P02788\|TRFL_HUMAN Lactotransferrin OS=Homo sapiens OX=9606 GN=LTF PE=1 SV=6 | 1270 | 20201113_gy_milk_1ng_glyco_hcd30_etd30_compound15.20514.20514.3 | controllerType=0 controllerNumber=1 scan=20514 | 46.2861 |
| 16206:4 | 1 | R.[+42.011]TAGWNVPIGTLRPFLN[+2018.708]WTGPPEPIEAAVAR.F | HexNAc(3)Hex(6)Fuc(1)NeuAc(1) | .[42], N[+2019] | 1323.6252 | 4 | 5291.4792 | 5291.4299 | 0 | 9.3 | 141 | Specific | 329.3 | 313.35 | 313.35 | 3.4 | 50 | >sp\|P02788\|TRFL_HUMAN Lactotransferrin OS=Homo sapiens OX=9606 GN=LTF PE=1 SV=6 | 1270 | 20201113_gy_milk_1ng_glyco_hcd30_etd30_compound15.19790.19790.4 | controllerType=0 controllerNumber=1 scan=19790 | 44.0747 |
| 16799:3 | 1 | R.TAGWNVPIGTLRPFLN[+2570.925]WTGPPEPIEAAVAR.F | HexNAc(5)Hex(6)Fuc(2)NeuAc(1) | N[+2571] | 1934.5592 | 3 | 5801.6629 | 5801.636 | 0 | 4.7 | 141 | Specific | 359.17 | 274.45 | 274.45 | 5.45 | 50 | >sp\|P02788\|TRFL_HUMAN Lactotransferrin OS=Homo sapiens OX=9606 GN=LTF PE=1 SV=6 | 1270 | 20201113_gy_milk_1ng_glyco_hcd30_etd30_compound15.20418.20418.3 | controllerType=0 controllerNumber=1 scan=20418 | 46.014 |
| 16788:4 | 1 | R.TAGWNVPIGTLRPFLN[+2716.983]WTGPPEPIEAAVAR.F | HexNAc(5)Hex(6)Fuc(3)NeuAc(1) | N[+2717] | 1487.6849 | 4 | 5947.7179 | 5947.6939 | 0 | 4 | 141 | Specific | 365.33 | 365.33 | 365.33 | 5.85 | 50 | >sp\|P02788\|TRFL_HUMAN Lactotransferrin OS=Homo sapiens OX=9606 GN=LTF PE=1 SV=6 | 1270 | 20201113_gy_milk_1ng_glyco_hcd30_etd30_compound15.20407.20407.4 | controllerType=0 controllerNumber=1 scan=20407 | 45.9907 |
| 16793:3 | 1 | R.TAGWNVPIGTLRPFLN[+2715.963]WTGPPEPIEAAVAR.F | HexNAc(5)Hex(6)Fuc(1)NeuAc(2) | N[+2716] | 1982.9101 | 3 | 5946.7158 | 5946.6735 | 0 | 7.1 | 141 | Specific | 379.31 | 256.57 | 256.57 | 5.05 | 50 | >sp\|P02788\|TRFL_HUMAN Lactotransferrin OS=Homo sapiens OX=9606 GN=LTF PE=1 SV=6 | 1270 | 20201113_gy_milk_1ng_glyco_hcd30_etd30_compound15.20412.20412.3 | controllerType=0 controllerNumber=1 scan=20412 | 45.9991 |
| 16215:4 | 1 | R.[+42.011]TAGWNVPIGTLRPFLN[+1872.651]WTGPPEPIEAAVAR.F | HexNAc(3)Hex(6)NeuAc(1) | .[42], N[+1873] | 1287.1106 | 4 | 5145.4206 | 5145.372 | 0 | 9.4 | 141 | Specific | 405.57 | 405.57 | 405.57 | 5.29 | 50 | >sp\|P02788\|TRFL_HUMAN Lactotransferrin OS=Homo sapiens OX=9606 GN=LTF PE=1 SV=6 | 1270 | 20201113_gy_milk_1ng_glyco_hcd30_etd30_compound15.19800.19800.4 | controllerType=0 controllerNumber=1 scan=19800 | 44.1182 |
| 16918:4 | 1 | R.TAGWNVPIGTLRPFLN[+2059.735]WTGPPEPIEAAVAR.F | HexNAc(4)Hex(5)Fuc(1)NeuAc(1) | N[+2060] | 1323.3711 | 4 | 5290.4626 | 5290.4459 | 0 | 3.2 | 141 | Specific | 418.32 | 418.32 | 418.32 | 6.14 | 50 | >sp\|P02788\|TRFL_HUMAN Lactotransferrin OS=Homo sapiens OX=9606 GN=LTF PE=1 SV=6 | 1270 | 20201113_gy_milk_1ng_glyco_hcd30_etd30_compound15.20545.20545.4 | controllerType=0 controllerNumber=1 scan=20545 | 46.3872 |
| 16225:3 | 1 | R.TAGWNVPIGTLRPFLN[+1768.640]WTGPPEPIEAAVAR.F | HexNAc(4)Hex(5)Fuc(1) | N[+1769] | 1667.1268 | 3 | 4999.3658 | 4999.3504 | 0 | 3.1 | 141 | Specific | 481.69 | 398.52 | 398.52 | 6.44 | 50 | >sp\|P02788\|TRFL_HUMAN Lactotransferrin OS=Homo sapiens OX=9606 GN=LTF PE=1 SV=6 | 1270 | 20201113_gy_milk_1ng_glyco_hcd30_etd30_compound15.19811.19811.3 | controllerType=0 controllerNumber=1 scan=19811 | 44.1611 |
| 16855:4 | 1 | R.TAGWNVPIGTLRPFLN[+2716.983]WTGPPEPIEAAVAR.F | HexNAc(5)Hex(6)Fuc(3)NeuAc(1) | N[+2717] | 1487.682 | 4 | 5947.7062 | 5947.6939 | 0 | 2.1 | 141 | Specific | 506.61 | 506.61 | 506.61 | 6.99 | 50 | >sp\|P02788\|TRFL_HUMAN Lactotransferrin OS=Homo sapiens OX=9606 GN=LTF PE=1 SV=6 | 1270 | 20201113_gy_milk_1ng_glyco_hcd30_etd30_compound15.20477.20477.4 | controllerType=0 controllerNumber=1 scan=20477 | 46.1522 |
| 16879:3 | 1 | R.TAGWNVPIGTLRPFLN[+2205.793]WTGPPEPIEAAVAR.F | HexNAc(4)Hex(5)Fuc(2)NeuAc(1) | N[+2206] | 1812.8419 | 3 | 5436.5111 | 5436.5038 | 0 | 1.3 | 141 | Specific | 596.42 | 490.74 | 490.74 | 9.2 | 50 | >sp\|P02788\|TRFL_HUMAN Lactotransferrin OS=Homo sapiens OX=9606 GN=LTF PE=1 SV=6 | 1270 | 20201113_gy_milk_1ng_glyco_hcd30_etd30_compound15.20503.20503.3 | controllerType=0 controllerNumber=1 scan=20503 | 46.2609 |
| 17956:4 | 1 | R.TAGWNVPIGTLRPFLN[+2350.830]WTGPPEPIEAAVAR.F | HexNAc(4)Hex(5)Fuc(1)NeuAc(2) | N[+2351] | 1396.1449 | 4 | 5581.5578 | 5581.5413 | 0 | 3 | 141 | Specific | 849.94 | 841.39 | 841.39 | 14.2 | 50 | >sp\|P02788\|TRFL_HUMAN Lactotransferrin OS=Homo sapiens OX=9606 GN=LTF PE=1 SV=6 | 1270 | 20201113_gy_milk_1ng_glyco_hcd30_etd30_compound15.21642.21642.4 | controllerType=0 controllerNumber=1 scan=21642 | 49.1701 |
| 16794:4 | 1 | R.TAGWNVPIGTLRPFLN[+2570.925]WTGPPEPIEAAVAR.F | HexNAc(5)Hex(6)Fuc(2)NeuAc(1) | N[+2571] | 1451.1716 | 4 | 5801.6647 | 5801.636 | 0 | 5 | 141 | Specific | 869.44 | 869.44 | 869.44 | 14.41 | 50 | >sp\|P02788\|TRFL_HUMAN Lactotransferrin OS=Homo sapiens OX=9606 GN=LTF PE=1 SV=6 | 1270 | 20201113_gy_milk_1ng_glyco_hcd30_etd30_compound15.20413.20413.4 | controllerType=0 controllerNumber=1 scan=20413 | 46.0041 |
| 15422:2 | 1 | R.TAGWNIPMGLLFN[+1216.423]QTGSC[+57.021]K.F | HexNAc(2)Hex(5) | C[+57], N[+1216] | 1656.2197 | 2 | 3311.4321 | 3311.4328 | 0 | -0.2 | 485 | Specific | 267.71 | 201.5 | 107.26 | 4.43 | 50 | >sp\|P02788\|TRFL_HUMAN Lactotransferrin OS=Homo sapiens OX=9606 GN=LTF PE=1 SV=6 | 1270 | 20201113_gy_milk_1ng_glyco_hcd30_etd30_compound15.18961.18961.2 | controllerType=0 controllerNumber=1 scan=18961 | 41.6127 |
| 15249:3 | 1 | R.TAGWNIPMGLLFN[+1606.587]QTGSC[+57.021]K.F | HexNAc(4)Hex(4)Fuc(1) | C[+57], N[+1607] | 1234.5412 | 3 | 3701.609 | 3701.5966 | 0 | 3.3 | 485 | Specific | 269.71 | 218.05 | 218.05 | 3.59 | 50 | >sp\|P02788\|TRFL_HUMAN Lactotransferrin OS=Homo sapiens OX=9606 GN=LTF PE=1 SV=6 | 1270 | 20201113_gy_milk_1ng_glyco_hcd30_etd30_compound15.18778.18778.3 | controllerType=0 controllerNumber=1 scan=18778 | 41.0417 |
| 16625:3 | 1 | R.TAGWNIPMGLLFN[+1751.624]QTGSC[+57.021]K.F | HexNAc(4)Hex(4)NeuAc(1) | C[+57], N[+1752] | 1282.8871 | 3 | 3846.6469 | 3846.6341 | 0 | 3.3 | 485 | Specific | 275.53 | 201.41 | 201.41 | 4.7 | 50 | >sp\|P02788\|TRFL_HUMAN Lactotransferrin OS=Homo sapiens OX=9606 GN=LTF PE=1 SV=6 | 1270 | 20201113_gy_milk_1ng_glyco_hcd30_etd30_compound15.20236.20236.3 | controllerType=0 controllerNumber=1 scan=20236 | 45.5473 |
| 18070:3 | 1 | R.TAGWNIPMGLLFN[+2350.830]QTGSC[+57.021]K.F | HexNAc(4)Hex(5)Fuc(1)NeuAc(2) | C[+57], N[+2351] | 1482.6198 | 3 | 4445.8449 | 4445.8403 | 0 | 1 | 485 | Specific | 285.65 | 169.64 | 169.64 | 4.46 | 50 | >sp\|P02788\|TRFL_HUMAN Lactotransferrin OS=Homo sapiens OX=9606 GN=LTF PE=1 SV=6 | 1270 | 20201113_gy_milk_1ng_glyco_hcd30_etd30_compound15.21761.21761.2 | controllerType=0 controllerNumber=1 scan=21761 | 49.4674 |
| 16414:3 | 1 | R.TAGWNIPMGLLFN[+2351.851]QTGSC[+57.021]K.F | HexNAc(4)Hex(5)Fuc(3)NeuAc(1) | C[+57], N[+2352] | 1482.9636 | 3 | 4446.8761 | 4446.8607 | 0 | 3.5 | 485 | Specific | 291.04 | 211.53 | 211.53 | 4.66 | 50 | >sp\|P02788\|TRFL_HUMAN Lactotransferrin OS=Homo sapiens OX=9606 GN=LTF PE=1 SV=6 | 1270 | 20201113_gy_milk_1ng_glyco_hcd30_etd30_compound15.20011.20011.3 | controllerType=0 controllerNumber=1 scan=20011 | 44.7709 |
| 15358:3 | 1 | R.TAGWNIPMGLLFN[+1378.476]QTGSC[+57.021]K.F | HexNAc(2)Hex(6) | C[+57], N[+1378] | 1158.503 | 3 | 3473.4944 | 3473.4856 | 0 | 2.5 | 485 | Specific | 298.44 | 277.32 | 149.05 | 5.78 | 50 | >sp\|P02788\|TRFL_HUMAN Lactotransferrin OS=Homo sapiens OX=9606 GN=LTF PE=1 SV=6 | 1270 | 20201113_gy_milk_1ng_glyco_hcd30_etd30_compound15.18893.18893.3 | controllerType=0 controllerNumber=1 scan=18893 | 41.4218 |
| 16577:3 | 1 | R.TAGWNIPMGLLFN[+1872.651]QTGSC[+57.021]K.F | HexNAc(3)Hex(6)NeuAc(1) | C[+57], N[+1873] | 1323.2298 | 3 | 3967.6748 | 3967.6604 | 0 | 3.6 | 485 | Specific | 350.42 | 251.56 | 226.07 | 4.71 | 50 | >sp\|P02788\|TRFL_HUMAN Lactotransferrin OS=Homo sapiens OX=9606 GN=LTF PE=1 SV=6 | 1270 | 20201113_gy_milk_1ng_glyco_hcd30_etd30_compound15.20185.20185.3 | controllerType=0 controllerNumber=1 scan=20185 | 45.3584 |
| 15405:3 | 1 | R.TAGWNIPMGLLFN[+1216.423]QTGSC[+57.021]K.F | HexNAc(2)Hex(5) | C[+57], N[+1216] | 1104.4851 | 3 | 3311.4409 | 3311.4328 | 0 | 2.4 | 485 | Specific | 353.06 | 301.39 | 205.92 | 5.76 | 50 | >sp\|P02788\|TRFL_HUMAN Lactotransferrin OS=Homo sapiens OX=9606 GN=LTF PE=1 SV=6 | 1270 | 20201113_gy_milk_1ng_glyco_hcd30_etd30_compound15.18943.18943.3 | controllerType=0 controllerNumber=1 scan=18943 | 41.5773 |
| 15264:3 | 1 | R.TAGWNIPMGLLFN[+1622.582]QTGSC[+57.021]K.F | HexNAc(4)Hex(5) | C[+57], N[+1623] | 1239.8716 | 3 | 3717.6004 | 3717.5916 | 0 | 2.4 | 485 | Specific | 365.72 | 298.13 | 233.51 | 5.99 | 50 | >sp\|P02788\|TRFL_HUMAN Lactotransferrin OS=Homo sapiens OX=9606 GN=LTF PE=1 SV=6 | 1270 | 20201113_gy_milk_1ng_glyco_hcd30_etd30_compound15.18794.18794.3 | controllerType=0 controllerNumber=1 scan=18794 | 41.0854 |
| 15129:3 | 1 | R.TAGWNIPMGLLFN[+1768.640]QTGSC[+57.021]K.F | HexNAc(4)Hex(5)Fuc(1) | C[+57], N[+1769] | 1288.5577 | 3 | 3863.6585 | 3863.6495 | 0 | 2.3 | 485 | Specific | 373.93 | 268.63 | 268.63 | 5.98 | 50 | >sp\|P02788\|TRFL_HUMAN Lactotransferrin OS=Homo sapiens OX=9606 GN=LTF PE=1 SV=6 | 1270 | 20201113_gy_milk_1ng_glyco_hcd30_etd30_compound15.18652.18652.3 | controllerType=0 controllerNumber=1 scan=18652 | 40.6953 |
| 16561:3 | 1 | R.TAGWNIPMGLLFN[+1872.651]QTGSC[+57.021]K.F | HexNAc(3)Hex(6)NeuAc(1) | C[+57], N[+1873] | 1323.2286 | 3 | 3967.6712 | 3967.6604 | 0 | 2.7 | 485 | Specific | 381.8 | 270.47 | 249.06 | 5.98 | 50 | >sp\|P02788\|TRFL_HUMAN Lactotransferrin OS=Homo sapiens OX=9606 GN=LTF PE=1 SV=6 | 1270 | 20201113_gy_milk_1ng_glyco_hcd30_etd30_compound15.20168.20168.3 | controllerType=0 controllerNumber=1 scan=20168 | 45.3095 |
| 18186:3 | 1 | R.TAGWNIPMGLLFN[+2204.772]QTGSC[+57.021]K.F | HexNAc(4)Hex(5)NeuAc(2) | C[+57], N[+2205] | 1433.9401 | 3 | 4299.8058 | 4299.7824 | 0 | 5.4 | 485 | Specific | 384.65 | 268.98 | 268.98 | 5.19 | 50 | >sp\|P02788\|TRFL_HUMAN Lactotransferrin OS=Homo sapiens OX=9606 GN=LTF PE=1 SV=6 | 1270 | 20201113_gy_milk_1ng_glyco_hcd30_etd30_compound15.21884.21884.3 | controllerType=0 controllerNumber=1 scan=21884 | 49.8256 |
| 18029:3 | 1 | R.TAGWNIPMGLLFN[+2350.830]QTGSC[+57.021]K.F | HexNAc(4)Hex(5)Fuc(1)NeuAc(2) | C[+57], N[+2351] | 1482.6196 | 3 | 4445.8441 | 4445.8403 | 0 | 0.9 | 485 | Specific | 413.2 | 273.35 | 273.35 | 4.96 | 50 | >sp\|P02788\|TRFL_HUMAN Lactotransferrin OS=Homo sapiens OX=9606 GN=LTF PE=1 SV=6 | 1270 | 20201113_gy_milk_1ng_glyco_hcd30_etd30_compound15.21719.21719.3 | controllerType=0 controllerNumber=1 scan=21719 | 49.3808 |
| 16521:3 | 1 | R.TAGWNIPMGLLFN[+1913.677]QTGSC[+57.021]K.F | HexNAc(4)Hex(5)NeuAc(1) | C[+57], N[+1914] | 1336.9033 | 3 | 4008.6952 | 4008.687 | 0 | 2.1 | 485 | Specific | 435.06 | 275.5 | 275.5 | 6.03 | 50 | >sp\|P02788\|TRFL_HUMAN Lactotransferrin OS=Homo sapiens OX=9606 GN=LTF PE=1 SV=6 | 1270 | 20201113_gy_milk_1ng_glyco_hcd30_etd30_compound15.20126.20126.3 | controllerType=0 controllerNumber=1 scan=20126 | 45.1656 |
| 16440:3 | 1 | R.TAGWNIPMGLLFN[+2059.735]QTGSC[+57.021]K.F | HexNAc(4)Hex(5)Fuc(1)NeuAc(1) | C[+57], N[+2060] | 1385.5957 | 3 | 4154.7725 | 4154.7449 | 0 | 6.7 | 485 | Specific | 447.24 | 308.6 | 308.6 | 5.01 | 50 | >sp\|P02788\|TRFL_HUMAN Lactotransferrin OS=Homo sapiens OX=9606 GN=LTF PE=1 SV=6 | 1270 | 20201113_gy_milk_1ng_glyco_hcd30_etd30_compound15.20038.20038.3 | controllerType=0 controllerNumber=1 scan=20038 | 44.8502 |
| 16467:3 | 1 | R.TAGWNIPMGLLFN[+2205.793]QTGSC[+57.021]K.F | HexNAc(4)Hex(5)Fuc(2)NeuAc(1) | C[+57], N[+2206] | 1434.2847 | 3 | 4300.8396 | 4300.8028 | 0 | 8.6 | 485 | Specific | 458.98 | 369.07 | 369.07 | 6.43 | 50 | >sp\|P02788\|TRFL_HUMAN Lactotransferrin OS=Homo sapiens OX=9606 GN=LTF PE=1 SV=6 | 1270 | 20201113_gy_milk_1ng_glyco_hcd30_etd30_compound15.20068.20068.3 | controllerType=0 controllerNumber=1 scan=20068 | 44.9533 |
| 15501:3 | 1 | R.TAGWNIPM[+15.995]GLLFN[+2059.735]QTGSC[+57.021]K.F | HexNAc(4)Hex(5)Fuc(1)NeuAc(1) | C[+57], M[+16], N[+2060] | 1390.9207 | 3 | 4170.7475 | 4170.7398 | 0 | 1.9 | 485 | Specific | 250.53 | 177.81 | 177.81 | 5.58 | 50 | >sp\|P02788\|TRFL_HUMAN Lactotransferrin OS=Homo sapiens OX=9606 GN=LTF PE=1 SV=6 | 1270 | 20201113_gy_milk_1ng_glyco_hcd30_etd30_compound15.19045.19045.3 | controllerType=0 controllerNumber=1 scan=19045 | 41.8451 |
| 15502:3 | 1 | R.TAGWNIPM[+15.995]GLLFN[+1913.677]QTGSC[+57.021]K.F | HexNAc(4)Hex(5)NeuAc(1) | C[+57], M[+16], N[+1914] | 1342.2328 | 3 | 4024.6838 | 4024.6819 | 0 | 0.5 | 485 | Specific | 262.21 | 160.65 | 160.65 | 4.51 | 50 | >sp\|P02788\|TRFL_HUMAN Lactotransferrin OS=Homo sapiens OX=9606 GN=LTF PE=1 SV=6 | 1270 | 20201113_gy_milk_1ng_glyco_hcd30_etd30_compound15.19046.19046.3 | controllerType=0 controllerNumber=1 scan=19046 | 41.8524 |
| 15960:3 | 1 | R.TAGWNIPMGLLFN[+1622.582]QTGSC[+57.021]KFDEYFSQSC[+57.021]APGSDPR.S | HexNAc(4)Hex(5) | C[+57]*2, N[+1623] | 1854.457 | 3 | 5561.3564 | 5561.3387 | 0 | 3.2 | 485 | Specific | 247.86 | 201.73 | 107.49 | 4.55 | 50 | >sp\|P02788\|TRFL_HUMAN Lactotransferrin OS=Homo sapiens OX=9606 GN=LTF PE=1 SV=6 | 1270 | 20201113_gy_milk_1ng_glyco_hcd30_etd30_compound15.19532.19532.3 | controllerType=0 controllerNumber=1 scan=19532 | 43.3874 |
| 16658:3 | 1 | R.TAGWNIPMGLLFN[+1913.677]QTGSC[+57.021]KFDEYFSQSC[+57.021]APGSDPR.S | HexNAc(4)Hex(5)NeuAc(1) | C[+57]*2, N[+1914] | 1951.4887 | 3 | 5852.4516 | 5852.4341 | 0 | 3 | 485 | Specific | 264.34 | 164.13 | 164.13 | 4.63 | 50 | >sp\|P02788\|TRFL_HUMAN Lactotransferrin OS=Homo sapiens OX=9606 GN=LTF PE=1 SV=6 | 1270 | 20201113_gy_milk_1ng_glyco_hcd30_etd30_compound15.20271.20271.3 | controllerType=0 controllerNumber=1 scan=20271 | 45.6569 |
| 16653:4 | 1 | R.TAGWNIPMGLLFN[+1913.677]QTGSC[+57.021]KFDEYFSQSC[+57.021]APGSDPR.S | HexNAc(4)Hex(5)NeuAc(1) | C[+57]*2, N[+1914] | 1463.8698 | 4 | 5852.4572 | 5852.4341 | 0 | 3.9 | 485 | Specific | 479.62 | 429.44 | 429.44 | 5.64 | 50 | >sp\|P02788\|TRFL_HUMAN Lactotransferrin OS=Homo sapiens OX=9606 GN=LTF PE=1 SV=6 | 1270 | 20201113_gy_milk_1ng_glyco_hcd30_etd30_compound15.20266.20266.4 | controllerType=0 controllerNumber=1 scan=20266 | 45.6397 |
| 15910:3 | 1 | R.TAGWNIPMGLLFN[+1768.640]QTGSC[+57.021]KFDEYFSQSC[+57.021]APGSDPR.S | HexNAc(4)Hex(5)Fuc(1) | C[+57]*2, N[+1769] | 1903.1466 | 3 | 5707.4252 | 5707.3966 | 0 | 5 | 485 | Specific | 487.92 | 393.33 | 393.33 | 6.7 | 50 | >sp\|P02788\|TRFL_HUMAN Lactotransferrin OS=Homo sapiens OX=9606 GN=LTF PE=1 SV=6 | 1270 | 20201113_gy_milk_1ng_glyco_hcd30_etd30_compound15.19479.19479.3 | controllerType=0 controllerNumber=1 scan=19479 | 43.2177 |
| 16604:4 | 1 | R.TAGWNIPMGLLFN[+2205.793]QTGSC[+57.021]KFDEYFSQSC[+57.021]APGSDPR.S | HexNAc(4)Hex(5)Fuc(2)NeuAc(1) | C[+57]*2, N[+2206] | 1536.8951 | 4 | 6144.5587 | 6144.5499 | 0 | 1.4 | 485 | Specific | 706.49 | 626.39 | 626.39 | 11.55 | 50 | >sp\|P02788\|TRFL_HUMAN Lactotransferrin OS=Homo sapiens OX=9606 GN=LTF PE=1 SV=6 | 1270 | 20201113_gy_milk_1ng_glyco_hcd30_etd30_compound15.20214.20214.4 | controllerType=0 controllerNumber=1 scan=20214 | 45.4634 |
| 16614:4 | 1 | R.TAGWNIPMGLLFN[+2059.735]QTGSC[+57.021]KFDEYFSQSC[+57.021]APGSDPR.S | HexNAc(4)Hex(5)Fuc(1)NeuAc(1) | C[+57]*2, N[+2060] | 1500.382 | 4 | 5998.506 | 5998.492 | 0 | 2.3 | 485 | Specific | 813.45 | 699.35 | 699.35 | 13.55 | 50 | >sp\|P02788\|TRFL_HUMAN Lactotransferrin OS=Homo sapiens OX=9606 GN=LTF PE=1 SV=6 | 1270 | 20201113_gy_milk_1ng_glyco_hcd30_etd30_compound15.20224.20224.4 | controllerType=0 controllerNumber=1 scan=20224 | 45.511 |
| 15909:4 | 1 | R.TAGWNIPMGLLFN[+1768.640]QTGSC[+57.021]KFDEYFSQSC[+57.021]APGSDPR.S | HexNAc(4)Hex(5)Fuc(1) | C[+57]*2, N[+1769] | 1427.6135 | 4 | 5707.4323 | 5707.3966 | 0 | 6.2 | 485 | Specific | 848 | 781.58 | 781.58 | 12.76 | 50 | >sp\|P02788\|TRFL_HUMAN Lactotransferrin OS=Homo sapiens OX=9606 GN=LTF PE=1 SV=6 | 1270 | 20201113_gy_milk_1ng_glyco_hcd30_etd30_compound15.19478.19478.4 | controllerType=0 controllerNumber=1 scan=19478 | 43.2126 |
| 08330:3 | 1 | R.N[+2205.793]GSDC[+57.021]PDKFC[+57.021]LFQSETK.N | HexNAc(4)Hex(5)Fuc(2)NeuAc(1) | C[+57]*2, N[+2206] | 1413.5627 | 3 | 4238.6735 | 4238.6667 | 0 | 1.6 | 642 | Specific | 275.6 | 180.84 | 180.84 | 5.49 | 50 | >sp\|P02788\|TRFL_HUMAN Lactotransferrin OS=Homo sapiens OX=9606 GN=LTF PE=1 SV=6 | 1270 | 20201113_gy_milk_1ng_glyco_hcd30_etd30_compound15.11521.11521.2 | controllerType=0 controllerNumber=1 scan=11521 | 26.5577 |
| 08345:3 | 1 | R.N[+2205.793]GSDC[+57.021]PDKFC[+57.021]LFQSETK.N | HexNAc(4)Hex(5)Fuc(2)NeuAc(1) | C[+57]*2, N[+2206] | 1413.5634 | 3 | 4238.6757 | 4238.6667 | 0 | 2.1 | 642 | Specific | 278.8 | 170.73 | 170.73 | 5.49 | 50 | >sp\|P02788\|TRFL_HUMAN Lactotransferrin OS=Homo sapiens OX=9606 GN=LTF PE=1 SV=6 | 1270 | 20201113_gy_milk_1ng_glyco_hcd30_etd30_compound15.11537.11537.3 | controllerType=0 controllerNumber=1 scan=11537 | 26.5973 |
| 08352:3 | 1 | R.N[+2205.793]GSDC[+57.021]PDKFC[+57.021]LFQSETK.N | HexNAc(4)Hex(5)Fuc(2)NeuAc(1) | C[+57]*2, N[+2206] | 1413.5641 | 3 | 4238.6779 | 4238.6667 | 0 | 2.6 | 642 | Specific | 281.87 | 168.24 | 168.24 | 5.44 | 50 | >sp\|P02788\|TRFL_HUMAN Lactotransferrin OS=Homo sapiens OX=9606 GN=LTF PE=1 SV=6 | 1270 | 20201113_gy_milk_1ng_glyco_hcd30_etd30_compound15.11545.11545.3 | controllerType=0 controllerNumber=1 scan=11545 | 26.6185 |
| 06804:3 | 3 | R.FSDGLESN[+1864.634]SSTQFEVK.K | HexNAc(2)Hex(9) | N[+1865] | 1213.8253 | 3 | 3639.4612 | 3639.4472 | 0 | 3.9 | 219 | Specific | 407.2 | 402.45 | 402.45 | 6.56 | 9 | >sp\|P0C0L4\|CO4A_HUMAN Complement C4-A OS=Homo sapiens OX=9606 GN=C4A PE=1 SV=2 | 1235 | 20201113_gy_milk_1ng_glyco_hcd30_etd30_compound15.9928.9928.3 | controllerType=0 controllerNumber=1 scan=9928 | 23.5352 |
| 04436:3 | 3 | R.FSDGLESN[+1864.634]SSTQFEVKK.Y | HexNAc(2)Hex(9) | N[+1865] | 1256.5242 | 3 | 3767.5581 | 3767.5421 | 0 | 4.2 | 219 | Specific | 221.74 | 156.77 | 156.77 | 4.1 | 9 | >sp\|P0C0L4\|CO4A_HUMAN Complement C4-A OS=Homo sapiens OX=9606 GN=C4A PE=1 SV=2 | 1235 | 20201113_gy_milk_1ng_glyco_hcd30_etd30_compound15.7476.7476.3 | controllerType=0 controllerNumber=1 scan=7476 | 19.2964 |
| 16571:3 | 4 | K.QIGLYPVLVIDSSGYVNPN[+2059.735]YTGR.I | HexNAc(4)Hex(5)Fuc(1)NeuAc(1) | N[+2060] | 1529.0248 | 3 | 4585.06 | 4585.0384 | 0 | 4.7 | 168 | Specific | 492.14 | 415.31 | 415.31 | 6.62 | 17 | >sp\|P01833\|PIGR_HUMAN Polymeric immunoglobulin receptor OS=Homo sapiens OX=9606 GN=PIGR PE=1 SV=4 | 902 | 20201113_gy_milk_1ng_glyco_hcd30_etd30_compound15.20179.20179.3 | controllerType=0 controllerNumber=1 scan=20179 | 45.3455 |
| 19777:3 | 4 | R.LSLLEEPGN[+2350.830]GTFTVILNQLTSR.D | HexNAc(4)Hex(5)Fuc(1)NeuAc(2) | N[+2351] | 1585.0522 | 3 | 4753.142 | 4753.1229 | 0 | 4 | 413 | Specific | 395.85 | 274.17 | 274.17 | 4.94 | 17 | >sp\|P01833\|PIGR_HUMAN Polymeric immunoglobulin receptor OS=Homo sapiens OX=9606 GN=PIGR PE=1 SV=4 | 902 | 20201113_gy_milk_1ng_glyco_hcd30_etd30_compound15.23555.23555.3 | controllerType=0 controllerNumber=1 scan=23555 | 53.2983 |
| 14505:3 | 4 | K.IIEGEPNLKVPGN[+2205.793]VTAVLGETLK.V | HexNAc(4)Hex(5)Fuc(2)NeuAc(1) | N[+2206] | 1533.0544 | 3 | 4597.1486 | 4597.1422 | 0 | 1.4 | 457 | Specific | 419.7 | 343.36 | 343.36 | 4.76 | 17 | >sp\|P01833\|PIGR_HUMAN Polymeric immunoglobulin receptor OS=Homo sapiens OX=9606 GN=PIGR PE=1 SV=4 | 902 | 20201113_gy_milk_1ng_glyco_hcd30_etd30_compound15.17993.17993.3 | controllerType=0 controllerNumber=1 scan=17993 | 39.1989 |
| 15329:4 | 4 | K.IIEGEPNLKVPGN[+2350.830]VTAVLGETLKVPC[+57.021]HFPC[+57.021]K.F | HexNAc(4)Hex(5)Fuc(1)NeuAc(2) | C[+57]*2, N[+2351] | 1442.6688 | 4 | 5767.6535 | 5767.6372 | 0 | 2.8 | 457 | Specific | 268.15 | 268.15 | 268.15 | 4.11 | 17 | >sp\|P01833\|PIGR_HUMAN Polymeric immunoglobulin receptor OS=Homo sapiens OX=9606 GN=PIGR PE=1 SV=4 | 902 | 20201113_gy_milk_1ng_glyco_hcd30_etd30_compound15.18863.18863.4 | controllerType=0 controllerNumber=1 scan=18863 | 41.3219 |
| 14114:4 | 4 | K.IIEGEPNLKVPGN[+2059.735]VTAVLGETLKVPC[+57.021]HFPC[+57.021]K.F | HexNAc(4)Hex(5)Fuc(1)NeuAc(1) | C[+57]*2, N[+2060] | 1369.8921 | 4 | 5476.5466 | 5476.5418 | 0 | 0.9 | 457 | Specific | 357.29 | 357.29 | 357.29 | 3.22 | 17 | >sp\|P01833\|PIGR_HUMAN Polymeric immunoglobulin receptor OS=Homo sapiens OX=9606 GN=PIGR PE=1 SV=4 | 902 | 20201113_gy_milk_1ng_glyco_hcd30_etd30_compound15.17562.17562.4 | controllerType=0 controllerNumber=1 scan=17562 | 38.268 |
| 14085:5 | 4 | K.IIEGEPNLKVPGN[+2059.735]VTAVLGETLKVPC[+57.021]HFPC[+57.021]K.F | HexNAc(4)Hex(5)Fuc(1)NeuAc(1) | C[+57]*2, N[+2060] | 1096.1229 | 5 | 5476.5855 | 5476.5418 | 0 | 8 | 457 | Specific | 365.29 | 365.29 | 365.29 | 4.91 | 17 | >sp\|P01833\|PIGR_HUMAN Polymeric immunoglobulin receptor OS=Homo sapiens OX=9606 GN=PIGR PE=1 SV=4 | 902 | 20201113_gy_milk_1ng_glyco_hcd30_etd30_compound15.17531.17531.3 | controllerType=0 controllerNumber=1 scan=17531 | 38.1991 |
| 12444:4 | 4 | K.VPGN[+2205.793]VTAVLGETLKVPC[+57.021]HFPC[+57.021]K.F | HexNAc(4)Hex(5)Fuc(2)NeuAc(1) | C[+57]*2, N[+2206] | 1158.0232 | 4 | 4629.0711 | 4629.0502 | 0 | 4.5 | 466 | Specific | 281.57 | 281.57 | 281.57 | 2.95 | 17 | >sp\|P01833\|PIGR_HUMAN Polymeric immunoglobulin receptor OS=Homo sapiens OX=9606 GN=PIGR PE=1 SV=4 | 902 | 20201113_gy_milk_1ng_glyco_hcd30_etd30_compound15.15821.15821.4 | controllerType=0 controllerNumber=1 scan=15821 | 34.649 |
| 12449:4 | 4 | K.VPGN[+2059.735]VTAVLGETLKVPC[+57.021]HFPC[+57.021]K.F | HexNAc(4)Hex(5)Fuc(1)NeuAc(1) | C[+57]*2, N[+2060] | 1121.5103 | 4 | 4483.0194 | 4482.9923 | 0 | 6 | 466 | Specific | 327.57 | 327.57 | 327.57 | 3.12 | 17 | >sp\|P01833\|PIGR_HUMAN Polymeric immunoglobulin receptor OS=Homo sapiens OX=9606 GN=PIGR PE=1 SV=4 | 902 | 20201113_gy_milk_1ng_glyco_hcd30_etd30_compound15.15826.15826.4 | controllerType=0 controllerNumber=1 scan=15826 | 34.6626 |
| 05231:3 | 4 | K.WN[+2059.735]NTGC[+57.021]QALPSQDEGPSK.A | HexNAc(4)Hex(5)Fuc(1)NeuAc(1) | C[+57], N[+2060] | 1350.2121 | 3 | 4048.6217 | 4048.6116 | 0 | 2.5 | 498 | Specific | 425.54 | 293.13 | 293.13 | 6.03 | 17 | >sp\|P01833\|PIGR_HUMAN Polymeric immunoglobulin receptor OS=Homo sapiens OX=9606 GN=PIGR PE=1 SV=4 | 902 | 20201113_gy_milk_1ng_glyco_hcd30_etd30_compound15.8299.8299.3 | controllerType=0 controllerNumber=1 scan=8299 | 20.6951 |
| 15745:3 | 5 | R.LSLHRPALEDLLLGSEAN[+1216.423]LTC[+57.021]TLTGLR.D | HexNAc(2)Hex(5) | C[+57], N[+1216] | 1394.014 | 3 | 4180.0274 | 4180.0211 | 0 | 1.5 | 127 | Specific | 398.21 | 398.21 | 398.21 | 7.41 | 17 | >sp\|P01876\|IGHA1_HUMAN Immunoglobulin heavy constant alpha 1 OS=Homo sapiens OX=9606 GN=IGHA1 PE=1 SV=2 | 255 | 20201113_gy_milk_1ng_glyco_hcd30_etd30_compound15.19304.19304.3 | controllerType=0 controllerNumber=1 scan=19304 | 42.6551 |
| 15700:3 | 5 | R.LSLHRPALEDLLLGSEAN[+1866.688]LTC[+57.021]TLTGLR.D | HexNAc(6)Hex(4) | C[+57], N[+1867] | 1610.7535 | 3 | 4830.2459 | 4830.2858 | 0 | -8.3 | 127 | Specific | 400.9 | 347.36 | 347.36 | 4.54 | 17 | >sp\|P01876\|IGHA1_HUMAN Immunoglobulin heavy constant alpha 1 OS=Homo sapiens OX=9606 GN=IGHA1 PE=1 SV=2 | 255 | 20201113_gy_milk_1ng_glyco_hcd30_etd30_compound15.19255.19255.3 | controllerType=0 controllerNumber=1 scan=19255 | 42.4373 |
| 16434:4 | 5 | R.LSLHRPALEDLLLGSEAN[+1955.724]LTC[+57.021]TLTGLR.D | HexNAc(5)Hex(4)Fuc(2) | C[+57], N[+1956] | 1230.5831 | 4 | 4919.3104 | 4919.3222 | 0 | -2.4 | 127 | Specific | 402.73 | 402.73 | 402.73 | 6.74 | 17 | >sp\|P01876\|IGHA1_HUMAN Immunoglobulin heavy constant alpha 1 OS=Homo sapiens OX=9606 GN=IGHA1 PE=1 SV=2 | 255 | 20201113_gy_milk_1ng_glyco_hcd30_etd30_compound15.20032.20032.4 | controllerType=0 controllerNumber=1 scan=20032 | 44.8247 |
| 16419:3 | 5 | R.LSLHRPALEDLLLGSEAN[+1751.624]LTC[+57.021]TLTGLR.D | HexNAc(4)Hex(4)NeuAc(1) | C[+57], N[+1752] | 1572.4177 | 3 | 4715.2384 | 4715.2225 | 0 | 3.4 | 127 | Specific | 471.1 | 424.57 | 424.57 | 6.8 | 17 | >sp\|P01876\|IGHA1_HUMAN Immunoglobulin heavy constant alpha 1 OS=Homo sapiens OX=9606 GN=IGHA1 PE=1 SV=2 | 255 | 20201113_gy_milk_1ng_glyco_hcd30_etd30_compound15.20016.20016.3 | controllerType=0 controllerNumber=1 scan=20016 | 44.7855 |
| 16018:4 | 5 | R.LSLHRPALEDLLLGSEAN[+1501.555]LTC[+57.021]TLTGLR.D | HexNAc(5)Hex(3) | C[+57], N[+1502] | 1117.05 | 4 | 4465.178 | 4465.1536 | 0 | 5.5 | 127 | Specific | 728.08 | 708.97 | 708.97 | 11.41 | 17 | >sp\|P01876\|IGHA1_HUMAN Immunoglobulin heavy constant alpha 1 OS=Homo sapiens OX=9606 GN=IGHA1 PE=1 SV=2 | 255 | 20201113_gy_milk_1ng_glyco_hcd30_etd30_compound15.19593.19593.4 | controllerType=0 controllerNumber=1 scan=19593 | 43.5343 |
| 15725:4 | 5 | R.LSLHRPALEDLLLGSEAN[+1540.529]LTC[+57.021]TLTGLR.D | HexNAc(2)Hex(7) | C[+57], N[+1541] | 1126.7915 | 4 | 4504.1444 | 4504.1268 | 0 | 3.9 | 127 | Specific | 905.68 | 846.73 | 846.73 | 15.4 | 17 | >sp\|P01876\|IGHA1_HUMAN Immunoglobulin heavy constant alpha 1 OS=Homo sapiens OX=9606 GN=IGHA1 PE=1 SV=2 | 255 | 20201113_gy_milk_1ng_glyco_hcd30_etd30_compound15.19282.19282.4 | controllerType=0 controllerNumber=1 scan=19282 | 42.5439 |
| 15681:4 | 5 | R.LSLHRPALEDLLLGSEAN[+1540.529]LTC[+57.021]TLTGLR.D | HexNAc(2)Hex(7) | C[+57], N[+1541] | 1126.788 | 4 | 4504.1302 | 4504.1268 | 0 | 0.8 | 127 | Specific | 993.96 | 986.09 | 986.09 | 16.12 | 17 | >sp\|P01876\|IGHA1_HUMAN Immunoglobulin heavy constant alpha 1 OS=Homo sapiens OX=9606 GN=IGHA1 PE=1 SV=2 | 255 | 20201113_gy_milk_1ng_glyco_hcd30_etd30_compound15.19235.19235.2 | controllerType=0 controllerNumber=1 scan=19235 | 42.3695 |
| 11964:3 | 5 | R.LAGKPTHVN[+1913.677]VSVVMAEVDGTC[+57.021]Y.- | HexNAc(4)Hex(5)NeuAc(1) | C[+57], N[+1914] | 1420.9455 | 3 | 4260.8219 | 4260.8191 | 0 | 0.7 | 332 | CRagged | 240.86 | 126.2 | 126.2 | 2.76 | 17 | >sp\|P01876\|IGHA1_HUMAN Immunoglobulin heavy constant alpha 1 OS=Homo sapiens OX=9606 GN=IGHA1 PE=1 SV=2 | 255 | 20201113_gy_milk_1ng_glyco_hcd30_etd30_compound15.15321.15321.3 | controllerType=0 controllerNumber=1 scan=15321 | 33.6289 |
| 10357:3 | 5 | R.LAGKPTHVN[+1702.581]VSVVMAEVDGTC[+57.021]Y.- | HexNAc(2)Hex(8) | C[+57], N[+1703] | 1350.5873 | 3 | 4049.7475 | 4049.7234 | 0 | 5.9 | 332 | CRagged | 241.52 | 194.28 | 184.64 | 3.08 | 17 | >sp\|P01876\|IGHA1_HUMAN Immunoglobulin heavy constant alpha 1 OS=Homo sapiens OX=9606 GN=IGHA1 PE=1 SV=2 | 255 | 20201113_gy_milk_1ng_glyco_hcd30_etd30_compound15.13627.13627.3 | controllerType=0 controllerNumber=1 scan=13627 | 30.3991 |
| 10468:3 | 5 | R.LAGKPTHVN[+1647.613]VSVVMAEVDGTC[+57.021]Y.- | HexNAc(5)Hex(3)Fuc(1) | C[+57], N[+1648] | 1332.2682 | 3 | 3994.7902 | 3994.7553 | 0 | 8.7 | 332 | CRagged | 249.09 | 166.28 | 166.28 | 2.85 | 17 | >sp\|P01876\|IGHA1_HUMAN Immunoglobulin heavy constant alpha 1 OS=Homo sapiens OX=9606 GN=IGHA1 PE=1 SV=2 | 255 | 20201113_gy_milk_1ng_glyco_hcd30_etd30_compound15.13743.13743.3 | controllerType=0 controllerNumber=1 scan=13743 | 30.679 |
| 10437:3 | 5 | R.LAGKPTHVN[+1809.666]VSVVMAEVDGTC[+57.021]Y.- | HexNAc(5)Hex(4)Fuc(1) | C[+57], N[+1810] | 1386.2742 | 3 | 4156.8081 | 4156.8081 | 0 | 0 | 332 | CRagged | 266.36 | 175.27 | 175.27 | 2.92 | 17 | >sp\|P01876\|IGHA1_HUMAN Immunoglobulin heavy constant alpha 1 OS=Homo sapiens OX=9606 GN=IGHA1 PE=1 SV=2 | 255 | 20201113_gy_milk_1ng_glyco_hcd30_etd30_compound15.13711.13711.3 | controllerType=0 controllerNumber=1 scan=13711 | 30.6108 |
| 10363:3 | 5 | R.LAGKPTHVN[+1540.529]VSVVMAEVDGTC[+57.021]Y.- | HexNAc(2)Hex(7) | C[+57], N[+1541] | 1296.57 | 3 | 3887.6955 | 3887.6706 | 0 | 6.4 | 332 | CRagged | 276.99 | 230.42 | 208.26 | 2.85 | 17 | >sp\|P01876\|IGHA1_HUMAN Immunoglobulin heavy constant alpha 1 OS=Homo sapiens OX=9606 GN=IGHA1 PE=1 SV=2 | 255 | 20201113_gy_milk_1ng_glyco_hcd30_etd30_compound15.13633.13633.3 | controllerType=0 controllerNumber=1 scan=13633 | 30.4181 |
| 12164:3 | 8 | R.IIVPLNNREN[+2204.772]ISDPTSPLR.T | HexNAc(4)Hex(5)NeuAc(2) | N[+2205] | 1451.6544 | 3 | 4352.9485 | 4352.9496 | 0 | -0.2 | 62 | Specific | 220.88 | 122.06 | 122.06 | 2.54 | 9 | >sp\|P01591\|IGJ_HUMAN Immunoglobulin J chain OS=Homo sapiens OX=9606 GN=JCHAIN PE=1 SV=4 | 1392 | 20201113_gy_milk_1ng_glyco_hcd30_etd30_compound15.15530.15530.3 | controllerType=0 controllerNumber=1 scan=15530 | 34.0917 |
| 10496:3 | 8 | R.IIVPLNNREN[+1694.603]ISDPTSPLR.T | HexNAc(3)Hex(4)Fuc(1)NeuAc(1) | N[+1695] | 1281.6016 | 3 | 3842.7903 | 3842.7799 | 0 | 2.7 | 62 | Specific | 240.27 | 161.32 | 161.32 | 4.01 | 9 | >sp\|P01591\|IGJ_HUMAN Immunoglobulin J chain OS=Homo sapiens OX=9606 GN=JCHAIN PE=1 SV=4 | 1392 | 20201113_gy_milk_1ng_glyco_hcd30_etd30_compound15.13773.13773.3 | controllerType=0 controllerNumber=1 scan=13773 | 30.7773 |
| 10433:3 | 8 | R.IIVPLNNREN[+1856.656]ISDPTSPLR.T | HexNAc(3)Hex(5)Fuc(1)NeuAc(1) | N[+1857] | 1335.6158 | 3 | 4004.8328 | 4004.8327 | 0 | 0 | 62 | Specific | 272.89 | 213.75 | 213.75 | 2.96 | 9 | >sp\|P01591\|IGJ_HUMAN Immunoglobulin J chain OS=Homo sapiens OX=9606 GN=JCHAIN PE=1 SV=4 | 1392 | 20201113_gy_milk_1ng_glyco_hcd30_etd30_compound15.13706.13706.3 | controllerType=0 controllerNumber=1 scan=13706 | 30.5989 |
| 10358:3 | 8 | R.IIVPLNNREN[+2059.735]ISDPTSPLR.T | HexNAc(4)Hex(5)Fuc(1)NeuAc(1) | N[+2060] | 1403.3174 | 3 | 4207.9378 | 4207.9121 | 0 | 6.1 | 62 | Specific | 274.85 | 234.27 | 234.27 | 3.27 | 9 | >sp\|P01591\|IGJ_HUMAN Immunoglobulin J chain OS=Homo sapiens OX=9606 GN=JCHAIN PE=1 SV=4 | 1392 | 20201113_gy_milk_1ng_glyco_hcd30_etd30_compound15.13628.13628.3 | controllerType=0 controllerNumber=1 scan=13628 | 30.406 |
| 11995:4 | 12 | K.KGN[+2205.793]TTLNSFVIPSESDVPTHC[+57.021]PSQWWPYAGHC[+57.021]YK.I | HexNAc(4)Hex(5)Fuc(2)NeuAc(1) | C[+57]*2, N[+2206] | 1543.161 | 4 | 6169.6222 | 6169.6146 | 0 | 1.2 | 342 | Specific | 258.44 | 258.44 | 258.44 | 4.49 | 6 | >sp\|P22897\|MRC1_HUMAN Macrophage mannose receptor 1 OS=Homo sapiens OX=9606 GN=MRC1 PE=1 SV=1 | 1049 | 20201113_gy_milk_1ng_glyco_hcd30_etd30_compound15.15354.15354.4 | controllerType=0 controllerNumber=1 scan=15354 | 33.7164 |
| 20960:3 | 25 | R.QLAHQSN[+2204.772]STNIFFSPVSIATAFAMLSLGTK.A | HexNAc(4)Hex(5)NeuAc(2) | N[+2205] | 1796.1399 | 3 | 5386.405 | 5386.4075 | 0 | -0.5 | 64 | Specific | 268.03 | 163.16 | 163.16 | 4.15 | 3 | >sp\|P01009\|A1AT_HUMAN Alpha-1-antitrypsin OS=Homo sapiens OX=9606 GN=SERPINA1 PE=1 SV=3 | 642 | 20201113_gy_milk_1ng_glyco_hcd30_etd30_compound15.24789.24789.3 | controllerType=0 controllerNumber=1 scan=24789 | 55.9025 |

Table S4. Cont.

| Query #:z | Protein Rank | Glycopeptide | Glycan | Modification Type(s) | Observed m/z | z | Observed (M+H) | Calc. mass (M+H) | Off-by-x error | Mass error (ppm) | StartPosition | Cleavage | Score | Delta | Delta Mod | \|Log Prob\| | # of unique peptides | Protein Name | Protein DB number | Comment | Scan # | Scan Time |
|---|---|---|---|---|---|---|---|---|---|---|---|---|---|---|---|---|---|---|---|---|---|---|
| 17460:4 | 25 | K.YLGN[+2204.772]ATAIFFLPDEGKLQHLENELTHDIITK.F | HexNAc(4)Hex(5)NeuAc(2) | N[+2205] | 1437.1608 | 4 | 5745.6212 | 5745.6098 | 0 | 2 | 268 | Specific | 295.67 | 291.96 | 291.96 | 4.64 | 3 | >sp\|P01009\|A1AT_HUMAN Alpha-1-antitrypsin OS=Homo sapiens OX=9606 GN=SERPINA1 PE=1 SV=3 | 642 | 20201113_gy_milk_1ng_glyco_hcd30_etd30_compound15.21117.21117.4 | controllerType=0 controllerNumber=1 scan=21117 | 47.6868 |
| 02683:3 | 36 | K.TPLTAN[+1809.666]ITK.S | HexNAc(5)Hex(4)Fuc(1) | N[+1810] | 923.4142 | 3 | 2768.2279 | 2768.2228 | 0 | 1.8 | 200 | Specific | 266.01 | 118.73 | 118.73 | 2.78 | 4 | >sp\|P01877\|IGHA2_HUMAN Immunoglobulin heavy constant alpha 2 OS=Homo sapiens OX=9606 GN=IGHA2 PE=1 SV=4 | 576 | 20201113_gy_milk_1ng_glyco_hcd30_etd30_compound15.5658.5658.3 | controllerType=0 controllerNumber=1 scan=5658 | 16.104 |
| 02708:2 | 36 | K.TPLTAN[+1647.613]ITK.S | HexNAc(5)Hex(3)Fuc(1) | N[+1648] | 1303.5916 | 2 | 2606.1759 | 2606.17 | 0 | 2.3 | 200 | Specific | 295.83 | 121.77 | 121.77 | 2.98 | 4 | >sp\|P01877\|IGHA2_HUMAN Immunoglobulin heavy constant alpha 2 OS=Homo sapiens OX=9606 GN=IGHA2 PE=1 SV=4 | 576 | 20201113_gy_milk_1ng_glyco_hcd30_etd30_compound15.5684.5684.2 | controllerType=0 controllerNumber=1 scan=5684 | 16.1655 |
| 07114:3 | 48 | R.LSPN[+2350.830]ASAEHLELR.W | HexNAc(4)Hex(5)Fuc(1)NeuAc(2) | N[+2351] | 1263.2006 | 3 | 3787.5873 | 3787.5796 | 0 | 2 | 52 | Specific | 363.19 | 233.58 | 233.58 | 4.78 | 3 | >sp\|Q13410\|BT1A1_HUMAN Butyrophilin subfamily 1 member A1 OS=Homo sapiens OX=9606 GN=BTN1A1 PE=1 SV=3 | 179 | 20201113_gy_milk_1ng_glyco_hcd30_etd30_compound15.10252.10252.3 | controllerType=0 controllerNumber=1 scan=10252 | 24.1481 |
| 15668:3 | 48 | K.N[+2059.735]VSC[+57.021]YIQNLLLGQEK.K | HexNAc(4)Hex(5)Fuc(1)NeuAc(1) | C[+57], N[+2060] | 1280.2225 | 3 | 3838.6529 | 3838.6455 | 0 | 1.9 | 215 | Specific | 205.58 | 58.46 | 58.46 | 3.88 | 3 | >sp\|Q13410\|BT1A1_HUMAN Butyrophilin subfamily 1 member A1 OS=Homo sapiens OX=9606 GN=BTN1A1 PE=1 SV=3 | 179 | 20201113_gy_milk_1ng_glyco_hcd30_etd30_compound15.19221.19221.3 | controllerType=0 controllerNumber=1 scan=19221 | 42.3209 |
| 18425:3 | 73 | K.MLN[+1913.677]TSSLLEQLNEQFNWVSR.L | HexNAc(4)Hex(5)NeuAc(1) | N[+1914] | 1441.6324 | 3 | 4322.8826 | 4322.8637 | 0 | 4.4 | 352 | Specific | 221.63 | 140.45 | 140.45 | 3.14 | 1 | >sp\|P10909\|CLUS_HUMAN Clusterin OS=Homo sapiens OX=9606 GN=CLU PE=1 SV=1 | 1467 | 20201113_gy_milk_1ng_glyco_hcd30_etd30_compound15.22133.22133.3 | controllerType=0 controllerNumber=1 scan=22133 | 50.3733 |
| 15752:4 | 1 | K.VFPLSLDSTPQDGNVVVAC[+57.021]LVQGFFPQEPLSVTWSESGQN[+1809.666]VTAI | HexNAc(5)Hex(4)Fuc(1) | C[+57], N[+1810] | 1647.2656 | 4 | 6586.0407 | 6586.0275 | 0 | 2 | 8 | Specific | 215.92 | 215.92 | 215.92 | 5.91 | 93 | >sp\|P01877\|IGHA2_HUMAN Immunoglobulin heavy constant alpha 2 OS=Homo sapiens OX=9606 GN=IGHA2 PE=1 SV=4 | 576 | 20201114_qwl_colo_milk_1ng_glyco_hcd30_etd30_compound15.19364.19364.4 | controllerType=0 controllerNumber=1 scan=19364 | 53.4513 |
| 16304:4 | 1 | K.VFPLSLDSTPQDGNVVVAC[+57.021]LVQGFFPQEPLSVTWSESGQN[+2262.814]VTAI | HexNAc(5)Hex(5)Fuc(1)NeuAc(1) | C[+57], N[+2263] | 1760.5578 | 4 | 7039.2095 | 7039.1757 | 0 | 4.8 | 8 | Specific | 319.17 | 319.17 | 319.17 | 6.5 | 93 | >sp\|P01877\|IGHA2_HUMAN Immunoglobulin heavy constant alpha 2 OS=Homo sapiens OX=9606 GN=IGHA2 PE=1 SV=4 | 576 | 20201114_qwl_colo_milk_1ng_glyco_hcd30_etd30_compound15.19956.19956.4 | controllerType=0 controllerNumber=1 scan=19956 | 55.4902 |
| 16239:4 | 1 | K.VFPLSLDSTPQDGNVVVAC[+57.021]LVQGFFPQEPLSVTWSESGQN[+2262.814]VTAI | HexNAc(5)Hex(5)Fuc(1)NeuAc(1) | C[+57], N[+2263] | 1760.5606 | 4 | 7039.2207 | 7039.1757 | 0 | 6.4 | 8 | Specific | 357.1 | 357.1 | 357.1 | 6.89 | 93 | >sp\|P01877\|IGHA2_HUMAN Immunoglobulin heavy constant alpha 2 OS=Homo sapiens OX=9606 GN=IGHA2 PE=1 SV=4 | 576 | 20201114_qwl_colo_milk_1ng_glyco_hcd30_etd30_compound15.19887.19887.4 | controllerType=0 controllerNumber=1 scan=19887 | 55.2516 |
| 15596:4 | 1 | K.VFPLSLDSTPQDGNVVVAC[+57.021]LVQGFFPQEPLSVTWSESGQN[+1971.719]VTAI | HexNAc(5)Hex(5)Fuc(1) | C[+57], N[+1972] | 1687.7887 | 4 | 6748.1328 | 6748.0803 | 0 | 7.8 | 8 | Specific | 365.41 | 365.41 | 365.41 | 7.78 | 93 | >sp\|P01877\|IGHA2_HUMAN Immunoglobulin heavy constant alpha 2 OS=Homo sapiens OX=9606 GN=IGHA2 PE=1 SV=4 | 576 | 20201114_qwl_colo_milk_1ng_glyco_hcd30_etd30_compound15.19196.19196.4 | controllerType=0 controllerNumber=1 scan=19196 | 52.8471 |
| 15770:4 | 1 | K.VFPLSLDSTPQDGNVVVAC[+57.021]LVQGFFPQEPLSVTWSESGQN[+1444.534]VTAI | HexNAc(4)Hex(3)Fuc(1) | C[+57], N[+1445] | 1555.9808 | 4 | 6220.9015 | 6220.8953 | 0 | 1 | 8 | Specific | 369.96 | 369.96 | 369.96 | 8.16 | 93 | >sp\|P01877\|IGHA2_HUMAN Immunoglobulin heavy constant alpha 2 OS=Homo sapiens OX=9606 GN=IGHA2 PE=1 SV=4 | 576 | 20201114_qwl_colo_milk_1ng_glyco_hcd30_etd30_compound15.19383.19383.4 | controllerType=0 controllerNumber=1 scan=19383 | 53.5225 |
| 15732:4 | 1 | K.VFPLSLDSTPQDGNVVVAC[+57.021]LVQGFFPQEPLSVTWSESGQN[+1444.534]VTAI | HexNAc(4)Hex(3)Fuc(1) | C[+57], N[+1445] | 1555.9833 | 4 | 6220.9113 | 6220.8953 | 0 | 2.6 | 8 | Specific | 390.16 | 390.16 | 390.16 | 8.37 | 93 | >sp\|P01877\|IGHA2_HUMAN Immunoglobulin heavy constant alpha 2 OS=Homo sapiens OX=9606 GN=IGHA2 PE=1 SV=4 | 576 | 20201114_qwl_colo_milk_1ng_glyco_hcd30_etd30_compound15.19343.19343.4 | controllerType=0 controllerNumber=1 scan=19343 | 53.3944 |
| 15365:4 | 1 | K.VFPLSLDSTPQDGNVVVAC[+57.021]LVQGFFPQEPLSVTWSESGQN[+1606.587]VTAI | HexNAc(4)Hex(4)Fuc(1) | C[+57], N[+1607] | 1596.5029 | 4 | 6382.9899 | 6382.9481 | 0 | 6.6 | 8 | Specific | 415.34 | 415.34 | 415.34 | 7.89 | 93 | >sp\|P01877\|IGHA2_HUMAN Immunoglobulin heavy constant alpha 2 OS=Homo sapiens OX=9606 GN=IGHA2 PE=1 SV=4 | 576 | 20201114_qwl_colo_milk_1ng_glyco_hcd30_etd30_compound15.18949.18949.4 | controllerType=0 controllerNumber=1 scan=18949 | 51.9798 |
| 15720:4 | 1 | K.VFPLSLDSTPQDGNVVVAC[+57.021]LVQGFFPQEPLSVTWSESGQN[+1809.666]VTAI | HexNAc(5)Hex(4)Fuc(1) | C[+57], N[+1810] | 1647.2742 | 4 | 6586.0749 | 6586.0275 | 0 | 7.2 | 8 | Specific | 497.39 | 497.39 | 497.39 | 8.02 | 93 | >sp\|P01877\|IGHA2_HUMAN Immunoglobulin heavy constant alpha 2 OS=Homo sapiens OX=9606 GN=IGHA2 PE=1 SV=4 | 576 | 20201114_qwl_colo_milk_1ng_glyco_hcd30_etd30_compound15.19330.19330.4 | controllerType=0 controllerNumber=1 scan=19330 | 53.3427 |
| 15600:4 | 1 | K.VFPLSLDSTPQDGNVVVAC[+57.021]LVQGFFPQEPLSVTWSESGQN[+1809.666]VTAI | HexNAc(5)Hex(4)Fuc(1) | C[+57], N[+1810] | 1647.2738 | 4 | 6586.0734 | 6586.0275 | 0 | 7 | 8 | Specific | 504.79 | 504.79 | 504.79 | 8.02 | 93 | >sp\|P01877\|IGHA2_HUMAN Immunoglobulin heavy constant alpha 2 OS=Homo sapiens OX=9606 GN=IGHA2 PE=1 SV=4 | 576 | 20201114_qwl_colo_milk_1ng_glyco_hcd30_etd30_compound15.19201.19201.4 | controllerType=0 controllerNumber=1 scan=19201 | 52.8654 |
| 15369:4 | 1 | K.VFPLSLDSTPQDGNVVVAC[+57.021]LVQGFFPQEPLSVTWSESGQN[+1444.534]VTAI | HexNAc(4)Hex(3)Fuc(1) | C[+57], N[+1445] | 1555.9858 | 4 | 6220.9215 | 6220.8953 | 0 | 4.2 | 8 | Specific | 572.57 | 572.57 | 572.57 | 9.1 | 93 | >sp\|P01877\|IGHA2_HUMAN Immunoglobulin heavy constant alpha 2 OS=Homo sapiens OX=9606 GN=IGHA2 PE=1 SV=4 | 576 | 20201114_qwl_colo_milk_1ng_glyco_hcd30_etd30_compound15.18953.18953.4 | controllerType=0 controllerNumber=1 scan=18953 | 51.9957 |
| 15352:4 | 1 | K.VFPLSLDSTPQDGNVVVAC[+57.021]LVQGFFPQEPLSVTWSESGQN[+1647.613]VTAI | HexNAc(5)Hex(3)Fuc(1) | C[+57], N[+1648] | 1606.7649 | 4 | 6424.0378 | 6423.9746 | 0 | 9.8 | 8 | Specific | 885.96 | 885.96 | 885.96 | 15.18 | 93 | >sp\|P01877\|IGHA2_HUMAN Immunoglobulin heavy constant alpha 2 OS=Homo sapiens OX=9606 GN=IGHA2 PE=1 SV=4 | 576 | 20201114_qwl_colo_milk_1ng_glyco_hcd30_etd30_compound15.18935.18935.4 | controllerType=0 controllerNumber=1 scan=18935 | 51.9163 |
| 01314:3 | 1 | K.HYTN[+1444.534]SSQDVTVPC[+57.021]R.V | HexNAc(4)Hex(3)Fuc(1) | C[+57], N[+1445] | 1036.7677 | 3 | 3108.2886 | 3108.2832 | 0 | 1.8 | 89 | Specific | 206.21 | 145.21 | 145.21 | 4.41 | 93 | >sp\|P01877\|IGHA2_HUMAN Immunoglobulin heavy constant alpha 2 OS=Homo sapiens OX=9606 GN=IGHA2 PE=1 SV=4 | 576 | 20201114_qwl_colo_milk_1ng_glyco_hcd30_etd30_compound15.4157.4157.3 | controllerType=0 controllerNumber=1 scan=4157 | 13.8906 |
| 00862:2 | 1 | K.HYTN[+1444.534]SSQDVTVPC[+57.021]R.V | HexNAc(4)Hex(3)Fuc(1) | C[+57], N[+1445] | 1554.6524 | 2 | 3108.2975 | 3108.2832 | 0 | 4.6 | 89 | Specific | 227.57 | 166.74 | 166.74 | 3.59 | 93 | >sp\|P01877\|IGHA2_HUMAN Immunoglobulin heavy constant alpha 2 OS=Homo sapiens OX=9606 GN=IGHA2 PE=1 SV=4 | 576 | 20201114_qwl_colo_milk_1ng_glyco_hcd30_etd30_compound15.3686.3686.2 | controllerType=0 controllerNumber=1 scan=3686 | 12.9294 |
| 01556:3 | 1 | K.HYTN[+2059.735]SSQDVTVPC[+57.021]R.V | HexNAc(4)Hex(5)Fuc(1)NeuAc(1) | C[+57], N[+2060] | 1241.836 | 3 | 3723.4934 | 3723.4842 | 0 | 2.5 | 89 | Specific | 316.11 | 200.76 | 200.76 | 5.52 | 93 | >sp\|P01877\|IGHA2_HUMAN Immunoglobulin heavy constant alpha 2 OS=Homo sapiens OX=9606 GN=IGHA2 PE=1 SV=4 | 576 | 20201114_qwl_colo_milk_1ng_glyco_hcd30_etd30_compound15.4408.4408.3 | controllerType=0 controllerNumber=1 scan=4408 | 14.3393 |
| 00805:3 | 1 | K.HYTN[+1403.507]SSQDVTVPC[+57.021]R.V | HexNAc(3)Hex(4)Fuc(1) | C[+57], N[+1404] | 1023.0991 | 3 | 3067.2827 | 3067.2566 | 0 | 8.5 | 89 | Specific | 329.49 | 249.7 | 249.7 | 5.45 | 93 | >sp\|P01877\|IGHA2_HUMAN Immunoglobulin heavy constant alpha 2 OS=Homo sapiens OX=9606 GN=IGHA2 PE=1 SV=4 | 576 | 20201114_qwl_colo_milk_1ng_glyco_hcd30_etd30_compound15.3627.3627.3 | controllerType=0 controllerNumber=1 scan=3627 | 12.7794 |
| 01598:3 | 1 | K.HYTN[+2262.814]SSQDVTVPC[+57.021]R.V | HexNAc(5)Hex(5)Fuc(1)NeuAc(1) | C[+57], N[+2263] | 1309.529 | 3 | 3926.5724 | 3926.5636 | 0 | 2.2 | 89 | Specific | 337.93 | 223.45 | 223.45 | 5.86 | 93 | >sp\|P01877\|IGHA2_HUMAN Immunoglobulin heavy constant alpha 2 OS=Homo sapiens OX=9606 GN=IGHA2 PE=1 SV=4 | 576 | 20201114_qwl_colo_milk_1ng_glyco_hcd30_etd30_compound15.4452.4452.3 | controllerType=0 controllerNumber=1 scan=4452 | 14.4331 |
| 00826:3 | 1 | K.HYTN[+1971.719]SSQDVTVPC[+57.021]R.V | HexNAc(5)Hex(5)Fuc(1) | C[+57], N[+1972] | 1212.4949 | 3 | 3635.4702 | 3635.4682 | 0 | 0.6 | 89 | Specific | 404.59 | 332.49 | 332.49 | 5.93 | 93 | >sp\|P01877\|IGHA2_HUMAN Immunoglobulin heavy constant alpha 2 OS=Homo sapiens OX=9606 GN=IGHA2 PE=1 SV=4 | 576 | 20201114_qwl_colo_milk_1ng_glyco_hcd30_etd30_compound15.3649.3649.3 | controllerType=0 controllerNumber=1 scan=3649 | 12.8325 |
| 00865:3 | 1 | K.HYTN[+1809.666]SSQDVTVPC[+57.021]R.V | HexNAc(5)Hex(4)Fuc(1) | C[+57], N[+1810] | 1158.4788 | 3 | 3473.4219 | 3473.4154 | 0 | 1.9 | 89 | Specific | 417.13 | 332.96 | 332.96 | 6.14 | 93 | >sp\|P01877\|IGHA2_HUMAN Immunoglobulin heavy constant alpha 2 OS=Homo sapiens OX=9606 GN=IGHA2 PE=1 SV=4 | 576 | 20201114_qwl_colo_milk_1ng_glyco_hcd30_etd30_compound15.3690.3690.3 | controllerType=0 controllerNumber=1 scan=3690 | 12.9443 |
| 00869:3 | 1 | K.HYTN[+1647.613]SSQDVTVPC[+57.021]R.V | HexNAc(5)Hex(3)Fuc(1) | C[+57], N[+1648] | 1104.4618 | 3 | 3311.3709 | 3311.3625 | 0 | 2.5 | 89 | Specific | 524.21 | 459.07 | 459.07 | 7.45 | 93 | >sp\|P01877\|IGHA2_HUMAN Immunoglobulin heavy constant alpha 2 OS=Homo sapiens OX=9606 GN=IGHA2 PE=1 SV=4 | 576 | 20201114_qwl_colo_milk_1ng_glyco_hcd30_etd30_compound15.3694.3694.3 | controllerType=0 controllerNumber=1 scan=3694 | 12.9668 |
| 00808:3 | 1 | K.HYTN[+1444.534]SSQDVTVPC[+57.021]R.V | HexNAc(4)Hex(3)Fuc(1) | C[+57], N[+1445] | 1036.7746 | 3 | 3108.3091 | 3108.2832 | 0 | 8.4 | 89 | Specific | 527.23 | 455.35 | 455.35 | 7 | 93 | >sp\|P01877\|IGHA2_HUMAN Immunoglobulin heavy constant alpha 2 OS=Homo sapiens OX=9606 GN=IGHA2 PE=1 SV=4 | 576 | 20201114_qwl_colo_milk_1ng_glyco_hcd30_etd30_compound15.3630.3630.3 | controllerType=0 controllerNumber=1 scan=3630 | 12.7907 |
| 00853:3 | 1 | K.HYTN[+1241.454]SSQDVTVPC[+57.021]R.V | HexNAc(3)Hex(3)Fuc(1) | C[+57], N[+1241] | 969.0772 | 3 | 2905.217 | 2905.2038 | 0 | 4.5 | 89 | Specific | 527.9 | 410.97 | 410.97 | 6.92 | 93 | >sp\|P01877\|IGHA2_HUMAN Immunoglobulin heavy constant alpha 2 OS=Homo sapiens OX=9606 GN=IGHA2 PE=1 SV=4 | 576 | 20201114_qwl_colo_milk_1ng_glyco_hcd30_etd30_compound15.3677.3677.3 | controllerType=0 controllerNumber=1 scan=3677 | 12.9028 |
| 00781:3 | 1 | K.HYTN[+1606.587]SSQDVTVPC[+57.021]R.V | HexNAc(4)Hex(4)Fuc(1) | C[+57], N[+1607] | 1090.7852 | 3 | 3270.341 | 3270.336 | 0 | 1.5 | 89 | Specific | 542.77 | 463.89 | 463.89 | 7.24 | 93 | >sp\|P01877\|IGHA2_HUMAN Immunoglobulin heavy constant alpha 2 OS=Homo sapiens OX=9606 GN=IGHA2 PE=1 SV=4 | 576 | 20201114_qwl_colo_milk_1ng_glyco_hcd30_etd30_compound15.3602.3602.3 | controllerType=0 controllerNumber=1 scan=3602 | 12.7285 |
| 00852:3 | 1 | K.HYTN[+1038.375]SSQDVTVPC[+57.021]R.V | HexNAc(2)Hex(3)Fuc(1) | C[+57], N[+1038] | 901.3836 | 3 | 2702.1362 | 2702.1244 | 0 | 4.3 | 89 | Specific | 552.07 | 451.3 | 451.3 | 6.92 | 93 | >sp\|P01877\|IGHA2_HUMAN Immunoglobulin heavy constant alpha 2 OS=Homo sapiens OX=9606 GN=IGHA2 PE=1 SV=4 | 576 | 20201114_qwl_colo_milk_1ng_glyco_hcd30_etd30_compound15.3676.3676.3 | controllerType=0 controllerNumber=1 scan=3676 | 12.8947 |
| 03045:5 | 1 | K.HYTN[+2262.814]SSQDVTVPC[+57.021]RVPPPPPC[+57.021]C[+57.021]HPR.L | HexNAc(5)Hex(5)Fuc(1)NeuAc(1) | C[+57]*3, N[+2263] | 1045.0477 | 5 | 5221.2096 | 5221.1699 | 0 | 7.6 | 89 | Specific | 319.14 | 274.35 | 274.35 | 3.79 | 93 | >sp\|P01877\|IGHA2_HUMAN Immunoglobulin heavy constant alpha 2 OS=Homo sapiens OX=9606 GN=IGHA2 PE=1 SV=4 | 576 | 20201114_qwl_colo_milk_1ng_glyco_hcd30_etd30_compound15.5966.5966.5 | controllerType=0 controllerNumber=1 scan=5966 | 17.778 |
| 02530:5 | 1 | K.HYTN[+1606.587]SSQDVTVPC[+57.021]RVPPPPPC[+57.021]C[+57.021]HPR.L | HexNAc(4)Hex(4)Fuc(1) | C[+57]*3, N[+1607] | 913.7958 | 5 | 4564.9499 | 4564.9423 | 0 | 1.7 | 89 | Specific | 344.09 | 344.09 | 344.09 | 4.39 | 93 | >sp\|P01877\|IGHA2_HUMAN Immunoglobulin heavy constant alpha 2 OS=Homo sapiens OX=9606 GN=IGHA2 PE=1 SV=4 | 576 | 20201114_qwl_colo_milk_1ng_glyco_hcd30_etd30_compound15.5427.5427.5 | controllerType=0 controllerNumber=1 scan=5427 | 16.5865 |
| 02490:4 | 1 | K.HYTN[+1647.613]SSQDVTVPC[+57.021]RVPPPPPC[+57.021]C[+57.021]HPR.L | HexNAc(5)Hex(3)Fuc(1) | C[+57]*3, N[+1648] | 1152.2486 | 4 | 4605.9725 | 4605.9689 | 0 | 0.8 | 89 | Specific | 363.68 | 358.89 | 358.89 | 4.16 | 93 | >sp\|P01877\|IGHA2_HUMAN Immunoglobulin heavy constant alpha 2 OS=Homo sapiens OX=9606 GN=IGHA2 PE=1 SV=4 | 576 | 20201114_qwl_colo_milk_1ng_glyco_hcd30_etd30_compound15.5385.5385.4 | controllerType=0 controllerNumber=1 scan=5385 | 16.483 |
| 02505:4 | 1 | K.HYTN[+1809.666]SSQDVTVPC[+57.021]RVPPPPPC[+57.021]C[+57.021]HPR.L | HexNAc(5)Hex(4)Fuc(1) | C[+57]*3, N[+1810] | 1192.7633 | 4 | 4768.0316 | 4768.0217 | 0 | 2.1 | 89 | Specific | 490.78 | 451.24 | 451.24 | 5.72 | 93 | >sp\|P01877\|IGHA2_HUMAN Immunoglobulin heavy constant alpha 2 OS=Homo sapiens OX=9606 GN=IGHA2 PE=1 SV=4 | 576 | 20201114_qwl_colo_milk_1ng_glyco_hcd30_etd30_compound15.5401.5401.4 | controllerType=0 controllerNumber=1 scan=5401 | 16.533 |
| 02524:5 | 1 | K.HYTN[+1647.613]SSQDVTVPC[+57.021]RVPPPPPC[+57.021]C[+57.021]HPR.L | HexNAc(5)Hex(3)Fuc(1) | C[+57]*3, N[+1648] | 922.0016 | 5 | 4605.9789 | 4605.9689 | 0 | 2.2 | 89 | Specific | 653.29 | 606.56 | 606.56 | 9.47 | 93 | >sp\|P01877\|IGHA2_HUMAN Immunoglobulin heavy constant alpha 2 OS=Homo sapiens OX=9606 GN=IGHA2 PE=1 SV=4 | 576 | 20201114_qwl_colo_milk_1ng_glyco_hcd30_etd30_compound15.5421.5421.5 | controllerType=0 controllerNumber=1 scan=5421 | 16.5731 |
| 02493:5 | 1 | K.HYTN[+1444.534]SSQDVTVPC[+57.021]RVPPPPPC[+57.021]C[+57.021]HPR.L | HexNAc(4)Hex(3)Fuc(1) | C[+57]*3, N[+1445] | 881.3848 | 5 | 4402.8948 | 4402.8895 | 0 | 1.2 | 89 | Specific | 676.66 | 641.22 | 641.22 | 8.89 | 93 | >sp\|P01877\|IGHA2_HUMAN Immunoglobulin heavy constant alpha 2 OS=Homo sapiens OX=9606 GN=IGHA2 PE=1 SV=4 | 576 | 20201114_qwl_colo_milk_1ng_glyco_hcd30_etd30_compound15.5388.5388.5 | controllerType=0 controllerNumber=1 scan=5388 | 16.4949 |
| 02489:4 | 1 | K.HYTN[+1444.534]SSQDVTVPC[+57.021]RVPPPPPC[+57.021]C[+57.021]HPR.L | HexNAc(4)Hex(3)Fuc(1) | C[+57]*3, N[+1445] | 1101.4782 | 4 | 4402.8909 | 4402.8895 | 0 | 0.3 | 89 | Specific | 764.95 | 698.59 | 698.59 | 11.82 | 93 | >sp\|P01877\|IGHA2_HUMAN Immunoglobulin heavy constant alpha 2 OS=Homo sapiens OX=9606 GN=IGHA2 PE=1 SV=4 | 576 | 20201114_qwl_colo_milk_1ng_glyco_hcd30_etd30_compound15.5384.5384.4 | controllerType=0 controllerNumber=1 scan=5384 | 16.4761 |
| 13273:3 | 1 | R.LSLHRPALEDLLLGSEAN[+1913.677]LTC[+57.021]TLTGLR.D | HexNAc(4)Hex(5)NeuAc(1) | C[+57], N[+1914] | 1626.4412 | 3 | 4877.3091 | 4877.2753 | 0 | 6.9 | 114 | Specific | 202.05 | 202.05 | 202.05 | 4.74 | 93 | >sp\|P01877\|IGHA2_HUMAN Immunoglobulin heavy constant alpha 2 OS=Homo sapiens OX=9606 GN=IGHA2 PE=1 SV=4 | 576 | 20201114_qwl_colo_milk_1ng_glyco_hcd30_etd30_compound15.16717.16717.3 | controllerType=0 controllerNumber=1 scan=16717 | 44.5971 |
| 12706:3 | 1 | R.LSLHRPALEDLLLGSEAN[+1054.370]LTC[+57.021]TLTGLR.D | HexNAc(2)Hex(4) | C[+57], N[+1054] | 1339.9971 | 3 | 4017.9768 | 4017.9683 | 0 | 2.1 | 114 | Specific | 287 | 287 | 287 | 6.56 | 93 | >sp\|P01877\|IGHA2_HUMAN Immunoglobulin heavy constant alpha 2 OS=Homo sapiens OX=9606 GN=IGHA2 PE=1 SV=4 | 576 | 20201114_qwl_colo_milk_1ng_glyco_hcd30_etd30_compound15.16115.16115.3 | controllerType=0 controllerNumber=1 scan=16115 | 42.6607 |
| 12848:4 | 1 | R.LSLHRPALEDLLLGSEAN[+1460.529]LTC[+57.021]TLTGLR.D | HexNAc(4)Hex(4) | C[+57], N[+1461] | 1106.7942 | 4 | 4424.1551 | 4424.1271 | 0 | 6.3 | 114 | Specific | 298.4 | 298.4 | 298.4 | 5.06 | 93 | >sp\|P01877\|IGHA2_HUMAN Immunoglobulin heavy constant alpha 2 OS=Homo sapiens OX=9606 GN=IGHA2 PE=1 SV=4 | 576 | 20201114_qwl_colo_milk_1ng_glyco_hcd30_etd30_compound15.16266.16266.4 | controllerType=0 controllerNumber=1 scan=16266 | 43.197 |
| 13325:3 | 1 | R.LSLHRPALEDLLLGSEAN[+1872.651]LTC[+57.021]TLTGLR.D | HexNAc(3)Hex(6)NeuAc(1) | C[+57], N[+1873] | 1612.7662 | 3 | 4836.284 | 4836.2487 | 0 | 7.3 | 114 | Specific | 326.18 | 286.52 | 286.52 | 5.23 | 93 | >sp\|P01877\|IGHA2_HUMAN Immunoglobulin heavy constant alpha 2 OS=Homo sapiens OX=9606 GN=IGHA2 PE=1 SV=4 | 576 | 20201114_qwl_colo_milk_1ng_glyco_hcd30_etd30_compound15.16773.16773.3 | controllerType=0 controllerNumber=1 scan=16773 | 44.8159 |
| 13262:4 | 1 | R.LSLHRPALEDLLLGSEAN[+1913.677]LTC[+57.021]TLTGLR.D | HexNAc(4)Hex(5)NeuAc(1) | C[+57], N[+1914] | 1220.0824 | 4 | 4877.3079 | 4877.2753 | 0 | 6.7 | 114 | Specific | 340.98 | 340.98 | 340.98 | 5.84 | 93 | >sp\|P01877\|IGHA2_HUMAN Immunoglobulin heavy constant alpha 2 OS=Homo sapiens OX=9606 GN=IGHA2 PE=1 SV=4 | 576 | 20201114_qwl_colo_milk_1ng_glyco_hcd30_etd30_compound15.16706.16706.4 | controllerType=0 controllerNumber=1 scan=16706 | 44.5619 |
| 15119:4 | 1 | R.LSLHRPALEDLLLGSEAN[+1501.555]LTC[+57.021]TLTGLR.D | HexNAc(5)Hex(3) | C[+57], N[+1502] | 1117.0536 | 4 | 4465.1927 | 4465.1536 | 0 | 8.8 | 114 | Specific | 397.35 | 397.35 | 397.35 | 7.86 | 93 | >sp\|P01877\|IGHA2_HUMAN Immunoglobulin heavy constant alpha 2 OS=Homo sapiens OX=9606 GN=IGHA2 PE=1 SV=4 | 576 | 20201114_qwl_colo_milk_1ng_glyco_hcd30_etd30_compound15.18687.18687.2 | controllerType=0 controllerNumber=1 scan=18687 | 51.166 |
| 12622:3 | 1 | R.LSLHRPALEDLLLGSEAN[+1702.581]LTC[+57.021]TLTGLR.D | HexNAc(2)Hex(8) | C[+57], N[+1703] | 1556.0764 | 3 | 4666.2145 | 4666.1796 | 0 | 7.5 | 114 | Specific | 412.72 | 409.85 | 409.85 | 7.4 | 93 | >sp\|P01877\|IGHA2_HUMAN Immunoglobulin heavy constant alpha 2 OS=Homo sapiens OX=9606 GN=IGHA2 PE=1 SV=4 | 576 | 20201114_qwl_colo_milk_1ng_glyco_hcd30_etd30_compound15.16024.16024.3 | controllerType=0 controllerNumber=1 scan=16024 | 42.287 |
| 12841:4 | 1 | R.LSLHRPALEDLLLGSEAN[+1444.534]LTC[+57.021]TLTGLR.D | HexNAc(4)Hex(3)Fuc(1) | C[+57], N[+1445] | 1102.7963 | 4 | 4408.1634 | 4408.1321 | 0 | 7.1 | 114 | Specific | 437.86 | 437.86 | 437.86 | 7.75 | 93 | >sp\|P01877\|IGHA2_HUMAN Immunoglobulin heavy constant alpha 2 OS=Homo sapiens OX=9606 GN=IGHA2 PE=1 SV=4 | 576 | 20201114_qwl_colo_milk_1ng_glyco_hcd30_etd30_compound15.16259.16259.4 | controllerType=0 controllerNumber=1 scan=16259 | 43.1802 |
| 12907:4 | 1 | R.LSLHRPALEDLLLGSEAN[+1663.608]LTC[+57.021]TLTGLR.D | HexNAc(5)Hex(4) | C[+57], N[+1664] | 1157.5669 | 4 | 4627.2458 | 4627.2064 | 0 | 8.5 | 114 | Specific | 437.91 | 437.91 | 437.91 | 6.61 | 93 | >sp\|P01877\|IGHA2_HUMAN Immunoglobulin heavy constant alpha 2 OS=Homo sapiens OX=9606 GN=IGHA2 PE=1 SV=4 | 576 | 20201114_qwl_colo_milk_1ng_glyco_hcd30_etd30_compound15.16328.16328.4 | controllerType=0 controllerNumber=1 scan=16328 | 43.4103 |
| 12599:3 | 1 | R.LSLHRPALEDLLLGSEAN[+1864.634]LTC[+57.021]TLTGLR.D | HexNAc(2)Hex(9) | C[+57], N[+1865] | 1610.0973 | 3 | 4828.2775 | 4828.2324 | 0 | 9.3 | 114 | Specific | 475.75 | 456.32 | 456.32 | 7.54 | 93 | >sp\|P01877\|IGHA2_HUMAN Immunoglobulin heavy constant alpha 2 OS=Homo sapiens OX=9606 GN=IGHA2 PE=1 SV=4 | 576 | 20201114_qwl_colo_milk_1ng_glyco_hcd30_etd30_compound15.15999.15999.3 | controllerType=0 controllerNumber=1 scan=15999 | 42.1851 |
| 12716:3 | 1 | R.LSLHRPALEDLLLGSEAN[+1501.555]LTC[+57.021]TLTGLR.D | HexNAc(5)Hex(3) | C[+57], N[+1502] | 1489.06 | 3 | 4465.1654 | 4465.1536 | 0 | 2.6 | 114 | Specific | 480.91 | 434.13 | 434.13 | 8.19 | 93 | >sp\|P01877\|IGHA2_HUMAN Immunoglobulin heavy constant alpha 2 OS=Homo sapiens OX=9606 GN=IGHA2 PE=1 SV=4 | 576 | 20201114_qwl_colo_milk_1ng_glyco_hcd30_etd30_compound15.16126.16126.3 | controllerType=0 controllerNumber=1 scan=16126 | 42.7044 |
| 24037:4 | 1 | R.LSLHRPALEDLLLGSEAN[+1501.555]LTC[+57.021]TLTGLR.D | HexNAc(5)Hex(3) | C[+57], N[+1502] | 1117.047 | 4 | 4465.1663 | 4465.1536 | 0 | 2.9 | 114 | Specific | 483.15 | 483.15 | 483.15 | 8.55 | 93 | >sp\|P01877\|IGHA2_HUMAN Immunoglobulin heavy constant alpha 2 OS=Homo sapiens OX=9606 GN=IGHA2 PE=1 SV=4 | 576 | 20201114_qwl_colo_milk_1ng_glyco_hcd30_etd30_compound15.32019.32019.4 | controllerType=0 controllerNumber=1 scan=32019 | 88.8063 |
| 13320:3 | 1 | R.LSLHRPALEDLLLGSEAN[+1751.624]LTC[+57.021]TLTGLR.D | HexNAc(4)Hex(4)NeuAc(1) | C[+57], N[+1752] | 1572.4247 | 3 | 4715.2597 | 4715.2225 | 0 | 7.9 | 114 | Specific | 494.38 | 454.75 | 454.75 | 7.61 | 93 | >sp\|P01877\|IGHA2_HUMAN Immunoglobulin heavy constant alpha 2 OS=Homo sapiens OX=9606 GN=IGHA2 PE=1 SV=4 | 576 | 20201114_qwl_colo_milk_1ng_glyco_hcd30_etd30_compound15.16768.16768.3 | controllerType=0 controllerNumber=1 scan=16768 | 44.7951 |
| 12637:3 | 1 | R.LSLHRPALEDLLLGSEAN[+1540.529]LTC[+57.021]TLTGLR.D | HexNAc(2)Hex(7) | C[+57], N[+1541] | 1502.0462 | 3 | 4504.124 | 4504.1268 | 0 | -0.6 | 114 | Specific | 509.26 | 477.65 | 477.65 | 7.62 | 93 | >sp\|P01877\|IGHA2_HUMAN Immunoglobulin heavy constant alpha 2 OS=Homo sapiens OX=9606 GN=IGHA2 PE=1 SV=4 | 576 | 20201114_qwl_colo_milk_1ng_glyco_hcd30_etd30_compound15.16040.16040.3 | controllerType=0 controllerNumber=1 scan=16040 | 42.3584 |
| 12665:3 | 1 | R.LSLHRPALEDLLLGSEAN[+1501.555]LTC[+57.021]TLTGLR.D | HexNAc(5)Hex(3) | C[+57], N[+1502] | 1489.0606 | 3 | 4465.1673 | 4465.1536 | 0 | 3.1 | 114 | Specific | 530.74 | 497.89 | 497.89 | 8.27 | 93 | >sp\|P01877\|IGHA2_HUMAN Immunoglobulin heavy constant alpha 2 OS=Homo sapiens OX=9606 GN=IGHA2 PE=1 SV=4 | 576 | 20201114_qwl_colo_milk_1ng_glyco_hcd30_etd30_compound15.16071.16071.3 | controllerType=0 controllerNumber=1 scan=16071 | 42.4845 |
| 12651:3 | 1 | R.LSLHRPALEDLLLGSEAN[+1460.529]LTC[+57.021]TLTGLR.D | HexNAc(4)Hex(4) | C[+57], N[+1461] | 1475.3848 | 3 | 4424.1398 | 4424.1271 | 0 | 2.9 | 114 | Specific | 531.16 | 470.82 | 470.82 | 8.27 | 93 | >sp\|P01877\|IGHA2_HUMAN Immunoglobulin heavy constant alpha 2 OS=Homo sapiens OX=9606 GN=IGHA2 PE=1 SV=4 | 576 | 20201114_qwl_colo_milk_1ng_glyco_hcd30_etd30_compound15.16055.16055.3 | controllerType=0 controllerNumber=1 scan=16055 | 42.416 |
| 12753:4 | 1 | R.LSLHRPALEDLLLGSEAN[+1257.449]LTC[+57.021]TLTGLR.D | HexNAc(3)Hex(4) | C[+57], N[+1257] | 1056.0247 | 4 | 4221.077 | 4221.0477 | 0 | 6.9 | 114 | Specific | 542.83 | 542.83 | 542.83 | 8.04 | 93 | >sp\|P01877\|IGHA2_HUMAN Immunoglobulin heavy constant alpha 2 OS=Homo sapiens OX=9606 GN=IGHA2 PE=1 SV=4 | 576 | 20201114_qwl_colo_milk_1ng_glyco_hcd30_etd30_compound15.16165.16165.4 | controllerType=0 controllerNumber=1 scan=16165 | 42.8431 |
| 12641:4 | 1 | R.LSLHRPALEDLLLGSEAN[+1378.476]LTC[+57.021]TLTGLR.D | HexNAc(2)Hex(6) | C[+57], N[+1378] | 1086.2741 | 4 | 4342.0745 | 4342.074 | 0 | 0.1 | 114 | Specific | 557.37 | 557.37 | 557.37 | 8.41 | 93 | >sp\|P01877\|IGHA2_HUMAN Immunoglobulin heavy constant alpha 2 OS=Homo sapiens OX=9606 GN=IGHA2 PE=1 SV=4 | 576 | 20201114_qwl_colo_milk_1ng_glyco_hcd30_etd30_compound15.16044.16044.4 | controllerType=0 controllerNumber=1 scan=16044 | 42.3716 |
| 12721:3 | 1 | R.LSLHRPALEDLLLGSEAN[+1054.370]LTC[+57.021]TLTGLR.D | HexNAc(2)Hex(4) | C[+57], N[+1054] | 1339.9972 | 3 | 4017.9772 | 4017.9683 | 0 | 2.2 | 114 | Specific | 563.11 | 536.7 | 536.7 | 9.21 | 93 | >sp\|P01877\|IGHA2_HUMAN Immunoglobulin heavy constant alpha 2 OS=Homo sapiens OX=9606 GN=IGHA2 PE=1 SV=4 | 576 | 20201114_qwl_colo_milk_1ng_glyco_hcd30_etd30_compound15.16131.16131.3 | controllerType=0 controllerNumber=1 scan=16131 | 42.726 |
| 12632:3 | 1 | R.LSLHRPALEDLLLGSEAN[+1216.423]LTC[+57.021]TLTGLR.D | HexNAc(2)Hex(5) | C[+57], N[+1216] | 1394.0201 | 3 | 4180.0457 | 4180.0211 | 0 | 5.9 | 114 | Specific | 566.69 | 557.18 | 557.18 | 8.7 | 93 | >sp\|P01877\|IGHA2_HUMAN Immunoglobulin heavy constant alpha 2 OS=Homo sapiens OX=9606 GN=IGHA2 PE=1 SV=4 | 576 | 20201114_qwl_colo_milk_1ng_glyco_hcd30_etd30_compound15.16034.16034.3 | controllerType=0 controllerNumber=1 scan=16034 | 42.3366 |
| 12682:3 | 1 | R.LSLHRPALEDLLLGSEAN[+203.079]LTC[+57.021]TLTGLR.D | HexNAc(1) | C[+57], N[+203] | 1056.2317 | 3 | 3166.6806 | 3166.6776 | 0 | 0.9 | 114 | Specific | 648.21 | 648.21 | 648.21 | 10 | 93 | >sp\|P01877\|IGHA2_HUMAN Immunoglobulin heavy constant alpha 2 OS=Homo sapiens OX=9606 GN=IGHA2 PE=1 SV=4 | 576 | 20201114_qwl_colo_milk_1ng_glyco_hcd30_etd30_compound15.16089.16089.4 | controllerType=0 controllerNumber=1 scan=16089 | 42.5472 |
| 13324:4 | 1 | R.LSLHRPALEDLLLGSEAN[+1872.651]LTC[+57.021]TLTGLR.D | HexNAc(3)Hex(6)NeuAc(1) | C[+57], N[+1873] | 1209.8268 | 4 | 4836.2855 | 4836.2487 | 0 | 7.6 | 114 | Specific | 693.72 | 622.51 | 622.51 | 10.98 | 93 | >sp\|P01877\|IGHA2_HUMAN Immunoglobulin heavy constant alpha 2 OS=Homo sapiens OX=9606 GN=IGHA2 PE=1 SV=4 | 576 | 20201114_qwl_colo_milk_1ng_glyco_hcd30_etd30_compound15.16772.16772.4 | controllerType=0 controllerNumber=1 scan=16772 | 44.8074 |
| 12642:4 | 1 | R.LSLHRPALEDLLLGSEAN[+1501.555]LTC[+57.021]TLTGLR.D | HexNAc(5)Hex(3) | C[+57], N[+1502] | 1117.0441 | 4 | 4465.1546 | 4465.1536 | 0 | 0.2 | 114 | Specific | 879.61 | 879.22 | 879.22 | 15.35 | 93 | >sp\|P01877\|IGHA2_HUMAN Immunoglobulin heavy constant alpha 2 OS=Homo sapiens OX=9606 GN=IGHA2 PE=1 SV=4 | 576 | 20201114_qwl_colo_milk_1ng_glyco_hcd30_etd30_compound15.16045.16045.4 | controllerType=0 controllerNumber=1 scan=16045 | 42.3796 |
| 12602:4 | 1 | R.LSLHRPALEDLLLGSEAN[+1864.634]LTC[+57.021]TLTGLR.D | HexNAc(2)Hex(9) | C[+57], N[+1865] | 1207.8227 | 4 | 4828.2689 | 4828.2324 | 0 | 7.5 | 114 | Specific | 885.15 | 864.34 | 864.34 | 14.98 | 93 | >sp\|P01877\|IGHA2_HUMAN Immunoglobulin heavy constant alpha 2 OS=Homo sapiens OX=9606 GN=IGHA2 PE=1 SV=4 | 576 | 20201114_qwl_colo_milk_1ng_glyco_hcd30_etd30_compound15.16002.16002.4 | controllerType=0 controllerNumber=1 scan=16002 | 42.2 |
| 12849:4 | 1 | R.LSLHRPALEDLLLGSEAN[+1622.582]LTC[+57.021]TLTGLR.D | HexNAc(4)Hex(5) | C[+57], N[+1623] | 1147.3064 | 4 | 4586.2039 | 4586.1799 | 0 | 5.2 | 114 | Specific | 892.8 | 829.86 | 829.86 | 15.06 | 93 | >sp\|P01877\|IGHA2_HUMAN Immunoglobulin heavy constant alpha 2 OS=Homo sapiens OX=9606 GN=IGHA2 PE=1 SV=4 | 576 | 20201114_qwl_colo_milk_1ng_glyco_hcd30_etd30_compound15.16267.16267.4 | controllerType=0 controllerNumber=1 scan=16267 | 43.2032 |
| 13295:4 | 1 | R.LSLHRPALEDLLLGSEAN[+1751.624]LTC[+57.021]TLTGLR.D | HexNAc(4)Hex(4)NeuAc(1) | C[+57], N[+1752] | 1179.5671 | 4 | 4715.2464 | 4715.2225 | 0 | 5.1 | 114 | Specific | 909.44 | 836.92 | 836.92 | 16.06 | 93 | >sp\|P01877\|IGHA2_HUMAN Immunoglobulin heavy constant alpha 2 OS=Homo sapiens OX=9606 GN=IGHA2 PE=1 SV=4 | 576 | 20201114_qwl_colo_milk_1ng_glyco_hcd30_etd30_compound15.16741.16741.4 | controllerType=0 controllerNumber=1 scan=16741 | 44.6883 |
| 12621:4 | 1 | R.LSLHRPALEDLLLGSEAN[+1702.581]LTC[+57.021]TLTGLR.D | HexNAc(2)Hex(8) | C[+57], N[+1703] | 1167.3064 | 4 | 4666.2039 | 4666.1796 | 0 | 5.2 | 114 | Specific | 973.34 | 964.26 | 964.26 | 17.06 | 93 | >sp\|P01877\|IGHA2_HUMAN Immunoglobulin heavy constant alpha 2 OS=Homo sapiens OX=9606 GN=IGHA2 PE=1 SV=4 | 576 | 20201114_qwl_colo_milk_1ng_glyco_hcd30_etd30_compound15.16023.16023.4 | controllerType=0 controllerNumber=1 scan=16023 | 42.2823 |
| 12725:4 | 1 | R.LSLHRPALEDLLLGSEAN[+892.317]LTC[+57.021]TLTGLR.D | HexNAc(2)Hex(3) | C[+57], N[+892] | 964.7358 | 4 | 3855.9212 | 3855.9155 | 0 | 1.5 | 114 | Specific | 1094.89 | 1075.57 | 1075.57 | 19.35 | 93 | >sp\|P01877\|IGHA2_HUMAN Immunoglobulin heavy constant alpha 2 OS=Homo sapiens OX=9606 GN=IGHA2 PE=1 SV=4 | 576 | 20201114_qwl_colo_milk_1ng_glyco_hcd30_etd30_compound15.16136.16136.4 | controllerType=0 controllerNumber=1 scan=16136 | 42.747 |
| 12710:4 | 1 | R.LSLHRPALEDLLLGSEAN[+1095.397]LTC[+57.021]TLTGLR.D | HexNAc(3)Hex(3) | C[+57], N[+1095] | 1015.5036 | 4 | 4058.9925 | 4058.9949 | 0 | -0.6 | 114 | Specific | 1094.93 | 1063.75 | 1063.75 | 18.99 | 93 | >sp\|P01877\|IGHA2_HUMAN Immunoglobulin heavy constant alpha 2 OS=Homo sapiens OX=9606 GN=IGHA2 PE=1 SV=4 | 576 | 20201114_qwl_colo_milk_1ng_glyco_hcd30_etd30_compound15.16119.16119.4 | controllerType=0 controllerNumber=1 scan=16119 | [illegible] |
| 12671:4 | 1 | R.LSLHRPALEDLLLGSEAN[+1501.555]LTC[+57.021]TLTGLR.D | HexNAc(5)Hex(3) | C[+57], N[+1502] | 1117.047 | 4 | 4465.1663 | 4465.1536 | 0 | 2.9 | 114 | Specific | 1114.83 | 1010.37 | 1010.37 | 20.56 | 93 | >sp\|P01877\|IGHA2_HUMAN Immunoglobulin heavy constant alpha 2 OS=Homo sapiens OX=9606 GN=IGHA2 PE=1 SV=4 | 576 | 20201114_qwl_colo_milk_1ng_glyco_hcd30_etd30_compound15.16077.16077.6 | controllerType=0 controllerNumber=1 scan=16077 | [illegible] |
| 12670:4 | 1 | R.LSLHRPALEDLLLGSEAN[+1298.476]LTC[+57.021]TLTGLR.D | HexNAc(4)Hex(3) | C[+57], N[+1298] | 1066.276 | 4 | 4262.0823 | 4262.0742 | 0 | 1.9 | 114 | Specific | 1285.14 | 1229.6 | 1229.6 | 23.56 | 93 | >sp\|P01877\|IGHA2_HUMAN Immunoglobulin heavy constant alpha 2 OS=Homo sapiens OX=9606 GN=IGHA2 PE=1 SV=4 | 576 | 20201114_qwl_colo_milk_1ng_glyco_hcd30_etd30_compound15.16076.16076.4 | controllerType=0 controllerNumber=1 scan=16076 | 42.4998 |
| 12664:4 | 1 | R.LSLHRPALEDLLLGSEAN[+1216.423]LTC[+57.021]TLTGLR.D | HexNAc(2)Hex(5) | C[+57], N[+1216] | 1045.7636 | 4 | 4180.0325 | 4180.0211 | 0 | 2.7 | 114 | Specific | 1316.78 | 1218.35 | 1218.35 | 24.56 | 93 | >sp\|P01877\|IGHA2_HUMAN Immunoglobulin heavy constant alpha 2 OS=Homo sapiens OX=9606 GN=IGHA2 PE=1 SV=4 | 576 | 20201114_qwl_colo_milk_1ng_glyco_hcd30_etd30_compound15.16070.16070.4 | controllerType=0 controllerNumber=1 scan=16070 | 42.4702 |
| 09531:6 | 1 | K.SAVQGPPERDLC[+57.021]GC[+57.021]YSVSSVLPGC[+57.021]AQPWNHGETFTC[+ | HexNAc(4)Hex(3)Fuc(1) | C[+57]*4, N[+1445] | 1210.0557 | 6 | 7255.2979 | 7255.2797 | 0 | 2.5 | 156 | Specific | 289.65 | 289.65 | 289.65 | 3.27 | 93 | >sp\|P01877\|IGHA2_HUMAN Immunoglobulin heavy constant alpha 2 OS=Homo sapiens OX=9606 GN=IGHA2 PE=1 SV=4 | 576 | 20201114_qwl_colo_milk_1ng_glyco_hcd30_etd30_compound15.12758.12758.3 | controllerType=0 controllerNumber=1 scan=12758 | 33.1129 |
| 03726:2 | 1 | K.TPLTAN[+2262.814]ITK.S | HexNAc(5)Hex(5)Fuc(1)NeuAc(1) | N[+2263] | 1611.1949 | 2 | 3221.3825 | 3221.3711 | 0 | 3.5 | 200 | Specific | 208.32 | 62.09 | 62.09 | 2.92 | 93 | >sp\|P01877\|IGHA2_HUMAN Immunoglobulin heavy constant alpha 2 OS=Homo sapiens OX=9606 GN=IGHA2 PE=1 SV=4 | 576 | 20201114_qwl_colo_milk_1ng_glyco_hcd30_etd30_compound15.6682.6682.2 | controllerType=0 controllerNumber=1 scan=6682 | 19.1745 |
| 02284:2 | 1 | K.TPLTAN[+1971.719]ITK.S | HexNAc(5)Hex(5)Fuc(1) | N[+1972] | 1465.647 | 2 | 2930.2868 | 2930.2757 | 0 | 3.8 | 200 | Specific | 249.03 | 82.86 | 82.86 | 2.82 | 93 | >sp\|P01877\|IGHA2_HUMAN Immunoglobulin heavy constant alpha 2 OS=Homo sapiens OX=9606 GN=IGHA2 PE=1 SV=4 | 576 | 20201114_qwl_colo_milk_1ng_glyco_hcd30_etd30_compound15.5165.5165.2 | controllerType=0 controllerNumber=1 scan=5165 | 15.7821 |
| 03715:3 | 1 | K.TPLTAN[+2262.814]ITK.S | HexNAc(5)Hex(5)Fuc(1)NeuAc(1) | N[+2263] | 1074.4658 | 3 | 3221.383 | 3221.3711 | 0 | 3.7 | 200 | Specific | 252.9 | 120.71 | 120.71 | 3.21 | 93 | >sp\|P01877\|IGHA2_HUMAN Immunoglobulin heavy constant alpha 2 OS=Homo sapiens OX=9606 GN=IGHA2 PE=1 SV=4 | 576 | 20201114_qwl_colo_milk_1ng_glyco_hcd30_etd30_compound15.6671.6671.3 | controllerType=0 controllerNumber=1 scan=6671 | 19.1492 |
| 02242:2 | 1 | K.TPLTAN[+1768.640]ITK.S | HexNAc(4)Hex(5)Fuc(1) | N[+1769] | 1364.104 | 2 | 2727.2008 | 2727.1963 | 0 | 1.7 | 200 | Specific | 255.7 | 81.82 | 81.82 | 2.95 | 93 | >sp\|P01877\|IGHA2_HUMAN Immunoglobulin heavy constant alpha 2 OS=Homo sapiens OX=9606 GN=IGHA2 PE=1 SV=4 | 576 | 20201114_qwl_colo_milk_1ng_glyco_hcd30_etd30_compound15.5122.5122.2 | controllerType=0 controllerNumber=1 scan=5122 | 15.6979 |
| 03683:3 | 1 | K.TPLTAN[+2059.735]ITK.S | HexNAc(4)Hex(5)Fuc(1)NeuAc(1) | N[+2060] | 1006.7723 | 3 | 3018.3024 | 3018.2917 | 0 | 3.5 | 200 | Specific | 256.17 | 114.73 | 114.73 | 3.21 | 93 | >sp\|P01877\|IGHA2_HUMAN Immunoglobulin heavy constant alpha 2 OS=Homo sapiens OX=9606 GN=IGHA2 PE=1 SV=4 | 576 | 20201114_qwl_colo_milk_1ng_glyco_hcd30_etd30_compound15.6636.6636.3 | controllerType=0 controllerNumber=1 scan=6636 | 19.0687 |
| 02344:3 | 1 | K.TPLTAN[+1809.666]ITK.S | HexNAc(5)Hex(4)Fuc(1) | N[+1810] | 923.4144 | 3 | 2768.2286 | 2768.2228 | 0 | 2.1 | 200 | Specific | 263.5 | 124.56 | 124.56 | 3.24 | 93 | >sp\|P01877\|IGHA2_HUMAN Immunoglobulin heavy constant alpha 2 OS=Homo sapiens OX=9606 GN=IGHA2 PE=1 SV=4 | 576 | 20201114_qwl_colo_milk_1ng_glyco_hcd30_etd30_compound15.5230.5230.3 | controllerType=0 controllerNumber=1 scan=5230 | 15.9595 |
| 02352:2 | 1 | K.TPLTAN[+1663.608]ITK.S | HexNAc(5)Hex(4) | N[+1664] | 1311.5885 | 2 | 2622.1698 | 2622.1649 | 0 | 1.9 | 200 | Specific | 271.38 | 88.64 | 88.64 | 2.91 | 93 | >sp\|P01877\|IGHA2_HUMAN Immunoglobulin heavy constant alpha 2 OS=Homo sapiens OX=9606 GN=IGHA2 PE=1 SV=4 | 576 | 20201114_qwl_colo_milk_1ng_glyco_hcd30_etd30_compound15.5238.5238.2 | controllerType=0 controllerNumber=1 scan=5238 | 15.9885 |
| 02315:2 | 1 | K.TPLTAN[+1809.666]ITK.S | HexNAc(5)Hex(4)Fuc(1) | N[+1810] | 1384.618 | 2 | 2768.2286 | 2768.2228 | 0 | 2.1 | 200 | Specific | 274.9 | 102.22 | 102.22 | 3.27 | 93 | >sp\|P01877\|IGHA2_HUMAN Immunoglobulin heavy constant alpha 2 OS=Homo sapiens OX=9606 GN=IGHA2 PE=1 SV=4 | 576 | 20201114_qwl_colo_milk_1ng_glyco_hcd30_etd30_compound15.5198.5198.2 | controllerType=0 controllerNumber=1 scan=5198 | 15.8619 |
| 02311:2 | 1 | K.TPLTAN[+1971.719]ITK.S | HexNAc(5)Hex(5)Fuc(1) | N[+1972] | 1465.6447 | 2 | 2930.2821 | 2930.2757 | 0 | 2.2 | 200 | Specific | 282.51 | 104.46 | 104.46 | 3.09 | 93 | >sp\|P01877\|IGHA2_HUMAN Immunoglobulin heavy constant alpha 2 OS=Homo sapiens OX=9606 GN=IGHA2 PE=1 SV=4 | 576 | 20201114_qwl_colo_milk_1ng_glyco_hcd30_etd30_compound15.5194.5194.2 | controllerType=0 controllerNumber=1 scan=5194 | 15.8496 |
| 02394:3 | 1 | K.TPLTAN[+1647.613]ITK.S | HexNAc(5)Hex(3)Fuc(1) | N[+1648] | 869.3969 | 3 | 2606.1763 | 2606.17 | 0 | 2.4 | 200 | Specific | 290.38 | 131.98 | 131.98 | 3.28 | 93 | >sp\|P01877\|IGHA2_HUMAN Immunoglobulin heavy constant alpha 2 OS=Homo sapiens OX=9606 GN=IGHA2 PE=1 SV=4 | 576 | 20201114_qwl_colo_milk_1ng_glyco_hcd30_etd30_compound15.5283.5283.3 | controllerType=0 controllerNumber=1 scan=5283 | 16.1532 |
| 02307:3 | 1 | K.TPLTAN[+1971.719]ITK.S | HexNAc(5)Hex(5)Fuc(1) | N[+1972] | 977.4329 | 3 | 2930.2841 | 2930.2757 | 0 | 2.9 | 200 | Specific | 297.92 | 135.71 | 135.71 | 3.37 | 93 | >sp\|P01877\|IGHA2_HUMAN Immunoglobulin heavy constant alpha 2 OS=Homo sapiens OX=9606 GN=IGHA2 PE=1 SV=4 | 576 | 20201114_qwl_colo_milk_1ng_glyco_hcd30_etd30_compound15.5189.5189.3 | controllerType=0 controllerNumber=1 scan=5189 | 15.8339 |
| 02384:2 | 1 | K.TPLTAN[+1647.613]ITK.S | HexNAc(5)Hex(3)Fuc(1) | N[+1648] | 1303.5921 | 2 | 2606.1769 | 2606.17 | 0 | 2.6 | 200 | Specific | 299.86 | 117.68 | 117.68 | 3.32 | 93 | >sp\|P01877\|IGHA2_HUMAN Immunoglobulin heavy constant alpha 2 OS=Homo sapiens OX=9606 GN=IGHA2 PE=1 SV=4 | 576 | 20201114_qwl_colo_milk_1ng_glyco_hcd30_etd30_compound15.5272.5272.2 | controllerType=0 controllerNumber=1 scan=5272 | 16.1146 |
| 02351:3 | 1 | K.TPLTAN[+1606.587]ITK.S | HexNAc(4)Hex(4)Fuc(1) | N[+1607] | 855.7215 | 3 | 2565.1498 | 2565.1435 | 0 | 2.5 | 200 | Specific | 307.44 | 132.77 | 132.77 | 3.37 | 93 | >sp\|P01877\|IGHA2_HUMAN Immunoglobulin heavy constant alpha 2 OS=Homo sapiens OX=9606 GN=IGHA2 PE=1 SV=4 | 576 | 20201114_qwl_colo_milk_1ng_glyco_hcd30_etd30_compound15.5237.5237.3 | controllerType=0 controllerNumber=1 scan=5237 | 15.9816 |
| 02318:2 | 1 | K.TPLTAN[+1606.587]ITK.S | HexNAc(4)Hex(4)Fuc(1) | N[+1607] | 1283.0778 | 2 | 2565.1483 | 2565.1435 | 0 | 1.9 | 200 | Specific | 318.28 | 125.47 | 125.47 | 3.09 | 93 | >sp\|P01877\|IGHA2_HUMAN Immunoglobulin heavy constant alpha 2 OS=Homo sapiens OX=9606 GN=IGHA2 PE=1 SV=4 | 576 | 20201114_qwl_colo_milk_1ng_glyco_hcd30_etd30_compound15.5201.5201.2 | controllerType=0 controllerNumber=1 scan=5201 | 15.8734 |
| 02416:2 | 1 | K.TPLTAN[+1038.375]ITK.S | HexNAc(2)Hex(3)Fuc(1) | N[+1038] | 998.9701 | 2 | 1996.933 | 1996.9319 | 0 | 0.5 | 200 | Specific | 320.22 | 101.35 | 101.35 | 3.07 | 93 | >sp\|P01877\|IGHA2_HUMAN Immunoglobulin heavy constant alpha 2 OS=Homo sapiens OX=9606 GN=IGHA2 PE=1 SV=4 | 576 | 20201114_qwl_colo_milk_1ng_glyco_hcd30_etd30_compound15.5307.5307.2 | controllerType=0 controllerNumber=1 scan=5307 | 16.254 |
| 02389:2 | 1 | K.TPLTAN[+1241.454]ITK.S | HexNAc(3)Hex(3)Fuc(1) | N[+1241] | 1100.5118 | 2 | 2200.0162 | 2200.0113 | 0 | 2.3 | 200 | Specific | 339.76 | 120.8 | 120.8 | 3.19 | 93 | >sp\|P01877\|IGHA2_HUMAN Immunoglobulin heavy constant alpha 2 OS=Homo sapiens OX=9606 GN=IGHA2 PE=1 SV=4 | 576 | 20201114_qwl_colo_milk_1ng_glyco_hcd30_etd30_compound15.5278.5278.2 | controllerType=0 controllerNumber=1 scan=5278 | 16.1391 |
| 02380:2 | 1 | K.TPLTAN[+1298.476]ITK.S | HexNAc(4)Hex(3) | N[+1298] | 1129.0231 | 2 | 2257.039 | 2257.0327 | 0 | 2.8 | 200 | Specific | 345.01 | 115.41 | 115.41 | 3.19 | 93 | >sp\|P01877\|IGHA2_HUMAN Immunoglobulin heavy constant alpha 2 OS=Homo sapiens OX=9606 GN=IGHA2 PE=1 SV=4 | 576 | 20201114_qwl_colo_milk_1ng_glyco_hcd30_etd30_compound15.5268.5268.2 | controllerType=0 controllerNumber=1 scan=5268 | 16.097 |
| 02360:2 | 1 | K.TPLTAN[+1444.534]ITK.S | HexNAc(4)Hex(3)Fuc(1) | N[+1445] | 1202.0511 | 2 | 2403.0949 | 2403.0906 | 0 | 1.8 | 200 | Specific | 350.14 | 121.84 | 121.84 | 3.19 | 93 | >sp\|P01877\|IGHA2_HUMAN Immunoglobulin heavy constant alpha 2 OS=Homo sapiens OX=9606 GN=IGHA2 PE=1 SV=4 | 576 | 20201114_qwl_colo_milk_1ng_glyco_hcd30_etd30_compound15.5247.5247.2 | controllerType=0 controllerNumber=1 scan=5247 | 16.0196 |
| 02335:2 | 1 | K.TPLTAN[+1403.507]ITK.S | HexNAc(3)Hex(4)Fuc(1) | N[+1404] | 1181.5377 | 2 | 2362.0681 | 2362.0641 | 0 | 1.7 | 200 | Specific | 355.49 | 128 | 128 | 3.19 | 93 | >sp\|P01877\|IGHA2_HUMAN Immunoglobulin heavy constant alpha 2 OS=Homo sapiens OX=9606 GN=IGHA2 PE=1 SV=4 | 576 | 20201114_qwl_colo_milk_1ng_glyco_hcd30_etd30_compound15.5219.5219.2 | controllerType=0 controllerNumber=1 scan=5219 | 15.92 |
| 02375:2 | 1 | K.TPLTAN[+1501.555]ITK.S | HexNAc(5)Hex(3) | N[+1502] | 1230.563 | 2 | 2460.1188 | 2460.1121 | 0 | 2.7 | 200 | Specific | 362.88 | 135.96 | 135.96 | 3.16 | 93 | >sp\|P01877\|IGHA2_HUMAN Immunoglobulin heavy constant alpha 2 OS=Homo sapiens OX=9606 GN=IGHA2 PE=1 SV=4 | 576 | 20201114_qwl_colo_milk_1ng_glyco_hcd30_etd30_compound15.5263.5263.2 | controllerType=0 controllerNumber=1 scan=5263 | 16.0765 |
| 02409:3 | 1 | K.TPLTAN[+1444.534]ITK.S | HexNAc(4)Hex(3)Fuc(1) | N[+1445] | 801.7036 | 3 | 2403.0964 | 2403.0906 | 0 | 2.4 | 200 | Specific | 469.12 | 279.18 | 279.18 | 4.36 | 93 | >sp\|P01877\|IGHA2_HUMAN Immunoglobulin heavy constant alpha 2 OS=Homo sapiens OX=9606 GN=IGHA2 PE=1 SV=4 | 576 | 20201114_qwl_colo_milk_1ng_glyco_hcd30_etd30_compound15.5299.5299.3 | controllerType=0 controllerNumber=1 scan=5299 | 16.2291 |
| 06442:3 | 1 | R.MAGKPTHIN[+1216.423]VSVVMAEADGTC[+57.021].Y | HexNAc(2)Hex(5) | C[+57], N[+1216] | 1135.4884 | 3 | 3404.4507 | 3404.4424 | 0 | 2.5 | 319 | CRagged | 272.03 | 234.4 | 196.5 | 5.11 | 93 | >sp\|P01877\|IGHA2_HUMAN Immunoglobulin heavy constant alpha 2 OS=Homo sapiens OX=9606 GN=IGHA2 PE=1 SV=4 | 576 | 20201114_qwl_colo_milk_1ng_glyco_hcd30_etd30_compound15.9510.9510.3 | controllerType=0 controllerNumber=1 scan=9510 | 24.8997 |
| 07831:3 | 1 | R.MAGKPTHIN[+1702.581]VSVVMAEADGTC[+57.021]Y.- | HexNAc(2)Hex(8) | C[+57], N[+1703] | 1351.9025 | 3 | 4053.693 | 4053.6642 | 0 | 7.1 | 319 | CRagged | 232.27 | 232.27 | 184.37 | 3.33 | 93 | >sp\|P01877\|IGHA2_HUMAN Immunoglobulin heavy constant alpha 2 OS=Homo sapiens OX=9606 GN=IGHA2 PE=1 SV=4 | 576 | 20201114_qwl_colo_milk_1ng_glyco_hcd30_etd30_compound15.10970.10970.3 | controllerType=0 controllerNumber=1 scan=10970 | 28.5347 |
| 07382:3 | 1 | R.MAGKPTHIN[+1378.476]VSVVMAEADGTC[+57.021]Y.- | HexNAc(2)Hex(6) | C[+57], N[+1378] | 1243.8687 | 3 | 3729.5916 | 3729.5585 | 0 | 8.9 | 319 | CRagged | 236.39 | 236.39 | 158.28 | 3.43 | 93 | >sp\|P01877\|IGHA2_HUMAN Immunoglobulin heavy constant alpha 2 OS=Homo sapiens OX=9606 GN=IGHA2 PE=1 SV=4 | 576 | 20201114_qwl_colo_milk_1ng_glyco_hcd30_etd30_compound15.10498.10498.3 | controllerType=0 controllerNumber=1 scan=10498 | 27.2723 |
| 07913:3 | 1 | R.MAGKPTHIN[+1501.555]VSVVMAEADGTC[+57.021]Y.- | HexNAc(5)Hex(3) | C[+57], N[+1502] | 1284.8862 | 3 | 3852.6439 | 3852.6382 | 0 | 1.5 | 319 | CRagged | 261.92 | 197.3 | 197.3 | 4.24 | 93 | >sp\|P01877\|IGHA2_HUMAN Immunoglobulin heavy constant alpha 2 OS=Homo sapiens OX=9606 GN=IGHA2 PE=1 SV=4 | 576 | 20201114_qwl_colo_milk_1ng_glyco_hcd30_etd30_compound15.11058.11058.3 | controllerType=0 controllerNumber=1 scan=11058 | 28.8182 |
| 07893:3 | 1 | R.MAGKPTHIN[+1216.423]VSVVMAEADGTC[+57.021]Y.- | HexNAc(2)Hex(5) | C[+57], N[+1216] | 1189.8473 | 3 | 3567.5275 | 3567.5057 | 0 | 6.1 | 319 | CRagged | 301.06 | 281 | 211.29 | 4.46 | 93 | >sp\|P01877\|IGHA2_HUMAN Immunoglobulin heavy constant alpha 2 OS=Homo sapiens OX=9606 GN=IGHA2 PE=1 SV=4 | 576 | 20201114_qwl_colo_milk_1ng_glyco_hcd30_etd30_compound15.11037.11037.3 | controllerType=0 controllerNumber=1 scan=11037 | 28.7576 |
| 07540:3 | 1 | R.M[+15.995]AGKPTHIN[+1378.476]VSVVMAEADGTC[+57.021]Y.- | HexNAc(2)Hex(6) | C[+57], M[+16], N[+1378] | 1249.1921 | 3 | 3745.5617 | 3745.5535 | 0 | 2.2 | 319 | CRagged | 211.41 | 149.84 | 117.76 | 3.66 | 93 | >sp\|P01877\|IGHA2_HUMAN Immunoglobulin heavy constant alpha 2 OS=Homo sapiens OX=9606 GN=IGHA2 PE=1 SV=4 | 576 | 20201114_qwl_colo_milk_1ng_glyco_hcd30_etd30_compound15.10664.10664.3 | controllerType=0 controllerNumber=1 scan=10664 | 27.708 |
| 12986:4 | 2 | R.RTAGWNVPIGTLRPFLN[+2059.735]WTGPPEPIEAAVAR.F | HexNAc(4)Hex(5)Fuc(1)NeuAc(1) | N[+2060] | 1362.3997 | 4 | 5446.5758 | 5446.547 | 0 | 5.5 | 140 | Specific | 458.36 | 458.36 | 458.36 | 7.36 | 46 | >sp\|P02788\|TRFL_HUMAN Lactotransferrin OS=Homo sapiens OX=9606 GN=LTF PE=1 SV=6 | 1270 | 20201114_qwl_colo_milk_1ng_glyco_hcd30_etd30_compound15.16413.16413.4 | controllerType=0 controllerNumber=1 scan=16413 | 43.6797 |
| 12869:4 | 2 | R.RTAGWNVPIGTLRPFLN[+2570.925]WTGPPEPIEAAVAR.F | HexNAc(5)Hex(6)Fuc(2)NeuAc(1) | N[+2571] | 1490.192 | 4 | 5957.7462 | 5957.7371 | 0 | 1.5 | 140 | Specific | 459.07 | 459.07 | 459.07 | 7.65 | 46 | >sp\|P02788\|TRFL_HUMAN Lactotransferrin OS=Homo sapiens OX=9606 GN=LTF PE=1 SV=6 | 1270 | 20201114_qwl_colo_milk_1ng_glyco_hcd30_etd30_compound15.16288.16288.4 | controllerType=0 controllerNumber=1 scan=16288 | 43.2688 |
| 12362:4 | 2 | R.RTAGWNVPIGTLRPFLN[+1768.640]WTGPPEPIEAAVAR.F | HexNAc(4)Hex(5)Fuc(1) | N[+1769] | 1289.6296 | 4 | 5155.4967 | 5155.4516 | 0 | 8.8 | 140 | Specific | 625.22 | 625.22 | 625.22 | 9.64 | 46 | >sp\|P02788\|TRFL_HUMAN Lactotransferrin OS=Homo sapiens OX=9606 GN=LTF PE=1 SV=6 | 1270 | 20201114_qwl_colo_milk_1ng_glyco_hcd30_etd30_compound15.15747.15747.4 | controllerType=0 controllerNumber=1 scan=15747 | 41.3866 |
| 13705:4 | 2 | R.TAGWNVPIGTLRPFLN[+2424.867]WTGPPEPIEAAVAR.F | HexNAc(5)Hex(6)Fuc(1)NeuAc(1) | N[+2425] | 1414.6545 | 4 | 5655.5963 | 5655.5781 | 0 | 3.2 | 141 | Specific | 256.7 | 256.7 | 256.7 | 5.39 | 46 | >sp\|P02788\|TRFL_HUMAN Lactotransferrin OS=Homo sapiens OX=9606 GN=LTF PE=1 SV=6 | 1270 | 20201114_qwl_colo_milk_1ng_glyco_hcd30_etd30_compound15.17178.17178.4 | controllerType=0 controllerNumber=1 scan=17178 | 46.1402 |
| 13121:3 | 2 | R.TAGWNVPIGTLRPFLN[+1768.640]WTGPPEPIEAAVAR.F | HexNAc(4)Hex(5)Fuc(1) | N[+1769] | 1667.1246 | 3 | 4999.3592 | 4999.3504 | 0 | 1.8 | 141 | Specific | 269.57 | 230.5 | 230.5 | 5.39 | 46 | >sp\|P02788\|TRFL_HUMAN Lactotransferrin OS=Homo sapiens OX=9606 GN=LTF PE=1 SV=6 | 1270 | 20201114_qwl_colo_milk_1ng_glyco_hcd30_etd30_compound15.16555.16555.3 | controllerType=0 controllerNumber=1 scan=16555 | 44.093 |
| 13745:4 | 2 | R.TAGWNVPIGTLRPFLN[+2205.793]WTGPPEPIEAAVAR.F | HexNAc(4)Hex(5)Fuc(2)NeuAc(1) | N[+2206] | 1359.8788 | 4 | 5436.4933 | 5436.5038 | 0 | -1.9 | 141 | Specific | 273.02 | 273.02 | 273.02 | 5.01 | 46 | >sp\|P02788\|TRFL_HUMAN Lactotransferrin OS=Homo sapiens OX=9606 GN=LTF PE=1 SV=6 | 1270 | 20201114_qwl_colo_milk_1ng_glyco_hcd30_etd30_compound15.17222.17222.2 | controllerType=0 controllerNumber=1 scan=17222 | 46.2726 |
| 13725:4 | 2 | R.[+42.011]TAGWNVPIGTLRPFLN[+2018.708]WTGPPEPIEAAVAR.F | HexNAc(3)Hex(6)Fuc(1)NeuAc(1) | .[42], N[+2019] | 1323.6167 | 4 | 5291.445 | 5291.4299 | 0 | 2.9 | 141 | Specific | 304.3 | 304.3 | 304.3 | 6.1 | 46 | >sp\|P02788\|TRFL_HUMAN Lactotransferrin OS=Homo sapiens OX=9606 GN=LTF PE=1 SV=6 | 1270 | 20201114_qwl_colo_milk_1ng_glyco_hcd30_etd30_compound15.17201.17201.4 | controllerType=0 controllerNumber=1 scan=17201 | 46.2141 |
| 14571:4 | 2 | R.[+42.011]TAGWNVPIGTLRPFLN[+2336.851]WTGPPEPIEAAVAR.F | HexNAc(6)Hex(6)Fuc(1) | .[42], N[+2337] | 1403.146 | 4 | 5609.5622 | 5609.5726 | 0 | -1.9 | 141 | Specific | 365.78 | 330.53 | 330.53 | 5.46 | 46 | >sp\|P02788\|TRFL_HUMAN Lactotransferrin OS=Homo sapiens OX=9606 GN=LTF PE=1 SV=6 | 1270 | 20201114_qwl_colo_milk_1ng_glyco_hcd30_etd30_compound15.18105.18105.4 | controllerType=0 controllerNumber=1 scan=18105 | 49.2109 |
| 13726:3 | 2 | R.TAGWNVPIGTLRPFLN[+2059.735]WTGPPEPIEAAVAR.F | HexNAc(4)Hex(5)Fuc(1)NeuAc(1) | N[+2060] | 1764.1578 | 3 | 5290.4588 | 5290.4459 | 0 | 2.5 | 141 | Specific | 408.73 | 332.77 | 332.77 | 7.31 | 46 | >sp\|P02788\|TRFL_HUMAN Lactotransferrin OS=Homo sapiens OX=9606 GN=LTF PE=1 SV=6 | 1270 | 20201114_qwl_colo_milk_1ng_glyco_hcd30_etd30_compound15.17202.17202.3 | controllerType=0 controllerNumber=1 scan=17202 | 46.2194 |
| 14586:4 | 2 | R.TAGWNVPIGTLRPFLN[+2496.888]WTGPPEPIEAAVAR.F | HexNAc(4)Hex(5)Fuc(2)NeuAc(2) | N[+2497] | 1432.6592 | 4 | 5727.6149 | 5727.5992 | 0 | 2.7 | 141 | Specific | 495.77 | 495.77 | 495.77 | 7.7 | 46 | >sp\|P02788\|TRFL_HUMAN Lactotransferrin OS=Homo sapiens OX=9606 GN=LTF PE=1 SV=6 | 1270 | 20201114_qwl_colo_milk_1ng_glyco_hcd30_etd30_compound15.18121.18121.4 | controllerType=0 controllerNumber=1 scan=18121 | 49.2675 |
| 13640:4 | 2 | R.TAGWNVPIGTLRPFLN[+2862.021]WTGPPEPIEAAVAR.F | HexNAc(5)Hex(6)Fuc(2)NeuAc(2) | N[+2862] | 1523.9418 | 4 | 6092.7453 | 6092.7314 | 0 | 2.3 | 141 | Specific | 719.46 | 705.67 | 705.67 | 12.99 | 46 | >sp\|P02788\|TRFL_HUMAN Lactotransferrin OS=Homo sapiens OX=9606 GN=LTF PE=1 SV=6 | 1270 | 20201114_qwl_colo_milk_1ng_glyco_hcd30_etd30_compound15.17108.17108.4 | controllerType=0 controllerNumber=1 scan=17108 | 45.8765 |
| 13749:4 | 2 | R.TAGWNVPIGTLRPFLN[+2059.735]WTGPPEPIEAAVAR.F | HexNAc(4)Hex(5)Fuc(1)NeuAc(1) | N[+2060] | 1323.365 | 4 | 5290.4381 | 5290.4459 | 0 | -1.5 | 141 | Specific | 867.5 | 809.68 | 809.68 | 15.41 | 46 | >sp\|P02788\|TRFL_HUMAN Lactotransferrin OS=Homo sapiens OX=9606 GN=LTF PE=1 SV=6 | 1270 | 20201114_qwl_colo_milk_1ng_glyco_hcd30_etd30_compound15.17226.17226.2 | controllerType=0 controllerNumber=1 scan=17226 | 46.2839 |
| 13319:3 | 2 | R.TAGWNIPMGLLFN[+2205.793]QTGSC[+57.021]K.F | HexNAc(4)Hex(5)Fuc(2)NeuAc(1) | C[+57], N[+2206] | 1434.284 | 3 | 4300.8374 | 4300.8028 | 0 | 8.1 | 485 | Specific | 211.54 | 141.04 | 141.04 | 4.25 | 46 | >sp\|P02788\|TRFL_HUMAN Lactotransferrin OS=Homo sapiens OX=9606 GN=LTF PE=1 SV=6 | 1270 | 20201114_qwl_colo_milk_1ng_glyco_hcd30_etd30_compound15.16767.16767.3 | controllerType=0 controllerNumber=1 scan=16767 | 44.7877 |
| 12166:3 | 2 | R.TAGWNIPMGLLFN[+1971.719]QTGSC[+57.021]K.F | HexNAc(5)Hex(5)Fuc(1) | C[+57], N[+1972] | 1356.2546 | 3 | 4066.7492 | 4066.7288 | 0 | 5 | 485 | Specific | 276.33 | 264.94 | 264.94 | 4.2 | 46 | >sp\|P02788\|TRFL_HUMAN Lactotransferrin OS=Homo sapiens OX=9606 GN=LTF PE=1 SV=6 | 1270 | 20201114_qwl_colo_milk_1ng_glyco_hcd30_etd30_compound15.15539.15539.3 | controllerType=0 controllerNumber=1 scan=15539 | 40.7107 |
| 12147:3 | 2 | R.TAGWNIPMGLLFN[+1768.640]QTGSC[+57.021]K.F | HexNAc(4)Hex(5)Fuc(1) | C[+57], N[+1769] | 1288.5602 | 3 | 3863.6662 | 3863.6495 | 0 | 4.3 | 485 | Specific | 321.89 | 253.27 | 253.27 | 4.53 | 46 | >sp\|P02788\|TRFL_HUMAN Lactotransferrin OS=Homo sapiens OX=9606 GN=LTF PE=1 SV=6 | 1270 | 20201114_qwl_colo_milk_1ng_glyco_hcd30_etd30_compound15.15520.15520.3 | controllerType=0 controllerNumber=1 scan=15520 | 40.6749 |
| 12022:3 | 2 | R.TAGWNIPMGLLFN[+2279.830]QTGSC[+57.021]K.F | HexNAc(5)Hex(6)Fuc(2) | C[+57], N[+2280] | 1458.9628 | 3 | 4374.8739 | 4374.8396 | 0 | 7.9 | 485 | Specific | 357.91 | 310.58 | 220.36 | 5.8 | 46 | >sp\|P02788\|TRFL_HUMAN Lactotransferrin OS=Homo sapiens OX=9606 GN=LTF PE=1 SV=6 | 1270 | 20201114_qwl_colo_milk_1ng_glyco_hcd30_etd30_compound15.15386.15386.3 | controllerType=0 controllerNumber=1 scan=15386 | 40.2635 |
| 12419:3 | 2 | R.TAGWNIPMGLLFN[+1216.423]QTGSC[+57.021]K.F | HexNAc(2)Hex(5) | C[+57], N[+1216] | 1104.4872 | 3 | 3311.4471 | 3311.4328 | 0 | 4.3 | 485 | Specific | 359.61 | 353.27 | 185.07 | 6.06 | 46 | >sp\|P02788\|TRFL_HUMAN Lactotransferrin OS=Homo sapiens OX=9606 GN=LTF PE=1 SV=6 | 1270 | 20201114_qwl_colo_milk_1ng_glyco_hcd30_etd30_compound15.15808.15808.3 | controllerType=0 controllerNumber=1 scan=15808 | 41.5625 |
| 12271:3 | 2 | R.TAGWNIPMGLLFN[+1622.582]QTGSC[+57.021]K.F | HexNAc(4)Hex(5) | C[+57], N[+1623] | 1239.8714 | 3 | 3717.5996 | 3717.5916 | 0 | 2.2 | 485 | Specific | 387.11 | 304.92 | 219.83 | 6.34 | 46 | >sp\|P02788\|TRFL_HUMAN Lactotransferrin OS=Homo sapiens OX=9606 GN=LTF PE=1 SV=6 | 1270 | 20201114_qwl_colo_milk_1ng_glyco_hcd30_etd30_compound15.15651.15651.3 | controllerType=0 controllerNumber=1 scan=15651 | 41.0668 |
| 13372:3 | 2 | R.TAGWNIPMGLLFN[+2059.735]QTGSC[+57.021]K.F | HexNAc(4)Hex(5)Fuc(1)NeuAc(1) | C[+57], N[+2060] | 1385.593 | 3 | 4154.7643 | 4154.7449 | 0 | 4.7 | 485 | Specific | 395.31 | 286.47 | 286.47 | 5.75 | 46 | >sp\|P02788\|TRFL_HUMAN Lactotransferrin OS=Homo sapiens OX=9606 GN=LTF PE=1 SV=6 | 1270 | 20201114_qwl_colo_milk_1ng_glyco_hcd30_etd30_compound15.16824.16824.3 | controllerType=0 controllerNumber=1 scan=16824 | 44.9854 |
| 13331:3 | 2 | R.TAGWNIPMGLLFN[+2059.735]QTGSC[+57.021]K.F | HexNAc(4)Hex(5)Fuc(1)NeuAc(1) | C[+57], N[+2060] | 1385.585 | 3 | 4154.7405 | 4154.7449 | 0 | -1.1 | 485 | Specific | 405.87 | 279.87 | 279.87 | 5.73 | 46 | >sp\|P02788\|TRFL_HUMAN Lactotransferrin OS=Homo sapiens OX=9606 GN=LTF PE=1 SV=6 | 1270 | 20201114_qwl_colo_milk_1ng_glyco_hcd30_etd30_compound15.16780.16780.3 | controllerType=0 controllerNumber=1 scan=16780 | 44.8491 |
| 14714:3 | 2 | R.TAGWNIPMGLLFN[+2204.772]QTGSC[+57.021]K.F | HexNAc(4)Hex(5)NeuAc(2) | C[+57], N[+2205] | 1433.9377 | 3 | 4299.7985 | 4299.7824 | 0 | 3.7 | 485 | Specific | 412.01 | 300.33 | 300.33 | 5.84 | 46 | >sp\|P02788\|TRFL_HUMAN Lactotransferrin OS=Homo sapiens OX=9606 GN=LTF PE=1 SV=6 | 1270 | 20201114_qwl_colo_milk_1ng_glyco_hcd30_etd30_compound15.18257.18257.3 | controllerType=0 controllerNumber=1 scan=18257 | 49.7228 |
| 14602:3 | 2 | R.TAGWNIPMGLLFN[+2350.830]QTGSC[+57.021]K.F | HexNAc(4)Hex(5)Fuc(1)NeuAc(2) | C[+57], N[+2351] | 1482.6226 | 3 | 4445.8533 | 4445.8403 | 0 | 2.9 | 485 | Specific | 472.62 | 341.63 | 341.63 | 6.57 | 46 | >sp\|P02788\|TRFL_HUMAN Lactotransferrin OS=Homo sapiens OX=9606 GN=LTF PE=1 SV=6 | 1270 | 20201114_qwl_colo_milk_1ng_glyco_hcd30_etd30_compound15.18138.18138.3 | controllerType=0 controllerNumber=1 scan=18138 | 49.3217 |
| 12385:3 | 2 | R.TAGWNIPMGLLFN[+1216.423]QTGSC[+57.021]K.F | HexNAc(2)Hex(5) | C[+57], N[+1216] | 1104.4909 | 3 | 3311.4581 | 3311.4328 | 0 | 7.6 | 485 | Specific | 501.87 | 467.87 | 260.38 | 7.21 | 46 | >sp\|P02788\|TRFL_HUMAN Lactotransferrin OS=Homo sapiens OX=9606 GN=LTF PE=1 SV=6 | 1270 | 20201114_qwl_colo_milk_1ng_glyco_hcd30_etd30_compound15.15772.15772.3 | controllerType=0 controllerNumber=1 scan=15772 | 41.4544 |
| 12390:2 | 2 | R.TAGWNIPMGLLFN[+1216.423]QTGSC[+57.021]K.F | HexNAc(2)Hex(5) | C[+57], N[+1216] | 1656.2352 | 2 | 3311.463 | 3311.4328 | 0 | 9.1 | 485 | Specific | 540.28 | 484.23 | 169.97 | 7 | 46 | >sp\|P02788\|TRFL_HUMAN Lactotransferrin OS=Homo sapiens OX=9606 GN=LTF PE=1 SV=6 | 1270 | 20201114_qwl_colo_milk_1ng_glyco_hcd30_etd30_compound15.15777.15777.2 | controllerType=0 controllerNumber=1 scan=15777 | 41.4776 |
| 12939:3 | 2 | R.TAGWNIPMGLLFN[+1216.423]QTGSC[+57.021]KFDEYFSQSC[+57.021]APGSDPR.S | HexNAc(2)Hex(5) | C[+57]*2, N[+1216] | 1719.0711 | 3 | 5155.1988 | 5155.18 | 0 | 3.7 | 485 | Specific | 252.81 | 222.37 | 111.08 | 4.28 | 46 | >sp\|P02788\|TRFL_HUMAN Lactotransferrin OS=Homo sapiens OX=9606 GN=LTF PE=1 SV=6 | 1270 | 20201114_qwl_colo_milk_1ng_glyco_hcd30_etd30_compound15.16363.16363.3 | controllerType=0 controllerNumber=1 scan=16363 | 43.5053 |
| 12860:4 | 2 | R.TAGWNIPMGLLFN[+1768.640]QTGSC[+57.021]KFDEYFSQSC[+57.021]APGSDPR.S | HexNAc(4)Hex(5)Fuc(1) | C[+57]*2, N[+1769] | 1427.6085 | 4 | 5707.4123 | 5707.3966 | 0 | 2.7 | 485 | Specific | 334.58 | 294.94 | 294.94 | 5.66 | 46 | >sp\|P02788\|TRFL_HUMAN Lactotransferrin OS=Homo sapiens OX=9606 GN=LTF PE=1 SV=6 | 1270 | 20201114_qwl_colo_milk_1ng_glyco_hcd30_etd30_compound15.16279.16279.4 | controllerType=0 controllerNumber=1 scan=16279 | 43.2391 |
| 14312:4 | 2 | R.TAGWNIPMGLLFN[+2350.830]QTGSC[+57.021]KFDEYFSQSC[+57.021]APGSDPR.S | HexNAc(4)Hex(5)Fuc(1)NeuAc(2) | C[+57]*2, N[+2351] | 1573.1516 | 4 | 6289.5846 | 6289.5875 | 0 | -0.4 | 485 | Specific | 686.53 | 642.68 | 642.68 | 10.43 | 46 | >sp\|P02788\|TRFL_HUMAN Lactotransferrin OS=Homo sapiens OX=9606 GN=LTF PE=1 SV=6 | 1270 | 20201114_qwl_colo_milk_1ng_glyco_hcd30_etd30_compound15.17829.17829.4 | controllerType=0 controllerNumber=1 scan=17829 | 48.2949 |
| 11286:3 | 3 | R.LN[+1378.476]YSLPTGQWVGVQLPR.N | HexNAc(2)Hex(6) | N[+1378] | 1102.8443 | 3 | 3306.5182 | 3306.5145 | 0 | 1.1 | 1017 | Specific | 239.33 | 239.33 | 239.33 | 3.67 | 24 | >sp\|P24821\|TENA_HUMAN Tenascin OS=Homo sapiens OX=9606 GN=TNC PE=1 SV=3 | 486 | 20201114_qwl_colo_milk_1ng_glyco_hcd30_etd30_compound15.14606.14606.2 | controllerType=0 controllerNumber=1 scan=14606 | 38.0134 |
| 11321:2 | 3 | R.LN[+1216.423]YSLPTGQWVGVQLPR.N | HexNAc(2)Hex(5) | N[+1216] | 1572.7377 | 2 | 3144.4682 | 3144.4617 | 0 | 2 | 1017 | Specific | 540.79 | 483.3 | 483.3 | 7.45 | 24 | >sp\|P24821\|TENA_HUMAN Tenascin OS=Homo sapiens OX=9606 GN=TNC PE=1 SV=3 | 486 | 20201114_qwl_colo_milk_1ng_glyco_hcd30_etd30_compound15.14643.14643.2 | controllerType=0 controllerNumber=1 scan=14643 | 38.1342 |
| 11316:3 | 3 | R.LN[+1216.423]YSLPTGQWVGVQLPR.N | HexNAc(2)Hex(5) | N[+1216] | 1048.8312 | 3 | 3144.4791 | 3144.4617 | 0 | 5.5 | 1017 | Specific | 648.64 | 583.78 | 583.78 | 9.31 | 24 | >sp\|P24821\|TENA_HUMAN Tenascin OS=Homo sapiens OX=9606 GN=TNC PE=1 SV=3 | 486 | 20201114_qwl_colo_milk_1ng_glyco_hcd30_etd30_compound15.14638.14638.3 | controllerType=0 controllerNumber=1 scan=14638 | 38.1109 |
| 14716:3 | 4 | K.QIGLYPVLVIDSSGYVNPN[+2350.830]YTGR.I | HexNAc(4)Hex(5)Fuc(1)NeuAc(2) | N[+2351] | 1626.0616 | 3 | 4876.1702 | 4876.1338 | 0 | 7.5 | 168 | Specific | 249.59 | 170.82 | 170.82 | 5.58 | 46 | >sp\|P01833\|PIGR_HUMAN Polymeric immunoglobulin receptor OS=Homo sapiens OX=9606 GN=PIGR PE=1 SV=4 | 902 | 20201114_qwl_colo_milk_1ng_glyco_hcd30_etd30_compound15.18259.18259.3 | controllerType=0 controllerNumber=1 scan=18259 | 49.7346 |
| 15881:3 | 4 | K.QIGLYPVLVIDSSGYVNPN[+2188.740]YTGR.I | HexNAc(2)Hex(11) | N[+2189] | 1572.0339 | 3 | 4714.0871 | 4714.0433 | 0 | 9.3 | 168 | Specific | 277.58 | 216.28 | 216.28 | 6 | 46 | >sp\|P01833\|PIGR_HUMAN Polymeric immunoglobulin receptor OS=Homo sapiens OX=9606 GN=PIGR PE=1 SV=4 | 902 | 20201114_qwl_colo_milk_1ng_glyco_hcd30_etd30_compound15.19501.19501.3 | controllerType=0 controllerNumber=1 scan=19501 | 53.9222 |
| 14846:3 | 4 | K.QIGLYPVLVIDSSGYVNPN[+2204.772]YTGR.I | HexNAc(4)Hex(5)NeuAc(2) | N[+2205] | 1577.3692 | 3 | 4730.093 | 4730.0759 | 0 | 3.6 | 168 | Specific | 287.14 | 183.36 | 183.36 | 5.92 | 46 | >sp\|P01833\|PIGR_HUMAN Polymeric immunoglobulin receptor OS=Homo sapiens OX=9606 GN=PIGR PE=1 SV=4 | 902 | 20201114_qwl_colo_milk_1ng_glyco_hcd30_etd30_compound15.18398.18398.3 | controllerType=0 controllerNumber=1 scan=18398 | 50.2355 |
| 19392:3 | 4 | K.QIGLYPVLVIDSSGYVNPN[+2188.740]YTGR.I | HexNAc(2)Hex(11) | N[+2189] | 1572.028 | 3 | 4714.0694 | 4714.0433 | 0 | 5.6 | 168 | Specific | 330.07 | 229.98 | 229.98 | 6.28 | 46 | >sp\|P01833\|PIGR_HUMAN Polymeric immunoglobulin receptor OS=Homo sapiens OX=9606 GN=PIGR PE=1 SV=4 | 902 | 20201114_qwl_colo_milk_1ng_glyco_hcd30_etd30_compound15.23298.23298.3 | controllerType=0 controllerNumber=1 scan=23298 | 63.9181 |
| 14702:3 | 4 | K.QIGLYPVLVIDSSGYVNPN[+2350.830]YTGR.I | HexNAc(4)Hex(5)Fuc(1)NeuAc(2) | N[+2351] | 1626.0522 | 3 | 4876.142 | 4876.1338 | 0 | 1.7 | 168 | Specific | 367.8 | 236.63 | 236.63 | 6.74 | 46 | >sp\|P01833\|PIGR_HUMAN Polymeric immunoglobulin receptor OS=Homo sapiens OX=9606 GN=PIGR PE=1 SV=4 | 902 | 20201114_qwl_colo_milk_1ng_glyco_hcd30_etd30_compound15.18244.18244.3 | controllerType=0 controllerNumber=1 scan=18244 | 49.6655 |
| 14421:4 | 4 | R.LSLLEEPGN[+2351.851]GTFTVILNQLTSR.D | HexNAc(4)Hex(5)Fuc(3)NeuAc(1) | N[+2352] | 1189.2957 | 4 | 4754.161 | 4754.1433 | 0 | 3.7 | 413 | Specific | 219.21 | 219.21 | 219.21 | 3.73 | 46 | >sp\|P01833\|PIGR_HUMAN Polymeric immunoglobulin receptor OS=Homo sapiens OX=9606 GN=PIGR PE=1 SV=4 | 902 | 20201114_qwl_colo_milk_1ng_glyco_hcd30_etd30_compound15.17945.17945.4 | controllerType=0 controllerNumber=1 scan=17945 | 48.6744 |
| 15751:3 | 4 | R.[+42.011]LSLLEEPGN[+2336.851]GTFTVILNQLTSR.D | HexNAc(6)Hex(6)Fuc(1) | .[42], N[+2337] | 1594.3832 | 3 | 4781.1352 | 4781.1542 | 0 | -4 | 413 | Specific | 267.5 | 175.3 | 175.3 | 3.86 | 46 | >sp\|P01833\|PIGR_HUMAN Polymeric immunoglobulin receptor OS=Homo sapiens OX=9606 GN=PIGR PE=1 SV=4 | 902 | 20201114_qwl_colo_milk_1ng_glyco_hcd30_etd30_compound15.19363.19363.3 | controllerType=0 controllerNumber=1 scan=19363 | 53.4453 |
| 16000:3 | 4 | R.LSLLEEPGN[+2350.830]GTFTVILNQLTSR.D | HexNAc(4)Hex(5)Fuc(1)NeuAc(2) | N[+2351] | 1585.0482 | 3 | 4753.13 | 4753.1229 | 0 | 1.5 | 413 | Specific | 324.9 | 195.76 | 195.76 | 6.4 | 46 | >sp\|P01833\|PIGR_HUMAN Polymeric immunoglobulin receptor OS=Homo sapiens OX=9606 GN=PIGR PE=1 SV=4 | 902 | 20201114_qwl_colo_milk_1ng_glyco_hcd30_etd30_compound15.19630.19630.3 | controllerType=0 controllerNumber=1 scan=19630 | 54.3627 |
| 17499:3 | 4 | R.LSLLEEPGN[+3153.116]GTFTVILNQLTSR.D | HexNAc(5)Hex(6)Fuc(2)NeuAc(3) | N[+3153] | 1852.4782 | 3 | 5555.4202 | 5555.4084 | 0 | 2.1 | 413 | Specific | 329.29 | 238.31 | 238.31 | 6.79 | 46 | >sp\|P01833\|PIGR_HUMAN Polymeric immunoglobulin receptor OS=Homo sapiens OX=9606 GN=PIGR PE=1 SV=4 | 902 | 20201114_qwl_colo_milk_1ng_glyco_hcd30_etd30_compound15.21232.21232.3 | controllerType=0 controllerNumber=1 scan=21232 | 59.0653 |

Table S4. Cont.

| Query #:z | Protein Rank | Glycopeptide | Glycan | Modification Type(s) | Observed m/z | z | Observed (M+H) | Calc. mass (M+H) | Off-by-x error | Mass error (ppm) | StartPosition | Cleavage | Score | Delta | Delta Mod | \|Log Prob\| | # of unique peptides | Protein Name | Protein DB number | Comment | Scan # | Scan Time |
|---|---|---|---|---|---|---|---|---|---|---|---|---|---|---|---|---|---|---|---|---|---|---|
| 14437:3 | 4 | R.LSLLEEPGN[+2205.793]GTFTVILNQLTSR.D | HexNAc(4)Hex(5)Fuc(2)NeuAc(1) | N[+2206] | 1536.7055 | 3 | 4608.102 | 4608.0854 | 0 | 3.6 | 413 | Specific | 413.29 | 281.76 | 281.76 | 6.28 | 46 | >sp\|P01833\|PIGR_HUMAN Polymeric immunoglobulin receptor OS=Homo sapiens OX=9606 GN=PIGR PE=1 SV=4 | 902 | 20201114_qwl_colo_milk_1ng_glyco_hcd30_etd30_compound15.17963.17963.3 | controllerType=0 controllerNumber=1 scan=17963 | 48.7457 |
| 14438:4 | 4 | R.LSLLEEPGN[+2205.793]GTFTVILNQLTSR.D | HexNAc(4)Hex(5)Fuc(2)NeuAc(1) | N[+2206] | 1152.779 | 4 | 4608.0941 | 4608.0854 | 0 | 1.9 | 413 | Specific | 505.48 | 462.96 | 462.96 | 7.02 | 46 | >sp\|P01833\|PIGR_HUMAN Polymeric immunoglobulin receptor OS=Homo sapiens OX=9606 GN=PIGR PE=1 SV=4 | 902 | 20201114_qwl_colo_milk_1ng_glyco_hcd30_etd30_compound15.17964.17964.4 | controllerType=0 controllerNumber=1 scan=17964 | 48.7487 |
| 15679:4 | 4 | R.LSLLEEPGN[+2350.830]GTFTVILNQLTSR.D | HexNAc(4)Hex(5)Fuc(1)NeuAc(2) | N[+2351] | 1189.0378 | 4 | 4753.1292 | 4753.1229 | 0 | 1.3 | 413 | Specific | 511.4 | 455.75 | 455.75 | 6.81 | 46 | >sp\|P01833\|PIGR_HUMAN Polymeric immunoglobulin receptor OS=Homo sapiens OX=9606 GN=PIGR PE=1 SV=4 | 902 | 20201114_qwl_colo_milk_1ng_glyco_hcd30_etd30_compound15.19286.19286.4 | controllerType=0 controllerNumber=1 scan=19286 | 53.1721 |
| 14512:3 | 4 | R.LSLLEEPGN[+2351.851]GTFTVILNQLTSR.D | HexNAc(4)Hex(5)Fuc(3)NeuAc(1) | N[+2352] | 1585.3908 | 3 | 4754.1579 | 4754.1433 | 0 | 3.1 | 413 | Specific | 530.53 | 430.54 | 430.54 | 8.38 | 46 | >sp\|P01833\|PIGR_HUMAN Polymeric immunoglobulin receptor OS=Homo sapiens OX=9606 GN=PIGR PE=1 SV=4 | 902 | 20201114_qwl_colo_milk_1ng_glyco_hcd30_etd30_compound15.18042.18042.3 | controllerType=0 controllerNumber=1 scan=18042 | 48.9856 |
| 14495:3 | 4 | R.LSLLEEPGN[+2351.851]GTFTVILNQLTSR.D | HexNAc(4)Hex(5)Fuc(3)NeuAc(1) | N[+2352] | 1585.3893 | 3 | 4754.1535 | 4754.1433 | 0 | 2.1 | 413 | Specific | 551.88 | 400.54 | 400.54 | 8.38 | 46 | >sp\|P01833\|PIGR_HUMAN Polymeric immunoglobulin receptor OS=Homo sapiens OX=9606 GN=PIGR PE=1 SV=4 | 902 | 20201114_qwl_colo_milk_1ng_glyco_hcd30_etd30_compound15.18024.18024.3 | controllerType=0 controllerNumber=1 scan=18024 | 48.9383 |
| 15715:3 | 4 | R.LSLLEEPGN[+2496.888]GTFTVILNQLTSR.D | HexNAc(4)Hex(5)Fuc(2)NeuAc(2) | N[+2497] | 1633.7326 | 3 | 4899.1832 | 4899.1808 | 0 | 0.5 | 413 | Specific | 618.79 | 483.13 | 483.13 | 10.11 | 46 | >sp\|P01833\|PIGR_HUMAN Polymeric immunoglobulin receptor OS=Homo sapiens OX=9606 GN=PIGR PE=1 SV=4 | 902 | 20201114_qwl_colo_milk_1ng_glyco_hcd30_etd30_compound15.19324.19324.3 | controllerType=0 controllerNumber=1 scan=19324 | 53.3268 |
| 15678:3 | 4 | R.LSLLEEPGN[+2350.830]GTFTVILNQLTSR.D | HexNAc(4)Hex(5)Fuc(1)NeuAc(2) | N[+2351] | 1585.0507 | 3 | 4753.1376 | 4753.1229 | 0 | 3.1 | 413 | Specific | 662.6 | 482.41 | 482.41 | 11.32 | 46 | >sp\|P01833\|PIGR_HUMAN Polymeric immunoglobulin receptor OS=Homo sapiens OX=9606 GN=PIGR PE=1 SV=4 | 902 | 20201114_qwl_colo_milk_1ng_glyco_hcd30_etd30_compound15.19285.19285.3 | controllerType=0 controllerNumber=1 scan=19285 | 53.1688 |
| 11595:3 | 4 | K.IIEGEPNLKVPGN[+2351.851]VTAVLGETLK.V | HexNAc(4)Hex(5)Fuc(3)NeuAc(1) | N[+2352] | 1581.7444 | 3 | 4743.2188 | 4743.2001 | 0 | 3.9 | 457 | Specific | 330.77 | 288.41 | 288.41 | 4.52 | 46 | >sp\|P01833\|PIGR_HUMAN Polymeric immunoglobulin receptor OS=Homo sapiens OX=9606 GN=PIGR PE=1 SV=4 | 902 | 20201114_qwl_colo_milk_1ng_glyco_hcd30_etd30_compound15.14937.14937.3 | controllerType=0 controllerNumber=1 scan=14937 | 39.0745 |
| 11668:3 | 4 | K.IIEGEPNLKVPGN[+2059.735]VTAVLGETLK.V | HexNAc(4)Hex(5)Fuc(1)NeuAc(1) | N[+2060] | 1484.3754 | 3 | 4451.1117 | 4451.0843 | 0 | 6.2 | 457 | Specific | 359.92 | 321.54 | 321.54 | 4.16 | 46 | >sp\|P01833\|PIGR_HUMAN Polymeric immunoglobulin receptor OS=Homo sapiens OX=9606 GN=PIGR PE=1 SV=4 | 902 | 20201114_qwl_colo_milk_1ng_glyco_hcd30_etd30_compound15.15014.15014.3 | controllerType=0 controllerNumber=1 scan=15014 | 39.2892 |
| 11418:5 | 4 | K.IIEGEPNLKVPGN[+2205.793]VTAVLGETLKVPC[+57.021]HFPC[+57.021]K.F | HexNAc(4)Hex(5)Fuc(2)NeuAc(1) | C[+57]*2, N[+2206] | 1125.3289 | 5 | 5622.6152 | 5622.5997 | 0 | 2.8 | 457 | Specific | 329.62 | 329.62 | 329.62 | 4.52 | 46 | >sp\|P01833\|PIGR_HUMAN Polymeric immunoglobulin receptor OS=Homo sapiens OX=9606 GN=PIGR PE=1 SV=4 | 902 | 20201114_qwl_colo_milk_1ng_glyco_hcd30_etd30_compound15.14748.14748.5 | controllerType=0 controllerNumber=1 scan=14748 | 38.4861 |
| 11310:5 | 4 | K.IIEGEPNLKVPGN[+2204.772]VTAVLGETLKVPC[+57.021]HFPC[+57.021]K.F | HexNAc(4)Hex(5)NeuAc(2) | C[+57]*2, N[+2205] | 1125.1285 | 5 | 5621.6136 | 5621.5793 | 0 | 6.1 | 457 | Specific | 364.82 | 364.82 | 364.82 | 4.98 | 46 | >sp\|P01833\|PIGR_HUMAN Polymeric immunoglobulin receptor OS=Homo sapiens OX=9606 GN=PIGR PE=1 SV=4 | 902 | 20201114_qwl_colo_milk_1ng_glyco_hcd30_etd30_compound15.14632.14632.5 | controllerType=0 controllerNumber=1 scan=14632 | 38.0951 |
| 11398:5 | 4 | K.IIEGEPNLKVPGN[+2059.735]VTAVLGETLKVPC[+57.021]HFPC[+57.021]K.F | HexNAc(4)Hex(5)Fuc(1)NeuAc(1) | C[+57]*2, N[+2060] | 1096.1191 | 5 | 5476.5664 | 5476.5418 | 0 | 4.5 | 457 | Specific | 367.35 | 367.35 | 367.35 | 4.84 | 46 | >sp\|P01833\|PIGR_HUMAN Polymeric immunoglobulin receptor OS=Homo sapiens OX=9606 GN=PIGR PE=1 SV=4 | 902 | 20201114_qwl_colo_milk_1ng_glyco_hcd30_etd30_compound15.14726.14726.5 | controllerType=0 controllerNumber=1 scan=14726 | 38.4303 |
| 12319:4 | 4 | K.IIEGEPNLKVPGN[+2350.830]VTAVLGETLKVPC[+57.021]HFPC[+57.021]K.F | HexNAc(4)Hex(5)Fuc(1)NeuAc(2) | C[+57]*2, N[+2351] | 1442.6781 | 4 | 5767.6906 | 5767.6372 | 0 | 9.3 | 457 | Specific | 451.25 | 451.25 | 451.25 | 5.18 | 46 | >sp\|P01833\|PIGR_HUMAN Polymeric immunoglobulin receptor OS=Homo sapiens OX=9606 GN=PIGR PE=1 SV=4 | 902 | 20201114_qwl_colo_milk_1ng_glyco_hcd30_etd30_compound15.15702.15702.4 | controllerType=0 controllerNumber=1 scan=15702 | 41.2221 |
| 12320:5 | 4 | K.IIEGEPNLKVPGN[+2350.830]VTAVLGETLKVPC[+57.021]HFPC[+57.021]K.F | HexNAc(4)Hex(5)Fuc(1)NeuAc(2) | C[+57]*2, N[+2351] | 1154.3413 | 5 | 5767.6774 | 5767.6372 | 0 | 7 | 457 | Specific | 457.35 | 457.35 | 457.35 | 4.82 | 46 | >sp\|P01833\|PIGR_HUMAN Polymeric immunoglobulin receptor OS=Homo sapiens OX=9606 GN=PIGR PE=1 SV=4 | 902 | 20201114_qwl_colo_milk_1ng_glyco_hcd30_etd30_compound15.15703.15703.5 | controllerType=0 controllerNumber=1 scan=15703 | 41.2261 |
| 11399:4 | 4 | K.IIEGEPNLKVPGN[+2059.735]VTAVLGETLKVPC[+57.021]HFPC[+57.021]K.F | HexNAc(4)Hex(5)Fuc(1)NeuAc(1) | C[+57]*2, N[+2060] | 1369.8971 | 4 | 5476.5666 | 5476.5418 | 0 | 4.5 | 457 | Specific | 485.6 | 485.6 | 485.6 | 5.21 | 46 | >sp\|P01833\|PIGR_HUMAN Polymeric immunoglobulin receptor OS=Homo sapiens OX=9606 GN=PIGR PE=1 SV=4 | 902 | 20201114_qwl_colo_milk_1ng_glyco_hcd30_etd30_compound15.14727.14727.4 | controllerType=0 controllerNumber=1 scan=14727 | 38.4386 |
| 11311:4 | 4 | K.IIEGEPNLKVPGN[+2205.793]VTAVLGETLKVPC[+57.021]HFPC[+57.021]K.F | HexNAc(4)Hex(5)Fuc(2)NeuAc(1) | C[+57]*2, N[+2206] | 1406.4072 | 4 | 5622.6071 | 5622.5997 | 0 | 1.3 | 457 | Specific | 520.12 | 520.12 | 520.12 | 5.73 | 46 | >sp\|P01833\|PIGR_HUMAN Polymeric immunoglobulin receptor OS=Homo sapiens OX=9606 GN=PIGR PE=1 SV=4 | 902 | 20201114_qwl_colo_milk_1ng_glyco_hcd30_etd30_compound15.14633.14633.4 | controllerType=0 controllerNumber=1 scan=14633 | 38.0992 |
| 11424:4 | 4 | K.IIEGEPNLKVPGN[+2205.793]VTAVLGETLKVPC[+57.021]HFPC[+57.021]K.F | HexNAc(4)Hex(5)Fuc(2)NeuAc(1) | C[+57]*2, N[+2206] | 1406.4105 | 4 | 5622.6203 | 5622.5997 | 0 | 3.7 | 457 | Specific | 539.33 | 539.33 | 539.33 | 5.41 | 46 | >sp\|P01833\|PIGR_HUMAN Polymeric immunoglobulin receptor OS=Homo sapiens OX=9606 GN=PIGR PE=1 SV=4 | 902 | 20201114_qwl_colo_milk_1ng_glyco_hcd30_etd30_compound15.14754.14754.4 | controllerType=0 controllerNumber=1 scan=14754 | 38.5136 |
| 11353:4 | 4 | K.VPGN[+2350.830]VTAVLGETLKVPC[+57.021]HFPC[+57.021]K.F | HexNAc(4)Hex(5)Fuc(1)NeuAc(2) | C[+57]*2, N[+2351] | 1194.2819 | 4 | 4774.1058 | 4774.0877 | 0 | 3.8 | 466 | Specific | 302.04 | 280.79 | 280.79 | 3.52 | 46 | >sp\|P01833\|PIGR_HUMAN Polymeric immunoglobulin receptor OS=Homo sapiens OX=9606 GN=PIGR PE=1 SV=4 | 902 | 20201114_qwl_colo_milk_1ng_glyco_hcd30_etd30_compound15.14677.14677.4 | controllerType=0 controllerNumber=1 scan=14677 | 38.252 |
| 08975:4 | 4 | K.[+42.011]VPGN[+2018.708]VTAVLGETLKVPC[+57.021]HFPC[+57.021]K.F | HexNAc(3)Hex(6)Fuc(1)NeuAc(1) | .[42], C[+57]*2, N[+2019] | 1121.7605 | 4 | 4484.0203 | 4483.9763 | 0 | 9.8 | 466 | Specific | 314.54 | 314.54 | 314.54 | 3.54 | 46 | >sp\|P01833\|PIGR_HUMAN Polymeric immunoglobulin receptor OS=Homo sapiens OX=9606 GN=PIGR PE=1 SV=4 | 902 | 20201114_qwl_colo_milk_1ng_glyco_hcd30_etd30_compound15.12177.12177.4 | controllerType=0 controllerNumber=1 scan=12177 | 31.7976 |
| 10046:4 | 4 | K.VPGN[+2205.793]VTAVLGETLKVPC[+57.021]HFPC[+57.021]K.F | HexNAc(4)Hex(5)Fuc(2)NeuAc(1) | C[+57]*2, N[+2206] | 1158.0201 | 4 | 4629.0584 | 4629.0502 | 0 | 1.8 | 466 | Specific | 429.53 | 397.83 | 397.83 | 5.75 | 46 | >sp\|P01833\|PIGR_HUMAN Polymeric immunoglobulin receptor OS=Homo sapiens OX=9606 GN=PIGR PE=1 SV=4 | 902 | 20201114_qwl_colo_milk_1ng_glyco_hcd30_etd30_compound15.13301.13301.4 | controllerType=0 controllerNumber=1 scan=13301 | 34.4898 |
| 06234:3 | 4 | K.WN[+2204.772]NTGC[+57.021]QALPSQDEGPSK.A | HexNAc(4)Hex(5)NeuAc(2) | C[+57], N[+2205] | 1398.5571 | 3 | 4193.6566 | 4193.6491 | 0 | 1.8 | 498 | Specific | 354.01 | 224.12 | 224.12 | 6.29 | 46 | >sp\|P01833\|PIGR_HUMAN Polymeric immunoglobulin receptor OS=Homo sapiens OX=9606 GN=PIGR PE=1 SV=4 | 902 | 20201114_qwl_colo_milk_1ng_glyco_hcd30_etd30_compound15.9291.9291.3 | controllerType=0 controllerNumber=1 scan=9291 | 24.3571 |
| 04469:3 | 4 | K.WN[+2059.735]NTGC[+57.021]QALPSQDEGPSK.A | HexNAc(4)Hex(5)Fuc(1)NeuAc(1) | C[+57], N[+2060] | 1350.2209 | 3 | 4048.6481 | 4048.6116 | 0 | 9 | 498 | Specific | 398.3 | 279.75 | 279.75 | 5.83 | 46 | >sp\|P01833\|PIGR_HUMAN Polymeric immunoglobulin receptor OS=Homo sapiens OX=9606 GN=PIGR PE=1 SV=4 | 902 | 20201114_qwl_colo_milk_1ng_glyco_hcd30_etd30_compound15.7455.7455.3 | controllerType=0 controllerNumber=1 scan=7455 | 20.6862 |
| 04425:3 | 4 | K.WN[+2205.793]NTGC[+57.021]QALPSQDEGPSK.A | HexNAc(4)Hex(5)Fuc(2)NeuAc(1) | C[+57], N[+2206] | 1398.8989 | 3 | 4194.682 | 4194.6695 | 0 | 3 | 498 | Specific | 418.47 | 286.77 | 286.77 | 6.31 | 46 | >sp\|P01833\|PIGR_HUMAN Polymeric immunoglobulin receptor OS=Homo sapiens OX=9606 GN=PIGR PE=1 SV=4 | 902 | 20201114_qwl_colo_milk_1ng_glyco_hcd30_etd30_compound15.7409.7409.3 | controllerType=0 controllerNumber=1 scan=7409 | 20.5854 |
| 04538:3 | 4 | K.WN[+1872.651]NTGC[+57.021]QALPSQDEGPSK.A | HexNAc(3)Hex(6)NeuAc(1) | C[+57], N[+1873] | 1287.8521 | 3 | 3861.5418 | 3861.5272 | 0 | 3.8 | 498 | Specific | 430.57 | 273.82 | 273.82 | 5.78 | 46 | >sp\|P01833\|PIGR_HUMAN Polymeric immunoglobulin receptor OS=Homo sapiens OX=9606 GN=PIGR PE=1 SV=4 | 902 | 20201114_qwl_colo_milk_1ng_glyco_hcd30_etd30_compound15.7527.7527.3 | controllerType=0 controllerNumber=1 scan=7527 | 20.8472 |
| 04524:3 | 4 | K.WN[+1913.677]NTGC[+57.021]QALPSQDEGPSK.A | HexNAc(4)Hex(5)NeuAc(1) | C[+57], N[+1914] | 1301.5265 | 3 | 3902.5651 | 3902.5537 | 0 | 2.9 | 498 | Specific | 488.51 | 391.42 | 391.42 | 7.8 | 46 | >sp\|P01833\|PIGR_HUMAN Polymeric immunoglobulin receptor OS=Homo sapiens OX=9606 GN=PIGR PE=1 SV=4 | 902 | 20201114_qwl_colo_milk_1ng_glyco_hcd30_etd30_compound15.7512.7512.3 | controllerType=0 controllerNumber=1 scan=7512 | 20.8053 |
| 03208:3 | 4 | K.WN[+1216.423]NTGC[+57.021]QALPSQDEGPSK.A | HexNAc(2)Hex(5) | C[+57], N[+1216] | 1069.1023 | 3 | 3205.2924 | 3205.2995 | 0 | -2.2 | 498 | Specific | 503.41 | 444.41 | 444.41 | 6.95 | 46 | >sp\|P01833\|PIGR_HUMAN Polymeric immunoglobulin receptor OS=Homo sapiens OX=9606 GN=PIGR PE=1 SV=4 | 902 | 20201114_qwl_colo_milk_1ng_glyco_hcd30_etd30_compound15.6139.6139.3 | controllerType=0 controllerNumber=1 scan=6139 | 18.1618 |
| 04593:3 | 4 | K.WN[+1710.598]NTGC[+57.021]QALPSQDEGPSK.A | HexNAc(3)Hex(5)NeuAc(1) | C[+57], N[+1711] | 1233.8326 | 3 | 3699.4832 | 3699.4743 | 0 | 2.4 | 498 | Specific | 516.42 | 373.42 | 373.42 | 7.8 | 46 | >sp\|P01833\|PIGR_HUMAN Polymeric immunoglobulin receptor OS=Homo sapiens OX=9606 GN=PIGR PE=1 SV=4 | 902 | 20201114_qwl_colo_milk_1ng_glyco_hcd30_etd30_compound15.7584.7584.3 | controllerType=0 controllerNumber=1 scan=7584 | 20.9509 |
| 07880:3 | 5 | R.LAGKPTHVN[+1647.613]VSVVMAEVDGTC[+57.021].Y | HexNAc(5)Hex(3)Fuc(1) | C[+57], N[+1648] | 1277.9108 | 3 | 3831.7179 | 3831.692 | 0 | 6.8 | 332 | CRagged | 220.85 | 164.69 | 164.69 | 2.93 | 46 | >sp\|P01876\|IGHA1_HUMAN Immunoglobulin heavy constant alpha 1 OS=Homo sapiens OX=9606 GN=IGHA1 PE=1 SV=2 | 255 | 20201114_qwl_colo_milk_1ng_glyco_hcd30_etd30_compound15.11023.11023.3 | controllerType=0 controllerNumber=1 scan=11023 | 28.7074 |
| 06883:3 | 5 | R.LAGKPTHVN[+1378.476]VSVVMAEVDGTC[+57.021].Y | HexNAc(2)Hex(6) | C[+57], N[+1378] | 1188.1934 | 3 | 3562.5657 | 3562.5544 | 0 | 3.2 | 332 | CRagged | 286.49 | 230.74 | 207.51 | 4.9 | 46 | >sp\|P01876\|IGHA1_HUMAN Immunoglobulin heavy constant alpha 1 OS=Homo sapiens OX=9606 GN=IGHA1 PE=1 SV=2 | 255 | 20201114_qwl_colo_milk_1ng_glyco_hcd30_etd30_compound15.9972.9972.3 | controllerType=0 controllerNumber=1 scan=9972 | 25.9759 |
| 06902:3 | 5 | R.LAGKPTHVN[+1216.423]VSVVMAEVDGTC[+57.021].Y | HexNAc(2)Hex(5) | C[+57], N[+1216] | 1134.1758 | 3 | 3400.5129 | 3400.5016 | 0 | 3.3 | 332 | CRagged | 319.16 | 257.71 | 217.79 | 4.42 | 46 | >sp\|P01876\|IGHA1_HUMAN Immunoglobulin heavy constant alpha 1 OS=Homo sapiens OX=9606 GN=IGHA1 PE=1 SV=2 | 255 | 20201114_qwl_colo_milk_1ng_glyco_hcd30_etd30_compound15.9992.9992.3 | controllerType=0 controllerNumber=1 scan=9992 | 26.0323 |
| 06965:3 | 5 | R.LAGKPTHVN[+1054.370]VSVVMAEVDGTC[+57.021].Y | HexNAc(2)Hex(4) | C[+57], N[+1054] | 1080.1575 | 3 | 3238.4579 | 3238.4488 | 0 | 2.8 | 332 | CRagged | 395.14 | 330.66 | 225.23 | 5.7 | 46 | >sp\|P01876\|IGHA1_HUMAN Immunoglobulin heavy constant alpha 1 OS=Homo sapiens OX=9606 GN=IGHA1 PE=1 SV=2 | 255 | 20201114_qwl_colo_milk_1ng_glyco_hcd30_etd30_compound15.10058.10058.3 | controllerType=0 controllerNumber=1 scan=10058 | 26.204 |
| 05135:3 | 5 | R.LAGKPTHVN[+1054.370]VSVVM[+15.995]AEVDGTC[+57.021].Y | HexNAc(2)Hex(4) | C[+57], M[+16], N[+1054] | 1085.4888 | 3 | 3254.4518 | 3254.4437 | 0 | 2.5 | 332 | CRagged | 202.11 | 188.13 | 188.13 | 3.36 | 46 | >sp\|P01876\|IGHA1_HUMAN Immunoglobulin heavy constant alpha 1 OS=Homo sapiens OX=9606 GN=IGHA1 PE=1 SV=2 | 255 | 20201114_qwl_colo_milk_1ng_glyco_hcd30_etd30_compound15.8147.8147.3 | controllerType=0 controllerNumber=1 scan=8147 | 22.062 |
| 05764:3 | 5 | R.LAGKPTHVN[+1216.423]VSVVM[+15.995]AEVDGTC[+57.021].Y | HexNAc(2)Hex(5) | C[+57], M[+16], N[+1216] | 1139.5093 | 3 | 3416.5134 | 3416.4965 | 0 | 4.9 | 332 | CRagged | 225.14 | 219.09 | 219.09 | 3.45 | 46 | >sp\|P01876\|IGHA1_HUMAN Immunoglobulin heavy constant alpha 1 OS=Homo sapiens OX=9606 GN=IGHA1 PE=1 SV=2 | 255 | 20201114_qwl_colo_milk_1ng_glyco_hcd30_etd30_compound15.8800.8800.3 | controllerType=0 controllerNumber=1 scan=8800 | 23.3061 |
| 05057:3 | 5 | R.LAGKPTHVN[+1216.423]VSVVM[+15.995]AEVDGTC[+57.021].Y | HexNAc(2)Hex(5) | C[+57], M[+16], N[+1216] | 1139.5061 | 3 | 3416.5038 | 3416.4965 | 0 | 2.1 | 332 | CRagged | 264.9 | 263.8 | 263.8 | 4.67 | 46 | >sp\|P01876\|IGHA1_HUMAN Immunoglobulin heavy constant alpha 1 OS=Homo sapiens OX=9606 GN=IGHA1 PE=1 SV=2 | 255 | 20201114_qwl_colo_milk_1ng_glyco_hcd30_etd30_compound15.8066.8066.3 | controllerType=0 controllerNumber=1 scan=8066 | 21.9103 |
| 08482:3 | 5 | R.LAGKPTHVN[+1971.719]VSVVMAEVDGTC[+57.021]Y.- | HexNAc(5)Hex(5)Fuc(1) | C[+57], N[+1972] | 1440.2922 | 3 | 4318.862 | 4318.861 | 0 | 0.2 | 332 | CRagged | 201.08 | 134.97 | 134.97 | 3.01 | 46 | >sp\|P01876\|IGHA1_HUMAN Immunoglobulin heavy constant alpha 1 OS=Homo sapiens OX=9606 GN=IGHA1 PE=1 SV=2 | 255 | 20201114_qwl_colo_milk_1ng_glyco_hcd30_etd30_compound15.11657.11657.3 | controllerType=0 controllerNumber=1 scan=11657 | 30.413 |
| 07802:3 | 5 | R.LAGKPTHVN[+1581.555]VSVVMAEVDGTC[+57.021]Y.- | HexNAc(3)Hex(6) | C[+57], N[+1582] | 1310.2502 | 3 | 3928.736 | 3928.6971 | 0 | 9.9 | 332 | CRagged | 201.44 | 188.65 | 158.39 | 3.3 | 46 | >sp\|P01876\|IGHA1_HUMAN Immunoglobulin heavy constant alpha 1 OS=Homo sapiens OX=9606 GN=IGHA1 PE=1 SV=2 | 255 | 20201114_qwl_colo_milk_1ng_glyco_hcd30_etd30_compound15.10939.10939.3 | controllerType=0 controllerNumber=1 scan=10939 | 28.3939 |
| 08626:3 | 5 | R.LAGKPTHVN[+1378.476]VSVVMAEVDGTC[+57.021]Y.- | HexNAc(2)Hex(6) | C[+57], N[+1378] | 1242.5463 | 3 | 3725.6244 | 3725.6178 | 0 | 1.8 | 332 | CRagged | 210.24 | 210.24 | 171.43 | 3.75 | 46 | >sp\|P01876\|IGHA1_HUMAN Immunoglobulin heavy constant alpha 1 OS=Homo sapiens OX=9606 GN=IGHA1 PE=1 SV=2 | 255 | 20201114_qwl_colo_milk_1ng_glyco_hcd30_etd30_compound15.11810.11810.3 | controllerType=0 controllerNumber=1 scan=11810 | 30.833 |
| 08496:3 | 5 | R.LAGKPTHVN[+1606.587]VSVVMAEVDGTC[+57.021]Y.- | HexNAc(4)Hex(4)Fuc(1) | C[+57], N[+1607] | 1318.5845 | 3 | 3953.739 | 3953.7288 | 0 | 2.6 | 332 | CRagged | 210.43 | 152.95 | 152.95 | 3.29 | 46 | >sp\|P01876\|IGHA1_HUMAN Immunoglobulin heavy constant alpha 1 OS=Homo sapiens OX=9606 GN=IGHA1 PE=1 SV=2 | 255 | 20201114_qwl_colo_milk_1ng_glyco_hcd30_etd30_compound15.11672.11672.3 | controllerType=0 controllerNumber=1 scan=11672 | 30.4578 |
| 08565:3 | 5 | R.LAGKPTHVN[+1501.555]VSVVMAEVDGTC[+57.021]Y.- | HexNAc(5)Hex(3) | C[+57], N[+1502] | 1283.5729 | 3 | 3848.7042 | 3848.6974 | 0 | 1.8 | 332 | CRagged | 211.84 | 146.96 | 146.96 | 3.29 | 46 | >sp\|P01876\|IGHA1_HUMAN Immunoglobulin heavy constant alpha 1 OS=Homo sapiens OX=9606 GN=IGHA1 PE=1 SV=2 | 255 | 20201114_qwl_colo_milk_1ng_glyco_hcd30_etd30_compound15.11746.11746.3 | controllerType=0 controllerNumber=1 scan=11746 | 30.656 |
| 08519:3 | 5 | R.LAGKPTHVN[+1378.476]VSVVMAEVDGTC[+57.021]Y.- | HexNAc(2)Hex(6) | C[+57], N[+1378] | 1242.5453 | 3 | 3725.6215 | 3725.6178 | 0 | 1 | 332 | CRagged | 213.84 | 160.24 | 160.24 | 3.4 | 46 | >sp\|P01876\|IGHA1_HUMAN Immunoglobulin heavy constant alpha 1 OS=Homo sapiens OX=9606 GN=IGHA1 PE=1 SV=2 | 255 | 20201114_qwl_colo_milk_1ng_glyco_hcd30_etd30_compound15.11697.11697.3 | controllerType=0 controllerNumber=1 scan=11697 | 30.5351 |
| 09687:3 | 5 | R.LAGKPTHVN[+2262.814]VSVVMAEVDGTC[+57.021]Y.- | HexNAc(5)Hex(5)Fuc(1)NeuAc(1) | C[+57], N[+2263] | 1537.3239 | 3 | 4609.9572 | 4609.9564 | 0 | 0.2 | 332 | CRagged | 216.32 | 68.02 | 68.02 | 2.95 | 46 | >sp\|P01876\|IGHA1_HUMAN Immunoglobulin heavy constant alpha 1 OS=Homo sapiens OX=9606 GN=IGHA1 PE=1 SV=2 | 255 | 20201114_qwl_colo_milk_1ng_glyco_hcd30_etd30_compound15.12923.12923.3 | controllerType=0 controllerNumber=1 scan=12923 | 33.5728 |
| 08540:3 | 5 | R.LAGKPTHVN[+1647.613]VSVVMAEVDGTC[+57.021]Y.- | HexNAc(5)Hex(3)Fuc(1) | C[+57], N[+1648] | 1332.2584 | 3 | 3994.7605 | 3994.7553 | 0 | 1.3 | 332 | CRagged | 218.85 | 147.8 | 147.8 | 3.15 | 46 | >sp\|P01876\|IGHA1_HUMAN Immunoglobulin heavy constant alpha 1 OS=Homo sapiens OX=9606 GN=IGHA1 PE=1 SV=2 | 255 | 20201114_qwl_colo_milk_1ng_glyco_hcd30_etd30_compound15.11719.11719.3 | controllerType=0 controllerNumber=1 scan=11719 | 30.5828 |
| 09703:4 | 5 | R.LAGKPTHVN[+2262.814]VSVVMAEVDGTC[+57.021]Y.- | HexNAc(5)Hex(5)Fuc(1)NeuAc(1) | C[+57], N[+2263] | 1153.2448 | 4 | 4609.9573 | 4609.9564 | 0 | 0.2 | 332 | CRagged | 219 | 189.47 | 189.47 | 3.95 | 46 | >sp\|P01876\|IGHA1_HUMAN Immunoglobulin heavy constant alpha 1 OS=Homo sapiens OX=9606 GN=IGHA1 PE=1 SV=2 | 255 | 20201114_qwl_colo_milk_1ng_glyco_hcd30_etd30_compound15.12939.12939.4 | controllerType=0 controllerNumber=1 scan=12939 | 33.6165 |
| 07872:3 | 5 | R.LAGKPTHVN[+1095.397]VSVVMAEVDGTC[+57.021]Y.- | HexNAc(3)Hex(3) | C[+57], N[+1095] | 1148.1942 | 3 | 3442.5682 | 3442.5387 | 0 | 8.6 | 332 | CRagged | 220.6 | 151.53 | 151.53 | 3.23 | 46 | >sp\|P01876\|IGHA1_HUMAN Immunoglobulin heavy constant alpha 1 OS=Homo sapiens OX=9606 GN=IGHA1 PE=1 SV=2 | 255 | 20201114_qwl_colo_milk_1ng_glyco_hcd30_etd30_compound15.11014.11014.3 | controllerType=0 controllerNumber=1 scan=11014 | 28.6737 |
| 07838:3 | 5 | R.LAGKPTHVN[+1501.555]VSVVMAEVDGTC[+57.021]Y.- | HexNAc(5)Hex(3) | C[+57], N[+1502] | 1283.5801 | 3 | 3848.7257 | 3848.6974 | 0 | 7.4 | 332 | CRagged | 223.61 | 160.54 | 160.54 | 2.97 | 46 | >sp\|P01876\|IGHA1_HUMAN Immunoglobulin heavy constant alpha 1 OS=Homo sapiens OX=9606 GN=IGHA1 PE=1 SV=2 | 255 | 20201114_qwl_colo_milk_1ng_glyco_hcd30_etd30_compound15.10978.10978.3 | controllerType=0 controllerNumber=1 scan=10978 | 28.5648 |
| 08453:3 | 5 | R.LAGKPTHVN[+1702.581]VSVVMAEVDGTC[+57.021]Y.- | HexNAc(2)Hex(8) | C[+57], N[+1703] | 1350.5843 | 3 | 4049.7383 | 4049.7234 | 0 | 3.7 | 332 | CRagged | 229.8 | 195.79 | 182.72 | 3.32 | 46 | >sp\|P01876\|IGHA1_HUMAN Immunoglobulin heavy constant alpha 1 OS=Homo sapiens OX=9606 GN=IGHA1 PE=1 SV=2 | 255 | 20201114_qwl_colo_milk_1ng_glyco_hcd30_etd30_compound15.11626.11626.3 | controllerType=0 controllerNumber=1 scan=11626 | 30.2684 |
| 07798:3 | 5 | R.LAGKPTHVN[+1663.608]VSVVMAEVDGTC[+57.021]Y.- | HexNAc(5)Hex(4) | C[+57], N[+1664] | 1337.601 | 3 | 4010.7885 | 4010.7502 | 0 | 9.5 | 332 | CRagged | 233.2 | 214.7 | 169.99 | 3.18 | 46 | >sp\|P01876\|IGHA1_HUMAN Immunoglobulin heavy constant alpha 1 OS=Homo sapiens OX=9606 GN=IGHA1 PE=1 SV=2 | 255 | 20201114_qwl_colo_milk_1ng_glyco_hcd30_etd30_compound15.10935.10935.3 | controllerType=0 controllerNumber=1 scan=10935 | 28.3828 |
| 09784:3 | 5 | R.LAGKPTHVN[+1872.651]VSVVMAEVDGTC[+57.021]Y.- | HexNAc(3)Hex(6)NeuAc(1) | C[+57], N[+1873] | 1407.2742 | 3 | 4219.8081 | 4219.7926 | 0 | 3.7 | 332 | CRagged | 241.3 | 163.79 | 147.16 | 3.94 | 46 | >sp\|P01876\|IGHA1_HUMAN Immunoglobulin heavy constant alpha 1 OS=Homo sapiens OX=9606 GN=IGHA1 PE=1 SV=2 | 255 | 20201114_qwl_colo_milk_1ng_glyco_hcd30_etd30_compound15.13024.13024.3 | controllerType=0 controllerNumber=1 scan=13024 | 33.7969 |
| 08513:3 | 5 | R.LAGKPTHVN[+1419.502]VSVVMAEVDGTC[+57.021]Y.- | HexNAc(3)Hex(5) | C[+57], N[+1420] | 1256.2211 | 3 | 3766.6487 | 3766.6443 | 0 | 1.2 | 332 | CRagged | 244.38 | 172.47 | 157.43 | 4.15 | 46 | >sp\|P01876\|IGHA1_HUMAN Immunoglobulin heavy constant alpha 1 OS=Homo sapiens OX=9606 GN=IGHA1 PE=1 SV=2 | 255 | 20201114_qwl_colo_milk_1ng_glyco_hcd30_etd30_compound15.11690.11690.3 | controllerType=0 controllerNumber=1 scan=11690 | 30.5148 |
| 07806:3 | 5 | R.LAGKPTHVN[+1419.502]VSVVMAEVDGTC[+57.021]Y.- | HexNAc(3)Hex(5) | C[+57], N[+1420] | 1256.2293 | 3 | 3766.6734 | 3766.6443 | 0 | 7.7 | 332 | CRagged | 246.75 | 208.03 | 188.72 | 3.89 | 46 | >sp\|P01876\|IGHA1_HUMAN Immunoglobulin heavy constant alpha 1 OS=Homo sapiens OX=9606 GN=IGHA1 PE=1 SV=2 | 255 | 20201114_qwl_colo_milk_1ng_glyco_hcd30_etd30_compound15.10943.10943.3 | controllerType=0 controllerNumber=1 scan=10943 | 28.419 |
| 08512:3 | 5 | R.LAGKPTHVN[+1622.582]VSVVMAEVDGTC[+57.021]Y.- | HexNAc(4)Hex(5) | C[+57], N[+1623] | 1323.9161 | 3 | 3969.7337 | 3969.7237 | 0 | 2.5 | 332 | CRagged | 251.31 | 164.51 | 164.51 | 4.45 | 46 | >sp\|P01876\|IGHA1_HUMAN Immunoglobulin heavy constant alpha 1 OS=Homo sapiens OX=9606 GN=IGHA1 PE=1 SV=2 | 255 | 20201114_qwl_colo_milk_1ng_glyco_hcd30_etd30_compound15.11689.11689.3 | controllerType=0 controllerNumber=1 scan=11689 | 30.5079 |
| 07784:3 | 5 | R.LAGKPTHVN[+1606.587]VSVVMAEVDGTC[+57.021]Y.- | HexNAc(4)Hex(4)Fuc(1) | C[+57], N[+1607] | 1318.5849 | 3 | 3953.7401 | 3953.7288 | 0 | 2.9 | 332 | CRagged | 253.37 | 168.79 | 168.79 | 4 | 46 | >sp\|P01876\|IGHA1_HUMAN Immunoglobulin heavy constant alpha 1 OS=Homo sapiens OX=9606 GN=IGHA1 PE=1 SV=2 | 255 | 20201114_qwl_colo_milk_1ng_glyco_hcd30_etd30_compound15.10920.10920.3 | controllerType=0 controllerNumber=1 scan=10920 | 28.3482 |
| 08477:3 | 5 | R.LAGKPTHVN[+1663.608]VSVVMAEVDGTC[+57.021]Y.- | HexNAc(5)Hex(4) | C[+57], N[+1664] | 1337.5879 | 3 | 4010.7493 | 4010.7502 | 0 | -0.2 | 332 | CRagged | 254.14 | 198.49 | 188.64 | 3.58 | 46 | >sp\|P01876\|IGHA1_HUMAN Immunoglobulin heavy constant alpha 1 OS=Homo sapiens OX=9606 GN=IGHA1 PE=1 SV=2 | 255 | 20201114_qwl_colo_milk_1ng_glyco_hcd30_etd30_compound15.11652.11652.3 | controllerType=0 controllerNumber=1 scan=11652 | 30.3843 |
| 08507:3 | 5 | R.LAGKPTHVN[+1460.529]VSVVMAEVDGTC[+57.021]Y.- | HexNAc(4)Hex(4) | C[+57], N[+1461] | 1269.8978 | 3 | 3807.6788 | 3807.6709 | 0 | 2.1 | 332 | CRagged | 257.62 | 195.33 | 191.65 | 4.55 | 46 | >sp\|P01876\|IGHA1_HUMAN Immunoglobulin heavy constant alpha 1 OS=Homo sapiens OX=9606 GN=IGHA1 PE=1 SV=2 | 255 | 20201114_qwl_colo_milk_1ng_glyco_hcd30_etd30_compound15.11684.11684.3 | controllerType=0 controllerNumber=1 scan=11684 | 30.4892 |
| 08239:3 | 5 | R.LAGKPTHVN[+1216.423]VSVVMAEVDGTC[+57.021]Y.- | HexNAc(2)Hex(5) | C[+57], N[+1216] | 1188.5285 | 3 | 3563.5709 | 3563.5649 | 0 | 1.7 | 332 | CRagged | 258.98 | 233.52 | 196.11 | 4.63 | 46 | >sp\|P01876\|IGHA1_HUMAN Immunoglobulin heavy constant alpha 1 OS=Homo sapiens OX=9606 GN=IGHA1 PE=1 SV=2 | 255 | 20201114_qwl_colo_milk_1ng_glyco_hcd30_etd30_compound15.11400.11400.3 | controllerType=0 controllerNumber=1 scan=11400 | 29.6906 |
| 09819:3 | 5 | R.LAGKPTHVN[+1751.624]VSVVMAEVDGTC[+57.021]Y.- | HexNAc(4)Hex(4)NeuAc(1) | C[+57], N[+1752] | 1366.9299 | 3 | 4098.7751 | 4098.7663 | 0 | 2.2 | 332 | CRagged | 262.85 | 152.84 | 152.84 | 4.35 | 46 | >sp\|P01876\|IGHA1_HUMAN Immunoglobulin heavy constant alpha 1 OS=Homo sapiens OX=9606 GN=IGHA1 PE=1 SV=2 | 255 | 20201114_qwl_colo_milk_1ng_glyco_hcd30_etd30_compound15.13061.13061.3 | controllerType=0 controllerNumber=1 scan=13061 | 33.8941 |
| 08602:3 | 5 | R.LAGKPTHVN[+1216.423]VSVVMAEVDGTC[+57.021]Y.- | HexNAc(2)Hex(5) | C[+57], N[+1216] | 1188.5254 | 3 | 3563.5618 | 3563.5649 | 0 | -0.9 | 332 | CRagged | 263.1 | 223.8 | 182.36 | 4.17 | 46 | >sp\|P01876\|IGHA1_HUMAN Immunoglobulin heavy constant alpha 1 OS=Homo sapiens OX=9606 GN=IGHA1 PE=1 SV=2 | 255 | 20201114_qwl_colo_milk_1ng_glyco_hcd30_etd30_compound15.11785.11785.3 | controllerType=0 controllerNumber=1 scan=11785 | 30.7655 |
| 09725:3 | 5 | R.LAGKPTHVN[+1872.651]VSVVMAEVDGTC[+57.021]Y.- | HexNAc(3)Hex(6)NeuAc(1) | C[+57], N[+1873] | 1407.2753 | 3 | 4219.8114 | 4219.7926 | 0 | 4.5 | 332 | CRagged | 268.77 | 127.5 | 127.5 | 3.69 | 46 | >sp\|P01876\|IGHA1_HUMAN Immunoglobulin heavy constant alpha 1 OS=Homo sapiens OX=9606 GN=IGHA1 PE=1 SV=2 | 255 | 20201114_qwl_colo_milk_1ng_glyco_hcd30_etd30_compound15.12962.12962.3 | controllerType=0 controllerNumber=1 scan=12962 | 33.6646 |
| 07777:3 | 5 | R.LAGKPTHVN[+1378.476]VSVVMAEVDGTC[+57.021]Y.- | HexNAc(2)Hex(6) | C[+57], N[+1378] | 1242.5477 | 3 | 3725.6284 | 3725.6178 | 0 | 2.9 | 332 | CRagged | 273.42 | 273.42 | 204.66 | 4.79 | 46 | >sp\|P01876\|IGHA1_HUMAN Immunoglobulin heavy constant alpha 1 OS=Homo sapiens OX=9606 GN=IGHA1 PE=1 SV=2 | 255 | 20201114_qwl_colo_milk_1ng_glyco_hcd30_etd30_compound15.10913.10913.3 | controllerType=0 controllerNumber=1 scan=10913 | 28.3317 |
| 08501:3 | 5 | R.LAGKPTHVN[+1809.666]VSVVMAEVDGTC[+57.021]Y.- | HexNAc(5)Hex(4)Fuc(1) | C[+57], N[+1810] | 1386.2735 | 3 | 4156.8059 | 4156.8081 | 0 | -0.5 | 332 | CRagged | 273.52 | 190.82 | 190.82 | 3.58 | 46 | >sp\|P01876\|IGHA1_HUMAN Immunoglobulin heavy constant alpha 1 OS=Homo sapiens OX=9606 GN=IGHA1 PE=1 SV=2 | 255 | 20201114_qwl_colo_milk_1ng_glyco_hcd30_etd30_compound15.11678.11678.3 | controllerType=0 controllerNumber=1 scan=11678 | 30.4746 |
| 08589:3 | 5 | R.LAGKPTHVN[+1095.397]VSVVMAEVDGTC[+57.021]Y.- | HexNAc(3)Hex(3) | C[+57], N[+1095] | 1148.1853 | 3 | 3442.5414 | 3442.5387 | 0 | 0.8 | 332 | CRagged | 279.21 | 191.13 | 191.13 | 4.24 | 46 | >sp\|P01876\|IGHA1_HUMAN Immunoglobulin heavy constant alpha 1 OS=Homo sapiens OX=9606 GN=IGHA1 PE=1 SV=2 | 255 | 20201114_qwl_colo_milk_1ng_glyco_hcd30_etd30_compound15.11771.11771.3 | controllerType=0 controllerNumber=1 scan=11771 | 30.7157 |
| 07780:3 | 5 | R.LAGKPTHVN[+1809.666]VSVVMAEVDGTC[+57.021]Y.- | HexNAc(5)Hex(4)Fuc(1) | C[+57], N[+1810] | 1386.278 | 3 | 4156.8195 | 4156.8081 | 0 | 2.7 | 332 | CRagged | 280.54 | 154.45 | 154.45 | 4.52 | 46 | >sp\|P01876\|IGHA1_HUMAN Immunoglobulin heavy constant alpha 1 OS=Homo sapiens OX=9606 GN=IGHA1 PE=1 SV=2 | 255 | 20201114_qwl_colo_milk_1ng_glyco_hcd30_etd30_compound15.10916.10916.3 | controllerType=0 controllerNumber=1 scan=10916 | 28.3381 |
| 08571:3 | 5 | R.LAGKPTHVN[+1444.534]VSVVMAEVDGTC[+57.021]Y.- | HexNAc(4)Hex(3)Fuc(1) | C[+57], N[+1445] | 1264.5659 | 3 | 3791.683 | 3791.6759 | 0 | 1.9 | 332 | CRagged | 286.28 | 182.91 | 182.91 | 4.63 | 46 | >sp\|P01876\|IGHA1_HUMAN Immunoglobulin heavy constant alpha 1 OS=Homo sapiens OX=9606 GN=IGHA1 PE=1 SV=2 | 255 | 20201114_qwl_colo_milk_1ng_glyco_hcd30_etd30_compound15.11752.11752.3 | controllerType=0 controllerNumber=1 scan=11752 | 30.6673 |
| 07811:3 | 5 | R.LAGKPTHVN[+1216.423]VSVVMAEVDGTC[+57.021]Y.- | HexNAc(2)Hex(5) | C[+57], N[+1216] | 1188.5284 | 3 | 3563.5706 | 3563.5649 | 0 | 1.6 | 332 | CRagged | 288.45 | 261.2 | 200.32 | 4.75 | 46 | >sp\|P01876\|IGHA1_HUMAN Immunoglobulin heavy constant alpha 1 OS=Homo sapiens OX=9606 GN=IGHA1 PE=1 SV=2 | 255 | 20201114_qwl_colo_milk_1ng_glyco_hcd30_etd30_compound15.10949.10949.3 | controllerType=0 controllerNumber=1 scan=10949 | 28.4566 |
| 09791:3 | 5 | R.LAGKPTHVN[+1710.598]VSVVMAEVDGTC[+57.021]Y.- | HexNAc(3)Hex(5)NeuAc(1) | C[+57], N[+1711] | 1353.2551 | 3 | 4057.7506 | 4057.7397 | 0 | 2.7 | 332 | CRagged | 315.66 | 231.2 | 186.75 | 4.9 | 46 | >sp\|P01876\|IGHA1_HUMAN Immunoglobulin heavy constant alpha 1 OS=Homo sapiens OX=9606 GN=IGHA1 PE=1 SV=2 | 255 | 20201114_qwl_colo_milk_1ng_glyco_hcd30_etd30_compound15.13031.13031.3 | controllerType=0 controllerNumber=1 scan=13031 | 33.8135 |
| 08567:3 | 5 | R.LAGKPTHVN[+1054.370]VSVVMAEVDGTC[+57.021]Y.- | HexNAc(2)Hex(4) | C[+57], N[+1054] | 1134.5104 | 3 | 3401.5167 | 3401.5121 | 0 | 1.3 | 332 | CRagged | 328.6 | 294.1 | 227.99 | 5.34 | 46 | >sp\|P01876\|IGHA1_HUMAN Immunoglobulin heavy constant alpha 1 OS=Homo sapiens OX=9606 GN=IGHA1 PE=1 SV=2 | 255 | 20201114_qwl_colo_milk_1ng_glyco_hcd30_etd30_compound15.11748.11748.3 | controllerType=0 controllerNumber=1 scan=11748 | [illegible] |
| 08609:3 | 5 | R.LAGKPTHVN[+892.317]VSVVMAEVDGTC[+57.021]Y.- | HexNAc(2)Hex(3) | C[+57], N[+892] | 1080.4917 | 3 | 3239.4606 | 3239.4593 | 0 | 0.4 | 332 | CRagged | 355.17 | 328.4 | 150.71 | 5.17 | 46 | >sp\|P01876\|IGHA1_HUMAN Immunoglobulin heavy constant alpha 1 OS=Homo sapiens OX=9606 GN=IGHA1 PE=1 SV=2 | 255 | 20201114_qwl_colo_milk_1ng_glyco_hcd30_etd30_compound15.11792.11792.3 | controllerType=0 controllerNumber=1 scan=11792 | 30.7794 |
| 08527:2 | 5 | R.LAGKPTHVN[+1216.423]VSVVMAEVDGTC[+57.021]Y.- | HexNAc(2)Hex(5) | C[+57], N[+1216] | 1782.2883 | 2 | 3563.5693 | 3563.5649 | 0 | 1.2 | 332 | CRagged | 367.7 | 361.47 | 148.05 | 6.05 | 46 | >sp\|P01876\|IGHA1_HUMAN Immunoglobulin heavy constant alpha 1 OS=Homo sapiens OX=9606 GN=IGHA1 PE=1 SV=2 | 255 | 20201114_qwl_colo_milk_1ng_glyco_hcd30_etd30_compound15.11706.11706.2 | controllerType=0 controllerNumber=1 scan=11706 | 30.5516 |
| 06849:3 | 5 | R.LAGKPTHVN[+1216.423]VSVVM[+15.995]AEVDGTC[+57.021]Y.- | HexNAc(2)Hex(5) | C[+57], M[+16], N[+1216] | 1193.8593 | 3 | 3579.5634 | 3579.5599 | 0 | 1 | 332 | CRagged | 242.57 | 210.04 | 210.04 | 4.24 | 46 | >sp\|P01876\|IGHA1_HUMAN Immunoglobulin heavy constant alpha 1 OS=Homo sapiens OX=9606 GN=IGHA1 PE=1 SV=2 | 255 | 20201114_qwl_colo_milk_1ng_glyco_hcd30_etd30_compound15.9936.9936.3 | controllerType=0 controllerNumber=1 scan=9936 | 25.8849 |
| 08536:3 | 9 | R.IIVPLNNREN[+1913.677]ISDPTSPLR.T | HexNAc(4)Hex(5)NeuAc(1) | N[+1914] | 1354.624 | 3 | 4061.8573 | 4061.8542 | 0 | 0.8 | 62 | Specific | 235.07 | 191.48 | 191.48 | 2.95 | 22 | >sp\|P01591\|IGJ_HUMAN Immunoglobulin J chain OS=Homo sapiens OX=9606 GN=JCHAIN PE=1 SV=4 | 1392 | 20201114_qwl_colo_milk_1ng_glyco_hcd30_etd30_compound15.11715.11715.3 | controllerType=0 controllerNumber=1 scan=11715 | 30.5726 |
| 09862:3 | 9 | R.IIVPLNNREN[+2204.772]ISDPTSPLR.T | HexNAc(4)Hex(5)NeuAc(2) | N[+2205] | 1451.6576 | 3 | 4352.9584 | 4352.9496 | 0 | 2 | 62 | Specific | 245.09 | 155.64 | 155.64 | 3.53 | 22 | >sp\|P01591\|IGJ_HUMAN Immunoglobulin J chain OS=Homo sapiens OX=9606 GN=JCHAIN PE=1 SV=4 | 1392 | 20201114_qwl_colo_milk_1ng_glyco_hcd30_etd30_compound15.13107.13107.3 | controllerType=0 controllerNumber=1 scan=13107 | 34.0001 |
| 08610:3 | 9 | R.IIVPLNNREN[+1548.545]ISDPTSPLR.T | HexNAc(3)Hex(4)NeuAc(1) | N[+1549] | 1232.9123 | 3 | 3696.7223 | 3696.722 | 0 | 0.1 | 62 | Specific | 253.11 | 223.72 | 223.72 | 3.57 | 22 | >sp\|P01591\|IGJ_HUMAN Immunoglobulin J chain OS=Homo sapiens OX=9606 GN=JCHAIN PE=1 SV=4 | 1392 | 20201114_qwl_colo_milk_1ng_glyco_hcd30_etd30_compound15.11793.11793.3 | controllerType=0 controllerNumber=1 scan=11793 | 30.7851 |
| 08486:3 | 9 | R.IIVPLNNREN[+1913.677]ISDPTSPLR.T | HexNAc(4)Hex(5)NeuAc(1) | N[+1914] | 1354.6263 | 3 | 4061.8643 | 4061.8542 | 0 | 2.5 | 62 | Specific | 257.27 | 205 | 205 | 3.6 | 22 | >sp\|P01591\|IGJ_HUMAN Immunoglobulin J chain OS=Homo sapiens OX=9606 GN=JCHAIN PE=1 SV=4 | 1392 | 20201114_qwl_colo_milk_1ng_glyco_hcd30_etd30_compound15.11662.11662.3 | controllerType=0 controllerNumber=1 scan=11662 | 30.424 |
| 09880:3 | 9 | R.IIVPLNNREN[+2204.772]ISDPTSPLR.T | HexNAc(4)Hex(5)NeuAc(2) | N[+2205] | 1451.6575 | 3 | 4352.958 | 4352.9496 | 0 | 1.9 | 62 | Specific | 276.45 | 196.75 | 196.75 | 3.62 | 22 | >sp\|P01591\|IGJ_HUMAN Immunoglobulin J chain OS=Homo sapiens OX=9606 GN=JCHAIN PE=1 SV=4 | 1392 | 20201114_qwl_colo_milk_1ng_glyco_hcd30_etd30_compound15.13126.13126.3 | controllerType=0 controllerNumber=1 scan=13126 | 34.0456 |
| 07510:3 | 9 | R.IIVPLNNREN[+1257.449]ISDPTSPLR.T | HexNAc(3)Hex(4) | N[+1257] | 1135.8836 | 3 | 3405.6362 | 3405.6266 | 0 | 2.8 | 62 | Specific | 295.3 | 248.5 | 248.5 | 5.19 | 22 | >sp\|P01591\|IGJ_HUMAN Immunoglobulin J chain OS=Homo sapiens OX=9606 GN=JCHAIN PE=1 SV=4 | 1392 | 20201114_qwl_colo_milk_1ng_glyco_hcd30_etd30_compound15.10632.10632.3 | controllerType=0 controllerNumber=1 scan=10632 | 27.6212 |
| 07522:3 | 9 | R.IIVPLNNREN[+1501.555]ISDPTSPLR.T | HexNAc(5)Hex(3) | N[+1502] | 1217.2559 | 3 | 3649.7532 | 3649.7325 | 0 | 5.7 | 62 | Specific | 300.32 | 252.61 | 252.61 | 3.53 | 22 | >sp\|P01591\|IGJ_HUMAN Immunoglobulin J chain OS=Homo sapiens OX=9606 GN=JCHAIN PE=1 SV=4 | 1392 | 20201114_qwl_colo_milk_1ng_glyco_hcd30_etd30_compound15.10645.10645.3 | controllerType=0 controllerNumber=1 scan=10645 | 27.6578 |
| 07460:3 | 9 | R.IIVPLNNREN[+1663.608]ISDPTSPLR.T | HexNAc(5)Hex(4) | N[+1664] | 1271.2691 | 3 | 3811.7927 | 3811.7853 | 0 | 2 | 62 | Specific | 304.64 | 225.91 | 225.91 | 3.92 | 22 | >sp\|P01591\|IGJ_HUMAN Immunoglobulin J chain OS=Homo sapiens OX=9606 GN=JCHAIN PE=1 SV=4 | 1392 | 20201114_qwl_colo_milk_1ng_glyco_hcd30_etd30_compound15.10580.10580.3 | controllerType=0 controllerNumber=1 scan=10580 | 27.5071 |
| 07443:3 | 9 | R.IIVPLNNREN[+1809.666]ISDPTSPLR.T | HexNAc(5)Hex(4)Fuc(1) | N[+1810] | 1319.9623 | 3 | 3957.8725 | 3957.8432 | 0 | 7.4 | 62 | Specific | 312.99 | 223.62 | 223.62 | 3.42 | 22 | >sp\|P01591\|IGJ_HUMAN Immunoglobulin J chain OS=Homo sapiens OX=9606 GN=JCHAIN PE=1 SV=4 | 1392 | 20201114_qwl_colo_milk_1ng_glyco_hcd30_etd30_compound15.10562.10562.3 | controllerType=0 controllerNumber=1 scan=10562 | 27.4429 |
| 07500:3 | 9 | R.IIVPLNNREN[+1768.640]ISDPTSPLR.T | HexNAc(4)Hex(5)Fuc(1) | N[+1769] | 1306.2817 | 3 | 3916.8305 | 3916.8167 | 0 | 3.5 | 62 | Specific | 313 | 231.01 | 231.01 | 3.46 | 22 | >sp\|P01591\|IGJ_HUMAN Immunoglobulin J chain OS=Homo sapiens OX=9606 GN=JCHAIN PE=1 SV=4 | 1392 | 20201114_qwl_colo_milk_1ng_glyco_hcd30_etd30_compound15.10622.10622.3 | controllerType=0 controllerNumber=1 scan=10622 | 27.5935 |
| 07509:3 | 9 | R.IIVPLNNREN[+1647.613]ISDPTSPLR.T | HexNAc(5)Hex(3)Fuc(1) | N[+1648] | 1265.9389 | 3 | 3795.8021 | 3795.7904 | 0 | 3.1 | 62 | Specific | 321.86 | 228.05 | 228.05 | 3.98 | 22 | >sp\|P01591\|IGJ_HUMAN Immunoglobulin J chain OS=Homo sapiens OX=9606 GN=JCHAIN PE=1 SV=4 | 1392 | 20201114_qwl_colo_milk_1ng_glyco_hcd30_etd30_compound15.10631.10631.3 | controllerType=0 controllerNumber=1 scan=10631 | 27.6164 |
| 08577:3 | 9 | R.IIVPLNNREN[+1694.603]ISDPTSPLR.T | HexNAc(3)Hex(4)Fuc(1)NeuAc(1) | N[+1695] | 1281.6005 | 3 | 3842.787 | 3842.7799 | 0 | 1.9 | 62 | Specific | 347.97 | 225.63 | 225.63 | 4.07 | 22 | >sp\|P01591\|IGJ_HUMAN Immunoglobulin J chain OS=Homo sapiens OX=9606 GN=JCHAIN PE=1 SV=4 | 1392 | 20201114_qwl_colo_milk_1ng_glyco_hcd30_etd30_compound15.11758.11758.3 | controllerType=0 controllerNumber=1 scan=11758 | 30.6844 |
| 07479:3 | 9 | R.IIVPLNNREN[+1460.529]ISDPTSPLR.T | HexNAc(4)Hex(4) | N[+1461] | 1203.5753 | 3 | 3608.7112 | 3608.7059 | 0 | 1.5 | 62 | Specific | 349.05 | 234.02 | 234.02 | 3.89 | 22 | >sp\|P01591\|IGJ_HUMAN Immunoglobulin J chain OS=Homo sapiens OX=9606 GN=JCHAIN PE=1 SV=4 | 1392 | 20201114_qwl_colo_milk_1ng_glyco_hcd30_etd30_compound15.10600.10600.3 | controllerType=0 controllerNumber=1 scan=10600 | 27.5481 |
| 08506:3 | 9 | R.IIVPLNNREN[+1856.656]ISDPTSPLR.T | HexNAc(3)Hex(5)Fuc(1)NeuAc(1) | N[+1857] | 1335.6103 | 3 | 4004.8163 | 4004.8327 | 0 | -4.1 | 62 | Specific | 377 | 292.54 | 292.54 | 3.19 | 22 | >sp\|P01591\|IGJ_HUMAN Immunoglobulin J chain OS=Homo sapiens OX=9606 GN=JCHAIN PE=1 SV=4 | 1392 | 20201114_qwl_colo_milk_1ng_glyco_hcd30_etd30_compound15.11683.11683.3 | controllerType=0 controllerNumber=1 scan=11683 | 30.4851 |
| 03924:3 | 9 | R.EN[+1663.608]ISDPTSPLR.T | HexNAc(5)Hex(4) | N[+1664] | 964.7478 | 3 | 2892.2289 | 2892.225 | 0 | 1.4 | 70 | Specific | 284.8 | 155.41 | 155.41 | 3.34 | 22 | >sp\|P01591\|IGJ_HUMAN Immunoglobulin J chain OS=Homo sapiens OX=9606 GN=JCHAIN PE=1 SV=4 | 1392 | 20201114_qwl_colo_milk_1ng_glyco_hcd30_etd30_compound15.6889.6889.3 | controllerType=0 controllerNumber=1 scan=6889 | 19.5931 |
| 16377:3 | 11 | R.QLAHQSN[+2204.772]STNIFFSPVSIATAFAMLSLGTK.A | HexNAc(4)Hex(5)NeuAc(2) | N[+2205] | 1796.1466 | 3 | 5386.4252 | 5386.4075 | 0 | 3.3 | 64 | Specific | 206.08 | 102.22 | 102.22 | 4.49 | 9 | >sp\|P01009\|A1AT_HUMAN Alpha-1-antitrypsin OS=Homo sapiens OX=9606 GN=SERPINA1 PE=1 SV=3 | 642 | 20201114_qwl_colo_milk_1ng_glyco_hcd30_etd30_compound15.20035.20035.3 | controllerType=0 controllerNumber=1 scan=20035 | 55.7831 |
| 16368:3 | 11 | R.QLAHQSN[+2350.830]STNIFFSPVSIATAFAMLSLGTK.A | HexNAc(4)Hex(5)Fuc(1)NeuAc(2) | N[+2351] | 1844.8187 | 3 | 5532.4415 | 5532.4654 | 0 | -4.3 | 64 | Specific | 206.33 | 107.56 | 107.56 | 4.26 | 9 | >sp\|P01009\|A1AT_HUMAN Alpha-1-antitrypsin OS=Homo sapiens OX=9606 GN=SERPINA1 PE=1 SV=3 | 642 | 20201114_qwl_colo_milk_1ng_glyco_hcd30_etd30_compound15.20025.20025.3 | controllerType=0 controllerNumber=1 scan=20025 | 55.7469 |
| 16455:3 | 11 | R.QLAHQSN[+2204.772]STNIFFSPVSIATAFAMLSLGTK.A | HexNAc(4)Hex(5)NeuAc(2) | N[+2205] | 1796.1353 | 3 | 5386.3915 | 5386.4075 | 0 | -3 | 64 | Specific | 213.1 | 117.03 | 117.03 | 3.39 | 9 | >sp\|P01009\|A1AT_HUMAN Alpha-1-antitrypsin OS=Homo sapiens OX=9606 GN=SERPINA1 PE=1 SV=3 | 642 | 20201114_qwl_colo_milk_1ng_glyco_hcd30_etd30_compound15.20118.20118.3 | controllerType=0 controllerNumber=1 scan=20118 | 56.0668 |
| 16378:4 | 11 | R.QLAHQSN[+2204.772]STNIFFSPVSIATAFAMLSLGTK.A | HexNAc(4)Hex(5)NeuAc(2) | N[+2205] | 1347.3615 | 4 | 5386.424 | 5386.4075 | 0 | 3.1 | 64 | Specific | 373.89 | 294.04 | 294.04 | 6.21 | 9 | >sp\|P01009\|A1AT_HUMAN Alpha-1-antitrypsin OS=Homo sapiens OX=9606 GN=SERPINA1 PE=1 SV=3 | 642 | 20201114_qwl_colo_milk_1ng_glyco_hcd30_etd30_compound15.20036.20036.4 | controllerType=0 controllerNumber=1 scan=20036 | 55.7875 |
| 16364:4 | 11 | R.QLAHQSN[+2350.830]STNIFFSPVSIATAFAMLSLGTK.A | HexNAc(4)Hex(5)Fuc(1)NeuAc(2) | N[+2351] | 1383.8689 | 4 | 5532.4538 | 5532.4654 | 0 | -2.1 | 64 | Specific | 652.77 | 561.36 | 561.36 | 11.14 | 9 | >sp\|P01009\|A1AT_HUMAN Alpha-1-antitrypsin OS=Homo sapiens OX=9606 GN=SERPINA1 PE=1 SV=3 | 642 | 20201114_qwl_colo_milk_1ng_glyco_hcd30_etd30_compound15.20021.20021.4 | controllerType=0 controllerNumber=1 scan=20021 | 55.7289 |
| 10320:4 | 12 | K.LKELPGVC[+57.021]N[+2205.793]ETMMALWEEC[+57.021]KPC[+57.021]LK.Q | HexNAc(4)Hex(5)Fuc(2)NeuAc(1) | C[+57]*3, N[+2206] | 1286.3043 | 4 | 5142.1955 | 5142.1776 | 0 | 3.5 | 95 | Specific | 408.54 | 319.67 | 319.67 | 4.03 | 15 | >sp\|P10909\|CLUS_HUMAN Clusterin OS=Homo sapiens OX=9606 GN=CLU PE=1 SV=1 | 1467 | 20201114_qwl_colo_milk_1ng_glyco_hcd30_etd30_compound15.13589.13589.4 | controllerType=0 controllerNumber=1 scan=13589 | 35.2994 |
| 10335:4 | 12 | K.LKELPGVC[+57.021]N[+2059.735]ETMMALWEEC[+57.021]KPC[+57.021]LK.Q | HexNAc(4)Hex(5)Fuc(1)NeuAc(1) | C[+57]*3, N[+2060] | 1249.7907 | 4 | 4996.1408 | 4996.1196 | 0 | 4.2 | 95 | Specific | 646.65 | 476.24 | 476.24 | 7.94 | 15 | >sp\|P10909\|CLUS_HUMAN Clusterin OS=Homo sapiens OX=9606 GN=CLU PE=1 SV=1 | 1467 | 20201114_qwl_colo_milk_1ng_glyco_hcd30_etd30_compound15.13605.13605.4 | controllerType=0 controllerNumber=1 scan=13605 | 35.3292 |
| 00134:4 | 12 | R.EIRHN[+2424.867]STGC[+57.021]LR.M | HexNAc(5)Hex(6)Fuc(1)NeuAc(1) | C[+57], N[+2425] | 942.6416 | 4 | 3767.5447 | 3767.5316 | 0 | 3.5 | 287 | Specific | 389.07 | 297.9 | 297.9 | 4.22 | 15 | >sp\|P10909\|CLUS_HUMAN Clusterin OS=Homo sapiens OX=9606 GN=CLU PE=1 SV=1 | 1467 | 20201114_qwl_colo_milk_1ng_glyco_hcd30_etd30_compound15.2919.2919.4 | controllerType=0 controllerNumber=1 scan=2919 | 11.3051 |
| 00118:4 | 12 | R.EIRHN[+2059.735]STGC[+57.021]LR.M | HexNAc(4)Hex(5)Fuc(1)NeuAc(1) | C[+57], N[+2060] | 851.357 | 4 | 3402.406 | 3402.3994 | 0 | 1.9 | 287 | Specific | 493.58 | 362.32 | 362.32 | 5.18 | 15 | >sp\|P10909\|CLUS_HUMAN Clusterin OS=Homo sapiens OX=9606 GN=CLU PE=1 SV=1 | 1467 | 20201114_qwl_colo_milk_1ng_glyco_hcd30_etd30_compound15.2901.2901.4 | controllerType=0 controllerNumber=1 scan=2901 | 11.2477 |
| 14710:3 | 12 | K.MLN[+2205.793]TSSLLEQLNEQFNWVSR.L | HexNAc(4)Hex(5)Fuc(2)NeuAc(1) | N[+2206] | 1539.0062 | 3 | 4615.0039 | 4614.9796 | 0 | 5.3 | 352 | Specific | 447.42 | 278.68 | 278.68 | 5.03 | 15 | >sp\|P10909\|CLUS_HUMAN Clusterin OS=Homo sapiens OX=9606 GN=CLU PE=1 SV=1 | 1467 | 20201114_qwl_colo_milk_1ng_glyco_hcd30_etd30_compound15.18253.18253.3 | controllerType=0 controllerNumber=1 scan=18253 | 49.7101 |
| 14724:3 | 12 | K.MLN[+2059.735]TSSLLEQLNEQFNWVSR.L | HexNAc(4)Hex(5)Fuc(1)NeuAc(1) | N[+2060] | 1490.3177 | 3 | 4468.9385 | 4468.9217 | 0 | 3.8 | 352 | Specific | 455.01 | 271.77 | 271.77 | 6.28 | 15 | >sp\|P10909\|CLUS_HUMAN Clusterin OS=Homo sapiens OX=9606 GN=CLU PE=1 SV=1 | 1467 | 20201114_qwl_colo_milk_1ng_glyco_hcd30_etd30_compound15.18268.18268.3 | controllerType=0 controllerNumber=1 scan=18268 | 49.7666 |
| 14676:3 | 12 | K.MLN[+2424.867]TSSLLEQLNEQFNWVSR.L | HexNAc(5)Hex(6)Fuc(1)NeuAc(1) | N[+2425] | 1612.0272 | 3 | 4834.0669 | 4834.0539 | 0 | 2.7 | 352 | Specific | 492.09 | 323.95 | 118.4 | 7.08 | 15 | >sp\|P10909\|CLUS_HUMAN Clusterin OS=Homo sapiens OX=9606 GN=CLU PE=1 SV=1 | 1467 | 20201114_qwl_colo_milk_1ng_glyco_hcd30_etd30_compound15.18216.18216.3 | controllerType=0 controllerNumber=1 scan=18216 | 49.5494 |
| 14767:3 | 12 | K.MLN[+1856.656]TSSLLEQLNEQFNWVSR.L | HexNAc(3)Hex(5)Fuc(1)NeuAc(1) | N[+1857] | 1422.6251 | 3 | 4265.8606 | 4265.8423 | 0 | 4.3 | 352 | Specific | 525.05 | 410.3 | 139.82 | 7.85 | 15 | >sp\|P10909\|CLUS_HUMAN Clusterin OS=Homo sapiens OX=9606 GN=CLU PE=1 SV=1 | 1467 | 20201114_qwl_colo_milk_1ng_glyco_hcd30_etd30_compound15.18313.18313.3 | controllerType=0 controllerNumber=1 scan=18313 | 49.9188 |
| 14867:3 | 12 | K.MLN[+1913.677]TSSLLEQLNEQFNWVSR.L | HexNAc(4)Hex(5)NeuAc(1) | N[+1914] | 1441.6365 | 3 | 4322.895 | 4322.8637 | 0 | 7.2 | 352 | Specific | 590.57 | 420.74 | 420.74 | 8.73 | 15 | >sp\|P10909\|CLUS_HUMAN Clusterin OS=Homo sapiens OX=9606 GN=CLU PE=1 SV=1 | 1467 | 20201114_qwl_colo_milk_1ng_glyco_hcd30_etd30_compound15.18420.18420.3 | controllerType=0 controllerNumber=1 scan=18420 | 50.2927 |
| 07747:3 | 12 | R.LAN[+2716.983]LTQGEDQYYLR.V | HexNAc(5)Hex(6)Fuc(3)NeuAc(1) | N[+2717] | 1467.6153 | 3 | 4400.8313 | 4400.8166 | 0 | 3.3 | 372 | Specific | 282.4 | 207.24 | 207.24 | 5.41 | 15 | >sp\|P10909\|CLUS_HUMAN Clusterin OS=Homo sapiens OX=9606 GN=CLU PE=1 SV=1 | 1467 | 20201114_qwl_colo_milk_1ng_glyco_hcd30_etd30_compound15.10881.10881.3 | controllerType=0 controllerNumber=1 scan=10881 | 28.2545 |
| 03030:2 | 16 | K.AAIPSALDTN[+1216.423]SSK.S | HexNAc(2)Hex(5) | N[+1216] | 1246.0467 | 2 | 2491.0861 | 2491.0815 | 0 | 1.8 | 542 | Specific | 263.17 | 89.64 | 89.64 | 2.84 | 5 | >sp\|Q08380\|LG3BP_HUMAN Galectin-3-binding protein OS=Homo sapiens OX=9606 GN=LGALS3BP PE=1 SV=1 | 1562 | 20201114_qwl_colo_milk_1ng_glyco_hcd30_etd30_compound15.5950.5950.2 | controllerType=0 controllerNumber=1 scan=5950 | 17.7339 |
| 01560:3 | 27 | R.LRN[+1216.423]VSWATGR.S | HexNAc(2)Hex(5) | N[+1216] | 792.6924 | 3 | 2376.0625 | 2376.0559 | 0 | 2.8 | 149 | Specific | 349.72 | 156.62 | 156.62 | 2.84 | 5 | >sp\|P08571\|CD14_HUMAN Monocyte differentiation antigen CD14 OS=Homo sapiens OX=9606 GN=CD14 PE=1 SV=2 | 1232 | 20201114_qwl_colo_milk_1ng_glyco_hcd30_etd30_compound15.4412.4412.3 | controllerType=0 controllerNumber=1 scan=4412 | 14.3518 |
| 01546:2 | 27 | R.N[+1216.423]VSWATGR.S | HexNAc(2)Hex(5) | N[+1216] | 1053.9421 | 2 | 2106.8768 | 2106.8708 | 0 | 2.9 | 151 | Specific | 226.05 | 103.78 | 103.78 | 2.89 | 5 | >sp\|P08571\|CD14_HUMAN Monocyte differentiation antigen CD14 OS=Homo sapiens OX=9606 GN=CD14 PE=1 SV=2 | 1232 | 20201114_qwl_colo_milk_1ng_glyco_hcd30_etd30_compound15.4397.4397.2 | controllerType=0 controllerNumber=1 scan=4397 | 14.3132 |
| 15793:4 | 1 | K.SC[+57.021]HTGLRRTAGWNVPIGTLRPFLN[+1540.529]WTGPPEPIEAAVAR.F | HexNAc(2)Hex(7) | C[+57], N[+1541] | 1435.4355 | 4 | 5738.7204 | 5738.7165 | 0 | 0.7 | 133 | Specific | 273.34 | 273.34 | 273.34 | 3.01 | 48 | >sp\|P02788\|TRFL_HUMAN Lactotransferrin OS=Homo sapiens OX=9606 GN=LTF PE=1 SV=6 | 1270 | 20201114_qwl_milk_1ng_glyco_hcd30_etd30_compound15.19298.19298.4 | controllerType=0 controllerNumber=1 scan=19298 | 43.5338 |
| 15735:4 | 1 | R.RTAGWNVPIGTLRPFLN[+2570.925]WTGPPEPIEAAVAR.F | HexNAc(5)Hex(6)Fuc(2)NeuAc(1) | N[+2571] | 1490.1945 | 4 | 5957.756 | 5957.7371 | 0 | 3.2 | 140 | Specific | 265.04 | 265.04 | 265.04 | 3.85 | 48 | >sp\|P02788\|TRFL_HUMAN Lactotransferrin OS=Homo sapiens OX=9606 GN=LTF PE=1 SV=6 | 1270 | 20201114_qwl_milk_1ng_glyco_hcd30_etd30_compound15.19237.19237.4 | controllerType=0 controllerNumber=1 scan=19237 | 43.3824 |
| 15773:4 | 1 | R.RTAGWNVPIGTLRPFLN[+2205.793]WTGPPEPIEAAVAR.F | HexNAc(4)Hex(5)Fuc(2)NeuAc(1) | N[+2206] | 1398.9086 | 4 | 5592.6125 | 5592.6049 | 0 | 1.4 | 140 | Specific | 314.66 | 314.66 | 314.66 | 4.92 | 48 | >sp\|P02788\|TRFL_HUMAN Lactotransferrin OS=Homo sapiens OX=9606 GN=LTF PE=1 SV=6 | 1270 | 20201114_qwl_milk_1ng_glyco_hcd30_etd30_compound15.19277.19277.4 | controllerType=0 controllerNumber=1 scan=19277 | 43.4811 |
| 15716:4 | 1 | R.RTAGWNVPIGTLRPFLN[+2716.983]WTGPPEPIEAAVAR.F | HexNAc(5)Hex(6)Fuc(3)NeuAc(1) | N[+2717] | 1526.7137 | 4 | 6103.8332 | 6103.795 | 0 | 6.3 | 140 | Specific | 441.4 | 441.4 | 441.4 | 6.82 | 48 | >sp\|P02788\|TRFL_HUMAN Lactotransferrin OS=Homo sapiens OX=9606 GN=LTF PE=1 SV=6 | 1270 | 20201114_qwl_milk_1ng_glyco_hcd30_etd30_compound15.19217.19217.4 | controllerType=0 controllerNumber=1 scan=19217 | 43.3238 |
| 15083:4 | 1 | R.RTAGWNVPIGTLRPFLN[+1768.640]WTGPPEPIEAAVAR.F | HexNAc(4)Hex(5)Fuc(1) | N[+1769] | 1289.6295 | 4 | 5155.4962 | 5155.4516 | 0 | 8.7 | 140 | Specific | 783.53 | 783.53 | 783.53 | 12.6 | 48 | >sp\|P02788\|TRFL_HUMAN Lactotransferrin OS=Homo sapiens OX=9606 GN=LTF PE=1 SV=6 | 1270 | 20201114_qwl_milk_1ng_glyco_hcd30_etd30_compound15.18546.18546.4 | controllerType=0 controllerNumber=1 scan=18546 | 41.4134 |
| 15810:4 | 1 | R.RTAGWNVPIGTLRPFLN[+2204.772]WTGPPEPIEAAVAR.F | HexNAc(4)Hex(5)NeuAc(2) | N[+2205] | 1398.6597 | 4 | 5591.6169 | 5591.5845 | 0 | 5.8 | 140 | Specific | 799.96 | 758.18 | 758.18 | 12.07 | 48 | >sp\|P02788\|TRFL_HUMAN Lactotransferrin OS=Homo sapiens OX=9606 GN=LTF PE=1 SV=6 | 1270 | 20201114_qwl_milk_1ng_glyco_hcd30_etd30_compound15.19316.19316.4 | controllerType=0 controllerNumber=1 scan=19316 | 43.5993 |
| 16631:3 | 1 | R.TAGWNVPIGTLRPFLN[+2351.851]WTGPPEPIEAAVAR.F | HexNAc(4)Hex(5)Fuc(3)NeuAc(1) | N[+2352] | 1861.5282 | 3 | 5582.5699 | 5582.5617 | 0 | 1.5 | 141 | Specific | 207.68 | 132.55 | 132.55 | 3.76 | 48 | >sp\|P02788\|TRFL_HUMAN Lactotransferrin OS=Homo sapiens OX=9606 GN=LTF PE=1 SV=6 | 1270 | 20201114_qwl_milk_1ng_glyco_hcd30_etd30_compound15.20187.20187.3 | controllerType=0 controllerNumber=1 scan=20187 | 46.2702 |
| 16543:3 | 1 | R.TAGWNVPIGTLRPFLN[+2716.983]WTGPPEPIEAAVAR.F | HexNAc(5)Hex(6)Fuc(3)NeuAc(1) | N[+2717] | 1983.2368 | 3 | 5947.6958 | 5947.6939 | 0 | 0.3 | 141 | Specific | 216.54 | 177.58 | 177.58 | 4.65 | 48 | >sp\|P02788\|TRFL_HUMAN Lactotransferrin OS=Homo sapiens OX=9606 GN=LTF PE=1 SV=6 | 1270 | 20201114_qwl_milk_1ng_glyco_hcd30_etd30_compound15.20093.20093.3 | controllerType=0 controllerNumber=1 scan=20093 | 45.9767 |
| 16689:3 | 1 | R.TAGWNVPIGTLRPFLN[+2205.793]WTGPPEPIEAAVAR.F | HexNAc(4)Hex(5)Fuc(2)NeuAc(1) | N[+2206] | 1812.8546 | 3 | 5436.5492 | 5436.5038 | 0 | 8.4 | 141 | Specific | 347.84 | 262.13 | 262.13 | 5.76 | 48 | >sp\|P02788\|TRFL_HUMAN Lactotransferrin OS=Homo sapiens OX=9606 GN=LTF PE=1 SV=6 | 1270 | 20201114_qwl_milk_1ng_glyco_hcd30_etd30_compound15.20247.20247.2 | controllerType=0 controllerNumber=1 scan=20247 | 46.3974 |
| 16603:3 | 1 | R.TAGWNVPIGTLRPFLN[+2570.925]WTGPPEPIEAAVAR.F | HexNAc(5)Hex(6)Fuc(2)NeuAc(1) | N[+2571] | 1934.5509 | 3 | 5801.6381 | 5801.636 | 0 | 0.4 | 141 | Specific | 377.69 | 308.06 | 308.06 | 5.87 | 48 | >sp\|P02788\|TRFL_HUMAN Lactotransferrin OS=Homo sapiens OX=9606 GN=LTF PE=1 SV=6 | 1270 | 20201114_qwl_milk_1ng_glyco_hcd30_etd30_compound15.20157.20157.3 | controllerType=0 controllerNumber=1 scan=20157 | 46.1799 |
| 15922:4 | 1 | R.TAGWNVPIGTLRPFLN[+2425.887]WTGPPEPIEAAVAR.F | HexNAc(5)Hex(6)Fuc(3) | N[+2426] | 1414.9153 | 4 | 5656.6394 | 5656.5985 | 0 | 7.2 | 141 | Specific | 388.43 | 298.79 | 298.79 | 3.72 | 48 | >sp\|P02788\|TRFL_HUMAN Lactotransferrin OS=Homo sapiens OX=9606 GN=LTF PE=1 SV=6 | 1270 | 20201114_qwl_milk_1ng_glyco_hcd30_etd30_compound15.19433.19433.4 | controllerType=0 controllerNumber=1 scan=19433 | 43.897 |
| 16650:4 | 1 | R.TAGWNVPIGTLRPFLN[+2205.793]WTGPPEPIEAAVAR.F | HexNAc(4)Hex(5)Fuc(2)NeuAc(1) | N[+2206] | 1359.8828 | 4 | 5436.5094 | 5436.5038 | 0 | 1 | 141 | Specific | 398.78 | 398.78 | 398.78 | 5.68 | 48 | >sp\|P02788\|TRFL_HUMAN Lactotransferrin OS=Homo sapiens OX=9606 GN=LTF PE=1 SV=6 | 1270 | 20201114_qwl_milk_1ng_glyco_hcd30_etd30_compound15.20207.20207.4 | controllerType=0 controllerNumber=1 scan=20207 | 46.3162 |
| 16630:4 | 1 | R.TAGWNVPIGTLRPFLN[+2351.851]WTGPPEPIEAAVAR.F | HexNAc(4)Hex(5)Fuc(3)NeuAc(1) | N[+2352] | 1396.3966 | 4 | 5582.5646 | 5582.5617 | 0 | 0.5 | 141 | Specific | 476.21 | 476.21 | 476.21 | 5.8 | 48 | >sp\|P02788\|TRFL_HUMAN Lactotransferrin OS=Homo sapiens OX=9606 GN=LTF PE=1 SV=6 | 1270 | 20201114_qwl_milk_1ng_glyco_hcd30_etd30_compound15.20186.20186.4 | controllerType=0 controllerNumber=1 scan=20186 | 46.2673 |
| 16635:3 | 1 | R.TAGWNVPIGTLRPFLN[+2059.735]WTGPPEPIEAAVAR.F | HexNAc(4)Hex(5)Fuc(1)NeuAc(1) | N[+2060] | 1764.1576 | 3 | 5290.4581 | 5290.4459 | 0 | 2.3 | 141 | Specific | 476.67 | 361.56 | 361.56 | 5.44 | 48 | >sp\|P02788\|TRFL_HUMAN Lactotransferrin OS=Homo sapiens OX=9606 GN=LTF PE=1 SV=6 | 1270 | 20201114_qwl_milk_1ng_glyco_hcd30_etd30_compound15.20191.20191.3 | controllerType=0 controllerNumber=1 scan=20191 | 46.2792 |
| 16674:3 | 1 | R.TAGWNVPIGTLRPFLN[+2059.735]WTGPPEPIEAAVAR.F | HexNAc(4)Hex(5)Fuc(1)NeuAc(1) | N[+2060] | 1764.1687 | 3 | 5290.4914 | 5290.4459 | 0 | 8.6 | 141 | Specific | 500.37 | 397.17 | 397.17 | 6.63 | 48 | >sp\|P02788\|TRFL_HUMAN Lactotransferrin OS=Homo sapiens OX=9606 GN=LTF PE=1 SV=6 | 1270 | 20201114_qwl_milk_1ng_glyco_hcd30_etd30_compound15.20232.20232.3 | controllerType=0 controllerNumber=1 scan=20232 | 46.365 |
| 16602:4 | 1 | R.TAGWNVPIGTLRPFLN[+2569.905]WTGPPEPIEAAVAR.F | HexNAc(5)Hex(6)NeuAc(2) | N[+2570] | 1450.9108 | 4 | 5800.6212 | 5800.6156 | 0 | 1 | 141 | Specific | 649.87 | 600.06 | 600.06 | 8.05 | 48 | >sp\|P02788\|TRFL_HUMAN Lactotransferrin OS=Homo sapiens OX=9606 GN=LTF PE=1 SV=6 | 1270 | 20201114_qwl_milk_1ng_glyco_hcd30_etd30_compound15.20156.20156.4 | controllerType=0 controllerNumber=1 scan=20156 | 46.173 |
| 15995:4 | 1 | R.TAGWNVPIGTLRPFLN[+1768.640]WTGPPEPIEAAVAR.F | HexNAc(4)Hex(5)Fuc(1) | N[+1769] | 1250.5945 | 4 | 4999.3561 | 4999.3504 | 0 | 1.1 | 141 | Specific | 668.13 | 668.13 | 668.13 | 9.86 | 48 | >sp\|P02788\|TRFL_HUMAN Lactotransferrin OS=Homo sapiens OX=9606 GN=LTF PE=1 SV=6 | 1270 | 20201114_qwl_milk_1ng_glyco_hcd30_etd30_compound15.19511.19511.4 | controllerType=0 controllerNumber=1 scan=19511 | 44.1358 |
| 14998:3 | 1 | R.TAGWNIPMGLLFN[+1622.582]QTGSC[+57.021]K.F | HexNAc(4)Hex(5) | C[+57], N[+1623] | 1239.873 | 3 | 3717.6044 | 3717.5916 | 0 | 3.5 | 485 | Specific | 234.61 | 154.24 | 154.24 | 3.02 | 48 | >sp\|P02788\|TRFL_HUMAN Lactotransferrin OS=Homo sapiens OX=9606 GN=LTF PE=1 SV=6 | 1270 | 20201114_qwl_milk_1ng_glyco_hcd30_etd30_compound15.18456.18456.3 | controllerType=0 controllerNumber=1 scan=18456 | 41.1413 |
| 14951:3 | 1 | R.TAGWNIPMGLLFN[+1606.587]QTGSC[+57.021]K.F | HexNAc(4)Hex(4)Fuc(1) | C[+57], N[+1607] | 1234.5425 | 3 | 3701.6131 | 3701.5966 | 0 | 4.4 | 485 | Specific | 289.49 | 268.43 | 268.43 | 3.48 | 48 | >sp\|P02788\|TRFL_HUMAN Lactotransferrin OS=Homo sapiens OX=9606 GN=LTF PE=1 SV=6 | 1270 | 20201114_qwl_milk_1ng_glyco_hcd30_etd30_compound15.18406.18406.3 | controllerType=0 controllerNumber=1 scan=18406 | 41.0041 |
| 14845:3 | 1 | R.TAGWNIPMGLLFN[+1768.640]QTGSC[+57.021]K.F | HexNAc(4)Hex(5)Fuc(1) | C[+57], N[+1769] | 1288.567 | 3 | 3863.6863 | 3863.6495 | 0 | 9.5 | 485 | Specific | 340.82 | 261.9 | 261.9 | 4.41 | 48 | >sp\|P02788\|TRFL_HUMAN Lactotransferrin OS=Homo sapiens OX=9606 GN=LTF PE=1 SV=6 | 1270 | 20201114_qwl_milk_1ng_glyco_hcd30_etd30_compound15.18295.18295.3 | controllerType=0 controllerNumber=1 scan=18295 | 40.735 |
| 16200:3 | 1 | R.TAGWNIPMGLLFN[+2205.793]QTGSC[+57.021]K.F | HexNAc(4)Hex(5)Fuc(2)NeuAc(1) | C[+57], N[+2206] | 1434.2848 | 3 | 4300.84 | 4300.8028 | 0 | 8.7 | 485 | Specific | 349.98 | 255.96 | 255.96 | 4.99 | 48 | >sp\|P02788\|TRFL_HUMAN Lactotransferrin OS=Homo sapiens OX=9606 GN=LTF PE=1 SV=6 | 1270 | 20201114_qwl_milk_1ng_glyco_hcd30_etd30_compound15.19730.19730.3 | controllerType=0 controllerNumber=1 scan=19730 | 44.8247 |
| 16301:3 | 1 | R.TAGWNIPMGLLFN[+1913.677]QTGSC[+57.021]K.F | HexNAc(4)Hex(5)NeuAc(1) | C[+57], N[+1914] | 1336.9018 | 3 | 4008.6908 | 4008.687 | 0 | 1 | 485 | Specific | 357.87 | 256.96 | 256.96 | 4.87 | 48 | >sp\|P02788\|TRFL_HUMAN Lactotransferrin OS=Homo sapiens OX=9606 GN=LTF PE=1 SV=6 | 1270 | 20201114_qwl_milk_1ng_glyco_hcd30_etd30_compound15.19838.19838.3 | controllerType=0 controllerNumber=1 scan=19838 | 45.2052 |
| 16209:3 | 1 | R.TAGWNIPMGLLFN[+2059.735]QTGSC[+57.021]K.F | HexNAc(4)Hex(5)Fuc(1)NeuAc(1) | C[+57], N[+2060] | 1385.5986 | 3 | 4154.7812 | 4154.7449 | 0 | 8.7 | 485 | Specific | 380.85 | 265.2 | 265.2 | 5.33 | 48 | >sp\|P02788\|TRFL_HUMAN Lactotransferrin OS=Homo sapiens OX=9606 GN=LTF PE=1 SV=6 | 1270 | 20201114_qwl_milk_1ng_glyco_hcd30_etd30_compound15.19739.19739.3 | controllerType=0 controllerNumber=1 scan=19739 | 44.8692 |
| 15723:4 | 1 | R.TAGWNIPMGLLFN[+1606.587]QTGSC[+57.021]KFDEYFSQSC[+57.021]APGSDPR.S | HexNAc(4)Hex(4)Fuc(1) | C[+57]*2, N[+1607] | 1387.0988 | 4 | 5545.3732 | 5545.3438 | 0 | 5.3 | 485 | Specific | 283.7 | 283.7 | 283.7 | 3.19 | 48 | >sp\|P02788\|TRFL_HUMAN Lactotransferrin OS=Homo sapiens OX=9606 GN=LTF PE=1 SV=6 | 1270 | 20201114_qwl_milk_1ng_glyco_hcd30_etd30_compound15.19224.19224.4 | controllerType=0 controllerNumber=1 scan=19224 | 43.3484 |
| 16493:4 | 1 | R.TAGWNIPMGLLFN[+2059.735]QTGSC[+57.021]KFDEYFSQSC[+57.021]APGSDPR.S | HexNAc(4)Hex(5)Fuc(1)NeuAc(1) | C[+57]*2, N[+2060] | 1500.3898 | 4 | 5998.5373 | 5998.492 | 0 | 7.5 | 485 | Specific | 312.91 | 290.37 | 290.37 | 3.81 | 48 | >sp\|P02788\|TRFL_HUMAN Lactotransferrin OS=Homo sapiens OX=9606 GN=LTF PE=1 SV=6 | 1270 | 20201114_qwl_milk_1ng_glyco_hcd30_etd30_compound15.20040.20040.2 | controllerType=0 controllerNumber=1 scan=20040 | 45.7874 |
| 16442:3 | 1 | R.TAGWNIPMGLLFN[+1913.677]QTGSC[+57.021]KFDEYFSQSC[+57.021]APGSDPR.S | HexNAc(4)Hex(5)NeuAc(1) | C[+57]*2, N[+1914] | 1951.5001 | 3 | 5852.4857 | 5852.4341 | 0 | 8.8 | 485 | Specific | 433.63 | 325.39 | 325.39 | 5.72 | 48 | >sp\|P02788\|TRFL_HUMAN Lactotransferrin OS=Homo sapiens OX=9606 GN=LTF PE=1 SV=6 | 1270 | 20201114_qwl_milk_1ng_glyco_hcd30_etd30_compound15.19986.19986.3 | controllerType=0 controllerNumber=1 scan=19986 | 45.6026 |
| 15711:4 | 1 | R.TAGWNIPMGLLFN[+1622.582]QTGSC[+57.021]KFDEYFSQSC[+57.021]APGSDPR.S | HexNAc(4)Hex(5) | C[+57]*2, N[+1623] | 1391.0994 | 4 | 5561.3756 | 5561.3387 | 0 | 6.6 | 485 | Specific | 467.33 | 370.96 | 149.32 | 5.12 | 48 | >sp\|P02788\|TRFL_HUMAN Lactotransferrin OS=Homo sapiens OX=9606 GN=LTF PE=1 SV=6 | 1270 | 20201114_qwl_milk_1ng_glyco_hcd30_etd30_compound15.19212.19212.4 | controllerType=0 controllerNumber=1 scan=19212 | 43.3116 |
| 15779:3 | 1 | R.TAGWNIPMGLLFN[+1216.423]QTGSC[+57.021]KFDEYFSQSC[+57.021]APGSDPR.S | HexNAc(2)Hex(5) | C[+57]*2, N[+1216] | 1719.0686 | 3 | 5155.1911 | 5155.18 | 0 | 2.2 | 485 | Specific | 468.22 | 387.44 | 162.1 | 7.23 | 48 | >sp\|P02788\|TRFL_HUMAN Lactotransferrin OS=Homo sapiens OX=9606 GN=LTF PE=1 SV=6 | 1270 | 20201114_qwl_milk_1ng_glyco_hcd30_etd30_compound15.19283.19283.3 | controllerType=0 controllerNumber=1 scan=19283 | 43.4975 |
| 15717:3 | 1 | R.TAGWNIPMGLLFN[+1606.587]QTGSC[+57.021]KFDEYFSQSC[+57.021]APGSDPR.S | HexNAc(4)Hex(4)Fuc(1) | C[+57]*2, N[+1607] | 1849.1296 | 3 | 5545.3743 | 5545.3438 | 0 | 5.5 | 485 | Specific | 510.72 | 435.33 | 435.33 | 6.83 | 48 | >sp\|P02788\|TRFL_HUMAN Lactotransferrin OS=Homo sapiens OX=9606 GN=LTF PE=1 SV=6 | 1270 | 20201114_qwl_milk_1ng_glyco_hcd30_etd30_compound15.19218.19218.3 | controllerType=0 controllerNumber=1 scan=19218 | 43.3309 |
| 15663:3 | 1 | R.TAGWNIPMGLLFN[+1768.640]QTGSC[+57.021]KFDEYFSQSC[+57.021]APGSDPR.S | HexNAc(4)Hex(5)Fuc(1) | C[+57]*2, N[+1769] | 1903.1547 | 3 | 5707.4497 | 5707.3966 | 0 | 9.3 | 485 | Specific | 564.58 | 459.35 | 459.35 | 8.52 | 48 | >sp\|P02788\|TRFL_HUMAN Lactotransferrin OS=Homo sapiens OX=9606 GN=LTF PE=1 SV=6 | 1270 | 20201114_qwl_milk_1ng_glyco_hcd30_etd30_compound15.19161.19161.3 | controllerType=0 controllerNumber=1 scan=19161 | 43.1682 |
| 16365:4 | 1 | R.TAGWNIPMGLLFN[+2205.793]QTGSC[+57.021]KFDEYFSQSC[+57.021]APGSDPR.S | HexNAc(4)Hex(5)Fuc(2)NeuAc(1) | C[+57]*2, N[+2206] | 1536.9065 | 4 | 6144.6042 | 6144.5499 | 0 | 8.8 | 485 | Specific | 735.21 | 707.98 | 707.98 | 10.9 | 48 | >sp\|P02788\|TRFL_HUMAN Lactotransferrin OS=Homo sapiens OX=9606 GN=LTF PE=1 SV=6 | 1270 | 20201114_qwl_milk_1ng_glyco_hcd30_etd30_compound15.19905.19905.4 | controllerType=0 controllerNumber=1 scan=19905 | 45.4068 |
| 16386:4 | 1 | R.TAGWNIPMGLLFN[+2059.735]QTGSC[+57.021]KFDEYFSQSC[+57.021]APGSDPR.S | HexNAc(4)Hex(5)Fuc(1)NeuAc(1) | C[+57]*2, N[+2060] | 1500.3814 | 4 | 5998.5036 | 5998.492 | 0 | 1.9 | 485 | Specific | 802.7 | 741.73 | 741.73 | 12.85 | 48 | >sp\|P02788\|TRFL_HUMAN Lactotransferrin OS=Homo sapiens OX=9606 GN=LTF PE=1 SV=6 | 1270 | 20201114_qwl_milk_1ng_glyco_hcd30_etd30_compound15.19928.19928.4 | controllerType=0 controllerNumber=1 scan=19928 | 45.4815 |
| 07351:3 | 1 | K.FGRN[+2205.793]GSDC[+57.021]PDKFC[+57.021]LFQSETK.N | HexNAc(4)Hex(5)Fuc(2)NeuAc(1) | C[+57]*2, N[+2206] | 1533.6287 | 3 | 4598.8716 | 4598.8577 | 0 | 3 | 639 | Specific | 255.87 | 183.34 | 183.34 | 3.46 | 48 | >sp\|P02788\|TRFL_HUMAN Lactotransferrin OS=Homo sapiens OX=9606 GN=LTF PE=1 SV=6 | 1270 | 20201114_qwl_milk_1ng_glyco_hcd30_etd30_compound15.10454.10454.3 | controllerType=0 controllerNumber=1 scan=10454 | 24.7405 |

Table S4. Cont.

| Query #:z | Protein Rank | Glycopeptide | Glycan | Modification Type(s) | Observed m/z | z | Observed (M+H) | Calc. mass (M+H) | Off-by-x error | Mass error (ppm) | StartPosition | Cleavage | Score | Delta | Delta Mod | \|Log Prob\| | # of unique peptides | Protein Name | Protein DB number | Comment | Scan # | Scan Time |
|---|---|---|---|---|---|---|---|---|---|---|---|---|---|---|---|---|---|---|---|---|---|---|
| 07431:4 | 1 | K.FGRN[+2059.735]GSDC[+57.021]PDKFC[+57.021]LFQSETK.N | HexNAc(4)Hex(5)Fuc(1)NeuAc(1) | C[+57]*2, N[+2060] | 1113.9587 | 4 | 4452.8128 | 4452.7998 | 0 | 2.9 | 639 | Specific | 569.13 | 461.27 | 461.27 | 8.22 | 48 | >sp\|P02788\|TRFL_HUMAN Lactotransferrin OS=Homo sapiens OX=9606 GN=LTF PE=1 SV=6 | 1270 | 20201114_qwl_milk_1ng_glyco_hcd30_etd30_compound15.10538.10538.4 | controllerType=0 controllerNumber=1 scan=10538 | 24.9005 |
| 07357:4 | 1 | K.FGRN[+2205.793]GSDC[+57.021]PDKFC[+57.021]LFQSETK.N | HexNAc(4)Hex(5)Fuc(2)NeuAc(1) | C[+57]*2, N[+2206] | 1150.4704 | 4 | 4598.8597 | 4598.8577 | 0 | 0.4 | 639 | Specific | 596.95 | 496.01 | 496.01 | 8.14 | 48 | >sp\|P02788\|TRFL_HUMAN Lactotransferrin OS=Homo sapiens OX=9606 GN=LTF PE=1 SV=6 | 1270 | 20201114_qwl_milk_1ng_glyco_hcd30_etd30_compound15.10461.10461.4 | controllerType=0 controllerNumber=1 scan=10461 | 24.7637 |
| 07340:4 | 1 | K.FGRN[+2205.793]GSDC[+57.021]PDKFC[+57.021]LFQSETK.N | HexNAc(4)Hex(5)Fuc(2)NeuAc(1) | C[+57]*2, N[+2206] | 1150.4726 | 4 | 4598.8685 | 4598.8577 | 0 | 2.3 | 639 | Specific | 619.43 | 450.98 | 450.98 | 9.22 | 48 | >sp\|P02788\|TRFL_HUMAN Lactotransferrin OS=Homo sapiens OX=9606 GN=LTF PE=1 SV=6 | 1270 | 20201114_qwl_milk_1ng_glyco_hcd30_etd30_compound15.10443.10443.4 | controllerType=0 controllerNumber=1 scan=10443 | 24.7195 |
| 18610:4 | 3 | K.VFPLSLDSTPQDGNVVVAC[+57.021]LVQGFFPQEPLSVTWSESGQN[+1809.666]VTAI | HexNAc(5)Hex(4)Fuc(1) | C[+57], N[+1810] | 1647.2577 | 4 | 6586.0089 | 6586.0275 | 0 | -2.8 | 8 | Specific | 371 | 371 | 371 | 5.11 | 26 | >sp\|P01877\|IGHA2_HUMAN Immunoglobulin heavy constant alpha 2 OS=Homo sapiens OX=9606 GN=IGHA2 PE=1 SV=4 | 576 | 20201114_qwl_milk_1ng_glyco_hcd30_etd30_compound15.22277.22277.4 | controllerType=0 controllerNumber=1 scan=22277 | 51.7457 |
| 18693:4 | 3 | K.VFPLSLDSTPQDGNVVVAC[+57.021]LVQGFFPQEPLSVTWSESGQN[+1809.666]VTAI | HexNAc(5)Hex(4)Fuc(1) | C[+57], N[+1810] | 1647.2766 | 4 | 6586.0846 | 6586.0275 | 0 | 8.7 | 8 | Specific | 487.46 | 487.46 | 487.46 | 6.53 | 26 | >sp\|P01877\|IGHA2_HUMAN Immunoglobulin heavy constant alpha 2 OS=Homo sapiens OX=9606 GN=IGHA2 PE=1 SV=4 | 576 | 20201114_qwl_milk_1ng_glyco_hcd30_etd30_compound15.22365.22365.4 | controllerType=0 controllerNumber=1 scan=22365 | 51.9868 |
| 18673:4 | 3 | K.VFPLSLDSTPQDGNVVVAC[+57.021]LVQGFFPQEPLSVTWSESGQN[+1647.613]VTAI | HexNAc(5)Hex(3)Fuc(1) | C[+57], N[+1648] | 1606.7515 | 4 | 6423.984 | 6423.9746 | 0 | 1.5 | 8 | Specific | 496.48 | 496.48 | 496.48 | 6.48 | 26 | >sp\|P01877\|IGHA2_HUMAN Immunoglobulin heavy constant alpha 2 OS=Homo sapiens OX=9606 GN=IGHA2 PE=1 SV=4 | 576 | 20201114_qwl_milk_1ng_glyco_hcd30_etd30_compound15.22344.22344.4 | controllerType=0 controllerNumber=1 scan=22344 | 51.9318 |
| 00948:3 | 3 | K.HYTN[+1809.666]SSQDVTVPC[+57.021]R.V | HexNAc(5)Hex(4)Fuc(1) | C[+57], N[+1810] | 1158.4792 | 3 | 3473.423 | 3473.4154 | 0 | 2.2 | 89 | Specific | 201.5 | 154.18 | 154.18 | 4.09 | 26 | >sp\|P01877\|IGHA2_HUMAN Immunoglobulin heavy constant alpha 2 OS=Homo sapiens OX=9606 GN=IGHA2 PE=1 SV=4 | 576 | 20201114_qwl_milk_1ng_glyco_hcd30_etd30_compound15.3802.3802.3 | controllerType=0 controllerNumber=1 scan=3802 | 12.9054 |
| 00963:3 | 3 | K.HYTN[+1647.613]SSQDVTVPC[+57.021]R.V | HexNAc(5)Hex(3)Fuc(1) | C[+57], N[+1648] | 1104.4598 | 3 | 3311.3648 | 3311.3625 | 0 | 0.7 | 89 | Specific | 434.79 | 375.87 | 375.87 | 6.28 | 26 | >sp\|P01877\|IGHA2_HUMAN Immunoglobulin heavy constant alpha 2 OS=Homo sapiens OX=9606 GN=IGHA2 PE=1 SV=4 | 576 | 20201114_qwl_milk_1ng_glyco_hcd30_etd30_compound15.3818.3818.3 | controllerType=0 controllerNumber=1 scan=3818 | 12.9475 |
| 02841:4 | 3 | K.HYTN[+1647.613]SSQDVTVPC[+57.021]RVPPPPPC[+57.021]C[+57.021]HPR.L | HexNAc(5)Hex(3)Fuc(1) | C[+57]*3, N[+1648] | 1152.2503 | 4 | 4605.9793 | 4605.9689 | 0 | 2.3 | 89 | Specific | 364.4 | 302.71 | 302.71 | 3.52 | 26 | >sp\|P01877\|IGHA2_HUMAN Immunoglobulin heavy constant alpha 2 OS=Homo sapiens OX=9606 GN=IGHA2 PE=1 SV=4 | 576 | 20201114_qwl_milk_1ng_glyco_hcd30_etd30_compound15.5770.5770.4 | controllerType=0 controllerNumber=1 scan=5770 | 16.4856 |
| 02844:5 | 3 | K.HYTN[+1647.613]SSQDVTVPC[+57.021]RVPPPPPC[+57.021]C[+57.021]HPR.L | HexNAc(5)Hex(3)Fuc(1) | C[+57]*3, N[+1648] | 922.0015 | 5 | 4605.9783 | 4605.9689 | 0 | 2.1 | 89 | Specific | 536.57 | 510 | 510 | 5.71 | 26 | >sp\|P01877\|IGHA2_HUMAN Immunoglobulin heavy constant alpha 2 OS=Homo sapiens OX=9606 GN=IGHA2 PE=1 SV=4 | 576 | 20201114_qwl_milk_1ng_glyco_hcd30_etd30_compound15.5773.5773.5 | controllerType=0 controllerNumber=1 scan=5773 | 16.4973 |
| 15454:3 | 3 | R.LSLHRPALEDLLLGSEAN[+1663.608]LTC[+57.021]TLTGLR.D | HexNAc(5)Hex(4) | C[+57], N[+1664] | 1543.0776 | 3 | 4627.2182 | 4627.2064 | 0 | 2.5 | 114 | Specific | 352.76 | 298.1 | 298.1 | 4.05 | 26 | >sp\|P01877\|IGHA2_HUMAN Immunoglobulin heavy constant alpha 2 OS=Homo sapiens OX=9606 GN=IGHA2 PE=1 SV=4 | 576 | 20201114_qwl_milk_1ng_glyco_hcd30_etd30_compound15.18939.18939.3 | controllerType=0 controllerNumber=1 scan=18939 | 42.5092 |
| 15448:4 | 3 | R.LSLHRPALEDLLLGSEAN[+1663.608]LTC[+57.021]TLTGLR.D | HexNAc(5)Hex(4) | C[+57], N[+1664] | 1157.5601 | 4 | 4627.2186 | 4627.2064 | 0 | 2.6 | 114 | Specific | 391.36 | 391.36 | 391.36 | 5.57 | 26 | >sp\|P01877\|IGHA2_HUMAN Immunoglobulin heavy constant alpha 2 OS=Homo sapiens OX=9606 GN=IGHA2 PE=1 SV=4 | 576 | 20201114_qwl_milk_1ng_glyco_hcd30_etd30_compound15.18933.18933.4 | controllerType=0 controllerNumber=1 scan=18933 | 42.49 |
| 15379:3 | 3 | R.LSLHRPALEDLLLGSEAN[+1702.581]LTC[+57.021]TLTGLR.D | HexNAc(2)Hex(8) | C[+57], N[+1703] | 1556.0684 | 3 | 4666.1907 | 4666.1796 | 0 | 2.4 | 114 | Specific | 467.45 | 435.65 | 435.65 | 6.33 | 26 | >sp\|P01877\|IGHA2_HUMAN Immunoglobulin heavy constant alpha 2 OS=Homo sapiens OX=9606 GN=IGHA2 PE=1 SV=4 | 576 | 20201114_qwl_milk_1ng_glyco_hcd30_etd30_compound15.18860.18860.3 | controllerType=0 controllerNumber=1 scan=18860 | 42.26 |
| 15449:3 | 3 | R.LSLHRPALEDLLLGSEAN[+1540.529]LTC[+57.021]TLTGLR.D | HexNAc(2)Hex(7) | C[+57], N[+1541] | 1502.0505 | 3 | 4504.1369 | 4504.1268 | 0 | 2.2 | 114 | Specific | 502.68 | 467.77 | 467.77 | 6.42 | 26 | >sp\|P01877\|IGHA2_HUMAN Immunoglobulin heavy constant alpha 2 OS=Homo sapiens OX=9606 GN=IGHA2 PE=1 SV=4 | 576 | 20201114_qwl_milk_1ng_glyco_hcd30_etd30_compound15.18934.18934.3 | controllerType=0 controllerNumber=1 scan=18934 | 42.493 |
| 16194:4 | 3 | R.LSLHRPALEDLLLGSEAN[+1710.598]LTC[+57.021]TLTGLR.D | HexNAc(3)Hex(5)NeuAc(1) | C[+57], N[+1711] | 1169.3072 | 4 | 4674.2069 | 4674.1959 | 0 | 2.3 | 114 | Specific | 800.52 | 776.16 | 776.16 | 13.11 | 26 | >sp\|P01877\|IGHA2_HUMAN Immunoglobulin heavy constant alpha 2 OS=Homo sapiens OX=9606 GN=IGHA2 PE=1 SV=4 | 576 | 20201114_qwl_milk_1ng_glyco_hcd30_etd30_compound15.19723.19723.4 | controllerType=0 controllerNumber=1 scan=19723 | 44.7809 |
| 15498:4 | 3 | R.LSLHRPALEDLLLGSEAN[+1298.476]LTC[+57.021]TLTGLR.D | HexNAc(4)Hex(3) | C[+57], N[+1298] | 1066.2841 | 4 | 4262.1146 | 4262.0742 | 0 | 9.5 | 114 | Specific | 889.26 | 889.26 | 889.26 | 14.15 | 26 | >sp\|P01877\|IGHA2_HUMAN Immunoglobulin heavy constant alpha 2 OS=Homo sapiens OX=9606 GN=IGHA2 PE=1 SV=4 | 576 | 20201114_qwl_milk_1ng_glyco_hcd30_etd30_compound15.18986.18986.4 | controllerType=0 controllerNumber=1 scan=18986 | 42.7049 |
| 15374:4 | 3 | R.LSLHRPALEDLLLGSEAN[+1702.581]LTC[+57.021]TLTGLR.D | HexNAc(2)Hex(8) | C[+57], N[+1703] | 1167.303 | 4 | 4666.1903 | 4666.1796 | 0 | 2.3 | 114 | Specific | 946.13 | 921.28 | 921.28 | 15.11 | 26 | >sp\|P01877\|IGHA2_HUMAN Immunoglobulin heavy constant alpha 2 OS=Homo sapiens OX=9606 GN=IGHA2 PE=1 SV=4 | 576 | 20201114_qwl_milk_1ng_glyco_hcd30_etd30_compound15.18855.18855.4 | controllerType=0 controllerNumber=1 scan=18855 | 42.24 |
| 15445:4 | 3 | R.LSLHRPALEDLLLGSEAN[+1540.529]LTC[+57.021]TLTGLR.D | HexNAc(2)Hex(7) | C[+57], N[+1541] | 1126.7895 | 4 | 4504.1361 | 4504.1268 | 0 | 2.1 | 114 | Specific | 1049 | 998.29 | 998.29 | 17.11 | 26 | >sp\|P01877\|IGHA2_HUMAN Immunoglobulin heavy constant alpha 2 OS=Homo sapiens OX=9606 GN=IGHA2 PE=1 SV=4 | 576 | 20201114_qwl_milk_1ng_glyco_hcd30_etd30_compound15.18930.18930.4 | controllerType=0 controllerNumber=1 scan=18930 | 42.4817 |
| 15513:4 | 3 | R.LSLHRPALEDLLLGSEAN[+1095.397]LTC[+57.021]TLTGLR.D | HexNAc(3)Hex(3) | C[+57], N[+1095] | 1015.5053 | 4 | 4058.9993 | 4058.9949 | 0 | 1.1 | 114 | Specific | 1060.62 | 1055.26 | 1055.26 | 18.04 | 26 | >sp\|P01877\|IGHA2_HUMAN Immunoglobulin heavy constant alpha 2 OS=Homo sapiens OX=9606 GN=IGHA2 PE=1 SV=4 | 576 | 20201114_qwl_milk_1ng_glyco_hcd30_etd30_compound15.19003.19003.4 | controllerType=0 controllerNumber=1 scan=19003 | 42.7376 |
| 15468:4 | 3 | R.LSLHRPALEDLLLGSEAN[+1216.423]LTC[+57.021]TLTGLR.D | HexNAc(2)Hex(5) | C[+57], N[+1216] | 1045.771 | 4 | 4180.0623 | 4180.0211 | 0 | 9.9 | 114 | Specific | 1187.3 | 1141.68 | 1141.68 | 20.15 | 26 | >sp\|P01877\|IGHA2_HUMAN Immunoglobulin heavy constant alpha 2 OS=Homo sapiens OX=9606 GN=IGHA2 PE=1 SV=4 | 576 | 20201114_qwl_milk_1ng_glyco_hcd30_etd30_compound15.18954.18954.4 | controllerType=0 controllerNumber=1 scan=18954 | 42.5497 |
| 11559:5 | 3 | K.SAVQGPPERDLC[+57.021]GC[+57.021]YSVSSVLPGC[+57.021]AQPWNHGETFTC[+ | HexNAc(5)Hex(3)Fuc(1) | C[+57]*4, N[+1648] | 1492.4795 | 5 | 7458.3686 | 7458.3591 | 0 | 1.3 | 156 | Specific | 240.15 | 240.15 | 240.15 | 2.56 | 26 | >sp\|P01877\|IGHA2_HUMAN Immunoglobulin heavy constant alpha 2 OS=Homo sapiens OX=9606 GN=IGHA2 PE=1 SV=4 | 576 | 20201114_qwl_milk_1ng_glyco_hcd30_etd30_compound15.14840.14840.5 | controllerType=0 controllerNumber=1 scan=14840 | 33.0439 |
| 11517:6 | 3 | K.SAVQGPPERDLC[+57.021]GC[+57.021]YSVSSVLPGC[+57.021]AQPWNHGETFTC[+ | HexNAc(5)Hex(4)Fuc(1) | C[+57]*4, N[+1810] | 1270.9136 | 6 | 7620.4454 | 7620.4119 | 0 | 4.4 | 156 | Specific | 738.14 | 738.14 | 738.14 | 10.03 | 26 | >sp\|P01877\|IGHA2_HUMAN Immunoglobulin heavy constant alpha 2 OS=Homo sapiens OX=9606 GN=IGHA2 PE=1 SV=4 | 576 | 20201114_qwl_milk_1ng_glyco_hcd30_etd30_compound15.14796.14796.6 | controllerType=0 controllerNumber=1 scan=14796 | 32.9391 |
| 12422:5 | 3 | R.DLC[+57.021]GC[+57.021]YSVSSVLPGC[+57.021]AQPWNHGETFTC[+57.021]TAAH | HexNAc(5)Hex(3)Fuc(1) | C[+57]*4, N[+1648] | 1308.1926 | 5 | 6536.934 | 6536.8923 | 0 | 6.4 | 165 | Specific | 203.32 | 203.32 | 203.32 | 2.71 | 26 | >sp\|P01877\|IGHA2_HUMAN Immunoglobulin heavy constant alpha 2 OS=Homo sapiens OX=9606 GN=IGHA2 PE=1 SV=4 | 576 | 20201114_qwl_milk_1ng_glyco_hcd30_etd30_compound15.15743.15743.5 | controllerType=0 controllerNumber=1 scan=15743 | 35.0228 |
| 02659:2 | 3 | K.TPLTAN[+1647.613]ITK.S | HexNAc(5)Hex(3)Fuc(1) | N[+1648] | 1303.5917 | 2 | 2606.1762 | 2606.17 | 0 | 2.4 | 200 | Specific | 223.4 | 74.63 | 74.63 | 2.86 | 26 | >sp\|P01877\|IGHA2_HUMAN Immunoglobulin heavy constant alpha 2 OS=Homo sapiens OX=9606 GN=IGHA2 PE=1 SV=4 | 576 | 20201114_qwl_milk_1ng_glyco_hcd30_etd30_compound15.5580.5580.2 | controllerType=0 controllerNumber=1 scan=5580 | 16.113 |
| 02677:2 | 3 | K.TPLTAN[+1501.555]ITK.S | HexNAc(5)Hex(3) | N[+1502] | 1230.5599 | 2 | 2460.1124 | 2460.1121 | 0 | 0.1 | 200 | Specific | 274.77 | 73.61 | 73.61 | 2.58 | 26 | >sp\|P01877\|IGHA2_HUMAN Immunoglobulin heavy constant alpha 2 OS=Homo sapiens OX=9606 GN=IGHA2 PE=1 SV=4 | 576 | 20201114_qwl_milk_1ng_glyco_hcd30_etd30_compound15.5599.5599.2 | controllerType=0 controllerNumber=1 scan=5599 | 16.1567 |
| 02615:2 | 3 | K.TPLTAN[+1809.666]ITK.S | HexNAc(5)Hex(4)Fuc(1) | N[+1810] | 1384.6182 | 2 | 2768.2291 | 2768.2228 | 0 | 2.3 | 200 | Specific | 291.43 | 110.98 | 110.98 | 2.86 | 26 | >sp\|P01877\|IGHA2_HUMAN Immunoglobulin heavy constant alpha 2 OS=Homo sapiens OX=9606 GN=IGHA2 PE=1 SV=4 | 576 | 20201114_qwl_milk_1ng_glyco_hcd30_etd30_compound15.5534.5534.2 | controllerType=0 controllerNumber=1 scan=5534 | 16.0068 |
| 09357:3 | 3 | R.MAGKPTHIN[+1702.581]VSVVMAEADGTC[+57.021]Y.- | HexNAc(2)Hex(8) | C[+57], N[+1703] | 1351.8956 | 3 | 4053.6721 | 4053.6642 | 0 | 2 | 319 | CRagged | 230.56 | 230.56 | 192.51 | 3.8 | 26 | >sp\|P01877\|IGHA2_HUMAN Immunoglobulin heavy constant alpha 2 OS=Homo sapiens OX=9606 GN=IGHA2 PE=1 SV=4 | 576 | 20201114_qwl_milk_1ng_glyco_hcd30_etd30_compound15.12541.12541.3 | controllerType=0 controllerNumber=1 scan=12541 | 28.4918 |
| 09379:3 | 3 | R.MAGKPTHIN[+1540.529]VSVVMAEADGTC[+57.021]Y.- | HexNAc(2)Hex(7) | C[+57], N[+1541] | 1297.8797 | 3 | 3891.6245 | 3891.6114 | 0 | 3.4 | 319 | CRagged | 236.11 | 236.11 | 171.66 | 3.38 | 26 | >sp\|P01877\|IGHA2_HUMAN Immunoglobulin heavy constant alpha 2 OS=Homo sapiens OX=9606 GN=IGHA2 PE=1 SV=4 | 576 | 20201114_qwl_milk_1ng_glyco_hcd30_etd30_compound15.12564.12564.3 | controllerType=0 controllerNumber=1 scan=12564 | 28.5462 |
| 09457:3 | 3 | R.MAGKPTHIN[+1216.423]VSVVMAEADGTC[+57.021]Y.- | HexNAc(2)Hex(5) | C[+57], N[+1216] | 1189.8428 | 3 | 3567.5139 | 3567.5057 | 0 | 2.3 | 319 | CRagged | 247.57 | 220.81 | 185.48 | 4.18 | 26 | >sp\|P01877\|IGHA2_HUMAN Immunoglobulin heavy constant alpha 2 OS=Homo sapiens OX=9606 GN=IGHA2 PE=1 SV=4 | 576 | 20201114_qwl_milk_1ng_glyco_hcd30_etd30_compound15.12645.12645.3 | controllerType=0 controllerNumber=1 scan=12645 | 28.7052 |
| 09396:3 | 3 | R.MAGKPTHIN[+1378.476]VSVVMAEADGTC[+57.021]Y.- | HexNAc(2)Hex(6) | C[+57], N[+1378] | 1243.8608 | 3 | 3729.5678 | 3729.5585 | 0 | 2.5 | 319 | CRagged | 277.18 | 277.18 | 207.79 | 4.5 | 26 | >sp\|P01877\|IGHA2_HUMAN Immunoglobulin heavy constant alpha 2 OS=Homo sapiens OX=9606 GN=IGHA2 PE=1 SV=4 | 576 | 20201114_qwl_milk_1ng_glyco_hcd30_etd30_compound15.12582.12582.3 | controllerType=0 controllerNumber=1 scan=12582 | 28.5939 |
| 10431:3 | 7 | R.LAGKPTHVN[+1647.613]VSVVMAEVDGTC[+57.021]Y.- | HexNAc(5)Hex(3)Fuc(1) | C[+57], N[+1648] | 1332.2579 | 3 | 3994.7591 | 3994.7553 | 0 | 0.9 | 332 | CRagged | 207.84 | 55.75 | 55.75 | 2.75 | 14 | >sp\|P01876\|IGHA1_HUMAN Immunoglobulin heavy constant alpha 1 OS=Homo sapiens OX=9606 GN=IGHA1 PE=1 SV=2 | 255 | 20201114_qwl_milk_1ng_glyco_hcd30_etd30_compound15.13659.13659.3 | controllerType=0 controllerNumber=1 scan=13659 | 30.699 |
| 10311:3 | 7 | R.LAGKPTHVN[+1540.529]VSVVMAEVDGTC[+57.021]Y.- | HexNAc(2)Hex(7) | C[+57], N[+1541] | 1296.5643 | 3 | 3887.6782 | 3887.6706 | 0 | 2 | 332 | CRagged | 237.21 | 224.44 | 182.51 | 3.8 | 14 | >sp\|P01876\|IGHA1_HUMAN Immunoglobulin heavy constant alpha 1 OS=Homo sapiens OX=9606 GN=IGHA1 PE=1 SV=2 | 255 | 20201114_qwl_milk_1ng_glyco_hcd30_etd30_compound15.13533.13533.3 | controllerType=0 controllerNumber=1 scan=13533 | 30.3566 |
| 10424:3 | 7 | R.LAGKPTHVN[+1501.555]VSVVMAEVDGTC[+57.021]Y.- | HexNAc(5)Hex(3) | C[+57], N[+1502] | 1283.574 | 3 | 3848.7075 | 3848.6974 | 0 | 2.6 | 332 | CRagged | 246.91 | 164.84 | 164.84 | 3.87 | 14 | >sp\|P01876\|IGHA1_HUMAN Immunoglobulin heavy constant alpha 1 OS=Homo sapiens OX=9606 GN=IGHA1 PE=1 SV=2 | 255 | 20201114_qwl_milk_1ng_glyco_hcd30_etd30_compound15.13651.13651.3 | controllerType=0 controllerNumber=1 scan=13651 | 30.6768 |
| 10353:3 | 7 | R.LAGKPTHVN[+1809.666]VSVVMAEVDGTC[+57.021]Y.- | HexNAc(5)Hex(4)Fuc(1) | C[+57], N[+1810] | 1386.2765 | 3 | 4156.8151 | 4156.8081 | 0 | 1.7 | 332 | CRagged | 262.16 | 155.69 | 155.69 | 3.87 | 14 | >sp\|P01876\|IGHA1_HUMAN Immunoglobulin heavy constant alpha 1 OS=Homo sapiens OX=9606 GN=IGHA1 PE=1 SV=2 | 255 | 20201114_qwl_milk_1ng_glyco_hcd30_etd30_compound15.13577.13577.3 | controllerType=0 controllerNumber=1 scan=13577 | 30.4743 |
| 10297:3 | 7 | R.LAGKPTHVN[+1702.581]VSVVMAEVDGTC[+57.021]Y.- | HexNAc(2)Hex(8) | C[+57], N[+1703] | 1350.5842 | 3 | 4049.7379 | 4049.7234 | 0 | 3.6 | 332 | CRagged | 263.2 | 250.92 | 190.75 | 3.72 | 14 | >sp\|P01876\|IGHA1_HUMAN Immunoglobulin heavy constant alpha 1 OS=Homo sapiens OX=9606 GN=IGHA1 PE=1 SV=2 | 255 | 20201114_qwl_milk_1ng_glyco_hcd30_etd30_compound15.13518.13518.3 | controllerType=0 controllerNumber=1 scan=13518 | 30.3265 |
| 10327:3 | 7 | R.LAGKPTHVN[+1378.476]VSVVMAEVDGTC[+57.021]Y.- | HexNAc(2)Hex(6) | C[+57], N[+1378] | 1242.5478 | 3 | 3725.6288 | 3725.6178 | 0 | 3 | 332 | CRagged | 286.91 | 225.09 | 206.62 | 4.19 | 14 | >sp\|P01876\|IGHA1_HUMAN Immunoglobulin heavy constant alpha 1 OS=Homo sapiens OX=9606 GN=IGHA1 PE=1 SV=2 | 255 | 20201114_qwl_milk_1ng_glyco_hcd30_etd30_compound15.13550.13550.3 | controllerType=0 controllerNumber=1 scan=13550 | 30.4158 |
| 10382:3 | 7 | R.LAGKPTHVN[+1216.423]VSVVMAEVDGTC[+57.021]Y.- | HexNAc(2)Hex(5) | C[+57], N[+1216] | 1188.5272 | 3 | 3563.5669 | 3563.5649 | 0 | 0.6 | 332 | CRagged | 312.02 | 281.87 | 205.45 | 4.33 | 14 | >sp\|P01876\|IGHA1_HUMAN Immunoglobulin heavy constant alpha 1 OS=Homo sapiens OX=9606 GN=IGHA1 PE=1 SV=2 | 255 | 20201114_qwl_milk_1ng_glyco_hcd30_etd30_compound15.13607.13607.3 | controllerType=0 controllerNumber=1 scan=13607 | 30.5695 |
| 06698:3 | 8 | R.FSDGLESN[+1864.634]SSTQFEVK.K | HexNAc(2)Hex(9) | N[+1865] | 1213.8229 | 3 | 3639.4542 | 3639.4472 | 0 | 1.9 | 219 | Specific | 244.04 | 244.04 | 244.04 | 4.11 | 8 | >sp\|P0C0L4\|CO4A_HUMAN Complement C4-A OS=Homo sapiens OX=9606 GN=C4A PE=1 SV=2 | 1235 | 20201114_qwl_milk_1ng_glyco_hcd30_etd30_compound15.9772.9772.3 | controllerType=0 controllerNumber=1 scan=9772 | 23.5396 |
| 04345:3 | 8 | R.FSDGLESN[+1864.634]SSTQFEVKK.Y | HexNAc(2)Hex(9) | N[+1865] | 1256.5229 | 3 | 3767.5541 | 3767.5421 | 0 | 3.2 | 219 | Specific | 203.58 | 118.14 | 118.14 | 3.71 | 8 | >sp\|P0C0L4\|CO4A_HUMAN Complement C4-A OS=Homo sapiens OX=9606 GN=C4A PE=1 SV=2 | 1235 | 20201114_qwl_milk_1ng_glyco_hcd30_etd30_compound15.7332.7332.3 | controllerType=0 controllerNumber=1 scan=7332 | 19.2811 |
| 19283:3 | 9 | R.LSLLEEPGN[+2351.851]GTFTVILNQLTSR.D | HexNAc(4)Hex(5)Fuc(3)NeuAc(1) | N[+2352] | 1585.3885 | 3 | 4754.1509 | 4754.1433 | 0 | 1.6 | 413 | Specific | 204.32 | 149.85 | 149.85 | 4 | 10 | >sp\|P01833\|PIGR_HUMAN Polymeric immunoglobulin receptor OS=Homo sapiens OX=9606 GN=PIGR PE=1 SV=4 | 902 | 20201114_qwl_milk_1ng_glyco_hcd30_etd30_compound15.22985.22985.3 | controllerType=0 controllerNumber=1 scan=22985 | 53.4451 |
| 17499:3 | 9 | R.LSLLEEPGN[+2205.793]GTFTVILNQLTSR.D | HexNAc(4)Hex(5)Fuc(2)NeuAc(1) | N[+2206] | 1536.704 | 3 | 4608.0976 | 4608.0854 | 0 | 2.6 | 413 | Specific | 519.96 | 375.89 | 375.89 | 6.84 | 10 | >sp\|P01833\|PIGR_HUMAN Polymeric immunoglobulin receptor OS=Homo sapiens OX=9606 GN=PIGR PE=1 SV=4 | 902 | 20201114_qwl_milk_1ng_glyco_hcd30_etd30_compound15.21106.21106.3 | controllerType=0 controllerNumber=1 scan=21106 | 48.8631 |
| 17446:3 | 9 | R.LSLLEEPGN[+2351.851]GTFTVILNQLTSR.D | HexNAc(4)Hex(5)Fuc(3)NeuAc(1) | N[+2352] | 1585.3838 | 3 | 4754.137 | 4754.1433 | 0 | -1.3 | 413 | Specific | 538.23 | 392.11 | 392.11 | 6.28 | 10 | >sp\|P01833\|PIGR_HUMAN Polymeric immunoglobulin receptor OS=Homo sapiens OX=9606 GN=PIGR PE=1 SV=4 | 902 | 20201114_qwl_milk_1ng_glyco_hcd30_etd30_compound15.21050.21050.3 | controllerType=0 controllerNumber=1 scan=21050 | 48.7141 |
| 14238:3 | 9 | K.IIEGEPNLKVPGN[+2205.793]VTAVLGETLK.V | HexNAc(4)Hex(5)Fuc(2)NeuAc(1) | N[+2206] | 1533.0543 | 3 | 4597.1482 | 4597.1422 | 0 | 1.3 | 457 | Specific | 359.75 | 246.63 | 246.63 | 3.61 | 10 | >sp\|P01833\|PIGR_HUMAN Polymeric immunoglobulin receptor OS=Homo sapiens OX=9606 GN=PIGR PE=1 SV=4 | 902 | 20201114_qwl_milk_1ng_glyco_hcd30_etd30_compound15.17655.17655.3 | controllerType=0 controllerNumber=1 scan=17655 | 39.255 |
| 15022:5 | 9 | K.IIEGEPNLKVPGN[+2350.830]VTAVLGETLKVPC[+57.021]HFPC[+57.021]K.F | HexNAc(4)Hex(5)Fuc(1)NeuAc(2) | C[+57]*2, N[+2351] | 1154.3447 | 5 | 5767.6945 | 5767.6372 | 0 | 9.9 | 457 | Specific | 267.07 | 267.07 | 267.07 | 2.95 | 10 | >sp\|P01833\|PIGR_HUMAN Polymeric immunoglobulin receptor OS=Homo sapiens OX=9606 GN=PIGR PE=1 SV=4 | 902 | 20201114_qwl_milk_1ng_glyco_hcd30_etd30_compound15.18481.18481.5 | controllerType=0 controllerNumber=1 scan=18481 | 41.2098 |
| 13838:4 | 9 | K.IIEGEPNLKVPGN[+2205.793]VTAVLGETLKVPC[+57.021]HFPC[+57.021]K.F | HexNAc(4)Hex(5)Fuc(2)NeuAc(1) | C[+57]*2, N[+2206] | 1406.4083 | 4 | 5622.6115 | 5622.5997 | 0 | 2.1 | 457 | Specific | 356.69 | 356.69 | 356.69 | 3.78 | 10 | >sp\|P01833\|PIGR_HUMAN Polymeric immunoglobulin receptor OS=Homo sapiens OX=9606 GN=PIGR PE=1 SV=4 | 902 | 20201114_qwl_milk_1ng_glyco_hcd30_etd30_compound15.17230.17230.4 | controllerType=0 controllerNumber=1 scan=17230 | 38.3345 |
| 13780:4 | 9 | K.IIEGEPNLKVPGN[+2205.793]VTAVLGETLKVPC[+57.021]HFPC[+57.021]K.F | HexNAc(4)Hex(5)Fuc(2)NeuAc(1) | C[+57]*2, N[+2206] | 1406.4087 | 4 | 5622.6129 | 5622.5997 | 0 | 2.4 | 457 | Specific | 457.98 | 457.98 | 457.98 | 5.2 | 10 | >sp\|P01833\|PIGR_HUMAN Polymeric immunoglobulin receptor OS=Homo sapiens OX=9606 GN=PIGR PE=1 SV=4 | 902 | 20201114_qwl_milk_1ng_glyco_hcd30_etd30_compound15.17169.17169.4 | controllerType=0 controllerNumber=1 scan=17169 | 38.1758 |
| 13837:5 | 9 | K.IIEGEPNLKVPGN[+2205.793]VTAVLGETLKVPC[+57.021]HFPC[+57.021]K.F | HexNAc(4)Hex(5)Fuc(2)NeuAc(1) | C[+57]*2, N[+2206] | 1125.327 | 5 | 5622.606 | 5622.5997 | 0 | 1.1 | 457 | Specific | 468.91 | 468.91 | 468.91 | 5.53 | 10 | >sp\|P01833\|PIGR_HUMAN Polymeric immunoglobulin receptor OS=Homo sapiens OX=9606 GN=PIGR PE=1 SV=4 | 902 | 20201114_qwl_milk_1ng_glyco_hcd30_etd30_compound15.17229.17229.5 | controllerType=0 controllerNumber=1 scan=17229 | 38.3218 |
| 09266:3 | 11 | K.TVLTPATNHMGN[+1378.476]VTFTIPANR.E | HexNAc(2)Hex(6) | N[+1378] | 1211.8842 | 3 | 3633.6381 | 3633.6358 | 0 | 0.6 | 74 | Specific | 444.39 | 347.65 | 161.95 | 6.24 | 5 | >sp\|P01024\|CO3_HUMAN Complement C3 OS=Homo sapiens OX=9606 GN=C3 PE=1 SV=2 | 492 | 20201114_qwl_milk_1ng_glyco_hcd30_etd30_compound15.12446.12446.3 | controllerType=0 controllerNumber=1 scan=12446 | 28.2812 |
| 11814:5 | 12 | K.KGN[+2205.793]TTLNSFVIPSESDVPTHC[+57.021]PSQWWPYAGHC[+57.021]YK.I | HexNAc(4)Hex(5)Fuc(2)NeuAc(1) | C[+57]*2, N[+2206] | 1234.733 | 5 | 6169.6361 | 6169.6146 | 0 | 3.5 | 342 | Specific | 361.58 | 354.91 | 354.91 | 4.47 | 5 | >sp\|P22897\|MRC1_HUMAN Macrophage mannose receptor 1 OS=Homo sapiens OX=9606 GN=MRC1 PE=1 SV=1 | 1049 | 20201114_qwl_milk_1ng_glyco_hcd30_etd30_compound15.15108.15108.5 | controllerType=0 controllerNumber=1 scan=15108 | 33.5819 |
| 12535:4 | 13 | K.LKELPGVC[+57.021]N[+2351.851]ETMMALWEEC[+57.021]KPC[+57.021]LK.Q | HexNAc(4)Hex(5)Fuc(3)NeuAc(1) | C[+57]*3, N[+2352] | 1322.8084 | 4 | 5288.2116 | 5288.2355 | 0 | -4.5 | 95 | Specific | 310.26 | 192.35 | 192.35 | 3.22 | 11 | >sp\|P10909\|CLUS_HUMAN Clusterin OS=Homo sapiens OX=9606 GN=CLU PE=1 SV=1 | 1467 | 20201114_qwl_milk_1ng_glyco_hcd30_etd30_compound15.15861.15861.4 | controllerType=0 controllerNumber=1 scan=15861 | 35.271 |
| 12595:4 | 13 | K.LKELPGVC[+57.021]N[+2059.735]ETMMALWEEC[+57.021]KPC[+57.021]LK.Q | HexNAc(4)Hex(5)Fuc(1)NeuAc(1) | C[+57]*3, N[+2060] | 1249.7874 | 4 | 4996.1276 | 4996.1196 | 0 | 1.6 | 95 | Specific | 384.51 | 269.51 | 269.51 | 3.93 | 11 | >sp\|P10909\|CLUS_HUMAN Clusterin OS=Homo sapiens OX=9606 GN=CLU PE=1 SV=1 | 1467 | 20201114_qwl_milk_1ng_glyco_hcd30_etd30_compound15.15924.15924.4 | controllerType=0 controllerNumber=1 scan=15924 | 35.4033 |
| 12507:4 | 13 | K.LKELPGVC[+57.021]N[+2351.851]ETMMALWEEC[+57.021]KPC[+57.021]LK.Q | HexNAc(4)Hex(5)Fuc(3)NeuAc(1) | C[+57]*3, N[+2352] | 1322.8263 | 4 | 5288.2834 | 5288.2355 | 0 | 9.1 | 95 | Specific | 432.8 | 313.72 | 313.72 | 3.59 | 11 | >sp\|P10909\|CLUS_HUMAN Clusterin OS=Homo sapiens OX=9606 GN=CLU PE=1 SV=1 | 1467 | 20201114_qwl_milk_1ng_glyco_hcd30_etd30_compound15.15832.15832.4 | controllerType=0 controllerNumber=1 scan=15832 | 35.1934 |
| 12532:4 | 13 | K.LKELPGVC[+57.021]N[+2205.793]ETMMALWEEC[+57.021]KPC[+57.021]LK.Q | HexNAc(4)Hex(5)Fuc(2)NeuAc(1) | C[+57]*3, N[+2206] | 1286.301 | 4 | 5142.1823 | 5142.1776 | 0 | 0.9 | 95 | Specific | 448.9 | 297.3 | 297.3 | 4.07 | 11 | >sp\|P10909\|CLUS_HUMAN Clusterin OS=Homo sapiens OX=9606 GN=CLU PE=1 SV=1 | 1467 | 20201114_qwl_milk_1ng_glyco_hcd30_etd30_compound15.15858.15858.4 | controllerType=0 controllerNumber=1 scan=15858 | 35.2615 |
| 00138:4 | 13 | R.EIRHN[+2059.735]STGC[+57.021]LR.M | HexNAc(4)Hex(5)Fuc(1)NeuAc(1) | C[+57], N[+2060] | 851.3576 | 4 | 3402.4084 | 3402.3994 | 0 | 2.7 | 287 | Specific | 605.38 | 466.15 | 466.15 | 8.74 | 11 | >sp\|P10909\|CLUS_HUMAN Clusterin OS=Homo sapiens OX=9606 GN=CLU PE=1 SV=1 | 1467 | 20201114_qwl_milk_1ng_glyco_hcd30_etd30_compound15.2957.2957.4 | controllerType=0 controllerNumber=1 scan=2957 | 11.2887 |
| 19662:3 | 13 | K.MLN[+2350.830]TSSLLEQLNEQFNWVSR.L | HexNAc(4)Hex(5)Fuc(1)NeuAc(2) | N[+2351] | 1587.3572 | 3 | 4760.0572 | 4760.0171 | 0 | 8.4 | 352 | Specific | 320.57 | 175.96 | 175.96 | 4.71 | 11 | >sp\|P10909\|CLUS_HUMAN Clusterin OS=Homo sapiens OX=9606 GN=CLU PE=1 SV=1 | 1467 | 20201114_qwl_milk_1ng_glyco_hcd30_etd30_compound15.23385.23385.3 | controllerType=0 controllerNumber=1 scan=23385 | 54.3843 |
| 17798:3 | 13 | K.MLN[+2059.735]TSSLLEQLNEQFNWVSR.L | HexNAc(4)Hex(5)Fuc(1)NeuAc(1) | N[+2060] | 1490.3183 | 3 | 4468.9403 | 4468.9217 | 0 | 4.2 | 352 | Specific | 467.54 | 306.58 | 306.58 | 5.07 | 11 | >sp\|P10909\|CLUS_HUMAN Clusterin OS=Homo sapiens OX=9606 GN=CLU PE=1 SV=1 | 1467 | 20201114_qwl_milk_1ng_glyco_hcd30_etd30_compound15.21422.21422.3 | controllerType=0 controllerNumber=1 scan=21422 | 49.7648 |
| 20204:3 | 14 | R.QLAHQSN[+2204.772]STNIFFSPVSIATAFAMLSLGTK.A | HexNAc(4)Hex(5)NeuAc(2) | N[+2205] | 1796.1447 | 3 | 5386.4197 | 5386.4075 | 0 | 2.3 | 64 | Specific | 482.19 | 346.38 | 346.38 | 6.32 | 3 | >sp\|P01009\|A1AT_HUMAN Alpha-1-antitrypsin OS=Homo sapiens OX=9606 GN=SERPINA1 PE=1 SV=3 | 642 | 20201114_qwl_milk_1ng_glyco_hcd30_etd30_compound15.23955.23955.3 | controllerType=0 controllerNumber=1 scan=23955 | 55.782 |
| 20203:4 | 14 | R.QLAHQSN[+2204.772]STNIFFSPVSIATAFAMLSLGTK.A | HexNAc(4)Hex(5)NeuAc(2) | N[+2205] | 1347.3608 | 4 | 5386.4215 | 5386.4075 | 0 | 2.6 | 64 | Specific | 701.46 | 594.86 | 340.31 | 10.93 | 3 | >sp\|P01009\|A1AT_HUMAN Alpha-1-antitrypsin OS=Homo sapiens OX=9606 GN=SERPINA1 PE=1 SV=3 | 642 | 20201114_qwl_milk_1ng_glyco_hcd30_etd30_compound15.23954.23954.4 | controllerType=0 controllerNumber=1 scan=23954 | 55.7743 |
| 10402:3 | 17 | R.IIVPLNNREN[+1856.656]ISDPTSPLR.T | HexNAc(3)Hex(5)Fuc(1)NeuAc(1) | N[+1857] | 1335.6181 | 3 | 4004.8397 | 4004.8327 | 0 | 1.8 | 62 | Specific | 237.08 | 215.51 | 215.51 | 3.64 | 8 | >sp\|P01591\|IGJ_HUMAN Immunoglobulin J chain OS=Homo sapiens OX=9606 GN=JCHAIN PE=1 SV=4 | 1392 | 20201114_qwl_milk_1ng_glyco_hcd30_etd30_compound15.13628.13628.3 | controllerType=0 controllerNumber=1 scan=13628 | 30.6245 |
| 08895:3 | 17 | R.IIVPLNNREN[+1663.608]ISDPTSPLR.T | HexNAc(5)Hex(4) | N[+1664] | 1271.268 | 3 | 3811.7895 | 3811.7853 | 0 | 1.1 | 62 | Specific | 267.75 | 190.27 | 190.27 | 3.92 | 8 | >sp\|P01591\|IGJ_HUMAN Immunoglobulin J chain OS=Homo sapiens OX=9606 GN=JCHAIN PE=1 SV=4 | 1392 | 20201114_qwl_milk_1ng_glyco_hcd30_etd30_compound15.12060.12060.3 | controllerType=0 controllerNumber=1 scan=12060 | 27.6137 |
| 08951:3 | 17 | R.IIVPLNNREN[+1501.555]ISDPTSPLR.T | HexNAc(5)Hex(3) | N[+1502] | 1217.2516 | 3 | 3649.7404 | 3649.7325 | 0 | 2.2 | 62 | Specific | 275.6 | 222.1 | 222.1 | 4.01 | 8 | >sp\|P01591\|IGJ_HUMAN Immunoglobulin J chain OS=Homo sapiens OX=9606 GN=JCHAIN PE=1 SV=4 | 1392 | 20201114_qwl_milk_1ng_glyco_hcd30_etd30_compound15.12118.12118.3 | controllerType=0 controllerNumber=1 scan=12118 | 27.7097 |
| 10314:3 | 17 | R.IIVPLNNREN[+1913.677]ISDPTSPLR.T | HexNAc(4)Hex(5)NeuAc(1) | N[+1914] | 1354.6252 | 3 | 4061.861 | 4061.8542 | 0 | 1.7 | 62 | Specific | 304.27 | 212.46 | 212.46 | 4.04 | 8 | >sp\|P01591\|IGJ_HUMAN Immunoglobulin J chain OS=Homo sapiens OX=9606 GN=JCHAIN PE=1 SV=4 | 1392 | 20201114_qwl_milk_1ng_glyco_hcd30_etd30_compound15.13536.13536.3 | controllerType=0 controllerNumber=1 scan=13536 | 30.3653 |
| 11971:3 | 17 | R.IIVPLNNREN[+2350.830]ISDPTSPLR.T | HexNAc(4)Hex(5)Fuc(1)NeuAc(2) | N[+2351] | 1500.3423 | 3 | 4499.0125 | 4499.0075 | 0 | 1.1 | 62 | Specific | 307.22 | 218.35 | 218.35 | 3.97 | 8 | >sp\|P01591\|IGJ_HUMAN Immunoglobulin J chain OS=Homo sapiens OX=9606 GN=JCHAIN PE=1 SV=4 | 1392 | 20201114_qwl_milk_1ng_glyco_hcd30_etd30_compound15.15272.15272.3 | controllerType=0 controllerNumber=1 scan=15272 | 33.9291 |
| 08861:3 | 17 | R.IIVPLNNREN[+1663.608]ISDPTSPLR.T | HexNAc(5)Hex(4) | N[+1664] | 1271.2684 | 3 | 3811.7905 | 3811.7853 | 0 | 1.4 | 62 | Specific | 319.46 | 247.26 | 247.26 | 4.02 | 8 | >sp\|P01591\|IGJ_HUMAN Immunoglobulin J chain OS=Homo sapiens OX=9606 GN=JCHAIN PE=1 SV=4 | 1392 | 20201114_qwl_milk_1ng_glyco_hcd30_etd30_compound15.12024.12024.3 | controllerType=0 controllerNumber=1 scan=12024 | 27.5347 |
| 00757:2 | 18 | R.SYN[+1038.375]DSVDPR.I | HexNAc(2)Hex(3)Fuc(1) | N[+1038] | 1045.9244 | 2 | 2090.8414 | 2090.8395 | 0 | 1 | 481 | Specific | 284.94 | 123.23 | 123.23 | 3.14 | 5 | >sp\|P05164\|PERM_HUMAN Myeloperoxidase OS=Homo sapiens OX=9606 GN=MPO PE=1 SV=1 | 12 | 20201114_qwl_milk_1ng_glyco_hcd30_etd30_compound15.3602.3602.2 | controllerType=0 controllerNumber=1 scan=3602 | [illegible] |
| 05093:3 | 20 | R.LSPN[+2351.851]ASAEHLELR.W | HexNAc(4)Hex(5)Fuc(3)NeuAc(1) | N[+2352] | 1263.5417 | 3 | 3788.6105 | 3788.6 | 0 | 2.8 | 52 | Specific | 227.45 | 137.56 | 137.56 | 3.17 | 7 | >sp\|Q13410\|BT1A1_HUMAN Butyrophilin subfamily 1 member A1 OS=Homo sapiens OX=9606 GN=BTN1A1 PE=1 SV=3 | 179 | 20201114_qwl_milk_1ng_glyco_hcd30_etd30_compound15.8107.8107.3 | controllerType=0 controllerNumber=1 scan=8107 | [illegible] |
| 05061:3 | 20 | R.LSPN[+2205.793]ASAEHLELR.W | HexNAc(4)Hex(5)Fuc(2)NeuAc(1) | N[+2206] | 1214.8542 | 3 | 3642.548 | 3642.5421 | 0 | 1.6 | 52 | Specific | 349.16 | 241.47 | 241.47 | 3.71 | 7 | >sp\|Q13410\|BT1A1_HUMAN Butyrophilin subfamily 1 member A1 OS=Homo sapiens OX=9606 GN=BTN1A1 PE=1 SV=3 | 179 | 20201114_qwl_milk_1ng_glyco_hcd30_etd30_compound15.8074.8074.3 | controllerType=0 controllerNumber=1 scan=8074 | 20.5654 |
| 07122:3 | 58 | K.IISKN[+2205.793]C[+57.021]TSYGVLDISK.C | HexNAc(4)Hex(5)Fuc(2)NeuAc(1) | C[+57], N[+2206] | 1335.2502 | 3 | 4003.736 | 4003.7344 | 0 | 0.4 | 317 | Specific | 233.38 | 154.29 | 154.29 | 2.84 | 2 | >sp\|P16671\|CD36_HUMAN Platelet glycoprotein 4 OS=Homo sapiens OX=9606 GN=CD36 PE=1 SV=2 | 812 | 20201114_qwl_milk_1ng_glyco_hcd30_etd30_compound15.10215.10215.3 | controllerType=0 controllerNumber=1 scan=10215 | 24.3143 |
| 15412:3 | 58 | R.NYIVPILWLN[+1864.634]ETGTIGDEKANMFR.S | HexNAc(2)Hex(9) | N[+1865] | 1553.6922 | 3 | 4659.062 | 4659.0574 | 0 | 1 | 408 | Specific | 583.4 | 547.97 | 547.97 | 7.86 | 2 | >sp\|P16671\|CD36_HUMAN Platelet glycoprotein 4 OS=Homo sapiens OX=9606 GN=CD36 PE=1 SV=2 | 812 | 20201114_qwl_milk_1ng_glyco_hcd30_etd30_compound15.18895.18895.3 | controllerType=0 controllerNumber=1 scan=18895 | 42.3647 |
| 00764:4 | 71 | K.TKPREEQFN[+1606.587]STFR.V | HexNAc(4)Hex(4)Fuc(1) | N[+1607] | 812.3581 | 4 | 3246.4106 | 3246.4054 | 0 | 1.6 | 168 | Specific | 334.52 | 306.91 | 306.91 | 3.73 | 2 | >sp\|P01859\|IGHG2_HUMAN Immunoglobulin heavy constant gamma 2 OS=Homo sapiens OX=9606 GN=IGHG2 PE=1 SV=2 | 387 | 20201114_qwl_milk_1ng_glyco_hcd30_etd30_compound15.3609.3609.4 | controllerType=0 controllerNumber=1 scan=3609 | 12.5316 |
| 00765:4 | 71 | K.TKPREEQFN[+1444.534]STFR.V | HexNAc(4)Hex(3)Fuc(1) | N[+1445] | 771.8446 | 4 | 3084.3567 | 3084.3526 | 0 | 1.3 | 168 | Specific | 393.82 | 356.76 | 356.76 | 4.14 | 2 | >sp\|P01859\|IGHG2_HUMAN Immunoglobulin heavy constant gamma 2 OS=Homo sapiens OX=9606 GN=IGHG2 PE=1 SV=2 | 387 | 20201114_qwl_milk_1ng_glyco_hcd30_etd30_compound15.3610.3610.4 | controllerType=0 controllerNumber=1 scan=3610 | 12.5392 |
| 17562:4 | 1 | R.RTAGWNVPIGTLRPFLN[+2350.830]WTGPPEPIEAAVAR.F | HexNAc(4)Hex(5)Fuc(1)NeuAc(2) | N[+2351] | 1435.1718 | 4 | 5737.6652 | 5737.6424 | 0 | 4 | 140 | Specific | 203.04 | 203.04 | 203.04 | 5.15 | 136 | >sp\|P02788\|TRFL_HUMAN Lactotransferrin OS=Homo sapiens OX=9606 GN=LTF PE=1 SV=6 | 1270 | 20201115_07413_1ng_glyco_hcd203040_compound15.21313.21313.4 | controllerType=0 controllerNumber=1 scan=21313 | 45.9698 |
| 17841:3 | 1 | R.[+42.011]RTAGWNVPIGTLRPFLN[+1864.634]WTGPPEPIEAAVAR.F | HexNAc(2)Hex(9) | .[42], N[+1865] | 1765.1597 | 3 | 5293.4645 | 5293.4568 | 0 | 1.5 | 140 | Specific | 319.15 | 319.15 | 319.15 | 6.6 | 136 | >sp\|P02788\|TRFL_HUMAN Lactotransferrin OS=Homo sapiens OX=9606 GN=LTF PE=1 SV=6 | 1270 | 20201115_07413_1ng_glyco_hcd203040_compound15.21611.21611.3 | controllerType=0 controllerNumber=1 scan=21611 | 46.5793 |
| 17844:3 | 1 | R.[+42.011]RTAGWNVPIGTLRPFLN[+1864.634]WTGPPEPIEAAVAR.F | HexNAc(2)Hex(9) | .[42], N[+1865] | 1765.1597 | 3 | 5293.4645 | 5293.4568 | 0 | 1.5 | 140 | Specific | 328.07 | 328.07 | 328.07 | 6.86 | 136 | >sp\|P02788\|TRFL_HUMAN Lactotransferrin OS=Homo sapiens OX=9606 GN=LTF PE=1 SV=6 | 1270 | 20201115_07413_1ng_glyco_hcd203040_compound15.21614.21614.3 | controllerType=0 controllerNumber=1 scan=21614 | 46.5852 |
| 17521:4 | 1 | R.TAGWNVPIGTLRPFLN[+2076.750]WTGPPEPIEAAVAR.F | HexNAc(4)Hex(6)Fuc(2) | N[+2077] | 1327.6155 | 4 | 5307.4401 | 5307.4612 | 0 | -4 | 141 | Specific | 202.99 | 198.51 | 198.51 | 2.26 | 136 | >sp\|P02788\|TRFL_HUMAN Lactotransferrin OS=Homo sapiens OX=9606 GN=LTF PE=1 SV=6 | 1270 | 20201115_07413_1ng_glyco_hcd203040_compound15.21271.21271.3 | controllerType=0 controllerNumber=1 scan=21271 | 45.8925 |
| 17524:4 | 1 | R.TAGWNVPIGTLRPFLN[+2076.750]WTGPPEPIEAAVAR.F | HexNAc(4)Hex(6)Fuc(2) | N[+2077] | 1327.6155 | 4 | 5307.4401 | 5307.4612 | 0 | -4 | 141 | Specific | 212.73 | 212.73 | 212.73 | 2.26 | 136 | >sp\|P02788\|TRFL_HUMAN Lactotransferrin OS=Homo sapiens OX=9606 GN=LTF PE=1 SV=6 | 1270 | 20201115_07413_1ng_glyco_hcd203040_compound15.21274.21274.3 | controllerType=0 controllerNumber=1 scan=21274 | 45.8979 |
| 16727:4 | 1 | R.[+42.011]TAGWNVPIGTLRPFLN[+1872.651]WTGPPEPIEAAVAR.F | HexNAc(3)Hex(6)NeuAc(1) | .[42], N[+1873] | 1287.1091 | 4 | 5145.4147 | 5145.372 | 0 | 8.3 | 141 | Specific | 241.2 | 197.92 | 197.92 | 5.41 | 136 | >sp\|P02788\|TRFL_HUMAN Lactotransferrin OS=Homo sapiens OX=9606 GN=LTF PE=1 SV=6 | 1270 | 20201115_07413_1ng_glyco_hcd203040_compound15.20437.20437.4 | controllerType=0 controllerNumber=1 scan=20437 | 44.0562 |
| 16719:3 | 1 | R.[+42.011]TAGWNVPIGTLRPFLN[+1872.651]WTGPPEPIEAAVAR.F | HexNAc(3)Hex(6)NeuAc(1) | .[42], N[+1873] | 1715.8104 | 3 | 5145.4166 | 5145.372 | 0 | 8.7 | 141 | Specific | 245.8 | 240.97 | 240.97 | 4.37 | 136 | >sp\|P02788\|TRFL_HUMAN Lactotransferrin OS=Homo sapiens OX=9606 GN=LTF PE=1 SV=6 | 1270 | 20201115_07413_1ng_glyco_hcd203040_compound15.20429.20429.3 | controllerType=0 controllerNumber=1 scan=20429 | 44.0392 |
| 16716:3 | 1 | R.[+42.011]TAGWNVPIGTLRPFLN[+1872.651]WTGPPEPIEAAVAR.F | HexNAc(3)Hex(6)NeuAc(1) | .[42], N[+1873] | 1715.8104 | 3 | 5145.4166 | 5145.372 | 0 | 8.7 | 141 | Specific | 257.02 | 223.19 | 223.19 | 5.82 | 136 | >sp\|P02788\|TRFL_HUMAN Lactotransferrin OS=Homo sapiens OX=9606 GN=LTF PE=1 SV=6 | 1270 | 20201115_07413_1ng_glyco_hcd203040_compound15.20426.20426.3 | controllerType=0 controllerNumber=1 scan=20426 | 44.0345 |
| 17579:3 | 1 | R.TAGWNVPIGTLRPFLN[+2716.983]WTGPPEPIEAAVAR.F | HexNAc(5)Hex(6)Fuc(3)NeuAc(1) | N[+2717] | 1983.2439 | 3 | 5947.717 | 5947.6939 | 0 | 3.9 | 141 | Specific | 271.09 | 265.12 | 265.12 | 6.76 | 136 | >sp\|P02788\|TRFL_HUMAN Lactotransferrin OS=Homo sapiens OX=9606 GN=LTF PE=1 SV=6 | 1270 | 20201115_07413_1ng_glyco_hcd203040_compound15.21331.21331.3 | controllerType=0 controllerNumber=1 scan=21331 | 46.0021 |
| 19012:4 | 1 | R.[+42.011]TAGWNVPIGTLRPFLN[+2336.851]WTGPPEPIEAAVAR.F | HexNAc(6)Hex(6)Fuc(1) | .[42], N[+2337] | 1403.1486 | 4 | 5609.5724 | 5609.5726 | 0 | 0 | 141 | Specific | 291.97 | 254.64 | 254.64 | 6.2 | 136 | >sp\|P02788\|TRFL_HUMAN Lactotransferrin OS=Homo sapiens OX=9606 GN=LTF PE=1 SV=6 | 1270 | 20201115_07413_1ng_glyco_hcd203040_compound15.22841.22841.4 | controllerType=0 controllerNumber=1 scan=22841 | 49.2899 |
| 18863:4 | 1 | R.TAGWNVPIGTLRPFLN[+2862.021]WTGPPEPIEAAVAR.F | HexNAc(5)Hex(6)Fuc(2)NeuAc(2) | N[+2862] | 1523.9522 | 4 | 6092.7868 | 6092.7314 | 0 | 9.1 | 141 | Specific | 327.69 | 327.54 | 327.54 | 6.71 | 136 | >sp\|P02788\|TRFL_HUMAN Lactotransferrin OS=Homo sapiens OX=9606 GN=LTF PE=1 SV=6 | 1270 | 20201115_07413_1ng_glyco_hcd203040_compound15.22684.22684.4 | controllerType=0 controllerNumber=1 scan=22684 | 48.9088 |
| 19009:4 | 1 | R.[+42.011]TAGWNVPIGTLRPFLN[+2336.851]WTGPPEPIEAAVAR.F | HexNAc(6)Hex(6)Fuc(1) | .[42], N[+2337] | 1403.1486 | 4 | 5609.5724 | 5609.5726 | 0 | 0 | 141 | Specific | 355.93 | 355.93 | 355.93 | 7.03 | 136 | >sp\|P02788\|TRFL_HUMAN Lactotransferrin OS=Homo sapiens OX=9606 GN=LTF PE=1 SV=6 | 1270 | 20201115_07413_1ng_glyco_hcd203040_compound15.22838.22838.4 | controllerType=0 controllerNumber=1 scan=22838 | 49.2847 |
| 17557:3 | 1 | R.TAGWNVPIGTLRPFLN[+2716.983]WTGPPEPIEAAVAR.F | HexNAc(5)Hex(6)Fuc(3)NeuAc(1) | N[+2717] | 1983.2435 | 3 | 5947.7159 | 5947.6939 | 0 | 3.7 | 141 | Specific | 391.98 | 348.59 | 348.59 | 7.69 | 136 | >sp\|P02788\|TRFL_HUMAN Lactotransferrin OS=Homo sapiens OX=9606 GN=LTF PE=1 SV=6 | 1270 | 20201115_07413_1ng_glyco_hcd203040_compound15.21308.21308.3 | controllerType=0 controllerNumber=1 scan=21308 | 45.9594 |
| 18996:3 | 1 | R.TAGWNVPIGTLRPFLN[+2350.830]WTGPPEPIEAAVAR.F | HexNAc(4)Hex(5)Fuc(1)NeuAc(2) | N[+2351] | 1861.1887 | 3 | 5581.5515 | 5581.5413 | 0 | 1.8 | 141 | Specific | 405.42 | 256.26 | 256.26 | 7.2 | 136 | >sp\|P02788\|TRFL_HUMAN Lactotransferrin OS=Homo sapiens OX=9606 GN=LTF PE=1 SV=6 | 1270 | 20201115_07413_1ng_glyco_hcd203040_compound15.22824.22824.2 | controllerType=0 controllerNumber=1 scan=22824 | 49.2547 |
| 18865:4 | 1 | R.TAGWNVPIGTLRPFLN[+2862.021]WTGPPEPIEAAVAR.F | HexNAc(5)Hex(6)Fuc(2)NeuAc(2) | N[+2862] | 1523.9522 | 4 | 6092.7868 | 6092.7314 | 0 | 9.1 | 141 | Specific | 417.21 | 309.57 | 309.57 | 7.02 | 136 | >sp\|P02788\|TRFL_HUMAN Lactotransferrin OS=Homo sapiens OX=9606 GN=LTF PE=1 SV=6 | 1270 | 20201115_07413_1ng_glyco_hcd203040_compound15.22686.22686.4 | controllerType=0 controllerNumber=1 scan=22686 | 48.9135 |
| 17713:5 | 1 | R.TAGWNVPIGTLRPFLN[+2059.735]WTGPPEPIEAAVAR.F | HexNAc(4)Hex(5)Fuc(1)NeuAc(1) | N[+2060] | 1058.9 | 5 | 5290.471 | 5290.4459 | 0 | 4.8 | 141 | Specific | 436.09 | 358.8 | 358.8 | 7.67 | 136 | >sp\|P02788\|TRFL_HUMAN Lactotransferrin OS=Homo sapiens OX=9606 GN=LTF PE=1 SV=6 | 1270 | 20201115_07413_1ng_glyco_hcd203040_compound15.21472.21472.5 | controllerType=0 controllerNumber=1 scan=21472 | 46.3289 |
| 17591:4 | 1 | R.TAGWNVPIGTLRPFLN[+2570.925]WTGPPEPIEAAVAR.F | HexNAc(5)Hex(6)Fuc(2)NeuAc(1) | N[+2571] | 1451.1671 | 4 | 5801.6466 | 5801.636 | 0 | 1.8 | 141 | Specific | 458.74 | 344.94 | 344.94 | 7.6 | 136 | >sp\|P02788\|TRFL_HUMAN Lactotransferrin OS=Homo sapiens OX=9606 GN=LTF PE=1 SV=6 | 1270 | 20201115_07413_1ng_glyco_hcd203040_compound15.21344.21344.4 | controllerType=0 controllerNumber=1 scan=21344 | 46.0284 |
| 17901:3 | 1 | R.TAGWNVPIGTLRPFLN[+2059.735]WTGPPEPIEAAVAR.F | HexNAc(4)Hex(5)Fuc(1)NeuAc(1) | N[+2060] | 1764.1584 | 3 | 5290.4607 | 5290.4459 | 0 | 2.8 | 141 | Specific | 476.66 | 420.11 | 420.11 | 9.27 | 136 | >sp\|P02788\|TRFL_HUMAN Lactotransferrin OS=Homo sapiens OX=9606 GN=LTF PE=1 SV=6 | 1270 | 20201115_07413_1ng_glyco_hcd203040_compound15.21675.21675.3 | controllerType=0 controllerNumber=1 scan=21675 | 46.7238 |
| 17564:4 | 1 | R.TAGWNVPIGTLRPFLN[+2715.963]WTGPPEPIEAAVAR.F | HexNAc(5)Hex(6)Fuc(1)NeuAc(2) | N[+2716] | 1487.4369 | 4 | 5946.7257 | 5946.6735 | 0 | 8.8 | 141 | Specific | 498.69 | 465.34 | 465.34 | 8.72 | 136 | >sp\|P02788\|TRFL_HUMAN Lactotransferrin OS=Homo sapiens OX=9606 GN=LTF PE=1 SV=6 | 1270 | 20201115_07413_1ng_glyco_hcd203040_compound15.21316.21316.4 | controllerType=0 controllerNumber=1 scan=21316 | 45.9757 |
| 16671:4 | 1 | R.TAGWNVPIGTLRPFLN[+2279.830]WTGPPEPIEAAVAR.F | HexNAc(5)Hex(6)Fuc(2) | N[+2280] | 1378.3992 | 4 | 5510.5749 | 5510.5406 | 0 | 6.2 | 141 | Specific | 531.85 | 509.21 | 509.21 | 8.84 | 136 | >sp\|P02788\|TRFL_HUMAN Lactotransferrin OS=Homo sapiens OX=9606 GN=LTF PE=1 SV=6 | 1270 | 20201115_07413_1ng_glyco_hcd203040_compound15.20378.20378.4 | controllerType=0 controllerNumber=1 scan=20378 | 43.9031 |
| 18242:4 | 1 | R.TAGWNVPIGTLRPFLN[+2059.735]WTGPPEPIEAAVAR.F | HexNAc(4)Hex(5)Fuc(1)NeuAc(1) | N[+2060] | 1323.3711 | 4 | 5290.4626 | 5290.4459 | 0 | 3.2 | 141 | Specific | 561.46 | 561.46 | 561.46 | 9.51 | 136 | >sp\|P02788\|TRFL_HUMAN Lactotransferrin OS=Homo sapiens OX=9606 GN=LTF PE=1 SV=6 | 1270 | 20201115_07413_1ng_glyco_hcd203040_compound15.22032.22032.2 | controllerType=0 controllerNumber=1 scan=22032 | 47.4564 |
| 18392:4 | 1 | R.TAGWNVPIGTLRPFLN[+2059.735]WTGPPEPIEAAVAR.F | HexNAc(4)Hex(5)Fuc(1)NeuAc(1) | N[+2060] | 1323.3665 | 4 | 5290.444 | 5290.4459 | 0 | -0.4 | 141 | Specific | 577.07 | 538.44 | 538.44 | 9.88 | 136 | >sp\|P02788\|TRFL_HUMAN Lactotransferrin OS=Homo sapiens OX=9606 GN=LTF PE=1 SV=6 | 1270 | 20201115_07413_1ng_glyco_hcd203040_compound15.22189.22189.4 | controllerType=0 controllerNumber=1 scan=22189 | 47.7852 |
| 18393:4 | 1 | R.TAGWNVPIGTLRPFLN[+2059.735]WTGPPEPIEAAVAR.F | HexNAc(4)Hex(5)Fuc(1)NeuAc(1) | N[+2060] | 1323.3665 | 4 | 5290.444 | 5290.4459 | 0 | -0.4 | 141 | Specific | 592.23 | 580.28 | 580.28 | 9.88 | 136 | >sp\|P02788\|TRFL_HUMAN Lactotransferrin OS=Homo sapiens OX=9606 GN=LTF PE=1 SV=6 | 1270 | 20201115_07413_1ng_glyco_hcd203040_compound15.22190.22190.4 | controllerType=0 controllerNumber=1 scan=22190 | 47.7892 |
| 17566:4 | 1 | R.TAGWNVPIGTLRPFLN[+2715.963]WTGPPEPIEAAVAR.F | HexNAc(5)Hex(6)Fuc(1)NeuAc(2) | N[+2716] | 1487.4369 | 4 | 5946.7257 | 5946.6735 | 0 | 8.8 | 141 | Specific | 605.32 | 493.9 | 493.9 | 11.19 | 136 | >sp\|P02788\|TRFL_HUMAN Lactotransferrin OS=Homo sapiens OX=9606 GN=LTF PE=1 SV=6 | 1270 | 20201115_07413_1ng_glyco_hcd203040_compound15.21318.21318.4 | controllerType=0 controllerNumber=1 scan=21318 | 45.9819 |
| 17559:3 | 1 | R.TAGWNVPIGTLRPFLN[+2716.983]WTGPPEPIEAAVAR.F | HexNAc(5)Hex(6)Fuc(3)NeuAc(1) | N[+2717] | 1983.2435 | 3 | 5947.7159 | 5947.6939 | 0 | 3.7 | 141 | Specific | 619.98 | 564.44 | 564.44 | 11.95 | 136 | >sp\|P02788\|TRFL_HUMAN Lactotransferrin OS=Homo sapiens OX=9606 GN=LTF PE=1 SV=6 | 1270 | 20201115_07413_1ng_glyco_hcd203040_compound15.21310.21310.3 | controllerType=0 controllerNumber=1 scan=21310 | 45.9652 |
| 17728:3 | 1 | R.TAGWNVPIGTLRPFLN[+2205.793]WTGPPEPIEAAVAR.F | HexNAc(4)Hex(5)Fuc(2)NeuAc(1) | N[+2206] | 1812.8429 | 3 | 5436.514 | 5436.5038 | 0 | 1.9 | 141 | Specific | 635.49 | 586.24 | 586.24 | 11.96 | 136 | >sp\|P02788\|TRFL_HUMAN Lactotransferrin OS=Homo sapiens OX=9606 GN=LTF PE=1 SV=6 | 1270 | 20201115_07413_1ng_glyco_hcd203040_compound15.21488.21488.2 | controllerType=0 controllerNumber=1 scan=21488 | 46.377 |
| 17711:5 | 1 | R.TAGWNVPIGTLRPFLN[+2059.735]WTGPPEPIEAAVAR.F | HexNAc(4)Hex(5)Fuc(1)NeuAc(1) | N[+2060] | 1058.9 | 5 | 5290.471 | 5290.4459 | 0 | 4.8 | 141 | Specific | 645.15 | 569.79 | 569.79 | 11.72 | 136 | >sp\|P02788\|TRFL_HUMAN Lactotransferrin OS=Homo sapiens OX=9606 GN=LTF PE=1 SV=6 | 1270 | 20201115_07413_1ng_glyco_hcd203040_compound15.21470.21470.5 | controllerType=0 controllerNumber=1 scan=21470 | 46.321 |
| 18994:3 | 1 | R.TAGWNVPIGTLRPFLN[+2350.830]WTGPPEPIEAAVAR.F | HexNAc(4)Hex(5)Fuc(1)NeuAc(2) | N[+2351] | 1861.1887 | 3 | 5581.5515 | 5581.5413 | 0 | 1.8 | 141 | Specific | 675.72 | 515.47 | 515.47 | 12.96 | 136 | >sp\|P02788\|TRFL_HUMAN Lactotransferrin OS=Homo sapiens OX=9606 GN=LTF PE=1 SV=6 | 1270 | 20201115_07413_1ng_glyco_hcd203040_compound15.22822.22822.2 | controllerType=0 controllerNumber=1 scan=22822 | 49.2453 |
| 16673:4 | 1 | R.TAGWNVPIGTLRPFLN[+2279.830]WTGPPEPIEAAVAR.F | HexNAc(5)Hex(6)Fuc(2) | N[+2280] | 1378.3992 | 4 | 5510.5749 | 5510.5406 | 0 | 6.2 | 141 | Specific | 706.36 | 623.87 | 623.87 | 13.19 | 136 | >sp\|P02788\|TRFL_HUMAN Lactotransferrin OS=Homo sapiens OX=9606 GN=LTF PE=1 SV=6 | 1270 | 20201115_07413_1ng_glyco_hcd203040_compound15.20380.20380.4 | controllerType=0 controllerNumber=1 scan=20380 | 43.9087 |
| 17731:3 | 1 | R.TAGWNVPIGTLRPFLN[+2205.793]WTGPPEPIEAAVAR.F | HexNAc(4)Hex(5)Fuc(2)NeuAc(1) | N[+2206] | 1812.8429 | 3 | 5436.514 | 5436.5038 | 0 | 1.9 | 141 | Specific | 757.79 | 635.53 | 635.53 | 14.96 | 136 | >sp\|P02788\|TRFL_HUMAN Lactotransferrin OS=Homo sapiens OX=9606 GN=LTF PE=1 SV=6 | 1270 | 20201115_07413_1ng_glyco_hcd203040_compound15.21491.21491.2 | controllerType=0 controllerNumber=1 scan=21491 | 46.3842 |
| 17594:4 | 1 | R.TAGWNVPIGTLRPFLN[+2570.925]WTGPPEPIEAAVAR.F | HexNAc(5)Hex(6)Fuc(2)NeuAc(1) | N[+2571] | 1451.1671 | 4 | 5801.6466 | 5801.636 | 0 | 1.8 | 141 | Specific | 874.5 | 777.92 | 777.92 | 16.69 | 136 | >sp\|P02788\|TRFL_HUMAN Lactotransferrin OS=Homo sapiens OX=9606 GN=LTF PE=1 SV=6 | 1270 | 20201115_07413_1ng_glyco_hcd203040_compound15.21347.21347.4 | controllerType=0 controllerNumber=1 scan=21347 | 46.0382 |
| 18244:4 | 1 | R.TAGWNVPIGTLRPFLN[+2059.735]WTGPPEPIEAAVAR.F | HexNAc(4)Hex(5)Fuc(1)NeuAc(1) | N[+2060] | 1323.3711 | 4 | 5290.4626 | 5290.4459 | 0 | 3.2 | 141 | Specific | 911.32 | 830.39 | 830.39 | 17.67 | 136 | >sp\|P02788\|TRFL_HUMAN Lactotransferrin OS=Homo sapiens OX=9606 GN=LTF PE=1 SV=6 | 1270 | 20201115_07413_1ng_glyco_hcd203040_compound15.22034.22034.2 | controllerType=0 controllerNumber=1 scan=22034 | 47.4606 |
| 15968:3 | 1 | R.[+42.011]PFLN[+2018.708]WTGPPEPIEAAVAR.F | HexNAc(3)Hex(6)Fuc(1)NeuAc(1) | .[42], N[+2019] | 1342.5887 | 3 | 4025.7515 | 4025.7418 | 0 | 2.4 | 153 | Specific | 297.06 | 214.38 | 214.38 | 5.11 | 136 | >sp\|P02788\|TRFL_HUMAN Lactotransferrin OS=Homo sapiens OX=9606 GN=LTF PE=1 SV=6 | 1270 | 20201115_07413_1ng_glyco_hcd203040_compound15.19643.19643.3 | controllerType=0 controllerNumber=1 scan=19643 | 42.3208 |
| 15966:3 | 1 | R.[+42.011]PFLN[+2018.708]WTGPPEPIEAAVAR.F | HexNAc(3)Hex(6)Fuc(1)NeuAc(1) | .[42], N[+2019] | 1342.5887 | 3 | 4025.7515 | 4025.7418 | 0 | 2.4 | 153 | Specific | 375.79 | 281 | 281 | 6.93 | 136 | >sp\|P02788\|TRFL_HUMAN Lactotransferrin OS=Homo sapiens OX=9606 GN=LTF PE=1 SV=6 | 1270 | 20201115_07413_1ng_glyco_hcd203040_compound15.19641.19641.3 | controllerType=0 controllerNumber=1 scan=19641 | 42.3162 |
| 14985:3 | 1 | R.TAGWNIPMGLLFN[+2278.809]QTGSC[+57.021]K.F | HexNAc(5)Hex(6)NeuAc(1) | C[+57], N[+2279] | 1458.6193 | 3 | 4373.8434 | 4373.8192 | 0 | 5.5 | 485 | Specific | 225.17 | 225.17 | 225.17 | 4.81 | 136 | >sp\|P02788\|TRFL_HUMAN Lactotransferrin OS=Homo sapiens OX=9606 GN=LTF PE=1 SV=6 | 1270 | 20201115_07413_1ng_glyco_hcd203040_compound15.18615.18615.3 | controllerType=0 controllerNumber=1 scan=18615 | 40.2601 |
| 15724:3 | 1 | R.TAGWNIPMGLLFN[+1095.397]QTGSC[+57.021]K.F | HexNAc(3)Hex(3) | C[+57], N[+1095] | 1064.1494 | 3 | 3190.4338 | 3190.4065 | 0 | 8.5 | 485 | Specific | 367.26 | 316.44 | 316.44 | 6.72 | 136 | >sp\|P02788\|TRFL_HUMAN Lactotransferrin OS=Homo sapiens OX=9606 GN=LTF PE=1 SV=6 | 1270 | 20201115_07413_1ng_glyco_hcd203040_compound15.19388.19388.2 | controllerType=0 controllerNumber=1 scan=19388 | 41.8046 |
| 14987:3 | 1 | R.TAGWNIPMGLLFN[+2278.809]QTGSC[+57.021]K.F | HexNAc(5)Hex(6)NeuAc(1) | C[+57], N[+2279] | 1458.6193 | 3 | 4373.8434 | 4373.8192 | 0 | 5.5 | 485 | Specific | 401.87 | 401.87 | 297.79 | 8.5 | 136 | >sp\|P02788\|TRFL_HUMAN Lactotransferrin OS=Homo sapiens OX=9606 GN=LTF PE=1 SV=6 | 1270 | 20201115_07413_1ng_glyco_hcd203040_compound15.18617.18617.3 | controllerType=0 controllerNumber=1 scan=18617 | 40.2636 |
| 15197:2 | 1 | R.TAGWNIPMGLLFN[+1768.640]QTGSC[+57.021]K.F | HexNAc(4)Hex(5)Fuc(1) | C[+57], N[+1769] | 1932.335 | 2 | 3863.6627 | 3863.6495 | 0 | 3.4 | 485 | Specific | 428.02 | 396.76 | 396.76 | 9.29 | 136 | >sp\|P02788\|TRFL_HUMAN Lactotransferrin OS=Homo sapiens OX=9606 GN=LTF PE=1 SV=6 | 1270 | 20201115_07413_1ng_glyco_hcd203040_compound15.18837.18837.2 | controllerType=0 controllerNumber=1 scan=18837 | 40.6696 |
| 15433:3 | 1 | R.TAGWNIPMGLLFN[+1581.555]QTGSC[+57.021]K.F | HexNAc(3)Hex(6) | C[+57], N[+1582] | 1226.1962 | 3 | 3676.5741 | 3676.565 | 0 | 2.5 | 485 | Specific | 476.1 | 476.1 | 203.82 | 9 | 136 | >sp\|P02788\|TRFL_HUMAN Lactotransferrin OS=Homo sapiens OX=9606 GN=LTF PE=1 SV=6 | 1270 | 20201115_07413_1ng_glyco_hcd203040_compound15.19082.19082.3 | controllerType=0 controllerNumber=1 scan=19082 | 41.1287 |
| 15329:3 | 1 | R.TAGWNIPMGLLFN[+1622.582]QTGSC[+57.021]K.F | HexNAc(4)Hex(5) | C[+57], N[+1623] | 1239.8729 | 3 | 3717.604 | 3717.5916 | 0 | 3.4 | 485 | Specific | 502.99 | 502.99 | 235.11 | 9.2 | 136 | >sp\|P02788\|TRFL_HUMAN Lactotransferrin OS=Homo sapiens OX=9606 GN=LTF PE=1 SV=6 | 1270 | 20201115_07413_1ng_glyco_hcd203040_compound15.18975.18975.3 | controllerType=0 controllerNumber=1 scan=18975 | 40.9472 |
| 17240:3 | 1 | R.TAGWNIPMGLLFN[+1694.603]QTGSC[+57.021]K.F | HexNAc(3)Hex(4)Fuc(1)NeuAc(1) | C[+57], N[+1695] | 1263.886 | 3 | 3789.6436 | 3789.6127 | 0 | 8.2 | 485 | Specific | 526.34 | 445.62 | 445.62 | 8.95 | 136 | >sp\|P02788\|TRFL_HUMAN Lactotransferrin OS=Homo sapiens OX=9606 GN=LTF PE=1 SV=6 | 1270 | 20201115_07413_1ng_glyco_hcd203040_compound15.20978.20978.3 | controllerType=0 controllerNumber=1 scan=20978 | 45.3 |
| 17321:3 | 1 | R.TAGWNIPMGLLFN[+1710.598]QTGSC[+57.021]K.F | HexNAc(3)Hex(5)NeuAc(1) | C[+57], N[+1711] | 1269.2149 | 3 | 3805.6301 | 3805.6076 | 0 | 5.9 | 485 | Specific | 539.2 | 462.14 | 264.45 | 8.04 | 136 | >sp\|P02788\|TRFL_HUMAN Lactotransferrin OS=Homo sapiens OX=9606 GN=LTF PE=1 SV=6 | 1270 | 20201115_07413_1ng_glyco_hcd203040_compound15.21062.21062.3 | controllerType=0 controllerNumber=1 scan=21062 | 45.4784 |
| 15453:3 | 1 | R.TAGWNIPMGLLFN[+1581.555]QTGSC[+57.021]K.F | HexNAc(3)Hex(6) | C[+57], N[+1582] | 1226.1972 | 3 | 3676.577 | 3676.565 | 0 | 3.3 | 485 | Specific | 571.06 | 571.06 | 332.25 | 10.67 | 136 | >sp\|P02788\|TRFL_HUMAN Lactotransferrin OS=Homo sapiens OX=9606 GN=LTF PE=1 SV=6 | 1270 | 20201115_07413_1ng_glyco_hcd203040_compound15.19103.19103.3 | controllerType=0 controllerNumber=1 scan=19103 | 41.1845 |
| 15524:3 | 1 | R.TAGWNIPMGLLFN[+1460.529]QTGSC[+57.021]K.F | HexNAc(4)Hex(4) | C[+57], N[+1461] | 1185.8555 | 3 | 3555.552 | 3555.5387 | 0 | 3.7 | 485 | Specific | 573.61 | 573.61 | 253.69 | 10.67 | 136 | >sp\|P02788\|TRFL_HUMAN Lactotransferrin OS=Homo sapiens OX=9606 GN=LTF PE=1 SV=6 | 1270 | 20201115_07413_1ng_glyco_hcd203040_compound15.19178.19178.3 | controllerType=0 controllerNumber=1 scan=19178 | 41.3515 |
| 17310:3 | 1 | R.TAGWNIPMGLLFN[+1751.624]QTGSC[+57.021]K.F | HexNAc(4)Hex(4)NeuAc(1) | C[+57], N[+1752] | 1282.8831 | 3 | 3846.6348 | 3846.6341 | 0 | 0.2 | 485 | Specific | 606.01 | 606.01 | 606.01 | 11.38 | 136 | >sp\|P02788\|TRFL_HUMAN Lactotransferrin OS=Homo sapiens OX=9606 GN=LTF PE=1 SV=6 | 1270 | 20201115_07413_1ng_glyco_hcd203040_compound15.21051.21051.3 | controllerType=0 controllerNumber=1 scan=21051 | 45.4444 |
| 17242:3 | 1 | R.TAGWNIPMGLLFN[+1694.603]QTGSC[+57.021]K.F | HexNAc(3)Hex(4)Fuc(1)NeuAc(1) | C[+57], N[+1695] | 1263.886 | 3 | 3789.6436 | 3789.6127 | 0 | 8.2 | 485 | Specific | 634.07 | 623.89 | 623.89 | 11.3 | 136 | >sp\|P02788\|TRFL_HUMAN Lactotransferrin OS=Homo sapiens OX=9606 GN=LTF PE=1 SV=6 | 1270 | 20201115_07413_1ng_glyco_hcd203040_compound15.20980.20980.3 | controllerType=0 controllerNumber=1 scan=20980 | 45.3041 |
| 17078:3 | 1 | R.TAGWNIPMGLLFN[+2059.735]QTGSC[+57.021]K.F | HexNAc(4)Hex(5)Fuc(1)NeuAc(1) | C[+57], N[+2060] | 1385.5851 | 3 | 4154.7409 | 4154.7449 | 0 | -1 | 485 | Specific | 646.27 | 642.73 | 642.73 | 10.88 | 136 | >sp\|P02788\|TRFL_HUMAN Lactotransferrin OS=Homo sapiens OX=9606 GN=LTF PE=1 SV=6 | 1270 | 20201115_07413_1ng_glyco_hcd203040_compound15.20807.20807.3 | controllerType=0 controllerNumber=1 scan=20807 | 44.8657 |
| 17117:3 | 1 | R.TAGWNIPMGLLFN[+2205.793]QTGSC[+57.021]K.F | HexNAc(4)Hex(5)Fuc(2)NeuAc(1) | C[+57], N[+2206] | 1434.285 | 3 | 4300.8404 | 4300.8028 | 0 | 8.7 | 485 | Specific | 648.81 | 644.35 | 644.35 | 11.18 | 136 | >sp\|P02788\|TRFL_HUMAN Lactotransferrin OS=Homo sapiens OX=9606 GN=LTF PE=1 SV=6 | 1270 | 20201115_07413_1ng_glyco_hcd203040_compound15.20847.20847.3 | controllerType=0 controllerNumber=1 scan=20847 | 44.9238 |
| 15199:2 | 1 | R.TAGWNIPMGLLFN[+1768.640]QTGSC[+57.021]K.F | HexNAc(4)Hex(5)Fuc(1) | C[+57], N[+1769] | 1932.335 | 2 | 3863.6627 | 3863.6495 | 0 | 3.4 | 485 | Specific | 661.21 | 631.58 | 631.58 | 12.94 | 136 | >sp\|P02788\|TRFL_HUMAN Lactotransferrin OS=Homo sapiens OX=9606 GN=LTF PE=1 SV=6 | 1270 | 20201115_07413_1ng_glyco_hcd203040_compound15.18839.18839.2 | controllerType=0 controllerNumber=1 scan=18839 | 40.6754 |
| 19099:3 | 1 | R.TAGWNIPMGLLFN[+2350.830]QTGSC[+57.021]K.F | HexNAc(4)Hex(5)Fuc(1)NeuAc(2) | C[+57], N[+2351] | 1482.6253 | 3 | 4445.8614 | 4445.8403 | 0 | 4.7 | 485 | Specific | 681.31 | 651.74 | 651.74 | 12.34 | 136 | >sp\|P02788\|TRFL_HUMAN Lactotransferrin OS=Homo sapiens OX=9606 GN=LTF PE=1 SV=6 | 1270 | 20201115_07413_1ng_glyco_hcd203040_compound15.22932.22932.3 | controllerType=0 controllerNumber=1 scan=22932 | 49.4957 |
| 17318:3 | 1 | R.TAGWNIPMGLLFN[+1710.598]QTGSC[+57.021]K.F | HexNAc(3)Hex(5)NeuAc(1) | C[+57], N[+1711] | 1269.2149 | 3 | 3805.6301 | 3805.6076 | 0 | 5.9 | 485 | Specific | 683.82 | 675.03 | 272.74 | 12.18 | 136 | >sp\|P02788\|TRFL_HUMAN Lactotransferrin OS=Homo sapiens OX=9606 GN=LTF PE=1 SV=6 | 1270 | 20201115_07413_1ng_glyco_hcd203040_compound15.21059.21059.3 | controllerType=0 controllerNumber=1 scan=21059 | 45.456 |
| 15276:3 | 1 | R.TAGWNIPMGLLFN[+1606.587]QTGSC[+57.021]K.F | HexNAc(4)Hex(4)Fuc(1) | C[+57], N[+1607] | 1234.5406 | 3 | 3701.6072 | 3701.5966 | 0 | 2.9 | 485 | Specific | 687.53 | 687.53 | 687.53 | 12.67 | 136 | >sp\|P02788\|TRFL_HUMAN Lactotransferrin OS=Homo sapiens OX=9606 GN=LTF PE=1 SV=6 | 1270 | 20201115_07413_1ng_glyco_hcd203040_compound15.18920.18920.3 | controllerType=0 controllerNumber=1 scan=18920 | 40.844 |
| 15451:3 | 1 | R.TAGWNIPMGLLFN[+1581.555]QTGSC[+57.021]K.F | HexNAc(3)Hex(6) | C[+57], N[+1582] | 1226.1972 | 3 | 3676.577 | 3676.565 | 0 | 3.3 | 485 | Specific | 719.65 | 719.65 | 296.72 | 13.67 | 136 | >sp\|P02788\|TRFL_HUMAN Lactotransferrin OS=Homo sapiens OX=9606 GN=LTF PE=1 SV=6 | 1270 | 20201115_07413_1ng_glyco_hcd203040_compound15.19101.19101.3 | controllerType=0 controllerNumber=1 scan=19101 | 41.1803 |
| 17225:3 | 1 | R.TAGWNIPMGLLFN[+1872.651]QTGSC[+57.021]K.F | HexNAc(3)Hex(6)NeuAc(1) | C[+57], N[+1873] | 1323.225 | 3 | 3967.6605 | 3967.6604 | 0 | 0 | 485 | Specific | 734.39 | 622.71 | 293.77 | 13.38 | 136 | >sp\|P02788\|TRFL_HUMAN Lactotransferrin OS=Homo sapiens OX=9606 GN=LTF PE=1 SV=6 | 1270 | 20201115_07413_1ng_glyco_hcd203040_compound15.20962.20962.3 | controllerType=0 controllerNumber=1 scan=20962 | 45.2648 |
| 19101:3 | 1 | R.TAGWNIPMGLLFN[+2350.830]QTGSC[+57.021]K.F | HexNAc(4)Hex(5)Fuc(1)NeuAc(2) | C[+57], N[+2351] | 1482.6253 | 3 | 4445.8614 | 4445.8403 | 0 | 4.7 | 485 | Specific | 765.62 | 742.12 | 742.12 | 14.34 | 136 | >sp\|P02788\|TRFL_HUMAN Lactotransferrin OS=Homo sapiens OX=9606 GN=LTF PE=1 SV=6 | 1270 | 20201115_07413_1ng_glyco_hcd203040_compound15.22934.22934.3 | controllerType=0 controllerNumber=1 scan=22934 | 49.5005 |
| 15311:3 | 1 | R.TAGWNIPMGLLFN[+1622.582]QTGSC[+57.021]K.F | HexNAc(4)Hex(5) | C[+57], N[+1623] | 1239.8764 | 3 | 3717.6146 | 3717.5916 | 0 | 6.2 | 485 | Specific | 779.16 | 653.51 | 341.56 | 14.18 | 136 | >sp\|P02788\|TRFL_HUMAN Lactotransferrin OS=Homo sapiens OX=9606 GN=LTF PE=1 SV=6 | 1270 | 20201115_07413_1ng_glyco_hcd203040_compound15.18956.18956.3 | controllerType=0 controllerNumber=1 scan=18956 | 40.9082 |
| 17313:3 | 1 | R.TAGWNIPMGLLFN[+1751.624]QTGSC[+57.021]K.F | HexNAc(4)Hex(4)NeuAc(1) | C[+57], N[+1752] | 1282.8831 | 3 | 3846.6348 | 3846.6341 | 0 | 0.2 | 485 | Specific | 783.84 | 768.46 | 768.46 | 14.38 | 136 | >sp\|P02788\|TRFL_HUMAN Lactotransferrin OS=Homo sapiens OX=9606 GN=LTF PE=1 SV=6 | 1270 | 20201115_07413_1ng_glyco_hcd203040_compound15.21054.21054.3 | controllerType=0 controllerNumber=1 scan=21054 | 45.4484 |
| 17080:3 | 1 | R.TAGWNIPMGLLFN[+2059.735]QTGSC[+57.021]K.F | HexNAc(4)Hex(5)Fuc(1)NeuAc(1) | C[+57], N[+2060] | 1385.5851 | 3 | 4154.7409 | 4154.7449 | 0 | -1 | 485 | Specific | 811.29 | 665.01 | 665.01 | 14.88 | 136 | >sp\|P02788\|TRFL_HUMAN Lactotransferrin OS=Homo sapiens OX=9606 GN=LTF PE=1 SV=6 | 1270 | 20201115_07413_1ng_glyco_hcd203040_compound15.20809.20809.3 | controllerType=0 controllerNumber=1 scan=20809 | 44.8695 |
| 15526:3 | 1 | R.TAGWNIPMGLLFN[+1460.529]QTGSC[+57.021]K.F | HexNAc(4)Hex(4) | C[+57], N[+1461] | 1185.8555 | 3 | 3555.552 | 3555.5387 | 0 | 3.7 | 485 | Specific | 820.12 | 645.95 | 365.36 | 15.67 | 136 | >sp\|P02788\|TRFL_HUMAN Lactotransferrin OS=Homo sapiens OX=9606 GN=LTF PE=1 SV=6 | 1270 | 20201115_07413_1ng_glyco_hcd203040_compound15.19180.19180.3 | controllerType=0 controllerNumber=1 scan=19180 | 41.3558 |
| 15515:3 | 1 | R.TAGWNIPMGLLFN[+1419.502]QTGSC[+57.021]K.F | HexNAc(3)Hex(5) | C[+57], N[+1420] | 1172.1746 | 3 | 3514.5093 | 3514.5122 | 0 | -0.8 | 485 | Specific | 840.56 | 755.06 | 395.24 | 14.88 | 136 | >sp\|P02788\|TRFL_HUMAN Lactotransferrin OS=Homo sapiens OX=9606 GN=LTF PE=1 SV=6 | 1270 | 20201115_07413_1ng_glyco_hcd203040_compound15.19168.19168.3 | controllerType=0 controllerNumber=1 scan=19168 | 41.3303 |
| 17227:3 | 1 | R.TAGWNIPMGLLFN[+1872.651]QTGSC[+57.021]K.F | HexNAc(3)Hex(6)NeuAc(1) | C[+57], N[+1873] | 1323.225 | 3 | 3967.6605 | 3967.6604 | 0 | 0 | 485 | Specific | 843.05 | 793.82 | 328.5 | 15.38 | 136 | >sp\|P02788\|TRFL_HUMAN Lactotransferrin OS=Homo sapiens OX=9606 GN=LTF PE=1 SV=6 | 1270 | 20201115_07413_1ng_glyco_hcd203040_compound15.20964.20964.3 | controllerType=0 controllerNumber=1 scan=20964 | 45.2697 |
| 15573:2 | 1 | R.TAGWNIPMGLLFN[+1216.423]QTGSC[+57.021]K.F | HexNAc(2)Hex(5) | C[+57], N[+1216] | 1656.2227 | 2 | 3311.4381 | 3311.4328 | 0 | 1.6 | 485 | Specific | 846.47 | 763.47 | 245.66 | 15.95 | 136 | >sp\|P02788\|TRFL_HUMAN Lactotransferrin OS=Homo sapiens OX=9606 GN=LTF PE=1 SV=6 | 1270 | 20201115_07413_1ng_glyco_hcd203040_compound15.19229.19229.2 | controllerType=0 controllerNumber=1 scan=19229 | 41.4667 |
| 15436:3 | 1 | R.TAGWNIPMGLLFN[+1581.555]QTGSC[+57.021]K.F | HexNAc(3)Hex(6) | C[+57], N[+1582] | 1226.1962 | 3 | 3676.5741 | 3676.565 | 0 | 2.5 | 485 | Specific | 847.34 | 808.02 | 396.6 | 15.68 | 136 | >sp\|P02788\|TRFL_HUMAN Lactotransferrin OS=Homo sapiens OX=9606 GN=LTF PE=1 SV=6 | 1270 | 20201115_07413_1ng_glyco_hcd203040_compound15.19085.19085.3 | controllerType=0 controllerNumber=1 scan=19085 | 41.1329 |
| 15480:3 | 1 | R.TAGWNIPMGLLFN[+1378.476]QTGSC[+57.021]K.F | HexNAc(2)Hex(6) | C[+57], N[+1378] | 1158.507 | 3 | 3473.5065 | 3473.4856 | 0 | 6 | 485 | Specific | 866.05 | 812.67 | 374.92 | 16.18 | 136 | >sp\|P02788\|TRFL_HUMAN Lactotransferrin OS=Homo sapiens OX=9606 GN=LTF PE=1 SV=6 | 1270 | 20201115_07413_1ng_glyco_hcd203040_compound15.19132.19132.3 | controllerType=0 controllerNumber=1 scan=19132 | 41.2477 |

Table S4. Cont.

| Query #:z | Protein Rank | Glycopeptide | Glycan | Modification Type(s) | Observed m/z | z | Observed (M+H) | Calc. mass (M+H) | Off-by-x error | Mass error (ppm) | StartPosition | Cleavage | Score | Delta | Delta Mod | \|Log Prob\| | # of unique peptides | Protein Name | Protein DB number | Comment | Scan # | Scan Time |
|---|---|---|---|---|---|---|---|---|---|---|---|---|---|---|---|---|---|---|---|---|---|---|
| 17035:3 | 1 | R.TAGWNIPMGLLFN[+2205.793]QTGSC[+57.021]K.F | HexNAc(4)Hex(5)Fuc(2)NeuAc(1) | C[+57], N[+2206] | 1434.2866 | 3 | 4300.8451 | 4300.8028 | 0 | 9.8 | 485 | Specific | 872.96 | 872.96 | 872.96 | 16.18 | 136 | >sp\|P02788\|TRFL_HUMAN Lactotransferrin OS=Homo sapiens OX=9606 GN=LTF PE=1 SV=6 | 1270 | 20201115_07413_1ng_glyco_hcd203040_compound15.20761.20761.3 | controllerType=0 controllerNumber=1 scan=20761 | 44.7509 |
| 17231:3 | 1 | R.TAGWNIPMGLLFN[+1913.677]QTGSC[+57.021]K.F | HexNAc(4)Hex(5)NeuAc(1) | C[+57], N[+1914] | 1336.9136 | 3 | 4008.7263 | 4008.687 | 0 | 9.8 | 485 | Specific | 874.67 | 849.66 | 849.66 | 16.18 | 136 | >sp\|P02788\|TRFL_HUMAN Lactotransferrin OS=Homo sapiens OX=9606 GN=LTF PE=1 SV=6 | 1270 | 20201115_07413_1ng_glyco_hcd203040_compound15.20969.20969.3 | controllerType=0 controllerNumber=1 scan=20969 | 45.2798 |
| 15513:3 | 1 | R.TAGWNIPMGLLFN[+1419.502]QTGSC[+57.021]K.F | HexNAc(3)Hex(5) | C[+57], N[+1420] | 1172.1746 | 3 | 3514.5093 | 3514.5122 | 0 | -0.8 | 485 | Specific | 880.99 | 818.02 | 370.41 | 15.88 | 136 | >sp\|P02788\|TRFL_HUMAN Lactotransferrin OS=Homo sapiens OX=9606 GN=LTF PE=1 SV=6 | 1270 | 20201115_07413_1ng_glyco_hcd203040_compound15.19166.19166.3 | controllerType=0 controllerNumber=1 scan=19166 | 41.3246 |
| 17203:3 | 1 | R.TAGWNIPMGLLFN[+1913.677]QTGSC[+57.021]K.F | HexNAc(4)Hex(5)NeuAc(1) | C[+57], N[+1914] | 1336.9113 | 3 | 4008.7194 | 4008.687 | 0 | 8.1 | 485 | Specific | 889.32 | 847.49 | 847.49 | 16.3 | 136 | >sp\|P02788\|TRFL_HUMAN Lactotransferrin OS=Homo sapiens OX=9606 GN=LTF PE=1 SV=6 | 1270 | 20201115_07413_1ng_glyco_hcd203040_compound15.20938.20938.3 | controllerType=0 controllerNumber=1 scan=20938 | 45.1695 |
| 15332:3 | 1 | R.TAGWNIPMGLLFN[+1622.582]QTGSC[+57.021]K.F | HexNAc(4)Hex(5) | C[+57], N[+1623] | 1239.8729 | 3 | 3717.604 | 3717.5916 | 0 | 3.4 | 485 | Specific | 902.81 | 902.81 | 434.08 | 17.67 | 136 | >sp\|P02788\|TRFL_HUMAN Lactotransferrin OS=Homo sapiens OX=9606 GN=LTF PE=1 SV=6 | 1270 | 20201115_07413_1ng_glyco_hcd203040_compound15.18978.18978.3 | controllerType=0 controllerNumber=1 scan=18978 | 40.9535 |
| 17200:3 | 1 | R.TAGWNIPMGLLFN[+1913.677]QTGSC[+57.021]K.F | HexNAc(4)Hex(5)NeuAc(1) | C[+57], N[+1914] | 1336.9113 | 3 | 4008.7194 | 4008.687 | 0 | 8.1 | 485 | Specific | 922.43 | 892.14 | 892.14 | 17.3 | 136 | >sp\|P02788\|TRFL_HUMAN Lactotransferrin OS=Homo sapiens OX=9606 GN=LTF PE=1 SV=6 | 1270 | 20201115_07413_1ng_glyco_hcd203040_compound15.20935.20935.3 | controllerType=0 controllerNumber=1 scan=20935 | 45.1394 |
| 15169:3 | 1 | R.TAGWNIPMGLLFN[+1768.640]QTGSC[+57.021]K.F | HexNAc(4)Hex(5)Fuc(1) | C[+57], N[+1769] | 1288.5597 | 3 | 3863.6647 | 3863.6495 | 0 | 3.9 | 485 | Specific | 923.1 | 923.1 | 923.1 | 17.67 | 136 | >sp\|P02788\|TRFL_HUMAN Lactotransferrin OS=Homo sapiens OX=9606 GN=LTF PE=1 SV=6 | 1270 | 20201115_07413_1ng_glyco_hcd203040_compound15.18808.18808.3 | controllerType=0 controllerNumber=1 scan=18808 | 40.6032 |
| 15279:3 | 1 | R.TAGWNIPMGLLFN[+1606.587]QTGSC[+57.021]K.F | HexNAc(4)Hex(4)Fuc(1) | C[+57], N[+1607] | 1234.5406 | 3 | 3701.6072 | 3701.5966 | 0 | 2.9 | 485 | Specific | 929.61 | 899.55 | 899.55 | 17.67 | 136 | >sp\|P02788\|TRFL_HUMAN Lactotransferrin OS=Homo sapiens OX=9606 GN=LTF PE=1 SV=6 | 1270 | 20201115_07413_1ng_glyco_hcd203040_compound15.18923.18923.3 | controllerType=0 controllerNumber=1 scan=18923 | 40.8492 |
| 15571:2 | 1 | R.TAGWNIPMGLLFN[+1216.423]QTGSC[+57.021]K.F | HexNAc(2)Hex(5) | C[+57], N[+1216] | 1656.2227 | 2 | 3311.4381 | 3311.4328 | 0 | 1.6 | 485 | Specific | 932.47 | 900.85 | 358.64 | 17.95 | 136 | >sp\|P02788\|TRFL_HUMAN Lactotransferrin OS=Homo sapiens OX=9606 GN=LTF PE=1 SV=6 | 1270 | 20201115_07413_1ng_glyco_hcd203040_compound15.19227.19227.2 | controllerType=0 controllerNumber=1 scan=19227 | 41.4552 |
| 17068:3 | 1 | R.TAGWNIPMGLLFN[+2059.735]QTGSC[+57.021]K.F | HexNAc(4)Hex(5)Fuc(1)NeuAc(1) | C[+57], N[+2060] | 1385.5856 | 3 | 4154.7423 | 4154.7449 | 0 | -0.6 | 485 | Specific | 942.46 | 937.83 | 937.83 | 16.88 | 136 | >sp\|P02788\|TRFL_HUMAN Lactotransferrin OS=Homo sapiens OX=9606 GN=LTF PE=1 SV=6 | 1270 | 20201115_07413_1ng_glyco_hcd203040_compound15.20796.20796.3 | controllerType=0 controllerNumber=1 scan=20796 | 44.8441 |
| 17040:3 | 1 | R.TAGWNIPMGLLFN[+2059.735]QTGSC[+57.021]K.F | HexNAc(4)Hex(5)Fuc(1)NeuAc(1) | C[+57], N[+2060] | 1385.592 | 3 | 4154.7614 | 4154.7449 | 0 | 4 | 485 | Specific | 951.17 | 925.05 | 925.05 | 18.67 | 136 | >sp\|P02788\|TRFL_HUMAN Lactotransferrin OS=Homo sapiens OX=9606 GN=LTF PE=1 SV=6 | 1270 | 20201115_07413_1ng_glyco_hcd203040_compound15.20767.20767.3 | controllerType=0 controllerNumber=1 scan=20767 | 44.7665 |
| 17066:3 | 1 | R.TAGWNIPMGLLFN[+2059.735]QTGSC[+57.021]K.F | HexNAc(4)Hex(5)Fuc(1)NeuAc(1) | C[+57], N[+2060] | 1385.5856 | 3 | 4154.7423 | 4154.7449 | 0 | -0.6 | 485 | Specific | 965.61 | 919.29 | 919.29 | 17.88 | 136 | >sp\|P02788\|TRFL_HUMAN Lactotransferrin OS=Homo sapiens OX=9606 GN=LTF PE=1 SV=6 | 1270 | 20201115_07413_1ng_glyco_hcd203040_compound15.20794.20794.3 | controllerType=0 controllerNumber=1 scan=20794 | 44.8394 |
| 17033:3 | 1 | R.TAGWNIPMGLLFN[+2205.793]QTGSC[+57.021]K.F | HexNAc(4)Hex(5)Fuc(2)NeuAc(1) | C[+57], N[+2206] | 1434.2866 | 3 | 4300.8451 | 4300.8028 | 0 | 9.8 | 485 | Specific | 976.34 | 883.68 | 883.68 | 18.18 | 136 | >sp\|P02788\|TRFL_HUMAN Lactotransferrin OS=Homo sapiens OX=9606 GN=LTF PE=1 SV=6 | 1270 | 20201115_07413_1ng_glyco_hcd203040_compound15.20759.20759.3 | controllerType=0 controllerNumber=1 scan=20759 | 44.7378 |
| 15568:3 | 1 | R.TAGWNIPMGLLFN[+1216.423]QTGSC[+57.021]K.F | HexNAc(2)Hex(5) | C[+57], N[+1216] | 1104.489 | 3 | 3311.4526 | 3311.4328 | 0 | 6 | 485 | Specific | 990.17 | 872.66 | 463.63 | 18.18 | 136 | >sp\|P02788\|TRFL_HUMAN Lactotransferrin OS=Homo sapiens OX=9606 GN=LTF PE=1 SV=6 | 1270 | 20201115_07413_1ng_glyco_hcd203040_compound15.19223.19223.3 | controllerType=0 controllerNumber=1 scan=19223 | 41.4463 |
| 15482:3 | 1 | R.TAGWNIPMGLLFN[+1378.476]QTGSC[+57.021]K.F | HexNAc(2)Hex(6) | C[+57], N[+1378] | 1158.507 | 3 | 3473.5065 | 3473.4856 | 0 | 6 | 485 | Specific | 993.4 | 827.04 | 474.53 | 18.18 | 136 | >sp\|P02788\|TRFL_HUMAN Lactotransferrin OS=Homo sapiens OX=9606 GN=LTF PE=1 SV=6 | 1270 | 20201115_07413_1ng_glyco_hcd203040_compound15.19134.19134.3 | controllerType=0 controllerNumber=1 scan=19134 | 41.2537 |
| 15565:3 | 1 | R.TAGWNIPMGLLFN[+1216.423]QTGSC[+57.021]K.F | HexNAc(2)Hex(5) | C[+57], N[+1216] | 1104.489 | 3 | 3311.4526 | 3311.4328 | 0 | 6 | 485 | Specific | 1019.49 | 926 | 474.66 | 19.18 | 136 | >sp\|P02788\|TRFL_HUMAN Lactotransferrin OS=Homo sapiens OX=9606 GN=LTF PE=1 SV=6 | 1270 | 20201115_07413_1ng_glyco_hcd203040_compound15.19220.19220.3 | controllerType=0 controllerNumber=1 scan=19220 | 41.4316 |
| 15041:3 | 1 | R.TAGWNIPM[+15.995]GLLFN[+1913.677]QTGSC[+57.021]K.F | HexNAc(4)Hex(5)NeuAc(1) | C[+57], M[+16], N[+1914] | 1342.2324 | 3 | 4024.6825 | 4024.6819 | 0 | 0.2 | 485 | Specific | 272.13 | 215.87 | 215.87 | 5.84 | 136 | >sp\|P02788\|TRFL_HUMAN Lactotransferrin OS=Homo sapiens OX=9606 GN=LTF PE=1 SV=6 | 1270 | 20201115_07413_1ng_glyco_hcd203040_compound15.18673.18673.3 | controllerType=0 controllerNumber=1 scan=18673 | 40.3708 |
| 15660:3 | 1 | R.TAGWNIPM[+15.995]GLLFN[+2059.735]QTGSC[+57.021]K.F | HexNAc(4)Hex(5)Fuc(1)NeuAc(1) | C[+57], M[+16], N[+2060] | 1390.9267 | 3 | 4170.7655 | 4170.7398 | 0 | 6.2 | 485 | Specific | 276.87 | 250.29 | 250.29 | 5.99 | 136 | >sp\|P02788\|TRFL_HUMAN Lactotransferrin OS=Homo sapiens OX=9606 GN=LTF PE=1 SV=6 | 1270 | 20201115_07413_1ng_glyco_hcd203040_compound15.19321.19321.3 | controllerType=0 controllerNumber=1 scan=19321 | 41.6698 |
| 15018:3 | 1 | R.TAGWNIPM[+15.995]GLLFN[+1913.677]QTGSC[+57.021]K.F | HexNAc(4)Hex(5)NeuAc(1) | C[+57], M[+16], N[+1914] | 1342.2357 | 3 | 4024.6924 | 4024.6819 | 0 | 2.6 | 485 | Specific | 356.46 | 356.46 | 356.46 | 7.32 | 136 | >sp\|P02788\|TRFL_HUMAN Lactotransferrin OS=Homo sapiens OX=9606 GN=LTF PE=1 SV=6 | 1270 | 20201115_07413_1ng_glyco_hcd203040_compound15.18649.18649.3 | controllerType=0 controllerNumber=1 scan=18649 | 40.3275 |
| 15043:3 | 1 | R.TAGWNIPM[+15.995]GLLFN[+1913.677]QTGSC[+57.021]K.F | HexNAc(4)Hex(5)NeuAc(1) | C[+57], M[+16], N[+1914] | 1342.2324 | 3 | 4024.6825 | 4024.6819 | 0 | 0.2 | 485 | Specific | 405.82 | 373.52 | 373.52 | 8.53 | 136 | >sp\|P02788\|TRFL_HUMAN Lactotransferrin OS=Homo sapiens OX=9606 GN=LTF PE=1 SV=6 | 1270 | 20201115_07413_1ng_glyco_hcd203040_compound15.18675.18675.3 | controllerType=0 controllerNumber=1 scan=18675 | 40.3743 |
| 15754:3 | 1 | R.TAGWNIPM[+15.995]GLLFN[+2059.735]QTGSC[+57.021]K.F | HexNAc(4)Hex(5)Fuc(1)NeuAc(1) | C[+57], M[+16], N[+2060] | 1390.9246 | 3 | 4170.7593 | 4170.7398 | 0 | 4.7 | 485 | Specific | 448.98 | 357.13 | 357.13 | 7.26 | 136 | >sp\|P02788\|TRFL_HUMAN Lactotransferrin OS=Homo sapiens OX=9606 GN=LTF PE=1 SV=6 | 1270 | 20201115_07413_1ng_glyco_hcd203040_compound15.19419.19419.3 | controllerType=0 controllerNumber=1 scan=19419 | 41.8748 |
| 15662:3 | 1 | R.TAGWNIPM[+15.995]GLLFN[+2059.735]QTGSC[+57.021]K.F | HexNAc(4)Hex(5)Fuc(1)NeuAc(1) | C[+57], M[+16], N[+2060] | 1390.9267 | 3 | 4170.7655 | 4170.7398 | 0 | 6.2 | 485 | Specific | 687.68 | 647.57 | 647.57 | 12.18 | 136 | >sp\|P02788\|TRFL_HUMAN Lactotransferrin OS=Homo sapiens OX=9606 GN=LTF PE=1 SV=6 | 1270 | 20201115_07413_1ng_glyco_hcd203040_compound15.19323.19323.3 | controllerType=0 controllerNumber=1 scan=19323 | 41.6735 |
| 15756:3 | 1 | R.TAGWNIPM[+15.995]GLLFN[+2059.735]QTGSC[+57.021]K.F | HexNAc(4)Hex(5)Fuc(1)NeuAc(1) | C[+57], M[+16], N[+2060] | 1390.9246 | 3 | 4170.7593 | 4170.7398 | 0 | 4.7 | 485 | Specific | 832.96 | 748.76 | 748.76 | 15.34 | 136 | >sp\|P02788\|TRFL_HUMAN Lactotransferrin OS=Homo sapiens OX=9606 GN=LTF PE=1 SV=6 | 1270 | 20201115_07413_1ng_glyco_hcd203040_compound15.19421.19421.3 | controllerType=0 controllerNumber=1 scan=19421 | 41.8791 |
| 16394:3 | 1 | R.TAGWNIPMGLLFN[+1606.587]QTGSC[+57.021]KFDEYFSQSC[+57.021]APGSDPR.S | HexNAc(4)Hex(4)Fuc(1) | C[+57]*2, N[+1607] | 1849.1284 | 3 | 5545.3706 | 5545.3438 | 0 | 4.8 | 485 | Specific | 233.83 | 226.72 | 226.72 | 5.98 | 136 | >sp\|P02788\|TRFL_HUMAN Lactotransferrin OS=Homo sapiens OX=9606 GN=LTF PE=1 SV=6 | 1270 | 20201115_07413_1ng_glyco_hcd203040_compound15.20090.20090.3 | controllerType=0 controllerNumber=1 scan=20090 | 43.3658 |
| 16392:3 | 1 | R.TAGWNIPMGLLFN[+1606.587]QTGSC[+57.021]KFDEYFSQSC[+57.021]APGSDPR.S | HexNAc(4)Hex(4)Fuc(1) | C[+57]*2, N[+1607] | 1849.1284 | 3 | 5545.3706 | 5545.3438 | 0 | 4.8 | 485 | Specific | 310.17 | 310.17 | 310.17 | 6.88 | 136 | >sp\|P02788\|TRFL_HUMAN Lactotransferrin OS=Homo sapiens OX=9606 GN=LTF PE=1 SV=6 | 1270 | 20201115_07413_1ng_glyco_hcd203040_compound15.20088.20088.3 | controllerType=0 controllerNumber=1 scan=20088 | 43.3617 |
| 17312:4 | 1 | R.TAGWNIPMGLLFN[+2205.793]QTGSC[+57.021]KFDEYFSQSC[+57.021]APGSDPR.S | HexNAc(4)Hex(5)Fuc(2)NeuAc(1) | C[+57]*2, N[+2206] | 1536.8918 | 4 | 6144.5456 | 6144.5499 | 0 | -0.7 | 485 | Specific | 459.53 | 450.54 | 450.54 | 8.2 | 136 | >sp\|P02788\|TRFL_HUMAN Lactotransferrin OS=Homo sapiens OX=9606 GN=LTF PE=1 SV=6 | 1270 | 20201115_07413_1ng_glyco_hcd203040_compound15.21053.21053.4 | controllerType=0 controllerNumber=1 scan=21053 | 45.4466 |
| 16381:4 | 1 | R.TAGWNIPMGLLFN[+1606.587]QTGSC[+57.021]KFDEYFSQSC[+57.021]APGSDPR.S | HexNAc(4)Hex(4)Fuc(1) | C[+57]*2, N[+1607] | 1387.0929 | 4 | 5545.3498 | 5545.3438 | 0 | 1.1 | 485 | Specific | 490.25 | 490.25 | 490.25 | 8.91 | 136 | >sp\|P02788\|TRFL_HUMAN Lactotransferrin OS=Homo sapiens OX=9606 GN=LTF PE=1 SV=6 | 1270 | 20201115_07413_1ng_glyco_hcd203040_compound15.20077.20077.4 | controllerType=0 controllerNumber=1 scan=20077 | 43.3377 |
| 16325:3 | 1 | R.TAGWNIPMGLLFN[+1768.640]QTGSC[+57.021]KFDEYFSQSC[+57.021]APGSDPR.S | HexNAc(4)Hex(5)Fuc(1) | C[+57]*2, N[+1769] | 1903.1395 | 3 | 5707.4039 | 5707.3966 | 0 | 1.3 | 485 | Specific | 566.06 | 436.29 | 436.29 | 9.79 | 136 | >sp\|P02788\|TRFL_HUMAN Lactotransferrin OS=Homo sapiens OX=9606 GN=LTF PE=1 SV=6 | 1270 | 20201115_07413_1ng_glyco_hcd203040_compound15.20019.20019.3 | controllerType=0 controllerNumber=1 scan=20019 | 43.224 |
| 17400:3 | 1 | R.TAGWNIPMGLLFN[+1913.677]QTGSC[+57.021]KFDEYFSQSC[+57.021]APGSDPR.S | HexNAc(4)Hex(5)NeuAc(1) | C[+57]*2, N[+1914] | 1951.4843 | 3 | 5852.4385 | 5852.4341 | 0 | 0.7 | 485 | Specific | 575.12 | 455.71 | 455.71 | 10.65 | 136 | >sp\|P02788\|TRFL_HUMAN Lactotransferrin OS=Homo sapiens OX=9606 GN=LTF PE=1 SV=6 | 1270 | 20201115_07413_1ng_glyco_hcd203040_compound15.21144.21144.3 | controllerType=0 controllerNumber=1 scan=21144 | 45.6162 |
| 16323:3 | 1 | R.TAGWNIPMGLLFN[+1768.640]QTGSC[+57.021]KFDEYFSQSC[+57.021]APGSDPR.S | HexNAc(4)Hex(5)Fuc(1) | C[+57]*2, N[+1769] | 1903.1395 | 3 | 5707.4039 | 5707.3966 | 0 | 1.3 | 485 | Specific | 601.8 | 535.81 | 535.81 | 11.95 | 136 | >sp\|P02788\|TRFL_HUMAN Lactotransferrin OS=Homo sapiens OX=9606 GN=LTF PE=1 SV=6 | 1270 | 20201115_07413_1ng_glyco_hcd203040_compound15.20017.20017.3 | controllerType=0 controllerNumber=1 scan=20017 | 43.2187 |
| 17394:4 | 1 | R.TAGWNIPMGLLFN[+1913.677]QTGSC[+57.021]KFDEYFSQSC[+57.021]APGSDPR.S | HexNAc(4)Hex(5)NeuAc(1) | C[+57]*2, N[+1914] | 1463.8644 | 4 | 5852.4357 | 5852.4341 | 0 | 0.3 | 485 | Specific | 673.31 | 590.22 | 590.22 | 12.38 | 136 | >sp\|P02788\|TRFL_HUMAN Lactotransferrin OS=Homo sapiens OX=9606 GN=LTF PE=1 SV=6 | 1270 | 20201115_07413_1ng_glyco_hcd203040_compound15.21138.21138.4 | controllerType=0 controllerNumber=1 scan=21138 | 45.6052 |
| 16383:4 | 1 | R.TAGWNIPMGLLFN[+1606.587]QTGSC[+57.021]KFDEYFSQSC[+57.021]APGSDPR.S | HexNAc(4)Hex(4)Fuc(1) | C[+57]*2, N[+1607] | 1387.0929 | 4 | 5545.3498 | 5545.3438 | 0 | 1.1 | 485 | Specific | 696.29 | 690.47 | 690.47 | 12.38 | 136 | >sp\|P02788\|TRFL_HUMAN Lactotransferrin OS=Homo sapiens OX=9606 GN=LTF PE=1 SV=6 | 1270 | 20201115_07413_1ng_glyco_hcd203040_compound15.20079.20079.4 | controllerType=0 controllerNumber=1 scan=20079 | 43.3414 |
| 17315:4 | 1 | R.TAGWNIPMGLLFN[+2205.793]QTGSC[+57.021]KFDEYFSQSC[+57.021]APGSDPR.S | HexNAc(4)Hex(5)Fuc(2)NeuAc(1) | C[+57]*2, N[+2206] | 1536.8918 | 4 | 6144.5456 | 6144.5499 | 0 | -0.7 | 485 | Specific | 713.3 | 677.44 | 677.44 | 12.88 | 136 | >sp\|P02788\|TRFL_HUMAN Lactotransferrin OS=Homo sapiens OX=9606 GN=LTF PE=1 SV=6 | 1270 | 20201115_07413_1ng_glyco_hcd203040_compound15.21056.21056.4 | controllerType=0 controllerNumber=1 scan=21056 | 45.4522 |
| 16322:4 | 1 | R.TAGWNIPMGLLFN[+1768.640]QTGSC[+57.021]KFDEYFSQSC[+57.021]APGSDPR.S | HexNAc(4)Hex(5)Fuc(1) | C[+57]*2, N[+1769] | 1427.6057 | 4 | 5707.401 | 5707.3966 | 0 | 0.8 | 485 | Specific | 741.83 | 719.19 | 719.19 | 13.38 | 136 | >sp\|P02788\|TRFL_HUMAN Lactotransferrin OS=Homo sapiens OX=9606 GN=LTF PE=1 SV=6 | 1270 | 20201115_07413_1ng_glyco_hcd203040_compound15.20016.20016.4 | controllerType=0 controllerNumber=1 scan=20016 | 43.2174 |
| 17307:4 | 1 | R.TAGWNIPMGLLFN[+2059.735]QTGSC[+57.021]KFDEYFSQSC[+57.021]APGSDPR.S | HexNAc(4)Hex(5)Fuc(1)NeuAc(1) | C[+57]*2, N[+2060] | 1500.3793 | 4 | 5998.4953 | 5998.492 | 0 | 0.5 | 485 | Specific | 803.13 | 763.93 | 763.93 | 15.38 | 136 | >sp\|P02788\|TRFL_HUMAN Lactotransferrin OS=Homo sapiens OX=9606 GN=LTF PE=1 SV=6 | 1270 | 20201115_07413_1ng_glyco_hcd203040_compound15.21048.21048.4 | controllerType=0 controllerNumber=1 scan=21048 | 45.4391 |
| 17398:3 | 1 | R.TAGWNIPMGLLFN[+1913.677]QTGSC[+57.021]KFDEYFSQSC[+57.021]APGSDPR.S | HexNAc(4)Hex(5)NeuAc(1) | C[+57]*2, N[+1914] | 1951.4843 | 3 | 5852.4385 | 5852.4341 | 0 | 0.7 | 485 | Specific | 808.69 | 683.87 | 683.87 | 15.65 | 136 | >sp\|P02788\|TRFL_HUMAN Lactotransferrin OS=Homo sapiens OX=9606 GN=LTF PE=1 SV=6 | 1270 | 20201115_07413_1ng_glyco_hcd203040_compound15.21142.21142.3 | controllerType=0 controllerNumber=1 scan=21142 | 45.6115 |
| 17309:4 | 1 | R.TAGWNIPMGLLFN[+2059.735]QTGSC[+57.021]KFDEYFSQSC[+57.021]APGSDPR.S | HexNAc(4)Hex(5)Fuc(1)NeuAc(1) | C[+57]*2, N[+2060] | 1500.3793 | 4 | 5998.4953 | 5998.492 | 0 | 0.5 | 485 | Specific | 846.19 | 798.25 | 798.25 | 15.38 | 136 | >sp\|P02788\|TRFL_HUMAN Lactotransferrin OS=Homo sapiens OX=9606 GN=LTF PE=1 SV=6 | 1270 | 20201115_07413_1ng_glyco_hcd203040_compound15.21050.21050.4 | controllerType=0 controllerNumber=1 scan=21050 | 45.4437 |
| 16324:4 | 1 | R.TAGWNIPMGLLFN[+1768.640]QTGSC[+57.021]KFDEYFSQSC[+57.021]APGSDPR.S | HexNAc(4)Hex(5)Fuc(1) | C[+57]*2, N[+1769] | 1427.6057 | 4 | 5707.401 | 5707.3966 | 0 | 0.8 | 485 | Specific | 859.59 | 796.57 | 796.57 | 16.38 | 136 | >sp\|P02788\|TRFL_HUMAN Lactotransferrin OS=Homo sapiens OX=9606 GN=LTF PE=1 SV=6 | 1270 | 20201115_07413_1ng_glyco_hcd203040_compound15.20018.20018.4 | controllerType=0 controllerNumber=1 scan=20018 | 43.2225 |
| 18620:4 | 1 | R.TAGWNIPMGLLFN[+2350.830]QTGSC[+57.021]KFDEYFSQSC[+57.021]APGSDPR.S | HexNAc(4)Hex(5)Fuc(1)NeuAc(2) | C[+57]*2, N[+2351] | 1573.1533 | 4 | 6289.5915 | 6289.5875 | 0 | 0.6 | 485 | Specific | 865.64 | 820.24 | 820.24 | 16.38 | 136 | >sp\|P02788\|TRFL_HUMAN Lactotransferrin OS=Homo sapiens OX=9606 GN=LTF PE=1 SV=6 | 1270 | 20201115_07413_1ng_glyco_hcd203040_compound15.22429.22429.4 | controllerType=0 controllerNumber=1 scan=22429 | 48.3319 |
| 17397:4 | 1 | R.TAGWNIPMGLLFN[+1913.677]QTGSC[+57.021]KFDEYFSQSC[+57.021]APGSDPR.S | HexNAc(4)Hex(5)NeuAc(1) | C[+57]*2, N[+1914] | 1463.8644 | 4 | 5852.4357 | 5852.4341 | 0 | 0.3 | 485 | Specific | 915.59 | 863.35 | 863.35 | 17.38 | 136 | >sp\|P02788\|TRFL_HUMAN Lactotransferrin OS=Homo sapiens OX=9606 GN=LTF PE=1 SV=6 | 1270 | 20201115_07413_1ng_glyco_hcd203040_compound15.21141.21141.4 | controllerType=0 controllerNumber=1 scan=21141 | 45.6106 |
| 18618:4 | 1 | R.TAGWNIPMGLLFN[+2350.830]QTGSC[+57.021]KFDEYFSQSC[+57.021]APGSDPR.S | HexNAc(4)Hex(5)Fuc(1)NeuAc(2) | C[+57]*2, N[+2351] | 1573.1533 | 4 | 6289.5915 | 6289.5875 | 0 | 0.6 | 485 | Specific | 951.71 | 912.45 | 912.45 | 18.38 | 136 | >sp\|P02788\|TRFL_HUMAN Lactotransferrin OS=Homo sapiens OX=9606 GN=LTF PE=1 SV=6 | 1270 | 20201115_07413_1ng_glyco_hcd203040_compound15.22427.22427.4 | controllerType=0 controllerNumber=1 scan=22427 | 48.3266 |
| 16129:4 | 1 | R.TAGWNIPM[+15.995]GLLFN[+2059.735]QTGSC[+57.021]KFDEYFSQSC[+57.021]A | HexNAc(4)Hex(5)Fuc(1)NeuAc(1) | C[+57]*2, M[+16], N[+2060] | 1504.3801 | 4 | 6014.4987 | 6014.487 | 0 | 2 | 485 | Specific | 803.91 | 576.09 | 576.09 | 15.68 | 136 | >sp\|P02788\|TRFL_HUMAN Lactotransferrin OS=Homo sapiens OX=9606 GN=LTF PE=1 SV=6 | 1270 | 20201115_07413_1ng_glyco_hcd203040_compound15.19813.19813.4 | controllerType=0 controllerNumber=1 scan=19813 | 42.762 |
| 07772:3 | 1 | R.N[+2059.735]GSDC[+57.021]PDKFC[+57.021]LFQSETK.N | HexNAc(4)Hex(5)Fuc(1)NeuAc(1) | C[+57]*2, N[+2060] | 1364.8777 | 3 | 4092.6187 | 4092.6088 | 0 | 2.4 | 642 | Specific | 226.67 | 150.03 | 150.03 | 3.88 | 136 | >sp\|P02788\|TRFL_HUMAN Lactotransferrin OS=Homo sapiens OX=9606 GN=LTF PE=1 SV=6 | 1270 | 20201115_07413_1ng_glyco_hcd203040_compound15.11026.11026.3 | controllerType=0 controllerNumber=1 scan=11026 | 26.5905 |
| 07770:3 | 1 | R.N[+2059.735]GSDC[+57.021]PDKFC[+57.021]LFQSETK.N | HexNAc(4)Hex(5)Fuc(1)NeuAc(1) | C[+57]*2, N[+2060] | 1364.8777 | 3 | 4092.6187 | 4092.6088 | 0 | 2.4 | 642 | Specific | 237.84 | 176.74 | 176.74 | 5.29 | 136 | >sp\|P02788\|TRFL_HUMAN Lactotransferrin OS=Homo sapiens OX=9606 GN=LTF PE=1 SV=6 | 1270 | 20201115_07413_1ng_glyco_hcd203040_compound15.11024.11024.3 | controllerType=0 controllerNumber=1 scan=11024 | 26.5869 |
| 10044:3 | 1 | R.N[+2350.830]GSDC[+57.021]PDKFC[+57.021]LFQSETK.N | HexNAc(4)Hex(5)Fuc(1)NeuAc(2) | C[+57]*2, N[+2351] | 1461.9112 | 3 | 4383.719 | 4383.7043 | 0 | 3.4 | 642 | Specific | 238.85 | 147.64 | 147.64 | 3.81 | 136 | >sp\|P02788\|TRFL_HUMAN Lactotransferrin OS=Homo sapiens OX=9606 GN=LTF PE=1 SV=6 | 1270 | 20201115_07413_1ng_glyco_hcd203040_compound15.13418.13418.3 | controllerType=0 controllerNumber=1 scan=13418 | 30.86 |
| 07703:3 | 1 | R.N[+2205.793]GSDC[+57.021]PDKFC[+57.021]LFQSETK.N | HexNAc(4)Hex(5)Fuc(2)NeuAc(1) | C[+57]*2, N[+2206] | 1413.5657 | 3 | 4238.6825 | 4238.6667 | 0 | 3.7 | 642 | Specific | 305.91 | 243.28 | 243.28 | 6.17 | 136 | >sp\|P02788\|TRFL_HUMAN Lactotransferrin OS=Homo sapiens OX=9606 GN=LTF PE=1 SV=6 | 1270 | 20201115_07413_1ng_glyco_hcd203040_compound15.10953.10953.3 | controllerType=0 controllerNumber=1 scan=10953 | 26.4617 |
| 07701:3 | 1 | R.N[+2205.793]GSDC[+57.021]PDKFC[+57.021]LFQSETK.N | HexNAc(4)Hex(5)Fuc(2)NeuAc(1) | C[+57]*2, N[+2206] | 1413.5657 | 3 | 4238.6825 | 4238.6667 | 0 | 3.7 | 642 | Specific | 326.8 | 219.84 | 219.84 | 6.43 | 136 | >sp\|P02788\|TRFL_HUMAN Lactotransferrin OS=Homo sapiens OX=9606 GN=LTF PE=1 SV=6 | 1270 | 20201115_07413_1ng_glyco_hcd203040_compound15.10951.10951.3 | controllerType=0 controllerNumber=1 scan=10951 | 26.4583 |
| 10008:3 | 1 | R.N[+2350.830]GSDC[+57.021]PDKFC[+57.021]LFQSETK.N | HexNAc(4)Hex(5)Fuc(1)NeuAc(2) | C[+57]*2, N[+2351] | 1461.9128 | 3 | 4383.7238 | 4383.7043 | 0 | 4.5 | 642 | Specific | 380.4 | 313.03 | 313.03 | 6.63 | 136 | >sp\|P02788\|TRFL_HUMAN Lactotransferrin OS=Homo sapiens OX=9606 GN=LTF PE=1 SV=6 | 1270 | 20201115_07413_1ng_glyco_hcd203040_compound15.13380.13380.3 | controllerType=0 controllerNumber=1 scan=13380 | 30.7889 |
| 10042:3 | 1 | R.N[+2350.830]GSDC[+57.021]PDKFC[+57.021]LFQSETK.N | HexNAc(4)Hex(5)Fuc(1)NeuAc(2) | C[+57]*2, N[+2351] | 1461.9112 | 3 | 4383.719 | 4383.7043 | 0 | 3.4 | 642 | Specific | 386.33 | 236.54 | 236.54 | 6.55 | 136 | >sp\|P02788\|TRFL_HUMAN Lactotransferrin OS=Homo sapiens OX=9606 GN=LTF PE=1 SV=6 | 1270 | 20201115_07413_1ng_glyco_hcd203040_compound15.13416.13416.3 | controllerType=0 controllerNumber=1 scan=13416 | 30.8561 |
| 07802:3 | 1 | R.N[+2059.735]GSDC[+57.021]PDKFC[+57.021]LFQSETK.N | HexNAc(4)Hex(5)Fuc(1)NeuAc(1) | C[+57]*2, N[+2060] | 1364.8791 | 3 | 4092.6227 | 4092.6088 | 0 | 3.4 | 642 | Specific | 417.36 | 285.35 | 285.35 | 7.05 | 136 | >sp\|P02788\|TRFL_HUMAN Lactotransferrin OS=Homo sapiens OX=9606 GN=LTF PE=1 SV=6 | 1270 | 20201115_07413_1ng_glyco_hcd203040_compound15.11057.11057.3 | controllerType=0 controllerNumber=1 scan=11057 | 26.6423 |
| 07804:3 | 1 | R.N[+2059.735]GSDC[+57.021]PDKFC[+57.021]LFQSETK.N | HexNAc(4)Hex(5)Fuc(1)NeuAc(1) | C[+57]*2, N[+2060] | 1364.8791 | 3 | 4092.6227 | 4092.6088 | 0 | 3.4 | 642 | Specific | 531.61 | 397.4 | 397.4 | 9.06 | 136 | >sp\|P02788\|TRFL_HUMAN Lactotransferrin OS=Homo sapiens OX=9606 GN=LTF PE=1 SV=6 | 1270 | 20201115_07413_1ng_glyco_hcd203040_compound15.11059.11059.3 | controllerType=0 controllerNumber=1 scan=11059 | 26.6463 |
| 10010:3 | 1 | R.N[+2350.830]GSDC[+57.021]PDKFC[+57.021]LFQSETK.N | HexNAc(4)Hex(5)Fuc(1)NeuAc(2) | C[+57]*2, N[+2351] | 1461.9128 | 3 | 4383.7238 | 4383.7043 | 0 | 4.5 | 642 | Specific | 559.72 | 456.74 | 456.74 | 8.92 | 136 | >sp\|P02788\|TRFL_HUMAN Lactotransferrin OS=Homo sapiens OX=9606 GN=LTF PE=1 SV=6 | 1270 | 20201115_07413_1ng_glyco_hcd203040_compound15.13382.13382.3 | controllerType=0 controllerNumber=1 scan=13382 | 30.7944 |
| 08652:4 | 2 | K.RQSGVN[+2059.735]ATLPEENQPVVFNHVYNIK.L | HexNAc(4)Hex(5)Fuc(1)NeuAc(1) | N[+2060] | 1229.0675 | 4 | 4913.2482 | 4913.1991 | 0 | 10 | 33 | Specific | 522.12 | 480.86 | 480.86 | 9.39 | 83 | >sp\|P24821\|TENA_HUMAN Tenascin OS=Homo sapiens OX=9606 GN=TNC PE=1 SV=3 | 486 | 20201115_07413_1ng_glyco_hcd203040_compound15.11954.11954.4 | controllerType=0 controllerNumber=1 scan=11954 | 28.1911 |
| 08654:4 | 2 | K.RQSGVN[+2059.735]ATLPEENQPVVFNHVYNIK.L | HexNAc(4)Hex(5)Fuc(1)NeuAc(1) | N[+2060] | 1229.0675 | 4 | 4913.2482 | 4913.1991 | 0 | 10 | 33 | Specific | 540.15 | 438.21 | 438.21 | 9.57 | 83 | >sp\|P24821\|TENA_HUMAN Tenascin OS=Homo sapiens OX=9606 GN=TNC PE=1 SV=3 | 486 | 20201115_07413_1ng_glyco_hcd203040_compound15.11956.11956.4 | controllerType=0 controllerNumber=1 scan=11956 | 28.1951 |
| 10438:3 | 2 | R.QSGVN[+2059.735]ATLPEENQPVVFNHVYNIK.L | HexNAc(4)Hex(5)Fuc(1)NeuAc(1) | N[+2060] | 1586.3771 | 3 | 4757.1168 | 4757.098 | 0 | 4 | 34 | Specific | 312.84 | 261.51 | 261.51 | 7.09 | 83 | >sp\|P24821\|TENA_HUMAN Tenascin OS=Homo sapiens OX=9606 GN=TNC PE=1 SV=3 | 486 | 20201115_07413_1ng_glyco_hcd203040_compound15.13829.13829.3 | controllerType=0 controllerNumber=1 scan=13829 | 31.5692 |
| 13833:3 | 2 | R.LN[+1378.476]YSLPTGQWVGVQLPR.N | HexNAc(2)Hex(6) | N[+1378] | 1102.8462 | 3 | 3306.5241 | 3306.5145 | 0 | 2.9 | 1017 | Specific | 536.32 | 536.32 | 536.32 | 9.22 | 83 | >sp\|P24821\|TENA_HUMAN Tenascin OS=Homo sapiens OX=9606 GN=TNC PE=1 SV=3 | 486 | 20201115_07413_1ng_glyco_hcd203040_compound15.17388.17388.3 | controllerType=0 controllerNumber=1 scan=17388 | 37.9887 |
| 13757:3 | 2 | R.LN[+1540.529]YSLPTGQWVGVQLPR.N | HexNAc(2)Hex(7) | N[+1541] | 1156.866 | 3 | 3468.5835 | 3468.5674 | 0 | 4.7 | 1017 | Specific | 845.43 | 845.43 | 845.43 | 15.25 | 83 | >sp\|P24821\|TENA_HUMAN Tenascin OS=Homo sapiens OX=9606 GN=TNC PE=1 SV=3 | 486 | 20201115_07413_1ng_glyco_hcd203040_compound15.17309.17309.3 | controllerType=0 controllerNumber=1 scan=17309 | 37.8307 |
| 13835:3 | 2 | R.LN[+1378.476]YSLPTGQWVGVQLPR.N | HexNAc(2)Hex(6) | N[+1378] | 1102.8462 | 3 | 3306.5241 | 3306.5145 | 0 | 2.9 | 1017 | Specific | 888.49 | 773.33 | 773.33 | 16.57 | 83 | >sp\|P24821\|TENA_HUMAN Tenascin OS=Homo sapiens OX=9606 GN=TNC PE=1 SV=3 | 486 | 20201115_07413_1ng_glyco_hcd203040_compound15.17390.17390.3 | controllerType=0 controllerNumber=1 scan=17390 | 37.9923 |
| 13758:3 | 2 | R.LN[+1540.529]YSLPTGQWVGVQLPR.N | HexNAc(2)Hex(7) | N[+1541] | 1156.866 | 3 | 3468.5835 | 3468.5674 | 0 | 4.7 | 1017 | Specific | 1050.31 | 1000.62 | 1000.62 | 20.25 | 83 | >sp\|P24821\|TENA_HUMAN Tenascin OS=Homo sapiens OX=9606 GN=TNC PE=1 SV=3 | 486 | 20201115_07413_1ng_glyco_hcd203040_compound15.17310.17310.3 | controllerType=0 controllerNumber=1 scan=17310 | 37.8353 |
| 08656:3 | 5 | K.TVLTPATNHMGN[+1540.529]VTFTIPANR.E | HexNAc(2)Hex(7) | N[+1541] | 1265.9132 | 3 | 3795.725 | 3795.6886 | 0 | 9.6 | 74 | Specific | 496.85 | 496.85 | 254.04 | 8.98 | 88 | >sp\|P01024\|CO3_HUMAN Complement C3 OS=Homo sapiens OX=9606 GN=C3 PE=1 SV=2 | 492 | 20201115_07413_1ng_glyco_hcd203040_compound15.11958.11958.3 | controllerType=0 controllerNumber=1 scan=11958 | 28.1985 |
| 08658:3 | 5 | K.TVLTPATNHMGN[+1540.529]VTFTIPANR.E | HexNAc(2)Hex(7) | N[+1541] | 1265.9132 | 3 | 3795.725 | 3795.6886 | 0 | 9.6 | 74 | Specific | 578.42 | 559.43 | 253.86 | 10.45 | 88 | >sp\|P01024\|CO3_HUMAN Complement C3 OS=Homo sapiens OX=9606 GN=C3 PE=1 SV=2 | 492 | 20201115_07413_1ng_glyco_hcd203040_compound15.11960.11960.3 | controllerType=0 controllerNumber=1 scan=11960 | 28.2026 |
| 17879:3 | 7 | R.GLSFDVSLEVSQGPGLLN[+2350.830]DTK.V | HexNAc(4)Hex(5)Fuc(1)NeuAc(2) | N[+2351] | 1509.653 | 3 | 4526.9445 | 4526.9436 | 0 | 0.2 | 118 | Specific | 393.15 | 254.33 | 254.33 | 6.62 | 62 | >sp\|P01833\|PIGR_HUMAN Polymeric immunoglobulin receptor OS=Homo sapiens OX=9606 GN=PIGR PE=1 SV=4 | 902 | 20201115_07413_1ng_glyco_hcd203040_compound15.21652.21652.3 | controllerType=0 controllerNumber=1 scan=21652 | 46.6712 |
| 17858:3 | 7 | R.GLSFDVSLEVSQGPGLLN[+2350.830]DTK.V | HexNAc(4)Hex(5)Fuc(1)NeuAc(2) | N[+2351] | 1509.6619 | 3 | 4526.9712 | 4526.9436 | 0 | 6.1 | 118 | Specific | 943.16 | 921.18 | 921.18 | 17.18 | 62 | >sp\|P01833\|PIGR_HUMAN Polymeric immunoglobulin receptor OS=Homo sapiens OX=9606 GN=PIGR PE=1 SV=4 | 902 | 20201115_07413_1ng_glyco_hcd203040_compound15.21630.21630.3 | controllerType=0 controllerNumber=1 scan=21630 | 46.6205 |
| 17881:3 | 7 | R.GLSFDVSLEVSQGPGLLN[+2350.830]DTK.V | HexNAc(4)Hex(5)Fuc(1)NeuAc(2) | N[+2351] | 1509.653 | 3 | 4526.9445 | 4526.9436 | 0 | 0.2 | 118 | Specific | 963.95 | 903.41 | 903.41 | 18.38 | 62 | >sp\|P01833\|PIGR_HUMAN Polymeric immunoglobulin receptor OS=Homo sapiens OX=9606 GN=PIGR PE=1 SV=4 | 902 | 20201115_07413_1ng_glyco_hcd203040_compound15.21654.21654.3 | controllerType=0 controllerNumber=1 scan=21654 | 46.6765 |
| 17311:3 | 7 | K.QIGLYPVLVIDSSGYVNPN[+1913.677]YTGR.I | HexNAc(4)Hex(5)NeuAc(1) | N[+1914] | 1480.3296 | 3 | 4438.9744 | 4438.9805 | 0 | -1.4 | 168 | Specific | 383.58 | 352.78 | 352.78 | 6.52 | 62 | >sp\|P01833\|PIGR_HUMAN Polymeric immunoglobulin receptor OS=Homo sapiens OX=9606 GN=PIGR PE=1 SV=4 | 902 | 20201115_07413_1ng_glyco_hcd203040_compound15.21052.21052.3 | controllerType=0 controllerNumber=1 scan=21052 | 45.4455 |
| 17314:3 | 7 | K.QIGLYPVLVIDSSGYVNPN[+1913.677]YTGR.I | HexNAc(4)Hex(5)NeuAc(1) | N[+1914] | 1480.3296 | 3 | 4438.9744 | 4438.9805 | 0 | -1.4 | 168 | Specific | 707.87 | 604.28 | 604.28 | 12.88 | 62 | >sp\|P01833\|PIGR_HUMAN Polymeric immunoglobulin receptor OS=Homo sapiens OX=9606 GN=PIGR PE=1 SV=4 | 902 | 20201115_07413_1ng_glyco_hcd203040_compound15.21055.21055.3 | controllerType=0 controllerNumber=1 scan=21055 | 45.4504 |
| 26906:3 | 7 | R.[+42.011]LSLLEEPGN[+2351.851]GTFTVILNQLTSR.D | HexNAc(4)Hex(5)Fuc(3)NeuAc(1) | .[42], N[+2352] | 1599.3948 | 3 | 4796.1699 | 4796.1539 | 0 | 3.3 | 413 | Specific | 267.2 | 245.33 | 245.33 | 6.38 | 62 | >sp\|P01833\|PIGR_HUMAN Polymeric immunoglobulin receptor OS=Homo sapiens OX=9606 GN=PIGR PE=1 SV=4 | 902 | 20201115_07413_1ng_glyco_hcd203040_compound15.31221.31221.3 | controllerType=0 controllerNumber=1 scan=31221 | 65.1214 |
| 18822:3 | 7 | R.LSLLEEPGN[+2076.750]GTFTVILNQLTSR.D | HexNAc(4)Hex(6)Fuc(2) | N[+2077] | 1493.7005 | 3 | 4479.0869 | 4479.0428 | 0 | 9.9 | 413 | Specific | 437.09 | 343.8 | 343.8 | 5.64 | 62 | >sp\|P01833\|PIGR_HUMAN Polymeric immunoglobulin receptor OS=Homo sapiens OX=9606 GN=PIGR PE=1 SV=4 | 902 | 20201115_07413_1ng_glyco_hcd203040_compound15.22641.22641.3 | controllerType=0 controllerNumber=1 scan=22641 | 48.7998 |
| 18656:3 | 7 | R.LSLLEEPGN[+2570.925]GTFTVILNQLTSR.D | HexNAc(5)Hex(6)Fuc(2)NeuAc(1) | N[+2571] | 1658.419 | 3 | 4973.2425 | 4973.2176 | 0 | 5 | 413 | Specific | 572.03 | 498.65 | 498.65 | 10.34 | 62 | >sp\|P01833\|PIGR_HUMAN Polymeric immunoglobulin receptor OS=Homo sapiens OX=9606 GN=PIGR PE=1 SV=4 | 902 | 20201115_07413_1ng_glyco_hcd203040_compound15.22466.22466.3 | controllerType=0 controllerNumber=1 scan=22466 | 48.4032 |
| 18654:3 | 7 | R.LSLLEEPGN[+2570.925]GTFTVILNQLTSR.D | HexNAc(5)Hex(6)Fuc(2)NeuAc(1) | N[+2571] | 1658.419 | 3 | 4973.2425 | 4973.2176 | 0 | 5 | 413 | Specific | 711.76 | 543.66 | 543.66 | 13.34 | 62 | >sp\|P01833\|PIGR_HUMAN Polymeric immunoglobulin receptor OS=Homo sapiens OX=9606 GN=PIGR PE=1 SV=4 | 902 | 20201115_07413_1ng_glyco_hcd203040_compound15.22464.22464.3 | controllerType=0 controllerNumber=1 scan=22464 | 48.3994 |
| 21014:3 | 7 | R.LSLLEEPGN[+2350.830]GTFTVILNQLTSR.D | HexNAc(4)Hex(5)Fuc(1)NeuAc(2) | N[+2351] | 1585.0566 | 3 | 4753.1552 | 4753.1229 | 0 | 6.8 | 413 | Specific | 894.56 | 672.42 | 672.42 | 16.18 | 62 | >sp\|P01833\|PIGR_HUMAN Polymeric immunoglobulin receptor OS=Homo sapiens OX=9606 GN=PIGR PE=1 SV=4 | 902 | 20201115_07413_1ng_glyco_hcd203040_compound15.24939.24939.2 | controllerType=0 controllerNumber=1 scan=24939 | 53.3933 |
| 14460:3 | 7 | K.[+42.011]IIEGEPNLKVPGN[+1872.651]VTAVLGETLK.V | HexNAc(3)Hex(6)NeuAc(1) | .[42], N[+1873] | 1436.0038 | 3 | 4305.9968 | 4306.0104 | 0 | -3.1 | 457 | Specific | 225.06 | 225.06 | 225.06 | 3.66 | 62 | >sp\|P01833\|PIGR_HUMAN Polymeric immunoglobulin receptor OS=Homo sapiens OX=9606 GN=PIGR PE=1 SV=4 | 902 | 20201115_07413_1ng_glyco_hcd203040_compound15.18061.18061.3 | controllerType=0 controllerNumber=1 scan=18061 | 39.2417 |
| 14391:3 | 7 | K.IIEGEPNLKVPGN[+2351.851]VTAVLGETLK.V | HexNAc(4)Hex(5)Fuc(3)NeuAc(1) | N[+2352] | 1581.7391 | 3 | 4743.2027 | 4743.2001 | 0 | 0.6 | 457 | Specific | 298.97 | 278.72 | 278.72 | 6.28 | 62 | >sp\|P01833\|PIGR_HUMAN Polymeric immunoglobulin receptor OS=Homo sapiens OX=9606 GN=PIGR PE=1 SV=4 | 902 | 20201115_07413_1ng_glyco_hcd203040_compound15.17987.17987.3 | controllerType=0 controllerNumber=1 scan=17987 | 39.1012 |
| 14389:3 | 7 | K.IIEGEPNLKVPGN[+2351.851]VTAVLGETLK.V | HexNAc(4)Hex(5)Fuc(3)NeuAc(1) | N[+2352] | 1581.7391 | 3 | 4743.2027 | 4743.2001 | 0 | 0.6 | 457 | Specific | 555.07 | 522.21 | 522.21 | 9.05 | 62 | >sp\|P01833\|PIGR_HUMAN Polymeric immunoglobulin receptor OS=Homo sapiens OX=9606 GN=PIGR PE=1 SV=4 | 902 | 20201115_07413_1ng_glyco_hcd203040_compound15.17985.17985.3 | controllerType=0 controllerNumber=1 scan=17985 | 39.0975 |
| 16296:4 | 7 | K.IIEGEPNLKVPGN[+2350.830]VTAVLGETLK.V | HexNAc(4)Hex(5)Fuc(1)NeuAc(2) | N[+2351] | 1186.3049 | 4 | 4742.1976 | 4742.1797 | 0 | 3.8 | 457 | Specific | 594.06 | 526.85 | 526.85 | 10.23 | 62 | >sp\|P01833\|PIGR_HUMAN Polymeric immunoglobulin receptor OS=Homo sapiens OX=9606 GN=PIGR PE=1 SV=4 | 902 | 20201115_07413_1ng_glyco_hcd203040_compound15.19989.19989.4 | controllerType=0 controllerNumber=1 scan=19989 | 43.1674 |
| 16299:4 | 7 | K.IIEGEPNLKVPGN[+2350.830]VTAVLGETLK.V | HexNAc(4)Hex(5)Fuc(1)NeuAc(2) | N[+2351] | 1186.3049 | 4 | 4742.1976 | 4742.1797 | 0 | 3.8 | 457 | Specific | 632.89 | 525.29 | 525.29 | 11.23 | 62 | >sp\|P01833\|PIGR_HUMAN Polymeric immunoglobulin receptor OS=Homo sapiens OX=9606 GN=PIGR PE=1 SV=4 | 902 | 20201115_07413_1ng_glyco_hcd203040_compound15.19992.19992.4 | controllerType=0 controllerNumber=1 scan=19992 | 43.1727 |
| 16264:3 | 7 | K.IIEGEPNLKVPGN[+2350.830]VTAVLGETLK.V | HexNAc(4)Hex(5)Fuc(1)NeuAc(2) | N[+2351] | 1581.4057 | 3 | 4742.2025 | 4742.1797 | 0 | 4.8 | 457 | Specific | 730.28 | 557.93 | 557.93 | 13.18 | 62 | >sp\|P01833\|PIGR_HUMAN Polymeric immunoglobulin receptor OS=Homo sapiens OX=9606 GN=PIGR PE=1 SV=4 | 902 | 20201115_07413_1ng_glyco_hcd203040_compound15.19955.19955.2 | controllerType=0 controllerNumber=1 scan=19955 | 43.1029 |
| 16261:3 | 7 | K.IIEGEPNLKVPGN[+2350.830]VTAVLGETLK.V | HexNAc(4)Hex(5)Fuc(1)NeuAc(2) | N[+2351] | 1581.4057 | 3 | 4742.2025 | 4742.1797 | 0 | 4.8 | 457 | Specific | 1035.25 | 959.88 | 959.88 | 19.18 | 62 | >sp\|P01833\|PIGR_HUMAN Polymeric immunoglobulin receptor OS=Homo sapiens OX=9606 GN=PIGR PE=1 SV=4 | 902 | 20201115_07413_1ng_glyco_hcd203040_compound15.19952.19952.2 | controllerType=0 controllerNumber=1 scan=19952 | 43.091 |
| 15439:4 | 7 | K.IIEGEPNLKVPGN[+2350.830]VTAVLGETLKVPC[+57.021]HFPC[+57.021]K.F | HexNAc(4)Hex(5)Fuc(1)NeuAc(2) | C[+57]*2, N[+2351] | 1442.6703 | 4 | 5767.6593 | 5767.6372 | 0 | 3.8 | 457 | Specific | 684.76 | 666.24 | 666.24 | 11.57 | 62 | >sp\|P01833\|PIGR_HUMAN Polymeric immunoglobulin receptor OS=Homo sapiens OX=9606 GN=PIGR PE=1 SV=4 | 902 | 20201115_07413_1ng_glyco_hcd203040_compound15.19089.19089.4 | controllerType=0 controllerNumber=1 scan=19089 | 41.1435 |
| 15440:5 | 7 | K.IIEGEPNLKVPGN[+2350.830]VTAVLGETLKVPC[+57.021]HFPC[+57.021]K.F | HexNAc(4)Hex(5)Fuc(1)NeuAc(2) | C[+57]*2, N[+2351] | 1154.3364 | 5 | 5767.653 | 5767.6372 | 0 | 2.7 | 457 | Specific | 941.95 | 687.89 | 687.89 | 17.19 | 62 | >sp\|P01833\|PIGR_HUMAN Polymeric immunoglobulin receptor OS=Homo sapiens OX=9606 GN=PIGR PE=1 SV=4 | 902 | 20201115_07413_1ng_glyco_hcd203040_compound15.19090.19090.5 | controllerType=0 controllerNumber=1 scan=19090 | 41.149 |
| 15437:4 | 7 | K.IIEGEPNLKVPGN[+2350.830]VTAVLGETLKVPC[+57.021]HFPC[+57.021]K.F | HexNAc(4)Hex(5)Fuc(1)NeuAc(2) | C[+57]*2, N[+2351] | 1442.6703 | 4 | 5767.6593 | 5767.6372 | 0 | 3.8 | 457 | Specific | 1020.68 | 929.26 | 929.26 | 18.57 | 62 | >sp\|P01833\|PIGR_HUMAN Polymeric immunoglobulin receptor OS=Homo sapiens OX=9606 GN=PIGR PE=1 SV=4 | 902 | 20201115_07413_1ng_glyco_hcd203040_compound15.19087.19087.4 | controllerType=0 controllerNumber=1 scan=19087 | 41.1376 |
| 15438:5 | 7 | K.IIEGEPNLKVPGN[+2350.830]VTAVLGETLKVPC[+57.021]HFPC[+57.021]K.F | HexNAc(4)Hex(5)Fuc(1)NeuAc(2) | C[+57]*2, N[+2351] | 1154.3364 | 5 | 5767.653 | 5767.6372 | 0 | 2.7 | 457 | Specific | 1112.71 | 893.14 | 893.14 | 21.19 | 62 | >sp\|P01833\|PIGR_HUMAN Polymeric immunoglobulin receptor OS=Homo sapiens OX=9606 GN=PIGR PE=1 SV=4 | 902 | 20201115_07413_1ng_glyco_hcd203040_compound15.19088.19088.5 | controllerType=0 controllerNumber=1 scan=19088 | 41.139 |
| 12074:3 | 7 | K.VPGN[+2059.735]VTAVLGETLK.V | HexNAc(4)Hex(5)Fuc(1)NeuAc(1) | N[+2060] | 1153.1843 | 3 | 3457.5383 | 3457.5348 | 0 | 1 | 466 | Specific | 766.26 | 496.36 | 496.36 | 14.28 | 62 | >sp\|P01833\|PIGR_HUMAN Polymeric immunoglobulin receptor OS=Homo sapiens OX=9606 GN=PIGR PE=1 SV=4 | 902 | 20201115_07413_1ng_glyco_hcd203040_compound15.15555.15555.3 | controllerType=0 controllerNumber=1 scan=15555 | 34.637 |
| 12072:3 | 7 | K.VPGN[+2059.735]VTAVLGETLK.V | HexNAc(4)Hex(5)Fuc(1)NeuAc(1) | N[+2060] | 1153.1843 | 3 | 3457.5383 | 3457.5348 | 0 | 1 | 466 | Specific | 775.65 | 537.5 | 537.5 | 14.28 | 62 | >sp\|P01833\|PIGR_HUMAN Polymeric immunoglobulin receptor OS=Homo sapiens OX=9606 GN=PIGR PE=1 SV=4 | 902 | 20201115_07413_1ng_glyco_hcd203040_compound15.15553.15553.3 | controllerType=0 controllerNumber=1 scan=15553 | 34.6282 |
| 12117:3 | 7 | K.VPGN[+2059.735]VTAVLGETLKVPC[+57.021]HFPC[+57.021]K.F | HexNAc(4)Hex(5)Fuc(1)NeuAc(1) | C[+57]*2, N[+2060] | 1495.0084 | 3 | 4483.0106 | 4482.9923 | 0 | 4.1 | 466 | Specific | 402.56 | 390.23 | 390.23 | 7.72 | 62 | >sp\|P01833\|PIGR_HUMAN Polymeric immunoglobulin receptor OS=Homo sapiens OX=9606 GN=PIGR PE=1 SV=4 | 902 | 20201115_07413_1ng_glyco_hcd203040_compound15.15600.15600.3 | controllerType=0 controllerNumber=1 scan=15600 | 34.7205 |
| 12032:4 | 7 | K.VPGN[+2205.793]VTAVLGETLKVPC[+57.021]HFPC[+57.021]K.F | HexNAc(4)Hex(5)Fuc(2)NeuAc(1) | C[+57]*2, N[+2206] | 1158.0254 | 4 | 4629.0799 | 4629.0502 | 0 | 6.4 | 466 | Specific | 670.83 | 670.83 | 670.83 | 12.02 | 62 | >sp\|P01833\|PIGR_HUMAN Polymeric immunoglobulin receptor OS=Homo sapiens OX=9606 GN=PIGR PE=1 SV=4 | 902 | 20201115_07413_1ng_glyco_hcd203040_compound15.15511.15511.3 | controllerType=0 controllerNumber=1 scan=15511 | 34.5288 |
| 12049:4 | 7 | K.VPGN[+2059.735]VTAVLGETLKVPC[+57.021]HFPC[+57.021]K.F | HexNAc(4)Hex(5)Fuc(1)NeuAc(1) | C[+57]*2, N[+2060] | 1121.5133 | 4 | 4483.0316 | 4482.9923 | 0 | 8.8 | 466 | Specific | 885.94 | 885.94 | 885.94 | 16.02 | 62 | >sp\|P01833\|PIGR_HUMAN Polymeric immunoglobulin receptor OS=Homo sapiens OX=9606 GN=PIGR PE=1 SV=4 | 902 | 20201115_07413_1ng_glyco_hcd203040_compound15.15528.15528.4 | controllerType=0 controllerNumber=1 scan=15528 | 34.563 |
| 12047:4 | 7 | K.VPGN[+2059.735]VTAVLGETLKVPC[+57.021]HFPC[+57.021]K.F | HexNAc(4)Hex(5)Fuc(1)NeuAc(1) | C[+57]*2, N[+2060] | 1121.5133 | 4 | 4483.0316 | 4482.9923 | 0 | 8.8 | 466 | Specific | 950.87 | 898.57 | 898.57 | 18.02 | 62 | >sp\|P01833\|PIGR_HUMAN Polymeric immunoglobulin receptor OS=Homo sapiens OX=9606 GN=PIGR PE=1 SV=4 | 902 | 20201115_07413_1ng_glyco_hcd203040_compound15.15526.15526.4 | controllerType=0 controllerNumber=1 scan=15526 | 34.5574 |
| 12030:4 | 7 | K.VPGN[+2205.793]VTAVLGETLKVPC[+57.021]HFPC[+57.021]K.F | HexNAc(4)Hex(5)Fuc(2)NeuAc(1) | C[+57]*2, N[+2206] | 1158.0254 | 4 | 4629.0799 | 4629.0502 | 0 | 6.4 | 466 | Specific | 970.24 | 650.99 | 650.99 | 18.02 | 62 | >sp\|P01833\|PIGR_HUMAN Polymeric immunoglobulin receptor OS=Homo sapiens OX=9606 GN=PIGR PE=1 SV=4 | 902 | 20201115_07413_1ng_glyco_hcd203040_compound15.15509.15509.3 | controllerType=0 controllerNumber=1 scan=15509 | 34.5209 |
| 12115:3 | 7 | K.VPGN[+2059.735]VTAVLGETLKVPC[+57.021]HFPC[+57.021]K.F | HexNAc(4)Hex(5)Fuc(1)NeuAc(1) | C[+57]*2, N[+2060] | 1495.0084 | 3 | 4483.0106 | 4482.9923 | 0 | 4.1 | 466 | Specific | 1025.54 | 899.43 | 899.43 | 18.56 | 62 | >sp\|P01833\|PIGR_HUMAN Polymeric immunoglobulin receptor OS=Homo sapiens OX=9606 GN=PIGR PE=1 SV=4 | 902 | 20201115_07413_1ng_glyco_hcd203040_compound15.15598.15598.3 | controllerType=0 controllerNumber=1 scan=15598 | 34.7154 |
| 04764:3 | 7 | K.WN[+1548.545]NTGC[+57.021]QALPSQDEGPSK.A | HexNAc(3)Hex(4)NeuAc(1) | C[+57], N[+1549] | 1179.817 | 3 | 3537.4364 | 3537.4215 | 0 | 4.2 | 498 | Specific | 202.53 | 194.65 | 194.65 | 4.46 | 62 | >sp\|P01833\|PIGR_HUMAN Polymeric immunoglobulin receptor OS=Homo sapiens OX=9606 GN=PIGR PE=1 SV=4 | 902 | 20201115_07413_1ng_glyco_hcd203040_compound15.7835.7835.3 | controllerType=0 controllerNumber=1 scan=7835 | 21.0077 |
| 04597:3 | 7 | K.WN[+1872.651]NTGC[+57.021]QALPSQDEGPSK.A | HexNAc(3)Hex(6)NeuAc(1) | C[+57], N[+1873] | 1287.8494 | 3 | 3861.5336 | 3861.5272 | 0 | 1.7 | 498 | Specific | 374.04 | 353.81 | 353.81 | 7.23 | 62 | >sp\|P01833\|PIGR_HUMAN Polymeric immunoglobulin receptor OS=Homo sapiens OX=9606 GN=PIGR PE=1 SV=4 | 902 | 20201115_07413_1ng_glyco_hcd203040_compound15.7659.7659.3 | controllerType=0 controllerNumber=1 scan=7659 | 20.6965 |
| 04599:3 | 7 | K.WN[+1872.651]NTGC[+57.021]QALPSQDEGPSK.A | HexNAc(3)Hex(6)NeuAc(1) | C[+57], N[+1873] | 1287.8494 | 3 | 3861.5336 | 3861.5272 | 0 | 1.7 | 498 | Specific | 450.7 | 418.73 | 418.73 | 8.91 | 62 | >sp\|P01833\|PIGR_HUMAN Polymeric immunoglobulin receptor OS=Homo sapiens OX=9606 GN=PIGR PE=1 SV=4 | 902 | 20201115_07413_1ng_glyco_hcd203040_compound15.7661.7661.3 | controllerType=0 controllerNumber=1 scan=7661 | 20.7006 |
| 06517:3 | 7 | K.WN[+2204.772]NTGC[+57.021]QALPSQDEGPSK.A | HexNAc(4)Hex(5)NeuAc(2) | C[+57], N[+2205] | 1398.5596 | 3 | 4193.6643 | 4193.6491 | 0 | 3.6 | 498 | Specific | 532.86 | 466.39 | 466.39 | 9.22 | 62 | >sp\|P01833\|PIGR_HUMAN Polymeric immunoglobulin receptor OS=Homo sapiens OX=9606 GN=PIGR PE=1 SV=4 | 902 | 20201115_07413_1ng_glyco_hcd203040_compound15.9690.9690.3 | controllerType=0 controllerNumber=1 scan=9690 | 24.2306 |
| 04766:3 | 7 | K.WN[+1548.545]NTGC[+57.021]QALPSQDEGPSK.A | HexNAc(3)Hex(4)NeuAc(1) | C[+57], N[+1549] | 1179.817 | 3 | 3537.4364 | 3537.4215 | 0 | 4.2 | 498 | Specific | 572.31 | 507.46 | 507.46 | 10.25 | 62 | >sp\|P01833\|PIGR_HUMAN Polymeric immunoglobulin receptor OS=Homo sapiens OX=9606 GN=PIGR PE=1 SV=4 | 902 | 20201115_07413_1ng_glyco_hcd203040_compound15.7837.7837.3 | controllerType=0 controllerNumber=1 scan=7837 | 21.0116 |
| 06519:3 | 7 | K.WN[+2204.772]NTGC[+57.021]QALPSQDEGPSK.A | HexNAc(4)Hex(5)NeuAc(2) | C[+57], N[+2205] | 1398.5596 | 3 | 4193.6643 | 4193.6491 | 0 | 3.6 | 498 | Specific | 592.45 | 504.9 | 504.9 | 10.57 | 62 | >sp\|P01833\|PIGR_HUMAN Polymeric immunoglobulin receptor OS=Homo sapiens OX=9606 GN=PIGR PE=1 SV=4 | 902 | 20201115_07413_1ng_glyco_hcd203040_compound15.9692.9692.3 | controllerType=0 controllerNumber=1 scan=9692 | 24.2357 |
| 04682:3 | 7 | K.WN[+1710.598]NTGC[+57.021]QALPSQDEGPSK.A | HexNAc(3)Hex(5)NeuAc(1) | C[+57], N[+1711] | 1233.8303 | 3 | 3699.4763 | 3699.4743 | 0 | 0.5 | 498 | Specific | 640.02 | 589.58 | 589.58 | 11.28 | 62 | >sp\|P01833\|PIGR_HUMAN Polymeric immunoglobulin receptor OS=Homo sapiens OX=9606 GN=PIGR PE=1 SV=4 | 902 | 20201115_07413_1ng_glyco_hcd203040_compound15.7748.7748.3 | controllerType=0 controllerNumber=1 scan=7748 | 20.8515 |
| 04684:3 | 7 | K.WN[+1710.598]NTGC[+57.021]QALPSQDEGPSK.A | HexNAc(3)Hex(5)NeuAc(1) | C[+57], N[+1711] | 1233.8303 | 3 | 3699.4763 | 3699.4743 | 0 | 0.5 | 498 | Specific | 702.12 | 626.19 | 626.19 | 13.28 | 62 | >sp\|P01833\|PIGR_HUMAN Polymeric immunoglobulin receptor OS=Homo sapiens OX=9606 GN=PIGR PE=1 SV=4 | 902 | 20201115_07413_1ng_glyco_hcd203040_compound15.7750.7750.3 | controllerType=0 controllerNumber=1 scan=7750 | 20.8549 |
| 11616:5 | 8 | K.KGN[+2059.735]TTLNSFVIPSESDVPTHC[+57.021]PSQWWPYAGHC[+57.021]YK.I | HexNAc(4)Hex(5)Fuc(1)NeuAc(1) | C[+57]*2, N[+2060] | 1205.52 | 5 | 6023.571 | 6023.5567 | 0 | 2.4 | 342 | Specific | 208.8 | 208.8 | 208.8 | 5.59 | 69 | >sp\|P22897\|MRC1_HUMAN Macrophage mannose receptor 1 OS=Homo sapiens OX=9606 GN=MRC1 PE=1 SV=1 | 1049 | 20201115_07413_1ng_glyco_hcd203040_compound15.15075.15075.3 | controllerType=0 controllerNumber=1 scan=15075 | 33.7562 |
| 15557:4 | 9 | K.RNIAAFGGDPNN[+1913.677]ITLFGESAGGASVSLQTLSPYNK.G | HexNAc(4)Hex(5)NeuAc(1) | N[+1914] | 1370.8796 | 4 | 5480.4967 | 5480.4644 | 0 | 5.9 | 196 | Specific | 431.42 | 405.82 | 405.82 | 8.96 | 49 | >sp\|P19835\|CEL_HUMAN Bile salt-activated lipase OS=Homo sapiens OX=9606 GN=CEL PE=1 SV=3 | 476 | 20201115_07413_1ng_glyco_hcd203040_compound15.19212.19212.4 | controllerType=0 controllerNumber=1 scan=19212 | 41.4163 |
| 15559:3 | 9 | K.RNIAAFGGDPNN[+1913.677]ITLFGESAGGASVSLQTLSPYNK.G | HexNAc(4)Hex(5)NeuAc(1) | N[+1914] | 1827.5039 | 3 | 5480.4971 | 5480.4644 | 0 | 6 | 196 | Specific | 455.89 | 368.63 | 368.63 | 7.8 | 49 | >sp\|P19835\|CEL_HUMAN Bile salt-activated lipase OS=Homo sapiens OX=9606 GN=CEL PE=1 SV=3 | 476 | 20201115_07413_1ng_glyco_hcd203040_compound15.19214.19214.3 | controllerType=0 controllerNumber=1 scan=19214 | 41.4186 |
| 15561:3 | 9 | K.RNIAAFGGDPNN[+1913.677]ITLFGESAGGASVSLQTLSPYNK.G | HexNAc(4)Hex(5)NeuAc(1) | N[+1914] | 1827.5039 | 3 | 5480.4971 | 5480.4644 | 0 | 6 | 196 | Specific | 774.93 | 702.9 | 702.9 | 14.89 | 49 | >sp\|P19835\|CEL_HUMAN Bile salt-activated lipase OS=Homo sapiens OX=9606 GN=CEL PE=1 SV=3 | 476 | 20201115_07413_1ng_glyco_hcd203040_compound15.19216.19216.3 | controllerType=0 controllerNumber=1 scan=19216 | 41.4263 |
| 15560:4 | 9 | K.RNIAAFGGDPNN[+1913.677]ITLFGESAGGASVSLQTLSPYNK.G | HexNAc(4)Hex(5)NeuAc(1) | N[+1914] | 1370.8796 | 4 | 5480.4967 | 5480.4644 | 0 | 5.9 | 196 | Specific | 990.23 | 921.09 | 921.09 | 18.62 | 49 | >sp\|P19835\|CEL_HUMAN Bile salt-activated lipase OS=Homo sapiens OX=9606 GN=CEL PE=1 SV=3 | 476 | 20201115_07413_1ng_glyco_hcd203040_compound15.19215.19215.4 | controllerType=0 controllerNumber=1 scan=19215 | 41.42 |
| 17235:4 | 9 | R.NIAAFGGDPNN[+1913.677]ITLFGESAGGASVSLQTLSPYNK.G | HexNAc(4)Hex(5)NeuAc(1) | N[+1914] | 1331.8562 | 4 | 5324.403 | 5324.3633 | 0 | 7.5 | 197 | Specific | 867.04 | 829.2 | 829.2 | 17.1 | 49 | >sp\|P19835\|CEL_HUMAN Bile salt-activated lipase OS=Homo sapiens OX=9606 GN=CEL PE=1 SV=3 | 476 | 20201115_07413_1ng_glyco_hcd203040_compound15.20973.20973.3 | controllerType=0 controllerNumber=1 scan=20973 | 45.2859 |
| 17238:4 | 9 | R.NIAAFGGDPNN[+1913.677]ITLFGESAGGASVSLQTLSPYNK.G | HexNAc(4)Hex(5)NeuAc(1) | N[+1914] | 1331.8562 | 4 | 5324.403 | 5324.3633 | 0 | 7.5 | 197 | Specific | 909.36 | 898.95 | 898.95 | 18.1 | 49 | >sp\|P19835\|CEL_HUMAN Bile salt-activated lipase OS=Homo sapiens OX=9606 GN=CEL PE=1 SV=3 | 476 | 20201115_07413_1ng_glyco_hcd203040_compound15.20976.20976.3 | controllerType=0 controllerNumber=1 scan=20976 | 45.2967 |
| 17263:3 | 9 | R.NIAAFGGDPNN[+1913.677]ITLFGESAGGASVSLQTLSPYNK.G | HexNAc(4)Hex(5)NeuAc(1) | N[+1914] | 1775.4588 | 3 | 5324.3619 | 5324.3633 | 0 | -0.3 | 197 | Specific | 943.93 | 875.82 | 875.82 | 17.81 | 49 | >sp\|P19835\|CEL_HUMAN Bile salt-activated lipase OS=Homo sapiens OX=9606 GN=CEL PE=1 SV=3 | 476 | 20201115_07413_1ng_glyco_hcd203040_compound15.21002.21002.3 | controllerType=0 controllerNumber=1 scan=21002 | 45.3522 |
| 17265:3 | 9 | R.NIAAFGGDPNN[+1913.677]ITLFGESAGGASVSLQTLSPYNK.G | HexNAc(4)Hex(5)NeuAc(1) | N[+1914] | 1775.4588 | 3 | 5324.3619 | 5324.3633 | 0 | -0.3 | 197 | Specific | 1072.6 | 1038.64 | 1038.64 | 20.81 | 49 | >sp\|P19835\|CEL_HUMAN Bile salt-activated lipase OS=Homo sapiens OX=9606 GN=CEL PE=1 SV=3 | 476 | 20201115_07413_1ng_glyco_hcd203040_compound15.21004.21004.3 | controllerType=0 controllerNumber=1 scan=21004 | 45.3627 |
| 16054:3 | 10 | R.LSLHRPALEDLLLGSEAN[+1702.581]LTC[+57.021]TLTGLR.D | HexNAc(2)Hex(8) | C[+57], N[+1703] | 1556.0796 | 3 | 4666.2242 | 4666.1796 | 0 | 9.6 | 127 | Specific | 205.3 | 205.3 | 205.3 | 4.34 | 52 | >sp\|P01876\|IGHA1_HUMAN Immunoglobulin heavy constant alpha 1 OS=Homo sapiens OX=9606 GN=IGHA1 PE=1 SV=2 | 255 | 20201115_07413_1ng_glyco_hcd203040_compound15.19733.19733.3 | controllerType=0 controllerNumber=1 scan=19733 | 42.5283 |
| 16045:3 | 10 | R.LSLHRPALEDLLLGSEAN[+1540.529]LTC[+57.021]TLTGLR.D | HexNAc(2)Hex(7) | C[+57], N[+1541] | 1502.0549 | 3 | 4504.15 | 4504.1268 | 0 | 5.2 | 127 | Specific | 260.19 | 260.19 | 260.19 | 5.75 | 52 | >sp\|P01876\|IGHA1_HUMAN Immunoglobulin heavy constant alpha 1 OS=Homo sapiens OX=9606 GN=IGHA1 PE=1 SV=2 | 255 | 20201115_07413_1ng_glyco_hcd203040_compound15.19724.19724.3 | controllerType=0 controllerNumber=1 scan=19724 | 42.4983 |
| 16990:3 | 10 | R.LSLHRPALEDLLLGSEAN[+2116.756]LTC[+57.021]TLTGLR.D | HexNAc(5)Hex(5)NeuAc(1) | C[+57], N[+2117] | 1694.1369 | 3 | 5080.3962 | 5080.3547 | 0 | 8.2 | 127 | Specific | 270.75 | 169.46 | 169.46 | 5.46 | 52 | >sp\|P01876\|IGHA1_HUMAN Immunoglobulin heavy constant alpha 1 OS=Homo sapiens OX=9606 GN=IGHA1 PE=1 SV=2 | 255 | 20201115_07413_1ng_glyco_hcd203040_compound15.20714.20714.3 | controllerType=0 controllerNumber=1 scan=20714 | 44.6068 |
| 18128:3 | 10 | R.LSLHRPALEDLLLGSEAN[+2204.772]LTC[+57.021]TLTGLR.D | HexNAc(4)Hex(5)NeuAc(2) | C[+57], N[+2205] | 1723.4718 | 3 | 5168.4008 | 5168.3707 | 0 | 5.8 | 127 | Specific | 381.79 | 304.66 | 304.66 | 6.46 | 52 | >sp\|P01876\|IGHA1_HUMAN Immunoglobulin heavy constant alpha 1 OS=Homo sapiens OX=9606 GN=IGHA1 PE=1 SV=2 | 255 | 20201115_07413_1ng_glyco_hcd203040_compound15.21913.21913.3 | controllerType=0 controllerNumber=1 scan=21913 | 47.2082 |
| 16137:3 | 10 | R.LSLHRPALEDLLLGSEAN[+1501.555]LTC[+57.021]TLTGLR.D | HexNAc(5)Hex(3) | C[+57], N[+1502] | 1489.0682 | 3 | 4465.19 | 4465.1536 | 0 | 8.1 | 127 | Specific | 414.46 | 414.46 | 414.46 | 8.39 | 52 | >sp\|P01876\|IGHA1_HUMAN Immunoglobulin heavy constant alpha 1 OS=Homo sapiens OX=9606 GN=IGHA1 PE=1 SV=2 | 255 | 20201115_07413_1ng_glyco_hcd203040_compound15.19822.19822.3 | controllerType=0 controllerNumber=1 scan=19822 | 42.7828 |
| 18121:4 | 10 | R.LSLHRPALEDLLLGSEAN[+2204.772]LTC[+57.021]TLTGLR.D | HexNAc(4)Hex(5)NeuAc(2) | C[+57], N[+2205] | 1292.8521 | 4 | 5168.3864 | 5168.3707 | 0 | 3 | 127 | Specific | 425.93 | 375.21 | 375.21 | 9.57 | 52 | >sp\|P01876\|IGHA1_HUMAN Immunoglobulin heavy constant alpha 1 OS=Homo sapiens OX=9606 GN=IGHA1 PE=1 SV=2 | 255 | 20201115_07413_1ng_glyco_hcd203040_compound15.21906.21906.4 | controllerType=0 controllerNumber=1 scan=21906 | 47.1934 |
| 16052:3 | 10 | R.LSLHRPALEDLLLGSEAN[+1702.581]LTC[+57.021]TLTGLR.D | HexNAc(2)Hex(8) | C[+57], N[+1703] | 1556.0796 | 3 | 4666.2242 | 4666.1796 | 0 | 9.6 | 127 | Specific | 446.03 | 446.03 | 446.03 | 8.47 | 52 | >sp\|P01876\|IGHA1_HUMAN Immunoglobulin heavy constant alpha 1 OS=Homo sapiens OX=9606 GN=IGHA1 PE=1 SV=2 | 255 | 20201115_07413_1ng_glyco_hcd203040_compound15.19731.19731.3 | controllerType=0 controllerNumber=1 scan=19731 | 42.5123 |
| 16135:3 | 10 | R.LSLHRPALEDLLLGSEAN[+1501.555]LTC[+57.021]TLTGLR.D | HexNAc(5)Hex(3) | C[+57], N[+1502] | 1489.0682 | 3 | 4465.19 | 4465.1536 | 0 | 8.1 | 127 | Specific | 452.51 | 452.51 | 452.51 | 8.56 | 52 | >sp\|P01876\|IGHA1_HUMAN Immunoglobulin heavy constant alpha 1 OS=Homo sapiens OX=9606 GN=IGHA1 PE=1 SV=2 | 255 | 20201115_07413_1ng_glyco_hcd203040_compound15.19820.19820.3 | controllerType=0 controllerNumber=1 scan=19820 | 42.7789 |
| 16982:4 | 10 | R.LSLHRPALEDLLLGSEAN[+1913.677]LTC[+57.021]TLTGLR.D | HexNAc(4)Hex(5)NeuAc(1) | C[+57], N[+1914] | 1220.0829 | 4 | 4877.3099 | 4877.2753 | 0 | 7.1 | 127 | Specific | 493.16 | 493.16 | 493.16 | 9.39 | 52 | >sp\|P01876\|IGHA1_HUMAN Immunoglobulin heavy constant alpha 1 OS=Homo sapiens OX=9606 GN=IGHA1 PE=1 SV=2 | 255 | 20201115_07413_1ng_glyco_hcd203040_compound15.20706.20706.4 | controllerType=0 controllerNumber=1 scan=20706 | 44.5943 |
| 17055:3 | 10 | R.LSLHRPALEDLLLGSEAN[+1751.624]LTC[+57.021]TLTGLR.D | HexNAc(4)Hex(4)NeuAc(1) | C[+57], N[+1752] | 1572.4124 | 3 | 4715.2227 | 4715.2225 | 0 | 0 | 127 | Specific | 517.32 | 498.86 | 498.86 | 8.97 | 52 | >sp\|P01876\|IGHA1_HUMAN Immunoglobulin heavy constant alpha 1 OS=Homo sapiens OX=9606 GN=IGHA1 PE=1 SV=2 | 255 | 20201115_07413_1ng_glyco_hcd203040_compound15.20783.20783.3 | controllerType=0 controllerNumber=1 scan=20783 | 44.82 |
| 18076:4 | 10 | R.LSLHRPALEDLLLGSEAN[+2407.852]LTC[+57.021]TLTGLR.D | HexNAc(5)Hex(5)NeuAc(2) | C[+57], N[+2408] | 1343.6205 | 4 | 5371.4601 | 5371.4501 | 0 | 1.9 | 127 | Specific | 577.3 | 506.75 | 506.75 | 11.24 | 52 | >sp\|P01876\|IGHA1_HUMAN Immunoglobulin heavy constant alpha 1 OS=Homo sapiens OX=9606 GN=IGHA1 PE=1 SV=2 | 255 | 20201115_07413_1ng_glyco_hcd203040_compound15.21859.21859.4 | controllerType=0 controllerNumber=1 scan=21859 | 47.099 |
| 16046:3 | 10 | R.LSLHRPALEDLLLGSEAN[+1663.608]LTC[+57.021]TLTGLR.D | HexNAc(5)Hex(4) | C[+57], N[+1664] | 1543.0811 | 3 | 4627.2288 | 4627.2064 | 0 | 4.8 | 127 | Specific | 583.76 | 556.02 | 556.02 | 10.28 | 52 | >sp\|P01876\|IGHA1_HUMAN Immunoglobulin heavy constant alpha 1 OS=Homo sapiens OX=9606 GN=IGHA1 PE=1 SV=2 | 255 | 20201115_07413_1ng_glyco_hcd203040_compound15.19725.19725.3 | controllerType=0 controllerNumber=1 scan=19725 | 42.4995 |
| 16049:3 | 10 | R.LSLHRPALEDLLLGSEAN[+1663.608]LTC[+57.021]TLTGLR.D | HexNAc(5)Hex(4) | C[+57], N[+1664] | 1543.0811 | 3 | 4627.2288 | 4627.2064 | 0 | 4.8 | 127 | Specific | 604.23 | 533.29 | 533.29 | 11.28 | 52 | >sp\|P01876\|IGHA1_HUMAN Immunoglobulin heavy constant alpha 1 OS=Homo sapiens OX=9606 GN=IGHA1 PE=1 SV=2 | 255 | 20201115_07413_1ng_glyco_hcd203040_compound15.19728.19728.3 | controllerType=0 controllerNumber=1 scan=19728 | 42.5079 |
| 16039:4 | 10 | R.LSLHRPALEDLLLGSEAN[+1540.529]LTC[+57.021]TLTGLR.D | HexNAc(2)Hex(7) | C[+57], N[+1541] | 1126.7915 | 4 | 4504.1444 | 4504.1268 | 0 | 3.9 | 127 | Specific | 613.53 | 613.53 | 613.53 | 12.22 | 52 | >sp\|P01876\|IGHA1_HUMAN Immunoglobulin heavy constant alpha 1 OS=Homo sapiens OX=9606 GN=IGHA1 PE=1 SV=2 | 255 | 20201115_07413_1ng_glyco_hcd203040_compound15.19718.19718.4 | controllerType=0 controllerNumber=1 scan=19718 | 42.4837 |
| 16992:3 | 10 | R.LSLHRPALEDLLLGSEAN[+2116.756]LTC[+57.021]TLTGLR.D | HexNAc(5)Hex(5)NeuAc(1) | C[+57], N[+2117] | 1694.1369 | 3 | 5080.3962 | 5080.3547 | 0 | 8.2 | 127 | Specific | 614.17 | 584.13 | 584.13 | 11.24 | 52 | >sp\|P01876\|IGHA1_HUMAN Immunoglobulin heavy constant alpha 1 OS=Homo sapiens OX=9606 GN=IGHA1 PE=1 SV=2 | 255 | 20201115_07413_1ng_glyco_hcd203040_compound15.20716.20716.3 | controllerType=0 controllerNumber=1 scan=20716 | 44.6114 |
| 17054:4 | 10 | R.LSLHRPALEDLLLGSEAN[+1751.624]LTC[+57.021]TLTGLR.D | HexNAc(4)Hex(4)NeuAc(1) | C[+57], N[+1752] | 1179.561 | 4 | 4715.222 | 4715.2225 | 0 | -0.1 | 127 | Specific | 630.82 | 582.82 | 582.82 | 11.94 | 52 | >sp\|P01876\|IGHA1_HUMAN Immunoglobulin heavy constant alpha 1 OS=Homo sapiens OX=9606 GN=IGHA1 PE=1 SV=2 | 255 | 20201115_07413_1ng_glyco_hcd203040_compound15.20782.20782.4 | controllerType=0 controllerNumber=1 scan=20782 | 44.8187 |
| 18078:4 | 10 | R.LSLHRPALEDLLLGSEAN[+2407.852]LTC[+57.021]TLTGLR.D | HexNAc(5)Hex(5)NeuAc(2) | C[+57], N[+2408] | 1343.6205 | 4 | 5371.4601 | 5371.4501 | 0 | 1.9 | 127 | Specific | 634.65 | 538.49 | 538.49 | 12.24 | 52 | >sp\|P01876\|IGHA1_HUMAN Immunoglobulin heavy constant alpha 1 OS=Homo sapiens OX=9606 GN=IGHA1 PE=1 SV=2 | 255 | 20201115_07413_1ng_glyco_hcd203040_compound15.21861.21861.4 | controllerType=0 controllerNumber=1 scan=21861 | 47.1041 |
| 16077:3 | 10 | R.LSLHRPALEDLLLGSEAN[+1216.423]LTC[+57.021]TLTGLR.D | HexNAc(2)Hex(5) | C[+57], N[+1216] | 1394.0251 | 3 | 4180.0607 | 4180.0211 | 0 | 9.5 | 127 | Specific | 645.05 | 645.05 | 645.05 | 11.12 | 52 | >sp\|P01876\|IGHA1_HUMAN Immunoglobulin heavy constant alpha 1 OS=Homo sapiens OX=9606 GN=IGHA1 PE=1 SV=2 | 255 | 20201115_07413_1ng_glyco_hcd203040_compound15.19758.19758.3 | controllerType=0 controllerNumber=1 scan=19758 | 42.6211 |
| 16048:3 | 10 | R.LSLHRPALEDLLLGSEAN[+1540.529]LTC[+57.021]TLTGLR.D | HexNAc(2)Hex(7) | C[+57], N[+1541] | 1502.0549 | 3 | 4504.15 | 4504.1268 | 0 | 5.2 | 127 | Specific | 681.55 | 629.94 | 629.94 | 12.28 | 52 | >sp\|P01876\|IGHA1_HUMAN Immunoglobulin heavy constant alpha 1 OS=Homo sapiens OX=9606 GN=IGHA1 PE=1 SV=2 | 255 | 20201115_07413_1ng_glyco_hcd203040_compound15.19727.19727.3 | controllerType=0 controllerNumber=1 scan=19727 | 42.506 |
| 16040:4 | 10 | R.LSLHRPALEDLLLGSEAN[+1663.608]LTC[+57.021]TLTGLR.D | HexNAc(5)Hex(4) | C[+57], N[+1664] | 1157.5627 | 4 | 4627.2288 | 4627.2064 | 0 | 4.8 | 127 | Specific | 703.76 | 703.76 | 703.76 | 13.9 | 52 | >sp\|P01876\|IGHA1_HUMAN Immunoglobulin heavy constant alpha 1 OS=Homo sapiens OX=9606 GN=IGHA1 PE=1 SV=2 | 255 | 20201115_07413_1ng_glyco_hcd203040_compound15.19719.19719.4 | controllerType=0 controllerNumber=1 scan=19719 | 42.4851 |
| 16050:3 | 10 | R.LSLHRPALEDLLLGSEAN[+1378.476]LTC[+57.021]TLTGLR.D | HexNAc(2)Hex(6) | C[+57], N[+1378] | 1448.0368 | 3 | 4342.0958 | 4342.074 | 0 | 5 | 127 | Specific | 745.45 | 675.59 | 675.59 | 13.28 | 52 | >sp\|P01876\|IGHA1_HUMAN Immunoglobulin heavy constant alpha 1 OS=Homo sapiens OX=9606 GN=IGHA1 PE=1 SV=2 | 255 | 20201115_07413_1ng_glyco_hcd203040_compound15.19729.19729.3 | controllerType=0 controllerNumber=1 scan=19729 | 42.5097 |
| 16073:3 | 10 | R.LSLHRPALEDLLLGSEAN[+1216.423]LTC[+57.021]TLTGLR.D | HexNAc(2)Hex(5) | C[+57], N[+1216] | 1394.0251 | 3 | 4180.0607 | 4180.0211 | 0 | 9.5 | 127 | Specific | 745.65 | 745.65 | 745.65 | 13.12 | 52 | >sp\|P01876\|IGHA1_HUMAN Immunoglobulin heavy constant alpha 1 OS=Homo sapiens OX=9606 GN=IGHA1 PE=1 SV=2 | 255 | 20201115_07413_1ng_glyco_hcd203040_compound15.19754.19754.3 | controllerType=0 controllerNumber=1 scan=19754 | 42.6011 |
| 16985:4 | 10 | R.LSLHRPALEDLLLGSEAN[+1913.677]LTC[+57.021]TLTGLR.D | HexNAc(4)Hex(5)NeuAc(1) | C[+57], N[+1914] | 1220.0829 | 4 | 4877.3099 | 4877.2753 | 0 | 7.1 | 127 | Specific | 752.08 | 716.53 | 716.53 | 14.86 | 52 | >sp\|P01876\|IGHA1_HUMAN Immunoglobulin heavy constant alpha 1 OS=Homo sapiens OX=9606 GN=IGHA1 PE=1 SV=2 | 255 | 20201115_07413_1ng_glyco_hcd203040_compound15.20709.20709.4 | controllerType=0 controllerNumber=1 scan=20709 | 44.601 |
| 18124:4 | 10 | R.LSLHRPALEDLLLGSEAN[+2204.772]LTC[+57.021]TLTGLR.D | HexNAc(4)Hex(5)NeuAc(2) | C[+57], N[+2205] | 1292.8521 | 4 | 5168.3864 | 5168.3707 | 0 | 3 | 127 | Specific | 797.59 | 775.02 | 775.02 | 15.22 | 52 | >sp\|P01876\|IGHA1_HUMAN Immunoglobulin heavy constant alpha 1 OS=Homo sapiens OX=9606 GN=IGHA1 PE=1 SV=2 | 255 | 20201115_07413_1ng_glyco_hcd203040_compound15.21909.21909.4 | controllerType=0 controllerNumber=1 scan=21909 | 47.2016 |
| 16053:3 | 10 | R.LSLHRPALEDLLLGSEAN[+1378.476]LTC[+57.021]TLTGLR.D | HexNAc(2)Hex(6) | C[+57], N[+1378] | 1448.0368 | 3 | 4342.0958 | 4342.074 | 0 | 5 | 127 | Specific | 806.34 | 801.43 | 801.43 | 15.28 | 52 | >sp\|P01876\|IGHA1_HUMAN Immunoglobulin heavy constant alpha 1 OS=Homo sapiens OX=9606 GN=IGHA1 PE=1 SV=2 | 255 | 20201115_07413_1ng_glyco_hcd203040_compound15.19732.19732.3 | controllerType=0 controllerNumber=1 scan=19732 | 42.5185 |
| 16042:4 | 10 | R.LSLHRPALEDLLLGSEAN[+1663.608]LTC[+57.021]TLTGLR.D | HexNAc(5)Hex(4) | C[+57], N[+1664] | 1157.5627 | 4 | 4627.2288 | 4627.2064 | 0 | 4.8 | 127 | Specific | 817.5 | 810.53 | 810.53 | 15.9 | 52 | >sp\|P01876\|IGHA1_HUMAN Immunoglobulin heavy constant alpha 1 OS=Homo sapiens OX=9606 GN=IGHA1 PE=1 SV=2 | 255 | 20201115_07413_1ng_glyco_hcd203040_compound15.19721.19721.4 | controllerType=0 controllerNumber=1 scan=19721 | 42.4942 |
| 16023:4 | 10 | R.LSLHRPALEDLLLGSEAN[+1864.634]LTC[+57.021]TLTGLR.D | HexNAc(2)Hex(9) | C[+57], N[+1865] | 1207.8238 | 4 | 4828.2733 | 4828.2324 | 0 | 8.5 | 127 | Specific | 825.09 | 825.09 | 825.09 | 15.86 | 52 | >sp\|P01876\|IGHA1_HUMAN Immunoglobulin heavy constant alpha 1 OS=Homo sapiens OX=9606 GN=IGHA1 PE=1 SV=2 | 255 | 20201115_07413_1ng_glyco_hcd203040_compound15.19701.19701.4 | controllerType=0 controllerNumber=1 scan=19701 | 42.4464 |
| 16044:4 | 10 | R.LSLHRPALEDLLLGSEAN[+1378.476]LTC[+57.021]TLTGLR.D | HexNAc(2)Hex(6) | C[+57], N[+1378] | 1086.279 | 4 | 4342.0941 | 4342.074 | 0 | 4.6 | 127 | Specific | 848.86 | 781.35 | 781.35 | 15.9 | 52 | >sp\|P01876\|IGHA1_HUMAN Immunoglobulin heavy constant alpha 1 OS=Homo sapiens OX=9606 GN=IGHA1 PE=1 SV=2 | 255 | 20201115_07413_1ng_glyco_hcd203040_compound15.19723.19723.4 | controllerType=0 controllerNumber=1 scan=19723 | 42.4971 |

Table S4. Cont.

| Query #:z | Protein Rank | Glycopeptide | Glycan | Modification Type(s) | Observed m/z | z | Observed (M+H) | Calc. mass (M+H) | Off-by-x error | Mass error (ppm) | StartPosition | Cleavage | Score | Delta | Delta Mod | \|Log Prob\| | # of unique peptides | Protein Name | Protein DB number | Comment | Scan # | Scan Time |
|---|---|---|---|---|---|---|---|---|---|---|---|---|---|---|---|---|---|---|---|---|---|---|
| 16003:4 | 10 | R.LSLHRPALEDLLLGSEAN[+1702.581]LTC[+57.021]TLTGLR.D | HexNAc(2)Hex(8) | C[+57], N[+1703] | 1167.3014 | 4 | 4666.1839 | 4666.1796 | 0 | 0.9 | 127 | Specific | 905.16 | 905.16 | 905.16 | 17.94 | 52 | >sp\|P01876\|IGHA1_HUMAN Immunoglobulin heavy constant alpha 1 OS=Homo sapiens OX=9606 GN=IGHA1 PE=1 SV=2 | 255 | 20201115_07413_1ng_glyco_hcd203040_compound15.19680.19680.4 | controllerType=0 controllerNumber=1 scan=19680 | 42.4032 |
| 16026:4 | 10 | R.LSLHRPALEDLLLGSEAN[+1864.634]LTC[+57.021]TLTGLR.D | HexNAc(2)Hex(9) | C[+57], N[+1865] | 1207.8238 | 4 | 4828.2733 | 4828.2324 | 0 | 8.5 | 127 | Specific | 906.85 | 835.9 | 835.9 | 17.86 | 52 | >sp\|P01876\|IGHA1_HUMAN Immunoglobulin heavy constant alpha 1 OS=Homo sapiens OX=9606 GN=IGHA1 PE=1 SV=2 | 255 | 20201115_07413_1ng_glyco_hcd203040_compound15.19704.19704.4 | controllerType=0 controllerNumber=1 scan=19704 | 42.4536 |
| 16074:4 | 10 | R.LSLHRPALEDLLLGSEAN[+1216.423]LTC[+57.021]TLTGLR.D | HexNAc(2)Hex(5) | C[+57], N[+1216] | 1045.7702 | 4 | 4180.0589 | 4180.0211 | 0 | 9 | 127 | Specific | 911.06 | 911.06 | 911.06 | 17.74 | 52 | >sp\|P01876\|IGHA1_HUMAN Immunoglobulin heavy constant alpha 1 OS=Homo sapiens OX=9606 GN=IGHA1 PE=1 SV=2 | 255 | 20201115_07413_1ng_glyco_hcd203040_compound15.19755.19755.4 | controllerType=0 controllerNumber=1 scan=19755 | 42.6026 |
| 16041:4 | 10 | R.LSLHRPALEDLLLGSEAN[+1540.529]LTC[+57.021]TLTGLR.D | HexNAc(2)Hex(7) | C[+57], N[+1541] | 1126.7915 | 4 | 4504.1444 | 4504.1268 | 0 | 3.9 | 127 | Specific | 931.32 | 886.38 | 886.38 | 18.22 | 52 | >sp\|P01876\|IGHA1_HUMAN Immunoglobulin heavy constant alpha 1 OS=Homo sapiens OX=9606 GN=IGHA1 PE=1 SV=2 | 255 | 20201115_07413_1ng_glyco_hcd203040_compound15.19720.19720.4 | controllerType=0 controllerNumber=1 scan=19720 | 42.4882 |
| 16084:4 | 10 | R.LSLHRPALEDLLLGSEAN[+1257.449]LTC[+57.021]TLTGLR.D | HexNAc(3)Hex(4) | C[+57], N[+1257] | 1056.0254 | 4 | 4221.0799 | 4221.0477 | 0 | 7.6 | 127 | Specific | 951.34 | 893.15 | 893.15 | 18.86 | 52 | >sp\|P01876\|IGHA1_HUMAN Immunoglobulin heavy constant alpha 1 OS=Homo sapiens OX=9606 GN=IGHA1 PE=1 SV=2 | 255 | 20201115_07413_1ng_glyco_hcd203040_compound15.19765.19765.4 | controllerType=0 controllerNumber=1 scan=19765 | 42.6351 |
| 16125:4 | 10 | R.LSLHRPALEDLLLGSEAN[+1095.397]LTC[+57.021]TLTGLR.D | HexNAc(3)Hex(3) | C[+57], N[+1095] | 1015.5083 | 4 | 4059.0113 | 4058.9949 | 0 | 4.1 | 127 | Specific | 992.2 | 992.2 | 992.2 | 19.22 | 52 | >sp\|P01876\|IGHA1_HUMAN Immunoglobulin heavy constant alpha 1 OS=Homo sapiens OX=9606 GN=IGHA1 PE=1 SV=2 | 255 | 20201115_07413_1ng_glyco_hcd203040_compound15.19809.19809.4 | controllerType=0 controllerNumber=1 scan=19809 | 42.752 |
| 16006:4 | 10 | R.LSLHRPALEDLLLGSEAN[+1702.581]LTC[+57.021]TLTGLR.D | HexNAc(2)Hex(8) | C[+57], N[+1703] | 1167.3014 | 4 | 4666.1839 | 4666.1796 | 0 | 0.9 | 127 | Specific | 1009.18 | 1009.18 | 1009.18 | 19.94 | 52 | >sp\|P01876\|IGHA1_HUMAN Immunoglobulin heavy constant alpha 1 OS=Homo sapiens OX=9606 GN=IGHA1 PE=1 SV=2 | 255 | 20201115_07413_1ng_glyco_hcd203040_compound15.19683.19683.4 | controllerType=0 controllerNumber=1 scan=19683 | 42.4103 |
| 16047:4 | 10 | R.LSLHRPALEDLLLGSEAN[+1378.476]LTC[+57.021]TLTGLR.D | HexNAc(2)Hex(6) | C[+57], N[+1378] | 1086.279 | 4 | 4342.0941 | 4342.074 | 0 | 4.6 | 127 | Specific | 1009.24 | 946.85 | 946.85 | 19.9 | 52 | >sp\|P01876\|IGHA1_HUMAN Immunoglobulin heavy constant alpha 1 OS=Homo sapiens OX=9606 GN=IGHA1 PE=1 SV=2 | 255 | 20201115_07413_1ng_glyco_hcd203040_compound15.19726.19726.4 | controllerType=0 controllerNumber=1 scan=19726 | 42.5027 |
| 16085:4 | 10 | R.LSLHRPALEDLLLGSEAN[+1257.449]LTC[+57.021]TLTGLR.D | HexNAc(3)Hex(4) | C[+57], N[+1257] | 1056.0254 | 4 | 4221.0799 | 4221.0477 | 0 | 7.6 | 127 | Specific | 1069.25 | 1069.25 | 1069.25 | 20.86 | 52 | >sp\|P01876\|IGHA1_HUMAN Immunoglobulin heavy constant alpha 1 OS=Homo sapiens OX=9606 GN=IGHA1 PE=1 SV=2 | 255 | 20201115_07413_1ng_glyco_hcd203040_compound15.19766.19766.4 | controllerType=0 controllerNumber=1 scan=19766 | 42.6425 |
| 10096:3 | 10 | R.LAGKPTHVN[+1809.666]VSVVMAEVDGTC[+57.021]Y.- | HexNAc(5)Hex(4)Fuc(1) | C[+57], N[+1810] | 1386.2872 | 3 | 4156.8469 | 4156.8081 | 0 | 9.3 | 332 | CRagged | 206.24 | 206.24 | 206.24 | 4.66 | 52 | >sp\|P01876\|IGHA1_HUMAN Immunoglobulin heavy constant alpha 1 OS=Homo sapiens OX=9606 GN=IGHA1 PE=1 SV=2 | 255 | 20201115_07413_1ng_glyco_hcd203040_compound15.13471.13471.3 | controllerType=0 controllerNumber=1 scan=13471 | 30.9499 |
| 11603:3 | 10 | R.LAGKPTHVN[+1913.677]VSVVMAEVDGTC[+57.021]Y.- | HexNAc(4)Hex(5)NeuAc(1) | C[+57], N[+1914] | 1420.9496 | 3 | 4260.8344 | 4260.8191 | 0 | 3.6 | 332 | CRagged | 265.27 | 242.69 | 242.69 | 6.65 | 52 | >sp\|P01876\|IGHA1_HUMAN Immunoglobulin heavy constant alpha 1 OS=Homo sapiens OX=9606 GN=IGHA1 PE=1 SV=2 | 255 | 20201115_07413_1ng_glyco_hcd203040_compound15.15062.15062.3 | controllerType=0 controllerNumber=1 scan=15062 | 33.7344 |
| 13461:3 | 10 | R.LAGKPTHVN[+2553.910]VSVVMAEVDGTC[+57.021]Y.- | HexNAc(5)Hex(5)Fuc(1)NeuAc(2) | C[+57], N[+2554] | 1634.3644 | 3 | 4901.0788 | 4901.0518 | 0 | 5.5 | 332 | CRagged | 316.76 | 274.03 | 274.03 | 6.48 | 52 | >sp\|P01876\|IGHA1_HUMAN Immunoglobulin heavy constant alpha 1 OS=Homo sapiens OX=9606 GN=IGHA1 PE=1 SV=2 | 255 | 20201115_07413_1ng_glyco_hcd203040_compound15.17003.17003.3 | controllerType=0 controllerNumber=1 scan=17003 | 37.2827 |
| 13463:4 | 10 | R.LAGKPTHVN[+2553.910]VSVVMAEVDGTC[+57.021]Y.- | HexNAc(5)Hex(5)Fuc(1)NeuAc(2) | C[+57], N[+2554] | 1226.0251 | 4 | 4901.0784 | 4901.0518 | 0 | 5.4 | 332 | CRagged | 361.14 | 289.11 | 289.11 | 6.69 | 52 | >sp\|P01876\|IGHA1_HUMAN Immunoglobulin heavy constant alpha 1 OS=Homo sapiens OX=9606 GN=IGHA1 PE=1 SV=2 | 255 | 20201115_07413_1ng_glyco_hcd203040_compound15.17005.17005.4 | controllerType=0 controllerNumber=1 scan=17005 | 37.2865 |
| 11704:3 | 10 | R.LAGKPTHVN[+1548.545]VSVVMAEVDGTC[+57.021]Y.- | HexNAc(3)Hex(4)NeuAc(1) | C[+57], N[+1549] | 1299.2361 | 3 | 3895.6939 | 3895.6869 | 0 | 1.8 | 332 | CRagged | 417.4 | 417.4 | 417.4 | 9.3 | 52 | >sp\|P01876\|IGHA1_HUMAN Immunoglobulin heavy constant alpha 1 OS=Homo sapiens OX=9606 GN=IGHA1 PE=1 SV=2 | 255 | 20201115_07413_1ng_glyco_hcd203040_compound15.15166.15166.3 | controllerType=0 controllerNumber=1 scan=15166 | 33.9111 |
| 09998:3 | 10 | R.LAGKPTHVN[+1216.423]VSVVMAEVDGTC[+57.021]Y.- | HexNAc(2)Hex(5) | C[+57], N[+1216] | 1188.5316 | 3 | 3563.5801 | 3563.5649 | 0 | 4.3 | 332 | CRagged | 452.87 | 408.81 | 359.32 | 8.94 | 52 | >sp\|P01876\|IGHA1_HUMAN Immunoglobulin heavy constant alpha 1 OS=Homo sapiens OX=9606 GN=IGHA1 PE=1 SV=2 | 255 | 20201115_07413_1ng_glyco_hcd203040_compound15.13370.13370.3 | controllerType=0 controllerNumber=1 scan=13370 | 30.7735 |
| 11621:3 | 10 | R.LAGKPTHVN[+1872.651]VSVVMAEVDGTC[+57.021]Y.- | HexNAc(3)Hex(6)NeuAc(1) | C[+57], N[+1873] | 1407.2707 | 3 | 4219.7975 | 4219.7926 | 0 | 1.2 | 332 | CRagged | 467.66 | 450.5 | 310.22 | 8.97 | 52 | >sp\|P01876\|IGHA1_HUMAN Immunoglobulin heavy constant alpha 1 OS=Homo sapiens OX=9606 GN=IGHA1 PE=1 SV=2 | 255 | 20201115_07413_1ng_glyco_hcd203040_compound15.15080.15080.3 | controllerType=0 controllerNumber=1 scan=15080 | 33.7662 |
| 09996:3 | 10 | R.LAGKPTHVN[+1216.423]VSVVMAEVDGTC[+57.021]Y.- | HexNAc(2)Hex(5) | C[+57], N[+1216] | 1188.5316 | 3 | 3563.5801 | 3563.5649 | 0 | 4.3 | 332 | CRagged | 499.47 | 499.47 | 286.5 | 9.15 | 52 | >sp\|P01876\|IGHA1_HUMAN Immunoglobulin heavy constant alpha 1 OS=Homo sapiens OX=9606 GN=IGHA1 PE=1 SV=2 | 255 | 20201115_07413_1ng_glyco_hcd203040_compound15.13368.13368.3 | controllerType=0 controllerNumber=1 scan=13368 | 30.7697 |
| 09940:3 | 10 | R.LAGKPTHVN[+1647.613]VSVVMAEVDGTC[+57.021]Y.- | HexNAc(5)Hex(3)Fuc(1) | C[+57], N[+1648] | 1332.2569 | 3 | 3994.7561 | 3994.7553 | 0 | 0.2 | 332 | CRagged | 532.54 | 494.86 | 494.86 | 9.3 | 52 | >sp\|P01876\|IGHA1_HUMAN Immunoglobulin heavy constant alpha 1 OS=Homo sapiens OX=9606 GN=IGHA1 PE=1 SV=2 | 255 | 20201115_07413_1ng_glyco_hcd203040_compound15.13307.13307.3 | controllerType=0 controllerNumber=1 scan=13307 | 30.6219 |
| 11619:3 | 10 | R.LAGKPTHVN[+1872.651]VSVVMAEVDGTC[+57.021]Y.- | HexNAc(3)Hex(6)NeuAc(1) | C[+57], N[+1873] | 1407.2707 | 3 | 4219.7975 | 4219.7926 | 0 | 1.2 | 332 | CRagged | 538.97 | 507.45 | 353.46 | 9.49 | 52 | >sp\|P01876\|IGHA1_HUMAN Immunoglobulin heavy constant alpha 1 OS=Homo sapiens OX=9606 GN=IGHA1 PE=1 SV=2 | 255 | 20201115_07413_1ng_glyco_hcd203040_compound15.15078.15078.3 | controllerType=0 controllerNumber=1 scan=15078 | 33.7618 |
| 11601:3 | 10 | R.LAGKPTHVN[+1913.677]VSVVMAEVDGTC[+57.021]Y.- | HexNAc(4)Hex(5)NeuAc(1) | C[+57], N[+1914] | 1420.9496 | 3 | 4260.8344 | 4260.8191 | 0 | 3.6 | 332 | CRagged | 543.25 | 439.76 | 439.76 | 9.77 | 52 | >sp\|P01876\|IGHA1_HUMAN Immunoglobulin heavy constant alpha 1 OS=Homo sapiens OX=9606 GN=IGHA1 PE=1 SV=2 | 255 | 20201115_07413_1ng_glyco_hcd203040_compound15.15060.15060.3 | controllerType=0 controllerNumber=1 scan=15060 | 33.7292 |
| 09857:3 | 10 | R.LAGKPTHVN[+1809.666]VSVVMAEVDGTC[+57.021]Y.- | HexNAc(5)Hex(4)Fuc(1) | C[+57], N[+1810] | 1386.2763 | 3 | 4156.8144 | 4156.8081 | 0 | 1.5 | 332 | CRagged | 554.27 | 554.27 | 369.39 | 9.79 | 52 | >sp\|P01876\|IGHA1_HUMAN Immunoglobulin heavy constant alpha 1 OS=Homo sapiens OX=9606 GN=IGHA1 PE=1 SV=2 | 255 | 20201115_07413_1ng_glyco_hcd203040_compound15.13221.13221.3 | controllerType=0 controllerNumber=1 scan=13221 | 30.4495 |
| 08782:3 | 10 | R.LAGKPTHVN[+1216.423]VSVVMAEVDGTC[+57.021]Y.- | HexNAc(2)Hex(5) | C[+57], N[+1216] | 1188.5279 | 3 | 3563.5691 | 3563.5649 | 0 | 1.2 | 332 | CRagged | 574.79 | 574.79 | 363.96 | 10.95 | 52 | >sp\|P01876\|IGHA1_HUMAN Immunoglobulin heavy constant alpha 1 OS=Homo sapiens OX=9606 GN=IGHA1 PE=1 SV=2 | 255 | 20201115_07413_1ng_glyco_hcd203040_compound15.12090.12090.3 | controllerType=0 controllerNumber=1 scan=12090 | 28.4318 |
| 11706:3 | 10 | R.LAGKPTHVN[+1548.545]VSVVMAEVDGTC[+57.021]Y.- | HexNAc(3)Hex(4)NeuAc(1) | C[+57], N[+1549] | 1299.2361 | 3 | 3895.6939 | 3895.6869 | 0 | 1.8 | 332 | CRagged | 634.73 | 605.49 | 605.49 | 11.95 | 52 | >sp\|P01876\|IGHA1_HUMAN Immunoglobulin heavy constant alpha 1 OS=Homo sapiens OX=9606 GN=IGHA1 PE=1 SV=2 | 255 | 20201115_07413_1ng_glyco_hcd203040_compound15.15168.15168.3 | controllerType=0 controllerNumber=1 scan=15168 | 33.9147 |
| 09956:3 | 10 | R.LAGKPTHVN[+1501.555]VSVVMAEVDGTC[+57.021]Y.- | HexNAc(5)Hex(3) | C[+57], N[+1502] | 1283.5803 | 3 | 3848.7262 | 3848.6974 | 0 | 7.5 | 332 | CRagged | 682.83 | 660.56 | 660.56 | 12.57 | 52 | >sp\|P01876\|IGHA1_HUMAN Immunoglobulin heavy constant alpha 1 OS=Homo sapiens OX=9606 GN=IGHA1 PE=1 SV=2 | 255 | 20201115_07413_1ng_glyco_hcd203040_compound15.13326.13326.3 | controllerType=0 controllerNumber=1 scan=13326 | 30.6767 |
| 13459:3 | 10 | R.LAGKPTHVN[+2553.910]VSVVMAEVDGTC[+57.021]Y.- | HexNAc(5)Hex(5)Fuc(1)NeuAc(2) | C[+57], N[+2554] | 1634.3644 | 3 | 4901.0788 | 4901.0518 | 0 | 5.5 | 332 | CRagged | 705.82 | 515.85 | 515.85 | 13.62 | 52 | >sp\|P01876\|IGHA1_HUMAN Immunoglobulin heavy constant alpha 1 OS=Homo sapiens OX=9606 GN=IGHA1 PE=1 SV=2 | 255 | 20201115_07413_1ng_glyco_hcd203040_compound15.17001.17001.3 | controllerType=0 controllerNumber=1 scan=17001 | 37.2773 |
| 09959:3 | 10 | R.LAGKPTHVN[+1501.555]VSVVMAEVDGTC[+57.021]Y.- | HexNAc(5)Hex(3) | C[+57], N[+1502] | 1283.5803 | 3 | 3848.7262 | 3848.6974 | 0 | 7.5 | 332 | CRagged | 721.84 | 680.92 | 680.92 | 13.57 | 52 | >sp\|P01876\|IGHA1_HUMAN Immunoglobulin heavy constant alpha 1 OS=Homo sapiens OX=9606 GN=IGHA1 PE=1 SV=2 | 255 | 20201115_07413_1ng_glyco_hcd203040_compound15.13329.13329.3 | controllerType=0 controllerNumber=1 scan=13329 | 30.6831 |
| 09896:3 | 10 | R.LAGKPTHVN[+1216.423]VSVVMAEVDGTC[+57.021]Y.- | HexNAc(2)Hex(5) | C[+57], N[+1216] | 1188.5273 | 3 | 3563.5673 | 3563.5649 | 0 | 0.7 | 332 | CRagged | 751.62 | 684.67 | 433.71 | 14.65 | 52 | >sp\|P01876\|IGHA1_HUMAN Immunoglobulin heavy constant alpha 1 OS=Homo sapiens OX=9606 GN=IGHA1 PE=1 SV=2 | 255 | 20201115_07413_1ng_glyco_hcd203040_compound15.13262.13262.3 | controllerType=0 controllerNumber=1 scan=13262 | 30.5364 |
| 09801:3 | 10 | R.LAGKPTHVN[+1540.529]VSVVMAEVDGTC[+57.021]Y.- | HexNAc(2)Hex(7) | C[+57], N[+1541] | 1296.5589 | 3 | 3887.6621 | 3887.6706 | 0 | -2.2 | 332 | CRagged | 757.79 | 589.67 | 459.86 | 13.5 | 52 | >sp\|P01876\|IGHA1_HUMAN Immunoglobulin heavy constant alpha 1 OS=Homo sapiens OX=9606 GN=IGHA1 PE=1 SV=2 | 255 | 20201115_07413_1ng_glyco_hcd203040_compound15.13163.13163.3 | controllerType=0 controllerNumber=1 scan=13163 | 30.332 |
| 09789:3 | 10 | R.LAGKPTHVN[+1702.581]VSVVMAEVDGTC[+57.021]Y.- | HexNAc(2)Hex(8) | C[+57], N[+1703] | 1350.5843 | 3 | 4049.7383 | 4049.7234 | 0 | 3.7 | 332 | CRagged | 861.43 | 842.53 | 525.86 | 16.94 | 52 | >sp\|P01876\|IGHA1_HUMAN Immunoglobulin heavy constant alpha 1 OS=Homo sapiens OX=9606 GN=IGHA1 PE=1 SV=2 | 255 | 20201115_07413_1ng_glyco_hcd203040_compound15.13150.13150.3 | controllerType=0 controllerNumber=1 scan=13150 | 30.301 |
| 09850:3 | 10 | R.LAGKPTHVN[+1378.476]VSVVMAEVDGTC[+57.021]Y.- | HexNAc(2)Hex(6) | C[+57], N[+1378] | 1242.5462 | 3 | 3725.624 | 3725.6178 | 0 | 1.7 | 332 | CRagged | 872.6 | 818.35 | 515.37 | 16.95 | 52 | >sp\|P01876\|IGHA1_HUMAN Immunoglobulin heavy constant alpha 1 OS=Homo sapiens OX=9606 GN=IGHA1 PE=1 SV=2 | 255 | 20201115_07413_1ng_glyco_hcd203040_compound15.13214.13214.3 | controllerType=0 controllerNumber=1 scan=13214 | 30.4364 |
| 09938:3 | 10 | R.LAGKPTHVN[+1647.613]VSVVMAEVDGTC[+57.021]Y.- | HexNAc(5)Hex(3)Fuc(1) | C[+57], N[+1648] | 1332.2569 | 3 | 3994.7561 | 3994.7553 | 0 | 0.2 | 332 | CRagged | 905.92 | 856.92 | 856.92 | 17.65 | 52 | >sp\|P01876\|IGHA1_HUMAN Immunoglobulin heavy constant alpha 1 OS=Homo sapiens OX=9606 GN=IGHA1 PE=1 SV=2 | 255 | 20201115_07413_1ng_glyco_hcd203040_compound15.13305.13305.3 | controllerType=0 controllerNumber=1 scan=13305 | 30.6139 |
| 09898:3 | 10 | R.LAGKPTHVN[+1216.423]VSVVMAEVDGTC[+57.021]Y.- | HexNAc(2)Hex(5) | C[+57], N[+1216] | 1188.5273 | 3 | 3563.5673 | 3563.5649 | 0 | 0.7 | 332 | CRagged | 915.8 | 902.18 | 550.77 | 17.65 | 52 | >sp\|P01876\|IGHA1_HUMAN Immunoglobulin heavy constant alpha 1 OS=Homo sapiens OX=9606 GN=IGHA1 PE=1 SV=2 | 255 | 20201115_07413_1ng_glyco_hcd203040_compound15.13264.13264.3 | controllerType=0 controllerNumber=1 scan=13264 | 30.5414 |
| 09848:3 | 10 | R.LAGKPTHVN[+1378.476]VSVVMAEVDGTC[+57.021]Y.- | HexNAc(2)Hex(6) | C[+57], N[+1378] | 1242.5462 | 3 | 3725.624 | 3725.6178 | 0 | 1.7 | 332 | CRagged | 919.57 | 919.57 | 575.55 | 17.95 | 52 | >sp\|P01876\|IGHA1_HUMAN Immunoglobulin heavy constant alpha 1 OS=Homo sapiens OX=9606 GN=IGHA1 PE=1 SV=2 | 255 | 20201115_07413_1ng_glyco_hcd203040_compound15.13212.13212.3 | controllerType=0 controllerNumber=1 scan=13212 | 30.4286 |
| 09854:3 | 10 | R.LAGKPTHVN[+1809.666]VSVVMAEVDGTC[+57.021]Y.- | HexNAc(5)Hex(4)Fuc(1) | C[+57], N[+1810] | 1386.2763 | 3 | 4156.8144 | 4156.8081 | 0 | 1.5 | 332 | CRagged | 927.47 | 776.34 | 627.42 | 17.95 | 52 | >sp\|P01876\|IGHA1_HUMAN Immunoglobulin heavy constant alpha 1 OS=Homo sapiens OX=9606 GN=IGHA1 PE=1 SV=2 | 255 | 20201115_07413_1ng_glyco_hcd203040_compound15.13218.13218.3 | controllerType=0 controllerNumber=1 scan=13218 | 30.4406 |
| 09799:3 | 10 | R.LAGKPTHVN[+1540.529]VSVVMAEVDGTC[+57.021]Y.- | HexNAc(2)Hex(7) | C[+57], N[+1541] | 1296.5589 | 3 | 3887.6621 | 3887.6706 | 0 | -2.2 | 332 | CRagged | 947.43 | 947.43 | 606.61 | 16.5 | 52 | >sp\|P01876\|IGHA1_HUMAN Immunoglobulin heavy constant alpha 1 OS=Homo sapiens OX=9606 GN=IGHA1 PE=1 SV=2 | 255 | 20201115_07413_1ng_glyco_hcd203040_compound15.13161.13161.3 | controllerType=0 controllerNumber=1 scan=13161 | 30.3223 |
| 09787:3 | 10 | R.LAGKPTHVN[+1702.581]VSVVMAEVDGTC[+57.021]Y.- | HexNAc(2)Hex(8) | C[+57], N[+1703] | 1350.5843 | 3 | 4049.7383 | 4049.7234 | 0 | 3.7 | 332 | CRagged | 949.6 | 949.6 | 607.68 | 17.94 | 52 | >sp\|P01876\|IGHA1_HUMAN Immunoglobulin heavy constant alpha 1 OS=Homo sapiens OX=9606 GN=IGHA1 PE=1 SV=2 | 255 | 20201115_07413_1ng_glyco_hcd203040_compound15.13148.13148.3 | controllerType=0 controllerNumber=1 scan=13148 | 30.2923 |
| 22129:3 | 13 | R.QLAHQSN[+2204.772]STNIFFSPVSIATAFAMLSLGTK.A | HexNAc(4)Hex(5)NeuAc(2) | N[+2205] | 1796.1468 | 3 | 5386.4259 | 5386.4075 | 0 | 3.4 | 64 | Specific | 414.87 | 414.87 | 414.87 | 9.1 | 39 | >sp\|P01009\|A1AT_HUMAN Alpha-1-antitrypsin OS=Homo sapiens OX=9606 GN=SERPINA1 PE=1 SV=3 | 642 | 20201115_07413_1ng_glyco_hcd203040_compound15.26124.26124.3 | controllerType=0 controllerNumber=1 scan=26124 | 55.6835 |
| 22155:3 | 13 | R.QLAHQSN[+2204.772]STNIFFSPVSIATAFAMLSLGTK.A | HexNAc(4)Hex(5)NeuAc(2) | N[+2205] | 1796.1393 | 3 | 5386.4033 | 5386.4075 | 0 | -0.8 | 64 | Specific | 526.41 | 470.97 | 448.26 | 8.81 | 39 | >sp\|P01009\|A1AT_HUMAN Alpha-1-antitrypsin OS=Homo sapiens OX=9606 GN=SERPINA1 PE=1 SV=3 | 642 | 20201115_07413_1ng_glyco_hcd203040_compound15.26151.26151.3 | controllerType=0 controllerNumber=1 scan=26151 | 55.7393 |
| 22179:3 | 13 | R.QLAHQSN[+2204.772]STNIFFSPVSIATAFAMLSLGTK.A | HexNAc(4)Hex(5)NeuAc(2) | N[+2205] | 1796.1504 | 3 | 5386.4365 | 5386.4075 | 0 | 5.4 | 64 | Specific | 585.18 | 566.59 | 566.59 | 10.62 | 39 | >sp\|P01009\|A1AT_HUMAN Alpha-1-antitrypsin OS=Homo sapiens OX=9606 GN=SERPINA1 PE=1 SV=3 | 642 | 20201115_07413_1ng_glyco_hcd203040_compound15.26177.26177.3 | controllerType=0 controllerNumber=1 scan=26177 | 55.7979 |
| 22130:4 | 13 | R.QLAHQSN[+2350.830]STNIFFSPVSIATAFAMLSLGTK.A | HexNAc(4)Hex(5)Fuc(1)NeuAc(2) | N[+2351] | 1383.876 | 4 | 5532.4821 | 5532.4654 | 0 | 3 | 64 | Specific | 590.45 | 557.17 | 557.17 | 10.67 | 39 | >sp\|P01009\|A1AT_HUMAN Alpha-1-antitrypsin OS=Homo sapiens OX=9606 GN=SERPINA1 PE=1 SV=3 | 642 | 20201115_07413_1ng_glyco_hcd203040_compound15.26125.26125.4 | controllerType=0 controllerNumber=1 scan=26125 | 55.6855 |
| 22177:3 | 13 | R.QLAHQSN[+2204.772]STNIFFSPVSIATAFAMLSLGTK.A | HexNAc(4)Hex(5)NeuAc(2) | N[+2205] | 1796.1504 | 3 | 5386.4365 | 5386.4075 | 0 | 5.4 | 64 | Specific | 659.65 | 615.18 | 615.18 | 12.62 | 39 | >sp\|P01009\|A1AT_HUMAN Alpha-1-antitrypsin OS=Homo sapiens OX=9606 GN=SERPINA1 PE=1 SV=3 | 642 | 20201115_07413_1ng_glyco_hcd203040_compound15.26175.26175.3 | controllerType=0 controllerNumber=1 scan=26175 | 55.7889 |
| 22178:4 | 13 | R.QLAHQSN[+2204.772]STNIFFSPVSIATAFAMLSLGTK.A | HexNAc(4)Hex(5)NeuAc(2) | N[+2205] | 1347.3635 | 4 | 5386.4323 | 5386.4075 | 0 | 4.6 | 64 | Specific | 722.77 | 677.85 | 677.85 | 13.35 | 39 | >sp\|P01009\|A1AT_HUMAN Alpha-1-antitrypsin OS=Homo sapiens OX=9606 GN=SERPINA1 PE=1 SV=3 | 642 | 20201115_07413_1ng_glyco_hcd203040_compound15.26176.26176.4 | controllerType=0 controllerNumber=1 scan=26176 | 55.796 |
| 22153:3 | 13 | R.QLAHQSN[+2204.772]STNIFFSPVSIATAFAMLSLGTK.A | HexNAc(4)Hex(5)NeuAc(2) | N[+2205] | 1796.1393 | 3 | 5386.4033 | 5386.4075 | 0 | -0.8 | 64 | Specific | 760.63 | 712.09 | 522.08 | 14.16 | 39 | >sp\|P01009\|A1AT_HUMAN Alpha-1-antitrypsin OS=Homo sapiens OX=9606 GN=SERPINA1 PE=1 SV=3 | 642 | 20201115_07413_1ng_glyco_hcd203040_compound15.26149.26149.3 | controllerType=0 controllerNumber=1 scan=26149 | 55.7328 |
| 22132:4 | 13 | R.QLAHQSN[+2350.830]STNIFFSPVSIATAFAMLSLGTK.A | HexNAc(4)Hex(5)Fuc(1)NeuAc(2) | N[+2351] | 1383.876 | 4 | 5532.4821 | 5532.4654 | 0 | 3 | 64 | Specific | 850.03 | 786.63 | 786.63 | 16.67 | 39 | >sp\|P01009\|A1AT_HUMAN Alpha-1-antitrypsin OS=Homo sapiens OX=9606 GN=SERPINA1 PE=1 SV=3 | 642 | 20201115_07413_1ng_glyco_hcd203040_compound15.26127.26127.4 | controllerType=0 controllerNumber=1 scan=26127 | 55.6908 |
| 22154:4 | 13 | R.QLAHQSN[+2204.772]STNIFFSPVSIATAFAMLSLGTK.A | HexNAc(4)Hex(5)NeuAc(2) | N[+2205] | 1347.3563 | 4 | 5386.4034 | 5386.4075 | 0 | -0.8 | 64 | Specific | 873.89 | 780.89 | 549.81 | 15.88 | 39 | >sp\|P01009\|A1AT_HUMAN Alpha-1-antitrypsin OS=Homo sapiens OX=9606 GN=SERPINA1 PE=1 SV=3 | 642 | 20201115_07413_1ng_glyco_hcd203040_compound15.26150.26150.4 | controllerType=0 controllerNumber=1 scan=26150 | 55.7377 |
| 22152:4 | 13 | R.QLAHQSN[+2204.772]STNIFFSPVSIATAFAMLSLGTK.A | HexNAc(4)Hex(5)NeuAc(2) | N[+2205] | 1347.3563 | 4 | 5386.4034 | 5386.4075 | 0 | -0.8 | 64 | Specific | 913.41 | 831.86 | 598.27 | 16.88 | 39 | >sp\|P01009\|A1AT_HUMAN Alpha-1-antitrypsin OS=Homo sapiens OX=9606 GN=SERPINA1 PE=1 SV=3 | 642 | 20201115_07413_1ng_glyco_hcd203040_compound15.26148.26148.4 | controllerType=0 controllerNumber=1 scan=26148 | 55.7314 |
| 22176:4 | 13 | R.QLAHQSN[+2204.772]STNIFFSPVSIATAFAMLSLGTK.A | HexNAc(4)Hex(5)NeuAc(2) | N[+2205] | 1347.3635 | 4 | 5386.4323 | 5386.4075 | 0 | 4.6 | 64 | Specific | 1002.72 | 950.52 | 950.52 | 19.35 | 39 | >sp\|P01009\|A1AT_HUMAN Alpha-1-antitrypsin OS=Homo sapiens OX=9606 GN=SERPINA1 PE=1 SV=3 | 642 | 20201115_07413_1ng_glyco_hcd203040_compound15.26174.26174.4 | controllerType=0 controllerNumber=1 scan=26174 | 55.7874 |
| 21062:4 | 13 | K.ADTHDEILEGLNFN[+2862.021]LTEIPEAQIHEGFQELLR.T | HexNAc(5)Hex(6)Fuc(2)NeuAc(2) | N[+2862] | 1639.2293 | 4 | 6553.8952 | 6553.8443 | 0 | 7.8 | 94 | Specific | 256.49 | 256.49 | 256.49 | 6.04 | 39 | >sp\|P01009\|A1AT_HUMAN Alpha-1-antitrypsin OS=Homo sapiens OX=9606 GN=SERPINA1 PE=1 SV=3 | 642 | 20201115_07413_1ng_glyco_hcd203040_compound15.24990.24990.2 | controllerType=0 controllerNumber=1 scan=24990 | 53.4971 |
| 20783:4 | 13 | K.ADTHDEILEGLNFN[+2861.000]LTEIPEAQIHEGFQELLR.T | HexNAc(5)Hex(6)NeuAc(3) | N[+2861] | 1638.9727 | 4 | 6552.8688 | 6552.8239 | 0 | 6.9 | 94 | Specific | 979.54 | 916.28 | 916.28 | 18.46 | 39 | >sp\|P01009\|A1AT_HUMAN Alpha-1-antitrypsin OS=Homo sapiens OX=9606 GN=SERPINA1 PE=1 SV=3 | 642 | 20201115_07413_1ng_glyco_hcd203040_compound15.24700.24700.4 | controllerType=0 controllerNumber=1 scan=24700 | 52.9278 |
| 18533:3 | 13 | K.YLGN[+2204.772]ATAIFFLPDEGK.L | HexNAc(4)Hex(5)NeuAc(2) | N[+2205] | 1320.8965 | 3 | 3960.6751 | 3960.6677 | 0 | 1.9 | 268 | Specific | 776.04 | 691.34 | 691.34 | 14.58 | 39 | >sp\|P01009\|A1AT_HUMAN Alpha-1-antitrypsin OS=Homo sapiens OX=9606 GN=SERPINA1 PE=1 SV=3 | 642 | 20201115_07413_1ng_glyco_hcd203040_compound15.22337.22337.3 | controllerType=0 controllerNumber=1 scan=22337 | 48.0918 |
| 18531:3 | 13 | K.YLGN[+2204.772]ATAIFFLPDEGK.L | HexNAc(4)Hex(5)NeuAc(2) | N[+2205] | 1320.8965 | 3 | 3960.6751 | 3960.6677 | 0 | 1.9 | 268 | Specific | 824.39 | 692.01 | 692.01 | 15.58 | 39 | >sp\|P01009\|A1AT_HUMAN Alpha-1-antitrypsin OS=Homo sapiens OX=9606 GN=SERPINA1 PE=1 SV=3 | 642 | 20201115_07413_1ng_glyco_hcd203040_compound15.22335.22335.3 | controllerType=0 controllerNumber=1 scan=22335 | 48.0868 |
| 14129:4 | 14 | K.LKELPGVC[+57.021]N[+2350.830]ETMMALWEEC[+57.021]KPC[+57.021]LK.Q | HexNAc(4)Hex(5)Fuc(1)NeuAc(2) | C[+57]*3, N[+2351] | 1322.5686 | 4 | 5287.2526 | 5287.2151 | 0 | 7.1 | 95 | Specific | 221.42 | 219.66 | 219.66 | 4.89 | 32 | >sp\|P10909\|CLUS_HUMAN Clusterin OS=Homo sapiens OX=9606 GN=CLU PE=1 SV=1 | 1467 | 20201115_07413_1ng_glyco_hcd203040_compound15.17697.17697.4 | controllerType=0 controllerNumber=1 scan=17697 | 38.5749 |
| 12401:3 | 14 | K.LKELPGVC[+57.021]N[+2204.772]ETMMALWEEC[+57.021]KPC[+57.021]LK.Q | HexNAc(4)Hex(5)NeuAc(2) | C[+57]*3, N[+2205] | 1714.4034 | 3 | 5141.1957 | 5141.1572 | 0 | 7.5 | 95 | Specific | 275.22 | 275.22 | 275.22 | 5.62 | 32 | >sp\|P10909\|CLUS_HUMAN Clusterin OS=Homo sapiens OX=9606 GN=CLU PE=1 SV=1 | 1467 | 20201115_07413_1ng_glyco_hcd203040_compound15.15896.15896.3 | controllerType=0 controllerNumber=1 scan=15896 | 35.2629 |
| 12357:4 | 14 | K.LKELPGVC[+57.021]N[+2424.867]ETMMALWEEC[+57.021]KPC[+57.021]LK.Q | HexNAc(5)Hex(6)Fuc(1)NeuAc(1) | C[+57]*3, N[+2425] | 1341.0811 | 4 | 5361.3024 | 5361.2518 | 0 | 9.4 | 95 | Specific | 325.74 | 256.97 | 213.04 | 6.42 | 32 | >sp\|P10909\|CLUS_HUMAN Clusterin OS=Homo sapiens OX=9606 GN=CLU PE=1 SV=1 | 1467 | 20201115_07413_1ng_glyco_hcd203040_compound15.15851.15851.4 | controllerType=0 controllerNumber=1 scan=15851 | 35.1845 |
| 12287:4 | 14 | K.LKELPGVC[+57.021]N[+2569.905]ETMMALWEEC[+57.021]KPC[+57.021]LK.Q | HexNAc(5)Hex(6)NeuAc(2) | C[+57]*3, N[+2570] | 1377.335 | 4 | 5506.318 | 5506.2893 | 0 | 5.2 | 95 | Specific | 601.97 | 580.27 | 291.87 | 11.47 | 32 | >sp\|P10909\|CLUS_HUMAN Clusterin OS=Homo sapiens OX=9606 GN=CLU PE=1 SV=1 | 1467 | 20201115_07413_1ng_glyco_hcd203040_compound15.15778.15778.4 | controllerType=0 controllerNumber=1 scan=15778 | 35.0608 |
| 12418:3 | 14 | K.LKELPGVC[+57.021]N[+2059.735]ETMMALWEEC[+57.021]KPC[+57.021]LK.Q | HexNAc(4)Hex(5)Fuc(1)NeuAc(1) | C[+57]*3, N[+2060] | 1666.0512 | 3 | 4996.1391 | 4996.1196 | 0 | 3.9 | 95 | Specific | 638.45 | 524.29 | 524.29 | 11.17 | 32 | >sp\|P10909\|CLUS_HUMAN Clusterin OS=Homo sapiens OX=9606 GN=CLU PE=1 SV=1 | 1467 | 20201115_07413_1ng_glyco_hcd203040_compound15.15914.15914.3 | controllerType=0 controllerNumber=1 scan=15914 | 35.3017 |
| 12360:4 | 14 | K.LKELPGVC[+57.021]N[+2424.867]ETMMALWEEC[+57.021]KPC[+57.021]LK.Q | HexNAc(5)Hex(6)Fuc(1)NeuAc(1) | C[+57]*3, N[+2425] | 1341.0811 | 4 | 5361.3024 | 5361.2518 | 0 | 9.4 | 95 | Specific | 706.92 | 529.72 | 484.18 | 13.3 | 32 | >sp\|P10909\|CLUS_HUMAN Clusterin OS=Homo sapiens OX=9606 GN=CLU PE=1 SV=1 | 1467 | 20201115_07413_1ng_glyco_hcd203040_compound15.15854.15854.4 | controllerType=0 controllerNumber=1 scan=15854 | 35.1887 |
| 12416:3 | 14 | K.LKELPGVC[+57.021]N[+2059.735]ETMMALWEEC[+57.021]KPC[+57.021]LK.Q | HexNAc(4)Hex(5)Fuc(1)NeuAc(1) | C[+57]*3, N[+2060] | 1666.0512 | 3 | 4996.1391 | 4996.1196 | 0 | 3.9 | 95 | Specific | 802.4 | 798.36 | 798.36 | 15.17 | 32 | >sp\|P10909\|CLUS_HUMAN Clusterin OS=Homo sapiens OX=9606 GN=CLU PE=1 SV=1 | 1467 | 20201115_07413_1ng_glyco_hcd203040_compound15.15912.15912.3 | controllerType=0 controllerNumber=1 scan=15912 | 35.2972 |
| 12390:4 | 14 | K.LKELPGVC[+57.021]N[+2205.793]ETMMALWEEC[+57.021]KPC[+57.021]LK.Q | HexNAc(4)Hex(5)Fuc(2)NeuAc(1) | C[+57]*3, N[+2206] | 1286.3058 | 4 | 5142.2013 | 5142.1776 | 0 | 4.6 | 95 | Specific | 831.03 | 794.02 | 794.02 | 15.47 | 32 | >sp\|P10909\|CLUS_HUMAN Clusterin OS=Homo sapiens OX=9606 GN=CLU PE=1 SV=1 | 1467 | 20201115_07413_1ng_glyco_hcd203040_compound15.15885.15885.4 | controllerType=0 controllerNumber=1 scan=15885 | 35.2452 |
| 12411:4 | 14 | K.LKELPGVC[+57.021]N[+2059.735]ETMMALWEEC[+57.021]KPC[+57.021]LK.Q | HexNAc(4)Hex(5)Fuc(1)NeuAc(1) | C[+57]*3, N[+2060] | 1249.7902 | 4 | 4996.1388 | 4996.1196 | 0 | 3.8 | 95 | Specific | 873.77 | 793.28 | 793.28 | 16.79 | 32 | >sp\|P10909\|CLUS_HUMAN Clusterin OS=Homo sapiens OX=9606 GN=CLU PE=1 SV=1 | 1467 | 20201115_07413_1ng_glyco_hcd203040_compound15.15907.15907.4 | controllerType=0 controllerNumber=1 scan=15907 | 35.2887 |
| 12409:4 | 14 | K.LKELPGVC[+57.021]N[+2059.735]ETMMALWEEC[+57.021]KPC[+57.021]LK.Q | HexNAc(4)Hex(5)Fuc(1)NeuAc(1) | C[+57]*3, N[+2060] | 1249.7902 | 4 | 4996.1388 | 4996.1196 | 0 | 3.8 | 95 | Specific | 1123.46 | 1065.97 | 1065.97 | 21.79 | 32 | >sp\|P10909\|CLUS_HUMAN Clusterin OS=Homo sapiens OX=9606 GN=CLU PE=1 SV=1 | 1467 | 20201115_07413_1ng_glyco_hcd203040_compound15.15905.15905.4 | controllerType=0 controllerNumber=1 scan=15905 | 35.2825 |
| 14226:4 | 14 | K.ELPGVC[+57.021]N[+2059.735]ETMMALWEEC[+57.021]KPC[+57.021]LK.Q | HexNAc(4)Hex(5)Fuc(1)NeuAc(1) | C[+57]*3, N[+2060] | 1189.4989 | 4 | 4754.9739 | 4754.9406 | 0 | 7 | 97 | Specific | 214.93 | 204.02 | 204.02 | 4.51 | 32 | >sp\|P10909\|CLUS_HUMAN Clusterin OS=Homo sapiens OX=9606 GN=CLU PE=1 SV=1 | 1467 | 20201115_07413_1ng_glyco_hcd203040_compound15.17798.17798.2 | controllerType=0 controllerNumber=1 scan=17798 | 38.7412 |
| 14228:4 | 14 | K.ELPGVC[+57.021]N[+2059.735]ETMMALWEEC[+57.021]KPC[+57.021]LK.Q | HexNAc(4)Hex(5)Fuc(1)NeuAc(1) | C[+57]*3, N[+2060] | 1189.4989 | 4 | 4754.9739 | 4754.9406 | 0 | 7 | 97 | Specific | 312.01 | 298.38 | 298.38 | 4.89 | 32 | >sp\|P10909\|CLUS_HUMAN Clusterin OS=Homo sapiens OX=9606 GN=CLU PE=1 SV=1 | 1467 | 20201115_07413_1ng_glyco_hcd203040_compound15.17800.17800.2 | controllerType=0 controllerNumber=1 scan=17800 | 38.745 |
| 18771:3 | 14 | K.M[+15.995]LN[+2059.735]TSSLLEQLNEQFNWVSR.L | HexNAc(4)Hex(5)Fuc(1)NeuAc(1) | M[+16], N[+2060] | 1495.6506 | 3 | 4484.9372 | 4484.9166 | 0 | 4.6 | 352 | Specific | 569.36 | 562.09 | 562.09 | 10.34 | 32 | >sp\|P10909\|CLUS_HUMAN Clusterin OS=Homo sapiens OX=9606 GN=CLU PE=1 SV=1 | 1467 | 20201115_07413_1ng_glyco_hcd203040_compound15.22587.22587.3 | controllerType=0 controllerNumber=1 scan=22587 | 48.6615 |
| 18773:3 | 14 | K.M[+15.995]LN[+2059.735]TSSLLEQLNEQFNWVSR.L | HexNAc(4)Hex(5)Fuc(1)NeuAc(1) | M[+16], N[+2060] | 1495.6506 | 3 | 4484.9372 | 4484.9166 | 0 | 4.6 | 352 | Specific | 896.4 | 816.15 | 816.15 | 16.34 | 32 | >sp\|P10909\|CLUS_HUMAN Clusterin OS=Homo sapiens OX=9606 GN=CLU PE=1 SV=1 | 1467 | 20201115_07413_1ng_glyco_hcd203040_compound15.22589.22589.3 | controllerType=0 controllerNumber=1 scan=22589 | 48.6652 |
| 08961:3 | 14 | R.LAN[+2205.793]LTQGEDQYYLR.V | HexNAc(4)Hex(5)Fuc(2)NeuAc(1) | N[+2206] | 1297.2266 | 3 | 3889.6652 | 3889.6265 | 0 | 10 | 372 | Specific | 287.65 | 287.65 | 287.65 | 6.09 | 32 | >sp\|P10909\|CLUS_HUMAN Clusterin OS=Homo sapiens OX=9606 GN=CLU PE=1 SV=1 | 1467 | 20201115_07413_1ng_glyco_hcd203040_compound15.12277.12277.3 | controllerType=0 controllerNumber=1 scan=12277 | 28.7596 |
| 08963:3 | 14 | R.LAN[+2205.793]LTQGEDQYYLR.V | HexNAc(4)Hex(5)Fuc(2)NeuAc(1) | N[+2206] | 1297.2266 | 3 | 3889.6652 | 3889.6265 | 0 | 10 | 372 | Specific | 340.64 | 324.41 | 324.41 | 6.6 | 32 | >sp\|P10909\|CLUS_HUMAN Clusterin OS=Homo sapiens OX=9606 GN=CLU PE=1 SV=1 | 1467 | 20201115_07413_1ng_glyco_hcd203040_compound15.12279.12279.3 | controllerType=0 controllerNumber=1 scan=12279 | 28.7637 |
| 12000:4 | 15 | R.FELLGC[+57.021]ELNGC[+57.021]ANPLGLKN[+1378.476]NSIPDKQITASSSYK.T | HexNAc(2)Hex(6) | C[+57]*2, N[+1378] | 1280.0947 | 4 | 5117.3571 | 5117.3223 | 0 | 6.8 | 220 | Specific | 206.81 | 206.81 | 206.81 | 4.35 | 31 | >sp\|Q08431\|MFGM_HUMAN Lactadherin OS=Homo sapiens OX=9606 GN=MFGE8 PE=1 SV=3 | 1064 | 20201115_07413_1ng_glyco_hcd203040_compound15.15477.15477.4 | controllerType=0 controllerNumber=1 scan=15477 | 34.4597 |
| 12002:4 | 15 | R.FELLGC[+57.021]ELNGC[+57.021]ANPLGLKN[+1378.476]NSIPDKQITASSSYK.T | HexNAc(2)Hex(6) | C[+57]*2, N[+1378] | 1280.0947 | 4 | 5117.3571 | 5117.3223 | 0 | 6.8 | 220 | Specific | 631.39 | 618.65 | 618.65 | 11.13 | 31 | >sp\|Q08431\|MFGM_HUMAN Lactadherin OS=Homo sapiens OX=9606 GN=MFGE8 PE=1 SV=3 | 1064 | 20201115_07413_1ng_glyco_hcd203040_compound15.15479.15479.4 | controllerType=0 controllerNumber=1 scan=15479 | 34.4638 |
| 08099:4 | 16 | K.QTDEIKDTRN[+2059.735]ESTQNC[+57.021]VVAEPEKMESSISSSSEEMSLSK.C | HexNAc(4)Hex(5)Fuc(1)NeuAc(1) | C[+57], N[+2060] | 1613.4379 | 4 | 6450.7296 | 6450.7156 | 0 | 2.2 | 60 | Specific | 443.1 | 341.43 | 341.43 | 6.87 | 31 | >sp\|P47710\|CASA1_HUMAN Alpha-S1-casein OS=Homo sapiens OX=9606 GN=CSN1S1 PE=1 SV=1 | 989 | 20201115_07413_1ng_glyco_hcd203040_compound15.11372.11372.4 | controllerType=0 controllerNumber=1 scan=11372 | 27.2084 |
| 08101:4 | 16 | K.QTDEIKDTRN[+2059.735]ESTQNC[+57.021]VVAEPEKMESSISSSSEEMSLSK.C | HexNAc(4)Hex(5)Fuc(1)NeuAc(1) | C[+57], N[+2060] | 1613.4379 | 4 | 6450.7296 | 6450.7156 | 0 | 2.2 | 60 | Specific | 668.65 | 444.04 | 444.04 | 11.93 | 31 | >sp\|P47710\|CASA1_HUMAN Alpha-S1-casein OS=Homo sapiens OX=9606 GN=CSN1S1 PE=1 SV=1 | 989 | 20201115_07413_1ng_glyco_hcd203040_compound15.11374.11374.4 | controllerType=0 controllerNumber=1 scan=11374 | 27.2123 |
| 06451:3 | 17 | R.LSPN[+2350.830]ASAEHLELR.W | HexNAc(4)Hex(5)Fuc(1)NeuAc(2) | N[+2351] | 1263.2083 | 3 | 3787.6103 | 3787.5796 | 0 | 8.1 | 52 | Specific | 450.13 | 407.78 | 407.78 | 8.51 | 32 | >sp\|Q13410\|BT1A1_HUMAN Butyrophilin subfamily 1 member A1 OS=Homo sapiens OX=9606 GN=BTN1A1 PE=1 SV=3 | 179 | 20201115_07413_1ng_glyco_hcd203040_compound15.9617.9617.3 | controllerType=0 controllerNumber=1 scan=9617 | 24.0968 |
| 04505:3 | 17 | R.LSPN[+2205.793]ASAEHLELR.W | HexNAc(4)Hex(5)Fuc(2)NeuAc(1) | N[+2206] | 1214.8554 | 3 | 3642.5517 | 3642.5421 | 0 | 2.6 | 52 | Specific | 519.61 | 491.27 | 491.27 | 9.21 | 32 | >sp\|Q13410\|BT1A1_HUMAN Butyrophilin subfamily 1 member A1 OS=Homo sapiens OX=9606 GN=BTN1A1 PE=1 SV=3 | 179 | 20201115_07413_1ng_glyco_hcd203040_compound15.7564.7564.3 | controllerType=0 controllerNumber=1 scan=7564 | 20.5338 |
| 06449:3 | 17 | R.LSPN[+2350.830]ASAEHLELR.W | HexNAc(4)Hex(5)Fuc(1)NeuAc(2) | N[+2351] | 1263.2083 | 3 | 3787.6103 | 3787.5796 | 0 | 8.1 | 52 | Specific | 634.18 | 602.83 | 602.83 | 11.19 | 32 | >sp\|Q13410\|BT1A1_HUMAN Butyrophilin subfamily 1 member A1 OS=Homo sapiens OX=9606 GN=BTN1A1 PE=1 SV=3 | 179 | 20201115_07413_1ng_glyco_hcd203040_compound15.9615.9615.3 | controllerType=0 controllerNumber=1 scan=9615 | 24.0929 |
| 16014:3 | 17 | K.N[+1913.677]VSC[+57.021]YIQNLLLGQEK.K | HexNAc(4)Hex(5)NeuAc(1) | C[+57], N[+1914] | 1231.5334 | 3 | 3692.5856 | 3692.5876 | 0 | -0.5 | 215 | Specific | 498.54 | 374.27 | 374.27 | 8.31 | 32 | >sp\|Q13410\|BT1A1_HUMAN Butyrophilin subfamily 1 member A1 OS=Homo sapiens OX=9606 GN=BTN1A1 PE=1 SV=3 | 179 | 20201115_07413_1ng_glyco_hcd203040_compound15.19691.19691.3 | controllerType=0 controllerNumber=1 scan=19691 | 42.4259 |
| 16012:3 | 17 | K.N[+1913.677]VSC[+57.021]YIQNLLLGQEK.K | HexNAc(4)Hex(5)NeuAc(1) | C[+57], N[+1914] | 1231.5334 | 3 | 3692.5856 | 3692.5876 | 0 | -0.5 | 215 | Specific | 691.41 | 611.05 | 611.05 | 11.78 | 32 | >sp\|Q13410\|BT1A1_HUMAN Butyrophilin subfamily 1 member A1 OS=Homo sapiens OX=9606 GN=BTN1A1 PE=1 SV=3 | 179 | 20201115_07413_1ng_glyco_hcd203040_compound15.19689.19689.3 | controllerType=0 controllerNumber=1 scan=19689 | 42.4208 |
| 07259:5 | 20 | K.SC[+57.021]QHN[+2059.735]GTMYQHGEIFSAHELFPSR.L | HexNAc(4)Hex(5)Fuc(1)NeuAc(1) | C[+57], N[+2060] | 979.4058 | 5 | 4892.9998 | 4892.9919 | 0 | 1.6 | 110 | Specific | 726.08 | 631.02 | 631.02 | 14.24 | 22 | >sp\|Q6WN34\|CRDL2_HUMAN Chordin-like protein 2 OS=Homo sapiens OX=9606 GN=CHRDL2 PE=1 SV=1 | 1189 | 20201115_07413_1ng_glyco_hcd203040_compound15.10481.10481.5 | controllerType=0 controllerNumber=1 scan=10481 | 25.6342 |
| 07261:5 | 20 | K.SC[+57.021]QHN[+2059.735]GTMYQHGEIFSAHELFPSR.L | HexNAc(4)Hex(5)Fuc(1)NeuAc(1) | C[+57], N[+2060] | 979.4058 | 5 | 4892.9998 | 4892.9919 | 0 | 1.6 | 110 | Specific | 757.28 | 757.28 | 757.28 | 15.24 | 22 | >sp\|Q6WN34\|CRDL2_HUMAN Chordin-like protein 2 OS=Homo sapiens OX=9606 GN=CHRDL2 PE=1 SV=1 | 1189 | 20201115_07413_1ng_glyco_hcd203040_compound15.10483.10483.5 | controllerType=0 controllerNumber=1 scan=10483 | 25.6383 |
| 24364:5 | 36 | K.VFPLSLDSTPQDGNVVVAC[+57.021]LVQGFFPQEPLSVTWSESGQN[+2553.910]VTAI | HexNAc(5)Hex(5)Fuc(1)NeuAc(2) | C[+57], N[+2554] | 1466.8682 | 5 | 7330.3119 | 7330.2711 | 0 | 5.6 | 123 | Specific | 222.3 | 222.3 | 222.3 | 5.62 | 21 | >sp\|P0DOX2\|IGA2_HUMAN Immunoglobulin alpha-2 heavy chain OS=Homo sapiens OX=9606 PE=1 SV=2 | 631 | 20201115_07413_1ng_glyco_hcd203040_compound15.28506.28506.5 | controllerType=0 controllerNumber=1 scan=28506 | 60.1484 |
| 20270:4 | 36 | K.VFPLSLDSTPQDGNVVVAC[+57.021]LVQGFFPQEPLSVTWSESGQN[+1809.666]VTAI | HexNAc(5)Hex(4)Fuc(1) | C[+57], N[+1810] | 1647.2786 | 4 | 6586.0926 | 6586.0275 | 0 | 9.9 | 123 | Specific | 379.96 | 379.96 | 379.96 | 9.21 | 21 | >sp\|P0DOX2\|IGA2_HUMAN Immunoglobulin alpha-2 heavy chain OS=Homo sapiens OX=9606 PE=1 SV=2 | 631 | 20201115_07413_1ng_glyco_hcd203040_compound15.24156.24156.4 | controllerType=0 controllerNumber=1 scan=24156 | 51.96 |
| 20463:4 | 36 | K.VFPLSLDSTPQDGNVVVAC[+57.021]LVQGFFPQEPLSVTWSESGQN[+1647.613]VTAI | HexNAc(5)Hex(3)Fuc(1) | C[+57], N[+1648] | 1606.7648 | 4 | 6424.0373 | 6423.9746 | 0 | 9.7 | 123 | Specific | 548.44 | 548.44 | 548.44 | 9.99 | 21 | >sp\|P0DOX2\|IGA2_HUMAN Immunoglobulin alpha-2 heavy chain OS=Homo sapiens OX=9606 PE=1 SV=2 | 631 | 20201115_07413_1ng_glyco_hcd203040_compound15.24362.24362.4 | controllerType=0 controllerNumber=1 scan=24362 | 52.3112 |
| 20268:4 | 36 | K.VFPLSLDSTPQDGNVVVAC[+57.021]LVQGFFPQEPLSVTWSESGQN[+1809.666]VTAI | HexNAc(5)Hex(4)Fuc(1) | C[+57], N[+1810] | 1647.2786 | 4 | 6586.0926 | 6586.0275 | 0 | 9.9 | 123 | Specific | 668.51 | 611.57 | 611.57 | 13.16 | 21 | >sp\|P0DOX2\|IGA2_HUMAN Immunoglobulin alpha-2 heavy chain OS=Homo sapiens OX=9606 PE=1 SV=2 | 631 | 20201115_07413_1ng_glyco_hcd203040_compound15.24154.24154.4 | controllerType=0 controllerNumber=1 scan=24154 | 51.9495 |
| 20466:4 | 36 | K.VFPLSLDSTPQDGNVVVAC[+57.021]LVQGFFPQEPLSVTWSESGQN[+1647.613]VTAI | HexNAc(5)Hex(3)Fuc(1) | C[+57], N[+1648] | 1606.7648 | 4 | 6424.0373 | 6423.9746 | 0 | 9.7 | 123 | Specific | 687.96 | 687.96 | 687.96 | 13.16 | 21 | >sp\|P0DOX2\|IGA2_HUMAN Immunoglobulin alpha-2 heavy chain OS=Homo sapiens OX=9606 PE=1 SV=2 | 631 | 20201115_07413_1ng_glyco_hcd203040_compound15.24365.24365.4 | controllerType=0 controllerNumber=1 scan=24365 | 52.3169 |
| 03728:3 | 36 | K.TPLTAN[+2262.814]ITK.S | HexNAc(5)Hex(5)Fuc(1)NeuAc(1) | N[+2263] | 1074.4641 | 3 | 3221.3779 | 3221.3711 | 0 | 2.1 | 315 | Specific | 202.06 | 13.21 | 13.21 | 3.17 | 21 | >sp\|P0DOX2\|IGA2_HUMAN Immunoglobulin alpha-2 heavy chain OS=Homo sapiens OX=9606 PE=1 SV=2 | 631 | 20201115_07413_1ng_glyco_hcd203040_compound15.6736.6736.3 | controllerType=0 controllerNumber=1 scan=6736 | 19.0502 |
| 02121:3 | 36 | K.TPLTAN[+1971.719]ITK.S | HexNAc(5)Hex(5)Fuc(1) | N[+1972] | 977.4322 | 3 | 2930.2819 | 2930.2757 | 0 | 2.1 | 315 | Specific | 223.27 | 213.41 | 213.41 | 4.51 | 21 | >sp\|P0DOX2\|IGA2_HUMAN Immunoglobulin alpha-2 heavy chain OS=Homo sapiens OX=9606 PE=1 SV=2 | 631 | 20201115_07413_1ng_glyco_hcd203040_compound15.5005.5005.3 | controllerType=0 controllerNumber=1 scan=5005 | 15.851 |
| 02122:2 | 36 | K.TPLTAN[+1971.719]ITK.S | HexNAc(5)Hex(5)Fuc(1) | N[+1972] | 1465.6446 | 2 | 2930.2819 | 2930.2757 | 0 | 2.1 | 315 | Specific | 264.45 | 90.89 | 90.89 | 4.51 | 21 | >sp\|P0DOX2\|IGA2_HUMAN Immunoglobulin alpha-2 heavy chain OS=Homo sapiens OX=9606 PE=1 SV=2 | 631 | 20201115_07413_1ng_glyco_hcd203040_compound15.5006.5006.2 | controllerType=0 controllerNumber=1 scan=5006 | 15.8525 |
| 02233:3 | 36 | K.TPLTAN[+1647.613]ITK.S | HexNAc(5)Hex(3)Fuc(1) | N[+1648] | 869.3967 | 3 | 2606.1755 | 2606.17 | 0 | 2.1 | 315 | Specific | 313.92 | 48.02 | 48.02 | 4.15 | 21 | >sp\|P0DOX2\|IGA2_HUMAN Immunoglobulin alpha-2 heavy chain OS=Homo sapiens OX=9606 PE=1 SV=2 | 631 | 20201115_07413_1ng_glyco_hcd203040_compound15.5121.5121.3 | controllerType=0 controllerNumber=1 scan=5121 | 16.0751 |
| 02123:3 | 36 | K.TPLTAN[+1971.719]ITK.S | HexNAc(5)Hex(5)Fuc(1) | N[+1972] | 977.4322 | 3 | 2930.2819 | 2930.2757 | 0 | 2.1 | 315 | Specific | 359.22 | 233.7 | 233.7 | 6.34 | 21 | >sp\|P0DOX2\|IGA2_HUMAN Immunoglobulin alpha-2 heavy chain OS=Homo sapiens OX=9606 PE=1 SV=2 | 631 | 20201115_07413_1ng_glyco_hcd203040_compound15.5007.5007.3 | controllerType=0 controllerNumber=1 scan=5007 | 15.8573 |
| 02124:2 | 36 | K.TPLTAN[+1971.719]ITK.S | HexNAc(5)Hex(5)Fuc(1) | N[+1972] | 1465.6446 | 2 | 2930.2819 | 2930.2757 | 0 | 2.1 | 315 | Specific | 399.51 | 267.9 | 267.9 | 6.81 | 21 | >sp\|P0DOX2\|IGA2_HUMAN Immunoglobulin alpha-2 heavy chain OS=Homo sapiens OX=9606 PE=1 SV=2 | 631 | 20201115_07413_1ng_glyco_hcd203040_compound15.5008.5008.2 | controllerType=0 controllerNumber=1 scan=5008 | 15.8592 |
| 02230:2 | 36 | K.TPLTAN[+1501.555]ITK.S | HexNAc(5)Hex(3) | N[+1502] | 1230.5628 | 2 | 2460.1183 | 2460.1121 | 0 | 2.5 | 315 | Specific | 436.08 | 367.1 | 367.1 | 7.51 | 21 | >sp\|P0DOX2\|IGA2_HUMAN Immunoglobulin alpha-2 heavy chain OS=Homo sapiens OX=9606 PE=1 SV=2 | 631 | 20201115_07413_1ng_glyco_hcd203040_compound15.5118.5118.2 | controllerType=0 controllerNumber=1 scan=5118 | 16.0695 |
| 03730:3 | 36 | K.TPLTAN[+2262.814]ITK.S | HexNAc(5)Hex(5)Fuc(1)NeuAc(1) | N[+2263] | 1074.4641 | 3 | 3221.3779 | 3221.3711 | 0 | 2.1 | 315 | Specific | 444.82 | 232.87 | 232.87 | 6.64 | 21 | >sp\|P0DOX2\|IGA2_HUMAN Immunoglobulin alpha-2 heavy chain OS=Homo sapiens OX=9606 PE=1 SV=2 | 631 | 20201115_07413_1ng_glyco_hcd203040_compound15.6738.6738.3 | controllerType=0 controllerNumber=1 scan=6738 | 19.054 |
| 02167:3 | 36 | K.TPLTAN[+1809.666]ITK.S | HexNAc(5)Hex(4)Fuc(1) | N[+1810] | 923.4145 | 3 | 2768.2288 | 2768.2228 | 0 | 2.2 | 315 | Specific | 459 | 176.96 | 176.96 | 6.12 | 21 | >sp\|P0DOX2\|IGA2_HUMAN Immunoglobulin alpha-2 heavy chain OS=Homo sapiens OX=9606 PE=1 SV=2 | 631 | 20201115_07413_1ng_glyco_hcd203040_compound15.5052.5052.3 | controllerType=0 controllerNumber=1 scan=5052 | 15.9423 |
| 02265:2 | 36 | K.TPLTAN[+1444.534]ITK.S | HexNAc(4)Hex(3)Fuc(1) | N[+1445] | 1202.0534 | 2 | 2403.0995 | 2403.0906 | 0 | 3.7 | 315 | Specific | 519.72 | 274.92 | 274.92 | 7.3 | 21 | >sp\|P0DOX2\|IGA2_HUMAN Immunoglobulin alpha-2 heavy chain OS=Homo sapiens OX=9606 PE=1 SV=2 | 631 | 20201115_07413_1ng_glyco_hcd203040_compound15.5154.5154.2 | controllerType=0 controllerNumber=1 scan=5154 | 16.1389 |
| 02169:3 | 36 | K.TPLTAN[+1809.666]ITK.S | HexNAc(5)Hex(4)Fuc(1) | N[+1810] | 923.4145 | 3 | 2768.2288 | 2768.2228 | 0 | 2.2 | 315 | Specific | 543.28 | 255.71 | 255.71 | 7.23 | 21 | >sp\|P0DOX2\|IGA2_HUMAN Immunoglobulin alpha-2 heavy chain OS=Homo sapiens OX=9606 PE=1 SV=2 | 631 | 20201115_07413_1ng_glyco_hcd203040_compound15.5054.5054.3 | controllerType=0 controllerNumber=1 scan=5054 | 15.9483 |
| 02149:2 | 36 | K.TPLTAN[+1809.666]ITK.S | HexNAc(5)Hex(4)Fuc(1) | N[+1810] | 1384.6185 | 2 | 2768.2296 | 2768.2228 | 0 | 2.5 | 315 | Specific | 549.04 | 302.84 | 302.84 | 7.7 | 21 | >sp\|P0DOX2\|IGA2_HUMAN Immunoglobulin alpha-2 heavy chain OS=Homo sapiens OX=9606 PE=1 SV=2 | 631 | 20201115_07413_1ng_glyco_hcd203040_compound15.5034.5034.2 | controllerType=0 controllerNumber=1 scan=5034 | 15.9103 |
| 02151:2 | 36 | K.TPLTAN[+1809.666]ITK.S | HexNAc(5)Hex(4)Fuc(1) | N[+1810] | 1384.6185 | 2 | 2768.2296 | 2768.2228 | 0 | 2.5 | 315 | Specific | 556.28 | 319.44 | 319.44 | 7.9 | 21 | >sp\|P0DOX2\|IGA2_HUMAN Immunoglobulin alpha-2 heavy chain OS=Homo sapiens OX=9606 PE=1 SV=2 | 631 | 20201115_07413_1ng_glyco_hcd203040_compound15.5036.5036.2 | controllerType=0 controllerNumber=1 scan=5036 | 15.9149 |
| 02235:3 | 36 | K.TPLTAN[+1647.613]ITK.S | HexNAc(5)Hex(3)Fuc(1) | N[+1648] | 869.3967 | 3 | 2606.1755 | 2606.17 | 0 | 2.1 | 315 | Specific | 573.49 | 292.07 | 292.07 | 8.59 | 21 | >sp\|P0DOX2\|IGA2_HUMAN Immunoglobulin alpha-2 heavy chain OS=Homo sapiens OX=9606 PE=1 SV=2 | 631 | 20201115_07413_1ng_glyco_hcd203040_compound15.5123.5123.3 | controllerType=0 controllerNumber=1 scan=5123 | 16.0798 |
| 02225:2 | 36 | K.TPLTAN[+1647.613]ITK.S | HexNAc(5)Hex(3)Fuc(1) | N[+1648] | 1303.5917 | 2 | 2606.1762 | 2606.17 | 0 | 2.4 | 315 | Specific | 594.41 | 292.57 | 292.57 | 8.86 | 21 | >sp\|P0DOX2\|IGA2_HUMAN Immunoglobulin alpha-2 heavy chain OS=Homo sapiens OX=9606 PE=1 SV=2 | 631 | 20201115_07413_1ng_glyco_hcd203040_compound15.5113.5113.2 | controllerType=0 controllerNumber=1 scan=5113 | 16.0615 |
| 02263:2 | 36 | K.TPLTAN[+1444.534]ITK.S | HexNAc(4)Hex(3)Fuc(1) | N[+1445] | 1202.0534 | 2 | 2403.0995 | 2403.0906 | 0 | 3.7 | 315 | Specific | 615.68 | 324.71 | 324.71 | 10.05 | 21 | >sp\|P0DOX2\|IGA2_HUMAN Immunoglobulin alpha-2 heavy chain OS=Homo sapiens OX=9606 PE=1 SV=2 | 631 | 20201115_07413_1ng_glyco_hcd203040_compound15.5152.5152.2 | controllerType=0 controllerNumber=1 scan=5152 | 16.1344 |
| 02223:2 | 36 | K.TPLTAN[+1647.613]ITK.S | HexNAc(5)Hex(3)Fuc(1) | N[+1648] | 1303.5917 | 2 | 2606.1762 | 2606.17 | 0 | 2.4 | 315 | Specific | 617.13 | 347.27 | 347.27 | 10.16 | 21 | >sp\|P0DOX2\|IGA2_HUMAN Immunoglobulin alpha-2 heavy chain OS=Homo sapiens OX=9606 PE=1 SV=2 | 631 | 20201115_07413_1ng_glyco_hcd203040_compound15.5111.5111.2 | controllerType=0 controllerNumber=1 scan=5111 | 16.0546 |
| 08328:3 | 40 | R.IIVPLNNREN[+1663.608]ISDPTSPLR.T | HexNAc(5)Hex(4) | N[+1664] | 1271.2714 | 3 | 3811.7997 | 3811.7853 | 0 | 3.8 | 62 | Specific | 278.79 | 278.79 | 278.79 | 6.52 | 18 | >sp\|P01591\|IGJ_HUMAN Immunoglobulin J chain OS=Homo sapiens OX=9606 GN=JCHAIN PE=1 SV=4 | 1392 | 20201115_07413_1ng_glyco_hcd203040_compound15.11610.11610.3 | controllerType=0 controllerNumber=1 scan=11610 | 27.5987 |
| 09824:3 | 40 | R.IIVPLNNREN[+1913.677]ISDPTSPLR.T | HexNAc(4)Hex(5)NeuAc(1) | N[+1914] | 1354.6249 | 3 | 4061.8603 | 4061.8542 | 0 | 1.5 | 62 | Specific | 325 | 323.53 | 323.53 | 7.04 | 18 | >sp\|P01591\|IGJ_HUMAN Immunoglobulin J chain OS=Homo sapiens OX=9606 GN=JCHAIN PE=1 SV=4 | 1392 | 20201115_07413_1ng_glyco_hcd203040_compound15.13187.13187.3 | controllerType=0 controllerNumber=1 scan=13187 | 30.3747 |
| 09777:3 | 40 | R.IIVPLNNREN[+2059.735]ISDPTSPLR.T | HexNAc(4)Hex(5)Fuc(1)NeuAc(1) | N[+2060] | 1403.3141 | 3 | 4207.9279 | 4207.9121 | 0 | 3.8 | 62 | Specific | 447.88 | 425.65 | 425.65 | 8.83 | 18 | >sp\|P01591\|IGJ_HUMAN Immunoglobulin J chain OS=Homo sapiens OX=9606 GN=JCHAIN PE=1 SV=4 | 1392 | 20201115_07413_1ng_glyco_hcd203040_compound15.13138.13138.3 | controllerType=0 controllerNumber=1 scan=13138 | 30.2713 |
| 09897:3 | 40 | R.IIVPLNNREN[+1856.656]ISDPTSPLR.T | HexNAc(3)Hex(5)Fuc(1)NeuAc(1) | N[+1857] | 1335.6169 | 3 | 4004.8361 | 4004.8327 | 0 | 0.8 | 62 | Specific | 466.33 | 466.33 | 466.33 | 8.54 | 18 | >sp\|P01591\|IGJ_HUMAN Immunoglobulin J chain OS=Homo sapiens OX=9606 GN=JCHAIN PE=1 SV=4 | 1392 | 20201115_07413_1ng_glyco_hcd203040_compound15.13263.13263.3 | controllerType=0 controllerNumber=1 scan=13263 | 30.5378 |
| 11686:3 | 40 | R.IIVPLNNREN[+2350.830]ISDPTSPLR.T | HexNAc(4)Hex(5)Fuc(1)NeuAc(2) | N[+2351] | 1500.3434 | 3 | 4499.0158 | 4499.0075 | 0 | 1.8 | 62 | Specific | 470.17 | 470.17 | 470.17 | 8.84 | 18 | >sp\|P01591\|IGJ_HUMAN Immunoglobulin J chain OS=Homo sapiens OX=9606 GN=JCHAIN PE=1 SV=4 | 1392 | 20201115_07413_1ng_glyco_hcd203040_compound15.15148.15148.3 | controllerType=0 controllerNumber=1 scan=15148 | 33.8829 |
| 11773:3 | 40 | R.IIVPLNNREN[+2205.793]ISDPTSPLR.T | HexNAc(4)Hex(5)Fuc(2)NeuAc(1) | N[+2206] | 1451.9915 | 3 | 4353.9599 | 4353.97 | 0 | -2.3 | 62 | Specific | 476.16 | 476.16 | 476.16 | 7.39 | 18 | >sp\|P01591\|IGJ_HUMAN Immunoglobulin J chain OS=Homo sapiens OX=9606 GN=JCHAIN PE=1 SV=4 | 1392 | 20201115_07413_1ng_glyco_hcd203040_compound15.15238.15238.3 | controllerType=0 controllerNumber=1 scan=15238 | 34.0367 |
| 09779:3 | 40 | R.IIVPLNNREN[+2059.735]ISDPTSPLR.T | HexNAc(4)Hex(5)Fuc(1)NeuAc(1) | N[+2060] | 1403.3141 | 3 | 4207.9279 | 4207.9121 | 0 | 3.8 | 62 | Specific | 496.44 | 454.52 | 454.52 | 9.04 | 18 | >sp\|P01591\|IGJ_HUMAN Immunoglobulin J chain OS=Homo sapiens OX=9606 GN=JCHAIN PE=1 SV=4 | 1392 | 20201115_07413_1ng_glyco_hcd203040_compound15.13140.13140.3 | controllerType=0 controllerNumber=1 scan=13140 | 30.2762 |
| 11771:3 | 40 | R.IIVPLNNREN[+2205.793]ISDPTSPLR.T | HexNAc(4)Hex(5)Fuc(2)NeuAc(1) | N[+2206] | 1451.9915 | 3 | 4353.9599 | 4353.97 | 0 | -2.3 | 62 | Specific | 501.48 | 496.36 | 496.36 | 7.59 | 18 | >sp\|P01591\|IGJ_HUMAN Immunoglobulin J chain OS=Homo sapiens OX=9606 GN=JCHAIN PE=1 SV=4 | 1392 | 20201115_07413_1ng_glyco_hcd203040_compound15.15236.15236.3 | controllerType=0 controllerNumber=1 scan=15236 | 34.0329 |
| 11684:3 | 40 | R.IIVPLNNREN[+2350.830]ISDPTSPLR.T | HexNAc(4)Hex(5)Fuc(1)NeuAc(2) | N[+2351] | 1500.3434 | 3 | 4499.0158 | 4499.0075 | 0 | 1.8 | 62 | Specific | 511.49 | 485.46 | 485.46 | 9.17 | 18 | >sp\|P01591\|IGJ_HUMAN Immunoglobulin J chain OS=Homo sapiens OX=9606 GN=JCHAIN PE=1 SV=4 | 1392 | 20201115_07413_1ng_glyco_hcd203040_compound15.15146.15146.3 | controllerType=0 controllerNumber=1 scan=15146 | 33.8782 |
| 09826:3 | 40 | R.IIVPLNNREN[+1913.677]ISDPTSPLR.T | HexNAc(4)Hex(5)NeuAc(1) | N[+1914] | 1354.6249 | 3 | 4061.8603 | 4061.8542 | 0 | 1.5 | 62 | Specific | 511.99 | 393.03 | 393.03 | 9.17 | 18 | >sp\|P01591\|IGJ_HUMAN Immunoglobulin J chain OS=Homo sapiens OX=9606 GN=JCHAIN PE=1 SV=4 | 1392 | 20201115_07413_1ng_glyco_hcd203040_compound15.13189.13189.3 | controllerType=0 controllerNumber=1 scan=13189 | 30.3787 |
| 11753:3 | 40 | R.IIVPLNNREN[+2204.772]ISDPTSPLR.T | HexNAc(4)Hex(5)NeuAc(2) | N[+2205] | 1451.6566 | 3 | 4352.9551 | 4352.9496 | 0 | 1.3 | 62 | Specific | 518.52 | 512.8 | 512.8 | 9.17 | 18 | >sp\|P01591\|IGJ_HUMAN Immunoglobulin J chain OS=Homo sapiens OX=9606 GN=JCHAIN PE=1 SV=4 | 1392 | 20201115_07413_1ng_glyco_hcd203040_compound15.15217.15217.3 | controllerType=0 controllerNumber=1 scan=15217 | 34.0006 |
| 09899:3 | 40 | R.IIVPLNNREN[+1856.656]ISDPTSPLR.T | HexNAc(3)Hex(5)Fuc(1)NeuAc(1) | N[+1857] | 1335.6169 | 3 | 4004.8361 | 4004.8327 | 0 | 0.8 | 62 | Specific | 564.64 | 564.64 | 564.64 | 9.05 | 18 | >sp\|P01591\|IGJ_HUMAN Immunoglobulin J chain OS=Homo sapiens OX=9606 GN=JCHAIN PE=1 SV=4 | 1392 | 20201115_07413_1ng_glyco_hcd203040_compound15.13265.13265.3 | controllerType=0 controllerNumber=1 scan=13265 | 30.543 |
| 11751:3 | 40 | R.IIVPLNNREN[+2204.772]ISDPTSPLR.T | HexNAc(4)Hex(5)NeuAc(2) | N[+2205] | 1451.6566 | 3 | 4352.9551 | 4352.9496 | 0 | 1.3 | 62 | Specific | 571.58 | 522 | 522 | 10.52 | 18 | >sp\|P01591\|IGJ_HUMAN Immunoglobulin J chain OS=Homo sapiens OX=9606 GN=JCHAIN PE=1 SV=4 | 1392 | 20201115_07413_1ng_glyco_hcd203040_compound15.15215.15215.3 | controllerType=0 controllerNumber=1 scan=15215 | 33.9955 |
| 14955:3 | 59 | R.NYIVPILWLN[+1864.634]ETGTIGDEKANM[+15.995]FR.S | HexNAc(2)Hex(9) | M[+16], N[+1865] | 1559.0193 | 3 | 4675.0435 | 4675.0523 | 0 | -1.9 | 408 | Specific | 325.5 | 325.5 | 325.5 | 5.87 | 11 | >sp\|P16671\|CD36_HUMAN Platelet glycoprotein 4 OS=Homo sapiens OX=9606 GN=CD36 PE=1 SV=2 | 812 | 20201115_07413_1ng_glyco_hcd203040_compound15.18584.18584.3 | controllerType=0 controllerNumber=1 scan=18584 | 40.2024 |
| 14953:3 | 59 | R.NYIVPILWLN[+1864.634]ETGTIGDEKANM[+15.995]FR.S | HexNAc(2)Hex(9) | M[+16], N[+1865] | 1559.0193 | 3 | 4675.0435 | 4675.0523 | 0 | -1.9 | 408 | Specific | 456.66 | 434.84 | 434.84 | 7.67 | 11 | >sp\|P16671\|CD36_HUMAN Platelet glycoprotein 4 OS=Homo sapiens OX=9606 GN=CD36 PE=1 SV=2 | 812 | 20201115_07413_1ng_glyco_hcd203040_compound15.18582.18582.3 | controllerType=0 controllerNumber=1 scan=18582 | 40.1988 |
| 14887:3 | 59 | R.NYIVPILWLN[+1864.634]ETGTIGDEKANM[+15.995]FR.S | HexNAc(2)Hex(9) | M[+16], N[+1865] | 1559.0262 | 3 | 4675.064 | 4675.0523 | 0 | 2.5 | 408 | Specific | 558.02 | 558.02 | 558.02 | 9.64 | 11 | >sp\|P16671\|CD36_HUMAN Platelet glycoprotein 4 OS=Homo sapiens OX=9606 GN=CD36 PE=1 SV=2 | 812 | 20201115_07413_1ng_glyco_hcd203040_compound15.18513.18513.3 | controllerType=0 controllerNumber=1 scan=18513 | 40.0712 |
| 14889:3 | 59 | R.NYIVPILWLN[+1864.634]ETGTIGDEKANM[+15.995]FR.S | HexNAc(2)Hex(9) | M[+16], N[+1865] | 1559.0262 | 3 | 4675.064 | 4675.0523 | 0 | 2.5 | 408 | Specific | 626.54 | 614.92 | 614.92 | 11.81 | 11 | >sp\|P16671\|CD36_HUMAN Platelet glycoprotein 4 OS=Homo sapiens OX=9606 GN=CD36 PE=1 SV=2 | 812 | 20201115_07413_1ng_glyco_hcd203040_compound15.18515.18515.3 | controllerType=0 controllerNumber=1 scan=18515 | 40.0752 |
| 15042:3 | 70 | K.AALAAFNAQNN[+2205.793]GSNFQLEEISR.A | HexNAc(4)Hex(5)Fuc(2)NeuAc(1) | N[+2206] | 1524.3168 | 3 | 4570.9358 | 4570.946 | 0 | -2.2 | 166 | Specific | 239.67 | 239.67 | 239.67 | 4.6 | 8 | >sp\|P02765\|FETUA_HUMAN Alpha-2-HS-glycoprotein OS=Homo sapiens OX=9606 GN=AHSG PE=1 SV=2 | 135 | 20201115_07413_1ng_glyco_hcd203040_compound15.18674.18674.3 | controllerType=0 controllerNumber=1 scan=18674 | 40.3722 |
| 15020:3 | 70 | K.AALAAFNAQNN[+2204.772]GSNFQLEEISR.A | HexNAc(4)Hex(5)NeuAc(2) | N[+2205] | 1523.9893 | 3 | 4569.9534 | 4569.9256 | 0 | 6.1 | 166 | Specific | 262.07 | 255.43 | 255.43 | 5.99 | 8 | >sp\|P02765\|FETUA_HUMAN Alpha-2-HS-glycoprotein OS=Homo sapiens OX=9606 GN=AHSG PE=1 SV=2 | 135 | 20201115_07413_1ng_glyco_hcd203040_compound15.18651.18651.3 | controllerType=0 controllerNumber=1 scan=18651 | 40.3309 |
| 15022:3 | 70 | K.AALAAFNAQNN[+2204.772]GSNFQLEEISR.A | HexNAc(4)Hex(5)NeuAc(2) | N[+2205] | 1523.9893 | 3 | 4569.9534 | 4569.9256 | 0 | 6.1 | 166 | Specific | 422.83 | 361.71 | 361.71 | 7.12 | 8 | >sp\|P02765\|FETUA_HUMAN Alpha-2-HS-glycoprotein OS=Homo sapiens OX=9606 GN=AHSG PE=1 SV=2 | 135 | 20201115_07413_1ng_glyco_hcd203040_compound15.18653.18653.3 | controllerType=0 controllerNumber=1 scan=18653 | 40.3356 |
| 17481:4 | 1 | R.RTAGWNVPIGTLRPFLN[+1768.640]WTGPPEPIEAAVAR.F | HexNAc(4)Hex(5)Fuc(1) | N[+1769] | 1289.6216 | 4 | 5155.4645 | 5155.4516 | 0 | 2.5 | 140 | Specific | 400.99 | 311.09 | 311.09 | 8.99 | 111 | >sp\|P02788\|TRFL_HUMAN Lactotransferrin OS=Homo sapiens OX=9606 GN=LTF PE=1 SV=6 | 1270 | 20201115_07413_1ng_glyco_hcd203040_compound15.20872.20872.4 | controllerType=0 controllerNumber=1 scan=20872 | 41.3934 |
| 18451:5 | 1 | R.RTAGWNVPIGTLRPFLN[+2205.793]WTGPPEPIEAAVAR.F | HexNAc(4)Hex(5)Fuc(2)NeuAc(1) | N[+2206] | 1119.3302 | 5 | 5592.6219 | 5592.6049 | 0 | 3 | 140 | Specific | 425.19 | 402.47 | 402.47 | 10.7 | 111 | >sp\|P02788\|TRFL_HUMAN Lactotransferrin OS=Homo sapiens OX=9606 GN=LTF PE=1 SV=6 | 1270 | 20201115_07433_1ng_glyco_hcd203040_compound15.21882.21882.5 | controllerType=0 controllerNumber=1 scan=21882 | 43.5601 |
| 18453:5 | 1 | R.RTAGWNVPIGTLRPFLN[+2205.793]WTGPPEPIEAAVAR.F | HexNAc(4)Hex(5)Fuc(2)NeuAc(1) | N[+2206] | 1119.3302 | 5 | 5592.6219 | 5592.6049 | 0 | 3 | 140 | Specific | 440.81 | 359.62 | 359.62 | 9.38 | 111 | >sp\|P02788\|TRFL_HUMAN Lactotransferrin OS=Homo sapiens OX=9606 GN=LTF PE=1 SV=6 | 1270 | 20201115_07433_1ng_glyco_hcd203040_compound15.21884.21884.5 | controllerType=0 controllerNumber=1 scan=21884 | 43.5735 |
| 18454:4 | 1 | R.RTAGWNVPIGTLRPFLN[+2059.735]WTGPPEPIEAAVAR.F | HexNAc(4)Hex(5)Fuc(1)NeuAc(1) | N[+2060] | 1362.3971 | 4 | 5446.5666 | 5446.547 | 0 | 3.6 | 140 | Specific | 546.82 | 546.82 | 546.82 | 11.31 | 111 | >sp\|P02788\|TRFL_HUMAN Lactotransferrin OS=Homo sapiens OX=9606 GN=LTF PE=1 SV=6 | 1270 | 20201115_07433_1ng_glyco_hcd203040_compound15.21886.21886.4 | controllerType=0 controllerNumber=1 scan=21886 | 43.578 |
| 18442:4 | 1 | R.RTAGWNVPIGTLRPFLN[+2205.793]WTGPPEPIEAAVAR.F | HexNAc(4)Hex(5)Fuc(2)NeuAc(1) | N[+2206] | 1398.9102 | 4 | 5592.6188 | 5592.6049 | 0 | 2.5 | 140 | Specific | 566.99 | 466.82 | 466.82 | 11.59 | 111 | >sp\|P02788\|TRFL_HUMAN Lactotransferrin OS=Homo sapiens OX=9606 GN=LTF PE=1 SV=6 | 1270 | 20201115_07433_1ng_glyco_hcd203040_compound15.21873.21873.4 | controllerType=0 controllerNumber=1 scan=21873 | 43.5431 |
| 18439:4 | 1 | R.RTAGWNVPIGTLRPFLN[+2205.793]WTGPPEPIEAAVAR.F | HexNAc(4)Hex(5)Fuc(2)NeuAc(1) | N[+2206] | 1398.9102 | 4 | 5592.6188 | 5592.6049 | 0 | 2.5 | 140 | Specific | 631.98 | 442.71 | 442.71 | 13.88 | 111 | >sp\|P02788\|TRFL_HUMAN Lactotransferrin OS=Homo sapiens OX=9606 GN=LTF PE=1 SV=6 | 1270 | 20201115_07433_1ng_glyco_hcd203040_compound15.21870.21870.4 | controllerType=0 controllerNumber=1 scan=21870 | 43.5322 |
| 17484:4 | 1 | R.RTAGWNVPIGTLRPFLN[+1768.640]WTGPPEPIEAAVAR.F | HexNAc(4)Hex(5)Fuc(1) | N[+1769] | 1289.6216 | 4 | 5155.4645 | 5155.4516 | 0 | 2.5 | 140 | Specific | 655.11 | 510 | 510 | 14.88 | 111 | >sp\|P02788\|TRFL_HUMAN Lactotransferrin OS=Homo sapiens OX=9606 GN=LTF PE=1 SV=6 | 1270 | 20201115_07433_1ng_glyco_hcd203040_compound15.20875.20875.4 | controllerType=0 controllerNumber=1 scan=20875 | 41.402 |
| 18456:4 | 1 | R.RTAGWNVPIGTLRPFLN[+2059.735]WTGPPEPIEAAVAR.F | HexNAc(4)Hex(5)Fuc(1)NeuAc(1) | N[+2060] | 1362.3971 | 4 | 5446.5666 | 5446.547 | 0 | 3.6 | 140 | Specific | 689.65 | 566.18 | 566.18 | 14.72 | 111 | >sp\|P02788\|TRFL_HUMAN Lactotransferrin OS=Homo sapiens OX=9606 GN=LTF PE=1 SV=6 | 1270 | 20201115_07433_1ng_glyco_hcd203040_compound15.21888.21888.4 | controllerType=0 controllerNumber=1 scan=21888 | 43.5828 |
| 19496:4 | 1 | R.TAGWNVPIGTLRPFLN[+2716.983]WTGPPEPIEAAVAR.F | HexNAc(5)Hex(6)Fuc(3)NeuAc(1) | N[+2717] | 1487.6852 | 4 | 5947.7189 | 5947.6939 | 0 | 4.2 | 141 | Specific | 268.11 | 268.11 | 268.11 | 7.77 | 111 | >sp\|P02788\|TRFL_HUMAN Lactotransferrin OS=Homo sapiens OX=9606 GN=LTF PE=1 SV=6 | 1270 | 20201115_07433_1ng_glyco_hcd203040_compound15.22975.22975.4 | controllerType=0 controllerNumber=1 scan=22975 | 45.9343 |
| 18650:4 | 1 | R.TAGWNVPIGTLRPFLN[+2279.830]WTGPPEPIEAAVAR.F | HexNAc(5)Hex(6)Fuc(2) | N[+2280] | 1378.3955 | 4 | 5510.5602 | 5510.5406 | 0 | 3.6 | 141 | Specific | 336.4 | 300.67 | 300.67 | 8.17 | 111 | >sp\|P02788\|TRFL_HUMAN Lactotransferrin OS=Homo sapiens OX=9606 GN=LTF PE=1 SV=6 | 1270 | 20201115_07433_1ng_glyco_hcd203040_compound15.22089.22089.4 | controllerType=0 controllerNumber=1 scan=22089 | 43.9723 |
| 19499:4 | 1 | R.TAGWNVPIGTLRPFLN[+2716.983]WTGPPEPIEAAVAR.F | HexNAc(5)Hex(6)Fuc(3)NeuAc(1) | N[+2717] | 1487.6852 | 4 | 5947.7189 | 5947.6939 | 0 | 4.2 | 141 | Specific | 417.69 | 399.63 | 399.63 | 10.4 | 111 | >sp\|P02788\|TRFL_HUMAN Lactotransferrin OS=Homo sapiens OX=9606 GN=LTF PE=1 SV=6 | 1270 | 20201115_07433_1ng_glyco_hcd203040_compound15.22978.22978.4 | controllerType=0 controllerNumber=1 scan=22978 | 45.939 |
| 21024:5 | 1 | R.TAGWNVPIGTLRPFLN[+2350.830]WTGPPEPIEAAVAR.F | HexNAc(4)Hex(5)Fuc(1)NeuAc(2) | N[+2351] | 1117.1183 | 5 | 5581.5623 | 5581.5413 | 0 | 3.8 | 141 | Specific | 542.7 | 434.25 | 434.25 | 11.9 | 111 | >sp\|P02788\|TRFL_HUMAN Lactotransferrin OS=Homo sapiens OX=9606 GN=LTF PE=1 SV=6 | 1270 | 20201115_07433_1ng_glyco_hcd203040_compound15.24570.24570.5 | controllerType=0 controllerNumber=1 scan=24570 | 49.1982 |
| 21011:3 | 1 | R.TAGWNVPIGTLRPFLN[+2350.830]WTGPPEPIEAAVAR.F | HexNAc(4)Hex(5)Fuc(1)NeuAc(2) | N[+2351] | 1861.1934 | 3 | 5581.5658 | 5581.5413 | 0 | 4.4 | 141 | Specific | 566.41 | 400.9 | 400.9 | 11.12 | 111 | >sp\|P02788\|TRFL_HUMAN Lactotransferrin OS=Homo sapiens OX=9606 GN=LTF PE=1 SV=6 | 1270 | 20201115_07433_1ng_glyco_hcd203040_compound15.24557.24557.3 | controllerType=0 controllerNumber=1 scan=24557 | 49.1736 |
| 21026:5 | 1 | R.TAGWNVPIGTLRPFLN[+2350.830]WTGPPEPIEAAVAR.F | HexNAc(4)Hex(5)Fuc(1)NeuAc(2) | N[+2351] | 1117.1183 | 5 | 5581.5623 | 5581.5413 | 0 | 3.8 | 141 | Specific | 599.86 | 372.88 | 372.88 | 11.85 | 111 | >sp\|P02788\|TRFL_HUMAN Lactotransferrin OS=Homo sapiens OX=9606 GN=LTF PE=1 SV=6 | 1270 | 20201115_07433_1ng_glyco_hcd203040_compound15.24572.24572.5 | controllerType=0 controllerNumber=1 scan=24572 | 49.2052 |
| 21014:3 | 1 | R.TAGWNVPIGTLRPFLN[+2350.830]WTGPPEPIEAAVAR.F | HexNAc(4)Hex(5)Fuc(1)NeuAc(2) | N[+2351] | 1861.1934 | 3 | 5581.5658 | 5581.5413 | 0 | 4.4 | 141 | Specific | 613.7 | 452.21 | 452.21 | 13.42 | 111 | >sp\|P02788\|TRFL_HUMAN Lactotransferrin OS=Homo sapiens OX=9606 GN=LTF PE=1 SV=6 | 1270 | 20201115_07433_1ng_glyco_hcd203040_compound15.24560.24560.3 | controllerType=0 controllerNumber=1 scan=24560 | 49.1781 |

Table S4. Cont.

| Query #:z | Protein Rank | Glycopeptide | Glycan | Modification Type(s) | Observed m/z | z | Observed (M+H) | Calc. mass (M+H) | Off-by-x error | Mass error (ppm) | StartPosition | Cleavage | Score | Delta | Delta Mod | \|Log Prob\| | # of unique peptides | Protein Name | Protein DB number | Comment | Scan # | Scan Time |
|---|---|---|---|---|---|---|---|---|---|---|---|---|---|---|---|---|---|---|---|---|---|---|
| 19625:3 | 1 | R.TAGWNVPIGTLRPFLN[+2205.793]WTGPPEPIEAAVAR.F | HexNAc(4)Hex(5)Fuc(2)NeuAc(1) | N[+2206] | 1812.8479 | 3 | 5436.529 | 5436.5038 | 0 | 4.6 | 141 | Specific | 635.34 | 567.72 | 567.72 | 13.42 | 111 | >sp\|P02788\|TRFL_HUMAN Lactotransferrin OS=Homo sapiens OX=9606 GN=LTF PE=1 SV=6 | 1270 | 20201115_07433_1ng_glyco_hcd203040_compound15.23109.23109.3 | controllerType=0 controllerNumber=1 scan=23109 | 46.196 |
| 21007:4 | 1 | R.TAGWNVPIGTLRPFLN[+2350.830]WTGPPEPIEAAVAR.F | HexNAc(4)Hex(5)Fuc(1)NeuAc(2) | N[+2351] | 1396.145 | 4 | 5581.5583 | 5581.5413 | 0 | 3 | 141 | Specific | 636.68 | 511.68 | 511.68 | 13.71 | 111 | >sp\|P02788\|TRFL_HUMAN Lactotransferrin OS=Homo sapiens OX=9606 GN=LTF PE=1 SV=6 | 1270 | 20201115_07433_1ng_glyco_hcd203040_compound15.24553.24553.4 | controllerType=0 controllerNumber=1 scan=24553 | 49.1672 |
| 19628:3 | 1 | R.TAGWNVPIGTLRPFLN[+2205.793]WTGPPEPIEAAVAR.F | HexNAc(4)Hex(5)Fuc(2)NeuAc(1) | N[+2206] | 1812.8479 | 3 | 5436.529 | 5436.5038 | 0 | 4.6 | 141 | Specific | 640.98 | 559.81 | 559.81 | 13.42 | 111 | >sp\|P02788\|TRFL_HUMAN Lactotransferrin OS=Homo sapiens OX=9606 GN=LTF PE=1 SV=6 | 1270 | 20201115_07433_1ng_glyco_hcd203040_compound15.23112.23112.3 | controllerType=0 controllerNumber=1 scan=23112 | 46.2017 |
| 19620:4 | 1 | R.TAGWNVPIGTLRPFLN[+2205.793]WTGPPEPIEAAVAR.F | HexNAc(4)Hex(5)Fuc(2)NeuAc(1) | N[+2206] | 1359.8878 | 4 | 5436.5295 | 5436.5038 | 0 | 4.7 | 141 | Specific | 679.3 | 615.66 | 615.66 | 14.41 | 111 | >sp\|P02788\|TRFL_HUMAN Lactotransferrin OS=Homo sapiens OX=9606 GN=LTF PE=1 SV=6 | 1270 | 20201115_07433_1ng_glyco_hcd203040_compound15.23104.23104.4 | controllerType=0 controllerNumber=1 scan=23104 | 46.1843 |
| 18705:3 | 1 | R.TAGWNVPIGTLRPFLN[+1768.640]WTGPPEPIEAAVAR.F | HexNAc(4)Hex(5)Fuc(1) | N[+1769] | 1667.1257 | 3 | 4999.3625 | 4999.3504 | 0 | 2.4 | 141 | Specific | 694.12 | 585.78 | 585.78 | 14.88 | 111 | >sp\|P02788\|TRFL_HUMAN Lactotransferrin OS=Homo sapiens OX=9606 GN=LTF PE=1 SV=6 | 1270 | 20201115_07433_1ng_glyco_hcd203040_compound15.22146.22146.3 | controllerType=0 controllerNumber=1 scan=22146 | 44.0862 |
| 18703:3 | 1 | R.TAGWNVPIGTLRPFLN[+1768.640]WTGPPEPIEAAVAR.F | HexNAc(4)Hex(5)Fuc(1) | N[+1769] | 1667.1257 | 3 | 4999.3625 | 4999.3504 | 0 | 2.4 | 141 | Specific | 698.43 | 568.26 | 568.26 | 14.88 | 111 | >sp\|P02788\|TRFL_HUMAN Lactotransferrin OS=Homo sapiens OX=9606 GN=LTF PE=1 SV=6 | 1270 | 20201115_07433_1ng_glyco_hcd203040_compound15.22144.22144.3 | controllerType=0 controllerNumber=1 scan=22144 | 44.0806 |
| 18696:4 | 1 | R.TAGWNVPIGTLRPFLN[+1768.640]WTGPPEPIEAAVAR.F | HexNAc(4)Hex(5)Fuc(1) | N[+1769] | 1250.5973 | 4 | 4999.3673 | 4999.3504 | 0 | 3.4 | 141 | Specific | 702.93 | 668.96 | 668.96 | 15.71 | 111 | >sp\|P02788\|TRFL_HUMAN Lactotransferrin OS=Homo sapiens OX=9606 GN=LTF PE=1 SV=6 | 1270 | 20201115_07433_1ng_glyco_hcd203040_compound15.22137.22137.4 | controllerType=0 controllerNumber=1 scan=22137 | 44.0653 |
| 18653:4 | 1 | R.TAGWNVPIGTLRPFLN[+2279.830]WTGPPEPIEAAVAR.F | HexNAc(5)Hex(6)Fuc(2) | N[+2280] | 1378.3955 | 4 | 5510.5602 | 5510.5406 | 0 | 3.6 | 141 | Specific | 711.9 | 674.85 | 674.85 | 15.71 | 111 | >sp\|P02788\|TRFL_HUMAN Lactotransferrin OS=Homo sapiens OX=9606 GN=LTF PE=1 SV=6 | 1270 | 20201115_07433_1ng_glyco_hcd203040_compound15.22092.22092.4 | controllerType=0 controllerNumber=1 scan=22092 | 43.9807 |
| 19639:5 | 1 | R.TAGWNVPIGTLRPFLN[+2205.793]WTGPPEPIEAAVAR.F | HexNAc(4)Hex(5)Fuc(2)NeuAc(1) | N[+2206] | 1088.1124 | 5 | 5436.533 | 5436.5038 | 0 | 5.4 | 141 | Specific | 712.34 | 578.97 | 578.97 | 15.58 | 111 | >sp\|P02788\|TRFL_HUMAN Lactotransferrin OS=Homo sapiens OX=9606 GN=LTF PE=1 SV=6 | 1270 | 20201115_07433_1ng_glyco_hcd203040_compound15.23123.23123.5 | controllerType=0 controllerNumber=1 scan=23123 | 46.223 |
| 21009:4 | 1 | R.TAGWNVPIGTLRPFLN[+2350.830]WTGPPEPIEAAVAR.F | HexNAc(4)Hex(5)Fuc(1)NeuAc(2) | N[+2351] | 1396.145 | 4 | 5581.5583 | 5581.5413 | 0 | 3 | 141 | Specific | 794.37 | 545.5 | 545.5 | 16.71 | 111 | >sp\|P02788\|TRFL_HUMAN Lactotransferrin OS=Homo sapiens OX=9606 GN=LTF PE=1 SV=6 | 1270 | 20201115_07433_1ng_glyco_hcd203040_compound15.24555.24555.4 | controllerType=0 controllerNumber=1 scan=24555 | 49.1716 |
| 19622:4 | 1 | R.TAGWNVPIGTLRPFLN[+2205.793]WTGPPEPIEAAVAR.F | HexNAc(4)Hex(5)Fuc(2)NeuAc(1) | N[+2206] | 1359.8878 | 4 | 5436.5295 | 5436.5038 | 0 | 4.7 | 141 | Specific | 852.47 | 714.01 | 714.01 | 18.41 | 111 | >sp\|P02788\|TRFL_HUMAN Lactotransferrin OS=Homo sapiens OX=9606 GN=LTF PE=1 SV=6 | 1270 | 20201115_07433_1ng_glyco_hcd203040_compound15.23106.23106.4 | controllerType=0 controllerNumber=1 scan=23106 | 46.1908 |
| 19642:5 | 1 | R.TAGWNVPIGTLRPFLN[+2205.793]WTGPPEPIEAAVAR.F | HexNAc(4)Hex(5)Fuc(2)NeuAc(1) | N[+2206] | 1088.1124 | 5 | 5436.533 | 5436.5038 | 0 | 5.4 | 141 | Specific | 903.48 | 698.13 | 698.13 | 19.58 | 111 | >sp\|P02788\|TRFL_HUMAN Lactotransferrin OS=Homo sapiens OX=9606 GN=LTF PE=1 SV=6 | 1270 | 20201115_07433_1ng_glyco_hcd203040_compound15.23126.23126.5 | controllerType=0 controllerNumber=1 scan=23126 | 46.23 |
| 18699:4 | 1 | R.TAGWNVPIGTLRPFLN[+1768.640]WTGPPEPIEAAVAR.F | HexNAc(4)Hex(5)Fuc(1) | N[+1769] | 1250.5973 | 4 | 4999.3673 | 4999.3504 | 0 | 3.4 | 141 | Specific | 910.66 | 815.58 | 815.58 | 19.71 | 111 | >sp\|P02788\|TRFL_HUMAN Lactotransferrin OS=Homo sapiens OX=9606 GN=LTF PE=1 SV=6 | 1270 | 20201115_07433_1ng_glyco_hcd203040_compound15.22140.22140.4 | controllerType=0 controllerNumber=1 scan=22140 | 44.0712 |
| 21173:3 | 1 | R.TAGWNIPMGLLFN[+2350.830]QTGSC[+57.021]K.F | HexNAc(4)Hex(5)Fuc(1)NeuAc(2) | C[+57], N[+2351] | 1482.6235 | 3 | 4445.8559 | 4445.8403 | 0 | 3.5 | 485 | Specific | 230.31 | 168.94 | 168.94 | 4.15 | 111 | >sp\|P02788\|TRFL_HUMAN Lactotransferrin OS=Homo sapiens OX=9606 GN=LTF PE=1 SV=6 | 1270 | 20201115_07433_1ng_glyco_hcd203040_compound15.24725.24725.2 | controllerType=0 controllerNumber=1 scan=24725 | 49.4967 |
| 17157:2 | 1 | R.TAGWNIPMGLLFN[+1768.640]QTGSC[+57.021]K.F | HexNAc(4)Hex(5)Fuc(1) | C[+57], N[+1769] | 1932.3333 | 2 | 3863.6594 | 3863.6495 | 0 | 2.6 | 485 | Specific | 353.81 | 330.55 | 330.55 | 8.15 | 111 | >sp\|P02788\|TRFL_HUMAN Lactotransferrin OS=Homo sapiens OX=9606 GN=LTF PE=1 SV=6 | 1270 | 20201115_07433_1ng_glyco_hcd203040_compound15.20534.20534.2 | controllerType=0 controllerNumber=1 scan=20534 | 40.6955 |
| 19084:3 | 1 | R.TAGWNIPMGLLFN[+2205.793]QTGSC[+57.021]K.F | HexNAc(4)Hex(5)Fuc(2)NeuAc(1) | C[+57], N[+2206] | 1434.2794 | 3 | 4300.8235 | 4300.8028 | 0 | 4.8 | 485 | Specific | 431.65 | 424 | 424 | 9.73 | 111 | >sp\|P02788\|TRFL_HUMAN Lactotransferrin OS=Homo sapiens OX=9606 GN=LTF PE=1 SV=6 | 1270 | 20201115_07433_1ng_glyco_hcd203040_compound15.22543.22543.3 | controllerType=0 controllerNumber=1 scan=22543 | 44.9477 |
| 17160:2 | 1 | R.TAGWNIPMGLLFN[+1768.640]QTGSC[+57.021]K.F | HexNAc(4)Hex(5)Fuc(1) | C[+57], N[+1769] | 1932.3333 | 2 | 3863.6594 | 3863.6495 | 0 | 2.6 | 485 | Specific | 599.32 | 516.57 | 516.57 | 12.21 | 111 | >sp\|P02788\|TRFL_HUMAN Lactotransferrin OS=Homo sapiens OX=9606 GN=LTF PE=1 SV=6 | 1270 | 20201115_07433_1ng_glyco_hcd203040_compound15.20537.20537.2 | controllerType=0 controllerNumber=1 scan=20537 | 40.7017 |
| 19192:4 | 1 | R.TAGWNIPMGLLFN[+1913.677]QTGSC[+57.021]K.F | HexNAc(4)Hex(5)NeuAc(1) | C[+57], N[+1914] | 1002.9313 | 4 | 4008.7034 | 4008.687 | 0 | 4.1 | 485 | Specific | 602.15 | 509.88 | 509.88 | 13.63 | 111 | >sp\|P02788\|TRFL_HUMAN Lactotransferrin OS=Homo sapiens OX=9606 GN=LTF PE=1 SV=6 | 1270 | 20201115_07433_1ng_glyco_hcd203040_compound15.22657.22657.4 | controllerType=0 controllerNumber=1 scan=22657 | 45.2769 |
| 17328:3 | 1 | R.TAGWNIPMGLLFN[+1622.582]QTGSC[+57.021]K.F | HexNAc(4)Hex(5) | C[+57], N[+1623] | 1239.8727 | 3 | 3717.6037 | 3717.5916 | 0 | 3.3 | 485 | Specific | 627.19 | 627.19 | 242.74 | 13.03 | 111 | >sp\|P02788\|TRFL_HUMAN Lactotransferrin OS=Homo sapiens OX=9606 GN=LTF PE=1 SV=6 | 1270 | 20201115_07433_1ng_glyco_hcd203040_compound15.20713.20713.3 | controllerType=0 controllerNumber=1 scan=20713 | 41.0737 |
| 19086:3 | 1 | R.TAGWNIPMGLLFN[+2205.793]QTGSC[+57.021]K.F | HexNAc(4)Hex(5)Fuc(2)NeuAc(1) | C[+57], N[+2206] | 1434.2794 | 3 | 4300.8235 | 4300.8028 | 0 | 4.8 | 485 | Specific | 630.43 | 500.48 | 500.48 | 12.73 | 111 | >sp\|P02788\|TRFL_HUMAN Lactotransferrin OS=Homo sapiens OX=9606 GN=LTF PE=1 SV=6 | 1270 | 20201115_07433_1ng_glyco_hcd203040_compound15.22545.22545.3 | controllerType=0 controllerNumber=1 scan=22545 | 44.9528 |
| 19195:4 | 1 | R.TAGWNIPMGLLFN[+1913.677]QTGSC[+57.021]K.F | HexNAc(4)Hex(5)NeuAc(1) | C[+57], N[+1914] | 1002.9313 | 4 | 4008.7034 | 4008.687 | 0 | 4.1 | 485 | Specific | 678.91 | 486.32 | 486.32 | 14.63 | 111 | >sp\|P02788\|TRFL_HUMAN Lactotransferrin OS=Homo sapiens OX=9606 GN=LTF PE=1 SV=6 | 1270 | 20201115_07433_1ng_glyco_hcd203040_compound15.22660.22660.4 | controllerType=0 controllerNumber=1 scan=22660 | 45.2828 |
| 17150:4 | 1 | R.TAGWNIPMGLLFN[+1768.640]QTGSC[+57.021]K.F | HexNAc(4)Hex(5)Fuc(1) | C[+57], N[+1769] | 966.6722 | 4 | 3863.6671 | 3863.6495 | 0 | 4.6 | 485 | Specific | 700.71 | 682.62 | 682.62 | 15.33 | 111 | >sp\|P02788\|TRFL_HUMAN Lactotransferrin OS=Homo sapiens OX=9606 GN=LTF PE=1 SV=6 | 1270 | 20201115_07433_1ng_glyco_hcd203040_compound15.20527.20527.4 | controllerType=0 controllerNumber=1 scan=20527 | 40.683 |
| 17249:3 | 1 | R.TAGWNIPMGLLFN[+1606.587]QTGSC[+57.021]K.F | HexNAc(4)Hex(4)Fuc(1) | C[+57], N[+1607] | 1234.5394 | 3 | 3701.6035 | 3701.5966 | 0 | 1.9 | 485 | Specific | 716.16 | 716.16 | 716.16 | 15.2 | 111 | >sp\|P02788\|TRFL_HUMAN Lactotransferrin OS=Homo sapiens OX=9606 GN=LTF PE=1 SV=6 | 1270 | 20201115_07433_1ng_glyco_hcd203040_compound15.20630.20630.3 | controllerType=0 controllerNumber=1 scan=20630 | 40.9018 |
| 17080:3 | 1 | R.TAGWNIPMGLLFN[+1768.640]QTGSC[+57.021]K.F | HexNAc(4)Hex(5)Fuc(1) | C[+57], N[+1769] | 1288.5583 | 3 | 3863.6603 | 3863.6495 | 0 | 2.8 | 485 | Specific | 767.22 | 765.55 | 765.55 | 16.2 | 111 | >sp\|P02788\|TRFL_HUMAN Lactotransferrin OS=Homo sapiens OX=9606 GN=LTF PE=1 SV=6 | 1270 | 20201115_07433_1ng_glyco_hcd203040_compound15.20454.20454.3 | controllerType=0 controllerNumber=1 scan=20454 | 40.5286 |
| 17251:3 | 1 | R.TAGWNIPMGLLFN[+1606.587]QTGSC[+57.021]K.F | HexNAc(4)Hex(4)Fuc(1) | C[+57], N[+1607] | 1234.5394 | 3 | 3701.6035 | 3701.5966 | 0 | 1.9 | 485 | Specific | 794.56 | 788.19 | 788.19 | 16.2 | 111 | >sp\|P02788\|TRFL_HUMAN Lactotransferrin OS=Homo sapiens OX=9606 GN=LTF PE=1 SV=6 | 1270 | 20201115_07433_1ng_glyco_hcd203040_compound15.20632.20632.3 | controllerType=0 controllerNumber=1 scan=20632 | 40.9059 |
| 19171:3 | 1 | R.TAGWNIPMGLLFN[+1913.677]QTGSC[+57.021]K.F | HexNAc(4)Hex(5)NeuAc(1) | C[+57], N[+1914] | 1336.9057 | 3 | 4008.7025 | 4008.687 | 0 | 3.9 | 485 | Specific | 806.56 | 795.5 | 795.5 | 17.03 | 111 | >sp\|P02788\|TRFL_HUMAN Lactotransferrin OS=Homo sapiens OX=9606 GN=LTF PE=1 SV=6 | 1270 | 20201115_07433_1ng_glyco_hcd203040_compound15.22636.22636.3 | controllerType=0 controllerNumber=1 scan=22636 | 45.2385 |
| 17153:4 | 1 | R.TAGWNIPMGLLFN[+1768.640]QTGSC[+57.021]K.F | HexNAc(4)Hex(5)Fuc(1) | C[+57], N[+1769] | 966.6722 | 4 | 3863.6671 | 3863.6495 | 0 | 4.6 | 485 | Specific | 839.88 | 610.07 | 610.07 | 17.33 | 111 | >sp\|P02788\|TRFL_HUMAN Lactotransferrin OS=Homo sapiens OX=9606 GN=LTF PE=1 SV=6 | 1270 | 20201115_07433_1ng_glyco_hcd203040_compound15.20530.20530.4 | controllerType=0 controllerNumber=1 scan=20530 | 40.6882 |
| 17102:3 | 1 | R.TAGWNIPMGLLFN[+1768.640]QTGSC[+57.021]K.F | HexNAc(4)Hex(5)Fuc(1) | C[+57], N[+1769] | 1288.559 | 3 | 3863.6625 | 3863.6495 | 0 | 3.4 | 485 | Specific | 881.18 | 881.18 | 881.18 | 18.03 | 111 | >sp\|P02788\|TRFL_HUMAN Lactotransferrin OS=Homo sapiens OX=9606 GN=LTF PE=1 SV=6 | 1270 | 20201115_07433_1ng_glyco_hcd203040_compound15.20477.20477.3 | controllerType=0 controllerNumber=1 scan=20477 | 40.5814 |
| 17330:3 | 1 | R.TAGWNIPMGLLFN[+1622.582]QTGSC[+57.021]K.F | HexNAc(4)Hex(5) | C[+57], N[+1623] | 1239.8727 | 3 | 3717.6037 | 3717.5916 | 0 | 3.3 | 485 | Specific | 885.29 | 863.91 | 422.49 | 18.03 | 111 | >sp\|P02788\|TRFL_HUMAN Lactotransferrin OS=Homo sapiens OX=9606 GN=LTF PE=1 SV=6 | 1270 | 20201115_07433_1ng_glyco_hcd203040_compound15.20715.20715.3 | controllerType=0 controllerNumber=1 scan=20715 | 41.0781 |
| 19174:3 | 1 | R.TAGWNIPMGLLFN[+1913.677]QTGSC[+57.021]K.F | HexNAc(4)Hex(5)NeuAc(1) | C[+57], N[+1914] | 1336.9057 | 3 | 4008.7025 | 4008.687 | 0 | 3.9 | 485 | Specific | 892.3 | 890.55 | 890.55 | 18.03 | 111 | >sp\|P02788\|TRFL_HUMAN Lactotransferrin OS=Homo sapiens OX=9606 GN=LTF PE=1 SV=6 | 1270 | 20201115_07433_1ng_glyco_hcd203040_compound15.22639.22639.3 | controllerType=0 controllerNumber=1 scan=22639 | 45.2452 |
| 19023:3 | 1 | R.TAGWNIPMGLLFN[+2059.735]QTGSC[+57.021]K.F | HexNAc(4)Hex(5)Fuc(1)NeuAc(1) | C[+57], N[+2060] | 1385.5842 | 3 | 4154.7379 | 4154.7449 | 0 | -1.7 | 485 | Specific | 911.24 | 834.37 | 834.37 | 17.67 | 111 | >sp\|P02788\|TRFL_HUMAN Lactotransferrin OS=Homo sapiens OX=9606 GN=LTF PE=1 SV=6 | 1270 | 20201115_07433_1ng_glyco_hcd203040_compound15.22479.22479.3 | controllerType=0 controllerNumber=1 scan=22479 | 44.8248 |
| 19020:3 | 1 | R.TAGWNIPMGLLFN[+2059.735]QTGSC[+57.021]K.F | HexNAc(4)Hex(5)Fuc(1)NeuAc(1) | C[+57], N[+2060] | 1385.5842 | 3 | 4154.7379 | 4154.7449 | 0 | -1.7 | 485 | Specific | 923.31 | 895.77 | 895.77 | 17.67 | 111 | >sp\|P02788\|TRFL_HUMAN Lactotransferrin OS=Homo sapiens OX=9606 GN=LTF PE=1 SV=6 | 1270 | 20201115_07433_1ng_glyco_hcd203040_compound15.22476.22476.3 | controllerType=0 controllerNumber=1 scan=22476 | 44.8154 |
| 17083:3 | 1 | R.TAGWNIPMGLLFN[+1768.640]QTGSC[+57.021]K.F | HexNAc(4)Hex(5)Fuc(1) | C[+57], N[+1769] | 1288.5583 | 3 | 3863.6603 | 3863.6495 | 0 | 2.8 | 485 | Specific | 961.55 | 935.38 | 935.38 | 20.2 | 111 | >sp\|P02788\|TRFL_HUMAN Lactotransferrin OS=Homo sapiens OX=9606 GN=LTF PE=1 SV=6 | 1270 | 20201115_07433_1ng_glyco_hcd203040_compound15.20457.20457.3 | controllerType=0 controllerNumber=1 scan=20457 | 40.5334 |
| 17099:3 | 1 | R.TAGWNIPMGLLFN[+1768.640]QTGSC[+57.021]K.F | HexNAc(4)Hex(5)Fuc(1) | C[+57], N[+1769] | 1288.559 | 3 | 3863.6625 | 3863.6495 | 0 | 3.4 | 485 | Specific | 995.29 | 995.29 | 995.29 | 20.03 | 111 | >sp\|P02788\|TRFL_HUMAN Lactotransferrin OS=Homo sapiens OX=9606 GN=LTF PE=1 SV=6 | 1270 | 20201115_07433_1ng_glyco_hcd203040_compound15.20474.20474.3 | controllerType=0 controllerNumber=1 scan=20474 | 40.574 |
| 18264:4 | 1 | R.[+42.011]TAGWNIPMGLLFN[+1872.651]QTGSC[+57.021]KFDEYFSQSC[+57.021]A | HexNAc(3)Hex(6)NeuAc(1) | .[42], C[+57]*2, N[+1873] | 1464.1213 | 4 | 5853.4635 | 5853.4181 | 0 | 7.8 | 485 | Specific | 231.13 | 219.1 | 219.1 | 3.21 | 111 | >sp\|P02788\|TRFL_HUMAN Lactotransferrin OS=Homo sapiens OX=9606 GN=LTF PE=1 SV=6 | 1270 | 20201115_07433_1ng_glyco_hcd203040_compound15.21688.21688.4 | controllerType=0 controllerNumber=1 scan=21688 | 43.1462 |
| 18270:3 | 1 | R.TAGWNIPMGLLFN[+1768.640]QTGSC[+57.021]KFDEYFSQSC[+57.021]APGSDPR.S | HexNAc(4)Hex(5)Fuc(1) | C[+57]*2, N[+1769] | 1903.1422 | 3 | 5707.412 | 5707.3966 | 0 | 2.7 | 485 | Specific | 661.1 | 575.73 | 575.73 | 14.25 | 111 | >sp\|P02788\|TRFL_HUMAN Lactotransferrin OS=Homo sapiens OX=9606 GN=LTF PE=1 SV=6 | 1270 | 20201115_07433_1ng_glyco_hcd203040_compound15.21694.21694.3 | controllerType=0 controllerNumber=1 scan=21694 | 43.155 |
| 18261:4 | 1 | R.TAGWNIPMGLLFN[+1768.640]QTGSC[+57.021]KFDEYFSQSC[+57.021]APGSDPR.S | HexNAc(4)Hex(5)Fuc(1) | C[+57]*2, N[+1769] | 1427.6086 | 4 | 5707.4128 | 5707.3966 | 0 | 2.8 | 485 | Specific | 704.68 | 660.25 | 660.25 | 15.24 | 111 | >sp\|P02788\|TRFL_HUMAN Lactotransferrin OS=Homo sapiens OX=9606 GN=LTF PE=1 SV=6 | 1270 | 20201115_07433_1ng_glyco_hcd203040_compound15.21685.21685.4 | controllerType=0 controllerNumber=1 scan=21685 | 43.1408 |
| 18349:4 | 1 | R.TAGWNIPMGLLFN[+1622.582]QTGSC[+57.021]KFDEYFSQSC[+57.021]APGSDPR.S | HexNAc(4)Hex(5) | C[+57]*2, N[+1623] | 1391.0955 | 4 | 5561.36 | 5561.3387 | 0 | 3.8 | 485 | Specific | 711.12 | 662.59 | 170.8 | 15.07 | 111 | >sp\|P02788\|TRFL_HUMAN Lactotransferrin OS=Homo sapiens OX=9606 GN=LTF PE=1 SV=6 | 1270 | 20201115_07433_1ng_glyco_hcd203040_compound15.21776.21776.4 | controllerType=0 controllerNumber=1 scan=21776 | 43.3095 |
| 18273:3 | 1 | R.TAGWNIPMGLLFN[+1768.640]QTGSC[+57.021]KFDEYFSQSC[+57.021]APGSDPR.S | HexNAc(4)Hex(5)Fuc(1) | C[+57]*2, N[+1769] | 1903.1422 | 3 | 5707.412 | 5707.3966 | 0 | 2.7 | 485 | Specific | 717.16 | 568.53 | 568.53 | 15.25 | 111 | >sp\|P02788\|TRFL_HUMAN Lactotransferrin OS=Homo sapiens OX=9606 GN=LTF PE=1 SV=6 | 1270 | 20201115_07433_1ng_glyco_hcd203040_compound15.21697.21697.3 | controllerType=0 controllerNumber=1 scan=21697 | 43.1589 |
| 20631:4 | 1 | R.TAGWNIPMGLLFN[+2350.830]QTGSC[+57.021]KFDEYFSQSC[+57.021]APGSDPR.S | HexNAc(4)Hex(5)Fuc(1)NeuAc(2) | C[+57]*2, N[+2351] | 1573.1585 | 4 | 6289.612 | 6289.5875 | 0 | 3.9 | 485 | Specific | 876.36 | 726.22 | 726.22 | 18.07 | 111 | >sp\|P02788\|TRFL_HUMAN Lactotransferrin OS=Homo sapiens OX=9606 GN=LTF PE=1 SV=6 | 1270 | 20201115_07433_1ng_glyco_hcd203040_compound15.24161.24161.4 | controllerType=0 controllerNumber=1 scan=24161 | 48.3311 |
| 20633:4 | 1 | R.TAGWNIPMGLLFN[+2350.830]QTGSC[+57.021]KFDEYFSQSC[+57.021]APGSDPR.S | HexNAc(4)Hex(5)Fuc(1)NeuAc(2) | C[+57]*2, N[+2351] | 1573.1585 | 4 | 6289.612 | 6289.5875 | 0 | 3.9 | 485 | Specific | 898.04 | 850.34 | 850.34 | 18.07 | 111 | >sp\|P02788\|TRFL_HUMAN Lactotransferrin OS=Homo sapiens OX=9606 GN=LTF PE=1 SV=6 | 1270 | 20201115_07433_1ng_glyco_hcd203040_compound15.24163.24163.4 | controllerType=0 controllerNumber=1 scan=24163 | 48.3358 |
| 18352:4 | 1 | R.TAGWNIPMGLLFN[+1622.582]QTGSC[+57.021]KFDEYFSQSC[+57.021]APGSDPR.S | HexNAc(4)Hex(5) | C[+57]*2, N[+1623] | 1391.0955 | 4 | 5561.36 | 5561.3387 | 0 | 3.8 | 485 | Specific | 933.13 | 837.55 | 327.14 | 19.07 | 111 | >sp\|P02788\|TRFL_HUMAN Lactotransferrin OS=Homo sapiens OX=9606 GN=LTF PE=1 SV=6 | 1270 | 20201115_07433_1ng_glyco_hcd203040_compound15.21779.21779.4 | controllerType=0 controllerNumber=1 scan=21779 | 43.3187 |
| 19336:4 | 1 | R.TAGWNIPMGLLFN[+1913.677]QTGSC[+57.021]KFDEYFSQSC[+57.021]APGSDPR.S | HexNAc(4)Hex(5)NeuAc(1) | C[+57]*2, N[+1914] | 1463.8683 | 4 | 5852.4513 | 5852.4341 | 0 | 2.9 | 485 | Specific | 948.68 | 899.55 | 899.55 | 19.24 | 111 | >sp\|P02788\|TRFL_HUMAN Lactotransferrin OS=Homo sapiens OX=9606 GN=LTF PE=1 SV=6 | 1270 | 20201115_07433_1ng_glyco_hcd203040_compound15.22808.22808.4 | controllerType=0 controllerNumber=1 scan=22808 | 45.5908 |
| 18263:4 | 1 | R.TAGWNIPMGLLFN[+1768.640]QTGSC[+57.021]KFDEYFSQSC[+57.021]APGSDPR.S | HexNAc(4)Hex(5)Fuc(1) | C[+57]*2, N[+1769] | 1427.6086 | 4 | 5707.4128 | 5707.3966 | 0 | 2.8 | 485 | Specific | 952.42 | 906.53 | 906.53 | 20.24 | 111 | >sp\|P02788\|TRFL_HUMAN Lactotransferrin OS=Homo sapiens OX=9606 GN=LTF PE=1 SV=6 | 1270 | 20201115_07433_1ng_glyco_hcd203040_compound15.21687.21687.4 | controllerType=0 controllerNumber=1 scan=21687 | 43.1452 |
| 19334:4 | 1 | R.TAGWNIPMGLLFN[+1913.677]QTGSC[+57.021]KFDEYFSQSC[+57.021]APGSDPR.S | HexNAc(4)Hex(5)NeuAc(1) | C[+57]*2, N[+1914] | 1463.8683 | 4 | 5852.4513 | 5852.4341 | 0 | 2.9 | 485 | Specific | 983.76 | 914.25 | 914.25 | 20.24 | 111 | >sp\|P02788\|TRFL_HUMAN Lactotransferrin OS=Homo sapiens OX=9606 GN=LTF PE=1 SV=6 | 1270 | 20201115_07433_1ng_glyco_hcd203040_compound15.22806.22806.4 | controllerType=0 controllerNumber=1 scan=22806 | 45.5847 |
| 09339:3 | 1 | R.N[+2205.793]GSDC[+57.021]PDKFC[+57.021]LFQSETK.N | HexNAc(4)Hex(5)Fuc(2)NeuAc(1) | C[+57]*2, N[+2206] | 1413.561 | 3 | 4238.6684 | 4238.6667 | 0 | 0.4 | 642 | Specific | 382.69 | 286.84 | 286.84 | 6.62 | 111 | >sp\|P02788\|TRFL_HUMAN Lactotransferrin OS=Homo sapiens OX=9606 GN=LTF PE=1 SV=6 | 1270 | 20201115_07433_1ng_glyco_hcd203040_compound15.12353.12353.3 | controllerType=0 controllerNumber=1 scan=12353 | 26.4579 |
| 09341:3 | 1 | R.N[+2205.793]GSDC[+57.021]PDKFC[+57.021]LFQSETK.N | HexNAc(4)Hex(5)Fuc(2)NeuAc(1) | C[+57]*2, N[+2206] | 1413.561 | 3 | 4238.6684 | 4238.6667 | 0 | 0.4 | 642 | Specific | 411.23 | 255.53 | 255.53 | 6.49 | 111 | >sp\|P02788\|TRFL_HUMAN Lactotransferrin OS=Homo sapiens OX=9606 GN=LTF PE=1 SV=6 | 1270 | 20201115_07433_1ng_glyco_hcd203040_compound15.12355.12355.3 | controllerType=0 controllerNumber=1 scan=12355 | 26.4616 |
| 09373:3 | 1 | R.N[+2205.793]GSDC[+57.021]PDKFC[+57.021]LFQSETK.N | HexNAc(4)Hex(5)Fuc(2)NeuAc(1) | C[+57]*2, N[+2206] | 1413.565 | 3 | 4238.6804 | 4238.6667 | 0 | 3.2 | 642 | Specific | 468.76 | 365.63 | 365.63 | 8.21 | 111 | >sp\|P02788\|TRFL_HUMAN Lactotransferrin OS=Homo sapiens OX=9606 GN=LTF PE=1 SV=6 | 1270 | 20201115_07433_1ng_glyco_hcd203040_compound15.12389.12389.3 | controllerType=0 controllerNumber=1 scan=12389 | 26.524 |
| 09465:3 | 1 | R.N[+2059.735]GSDC[+57.021]PDKFC[+57.021]LFQSETK.N | HexNAc(4)Hex(5)Fuc(1)NeuAc(1) | C[+57]*2, N[+2060] | 1364.8804 | 3 | 4092.6267 | 4092.6088 | 0 | 4.4 | 642 | Specific | 496.47 | 394.24 | 394.24 | 9.57 | 111 | >sp\|P02788\|TRFL_HUMAN Lactotransferrin OS=Homo sapiens OX=9606 GN=LTF PE=1 SV=6 | 1270 | 20201115_07433_1ng_glyco_hcd203040_compound15.12484.12484.3 | controllerType=0 controllerNumber=1 scan=12484 | 26.677 |
| 09375:3 | 1 | R.N[+2205.793]GSDC[+57.021]PDKFC[+57.021]LFQSETK.N | HexNAc(4)Hex(5)Fuc(2)NeuAc(1) | C[+57]*2, N[+2206] | 1413.565 | 3 | 4238.6804 | 4238.6667 | 0 | 3.2 | 642 | Specific | 510.78 | 405.99 | 405.99 | 9.95 | 111 | >sp\|P02788\|TRFL_HUMAN Lactotransferrin OS=Homo sapiens OX=9606 GN=LTF PE=1 SV=6 | 1270 | 20201115_07433_1ng_glyco_hcd203040_compound15.12391.12391.3 | controllerType=0 controllerNumber=1 scan=12391 | 26.5282 |
| 09467:3 | 1 | R.N[+2059.735]GSDC[+57.021]PDKFC[+57.021]LFQSETK.N | HexNAc(4)Hex(5)Fuc(1)NeuAc(1) | C[+57]*2, N[+2060] | 1364.8804 | 3 | 4092.6267 | 4092.6088 | 0 | 4.4 | 642 | Specific | 585.86 | 465.16 | 465.16 | 11.24 | 111 | >sp\|P02788\|TRFL_HUMAN Lactotransferrin OS=Homo sapiens OX=9606 GN=LTF PE=1 SV=6 | 1270 | 20201115_07433_1ng_glyco_hcd203040_compound15.12486.12486.3 | controllerType=0 controllerNumber=1 scan=12486 | 26.681 |
| 07902:3 | 4 | K.WEC[+57.021]KN[+2205.793]DTLLGIK.G | HexNAc(4)Hex(5)Fuc(2)NeuAc(1) | C[+57], N[+2206] | 1228.1905 | 3 | 3682.5569 | 3682.5444 | 0 | 3.4 | 100 | Specific | 313.95 | 50.42 | 50.42 | 4.67 | 92 | >sp\|P22897\|MRC1_HUMAN Macrophage mannose receptor 1 OS=Homo sapiens OX=9606 GN=MRC1 PE=1 SV=1 | 1049 | 20201115_07433_1ng_glyco_hcd203040_compound15.10841.10841.3 | controllerType=0 controllerNumber=1 scan=10841 | 23.9149 |
| 13470:5 | 4 | K.KGN[+2059.735]TTLNSFVIPSESDVPTHC[+57.021]PSQWWPYAGHC[+57.021]YK.I | HexNAc(4)Hex(5)Fuc(1)NeuAc(1) | C[+57]*2, N[+2060] | 1205.5223 | 5 | 6023.5826 | 6023.5567 | 0 | 4.3 | 342 | Specific | 411.96 | 411.96 | 411.96 | 10.25 | 92 | >sp\|P22897\|MRC1_HUMAN Macrophage mannose receptor 1 OS=Homo sapiens OX=9606 GN=MRC1 PE=1 SV=1 | 1049 | 20201115_07433_1ng_glyco_hcd203040_compound15.16657.16657.5 | controllerType=0 controllerNumber=1 scan=16657 | 33.6512 |
| 13468:5 | 4 | K.KGN[+2059.735]TTLNSFVIPSESDVPTHC[+57.021]PSQWWPYAGHC[+57.021]YK.I | HexNAc(4)Hex(5)Fuc(1)NeuAc(1) | C[+57]*2, N[+2060] | 1205.5223 | 5 | 6023.5826 | 6023.5567 | 0 | 4.3 | 342 | Specific | 510.64 | 510.64 | 510.64 | 10.67 | 92 | >sp\|P22897\|MRC1_HUMAN Macrophage mannose receptor 1 OS=Homo sapiens OX=9606 GN=MRC1 PE=1 SV=1 | 1049 | 20201115_07433_1ng_glyco_hcd203040_compound15.16655.16655.5 | controllerType=0 controllerNumber=1 scan=16655 | 33.6456 |
| 16838:3 | 4 | R.VWIALNSN[+2205.793]LTDNQYTWTDKWR.V | HexNAc(4)Hex(5)Fuc(2)NeuAc(1) | N[+2206] | 1610.6926 | 3 | 4830.0631 | 4830.0821 | 0 | -3.9 | 1153 | Specific | 283.89 | 167.17 | 167.17 | 4.68 | 92 | >sp\|P22897\|MRC1_HUMAN Macrophage mannose receptor 1 OS=Homo sapiens OX=9606 GN=MRC1 PE=1 SV=1 | 1049 | 20201115_07433_1ng_glyco_hcd203040_compound15.20203.20203.3 | controllerType=0 controllerNumber=1 scan=20203 | 40.0765 |
| 08733:3 | 4 | K.TAHC[+57.021]N[+2059.735]ESFYFLC[+57.021]K.R | HexNAc(4)Hex(5)Fuc(1)NeuAc(1) | C[+57]*2, N[+2060] | 1246.158 | 3 | 3736.4595 | 3736.4545 | 0 | 1.3 | 1201 | Specific | 806.85 | 731.11 | 731.11 | 16.58 | 92 | >sp\|P22897\|MRC1_HUMAN Macrophage mannose receptor 1 OS=Homo sapiens OX=9606 GN=MRC1 PE=1 SV=1 | 1049 | 20201115_07433_1ng_glyco_hcd203040_compound15.11714.11714.3 | controllerType=0 controllerNumber=1 scan=11714 | 25.3841 |
| 08735:3 | 4 | K.TAHC[+57.021]N[+2059.735]ESFYFLC[+57.021]K.R | HexNAc(4)Hex(5)Fuc(1)NeuAc(1) | C[+57]*2, N[+2060] | 1246.158 | 3 | 3736.4595 | 3736.4545 | 0 | 1.3 | 1201 | Specific | 827.56 | 722.07 | 722.07 | 16.58 | 92 | >sp\|P22897\|MRC1_HUMAN Macrophage mannose receptor 1 OS=Homo sapiens OX=9606 GN=MRC1 PE=1 SV=1 | 1049 | 20201115_07433_1ng_glyco_hcd203040_compound15.11716.11716.3 | controllerType=0 controllerNumber=1 scan=11716 | 25.3903 |
| 17465:4 | 5 | K.RNIAAFGGDPNN[+1913.677]ITLFGESAGGASVSLQTLSPYNK.G | HexNAc(4)Hex(5)NeuAc(1) | N[+1914] | 1370.8737 | 4 | 5480.4728 | 5480.4644 | 0 | 1.5 | 196 | Specific | 980.67 | 966.36 | 966.36 | 20.32 | 67 | >sp\|P19835\|CEL_HUMAN Bile salt-activated lipase OS=Homo sapiens OX=9606 GN=CEL PE=1 SV=3 | 476 | 20201115_07433_1ng_glyco_hcd203040_compound15.20855.20855.4 | controllerType=0 controllerNumber=1 scan=20855 | 41.3541 |
| 17467:4 | 5 | K.RNIAAFGGDPNN[+1913.677]ITLFGESAGGASVSLQTLSPYNK.G | HexNAc(4)Hex(5)NeuAc(1) | N[+1914] | 1370.8737 | 4 | 5480.4728 | 5480.4644 | 0 | 1.5 | 196 | Specific | 1006.65 | 927.7 | 927.7 | 21.32 | 67 | >sp\|P19835\|CEL_HUMAN Bile salt-activated lipase OS=Homo sapiens OX=9606 GN=CEL PE=1 SV=3 | 476 | 20201115_07433_1ng_glyco_hcd203040_compound15.20857.20857.4 | controllerType=0 controllerNumber=1 scan=20857 | 41.3653 |
| 19244:4 | 5 | R.NIAAFGGDPNN[+2059.735]ITLFGESAGGASVSLQTLSPYNK.G | HexNAc(4)Hex(5)Fuc(1)NeuAc(1) | N[+2060] | 1368.3633 | 4 | 5470.4313 | 5470.4212 | 0 | 1.8 | 197 | Specific | 519.94 | 495.13 | 495.13 | 11.32 | 67 | >sp\|P19835\|CEL_HUMAN Bile salt-activated lipase OS=Homo sapiens OX=9606 GN=CEL PE=1 SV=3 | 476 | 20201115_07433_1ng_glyco_hcd203040_compound15.22711.22711.4 | controllerType=0 controllerNumber=1 scan=22711 | 45.3897 |
| 18065:3 | 5 | R.NIAAFGGDPNN[+1622.582]ITLFGESAGGASVSLQTLSPYNK.G | HexNAc(4)Hex(5) | N[+1623] | 1678.4316 | 3 | 5033.2803 | 5033.2679 | 0 | 2.5 | 197 | Specific | 640.28 | 640.28 | 640.28 | 13.72 | 67 | >sp\|P19835\|CEL_HUMAN Bile salt-activated lipase OS=Homo sapiens OX=9606 GN=CEL PE=1 SV=3 | 476 | 20201115_07433_1ng_glyco_hcd203040_compound15.21480.21480.3 | controllerType=0 controllerNumber=1 scan=21480 | 42.6976 |
| 18068:3 | 5 | R.NIAAFGGDPNN[+1622.582]ITLFGESAGGASVSLQTLSPYNK.G | HexNAc(4)Hex(5) | N[+1623] | 1678.4316 | 3 | 5033.2803 | 5033.2679 | 0 | 2.5 | 197 | Specific | 722.7 | 679.6 | 679.6 | 15.72 | 67 | >sp\|P19835\|CEL_HUMAN Bile salt-activated lipase OS=Homo sapiens OX=9606 GN=CEL PE=1 SV=3 | 476 | 20201115_07433_1ng_glyco_hcd203040_compound15.21483.21483.3 | controllerType=0 controllerNumber=1 scan=21483 | 42.7118 |
| 19212:4 | 5 | R.NIAAFGGDPNN[+1913.677]ITLFGESAGGASVSLQTLSPYNK.G | HexNAc(4)Hex(5)NeuAc(1) | N[+1914] | 1331.8492 | 4 | 5324.3752 | 5324.3633 | 0 | 2.2 | 197 | Specific | 998.43 | 971.53 | 971.53 | 21.31 | 67 | >sp\|P19835\|CEL_HUMAN Bile salt-activated lipase OS=Homo sapiens OX=9606 GN=CEL PE=1 SV=3 | 476 | 20201115_07433_1ng_glyco_hcd203040_compound15.22678.22678.4 | controllerType=0 controllerNumber=1 scan=22678 | 45.3315 |
| 19210:4 | 5 | R.NIAAFGGDPNN[+1913.677]ITLFGESAGGASVSLQTLSPYNK.G | HexNAc(4)Hex(5)NeuAc(1) | N[+1914] | 1331.8492 | 4 | 5324.3752 | 5324.3633 | 0 | 2.2 | 197 | Specific | 1064.91 | 1020.65 | 1020.65 | 23.31 | 67 | >sp\|P19835\|CEL_HUMAN Bile salt-activated lipase OS=Homo sapiens OX=9606 GN=CEL PE=1 SV=3 | 476 | 20201115_07433_1ng_glyco_hcd203040_compound15.22676.22676.4 | controllerType=0 controllerNumber=1 scan=22676 | 45.3194 |
| 19883:4 | 7 | R.GLSFDVSLEVSQGPGLLN[+2351.851]DTK.V | HexNAc(4)Hex(5)Fuc(3)NeuAc(1) | N[+2352] | 1132.7435 | 4 | 4527.952 | 4527.964 | 0 | -2.6 | 118 | Specific | 297.8 | 97.77 | 97.77 | 4.88 | 64 | >sp\|P01833\|PIGR_HUMAN Polymeric immunoglobulin receptor OS=Homo sapiens OX=9606 GN=PIGR PE=1 SV=4 | 902 | 20201115_07433_1ng_glyco_hcd203040_compound15.23379.23379.4 | controllerType=0 controllerNumber=1 scan=23379 | [illegible] |
| 19886:4 | 7 | R.GLSFDVSLEVSQGPGLLN[+2351.851]DTK.V | HexNAc(4)Hex(5)Fuc(3)NeuAc(1) | N[+2352] | 1132.7435 | 4 | 4527.952 | 4527.964 | 0 | -2.6 | 118 | Specific | 715.72 | 475.71 | 475.71 | 14.26 | 64 | >sp\|P01833\|PIGR_HUMAN Polymeric immunoglobulin receptor OS=Homo sapiens OX=9606 GN=PIGR PE=1 SV=4 | 902 | 20201115_07433_1ng_glyco_hcd203040_compound15.23382.23382.4 | controllerType=0 controllerNumber=1 scan=23382 | [illegible] |
| 17395:3 | 7 | K.QIGLYPVLVIDSSGYVNPN[+1913.677]YTGR.I | HexNAc(4)Hex(5)NeuAc(1) | N[+1914] | 1480.3454 | 3 | 4439.0216 | 4438.9805 | 0 | 9.3 | 168 | Specific | 259.96 | 259.96 | 259.96 | 4.48 | 64 | >sp\|P01833\|PIGR_HUMAN Polymeric immunoglobulin receptor OS=Homo sapiens OX=9606 GN=PIGR PE=1 SV=4 | 902 | 20201115_07433_1ng_glyco_hcd203040_compound15.20782.20782.3 | controllerType=0 controllerNumber=1 scan=20782 | 41.2104 |
| 19223:3 | 7 | K.QIGLYPVLVIDSSGYVNPN[+2059.735]YTGR.I | HexNAc(4)Hex(5)Fuc(1)NeuAc(1) | N[+2060] | 1529.0229 | 3 | 4585.0541 | 4585.0384 | 0 | 3.4 | 168 | Specific | 287.64 | 287.64 | 287.64 | 7.51 | 64 | >sp\|P01833\|PIGR_HUMAN Polymeric immunoglobulin receptor OS=Homo sapiens OX=9606 GN=PIGR PE=1 SV=4 | 902 | 20201115_07433_1ng_glyco_hcd203040_compound15.22690.22690.3 | controllerType=0 controllerNumber=1 scan=22690 | 45.3513 |
| 19264:3 | 7 | K.QIGLYPVLVIDSSGYVNPN[+2059.735]YTGR.I | HexNAc(4)Hex(5)Fuc(1)NeuAc(1) | N[+2060] | 1529.0178 | 3 | 4585.0387 | 4585.0384 | 0 | 0.1 | 168 | Specific | 384.24 | 343.01 | 343.01 | 7.63 | 64 | >sp\|P01833\|PIGR_HUMAN Polymeric immunoglobulin receptor OS=Homo sapiens OX=9606 GN=PIGR PE=1 SV=4 | 902 | 20201115_07433_1ng_glyco_hcd203040_compound15.22732.22732.3 | controllerType=0 controllerNumber=1 scan=22732 | 45.4288 |
| 17397:3 | 7 | K.QIGLYPVLVIDSSGYVNPN[+1913.677]YTGR.I | HexNAc(4)Hex(5)NeuAc(1) | N[+1914] | 1480.3454 | 3 | 4439.0216 | 4438.9805 | 0 | 9.3 | 168 | Specific | 416.66 | 402.34 | 402.34 | 7.04 | 64 | >sp\|P01833\|PIGR_HUMAN Polymeric immunoglobulin receptor OS=Homo sapiens OX=9606 GN=PIGR PE=1 SV=4 | 902 | 20201115_07433_1ng_glyco_hcd203040_compound15.20784.20784.3 | controllerType=0 controllerNumber=1 scan=20784 | 41.2154 |
| 19267:3 | 7 | K.QIGLYPVLVIDSSGYVNPN[+2059.735]YTGR.I | HexNAc(4)Hex(5)Fuc(1)NeuAc(1) | N[+2060] | 1529.0178 | 3 | 4585.0387 | 4585.0384 | 0 | 0.1 | 168 | Specific | 601.4 | 448.05 | 448.05 | 12.57 | 64 | >sp\|P01833\|PIGR_HUMAN Polymeric immunoglobulin receptor OS=Homo sapiens OX=9606 GN=PIGR PE=1 SV=4 | 902 | 20201115_07433_1ng_glyco_hcd203040_compound15.22735.22735.3 | controllerType=0 controllerNumber=1 scan=22735 | 45.4341 |
| 19227:4 | 7 | K.QIGLYPVLVIDSSGYVNPN[+2059.735]YTGR.I | HexNAc(4)Hex(5)Fuc(1)NeuAc(1) | N[+2060] | 1147.0188 | 4 | 4585.0535 | 4585.0384 | 0 | 3.3 | 168 | Specific | 720.84 | 656.22 | 656.22 | 15.86 | 64 | >sp\|P01833\|PIGR_HUMAN Polymeric immunoglobulin receptor OS=Homo sapiens OX=9606 GN=PIGR PE=1 SV=4 | 902 | 20201115_07433_1ng_glyco_hcd203040_compound15.22694.22694.4 | controllerType=0 controllerNumber=1 scan=22694 | 45.3572 |
| 19226:3 | 7 | K.QIGLYPVLVIDSSGYVNPN[+2059.735]YTGR.I | HexNAc(4)Hex(5)Fuc(1)NeuAc(1) | N[+2060] | 1529.0229 | 3 | 4585.0541 | 4585.0384 | 0 | 3.4 | 168 | Specific | 855.12 | 599.74 | 599.74 | 18.27 | 64 | >sp\|P01833\|PIGR_HUMAN Polymeric immunoglobulin receptor OS=Homo sapiens OX=9606 GN=PIGR PE=1 SV=4 | 902 | 20201115_07433_1ng_glyco_hcd203040_compound15.22693.22693.3 | controllerType=0 controllerNumber=1 scan=22693 | 45.3548 |
| 29418:3 | 7 | R.[+42.011]LSLLEEPGN[+2351.851]GTFTVILNQLTSR.D | HexNAc(4)Hex(5)Fuc(3)NeuAc(1) | .[42], N[+2352] | 1599.3848 | 3 | 4796.1399 | 4796.1539 | 0 | -2.9 | 413 | Specific | 207.83 | 190.31 | 190.31 | 4.05 | 64 | >sp\|P01833\|PIGR_HUMAN Polymeric immunoglobulin receptor OS=Homo sapiens OX=9606 GN=PIGR PE=1 SV=4 | 902 | 20201115_07433_1ng_glyco_hcd203040_compound15.33335.33335.3 | controllerType=0 controllerNumber=1 scan=33335 | 65.2009 |
| 29420:3 | 7 | R.[+42.011]LSLLEEPGN[+2351.851]GTFTVILNQLTSR.D | HexNAc(4)Hex(5)Fuc(3)NeuAc(1) | .[42], N[+2352] | 1599.3848 | 3 | 4796.1399 | 4796.1539 | 0 | -2.9 | 413 | Specific | 230.79 | 215.03 | 215.03 | 5.16 | 64 | >sp\|P01833\|PIGR_HUMAN Polymeric immunoglobulin receptor OS=Homo sapiens OX=9606 GN=PIGR PE=1 SV=4 | 902 | 20201115_07433_1ng_glyco_hcd203040_compound15.33337.33337.3 | controllerType=0 controllerNumber=1 scan=33337 | 65.2049 |
| 23275:3 | 7 | R.LSLLEEPGN[+2351.851]GTFTVILNQLTSR.D | HexNAc(4)Hex(5)Fuc(3)NeuAc(1) | N[+2352] | 1585.387 | 3 | 4754.1464 | 4754.1433 | 0 | 0.7 | 413 | Specific | 326.98 | 289.61 | 289.61 | 6.8 | 64 | >sp\|P01833\|PIGR_HUMAN Polymeric immunoglobulin receptor OS=Homo sapiens OX=9606 GN=PIGR PE=1 SV=4 | 902 | 20201115_07433_1ng_glyco_hcd203040_compound15.26929.26929.3 | controllerType=0 controllerNumber=1 scan=26929 | 53.4994 |
| 20874:4 | 7 | R.LSLLEEPGN[+2076.750]GTFTVILNQLTSR.D | HexNAc(4)Hex(6)Fuc(2) | N[+2077] | 1120.522 | 4 | 4479.0662 | 4479.0428 | 0 | 5.2 | 413 | Specific | 433.84 | 338.49 | 338.49 | 8.52 | 64 | >sp\|P01833\|PIGR_HUMAN Polymeric immunoglobulin receptor OS=Homo sapiens OX=9606 GN=PIGR PE=1 SV=4 | 902 | 20201115_07433_1ng_glyco_hcd203040_compound15.24414.24414.4 | controllerType=0 controllerNumber=1 scan=24414 | 48.8597 |
| 23245:3 | 7 | R.LSLLEEPGN[+2350.830]GTFTVILNQLTSR.D | HexNAc(4)Hex(5)Fuc(1)NeuAc(2) | N[+2351] | 1585.0502 | 3 | 4753.1361 | 4753.1229 | 0 | 2.8 | 413 | Specific | 449.06 | 410 | 410 | 10.32 | 64 | >sp\|P01833\|PIGR_HUMAN Polymeric immunoglobulin receptor OS=Homo sapiens OX=9606 GN=PIGR PE=1 SV=4 | 902 | 20201115_07433_1ng_glyco_hcd203040_compound15.26898.26898.3 | controllerType=0 controllerNumber=1 scan=26898 | 53.4341 |
| 23278:3 | 7 | R.LSLLEEPGN[+2351.851]GTFTVILNQLTSR.D | HexNAc(4)Hex(5)Fuc(3)NeuAc(1) | N[+2352] | 1585.387 | 3 | 4754.1464 | 4754.1433 | 0 | 0.7 | 413 | Specific | 600.49 | 398.05 | 398.05 | 12.34 | 64 | >sp\|P01833\|PIGR_HUMAN Polymeric immunoglobulin receptor OS=Homo sapiens OX=9606 GN=PIGR PE=1 SV=4 | 902 | 20201115_07433_1ng_glyco_hcd203040_compound15.26932.26932.3 | controllerType=0 controllerNumber=1 scan=26932 | 53.5036 |
| 23248:3 | 7 | R.LSLLEEPGN[+2350.830]GTFTVILNQLTSR.D | HexNAc(4)Hex(5)Fuc(1)NeuAc(2) | N[+2351] | 1585.0502 | 3 | 4753.1361 | 4753.1229 | 0 | 2.8 | 413 | Specific | 762.18 | 514.88 | 514.88 | 16.2 | 64 | >sp\|P01833\|PIGR_HUMAN Polymeric immunoglobulin receptor OS=Homo sapiens OX=9606 GN=PIGR PE=1 SV=4 | 902 | 20201115_07433_1ng_glyco_hcd203040_compound15.26901.26901.3 | controllerType=0 controllerNumber=1 scan=26901 | 53.438 |
| 20848:4 | 7 | R.LSLLEEPGN[+2205.793]GTFTVILNQLTSR.D | HexNAc(4)Hex(5)Fuc(2)NeuAc(1) | N[+2206] | 1152.7832 | 4 | 4608.1112 | 4608.0854 | 0 | 5.6 | 413 | Specific | 789.61 | 603.98 | 603.98 | 15.91 | 64 | >sp\|P01833\|PIGR_HUMAN Polymeric immunoglobulin receptor OS=Homo sapiens OX=9606 GN=PIGR PE=1 SV=4 | 902 | 20201115_07433_1ng_glyco_hcd203040_compound15.24387.24387.4 | controllerType=0 controllerNumber=1 scan=24387 | 48.8015 |
| 20818:4 | 7 | R.LSLLEEPGN[+2205.793]GTFTVILNQLTSR.D | HexNAc(4)Hex(5)Fuc(2)NeuAc(1) | N[+2206] | 1152.783 | 4 | 4608.1102 | 4608.0854 | 0 | 5.4 | 413 | Specific | 822.66 | 639.56 | 639.56 | 16.91 | 64 | >sp\|P01833\|PIGR_HUMAN Polymeric immunoglobulin receptor OS=Homo sapiens OX=9606 GN=PIGR PE=1 SV=4 | 902 | 20201115_07433_1ng_glyco_hcd203040_compound15.24355.24355.3 | controllerType=0 controllerNumber=1 scan=24355 | 48.7293 |
| 20845:3 | 7 | R.LSLLEEPGN[+2205.793]GTFTVILNQLTSR.D | HexNAc(4)Hex(5)Fuc(2)NeuAc(1) | N[+2206] | 1536.7057 | 3 | 4608.1027 | 4608.0854 | 0 | 3.7 | 413 | Specific | 870.16 | 624.52 | 624.52 | 18.03 | 64 | >sp\|P01833\|PIGR_HUMAN Polymeric immunoglobulin receptor OS=Homo sapiens OX=9606 GN=PIGR PE=1 SV=4 | 902 | 20201115_07433_1ng_glyco_hcd203040_compound15.24384.24384.3 | controllerType=0 controllerNumber=1 scan=24384 | 48.7959 |
| 20821:4 | 7 | R.LSLLEEPGN[+2205.793]GTFTVILNQLTSR.D | HexNAc(4)Hex(5)Fuc(2)NeuAc(1) | N[+2206] | 1152.783 | 4 | 4608.1102 | 4608.0854 | 0 | 5.4 | 413 | Specific | 877.85 | 685.13 | 685.13 | 17.91 | 64 | >sp\|P01833\|PIGR_HUMAN Polymeric immunoglobulin receptor OS=Homo sapiens OX=9606 GN=PIGR PE=1 SV=4 | 902 | 20201115_07433_1ng_glyco_hcd203040_compound15.24358.24358.3 | controllerType=0 controllerNumber=1 scan=24358 | 48.7379 |
| 20847:3 | 7 | R.LSLLEEPGN[+2205.793]GTFTVILNQLTSR.D | HexNAc(4)Hex(5)Fuc(2)NeuAc(1) | N[+2206] | 1536.7057 | 3 | 4608.1027 | 4608.0854 | 0 | 3.7 | 413 | Specific | 943.81 | 672.75 | 672.75 | 19.03 | 64 | >sp\|P01833\|PIGR_HUMAN Polymeric immunoglobulin receptor OS=Homo sapiens OX=9606 GN=PIGR PE=1 SV=4 | 902 | 20201115_07433_1ng_glyco_hcd203040_compound15.24386.24386.3 | controllerType=0 controllerNumber=1 scan=24386 | 48.8002 |
| 20851:4 | 7 | R.LSLLEEPGN[+2205.793]GTFTVILNQLTSR.D | HexNAc(4)Hex(5)Fuc(2)NeuAc(1) | N[+2206] | 1152.7832 | 4 | 4608.1112 | 4608.0854 | 0 | 5.6 | 413 | Specific | 980.41 | 775.8 | 775.8 | 19.91 | 64 | >sp\|P01833\|PIGR_HUMAN Polymeric immunoglobulin receptor OS=Homo sapiens OX=9606 GN=PIGR PE=1 SV=4 | 902 | 20201115_07433_1ng_glyco_hcd203040_compound15.24390.24390.4 | controllerType=0 controllerNumber=1 scan=24390 | 48.8061 |
| 16344:4 | 7 | K.IIEGEPNLKVPGN[+2204.772]VTAVLGETLK.V | HexNAc(4)Hex(5)NeuAc(2) | N[+2205] | 1149.7896 | 4 | 4596.1365 | 4596.1218 | 0 | 3.2 | 457 | Specific | 676.9 | 676.9 | 676.9 | 13.79 | 64 | >sp\|P01833\|PIGR_HUMAN Polymeric immunoglobulin receptor OS=Homo sapiens OX=9606 GN=PIGR PE=1 SV=4 | 902 | 20201115_07433_1ng_glyco_hcd203040_compound15.19687.19687.4 | controllerType=0 controllerNumber=1 scan=19687 | 39.2036 |
| 16374:4 | 7 | K.IIEGEPNLKVPGN[+2205.793]VTAVLGETLK.V | HexNAc(4)Hex(5)Fuc(2)NeuAc(1) | N[+2206] | 1150.0436 | 4 | 4597.1527 | 4597.1422 | 0 | 2.3 | 457 | Specific | 684.4 | 652.34 | 652.34 | 13.96 | 64 | >sp\|P01833\|PIGR_HUMAN Polymeric immunoglobulin receptor OS=Homo sapiens OX=9606 GN=PIGR PE=1 SV=4 | 902 | 20201115_07433_1ng_glyco_hcd203040_compound15.19718.19718.4 | controllerType=0 controllerNumber=1 scan=19718 | 39.2617 |
| 16371:4 | 7 | K.IIEGEPNLKVPGN[+2205.793]VTAVLGETLK.V | HexNAc(4)Hex(5)Fuc(2)NeuAc(1) | N[+2206] | 1150.0436 | 4 | 4597.1527 | 4597.1422 | 0 | 2.3 | 457 | Specific | 702.14 | 702.14 | 702.14 | 14.96 | 64 | >sp\|P01833\|PIGR_HUMAN Polymeric immunoglobulin receptor OS=Homo sapiens OX=9606 GN=PIGR PE=1 SV=4 | 902 | 20201115_07433_1ng_glyco_hcd203040_compound15.19715.19715.4 | controllerType=0 controllerNumber=1 scan=19715 | 39.2565 |
| 16395:4 | 7 | K.IIEGEPNLKVPGN[+2059.735]VTAVLGETLK.V | HexNAc(4)Hex(5)Fuc(1)NeuAc(1) | N[+2060] | 1113.5306 | 4 | 4451.1004 | 4451.0843 | 0 | 3.6 | 457 | Specific | 703.61 | 703.61 | 703.61 | 14.79 | 64 | >sp\|P01833\|PIGR_HUMAN Polymeric immunoglobulin receptor OS=Homo sapiens OX=9606 GN=PIGR PE=1 SV=4 | 902 | 20201115_07433_1ng_glyco_hcd203040_compound15.19740.19740.4 | controllerType=0 controllerNumber=1 scan=19740 | 39.3108 |
| 16347:4 | 7 | K.IIEGEPNLKVPGN[+2204.772]VTAVLGETLK.V | HexNAc(4)Hex(5)NeuAc(2) | N[+2205] | 1149.7896 | 4 | 4596.1365 | 4596.1218 | 0 | 3.2 | 457 | Specific | 709.69 | 585.85 | 585.85 | 14.79 | 64 | >sp\|P01833\|PIGR_HUMAN Polymeric immunoglobulin receptor OS=Homo sapiens OX=9606 GN=PIGR PE=1 SV=4 | 902 | 20201115_07433_1ng_glyco_hcd203040_compound15.19690.19690.4 | controllerType=0 controllerNumber=1 scan=19690 | 39.2091 |
| 16397:4 | 7 | K.IIEGEPNLKVPGN[+2059.735]VTAVLGETLK.V | HexNAc(4)Hex(5)Fuc(1)NeuAc(1) | N[+2060] | 1113.5306 | 4 | 4451.1004 | 4451.0843 | 0 | 3.6 | 457 | Specific | 713.53 | 643.67 | 643.67 | 14.79 | 64 | >sp\|P01833\|PIGR_HUMAN Polymeric immunoglobulin receptor OS=Homo sapiens OX=9606 GN=PIGR PE=1 SV=4 | 902 | 20201115_07433_1ng_glyco_hcd203040_compound15.19742.19742.4 | controllerType=0 controllerNumber=1 scan=19742 | 39.3152 |
| 16323:3 | 7 | K.IIEGEPNLKVPGN[+2205.793]VTAVLGETLK.V | HexNAc(4)Hex(5)Fuc(2)NeuAc(1) | N[+2206] | 1533.0513 | 3 | 4597.1394 | 4597.1422 | 0 | -0.6 | 457 | Specific | 829.94 | 800.25 | 800.25 | 16.1 | 64 | >sp\|P01833\|PIGR_HUMAN Polymeric immunoglobulin receptor OS=Homo sapiens OX=9606 GN=PIGR PE=1 SV=4 | 902 | 20201115_07433_1ng_glyco_hcd203040_compound15.19664.19664.3 | controllerType=0 controllerNumber=1 scan=19664 | 39.1603 |
| 16390:3 | 7 | K.IIEGEPNLKVPGN[+2059.735]VTAVLGETLK.V | HexNAc(4)Hex(5)Fuc(1)NeuAc(1) | N[+2060] | 1484.3716 | 3 | 4451.1004 | 4451.0843 | 0 | 3.6 | 457 | Specific | 830.85 | 642.94 | 642.94 | 16.8 | 64 | >sp\|P01833\|PIGR_HUMAN Polymeric immunoglobulin receptor OS=Homo sapiens OX=9606 GN=PIGR PE=1 SV=4 | 902 | 20201115_07433_1ng_glyco_hcd203040_compound15.19735.19735.3 | controllerType=0 controllerNumber=1 scan=19735 | 39.3001 |
| 16320:3 | 7 | K.IIEGEPNLKVPGN[+2205.793]VTAVLGETLK.V | HexNAc(4)Hex(5)Fuc(2)NeuAc(1) | N[+2206] | 1533.0513 | 3 | 4597.1394 | 4597.1422 | 0 | -0.6 | 457 | Specific | 958.33 | 901.86 | 901.86 | 19.1 | 64 | >sp\|P01833\|PIGR_HUMAN Polymeric immunoglobulin receptor OS=Homo sapiens OX=9606 GN=PIGR PE=1 SV=4 | 902 | 20201115_07433_1ng_glyco_hcd203040_compound15.19661.19661.3 | controllerType=0 controllerNumber=1 scan=19661 | 39.1562 |
| 17494:5 | 7 | K.IIEGEPNLKVPGN[+2350.830]VTAVLGETLKVPC[+57.021]HFPC[+57.021]K.F | HexNAc(4)Hex(5)Fuc(1)NeuAc(2) | C[+57]*2, N[+2351] | 1154.3325 | 5 | 5767.6335 | 5767.6372 | 0 | -0.6 | 457 | Specific | 295.99 | 295.99 | 295.99 | 6.19 | 64 | >sp\|P01833\|PIGR_HUMAN Polymeric immunoglobulin receptor OS=Homo sapiens OX=9606 GN=PIGR PE=1 SV=4 | 902 | 20201115_07433_1ng_glyco_hcd203040_compound15.20885.20885.5 | controllerType=0 controllerNumber=1 scan=20885 | 41.4221 |
| 17496:5 | 7 | K.IIEGEPNLKVPGN[+2350.830]VTAVLGETLKVPC[+57.021]HFPC[+57.021]K.F | HexNAc(4)Hex(5)Fuc(1)NeuAc(2) | C[+57]*2, N[+2351] | 1154.3325 | 5 | 5767.6335 | 5767.6372 | 0 | -0.6 | 457 | Specific | 414.24 | 329.08 | 329.08 | 7.09 | 64 | >sp\|P01833\|PIGR_HUMAN Polymeric immunoglobulin receptor OS=Homo sapiens OX=9606 GN=PIGR PE=1 SV=4 | 902 | 20201115_07433_1ng_glyco_hcd203040_compound15.20887.20887.5 | controllerType=0 controllerNumber=1 scan=20887 | 41.4272 |
| 15851:5 | 7 | K.IIEGEPNLKVPGN[+2059.735]VTAVLGETLKVPC[+57.021]HFPC[+57.021]K.F | HexNAc(4)Hex(5)Fuc(1)NeuAc(1) | C[+57]*2, N[+2060] | 1096.1189 | 5 | 5476.5654 | 5476.5418 | 0 | 4.3 | 457 | Specific | 703.69 | 610.18 | 610.18 | 14.75 | 64 | >sp\|P01833\|PIGR_HUMAN Polymeric immunoglobulin receptor OS=Homo sapiens OX=9606 GN=PIGR PE=1 SV=4 | 902 | 20201115_07433_1ng_glyco_hcd203040_compound15.19124.19124.3 | controllerType=0 controllerNumber=1 scan=19124 | 38.1075 |
| 17357:4 | 7 | K.IIEGEPNLKVPGN[+2350.830]VTAVLGETLKVPC[+57.021]HFPC[+57.021]K.F | HexNAc(4)Hex(5)Fuc(1)NeuAc(2) | C[+57]*2, N[+2351] | 1442.6708 | 4 | 5767.6613 | 5767.6372 | 0 | 4.2 | 457 | Specific | 707.85 | 587.7 | 587.7 | 14.21 | 64 | >sp\|P01833\|PIGR_HUMAN Polymeric immunoglobulin receptor OS=Homo sapiens OX=9606 GN=PIGR PE=1 SV=4 | 902 | 20201115_07433_1ng_glyco_hcd203040_compound15.20743.20743.4 | controllerType=0 controllerNumber=1 scan=20743 | 41.1365 |
| 15859:5 | 7 | K.IIEGEPNLKVPGN[+2204.772]VTAVLGETLKVPC[+57.021]HFPC[+57.021]K.F | HexNAc(4)Hex(5)NeuAc(2) | C[+57]*2, N[+2205] | 1125.1283 | 5 | 5621.6124 | 5621.5793 | 0 | 5.9 | 457 | Specific | 756.23 | 572.5 | 572.5 | 15.33 | 64 | >sp\|P01833\|PIGR_HUMAN Polymeric immunoglobulin receptor OS=Homo sapiens OX=9606 GN=PIGR PE=1 SV=4 | 902 | 20201115_07433_1ng_glyco_hcd203040_compound15.19133.19133.5 | controllerType=0 controllerNumber=1 scan=19133 | 38.1299 |
| 15869:4 | 7 | K.IIEGEPNLKVPGN[+2205.793]VTAVLGETLKVPC[+57.021]HFPC[+57.021]K.F | HexNAc(4)Hex(5)Fuc(2)NeuAc(1) | C[+57]*2, N[+2206] | 1406.4085 | 4 | 5622.612 | 5622.5997 | 0 | 2.2 | 457 | Specific | 793.97 | 793.97 | 793.97 | 15.68 | 64 | >sp\|P01833\|PIGR_HUMAN Polymeric immunoglobulin receptor OS=Homo sapiens OX=9606 GN=PIGR PE=1 SV=4 | 902 | 20201115_07433_1ng_glyco_hcd203040_compound15.19143.19143.4 | controllerType=0 controllerNumber=1 scan=19143 | 38.1519 |
| 15931:4 | 7 | K.IIEGEPNLKVPGN[+2059.735]VTAVLGETLKVPC[+57.021]HFPC[+57.021]K.F | HexNAc(4)Hex(5)Fuc(1)NeuAc(1) | C[+57]*2, N[+2060] | 1369.8927 | 4 | 5476.549 | 5476.5418 | 0 | 1.3 | 457 | Specific | 855.22 | 816.08 | 816.08 | 17.27 | 64 | >sp\|P01833\|PIGR_HUMAN Polymeric immunoglobulin receptor OS=Homo sapiens OX=9606 GN=PIGR PE=1 SV=4 | 902 | 20201115_07433_1ng_glyco_hcd203040_compound15.19208.19208.4 | controllerType=0 controllerNumber=1 scan=19208 | 38.2813 |
| 17349:5 | 7 | K.IIEGEPNLKVPGN[+2350.830]VTAVLGETLKVPC[+57.021]HFPC[+57.021]K.F | HexNAc(4)Hex(5)Fuc(1)NeuAc(2) | C[+57]*2, N[+2351] | 1154.3358 | 5 | 5767.65 | 5767.6372 | 0 | 2.2 | 457 | Specific | 861.98 | 861.98 | 861.98 | 18.22 | 64 | >sp\|P01833\|PIGR_HUMAN Polymeric immunoglobulin receptor OS=Homo sapiens OX=9606 GN=PIGR PE=1 SV=4 | 902 | 20201115_07433_1ng_glyco_hcd203040_compound15.20734.20734.5 | controllerType=0 controllerNumber=1 scan=20734 | 41.1193 |
| 15930:5 | 7 | K.IIEGEPNLKVPGN[+2059.735]VTAVLGETLKVPC[+57.021]HFPC[+57.021]K.F | HexNAc(4)Hex(5)Fuc(1)NeuAc(1) | C[+57]*2, N[+2060] | 1096.1172 | 5 | 5476.5568 | 5476.5418 | 0 | 2.7 | 457 | Specific | 892.49 | 813.64 | 813.64 | 18.22 | 64 | >sp\|P01833\|PIGR_HUMAN Polymeric immunoglobulin receptor OS=Homo sapiens OX=9606 GN=PIGR PE=1 SV=4 | 902 | 20201115_07433_1ng_glyco_hcd203040_compound15.19207.19207.5 | controllerType=0 controllerNumber=1 scan=19207 | 38.2793 |
| 15853:5 | 7 | K.IIEGEPNLKVPGN[+2059.735]VTAVLGETLKVPC[+57.021]HFPC[+57.021]K.F | HexNAc(4)Hex(5)Fuc(1)NeuAc(1) | C[+57]*2, N[+2060] | 1096.1189 | 5 | 5476.5654 | 5476.5418 | 0 | 4.3 | 457 | Specific | 893.53 | 810.2 | 810.2 | 17.75 | 64 | >sp\|P01833\|PIGR_HUMAN Polymeric immunoglobulin receptor OS=Homo sapiens OX=9606 GN=PIGR PE=1 SV=4 | 902 | 20201115_07433_1ng_glyco_hcd203040_compound15.19126.19126.3 | controllerType=0 controllerNumber=1 scan=19126 | 38.1127 |
| 15929:4 | 7 | K.IIEGEPNLKVPGN[+2059.735]VTAVLGETLKVPC[+57.021]HFPC[+57.021]K.F | HexNAc(4)Hex(5)Fuc(1)NeuAc(1) | C[+57]*2, N[+2060] | 1369.8927 | 4 | 5476.549 | 5476.5418 | 0 | 1.3 | 457 | Specific | 920.93 | 858.43 | 858.43 | 18.27 | 64 | >sp\|P01833\|PIGR_HUMAN Polymeric immunoglobulin receptor OS=Homo sapiens OX=9606 GN=PIGR PE=1 SV=4 | 902 | 20201115_07433_1ng_glyco_hcd203040_compound15.19206.19206.4 | controllerType=0 controllerNumber=1 scan=19206 | 38.2756 |
| 17347:5 | 7 | K.IIEGEPNLKVPGN[+2350.830]VTAVLGETLKVPC[+57.021]HFPC[+57.021]K.F | HexNAc(4)Hex(5)Fuc(1)NeuAc(2) | C[+57]*2, N[+2351] | 1154.3358 | 5 | 5767.65 | 5767.6372 | 0 | 2.2 | 457 | Specific | 982.89 | 760.71 | 760.71 | 20.22 | 64 | >sp\|P01833\|PIGR_HUMAN Polymeric immunoglobulin receptor OS=Homo sapiens OX=9606 GN=PIGR PE=1 SV=4 | 902 | 20201115_07433_1ng_glyco_hcd203040_compound15.20732.20732.5 | controllerType=0 controllerNumber=1 scan=20732 | 41.1145 |
| 15867:4 | 7 | K.IIEGEPNLKVPGN[+2205.793]VTAVLGETLKVPC[+57.021]HFPC[+57.021]K.F | HexNAc(4)Hex(5)Fuc(2)NeuAc(1) | C[+57]*2, N[+2206] | 1406.4085 | 4 | 5622.612 | 5622.5997 | 0 | 2.2 | 457 | Specific | 1012.25 | 963.64 | 963.64 | 20.68 | 64 | >sp\|P01833\|PIGR_HUMAN Polymeric immunoglobulin receptor OS=Homo sapiens OX=9606 GN=PIGR PE=1 SV=4 | 902 | 20201115_07433_1ng_glyco_hcd203040_compound15.19141.19141.4 | controllerType=0 controllerNumber=1 scan=19141 | 38.1464 |
| 17355:4 | 7 | K.IIEGEPNLKVPGN[+2350.830]VTAVLGETLKVPC[+57.021]HFPC[+57.021]K.F | HexNAc(4)Hex(5)Fuc(1)NeuAc(2) | C[+57]*2, N[+2351] | 1442.6708 | 4 | 5767.6613 | 5767.6372 | 0 | 4.2 | 457 | Specific | 1023.94 | 794.23 | 794.23 | 20.21 | 64 | >sp\|P01833\|PIGR_HUMAN Polymeric immunoglobulin receptor OS=Homo sapiens OX=9606 GN=PIGR PE=1 SV=4 | 902 | 20201115_07433_1ng_glyco_hcd203040_compound15.20741.20741.4 | controllerType=0 controllerNumber=1 scan=20741 | 41.1313 |
| 15928:5 | 7 | K.IIEGEPNLKVPGN[+2059.735]VTAVLGETLKVPC[+57.021]HFPC[+57.021]K.F | HexNAc(4)Hex(5)Fuc(1)NeuAc(1) | C[+57]*2, N[+2060] | 1096.1172 | 5 | 5476.5568 | 5476.5418 | 0 | 2.7 | 457 | Specific | 1028.28 | 883.15 | 883.15 | 21.22 | 64 | >sp\|P01833\|PIGR_HUMAN Polymeric immunoglobulin receptor OS=Homo sapiens OX=9606 GN=PIGR PE=1 SV=4 | 902 | 20201115_07433_1ng_glyco_hcd203040_compound15.19205.19205.5 | controllerType=0 controllerNumber=1 scan=19205 | 38.2742 |
| 15857:5 | 7 | K.IIEGEPNLKVPGN[+2204.772]VTAVLGETLKVPC[+57.021]HFPC[+57.021]K.F | HexNAc(4)Hex(5)NeuAc(2) | C[+57]*2, N[+2205] | 1125.1283 | 5 | 5621.6124 | 5621.5793 | 0 | 5.9 | 457 | Specific | 1036.08 | 797.88 | 797.88 | 20.33 | 64 | >sp\|P01833\|PIGR_HUMAN Polymeric immunoglobulin receptor OS=Homo sapiens OX=9606 GN=PIGR PE=1 SV=4 | 902 | 20201115_07433_1ng_glyco_hcd203040_compound15.19131.19131.5 | controllerType=0 controllerNumber=1 scan=19131 | 38.1234 |
| 13919:3 | 7 | K.VPGN[+2205.793]VTAVLGETLK.V | HexNAc(4)Hex(5)Fuc(2)NeuAc(1) | N[+2206] | 1201.8732 | 3 | 3603.6051 | 3603.5927 | 0 | 3.4 | 466 | Specific | 444.43 | 444.43 | 444.43 | 9.93 | 64 | >sp\|P01833\|PIGR_HUMAN Polymeric immunoglobulin receptor OS=Homo sapiens OX=9606 GN=PIGR PE=1 SV=4 | 902 | 20201115_07433_1ng_glyco_hcd203040_compound15.17121.17121.3 | controllerType=0 controllerNumber=1 scan=17121 | 34.427 |
| 13921:3 | 7 | K.VPGN[+2205.793]VTAVLGETLK.V | HexNAc(4)Hex(5)Fuc(2)NeuAc(1) | N[+2206] | 1201.8732 | 3 | 3603.6051 | 3603.5927 | 0 | 3.4 | 466 | Specific | 532.6 | 532.6 | 532.6 | 10.41 | 64 | >sp\|P01833\|PIGR_HUMAN Polymeric immunoglobulin receptor OS=Homo sapiens OX=9606 GN=PIGR PE=1 SV=4 | 902 | 20201115_07433_1ng_glyco_hcd203040_compound15.17123.17123.3 | controllerType=0 controllerNumber=1 scan=17123 | 34.4307 |
| 13960:4 | 7 | K.VPGN[+2204.772]VTAVLGETLKVPC[+57.021]HFPC[+57.021]K.F | HexNAc(4)Hex(5)NeuAc(2) | C[+57]*2, N[+2205] | 1157.7576 | 4 | 4628.0086 | 4628.0298 | 0 | -4.6 | 466 | Specific | 468.37 | 350.11 | 350.11 | 6.51 | 64 | >sp\|P01833\|PIGR_HUMAN Polymeric immunoglobulin receptor OS=Homo sapiens OX=9606 GN=PIGR PE=1 SV=4 | 902 | 20201115_07433_1ng_glyco_hcd203040_compound15.17164.17164.3 | controllerType=0 controllerNumber=1 scan=17164 | 34.5194 |
| 14105:4 | 7 | K.VPGN[+2059.735]VTAVLGETLKVPC[+57.021]HFPC[+57.021]K.F | HexNAc(4)Hex(5)Fuc(1)NeuAc(1) | C[+57]*2, N[+2060] | 1121.5074 | 4 | 4483.0076 | 4482.9923 | 0 | 3.4 | 466 | Specific | 567.32 | 487.52 | 487.52 | 10.27 | 64 | >sp\|P01833\|PIGR_HUMAN Polymeric immunoglobulin receptor OS=Homo sapiens OX=9606 GN=PIGR PE=1 SV=4 | 902 | 20201115_07433_1ng_glyco_hcd203040_compound15.17315.17315.4 | controllerType=0 controllerNumber=1 scan=17315 | 34.821 |
| 13958:4 | 7 | K.VPGN[+2204.772]VTAVLGETLKVPC[+57.021]HFPC[+57.021]K.F | HexNAc(4)Hex(5)NeuAc(2) | C[+57]*2, N[+2205] | 1157.7576 | 4 | 4628.0086 | 4628.0298 | 0 | -4.6 | 466 | Specific | 597.84 | 445.86 | 445.86 | 10.18 | 64 | >sp\|P01833\|PIGR_HUMAN Polymeric immunoglobulin receptor OS=Homo sapiens OX=9606 GN=PIGR PE=1 SV=4 | 902 | 20201115_07433_1ng_glyco_hcd203040_compound15.17162.17162.3 | controllerType=0 controllerNumber=1 scan=17162 | 34.509 |
| 14107:4 | 7 | K.VPGN[+2059.735]VTAVLGETLKVPC[+57.021]HFPC[+57.021]K.F | HexNAc(4)Hex(5)Fuc(1)NeuAc(1) | C[+57]*2, N[+2060] | 1121.5074 | 4 | 4483.0076 | 4482.9923 | 0 | 3.4 | 466 | Specific | 658.01 | 478.22 | 478.22 | 13.56 | 64 | >sp\|P01833\|PIGR_HUMAN Polymeric immunoglobulin receptor OS=Homo sapiens OX=9606 GN=PIGR PE=1 SV=4 | 902 | 20201115_07433_1ng_glyco_hcd203040_compound15.17317.17317.4 | controllerType=0 controllerNumber=1 scan=17317 | 34.8249 |
| 14032:4 | 7 | K.VPGN[+2205.793]VTAVLGETLKVPC[+57.021]HFPC[+57.021]K.F | HexNAc(4)Hex(5)Fuc(2)NeuAc(1) | C[+57]*2, N[+2206] | 1158.0218 | 4 | 4629.0653 | 4629.0502 | 0 | 3.2 | 466 | Specific | 699.52 | 657.89 | 657.89 | 13.56 | 64 | >sp\|P01833\|PIGR_HUMAN Polymeric immunoglobulin receptor OS=Homo sapiens OX=9606 GN=PIGR PE=1 SV=4 | 902 | 20201115_07433_1ng_glyco_hcd203040_compound15.17239.17239.4 | controllerType=0 controllerNumber=1 scan=17239 | 34.6707 |
| 14030:4 | 7 | K.VPGN[+2205.793]VTAVLGETLKVPC[+57.021]HFPC[+57.021]K.F | HexNAc(4)Hex(5)Fuc(2)NeuAc(1) | C[+57]*2, N[+2206] | 1158.0218 | 4 | 4629.0653 | 4629.0502 | 0 | 3.2 | 466 | Specific | 735.02 | 474.08 | 474.08 | 14.56 | 64 | >sp\|P01833\|PIGR_HUMAN Polymeric immunoglobulin receptor OS=Homo sapiens OX=9606 GN=PIGR PE=1 SV=4 | 902 | 20201115_07433_1ng_glyco_hcd203040_compound15.17237.17237.4 | controllerType=0 controllerNumber=1 scan=17237 | 34.6652 |
| 05973:3 | 7 | K.WN[+2059.735]NTGC[+57.021]QALPSQDEGPSK.A | HexNAc(4)Hex(5)Fuc(1)NeuAc(1) | C[+57], N[+2060] | 1350.211 | 3 | 4048.6184 | 4048.6116 | 0 | 1.7 | 498 | Specific | 614.6 | 544.11 | 544.11 | 12.77 | 64 | >sp\|P01833\|PIGR_HUMAN Polymeric immunoglobulin receptor OS=Homo sapiens OX=9606 GN=PIGR PE=1 SV=4 | 902 | 20201115_07433_1ng_glyco_hcd203040_compound15.8844.8844.3 | controllerType=0 controllerNumber=1 scan=8844 | 20.6375 |
| 05975:3 | 7 | K.WN[+2059.735]NTGC[+57.021]QALPSQDEGPSK.A | HexNAc(4)Hex(5)Fuc(1)NeuAc(1) | C[+57], N[+2060] | 1350.211 | 3 | 4048.6184 | 4048.6116 | 0 | 1.7 | 498 | Specific | 671.56 | 595.6 | 595.6 | 13.77 | 64 | >sp\|P01833\|PIGR_HUMAN Polymeric immunoglobulin receptor OS=Homo sapiens OX=9606 GN=PIGR PE=1 SV=4 | 902 | 20201115_07433_1ng_glyco_hcd203040_compound15.8846.8846.3 | controllerType=0 controllerNumber=1 scan=8846 | 20.6418 |
| 06010:3 | 7 | K.WN[+1913.677]NTGC[+57.021]QALPSQDEGPSK.A | HexNAc(4)Hex(5)NeuAc(1) | C[+57], N[+1914] | 1301.5283 | 3 | 3902.5702 | 3902.5537 | 0 | 4.2 | 498 | Specific | 760.74 | 708.06 | 708.06 | 15.71 | 64 | >sp\|P01833\|PIGR_HUMAN Polymeric immunoglobulin receptor OS=Homo sapiens OX=9606 GN=PIGR PE=1 SV=4 | 902 | 20201115_07433_1ng_glyco_hcd203040_compound15.8882.8882.3 | controllerType=0 controllerNumber=1 scan=8882 | 20.702 |
| 06008:3 | 7 | K.WN[+1913.677]NTGC[+57.021]QALPSQDEGPSK.A | HexNAc(4)Hex(5)NeuAc(1) | C[+57], N[+1914] | 1301.5283 | 3 | 3902.5702 | 3902.5537 | 0 | 4.2 | 498 | Specific | 843.92 | 788.82 | 788.82 | 16.71 | 64 | >sp\|P01833\|PIGR_HUMAN Polymeric immunoglobulin receptor OS=Homo sapiens OX=9606 GN=PIGR PE=1 SV=4 | 902 | 20201115_07433_1ng_glyco_hcd203040_compound15.8880.8880.3 | controllerType=0 controllerNumber=1 scan=8880 | 20.6976 |
| 18038:3 | 11 | R.LSLHRPALEDLLLGSEAN[+1378.476]LTC[+57.021]TLTGLR.D | HexNAc(2)Hex(6) | C[+57], N[+1378] | 1448.0323 | 3 | 4342.0823 | 4342.074 | 0 | 1.9 | 127 | Specific | 229.41 | 229.41 | 229.41 | 4.45 | 44 | >sp\|P01876\|IGHA1_HUMAN Immunoglobulin heavy constant alpha 1 OS=Homo sapiens OX=9606 GN=IGHA1 PE=1 SV=2 | 255 | 20201115_07433_1ng_glyco_hcd203040_compound15.21452.21452.3 | controllerType=0 controllerNumber=1 scan=21452 | 42.6264 |
| 18135:3 | 11 | R.LSLHRPALEDLLLGSEAN[+1095.397]LTC[+57.021]TLTGLR.D | HexNAc(3)Hex(3) | C[+57], N[+1095] | 1353.6742 | 3 | 4059.008 | 4058.9949 | 0 | 3.2 | 127 | Specific | 357.57 | 357.57 | 357.57 | 7.88 | 44 | >sp\|P01876\|IGHA1_HUMAN Immunoglobulin heavy constant alpha 1 OS=Homo sapiens OX=9606 GN=IGHA1 PE=1 SV=2 | 255 | 20201115_07433_1ng_glyco_hcd203040_compound15.21553.21553.3 | controllerType=0 controllerNumber=1 scan=21553 | 42.8572 |
| 18075:3 | 11 | R.LSLHRPALEDLLLGSEAN[+1216.423]LTC[+57.021]TLTGLR.D | HexNAc(2)Hex(5) | C[+57], N[+1216] | 1394.0157 | 3 | 4180.0325 | 4180.0211 | 0 | 2.7 | 127 | Specific | 392.68 | 392.68 | 392.68 | 9.66 | 44 | >sp\|P01876\|IGHA1_HUMAN Immunoglobulin heavy constant alpha 1 OS=Homo sapiens OX=9606 GN=IGHA1 PE=1 SV=2 | 255 | 20201115_07433_1ng_glyco_hcd203040_compound15.21491.21491.3 | controllerType=0 controllerNumber=1 scan=21491 | 42.7267 |
| 18125:4 | 11 | R.LSLHRPALEDLLLGSEAN[+1095.397]LTC[+57.021]TLTGLR.D | HexNAc(3)Hex(3) | C[+57], N[+1095] | 1015.5073 | 4 | 4059.0074 | 4058.9949 | 0 | 3.1 | 127 | Specific | 394.22 | 391.25 | 391.25 | 10.03 | 44 | >sp\|P01876\|IGHA1_HUMAN Immunoglobulin heavy constant alpha 1 OS=Homo sapiens OX=9606 GN=IGHA1 PE=1 SV=2 | 255 | 20201115_07433_1ng_glyco_hcd203040_compound15.21543.21543.4 | controllerType=0 controllerNumber=1 scan=21543 | 42.8362 |
| 18035:3 | 11 | R.LSLHRPALEDLLLGSEAN[+1378.476]LTC[+57.021]TLTGLR.D | HexNAc(2)Hex(6) | C[+57], N[+1378] | 1448.0323 | 3 | 4342.0823 | 4342.074 | 0 | 1.9 | 127 | Specific | 398.47 | 398.47 | 398.47 | 9.66 | 44 | >sp\|P01876\|IGHA1_HUMAN Immunoglobulin heavy constant alpha 1 OS=Homo sapiens OX=9606 GN=IGHA1 PE=1 SV=2 | 255 | 20201115_07433_1ng_glyco_hcd203040_compound15.21449.21449.3 | controllerType=0 controllerNumber=1 scan=21449 | 42.6202 |
| 18047:3 | 11 | R.LSLHRPALEDLLLGSEAN[+1460.529]LTC[+57.021]TLTGLR.D | HexNAc(4)Hex(4) | C[+57], N[+1461] | 1475.3841 | 3 | 4424.1377 | 4424.1271 | 0 | 2.4 | 127 | Specific | 544.44 | 538 | 538 | 10.36 | 44 | >sp\|P01876\|IGHA1_HUMAN Immunoglobulin heavy constant alpha 1 OS=Homo sapiens OX=9606 GN=IGHA1 PE=1 SV=2 | 255 | 20201115_07433_1ng_glyco_hcd203040_compound15.21461.21461.3 | controllerType=0 controllerNumber=1 scan=21461 | 42.6492 |
| 18122:4 | 11 | R.LSLHRPALEDLLLGSEAN[+1095.397]LTC[+57.021]TLTGLR.D | HexNAc(3)Hex(3) | C[+57], N[+1095] | 1015.5073 | 4 | 4059.0074 | 4058.9949 | 0 | 3.1 | 127 | Specific | 548.61 | 548.61 | 548.61 | 10.73 | 44 | >sp\|P01876\|IGHA1_HUMAN Immunoglobulin heavy constant alpha 1 OS=Homo sapiens OX=9606 GN=IGHA1 PE=1 SV=2 | 255 | 20201115_07433_1ng_glyco_hcd203040_compound15.21540.21540.4 | controllerType=0 controllerNumber=1 scan=21540 | 42.8308 |
| 17991:3 | 11 | R.LSLHRPALEDLLLGSEAN[+1663.608]LTC[+57.021]TLTGLR.D | HexNAc(5)Hex(4) | C[+57], N[+1664] | 1543.0779 | 3 | 4627.2193 | 4627.2064 | 0 | 2.8 | 127 | Specific | 587.46 | 546.49 | 546.49 | 11.76 | 44 | >sp\|P01876\|IGHA1_HUMAN Immunoglobulin heavy constant alpha 1 OS=Homo sapiens OX=9606 GN=IGHA1 PE=1 SV=2 | 255 | 20201115_07433_1ng_glyco_hcd203040_compound15.21403.21403.3 | controllerType=0 controllerNumber=1 scan=21403 | 42.5146 |
| 18078:3 | 11 | R.LSLHRPALEDLLLGSEAN[+1216.423]LTC[+57.021]TLTGLR.D | HexNAc(2)Hex(5) | C[+57], N[+1216] | 1394.0157 | 3 | 4180.0325 | 4180.0211 | 0 | 2.7 | 127 | Specific | 588.44 | 563.25 | 563.25 | 11.76 | 44 | >sp\|P01876\|IGHA1_HUMAN Immunoglobulin heavy constant alpha 1 OS=Homo sapiens OX=9606 GN=IGHA1 PE=1 SV=2 | 255 | 20201115_07433_1ng_glyco_hcd203040_compound15.21494.21494.3 | controllerType=0 controllerNumber=1 scan=21494 | 42.7371 |
| 17981:4 | 11 | R.LSLHRPALEDLLLGSEAN[+1622.582]LTC[+57.021]TLTGLR.D | HexNAc(4)Hex(5) | C[+57], N[+1623] | 1147.3038 | 4 | 4586.1932 | 4586.1799 | 0 | 2.9 | 127 | Specific | 592.86 | 513.44 | 513.44 | 12.13 | 44 | >sp\|P01876\|IGHA1_HUMAN Immunoglobulin heavy constant alpha 1 OS=Homo sapiens OX=9606 GN=IGHA1 PE=1 SV=2 | 255 | 20201115_07433_1ng_glyco_hcd203040_compound15.21392.21392.2 | controllerType=0 controllerNumber=1 scan=21392 | 42.4901 |
| 17985:4 | 11 | R.LSLHRPALEDLLLGSEAN[+1663.608]LTC[+57.021]TLTGLR.D | HexNAc(5)Hex(4) | C[+57], N[+1664] | 1157.5601 | 4 | 4627.2186 | 4627.2064 | 0 | 2.6 | 127 | Specific | 597.52 | 597.52 | 597.52 | 12.3 | 44 | >sp\|P01876\|IGHA1_HUMAN Immunoglobulin heavy constant alpha 1 OS=Homo sapiens OX=9606 GN=IGHA1 PE=1 SV=2 | 255 | 20201115_07433_1ng_glyco_hcd203040_compound15.21397.21397.4 | controllerType=0 controllerNumber=1 scan=21397 | 42.5023 |
| 18002:3 | 11 | R.LSLHRPALEDLLLGSEAN[+1622.582]LTC[+57.021]TLTGLR.D | HexNAc(4)Hex(5) | C[+57], N[+1623] | 1529.4022 | 3 | 4586.1919 | 4586.1799 | 0 | 2.6 | 127 | Specific | 598.7 | 595.6 | 595.6 | 11.76 | 44 | >sp\|P01876\|IGHA1_HUMAN Immunoglobulin heavy constant alpha 1 OS=Homo sapiens OX=9606 GN=IGHA1 PE=1 SV=2 | 255 | 20201115_07433_1ng_glyco_hcd203040_compound15.21414.21414.3 | controllerType=0 controllerNumber=1 scan=21414 | 42.5358 |

Table S4. Cont.

| Query #:z | Protein Rank | Glycopeptide | Glycan | Modification Type(s) | Observed m/z | z | Observed (M+H) | Calc. mass (M+H) | Off-by-x error | Mass error (ppm) | StartPosition | Cleavage | Score | Delta | Delta Mod | \|Log Prob\| | # of unique peptides | Protein Name | Protein DB number | Comment | Scan # | Scan Time |
|---|---|---|---|---|---|---|---|---|---|---|---|---|---|---|---|---|---|---|---|---|---|---|
| 18049:3 | 11 | R.LSLHRPALEDLLLGSEAN[+1460.529]LTC[+57.021]TLTGLR.D | HexNAc(4)Hex(4) | C[+57], N[+1461] | 1475.3841 | 3 | 4424.1377 | 4424.1271 | 0 | 2.4 | 127 | Specific | 600.71 | 519.36 | 519.36 | 12.76 | 44 | >sp\|P01876\|IGHA1_HUMAN Immunoglobulin heavy constant alpha 1 OS=Homo sapiens OX=9606 GN=IGHA1 PE=1 SV=2 | 255 | 20201115_07433_1ng_glyco_hcd203040_compound15.21463.21463.3 | controllerType=0 controllerNumber=1 scan=21463 | 42.6575 |
| 18005:3 | 11 | R.LSLHRPALEDLLLGSEAN[+1622.582]LTC[+57.021]TLTGLR.D | HexNAc(4)Hex(5) | C[+57], N[+1623] | 1529.4022 | 3 | 4586.1919 | 4586.1799 | 0 | 2.6 | 127 | Specific | 604.78 | 523 | 523 | 12.76 | 44 | >sp\|P01876\|IGHA1_HUMAN Immunoglobulin heavy constant alpha 1 OS=Homo sapiens OX=9606 GN=IGHA1 PE=1 SV=2 | 255 | 20201115_07433_1ng_glyco_hcd203040_compound15.21417.21417.3 | controllerType=0 controllerNumber=1 scan=21417 | 42.5448 |
| 17994:3 | 11 | R.LSLHRPALEDLLLGSEAN[+1663.608]LTC[+57.021]TLTGLR.D | HexNAc(5)Hex(4) | C[+57], N[+1664] | 1543.0779 | 3 | 4627.2193 | 4627.2064 | 0 | 2.8 | 127 | Specific | 627.14 | 565.42 | 565.42 | 12.76 | 44 | >sp\|P01876\|IGHA1_HUMAN Immunoglobulin heavy constant alpha 1 OS=Homo sapiens OX=9606 GN=IGHA1 PE=1 SV=2 | 255 | 20201115_07433_1ng_glyco_hcd203040_compound15.21406.21406.3 | controllerType=0 controllerNumber=1 scan=21406 | 42.5232 |
| 18010:4 | 11 | R.LSLHRPALEDLLLGSEAN[+1419.502]LTC[+57.021]TLTGLR.D | HexNAc(3)Hex(5) | C[+57], N[+1420] | 1096.5351 | 4 | 4383.1185 | 4383.1005 | 0 | 4.1 | 127 | Specific | 631.37 | 631.37 | 631.37 | 13.13 | 44 | >sp\|P01876\|IGHA1_HUMAN Immunoglobulin heavy constant alpha 1 OS=Homo sapiens OX=9606 GN=IGHA1 PE=1 SV=2 | 255 | 20201115_07433_1ng_glyco_hcd203040_compound15.21423.21423.4 | controllerType=0 controllerNumber=1 scan=21423 | 42.5584 |
| 18016:3 | 11 | R.LSLHRPALEDLLLGSEAN[+1501.555]LTC[+57.021]TLTGLR.D | HexNAc(5)Hex(3) | C[+57], N[+1502] | 1489.061 | 3 | 4465.1684 | 4465.1536 | 0 | 3.3 | 127 | Specific | 634.93 | 634.93 | 634.93 | 12.6 | 44 | >sp\|P01876\|IGHA1_HUMAN Immunoglobulin heavy constant alpha 1 OS=Homo sapiens OX=9606 GN=IGHA1 PE=1 SV=2 | 255 | 20201115_07433_1ng_glyco_hcd203040_compound15.21429.21429.3 | controllerType=0 controllerNumber=1 scan=21429 | 42.5693 |
| 18019:3 | 11 | R.LSLHRPALEDLLLGSEAN[+1501.555]LTC[+57.021]TLTGLR.D | HexNAc(5)Hex(3) | C[+57], N[+1502] | 1489.061 | 3 | 4465.1684 | 4465.1536 | 0 | 3.3 | 127 | Specific | 651.78 | 595.87 | 595.87 | 13.6 | 44 | >sp\|P01876\|IGHA1_HUMAN Immunoglobulin heavy constant alpha 1 OS=Homo sapiens OX=9606 GN=IGHA1 PE=1 SV=2 | 255 | 20201115_07433_1ng_glyco_hcd203040_compound15.21432.21432.3 | controllerType=0 controllerNumber=1 scan=21432 | 42.58 |
| 17955:3 | 11 | R.LSLHRPALEDLLLGSEAN[+1540.529]LTC[+57.021]TLTGLR.D | HexNAc(2)Hex(7) | C[+57], N[+1541] | 1502.053 | 3 | 4504.1446 | 4504.1268 | 0 | 3.9 | 127 | Specific | 661.34 | 580.72 | 580.72 | 13.6 | 44 | >sp\|P01876\|IGHA1_HUMAN Immunoglobulin heavy constant alpha 1 OS=Homo sapiens OX=9606 GN=IGHA1 PE=1 SV=2 | 255 | 20201115_07433_1ng_glyco_hcd203040_compound15.21365.21365.3 | controllerType=0 controllerNumber=1 scan=21365 | 42.4141 |
| 18107:4 | 11 | R.LSLHRPALEDLLLGSEAN[+1054.370]LTC[+57.021]TLTGLR.D | HexNAc(2)Hex(4) | C[+57], N[+1054] | 1005.2512 | 4 | 4017.983 | 4017.9683 | 0 | 3.7 | 127 | Specific | 667.49 | 646.46 | 646.46 | 14.13 | 44 | >sp\|P01876\|IGHA1_HUMAN Immunoglobulin heavy constant alpha 1 OS=Homo sapiens OX=9606 GN=IGHA1 PE=1 SV=2 | 255 | 20201115_07433_1ng_glyco_hcd203040_compound15.21524.21524.4 | controllerType=0 controllerNumber=1 scan=21524 | 42.7978 |
| 17919:3 | 11 | R.LSLHRPALEDLLLGSEAN[+1702.581]LTC[+57.021]TLTGLR.D | HexNAc(2)Hex(8) | C[+57], N[+1703] | 1556.0678 | 3 | 4666.1889 | 4666.1796 | 0 | 2 | 127 | Specific | 682 | 662.25 | 662.25 | 13.76 | 44 | >sp\|P01876\|IGHA1_HUMAN Immunoglobulin heavy constant alpha 1 OS=Homo sapiens OX=9606 GN=IGHA1 PE=1 SV=2 | 255 | 20201115_07433_1ng_glyco_hcd203040_compound15.21327.21327.3 | controllerType=0 controllerNumber=1 scan=21327 | 42.3231 |
| 18031:4 | 11 | R.LSLHRPALEDLLLGSEAN[+1460.529]LTC[+57.021]TLTGLR.D | HexNAc(4)Hex(4) | C[+57], N[+1461] | 1106.7911 | 4 | 4424.1424 | 4424.1271 | 0 | 3.5 | 127 | Specific | 693.86 | 681.85 | 681.85 | 14.13 | 44 | >sp\|P01876\|IGHA1_HUMAN Immunoglobulin heavy constant alpha 1 OS=Homo sapiens OX=9606 GN=IGHA1 PE=1 SV=2 | 255 | 20201115_07433_1ng_glyco_hcd203040_compound15.21445.21445.4 | controllerType=0 controllerNumber=1 scan=21445 | 42.6124 |
| 17917:3 | 11 | R.LSLHRPALEDLLLGSEAN[+1702.581]LTC[+57.021]TLTGLR.D | HexNAc(2)Hex(8) | C[+57], N[+1703] | 1556.0678 | 3 | 4666.1889 | 4666.1796 | 0 | 2 | 127 | Specific | 703.16 | 703.16 | 703.16 | 14.76 | 44 | >sp\|P01876\|IGHA1_HUMAN Immunoglobulin heavy constant alpha 1 OS=Homo sapiens OX=9606 GN=IGHA1 PE=1 SV=2 | 255 | 20201115_07433_1ng_glyco_hcd203040_compound15.21325.21325.3 | controllerType=0 controllerNumber=1 scan=21325 | 42.317 |
| 18009:4 | 11 | R.LSLHRPALEDLLLGSEAN[+1501.555]LTC[+57.021]TLTGLR.D | HexNAc(5)Hex(3) | C[+57], N[+1502] | 1117.0474 | 4 | 4465.1678 | 4465.1536 | 0 | 3.2 | 127 | Specific | 711.93 | 711.93 | 711.93 | 15.13 | 44 | >sp\|P01876\|IGHA1_HUMAN Immunoglobulin heavy constant alpha 1 OS=Homo sapiens OX=9606 GN=IGHA1 PE=1 SV=2 | 255 | 20201115_07433_1ng_glyco_hcd203040_compound15.21422.21422.4 | controllerType=0 controllerNumber=1 scan=21422 | 42.557 |
| 17953:3 | 11 | R.LSLHRPALEDLLLGSEAN[+1540.529]LTC[+57.021]TLTGLR.D | HexNAc(2)Hex(7) | C[+57], N[+1541] | 1502.053 | 3 | 4504.1446 | 4504.1268 | 0 | 3.9 | 127 | Specific | 715.68 | 682.25 | 682.25 | 14.6 | 44 | >sp\|P01876\|IGHA1_HUMAN Immunoglobulin heavy constant alpha 1 OS=Homo sapiens OX=9606 GN=IGHA1 PE=1 SV=2 | 255 | 20201115_07433_1ng_glyco_hcd203040_compound15.21363.21363.3 | controllerType=0 controllerNumber=1 scan=21363 | 42.4077 |
| 17987:4 | 11 | R.LSLHRPALEDLLLGSEAN[+1663.608]LTC[+57.021]TLTGLR.D | HexNAc(5)Hex(4) | C[+57], N[+1664] | 1157.5601 | 4 | 4627.2186 | 4627.2064 | 0 | 2.6 | 127 | Specific | 729.4 | 680.96 | 680.96 | 15.3 | 44 | >sp\|P01876\|IGHA1_HUMAN Immunoglobulin heavy constant alpha 1 OS=Homo sapiens OX=9606 GN=IGHA1 PE=1 SV=2 | 255 | 20201115_07433_1ng_glyco_hcd203040_compound15.21399.21399.4 | controllerType=0 controllerNumber=1 scan=21399 | 42.5071 |
| 17983:4 | 11 | R.LSLHRPALEDLLLGSEAN[+1622.582]LTC[+57.021]TLTGLR.D | HexNAc(4)Hex(5) | C[+57], N[+1623] | 1147.3038 | 4 | 4586.1932 | 4586.1799 | 0 | 2.9 | 127 | Specific | 773.86 | 545.48 | 545.48 | 16.13 | 44 | >sp\|P01876\|IGHA1_HUMAN Immunoglobulin heavy constant alpha 1 OS=Homo sapiens OX=9606 GN=IGHA1 PE=1 SV=2 | 255 | 20201115_07433_1ng_glyco_hcd203040_compound15.21394.21394.2 | controllerType=0 controllerNumber=1 scan=21394 | 42.4961 |
| 18011:4 | 11 | R.LSLHRPALEDLLLGSEAN[+1501.555]LTC[+57.021]TLTGLR.D | HexNAc(5)Hex(3) | C[+57], N[+1502] | 1117.0474 | 4 | 4465.1678 | 4465.1536 | 0 | 3.2 | 127 | Specific | 802.05 | 800.66 | 800.66 | 17.13 | 44 | >sp\|P01876\|IGHA1_HUMAN Immunoglobulin heavy constant alpha 1 OS=Homo sapiens OX=9606 GN=IGHA1 PE=1 SV=2 | 255 | 20201115_07433_1ng_glyco_hcd203040_compound15.21424.21424.4 | controllerType=0 controllerNumber=1 scan=21424 | 42.5629 |
| 18033:4 | 11 | R.LSLHRPALEDLLLGSEAN[+1460.529]LTC[+57.021]TLTGLR.D | HexNAc(4)Hex(4) | C[+57], N[+1461] | 1106.7911 | 4 | 4424.1424 | 4424.1271 | 0 | 3.5 | 127 | Specific | 842.38 | 701.13 | 701.13 | 17.13 | 44 | >sp\|P01876\|IGHA1_HUMAN Immunoglobulin heavy constant alpha 1 OS=Homo sapiens OX=9606 GN=IGHA1 PE=1 SV=2 | 255 | 20201115_07433_1ng_glyco_hcd203040_compound15.21447.21447.4 | controllerType=0 controllerNumber=1 scan=21447 | 42.6179 |
| 18110:4 | 11 | R.LSLHRPALEDLLLGSEAN[+1054.370]LTC[+57.021]TLTGLR.D | HexNAc(2)Hex(4) | C[+57], N[+1054] | 1005.2512 | 4 | 4017.983 | 4017.9683 | 0 | 3.7 | 127 | Specific | 873.03 | 821.95 | 821.95 | 18.13 | 44 | >sp\|P01876\|IGHA1_HUMAN Immunoglobulin heavy constant alpha 1 OS=Homo sapiens OX=9606 GN=IGHA1 PE=1 SV=2 | 255 | 20201115_07433_1ng_glyco_hcd203040_compound15.21527.21527.4 | controllerType=0 controllerNumber=1 scan=21527 | 42.8057 |
| 18026:4 | 11 | R.LSLHRPALEDLLLGSEAN[+1216.423]LTC[+57.021]TLTGLR.D | HexNAc(2)Hex(5) | C[+57], N[+1216] | 1045.7641 | 4 | 4180.0345 | 4180.0211 | 0 | 3.2 | 127 | Specific | 893.34 | 874.38 | 874.38 | 18.13 | 44 | >sp\|P01876\|IGHA1_HUMAN Immunoglobulin heavy constant alpha 1 OS=Homo sapiens OX=9606 GN=IGHA1 PE=1 SV=2 | 255 | 20201115_07433_1ng_glyco_hcd203040_compound15.21439.21439.4 | controllerType=0 controllerNumber=1 scan=21439 | 42.5965 |
| 18012:4 | 11 | R.LSLHRPALEDLLLGSEAN[+1419.502]LTC[+57.021]TLTGLR.D | HexNAc(3)Hex(5) | C[+57], N[+1420] | 1096.5351 | 4 | 4383.1185 | 4383.1005 | 0 | 4.1 | 127 | Specific | 905.65 | 870.23 | 870.23 | 19.13 | 44 | >sp\|P01876\|IGHA1_HUMAN Immunoglobulin heavy constant alpha 1 OS=Homo sapiens OX=9606 GN=IGHA1 PE=1 SV=2 | 255 | 20201115_07433_1ng_glyco_hcd203040_compound15.21425.21425.4 | controllerType=0 controllerNumber=1 scan=21425 | 42.5645 |
| 18055:4 | 11 | R.LSLHRPALEDLLLGSEAN[+1257.449]LTC[+57.021]TLTGLR.D | HexNAc(3)Hex(4) | C[+57], N[+1257] | 1056.0196 | 4 | 4221.0565 | 4221.0477 | 0 | 2.1 | 127 | Specific | 926.56 | 856.06 | 856.06 | 19.3 | 44 | >sp\|P01876\|IGHA1_HUMAN Immunoglobulin heavy constant alpha 1 OS=Homo sapiens OX=9606 GN=IGHA1 PE=1 SV=2 | 255 | 20201115_07433_1ng_glyco_hcd203040_compound15.21470.21470.4 | controllerType=0 controllerNumber=1 scan=21470 | 42.6752 |
| 17935:4 | 11 | R.LSLHRPALEDLLLGSEAN[+1540.529]LTC[+57.021]TLTGLR.D | HexNAc(2)Hex(7) | C[+57], N[+1541] | 1126.7915 | 4 | 4504.1444 | 4504.1268 | 0 | 3.9 | 127 | Specific | 948.8 | 869.86 | 869.86 | 19.13 | 44 | >sp\|P01876\|IGHA1_HUMAN Immunoglobulin heavy constant alpha 1 OS=Homo sapiens OX=9606 GN=IGHA1 PE=1 SV=2 | 255 | 20201115_07433_1ng_glyco_hcd203040_compound15.21344.21344.4 | controllerType=0 controllerNumber=1 scan=21344 | 42.3548 |
| 18028:4 | 11 | R.LSLHRPALEDLLLGSEAN[+1216.423]LTC[+57.021]TLTGLR.D | HexNAc(2)Hex(5) | C[+57], N[+1216] | 1045.7641 | 4 | 4180.0345 | 4180.0211 | 0 | 3.2 | 127 | Specific | 1002.78 | 1002.78 | 1002.78 | 21.13 | 44 | >sp\|P01876\|IGHA1_HUMAN Immunoglobulin heavy constant alpha 1 OS=Homo sapiens OX=9606 GN=IGHA1 PE=1 SV=2 | 255 | 20201115_07433_1ng_glyco_hcd203040_compound15.21441.21441.4 | controllerType=0 controllerNumber=1 scan=21441 | 42.6046 |
| 17937:4 | 11 | R.LSLHRPALEDLLLGSEAN[+1540.529]LTC[+57.021]TLTGLR.D | HexNAc(2)Hex(7) | C[+57], N[+1541] | 1126.7915 | 4 | 4504.1444 | 4504.1268 | 0 | 3.9 | 127 | Specific | 1023.02 | 986.46 | 986.46 | 21.13 | 44 | >sp\|P01876\|IGHA1_HUMAN Immunoglobulin heavy constant alpha 1 OS=Homo sapiens OX=9606 GN=IGHA1 PE=1 SV=2 | 255 | 20201115_07433_1ng_glyco_hcd203040_compound15.21346.21346.4 | controllerType=0 controllerNumber=1 scan=21346 | 42.3647 |
| 18058:4 | 11 | R.LSLHRPALEDLLLGSEAN[+1257.449]LTC[+57.021]TLTGLR.D | HexNAc(3)Hex(4) | C[+57], N[+1257] | 1056.0196 | 4 | 4221.0565 | 4221.0477 | 0 | 2.1 | 127 | Specific | 1084.19 | 1084.19 | 1084.19 | 22.3 | 44 | >sp\|P01876\|IGHA1_HUMAN Immunoglobulin heavy constant alpha 1 OS=Homo sapiens OX=9606 GN=IGHA1 PE=1 SV=2 | 255 | 20201115_07433_1ng_glyco_hcd203040_compound15.21473.21473.4 | controllerType=0 controllerNumber=1 scan=21473 | 42.6835 |
| 15431:4 | 11 | R.LAGKPTHVN[+2553.910]VSVVMAEVDGTC[+57.021]Y.- | HexNAc(5)Hex(5)Fuc(1)NeuAc(2) | C[+57], N[+2554] | 1226.0237 | 4 | 4901.0731 | 4901.0518 | 0 | 4.3 | 332 | CRagged | 220.11 | 220.11 | 220.11 | 5.12 | 44 | >sp\|P01876\|IGHA1_HUMAN Immunoglobulin heavy constant alpha 1 OS=Homo sapiens OX=9606 GN=IGHA1 PE=1 SV=2 | 255 | 20201115_07433_1ng_glyco_hcd203040_compound15.18689.18689.4 | controllerType=0 controllerNumber=1 scan=18689 | 37.3005 |
| 11837:3 | 11 | R.LAGKPTHVN[+1501.555]VSVVMAEVDGTC[+57.021]Y.- | HexNAc(5)Hex(3) | C[+57], N[+1502] | 1283.5728 | 3 | 3848.7039 | 3848.6974 | 0 | 1.7 | 332 | CRagged | 233.66 | 114.58 | 114.58 | 3.62 | 44 | >sp\|P01876\|IGHA1_HUMAN Immunoglobulin heavy constant alpha 1 OS=Homo sapiens OX=9606 GN=IGHA1 PE=1 SV=2 | 255 | 20201115_07433_1ng_glyco_hcd203040_compound15.14937.14937.3 | controllerType=0 controllerNumber=1 scan=14937 | 30.7728 |
| 15420:3 | 11 | R.LAGKPTHVN[+2350.830]VSVVMAEVDGTC[+57.021]Y.- | HexNAc(4)Hex(5)Fuc(1)NeuAc(2) | C[+57], N[+2351] | 1566.6679 | 3 | 4697.9892 | 4697.9724 | 0 | 3.6 | 332 | CRagged | 235.44 | 233.26 | 233.26 | 6.53 | 44 | >sp\|P01876\|IGHA1_HUMAN Immunoglobulin heavy constant alpha 1 OS=Homo sapiens OX=9606 GN=IGHA1 PE=1 SV=2 | 255 | 20201115_07433_1ng_glyco_hcd203040_compound15.18677.18677.3 | controllerType=0 controllerNumber=1 scan=18677 | 37.2749 |
| 15433:4 | 11 | R.LAGKPTHVN[+2553.910]VSVVMAEVDGTC[+57.021]Y.- | HexNAc(5)Hex(5)Fuc(1)NeuAc(2) | C[+57], N[+2554] | 1226.0237 | 4 | 4901.0731 | 4901.0518 | 0 | 4.3 | 332 | CRagged | 305.49 | 258.91 | 258.91 | 6.88 | 44 | >sp\|P01876\|IGHA1_HUMAN Immunoglobulin heavy constant alpha 1 OS=Homo sapiens OX=9606 GN=IGHA1 PE=1 SV=2 | 255 | 20201115_07433_1ng_glyco_hcd203040_compound15.18691.18691.4 | controllerType=0 controllerNumber=1 scan=18691 | 37.3044 |
| 11734:4 | 11 | R.LAGKPTHVN[+1809.666]VSVVMAEVDGTC[+57.021]Y.- | HexNAc(5)Hex(4)Fuc(1) | C[+57], N[+1810] | 1039.9572 | 4 | 4156.807 | 4156.8081 | 0 | -0.3 | 332 | CRagged | 311.79 | 160.09 | 160.09 | 3.61 | 44 | >sp\|P01876\|IGHA1_HUMAN Immunoglobulin heavy constant alpha 1 OS=Homo sapiens OX=9606 GN=IGHA1 PE=1 SV=2 | 255 | 20201115_07433_1ng_glyco_hcd203040_compound15.14830.14830.4 | controllerType=0 controllerNumber=1 scan=14830 | 30.5736 |
| 11812:4 | 11 | R.LAGKPTHVN[+1647.613]VSVVMAEVDGTC[+57.021]Y.- | HexNAc(5)Hex(3)Fuc(1) | C[+57], N[+1648] | 999.4482 | 4 | 3994.7711 | 3994.7553 | 0 | 3.9 | 332 | CRagged | 338.15 | 338.15 | 338.15 | 7.78 | 44 | >sp\|P01876\|IGHA1_HUMAN Immunoglobulin heavy constant alpha 1 OS=Homo sapiens OX=9606 GN=IGHA1 PE=1 SV=2 | 255 | 20201115_07433_1ng_glyco_hcd203040_compound15.14911.14911.4 | controllerType=0 controllerNumber=1 scan=14911 | 30.727 |
| 11731:4 | 11 | R.LAGKPTHVN[+1809.666]VSVVMAEVDGTC[+57.021]Y.- | HexNAc(5)Hex(4)Fuc(1) | C[+57], N[+1810] | 1039.9572 | 4 | 4156.807 | 4156.8081 | 0 | -0.3 | 332 | CRagged | 346.29 | 300.61 | 300.61 | 7 | 44 | >sp\|P01876\|IGHA1_HUMAN Immunoglobulin heavy constant alpha 1 OS=Homo sapiens OX=9606 GN=IGHA1 PE=1 SV=2 | 255 | 20201115_07433_1ng_glyco_hcd203040_compound15.14827.14827.4 | controllerType=0 controllerNumber=1 scan=14827 | 30.5642 |
| 11815:4 | 11 | R.LAGKPTHVN[+1647.613]VSVVMAEVDGTC[+57.021]Y.- | HexNAc(5)Hex(3)Fuc(1) | C[+57], N[+1648] | 999.4482 | 4 | 3994.7711 | 3994.7553 | 0 | 3.9 | 332 | CRagged | 433.28 | 314.04 | 314.04 | 6.67 | 44 | >sp\|P01876\|IGHA1_HUMAN Immunoglobulin heavy constant alpha 1 OS=Homo sapiens OX=9606 GN=IGHA1 PE=1 SV=2 | 255 | 20201115_07433_1ng_glyco_hcd203040_compound15.14914.14914.4 | controllerType=0 controllerNumber=1 scan=14914 | 30.7314 |
| 11717:3 | 11 | R.LAGKPTHVN[+1501.555]VSVVMAEVDGTC[+57.021]Y.- | HexNAc(5)Hex(3) | C[+57], N[+1502] | 1283.5695 | 3 | 3848.694 | 3848.6974 | 0 | -0.9 | 332 | CRagged | 526.66 | 492.76 | 492.76 | 9.28 | 44 | >sp\|P01876\|IGHA1_HUMAN Immunoglobulin heavy constant alpha 1 OS=Homo sapiens OX=9606 GN=IGHA1 PE=1 SV=2 | 255 | 20201115_07433_1ng_glyco_hcd203040_compound15.14813.14813.3 | controllerType=0 controllerNumber=1 scan=14813 | 30.5413 |
| 11719:3 | 11 | R.LAGKPTHVN[+1501.555]VSVVMAEVDGTC[+57.021]Y.- | HexNAc(5)Hex(3) | C[+57], N[+1502] | 1283.5695 | 3 | 3848.694 | 3848.6974 | 0 | -0.9 | 332 | CRagged | 617.58 | 583.55 | 583.55 | 11.87 | 44 | >sp\|P01876\|IGHA1_HUMAN Immunoglobulin heavy constant alpha 1 OS=Homo sapiens OX=9606 GN=IGHA1 PE=1 SV=2 | 255 | 20201115_07433_1ng_glyco_hcd203040_compound15.14815.14815.3 | controllerType=0 controllerNumber=1 scan=14815 | 30.5459 |
| 11679:3 | 11 | R.LAGKPTHVN[+1663.608]VSVVMAEVDGTC[+57.021]Y.- | HexNAc(5)Hex(4) | C[+57], N[+1664] | 1337.5911 | 3 | 4010.7588 | 4010.7502 | 0 | 2.1 | 332 | CRagged | 708.91 | 708.91 | 441.87 | 15.21 | 44 | >sp\|P01876\|IGHA1_HUMAN Immunoglobulin heavy constant alpha 1 OS=Homo sapiens OX=9606 GN=IGHA1 PE=1 SV=2 | 255 | 20201115_07433_1ng_glyco_hcd203040_compound15.14773.14773.3 | controllerType=0 controllerNumber=1 scan=14773 | 30.4507 |
| 11634:3 | 11 | R.LAGKPTHVN[+1540.529]VSVVMAEVDGTC[+57.021]Y.- | HexNAc(2)Hex(7) | C[+57], N[+1541] | 1296.5665 | 3 | 3887.6848 | 3887.6706 | 0 | 3.7 | 332 | CRagged | 766.46 | 766.46 | 429.28 | 16.04 | 44 | >sp\|P01876\|IGHA1_HUMAN Immunoglobulin heavy constant alpha 1 OS=Homo sapiens OX=9606 GN=IGHA1 PE=1 SV=2 | 255 | 20201115_07433_1ng_glyco_hcd203040_compound15.14726.14726.3 | controllerType=0 controllerNumber=1 scan=14726 | 30.3433 |
| 11769:3 | 11 | R.LAGKPTHVN[+1647.613]VSVVMAEVDGTC[+57.021]Y.- | HexNAc(5)Hex(3)Fuc(1) | C[+57], N[+1648] | 1332.2613 | 3 | 3994.7693 | 3994.7553 | 0 | 3.5 | 332 | CRagged | 766.93 | 610.61 | 610.61 | 16.04 | 44 | >sp\|P01876\|IGHA1_HUMAN Immunoglobulin heavy constant alpha 1 OS=Homo sapiens OX=9606 GN=IGHA1 PE=1 SV=2 | 255 | 20201115_07433_1ng_glyco_hcd203040_compound15.14866.14866.3 | controllerType=0 controllerNumber=1 scan=14866 | 30.645 |
| 11699:3 | 11 | R.LAGKPTHVN[+1809.666]VSVVMAEVDGTC[+57.021]Y.- | HexNAc(5)Hex(4)Fuc(1) | C[+57], N[+1810] | 1386.277 | 3 | 4156.8166 | 4156.8081 | 0 | 2 | 332 | CRagged | 781.58 | 781.58 | 503 | 16.21 | 44 | >sp\|P01876\|IGHA1_HUMAN Immunoglobulin heavy constant alpha 1 OS=Homo sapiens OX=9606 GN=IGHA1 PE=1 SV=2 | 255 | 20201115_07433_1ng_glyco_hcd203040_compound15.14794.14794.3 | controllerType=0 controllerNumber=1 scan=14794 | 30.4947 |
| 11702:3 | 11 | R.LAGKPTHVN[+1809.666]VSVVMAEVDGTC[+57.021]Y.- | HexNAc(5)Hex(4)Fuc(1) | C[+57], N[+1810] | 1386.277 | 3 | 4156.8166 | 4156.8081 | 0 | 2 | 332 | CRagged | 787.15 | 633.69 | 518.73 | 16.21 | 44 | >sp\|P01876\|IGHA1_HUMAN Immunoglobulin heavy constant alpha 1 OS=Homo sapiens OX=9606 GN=IGHA1 PE=1 SV=2 | 255 | 20201115_07433_1ng_glyco_hcd203040_compound15.14797.14797.3 | controllerType=0 controllerNumber=1 scan=14797 | 30.5055 |
| 11636:3 | 11 | R.LAGKPTHVN[+1540.529]VSVVMAEVDGTC[+57.021]Y.- | HexNAc(2)Hex(7) | C[+57], N[+1541] | 1296.5665 | 3 | 3887.6848 | 3887.6706 | 0 | 3.7 | 332 | CRagged | 789.01 | 787.75 | 454.56 | 16.04 | 44 | >sp\|P01876\|IGHA1_HUMAN Immunoglobulin heavy constant alpha 1 OS=Homo sapiens OX=9606 GN=IGHA1 PE=1 SV=2 | 255 | 20201115_07433_1ng_glyco_hcd203040_compound15.14728.14728.3 | controllerType=0 controllerNumber=1 scan=14728 | 30.3558 |
| 11682:3 | 11 | R.LAGKPTHVN[+1663.608]VSVVMAEVDGTC[+57.021]Y.- | HexNAc(5)Hex(4) | C[+57], N[+1664] | 1337.5911 | 3 | 4010.7588 | 4010.7502 | 0 | 2.1 | 332 | CRagged | 802.57 | 742.16 | 478.97 | 17.21 | 44 | >sp\|P01876\|IGHA1_HUMAN Immunoglobulin heavy constant alpha 1 OS=Homo sapiens OX=9606 GN=IGHA1 PE=1 SV=2 | 255 | 20201115_07433_1ng_glyco_hcd203040_compound15.14776.14776.3 | controllerType=0 controllerNumber=1 scan=14776 | 30.456 |
| 11612:3 | 11 | R.LAGKPTHVN[+1702.581]VSVVMAEVDGTC[+57.021]Y.- | HexNAc(2)Hex(8) | C[+57], N[+1703] | 1350.5845 | 3 | 4049.739 | 4049.7234 | 0 | 3.9 | 332 | CRagged | 875.07 | 875.07 | 586.06 | 18.04 | 44 | >sp\|P01876\|IGHA1_HUMAN Immunoglobulin heavy constant alpha 1 OS=Homo sapiens OX=9606 GN=IGHA1 PE=1 SV=2 | 255 | 20201115_07433_1ng_glyco_hcd203040_compound15.14703.14703.3 | controllerType=0 controllerNumber=1 scan=14703 | 30.283 |
| 11614:3 | 11 | R.LAGKPTHVN[+1702.581]VSVVMAEVDGTC[+57.021]Y.- | HexNAc(2)Hex(8) | C[+57], N[+1703] | 1350.5845 | 3 | 4049.739 | 4049.7234 | 0 | 3.9 | 332 | CRagged | 875.28 | 875.28 | 517.16 | 18.04 | 44 | >sp\|P01876\|IGHA1_HUMAN Immunoglobulin heavy constant alpha 1 OS=Homo sapiens OX=9606 GN=IGHA1 PE=1 SV=2 | 255 | 20201115_07433_1ng_glyco_hcd203040_compound15.14705.14705.3 | controllerType=0 controllerNumber=1 scan=14705 | 30.2888 |
| 11767:3 | 11 | R.LAGKPTHVN[+1647.613]VSVVMAEVDGTC[+57.021]Y.- | HexNAc(5)Hex(3)Fuc(1) | C[+57], N[+1648] | 1332.2613 | 3 | 3994.7693 | 3994.7553 | 0 | 3.5 | 332 | CRagged | 883.25 | 883.25 | 883.25 | 18.04 | 44 | >sp\|P01876\|IGHA1_HUMAN Immunoglobulin heavy constant alpha 1 OS=Homo sapiens OX=9606 GN=IGHA1 PE=1 SV=2 | 255 | 20201115_07433_1ng_glyco_hcd203040_compound15.14864.14864.3 | controllerType=0 controllerNumber=1 scan=14864 | 30.6356 |
| 11705:3 | 11 | R.LAGKPTHVN[+1216.423]VSVVMAEVDGTC[+57.021]Y.- | HexNAc(2)Hex(5) | C[+57], N[+1216] | 1188.5287 | 3 | 3563.5717 | 3563.5649 | 0 | 1.9 | 332 | CRagged | 886.76 | 886.76 | 517.2 | 18.21 | 44 | >sp\|P01876\|IGHA1_HUMAN Immunoglobulin heavy constant alpha 1 OS=Homo sapiens OX=9606 GN=IGHA1 PE=1 SV=2 | 255 | 20201115_07433_1ng_glyco_hcd203040_compound15.14800.14800.3 | controllerType=0 controllerNumber=1 scan=14800 | 30.5096 |
| 11667:3 | 11 | R.LAGKPTHVN[+1378.476]VSVVMAEVDGTC[+57.021]Y.- | HexNAc(2)Hex(6) | C[+57], N[+1378] | 1242.5477 | 3 | 3725.6284 | 3725.6178 | 0 | 2.9 | 332 | CRagged | 896.99 | 832.52 | 550.56 | 18.21 | 44 | >sp\|P01876\|IGHA1_HUMAN Immunoglobulin heavy constant alpha 1 OS=Homo sapiens OX=9606 GN=IGHA1 PE=1 SV=2 | 255 | 20201115_07433_1ng_glyco_hcd203040_compound15.14761.14761.3 | controllerType=0 controllerNumber=1 scan=14761 | 30.436 |
| 11708:3 | 11 | R.LAGKPTHVN[+1216.423]VSVVMAEVDGTC[+57.021]Y.- | HexNAc(2)Hex(5) | C[+57], N[+1216] | 1188.5287 | 3 | 3563.5717 | 3563.5649 | 0 | 1.9 | 332 | CRagged | 917.24 | 905.3 | 550.48 | 19.21 | 44 | >sp\|P01876\|IGHA1_HUMAN Immunoglobulin heavy constant alpha 1 OS=Homo sapiens OX=9606 GN=IGHA1 PE=1 SV=2 | 255 | 20201115_07433_1ng_glyco_hcd203040_compound15.14803.14803.3 | controllerType=0 controllerNumber=1 scan=14803 | 30.5246 |
| 11664:3 | 11 | R.LAGKPTHVN[+1378.476]VSVVMAEVDGTC[+57.021]Y.- | HexNAc(2)Hex(6) | C[+57], N[+1378] | 1242.5477 | 3 | 3725.6284 | 3725.6178 | 0 | 2.9 | 332 | CRagged | 941.59 | 941.59 | 566.88 | 19.21 | 44 | >sp\|P01876\|IGHA1_HUMAN Immunoglobulin heavy constant alpha 1 OS=Homo sapiens OX=9606 GN=IGHA1 PE=1 SV=2 | 255 | 20201115_07433_1ng_glyco_hcd203040_compound15.14758.14758.3 | controllerType=0 controllerNumber=1 scan=14758 | 30.4295 |
| 05909:3 | 12 | R.LSPN[+2351.851]ASAEHLELR.W | HexNAc(4)Hex(5)Fuc(3)NeuAc(1) | N[+2352] | 1263.5419 | 3 | 3788.6112 | 3788.6 | 0 | 3 | 52 | Specific | 295.35 | 151.68 | 151.68 | 6.22 | 36 | >sp\|Q13410\|BT1A1_HUMAN Butyrophilin subfamily 1 member A1 OS=Homo sapiens OX=9606 GN=BTN1A1 PE=1 SV=3 | 179 | 20201115_07433_1ng_glyco_hcd203040_compound15.8777.8777.3 | controllerType=0 controllerNumber=1 scan=8777 | 20.5221 |
| 05907:3 | 12 | R.LSPN[+2351.851]ASAEHLELR.W | HexNAc(4)Hex(5)Fuc(3)NeuAc(1) | N[+2352] | 1263.5419 | 3 | 3788.6112 | 3788.6 | 0 | 3 | 52 | Specific | 336.33 | 251.95 | 251.95 | 7.12 | 36 | >sp\|Q13410\|BT1A1_HUMAN Butyrophilin subfamily 1 member A1 OS=Homo sapiens OX=9606 GN=BTN1A1 PE=1 SV=3 | 179 | 20201115_07433_1ng_glyco_hcd203040_compound15.8775.8775.3 | controllerType=0 controllerNumber=1 scan=8775 | 20.5182 |
| 08011:3 | 12 | R.LSPN[+2350.830]ASAEHLELR.W | HexNAc(4)Hex(5)Fuc(1)NeuAc(2) | N[+2351] | 1263.1965 | 3 | 3787.5749 | 3787.5796 | 0 | -1.2 | 52 | Specific | 443.05 | 224.01 | 224.01 | 6.22 | 36 | >sp\|Q13410\|BT1A1_HUMAN Butyrophilin subfamily 1 member A1 OS=Homo sapiens OX=9606 GN=BTN1A1 PE=1 SV=3 | 179 | 20201115_07433_1ng_glyco_hcd203040_compound15.10954.10954.3 | controllerType=0 controllerNumber=1 scan=10954 | 24.1001 |
| 08013:3 | 12 | R.LSPN[+2350.830]ASAEHLELR.W | HexNAc(4)Hex(5)Fuc(1)NeuAc(2) | N[+2351] | 1263.1965 | 3 | 3787.5749 | 3787.5796 | 0 | -1.2 | 52 | Specific | 456.27 | 290.5 | 290.5 | 6.69 | 36 | >sp\|Q13410\|BT1A1_HUMAN Butyrophilin subfamily 1 member A1 OS=Homo sapiens OX=9606 GN=BTN1A1 PE=1 SV=3 | 179 | 20201115_07433_1ng_glyco_hcd203040_compound15.10956.10956.3 | controllerType=0 controllerNumber=1 scan=10956 | 24.1045 |
| 05897:3 | 12 | R.LSPN[+2205.793]ASAEHLELR.W | HexNAc(4)Hex(5)Fuc(2)NeuAc(1) | N[+2206] | 1214.8571 | 3 | 3642.5568 | 3642.5421 | 0 | 4 | 52 | Specific | 662.85 | 570.73 | 570.73 | 13.82 | 36 | >sp\|Q13410\|BT1A1_HUMAN Butyrophilin subfamily 1 member A1 OS=Homo sapiens OX=9606 GN=BTN1A1 PE=1 SV=3 | 179 | 20201115_07433_1ng_glyco_hcd203040_compound15.8765.8765.3 | controllerType=0 controllerNumber=1 scan=8765 | 20.5021 |
| 05895:3 | 12 | R.LSPN[+2205.793]ASAEHLELR.W | HexNAc(4)Hex(5)Fuc(2)NeuAc(1) | N[+2206] | 1214.8571 | 3 | 3642.5568 | 3642.5421 | 0 | 4 | 52 | Specific | 695.84 | 398.13 | 398.13 | 13.82 | 36 | >sp\|Q13410\|BT1A1_HUMAN Butyrophilin subfamily 1 member A1 OS=Homo sapiens OX=9606 GN=BTN1A1 PE=1 SV=3 | 179 | 20201115_07433_1ng_glyco_hcd203040_compound15.8763.8763.3 | controllerType=0 controllerNumber=1 scan=8763 | 20.4975 |
| 17977:3 | 12 | K.N[+1913.677]VSC[+57.021]YIQNLLLGQEK.K | HexNAc(4)Hex(5)NeuAc(1) | C[+57], N[+1914] | 1231.5373 | 3 | 3692.5973 | 3692.5876 | 0 | 2.6 | 215 | Specific | 570.65 | 507.92 | 507.92 | 10.68 | 36 | >sp\|Q13410\|BT1A1_HUMAN Butyrophilin subfamily 1 member A1 OS=Homo sapiens OX=9606 GN=BTN1A1 PE=1 SV=3 | 179 | 20201115_07433_1ng_glyco_hcd203040_compound15.21388.21388.3 | controllerType=0 controllerNumber=1 scan=21388 | 42.4695 |
| 17980:3 | 12 | K.N[+1913.677]VSC[+57.021]YIQNLLLGQEK.K | HexNAc(4)Hex(5)NeuAc(1) | C[+57], N[+1914] | 1231.5373 | 3 | 3692.5973 | 3692.5876 | 0 | 2.6 | 215 | Specific | 680.82 | 490.73 | 490.73 | 13.98 | 36 | >sp\|Q13410\|BT1A1_HUMAN Butyrophilin subfamily 1 member A1 OS=Homo sapiens OX=9606 GN=BTN1A1 PE=1 SV=3 | 179 | 20201115_07433_1ng_glyco_hcd203040_compound15.21391.21391.3 | controllerType=0 controllerNumber=1 scan=21391 | 42.4882 |
| 02830:4 | 13 | K.QTDEIKDTRN[+2059.735]ESTQNC[+57.021]VVAEPEK.M | HexNAc(4)Hex(5)Fuc(1)NeuAc(1) | C[+57], N[+2060] | 1188.5072 | 4 | 4751.0072 | 4750.9876 | 0 | 4.1 | 60 | Specific | 254.94 | 254.94 | 254.94 | 6.69 | 33 | >sp\|P47710\|CASA1_HUMAN Alpha-S1-casein OS=Homo sapiens OX=9606 GN=CSN1S1 PE=1 SV=1 | 989 | 20201115_07433_1ng_glyco_hcd203040_compound15.5592.5592.4 | controllerType=0 controllerNumber=1 scan=5592 | 15.2358 |
| 02828:4 | 13 | K.QTDEIKDTRN[+2059.735]ESTQNC[+57.021]VVAEPEK.M | HexNAc(4)Hex(5)Fuc(1)NeuAc(1) | C[+57], N[+2060] | 1188.5072 | 4 | 4751.0072 | 4750.9876 | 0 | 4.1 | 60 | Specific | 319.52 | 148.31 | 148.31 | 6.16 | 33 | >sp\|P47710\|CASA1_HUMAN Alpha-S1-casein OS=Homo sapiens OX=9606 GN=CSN1S1 PE=1 SV=1 | 989 | 20201115_07433_1ng_glyco_hcd203040_compound15.5590.5590.4 | controllerType=0 controllerNumber=1 scan=5590 | 15.2321 |
| 09793:4 | 13 | K.QTDEIKDTRN[+2059.735]ESTQNC[+57.021]VVAEPEKMESSISSSSEEMSLSK.C | HexNAc(4)Hex(5)Fuc(1)NeuAc(1) | C[+57], N[+2060] | 1613.4353 | 4 | 6450.7194 | 6450.7156 | 0 | 0.6 | 60 | Specific | 726.69 | 542.45 | 542.45 | 12.33 | 33 | >sp\|P47710\|CASA1_HUMAN Alpha-S1-casein OS=Homo sapiens OX=9606 GN=CSN1S1 PE=1 SV=1 | 989 | 20201115_07433_1ng_glyco_hcd203040_compound15.12823.12823.4 | controllerType=0 controllerNumber=1 scan=12823 | 27.2295 |
| 09791:4 | 13 | K.QTDEIKDTRN[+2059.735]ESTQNC[+57.021]VVAEPEKMESSISSSSEEMSLSK.C | HexNAc(4)Hex(5)Fuc(1)NeuAc(1) | C[+57], N[+2060] | 1613.4353 | 4 | 6450.7194 | 6450.7156 | 0 | 0.6 | 60 | Specific | 887.26 | 680.51 | 680.51 | 15.33 | 33 | >sp\|P47710\|CASA1_HUMAN Alpha-S1-casein OS=Homo sapiens OX=9606 GN=CSN1S1 PE=1 SV=1 | 989 | 20201115_07433_1ng_glyco_hcd203040_compound15.12821.12821.4 | controllerType=0 controllerNumber=1 scan=12821 | 27.2238 |
| 27321:4 | 14 | R.QLAHQSN[+2861.000]STNIFFSPVSIATAFAMLSLGTK.A | HexNAc(5)Hex(6)NeuAc(3) | N[+2861] | 1511.4147 | 4 | 6042.6369 | 6042.6351 | 0 | 0.3 | 64 | Specific | 237.99 | 226.31 | 226.31 | 6.5 | 37 | >sp\|P01009\|A1AT_HUMAN Alpha-1-antitrypsin OS=Homo sapiens OX=9606 GN=SERPINA1 PE=1 SV=3 | 642 | 20201115_07433_1ng_glyco_hcd203040_compound15.31138.31138.2 | controllerType=0 controllerNumber=1 scan=31138 | 61.1829 |
| 24491:5 | 14 | R.QLAHQSN[+2204.772]STNIFFSPVSIATAFAMLSLGTK.A | HexNAc(4)Hex(5)NeuAc(2) | N[+2205] | 1078.0936 | 5 | 5386.439 | 5386.4075 | 0 | 5.9 | 64 | Specific | 426.28 | 412.63 | 412.63 | 10.57 | 37 | >sp\|P01009\|A1AT_HUMAN Alpha-1-antitrypsin OS=Homo sapiens OX=9606 GN=SERPINA1 PE=1 SV=3 | 642 | 20201115_07433_1ng_glyco_hcd203040_compound15.28195.28195.5 | controllerType=0 controllerNumber=1 scan=28195 | 55.9081 |
| 24489:5 | 14 | R.QLAHQSN[+2204.772]STNIFFSPVSIATAFAMLSLGTK.A | HexNAc(4)Hex(5)NeuAc(2) | N[+2205] | 1078.0936 | 5 | 5386.439 | 5386.4075 | 0 | 5.9 | 64 | Specific | 472.33 | 472.33 | 472.33 | 10.7 | 37 | >sp\|P01009\|A1AT_HUMAN Alpha-1-antitrypsin OS=Homo sapiens OX=9606 GN=SERPINA1 PE=1 SV=3 | 642 | 20201115_07433_1ng_glyco_hcd203040_compound15.28193.28193.5 | controllerType=0 controllerNumber=1 scan=28193 | 55.904 |
| 24424:4 | 14 | R.QLAHQSN[+2350.830]STNIFFSPVSIATAFAMLSLGTK.A | HexNAc(4)Hex(5)Fuc(1)NeuAc(2) | N[+2351] | 1383.8713 | 4 | 5532.4635 | 5532.4654 | 0 | -0.3 | 64 | Specific | 492.07 | 443.07 | 443.07 | 10.35 | 37 | >sp\|P01009\|A1AT_HUMAN Alpha-1-antitrypsin OS=Homo sapiens OX=9606 GN=SERPINA1 PE=1 SV=3 | 642 | 20201115_07433_1ng_glyco_hcd203040_compound15.28125.28125.4 | controllerType=0 controllerNumber=1 scan=28125 | 55.7669 |
| 24426:4 | 14 | R.QLAHQSN[+2350.830]STNIFFSPVSIATAFAMLSLGTK.A | HexNAc(4)Hex(5)Fuc(1)NeuAc(2) | N[+2351] | 1383.8713 | 4 | 5532.4635 | 5532.4654 | 0 | -0.3 | 64 | Specific | 627.94 | 551.77 | 551.77 | 13.02 | 37 | >sp\|P01009\|A1AT_HUMAN Alpha-1-antitrypsin OS=Homo sapiens OX=9606 GN=SERPINA1 PE=1 SV=3 | 642 | 20201115_07433_1ng_glyco_hcd203040_compound15.28127.28127.4 | controllerType=0 controllerNumber=1 scan=28127 | 55.7723 |
| 24480:3 | 14 | R.QLAHQSN[+2204.772]STNIFFSPVSIATAFAMLSLGTK.A | HexNAc(4)Hex(5)NeuAc(2) | N[+2205] | 1796.1494 | 3 | 5386.4336 | 5386.4075 | 0 | 4.8 | 64 | Specific | 692.06 | 690.3 | 690.3 | 14.42 | 37 | >sp\|P01009\|A1AT_HUMAN Alpha-1-antitrypsin OS=Homo sapiens OX=9606 GN=SERPINA1 PE=1 SV=3 | 642 | 20201115_07433_1ng_glyco_hcd203040_compound15.28184.28184.3 | controllerType=0 controllerNumber=1 scan=28184 | 55.885 |
| 24482:3 | 14 | R.QLAHQSN[+2204.772]STNIFFSPVSIATAFAMLSLGTK.A | HexNAc(4)Hex(5)NeuAc(2) | N[+2205] | 1796.1494 | 3 | 5386.4336 | 5386.4075 | 0 | 4.8 | 64 | Specific | 704.9 | 599.22 | 599.22 | 15.42 | 37 | >sp\|P01009\|A1AT_HUMAN Alpha-1-antitrypsin OS=Homo sapiens OX=9606 GN=SERPINA1 PE=1 SV=3 | 642 | 20201115_07433_1ng_glyco_hcd203040_compound15.28186.28186.3 | controllerType=0 controllerNumber=1 scan=28186 | 55.8891 |
| 24476:4 | 14 | R.QLAHQSN[+2204.772]STNIFFSPVSIATAFAMLSLGTK.A | HexNAc(4)Hex(5)NeuAc(2) | N[+2205] | 1347.3627 | 4 | 5386.4289 | 5386.4075 | 0 | 4 | 64 | Specific | 874.28 | 849.19 | 849.19 | 18.71 | 37 | >sp\|P01009\|A1AT_HUMAN Alpha-1-antitrypsin OS=Homo sapiens OX=9606 GN=SERPINA1 PE=1 SV=3 | 642 | 20201115_07433_1ng_glyco_hcd203040_compound15.28180.28180.4 | controllerType=0 controllerNumber=1 scan=28180 | 55.8787 |
| 24474:4 | 14 | R.QLAHQSN[+2204.772]STNIFFSPVSIATAFAMLSLGTK.A | HexNAc(4)Hex(5)NeuAc(2) | N[+2205] | 1347.3627 | 4 | 5386.4289 | 5386.4075 | 0 | 4 | 64 | Specific | 942.33 | 904.63 | 904.63 | 19.71 | 37 | >sp\|P01009\|A1AT_HUMAN Alpha-1-antitrypsin OS=Homo sapiens OX=9606 GN=SERPINA1 PE=1 SV=3 | 642 | 20201115_07433_1ng_glyco_hcd203040_compound15.28178.28178.4 | controllerType=0 controllerNumber=1 scan=28178 | 55.8709 |
| 20601:3 | 14 | K.YLGN[+2205.793]ATAIFFLPDEGK.L | HexNAc(4)Hex(5)Fuc(2)NeuAc(1) | N[+2206] | 1321.2253 | 3 | 3961.6613 | 3961.6881 | 0 | -6.7 | 268 | Specific | 225.6 | 172.63 | 172.63 | 2.32 | 37 | >sp\|P01009\|A1AT_HUMAN Alpha-1-antitrypsin OS=Homo sapiens OX=9606 GN=SERPINA1 PE=1 SV=3 | 642 | 20201115_07433_1ng_glyco_hcd203040_compound15.24128.24128.3 | controllerType=0 controllerNumber=1 scan=24128 | 48.2396 |
| 20361:4 | 14 | K.YLGN[+2204.772]ATAIFFLPDEGKLQHLENELTHDIITK.F | HexNAc(4)Hex(5)NeuAc(2) | N[+2205] | 1437.1627 | 4 | 5745.6291 | 5745.6098 | 0 | 3.4 | 268 | Specific | 562.56 | 478.44 | 478.44 | 10.41 | 37 | >sp\|P01009\|A1AT_HUMAN Alpha-1-antitrypsin OS=Homo sapiens OX=9606 GN=SERPINA1 PE=1 SV=3 | 642 | 20201115_07433_1ng_glyco_hcd203040_compound15.23878.23878.4 | controllerType=0 controllerNumber=1 scan=23878 | 47.7096 |
| 20364:5 | 14 | K.YLGN[+2204.772]ATAIFFLPDEGKLQHLENELTHDIITK.F | HexNAc(4)Hex(5)NeuAc(2) | N[+2205] | 1149.9309 | 5 | 5745.6254 | 5745.6098 | 0 | 2.7 | 268 | Specific | 702.1 | 460.68 | 460.68 | 15.41 | 37 | >sp\|P01009\|A1AT_HUMAN Alpha-1-antitrypsin OS=Homo sapiens OX=9606 GN=SERPINA1 PE=1 SV=3 | 642 | 20201115_07433_1ng_glyco_hcd203040_compound15.23881.23881.5 | controllerType=0 controllerNumber=1 scan=23881 | 47.7156 |
| 20363:4 | 14 | K.YLGN[+2204.772]ATAIFFLPDEGKLQHLENELTHDIITK.F | HexNAc(4)Hex(5)NeuAc(2) | N[+2205] | 1437.1627 | 4 | 5745.6291 | 5745.6098 | 0 | 3.4 | 268 | Specific | 941.95 | 749.72 | 749.72 | 18.7 | 37 | >sp\|P01009\|A1AT_HUMAN Alpha-1-antitrypsin OS=Homo sapiens OX=9606 GN=SERPINA1 PE=1 SV=3 | 642 | 20201115_07433_1ng_glyco_hcd203040_compound15.23880.23880.4 | controllerType=0 controllerNumber=1 scan=23880 | 47.7137 |
| 23717:4 | 28 | K.MLN[+2350.830]TSSLLEQLNEQFNWVSR.L | HexNAc(4)Hex(5)Fuc(1)NeuAc(2) | N[+2351] | 1190.7687 | 4 | 4760.053 | 4760.0171 | 0 | 7.6 | 352 | Specific | 514.12 | 427.17 | 427.17 | 10.06 | 23 | >sp\|P10909\|CLUS_HUMAN Clusterin OS=Homo sapiens OX=9606 GN=CLU PE=1 SV=1 | 1467 | 20201115_07433_1ng_glyco_hcd203040_compound15.27389.27389.4 | controllerType=0 controllerNumber=1 scan=27389 | 54.4107 |
| 23719:4 | 28 | K.MLN[+2350.830]TSSLLEQLNEQFNWVSR.L | HexNAc(4)Hex(5)Fuc(1)NeuAc(2) | N[+2351] | 1190.7687 | 4 | 4760.053 | 4760.0171 | 0 | 7.6 | 352 | Specific | 545.45 | 429.1 | 429.1 | 10.24 | 23 | >sp\|P10909\|CLUS_HUMAN Clusterin OS=Homo sapiens OX=9606 GN=CLU PE=1 SV=1 | 1467 | 20201115_07433_1ng_glyco_hcd203040_compound15.27391.27391.4 | controllerType=0 controllerNumber=1 scan=27391 | 54.416 |
| 23735:4 | 28 | K.M[+15.995]LN[+2336.851]TSSLLEQLNEQFNWVSR.L | HexNAc(6)Hex(6)Fuc(1) | M[+16], N[+2337] | 1191.27 | 4 | 4762.0582 | 4762.0327 | 0 | 5.3 | 352 | Specific | 205.99 | 201.95 | 201.95 | 3.28 | 23 | >sp\|P10909\|CLUS_HUMAN Clusterin OS=Homo sapiens OX=9606 GN=CLU PE=1 SV=1 | 1467 | 20201115_07433_1ng_glyco_hcd203040_compound15.27408.27408.4 | controllerType=0 controllerNumber=1 scan=27408 | 54.4548 |
| 23733:4 | 28 | K.M[+15.995]LN[+2336.851]TSSLLEQLNEQFNWVSR.L | HexNAc(6)Hex(6)Fuc(1) | M[+16], N[+2337] | 1191.27 | 4 | 4762.0582 | 4762.0327 | 0 | 5.3 | 352 | Specific | 211.02 | 152.62 | 152.62 | 3.25 | 23 | >sp\|P10909\|CLUS_HUMAN Clusterin OS=Homo sapiens OX=9606 GN=CLU PE=1 SV=1 | 1467 | 20201115_07433_1ng_glyco_hcd203040_compound15.27406.27406.4 | controllerType=0 controllerNumber=1 scan=27406 | 54.4481 |
| 24096:5 | 51 | K.VFPLSLDSTPQDGNVVVAC[+57.021]LVQGFFPQEPLSVTWSESGQN[+2262.814]VTAI | HexNAc(5)Hex(5)Fuc(1)NeuAc(1) | C[+57], N[+2263] | 1408.6526 | 5 | 7039.2341 | 7039.1757 | 0 | 8.3 | 123 | Specific | 321.54 | 321.54 | 321.54 | 7.88 | 14 | >sp\|P0DOX2\|IGA2_HUMAN Immunoglobulin alpha-2 heavy chain OS=Homo sapiens OX=9606 PE=1 SV=2 | 631 | 20201115_07433_1ng_glyco_hcd203040_compound15.27782.27782.5 | controllerType=0 controllerNumber=1 scan=27782 | 55.1233 |
| 24098:5 | 51 | K.VFPLSLDSTPQDGNVVVAC[+57.021]LVQGFFPQEPLSVTWSESGQN[+2262.814]VTAI | HexNAc(5)Hex(5)Fuc(1)NeuAc(1) | C[+57], N[+2263] | 1408.6526 | 5 | 7039.2341 | 7039.1757 | 0 | 8.3 | 123 | Specific | 436.63 | 436.63 | 436.63 | 10.24 | 14 | >sp\|P0DOX2\|IGA2_HUMAN Immunoglobulin alpha-2 heavy chain OS=Homo sapiens OX=9606 PE=1 SV=2 | 631 | 20201115_07433_1ng_glyco_hcd203040_compound15.27784.27784.5 | controllerType=0 controllerNumber=1 scan=27784 | 55.1291 |
| 03179:3 | 51 | K.TPLTAN[+1971.719]ITK.S | HexNAc(5)Hex(5)Fuc(1) | N[+1972] | 977.4324 | 3 | 2930.2827 | 2930.2757 | 0 | 2.4 | 315 | Specific | 218.37 | 142.85 | 142.85 | 3.22 | 14 | >sp\|P0DOX2\|IGA2_HUMAN Immunoglobulin alpha-2 heavy chain OS=Homo sapiens OX=9606 PE=1 SV=2 | 631 | 20201115_07433_1ng_glyco_hcd203040_compound15.5953.5953.3 | controllerType=0 controllerNumber=1 scan=5953 | 15.8588 |
| 03289:2 | 51 | K.TPLTAN[+1501.555]ITK.S | HexNAc(5)Hex(3) | N[+1502] | 1230.5655 | 2 | 2460.1237 | 2460.1121 | 0 | 4.7 | 315 | Specific | 256.19 | 68.55 | 68.55 | 4.54 | 14 | >sp\|P0DOX2\|IGA2_HUMAN Immunoglobulin alpha-2 heavy chain OS=Homo sapiens OX=9606 PE=1 SV=2 | 631 | 20201115_07433_1ng_glyco_hcd203040_compound15.6067.6067.2 | controllerType=0 controllerNumber=1 scan=6067 | 16.0703 |
| 03291:2 | 51 | K.TPLTAN[+1501.555]ITK.S | HexNAc(5)Hex(3) | N[+1502] | 1230.5655 | 2 | 2460.1237 | 2460.1121 | 0 | 4.7 | 315 | Specific | 441 | 242.6 | 242.6 | 7.14 | 14 | >sp\|P0DOX2\|IGA2_HUMAN Immunoglobulin alpha-2 heavy chain OS=Homo sapiens OX=9606 PE=1 SV=2 | 631 | 20201115_07433_1ng_glyco_hcd203040_compound15.6069.6069.2 | controllerType=0 controllerNumber=1 scan=6069 | 16.0744 |
| 03284:3 | 51 | K.TPLTAN[+1647.613]ITK.S | HexNAc(5)Hex(3)Fuc(1) | N[+1648] | 869.398 | 3 | 2606.1794 | 2606.17 | 0 | 3.6 | 315 | Specific | 485.38 | 204.63 | 204.63 | 7.29 | 14 | >sp\|P0DOX2\|IGA2_HUMAN Immunoglobulin alpha-2 heavy chain OS=Homo sapiens OX=9606 PE=1 SV=2 | 631 | 20201115_07433_1ng_glyco_hcd203040_compound15.6062.6062.3 | controllerType=0 controllerNumber=1 scan=6062 | 16.0598 |
| 03226:3 | 51 | K.TPLTAN[+1809.666]ITK.S | HexNAc(5)Hex(4)Fuc(1) | N[+1810] | 923.4152 | 3 | 2768.231 | 2768.2228 | 0 | 3 | 315 | Specific | 493.44 | 187.93 | 187.93 | 7.39 | 14 | >sp\|P0DOX2\|IGA2_HUMAN Immunoglobulin alpha-2 heavy chain OS=Homo sapiens OX=9606 PE=1 SV=2 | 631 | 20201115_07433_1ng_glyco_hcd203040_compound15.6002.6002.3 | controllerType=0 controllerNumber=1 scan=6002 | 15.9471 |
| 03228:3 | 51 | K.TPLTAN[+1809.666]ITK.S | HexNAc(5)Hex(4)Fuc(1) | N[+1810] | 923.4152 | 3 | 2768.231 | 2768.2228 | 0 | 3 | 315 | Specific | 533.01 | 304.36 | 304.36 | 8.24 | 14 | >sp\|P0DOX2\|IGA2_HUMAN Immunoglobulin alpha-2 heavy chain OS=Homo sapiens OX=9606 PE=1 SV=2 | 631 | 20201115_07433_1ng_glyco_hcd203040_compound15.6004.6004.3 | controllerType=0 controllerNumber=1 scan=6004 | 15.9512 |
| 03286:3 | 51 | K.TPLTAN[+1647.613]ITK.S | HexNAc(5)Hex(3)Fuc(1) | N[+1648] | 869.398 | 3 | 2606.1794 | 2606.17 | 0 | 3.6 | 315 | Specific | 573.85 | 280.84 | 280.84 | 8.29 | 14 | >sp\|P0DOX2\|IGA2_HUMAN Immunoglobulin alpha-2 heavy chain OS=Homo sapiens OX=9606 PE=1 SV=2 | 631 | 20201115_07433_1ng_glyco_hcd203040_compound15.6064.6064.3 | controllerType=0 controllerNumber=1 scan=6064 | 16.0655 |
| 03279:2 | 51 | K.TPLTAN[+1647.613]ITK.S | HexNAc(5)Hex(3)Fuc(1) | N[+1648] | 1303.594 | 2 | 2606.1808 | 2606.17 | 0 | 4.1 | 315 | Specific | 587.69 | 317 | 317 | 9.48 | 14 | >sp\|P0DOX2\|IGA2_HUMAN Immunoglobulin alpha-2 heavy chain OS=Homo sapiens OX=9606 PE=1 SV=2 | 631 | 20201115_07433_1ng_glyco_hcd203040_compound15.6057.6057.2 | controllerType=0 controllerNumber=1 scan=6057 | 16.0519 |
| 03234:2 | 51 | K.TPLTAN[+1809.666]ITK.S | HexNAc(5)Hex(4)Fuc(1) | N[+1810] | 1384.6193 | 2 | 2768.2313 | 2768.2228 | 0 | 3.1 | 315 | Specific | 601.26 | 294.67 | 294.67 | 10.48 | 14 | >sp\|P0DOX2\|IGA2_HUMAN Immunoglobulin alpha-2 heavy chain OS=Homo sapiens OX=9606 PE=1 SV=2 | 631 | 20201115_07433_1ng_glyco_hcd203040_compound15.6010.6010.2 | controllerType=0 controllerNumber=1 scan=6010 | 15.962 |
| 03277:2 | 51 | K.TPLTAN[+1647.613]ITK.S | HexNAc(5)Hex(3)Fuc(1) | N[+1648] | 1303.594 | 2 | 2606.1808 | 2606.17 | 0 | 4.1 | 315 | Specific | 604.16 | 331.08 | 331.08 | 10.46 | 14 | >sp\|P0DOX2\|IGA2_HUMAN Immunoglobulin alpha-2 heavy chain OS=Homo sapiens OX=9606 PE=1 SV=2 | 631 | 20201115_07433_1ng_glyco_hcd203040_compound15.6055.6055.2 | controllerType=0 controllerNumber=1 scan=6055 | 16.0451 |
| 03232:2 | 51 | K.TPLTAN[+1809.666]ITK.S | HexNAc(5)Hex(4)Fuc(1) | N[+1810] | 1384.6193 | 2 | 2768.2313 | 2768.2228 | 0 | 3.1 | 315 | Specific | 606.52 | 330.4 | 330.4 | 10.76 | 14 | >sp\|P0DOX2\|IGA2_HUMAN Immunoglobulin alpha-2 heavy chain OS=Homo sapiens OX=9606 PE=1 SV=2 | 631 | 20201115_07433_1ng_glyco_hcd203040_compound15.6008.6008.2 | controllerType=0 controllerNumber=1 scan=6008 | 15.9575 |
| 03321:2 | 51 | K.TPLTAN[+1444.534]ITK.S | HexNAc(4)Hex(3)Fuc(1) | N[+1445] | 1202.0523 | 2 | 2403.0973 | 2403.0906 | 0 | 2.8 | 315 | Specific | 621.94 | 303.8 | 303.8 | 10.65 | 14 | >sp\|P0DOX2\|IGA2_HUMAN Immunoglobulin alpha-2 heavy chain OS=Homo sapiens OX=9606 PE=1 SV=2 | 631 | 20201115_07433_1ng_glyco_hcd203040_compound15.6100.6100.2 | controllerType=0 controllerNumber=1 scan=6100 | 16.1441 |
| 03319:2 | 51 | K.TPLTAN[+1444.534]ITK.S | HexNAc(4)Hex(3)Fuc(1) | N[+1445] | 1202.0523 | 2 | 2403.0973 | 2403.0906 | 0 | 2.8 | 315 | Specific | 647.88 | 376.75 | 376.75 | 12.73 | 14 | >sp\|P0DOX2\|IGA2_HUMAN Immunoglobulin alpha-2 heavy chain OS=Homo sapiens OX=9606 PE=1 SV=2 | 631 | 20201115_07433_1ng_glyco_hcd203040_compound15.6098.6098.2 | controllerType=0 controllerNumber=1 scan=6098 | 16.1369 |
| 13617:3 | 61 | R.IIVPLNNREN[+2350.830]ISDPTSPLR.T | HexNAc(4)Hex(5)Fuc(1)NeuAc(2) | N[+2351] | 1500.345 | 3 | 4499.0205 | 4499.0075 | 0 | 2.9 | 62 | Specific | 278.28 | 278.28 | 278.28 | 7.09 | 13 | >sp\|P01591\|IGJ_HUMAN Immunoglobulin J chain OS=Homo sapiens OX=9606 GN=JCHAIN PE=1 SV=4 | 1392 | 20201115_07433_1ng_glyco_hcd203040_compound15.16809.16809.3 | controllerType=0 controllerNumber=1 scan=16809 | 33.9066 |
| 13620:3 | 61 | R.IIVPLNNREN[+2350.830]ISDPTSPLR.T | HexNAc(4)Hex(5)Fuc(1)NeuAc(2) | N[+2351] | 1500.345 | 3 | 4499.0205 | 4499.0075 | 0 | 2.9 | 62 | Specific | 288.56 | 265.94 | 265.94 | 7.09 | 13 | >sp\|P01591\|IGJ_HUMAN Immunoglobulin J chain OS=Homo sapiens OX=9606 GN=JCHAIN PE=1 SV=4 | 1392 | 20201115_07433_1ng_glyco_hcd203040_compound15.16812.16812.3 | controllerType=0 controllerNumber=1 scan=16812 | 33.9109 |
| 09975:3 | 61 | R.IIVPLNNREN[+1663.608]ISDPTSPLR.T | HexNAc(5)Hex(4) | N[+1664] | 1271.2656 | 3 | 3811.7821 | 3811.7853 | 0 | -0.8 | 62 | Specific | 338.86 | 316.71 | 316.71 | 6.16 | 13 | >sp\|P01591\|IGJ_HUMAN Immunoglobulin J chain OS=Homo sapiens OX=9606 GN=JCHAIN PE=1 SV=4 | 1392 | 20201115_07433_1ng_glyco_hcd203040_compound15.13011.13011.3 | controllerType=0 controllerNumber=1 scan=13011 | 27.5325 |
| 09977:3 | 61 | R.IIVPLNNREN[+1663.608]ISDPTSPLR.T | HexNAc(5)Hex(4) | N[+1664] | 1271.2656 | 3 | 3811.7821 | 3811.7853 | 0 | -0.8 | 62 | Specific | 349.41 | 288.14 | 288.14 | 6.06 | 13 | >sp\|P01591\|IGJ_HUMAN Immunoglobulin J chain OS=Homo sapiens OX=9606 GN=JCHAIN PE=1 SV=4 | 1392 | 20201115_07433_1ng_glyco_hcd203040_compound15.13013.13013.3 | controllerType=0 controllerNumber=1 scan=13013 | 27.5362 |
| 06938:3 | 345 | R.[+42.011]VEDAYILTC[+57.021]N[+730.264]VSLEYEK.T | HexNAc(2)Hex(2) | .[42], C[+57], N[+730] | 940.0935 | 3 | 2818.2661 | 2818.2486 | 0 | 6.2 | 225 | Specific | 263.27 | 184.44 | 184.44 | 3.25 | 3 | >sp\|P40227\|TCPZ_HUMAN T-complex protein 1 subunit zeta OS=Homo sapiens OX=9606 GN=CCT6A PE=1 SV=3 | 1115 | 20201115_07433_1ng_glyco_hcd203040_compound15.9843.9843.3 | controllerType=0 controllerNumber=1 scan=9843 | 22.2825 |
| 06910:3 | 345 | R.[+42.011]VEDAYILTC[+57.021]N[+730.264]VSLEYEK.T | HexNAc(2)Hex(2) | .[42], C[+57], N[+730] | 940.0946 | 3 | 2818.2692 | 2818.2486 | 0 | 7.3 | 225 | Specific | 333.89 | 198.86 | 198.86 | 3.49 | 3 | >sp\|P40227\|TCPZ_HUMAN T-complex protein 1 subunit zeta OS=Homo sapiens OX=9606 GN=CCT6A PE=1 SV=3 | 1115 | 20201115_07433_1ng_glyco_hcd203040_compound15.9814.9814.3 | controllerType=0 controllerNumber=1 scan=9814 | 22.236 |
| 17822:4 | 1 | R.[+42.011]RTAGWNVPIGTLRPFLN[+2336.851]WTGPPEPIEAAVAR.F | HexNAc(6)Hex(6)Fuc(1) | .[42], N[+2337] | 1442.1704 | 4 | 5765.6598 | 5765.6737 | 0 | -2.4 | 140 | Specific | 211.87 | 211.87 | 211.87 | 3.22 | 161 | >sp\|P02788\|TRFL_HUMAN Lactotransferrin OS=Homo sapiens OX=9606 GN=LTF PE=1 SV=6 | 1270 | 20201117_08512_1ng_glyco_hcd203040_compound15.21409.21409.4 | controllerType=0 controllerNumber=1 scan=21409 | 45.8467 |
| 16927:5 | 1 | R.RTAGWNVPIGTLRPFLN[+2059.735]WTGPPEPIEAAVAR.F | HexNAc(4)Hex(5)Fuc(1)NeuAc(1) | N[+2060] | 1090.1184 | 5 | 5446.5629 | 5446.547 | 0 | 2.9 | 140 | Specific | 243.88 | 243.88 | 243.88 | 6.26 | 161 | >sp\|P02788\|TRFL_HUMAN Lactotransferrin OS=Homo sapiens OX=9606 GN=LTF PE=1 SV=6 | 1270 | 20201117_08512_1ng_glyco_hcd203040_compound15.20473.20473.5 | controllerType=0 controllerNumber=1 scan=20473 | 43.709 |
| 17865:4 | 1 | R.RTAGWNVPIGTLRPFLN[+2350.830]WTGPPEPIEAAVAR.F | HexNAc(4)Hex(5)Fuc(1)NeuAc(2) | N[+2351] | 1435.1688 | 4 | 5737.6535 | 5737.6424 | 0 | 1.9 | 140 | Specific | 249.62 | 249.62 | 249.62 | 6.8 | 161 | >sp\|P02788\|TRFL_HUMAN Lactotransferrin OS=Homo sapiens OX=9606 GN=LTF PE=1 SV=6 | 1270 | 20201117_08512_1ng_glyco_hcd203040_compound15.21454.21454.4 | controllerType=0 controllerNumber=1 scan=21454 | 45.9385 |
| 16769:5 | 1 | R.RTAGWNVPIGTLRPFLN[+2570.925]WTGPPEPIEAAVAR.F | HexNAc(5)Hex(6)Fuc(2)NeuAc(1) | N[+2571] | 1192.3583 | 5 | 5957.7623 | 5957.7371 | 0 | 4.2 | 140 | Specific | 281.5 | 249.78 | 249.78 | 6.48 | 161 | >sp\|P02788\|TRFL_HUMAN Lactotransferrin OS=Homo sapiens OX=9606 GN=LTF PE=1 SV=6 | 1270 | 20201117_08512_1ng_glyco_hcd203040_compound15.20308.20308.5 | controllerType=0 controllerNumber=1 scan=20308 | 43.3328 |
| 16919:4 | 1 | R.RTAGWNVPIGTLRPFLN[+2059.735]WTGPPEPIEAAVAR.F | HexNAc(4)Hex(5)Fuc(1)NeuAc(1) | N[+2060] | 1362.3958 | 4 | 5446.5612 | 5446.547 | 0 | 2.6 | 140 | Specific | 353.24 | 311.71 | 311.71 | 7.63 | 161 | >sp\|P02788\|TRFL_HUMAN Lactotransferrin OS=Homo sapiens OX=9606 GN=LTF PE=1 SV=6 | 1270 | 20201117_08512_1ng_glyco_hcd203040_compound15.20465.20465.4 | controllerType=0 controllerNumber=1 scan=20465 | 43.6958 |
| 17905:4 | 1 | R.RTAGWNVPIGTLRPFLN[+2350.830]WTGPPEPIEAAVAR.F | HexNAc(4)Hex(5)Fuc(1)NeuAc(2) | N[+2351] | 1435.165 | 4 | 5737.6383 | 5737.6424 | 0 | -0.7 | 140 | Specific | 398.58 | 398.58 | 398.58 | 8.62 | 161 | >sp\|P02788\|TRFL_HUMAN Lactotransferrin OS=Homo sapiens OX=9606 GN=LTF PE=1 SV=6 | 1270 | 20201117_08512_1ng_glyco_hcd203040_compound15.21495.21495.4 | controllerType=0 controllerNumber=1 scan=21495 | 46.0204 |
| 16772:5 | 1 | R.RTAGWNVPIGTLRPFLN[+2570.925]WTGPPEPIEAAVAR.F | HexNAc(5)Hex(6)Fuc(2)NeuAc(1) | N[+2571] | 1192.3583 | 5 | 5957.7623 | 5957.7371 | 0 | 4.2 | 140 | Specific | 425.99 | 425.99 | 425.99 | 9.5 | 161 | >sp\|P02788\|TRFL_HUMAN Lactotransferrin OS=Homo sapiens OX=9606 GN=LTF PE=1 SV=6 | 1270 | 20201117_08512_1ng_glyco_hcd203040_compound15.20311.20311.5 | controllerType=0 controllerNumber=1 scan=20311 | 43.3511 |
| 16858:4 | 1 | R.RTAGWNVPIGTLRPFLN[+2351.851]WTGPPEPIEAAVAR.F | HexNAc(4)Hex(5)Fuc(3)NeuAc(1) | N[+2352] | 1435.4246 | 4 | 5738.6764 | 5738.6628 | 0 | 2.4 | 140 | Specific | 492.27 | 405.55 | 405.55 | 10.01 | 161 | >sp\|P02788\|TRFL_HUMAN Lactotransferrin OS=Homo sapiens OX=9606 GN=LTF PE=1 SV=6 | 1270 | 20201117_08512_1ng_glyco_hcd203040_compound15.20401.20401.4 | controllerType=0 controllerNumber=1 scan=20401 | 43.5468 |
| 16930:5 | 1 | R.RTAGWNVPIGTLRPFLN[+2059.735]WTGPPEPIEAAVAR.F | HexNAc(4)Hex(5)Fuc(1)NeuAc(1) | N[+2060] | 1090.1184 | 5 | 5446.5629 | 5446.547 | 0 | 2.9 | 140 | Specific | 492.37 | 465.07 | 465.07 | 9.83 | 161 | >sp\|P02788\|TRFL_HUMAN Lactotransferrin OS=Homo sapiens OX=9606 GN=LTF PE=1 SV=6 | 1270 | 20201117_08512_1ng_glyco_hcd203040_compound15.20476.20476.5 | controllerType=0 controllerNumber=1 scan=20476 | 43.7137 |
| 15943:4 | 1 | R.RTAGWNVPIGTLRPFLN[+1768.640]WTGPPEPIEAAVAR.F | HexNAc(4)Hex(5)Fuc(1) | N[+1769] | 1289.6199 | 4 | 5155.4577 | 5155.4516 | 0 | 1.2 | 140 | Specific | 535.14 | 470.96 | 470.96 | 10.34 | 161 | >sp\|P02788\|TRFL_HUMAN Lactotransferrin OS=Homo sapiens OX=9606 GN=LTF PE=1 SV=6 | 1270 | 20201117_08512_1ng_glyco_hcd203040_compound15.19446.19446.4 | controllerType=0 controllerNumber=1 scan=19446 | 41.3757 |
| 16861:4 | 1 | R.RTAGWNVPIGTLRPFLN[+2351.851]WTGPPEPIEAAVAR.F | HexNAc(4)Hex(5)Fuc(3)NeuAc(1) | N[+2352] | 1435.4246 | 4 | 5738.6764 | 5738.6628 | 0 | 2.4 | 140 | Specific | 538.01 | 376.03 | 376.03 | 8.77 | 161 | >sp\|P02788\|TRFL_HUMAN Lactotransferrin OS=Homo sapiens OX=9606 GN=LTF PE=1 SV=6 | 1270 | 20201117_08512_1ng_glyco_hcd203040_compound15.20404.20404.4 | controllerType=0 controllerNumber=1 scan=20404 | 43.5524 |
| 16921:4 | 1 | R.RTAGWNVPIGTLRPFLN[+2059.735]WTGPPEPIEAAVAR.F | HexNAc(4)Hex(5)Fuc(1)NeuAc(1) | N[+2060] | 1362.3958 | 4 | 5446.5612 | 5446.547 | 0 | 2.6 | 140 | Specific | 544.17 | 431.39 | 431.39 | 10.23 | 161 | >sp\|P02788\|TRFL_HUMAN Lactotransferrin OS=Homo sapiens OX=9606 GN=LTF PE=1 SV=6 | 1270 | 20201117_08512_1ng_glyco_hcd203040_compound15.20467.20467.4 | controllerType=0 controllerNumber=1 scan=20467 | 43.6998 |
| 17867:4 | 1 | R.RTAGWNVPIGTLRPFLN[+2350.830]WTGPPEPIEAAVAR.F | HexNAc(4)Hex(5)Fuc(1)NeuAc(2) | N[+2351] | 1435.1688 | 4 | 5737.6535 | 5737.6424 | 0 | 1.9 | 140 | Specific | 626.22 | 498.49 | 498.49 | 12.81 | 161 | >sp\|P02788\|TRFL_HUMAN Lactotransferrin OS=Homo sapiens OX=9606 GN=LTF PE=1 SV=6 | 1270 | 20201117_08512_1ng_glyco_hcd203040_compound15.21456.21456.4 | controllerType=0 controllerNumber=1 scan=21456 | 45.9419 |
| 15945:4 | 1 | R.RTAGWNVPIGTLRPFLN[+1768.640]WTGPPEPIEAAVAR.F | HexNAc(4)Hex(5)Fuc(1) | N[+1769] | 1289.6199 | 4 | 5155.4577 | 5155.4516 | 0 | 1.2 | 140 | Specific | 685.07 | 533.9 | 533.9 | 13.81 | 161 | >sp\|P02788\|TRFL_HUMAN Lactotransferrin OS=Homo sapiens OX=9606 GN=LTF PE=1 SV=6 | 1270 | 20201117_08512_1ng_glyco_hcd203040_compound15.19448.19448.4 | controllerType=0 controllerNumber=1 scan=19448 | 41.381 |
| 17089:4 | 1 | R.TAGWNVPIGTLRPFLN[+2279.830]WTGPPEPIEAAVAR.F | HexNAc(5)Hex(6)Fuc(2) | N[+2280] | 1378.3945 | 4 | 5510.5562 | 5510.5406 | 0 | 2.8 | 141 | Specific | 208.85 | 208.85 | 208.85 | 4.82 | 161 | >sp\|P02788\|TRFL_HUMAN Lactotransferrin OS=Homo sapiens OX=9606 GN=LTF PE=1 SV=6 | 1270 | 20201117_08512_1ng_glyco_hcd203040_compound15.20641.20641.4 | controllerType=0 controllerNumber=1 scan=20641 | 44.0319 |
| 18116:3 | 1 | R.TAGWNVPIGTLRPFLN[+2059.735]WTGPPEPIEAAVAR.F | HexNAc(4)Hex(5)Fuc(1)NeuAc(1) | N[+2060] | 1764.1554 | 3 | 5290.4515 | 5290.4459 | 0 | 1.1 | 141 | Specific | 251.31 | 251.31 | 251.31 | 6.8 | 161 | >sp\|P02788\|TRFL_HUMAN Lactotransferrin OS=Homo sapiens OX=9606 GN=LTF PE=1 SV=6 | 1270 | 20201117_08512_1ng_glyco_hcd203040_compound15.21715.21715.2 | controllerType=0 controllerNumber=1 scan=21715 | 46.5017 |
| 19335:5 | 1 | R.[+42.011]TAGWNVPIGTLRPFLN[+2336.851]WTGPPEPIEAAVAR.F | HexNAc(6)Hex(6)Fuc(1) | .[42], N[+2337] | 1122.7216 | 5 | 5609.5787 | 5609.5726 | 0 | 1.1 | 141 | Specific | 261.31 | 213.82 | 213.82 | 5.7 | 161 | >sp\|P02788\|TRFL_HUMAN Lactotransferrin OS=Homo sapiens OX=9606 GN=LTF PE=1 SV=6 | 1270 | 20201117_08512_1ng_glyco_hcd203040_compound15.22989.22989.5 | controllerType=0 controllerNumber=1 scan=22989 | 49.3836 |
| 17094:4 | 1 | R.[+42.011]TAGWNVPIGTLRPFLN[+1872.651]WTGPPEPIEAAVAR.F | HexNAc(3)Hex(6)NeuAc(1) | .[42], N[+1873] | 1287.111 | 4 | 5145.4222 | 5145.372 | 0 | 9.8 | 141 | Specific | 262.8 | 262.8 | 262.8 | 5.08 | 161 | >sp\|P02788\|TRFL_HUMAN Lactotransferrin OS=Homo sapiens OX=9606 GN=LTF PE=1 SV=6 | 1270 | 20201117_08512_1ng_glyco_hcd203040_compound15.20646.20646.4 | controllerType=0 controllerNumber=1 scan=20646 | 44.0418 |
| 17982:4 | 1 | R.TAGWNVPIGTLRPFLN[+2715.963]WTGPPEPIEAAVAR.F | HexNAc(5)Hex(6)Fuc(1)NeuAc(2) | N[+2716] | 1487.4308 | 4 | 5946.7013 | 5946.6735 | 0 | 4.7 | 141 | Specific | 266.19 | 163.52 | 163.52 | 5.26 | 161 | >sp\|P02788\|TRFL_HUMAN Lactotransferrin OS=Homo sapiens OX=9606 GN=LTF PE=1 SV=6 | 1270 | 20201117_08512_1ng_glyco_hcd203040_compound15.21575.21575.4 | controllerType=0 controllerNumber=1 scan=21575 | 46.1833 |
| 17095:3 | 1 | R.[+42.011]TAGWNVPIGTLRPFLN[+1872.651]WTGPPEPIEAAVAR.F | HexNAc(3)Hex(6)NeuAc(1) | .[42], N[+1873] | 1715.8123 | 3 | 5145.4223 | 5145.372 | 0 | 9.8 | 141 | Specific | 270.05 | 167.4 | 167.4 | 4.74 | 161 | >sp\|P02788\|TRFL_HUMAN Lactotransferrin OS=Homo sapiens OX=9606 GN=LTF PE=1 SV=6 | 1270 | 20201117_08512_1ng_glyco_hcd203040_compound15.20647.20647.3 | controllerType=0 controllerNumber=1 scan=20647 | 44.043 |
| 17057:4 | 1 | R.TAGWNVPIGTLRPFLN[+2498.904]WTGPPEPIEAAVAR.F | HexNAc(6)Hex(7)Fuc(1) | N[+2499] | 1433.162 | 4 | 5729.6261 | 5729.6148 | 0 | 2 | 141 | Specific | 278.39 | 270.57 | 270.57 | 6.83 | 161 | >sp\|P02788\|TRFL_HUMAN Lactotransferrin OS=Homo sapiens OX=9606 GN=LTF PE=1 SV=6 | 1270 | 20201117_08512_1ng_glyco_hcd203040_compound15.20607.20607.4 | controllerType=0 controllerNumber=1 scan=20607 | 43.9596 |
| 17098:3 | 1 | R.[+42.011]TAGWNVPIGTLRPFLN[+1872.651]WTGPPEPIEAAVAR.F | HexNAc(3)Hex(6)NeuAc(1) | .[42], N[+1873] | 1715.8123 | 3 | 5145.4223 | 5145.372 | 0 | 9.8 | 141 | Specific | 284.51 | 211.56 | 211.56 | 3.48 | 161 | >sp\|P02788\|TRFL_HUMAN Lactotransferrin OS=Homo sapiens OX=9606 GN=LTF PE=1 SV=6 | 1270 | 20201117_08512_1ng_glyco_hcd203040_compound15.20650.20650.3 | controllerType=0 controllerNumber=1 scan=20650 | 44.0494 |
| 17105:3 | 1 | R.TAGWNVPIGTLRPFLN[+2279.830]WTGPPEPIEAAVAR.F | HexNAc(5)Hex(6)Fuc(2) | N[+2280] | 1837.5237 | 3 | 5510.5565 | 5510.5406 | 0 | 2.9 | 141 | Specific | 284.67 | 222.93 | 222.93 | 6.65 | 161 | >sp\|P02788\|TRFL_HUMAN Lactotransferrin OS=Homo sapiens OX=9606 GN=LTF PE=1 SV=6 | 1270 | 20201117_08512_1ng_glyco_hcd203040_compound15.20657.20657.3 | controllerType=0 controllerNumber=1 scan=20657 | 44.0633 |
| 18021:4 | 1 | R.TAGWNVPIGTLRPFLN[+2076.750]WTGPPEPIEAAVAR.F | HexNAc(4)Hex(6)Fuc(2) | N[+2077] | 1327.624 | 4 | 5307.4743 | 5307.4612 | 0 | 2.5 | 141 | Specific | 298.94 | 298.94 | 298.94 | 7.2 | 161 | >sp\|P02788\|TRFL_HUMAN Lactotransferrin OS=Homo sapiens OX=9606 GN=LTF PE=1 SV=6 | 1270 | 20201117_08512_1ng_glyco_hcd203040_compound15.21616.21616.4 | controllerType=0 controllerNumber=1 scan=21616 | 46.2951 |
| 19332:5 | 1 | R.[+42.011]TAGWNVPIGTLRPFLN[+2336.851]WTGPPEPIEAAVAR.F | HexNAc(6)Hex(6)Fuc(1) | .[42], N[+2337] | 1122.7216 | 5 | 5609.5787 | 5609.5726 | 0 | 1.1 | 141 | Specific | 299.5 | 280.45 | 280.45 | 7.61 | 161 | >sp\|P02788\|TRFL_HUMAN Lactotransferrin OS=Homo sapiens OX=9606 GN=LTF PE=1 SV=6 | 1270 | 20201117_08512_1ng_glyco_hcd203040_compound15.22986.22986.5 | controllerType=0 controllerNumber=1 scan=22986 | 49.3689 |
| 18042:3 | 1 | R.TAGWNVPIGTLRPFLN[+2205.793]WTGPPEPIEAAVAR.F | HexNAc(4)Hex(5)Fuc(2)NeuAc(1) | N[+2206] | 1812.849 | 3 | 5436.5323 | 5436.5038 | 0 | 5.3 | 141 | Specific | 301.12 | 287.68 | 287.68 | 6.74 | 161 | >sp\|P02788\|TRFL_HUMAN Lactotransferrin OS=Homo sapiens OX=9606 GN=LTF PE=1 SV=6 | 1270 | 20201117_08512_1ng_glyco_hcd203040_compound15.21638.21638.2 | controllerType=0 controllerNumber=1 scan=21638 | 46.3458 |
| 19329:4 | 1 | R.[+42.011]TAGWNVPIGTLRPFLN[+2336.851]WTGPPEPIEAAVAR.F | HexNAc(6)Hex(6)Fuc(1) | .[42], N[+2337] | 1403.1512 | 4 | 5609.5832 | 5609.5726 | 0 | 1.9 | 141 | Specific | 303.29 | 303.29 | 303.29 | 7.31 | 161 | >sp\|P02788\|TRFL_HUMAN Lactotransferrin OS=Homo sapiens OX=9606 GN=LTF PE=1 SV=6 | 1270 | 20201117_08512_1ng_glyco_hcd203040_compound15.22983.22983.4 | controllerType=0 controllerNumber=1 scan=22983 | 49.3648 |
| 17097:4 | 1 | R.[+42.011]TAGWNVPIGTLRPFLN[+1872.651]WTGPPEPIEAAVAR.F | HexNAc(3)Hex(6)NeuAc(1) | .[42], N[+1873] | 1287.111 | 4 | 5145.4222 | 5145.372 | 0 | 9.8 | 141 | Specific | 313.81 | 274.55 | 274.55 | 4.12 | 161 | >sp\|P02788\|TRFL_HUMAN Lactotransferrin OS=Homo sapiens OX=9606 GN=LTF PE=1 SV=6 | 1270 | 20201117_08512_1ng_glyco_hcd203040_compound15.20649.20649.4 | controllerType=0 controllerNumber=1 scan=20649 | 44.0463 |
| 19327:4 | 1 | R.[+42.011]TAGWNVPIGTLRPFLN[+2336.851]WTGPPEPIEAAVAR.F | HexNAc(6)Hex(6)Fuc(1) | .[42], N[+2337] | 1403.1512 | 4 | 5609.5832 | 5609.5726 | 0 | 1.9 | 141 | Specific | 324.11 | 324.11 | 324.11 | 7.5 | 161 | >sp\|P02788\|TRFL_HUMAN Lactotransferrin OS=Homo sapiens OX=9606 GN=LTF PE=1 SV=6 | 1270 | 20201117_08512_1ng_glyco_hcd203040_compound15.22981.22981.4 | controllerType=0 controllerNumber=1 scan=22981 | 49.3566 |
| 19263:3 | 1 | R.TAGWNVPIGTLRPFLN[+2350.830]WTGPPEPIEAAVAR.F | HexNAc(4)Hex(5)Fuc(1)NeuAc(2) | N[+2351] | 1861.1915 | 3 | 5581.5599 | 5581.5413 | 0 | 3.3 | 141 | Specific | 325.5 | 140.23 | 140.23 | 6.08 | 161 | >sp\|P02788\|TRFL_HUMAN Lactotransferrin OS=Homo sapiens OX=9606 GN=LTF PE=1 SV=6 | 1270 | 20201117_08512_1ng_glyco_hcd203040_compound15.22914.22914.3 | controllerType=0 controllerNumber=1 scan=22914 | 49.1968 |
| 18028:4 | 1 | R.TAGWNVPIGTLRPFLN[+2351.851]WTGPPEPIEAAVAR.F | HexNAc(4)Hex(5)Fuc(3)NeuAc(1) | N[+2352] | 1396.4028 | 4 | 5582.5895 | 5582.5617 | 0 | 5 | 141 | Specific | 339.16 | 339.16 | 339.16 | 6.78 | 161 | >sp\|P02788\|TRFL_HUMAN Lactotransferrin OS=Homo sapiens OX=9606 GN=LTF PE=1 SV=6 | 1270 | 20201117_08512_1ng_glyco_hcd203040_compound15.21624.21624.4 | controllerType=0 controllerNumber=1 scan=21624 | 46.3146 |
| 17059:4 | 1 | R.TAGWNVPIGTLRPFLN[+2498.904]WTGPPEPIEAAVAR.F | HexNAc(6)Hex(7)Fuc(1) | N[+2499] | 1433.162 | 4 | 5729.6261 | 5729.6148 | 0 | 2 | 141 | Specific | 387.27 | 297.77 | 297.77 | 7.59 | 161 | >sp\|P02788\|TRFL_HUMAN Lactotransferrin OS=Homo sapiens OX=9606 GN=LTF PE=1 SV=6 | 1270 | 20201117_08512_1ng_glyco_hcd203040_compound15.20609.20609.4 | controllerType=0 controllerNumber=1 scan=20609 | 43.9633 |
| 18417:4 | 1 | R.TAGWNVPIGTLRPFLN[+2059.735]WTGPPEPIEAAVAR.F | HexNAc(4)Hex(5)Fuc(1)NeuAc(1) | N[+2060] | 1323.3682 | 4 | 5290.4508 | 5290.4459 | 0 | 0.9 | 141 | Specific | 407.68 | 382.78 | 382.78 | 9.51 | 161 | >sp\|P02788\|TRFL_HUMAN Lactotransferrin OS=Homo sapiens OX=9606 GN=LTF PE=1 SV=6 | 1270 | 20201117_08512_1ng_glyco_hcd203040_compound15.22029.22029.4 | controllerType=0 controllerNumber=1 scan=22029 | 47.176 |
| 17917:4 | 1 | R.TAGWNVPIGTLRPFLN[+2716.983]WTGPPEPIEAAVAR.F | HexNAc(5)Hex(6)Fuc(3)NeuAc(1) | N[+2717] | 1487.6815 | 4 | 5947.7043 | 5947.6939 | 0 | 1.7 | 141 | Specific | 408.23 | 408.23 | 408.23 | 9.51 | 161 | >sp\|P02788\|TRFL_HUMAN Lactotransferrin OS=Homo sapiens OX=9606 GN=LTF PE=1 SV=6 | 1270 | 20201117_08512_1ng_glyco_hcd203040_compound15.21508.21508.4 | controllerType=0 controllerNumber=1 scan=21508 | 46.0442 |
| 17903:3 | 1 | R.TAGWNVPIGTLRPFLN[+2424.867]WTGPPEPIEAAVAR.F | HexNAc(5)Hex(6)Fuc(1)NeuAc(1) | N[+2425] | 1885.8692 | 3 | 5655.5931 | 5655.5781 | 0 | 2.7 | 141 | Specific | 410.15 | 290.47 | 290.47 | 7.73 | 161 | >sp\|P02788\|TRFL_HUMAN Lactotransferrin OS=Homo sapiens OX=9606 GN=LTF PE=1 SV=6 | 1270 | 20201117_08512_1ng_glyco_hcd203040_compound15.21493.21493.3 | controllerType=0 controllerNumber=1 scan=21493 | 46.0174 |
| 18018:4 | 1 | R.TAGWNVPIGTLRPFLN[+2076.750]WTGPPEPIEAAVAR.F | HexNAc(4)Hex(6)Fuc(2) | N[+2077] | 1327.624 | 4 | 5307.4743 | 5307.4612 | 0 | 2.5 | 141 | Specific | 438.84 | 438.84 | 438.84 | 9.64 | 161 | >sp\|P02788\|TRFL_HUMAN Lactotransferrin OS=Homo sapiens OX=9606 GN=LTF PE=1 SV=6 | 1270 | 20201117_08512_1ng_glyco_hcd203040_compound15.21613.21613.4 | controllerType=0 controllerNumber=1 scan=21613 | 46.2895 |

Table S4. Cont.

| Query #:z | Protein Rank | Glycopeptide | Glycan | Modification Type(s) | Observed m/z | z | Observed (M+H) | Calc. mass (M+H) | Off-by-x error | Mass error (ppm) | StartPosition | Cleavage | Score | Delta | Delta Mod | \|Log Prob\| | # of unique peptides | Protein Name | Protein DB number | Comment | Scan # | Scan Time |
|---|---|---|---|---|---|---|---|---|---|---|---|---|---|---|---|---|---|---|---|---|---|---|
| 18047:5 | 1 | R.TAGWNVPIGTLRPFLN[+2350.830]WTGPPEPIEAAVAR.F | HexNAc(4)Hex(5)Fuc(1)NeuAc(2) | N[+2351] | 1117.1213 | 5 | 5581.5776 | 5581.5413 | 0 | 6.5 | 141 | Specific | 471.6 | 471.6 | 471.6 | 9.36 | 161 | >sp\|P02788\|TRFL_HUMAN Lactotransferrin OS=Homo sapiens OX=9606 GN=LTF PE=1 SV=6 | 1270 | 20201117_08512_1ng_glyco_hcd203040_compound15.21643.21643.5 | controllerType=0 controllerNumber=1 scan=21643 | 46.3626 |
| 17144:3 | 1 | R.TAGWNVPIGTLRPFLN[+1768.640]WTGPPEPIEAAVAR.F | HexNAc(4)Hex(5)Fuc(1) | N[+1769] | 1667.1259 | 3 | 4999.3633 | 4999.3504 | 0 | 2.6 | 141 | Specific | 484.6 | 373.93 | 373.93 | 8.54 | 161 | >sp\|P02788\|TRFL_HUMAN Lactotransferrin OS=Homo sapiens OX=9606 GN=LTF PE=1 SV=6 | 1270 | 20201117_08512_1ng_glyco_hcd203040_compound15.20700.20700.3 | controllerType=0 controllerNumber=1 scan=20700 | 44.1924 |
| 17984:4 | 1 | R.TAGWNVPIGTLRPFLN[+2715.963]WTGPPEPIEAAVAR.F | HexNAc(5)Hex(6)Fuc(1)NeuAc(2) | N[+2716] | 1487.4308 | 4 | 5946.7013 | 5946.6735 | 0 | 4.7 | 141 | Specific | 503 | 481.52 | 481.52 | 9.19 | 161 | >sp\|P02788\|TRFL_HUMAN Lactotransferrin OS=Homo sapiens OX=9606 GN=LTF PE=1 SV=6 | 1270 | 20201117_08512_1ng_glyco_hcd203040_compound15.21577.21577.4 | controllerType=0 controllerNumber=1 scan=21577 | 46.1866 |
| 17993:4 | 1 | R.TAGWNVPIGTLRPFLN[+2424.867]WTGPPEPIEAAVAR.F | HexNAc(5)Hex(6)Fuc(1)NeuAc(1) | N[+2425] | 1414.6558 | 4 | 5655.6012 | 5655.5781 | 0 | 4.1 | 141 | Specific | 546.98 | 457.08 | 457.08 | 9.97 | 161 | >sp\|P02788\|TRFL_HUMAN Lactotransferrin OS=Homo sapiens OX=9606 GN=LTF PE=1 SV=6 | 1270 | 20201117_08512_1ng_glyco_hcd203040_compound15.21587.21587.4 | controllerType=0 controllerNumber=1 scan=21587 | 46.211 |
| 19265:3 | 1 | R.TAGWNVPIGTLRPFLN[+2350.830]WTGPPEPIEAAVAR.F | HexNAc(4)Hex(5)Fuc(1)NeuAc(2) | N[+2351] | 1861.1915 | 3 | 5581.5599 | 5581.5413 | 0 | 3.3 | 141 | Specific | 555.24 | 355.73 | 355.73 | 8.91 | 161 | >sp\|P02788\|TRFL_HUMAN Lactotransferrin OS=Homo sapiens OX=9606 GN=LTF PE=1 SV=6 | 1270 | 20201117_08512_1ng_glyco_hcd203040_compound15.22916.22916.3 | controllerType=0 controllerNumber=1 scan=22916 | 49.2014 |
| 17889:4 | 1 | R.TAGWNVPIGTLRPFLN[+2936.057]WTGPPEPIEAAVAR.F | HexNAc(6)Hex(7)Fuc(2)NeuAc(1) | N[+2936] | 1542.4524 | 4 | 6166.7878 | 6166.7682 | 0 | 3.2 | 141 | Specific | 564.18 | 500.45 | 500.45 | 10.2 | 161 | >sp\|P02788\|TRFL_HUMAN Lactotransferrin OS=Homo sapiens OX=9606 GN=LTF PE=1 SV=6 | 1270 | 20201117_08512_1ng_glyco_hcd203040_compound15.21478.21478.4 | controllerType=0 controllerNumber=1 scan=21478 | 45.9851 |
| 17906:3 | 1 | R.TAGWNVPIGTLRPFLN[+2424.867]WTGPPEPIEAAVAR.F | HexNAc(5)Hex(6)Fuc(1)NeuAc(1) | N[+2425] | 1885.8692 | 3 | 5655.5931 | 5655.5781 | 0 | 2.7 | 141 | Specific | 567.59 | 420.27 | 420.27 | 10.46 | 161 | >sp\|P02788\|TRFL_HUMAN Lactotransferrin OS=Homo sapiens OX=9606 GN=LTF PE=1 SV=6 | 1270 | 20201117_08512_1ng_glyco_hcd203040_compound15.21496.21496.3 | controllerType=0 controllerNumber=1 scan=21496 | 46.0223 |
| 18013:3 | 1 | R.TAGWNVPIGTLRPFLN[+2205.793]WTGPPEPIEAAVAR.F | HexNAc(4)Hex(5)Fuc(2)NeuAc(1) | N[+2206] | 1812.8438 | 3 | 5436.517 | 5436.5038 | 0 | 2.4 | 141 | Specific | 618.02 | 565.2 | 565.2 | 12.71 | 161 | >sp\|P02788\|TRFL_HUMAN Lactotransferrin OS=Homo sapiens OX=9606 GN=LTF PE=1 SV=6 | 1270 | 20201117_08512_1ng_glyco_hcd203040_compound15.21608.21608.3 | controllerType=0 controllerNumber=1 scan=21608 | 46.2769 |
| 17141:4 | 1 | R.TAGWNVPIGTLRPFLN[+1768.640]WTGPPEPIEAAVAR.F | HexNAc(4)Hex(5)Fuc(1) | N[+1769] | 1250.5957 | 4 | 4999.361 | 4999.3504 | 0 | 2.1 | 141 | Specific | 626.69 | 582.83 | 582.83 | 12.54 | 161 | >sp\|P02788\|TRFL_HUMAN Lactotransferrin OS=Homo sapiens OX=9606 GN=LTF PE=1 SV=6 | 1270 | 20201117_08512_1ng_glyco_hcd203040_compound15.20697.20697.4 | controllerType=0 controllerNumber=1 scan=20697 | 44.1865 |
| 17147:3 | 1 | R.TAGWNVPIGTLRPFLN[+1768.640]WTGPPEPIEAAVAR.F | HexNAc(4)Hex(5)Fuc(1) | N[+1769] | 1667.1259 | 3 | 4999.3633 | 4999.3504 | 0 | 2.6 | 141 | Specific | 627.44 | 585.36 | 585.36 | 12.71 | 161 | >sp\|P02788\|TRFL_HUMAN Lactotransferrin OS=Homo sapiens OX=9606 GN=LTF PE=1 SV=6 | 1270 | 20201117_08512_1ng_glyco_hcd203040_compound15.20703.20703.3 | controllerType=0 controllerNumber=1 scan=20703 | 44.197 |
| 18119:3 | 1 | R.TAGWNVPIGTLRPFLN[+2059.735]WTGPPEPIEAAVAR.F | HexNAc(4)Hex(5)Fuc(1)NeuAc(1) | N[+2060] | 1764.1554 | 3 | 5290.4515 | 5290.4459 | 0 | 1.1 | 141 | Specific | 637.4 | 544.66 | 544.66 | 12.81 | 161 | >sp\|P02788\|TRFL_HUMAN Lactotransferrin OS=Homo sapiens OX=9606 GN=LTF PE=1 SV=6 | 1270 | 20201117_08512_1ng_glyco_hcd203040_compound15.21718.21718.2 | controllerType=0 controllerNumber=1 scan=21718 | 46.5075 |
| 18035:4 | 1 | R.TAGWNVPIGTLRPFLN[+2076.750]WTGPPEPIEAAVAR.F | HexNAc(4)Hex(6)Fuc(2) | N[+2077] | 1327.6256 | 4 | 5307.4806 | 5307.4612 | 0 | 3.7 | 141 | Specific | 638.06 | 625.53 | 625.53 | 12.44 | 161 | >sp\|P02788\|TRFL_HUMAN Lactotransferrin OS=Homo sapiens OX=9606 GN=LTF PE=1 SV=6 | 1270 | 20201117_08512_1ng_glyco_hcd203040_compound15.21631.21631.4 | controllerType=0 controllerNumber=1 scan=21631 | 46.3325 |
| 18032:4 | 1 | R.TAGWNVPIGTLRPFLN[+2076.750]WTGPPEPIEAAVAR.F | HexNAc(4)Hex(6)Fuc(2) | N[+2077] | 1327.6256 | 4 | 5307.4806 | 5307.4612 | 0 | 3.7 | 141 | Specific | 644.48 | 617.57 | 617.57 | 12.44 | 161 | >sp\|P02788\|TRFL_HUMAN Lactotransferrin OS=Homo sapiens OX=9606 GN=LTF PE=1 SV=6 | 1270 | 20201117_08512_1ng_glyco_hcd203040_compound15.21628.21628.4 | controllerType=0 controllerNumber=1 scan=21628 | 46.3241 |
| 18008:4 | 1 | R.TAGWNVPIGTLRPFLN[+2205.793]WTGPPEPIEAAVAR.F | HexNAc(4)Hex(5)Fuc(2)NeuAc(1) | N[+2206] | 1359.8843 | 4 | 5436.5153 | 5436.5038 | 0 | 2.1 | 141 | Specific | 661.8 | 590.57 | 590.57 | 13.54 | 161 | >sp\|P02788\|TRFL_HUMAN Lactotransferrin OS=Homo sapiens OX=9606 GN=LTF PE=1 SV=6 | 1270 | 20201117_08512_1ng_glyco_hcd203040_compound15.21603.21603.4 | controllerType=0 controllerNumber=1 scan=21603 | 46.2632 |
| 18016:3 | 1 | R.TAGWNVPIGTLRPFLN[+2205.793]WTGPPEPIEAAVAR.F | HexNAc(4)Hex(5)Fuc(2)NeuAc(1) | N[+2206] | 1812.8438 | 3 | 5436.517 | 5436.5038 | 0 | 2.4 | 141 | Specific | 664.32 | 608.93 | 608.93 | 13.71 | 161 | >sp\|P02788\|TRFL_HUMAN Lactotransferrin OS=Homo sapiens OX=9606 GN=LTF PE=1 SV=6 | 1270 | 20201117_08512_1ng_glyco_hcd203040_compound15.21611.21611.3 | controllerType=0 controllerNumber=1 scan=21611 | 46.2873 |
| 19262:4 | 1 | R.TAGWNVPIGTLRPFLN[+2350.830]WTGPPEPIEAAVAR.F | HexNAc(4)Hex(5)Fuc(1)NeuAc(2) | N[+2351] | 1396.1455 | 4 | 5581.5602 | 5581.5413 | 0 | 3.4 | 141 | Specific | 697.25 | 561.12 | 561.12 | 13.44 | 161 | >sp\|P02788\|TRFL_HUMAN Lactotransferrin OS=Homo sapiens OX=9606 GN=LTF PE=1 SV=6 | 1270 | 20201117_08512_1ng_glyco_hcd203040_compound15.22913.22913.4 | controllerType=0 controllerNumber=1 scan=22913 | 49.1954 |
| 18017:3 | 1 | R.TAGWNVPIGTLRPFLN[+2059.735]WTGPPEPIEAAVAR.F | HexNAc(4)Hex(5)Fuc(1)NeuAc(1) | N[+2060] | 1764.1584 | 3 | 5290.4607 | 5290.4459 | 0 | 2.8 | 141 | Specific | 715.13 | 576.48 | 576.48 | 14.71 | 161 | >sp\|P02788\|TRFL_HUMAN Lactotransferrin OS=Homo sapiens OX=9606 GN=LTF PE=1 SV=6 | 1270 | 20201117_08512_1ng_glyco_hcd203040_compound15.21612.21612.3 | controllerType=0 controllerNumber=1 scan=21612 | 46.2883 |
| 18014:3 | 1 | R.TAGWNVPIGTLRPFLN[+2059.735]WTGPPEPIEAAVAR.F | HexNAc(4)Hex(5)Fuc(1)NeuAc(1) | N[+2060] | 1764.1584 | 3 | 5290.4607 | 5290.4459 | 0 | 2.8 | 141 | Specific | 721.56 | 636.52 | 636.52 | 14.71 | 161 | >sp\|P02788\|TRFL_HUMAN Lactotransferrin OS=Homo sapiens OX=9606 GN=LTF PE=1 SV=6 | 1270 | 20201117_08512_1ng_glyco_hcd203040_compound15.21609.21609.3 | controllerType=0 controllerNumber=1 scan=21609 | 46.278 |
| 18049:5 | 1 | R.TAGWNVPIGTLRPFLN[+2059.735]WTGPPEPIEAAVAR.F | HexNAc(4)Hex(5)Fuc(1)NeuAc(1) | N[+2060] | 1058.8983 | 5 | 5290.4625 | 5290.4459 | 0 | 3.1 | 141 | Specific | 725.31 | 691.84 | 691.84 | 15.07 | 161 | >sp\|P02788\|TRFL_HUMAN Lactotransferrin OS=Homo sapiens OX=9606 GN=LTF PE=1 SV=6 | 1270 | 20201117_08512_1ng_glyco_hcd203040_compound15.21646.21646.5 | controllerType=0 controllerNumber=1 scan=21646 | 46.371 |
| 18009:4 | 1 | R.TAGWNVPIGTLRPFLN[+2059.735]WTGPPEPIEAAVAR.F | HexNAc(4)Hex(5)Fuc(1)NeuAc(1) | N[+2060] | 1323.3716 | 4 | 5290.4645 | 5290.4459 | 0 | 3.5 | 141 | Specific | 738.74 | 670.31 | 670.31 | 14.44 | 161 | >sp\|P02788\|TRFL_HUMAN Lactotransferrin OS=Homo sapiens OX=9606 GN=LTF PE=1 SV=6 | 1270 | 20201117_08512_1ng_glyco_hcd203040_compound15.21604.21604.4 | controllerType=0 controllerNumber=1 scan=21604 | 46.2646 |
| 17995:4 | 1 | R.TAGWNVPIGTLRPFLN[+2424.867]WTGPPEPIEAAVAR.F | HexNAc(5)Hex(6)Fuc(1)NeuAc(1) | N[+2425] | 1414.6558 | 4 | 5655.6012 | 5655.5781 | 0 | 4.1 | 141 | Specific | 747.44 | 544.34 | 544.34 | 14.44 | 161 | >sp\|P02788\|TRFL_HUMAN Lactotransferrin OS=Homo sapiens OX=9606 GN=LTF PE=1 SV=6 | 1270 | 20201117_08512_1ng_glyco_hcd203040_compound15.21589.21589.4 | controllerType=0 controllerNumber=1 scan=21589 | 46.2208 |
| 17919:4 | 1 | R.TAGWNVPIGTLRPFLN[+2716.983]WTGPPEPIEAAVAR.F | HexNAc(5)Hex(6)Fuc(3)NeuAc(1) | N[+2717] | 1487.6815 | 4 | 5947.7043 | 5947.6939 | 0 | 1.7 | 141 | Specific | 753.91 | 630.37 | 630.37 | 15.64 | 161 | >sp\|P02788\|TRFL_HUMAN Lactotransferrin OS=Homo sapiens OX=9606 GN=LTF PE=1 SV=6 | 1270 | 20201117_08512_1ng_glyco_hcd203040_compound15.21510.21510.4 | controllerType=0 controllerNumber=1 scan=21510 | 46.0479 |
| 17092:4 | 1 | R.TAGWNVPIGTLRPFLN[+2279.830]WTGPPEPIEAAVAR.F | HexNAc(5)Hex(6)Fuc(2) | N[+2280] | 1378.3945 | 4 | 5510.5562 | 5510.5406 | 0 | 2.8 | 141 | Specific | 773.05 | 661.76 | 661.76 | 15.54 | 161 | >sp\|P02788\|TRFL_HUMAN Lactotransferrin OS=Homo sapiens OX=9606 GN=LTF PE=1 SV=6 | 1270 | 20201117_08512_1ng_glyco_hcd203040_compound15.20644.20644.4 | controllerType=0 controllerNumber=1 scan=20644 | 44.0381 |
| 19266:5 | 1 | R.TAGWNVPIGTLRPFLN[+2350.830]WTGPPEPIEAAVAR.F | HexNAc(4)Hex(5)Fuc(1)NeuAc(2) | N[+2351] | 1117.1178 | 5 | 5581.5599 | 5581.5413 | 0 | 3.3 | 141 | Specific | 785.61 | 745.21 | 745.21 | 16.07 | 161 | >sp\|P02788\|TRFL_HUMAN Lactotransferrin OS=Homo sapiens OX=9606 GN=LTF PE=1 SV=6 | 1270 | 20201117_08512_1ng_glyco_hcd203040_compound15.22917.22917.5 | controllerType=0 controllerNumber=1 scan=22917 | 49.2025 |
| 19264:4 | 1 | R.TAGWNVPIGTLRPFLN[+2350.830]WTGPPEPIEAAVAR.F | HexNAc(4)Hex(5)Fuc(1)NeuAc(2) | N[+2351] | 1396.1455 | 4 | 5581.5602 | 5581.5413 | 0 | 3.4 | 141 | Specific | 814.17 | 636.87 | 636.87 | 16.44 | 161 | >sp\|P02788\|TRFL_HUMAN Lactotransferrin OS=Homo sapiens OX=9606 GN=LTF PE=1 SV=6 | 1270 | 20201117_08512_1ng_glyco_hcd203040_compound15.22915.22915.4 | controllerType=0 controllerNumber=1 scan=22915 | 49.2003 |
| 18011:4 | 1 | R.TAGWNVPIGTLRPFLN[+2059.735]WTGPPEPIEAAVAR.F | HexNAc(4)Hex(5)Fuc(1)NeuAc(1) | N[+2060] | 1323.3716 | 4 | 5290.4645 | 5290.4459 | 0 | 3.5 | 141 | Specific | 830.77 | 721.72 | 721.72 | 16.44 | 161 | >sp\|P02788\|TRFL_HUMAN Lactotransferrin OS=Homo sapiens OX=9606 GN=LTF PE=1 SV=6 | 1270 | 20201117_08512_1ng_glyco_hcd203040_compound15.21606.21606.4 | controllerType=0 controllerNumber=1 scan=21606 | 46.2746 |
| 18010:4 | 1 | R.TAGWNVPIGTLRPFLN[+2205.793]WTGPPEPIEAAVAR.F | HexNAc(4)Hex(5)Fuc(2)NeuAc(1) | N[+2206] | 1359.8843 | 4 | 5436.5153 | 5436.5038 | 0 | 2.1 | 141 | Specific | 839.26 | 723.11 | 723.11 | 16.54 | 161 | >sp\|P02788\|TRFL_HUMAN Lactotransferrin OS=Homo sapiens OX=9606 GN=LTF PE=1 SV=6 | 1270 | 20201117_08512_1ng_glyco_hcd203040_compound15.21605.21605.4 | controllerType=0 controllerNumber=1 scan=21605 | 46.2691 |
| 18030:4 | 1 | R.TAGWNVPIGTLRPFLN[+2351.851]WTGPPEPIEAAVAR.F | HexNAc(4)Hex(5)Fuc(3)NeuAc(1) | N[+2352] | 1396.4028 | 4 | 5582.5895 | 5582.5617 | 0 | 5 | 141 | Specific | 849.22 | 733.47 | 733.47 | 15.9 | 161 | >sp\|P02788\|TRFL_HUMAN Lactotransferrin OS=Homo sapiens OX=9606 GN=LTF PE=1 SV=6 | 1270 | 20201117_08512_1ng_glyco_hcd203040_compound15.21626.21626.4 | controllerType=0 controllerNumber=1 scan=21626 | 46.3185 |
| 17143:4 | 1 | R.TAGWNVPIGTLRPFLN[+1768.640]WTGPPEPIEAAVAR.F | HexNAc(4)Hex(5)Fuc(1) | N[+1769] | 1250.5957 | 4 | 4999.361 | 4999.3504 | 0 | 2.1 | 141 | Specific | 872.8 | 773.31 | 773.31 | 17.54 | 161 | >sp\|P02788\|TRFL_HUMAN Lactotransferrin OS=Homo sapiens OX=9606 GN=LTF PE=1 SV=6 | 1270 | 20201117_08512_1ng_glyco_hcd203040_compound15.20699.20699.4 | controllerType=0 controllerNumber=1 scan=20699 | 44.191 |
| 19269:5 | 1 | R.TAGWNVPIGTLRPFLN[+2350.830]WTGPPEPIEAAVAR.F | HexNAc(4)Hex(5)Fuc(1)NeuAc(2) | N[+2351] | 1117.1178 | 5 | 5581.5599 | 5581.5413 | 0 | 3.3 | 141 | Specific | 874.42 | 750.08 | 750.08 | 18.07 | 161 | >sp\|P02788\|TRFL_HUMAN Lactotransferrin OS=Homo sapiens OX=9606 GN=LTF PE=1 SV=6 | 1270 | 20201117_08512_1ng_glyco_hcd203040_compound15.22920.22920.5 | controllerType=0 controllerNumber=1 scan=22920 | 49.2088 |
| 18419:4 | 1 | R.TAGWNVPIGTLRPFLN[+2059.735]WTGPPEPIEAAVAR.F | HexNAc(4)Hex(5)Fuc(1)NeuAc(1) | N[+2060] | 1323.3682 | 4 | 5290.4508 | 5290.4459 | 0 | 0.9 | 141 | Specific | 928.85 | 771.25 | 771.25 | 18.64 | 161 | >sp\|P02788\|TRFL_HUMAN Lactotransferrin OS=Homo sapiens OX=9606 GN=LTF PE=1 SV=6 | 1270 | 20201117_08512_1ng_glyco_hcd203040_compound15.22031.22031.4 | controllerType=0 controllerNumber=1 scan=22031 | 47.1805 |
| 18051:5 | 1 | R.TAGWNVPIGTLRPFLN[+2059.735]WTGPPEPIEAAVAR.F | HexNAc(4)Hex(5)Fuc(1)NeuAc(1) | N[+2060] | 1058.8983 | 5 | 5290.4625 | 5290.4459 | 0 | 3.1 | 141 | Specific | 936.99 | 787.02 | 787.02 | 19.07 | 161 | >sp\|P02788\|TRFL_HUMAN Lactotransferrin OS=Homo sapiens OX=9606 GN=LTF PE=1 SV=6 | 1270 | 20201117_08512_1ng_glyco_hcd203040_compound15.21648.21648.5 | controllerType=0 controllerNumber=1 scan=21648 | 46.3757 |
| 16100:3 | 1 | R.TAGWNIPMGLLFN[+1216.423]QTGSC[+57.021]K.F | HexNAc(2)Hex(5) | C[+57], N[+1216] | 1104.4859 | 3 | 3311.4431 | 3311.4328 | 0 | 3.1 | 485 | Specific | 223.56 | 223.56 | 105.75 | 4.79 | 161 | >sp\|P02788\|TRFL_HUMAN Lactotransferrin OS=Homo sapiens OX=9606 GN=LTF PE=1 SV=6 | 1270 | 20201117_08512_1ng_glyco_hcd203040_compound15.19609.19609.2 | controllerType=0 controllerNumber=1 scan=19609 | 41.7255 |
| 19401:3 | 1 | R.[+42.011]TAGWNIPMGLLFN[+2336.851]QTGSC[+57.021]K.F | HexNAc(6)Hex(6)Fuc(1) | .[42], C[+57], N[+2337] | 1491.9515 | 3 | 4473.8399 | 4473.8716 | 0 | -7.1 | 485 | Specific | 272 | 220.19 | 220.19 | 4.13 | 161 | >sp\|P02788\|TRFL_HUMAN Lactotransferrin OS=Homo sapiens OX=9606 GN=LTF PE=1 SV=6 | 1270 | 20201117_08512_1ng_glyco_hcd203040_compound15.23058.23058.3 | controllerType=0 controllerNumber=1 scan=23058 | 49.5545 |
| 15851:3 | 1 | R.TAGWNIPMGLLFN[+1622.582]QTGSC[+57.021]K.F | HexNAc(4)Hex(5) | C[+57], N[+1623] | 1239.871 | 3 | 3717.5985 | 3717.5916 | 0 | 1.9 | 485 | Specific | 476.63 | 476.63 | 195.64 | 9.73 | 161 | >sp\|P02788\|TRFL_HUMAN Lactotransferrin OS=Homo sapiens OX=9606 GN=LTF PE=1 SV=6 | 1270 | 20201117_08512_1ng_glyco_hcd203040_compound15.19349.19349.3 | controllerType=0 controllerNumber=1 scan=19349 | 41.1484 |
| 19363:4 | 1 | R.TAGWNIPMGLLFN[+2350.830]QTGSC[+57.021]K.F | HexNAc(4)Hex(5)Fuc(1)NeuAc(2) | C[+57], N[+2351] | 1112.219 | 4 | 4445.8543 | 4445.8403 | 0 | 3.2 | 485 | Specific | 519.25 | 376.29 | 376.29 | 8.91 | 161 | >sp\|P02788\|TRFL_HUMAN Lactotransferrin OS=Homo sapiens OX=9606 GN=LTF PE=1 SV=6 | 1270 | 20201117_08512_1ng_glyco_hcd203040_compound15.23019.23019.4 | controllerType=0 controllerNumber=1 scan=23019 | 49.4639 |
| 17578:3 | 1 | R.TAGWNIPMGLLFN[+1913.677]QTGSC[+57.021]K.F | HexNAc(4)Hex(5)NeuAc(1) | C[+57], N[+1914] | 1336.9058 | 3 | 4008.7029 | 4008.687 | 0 | 4 | 485 | Specific | 552.12 | 499.03 | 499.03 | 9.88 | 161 | >sp\|P02788\|TRFL_HUMAN Lactotransferrin OS=Homo sapiens OX=9606 GN=LTF PE=1 SV=6 | 1270 | 20201117_08512_1ng_glyco_hcd203040_compound15.21155.21155.3 | controllerType=0 controllerNumber=1 scan=21155 | 45.2876 |
| 16124:3 | 1 | R.TAGWNIPMGLLFN[+1216.423]QTGSC[+57.021]K.F | HexNAc(2)Hex(5) | C[+57], N[+1216] | 1104.4856 | 3 | 3311.4423 | 3311.4328 | 0 | 2.9 | 485 | Specific | 563.84 | 468.56 | 173.05 | 10.2 | 161 | >sp\|P02788\|TRFL_HUMAN Lactotransferrin OS=Homo sapiens OX=9606 GN=LTF PE=1 SV=6 | 1270 | 20201117_08512_1ng_glyco_hcd203040_compound15.19634.19634.3 | controllerType=0 controllerNumber=1 scan=19634 | 41.7856 |
| 17585:4 | 1 | R.TAGWNIPMGLLFN[+1913.677]QTGSC[+57.021]K.F | HexNAc(4)Hex(5)NeuAc(1) | C[+57], N[+1914] | 1002.9312 | 4 | 4008.703 | 4008.687 | 0 | 4 | 485 | Specific | 603.09 | 603.09 | 603.09 | 12.98 | 161 | >sp\|P02788\|TRFL_HUMAN Lactotransferrin OS=Homo sapiens OX=9606 GN=LTF PE=1 SV=6 | 1270 | 20201117_08512_1ng_glyco_hcd203040_compound15.21162.21162.4 | controllerType=0 controllerNumber=1 scan=21162 | 45.2991 |
| 15854:3 | 1 | R.TAGWNIPMGLLFN[+1622.582]QTGSC[+57.021]K.F | HexNAc(4)Hex(5) | C[+57], N[+1623] | 1239.871 | 3 | 3717.5985 | 3717.5916 | 0 | 1.9 | 485 | Specific | 688.64 | 626.37 | 339.89 | 13.55 | 161 | >sp\|P02788\|TRFL_HUMAN Lactotransferrin OS=Homo sapiens OX=9606 GN=LTF PE=1 SV=6 | 1270 | 20201117_08512_1ng_glyco_hcd203040_compound15.19352.19352.3 | controllerType=0 controllerNumber=1 scan=19352 | 41.1527 |
| 19339:4 | 1 | R.TAGWNIPMGLLFN[+2350.830]QTGSC[+57.021]K.F | HexNAc(4)Hex(5)Fuc(1)NeuAc(2) | C[+57], N[+2351] | 1112.222 | 4 | 4445.866 | 4445.8403 | 0 | 5.8 | 485 | Specific | 727.31 | 669.73 | 669.73 | 14.09 | 161 | >sp\|P02788\|TRFL_HUMAN Lactotransferrin OS=Homo sapiens OX=9606 GN=LTF PE=1 SV=6 | 1270 | 20201117_08512_1ng_glyco_hcd203040_compound15.22993.22993.4 | controllerType=0 controllerNumber=1 scan=22993 | 49.3935 |
| 16102:3 | 1 | R.TAGWNIPMGLLFN[+1216.423]QTGSC[+57.021]K.F | HexNAc(2)Hex(5) | C[+57], N[+1216] | 1104.4859 | 3 | 3311.4431 | 3311.4328 | 0 | 3.1 | 485 | Specific | 748.78 | 638.91 | 294.54 | 14.45 | 161 | >sp\|P02788\|TRFL_HUMAN Lactotransferrin OS=Homo sapiens OX=9606 GN=LTF PE=1 SV=6 | 1270 | 20201117_08512_1ng_glyco_hcd203040_compound15.19611.19611.2 | controllerType=0 controllerNumber=1 scan=19611 | 41.7287 |
| 17405:3 | 1 | R.TAGWNIPMGLLFN[+2205.793]QTGSC[+57.021]K.F | HexNAc(4)Hex(5)Fuc(2)NeuAc(1) | C[+57], N[+2206] | 1434.2762 | 3 | 4300.814 | 4300.8028 | 0 | 2.6 | 485 | Specific | 771.15 | 751.89 | 751.89 | 15.45 | 161 | >sp\|P02788\|TRFL_HUMAN Lactotransferrin OS=Homo sapiens OX=9606 GN=LTF PE=1 SV=6 | 1270 | 20201117_08512_1ng_glyco_hcd203040_compound15.20972.20972.3 | controllerType=0 controllerNumber=1 scan=20972 | 44.8254 |
| 19365:4 | 1 | R.TAGWNIPMGLLFN[+2350.830]QTGSC[+57.021]K.F | HexNAc(4)Hex(5)Fuc(1)NeuAc(2) | C[+57], N[+2351] | 1112.219 | 4 | 4445.8543 | 4445.8403 | 0 | 3.2 | 485 | Specific | 786.29 | 560.27 | 560.27 | 15.98 | 161 | >sp\|P02788\|TRFL_HUMAN Lactotransferrin OS=Homo sapiens OX=9606 GN=LTF PE=1 SV=6 | 1270 | 20201117_08512_1ng_glyco_hcd203040_compound15.23021.23021.4 | controllerType=0 controllerNumber=1 scan=23021 | 49.4686 |
| 19342:4 | 1 | R.TAGWNIPMGLLFN[+2350.830]QTGSC[+57.021]K.F | HexNAc(4)Hex(5)Fuc(1)NeuAc(2) | C[+57], N[+2351] | 1112.222 | 4 | 4445.866 | 4445.8403 | 0 | 5.8 | 485 | Specific | 806.88 | 601.47 | 601.47 | 16.09 | 161 | >sp\|P02788\|TRFL_HUMAN Lactotransferrin OS=Homo sapiens OX=9606 GN=LTF PE=1 SV=6 | 1270 | 20201117_08512_1ng_glyco_hcd203040_compound15.22996.22996.4 | controllerType=0 controllerNumber=1 scan=22996 | 49.3996 |
| 16126:3 | 1 | R.TAGWNIPMGLLFN[+1216.423]QTGSC[+57.021]K.F | HexNAc(2)Hex(5) | C[+57], N[+1216] | 1104.4856 | 3 | 3311.4423 | 3311.4328 | 0 | 2.9 | 485 | Specific | 820.8 | 765.14 | 376.61 | 16.45 | 161 | >sp\|P02788\|TRFL_HUMAN Lactotransferrin OS=Homo sapiens OX=9606 GN=LTF PE=1 SV=6 | 1270 | 20201117_08512_1ng_glyco_hcd203040_compound15.19636.19636.3 | controllerType=0 controllerNumber=1 scan=19636 | 41.791 |
| 17534:4 | 1 | R.TAGWNIPMGLLFN[+1913.677]QTGSC[+57.021]K.F | HexNAc(4)Hex(5)NeuAc(1) | C[+57], N[+1914] | 1002.9307 | 4 | 4008.701 | 4008.687 | 0 | 3.5 | 485 | Specific | 870.83 | 790.74 | 790.74 | 17.98 | 161 | >sp\|P02788\|TRFL_HUMAN Lactotransferrin OS=Homo sapiens OX=9606 GN=LTF PE=1 SV=6 | 1270 | 20201117_08512_1ng_glyco_hcd203040_compound15.21108.21108.2 | controllerType=0 controllerNumber=1 scan=21108 | 45.1583 |
| 17407:3 | 1 | R.TAGWNIPMGLLFN[+2205.793]QTGSC[+57.021]K.F | HexNAc(4)Hex(5)Fuc(2)NeuAc(1) | C[+57], N[+2206] | 1434.2762 | 3 | 4300.814 | 4300.8028 | 0 | 2.6 | 485 | Specific | 872.38 | 759.78 | 759.78 | 17.45 | 161 | >sp\|P02788\|TRFL_HUMAN Lactotransferrin OS=Homo sapiens OX=9606 GN=LTF PE=1 SV=6 | 1270 | 20201117_08512_1ng_glyco_hcd203040_compound15.20974.20974.3 | controllerType=0 controllerNumber=1 scan=20974 | 44.8298 |
| 19334:3 | 1 | R.TAGWNIPMGLLFN[+2350.830]QTGSC[+57.021]K.F | HexNAc(4)Hex(5)Fuc(1)NeuAc(2) | C[+57], N[+2351] | 1482.6327 | 3 | 4445.8837 | 4445.8403 | 0 | 9.8 | 485 | Specific | 883.85 | 883.85 | 883.85 | 15.85 | 161 | >sp\|P02788\|TRFL_HUMAN Lactotransferrin OS=Homo sapiens OX=9606 GN=LTF PE=1 SV=6 | 1270 | 20201117_08512_1ng_glyco_hcd203040_compound15.22988.22988.3 | controllerType=0 controllerNumber=1 scan=22988 | 49.3818 |
| 17588:4 | 1 | R.TAGWNIPMGLLFN[+1913.677]QTGSC[+57.021]K.F | HexNAc(4)Hex(5)NeuAc(1) | C[+57], N[+1914] | 1002.9312 | 4 | 4008.703 | 4008.687 | 0 | 4 | 485 | Specific | 894.76 | 856.41 | 856.41 | 17.98 | 161 | >sp\|P02788\|TRFL_HUMAN Lactotransferrin OS=Homo sapiens OX=9606 GN=LTF PE=1 SV=6 | 1270 | 20201117_08512_1ng_glyco_hcd203040_compound15.21165.21165.4 | controllerType=0 controllerNumber=1 scan=21165 | 45.3103 |
| 17440:3 | 1 | R.TAGWNIPMGLLFN[+2059.735]QTGSC[+57.021]K.F | HexNAc(4)Hex(5)Fuc(1)NeuAc(1) | C[+57], N[+2060] | 1385.5901 | 3 | 4154.7559 | 4154.7449 | 0 | 2.6 | 485 | Specific | 895.34 | 892.18 | 892.18 | 17.45 | 161 | >sp\|P02788\|TRFL_HUMAN Lactotransferrin OS=Homo sapiens OX=9606 GN=LTF PE=1 SV=6 | 1270 | 20201117_08512_1ng_glyco_hcd203040_compound15.21009.21009.3 | controllerType=0 controllerNumber=1 scan=21009 | 44.9056 |
| 19331:3 | 1 | R.TAGWNIPMGLLFN[+2350.830]QTGSC[+57.021]K.F | HexNAc(4)Hex(5)Fuc(1)NeuAc(2) | C[+57], N[+2351] | 1482.6327 | 3 | 4445.8837 | 4445.8403 | 0 | 9.8 | 485 | Specific | 898.4 | 898.4 | 898.4 | 15.85 | 161 | >sp\|P02788\|TRFL_HUMAN Lactotransferrin OS=Homo sapiens OX=9606 GN=LTF PE=1 SV=6 | 1270 | 20201117_08512_1ng_glyco_hcd203040_compound15.22985.22985.3 | controllerType=0 controllerNumber=1 scan=22985 | 49.3678 |
| 17537:4 | 1 | R.TAGWNIPMGLLFN[+1913.677]QTGSC[+57.021]K.F | HexNAc(4)Hex(5)NeuAc(1) | C[+57], N[+1914] | 1002.9307 | 4 | 4008.701 | 4008.687 | 0 | 3.5 | 485 | Specific | 902.71 | 855.35 | 855.35 | 18.98 | 161 | >sp\|P02788\|TRFL_HUMAN Lactotransferrin OS=Homo sapiens OX=9606 GN=LTF PE=1 SV=6 | 1270 | 20201117_08512_1ng_glyco_hcd203040_compound15.21111.21111.2 | controllerType=0 controllerNumber=1 scan=21111 | 45.1676 |
| 15630:3 | 1 | R.TAGWNIPMGLLFN[+1768.640]QTGSC[+57.021]K.F | HexNAc(4)Hex(5)Fuc(1) | C[+57], N[+1769] | 1288.5566 | 3 | 3863.6552 | 3863.6495 | 0 | 1.5 | 485 | Specific | 908.3 | 749.78 | 749.78 | 18.55 | 161 | >sp\|P02788\|TRFL_HUMAN Lactotransferrin OS=Homo sapiens OX=9606 GN=LTF PE=1 SV=6 | 1270 | 20201117_08512_1ng_glyco_hcd203040_compound15.19119.19119.3 | controllerType=0 controllerNumber=1 scan=19119 | 40.6541 |
| 16021:2 | 1 | R.TAGWNIPMGLLFN[+1216.423]QTGSC[+57.021]K.F | HexNAc(2)Hex(5) | C[+57], N[+1216] | 1656.2234 | 2 | 3311.4396 | 3311.4328 | 0 | 2 | 485 | Specific | 909.84 | 792.41 | 297.36 | 18.63 | 161 | >sp\|P02788\|TRFL_HUMAN Lactotransferrin OS=Homo sapiens OX=9606 GN=LTF PE=1 SV=6 | 1270 | 20201117_08512_1ng_glyco_hcd203040_compound15.19527.19527.2 | controllerType=0 controllerNumber=1 scan=19527 | 41.551 |
| 17580:3 | 1 | R.TAGWNIPMGLLFN[+1913.677]QTGSC[+57.021]K.F | HexNAc(4)Hex(5)NeuAc(1) | C[+57], N[+1914] | 1336.9058 | 3 | 4008.7029 | 4008.687 | 0 | 4 | 485 | Specific | 914.69 | 874 | 874 | 18.36 | 161 | >sp\|P02788\|TRFL_HUMAN Lactotransferrin OS=Homo sapiens OX=9606 GN=LTF PE=1 SV=6 | 1270 | 20201117_08512_1ng_glyco_hcd203040_compound15.21157.21157.3 | controllerType=0 controllerNumber=1 scan=21157 | 45.2919 |
| 16018:2 | 1 | R.TAGWNIPMGLLFN[+1216.423]QTGSC[+57.021]K.F | HexNAc(2)Hex(5) | C[+57], N[+1216] | 1656.2234 | 2 | 3311.4396 | 3311.4328 | 0 | 2 | 485 | Specific | 924.2 | 903.48 | 341.85 | 18.63 | 161 | >sp\|P02788\|TRFL_HUMAN Lactotransferrin OS=Homo sapiens OX=9606 GN=LTF PE=1 SV=6 | 1270 | 20201117_08512_1ng_glyco_hcd203040_compound15.19524.19524.2 | controllerType=0 controllerNumber=1 scan=19524 | 41.5449 |
| 15628:3 | 1 | R.TAGWNIPMGLLFN[+1768.640]QTGSC[+57.021]K.F | HexNAc(4)Hex(5)Fuc(1) | C[+57], N[+1769] | 1288.5566 | 3 | 3863.6552 | 3863.6495 | 0 | 1.5 | 485 | Specific | 924.82 | 924.82 | 924.82 | 18.55 | 161 | >sp\|P02788\|TRFL_HUMAN Lactotransferrin OS=Homo sapiens OX=9606 GN=LTF PE=1 SV=6 | 1270 | 20201117_08512_1ng_glyco_hcd203040_compound15.19117.19117.3 | controllerType=0 controllerNumber=1 scan=19117 | 40.6486 |
| 15991:3 | 1 | R.TAGWNIPMGLLFN[+1216.423]QTGSC[+57.021]K.F | HexNAc(2)Hex(5) | C[+57], N[+1216] | 1104.4848 | 3 | 3311.4398 | 3311.4328 | 0 | 2.1 | 485 | Specific | 950.93 | 950.93 | 405.39 | 19.45 | 161 | >sp\|P02788\|TRFL_HUMAN Lactotransferrin OS=Homo sapiens OX=9606 GN=LTF PE=1 SV=6 | 1270 | 20201117_08512_1ng_glyco_hcd203040_compound15.19496.19496.3 | controllerType=0 controllerNumber=1 scan=19496 | 41.4878 |
| 17438:3 | 1 | R.TAGWNIPMGLLFN[+2059.735]QTGSC[+57.021]K.F | HexNAc(4)Hex(5)Fuc(1)NeuAc(1) | C[+57], N[+2060] | 1385.5901 | 3 | 4154.7559 | 4154.7449 | 0 | 2.6 | 485 | Specific | 950.99 | 853.46 | 853.46 | 19.45 | 161 | >sp\|P02788\|TRFL_HUMAN Lactotransferrin OS=Homo sapiens OX=9606 GN=LTF PE=1 SV=6 | 1270 | 20201117_08512_1ng_glyco_hcd203040_compound15.21007.21007.3 | controllerType=0 controllerNumber=1 scan=21007 | 44.9006 |
| 15994:3 | 1 | R.TAGWNIPMGLLFN[+1216.423]QTGSC[+57.021]K.F | HexNAc(2)Hex(5) | C[+57], N[+1216] | 1104.4848 | 3 | 3311.4398 | 3311.4328 | 0 | 2.1 | 485 | Specific | 1040.95 | 885.51 | 499.12 | 20.45 | 161 | >sp\|P02788\|TRFL_HUMAN Lactotransferrin OS=Homo sapiens OX=9606 GN=LTF PE=1 SV=6 | 1270 | 20201117_08512_1ng_glyco_hcd203040_compound15.19499.19499.3 | controllerType=0 controllerNumber=1 scan=19499 | 41.4942 |
| 16133:3 | 1 | R.TAGWNIPM[+15.995]GLLFN[+2059.735]QTGSC[+57.021]K.F | HexNAc(4)Hex(5)Fuc(1)NeuAc(1) | C[+57], M[+16], N[+2060] | 1390.9266 | 3 | 4170.7652 | 4170.7398 | 0 | 6.1 | 485 | Specific | 524.87 | 503.13 | 503.13 | 8.86 | 161 | >sp\|P02788\|TRFL_HUMAN Lactotransferrin OS=Homo sapiens OX=9606 GN=LTF PE=1 SV=6 | 1270 | 20201117_08512_1ng_glyco_hcd203040_compound15.19644.19644.3 | controllerType=0 controllerNumber=1 scan=19644 | 41.8093 |
| 16135:3 | 1 | R.TAGWNIPM[+15.995]GLLFN[+2059.735]QTGSC[+57.021]K.F | HexNAc(4)Hex(5)Fuc(1)NeuAc(1) | C[+57], M[+16], N[+2060] | 1390.9266 | 3 | 4170.7652 | 4170.7398 | 0 | 6.1 | 485 | Specific | 662.85 | 551.86 | 551.86 | 12.47 | 161 | >sp\|P02788\|TRFL_HUMAN Lactotransferrin OS=Homo sapiens OX=9606 GN=LTF PE=1 SV=6 | 1270 | 20201117_08512_1ng_glyco_hcd203040_compound15.19646.19646.3 | controllerType=0 controllerNumber=1 scan=19646 | 41.8161 |
| 17629:4 | 1 | R.TAGWNIPMGLLFN[+2076.750]QTGSC[+57.021]KFDEYFSQSC[+57.021]APGSDPR.S | HexNAc(4)Hex(6)Fuc(2) | C[+57]*2, N[+2077] | 1504.6403 | 4 | 6015.5392 | 6015.5074 | 0 | 5.3 | 485 | Specific | 359.11 | 359.11 | 359.11 | 6.67 | 161 | >sp\|P02788\|TRFL_HUMAN Lactotransferrin OS=Homo sapiens OX=9606 GN=LTF PE=1 SV=6 | 1270 | 20201117_08512_1ng_glyco_hcd203040_compound15.21208.21208.4 | controllerType=0 controllerNumber=1 scan=21208 | 45.4352 |
| 17636:4 | 1 | R.TAGWNIPMGLLFN[+2076.750]QTGSC[+57.021]KFDEYFSQSC[+57.021]APGSDPR.S | HexNAc(4)Hex(6)Fuc(2) | C[+57]*2, N[+2077] | 1504.6431 | 4 | 6015.5504 | 6015.5074 | 0 | 7.2 | 485 | Specific | 369.03 | 369.03 | 369.03 | 6.32 | 161 | >sp\|P02788\|TRFL_HUMAN Lactotransferrin OS=Homo sapiens OX=9606 GN=LTF PE=1 SV=6 | 1270 | 20201117_08512_1ng_glyco_hcd203040_compound15.21216.21216.4 | controllerType=0 controllerNumber=1 scan=21216 | 45.4512 |
| 16875:4 | 1 | R.TAGWNIPMGLLFN[+1216.423]QTGSC[+57.021]KFDEYFSQSC[+57.021]APGSDPR.S | HexNAc(2)Hex(5) | C[+57]*2, N[+1216] | 1289.5527 | 4 | 5155.1891 | 5155.18 | 0 | 1.8 | 485 | Specific | 435.57 | 417.47 | 91.65 | 9.25 | 161 | >sp\|P02788\|TRFL_HUMAN Lactotransferrin OS=Homo sapiens OX=9606 GN=LTF PE=1 SV=6 | 1270 | 20201117_08512_1ng_glyco_hcd203040_compound15.20419.20419.4 | controllerType=0 controllerNumber=1 scan=20419 | 43.5883 |
| 17797:3 | 1 | R.TAGWNIPMGLLFN[+1913.677]QTGSC[+57.021]KFDEYFSQSC[+57.021]APGSDPR.S | HexNAc(4)Hex(5)NeuAc(1) | C[+57]*2, N[+1914] | 1951.4818 | 3 | 5852.4308 | 5852.4341 | 0 | -0.6 | 485 | Specific | 439.66 | 334.87 | 334.87 | 6.56 | 161 | >sp\|P02788\|TRFL_HUMAN Lactotransferrin OS=Homo sapiens OX=9606 GN=LTF PE=1 SV=6 | 1270 | 20201117_08512_1ng_glyco_hcd203040_compound15.21383.21383.3 | controllerType=0 controllerNumber=1 scan=21383 | 45.7912 |
| 17765:5 | 1 | R.TAGWNIPMGLLFN[+1913.677]QTGSC[+57.021]KFDEYFSQSC[+57.021]APGSDPR.S | HexNAc(4)Hex(5)NeuAc(1) | C[+57]*2, N[+1914] | 1171.2952 | 5 | 5852.4467 | 5852.4341 | 0 | 2.2 | 485 | Specific | 440.71 | 440.71 | 440.71 | 9.77 | 161 | >sp\|P02788\|TRFL_HUMAN Lactotransferrin OS=Homo sapiens OX=9606 GN=LTF PE=1 SV=6 | 1270 | 20201117_08512_1ng_glyco_hcd203040_compound15.21350.21350.5 | controllerType=0 controllerNumber=1 scan=21350 | 45.7191 |
| 19042:4 | 1 | R.TAGWNIPMGLLFN[+2204.772]QTGSC[+57.021]KFDEYFSQSC[+57.021]APGSDPR.S | HexNAc(4)Hex(5)NeuAc(2) | C[+57]*2, N[+2205] | 1536.6366 | 4 | 6143.5246 | 6143.5295 | 0 | -0.8 | 485 | Specific | 442.36 | 427.67 | 192.42 | 8.18 | 161 | >sp\|P02788\|TRFL_HUMAN Lactotransferrin OS=Homo sapiens OX=9606 GN=LTF PE=1 SV=6 | 1270 | 20201117_08512_1ng_glyco_hcd203040_compound15.22682.22682.4 | controllerType=0 controllerNumber=1 scan=22682 | 48.6573 |
| 17757:5 | 1 | R.TAGWNIPMGLLFN[+1913.677]QTGSC[+57.021]KFDEYFSQSC[+57.021]APGSDPR.S | HexNAc(4)Hex(5)NeuAc(1) | C[+57]*2, N[+1914] | 1171.2939 | 5 | 5852.4406 | 5852.4341 | 0 | 1.1 | 485 | Specific | 443.6 | 443.6 | 443.6 | 9.87 | 161 | >sp\|P02788\|TRFL_HUMAN Lactotransferrin OS=Homo sapiens OX=9606 GN=LTF PE=1 SV=6 | 1270 | 20201117_08512_1ng_glyco_hcd203040_compound15.21341.21341.5 | controllerType=0 controllerNumber=1 scan=21341 | 45.7046 |
| 17788:4 | 1 | R.TAGWNIPMGLLFN[+1913.677]QTGSC[+57.021]KFDEYFSQSC[+57.021]APGSDPR.S | HexNAc(4)Hex(5)NeuAc(1) | C[+57]*2, N[+1914] | 1463.8672 | 4 | 5852.4469 | 5852.4341 | 0 | 2.2 | 485 | Specific | 456.12 | 397.65 | 397.65 | 9.15 | 161 | >sp\|P02788\|TRFL_HUMAN Lactotransferrin OS=Homo sapiens OX=9606 GN=LTF PE=1 SV=6 | 1270 | 20201117_08512_1ng_glyco_hcd203040_compound15.21374.21374.4 | controllerType=0 controllerNumber=1 scan=21374 | 45.7751 |
| 16709:4 | 1 | R.TAGWNIPMGLLFN[+1971.719]QTGSC[+57.021]KFDEYFSQSC[+57.021]APGSDPR.S | HexNAc(5)Hex(5)Fuc(1) | C[+57]*2, N[+1972] | 1478.3744 | 4 | 5910.4757 | 5910.476 | 0 | 0 | 485 | Specific | 459.86 | 416.07 | 23.47 | 8.68 | 161 | >sp\|P02788\|TRFL_HUMAN Lactotransferrin OS=Homo sapiens OX=9606 GN=LTF PE=1 SV=6 | 1270 | 20201117_08512_1ng_glyco_hcd203040_compound15.20245.20245.2 | controllerType=0 controllerNumber=1 scan=20245 | 43.1969 |
| 16715:3 | 1 | R.TAGWNIPMGLLFN[+1768.640]QTGSC[+57.021]KFDEYFSQSC[+57.021]APGSDPR.S | HexNAc(4)Hex(5)Fuc(1) | C[+57]*2, N[+1769] | 1903.144 | 3 | 5707.4175 | 5707.3966 | 0 | 3.7 | 485 | Specific | 528.55 | 457.59 | 457.59 | 9.52 | 161 | >sp\|P02788\|TRFL_HUMAN Lactotransferrin OS=Homo sapiens OX=9606 GN=LTF PE=1 SV=6 | 1270 | 20201117_08512_1ng_glyco_hcd203040_compound15.20252.20252.3 | controllerType=0 controllerNumber=1 scan=20252 | 43.2122 |
| 16877:4 | 1 | R.TAGWNIPMGLLFN[+1216.423]QTGSC[+57.021]KFDEYFSQSC[+57.021]APGSDPR.S | HexNAc(2)Hex(5) | C[+57]*2, N[+1216] | 1289.5527 | 4 | 5155.1891 | 5155.18 | 0 | 1.8 | 485 | Specific | 545.91 | 545.91 | 230.4 | 9.67 | 161 | >sp\|P02788\|TRFL_HUMAN Lactotransferrin OS=Homo sapiens OX=9606 GN=LTF PE=1 SV=6 | 1270 | 20201117_08512_1ng_glyco_hcd203040_compound15.20421.20421.4 | controllerType=0 controllerNumber=1 scan=20421 | 43.5935 |
| 16727:4 | 1 | R.TAGWNIPMGLLFN[+1971.719]QTGSC[+57.021]KFDEYFSQSC[+57.021]APGSDPR.S | HexNAc(5)Hex(5)Fuc(1) | C[+57]*2, N[+1972] | 1478.382 | 4 | 5910.506 | 5910.476 | 0 | 5.1 | 485 | Specific | 560.08 | 560.08 | 165.07 | 9.16 | 161 | >sp\|P02788\|TRFL_HUMAN Lactotransferrin OS=Homo sapiens OX=9606 GN=LTF PE=1 SV=6 | 1270 | 20201117_08512_1ng_glyco_hcd203040_compound15.20264.20264.4 | controllerType=0 controllerNumber=1 scan=20264 | 43.2341 |
| 17637:4 | 1 | R.TAGWNIPMGLLFN[+2351.851]QTGSC[+57.021]KFDEYFSQSC[+57.021]APGSDPR.S | HexNAc(4)Hex(5)Fuc(3)NeuAc(1) | C[+57]*2, N[+2352] | 1573.4213 | 4 | 6290.6632 | 6290.6079 | 0 | 8.8 | 485 | Specific | 571.41 | 526.75 | 526.75 | 8.27 | 161 | >sp\|P02788\|TRFL_HUMAN Lactotransferrin OS=Homo sapiens OX=9606 GN=LTF PE=1 SV=6 | 1270 | 20201117_08512_1ng_glyco_hcd203040_compound15.21217.21217.4 | controllerType=0 controllerNumber=1 scan=21217 | 45.4526 |
| 16697:3 | 1 | R.TAGWNIPMGLLFN[+1768.640]QTGSC[+57.021]KFDEYFSQSC[+57.021]APGSDPR.S | HexNAc(4)Hex(5)Fuc(1) | C[+57]*2, N[+1769] | 1903.1417 | 3 | 5707.4105 | 5707.3966 | 0 | 2.4 | 485 | Specific | 612.37 | 549.22 | 549.22 | 12.22 | 161 | >sp\|P02788\|TRFL_HUMAN Lactotransferrin OS=Homo sapiens OX=9606 GN=LTF PE=1 SV=6 | 1270 | 20201117_08512_1ng_glyco_hcd203040_compound15.20233.20233.3 | controllerType=0 controllerNumber=1 scan=20233 | 43.1742 |
| 19044:4 | 1 | R.TAGWNIPMGLLFN[+2204.772]QTGSC[+57.021]KFDEYFSQSC[+57.021]APGSDPR.S | HexNAc(4)Hex(5)NeuAc(2) | C[+57]*2, N[+2205] | 1536.6366 | 4 | 6143.5246 | 6143.5295 | 0 | -0.8 | 485 | Specific | 650.5 | 609.63 | 374.65 | 12.08 | 161 | >sp\|P02788\|TRFL_HUMAN Lactotransferrin OS=Homo sapiens OX=9606 GN=LTF PE=1 SV=6 | 1270 | 20201117_08512_1ng_glyco_hcd203040_compound15.22684.22684.4 | controllerType=0 controllerNumber=1 scan=22684 | 48.6636 |
| 17733:3 | 1 | R.TAGWNIPMGLLFN[+1913.677]QTGSC[+57.021]KFDEYFSQSC[+57.021]APGSDPR.S | HexNAc(4)Hex(5)NeuAc(1) | C[+57]*2, N[+1914] | 1951.4839 | 3 | 5852.437 | 5852.4341 | 0 | 0.5 | 485 | Specific | 652.85 | 544.47 | 544.47 | 12.67 | 161 | >sp\|P02788\|TRFL_HUMAN Lactotransferrin OS=Homo sapiens OX=9606 GN=LTF PE=1 SV=6 | 1270 | 20201117_08512_1ng_glyco_hcd203040_compound15.21317.21317.3 | controllerType=0 controllerNumber=1 scan=21317 | 45.6665 |
| 16699:3 | 1 | R.TAGWNIPMGLLFN[+1768.640]QTGSC[+57.021]KFDEYFSQSC[+57.021]APGSDPR.S | HexNAc(4)Hex(5)Fuc(1) | C[+57]*2, N[+1769] | 1903.1417 | 3 | 5707.4105 | 5707.3966 | 0 | 2.4 | 485 | Specific | 661.98 | 519.35 | 519.35 | 13.22 | 161 | >sp\|P02788\|TRFL_HUMAN Lactotransferrin OS=Homo sapiens OX=9606 GN=LTF PE=1 SV=6 | 1270 | 20201117_08512_1ng_glyco_hcd203040_compound15.20235.20235.3 | controllerType=0 controllerNumber=1 scan=20235 | 43.1787 |
| 16729:4 | 1 | R.TAGWNIPMGLLFN[+1971.719]QTGSC[+57.021]KFDEYFSQSC[+57.021]APGSDPR.S | HexNAc(5)Hex(5)Fuc(1) | C[+57]*2, N[+1972] | 1478.382 | 4 | 5910.506 | 5910.476 | 0 | 5.1 | 485 | Specific | 684.62 | 679.52 | 319.91 | 12.41 | 161 | >sp\|P02788\|TRFL_HUMAN Lactotransferrin OS=Homo sapiens OX=9606 GN=LTF PE=1 SV=6 | 1270 | 20201117_08512_1ng_glyco_hcd203040_compound15.20266.20266.4 | controllerType=0 controllerNumber=1 scan=20266 | 43.2378 |
| 17638:4 | 1 | R.TAGWNIPMGLLFN[+2076.750]QTGSC[+57.021]KFDEYFSQSC[+57.021]APGSDPR.S | HexNAc(4)Hex(6)Fuc(2) | C[+57]*2, N[+2077] | 1504.6431 | 4 | 6015.5504 | 6015.5074 | 0 | 7.2 | 485 | Specific | 687.96 | 687.96 | 687.96 | 11.97 | 161 | >sp\|P02788\|TRFL_HUMAN Lactotransferrin OS=Homo sapiens OX=9606 GN=LTF PE=1 SV=6 | 1270 | 20201117_08512_1ng_glyco_hcd203040_compound15.21218.21218.4 | controllerType=0 controllerNumber=1 scan=21218 | 45.4546 |
| 16712:4 | 1 | R.TAGWNIPMGLLFN[+1768.640]QTGSC[+57.021]KFDEYFSQSC[+57.021]APGSDPR.S | HexNAc(4)Hex(5)Fuc(1) | C[+57]*2, N[+1769] | 1427.6105 | 4 | 5707.4201 | 5707.3966 | 0 | 4.1 | 485 | Specific | 707.5 | 653.5 | 653.5 | 13.95 | 161 | >sp\|P02788\|TRFL_HUMAN Lactotransferrin OS=Homo sapiens OX=9606 GN=LTF PE=1 SV=6 | 1270 | 20201117_08512_1ng_glyco_hcd203040_compound15.20249.20249.4 | controllerType=0 controllerNumber=1 scan=20249 | 43.2066 |
| 16711:4 | 1 | R.TAGWNIPMGLLFN[+1971.719]QTGSC[+57.021]KFDEYFSQSC[+57.021]APGSDPR.S | HexNAc(5)Hex(5)Fuc(1) | C[+57]*2, N[+1972] | 1478.3744 | 4 | 5910.4757 | 5910.476 | 0 | 0 | 485 | Specific | 725.22 | 675.24 | 281.5 | 13.49 | 161 | >sp\|P02788\|TRFL_HUMAN Lactotransferrin OS=Homo sapiens OX=9606 GN=LTF PE=1 SV=6 | 1270 | 20201117_08512_1ng_glyco_hcd203040_compound15.20247.20247.2 | controllerType=0 controllerNumber=1 scan=20247 | 43.2021 |
| 17847:4 | 1 | R.TAGWNIPMGLLFN[+1548.545]QTGSC[+57.021]KFDEYFSQSC[+57.021]APGSDPR.S | HexNAc(3)Hex(4)NeuAc(1) | C[+57]*2, N[+1549] | 1372.5862 | 4 | 5487.3229 | 5487.3019 | 0 | 3.8 | 485 | Specific | 740.92 | 717.05 | 717.05 | 13.95 | 161 | >sp\|P02788\|TRFL_HUMAN Lactotransferrin OS=Homo sapiens OX=9606 GN=LTF PE=1 SV=6 | 1270 | 20201117_08512_1ng_glyco_hcd203040_compound15.21435.21435.4 | controllerType=0 controllerNumber=1 scan=21435 | 45.8919 |
| 16718:3 | 1 | R.TAGWNIPMGLLFN[+1768.640]QTGSC[+57.021]KFDEYFSQSC[+57.021]APGSDPR.S | HexNAc(4)Hex(5)Fuc(1) | C[+57]*2, N[+1769] | 1903.144 | 3 | 5707.4175 | 5707.3966 | 0 | 3.7 | 485 | Specific | 747.02 | 619.05 | 619.05 | 14.13 | 161 | >sp\|P02788\|TRFL_HUMAN Lactotransferrin OS=Homo sapiens OX=9606 GN=LTF PE=1 SV=6 | 1270 | 20201117_08512_1ng_glyco_hcd203040_compound15.20255.20255.3 | controllerType=0 controllerNumber=1 scan=20255 | 43.216 |
| 17736:3 | 1 | R.TAGWNIPMGLLFN[+1913.677]QTGSC[+57.021]KFDEYFSQSC[+57.021]APGSDPR.S | HexNAc(4)Hex(5)NeuAc(1) | C[+57]*2, N[+1914] | 1951.4839 | 3 | 5852.437 | 5852.4341 | 0 | 0.5 | 485 | Specific | 758.81 | 601.34 | 601.34 | 14.67 | 161 | >sp\|P02788\|TRFL_HUMAN Lactotransferrin OS=Homo sapiens OX=9606 GN=LTF PE=1 SV=6 | 1270 | 20201117_08512_1ng_glyco_hcd203040_compound15.21320.21320.3 | controllerType=0 controllerNumber=1 scan=21320 | 45.6704 |
| 17624:4 | 1 | R.TAGWNIPMGLLFN[+2351.851]QTGSC[+57.021]KFDEYFSQSC[+57.021]APGSDPR.S | HexNAc(4)Hex(5)Fuc(3)NeuAc(1) | C[+57]*2, N[+2352] | 1573.4128 | 4 | 6290.6296 | 6290.6079 | 0 | 3.4 | 485 | Specific | 838.18 | 825.01 | 825.01 | 15.95 | 161 | >sp\|P02788\|TRFL_HUMAN Lactotransferrin OS=Homo sapiens OX=9606 GN=LTF PE=1 SV=6 | 1270 | 20201117_08512_1ng_glyco_hcd203040_compound15.21203.21203.4 | controllerType=0 controllerNumber=1 scan=21203 | 45.4214 |
| 17676:4 | 1 | R.TAGWNIPMGLLFN[+2059.735]QTGSC[+57.021]KFDEYFSQSC[+57.021]APGSDPR.S | HexNAc(4)Hex(5)Fuc(1)NeuAc(1) | C[+57]*2, N[+2060] | 1500.3827 | 4 | 5998.5089 | 5998.492 | 0 | 2.8 | 485 | Specific | 869.78 | 832.62 | 832.62 | 17.04 | 161 | >sp\|P02788\|TRFL_HUMAN Lactotransferrin OS=Homo sapiens OX=9606 GN=LTF PE=1 SV=6 | 1270 | 20201117_08512_1ng_glyco_hcd203040_compound15.21258.21258.4 | controllerType=0 controllerNumber=1 scan=21258 | 45.557 |
| 17760:5 | 1 | R.TAGWNIPMGLLFN[+1913.677]QTGSC[+57.021]KFDEYFSQSC[+57.021]APGSDPR.S | HexNAc(4)Hex(5)NeuAc(1) | C[+57]*2, N[+1914] | 1171.2939 | 5 | 5852.4406 | 5852.4341 | 0 | 1.1 | 485 | Specific | 870.89 | 801.1 | 801.1 | 17.77 | 161 | >sp\|P02788\|TRFL_HUMAN Lactotransferrin OS=Homo sapiens OX=9606 GN=LTF PE=1 SV=6 | 1270 | 20201117_08512_1ng_glyco_hcd203040_compound15.21344.21344.5 | controllerType=0 controllerNumber=1 scan=21344 | 45.7099 |
| 17639:4 | 1 | R.TAGWNIPMGLLFN[+2351.851]QTGSC[+57.021]KFDEYFSQSC[+57.021]APGSDPR.S | HexNAc(4)Hex(5)Fuc(3)NeuAc(1) | C[+57]*2, N[+2352] | 1573.4213 | 4 | 6290.6632 | 6290.6079 | 0 | 8.8 | 485 | Specific | 878.25 | 868.05 | 868.05 | 15.52 | 161 | >sp\|P02788\|TRFL_HUMAN Lactotransferrin OS=Homo sapiens OX=9606 GN=LTF PE=1 SV=6 | 1270 | 20201117_08512_1ng_glyco_hcd203040_compound15.21219.21219.4 | controllerType=0 controllerNumber=1 scan=21219 | 45.457 |
| 16714:4 | 1 | R.TAGWNIPMGLLFN[+1768.640]QTGSC[+57.021]KFDEYFSQSC[+57.021]APGSDPR.S | HexNAc(4)Hex(5)Fuc(1) | C[+57]*2, N[+1769] | 1427.6105 | 4 | 5707.4201 | 5707.3966 | 0 | 4.1 | 485 | Specific | 900.66 | 867.25 | 867.25 | 17.95 | 161 | >sp\|P02788\|TRFL_HUMAN Lactotransferrin OS=Homo sapiens OX=9606 GN=LTF PE=1 SV=6 | 1270 | 20201117_08512_1ng_glyco_hcd203040_compound15.20251.20251.4 | controllerType=0 controllerNumber=1 scan=20251 | 43.2112 |
| 16840:3 | 1 | R.TAGWNIPMGLLFN[+1216.423]QTGSC[+57.021]KFDEYFSQSC[+57.021]APGSDPR.S | HexNAc(2)Hex(5) | C[+57]*2, N[+1216] | 1719.0694 | 3 | 5155.1937 | 5155.18 | 0 | 2.7 | 485 | Specific | 907.65 | 810.45 | 258.35 | 18.22 | 161 | >sp\|P02788\|TRFL_HUMAN Lactotransferrin OS=Homo sapiens OX=9606 GN=LTF PE=1 SV=6 | 1270 | 20201117_08512_1ng_glyco_hcd203040_compound15.20382.20382.3 | controllerType=0 controllerNumber=1 scan=20382 | 43.5021 |
| 17791:4 | 1 | R.TAGWNIPMGLLFN[+1913.677]QTGSC[+57.021]KFDEYFSQSC[+57.021]APGSDPR.S | HexNAc(4)Hex(5)NeuAc(1) | C[+57]*2, N[+1914] | 1463.8672 | 4 | 5852.4469 | 5852.4341 | 0 | 2.2 | 485 | Specific | 915.78 | 825.22 | 825.22 | 18.04 | 161 | >sp\|P02788\|TRFL_HUMAN Lactotransferrin OS=Homo sapiens OX=9606 GN=LTF PE=1 SV=6 | 1270 | 20201117_08512_1ng_glyco_hcd203040_compound15.21377.21377.4 | controllerType=0 controllerNumber=1 scan=21377 | 45.7783 |
| 17627:4 | 1 | R.TAGWNIPMGLLFN[+2351.851]QTGSC[+57.021]KFDEYFSQSC[+57.021]APGSDPR.S | HexNAc(4)Hex(5)Fuc(3)NeuAc(1) | C[+57]*2, N[+2352] | 1573.4128 | 4 | 6290.6296 | 6290.6079 | 0 | 3.4 | 485 | Specific | 919.82 | 883.23 | 883.23 | 17.95 | 161 | >sp\|P02788\|TRFL_HUMAN Lactotransferrin OS=Homo sapiens OX=9606 GN=LTF PE=1 SV=6 | 1270 | 20201117_08512_1ng_glyco_hcd203040_compound15.21206.21206.4 | controllerType=0 controllerNumber=1 scan=21206 | 45.4317 |
| 17768:5 | 1 | R.TAGWNIPMGLLFN[+1913.677]QTGSC[+57.021]KFDEYFSQSC[+57.021]APGSDPR.S | HexNAc(4)Hex(5)NeuAc(1) | C[+57]*2, N[+1914] | 1171.2952 | 5 | 5852.4467 | 5852.4341 | 0 | 2.2 | 485 | Specific | 939.74 | 893.81 | 893.81 | 18.67 | 161 | >sp\|P02788\|TRFL_HUMAN Lactotransferrin OS=Homo sapiens OX=9606 GN=LTF PE=1 SV=6 | 1270 | 20201117_08512_1ng_glyco_hcd203040_compound15.21353.21353.5 | controllerType=0 controllerNumber=1 scan=21353 | 45.7228 |
| 17660:4 | 1 | R.TAGWNIPMGLLFN[+2059.735]QTGSC[+57.021]KFDEYFSQSC[+57.021]APGSDPR.S | HexNAc(4)Hex(5)Fuc(1)NeuAc(1) | C[+57]*2, N[+2060] | 1500.3844 | 4 | 5998.5158 | 5998.492 | 0 | 4 | 485 | Specific | 952.79 | 863.57 | 863.57 | 18.95 | 161 | >sp\|P02788\|TRFL_HUMAN Lactotransferrin OS=Homo sapiens OX=9606 GN=LTF PE=1 SV=6 | 1270 | 20201117_08512_1ng_glyco_hcd203040_compound15.21241.21241.4 | controllerType=0 controllerNumber=1 scan=21241 | 45.5023 |
| 17663:4 | 1 | R.TAGWNIPMGLLFN[+2059.735]QTGSC[+57.021]KFDEYFSQSC[+57.021]APGSDPR.S | HexNAc(4)Hex(5)Fuc(1)NeuAc(1) | C[+57]*2, N[+2060] | 1500.3844 | 4 | 5998.5158 | 5998.492 | 0 | 4 | 485 | Specific | 959.25 | 867.97 | 867.97 | 18.95 | 161 | >sp\|P02788\|TRFL_HUMAN Lactotransferrin OS=Homo sapiens OX=9606 GN=LTF PE=1 SV=6 | 1270 | 20201117_08512_1ng_glyco_hcd203040_compound15.21244.21244.4 | controllerType=0 controllerNumber=1 scan=21244 | 45.5138 |
| 17678:4 | 1 | R.TAGWNIPMGLLFN[+2059.735]QTGSC[+57.021]KFDEYFSQSC[+57.021]APGSDPR.S | HexNAc(4)Hex(5)Fuc(1)NeuAc(1) | C[+57]*2, N[+2060] | 1500.3827 | 4 | 5998.5089 | 5998.492 | 0 | 2.8 | 485 | Specific | 980.7 | 910.39 | 910.39 | 19.04 | 161 | >sp\|P02788\|TRFL_HUMAN Lactotransferrin OS=Homo sapiens OX=9606 GN=LTF PE=1 SV=6 | 1270 | 20201117_08512_1ng_glyco_hcd203040_compound15.21260.21260.4 | controllerType=0 controllerNumber=1 scan=21260 | 45.5623 |
| 17850:4 | 1 | R.TAGWNIPMGLLFN[+1548.545]QTGSC[+57.021]KFDEYFSQSC[+57.021]APGSDPR.S | HexNAc(3)Hex(4)NeuAc(1) | C[+57]*2, N[+1549] | 1372.5862 | 4 | 5487.3229 | 5487.3019 | 0 | 3.8 | 485 | Specific | 1016.98 | 962.5 | 962.5 | 19.95 | 161 | >sp\|P02788\|TRFL_HUMAN Lactotransferrin OS=Homo sapiens OX=9606 GN=LTF PE=1 SV=6 | 1270 | 20201117_08512_1ng_glyco_hcd203040_compound15.21438.21438.4 | controllerType=0 controllerNumber=1 scan=21438 | 45.9074 |
| 16838:3 | 1 | R.TAGWNIPMGLLFN[+1216.423]QTGSC[+57.021]KFDEYFSQSC[+57.021]APGSDPR.S | HexNAc(2)Hex(5) | C[+57]*2, N[+1216] | 1719.0694 | 3 | 5155.1937 | 5155.18 | 0 | 2.7 | 485 | Specific | 1109.88 | 1019.64 | 282.17 | 22.22 | 161 | >sp\|P02788\|TRFL_HUMAN Lactotransferrin OS=Homo sapiens OX=9606 GN=LTF PE=1 SV=6 | 1270 | 20201117_08512_1ng_glyco_hcd203040_compound15.20380.20380.3 | controllerType=0 controllerNumber=1 scan=20380 | 43.4955 |
| 17709:4 | 1 | R.TAGWNIPM[+15.995]GLLFN[+2190.793]QTGSC[+57.021]KFDEYFSQSC[+57.021]A | HexNAc(6)Hex(6) | C[+57]*2, M[+16], N[+2191] | 1537.1479 | 4 | 6145.5698 | 6145.5452 | 0 | 4 | 485 | Specific | 202.02 | 202.02 | 202.02 | 4.25 | 161 | >sp\|P02788\|TRFL_HUMAN Lactotransferrin OS=Homo sapiens OX=9606 GN=LTF PE=1 SV=6 | 1270 | 20201117_08512_1ng_glyco_hcd203040_compound15.21292.21292.4 | controllerType=0 controllerNumber=1 scan=21292 | 45.6243 |
| 17722:4 | 1 | R.TAGWNIPM[+15.995]GLLFN[+2116.756]QTGSC[+57.021]KFDEYFSQSC[+57.021]A | HexNAc(5)Hex(5)NeuAc(1) | C[+57]*2, M[+16], N[+2117] | 1518.621 | 4 | 6071.4622 | 6071.5084 | 0 | -7.6 | 485 | Specific | 259.35 | 208.67 | 208.67 | 3.75 | 161 | >sp\|P02788\|TRFL_HUMAN Lactotransferrin OS=Homo sapiens OX=9606 GN=LTF PE=1 SV=6 | 1270 | 20201117_08512_1ng_glyco_hcd203040_compound15.21305.21305.4 | controllerType=0 controllerNumber=1 scan=21305 | 45.6454 |
| 17711:4 | 1 | R.TAGWNIPM[+15.995]GLLFN[+2190.793]QTGSC[+57.021]KFDEYFSQSC[+57.021]A | HexNAc(6)Hex(6) | C[+57]*2, M[+16], N[+2191] | 1537.1479 | 4 | 6145.5698 | 6145.5452 | 0 | 4 | 485 | Specific | 270.91 | 270.91 | 270.91 | 6.14 | 161 | >sp\|P02788\|TRFL_HUMAN Lactotransferrin OS=Homo sapiens OX=9606 GN=LTF PE=1 SV=6 | 1270 | 20201117_08512_1ng_glyco_hcd203040_compound15.21294.21294.4 | controllerType=0 controllerNumber=1 scan=21294 | 45.6277 |
| 16708:4 | 1 | R.TAGWNIPM[+15.995]GLLFN[+2059.735]QTGSC[+57.021]KFDEYFSQSC[+57.021]A | HexNAc(4)Hex(5)Fuc(1)NeuAc(1) | C[+57]*2, M[+16], N[+2060] | 1504.3805 | 4 | 6014.5002 | 6014.487 | 0 | 2.2 | 485 | Specific | 503.28 | 427.11 | 427.11 | 9.34 | 161 | >sp\|P02788\|TRFL_HUMAN Lactotransferrin OS=Homo sapiens OX=9606 GN=LTF PE=1 SV=6 | 1270 | 20201117_08512_1ng_glyco_hcd203040_compound15.20244.20244.2 | controllerType=0 controllerNumber=1 scan=20244 | 43.1958 |
| 16742:4 | 1 | R.TAGWNIPM[+15.995]GLLFN[+2059.735]QTGSC[+57.021]KFDEYFSQSC[+57.021]A | HexNAc(4)Hex(5)Fuc(1)NeuAc(1) | C[+57]*2, M[+16], N[+2060] | 1504.3818 | 4 | 6014.5055 | 6014.487 | 0 | 3.1 | 485 | Specific | 649.46 | 589.85 | 589.85 | 12.04 | 161 | >sp\|P02788\|TRFL_HUMAN Lactotransferrin OS=Homo sapiens OX=9606 GN=LTF PE=1 SV=6 | 1270 | 20201117_08512_1ng_glyco_hcd203040_compound15.20280.20280.4 | controllerType=0 controllerNumber=1 scan=20280 | 43.2713 |
| 16710:4 | 1 | R.TAGWNIPM[+15.995]GLLFN[+2059.735]QTGSC[+57.021]KFDEYFSQSC[+57.021]A | HexNAc(4)Hex(5)Fuc(1)NeuAc(1) | C[+57]*2, M[+16], N[+2060] | 1504.3805 | 4 | 6014.5002 | 6014.487 | 0 | 2.2 | 485 | Specific | 839.83 | 731.1 | 731.1 | 16.04 | 161 | >sp\|P02788\|TRFL_HUMAN Lactotransferrin OS=Homo sapiens OX=9606 GN=LTF PE=1 SV=6 | 1270 | 20201117_08512_1ng_glyco_hcd203040_compound15.20246.20246.2 | controllerType=0 controllerNumber=1 scan=20246 | 43.1994 |
| 16745:4 | 1 | R.TAGWNIPM[+15.995]GLLFN[+2059.735]QTGSC[+57.021]KFDEYFSQSC[+57.021]A | HexNAc(4)Hex(5)Fuc(1)NeuAc(1) | C[+57]*2, M[+16], N[+2060] | 1504.3818 | 4 | 6014.5055 | 6014.487 | 0 | 3.1 | 485 | Specific | 929.25 | 845.15 | 845.15 | 18.04 | 161 | >sp\|P02788\|TRFL_HUMAN Lactotransferrin OS=Homo sapiens OX=9606 GN=LTF PE=1 SV=6 | 1270 | 20201117_08512_1ng_glyco_hcd203040_compound15.20283.20283.4 | controllerType=0 controllerNumber=1 scan=20283 | 43.2786 |
| 07792:3 | 1 | K.FGRN[+2059.735]GSDC[+57.021]PDKFC[+57.021]LFQSETK.N | HexNAc(4)Hex(5)Fuc(1)NeuAc(1) | C[+57]*2, N[+2060] | 1484.9421 | 3 | 4452.8117 | 4452.7998 | 0 | 2.7 | 639 | Specific | 516.09 | 516.09 | 516.09 | 9.16 | 161 | >sp\|P02788\|TRFL_HUMAN Lactotransferrin OS=Homo sapiens OX=9606 GN=LTF PE=1 SV=6 | 1270 | 20201117_08512_1ng_glyco_hcd203040_compound15.10949.10949.3 | controllerType=0 controllerNumber=1 scan=10949 | 24.9187 |
| 07794:3 | 1 | K.FGRN[+2059.735]GSDC[+57.021]PDKFC[+57.021]LFQSETK.N | HexNAc(4)Hex(5)Fuc(1)NeuAc(1) | C[+57]*2, N[+2060] | 1484.9421 | 3 | 4452.8117 | 4452.7998 | 0 | 2.7 | 639 | Specific | 634.43 | 533.7 | 533.7 | 11.77 | 161 | >sp\|P02788\|TRFL_HUMAN Lactotransferrin OS=Homo sapiens OX=9606 GN=LTF PE=1 SV=6 | 1270 | 20201117_08512_1ng_glyco_hcd203040_compound15.10951.10951.3 | controllerType=0 controllerNumber=1 scan=10951 | 24.9228 |
| 07789:4 | 1 | K.FGRN[+2059.735]GSDC[+57.021]PDKFC[+57.021]LFQSETK.N | HexNAc(4)Hex(5)Fuc(1)NeuAc(1) | C[+57]*2, N[+2060] | 1113.9572 | 4 | 4452.807 | 4452.7998 | 0 | 1.6 | 639 | Specific | 650.68 | 650.68 | 650.68 | 13.34 | 161 | >sp\|P02788\|TRFL_HUMAN Lactotransferrin OS=Homo sapiens OX=9606 GN=LTF PE=1 SV=6 | 1270 | 20201117_08512_1ng_glyco_hcd203040_compound15.10946.10946.4 | controllerType=0 controllerNumber=1 scan=10946 | 24.9118 |
| 07791:4 | 1 | K.FGRN[+2059.735]GSDC[+57.021]PDKFC[+57.021]LFQSETK.N | HexNAc(4)Hex(5)Fuc(1)NeuAc(1) | C[+57]*2, N[+2060] | 1113.9572 | 4 | 4452.807 | 4452.7998 | 0 | 1.6 | 639 | Specific | 682.5 | 673.96 | 673.96 | 13.34 | 161 | >sp\|P02788\|TRFL_HUMAN Lactotransferrin OS=Homo sapiens OX=9606 GN=LTF PE=1 SV=6 | 1270 | 20201117_08512_1ng_glyco_hcd203040_compound15.10948.10948.4 | controllerType=0 controllerNumber=1 scan=10948 | 24.9168 |
| 10919:3 | 1 | R.N[+2350.830]GSDC[+57.021]PDKFC[+57.021]LFQSETK.N | HexNAc(4)Hex(5)Fuc(1)NeuAc(2) | C[+57]*2, N[+2351] | 1461.9107 | 3 | 4383.7176 | 4383.7043 | 0 | 3 | 642 | Specific | 205.22 | 150.82 | 150.82 | 3.82 | 161 | >sp\|P02788\|TRFL_HUMAN Lactotransferrin OS=Homo sapiens OX=9606 GN=LTF PE=1 SV=6 | 1270 | 20201117_08512_1ng_glyco_hcd203040_compound15.14195.14195.3 | controllerType=0 controllerNumber=1 scan=14195 | 31.0114 |
| 10917:3 | 1 | R.N[+2350.830]GSDC[+57.021]PDKFC[+57.021]LFQSETK.N | HexNAc(4)Hex(5)Fuc(1)NeuAc(2) | C[+57]*2, N[+2351] | 1461.9107 | 3 | 4383.7176 | 4383.7043 | 0 | 3 | 642 | Specific | 231.53 | 169.46 | 169.46 | 5.06 | 161 | >sp\|P02788\|TRFL_HUMAN Lactotransferrin OS=Homo sapiens OX=9606 GN=LTF PE=1 SV=6 | 1270 | 20201117_08512_1ng_glyco_hcd203040_compound15.14193.14193.3 | controllerType=0 controllerNumber=1 scan=14193 | 31.0039 |
| 08546:3 | 1 | R.N[+2571.946]GSDC[+57.021]PDKFC[+57.021]LFQSETK.N | HexNAc(5)Hex(6)Fuc(4) | C[+57]*2, N[+2572] | 1535.6041 | 3 | 4604.7977 | 4604.8193 | 0 | -4.7 | 642 | Specific | 237.51 | 173.57 | 173.57 | 3.91 | 161 | >sp\|P02788\|TRFL_HUMAN Lactotransferrin OS=Homo sapiens OX=9606 GN=LTF PE=1 SV=6 | 1270 | 20201117_08512_1ng_glyco_hcd203040_compound15.11730.11730.3 | controllerType=0 controllerNumber=1 scan=11730 | 26.3161 |
| 08544:3 | 1 | R.N[+2571.946]GSDC[+57.021]PDKFC[+57.021]LFQSETK.N | HexNAc(5)Hex(6)Fuc(4) | C[+57]*2, N[+2572] | 1535.6041 | 3 | 4604.7977 | 4604.8193 | 0 | -4.7 | 642 | Specific | 289.46 | 264.35 | 264.35 | 4.69 | 161 | >sp\|P02788\|TRFL_HUMAN Lactotransferrin OS=Homo sapiens OX=9606 GN=LTF PE=1 SV=6 | 1270 | 20201117_08512_1ng_glyco_hcd203040_compound15.11728.11728.3 | controllerType=0 controllerNumber=1 scan=11728 | 26.3126 |
| 08746:3 | 1 | R.N[+2059.735]GSDC[+57.021]PDKFC[+57.021]LFQSETK.N | HexNAc(4)Hex(5)Fuc(1)NeuAc(1) | C[+57]*2, N[+2060] | 1364.8777 | 3 | 4092.6187 | 4092.6088 | 0 | 2.4 | 642 | Specific | 500.47 | 340.29 | 340.29 | 7.29 | 161 | >sp\|P02788\|TRFL_HUMAN Lactotransferrin OS=Homo sapiens OX=9606 GN=LTF PE=1 SV=6 | 1270 | 20201117_08512_1ng_glyco_hcd203040_compound15.11939.11939.3 | controllerType=0 controllerNumber=1 scan=11939 | 26.7159 |
| 10865:3 | 1 | R.N[+2350.830]GSDC[+57.021]PDKFC[+57.021]LFQSETK.N | HexNAc(4)Hex(5)Fuc(1)NeuAc(2) | C[+57]*2, N[+2351] | 1461.9101 | 3 | 4383.7157 | 4383.7043 | 0 | 2.6 | 642 | Specific | 517.3 | 394.8 | 394.8 | 9.18 | 161 | >sp\|P02788\|TRFL_HUMAN Lactotransferrin OS=Homo sapiens OX=9606 GN=LTF PE=1 SV=6 | 1270 | 20201117_08512_1ng_glyco_hcd203040_compound15.14139.14139.3 | controllerType=0 controllerNumber=1 scan=14139 | 30.8845 |
| 10862:3 | 1 | R.N[+2350.830]GSDC[+57.021]PDKFC[+57.021]LFQSETK.N | HexNAc(4)Hex(5)Fuc(1)NeuAc(2) | C[+57]*2, N[+2351] | 1461.9101 | 3 | 4383.7157 | 4383.7043 | 0 | 2.6 | 642 | Specific | 561.72 | 457.59 | 457.59 | 9.54 | 161 | >sp\|P02788\|TRFL_HUMAN Lactotransferrin OS=Homo sapiens OX=9606 GN=LTF PE=1 SV=6 | 1270 | 20201117_08512_1ng_glyco_hcd203040_compound15.14136.14136.3 | controllerType=0 controllerNumber=1 scan=14136 | 30.8793 |
| 08743:3 | 1 | R.N[+2059.735]GSDC[+57.021]PDKFC[+57.021]LFQSETK.N | HexNAc(4)Hex(5)Fuc(1)NeuAc(1) | C[+57]*2, N[+2060] | 1364.8777 | 3 | 4092.6187 | 4092.6088 | 0 | 2.4 | 642 | Specific | 671.83 | 505.43 | 505.43 | 12.79 | 161 | >sp\|P02788\|TRFL_HUMAN Lactotransferrin OS=Homo sapiens OX=9606 GN=LTF PE=1 SV=6 | 1270 | 20201117_08512_1ng_glyco_hcd203040_compound15.11936.11936.3 | controllerType=0 controllerNumber=1 scan=11936 | 26.7089 |
| 07811:3 | 2 | R.GN[+2059.735]FSTEGC[+57.021]GC[+57.021]VC[+57.021]EPGWK.G | HexNAc(4)Hex(5)Fuc(1)NeuAc(1) | C[+57]*3, N[+2060] | 1335.5077 | 3 | 4004.5087 | 4004.5023 | 0 | 1.6 | 165 | Specific | 240.27 | 176.68 | 176.68 | 5.36 | 146 | >sp\|P24821\|TENA_HUMAN Tenascin OS=Homo sapiens OX=9606 GN=TNC PE=1 SV=3 | 486 | 20201117_08512_1ng_glyco_hcd203040_compound15.10968.10968.3 | controllerType=0 controllerNumber=1 scan=10968 | 24.9547 |
| 07809:3 | 2 | R.GN[+2059.735]FSTEGC[+57.021]GC[+57.021]VC[+57.021]EPGWK.G | HexNAc(4)Hex(5)Fuc(1)NeuAc(1) | C[+57]*3, N[+2060] | 1335.5077 | 3 | 4004.5087 | 4004.5023 | 0 | 1.6 | 165 | Specific | 242.44 | 174.79 | 174.79 | 5.36 | 146 | >sp\|P24821\|TENA_HUMAN Tenascin OS=Homo sapiens OX=9606 GN=TNC PE=1 SV=3 | 486 | 20201117_08512_1ng_glyco_hcd203040_compound15.10966.10966.3 | controllerType=0 controllerNumber=1 scan=10966 | 24.9494 |
| 01531:3 | 2 | K.GPN[+2059.735]C[+57.021]SEPEC[+57.021]PGNC[+57.021]HLR.G | HexNAc(4)Hex(5)Fuc(1)NeuAc(1) | C[+57]*3, N[+2060] | 1315.1717 | 3 | 3943.5005 | 3943.4931 | 0 | 1.9 | 182 | Specific | 233.32 | 200.89 | 200.89 | 5.52 | 146 | >sp\|P24821\|TENA_HUMAN Tenascin OS=Homo sapiens OX=9606 GN=TNC PE=1 SV=3 | 486 | 20201117_08512_1ng_glyco_hcd203040_compound15.4462.4462.3 | controllerType=0 controllerNumber=1 scan=4462 | 13.8048 |
| 01533:3 | 2 | K.GPN[+2059.735]C[+57.021]SEPEC[+57.021]PGNC[+57.021]HLR.G | HexNAc(4)Hex(5)Fuc(1)NeuAc(1) | C[+57]*3, N[+2060] | 1315.1717 | 3 | 3943.5005 | 3943.4931 | 0 | 1.9 | 182 | Specific | 257.91 | 209.08 | 209.08 | 5.78 | 146 | >sp\|P24821\|TENA_HUMAN Tenascin OS=Homo sapiens OX=9606 GN=TNC PE=1 SV=3 | 486 | 20201117_08512_1ng_glyco_hcd203040_compound15.4464.4464.3 | controllerType=0 controllerNumber=1 scan=4464 | 13.8087 |
| 03722:5 | 2 | R.C[+57.021]IN[+2059.735]GTC[+57.021]YC[+57.021]EEGFTGEDC[+57.021]GKPTC[ | HexNAc(4)Hex(5)Fuc(1)NeuAc(1) | C[+57]*6, N[+2060] | 1144.6407 | 5 | 5719.1746 | 5719.1633 | 0 | 2 | 325 | Specific | 578.79 | 578.79 | 578.79 | 11.71 | 146 | >sp\|P24821\|TENA_HUMAN Tenascin OS=Homo sapiens OX=9606 GN=TNC PE=1 SV=3 | 486 | 20201117_08512_1ng_glyco_hcd203040_compound15.6735.6735.5 | controllerType=0 controllerNumber=1 scan=6735 | 17.7716 |
| 03724:5 | 2 | R.C[+57.021]IN[+2059.735]GTC[+57.021]YC[+57.021]EEGFTGEDC[+57.021]GKPTC[ | HexNAc(4)Hex(5)Fuc(1)NeuAc(1) | C[+57]*6, N[+2060] | 1144.6407 | 5 | 5719.1746 | 5719.1633 | 0 | 2 | 325 | Specific | 663.51 | 663.51 | 663.51 | 13.71 | 146 | >sp\|P24821\|TENA_HUMAN Tenascin OS=Homo sapiens OX=9606 GN=TNC PE=1 SV=3 | 486 | 20201117_08512_1ng_glyco_hcd203040_compound15.6737.6737.5 | controllerType=0 controllerNumber=1 scan=6737 | 17.7757 |
| 08660:4 | 2 | R.QTGLAPGQEYEISLHIVKN[+2205.793]NTR.G | HexNAc(4)Hex(5)Fuc(2)NeuAc(1) | N[+2206] | 1169.2774 | 4 | 4674.0877 | 4674.0821 | 0 | 1.2 | 770 | Specific | 281.05 | 215.86 | 215.86 | 5.22 | 146 | >sp\|P24821\|TENA_HUMAN Tenascin OS=Homo sapiens OX=9606 GN=TNC PE=1 SV=3 | 486 | 20201117_08512_1ng_glyco_hcd203040_compound15.11849.11849.2 | controllerType=0 controllerNumber=1 scan=11849 | 26.5445 |
| 08687:4 | 2 | R.QTGLAPGQEYEISLHIVKN[+2059.735]NTR.G | HexNAc(4)Hex(5)Fuc(1)NeuAc(1) | N[+2060] | 1132.7636 | 4 | 4528.0325 | 4528.0241 | 0 | 1.9 | 770 | Specific | 321.44 | 220.54 | 220.54 | 6.83 | 146 | >sp\|P24821\|TENA_HUMAN Tenascin OS=Homo sapiens OX=9606 GN=TNC PE=1 SV=3 | 486 | 20201117_08512_1ng_glyco_hcd203040_compound15.11878.11878.4 | controllerType=0 controllerNumber=1 scan=11878 | 26.6019 |
| 08767:3 | 2 | R.QTGLAPGQEYEISLHIVKN[+1913.677]NTR.G | HexNAc(4)Hex(5)NeuAc(1) | N[+1914] | 1461.3298 | 3 | 4381.9748 | 4381.9662 | 0 | 1.9 | 770 | Specific | 361.94 | 189.34 | 189.34 | 6.14 | 146 | >sp\|P24821\|TENA_HUMAN Tenascin OS=Homo sapiens OX=9606 GN=TNC PE=1 SV=3 | 486 | 20201117_08512_1ng_glyco_hcd203040_compound15.11961.11961.3 | controllerType=0 controllerNumber=1 scan=11961 | 26.7577 |
| 08688:3 | 2 | R.QTGLAPGQEYEISLHIVKN[+2059.735]NTR.G | HexNAc(4)Hex(5)Fuc(1)NeuAc(1) | N[+2060] | 1510.0153 | 3 | 4528.0314 | 4528.0241 | 0 | 1.6 | 770 | Specific | 372.37 | 253.14 | 253.14 | 6.59 | 146 | >sp\|P24821\|TENA_HUMAN Tenascin OS=Homo sapiens OX=9606 GN=TNC PE=1 SV=3 | 486 | 20201117_08512_1ng_glyco_hcd203040_compound15.11879.11879.3 | controllerType=0 controllerNumber=1 scan=11879 | 26.6033 |
| 08689:4 | 2 | R.QTGLAPGQEYEISLHIVKN[+2059.735]NTR.G | HexNAc(4)Hex(5)Fuc(1)NeuAc(1) | N[+2060] | 1132.7636 | 4 | 4528.0325 | 4528.0241 | 0 | 1.9 | 770 | Specific | 467.51 | 293.02 | 293.02 | 7.91 | 146 | >sp\|P24821\|TENA_HUMAN Tenascin OS=Homo sapiens OX=9606 GN=TNC PE=1 SV=3 | 486 | 20201117_08512_1ng_glyco_hcd203040_compound15.11880.11880.4 | controllerType=0 controllerNumber=1 scan=11880 | 26.6066 |
| 08690:3 | 2 | R.QTGLAPGQEYEISLHIVKN[+2059.735]NTR.G | HexNAc(4)Hex(5)Fuc(1)NeuAc(1) | N[+2060] | 1510.0153 | 3 | 4528.0314 | 4528.0241 | 0 | 1.6 | 770 | Specific | 602.07 | 358.5 | 358.5 | 10.65 | 146 | >sp\|P24821\|TENA_HUMAN Tenascin OS=Homo sapiens OX=9606 GN=TNC PE=1 SV=3 | 486 | 20201117_08512_1ng_glyco_hcd203040_compound15.11881.11881.3 | controllerType=0 controllerNumber=1 scan=11881 | 26.6083 |
| 14347:2 | 2 | R.LN[+1378.476]YSLPTGQWVGVQLPR.N | HexNAc(2)Hex(6) | N[+1378] | 1653.7615 | 2 | 3306.5158 | 3306.5145 | 0 | 0.4 | 1017 | Specific | 603.01 | 463.22 | 463.22 | 11.53 | 146 | >sp\|P24821\|TENA_HUMAN Tenascin OS=Homo sapiens OX=9606 GN=TNC PE=1 SV=3 | 486 | 20201117_08512_1ng_glyco_hcd203040_compound15.17765.17765.2 | controllerType=0 controllerNumber=1 scan=17765 | 38.0407 |
| 14281:2 | 2 | R.LN[+1540.529]YSLPTGQWVGVQLPR.N | HexNAc(2)Hex(7) | N[+1541] | 1734.7917 | 2 | 3468.5761 | 3468.5674 | 0 | 2.5 | 1017 | Specific | 648.13 | 496.77 | 496.77 | 12.08 | 146 | >sp\|P24821\|TENA_HUMAN Tenascin OS=Homo sapiens OX=9606 GN=TNC PE=1 SV=3 | 486 | 20201117_08512_1ng_glyco_hcd203040_compound15.17696.17696.2 | controllerType=0 controllerNumber=1 scan=17696 | 37.8908 |
| 14344:2 | 2 | R.LN[+1378.476]YSLPTGQWVGVQLPR.N | HexNAc(2)Hex(6) | N[+1378] | 1653.7615 | 2 | 3306.5158 | 3306.5145 | 0 | 0.4 | 1017 | Specific | 703.21 | 608.85 | 608.85 | 13.53 | 146 | >sp\|P24821\|TENA_HUMAN Tenascin OS=Homo sapiens OX=9606 GN=TNC PE=1 SV=3 | 486 | 20201117_08512_1ng_glyco_hcd203040_compound15.17762.17762.2 | controllerType=0 controllerNumber=1 scan=17762 | 38.0313 |

Table S4. Cont.

| Query #:z | Protein Rank | Glycopeptide | Glycan | Modification Type(s) | Observed m/z | z | Observed (M+H) | Calc. mass (M+H) | Off-by-x error | Mass error (ppm) | StartPosition | Cleavage | Score | Delta | Delta Mod | \|Log Prob\| | # of unique peptides | Protein Name | Protein DB number | Comment | Scan # | Scan Time |
|---|---|---|---|---|---|---|---|---|---|---|---|---|---|---|---|---|---|---|---|---|---|---|
| 14279:2 | 2 | R.LN[+1540.529]YSLPTGQWVGVQLPR.N | HexNAc(2)Hex(7) | N[+1541] | 1734.7917 | 2 | 3468.5761 | 3468.5674 | 0 | 2.5 | 1017 | Specific | 723.92 | 618.97 | 618.97 | 14.08 | 146 | >sp\|P24821\|TENA_HUMAN Tenascin OS=Homo sapiens OX=9606 GN=TNC PE=1 SV=3 | 486 | 20201117_08512_1ng_glyco_hcd203040_compound15.17694.17694.2 | controllerType=0 controllerNumber=1 scan=17694 | 37.8874 |
| 14383:2 | 2 | R.LN[+1216.423]YSLPTGQWVGVQLPR.N | HexNAc(2)Hex(5) | N[+1216] | 1572.7372 | 2 | 3144.4672 | 3144.4617 | 0 | 1.7 | 1017 | Specific | 839.16 | 786.95 | 786.95 | 16.18 | 146 | >sp\|P24821\|TENA_HUMAN Tenascin OS=Homo sapiens OX=9606 GN=TNC PE=1 SV=3 | 486 | 20201117_08512_1ng_glyco_hcd203040_compound15.17803.17803.2 | controllerType=0 controllerNumber=1 scan=17803 | 38.1339 |
| 14385:2 | 2 | R.LN[+1216.423]YSLPTGQWVGVQLPR.N | HexNAc(2)Hex(5) | N[+1216] | 1572.7372 | 2 | 3144.4672 | 3144.4617 | 0 | 1.7 | 1017 | Specific | 859.98 | 686.83 | 686.83 | 17.18 | 146 | >sp\|P24821\|TENA_HUMAN Tenascin OS=Homo sapiens OX=9606 GN=TNC PE=1 SV=3 | 486 | 20201117_08512_1ng_glyco_hcd203040_compound15.17805.17805.2 | controllerType=0 controllerNumber=1 scan=17805 | 38.1384 |
| 14194:3 | 2 | R.LN[+1702.581]YSLPTGQWVGVQLPR.N | HexNAc(2)Hex(8) | N[+1703] | 1210.8806 | 3 | 3630.6271 | 3630.6202 | 0 | 1.9 | 1017 | Specific | 899.84 | 836.05 | 836.05 | 17 | 146 | >sp\|P24821\|TENA_HUMAN Tenascin OS=Homo sapiens OX=9606 GN=TNC PE=1 SV=3 | 486 | 20201117_08512_1ng_glyco_hcd203040_compound15.17606.17606.3 | controllerType=0 controllerNumber=1 scan=17606 | 37.6996 |
| 14311:3 | 2 | R.LN[+1378.476]YSLPTGQWVGVQLPR.N | HexNAc(2)Hex(6) | N[+1378] | 1102.8461 | 3 | 3306.5237 | 3306.5145 | 0 | 2.8 | 1017 | Specific | 939.85 | 939.85 | 939.85 | 17.9 | 146 | >sp\|P24821\|TENA_HUMAN Tenascin OS=Homo sapiens OX=9606 GN=TNC PE=1 SV=3 | 486 | 20201117_08512_1ng_glyco_hcd203040_compound15.17728.17728.3 | controllerType=0 controllerNumber=1 scan=17728 | 37.9579 |
| 14264:3 | 2 | R.LN[+1540.529]YSLPTGQWVGVQLPR.N | HexNAc(2)Hex(7) | N[+1541] | 1156.8628 | 3 | 3468.574 | 3468.5674 | 0 | 1.9 | 1017 | Specific | 973.15 | 962.03 | 962.03 | 19 | 146 | >sp\|P24821\|TENA_HUMAN Tenascin OS=Homo sapiens OX=9606 GN=TNC PE=1 SV=3 | 486 | 20201117_08512_1ng_glyco_hcd203040_compound15.17679.17679.3 | controllerType=0 controllerNumber=1 scan=17679 | 37.8527 |
| 14197:3 | 2 | R.LN[+1702.581]YSLPTGQWVGVQLPR.N | HexNAc(2)Hex(8) | N[+1703] | 1210.8806 | 3 | 3630.6271 | 3630.6202 | 0 | 1.9 | 1017 | Specific | 1055.15 | 871.12 | 871.12 | 21 | 146 | >sp\|P24821\|TENA_HUMAN Tenascin OS=Homo sapiens OX=9606 GN=TNC PE=1 SV=3 | 486 | 20201117_08512_1ng_glyco_hcd203040_compound15.17609.17609.3 | controllerType=0 controllerNumber=1 scan=17609 | 37.7042 |
| 14358:3 | 2 | R.LN[+1216.423]YSLPTGQWVGVQLPR.N | HexNAc(2)Hex(5) | N[+1216] | 1048.8295 | 3 | 3144.4739 | 3144.4617 | 0 | 3.9 | 1017 | Specific | 1059.06 | 1059.06 | 1059.06 | 20.81 | 146 | >sp\|P24821\|TENA_HUMAN Tenascin OS=Homo sapiens OX=9606 GN=TNC PE=1 SV=3 | 486 | 20201117_08512_1ng_glyco_hcd203040_compound15.17777.17777.3 | controllerType=0 controllerNumber=1 scan=17777 | 38.084 |
| 14313:3 | 2 | R.LN[+1378.476]YSLPTGQWVGVQLPR.N | HexNAc(2)Hex(6) | N[+1378] | 1102.8461 | 3 | 3306.5237 | 3306.5145 | 0 | 2.8 | 1017 | Specific | 1073.63 | 890.61 | 890.61 | 20.9 | 146 | >sp\|P24821\|TENA_HUMAN Tenascin OS=Homo sapiens OX=9606 GN=TNC PE=1 SV=3 | 486 | 20201117_08512_1ng_glyco_hcd203040_compound15.17730.17730.3 | controllerType=0 controllerNumber=1 scan=17730 | 37.9627 |
| 14266:3 | 2 | R.LN[+1540.529]YSLPTGQWVGVQLPR.N | HexNAc(2)Hex(7) | N[+1541] | 1156.8628 | 3 | 3468.574 | 3468.5674 | 0 | 1.9 | 1017 | Specific | 1086.71 | 919.23 | 919.23 | 21 | 146 | >sp\|P24821\|TENA_HUMAN Tenascin OS=Homo sapiens OX=9606 GN=TNC PE=1 SV=3 | 486 | 20201117_08512_1ng_glyco_hcd203040_compound15.17681.17681.3 | controllerType=0 controllerNumber=1 scan=17681 | 37.8582 |
| 14361:3 | 2 | R.LN[+1216.423]YSLPTGQWVGVQLPR.N | HexNAc(2)Hex(5) | N[+1216] | 1048.8295 | 3 | 3144.4739 | 3144.4617 | 0 | 3.9 | 1017 | Specific | 1092.28 | 930.06 | 930.06 | 20.81 | 146 | >sp\|P24821\|TENA_HUMAN Tenascin OS=Homo sapiens OX=9606 GN=TNC PE=1 SV=3 | 486 | 20201117_08512_1ng_glyco_hcd203040_compound15.17780.17780.3 | controllerType=0 controllerNumber=1 scan=17780 | 38.092 |
| 03687:2 | 2 | R.N[+2059.735]TTSYVLR.G | HexNAc(4)Hex(5)Fuc(1)NeuAc(1) | N[+2060] | 1507.1282 | 2 | 3013.2492 | 3013.24 | 0 | 3 | 1034 | Specific | 213.52 | 100.09 | 100.09 | 3.38 | 146 | >sp\|P24821\|TENA_HUMAN Tenascin OS=Homo sapiens OX=9606 GN=TNC PE=1 SV=3 | 486 | 20201117_08512_1ng_glyco_hcd203040_compound15.6698.6698.2 | controllerType=0 controllerNumber=1 scan=6698 | 17.6985 |
| 03679:3 | 2 | R.N[+2059.735]TTSYVLR.G | HexNAc(4)Hex(5)Fuc(1)NeuAc(1) | N[+2060] | 1005.0883 | 3 | 3013.2505 | 3013.24 | 0 | 3.5 | 1034 | Specific | 219.23 | 137.01 | 137.01 | 3.53 | 146 | >sp\|P24821\|TENA_HUMAN Tenascin OS=Homo sapiens OX=9606 GN=TNC PE=1 SV=3 | 486 | 20201117_08512_1ng_glyco_hcd203040_compound15.6690.6690.3 | controllerType=0 controllerNumber=1 scan=6690 | 17.6794 |
| 03681:3 | 2 | R.N[+2059.735]TTSYVLR.G | HexNAc(4)Hex(5)Fuc(1)NeuAc(1) | N[+2060] | 1005.0883 | 3 | 3013.2505 | 3013.24 | 0 | 3.5 | 1034 | Specific | 287.34 | 181.74 | 181.74 | 5.27 | 146 | >sp\|P24821\|TENA_HUMAN Tenascin OS=Homo sapiens OX=9606 GN=TNC PE=1 SV=3 | 486 | 20201117_08512_1ng_glyco_hcd203040_compound15.6692.6692.3 | controllerType=0 controllerNumber=1 scan=6692 | 17.6894 |
| 03689:2 | 2 | R.N[+2059.735]TTSYVLR.G | HexNAc(4)Hex(5)Fuc(1)NeuAc(1) | N[+2060] | 1507.1282 | 2 | 3013.2492 | 3013.24 | 0 | 3 | 1034 | Specific | 349.74 | 203.98 | 203.98 | 5.88 | 146 | >sp\|P24821\|TENA_HUMAN Tenascin OS=Homo sapiens OX=9606 GN=TNC PE=1 SV=3 | 486 | 20201117_08512_1ng_glyco_hcd203040_compound15.6700.6700.2 | controllerType=0 controllerNumber=1 scan=6700 | 17.7049 |
| 22539:4 | 2 | R.GVTQDFSTTPLSVEVLTEEVPDMGN[+2350.830]LTVTEVSWDALR.L | HexNAc(4)Hex(5)Fuc(1)NeuAc(2) | N[+2351] | 1597.4569 | 4 | 6386.8058 | 6386.8047 | 0 | 0.2 | 1237 | Specific | 334.31 | 334.31 | 334.31 | 7.87 | 146 | >sp\|P24821\|TENA_HUMAN Tenascin OS=Homo sapiens OX=9606 GN=TNC PE=1 SV=3 | 486 | 20201117_08512_1ng_glyco_hcd203040_compound15.26335.26335.4 | controllerType=0 controllerNumber=1 scan=26335 | 56.5418 |
| 22541:4 | 2 | R.GVTQDFSTTPLSVEVLTEEVPDMGN[+2350.830]LTVTEVSWDALR.L | HexNAc(4)Hex(5)Fuc(1)NeuAc(2) | N[+2351] | 1597.4569 | 4 | 6386.8058 | 6386.8047 | 0 | 0.2 | 1237 | Specific | 409.39 | 409.39 | 409.39 | 9.85 | 146 | >sp\|P24821\|TENA_HUMAN Tenascin OS=Homo sapiens OX=9606 GN=TNC PE=1 SV=3 | 486 | 20201117_08512_1ng_glyco_hcd203040_compound15.26337.26337.4 | controllerType=0 controllerNumber=1 scan=26337 | 56.5464 |
| 09550:3 | 3 | K.TVLTPATNHMGN[+1540.529]VTFTIPANR.E | HexNAc(2)Hex(7) | N[+1541] | 1265.9066 | 3 | 3795.7051 | 3795.6886 | 0 | 4.3 | 74 | Specific | 636.52 | 601.25 | 287.05 | 11.99 | 128 | >sp\|P01024\|CO3_HUMAN Complement C3 OS=Homo sapiens OX=9606 GN=C3 PE=1 SV=2 | 492 | 20201117_08512_1ng_glyco_hcd203040_compound15.12774.12774.3 | controllerType=0 controllerNumber=1 scan=12774 | 28.2658 |
| 09548:3 | 3 | K.TVLTPATNHMGN[+1540.529]VTFTIPANR.E | HexNAc(2)Hex(7) | N[+1541] | 1265.9066 | 3 | 3795.7051 | 3795.6886 | 0 | 4.3 | 74 | Specific | 685.03 | 685.03 | 401 | 12.99 | 128 | >sp\|P01024\|CO3_HUMAN Complement C3 OS=Homo sapiens OX=9606 GN=C3 PE=1 SV=2 | 492 | 20201117_08512_1ng_glyco_hcd203040_compound15.12772.12772.3 | controllerType=0 controllerNumber=1 scan=12772 | 28.2618 |
| 09578:3 | 3 | K.TVLTPATNHMGN[+1378.476]VTFTIPANR.E | HexNAc(2)Hex(6) | N[+1378] | 1211.8868 | 3 | 3633.6458 | 3633.6358 | 0 | 2.7 | 74 | Specific | 723.48 | 723.48 | 351.36 | 14.63 | 128 | >sp\|P01024\|CO3_HUMAN Complement C3 OS=Homo sapiens OX=9606 GN=C3 PE=1 SV=2 | 492 | 20201117_08512_1ng_glyco_hcd203040_compound15.12803.12803.3 | controllerType=0 controllerNumber=1 scan=12803 | 28.3174 |
| 09576:3 | 3 | K.TVLTPATNHMGN[+1378.476]VTFTIPANR.E | HexNAc(2)Hex(6) | N[+1378] | 1211.8868 | 3 | 3633.6458 | 3633.6358 | 0 | 2.7 | 74 | Specific | 839.34 | 832.21 | 461.42 | 16.63 | 128 | >sp\|P01024\|CO3_HUMAN Complement C3 OS=Homo sapiens OX=9606 GN=C3 PE=1 SV=2 | 492 | 20201117_08512_1ng_glyco_hcd203040_compound15.12801.12801.3 | controllerType=0 controllerNumber=1 scan=12801 | 28.3128 |
| 18280:3 | 5 | R.[+42.011]GLSFDVSLEVSQGPGLLN[+2336.851]DTK.V | HexNAc(6)Hex(6)Fuc(1) | .[42], N[+2337] | 1518.99 | 3 | 4554.9556 | 4554.9749 | 0 | -4.2 | 118 | Specific | 275.41 | 218.94 | 218.94 | 3.94 | 91 | >sp\|P01833\|PIGR_HUMAN Polymeric immunoglobulin receptor OS=Homo sapiens OX=9606 GN=PIGR PE=1 SV=4 | 902 | 20201117_08512_1ng_glyco_hcd203040_compound15.21886.21886.3 | controllerType=0 controllerNumber=1 scan=21886 | 46.8834 |
| 16322:3 | 5 | R.GLSFDVSLEVSQGPGLLN[+2570.925]DTK.V | HexNAc(5)Hex(6)Fuc(2)NeuAc(1) | N[+2571] | 1583.0259 | 3 | 4747.0633 | 4747.0382 | 0 | 5.3 | 118 | Specific | 276.49 | 276.49 | 276.49 | 6 | 91 | >sp\|P01833\|PIGR_HUMAN Polymeric immunoglobulin receptor OS=Homo sapiens OX=9606 GN=PIGR PE=1 SV=4 | 902 | 20201117_08512_1ng_glyco_hcd203040_compound15.19841.19841.3 | controllerType=0 controllerNumber=1 scan=19841 | 42.2543 |
| 18277:3 | 5 | R.[+42.011]GLSFDVSLEVSQGPGLLN[+2336.851]DTK.V | HexNAc(6)Hex(6)Fuc(1) | .[42], N[+2337] | 1518.99 | 3 | 4554.9556 | 4554.9749 | 0 | -4.2 | 118 | Specific | 377.3 | 350.25 | 350.25 | 6.17 | 91 | >sp\|P01833\|PIGR_HUMAN Polymeric immunoglobulin receptor OS=Homo sapiens OX=9606 GN=PIGR PE=1 SV=4 | 902 | 20201117_08512_1ng_glyco_hcd203040_compound15.21883.21883.3 | controllerType=0 controllerNumber=1 scan=21883 | 46.8765 |
| 16230:3 | 5 | R.GLSFDVSLEVSQGPGLLN[+2716.983]DTK.V | HexNAc(5)Hex(6)Fuc(3)NeuAc(1) | N[+2717] | 1631.7067 | 3 | 4893.1056 | 4893.0962 | 0 | 1.9 | 118 | Specific | 449.59 | 363.33 | 363.33 | 8.19 | 91 | >sp\|P01833\|PIGR_HUMAN Polymeric immunoglobulin receptor OS=Homo sapiens OX=9606 GN=PIGR PE=1 SV=4 | 902 | 20201117_08512_1ng_glyco_hcd203040_compound15.19745.19745.3 | controllerType=0 controllerNumber=1 scan=19745 | 42.0457 |
| 16325:3 | 5 | R.GLSFDVSLEVSQGPGLLN[+2570.925]DTK.V | HexNAc(5)Hex(6)Fuc(2)NeuAc(1) | N[+2571] | 1583.0259 | 3 | 4747.0633 | 4747.0382 | 0 | 5.3 | 118 | Specific | 604.6 | 380.06 | 380.06 | 11.81 | 91 | >sp\|P01833\|PIGR_HUMAN Polymeric immunoglobulin receptor OS=Homo sapiens OX=9606 GN=PIGR PE=1 SV=4 | 902 | 20201117_08512_1ng_glyco_hcd203040_compound15.19844.19844.3 | controllerType=0 controllerNumber=1 scan=19844 | 42.2613 |
| 16232:3 | 5 | R.GLSFDVSLEVSQGPGLLN[+2716.983]DTK.V | HexNAc(5)Hex(6)Fuc(3)NeuAc(1) | N[+2717] | 1631.7067 | 3 | 4893.1056 | 4893.0962 | 0 | 1.9 | 118 | Specific | 621.25 | 615.81 | 615.81 | 12.55 | 91 | >sp\|P01833\|PIGR_HUMAN Polymeric immunoglobulin receptor OS=Homo sapiens OX=9606 GN=PIGR PE=1 SV=4 | 902 | 20201117_08512_1ng_glyco_hcd203040_compound15.19747.19747.3 | controllerType=0 controllerNumber=1 scan=19747 | 42.0507 |
| 18185:3 | 5 | R.GLSFDVSLEVSQGPGLLN[+2350.830]DTK.V | HexNAc(4)Hex(5)Fuc(1)NeuAc(2) | N[+2351] | 1509.6564 | 3 | 4526.9547 | 4526.9436 | 0 | 2.5 | 118 | Specific | 956.17 | 878.08 | 878.08 | 19.45 | 91 | >sp\|P01833\|PIGR_HUMAN Polymeric immunoglobulin receptor OS=Homo sapiens OX=9606 GN=PIGR PE=1 SV=4 | 902 | 20201117_08512_1ng_glyco_hcd203040_compound15.21787.21787.3 | controllerType=0 controllerNumber=1 scan=21787 | 46.6512 |
| 18188:3 | 5 | R.GLSFDVSLEVSQGPGLLN[+2350.830]DTK.V | HexNAc(4)Hex(5)Fuc(1)NeuAc(2) | N[+2351] | 1509.6564 | 3 | 4526.9547 | 4526.9436 | 0 | 2.5 | 118 | Specific | 1010.71 | 985.34 | 985.34 | 20.45 | 91 | >sp\|P01833\|PIGR_HUMAN Polymeric immunoglobulin receptor OS=Homo sapiens OX=9606 GN=PIGR PE=1 SV=4 | 902 | 20201117_08512_1ng_glyco_hcd203040_compound15.21790.21790.3 | controllerType=0 controllerNumber=1 scan=21790 | 46.6574 |
| 18465:3 | 5 | R.GLSFDVSLEVSQGPGLLN[+2350.830]DTKVYTVDLGR.T | HexNAc(4)Hex(5)Fuc(1)NeuAc(2) | N[+2351] | 1810.8221 | 3 | 5430.4518 | 5430.425 | 0 | 4.9 | 118 | Specific | 372.42 | 372.42 | 372.42 | 6.56 | 91 | >sp\|P01833\|PIGR_HUMAN Polymeric immunoglobulin receptor OS=Homo sapiens OX=9606 GN=PIGR PE=1 SV=4 | 902 | 20201117_08512_1ng_glyco_hcd203040_compound15.22079.22079.3 | controllerType=0 controllerNumber=1 scan=22079 | 47.2799 |
| 18468:3 | 5 | R.GLSFDVSLEVSQGPGLLN[+2350.830]DTKVYTVDLGR.T | HexNAc(4)Hex(5)Fuc(1)NeuAc(2) | N[+2351] | 1810.8221 | 3 | 5430.4518 | 5430.425 | 0 | 4.9 | 118 | Specific | 450.61 | 415.82 | 415.82 | 8.31 | 91 | >sp\|P01833\|PIGR_HUMAN Polymeric immunoglobulin receptor OS=Homo sapiens OX=9606 GN=PIGR PE=1 SV=4 | 902 | 20201117_08512_1ng_glyco_hcd203040_compound15.22082.22082.3 | controllerType=0 controllerNumber=1 scan=22082 | 47.293 |
| 18458:4 | 5 | R.GLSFDVSLEVSQGPGLLN[+2350.830]DTKVYTVDLGR.T | HexNAc(4)Hex(5)Fuc(1)NeuAc(2) | N[+2351] | 1358.3662 | 4 | 5430.443 | 5430.425 | 0 | 3.3 | 118 | Specific | 631.35 | 468.64 | 468.64 | 11.58 | 91 | >sp\|P01833\|PIGR_HUMAN Polymeric immunoglobulin receptor OS=Homo sapiens OX=9606 GN=PIGR PE=1 SV=4 | 902 | 20201117_08512_1ng_glyco_hcd203040_compound15.22072.22072.4 | controllerType=0 controllerNumber=1 scan=22072 | 47.267 |
| 18460:4 | 5 | R.GLSFDVSLEVSQGPGLLN[+2350.830]DTKVYTVDLGR.T | HexNAc(4)Hex(5)Fuc(1)NeuAc(2) | N[+2351] | 1358.3662 | 4 | 5430.443 | 5430.425 | 0 | 3.3 | 118 | Specific | 806.05 | 779.82 | 779.82 | 15.58 | 91 | >sp\|P01833\|PIGR_HUMAN Polymeric immunoglobulin receptor OS=Homo sapiens OX=9606 GN=PIGR PE=1 SV=4 | 902 | 20201117_08512_1ng_glyco_hcd203040_compound15.22074.22074.4 | controllerType=0 controllerNumber=1 scan=22074 | 47.2729 |
| 19453:3 | 5 | K.QIGLYPVLVIDSSGYVNPN[+2350.830]YTGR.I | HexNAc(4)Hex(5)Fuc(1)NeuAc(2) | N[+2351] | 1626.0571 | 3 | 4876.1566 | 4876.1338 | 0 | 4.7 | 168 | Specific | 538.12 | 459.44 | 459.44 | 9.47 | 91 | >sp\|P01833\|PIGR_HUMAN Polymeric immunoglobulin receptor OS=Homo sapiens OX=9606 GN=PIGR PE=1 SV=4 | 902 | 20201117_08512_1ng_glyco_hcd203040_compound15.23113.23113.3 | controllerType=0 controllerNumber=1 scan=23113 | 49.6975 |
| 19511:5 | 5 | K.QIGLYPVLVIDSSGYVNPN[+2204.772]YTGR.I | HexNAc(4)Hex(5)NeuAc(2) | N[+2205] | 946.8243 | 5 | 4730.0924 | 4730.0759 | 0 | 3.5 | 168 | Specific | 599.66 | 521.66 | 521.66 | 11.08 | 91 | >sp\|P01833\|PIGR_HUMAN Polymeric immunoglobulin receptor OS=Homo sapiens OX=9606 GN=PIGR PE=1 SV=4 | 902 | 20201117_08512_1ng_glyco_hcd203040_compound15.23174.23174.5 | controllerType=0 controllerNumber=1 scan=23174 | 49.8707 |
| 19513:5 | 5 | K.QIGLYPVLVIDSSGYVNPN[+2204.772]YTGR.I | HexNAc(4)Hex(5)NeuAc(2) | N[+2205] | 946.8243 | 5 | 4730.0924 | 4730.0759 | 0 | 3.5 | 168 | Specific | 616.57 | 596.45 | 596.45 | 12.08 | 91 | >sp\|P01833\|PIGR_HUMAN Polymeric immunoglobulin receptor OS=Homo sapiens OX=9606 GN=PIGR PE=1 SV=4 | 902 | 20201117_08512_1ng_glyco_hcd203040_compound15.23176.23176.5 | controllerType=0 controllerNumber=1 scan=23176 | 49.8763 |
| 19455:3 | 5 | K.QIGLYPVLVIDSSGYVNPN[+2350.830]YTGR.I | HexNAc(4)Hex(5)Fuc(1)NeuAc(2) | N[+2351] | 1626.0571 | 3 | 4876.1566 | 4876.1338 | 0 | 4.7 | 168 | Specific | 741.43 | 508.88 | 508.88 | 13.95 | 91 | >sp\|P01833\|PIGR_HUMAN Polymeric immunoglobulin receptor OS=Homo sapiens OX=9606 GN=PIGR PE=1 SV=4 | 902 | 20201117_08512_1ng_glyco_hcd203040_compound15.23115.23115.3 | controllerType=0 controllerNumber=1 scan=23115 | 49.7103 |
| 17591:4 | 5 | K.QIGLYPVLVIDSSGYVNPN[+2059.735]YTGR.I | HexNAc(4)Hex(5)Fuc(1)NeuAc(1) | N[+2060] | 1147.0198 | 4 | 4585.0574 | 4585.0384 | 0 | 4.2 | 168 | Specific | 784.76 | 779 | 779 | 16.12 | 91 | >sp\|P01833\|PIGR_HUMAN Polymeric immunoglobulin receptor OS=Homo sapiens OX=9606 GN=PIGR PE=1 SV=4 | 902 | 20201117_08512_1ng_glyco_hcd203040_compound15.21168.21168.4 | controllerType=0 controllerNumber=1 scan=21168 | 45.3145 |
| 17579:3 | 5 | K.QIGLYPVLVIDSSGYVNPN[+2059.735]YTGR.I | HexNAc(4)Hex(5)Fuc(1)NeuAc(1) | N[+2060] | 1529.0252 | 3 | 4585.0611 | 4585.0384 | 0 | 4.9 | 168 | Specific | 839.74 | 790.97 | 790.97 | 15.95 | 91 | >sp\|P01833\|PIGR_HUMAN Polymeric immunoglobulin receptor OS=Homo sapiens OX=9606 GN=PIGR PE=1 SV=4 | 902 | 20201117_08512_1ng_glyco_hcd203040_compound15.21156.21156.3 | controllerType=0 controllerNumber=1 scan=21156 | 45.2891 |
| 17594:4 | 5 | K.QIGLYPVLVIDSSGYVNPN[+2059.735]YTGR.I | HexNAc(4)Hex(5)Fuc(1)NeuAc(1) | N[+2060] | 1147.0198 | 4 | 4585.0574 | 4585.0384 | 0 | 4.2 | 168 | Specific | 854.76 | 834.23 | 834.23 | 18.12 | 91 | >sp\|P01833\|PIGR_HUMAN Polymeric immunoglobulin receptor OS=Homo sapiens OX=9606 GN=PIGR PE=1 SV=4 | 902 | 20201117_08512_1ng_glyco_hcd203040_compound15.21171.21171.4 | controllerType=0 controllerNumber=1 scan=21171 | 45.3344 |
| 17570:3 | 5 | K.QIGLYPVLVIDSSGYVNPN[+2059.735]YTGR.I | HexNAc(4)Hex(5)Fuc(1)NeuAc(1) | N[+2060] | 1529.0257 | 3 | 4585.0625 | 4585.0384 | 0 | 5.3 | 168 | Specific | 873.99 | 797.99 | 797.99 | 16.95 | 91 | >sp\|P01833\|PIGR_HUMAN Polymeric immunoglobulin receptor OS=Homo sapiens OX=9606 GN=PIGR PE=1 SV=4 | 902 | 20201117_08512_1ng_glyco_hcd203040_compound15.21146.21146.3 | controllerType=0 controllerNumber=1 scan=21146 | 45.2672 |
| 17568:3 | 5 | K.QIGLYPVLVIDSSGYVNPN[+2059.735]YTGR.I | HexNAc(4)Hex(5)Fuc(1)NeuAc(1) | N[+2060] | 1529.0257 | 3 | 4585.0625 | 4585.0384 | 0 | 5.3 | 168 | Specific | 926.71 | 794.31 | 794.31 | 17.95 | 91 | >sp\|P01833\|PIGR_HUMAN Polymeric immunoglobulin receptor OS=Homo sapiens OX=9606 GN=PIGR PE=1 SV=4 | 902 | 20201117_08512_1ng_glyco_hcd203040_compound15.21144.21144.3 | controllerType=0 controllerNumber=1 scan=21144 | 45.2589 |
| 17581:3 | 5 | K.QIGLYPVLVIDSSGYVNPN[+2059.735]YTGR.I | HexNAc(4)Hex(5)Fuc(1)NeuAc(1) | N[+2060] | 1529.0252 | 3 | 4585.0611 | 4585.0384 | 0 | 4.9 | 168 | Specific | 928.73 | 854.25 | 854.25 | 17.95 | 91 | >sp\|P01833\|PIGR_HUMAN Polymeric immunoglobulin receptor OS=Homo sapiens OX=9606 GN=PIGR PE=1 SV=4 | 902 | 20201117_08512_1ng_glyco_hcd203040_compound15.21158.21158.3 | controllerType=0 controllerNumber=1 scan=21158 | 45.2944 |
| 21130:5 | 5 | R.[+42.011]C[+57.021]PLLVDSEGWVKAQYEGRLSLLEEPGN[+1403.507]GTFTVILNQL | HexNAc(3)Hex(4)Fuc(1) | .[42], C[+57], N[+1404] | 1187.9722 | 5 | 5935.8317 | 5935.8203 | 0 | 1.9 | 395 | Specific | 264.32 | 264.32 | 264.32 | 6.52 | 91 | >sp\|P01833\|PIGR_HUMAN Polymeric immunoglobulin receptor OS=Homo sapiens OX=9606 GN=PIGR PE=1 SV=4 | 902 | 20201117_08512_1ng_glyco_hcd203040_compound15.24866.24866.4 | controllerType=0 controllerNumber=1 scan=24866 | 53.5022 |
| 21133:5 | 5 | R.[+42.011]C[+57.021]PLLVDSEGWVKAQYEGRLSLLEEPGN[+1403.507]GTFTVILNQL | HexNAc(3)Hex(4)Fuc(1) | .[42], C[+57], N[+1404] | 1187.9722 | 5 | 5935.8317 | 5935.8203 | 0 | 1.9 | 395 | Specific | 309.18 | 309.18 | 309.18 | 7.09 | 91 | >sp\|P01833\|PIGR_HUMAN Polymeric immunoglobulin receptor OS=Homo sapiens OX=9606 GN=PIGR PE=1 SV=4 | 902 | 20201117_08512_1ng_glyco_hcd203040_compound15.24869.24869.4 | controllerType=0 controllerNumber=1 scan=24869 | 53.5094 |
| 24148:4 | 5 | R.LSLLEEPGN[+3007.058]GTFTVILNQLTSR.D | HexNAc(5)Hex(6)Fuc(1)NeuAc(3) | N[+3007] | 1353.098 | 4 | 5409.3703 | 5409.3505 | 0 | 3.6 | 413 | Specific | 215.46 | 141.07 | 141.07 | 4.89 | 91 | >sp\|P01833\|PIGR_HUMAN Polymeric immunoglobulin receptor OS=Homo sapiens OX=9606 GN=PIGR PE=1 SV=4 | 902 | 20201117_08512_1ng_glyco_hcd203040_compound15.28006.28006.4 | controllerType=0 controllerNumber=1 scan=28006 | 59.6057 |
| 27172:4 | 5 | R.[+42.011]LSLLEEPGN[+2351.851]GTFTVILNQLTSR.D | HexNAc(4)Hex(5)Fuc(3)NeuAc(1) | .[42], N[+2352] | 1199.7922 | 4 | 4796.1468 | 4796.1539 | 0 | -1.5 | 413 | Specific | 276.52 | 276.52 | 276.52 | 6.43 | 91 | >sp\|P01833\|PIGR_HUMAN Polymeric immunoglobulin receptor OS=Homo sapiens OX=9606 GN=PIGR PE=1 SV=4 | 902 | 20201117_08512_1ng_glyco_hcd203040_compound15.31156.31156.4 | controllerType=0 controllerNumber=1 scan=31156 | 65.2916 |
| 20839:4 | 5 | R.LSLLEEPGN[+2862.021]GTFTVILNQLTSR.D | HexNAc(5)Hex(6)Fuc(2)NeuAc(2) | N[+2862] | 1316.8386 | 4 | 5264.3327 | 5264.313 | 0 | 3.7 | 413 | Specific | 344.3 | 344.3 | 344.3 | 8.05 | 91 | >sp\|P01833\|PIGR_HUMAN Polymeric immunoglobulin receptor OS=Homo sapiens OX=9606 GN=PIGR PE=1 SV=4 | 902 | 20201117_08512_1ng_glyco_hcd203040_compound15.24562.24562.4 | controllerType=0 controllerNumber=1 scan=24562 | 52.8462 |
| 24091:4 | 5 | R.LSLLEEPGN[+3007.058]GTFTVILNQLTSR.D | HexNAc(5)Hex(6)Fuc(1)NeuAc(3) | N[+3007] | 1353.0949 | 4 | 5409.3576 | 5409.3505 | 0 | 1.3 | 413 | Specific | 353.68 | 292.21 | 292.21 | 8.18 | 91 | >sp\|P01833\|PIGR_HUMAN Polymeric immunoglobulin receptor OS=Homo sapiens OX=9606 GN=PIGR PE=1 SV=4 | 902 | 20201117_08512_1ng_glyco_hcd203040_compound15.27947.27947.4 | controllerType=0 controllerNumber=1 scan=27947 | 59.5027 |
| 27171:3 | 5 | R.[+42.011]LSLLEEPGN[+2351.851]GTFTVILNQLTSR.D | HexNAc(4)Hex(5)Fuc(3)NeuAc(1) | .[42], N[+2352] | 1599.3838 | 3 | 4796.137 | 4796.1539 | 0 | -3.5 | 413 | Specific | 355.08 | 297.95 | 297.95 | 6.01 | 91 | >sp\|P01833\|PIGR_HUMAN Polymeric immunoglobulin receptor OS=Homo sapiens OX=9606 GN=PIGR PE=1 SV=4 | 902 | 20201117_08512_1ng_glyco_hcd203040_compound15.31155.31155.3 | controllerType=0 controllerNumber=1 scan=31155 | 65.289 |
| 21084:3 | 5 | R.[+42.011]LSLLEEPGN[+2336.851]GTFTVILNQLTSR.D | HexNAc(6)Hex(6)Fuc(1) | .[42], N[+2337] | 1594.3826 | 3 | 4781.1333 | 4781.1542 | 0 | -4.4 | 413 | Specific | 369.64 | 369.64 | 369.64 | 6.63 | 91 | >sp\|P01833\|PIGR_HUMAN Polymeric immunoglobulin receptor OS=Homo sapiens OX=9606 GN=PIGR PE=1 SV=4 | 902 | 20201117_08512_1ng_glyco_hcd203040_compound15.24818.24818.3 | controllerType=0 controllerNumber=1 scan=24818 | 53.3945 |
| 27173:3 | 5 | R.[+42.011]LSLLEEPGN[+2351.851]GTFTVILNQLTSR.D | HexNAc(4)Hex(5)Fuc(3)NeuAc(1) | .[42], N[+2352] | 1599.3838 | 3 | 4796.137 | 4796.1539 | 0 | -3.5 | 413 | Specific | 376.52 | 301.11 | 301.11 | 6.09 | 91 | >sp\|P01833\|PIGR_HUMAN Polymeric immunoglobulin receptor OS=Homo sapiens OX=9606 GN=PIGR PE=1 SV=4 | 902 | 20201117_08512_1ng_glyco_hcd203040_compound15.31157.31157.3 | controllerType=0 controllerNumber=1 scan=31157 | 65.294 |
| 24146:4 | 5 | R.LSLLEEPGN[+3007.058]GTFTVILNQLTSR.D | HexNAc(5)Hex(6)Fuc(1)NeuAc(3) | N[+3007] | 1353.098 | 4 | 5409.3703 | 5409.3505 | 0 | 3.6 | 413 | Specific | 381.92 | 380.31 | 380.31 | 9.92 | 91 | >sp\|P01833\|PIGR_HUMAN Polymeric immunoglobulin receptor OS=Homo sapiens OX=9606 GN=PIGR PE=1 SV=4 | 902 | 20201117_08512_1ng_glyco_hcd203040_compound15.28004.28004.4 | controllerType=0 controllerNumber=1 scan=28004 | 59.6019 |
| 21082:3 | 5 | R.[+42.011]LSLLEEPGN[+2336.851]GTFTVILNQLTSR.D | HexNAc(6)Hex(6)Fuc(1) | .[42], N[+2337] | 1594.3826 | 3 | 4781.1333 | 4781.1542 | 0 | -4.4 | 413 | Specific | 410.67 | 402.47 | 402.47 | 8.16 | 91 | >sp\|P01833\|PIGR_HUMAN Polymeric immunoglobulin receptor OS=Homo sapiens OX=9606 GN=PIGR PE=1 SV=4 | 902 | 20201117_08512_1ng_glyco_hcd203040_compound15.24816.24816.3 | controllerType=0 controllerNumber=1 scan=24816 | 53.3905 |
| 24093:4 | 5 | R.LSLLEEPGN[+3007.058]GTFTVILNQLTSR.D | HexNAc(5)Hex(6)Fuc(1)NeuAc(3) | N[+3007] | 1353.0949 | 4 | 5409.3576 | 5409.3505 | 0 | 1.3 | 413 | Specific | 456.93 | 377.44 | 377.44 | 10.41 | 91 | >sp\|P01833\|PIGR_HUMAN Polymeric immunoglobulin receptor OS=Homo sapiens OX=9606 GN=PIGR PE=1 SV=4 | 902 | 20201117_08512_1ng_glyco_hcd203040_compound15.27949.27949.4 | controllerType=0 controllerNumber=1 scan=27949 | 59.5067 |
| 21077:3 | 5 | R.LSLLEEPGN[+2553.910]GTFTVILNQLTSR.D | HexNAc(5)Hex(5)Fuc(1)NeuAc(2) | N[+2554] | 1652.748 | 3 | 4956.2295 | 4956.2023 | 0 | 5.5 | 413 | Specific | 466.79 | 437.45 | 437.45 | 8.78 | 91 | >sp\|P01833\|PIGR_HUMAN Polymeric immunoglobulin receptor OS=Homo sapiens OX=9606 GN=PIGR PE=1 SV=4 | 902 | 20201117_08512_1ng_glyco_hcd203040_compound15.24810.24810.3 | controllerType=0 controllerNumber=1 scan=24810 | 53.3787 |
| 21074:3 | 5 | R.LSLLEEPGN[+2553.910]GTFTVILNQLTSR.D | HexNAc(5)Hex(5)Fuc(1)NeuAc(2) | N[+2554] | 1652.748 | 3 | 4956.2295 | 4956.2023 | 0 | 5.5 | 413 | Specific | 489.92 | 465.66 | 465.66 | 8.9 | 91 | >sp\|P01833\|PIGR_HUMAN Polymeric immunoglobulin receptor OS=Homo sapiens OX=9606 GN=PIGR PE=1 SV=4 | 902 | 20201117_08512_1ng_glyco_hcd203040_compound15.24807.24807.3 | controllerType=0 controllerNumber=1 scan=24807 | 53.3738 |
| 20842:4 | 5 | R.LSLLEEPGN[+2862.021]GTFTVILNQLTSR.D | HexNAc(5)Hex(6)Fuc(2)NeuAc(2) | N[+2862] | 1316.8386 | 4 | 5264.3327 | 5264.313 | 0 | 3.7 | 413 | Specific | 504.93 | 504.93 | 504.93 | 10.41 | 91 | >sp\|P01833\|PIGR_HUMAN Polymeric immunoglobulin receptor OS=Homo sapiens OX=9606 GN=PIGR PE=1 SV=4 | 902 | 20201117_08512_1ng_glyco_hcd203040_compound15.24565.24565.4 | controllerType=0 controllerNumber=1 scan=24565 | 52.8546 |
| 18971:3 | 5 | R.LSLLEEPGN[+2715.963]GTFTVILNQLTSR.D | HexNAc(5)Hex(6)Fuc(1)NeuAc(2) | N[+2716] | 1706.7689 | 3 | 5118.2921 | 5118.2551 | 0 | 7.2 | 413 | Specific | 544.52 | 390.57 | 390.57 | 9.03 | 91 | >sp\|P01833\|PIGR_HUMAN Polymeric immunoglobulin receptor OS=Homo sapiens OX=9606 GN=PIGR PE=1 SV=4 | 902 | 20201117_08512_1ng_glyco_hcd203040_compound15.22607.22607.3 | controllerType=0 controllerNumber=1 scan=22607 | 48.4859 |
| 18973:3 | 5 | R.LSLLEEPGN[+2715.963]GTFTVILNQLTSR.D | HexNAc(5)Hex(6)Fuc(1)NeuAc(2) | N[+2716] | 1706.7689 | 3 | 5118.2921 | 5118.2551 | 0 | 7.2 | 413 | Specific | 575.58 | 350.88 | 350.88 | 7.47 | 91 | >sp\|P01833\|PIGR_HUMAN Polymeric immunoglobulin receptor OS=Homo sapiens OX=9606 GN=PIGR PE=1 SV=4 | 902 | 20201117_08512_1ng_glyco_hcd203040_compound15.22609.22609.3 | controllerType=0 controllerNumber=1 scan=22609 | 48.4895 |
| 19138:3 | 5 | R.LSLLEEPGN[+2059.735]GTFTVILNQLTSR.D | HexNAc(4)Hex(5)Fuc(1)NeuAc(1) | N[+2060] | 1488.0222 | 3 | 4462.0519 | 4462.0275 | 0 | 5.5 | 413 | Specific | 907.82 | 702.55 | 702.55 | 17.6 | 91 | >sp\|P01833\|PIGR_HUMAN Polymeric immunoglobulin receptor OS=Homo sapiens OX=9606 GN=PIGR PE=1 SV=4 | 902 | 20201117_08512_1ng_glyco_hcd203040_compound15.22782.22782.3 | controllerType=0 controllerNumber=1 scan=22782 | 48.867 |
| 19141:4 | 5 | R.LSLLEEPGN[+2205.793]GTFTVILNQLTSR.D | HexNAc(4)Hex(5)Fuc(2)NeuAc(1) | N[+2206] | 1152.7811 | 4 | 4608.1026 | 4608.0854 | 0 | 3.7 | 413 | Specific | 915.17 | 704.75 | 704.75 | 19.12 | 91 | >sp\|P01833\|PIGR_HUMAN Polymeric immunoglobulin receptor OS=Homo sapiens OX=9606 GN=PIGR PE=1 SV=4 | 902 | 20201117_08512_1ng_glyco_hcd203040_compound15.22785.22785.4 | controllerType=0 controllerNumber=1 scan=22785 | 48.8729 |
| 19094:3 | 5 | R.LSLLEEPGN[+2205.793]GTFTVILNQLTSR.D | HexNAc(4)Hex(5)Fuc(2)NeuAc(1) | N[+2206] | 1536.7067 | 3 | 4608.1056 | 4608.0854 | 0 | 4.4 | 413 | Specific | 915.4 | 640.35 | 640.35 | 17.95 | 91 | >sp\|P01833\|PIGR_HUMAN Polymeric immunoglobulin receptor OS=Homo sapiens OX=9606 GN=PIGR PE=1 SV=4 | 902 | 20201117_08512_1ng_glyco_hcd203040_compound15.22736.22736.3 | controllerType=0 controllerNumber=1 scan=22736 | 48.7716 |
| 19106:4 | 5 | R.LSLLEEPGN[+2205.793]GTFTVILNQLTSR.D | HexNAc(4)Hex(5)Fuc(2)NeuAc(1) | N[+2206] | 1152.7834 | 4 | 4608.1116 | 4608.0854 | 0 | 5.7 | 413 | Specific | 940.31 | 768.47 | 768.47 | 18.22 | 91 | >sp\|P01833\|PIGR_HUMAN Polymeric immunoglobulin receptor OS=Homo sapiens OX=9606 GN=PIGR PE=1 SV=4 | 902 | 20201117_08512_1ng_glyco_hcd203040_compound15.22748.22748.4 | controllerType=0 controllerNumber=1 scan=22748 | 48.7[illegible]38 |
| 19140:3 | 5 | R.LSLLEEPGN[+2059.735]GTFTVILNQLTSR.D | HexNAc(4)Hex(5)Fuc(1)NeuAc(1) | N[+2060] | 1488.0222 | 3 | 4462.0519 | 4462.0275 | 0 | 5.5 | 413 | Specific | 977.71 | 731.33 | 731.33 | 18.6 | 91 | >sp\|P01833\|PIGR_HUMAN Polymeric immunoglobulin receptor OS=Homo sapiens OX=9606 GN=PIGR PE=1 SV=4 | 902 | 20201117_08512_1ng_glyco_hcd203040_compound15.22784.22784.3 | controllerType=0 controllerNumber=1 scan=22784 | 48.[illegible]07 |
| 19092:3 | 5 | R.LSLLEEPGN[+2205.793]GTFTVILNQLTSR.D | HexNAc(4)Hex(5)Fuc(2)NeuAc(1) | N[+2206] | 1536.7067 | 3 | 4608.1056 | 4608.0854 | 0 | 4.4 | 413 | Specific | 1064.44 | 827.98 | 827.98 | 20.95 | 91 | >sp\|P01833\|PIGR_HUMAN Polymeric immunoglobulin receptor OS=Homo sapiens OX=9606 GN=PIGR PE=1 SV=4 | 902 | 20201117_08512_1ng_glyco_hcd203040_compound15.22734.22734.3 | controllerType=0 controllerNumber=1 scan=22734 | 48.[illegible]41 |
| 19103:4 | 5 | R.LSLLEEPGN[+2205.793]GTFTVILNQLTSR.D | HexNAc(4)Hex(5)Fuc(2)NeuAc(1) | N[+2206] | 1152.7834 | 4 | 4608.1116 | 4608.0854 | 0 | 5.7 | 413 | Specific | 1100.19 | 911.46 | 911.46 | 22.22 | 91 | >sp\|P01833\|PIGR_HUMAN Polymeric immunoglobulin receptor OS=Homo sapiens OX=9606 GN=PIGR PE=1 SV=4 | 902 | 20201117_08512_1ng_glyco_hcd203040_compound15.22745.22745.4 | controllerType=0 controllerNumber=1 scan=22745 | 48.7861 |
| 14916:3 | 5 | K.IIEGEPNLKVPGN[+2351.851]VTAVLGETLK.V | HexNAc(4)Hex(5)Fuc(3)NeuAc(1) | N[+2352] | 1581.7404 | 3 | 4743.2067 | 4743.2001 | 0 | 1.4 | 457 | Specific | 547.13 | 523.12 | 523.12 | 10.1 | 91 | >sp\|P01833\|PIGR_HUMAN Polymeric immunoglobulin receptor OS=Homo sapiens OX=9606 GN=PIGR PE=1 SV=4 | 902 | 20201117_08512_1ng_glyco_hcd203040_compound15.18373.18373.3 | controllerType=0 controllerNumber=1 scan=18373 | 39.2019 |
| 14965:4 | 5 | K.IIEGEPNLKVPGN[+2205.793]VTAVLGETLK.V | HexNAc(4)Hex(5)Fuc(2)NeuAc(1) | N[+2206] | 1150.0445 | 4 | 4597.1561 | 4597.1422 | 0 | 3 | 457 | Specific | 576.59 | 556.94 | 556.94 | 11.3 | 91 | >sp\|P01833\|PIGR_HUMAN Polymeric immunoglobulin receptor OS=Homo sapiens OX=9606 GN=PIGR PE=1 SV=4 | 902 | 20201117_08512_1ng_glyco_hcd203040_compound15.18424.18424.4 | controllerType=0 controllerNumber=1 scan=18424 | 39.3046 |
| 14914:3 | 5 | K.IIEGEPNLKVPGN[+2351.851]VTAVLGETLK.V | HexNAc(4)Hex(5)Fuc(3)NeuAc(1) | N[+2352] | 1581.7404 | 3 | 4743.2067 | 4743.2001 | 0 | 1.4 | 457 | Specific | 614.38 | 596.83 | 596.83 | 12.58 | 91 | >sp\|P01833\|PIGR_HUMAN Polymeric immunoglobulin receptor OS=Homo sapiens OX=9606 GN=PIGR PE=1 SV=4 | 902 | 20201117_08512_1ng_glyco_hcd203040_compound15.18371.18371.3 | controllerType=0 controllerNumber=1 scan=18371 | 39.1984 |
| 14973:4 | 5 | K.IIEGEPNLKVPGN[+2059.735]VTAVLGETLK.V | HexNAc(4)Hex(5)Fuc(1)NeuAc(1) | N[+2060] | 1113.5286 | 4 | 4451.0926 | 4451.0843 | 0 | 1.9 | 457 | Specific | 620.91 | 599.61 | 599.61 | 12.4 | 91 | >sp\|P01833\|PIGR_HUMAN Polymeric immunoglobulin receptor OS=Homo sapiens OX=9606 GN=PIGR PE=1 SV=4 | 902 | 20201117_08512_1ng_glyco_hcd203040_compound15.18433.18433.4 | controllerType=0 controllerNumber=1 scan=18433 | 39.3232 |
| 14963:4 | 5 | K.IIEGEPNLKVPGN[+2205.793]VTAVLGETLK.V | HexNAc(4)Hex(5)Fuc(2)NeuAc(1) | N[+2206] | 1150.0445 | 4 | 4597.1561 | 4597.1422 | 0 | 3 | 457 | Specific | 648.27 | 570.08 | 570.08 | 12.3 | 91 | >sp\|P01833\|PIGR_HUMAN Polymeric immunoglobulin receptor OS=Homo sapiens OX=9606 GN=PIGR PE=1 SV=4 | 902 | 20201117_08512_1ng_glyco_hcd203040_compound15.18422.18422.4 | controllerType=0 controllerNumber=1 scan=18422 | 39.2999 |
| 14894:3 | 5 | K.IIEGEPNLKVPGN[+2205.793]VTAVLGETLK.V | HexNAc(4)Hex(5)Fuc(2)NeuAc(1) | N[+2206] | 1533.0525 | 3 | 4597.1431 | 4597.1422 | 0 | 0.2 | 457 | Specific | 772.89 | 700.81 | 700.81 | 14.93 | 91 | >sp\|P01833\|PIGR_HUMAN Polymeric immunoglobulin receptor OS=Homo sapiens OX=9606 GN=PIGR PE=1 SV=4 | 902 | 20201117_08512_1ng_glyco_hcd203040_compound15.18350.18350.3 | controllerType=0 controllerNumber=1 scan=18350 | 39.1624 |
| 14976:4 | 5 | K.IIEGEPNLKVPGN[+2059.735]VTAVLGETLK.V | HexNAc(4)Hex(5)Fuc(1)NeuAc(1) | N[+2060] | 1113.5286 | 4 | 4451.0926 | 4451.0843 | 0 | 1.9 | 457 | Specific | 823.7 | 708.61 | 708.61 | 16.4 | 91 | >sp\|P01833\|PIGR_HUMAN Polymeric immunoglobulin receptor OS=Homo sapiens OX=9606 GN=PIGR PE=1 SV=4 | 902 | 20201117_08512_1ng_glyco_hcd203040_compound15.18436.18436.4 | controllerType=0 controllerNumber=1 scan=18436 | 39.3278 |
| 14934:3 | 5 | K.IIEGEPNLKVPGN[+2059.735]VTAVLGETLK.V | HexNAc(4)Hex(5)Fuc(1)NeuAc(1) | N[+2060] | 1484.3685 | 3 | 4451.0908 | 4451.0843 | 0 | 1.5 | 457 | Specific | 864.39 | 663.25 | 663.25 | 17.58 | 91 | >sp\|P01833\|PIGR_HUMAN Polymeric immunoglobulin receptor OS=Homo sapiens OX=9606 GN=PIGR PE=1 SV=4 | 902 | 20201117_08512_1ng_glyco_hcd203040_compound15.18392.18392.3 | controllerType=0 controllerNumber=1 scan=18392 | 39.2399 |
| 14892:3 | 5 | K.IIEGEPNLKVPGN[+2205.793]VTAVLGETLK.V | HexNAc(4)Hex(5)Fuc(2)NeuAc(1) | N[+2206] | 1533.0525 | 3 | 4597.1431 | 4597.1422 | 0 | 0.2 | 457 | Specific | 997.17 | 911.26 | 911.26 | 18.93 | 91 | >sp\|P01833\|PIGR_HUMAN Polymeric immunoglobulin receptor OS=Homo sapiens OX=9606 GN=PIGR PE=1 SV=4 | 902 | 20201117_08512_1ng_glyco_hcd203040_compound15.18348.18348.3 | controllerType=0 controllerNumber=1 scan=18348 | 39.1579 |
| 14931:3 | 5 | K.IIEGEPNLKVPGN[+2059.735]VTAVLGETLK.V | HexNAc(4)Hex(5)Fuc(1)NeuAc(1) | N[+2060] | 1484.3685 | 3 | 4451.0908 | 4451.0843 | 0 | 1.5 | 457 | Specific | 1063.19 | 960.46 | 960.46 | 21.58 | 91 | >sp\|P01833\|PIGR_HUMAN Polymeric immunoglobulin receptor OS=Homo sapiens OX=9606 GN=PIGR PE=1 SV=4 | 902 | 20201117_08512_1ng_glyco_hcd203040_compound15.18389.18389.3 | controllerType=0 controllerNumber=1 scan=18389 | 39.233 |
| 16125:5 | 5 | K.IIEGEPNLKVPGN[+2350.830]VTAVLGETLKVPC[+57.021]HFPC[+57.021]K.F | HexNAc(4)Hex(5)Fuc(1)NeuAc(2) | C[+57]*2, N[+2351] | 1154.3364 | 5 | 5767.653 | 5767.6372 | 0 | 2.7 | 457 | Specific | 239.36 | 239.36 | 239.36 | 4.35 | 91 | >sp\|P01833\|PIGR_HUMAN Polymeric immunoglobulin receptor OS=Homo sapiens OX=9606 GN=PIGR PE=1 SV=4 | 902 | 20201117_08512_1ng_glyco_hcd203040_compound15.19635.19635.3 | controllerType=0 controllerNumber=1 scan=19635 | 41.7877 |
| 15778:4 | 5 | K.IIEGEPNLKVPGN[+2351.851]VTAVLGETLKVPC[+57.021]HFPC[+57.021]K.F | HexNAc(4)Hex(5)Fuc(3)NeuAc(1) | C[+57]*2, N[+2352] | 1442.9191 | 4 | 5768.6545 | 5768.6576 | 0 | -0.6 | 457 | Specific | 240.75 | 240.75 | 240.75 | 4.63 | 91 | >sp\|P01833\|PIGR_HUMAN Polymeric immunoglobulin receptor OS=Homo sapiens OX=9606 GN=PIGR PE=1 SV=4 | 902 | 20201117_08512_1ng_glyco_hcd203040_compound15.19273.19273.3 | controllerType=0 controllerNumber=1 scan=19273 | 40.9797 |
| 14665:5 | 5 | K.IIEGEPNLKVPGN[+2059.735]VTAVLGETLKVPC[+57.021]HFPC[+57.021]K.F | HexNAc(4)Hex(5)Fuc(1)NeuAc(1) | C[+57]*2, N[+2060] | 1096.1165 | 5 | 5476.5532 | 5476.5418 | 0 | 2.1 | 457 | Specific | 264.94 | 264.94 | 264.94 | 5.21 | 91 | >sp\|P01833\|PIGR_HUMAN Polymeric immunoglobulin receptor OS=Homo sapiens OX=9606 GN=PIGR PE=1 SV=4 | 902 | 20201117_08512_1ng_glyco_hcd203040_compound15.18096.18096.3 | controllerType=0 controllerNumber=1 scan=18096 | 38.6946 |
| 13456:5 | 5 | K.IIEGEPNLKVPGN[+2205.793]VTAVLGETLKVPC[+57.021]HFPC[+57.021]K.F | HexNAc(4)Hex(5)Fuc(2)NeuAc(1) | C[+57]*2, N[+2206] | 1125.3307 | 5 | 5622.6243 | 5622.5997 | 0 | 4.4 | 457 | Specific | 267.14 | 267.14 | 267.14 | 5.83 | 91 | >sp\|P01833\|PIGR_HUMAN Polymeric immunoglobulin receptor OS=Homo sapiens OX=9606 GN=PIGR PE=1 SV=4 | 902 | 20201117_08512_1ng_glyco_hcd203040_compound15.16837.16837.5 | controllerType=0 controllerNumber=1 scan=16837 | 36.1611 |
| 15780:4 | 5 | K.IIEGEPNLKVPGN[+2351.851]VTAVLGETLKVPC[+57.021]HFPC[+57.021]K.F | HexNAc(4)Hex(5)Fuc(3)NeuAc(1) | C[+57]*2, N[+2352] | 1442.9191 | 4 | 5768.6545 | 5768.6576 | 0 | -0.6 | 457 | Specific | 271.83 | 271.83 | 271.83 | 4.19 | 91 | >sp\|P01833\|PIGR_HUMAN Polymeric immunoglobulin receptor OS=Homo sapiens OX=9606 GN=PIGR PE=1 SV=4 | 902 | 20201117_08512_1ng_glyco_hcd203040_compound15.19275.19275.3 | controllerType=0 controllerNumber=1 scan=19275 | 40.988 |
| 14668:5 | 5 | K.IIEGEPNLKVPGN[+2059.735]VTAVLGETLKVPC[+57.021]HFPC[+57.021]K.F | HexNAc(4)Hex(5)Fuc(1)NeuAc(1) | C[+57]*2, N[+2060] | 1096.1165 | 5 | 5476.5532 | 5476.5418 | 0 | 2.1 | 457 | Specific | 287.87 | 287.87 | 287.87 | 4.88 | 91 | >sp\|P01833\|PIGR_HUMAN Polymeric immunoglobulin receptor OS=Homo sapiens OX=9606 GN=PIGR PE=1 SV=4 | 902 | 20201117_08512_1ng_glyco_hcd203040_compound15.18099.18099.3 | controllerType=0 controllerNumber=1 scan=18099 | 38.7019 |
| 13454:5 | 5 | K.IIEGEPNLKVPGN[+2205.793]VTAVLGETLKVPC[+57.021]HFPC[+57.021]K.F | HexNAc(4)Hex(5)Fuc(2)NeuAc(1) | C[+57]*2, N[+2206] | 1125.3307 | 5 | 5622.6243 | 5622.5997 | 0 | 4.4 | 457 | Specific | 318.67 | 310.36 | 310.36 | 6.55 | 91 | >sp\|P01833\|PIGR_HUMAN Polymeric immunoglobulin receptor OS=Homo sapiens OX=9606 GN=PIGR PE=1 SV=4 | 902 | 20201117_08512_1ng_glyco_hcd203040_compound15.16835.16835.5 | controllerType=0 controllerNumber=1 scan=16835 | 36.1575 |
| 15898:4 | 5 | K.IIEGEPNLKVPGN[+2204.772]VTAVLGETLKVPC[+57.021]HFPC[+57.021]K.F | HexNAc(4)Hex(5)NeuAc(2) | C[+57]*2, N[+2205] | 1406.1543 | 4 | 5621.5954 | 5621.5793 | 0 | 2.9 | 457 | Specific | 502.45 | 435.62 | 435.62 | 9.15 | 91 | >sp\|P01833\|PIGR_HUMAN Polymeric immunoglobulin receptor OS=Homo sapiens OX=9606 GN=PIGR PE=1 SV=4 | 902 | 20201117_08512_1ng_glyco_hcd203040_compound15.19398.19398.4 | controllerType=0 controllerNumber=1 scan=19398 | 41.2594 |
| 14309:5 | 5 | K.IIEGEPNLKVPGN[+2570.925]VTAVLGETLKVPC[+57.021]HFPC[+57.021]K.F | HexNAc(5)Hex(6)Fuc(2)NeuAc(1) | C[+57]*2, N[+2571] | 1198.3555 | 5 | 5987.7483 | 5987.7319 | 0 | 2.7 | 457 | Specific | 522.29 | 457.53 | 457.53 | 9.72 | 91 | >sp\|P01833\|PIGR_HUMAN Polymeric immunoglobulin receptor OS=Homo sapiens OX=9606 GN=PIGR PE=1 SV=4 | 902 | 20201117_08512_1ng_glyco_hcd203040_compound15.17725.17725.5 | controllerType=0 controllerNumber=1 scan=17725 | 37.9518 |
| 13396:5 | 5 | K.IIEGEPNLKVPGN[+2205.793]VTAVLGETLKVPC[+57.021]HFPC[+57.021]K.F | HexNAc(4)Hex(5)Fuc(2)NeuAc(1) | C[+57]*2, N[+2206] | 1125.3344 | 5 | 5622.6427 | 5622.5997 | 0 | 7.6 | 457 | Specific | 527.46 | 527.46 | 527.46 | 8.64 | 91 | >sp\|P01833\|PIGR_HUMAN Polymeric immunoglobulin receptor OS=Homo sapiens OX=9606 GN=PIGR PE=1 SV=4 | 902 | 20201117_08512_1ng_glyco_hcd203040_compound15.16775.16775.5 | controllerType=0 controllerNumber=1 scan=16775 | 36.031 |
| 15860:6 | 5 | K.IIEGEPNLKVPGN[+2350.830]VTAVLGETLKVPC[+57.021]HFPC[+57.021]K.F | HexNAc(4)Hex(5)Fuc(1)NeuAc(2) | C[+57]*2, N[+2351] | 962.1154 | 6 | 5767.656 | 5767.6372 | 0 | 3.3 | 457 | Specific | 555.78 | 478.12 | 478.12 | 9.81 | 91 | >sp\|P01833\|PIGR_HUMAN Polymeric immunoglobulin receptor OS=Homo sapiens OX=9606 GN=PIGR PE=1 SV=4 | 902 | 20201117_08512_1ng_glyco_hcd203040_compound15.19359.19359.6 | controllerType=0 controllerNumber=1 scan=19359 | 41.1684 |
| 15863:6 | 5 | K.IIEGEPNLKVPGN[+2350.830]VTAVLGETLKVPC[+57.021]HFPC[+57.021]K.F | HexNAc(4)Hex(5)Fuc(1)NeuAc(2) | C[+57]*2, N[+2351] | 962.1154 | 6 | 5767.656 | 5767.6372 | 0 | 3.3 | 457 | Specific | 566.12 | 441.32 | 441.32 | 9.81 | 91 | >sp\|P01833\|PIGR_HUMAN Polymeric immunoglobulin receptor OS=Homo sapiens OX=9606 GN=PIGR PE=1 SV=4 | 902 | 20201117_08512_1ng_glyco_hcd203040_compound15.19362.19362.6 | controllerType=0 controllerNumber=1 scan=19362 | 41.1761 |
| 14457:5 | 5 | K.IIEGEPNLKVPGN[+2351.851]VTAVLGETLKVPC[+57.021]HFPC[+57.021]K.F | HexNAc(4)Hex(5)Fuc(3)NeuAc(1) | C[+57]*2, N[+2352] | 1154.5408 | 5 | 5768.6748 | 5768.6576 | 0 | 3 | 457 | Specific | 573.98 | 573.98 | 573.98 | 10.08 | 91 | >sp\|P01833\|PIGR_HUMAN Polymeric immunoglobulin receptor OS=Homo sapiens OX=9606 GN=PIGR PE=1 SV=4 | 902 | 20201117_08512_1ng_glyco_hcd203040_compound15.17880.17880.5 | controllerType=0 controllerNumber=1 scan=17880 | 38.2877 |
| 13398:5 | 5 | K.IIEGEPNLKVPGN[+2205.793]VTAVLGETLKVPC[+57.021]HFPC[+57.021]K.F | HexNAc(4)Hex(5)Fuc(2)NeuAc(1) | C[+57]*2, N[+2206] | 1125.3344 | 5 | 5622.6427 | 5622.5997 | 0 | 7.6 | 457 | Specific | 613.66 | 613.66 | 613.66 | 11.25 | 91 | >sp\|P01833\|PIGR_HUMAN Polymeric immunoglobulin receptor OS=Homo sapiens OX=9606 GN=PIGR PE=1 SV=4 | 902 | 20201117_08512_1ng_glyco_hcd203040_compound15.16777.16777.5 | controllerType=0 controllerNumber=1 scan=16777 | 36.0353 |
| 15995:4 | 5 | K.IIEGEPNLKVPGN[+2350.830]VTAVLGETLKVPC[+57.021]HFPC[+57.021]K.F | HexNAc(4)Hex(5)Fuc(1)NeuAc(2) | C[+57]*2, N[+2351] | 1442.672 | 4 | 5767.6662 | 5767.6372 | 0 | 5 | 457 | Specific | 634.67 | 498.7 | 498.7 | 11.22 | 91 | >sp\|P01833\|PIGR_HUMAN Polymeric immunoglobulin receptor OS=Homo sapiens OX=9606 GN=PIGR PE=1 SV=4 | 902 | 20201117_08512_1ng_glyco_hcd203040_compound15.19500.19500.4 | controllerType=0 controllerNumber=1 scan=19500 | 41.4968 |
| 14534:4 | 5 | K.IIEGEPNLKVPGN[+2205.793]VTAVLGETLKVPC[+57.021]HFPC[+57.021]K.F | HexNAc(4)Hex(5)Fuc(2)NeuAc(1) | C[+57]*2, N[+2206] | 1406.4083 | 4 | 5622.6115 | 5622.5997 | 0 | 2.1 | 457 | Specific | 673.35 | 637.72 | 637.72 | 12.86 | 91 | >sp\|P01833\|PIGR_HUMAN Polymeric immunoglobulin receptor OS=Homo sapiens OX=9606 GN=PIGR PE=1 SV=4 | 902 | 20201117_08512_1ng_glyco_hcd203040_compound15.17960.17960.4 | controllerType=0 controllerNumber=1 scan=17960 | 38.4222 |
| 15838:4 | 5 | K.IIEGEPNLKVPGN[+2350.830]VTAVLGETLKVPC[+57.021]HFPC[+57.021]K.F | HexNAc(4)Hex(5)Fuc(1)NeuAc(2) | C[+57]*2, N[+2351] | 1442.6689 | 4 | 5767.654 | 5767.6372 | 0 | 2.9 | 457 | Specific | 719.35 | 719.35 | 719.35 | 13.86 | 91 | >sp\|P01833\|PIGR_HUMAN Polymeric immunoglobulin receptor OS=Homo sapiens OX=9606 GN=PIGR PE=1 SV=4 | 902 | 20201117_08512_1ng_glyco_hcd203040_compound15.19336.19336.4 | controllerType=0 controllerNumber=1 scan=19336 | 41.118 |
| 15993:4 | 5 | K.IIEGEPNLKVPGN[+2350.830]VTAVLGETLKVPC[+57.021]HFPC[+57.021]K.F | HexNAc(4)Hex(5)Fuc(1)NeuAc(2) | C[+57]*2, N[+2351] | 1442.672 | 4 | 5767.6662 | 5767.6372 | 0 | 5 | 457 | Specific | 728.61 | 566.77 | 566.77 | 13.22 | 91 | >sp\|P01833\|PIGR_HUMAN Polymeric immunoglobulin receptor OS=Homo sapiens OX=9606 GN=PIGR PE=1 SV=4 | 902 | 20201117_08512_1ng_glyco_hcd203040_compound15.19498.19498.4 | controllerType=0 controllerNumber=1 scan=19498 | 41.49 |
| 14460:5 | 5 | K.IIEGEPNLKVPGN[+2351.851]VTAVLGETLKVPC[+57.021]HFPC[+57.021]K.F | HexNAc(4)Hex(5)Fuc(3)NeuAc(1) | C[+57]*2, N[+2352] | 1154.5408 | 5 | 5768.6748 | 5768.6576 | 0 | 3 | 457 | Specific | 733.56 | 605.01 | 605.01 | 14.33 | 91 | >sp\|P01833\|PIGR_HUMAN Polymeric immunoglobulin receptor OS=Homo sapiens OX=9606 GN=PIGR PE=1 SV=4 | 902 | 20201117_08512_1ng_glyco_hcd203040_compound15.17883.17883.5 | controllerType=0 controllerNumber=1 scan=17883 | 38.292 |
| 14537:4 | 5 | K.IIEGEPNLKVPGN[+2205.793]VTAVLGETLKVPC[+57.021]HFPC[+57.021]K.F | HexNAc(4)Hex(5)Fuc(2)NeuAc(1) | C[+57]*2, N[+2206] | 1406.4083 | 4 | 5622.6115 | 5622.5997 | 0 | 2.1 | 457 | Specific | 755.72 | 755.72 | 755.72 | 14.86 | 91 | >sp\|P01833\|PIGR_HUMAN Polymeric immunoglobulin receptor OS=Homo sapiens OX=9606 GN=PIGR PE=1 SV=4 | 902 | 20201117_08512_1ng_glyco_hcd203040_compound15.17963.17963.4 | controllerType=0 controllerNumber=1 scan=17963 | 38.4264 |
| 14610:4 | 5 | K.IIEGEPNLKVPGN[+2059.735]VTAVLGETLKVPC[+57.021]HFPC[+57.021]K.F | HexNAc(4)Hex(5)Fuc(1)NeuAc(1) | C[+57]*2, N[+2060] | 1369.8918 | 4 | 5476.5456 | 5476.5418 | 0 | 0.7 | 457 | Specific | 769.07 | 647.07 | 647.07 | 14.31 | 91 | >sp\|P01833\|PIGR_HUMAN Polymeric immunoglobulin receptor OS=Homo sapiens OX=9606 GN=PIGR PE=1 SV=4 | 902 | 20201117_08512_1ng_glyco_hcd203040_compound15.18039.18039.4 | controllerType=0 controllerNumber=1 scan=18039 | 38.5799 |
| 14456:4 | 5 | K.IIEGEPNLKVPGN[+2059.735]VTAVLGETLKVPC[+57.021]HFPC[+57.021]K.F | HexNAc(4)Hex(5)Fuc(1)NeuAc(1) | C[+57]*2, N[+2060] | 1369.8931 | 4 | 5476.5504 | 5476.5418 | 0 | 1.6 | 457 | Specific | 775.67 | 738.74 | 738.74 | 14.96 | 91 | >sp\|P01833\|PIGR_HUMAN Polymeric immunoglobulin receptor OS=Homo sapiens OX=9606 GN=PIGR PE=1 SV=4 | 902 | 20201117_08512_1ng_glyco_hcd203040_compound15.17879.17879.4 | controllerType=0 controllerNumber=1 scan=17879 | 38.2866 |
| 14386:4 | 5 | K.IIEGEPNLKVPGN[+2205.793]VTAVLGETLKVPC[+57.021]HFPC[+57.021]K.F | HexNAc(4)Hex(5)Fuc(2)NeuAc(1) | C[+57]*2, N[+2206] | 1406.4074 | 4 | 5622.6076 | 5622.5997 | 0 | 1.4 | 457 | Specific | 787.79 | 787.79 | 787.79 | 14.96 | 91 | >sp\|P01833\|PIGR_HUMAN Polymeric immunoglobulin receptor OS=Homo sapiens OX=9606 GN=PIGR PE=1 SV=4 | 902 | 20201117_08512_1ng_glyco_hcd203040_compound15.17806.17806.4 | controllerType=0 controllerNumber=1 scan=17806 | 38.1431 |
| 13434:5 | 5 | K.IIEGEPNLKVPGN[+1768.640]VTAVLGETLKVPC[+57.021]HFPC[+57.021]K.F | HexNAc(4)Hex(5)Fuc(1) | C[+57]*2, N[+1769] | 1037.8965 | 5 | 5185.4533 | 5185.4464 | 0 | 1.3 | 457 | Specific | 813.97 | 680.21 | 680.21 | 16.43 | 91 | >sp\|P01833\|PIGR_HUMAN Polymeric immunoglobulin receptor OS=Homo sapiens OX=9606 GN=PIGR PE=1 SV=4 | 902 | 20201117_08512_1ng_glyco_hcd203040_compound15.16814.16814.5 | controllerType=0 controllerNumber=1 scan=16814 | 36.1158 |
| 14650:5 | 5 | K.IIEGEPNLKVPGN[+2059.735]VTAVLGETLKVPC[+57.021]HFPC[+57.021]K.F | HexNAc(4)Hex(5)Fuc(1)NeuAc(1) | C[+57]*2, N[+2060] | 1096.1165 | 5 | 5476.5532 | 5476.5418 | 0 | 2.1 | 457 | Specific | 818.71 | 714.49 | 714.49 | 16.33 | 91 | >sp\|P01833\|PIGR_HUMAN Polymeric immunoglobulin receptor OS=Homo sapiens OX=9606 GN=PIGR PE=1 SV=4 | 902 | 20201117_08512_1ng_glyco_hcd203040_compound15.18081.18081.5 | controllerType=0 controllerNumber=1 scan=18081 | 38.6561 |
| 14454:5 | 5 | K.IIEGEPNLKVPGN[+2059.735]VTAVLGETLKVPC[+57.021]HFPC[+57.021]K.F | HexNAc(4)Hex(5)Fuc(1)NeuAc(1) | C[+57]*2, N[+2060] | 1096.1166 | 5 | 5476.5538 | 5476.5418 | 0 | 2.2 | 457 | Specific | 832.12 | 748.75 | 748.75 | 16.33 | 91 | >sp\|P01833\|PIGR_HUMAN Polymeric immunoglobulin receptor OS=Homo sapiens OX=9606 GN=PIGR PE=1 SV=4 | 902 | 20201117_08512_1ng_glyco_hcd203040_compound15.17877.17877.5 | controllerType=0 controllerNumber=1 scan=17877 | 38.2826 |
| 15837:5 | 5 | K.IIEGEPNLKVPGN[+2350.830]VTAVLGETLKVPC[+57.021]HFPC[+57.021]K.F | HexNAc(4)Hex(5)Fuc(1)NeuAc(2) | C[+57]*2, N[+2351] | 1154.3358 | 5 | 5767.65 | 5767.6372 | 0 | 2.2 | 457 | Specific | 869.15 | 691.14 | 691.14 | 17.33 | 91 | >sp\|P01833\|PIGR_HUMAN Polymeric immunoglobulin receptor OS=Homo sapiens OX=9606 GN=PIGR PE=1 SV=4 | 902 | 20201117_08512_1ng_glyco_hcd203040_compound15.19335.19335.5 | controllerType=0 controllerNumber=1 scan=19335 | 41.1158 |
| 14653:5 | 5 | K.IIEGEPNLKVPGN[+2059.735]VTAVLGETLKVPC[+57.021]HFPC[+57.021]K.F | HexNAc(4)Hex(5)Fuc(1)NeuAc(1) | C[+57]*2, N[+2060] | 1096.1165 | 5 | 5476.5532 | 5476.5418 | 0 | 2.1 | 457 | Specific | 913.58 | 787.27 | 787.27 | 18.33 | 91 | >sp\|P01833\|PIGR_HUMAN Polymeric immunoglobulin receptor OS=Homo sapiens OX=9606 GN=PIGR PE=1 SV=4 | 902 | 20201117_08512_1ng_glyco_hcd203040_compound15.18084.18084.5 | controllerType=0 controllerNumber=1 scan=18084 | 38.6609 |
| 15835:5 | 5 | K.IIEGEPNLKVPGN[+2350.830]VTAVLGETLKVPC[+57.021]HFPC[+57.021]K.F | HexNAc(4)Hex(5)Fuc(1)NeuAc(2) | C[+57]*2, N[+2351] | 1154.3358 | 5 | 5767.65 | 5767.6372 | 0 | 2.2 | 457 | Specific | 961.91 | 855.49 | 855.49 | 19.33 | 91 | >sp\|P01833\|PIGR_HUMAN Polymeric immunoglobulin receptor OS=Homo sapiens OX=9606 GN=PIGR PE=1 SV=4 | 902 | 20201117_08512_1ng_glyco_hcd203040_compound15.19333.19333.5 | controllerType=0 controllerNumber=1 scan=19333 | 41.1104 |
| 15836:4 | 5 | K.IIEGEPNLKVPGN[+2350.830]VTAVLGETLKVPC[+57.021]HFPC[+57.021]K.F | HexNAc(4)Hex(5)Fuc(1)NeuAc(2) | C[+57]*2, N[+2351] | 1442.6689 | 4 | 5767.654 | 5767.6372 | 0 | 2.9 | 457 | Specific | 992.44 | 775.41 | 775.41 | 18.86 | 91 | >sp\|P01833\|PIGR_HUMAN Polymeric immunoglobulin receptor OS=Homo sapiens OX=9606 GN=PIGR PE=1 SV=4 | 902 | 20201117_08512_1ng_glyco_hcd203040_compound15.19334.19334.4 | controllerType=0 controllerNumber=1 scan=19334 | 41.1118 |
| 13432:5 | 5 | K.IIEGEPNLKVPGN[+1768.640]VTAVLGETLKVPC[+57.021]HFPC[+57.021]K.F | HexNAc(4)Hex(5)Fuc(1) | C[+57]*2, N[+1769] | 1037.8965 | 5 | 5185.4533 | 5185.4464 | 0 | 1.3 | 457 | Specific | 1014.34 | 958.4 | 958.4 | 20.43 | 91 | >sp\|P01833\|PIGR_HUMAN Polymeric immunoglobulin receptor OS=Homo sapiens OX=9606 GN=PIGR PE=1 SV=4 | 902 | 20201117_08512_1ng_glyco_hcd203040_compound15.16812.16812.5 | controllerType=0 controllerNumber=1 scan=16812 | 36.1087 |
| 14451:5 | 5 | K.IIEGEPNLKVPGN[+2059.735]VTAVLGETLKVPC[+57.021]HFPC[+57.021]K.F | HexNAc(4)Hex(5)Fuc(1)NeuAc(1) | C[+57]*2, N[+2060] | 1096.1166 | 5 | 5476.5538 | 5476.5418 | 0 | 2.2 | 457 | Specific | 1046.45 | 944.22 | 944.22 | 20.33 | 91 | >sp\|P01833\|PIGR_HUMAN Polymeric immunoglobulin receptor OS=Homo sapiens OX=9606 GN=PIGR PE=1 SV=4 | 902 | 20201117_08512_1ng_glyco_hcd203040_compound15.17874.17874.5 | controllerType=0 controllerNumber=1 scan=17874 | 38.2781 |
| 14607:4 | 5 | K.IIEGEPNLKVPGN[+2059.735]VTAVLGETLKVPC[+57.021]HFPC[+57.021]K.F | HexNAc(4)Hex(5)Fuc(1)NeuAc(1) | C[+57]*2, N[+2060] | 1369.8918 | 4 | 5476.5456 | 5476.5418 | 0 | 0.7 | 457 | Specific | 1060.08 | 939.21 | 939.21 | 20.31 | 91 | >sp\|P01833\|PIGR_HUMAN Polymeric immunoglobulin receptor OS=Homo sapiens OX=9606 GN=PIGR PE=1 SV=4 | 902 | 20201117_08512_1ng_glyco_hcd203040_compound15.18036.18036.4 | controllerType=0 controllerNumber=1 scan=18036 | 38.5728 |
| 14453:4 | 5 | K.IIEGEPNLKVPGN[+2059.735]VTAVLGETLKVPC[+57.021]HFPC[+57.021]K.F | HexNAc(4)Hex(5)Fuc(1)NeuAc(1) | C[+57]*2, N[+2060] | 1369.8931 | 4 | 5476.5504 | 5476.5418 | 0 | 1.6 | 457 | Specific | 1095.99 | 1005.81 | 1005.81 | 20.96 | 91 | >sp\|P01833\|PIGR_HUMAN Polymeric immunoglobulin receptor OS=Homo sapiens OX=9606 GN=PIGR PE=1 SV=4 | 902 | 20201117_08512_1ng_glyco_hcd203040_compound15.17876.17876.4 | controllerType=0 controllerNumber=1 scan=17876 | 38.2805 |
| 14384:4 | 5 | K.IIEGEPNLKVPGN[+2205.793]VTAVLGETLKVPC[+57.021]HFPC[+57.021]K.F | HexNAc(4)Hex(5)Fuc(2)NeuAc(1) | C[+57]*2, N[+2206] | 1406.4074 | 4 | 5622.6076 | 5622.5997 | 0 | 1.4 | 457 | Specific | 1109.22 | 1020.05 | 1020.05 | 21.96 | 91 | >sp\|P01833\|PIGR_HUMAN Polymeric immunoglobulin receptor OS=Homo sapiens OX=9606 GN=PIGR PE=1 SV=4 | 902 | 20201117_08512_1ng_glyco_hcd203040_compound15.17804.17804.4 | controllerType=0 controllerNumber=1 scan=17804 | 38.1353 |
| 14458:4 | 5 | K.VPGN[+2350.830]VTAVLGETLKVPC[+57.021]HFPC[+57.021]K.F | HexNAc(4)Hex(5)Fuc(1)NeuAc(2) | C[+57]*2, N[+2351] | 1194.2818 | 4 | 4774.1053 | 4774.0877 | 0 | 3.7 | 466 | Specific | 378.32 | 378.32 | 378.32 | 9.19 | 91 | >sp\|P01833\|PIGR_HUMAN Polymeric immunoglobulin receptor OS=Homo sapiens OX=9606 GN=PIGR PE=1 SV=4 | 902 | 20201117_08512_1ng_glyco_hcd203040_compound15.17881.17881.4 | controllerType=0 controllerNumber=1 scan=17881 | 38.2888 |
| 12773:4 | 5 | K.VPGN[+2205.793]VTAVLGETLKVPC[+57.021]HFPC[+57.021]K.F | HexNAc(4)Hex(5)Fuc(2)NeuAc(1) | C[+57]*2, N[+2206] | 1158.0234 | 4 | 4629.0716 | 4629.0502 | 0 | 4.6 | 466 | Specific | 611.33 | 453.06 | 453.06 | 11.84 | 91 | >sp\|P01833\|PIGR_HUMAN Polymeric immunoglobulin receptor OS=Homo sapiens OX=9606 GN=PIGR PE=1 SV=4 | 902 | 20201117_08512_1ng_glyco_hcd203040_compound15.16125.16125.3 | controllerType=0 controllerNumber=1 scan=16125 | 34.6829 |
| 12724:4 | 5 | K.VPGN[+2205.793]VTAVLGETLKVPC[+57.021]HFPC[+57.021]K.F | HexNAc(4)Hex(5)Fuc(2)NeuAc(1) | C[+57]*2, N[+2206] | 1158.022 | 4 | 4629.0662 | 4629.0502 | 0 | 3.5 | 466 | Specific | 630.06 | 557.29 | 557.29 | 12.39 | 91 | >sp\|P01833\|PIGR_HUMAN Polymeric immunoglobulin receptor OS=Homo sapiens OX=9606 GN=PIGR PE=1 SV=4 | 902 | 20201117_08512_1ng_glyco_hcd203040_compound15.16074.16074.4 | controllerType=0 controllerNumber=1 scan=16074 | 34.5524 |
| 12726:4 | 5 | K.VPGN[+2205.793]VTAVLGETLKVPC[+57.021]HFPC[+57.021]K.F | HexNAc(4)Hex(5)Fuc(2)NeuAc(1) | C[+57]*2, N[+2206] | 1158.022 | 4 | 4629.0662 | 4629.0502 | 0 | 3.5 | 466 | Specific | 679.07 | 418.82 | 418.82 | 13.39 | 91 | >sp\|P01833\|PIGR_HUMAN Polymeric immunoglobulin receptor OS=Homo sapiens OX=9606 GN=PIGR PE=1 SV=4 | 902 | 20201117_08512_1ng_glyco_hcd203040_compound15.16076.16076.4 | controllerType=0 controllerNumber=1 scan=16076 | 34.558 |
| 14461:4 | 5 | K.VPGN[+2350.830]VTAVLGETLKVPC[+57.021]HFPC[+57.021]K.F | HexNAc(4)Hex(5)Fuc(1)NeuAc(2) | C[+57]*2, N[+2351] | 1194.2818 | 4 | 4774.1053 | 4774.0877 | 0 | 3.7 | 466 | Specific | 717.35 | 702.1 | 702.1 | 14.39 | 91 | >sp\|P01833\|PIGR_HUMAN Polymeric immunoglobulin receptor OS=Homo sapiens OX=9606 GN=PIGR PE=1 SV=4 | 902 | 20201117_08512_1ng_glyco_hcd203040_compound15.17884.17884.4 | controllerType=0 controllerNumber=1 scan=17884 | 38.2939 |
| 12770:4 | 5 | K.VPGN[+2205.793]VTAVLGETLKVPC[+57.021]HFPC[+57.021]K.F | HexNAc(4)Hex(5)Fuc(2)NeuAc(1) | C[+57]*2, N[+2206] | 1158.0234 | 4 | 4629.0716 | 4629.0502 | 0 | 4.6 | 466 | Specific | 756.61 | 551.57 | 551.57 | 14.84 | 91 | >sp\|P01833\|PIGR_HUMAN Polymeric immunoglobulin receptor OS=Homo sapiens OX=9606 GN=PIGR PE=1 SV=4 | 902 | 20201117_08512_1ng_glyco_hcd203040_compound15.16122.16122.3 | controllerType=0 controllerNumber=1 scan=16122 | 34.6764 |
| 12791:4 | 5 | K.VPGN[+2059.735]VTAVLGETLKVPC[+57.021]HFPC[+57.021]K.F | HexNAc(4)Hex(5)Fuc(1)NeuAc(1) | C[+57]*2, N[+2060] | 1121.5069 | 4 | 4483.0057 | 4482.9923 | 0 | 3 | 466 | Specific | 924.59 | 795.16 | 795.16 | 18.48 | 91 | >sp\|P01833\|PIGR_HUMAN Polymeric immunoglobulin receptor OS=Homo sapiens OX=9606 GN=PIGR PE=1 SV=4 | 902 | 20201117_08512_1ng_glyco_hcd203040_compound15.16144.16144.4 | controllerType=0 controllerNumber=1 scan=16144 | 34.7187 |
| 07502:3 | 5 | K.WN[+2204.772]NTGC[+57.021]QALPSQDEGPSK.A | HexNAc(4)Hex(5)NeuAc(2) | C[+57], N[+2205] | 1398.5591 | 3 | 4193.6627 | 4193.6491 | 0 | 3.2 | 498 | Specific | 270.26 | 208.19 | 208.19 | 5.41 | 91 | >sp\|P01833\|PIGR_HUMAN Polymeric immunoglobulin receptor OS=Homo sapiens OX=9606 GN=PIGR PE=1 SV=4 | 902 | 20201117_08512_1ng_glyco_hcd203040_compound15.10646.10646.3 | controllerType=0 controllerNumber=1 scan=10646 | 24.3676 |
| 07504:3 | 5 | K.WN[+2204.772]NTGC[+57.021]QALPSQDEGPSK.A | HexNAc(4)Hex(5)NeuAc(2) | C[+57], N[+2205] | 1398.5591 | 3 | 4193.6627 | 4193.6491 | 0 | 3.2 | 498 | Specific | 324.02 | 290.93 | 290.93 | 6.63 | 91 | >sp\|P01833\|PIGR_HUMAN Polymeric immunoglobulin receptor OS=Homo sapiens OX=9606 GN=PIGR PE=1 SV=4 | 902 | 20201117_08512_1ng_glyco_hcd203040_compound15.10648.10648.3 | controllerType=0 controllerNumber=1 scan=10648 | 24.3714 |
| 05657:3 | 5 | K.WN[+1548.545]NTGC[+57.021]QALPSQDEGPSK.A | HexNAc(3)Hex(4)NeuAc(1) | C[+57], N[+1549] | 1179.8144 | 3 | 3537.4286 | 3537.4215 | 0 | 2 | 498 | Specific | 422.24 | 367.65 | 367.65 | 7.5 | 91 | >sp\|P01833\|PIGR_HUMAN Polymeric immunoglobulin receptor OS=Homo sapiens OX=9606 GN=PIGR PE=1 SV=4 | 902 | 20201117_08512_1ng_glyco_hcd203040_compound15.8736.8736.3 | controllerType=0 controllerNumber=1 scan=8736 | 21.0951 |
| 05379:3 | 5 | K.WN[+2059.735]NTGC[+57.021]QALPSQDEGPSK.A | HexNAc(4)Hex(5)Fuc(1)NeuAc(1) | C[+57], N[+2060] | 1350.2129 | 3 | 4048.6243 | 4048.6116 | 0 | 3.1 | 498 | Specific | 541.18 | 501.31 | 501.31 | 9.33 | 91 | >sp\|P01833\|PIGR_HUMAN Polymeric immunoglobulin receptor OS=Homo sapiens OX=9606 GN=PIGR PE=1 SV=4 | 902 | 20201117_08512_1ng_glyco_hcd203040_compound15.8449.8449.3 | controllerType=0 controllerNumber=1 scan=8449 | 20.6255 |
| 05381:3 | 5 | K.WN[+2059.735]NTGC[+57.021]QALPSQDEGPSK.A | HexNAc(4)Hex(5)Fuc(1)NeuAc(1) | C[+57], N[+2060] | 1350.2129 | 3 | 4048.6243 | 4048.6116 | 0 | 3.1 | 498 | Specific | 561.16 | 511.93 | 511.93 | 9.56 | 91 | >sp\|P01833\|PIGR_HUMAN Polymeric immunoglobulin receptor OS=Homo sapiens OX=9606 GN=PIGR PE=1 SV=4 | 902 | 20201117_08512_1ng_glyco_hcd203040_compound15.8451.8451.3 | controllerType=0 controllerNumber=1 scan=8451 | 20.629 |
| 05659:3 | 5 | K.WN[+1548.545]NTGC[+57.021]QALPSQDEGPSK.A | HexNAc(3)Hex(4)NeuAc(1) | C[+57], N[+1549] | 1179.8144 | 3 | 3537.4286 | 3537.4215 | 0 | 2 | 498 | Specific | 676.2 | 556.51 | 556.51 | 12.9 | 91 | >sp\|P01833\|PIGR_HUMAN Polymeric immunoglobulin receptor OS=Homo sapiens OX=9606 GN=PIGR PE=1 SV=4 | 902 | 20201117_08512_1ng_glyco_hcd203040_compound15.8738.8738.3 | controllerType=0 controllerNumber=1 scan=8738 | 21.0987 |
| 05442:3 | 5 | K.WN[+1872.651]NTGC[+57.021]QALPSQDEGPSK.A | HexNAc(3)Hex(6)NeuAc(1) | C[+57], N[+1873] | 1287.8498 | 3 | 3861.5348 | 3861.5272 | 0 | 2 | 498 | Specific | 816.32 | 729.64 | 729.64 | 15.9 | 91 | >sp\|P01833\|PIGR_HUMAN Polymeric immunoglobulin receptor OS=Homo sapiens OX=9606 GN=PIGR PE=1 SV=4 | 902 | 20201117_08512_1ng_glyco_hcd203040_compound15.8514.8514.3 | controllerType=0 controllerNumber=1 scan=8514 | 20.7304 |
| 05447:3 | 5 | K.WN[+1913.677]NTGC[+57.021]QALPSQDEGPSK.A | HexNAc(4)Hex(5)NeuAc(1) | C[+57], N[+1914] | 1301.5245 | 3 | 3902.5589 | 3902.5537 | 0 | 1.3 | 498 | Specific | 824.46 | 722.57 | 722.57 | 16 | 91 | >sp\|P01833\|PIGR_HUMAN Polymeric immunoglobulin receptor OS=Homo sapiens OX=9606 GN=PIGR PE=1 SV=4 | 902 | 20201117_08512_1ng_glyco_hcd203040_compound15.8519.8519.3 | controllerType=0 controllerNumber=1 scan=8519 | 20.7404 |
| 05440:3 | 5 | K.WN[+1872.651]NTGC[+57.021]QALPSQDEGPSK.A | HexNAc(3)Hex(6)NeuAc(1) | C[+57], N[+1873] | 1287.8498 | 3 | 3861.5348 | 3861.5272 | 0 | 2 | 498 | Specific | 846.79 | 767.9 | 767.9 | 15.9 | 91 | >sp\|P01833\|PIGR_HUMAN Polymeric immunoglobulin receptor OS=Homo sapiens OX=9606 GN=PIGR PE=1 SV=4 | 902 | 20201117_08512_1ng_glyco_hcd203040_compound15.8512.8512.3 | controllerType=0 controllerNumber=1 scan=8512 | 20.7262 |
| 05531:3 | 5 | K.WN[+1710.598]NTGC[+57.021]QALPSQDEGPSK.A | HexNAc(3)Hex(5)NeuAc(1) | C[+57], N[+1711] | 1233.8339 | 3 | 3699.4872 | 3699.4743 | 0 | 3.5 | 498 | Specific | 857.51 | 792.86 | 792.86 | 16.81 | 91 | >sp\|P01833\|PIGR_HUMAN Polymeric immunoglobulin receptor OS=Homo sapiens OX=9606 GN=PIGR PE=1 SV=4 | 902 | 20201117_08512_1ng_glyco_hcd203040_compound15.8606.8606.3 | controllerType=0 controllerNumber=1 scan=8606 | 20.8787 |
| 05533:3 | 5 | K.WN[+1710.598]NTGC[+57.021]QALPSQDEGPSK.A | HexNAc(3)Hex(5)NeuAc(1) | C[+57], N[+1711] | 1233.8339 | 3 | 3699.4872 | 3699.4743 | 0 | 3.5 | 498 | Specific | 860.51 | 762.56 | 762.56 | 16.81 | 91 | >sp\|P01833\|PIGR_HUMAN Polymeric immunoglobulin receptor OS=Homo sapiens OX=9606 GN=PIGR PE=1 SV=4 | 902 | 20201117_08512_1ng_glyco_hcd203040_compound15.8608.8608.3 | controllerType=0 controllerNumber=1 scan=8608 | 20.8833 |
| 05445:3 | 5 | K.WN[+1913.677]NTGC[+57.021]QALPSQDEGPSK.A | HexNAc(4)Hex(5)NeuAc(1) | C[+57], N[+1914] | 1301.5245 | 3 | 3902.5589 | 3902.5537 | 0 | 1.3 | 498 | Specific | 917.37 | 852.28 | 852.28 | 18 | 91 | >sp\|P01833\|PIGR_HUMAN Polymeric immunoglobulin receptor OS=Homo sapiens OX=9606 GN=PIGR PE=1 SV=4 | 902 | 20201117_08512_1ng_glyco_hcd203040_compound15.8517.8517.3 | controllerType=0 controllerNumber=1 scan=8517 | 20.7353 |
| 16552:3 | 7 | R.LSLHRPALEDLLLGSEAN[+1501.555]LTC[+57.021]TLTGLR.D | HexNAc(5)Hex(3) | C[+57], N[+1502] | 1489.0602 | 3 | 4465.1662 | 4465.1536 | 0 | 2.8 | 127 | Specific | 294.45 | 294.45 | 294.45 | 6.33 | 66 | >sp\|P01876\|IGHA1_HUMAN Immunoglobulin heavy constant alpha 1 OS=Homo sapiens OX=9606 GN=IGHA1 PE=1 SV=2 | 255 | 20201117_08512_1ng_glyco_hcd203040_compound15.20082.20082.6 | controllerType=0 controllerNumber=1 scan=20082 | 42.8415 |
| 16675:3 | 7 | R.LSLHRPALEDLLLGSEAN[+1501.555]LTC[+57.021]TLTGLR.D | HexNAc(5)Hex(3) | C[+57], N[+1502] | 1489.0599 | 3 | 4465.1651 | 4465.1536 | 0 | 2.6 | 127 | Specific | 364.97 | 338.37 | 338.37 | 6.68 | 66 | >sp\|P01876\|IGHA1_HUMAN Immunoglobulin heavy constant alpha 1 OS=Homo sapiens OX=9606 GN=IGHA1 PE=1 SV=2 | 255 | 20201117_08512_1ng_glyco_hcd203040_compound15.20210.20210.3 | controllerType=0 controllerNumber=1 scan=20210 | 43.1315 |
| 16562:5 | 7 | R.LSLHRPALEDLLLGSEAN[+1524.534]LTC[+57.021]TLTGLR.D | HexNAc(2)Hex(6)Fuc(1) | C[+57], N[+1525] | 898.4298 | 5 | 4488.1197 | 4488.1319 | 0 | -2.7 | 127 | Specific | 379.83 | 379.83 | 379.83 | 7.32 | 66 | >sp\|P01876\|IGHA1_HUMAN Immunoglobulin heavy constant alpha 1 OS=Homo sapiens OX=9606 GN=IGHA1 PE=1 SV=2 | 255 | 20201117_08512_1ng_glyco_hcd203040_compound15.20092.20092.2 | controllerType=0 controllerNumber=1 scan=20092 | 42.8582 |
| 17452:3 | 7 | R.LSLHRPALEDLLLGSEAN[+1710.598]LTC[+57.021]TLTGLR.D | HexNAc(3)Hex(5)NeuAc(1) | C[+57], N[+1711] | 1558.7413 | 3 | 4674.2093 | 4674.1959 | 0 | 2.9 | 127 | Specific | 383.59 | 383.29 | 383.29 | 8.47 | 66 | >sp\|P01876\|IGHA1_HUMAN Immunoglobulin heavy constant alpha 1 OS=Homo sapiens OX=9606 GN=IGHA1 PE=1 SV=2 | 255 | 20201117_08512_1ng_glyco_hcd203040_compound15.21021.21021.3 | controllerType=0 controllerNumber=1 scan=21021 | 44.9268 |

Table S4. Cont.

| Query #:z | Protein Rank | Glycopeptide | Glycan | Modification Type(s) | Observed m/z | z | Observed (M+H) | Calc. mass (M+H) | Off-by-x error | Mass error (ppm) | StartPosition | Cleavage | Score | Delta | Delta Mod | \|Log Prob\| | # of unique peptides | Protein Name | Protein DB number | Comment | Scan # | Scan Time |
|---|---|---|---|---|---|---|---|---|---|---|---|---|---|---|---|---|---|---|---|---|---|---|
| 17372:3 | 7 | R.LSLHRPALEDLLLGSEAN[+2116.756]LTC[+57.021]TLTGLR.D | HexNAc(5)Hex(5)NeuAc(1) | C[+57], N[+2117] | 1694.1289 | 3 | 5080.3721 | 5080.3547 | 0 | 3.4 | 127 | Specific | 394.23 | 394.17 | 394.17 | 8.44 | 66 | >sp\|P01876\|IGHA1_HUMAN Immunoglobulin heavy constant alpha 1 OS=Homo sapiens OX=9606 GN=IGHA1 PE=1 SV=2 | 255 | 20201117_08512_1ng_glyco_hcd203040_compound15.20938.20938.3 | controllerType=0 controllerNumber=1 scan=20938 | 44.7396 |
| 16608:4 | 7 | R.LSLHRPALEDLLLGSEAN[+1460.529]LTC[+57.021]TLTGLR.D | HexNAc(4)Hex(4) | C[+57], N[+1461] | 1106.7913 | 4 | 4424.1434 | 4424.1271 | 0 | 3.7 | 127 | Specific | 412.98 | 412.98 | 412.98 | 9.1 | 66 | >sp\|P01876\|IGHA1_HUMAN Immunoglobulin heavy constant alpha 1 OS=Homo sapiens OX=9606 GN=IGHA1 PE=1 SV=2 | 255 | 20201117_08512_1ng_glyco_hcd203040_compound15.20140.20140.4 | controllerType=0 controllerNumber=1 scan=20140 | 42.9855 |
| 16454:4 | 7 | R.LSLHRPALEDLLLGSEAN[+1825.661]LTC[+57.021]TLTGLR.D | HexNAc(5)Hex(5) | C[+57], N[+1826] | 1198.075 | 4 | 4789.2781 | 4789.2592 | 0 | 3.9 | 127 | Specific | 456.87 | 456.87 | 456.87 | 9.14 | 66 | >sp\|P01876\|IGHA1_HUMAN Immunoglobulin heavy constant alpha 1 OS=Homo sapiens OX=9606 GN=IGHA1 PE=1 SV=2 | 255 | 20201117_08512_1ng_glyco_hcd203040_compound15.19979.19979.4 | controllerType=0 controllerNumber=1 scan=19979 | 42.561 |
| 17454:3 | 7 | R.LSLHRPALEDLLLGSEAN[+1710.598]LTC[+57.021]TLTGLR.D | HexNAc(3)Hex(5)NeuAc(1) | C[+57], N[+1711] | 1558.7413 | 3 | 4674.2093 | 4674.1959 | 0 | 2.9 | 127 | Specific | 486.46 | 402.05 | 402.05 | 8.96 | 66 | >sp\|P01876\|IGHA1_HUMAN Immunoglobulin heavy constant alpha 1 OS=Homo sapiens OX=9606 GN=IGHA1 PE=1 SV=2 | 255 | 20201117_08512_1ng_glyco_hcd203040_compound15.21023.21023.3 | controllerType=0 controllerNumber=1 scan=21023 | 44.933 |
| 16672:3 | 7 | R.LSLHRPALEDLLLGSEAN[+1501.555]LTC[+57.021]TLTGLR.D | HexNAc(5)Hex(3) | C[+57], N[+1502] | 1489.0599 | 3 | 4465.1651 | 4465.1536 | 0 | 2.6 | 127 | Specific | 525.21 | 499.9 | 499.9 | 9.05 | 66 | >sp\|P01876\|IGHA1_HUMAN Immunoglobulin heavy constant alpha 1 OS=Homo sapiens OX=9606 GN=IGHA1 PE=1 SV=2 | 255 | 20201117_08512_1ng_glyco_hcd203040_compound15.20207.20207.3 | controllerType=0 controllerNumber=1 scan=20207 | 43.1246 |
| 16565:5 | 7 | R.LSLHRPALEDLLLGSEAN[+1524.534]LTC[+57.021]TLTGLR.D | HexNAc(2)Hex(6)Fuc(1) | C[+57], N[+1525] | 898.4298 | 5 | 4488.1197 | 4488.1319 | 0 | -2.7 | 127 | Specific | 551.41 | 551.41 | 551.41 | 8.03 | 66 | >sp\|P01876\|IGHA1_HUMAN Immunoglobulin heavy constant alpha 1 OS=Homo sapiens OX=9606 GN=IGHA1 PE=1 SV=2 | 255 | 20201117_08512_1ng_glyco_hcd203040_compound15.20095.20095.2 | controllerType=0 controllerNumber=1 scan=20095 | 42.8654 |
| 16543:5 | 7 | R.LSLHRPALEDLLLGSEAN[+1501.555]LTC[+57.021]TLTGLR.D | HexNAc(5)Hex(3) | C[+57], N[+1502] | 893.8391 | 5 | 4465.1663 | 4465.1536 | 0 | 2.8 | 127 | Specific | 552.41 | 552.41 | 552.41 | 9.48 | 66 | >sp\|P01876\|IGHA1_HUMAN Immunoglobulin heavy constant alpha 1 OS=Homo sapiens OX=9606 GN=IGHA1 PE=1 SV=2 | 255 | 20201117_08512_1ng_glyco_hcd203040_compound15.20073.20073.5 | controllerType=0 controllerNumber=1 scan=20073 | 42.8279 |
| 17399:3 | 7 | R.LSLHRPALEDLLLGSEAN[+1872.651]LTC[+57.021]TLTGLR.D | HexNAc(3)Hex(6)NeuAc(1) | C[+57], N[+1873] | 1612.7595 | 3 | 4836.2638 | 4836.2487 | 0 | 3.1 | 127 | Specific | 564.42 | 564.42 | 564.42 | 9.32 | 66 | >sp\|P01876\|IGHA1_HUMAN Immunoglobulin heavy constant alpha 1 OS=Homo sapiens OX=9606 GN=IGHA1 PE=1 SV=2 | 255 | 20201117_08512_1ng_glyco_hcd203040_compound15.20966.20966.3 | controllerType=0 controllerNumber=1 scan=20966 | 44.8101 |
| 16528:3 | 7 | R.LSLHRPALEDLLLGSEAN[+1501.555]LTC[+57.021]TLTGLR.D | HexNAc(5)Hex(3) | C[+57], N[+1502] | 1489.0593 | 3 | 4465.1632 | 4465.1536 | 0 | 2.2 | 127 | Specific | 589.05 | 550.32 | 550.32 | 10.66 | 66 | >sp\|P01876\|IGHA1_HUMAN Immunoglobulin heavy constant alpha 1 OS=Homo sapiens OX=9606 GN=IGHA1 PE=1 SV=2 | 255 | 20201117_08512_1ng_glyco_hcd203040_compound15.20057.20057.2 | controllerType=0 controllerNumber=1 scan=20057 | 42.7908 |
| 16525:3 | 7 | R.LSLHRPALEDLLLGSEAN[+1501.555]LTC[+57.021]TLTGLR.D | HexNAc(5)Hex(3) | C[+57], N[+1502] | 1489.0593 | 3 | 4465.1632 | 4465.1536 | 0 | 2.2 | 127 | Specific | 589.94 | 589.37 | 589.37 | 10.66 | 66 | >sp\|P01876\|IGHA1_HUMAN Immunoglobulin heavy constant alpha 1 OS=Homo sapiens OX=9606 GN=IGHA1 PE=1 SV=2 | 255 | 20201117_08512_1ng_glyco_hcd203040_compound15.20054.20054.2 | controllerType=0 controllerNumber=1 scan=20054 | 42.7816 |
| 16446:5 | 7 | R.LSLHRPALEDLLLGSEAN[+1864.634]LTC[+57.021]TLTGLR.D | HexNAc(2)Hex(9) | C[+57], N[+1865] | 966.4535 | 5 | 4828.2385 | 4828.2324 | 0 | 1.3 | 127 | Specific | 595.41 | 595.41 | 595.41 | 11.06 | 66 | >sp\|P01876\|IGHA1_HUMAN Immunoglobulin heavy constant alpha 1 OS=Homo sapiens OX=9606 GN=IGHA1 PE=1 SV=2 | 255 | 20201117_08512_1ng_glyco_hcd203040_compound15.19970.19970.5 | controllerType=0 controllerNumber=1 scan=19970 | 42.5358 |
| 17373:3 | 7 | R.LSLHRPALEDLLLGSEAN[+1913.677]LTC[+57.021]TLTGLR.D | HexNAc(4)Hex(5)NeuAc(1) | C[+57], N[+1914] | 1626.4354 | 3 | 4877.2915 | 4877.2753 | 0 | 3.3 | 127 | Specific | 599.45 | 599.45 | 599.45 | 10.57 | 66 | >sp\|P01876\|IGHA1_HUMAN Immunoglobulin heavy constant alpha 1 OS=Homo sapiens OX=9606 GN=IGHA1 PE=1 SV=2 | 255 | 20201117_08512_1ng_glyco_hcd203040_compound15.20939.20939.3 | controllerType=0 controllerNumber=1 scan=20939 | 44.7407 |
| 17442:4 | 7 | R.LSLHRPALEDLLLGSEAN[+1710.598]LTC[+57.021]TLTGLR.D | HexNAc(3)Hex(5)NeuAc(1) | C[+57], N[+1711] | 1169.3074 | 4 | 4674.2078 | 4674.1959 | 0 | 2.5 | 127 | Specific | 613.83 | 613.83 | 613.83 | 12.13 | 66 | >sp\|P01876\|IGHA1_HUMAN Immunoglobulin heavy constant alpha 1 OS=Homo sapiens OX=9606 GN=IGHA1 PE=1 SV=2 | 255 | 20201117_08512_1ng_glyco_hcd203040_compound15.21011.21011.4 | controllerType=0 controllerNumber=1 scan=21011 | 44.908 |
| 16461:3 | 7 | R.LSLHRPALEDLLLGSEAN[+1663.608]LTC[+57.021]TLTGLR.D | HexNAc(5)Hex(4) | C[+57], N[+1664] | 1543.0803 | 3 | 4627.2262 | 4627.2064 | 0 | 4.3 | 127 | Specific | 620.14 | 580.11 | 580.11 | 11.57 | 66 | >sp\|P01876\|IGHA1_HUMAN Immunoglobulin heavy constant alpha 1 OS=Homo sapiens OX=9606 GN=IGHA1 PE=1 SV=2 | 255 | 20201117_08512_1ng_glyco_hcd203040_compound15.19986.19986.3 | controllerType=0 controllerNumber=1 scan=19986 | 42.5785 |
| 16555:3 | 7 | R.LSLHRPALEDLLLGSEAN[+1501.555]LTC[+57.021]TLTGLR.D | HexNAc(5)Hex(3) | C[+57], N[+1502] | 1489.0602 | 3 | 4465.1662 | 4465.1536 | 0 | 2.8 | 127 | Specific | 620.45 | 501.82 | 501.82 | 11.66 | 66 | >sp\|P01876\|IGHA1_HUMAN Immunoglobulin heavy constant alpha 1 OS=Homo sapiens OX=9606 GN=IGHA1 PE=1 SV=2 | 255 | 20201117_08512_1ng_glyco_hcd203040_compound15.20085.20085.6 | controllerType=0 controllerNumber=1 scan=20085 | 42.8484 |
| 17375:3 | 7 | R.LSLHRPALEDLLLGSEAN[+2116.756]LTC[+57.021]TLTGLR.D | HexNAc(5)Hex(5)NeuAc(1) | C[+57], N[+2117] | 1694.1289 | 3 | 5080.3721 | 5080.3547 | 0 | 3.4 | 127 | Specific | 622.23 | 536.75 | 536.75 | 11.57 | 66 | >sp\|P01876\|IGHA1_HUMAN Immunoglobulin heavy constant alpha 1 OS=Homo sapiens OX=9606 GN=IGHA1 PE=1 SV=2 | 255 | 20201117_08512_1ng_glyco_hcd203040_compound15.20941.20941.3 | controllerType=0 controllerNumber=1 scan=20941 | 44.7436 |
| 16383:3 | 7 | R.LSLHRPALEDLLLGSEAN[+1864.634]LTC[+57.021]TLTGLR.D | HexNAc(2)Hex(9) | C[+57], N[+1865] | 1610.0848 | 3 | 4828.2398 | 4828.2324 | 0 | 1.5 | 127 | Specific | 633.02 | 601.58 | 601.58 | 11.77 | 66 | >sp\|P01876\|IGHA1_HUMAN Immunoglobulin heavy constant alpha 1 OS=Homo sapiens OX=9606 GN=IGHA1 PE=1 SV=2 | 255 | 20201117_08512_1ng_glyco_hcd203040_compound15.19905.19905.3 | controllerType=0 controllerNumber=1 scan=19905 | 42.3904 |
| 16541:5 | 7 | R.LSLHRPALEDLLLGSEAN[+1501.555]LTC[+57.021]TLTGLR.D | HexNAc(5)Hex(3) | C[+57], N[+1502] | 893.8391 | 5 | 4465.1663 | 4465.1536 | 0 | 2.8 | 127 | Specific | 634.81 | 634.81 | 634.81 | 11.96 | 66 | >sp\|P01876\|IGHA1_HUMAN Immunoglobulin heavy constant alpha 1 OS=Homo sapiens OX=9606 GN=IGHA1 PE=1 SV=2 | 255 | 20201117_08512_1ng_glyco_hcd203040_compound15.20071.20071.5 | controllerType=0 controllerNumber=1 scan=20071 | 42.8228 |
| 17402:3 | 7 | R.LSLHRPALEDLLLGSEAN[+1872.651]LTC[+57.021]TLTGLR.D | HexNAc(3)Hex(6)NeuAc(1) | C[+57], N[+1873] | 1612.7595 | 3 | 4836.2638 | 4836.2487 | 0 | 3.1 | 127 | Specific | 635.32 | 570.11 | 570.11 | 11.57 | 66 | >sp\|P01876\|IGHA1_HUMAN Immunoglobulin heavy constant alpha 1 OS=Homo sapiens OX=9606 GN=IGHA1 PE=1 SV=2 | 255 | 20201117_08512_1ng_glyco_hcd203040_compound15.20969.20969.3 | controllerType=0 controllerNumber=1 scan=20969 | 44.8149 |
| 16494:3 | 7 | R.LSLHRPALEDLLLGSEAN[+1501.555]LTC[+57.021]TLTGLR.D | HexNAc(5)Hex(3) | C[+57], N[+1502] | 1489.0693 | 3 | 4465.1933 | 4465.1536 | 0 | 8.9 | 127 | Specific | 640.29 | 623.81 | 623.81 | 10.07 | 66 | >sp\|P01876\|IGHA1_HUMAN Immunoglobulin heavy constant alpha 1 OS=Homo sapiens OX=9606 GN=IGHA1 PE=1 SV=2 | 255 | 20201117_08512_1ng_glyco_hcd203040_compound15.20021.20021.3 | controllerType=0 controllerNumber=1 scan=20021 | 42.6812 |
| 16491:3 | 7 | R.LSLHRPALEDLLLGSEAN[+1501.555]LTC[+57.021]TLTGLR.D | HexNAc(5)Hex(3) | C[+57], N[+1502] | 1489.0693 | 3 | 4465.1933 | 4465.1536 | 0 | 8.9 | 127 | Specific | 647.36 | 647.36 | 647.36 | 10.07 | 66 | >sp\|P01876\|IGHA1_HUMAN Immunoglobulin heavy constant alpha 1 OS=Homo sapiens OX=9606 GN=IGHA1 PE=1 SV=2 | 255 | 20201117_08512_1ng_glyco_hcd203040_compound15.20018.20018.3 | controllerType=0 controllerNumber=1 scan=20018 | 42.674 |
| 17367:4 | 7 | R.LSLHRPALEDLLLGSEAN[+2116.756]LTC[+57.021]TLTGLR.D | HexNAc(5)Hex(5)NeuAc(1) | C[+57], N[+2117] | 1270.8473 | 4 | 5080.3673 | 5080.3547 | 0 | 2.5 | 127 | Specific | 648.65 | 648.65 | 648.65 | 12.13 | 66 | >sp\|P01876\|IGHA1_HUMAN Immunoglobulin heavy constant alpha 1 OS=Homo sapiens OX=9606 GN=IGHA1 PE=1 SV=2 | 255 | 20201117_08512_1ng_glyco_hcd203040_compound15.20933.20933.4 | controllerType=0 controllerNumber=1 scan=20933 | 44.7266 |
| 16464:3 | 7 | R.LSLHRPALEDLLLGSEAN[+1663.608]LTC[+57.021]TLTGLR.D | HexNAc(5)Hex(4) | C[+57], N[+1664] | 1543.0803 | 3 | 4627.2262 | 4627.2064 | 0 | 4.3 | 127 | Specific | 654.09 | 597.16 | 597.16 | 12.57 | 66 | >sp\|P01876\|IGHA1_HUMAN Immunoglobulin heavy constant alpha 1 OS=Homo sapiens OX=9606 GN=IGHA1 PE=1 SV=2 | 255 | 20201117_08512_1ng_glyco_hcd203040_compound15.19989.19989.3 | controllerType=0 controllerNumber=1 scan=19989 | 42.5853 |
| 17376:3 | 7 | R.LSLHRPALEDLLLGSEAN[+1913.677]LTC[+57.021]TLTGLR.D | HexNAc(4)Hex(5)NeuAc(1) | C[+57], N[+1914] | 1626.4354 | 3 | 4877.2915 | 4877.2753 | 0 | 3.3 | 127 | Specific | 657.15 | 589.12 | 589.12 | 12.57 | 66 | >sp\|P01876\|IGHA1_HUMAN Immunoglobulin heavy constant alpha 1 OS=Homo sapiens OX=9606 GN=IGHA1 PE=1 SV=2 | 255 | 20201117_08512_1ng_glyco_hcd203040_compound15.20942.20942.3 | controllerType=0 controllerNumber=1 scan=20942 | 44.7472 |
| 16442:3 | 7 | R.LSLHRPALEDLLLGSEAN[+1540.529]LTC[+57.021]TLTGLR.D | HexNAc(2)Hex(7) | C[+57], N[+1541] | 1502.0499 | 3 | 4504.135 | 4504.1268 | 0 | 1.8 | 127 | Specific | 661.63 | 607.38 | 607.38 | 12.77 | 66 | >sp\|P01876\|IGHA1_HUMAN Immunoglobulin heavy constant alpha 1 OS=Homo sapiens OX=9606 GN=IGHA1 PE=1 SV=2 | 255 | 20201117_08512_1ng_glyco_hcd203040_compound15.19966.19966.3 | controllerType=0 controllerNumber=1 scan=19966 | 42.5307 |
| 16542:4 | 7 | R.LSLHRPALEDLLLGSEAN[+1298.476]LTC[+57.021]TLTGLR.D | HexNAc(4)Hex(3) | C[+57], N[+1298] | 1066.2762 | 4 | 4262.0828 | 4262.0742 | 0 | 2 | 127 | Specific | 662.18 | 661.11 | 661.11 | 13.13 | 66 | >sp\|P01876\|IGHA1_HUMAN Immunoglobulin heavy constant alpha 1 OS=Homo sapiens OX=9606 GN=IGHA1 PE=1 SV=2 | 255 | 20201117_08512_1ng_glyco_hcd203040_compound15.20072.20072.4 | controllerType=0 controllerNumber=1 scan=20072 | 42.8243 |
| 16488:3 | 7 | R.LSLHRPALEDLLLGSEAN[+1216.423]LTC[+57.021]TLTGLR.D | HexNAc(2)Hex(5) | C[+57], N[+1216] | 1394.0219 | 3 | 4180.0512 | 4180.0211 | 0 | 7.2 | 127 | Specific | 665.92 | 644.84 | 644.84 | 11.59 | 66 | >sp\|P01876\|IGHA1_HUMAN Immunoglobulin heavy constant alpha 1 OS=Homo sapiens OX=9606 GN=IGHA1 PE=1 SV=2 | 255 | 20201117_08512_1ng_glyco_hcd203040_compound15.20015.20015.3 | controllerType=0 controllerNumber=1 scan=20015 | 42.662 |
| 16627:4 | 7 | R.LSLHRPALEDLLLGSEAN[+1378.476]LTC[+57.021]TLTGLR.D | HexNAc(2)Hex(6) | C[+57], N[+1378] | 1086.2748 | 4 | 4342.0775 | 4342.074 | 0 | 0.8 | 127 | Specific | 672.64 | 672.64 | 672.64 | 13.23 | 66 | >sp\|P01876\|IGHA1_HUMAN Immunoglobulin heavy constant alpha 1 OS=Homo sapiens OX=9606 GN=IGHA1 PE=1 SV=2 | 255 | 20201117_08512_1ng_glyco_hcd203040_compound15.20160.20160.2 | controllerType=0 controllerNumber=1 scan=20160 | 43.0248 |
| 16490:3 | 7 | R.LSLHRPALEDLLLGSEAN[+1216.423]LTC[+57.021]TLTGLR.D | HexNAc(2)Hex(5) | C[+57], N[+1216] | 1394.0219 | 3 | 4180.0512 | 4180.0211 | 0 | 7.2 | 127 | Specific | 689.57 | 637.49 | 637.49 | 11.59 | 66 | >sp\|P01876\|IGHA1_HUMAN Immunoglobulin heavy constant alpha 1 OS=Homo sapiens OX=9606 GN=IGHA1 PE=1 SV=2 | 255 | 20201117_08512_1ng_glyco_hcd203040_compound15.20017.20017.3 | controllerType=0 controllerNumber=1 scan=20017 | 42.6721 |
| 16381:3 | 7 | R.LSLHRPALEDLLLGSEAN[+1864.634]LTC[+57.021]TLTGLR.D | HexNAc(2)Hex(9) | C[+57], N[+1865] | 1610.0848 | 3 | 4828.2398 | 4828.2324 | 0 | 1.5 | 127 | Specific | 699.09 | 697.74 | 697.74 | 12.77 | 66 | >sp\|P01876\|IGHA1_HUMAN Immunoglobulin heavy constant alpha 1 OS=Homo sapiens OX=9606 GN=IGHA1 PE=1 SV=2 | 255 | 20201117_08512_1ng_glyco_hcd203040_compound15.19903.19903.3 | controllerType=0 controllerNumber=1 scan=19903 | 42.382 |
| 16409:3 | 7 | R.LSLHRPALEDLLLGSEAN[+1702.581]LTC[+57.021]TLTGLR.D | HexNAc(2)Hex(8) | C[+57], N[+1703] | 1556.0687 | 3 | 4666.1914 | 4666.1796 | 0 | 2.5 | 127 | Specific | 701.86 | 669.69 | 669.69 | 13.66 | 66 | >sp\|P01876\|IGHA1_HUMAN Immunoglobulin heavy constant alpha 1 OS=Homo sapiens OX=9606 GN=IGHA1 PE=1 SV=2 | 255 | 20201117_08512_1ng_glyco_hcd203040_compound15.19932.19932.3 | controllerType=0 controllerNumber=1 scan=19932 | 42.4519 |
| 16479:3 | 7 | R.LSLHRPALEDLLLGSEAN[+1378.476]LTC[+57.021]TLTGLR.D | HexNAc(2)Hex(6) | C[+57], N[+1378] | 1448.0345 | 3 | 4342.0889 | 4342.074 | 0 | 3.4 | 127 | Specific | 714.45 | 679.59 | 679.59 | 13.57 | 66 | >sp\|P01876\|IGHA1_HUMAN Immunoglobulin heavy constant alpha 1 OS=Homo sapiens OX=9606 GN=IGHA1 PE=1 SV=2 | 255 | 20201117_08512_1ng_glyco_hcd203040_compound15.20005.20005.3 | controllerType=0 controllerNumber=1 scan=20005 | 42.6366 |
| 17351:4 | 7 | R.LSLHRPALEDLLLGSEAN[+1913.677]LTC[+57.021]TLTGLR.D | HexNAc(4)Hex(5)NeuAc(1) | C[+57], N[+1914] | 1220.0793 | 4 | 4877.2952 | 4877.2753 | 0 | 4.1 | 127 | Specific | 725.39 | 725.39 | 725.39 | 14.04 | 66 | >sp\|P01876\|IGHA1_HUMAN Immunoglobulin heavy constant alpha 1 OS=Homo sapiens OX=9606 GN=IGHA1 PE=1 SV=2 | 255 | 20201117_08512_1ng_glyco_hcd203040_compound15.20916.20916.4 | controllerType=0 controllerNumber=1 scan=20916 | 44.6858 |
| 16451:4 | 7 | R.LSLHRPALEDLLLGSEAN[+1663.608]LTC[+57.021]TLTGLR.D | HexNAc(5)Hex(4) | C[+57], N[+1664] | 1157.5621 | 4 | 4627.2264 | 4627.2064 | 0 | 4.3 | 127 | Specific | 733.81 | 733.81 | 733.81 | 13.49 | 66 | >sp\|P01876\|IGHA1_HUMAN Immunoglobulin heavy constant alpha 1 OS=Homo sapiens OX=9606 GN=IGHA1 PE=1 SV=2 | 255 | 20201117_08512_1ng_glyco_hcd203040_compound15.19976.19976.4 | controllerType=0 controllerNumber=1 scan=19976 | 42.5498 |
| 17383:4 | 7 | R.LSLHRPALEDLLLGSEAN[+1872.651]LTC[+57.021]TLTGLR.D | HexNAc(3)Hex(6)NeuAc(1) | C[+57], N[+1873] | 1209.8207 | 4 | 4836.2611 | 4836.2487 | 0 | 2.5 | 127 | Specific | 735.39 | 683.83 | 683.83 | 14.13 | 66 | >sp\|P01876\|IGHA1_HUMAN Immunoglobulin heavy constant alpha 1 OS=Homo sapiens OX=9606 GN=IGHA1 PE=1 SV=2 | 255 | 20201117_08512_1ng_glyco_hcd203040_compound15.20949.20949.4 | controllerType=0 controllerNumber=1 scan=20949 | 44.7633 |
| 16507:4 | 7 | R.LSLHRPALEDLLLGSEAN[+1501.555]LTC[+57.021]TLTGLR.D | HexNAc(5)Hex(3) | C[+57], N[+1502] | 1117.0451 | 4 | 4465.1585 | 4465.1536 | 0 | 1.1 | 127 | Specific | 743.23 | 743.23 | 743.23 | 14.23 | 66 | >sp\|P01876\|IGHA1_HUMAN Immunoglobulin heavy constant alpha 1 OS=Homo sapiens OX=9606 GN=IGHA1 PE=1 SV=2 | 255 | 20201117_08512_1ng_glyco_hcd203040_compound15.20035.20035.4 | controllerType=0 controllerNumber=1 scan=20035 | 42.7172 |
| 16457:4 | 7 | R.LSLHRPALEDLLLGSEAN[+1825.661]LTC[+57.021]TLTGLR.D | HexNAc(5)Hex(5) | C[+57], N[+1826] | 1198.075 | 4 | 4789.2781 | 4789.2592 | 0 | 3.9 | 127 | Specific | 762.96 | 597.05 | 597.05 | 15.04 | 66 | >sp\|P01876\|IGHA1_HUMAN Immunoglobulin heavy constant alpha 1 OS=Homo sapiens OX=9606 GN=IGHA1 PE=1 SV=2 | 255 | 20201117_08512_1ng_glyco_hcd203040_compound15.19982.19982.4 | controllerType=0 controllerNumber=1 scan=19982 | 42.5653 |
| 16406:3 | 7 | R.LSLHRPALEDLLLGSEAN[+1702.581]LTC[+57.021]TLTGLR.D | HexNAc(2)Hex(8) | C[+57], N[+1703] | 1556.0687 | 3 | 4666.1914 | 4666.1796 | 0 | 2.5 | 127 | Specific | 765.31 | 754.56 | 754.56 | 14.66 | 66 | >sp\|P01876\|IGHA1_HUMAN Immunoglobulin heavy constant alpha 1 OS=Homo sapiens OX=9606 GN=IGHA1 PE=1 SV=2 | 255 | 20201117_08512_1ng_glyco_hcd203040_compound15.19929.19929.3 | controllerType=0 controllerNumber=1 scan=19929 | 42.4419 |
| 16449:5 | 7 | R.LSLHRPALEDLLLGSEAN[+1864.634]LTC[+57.021]TLTGLR.D | HexNAc(2)Hex(9) | C[+57], N[+1865] | 966.4535 | 5 | 4828.2385 | 4828.2324 | 0 | 1.3 | 127 | Specific | 769.43 | 622.33 | 622.33 | 15.06 | 66 | >sp\|P01876\|IGHA1_HUMAN Immunoglobulin heavy constant alpha 1 OS=Homo sapiens OX=9606 GN=IGHA1 PE=1 SV=2 | 255 | 20201117_08512_1ng_glyco_hcd203040_compound15.19973.19973.5 | controllerType=0 controllerNumber=1 scan=19973 | 42.5439 |
| 16520:4 | 7 | R.LSLHRPALEDLLLGSEAN[+1298.476]LTC[+57.021]TLTGLR.D | HexNAc(4)Hex(3) | C[+57], N[+1298] | 1066.276 | 4 | 4262.0823 | 4262.0742 | 0 | 1.9 | 127 | Specific | 770.37 | 770.37 | 770.37 | 15.23 | 66 | >sp\|P01876\|IGHA1_HUMAN Immunoglobulin heavy constant alpha 1 OS=Homo sapiens OX=9606 GN=IGHA1 PE=1 SV=2 | 255 | 20201117_08512_1ng_glyco_hcd203040_compound15.20049.20049.4 | controllerType=0 controllerNumber=1 scan=20049 | 42.77 |
| 16439:3 | 7 | R.LSLHRPALEDLLLGSEAN[+1540.529]LTC[+57.021]TLTGLR.D | HexNAc(2)Hex(7) | C[+57], N[+1541] | 1502.0499 | 3 | 4504.135 | 4504.1268 | 0 | 1.8 | 127 | Specific | 770.97 | 699.88 | 699.88 | 14.77 | 66 | >sp\|P01876\|IGHA1_HUMAN Immunoglobulin heavy constant alpha 1 OS=Homo sapiens OX=9606 GN=IGHA1 PE=1 SV=2 | 255 | 20201117_08512_1ng_glyco_hcd203040_compound15.19963.19963.3 | controllerType=0 controllerNumber=1 scan=19963 | 42.5135 |
| 17369:4 | 7 | R.LSLHRPALEDLLLGSEAN[+2116.756]LTC[+57.021]TLTGLR.D | HexNAc(5)Hex(5)NeuAc(1) | C[+57], N[+2117] | 1270.8473 | 4 | 5080.3673 | 5080.3547 | 0 | 2.5 | 127 | Specific | 775.41 | 718.5 | 718.5 | 15.13 | 66 | >sp\|P01876\|IGHA1_HUMAN Immunoglobulin heavy constant alpha 1 OS=Homo sapiens OX=9606 GN=IGHA1 PE=1 SV=2 | 255 | 20201117_08512_1ng_glyco_hcd203040_compound15.20935.20935.4 | controllerType=0 controllerNumber=1 scan=20935 | 44.7318 |
| 16476:3 | 7 | R.LSLHRPALEDLLLGSEAN[+1378.476]LTC[+57.021]TLTGLR.D | HexNAc(2)Hex(6) | C[+57], N[+1378] | 1448.0345 | 3 | 4342.0889 | 4342.074 | 0 | 3.4 | 127 | Specific | 781.51 | 706.6 | 706.6 | 14.57 | 66 | >sp\|P01876\|IGHA1_HUMAN Immunoglobulin heavy constant alpha 1 OS=Homo sapiens OX=9606 GN=IGHA1 PE=1 SV=2 | 255 | 20201117_08512_1ng_glyco_hcd203040_compound15.20002.20002.3 | controllerType=0 controllerNumber=1 scan=20002 | 42.6294 |
| 16453:4 | 7 | R.LSLHRPALEDLLLGSEAN[+1663.608]LTC[+57.021]TLTGLR.D | HexNAc(5)Hex(4) | C[+57], N[+1664] | 1157.5621 | 4 | 4627.2264 | 4627.2064 | 0 | 4.3 | 127 | Specific | 782.32 | 782.32 | 782.32 | 14.49 | 66 | >sp\|P01876\|IGHA1_HUMAN Immunoglobulin heavy constant alpha 1 OS=Homo sapiens OX=9606 GN=IGHA1 PE=1 SV=2 | 255 | 20201117_08512_1ng_glyco_hcd203040_compound15.19978.19978.4 | controllerType=0 controllerNumber=1 scan=19978 | 42.5599 |
| 16473:4 | 7 | R.LSLHRPALEDLLLGSEAN[+1419.502]LTC[+57.021]TLTGLR.D | HexNAc(3)Hex(5) | C[+57], N[+1420] | 1096.5336 | 4 | 4383.1126 | 4383.1005 | 0 | 2.8 | 127 | Specific | 797.43 | 751.64 | 751.64 | 15.13 | 66 | >sp\|P01876\|IGHA1_HUMAN Immunoglobulin heavy constant alpha 1 OS=Homo sapiens OX=9606 GN=IGHA1 PE=1 SV=2 | 255 | 20201117_08512_1ng_glyco_hcd203040_compound15.19999.19999.4 | controllerType=0 controllerNumber=1 scan=19999 | 42.6113 |
| 16544:4 | 7 | R.LSLHRPALEDLLLGSEAN[+1298.476]LTC[+57.021]TLTGLR.D | HexNAc(4)Hex(3) | C[+57], N[+1298] | 1066.2762 | 4 | 4262.0828 | 4262.0742 | 0 | 2 | 127 | Specific | 797.53 | 653.63 | 653.63 | 15.13 | 66 | >sp\|P01876\|IGHA1_HUMAN Immunoglobulin heavy constant alpha 1 OS=Homo sapiens OX=9606 GN=IGHA1 PE=1 SV=2 | 255 | 20201117_08512_1ng_glyco_hcd203040_compound15.20074.20074.4 | controllerType=0 controllerNumber=1 scan=20074 | 42.8306 |
| 16510:4 | 7 | R.LSLHRPALEDLLLGSEAN[+1501.555]LTC[+57.021]TLTGLR.D | HexNAc(5)Hex(3) | C[+57], N[+1502] | 1117.0451 | 4 | 4465.1585 | 4465.1536 | 0 | 1.1 | 127 | Specific | 800.06 | 739.86 | 739.86 | 16.23 | 66 | >sp\|P01876\|IGHA1_HUMAN Immunoglobulin heavy constant alpha 1 OS=Homo sapiens OX=9606 GN=IGHA1 PE=1 SV=2 | 255 | 20201117_08512_1ng_glyco_hcd203040_compound15.20038.20038.4 | controllerType=0 controllerNumber=1 scan=20038 | 42.7313 |
| 17353:4 | 7 | R.LSLHRPALEDLLLGSEAN[+1913.677]LTC[+57.021]TLTGLR.D | HexNAc(4)Hex(5)NeuAc(1) | C[+57], N[+1914] | 1220.0793 | 4 | 4877.2952 | 4877.2753 | 0 | 4.1 | 127 | Specific | 804.24 | 762.01 | 762.01 | 16.04 | 66 | >sp\|P01876\|IGHA1_HUMAN Immunoglobulin heavy constant alpha 1 OS=Homo sapiens OX=9606 GN=IGHA1 PE=1 SV=2 | 255 | 20201117_08512_1ng_glyco_hcd203040_compound15.20918.20918.4 | controllerType=0 controllerNumber=1 scan=20918 | 44.6933 |
| 17386:4 | 7 | R.LSLHRPALEDLLLGSEAN[+1872.651]LTC[+57.021]TLTGLR.D | HexNAc(3)Hex(6)NeuAc(1) | C[+57], N[+1873] | 1209.8207 | 4 | 4836.2611 | 4836.2487 | 0 | 2.5 | 127 | Specific | 823.1 | 823.1 | 823.1 | 16.13 | 66 | >sp\|P01876\|IGHA1_HUMAN Immunoglobulin heavy constant alpha 1 OS=Homo sapiens OX=9606 GN=IGHA1 PE=1 SV=2 | 255 | 20201117_08512_1ng_glyco_hcd203040_compound15.20952.20952.4 | controllerType=0 controllerNumber=1 scan=20952 | 44.7778 |
| 16570:4 | 7 | R.LSLHRPALEDLLLGSEAN[+1054.370]LTC[+57.021]TLTGLR.D | HexNAc(2)Hex(4) | C[+57], N[+1054] | 1005.2503 | 4 | 4017.9796 | 4017.9683 | 0 | 2.8 | 127 | Specific | 825.06 | 790.14 | 790.14 | 16.13 | 66 | >sp\|P01876\|IGHA1_HUMAN Immunoglobulin heavy constant alpha 1 OS=Homo sapiens OX=9606 GN=IGHA1 PE=1 SV=2 | 255 | 20201117_08512_1ng_glyco_hcd203040_compound15.20101.20101.4 | controllerType=0 controllerNumber=1 scan=20101 | 42.8803 |
| 16522:4 | 7 | R.LSLHRPALEDLLLGSEAN[+1298.476]LTC[+57.021]TLTGLR.D | HexNAc(4)Hex(3) | C[+57], N[+1298] | 1066.276 | 4 | 4262.0823 | 4262.0742 | 0 | 1.9 | 127 | Specific | 856.06 | 796.21 | 796.21 | 17.23 | 66 | >sp\|P01876\|IGHA1_HUMAN Immunoglobulin heavy constant alpha 1 OS=Homo sapiens OX=9606 GN=IGHA1 PE=1 SV=2 | 255 | 20201117_08512_1ng_glyco_hcd203040_compound15.20051.20051.4 | controllerType=0 controllerNumber=1 scan=20051 | 42.7781 |
| 17445:4 | 7 | R.LSLHRPALEDLLLGSEAN[+1710.598]LTC[+57.021]TLTGLR.D | HexNAc(3)Hex(5)NeuAc(1) | C[+57], N[+1711] | 1169.3074 | 4 | 4674.2078 | 4674.1959 | 0 | 2.5 | 127 | Specific | 868.23 | 839.76 | 839.76 | 17.13 | 66 | >sp\|P01876\|IGHA1_HUMAN Immunoglobulin heavy constant alpha 1 OS=Homo sapiens OX=9606 GN=IGHA1 PE=1 SV=2 | 255 | 20201117_08512_1ng_glyco_hcd203040_compound15.21014.21014.4 | controllerType=0 controllerNumber=1 scan=21014 | 44.9124 |
| 16380:4 | 7 | R.LSLHRPALEDLLLGSEAN[+1864.634]LTC[+57.021]TLTGLR.D | HexNAc(2)Hex(9) | C[+57], N[+1865] | 1207.8151 | 4 | 4828.2386 | 4828.2324 | 0 | 1.3 | 127 | Specific | 870.25 | 870.25 | 870.25 | 17.23 | 66 | >sp\|P01876\|IGHA1_HUMAN Immunoglobulin heavy constant alpha 1 OS=Homo sapiens OX=9606 GN=IGHA1 PE=1 SV=2 | 255 | 20201117_08512_1ng_glyco_hcd203040_compound15.19902.19902.4 | controllerType=0 controllerNumber=1 scan=19902 | 42.3806 |
| 16611:4 | 7 | R.LSLHRPALEDLLLGSEAN[+1460.529]LTC[+57.021]TLTGLR.D | HexNAc(4)Hex(4) | C[+57], N[+1461] | 1106.7913 | 4 | 4424.1434 | 4424.1271 | 0 | 3.7 | 127 | Specific | 870.94 | 759.21 | 759.21 | 17.04 | 66 | >sp\|P01876\|IGHA1_HUMAN Immunoglobulin heavy constant alpha 1 OS=Homo sapiens OX=9606 GN=IGHA1 PE=1 SV=2 | 255 | 20201117_08512_1ng_glyco_hcd203040_compound15.20143.20143.4 | controllerType=0 controllerNumber=1 scan=20143 | 42.9917 |
| 16402:4 | 7 | R.LSLHRPALEDLLLGSEAN[+1702.581]LTC[+57.021]TLTGLR.D | HexNAc(2)Hex(8) | C[+57], N[+1703] | 1167.3029 | 4 | 4666.1898 | 4666.1796 | 0 | 2.2 | 127 | Specific | 872.69 | 872.69 | 872.69 | 17.13 | 66 | >sp\|P01876\|IGHA1_HUMAN Immunoglobulin heavy constant alpha 1 OS=Homo sapiens OX=9606 GN=IGHA1 PE=1 SV=2 | 255 | 20201117_08512_1ng_glyco_hcd203040_compound15.19925.19925.4 | controllerType=0 controllerNumber=1 scan=19925 | 42.4337 |
| 16472:4 | 7 | R.LSLHRPALEDLLLGSEAN[+1378.476]LTC[+57.021]TLTGLR.D | HexNAc(2)Hex(6) | C[+57], N[+1378] | 1086.2775 | 4 | 4342.0882 | 4342.074 | 0 | 3.3 | 127 | Specific | 883.28 | 841.62 | 841.62 | 17.04 | 66 | >sp\|P01876\|IGHA1_HUMAN Immunoglobulin heavy constant alpha 1 OS=Homo sapiens OX=9606 GN=IGHA1 PE=1 SV=2 | 255 | 20201117_08512_1ng_glyco_hcd203040_compound15.19998.19998.4 | controllerType=0 controllerNumber=1 scan=19998 | 42.6099 |
| 16487:4 | 7 | R.LSLHRPALEDLLLGSEAN[+1216.423]LTC[+57.021]TLTGLR.D | HexNAc(2)Hex(5) | C[+57], N[+1216] | 1045.769 | 4 | 4180.054 | 4180.0211 | 0 | 7.9 | 127 | Specific | 913.76 | 913.76 | 913.76 | 16.61 | 66 | >sp\|P01876\|IGHA1_HUMAN Immunoglobulin heavy constant alpha 1 OS=Homo sapiens OX=9606 GN=IGHA1 PE=1 SV=2 | 255 | 20201117_08512_1ng_glyco_hcd203040_compound15.20014.20014.4 | controllerType=0 controllerNumber=1 scan=20014 | 42.6607 |
| 16537:4 | 7 | R.LSLHRPALEDLLLGSEAN[+1054.370]LTC[+57.021]TLTGLR.D | HexNAc(2)Hex(4) | C[+57], N[+1054] | 1005.2496 | 4 | 4017.9764 | 4017.9683 | 0 | 2 | 127 | Specific | 917.64 | 895.83 | 895.83 | 18.13 | 66 | >sp\|P01876\|IGHA1_HUMAN Immunoglobulin heavy constant alpha 1 OS=Homo sapiens OX=9606 GN=IGHA1 PE=1 SV=2 | 255 | 20201117_08512_1ng_glyco_hcd203040_compound15.20066.20066.2 | controllerType=0 controllerNumber=1 scan=20066 | 42.8052 |
| 16428:4 | 7 | R.LSLHRPALEDLLLGSEAN[+1540.529]LTC[+57.021]TLTGLR.D | HexNAc(2)Hex(7) | C[+57], N[+1541] | 1126.789 | 4 | 4504.1341 | 4504.1268 | 0 | 1.6 | 127 | Specific | 918.72 | 842.34 | 842.34 | 18.23 | 66 | >sp\|P01876\|IGHA1_HUMAN Immunoglobulin heavy constant alpha 1 OS=Homo sapiens OX=9606 GN=IGHA1 PE=1 SV=2 | 255 | 20201117_08512_1ng_glyco_hcd203040_compound15.19952.19952.4 | controllerType=0 controllerNumber=1 scan=19952 | 42.4911 |
| 16475:4 | 7 | R.LSLHRPALEDLLLGSEAN[+1419.502]LTC[+57.021]TLTGLR.D | HexNAc(3)Hex(5) | C[+57], N[+1420] | 1096.5336 | 4 | 4383.1126 | 4383.1005 | 0 | 2.8 | 127 | Specific | 923.03 | 923.03 | 923.03 | 18.13 | 66 | >sp\|P01876\|IGHA1_HUMAN Immunoglobulin heavy constant alpha 1 OS=Homo sapiens OX=9606 GN=IGHA1 PE=1 SV=2 | 255 | 20201117_08512_1ng_glyco_hcd203040_compound15.20001.20001.4 | controllerType=0 controllerNumber=1 scan=20001 | 42.6283 |
| 16489:4 | 7 | R.LSLHRPALEDLLLGSEAN[+1216.423]LTC[+57.021]TLTGLR.D | HexNAc(2)Hex(5) | C[+57], N[+1216] | 1045.769 | 4 | 4180.054 | 4180.0211 | 0 | 7.9 | 127 | Specific | 930.52 | 930.52 | 930.52 | 16.61 | 66 | >sp\|P01876\|IGHA1_HUMAN Immunoglobulin heavy constant alpha 1 OS=Homo sapiens OX=9606 GN=IGHA1 PE=1 SV=2 | 255 | 20201117_08512_1ng_glyco_hcd203040_compound15.20016.20016.4 | controllerType=0 controllerNumber=1 scan=20016 | 42.6693 |
| 16404:4 | 7 | R.LSLHRPALEDLLLGSEAN[+1702.581]LTC[+57.021]TLTGLR.D | HexNAc(2)Hex(8) | C[+57], N[+1703] | 1167.3029 | 4 | 4666.1898 | 4666.1796 | 0 | 2.2 | 127 | Specific | 975.61 | 936.51 | 936.51 | 19.13 | 66 | >sp\|P01876\|IGHA1_HUMAN Immunoglobulin heavy constant alpha 1 OS=Homo sapiens OX=9606 GN=IGHA1 PE=1 SV=2 | 255 | 20201117_08512_1ng_glyco_hcd203040_compound15.19927.19927.4 | controllerType=0 controllerNumber=1 scan=19927 | 42.4399 |
| 16430:4 | 7 | R.LSLHRPALEDLLLGSEAN[+1540.529]LTC[+57.021]TLTGLR.D | HexNAc(2)Hex(7) | C[+57], N[+1541] | 1126.789 | 4 | 4504.1341 | 4504.1268 | 0 | 1.6 | 127 | Specific | 976.17 | 976.17 | 976.17 | 19.23 | 66 | >sp\|P01876\|IGHA1_HUMAN Immunoglobulin heavy constant alpha 1 OS=Homo sapiens OX=9606 GN=IGHA1 PE=1 SV=2 | 255 | 20201117_08512_1ng_glyco_hcd203040_compound15.19954.19954.4 | controllerType=0 controllerNumber=1 scan=19954 | 42.4985 |
| 16382:4 | 7 | R.LSLHRPALEDLLLGSEAN[+1864.634]LTC[+57.021]TLTGLR.D | HexNAc(2)Hex(9) | C[+57], N[+1865] | 1207.8151 | 4 | 4828.2386 | 4828.2324 | 0 | 1.3 | 127 | Specific | 989.41 | 989.41 | 989.41 | 19.23 | 66 | >sp\|P01876\|IGHA1_HUMAN Immunoglobulin heavy constant alpha 1 OS=Homo sapiens OX=9606 GN=IGHA1 PE=1 SV=2 | 255 | 20201117_08512_1ng_glyco_hcd203040_compound15.19904.19904.4 | controllerType=0 controllerNumber=1 scan=19904 | 42.387 |
| 16572:4 | 7 | R.LSLHRPALEDLLLGSEAN[+1054.370]LTC[+57.021]TLTGLR.D | HexNAc(2)Hex(4) | C[+57], N[+1054] | 1005.2503 | 4 | 4017.9796 | 4017.9683 | 0 | 2.8 | 127 | Specific | 994.54 | 994.54 | 994.54 | 19.13 | 66 | >sp\|P01876\|IGHA1_HUMAN Immunoglobulin heavy constant alpha 1 OS=Homo sapiens OX=9606 GN=IGHA1 PE=1 SV=2 | 255 | 20201117_08512_1ng_glyco_hcd203040_compound15.20103.20103.4 | controllerType=0 controllerNumber=1 scan=20103 | 42.8863 |
| 16630:4 | 7 | R.LSLHRPALEDLLLGSEAN[+1378.476]LTC[+57.021]TLTGLR.D | HexNAc(2)Hex(6) | C[+57], N[+1378] | 1086.2748 | 4 | 4342.0775 | 4342.074 | 0 | 0.8 | 127 | Specific | 1014.21 | 953.97 | 953.97 | 20.23 | 66 | >sp\|P01876\|IGHA1_HUMAN Immunoglobulin heavy constant alpha 1 OS=Homo sapiens OX=9606 GN=IGHA1 PE=1 SV=2 | 255 | 20201117_08512_1ng_glyco_hcd203040_compound15.20163.20163.2 | controllerType=0 controllerNumber=1 scan=20163 | 43.0337 |
| 16474:4 | 7 | R.LSLHRPALEDLLLGSEAN[+1378.476]LTC[+57.021]TLTGLR.D | HexNAc(2)Hex(6) | C[+57], N[+1378] | 1086.2775 | 4 | 4342.0882 | 4342.074 | 0 | 3.3 | 127 | Specific | 1020.57 | 1020.57 | 1020.57 | 20.04 | 66 | >sp\|P01876\|IGHA1_HUMAN Immunoglobulin heavy constant alpha 1 OS=Homo sapiens OX=9606 GN=IGHA1 PE=1 SV=2 | 255 | 20201117_08512_1ng_glyco_hcd203040_compound15.20000.20000.4 | controllerType=0 controllerNumber=1 scan=20000 | 42.6244 |
| 16540:4 | 7 | R.LSLHRPALEDLLLGSEAN[+1054.370]LTC[+57.021]TLTGLR.D | HexNAc(2)Hex(4) | C[+57], N[+1054] | 1005.2496 | 4 | 4017.9764 | 4017.9683 | 0 | 2 | 127 | Specific | 1021.99 | 1021.99 | 1021.99 | 20.13 | 66 | >sp\|P01876\|IGHA1_HUMAN Immunoglobulin heavy constant alpha 1 OS=Homo sapiens OX=9606 GN=IGHA1 PE=1 SV=2 | 255 | 20201117_08512_1ng_glyco_hcd203040_compound15.20069.20069.2 | controllerType=0 controllerNumber=1 scan=20069 | 42.8184 |
| 10713:3 | 7 | R.LAGKPTHVN[+2133.772]VSVVMAEVDGTC[+57.021].Y | HexNAc(5)Hex(6)Fuc(1) | C[+57], N[+2134] | 1439.9623 | 3 | 4317.8725 | 4317.8505 | 0 | 5.1 | 332 | CRagged | 222.96 | 144.76 | 1.82 | 2.06 | 66 | >sp\|P01876\|IGHA1_HUMAN Immunoglobulin heavy constant alpha 1 OS=Homo sapiens OX=9606 GN=IGHA1 PE=1 SV=2 | 255 | 20201117_08512_1ng_glyco_hcd203040_compound15.13981.13981.3 | controllerType=0 controllerNumber=1 scan=13981 | 30.5586 |
| 12300:3 | 7 | R.LAGKPTHVN[+2263.835]VSVVMAEVDGTC[+57.021].Y | HexNAc(5)Hex(5)Fuc(3) | C[+57], N[+2264] | 1483.3101 | 3 | 4447.9158 | 4447.9135 | 0 | 0.5 | 332 | CRagged | 296.43 | 269.4 | 269.4 | 6.27 | 66 | >sp\|P01876\|IGHA1_HUMAN Immunoglobulin heavy constant alpha 1 OS=Homo sapiens OX=9606 GN=IGHA1 PE=1 SV=2 | 255 | 20201117_08512_1ng_glyco_hcd203040_compound15.15633.15633.3 | controllerType=0 controllerNumber=1 scan=15633 | 33.6754 |
| 10711:3 | 7 | R.LAGKPTHVN[+2133.772]VSVVMAEVDGTC[+57.021].Y | HexNAc(5)Hex(6)Fuc(1) | C[+57], N[+2134] | 1439.9623 | 3 | 4317.8725 | 4317.8505 | 0 | 5.1 | 332 | CRagged | 321.67 | 207.13 | 94.78 | 5.79 | 66 | >sp\|P01876\|IGHA1_HUMAN Immunoglobulin heavy constant alpha 1 OS=Homo sapiens OX=9606 GN=IGHA1 PE=1 SV=2 | 255 | 20201117_08512_1ng_glyco_hcd203040_compound15.13979.13979.3 | controllerType=0 controllerNumber=1 scan=13979 | 30.5543 |
| 10823:3 | 7 | R.LAGKPTHVN[+1419.502]VSVVMAEVDGTC[+57.021]Y.- | HexNAc(3)Hex(5) | C[+57], N[+1420] | 1256.2219 | 3 | 3766.651 | 3766.6443 | 0 | 1.8 | 332 | CRagged | 262.09 | 200.15 | 200.15 | 5.54 | 66 | >sp\|P01876\|IGHA1_HUMAN Immunoglobulin heavy constant alpha 1 OS=Homo sapiens OX=9606 GN=IGHA1 PE=1 SV=2 | 255 | 20201117_08512_1ng_glyco_hcd203040_compound15.14095.14095.3 | controllerType=0 controllerNumber=1 scan=14095 | 30.7963 |
| 10851:4 | 7 | R.LAGKPTHVN[+1647.613]VSVVMAEVDGTC[+57.021]Y.- | HexNAc(5)Hex(3)Fuc(1) | C[+57], N[+1648] | 999.4461 | 4 | 3994.7625 | 3994.7553 | 0 | 1.8 | 332 | CRagged | 265.74 | 236.08 | 236.08 | 6.58 | 66 | >sp\|P01876\|IGHA1_HUMAN Immunoglobulin heavy constant alpha 1 OS=Homo sapiens OX=9606 GN=IGHA1 PE=1 SV=2 | 255 | 20201117_08512_1ng_glyco_hcd203040_compound15.14125.14125.4 | controllerType=0 controllerNumber=1 scan=14125 | 30.8636 |
| 14161:3 | 7 | R.LAGKPTHVN[+2204.772]VSVVMAEVDGTC[+57.021]Y.- | HexNAc(4)Hex(5)NeuAc(2) | C[+57], N[+2205] | 1517.9811 | 3 | 4551.9289 | 4551.9145 | 0 | 3.1 | 332 | CRagged | 280.16 | 251.64 | 251.64 | 6.79 | 66 | >sp\|P01876\|IGHA1_HUMAN Immunoglobulin heavy constant alpha 1 OS=Homo sapiens OX=9606 GN=IGHA1 PE=1 SV=2 | 255 | 20201117_08512_1ng_glyco_hcd203040_compound15.17571.17571.3 | controllerType=0 controllerNumber=1 scan=17571 | 37.6186 |
| 14055:4 | 7 | R.LAGKPTHVN[+2350.830]VSVVMAEVDGTC[+57.021]Y.- | HexNAc(4)Hex(5)Fuc(1)NeuAc(2) | C[+57], N[+2351] | 1175.2525 | 4 | 4697.9881 | 4697.9724 | 0 | 3.3 | 332 | CRagged | 296.55 | 251.27 | 251.27 | 5.49 | 66 | >sp\|P01876\|IGHA1_HUMAN Immunoglobulin heavy constant alpha 1 OS=Homo sapiens OX=9606 GN=IGHA1 PE=1 SV=2 | 255 | 20201117_08512_1ng_glyco_hcd203040_compound15.17460.17460.4 | controllerType=0 controllerNumber=1 scan=17460 | 37.373 |
| 14159:3 | 7 | R.LAGKPTHVN[+2204.772]VSVVMAEVDGTC[+57.021]Y.- | HexNAc(4)Hex(5)NeuAc(2) | C[+57], N[+2205] | 1517.9811 | 3 | 4551.9289 | 4551.9145 | 0 | 3.1 | 332 | CRagged | 305.34 | 260.52 | 260.52 | 7.11 | 66 | >sp\|P01876\|IGHA1_HUMAN Immunoglobulin heavy constant alpha 1 OS=Homo sapiens OX=9606 GN=IGHA1 PE=1 SV=2 | 255 | 20201117_08512_1ng_glyco_hcd203040_compound15.17569.17569.3 | controllerType=0 controllerNumber=1 scan=17569 | 37.6056 |
| 10747:4 | 7 | R.LAGKPTHVN[+1663.608]VSVVMAEVDGTC[+57.021]Y.- | HexNAc(5)Hex(4) | C[+57], N[+1664] | 1003.4409 | 4 | 4010.7418 | 4010.7502 | 0 | -2.1 | 332 | CRagged | 307.98 | 272.49 | 254.49 | 3.98 | 66 | >sp\|P01876\|IGHA1_HUMAN Immunoglobulin heavy constant alpha 1 OS=Homo sapiens OX=9606 GN=IGHA1 PE=1 SV=2 | 255 | 20201117_08512_1ng_glyco_hcd203040_compound15.14016.14016.2 | controllerType=0 controllerNumber=1 scan=14016 | 30.635 |
| 14053:4 | 7 | R.LAGKPTHVN[+2350.830]VSVVMAEVDGTC[+57.021]Y.- | HexNAc(4)Hex(5)Fuc(1)NeuAc(2) | C[+57], N[+2351] | 1175.2525 | 4 | 4697.9881 | 4697.9724 | 0 | 3.3 | 332 | CRagged | 327.65 | 327.65 | 327.65 | 7.36 | 66 | >sp\|P01876\|IGHA1_HUMAN Immunoglobulin heavy constant alpha 1 OS=Homo sapiens OX=9606 GN=IGHA1 PE=1 SV=2 | 255 | 20201117_08512_1ng_glyco_hcd203040_compound15.17458.17458.4 | controllerType=0 controllerNumber=1 scan=17458 | 37.3691 |
| 09743:3 | 7 | R.LAGKPTHVN[+1647.613]VSVVMAEVDGTC[+57.021]Y.- | HexNAc(5)Hex(3)Fuc(1) | C[+57], N[+1648] | 1332.2587 | 3 | 3994.7616 | 3994.7553 | 0 | 1.6 | 332 | CRagged | 394.07 | 394.07 | 394.07 | 9.73 | 66 | >sp\|P01876\|IGHA1_HUMAN Immunoglobulin heavy constant alpha 1 OS=Homo sapiens OX=9606 GN=IGHA1 PE=1 SV=2 | 255 | 20201117_08512_1ng_glyco_hcd203040_compound15.12974.12974.3 | controllerType=0 controllerNumber=1 scan=12974 | 28.6374 |
| 12378:4 | 7 | R.LAGKPTHVN[+2262.814]VSVVMAEVDGTC[+57.021]Y.- | HexNAc(5)Hex(5)Fuc(1)NeuAc(1) | C[+57], N[+2263] | 1153.2467 | 4 | 4609.9652 | 4609.9564 | 0 | 1.9 | 332 | CRagged | 411.24 | 411.24 | 411.24 | 9.56 | 66 | >sp\|P01876\|IGHA1_HUMAN Immunoglobulin heavy constant alpha 1 OS=Homo sapiens OX=9606 GN=IGHA1 PE=1 SV=2 | 255 | 20201117_08512_1ng_glyco_hcd203040_compound15.15714.15714.4 | controllerType=0 controllerNumber=1 scan=15714 | 33.8343 |
| 10797:2 | 7 | R.LAGKPTHVN[+1216.423]VSVVMAEVDGTC[+57.021]Y.- | HexNAc(2)Hex(5) | C[+57], N[+1216] | 1782.2871 | 2 | 3563.5669 | 3563.5649 | 0 | 0.5 | 332 | CRagged | 417.39 | 417.39 | 246.96 | 8.8 | 66 | >sp\|P01876\|IGHA1_HUMAN Immunoglobulin heavy constant alpha 1 OS=Homo sapiens OX=9606 GN=IGHA1 PE=1 SV=2 | 255 | 20201117_08512_1ng_glyco_hcd203040_compound15.14068.14068.2 | controllerType=0 controllerNumber=1 scan=14068 | 30.7441 |
| 12377:3 | 7 | R.LAGKPTHVN[+2262.814]VSVVMAEVDGTC[+57.021]Y.- | HexNAc(5)Hex(5)Fuc(1)NeuAc(1) | C[+57], N[+2263] | 1537.3272 | 3 | 4609.9671 | 4609.9564 | 0 | 2.3 | 332 | CRagged | 449.22 | 439.2 | 439.2 | 9.86 | 66 | >sp\|P01876\|IGHA1_HUMAN Immunoglobulin heavy constant alpha 1 OS=Homo sapiens OX=9606 GN=IGHA1 PE=1 SV=2 | 255 | 20201117_08512_1ng_glyco_hcd203040_compound15.15713.15713.3 | controllerType=0 controllerNumber=1 scan=15713 | 33.8329 |
| 14150:3 | 7 | R.LAGKPTHVN[+2204.772]VSVVMAEVDGTC[+57.021]Y.- | HexNAc(4)Hex(5)NeuAc(2) | C[+57], N[+2205] | 1517.9778 | 3 | 4551.919 | 4551.9145 | 0 | 1 | 332 | CRagged | 460.15 | 439.18 | 425.51 | 10.05 | 66 | >sp\|P01876\|IGHA1_HUMAN Immunoglobulin heavy constant alpha 1 OS=Homo sapiens OX=9606 GN=IGHA1 PE=1 SV=2 | 255 | 20201117_08512_1ng_glyco_hcd203040_compound15.17559.17559.3 | controllerType=0 controllerNumber=1 scan=17559 | 37.5847 |
| 09746:3 | 7 | R.LAGKPTHVN[+1647.613]VSVVMAEVDGTC[+57.021]Y.- | HexNAc(5)Hex(3)Fuc(1) | C[+57], N[+1648] | 1332.2587 | 3 | 3994.7616 | 3994.7553 | 0 | 1.6 | 332 | CRagged | 463.3 | 343.14 | 343.14 | 8.25 | 66 | >sp\|P01876\|IGHA1_HUMAN Immunoglobulin heavy constant alpha 1 OS=Homo sapiens OX=9606 GN=IGHA1 PE=1 SV=2 | 255 | 20201117_08512_1ng_glyco_hcd203040_compound15.12977.12977.3 | controllerType=0 controllerNumber=1 scan=12977 | 28.644 |
| 14030:3 | 7 | R.LAGKPTHVN[+2350.830]VSVVMAEVDGTC[+57.021]Y.- | HexNAc(4)Hex(5)Fuc(1)NeuAc(2) | C[+57], N[+2351] | 1566.6684 | 3 | 4697.9906 | 4697.9724 | 0 | 3.9 | 332 | CRagged | 493.98 | 352.35 | 352.35 | 8.17 | 66 | >sp\|P01876\|IGHA1_HUMAN Immunoglobulin heavy constant alpha 1 OS=Homo sapiens OX=9606 GN=IGHA1 PE=1 SV=2 | 255 | 20201117_08512_1ng_glyco_hcd203040_compound15.17434.17434.3 | controllerType=0 controllerNumber=1 scan=17434 | 37.3192 |
| 09661:3 | 7 | R.LAGKPTHVN[+1809.666]VSVVMAEVDGTC[+57.021]Y.- | HexNAc(5)Hex(4)Fuc(1) | C[+57], N[+1810] | 1386.2784 | 3 | 4156.8206 | 4156.8081 | 0 | 3 | 332 | CRagged | 509.76 | 509.76 | 357.64 | 10.15 | 66 | >sp\|P01876\|IGHA1_HUMAN Immunoglobulin heavy constant alpha 1 OS=Homo sapiens OX=9606 GN=IGHA1 PE=1 SV=2 | 255 | 20201117_08512_1ng_glyco_hcd203040_compound15.12889.12889.3 | controllerType=0 controllerNumber=1 scan=12889 | 28.4753 |
| 10798:2 | 7 | R.LAGKPTHVN[+1216.423]VSVVMAEVDGTC[+57.021]Y.- | HexNAc(2)Hex(5) | C[+57], N[+1216] | 1782.2871 | 2 | 3563.5669 | 3563.5649 | 0 | 0.5 | 332 | CRagged | 511.27 | 428.61 | 202.72 | 7.93 | 66 | >sp\|P01876\|IGHA1_HUMAN Immunoglobulin heavy constant alpha 1 OS=Homo sapiens OX=9606 GN=IGHA1 PE=1 SV=2 | 255 | 20201117_08512_1ng_glyco_hcd203040_compound15.14069.14069.2 | controllerType=0 controllerNumber=1 scan=14069 | 30.7473 |
| 12308:3 | 7 | R.LAGKPTHVN[+2116.756]VSVVMAEVDGTC[+57.021]Y.- | HexNAc(5)Hex(5)NeuAc(1) | C[+57], N[+2117] | 1488.642 | 3 | 4463.9115 | 4463.8985 | 0 | 2.9 | 332 | CRagged | 529.95 | 450.35 | 450.35 | 10.28 | 66 | >sp\|P01876\|IGHA1_HUMAN Immunoglobulin heavy constant alpha 1 OS=Homo sapiens OX=9606 GN=IGHA1 PE=1 SV=2 | 255 | 20201117_08512_1ng_glyco_hcd203040_compound15.15641.15641.3 | controllerType=0 controllerNumber=1 scan=15641 | 33.6903 |
| 10853:4 | 7 | R.LAGKPTHVN[+1647.613]VSVVMAEVDGTC[+57.021]Y.- | HexNAc(5)Hex(3)Fuc(1) | C[+57], N[+1648] | 999.4461 | 4 | 3994.7625 | 3994.7553 | 0 | 1.8 | 332 | CRagged | 530.79 | 442.26 | 442.26 | 10.21 | 66 | >sp\|P01876\|IGHA1_HUMAN Immunoglobulin heavy constant alpha 1 OS=Homo sapiens OX=9606 GN=IGHA1 PE=1 SV=2 | 255 | 20201117_08512_1ng_glyco_hcd203040_compound15.14127.14127.4 | controllerType=0 controllerNumber=1 scan=14127 | 30.8667 |
| 12443:3 | 7 | R.LAGKPTHVN[+1710.598]VSVVMAEVDGTC[+57.021]Y.- | HexNAc(3)Hex(5)NeuAc(1) | C[+57], N[+1711] | 1353.2546 | 3 | 4057.7492 | 4057.7397 | 0 | 2.3 | 332 | CRagged | 536.19 | 536.19 | 358.75 | 10.28 | 66 | >sp\|P01876\|IGHA1_HUMAN Immunoglobulin heavy constant alpha 1 OS=Homo sapiens OX=9606 GN=IGHA1 PE=1 SV=2 | 255 | 20201117_08512_1ng_glyco_hcd203040_compound15.15781.15781.3 | controllerType=0 controllerNumber=1 scan=15781 | 33.9602 |
| 09663:3 | 7 | R.LAGKPTHVN[+1809.666]VSVVMAEVDGTC[+57.021]Y.- | HexNAc(5)Hex(4)Fuc(1) | C[+57], N[+1810] | 1386.2784 | 3 | 4156.8206 | 4156.8081 | 0 | 3 | 332 | CRagged | 551.54 | 258.32 | 258.32 | 7.96 | 66 | >sp\|P01876\|IGHA1_HUMAN Immunoglobulin heavy constant alpha 1 OS=Homo sapiens OX=9606 GN=IGHA1 PE=1 SV=2 | 255 | 20201117_08512_1ng_glyco_hcd203040_compound15.12891.12891.3 | controllerType=0 controllerNumber=1 scan=12891 | 28.481 |
| 14148:3 | 7 | R.LAGKPTHVN[+2204.772]VSVVMAEVDGTC[+57.021]Y.- | HexNAc(4)Hex(5)NeuAc(2) | C[+57], N[+2205] | 1517.9778 | 3 | 4551.919 | 4551.9145 | 0 | 1 | 332 | CRagged | 556.31 | 521.6 | 478.32 | 10.61 | 66 | >sp\|P01876\|IGHA1_HUMAN Immunoglobulin heavy constant alpha 1 OS=Homo sapiens OX=9606 GN=IGHA1 PE=1 SV=2 | 255 | 20201117_08512_1ng_glyco_hcd203040_compound15.17557.17557.3 | controllerType=0 controllerNumber=1 scan=17557 | 37.5752 |
| 10752:3 | 7 | R.LAGKPTHVN[+1663.608]VSVVMAEVDGTC[+57.021]Y.- | HexNAc(5)Hex(4) | C[+57], N[+1664] | 1337.5898 | 3 | 4010.7548 | 4010.7502 | 0 | 1.1 | 332 | CRagged | 557.17 | 557.17 | 357.27 | 10.61 | 66 | >sp\|P01876\|IGHA1_HUMAN Immunoglobulin heavy constant alpha 1 OS=Homo sapiens OX=9606 GN=IGHA1 PE=1 SV=2 | 255 | 20201117_08512_1ng_glyco_hcd203040_compound15.14022.14022.3 | controllerType=0 controllerNumber=1 scan=14022 | 30.6473 |
| 14032:3 | 7 | R.LAGKPTHVN[+2350.830]VSVVMAEVDGTC[+57.021]Y.- | HexNAc(4)Hex(5)Fuc(1)NeuAc(2) | C[+57], N[+2351] | 1566.6684 | 3 | 4697.9906 | 4697.9724 | 0 | 3.9 | 332 | CRagged | 559.29 | 559.29 | 559.29 | 10.42 | 66 | >sp\|P01876\|IGHA1_HUMAN Immunoglobulin heavy constant alpha 1 OS=Homo sapiens OX=9606 GN=IGHA1 PE=1 SV=2 | 255 | 20201117_08512_1ng_glyco_hcd203040_compound15.17436.17436.3 | controllerType=0 controllerNumber=1 scan=17436 | 37.3232 |
| 14049:3 | 7 | R.LAGKPTHVN[+2553.910]VSVVMAEVDGTC[+57.021]Y.- | HexNAc(5)Hex(5)Fuc(1)NeuAc(2) | C[+57], N[+2554] | 1634.3597 | 3 | 4901.0645 | 4901.0518 | 0 | 2.6 | 332 | CRagged | 603.1 | 415.15 | 415.15 | 12.76 | 66 | >sp\|P01876\|IGHA1_HUMAN Immunoglobulin heavy constant alpha 1 OS=Homo sapiens OX=9606 GN=IGHA1 PE=1 SV=2 | 255 | 20201117_08512_1ng_glyco_hcd203040_compound15.17454.17454.3 | controllerType=0 controllerNumber=1 scan=17454 | 37.362 |
| 10947:3 | 7 | R.LAGKPTHVN[+1216.423]VSVVMAEVDGTC[+57.021]Y.- | HexNAc(2)Hex(5) | C[+57], N[+1216] | 1188.5267 | 3 | 3563.5655 | 3563.5649 | 0 | 0.1 | 332 | CRagged | 624.94 | 624.94 | 326.64 | 12.21 | 66 | >sp\|P01876\|IGHA1_HUMAN Immunoglobulin heavy constant alpha 1 OS=Homo sapiens OX=9606 GN=IGHA1 PE=1 SV=2 | 255 | 20201117_08512_1ng_glyco_hcd203040_compound15.14224.14224.3 | controllerType=0 controllerNumber=1 scan=14224 | 31.0656 |
| 14041:4 | 7 | R.LAGKPTHVN[+2553.910]VSVVMAEVDGTC[+57.021]Y.- | HexNAc(5)Hex(5)Fuc(1)NeuAc(2) | C[+57], N[+2554] | 1226.0221 | 4 | 4901.0667 | 4901.0518 | 0 | 3 | 332 | CRagged | 627.72 | 467.92 | 467.92 | 12.58 | 66 | >sp\|P01876\|IGHA1_HUMAN Immunoglobulin heavy constant alpha 1 OS=Homo sapiens OX=9606 GN=IGHA1 PE=1 SV=2 | 255 | 20201117_08512_1ng_glyco_hcd203040_compound15.17446.17446.4 | controllerType=0 controllerNumber=1 scan=17446 | 37.3442 |
| 14051:3 | 7 | R.LAGKPTHVN[+2553.910]VSVVMAEVDGTC[+57.021]Y.- | HexNAc(5)Hex(5)Fuc(1)NeuAc(2) | C[+57], N[+2554] | 1634.3597 | 3 | 4901.0645 | 4901.0518 | 0 | 2.6 | 332 | CRagged | 632.33 | 492.29 | 492.29 | 12.76 | 66 | >sp\|P01876\|IGHA1_HUMAN Immunoglobulin heavy constant alpha 1 OS=Homo sapiens OX=9606 GN=IGHA1 PE=1 SV=2 | 255 | 20201117_08512_1ng_glyco_hcd203040_compound15.17456.17456.3 | controllerType=0 controllerNumber=1 scan=17456 | 37.3661 |
| 10767:4 | 7 | R.LAGKPTHVN[+1809.666]VSVVMAEVDGTC[+57.021]Y.- | HexNAc(5)Hex(4)Fuc(1) | C[+57], N[+1810] | 1039.9589 | 4 | 4156.8138 | 4156.8081 | 0 | 1.4 | 332 | CRagged | 642.79 | 554.95 | 497.63 | 12.69 | 66 | >sp\|P01876\|IGHA1_HUMAN Immunoglobulin heavy constant alpha 1 OS=Homo sapiens OX=9606 GN=IGHA1 PE=1 SV=2 | 255 | 20201117_08512_1ng_glyco_hcd203040_compound15.14037.14037.4 | controllerType=0 controllerNumber=1 scan=14037 | 30.6778 |
| 12380:4 | 7 | R.LAGKPTHVN[+2262.814]VSVVMAEVDGTC[+57.021]Y.- | HexNAc(5)Hex(5)Fuc(1)NeuAc(1) | C[+57], N[+2263] | 1153.2467 | 4 | 4609.9652 | 4609.9564 | 0 | 1.9 | 332 | CRagged | 649.65 | 546.16 | 546.16 | 12.69 | 66 | >sp\|P01876\|IGHA1_HUMAN Immunoglobulin heavy constant alpha 1 OS=Homo sapiens OX=9606 GN=IGHA1 PE=1 SV=2 | 255 | 20201117_08512_1ng_glyco_hcd203040_compound15.15716.15716.4 | controllerType=0 controllerNumber=1 scan=15716 | 33.8386 |
| 10949:3 | 7 | R.LAGKPTHVN[+1216.423]VSVVMAEVDGTC[+57.021]Y.- | HexNAc(2)Hex(5) | C[+57], N[+1216] | 1188.5267 | 3 | 3563.5655 | 3563.5649 | 0 | 0.1 | 332 | CRagged | 657.31 | 517.42 | 377.61 | 13.21 | 66 | >sp\|P01876\|IGHA1_HUMAN Immunoglobulin heavy constant alpha 1 OS=Homo sapiens OX=9606 GN=IGHA1 PE=1 SV=2 | 255 | 20201117_08512_1ng_glyco_hcd203040_compound15.14226.14226.3 | controllerType=0 controllerNumber=1 scan=14226 | 31.0699 |
| 14044:4 | 7 | R.LAGKPTHVN[+2553.910]VSVVMAEVDGTC[+57.021]Y.- | HexNAc(5)Hex(5)Fuc(1)NeuAc(2) | C[+57], N[+2554] | 1226.0221 | 4 | 4901.0667 | 4901.0518 | 0 | 3 | 332 | CRagged | 677.6 | 535.09 | 535.09 | 13.58 | 66 | >sp\|P01876\|IGHA1_HUMAN Immunoglobulin heavy constant alpha 1 OS=Homo sapiens OX=9606 GN=IGHA1 PE=1 SV=2 | 255 | 20201117_08512_1ng_glyco_hcd203040_compound15.17449.17449.4 | controllerType=0 controllerNumber=1 scan=17449 | 37.3521 |
| 10770:4 | 7 | R.LAGKPTHVN[+1809.666]VSVVMAEVDGTC[+57.021]Y.- | HexNAc(5)Hex(4)Fuc(1) | C[+57], N[+1810] | 1039.9589 | 4 | 4156.8138 | 4156.8081 | 0 | 1.4 | 332 | CRagged | 678.68 | 674.96 | 454.59 | 13.69 | 66 | >sp\|P01876\|IGHA1_HUMAN Immunoglobulin heavy constant alpha 1 OS=Homo sapiens OX=9606 GN=IGHA1 PE=1 SV=2 | 255 | 20201117_08512_1ng_glyco_hcd203040_compound15.14040.14040.4 | controllerType=0 controllerNumber=1 scan=14040 | 30.6862 |
| 12358:3 | 7 | R.LAGKPTHVN[+1872.651]VSVVMAEVDGTC[+57.021]Y.- | HexNAc(3)Hex(6)NeuAc(1) | C[+57], N[+1873] | 1407.2722 | 3 | 4219.8019 | 4219.7926 | 0 | 2.2 | 332 | CRagged | 683.93 | 613.89 | 373.72 | 13.76 | 66 | >sp\|P01876\|IGHA1_HUMAN Immunoglobulin heavy constant alpha 1 OS=Homo sapiens OX=9606 GN=IGHA1 PE=1 SV=2 | 255 | 20201117_08512_1ng_glyco_hcd203040_compound15.15693.15693.3 | controllerType=0 controllerNumber=1 scan=15693 | 33.7905 |
| 12379:3 | 7 | R.LAGKPTHVN[+2262.814]VSVVMAEVDGTC[+57.021]Y.- | HexNAc(5)Hex(5)Fuc(1)NeuAc(1) | C[+57], N[+2263] | 1537.3272 | 3 | 4609.9671 | 4609.9564 | 0 | 2.3 | 332 | CRagged | 694.53 | 565.73 | 565.73 | 13.76 | 66 | >sp\|P01876\|IGHA1_HUMAN Immunoglobulin heavy constant alpha 1 OS=Homo sapiens OX=9606 GN=IGHA1 PE=1 SV=2 | 255 | 20201117_08512_1ng_glyco_hcd203040_compound15.15715.15715.3 | controllerType=0 controllerNumber=1 scan=15715 | 33.8367 |
| 12489:3 | 7 | R.LAGKPTHVN[+1548.545]VSVVMAEVDGTC[+57.021]Y.- | HexNAc(3)Hex(4)NeuAc(1) | C[+57], N[+1549] | 1299.2372 | 3 | 3895.6972 | 3895.6869 | 0 | 2.6 | 332 | CRagged | 705.93 | 696.37 | 696.37 | 14.76 | 66 | >sp\|P01876\|IGHA1_HUMAN Immunoglobulin heavy constant alpha 1 OS=Homo sapiens OX=9606 GN=IGHA1 PE=1 SV=2 | 255 | 20201117_08512_1ng_glyco_hcd203040_compound15.15829.15829.3 | controllerType=0 controllerNumber=1 scan=15829 | 34.0421 |
| 12311:3 | 7 | R.LAGKPTHVN[+2116.756]VSVVMAEVDGTC[+57.021]Y.- | HexNAc(5)Hex(5)NeuAc(1) | C[+57], N[+2117] | 1488.642 | 3 | 4463.9115 | 4463.8985 | 0 | 2.9 | 332 | CRagged | 723.89 | 521.71 | 521.71 | 14.76 | 66 | >sp\|P01876\|IGHA1_HUMAN Immunoglobulin heavy constant alpha 1 OS=Homo sapiens OX=9606 GN=IGHA1 PE=1 SV=2 | 255 | 20201117_08512_1ng_glyco_hcd203040_compound15.15644.15644.3 | controllerType=0 controllerNumber=1 scan=15644 | 33.6944 |
| 10681:3 | 7 | R.LAGKPTHVN[+1825.661]VSVVMAEVDGTC[+57.021]Y.- | HexNAc(5)Hex(5) | C[+57], N[+1826] | 1391.6067 | 3 | 4172.8057 | 4172.8031 | 0 | 0.6 | 332 | CRagged | 730.99 | 718.54 | 462.4 | 14.21 | 66 | >sp\|P01876\|IGHA1_HUMAN Immunoglobulin heavy constant alpha 1 OS=Homo sapiens OX=9606 GN=IGHA1 PE=1 SV=2 | 255 | 20201117_08512_1ng_glyco_hcd203040_compound15.13948.13948.3 | controllerType=0 controllerNumber=1 scan=13948 | 30.4904 |
| 12491:3 | 7 | R.LAGKPTHVN[+1548.545]VSVVMAEVDGTC[+57.021]Y.- | HexNAc(3)Hex(4)NeuAc(1) | C[+57], N[+1549] | 1299.2372 | 3 | 3895.6972 | 3895.6869 | 0 | 2.6 | 332 | CRagged | 743.28 | 713.88 | 713.88 | 14.76 | 66 | >sp\|P01876\|IGHA1_HUMAN Immunoglobulin heavy constant alpha 1 OS=Homo sapiens OX=9606 GN=IGHA1 PE=1 SV=2 | 255 | 20201117_08512_1ng_glyco_hcd203040_compound15.15831.15831.3 | controllerType=0 controllerNumber=1 scan=15831 | 34.0469 |
| 12445:3 | 7 | R.LAGKPTHVN[+1710.598]VSVVMAEVDGTC[+57.021]Y.- | HexNAc(3)Hex(5)NeuAc(1) | C[+57], N[+1711] | 1353.2546 | 3 | 4057.7492 | 4057.7397 | 0 | 2.3 | 332 | CRagged | 745.63 | 633.73 | 428.93 | 14.76 | 66 | >sp\|P01876\|IGHA1_HUMAN Immunoglobulin heavy constant alpha 1 OS=Homo sapiens OX=9606 GN=IGHA1 PE=1 SV=2 | 255 | 20201117_08512_1ng_glyco_hcd203040_compound15.15783.15783.3 | controllerType=0 controllerNumber=1 scan=15783 | 33.9643 |
| 09636:3 | 7 | R.LAGKPTHVN[+1378.476]VSVVMAEVDGTC[+57.021]Y.- | HexNAc(2)Hex(6) | C[+57], N[+1378] | 1242.5464 | 3 | 3725.6248 | 3725.6178 | 0 | 1.9 | 332 | CRagged | 750.05 | 552.01 | 461.86 | 15.86 | 66 | >sp\|P01876\|IGHA1_HUMAN Immunoglobulin heavy constant alpha 1 OS=Homo sapiens OX=9606 GN=IGHA1 PE=1 SV=2 | 255 | 20201117_08512_1ng_glyco_hcd203040_compound15.12863.12863.3 | controllerType=0 controllerNumber=1 scan=12863 | 28.4268 |
| 09634:3 | 7 | R.LAGKPTHVN[+1378.476]VSVVMAEVDGTC[+57.021]Y.- | HexNAc(2)Hex(6) | C[+57], N[+1378] | 1242.5464 | 3 | 3725.6248 | 3725.6178 | 0 | 1.9 | 332 | CRagged | 755.56 | 755.56 | 415.94 | 15.86 | 66 | >sp\|P01876\|IGHA1_HUMAN Immunoglobulin heavy constant alpha 1 OS=Homo sapiens OX=9606 GN=IGHA1 PE=1 SV=2 | 255 | 20201117_08512_1ng_glyco_hcd203040_compound15.12861.12861.3 | controllerType=0 controllerNumber=1 scan=12861 | 28.4218 |
| 10800:3 | 7 | R.LAGKPTHVN[+1501.555]VSVVMAEVDGTC[+57.021]Y.- | HexNAc(5)Hex(3) | C[+57], N[+1502] | 1283.5734 | 3 | 3848.7057 | 3848.6974 | 0 | 2.2 | 332 | CRagged | 760.17 | 760.17 | 760.17 | 15.76 | 66 | >sp\|P01876\|IGHA1_HUMAN Immunoglobulin heavy constant alpha 1 OS=Homo sapiens OX=9606 GN=IGHA1 PE=1 SV=2 | 255 | 20201117_08512_1ng_glyco_hcd203040_compound15.14072.14072.3 | controllerType=0 controllerNumber=1 scan=14072 | 30.7532 |
| 10816:3 | 7 | R.LAGKPTHVN[+1647.613]VSVVMAEVDGTC[+57.021]Y.- | HexNAc(5)Hex(3)Fuc(1) | C[+57], N[+1648] | 1332.2593 | 3 | 3994.7635 | 3994.7553 | 0 | 2 | 332 | CRagged | 762.69 | 739.3 | 739.3 | 15.76 | 66 | >sp\|P01876\|IGHA1_HUMAN Immunoglobulin heavy constant alpha 1 OS=Homo sapiens OX=9606 GN=IGHA1 PE=1 SV=2 | 255 | 20201117_08512_1ng_glyco_hcd203040_compound15.14088.14088.3 | controllerType=0 controllerNumber=1 scan=14088 | 30.7835 |
| 09681:3 | 7 | R.LAGKPTHVN[+1216.423]VSVVMAEVDGTC[+57.021]Y.- | HexNAc(2)Hex(5) | C[+57], N[+1216] | 1188.5302 | 3 | 3563.5761 | 3563.5649 | 0 | 3.1 | 332 | CRagged | 770.6 | 770.6 | 420.61 | 15.67 | 66 | >sp\|P01876\|IGHA1_HUMAN Immunoglobulin heavy constant alpha 1 OS=Homo sapiens OX=9606 GN=IGHA1 PE=1 SV=2 | 255 | 20201117_08512_1ng_glyco_hcd203040_compound15.12910.12910.3 | controllerType=0 controllerNumber=1 scan=12910 | 28.5147 |
| 10750:3 | 7 | R.LAGKPTHVN[+1809.666]VSVVMAEVDGTC[+57.021]Y.- | HexNAc(5)Hex(4)Fuc(1) | C[+57], N[+1810] | 1386.2758 | 3 | 4156.8129 | 4156.8081 | 0 | 1.1 | 332 | CRagged | 781.53 | 781.53 | 487.9 | 15.86 | 66 | >sp\|P01876\|IGHA1_HUMAN Immunoglobulin heavy constant alpha 1 OS=Homo sapiens OX=9606 GN=IGHA1 PE=1 SV=2 | 255 | 20201117_08512_1ng_glyco_hcd203040_compound15.14020.14020.3 | controllerType=0 controllerNumber=1 scan=14020 | 30.645 |
| 10679:3 | 7 | R.LAGKPTHVN[+1825.661]VSVVMAEVDGTC[+57.021]Y.- | HexNAc(5)Hex(5) | C[+57], N[+1826] | 1391.6067 | 3 | 4172.8057 | 4172.8031 | 0 | 0.6 | 332 | CRagged | 783.47 | 772.91 | 480.08 | 15.21 | 66 | >sp\|P01876\|IGHA1_HUMAN Immunoglobulin heavy constant alpha 1 OS=Homo sapiens OX=9606 GN=IGHA1 PE=1 SV=2 | 255 | 20201117_08512_1ng_glyco_hcd203040_compound15.13946.13946.3 | controllerType=0 controllerNumber=1 scan=13946 | 30.4826 |
| 10802:3 | 7 | R.LAGKPTHVN[+1501.555]VSVVMAEVDGTC[+57.021]Y.- | HexNAc(5)Hex(3) | C[+57], N[+1502] | 1283.5734 | 3 | 3848.7057 | 3848.6974 | 0 | 2.2 | 332 | CRagged | 784.41 | 755.98 | 755.98 | 15.76 | 66 | >sp\|P01876\|IGHA1_HUMAN Immunoglobulin heavy constant alpha 1 OS=Homo sapiens OX=9606 GN=IGHA1 PE=1 SV=2 | 255 | 20201117_08512_1ng_glyco_hcd203040_compound15.14074.14074.3 | controllerType=0 controllerNumber=1 scan=14074 | 30.7586 |
| 10721:3 | 7 | R.LAGKPTHVN[+1809.666]VSVVMAEVDGTC[+57.021]Y.- | HexNAc(5)Hex(4)Fuc(1) | C[+57], N[+1810] | 1386.2751 | 3 | 4156.8107 | 4156.8081 | 0 | 0.6 | 332 | CRagged | 806.37 | 745.74 | 524.7 | 16.21 | 66 | >sp\|P01876\|IGHA1_HUMAN Immunoglobulin heavy constant alpha 1 OS=Homo sapiens OX=9606 GN=IGHA1 PE=1 SV=2 | 255 | 20201117_08512_1ng_glyco_hcd203040_compound15.13989.13989.3 | controllerType=0 controllerNumber=1 scan=13989 | 30.5779 |
| 10753:3 | 7 | R.LAGKPTHVN[+1809.666]VSVVMAEVDGTC[+57.021]Y.- | HexNAc(5)Hex(4)Fuc(1) | C[+57], N[+1810] | 1386.2758 | 3 | 4156.8129 | 4156.8081 | 0 | 1.1 | 332 | CRagged | 816.8 | 737.13 | 537.78 | 16.86 | 66 | >sp\|P01876\|IGHA1_HUMAN Immunoglobulin heavy constant alpha 1 OS=Homo sapiens OX=9606 GN=IGHA1 PE=1 SV=2 | 255 | 20201117_08512_1ng_glyco_hcd203040_compound15.14023.14023.3 | controllerType=0 controllerNumber=1 scan=14023 | 30.6501 |
| 12355:3 | 7 | R.LAGKPTHVN[+1872.651]VSVVMAEVDGTC[+57.021]Y.- | HexNAc(3)Hex(6)NeuAc(1) | C[+57], N[+1873] | 1407.2722 | 3 | 4219.8019 | 4219.7926 | 0 | 2.2 | 332 | CRagged | 817.67 | 676.55 | 473.09 | 16.76 | 66 | >sp\|P01876\|IGHA1_HUMAN Immunoglobulin heavy constant alpha 1 OS=Homo sapiens OX=9606 GN=IGHA1 PE=1 SV=2 | 255 | 20201117_08512_1ng_glyco_hcd203040_compound15.15690.15690.3 | controllerType=0 controllerNumber=1 scan=15690 | 33.7847 |
| 10754:3 | 7 | R.LAGKPTHVN[+1663.608]VSVVMAEVDGTC[+57.021]Y.- | HexNAc(5)Hex(4) | C[+57], N[+1664] | 1337.5898 | 3 | 4010.7548 | 4010.7502 | 0 | 1.1 | 332 | CRagged | 819.27 | 819.27 | 502.47 | 16.86 | 66 | >sp\|P01876\|IGHA1_HUMAN Immunoglobulin heavy constant alpha 1 OS=Homo sapiens OX=9606 GN=IGHA1 PE=1 SV=2 | 255 | 20201117_08512_1ng_glyco_hcd203040_compound15.14024.14024.3 | controllerType=0 controllerNumber=1 scan=14024 | 30.6519 |
| 10662:3 | 7 | R.LAGKPTHVN[+1702.581]VSVVMAEVDGTC[+57.021]Y.- | HexNAc(2)Hex(8) | C[+57], N[+1703] | 1350.5815 | 3 | 4049.7299 | 4049.7234 | 0 | 1.6 | 332 | CRagged | 819.61 | 762.85 | 486.75 | 16.86 | 66 | >sp\|P01876\|IGHA1_HUMAN Immunoglobulin heavy constant alpha 1 OS=Homo sapiens OX=9606 GN=IGHA1 PE=1 SV=2 | 255 | 20201117_08512_1ng_glyco_hcd203040_compound15.13928.13928.3 | controllerType=0 controllerNumber=1 scan=13928 | 30.4405 |
| 10683:3 | 7 | R.LAGKPTHVN[+1540.529]VSVVMAEVDGTC[+57.021]Y.- | HexNAc(2)Hex(7) | C[+57], N[+1541] | 1296.5633 | 3 | 3887.6753 | 3887.6706 | 0 | 1.2 | 332 | CRagged | 824.23 | 815.37 | 495.84 | 16.86 | 66 | >sp\|P01876\|IGHA1_HUMAN Immunoglobulin heavy constant alpha 1 OS=Homo sapiens OX=9606 GN=IGHA1 PE=1 SV=2 | 255 | 20201117_08512_1ng_glyco_hcd203040_compound15.13950.13950.3 | controllerType=0 controllerNumber=1 scan=13950 | 30.4935 |
| 10814:3 | 7 | R.LAGKPTHVN[+1647.613]VSVVMAEVDGTC[+57.021]Y.- | HexNAc(5)Hex(3)Fuc(1) | C[+57], N[+1648] | 1332.2593 | 3 | 3994.7635 | 3994.7553 | 0 | 2 | 332 | CRagged | 866.49 | 866.49 | 866.49 | 17.76 | 66 | >sp\|P01876\|IGHA1_HUMAN Immunoglobulin heavy constant alpha 1 OS=Homo sapiens OX=9606 GN=IGHA1 PE=1 SV=2 | 255 | 20201117_08512_1ng_glyco_hcd203040_compound15.14086.14086.3 | controllerType=0 controllerNumber=1 scan=14086 | 30.7758 |
| 09684:3 | 7 | R.LAGKPTHVN[+1216.423]VSVVMAEVDGTC[+57.021]Y.- | HexNAc(2)Hex(5) | C[+57], N[+1216] | 1188.5302 | 3 | 3563.5761 | 3563.5649 | 0 | 3.1 | 332 | CRagged | 869.25 | 829.97 | 484.84 | 17.67 | 66 | >sp\|P01876\|IGHA1_HUMAN Immunoglobulin heavy constant alpha 1 OS=Homo sapiens OX=9606 GN=IGHA1 PE=1 SV=2 | 255 | 20201117_08512_1ng_glyco_hcd203040_compound15.12913.12913.3 | controllerType=0 controllerNumber=1 scan=12913 | 28.5231 |
| 10760:3 | 7 | R.LAGKPTHVN[+1216.423]VSVVMAEVDGTC[+57.021]Y.- | HexNAc(2)Hex(5) | C[+57], N[+1216] | 1188.528 | 3 | 3563.5695 | 3563.5649 | 0 | 1.3 | 332 | CRagged | 872.9 | 872.9 | 500.74 | 17.86 | 66 | >sp\|P01876\|IGHA1_HUMAN Immunoglobulin heavy constant alpha 1 OS=Homo sapiens OX=9606 GN=IGHA1 PE=1 SV=2 | 255 | 20201117_08512_1ng_glyco_hcd203040_compound15.14030.14030.3 | controllerType=0 controllerNumber=1 scan=14030 | 30.6623 |
| 10659:3 | 7 | R.LAGKPTHVN[+1702.581]VSVVMAEVDGTC[+57.021]Y.- | HexNAc(2)Hex(8) | C[+57], N[+1703] | 1350.5815 | 3 | 4049.7299 | 4049.7234 | 0 | 1.6 | 332 | CRagged | 874.92 | 874.92 | 552.73 | 17.86 | 66 | >sp\|P01876\|IGHA1_HUMAN Immunoglobulin heavy constant alpha 1 OS=Homo sapiens OX=9606 GN=IGHA1 PE=1 SV=2 | 255 | 20201117_08512_1ng_glyco_hcd203040_compound15.13925.13925.3 | controllerType=0 controllerNumber=1 scan=13925 | 30.4311 |
| 10686:3 | 7 | R.LAGKPTHVN[+1540.529]VSVVMAEVDGTC[+57.021]Y.- | HexNAc(2)Hex(7) | C[+57], N[+1541] | 1296.5633 | 3 | 3887.6753 | 3887.6706 | 0 | 1.2 | 332 | CRagged | 879.31 | 786.53 | 519.18 | 17.86 | 66 | >sp\|P01876\|IGHA1_HUMAN Immunoglobulin heavy constant alpha 1 OS=Homo sapiens OX=9606 GN=IGHA1 PE=1 SV=2 | 255 | 20201117_08512_1ng_glyco_hcd203040_compound15.13953.13953.3 | controllerType=0 controllerNumber=1 scan=13953 | 30.505 |
| 10703:3 | 7 | R.LAGKPTHVN[+1378.476]VSVVMAEVDGTC[+57.021]Y.- | HexNAc(2)Hex(6) | C[+57], N[+1378] | 1242.5463 | 3 | 3725.6244 | 3725.6178 | 0 | 1.8 | 332 | CRagged | 891.05 | 873.02 | 543.12 | 17.86 | 66 | >sp\|P01876\|IGHA1_HUMAN Immunoglobulin heavy constant alpha 1 OS=Homo sapiens OX=9606 GN=IGHA1 PE=1 SV=2 | 255 | 20201117_08512_1ng_glyco_hcd203040_compound15.13971.13971.3 | controllerType=0 controllerNumber=1 scan=13971 | 30.5435 |
| 10700:3 | 7 | R.LAGKPTHVN[+1378.476]VSVVMAEVDGTC[+57.021]Y.- | HexNAc(2)Hex(6) | C[+57], N[+1378] | 1242.5463 | 3 | 3725.6244 | 3725.6178 | 0 | 1.8 | 332 | CRagged | 904.58 | 904.58 | 539.85 | 18.86 | 66 | >sp\|P01876\|IGHA1_HUMAN Immunoglobulin heavy constant alpha 1 OS=Homo sapiens OX=9606 GN=IGHA1 PE=1 SV=2 | 255 | 20201117_08512_1ng_glyco_hcd203040_compound15.13968.13968.3 | controllerType=0 controllerNumber=1 scan=13968 | 30.5363 |
| 10719:3 | 7 | R.LAGKPTHVN[+1809.666]VSVVMAEVDGTC[+57.021]Y.- | HexNAc(5)Hex(4)Fuc(1) | C[+57], N[+1810] | 1386.2751 | 3 | 4156.8107 | 4156.8081 | 0 | 0.6 | 332 | CRagged | 910.02 | 910.02 | 598.31 | 18.21 | 66 | >sp\|P01876\|IGHA1_HUMAN Immunoglobulin heavy constant alpha 1 OS=Homo sapiens OX=9606 GN=IGHA1 PE=1 SV=2 | 255 | 20201117_08512_1ng_glyco_hcd203040_compound15.13987.13987.3 | controllerType=0 controllerNumber=1 scan=13987 | 30.5694 |
| 10763:3 | 7 | R.LAGKPTHVN[+1216.423]VSVVMAEVDGTC[+57.021]Y.- | HexNAc(2)Hex(5) | C[+57], N[+1216] | 1188.528 | 3 | 3563.5695 | 3563.5649 | 0 | 1.3 | 332 | CRagged | 954.93 | 889.98 | 578.34 | 19.86 | 66 | >sp\|P01876\|IGHA1_HUMAN Immunoglobulin heavy constant alpha 1 OS=Homo sapiens OX=9606 GN=IGHA1 PE=1 SV=2 | 255 | 20201117_08512_1ng_glyco_hcd203040_compound15.14033.14033.3 | controllerType=0 controllerNumber=1 scan=14033 | 30.6704 |
| 08862:3 | 7 | R.LAGKPTHVN[+1378.476]VSVVM[+15.995]AEVDGTC[+57.021]Y.- | HexNAc(2)Hex(6) | C[+57], M[+16], N[+1378] | 1247.8812 | 3 | 3741.6289 | 3741.6127 | 0 | 4.3 | 332 | CRagged | 251.55 | 251.55 | 251.55 | 6.11 | 66 | >sp\|P01876\|IGHA1_HUMAN Immunoglobulin heavy constant alpha 1 OS=Homo sapiens OX=9606 GN=IGHA1 PE=1 SV=2 | 255 | 20201117_08512_1ng_glyco_hcd203040_compound15.12059.12059.3 | controllerType=0 controllerNumber=1 scan=12059 | 26.939 |
| 08864:3 | 7 | R.LAGKPTHVN[+1378.476]VSVVM[+15.995]AEVDGTC[+57.021]Y.- | HexNAc(2)Hex(6) | C[+57], M[+16], N[+1378] | 1247.8812 | 3 | 3741.6289 | 3741.6127 | 0 | 4.3 | 332 | CRagged | 326.6 | 311.37 | 311.37 | 6.99 | 66 | >sp\|P01876\|IGHA1_HUMAN Immunoglobulin heavy constant alpha 1 OS=Homo sapiens OX=9606 GN=IGHA1 PE=1 SV=2 | 255 | 20201117_08512_1ng_glyco_hcd203040_compound15.12061.12061.3 | controllerType=0 controllerNumber=1 scan=12061 | 26.9425 |

Table S4. Cont.

| Query #:z | Protein Rank | Glycopeptide | Glycan | Modification Type(s) | Observed m/z | z | Observed (M+H) | Calc. mass (M+H) | Off-by-x error | Mass error (ppm) | StartPosition | Cleavage | Score | Delta | Delta Mod | \|Log Prob\| | # of unique peptides | Protein Name | Protein DB number | Comment | Scan # | Scan Time |
|---|---|---|---|---|---|---|---|---|---|---|---|---|---|---|---|---|---|---|---|---|---|---|
| 08372:3 | 7 | R.LAGKPTHVN[+1216.423]VSVVM[+15.995]AEVDGTC[+57.021]Y.- | HexNAc(2)Hex(5) | C[+57], M[+16], N[+1216] | 1193.861 | 3 | 3579.5685 | 3579.5599 | 0 | 2.4 | 332 | CRagged | 526.95 | 526.95 | 526.95 | 10.15 | 66 | >sp\|P01876\|IGHA1_HUMAN Immunoglobulin heavy constant alpha 1 OS=Homo sapiens OX=9606 GN=IGHA1 PE=1 SV=2 | 255 | 20201117_08512_1ng_glyco_hcd203040_compound15.11550.11550.3 | controllerType=0 controllerNumber=1 scan=11550 | 25.9945 |
| 08374:3 | 7 | R.LAGKPTHVN[+1216.423]VSVVM[+15.995]AEVDGTC[+57.021]Y.- | HexNAc(2)Hex(5) | C[+57], M[+16], N[+1216] | 1193.861 | 3 | 3579.5685 | 3579.5599 | 0 | 2.4 | 332 | CRagged | 592.38 | 592.38 | 592.38 | 11.76 | 66 | >sp\|P01876\|IGHA1_HUMAN Immunoglobulin heavy constant alpha 1 OS=Homo sapiens OX=9606 GN=IGHA1 PE=1 SV=2 | 255 | 20201117_08512_1ng_glyco_hcd203040_compound15.11552.11552.3 | controllerType=0 controllerNumber=1 scan=11552 | 26 |
| 21865:4 | 10 | K.VFPLSLDSTPQDGNVVVAC[+57.021]LVQGFFPQEPLSVTWSESGQN[+2279.830]VTAI | HexNAc(5)Hex(6)Fuc(2) | C[+57], N[+2280] | 1764.8128 | 4 | 7056.2295 | 7056.191 | 0 | 5.5 | 8 | Specific | 249.62 | 202.97 | 202.97 | 6.41 | 62 | >sp\|P01877\|IGHA2_HUMAN Immunoglobulin heavy constant alpha 2 OS=Homo sapiens OX=9606 GN=IGHA2 PE=1 SV=4 | 576 | 20201117_08512_1ng_glyco_hcd203040_compound15.25633.25633.4 | controllerType=0 controllerNumber=1 scan=25633 | 55.1398 |
| 21794:4 | 10 | K.VFPLSLDSTPQDGNVVVAC[+57.021]LVQGFFPQEPLSVTWSESGQN[+2262.814]VTAI | HexNAc(5)Hex(5)Fuc(1)NeuAc(1) | C[+57], N[+2263] | 1760.5531 | 4 | 7039.1904 | 7039.1757 | 0 | 2.1 | 8 | Specific | 334.78 | 316.09 | 316.09 | 8.66 | 62 | >sp\|P01877\|IGHA2_HUMAN Immunoglobulin heavy constant alpha 2 OS=Homo sapiens OX=9606 GN=IGHA2 PE=1 SV=4 | 576 | 20201117_08512_1ng_glyco_hcd203040_compound15.25559.25559.2 | controllerType=0 controllerNumber=1 scan=25559 | 54.9918 |
| 21853:6 | 10 | K.VFPLSLDSTPQDGNVVVAC[+57.021]LVQGFFPQEPLSVTWSESGQN[+2262.814]VTAI | HexNAc(5)Hex(5)Fuc(1)NeuAc(1) | C[+57], N[+2263] | 1174.0363 | 6 | 7039.1815 | 7039.1757 | 0 | 0.8 | 8 | Specific | 354 | 354 | 354 | 8.13 | 62 | >sp\|P01877\|IGHA2_HUMAN Immunoglobulin heavy constant alpha 2 OS=Homo sapiens OX=9606 GN=IGHA2 PE=1 SV=4 | 576 | 20201117_08512_1ng_glyco_hcd203040_compound15.25620.25620.6 | controllerType=0 controllerNumber=1 scan=25620 | 55.1113 |
| 20654:4 | 10 | K.VFPLSLDSTPQDGNVVVAC[+57.021]LVQGFFPQEPLSVTWSESGQN[+1647.613]VTAI | HexNAc(5)Hex(3)Fuc(1) | C[+57], N[+1648] | 1606.7563 | 4 | 6424.0034 | 6423.9746 | 0 | 4.5 | 8 | Specific | 359.33 | 359.33 | 359.33 | 8.43 | 62 | >sp\|P01877\|IGHA2_HUMAN Immunoglobulin heavy constant alpha 2 OS=Homo sapiens OX=9606 GN=IGHA2 PE=1 SV=4 | 576 | 20201117_08512_1ng_glyco_hcd203040_compound15.24369.24369.4 | controllerType=0 controllerNumber=1 scan=24369 | 52.4487 |
| 20304:4 | 10 | K.VFPLSLDSTPQDGNVVVAC[+57.021]LVQGFFPQEPLSVTWSESGQN[+1809.666]VTAI | HexNAc(5)Hex(4)Fuc(1) | C[+57], N[+1810] | 1647.2699 | 4 | 6586.0578 | 6586.0275 | 0 | 4.6 | 8 | Specific | 380.16 | 380.16 | 380.16 | 9.97 | 62 | >sp\|P01877\|IGHA2_HUMAN Immunoglobulin heavy constant alpha 2 OS=Homo sapiens OX=9606 GN=IGHA2 PE=1 SV=4 | 576 | 20201117_08512_1ng_glyco_hcd203040_compound15.24003.24003.2 | controllerType=0 controllerNumber=1 scan=24003 | 51.7269 |
| 21855:6 | 10 | K.VFPLSLDSTPQDGNVVVAC[+57.021]LVQGFFPQEPLSVTWSESGQN[+2262.814]VTAI | HexNAc(5)Hex(5)Fuc(1)NeuAc(1) | C[+57], N[+2263] | 1174.0363 | 6 | 7039.1815 | 7039.1757 | 0 | 0.8 | 8 | Specific | 404.75 | 404.75 | 404.75 | 9.74 | 62 | >sp\|P01877\|IGHA2_HUMAN Immunoglobulin heavy constant alpha 2 OS=Homo sapiens OX=9606 GN=IGHA2 PE=1 SV=4 | 576 | 20201117_08512_1ng_glyco_hcd203040_compound15.25622.25622.6 | controllerType=0 controllerNumber=1 scan=25622 | 55.1161 |
| 21992:5 | 10 | K.VFPLSLDSTPQDGNVVVAC[+57.021]LVQGFFPQEPLSVTWSESGQN[+2262.814]VTAI | HexNAc(5)Hex(5)Fuc(1)NeuAc(1) | C[+57], N[+2263] | 1408.6425 | 5 | 7039.1834 | 7039.1757 | 0 | 1.1 | 8 | Specific | 467.17 | 467.17 | 467.17 | 10.05 | 62 | >sp\|P01877\|IGHA2_HUMAN Immunoglobulin heavy constant alpha 2 OS=Homo sapiens OX=9606 GN=IGHA2 PE=1 SV=4 | 576 | 20201117_08512_1ng_glyco_hcd203040_compound15.25765.25765.5 | controllerType=0 controllerNumber=1 scan=25765 | 55.4066 |
| 21842:4 | 10 | K.VFPLSLDSTPQDGNVVVAC[+57.021]LVQGFFPQEPLSVTWSESGQN[+2262.814]VTAI | HexNAc(5)Hex(5)Fuc(1)NeuAc(1) | C[+57], N[+2263] | 1760.5581 | 4 | 7039.2104 | 7039.1757 | 0 | 4.9 | 8 | Specific | 492.1 | 484.71 | 484.71 | 10.45 | 62 | >sp\|P01877\|IGHA2_HUMAN Immunoglobulin heavy constant alpha 2 OS=Homo sapiens OX=9606 GN=IGHA2 PE=1 SV=4 | 576 | 20201117_08512_1ng_glyco_hcd203040_compound15.25609.25609.4 | controllerType=0 controllerNumber=1 scan=25609 | 55.0853 |
| 20306:4 | 10 | K.VFPLSLDSTPQDGNVVVAC[+57.021]LVQGFFPQEPLSVTWSESGQN[+1809.666]VTAI | HexNAc(5)Hex(4)Fuc(1) | C[+57], N[+1810] | 1647.2699 | 4 | 6586.0578 | 6586.0275 | 0 | 4.6 | 8 | Specific | 549.52 | 514.24 | 514.24 | 10.68 | 62 | >sp\|P01877\|IGHA2_HUMAN Immunoglobulin heavy constant alpha 2 OS=Homo sapiens OX=9606 GN=IGHA2 PE=1 SV=4 | 576 | 20201117_08512_1ng_glyco_hcd203040_compound15.24005.24005.2 | controllerType=0 controllerNumber=1 scan=24005 | 51.7318 |
| 21841:5 | 10 | K.VFPLSLDSTPQDGNVVVAC[+57.021]LVQGFFPQEPLSVTWSESGQN[+2262.814]VTAI | HexNAc(5)Hex(5)Fuc(1)NeuAc(1) | C[+57], N[+2263] | 1408.6445 | 5 | 7039.1932 | 7039.1757 | 0 | 2.5 | 8 | Specific | 571.66 | 571 | 571 | 10.52 | 62 | >sp\|P01877\|IGHA2_HUMAN Immunoglobulin heavy constant alpha 2 OS=Homo sapiens OX=9606 GN=IGHA2 PE=1 SV=4 | 576 | 20201117_08512_1ng_glyco_hcd203040_compound15.25608.25608.5 | controllerType=0 controllerNumber=1 scan=25608 | 55.0832 |
| 20656:4 | 10 | K.VFPLSLDSTPQDGNVVVAC[+57.021]LVQGFFPQEPLSVTWSESGQN[+1647.613]VTAI | HexNAc(5)Hex(3)Fuc(1) | C[+57], N[+1648] | 1606.7563 | 4 | 6424.0034 | 6423.9746 | 0 | 4.5 | 8 | Specific | 670.35 | 637.42 | 637.42 | 14.16 | 62 | >sp\|P01877\|IGHA2_HUMAN Immunoglobulin heavy constant alpha 2 OS=Homo sapiens OX=9606 GN=IGHA2 PE=1 SV=4 | 576 | 20201117_08512_1ng_glyco_hcd203040_compound15.24371.24371.4 | controllerType=0 controllerNumber=1 scan=24371 | 52.4523 |
| 20386:4 | 10 | K.VFPLSLDSTPQDGNVVVAC[+57.021]LVQGFFPQEPLSVTWSESGQN[+1809.666]VTAI | HexNAc(5)Hex(4)Fuc(1) | C[+57], N[+1810] | 1647.271 | 4 | 6586.0622 | 6586.0275 | 0 | 5.3 | 8 | Specific | 686.3 | 674.57 | 674.57 | 14.16 | 62 | >sp\|P01877\|IGHA2_HUMAN Immunoglobulin heavy constant alpha 2 OS=Homo sapiens OX=9606 GN=IGHA2 PE=1 SV=4 | 576 | 20201117_08512_1ng_glyco_hcd203040_compound15.24089.24089.4 | controllerType=0 controllerNumber=1 scan=24089 | 51.9106 |
| 21844:4 | 10 | K.VFPLSLDSTPQDGNVVVAC[+57.021]LVQGFFPQEPLSVTWSESGQN[+2262.814]VTAI | HexNAc(5)Hex(5)Fuc(1)NeuAc(1) | C[+57], N[+2263] | 1760.5581 | 4 | 7039.2104 | 7039.1757 | 0 | 4.9 | 8 | Specific | 705.28 | 698.66 | 698.66 | 15.16 | 62 | >sp\|P01877\|IGHA2_HUMAN Immunoglobulin heavy constant alpha 2 OS=Homo sapiens OX=9606 GN=IGHA2 PE=1 SV=4 | 576 | 20201117_08512_1ng_glyco_hcd203040_compound15.25611.25611.4 | controllerType=0 controllerNumber=1 scan=25611 | 55.0913 |
| 21994:5 | 10 | K.VFPLSLDSTPQDGNVVVAC[+57.021]LVQGFFPQEPLSVTWSESGQN[+2262.814]VTAI | HexNAc(5)Hex(5)Fuc(1)NeuAc(1) | C[+57], N[+2263] | 1408.6425 | 5 | 7039.1834 | 7039.1757 | 0 | 1.1 | 8 | Specific | 745.31 | 692.9 | 692.9 | 14.87 | 62 | >sp\|P01877\|IGHA2_HUMAN Immunoglobulin heavy constant alpha 2 OS=Homo sapiens OX=9606 GN=IGHA2 PE=1 SV=4 | 576 | 20201117_08512_1ng_glyco_hcd203040_compound15.25767.25767.5 | controllerType=0 controllerNumber=1 scan=25767 | 55.4127 |
| 21843:5 | 10 | K.VFPLSLDSTPQDGNVVVAC[+57.021]LVQGFFPQEPLSVTWSESGQN[+2262.814]VTAI | HexNAc(5)Hex(5)Fuc(1)NeuAc(1) | C[+57], N[+2263] | 1408.6445 | 5 | 7039.1932 | 7039.1757 | 0 | 2.5 | 8 | Specific | 781.78 | 755.39 | 755.39 | 15.76 | 62 | >sp\|P01877\|IGHA2_HUMAN Immunoglobulin heavy constant alpha 2 OS=Homo sapiens OX=9606 GN=IGHA2 PE=1 SV=4 | 576 | 20201117_08512_1ng_glyco_hcd203040_compound15.25610.25610.5 | controllerType=0 controllerNumber=1 scan=25610 | 55.0893 |
| 20388:4 | 10 | K.VFPLSLDSTPQDGNVVVAC[+57.021]LVQGFFPQEPLSVTWSESGQN[+1809.666]VTAI | HexNAc(5)Hex(4)Fuc(1) | C[+57], N[+1810] | 1647.271 | 4 | 6586.0622 | 6586.0275 | 0 | 5.3 | 8 | Specific | 849.43 | 820.03 | 820.03 | 17.16 | 62 | >sp\|P01877\|IGHA2_HUMAN Immunoglobulin heavy constant alpha 2 OS=Homo sapiens OX=9606 GN=IGHA2 PE=1 SV=4 | 576 | 20201117_08512_1ng_glyco_hcd203040_compound15.24091.24091.4 | controllerType=0 controllerNumber=1 scan=24091 | 51.917 |
| 01080:2 | 10 | K.HYTN[+1647.613]SSQDVTVPC[+57.021]R.V | HexNAc(5)Hex(3)Fuc(1) | C[+57], N[+1648] | 1656.1808 | 2 | 3311.3543 | 3311.3625 | 0 | -2.5 | 89 | Specific | 208.02 | 182.72 | 182.72 | 2.39 | 62 | >sp\|P01877\|IGHA2_HUMAN Immunoglobulin heavy constant alpha 2 OS=Homo sapiens OX=9606 GN=IGHA2 PE=1 SV=4 | 576 | 20201117_08512_1ng_glyco_hcd203040_compound15.3995.3995.2 | controllerType=0 controllerNumber=1 scan=3995 | 13.0001 |
| 01086:3 | 10 | K.HYTN[+1444.534]SSQDVTVPC[+57.021]R.V | HexNAc(4)Hex(3)Fuc(1) | C[+57], N[+1445] | 1036.7665 | 3 | 3108.285 | 3108.2832 | 0 | 0.6 | 89 | Specific | 365.4 | 295.33 | 295.33 | 6.49 | 62 | >sp\|P01877\|IGHA2_HUMAN Immunoglobulin heavy constant alpha 2 OS=Homo sapiens OX=9606 GN=IGHA2 PE=1 SV=4 | 576 | 20201117_08512_1ng_glyco_hcd203040_compound15.4001.4001.3 | controllerType=0 controllerNumber=1 scan=4001 | 13.0113 |
| 00965:3 | 10 | K.HYTN[+1971.719]SSQDVTVPC[+57.021]R.V | HexNAc(5)Hex(5)Fuc(1) | C[+57], N[+1972] | 1212.4974 | 3 | 3635.4776 | 3635.4682 | 0 | 2.6 | 89 | Specific | 376.21 | 202.43 | 202.43 | 6.18 | 62 | >sp\|P01877\|IGHA2_HUMAN Immunoglobulin heavy constant alpha 2 OS=Homo sapiens OX=9606 GN=IGHA2 PE=1 SV=4 | 576 | 20201117_08512_1ng_glyco_hcd203040_compound15.3876.3876.3 | controllerType=0 controllerNumber=1 scan=3876 | 12.7945 |
| 00986:3 | 10 | K.HYTN[+1606.587]SSQDVTVPC[+57.021]R.V | HexNAc(4)Hex(4)Fuc(1) | C[+57], N[+1607] | 1090.7881 | 3 | 3270.3498 | 3270.336 | 0 | 4.2 | 89 | Specific | 392.86 | 277.98 | 277.98 | 6.69 | 62 | >sp\|P01877\|IGHA2_HUMAN Immunoglobulin heavy constant alpha 2 OS=Homo sapiens OX=9606 GN=IGHA2 PE=1 SV=4 | 576 | 20201117_08512_1ng_glyco_hcd203040_compound15.3898.3898.3 | controllerType=0 controllerNumber=1 scan=3898 | 12.8336 |
| 00988:3 | 10 | K.HYTN[+1606.587]SSQDVTVPC[+57.021]R.V | HexNAc(4)Hex(4)Fuc(1) | C[+57], N[+1607] | 1090.7881 | 3 | 3270.3498 | 3270.336 | 0 | 4.2 | 89 | Specific | 404.62 | 268.5 | 268.5 | 6.69 | 62 | >sp\|P01877\|IGHA2_HUMAN Immunoglobulin heavy constant alpha 2 OS=Homo sapiens OX=9606 GN=IGHA2 PE=1 SV=4 | 576 | 20201117_08512_1ng_glyco_hcd203040_compound15.3900.3900.3 | controllerType=0 controllerNumber=1 scan=3900 | 12.8377 |
| 01084:3 | 10 | K.HYTN[+1444.534]SSQDVTVPC[+57.021]R.V | HexNAc(4)Hex(3)Fuc(1) | C[+57], N[+1445] | 1036.7665 | 3 | 3108.285 | 3108.2832 | 0 | 0.6 | 89 | Specific | 417.81 | 299.21 | 299.21 | 6.74 | 62 | >sp\|P01877\|IGHA2_HUMAN Immunoglobulin heavy constant alpha 2 OS=Homo sapiens OX=9606 GN=IGHA2 PE=1 SV=4 | 576 | 20201117_08512_1ng_glyco_hcd203040_compound15.3999.3999.3 | controllerType=0 controllerNumber=1 scan=3999 | 13.0066 |
| 00963:3 | 10 | K.HYTN[+1971.719]SSQDVTVPC[+57.021]R.V | HexNAc(5)Hex(5)Fuc(1) | C[+57], N[+1972] | 1212.4974 | 3 | 3635.4776 | 3635.4682 | 0 | 2.6 | 89 | Specific | 420.02 | 216.07 | 216.07 | 6.72 | 62 | >sp\|P01877\|IGHA2_HUMAN Immunoglobulin heavy constant alpha 2 OS=Homo sapiens OX=9606 GN=IGHA2 PE=1 SV=4 | 576 | 20201117_08512_1ng_glyco_hcd203040_compound15.3874.3874.3 | controllerType=0 controllerNumber=1 scan=3874 | 12.791 |
| 00979:3 | 10 | K.HYTN[+1971.719]SSQDVTVPC[+57.021]R.V | HexNAc(5)Hex(5)Fuc(1) | C[+57], N[+1972] | 1212.498 | 3 | 3635.4794 | 3635.4682 | 0 | 3.1 | 89 | Specific | 463.23 | 264.27 | 264.27 | 7.1 | 62 | >sp\|P01877\|IGHA2_HUMAN Immunoglobulin heavy constant alpha 2 OS=Homo sapiens OX=9606 GN=IGHA2 PE=1 SV=4 | 576 | 20201117_08512_1ng_glyco_hcd203040_compound15.3891.3891.3 | controllerType=0 controllerNumber=1 scan=3891 | 12.8219 |
| 01002:3 | 10 | K.HYTN[+1444.534]SSQDVTVPC[+57.021]R.V | HexNAc(4)Hex(3)Fuc(1) | C[+57], N[+1445] | 1036.7685 | 3 | 3108.2908 | 3108.2832 | 0 | 2.5 | 89 | Specific | 467.87 | 435.75 | 435.75 | 9.27 | 62 | >sp\|P01877\|IGHA2_HUMAN Immunoglobulin heavy constant alpha 2 OS=Homo sapiens OX=9606 GN=IGHA2 PE=1 SV=4 | 576 | 20201117_08512_1ng_glyco_hcd203040_compound15.3914.3914.3 | controllerType=0 controllerNumber=1 scan=3914 | 12.8614 |
| 01089:3 | 10 | K.HYTN[+1647.613]SSQDVTVPC[+57.021]R.V | HexNAc(5)Hex(3)Fuc(1) | C[+57], N[+1648] | 1104.4613 | 3 | 3311.3694 | 3311.3625 | 0 | 2.1 | 89 | Specific | 524.52 | 327.25 | 327.25 | 7.66 | 62 | >sp\|P01877\|IGHA2_HUMAN Immunoglobulin heavy constant alpha 2 OS=Homo sapiens OX=9606 GN=IGHA2 PE=1 SV=4 | 576 | 20201117_08512_1ng_glyco_hcd203040_compound15.4004.4004.2 | controllerType=0 controllerNumber=1 scan=4004 | 13.0181 |
| 01894:3 | 10 | K.HYTN[+2262.814]SSQDVTVPC[+57.021]R.V | HexNAc(5)Hex(5)Fuc(1)NeuAc(1) | C[+57], N[+2263] | 1309.5302 | 3 | 3926.5761 | 3926.5636 | 0 | 3.2 | 89 | Specific | 540.93 | 317.34 | 317.34 | 7.7 | 62 | >sp\|P01877\|IGHA2_HUMAN Immunoglobulin heavy constant alpha 2 OS=Homo sapiens OX=9606 GN=IGHA2 PE=1 SV=4 | 576 | 20201117_08512_1ng_glyco_hcd203040_compound15.4841.4841.3 | controllerType=0 controllerNumber=1 scan=4841 | 14.4478 |
| 01087:3 | 10 | K.HYTN[+1647.613]SSQDVTVPC[+57.021]R.V | HexNAc(5)Hex(3)Fuc(1) | C[+57], N[+1648] | 1104.4613 | 3 | 3311.3694 | 3311.3625 | 0 | 2.1 | 89 | Specific | 561.51 | 345.98 | 345.98 | 8.04 | 62 | >sp\|P01877\|IGHA2_HUMAN Immunoglobulin heavy constant alpha 2 OS=Homo sapiens OX=9606 GN=IGHA2 PE=1 SV=4 | 576 | 20201117_08512_1ng_glyco_hcd203040_compound15.4002.4002.2 | controllerType=0 controllerNumber=1 scan=4002 | 13.0133 |
| 01017:3 | 10 | K.HYTN[+1444.534]SSQDVTVPC[+57.021]R.V | HexNAc(4)Hex(3)Fuc(1) | C[+57], N[+1445] | 1036.7697 | 3 | 3108.2945 | 3108.2832 | 0 | 3.6 | 89 | Specific | 570.25 | 415.02 | 415.02 | 9.75 | 62 | >sp\|P01877\|IGHA2_HUMAN Immunoglobulin heavy constant alpha 2 OS=Homo sapiens OX=9606 GN=IGHA2 PE=1 SV=4 | 576 | 20201117_08512_1ng_glyco_hcd203040_compound15.3930.3930.3 | controllerType=0 controllerNumber=1 scan=3930 | 12.8901 |
| 00981:3 | 10 | K.HYTN[+1971.719]SSQDVTVPC[+57.021]R.V | HexNAc(5)Hex(5)Fuc(1) | C[+57], N[+1972] | 1212.498 | 3 | 3635.4794 | 3635.4682 | 0 | 3.1 | 89 | Specific | 580.58 | 404.81 | 404.81 | 11.09 | 62 | >sp\|P01877\|IGHA2_HUMAN Immunoglobulin heavy constant alpha 2 OS=Homo sapiens OX=9606 GN=IGHA2 PE=1 SV=4 | 576 | 20201117_08512_1ng_glyco_hcd203040_compound15.3893.3893.3 | controllerType=0 controllerNumber=1 scan=3893 | 12.8257 |
| 01896:3 | 10 | K.HYTN[+2262.814]SSQDVTVPC[+57.021]R.V | HexNAc(5)Hex(5)Fuc(1)NeuAc(1) | C[+57], N[+2263] | 1309.5302 | 3 | 3926.5761 | 3926.5636 | 0 | 3.2 | 89 | Specific | 595.33 | 429.72 | 429.72 | 10.99 | 62 | >sp\|P01877\|IGHA2_HUMAN Immunoglobulin heavy constant alpha 2 OS=Homo sapiens OX=9606 GN=IGHA2 PE=1 SV=4 | 576 | 20201117_08512_1ng_glyco_hcd203040_compound15.4843.4843.3 | controllerType=0 controllerNumber=1 scan=4843 | 14.4523 |
| 01019:3 | 10 | K.HYTN[+1444.534]SSQDVTVPC[+57.021]R.V | HexNAc(4)Hex(3)Fuc(1) | C[+57], N[+1445] | 1036.7697 | 3 | 3108.2945 | 3108.2832 | 0 | 3.6 | 89 | Specific | 622.61 | 604.06 | 604.06 | 11.99 | 62 | >sp\|P01877\|IGHA2_HUMAN Immunoglobulin heavy constant alpha 2 OS=Homo sapiens OX=9606 GN=IGHA2 PE=1 SV=4 | 576 | 20201117_08512_1ng_glyco_hcd203040_compound15.3932.3932.3 | controllerType=0 controllerNumber=1 scan=3932 | 12.8937 |
| 00989:3 | 10 | K.HYTN[+1809.666]SSQDVTVPC[+57.021]R.V | HexNAc(5)Hex(4)Fuc(1) | C[+57], N[+1810] | 1158.4794 | 3 | 3473.4237 | 3473.4154 | 0 | 2.4 | 89 | Specific | 639.94 | 527.29 | 527.29 | 12.09 | 62 | >sp\|P01877\|IGHA2_HUMAN Immunoglobulin heavy constant alpha 2 OS=Homo sapiens OX=9606 GN=IGHA2 PE=1 SV=4 | 576 | 20201117_08512_1ng_glyco_hcd203040_compound15.3901.3901.3 | controllerType=0 controllerNumber=1 scan=3901 | 12.8409 |
| 01003:3 | 10 | K.HYTN[+1444.534]SSQDVTVPC[+57.021]R.V | HexNAc(4)Hex(3)Fuc(1) | C[+57], N[+1445] | 1036.7685 | 3 | 3108.2908 | 3108.2832 | 0 | 2.5 | 89 | Specific | 673.07 | 520.65 | 520.65 | 13.09 | 62 | >sp\|P01877\|IGHA2_HUMAN Immunoglobulin heavy constant alpha 2 OS=Homo sapiens OX=9606 GN=IGHA2 PE=1 SV=4 | 576 | 20201117_08512_1ng_glyco_hcd203040_compound15.3915.3915.3 | controllerType=0 controllerNumber=1 scan=3915 | 12.8655 |
| 00987:3 | 10 | K.HYTN[+1809.666]SSQDVTVPC[+57.021]R.V | HexNAc(5)Hex(4)Fuc(1) | C[+57], N[+1810] | 1158.4794 | 3 | 3473.4237 | 3473.4154 | 0 | 2.4 | 89 | Specific | 695.79 | 479.19 | 479.19 | 13.09 | 62 | >sp\|P01877\|IGHA2_HUMAN Immunoglobulin heavy constant alpha 2 OS=Homo sapiens OX=9606 GN=IGHA2 PE=1 SV=4 | 576 | 20201117_08512_1ng_glyco_hcd203040_compound15.3899.3899.3 | controllerType=0 controllerNumber=1 scan=3899 | 12.8352 |
| 01037:3 | 10 | K.HYTN[+1647.613]SSQDVTVPC[+57.021]R.V | HexNAc(5)Hex(3)Fuc(1) | C[+57], N[+1648] | 1104.4615 | 3 | 3311.3698 | 3311.3625 | 0 | 2.2 | 89 | Specific | 772.89 | 721.93 | 721.93 | 15.09 | 62 | >sp\|P01877\|IGHA2_HUMAN Immunoglobulin heavy constant alpha 2 OS=Homo sapiens OX=9606 GN=IGHA2 PE=1 SV=4 | 576 | 20201117_08512_1ng_glyco_hcd203040_compound15.3951.3951.3 | controllerType=0 controllerNumber=1 scan=3951 | 12.9292 |
| 01035:3 | 10 | K.HYTN[+1647.613]SSQDVTVPC[+57.021]R.V | HexNAc(5)Hex(3)Fuc(1) | C[+57], N[+1648] | 1104.4615 | 3 | 3311.3698 | 3311.3625 | 0 | 2.2 | 89 | Specific | 774.59 | 580.25 | 580.25 | 15.09 | 62 | >sp\|P01877\|IGHA2_HUMAN Immunoglobulin heavy constant alpha 2 OS=Homo sapiens OX=9606 GN=IGHA2 PE=1 SV=4 | 576 | 20201117_08512_1ng_glyco_hcd203040_compound15.3949.3949.3 | controllerType=0 controllerNumber=1 scan=3949 | 12.9233 |
| 03882:4 | 10 | K.HYTN[+2263.835]SSQDVTVPC[+57.021]RVPPPPPC[+57.021]C[+57.021]HPR.L | HexNAc(5)Hex(5)Fuc(3) | C[+57]*3, N[+2264] | 1306.2986 | 4 | 5222.1725 | 5222.1903 | 0 | -3.4 | 89 | Specific | 255.02 | 255.02 | 255.02 | 4.56 | 62 | >sp\|P01877\|IGHA2_HUMAN Immunoglobulin heavy constant alpha 2 OS=Homo sapiens OX=9606 GN=IGHA2 PE=1 SV=4 | 576 | 20201117_08512_1ng_glyco_hcd203040_compound15.6900.6900.2 | controllerType=0 controllerNumber=1 scan=6900 | 18.0563 |
| 03059:3 | 10 | K.HYTN[+1647.613]SSQDVTVPC[+57.021]RVPPPPPC[+57.021]C[+57.021]HPR.L | HexNAc(5)Hex(3)Fuc(1) | C[+57]*3, N[+1648] | 1535.9986 | 3 | 4605.9812 | 4605.9689 | 0 | 2.7 | 89 | Specific | 305.42 | 233.13 | 233.13 | 5.57 | 62 | >sp\|P01877\|IGHA2_HUMAN Immunoglobulin heavy constant alpha 2 OS=Homo sapiens OX=9606 GN=IGHA2 PE=1 SV=4 | 576 | 20201117_08512_1ng_glyco_hcd203040_compound15.6047.6047.3 | controllerType=0 controllerNumber=1 scan=6047 | 16.5769 |
| 03092:5 | 10 | K.HYTN[+1444.534]SSQDVTVPC[+57.021]RVPPPPPC[+57.021]C[+57.021]HPR.L | HexNAc(4)Hex(3)Fuc(1) | C[+57]*3, N[+1445] | 881.3855 | 5 | 4402.8985 | 4402.8895 | 0 | 2 | 89 | Specific | 377.22 | 377.22 | 377.22 | 9.28 | 62 | >sp\|P01877\|IGHA2_HUMAN Immunoglobulin heavy constant alpha 2 OS=Homo sapiens OX=9606 GN=IGHA2 PE=1 SV=4 | 576 | 20201117_08512_1ng_glyco_hcd203040_compound15.6081.6081.5 | controllerType=0 controllerNumber=1 scan=6081 | 16.6335 |
| 03061:3 | 10 | K.HYTN[+1647.613]SSQDVTVPC[+57.021]RVPPPPPC[+57.021]C[+57.021]HPR.L | HexNAc(5)Hex(3)Fuc(1) | C[+57]*3, N[+1648] | 1535.9986 | 3 | 4605.9812 | 4605.9689 | 0 | 2.7 | 89 | Specific | 389.24 | 245.53 | 245.53 | 4.81 | 62 | >sp\|P01877\|IGHA2_HUMAN Immunoglobulin heavy constant alpha 2 OS=Homo sapiens OX=9606 GN=IGHA2 PE=1 SV=4 | 576 | 20201117_08512_1ng_glyco_hcd203040_compound15.6049.6049.3 | controllerType=0 controllerNumber=1 scan=6049 | 16.5804 |
| 03587:5 | 10 | K.HYTN[+1647.613]SSQDVTVPC[+57.021]RVPPPPPC[+57.021]C[+57.021]HPR.L | HexNAc(5)Hex(3)Fuc(1) | C[+57]*3, N[+1648] | 922.0003 | 5 | 4605.9725 | 4605.9689 | 0 | 0.8 | 89 | Specific | 396.73 | 393.64 | 393.64 | 9.45 | 62 | >sp\|P01877\|IGHA2_HUMAN Immunoglobulin heavy constant alpha 2 OS=Homo sapiens OX=9606 GN=IGHA2 PE=1 SV=4 | 576 | 20201117_08512_1ng_glyco_hcd203040_compound15.6595.6595.5 | controllerType=0 controllerNumber=1 scan=6595 | 17.5133 |
| 03133:5 | 10 | K.HYTN[+1444.534]SSQDVTVPC[+57.021]RVPPPPPC[+57.021]C[+57.021]HPR.L | HexNAc(4)Hex(3)Fuc(1) | C[+57]*3, N[+1445] | 881.3847 | 5 | 4402.8942 | 4402.8895 | 0 | 1.1 | 89 | Specific | 447.34 | 447.34 | 447.34 | 9.68 | 62 | >sp\|P01877\|IGHA2_HUMAN Immunoglobulin heavy constant alpha 2 OS=Homo sapiens OX=9606 GN=IGHA2 PE=1 SV=4 | 576 | 20201117_08512_1ng_glyco_hcd203040_compound15.6123.6123.5 | controllerType=0 controllerNumber=1 scan=6123 | 16.7072 |
| 03589:5 | 10 | K.HYTN[+1647.613]SSQDVTVPC[+57.021]RVPPPPPC[+57.021]C[+57.021]HPR.L | HexNAc(5)Hex(3)Fuc(1) | C[+57]*3, N[+1648] | 922.0003 | 5 | 4605.9725 | 4605.9689 | 0 | 0.8 | 89 | Specific | 476.1 | 404.11 | 404.11 | 9.76 | 62 | >sp\|P01877\|IGHA2_HUMAN Immunoglobulin heavy constant alpha 2 OS=Homo sapiens OX=9606 GN=IGHA2 PE=1 SV=4 | 576 | 20201117_08512_1ng_glyco_hcd203040_compound15.6597.6597.5 | controllerType=0 controllerNumber=1 scan=6597 | 17.5169 |
| 02972:5 | 10 | K.HYTN[+1809.666]SSQDVTVPC[+57.021]RVPPPPPC[+57.021]C[+57.021]HPR.L | HexNAc(5)Hex(4)Fuc(1) | C[+57]*3, N[+1810] | 954.413 | 5 | 4768.0358 | 4768.0217 | 0 | 3 | 89 | Specific | 513.53 | 433.56 | 433.56 | 9.87 | 62 | >sp\|P01877\|IGHA2_HUMAN Immunoglobulin heavy constant alpha 2 OS=Homo sapiens OX=9606 GN=IGHA2 PE=1 SV=4 | 576 | 20201117_08512_1ng_glyco_hcd203040_compound15.5957.5957.5 | controllerType=0 controllerNumber=1 scan=5957 | 16.4088 |
| 02975:5 | 10 | K.HYTN[+1809.666]SSQDVTVPC[+57.021]RVPPPPPC[+57.021]C[+57.021]HPR.L | HexNAc(5)Hex(4)Fuc(1) | C[+57]*3, N[+1810] | 954.413 | 5 | 4768.0358 | 4768.0217 | 0 | 3 | 89 | Specific | 535.75 | 404.43 | 404.43 | 10 | 62 | >sp\|P01877\|IGHA2_HUMAN Immunoglobulin heavy constant alpha 2 OS=Homo sapiens OX=9606 GN=IGHA2 PE=1 SV=4 | 576 | 20201117_08512_1ng_glyco_hcd203040_compound15.5960.5960.5 | controllerType=0 controllerNumber=1 scan=5960 | 16.4161 |
| 02983:4 | 10 | K.HYTN[+1971.719]SSQDVTVPC[+57.021]RVPPPPPC[+57.021]C[+57.021]HPR.L | HexNAc(5)Hex(5)Fuc(1) | C[+57]*3, N[+1972] | 1233.2771 | 4 | 4930.0867 | 4930.0745 | 0 | 2.5 | 89 | Specific | 536.56 | 367.87 | 367.87 | 8.07 | 62 | >sp\|P01877\|IGHA2_HUMAN Immunoglobulin heavy constant alpha 2 OS=Homo sapiens OX=9606 GN=IGHA2 PE=1 SV=4 | 576 | 20201117_08512_1ng_glyco_hcd203040_compound15.5968.5968.4 | controllerType=0 controllerNumber=1 scan=5968 | 16.4325 |
| 02981:4 | 10 | K.HYTN[+1971.719]SSQDVTVPC[+57.021]RVPPPPPC[+57.021]C[+57.021]HPR.L | HexNAc(5)Hex(5)Fuc(1) | C[+57]*3, N[+1972] | 1233.2771 | 4 | 4930.0867 | 4930.0745 | 0 | 2.5 | 89 | Specific | 562.44 | 507.07 | 507.07 | 9.76 | 62 | >sp\|P01877\|IGHA2_HUMAN Immunoglobulin heavy constant alpha 2 OS=Homo sapiens OX=9606 GN=IGHA2 PE=1 SV=4 | 576 | 20201117_08512_1ng_glyco_hcd203040_compound15.5966.5966.4 | controllerType=0 controllerNumber=1 scan=5966 | 16.4268 |
| 03696:5 | 10 | K.HYTN[+2262.814]SSQDVTVPC[+57.021]RVPPPPPC[+57.021]C[+57.021]HPR.L | HexNAc(5)Hex(5)Fuc(1)NeuAc(1) | C[+57]*3, N[+2263] | 1045.0416 | 5 | 5221.179 | 5221.1699 | 0 | 1.7 | 89 | Specific | 621.91 | 388.41 | 388.41 | 12.58 | 62 | >sp\|P01877\|IGHA2_HUMAN Immunoglobulin heavy constant alpha 2 OS=Homo sapiens OX=9606 GN=IGHA2 PE=1 SV=4 | 576 | 20201117_08512_1ng_glyco_hcd203040_compound15.6708.6708.5 | controllerType=0 controllerNumber=1 scan=6708 | 17.7231 |
| 03695:4 | 10 | K.HYTN[+2262.814]SSQDVTVPC[+57.021]RVPPPPPC[+57.021]C[+57.021]HPR.L | HexNAc(5)Hex(5)Fuc(1)NeuAc(1) | C[+57]*3, N[+2263] | 1306.0516 | 4 | 5221.1847 | 5221.1699 | 0 | 2.8 | 89 | Specific | 654.1 | 479.02 | 479.02 | 13.01 | 62 | >sp\|P01877\|IGHA2_HUMAN Immunoglobulin heavy constant alpha 2 OS=Homo sapiens OX=9606 GN=IGHA2 PE=1 SV=4 | 576 | 20201117_08512_1ng_glyco_hcd203040_compound15.6707.6707.4 | controllerType=0 controllerNumber=1 scan=6707 | 17.7215 |
| 03094:5 | 10 | K.HYTN[+1444.534]SSQDVTVPC[+57.021]RVPPPPPC[+57.021]C[+57.021]HPR.L | HexNAc(4)Hex(3)Fuc(1) | C[+57]*3, N[+1445] | 881.3855 | 5 | 4402.8985 | 4402.8895 | 0 | 2 | 89 | Specific | 655.04 | 581.54 | 581.54 | 13.48 | 62 | >sp\|P01877\|IGHA2_HUMAN Immunoglobulin heavy constant alpha 2 OS=Homo sapiens OX=9606 GN=IGHA2 PE=1 SV=4 | 576 | 20201117_08512_1ng_glyco_hcd203040_compound15.6083.6083.5 | controllerType=0 controllerNumber=1 scan=6083 | 16.6373 |
| 03697:4 | 10 | K.HYTN[+2262.814]SSQDVTVPC[+57.021]RVPPPPPC[+57.021]C[+57.021]HPR.L | HexNAc(5)Hex(5)Fuc(1)NeuAc(1) | C[+57]*3, N[+2263] | 1306.0516 | 4 | 5221.1847 | 5221.1699 | 0 | 2.8 | 89 | Specific | 657.78 | 394.85 | 394.85 | 13.01 | 62 | >sp\|P01877\|IGHA2_HUMAN Immunoglobulin heavy constant alpha 2 OS=Homo sapiens OX=9606 GN=IGHA2 PE=1 SV=4 | 576 | 20201117_08512_1ng_glyco_hcd203040_compound15.6709.6709.4 | controllerType=0 controllerNumber=1 scan=6709 | 17.7258 |
| 02991:5 | 10 | K.HYTN[+1647.613]SSQDVTVPC[+57.021]RVPPPPPC[+57.021]C[+57.021]HPR.L | HexNAc(5)Hex(3)Fuc(1) | C[+57]*3, N[+1648] | 922.0017 | 5 | 4605.9795 | 4605.9689 | 0 | 2.3 | 89 | Specific | 661.43 | 452.06 | 452.06 | 13.48 | 62 | >sp\|P01877\|IGHA2_HUMAN Immunoglobulin heavy constant alpha 2 OS=Homo sapiens OX=9606 GN=IGHA2 PE=1 SV=4 | 576 | 20201117_08512_1ng_glyco_hcd203040_compound15.5976.5976.5 | controllerType=0 controllerNumber=1 scan=5976 | 16.4448 |
| 02993:5 | 10 | K.HYTN[+1647.613]SSQDVTVPC[+57.021]RVPPPPPC[+57.021]C[+57.021]HPR.L | HexNAc(5)Hex(3)Fuc(1) | C[+57]*3, N[+1648] | 922.0017 | 5 | 4605.9795 | 4605.9689 | 0 | 2.3 | 89 | Specific | 716.23 | 547.67 | 547.67 | 14.48 | 62 | >sp\|P01877\|IGHA2_HUMAN Immunoglobulin heavy constant alpha 2 OS=Homo sapiens OX=9606 GN=IGHA2 PE=1 SV=4 | 576 | 20201117_08512_1ng_glyco_hcd203040_compound15.5978.5978.5 | controllerType=0 controllerNumber=1 scan=5978 | 16.4508 |
| 02977:5 | 10 | K.HYTN[+1971.719]SSQDVTVPC[+57.021]RVPPPPPC[+57.021]C[+57.021]HPR.L | HexNAc(5)Hex(5)Fuc(1) | C[+57]*3, N[+1972] | 986.823 | 5 | 4930.0857 | 4930.0745 | 0 | 2.3 | 89 | Specific | 740.77 | 569.2 | 569.2 | 14.48 | 62 | >sp\|P01877\|IGHA2_HUMAN Immunoglobulin heavy constant alpha 2 OS=Homo sapiens OX=9606 GN=IGHA2 PE=1 SV=4 | 576 | 20201117_08512_1ng_glyco_hcd203040_compound15.5962.5962.5 | controllerType=0 controllerNumber=1 scan=5962 | 16.4188 |
| 03698:5 | 10 | K.HYTN[+2262.814]SSQDVTVPC[+57.021]RVPPPPPC[+57.021]C[+57.021]HPR.L | HexNAc(5)Hex(5)Fuc(1)NeuAc(1) | C[+57]*3, N[+2263] | 1045.0416 | 5 | 5221.179 | 5221.1699 | 0 | 1.7 | 89 | Specific | 801.83 | 603.08 | 603.08 | 16.58 | 62 | >sp\|P01877\|IGHA2_HUMAN Immunoglobulin heavy constant alpha 2 OS=Homo sapiens OX=9606 GN=IGHA2 PE=1 SV=4 | 576 | 20201117_08512_1ng_glyco_hcd203040_compound15.6710.6710.5 | controllerType=0 controllerNumber=1 scan=6710 | 17.7279 |
| 03033:4 | 10 | K.HYTN[+1444.534]SSQDVTVPC[+57.021]RVPPPPPC[+57.021]C[+57.021]HPR.L | HexNAc(4)Hex(3)Fuc(1) | C[+57]*3, N[+1445] | 1101.4798 | 4 | 4402.8973 | 4402.8895 | 0 | 1.8 | 89 | Specific | 847.34 | 634.41 | 634.41 | 16.11 | 62 | >sp\|P01877\|IGHA2_HUMAN Immunoglobulin heavy constant alpha 2 OS=Homo sapiens OX=9606 GN=IGHA2 PE=1 SV=4 | 576 | 20201117_08512_1ng_glyco_hcd203040_compound15.6020.6020.4 | controllerType=0 controllerNumber=1 scan=6020 | 16.5286 |
| 02979:5 | 10 | K.HYTN[+1971.719]SSQDVTVPC[+57.021]RVPPPPPC[+57.021]C[+57.021]HPR.L | HexNAc(5)Hex(5)Fuc(1) | C[+57]*3, N[+1972] | 986.823 | 5 | 4930.0857 | 4930.0745 | 0 | 2.3 | 89 | Specific | 852.07 | 608.87 | 608.87 | 17.48 | 62 | >sp\|P01877\|IGHA2_HUMAN Immunoglobulin heavy constant alpha 2 OS=Homo sapiens OX=9606 GN=IGHA2 PE=1 SV=4 | 576 | 20201117_08512_1ng_glyco_hcd203040_compound15.5964.5964.5 | controllerType=0 controllerNumber=1 scan=5964 | 16.4234 |
| 03035:4 | 10 | K.HYTN[+1444.534]SSQDVTVPC[+57.021]RVPPPPPC[+57.021]C[+57.021]HPR.L | HexNAc(4)Hex(3)Fuc(1) | C[+57]*3, N[+1445] | 1101.4798 | 4 | 4402.8973 | 4402.8895 | 0 | 1.8 | 89 | Specific | 875.97 | 571.48 | 571.48 | 17.11 | 62 | >sp\|P01877\|IGHA2_HUMAN Immunoglobulin heavy constant alpha 2 OS=Homo sapiens OX=9606 GN=IGHA2 PE=1 SV=4 | 576 | 20201117_08512_1ng_glyco_hcd203040_compound15.6022.6022.4 | controllerType=0 controllerNumber=1 scan=6022 | 16.533 |
| 02982:4 | 10 | K.HYTN[+1809.666]SSQDVTVPC[+57.021]RVPPPPPC[+57.021]C[+57.021]HPR.L | HexNAc(5)Hex(4)Fuc(1) | C[+57]*3, N[+1810] | 1192.7647 | 4 | 4768.0369 | 4768.0217 | 0 | 3.2 | 89 | Specific | 887.7 | 684.45 | 684.45 | 16.92 | 62 | >sp\|P01877\|IGHA2_HUMAN Immunoglobulin heavy constant alpha 2 OS=Homo sapiens OX=9606 GN=IGHA2 PE=1 SV=4 | 576 | 20201117_08512_1ng_glyco_hcd203040_compound15.5967.5967.4 | controllerType=0 controllerNumber=1 scan=5967 | 16.4306 |
| 02980:4 | 10 | K.HYTN[+1809.666]SSQDVTVPC[+57.021]RVPPPPPC[+57.021]C[+57.021]HPR.L | HexNAc(5)Hex(4)Fuc(1) | C[+57]*3, N[+1810] | 1192.7647 | 4 | 4768.0369 | 4768.0217 | 0 | 3.2 | 89 | Specific | 901.88 | 651.69 | 651.69 | 17.92 | 62 | >sp\|P01877\|IGHA2_HUMAN Immunoglobulin heavy constant alpha 2 OS=Homo sapiens OX=9606 GN=IGHA2 PE=1 SV=4 | 576 | 20201117_08512_1ng_glyco_hcd203040_compound15.5965.5965.4 | controllerType=0 controllerNumber=1 scan=5965 | 16.4253 |
| 03001:4 | 10 | K.HYTN[+1647.613]SSQDVTVPC[+57.021]RVPPPPPC[+57.021]C[+57.021]HPR.L | HexNAc(5)Hex(3)Fuc(1) | C[+57]*3, N[+1648] | 1152.2499 | 4 | 4605.9779 | 4605.9689 | 0 | 2 | 89 | Specific | 905.47 | 664.17 | 664.17 | 18.01 | 62 | >sp\|P01877\|IGHA2_HUMAN Immunoglobulin heavy constant alpha 2 OS=Homo sapiens OX=9606 GN=IGHA2 PE=1 SV=4 | 576 | 20201117_08512_1ng_glyco_hcd203040_compound15.5987.5987.4 | controllerType=0 controllerNumber=1 scan=5987 | [illegible] |
| 02999:4 | 10 | K.HYTN[+1647.613]SSQDVTVPC[+57.021]RVPPPPPC[+57.021]C[+57.021]HPR.L | HexNAc(5)Hex(3)Fuc(1) | C[+57]*3, N[+1648] | 1152.2499 | 4 | 4605.9779 | 4605.9689 | 0 | 2 | 89 | Specific | 945.65 | 761.76 | 761.76 | 18.01 | 62 | >sp\|P01877\|IGHA2_HUMAN Immunoglobulin heavy constant alpha 2 OS=Homo sapiens OX=9606 GN=IGHA2 PE=1 SV=4 | 576 | 20201117_08512_1ng_glyco_hcd203040_compound15.5985.5985.4 | controllerType=0 controllerNumber=1 scan=5985 | [illegible] |
| 04470:3 | 10 | K.TPLTAN[+2059.735]ITK.S | HexNAc(4)Hex(5)Fuc(1)NeuAc(1) | N[+2060] | 1006.7705 | 3 | 3018.2969 | 3018.2917 | 0 | 1.7 | 200 | Specific | 240.41 | 181.05 | 181.05 | 5.16 | 62 | >sp\|P01877\|IGHA2_HUMAN Immunoglobulin heavy constant alpha 2 OS=Homo sapiens OX=9606 GN=IGHA2 PE=1 SV=4 | 576 | 20201117_08512_1ng_glyco_hcd203040_compound15.7509.7509.3 | controllerType=0 controllerNumber=1 scan=7509 | [illegible] |
| 02672:2 | 10 | K.TPLTAN[+1971.719]ITK.S | HexNAc(5)Hex(5)Fuc(1) | N[+1972] | 1465.6459 | 2 | 2930.2846 | 2930.2757 | 0 | 3 | 200 | Specific | 337.58 | 160.49 | 160.49 | 5.6 | 62 | >sp\|P01877\|IGHA2_HUMAN Immunoglobulin heavy constant alpha 2 OS=Homo sapiens OX=9606 GN=IGHA2 PE=1 SV=4 | 576 | 20201117_08512_1ng_glyco_hcd203040_compound15.5646.5646.2 | controllerType=0 controllerNumber=1 scan=5646 | 15.8338 |
| 04477:3 | 10 | K.TPLTAN[+2262.814]ITK.S | HexNAc(5)Hex(5)Fuc(1)NeuAc(1) | N[+2263] | 1074.464 | 3 | 3221.3775 | 3221.3711 | 0 | 2 | 200 | Specific | 347.18 | 125.26 | 125.26 | 5.2 | 62 | >sp\|P01877\|IGHA2_HUMAN Immunoglobulin heavy constant alpha 2 OS=Homo sapiens OX=9606 GN=IGHA2 PE=1 SV=4 | 576 | 20201117_08512_1ng_glyco_hcd203040_compound15.7517.7517.3 | controllerType=0 controllerNumber=1 scan=7517 | 19.0669 |
| 02644:3 | 10 | K.TPLTAN[+1971.719]ITK.S | HexNAc(5)Hex(5)Fuc(1) | N[+1972] | 977.4326 | 3 | 2930.2834 | 2930.2757 | 0 | 2.6 | 200 | Specific | 386.54 | 128.25 | 128.25 | 5.28 | 62 | >sp\|P01877\|IGHA2_HUMAN Immunoglobulin heavy constant alpha 2 OS=Homo sapiens OX=9606 GN=IGHA2 PE=1 SV=4 | 576 | 20201117_08512_1ng_glyco_hcd203040_compound15.5617.5617.3 | controllerType=0 controllerNumber=1 scan=5617 | 15.7806 |
| 02738:2 | 10 | K.TPLTAN[+1606.587]ITK.S | HexNAc(4)Hex(4)Fuc(1) | N[+1607] | 1283.079 | 2 | 2565.1508 | 2565.1435 | 0 | 2.8 | 200 | Specific | 427.96 | 197.09 | 197.09 | 6.22 | 62 | >sp\|P01877\|IGHA2_HUMAN Immunoglobulin heavy constant alpha 2 OS=Homo sapiens OX=9606 GN=IGHA2 PE=1 SV=4 | 576 | 20201117_08512_1ng_glyco_hcd203040_compound15.5714.5714.2 | controllerType=0 controllerNumber=1 scan=5714 | 15.975 |
| 02773:3 | 10 | K.TPLTAN[+1647.613]ITK.S | HexNAc(5)Hex(3)Fuc(1) | N[+1648] | 869.3979 | 3 | 2606.179 | 2606.17 | 0 | 3.5 | 200 | Specific | 432.64 | 159.91 | 159.91 | 5.71 | 62 | >sp\|P01877\|IGHA2_HUMAN Immunoglobulin heavy constant alpha 2 OS=Homo sapiens OX=9606 GN=IGHA2 PE=1 SV=4 | 576 | 20201117_08512_1ng_glyco_hcd203040_compound15.5751.5751.3 | controllerType=0 controllerNumber=1 scan=5751 | 16.0418 |
| 02740:2 | 10 | K.TPLTAN[+1606.587]ITK.S | HexNAc(4)Hex(4)Fuc(1) | N[+1607] | 1283.079 | 2 | 2565.1508 | 2565.1435 | 0 | 2.8 | 200 | Specific | 443.58 | 291.09 | 291.09 | 7.12 | 62 | >sp\|P01877\|IGHA2_HUMAN Immunoglobulin heavy constant alpha 2 OS=Homo sapiens OX=9606 GN=IGHA2 PE=1 SV=4 | 576 | 20201117_08512_1ng_glyco_hcd203040_compound15.5716.5716.2 | controllerType=0 controllerNumber=1 scan=5716 | 15.9786 |
| 02779:2 | 10 | K.TPLTAN[+1501.555]ITK.S | HexNAc(5)Hex(3) | N[+1502] | 1230.5644 | 2 | 2460.1215 | 2460.1121 | 0 | 3.8 | 200 | Specific | 455.17 | 284.62 | 284.62 | 7.03 | 62 | >sp\|P01877\|IGHA2_HUMAN Immunoglobulin heavy constant alpha 2 OS=Homo sapiens OX=9606 GN=IGHA2 PE=1 SV=4 | 576 | 20201117_08512_1ng_glyco_hcd203040_compound15.5757.5757.2 | controllerType=0 controllerNumber=1 scan=5757 | 16.0568 |
| 02674:2 | 10 | K.TPLTAN[+1971.719]ITK.S | HexNAc(5)Hex(5)Fuc(1) | N[+1972] | 1465.6459 | 2 | 2930.2846 | 2930.2757 | 0 | 3 | 200 | Specific | 478.5 | 301.15 | 301.15 | 7.2 | 62 | >sp\|P01877\|IGHA2_HUMAN Immunoglobulin heavy constant alpha 2 OS=Homo sapiens OX=9606 GN=IGHA2 PE=1 SV=4 | 576 | 20201117_08512_1ng_glyco_hcd203040_compound15.5648.5648.2 | controllerType=0 controllerNumber=1 scan=5648 | 15.8374 |
| 04479:3 | 10 | K.TPLTAN[+2262.814]ITK.S | HexNAc(5)Hex(5)Fuc(1)NeuAc(1) | N[+2263] | 1074.464 | 3 | 3221.3775 | 3221.3711 | 0 | 2 | 200 | Specific | 497.75 | 267.51 | 267.51 | 6.82 | 62 | >sp\|P01877\|IGHA2_HUMAN Immunoglobulin heavy constant alpha 2 OS=Homo sapiens OX=9606 GN=IGHA2 PE=1 SV=4 | 576 | 20201117_08512_1ng_glyco_hcd203040_compound15.7519.7519.3 | controllerType=0 controllerNumber=1 scan=7519 | 19.0713 |
| 02781:2 | 10 | K.TPLTAN[+1501.555]ITK.S | HexNAc(5)Hex(3) | N[+1502] | 1230.5644 | 2 | 2460.1215 | 2460.1121 | 0 | 3.8 | 200 | Specific | 500.76 | 301.23 | 301.23 | 7.22 | 62 | >sp\|P01877\|IGHA2_HUMAN Immunoglobulin heavy constant alpha 2 OS=Homo sapiens OX=9606 GN=IGHA2 PE=1 SV=4 | 576 | 20201117_08512_1ng_glyco_hcd203040_compound15.5759.5759.2 | controllerType=0 controllerNumber=1 scan=5759 | 16.0604 |
| 02702:3 | 10 | K.TPLTAN[+1809.666]ITK.S | HexNAc(5)Hex(4)Fuc(1) | N[+1810] | 923.4144 | 3 | 2768.2286 | 2768.2228 | 0 | 2.1 | 200 | Specific | 504.02 | 218.12 | 218.12 | 6.57 | 62 | >sp\|P01877\|IGHA2_HUMAN Immunoglobulin heavy constant alpha 2 OS=Homo sapiens OX=9606 GN=IGHA2 PE=1 SV=4 | 576 | 20201117_08512_1ng_glyco_hcd203040_compound15.5677.5677.3 | controllerType=0 controllerNumber=1 scan=5677 | 15.888 |
| 02646:3 | 10 | K.TPLTAN[+1971.719]ITK.S | HexNAc(5)Hex(5)Fuc(1) | N[+1972] | 977.4326 | 3 | 2930.2834 | 2930.2757 | 0 | 2.6 | 200 | Specific | 509.59 | 255.36 | 255.36 | 6.76 | 62 | >sp\|P01877\|IGHA2_HUMAN Immunoglobulin heavy constant alpha 2 OS=Homo sapiens OX=9606 GN=IGHA2 PE=1 SV=4 | 576 | 20201117_08512_1ng_glyco_hcd203040_compound15.5619.5619.3 | controllerType=0 controllerNumber=1 scan=5619 | 15.7849 |
| 02704:3 | 10 | K.TPLTAN[+1809.666]ITK.S | HexNAc(5)Hex(4)Fuc(1) | N[+1810] | 923.4144 | 3 | 2768.2286 | 2768.2228 | 0 | 2.1 | 200 | Specific | 517.75 | 263.09 | 263.09 | 6.92 | 62 | >sp\|P01877\|IGHA2_HUMAN Immunoglobulin heavy constant alpha 2 OS=Homo sapiens OX=9606 GN=IGHA2 PE=1 SV=4 | 576 | 20201117_08512_1ng_glyco_hcd203040_compound15.5679.5679.3 | controllerType=0 controllerNumber=1 scan=5679 | 15.8928 |
| 02775:3 | 10 | K.TPLTAN[+1647.613]ITK.S | HexNAc(5)Hex(3)Fuc(1) | N[+1648] | 869.3979 | 3 | 2606.179 | 2606.17 | 0 | 3.5 | 200 | Specific | 559.92 | 265.34 | 265.34 | 7.19 | 62 | >sp\|P01877\|IGHA2_HUMAN Immunoglobulin heavy constant alpha 2 OS=Homo sapiens OX=9606 GN=IGHA2 PE=1 SV=4 | 576 | 20201117_08512_1ng_glyco_hcd203040_compound15.5753.5753.3 | controllerType=0 controllerNumber=1 scan=5753 | 16.0505 |
| 02741:2 | 10 | K.TPLTAN[+1444.534]ITK.S | HexNAc(4)Hex(3)Fuc(1) | N[+1445] | 1202.0535 | 2 | 2403.0997 | 2403.0906 | 0 | 3.8 | 200 | Specific | 578.52 | 334.13 | 334.13 | 8.99 | 62 | >sp\|P01877\|IGHA2_HUMAN Immunoglobulin heavy constant alpha 2 OS=Homo sapiens OX=9606 GN=IGHA2 PE=1 SV=4 | 576 | 20201117_08512_1ng_glyco_hcd203040_compound15.5717.5717.2 | controllerType=0 controllerNumber=1 scan=5717 | 15.9806 |
| 02743:2 | 10 | K.TPLTAN[+1444.534]ITK.S | HexNAc(4)Hex(3)Fuc(1) | N[+1445] | 1202.0535 | 2 | 2403.0997 | 2403.0906 | 0 | 3.8 | 200 | Specific | 602.21 | 340.13 | 340.13 | 9.99 | 62 | >sp\|P01877\|IGHA2_HUMAN Immunoglobulin heavy constant alpha 2 OS=Homo sapiens OX=9606 GN=IGHA2 PE=1 SV=4 | 576 | 20201117_08512_1ng_glyco_hcd203040_compound15.5719.5719.2 | controllerType=0 controllerNumber=1 scan=5719 | 15.9845 |
| 02774:2 | 10 | K.TPLTAN[+1647.613]ITK.S | HexNAc(5)Hex(3)Fuc(1) | N[+1648] | 1303.5933 | 2 | 2606.1793 | 2606.17 | 0 | 3.6 | 200 | Specific | 607.38 | 326.19 | 326.19 | 9.97 | 62 | >sp\|P01877\|IGHA2_HUMAN Immunoglobulin heavy constant alpha 2 OS=Homo sapiens OX=9606 GN=IGHA2 PE=1 SV=4 | 576 | 20201117_08512_1ng_glyco_hcd203040_compound15.5752.5752.2 | controllerType=0 controllerNumber=1 scan=5752 | 16.0475 |
| 02772:2 | 10 | K.TPLTAN[+1647.613]ITK.S | HexNAc(5)Hex(3)Fuc(1) | N[+1648] | 1303.5933 | 2 | 2606.1793 | 2606.17 | 0 | 3.6 | 200 | Specific | 610.55 | 352.45 | 352.45 | 9.99 | 62 | >sp\|P01877\|IGHA2_HUMAN Immunoglobulin heavy constant alpha 2 OS=Homo sapiens OX=9606 GN=IGHA2 PE=1 SV=4 | 576 | 20201117_08512_1ng_glyco_hcd203040_compound15.5750.5750.2 | controllerType=0 controllerNumber=1 scan=5750 | 16.0404 |
| 02717:2 | 10 | K.TPLTAN[+1809.666]ITK.S | HexNAc(5)Hex(4)Fuc(1) | N[+1810] | 1384.6187 | 2 | 2768.2301 | 2768.2228 | 0 | 2.6 | 200 | Specific | 675.39 | 328.53 | 328.53 | 11.06 | 62 | >sp\|P01877\|IGHA2_HUMAN Immunoglobulin heavy constant alpha 2 OS=Homo sapiens OX=9606 GN=IGHA2 PE=1 SV=4 | 576 | 20201117_08512_1ng_glyco_hcd203040_compound15.5692.5692.2 | controllerType=0 controllerNumber=1 scan=5692 | 15.9324 |
| 02715:2 | 10 | K.TPLTAN[+1809.666]ITK.S | HexNAc(5)Hex(4)Fuc(1) | N[+1810] | 1384.6187 | 2 | 2768.2301 | 2768.2228 | 0 | 2.6 | 200 | Specific | 676.26 | 352.21 | 352.21 | 11.08 | 62 | >sp\|P01877\|IGHA2_HUMAN Immunoglobulin heavy constant alpha 2 OS=Homo sapiens OX=9606 GN=IGHA2 PE=1 SV=4 | 576 | 20201117_08512_1ng_glyco_hcd203040_compound15.5690.5690.2 | controllerType=0 controllerNumber=1 scan=5690 | 15.9233 |
| 09733:3 | 10 | R.MAGKPTHIN[+1540.529]VSVVMAEADGTC[+57.021]Y.- | HexNAc(2)Hex(7) | C[+57], N[+1541] | 1297.8782 | 3 | 3891.6201 | 3891.6114 | 0 | 2.3 | 319 | CRagged | 205.35 | 205.35 | 127.8 | 3.9 | 62 | >sp\|P01877\|IGHA2_HUMAN Immunoglobulin heavy constant alpha 2 OS=Homo sapiens OX=9606 GN=IGHA2 PE=1 SV=4 | 576 | 20201117_08512_1ng_glyco_hcd203040_compound15.12964.12964.3 | controllerType=0 controllerNumber=1 scan=12964 | 28.6202 |
| 11331:3 | 10 | R.MAGKPTHIN[+1913.677]VSVVMAEADGTC[+57.021]Y.- | HexNAc(4)Hex(5)NeuAc(1) | C[+57], N[+1914] | 1422.2664 | 3 | 4264.7847 | 4264.7599 | 0 | 5.8 | 319 | CRagged | 346.16 | 346.16 | 346.16 | 6.71 | 62 | >sp\|P01877\|IGHA2_HUMAN Immunoglobulin heavy constant alpha 2 OS=Homo sapiens OX=9606 GN=IGHA2 PE=1 SV=4 | 576 | 20201117_08512_1ng_glyco_hcd203040_compound15.14622.14622.3 | controllerType=0 controllerNumber=1 scan=14622 | 31.8163 |
| 11329:3 | 10 | R.MAGKPTHIN[+1913.677]VSVVMAEADGTC[+57.021]Y.- | HexNAc(4)Hex(5)NeuAc(1) | C[+57], N[+1914] | 1422.2664 | 3 | 4264.7847 | 4264.7599 | 0 | 5.8 | 319 | CRagged | 353.83 | 352.51 | 352.51 | 6.71 | 62 | >sp\|P01877\|IGHA2_HUMAN Immunoglobulin heavy constant alpha 2 OS=Homo sapiens OX=9606 GN=IGHA2 PE=1 SV=4 | 576 | 20201117_08512_1ng_glyco_hcd203040_compound15.14620.14620.3 | controllerType=0 controllerNumber=1 scan=14620 | 31.8123 |
| 09889:3 | 10 | R.MAGKPTHIN[+1647.613]VSVVMAEADGTC[+57.021]Y.- | HexNAc(5)Hex(3)Fuc(1) | C[+57], N[+1648] | 1333.5735 | 3 | 3998.7061 | 3998.6961 | 0 | 2.5 | 319 | CRagged | 473.12 | 473.12 | 473.12 | 9.94 | 62 | >sp\|P01877\|IGHA2_HUMAN Immunoglobulin heavy constant alpha 2 OS=Homo sapiens OX=9606 GN=IGHA2 PE=1 SV=4 | 576 | 20201117_08512_1ng_glyco_hcd203040_compound15.13126.13126.3 | controllerType=0 controllerNumber=1 scan=13126 | 28.9674 |
| 11411:3 | 10 | R.MAGKPTHIN[+1710.598]VSVVMAEADGTC[+57.021]Y.- | HexNAc(3)Hex(5)NeuAc(1) | C[+57], N[+1711] | 1354.5672 | 3 | 4061.687 | 4061.6805 | 0 | 1.6 | 319 | CRagged | 504.86 | 435.68 | 314.23 | 10.16 | 62 | >sp\|P01877\|IGHA2_HUMAN Immunoglobulin heavy constant alpha 2 OS=Homo sapiens OX=9606 GN=IGHA2 PE=1 SV=4 | 576 | 20201117_08512_1ng_glyco_hcd203040_compound15.14705.14705.3 | controllerType=0 controllerNumber=1 scan=14705 | 31.9699 |
| 09890:3 | 10 | R.MAGKPTHIN[+1647.613]VSVVMAEADGTC[+57.021]Y.- | HexNAc(5)Hex(3)Fuc(1) | C[+57], N[+1648] | 1333.5735 | 3 | 3998.7061 | 3998.6961 | 0 | 2.5 | 319 | CRagged | 589.8 | 395.31 | 395.31 | 11.76 | 62 | >sp\|P01877\|IGHA2_HUMAN Immunoglobulin heavy constant alpha 2 OS=Homo sapiens OX=9606 GN=IGHA2 PE=1 SV=4 | 576 | 20201117_08512_1ng_glyco_hcd203040_compound15.13127.13127.3 | controllerType=0 controllerNumber=1 scan=13127 | 28.9708 |
| 11413:3 | 10 | R.MAGKPTHIN[+1710.598]VSVVMAEADGTC[+57.021]Y.- | HexNAc(3)Hex(5)NeuAc(1) | C[+57], N[+1711] | 1354.5672 | 3 | 4061.687 | 4061.6805 | 0 | 1.6 | 319 | CRagged | 637.41 | 580.25 | 368.33 | 12.86 | 62 | >sp\|P01877\|IGHA2_HUMAN Immunoglobulin heavy constant alpha 2 OS=Homo sapiens OX=9606 GN=IGHA2 PE=1 SV=4 | 576 | 20201117_08512_1ng_glyco_hcd203040_compound15.14707.14707.3 | controllerType=0 controllerNumber=1 scan=14707 | 31.9774 |
| 09797:3 | 10 | R.MAGKPTHIN[+1809.666]VSVVMAEADGTC[+57.021]Y.- | HexNAc(5)Hex(4)Fuc(1) | C[+57], N[+1810] | 1387.5933 | 3 | 4160.7654 | 4160.7489 | 0 | 4 | 319 | CRagged | 686.44 | 686.44 | 415.4 | 13.67 | 62 | >sp\|P01877\|IGHA2_HUMAN Immunoglobulin heavy constant alpha 2 OS=Homo sapiens OX=9606 GN=IGHA2 PE=1 SV=4 | 576 | 20201117_08512_1ng_glyco_hcd203040_compound15.13030.13030.3 | controllerType=0 controllerNumber=1 scan=13030 | 28.7545 |
| 09872:3 | 10 | R.MAGKPTHIN[+1501.555]VSVVMAEADGTC[+57.021]Y.- | HexNAc(5)Hex(3) | C[+57], N[+1502] | 1284.8881 | 3 | 3852.6498 | 3852.6382 | 0 | 3 | 319 | CRagged | 692.39 | 662.86 | 662.86 | 13.76 | 62 | >sp\|P01877\|IGHA2_HUMAN Immunoglobulin heavy constant alpha 2 OS=Homo sapiens OX=9606 GN=IGHA2 PE=1 SV=4 | 576 | 20201117_08512_1ng_glyco_hcd203040_compound15.13109.13109.3 | controllerType=0 controllerNumber=1 scan=13109 | 28.9345 |
| 09870:3 | 10 | R.MAGKPTHIN[+1501.555]VSVVMAEADGTC[+57.021]Y.- | HexNAc(5)Hex(3) | C[+57], N[+1502] | 1284.8881 | 3 | 3852.6498 | 3852.6382 | 0 | 3 | 319 | CRagged | 712.78 | 615.88 | 615.88 | 14.76 | 62 | >sp\|P01877\|IGHA2_HUMAN Immunoglobulin heavy constant alpha 2 OS=Homo sapiens OX=9606 GN=IGHA2 PE=1 SV=4 | 576 | 20201117_08512_1ng_glyco_hcd203040_compound15.13107.13107.3 | controllerType=0 controllerNumber=1 scan=13107 | 28.9304 |
| 09786:3 | 10 | R.MAGKPTHIN[+1663.608]VSVVMAEADGTC[+57.021]Y.- | HexNAc(5)Hex(4) | C[+57], N[+1664] | 1338.9068 | 3 | 4014.7058 | 4014.691 | 0 | 3.7 | 319 | CRagged | 741.16 | 689.95 | 411.75 | 14.67 | 62 | >sp\|P01877\|IGHA2_HUMAN Immunoglobulin heavy constant alpha 2 OS=Homo sapiens OX=9606 GN=IGHA2 PE=1 SV=4 | 576 | 20201117_08512_1ng_glyco_hcd203040_compound15.13019.13019.3 | controllerType=0 controllerNumber=1 scan=13019 | 28.7363 |
| 09799:3 | 10 | R.MAGKPTHIN[+1809.666]VSVVMAEADGTC[+57.021]Y.- | HexNAc(5)Hex(4)Fuc(1) | C[+57], N[+1810] | 1387.5933 | 3 | 4160.7654 | 4160.7489 | 0 | 4 | 319 | CRagged | 771.61 | 705.82 | 490.61 | 15.67 | 62 | >sp\|P01877\|IGHA2_HUMAN Immunoglobulin heavy constant alpha 2 OS=Homo sapiens OX=9606 GN=IGHA2 PE=1 SV=4 | 576 | 20201117_08512_1ng_glyco_hcd203040_compound15.13032.13032.3 | controllerType=0 controllerNumber=1 scan=13032 | 28.7588 |
| 09784:3 | 10 | R.MAGKPTHIN[+1663.608]VSVVMAEADGTC[+57.021]Y.- | HexNAc(5)Hex(4) | C[+57], N[+1664] | 1338.9068 | 3 | 4014.7058 | 4014.691 | 0 | 3.7 | 319 | CRagged | 815.05 | 815.05 | 474.14 | 16.67 | 62 | >sp\|P01877\|IGHA2_HUMAN Immunoglobulin heavy constant alpha 2 OS=Homo sapiens OX=9606 GN=IGHA2 PE=1 SV=4 | 576 | 20201117_08512_1ng_glyco_hcd203040_compound15.13017.13017.3 | controllerType=0 controllerNumber=1 scan=13017 | 28.7295 |
| 09759:3 | 10 | R.MAGKPTHIN[+1378.476]VSVVMAEADGTC[+57.021]Y.- | HexNAc(2)Hex(6) | C[+57], N[+1378] | 1243.8609 | 3 | 3729.5681 | 3729.5585 | 0 | 2.6 | 319 | CRagged | 924.64 | 828.1 | 544.18 | 18.76 | 62 | >sp\|P01877\|IGHA2_HUMAN Immunoglobulin heavy constant alpha 2 OS=Homo sapiens OX=9606 GN=IGHA2 PE=1 SV=4 | 576 | 20201117_08512_1ng_glyco_hcd203040_compound15.12991.12991.3 | controllerType=0 controllerNumber=1 scan=12991 | 28.6778 |
| 09809:3 | 10 | R.MAGKPTHIN[+1216.423]VSVVMAEADGTC[+57.021]Y.- | HexNAc(2)Hex(5) | C[+57], N[+1216] | 1189.8423 | 3 | 3567.5125 | 3567.5057 | 0 | 1.9 | 319 | CRagged | 929.54 | 795.86 | 545.98 | 18.86 | 62 | >sp\|P01877\|IGHA2_HUMAN Immunoglobulin heavy constant alpha 2 OS=Homo sapiens OX=9606 GN=IGHA2 PE=1 SV=4 | 576 | 20201117_08512_1ng_glyco_hcd203040_compound15.13043.13043.3 | controllerType=0 controllerNumber=1 scan=13043 | 28.7856 |
| 09807:3 | 10 | R.MAGKPTHIN[+1216.423]VSVVMAEADGTC[+57.021]Y.- | HexNAc(2)Hex(5) | C[+57], N[+1216] | 1189.8423 | 3 | 3567.5125 | 3567.5057 | 0 | 1.9 | 319 | CRagged | 949.74 | 783.27 | 555.78 | 18.86 | 62 | >sp\|P01877\|IGHA2_HUMAN Immunoglobulin heavy constant alpha 2 OS=Homo sapiens OX=9606 GN=IGHA2 PE=1 SV=4 | 576 | 20201117_08512_1ng_glyco_hcd203040_compound15.13041.13041.3 | controllerType=0 controllerNumber=1 scan=13041 | 28.7781 |
| 09757:3 | 10 | R.MAGKPTHIN[+1378.476]VSVVMAEADGTC[+57.021]Y.- | HexNAc(2)Hex(6) | C[+57], N[+1378] | 1243.8609 | 3 | 3729.5681 | 3729.5585 | 0 | 2.6 | 319 | CRagged | 1005.88 | 945.96 | 599.37 | 20.76 | 62 | >sp\|P01877\|IGHA2_HUMAN Immunoglobulin heavy constant alpha 2 OS=Homo sapiens OX=9606 GN=IGHA2 PE=1 SV=4 | 576 | 20201117_08512_1ng_glyco_hcd203040_compound15.12989.12989.3 | controllerType=0 controllerNumber=1 scan=12989 | 28.6699 |
| 09295:3 | 10 | R.M[+15.995]AGKPTHIN[+1378.476]VSVVMAEADGTC[+57.021]Y.- | HexNAc(2)Hex(6) | C[+57], M[+16], N[+1378] | 1249.1906 | 3 | 3745.5573 | 3745.5535 | 0 | 1 | 319 | CRagged | 668.97 | 668.97 | 210.46 | 13.86 | 62 | >sp\|P01877\|IGHA2_HUMAN Immunoglobulin heavy constant alpha 2 OS=Homo sapiens OX=9606 GN=IGHA2 PE=1 SV=4 | 576 | 20201117_08512_1ng_glyco_hcd203040_compound15.12509.12509.3 | controllerType=0 controllerNumber=1 scan=12509 | 27.7807 |
| 09372:3 | 10 | R.M[+15.995]AGKPTHIN[+1216.423]VSVVMAEADGTC[+57.021]Y.- | HexNAc(2)Hex(5) | C[+57], M[+16], N[+1216] | 1195.175 | 3 | 3583.5104 | 3583.5006 | 0 | 2.7 | 319 | CRagged | 672.43 | 662.54 | 249.86 | 13.76 | 62 | >sp\|P01877\|IGHA2_HUMAN Immunoglobulin heavy constant alpha 2 OS=Homo sapiens OX=9606 GN=IGHA2 PE=1 SV=4 | 576 | 20201117_08512_1ng_glyco_hcd203040_compound15.12589.12589.3 | controllerType=0 controllerNumber=1 scan=12589 | 27.9208 |
| 09292:3 | 10 | R.M[+15.995]AGKPTHIN[+1378.476]VSVVMAEADGTC[+57.021]Y.- | HexNAc(2)Hex(6) | C[+57], M[+16], N[+1378] | 1249.1906 | 3 | 3745.5573 | 3745.5535 | 0 | 1 | 319 | CRagged | 673.33 | 637.02 | 289.22 | 13.86 | 62 | >sp\|P01877\|IGHA2_HUMAN Immunoglobulin heavy constant alpha 2 OS=Homo sapiens OX=9606 GN=IGHA2 PE=1 SV=4 | 576 | 20201117_08512_1ng_glyco_hcd203040_compound15.12506.12506.3 | controllerType=0 controllerNumber=1 scan=12506 | 27.7751 |
| 09370:3 | 10 | R.M[+15.995]AGKPTHIN[+1216.423]VSVVMAEADGTC[+57.021]Y.- | HexNAc(2)Hex(5) | C[+57], M[+16], N[+1216] | 1195.175 | 3 | 3583.5104 | 3583.5006 | 0 | 2.7 | 319 | CRagged | 677.24 | 677.24 | 264.29 | 13.76 | 62 | >sp\|P01877\|IGHA2_HUMAN Immunoglobulin heavy constant alpha 2 OS=Homo sapiens OX=9606 GN=IGHA2 PE=1 SV=4 | 576 | 20201117_08512_1ng_glyco_hcd203040_compound15.12587.12587.3 | controllerType=0 controllerNumber=1 scan=12587 | 27.9163 |
| 22857:4 | 13 | K.APDKNVIFSPLSISTALAFLSLGAHN[+2862.021]TTLTEILK.G | HexNAc(5)Hex(6)Fuc(2)NeuAc(2) | N[+2862] | 1612.0061 | 4 | 6445.0026 | 6444.9986 | 0 | 0.6 | 68 | Specific | 279.97 | 279.97 | 279.97 | 5.86 | 42 | >sp\|P01011\|AACT_HUMAN Alpha-1-antichymotrypsin OS=Homo sapiens OX=9606 GN=SERPINA3 PE=1 SV=2 | 1314 | 20201117_08512_1ng_glyco_hcd203040_compound15.26665.26665.4 | controllerType=0 controllerNumber=1 scan=26665 | 57.1661 |
| 19835:4 | 13 | K.APDKNVIFSPLSISTALAFLSLGAHN[+2351.851]TTLTEILK.G | HexNAc(4)Hex(5)Fuc(3)NeuAc(1) | N[+2352] | 1484.4735 | 4 | 5934.8722 | 5934.8289 | 0 | 7.3 | 68 | Specific | 311.33 | 311.33 | 311.33 | 5.86 | 42 | >sp\|P01011\|AACT_HUMAN Alpha-1-antichymotrypsin OS=Homo sapiens OX=9606 GN=SERPINA3 PE=1 SV=2 | 1314 | 20201117_08512_1ng_glyco_hcd203040_compound15.23513.23513.4 | controllerType=0 controllerNumber=1 scan=23513 | 50.6511 |
| 19838:4 | 13 | K.APDKNVIFSPLSISTALAFLSLGAHN[+2351.851]TTLTEILK.G | HexNAc(4)Hex(5)Fuc(3)NeuAc(1) | N[+2352] | 1484.4735 | 4 | 5934.8722 | 5934.8289 | 0 | 7.3 | 68 | Specific | 338.21 | 338.21 | 338.21 | 6.03 | 42 | >sp\|P01011\|AACT_HUMAN Alpha-1-antichymotrypsin OS=Homo sapiens OX=9606 GN=SERPINA3 PE=1 SV=2 | 1314 | 20201117_08512_1ng_glyco_hcd203040_compound15.23516.23516.4 | controllerType=0 controllerNumber=1 scan=23516 | 50.6583 |
| 22855:4 | 13 | K.APDKNVIFSPLSISTALAFLSLGAHN[+2862.021]TTLTEILK.G | HexNAc(5)Hex(6)Fuc(2)NeuAc(2) | N[+2862] | 1612.0061 | 4 | 6445.0026 | 6444.9986 | 0 | 0.6 | 68 | Specific | 380.7 | 380.7 | 380.7 | 8.48 | 42 | >sp\|P01011\|AACT_HUMAN Alpha-1-antichymotrypsin OS=Homo sapiens OX=9606 GN=SERPINA3 PE=1 SV=2 | 1314 | 20201117_08512_1ng_glyco_hcd203040_compound15.26663.26663.4 | controllerType=0 controllerNumber=1 scan=26663 | 57.1626 |
| 19754:4 | 13 | K.APDKNVIFSPLSISTALAFLSLGAHN[+2570.925]TTLTEILK.G | HexNAc(5)Hex(6)Fuc(2)NeuAc(1) | N[+2571] | 1539.2329 | 4 | 6153.9098 | 6153.9032 | 0 | 1.1 | 68 | Specific | 616.41 | 592.62 | 592.62 | 12.32 | 42 | >sp\|P01011\|AACT_HUMAN Alpha-1-antichymotrypsin OS=Homo sapiens OX=9606 GN=SERPINA3 PE=1 SV=2 | 1314 | 20201117_08512_1ng_glyco_hcd203040_compound15.23429.23429.4 | controllerType=0 controllerNumber=1 scan=23429 | 50.4713 |
| 19751:4 | 13 | K.APDKNVIFSPLSISTALAFLSLGAHN[+2570.925]TTLTEILK.G | HexNAc(5)Hex(6)Fuc(2)NeuAc(1) | N[+2571] | 1539.2329 | 4 | 6153.9098 | 6153.9032 | 0 | 1.1 | 68 | Specific | 718.77 | 712.02 | 712.02 | 14.32 | 42 | >sp\|P01011\|AACT_HUMAN Alpha-1-antichymotrypsin OS=Homo sapiens OX=9606 GN=SERPINA3 PE=1 SV=2 | 1314 | 20201117_08512_1ng_glyco_hcd203040_compound15.23426.23426.4 | controllerType=0 controllerNumber=1 scan=23426 | 50.4658 |
| 21152:4 | 13 | K.APDKNVIFSPLSISTALAFLSLGAHN[+2350.830]TTLTEILK.G | HexNAc(4)Hex(5)Fuc(1)NeuAc(2) | N[+2351] | 1484.2141 | 4 | 5933.8346 | 5933.8085 | 0 | 4.4 | 68 | Specific | 760.86 | 760.86 | 760.86 | 14.58 | 42 | >sp\|P01011\|AACT_HUMAN Alpha-1-antichymotrypsin OS=Homo sapiens OX=9606 GN=SERPINA3 PE=1 SV=2 | 1314 | 20201117_08512_1ng_glyco_hcd203040_compound15.24889.24889.4 | controllerType=0 controllerNumber=1 scan=24889 | 53.5497 |
| 19846:4 | 13 | K.APDKNVIFSPLSISTALAFLSLGAHN[+2059.735]TTLTEILK.G | HexNAc(4)Hex(5)Fuc(1)NeuAc(1) | N[+2060] | 1411.4382 | 4 | 5642.7311 | 5642.7131 | 0 | 3.2 | 68 | Specific | 937.55 | 924.09 | 924.09 | 18.13 | 42 | >sp\|P01011\|AACT_HUMAN Alpha-1-antichymotrypsin OS=Homo sapiens OX=9606 GN=SERPINA3 PE=1 SV=2 | 1314 | 20201117_08512_1ng_glyco_hcd203040_compound15.23525.23525.4 | controllerType=0 controllerNumber=1 scan=23525 | 50.6966 |
| 21150:4 | 13 | K.APDKNVIFSPLSISTALAFLSLGAHN[+2350.830]TTLTEILK.G | HexNAc(4)Hex(5)Fuc(1)NeuAc(2) | N[+2351] | 1484.2141 | 4 | 5933.8346 | 5933.8085 | 0 | 4.4 | 68 | Specific | 1050.95 | 790.67 | 790.67 | 20.58 | 42 | >sp\|P01011\|AACT_HUMAN Alpha-1-antichymotrypsin OS=Homo sapiens OX=9606 GN=SERPINA3 PE=1 SV=2 | 1314 | 20201117_08512_1ng_glyco_hcd203040_compound15.24887.24887.4 | controllerType=0 controllerNumber=1 scan=24887 | 53.5443 |
| 19844:4 | 13 | K.APDKNVIFSPLSISTALAFLSLGAHN[+2059.735]TTLTEILK.G | HexNAc(4)Hex(5)Fuc(1)NeuAc(1) | N[+2060] | 1411.4382 | 4 | 5642.7311 | 5642.7131 | 0 | 3.2 | 68 | Specific | 1153.36 | 1138.46 | 1138.46 | 23.13 | 42 | >sp\|P01011\|AACT_HUMAN Alpha-1-antichymotrypsin OS=Homo sapiens OX=9606 GN=SERPINA3 PE=1 SV=2 | 1314 | 20201117_08512_1ng_glyco_hcd203040_compound15.23523.23523.4 | controllerType=0 controllerNumber=1 scan=23523 | 50.6902 |
| 07345:3 | 14 | R.LSPN[+2350.830]ASAEHLELR.W | HexNAc(4)Hex(5)Fuc(1)NeuAc(2) | N[+2351] | 1263.201 | 3 | 3787.5884 | 3787.5796 | 0 | 2.3 | 52 | Specific | 588.83 | 525.8 | 525.8 | 11.09 | 37 | >sp\|Q13410\|BT1A1_HUMAN Butyrophilin subfamily 1 member A1 OS=Homo sapiens OX=9606 GN=BTN1A1 PE=1 SV=3 | 179 | 20201117_08512_1ng_glyco_hcd203040_compound15.10483.10483.3 | controllerType=0 controllerNumber=1 scan=10483 | 24.0697 |
| 07343:3 | 14 | R.LSPN[+2350.830]ASAEHLELR.W | HexNAc(4)Hex(5)Fuc(1)NeuAc(2) | N[+2351] | 1263.201 | 3 | 3787.5884 | 3787.5796 | 0 | 2.3 | 52 | Specific | 601.08 | 502.68 | 502.68 | 12.09 | 37 | >sp\|Q13410\|BT1A1_HUMAN Butyrophilin subfamily 1 member A1 OS=Homo sapiens OX=9606 GN=BTN1A1 PE=1 SV=3 | 179 | 20201117_08512_1ng_glyco_hcd203040_compound15.10481.10481.3 | controllerType=0 controllerNumber=1 scan=10481 | 24.0648 |
| 05350:3 | 14 | R.LSPN[+2205.793]ASAEHLELR.W | HexNAc(4)Hex(5)Fuc(2)NeuAc(1) | N[+2206] | 1214.8543 | 3 | 3642.5484 | 3642.5421 | 0 | 1.7 | 52 | Specific | 610 | 337.74 | 337.74 | 10.39 | 37 | >sp\|Q13410\|BT1A1_HUMAN Butyrophilin subfamily 1 member A1 OS=Homo sapiens OX=9606 GN=BTN1A1 PE=1 SV=3 | 179 | 20201117_08512_1ng_glyco_hcd203040_compound15.8419.8419.3 | controllerType=0 controllerNumber=1 scan=8419 | 20.5728 |
| 05352:3 | 14 | R.LSPN[+2205.793]ASAEHLELR.W | HexNAc(4)Hex(5)Fuc(2)NeuAc(1) | N[+2206] | 1214.8543 | 3 | 3642.5484 | 3642.5421 | 0 | 1.7 | 52 | Specific | 638.61 | 519.72 | 519.72 | 12.19 | 37 | >sp\|Q13410\|BT1A1_HUMAN Butyrophilin subfamily 1 member A1 OS=Homo sapiens OX=9606 GN=BTN1A1 PE=1 SV=3 | 179 | 20201117_08512_1ng_glyco_hcd203040_compound15.8421.8421.3 | controllerType=0 controllerNumber=1 scan=8421 | 20.5769 |
| 12970:4 | 16 | K.LKELPGVC[+57.021]N[+2570.925]ETMMALWEEC[+57.021]KPC[+57.021]LK.Q | HexNAc(5)Hex(6)Fuc(2)NeuAc(1) | C[+57]*3, N[+2571] | 1377.5834 | 4 | 5507.3117 | 5507.3097 | 0 | 0.4 | 95 | Specific | 529.39 | 506.1 | 506.1 | 9.32 | 36 | >sp\|P10909\|CLUS_HUMAN Clusterin OS=Homo sapiens OX=9606 GN=CLU PE=1 SV=1 | 1467 | 20201117_08512_1ng_glyco_hcd203040_compound15.16330.16330.4 | controllerType=0 controllerNumber=1 scan=16330 | 35.0969 |
| 12968:4 | 16 | K.LKELPGVC[+57.021]N[+2570.925]ETMMALWEEC[+57.021]KPC[+57.021]LK.Q | HexNAc(5)Hex(6)Fuc(2)NeuAc(1) | C[+57]*3, N[+2571] | 1377.5834 | 4 | 5507.3117 | 5507.3097 | 0 | 0.4 | 95 | Specific | 584.79 | 584.79 | 584.79 | 10.93 | 36 | >sp\|P10909\|CLUS_HUMAN Clusterin OS=Homo sapiens OX=9606 GN=CLU PE=1 SV=1 | 1467 | 20201117_08512_1ng_glyco_hcd203040_compound15.16328.16328.4 | controllerType=0 controllerNumber=1 scan=16328 | 35.0919 |
| 12955:4 | 16 | K.LKELPGVC[+57.021]N[+2570.925]ETMMALWEEC[+57.021]KPC[+57.021]LK.Q | HexNAc(5)Hex(6)Fuc(2)NeuAc(1) | C[+57]*3, N[+2571] | 1377.5872 | 4 | 5507.3268 | 5507.3097 | 0 | 3.1 | 95 | Specific | 587.04 | 504.57 | 504.57 | 11.48 | 36 | >sp\|P10909\|CLUS_HUMAN Clusterin OS=Homo sapiens OX=9606 GN=CLU PE=1 SV=1 | 1467 | 20201117_08512_1ng_glyco_hcd203040_compound15.16314.16314.4 | controllerType=0 controllerNumber=1 scan=16314 | 35.0685 |
| 13118:3 | 16 | K.LKELPGVC[+57.021]N[+2059.735]ETMMALWEEC[+57.021]KPC[+57.021]LK.Q | HexNAc(4)Hex(5)Fuc(1)NeuAc(1) | C[+57]*3, N[+2060] | 1666.0504 | 3 | 4996.1366 | 4996.1196 | 0 | 3.4 | 95 | Specific | 626.89 | 598.71 | 598.71 | 11.92 | 36 | >sp\|P10909\|CLUS_HUMAN Clusterin OS=Homo sapiens OX=9606 GN=CLU PE=1 SV=1 | 1467 | 20201117_08512_1ng_glyco_hcd203040_compound15.16485.16485.3 | controllerType=0 controllerNumber=1 scan=16485 | 35.4201 |
| 12952:4 | 16 | K.LKELPGVC[+57.021]N[+2570.925]ETMMALWEEC[+57.021]KPC[+57.021]LK.Q | HexNAc(5)Hex(6)Fuc(2)NeuAc(1) | C[+57]*3, N[+2571] | 1377.5872 | 4 | 5507.3268 | 5507.3097 | 0 | 3.1 | 95 | Specific | 716.31 | 716.31 | 716.31 | 14.48 | 36 | >sp\|P10909\|CLUS_HUMAN Clusterin OS=Homo sapiens OX=9606 GN=CLU PE=1 SV=1 | 1467 | 20201117_08512_1ng_glyco_hcd203040_compound15.16311.16311.4 | controllerType=0 controllerNumber=1 scan=16311 | 35.0604 |
| 13100:4 | 16 | K.LKELPGVC[+57.021]N[+2059.735]ETMMALWEEC[+57.021]KPC[+57.021]LK.Q | HexNAc(4)Hex(5)Fuc(1)NeuAc(1) | C[+57]*3, N[+2060] | 1249.7894 | 4 | 4996.1359 | 4996.1196 | 0 | 3.3 | 95 | Specific | 843.08 | 766.79 | 766.79 | 16.39 | 36 | >sp\|P10909\|CLUS_HUMAN Clusterin OS=Homo sapiens OX=9606 GN=CLU PE=1 SV=1 | 1467 | 20201117_08512_1ng_glyco_hcd203040_compound15.16467.16467.4 | controllerType=0 controllerNumber=1 scan=16467 | 35.389 |
| 13115:3 | 16 | K.LKELPGVC[+57.021]N[+2059.735]ETMMALWEEC[+57.021]KPC[+57.021]LK.Q | HexNAc(4)Hex(5)Fuc(1)NeuAc(1) | C[+57]*3, N[+2060] | 1666.0504 | 3 | 4996.1366 | 4996.1196 | 0 | 3.4 | 95 | Specific | 851.37 | 851.37 | 851.37 | 16.92 | 36 | >sp\|P10909\|CLUS_HUMAN Clusterin OS=Homo sapiens OX=9606 GN=CLU PE=1 SV=1 | 1467 | 20201117_08512_1ng_glyco_hcd203040_compound15.16482.16482.3 | controllerType=0 controllerNumber=1 scan=16482 | 35.4146 |
| 13098:4 | 16 | K.LKELPGVC[+57.021]N[+2059.735]ETMMALWEEC[+57.021]KPC[+57.021]LK.Q | HexNAc(4)Hex(5)Fuc(1)NeuAc(1) | C[+57]*3, N[+2060] | 1249.7894 | 4 | 4996.1359 | 4996.1196 | 0 | 3.3 | 95 | Specific | 1149.42 | 1088.93 | 1088.93 | 22.39 | 36 | >sp\|P10909\|CLUS_HUMAN Clusterin OS=Homo sapiens OX=9606 GN=CLU PE=1 SV=1 | 1467 | 20201117_08512_1ng_glyco_hcd203040_compound15.16465.16465.4 | controllerType=0 controllerNumber=1 scan=16465 | 35.3836 |
| 14605:4 | 16 | K.LKELPGVC[+57.021]N[+2336.851]ETM[+15.995]MALWEEC[+57.021]KPC[+57.021 | HexNAc(6)Hex(6)Fuc(1) | C[+57]*3, M[+16], N[+2337] | 1323.059 | 4 | 5289.214 | 5289.2307 | 0 | -3.2 | 95 | Specific | 250.48 | 208.39 | 6.5 | 4.64 | 36 | >sp\|P10909\|CLUS_HUMAN Clusterin OS=Homo sapiens OX=9606 GN=CLU PE=1 SV=1 | 1467 | 20201117_08512_1ng_glyco_hcd203040_compound15.18034.18034.4 | controllerType=0 controllerNumber=1 scan=18034 | 38.5705 |
| 14608:4 | 16 | K.LKELPGVC[+57.021]N[+2336.851]ETM[+15.995]MALWEEC[+57.021]KPC[+57.021 | HexNAc(6)Hex(6)Fuc(1) | C[+57]*3, M[+16], N[+2337] | 1323.059 | 4 | 5289.214 | 5289.2307 | 0 | -3.2 | 95 | Specific | 281.75 | 220.24 | 5.29 | 3.79 | 36 | >sp\|P10909\|CLUS_HUMAN Clusterin OS=Homo sapiens OX=9606 GN=CLU PE=1 SV=1 | 1467 | 20201117_08512_1ng_glyco_hcd203040_compound15.18037.18037.4 | controllerType=0 controllerNumber=1 scan=18037 | 38.5751 |
| 14726:4 | 16 | K.ELPGVC[+57.021]N[+2059.735]ETMMALWEEC[+57.021]KPC[+57.021]LK.Q | HexNAc(4)Hex(5)Fuc(1)NeuAc(1) | C[+57]*3, N[+2060] | 1189.4935 | 4 | 4754.952 | 4754.9406 | 0 | 2.4 | 97 | Specific | 353.48 | 351.01 | 351.01 | 7.52 | 36 | >sp\|P10909\|CLUS_HUMAN Clusterin OS=Homo sapiens OX=9606 GN=CLU PE=1 SV=1 | 1467 | 20201117_08512_1ng_glyco_hcd203040_compound15.18160.18160.4 | controllerType=0 controllerNumber=1 scan=18160 | 38.8168 |
| 14728:4 | 16 | K.ELPGVC[+57.021]N[+2059.735]ETMMALWEEC[+57.021]KPC[+57.021]LK.Q | HexNAc(4)Hex(5)Fuc(1)NeuAc(1) | C[+57]*3, N[+2060] | 1189.4935 | 4 | 4754.952 | 4754.9406 | 0 | 2.4 | 97 | Specific | 553.53 | 480.32 | 480.32 | 10.34 | 36 | >sp\|P10909\|CLUS_HUMAN Clusterin OS=Homo sapiens OX=9606 GN=CLU PE=1 SV=1 | 1467 | 20201117_08512_1ng_glyco_hcd203040_compound15.18162.18162.4 | controllerType=0 controllerNumber=1 scan=18162 | 38.823 |
| 21599:4 | 16 | K.[+42.011]MLN[+2336.851]TSSLLEQLNEQFNWVSR.L | HexNAc(6)Hex(6)Fuc(1) | .[42], N[+2337] | 1197.764 | 4 | 4788.034 | 4788.0484 | 0 | -3 | 352 | Specific | 311.29 | 248.98 | 248.98 | 5.82 | 36 | >sp\|P10909\|CLUS_HUMAN Clusterin OS=Homo sapiens OX=9606 GN=CLU PE=1 SV=1 | 1467 | 20201117_08512_1ng_glyco_hcd203040_compound15.25356.25356.4 | controllerType=0 controllerNumber=1 scan=25356 | 54.5739 |
| 21597:4 | 16 | K.[+42.011]MLN[+2336.851]TSSLLEQLNEQFNWVSR.L | HexNAc(6)Hex(6)Fuc(1) | .[42], N[+2337] | 1197.764 | 4 | 4788.034 | 4788.0484 | 0 | -3 | 352 | Specific | 373.73 | 270.9 | 270.9 | 6.26 | 36 | >sp\|P10909\|CLUS_HUMAN Clusterin OS=Homo sapiens OX=9606 GN=CLU PE=1 SV=1 | 1467 | 20201117_08512_1ng_glyco_hcd203040_compound15.25354.25354.4 | controllerType=0 controllerNumber=1 scan=25354 | 54.57 |

Table S4. Cont.

| Query #:z | Protein Rank | Glycopeptide | Glycan | Modification Type(s) | Observed m/z | z | Observed (M+H) | Calc. mass (M+H) | Off-by-x error | Mass error (ppm) | StartPosition | Cleavage | Score | Delta | Delta Mod | \|Log Prob\| | # of unique peptides | Protein Name | Protein DB number | Comment | Scan # | Scan Time |
|---|---|---|---|---|---|---|---|---|---|---|---|---|---|---|---|---|---|---|---|---|---|---|
| 21525:4 | 16 | K.MLN[+2350.830]TSSLLEQLNEQFNWVSR.L | HexNAc(4)Hex(5)Fuc(1)NeuAc(2) | N[+2351] | 1190.7646 | 4 | 4760.0364 | 4760.0171 | 0 | 4.1 | 352 | Specific | 879.3 | 680.41 | 680.41 | 17.98 | 36 | >sp\|P10909\|CLUS_HUMAN Clusterin OS=Homo sapiens OX=9606 GN=CLU PE=1 SV=1 | 1467 | 20201117_08512_1ng_glyco_hcd203040_compound15.25278.25278.4 | controllerType=0 controllerNumber=1 scan=25278 | 54.4098 |
| 21516:4 | 16 | K.MLN[+2350.830]TSSLLEQLNEQFNWVSR.L | HexNAc(4)Hex(5)Fuc(1)NeuAc(2) | N[+2351] | 1190.7643 | 4 | 4760.0355 | 4760.0171 | 0 | 3.9 | 352 | Specific | 879.92 | 640.79 | 640.79 | 17.98 | 36 | >sp\|P10909\|CLUS_HUMAN Clusterin OS=Homo sapiens OX=9606 GN=CLU PE=1 SV=1 | 1467 | 20201117_08512_1ng_glyco_hcd203040_compound15.25268.25268.4 | controllerType=0 controllerNumber=1 scan=25268 | 54.3883 |
| 21528:4 | 16 | K.MLN[+2350.830]TSSLLEQLNEQFNWVSR.L | HexNAc(4)Hex(5)Fuc(1)NeuAc(2) | N[+2351] | 1190.7646 | 4 | 4760.0364 | 4760.0171 | 0 | 4.1 | 352 | Specific | 886.41 | 568.37 | 568.37 | 17.98 | 36 | >sp\|P10909\|CLUS_HUMAN Clusterin OS=Homo sapiens OX=9606 GN=CLU PE=1 SV=1 | 1467 | 20201117_08512_1ng_glyco_hcd203040_compound15.25281.25281.4 | controllerType=0 controllerNumber=1 scan=25281 | 54.4161 |
| 21523:3 | 16 | K.MLN[+2350.830]TSSLLEQLNEQFNWVSR.L | HexNAc(4)Hex(5)Fuc(1)NeuAc(2) | N[+2351] | 1587.3504 | 3 | 4760.0366 | 4760.0171 | 0 | 4.1 | 352 | Specific | 943.47 | 644.35 | 644.35 | 18.36 | 36 | >sp\|P10909\|CLUS_HUMAN Clusterin OS=Homo sapiens OX=9606 GN=CLU PE=1 SV=1 | 1467 | 20201117_08512_1ng_glyco_hcd203040_compound15.25276.25276.3 | controllerType=0 controllerNumber=1 scan=25276 | 54.4068 |
| 21514:4 | 16 | K.MLN[+2350.830]TSSLLEQLNEQFNWVSR.L | HexNAc(4)Hex(5)Fuc(1)NeuAc(2) | N[+2351] | 1190.7643 | 4 | 4760.0355 | 4760.0171 | 0 | 3.9 | 352 | Specific | 982.78 | 712.06 | 712.06 | 19.98 | 36 | >sp\|P10909\|CLUS_HUMAN Clusterin OS=Homo sapiens OX=9606 GN=CLU PE=1 SV=1 | 1467 | 20201117_08512_1ng_glyco_hcd203040_compound15.25266.25266.4 | controllerType=0 controllerNumber=1 scan=25266 | 54.3825 |
| 21521:3 | 16 | K.MLN[+2350.830]TSSLLEQLNEQFNWVSR.L | HexNAc(4)Hex(5)Fuc(1)NeuAc(2) | N[+2351] | 1587.3504 | 3 | 4760.0366 | 4760.0171 | 0 | 4.1 | 352 | Specific | 1094.44 | 828.16 | 828.16 | 21.36 | 36 | >sp\|P10909\|CLUS_HUMAN Clusterin OS=Homo sapiens OX=9606 GN=CLU PE=1 SV=1 | 1467 | 20201117_08512_1ng_glyco_hcd203040_compound15.25274.25274.3 | controllerType=0 controllerNumber=1 scan=25274 | 54.4017 |
| 12276:3 | 16 | R.LAN[+2350.830]LTQGEDQYYLR.V | HexNAc(4)Hex(5)Fuc(1)NeuAc(2) | N[+2351] | 1345.5685 | 3 | 4034.6911 | 4034.664 | 0 | 6.7 | 372 | Specific | 290.17 | 197.27 | 197.27 | 4.49 | 36 | >sp\|P10909\|CLUS_HUMAN Clusterin OS=Homo sapiens OX=9606 GN=CLU PE=1 SV=1 | 1467 | 20201117_08512_1ng_glyco_hcd203040_compound15.15608.15608.3 | controllerType=0 controllerNumber=1 scan=15608 | 33.6302 |
| 09609:3 | 16 | R.LAN[+2570.925]LTQGEDQYYLR.V | HexNAc(5)Hex(6)Fuc(2)NeuAc(1) | N[+2571] | 1418.929 | 3 | 4254.7725 | 4254.7587 | 0 | 3.2 | 372 | Specific | 409.67 | 374.04 | 374.04 | 7.23 | 36 | >sp\|P10909\|CLUS_HUMAN Clusterin OS=Homo sapiens OX=9606 GN=CLU PE=1 SV=1 | 1467 | 20201117_08512_1ng_glyco_hcd203040_compound15.12835.12835.3 | controllerType=0 controllerNumber=1 scan=12835 | 28.3726 |
| 09611:3 | 16 | R.LAN[+2570.925]LTQGEDQYYLR.V | HexNAc(5)Hex(6)Fuc(2)NeuAc(1) | N[+2571] | 1418.929 | 3 | 4254.7725 | 4254.7587 | 0 | 3.2 | 372 | Specific | 524.55 | 461.78 | 461.78 | 9.21 | 36 | >sp\|P10909\|CLUS_HUMAN Clusterin OS=Homo sapiens OX=9606 GN=CLU PE=1 SV=1 | 1467 | 20201117_08512_1ng_glyco_hcd203040_compound15.12837.12837.3 | controllerType=0 controllerNumber=1 scan=12837 | 28.3762 |
| 12231:3 | 16 | R.LAN[+2350.830]LTQGEDQYYLR.V | HexNAc(4)Hex(5)Fuc(1)NeuAc(2) | N[+2351] | 1345.5615 | 3 | 4034.6698 | 4034.664 | 0 | 1.4 | 372 | Specific | 552.06 | 464.65 | 464.65 | 9.53 | 36 | >sp\|P10909\|CLUS_HUMAN Clusterin OS=Homo sapiens OX=9606 GN=CLU PE=1 SV=1 | 1467 | 20201117_08512_1ng_glyco_hcd203040_compound15.15561.15561.3 | controllerType=0 controllerNumber=1 scan=15561 | 33.54 |
| 12234:3 | 16 | R.LAN[+2350.830]LTQGEDQYYLR.V | HexNAc(4)Hex(5)Fuc(1)NeuAc(2) | N[+2351] | 1345.5615 | 3 | 4034.6698 | 4034.664 | 0 | 1.4 | 372 | Specific | 653.85 | 579.36 | 579.36 | 13.01 | 36 | >sp\|P10909\|CLUS_HUMAN Clusterin OS=Homo sapiens OX=9606 GN=CLU PE=1 SV=1 | 1467 | 20201117_08512_1ng_glyco_hcd203040_compound15.15564.15564.3 | controllerType=0 controllerNumber=1 scan=15564 | 33.545 |
| 22212:4 | 17 | R.QLAHQSN[+2350.830]STNIFFSPVSIATAFAMLSLGTK.A | HexNAc(4)Hex(5)Fuc(1)NeuAc(2) | N[+2351] | 1383.8721 | 4 | 5532.4665 | 5532.4654 | 0 | 0.2 | 64 | Specific | 456 | 422.22 | 422.22 | 9.09 | 35 | >sp\|P01009\|A1AT_HUMAN Alpha-1-antitrypsin OS=Homo sapiens OX=9606 GN=SERPINA1 PE=1 SV=3 | 642 | 20201117_08512_1ng_glyco_hcd203040_compound15.25994.25994.4 | controllerType=0 controllerNumber=1 scan=25994 | 55.8793 |
| 22214:4 | 17 | R.QLAHQSN[+2350.830]STNIFFSPVSIATAFAMLSLGTK.A | HexNAc(4)Hex(5)Fuc(1)NeuAc(2) | N[+2351] | 1383.8721 | 4 | 5532.4665 | 5532.4654 | 0 | 0.2 | 64 | Specific | 487.38 | 463.85 | 463.85 | 9.28 | 35 | >sp\|P01009\|A1AT_HUMAN Alpha-1-antitrypsin OS=Homo sapiens OX=9606 GN=SERPINA1 PE=1 SV=3 | 642 | 20201117_08512_1ng_glyco_hcd203040_compound15.25996.25996.4 | controllerType=0 controllerNumber=1 scan=25996 | 55.8829 |
| 22219:5 | 17 | R.QLAHQSN[+2204.772]STNIFFSPVSIATAFAMLSLGTK.A | HexNAc(4)Hex(5)NeuAc(2) | N[+2205] | 1078.0897 | 5 | 5386.4195 | 5386.4075 | 0 | 2.2 | 64 | Specific | 554.51 | 495.98 | 495.98 | 10.91 | 35 | >sp\|P01009\|A1AT_HUMAN Alpha-1-antitrypsin OS=Homo sapiens OX=9606 GN=SERPINA1 PE=1 SV=3 | 642 | 20201117_08512_1ng_glyco_hcd203040_compound15.26001.26001.5 | controllerType=0 controllerNumber=1 scan=26001 | 55.893 |
| 22217:5 | 17 | R.QLAHQSN[+2204.772]STNIFFSPVSIATAFAMLSLGTK.A | HexNAc(4)Hex(5)NeuAc(2) | N[+2205] | 1078.0897 | 5 | 5386.4195 | 5386.4075 | 0 | 2.2 | 64 | Specific | 611.46 | 578.13 | 543.08 | 13.16 | 35 | >sp\|P01009\|A1AT_HUMAN Alpha-1-antitrypsin OS=Homo sapiens OX=9606 GN=SERPINA1 PE=1 SV=3 | 642 | 20201117_08512_1ng_glyco_hcd203040_compound15.25999.25999.5 | controllerType=0 controllerNumber=1 scan=25999 | 55.8886 |
| 22209:3 | 17 | R.QLAHQSN[+2204.772]STNIFFSPVSIATAFAMLSLGTK.A | HexNAc(4)Hex(5)NeuAc(2) | N[+2205] | 1796.1466 | 3 | 5386.4252 | 5386.4075 | 0 | 3.3 | 64 | Specific | 754.23 | 736.55 | 736.55 | 15.62 | 35 | >sp\|P01009\|A1AT_HUMAN Alpha-1-antitrypsin OS=Homo sapiens OX=9606 GN=SERPINA1 PE=1 SV=3 | 642 | 20201117_08512_1ng_glyco_hcd203040_compound15.25991.25991.3 | controllerType=0 controllerNumber=1 scan=25991 | 55.8718 |
| 22211:3 | 17 | R.QLAHQSN[+2204.772]STNIFFSPVSIATAFAMLSLGTK.A | HexNAc(4)Hex(5)NeuAc(2) | N[+2205] | 1796.1466 | 3 | 5386.4252 | 5386.4075 | 0 | 3.3 | 64 | Specific | 784.4 | 693.55 | 693.55 | 15.62 | 35 | >sp\|P01009\|A1AT_HUMAN Alpha-1-antitrypsin OS=Homo sapiens OX=9606 GN=SERPINA1 PE=1 SV=3 | 642 | 20201117_08512_1ng_glyco_hcd203040_compound15.25993.25993.3 | controllerType=0 controllerNumber=1 scan=25993 | 55.8774 |
| 22187:4 | 17 | R.QLAHQSN[+2204.772]STNIFFSPVSIATAFAMLSLGTK.A | HexNAc(4)Hex(5)NeuAc(2) | N[+2205] | 1347.3604 | 4 | 5386.4196 | 5386.4075 | 0 | 2.2 | 64 | Specific | 863.26 | 795.14 | 521.62 | 17.54 | 35 | >sp\|P01009\|A1AT_HUMAN Alpha-1-antitrypsin OS=Homo sapiens OX=9606 GN=SERPINA1 PE=1 SV=3 | 642 | 20201117_08512_1ng_glyco_hcd203040_compound15.25967.25967.4 | controllerType=0 controllerNumber=1 scan=25967 | 55.8009 |
| 22189:4 | 17 | R.QLAHQSN[+2204.772]STNIFFSPVSIATAFAMLSLGTK.A | HexNAc(4)Hex(5)NeuAc(2) | N[+2205] | 1347.3604 | 4 | 5386.4196 | 5386.4075 | 0 | 2.2 | 64 | Specific | 876.82 | 802.84 | 569.99 | 17.54 | 35 | >sp\|P01009\|A1AT_HUMAN Alpha-1-antitrypsin OS=Homo sapiens OX=9606 GN=SERPINA1 PE=1 SV=3 | 642 | 20201117_08512_1ng_glyco_hcd203040_compound15.25969.25969.4 | controllerType=0 controllerNumber=1 scan=25969 | 55.8063 |
| 22203:4 | 17 | R.QLAHQSN[+2204.772]STNIFFSPVSIATAFAMLSLGTK.A | HexNAc(4)Hex(5)NeuAc(2) | N[+2205] | 1347.36 | 4 | 5386.4181 | 5386.4075 | 0 | 2 | 64 | Specific | 954.4 | 902.77 | 616.62 | 19.64 | 35 | >sp\|P01009\|A1AT_HUMAN Alpha-1-antitrypsin OS=Homo sapiens OX=9606 GN=SERPINA1 PE=1 SV=3 | 642 | 20201117_08512_1ng_glyco_hcd203040_compound15.25984.25984.4 | controllerType=0 controllerNumber=1 scan=25984 | 55.8525 |
| 22201:4 | 17 | R.QLAHQSN[+2204.772]STNIFFSPVSIATAFAMLSLGTK.A | HexNAc(4)Hex(5)NeuAc(2) | N[+2205] | 1347.36 | 4 | 5386.4181 | 5386.4075 | 0 | 2 | 64 | Specific | 975.61 | 926.53 | 576.03 | 19.64 | 35 | >sp\|P01009\|A1AT_HUMAN Alpha-1-antitrypsin OS=Homo sapiens OX=9606 GN=SERPINA1 PE=1 SV=3 | 642 | 20201117_08512_1ng_glyco_hcd203040_compound15.25982.25982.4 | controllerType=0 controllerNumber=1 scan=25982 | 55.8323 |
| 12277:4 | 18 | R.FELLGC[+57.021]ELNGC[+57.021]ANPLGLKN[+1216.423]NSIPDKQITASSSYK.T | HexNAc(2)Hex(5) | C[+57]*2, N[+1216] | 1239.5752 | 4 | 4955.279 | 4955.2694 | 0 | 1.9 | 220 | Specific | 876.18 | 761.53 | 761.53 | 17.22 | 31 | >sp\|Q08431\|MFGM_HUMAN Lactadherin OS=Homo sapiens OX=9606 GN=MFGE8 PE=1 SV=3 | 1064 | 20201117_08512_1ng_glyco_hcd203040_compound15.15609.15609.4 | controllerType=0 controllerNumber=1 scan=15609 | 33.6338 |
| 12652:4 | 18 | R.FELLGC[+57.021]ELNGC[+57.021]ANPLGLKN[+1540.529]NSIPDKQITASSSYK.T | HexNAc(2)Hex(7) | C[+57]*2, N[+1541] | 1320.5977 | 4 | 5279.3688 | 5279.3751 | 0 | -1.2 | 220 | Specific | 936.76 | 677.89 | 677.89 | 17.15 | 31 | >sp\|Q08431\|MFGM_HUMAN Lactadherin OS=Homo sapiens OX=9606 GN=MFGE8 PE=1 SV=3 | 1064 | 20201117_08512_1ng_glyco_hcd203040_compound15.15998.15998.4 | controllerType=0 controllerNumber=1 scan=15998 | 34.3951 |
| 12275:4 | 18 | R.FELLGC[+57.021]ELNGC[+57.021]ANPLGLKN[+1216.423]NSIPDKQITASSSYK.T | HexNAc(2)Hex(5) | C[+57]*2, N[+1216] | 1239.5752 | 4 | 4955.279 | 4955.2694 | 0 | 1.9 | 220 | Specific | 951.33 | 889.61 | 889.61 | 19.22 | 31 | >sp\|Q08431\|MFGM_HUMAN Lactadherin OS=Homo sapiens OX=9606 GN=MFGE8 PE=1 SV=3 | 1064 | 20201117_08512_1ng_glyco_hcd203040_compound15.15607.15607.4 | controllerType=0 controllerNumber=1 scan=15607 | 33.6278 |
| 12651:4 | 18 | R.FELLGC[+57.021]ELNGC[+57.021]ANPLGLKN[+1540.529]NSIPDKQITASSSYK.T | HexNAc(2)Hex(7) | C[+57]*2, N[+1541] | 1320.5977 | 4 | 5279.3688 | 5279.3751 | 0 | -1.2 | 220 | Specific | 1172.2 | 951.19 | 951.19 | 22.15 | 31 | >sp\|Q08431\|MFGM_HUMAN Lactadherin OS=Homo sapiens OX=9606 GN=MFGE8 PE=1 SV=3 | 1064 | 20201117_08512_1ng_glyco_hcd203040_compound15.15997.15997.4 | controllerType=0 controllerNumber=1 scan=15997 | 34.3897 |
| 01954:3 | 18 | K.N[+1216.423]NSIPDKQITASSSYK.T | HexNAc(2)Hex(5) | N[+1216] | 990.4408 | 3 | 2969.3079 | 2969.2991 | 0 | 3 | 238 | Specific | 308.14 | 144.65 | 144.65 | 5.36 | 31 | >sp\|Q08431\|MFGM_HUMAN Lactadherin OS=Homo sapiens OX=9606 GN=MFGE8 PE=1 SV=3 | 1064 | 20201117_08512_1ng_glyco_hcd203040_compound15.4903.4903.3 | controllerType=0 controllerNumber=1 scan=4903 | 14.5597 |
| 01952:3 | 18 | K.N[+1216.423]NSIPDKQITASSSYK.T | HexNAc(2)Hex(5) | N[+1216] | 990.4408 | 3 | 2969.3079 | 2969.2991 | 0 | 3 | 238 | Specific | 333.06 | 179.5 | 179.5 | 5.77 | 31 | >sp\|Q08431\|MFGM_HUMAN Lactadherin OS=Homo sapiens OX=9606 GN=MFGE8 PE=1 SV=3 | 1064 | 20201117_08512_1ng_glyco_hcd203040_compound15.4901.4901.3 | controllerType=0 controllerNumber=1 scan=4901 | 14.5556 |
| 08216:5 | 26 | K.SC[+57.021]QHN[+2059.735]GTMYQHGEIFSAHELFPSR.L | HexNAc(4)Hex(5)Fuc(1)NeuAc(1) | C[+57], N[+2060] | 979.4071 | 5 | 4893.0062 | 4892.9919 | 0 | 2.9 | 110 | Specific | 738.87 | 738.87 | 738.87 | 14.76 | 21 | >sp\|Q6WN34\|CRDL2_HUMAN Chordin-like protein 2 OS=Homo sapiens OX=9606 GN=CHRDL2 PE=1 SV=1 | 1189 | 20201117_08512_1ng_glyco_hcd203040_compound15.11388.11388.5 | controllerType=0 controllerNumber=1 scan=11388 | 25.6944 |
| 08218:5 | 26 | K.SC[+57.021]QHN[+2059.735]GTMYQHGEIFSAHELFPSR.L | HexNAc(4)Hex(5)Fuc(1)NeuAc(1) | C[+57], N[+2060] | 979.4071 | 5 | 4893.0062 | 4892.9919 | 0 | 2.9 | 110 | Specific | 865.52 | 865.52 | 865.52 | 17.76 | 21 | >sp\|Q6WN34\|CRDL2_HUMAN Chordin-like protein 2 OS=Homo sapiens OX=9606 GN=CHRDL2 PE=1 SV=1 | 1189 | 20201117_08512_1ng_glyco_hcd203040_compound15.11390.11390.5 | controllerType=0 controllerNumber=1 scan=11390 | 25.6988 |
| 13298:2 | 29 | K.LC[+57.021]DN[+1866.688]LSTK.N | HexNAc(6)Hex(4) | C[+57], N[+1867] | 1409.0715 | 2 | 2817.1356 | 2817.1487 | 0 | -4.6 | 285 | Specific | 329.79 | 188.53 | 188.53 | 3.24 | 24 | >sp\|P02774\|VTDB_HUMAN Vitamin D-binding protein OS=Homo sapiens OX=9606 GN=GC PE=1 SV=2 | 1103 | 20201117_08512_1ng_glyco_hcd203040_compound15.16673.16673.2 | controllerType=0 controllerNumber=1 scan=16673 | 35.8247 |
| 12562:3 | 31 | R.IIVPLNNREN[+2205.793]ISDPTSPLR.T | HexNAc(4)Hex(5)Fuc(2)NeuAc(1) | N[+2206] | 1451.9932 | 3 | 4353.965 | 4353.97 | 0 | -1.1 | 62 | Specific | 205.93 | 205.93 | 205.93 | 3.74 | 24 | >sp\|P01591\|IGJ_HUMAN Immunoglobulin J chain OS=Homo sapiens OX=9606 GN=JCHAIN PE=1 SV=4 | 1392 | 20201117_08512_1ng_glyco_hcd203040_compound15.15905.15905.3 | controllerType=0 controllerNumber=1 scan=15905 | 34.2064 |
| 09185:4 | 31 | R.IIVPLNNREN[+1663.608]ISDPTSPLR.T | HexNAc(5)Hex(4) | N[+1664] | 953.7036 | 4 | 3811.7926 | 3811.7853 | 0 | 1.9 | 62 | Specific | 218.11 | 218.11 | 218.11 | 4.84 | 24 | >sp\|P01591\|IGJ_HUMAN Immunoglobulin J chain OS=Homo sapiens OX=9606 GN=JCHAIN PE=1 SV=4 | 1392 | 20201117_08512_1ng_glyco_hcd203040_compound15.12395.12395.4 | controllerType=0 controllerNumber=1 scan=12395 | 27.5682 |
| 12565:3 | 31 | R.IIVPLNNREN[+2205.793]ISDPTSPLR.T | HexNAc(4)Hex(5)Fuc(2)NeuAc(1) | N[+2206] | 1451.9932 | 3 | 4353.965 | 4353.97 | 0 | -1.1 | 62 | Specific | 261.15 | 261.15 | 261.15 | 5.52 | 24 | >sp\|P01591\|IGJ_HUMAN Immunoglobulin J chain OS=Homo sapiens OX=9606 GN=JCHAIN PE=1 SV=4 | 1392 | 20201117_08512_1ng_glyco_hcd203040_compound15.15908.15908.3 | controllerType=0 controllerNumber=1 scan=15908 | 34.2109 |
| 12504:4 | 31 | R.IIVPLNNREN[+2204.772]ISDPTSPLR.T | HexNAc(4)Hex(5)NeuAc(2) | N[+2205] | 1088.9945 | 4 | 4352.9564 | 4352.9496 | 0 | 1.6 | 62 | Specific | 336.56 | 336.56 | 336.56 | 7.15 | 24 | >sp\|P01591\|IGJ_HUMAN Immunoglobulin J chain OS=Homo sapiens OX=9606 GN=JCHAIN PE=1 SV=4 | 1392 | 20201117_08512_1ng_glyco_hcd203040_compound15.15844.15844.4 | controllerType=0 controllerNumber=1 scan=15844 | 34.0767 |
| 12430:3 | 31 | R.IIVPLNNREN[+2350.830]ISDPTSPLR.T | HexNAc(4)Hex(5)Fuc(1)NeuAc(2) | N[+2351] | 1500.3443 | 3 | 4499.0183 | 4499.0075 | 0 | 2.4 | 62 | Specific | 365.14 | 365.14 | 365.14 | 7.69 | 24 | >sp\|P01591\|IGJ_HUMAN Immunoglobulin J chain OS=Homo sapiens OX=9606 GN=JCHAIN PE=1 SV=4 | 1392 | 20201117_08512_1ng_glyco_hcd203040_compound15.15768.15768.3 | controllerType=0 controllerNumber=1 scan=15768 | 33.9342 |
| 09203:3 | 31 | R.IIVPLNNREN[+1768.640]ISDPTSPLR.T | HexNAc(4)Hex(5)Fuc(1) | N[+1769] | 1306.2775 | 3 | 3916.818 | 3916.8167 | 0 | 0.3 | 62 | Specific | 400.59 | 350.92 | 350.92 | 6.87 | 24 | >sp\|P01591\|IGJ_HUMAN Immunoglobulin J chain OS=Homo sapiens OX=9606 GN=JCHAIN PE=1 SV=4 | 1392 | 20201117_08512_1ng_glyco_hcd203040_compound15.12414.12414.3 | controllerType=0 controllerNumber=1 scan=12414 | 27.6035 |
| 12506:4 | 31 | R.IIVPLNNREN[+2204.772]ISDPTSPLR.T | HexNAc(4)Hex(5)NeuAc(2) | N[+2205] | 1088.9945 | 4 | 4352.9564 | 4352.9496 | 0 | 1.6 | 62 | Specific | 410.06 | 410.06 | 410.06 | 9.14 | 24 | >sp\|P01591\|IGJ_HUMAN Immunoglobulin J chain OS=Homo sapiens OX=9606 GN=JCHAIN PE=1 SV=4 | 1392 | 20201117_08512_1ng_glyco_hcd203040_compound15.15846.15846.4 | controllerType=0 controllerNumber=1 scan=15846 | 34.0809 |
| 09205:3 | 31 | R.IIVPLNNREN[+1768.640]ISDPTSPLR.T | HexNAc(4)Hex(5)Fuc(1) | N[+1769] | 1306.2775 | 3 | 3916.818 | 3916.8167 | 0 | 0.3 | 62 | Specific | 423.71 | 343.03 | 343.03 | 7.06 | 24 | >sp\|P01591\|IGJ_HUMAN Immunoglobulin J chain OS=Homo sapiens OX=9606 GN=JCHAIN PE=1 SV=4 | 1392 | 20201117_08512_1ng_glyco_hcd203040_compound15.12416.12416.3 | controllerType=0 controllerNumber=1 scan=12416 | 27.6088 |
| 12432:3 | 31 | R.IIVPLNNREN[+2350.830]ISDPTSPLR.T | HexNAc(4)Hex(5)Fuc(1)NeuAc(2) | N[+2351] | 1500.3443 | 3 | 4499.0183 | 4499.0075 | 0 | 2.4 | 62 | Specific | 485.53 | 414.71 | 414.71 | 9.63 | 24 | >sp\|P01591\|IGJ_HUMAN Immunoglobulin J chain OS=Homo sapiens OX=9606 GN=JCHAIN PE=1 SV=4 | 1392 | 20201117_08512_1ng_glyco_hcd203040_compound15.15770.15770.3 | controllerType=0 controllerNumber=1 scan=15770 | 33.9387 |
| 09170:3 | 31 | R.IIVPLNNREN[+1663.608]ISDPTSPLR.T | HexNAc(5)Hex(4) | N[+1664] | 1271.2679 | 3 | 3811.7891 | 3811.7853 | 0 | 1 | 62 | Specific | 486.93 | 460.31 | 460.31 | 9.74 | 24 | >sp\|P01591\|IGJ_HUMAN Immunoglobulin J chain OS=Homo sapiens OX=9606 GN=JCHAIN PE=1 SV=4 | 1392 | 20201117_08512_1ng_glyco_hcd203040_compound15.12380.12380.3 | controllerType=0 controllerNumber=1 scan=12380 | 27.5425 |
| 09129:3 | 31 | R.IIVPLNNREN[+1809.666]ISDPTSPLR.T | HexNAc(5)Hex(4)Fuc(1) | N[+1810] | 1319.9562 | 3 | 3957.8541 | 3957.8432 | 0 | 2.8 | 62 | Specific | 518.47 | 481.26 | 481.26 | 9.73 | 24 | >sp\|P01591\|IGJ_HUMAN Immunoglobulin J chain OS=Homo sapiens OX=9606 GN=JCHAIN PE=1 SV=4 | 1392 | 20201117_08512_1ng_glyco_hcd203040_compound15.12337.12337.3 | controllerType=0 controllerNumber=1 scan=12337 | 27.4494 |
| 09131:3 | 31 | R.IIVPLNNREN[+1809.666]ISDPTSPLR.T | HexNAc(5)Hex(4)Fuc(1) | N[+1810] | 1319.9562 | 3 | 3957.8541 | 3957.8432 | 0 | 2.8 | 62 | Specific | 526.09 | 384.52 | 384.52 | 9.73 | 24 | >sp\|P01591\|IGJ_HUMAN Immunoglobulin J chain OS=Homo sapiens OX=9606 GN=JCHAIN PE=1 SV=4 | 1392 | 20201117_08512_1ng_glyco_hcd203040_compound15.12339.12339.3 | controllerType=0 controllerNumber=1 scan=12339 | 27.4545 |
| 10620:3 | 31 | R.IIVPLNNREN[+1913.677]ISDPTSPLR.T | HexNAc(4)Hex(5)NeuAc(1) | N[+1914] | 1354.6265 | 3 | 4061.865 | 4061.8542 | 0 | 2.7 | 62 | Specific | 527.7 | 496.07 | 496.07 | 9.73 | 24 | >sp\|P01591\|IGJ_HUMAN Immunoglobulin J chain OS=Homo sapiens OX=9606 GN=JCHAIN PE=1 SV=4 | 1392 | 20201117_08512_1ng_glyco_hcd203040_compound15.13884.13884.3 | controllerType=0 controllerNumber=1 scan=13884 | 30.3366 |
| 10587:3 | 31 | R.IIVPLNNREN[+2059.735]ISDPTSPLR.T | HexNAc(4)Hex(5)Fuc(1)NeuAc(1) | N[+2060] | 1403.313 | 3 | 4207.9246 | 4207.9121 | 0 | 3 | 62 | Specific | 530.11 | 495.69 | 495.69 | 9.86 | 24 | >sp\|P01591\|IGJ_HUMAN Immunoglobulin J chain OS=Homo sapiens OX=9606 GN=JCHAIN PE=1 SV=4 | 1392 | 20201117_08512_1ng_glyco_hcd203040_compound15.13849.13849.3 | controllerType=0 controllerNumber=1 scan=13849 | 30.2602 |
| 12481:3 | 31 | R.IIVPLNNREN[+2204.772]ISDPTSPLR.T | HexNAc(4)Hex(5)NeuAc(2) | N[+2205] | 1451.6572 | 3 | 4352.9569 | 4352.9496 | 0 | 1.7 | 62 | Specific | 535.25 | 517.34 | 517.34 | 9.96 | 24 | >sp\|P01591\|IGJ_HUMAN Immunoglobulin J chain OS=Homo sapiens OX=9606 GN=JCHAIN PE=1 SV=4 | 1392 | 20201117_08512_1ng_glyco_hcd203040_compound15.15820.15820.3 | controllerType=0 controllerNumber=1 scan=15820 | 34.0261 |
| 12479:3 | 31 | R.IIVPLNNREN[+2204.772]ISDPTSPLR.T | HexNAc(4)Hex(5)NeuAc(2) | N[+2205] | 1451.6572 | 3 | 4352.9569 | 4352.9496 | 0 | 1.7 | 62 | Specific | 536.25 | 500.88 | 500.88 | 9.96 | 24 | >sp\|P01591\|IGJ_HUMAN Immunoglobulin J chain OS=Homo sapiens OX=9606 GN=JCHAIN PE=1 SV=4 | 1392 | 20201117_08512_1ng_glyco_hcd203040_compound15.15818.15818.3 | controllerType=0 controllerNumber=1 scan=15818 | 34.0212 |
| 10622:3 | 31 | R.IIVPLNNREN[+1913.677]ISDPTSPLR.T | HexNAc(4)Hex(5)NeuAc(1) | N[+1914] | 1354.6265 | 3 | 4061.865 | 4061.8542 | 0 | 2.7 | 62 | Specific | 542.01 | 542.01 | 542.01 | 9.86 | 24 | >sp\|P01591\|IGJ_HUMAN Immunoglobulin J chain OS=Homo sapiens OX=9606 GN=JCHAIN PE=1 SV=4 | 1392 | 20201117_08512_1ng_glyco_hcd203040_compound15.13886.13886.3 | controllerType=0 controllerNumber=1 scan=13886 | 30.3429 |
| 10589:3 | 31 | R.IIVPLNNREN[+2059.735]ISDPTSPLR.T | HexNAc(4)Hex(5)Fuc(1)NeuAc(1) | N[+2060] | 1403.313 | 3 | 4207.9246 | 4207.9121 | 0 | 3 | 62 | Specific | 546.1 | 455.21 | 455.21 | 9.86 | 24 | >sp\|P01591\|IGJ_HUMAN Immunoglobulin J chain OS=Homo sapiens OX=9606 GN=JCHAIN PE=1 SV=4 | 1392 | 20201117_08512_1ng_glyco_hcd203040_compound15.13851.13851.3 | controllerType=0 controllerNumber=1 scan=13851 | 30.2655 |
| 10685:3 | 31 | R.IIVPLNNREN[+1856.656]ISDPTSPLR.T | HexNAc(3)Hex(5)Fuc(1)NeuAc(1) | N[+1857] | 1335.618 | 3 | 4004.8394 | 4004.8327 | 0 | 1.7 | 62 | Specific | 546.47 | 546.47 | 546.47 | 9.96 | 24 | >sp\|P01591\|IGJ_HUMAN Immunoglobulin J chain OS=Homo sapiens OX=9606 GN=JCHAIN PE=1 SV=4 | 1392 | 20201117_08512_1ng_glyco_hcd203040_compound15.13952.13952.3 | controllerType=0 controllerNumber=1 scan=13952 | 30.496 |
| 09172:3 | 31 | R.IIVPLNNREN[+1663.608]ISDPTSPLR.T | HexNAc(5)Hex(4) | N[+1664] | 1271.2679 | 3 | 3811.7891 | 3811.7853 | 0 | 1 | 62 | Specific | 570.43 | 564.72 | 564.72 | 10.19 | 24 | >sp\|P01591\|IGJ_HUMAN Immunoglobulin J chain OS=Homo sapiens OX=9606 GN=JCHAIN PE=1 SV=4 | 1392 | 20201117_08512_1ng_glyco_hcd203040_compound15.12382.12382.3 | controllerType=0 controllerNumber=1 scan=12382 | 27.5469 |
| 10687:3 | 31 | R.IIVPLNNREN[+1856.656]ISDPTSPLR.T | HexNAc(3)Hex(5)Fuc(1)NeuAc(1) | N[+1857] | 1335.618 | 3 | 4004.8394 | 4004.8327 | 0 | 1.7 | 62 | Specific | 574.3 | 574.3 | 574.3 | 10.19 | 24 | >sp\|P01591\|IGJ_HUMAN Immunoglobulin J chain OS=Homo sapiens OX=9606 GN=JCHAIN PE=1 SV=4 | 1392 | 20201117_08512_1ng_glyco_hcd203040_compound15.13954.13954.3 | controllerType=0 controllerNumber=1 scan=13954 | 30.5098 |
| 09146:3 | 31 | R.IIVPLNNREN[+1663.608]ISDPTSPLR.T | HexNAc(5)Hex(4) | N[+1664] | 1271.2696 | 3 | 3811.7942 | 3811.7853 | 0 | 2.3 | 62 | Specific | 592.13 | 567.86 | 567.86 | 11.34 | 24 | >sp\|P01591\|IGJ_HUMAN Immunoglobulin J chain OS=Homo sapiens OX=9606 GN=JCHAIN PE=1 SV=4 | 1392 | 20201117_08512_1ng_glyco_hcd203040_compound15.12354.12354.3 | controllerType=0 controllerNumber=1 scan=12354 | 27.4859 |
| 09144:3 | 31 | R.IIVPLNNREN[+1663.608]ISDPTSPLR.T | HexNAc(5)Hex(4) | N[+1664] | 1271.2696 | 3 | 3811.7942 | 3811.7853 | 0 | 2.3 | 62 | Specific | 622.1 | 587.59 | 587.59 | 12.34 | 24 | >sp\|P01591\|IGJ_HUMAN Immunoglobulin J chain OS=Homo sapiens OX=9606 GN=JCHAIN PE=1 SV=4 | 1392 | 20201117_08512_1ng_glyco_hcd203040_compound15.12352.12352.3 | controllerType=0 controllerNumber=1 scan=12352 | 27.4793 |
| 14730:3 | 44 | R.KLPPGLLAN[+2059.735]FTLLR.T | HexNAc(4)Hex(5)Fuc(1)NeuAc(1) | N[+2060] | 1204.9037 | 3 | 3612.6967 | 3612.6923 | 0 | 1.2 | 178 | Specific | 418.31 | 418.31 | 418.31 | 9.25 | 14 | >sp\|P02750\|A2GL_HUMAN Leucine-rich alpha-2-glycoprotein OS=Homo sapiens OX=9606 GN=LRG1 PE=1 SV=2 | 518 | 20201117_08512_1ng_glyco_hcd203040_compound15.18164.18164.3 | controllerType=0 controllerNumber=1 scan=18164 | 38.826 |
| 14732:3 | 44 | R.KLPPGLLAN[+2059.735]FTLLR.T | HexNAc(4)Hex(5)Fuc(1)NeuAc(1) | N[+2060] | 1204.9037 | 3 | 3612.6967 | 3612.6923 | 0 | 1.2 | 178 | Specific | 493.83 | 423.22 | 423.22 | 9.48 | 14 | >sp\|P02750\|A2GL_HUMAN Leucine-rich alpha-2-glycoprotein OS=Homo sapiens OX=9606 GN=LRG1 PE=1 SV=2 | 518 | 20201117_08512_1ng_glyco_hcd203040_compound15.18166.18166.3 | controllerType=0 controllerNumber=1 scan=18166 | 38.8317 |
| 15195:4 | 74 | R.FSLLGHASISC[+57.021]TVEN[+2350.830]ETIGVWRPSPPTC[+57.021]EK.I | HexNAc(4)Hex(5)Fuc(1)NeuAc(2) | C[+57]*2, N[+2351] | 1431.6193 | 4 | 5723.4552 | 5723.4655 | 0 | -1.8 | 207 | Specific | 223.31 | 176.91 | 176.91 | 3.81 | 10 | >sp\|P04003\|C4BPA_HUMAN C4b-binding protein alpha chain OS=Homo sapiens OX=9606 GN=C4BPA PE=1 SV=2 | 218 | 20201117_08512_1ng_glyco_hcd203040_compound15.18666.18666.4 | controllerType=0 controllerNumber=1 scan=18666 | 39.7495 |
| 00802:2 | 86 | R.SYN[+1038.375]DSVDPR.I | HexNAc(2)Hex(3)Fuc(1) | N[+1038] | 1045.9255 | 2 | 2090.8436 | 2090.8395 | 0 | 2 | 481 | Specific | 203.15 | 51.56 | 51.56 | 2.61 | 11 | >sp\|P05164\|PERM_HUMAN Myeloperoxidase OS=Homo sapiens OX=9606 GN=MPO PE=1 SV=1 | 12 | 20201117_08512_1ng_glyco_hcd203040_compound15.3707.3707.2 | controllerType=0 controllerNumber=1 scan=3707 | 12.4926 |
| 00800:2 | 86 | R.SYN[+1038.375]DSVDPR.I | HexNAc(2)Hex(3)Fuc(1) | N[+1038] | 1045.9255 | 2 | 2090.8436 | 2090.8395 | 0 | 2 | 481 | Specific | 216.32 | 178.33 | 178.33 | 4.2 | 11 | >sp\|P05164\|PERM_HUMAN Myeloperoxidase OS=Homo sapiens OX=9606 GN=MPO PE=1 SV=1 | 12 | 20201117_08512_1ng_glyco_hcd203040_compound15.3705.3705.2 | controllerType=0 controllerNumber=1 scan=3705 | 12.4882 |
| 22910:3 | 126 | R.AVN[+2204.772]ITSENLIDDVVSLIR.G | HexNAc(4)Hex(5)NeuAc(2) | N[+2205] | 1392.6265 | 3 | 4175.865 | 4175.8481 | 0 | 4 | 413 | Specific | 339.23 | 295.55 | 295.55 | 6.63 | 7 | >sp\|P02748\|CO9_HUMAN Complement component C9 OS=Homo sapiens OX=9606 GN=C9 PE=1 SV=2 | 1547 | 20201117_08512_1ng_glyco_hcd203040_compound15.26720.26720.3 | controllerType=0 controllerNumber=1 scan=26720 | 57.27 |
| 08549:3 | 230 | R.FVN[+876.322]VTVTPEDQC[+57.021]RPNNVC[+57.021]TGVLTR.R | HexNAc(2)Hex(2)Fuc(1) | C[+57]*2, N[+876] | 1218.2303 | 3 | 3652.6763 | 3652.6728 | 0 | 1 | 169 | Specific | 490.9 | 490.9 | 490.9 | 10.16 | 3 | >sp\|P20160\|CAP7_HUMAN Azurocidin OS=Homo sapiens OX=9606 GN=AZU1 PE=1 SV=3 | 850 | 20201117_08512_1ng_glyco_hcd203040_compound15.11733.11733.3 | controllerType=0 controllerNumber=1 scan=11733 | 26.3208 |
| 08551:3 | 230 | R.FVN[+876.322]VTVTPEDQC[+57.021]RPNNVC[+57.021]TGVLTR.R | HexNAc(2)Hex(2)Fuc(1) | C[+57]*2, N[+876] | 1218.2303 | 3 | 3652.6763 | 3652.6728 | 0 | 1 | 169 | Specific | 569.19 | 569.19 | 569.19 | 10.61 | 3 | >sp\|P20160\|CAP7_HUMAN Azurocidin OS=Homo sapiens OX=9606 GN=AZU1 PE=1 SV=3 | 850 | 20201117_08512_1ng_glyco_hcd203040_compound15.11735.11735.3 | controllerType=0 controllerNumber=1 scan=11735 | 26.3243 |
| 18818:4 | 1 | R.RTAGWNVPIGTLRPFLN[+2350.830]WTGPPEPIEAAVAR.F | HexNAc(4)Hex(5)Fuc(1)NeuAc(2) | N[+2351] | 1435.1704 | 4 | 5737.6598 | 5737.6424 | 0 | 3 | 140 | Specific | 219.54 | 219.54 | 219.54 | 5.06 | 130 | >sp\|P02788\|TRFL_HUMAN Lactotransferrin OS=Homo sapiens OX=9606 GN=LTF PE=1 SV=6 | 1270 | 20201119_08533_1ng_glyco_hcd203040_compound15.22534.22534.4 | controllerType=0 controllerNumber=1 scan=22534 | 45.9277 |
| 16880:4 | 1 | R.RTAGWNVPIGTLRPFLN[+1768.640]WTGPPEPIEAAVAR.F | HexNAc(4)Hex(5)Fuc(1) | N[+1769] | 1289.6219 | 4 | 5155.4658 | 5155.4516 | 0 | 2.8 | 140 | Specific | 279.32 | 233.37 | 233.37 | 6.37 | 130 | >sp\|P02788\|TRFL_HUMAN Lactotransferrin OS=Homo sapiens OX=9606 GN=LTF PE=1 SV=6 | 1270 | 20201119_08533_1ng_glyco_hcd203040_compound15.20511.20511.4 | controllerType=0 controllerNumber=1 scan=20511 | 41.4857 |
| 17793:5 | 1 | R.RTAGWNVPIGTLRPFLN[+2059.735]WTGPPEPIEAAVAR.F | HexNAc(4)Hex(5)Fuc(1)NeuAc(1) | N[+2060] | 1090.1179 | 5 | 5446.5605 | 5446.547 | 0 | 2.5 | 140 | Specific | 306.03 | 306.03 | 306.03 | 6.87 | 130 | >sp\|P02788\|TRFL_HUMAN Lactotransferrin OS=Homo sapiens OX=9606 GN=LTF PE=1 SV=6 | 1270 | 20201119_08533_1ng_glyco_hcd203040_compound15.21465.21465.5 | controllerType=0 controllerNumber=1 scan=21465 | 43.6045 |
| 17797:3 | 1 | R.RTAGWNVPIGTLRPFLN[+2059.735]WTGPPEPIEAAVAR.F | HexNAc(4)Hex(5)Fuc(1)NeuAc(1) | N[+2060] | 1816.1958 | 3 | 5446.5727 | 5446.547 | 0 | 4.7 | 140 | Specific | 318.48 | 318.48 | 318.48 | 5.48 | 130 | >sp\|P02788\|TRFL_HUMAN Lactotransferrin OS=Homo sapiens OX=9606 GN=LTF PE=1 SV=6 | 1270 | 20201119_08533_1ng_glyco_hcd203040_compound15.21469.21469.3 | controllerType=0 controllerNumber=1 scan=21469 | 43.6108 |
| 17788:4 | 1 | R.RTAGWNVPIGTLRPFLN[+2059.735]WTGPPEPIEAAVAR.F | HexNAc(4)Hex(5)Fuc(1)NeuAc(1) | N[+2060] | 1362.3977 | 4 | 5446.569 | 5446.547 | 0 | 4 | 140 | Specific | 328.58 | 328.58 | 328.58 | 7.31 | 130 | >sp\|P02788\|TRFL_HUMAN Lactotransferrin OS=Homo sapiens OX=9606 GN=LTF PE=1 SV=6 | 1270 | 20201119_08533_1ng_glyco_hcd203040_compound15.21460.21460.4 | controllerType=0 controllerNumber=1 scan=21460 | [illegible] |
| 16882:4 | 1 | R.RTAGWNVPIGTLRPFLN[+1768.640]WTGPPEPIEAAVAR.F | HexNAc(4)Hex(5)Fuc(1) | N[+1769] | 1289.6219 | 4 | 5155.4658 | 5155.4516 | 0 | 2.8 | 140 | Specific | 504.65 | 504.65 | 504.65 | 9.83 | 130 | >sp\|P02788\|TRFL_HUMAN Lactotransferrin OS=Homo sapiens OX=9606 GN=LTF PE=1 SV=6 | 1270 | 20201119_08533_1ng_glyco_hcd203040_compound15.20513.20513.4 | controllerType=0 controllerNumber=1 scan=20513 | 41.4904 |
| 17663:4 | 1 | R.RTAGWNVPIGTLRPFLN[+2570.925]WTGPPEPIEAAVAR.F | HexNAc(5)Hex(6)Fuc(2)NeuAc(1) | N[+2571] | 1490.1948 | 4 | 5957.7575 | 5957.7371 | 0 | 3.4 | 140 | Specific | 515.39 | 476.94 | 476.94 | 9.83 | 130 | >sp\|P02788\|TRFL_HUMAN Lactotransferrin OS=Homo sapiens OX=9606 GN=LTF PE=1 SV=6 | 1270 | 20201119_08533_1ng_glyco_hcd203040_compound15.21330.21330.4 | controllerType=0 controllerNumber=1 scan=21330 | 43.3282 |
| 17665:4 | 1 | R.RTAGWNVPIGTLRPFLN[+2570.925]WTGPPEPIEAAVAR.F | HexNAc(5)Hex(6)Fuc(2)NeuAc(1) | N[+2571] | 1490.1948 | 4 | 5957.7575 | 5957.7371 | 0 | 3.4 | 140 | Specific | 519.86 | 391.09 | 391.09 | 9.83 | 130 | >sp\|P02788\|TRFL_HUMAN Lactotransferrin OS=Homo sapiens OX=9606 GN=LTF PE=1 SV=6 | 1270 | 20201119_08533_1ng_glyco_hcd203040_compound15.21332.21332.4 | controllerType=0 controllerNumber=1 scan=21332 | 43.3332 |
| 17796:5 | 1 | R.RTAGWNVPIGTLRPFLN[+2059.735]WTGPPEPIEAAVAR.F | HexNAc(4)Hex(5)Fuc(1)NeuAc(1) | N[+2060] | 1090.1179 | 5 | 5446.5605 | 5446.547 | 0 | 2.5 | 140 | Specific | 558.4 | 558.4 | 558.4 | 9.94 | 130 | >sp\|P02788\|TRFL_HUMAN Lactotransferrin OS=Homo sapiens OX=9606 GN=LTF PE=1 SV=6 | 1270 | 20201119_08533_1ng_glyco_hcd203040_compound15.21468.21468.5 | controllerType=0 controllerNumber=1 scan=21468 | 43.6084 |
| 18820:4 | 1 | R.RTAGWNVPIGTLRPFLN[+2350.830]WTGPPEPIEAAVAR.F | HexNAc(4)Hex(5)Fuc(1)NeuAc(2) | N[+2351] | 1435.1704 | 4 | 5737.6598 | 5737.6424 | 0 | 3 | 140 | Specific | 598.81 | 491.18 | 491.18 | 11.37 | 130 | >sp\|P02788\|TRFL_HUMAN Lactotransferrin OS=Homo sapiens OX=9606 GN=LTF PE=1 SV=6 | 1270 | 20201119_08533_1ng_glyco_hcd203040_compound15.22536.22536.4 | controllerType=0 controllerNumber=1 scan=22536 | 45.9315 |
| 17790:4 | 1 | R.RTAGWNVPIGTLRPFLN[+2059.735]WTGPPEPIEAAVAR.F | HexNAc(4)Hex(5)Fuc(1)NeuAc(1) | N[+2060] | 1362.3977 | 4 | 5446.569 | 5446.547 | 0 | 4 | 140 | Specific | 613.86 | 499.73 | 499.73 | 12.11 | 130 | >sp\|P02788\|TRFL_HUMAN Lactotransferrin OS=Homo sapiens OX=9606 GN=LTF PE=1 SV=6 | 1270 | 20201119_08533_1ng_glyco_hcd203040_compound15.21462.21462.4 | controllerType=0 controllerNumber=1 scan=21462 | 43.5968 |
| 17980:3 | 1 | R.TAGWNVPIGTLRPFLN[+2279.830]WTGPPEPIEAAVAR.F | HexNAc(5)Hex(6)Fuc(2) | N[+2280] | 1837.5246 | 3 | 5510.5593 | 5510.5406 | 0 | 3.4 | 141 | Specific | 221.56 | 221.56 | 221.56 | 3.98 | 130 | >sp\|P02788\|TRFL_HUMAN Lactotransferrin OS=Homo sapiens OX=9606 GN=LTF PE=1 SV=6 | 1270 | 20201119_08533_1ng_glyco_hcd203040_compound15.21659.21659.3 | controllerType=0 controllerNumber=1 scan=21659 | 43.9869 |
| 18992:4 | 1 | R.TAGWNVPIGTLRPFLN[+2205.793]WTGPPEPIEAAVAR.F | HexNAc(4)Hex(5)Fuc(2)NeuAc(1) | N[+2206] | 1359.8831 | 4 | 5436.5104 | 5436.5038 | 0 | 1.2 | 141 | Specific | 423.83 | 347 | 347 | 7.23 | 130 | >sp\|P02788\|TRFL_HUMAN Lactotransferrin OS=Homo sapiens OX=9606 GN=LTF PE=1 SV=6 | 1270 | 20201119_08533_1ng_glyco_hcd203040_compound15.22716.22716.4 | controllerType=0 controllerNumber=1 scan=22716 | 46.3264 |
| 19016:3 | 1 | R.TAGWNVPIGTLRPFLN[+2059.735]WTGPPEPIEAAVAR.F | HexNAc(4)Hex(5)Fuc(1)NeuAc(1) | N[+2060] | 1764.1585 | 3 | 5290.461 | 5290.4459 | 0 | 2.9 | 141 | Specific | 442.11 | 404.03 | 404.03 | 9.59 | 130 | >sp\|P02788\|TRFL_HUMAN Lactotransferrin OS=Homo sapiens OX=9606 GN=LTF PE=1 SV=6 | 1270 | 20201119_08533_1ng_glyco_hcd203040_compound15.22741.22741.2 | controllerType=0 controllerNumber=1 scan=22741 | 46.383 |
| 18903:4 | 1 | R.TAGWNVPIGTLRPFLN[+2424.867]WTGPPEPIEAAVAR.F | HexNAc(5)Hex(6)Fuc(1)NeuAc(1) | N[+2425] | 1414.6543 | 4 | 5655.5954 | 5655.5781 | 0 | 3.1 | 141 | Specific | 495.52 | 380.92 | 380.92 | 9.58 | 130 | >sp\|P02788\|TRFL_HUMAN Lactotransferrin OS=Homo sapiens OX=9606 GN=LTF PE=1 SV=6 | 1270 | 20201119_08533_1ng_glyco_hcd203040_compound15.22623.22623.4 | controllerType=0 controllerNumber=1 scan=22623 | 46.1289 |
| 18847:3 | 1 | R.TAGWNVPIGTLRPFLN[+2570.925]WTGPPEPIEAAVAR.F | HexNAc(5)Hex(6)Fuc(2)NeuAc(1) | N[+2571] | 1934.5556 | 3 | 5801.6523 | 5801.636 | 0 | 2.8 | 141 | Specific | 538.88 | 406.31 | 406.31 | 10 | 130 | >sp\|P02788\|TRFL_HUMAN Lactotransferrin OS=Homo sapiens OX=9606 GN=LTF PE=1 SV=6 | 1270 | 20201119_08533_1ng_glyco_hcd203040_compound15.22564.22564.3 | controllerType=0 controllerNumber=1 scan=22564 | 45.9848 |
| 18849:3 | 1 | R.TAGWNVPIGTLRPFLN[+2570.925]WTGPPEPIEAAVAR.F | HexNAc(5)Hex(6)Fuc(2)NeuAc(1) | N[+2571] | 1934.5556 | 3 | 5801.6523 | 5801.636 | 0 | 2.8 | 141 | Specific | 548.03 | 426.21 | 426.21 | 10.21 | 130 | >sp\|P02788\|TRFL_HUMAN Lactotransferrin OS=Homo sapiens OX=9606 GN=LTF PE=1 SV=6 | 1270 | 20201119_08533_1ng_glyco_hcd203040_compound15.22566.22566.3 | controllerType=0 controllerNumber=1 scan=22566 | 45.9902 |
| 20308:3 | 1 | R.TAGWNVPIGTLRPFLN[+2350.830]WTGPPEPIEAAVAR.F | HexNAc(4)Hex(5)Fuc(1)NeuAc(2) | N[+2351] | 1861.1882 | 3 | 5581.55 | 5581.5413 | 0 | 1.6 | 141 | Specific | 632.35 | 444.35 | 444.35 | 12.34 | 130 | >sp\|P02788\|TRFL_HUMAN Lactotransferrin OS=Homo sapiens OX=9606 GN=LTF PE=1 SV=6 | 1270 | 20201119_08533_1ng_glyco_hcd203040_compound15.24094.24094.3 | controllerType=0 controllerNumber=1 scan=24094 | 49.1964 |
| 18993:3 | 1 | R.TAGWNVPIGTLRPFLN[+2205.793]WTGPPEPIEAAVAR.F | HexNAc(4)Hex(5)Fuc(2)NeuAc(1) | N[+2206] | 1812.8455 | 3 | 5436.5221 | 5436.5038 | 0 | 3.4 | 141 | Specific | 638.24 | 585.73 | 585.73 | 12.37 | 130 | >sp\|P02788\|TRFL_HUMAN Lactotransferrin OS=Homo sapiens OX=9606 GN=LTF PE=1 SV=6 | 1270 | 20201119_08533_1ng_glyco_hcd203040_compound15.22717.22717.3 | controllerType=0 controllerNumber=1 scan=22717 | 46.3281 |
| 20306:3 | 1 | R.TAGWNVPIGTLRPFLN[+2350.830]WTGPPEPIEAAVAR.F | HexNAc(4)Hex(5)Fuc(1)NeuAc(2) | N[+2351] | 1861.1882 | 3 | 5581.55 | 5581.5413 | 0 | 1.6 | 141 | Specific | 640.48 | 465.13 | 465.13 | 12.34 | 130 | >sp\|P02788\|TRFL_HUMAN Lactotransferrin OS=Homo sapiens OX=9606 GN=LTF PE=1 SV=6 | 1270 | 20201119_08533_1ng_glyco_hcd203040_compound15.24092.24092.3 | controllerType=0 controllerNumber=1 scan=24092 | 49.1896 |
| 18985:5 | 1 | R.TAGWNVPIGTLRPFLN[+2059.735]WTGPPEPIEAAVAR.F | HexNAc(4)Hex(5)Fuc(1)NeuAc(1) | N[+2060] | 1058.9003 | 5 | 5290.4722 | 5290.4459 | 0 | 5 | 141 | Specific | 661.88 | 632.51 | 632.51 | 13.51 | 130 | >sp\|P02788\|TRFL_HUMAN Lactotransferrin OS=Homo sapiens OX=9606 GN=LTF PE=1 SV=6 | 1270 | 20201119_08533_1ng_glyco_hcd203040_compound15.22708.22708.5 | controllerType=0 controllerNumber=1 scan=22708 | 46.305 |
| 18971:3 | 1 | R.TAGWNVPIGTLRPFLN[+2059.735]WTGPPEPIEAAVAR.F | HexNAc(4)Hex(5)Fuc(1)NeuAc(1) | N[+2060] | 1764.1626 | 3 | 5290.4731 | 5290.4459 | 0 | 5.2 | 141 | Specific | 663.5 | 602.32 | 602.32 | 13.11 | 130 | >sp\|P02788\|TRFL_HUMAN Lactotransferrin OS=Homo sapiens OX=9606 GN=LTF PE=1 SV=6 | 1270 | 20201119_08533_1ng_glyco_hcd203040_compound15.22694.22694.3 | controllerType=0 controllerNumber=1 scan=22694 | 46.2749 |
| 20292:4 | 1 | R.TAGWNVPIGTLRPFLN[+2350.830]WTGPPEPIEAAVAR.F | HexNAc(4)Hex(5)Fuc(1)NeuAc(2) | N[+2351] | 1396.1445 | 4 | 5581.5563 | 5581.5413 | 0 | 2.7 | 141 | Specific | 682.96 | 598.78 | 598.78 | 13.09 | 130 | >sp\|P02788\|TRFL_HUMAN Lactotransferrin OS=Homo sapiens OX=9606 GN=LTF PE=1 SV=6 | 1270 | 20201119_08533_1ng_glyco_hcd203040_compound15.24078.24078.4 | controllerType=0 controllerNumber=1 scan=24078 | 49.1468 |
| 18032:3 | 1 | R.TAGWNVPIGTLRPFLN[+1768.640]WTGPPEPIEAAVAR.F | HexNAc(4)Hex(5)Fuc(1) | N[+1769] | 1667.1263 | 3 | 4999.3644 | 4999.3504 | 0 | 2.8 | 141 | Specific | 689.39 | 558.65 | 558.65 | 13.37 | 130 | >sp\|P02788\|TRFL_HUMAN Lactotransferrin OS=Homo sapiens OX=9606 GN=LTF PE=1 SV=6 | 1270 | 20201119_08533_1ng_glyco_hcd203040_compound15.21714.21714.3 | controllerType=0 controllerNumber=1 scan=21714 | 44.1189 |
| 18988:5 | 1 | R.TAGWNVPIGTLRPFLN[+2059.735]WTGPPEPIEAAVAR.F | HexNAc(4)Hex(5)Fuc(1)NeuAc(1) | N[+2060] | 1058.9003 | 5 | 5290.4722 | 5290.4459 | 0 | 5 | 141 | Specific | 691.79 | 576.45 | 576.45 | 13.51 | 130 | >sp\|P02788\|TRFL_HUMAN Lactotransferrin OS=Homo sapiens OX=9606 GN=LTF PE=1 SV=6 | 1270 | 20201119_08533_1ng_glyco_hcd203040_compound15.22711.22711.5 | controllerType=0 controllerNumber=1 scan=22711 | 46.3175 |
| 18970:4 | 1 | R.TAGWNVPIGTLRPFLN[+2059.735]WTGPPEPIEAAVAR.F | HexNAc(4)Hex(5)Fuc(1)NeuAc(1) | N[+2060] | 1323.3744 | 4 | 5290.4757 | 5290.4459 | 0 | 5.6 | 141 | Specific | 691.89 | 644.77 | 644.77 | 12.35 | 130 | >sp\|P02788\|TRFL_HUMAN Lactotransferrin OS=Homo sapiens OX=9606 GN=LTF PE=1 SV=6 | 1270 | 20201119_08533_1ng_glyco_hcd203040_compound15.22693.22693.4 | controllerType=0 controllerNumber=1 scan=22693 | 46.2735 |
| 18940:4 | 1 | R.TAGWNVPIGTLRPFLN[+2351.851]WTGPPEPIEAAVAR.F | HexNAc(4)Hex(5)Fuc(3)NeuAc(1) | N[+2352] | 1396.3984 | 4 | 5582.5719 | 5582.5617 | 0 | 1.8 | 141 | Specific | 698.35 | 630.9 | 630.9 | 13.09 | 130 | >sp\|P02788\|TRFL_HUMAN Lactotransferrin OS=Homo sapiens OX=9606 GN=LTF PE=1 SV=6 | 1270 | 20201119_08533_1ng_glyco_hcd203040_compound15.22661.22661.4 | controllerType=0 controllerNumber=1 scan=22661 | 46.2107 |
| 18906:4 | 1 | R.TAGWNVPIGTLRPFLN[+2424.867]WTGPPEPIEAAVAR.F | HexNAc(5)Hex(6)Fuc(1)NeuAc(1) | N[+2425] | 1414.6543 | 4 | 5655.5954 | 5655.5781 | 0 | 3.1 | 141 | Specific | 705.21 | 576.98 | 576.98 | 14.12 | 130 | >sp\|P02788\|TRFL_HUMAN Lactotransferrin OS=Homo sapiens OX=9606 GN=LTF PE=1 SV=6 | 1270 | 20201119_08533_1ng_glyco_hcd203040_compound15.22626.22626.4 | controllerType=0 controllerNumber=1 scan=22626 | 46.1339 |
| 18031:4 | 1 | R.TAGWNVPIGTLRPFLN[+1768.640]WTGPPEPIEAAVAR.F | HexNAc(4)Hex(5)Fuc(1) | N[+1769] | 1250.5956 | 4 | 4999.3605 | 4999.3504 | 0 | 2 | 141 | Specific | 714.99 | 689.7 | 689.7 | 14.09 | 130 | >sp\|P02788\|TRFL_HUMAN Lactotransferrin OS=Homo sapiens OX=9606 GN=LTF PE=1 SV=6 | 1270 | 20201119_08533_1ng_glyco_hcd203040_compound15.21713.21713.4 | controllerType=0 controllerNumber=1 scan=21713 | 44.1155 |
| 18907:4 | 1 | R.TAGWNVPIGTLRPFLN[+2716.983]WTGPPEPIEAAVAR.F | HexNAc(5)Hex(6)Fuc(3)NeuAc(1) | N[+2717] | 1487.6846 | 4 | 5947.7165 | 5947.6939 | 0 | 3.8 | 141 | Specific | 721.51 | 607.85 | 607.85 | 14.12 | 130 | >sp\|P02788\|TRFL_HUMAN Lactotransferrin OS=Homo sapiens OX=9606 GN=LTF PE=1 SV=6 | 1270 | 20201119_08533_1ng_glyco_hcd203040_compound15.22627.22627.4 | controllerType=0 controllerNumber=1 scan=22627 | 46.1376 |
| 18030:3 | 1 | R.TAGWNVPIGTLRPFLN[+1768.640]WTGPPEPIEAAVAR.F | HexNAc(4)Hex(5)Fuc(1) | N[+1769] | 1667.1263 | 3 | 4999.3644 | 4999.3504 | 0 | 2.8 | 141 | Specific | 737.3 | 627.53 | 627.53 | 14.37 | 130 | >sp\|P02788\|TRFL_HUMAN Lactotransferrin OS=Homo sapiens OX=9606 GN=LTF PE=1 SV=6 | 1270 | 20201119_08533_1ng_glyco_hcd203040_compound15.21712.21712.3 | controllerType=0 controllerNumber=1 scan=21712 | 44.1141 |
| 20313:5 | 1 | R.TAGWNVPIGTLRPFLN[+2350.830]WTGPPEPIEAAVAR.F | HexNAc(4)Hex(5)Fuc(1)NeuAc(2) | N[+2351] | 1117.1183 | 5 | 5581.5623 | 5581.5413 | 0 | 3.8 | 141 | Specific | 744.45 | 658.45 | 658.45 | 14.76 | 130 | >sp\|P02788\|TRFL_HUMAN Lactotransferrin OS=Homo sapiens OX=9606 GN=LTF PE=1 SV=6 | 1270 | 20201119_08533_1ng_glyco_hcd203040_compound15.24100.24100.5 | controllerType=0 controllerNumber=1 scan=24100 | 49.2051 |
| 18996:3 | 1 | R.TAGWNVPIGTLRPFLN[+2205.793]WTGPPEPIEAAVAR.F | HexNAc(4)Hex(5)Fuc(2)NeuAc(1) | N[+2206] | 1812.8455 | 3 | 5436.5221 | 5436.5038 | 0 | 3.4 | 141 | Specific | 757.24 | 697.01 | 697.01 | 15.37 | 130 | >sp\|P02788\|TRFL_HUMAN Lactotransferrin OS=Homo sapiens OX=9606 GN=LTF PE=1 SV=6 | 1270 | 20201119_08533_1ng_glyco_hcd203040_compound15.22720.22720.3 | controllerType=0 controllerNumber=1 scan=22720 | 46.3365 |
| 18973:3 | 1 | R.TAGWNVPIGTLRPFLN[+2059.735]WTGPPEPIEAAVAR.F | HexNAc(4)Hex(5)Fuc(1)NeuAc(1) | N[+2060] | 1764.1626 | 3 | 5290.4731 | 5290.4459 | 0 | 5.2 | 141 | Specific | 778.71 | 621.35 | 621.35 | 15.11 | 130 | >sp\|P02788\|TRFL_HUMAN Lactotransferrin OS=Homo sapiens OX=9606 GN=LTF PE=1 SV=6 | 1270 | 20201119_08533_1ng_glyco_hcd203040_compound15.22696.22696.3 | controllerType=0 controllerNumber=1 scan=22696 | 46.2821 |
| 19019:3 | 1 | R.TAGWNVPIGTLRPFLN[+2059.735]WTGPPEPIEAAVAR.F | HexNAc(4)Hex(5)Fuc(1)NeuAc(1) | N[+2060] | 1764.1585 | 3 | 5290.461 | 5290.4459 | 0 | 2.9 | 141 | Specific | 779.67 | 654.35 | 654.35 | 15.37 | 130 | >sp\|P02788\|TRFL_HUMAN Lactotransferrin OS=Homo sapiens OX=9606 GN=LTF PE=1 SV=6 | 1270 | 20201119_08533_1ng_glyco_hcd203040_compound15.22744.22744.2 | controllerType=0 controllerNumber=1 scan=22744 | 46.3942 |
| 18943:4 | 1 | R.TAGWNVPIGTLRPFLN[+2351.851]WTGPPEPIEAAVAR.F | HexNAc(4)Hex(5)Fuc(3)NeuAc(1) | N[+2352] | 1396.3984 | 4 | 5582.5719 | 5582.5617 | 0 | 1.8 | 141 | Specific | 783.2 | 634.88 | 634.88 | 15.09 | 130 | >sp\|P02788\|TRFL_HUMAN Lactotransferrin OS=Homo sapiens OX=9606 GN=LTF PE=1 SV=6 | 1270 | 20201119_08533_1ng_glyco_hcd203040_compound15.22664.22664.4 | controllerType=0 controllerNumber=1 scan=22664 | 46.2185 |
| 18995:4 | 1 | R.TAGWNVPIGTLRPFLN[+2205.793]WTGPPEPIEAAVAR.F | HexNAc(4)Hex(5)Fuc(2)NeuAc(1) | N[+2206] | 1359.8831 | 4 | 5436.5104 | 5436.5038 | 0 | 1.2 | 141 | Specific | 818.94 | 686 | 686 | 15.54 | 130 | >sp\|P02788\|TRFL_HUMAN Lactotransferrin OS=Homo sapiens OX=9606 GN=LTF PE=1 SV=6 | 1270 | 20201119_08533_1ng_glyco_hcd203040_compound15.22719.22719.4 | controllerType=0 controllerNumber=1 scan=22719 | 46.332 |
| 18972:4 | 1 | R.TAGWNVPIGTLRPFLN[+2059.735]WTGPPEPIEAAVAR.F | HexNAc(4)Hex(5)Fuc(1)NeuAc(1) | N[+2060] | 1323.3744 | 4 | 5290.4757 | 5290.4459 | 0 | 5.6 | 141 | Specific | 849.79 | 703.91 | 703.91 | 15.35 | 130 | >sp\|P02788\|TRFL_HUMAN Lactotransferrin OS=Homo sapiens OX=9606 GN=LTF PE=1 SV=6 | 1270 | 20201119_08533_1ng_glyco_hcd203040_compound15.22695.22695.4 | controllerType=0 controllerNumber=1 scan=22695 | 46.2799 |
| 18033:4 | 1 | R.TAGWNVPIGTLRPFLN[+1768.640]WTGPPEPIEAAVAR.F | HexNAc(4)Hex(5)Fuc(1) | N[+1769] | 1250.5956 | 4 | 4999.3605 | 4999.3504 | 0 | 2 | 141 | Specific | 880.95 | 780.29 | 780.29 | 17.09 | 130 | >sp\|P02788\|TRFL_HUMAN Lactotransferrin OS=Homo sapiens OX=9606 GN=LTF PE=1 SV=6 | 1270 | 20201119_08533_1ng_glyco_hcd203040_compound15.21715.21715.4 | controllerType=0 controllerNumber=1 scan=21715 | 44.1208 |
| 20316:5 | 1 | R.TAGWNVPIGTLRPFLN[+2350.830]WTGPPEPIEAAVAR.F | HexNAc(4)Hex(5)Fuc(1)NeuAc(2) | N[+2351] | 1117.1183 | 5 | 5581.5623 | 5581.5413 | 0 | 3.8 | 141 | Specific | 917.69 | 628.12 | 628.12 | 18.76 | 130 | >sp\|P02788\|TRFL_HUMAN Lactotransferrin OS=Homo sapiens OX=9606 GN=LTF PE=1 SV=6 | 1270 | 20201119_08533_1ng_glyco_hcd203040_compound15.24103.24103.5 | controllerType=0 controllerNumber=1 scan=24103 | 49.2166 |
| 20598:3 | 1 | R.TAGWNIPMGLLFN[+2205.793]QTGSC[+57.021]K.F | HexNAc(4)Hex(5)Fuc(2)NeuAc(1) | C[+57], N[+2206] | 1434.2677 | 3 | 4300.7885 | 4300.8028 | 0 | -3.3 | 485 | Specific | 451.2 | 357.76 | 357.76 | 6.17 | 130 | >sp\|P02788\|TRFL_HUMAN Lactotransferrin OS=Homo sapiens OX=9606 GN=LTF PE=1 SV=6 | 1270 | 20201119_08533_1ng_glyco_hcd203040_compound15.24398.24398.3 | controllerType=0 controllerNumber=1 scan=24398 | 49.8767 |
| 20600:3 | 1 | R.TAGWNIPMGLLFN[+2205.793]QTGSC[+57.021]K.F | HexNAc(4)Hex(5)Fuc(2)NeuAc(1) | C[+57], N[+2206] | 1434.2677 | 3 | 4300.7885 | 4300.8028 | 0 | -3.3 | 485 | Specific | 471.41 | 414.26 | 414.26 | 7.94 | 130 | >sp\|P02788\|TRFL_HUMAN Lactotransferrin OS=Homo sapiens OX=9606 GN=LTF PE=1 SV=6 | 1270 | 20201119_08533_1ng_glyco_hcd203040_compound15.24400.24400.3 | controllerType=0 controllerNumber=1 scan=24400 | 49.8818 |
| 16879:3 | 1 | R.TAGWNIPMGLLFN[+1216.423]QTGSC[+57.021]K.F | HexNAc(2)Hex(5) | C[+57], N[+1216] | 1104.4878 | 3 | 3311.4488 | 3311.4328 | 0 | 4.8 | 485 | Specific | 522.93 | 522.93 | 150.34 | 9.52 | 130 | >sp\|P02788\|TRFL_HUMAN Lactotransferrin OS=Homo sapiens OX=9606 GN=LTF PE=1 SV=6 | 1270 | 20201119_08533_1ng_glyco_hcd203040_compound15.20510.20510.3 | controllerType=0 controllerNumber=1 scan=20510 | 41.4841 |
| 16724:3 | 1 | R.TAGWNIPMGLLFN[+1622.582]QTGSC[+57.021]K.F | HexNAc(4)Hex(5) | C[+57], N[+1623] | 1239.8697 | 3 | 3717.5945 | 3717.5916 | 0 | 0.8 | 485 | Specific | 523.46 | 523.46 | 205.96 | 9.19 | 130 | >sp\|P02788\|TRFL_HUMAN Lactotransferrin OS=Homo sapiens OX=9606 GN=LTF PE=1 SV=6 | 1270 | 20201119_08533_1ng_glyco_hcd203040_compound15.20348.20348.3 | controllerType=0 controllerNumber=1 scan=20348 | 41.1233 |
| 18319:3 | 1 | R.TAGWNIPMGLLFN[+2205.793]QTGSC[+57.021]K.F | HexNAc(4)Hex(5)Fuc(2)NeuAc(1) | C[+57], N[+2206] | 1434.2751 | 3 | 4300.8107 | 4300.8028 | 0 | 1.8 | 485 | Specific | 626.35 | 624.58 | 624.58 | 12.12 | 130 | >sp\|P02788\|TRFL_HUMAN Lactotransferrin OS=Homo sapiens OX=9606 GN=LTF PE=1 SV=6 | 1270 | 20201119_08533_1ng_glyco_hcd203040_compound15.22014.22014.3 | controllerType=0 controllerNumber=1 scan=22014 | 44.795 |
| 20438:3 | 1 | R.TAGWNIPMGLLFN[+2350.830]QTGSC[+57.021]K.F | HexNAc(4)Hex(5)Fuc(1)NeuAc(2) | C[+57], N[+2351] | 1482.6236 | 3 | 4445.8562 | 4445.8403 | 0 | 3.6 | 485 | Specific | 710.11 | 612.43 | 612.43 | 14.14 | 130 | >sp\|P02788\|TRFL_HUMAN Lactotransferrin OS=Homo sapiens OX=9606 GN=LTF PE=1 SV=6 | 1270 | 20201119_08533_1ng_glyco_hcd203040_compound15.24230.24230.3 | controllerType=0 controllerNumber=1 scan=24230 | 49.4644 |
| 16726:3 | 1 | R.TAGWNIPMGLLFN[+1622.582]QTGSC[+57.021]K.F | HexNAc(4)Hex(5) | C[+57], N[+1623] | 1239.8697 | 3 | 3717.5945 | 3717.5916 | 0 | 0.8 | 485 | Specific | 713.96 | 571.65 | 349.76 | 13.56 | 130 | >sp\|P02788\|TRFL_HUMAN Lactotransferrin OS=Homo sapiens OX=9606 GN=LTF PE=1 SV=6 | 1270 | 20201119_08533_1ng_glyco_hcd203040_compound15.20350.20350.3 | controllerType=0 controllerNumber=1 scan=20350 | 41.1272 |
| 16446:3 | 1 | R.TAGWNIPMGLLFN[+1768.640]QTGSC[+57.021]K.F | HexNAc(4)Hex(5)Fuc(1) | C[+57], N[+1769] | 1288.5639 | 3 | 3863.6771 | 3863.6495 | 0 | 7.2 | 485 | Specific | 855.81 | 788.77 | 788.77 | 15.94 | 130 | >sp\|P02788\|TRFL_HUMAN Lactotransferrin OS=Homo sapiens OX=9606 GN=LTF PE=1 SV=6 | 1270 | 20201119_08533_1ng_glyco_hcd203040_compound15.20058.20058.3 | controllerType=0 controllerNumber=1 scan=20058 | 40.5788 |
| 18321:3 | 1 | R.TAGWNIPMGLLFN[+2205.793]QTGSC[+57.021]K.F | HexNAc(4)Hex(5)Fuc(2)NeuAc(1) | C[+57], N[+2206] | 1434.2751 | 3 | 4300.8107 | 4300.8028 | 0 | 1.8 | 485 | Specific | 895.21 | 798.29 | 798.29 | 17.12 | 130 | >sp\|P02788\|TRFL_HUMAN Lactotransferrin OS=Homo sapiens OX=9606 GN=LTF PE=1 SV=6 | 1270 | 20201119_08533_1ng_glyco_hcd203040_compound15.22016.22016.3 | controllerType=0 controllerNumber=1 scan=22016 | 44.7986 |
| 18508:3 | 1 | R.TAGWNIPMGLLFN[+1913.677]QTGSC[+57.021]K.F | HexNAc(4)Hex(5)NeuAc(1) | C[+57], N[+1914] | 1336.904 | 3 | 4008.6974 | 4008.687 | 0 | 2.6 | 485 | Specific | 896.17 | 896.17 | 896.17 | 17.12 | 130 | >sp\|P02788\|TRFL_HUMAN Lactotransferrin OS=Homo sapiens OX=9606 GN=LTF PE=1 SV=6 | 1270 | 20201119_08533_1ng_glyco_hcd203040_compound15.22212.22212.3 | controllerType=0 controllerNumber=1 scan=22212 | 45.2756 |
| 16448:3 | 1 | R.TAGWNIPMGLLFN[+1768.640]QTGSC[+57.021]K.F | HexNAc(4)Hex(5)Fuc(1) | C[+57], N[+1769] | 1288.5639 | 3 | 3863.6771 | 3863.6495 | 0 | 7.2 | 485 | Specific | 907.34 | 696.63 | 696.63 | 16.94 | 130 | >sp\|P02788\|TRFL_HUMAN Lactotransferrin OS=Homo sapiens OX=9606 GN=LTF PE=1 SV=6 | 1270 | 20201119_08533_1ng_glyco_hcd203040_compound15.20060.20060.3 | controllerType=0 controllerNumber=1 scan=20060 | 40.5887 |
| 20440:3 | 1 | R.TAGWNIPMGLLFN[+2350.830]QTGSC[+57.021]K.F | HexNAc(4)Hex(5)Fuc(1)NeuAc(2) | C[+57], N[+2351] | 1482.6236 | 3 | 4445.8562 | 4445.8403 | 0 | 3.6 | 485 | Specific | 911.7 | 884.86 | 884.86 | 18.14 | 130 | >sp\|P02788\|TRFL_HUMAN Lactotransferrin OS=Homo sapiens OX=9606 GN=LTF PE=1 SV=6 | 1270 | 20201119_08533_1ng_glyco_hcd203040_compound15.24232.24232.3 | controllerType=0 controllerNumber=1 scan=24232 | 49.4692 |
| 18510:3 | 1 | R.TAGWNIPMGLLFN[+1913.677]QTGSC[+57.021]K.F | HexNAc(4)Hex(5)NeuAc(1) | C[+57], N[+1914] | 1336.904 | 3 | 4008.6974 | 4008.687 | 0 | 2.6 | 485 | Specific | 921.86 | 842.73 | 842.73 | 18.12 | 130 | >sp\|P02788\|TRFL_HUMAN Lactotransferrin OS=Homo sapiens OX=9606 GN=LTF PE=1 SV=6 | 1270 | 20201119_08533_1ng_glyco_hcd203040_compound15.22214.22214.3 | controllerType=0 controllerNumber=1 scan=22214 | 45.2799 |
| 18379:3 | 1 | R.TAGWNIPMGLLFN[+2059.735]QTGSC[+57.021]K.F | HexNAc(4)Hex(5)Fuc(1)NeuAc(1) | C[+57], N[+2060] | 1385.591 | 3 | 4154.7584 | 4154.7449 | 0 | 3.3 | 485 | Specific | 932.35 | 915.96 | 915.96 | 18.14 | 130 | >sp\|P02788\|TRFL_HUMAN Lactotransferrin OS=Homo sapiens OX=9606 GN=LTF PE=1 SV=6 | 1270 | 20201119_08533_1ng_glyco_hcd203040_compound15.22076.22076.3 | controllerType=0 controllerNumber=1 scan=22076 | 44.9038 |
| 16881:3 | 1 | R.TAGWNIPMGLLFN[+1216.423]QTGSC[+57.021]K.F | HexNAc(2)Hex(5) | C[+57], N[+1216] | 1104.4878 | 3 | 3311.4488 | 3311.4328 | 0 | 4.8 | 485 | Specific | 957.02 | 957.02 | 442.44 | 18.89 | 130 | >sp\|P02788\|TRFL_HUMAN Lactotransferrin OS=Homo sapiens OX=9606 GN=LTF PE=1 SV=6 | 1270 | 20201119_08533_1ng_glyco_hcd203040_compound15.20512.20512.3 | controllerType=0 controllerNumber=1 scan=20512 | 41.4878 |
| 18377:3 | 1 | R.TAGWNIPMGLLFN[+2059.735]QTGSC[+57.021]K.F | HexNAc(4)Hex(5)Fuc(1)NeuAc(1) | C[+57], N[+2060] | 1385.591 | 3 | 4154.7584 | 4154.7449 | 0 | 3.3 | 485 | Specific | 979.65 | 950.92 | 950.92 | 19.14 | 130 | >sp\|P02788\|TRFL_HUMAN Lactotransferrin OS=Homo sapiens OX=9606 GN=LTF PE=1 SV=6 | 1270 | 20201119_08533_1ng_glyco_hcd203040_compound15.22074.22074.3 | controllerType=0 controllerNumber=1 scan=22074 | 44.8986 |
| 16648:3 | 1 | R.TAGWNIPM[+15.995]GLLFN[+2059.735]QTGSC[+57.021]K.F | HexNAc(4)Hex(5)Fuc(1)NeuAc(1) | C[+57], M[+16], N[+2060] | 1390.9225 | 3 | 4170.7531 | 4170.7398 | 0 | 3.2 | 485 | Specific | 286.59 | 223.73 | 223.73 | 6.24 | 130 | >sp\|P02788\|TRFL_HUMAN Lactotransferrin OS=Homo sapiens OX=9606 GN=LTF PE=1 SV=6 | 1270 | 20201119_08533_1ng_glyco_hcd203040_compound15.20269.20269.3 | controllerType=0 controllerNumber=1 scan=20269 | 40.9792 |
| 16558:3 | 1 | R.TAGWNIPM[+15.995]GLLFN[+1751.624]QTGSC[+57.021]K.F | HexNAc(4)Hex(4)NeuAc(1) | C[+57], M[+16], N[+1752] | 1288.2205 | 3 | 3862.647 | 3862.6291 | 0 | 4.6 | 485 | Specific | 339.43 | 249.78 | 249.78 | 5.48 | 130 | >sp\|P02788\|TRFL_HUMAN Lactotransferrin OS=Homo sapiens OX=9606 GN=LTF PE=1 SV=6 | 1270 | 20201119_08533_1ng_glyco_hcd203040_compound15.20175.20175.3 | controllerType=0 controllerNumber=1 scan=20175 | 40.8008 |
| 16650:3 | 1 | R.TAGWNIPM[+15.995]GLLFN[+2059.735]QTGSC[+57.021]K.F | HexNAc(4)Hex(5)Fuc(1)NeuAc(1) | C[+57], M[+16], N[+2060] | 1390.9225 | 3 | 4170.7531 | 4170.7398 | 0 | 3.2 | 485 | Specific | 572.44 | 506.11 | 506.11 | 9.99 | 130 | >sp\|P02788\|TRFL_HUMAN Lactotransferrin OS=Homo sapiens OX=9606 GN=LTF PE=1 SV=6 | 1270 | 20201119_08533_1ng_glyco_hcd203040_compound15.20271.20271.3 | controllerType=0 controllerNumber=1 scan=20271 | 40.985 |
| 20123:4 | 1 | R.TAGWNIPMGLLFN[+2350.830]QTGSC[+57.021]KFDEYFSQSC[+57.021]APGSDPR.S | HexNAc(4)Hex(5)Fuc(1)NeuAc(2) | C[+57]*2, N[+2351] | 1573.1556 | 4 | 6289.6007 | 6289.5875 | 0 | 2.1 | 485 | Specific | 273.23 | 273.23 | 273.23 | 6.39 | 130 | >sp\|P02788\|TRFL_HUMAN Lactotransferrin OS=Homo sapiens OX=9606 GN=LTF PE=1 SV=6 | 1270 | 20201119_08533_1ng_glyco_hcd203040_compound15.23900.23900.4 | controllerType=0 controllerNumber=1 scan=23900 | 48.7189 |
| 20093:4 | 1 | R.TAGWNIPMGLLFN[+2350.830]QTGSC[+57.021]KFDEYFSQSC[+57.021]APGSDPR.S | HexNAc(4)Hex(5)Fuc(1)NeuAc(2) | C[+57]*2, N[+2351] | 1573.1594 | 4 | 6289.6159 | 6289.5875 | 0 | 4.5 | 485 | Specific | 275.63 | 250.63 | 250.63 | 5.95 | 130 | >sp\|P02788\|TRFL_HUMAN Lactotransferrin OS=Homo sapiens OX=9606 GN=LTF PE=1 SV=6 | 1270 | 20201119_08533_1ng_glyco_hcd203040_compound15.23869.23869.4 | controllerType=0 controllerNumber=1 scan=23869 | 48.6642 |
| 18730:4 | 1 | R.TAGWNIPMGLLFN[+1913.677]QTGSC[+57.021]KFDEYFSQSC[+57.021]APGSDPR.S | HexNAc(4)Hex(5)NeuAc(1) | C[+57]*2, N[+1914] | 1463.8677 | 4 | 5852.4489 | 5852.4341 | 0 | 2.5 | 485 | Specific | 406.94 | 348.24 | 348.24 | 7.31 | 130 | >sp\|P02788\|TRFL_HUMAN Lactotransferrin OS=Homo sapiens OX=9606 GN=LTF PE=1 SV=6 | 1270 | 20201119_08533_1ng_glyco_hcd203040_compound15.22443.22443.4 | controllerType=0 controllerNumber=1 scan=22443 | 45.771 |
| 20125:4 | 1 | R.TAGWNIPMGLLFN[+2350.830]QTGSC[+57.021]KFDEYFSQSC[+57.021]APGSDPR.S | HexNAc(4)Hex(5)Fuc(1)NeuAc(2) | C[+57]*2, N[+2351] | 1573.1556 | 4 | 6289.6007 | 6289.5875 | 0 | 2.1 | 485 | Specific | 520.87 | 511.46 | 511.46 | 9.58 | 130 | >sp\|P02788\|TRFL_HUMAN Lactotransferrin OS=Homo sapiens OX=9606 GN=LTF PE=1 SV=6 | 1270 | 20201119_08533_1ng_glyco_hcd203040_compound15.23902.23902.4 | controllerType=0 controllerNumber=1 scan=23902 | 48.7235 |
| 20095:4 | 1 | R.TAGWNIPMGLLFN[+2350.830]QTGSC[+57.021]KFDEYFSQSC[+57.021]APGSDPR.S | HexNAc(4)Hex(5)Fuc(1)NeuAc(2) | C[+57]*2, N[+2351] | 1573.1594 | 4 | 6289.6159 | 6289.5875 | 0 | 4.5 | 485 | Specific | 538.84 | 538.84 | 538.84 | 9.35 | 130 | >sp\|P02788\|TRFL_HUMAN Lactotransferrin OS=Homo sapiens OX=9606 GN=LTF PE=1 SV=6 | 1270 | 20201119_08533_1ng_glyco_hcd203040_compound15.23871.23871.4 | controllerType=0 controllerNumber=1 scan=23871 | 48.6682 |
| 18679:4 | 1 | R.TAGWNIPMGLLFN[+1913.677]QTGSC[+57.021]KFDEYFSQSC[+57.021]APGSDPR.S | HexNAc(4)Hex(5)NeuAc(1) | C[+57]*2, N[+1914] | 1463.8656 | 4 | 5852.4406 | 5852.4341 | 0 | 1.1 | 485 | Specific | 617.66 | 562.4 | 562.4 | 11.39 | 130 | >sp\|P02788\|TRFL_HUMAN Lactotransferrin OS=Homo sapiens OX=9606 GN=LTF PE=1 SV=6 | 1270 | 20201119_08533_1ng_glyco_hcd203040_compound15.22390.22390.4 | controllerType=0 controllerNumber=1 scan=22390 | 45.6766 |
| 18682:4 | 1 | R.TAGWNIPMGLLFN[+1913.677]QTGSC[+57.021]KFDEYFSQSC[+57.021]APGSDPR.S | HexNAc(4)Hex(5)NeuAc(1) | C[+57]*2, N[+1914] | 1463.8656 | 4 | 5852.4406 | 5852.4341 | 0 | 1.1 | 485 | Specific | 690.67 | 636.07 | 636.07 | 12.39 | 130 | >sp\|P02788\|TRFL_HUMAN Lactotransferrin OS=Homo sapiens OX=9606 GN=LTF PE=1 SV=6 | 1270 | 20201119_08533_1ng_glyco_hcd203040_compound15.22393.22393.4 | controllerType=0 controllerNumber=1 scan=22393 | 45.6845 |
| 17604:3 | 1 | R.TAGWNIPMGLLFN[+1768.640]QTGSC[+57.021]KFDEYFSQSC[+57.021]APGSDPR.S | HexNAc(4)Hex(5)Fuc(1) | C[+57]*2, N[+1769] | 1903.1421 | 3 | 5707.4116 | 5707.3966 | 0 | 2.6 | 485 | Specific | 730.23 | 664.58 | 664.58 | 14.22 | 130 | >sp\|P02788\|TRFL_HUMAN Lactotransferrin OS=Homo sapiens OX=9606 GN=LTF PE=1 SV=6 | 1270 | 20201119_08533_1ng_glyco_hcd203040_compound15.21268.21268.3 | controllerType=0 controllerNumber=1 scan=21268 | 43.1826 |
| 17658:4 | 1 | R.TAGWNIPMGLLFN[+1622.582]QTGSC[+57.021]KFDEYFSQSC[+57.021]APGSDPR.S | HexNAc(4)Hex(5) | C[+57]*2, N[+1623] | 1391.0944 | 4 | 5561.3556 | 5561.3387 | 0 | 3 | 485 | Specific | 734.17 | 686.43 | 151.13 | 13.97 | 130 | >sp\|P02788\|TRFL_HUMAN Lactotransferrin OS=Homo sapiens OX=9606 GN=LTF PE=1 SV=6 | 1270 | 20201119_08533_1ng_glyco_hcd203040_compound15.21324.21324.2 | controllerType=0 controllerNumber=1 scan=21324 | 43.3123 |
| 17606:3 | 1 | R.TAGWNIPMGLLFN[+1768.640]QTGSC[+57.021]KFDEYFSQSC[+57.021]APGSDPR.S | HexNAc(4)Hex(5)Fuc(1) | C[+57]*2, N[+1769] | 1903.1421 | 3 | 5707.4116 | 5707.3966 | 0 | 2.6 | 485 | Specific | 784.48 | 632.86 | 632.86 | 15.22 | 130 | >sp\|P02788\|TRFL_HUMAN Lactotransferrin OS=Homo sapiens OX=9606 GN=LTF PE=1 SV=6 | 1270 | 20201119_08533_1ng_glyco_hcd203040_compound15.21270.21270.3 | controllerType=0 controllerNumber=1 scan=21270 | 43.1901 |
| 18733:4 | 1 | R.TAGWNIPMGLLFN[+1913.677]QTGSC[+57.021]KFDEYFSQSC[+57.021]APGSDPR.S | HexNAc(4)Hex(5)NeuAc(1) | C[+57]*2, N[+1914] | 1463.8677 | 4 | 5852.4489 | 5852.4341 | 0 | 2.5 | 485 | Specific | 818.32 | 763.81 | 763.81 | 15.95 | 130 | >sp\|P02788\|TRFL_HUMAN Lactotransferrin OS=Homo sapiens OX=9606 GN=LTF PE=1 SV=6 | 1270 | 20201119_08533_1ng_glyco_hcd203040_compound15.22446.22446.4 | controllerType=0 controllerNumber=1 scan=22446 | 45.7746 |
| 18611:4 | 1 | R.TAGWNIPMGLLFN[+2059.735]QTGSC[+57.021]KFDEYFSQSC[+57.021]APGSDPR.S | HexNAc(4)Hex(5)Fuc(1)NeuAc(1) | C[+57]*2, N[+2060] | 1500.3854 | 4 | 5998.5197 | 5998.492 | 0 | 4.6 | 485 | Specific | 855.8 | 806.71 | 806.71 | 16.72 | 130 | >sp\|P02788\|TRFL_HUMAN Lactotransferrin OS=Homo sapiens OX=9606 GN=LTF PE=1 SV=6 | 1270 | 20201119_08533_1ng_glyco_hcd203040_compound15.22320.22320.4 | controllerType=0 controllerNumber=1 scan=22320 | 45.5332 |
| 17592:4 | 1 | R.TAGWNIPMGLLFN[+1768.640]QTGSC[+57.021]KFDEYFSQSC[+57.021]APGSDPR.S | HexNAc(4)Hex(5)Fuc(1) | C[+57]*2, N[+1769] | 1427.6082 | 4 | 5707.4108 | 5707.3966 | 0 | 2.5 | 485 | Specific | 922.88 | 839.24 | 839.24 | 17.95 | 130 | >sp\|P02788\|TRFL_HUMAN Lactotransferrin OS=Homo sapiens OX=9606 GN=LTF PE=1 SV=6 | 1270 | 20201119_08533_1ng_glyco_hcd203040_compound15.21256.21256.4 | controllerType=0 controllerNumber=1 scan=21256 | 43.1595 |
| 17661:4 | 1 | R.TAGWNIPMGLLFN[+1622.582]QTGSC[+57.021]KFDEYFSQSC[+57.021]APGSDPR.S | HexNAc(4)Hex(5) | C[+57]*2, N[+1623] | 1391.0944 | 4 | 5561.3556 | 5561.3387 | 0 | 3 | 485 | Specific | 957.52 | 932.54 | 344.1 | 18.97 | 130 | >sp\|P02788\|TRFL_HUMAN Lactotransferrin OS=Homo sapiens OX=9606 GN=LTF PE=1 SV=6 | 1270 | 20201119_08533_1ng_glyco_hcd203040_compound15.21327.21327.2 | controllerType=0 controllerNumber=1 scan=21327 | 43.3222 |
| 18613:4 | 1 | R.TAGWNIPMGLLFN[+2059.735]QTGSC[+57.021]KFDEYFSQSC[+57.021]APGSDPR.S | HexNAc(4)Hex(5)Fuc(1)NeuAc(1) | C[+57]*2, N[+2060] | 1500.3854 | 4 | 5998.5197 | 5998.492 | 0 | 4.6 | 485 | Specific | 968.4 | 923.09 | 923.09 | 18.72 | 130 | >sp\|P02788\|TRFL_HUMAN Lactotransferrin OS=Homo sapiens OX=9606 GN=LTF PE=1 SV=6 | 1270 | 20201119_08533_1ng_glyco_hcd203040_compound15.22322.22322.4 | controllerType=0 controllerNumber=1 scan=22322 | 45.538 |

Table S4. Cont.

| Query #:z | Protein Rank | Glycopeptide | Glycan | Modification Type(s) | Observed m/z | z | Observed (M+H) | Calc. mass (M+H) | Off-by-x error | Mass error (ppm) | StartPosition | Cleavage | Score | Delta | Delta Mod | \|Log Prob\| | # of unique peptides | Protein Name | Protein DB number | Comment | Scan # | Scan Time |
|---|---|---|---|---|---|---|---|---|---|---|---|---|---|---|---|---|---|---|---|---|---|---|
| 17595:4 | 1 | R.TAGWNIPMGLLFN[+1768.640]QTGSC[+57.021]KFDEYFSQSC[+57.021]APGSDPR.S | HexNAc(4)Hex(5)Fuc(1) | C[+57]*2, N[+1769] | 1427.6082 | 4 | 5707.4108 | 5707.3966 | 0 | 2.5 | 485 | Specific | 1042.15 | 975.47 | 975.47 | 19.95 | 130 | >sp\|P02788\|TRFL_HUMAN Lactotransferrin OS=Homo sapiens OX=9606 GN=LTF PE=1 SV=6 | 1270 | 20201119_08533_1ng_glyco_hcd203040_compound15.21259.21259.4 | controllerType=0 controllerNumber=1 scan=21259 | 43.165 |
| 17807:4 | 1 | R.TAGWNIPM[+15.995]GLLFN[+2059.735]QTGSC[+57.021]KFDEYFSQSC[+57.021]A | HexNAc(4)Hex(5)Fuc(1)NeuAc(1) | C[+57]*2, M[+16], N[+2060] | 1504.3848 | 4 | 6014.5172 | 6014.487 | 0 | 5 | 485 | Specific | 514.58 | 448.8 | 448.8 | 9.18 | 130 | >sp\|P02788\|TRFL_HUMAN Lactotransferrin OS=Homo sapiens OX=9606 GN=LTF PE=1 SV=6 | 1270 | 20201119_08533_1ng_glyco_hcd203040_compound15.21479.21479.4 | controllerType=0 controllerNumber=1 scan=21479 | 43.632 |
| 17646:4 | 1 | R.TAGWNIPM[+15.995]GLLFN[+2059.735]QTGSC[+57.021]KFDEYFSQSC[+57.021]A | HexNAc(4)Hex(5)Fuc(1)NeuAc(1) | C[+57]*2, M[+16], N[+2060] | 1504.3832 | 4 | 6014.5109 | 6014.487 | 0 | 4 | 485 | Specific | 589.49 | 533.31 | 533.31 | 10.97 | 130 | >sp\|P02788\|TRFL_HUMAN Lactotransferrin OS=Homo sapiens OX=9606 GN=LTF PE=1 SV=6 | 1270 | 20201119_08533_1ng_glyco_hcd203040_compound15.21312.21312.4 | controllerType=0 controllerNumber=1 scan=21312 | 43.2866 |
| 17648:4 | 1 | R.TAGWNIPM[+15.995]GLLFN[+2059.735]QTGSC[+57.021]KFDEYFSQSC[+57.021]A | HexNAc(4)Hex(5)Fuc(1)NeuAc(1) | C[+57]*2, M[+16], N[+2060] | 1504.3832 | 4 | 6014.5109 | 6014.487 | 0 | 4 | 485 | Specific | 808.41 | 738.4 | 738.4 | 15.97 | 130 | >sp\|P02788\|TRFL_HUMAN Lactotransferrin OS=Homo sapiens OX=9606 GN=LTF PE=1 SV=6 | 1270 | 20201119_08533_1ng_glyco_hcd203040_compound15.21314.21314.4 | controllerType=0 controllerNumber=1 scan=21314 | 43.29 |
| 08666:3 | 1 | R.N[+2205.793]GSDC[+57.021]PDKFC[+57.021]LFQSETK.N | HexNAc(4)Hex(5)Fuc(2)NeuAc(1) | C[+57]*2, N[+2206] | 1413.5684 | 3 | 4238.6907 | 4238.6667 | 0 | 5.7 | 642 | Specific | 283 | 237.34 | 237.34 | 5.33 | 130 | >sp\|P02788\|TRFL_HUMAN Lactotransferrin OS=Homo sapiens OX=9606 GN=LTF PE=1 SV=6 | 1270 | 20201119_08533_1ng_glyco_hcd203040_compound15.11905.11905.3 | controllerType=0 controllerNumber=1 scan=11905 | 26.4904 |
| 08668:3 | 1 | R.N[+2205.793]GSDC[+57.021]PDKFC[+57.021]LFQSETK.N | HexNAc(4)Hex(5)Fuc(2)NeuAc(1) | C[+57]*2, N[+2206] | 1413.5684 | 3 | 4238.6907 | 4238.6667 | 0 | 5.7 | 642 | Specific | 306.05 | 161.15 | 161.15 | 4.63 | 130 | >sp\|P02788\|TRFL_HUMAN Lactotransferrin OS=Homo sapiens OX=9606 GN=LTF PE=1 SV=6 | 1270 | 20201119_08533_1ng_glyco_hcd203040_compound15.11907.11907.3 | controllerType=0 controllerNumber=1 scan=11907 | 26.494 |
| 08773:3 | 1 | R.N[+2059.735]GSDC[+57.021]PDKFC[+57.021]LFQSETK.N | HexNAc(4)Hex(5)Fuc(1)NeuAc(1) | C[+57]*2, N[+2060] | 1364.8825 | 3 | 4092.633 | 4092.6088 | 0 | 5.9 | 642 | Specific | 387.1 | 330.08 | 330.08 | 6.68 | 130 | >sp\|P02788\|TRFL_HUMAN Lactotransferrin OS=Homo sapiens OX=9606 GN=LTF PE=1 SV=6 | 1270 | 20201119_08533_1ng_glyco_hcd203040_compound15.12016.12016.3 | controllerType=0 controllerNumber=1 scan=12016 | 26.6799 |
| 11187:3 | 1 | R.N[+2350.830]GSDC[+57.021]PDKFC[+57.021]LFQSETK.N | HexNAc(4)Hex(5)Fuc(1)NeuAc(2) | C[+57]*2, N[+2351] | 1461.9085 | 3 | 4383.711 | 4383.7043 | 0 | 1.5 | 642 | Specific | 388.97 | 281.39 | 281.39 | 7.03 | 130 | >sp\|P02788\|TRFL_HUMAN Lactotransferrin OS=Homo sapiens OX=9606 GN=LTF PE=1 SV=6 | 1270 | 20201119_08533_1ng_glyco_hcd203040_compound15.14516.14516.3 | controllerType=0 controllerNumber=1 scan=14516 | 30.9165 |
| 11185:3 | 1 | R.N[+2350.830]GSDC[+57.021]PDKFC[+57.021]LFQSETK.N | HexNAc(4)Hex(5)Fuc(1)NeuAc(2) | C[+57]*2, N[+2351] | 1461.9085 | 3 | 4383.711 | 4383.7043 | 0 | 1.5 | 642 | Specific | 392.9 | 332.05 | 332.05 | 7.42 | 130 | >sp\|P02788\|TRFL_HUMAN Lactotransferrin OS=Homo sapiens OX=9606 GN=LTF PE=1 SV=6 | 1270 | 20201119_08533_1ng_glyco_hcd203040_compound15.14514.14514.3 | controllerType=0 controllerNumber=1 scan=14514 | 30.9128 |
| 08775:3 | 1 | R.N[+2059.735]GSDC[+57.021]PDKFC[+57.021]LFQSETK.N | HexNAc(4)Hex(5)Fuc(1)NeuAc(1) | C[+57]*2, N[+2060] | 1364.8825 | 3 | 4092.633 | 4092.6088 | 0 | 5.9 | 642 | Specific | 441.41 | 335.03 | 335.03 | 6.91 | 130 | >sp\|P02788\|TRFL_HUMAN Lactotransferrin OS=Homo sapiens OX=9606 GN=LTF PE=1 SV=6 | 1270 | 20201119_08533_1ng_glyco_hcd203040_compound15.12018.12018.3 | controllerType=0 controllerNumber=1 scan=12018 | 26.6843 |
| 06953:3 | 4 | R.FSDGLESN[+1864.634]SSTQFEVK.K | HexNAc(2)Hex(9) | N[+1865] | 1213.826 | 3 | 3639.4634 | 3639.4472 | 0 | 4.5 | 219 | Specific | 462.39 | 435.32 | 435.32 | 8.85 | 86 | >sp\|P0C0L5\|CO4B_HUMAN Complement C4-B OS=Homo sapiens OX=9606 GN=C4B PE=1 SV=2 | 1424 | 20201119_08533_1ng_glyco_hcd203040_compound15.10098.10098.3 | controllerType=0 controllerNumber=1 scan=10098 | 23.4369 |
| 06951:3 | 4 | R.FSDGLESN[+1864.634]SSTQFEVK.K | HexNAc(2)Hex(9) | N[+1865] | 1213.826 | 3 | 3639.4634 | 3639.4472 | 0 | 4.5 | 219 | Specific | 469.11 | 469.11 | 469.11 | 9.01 | 86 | >sp\|P0C0L5\|CO4B_HUMAN Complement C4-B OS=Homo sapiens OX=9606 GN=C4B PE=1 SV=2 | 1424 | 20201119_08533_1ng_glyco_hcd203040_compound15.10096.10096.3 | controllerType=0 controllerNumber=1 scan=10096 | 23.433 |
| 04441:3 | 4 | R.FSDGLESN[+1864.634]SSTQFEVKK.Y | HexNAc(2)Hex(9) | N[+1865] | 1256.5211 | 3 | 3767.5486 | 3767.5421 | 0 | 1.7 | 219 | Specific | 429.97 | 429.97 | 429.97 | 9.21 | 86 | >sp\|P0C0L5\|CO4B_HUMAN Complement C4-B OS=Homo sapiens OX=9606 GN=C4B PE=1 SV=2 | 1424 | 20201119_08533_1ng_glyco_hcd203040_compound15.7500.7500.3 | controllerType=0 controllerNumber=1 scan=7500 | 19.1273 |
| 04439:3 | 4 | R.FSDGLESN[+1864.634]SSTQFEVKK.Y | HexNAc(2)Hex(9) | N[+1865] | 1256.5211 | 3 | 3767.5486 | 3767.5421 | 0 | 1.7 | 219 | Specific | 590.6 | 381.92 | 381.92 | 10.97 | 86 | >sp\|P0C0L5\|CO4B_HUMAN Complement C4-B OS=Homo sapiens OX=9606 GN=C4B PE=1 SV=2 | 1424 | 20201119_08533_1ng_glyco_hcd203040_compound15.7498.7498.3 | controllerType=0 controllerNumber=1 scan=7498 | 19.1235 |
| 14002:4 | 6 | K.KGN[+2350.830]TTLNSFVIPSESDVPTHC[+57.021]PSQWWPYAGHC[+57.021]YK.I | HexNAc(4)Hex(5)Fuc(1)NeuAc(2) | C[+57]*2, N[+2351] | 1579.4253 | 4 | 6314.6794 | 6314.6521 | 0 | 4.3 | 342 | Specific | 330.12 | 217.94 | 217.94 | 6.2 | 85 | >sp\|P22897\|MRC1_HUMAN Macrophage mannose receptor 1 OS=Homo sapiens OX=9606 GN=MRC1 PE=1 SV=1 | 1049 | 20201119_08533_1ng_glyco_hcd203040_compound15.17467.17467.4 | controllerType=0 controllerNumber=1 scan=17467 | 36.0123 |
| 12739:5 | 6 | K.KGN[+2205.793]TTLNSFVIPSESDVPTHC[+57.021]PSQWWPYAGHC[+57.021]YK.I | HexNAc(4)Hex(5)Fuc(2)NeuAc(1) | C[+57]*2, N[+2206] | 1234.7303 | 5 | 6169.6226 | 6169.6146 | 0 | 1.3 | 342 | Specific | 426.33 | 426.33 | 426.33 | 9.27 | 85 | >sp\|P22897\|MRC1_HUMAN Macrophage mannose receptor 1 OS=Homo sapiens OX=9606 GN=MRC1 PE=1 SV=1 | 1049 | 20201119_08533_1ng_glyco_hcd203040_compound15.16158.16158.5 | controllerType=0 controllerNumber=1 scan=16158 | 33.6908 |
| 12737:5 | 6 | K.KGN[+2205.793]TTLNSFVIPSESDVPTHC[+57.021]PSQWWPYAGHC[+57.021]YK.I | HexNAc(4)Hex(5)Fuc(2)NeuAc(1) | C[+57]*2, N[+2206] | 1234.7303 | 5 | 6169.6226 | 6169.6146 | 0 | 1.3 | 342 | Specific | 462.62 | 388.13 | 388.13 | 9.26 | 85 | >sp\|P22897\|MRC1_HUMAN Macrophage mannose receptor 1 OS=Homo sapiens OX=9606 GN=MRC1 PE=1 SV=1 | 1049 | 20201119_08533_1ng_glyco_hcd203040_compound15.16156.16156.5 | controllerType=0 controllerNumber=1 scan=16156 | 33.6874 |
| 14000:4 | 6 | K.KGN[+2350.830]TTLNSFVIPSESDVPTHC[+57.021]PSQWWPYAGHC[+57.021]YK.I | HexNAc(4)Hex(5)Fuc(1)NeuAc(2) | C[+57]*2, N[+2351] | 1579.4253 | 4 | 6314.6794 | 6314.6521 | 0 | 4.3 | 342 | Specific | 466.13 | 466.13 | 466.13 | 9.04 | 85 | >sp\|P22897\|MRC1_HUMAN Macrophage mannose receptor 1 OS=Homo sapiens OX=9606 GN=MRC1 PE=1 SV=1 | 1049 | 20201119_08533_1ng_glyco_hcd203040_compound15.17465.17465.4 | controllerType=0 controllerNumber=1 scan=17465 | 36.0088 |
| 12746:5 | 6 | K.KGN[+2059.735]TTLNSFVIPSESDVPTHC[+57.021]PSQWWPYAGHC[+57.021]YK.I | HexNAc(4)Hex(5)Fuc(1)NeuAc(1) | C[+57]*2, N[+2060] | 1205.5192 | 5 | 6023.5667 | 6023.5567 | 0 | 1.7 | 342 | Specific | 551.9 | 551.9 | 551.9 | 10.44 | 85 | >sp\|P22897\|MRC1_HUMAN Macrophage mannose receptor 1 OS=Homo sapiens OX=9606 GN=MRC1 PE=1 SV=1 | 1049 | 20201119_08533_1ng_glyco_hcd203040_compound15.16165.16165.5 | controllerType=0 controllerNumber=1 scan=16165 | 33.7029 |
| 12744:5 | 6 | K.KGN[+2059.735]TTLNSFVIPSESDVPTHC[+57.021]PSQWWPYAGHC[+57.021]YK.I | HexNAc(4)Hex(5)Fuc(1)NeuAc(1) | C[+57]*2, N[+2060] | 1205.5192 | 5 | 6023.5667 | 6023.5567 | 0 | 1.7 | 342 | Specific | 620.92 | 530.79 | 530.79 | 12.59 | 85 | >sp\|P22897\|MRC1_HUMAN Macrophage mannose receptor 1 OS=Homo sapiens OX=9606 GN=MRC1 PE=1 SV=1 | 1049 | 20201119_08533_1ng_glyco_hcd203040_compound15.16163.16163.5 | controllerType=0 controllerNumber=1 scan=16163 | 33.6982 |
| 16124:3 | 6 | R.VWIALNSN[+2204.772]LTDNQYTWTDKWR.V | HexNAc(4)Hex(5)NeuAc(2) | N[+2205] | 1610.3648 | 3 | 4829.0799 | 4829.0617 | 0 | 3.8 | 1153 | Specific | 549.54 | 421.12 | 421.12 | 10.23 | 85 | >sp\|P22897\|MRC1_HUMAN Macrophage mannose receptor 1 OS=Homo sapiens OX=9606 GN=MRC1 PE=1 SV=1 | 1049 | 20201119_08533_1ng_glyco_hcd203040_compound15.19722.19722.3 | controllerType=0 controllerNumber=1 scan=19722 | 39.9887 |
| 16122:3 | 6 | R.VWIALNSN[+2204.772]LTDNQYTWTDKWR.V | HexNAc(4)Hex(5)NeuAc(2) | N[+2205] | 1610.3648 | 3 | 4829.0799 | 4829.0617 | 0 | 3.8 | 1153 | Specific | 586.99 | 439.14 | 439.14 | 11.38 | 85 | >sp\|P22897\|MRC1_HUMAN Macrophage mannose receptor 1 OS=Homo sapiens OX=9606 GN=MRC1 PE=1 SV=1 | 1049 | 20201119_08533_1ng_glyco_hcd203040_compound15.19720.19720.3 | controllerType=0 controllerNumber=1 scan=19720 | 39.9849 |
| 16102:3 | 6 | R.VWIALNSN[+2205.793]LTDNQYTWTDKWR.V | HexNAc(4)Hex(5)Fuc(2)NeuAc(1) | N[+2206] | 1610.705 | 3 | 4830.1005 | 4830.0821 | 0 | 3.8 | 1153 | Specific | 592.99 | 456.93 | 456.93 | 11.38 | 85 | >sp\|P22897\|MRC1_HUMAN Macrophage mannose receptor 1 OS=Homo sapiens OX=9606 GN=MRC1 PE=1 SV=1 | 1049 | 20201119_08533_1ng_glyco_hcd203040_compound15.19699.19699.3 | controllerType=0 controllerNumber=1 scan=19699 | 39.9504 |
| 16100:3 | 6 | R.VWIALNSN[+2205.793]LTDNQYTWTDKWR.V | HexNAc(4)Hex(5)Fuc(2)NeuAc(1) | N[+2206] | 1610.705 | 3 | 4830.1005 | 4830.0821 | 0 | 3.8 | 1153 | Specific | 741.84 | 593.25 | 593.25 | 14.38 | 85 | >sp\|P22897\|MRC1_HUMAN Macrophage mannose receptor 1 OS=Homo sapiens OX=9606 GN=MRC1 PE=1 SV=1 | 1049 | 20201119_08533_1ng_glyco_hcd203040_compound15.19697.19697.3 | controllerType=0 controllerNumber=1 scan=19697 | 39.9451 |
| 08056:3 | 6 | K.TAHC[+57.021]N[+2059.735]ESFYFLC[+57.021]K.R | HexNAc(4)Hex(5)Fuc(1)NeuAc(1) | C[+57]*2, N[+2060] | 1246.1592 | 3 | 3736.4632 | 3736.4545 | 0 | 2.3 | 1201 | Specific | 732.1 | 621.42 | 621.42 | 14.11 | 85 | >sp\|P22897\|MRC1_HUMAN Macrophage mannose receptor 1 OS=Homo sapiens OX=9606 GN=MRC1 PE=1 SV=1 | 1049 | 20201119_08533_1ng_glyco_hcd203040_compound15.11251.11251.3 | controllerType=0 controllerNumber=1 scan=11251 | 25.3637 |
| 08054:3 | 6 | K.TAHC[+57.021]N[+2059.735]ESFYFLC[+57.021]K.R | HexNAc(4)Hex(5)Fuc(1)NeuAc(1) | C[+57]*2, N[+2060] | 1246.1592 | 3 | 3736.4632 | 3736.4545 | 0 | 2.3 | 1201 | Specific | 739.49 | 542.51 | 542.51 | 14.11 | 85 | >sp\|P22897\|MRC1_HUMAN Macrophage mannose receptor 1 OS=Homo sapiens OX=9606 GN=MRC1 PE=1 SV=1 | 1049 | 20201119_08533_1ng_glyco_hcd203040_compound15.11249.11249.3 | controllerType=0 controllerNumber=1 scan=11249 | 25.3597 |
| 09692:3 | 7 | K.TVLTPATNHMGN[+1378.476]VTFTIPANR.E | HexNAc(2)Hex(6) | N[+1378] | 1211.8867 | 3 | 3633.6454 | 3633.6358 | 0 | 2.6 | 74 | Specific | 621.78 | 597.92 | 322.97 | 12.78 | 92 | >sp\|P01024\|CO3_HUMAN Complement C3 OS=Homo sapiens OX=9606 GN=C3 PE=1 SV=2 | 492 | 20201119_08533_1ng_glyco_hcd203040_compound15.12969.12969.3 | controllerType=0 controllerNumber=1 scan=12969 | 28.2838 |
| 09690:3 | 7 | K.TVLTPATNHMGN[+1378.476]VTFTIPANR.E | HexNAc(2)Hex(6) | N[+1378] | 1211.8867 | 3 | 3633.6454 | 3633.6358 | 0 | 2.6 | 74 | Specific | 654.58 | 517.08 | 372.13 | 13.78 | 92 | >sp\|P01024\|CO3_HUMAN Complement C3 OS=Homo sapiens OX=9606 GN=C3 PE=1 SV=2 | 492 | 20201119_08533_1ng_glyco_hcd203040_compound15.12967.12967.3 | controllerType=0 controllerNumber=1 scan=12967 | 28.28 |
| 18595:3 | 8 | K.QIGLYPVLVIDSSGYVNPN[+1913.677]YTGR.I | HexNAc(4)Hex(5)NeuAc(1) | N[+1914] | 1480.3392 | 3 | 4439.003 | 4438.9805 | 0 | 5.1 | 168 | Specific | 590.25 | 560.04 | 560.04 | 11.27 | 58 | >sp\|P01833\|PIGR_HUMAN Polymeric immunoglobulin receptor OS=Homo sapiens OX=9606 GN=PIGR PE=1 SV=4 | 902 | 20201119_08533_1ng_glyco_hcd203040_compound15.22302.22302.3 | controllerType=0 controllerNumber=1 scan=22302 | 45.4717 |
| 18598:3 | 8 | K.QIGLYPVLVIDSSGYVNPN[+1913.677]YTGR.I | HexNAc(4)Hex(5)NeuAc(1) | N[+1914] | 1480.3392 | 3 | 4439.003 | 4438.9805 | 0 | 5.1 | 168 | Specific | 643.77 | 599.35 | 599.35 | 12.27 | 58 | >sp\|P01833\|PIGR_HUMAN Polymeric immunoglobulin receptor OS=Homo sapiens OX=9606 GN=PIGR PE=1 SV=4 | 902 | 20201119_08533_1ng_glyco_hcd203040_compound15.22305.22305.3 | controllerType=0 controllerNumber=1 scan=22305 | 45.4777 |
| 18519:3 | 8 | K.QIGLYPVLVIDSSGYVNPN[+2059.735]YTGR.I | HexNAc(4)Hex(5)Fuc(1)NeuAc(1) | N[+2060] | 1529.022 | 3 | 4585.0515 | 4585.0384 | 0 | 2.9 | 168 | Specific | 870.32 | 724.3 | 724.3 | 17.53 | 58 | >sp\|P01833\|PIGR_HUMAN Polymeric immunoglobulin receptor OS=Homo sapiens OX=9606 GN=PIGR PE=1 SV=4 | 902 | 20201119_08533_1ng_glyco_hcd203040_compound15.22223.22223.3 | controllerType=0 controllerNumber=1 scan=22223 | 45.2998 |
| 18522:3 | 8 | K.QIGLYPVLVIDSSGYVNPN[+2059.735]YTGR.I | HexNAc(4)Hex(5)Fuc(1)NeuAc(1) | N[+2060] | 1529.022 | 3 | 4585.0515 | 4585.0384 | 0 | 2.9 | 168 | Specific | 922.6 | 769.54 | 769.54 | 18.53 | 58 | >sp\|P01833\|PIGR_HUMAN Polymeric immunoglobulin receptor OS=Homo sapiens OX=9606 GN=PIGR PE=1 SV=4 | 902 | 20201119_08533_1ng_glyco_hcd203040_compound15.22226.22226.3 | controllerType=0 controllerNumber=1 scan=22226 | 45.3054 |
| 20192:3 | 8 | R.[+42.011]LSLLEEPGN[+2190.793]GTFTVILNQLTSR.D | HexNAc(6)Hex(6) | .[42], N[+2191] | 1545.7089 | 3 | 4635.1122 | 4635.0963 | 0 | 3.4 | 413 | Specific | 373.14 | 326.39 | 326.39 | 6.59 | 58 | >sp\|P01833\|PIGR_HUMAN Polymeric immunoglobulin receptor OS=Homo sapiens OX=9606 GN=PIGR PE=1 SV=4 | 902 | 20201119_08533_1ng_glyco_hcd203040_compound15.23973.23973.3 | controllerType=0 controllerNumber=1 scan=23973 | 48.8989 |
| 20230:4 | 8 | R.LSLLEEPGN[+2059.735]GTFTVILNQLTSR.D | HexNAc(4)Hex(5)Fuc(1)NeuAc(1) | N[+2060] | 1116.2691 | 4 | 4462.0545 | 4462.0275 | 0 | 6.1 | 413 | Specific | 405.23 | 346.8 | 346.8 | 7.85 | 58 | >sp\|P01833\|PIGR_HUMAN Polymeric immunoglobulin receptor OS=Homo sapiens OX=9606 GN=PIGR PE=1 SV=4 | 902 | 20201119_08533_1ng_glyco_hcd203040_compound15.24013.24013.4 | controllerType=0 controllerNumber=1 scan=24013 | 48.9985 |
| 20190:3 | 8 | R.[+42.011]LSLLEEPGN[+2190.793]GTFTVILNQLTSR.D | HexNAc(6)Hex(6) | .[42], N[+2191] | 1545.7089 | 3 | 4635.1122 | 4635.0963 | 0 | 3.4 | 413 | Specific | 436.86 | 326.23 | 326.23 | 8.22 | 58 | >sp\|P01833\|PIGR_HUMAN Polymeric immunoglobulin receptor OS=Homo sapiens OX=9606 GN=PIGR PE=1 SV=4 | 902 | 20201119_08533_1ng_glyco_hcd203040_compound15.23971.23971.3 | controllerType=0 controllerNumber=1 scan=23971 | 48.8941 |
| 22525:4 | 8 | R.LSLLEEPGN[+2350.830]GTFTVILNQLTSR.D | HexNAc(4)Hex(5)Fuc(1)NeuAc(2) | N[+2351] | 1189.0409 | 4 | 4753.1419 | 4753.1229 | 0 | 4 | 413 | Specific | 585.44 | 468.17 | 468.17 | 11.92 | 58 | >sp\|P01833\|PIGR_HUMAN Polymeric immunoglobulin receptor OS=Homo sapiens OX=9606 GN=PIGR PE=1 SV=4 | 902 | 20201119_08533_1ng_glyco_hcd203040_compound15.26410.26410.5 | controllerType=0 controllerNumber=1 scan=26410 | 53.5694 |
| 20233:4 | 8 | R.LSLLEEPGN[+2059.735]GTFTVILNQLTSR.D | HexNAc(4)Hex(5)Fuc(1)NeuAc(1) | N[+2060] | 1116.2691 | 4 | 4462.0545 | 4462.0275 | 0 | 6.1 | 413 | Specific | 712.08 | 585.29 | 585.29 | 14.41 | 58 | >sp\|P01833\|PIGR_HUMAN Polymeric immunoglobulin receptor OS=Homo sapiens OX=9606 GN=PIGR PE=1 SV=4 | 902 | 20201119_08533_1ng_glyco_hcd203040_compound15.24016.24016.4 | controllerType=0 controllerNumber=1 scan=24016 | 49.0028 |
| 22527:4 | 8 | R.LSLLEEPGN[+2350.830]GTFTVILNQLTSR.D | HexNAc(4)Hex(5)Fuc(1)NeuAc(2) | N[+2351] | 1189.0409 | 4 | 4753.1419 | 4753.1229 | 0 | 4 | 413 | Specific | 723.36 | 561.06 | 561.06 | 14.92 | 58 | >sp\|P01833\|PIGR_HUMAN Polymeric immunoglobulin receptor OS=Homo sapiens OX=9606 GN=PIGR PE=1 SV=4 | 902 | 20201119_08533_1ng_glyco_hcd203040_compound15.26412.26412.5 | controllerType=0 controllerNumber=1 scan=26412 | 53.5742 |
| 22453:3 | 8 | R.LSLLEEPGN[+2350.830]GTFTVILNQLTSR.D | HexNAc(4)Hex(5)Fuc(1)NeuAc(2) | N[+2351] | 1585.053 | 3 | 4753.1446 | 4753.1229 | 0 | 4.6 | 413 | Specific | 916.58 | 698.23 | 698.23 | 18.27 | 58 | >sp\|P01833\|PIGR_HUMAN Polymeric immunoglobulin receptor OS=Homo sapiens OX=9606 GN=PIGR PE=1 SV=4 | 902 | 20201119_08533_1ng_glyco_hcd203040_compound15.26335.26335.3 | controllerType=0 controllerNumber=1 scan=26335 | 53.4075 |
| 22456:3 | 8 | R.LSLLEEPGN[+2350.830]GTFTVILNQLTSR.D | HexNAc(4)Hex(5)Fuc(1)NeuAc(2) | N[+2351] | 1585.053 | 3 | 4753.1446 | 4753.1229 | 0 | 4.6 | 413 | Specific | 976.29 | 694.94 | 694.94 | 19.27 | 58 | >sp\|P01833\|PIGR_HUMAN Polymeric immunoglobulin receptor OS=Homo sapiens OX=9606 GN=PIGR PE=1 SV=4 | 902 | 20201119_08533_1ng_glyco_hcd203040_compound15.26338.26338.3 | controllerType=0 controllerNumber=1 scan=26338 | 53.4118 |
| 20160:3 | 8 | R.LSLLEEPGN[+2059.735]GTFTVILNQLTSR.D | HexNAc(4)Hex(5)Fuc(1)NeuAc(1) | N[+2060] | 1488.0191 | 3 | 4462.0427 | 4462.0275 | 0 | 3.4 | 413 | Specific | 984.7 | 724.01 | 724.01 | 19.53 | 58 | >sp\|P01833\|PIGR_HUMAN Polymeric immunoglobulin receptor OS=Homo sapiens OX=9606 GN=PIGR PE=1 SV=4 | 902 | 20201119_08533_1ng_glyco_hcd203040_compound15.23940.23940.3 | controllerType=0 controllerNumber=1 scan=23940 | 48.8114 |
| 20158:3 | 8 | R.LSLLEEPGN[+2059.735]GTFTVILNQLTSR.D | HexNAc(4)Hex(5)Fuc(1)NeuAc(1) | N[+2060] | 1488.0191 | 3 | 4462.0427 | 4462.0275 | 0 | 3.4 | 413 | Specific | 1107.54 | 865.68 | 865.68 | 22.53 | 58 | >sp\|P01833\|PIGR_HUMAN Polymeric immunoglobulin receptor OS=Homo sapiens OX=9606 GN=PIGR PE=1 SV=4 | 902 | 20201119_08533_1ng_glyco_hcd203040_compound15.23938.23938.3 | controllerType=0 controllerNumber=1 scan=23938 | 48.8015 |
| 15735:3 | 8 | K.IIEGEPNLKVPGN[+2059.735]VTAVLGETLK.V | HexNAc(4)Hex(5)Fuc(1)NeuAc(1) | N[+2060] | 1484.3702 | 3 | 4451.096 | 4451.0843 | 0 | 2.6 | 457 | Specific | 597.53 | 597.53 | 597.53 | 11.36 | 58 | >sp\|P01833\|PIGR_HUMAN Polymeric immunoglobulin receptor OS=Homo sapiens OX=9606 GN=PIGR PE=1 SV=4 | 902 | 20201119_08533_1ng_glyco_hcd203040_compound15.19314.19314.3 | controllerType=0 controllerNumber=1 scan=19314 | 39.2904 |
| 15681:3 | 8 | K.IIEGEPNLKVPGN[+2205.793]VTAVLGETLK.V | HexNAc(4)Hex(5)Fuc(2)NeuAc(1) | N[+2206] | 1533.0547 | 3 | 4597.1497 | 4597.1422 | 0 | 1.6 | 457 | Specific | 763.24 | 763.24 | 763.24 | 15.36 | 58 | >sp\|P01833\|PIGR_HUMAN Polymeric immunoglobulin receptor OS=Homo sapiens OX=9606 GN=PIGR PE=1 SV=4 | 902 | 20201119_08533_1ng_glyco_hcd203040_compound15.19256.19256.3 | controllerType=0 controllerNumber=1 scan=19256 | 39.1782 |
| 15679:3 | 8 | K.IIEGEPNLKVPGN[+2205.793]VTAVLGETLK.V | HexNAc(4)Hex(5)Fuc(2)NeuAc(1) | N[+2206] | 1533.0547 | 3 | 4597.1497 | 4597.1422 | 0 | 1.6 | 457 | Specific | 991.81 | 896.84 | 896.84 | 19.36 | 58 | >sp\|P01833\|PIGR_HUMAN Polymeric immunoglobulin receptor OS=Homo sapiens OX=9606 GN=PIGR PE=1 SV=4 | 902 | 20201119_08533_1ng_glyco_hcd203040_compound15.19254.19254.3 | controllerType=0 controllerNumber=1 scan=19254 | 39.1728 |
| 15733:3 | 8 | K.IIEGEPNLKVPGN[+2059.735]VTAVLGETLK.V | HexNAc(4)Hex(5)Fuc(1)NeuAc(1) | N[+2060] | 1484.3702 | 3 | 4451.096 | 4451.0843 | 0 | 2.6 | 457 | Specific | 1037.97 | 981.79 | 981.79 | 20.36 | 58 | >sp\|P01833\|PIGR_HUMAN Polymeric immunoglobulin receptor OS=Homo sapiens OX=9606 GN=PIGR PE=1 SV=4 | 902 | 20201119_08533_1ng_glyco_hcd203040_compound15.19312.19312.3 | controllerType=0 controllerNumber=1 scan=19312 | 39.2823 |
| 16868:4 | 8 | K.IIEGEPNLKVPGN[+2350.830]VTAVLGETLKVPC[+57.021]HFPC[+57.021]K.F | HexNAc(4)Hex(5)Fuc(1)NeuAc(2) | C[+57]*2, N[+2351] | 1442.6721 | 4 | 5767.6667 | 5767.6372 | 0 | 5.1 | 457 | Specific | 455.76 | 455.76 | 455.76 | 8.35 | 58 | >sp\|P01833\|PIGR_HUMAN Polymeric immunoglobulin receptor OS=Homo sapiens OX=9606 GN=PIGR PE=1 SV=4 | 902 | 20201119_08533_1ng_glyco_hcd203040_compound15.20498.20498.4 | controllerType=0 controllerNumber=1 scan=20498 | 41.4524 |
| 15337:4 | 8 | K.IIEGEPNLKVPGN[+2059.735]VTAVLGETLKVPC[+57.021]HFPC[+57.021]K.F | HexNAc(4)Hex(5)Fuc(1)NeuAc(1) | C[+57]*2, N[+2060] | 1369.8912 | 4 | 5476.5431 | 5476.5418 | 0 | 0.2 | 457 | Specific | 568.47 | 523.55 | 523.55 | 8.65 | 58 | >sp\|P01833\|PIGR_HUMAN Polymeric immunoglobulin receptor OS=Homo sapiens OX=9606 GN=PIGR PE=1 SV=4 | 902 | 20201119_08533_1ng_glyco_hcd203040_compound15.18855.18855.4 | controllerType=0 controllerNumber=1 scan=18855 | 38.4077 |
| 16866:4 | 8 | K.IIEGEPNLKVPGN[+2350.830]VTAVLGETLKVPC[+57.021]HFPC[+57.021]K.F | HexNAc(4)Hex(5)Fuc(1)NeuAc(2) | C[+57]*2, N[+2351] | 1442.6721 | 4 | 5767.6667 | 5767.6372 | 0 | 5.1 | 457 | Specific | 598.16 | 466.07 | 466.07 | 10.13 | 58 | >sp\|P01833\|PIGR_HUMAN Polymeric immunoglobulin receptor OS=Homo sapiens OX=9606 GN=PIGR PE=1 SV=4 | 902 | 20201119_08533_1ng_glyco_hcd203040_compound15.20496.20496.4 | controllerType=0 controllerNumber=1 scan=20496 | 41.4485 |
| 15236:4 | 8 | K.IIEGEPNLKVPGN[+2204.772]VTAVLGETLKVPC[+57.021]HFPC[+57.021]K.F | HexNAc(4)Hex(5)NeuAc(2) | C[+57]*2, N[+2205] | 1406.158 | 4 | 5621.61 | 5621.5793 | 0 | 5.5 | 457 | Specific | 717.1 | 610 | 610 | 12.62 | 58 | >sp\|P01833\|PIGR_HUMAN Polymeric immunoglobulin receptor OS=Homo sapiens OX=9606 GN=PIGR PE=1 SV=4 | 902 | 20201119_08533_1ng_glyco_hcd203040_compound15.18747.18747.4 | controllerType=0 controllerNumber=1 scan=18747 | 38.215 |
| 15239:4 | 8 | K.IIEGEPNLKVPGN[+2204.772]VTAVLGETLKVPC[+57.021]HFPC[+57.021]K.F | HexNAc(4)Hex(5)NeuAc(2) | C[+57]*2, N[+2205] | 1406.158 | 4 | 5621.61 | 5621.5793 | 0 | 5.5 | 457 | Specific | 743.89 | 536.59 | 536.59 | 12.62 | 58 | >sp\|P01833\|PIGR_HUMAN Polymeric immunoglobulin receptor OS=Homo sapiens OX=9606 GN=PIGR PE=1 SV=4 | 902 | 20201119_08533_1ng_glyco_hcd203040_compound15.18750.18750.4 | controllerType=0 controllerNumber=1 scan=18750 | 38.2191 |
| 16837:5 | 8 | K.IIEGEPNLKVPGN[+2350.830]VTAVLGETLKVPC[+57.021]HFPC[+57.021]K.F | HexNAc(4)Hex(5)Fuc(1)NeuAc(2) | C[+57]*2, N[+2351] | 1154.3361 | 5 | 5767.6512 | 5767.6372 | 0 | 2.4 | 457 | Specific | 920.72 | 710.73 | 710.73 | 17.9 | 58 | >sp\|P01833\|PIGR_HUMAN Polymeric immunoglobulin receptor OS=Homo sapiens OX=9606 GN=PIGR PE=1 SV=4 | 902 | 20201119_08533_1ng_glyco_hcd203040_compound15.20466.20466.5 | controllerType=0 controllerNumber=1 scan=20466 | 41.3923 |
| 16835:5 | 8 | K.IIEGEPNLKVPGN[+2350.830]VTAVLGETLKVPC[+57.021]HFPC[+57.021]K.F | HexNAc(4)Hex(5)Fuc(1)NeuAc(2) | C[+57]*2, N[+2351] | 1154.3361 | 5 | 5767.6512 | 5767.6372 | 0 | 2.4 | 457 | Specific | 1012.91 | 787.24 | 787.24 | 19.9 | 58 | >sp\|P01833\|PIGR_HUMAN Polymeric immunoglobulin receptor OS=Homo sapiens OX=9606 GN=PIGR PE=1 SV=4 | 902 | 20201119_08533_1ng_glyco_hcd203040_compound15.20464.20464.5 | controllerType=0 controllerNumber=1 scan=20464 | 41.3869 |
| 15335:4 | 8 | K.IIEGEPNLKVPGN[+2059.735]VTAVLGETLKVPC[+57.021]HFPC[+57.021]K.F | HexNAc(4)Hex(5)Fuc(1)NeuAc(1) | C[+57]*2, N[+2060] | 1369.8912 | 4 | 5476.5431 | 5476.5418 | 0 | 0.2 | 457 | Specific | 1125.33 | 964.68 | 964.68 | 20.8 | 58 | >sp\|P01833\|PIGR_HUMAN Polymeric immunoglobulin receptor OS=Homo sapiens OX=9606 GN=PIGR PE=1 SV=4 | 902 | 20201119_08533_1ng_glyco_hcd203040_compound15.18853.18853.4 | controllerType=0 controllerNumber=1 scan=18853 | 38.4016 |
| 13278:4 | 8 | K.VPGN[+2205.793]VTAVLGETLKVPC[+57.021]HFPC[+57.021]K.F | HexNAc(4)Hex(5)Fuc(2)NeuAc(1) | C[+57]*2, N[+2206] | 1158.022 | 4 | 4629.0662 | 4629.0502 | 0 | 3.5 | 466 | Specific | 505.64 | 373.27 | 373.27 | 8.22 | 58 | >sp\|P01833\|PIGR_HUMAN Polymeric immunoglobulin receptor OS=Homo sapiens OX=9606 GN=PIGR PE=1 SV=4 | 902 | 20201119_08533_1ng_glyco_hcd203040_compound15.16716.16716.3 | controllerType=0 controllerNumber=1 scan=16716 | 34.6751 |
| 13276:4 | 8 | K.VPGN[+2205.793]VTAVLGETLKVPC[+57.021]HFPC[+57.021]K.F | HexNAc(4)Hex(5)Fuc(2)NeuAc(1) | C[+57]*2, N[+2206] | 1158.022 | 4 | 4629.0662 | 4629.0502 | 0 | 3.5 | 466 | Specific | 510.54 | 398.13 | 398.13 | 9.85 | 58 | >sp\|P01833\|PIGR_HUMAN Polymeric immunoglobulin receptor OS=Homo sapiens OX=9606 GN=PIGR PE=1 SV=4 | 902 | 20201119_08533_1ng_glyco_hcd203040_compound15.16714.16714.3 | controllerType=0 controllerNumber=1 scan=16714 | 34.6708 |
| 05570:3 | 8 | K.WN[+1548.545]NTGC[+57.021]QALPSQDEGPSK.A | HexNAc(3)Hex(4)NeuAc(1) | C[+57], N[+1549] | 1179.8159 | 3 | 3537.433 | 3537.4215 | 0 | 3.3 | 498 | Specific | 244.87 | 211.57 | 211.57 | 6.07 | 58 | >sp\|P01833\|PIGR_HUMAN Polymeric immunoglobulin receptor OS=Homo sapiens OX=9606 GN=PIGR PE=1 SV=4 | 902 | 20201119_08533_1ng_glyco_hcd203040_compound15.8667.8667.3 | controllerType=0 controllerNumber=1 scan=8667 | 21.0795 |
| 05572:3 | 8 | K.WN[+1548.545]NTGC[+57.021]QALPSQDEGPSK.A | HexNAc(3)Hex(4)NeuAc(1) | C[+57], N[+1549] | 1179.8159 | 3 | 3537.433 | 3537.4215 | 0 | 3.3 | 498 | Specific | 264.45 | 209.49 | 209.49 | 6.19 | 58 | >sp\|P01833\|PIGR_HUMAN Polymeric immunoglobulin receptor OS=Homo sapiens OX=9606 GN=PIGR PE=1 SV=4 | 902 | 20201119_08533_1ng_glyco_hcd203040_compound15.8669.8669.3 | controllerType=0 controllerNumber=1 scan=8669 | 21.0834 |
| 05390:3 | 8 | K.WN[+1872.651]NTGC[+57.021]QALPSQDEGPSK.A | HexNAc(3)Hex(6)NeuAc(1) | C[+57], N[+1873] | 1287.8536 | 3 | 3861.5462 | 3861.5272 | 0 | 4.9 | 498 | Specific | 279.68 | 263.69 | 263.69 | 6.12 | 58 | >sp\|P01833\|PIGR_HUMAN Polymeric immunoglobulin receptor OS=Homo sapiens OX=9606 GN=PIGR PE=1 SV=4 | 902 | 20201119_08533_1ng_glyco_hcd203040_compound15.8481.8481.3 | controllerType=0 controllerNumber=1 scan=8481 | 20.7671 |
| 05392:3 | 8 | K.WN[+1872.651]NTGC[+57.021]QALPSQDEGPSK.A | HexNAc(3)Hex(6)NeuAc(1) | C[+57], N[+1873] | 1287.8536 | 3 | 3861.5462 | 3861.5272 | 0 | 4.9 | 498 | Specific | 323.45 | 297.73 | 297.73 | 6.94 | 58 | >sp\|P01833\|PIGR_HUMAN Polymeric immunoglobulin receptor OS=Homo sapiens OX=9606 GN=PIGR PE=1 SV=4 | 902 | 20201119_08533_1ng_glyco_hcd203040_compound15.8483.8483.3 | controllerType=0 controllerNumber=1 scan=8483 | 20.7712 |
| 05328:3 | 8 | K.WN[+2059.735]NTGC[+57.021]QALPSQDEGPSK.A | HexNAc(4)Hex(5)Fuc(1)NeuAc(1) | C[+57], N[+2060] | 1350.2144 | 3 | 4048.6287 | 4048.6116 | 0 | 4.2 | 498 | Specific | 379.41 | 335.36 | 335.36 | 7.28 | 58 | >sp\|P01833\|PIGR_HUMAN Polymeric immunoglobulin receptor OS=Homo sapiens OX=9606 GN=PIGR PE=1 SV=4 | 902 | 20201119_08533_1ng_glyco_hcd203040_compound15.8417.8417.3 | controllerType=0 controllerNumber=1 scan=8417 | 20.6619 |
| 05326:3 | 8 | K.WN[+2059.735]NTGC[+57.021]QALPSQDEGPSK.A | HexNAc(4)Hex(5)Fuc(1)NeuAc(1) | C[+57], N[+2060] | 1350.2144 | 3 | 4048.6287 | 4048.6116 | 0 | 4.2 | 498 | Specific | 457.49 | 405.24 | 405.24 | 9.11 | 58 | >sp\|P01833\|PIGR_HUMAN Polymeric immunoglobulin receptor OS=Homo sapiens OX=9606 GN=PIGR PE=1 SV=4 | 902 | 20201119_08533_1ng_glyco_hcd203040_compound15.8415.8415.3 | controllerType=0 controllerNumber=1 scan=8415 | 20.6578 |
| 05448:3 | 8 | K.WN[+1710.598]NTGC[+57.021]QALPSQDEGPSK.A | HexNAc(3)Hex(5)NeuAc(1) | C[+57], N[+1711] | 1233.8314 | 3 | 3699.4795 | 3699.4743 | 0 | 1.4 | 498 | Specific | 574.06 | 515.28 | 515.28 | 11.12 | 58 | >sp\|P01833\|PIGR_HUMAN Polymeric immunoglobulin receptor OS=Homo sapiens OX=9606 GN=PIGR PE=1 SV=4 | 902 | 20201119_08533_1ng_glyco_hcd203040_compound15.8541.8541.3 | controllerType=0 controllerNumber=1 scan=8541 | 20.8691 |
| 05450:3 | 8 | K.WN[+1710.598]NTGC[+57.021]QALPSQDEGPSK.A | HexNAc(3)Hex(5)NeuAc(1) | C[+57], N[+1711] | 1233.8314 | 3 | 3699.4795 | 3699.4743 | 0 | 1.4 | 498 | Specific | 640.88 | 578.25 | 578.25 | 12.12 | 58 | >sp\|P01833\|PIGR_HUMAN Polymeric immunoglobulin receptor OS=Homo sapiens OX=9606 GN=PIGR PE=1 SV=4 | 902 | 20201119_08533_1ng_glyco_hcd203040_compound15.8543.8543.3 | controllerType=0 controllerNumber=1 scan=8543 | 20.8727 |
| 17535:3 | 11 | R.LSLHRPALEDLLLGSEAN[+1095.397]LTC[+57.021]TLTGLR.D | HexNAc(3)Hex(3) | C[+57], N[+1095] | 1353.6802 | 3 | 4059.026 | 4058.9949 | 0 | 7.7 | 127 | Specific | 281.43 | 281.43 | 281.43 | 5.28 | 48 | >sp\|P01876\|IGHA1_HUMAN Immunoglobulin heavy constant alpha 1 OS=Homo sapiens OX=9606 GN=IGHA1 PE=1 SV=2 | 255 | 20201119_08533_1ng_glyco_hcd203040_compound15.21196.21196.3 | controllerType=0 controllerNumber=1 scan=21196 | 43.0106 |
| 17433:4 | 11 | R.LSLHRPALEDLLLGSEAN[+1581.555]LTC[+57.021]TLTGLR.D | HexNAc(3)Hex(6) | C[+57], N[+1582] | 1137.0538 | 4 | 4545.1932 | 4545.1533 | 0 | 8.8 | 127 | Specific | 321.95 | 321.95 | 321.95 | 4.83 | 48 | >sp\|P01876\|IGHA1_HUMAN Immunoglobulin heavy constant alpha 1 OS=Homo sapiens OX=9606 GN=IGHA1 PE=1 SV=2 | 255 | 20201119_08533_1ng_glyco_hcd203040_compound15.21088.21088.4 | controllerType=0 controllerNumber=1 scan=21088 | 42.[illegible]017 |
| 17537:3 | 11 | R.LSLHRPALEDLLLGSEAN[+1095.397]LTC[+57.021]TLTGLR.D | HexNAc(3)Hex(3) | C[+57], N[+1095] | 1353.6802 | 3 | 4059.026 | 4058.9949 | 0 | 7.7 | 127 | Specific | 325.46 | 290.45 | 290.45 | 4.74 | 48 | >sp\|P01876\|IGHA1_HUMAN Immunoglobulin heavy constant alpha 1 OS=Homo sapiens OX=9606 GN=IGHA1 PE=1 SV=2 | 255 | 20201119_08533_1ng_glyco_hcd203040_compound15.21198.21198.3 | controllerType=0 controllerNumber=1 scan=21198 | 43.014 |
| 17460:3 | 11 | R.LSLHRPALEDLLLGSEAN[+1501.555]LTC[+57.021]TLTGLR.D | HexNAc(5)Hex(3) | C[+57], N[+1502] | 1489.0617 | 3 | 4465.1706 | 4465.1536 | 0 | 3.8 | 127 | Specific | 528.07 | 494.94 | 494.94 | 9.57 | 48 | >sp\|P01876\|IGHA1_HUMAN Immunoglobulin heavy constant alpha 1 OS=Homo sapiens OX=9606 GN=IGHA1 PE=1 SV=2 | 255 | 20201119_08533_1ng_glyco_hcd203040_compound15.21117.21117.3 | controllerType=0 controllerNumber=1 scan=21117 | 42.7911 |
| 17449:3 | 11 | R.LSLHRPALEDLLLGSEAN[+1540.529]LTC[+57.021]TLTGLR.D | HexNAc(2)Hex(7) | C[+57], N[+1541] | 1502.0614 | 3 | 4504.1696 | 4504.1268 | 0 | 9.5 | 127 | Specific | 557.11 | 557.11 | 557.11 | 7.91 | 48 | >sp\|P01876\|IGHA1_HUMAN Immunoglobulin heavy constant alpha 1 OS=Homo sapiens OX=9606 GN=IGHA1 PE=1 SV=2 | 255 | 20201119_08533_1ng_glyco_hcd203040_compound15.21105.21105.3 | controllerType=0 controllerNumber=1 scan=21105 | 42.7487 |
| 17319:4 | 11 | R.LSLHRPALEDLLLGSEAN[+1864.634]LTC[+57.021]TLTGLR.D | HexNAc(2)Hex(9) | C[+57], N[+1865] | 1207.8182 | 4 | 4828.2508 | 4828.2324 | 0 | 3.8 | 127 | Specific | 564.54 | 564.54 | 564.54 | 10.32 | 48 | >sp\|P01876\|IGHA1_HUMAN Immunoglobulin heavy constant alpha 1 OS=Homo sapiens OX=9606 GN=IGHA1 PE=1 SV=2 | 255 | 20201119_08533_1ng_glyco_hcd203040_compound15.20969.20969.4 | controllerType=0 controllerNumber=1 scan=20969 | 42.3955 |
| 17382:3 | 11 | R.LSLHRPALEDLLLGSEAN[+1702.581]LTC[+57.021]TLTGLR.D | HexNAc(2)Hex(8) | C[+57], N[+1703] | 1556.07 | 3 | 4666.1955 | 4666.1796 | 0 | 3.4 | 127 | Specific | 597.93 | 567.06 | 567.06 | 10.94 | 48 | >sp\|P01876\|IGHA1_HUMAN Immunoglobulin heavy constant alpha 1 OS=Homo sapiens OX=9606 GN=IGHA1 PE=1 SV=2 | 255 | 20201119_08533_1ng_glyco_hcd203040_compound15.21035.21035.3 | controllerType=0 controllerNumber=1 scan=21035 | 42.573 |
| 17447:3 | 11 | R.LSLHRPALEDLLLGSEAN[+1540.529]LTC[+57.021]TLTGLR.D | HexNAc(2)Hex(7) | C[+57], N[+1541] | 1502.0614 | 3 | 4504.1696 | 4504.1268 | 0 | 9.5 | 127 | Specific | 616.56 | 607.44 | 607.44 | 10.06 | 48 | >sp\|P01876\|IGHA1_HUMAN Immunoglobulin heavy constant alpha 1 OS=Homo sapiens OX=9606 GN=IGHA1 PE=1 SV=2 | 255 | 20201119_08533_1ng_glyco_hcd203040_compound15.21103.21103.3 | controllerType=0 controllerNumber=1 scan=21103 | 42.7394 |
| 17452:3 | 11 | R.LSLHRPALEDLLLGSEAN[+1460.529]LTC[+57.021]TLTGLR.D | HexNAc(4)Hex(4) | C[+57], N[+1461] | 1475.388 | 3 | 4424.1494 | 4424.1271 | 0 | 5.1 | 127 | Specific | 641.81 | 565.67 | 565.67 | 11.68 | 48 | >sp\|P01876\|IGHA1_HUMAN Immunoglobulin heavy constant alpha 1 OS=Homo sapiens OX=9606 GN=IGHA1 PE=1 SV=2 | 255 | 20201119_08533_1ng_glyco_hcd203040_compound15.21108.21108.3 | controllerType=0 controllerNumber=1 scan=21108 | 42.7574 |
| 17458:3 | 11 | R.LSLHRPALEDLLLGSEAN[+1501.555]LTC[+57.021]TLTGLR.D | HexNAc(5)Hex(3) | C[+57], N[+1502] | 1489.0617 | 3 | 4465.1706 | 4465.1536 | 0 | 3.8 | 127 | Specific | 645.3 | 645.3 | 645.3 | 11.94 | 48 | >sp\|P01876\|IGHA1_HUMAN Immunoglobulin heavy constant alpha 1 OS=Homo sapiens OX=9606 GN=IGHA1 PE=1 SV=2 | 255 | 20201119_08533_1ng_glyco_hcd203040_compound15.21115.21115.3 | controllerType=0 controllerNumber=1 scan=21115 | 42.7809 |
| 17450:3 | 11 | R.LSLHRPALEDLLLGSEAN[+1460.529]LTC[+57.021]TLTGLR.D | HexNAc(4)Hex(4) | C[+57], N[+1461] | 1475.388 | 3 | 4424.1494 | 4424.1271 | 0 | 5.1 | 127 | Specific | 658.32 | 628.77 | 628.77 | 12.68 | 48 | >sp\|P01876\|IGHA1_HUMAN Immunoglobulin heavy constant alpha 1 OS=Homo sapiens OX=9606 GN=IGHA1 PE=1 SV=2 | 255 | 20201119_08533_1ng_glyco_hcd203040_compound15.21106.21106.3 | controllerType=0 controllerNumber=1 scan=21106 | 42.7505 |
| 17399:3 | 11 | R.LSLHRPALEDLLLGSEAN[+1378.476]LTC[+57.021]TLTGLR.D | HexNAc(2)Hex(6) | C[+57], N[+1378] | 1448.0361 | 3 | 4342.0937 | 4342.074 | 0 | 4.5 | 127 | Specific | 663.99 | 622.79 | 622.79 | 12.68 | 48 | >sp\|P01876\|IGHA1_HUMAN Immunoglobulin heavy constant alpha 1 OS=Homo sapiens OX=9606 GN=IGHA1 PE=1 SV=2 | 255 | 20201119_08533_1ng_glyco_hcd203040_compound15.21053.21053.3 | controllerType=0 controllerNumber=1 scan=21053 | 42.6303 |
| 17495:4 | 11 | R.LSLHRPALEDLLLGSEAN[+1298.476]LTC[+57.021]TLTGLR.D | HexNAc(4)Hex(3) | C[+57], N[+1298] | 1066.2787 | 4 | 4262.0931 | 4262.0742 | 0 | 4.4 | 127 | Specific | 689 | 689 | 689 | 13.22 | 48 | >sp\|P01876\|IGHA1_HUMAN Immunoglobulin heavy constant alpha 1 OS=Homo sapiens OX=9606 GN=IGHA1 PE=1 SV=2 | 255 | 20201119_08533_1ng_glyco_hcd203040_compound15.21154.21154.4 | controllerType=0 controllerNumber=1 scan=21154 | 42.9037 |
| 17380:3 | 11 | R.LSLHRPALEDLLLGSEAN[+1702.581]LTC[+57.021]TLTGLR.D | HexNAc(2)Hex(8) | C[+57], N[+1703] | 1556.07 | 3 | 4666.1955 | 4666.1796 | 0 | 3.4 | 127 | Specific | 689.33 | 681.65 | 681.65 | 12.94 | 48 | >sp\|P01876\|IGHA1_HUMAN Immunoglobulin heavy constant alpha 1 OS=Homo sapiens OX=9606 GN=IGHA1 PE=1 SV=2 | 255 | 20201119_08533_1ng_glyco_hcd203040_compound15.21033.21033.3 | controllerType=0 controllerNumber=1 scan=21033 | 42.5675 |
| 17490:4 | 11 | R.LSLHRPALEDLLLGSEAN[+1501.555]LTC[+57.021]TLTGLR.D | HexNAc(5)Hex(3) | C[+57], N[+1502] | 1117.0496 | 4 | 4465.1766 | 4465.1536 | 0 | 5.1 | 127 | Specific | 716.21 | 716.21 | 716.21 | 14.22 | 48 | >sp\|P01876\|IGHA1_HUMAN Immunoglobulin heavy constant alpha 1 OS=Homo sapiens OX=9606 GN=IGHA1 PE=1 SV=2 | 255 | 20201119_08533_1ng_glyco_hcd203040_compound15.21149.21149.4 | controllerType=0 controllerNumber=1 scan=21149 | 42.8883 |
| 17323:3 | 11 | R.LSLHRPALEDLLLGSEAN[+1864.634]LTC[+57.021]TLTGLR.D | HexNAc(2)Hex(9) | C[+57], N[+1865] | 1610.0903 | 3 | 4828.2562 | 4828.2324 | 0 | 4.9 | 127 | Specific | 717.66 | 706.73 | 706.73 | 13.68 | 48 | >sp\|P01876\|IGHA1_HUMAN Immunoglobulin heavy constant alpha 1 OS=Homo sapiens OX=9606 GN=IGHA1 PE=1 SV=2 | 255 | 20201119_08533_1ng_glyco_hcd203040_compound15.20973.20973.3 | controllerType=0 controllerNumber=1 scan=20973 | 42.4056 |
| 17456:3 | 11 | R.LSLHRPALEDLLLGSEAN[+1216.423]LTC[+57.021]TLTGLR.D | HexNAc(2)Hex(5) | C[+57], N[+1216] | 1394.0162 | 3 | 4180.034 | 4180.0211 | 0 | 3.1 | 127 | Specific | 748.71 | 716.83 | 716.83 | 13.94 | 48 | >sp\|P01876\|IGHA1_HUMAN Immunoglobulin heavy constant alpha 1 OS=Homo sapiens OX=9606 GN=IGHA1 PE=1 SV=2 | 255 | 20201119_08533_1ng_glyco_hcd203040_compound15.21112.21112.3 | controllerType=0 controllerNumber=1 scan=21112 | 42.7706 |
| 17492:4 | 11 | R.LSLHRPALEDLLLGSEAN[+1501.555]LTC[+57.021]TLTGLR.D | HexNAc(5)Hex(3) | C[+57], N[+1502] | 1117.0496 | 4 | 4465.1766 | 4465.1536 | 0 | 5.1 | 127 | Specific | 749.7 | 731.01 | 731.01 | 14.22 | 48 | >sp\|P01876\|IGHA1_HUMAN Immunoglobulin heavy constant alpha 1 OS=Homo sapiens OX=9606 GN=IGHA1 PE=1 SV=2 | 255 | 20201119_08533_1ng_glyco_hcd203040_compound15.21151.21151.4 | controllerType=0 controllerNumber=1 scan=21151 | 42.895 |
| 17454:3 | 11 | R.LSLHRPALEDLLLGSEAN[+1216.423]LTC[+57.021]TLTGLR.D | HexNAc(2)Hex(5) | C[+57], N[+1216] | 1394.0162 | 3 | 4180.034 | 4180.0211 | 0 | 3.1 | 127 | Specific | 754.82 | 708.57 | 708.57 | 14.94 | 48 | >sp\|P01876\|IGHA1_HUMAN Immunoglobulin heavy constant alpha 1 OS=Homo sapiens OX=9606 GN=IGHA1 PE=1 SV=2 | 255 | 20201119_08533_1ng_glyco_hcd203040_compound15.21110.21110.3 | controllerType=0 controllerNumber=1 scan=21110 | 42.7614 |
| 17446:4 | 11 | R.LSLHRPALEDLLLGSEAN[+1460.529]LTC[+57.021]TLTGLR.D | HexNAc(4)Hex(4) | C[+57], N[+1461] | 1106.7918 | 4 | 4424.1453 | 4424.1271 | 0 | 4.1 | 127 | Specific | 758.23 | 750.47 | 750.47 | 15.22 | 48 | >sp\|P01876\|IGHA1_HUMAN Immunoglobulin heavy constant alpha 1 OS=Homo sapiens OX=9606 GN=IGHA1 PE=1 SV=2 | 255 | 20201119_08533_1ng_glyco_hcd203040_compound15.21102.21102.4 | controllerType=0 controllerNumber=1 scan=21102 | 42.738 |
| 17397:3 | 11 | R.LSLHRPALEDLLLGSEAN[+1378.476]LTC[+57.021]TLTGLR.D | HexNAc(2)Hex(6) | C[+57], N[+1378] | 1448.0361 | 3 | 4342.0937 | 4342.074 | 0 | 4.5 | 127 | Specific | 759.87 | 698.71 | 698.71 | 14.68 | 48 | >sp\|P01876\|IGHA1_HUMAN Immunoglobulin heavy constant alpha 1 OS=Homo sapiens OX=9606 GN=IGHA1 PE=1 SV=2 | 255 | 20201119_08533_1ng_glyco_hcd203040_compound15.21051.21051.3 | controllerType=0 controllerNumber=1 scan=21051 | 42.6234 |
| 17497:4 | 11 | R.LSLHRPALEDLLLGSEAN[+1298.476]LTC[+57.021]TLTGLR.D | HexNAc(4)Hex(3) | C[+57], N[+1298] | 1066.2787 | 4 | 4262.0931 | 4262.0742 | 0 | 4.4 | 127 | Specific | 787.66 | 780.01 | 780.01 | 15.22 | 48 | >sp\|P01876\|IGHA1_HUMAN Immunoglobulin heavy constant alpha 1 OS=Homo sapiens OX=9606 GN=IGHA1 PE=1 SV=2 | 255 | 20201119_08533_1ng_glyco_hcd203040_compound15.21156.21156.4 | controllerType=0 controllerNumber=1 scan=21156 | 42.9101 |
| 17500:4 | 11 | R.LSLHRPALEDLLLGSEAN[+1095.397]LTC[+57.021]TLTGLR.D | HexNAc(3)Hex(3) | C[+57], N[+1095] | 1015.5085 | 4 | 4059.0123 | 4058.9949 | 0 | 4.3 | 127 | Specific | 811.67 | 811.67 | 811.67 | 16.22 | 48 | >sp\|P01876\|IGHA1_HUMAN Immunoglobulin heavy constant alpha 1 OS=Homo sapiens OX=9606 GN=IGHA1 PE=1 SV=2 | 255 | 20201119_08533_1ng_glyco_hcd203040_compound15.21159.21159.4 | controllerType=0 controllerNumber=1 scan=21159 | 42.9157 |
| 17326:3 | 11 | R.LSLHRPALEDLLLGSEAN[+1864.634]LTC[+57.021]TLTGLR.D | HexNAc(2)Hex(9) | C[+57], N[+1865] | 1610.0903 | 3 | 4828.2562 | 4828.2324 | 0 | 4.9 | 127 | Specific | 813.94 | 774.88 | 774.88 | 15.68 | 48 | >sp\|P01876\|IGHA1_HUMAN Immunoglobulin heavy constant alpha 1 OS=Homo sapiens OX=9606 GN=IGHA1 PE=1 SV=2 | 255 | 20201119_08533_1ng_glyco_hcd203040_compound15.20976.20976.3 | controllerType=0 controllerNumber=1 scan=20976 | 42.4132 |
| 17448:4 | 11 | R.LSLHRPALEDLLLGSEAN[+1460.529]LTC[+57.021]TLTGLR.D | HexNAc(4)Hex(4) | C[+57], N[+1461] | 1106.7918 | 4 | 4424.1453 | 4424.1271 | 0 | 4.1 | 127 | Specific | 824.08 | 709.01 | 709.01 | 16.22 | 48 | >sp\|P01876\|IGHA1_HUMAN Immunoglobulin heavy constant alpha 1 OS=Homo sapiens OX=9606 GN=IGHA1 PE=1 SV=2 | 255 | 20201119_08533_1ng_glyco_hcd203040_compound15.21104.21104.4 | controllerType=0 controllerNumber=1 scan=21104 | 42.7467 |
| 17376:4 | 11 | R.LSLHRPALEDLLLGSEAN[+1702.581]LTC[+57.021]TLTGLR.D | HexNAc(2)Hex(8) | C[+57], N[+1703] | 1167.3044 | 4 | 4666.1956 | 4666.1796 | 0 | 3.4 | 127 | Specific | 889.98 | 889.98 | 889.98 | 17.48 | 48 | >sp\|P01876\|IGHA1_HUMAN Immunoglobulin heavy constant alpha 1 OS=Homo sapiens OX=9606 GN=IGHA1 PE=1 SV=2 | 255 | 20201119_08533_1ng_glyco_hcd203040_compound15.21029.21029.4 | controllerType=0 controllerNumber=1 scan=21029 | 42.5534 |
| 17503:4 | 11 | R.LSLHRPALEDLLLGSEAN[+1054.370]LTC[+57.021]TLTGLR.D | HexNAc(2)Hex(4) | C[+57], N[+1054] | 1005.2517 | 4 | 4017.9849 | 4017.9683 | 0 | 4.1 | 127 | Specific | 921.82 | 921.82 | 921.82 | 18.22 | 48 | >sp\|P01876\|IGHA1_HUMAN Immunoglobulin heavy constant alpha 1 OS=Homo sapiens OX=9606 GN=IGHA1 PE=1 SV=2 | 255 | 20201119_08533_1ng_glyco_hcd203040_compound15.21162.21162.4 | controllerType=0 controllerNumber=1 scan=21162 | 42.9265 |
| 17443:4 | 11 | R.LSLHRPALEDLLLGSEAN[+1216.423]LTC[+57.021]TLTGLR.D | HexNAc(2)Hex(5) | C[+57], N[+1216] | 1045.7638 | 4 | 4180.0335 | 4180.0211 | 0 | 3 | 127 | Specific | 930.01 | 869.44 | 869.44 | 18.48 | 48 | >sp\|P01876\|IGHA1_HUMAN Immunoglobulin heavy constant alpha 1 OS=Homo sapiens OX=9606 GN=IGHA1 PE=1 SV=2 | 255 | 20201119_08533_1ng_glyco_hcd203040_compound15.21099.21099.4 | controllerType=0 controllerNumber=1 scan=21099 | 42.7236 |
| 17501:4 | 11 | R.LSLHRPALEDLLLGSEAN[+1054.370]LTC[+57.021]TLTGLR.D | HexNAc(2)Hex(4) | C[+57], N[+1054] | 1005.2517 | 4 | 4017.9849 | 4017.9683 | 0 | 4.1 | 127 | Specific | 935.63 | 901.88 | 901.88 | 18.22 | 48 | >sp\|P01876\|IGHA1_HUMAN Immunoglobulin heavy constant alpha 1 OS=Homo sapiens OX=9606 GN=IGHA1 PE=1 SV=2 | 255 | 20201119_08533_1ng_glyco_hcd203040_compound15.21160.21160.4 | controllerType=0 controllerNumber=1 scan=21160 | 42.9172 |
| 17445:4 | 11 | R.LSLHRPALEDLLLGSEAN[+1216.423]LTC[+57.021]TLTGLR.D | HexNAc(2)Hex(5) | C[+57], N[+1216] | 1045.7638 | 4 | 4180.0335 | 4180.0211 | 0 | 3 | 127 | Specific | 1000.59 | 1000.59 | 1000.59 | 20.48 | 48 | >sp\|P01876\|IGHA1_HUMAN Immunoglobulin heavy constant alpha 1 OS=Homo sapiens OX=9606 GN=IGHA1 PE=1 SV=2 | 255 | 20201119_08533_1ng_glyco_hcd203040_compound15.21101.21101.4 | controllerType=0 controllerNumber=1 scan=21101 | 42.7363 |
| 17321:4 | 11 | R.LSLHRPALEDLLLGSEAN[+1864.634]LTC[+57.021]TLTGLR.D | HexNAc(2)Hex(9) | C[+57], N[+1865] | 1207.8182 | 4 | 4828.2508 | 4828.2324 | 0 | 3.8 | 127 | Specific | 1007.82 | 1007.82 | 1007.82 | 20.48 | 48 | >sp\|P01876\|IGHA1_HUMAN Immunoglobulin heavy constant alpha 1 OS=Homo sapiens OX=9606 GN=IGHA1 PE=1 SV=2 | 255 | 20201119_08533_1ng_glyco_hcd203040_compound15.20971.20971.4 | controllerType=0 controllerNumber=1 scan=20971 | 42.399 |
| 17378:4 | 11 | R.LSLHRPALEDLLLGSEAN[+1702.581]LTC[+57.021]TLTGLR.D | HexNAc(2)Hex(8) | C[+57], N[+1703] | 1167.3044 | 4 | 4666.1956 | 4666.1796 | 0 | 3.4 | 127 | Specific | 1014.77 | 953.33 | 953.33 | 20.48 | 48 | >sp\|P01876\|IGHA1_HUMAN Immunoglobulin heavy constant alpha 1 OS=Homo sapiens OX=9606 GN=IGHA1 PE=1 SV=2 | 255 | 20201119_08533_1ng_glyco_hcd203040_compound15.21031.21031.4 | controllerType=0 controllerNumber=1 scan=21031 | 42.5617 |
| 17502:4 | 11 | R.LSLHRPALEDLLLGSEAN[+1095.397]LTC[+57.021]TLTGLR.D | HexNAc(3)Hex(3) | C[+57], N[+1095] | 1015.5085 | 4 | 4059.0123 | 4058.9949 | 0 | 4.3 | 127 | Specific | 1056.99 | 1035.45 | 1035.45 | 21.22 | 48 | >sp\|P01876\|IGHA1_HUMAN Immunoglobulin heavy constant alpha 1 OS=Homo sapiens OX=9606 GN=IGHA1 PE=1 SV=2 | 255 | 20201119_08533_1ng_glyco_hcd203040_compound15.21161.21161.4 | controllerType=0 controllerNumber=1 scan=21161 | 42.9225 |
| 09878:3 | 11 | R.LAGKPTHVN[+1216.423]VSVVMAEVDGTC[+57.021]Y.- | HexNAc(2)Hex(5) | C[+57], N[+1216] | 1188.5286 | 3 | 3563.5713 | 3563.5649 | 0 | 1.8 | 332 | CRagged | 205.38 | 205.38 | 205.2 | 5.48 | 48 | >sp\|P01876\|IGHA1_HUMAN Immunoglobulin heavy constant alpha 1 OS=Homo sapiens OX=9606 GN=IGHA1 PE=1 SV=2 | 255 | 20201119_08533_1ng_glyco_hcd203040_compound15.13161.13161.3 | controllerType=0 controllerNumber=1 scan=13161 | 28.598 |
| 11089:4 | 11 | R.LAGKPTHVN[+1809.666]VSVVMAEVDGTC[+57.021]Y.- | HexNAc(5)Hex(4)Fuc(1) | C[+57], N[+1810] | 1039.9645 | 4 | 4156.8362 | 4156.8081 | 0 | 6.7 | 332 | CRagged | 251.21 | 251.21 | 251.21 | 5.44 | 48 | >sp\|P01876\|IGHA1_HUMAN Immunoglobulin heavy constant alpha 1 OS=Homo sapiens OX=9606 GN=IGHA1 PE=1 SV=2 | 255 | 20201119_08533_1ng_glyco_hcd203040_compound15.14414.14414.4 | controllerType=0 controllerNumber=1 scan=14414 | 30.7417 |
| 12905:3 | 11 | R.LAGKPTHVN[+1710.598]VSVVMAEVDGTC[+57.021]Y.- | HexNAc(3)Hex(5)NeuAc(1) | C[+57], N[+1711] | 1353.2604 | 3 | 4057.7667 | 4057.7397 | 0 | 6.7 | 332 | CRagged | 265.05 | 265.05 | 210.73 | 5.8 | 48 | >sp\|P01876\|IGHA1_HUMAN Immunoglobulin heavy constant alpha 1 OS=Homo sapiens OX=9606 GN=IGHA1 PE=1 SV=2 | 255 | 20201119_08533_1ng_glyco_hcd203040_compound15.16330.16330.3 | controllerType=0 controllerNumber=1 scan=16330 | 33.9862 |
| 12907:3 | 11 | R.LAGKPTHVN[+1710.598]VSVVMAEVDGTC[+57.021]Y.- | HexNAc(3)Hex(5)NeuAc(1) | C[+57], N[+1711] | 1353.2604 | 3 | 4057.7667 | 4057.7397 | 0 | 6.7 | 332 | CRagged | 310.92 | 304.85 | 279.53 | 6.63 | 48 | >sp\|P01876\|IGHA1_HUMAN Immunoglobulin heavy constant alpha 1 OS=Homo sapiens OX=9606 GN=IGHA1 PE=1 SV=2 | 255 | 20201119_08533_1ng_glyco_hcd203040_compound15.16332.16332.3 | controllerType=0 controllerNumber=1 scan=16332 | 33.9903 |
| 11103:3 | 11 | R.LAGKPTHVN[+1378.476]VSVVMAEVDGTC[+57.021]Y.- | HexNAc(2)Hex(6) | C[+57], N[+1378] | 1242.5484 | 3 | 3725.6306 | 3725.6178 | 0 | 3.5 | 332 | CRagged | 386.23 | 161.26 | 161.26 | 6.63 | 48 | >sp\|P01876\|IGHA1_HUMAN Immunoglobulin heavy constant alpha 1 OS=Homo sapiens OX=9606 GN=IGHA1 PE=1 SV=2 | 255 | 20201119_08533_1ng_glyco_hcd203040_compound15.14429.14429.3 | controllerType=0 controllerNumber=1 scan=14429 | 30.7678 |
| 12786:3 | 11 | R.LAGKPTHVN[+2059.735]VSVVMAEVDGTC[+57.021]Y.- | HexNAc(4)Hex(5)Fuc(1)NeuAc(1) | C[+57], N[+2060] | 1469.6347 | 3 | 4406.8895 | 4406.877 | 0 | 2.8 | 332 | CRagged | 441.57 | 441.57 | 441.57 | 10 | 48 | >sp\|P01876\|IGHA1_HUMAN Immunoglobulin heavy constant alpha 1 OS=Homo sapiens OX=9606 GN=IGHA1 PE=1 SV=2 | 255 | 20201119_08533_1ng_glyco_hcd203040_compound15.16207.16207.3 | controllerType=0 controllerNumber=1 scan=16207 | 33.7759 |
| 12888:3 | 11 | R.LAGKPTHVN[+1751.624]VSVVMAEVDGTC[+57.021]Y.- | HexNAc(4)Hex(4)NeuAc(1) | C[+57], N[+1752] | 1366.929 | 3 | 4098.7725 | 4098.7663 | 0 | 1.5 | 332 | CRagged | 484.71 | 375.22 | 375.22 | 10.13 | 48 | >sp\|P01876\|IGHA1_HUMAN Immunoglobulin heavy constant alpha 1 OS=Homo sapiens OX=9606 GN=IGHA1 PE=1 SV=2 | 255 | 20201119_08533_1ng_glyco_hcd203040_compound15.16312.16312.3 | controllerType=0 controllerNumber=1 scan=16312 | 33.9559 |
| 11051:3 | 11 | R.LAGKPTHVN[+1663.608]VSVVMAEVDGTC[+57.021]Y.- | HexNAc(5)Hex(4) | C[+57], N[+1664] | 1337.5965 | 3 | 4010.7749 | 4010.7502 | 0 | 6.2 | 332 | CRagged | 505.18 | 496.1 | 317.46 | 9.47 | 48 | >sp\|P01876\|IGHA1_HUMAN Immunoglobulin heavy constant alpha 1 OS=Homo sapiens OX=9606 GN=IGHA1 PE=1 SV=2 | 255 | 20201119_08533_1ng_glyco_hcd203040_compound15.14375.14375.3 | controllerType=0 controllerNumber=1 scan=14375 | 30.6689 |
| 12788:3 | 11 | R.LAGKPTHVN[+2059.735]VSVVMAEVDGTC[+57.021]Y.- | HexNAc(4)Hex(5)Fuc(1)NeuAc(1) | C[+57], N[+2060] | 1469.6347 | 3 | 4406.8895 | 4406.877 | 0 | 2.8 | 332 | CRagged | 518.24 | 517.69 | 517.69 | 10.24 | 48 | >sp\|P01876\|IGHA1_HUMAN Immunoglobulin heavy constant alpha 1 OS=Homo sapiens OX=9606 GN=IGHA1 PE=1 SV=2 | 255 | 20201119_08533_1ng_glyco_hcd203040_compound15.16209.16209.3 | controllerType=0 controllerNumber=1 scan=16209 | 33.7813 |
| 12886:3 | 11 | R.LAGKPTHVN[+1751.624]VSVVMAEVDGTC[+57.021]Y.- | HexNAc(4)Hex(4)NeuAc(1) | C[+57], N[+1752] | 1366.929 | 3 | 4098.7725 | 4098.7663 | 0 | 1.5 | 332 | CRagged | 526.56 | 526.56 | 526.56 | 10.38 | 48 | >sp\|P01876\|IGHA1_HUMAN Immunoglobulin heavy constant alpha 1 OS=Homo sapiens OX=9606 GN=IGHA1 PE=1 SV=2 | 255 | 20201119_08533_1ng_glyco_hcd203040_compound15.16310.16310.3 | controllerType=0 controllerNumber=1 scan=16310 | 33.952 |
| 11053:3 | 11 | R.LAGKPTHVN[+1663.608]VSVVMAEVDGTC[+57.021]Y.- | HexNAc(5)Hex(4) | C[+57], N[+1664] | 1337.5965 | 3 | 4010.7749 | 4010.7502 | 0 | 6.2 | 332 | CRagged | 614.8 | 601.41 | 410.12 | 12.01 | 48 | >sp\|P01876\|IGHA1_HUMAN Immunoglobulin heavy constant alpha 1 OS=Homo sapiens OX=9606 GN=IGHA1 PE=1 SV=2 | 255 | 20201119_08533_1ng_glyco_hcd203040_compound15.14377.14377.3 | controllerType=0 controllerNumber=1 scan=14377 | 30.672 |
| 11018:3 | 11 | R.LAGKPTHVN[+1540.529]VSVVMAEVDGTC[+57.021]Y.- | HexNAc(2)Hex(7) | C[+57], N[+1541] | 1296.565 | 3 | 3887.6804 | 3887.6706 | 0 | 2.5 | 332 | CRagged | 629.17 | 594.09 | 410.07 | 12.75 | 48 | >sp\|P01876\|IGHA1_HUMAN Immunoglobulin heavy constant alpha 1 OS=Homo sapiens OX=9606 GN=IGHA1 PE=1 SV=2 | 255 | 20201119_08533_1ng_glyco_hcd203040_compound15.14340.14340.3 | controllerType=0 controllerNumber=1 scan=14340 | 30.5887 |
| 11025:3 | 11 | R.LAGKPTHVN[+1378.476]VSVVMAEVDGTC[+57.021]Y.- | HexNAc(2)Hex(6) | C[+57], N[+1378] | 1242.5474 | 3 | 3725.6277 | 3725.6178 | 0 | 2.7 | 332 | CRagged | 687.62 | 685.51 | 416.08 | 13.78 | 48 | >sp\|P01876\|IGHA1_HUMAN Immunoglobulin heavy constant alpha 1 OS=Homo sapiens OX=9606 GN=IGHA1 PE=1 SV=2 | 255 | 20201119_08533_1ng_glyco_hcd203040_compound15.14348.14348.3 | controllerType=0 controllerNumber=1 scan=14348 | 30.6055 |
| 10977:3 | 11 | R.LAGKPTHVN[+1540.529]VSVVMAEVDGTC[+57.021]Y.- | HexNAc(2)Hex(7) | C[+57], N[+1541] | 1296.5588 | 3 | 3887.6618 | 3887.6706 | 0 | -2.3 | 332 | CRagged | 694.88 | 694.88 | 443.99 | 12.2 | 48 | >sp\|P01876\|IGHA1_HUMAN Immunoglobulin heavy constant alpha 1 OS=Homo sapiens OX=9606 GN=IGHA1 PE=1 SV=2 | 255 | 20201119_08533_1ng_glyco_hcd203040_compound15.14298.14298.3 | controllerType=0 controllerNumber=1 scan=14298 | 30.5126 |
| 10926:3 | 11 | R.LAGKPTHVN[+1702.581]VSVVMAEVDGTC[+57.021]Y.- | HexNAc(2)Hex(8) | C[+57], N[+1703] | 1350.5828 | 3 | 4049.7339 | 4049.7234 | 0 | 2.6 | 332 | CRagged | 725.55 | 725.55 | 490 | 14.75 | 48 | >sp\|P01876\|IGHA1_HUMAN Immunoglobulin heavy constant alpha 1 OS=Homo sapiens OX=9606 GN=IGHA1 PE=1 SV=2 | 255 | 20201119_08533_1ng_glyco_hcd203040_compound15.14245.14245.3 | controllerType=0 controllerNumber=1 scan=14245 | 30.4112 |
| 11012:3 | 11 | R.LAGKPTHVN[+1663.608]VSVVMAEVDGTC[+57.021]Y.- | HexNAc(5)Hex(4) | C[+57], N[+1664] | 1337.5917 | 3 | 4010.7606 | 4010.7502 | 0 | 2.6 | 332 | CRagged | 739.5 | 739.5 | 496.59 | 14.75 | 48 | >sp\|P01876\|IGHA1_HUMAN Immunoglobulin heavy constant alpha 1 OS=Homo sapiens OX=9606 GN=IGHA1 PE=1 SV=2 | 255 | 20201119_08533_1ng_glyco_hcd203040_compound15.14334.14334.3 | controllerType=0 controllerNumber=1 scan=14334 | 30.5776 |
| 10980:3 | 11 | R.LAGKPTHVN[+1540.529]VSVVMAEVDGTC[+57.021]Y.- | HexNAc(2)Hex(7) | C[+57], N[+1541] | 1296.5588 | 3 | 3887.6618 | 3887.6706 | 0 | -2.3 | 332 | CRagged | 787.25 | 723.63 | 472.38 | 14.2 | 48 | >sp\|P01876\|IGHA1_HUMAN Immunoglobulin heavy constant alpha 1 OS=Homo sapiens OX=9606 GN=IGHA1 PE=1 SV=2 | 255 | 20201119_08533_1ng_glyco_hcd203040_compound15.14301.14301.3 | controllerType=0 controllerNumber=1 scan=14301 | 30.5249 |
| 10952:3 | 11 | R.LAGKPTHVN[+1540.529]VSVVMAEVDGTC[+57.021]Y.- | HexNAc(2)Hex(7) | C[+57], N[+1541] | 1296.5646 | 3 | 3887.6793 | 3887.6706 | 0 | 2.3 | 332 | CRagged | 797.14 | 781.32 | 489.81 | 15.75 | 48 | >sp\|P01876\|IGHA1_HUMAN Immunoglobulin heavy constant alpha 1 OS=Homo sapiens OX=9606 GN=IGHA1 PE=1 SV=2 | 255 | 20201119_08533_1ng_glyco_hcd203040_compound15.14271.14271.3 | controllerType=0 controllerNumber=1 scan=14271 | 30.4539 |
| 11028:3 | 11 | R.LAGKPTHVN[+1809.666]VSVVMAEVDGTC[+57.021]Y.- | HexNAc(5)Hex(4)Fuc(1) | C[+57], N[+1810] | 1386.2764 | 3 | 4156.8147 | 4156.8081 | 0 | 1.6 | 332 | CRagged | 797.55 | 665.68 | 494.57 | 15.75 | 48 | >sp\|P01876\|IGHA1_HUMAN Immunoglobulin heavy constant alpha 1 OS=Homo sapiens OX=9606 GN=IGHA1 PE=1 SV=2 | 255 | 20201119_08533_1ng_glyco_hcd203040_compound15.14351.14351.3 | controllerType=0 controllerNumber=1 scan=14351 | 30.6138 |
| 11060:3 | 11 | R.LAGKPTHVN[+1501.555]VSVVMAEVDGTC[+57.021]Y.- | HexNAc(5)Hex(3) | C[+57], N[+1502] | 1283.5793 | 3 | 3848.7233 | 3848.6974 | 0 | 6.7 | 332 | CRagged | 798.33 | 654.94 | 654.94 | 14.57 | 48 | >sp\|P01876\|IGHA1_HUMAN Immunoglobulin heavy constant alpha 1 OS=Homo sapiens OX=9606 GN=IGHA1 PE=1 SV=2 | 255 | 20201119_08533_1ng_glyco_hcd203040_compound15.14384.14384.3 | controllerType=0 controllerNumber=1 scan=14384 | 30.6874 |
| 10929:3 | 11 | R.LAGKPTHVN[+1702.581]VSVVMAEVDGTC[+57.021]Y.- | HexNAc(2)Hex(8) | C[+57], N[+1703] | 1350.5828 | 3 | 4049.7339 | 4049.7234 | 0 | 2.6 | 332 | CRagged | 829.93 | 810.25 | 494.49 | 16.75 | 48 | >sp\|P01876\|IGHA1_HUMAN Immunoglobulin heavy constant alpha 1 OS=Homo sapiens OX=9606 GN=IGHA1 PE=1 SV=2 | 255 | 20201119_08533_1ng_glyco_hcd203040_compound15.14248.14248.3 | controllerType=0 controllerNumber=1 scan=14248 | 30.4152 |
| 11058:3 | 11 | R.LAGKPTHVN[+1501.555]VSVVMAEVDGTC[+57.021]Y.- | HexNAc(5)Hex(3) | C[+57], N[+1502] | 1283.5793 | 3 | 3848.7233 | 3848.6974 | 0 | 6.7 | 332 | CRagged | 835.26 | 835.26 | 835.26 | 15.57 | 48 | >sp\|P01876\|IGHA1_HUMAN Immunoglobulin heavy constant alpha 1 OS=Homo sapiens OX=9606 GN=IGHA1 PE=1 SV=2 | 255 | 20201119_08533_1ng_glyco_hcd203040_compound15.14382.14382.3 | controllerType=0 controllerNumber=1 scan=14382 | 30.6799 |
| 10949:3 | 11 | R.LAGKPTHVN[+1540.529]VSVVMAEVDGTC[+57.021]Y.- | HexNAc(2)Hex(7) | C[+57], N[+1541] | 1296.5646 | 3 | 3887.6793 | 3887.6706 | 0 | 2.3 | 332 | CRagged | 851.93 | 851.93 | 515.43 | 17.75 | 48 | >sp\|P01876\|IGHA1_HUMAN Immunoglobulin heavy constant alpha 1 OS=Homo sapiens OX=9606 GN=IGHA1 PE=1 SV=2 | 255 | 20201119_08533_1ng_glyco_hcd203040_compound15.14268.14268.3 | controllerType=0 controllerNumber=1 scan=14268 | 30.4462 |
| 11030:3 | 11 | R.LAGKPTHVN[+1216.423]VSVVMAEVDGTC[+57.021]Y.- | HexNAc(2)Hex(5) | C[+57], N[+1216] | 1188.5291 | 3 | 3563.5728 | 3563.5649 | 0 | 2.2 | 332 | CRagged | 860.84 | 860.84 | 505.98 | 17.75 | 48 | >sp\|P01876\|IGHA1_HUMAN Immunoglobulin heavy constant alpha 1 OS=Homo sapiens OX=9606 GN=IGHA1 PE=1 SV=2 | 255 | 20201119_08533_1ng_glyco_hcd203040_compound15.14353.14353.3 | controllerType=0 controllerNumber=1 scan=14353 | 30.6165 |
| 11074:3 | 11 | R.LAGKPTHVN[+1647.613]VSVVMAEVDGTC[+57.021]Y.- | HexNAc(5)Hex(3)Fuc(1) | C[+57], N[+1648] | 1332.2571 | 3 | 3994.7569 | 3994.7553 | 0 | 0.4 | 332 | CRagged | 879.06 | 740.2 | 740.2 | 17.2 | 48 | >sp\|P01876\|IGHA1_HUMAN Immunoglobulin heavy constant alpha 1 OS=Homo sapiens OX=9606 GN=IGHA1 PE=1 SV=2 | 255 | 20201119_08533_1ng_glyco_hcd203040_compound15.14399.14399.3 | controllerType=0 controllerNumber=1 scan=14399 | 30.7183 |
| 11026:3 | 11 | R.LAGKPTHVN[+1809.666]VSVVMAEVDGTC[+57.021]Y.- | HexNAc(5)Hex(4)Fuc(1) | C[+57], N[+1810] | 1386.2764 | 3 | 4156.8147 | 4156.8081 | 0 | 1.6 | 332 | CRagged | 923.46 | 923.46 | 587.31 | 18.75 | 48 | >sp\|P01876\|IGHA1_HUMAN Immunoglobulin heavy constant alpha 1 OS=Homo sapiens OX=9606 GN=IGHA1 PE=1 SV=2 | 255 | 20201119_08533_1ng_glyco_hcd203040_compound15.14349.14349.3 | controllerType=0 controllerNumber=1 scan=14349 | 30.607 |
| 11072:3 | 11 | R.LAGKPTHVN[+1647.613]VSVVMAEVDGTC[+57.021]Y.- | HexNAc(5)Hex(3)Fuc(1) | C[+57], N[+1648] | 1332.2571 | 3 | 3994.7569 | 3994.7553 | 0 | 0.4 | 332 | CRagged | 927.95 | 922.69 | 922.69 | 18.2 | 48 | >sp\|P01876\|IGHA1_HUMAN Immunoglobulin heavy constant alpha 1 OS=Homo sapiens OX=9606 GN=IGHA1 PE=1 SV=2 | 255 | 20201119_08533_1ng_glyco_hcd203040_compound15.14397.14397.3 | controllerType=0 controllerNumber=1 scan=14397 | 30.7106 |
| 11027:3 | 11 | R.LAGKPTHVN[+1378.476]VSVVMAEVDGTC[+57.021]Y.- | HexNAc(2)Hex(6) | C[+57], N[+1378] | 1242.5474 | 3 | 3725.6277 | 3725.6178 | 0 | 2.7 | 332 | CRagged | 929.92 | 929.92 | 549.98 | 18.78 | 48 | >sp\|P01876\|IGHA1_HUMAN Immunoglobulin heavy constant alpha 1 OS=Homo sapiens OX=9606 GN=IGHA1 PE=1 SV=2 | 255 | 20201119_08533_1ng_glyco_hcd203040_compound15.14350.14350.3 | controllerType=0 controllerNumber=1 scan=14350 | 30.6101 |
| 11033:3 | 11 | R.LAGKPTHVN[+1216.423]VSVVMAEVDGTC[+57.021]Y.- | HexNAc(2)Hex(5) | C[+57], N[+1216] | 1188.5291 | 3 | 3563.5728 | 3563.5649 | 0 | 2.2 | 332 | CRagged | 931.52 | 931.52 | 554.87 | 18.75 | 48 | >sp\|P01876\|IGHA1_HUMAN Immunoglobulin heavy constant alpha 1 OS=Homo sapiens OX=9606 GN=IGHA1 PE=1 SV=2 | 255 | 20201119_08533_1ng_glyco_hcd203040_compound15.14356.14356.3 | controllerType=0 controllerNumber=1 scan=14356 | 30.6256 |
| 23527:4 | 13 | K.[+42.011]VFPLSLDSTPQDGNVVVAC[+57.021]LVQGFFPQEPLSVTWSESGQN[+2059 | HexNAc(4)Hex(5)Fuc(1)NeuAc(1) | .[42], C[+57], N[+2060] | 1720.2903 | 4 | 6878.1394 | 6878.1069 | 0 | 4.7 | 8 | Specific | 251.04 | 250.22 | 250.22 | 6.64 | 45 | >sp\|P01877\|IGHA2_HUMAN Immunoglobulin heavy constant alpha 2 OS=Homo sapiens OX=9606 GN=IGHA2 PE=1 SV=4 | 576 | 20201119_08533_1ng_glyco_hcd203040_compound15.27453.27453.4 | controllerType=0 controllerNumber=1 scan=27453 | 55.6035 |
| 23525:4 | 13 | K.[+42.011]VFPLSLDSTPQDGNVVVAC[+57.021]LVQGFFPQEPLSVTWSESGQN[+2059 | HexNAc(4)Hex(5)Fuc(1)NeuAc(1) | .[42], C[+57], N[+2060] | 1720.2903 | 4 | 6878.1394 | 6878.1069 | 0 | 4.7 | 8 | Specific | 291.75 | 291.75 | 291.75 | 7.89 | 45 | >sp\|P01877\|IGHA2_HUMAN Immunoglobulin heavy constant alpha 2 OS=Homo sapiens OX=9606 GN=IGHA2 PE=1 SV=4 | 576 | 20201119_08533_1ng_glyco_hcd203040_compound15.27451.27451.4 | controllerType=0 controllerNumber=1 scan=27451 | 55.5991 |
| 21619:4 | 13 | K.[+42.011]VFPLSLDSTPQDGNVVVAC[+57.021]LVQGFFPQEPLSVTWSESGQN[+1768 | HexNAc(4)Hex(5)Fuc(1) | .[42], C[+57], N[+1769] | 1647.5134 | 4 | 6587.0319 | 6587.0115 | 0 | 3.1 | 8 | Specific | 321.85 | 250.95 | 250.95 | 6.45 | 45 | >sp\|P01877\|IGHA2_HUMAN Immunoglobulin heavy constant alpha 2 OS=Homo sapiens OX=9606 GN=IGHA2 PE=1 SV=4 | 576 | 20201119_08533_1ng_glyco_hcd203040_compound15.25455.25455.4 | controllerType=0 controllerNumber=1 scan=25455 | 51.7997 |
| 21617:4 | 13 | K.[+42.011]VFPLSLDSTPQDGNVVVAC[+57.021]LVQGFFPQEPLSVTWSESGQN[+1768 | HexNAc(4)Hex(5)Fuc(1) | .[42], C[+57], N[+1769] | 1647.5134 | 4 | 6587.0319 | 6587.0115 | 0 | 3.1 | 8 | Specific | 428.43 | 410.39 | 410.39 | 10.85 | 45 | >sp\|P01877\|IGHA2_HUMAN Immunoglobulin heavy constant alpha 2 OS=Homo sapiens OX=9606 GN=IGHA2 PE=1 SV=4 | 576 | 20201119_08533_1ng_glyco_hcd203040_compound15.25453.25453.4 | controllerType=0 controllerNumber=1 scan=25453 | 51.794 |
| 21626:4 | 13 | K.VFPLSLDSTPQDGNVVVAC[+57.021]LVQGFFPQEPLSVTWSESGQN[+1809.666]VTAI | HexNAc(5)Hex(4)Fuc(1) | C[+57], N[+1810] | 1647.2786 | 4 | 6586.0924 | 6586.0275 | 0 | 9.9 | 8 | Specific | 570.06 | 567.12 | 567.12 | 9.58 | 45 | >sp\|P01877\|IGHA2_HUMAN Immunoglobulin heavy constant alpha 2 OS=Homo sapiens OX=9606 GN=IGHA2 PE=1 SV=4 | 576 | 20201119_08533_1ng_glyco_hcd203040_compound15.25463.25463.4 | controllerType=0 controllerNumber=1 scan=25463 | 51.8153 |
| 21628:4 | 13 | K.VFPLSLDSTPQDGNVVVAC[+57.021]LVQGFFPQEPLSVTWSESGQN[+1809.666]VTAI | HexNAc(5)Hex(4)Fuc(1) | C[+57], N[+1810] | 1647.2786 | 4 | 6586.0924 | 6586.0275 | 0 | 9.9 | 8 | Specific | 788.75 | 730.22 | 730.22 | 14.73 | 45 | >sp\|P01877\|IGHA2_HUMAN Immunoglobulin heavy constant alpha 2 OS=Homo sapiens OX=9606 GN=IGHA2 PE=1 SV=4 | 576 | 20201119_08533_1ng_glyco_hcd203040_compound15.25465.25465.4 | controllerType=0 controllerNumber=1 scan=25465 | 51.8201 |
| 21669:4 | 13 | K.VFPLSLDSTPQDGNVVVAC[+57.021]LVQGFFPQEPLSVTWSESGQN[+1647.613]VTAI | HexNAc(5)Hex(3)Fuc(1) | C[+57], N[+1648] | 1606.7561 | 4 | 6424.0026 | 6423.9746 | 0 | 4.4 | 8 | Specific | 877.98 | 807.25 | 807.25 | 18.36 | 45 | >sp\|P01877\|IGHA2_HUMAN Immunoglobulin heavy constant alpha 2 OS=Homo sapiens OX=9606 GN=IGHA2 PE=1 SV=4 | 576 | 20201119_08533_1ng_glyco_hcd203040_compound15.25507.25507.4 | controllerType=0 controllerNumber=1 scan=25507 | 51.9128 |
| 21671:4 | 13 | K.VFPLSLDSTPQDGNVVVAC[+57.021]LVQGFFPQEPLSVTWSESGQN[+1647.613]VTAI | HexNAc(5)Hex(3)Fuc(1) | C[+57], N[+1648] | 1606.7561 | 4 | 6424.0026 | 6423.9746 | 0 | 4.4 | 8 | Specific | 912.04 | 803.63 | 803.63 | 19.36 | 45 | >sp\|P01877\|IGHA2_HUMAN Immunoglobulin heavy constant alpha 2 OS=Homo sapiens OX=9606 GN=IGHA2 PE=1 SV=4 | 576 | 20201119_08533_1ng_glyco_hcd203040_compound15.25509.25509.4 | controllerType=0 controllerNumber=1 scan=25509 | 51.9275 |
| 01811:3 | 13 | K.HYTN[+2262.814]SSQDVTVPC[+57.021]R.V | HexNAc(5)Hex(5)Fuc(1)NeuAc(1) | C[+57], N[+2263] | 1309.5312 | 3 | 3926.579 | 3926.5636 | 0 | 3.9 | 89 | Specific | 208.2 | 133.44 | 133.44 | 3.86 | 45 | >sp\|P01877\|IGHA2_HUMAN Immunoglobulin heavy constant alpha 2 OS=Homo sapiens OX=9606 GN=IGHA2 PE=1 SV=4 | 576 | 20201119_08533_1ng_glyco_hcd203040_compound15.4771.4771.3 | controllerType=0 controllerNumber=1 scan=4771 | 14.4465 |

Table S4. Cont.

| Query #:z | Protein Rank | Glycopeptide | Glycan | Modification Type(s) | Observed m/z | z | Observed (M+H) | Calc. mass (M+H) | Off-by-x error | Mass error (ppm) | StartPosition | Cleavage | Score | Delta | Delta Mod | \|Log Prob\| | # of unique peptides | Protein Name | Protein DB number | Comment | Scan # | Scan Time |
|---|---|---|---|---|---|---|---|---|---|---|---|---|---|---|---|---|---|---|---|---|---|---|
| 01000:2 | 13 | K.HYTN[+1647.613]SSQDVTVPC[+57.021]R.V | HexNAc(5)Hex(3)Fuc(1) | C[+57], N[+1648] | 1656.1884 | 2 | 3311.3695 | 3311.3625 | 0 | 2.1 | 89 | Specific | 234.38 | 171.21 | 171.21 | 3.95 | 45 | >sp\|P01877\|IGHA2_HUMAN Immunoglobulin heavy constant alpha 2 OS=Homo sapiens OX=9606 GN=IGHA2 PE=1 SV=4 | 576 | 20201119_08533_1ng_glyco_hcd203040_compound15.3929.3929.2 | controllerType=0 controllerNumber=1 scan=3929 | 12.9894 |
| 01289:3 | 13 | K.HYTN[+1647.613]SSQDVTVPC[+57.021]R.V | HexNAc(5)Hex(3)Fuc(1) | C[+57], N[+1648] | 1104.464 | 3 | 3311.3775 | 3311.3625 | 0 | 4.5 | 89 | Specific | 266.46 | 266.46 | 266.46 | 6.11 | 45 | >sp\|P01877\|IGHA2_HUMAN Immunoglobulin heavy constant alpha 2 OS=Homo sapiens OX=9606 GN=IGHA2 PE=1 SV=4 | 576 | 20201119_08533_1ng_glyco_hcd203040_compound15.4228.4228.3 | controllerType=0 controllerNumber=1 scan=4228 | 13.5114 |
| 00930:3 | 13 | K.HYTN[+1971.719]SSQDVTVPC[+57.021]R.V | HexNAc(5)Hex(5)Fuc(1) | C[+57], N[+1972] | 1212.5005 | 3 | 3635.4871 | 3635.4682 | 0 | 5.2 | 89 | Specific | 267.38 | 267.38 | 267.38 | 6.11 | 45 | >sp\|P01877\|IGHA2_HUMAN Immunoglobulin heavy constant alpha 2 OS=Homo sapiens OX=9606 GN=IGHA2 PE=1 SV=4 | 576 | 20201119_08533_1ng_glyco_hcd203040_compound15.3857.3857.3 | controllerType=0 controllerNumber=1 scan=3857 | 12.8651 |
| 00928:3 | 13 | K.HYTN[+1971.719]SSQDVTVPC[+57.021]R.V | HexNAc(5)Hex(5)Fuc(1) | C[+57], N[+1972] | 1212.5005 | 3 | 3635.4871 | 3635.4682 | 0 | 5.2 | 89 | Specific | 297.65 | 297.65 | 297.65 | 6.65 | 45 | >sp\|P01877\|IGHA2_HUMAN Immunoglobulin heavy constant alpha 2 OS=Homo sapiens OX=9606 GN=IGHA2 PE=1 SV=4 | 576 | 20201119_08533_1ng_glyco_hcd203040_compound15.3855.3855.3 | controllerType=0 controllerNumber=1 scan=3855 | 12.862 |
| 00887:3 | 13 | K.HYTN[+1809.666]SSQDVTVPC[+57.021]R.V | HexNAc(5)Hex(4)Fuc(1) | C[+57], N[+1810] | 1158.4798 | 3 | 3473.4248 | 3473.4154 | 0 | 2.7 | 89 | Specific | 391.86 | 337.41 | 337.41 | 7.58 | 45 | >sp\|P01877\|IGHA2_HUMAN Immunoglobulin heavy constant alpha 2 OS=Homo sapiens OX=9606 GN=IGHA2 PE=1 SV=4 | 576 | 20201119_08533_1ng_glyco_hcd203040_compound15.3812.3812.3 | controllerType=0 controllerNumber=1 scan=3812 | 12.7811 |
| 00889:3 | 13 | K.HYTN[+1809.666]SSQDVTVPC[+57.021]R.V | HexNAc(5)Hex(4)Fuc(1) | C[+57], N[+1810] | 1158.4798 | 3 | 3473.4248 | 3473.4154 | 0 | 2.7 | 89 | Specific | 523.03 | 318.35 | 318.35 | 8.07 | 45 | >sp\|P01877\|IGHA2_HUMAN Immunoglobulin heavy constant alpha 2 OS=Homo sapiens OX=9606 GN=IGHA2 PE=1 SV=4 | 576 | 20201119_08533_1ng_glyco_hcd203040_compound15.3814.3814.3 | controllerType=0 controllerNumber=1 scan=3814 | 12.7856 |
| 00956:3 | 13 | K.HYTN[+1809.666]SSQDVTVPC[+57.021]R.V | HexNAc(5)Hex(4)Fuc(1) | C[+57], N[+1810] | 1158.4781 | 3 | 3473.4197 | 3473.4154 | 0 | 1.2 | 89 | Specific | 572.69 | 481 | 481 | 9.4 | 45 | >sp\|P01877\|IGHA2_HUMAN Immunoglobulin heavy constant alpha 2 OS=Homo sapiens OX=9606 GN=IGHA2 PE=1 SV=4 | 576 | 20201119_08533_1ng_glyco_hcd203040_compound15.3884.3884.3 | controllerType=0 controllerNumber=1 scan=3884 | 12.9123 |
| 00927:3 | 13 | K.HYTN[+1647.613]SSQDVTVPC[+57.021]R.V | HexNAc(5)Hex(3)Fuc(1) | C[+57], N[+1648] | 1104.4613 | 3 | 3311.3694 | 3311.3625 | 0 | 2.1 | 89 | Specific | 627.62 | 612.05 | 612.05 | 12.11 | 45 | >sp\|P01877\|IGHA2_HUMAN Immunoglobulin heavy constant alpha 2 OS=Homo sapiens OX=9606 GN=IGHA2 PE=1 SV=4 | 576 | 20201119_08533_1ng_glyco_hcd203040_compound15.3854.3854.3 | controllerType=0 controllerNumber=1 scan=3854 | 12.8601 |
| 00958:3 | 13 | K.HYTN[+1809.666]SSQDVTVPC[+57.021]R.V | HexNAc(5)Hex(4)Fuc(1) | C[+57], N[+1810] | 1158.4781 | 3 | 3473.4197 | 3473.4154 | 0 | 1.2 | 89 | Specific | 646.5 | 527.91 | 527.91 | 11.56 | 45 | >sp\|P01877\|IGHA2_HUMAN Immunoglobulin heavy constant alpha 2 OS=Homo sapiens OX=9606 GN=IGHA2 PE=1 SV=4 | 576 | 20201119_08533_1ng_glyco_hcd203040_compound15.3886.3886.3 | controllerType=0 controllerNumber=1 scan=3886 | 12.9161 |
| 00925:3 | 13 | K.HYTN[+1647.613]SSQDVTVPC[+57.021]R.V | HexNAc(5)Hex(3)Fuc(1) | C[+57], N[+1648] | 1104.4613 | 3 | 3311.3694 | 3311.3625 | 0 | 2.1 | 89 | Specific | 682.04 | 592.83 | 592.83 | 13.11 | 45 | >sp\|P01877\|IGHA2_HUMAN Immunoglobulin heavy constant alpha 2 OS=Homo sapiens OX=9606 GN=IGHA2 PE=1 SV=4 | 576 | 20201119_08533_1ng_glyco_hcd203040_compound15.3852.3852.3 | controllerType=0 controllerNumber=1 scan=3852 | 12.8552 |
| 03223:5 | 13 | K.HYTN[+1647.613]SSQDVTVPC[+57.021]RVPPPPPC[+57.021]C[+57.021]HPR.L | HexNAc(5)Hex(3)Fuc(1) | C[+57]*3, N[+1648] | 921.9999 | 5 | 4605.9704 | 4605.9689 | 0 | 0.3 | 89 | Specific | 353.34 | 353.34 | 353.34 | 6.76 | 45 | >sp\|P01877\|IGHA2_HUMAN Immunoglobulin heavy constant alpha 2 OS=Homo sapiens OX=9606 GN=IGHA2 PE=1 SV=4 | 576 | 20201119_08533_1ng_glyco_hcd203040_compound15.6236.6236.5 | controllerType=0 controllerNumber=1 scan=6236 | 17.0322 |
| 03225:5 | 13 | K.HYTN[+1647.613]SSQDVTVPC[+57.021]RVPPPPPC[+57.021]C[+57.021]HPR.L | HexNAc(5)Hex(3)Fuc(1) | C[+57]*3, N[+1648] | 921.9999 | 5 | 4605.9704 | 4605.9689 | 0 | 0.3 | 89 | Specific | 386.34 | 306.9 | 306.9 | 6.75 | 45 | >sp\|P01877\|IGHA2_HUMAN Immunoglobulin heavy constant alpha 2 OS=Homo sapiens OX=9606 GN=IGHA2 PE=1 SV=4 | 576 | 20201119_08533_1ng_glyco_hcd203040_compound15.6238.6238.5 | controllerType=0 controllerNumber=1 scan=6238 | 17.0359 |
| 02886:4 | 13 | K.HYTN[+1809.666]SSQDVTVPC[+57.021]RVPPPPPC[+57.021]C[+57.021]HPR.L | HexNAc(5)Hex(4)Fuc(1) | C[+57]*3, N[+1810] | 1192.7636 | 4 | 4768.0325 | 4768.0217 | 0 | 2.3 | 89 | Specific | 396.64 | 272.48 | 272.48 | 6.58 | 45 | >sp\|P01877\|IGHA2_HUMAN Immunoglobulin heavy constant alpha 2 OS=Homo sapiens OX=9606 GN=IGHA2 PE=1 SV=4 | 576 | 20201119_08533_1ng_glyco_hcd203040_compound15.5885.5885.4 | controllerType=0 controllerNumber=1 scan=5885 | 16.4219 |
| 02884:4 | 13 | K.HYTN[+1809.666]SSQDVTVPC[+57.021]RVPPPPPC[+57.021]C[+57.021]HPR.L | HexNAc(5)Hex(4)Fuc(1) | C[+57]*3, N[+1810] | 1192.7636 | 4 | 4768.0325 | 4768.0217 | 0 | 2.3 | 89 | Specific | 399.96 | 390.72 | 390.72 | 8.5 | 45 | >sp\|P01877\|IGHA2_HUMAN Immunoglobulin heavy constant alpha 2 OS=Homo sapiens OX=9606 GN=IGHA2 PE=1 SV=4 | 576 | 20201119_08533_1ng_glyco_hcd203040_compound15.5883.5883.4 | controllerType=0 controllerNumber=1 scan=5883 | 16.4184 |
| 02904:4 | 13 | K.HYTN[+1809.666]SSQDVTVPC[+57.021]RVPPPPPC[+57.021]C[+57.021]HPR.L | HexNAc(5)Hex(4)Fuc(1) | C[+57]*3, N[+1810] | 1192.7629 | 4 | 4768.0296 | 4768.0217 | 0 | 1.7 | 89 | Specific | 464.48 | 315.62 | 315.62 | 7.05 | 45 | >sp\|P01877\|IGHA2_HUMAN Immunoglobulin heavy constant alpha 2 OS=Homo sapiens OX=9606 GN=IGHA2 PE=1 SV=4 | 576 | 20201119_08533_1ng_glyco_hcd203040_compound15.5904.5904.4 | controllerType=0 controllerNumber=1 scan=5904 | 16.458 |
| 02902:4 | 13 | K.HYTN[+1809.666]SSQDVTVPC[+57.021]RVPPPPPC[+57.021]C[+57.021]HPR.L | HexNAc(5)Hex(4)Fuc(1) | C[+57]*3, N[+1810] | 1192.7629 | 4 | 4768.0296 | 4768.0217 | 0 | 1.7 | 89 | Specific | 492.2 | 492.2 | 492.2 | 8.9 | 45 | >sp\|P01877\|IGHA2_HUMAN Immunoglobulin heavy constant alpha 2 OS=Homo sapiens OX=9606 GN=IGHA2 PE=1 SV=4 | 576 | 20201119_08533_1ng_glyco_hcd203040_compound15.5902.5902.4 | controllerType=0 controllerNumber=1 scan=5902 | 16.4545 |
| 02896:4 | 13 | K.HYTN[+1647.613]SSQDVTVPC[+57.021]RVPPPPPC[+57.021]C[+57.021]HPR.L | HexNAc(5)Hex(3)Fuc(1) | C[+57]*3, N[+1648] | 1152.2498 | 4 | 4605.9774 | 4605.9689 | 0 | 1.8 | 89 | Specific | 683.04 | 465.26 | 465.26 | 12.52 | 45 | >sp\|P01877\|IGHA2_HUMAN Immunoglobulin heavy constant alpha 2 OS=Homo sapiens OX=9606 GN=IGHA2 PE=1 SV=4 | 576 | 20201119_08533_1ng_glyco_hcd203040_compound15.5896.5896.4 | controllerType=0 controllerNumber=1 scan=5896 | 16.443 |
| 02895:5 | 13 | K.HYTN[+1647.613]SSQDVTVPC[+57.021]RVPPPPPC[+57.021]C[+57.021]HPR.L | HexNAc(5)Hex(3)Fuc(1) | C[+57]*3, N[+1648] | 922.0011 | 5 | 4605.9765 | 4605.9689 | 0 | 1.7 | 89 | Specific | 758.55 | 511.76 | 511.76 | 15.06 | 45 | >sp\|P01877\|IGHA2_HUMAN Immunoglobulin heavy constant alpha 2 OS=Homo sapiens OX=9606 GN=IGHA2 PE=1 SV=4 | 576 | 20201119_08533_1ng_glyco_hcd203040_compound15.5895.5895.5 | controllerType=0 controllerNumber=1 scan=5895 | 16.441 |
| 02898:4 | 13 | K.HYTN[+1647.613]SSQDVTVPC[+57.021]RVPPPPPC[+57.021]C[+57.021]HPR.L | HexNAc(5)Hex(3)Fuc(1) | C[+57]*3, N[+1648] | 1152.2498 | 4 | 4605.9774 | 4605.9689 | 0 | 1.8 | 89 | Specific | 772.06 | 523.22 | 523.22 | 14.52 | 45 | >sp\|P01877\|IGHA2_HUMAN Immunoglobulin heavy constant alpha 2 OS=Homo sapiens OX=9606 GN=IGHA2 PE=1 SV=4 | 576 | 20201119_08533_1ng_glyco_hcd203040_compound15.5898.5898.4 | controllerType=0 controllerNumber=1 scan=5898 | 16.4481 |
| 02897:5 | 13 | K.HYTN[+1647.613]SSQDVTVPC[+57.021]RVPPPPPC[+57.021]C[+57.021]HPR.L | HexNAc(5)Hex(3)Fuc(1) | C[+57]*3, N[+1648] | 922.0011 | 5 | 4605.9765 | 4605.9689 | 0 | 1.7 | 89 | Specific | 845.53 | 632.19 | 632.19 | 16.06 | 45 | >sp\|P01877\|IGHA2_HUMAN Immunoglobulin heavy constant alpha 2 OS=Homo sapiens OX=9606 GN=IGHA2 PE=1 SV=4 | 576 | 20201119_08533_1ng_glyco_hcd203040_compound15.5897.5897.5 | controllerType=0 controllerNumber=1 scan=5897 | 16.4462 |
| 04406:3 | 13 | K.TPLTAN[+2262.814]ITK.S | HexNAc(5)Hex(5)Fuc(1)NeuAc(1) | N[+2263] | 1074.4634 | 3 | 3221.3757 | 3221.3711 | 0 | 1.4 | 200 | Specific | 221.45 | 94.7 | 94.7 | 3.29 | 45 | >sp\|P01877\|IGHA2_HUMAN Immunoglobulin heavy constant alpha 2 OS=Homo sapiens OX=9606 GN=IGHA2 PE=1 SV=4 | 576 | 20201119_08533_1ng_glyco_hcd203040_compound15.7463.7463.3 | controllerType=0 controllerNumber=1 scan=7463 | 19.0645 |
| 02574:3 | 13 | K.TPLTAN[+1971.719]ITK.S | HexNAc(5)Hex(5)Fuc(1) | N[+1972] | 977.4328 | 3 | 2930.2839 | 2930.2757 | 0 | 2.8 | 200 | Specific | 276.13 | 215.15 | 215.15 | 5.57 | 45 | >sp\|P01877\|IGHA2_HUMAN Immunoglobulin heavy constant alpha 2 OS=Homo sapiens OX=9606 GN=IGHA2 PE=1 SV=4 | 576 | 20201119_08533_1ng_glyco_hcd203040_compound15.5562.5562.3 | controllerType=0 controllerNumber=1 scan=5562 | 15.8333 |
| 02623:3 | 13 | K.TPLTAN[+1809.666]ITK.S | HexNAc(5)Hex(4)Fuc(1) | N[+1810] | 923.416 | 3 | 2768.2334 | 2768.2228 | 0 | 3.8 | 200 | Specific | 431.94 | 171.63 | 171.63 | 6 | 45 | >sp\|P01877\|IGHA2_HUMAN Immunoglobulin heavy constant alpha 2 OS=Homo sapiens OX=9606 GN=IGHA2 PE=1 SV=4 | 576 | 20201119_08533_1ng_glyco_hcd203040_compound15.5613.5613.3 | controllerType=0 controllerNumber=1 scan=5613 | 15.9257 |
| 02609:2 | 13 | K.TPLTAN[+1444.534]ITK.S | HexNAc(4)Hex(3)Fuc(1) | N[+1445] | 1202.0519 | 2 | 2403.0966 | 2403.0906 | 0 | 2.5 | 200 | Specific | 436.73 | 217.07 | 217.07 | 6.66 | 45 | >sp\|P01877\|IGHA2_HUMAN Immunoglobulin heavy constant alpha 2 OS=Homo sapiens OX=9606 GN=IGHA2 PE=1 SV=4 | 576 | 20201119_08533_1ng_glyco_hcd203040_compound15.5598.5598.2 | controllerType=0 controllerNumber=1 scan=5598 | 15.8973 |
| 02661:2 | 13 | K.TPLTAN[+1501.555]ITK.S | HexNAc(5)Hex(3) | N[+1502] | 1230.5639 | 2 | 2460.1205 | 2460.1121 | 0 | 3.4 | 200 | Specific | 452.04 | 275.82 | 275.82 | 7.11 | 45 | >sp\|P01877\|IGHA2_HUMAN Immunoglobulin heavy constant alpha 2 OS=Homo sapiens OX=9606 GN=IGHA2 PE=1 SV=4 | 576 | 20201119_08533_1ng_glyco_hcd203040_compound15.5652.5652.2 | controllerType=0 controllerNumber=1 scan=5652 | 15.995 |
| 02663:2 | 13 | K.TPLTAN[+1501.555]ITK.S | HexNAc(5)Hex(3) | N[+1502] | 1230.5639 | 2 | 2460.1205 | 2460.1121 | 0 | 3.4 | 200 | Specific | 473.59 | 282.77 | 282.77 | 7.26 | 45 | >sp\|P01877\|IGHA2_HUMAN Immunoglobulin heavy constant alpha 2 OS=Homo sapiens OX=9606 GN=IGHA2 PE=1 SV=4 | 576 | 20201119_08533_1ng_glyco_hcd203040_compound15.5654.5654.2 | controllerType=0 controllerNumber=1 scan=5654 | 15.9987 |
| 02680:3 | 13 | K.TPLTAN[+1647.613]ITK.S | HexNAc(5)Hex(3)Fuc(1) | N[+1648] | 869.3975 | 3 | 2606.1779 | 2606.17 | 0 | 3 | 200 | Specific | 478.21 | 190.08 | 190.08 | 6.61 | 45 | >sp\|P01877\|IGHA2_HUMAN Immunoglobulin heavy constant alpha 2 OS=Homo sapiens OX=9606 GN=IGHA2 PE=1 SV=4 | 576 | 20201119_08533_1ng_glyco_hcd203040_compound15.5672.5672.3 | controllerType=0 controllerNumber=1 scan=5672 | 16.0312 |
| 02625:3 | 13 | K.TPLTAN[+1809.666]ITK.S | HexNAc(5)Hex(4)Fuc(1) | N[+1810] | 923.416 | 3 | 2768.2334 | 2768.2228 | 0 | 3.8 | 200 | Specific | 512.77 | 270.55 | 270.55 | 7.1 | 45 | >sp\|P01877\|IGHA2_HUMAN Immunoglobulin heavy constant alpha 2 OS=Homo sapiens OX=9606 GN=IGHA2 PE=1 SV=4 | 576 | 20201119_08533_1ng_glyco_hcd203040_compound15.5615.5615.3 | controllerType=0 controllerNumber=1 scan=5615 | 15.9303 |
| 02611:2 | 13 | K.TPLTAN[+1444.534]ITK.S | HexNAc(4)Hex(3)Fuc(1) | N[+1445] | 1202.0519 | 2 | 2403.0966 | 2403.0906 | 0 | 2.5 | 200 | Specific | 514.89 | 268.41 | 268.41 | 7.32 | 45 | >sp\|P01877\|IGHA2_HUMAN Immunoglobulin heavy constant alpha 2 OS=Homo sapiens OX=9606 GN=IGHA2 PE=1 SV=4 | 576 | 20201119_08533_1ng_glyco_hcd203040_compound15.5600.5600.2 | controllerType=0 controllerNumber=1 scan=5600 | 15.901 |
| 02682:3 | 13 | K.TPLTAN[+1647.613]ITK.S | HexNAc(5)Hex(3)Fuc(1) | N[+1648] | 869.3975 | 3 | 2606.1779 | 2606.17 | 0 | 3 | 200 | Specific | 548.75 | 265.86 | 265.86 | 7.26 | 45 | >sp\|P01877\|IGHA2_HUMAN Immunoglobulin heavy constant alpha 2 OS=Homo sapiens OX=9606 GN=IGHA2 PE=1 SV=4 | 576 | 20201119_08533_1ng_glyco_hcd203040_compound15.5674.5674.3 | controllerType=0 controllerNumber=1 scan=5674 | 16.0364 |
| 02606:2 | 13 | K.TPLTAN[+1809.666]ITK.S | HexNAc(5)Hex(4)Fuc(1) | N[+1810] | 1384.6172 | 2 | 2768.2272 | 2768.2228 | 0 | 1.6 | 200 | Specific | 592.93 | 332.14 | 332.14 | 9.24 | 45 | >sp\|P01877\|IGHA2_HUMAN Immunoglobulin heavy constant alpha 2 OS=Homo sapiens OX=9606 GN=IGHA2 PE=1 SV=4 | 576 | 20201119_08533_1ng_glyco_hcd203040_compound15.5595.5595.2 | controllerType=0 controllerNumber=1 scan=5595 | 15.8925 |
| 02604:2 | 13 | K.TPLTAN[+1809.666]ITK.S | HexNAc(5)Hex(4)Fuc(1) | N[+1810] | 1384.6172 | 2 | 2768.2272 | 2768.2228 | 0 | 1.6 | 200 | Specific | 593.85 | 342.86 | 342.86 | 9.24 | 45 | >sp\|P01877\|IGHA2_HUMAN Immunoglobulin heavy constant alpha 2 OS=Homo sapiens OX=9606 GN=IGHA2 PE=1 SV=4 | 576 | 20201119_08533_1ng_glyco_hcd203040_compound15.5593.5593.2 | controllerType=0 controllerNumber=1 scan=5593 | 15.8883 |
| 02653:2 | 13 | K.TPLTAN[+1647.613]ITK.S | HexNAc(5)Hex(3)Fuc(1) | N[+1648] | 1303.5935 | 2 | 2606.1798 | 2606.17 | 0 | 3.8 | 200 | Specific | 614 | 373.58 | 373.58 | 11.81 | 45 | >sp\|P01877\|IGHA2_HUMAN Immunoglobulin heavy constant alpha 2 OS=Homo sapiens OX=9606 GN=IGHA2 PE=1 SV=4 | 576 | 20201119_08533_1ng_glyco_hcd203040_compound15.5644.5644.2 | controllerType=0 controllerNumber=1 scan=5644 | 15.9803 |
| 02655:2 | 13 | K.TPLTAN[+1647.613]ITK.S | HexNAc(5)Hex(3)Fuc(1) | N[+1648] | 1303.5935 | 2 | 2606.1798 | 2606.17 | 0 | 3.8 | 200 | Specific | 622.78 | 325.01 | 325.01 | 10.27 | 45 | >sp\|P01877\|IGHA2_HUMAN Immunoglobulin heavy constant alpha 2 OS=Homo sapiens OX=9606 GN=IGHA2 PE=1 SV=4 | 576 | 20201119_08533_1ng_glyco_hcd203040_compound15.5646.5646.2 | controllerType=0 controllerNumber=1 scan=5646 | 15.9861 |
| 10124:3 | 13 | R.MAGKPTHIN[+1095.397]VSVVMAEADGTC[+57.021]Y.- | HexNAc(3)Hex(3) | C[+57], N[+1095] | 1149.5011 | 3 | 3446.4887 | 3446.4794 | 0 | 2.7 | 319 | CRagged | 280.18 | 280.18 | 280.18 | 7.31 | 45 | >sp\|P01877\|IGHA2_HUMAN Immunoglobulin heavy constant alpha 2 OS=Homo sapiens OX=9606 GN=IGHA2 PE=1 SV=4 | 576 | 20201119_08533_1ng_glyco_hcd203040_compound15.13416.13416.3 | controllerType=0 controllerNumber=1 scan=13416 | 29.0528 |
| 09980:3 | 13 | R.MAGKPTHIN[+1663.608]VSVVMAEADGTC[+57.021]Y.- | HexNAc(5)Hex(4) | C[+57], N[+1664] | 1338.903 | 3 | 4014.6945 | 4014.691 | 0 | 0.9 | 319 | CRagged | 289.89 | 289.89 | 272.13 | 6.73 | 45 | >sp\|P01877\|IGHA2_HUMAN Immunoglobulin heavy constant alpha 2 OS=Homo sapiens OX=9606 GN=IGHA2 PE=1 SV=4 | 576 | 20201119_08533_1ng_glyco_hcd203040_compound15.13267.13267.3 | controllerType=0 controllerNumber=1 scan=13267 | 28.7943 |
| 11867:3 | 13 | R.MAGKPTHIN[+1548.545]VSVVMAEADGTC[+57.021]Y.- | HexNAc(3)Hex(4)NeuAc(1) | C[+57], N[+1549] | 1300.55 | 3 | 3899.6354 | 3899.6277 | 0 | 2 | 319 | CRagged | 303.81 | 281.7 | 281.7 | 7.29 | 45 | >sp\|P01877\|IGHA2_HUMAN Immunoglobulin heavy constant alpha 2 OS=Homo sapiens OX=9606 GN=IGHA2 PE=1 SV=4 | 576 | 20201119_08533_1ng_glyco_hcd203040_compound15.15225.15225.3 | controllerType=0 controllerNumber=1 scan=15225 | 32.115 |
| 10086:3 | 13 | R.MAGKPTHIN[+1647.613]VSVVMAEADGTC[+57.021]Y.- | HexNAc(5)Hex(3)Fuc(1) | C[+57], N[+1648] | 1333.5735 | 3 | 3998.7061 | 3998.6961 | 0 | 2.5 | 319 | CRagged | 510.52 | 510.52 | 510.52 | 10.21 | 45 | >sp\|P01877\|IGHA2_HUMAN Immunoglobulin heavy constant alpha 2 OS=Homo sapiens OX=9606 GN=IGHA2 PE=1 SV=4 | 576 | 20201119_08533_1ng_glyco_hcd203040_compound15.13377.13377.3 | controllerType=0 controllerNumber=1 scan=13377 | 28.9888 |
| 10084:3 | 13 | R.MAGKPTHIN[+1647.613]VSVVMAEADGTC[+57.021]Y.- | HexNAc(5)Hex(3)Fuc(1) | C[+57], N[+1648] | 1333.5735 | 3 | 3998.7061 | 3998.6961 | 0 | 2.5 | 319 | CRagged | 512 | 512 | 512 | 10.21 | 45 | >sp\|P01877\|IGHA2_HUMAN Immunoglobulin heavy constant alpha 2 OS=Homo sapiens OX=9606 GN=IGHA2 PE=1 SV=4 | 576 | 20201119_08533_1ng_glyco_hcd203040_compound15.13375.13375.3 | controllerType=0 controllerNumber=1 scan=13375 | 28.9852 |
| 09892:3 | 13 | R.MAGKPTHIN[+1540.529]VSVVMAEADGTC[+57.021]Y.- | HexNAc(2)Hex(7) | C[+57], N[+1541] | 1297.8807 | 3 | 3891.6275 | 3891.6114 | 0 | 4.1 | 319 | CRagged | 586.78 | 512.47 | 320.85 | 11.52 | 45 | >sp\|P01877\|IGHA2_HUMAN Immunoglobulin heavy constant alpha 2 OS=Homo sapiens OX=9606 GN=IGHA2 PE=1 SV=4 | 576 | 20201119_08533_1ng_glyco_hcd203040_compound15.13176.13176.3 | controllerType=0 controllerNumber=1 scan=13176 | 28.6257 |
| 10031:3 | 13 | R.MAGKPTHIN[+1501.555]VSVVMAEADGTC[+57.021]Y.- | HexNAc(5)Hex(3) | C[+57], N[+1502] | 1284.8915 | 3 | 3852.6601 | 3852.6382 | 0 | 5.7 | 319 | CRagged | 714.63 | 714.63 | 714.63 | 14.01 | 45 | >sp\|P01877\|IGHA2_HUMAN Immunoglobulin heavy constant alpha 2 OS=Homo sapiens OX=9606 GN=IGHA2 PE=1 SV=4 | 576 | 20201119_08533_1ng_glyco_hcd203040_compound15.13320.13320.3 | controllerType=0 controllerNumber=1 scan=13320 | 28.888 |
| 10018:3 | 13 | R.MAGKPTHIN[+1501.555]VSVVMAEADGTC[+57.021]Y.- | HexNAc(5)Hex(3) | C[+57], N[+1502] | 1284.8901 | 3 | 3852.6557 | 3852.6382 | 0 | 4.5 | 319 | CRagged | 720.49 | 720.49 | 720.49 | 14.52 | 45 | >sp\|P01877\|IGHA2_HUMAN Immunoglobulin heavy constant alpha 2 OS=Homo sapiens OX=9606 GN=IGHA2 PE=1 SV=4 | 576 | 20201119_08533_1ng_glyco_hcd203040_compound15.13306.13306.3 | controllerType=0 controllerNumber=1 scan=13306 | 28.8634 |
| 10016:3 | 13 | R.MAGKPTHIN[+1501.555]VSVVMAEADGTC[+57.021]Y.- | HexNAc(5)Hex(3) | C[+57], N[+1502] | 1284.8901 | 3 | 3852.6557 | 3852.6382 | 0 | 4.5 | 319 | CRagged | 730.33 | 730.33 | 730.33 | 14.52 | 45 | >sp\|P01877\|IGHA2_HUMAN Immunoglobulin heavy constant alpha 2 OS=Homo sapiens OX=9606 GN=IGHA2 PE=1 SV=4 | 576 | 20201119_08533_1ng_glyco_hcd203040_compound15.13304.13304.3 | controllerType=0 controllerNumber=1 scan=13304 | 28.8585 |
| 10033:3 | 13 | R.MAGKPTHIN[+1501.555]VSVVMAEADGTC[+57.021]Y.- | HexNAc(5)Hex(3) | C[+57], N[+1502] | 1284.8915 | 3 | 3852.6601 | 3852.6382 | 0 | 5.7 | 319 | CRagged | 760.81 | 760.81 | 760.81 | 15.01 | 45 | >sp\|P01877\|IGHA2_HUMAN Immunoglobulin heavy constant alpha 2 OS=Homo sapiens OX=9606 GN=IGHA2 PE=1 SV=4 | 576 | 20201119_08533_1ng_glyco_hcd203040_compound15.13322.13322.3 | controllerType=0 controllerNumber=1 scan=13322 | 28.8922 |
| 09836:3 | 13 | R.MAGKPTHIN[+1702.581]VSVVMAEADGTC[+57.021]Y.- | HexNAc(2)Hex(8) | C[+57], N[+1703] | 1351.898 | 3 | 4053.6795 | 4053.6642 | 0 | 3.8 | 319 | CRagged | 829.41 | 829.41 | 462.98 | 16.78 | 45 | >sp\|P01877\|IGHA2_HUMAN Immunoglobulin heavy constant alpha 2 OS=Homo sapiens OX=9606 GN=IGHA2 PE=1 SV=4 | 576 | 20201119_08533_1ng_glyco_hcd203040_compound15.13118.13118.3 | controllerType=0 controllerNumber=1 scan=13118 | 28.5264 |
| 09975:3 | 13 | R.MAGKPTHIN[+1216.423]VSVVMAEADGTC[+57.021]Y.- | HexNAc(2)Hex(5) | C[+57], N[+1216] | 1189.8427 | 3 | 3567.5136 | 3567.5057 | 0 | 2.2 | 319 | CRagged | 862.45 | 862.45 | 477.82 | 17.75 | 45 | >sp\|P01877\|IGHA2_HUMAN Immunoglobulin heavy constant alpha 2 OS=Homo sapiens OX=9606 GN=IGHA2 PE=1 SV=4 | 576 | 20201119_08533_1ng_glyco_hcd203040_compound15.13262.13262.3 | controllerType=0 controllerNumber=1 scan=13262 | 28.7815 |
| 09894:3 | 13 | R.MAGKPTHIN[+1540.529]VSVVMAEADGTC[+57.021]Y.- | HexNAc(2)Hex(7) | C[+57], N[+1541] | 1297.8807 | 3 | 3891.6275 | 3891.6114 | 0 | 4.1 | 319 | CRagged | 868.86 | 868.86 | 501.39 | 17.52 | 45 | >sp\|P01877\|IGHA2_HUMAN Immunoglobulin heavy constant alpha 2 OS=Homo sapiens OX=9606 GN=IGHA2 PE=1 SV=4 | 576 | 20201119_08533_1ng_glyco_hcd203040_compound15.13178.13178.3 | controllerType=0 controllerNumber=1 scan=13178 | 28.629 |
| 09904:3 | 13 | R.MAGKPTHIN[+1378.476]VSVVMAEADGTC[+57.021]Y.- | HexNAc(2)Hex(6) | C[+57], N[+1378] | 1243.8631 | 3 | 3729.5747 | 3729.5585 | 0 | 4.3 | 319 | CRagged | 893.14 | 893.14 | 514.31 | 17.52 | 45 | >sp\|P01877\|IGHA2_HUMAN Immunoglobulin heavy constant alpha 2 OS=Homo sapiens OX=9606 GN=IGHA2 PE=1 SV=4 | 576 | 20201119_08533_1ng_glyco_hcd203040_compound15.13188.13188.3 | controllerType=0 controllerNumber=1 scan=13188 | 28.6459 |
| 09977:3 | 13 | R.MAGKPTHIN[+1216.423]VSVVMAEADGTC[+57.021]Y.- | HexNAc(2)Hex(5) | C[+57], N[+1216] | 1189.8427 | 3 | 3567.5136 | 3567.5057 | 0 | 2.2 | 319 | CRagged | 905.76 | 905.76 | 495.19 | 18.75 | 45 | >sp\|P01877\|IGHA2_HUMAN Immunoglobulin heavy constant alpha 2 OS=Homo sapiens OX=9606 GN=IGHA2 PE=1 SV=4 | 576 | 20201119_08533_1ng_glyco_hcd203040_compound15.13264.13264.3 | controllerType=0 controllerNumber=1 scan=13264 | 28.7885 |
| 09906:3 | 13 | R.MAGKPTHIN[+1378.476]VSVVMAEADGTC[+57.021]Y.- | HexNAc(2)Hex(6) | C[+57], N[+1378] | 1243.8631 | 3 | 3729.5747 | 3729.5585 | 0 | 4.3 | 319 | CRagged | 970.73 | 861.05 | 595.82 | 19.52 | 45 | >sp\|P01877\|IGHA2_HUMAN Immunoglobulin heavy constant alpha 2 OS=Homo sapiens OX=9606 GN=IGHA2 PE=1 SV=4 | 576 | 20201119_08533_1ng_glyco_hcd203040_compound15.13190.13190.3 | controllerType=0 controllerNumber=1 scan=13190 | 28.6516 |
| 09475:3 | 13 | R.M[+15.995]AGKPTHIN[+1378.476]VSVVMAEADGTC[+57.021]Y.- | HexNAc(2)Hex(6) | C[+57], M[+16], N[+1378] | 1249.1944 | 3 | 3745.5686 | 3745.5535 | 0 | 4.1 | 319 | CRagged | 267.61 | 267.61 | 92.53 | 6.75 | 45 | >sp\|P01877\|IGHA2_HUMAN Immunoglobulin heavy constant alpha 2 OS=Homo sapiens OX=9606 GN=IGHA2 PE=1 SV=4 | 576 | 20201119_08533_1ng_glyco_hcd203040_compound15.12744.12744.3 | controllerType=0 controllerNumber=1 scan=12744 | 27.8903 |
| 09477:3 | 13 | R.M[+15.995]AGKPTHIN[+1378.476]VSVVMAEADGTC[+57.021]Y.- | HexNAc(2)Hex(6) | C[+57], M[+16], N[+1378] | 1249.1944 | 3 | 3745.5686 | 3745.5535 | 0 | 4.1 | 319 | CRagged | 306.2 | 306.2 | 40.7 | 7.29 | 45 | >sp\|P01877\|IGHA2_HUMAN Immunoglobulin heavy constant alpha 2 OS=Homo sapiens OX=9606 GN=IGHA2 PE=1 SV=4 | 576 | 20201119_08533_1ng_glyco_hcd203040_compound15.12746.12746.3 | controllerType=0 controllerNumber=1 scan=12746 | 27.9086 |
| 05296:3 | 14 | R.LSPN[+2205.793]ASAEHLELR.W | HexNAc(4)Hex(5)Fuc(2)NeuAc(1) | N[+2206] | 1214.8548 | 3 | 3642.5498 | 3642.5421 | 0 | 2.1 | 52 | Specific | 397.19 | 347.46 | 347.46 | 7.47 | 38 | >sp\|Q13410\|BT1A1_HUMAN Butyrophilin subfamily 1 member A1 OS=Homo sapiens OX=9606 GN=BTN1A1 PE=1 SV=3 | 179 | 20201119_08533_1ng_glyco_hcd203040_compound15.8384.8384.3 | controllerType=0 controllerNumber=1 scan=8384 | 20.6078 |
| 05294:3 | 14 | R.LSPN[+2205.793]ASAEHLELR.W | HexNAc(4)Hex(5)Fuc(2)NeuAc(1) | N[+2206] | 1214.8548 | 3 | 3642.5498 | 3642.5421 | 0 | 2.1 | 52 | Specific | 464.86 | 286.5 | 286.5 | 7.41 | 38 | >sp\|Q13410\|BT1A1_HUMAN Butyrophilin subfamily 1 member A1 OS=Homo sapiens OX=9606 GN=BTN1A1 PE=1 SV=3 | 179 | 20201119_08533_1ng_glyco_hcd203040_compound15.8382.8382.3 | controllerType=0 controllerNumber=1 scan=8382 | 20.6016 |
| 07306:3 | 14 | R.LSPN[+2350.830]ASAEHLELR.W | HexNAc(4)Hex(5)Fuc(1)NeuAc(2) | N[+2351] | 1263.2024 | 3 | 3787.5928 | 3787.5796 | 0 | 3.5 | 52 | Specific | 520.16 | 359.44 | 359.44 | 8.14 | 38 | >sp\|Q13410\|BT1A1_HUMAN Butyrophilin subfamily 1 member A1 OS=Homo sapiens OX=9606 GN=BTN1A1 PE=1 SV=3 | 179 | 20201119_08533_1ng_glyco_hcd203040_compound15.10463.10463.3 | controllerType=0 controllerNumber=1 scan=10463 | 24.034 |
| 07308:3 | 14 | R.LSPN[+2350.830]ASAEHLELR.W | HexNAc(4)Hex(5)Fuc(1)NeuAc(2) | N[+2351] | 1263.2024 | 3 | 3787.5928 | 3787.5796 | 0 | 3.5 | 52 | Specific | 547.75 | 516.13 | 516.13 | 9.98 | 38 | >sp\|Q13410\|BT1A1_HUMAN Butyrophilin subfamily 1 member A1 OS=Homo sapiens OX=9606 GN=BTN1A1 PE=1 SV=3 | 179 | 20201119_08533_1ng_glyco_hcd203040_compound15.10465.10465.3 | controllerType=0 controllerNumber=1 scan=10465 | 24.0382 |
| 22488:4 | 15 | K.ADTHDEILEGLNFN[+2861.000]LTEIPEAQIHEGFQELLR.T | HexNAc(5)Hex(6)NeuAc(3) | N[+2861] | 1638.9664 | 4 | 6552.8439 | 6552.8239 | 0 | 3.1 | 94 | Specific | 298.78 | 157.32 | 157.32 | 5.75 | 41 | >sp\|P01009\|A1AT_HUMAN Alpha-1-antitrypsin OS=Homo sapiens OX=9606 GN=SERPINA1 PE=1 SV=3 | 642 | 20201119_08533_1ng_glyco_hcd203040_compound15.26371.26371.4 | controllerType=0 controllerNumber=1 scan=26371 | 53.4865 |
| 22490:4 | 15 | K.ADTHDEILEGLNFN[+2861.000]LTEIPEAQIHEGFQELLR.T | HexNAc(5)Hex(6)NeuAc(3) | N[+2861] | 1638.9664 | 4 | 6552.8439 | 6552.8239 | 0 | 3.1 | 94 | Specific | 403.53 | 345.24 | 345.24 | 7.81 | 41 | >sp\|P01009\|A1AT_HUMAN Alpha-1-antitrypsin OS=Homo sapiens OX=9606 GN=SERPINA1 PE=1 SV=3 | 642 | 20201119_08533_1ng_glyco_hcd203040_compound15.26373.26373.4 | controllerType=0 controllerNumber=1 scan=26373 | 53.49 |
| 22430:4 | 15 | K.ADTHDEILEGLNFN[+2861.000]LTEIPEAQIHEGFQELLR.T | HexNAc(5)Hex(6)NeuAc(3) | N[+2861] | 1638.9716 | 4 | 6552.8644 | 6552.8239 | 0 | 6.2 | 94 | Specific | 863.65 | 606.6 | 606.6 | 16.6 | 41 | >sp\|P01009\|A1AT_HUMAN Alpha-1-antitrypsin OS=Homo sapiens OX=9606 GN=SERPINA1 PE=1 SV=3 | 642 | 20201119_08533_1ng_glyco_hcd203040_compound15.26311.26311.4 | controllerType=0 controllerNumber=1 scan=26311 | 53.3566 |
| 22428:4 | 15 | K.ADTHDEILEGLNFN[+2861.000]LTEIPEAQIHEGFQELLR.T | HexNAc(5)Hex(6)NeuAc(3) | N[+2861] | 1638.9716 | 4 | 6552.8644 | 6552.8239 | 0 | 6.2 | 94 | Specific | 970.72 | 838.29 | 838.29 | 18.6 | 41 | >sp\|P01009\|A1AT_HUMAN Alpha-1-antitrypsin OS=Homo sapiens OX=9606 GN=SERPINA1 PE=1 SV=3 | 642 | 20201119_08533_1ng_glyco_hcd203040_compound15.26309.26309.4 | controllerType=0 controllerNumber=1 scan=26309 | 53.3517 |
| 24687:4 | 17 | K.DIDGYGGIALPELIC[+57.021]TMFHTSGYDTQAIVENN[+2350.830]ESTEYGLFQISNK. | HexNAc(4)Hex(5)Fuc(1)NeuAc(2) | C[+57], N[+2351] | 1841.3083 | 4 | 7362.2114 | 7362.1592 | 0 | 7.1 | 33 | Specific | 341.82 | 311.78 | 311.78 | 6.75 | 27 | >sp\|P00709\|LALBA_HUMAN Alpha-lactalbumin OS=Homo sapiens OX=9606 GN=LALBA PE=1 SV=1 | 468 | 20201119_08533_1ng_glyco_hcd203040_compound15.28660.28660.4 | controllerType=0 controllerNumber=1 scan=28660 | 57.8499 |
| 22363:4 | 17 | K.DIDGYGGIALPELIC[+57.021]TMFHTSGYDTQAIVENN[+1686.586]ESTEYGLFQISNK. | HexNAc(2)Hex(7)Fuc(1) | C[+57], N[+1687] | 1675.2347 | 4 | 6697.917 | 6697.9153 | 0 | 0.3 | 33 | Specific | 380.01 | 263.46 | 263.46 | 6.98 | 27 | >sp\|P00709\|LALBA_HUMAN Alpha-lactalbumin OS=Homo sapiens OX=9606 GN=LALBA PE=1 SV=1 | 468 | 20201119_08533_1ng_glyco_hcd203040_compound15.26241.26241.4 | controllerType=0 controllerNumber=1 scan=26241 | 53.2118 |
| 22365:4 | 17 | K.DIDGYGGIALPELIC[+57.021]TMFHTSGYDTQAIVENN[+1686.586]ESTEYGLFQISNK. | HexNAc(2)Hex(7)Fuc(1) | C[+57], N[+1687] | 1675.2347 | 4 | 6697.917 | 6697.9153 | 0 | 0.3 | 33 | Specific | 424.11 | 424.11 | 424.11 | 9.42 | 27 | >sp\|P00709\|LALBA_HUMAN Alpha-lactalbumin OS=Homo sapiens OX=9606 GN=LALBA PE=1 SV=1 | 468 | 20201119_08533_1ng_glyco_hcd203040_compound15.26243.26243.4 | controllerType=0 controllerNumber=1 scan=26243 | 53.2161 |
| 09056:5 | 19 | K.[+42.011]QTDEIKDTRN[+2018.708]ESTQNC[+57.021]VVAEPEKMESSISSSSEEMSLS | HexNAc(3)Hex(6)Fuc(1)NeuAc(1) | .[42], C[+57], N[+2019] | 1291.1539 | 5 | 6451.7406 | 6451.6996 | 0 | 6.3 | 60 | Specific | 203.69 | 203.69 | 203.69 | 3.27 | 33 | >sp\|P47710\|CASA1_HUMAN Alpha-S1-casein OS=Homo sapiens OX=9606 GN=CSN1S1 PE=1 SV=1 | 989 | 20201119_08533_1ng_glyco_hcd203040_compound15.12309.12309.3 | controllerType=0 controllerNumber=1 scan=12309 | 27.1609 |
| 09166:5 | 19 | K.QTDEIKDTRN[+2059.735]ESTQNC[+57.021]VVAEPEKMESSISSSSEEMSLSK.C | HexNAc(4)Hex(5)Fuc(1)NeuAc(1) | C[+57], N[+2060] | 1290.9542 | 5 | 6450.742 | 6450.7156 | 0 | 4.1 | 60 | Specific | 211.86 | 160.95 | 160.95 | 3.92 | 33 | >sp\|P47710\|CASA1_HUMAN Alpha-S1-casein OS=Homo sapiens OX=9606 GN=CSN1S1 PE=1 SV=1 | 989 | 20201119_08533_1ng_glyco_hcd203040_compound15.12424.12424.5 | controllerType=0 controllerNumber=1 scan=12424 | 27.3501 |
| 09142:5 | 19 | K.[+42.011]QTDEIKDTRN[+2018.708]ESTQNC[+57.021]VVAEPEKMESSISSSSEEMSLS | HexNAc(3)Hex(6)Fuc(1)NeuAc(1) | .[42], C[+57], N[+2019] | 1291.1514 | 5 | 6451.7277 | 6451.6996 | 0 | 4.4 | 60 | Specific | 268.64 | 263.36 | 263.36 | 5.29 | 33 | >sp\|P47710\|CASA1_HUMAN Alpha-S1-casein OS=Homo sapiens OX=9606 GN=CSN1S1 PE=1 SV=1 | 989 | 20201119_08533_1ng_glyco_hcd203040_compound15.12399.12399.5 | controllerType=0 controllerNumber=1 scan=12399 | [illegible] |
| 09144:5 | 19 | K.[+42.011]QTDEIKDTRN[+2018.708]ESTQNC[+57.021]VVAEPEKMESSISSSSEEMSLS | HexNAc(3)Hex(6)Fuc(1)NeuAc(1) | .[42], C[+57], N[+2019] | 1291.1514 | 5 | 6451.7277 | 6451.6996 | 0 | 4.4 | 60 | Specific | 272.8 | 238.27 | 238.27 | 4.75 | 33 | >sp\|P47710\|CASA1_HUMAN Alpha-S1-casein OS=Homo sapiens OX=9606 GN=CSN1S1 PE=1 SV=1 | 989 | 20201119_08533_1ng_glyco_hcd203040_compound15.12401.12401.5 | controllerType=0 controllerNumber=1 scan=12401 | 27.3119 |
| 09164:5 | 19 | K.QTDEIKDTRN[+2059.735]ESTQNC[+57.021]VVAEPEKMESSISSSSEEMSLSK.C | HexNAc(4)Hex(5)Fuc(1)NeuAc(1) | C[+57], N[+2060] | 1290.9542 | 5 | 6450.742 | 6450.7156 | 0 | 4.1 | 60 | Specific | 299.01 | 218.7 | 218.7 | 5.97 | 33 | >sp\|P47710\|CASA1_HUMAN Alpha-S1-casein OS=Homo sapiens OX=9606 GN=CSN1S1 PE=1 SV=1 | 989 | 20201119_08533_1ng_glyco_hcd203040_compound15.12422.12422.5 | controllerType=0 controllerNumber=1 scan=12422 | 27.3465 |
| 09099:4 | 19 | K.QTDEIKDTRN[+2059.735]ESTQNC[+57.021]VVAEPEKMESSISSSSEEMSLSK.C | HexNAc(4)Hex(5)Fuc(1)NeuAc(1) | C[+57], N[+2060] | 1613.4364 | 4 | 6450.7238 | 6450.7156 | 0 | 1.3 | 60 | Specific | 391.37 | 290.42 | 290.42 | 6.07 | 33 | >sp\|P47710\|CASA1_HUMAN Alpha-S1-casein OS=Homo sapiens OX=9606 GN=CSN1S1 PE=1 SV=1 | 989 | 20201119_08533_1ng_glyco_hcd203040_compound15.12354.12354.4 | controllerType=0 controllerNumber=1 scan=12354 | 27.2362 |
| 09097:4 | 19 | K.QTDEIKDTRN[+2059.735]ESTQNC[+57.021]VVAEPEKMESSISSSSEEMSLSK.C | HexNAc(4)Hex(5)Fuc(1)NeuAc(1) | C[+57], N[+2060] | 1613.4364 | 4 | 6450.7238 | 6450.7156 | 0 | 1.3 | 60 | Specific | 428.54 | 304.19 | 304.19 | 6.55 | 33 | >sp\|P47710\|CASA1_HUMAN Alpha-S1-casein OS=Homo sapiens OX=9606 GN=CSN1S1 PE=1 SV=1 | 989 | 20201119_08533_1ng_glyco_hcd203040_compound15.12352.12352.4 | controllerType=0 controllerNumber=1 scan=12352 | 27.2321 |
| 13631:4 | 21 | K.LKELPGVC[+57.021]N[+2205.793]ETMMALWEEC[+57.021]KPC[+57.021]LK.Q | HexNAc(4)Hex(5)Fuc(2)NeuAc(1) | C[+57]*3, N[+2206] | 1286.3003 | 4 | 5142.1794 | 5142.1776 | 0 | 0.4 | 95 | Specific | 483.71 | 483.71 | 483.71 | 9.18 | 27 | >sp\|P10909\|CLUS_HUMAN Clusterin OS=Homo sapiens OX=9606 GN=CLU PE=1 SV=1 | 1467 | 20201119_08533_1ng_glyco_hcd203040_compound15.17083.17083.4 | controllerType=0 controllerNumber=1 scan=17083 | 35.3366 |
| 13633:4 | 21 | K.LKELPGVC[+57.021]N[+2205.793]ETMMALWEEC[+57.021]KPC[+57.021]LK.Q | HexNAc(4)Hex(5)Fuc(2)NeuAc(1) | C[+57]*3, N[+2206] | 1286.3003 | 4 | 5142.1794 | 5142.1776 | 0 | 0.4 | 95 | Specific | 554.91 | 554.91 | 554.91 | 9.65 | 27 | >sp\|P10909\|CLUS_HUMAN Clusterin OS=Homo sapiens OX=9606 GN=CLU PE=1 SV=1 | 1467 | 20201119_08533_1ng_glyco_hcd203040_compound15.17085.17085.4 | controllerType=0 controllerNumber=1 scan=17085 | 35.3405 |
| 13588:4 | 21 | K.LKELPGVC[+57.021]N[+2205.793]ETMMALWEEC[+57.021]KPC[+57.021]LK.Q | HexNAc(4)Hex(5)Fuc(2)NeuAc(1) | C[+57]*3, N[+2206] | 1286.3038 | 4 | 5142.1935 | 5142.1776 | 0 | 3.1 | 95 | Specific | 696.63 | 655.67 | 655.67 | 13.38 | 27 | >sp\|P10909\|CLUS_HUMAN Clusterin OS=Homo sapiens OX=9606 GN=CLU PE=1 SV=1 | 1467 | 20201119_08533_1ng_glyco_hcd203040_compound15.17038.17038.4 | controllerType=0 controllerNumber=1 scan=17038 | 35.2566 |
| 13590:4 | 21 | K.LKELPGVC[+57.021]N[+2205.793]ETMMALWEEC[+57.021]KPC[+57.021]LK.Q | HexNAc(4)Hex(5)Fuc(2)NeuAc(1) | C[+57]*3, N[+2206] | 1286.3038 | 4 | 5142.1935 | 5142.1776 | 0 | 3.1 | 95 | Specific | 712.92 | 695.96 | 695.96 | 14.38 | 27 | >sp\|P10909\|CLUS_HUMAN Clusterin OS=Homo sapiens OX=9606 GN=CLU PE=1 SV=1 | 1467 | 20201119_08533_1ng_glyco_hcd203040_compound15.17040.17040.4 | controllerType=0 controllerNumber=1 scan=17040 | 35.2608 |
| 23010:3 | 21 | K.[+42.011]MLN[+2336.851]TSSLLEQLNEQFNWVSR.L | HexNAc(6)Hex(6)Fuc(1) | .[42], N[+2337] | 1596.6941 | 3 | 4788.0679 | 4788.0484 | 0 | 4.1 | 352 | Specific | 313.14 | 12.78 | 12.78 | 4.21 | 27 | >sp\|P10909\|CLUS_HUMAN Clusterin OS=Homo sapiens OX=9606 GN=CLU PE=1 SV=1 | 1467 | 20201119_08533_1ng_glyco_hcd203040_compound15.26914.26914.3 | controllerType=0 controllerNumber=1 scan=26914 | 54.5865 |
| 22979:4 | 21 | K.MLN[+2350.830]TSSLLEQLNEQFNWVSR.L | HexNAc(4)Hex(5)Fuc(1)NeuAc(2) | N[+2351] | 1190.7636 | 4 | 4760.0325 | 4760.0171 | 0 | 3.2 | 352 | Specific | 611.81 | 477.01 | 477.01 | 12.79 | 27 | >sp\|P10909\|CLUS_HUMAN Clusterin OS=Homo sapiens OX=9606 GN=CLU PE=1 SV=1 | 1467 | 20201119_08533_1ng_glyco_hcd203040_compound15.26882.26882.4 | controllerType=0 controllerNumber=1 scan=26882 | 54.5121 |
| 22981:4 | 21 | K.MLN[+2350.830]TSSLLEQLNEQFNWVSR.L | HexNAc(4)Hex(5)Fuc(1)NeuAc(2) | N[+2351] | 1190.7636 | 4 | 4760.0325 | 4760.0171 | 0 | 3.2 | 352 | Specific | 841 | 640.67 | 640.67 | 16.79 | 27 | >sp\|P10909\|CLUS_HUMAN Clusterin OS=Homo sapiens OX=9606 GN=CLU PE=1 SV=1 | 1467 | 20201119_08533_1ng_glyco_hcd203040_compound15.26884.26884.4 | controllerType=0 controllerNumber=1 scan=26884 | 54.5167 |
| 22930:3 | 21 | K.MLN[+2350.830]TSSLLEQLNEQFNWVSR.L | HexNAc(4)Hex(5)Fuc(1)NeuAc(2) | N[+2351] | 1587.3476 | 3 | 4760.0282 | 4760.0171 | 0 | 2.3 | 352 | Specific | 843.65 | 577.34 | 577.34 | 16.12 | 27 | >sp\|P10909\|CLUS_HUMAN Clusterin OS=Homo sapiens OX=9606 GN=CLU PE=1 SV=1 | 1467 | 20201119_08533_1ng_glyco_hcd203040_compound15.26831.26831.3 | controllerType=0 controllerNumber=1 scan=26831 | 54.4099 |
| 22954:3 | 21 | K.MLN[+2350.830]TSSLLEQLNEQFNWVSR.L | HexNAc(4)Hex(5)Fuc(1)NeuAc(2) | N[+2351] | 1587.3498 | 3 | 4760.0348 | 4760.0171 | 0 | 3.7 | 352 | Specific | 888.9 | 617.75 | 617.75 | 17.14 | 27 | >sp\|P10909\|CLUS_HUMAN Clusterin OS=Homo sapiens OX=9606 GN=CLU PE=1 SV=1 | 1467 | 20201119_08533_1ng_glyco_hcd203040_compound15.26856.26856.3 | controllerType=0 controllerNumber=1 scan=26856 | 54.463 |
| 22952:3 | 21 | K.MLN[+2350.830]TSSLLEQLNEQFNWVSR.L | HexNAc(4)Hex(5)Fuc(1)NeuAc(2) | N[+2351] | 1587.3498 | 3 | 4760.0348 | 4760.0171 | 0 | 3.7 | 352 | Specific | 908.4 | 676.66 | 676.66 | 18.14 | 27 | >sp\|P10909\|CLUS_HUMAN Clusterin OS=Homo sapiens OX=9606 GN=CLU PE=1 SV=1 | 1467 | 20201119_08533_1ng_glyco_hcd203040_compound15.26854.26854.3 | controllerType=0 controllerNumber=1 scan=26854 | 54.4585 |
| 22932:3 | 21 | K.MLN[+2350.830]TSSLLEQLNEQFNWVSR.L | HexNAc(4)Hex(5)Fuc(1)NeuAc(2) | N[+2351] | 1587.3476 | 3 | 4760.0282 | 4760.0171 | 0 | 2.3 | 352 | Specific | 924.88 | 643.26 | 643.26 | 18.12 | 27 | >sp\|P10909\|CLUS_HUMAN Clusterin OS=Homo sapiens OX=9606 GN=CLU PE=1 SV=1 | 1467 | 20201119_08533_1ng_glyco_hcd203040_compound15.26833.26833.3 | controllerType=0 controllerNumber=1 scan=26833 | 54.4143 |
| 12655:3 | 21 | R.LAN[+2350.830]LTQGEDQYYLR.V | HexNAc(4)Hex(5)Fuc(1)NeuAc(2) | N[+2351] | 1345.5627 | 3 | 4034.6735 | 4034.664 | 0 | 2.3 | 372 | Specific | 342.61 | 323.63 | 323.63 | 7.2 | 27 | >sp\|P10909\|CLUS_HUMAN Clusterin OS=Homo sapiens OX=9606 GN=CLU PE=1 SV=1 | 1467 | 20201119_08533_1ng_glyco_hcd203040_compound15.16071.16071.3 | controllerType=0 controllerNumber=1 scan=16071 | 33.5462 |
| 12658:3 | 21 | R.LAN[+2350.830]LTQGEDQYYLR.V | HexNAc(4)Hex(5)Fuc(1)NeuAc(2) | N[+2351] | 1345.5627 | 3 | 4034.6735 | 4034.664 | 0 | 2.3 | 372 | Specific | 555.98 | 515.76 | 515.76 | 9.71 | 27 | >sp\|P10909\|CLUS_HUMAN Clusterin OS=Homo sapiens OX=9606 GN=CLU PE=1 SV=1 | 1467 | 20201119_08533_1ng_glyco_hcd203040_compound15.16074.16074.3 | controllerType=0 controllerNumber=1 scan=16074 | 33.55 |
| 09662:4 | 25 | K.SC[+57.021]QHN[+2350.830]GTMYQHGEIFSAHELFPSR.L | HexNAc(4)Hex(5)Fuc(1)NeuAc(2) | C[+57], N[+2351] | 1296.7881 | 4 | 5184.1305 | 5184.0873 | 0 | 8.3 | 110 | Specific | 272.88 | 272.88 | 272.88 | 5.16 | 22 | >sp\|Q6WN34\|CRDL2_HUMAN Chordin-like protein 2 OS=Homo sapiens OX=9606 GN=CHRDL2 PE=1 SV=1 | 1189 | 20201119_08533_1ng_glyco_hcd203040_compound15.12938.12938.2 | controllerType=0 controllerNumber=1 scan=12938 | 28.2324 |
| 09664:4 | 25 | K.SC[+57.021]QHN[+2350.830]GTMYQHGEIFSAHELFPSR.L | HexNAc(4)Hex(5)Fuc(1)NeuAc(2) | C[+57], N[+2351] | 1296.7881 | 4 | 5184.1305 | 5184.0873 | 0 | 8.3 | 110 | Specific | 291.79 | 291.79 | 291.79 | 5.25 | 22 | >sp\|Q6WN34\|CRDL2_HUMAN Chordin-like protein 2 OS=Homo sapiens OX=9606 GN=CHRDL2 PE=1 SV=1 | 1189 | 20201119_08533_1ng_glyco_hcd203040_compound15.12940.12940.2 | controllerType=0 controllerNumber=1 scan=12940 | 28.2366 |
| 13984:3 | 26 | K.VVLHPN[+2204.772]YSQVDIGLIK.L | HexNAc(4)Hex(5)NeuAc(2) | N[+2205] | 1333.9368 | 3 | 3999.7959 | 3999.7837 | 0 | 3.1 | 236 | Specific | 468.41 | 468.41 | 468.41 | 9.51 | 22 | >sp\|P00738\|HPT_HUMAN Haptoglobin OS=Homo sapiens OX=9606 GN=HP PE=1 SV=1 | 407 | 20201119_08533_1ng_glyco_hcd203040_compound15.17449.17449.3 | controllerType=0 controllerNumber=1 scan=17449 | 35.9816 |
| 13981:3 | 26 | K.VVLHPN[+2204.772]YSQVDIGLIK.L | HexNAc(4)Hex(5)NeuAc(2) | N[+2205] | 1333.9368 | 3 | 3999.7959 | 3999.7837 | 0 | 3.1 | 236 | Specific | 513.86 | 481.13 | 481.13 | 9.6 | 22 | >sp\|P00738\|HPT_HUMAN Haptoglobin OS=Homo sapiens OX=9606 GN=HP PE=1 SV=1 | 407 | 20201119_08533_1ng_glyco_hcd203040_compound15.17446.17446.3 | controllerType=0 controllerNumber=1 scan=17446 | 35.9774 |
| 18334:3 | 28 | R.C[+57.021]MWSSALNSLN[+1864.634]LSFAGLEQVPK.G | HexNAc(2)Hex(9) | C[+57], N[+1865] | 1439.6159 | 3 | 4316.8332 | 4316.8341 | 0 | -0.2 | 272 | Specific | 647.62 | 577.72 | 577.72 | 11.39 | 20 | >sp\|P08571\|CD14_HUMAN Monocyte differentiation antigen CD14 OS=Homo sapiens OX=9606 GN=CD14 PE=1 SV=2 | 1232 | 20201119_08533_1ng_glyco_hcd203040_compound15.22029.22029.3 | controllerType=0 controllerNumber=1 scan=22029 | 44.8202 |
| 09366:3 | 39 | R.IIVPLNNREN[+1647.613]ISDPTSPLR.T | HexNAc(5)Hex(3)Fuc(1) | N[+1648] | 1265.9407 | 3 | 3795.8075 | 3795.7904 | 0 | 4.5 | 62 | Specific | 229.04 | 165.96 | 165.96 | 5.2 | 22 | >sp\|P01591\|IGJ_HUMAN Immunoglobulin J chain OS=Homo sapiens OX=9606 GN=JCHAIN PE=1 SV=4 | 1392 | 20201119_08533_1ng_glyco_hcd203040_compound15.12631.12631.3 | controllerType=0 controllerNumber=1 scan=12631 | 27.698 |
| 09232:3 | 39 | R.IIVPLNNREN[+1606.587]ISDPTSPLR.T | HexNAc(4)Hex(4)Fuc(1) | N[+1607] | 1252.2651 | 3 | 3754.7807 | 3754.7638 | 0 | 4.5 | 62 | Specific | 242.92 | 172.95 | 172.95 | 5.2 | 22 | >sp\|P01591\|IGJ_HUMAN Immunoglobulin J chain OS=Homo sapiens OX=9606 GN=JCHAIN PE=1 SV=4 | 1392 | 20201119_08533_1ng_glyco_hcd203040_compound15.12492.12492.3 | controllerType=0 controllerNumber=1 scan=12492 | 27.4603 |
| 09272:3 | 39 | R.IIVPLNNREN[+1460.529]ISDPTSPLR.T | HexNAc(4)Hex(4) | N[+1461] | 1203.5759 | 3 | 3608.713 | 3608.7059 | 0 | 2 | 62 | Specific | 253.68 | 247.4 | 247.4 | 6.2 | 22 | >sp\|P01591\|IGJ_HUMAN Immunoglobulin J chain OS=Homo sapiens OX=9606 GN=JCHAIN PE=1 SV=4 | 1392 | 20201119_08533_1ng_glyco_hcd203040_compound15.12534.12534.3 | controllerType=0 controllerNumber=1 scan=12534 | 27.5346 |
| 09287:3 | 39 | R.IIVPLNNREN[+1768.640]ISDPTSPLR.T | HexNAc(4)Hex(5)Fuc(1) | N[+1769] | 1306.2779 | 3 | 3916.8191 | 3916.8167 | 0 | 0.6 | 62 | Specific | 262.15 | 258.26 | 258.26 | 5.65 | 22 | >sp\|P01591\|IGJ_HUMAN Immunoglobulin J chain OS=Homo sapiens OX=9606 GN=JCHAIN PE=1 SV=4 | 1392 | 20201119_08533_1ng_glyco_hcd203040_compound15.12549.12549.3 | controllerType=0 controllerNumber=1 scan=12549 | 27.5584 |
| 09289:3 | 39 | R.IIVPLNNREN[+1768.640]ISDPTSPLR.T | HexNAc(4)Hex(5)Fuc(1) | N[+1769] | 1306.2779 | 3 | 3916.8191 | 3916.8167 | 0 | 0.6 | 62 | Specific | 290.23 | 237.96 | 237.96 | 4.71 | 22 | >sp\|P01591\|IGJ_HUMAN Immunoglobulin J chain OS=Homo sapiens OX=9606 GN=JCHAIN PE=1 SV=4 | 1392 | 20201119_08533_1ng_glyco_hcd203040_compound15.12551.12551.3 | controllerType=0 controllerNumber=1 scan=12551 | 27.5619 |
| 09274:3 | 39 | R.IIVPLNNREN[+1460.529]ISDPTSPLR.T | HexNAc(4)Hex(4) | N[+1461] | 1203.5759 | 3 | 3608.713 | 3608.7059 | 0 | 2 | 62 | Specific | 301.58 | 250.9 | 250.9 | 6.3 | 22 | >sp\|P01591\|IGJ_HUMAN Immunoglobulin J chain OS=Homo sapiens OX=9606 GN=JCHAIN PE=1 SV=4 | 1392 | 20201119_08533_1ng_glyco_hcd203040_compound15.12536.12536.3 | controllerType=0 controllerNumber=1 scan=12536 | 27.5388 |
| 09234:3 | 39 | R.IIVPLNNREN[+1606.587]ISDPTSPLR.T | HexNAc(4)Hex(4)Fuc(1) | N[+1607] | 1252.2651 | 3 | 3754.7807 | 3754.7638 | 0 | 4.5 | 62 | Specific | 314.09 | 229.6 | 229.6 | 6.12 | 22 | >sp\|P01591\|IGJ_HUMAN Immunoglobulin J chain OS=Homo sapiens OX=9606 GN=JCHAIN PE=1 SV=4 | 1392 | 20201119_08533_1ng_glyco_hcd203040_compound15.12494.12494.3 | controllerType=0 controllerNumber=1 scan=12494 | 27.4638 |
| 09364:3 | 39 | R.IIVPLNNREN[+1647.613]ISDPTSPLR.T | HexNAc(5)Hex(3)Fuc(1) | N[+1648] | 1265.9407 | 3 | 3795.8075 | 3795.7904 | 0 | 4.5 | 62 | Specific | 319.26 | 257.46 | 257.46 | 6.35 | 22 | >sp\|P01591\|IGJ_HUMAN Immunoglobulin J chain OS=Homo sapiens OX=9606 GN=JCHAIN PE=1 SV=4 | 1392 | 20201119_08533_1ng_glyco_hcd203040_compound15.12629.12629.3 | controllerType=0 controllerNumber=1 scan=12629 | 27.6944 |
| 09212:3 | 39 | R.IIVPLNNREN[+1460.529]ISDPTSPLR.T | HexNAc(4)Hex(4) | N[+1461] | 1203.5781 | 3 | 3608.7196 | 3608.7059 | 0 | 3.8 | 62 | Specific | 321.8 | 302.39 | 302.39 | 7.05 | 22 | >sp\|P01591\|IGJ_HUMAN Immunoglobulin J chain OS=Homo sapiens OX=9606 GN=JCHAIN PE=1 SV=4 | 1392 | 20201119_08533_1ng_glyco_hcd203040_compound15.12472.12472.3 | controllerType=0 controllerNumber=1 scan=12472 | 27.4285 |
| 09214:3 | 39 | R.IIVPLNNREN[+1460.529]ISDPTSPLR.T | HexNAc(4)Hex(4) | N[+1461] | 1203.5781 | 3 | 3608.7196 | 3608.7059 | 0 | 3.8 | 62 | Specific | 322.21 | 246.85 | 246.85 | 6.6 | 22 | >sp\|P01591\|IGJ_HUMAN Immunoglobulin J chain OS=Homo sapiens OX=9606 GN=JCHAIN PE=1 SV=4 | 1392 | 20201119_08533_1ng_glyco_hcd203040_compound15.12474.12474.3 | controllerType=0 controllerNumber=1 scan=12474 | 27.4321 |
| 12861:3 | 39 | R.IIVPLNNREN[+2350.830]ISDPTSPLR.T | HexNAc(4)Hex(5)Fuc(1)NeuAc(2) | N[+2351] | 1500.3475 | 3 | 4499.0279 | 4499.0075 | 0 | 4.5 | 62 | Specific | 337.28 | 337.28 | 337.28 | 7.08 | 22 | >sp\|P01591\|IGJ_HUMAN Immunoglobulin J chain OS=Homo sapiens OX=9606 GN=JCHAIN PE=1 SV=4 | 1392 | 20201119_08533_1ng_glyco_hcd203040_compound15.16284.16284.3 | controllerType=0 controllerNumber=1 scan=16284 | 33.909 |
| 11016:3 | 39 | R.IIVPLNNREN[+1856.656]ISDPTSPLR.T | HexNAc(3)Hex(5)Fuc(1)NeuAc(1) | N[+1857] | 1335.6208 | 3 | 4004.8478 | 4004.8327 | 0 | 3.8 | 62 | Specific | 378.51 | 288.64 | 288.64 | 7.01 | 22 | >sp\|P01591\|IGJ_HUMAN Immunoglobulin J chain OS=Homo sapiens OX=9606 GN=JCHAIN PE=1 SV=4 | 1392 | 20201119_08533_1ng_glyco_hcd203040_compound15.14338.14338.3 | controllerType=0 controllerNumber=1 scan=14338 | 30.584 |
| 11019:3 | 39 | R.IIVPLNNREN[+1856.656]ISDPTSPLR.T | HexNAc(3)Hex(5)Fuc(1)NeuAc(1) | N[+1857] | 1335.6208 | 3 | 4004.8478 | 4004.8327 | 0 | 3.8 | 62 | Specific | 384.58 | 384.58 | 384.58 | 8.94 | 22 | >sp\|P01591\|IGJ_HUMAN Immunoglobulin J chain OS=Homo sapiens OX=9606 GN=JCHAIN PE=1 SV=4 | 1392 | 20201119_08533_1ng_glyco_hcd203040_compound15.14341.14341.3 | controllerType=0 controllerNumber=1 scan=14341 | 30.5906 |
| 09205:3 | 39 | R.IIVPLNNREN[+1809.666]ISDPTSPLR.T | HexNAc(5)Hex(4)Fuc(1) | N[+1810] | 1319.9578 | 3 | 3957.8589 | 3957.8432 | 0 | 4 | 62 | Specific | 412.55 | 336.59 | 336.59 | 7.44 | 22 | >sp\|P01591\|IGJ_HUMAN Immunoglobulin J chain OS=Homo sapiens OX=9606 GN=JCHAIN PE=1 SV=4 | 1392 | 20201119_08533_1ng_glyco_hcd203040_compound15.12464.12464.3 | controllerType=0 controllerNumber=1 scan=12464 | 27.4133 |
| 09365:3 | 39 | R.IIVPLNNREN[+1501.555]ISDPTSPLR.T | HexNAc(5)Hex(3) | N[+1502] | 1217.2545 | 3 | 3649.7488 | 3649.7325 | 0 | 4.5 | 62 | Specific | 415.88 | 370.49 | 370.49 | 7.36 | 22 | >sp\|P01591\|IGJ_HUMAN Immunoglobulin J chain OS=Homo sapiens OX=9606 GN=JCHAIN PE=1 SV=4 | 1392 | 20201119_08533_1ng_glyco_hcd203040_compound15.12630.12630.3 | controllerType=0 controllerNumber=1 scan=12630 | 27.6959 |
| 09367:3 | 39 | R.IIVPLNNREN[+1501.555]ISDPTSPLR.T | HexNAc(5)Hex(3) | N[+1502] | 1217.2545 | 3 | 3649.7488 | 3649.7325 | 0 | 4.5 | 62 | Specific | 426.4 | 372.69 | 372.69 | 7.36 | 22 | >sp\|P01591\|IGJ_HUMAN Immunoglobulin J chain OS=Homo sapiens OX=9606 GN=JCHAIN PE=1 SV=4 | 1392 | 20201119_08533_1ng_glyco_hcd203040_compound15.12632.12632.3 | controllerType=0 controllerNumber=1 scan=12632 | 27.7011 |
| 09241:3 | 39 | R.IIVPLNNREN[+1809.666]ISDPTSPLR.T | HexNAc(5)Hex(4)Fuc(1) | N[+1810] | 1319.9604 | 3 | 3957.8666 | 3957.8432 | 0 | 5.9 | 62 | Specific | 428.4 | 308.38 | 308.38 | 6.78 | 22 | >sp\|P01591\|IGJ_HUMAN Immunoglobulin J chain OS=Homo sapiens OX=9606 GN=JCHAIN PE=1 SV=4 | 1392 | 20201119_08533_1ng_glyco_hcd203040_compound15.12502.12502.3 | controllerType=0 controllerNumber=1 scan=12502 | 27.4807 |
| 12863:3 | 39 | R.IIVPLNNREN[+2350.830]ISDPTSPLR.T | HexNAc(4)Hex(5)Fuc(1)NeuAc(2) | N[+2351] | 1500.3475 | 3 | 4499.0279 | 4499.0075 | 0 | 4.5 | 62 | Specific | 430.74 | 335.93 | 335.93 | 7.44 | 22 | >sp\|P01591\|IGJ_HUMAN Immunoglobulin J chain OS=Homo sapiens OX=9606 GN=JCHAIN PE=1 SV=4 | 1392 | 20201119_08533_1ng_glyco_hcd203040_compound15.16286.16286.3 | controllerType=0 controllerNumber=1 scan=16286 | 33.9128 |
| 09243:3 | 39 | R.IIVPLNNREN[+1809.666]ISDPTSPLR.T | HexNAc(5)Hex(4)Fuc(1) | N[+1810] | 1319.9604 | 3 | 3957.8666 | 3957.8432 | 0 | 5.9 | 62 | Specific | 456.35 | 357.43 | 357.43 | 6.83 | 22 | >sp\|P01591\|IGJ_HUMAN Immunoglobulin J chain OS=Homo sapiens OX=9606 GN=JCHAIN PE=1 SV=4 | 1392 | 20201119_08533_1ng_glyco_hcd203040_compound15.12504.12504.3 | controllerType=0 controllerNumber=1 scan=12504 | 27.485 |
| 10849:3 | 39 | R.IIVPLNNREN[+2059.735]ISDPTSPLR.T | HexNAc(4)Hex(5)Fuc(1)NeuAc(1) | N[+2060] | 1403.3174 | 3 | 4207.9378 | 4207.9121 | 0 | 6.1 | 62 | Specific | 480.26 | 448.35 | 448.35 | 8.61 | 22 | >sp\|P01591\|IGJ_HUMAN Immunoglobulin J chain OS=Homo sapiens OX=9606 GN=JCHAIN PE=1 SV=4 | 1392 | 20201119_08533_1ng_glyco_hcd203040_compound15.14165.14165.3 | controllerType=0 controllerNumber=1 scan=14165 | 30.2737 |
| 09207:3 | 39 | R.IIVPLNNREN[+1809.666]ISDPTSPLR.T | HexNAc(5)Hex(4)Fuc(1) | N[+1810] | 1319.9578 | 3 | 3957.8589 | 3957.8432 | 0 | 4 | 62 | Specific | 512.94 | 433.54 | 433.54 | 9.46 | 22 | >sp\|P01591\|IGJ_HUMAN Immunoglobulin J chain OS=Homo sapiens OX=9606 GN=JCHAIN PE=1 SV=4 | 1392 | 20201119_08533_1ng_glyco_hcd203040_compound15.12466.12466.3 | controllerType=0 controllerNumber=1 scan=12466 | 27.4179 |
| 10851:3 | 39 | R.IIVPLNNREN[+2059.735]ISDPTSPLR.T | HexNAc(4)Hex(5)Fuc(1)NeuAc(1) | N[+2060] | 1403.3174 | 3 | 4207.9378 | 4207.9121 | 0 | 6.1 | 62 | Specific | 515.6 | 469.17 | 469.17 | 8.7 | 22 | >sp\|P01591\|IGJ_HUMAN Immunoglobulin J chain OS=Homo sapiens OX=9606 GN=JCHAIN PE=1 SV=4 | 1392 | 20201119_08533_1ng_glyco_hcd203040_compound15.14167.14167.3 | controllerType=0 controllerNumber=1 scan=14167 | 30.2782 |
| 10890:3 | 39 | R.IIVPLNNREN[+1913.677]ISDPTSPLR.T | HexNAc(4)Hex(5)NeuAc(1) | N[+1914] | 1354.6326 | 3 | 4061.8833 | 4061.8542 | 0 | 7.2 | 62 | Specific | 571.53 | 571.53 | 571.53 | 8.64 | 22 | >sp\|P01591\|IGJ_HUMAN Immunoglobulin J chain OS=Homo sapiens OX=9606 GN=JCHAIN PE=1 SV=4 | 1392 | 20201119_08533_1ng_glyco_hcd203040_compound15.14207.14207.3 | controllerType=0 controllerNumber=1 scan=14207 | 30.348 |
| 10888:3 | 39 | R.IIVPLNNREN[+1913.677]ISDPTSPLR.T | HexNAc(4)Hex(5)NeuAc(1) | N[+1914] | 1354.6326 | 3 | 4061.8833 | 4061.8542 | 0 | 7.2 | 62 | Specific | 615.39 | 592.35 | 592.35 | 10.8 | 22 | >sp\|P01591\|IGJ_HUMAN Immunoglobulin J chain OS=Homo sapiens OX=9606 GN=JCHAIN PE=1 SV=4 | 1392 | 20201119_08533_1ng_glyco_hcd203040_compound15.14205.14205.3 | controllerType=0 controllerNumber=1 scan=14205 | 30.3431 |
| 15524:3 | 61 | R.KLPPGLLAN[+2059.735]FTLLR.T | HexNAc(4)Hex(5)Fuc(1)NeuAc(1) | N[+2060] | 1204.9103 | 3 | 3612.7165 | 3612.6923 | 0 | 6.7 | 178 | Specific | 328.13 | 284.38 | 284.38 | 4.44 | 13 | >sp\|P02750\|A2GL_HUMAN Leucine-rich alpha-2-glycoprotein OS=Homo sapiens OX=9606 GN=LRG1 PE=1 SV=2 | 518 | 20201119_08533_1ng_glyco_hcd203040_compound15.19050.19050.3 | controllerType=0 controllerNumber=1 scan=19050 | 38.7549 |
| 24749:4 | 61 | R.QLDMLDLSN[+2204.772]NSLASVPEGLWASLGQPNWDMR.D | HexNAc(4)Hex(5)NeuAc(2) | N[+2205] | 1416.3713 | 4 | 5662.4635 | 5662.4239 | 0 | 7 | 261 | Specific | 338.28 | 338.28 | 338.28 | 6.9 | 13 | >sp\|P02750\|A2GL_HUMAN Leucine-rich alpha-2-glycoprotein OS=Homo sapiens OX=9606 GN=LRG1 PE=1 SV=2 | 518 | 20201119_08533_1ng_glyco_hcd203040_compound15.28724.28724.4 | controllerType=0 controllerNumber=1 scan=28724 | 57.9614 |
| 24751:4 | 61 | R.QLDMLDLSN[+2204.772]NSLASVPEGLWASLGQPNWDMR.D | HexNAc(4)Hex(5)NeuAc(2) | N[+2205] | 1416.3713 | 4 | 5662.4635 | 5662.4239 | 0 | 7 | 261 | Specific | 428.05 | 428.05 | 428.05 | 8.8 | 13 | >sp\|P02750\|A2GL_HUMAN Leucine-rich alpha-2-glycoprotein OS=Homo sapiens OX=9606 GN=LRG1 PE=1 SV=2 | 518 | 20201119_08533_1ng_glyco_hcd203040_compound15.28726.28726.4 | controllerType=0 controllerNumber=1 scan=28726 | 57.9656 |
| 16195:3 | 74 | R.NYIVPILWLN[+1864.634]ETGTIGDEKANM[+15.995]FR.S | HexNAc(2)Hex(9) | M[+16], N[+1865] | 1559.0193 | 3 | 4675.0433 | 4675.0523 | 0 | -1.9 | 408 | Specific | 454.18 | 440.1 | 440.1 | 8.03 | 11 | >sp\|P16671\|CD36_HUMAN Platelet glycoprotein 4 OS=Homo sapiens OX=9606 GN=CD36 PE=1 SV=2 | 812 | 20201119_08533_1ng_glyco_hcd203040_compound15.19796.19796.3 | controllerType=0 controllerNumber=1 scan=19796 | 40.1339 |
| 16198:3 | 74 | R.NYIVPILWLN[+1864.634]ETGTIGDEKANM[+15.995]FR.S | HexNAc(2)Hex(9) | M[+16], N[+1865] | 1559.0193 | 3 | 4675.0433 | 4675.0523 | 0 | -1.9 | 408 | Specific | 610.77 | 589.36 | 589.36 | 10.81 | 11 | >sp\|P16671\|CD36_HUMAN Platelet glycoprotein 4 OS=Homo sapiens OX=9606 GN=CD36 PE=1 SV=2 | 812 | 20201119_08533_1ng_glyco_hcd203040_compound15.19799.19799.3 | controllerType=0 controllerNumber=1 scan=19799 | 40.1398 |
| 24389:3 | 103 | R.AVN[+2204.772]ITSENLIDDVVSLIR.G | HexNAc(4)Hex(5)NeuAc(2) | N[+2205] | 1392.6271 | 3 | 4175.8667 | 4175.8481 | 0 | 4.5 | 413 | Specific | 249.49 | 240.21 | 240.21 | 5.77 | 10 | >sp\|P02748\|CO9_HUMAN Complement component C9 OS=Homo sapiens OX=9606 GN=C9 PE=1 SV=2 | 1547 | 20201119_08533_1ng_glyco_hcd203040_compound15.28350.28350.3 | controllerType=0 controllerNumber=1 scan=28350 | 57.2905 |
| 18467:3 | 1 | R.RTAGWNVPIGTLRPFLN[+2059.735]WTGPPEPIEAAVAR.F | HexNAc(4)Hex(5)Fuc(1)NeuAc(1) | N[+2060] | 1816.1921 | 3 | 5446.5618 | 5446.547 | 0 | 2.7 | 140 | Specific | 233.66 | 233.66 | 233.66 | 4.95 | 114 | >sp\|P02788\|TRFL_HUMAN Lactotransferrin OS=Homo sapiens OX=9606 GN=LTF PE=1 SV=6 | 1270 | 20201115_10113_1ng_glyco_hcd203040_compound15.21707.21707.3 | controllerType=0 controllerNumber=1 scan=21707 | 43.5377 |
| 18592:4 | 1 | R.[+42.011]RTAGWNVPIGTLRPFLN[+2018.708]WTGPPEPIEAAVAR.F | HexNAc(3)Hex(6)Fuc(1)NeuAc(1) | .[42], N[+2019] | 1362.6445 | 4 | 5447.5562 | 5447.531 | 0 | 4.6 | 140 | Specific | 238.15 | 20.56 | 20.56 | 3.96 | 114 | >sp\|P02788\|TRFL_HUMAN Lactotransferrin OS=Homo sapiens OX=9606 GN=LTF PE=1 SV=6 | 1270 | 20201115_10113_1ng_glyco_hcd203040_compound15.21836.21836.4 | controllerType=0 controllerNumber=1 scan=21836 | 43.7877 |
| 18471:5 | 1 | R.RTAGWNVPIGTLRPFLN[+2059.735]WTGPPEPIEAAVAR.F | HexNAc(4)Hex(5)Fuc(1)NeuAc(1) | N[+2060] | 1090.1176 | 5 | 5446.5587 | 5446.547 | 0 | 2.1 | 140 | Specific | 261.23 | 261.23 | 261.23 | 6.86 | 114 | >sp\|P02788\|TRFL_HUMAN Lactotransferrin OS=Homo sapiens OX=9606 GN=LTF PE=1 SV=6 | 1270 | 20201115_10113_1ng_glyco_hcd203040_compound15.21711.21711.5 | controllerType=0 controllerNumber=1 scan=21711 | 43.5422 |
| 18464:4 | 1 | R.RTAGWNVPIGTLRPFLN[+2059.735]WTGPPEPIEAAVAR.F | HexNAc(4)Hex(5)Fuc(1)NeuAc(1) | N[+2060] | 1362.3954 | 4 | 5446.5597 | 5446.547 | 0 | 2.3 | 140 | Specific | 321.89 | 297.89 | 297.89 | 7.46 | 114 | >sp\|P02788\|TRFL_HUMAN Lactotransferrin OS=Homo sapiens OX=9606 GN=LTF PE=1 SV=6 | 1270 | 20201115_10113_1ng_glyco_hcd203040_compound15.21704.21704.4 | controllerType=0 controllerNumber=1 scan=21704 | 43.5308 |
| 17552:4 | 1 | R.RTAGWNVPIGTLRPFLN[+1768.640]WTGPPEPIEAAVAR.F | HexNAc(4)Hex(5)Fuc(1) | N[+1769] | 1289.6213 | 4 | 5155.4635 | 5155.4516 | 0 | 2.3 | 140 | Specific | 391 | 337.67 | 337.67 | 8.13 | 114 | >sp\|P02788\|TRFL_HUMAN Lactotransferrin OS=Homo sapiens OX=9606 GN=LTF PE=1 SV=6 | 1270 | 20201115_10113_1ng_glyco_hcd203040_compound15.20751.20751.4 | controllerType=0 controllerNumber=1 scan=20751 | 41.4108 |
| 18431:5 | 1 | R.RTAGWNVPIGTLRPFLN[+2205.793]WTGPPEPIEAAVAR.F | HexNAc(4)Hex(5)Fuc(2)NeuAc(1) | N[+2206] | 1119.3348 | 5 | 5592.6451 | 5592.6049 | 0 | 7.2 | 140 | Specific | 440.44 | 440.44 | 440.44 | 8.84 | 114 | >sp\|P02788\|TRFL_HUMAN Lactotransferrin OS=Homo sapiens OX=9606 GN=LTF PE=1 SV=6 | 1270 | 20201115_10113_1ng_glyco_hcd203040_compound15.21669.21669.3 | controllerType=0 controllerNumber=1 scan=21669 | 43.4435 |
| 18469:4 | 1 | R.RTAGWNVPIGTLRPFLN[+2205.793]WTGPPEPIEAAVAR.F | HexNAc(4)Hex(5)Fuc(2)NeuAc(1) | N[+2206] | 1398.9075 | 4 | 5592.6081 | 5592.6049 | 0 | 0.6 | 140 | Specific | 449.65 | 449.65 | 449.65 | 9.23 | 114 | >sp\|P02788\|TRFL_HUMAN Lactotransferrin OS=Homo sapiens OX=9606 GN=LTF PE=1 SV=6 | 1270 | 20201115_10113_1ng_glyco_hcd203040_compound15.21709.21709.4 | controllerType=0 controllerNumber=1 scan=21709 | 43.5399 |
| 18360:4 | 1 | R.RTAGWNVPIGTLRPFLN[+2570.925]WTGPPEPIEAAVAR.F | HexNAc(5)Hex(6)Fuc(2)NeuAc(1) | N[+2571] | 1490.1948 | 4 | 5957.7575 | 5957.7371 | 0 | 3.4 | 140 | Specific | 516.18 | 484.66 | 484.66 | 10.38 | 114 | >sp\|P02788\|TRFL_HUMAN Lactotransferrin OS=Homo sapiens OX=9606 GN=LTF PE=1 SV=6 | 1270 | 20201115_10113_1ng_glyco_hcd203040_compound15.21594.21594.4 | controllerType=0 controllerNumber=1 scan=21594 | 43.2537 |
| 18473:5 | 1 | R.RTAGWNVPIGTLRPFLN[+2059.735]WTGPPEPIEAAVAR.F | HexNAc(4)Hex(5)Fuc(1)NeuAc(1) | N[+2060] | 1090.1176 | 5 | 5446.5587 | 5446.547 | 0 | 2.1 | 140 | Specific | 549.84 | 525.9 | 525.9 | 10.49 | 114 | >sp\|P02788\|TRFL_HUMAN Lactotransferrin OS=Homo sapiens OX=9606 GN=LTF PE=1 SV=6 | 1270 | 20201115_10113_1ng_glyco_hcd203040_compound15.21713.21713.5 | controllerType=0 controllerNumber=1 scan=21713 | 43.546 |
| 18363:4 | 1 | R.RTAGWNVPIGTLRPFLN[+2570.925]WTGPPEPIEAAVAR.F | HexNAc(5)Hex(6)Fuc(2)NeuAc(1) | N[+2571] | 1490.1948 | 4 | 5957.7575 | 5957.7371 | 0 | 3.4 | 140 | Specific | 559.4 | 474.9 | 474.9 | 10.69 | 114 | >sp\|P02788\|TRFL_HUMAN Lactotransferrin OS=Homo sapiens OX=9606 GN=LTF PE=1 SV=6 | 1270 | 20201115_10113_1ng_glyco_hcd203040_compound15.21597.21597.4 | controllerType=0 controllerNumber=1 scan=21597 | 43.26 |
| 18466:4 | 1 | R.RTAGWNVPIGTLRPFLN[+2059.735]WTGPPEPIEAAVAR.F | HexNAc(4)Hex(5)Fuc(1)NeuAc(1) | N[+2060] | 1362.3954 | 4 | 5446.5597 | 5446.547 | 0 | 2.3 | 140 | Specific | 578.31 | 464.03 | 464.03 | 10.71 | 114 | >sp\|P02788\|TRFL_HUMAN Lactotransferrin OS=Homo sapiens OX=9606 GN=LTF PE=1 SV=6 | 1270 | 20201115_10113_1ng_glyco_hcd203040_compound15.21706.21706.4 | controllerType=0 controllerNumber=1 scan=21706 | 43.5349 |
| 18434:5 | 1 | R.RTAGWNVPIGTLRPFLN[+2205.793]WTGPPEPIEAAVAR.F | HexNAc(4)Hex(5)Fuc(2)NeuAc(1) | N[+2206] | 1119.3348 | 5 | 5592.6451 | 5592.6049 | 0 | 7.2 | 140 | Specific | 630.55 | 528.44 | 528.44 | 11.7 | 114 | >sp\|P02788\|TRFL_HUMAN Lactotransferrin OS=Homo sapiens OX=9606 GN=LTF PE=1 SV=6 | 1270 | 20201115_10113_1ng_glyco_hcd203040_compound15.21672.21672.3 | controllerType=0 controllerNumber=1 scan=21672 | 43.4497 |
| 18472:4 | 1 | R.RTAGWNVPIGTLRPFLN[+2205.793]WTGPPEPIEAAVAR.F | HexNAc(4)Hex(5)Fuc(2)NeuAc(1) | N[+2206] | 1398.9075 | 4 | 5592.6081 | 5592.6049 | 0 | 0.6 | 140 | Specific | 637.39 | 442.63 | 442.63 | 12.09 | 114 | >sp\|P02788\|TRFL_HUMAN Lactotransferrin OS=Homo sapiens OX=9606 GN=LTF PE=1 SV=6 | 1270 | 20201115_10113_1ng_glyco_hcd203040_compound15.21712.21712.4 | controllerType=0 controllerNumber=1 scan=21712 | 43.5443 |
| 17554:4 | 1 | R.RTAGWNVPIGTLRPFLN[+1768.640]WTGPPEPIEAAVAR.F | HexNAc(4)Hex(5)Fuc(1) | N[+1769] | 1289.6213 | 4 | 5155.4635 | 5155.4516 | 0 | 2.3 | 140 | Specific | 658.48 | 553.87 | 553.87 | 13.77 | 114 | >sp\|P02788\|TRFL_HUMAN Lactotransferrin OS=Homo sapiens OX=9606 GN=LTF PE=1 SV=6 | 1270 | 20201115_10113_1ng_glyco_hcd203040_compound15.20753.20753.4 | controllerType=0 controllerNumber=1 scan=20753 | 41.4156 |
| 18718:4 | 1 | R.[+42.011]TAGWNVPIGTLRPFLN[+1872.651]WTGPPEPIEAAVAR.F | HexNAc(3)Hex(6)NeuAc(1) | .[42], N[+1873] | 1287.11 | 4 | 5145.4181 | 5145.372 | 0 | 9 | 141 | Specific | 230.11 | 173.24 | 173.24 | 3.72 | 114 | >sp\|P02788\|TRFL_HUMAN Lactotransferrin OS=Homo sapiens OX=9606 GN=LTF PE=1 SV=6 | 1270 | 20201115_10113_1ng_glyco_hcd203040_compound15.21967.21967.4 | controllerType=0 controllerNumber=1 scan=21967 | 44.0278 |
| 18717:3 | 1 | R.[+42.011]TAGWNVPIGTLRPFLN[+1872.651]WTGPPEPIEAAVAR.F | HexNAc(3)Hex(6)NeuAc(1) | .[42], N[+1873] | 1715.8111 | 3 | 5145.4188 | 5145.372 | 0 | 9.1 | 141 | Specific | 240.58 | 160.95 | 160.95 | 5.17 | 114 | >sp\|P02788\|TRFL_HUMAN Lactotransferrin OS=Homo sapiens OX=9606 GN=LTF PE=1 SV=6 | 1270 | 20201115_10113_1ng_glyco_hcd203040_compound15.21966.21966.3 | controllerType=0 controllerNumber=1 scan=21966 | 44.0254 |
| 18716:4 | 1 | R.[+42.011]TAGWNVPIGTLRPFLN[+1872.651]WTGPPEPIEAAVAR.F | HexNAc(3)Hex(6)NeuAc(1) | .[42], N[+1873] | 1287.11 | 4 | 5145.4181 | 5145.372 | 0 | 9 | 141 | Specific | 247.12 | 207.97 | 207.97 | 5.63 | 114 | >sp\|P02788\|TRFL_HUMAN Lactotransferrin OS=Homo sapiens OX=9606 GN=LTF PE=1 SV=6 | 1270 | 20201115_10113_1ng_glyco_hcd203040_compound15.21965.21965.4 | controllerType=0 controllerNumber=1 scan=21965 | 44.0224 |

Table S4. Cont.

| Query #:z | Protein Rank | Glycopeptide | Glycan | Modification Type(s) | Observed m/z | z | Observed (M+H) | Calc. mass (M+H) | Off-by-x error | Mass error (ppm) | StartPosition | Cleavage | Score | Delta | Delta Mod | \|Log Prob\| | # of unique peptides | Protein Name | Protein DB number | Comment | Scan # | Scan Time |
|---|---|---|---|---|---|---|---|---|---|---|---|---|---|---|---|---|---|---|---|---|---|---|
| 19714:5 | 1 | R.TAGWNVPIGTLRPFLN[+2205.793]WTGPPEPIEAAVAR.F | HexNAc(4)Hex(5)Fuc(2)NeuAc(1) | N[+2206] | 1088.1057 | 5 | 5436.4995 | 5436.5038 | 0 | -0.8 | 141 | Specific | 259.08 | 259.08 | 259.08 | 5.95 | 114 | >sp\|P02788\|TRFL_HUMAN Lactotransferrin OS=Homo sapiens OX=9606 GN=LTF PE=1 SV=6 | 1270 | 20201115_10113_1ng_glyco_hcd203040_compound15.23006.23006.5 | controllerType=0 controllerNumber=1 scan=23006 | 46.3856 |
| 18730:3 | 1 | R.[+42.011]TAGWNVPIGTLRPFLN[+2018.708]WTGPPEPIEAAVAR.F | HexNAc(3)Hex(6)Fuc(1)NeuAc(1) | .[42], N[+2019] | 1764.4983 | 3 | 5291.4802 | 5291.4299 | 0 | 9.5 | 141 | Specific | 272.6 | 214.29 | 214.29 | 6.08 | 114 | >sp\|P02788\|TRFL_HUMAN Lactotransferrin OS=Homo sapiens OX=9606 GN=LTF PE=1 SV=6 | 1270 | 20201115_10113_1ng_glyco_hcd203040_compound15.21979.21979.3 | controllerType=0 controllerNumber=1 scan=21979 | 44.0446 |
| 18684:4 | 1 | R.TAGWNVPIGTLRPFLN[+2279.830]WTGPPEPIEAAVAR.F | HexNAc(5)Hex(6)Fuc(2) | N[+2280] | 1378.3961 | 4 | 5510.5627 | 5510.5406 | 0 | 4 | 141 | Specific | 281.65 | 281.65 | 281.65 | 6.59 | 114 | >sp\|P02788\|TRFL_HUMAN Lactotransferrin OS=Homo sapiens OX=9606 GN=LTF PE=1 SV=6 | 1270 | 20201115_10113_1ng_glyco_hcd203040_compound15.21932.21932.4 | controllerType=0 controllerNumber=1 scan=21932 | 43.9611 |
| 19527:4 | 1 | R.TAGWNVPIGTLRPFLN[+2936.057]WTGPPEPIEAAVAR.F | HexNAc(6)Hex(7)Fuc(2)NeuAc(1) | N[+2936] | 1542.4489 | 4 | 6166.7736 | 6166.7682 | 0 | 0.9 | 141 | Specific | 282.88 | 246.06 | 246.06 | 6.82 | 114 | >sp\|P02788\|TRFL_HUMAN Lactotransferrin OS=Homo sapiens OX=9606 GN=LTF PE=1 SV=6 | 1270 | 20201115_10113_1ng_glyco_hcd203040_compound15.22810.22810.4 | controllerType=0 controllerNumber=1 scan=22810 | 45.9083 |
| 18719:3 | 1 | R.[+42.011]TAGWNVPIGTLRPFLN[+1872.651]WTGPPEPIEAAVAR.F | HexNAc(3)Hex(6)NeuAc(1) | .[42], N[+1873] | 1715.8111 | 3 | 5145.4188 | 5145.372 | 0 | 9.1 | 141 | Specific | 305.75 | 180.26 | 180.26 | 4.68 | 114 | >sp\|P02788\|TRFL_HUMAN Lactotransferrin OS=Homo sapiens OX=9606 GN=LTF PE=1 SV=6 | 1270 | 20201115_10113_1ng_glyco_hcd203040_compound15.21968.21968.3 | controllerType=0 controllerNumber=1 scan=21968 | 44.0292 |
| 19817:3 | 1 | R.TAGWNVPIGTLRPFLN[+2059.735]WTGPPEPIEAAVAR.F | HexNAc(4)Hex(5)Fuc(1)NeuAc(1) | N[+2060] | 1764.1679 | 3 | 5290.4892 | 5290.4459 | 0 | 8.2 | 141 | Specific | 329.53 | 324.72 | 324.72 | 6.75 | 114 | >sp\|P02788\|TRFL_HUMAN Lactotransferrin OS=Homo sapiens OX=9606 GN=LTF PE=1 SV=6 | 1270 | 20201115_10113_1ng_glyco_hcd203040_compound15.23114.23114.3 | controllerType=0 controllerNumber=1 scan=23114 | 46.6166 |
| 19677:3 | 1 | R.TAGWNVPIGTLRPFLN[+2204.772]WTGPPEPIEAAVAR.F | HexNAc(4)Hex(5)NeuAc(2) | N[+2205] | 1812.5109 | 3 | 5435.5183 | 5435.4834 | 0 | 6.4 | 141 | Specific | 337.93 | 177.6 | 177.6 | 6.08 | 114 | >sp\|P02788\|TRFL_HUMAN Lactotransferrin OS=Homo sapiens OX=9606 GN=LTF PE=1 SV=6 | 1270 | 20201115_10113_1ng_glyco_hcd203040_compound15.22968.22968.3 | controllerType=0 controllerNumber=1 scan=22968 | 46.3137 |
| 19525:4 | 1 | R.TAGWNVPIGTLRPFLN[+2936.057]WTGPPEPIEAAVAR.F | HexNAc(6)Hex(7)Fuc(2)NeuAc(1) | N[+2936] | 1542.4489 | 4 | 6166.7736 | 6166.7682 | 0 | 0.9 | 141 | Specific | 346.58 | 346.58 | 346.58 | 7.63 | 114 | >sp\|P02788\|TRFL_HUMAN Lactotransferrin OS=Homo sapiens OX=9606 GN=LTF PE=1 SV=6 | 1270 | 20201115_10113_1ng_glyco_hcd203040_compound15.22808.22808.4 | controllerType=0 controllerNumber=1 scan=22808 | 45.9049 |
| 18626:4 | 1 | R.TAGWNVPIGTLRPFLN[+2498.904]WTGPPEPIEAAVAR.F | HexNAc(6)Hex(7)Fuc(1) | N[+2499] | 1433.1608 | 4 | 5729.6212 | 5729.6148 | 0 | 1.1 | 141 | Specific | 356.27 | 356.27 | 356.27 | 7.72 | 114 | >sp\|P02788\|TRFL_HUMAN Lactotransferrin OS=Homo sapiens OX=9606 GN=LTF PE=1 SV=6 | 1270 | 20201115_10113_1ng_glyco_hcd203040_compound15.21872.21872.4 | controllerType=0 controllerNumber=1 scan=21872 | 43.8615 |
| 19676:4 | 1 | R.TAGWNVPIGTLRPFLN[+2205.793]WTGPPEPIEAAVAR.F | HexNAc(4)Hex(5)Fuc(2)NeuAc(1) | N[+2206] | 1359.8849 | 4 | 5436.5177 | 5436.5038 | 0 | 2.6 | 141 | Specific | 371.81 | 371.81 | 371.81 | 8.35 | 114 | >sp\|P02788\|TRFL_HUMAN Lactotransferrin OS=Homo sapiens OX=9606 GN=LTF PE=1 SV=6 | 1270 | 20201115_10113_1ng_glyco_hcd203040_compound15.22967.22967.4 | controllerType=0 controllerNumber=1 scan=22967 | 46.3123 |
| 19570:3 | 1 | R.TAGWNVPIGTLRPFLN[+2570.925]WTGPPEPIEAAVAR.F | HexNAc(5)Hex(6)Fuc(2)NeuAc(1) | N[+2571] | 1934.5571 | 3 | 5801.6567 | 5801.636 | 0 | 3.6 | 141 | Specific | 464.15 | 331.59 | 331.59 | 8.37 | 114 | >sp\|P02788\|TRFL_HUMAN Lactotransferrin OS=Homo sapiens OX=9606 GN=LTF PE=1 SV=6 | 1270 | 20201115_10113_1ng_glyco_hcd203040_compound15.22856.22856.3 | controllerType=0 controllerNumber=1 scan=22856 | 46.0371 |
| 18696:5 | 1 | R.TAGWNVPIGTLRPFLN[+2279.830]WTGPPEPIEAAVAR.F | HexNAc(5)Hex(6)Fuc(2) | N[+2280] | 1102.9192 | 5 | 5510.5669 | 5510.5406 | 0 | 4.8 | 141 | Specific | 533.13 | 455.52 | 455.52 | 9.91 | 114 | >sp\|P02788\|TRFL_HUMAN Lactotransferrin OS=Homo sapiens OX=9606 GN=LTF PE=1 SV=6 | 1270 | 20201115_10113_1ng_glyco_hcd203040_compound15.21944.21944.5 | controllerType=0 controllerNumber=1 scan=21944 | 43.9798 |
| 19680:5 | 1 | R.TAGWNVPIGTLRPFLN[+2204.772]WTGPPEPIEAAVAR.F | HexNAc(4)Hex(5)NeuAc(2) | N[+2205] | 1087.908 | 5 | 5435.5107 | 5435.4834 | 0 | 5 | 141 | Specific | 546.2 | 480.65 | 480.65 | 10.1 | 114 | >sp\|P02788\|TRFL_HUMAN Lactotransferrin OS=Homo sapiens OX=9606 GN=LTF PE=1 SV=6 | 1270 | 20201115_10113_1ng_glyco_hcd203040_compound15.22971.22971.5 | controllerType=0 controllerNumber=1 scan=22971 | 46.3208 |
| 18629:4 | 1 | R.TAGWNVPIGTLRPFLN[+2498.904]WTGPPEPIEAAVAR.F | HexNAc(6)Hex(7)Fuc(1) | N[+2499] | 1433.1608 | 4 | 5729.6212 | 5729.6148 | 0 | 1.1 | 141 | Specific | 546.78 | 501.28 | 501.28 | 10.43 | 114 | >sp\|P02788\|TRFL_HUMAN Lactotransferrin OS=Homo sapiens OX=9606 GN=LTF PE=1 SV=6 | 1270 | 20201115_10113_1ng_glyco_hcd203040_compound15.21875.21875.4 | controllerType=0 controllerNumber=1 scan=21875 | 43.8663 |
| 19452:4 | 1 | R.TAGWNVPIGTLRPFLN[+2936.057]WTGPPEPIEAAVAR.F | HexNAc(6)Hex(7)Fuc(2)NeuAc(1) | N[+2936] | 1542.4481 | 4 | 6166.7707 | 6166.7682 | 0 | 0.4 | 141 | Specific | 548.91 | 517.9 | 517.9 | 9.97 | 114 | >sp\|P02788\|TRFL_HUMAN Lactotransferrin OS=Homo sapiens OX=9606 GN=LTF PE=1 SV=6 | 1270 | 20201115_10113_1ng_glyco_hcd203040_compound15.22732.22732.2 | controllerType=0 controllerNumber=1 scan=22732 | 45.7581 |
| 19820:3 | 1 | R.TAGWNVPIGTLRPFLN[+2059.735]WTGPPEPIEAAVAR.F | HexNAc(4)Hex(5)Fuc(1)NeuAc(1) | N[+2060] | 1764.1679 | 3 | 5290.4892 | 5290.4459 | 0 | 8.2 | 141 | Specific | 557.01 | 486.83 | 486.83 | 9.73 | 114 | >sp\|P02788\|TRFL_HUMAN Lactotransferrin OS=Homo sapiens OX=9606 GN=LTF PE=1 SV=6 | 1270 | 20201115_10113_1ng_glyco_hcd203040_compound15.23117.23117.3 | controllerType=0 controllerNumber=1 scan=23117 | 46.6208 |
| 18704:4 | 1 | R.TAGWNVPIGTLRPFLN[+2133.772]WTGPPEPIEAAVAR.F | HexNAc(5)Hex(6)Fuc(1) | N[+2134] | 1341.8811 | 4 | 5364.5026 | 5364.4826 | 0 | 3.7 | 141 | Specific | 571.07 | 571.07 | 571.07 | 10.34 | 114 | >sp\|P02788\|TRFL_HUMAN Lactotransferrin OS=Homo sapiens OX=9606 GN=LTF PE=1 SV=6 | 1270 | 20201115_10113_1ng_glyco_hcd203040_compound15.21952.21952.4 | controllerType=0 controllerNumber=1 scan=21952 | 43.9951 |
| 18649:3 | 1 | R.TAGWNVPIGTLRPFLN[+2279.830]WTGPPEPIEAAVAR.F | HexNAc(5)Hex(6)Fuc(2) | N[+2280] | 1837.5224 | 3 | 5510.5527 | 5510.5406 | 0 | 2.2 | 141 | Specific | 578.2 | 476.15 | 476.15 | 10.86 | 114 | >sp\|P02788\|TRFL_HUMAN Lactotransferrin OS=Homo sapiens OX=9606 GN=LTF PE=1 SV=6 | 1270 | 20201115_10113_1ng_glyco_hcd203040_compound15.21895.21895.3 | controllerType=0 controllerNumber=1 scan=21895 | 43.8954 |
| 18652:3 | 1 | R.TAGWNVPIGTLRPFLN[+2279.830]WTGPPEPIEAAVAR.F | HexNAc(5)Hex(6)Fuc(2) | N[+2280] | 1837.5224 | 3 | 5510.5527 | 5510.5406 | 0 | 2.2 | 141 | Specific | 587.72 | 467.02 | 467.02 | 11.92 | 114 | >sp\|P02788\|TRFL_HUMAN Lactotransferrin OS=Homo sapiens OX=9606 GN=LTF PE=1 SV=6 | 1270 | 20201115_10113_1ng_glyco_hcd203040_compound15.21898.21898.3 | controllerType=0 controllerNumber=1 scan=21898 | 43.9001 |
| 20011:4 | 1 | R.TAGWNVPIGTLRPFLN[+2059.735]WTGPPEPIEAAVAR.F | HexNAc(4)Hex(5)Fuc(1)NeuAc(1) | N[+2060] | 1323.3795 | 4 | 5290.4962 | 5290.4459 | 0 | 9.5 | 141 | Specific | 600.91 | 554.09 | 554.09 | 11.89 | 114 | >sp\|P02788\|TRFL_HUMAN Lactotransferrin OS=Homo sapiens OX=9606 GN=LTF PE=1 SV=6 | 1270 | 20201115_10113_1ng_glyco_hcd203040_compound15.23317.23317.4 | controllerType=0 controllerNumber=1 scan=23317 | 47.0539 |
| 19679:3 | 1 | R.TAGWNVPIGTLRPFLN[+2204.772]WTGPPEPIEAAVAR.F | HexNAc(4)Hex(5)NeuAc(2) | N[+2205] | 1812.5109 | 3 | 5435.5183 | 5435.4834 | 0 | 6.4 | 141 | Specific | 624.1 | 421.17 | 421.17 | 12.05 | 114 | >sp\|P02788\|TRFL_HUMAN Lactotransferrin OS=Homo sapiens OX=9606 GN=LTF PE=1 SV=6 | 1270 | 20201115_10113_1ng_glyco_hcd203040_compound15.22970.22970.3 | controllerType=0 controllerNumber=1 scan=22970 | 46.3196 |
| 18707:4 | 1 | R.TAGWNVPIGTLRPFLN[+2133.772]WTGPPEPIEAAVAR.F | HexNAc(5)Hex(6)Fuc(1) | N[+2134] | 1341.8811 | 4 | 5364.5026 | 5364.4826 | 0 | 3.7 | 141 | Specific | 664.02 | 580.38 | 580.38 | 13.4 | 114 | >sp\|P02788\|TRFL_HUMAN Lactotransferrin OS=Homo sapiens OX=9606 GN=LTF PE=1 SV=6 | 1270 | 20201115_10113_1ng_glyco_hcd203040_compound15.21955.21955.4 | controllerType=0 controllerNumber=1 scan=21955 | 43.9998 |
| 18737:3 | 1 | R.TAGWNVPIGTLRPFLN[+1768.640]WTGPPEPIEAAVAR.F | HexNAc(4)Hex(5)Fuc(1) | N[+1769] | 1667.1247 | 3 | 4999.3596 | 4999.3504 | 0 | 1.8 | 141 | Specific | 685.87 | 594.92 | 594.92 | 13.92 | 114 | >sp\|P02788\|TRFL_HUMAN Lactotransferrin OS=Homo sapiens OX=9606 GN=LTF PE=1 SV=6 | 1270 | 20201115_10113_1ng_glyco_hcd203040_compound15.21986.21986.3 | controllerType=0 controllerNumber=1 scan=21986 | 44.0638 |
| 18723:4 | 1 | R.TAGWNVPIGTLRPFLN[+2059.735]WTGPPEPIEAAVAR.F | HexNAc(4)Hex(5)Fuc(1)NeuAc(1) | N[+2060] | 1323.3729 | 4 | 5290.4699 | 5290.4459 | 0 | 4.5 | 141 | Specific | 699.14 | 584.48 | 584.48 | 13.4 | 114 | >sp\|P02788\|TRFL_HUMAN Lactotransferrin OS=Homo sapiens OX=9606 GN=LTF PE=1 SV=6 | 1270 | 20201115_10113_1ng_glyco_hcd203040_compound15.21972.21972.4 | controllerType=0 controllerNumber=1 scan=21972 | 44.0349 |
| 21003:4 | 1 | R.TAGWNVPIGTLRPFLN[+2350.830]WTGPPEPIEAAVAR.F | HexNAc(4)Hex(5)Fuc(1)NeuAc(2) | N[+2351] | 1396.1414 | 4 | 5581.5436 | 5581.5413 | 0 | 0.4 | 141 | Specific | 699.85 | 549.68 | 549.68 | 13.15 | 114 | >sp\|P02788\|TRFL_HUMAN Lactotransferrin OS=Homo sapiens OX=9606 GN=LTF PE=1 SV=6 | 1270 | 20201115_10113_1ng_glyco_hcd203040_compound15.24355.24355.4 | controllerType=0 controllerNumber=1 scan=24355 | 49.4072 |
| 18732:4 | 1 | R.TAGWNVPIGTLRPFLN[+1768.640]WTGPPEPIEAAVAR.F | HexNAc(4)Hex(5)Fuc(1) | N[+1769] | 1250.5956 | 4 | 4999.3605 | 4999.3504 | 0 | 2 | 141 | Specific | 717.33 | 693.68 | 693.68 | 14.83 | 114 | >sp\|P02788\|TRFL_HUMAN Lactotransferrin OS=Homo sapiens OX=9606 GN=LTF PE=1 SV=6 | 1270 | 20201115_10113_1ng_glyco_hcd203040_compound15.21981.21981.4 | controllerType=0 controllerNumber=1 scan=21981 | 44.0506 |
| 18740:3 | 1 | R.TAGWNVPIGTLRPFLN[+1768.640]WTGPPEPIEAAVAR.F | HexNAc(4)Hex(5)Fuc(1) | N[+1769] | 1667.1247 | 3 | 4999.3596 | 4999.3504 | 0 | 1.8 | 141 | Specific | 730.76 | 594.47 | 594.47 | 14.92 | 114 | >sp\|P02788\|TRFL_HUMAN Lactotransferrin OS=Homo sapiens OX=9606 GN=LTF PE=1 SV=6 | 1270 | 20201115_10113_1ng_glyco_hcd203040_compound15.21989.21989.3 | controllerType=0 controllerNumber=1 scan=21989 | 44.0696 |
| 18645:4 | 1 | R.TAGWNVPIGTLRPFLN[+2279.830]WTGPPEPIEAAVAR.F | HexNAc(5)Hex(6)Fuc(2) | N[+2280] | 1378.3938 | 4 | 5510.5534 | 5510.5406 | 0 | 2.3 | 141 | Specific | 732.67 | 645.4 | 645.4 | 14.83 | 114 | >sp\|P02788\|TRFL_HUMAN Lactotransferrin OS=Homo sapiens OX=9606 GN=LTF PE=1 SV=6 | 1270 | 20201115_10113_1ng_glyco_hcd203040_compound15.21891.21891.2 | controllerType=0 controllerNumber=1 scan=21891 | 43.8877 |
| 18686:4 | 1 | R.TAGWNVPIGTLRPFLN[+2279.830]WTGPPEPIEAAVAR.F | HexNAc(5)Hex(6)Fuc(2) | N[+2280] | 1378.3961 | 4 | 5510.5627 | 5510.5406 | 0 | 4 | 141 | Specific | 736.46 | 678.9 | 678.9 | 14.4 | 114 | >sp\|P02788\|TRFL_HUMAN Lactotransferrin OS=Homo sapiens OX=9606 GN=LTF PE=1 SV=6 | 1270 | 20201115_10113_1ng_glyco_hcd203040_compound15.21934.21934.4 | controllerType=0 controllerNumber=1 scan=21934 | 43.9651 |
| 20013:4 | 1 | R.TAGWNVPIGTLRPFLN[+2059.735]WTGPPEPIEAAVAR.F | HexNAc(4)Hex(5)Fuc(1)NeuAc(1) | N[+2060] | 1323.3795 | 4 | 5290.4962 | 5290.4459 | 0 | 9.5 | 141 | Specific | 769.21 | 610.81 | 610.81 | 14.89 | 114 | >sp\|P02788\|TRFL_HUMAN Lactotransferrin OS=Homo sapiens OX=9606 GN=LTF PE=1 SV=6 | 1270 | 20201115_10113_1ng_glyco_hcd203040_compound15.23319.23319.4 | controllerType=0 controllerNumber=1 scan=23319 | 47.058 |
| 19717:5 | 1 | R.TAGWNVPIGTLRPFLN[+2205.793]WTGPPEPIEAAVAR.F | HexNAc(4)Hex(5)Fuc(2)NeuAc(1) | N[+2206] | 1088.1057 | 5 | 5436.4995 | 5436.5038 | 0 | -0.8 | 141 | Specific | 791.31 | 604.76 | 604.76 | 14.76 | 114 | >sp\|P02788\|TRFL_HUMAN Lactotransferrin OS=Homo sapiens OX=9606 GN=LTF PE=1 SV=6 | 1270 | 20201115_10113_1ng_glyco_hcd203040_compound15.23009.23009.5 | controllerType=0 controllerNumber=1 scan=23009 | 46.3904 |
| 19683:5 | 1 | R.TAGWNVPIGTLRPFLN[+2204.772]WTGPPEPIEAAVAR.F | HexNAc(4)Hex(5)NeuAc(2) | N[+2205] | 1087.908 | 5 | 5435.5107 | 5435.4834 | 0 | 5 | 141 | Specific | 795.29 | 569.85 | 569.85 | 15.28 | 114 | >sp\|P02788\|TRFL_HUMAN Lactotransferrin OS=Homo sapiens OX=9606 GN=LTF PE=1 SV=6 | 1270 | 20201115_10113_1ng_glyco_hcd203040_compound15.22974.22974.5 | controllerType=0 controllerNumber=1 scan=22974 | 46.3262 |
| 18648:4 | 1 | R.TAGWNVPIGTLRPFLN[+2279.830]WTGPPEPIEAAVAR.F | HexNAc(5)Hex(6)Fuc(2) | N[+2280] | 1378.3938 | 4 | 5510.5534 | 5510.5406 | 0 | 2.3 | 141 | Specific | 828.16 | 720.06 | 720.06 | 16.83 | 114 | >sp\|P02788\|TRFL_HUMAN Lactotransferrin OS=Homo sapiens OX=9606 GN=LTF PE=1 SV=6 | 1270 | 20201115_10113_1ng_glyco_hcd203040_compound15.21894.21894.2 | controllerType=0 controllerNumber=1 scan=21894 | 43.8942 |
| 19454:4 | 1 | R.TAGWNVPIGTLRPFLN[+2936.057]WTGPPEPIEAAVAR.F | HexNAc(6)Hex(7)Fuc(2)NeuAc(1) | N[+2936] | 1542.4481 | 4 | 6166.7707 | 6166.7682 | 0 | 0.4 | 141 | Specific | 851.83 | 736.18 | 736.18 | 17.15 | 114 | >sp\|P02788\|TRFL_HUMAN Lactotransferrin OS=Homo sapiens OX=9606 GN=LTF PE=1 SV=6 | 1270 | 20201115_10113_1ng_glyco_hcd203040_compound15.22734.22734.2 | controllerType=0 controllerNumber=1 scan=22734 | 45.7628 |
| 19678:4 | 1 | R.TAGWNVPIGTLRPFLN[+2205.793]WTGPPEPIEAAVAR.F | HexNAc(4)Hex(5)Fuc(2)NeuAc(1) | N[+2206] | 1359.8849 | 4 | 5436.5177 | 5436.5038 | 0 | 2.6 | 141 | Specific | 866.58 | 709.72 | 709.72 | 17.83 | 114 | >sp\|P02788\|TRFL_HUMAN Lactotransferrin OS=Homo sapiens OX=9606 GN=LTF PE=1 SV=6 | 1270 | 20201115_10113_1ng_glyco_hcd203040_compound15.22969.22969.4 | controllerType=0 controllerNumber=1 scan=22969 | 46.3184 |
| 18735:4 | 1 | R.TAGWNVPIGTLRPFLN[+1768.640]WTGPPEPIEAAVAR.F | HexNAc(4)Hex(5)Fuc(1) | N[+1769] | 1250.5956 | 4 | 4999.3605 | 4999.3504 | 0 | 2 | 141 | Specific | 906.64 | 823.29 | 823.29 | 18.83 | 114 | >sp\|P02788\|TRFL_HUMAN Lactotransferrin OS=Homo sapiens OX=9606 GN=LTF PE=1 SV=6 | 1270 | 20201115_10113_1ng_glyco_hcd203040_compound15.21984.21984.4 | controllerType=0 controllerNumber=1 scan=21984 | 44.0583 |
| 21039:3 | 1 | R.TAGWNIPMGLLFN[+2350.830]QTGSC[+57.021]K.F | HexNAc(4)Hex(5)Fuc(1)NeuAc(2) | C[+57], N[+2351] | 1482.6244 | 3 | 4445.8588 | 4445.8403 | 0 | 4.2 | 485 | Specific | 410.52 | 410.52 | 410.52 | 9.23 | 114 | >sp\|P02788\|TRFL_HUMAN Lactotransferrin OS=Homo sapiens OX=9606 GN=LTF PE=1 SV=6 | 1270 | 20201115_10113_1ng_glyco_hcd203040_compound15.24393.24393.3 | controllerType=0 controllerNumber=1 scan=24393 | 49.4979 |
| 17325:3 | 1 | R.TAGWNIPMGLLFN[+1606.587]QTGSC[+57.021]K.F | HexNAc(4)Hex(4)Fuc(1) | C[+57], N[+1607] | 1234.5392 | 3 | 3701.6032 | 3701.5966 | 0 | 1.8 | 485 | Specific | 419.54 | 419.54 | 419.54 | 9.71 | 114 | >sp\|P02788\|TRFL_HUMAN Lactotransferrin OS=Homo sapiens OX=9606 GN=LTF PE=1 SV=6 | 1270 | 20201115_10113_1ng_glyco_hcd203040_compound15.20515.20515.3 | controllerType=0 controllerNumber=1 scan=20515 | 40.948 |
| 17214:2 | 1 | R.TAGWNIPMGLLFN[+1768.640]QTGSC[+57.021]K.F | HexNAc(4)Hex(5)Fuc(1) | C[+57], N[+1769] | 1932.3341 | 2 | 3863.6609 | 3863.6495 | 0 | 3 | 485 | Specific | 452.95 | 385.75 | 385.75 | 10.02 | 114 | >sp\|P02788\|TRFL_HUMAN Lactotransferrin OS=Homo sapiens OX=9606 GN=LTF PE=1 SV=6 | 1270 | 20201115_10113_1ng_glyco_hcd203040_compound15.20399.20399.2 | controllerType=0 controllerNumber=1 scan=20399 | 40.7088 |
| 16978:3 | 1 | R.TAGWNIPMGLLFN[+2279.830]QTGSC[+57.021]K.F | HexNAc(5)Hex(6)Fuc(2) | C[+57], N[+2280] | 1458.9612 | 3 | 4374.8692 | 4374.8396 | 0 | 6.8 | 485 | Specific | 678.11 | 542.37 | 334.38 | 12.68 | 114 | >sp\|P02788\|TRFL_HUMAN Lactotransferrin OS=Homo sapiens OX=9606 GN=LTF PE=1 SV=6 | 1270 | 20201115_10113_1ng_glyco_hcd203040_compound15.20154.20154.3 | controllerType=0 controllerNumber=1 scan=20154 | 40.2257 |
| 17217:2 | 1 | R.TAGWNIPMGLLFN[+1768.640]QTGSC[+57.021]K.F | HexNAc(4)Hex(5)Fuc(1) | C[+57], N[+1769] | 1932.3341 | 2 | 3863.6609 | 3863.6495 | 0 | 3 | 485 | Specific | 680.64 | 579.05 | 579.05 | 13.62 | 114 | >sp\|P02788\|TRFL_HUMAN Lactotransferrin OS=Homo sapiens OX=9606 GN=LTF PE=1 SV=6 | 1270 | 20201115_10113_1ng_glyco_hcd203040_compound15.20402.20402.2 | controllerType=0 controllerNumber=1 scan=20402 | 40.715 |
| 19051:4 | 1 | R.TAGWNIPMGLLFN[+2205.793]QTGSC[+57.021]K.F | HexNAc(4)Hex(5)Fuc(2)NeuAc(1) | C[+57], N[+2206] | 1075.9612 | 4 | 4300.8231 | 4300.8028 | 0 | 4.7 | 485 | Specific | 696.8 | 636.29 | 636.29 | 13 | 114 | >sp\|P02788\|TRFL_HUMAN Lactotransferrin OS=Homo sapiens OX=9606 GN=LTF PE=1 SV=6 | 1270 | 20201115_10113_1ng_glyco_hcd203040_compound15.22314.22314.4 | controllerType=0 controllerNumber=1 scan=22314 | 44.7945 |
| 17205:4 | 1 | R.TAGWNIPMGLLFN[+1768.640]QTGSC[+57.021]K.F | HexNAc(4)Hex(5)Fuc(1) | C[+57], N[+1769] | 966.6708 | 4 | 3863.6612 | 3863.6495 | 0 | 3 | 485 | Specific | 744.42 | 624.65 | 624.65 | 14.57 | 114 | >sp\|P02788\|TRFL_HUMAN Lactotransferrin OS=Homo sapiens OX=9606 GN=LTF PE=1 SV=6 | 1270 | 20201115_10113_1ng_glyco_hcd203040_compound15.20390.20390.4 | controllerType=0 controllerNumber=1 scan=20390 | 40.6917 |
| 19018:3 | 1 | R.TAGWNIPMGLLFN[+2205.793]QTGSC[+57.021]K.F | HexNAc(4)Hex(5)Fuc(2)NeuAc(1) | C[+57], N[+2206] | 1434.2768 | 3 | 4300.8158 | 4300.8028 | 0 | 3 | 485 | Specific | 771.38 | 686.67 | 686.67 | 15.53 | 114 | >sp\|P02788\|TRFL_HUMAN Lactotransferrin OS=Homo sapiens OX=9606 GN=LTF PE=1 SV=6 | 1270 | 20201115_10113_1ng_glyco_hcd203040_compound15.22280.22280.3 | controllerType=0 controllerNumber=1 scan=22280 | 44.7242 |
| 16980:3 | 1 | R.TAGWNIPMGLLFN[+2279.830]QTGSC[+57.021]K.F | HexNAc(5)Hex(6)Fuc(2) | C[+57], N[+2280] | 1458.9612 | 3 | 4374.8692 | 4374.8396 | 0 | 6.8 | 485 | Specific | 772.51 | 722.86 | 430.82 | 14.68 | 114 | >sp\|P02788\|TRFL_HUMAN Lactotransferrin OS=Homo sapiens OX=9606 GN=LTF PE=1 SV=6 | 1270 | 20201115_10113_1ng_glyco_hcd203040_compound15.20156.20156.3 | controllerType=0 controllerNumber=1 scan=20156 | 40.2317 |
| 17608:2 | 1 | R.TAGWNIPMGLLFN[+1216.423]QTGSC[+57.021]K.F | HexNAc(2)Hex(5) | C[+57], N[+1216] | 1656.2258 | 2 | 3311.4442 | 3311.4328 | 0 | 3.4 | 485 | Specific | 777.92 | 688.45 | 229.52 | 15.62 | 114 | >sp\|P02788\|TRFL_HUMAN Lactotransferrin OS=Homo sapiens OX=9606 GN=LTF PE=1 SV=6 | 1270 | 20201115_10113_1ng_glyco_hcd203040_compound15.20809.20809.2 | controllerType=0 controllerNumber=1 scan=20809 | 41.549 |
| 17388:3 | 1 | R.TAGWNIPMGLLFN[+1606.587]QTGSC[+57.021]K.F | HexNAc(4)Hex(4)Fuc(1) | C[+57], N[+1607] | 1234.5412 | 3 | 3701.609 | 3701.5966 | 0 | 3.3 | 485 | Specific | 778.11 | 778.11 | 778.11 | 15.53 | 114 | >sp\|P02788\|TRFL_HUMAN Lactotransferrin OS=Homo sapiens OX=9606 GN=LTF PE=1 SV=6 | 1270 | 20201115_10113_1ng_glyco_hcd203040_compound15.20580.20580.3 | controllerType=0 controllerNumber=1 scan=20580 | 41.0734 |
| 21041:3 | 1 | R.TAGWNIPMGLLFN[+2350.830]QTGSC[+57.021]K.F | HexNAc(4)Hex(5)Fuc(1)NeuAc(2) | C[+57], N[+2351] | 1482.6244 | 3 | 4445.8588 | 4445.8403 | 0 | 4.2 | 485 | Specific | 825.53 | 741.25 | 741.25 | 16.12 | 114 | >sp\|P02788\|TRFL_HUMAN Lactotransferrin OS=Homo sapiens OX=9606 GN=LTF PE=1 SV=6 | 1270 | 20201115_10113_1ng_glyco_hcd203040_compound15.24395.24395.3 | controllerType=0 controllerNumber=1 scan=24395 | 49.5068 |
| 17170:3 | 1 | R.TAGWNIPMGLLFN[+1768.640]QTGSC[+57.021]K.F | HexNAc(4)Hex(5)Fuc(1) | C[+57], N[+1769] | 1288.5584 | 3 | 3863.6607 | 3863.6495 | 0 | 2.9 | 485 | Specific | 825.94 | 825.94 | 825.94 | 16.53 | 114 | >sp\|P02788\|TRFL_HUMAN Lactotransferrin OS=Homo sapiens OX=9606 GN=LTF PE=1 SV=6 | 1270 | 20201115_10113_1ng_glyco_hcd203040_compound15.20354.20354.3 | controllerType=0 controllerNumber=1 scan=20354 | 40.6292 |
| 20990:4 | 1 | R.TAGWNIPMGLLFN[+2350.830]QTGSC[+57.021]K.F | HexNAc(4)Hex(5)Fuc(1)NeuAc(2) | C[+57], N[+2351] | 1112.2188 | 4 | 4445.8533 | 4445.8403 | 0 | 2.9 | 485 | Specific | 830.77 | 657.51 | 657.51 | 16.57 | 114 | >sp\|P02788\|TRFL_HUMAN Lactotransferrin OS=Homo sapiens OX=9606 GN=LTF PE=1 SV=6 | 1270 | 20201115_10113_1ng_glyco_hcd203040_compound15.24341.24341.4 | controllerType=0 controllerNumber=1 scan=24341 | 49.3798 |
| 17328:3 | 1 | R.TAGWNIPMGLLFN[+1606.587]QTGSC[+57.021]K.F | HexNAc(4)Hex(4)Fuc(1) | C[+57], N[+1607] | 1234.5392 | 3 | 3701.6032 | 3701.5966 | 0 | 1.8 | 485 | Specific | 851.82 | 726 | 726 | 17.55 | 114 | >sp\|P02788\|TRFL_HUMAN Lactotransferrin OS=Homo sapiens OX=9606 GN=LTF PE=1 SV=6 | 1270 | 20201115_10113_1ng_glyco_hcd203040_compound15.20518.20518.3 | controllerType=0 controllerNumber=1 scan=20518 | 40.9524 |
| 17173:3 | 1 | R.TAGWNIPMGLLFN[+1768.640]QTGSC[+57.021]K.F | HexNAc(4)Hex(5)Fuc(1) | C[+57], N[+1769] | 1288.5584 | 3 | 3863.6607 | 3863.6495 | 0 | 2.9 | 485 | Specific | 858.21 | 690.42 | 690.42 | 17.53 | 114 | >sp\|P02788\|TRFL_HUMAN Lactotransferrin OS=Homo sapiens OX=9606 GN=LTF PE=1 SV=6 | 1270 | 20201115_10113_1ng_glyco_hcd203040_compound15.20357.20357.3 | controllerType=0 controllerNumber=1 scan=20357 | 40.6331 |
| 19053:4 | 1 | R.TAGWNIPMGLLFN[+2205.793]QTGSC[+57.021]K.F | HexNAc(4)Hex(5)Fuc(2)NeuAc(1) | C[+57], N[+2206] | 1075.9612 | 4 | 4300.8231 | 4300.8028 | 0 | 4.7 | 485 | Specific | 860.32 | 744.46 | 744.46 | 17 | 114 | >sp\|P02788\|TRFL_HUMAN Lactotransferrin OS=Homo sapiens OX=9606 GN=LTF PE=1 SV=6 | 1270 | 20201115_10113_1ng_glyco_hcd203040_compound15.22316.22316.4 | controllerType=0 controllerNumber=1 scan=22316 | 44.7996 |
| 20931:3 | 1 | R.TAGWNIPMGLLFN[+2350.830]QTGSC[+57.021]K.F | HexNAc(4)Hex(5)Fuc(1)NeuAc(2) | C[+57], N[+2351] | 1482.6201 | 3 | 4445.8456 | 4445.8403 | 0 | 1.2 | 485 | Specific | 872.8 | 754.4 | 754.4 | 17.34 | 114 | >sp\|P02788\|TRFL_HUMAN Lactotransferrin OS=Homo sapiens OX=9606 GN=LTF PE=1 SV=6 | 1270 | 20201115_10113_1ng_glyco_hcd203040_compound15.24279.24279.3 | controllerType=0 controllerNumber=1 scan=24279 | 49.2571 |
| 17208:4 | 1 | R.TAGWNIPMGLLFN[+1768.640]QTGSC[+57.021]K.F | HexNAc(4)Hex(5)Fuc(1) | C[+57], N[+1769] | 966.6708 | 4 | 3863.6612 | 3863.6495 | 0 | 3 | 485 | Specific | 887.48 | 671.74 | 671.74 | 17.57 | 114 | >sp\|P02788\|TRFL_HUMAN Lactotransferrin OS=Homo sapiens OX=9606 GN=LTF PE=1 SV=6 | 1270 | 20201115_10113_1ng_glyco_hcd203040_compound15.20393.20393.4 | controllerType=0 controllerNumber=1 scan=20393 | 40.6967 |
| 20987:4 | 1 | R.TAGWNIPMGLLFN[+2350.830]QTGSC[+57.021]K.F | HexNAc(4)Hex(5)Fuc(1)NeuAc(2) | C[+57], N[+2351] | 1112.2188 | 4 | 4445.8533 | 4445.8403 | 0 | 2.9 | 485 | Specific | 897.44 | 737.7 | 737.7 | 17.57 | 114 | >sp\|P02788\|TRFL_HUMAN Lactotransferrin OS=Homo sapiens OX=9606 GN=LTF PE=1 SV=6 | 1270 | 20201115_10113_1ng_glyco_hcd203040_compound15.24338.24338.4 | controllerType=0 controllerNumber=1 scan=24338 | 49.3683 |
| 17581:3 | 1 | R.TAGWNIPMGLLFN[+1216.423]QTGSC[+57.021]K.F | HexNAc(2)Hex(5) | C[+57], N[+1216] | 1104.4838 | 3 | 3311.4368 | 3311.4328 | 0 | 1.2 | 485 | Specific | 903.95 | 903.95 | 347.51 | 18.34 | 114 | >sp\|P02788\|TRFL_HUMAN Lactotransferrin OS=Homo sapiens OX=9606 GN=LTF PE=1 SV=6 | 1270 | 20201115_10113_1ng_glyco_hcd203040_compound15.20781.20781.3 | controllerType=0 controllerNumber=1 scan=20781 | 41.4632 |
| 19183:3 | 1 | R.TAGWNIPMGLLFN[+1913.677]QTGSC[+57.021]K.F | HexNAc(4)Hex(5)NeuAc(1) | C[+57], N[+1914] | 1336.9074 | 3 | 4008.7077 | 4008.687 | 0 | 5.2 | 485 | Specific | 905.85 | 905.85 | 905.85 | 17.97 | 114 | >sp\|P02788\|TRFL_HUMAN Lactotransferrin OS=Homo sapiens OX=9606 GN=LTF PE=1 SV=6 | 1270 | 20201115_10113_1ng_glyco_hcd203040_compound15.22452.22452.3 | controllerType=0 controllerNumber=1 scan=22452 | 45.1321 |
| 17382:3 | 1 | R.TAGWNIPMGLLFN[+1622.582]QTGSC[+57.021]K.F | HexNAc(4)Hex(5) | C[+57], N[+1623] | 1239.8715 | 3 | 3717.6 | 3717.5916 | 0 | 2.3 | 485 | Specific | 910.7 | 886.02 | 428.47 | 18.55 | 114 | >sp\|P02788\|TRFL_HUMAN Lactotransferrin OS=Homo sapiens OX=9606 GN=LTF PE=1 SV=6 | 1270 | 20201115_10113_1ng_glyco_hcd203040_compound15.20574.20574.3 | controllerType=0 controllerNumber=1 scan=20574 | 41.0628 |
| 20947:3 | 1 | R.TAGWNIPMGLLFN[+2350.830]QTGSC[+57.021]K.F | HexNAc(4)Hex(5)Fuc(1)NeuAc(2) | C[+57], N[+2351] | 1482.6202 | 3 | 4445.846 | 4445.8403 | 0 | 1.3 | 485 | Specific | 916.83 | 779.76 | 779.76 | 18.34 | 114 | >sp\|P02788\|TRFL_HUMAN Lactotransferrin OS=Homo sapiens OX=9606 GN=LTF PE=1 SV=6 | 1270 | 20201115_10113_1ng_glyco_hcd203040_compound15.24296.24296.3 | controllerType=0 controllerNumber=1 scan=24296 | 49.2877 |
| 19021:3 | 1 | R.TAGWNIPMGLLFN[+2205.793]QTGSC[+57.021]K.F | HexNAc(4)Hex(5)Fuc(2)NeuAc(1) | C[+57], N[+2206] | 1434.2768 | 3 | 4300.8158 | 4300.8028 | 0 | 3 | 485 | Specific | 919.06 | 902.39 | 902.39 | 18.53 | 114 | >sp\|P02788\|TRFL_HUMAN Lactotransferrin OS=Homo sapiens OX=9606 GN=LTF PE=1 SV=6 | 1270 | 20201115_10113_1ng_glyco_hcd203040_compound15.22283.22283.3 | controllerType=0 controllerNumber=1 scan=22283 | 44.7291 |
| 17605:2 | 1 | R.TAGWNIPMGLLFN[+1216.423]QTGSC[+57.021]K.F | HexNAc(2)Hex(5) | C[+57], N[+1216] | 1656.2258 | 2 | 3311.4442 | 3311.4328 | 0 | 3.4 | 485 | Specific | 924.13 | 890.75 | 354.18 | 18.62 | 114 | >sp\|P02788\|TRFL_HUMAN Lactotransferrin OS=Homo sapiens OX=9606 GN=LTF PE=1 SV=6 | 1270 | 20201115_10113_1ng_glyco_hcd203040_compound15.20806.20806.2 | controllerType=0 controllerNumber=1 scan=20806 | 41.5425 |
| 17380:3 | 1 | R.TAGWNIPMGLLFN[+1622.582]QTGSC[+57.021]K.F | HexNAc(4)Hex(5) | C[+57], N[+1623] | 1239.8715 | 3 | 3717.6 | 3717.5916 | 0 | 2.3 | 485 | Specific | 928.99 | 900.26 | 445.75 | 18.55 | 114 | >sp\|P02788\|TRFL_HUMAN Lactotransferrin OS=Homo sapiens OX=9606 GN=LTF PE=1 SV=6 | 1270 | 20201115_10113_1ng_glyco_hcd203040_compound15.20572.20572.3 | controllerType=0 controllerNumber=1 scan=20572 | 41.057 |
| 19180:3 | 1 | R.TAGWNIPMGLLFN[+1913.677]QTGSC[+57.021]K.F | HexNAc(4)Hex(5)NeuAc(1) | C[+57], N[+1914] | 1336.9074 | 3 | 4008.7077 | 4008.687 | 0 | 5.2 | 485 | Specific | 937.25 | 937.25 | 937.25 | 17.97 | 114 | >sp\|P02788\|TRFL_HUMAN Lactotransferrin OS=Homo sapiens OX=9606 GN=LTF PE=1 SV=6 | 1270 | 20201115_10113_1ng_glyco_hcd203040_compound15.22449.22449.3 | controllerType=0 controllerNumber=1 scan=22449 | 45.1262 |
| 20928:3 | 1 | R.TAGWNIPMGLLFN[+2350.830]QTGSC[+57.021]K.F | HexNAc(4)Hex(5)Fuc(1)NeuAc(2) | C[+57], N[+2351] | 1482.6201 | 3 | 4445.8456 | 4445.8403 | 0 | 1.2 | 485 | Specific | 962.43 | 930.77 | 930.77 | 19.34 | 114 | >sp\|P02788\|TRFL_HUMAN Lactotransferrin OS=Homo sapiens OX=9606 GN=LTF PE=1 SV=6 | 1270 | 20201115_10113_1ng_glyco_hcd203040_compound15.24276.24276.3 | controllerType=0 controllerNumber=1 scan=24276 | 49.2464 |
| 19042:3 | 1 | R.TAGWNIPMGLLFN[+2059.735]QTGSC[+57.021]K.F | HexNAc(4)Hex(5)Fuc(1)NeuAc(1) | C[+57], N[+2060] | 1385.5927 | 3 | 4154.7636 | 4154.7449 | 0 | 4.5 | 485 | Specific | 968.91 | 949.31 | 949.31 | 19.12 | 114 | >sp\|P02788\|TRFL_HUMAN Lactotransferrin OS=Homo sapiens OX=9606 GN=LTF PE=1 SV=6 | 1270 | 20201115_10113_1ng_glyco_hcd203040_compound15.22305.22305.3 | controllerType=0 controllerNumber=1 scan=22305 | 44.7803 |
| 20944:3 | 1 | R.TAGWNIPMGLLFN[+2350.830]QTGSC[+57.021]K.F | HexNAc(4)Hex(5)Fuc(1)NeuAc(2) | C[+57], N[+2351] | 1482.6202 | 3 | 4445.846 | 4445.8403 | 0 | 1.3 | 485 | Specific | 973.03 | 796.03 | 796.03 | 19.34 | 114 | >sp\|P02788\|TRFL_HUMAN Lactotransferrin OS=Homo sapiens OX=9606 GN=LTF PE=1 SV=6 | 1270 | 20201115_10113_1ng_glyco_hcd203040_compound15.24293.24293.3 | controllerType=0 controllerNumber=1 scan=24293 | 49.283 |
| 19040:3 | 1 | R.TAGWNIPMGLLFN[+2059.735]QTGSC[+57.021]K.F | HexNAc(4)Hex(5)Fuc(1)NeuAc(1) | C[+57], N[+2060] | 1385.5927 | 3 | 4154.7636 | 4154.7449 | 0 | 4.5 | 485 | Specific | 980.8 | 898.99 | 898.99 | 19.12 | 114 | >sp\|P02788\|TRFL_HUMAN Lactotransferrin OS=Homo sapiens OX=9606 GN=LTF PE=1 SV=6 | 1270 | 20201115_10113_1ng_glyco_hcd203040_compound15.22303.22303.3 | controllerType=0 controllerNumber=1 scan=22303 | 44.7737 |
| 19217:4 | 1 | R.TAGWNIPMGLLFN[+1913.677]QTGSC[+57.021]K.F | HexNAc(4)Hex(5)NeuAc(1) | C[+57], N[+1914] | 1002.9299 | 4 | 4008.6976 | 4008.687 | 0 | 2.6 | 485 | Specific | 987.68 | 865.31 | 865.31 | 19.58 | 114 | >sp\|P02788\|TRFL_HUMAN Lactotransferrin OS=Homo sapiens OX=9606 GN=LTF PE=1 SV=6 | 1270 | 20201115_10113_1ng_glyco_hcd203040_compound15.22487.22487.4 | controllerType=0 controllerNumber=1 scan=22487 | 45.2006 |
| 18319:5 | 1 | R.TAGWNIPMGLLFN[+1768.640]QTGSC[+57.021]KFDEYFSQSC[+57.021]APGSDPR.S | HexNAc(4)Hex(5)Fuc(1) | C[+57]*2, N[+1769] | 1142.2893 | 5 | 5707.4174 | 5707.3966 | 0 | 3.6 | 485 | Specific | 331.59 | 295.51 | 295.51 | 5.65 | 114 | >sp\|P02788\|TRFL_HUMAN Lactotransferrin OS=Homo sapiens OX=9606 GN=LTF PE=1 SV=6 | 1270 | 20201115_10113_1ng_glyco_hcd203040_compound15.21552.21552.5 | controllerType=0 controllerNumber=1 scan=21552 | 43.177 |
| 18321:5 | 1 | R.TAGWNIPMGLLFN[+1768.640]QTGSC[+57.021]KFDEYFSQSC[+57.021]APGSDPR.S | HexNAc(4)Hex(5)Fuc(1) | C[+57]*2, N[+1769] | 1142.2893 | 5 | 5707.4174 | 5707.3966 | 0 | 3.6 | 485 | Specific | 401.13 | 287.61 | 287.61 | 5.8 | 114 | >sp\|P02788\|TRFL_HUMAN Lactotransferrin OS=Homo sapiens OX=9606 GN=LTF PE=1 SV=6 | 1270 | 20201115_10113_1ng_glyco_hcd203040_compound15.21554.21554.5 | controllerType=0 controllerNumber=1 scan=21554 | 43.1829 |
| 18314:3 | 1 | R.TAGWNIPMGLLFN[+1768.640]QTGSC[+57.021]KFDEYFSQSC[+57.021]APGSDPR.S | HexNAc(4)Hex(5)Fuc(1) | C[+57]*2, N[+1769] | 1903.1447 | 3 | 5707.4197 | 5707.3966 | 0 | 4 | 485 | Specific | 587.09 | 525.17 | 525.17 | 10.86 | 114 | >sp\|P02788\|TRFL_HUMAN Lactotransferrin OS=Homo sapiens OX=9606 GN=LTF PE=1 SV=6 | 1270 | 20201115_10113_1ng_glyco_hcd203040_compound15.21547.21547.3 | controllerType=0 controllerNumber=1 scan=21547 | 43.17 |
| 18316:3 | 1 | R.TAGWNIPMGLLFN[+1768.640]QTGSC[+57.021]KFDEYFSQSC[+57.021]APGSDPR.S | HexNAc(4)Hex(5)Fuc(1) | C[+57]*2, N[+1769] | 1903.1447 | 3 | 5707.4197 | 5707.3966 | 0 | 4 | 485 | Specific | 722.33 | 586.16 | 586.16 | 13.86 | 114 | >sp\|P02788\|TRFL_HUMAN Lactotransferrin OS=Homo sapiens OX=9606 GN=LTF PE=1 SV=6 | 1270 | 20201115_10113_1ng_glyco_hcd203040_compound15.21549.21549.3 | controllerType=0 controllerNumber=1 scan=21549 | 43.1734 |
| 20542:4 | 1 | R.TAGWNIPMGLLFN[+2350.830]QTGSC[+57.021]KFDEYFSQSC[+57.021]APGSDPR.S | HexNAc(4)Hex(5)Fuc(1)NeuAc(2) | C[+57]*2, N[+2351] | 1573.1576 | 4 | 6289.6086 | 6289.5875 | 0 | 3.4 | 485 | Specific | 780.98 | 628.82 | 628.82 | 15.18 | 114 | >sp\|P02788\|TRFL_HUMAN Lactotransferrin OS=Homo sapiens OX=9606 GN=LTF PE=1 SV=6 | 1270 | 20201115_10113_1ng_glyco_hcd203040_compound15.23873.23873.4 | controllerType=0 controllerNumber=1 scan=23873 | 48.3411 |
| 18310:4 | 1 | R.TAGWNIPMGLLFN[+1768.640]QTGSC[+57.021]KFDEYFSQSC[+57.021]APGSDPR.S | HexNAc(4)Hex(5)Fuc(1) | C[+57]*2, N[+1769] | 1427.6108 | 4 | 5707.4215 | 5707.3966 | 0 | 4.4 | 485 | Specific | 933.17 | 905.41 | 905.41 | 17.77 | 114 | >sp\|P02788\|TRFL_HUMAN Lactotransferrin OS=Homo sapiens OX=9606 GN=LTF PE=1 SV=6 | 1270 | 20201115_10113_1ng_glyco_hcd203040_compound15.21542.21542.4 | controllerType=0 controllerNumber=1 scan=21542 | 43.1579 |
| 18312:4 | 1 | R.TAGWNIPMGLLFN[+1768.640]QTGSC[+57.021]KFDEYFSQSC[+57.021]APGSDPR.S | HexNAc(4)Hex(5)Fuc(1) | C[+57]*2, N[+1769] | 1427.6108 | 4 | 5707.4215 | 5707.3966 | 0 | 4.4 | 485 | Specific | 974.98 | 909.87 | 909.87 | 18.77 | 114 | >sp\|P02788\|TRFL_HUMAN Lactotransferrin OS=Homo sapiens OX=9606 GN=LTF PE=1 SV=6 | 1270 | 20201115_10113_1ng_glyco_hcd203040_compound15.21544.21544.4 | controllerType=0 controllerNumber=1 scan=21544 | 43.165 |
| 09500:3 | 1 | R.N[+2570.925]GSDC[+57.021]PDKFC[+57.021]LFQSETK.N | HexNAc(5)Hex(6)Fuc(2)NeuAc(1) | C[+57]*2, N[+2571] | 1535.2706 | 3 | 4603.7971 | 4603.7989 | 0 | -0.4 | 642 | Specific | 405.14 | 362.9 | 362.9 | 7.06 | 114 | >sp\|P02788\|TRFL_HUMAN Lactotransferrin OS=Homo sapiens OX=9606 GN=LTF PE=1 SV=6 | 1270 | 20201115_10113_1ng_glyco_hcd203040_compound15.12375.12375.3 | controllerType=0 controllerNumber=1 scan=12375 | 26.1974 |
| 09674:3 | 1 | R.N[+2205.793]GSDC[+57.021]PDKFC[+57.021]LFQSETK.N | HexNAc(4)Hex(5)Fuc(2)NeuAc(1) | C[+57]*2, N[+2206] | 1413.5656 | 3 | 4238.6823 | 4238.6667 | 0 | 3.7 | 642 | Specific | 507.32 | 428.22 | 428.22 | 9.62 | 114 | >sp\|P02788\|TRFL_HUMAN Lactotransferrin OS=Homo sapiens OX=9606 GN=LTF PE=1 SV=6 | 1270 | 20201115_10113_1ng_glyco_hcd203040_compound15.12562.12562.3 | controllerType=0 controllerNumber=1 scan=12562 | 26.525 |
| 09677:3 | 1 | R.N[+2205.793]GSDC[+57.021]PDKFC[+57.021]LFQSETK.N | HexNAc(4)Hex(5)Fuc(2)NeuAc(1) | C[+57]*2, N[+2206] | 1413.5656 | 3 | 4238.6823 | 4238.6667 | 0 | 3.7 | 642 | Specific | 520.49 | 418.12 | 418.12 | 9.75 | 114 | >sp\|P02788\|TRFL_HUMAN Lactotransferrin OS=Homo sapiens OX=9606 GN=LTF PE=1 SV=6 | 1270 | 20201115_10113_1ng_glyco_hcd203040_compound15.12565.12565.3 | controllerType=0 controllerNumber=1 scan=12565 | 26.5293 |
| 09754:3 | 1 | R.N[+2059.735]GSDC[+57.021]PDKFC[+57.021]LFQSETK.N | HexNAc(4)Hex(5)Fuc(1)NeuAc(1) | C[+57]*2, N[+2060] | 1364.8766 | 3 | 4092.6154 | 4092.6088 | 0 | 1.6 | 642 | Specific | 597.64 | 376.1 | 376.1 | 9.42 | 114 | >sp\|P02788\|TRFL_HUMAN Lactotransferrin OS=Homo sapiens OX=9606 GN=LTF PE=1 SV=6 | 1270 | 20201115_10113_1ng_glyco_hcd203040_compound15.12645.12645.3 | controllerType=0 controllerNumber=1 scan=12645 | 26.6739 |
| 09752:3 | 1 | R.N[+2059.735]GSDC[+57.021]PDKFC[+57.021]LFQSETK.N | HexNAc(4)Hex(5)Fuc(1)NeuAc(1) | C[+57]*2, N[+2060] | 1364.8766 | 3 | 4092.6154 | 4092.6088 | 0 | 1.6 | 642 | Specific | 714.21 | 546.8 | 546.8 | 13.93 | 114 | >sp\|P02788\|TRFL_HUMAN Lactotransferrin OS=Homo sapiens OX=9606 GN=LTF PE=1 SV=6 | 1270 | 20201115_10113_1ng_glyco_hcd203040_compound15.12643.12643.3 | controllerType=0 controllerNumber=1 scan=12643 | 26.6693 |
| 15742:3 | 2 | R.LN[+1702.581]YSLPTGQWVGVQLPR.N | HexNAc(2)Hex(8) | N[+1703] | 1210.878 | 3 | 3630.6194 | 3630.6202 | 0 | -0.2 | 1017 | Specific | 724.44 | 724.44 | 724.44 | 13.76 | 104 | >sp\|P24821\|TENA_HUMAN Tenascin OS=Homo sapiens OX=9606 GN=TNC PE=1 SV=3 | 486 | 20201115_10113_1ng_glyco_hcd203040_compound15.18861.18861.3 | controllerType=0 controllerNumber=1 scan=18861 | 37.7562 |
| 15720:3 | 2 | R.LN[+1702.581]YSLPTGQWVGVQLPR.N | HexNAc(2)Hex(8) | N[+1703] | 1210.8791 | 3 | 3630.6227 | 3630.6202 | 0 | 0.7 | 1017 | Specific | 818.71 | 809.74 | 809.74 | 16.23 | 104 | >sp\|P24821\|TENA_HUMAN Tenascin OS=Homo sapiens OX=9606 GN=TNC PE=1 SV=3 | 486 | 20201115_10113_1ng_glyco_hcd203040_compound15.18838.18838.2 | controllerType=0 controllerNumber=1 scan=18838 | 37.7117 |
| 15843:3 | 2 | R.LN[+1378.476]YSLPTGQWVGVQLPR.N | HexNAc(2)Hex(6) | N[+1378] | 1102.8533 | 3 | 3306.5453 | 3306.5145 | 0 | 9.3 | 1017 | Specific | 850.94 | 845.78 | 845.78 | 16.5 | 104 | >sp\|P24821\|TENA_HUMAN Tenascin OS=Homo sapiens OX=9606 GN=TNC PE=1 SV=3 | 486 | 20201115_10113_1ng_glyco_hcd203040_compound15.18966.18966.3 | controllerType=0 controllerNumber=1 scan=18966 | 37.996 |
| 15845:3 | 2 | R.LN[+1378.476]YSLPTGQWVGVQLPR.N | HexNAc(2)Hex(6) | N[+1378] | 1102.8533 | 3 | 3306.5453 | 3306.5145 | 0 | 9.3 | 1017 | Specific | 899.98 | 728.16 | 728.16 | 16.5 | 104 | >sp\|P24821\|TENA_HUMAN Tenascin OS=Homo sapiens OX=9606 GN=TNC PE=1 SV=3 | 486 | 20201115_10113_1ng_glyco_hcd203040_compound15.18968.18968.3 | controllerType=0 controllerNumber=1 scan=18968 | 38 |
| 15745:3 | 2 | R.LN[+1702.581]YSLPTGQWVGVQLPR.N | HexNAc(2)Hex(8) | N[+1703] | 1210.878 | 3 | 3630.6194 | 3630.6202 | 0 | -0.2 | 1017 | Specific | 921.86 | 785.24 | 785.24 | 17.76 | 104 | >sp\|P24821\|TENA_HUMAN Tenascin OS=Homo sapiens OX=9606 GN=TNC PE=1 SV=3 | 486 | 20201115_10113_1ng_glyco_hcd203040_compound15.18864.18864.3 | controllerType=0 controllerNumber=1 scan=18864 | 37.7608 |
| 15722:3 | 2 | R.LN[+1702.581]YSLPTGQWVGVQLPR.N | HexNAc(2)Hex(8) | N[+1703] | 1210.8791 | 3 | 3630.6227 | 3630.6202 | 0 | 0.7 | 1017 | Specific | 976.73 | 818.91 | 818.91 | 19.23 | 104 | >sp\|P24821\|TENA_HUMAN Tenascin OS=Homo sapiens OX=9606 GN=TNC PE=1 SV=3 | 486 | 20201115_10113_1ng_glyco_hcd203040_compound15.18840.18840.2 | controllerType=0 controllerNumber=1 scan=18840 | 37.7164 |
| 15890:3 | 2 | R.LN[+1216.423]YSLPTGQWVGVQLPR.N | HexNAc(2)Hex(5) | N[+1216] | 1048.8283 | 3 | 3144.4703 | 3144.4617 | 0 | 2.7 | 1017 | Specific | 1039.39 | 1039.39 | 1039.39 | 20.42 | 104 | >sp\|P24821\|TENA_HUMAN Tenascin OS=Homo sapiens OX=9606 GN=TNC PE=1 SV=3 | 486 | 20201115_10113_1ng_glyco_hcd203040_compound15.19015.19015.3 | controllerType=0 controllerNumber=1 scan=19015 | 38.1061 |
| 15887:3 | 2 | R.LN[+1216.423]YSLPTGQWVGVQLPR.N | HexNAc(2)Hex(5) | N[+1216] | 1048.8283 | 3 | 3144.4703 | 3144.4617 | 0 | 2.7 | 1017 | Specific | 1054.83 | 1054.83 | 1054.83 | 21.42 | 104 | >sp\|P24821\|TENA_HUMAN Tenascin OS=Homo sapiens OX=9606 GN=TNC PE=1 SV=3 | 486 | 20201115_10113_1ng_glyco_hcd203040_compound15.19012.19012.3 | controllerType=0 controllerNumber=1 scan=19012 | 38.1008 |
| 04457:3 | 2 | R.N[+2059.735]TTSYVLR.G | HexNAc(4)Hex(5)Fuc(1)NeuAc(1) | N[+2060] | 1005.0872 | 3 | 3013.247 | 3013.24 | 0 | 2.3 | 1034 | Specific | 273.37 | 186 | 186 | 5.91 | 104 | >sp\|P24821\|TENA_HUMAN Tenascin OS=Homo sapiens OX=9606 GN=TNC PE=1 SV=3 | 486 | 20201115_10113_1ng_glyco_hcd203040_compound15.7148.7148.3 | controllerType=0 controllerNumber=1 scan=7148 | 17.7123 |
| 10653:3 | 3 | K.TVLTPATNHMGN[+1540.529]VTFTIPANR.E | HexNAc(2)Hex(7) | N[+1541] | 1265.9052 | 3 | 3795.7011 | 3795.6886 | 0 | 3.3 | 74 | Specific | 738.9 | 716.19 | 342.24 | 14.99 | 113 | >sp\|P01024\|CO3_HUMAN Complement C3 OS=Homo sapiens OX=9606 GN=C3 PE=1 SV=2 | 492 | 20201115_10113_1ng_glyco_hcd203040_compound15.13576.13576.3 | controllerType=0 controllerNumber=1 scan=13576 | 28.2079 |
| 10650:3 | 3 | K.TVLTPATNHMGN[+1540.529]VTFTIPANR.E | HexNAc(2)Hex(7) | N[+1541] | 1265.9052 | 3 | 3795.7011 | 3795.6886 | 0 | 3.3 | 74 | Specific | 824.3 | 642.47 | 415.31 | 16.99 | 113 | >sp\|P01024\|CO3_HUMAN Complement C3 OS=Homo sapiens OX=9606 GN=C3 PE=1 SV=2 | 492 | 20201115_10113_1ng_glyco_hcd203040_compound15.13573.13573.3 | controllerType=0 controllerNumber=1 scan=13573 | 28.2022 |
| 10687:3 | 3 | K.TVLTPATNHMGN[+1378.476]VTFTIPANR.E | HexNAc(2)Hex(6) | N[+1378] | 1211.8851 | 3 | 3633.6406 | 3633.6358 | 0 | 1.3 | 74 | Specific | 899.76 | 884.57 | 455.27 | 17.8 | 113 | >sp\|P01024\|CO3_HUMAN Complement C3 OS=Homo sapiens OX=9606 GN=C3 PE=1 SV=2 | 492 | 20201115_10113_1ng_glyco_hcd203040_compound15.13611.13611.3 | controllerType=0 controllerNumber=1 scan=13611 | 28.2846 |
| 10685:3 | 3 | K.TVLTPATNHMGN[+1378.476]VTFTIPANR.E | HexNAc(2)Hex(6) | N[+1378] | 1211.8851 | 3 | 3633.6406 | 3633.6358 | 0 | 1.3 | 74 | Specific | 964.5 | 964.5 | 523.39 | 19.8 | 113 | >sp\|P01024\|CO3_HUMAN Complement C3 OS=Homo sapiens OX=9606 GN=C3 PE=1 SV=2 | 492 | 20201115_10113_1ng_glyco_hcd203040_compound15.13609.13609.3 | controllerType=0 controllerNumber=1 scan=13609 | 28.2729 |
| 10632:3 | 3 | K.TVLTPATNHMGN[+1540.529]VTFTIPANR.E | HexNAc(2)Hex(7) | N[+1541] | 1265.9026 | 3 | 3795.6934 | 3795.6886 | 0 | 1.3 | 74 | Specific | 969.88 | 875.92 | 432.52 | 19.8 | 113 | >sp\|P01024\|CO3_HUMAN Complement C3 OS=Homo sapiens OX=9606 GN=C3 PE=1 SV=2 | 492 | 20201115_10113_1ng_glyco_hcd203040_compound15.13554.13554.3 | controllerType=0 controllerNumber=1 scan=13554 | 28.1711 |
| 10630:3 | 3 | K.TVLTPATNHMGN[+1540.529]VTFTIPANR.E | HexNAc(2)Hex(7) | N[+1541] | 1265.9026 | 3 | 3795.6934 | 3795.6886 | 0 | 1.3 | 74 | Specific | 1023.15 | 915.98 | 522.53 | 20.8 | 113 | >sp\|P01024\|CO3_HUMAN Complement C3 OS=Homo sapiens OX=9606 GN=C3 PE=1 SV=2 | 492 | 20201115_10113_1ng_glyco_hcd203040_compound15.13552.13552.3 | controllerType=0 controllerNumber=1 scan=13552 | 28.1615 |
| 07949:3 | 5 | R.FSDGLESN[+1864.634]SSTQFEVK.K | HexNAc(2)Hex(9) | N[+1865] | 1213.8237 | 3 | 3639.4565 | 3639.4472 | 0 | 2.6 | 219 | Specific | 241.05 | 241.05 | 241.05 | 6.5 | 93 | >sp\|P0C0L4\|CO4A_HUMAN Complement C4-A OS=Homo sapiens OX=9606 GN=C4A PE=1 SV=2 | 1235 | 20201115_10113_1ng_glyco_hcd203040_compound15.10762.10762.3 | controllerType=0 controllerNumber=1 scan=10762 | 23.5013 |
| 07952:3 | 5 | R.FSDGLESN[+1864.634]SSTQFEVK.K | HexNAc(2)Hex(9) | N[+1865] | 1213.8237 | 3 | 3639.4565 | 3639.4472 | 0 | 2.6 | 219 | Specific | 511.96 | 511.96 | 511.96 | 9.94 | 93 | >sp\|P0C0L4\|CO4A_HUMAN Complement C4-A OS=Homo sapiens OX=9606 GN=C4A PE=1 SV=2 | 1235 | 20201115_10113_1ng_glyco_hcd203040_compound15.10765.10765.3 | controllerType=0 controllerNumber=1 scan=10765 | 23.5054 |
| 05350:3 | 5 | R.FSDGLESN[+1864.634]SSTQFEVKK.Y | HexNAc(2)Hex(9) | N[+1865] | 1256.5237 | 3 | 3767.5567 | 3767.5421 | 0 | 3.9 | 219 | Specific | 632.17 | 529.46 | 529.46 | 11.71 | 93 | >sp\|P0C0L4\|CO4A_HUMAN Complement C4-A OS=Homo sapiens OX=9606 GN=C4A PE=1 SV=2 | 1235 | 20201115_10113_1ng_glyco_hcd203040_compound15.8072.8072.3 | controllerType=0 controllerNumber=1 scan=8072 | 19.1718 |
| 05347:3 | 5 | R.FSDGLESN[+1864.634]SSTQFEVKK.Y | HexNAc(2)Hex(9) | N[+1865] | 1256.5237 | 3 | 3767.5567 | 3767.5421 | 0 | 3.9 | 219 | Specific | 843.45 | 592.01 | 592.01 | 15.71 | 93 | >sp\|P0C0L4\|CO4A_HUMAN Complement C4-A OS=Homo sapiens OX=9606 GN=C4A PE=1 SV=2 | 1235 | 20201115_10113_1ng_glyco_hcd203040_compound15.8069.8069.3 | controllerType=0 controllerNumber=1 scan=8069 | 19.1673 |
| 13929:3 | 5 | R.TYNVLDMKN[+2059.735]TTC[+57.021]QDLQIEVTVK.G | HexNAc(4)Hex(5)Fuc(1)NeuAc(1) | C[+57], N[+2060] | 1558.3511 | 3 | 4673.0388 | 4673.0248 | 0 | 3 | 1383 | Specific | 218.27 | 183.36 | 183.36 | 4.96 | 93 | >sp\|P0C0L4\|CO4A_HUMAN Complement C4-A OS=Homo sapiens OX=9606 GN=C4A PE=1 SV=2 | 1235 | 20201115_10113_1ng_glyco_hcd203040_compound15.16979.16979.3 | controllerType=0 controllerNumber=1 scan=16979 | 34.1776 |
| 17788:3 | 7 | R.GLSFDVSLEVSQGPGLLN[+2863.041]DTK.V | HexNAc(5)Hex(6)Fuc(4)NeuAc(1) | N[+2863] | 1680.3928 | 3 | 5039.1638 | 5039.1541 | 0 | 1.9 | 118 | Specific | 328.67 | 315.46 | 315.46 | 7.67 | 74 | >sp\|P01833\|PIGR_HUMAN Polymeric immunoglobulin receptor OS=Homo sapiens OX=9606 GN=PIGR PE=1 SV=4 | 902 | 20201115_10113_1ng_glyco_hcd203040_compound15.20997.20997.3 | controllerType=0 controllerNumber=1 scan=20997 | 41.9147 |
| 17786:3 | 7 | R.GLSFDVSLEVSQGPGLLN[+2863.041]DTK.V | HexNAc(5)Hex(6)Fuc(4)NeuAc(1) | N[+2863] | 1680.3928 | 3 | 5039.1638 | 5039.1541 | 0 | 1.9 | 118 | Specific | 657.66 | 634.97 | 634.97 | 13.92 | 74 | >sp\|P01833\|PIGR_HUMAN Polymeric immunoglobulin receptor OS=Homo sapiens OX=9606 GN=PIGR PE=1 SV=4 | 902 | 20201115_10113_1ng_glyco_hcd203040_compound15.20995.20995.3 | controllerType=0 controllerNumber=1 scan=20995 | 41.909 |
| 18102:3 | 7 | R.GLSFDVSLEVSQGPGLLN[+2059.735]DTK.V | HexNAc(4)Hex(5)Fuc(1)NeuAc(1) | N[+2060] | 1412.6276 | 3 | 4235.8683 | 4235.8481 | 0 | 4.8 | 118 | Specific | 712.96 | 696.28 | 696.28 | 14.49 | 74 | >sp\|P01833\|PIGR_HUMAN Polymeric immunoglobulin receptor OS=Homo sapiens OX=9606 GN=PIGR PE=1 SV=4 | 902 | 20201115_10113_1ng_glyco_hcd203040_compound15.21326.21326.3 | controllerType=0 controllerNumber=1 scan=21326 | 42.6865 |
| 17863:3 | 7 | R.GLSFDVSLEVSQGPGLLN[+2716.983]DTK.V | HexNAc(5)Hex(6)Fuc(3)NeuAc(1) | N[+2717] | 1631.7077 | 3 | 4893.1085 | 4893.0962 | 0 | 2.5 | 118 | Specific | 798.3 | 707.78 | 707.78 | 15.92 | 74 | >sp\|P01833\|PIGR_HUMAN Polymeric immunoglobulin receptor OS=Homo sapiens OX=9606 GN=PIGR PE=1 SV=4 | 902 | 20201115_10113_1ng_glyco_hcd203040_compound15.21075.21075.3 | controllerType=0 controllerNumber=1 scan=21075 | 42.0625 |
| 19811:4 | 7 | R.GLSFDVSLEVSQGPGLLN[+2350.830]DTK.V | HexNAc(4)Hex(5)Fuc(1)NeuAc(2) | N[+2351] | 1132.5005 | 4 | 4526.9803 | 4526.9436 | 0 | 8.1 | 118 | Specific | 803.3 | 772.18 | 772.18 | 15.95 | 74 | >sp\|P01833\|PIGR_HUMAN Polymeric immunoglobulin receptor OS=Homo sapiens OX=9606 GN=PIGR PE=1 SV=4 | 902 | 20201115_10113_1ng_glyco_hcd203040_compound15.23108.23108.4 | controllerType=0 controllerNumber=1 scan=23108 | 46.6045 |
| 17865:3 | 7 | R.GLSFDVSLEVSQGPGLLN[+2716.983]DTK.V | HexNAc(5)Hex(6)Fuc(3)NeuAc(1) | N[+2717] | 1631.7077 | 3 | 4893.1085 | 4893.0962 | 0 | 2.5 | 118 | Specific | 841.51 | 821.05 | 821.05 | 16.92 | 74 | >sp\|P01833\|PIGR_HUMAN Polymeric immunoglobulin receptor OS=Homo sapiens OX=9606 GN=PIGR PE=1 SV=4 | 902 | 20201115_10113_1ng_glyco_hcd203040_compound15.21077.21077.3 | controllerType=0 controllerNumber=1 scan=21077 | 42.072 |
| 19814:4 | 7 | R.GLSFDVSLEVSQGPGLLN[+2350.830]DTK.V | HexNAc(4)Hex(5)Fuc(1)NeuAc(2) | N[+2351] | 1132.5005 | 4 | 4526.9803 | 4526.9436 | 0 | 8.1 | 118 | Specific | 906.11 | 846.19 | 846.19 | 17.95 | 74 | >sp\|P01833\|PIGR_HUMAN Polymeric immunoglobulin receptor OS=Homo sapiens OX=9606 GN=PIGR PE=1 SV=4 | 902 | 20201115_10113_1ng_glyco_hcd203040_compound15.23111.23111.4 | controllerType=0 controllerNumber=1 scan=23111 | 46.6094 |
| 19805:3 | 7 | R.GLSFDVSLEVSQGPGLLN[+2350.830]DTK.V | HexNAc(4)Hex(5)Fuc(1)NeuAc(2) | N[+2351] | 1509.6653 | 3 | 4526.9815 | 4526.9436 | 0 | 8.4 | 118 | Specific | 994.66 | 953.77 | 953.77 | 18.92 | 74 | >sp\|P01833\|PIGR_HUMAN Polymeric immunoglobulin receptor OS=Homo sapiens OX=9606 GN=PIGR PE=1 SV=4 | 902 | 20201115_10113_1ng_glyco_hcd203040_compound15.23102.23102.3 | controllerType=0 controllerNumber=1 scan=23102 | 46.5897 |
| 21066:4 | 7 | K.QIGLYPVLVIDSSGYVNPN[+2350.830]YTGR.I | HexNAc(4)Hex(5)Fuc(1)NeuAc(2) | N[+2351] | 1219.7898 | 4 | 4876.1375 | 4876.1338 | 0 | 0.8 | 168 | Specific | 531.97 | 432.37 | 432.37 | 10.37 | 74 | >sp\|P01833\|PIGR_HUMAN Polymeric immunoglobulin receptor OS=Homo sapiens OX=9606 GN=PIGR PE=1 SV=4 | 902 | 20201115_10113_1ng_glyco_hcd203040_compound15.24421.24421.4 | controllerType=0 controllerNumber=1 scan=24421 | 49.5601 |
| 21063:4 | 7 | K.QIGLYPVLVIDSSGYVNPN[+2350.830]YTGR.I | HexNAc(4)Hex(5)Fuc(1)NeuAc(2) | N[+2351] | 1219.7898 | 4 | 4876.1375 | 4876.1338 | 0 | 0.8 | 168 | Specific | 580.31 | 491.37 | 491.37 | 11.74 | 74 | >sp\|P01833\|PIGR_HUMAN Polymeric immunoglobulin receptor OS=Homo sapiens OX=9606 GN=PIGR PE=1 SV=4 | 902 | 20201115_10113_1ng_glyco_hcd203040_compound15.24418.24418.4 | controllerType=0 controllerNumber=1 scan=24418 | 49.551 |
| 19240:4 | 7 | K.QIGLYPVLVIDSSGYVNPN[+2059.735]YTGR.I | HexNAc(4)Hex(5)Fuc(1)NeuAc(1) | N[+2060] | 1147.0263 | 4 | 4585.0833 | 4585.0384 | 0 | 9.8 | 168 | Specific | 822.15 | 769.77 | 769.77 | 16.01 | 74 | >sp\|P01833\|PIGR_HUMAN Polymeric immunoglobulin receptor OS=Homo sapiens OX=9606 GN=PIGR PE=1 SV=4 | 902 | 20201115_10113_1ng_glyco_hcd203040_compound15.22511.22511.4 | controllerType=0 controllerNumber=1 scan=22511 | 45.2598 |
| 19243:4 | 7 | K.QIGLYPVLVIDSSGYVNPN[+2059.735]YTGR.I | HexNAc(4)Hex(5)Fuc(1)NeuAc(1) | N[+2060] | 1147.0263 | 4 | 4585.0833 | 4585.0384 | 0 | 9.8 | 168 | Specific | 864.79 | 833.66 | 833.66 | 17.01 | 74 | >sp\|P01833\|PIGR_HUMAN Polymeric immunoglobulin receptor OS=Homo sapiens OX=9606 GN=PIGR PE=1 SV=4 | 902 | 20201115_10113_1ng_glyco_hcd203040_compound15.22514.22514.4 | controllerType=0 controllerNumber=1 scan=22514 | 45.2648 |
| 21145:4 | 7 | K.QIGLYPVLVIDSSGYVNPN[+2204.772]YTGR.I | HexNAc(4)Hex(5)NeuAc(2) | N[+2205] | 1183.2776 | 4 | 4730.0887 | 4730.0759 | 0 | 2.7 | 168 | Specific | 867.96 | 746.81 | 746.81 | 17.94 | 74 | >sp\|P01833\|PIGR_HUMAN Polymeric immunoglobulin receptor OS=Homo sapiens OX=9606 GN=PIGR PE=1 SV=4 | 902 | 20201115_10113_1ng_glyco_hcd203040_compound15.24504.24504.4 | controllerType=0 controllerNumber=1 scan=24504 | 49.7513 |
| 21142:4 | 7 | K.QIGLYPVLVIDSSGYVNPN[+2204.772]YTGR.I | HexNAc(4)Hex(5)NeuAc(2) | N[+2205] | 1183.2776 | 4 | 4730.0887 | 4730.0759 | 0 | 2.7 | 168 | Specific | 889.43 | 754.78 | 754.78 | 17.94 | 74 | >sp\|P01833\|PIGR_HUMAN Polymeric immunoglobulin receptor OS=Homo sapiens OX=9606 GN=PIGR PE=1 SV=4 | 902 | 20201115_10113_1ng_glyco_hcd203040_compound15.24501.24501.4 | controllerType=0 controllerNumber=1 scan=24501 | 49.7441 |
| 22678:4 | 7 | K.AQYEGRLSLLEEPGN[+1663.608]GTFTVILNQLTSR.D | HexNAc(5)Hex(4) | N[+1664] | 1193.3068 | 4 | 4770.2054 | 4770.2249 | 0 | -4.1 | 407 | Specific | 262.62 | 262.62 | 262.62 | 3.47 | 74 | >sp\|P01833\|PIGR_HUMAN Polymeric immunoglobulin receptor OS=Homo sapiens OX=9606 GN=PIGR PE=1 SV=4 | 902 | 20201115_10113_1ng_glyco_hcd203040_compound15.26111.26111.4 | controllerType=0 controllerNumber=1 scan=26111 | 53.1719 |
| 28535:3 | 7 | R.[+42.011]LSLLEEPGN[+2351.851]GTFTVILNQLTSR.D | HexNAc(4)Hex(5)Fuc(3)NeuAc(1) | .[42], N[+2352] | 1599.3837 | 3 | 4796.1366 | 4796.1539 | 0 | -3.6 | 413 | Specific | 208.75 | 185.24 | 185.24 | 3.83 | 74 | >sp\|P01833\|PIGR_HUMAN Polymeric immunoglobulin receptor OS=Homo sapiens OX=9606 GN=PIGR PE=1 SV=4 | 902 | 20201115_10113_1ng_glyco_hcd203040_compound15.32252.32252.3 | controllerType=0 controllerNumber=1 scan=32252 | 65.1254 |
| 28537:3 | 7 | R.[+42.011]LSLLEEPGN[+2351.851]GTFTVILNQLTSR.D | HexNAc(4)Hex(5)Fuc(3)NeuAc(1) | .[42], N[+2352] | 1599.3837 | 3 | 4796.1366 | 4796.1539 | 0 | -3.6 | 413 | Specific | 215.16 | 190.58 | 190.58 | 3.83 | 74 | >sp\|P01833\|PIGR_HUMAN Polymeric immunoglobulin receptor OS=Homo sapiens OX=9606 GN=PIGR PE=1 SV=4 | 902 | 20201115_10113_1ng_glyco_hcd203040_compound15.32254.32254.3 | controllerType=0 controllerNumber=1 scan=32254 | 65.1291 |
| 28613:3 | 7 | R.[+42.011]LSLLEEPGN[+2351.851]GTFTVILNQLTSR.D | HexNAc(4)Hex(5)Fuc(3)NeuAc(1) | .[42], N[+2352] | 1599.3863 | 3 | 4796.1443 | 4796.1539 | 0 | -2 | 413 | Specific | 239.28 | 228.53 | 228.53 | 5.4 | 74 | >sp\|P01833\|PIGR_HUMAN Polymeric immunoglobulin receptor OS=Homo sapiens OX=9606 GN=PIGR PE=1 SV=4 | 902 | 20201115_10113_1ng_glyco_hcd203040_compound15.32333.32333.3 | controllerType=0 controllerNumber=1 scan=32333 | 65.2741 |
| 22762:3 | 7 | R.LSLLEEPGN[+2350.830]GTFTVILNQLTSR.D | HexNAc(4)Hex(5)Fuc(1)NeuAc(2) | N[+2351] | 1585.0514 | 3 | 4753.1398 | 4753.1229 | 0 | 3.6 | 413 | Specific | 454.53 | 364.91 | 364.91 | 8.79 | 74 | >sp\|P01833\|PIGR_HUMAN Polymeric immunoglobulin receptor OS=Homo sapiens OX=9606 GN=PIGR PE=1 SV=4 | 902 | 20201115_10113_1ng_glyco_hcd203040_compound15.26199.26199.3 | controllerType=0 controllerNumber=1 scan=26199 | 53.3795 |
| 22765:3 | 7 | R.LSLLEEPGN[+2350.830]GTFTVILNQLTSR.D | HexNAc(4)Hex(5)Fuc(1)NeuAc(2) | N[+2351] | 1585.0514 | 3 | 4753.1398 | 4753.1229 | 0 | 3.6 | 413 | Specific | 740.75 | 544.3 | 544.3 | 14.9 | 74 | >sp\|P01833\|PIGR_HUMAN Polymeric immunoglobulin receptor OS=Homo sapiens OX=9606 GN=PIGR PE=1 SV=4 | 902 | 20201115_10113_1ng_glyco_hcd203040_compound15.26202.26202.3 | controllerType=0 controllerNumber=1 scan=26202 | 53.3837 |
| 20783:4 | 7 | R.LSLLEEPGN[+2059.735]GTFTVILNQLTSR.D | HexNAc(4)Hex(5)Fuc(1)NeuAc(1) | N[+2060] | 1116.2638 | 4 | 4462.0335 | 4462.0275 | 0 | 1.3 | 413 | Specific | 810.68 | 610.92 | 610.92 | 16.74 | 74 | >sp\|P01833\|PIGR_HUMAN Polymeric immunoglobulin receptor OS=Homo sapiens OX=9606 GN=PIGR PE=1 SV=4 | 902 | 20201115_10113_1ng_glyco_hcd203040_compound15.24124.24124.4 | controllerType=0 controllerNumber=1 scan=24124 | 48.8772 |
| 14781:3 | 7 | K.IIEGEPNLKVPGN[+1913.677]VTAVLGETLK.V | HexNAc(4)Hex(5)NeuAc(1) | N[+1914] | 1435.6906 | 3 | 4305.0573 | 4305.0264 | 0 | 7.2 | 457 | Specific | 399.95 | 399.95 | 399.95 | 8.75 | 74 | >sp\|P01833\|PIGR_HUMAN Polymeric immunoglobulin receptor OS=Homo sapiens OX=9606 GN=PIGR PE=1 SV=4 | 902 | 20201115_10113_1ng_glyco_hcd203040_compound15.17865.17865.3 | controllerType=0 controllerNumber=1 scan=17865 | 35.9514 |
| 14784:3 | 7 | K.IIEGEPNLKVPGN[+1913.677]VTAVLGETLK.V | HexNAc(4)Hex(5)NeuAc(1) | N[+1914] | 1435.6906 | 3 | 4305.0573 | 4305.0264 | 0 | 7.2 | 457 | Specific | 438.43 | 417.68 | 417.68 | 8.78 | 74 | >sp\|P01833\|PIGR_HUMAN Polymeric immunoglobulin receptor OS=Homo sapiens OX=9606 GN=PIGR PE=1 SV=4 | 902 | 20201115_10113_1ng_glyco_hcd203040_compound15.17868.17868.3 | controllerType=0 controllerNumber=1 scan=17868 | 35.9561 |
| 16534:4 | 7 | K.IIEGEPNLKVPGN[+2059.735]VTAVLGETLK.V | HexNAc(4)Hex(5)Fuc(1)NeuAc(1) | N[+2060] | 1113.5301 | 4 | 4451.0985 | 4451.0843 | 0 | 3.2 | 457 | Specific | 701.08 | 495.67 | 495.67 | 14.51 | 74 | >sp\|P01833\|PIGR_HUMAN Polymeric immunoglobulin receptor OS=Homo sapiens OX=9606 GN=PIGR PE=1 SV=4 | 902 | 20201115_10113_1ng_glyco_hcd203040_compound15.19691.19691.4 | controllerType=0 controllerNumber=1 scan=19691 | 39.3396 |
| 18320:3 | 7 | K.IIEGEPNLKVPGN[+2350.830]VTAVLGETLK.V | HexNAc(4)Hex(5)Fuc(1)NeuAc(2) | N[+2351] | 1581.4029 | 3 | 4742.1941 | 4742.1797 | 0 | 3 | 457 | Specific | 707.34 | 665.17 | 665.17 | 14.6 | 74 | >sp\|P01833\|PIGR_HUMAN Polymeric immunoglobulin receptor OS=Homo sapiens OX=9606 GN=PIGR PE=1 SV=4 | 902 | 20201115_10113_1ng_glyco_hcd203040_compound15.21553.21553.3 | controllerType=0 controllerNumber=1 scan=21553 | 43.1806 |
| 16463:4 | 7 | K.IIEGEPNLKVPGN[+2204.772]VTAVLGETLK.V | HexNAc(4)Hex(5)NeuAc(2) | N[+2205] | 1149.7939 | 4 | 4596.1536 | 4596.1218 | 0 | 6.9 | 457 | Specific | 756.1 | 728.34 | 728.34 | 14.55 | 74 | >sp\|P01833\|PIGR_HUMAN Polymeric immunoglobulin receptor OS=Homo sapiens OX=9606 GN=PIGR PE=1 SV=4 | 902 | 20201115_10113_1ng_glyco_hcd203040_compound15.19617.19617.4 | controllerType=0 controllerNumber=1 scan=19617 | 39.1812 |
| 16531:4 | 7 | K.IIEGEPNLKVPGN[+2059.735]VTAVLGETLK.V | HexNAc(4)Hex(5)Fuc(1)NeuAc(1) | N[+2060] | 1113.5301 | 4 | 4451.0985 | 4451.0843 | 0 | 3.2 | 457 | Specific | 763.88 | 589.72 | 589.72 | 15.51 | 74 | >sp\|P01833\|PIGR_HUMAN Polymeric immunoglobulin receptor OS=Homo sapiens OX=9606 GN=PIGR PE=1 SV=4 | 902 | 20201115_10113_1ng_glyco_hcd203040_compound15.19688.19688.4 | controllerType=0 controllerNumber=1 scan=19688 | 39.3344 |
| 16460:4 | 7 | K.IIEGEPNLKVPGN[+2204.772]VTAVLGETLK.V | HexNAc(4)Hex(5)NeuAc(2) | N[+2205] | 1149.7939 | 4 | 4596.1536 | 4596.1218 | 0 | 6.9 | 457 | Specific | 815.87 | 815.87 | 815.87 | 15.55 | 74 | >sp\|P01833\|PIGR_HUMAN Polymeric immunoglobulin receptor OS=Homo sapiens OX=9606 GN=PIGR PE=1 SV=4 | 902 | 20201115_10113_1ng_glyco_hcd203040_compound15.19614.19614.4 | controllerType=0 controllerNumber=1 scan=19614 | 39.1756 |
| 16477:3 | 7 | K.IIEGEPNLKVPGN[+2205.793]VTAVLGETLK.V | HexNAc(4)Hex(5)Fuc(2)NeuAc(1) | N[+2206] | 1533.056 | 3 | 4597.1533 | 4597.1422 | 0 | 2.4 | 457 | Specific | 845.04 | 737.16 | 737.16 | 16.61 | 74 | >sp\|P01833\|PIGR_HUMAN Polymeric immunoglobulin receptor OS=Homo sapiens OX=9606 GN=PIGR PE=1 SV=4 | 902 | 20201115_10113_1ng_glyco_hcd203040_compound15.19632.19632.3 | controllerType=0 controllerNumber=1 scan=19632 | 39.2122 |
| 16475:3 | 7 | K.IIEGEPNLKVPGN[+2205.793]VTAVLGETLK.V | HexNAc(4)Hex(5)Fuc(2)NeuAc(1) | N[+2206] | 1533.056 | 3 | 4597.1533 | 4597.1422 | 0 | 2.4 | 457 | Specific | 884.67 | 817.42 | 817.42 | 17.61 | 74 | >sp\|P01833\|PIGR_HUMAN Polymeric immunoglobulin receptor OS=Homo sapiens OX=9606 GN=PIGR PE=1 SV=4 | 902 | 20201115_10113_1ng_glyco_hcd203040_compound15.19630.19630.3 | controllerType=0 controllerNumber=1 scan=19630 | 39.2077 |
| 18317:3 | 7 | K.IIEGEPNLKVPGN[+2350.830]VTAVLGETLK.V | HexNAc(4)Hex(5)Fuc(1)NeuAc(2) | N[+2351] | 1581.4029 | 3 | 4742.1941 | 4742.1797 | 0 | 3 | 457 | Specific | 914.13 | 815.04 | 815.04 | 18.6 | 74 | >sp\|P01833\|PIGR_HUMAN Polymeric immunoglobulin receptor OS=Homo sapiens OX=9606 GN=PIGR PE=1 SV=4 | 902 | 20201115_10113_1ng_glyco_hcd203040_compound15.21550.21550.3 | controllerType=0 controllerNumber=1 scan=21550 | 43.1748 |
| 17565:4 | 7 | K.IIEGEPNLKVPGN[+2350.830]VTAVLGETLKVPC[+57.021]HFPC[+57.021]K.F | HexNAc(4)Hex(5)Fuc(1)NeuAc(2) | C[+57]*2, N[+2351] | 1442.6693 | 4 | 5767.6554 | 5767.6372 | 0 | 3.2 | 457 | Specific | 404.37 | 404.37 | 404.37 | 8.52 | 74 | >sp\|P01833\|PIGR_HUMAN Polymeric immunoglobulin receptor OS=Homo sapiens OX=9606 GN=PIGR PE=1 SV=4 | 902 | 20201115_10113_1ng_glyco_hcd203040_compound15.20764.20764.4 | controllerType=0 controllerNumber=1 scan=20764 | 41.4324 |
| 17563:4 | 7 | K.IIEGEPNLKVPGN[+2350.830]VTAVLGETLKVPC[+57.021]HFPC[+57.021]K.F | HexNAc(4)Hex(5)Fuc(1)NeuAc(2) | C[+57]*2, N[+2351] | 1442.6693 | 4 | 5767.6554 | 5767.6372 | 0 | 3.2 | 457 | Specific | 523.13 | 372.69 | 372.69 | 7.53 | 74 | >sp\|P01833\|PIGR_HUMAN Polymeric immunoglobulin receptor OS=Homo sapiens OX=9606 GN=PIGR PE=1 SV=4 | 902 | 20201115_10113_1ng_glyco_hcd203040_compound15.20762.20762.4 | controllerType=0 controllerNumber=1 scan=20762 | 41.4267 |
| 15962:4 | 7 | K.IIEGEPNLKVPGN[+2059.735]VTAVLGETLKVPC[+57.021]HFPC[+57.021]K.F | HexNAc(4)Hex(5)Fuc(1)NeuAc(1) | C[+57]*2, N[+2060] | 1369.8925 | 4 | 5476.548 | 5476.5418 | 0 | 1.1 | 457 | Specific | 642.24 | 605.32 | 605.32 | 11.22 | 74 | >sp\|P01833\|PIGR_HUMAN Polymeric immunoglobulin receptor OS=Homo sapiens OX=9606 GN=PIGR PE=1 SV=4 | 902 | 20201115_10113_1ng_glyco_hcd203040_compound15.19090.19090.4 | controllerType=0 controllerNumber=1 scan=19090 | 38.2526 |

Table S4. Cont.

| Query #:z | Protein Rank | Glycopeptide | Glycan | Modification Type(s) | Observed m/z | z | Observed (M+H) | Calc. mass (M+H) | Off-by-x error | Mass error (ppm) | StartPosition | Cleavage | Score | Delta | Delta Mod | |Log Prob| | # of unique peptides | Protein Name | Protein DB number | Comment | Scan # | Scan Time |
|---|---|---|---|---|---|---|---|---|---|---|---|---|---|---|---|---|---|---|---|---|---|---|
| 17413:4 | 7 | K.IIEGEPNLKVPGN[+2350.830]VTAVLGETLKVPC[+57.021]HFPC[+57.021]K.F | HexNAc(4)Hex(5)Fuc(1)NeuAc(2) | C[+57]*2, N[+2351] | 1442.6705 | 4 | 5767.6603 | 5767.6372 | 0 | 4 | 457 | Specific | 711.36 | 678.45 | 678.45 | 13 | 74 | >sp|P01833|PIGR_HUMAN Polymeric immunoglobulin receptor OS=Homo sapiens OX=9606 GN=PIGR PE=1 SV=4 | 902 | 20201115_10113_1ng_glyco_hcd203040_compound15.20606.20606.4 | controllerType=0 controllerNumber=1 scan=20606 | 41.1254 |
| 17407:5 | 7 | K.IIEGEPNLKVPGN[+2350.830]VTAVLGETLKVPC[+57.021]HFPC[+57.021]K.F | HexNAc(4)Hex(5)Fuc(1)NeuAc(2) | C[+57]*2, N[+2351] | 1154.3364 | 5 | 5767.653 | 5767.6372 | 0 | 2.7 | 457 | Specific | 805.04 | 603.63 | 603.63 | 16 | 74 | >sp|P01833|PIGR_HUMAN Polymeric immunoglobulin receptor OS=Homo sapiens OX=9606 GN=PIGR PE=1 SV=4 | 902 | 20201115_10113_1ng_glyco_hcd203040_compound15.20600.20600.5 | controllerType=0 controllerNumber=1 scan=20600 | 41.1114 |
| 15974:4 | 7 | K.IIEGEPNLKVPGN[+2205.793]VTAVLGETLKVPC[+57.021]HFPC[+57.021]K.F | HexNAc(4)Hex(5)Fuc(2)NeuAc(1) | C[+57]*2, N[+2206] | 1406.4105 | 4 | 5622.6203 | 5622.5997 | 0 | 3.7 | 457 | Specific | 807.34 | 807.34 | 807.34 | 15.41 | 74 | >sp|P01833|PIGR_HUMAN Polymeric immunoglobulin receptor OS=Homo sapiens OX=9606 GN=PIGR PE=1 SV=4 | 902 | 20201115_10113_1ng_glyco_hcd203040_compound15.19103.19103.4 | controllerType=0 controllerNumber=1 scan=19103 | 38.2807 |
| 16005:4 | 7 | K.IIEGEPNLKVPGN[+2059.735]VTAVLGETLKVPC[+57.021]HFPC[+57.021]K.F | HexNAc(4)Hex(5)Fuc(1)NeuAc(1) | C[+57]*2, N[+2060] | 1369.8914 | 4 | 5476.5436 | 5476.5418 | 0 | 0.3 | 457 | Specific | 814.09 | 768.25 | 768.25 | 14.75 | 74 | >sp|P01833|PIGR_HUMAN Polymeric immunoglobulin receptor OS=Homo sapiens OX=9606 GN=PIGR PE=1 SV=4 | 902 | 20201115_10113_1ng_glyco_hcd203040_compound15.19135.19135.4 | controllerType=0 controllerNumber=1 scan=19135 | 38.3365 |
| 16001:5 | 7 | K.IIEGEPNLKVPGN[+2059.735]VTAVLGETLKVPC[+57.021]HFPC[+57.021]K.F | HexNAc(4)Hex(5)Fuc(1)NeuAc(1) | C[+57]*2, N[+2060] | 1096.1157 | 5 | 5476.5495 | 5476.5418 | 0 | 1.4 | 457 | Specific | 823.87 | 774.09 | 774.09 | 15.8 | 74 | >sp|P01833|PIGR_HUMAN Polymeric immunoglobulin receptor OS=Homo sapiens OX=9606 GN=PIGR PE=1 SV=4 | 902 | 20201115_10113_1ng_glyco_hcd203040_compound15.19131.19131.5 | controllerType=0 controllerNumber=1 scan=19131 | 38.3303 |
| 15972:4 | 7 | K.IIEGEPNLKVPGN[+2205.793]VTAVLGETLKVPC[+57.021]HFPC[+57.021]K.F | HexNAc(4)Hex(5)Fuc(2)NeuAc(1) | C[+57]*2, N[+2206] | 1406.4105 | 4 | 5622.6203 | 5622.5997 | 0 | 3.7 | 457 | Specific | 868.13 | 837.55 | 837.55 | 16.41 | 74 | >sp|P01833|PIGR_HUMAN Polymeric immunoglobulin receptor OS=Homo sapiens OX=9606 GN=PIGR PE=1 SV=4 | 902 | 20201115_10113_1ng_glyco_hcd203040_compound15.19101.19101.4 | controllerType=0 controllerNumber=1 scan=19101 | 38.276 |
| 16002:4 | 7 | K.IIEGEPNLKVPGN[+2059.735]VTAVLGETLKVPC[+57.021]HFPC[+57.021]K.F | HexNAc(4)Hex(5)Fuc(1)NeuAc(1) | C[+57]*2, N[+2060] | 1369.8914 | 4 | 5476.5436 | 5476.5418 | 0 | 0.3 | 457 | Specific | 895.58 | 816.16 | 816.16 | 15.75 | 74 | >sp|P01833|PIGR_HUMAN Polymeric immunoglobulin receptor OS=Homo sapiens OX=9606 GN=PIGR PE=1 SV=4 | 902 | 20201115_10113_1ng_glyco_hcd203040_compound15.19132.19132.4 | controllerType=0 controllerNumber=1 scan=19132 | 38.331 |
| 15947:5 | 7 | K.IIEGEPNLKVPGN[+2205.793]VTAVLGETLKVPC[+57.021]HFPC[+57.021]K.F | HexNAc(4)Hex(5)Fuc(2)NeuAc(1) | C[+57]*2, N[+2206] | 1125.3285 | 5 | 5622.6134 | 5622.5997 | 0 | 2.4 | 457 | Specific | 902.28 | 842.42 | 842.42 | 18.02 | 74 | >sp|P01833|PIGR_HUMAN Polymeric immunoglobulin receptor OS=Homo sapiens OX=9606 GN=PIGR PE=1 SV=4 | 902 | 20201115_10113_1ng_glyco_hcd203040_compound15.19075.19075.5 | controllerType=0 controllerNumber=1 scan=19075 | 38.2287 |
| 16004:5 | 7 | K.IIEGEPNLKVPGN[+2059.735]VTAVLGETLKVPC[+57.021]HFPC[+57.021]K.F | HexNAc(4)Hex(5)Fuc(1)NeuAc(1) | C[+57]*2, N[+2060] | 1096.1157 | 5 | 5476.5495 | 5476.5418 | 0 | 1.4 | 457 | Specific | 908.59 | 779.84 | 779.84 | 17.8 | 74 | >sp|P01833|PIGR_HUMAN Polymeric immunoglobulin receptor OS=Homo sapiens OX=9606 GN=PIGR PE=1 SV=4 | 902 | 20201115_10113_1ng_glyco_hcd203040_compound15.19134.19134.5 | controllerType=0 controllerNumber=1 scan=19134 | 38.3348 |
| 17410:4 | 7 | K.IIEGEPNLKVPGN[+2350.830]VTAVLGETLKVPC[+57.021]HFPC[+57.021]K.F | HexNAc(4)Hex(5)Fuc(1)NeuAc(2) | C[+57]*2, N[+2351] | 1442.6705 | 4 | 5767.6603 | 5767.6372 | 0 | 4 | 457 | Specific | 931.14 | 710.66 | 710.66 | 17 | 74 | >sp|P01833|PIGR_HUMAN Polymeric immunoglobulin receptor OS=Homo sapiens OX=9606 GN=PIGR PE=1 SV=4 | 902 | 20201115_10113_1ng_glyco_hcd203040_compound15.20603.20603.4 | controllerType=0 controllerNumber=1 scan=20603 | 41.1198 |
| 15945:5 | 7 | K.IIEGEPNLKVPGN[+2205.793]VTAVLGETLKVPC[+57.021]HFPC[+57.021]K.F | HexNAc(4)Hex(5)Fuc(2)NeuAc(1) | C[+57]*2, N[+2206] | 1125.3285 | 5 | 5622.6134 | 5622.5997 | 0 | 2.4 | 457 | Specific | 1029.93 | 919.77 | 919.77 | 20.02 | 74 | >sp|P01833|PIGR_HUMAN Polymeric immunoglobulin receptor OS=Homo sapiens OX=9606 GN=PIGR PE=1 SV=4 | 902 | 20201115_10113_1ng_glyco_hcd203040_compound15.19073.19073.5 | controllerType=0 controllerNumber=1 scan=19073 | 38.2233 |
| 17404:5 | 7 | K.IIEGEPNLKVPGN[+2350.830]VTAVLGETLKVPC[+57.021]HFPC[+57.021]K.F | HexNAc(4)Hex(5)Fuc(1)NeuAc(2) | C[+57]*2, N[+2351] | 1154.3364 | 5 | 5767.653 | 5767.6372 | 0 | 2.7 | 457 | Specific | 1064.52 | 858.51 | 858.51 | 21 | 74 | >sp|P01833|PIGR_HUMAN Polymeric immunoglobulin receptor OS=Homo sapiens OX=9606 GN=PIGR PE=1 SV=4 | 902 | 20201115_10113_1ng_glyco_hcd203040_compound15.20597.20597.5 | controllerType=0 controllerNumber=1 scan=20597 | 41.1058 |
| 15959:4 | 7 | K.IIEGEPNLKVPGN[+2059.735]VTAVLGETLKVPC[+57.021]HFPC[+57.021]K.F | HexNAc(4)Hex(5)Fuc(1)NeuAc(1) | C[+57]*2, N[+2060] | 1369.8925 | 4 | 5476.548 | 5476.5418 | 0 | 1.1 | 457 | Specific | 1139.09 | 1139.09 | 1139.09 | 21.22 | 74 | >sp|P01833|PIGR_HUMAN Polymeric immunoglobulin receptor OS=Homo sapiens OX=9606 GN=PIGR PE=1 SV=4 | 902 | 20201115_10113_1ng_glyco_hcd203040_compound15.19087.19087.4 | controllerType=0 controllerNumber=1 scan=19087 | 38.2466 |
| 17080:3 | 7 | K.VPGN[+2351.851]VTAVLGETLK.V | HexNAc(4)Hex(5)Fuc(3)NeuAc(1) | N[+2352] | 1250.5501 | 3 | 3749.6358 | 3749.6506 | 0 | -4 | 466 | Specific | 239.95 | 230.52 | 230.52 | 3.25 | 74 | >sp|P01833|PIGR_HUMAN Polymeric immunoglobulin receptor OS=Homo sapiens OX=9606 GN=PIGR PE=1 SV=4 | 902 | 20201115_10113_1ng_glyco_hcd203040_compound15.20260.20260.3 | controllerType=0 controllerNumber=1 scan=20260 | 40.437 |
| 17078:3 | 7 | K.VPGN[+2351.851]VTAVLGETLK.V | HexNAc(4)Hex(5)Fuc(3)NeuAc(1) | N[+2352] | 1250.5501 | 3 | 3749.6358 | 3749.6506 | 0 | -4 | 466 | Specific | 302.77 | 302.77 | 302.77 | 5.37 | 74 | >sp|P01833|PIGR_HUMAN Polymeric immunoglobulin receptor OS=Homo sapiens OX=9606 GN=PIGR PE=1 SV=4 | 902 | 20201115_10113_1ng_glyco_hcd203040_compound15.20258.20258.3 | controllerType=0 controllerNumber=1 scan=20258 | 40.4335 |
| 14048:3 | 7 | K.VPGN[+2205.793]VTAVLGETLK.V | HexNAc(4)Hex(5)Fuc(2)NeuAc(1) | N[+2206] | 1201.8721 | 3 | 3603.6018 | 3603.5927 | 0 | 2.5 | 466 | Specific | 597.31 | 385.21 | 385.21 | 11.42 | 74 | >sp|P01833|PIGR_HUMAN Polymeric immunoglobulin receptor OS=Homo sapiens OX=9606 GN=PIGR PE=1 SV=4 | 902 | 20201115_10113_1ng_glyco_hcd203040_compound15.17104.17104.3 | controllerType=0 controllerNumber=1 scan=17104 | 34.4782 |
| 14046:3 | 7 | K.VPGN[+2205.793]VTAVLGETLK.V | HexNAc(4)Hex(5)Fuc(2)NeuAc(1) | N[+2206] | 1201.8721 | 3 | 3603.6018 | 3603.5927 | 0 | 2.5 | 466 | Specific | 608.41 | 255.77 | 255.77 | 10.15 | 74 | >sp|P01833|PIGR_HUMAN Polymeric immunoglobulin receptor OS=Homo sapiens OX=9606 GN=PIGR PE=1 SV=4 | 902 | 20201115_10113_1ng_glyco_hcd203040_compound15.17102.17102.3 | controllerType=0 controllerNumber=1 scan=17102 | 34.4728 |
| 14078:4 | 7 | K.VPGN[+2205.793]VTAVLGETLKVPC[+57.021]HFPC[+57.021]K.F | HexNAc(4)Hex(5)Fuc(2)NeuAc(1) | C[+57]*2, N[+2206] | 1158.0206 | 4 | 4629.0604 | 4629.0502 | 0 | 2.2 | 466 | Specific | 443.1 | 424.56 | 424.56 | 9.76 | 74 | >sp|P01833|PIGR_HUMAN Polymeric immunoglobulin receptor OS=Homo sapiens OX=9606 GN=PIGR PE=1 SV=4 | 902 | 20201115_10113_1ng_glyco_hcd203040_compound15.17135.17135.3 | controllerType=0 controllerNumber=1 scan=17135 | 34.5329 |
| 15978:4 | 7 | K.VPGN[+2350.830]VTAVLGETLKVPC[+57.021]HFPC[+57.021]K.F | HexNAc(4)Hex(5)Fuc(1)NeuAc(2) | C[+57]*2, N[+2351] | 1194.2808 | 4 | 4774.1014 | 4774.0877 | 0 | 2.9 | 466 | Specific | 495.12 | 495.12 | 495.12 | 10.1 | 74 | >sp|P01833|PIGR_HUMAN Polymeric immunoglobulin receptor OS=Homo sapiens OX=9606 GN=PIGR PE=1 SV=4 | 902 | 20201115_10113_1ng_glyco_hcd203040_compound15.19107.19107.4 | controllerType=0 controllerNumber=1 scan=19107 | 38.2862 |
| 14090:4 | 7 | K.VPGN[+2205.793]VTAVLGETLKVPC[+57.021]HFPC[+57.021]K.F | HexNAc(4)Hex(5)Fuc(2)NeuAc(1) | C[+57]*2, N[+2206] | 1158.0213 | 4 | 4629.0633 | 4629.0502 | 0 | 2.8 | 466 | Specific | 581.72 | 466.46 | 466.46 | 11.6 | 74 | >sp|P01833|PIGR_HUMAN Polymeric immunoglobulin receptor OS=Homo sapiens OX=9606 GN=PIGR PE=1 SV=4 | 902 | 20201115_10113_1ng_glyco_hcd203040_compound15.17148.17148.4 | controllerType=0 controllerNumber=1 scan=17148 | 34.5693 |
| 15981:4 | 7 | K.VPGN[+2350.830]VTAVLGETLKVPC[+57.021]HFPC[+57.021]K.F | HexNAc(4)Hex(5)Fuc(1)NeuAc(2) | C[+57]*2, N[+2351] | 1194.2808 | 4 | 4774.1014 | 4774.0877 | 0 | 2.9 | 466 | Specific | 624.96 | 615.9 | 615.9 | 12.6 | 74 | >sp|P01833|PIGR_HUMAN Polymeric immunoglobulin receptor OS=Homo sapiens OX=9606 GN=PIGR PE=1 SV=4 | 902 | 20201115_10113_1ng_glyco_hcd203040_compound15.19110.19110.4 | controllerType=0 controllerNumber=1 scan=19110 | 38.2907 |
| 14092:4 | 7 | K.VPGN[+2205.793]VTAVLGETLKVPC[+57.021]HFPC[+57.021]K.F | HexNAc(4)Hex(5)Fuc(2)NeuAc(1) | C[+57]*2, N[+2206] | 1158.0213 | 4 | 4629.0633 | 4629.0502 | 0 | 2.8 | 466 | Specific | 654.92 | 646.85 | 646.85 | 13.6 | 74 | >sp|P01833|PIGR_HUMAN Polymeric immunoglobulin receptor OS=Homo sapiens OX=9606 GN=PIGR PE=1 SV=4 | 902 | 20201115_10113_1ng_glyco_hcd203040_compound15.17150.17150.4 | controllerType=0 controllerNumber=1 scan=17150 | 34.5768 |
| 14081:4 | 7 | K.VPGN[+2205.793]VTAVLGETLKVPC[+57.021]HFPC[+57.021]K.F | HexNAc(4)Hex(5)Fuc(2)NeuAc(1) | C[+57]*2, N[+2206] | 1158.0206 | 4 | 4629.0604 | 4629.0502 | 0 | 2.2 | 466 | Specific | 681.14 | 383.06 | 383.06 | 13.61 | 74 | >sp|P01833|PIGR_HUMAN Polymeric immunoglobulin receptor OS=Homo sapiens OX=9606 GN=PIGR PE=1 SV=4 | 902 | 20201115_10113_1ng_glyco_hcd203040_compound15.17138.17138.3 | controllerType=0 controllerNumber=1 scan=17138 | 34.5366 |
| 15953:4 | 7 | K.VPGN[+2350.830]VTAVLGETLKVPC[+57.021]HFPC[+57.021]K.F | HexNAc(4)Hex(5)Fuc(1)NeuAc(2) | C[+57]*2, N[+2351] | 1194.2802 | 4 | 4774.0989 | 4774.0877 | 0 | 2.3 | 466 | Specific | 705.56 | 634.28 | 634.28 | 14.61 | 74 | >sp|P01833|PIGR_HUMAN Polymeric immunoglobulin receptor OS=Homo sapiens OX=9606 GN=PIGR PE=1 SV=4 | 902 | 20201115_10113_1ng_glyco_hcd203040_compound15.19081.19081.4 | controllerType=0 controllerNumber=1 scan=19081 | 38.2359 |
| 14154:4 | 7 | K.VPGN[+2059.735]VTAVLGETLKVPC[+57.021]HFPC[+57.021]K.F | HexNAc(4)Hex(5)Fuc(1)NeuAc(1) | C[+57]*2, N[+2060] | 1121.5053 | 4 | 4482.9993 | 4482.9923 | 0 | 1.6 | 466 | Specific | 743.04 | 454.37 | 454.37 | 14.61 | 74 | >sp|P01833|PIGR_HUMAN Polymeric immunoglobulin receptor OS=Homo sapiens OX=9606 GN=PIGR PE=1 SV=4 | 902 | 20201115_10113_1ng_glyco_hcd203040_compound15.17215.17215.4 | controllerType=0 controllerNumber=1 scan=17215 | 34.7271 |
| 14156:4 | 7 | K.VPGN[+2059.735]VTAVLGETLKVPC[+57.021]HFPC[+57.021]K.F | HexNAc(4)Hex(5)Fuc(1)NeuAc(1) | C[+57]*2, N[+2060] | 1121.5053 | 4 | 4482.9993 | 4482.9923 | 0 | 1.6 | 466 | Specific | 822.49 | 409.56 | 409.56 | 16.61 | 74 | >sp|P01833|PIGR_HUMAN Polymeric immunoglobulin receptor OS=Homo sapiens OX=9606 GN=PIGR PE=1 SV=4 | 902 | 20201115_10113_1ng_glyco_hcd203040_compound15.17217.17217.4 | controllerType=0 controllerNumber=1 scan=17217 | 34.7368 |
| 15956:4 | 7 | K.VPGN[+2350.830]VTAVLGETLKVPC[+57.021]HFPC[+57.021]K.F | HexNAc(4)Hex(5)Fuc(1)NeuAc(2) | C[+57]*2, N[+2351] | 1194.2802 | 4 | 4774.0989 | 4774.0877 | 0 | 2.3 | 466 | Specific | 824.13 | 456.18 | 456.18 | 16.61 | 74 | >sp|P01833|PIGR_HUMAN Polymeric immunoglobulin receptor OS=Homo sapiens OX=9606 GN=PIGR PE=1 SV=4 | 902 | 20201115_10113_1ng_glyco_hcd203040_compound15.19084.19084.4 | controllerType=0 controllerNumber=1 scan=19084 | 38.2413 |
| 06430:3 | 7 | K.WN[+1710.598]NTGC[+57.021]QALPSQDEGPSK.A | HexNAc(3)Hex(5)NeuAc(1) | C[+57], N[+1711] | 1233.8323 | 3 | 3699.4824 | 3699.4743 | 0 | 2.2 | 498 | Specific | 621.11 | 558.72 | 558.72 | 12.55 | 74 | >sp|P01833|PIGR_HUMAN Polymeric immunoglobulin receptor OS=Homo sapiens OX=9606 GN=PIGR PE=1 SV=4 | 902 | 20201115_10113_1ng_glyco_hcd203040_compound15.9187.9187.3 | controllerType=0 controllerNumber=1 scan=9187 | 20.9506 |
| 06307:3 | 7 | K.WN[+1872.651]NTGC[+57.021]QALPSQDEGPSK.A | HexNAc(3)Hex(6)NeuAc(1) | C[+57], N[+1873] | 1287.8509 | 3 | 3861.5381 | 3861.5272 | 0 | 2.8 | 498 | Specific | 626.05 | 597.62 | 597.62 | 12.53 | 74 | >sp|P01833|PIGR_HUMAN Polymeric immunoglobulin receptor OS=Homo sapiens OX=9606 GN=PIGR PE=1 SV=4 | 902 | 20201115_10113_1ng_glyco_hcd203040_compound15.9060.9060.3 | controllerType=0 controllerNumber=1 scan=9060 | 20.7474 |
| 06236:3 | 7 | K.WN[+2059.735]NTGC[+57.021]QALPSQDEGPSK.A | HexNAc(4)Hex(5)Fuc(1)NeuAc(1) | C[+57], N[+2060] | 1350.2097 | 3 | 4048.6144 | 4048.6116 | 0 | 0.7 | 498 | Specific | 657.04 | 584.56 | 584.56 | 13.34 | 74 | >sp|P01833|PIGR_HUMAN Polymeric immunoglobulin receptor OS=Homo sapiens OX=9606 GN=PIGR PE=1 SV=4 | 902 | 20201115_10113_1ng_glyco_hcd203040_compound15.8987.8987.3 | controllerType=0 controllerNumber=1 scan=8987 | 20.637 |
| 06234:3 | 7 | K.WN[+2059.735]NTGC[+57.021]QALPSQDEGPSK.A | HexNAc(4)Hex(5)Fuc(1)NeuAc(1) | C[+57], N[+2060] | 1350.2097 | 3 | 4048.6144 | 4048.6116 | 0 | 0.7 | 498 | Specific | 670.26 | 589.99 | 589.99 | 13.34 | 74 | >sp|P01833|PIGR_HUMAN Polymeric immunoglobulin receptor OS=Homo sapiens OX=9606 GN=PIGR PE=1 SV=4 | 902 | 20201115_10113_1ng_glyco_hcd203040_compound15.8985.8985.3 | controllerType=0 controllerNumber=1 scan=8985 | 20.6327 |
| 06428:3 | 7 | K.WN[+1710.598]NTGC[+57.021]QALPSQDEGPSK.A | HexNAc(3)Hex(5)NeuAc(1) | C[+57], N[+1711] | 1233.8323 | 3 | 3699.4824 | 3699.4743 | 0 | 2.2 | 498 | Specific | 678.32 | 603.85 | 603.85 | 13.55 | 74 | >sp|P01833|PIGR_HUMAN Polymeric immunoglobulin receptor OS=Homo sapiens OX=9606 GN=PIGR PE=1 SV=4 | 902 | 20201115_10113_1ng_glyco_hcd203040_compound15.9185.9185.3 | controllerType=0 controllerNumber=1 scan=9185 | 20.9469 |
| 06309:3 | 7 | K.WN[+1872.651]NTGC[+57.021]QALPSQDEGPSK.A | HexNAc(3)Hex(6)NeuAc(1) | C[+57], N[+1873] | 1287.8509 | 3 | 3861.5381 | 3861.5272 | 0 | 2.8 | 498 | Specific | 699.99 | 644.38 | 644.38 | 13.53 | 74 | >sp|P01833|PIGR_HUMAN Polymeric immunoglobulin receptor OS=Homo sapiens OX=9606 GN=PIGR PE=1 SV=4 | 902 | 20201115_10113_1ng_glyco_hcd203040_compound15.9062.9062.3 | controllerType=0 controllerNumber=1 scan=9062 | 20.7512 |
| 06329:3 | 7 | K.WN[+1913.677]NTGC[+57.021]QALPSQDEGPSK.A | HexNAc(4)Hex(5)NeuAc(1) | C[+57], N[+1914] | 1301.5242 | 3 | 3902.5581 | 3902.5537 | 0 | 1.1 | 498 | Specific | 749.96 | 660.67 | 660.67 | 14.34 | 74 | >sp|P01833|PIGR_HUMAN Polymeric immunoglobulin receptor OS=Homo sapiens OX=9606 GN=PIGR PE=1 SV=4 | 902 | 20201115_10113_1ng_glyco_hcd203040_compound15.9083.9083.3 | controllerType=0 controllerNumber=1 scan=9083 | 20.7876 |
| 06331:3 | 7 | K.WN[+1913.677]NTGC[+57.021]QALPSQDEGPSK.A | HexNAc(4)Hex(5)NeuAc(1) | C[+57], N[+1914] | 1301.5242 | 3 | 3902.5581 | 3902.5537 | 0 | 1.1 | 498 | Specific | 866.74 | 736.96 | 736.96 | 17.34 | 74 | >sp|P01833|PIGR_HUMAN Polymeric immunoglobulin receptor OS=Homo sapiens OX=9606 GN=PIGR PE=1 SV=4 | 902 | 20201115_10113_1ng_glyco_hcd203040_compound15.9085.9085.3 | controllerType=0 controllerNumber=1 scan=9085 | 20.7919 |
| 18990:3 | 8 | R.LSLHRPALEDLLLGSEAN[+2116.756]LTC[+57.021]TLTGLR.D | HexNAc(5)Hex(5)NeuAc(1) | C[+57], N[+2117] | 1694.1239 | 3 | 5080.357 | 5080.3547 | 0 | 0.5 | 127 | Specific | 260.53 | 260.53 | 260.53 | 5.78 | 67 | >sp|P01876|IGHA1_HUMAN Immunoglobulin heavy constant alpha 1 OS=Homo sapiens OX=9606 GN=IGHA1 PE=1 SV=2 | 255 | 20201115_10113_1ng_glyco_hcd203040_compound15.22250.22250.3 | controllerType=0 controllerNumber=1 scan=22250 | 44.6531 |
| 18163:3 | 8 | R.LSLHRPALEDLLLGSEAN[+1054.370]LTC[+57.021]TLTGLR.D | HexNAc(2)Hex(4) | C[+57], N[+1054] | 1339.9981 | 3 | 4017.9798 | 4017.9683 | 0 | 2.9 | 127 | Specific | 300.37 | 300.37 | 300.37 | 6.79 | 67 | >sp|P01876|IGHA1_HUMAN Immunoglobulin heavy constant alpha 1 OS=Homo sapiens OX=9606 GN=IGHA1 PE=1 SV=2 | 255 | 20201115_10113_1ng_glyco_hcd203040_compound15.21389.21389.3 | controllerType=0 controllerNumber=1 scan=21389 | 42.8139 |
| 19087:4 | 8 | R.LSLHRPALEDLLLGSEAN[+1913.677]LTC[+57.021]TLTGLR.D | HexNAc(4)Hex(5)NeuAc(1) | C[+57], N[+1914] | 1220.0856 | 4 | 4877.3206 | 4877.2753 | 0 | 9.3 | 127 | Specific | 305.57 | 305.57 | 305.57 | 6.61 | 67 | >sp|P01876|IGHA1_HUMAN Immunoglobulin heavy constant alpha 1 OS=Homo sapiens OX=9606 GN=IGHA1 PE=1 SV=2 | 255 | 20201115_10113_1ng_glyco_hcd203040_compound15.22351.22351.4 | controllerType=0 controllerNumber=1 scan=22351 | 44.8644 |
| 18138:4 | 8 | R.LSLHRPALEDLLLGSEAN[+1647.613]LTC[+57.021]TLTGLR.D | HexNAc(5)Hex(3)Fuc(1) | C[+57], N[+1648] | 1153.5574 | 4 | 4611.2078 | 4611.2115 | 0 | -0.8 | 127 | Specific | 306.14 | 304.57 | 304.57 | 4.75 | 67 | >sp|P01876|IGHA1_HUMAN Immunoglobulin heavy constant alpha 1 OS=Homo sapiens OX=9606 GN=IGHA1 PE=1 SV=2 | 255 | 20201115_10113_1ng_glyco_hcd203040_compound15.21363.21363.4 | controllerType=0 controllerNumber=1 scan=21363 | 42.7596 |
| 18166:3 | 8 | R.LSLHRPALEDLLLGSEAN[+1054.370]LTC[+57.021]TLTGLR.D | HexNAc(2)Hex(4) | C[+57], N[+1054] | 1339.9981 | 3 | 4017.9798 | 4017.9683 | 0 | 2.9 | 127 | Specific | 322.41 | 293.94 | 293.94 | 4.94 | 67 | >sp|P01876|IGHA1_HUMAN Immunoglobulin heavy constant alpha 1 OS=Homo sapiens OX=9606 GN=IGHA1 PE=1 SV=2 | 255 | 20201115_10113_1ng_glyco_hcd203040_compound15.21392.21392.3 | controllerType=0 controllerNumber=1 scan=21392 | 42.8183 |
| 18041:4 | 8 | R.LSLHRPALEDLLLGSEAN[+1419.502]LTC[+57.021]TLTGLR.D | HexNAc(3)Hex(5) | C[+57], N[+1420] | 1096.5317 | 4 | 4383.1048 | 4383.1005 | 0 | 1 | 127 | Specific | 346.51 | 346.51 | 346.51 | 7.66 | 67 | >sp|P01876|IGHA1_HUMAN Immunoglobulin heavy constant alpha 1 OS=Homo sapiens OX=9606 GN=IGHA1 PE=1 SV=2 | 255 | 20201115_10113_1ng_glyco_hcd203040_compound15.21262.21262.4 | controllerType=0 controllerNumber=1 scan=21262 | 42.5499 |
| 18010:4 | 8 | R.LSLHRPALEDLLLGSEAN[+1540.529]LTC[+57.021]TLTGLR.D | HexNAc(2)Hex(7) | C[+57], N[+1541] | 1126.7896 | 4 | 4504.1365 | 4504.1268 | 0 | 2.2 | 127 | Specific | 409.47 | 409.47 | 409.47 | 9.96 | 67 | >sp|P01876|IGHA1_HUMAN Immunoglobulin heavy constant alpha 1 OS=Homo sapiens OX=9606 GN=IGHA1 PE=1 SV=2 | 255 | 20201115_10113_1ng_glyco_hcd203040_compound15.21230.21230.4 | controllerType=0 controllerNumber=1 scan=21230 | 42.4506 |
| 18024:4 | 8 | R.LSLHRPALEDLLLGSEAN[+1663.608]LTC[+57.021]TLTGLR.D | HexNAc(5)Hex(4) | C[+57], N[+1664] | 1157.5589 | 4 | 4627.2137 | 4627.2064 | 0 | 1.6 | 127 | Specific | 419.05 | 419.05 | 419.05 | 9.8 | 67 | >sp|P01876|IGHA1_HUMAN Immunoglobulin heavy constant alpha 1 OS=Homo sapiens OX=9606 GN=IGHA1 PE=1 SV=2 | 255 | 20201115_10113_1ng_glyco_hcd203040_compound15.21245.21245.4 | controllerType=0 controllerNumber=1 scan=21245 | 42.5129 |
| 19069:4 | 8 | R.LSLHRPALEDLLLGSEAN[+1710.598]LTC[+57.021]TLTGLR.D | HexNAc(3)Hex(5)NeuAc(1) | C[+57], N[+1711] | 1169.3135 | 4 | 4674.2322 | 4674.1959 | 0 | 7.8 | 127 | Specific | 445.39 | 445.39 | 445.39 | 9.02 | 67 | >sp|P01876|IGHA1_HUMAN Immunoglobulin heavy constant alpha 1 OS=Homo sapiens OX=9606 GN=IGHA1 PE=1 SV=2 | 255 | 20201115_10113_1ng_glyco_hcd203040_compound15.22333.22333.4 | controllerType=0 controllerNumber=1 scan=22333 | 44.8298 |
| 18993:3 | 8 | R.LSLHRPALEDLLLGSEAN[+2116.756]LTC[+57.021]TLTGLR.D | HexNAc(5)Hex(5)NeuAc(1) | C[+57], N[+2117] | 1694.1239 | 3 | 5080.357 | 5080.3547 | 0 | 0.5 | 127 | Specific | 466.98 | 424.96 | 424.96 | 8.99 | 67 | >sp|P01876|IGHA1_HUMAN Immunoglobulin heavy constant alpha 1 OS=Homo sapiens OX=9606 GN=IGHA1 PE=1 SV=2 | 255 | 20201115_10113_1ng_glyco_hcd203040_compound15.22253.22253.3 | controllerType=0 controllerNumber=1 scan=22253 | 44.657 |
| 19045:3 | 8 | R.LSLHRPALEDLLLGSEAN[+1751.624]LTC[+57.021]TLTGLR.D | HexNAc(4)Hex(4)NeuAc(1) | C[+57], N[+1752] | 1572.4194 | 3 | 4715.2436 | 4715.2225 | 0 | 4.5 | 127 | Specific | 493.6 | 493.6 | 493.6 | 9.22 | 67 | >sp|P01876|IGHA1_HUMAN Immunoglobulin heavy constant alpha 1 OS=Homo sapiens OX=9606 GN=IGHA1 PE=1 SV=2 | 255 | 20201115_10113_1ng_glyco_hcd203040_compound15.22308.22308.3 | controllerType=0 controllerNumber=1 scan=22308 | 44.7837 |
| 18028:3 | 8 | R.LSLHRPALEDLLLGSEAN[+1663.608]LTC[+57.021]TLTGLR.D | HexNAc(5)Hex(4) | C[+57], N[+1664] | 1543.077 | 3 | 4627.2163 | 4627.2064 | 0 | 2.1 | 127 | Specific | 521.88 | 521.88 | 521.88 | 9.9 | 67 | >sp|P01876|IGHA1_HUMAN Immunoglobulin heavy constant alpha 1 OS=Homo sapiens OX=9606 GN=IGHA1 PE=1 SV=2 | 255 | 20201115_10113_1ng_glyco_hcd203040_compound15.21249.21249.3 | controllerType=0 controllerNumber=1 scan=21249 | 42.5206 |
| 18141:4 | 8 | R.LSLHRPALEDLLLGSEAN[+1647.613]LTC[+57.021]TLTGLR.D | HexNAc(5)Hex(3)Fuc(1) | C[+57], N[+1648] | 1153.5574 | 4 | 4611.2078 | 4611.2115 | 0 | -0.8 | 127 | Specific | 524.75 | 372.38 | 372.38 | 6.59 | 67 | >sp|P01876|IGHA1_HUMAN Immunoglobulin heavy constant alpha 1 OS=Homo sapiens OX=9606 GN=IGHA1 PE=1 SV=2 | 255 | 20201115_10113_1ng_glyco_hcd203040_compound15.21366.21366.4 | controllerType=0 controllerNumber=1 scan=21366 | 42.7653 |
| 19048:3 | 8 | R.LSLHRPALEDLLLGSEAN[+1751.624]LTC[+57.021]TLTGLR.D | HexNAc(4)Hex(4)NeuAc(1) | C[+57], N[+1752] | 1572.4194 | 3 | 4715.2436 | 4715.2225 | 0 | 4.5 | 127 | Specific | 561.71 | 475.75 | 475.75 | 9.78 | 67 | >sp|P01876|IGHA1_HUMAN Immunoglobulin heavy constant alpha 1 OS=Homo sapiens OX=9606 GN=IGHA1 PE=1 SV=2 | 255 | 20201115_10113_1ng_glyco_hcd203040_compound15.22311.22311.3 | controllerType=0 controllerNumber=1 scan=22311 | 44.7908 |
| 18050:3 | 8 | R.LSLHRPALEDLLLGSEAN[+1622.582]LTC[+57.021]TLTGLR.D | HexNAc(4)Hex(5) | C[+57], N[+1623] | 1529.4001 | 3 | 4586.1857 | 4586.1799 | 0 | 1.3 | 127 | Specific | 568.92 | 536.97 | 536.97 | 9.99 | 67 | >sp|P01876|IGHA1_HUMAN Immunoglobulin heavy constant alpha 1 OS=Homo sapiens OX=9606 GN=IGHA1 PE=1 SV=2 | 255 | 20201115_10113_1ng_glyco_hcd203040_compound15.21272.21272.3 | controllerType=0 controllerNumber=1 scan=21272 | 42.5741 |
| 17992:4 | 8 | R.LSLHRPALEDLLLGSEAN[+1825.661]LTC[+57.021]TLTGLR.D | HexNAc(5)Hex(5) | C[+57], N[+1826] | 1198.0818 | 4 | 4789.3055 | 4789.2592 | 0 | 9.7 | 127 | Specific | 574.43 | 574.43 | 574.43 | 9.85 | 67 | >sp|P01876|IGHA1_HUMAN Immunoglobulin heavy constant alpha 1 OS=Homo sapiens OX=9606 GN=IGHA1 PE=1 SV=2 | 255 | 20201115_10113_1ng_glyco_hcd203040_compound15.21211.21211.4 | controllerType=0 controllerNumber=1 scan=21211 | 42.4073 |
| 17986:4 | 8 | R.LSLHRPALEDLLLGSEAN[+1864.634]LTC[+57.021]TLTGLR.D | HexNAc(2)Hex(9) | C[+57], N[+1865] | 1207.8227 | 4 | 4828.2689 | 4828.2324 | 0 | 7.5 | 127 | Specific | 589.51 | 589.51 | 589.51 | 10.87 | 67 | >sp|P01876|IGHA1_HUMAN Immunoglobulin heavy constant alpha 1 OS=Homo sapiens OX=9606 GN=IGHA1 PE=1 SV=2 | 255 | 20201115_10113_1ng_glyco_hcd203040_compound15.21205.21205.4 | controllerType=0 controllerNumber=1 scan=21205 | 42.3981 |
| 19023:3 | 8 | R.LSLHRPALEDLLLGSEAN[+1710.598]LTC[+57.021]TLTGLR.D | HexNAc(3)Hex(5)NeuAc(1) | C[+57], N[+1711] | 1558.7414 | 3 | 4674.2096 | 4674.1959 | 0 | 2.9 | 127 | Specific | 591.85 | 528.98 | 528.98 | 11.25 | 67 | >sp|P01876|IGHA1_HUMAN Immunoglobulin heavy constant alpha 1 OS=Homo sapiens OX=9606 GN=IGHA1 PE=1 SV=2 | 255 | 20201115_10113_1ng_glyco_hcd203040_compound15.22285.22285.3 | controllerType=0 controllerNumber=1 scan=22285 | 44.7353 |
| 18060:3 | 8 | R.LSLHRPALEDLLLGSEAN[+1298.476]LTC[+57.021]TLTGLR.D | HexNAc(4)Hex(3) | C[+57], N[+1298] | 1421.3655 | 3 | 4262.0819 | 4262.0742 | 0 | 1.8 | 127 | Specific | 607.98 | 577.66 | 577.66 | 12.26 | 67 | >sp|P01876|IGHA1_HUMAN Immunoglobulin heavy constant alpha 1 OS=Homo sapiens OX=9606 GN=IGHA1 PE=1 SV=2 | 255 | 20201115_10113_1ng_glyco_hcd203040_compound15.21282.21282.3 | controllerType=0 controllerNumber=1 scan=21282 | 42.5956 |
| 18959:4 | 8 | R.LSLHRPALEDLLLGSEAN[+2116.756]LTC[+57.021]TLTGLR.D | HexNAc(5)Hex(5)NeuAc(1) | C[+57], N[+2117] | 1270.8425 | 4 | 5080.3483 | 5080.3547 | 0 | -1.3 | 127 | Specific | 608.9 | 604.78 | 604.78 | 11.75 | 67 | >sp|P01876|IGHA1_HUMAN Immunoglobulin heavy constant alpha 1 OS=Homo sapiens OX=9606 GN=IGHA1 PE=1 SV=2 | 255 | 20201115_10113_1ng_glyco_hcd203040_compound15.22218.22218.4 | controllerType=0 controllerNumber=1 scan=22218 | 44.5894 |
| 18063:3 | 8 | R.LSLHRPALEDLLLGSEAN[+1298.476]LTC[+57.021]TLTGLR.D | HexNAc(4)Hex(3) | C[+57], N[+1298] | 1421.3655 | 3 | 4262.0819 | 4262.0742 | 0 | 1.8 | 127 | Specific | 617.79 | 595.57 | 595.57 | 12.26 | 67 | >sp|P01876|IGHA1_HUMAN Immunoglobulin heavy constant alpha 1 OS=Homo sapiens OX=9606 GN=IGHA1 PE=1 SV=2 | 255 | 20201115_10113_1ng_glyco_hcd203040_compound15.21285.21285.3 | controllerType=0 controllerNumber=1 scan=21285 | 42.6005 |
| 19026:3 | 8 | R.LSLHRPALEDLLLGSEAN[+1710.598]LTC[+57.021]TLTGLR.D | HexNAc(3)Hex(5)NeuAc(1) | C[+57], N[+1711] | 1558.7414 | 3 | 4674.2096 | 4674.1959 | 0 | 2.9 | 127 | Specific | 618.15 | 555.07 | 555.07 | 12.25 | 67 | >sp|P01876|IGHA1_HUMAN Immunoglobulin heavy constant alpha 1 OS=Homo sapiens OX=9606 GN=IGHA1 PE=1 SV=2 | 255 | 20201115_10113_1ng_glyco_hcd203040_compound15.22288.22288.3 | controllerType=0 controllerNumber=1 scan=22288 | 44.7402 |
| 18052:3 | 8 | R.LSLHRPALEDLLLGSEAN[+1622.582]LTC[+57.021]TLTGLR.D | HexNAc(4)Hex(5) | C[+57], N[+1623] | 1529.4001 | 3 | 4586.1857 | 4586.1799 | 0 | 1.3 | 127 | Specific | 638.02 | 544.71 | 544.71 | 12.05 | 67 | >sp|P01876|IGHA1_HUMAN Immunoglobulin heavy constant alpha 1 OS=Homo sapiens OX=9606 GN=IGHA1 PE=1 SV=2 | 255 | 20201115_10113_1ng_glyco_hcd203040_compound15.21274.21274.3 | controllerType=0 controllerNumber=1 scan=21274 | 42.5784 |
| 18045:3 | 8 | R.LSLHRPALEDLLLGSEAN[+1501.555]LTC[+57.021]TLTGLR.D | HexNAc(5)Hex(3) | C[+57], N[+1502] | 1489.0585 | 3 | 4465.161 | 4465.1536 | 0 | 1.7 | 127 | Specific | 649.16 | 577.79 | 577.79 | 12.26 | 67 | >sp|P01876|IGHA1_HUMAN Immunoglobulin heavy constant alpha 1 OS=Homo sapiens OX=9606 GN=IGHA1 PE=1 SV=2 | 255 | 20201115_10113_1ng_glyco_hcd203040_compound15.21266.21266.3 | controllerType=0 controllerNumber=1 scan=21266 | 42.5576 |
| 18058:3 | 8 | R.LSLHRPALEDLLLGSEAN[+1460.529]LTC[+57.021]TLTGLR.D | HexNAc(4)Hex(4) | C[+57], N[+1461] | 1475.3785 | 3 | 4424.1209 | 4424.1271 | 0 | -1.4 | 127 | Specific | 651.65 | 581.57 | 581.57 | 12.16 | 67 | >sp|P01876|IGHA1_HUMAN Immunoglobulin heavy constant alpha 1 OS=Homo sapiens OX=9606 GN=IGHA1 PE=1 SV=2 | 255 | 20201115_10113_1ng_glyco_hcd203040_compound15.21280.21280.3 | controllerType=0 controllerNumber=1 scan=21280 | 42.5932 |
| 19090:4 | 8 | R.LSLHRPALEDLLLGSEAN[+1548.545]LTC[+57.021]TLTGLR.D | HexNAc(3)Hex(4)NeuAc(1) | C[+57], N[+1549] | 1128.7995 | 4 | 4512.1761 | 4512.1431 | 0 | 7.3 | 127 | Specific | 652.2 | 652.2 | 652.2 | 12.87 | 67 | >sp|P01876|IGHA1_HUMAN Immunoglobulin heavy constant alpha 1 OS=Homo sapiens OX=9606 GN=IGHA1 PE=1 SV=2 | 255 | 20201115_10113_1ng_glyco_hcd203040_compound15.22354.22354.3 | controllerType=0 controllerNumber=1 scan=22354 | 44.8704 |
| 18031:3 | 8 | R.LSLHRPALEDLLLGSEAN[+1663.608]LTC[+57.021]TLTGLR.D | HexNAc(5)Hex(4) | C[+57], N[+1664] | 1543.077 | 3 | 4627.2163 | 4627.2064 | 0 | 2.1 | 127 | Specific | 663.38 | 621.47 | 621.47 | 13.26 | 67 | >sp|P01876|IGHA1_HUMAN Immunoglobulin heavy constant alpha 1 OS=Homo sapiens OX=9606 GN=IGHA1 PE=1 SV=2 | 255 | 20201115_10113_1ng_glyco_hcd203040_compound15.21252.21252.3 | controllerType=0 controllerNumber=1 scan=21252 | 42.5247 |
| 17995:4 | 8 | R.LSLHRPALEDLLLGSEAN[+1825.661]LTC[+57.021]TLTGLR.D | HexNAc(5)Hex(5) | C[+57], N[+1826] | 1198.0818 | 4 | 4789.3055 | 4789.2592 | 0 | 9.7 | 127 | Specific | 678.84 | 621.2 | 621.2 | 12.91 | 67 | >sp|P01876|IGHA1_HUMAN Immunoglobulin heavy constant alpha 1 OS=Homo sapiens OX=9606 GN=IGHA1 PE=1 SV=2 | 255 | 20201115_10113_1ng_glyco_hcd203040_compound15.21214.21214.4 | controllerType=0 controllerNumber=1 scan=21214 | 42.4159 |
| 18018:3 | 8 | R.LSLHRPALEDLLLGSEAN[+1540.529]LTC[+57.021]TLTGLR.D | HexNAc(2)Hex(7) | C[+57], N[+1541] | 1502.051 | 3 | 4504.1383 | 4504.1268 | 0 | 2.6 | 127 | Specific | 685.84 | 685.84 | 685.84 | 13.26 | 67 | >sp|P01876|IGHA1_HUMAN Immunoglobulin heavy constant alpha 1 OS=Homo sapiens OX=9606 GN=IGHA1 PE=1 SV=2 | 255 | 20201115_10113_1ng_glyco_hcd203040_compound15.21238.21238.3 | controllerType=0 controllerNumber=1 scan=21238 | 42.4809 |
| 18961:4 | 8 | R.LSLHRPALEDLLLGSEAN[+2116.756]LTC[+57.021]TLTGLR.D | HexNAc(5)Hex(5)NeuAc(1) | C[+57], N[+2117] | 1270.8425 | 4 | 5080.3483 | 5080.3547 | 0 | -1.3 | 127 | Specific | 698.21 | 620.5 | 620.5 | 12.75 | 67 | >sp|P01876|IGHA1_HUMAN Immunoglobulin heavy constant alpha 1 OS=Homo sapiens OX=9606 GN=IGHA1 PE=1 SV=2 | 255 | 20201115_10113_1ng_glyco_hcd203040_compound15.22220.22220.4 | controllerType=0 controllerNumber=1 scan=22220 | [illegible] |
| 19093:4 | 8 | R.LSLHRPALEDLLLGSEAN[+1548.545]LTC[+57.021]TLTGLR.D | HexNAc(3)Hex(4)NeuAc(1) | C[+57], N[+1549] | 1128.7995 | 4 | 4512.1761 | 4512.1431 | 0 | 7.3 | 127 | Specific | 700.23 | 700.23 | 700.23 | 13.87 | 67 | >sp|P01876|IGHA1_HUMAN Immunoglobulin heavy constant alpha 1 OS=Homo sapiens OX=9606 GN=IGHA1 PE=1 SV=2 | 255 | 20201115_10113_1ng_glyco_hcd203040_compound15.22357.22357.3 | controllerType=0 controllerNumber=1 scan=22357 | [illegible] |
| 17999:4 | 8 | R.LSLHRPALEDLLLGSEAN[+1581.555]LTC[+57.021]TLTGLR.D | HexNAc(3)Hex(6) | C[+57], N[+1582] | 1137.0536 | 4 | 4545.1927 | 4545.1533 | 0 | 8.7 | 127 | Specific | 707.35 | 707.35 | 707.35 | 13.85 | 67 | >sp|P01876|IGHA1_HUMAN Immunoglobulin heavy constant alpha 1 OS=Homo sapiens OX=9606 GN=IGHA1 PE=1 SV=2 | 255 | 20201115_10113_1ng_glyco_hcd203040_compound15.21218.21218.2 | controllerType=0 controllerNumber=1 scan=21218 | 42.422 |
| 18016:3 | 8 | R.LSLHRPALEDLLLGSEAN[+1540.529]LTC[+57.021]TLTGLR.D | HexNAc(2)Hex(7) | C[+57], N[+1541] | 1502.051 | 3 | 4504.1383 | 4504.1268 | 0 | 2.6 | 127 | Specific | 713.83 | 640.24 | 640.24 | 14.26 | 67 | >sp|P01876|IGHA1_HUMAN Immunoglobulin heavy constant alpha 1 OS=Homo sapiens OX=9606 GN=IGHA1 PE=1 SV=2 | 255 | 20201115_10113_1ng_glyco_hcd203040_compound15.21236.21236.3 | controllerType=0 controllerNumber=1 scan=21236 | 42.4699 |
| 19028:4 | 8 | R.LSLHRPALEDLLLGSEAN[+1751.624]LTC[+57.021]TLTGLR.D | HexNAc(4)Hex(4)NeuAc(1) | C[+57], N[+1752] | 1179.5645 | 4 | 4715.2362 | 4715.2225 | 0 | 2.9 | 127 | Specific | 716.58 | 716.58 | 716.58 | 14.83 | 67 | >sp|P01876|IGHA1_HUMAN Immunoglobulin heavy constant alpha 1 OS=Homo sapiens OX=9606 GN=IGHA1 PE=1 SV=2 | 255 | 20201115_10113_1ng_glyco_hcd203040_compound15.22290.22290.4 | controllerType=0 controllerNumber=1 scan=22290 | 44.7447 |
| 18047:3 | 8 | R.LSLHRPALEDLLLGSEAN[+1501.555]LTC[+57.021]TLTGLR.D | HexNAc(5)Hex(3) | C[+57], N[+1502] | 1489.0585 | 3 | 4465.161 | 4465.1536 | 0 | 1.7 | 127 | Specific | 726.05 | 639.31 | 639.31 | 14.26 | 67 | >sp|P01876|IGHA1_HUMAN Immunoglobulin heavy constant alpha 1 OS=Homo sapiens OX=9606 GN=IGHA1 PE=1 SV=2 | 255 | 20201115_10113_1ng_glyco_hcd203040_compound15.21268.21268.3 | controllerType=0 controllerNumber=1 scan=21268 | 42.5664 |
| 18040:4 | 8 | R.LSLHRPALEDLLLGSEAN[+1460.529]LTC[+57.021]TLTGLR.D | HexNAc(4)Hex(4) | C[+57], N[+1461] | 1106.7893 | 4 | 4424.1356 | 4424.1271 | 0 | 1.9 | 127 | Specific | 741.49 | 737.27 | 737.27 | 14.85 | 67 | >sp|P01876|IGHA1_HUMAN Immunoglobulin heavy constant alpha 1 OS=Homo sapiens OX=9606 GN=IGHA1 PE=1 SV=2 | 255 | 20201115_10113_1ng_glyco_hcd203040_compound15.21261.21261.4 | controllerType=0 controllerNumber=1 scan=21261 | 42.5488 |
| 18029:4 | 8 | R.LSLHRPALEDLLLGSEAN[+1501.555]LTC[+57.021]TLTGLR.D | HexNAc(5)Hex(3) | C[+57], N[+1502] | 1117.0458 | 4 | 4465.1614 | 4465.1536 | 0 | 1.8 | 127 | Specific | 742.24 | 742.24 | 742.24 | 14.85 | 67 | >sp|P01876|IGHA1_HUMAN Immunoglobulin heavy constant alpha 1 OS=Homo sapiens OX=9606 GN=IGHA1 PE=1 SV=2 | 255 | 20201115_10113_1ng_glyco_hcd203040_compound15.21250.21250.4 | controllerType=0 controllerNumber=1 scan=21250 | 42.5217 |
| 18044:4 | 8 | R.LSLHRPALEDLLLGSEAN[+1419.502]LTC[+57.021]TLTGLR.D | HexNAc(3)Hex(5) | C[+57], N[+1420] | 1096.5317 | 4 | 4383.1048 | 4383.1005 | 0 | 1 | 127 | Specific | 788.36 | 563.75 | 563.75 | 15.64 | 67 | >sp|P01876|IGHA1_HUMAN Immunoglobulin heavy constant alpha 1 OS=Homo sapiens OX=9606 GN=IGHA1 PE=1 SV=2 | 255 | 20201115_10113_1ng_glyco_hcd203040_compound15.21265.21265.4 | controllerType=0 controllerNumber=1 scan=21265 | 42.5565 |
| 18026:4 | 8 | R.LSLHRPALEDLLLGSEAN[+1663.608]LTC[+57.021]TLTGLR.D | HexNAc(5)Hex(4) | C[+57], N[+1664] | 1157.5589 | 4 | 4627.2137 | 4627.2064 | 0 | 1.6 | 127 | Specific | 804.6 | 741.28 | 741.28 | 16.64 | 67 | >sp|P01876|IGHA1_HUMAN Immunoglobulin heavy constant alpha 1 OS=Homo sapiens OX=9606 GN=IGHA1 PE=1 SV=2 | 255 | 20201115_10113_1ng_glyco_hcd203040_compound15.21247.21247.4 | controllerType=0 controllerNumber=1 scan=21247 | 42.5169 |
| 18032:4 | 8 | R.LSLHRPALEDLLLGSEAN[+1501.555]LTC[+57.021]TLTGLR.D | HexNAc(5)Hex(3) | C[+57], N[+1502] | 1117.0458 | 4 | 4465.1614 | 4465.1536 | 0 | 1.8 | 127 | Specific | 805.94 | 754.76 | 754.76 | 16.85 | 67 | >sp|P01876|IGHA1_HUMAN Immunoglobulin heavy constant alpha 1 OS=Homo sapiens OX=9606 GN=IGHA1 PE=1 SV=2 | 255 | 20201115_10113_1ng_glyco_hcd203040_compound15.21253.21253.4 | controllerType=0 controllerNumber=1 scan=21253 | 42.5297 |
| 18998:4 | 8 | R.LSLHRPALEDLLLGSEAN[+1872.651]LTC[+57.021]TLTGLR.D | HexNAc(3)Hex(6)NeuAc(1) | C[+57], N[+1873] | 1209.8182 | 4 | 4836.2508 | 4836.2487 | 0 | 0.4 | 127 | Specific | 807.45 | 749.5 | 749.5 | 16.17 | 67 | >sp|P01876|IGHA1_HUMAN Immunoglobulin heavy constant alpha 1 OS=Homo sapiens OX=9606 GN=IGHA1 PE=1 SV=2 | 255 | 20201115_10113_1ng_glyco_hcd203040_compound15.22259.22259.4 | controllerType=0 controllerNumber=1 scan=22259 | 44.6684 |
| 19031:4 | 8 | R.LSLHRPALEDLLLGSEAN[+1751.624]LTC[+57.021]TLTGLR.D | HexNAc(4)Hex(4)NeuAc(1) | C[+57], N[+1752] | 1179.5645 | 4 | 4715.2362 | 4715.2225 | 0 | 2.9 | 127 | Specific | 814.09 | 741.86 | 741.86 | 16.83 | 67 | >sp|P01876|IGHA1_HUMAN Immunoglobulin heavy constant alpha 1 OS=Homo sapiens OX=9606 GN=IGHA1 PE=1 SV=2 | 255 | 20201115_10113_1ng_glyco_hcd203040_compound15.22293.22293.4 | controllerType=0 controllerNumber=1 scan=22293 | 44.751 |
| 18107:4 | 8 | R.LSLHRPALEDLLLGSEAN[+1095.397]LTC[+57.021]TLTGLR.D | HexNAc(3)Hex(3) | C[+57], N[+1095] | 1015.5083 | 4 | 4059.0115 | 4058.9949 | 0 | 4.1 | 127 | Specific | 822.72 | 822.72 | 822.72 | 16.42 | 67 | >sp|P01876|IGHA1_HUMAN Immunoglobulin heavy constant alpha 1 OS=Homo sapiens OX=9606 GN=IGHA1 PE=1 SV=2 | 255 | 20201115_10113_1ng_glyco_hcd203040_compound15.21331.21331.4 | controllerType=0 controllerNumber=1 scan=21331 | 42.6969 |
| 19071:4 | 8 | R.LSLHRPALEDLLLGSEAN[+1710.598]LTC[+57.021]TLTGLR.D | HexNAc(3)Hex(5)NeuAc(1) | C[+57], N[+1711] | 1169.3135 | 4 | 4674.2322 | 4674.1959 | 0 | 7.8 | 127 | Specific | 829.64 | 784.99 | 784.99 | 15.87 | 67 | >sp|P01876|IGHA1_HUMAN Immunoglobulin heavy constant alpha 1 OS=Homo sapiens OX=9606 GN=IGHA1 PE=1 SV=2 | 255 | 20201115_10113_1ng_glyco_hcd203040_compound15.22335.22335.4 | controllerType=0 controllerNumber=1 scan=22335 | 44.8332 |
| 18025:4 | 8 | R.LSLHRPALEDLLLGSEAN[+1378.476]LTC[+57.021]TLTGLR.D | HexNAc(2)Hex(6) | C[+57], N[+1378] | 1086.2756 | 4 | 4342.0804 | 4342.074 | 0 | 1.5 | 127 | Specific | 863.96 | 732.12 | 732.12 | 17.64 | 67 | >sp|P01876|IGHA1_HUMAN Immunoglobulin heavy constant alpha 1 OS=Homo sapiens OX=9606 GN=IGHA1 PE=1 SV=2 | 255 | 20201115_10113_1ng_glyco_hcd203040_compound15.21246.21246.4 | controllerType=0 controllerNumber=1 scan=21246 | 42.5143 |
| 19001:4 | 8 | R.LSLHRPALEDLLLGSEAN[+1872.651]LTC[+57.021]TLTGLR.D | HexNAc(3)Hex(6)NeuAc(1) | C[+57], N[+1873] | 1209.8182 | 4 | 4836.2508 | 4836.2487 | 0 | 0.4 | 127 | Specific | 865.07 | 865.07 | 865.07 | 17.17 | 67 | >sp|P01876|IGHA1_HUMAN Immunoglobulin heavy constant alpha 1 OS=Homo sapiens OX=9606 GN=IGHA1 PE=1 SV=2 | 255 | 20201115_10113_1ng_glyco_hcd203040_compound15.22262.22262.4 | controllerType=0 controllerNumber=1 scan=22262 | 44.6762 |
| 18043:4 | 8 | R.LSLHRPALEDLLLGSEAN[+1460.529]LTC[+57.021]TLTGLR.D | HexNAc(4)Hex(4) | C[+57], N[+1461] | 1106.7893 | 4 | 4424.1356 | 4424.1271 | 0 | 1.9 | 127 | Specific | 877.56 | 817.66 | 817.66 | 17.85 | 67 | >sp|P01876|IGHA1_HUMAN Immunoglobulin heavy constant alpha 1 OS=Homo sapiens OX=9606 GN=IGHA1 PE=1 SV=2 | 255 | 20201115_10113_1ng_glyco_hcd203040_compound15.21264.21264.4 | controllerType=0 controllerNumber=1 scan=21264 | 42.5555 |
| 18131:4 | 8 | R.LSLHRPALEDLLLGSEAN[+1054.370]LTC[+57.021]TLTGLR.D | HexNAc(2)Hex(4) | C[+57], N[+1054] | 1005.2497 | 4 | 4017.9769 | 4017.9683 | 0 | 2.1 | 127 | Specific | 882.47 | 882.47 | 882.47 | 17.85 | 67 | >sp|P01876|IGHA1_HUMAN Immunoglobulin heavy constant alpha 1 OS=Homo sapiens OX=9606 GN=IGHA1 PE=1 SV=2 | 255 | 20201115_10113_1ng_glyco_hcd203040_compound15.21356.21356.4 | controllerType=0 controllerNumber=1 scan=21356 | 42.7439 |
| 18002:4 | 8 | R.LSLHRPALEDLLLGSEAN[+1581.555]LTC[+57.021]TLTGLR.D | HexNAc(3)Hex(6) | C[+57], N[+1582] | 1137.0536 | 4 | 4545.1927 | 4545.1533 | 0 | 8.7 | 127 | Specific | 897.15 | 874.55 | 874.55 | 16.85 | 67 | >sp|P01876|IGHA1_HUMAN Immunoglobulin heavy constant alpha 1 OS=Homo sapiens OX=9606 GN=IGHA1 PE=1 SV=2 | 255 | 20201115_10113_1ng_glyco_hcd203040_compound15.21221.21221.2 | controllerType=0 controllerNumber=1 scan=21221 | 42.4328 |
| 18034:4 | 8 | R.LSLHRPALEDLLLGSEAN[+1216.423]LTC[+57.021]TLTGLR.D | HexNAc(2)Hex(5) | C[+57], N[+1216] | 1045.7624 | 4 | 4180.0277 | 4180.0211 | 0 | 1.6 | 127 | Specific | 902.41 | 869.43 | 869.43 | 18.85 | 67 | >sp|P01876|IGHA1_HUMAN Immunoglobulin heavy constant alpha 1 OS=Homo sapiens OX=9606 GN=IGHA1 PE=1 SV=2 | 255 | 20201115_10113_1ng_glyco_hcd203040_compound15.21255.21255.4 | controllerType=0 controllerNumber=1 scan=21255 | 42.5319 |
| 18027:4 | 8 | R.LSLHRPALEDLLLGSEAN[+1378.476]LTC[+57.021]TLTGLR.D | HexNAc(2)Hex(6) | C[+57], N[+1378] | 1086.2756 | 4 | 4342.0804 | 4342.074 | 0 | 1.5 | 127 | Specific | 980.21 | 921.25 | 921.25 | 19.64 | 67 | >sp|P01876|IGHA1_HUMAN Immunoglobulin heavy constant alpha 1 OS=Homo sapiens OX=9606 GN=IGHA1 PE=1 SV=2 | 255 | 20201115_10113_1ng_glyco_hcd203040_compound15.21248.21248.4 | controllerType=0 controllerNumber=1 scan=21248 | 42.5195 |
| 18037:4 | 8 | R.LSLHRPALEDLLLGSEAN[+1216.423]LTC[+57.021]TLTGLR.D | HexNAc(2)Hex(5) | C[+57], N[+1216] | 1045.7624 | 4 | 4180.0277 | 4180.0211 | 0 | 1.6 | 127 | Specific | 981.57 | 815.03 | 815.03 | 19.85 | 67 | >sp|P01876|IGHA1_HUMAN Immunoglobulin heavy constant alpha 1 OS=Homo sapiens OX=9606 GN=IGHA1 PE=1 SV=2 | 255 | 20201115_10113_1ng_glyco_hcd203040_compound15.21258.21258.4 | controllerType=0 controllerNumber=1 scan=21258 | 42.5458 |
| 18110:4 | 8 | R.LSLHRPALEDLLLGSEAN[+1095.397]LTC[+57.021]TLTGLR.D | HexNAc(3)Hex(3) | C[+57], N[+1095] | 1015.5083 | 4 | 4059.0115 | 4058.9949 | 0 | 4.1 | 127 | Specific | 991.83 | 991.83 | 991.83 | 19.42 | 67 | >sp|P01876|IGHA1_HUMAN Immunoglobulin heavy constant alpha 1 OS=Homo sapiens OX=9606 GN=IGHA1 PE=1 SV=2 | 255 | 20201115_10113_1ng_glyco_hcd203040_compound15.21334.21334.4 | controllerType=0 controllerNumber=1 scan=21334 | 42.7023 |
| 18134:4 | 8 | R.LSLHRPALEDLLLGSEAN[+1054.370]LTC[+57.021]TLTGLR.D | HexNAc(2)Hex(4) | C[+57], N[+1054] | 1005.2497 | 4 | 4017.9769 | 4017.9683 | 0 | 2.1 | 127 | Specific | 996.45 | 996.45 | 996.45 | 19.85 | 67 | >sp|P01876|IGHA1_HUMAN Immunoglobulin heavy constant alpha 1 OS=Homo sapiens OX=9606 GN=IGHA1 PE=1 SV=2 | 255 | 20201115_10113_1ng_glyco_hcd203040_compound15.21359.21359.4 | controllerType=0 controllerNumber=1 scan=21359 | 42.7498 |
| 18012:4 | 8 | R.LSLHRPALEDLLLGSEAN[+1540.529]LTC[+57.021]TLTGLR.D | HexNAc(2)Hex(7) | C[+57], N[+1541] | 1126.7896 | 4 | 4504.1365 | 4504.1268 | 0 | 2.2 | 127 | Specific | 1022.54 | 941.02 | 941.02 | 20.85 | 67 | >sp|P01876|IGHA1_HUMAN Immunoglobulin heavy constant alpha 1 OS=Homo sapiens OX=9606 GN=IGHA1 PE=1 SV=2 | 255 | 20201115_10113_1ng_glyco_hcd203040_compound15.21232.21232.4 | controllerType=0 controllerNumber=1 scan=21232 | 42.4541 |
| 17988:4 | 8 | R.LSLHRPALEDLLLGSEAN[+1864.634]LTC[+57.021]TLTGLR.D | HexNAc(2)Hex(9) | C[+57], N[+1865] | 1207.8227 | 4 | 4828.2689 | 4828.2324 | 0 | 7.5 | 127 | Specific | 1031.67 | 1031.67 | 1031.67 | 19.87 | 67 | >sp|P01876|IGHA1_HUMAN Immunoglobulin heavy constant alpha 1 OS=Homo sapiens OX=9606 GN=IGHA1 PE=1 SV=2 | 255 | 20201115_10113_1ng_glyco_hcd203040_compound15.21207.21207.4 | controllerType=0 controllerNumber=1 scan=21207 | 42.402 |
| 13614:3 | 8 | R.LAGKPTHVN[+2263.835]VSVVMAEVDGTC[+57.021].Y | HexNAc(5)Hex(5)Fuc(3) | C[+57], N[+2264] | 1483.3094 | 3 | 4447.9136 | 4447.9135 | 0 | 0 | 332 | CRagged | 278.8 | 264.72 | 264.72 | 6.31 | 67 | >sp|P01876|IGHA1_HUMAN Immunoglobulin heavy constant alpha 1 OS=Homo sapiens OX=9606 GN=IGHA1 PE=1 SV=2 | 255 | 20201115_10113_1ng_glyco_hcd203040_compound15.16653.16653.3 | controllerType=0 controllerNumber=1 scan=16653 | 33.5738 |
| 10737:3 | 8 | R.LAGKPTHVN[+1809.666]VSVVMAEVDGTC[+57.021]Y.- | HexNAc(5)Hex(4)Fuc(1) | C[+57], N[+1810] | 1386.2758 | 3 | 4156.8129 | 4156.8081 | 0 | 1.1 | 332 | CRagged | 271.7 | 271.7 | 271.7 | 6.98 | 67 | >sp|P01876|IGHA1_HUMAN Immunoglobulin heavy constant alpha 1 OS=Homo sapiens OX=9606 GN=IGHA1 PE=1 SV=2 | 255 | 20201115_10113_1ng_glyco_hcd203040_compound15.13663.13663.3 | controllerType=0 controllerNumber=1 scan=13663 | 28.3875 |
| 10739:3 | 8 | R.LAGKPTHVN[+1809.666]VSVVMAEVDGTC[+57.021]Y.- | HexNAc(5)Hex(4)Fuc(1) | C[+57], N[+1810] | 1386.2758 | 3 | 4156.8129 | 4156.8081 | 0 | 1.1 | 332 | CRagged | 282.66 | 282.66 | 225.8 | 6.99 | 67 | >sp|P01876|IGHA1_HUMAN Immunoglobulin heavy constant alpha 1 OS=Homo sapiens OX=9606 GN=IGHA1 PE=1 SV=2 | 255 | 20201115_10113_1ng_glyco_hcd203040_compound15.13665.13665.3 | controllerType=0 controllerNumber=1 scan=13665 | 28.3916 |
| 15527:3 | 8 | R.LAGKPTHVN[+2553.910]VSVVMAEVDGTC[+57.021]Y.- | HexNAc(5)Hex(5)Fuc(1)NeuAc(2) | C[+57], N[+2554] | 1634.3544 | 3 | 4901.0487 | 4901.0518 | 0 | -0.6 | 332 | CRagged | 311.58 | 294.78 | 294.78 | 6.36 | 67 | >sp|P01876|IGHA1_HUMAN Immunoglobulin heavy constant alpha 1 OS=Homo sapiens OX=9606 GN=IGHA1 PE=1 SV=2 | 255 | 20201115_10113_1ng_glyco_hcd203040_compound15.18638.18638.3 | controllerType=0 controllerNumber=1 scan=18638 | 37.3327 |
| 10768:3 | 8 | R.LAGKPTHVN[+1378.476]VSVVMAEVDGTC[+57.021]Y.- | HexNAc(2)Hex(6) | C[+57], N[+1378] | 1242.5475 | 3 | 3725.6281 | 3725.6178 | 0 | 2.8 | 332 | CRagged | 354.5 | 354.5 | 279.2 | 8.1 | 67 | >sp|P01876|IGHA1_HUMAN Immunoglobulin heavy constant alpha 1 OS=Homo sapiens OX=9606 GN=IGHA1 PE=1 SV=2 | 255 | 20201115_10113_1ng_glyco_hcd203040_compound15.13695.13695.3 | controllerType=0 controllerNumber=1 scan=13695 | 28.4467 |
| 13780:3 | 8 | R.LAGKPTHVN[+1710.598]VSVVMAEVDGTC[+57.021]Y.- | HexNAc(3)Hex(5)NeuAc(1) | C[+57], N[+1711] | 1353.2552 | 3 | 4057.751 | 4057.7397 | 0 | 2.8 | 332 | CRagged | 442.64 | 442.64 | 299.26 | 10.13 | 67 | >sp|P01876|IGHA1_HUMAN Immunoglobulin heavy constant alpha 1 OS=Homo sapiens OX=9606 GN=IGHA1 PE=1 SV=2 | 255 | 20201115_10113_1ng_glyco_hcd203040_compound15.16825.16825.3 | controllerType=0 controllerNumber=1 scan=16825 | 33.8902 |
| 15524:3 | 8 | R.LAGKPTHVN[+2553.910]VSVVMAEVDGTC[+57.021]Y.- | HexNAc(5)Hex(5)Fuc(1)NeuAc(2) | C[+57], N[+2554] | 1634.3544 | 3 | 4901.0487 | 4901.0518 | 0 | -0.6 | 332 | CRagged | 455.56 | 455.56 | 455.56 | 9.3 | 67 | >sp|P01876|IGHA1_HUMAN Immunoglobulin heavy constant alpha 1 OS=Homo sapiens OX=9606 GN=IGHA1 PE=1 SV=2 | 255 | 20201115_10113_1ng_glyco_hcd203040_compound15.18635.18635.3 | controllerType=0 controllerNumber=1 scan=18635 | 37.3286 |
| 11974:3 | 8 | R.LAGKPTHVN[+1501.555]VSVVMAEVDGTC[+57.021]Y.- | HexNAc(5)Hex(3) | C[+57], N[+1502] | 1283.5723 | 3 | 3848.7024 | 3848.6974 | 0 | 1.3 | 332 | CRagged | 460.42 | 367.65 | 367.65 | 8.69 | 67 | >sp|P01876|IGHA1_HUMAN Immunoglobulin heavy constant alpha 1 OS=Homo sapiens OX=9606 GN=IGHA1 PE=1 SV=2 | 255 | 20201115_10113_1ng_glyco_hcd203040_compound15.14947.14947.3 | controllerType=0 controllerNumber=1 scan=14947 | 30.6954 |
| 13673:3 | 8 | R.LAGKPTHVN[+1913.677]VSVVMAEVDGTC[+57.021]Y.- | HexNAc(4)Hex(5)NeuAc(1) | C[+57], N[+1914] | 1420.9485 | 3 | 4260.8309 | 4260.8191 | 0 | 2.8 | 332 | CRagged | 464.45 | 396.31 | 396.31 | 10.39 | 67 | >sp|P01876|IGHA1_HUMAN Immunoglobulin heavy constant alpha 1 OS=Homo sapiens OX=9606 GN=IGHA1 PE=1 SV=2 | 255 | 20201115_10113_1ng_glyco_hcd203040_compound15.16714.16714.3 | controllerType=0 controllerNumber=1 scan=16714 | 33.682 |
| 13721:3 | 8 | R.LAGKPTHVN[+1872.651]VSVVMAEVDGTC[+57.021]Y.- | HexNAc(3)Hex(6)NeuAc(1) | C[+57], N[+1873] | 1407.2754 | 3 | 4219.8118 | 4219.7926 | 0 | 4.6 | 332 | CRagged | 473.75 | 473.75 | 327.23 | 9.96 | 67 | >sp|P01876|IGHA1_HUMAN Immunoglobulin heavy constant alpha 1 OS=Homo sapiens OX=9606 GN=IGHA1 PE=1 SV=2 | 255 | 20201115_10113_1ng_glyco_hcd203040_compound15.16763.16763.3 | controllerType=0 controllerNumber=1 scan=16763 | 33.7677 |
| 10770:3 | 8 | R.LAGKPTHVN[+1378.476]VSVVMAEVDGTC[+57.021]Y.- | HexNAc(2)Hex(6) | C[+57], N[+1378] | 1242.5475 | 3 | 3725.6281 | 3725.6178 | 0 | 2.8 | 332 | CRagged | 483.99 | 483.99 | 353.68 | 10.37 | 67 | >sp|P01876|IGHA1_HUMAN Immunoglobulin heavy constant alpha 1 OS=Homo sapiens OX=9606 GN=IGHA1 PE=1 SV=2 | 255 | 20201115_10113_1ng_glyco_hcd203040_compound15.13697.13697.3 | controllerType=0 controllerNumber=1 scan=13697 | 28.4524 |
| 13783:3 | 8 | R.LAGKPTHVN[+1710.598]VSVVMAEVDGTC[+57.021]Y.- | HexNAc(3)Hex(5)NeuAc(1) | C[+57], N[+1711] | 1353.2552 | 3 | 4057.751 | 4057.7397 | 0 | 2.8 | 332 | CRagged | 528.03 | 528.03 | 391.8 | 10.62 | 67 | >sp|P01876|IGHA1_HUMAN Immunoglobulin heavy constant alpha 1 OS=Homo sapiens OX=9606 GN=IGHA1 PE=1 SV=2 | 255 | 20201115_10113_1ng_glyco_hcd203040_compound15.16828.16828.3 | controllerType=0 controllerNumber=1 scan=16828 | 33.8949 |
| 15496:3 | 8 | R.LAGKPTHVN[+2553.910]VSVVMAEVDGTC[+57.021]Y.- | HexNAc(5)Hex(5)Fuc(1)NeuAc(2) | C[+57], N[+2554] | 1634.3604 | 3 | 4901.0667 | 4901.0518 | 0 | 3 | 332 | CRagged | 547.33 | 395.35 | 395.35 | 10.81 | 67 | >sp|P01876|IGHA1_HUMAN Immunoglobulin heavy constant alpha 1 OS=Homo sapiens OX=9606 GN=IGHA1 PE=1 SV=2 | 255 | 20201115_10113_1ng_glyco_hcd203040_compound15.18606.18606.3 | controllerType=0 controllerNumber=1 scan=18606 | 37.2777 |
| 11981:4 | 8 | R.LAGKPTHVN[+1647.613]VSVVMAEVDGTC[+57.021]Y.- | HexNAc(5)Hex(3)Fuc(1) | C[+57], N[+1648] | 999.4458 | 4 | 3994.7615 | 3994.7553 | 0 | 1.6 | 332 | CRagged | 554.58 | 310.42 | 310.42 | 8.51 | 67 | >sp|P01876|IGHA1_HUMAN Immunoglobulin heavy constant alpha 1 OS=Homo sapiens OX=9606 GN=IGHA1 PE=1 SV=2 | 255 | 20201115_10113_1ng_glyco_hcd203040_compound15.14954.14954.4 | controllerType=0 controllerNumber=1 scan=14954 | 30.7045 |
| 13621:4 | 8 | R.LAGKPTHVN[+2262.814]VSVVMAEVDGTC[+57.021]Y.- | HexNAc(5)Hex(5)Fuc(1)NeuAc(1) | C[+57], N[+2263] | 1153.2476 | 4 | 4609.9686 | 4609.9564 | 0 | 2.6 | 332 | CRagged | 560 | 492.57 | 492.57 | 10.86 | 67 | >sp|P01876|IGHA1_HUMAN Immunoglobulin heavy constant alpha 1 OS=Homo sapiens OX=9606 GN=IGHA1 PE=1 SV=2 | 255 | 20201115_10113_1ng_glyco_hcd203040_compound15.16660.16660.4 | controllerType=0 controllerNumber=1 scan=16660 | 33.5849 |
| 13624:4 | 8 | R.LAGKPTHVN[+2262.814]VSVVMAEVDGTC[+57.021]Y.- | HexNAc(5)Hex(5)Fuc(1)NeuAc(1) | C[+57], N[+2263] | 1153.2476 | 4 | 4609.9686 | 4609.9564 | 0 | 2.6 | 332 | CRagged | 592.33 | 576.47 | 576.47 | 11.92 | 67 | >sp|P01876|IGHA1_HUMAN Immunoglobulin heavy constant alpha 1 OS=Homo sapiens OX=9606 GN=IGHA1 PE=1 SV=2 | 255 | 20201115_10113_1ng_glyco_hcd203040_compound15.16663.16663.4 | controllerType=0 controllerNumber=1 scan=16663 | 33.5939 |
| 15499:3 | 8 | R.LAGKPTHVN[+2553.910]VSVVMAEVDGTC[+57.021]Y.- | HexNAc(5)Hex(5)Fuc(1)NeuAc(2) | C[+57], N[+2554] | 1634.3604 | 3 | 4901.0667 | 4901.0518 | 0 | 3 | 332 | CRagged | 651.7 | 483.45 | 483.45 | 13.99 | 67 | >sp|P01876|IGHA1_HUMAN Immunoglobulin heavy constant alpha 1 OS=Homo sapiens OX=9606 GN=IGHA1 PE=1 SV=2 | 255 | 20201115_10113_1ng_glyco_hcd203040_compound15.18609.18609.3 | controllerType=0 controllerNumber=1 scan=18609 | 37.2825 |
| 11929:3 | 8 | R.LAGKPTHVN[+1501.555]VSVVMAEVDGTC[+57.021]Y.- | HexNAc(5)Hex(3) | C[+57], N[+1502] | 1283.5722 | 3 | 3848.702 | 3848.6974 | 0 | 1.2 | 332 | CRagged | 667.65 | 644.09 | 644.09 | 13.8 | 67 | >sp|P01876|IGHA1_HUMAN Immunoglobulin heavy constant alpha 1 OS=Homo sapiens OX=9606 GN=IGHA1 PE=1 SV=2 | 255 | 20201115_10113_1ng_glyco_hcd203040_compound15.14900.14900.3 | controllerType=0 controllerNumber=1 scan=14900 | 30.593 |
| 13723:3 | 8 | R.LAGKPTHVN[+1872.651]VSVVMAEVDGTC[+57.021]Y.- | HexNAc(3)Hex(6)NeuAc(1) | C[+57], N[+1873] | 1407.2754 | 3 | 4219.8118 | 4219.7926 | 0 | 4.6 | 332 | CRagged | 673.96 | 624.72 | 362.29 | 13.58 | 67 | >sp|P01876|IGHA1_HUMAN Immunoglobulin heavy constant alpha 1 OS=Homo sapiens OX=9606 GN=IGHA1 PE=1 SV=2 | 255 | 20201115_10113_1ng_glyco_hcd203040_compound15.16765.16765.3 | controllerType=0 controllerNumber=1 scan=16765 | 33.7711 |
| 11970:3 | 8 | R.LAGKPTHVN[+1647.613]VSVVMAEVDGTC[+57.021]Y.- | HexNAc(5)Hex(3)Fuc(1) | C[+57], N[+1648] | 1332.258 | 3 | 3994.7594 | 3994.7553 | 0 | 1 | 332 | CRagged | 696.55 | 594.84 | 594.84 | 13.8 | 67 | >sp|P01876|IGHA1_HUMAN Immunoglobulin heavy constant alpha 1 OS=Homo sapiens OX=9606 GN=IGHA1 PE=1 SV=2 | 255 | 20201115_10113_1ng_glyco_hcd203040_compound15.14943.14943.3 | controllerType=0 controllerNumber=1 scan=14943 | 30.6894 |
| 13676:3 | 8 | R.LAGKPTHVN[+1913.677]VSVVMAEVDGTC[+57.021]Y.- | HexNAc(4)Hex(5)NeuAc(1) | C[+57], N[+1914] | 1420.9485 | 3 | 4260.8309 | 4260.8191 | 0 | 2.8 | 332 | CRagged | 710.86 | 508.18 | 508.18 | 14.99 | 67 | >sp|P01876|IGHA1_HUMAN Immunoglobulin heavy constant alpha 1 OS=Homo sapiens OX=9606 GN=IGHA1 PE=1 SV=2 | 255 | 20201115_10113_1ng_glyco_hcd203040_compound15.16717.16717.3 | controllerType=0 controllerNumber=1 scan=16717 | 33.6856 |
| 13611:3 | 8 | R.LAGKPTHVN[+2262.814]VSVVMAEVDGTC[+57.021]Y.- | HexNAc(5)Hex(5)Fuc(1)NeuAc(1) | C[+57], N[+2263] | 1537.3298 | 3 | 4609.9748 | 4609.9564 | 0 | 4 | 332 | CRagged | 715.75 | 551.06 | 551.06 | 14.58 | 67 | >sp|P01876|IGHA1_HUMAN Immunoglobulin heavy constant alpha 1 OS=Homo sapiens OX=9606 GN=IGHA1 PE=1 SV=2 | 255 | 20201115_10113_1ng_glyco_hcd203040_compound15.16649.16649.3 | controllerType=0 controllerNumber=1 scan=16649 | 33.5524 |
| 11932:3 | 8 | R.LAGKPTHVN[+1501.555]VSVVMAEVDGTC[+57.021]Y.- | HexNAc(5)Hex(3) | C[+57], N[+1502] | 1283.5722 | 3 | 3848.702 | 3848.6974 | 0 | 1.2 | 332 | CRagged | 763.4 | 745.03 | 745.03 | 15.8 | 67 | >sp|P01876|IGHA1_HUMAN Immunoglobulin heavy constant alpha 1 OS=Homo sapiens OX=9606 GN=IGHA1 PE=1 SV=2 | 255 | 20201115_10113_1ng_glyco_hcd203040_compound15.14903.14903.3 | controllerType=0 controllerNumber=1 scan=14903 | 30.5991 |
| 10769:3 | 8 | R.LAGKPTHVN[+1216.423]VSVVMAEVDGTC[+57.021]Y.- | HexNAc(2)Hex(5) | C[+57], N[+1216] | 1188.5297 | 3 | 3563.5746 | 3563.5649 | 0 | 2.7 | 332 | CRagged | 764.09 | 693.77 | 447.04 | 15.99 | 67 | >sp|P01876|IGHA1_HUMAN Immunoglobulin heavy constant alpha 1 OS=Homo sapiens OX=9606 GN=IGHA1 PE=1 SV=2 | 255 | 20201115_10113_1ng_glyco_hcd203040_compound15.13696.13696.3 | controllerType=0 controllerNumber=1 scan=13696 | 28.4505 |
| 11829:3 | 8 | R.LAGKPTHVN[+1663.608]VSVVMAEVDGTC[+57.021]Y.- | HexNAc(5)Hex(4) | C[+57], N[+1664] | 1337.5915 | 3 | 4010.7599 | 4010.7502 | 0 | 2.4 | 332 | CRagged | 767.65 | 759.99 | 485.97 | 16.01 | 67 | >sp|P01876|IGHA1_HUMAN Immunoglobulin heavy constant alpha 1 OS=Homo sapiens OX=9606 GN=IGHA1 PE=1 SV=2 | 255 | 20201115_10113_1ng_glyco_hcd203040_compound15.14795.14795.3 | controllerType=0 controllerNumber=1 scan=14795 | 30.387 |
| 11834:3 | 8 | R.LAGKPTHVN[+1540.529]VSVVMAEVDGTC[+57.021]Y.- | HexNAc(2)Hex(7) | C[+57], N[+1541] | 1296.5689 | 3 | 3887.6922 | 3887.6706 | 0 | 5.5 | 332 | CRagged | 773.39 | 641.97 | 485.94 | 15.43 | 67 | >sp|P01876|IGHA1_HUMAN Immunoglobulin heavy constant alpha 1 OS=Homo sapiens OX=9606 GN=IGHA1 PE=1 SV=2 | 255 | 20201115_10113_1ng_glyco_hcd203040_compound15.14801.14801.3 | controllerType=0 controllerNumber=1 scan=14801 | 30.3985 |
| 10767:3 | 8 | R.LAGKPTHVN[+1216.423]VSVVMAEVDGTC[+57.021]Y.- | HexNAc(2)Hex(5) | C[+57], N[+1216] | 1188.5297 | 3 | 3563.5746 | 3563.5649 | 0 | 2.7 | 332 | CRagged | 773.4 | 710.81 | 427.7 | 15.99 | 67 | >sp|P01876|IGHA1_HUMAN Immunoglobulin heavy constant alpha 1 OS=Homo sapiens OX=9606 GN=IGHA1 PE=1 SV=2 | 255 | 20201115_10113_1ng_glyco_hcd203040_compound15.13694.13694.3 | controllerType=0 controllerNumber=1 scan=13694 | 28.4452 |
| 11977:3 | 8 | R.LAGKPTHVN[+1501.555]VSVVMAEVDGTC[+57.021]Y.- | HexNAc(5)Hex(3) | C[+57], N[+1502] | 1283.5723 | 3 | 3848.7024 | 3848.6974 | 0 | 1.3 | 332 | CRagged | 773.95 | 596.74 | 596.74 | 15.8 | 67 | >sp|P01876|IGHA1_HUMAN Immunoglobulin heavy constant alpha 1 OS=Homo sapiens OX=9606 GN=IGHA1 PE=1 SV=2 | 255 | 20201115_10113_1ng_glyco_hcd203040_compound15.14950.14950.3 | controllerType=0 controllerNumber=1 scan=14950 | 30.6992 |
| 11812:3 | 8 | R.LAGKPTHVN[+1702.581]VSVVMAEVDGTC[+57.021]Y.- | HexNAc(2)Hex(8) | C[+57], N[+1703] | 1350.5821 | 3 | 4049.7317 | 4049.7234 | 0 | 2 | 332 | CRagged | 813.14 | 813.14 | 524.38 | 17.01 | 67 | >sp|P01876|IGHA1_HUMAN Immunoglobulin heavy constant alpha 1 OS=Homo sapiens OX=9606 GN=IGHA1 PE=1 SV=2 | 255 | 20201115_10113_1ng_glyco_hcd203040_compound15.14778.14778.3 | controllerType=0 controllerNumber=1 scan=14778 | 30.3451 |
| 11843:3 | 8 | R.LAGKPTHVN[+1809.666]VSVVMAEVDGTC[+57.021]Y.- | HexNAc(5)Hex(4)Fuc(1) | C[+57], N[+1810] | 1386.2822 | 3 | 4156.8319 | 4156.8081 | 0 | 5.7 | 332 | CRagged | 814.32 | 661.25 | 518.14 | 16.43 | 67 | >sp|P01876|IGHA1_HUMAN Immunoglobulin heavy constant alpha 1 OS=Homo sapiens OX=9606 GN=IGHA1 PE=1 SV=2 | 255 | 20201115_10113_1ng_glyco_hcd203040_compound15.14810.14810.3 | controllerType=0 controllerNumber=1 scan=14810 | 30.4157 |
| 13626:3 | 8 | R.LAGKPTHVN[+2262.814]VSVVMAEVDGTC[+57.021]Y.- | HexNAc(5)Hex(5)Fuc(1)NeuAc(1) | C[+57], N[+2263] | 1537.3274 | 3 | 4609.9678 | 4609.9564 | 0 | 2.5 | 332 | CRagged | 835.56 | 526.13 | 526.13 | 17.01 | 67 | >sp|P01876|IGHA1_HUMAN Immunoglobulin heavy constant alpha 1 OS=Homo sapiens OX=9606 GN=IGHA1 PE=1 SV=2 | 255 | 20201115_10113_1ng_glyco_hcd203040_compound15.16665.16665.3 | controllerType=0 controllerNumber=1 scan=16665 | 33.5971 |
| 11972:3 | 8 | R.LAGKPTHVN[+1647.613]VSVVMAEVDGTC[+57.021]Y.- | HexNAc(5)Hex(3)Fuc(1) | C[+57], N[+1648] | 1332.258 | 3 | 3994.7594 | 3994.7553 | 0 | 1 | 332 | CRagged | 844.78 | 732.47 | 732.47 | 16.8 | 67 | >sp|P01876|IGHA1_HUMAN Immunoglobulin heavy constant alpha 1 OS=Homo sapiens OX=9606 GN=IGHA1 PE=1 SV=2 | 255 | 20201115_10113_1ng_glyco_hcd203040_compound15.14945.14945.3 | controllerType=0 controllerNumber=1 scan=14945 | 30.6935 |
| 11939:3 | 8 | R.LAGKPTHVN[+1647.613]VSVVMAEVDGTC[+57.021]Y.- | HexNAc(5)Hex(3)Fuc(1) | C[+57], N[+1648] | 1332.2595 | 3 | 3994.7638 | 3994.7553 | 0 | 2.1 | 332 | CRagged | 857.43 | 678.02 | 678.02 | 18.01 | 67 | >sp|P01876|IGHA1_HUMAN Immunoglobulin heavy constant alpha 1 OS=Homo sapiens OX=9606 GN=IGHA1 PE=1 SV=2 | 255 | 20201115_10113_1ng_glyco_hcd203040_compound15.14910.14910.3 | controllerType=0 controllerNumber=1 scan=14910 | 30.6101 |
| 13609:3 | 8 | R.LAGKPTHVN[+2262.814]VSVVMAEVDGTC[+57.021]Y.- | HexNAc(5)Hex(5)Fuc(1)NeuAc(1) | C[+57], N[+2263] | 1537.3298 | 3 | 4609.9748 | 4609.9564 | 0 | 4 | 332 | CRagged | 860.75 | 588.87 | 588.87 | 17.58 | 67 | >sp|P01876|IGHA1_HUMAN Immunoglobulin heavy constant alpha 1 OS=Homo sapiens OX=9606 GN=IGHA1 PE=1 SV=2 | 255 | 20201115_10113_1ng_glyco_hcd203040_compound15.16647.16647.3 | controllerType=0 controllerNumber=1 scan=16647 | 33.5376 |
| 11814:3 | 8 | R.LAGKPTHVN[+1702.581]VSVVMAEVDGTC[+57.021]Y.- | HexNAc(2)Hex(8) | C[+57], N[+1703] | 1350.5821 | 3 | 4049.7317 | 4049.7234 | 0 | 2 | 332 | CRagged | 864.06 | 822.14 | 529.93 | 18.01 | 67 | >sp|P01876|IGHA1_HUMAN Immunoglobulin heavy constant alpha 1 OS=Homo sapiens OX=9606 GN=IGHA1 PE=1 SV=2 | 255 | 20201115_10113_1ng_glyco_hcd203040_compound15.14780.14780.3 | controllerType=0 controllerNumber=1 scan=14780 | 30.3502 |
| 11775:3 | 8 | R.LAGKPTHVN[+1702.581]VSVVMAEVDGTC[+57.021]Y.- | HexNAc(2)Hex(8) | C[+57], N[+1703] | 1350.5821 | 3 | 4049.7317 | 4049.7234 | 0 | 2 | 332 | CRagged | 872.13 | 828.06 | 518.11 | 18.01 | 67 | >sp|P01876|IGHA1_HUMAN Immunoglobulin heavy constant alpha 1 OS=Homo sapiens OX=9606 GN=IGHA1 PE=1 SV=2 | 255 | 20201115_10113_1ng_glyco_hcd203040_compound15.14739.14739.3 | controllerType=0 controllerNumber=1 scan=14739 | 30.2631 |
| 11823:3 | 8 | R.LAGKPTHVN[+1378.476]VSVVMAEVDGTC[+57.021]Y.- | HexNAc(2)Hex(6) | C[+57], N[+1378] | 1242.5474 | 3 | 3725.6277 | 3725.6178 | 0 | 2.7 | 332 | CRagged | 876.27 | 829.69 | 527.3 | 17.99 | 67 | >sp|P01876|IGHA1_HUMAN Immunoglobulin heavy constant alpha 1 OS=Homo sapiens OX=9606 GN=IGHA1 PE=1 SV=2 | 255 | 20201115_10113_1ng_glyco_hcd203040_compound15.14789.14789.3 | controllerType=0 controllerNumber=1 scan=14789 | 30.3722 |
| 11840:3 | 8 | R.LAGKPTHVN[+1809.666]VSVVMAEVDGTC[+57.021]Y.- | HexNAc(5)Hex(4)Fuc(1) | C[+57], N[+1810] | 1386.2822 | 3 | 4156.8319 | 4156.8081 | 0 | 5.7 | 332 | CRagged | 877.75 | 726.1 | 594.73 | 17.43 | 67 | >sp|P01876|IGHA1_HUMAN Immunoglobulin heavy constant alpha 1 OS=Homo sapiens OX=9606 GN=IGHA1 PE=1 SV=2 | 255 | 20201115_10113_1ng_glyco_hcd203040_compound15.14807.14807.3 | controllerType=0 controllerNumber=1 scan=14807 | 30.4088 |
| 11836:3 | 8 | R.LAGKPTHVN[+1540.529]VSVVMAEVDGTC[+57.021]Y.- | HexNAc(2)Hex(7) | C[+57], N[+1541] | 1296.5689 | 3 | 3887.6922 | 3887.6706 | 0 | 5.5 | 332 | CRagged | 880.68 | 871.65 | 539.69 | 17.43 | 67 | >sp|P01876|IGHA1_HUMAN Immunoglobulin heavy constant alpha 1 OS=Homo sapiens OX=9606 GN=IGHA1 PE=1 SV=2 | 255 | 20201115_10113_1ng_glyco_hcd203040_compound15.14803.14803.3 | controllerType=0 controllerNumber=1 scan=14803 | 30.4034 |
| 11942:3 | 8 | R.LAGKPTHVN[+1647.613]VSVVMAEVDGTC[+57.021]Y.- | HexNAc(5)Hex(3)Fuc(1) | C[+57], N[+1648] | 1332.2595 | 3 | 3994.7638 | 3994.7553 | 0 | 2.1 | 332 | CRagged | 890.31 | 624.95 | 624.95 | 18.01 | 67 | >sp|P01876|IGHA1_HUMAN Immunoglobulin heavy constant alpha 1 OS=Homo sapiens OX=9606 GN=IGHA1 PE=1 SV=2 | 255 | 20201115_10113_1ng_glyco_hcd203040_compound15.14913.14913.3 | controllerType=0 controllerNumber=1 scan=14913 | 30.624 |
| 11826:3 | 8 | R.LAGKPTHVN[+1663.608]VSVVMAEVDGTC[+57.021]Y.- | HexNAc(5)Hex(4) | C[+57], N[+1664] | 1337.5915 | 3 | 4010.7599 | 4010.7502 | 0 | 2.4 | 332 | CRagged | 910.54 | 910.54 | 597.56 | 19.01 | 67 | >sp|P01876|IGHA1_HUMAN Immunoglobulin heavy constant alpha 1 OS=Homo sapiens OX=9606 GN=IGHA1 PE=1 SV=2 | 255 | 20201115_10113_1ng_glyco_hcd203040_compound15.14792.14792.3 | controllerType=0 controllerNumber=1 scan=14792 | 30.3759 |
| 11858:3 | 8 | R.LAGKPTHVN[+1216.423]VSVVMAEVDGTC[+57.021]Y.- | HexNAc(2)Hex(5) | C[+57], N[+1216] | 1188.5281 | 3 | 3563.5698 | 3563.5649 | 0 | 1.4 | 332 | CRagged | 915.43 | 915.43 | 525.12 | 18.8 | 67 | >sp|P01876|IGHA1_HUMAN Immunoglobulin heavy constant alpha 1 OS=Homo sapiens OX=9606 GN=IGHA1 PE=1 SV=2 | 255 | 20201115_10113_1ng_glyco_hcd203040_compound15.14826.14826.3 | controllerType=0 controllerNumber=1 scan=14826 | 30.4548 |
| 11820:3 | 8 | R.LAGKPTHVN[+1378.476]VSVVMAEVDGTC[+57.021]Y.- | HexNAc(2)Hex(6) | C[+57], N[+1378] | 1242.5474 | 3 | 3725.6277 | 3725.6178 | 0 | 2.7 | 332 | CRagged | 916.54 | 916.54 | 571.34 | 18.99 | 67 | >sp|P01876|IGHA1_HUMAN Immunoglobulin heavy constant alpha 1 OS=Homo sapiens OX=9606 GN=IGHA1 PE=1 SV=2 | 255 | 20201115_10113_1ng_glyco_hcd203040_compound15.14786.14786.3 | controllerType=0 controllerNumber=1 scan=14786 | 30.3626 |
| 11772:3 | 8 | R.LAGKPTHVN[+1702.581]VSVVMAEVDGTC[+57.021]Y.- | HexNAc(2)Hex(8) | C[+57], N[+1703] | 1350.5821 | 3 | 4049.7317 | 4049.7234 | 0 | 2 | 332 | CRagged | 928.93 | 928.93 | 592.34 | 19.01 | 67 | >sp|P01876|IGHA1_HUMAN Immunoglobulin heavy constant alpha 1 OS=Homo sapiens OX=9606 GN=IGHA1 PE=1 SV=2 | 255 | 20201115_10113_1ng_glyco_hcd203040_compound15.14736.14736.3 | controllerType=0 controllerNumber=1 scan=14736 | 30.255 |

Table S4. Cont.

| Query #:z | Protein Rank | Glycopeptide | Glycan | Modification Type(s) | Observed m/z | z | Observed (M+H) | Calc. mass (M+H) | Off-by-x error | Mass error (ppm) | StartPosition | Cleavage | Score | Delta | Delta Mod | |Log Prob| | # of unique peptides | Protein Name | Protein DB number | Comment | Scan # | Scan Time |
|---|---|---|---|---|---|---|---|---|---|---|---|---|---|---|---|---|---|---|---|---|---|---|
| 11862:3 | 8 | R.LAGKPTHVN[+1216.423]VSVVMAEVDGTC[+57.021]Y.- | HexNAc(2)Hex(5) | C[+57], N[+1216] | 1188.5281 | 3 | 3563.5698 | 3563.5649 | 0 | 1.4 | 332 | CRagged | 939.31 | 924.76 | 563.28 | 18.8 | 67 | >sp|P01876|IGHA1_HUMAN Immunoglobulin heavy constant alpha 1 OS=Homo sapiens OX=9606 GN=IGHA1 PE=1 SV=2 | 255 | 20201115_10113_1ng_glyco_hcd203040_compound15.14830.14830.3 | controllerType=0 controllerNumber=1 scan=14830 | 30.4681 |
| 09912:3 | 8 | R.LAGKPTHVN[+1216.423]VSVVM[+15.995]AEVDGTC[+57.021]Y.- | HexNAc(2)Hex(5) | C[+57], M[+16], N[+1216] | 1193.8609 | 3 | 3579.5681 | 3579.5599 | 0 | 2.3 | 332 | CRagged | 292.41 | 292.41 | 292.41 | 7.31 | 67 | >sp|P01876|IGHA1_HUMAN Immunoglobulin heavy constant alpha 1 OS=Homo sapiens OX=9606 GN=IGHA1 PE=1 SV=2 | 255 | 20201115_10113_1ng_glyco_hcd203040_compound15.12808.12808.3 | controllerType=0 controllerNumber=1 scan=12808 | 26.9377 |
| 09914:3 | 8 | R.LAGKPTHVN[+1216.423]VSVVM[+15.995]AEVDGTC[+57.021]Y.- | HexNAc(2)Hex(5) | C[+57], M[+16], N[+1216] | 1193.8609 | 3 | 3579.5681 | 3579.5599 | 0 | 2.3 | 332 | CRagged | 331.1 | 331.1 | 331.1 | 7.84 | 67 | >sp|P01876|IGHA1_HUMAN Immunoglobulin heavy constant alpha 1 OS=Homo sapiens OX=9606 GN=IGHA1 PE=1 SV=2 | 255 | 20201115_10113_1ng_glyco_hcd203040_compound15.12810.12810.3 | controllerType=0 controllerNumber=1 scan=12810 | 26.9413 |
| 24345:4 | 13 | K.APDKNVIFSPLSISTALAFLSLGAHN[+3007.058]TTLTEILK.G | HexNAc(5)Hex(6)Fuc(1)NeuAc(3) | N[+3007] | 1648.2725 | 4 | 6590.068 | 6590.0361 | 0 | 4.8 | 68 | Specific | 237.66 | 237.66 | 237.66 | 5.85 | 46 | >sp|P01011|AACT_HUMAN Alpha-1-antichymotrypsin OS=Homo sapiens OX=9606 GN=SERPINA3 PE=1 SV=2 | 1314 | 20201115_10113_1ng_glyco_hcd203040_compound15.27860.27860.3 | controllerType=0 controllerNumber=1 scan=27860 | 56.9857 |
| 24343:4 | 13 | K.APDKNVIFSPLSISTALAFLSLGAHN[+3007.058]TTLTEILK.G | HexNAc(5)Hex(6)Fuc(1)NeuAc(3) | N[+3007] | 1648.2725 | 4 | 6590.068 | 6590.0361 | 0 | 4.8 | 68 | Specific | 254.26 | 254.26 | 254.26 | 5.98 | 46 | >sp|P01011|AACT_HUMAN Alpha-1-antichymotrypsin OS=Homo sapiens OX=9606 GN=SERPINA3 PE=1 SV=2 | 1314 | 20201115_10113_1ng_glyco_hcd203040_compound15.27858.27858.3 | controllerType=0 controllerNumber=1 scan=27858 | 56.9818 |
| 21498:5 | 13 | K.APDKNVIFSPLSISTALAFLSLGAHN[+2424.867]TTLTEILK.G | HexNAc(5)Hex(6)Fuc(1)NeuAc(1) | N[+2425] | 1202.3793 | 5 | 6007.8673 | 6007.8453 | 0 | 3.7 | 68 | Specific | 472.02 | 214.45 | 214.45 | 7.33 | 46 | >sp|P01011|AACT_HUMAN Alpha-1-antichymotrypsin OS=Homo sapiens OX=9606 GN=SERPINA3 PE=1 SV=2 | 1314 | 20201115_10113_1ng_glyco_hcd203040_compound15.24873.24873.5 | controllerType=0 controllerNumber=1 scan=24873 | 50.4638 |
| 21500:5 | 13 | K.APDKNVIFSPLSISTALAFLSLGAHN[+2424.867]TTLTEILK.G | HexNAc(5)Hex(6)Fuc(1)NeuAc(1) | N[+2425] | 1202.3793 | 5 | 6007.8673 | 6007.8453 | 0 | 3.7 | 68 | Specific | 475.67 | 400.12 | 400.12 | 9.65 | 46 | >sp|P01011|AACT_HUMAN Alpha-1-antichymotrypsin OS=Homo sapiens OX=9606 GN=SERPINA3 PE=1 SV=2 | 1314 | 20201115_10113_1ng_glyco_hcd203040_compound15.24875.24875.5 | controllerType=0 controllerNumber=1 scan=24875 | 50.4685 |
| 21609:5 | 13 | K.APDKNVIFSPLSISTALAFLSLGAHN[+2059.735]TTLTEILK.G | HexNAc(4)Hex(5)Fuc(1)NeuAc(1) | N[+2060] | 1129.3543 | 5 | 5642.7421 | 5642.7131 | 0 | 5.1 | 68 | Specific | 576.67 | 576.67 | 576.67 | 9.64 | 46 | >sp|P01011|AACT_HUMAN Alpha-1-antichymotrypsin OS=Homo sapiens OX=9606 GN=SERPINA3 PE=1 SV=2 | 1314 | 20201115_10113_1ng_glyco_hcd203040_compound15.24989.24989.2 | controllerType=0 controllerNumber=1 scan=24989 | 50.7172 |
| 22785:5 | 13 | K.APDKNVIFSPLSISTALAFLSLGAHN[+2350.830]TTLTEILK.G | HexNAc(4)Hex(5)Fuc(1)NeuAc(2) | N[+2351] | 1187.5732 | 5 | 5933.8371 | 5933.8085 | 0 | 4.8 | 68 | Specific | 746.04 | 746.04 | 746.04 | 13.7 | 46 | >sp|P01011|AACT_HUMAN Alpha-1-antichymotrypsin OS=Homo sapiens OX=9606 GN=SERPINA3 PE=1 SV=2 | 1314 | 20201115_10113_1ng_glyco_hcd203040_compound15.26223.26223.5 | controllerType=0 controllerNumber=1 scan=26223 | 53.4364 |
| 22784:4 | 13 | K.APDKNVIFSPLSISTALAFLSLGAHN[+2350.830]TTLTEILK.G | HexNAc(4)Hex(5)Fuc(1)NeuAc(2) | N[+2351] | 1484.215 | 4 | 5933.838 | 5933.8085 | 0 | 5 | 68 | Specific | 786.03 | 775.21 | 775.21 | 14.79 | 46 | >sp|P01011|AACT_HUMAN Alpha-1-antichymotrypsin OS=Homo sapiens OX=9606 GN=SERPINA3 PE=1 SV=2 | 1314 | 20201115_10113_1ng_glyco_hcd203040_compound15.26222.26222.4 | controllerType=0 controllerNumber=1 scan=26222 | 53.4341 |
| 21607:5 | 13 | K.APDKNVIFSPLSISTALAFLSLGAHN[+2059.735]TTLTEILK.G | HexNAc(4)Hex(5)Fuc(1)NeuAc(1) | N[+2060] | 1129.3543 | 5 | 5642.7421 | 5642.7131 | 0 | 5.1 | 68 | Specific | 831.84 | 822.14 | 822.14 | 15.7 | 46 | >sp|P01011|AACT_HUMAN Alpha-1-antichymotrypsin OS=Homo sapiens OX=9606 GN=SERPINA3 PE=1 SV=2 | 1314 | 20201115_10113_1ng_glyco_hcd203040_compound15.24987.24987.2 | controllerType=0 controllerNumber=1 scan=24987 | 50.7108 |
| 21459:4 | 13 | K.APDKNVIFSPLSISTALAFLSLGAHN[+2570.925]TTLTEILK.G | HexNAc(5)Hex(6)Fuc(2)NeuAc(1) | N[+2571] | 1539.2385 | 4 | 6153.9322 | 6153.9032 | 0 | 4.7 | 68 | Specific | 832.08 | 832.08 | 832.08 | 15.95 | 46 | >sp|P01011|AACT_HUMAN Alpha-1-antichymotrypsin OS=Homo sapiens OX=9606 GN=SERPINA3 PE=1 SV=2 | 1314 | 20201115_10113_1ng_glyco_hcd203040_compound15.24832.24832.4 | controllerType=0 controllerNumber=1 scan=24832 | 50.3749 |
| 21457:4 | 13 | K.APDKNVIFSPLSISTALAFLSLGAHN[+2570.925]TTLTEILK.G | HexNAc(5)Hex(6)Fuc(2)NeuAc(1) | N[+2571] | 1539.2385 | 4 | 6153.9322 | 6153.9032 | 0 | 4.7 | 68 | Specific | 909.98 | 909.98 | 909.98 | 17.95 | 46 | >sp|P01011|AACT_HUMAN Alpha-1-antichymotrypsin OS=Homo sapiens OX=9606 GN=SERPINA3 PE=1 SV=2 | 1314 | 20201115_10113_1ng_glyco_hcd203040_compound15.24830.24830.4 | controllerType=0 controllerNumber=1 scan=24830 | 50.3657 |
| 22783:5 | 13 | K.APDKNVIFSPLSISTALAFLSLGAHN[+2350.830]TTLTEILK.G | HexNAc(4)Hex(5)Fuc(1)NeuAc(2) | N[+2351] | 1187.5732 | 5 | 5933.8371 | 5933.8085 | 0 | 4.8 | 68 | Specific | 935.7 | 935.7 | 935.7 | 17.7 | 46 | >sp|P01011|AACT_HUMAN Alpha-1-antichymotrypsin OS=Homo sapiens OX=9606 GN=SERPINA3 PE=1 SV=2 | 1314 | 20201115_10113_1ng_glyco_hcd203040_compound15.26221.26221.5 | controllerType=0 controllerNumber=1 scan=26221 | 53.43 |
| 22782:4 | 13 | K.APDKNVIFSPLSISTALAFLSLGAHN[+2350.830]TTLTEILK.G | HexNAc(4)Hex(5)Fuc(1)NeuAc(2) | N[+2351] | 1484.215 | 4 | 5933.838 | 5933.8085 | 0 | 5 | 68 | Specific | 1129.33 | 1079.4 | 1079.4 | 21.79 | 46 | >sp|P01011|AACT_HUMAN Alpha-1-antichymotrypsin OS=Homo sapiens OX=9606 GN=SERPINA3 PE=1 SV=2 | 1314 | 20201115_10113_1ng_glyco_hcd203040_compound15.26220.26220.4 | controllerType=0 controllerNumber=1 scan=26220 | 53.4283 |
| 15784:4 | 13 | K.GLKFN[+2350.830]LTETSEAEIHQSFQHLLR.T | HexNAc(4)Hex(5)Fuc(1)NeuAc(2) | N[+2351] | 1263.0652 | 4 | 5049.2389 | 5049.2251 | 0 | 2.7 | 102 | Specific | 831.39 | 533.94 | 533.94 | 16.01 | 46 | >sp|P01011|AACT_HUMAN Alpha-1-antichymotrypsin OS=Homo sapiens OX=9606 GN=SERPINA3 PE=1 SV=2 | 1314 | 20201115_10113_1ng_glyco_hcd203040_compound15.18905.18905.4 | controllerType=0 controllerNumber=1 scan=18905 | 37.8651 |
| 15782:4 | 13 | K.GLKFN[+2350.830]LTETSEAEIHQSFQHLLR.T | HexNAc(4)Hex(5)Fuc(1)NeuAc(2) | N[+2351] | 1263.0652 | 4 | 5049.2389 | 5049.2251 | 0 | 2.7 | 102 | Specific | 1179.38 | 871.82 | 871.82 | 23.01 | 46 | >sp|P01011|AACT_HUMAN Alpha-1-antichymotrypsin OS=Homo sapiens OX=9606 GN=SERPINA3 PE=1 SV=2 | 1314 | 20201115_10113_1ng_glyco_hcd203040_compound15.18903.18903.4 | controllerType=0 controllerNumber=1 scan=18903 | 37.8487 |
| 16182:3 | 13 | R.[+42.011]TLN[+2018.708]QSSDELQLSMGNAMFVK.E | HexNAc(3)Hex(6)Fuc(1)NeuAc(1) | .[42], N[+2019] | 1425.2682 | 3 | 4273.79 | 4273.7766 | 0 | 3.1 | 125 | Specific | 309.37 | 197.82 | 197.82 | 6.59 | 46 | >sp|P01011|AACT_HUMAN Alpha-1-antichymotrypsin OS=Homo sapiens OX=9606 GN=SERPINA3 PE=1 SV=2 | 1314 | 20201115_10113_1ng_glyco_hcd203040_compound15.19318.19318.3 | controllerType=0 controllerNumber=1 scan=19318 | 38.6471 |
| 16180:3 | 13 | R.[+42.011]TLN[+2018.708]QSSDELQLSMGNAMFVK.E | HexNAc(3)Hex(6)Fuc(1)NeuAc(1) | .[42], N[+2019] | 1425.2682 | 3 | 4273.79 | 4273.7766 | 0 | 3.1 | 125 | Specific | 379.9 | 326.49 | 326.49 | 7.64 | 46 | >sp|P01011|AACT_HUMAN Alpha-1-antichymotrypsin OS=Homo sapiens OX=9606 GN=SERPINA3 PE=1 SV=2 | 1314 | 20201115_10113_1ng_glyco_hcd203040_compound15.19316.19316.3 | controllerType=0 controllerNumber=1 scan=19316 | 38.642 |
| 16130:3 | 13 | R.[+42.011]TLN[+2018.708]QSSDELQLSMGNAMFVK.E | HexNAc(3)Hex(6)Fuc(1)NeuAc(1) | .[42], N[+2019] | 1425.2746 | 3 | 4273.8092 | 4273.7766 | 0 | 7.6 | 125 | Specific | 418.84 | 328.35 | 328.35 | 6.8 | 46 | >sp|P01011|AACT_HUMAN Alpha-1-antichymotrypsin OS=Homo sapiens OX=9606 GN=SERPINA3 PE=1 SV=2 | 1314 | 20201115_10113_1ng_glyco_hcd203040_compound15.19264.19264.3 | controllerType=0 controllerNumber=1 scan=19264 | 38.5572 |
| 16127:3 | 13 | R.[+42.011]TLN[+2018.708]QSSDELQLSMGNAMFVK.E | HexNAc(3)Hex(6)Fuc(1)NeuAc(1) | .[42], N[+2019] | 1425.2746 | 3 | 4273.8092 | 4273.7766 | 0 | 7.6 | 125 | Specific | 425.69 | 368.84 | 368.84 | 7.22 | 46 | >sp|P01011|AACT_HUMAN Alpha-1-antichymotrypsin OS=Homo sapiens OX=9606 GN=SERPINA3 PE=1 SV=2 | 1314 | 20201115_10113_1ng_glyco_hcd203040_compound15.19261.19261.3 | controllerType=0 controllerNumber=1 scan=19261 | 38.5524 |
| 25887:5 | 14 | K.VFPLSLDSTPQDGNVVVAC[+57.021]LVQGFFPQEPLSVTWSESGQN[+2553.910]VTAI | HexNAc(5)Hex(5)Fuc(1)NeuAc(2) | C[+57], N[+2554] | 1466.8666 | 5 | 7330.304 | 7330.2711 | 0 | 4.5 | 8 | Specific | 378.88 | 378.88 | 378.88 | 8.7 | 40 | >sp|P01877|IGHA2_HUMAN Immunoglobulin heavy constant alpha 2 OS=Homo sapiens OX=9606 GN=IGHA2 PE=1 SV=4 | 576 | 20201115_10113_1ng_glyco_hcd203040_compound15.29469.29469.5 | controllerType=0 controllerNumber=1 scan=29469 | 60.0317 |
| 25889:5 | 14 | K.VFPLSLDSTPQDGNVVVAC[+57.021]LVQGFFPQEPLSVTWSESGQN[+2553.910]VTAI | HexNAc(5)Hex(5)Fuc(1)NeuAc(2) | C[+57], N[+2554] | 1466.8666 | 5 | 7330.304 | 7330.2711 | 0 | 4.5 | 8 | Specific | 393.4 | 393.4 | 393.4 | 10.28 | 40 | >sp|P01877|IGHA2_HUMAN Immunoglobulin heavy constant alpha 2 OS=Homo sapiens OX=9606 GN=IGHA2 PE=1 SV=4 | 576 | 20201115_10113_1ng_glyco_hcd203040_compound15.29471.29471.5 | controllerType=0 controllerNumber=1 scan=29471 | 60.0371 |
| 22088:4 | 14 | K.VFPLSLDSTPQDGNVVVAC[+57.021]LVQGFFPQEPLSVTWSESGQN[+1809.666]VTAI | HexNAc(5)Hex(4)Fuc(1) | C[+57], N[+1810] | 1647.2605 | 4 | 6586.0202 | 6586.0275 | 0 | -1.1 | 8 | Specific | 715 | 702.42 | 702.42 | 14.5 | 40 | >sp|P01877|IGHA2_HUMAN Immunoglobulin heavy constant alpha 2 OS=Homo sapiens OX=9606 GN=IGHA2 PE=1 SV=4 | 576 | 20201115_10113_1ng_glyco_hcd203040_compound15.25490.25490.4 | controllerType=0 controllerNumber=1 scan=25490 | 51.8503 |
| 22113:4 | 14 | K.VFPLSLDSTPQDGNVVVAC[+57.021]LVQGFFPQEPLSVTWSESGQN[+1647.613]VTAI | HexNAc(5)Hex(3)Fuc(1) | C[+57], N[+1648] | 1606.7616 | 4 | 6424.0246 | 6423.9746 | 0 | 7.8 | 8 | Specific | 729.75 | 675.56 | 675.56 | 14.63 | 40 | >sp|P01877|IGHA2_HUMAN Immunoglobulin heavy constant alpha 2 OS=Homo sapiens OX=9606 GN=IGHA2 PE=1 SV=4 | 576 | 20201115_10113_1ng_glyco_hcd203040_compound15.25516.25516.4 | controllerType=0 controllerNumber=1 scan=25516 | 51.9027 |
| 22078:4 | 14 | K.VFPLSLDSTPQDGNVVVAC[+57.021]LVQGFFPQEPLSVTWSESGQN[+1647.613]VTAI | HexNAc(5)Hex(3)Fuc(1) | C[+57], N[+1648] | 1606.7539 | 4 | 6423.9938 | 6423.9746 | 0 | 3 | 8 | Specific | 862.02 | 789.82 | 789.82 | 18.59 | 40 | >sp|P01877|IGHA2_HUMAN Immunoglobulin heavy constant alpha 2 OS=Homo sapiens OX=9606 GN=IGHA2 PE=1 SV=4 | 576 | 20201115_10113_1ng_glyco_hcd203040_compound15.25479.25479.4 | controllerType=0 controllerNumber=1 scan=25479 | 51.8114 |
| 22091:4 | 14 | K.VFPLSLDSTPQDGNVVVAC[+57.021]LVQGFFPQEPLSVTWSESGQN[+1809.666]VTAI | HexNAc(5)Hex(4)Fuc(1) | C[+57], N[+1810] | 1647.2605 | 4 | 6586.0202 | 6586.0275 | 0 | -1.1 | 8 | Specific | 898.76 | 865.84 | 865.84 | 17.5 | 40 | >sp|P01877|IGHA2_HUMAN Immunoglobulin heavy constant alpha 2 OS=Homo sapiens OX=9606 GN=IGHA2 PE=1 SV=4 | 576 | 20201115_10113_1ng_glyco_hcd203040_compound15.25493.25493.4 | controllerType=0 controllerNumber=1 scan=25493 | 51.8566 |
| 22115:4 | 14 | K.VFPLSLDSTPQDGNVVVAC[+57.021]LVQGFFPQEPLSVTWSESGQN[+1647.613]VTAI | HexNAc(5)Hex(3)Fuc(1) | C[+57], N[+1648] | 1606.7616 | 4 | 6424.0246 | 6423.9746 | 0 | 7.8 | 8 | Specific | 961.64 | 961.64 | 961.64 | 19.63 | 40 | >sp|P01877|IGHA2_HUMAN Immunoglobulin heavy constant alpha 2 OS=Homo sapiens OX=9606 GN=IGHA2 PE=1 SV=4 | 576 | 20201115_10113_1ng_glyco_hcd203040_compound15.25518.25518.4 | controllerType=0 controllerNumber=1 scan=25518 | 51.911 |
| 01696:3 | 14 | K.HYTN[+1971.719]SSQDVTVPC[+57.021]R.V | HexNAc(5)Hex(5)Fuc(1) | C[+57], N[+1972] | 1212.4976 | 3 | 3635.4783 | 3635.4682 | 0 | 2.8 | 89 | Specific | 215.33 | 176.77 | 176.77 | 4.85 | 40 | >sp|P01877|IGHA2_HUMAN Immunoglobulin heavy constant alpha 2 OS=Homo sapiens OX=9606 GN=IGHA2 PE=1 SV=4 | 576 | 20201115_10113_1ng_glyco_hcd203040_compound15.4288.4288.3 | controllerType=0 controllerNumber=1 scan=4288 | 12.889 |
| 02608:4 | 14 | K.HYTN[+2262.814]SSQDVTVPC[+57.021]R.V | HexNAc(5)Hex(5)Fuc(1)NeuAc(1) | C[+57], N[+2263] | 982.3996 | 4 | 3926.5765 | 3926.5636 | 0 | 3.3 | 89 | Specific | 237.62 | 154.49 | 154.49 | 5.59 | 40 | >sp|P01877|IGHA2_HUMAN Immunoglobulin heavy constant alpha 2 OS=Homo sapiens OX=9606 GN=IGHA2 PE=1 SV=4 | 576 | 20201115_10113_1ng_glyco_hcd203040_compound15.5234.5234.4 | controllerType=0 controllerNumber=1 scan=5234 | 14.4539 |
| 02610:4 | 14 | K.HYTN[+2262.814]SSQDVTVPC[+57.021]R.V | HexNAc(5)Hex(5)Fuc(1)NeuAc(1) | C[+57], N[+2263] | 982.3996 | 4 | 3926.5765 | 3926.5636 | 0 | 3.3 | 89 | Specific | 367.59 | 299.53 | 299.53 | 7.24 | 40 | >sp|P01877|IGHA2_HUMAN Immunoglobulin heavy constant alpha 2 OS=Homo sapiens OX=9606 GN=IGHA2 PE=1 SV=4 | 576 | 20201115_10113_1ng_glyco_hcd203040_compound15.5236.5236.4 | controllerType=0 controllerNumber=1 scan=5236 | 14.4579 |
| 01646:3 | 14 | K.HYTN[+1971.719]SSQDVTVPC[+57.021]R.V | HexNAc(5)Hex(5)Fuc(1) | C[+57], N[+1972] | 1212.4974 | 3 | 3635.4776 | 3635.4682 | 0 | 2.6 | 89 | Specific | 375.04 | 290.81 | 290.81 | 7.29 | 40 | >sp|P01877|IGHA2_HUMAN Immunoglobulin heavy constant alpha 2 OS=Homo sapiens OX=9606 GN=IGHA2 PE=1 SV=4 | 576 | 20201115_10113_1ng_glyco_hcd203040_compound15.4236.4236.3 | controllerType=0 controllerNumber=1 scan=4236 | 12.797 |
| 01648:3 | 14 | K.HYTN[+1971.719]SSQDVTVPC[+57.021]R.V | HexNAc(5)Hex(5)Fuc(1) | C[+57], N[+1972] | 1212.4974 | 3 | 3635.4776 | 3635.4682 | 0 | 2.6 | 89 | Specific | 415 | 244.93 | 244.93 | 7.43 | 40 | >sp|P01877|IGHA2_HUMAN Immunoglobulin heavy constant alpha 2 OS=Homo sapiens OX=9606 GN=IGHA2 PE=1 SV=4 | 576 | 20201115_10113_1ng_glyco_hcd203040_compound15.4238.4238.3 | controllerType=0 controllerNumber=1 scan=4238 | 12.8004 |
| 02579:3 | 14 | K.HYTN[+2262.814]SSQDVTVPC[+57.021]R.V | HexNAc(5)Hex(5)Fuc(1)NeuAc(1) | C[+57], N[+2263] | 1309.5294 | 3 | 3926.5735 | 3926.5636 | 0 | 2.5 | 89 | Specific | 570.13 | 418.07 | 418.07 | 10.45 | 40 | >sp|P01877|IGHA2_HUMAN Immunoglobulin heavy constant alpha 2 OS=Homo sapiens OX=9606 GN=IGHA2 PE=1 SV=4 | 576 | 20201115_10113_1ng_glyco_hcd203040_compound15.5204.5204.3 | controllerType=0 controllerNumber=1 scan=5204 | 14.4035 |
| 02577:3 | 14 | K.HYTN[+2262.814]SSQDVTVPC[+57.021]R.V | HexNAc(5)Hex(5)Fuc(1)NeuAc(1) | C[+57], N[+2263] | 1309.5294 | 3 | 3926.5735 | 3926.5636 | 0 | 2.5 | 89 | Specific | 584.44 | 373.53 | 373.53 | 10 | 40 | >sp|P01877|IGHA2_HUMAN Immunoglobulin heavy constant alpha 2 OS=Homo sapiens OX=9606 GN=IGHA2 PE=1 SV=4 | 576 | 20201115_10113_1ng_glyco_hcd203040_compound15.5202.5202.3 | controllerType=0 controllerNumber=1 scan=5202 | 14.3997 |
| 01714:3 | 14 | K.HYTN[+1647.613]SSQDVTVPC[+57.021]R.V | HexNAc(5)Hex(3)Fuc(1) | C[+57], N[+1648] | 1104.4611 | 3 | 3311.3687 | 3311.3625 | 0 | 1.9 | 89 | Specific | 650.75 | 549.92 | 549.92 | 13.51 | 40 | >sp|P01877|IGHA2_HUMAN Immunoglobulin heavy constant alpha 2 OS=Homo sapiens OX=9606 GN=IGHA2 PE=1 SV=4 | 576 | 20201115_10113_1ng_glyco_hcd203040_compound15.4307.4307.3 | controllerType=0 controllerNumber=1 scan=4307 | 12.9226 |
| 01716:3 | 14 | K.HYTN[+1647.613]SSQDVTVPC[+57.021]R.V | HexNAc(5)Hex(3)Fuc(1) | C[+57], N[+1648] | 1104.4611 | 3 | 3311.3687 | 3311.3625 | 0 | 1.9 | 89 | Specific | 669.6 | 590.64 | 590.64 | 13.51 | 40 | >sp|P01877|IGHA2_HUMAN Immunoglobulin heavy constant alpha 2 OS=Homo sapiens OX=9606 GN=IGHA2 PE=1 SV=4 | 576 | 20201115_10113_1ng_glyco_hcd203040_compound15.4309.4309.3 | controllerType=0 controllerNumber=1 scan=4309 | 12.9268 |
| 01665:3 | 14 | K.HYTN[+1809.666]SSQDVTVPC[+57.021]R.V | HexNAc(5)Hex(4)Fuc(1) | C[+57], N[+1810] | 1158.4785 | 3 | 3473.4208 | 3473.4154 | 0 | 1.6 | 89 | Specific | 700.92 | 523.67 | 523.67 | 14.51 | 40 | >sp|P01877|IGHA2_HUMAN Immunoglobulin heavy constant alpha 2 OS=Homo sapiens OX=9606 GN=IGHA2 PE=1 SV=4 | 576 | 20201115_10113_1ng_glyco_hcd203040_compound15.4256.4256.3 | controllerType=0 controllerNumber=1 scan=4256 | 12.8296 |
| 01668:3 | 14 | K.HYTN[+1809.666]SSQDVTVPC[+57.021]R.V | HexNAc(5)Hex(4)Fuc(1) | C[+57], N[+1810] | 1158.4785 | 3 | 3473.4208 | 3473.4154 | 0 | 1.6 | 89 | Specific | 744.21 | 589.46 | 589.46 | 14.51 | 40 | >sp|P01877|IGHA2_HUMAN Immunoglobulin heavy constant alpha 2 OS=Homo sapiens OX=9606 GN=IGHA2 PE=1 SV=4 | 576 | 20201115_10113_1ng_glyco_hcd203040_compound15.4259.4259.3 | controllerType=0 controllerNumber=1 scan=4259 | 12.8351 |
| 03736:5 | 14 | K.HYTN[+1647.613]SSQDVTVPC[+57.021]RVPPPPPC[+57.021]C[+57.021]HPR.L | HexNAc(5)Hex(3)Fuc(1) | C[+57]*3, N[+1648] | 922.0012 | 5 | 4605.9771 | 4605.9689 | 0 | 1.8 | 89 | Specific | 748.53 | 587.29 | 587.29 | 14.46 | 40 | >sp|P01877|IGHA2_HUMAN Immunoglobulin heavy constant alpha 2 OS=Homo sapiens OX=9606 GN=IGHA2 PE=1 SV=4 | 576 | 20201115_10113_1ng_glyco_hcd203040_compound15.6402.6402.5 | controllerType=0 controllerNumber=1 scan=6402 | 16.4905 |
| 03739:5 | 14 | K.HYTN[+1647.613]SSQDVTVPC[+57.021]RVPPPPPC[+57.021]C[+57.021]HPR.L | HexNAc(5)Hex(3)Fuc(1) | C[+57]*3, N[+1648] | 922.0012 | 5 | 4605.9771 | 4605.9689 | 0 | 1.8 | 89 | Specific | 854.08 | 575.05 | 575.05 | 17.46 | 40 | >sp|P01877|IGHA2_HUMAN Immunoglobulin heavy constant alpha 2 OS=Homo sapiens OX=9606 GN=IGHA2 PE=1 SV=4 | 576 | 20201115_10113_1ng_glyco_hcd203040_compound15.6405.6405.5 | controllerType=0 controllerNumber=1 scan=6405 | 16.497 |
| 03737:4 | 14 | K.HYTN[+1647.613]SSQDVTVPC[+57.021]RVPPPPPC[+57.021]C[+57.021]HPR.L | HexNAc(5)Hex(3)Fuc(1) | C[+57]*3, N[+1648] | 1152.2497 | 4 | 4605.9769 | 4605.9689 | 0 | 1.7 | 89 | Specific | 943.23 | 707.75 | 707.75 | 17.87 | 40 | >sp|P01877|IGHA2_HUMAN Immunoglobulin heavy constant alpha 2 OS=Homo sapiens OX=9606 GN=IGHA2 PE=1 SV=4 | 576 | 20201115_10113_1ng_glyco_hcd203040_compound15.6403.6403.4 | controllerType=0 controllerNumber=1 scan=6403 | 16.4916 |
| 03740:4 | 14 | K.HYTN[+1647.613]SSQDVTVPC[+57.021]RVPPPPPC[+57.021]C[+57.021]HPR.L | HexNAc(5)Hex(3)Fuc(1) | C[+57]*3, N[+1648] | 1152.2497 | 4 | 4605.9769 | 4605.9689 | 0 | 1.7 | 89 | Specific | 946.69 | 696.13 | 696.13 | 17.87 | 40 | >sp|P01877|IGHA2_HUMAN Immunoglobulin heavy constant alpha 2 OS=Homo sapiens OX=9606 GN=IGHA2 PE=1 SV=4 | 576 | 20201115_10113_1ng_glyco_hcd203040_compound15.6406.6406.4 | controllerType=0 controllerNumber=1 scan=6406 | 16.4989 |
| 03436:2 | 14 | K.TPLTAN[+1971.719]ITK.S | HexNAc(5)Hex(5)Fuc(1) | N[+1972] | 1465.6452 | 2 | 2930.2831 | 2930.2757 | 0 | 2.5 | 200 | Specific | 256.54 | 193.8 | 193.8 | 6.1 | 40 | >sp|P01877|IGHA2_HUMAN Immunoglobulin heavy constant alpha 2 OS=Homo sapiens OX=9606 GN=IGHA2 PE=1 SV=4 | 576 | 20201115_10113_1ng_glyco_hcd203040_compound15.6090.6090.2 | controllerType=0 controllerNumber=1 scan=6090 | 15.8976 |
| 03438:2 | 14 | K.TPLTAN[+1971.719]ITK.S | HexNAc(5)Hex(5)Fuc(1) | N[+1972] | 1465.6452 | 2 | 2930.2831 | 2930.2757 | 0 | 2.5 | 200 | Specific | 300.33 | 183.02 | 183.02 | 6.12 | 40 | >sp|P01877|IGHA2_HUMAN Immunoglobulin heavy constant alpha 2 OS=Homo sapiens OX=9606 GN=IGHA2 PE=1 SV=4 | 576 | 20201115_10113_1ng_glyco_hcd203040_compound15.6092.6092.2 | controllerType=0 controllerNumber=1 scan=6092 | 15.9016 |
| 03419:3 | 14 | K.TPLTAN[+1971.719]ITK.S | HexNAc(5)Hex(5)Fuc(1) | N[+1972] | 977.4328 | 3 | 2930.2838 | 2930.2757 | 0 | 2.8 | 200 | Specific | 331.02 | 331.02 | 331.02 | 7.03 | 40 | >sp|P01877|IGHA2_HUMAN Immunoglobulin heavy constant alpha 2 OS=Homo sapiens OX=9606 GN=IGHA2 PE=1 SV=4 | 576 | 20201115_10113_1ng_glyco_hcd203040_compound15.6073.6073.3 | controllerType=0 controllerNumber=1 scan=6073 | 15.8647 |
| 03452:3 | 14 | K.TPLTAN[+1809.666]ITK.S | HexNAc(5)Hex(4)Fuc(1) | N[+1810] | 923.4145 | 3 | 2768.2288 | 2768.2228 | 0 | 2.2 | 200 | Specific | 473.18 | 197.15 | 197.15 | 6.96 | 40 | >sp|P01877|IGHA2_HUMAN Immunoglobulin heavy constant alpha 2 OS=Homo sapiens OX=9606 GN=IGHA2 PE=1 SV=4 | 576 | 20201115_10113_1ng_glyco_hcd203040_compound15.6107.6107.3 | controllerType=0 controllerNumber=1 scan=6107 | 15.9268 |
| 03522:3 | 14 | K.TPLTAN[+1647.613]ITK.S | HexNAc(5)Hex(3)Fuc(1) | N[+1648] | 869.397 | 3 | 2606.1765 | 2606.17 | 0 | 2.5 | 200 | Specific | 473.23 | 195.63 | 195.63 | 6.96 | 40 | >sp|P01877|IGHA2_HUMAN Immunoglobulin heavy constant alpha 2 OS=Homo sapiens OX=9606 GN=IGHA2 PE=1 SV=4 | 576 | 20201115_10113_1ng_glyco_hcd203040_compound15.6180.6180.3 | controllerType=0 controllerNumber=1 scan=6180 | 16.0834 |
| 03421:3 | 14 | K.TPLTAN[+1971.719]ITK.S | HexNAc(5)Hex(5)Fuc(1) | N[+1972] | 977.4328 | 3 | 2930.2838 | 2930.2757 | 0 | 2.8 | 200 | Specific | 475.27 | 259.74 | 259.74 | 7.31 | 40 | >sp|P01877|IGHA2_HUMAN Immunoglobulin heavy constant alpha 2 OS=Homo sapiens OX=9606 GN=IGHA2 PE=1 SV=4 | 576 | 20201115_10113_1ng_glyco_hcd203040_compound15.6075.6075.3 | controllerType=0 controllerNumber=1 scan=6075 | 15.8692 |
| 03534:2 | 14 | K.TPLTAN[+1501.555]ITK.S | HexNAc(5)Hex(3) | N[+1502] | 1230.563 | 2 | 2460.1188 | 2460.1121 | 0 | 2.7 | 200 | Specific | 527.55 | 334.58 | 334.58 | 7.99 | 40 | >sp|P01877|IGHA2_HUMAN Immunoglobulin heavy constant alpha 2 OS=Homo sapiens OX=9606 GN=IGHA2 PE=1 SV=4 | 576 | 20201115_10113_1ng_glyco_hcd203040_compound15.6192.6192.2 | controllerType=0 controllerNumber=1 scan=6192 | 16.1103 |
| 03454:3 | 14 | K.TPLTAN[+1809.666]ITK.S | HexNAc(5)Hex(4)Fuc(1) | N[+1810] | 923.4145 | 3 | 2768.2288 | 2768.2228 | 0 | 2.2 | 200 | Specific | 543.92 | 284.23 | 284.23 | 7.77 | 40 | >sp|P01877|IGHA2_HUMAN Immunoglobulin heavy constant alpha 2 OS=Homo sapiens OX=9606 GN=IGHA2 PE=1 SV=4 | 576 | 20201115_10113_1ng_glyco_hcd203040_compound15.6109.6109.3 | controllerType=0 controllerNumber=1 scan=6109 | 15.9322 |
| 03536:2 | 14 | K.TPLTAN[+1501.555]ITK.S | HexNAc(5)Hex(3) | N[+1502] | 1230.563 | 2 | 2460.1188 | 2460.1121 | 0 | 2.7 | 200 | Specific | 569.56 | 306.84 | 306.84 | 8.21 | 40 | >sp|P01877|IGHA2_HUMAN Immunoglobulin heavy constant alpha 2 OS=Homo sapiens OX=9606 GN=IGHA2 PE=1 SV=4 | 576 | 20201115_10113_1ng_glyco_hcd203040_compound15.6194.6194.2 | controllerType=0 controllerNumber=1 scan=6194 | 16.1142 |
| 03525:3 | 14 | K.TPLTAN[+1647.613]ITK.S | HexNAc(5)Hex(3)Fuc(1) | N[+1648] | 869.397 | 3 | 2606.1765 | 2606.17 | 0 | 2.5 | 200 | Specific | 576.91 | 281.68 | 281.68 | 7.9 | 40 | >sp|P01877|IGHA2_HUMAN Immunoglobulin heavy constant alpha 2 OS=Homo sapiens OX=9606 GN=IGHA2 PE=1 SV=4 | 576 | 20201115_10113_1ng_glyco_hcd203040_compound15.6183.6183.3 | controllerType=0 controllerNumber=1 scan=6183 | 16.0911 |
| 03461:2 | 14 | K.TPLTAN[+1809.666]ITK.S | HexNAc(5)Hex(4)Fuc(1) | N[+1810] | 1384.6176 | 2 | 2768.2279 | 2768.2228 | 0 | 1.8 | 200 | Specific | 607.1 | 355.37 | 355.37 | 10.56 | 40 | >sp|P01877|IGHA2_HUMAN Immunoglobulin heavy constant alpha 2 OS=Homo sapiens OX=9606 GN=IGHA2 PE=1 SV=4 | 576 | 20201115_10113_1ng_glyco_hcd203040_compound15.6116.6116.2 | controllerType=0 controllerNumber=1 scan=6116 | 15.9498 |
| 03517:2 | 14 | K.TPLTAN[+1647.613]ITK.S | HexNAc(5)Hex(3)Fuc(1) | N[+1648] | 1303.5923 | 2 | 2606.1774 | 2606.17 | 0 | 2.8 | 200 | Specific | 626.82 | 354.58 | 354.58 | 10.54 | 40 | >sp|P01877|IGHA2_HUMAN Immunoglobulin heavy constant alpha 2 OS=Homo sapiens OX=9606 GN=IGHA2 PE=1 SV=4 | 576 | 20201115_10113_1ng_glyco_hcd203040_compound15.6175.6175.2 | controllerType=0 controllerNumber=1 scan=6175 | 16.0716 |
| 03459:2 | 14 | K.TPLTAN[+1809.666]ITK.S | HexNAc(5)Hex(4)Fuc(1) | N[+1810] | 1384.6176 | 2 | 2768.2279 | 2768.2228 | 0 | 1.8 | 200 | Specific | 632.94 | 365.6 | 365.6 | 10.79 | 40 | >sp|P01877|IGHA2_HUMAN Immunoglobulin heavy constant alpha 2 OS=Homo sapiens OX=9606 GN=IGHA2 PE=1 SV=4 | 576 | 20201115_10113_1ng_glyco_hcd203040_compound15.6114.6114.2 | controllerType=0 controllerNumber=1 scan=6114 | 15.9425 |
| 03513:2 | 14 | K.TPLTAN[+1444.534]ITK.S | HexNAc(4)Hex(3)Fuc(1) | N[+1445] | 1202.0523 | 2 | 2403.0973 | 2403.0906 | 0 | 2.8 | 200 | Specific | 646.03 | 361.04 | 361.04 | 10.77 | 40 | >sp|P01877|IGHA2_HUMAN Immunoglobulin heavy constant alpha 2 OS=Homo sapiens OX=9606 GN=IGHA2 PE=1 SV=4 | 576 | 20201115_10113_1ng_glyco_hcd203040_compound15.6170.6170.2 | controllerType=0 controllerNumber=1 scan=6170 | 16.0598 |
| 03519:2 | 14 | K.TPLTAN[+1647.613]ITK.S | HexNAc(5)Hex(3)Fuc(1) | N[+1648] | 1303.5923 | 2 | 2606.1774 | 2606.17 | 0 | 2.8 | 200 | Specific | 648.16 | 341.84 | 341.84 | 10.54 | 40 | >sp|P01877|IGHA2_HUMAN Immunoglobulin heavy constant alpha 2 OS=Homo sapiens OX=9606 GN=IGHA2 PE=1 SV=4 | 576 | 20201115_10113_1ng_glyco_hcd203040_compound15.6177.6177.2 | controllerType=0 controllerNumber=1 scan=6177 | 16.0784 |
| 03515:2 | 14 | K.TPLTAN[+1444.534]ITK.S | HexNAc(4)Hex(3)Fuc(1) | N[+1445] | 1202.0523 | 2 | 2403.0973 | 2403.0906 | 0 | 2.8 | 200 | Specific | 649.71 | 286.64 | 286.64 | 10.03 | 40 | >sp|P01877|IGHA2_HUMAN Immunoglobulin heavy constant alpha 2 OS=Homo sapiens OX=9606 GN=IGHA2 PE=1 SV=4 | 576 | 20201115_10113_1ng_glyco_hcd203040_compound15.6172.6172.2 | controllerType=0 controllerNumber=1 scan=6172 | 16.0655 |
| 10866:3 | 14 | R.MAGKPTHIN[+1663.608]VSVVMAEADGTC[+57.021]Y.- | HexNAc(5)Hex(4) | C[+57], N[+1664] | 1338.9023 | 3 | 4014.6923 | 4014.691 | 0 | 0.3 | 319 | CRagged | 305.57 | 305.57 | 238.72 | 7.03 | 40 | >sp|P01877|IGHA2_HUMAN Immunoglobulin heavy constant alpha 2 OS=Homo sapiens OX=9606 GN=IGHA2 PE=1 SV=4 | 576 | 20201115_10113_1ng_glyco_hcd203040_compound15.13797.13797.3 | controllerType=0 controllerNumber=1 scan=13797 | 28.6547 |
| 10873:3 | 14 | R.MAGKPTHIN[+1809.666]VSVVMAEADGTC[+57.021]Y.- | HexNAc(5)Hex(4)Fuc(1) | C[+57], N[+1810] | 1387.5882 | 3 | 4160.75 | 4160.7489 | 0 | 0.3 | 319 | CRagged | 384.03 | 384.03 | 292.9 | 9.36 | 40 | >sp|P01877|IGHA2_HUMAN Immunoglobulin heavy constant alpha 2 OS=Homo sapiens OX=9606 GN=IGHA2 PE=1 SV=4 | 576 | 20201115_10113_1ng_glyco_hcd203040_compound15.13804.13804.3 | controllerType=0 controllerNumber=1 scan=13804 | 28.6661 |
| 10869:3 | 14 | R.MAGKPTHIN[+1663.608]VSVVMAEADGTC[+57.021]Y.- | HexNAc(5)Hex(4) | C[+57], N[+1664] | 1338.9023 | 3 | 4014.6923 | 4014.691 | 0 | 0.3 | 319 | CRagged | 410.59 | 252.48 | 252.48 | 7.25 | 40 | >sp|P01877|IGHA2_HUMAN Immunoglobulin heavy constant alpha 2 OS=Homo sapiens OX=9606 GN=IGHA2 PE=1 SV=4 | 576 | 20201115_10113_1ng_glyco_hcd203040_compound15.13800.13800.3 | controllerType=0 controllerNumber=1 scan=13800 | 28.6592 |
| 10871:3 | 14 | R.MAGKPTHIN[+1809.666]VSVVMAEADGTC[+57.021]Y.- | HexNAc(5)Hex(4)Fuc(1) | C[+57], N[+1810] | 1387.5882 | 3 | 4160.75 | 4160.7489 | 0 | 0.3 | 319 | CRagged | 456.6 | 456.6 | 312.22 | 9.73 | 40 | >sp|P01877|IGHA2_HUMAN Immunoglobulin heavy constant alpha 2 OS=Homo sapiens OX=9606 GN=IGHA2 PE=1 SV=4 | 576 | 20201115_10113_1ng_glyco_hcd203040_compound15.13802.13802.3 | controllerType=0 controllerNumber=1 scan=13802 | 28.6626 |
| 10953:3 | 14 | R.MAGKPTHIN[+1501.555]VSVVMAEADGTC[+57.021]Y.- | HexNAc(5)Hex(3) | C[+57], N[+1502] | 1284.8868 | 3 | 3852.6458 | 3852.6382 | 0 | 2 | 319 | CRagged | 495.27 | 495.27 | 495.27 | 10.5 | 40 | >sp|P01877|IGHA2_HUMAN Immunoglobulin heavy constant alpha 2 OS=Homo sapiens OX=9606 GN=IGHA2 PE=1 SV=4 | 576 | 20201115_10113_1ng_glyco_hcd203040_compound15.13887.13887.3 | controllerType=0 controllerNumber=1 scan=13887 | 28.8121 |
| 10955:3 | 14 | R.MAGKPTHIN[+1501.555]VSVVMAEADGTC[+57.021]Y.- | HexNAc(5)Hex(3) | C[+57], N[+1502] | 1284.8868 | 3 | 3852.6458 | 3852.6382 | 0 | 2 | 319 | CRagged | 511.07 | 511.07 | 511.07 | 10.5 | 40 | >sp|P01877|IGHA2_HUMAN Immunoglobulin heavy constant alpha 2 OS=Homo sapiens OX=9606 GN=IGHA2 PE=1 SV=4 | 576 | 20201115_10113_1ng_glyco_hcd203040_compound15.13889.13889.3 | controllerType=0 controllerNumber=1 scan=13889 | [illegible] |
| 10890:3 | 14 | R.MAGKPTHIN[+1216.423]VSVVMAEADGTC[+57.021]Y.- | HexNAc(2)Hex(5) | C[+57], N[+1216] | 1189.8421 | 3 | 3567.5117 | 3567.5057 | 0 | 1.7 | 319 | CRagged | 611.27 | 611.27 | 317.74 | 13.01 | 40 | >sp|P01877|IGHA2_HUMAN Immunoglobulin heavy constant alpha 2 OS=Homo sapiens OX=9606 GN=IGHA2 PE=1 SV=4 | 576 | 20201115_10113_1ng_glyco_hcd203040_compound15.13822.13822.3 | controllerType=0 controllerNumber=1 scan=13822 | [illegible] |
| 10833:3 | 14 | R.MAGKPTHIN[+1378.476]VSVVMAEADGTC[+57.021]Y.- | HexNAc(2)Hex(6) | C[+57], N[+1378] | 1243.8611 | 3 | 3729.5689 | 3729.5585 | 0 | 2.8 | 319 | CRagged | 877.44 | 783.01 | 506.67 | 17.99 | 40 | >sp|P01877|IGHA2_HUMAN Immunoglobulin heavy constant alpha 2 OS=Homo sapiens OX=9606 GN=IGHA2 PE=1 SV=4 | 576 | 20201115_10113_1ng_glyco_hcd203040_compound15.13763.13763.3 | controllerType=0 controllerNumber=1 scan=13763 | 28.5875 |
| 10787:3 | 14 | R.MAGKPTHIN[+1540.529]VSVVMAEADGTC[+57.021]Y.- | HexNAc(2)Hex(7) | C[+57], N[+1541] | 1297.8779 | 3 | 3891.619 | 3891.6114 | 0 | 2 | 319 | CRagged | 879.56 | 879.56 | 500.7 | 18.01 | 40 | >sp|P01877|IGHA2_HUMAN Immunoglobulin heavy constant alpha 2 OS=Homo sapiens OX=9606 GN=IGHA2 PE=1 SV=4 | 576 | 20201115_10113_1ng_glyco_hcd203040_compound15.13715.13715.3 | controllerType=0 controllerNumber=1 scan=13715 | 28.4914 |
| 10790:3 | 14 | R.MAGKPTHIN[+1540.529]VSVVMAEADGTC[+57.021]Y.- | HexNAc(2)Hex(7) | C[+57], N[+1541] | 1297.8779 | 3 | 3891.619 | 3891.6114 | 0 | 2 | 319 | CRagged | 881.59 | 881.59 | 506.25 | 18.01 | 40 | >sp|P01877|IGHA2_HUMAN Immunoglobulin heavy constant alpha 2 OS=Homo sapiens OX=9606 GN=IGHA2 PE=1 SV=4 | 576 | 20201115_10113_1ng_glyco_hcd203040_compound15.13718.13718.3 | controllerType=0 controllerNumber=1 scan=13718 | 28.4981 |
| 10774:3 | 14 | R.MAGKPTHIN[+1702.581]VSVVMAEADGTC[+57.021]Y.- | HexNAc(2)Hex(8) | C[+57], N[+1703] | 1351.8959 | 3 | 4053.6732 | 4053.6642 | 0 | 2.2 | 319 | CRagged | 884.62 | 715.94 | 502.7 | 18.01 | 40 | >sp|P01877|IGHA2_HUMAN Immunoglobulin heavy constant alpha 2 OS=Homo sapiens OX=9606 GN=IGHA2 PE=1 SV=4 | 576 | 20201115_10113_1ng_glyco_hcd203040_compound15.13701.13701.3 | controllerType=0 controllerNumber=1 scan=13701 | 28.4627 |
| 10772:3 | 14 | R.MAGKPTHIN[+1702.581]VSVVMAEADGTC[+57.021]Y.- | HexNAc(2)Hex(8) | C[+57], N[+1703] | 1351.8959 | 3 | 4053.6732 | 4053.6642 | 0 | 2.2 | 319 | CRagged | 921.17 | 921.17 | 526.82 | 19.01 | 40 | >sp|P01877|IGHA2_HUMAN Immunoglobulin heavy constant alpha 2 OS=Homo sapiens OX=9606 GN=IGHA2 PE=1 SV=4 | 576 | 20201115_10113_1ng_glyco_hcd203040_compound15.13699.13699.3 | controllerType=0 controllerNumber=1 scan=13699 | 28.4557 |
| 10887:3 | 14 | R.MAGKPTHIN[+1216.423]VSVVMAEADGTC[+57.021]Y.- | HexNAc(2)Hex(5) | C[+57], N[+1216] | 1189.8421 | 3 | 3567.5117 | 3567.5057 | 0 | 1.7 | 319 | CRagged | 942.4 | 788.81 | 532.48 | 19.01 | 40 | >sp|P01877|IGHA2_HUMAN Immunoglobulin heavy constant alpha 2 OS=Homo sapiens OX=9606 GN=IGHA2 PE=1 SV=4 | 576 | 20201115_10113_1ng_glyco_hcd203040_compound15.13819.13819.3 | controllerType=0 controllerNumber=1 scan=13819 | 28.6939 |
| 10831:3 | 14 | R.MAGKPTHIN[+1378.476]VSVVMAEADGTC[+57.021]Y.- | HexNAc(2)Hex(6) | C[+57], N[+1378] | 1243.8611 | 3 | 3729.5689 | 3729.5585 | 0 | 2.8 | 319 | CRagged | 1025.25 | 1006.84 | 605.88 | 20.99 | 40 | >sp|P01877|IGHA2_HUMAN Immunoglobulin heavy constant alpha 2 OS=Homo sapiens OX=9606 GN=IGHA2 PE=1 SV=4 | 576 | 20201115_10113_1ng_glyco_hcd203040_compound15.13761.13761.3 | controllerType=0 controllerNumber=1 scan=13761 | 28.581 |
| 08290:3 | 15 | R.LSPN[+2350.830]ASAEHLELR.W | HexNAc(4)Hex(5)Fuc(1)NeuAc(2) | N[+2351] | 1263.2006 | 3 | 3787.5873 | 3787.5796 | 0 | 2 | 52 | Specific | 563.95 | 499.33 | 499.33 | 10.45 | 37 | >sp|Q13410|BT1A1_HUMAN Butyrophilin subfamily 1 member A1 OS=Homo sapiens OX=9606 GN=BTN1A1 PE=1 SV=3 | 179 | 20201115_10113_1ng_glyco_hcd203040_compound15.11114.11114.3 | controllerType=0 controllerNumber=1 scan=11114 | 24.0784 |
| 08289:3 | 15 | R.LSPN[+2350.830]ASAEHLELR.W | HexNAc(4)Hex(5)Fuc(1)NeuAc(2) | N[+2351] | 1263.2006 | 3 | 3787.5873 | 3787.5796 | 0 | 2 | 52 | Specific | 570.6 | 367.52 | 367.52 | 8.94 | 37 | >sp|Q13410|BT1A1_HUMAN Butyrophilin subfamily 1 member A1 OS=Homo sapiens OX=9606 GN=BTN1A1 PE=1 SV=3 | 179 | 20201115_10113_1ng_glyco_hcd203040_compound15.11113.11113.3 | controllerType=0 controllerNumber=1 scan=11113 | 24.0737 |
| 18211:3 | 15 | R.DTSAKN[+2059.735]VSC[+57.021]YIQNLLLGQEK.K | HexNAc(4)Hex(5)Fuc(1)NeuAc(1) | C[+57], N[+2060] | 1447.6354 | 3 | 4340.8917 | 4340.8842 | 0 | 1.7 | 210 | Specific | 492.61 | 418.8 | 418.8 | 8.54 | 37 | >sp|Q13410|BT1A1_HUMAN Butyrophilin subfamily 1 member A1 OS=Homo sapiens OX=9606 GN=BTN1A1 PE=1 SV=3 | 179 | 20201115_10113_1ng_glyco_hcd203040_compound15.21439.21439.4 | controllerType=0 controllerNumber=1 scan=21439 | 42.9294 |
| 16135:4 | 16 | K.LKELPGVC[+57.021]N[+2351.851]ETMMALWEEC[+57.021]KPC[+57.021]LK.Q | HexNAc(4)Hex(5)Fuc(3)NeuAc(1) | C[+57]*3, N[+2352] | 1322.8105 | 4 | 5288.2202 | 5288.2355 | 0 | -2.9 | 95 | Specific | 553.38 | 527.11 | 527.11 | 8.94 | 39 | >sp|P10909|CLUS_HUMAN Clusterin OS=Homo sapiens OX=9606 GN=CLU PE=1 SV=1 | 1467 | 20201115_10113_1ng_glyco_hcd203040_compound15.19269.19269.4 | controllerType=0 controllerNumber=1 scan=19269 | 38.5644 |
| 14447:4 | 16 | K.LKELPGVC[+57.021]N[+2205.793]ETMMALWEEC[+57.021]KPC[+57.021]LK.Q | HexNAc(4)Hex(5)Fuc(2)NeuAc(1) | C[+57]*3, N[+2206] | 1286.304 | 4 | 5142.194 | 5142.1776 | 0 | 3.2 | 95 | Specific | 600.87 | 484.31 | 484.31 | 12.6 | 39 | >sp|P10909|CLUS_HUMAN Clusterin OS=Homo sapiens OX=9606 GN=CLU PE=1 SV=1 | 1467 | 20201115_10113_1ng_glyco_hcd203040_compound15.17519.17519.4 | controllerType=0 controllerNumber=1 scan=17519 | 35.3144 |
| 14445:4 | 16 | K.LKELPGVC[+57.021]N[+2059.735]ETMMALWEEC[+57.021]KPC[+57.021]LK.Q | HexNAc(4)Hex(5)Fuc(1)NeuAc(1) | C[+57]*3, N[+2060] | 1249.7888 | 4 | 4996.1335 | 4996.1196 | 0 | 2.8 | 95 | Specific | 717.56 | 694.1 | 694.1 | 14.6 | 39 | >sp|P10909|CLUS_HUMAN Clusterin OS=Homo sapiens OX=9606 GN=CLU PE=1 SV=1 | 1467 | 20201115_10113_1ng_glyco_hcd203040_compound15.17517.17517.4 | controllerType=0 controllerNumber=1 scan=17517 | 35.3074 |
| 14449:4 | 16 | K.LKELPGVC[+57.021]N[+2205.793]ETMMALWEEC[+57.021]KPC[+57.021]LK.Q | HexNAc(4)Hex(5)Fuc(2)NeuAc(1) | C[+57]*3, N[+2206] | 1286.304 | 4 | 5142.194 | 5142.1776 | 0 | 3.2 | 95 | Specific | 749.08 | 700.8 | 700.8 | 14.6 | 39 | >sp|P10909|CLUS_HUMAN Clusterin OS=Homo sapiens OX=9606 GN=CLU PE=1 SV=1 | 1467 | 20201115_10113_1ng_glyco_hcd203040_compound15.17521.17521.4 | controllerType=0 controllerNumber=1 scan=17521 | 35.318 |
| 14443:4 | 16 | K.LKELPGVC[+57.021]N[+2059.735]ETMMALWEEC[+57.021]KPC[+57.021]LK.Q | HexNAc(4)Hex(5)Fuc(1)NeuAc(1) | C[+57]*3, N[+2060] | 1249.7888 | 4 | 4996.1335 | 4996.1196 | 0 | 2.8 | 95 | Specific | 1059.99 | 994.69 | 994.69 | 21.6 | 39 | >sp|P10909|CLUS_HUMAN Clusterin OS=Homo sapiens OX=9606 GN=CLU PE=1 SV=1 | 1467 | 20201115_10113_1ng_glyco_hcd203040_compound15.17515.17515.4 | controllerType=0 controllerNumber=1 scan=17515 | 35.3023 |
| 16239:3 | 16 | K.ELPGVC[+57.021]N[+2059.735]ETMMALWEEC[+57.021]KPC[+57.021]LK.Q | HexNAc(4)Hex(5)Fuc(1)NeuAc(1) | C[+57]*3, N[+2060] | 1585.647 | 3 | 4754.9265 | 4754.9406 | 0 | -3 | 97 | Specific | 428.12 | 399.45 | 399.45 | 8.68 | 39 | >sp|P10909|CLUS_HUMAN Clusterin OS=Homo sapiens OX=9606 GN=CLU PE=1 SV=1 | 1467 | 20201115_10113_1ng_glyco_hcd203040_compound15.19377.19377.3 | controllerType=0 controllerNumber=1 scan=19377 | 38.7553 |
| 16236:3 | 16 | K.ELPGVC[+57.021]N[+2059.735]ETMMALWEEC[+57.021]KPC[+57.021]LK.Q | HexNAc(4)Hex(5)Fuc(1)NeuAc(1) | C[+57]*3, N[+2060] | 1585.647 | 3 | 4754.9265 | 4754.9406 | 0 | -3 | 97 | Specific | 570.45 | 359.59 | 359.59 | 7.95 | 39 | >sp|P10909|CLUS_HUMAN Clusterin OS=Homo sapiens OX=9606 GN=CLU PE=1 SV=1 | 1467 | 20201115_10113_1ng_glyco_hcd203040_compound15.19374.19374.3 | controllerType=0 controllerNumber=1 scan=19374 | 38.7484 |
| 00797:3 | 16 | R.EIRHN[+2059.735]STGC[+57.021]LR.M | HexNAc(4)Hex(5)Fuc(1)NeuAc(1) | C[+57], N[+2060] | 1134.809 | 3 | 3402.4124 | 3402.3994 | 0 | 3.8 | 287 | Specific | 244.4 | 130.39 | 130.39 | 4.15 | 39 | >sp|P10909|CLUS_HUMAN Clusterin OS=Homo sapiens OX=9606 GN=CLU PE=1 SV=1 | 1467 | 20201115_10113_1ng_glyco_hcd203040_compound15.3351.3351.3 | controllerType=0 controllerNumber=1 scan=3351 | 11.237 |
| 00814:3 | 16 | R.EIRHN[+2351.851]STGC[+57.021]LR.M | HexNAc(4)Hex(5)Fuc(3)NeuAc(1) | C[+57], N[+2352] | 1232.1775 | 3 | 3694.5181 | 3694.5152 | 0 | 0.8 | 287 | Specific | 249.4 | 170.51 | 170.51 | 4.67 | 39 | >sp|P10909|CLUS_HUMAN Clusterin OS=Homo sapiens OX=9606 GN=CLU PE=1 SV=1 | 1467 | 20201115_10113_1ng_glyco_hcd203040_compound15.3370.3370.3 | controllerType=0 controllerNumber=1 scan=3370 | 11.277 |
| 23152:4 | 16 | K.MLN[+2350.830]TSSLLEQLNEQFNWVSR.L | HexNAc(4)Hex(5)Fuc(1)NeuAc(2) | N[+2351] | 1190.7645 | 4 | 4760.0362 | 4760.0171 | 0 | 4 | 352 | Specific | 411.49 | 396.8 | 396.8 | 9.31 | 39 | >sp|P10909|CLUS_HUMAN Clusterin OS=Homo sapiens OX=9606 GN=CLU PE=1 SV=1 | 1467 | 20201115_10113_1ng_glyco_hcd203040_compound15.26608.26608.4 | controllerType=0 controllerNumber=1 scan=26608 | 54.327 |
| 23153:3 | 16 | K.MLN[+2350.830]TSSLLEQLNEQFNWVSR.L | HexNAc(4)Hex(5)Fuc(1)NeuAc(2) | N[+2351] | 1587.3503 | 3 | 4760.0363 | 4760.0171 | 0 | 4 | 352 | Specific | 850.15 | 590.59 | 590.59 | 17.12 | 39 | >sp|P10909|CLUS_HUMAN Clusterin OS=Homo sapiens OX=9606 GN=CLU PE=1 SV=1 | 1467 | 20201115_10113_1ng_glyco_hcd203040_compound15.26609.26609.3 | controllerType=0 controllerNumber=1 scan=26609 | 54.3297 |
| 23154:4 | 16 | K.MLN[+2350.830]TSSLLEQLNEQFNWVSR.L | HexNAc(4)Hex(5)Fuc(1)NeuAc(2) | N[+2351] | 1190.7645 | 4 | 4760.0362 | 4760.0171 | 0 | 4 | 352 | Specific | 930.21 | 677.86 | 677.86 | 18.16 | 39 | >sp|P10909|CLUS_HUMAN Clusterin OS=Homo sapiens OX=9606 GN=CLU PE=1 SV=1 | 1467 | 20201115_10113_1ng_glyco_hcd203040_compound15.26610.26610.4 | controllerType=0 controllerNumber=1 scan=26610 | 54.3323 |
| 21123:3 | 16 | K.MLN[+2059.735]TSSLLEQLNEQFNWVSR.L | HexNAc(4)Hex(5)Fuc(1)NeuAc(1) | N[+2060] | 1490.3168 | 3 | 4468.9359 | 4468.9217 | 0 | 3.2 | 352 | Specific | 958.05 | 708.69 | 708.69 | 19.53 | 39 | >sp|P10909|CLUS_HUMAN Clusterin OS=Homo sapiens OX=9606 GN=CLU PE=1 SV=1 | 1467 | 20201115_10113_1ng_glyco_hcd203040_compound15.24481.24481.3 | controllerType=0 controllerNumber=1 scan=24481 | 49.6916 |
| 23150:3 | 16 | K.MLN[+2350.830]TSSLLEQLNEQFNWVSR.L | HexNAc(4)Hex(5)Fuc(1)NeuAc(2) | N[+2351] | 1587.3503 | 3 | 4760.0363 | 4760.0171 | 0 | 4 | 352 | Specific | 1031.85 | 766.29 | 766.29 | 20.12 | 39 | >sp|P10909|CLUS_HUMAN Clusterin OS=Homo sapiens OX=9606 GN=CLU PE=1 SV=1 | 1467 | 20201115_10113_1ng_glyco_hcd203040_compound15.26606.26606.3 | controllerType=0 controllerNumber=1 scan=26606 | 54.3247 |
| 21120:3 | 16 | K.MLN[+2059.735]TSSLLEQLNEQFNWVSR.L | HexNAc(4)Hex(5)Fuc(1)NeuAc(1) | N[+2060] | 1490.3168 | 3 | 4468.9359 | 4468.9217 | 0 | 3.2 | 352 | Specific | 1091.46 | 826.21 | 826.21 | 21.53 | 39 | >sp|P10909|CLUS_HUMAN Clusterin OS=Homo sapiens OX=9606 GN=CLU PE=1 SV=1 | 1467 | 20201115_10113_1ng_glyco_hcd203040_compound15.24478.24478.3 | controllerType=0 controllerNumber=1 scan=24478 | 49.6864 |
| 20690:3 | 16 | K.M[+15.995]LN[+2059.735]TSSLLEQLNEQFNWVSR.L | HexNAc(4)Hex(5)Fuc(1)NeuAc(1) | M[+16], N[+2060] | 1495.6463 | 3 | 4484.9243 | 4484.9166 | 0 | 1.7 | 352 | Specific | 233.2 | 199.47 | 199.47 | 6.2 | 39 | >sp|P10909|CLUS_HUMAN Clusterin OS=Homo sapiens OX=9606 GN=CLU PE=1 SV=1 | 1467 | 20201115_10113_1ng_glyco_hcd203040_compound15.24027.24027.3 | controllerType=0 controllerNumber=1 scan=24027 | 48.6528 |
| 20693:3 | 16 | K.M[+15.995]LN[+2059.735]TSSLLEQLNEQFNWVSR.L | HexNAc(4)Hex(5)Fuc(1)NeuAc(1) | M[+16], N[+2060] | 1495.6463 | 3 | 4484.9243 | 4484.9166 | 0 | 1.7 | 352 | Specific | 578.44 | 403.49 | 403.49 | 10.49 | 39 | >sp|P10909|CLUS_HUMAN Clusterin OS=Homo sapiens OX=9606 GN=CLU PE=1 SV=1 | 1467 | 20201115_10113_1ng_glyco_hcd203040_compound15.24030.24030.3 | controllerType=0 controllerNumber=1 scan=24030 | 48.6611 |
| 13579:3 | 16 | R.LAN[+2350.830]LTQGEDQYYLR.V | HexNAc(4)Hex(5)Fuc(1)NeuAc(2) | N[+2351] | 1345.5618 | 3 | 4034.6709 | 4034.664 | 0 | 1.7 | 372 | Specific | 298.21 | 249.42 | 249.42 | 6.73 | 39 | >sp|P10909|CLUS_HUMAN Clusterin OS=Homo sapiens OX=9606 GN=CLU PE=1 SV=1 | 1467 | 20201115_10113_1ng_glyco_hcd203040_compound15.16616.16616.3 | controllerType=0 controllerNumber=1 scan=16616 | 33.4907 |
| 10665:3 | 16 | R.LAN[+2570.925]LTQGEDQYYLR.V | HexNAc(5)Hex(6)Fuc(2)NeuAc(1) | N[+2571] | 1418.9286 | 3 | 4254.7714 | 4254.7587 | 0 | 3 | 372 | Specific | 336.64 | 297.23 | 297.23 | 7.15 | 39 | >sp|P10909|CLUS_HUMAN Clusterin OS=Homo sapiens OX=9606 GN=CLU PE=1 SV=1 | 1467 | 20201115_10113_1ng_glyco_hcd203040_compound15.13588.13588.3 | controllerType=0 controllerNumber=1 scan=13588 | 28.232 |
| 10667:3 | 16 | R.LAN[+2570.925]LTQGEDQYYLR.V | HexNAc(5)Hex(6)Fuc(2)NeuAc(1) | N[+2571] | 1418.9286 | 3 | 4254.7714 | 4254.7587 | 0 | 3 | 372 | Specific | 466.82 | 365.99 | 365.99 | 8.29 | 39 | >sp|P10909|CLUS_HUMAN Clusterin OS=Homo sapiens OX=9606 GN=CLU PE=1 SV=1 | 1467 | 20201115_10113_1ng_glyco_hcd203040_compound15.13590.13590.3 | controllerType=0 controllerNumber=1 scan=13590 | 28.2356 |
| 13582:3 | 16 | R.LAN[+2350.830]LTQGEDQYYLR.V | HexNAc(4)Hex(5)Fuc(1)NeuAc(2) | N[+2351] | 1345.5618 | 3 | 4034.6709 | 4034.664 | 0 | 1.7 | 372 | Specific | 475.95 | 442.09 | 442.09 | 9.8 | 39 | >sp|P10909|CLUS_HUMAN Clusterin OS=Homo sapiens OX=9606 GN=CLU PE=1 SV=1 | 1467 | 20201115_10113_1ng_glyco_hcd203040_compound15.16619.16619.3 | controllerType=0 controllerNumber=1 scan=16619 | 33.495 |
| 02638:3 | 17 | K.N[+1216.423]NSIPDKQITASSSYK.T | HexNAc(2)Hex(5) | N[+1216] | 990.441 | 3 | 2969.3085 | 2969.2991 | 0 | 3.1 | 238 | Specific | 468.59 | 303.55 | 303.55 | 7.51 | 36 | >sp|Q08431|MFGM_HUMAN Lactadherin OS=Homo sapiens OX=9606 GN=MFGE8 PE=1 SV=3 | 1064 | 20201115_10113_1ng_glyco_hcd203040_compound15.5265.5265.3 | controllerType=0 controllerNumber=1 scan=5265 | 14.5095 |
| 02640:3 | 17 | K.N[+1216.423]NSIPDKQITASSSYK.T | HexNAc(2)Hex(5) | N[+1216] | 990.441 | 3 | 2969.3085 | 2969.2991 | 0 | 3.1 | 238 | Specific | 473.14 | 251.72 | 251.72 | 7.28 | 36 | >sp|Q08431|MFGM_HUMAN Lactadherin OS=Homo sapiens OX=9606 GN=MFGE8 PE=1 SV=3 | 1064 | 20201115_10113_1ng_glyco_hcd203040_compound15.5267.5267.3 | controllerType=0 controllerNumber=1 scan=5267 | 14.5131 |
| 23784:3 | 18 | R.QLAHQSN[+2350.830]STNIFFSPVSIATAFAMLSLGTK.A | HexNAc(4)Hex(5)Fuc(1)NeuAc(2) | N[+2351] | 1844.8305 | 3 | 5532.477 | 5532.4654 | 0 | 2.1 | 64 | Specific | 215.33 | 192.15 | 192.15 | 5.36 | 35 | >sp|P01009|A1AT_HUMAN Alpha-1-antitrypsin OS=Homo sapiens OX=9606 GN=SERPINA1 PE=1 SV=3 | 642 | 20201115_10113_1ng_glyco_hcd203040_compound15.27271.27271.3 | controllerType=0 controllerNumber=1 scan=27271 | 55.7617 |
| 23900:4 | 18 | R.[+42.011]QLAHQSN[+2190.793]STNIFFSPVSIATAFAMLSLGTK.A | HexNAc(6)Hex(6) | .[42], N[+2191] | 1354.3669 | 4 | 5414.446 | 5414.4388 | 0 | 1.3 | 64 | Specific | 230.15 | 205.04 | 205.04 | 6.26 | 35 | >sp|P01009|A1AT_HUMAN Alpha-1-antitrypsin OS=Homo sapiens OX=9606 GN=SERPINA1 PE=1 SV=3 | 642 | 20201115_10113_1ng_glyco_hcd203040_compound15.27393.27393.4 | controllerType=0 controllerNumber=1 scan=27393 | 56.0365 |
| 23898:4 | 18 | R.[+42.011]QLAHQSN[+2190.793]STNIFFSPVSIATAFAMLSLGTK.A | HexNAc(6)Hex(6) | .[42], N[+2191] | 1354.3669 | 4 | 5414.446 | 5414.4388 | 0 | 1.3 | 64 | Specific | 232.68 | 232.68 | 232.68 | 6.58 | 35 | >sp|P01009|A1AT_HUMAN Alpha-1-antitrypsin OS=Homo sapiens OX=9606 GN=SERPINA1 PE=1 SV=3 | 642 | 20201115_10113_1ng_glyco_hcd203040_compound15.27391.27391.4 | controllerType=0 controllerNumber=1 scan=27391 | 56.0331 |
| 23855:4 | 18 | R.[+42.011]QLAHQSN[+2190.793]STNIFFSPVSIATAFAMLSLGTK.A | HexNAc(6)Hex(6) | .[42], N[+2191] | 1354.3638 | 4 | 5414.4333 | 5414.4388 | 0 | -1 | 64 | Specific | 345.5 | 345.5 | 345.5 | 6.74 | 35 | >sp|P01009|A1AT_HUMAN Alpha-1-antitrypsin OS=Homo sapiens OX=9606 GN=SERPINA1 PE=1 SV=3 | 642 | 20201115_10113_1ng_glyco_hcd203040_compound15.27346.27346.4 | controllerType=0 controllerNumber=1 scan=27346 | 55.9188 |
| 23776:4 | 18 | R.QLAHQSN[+2350.830]STNIFFSPVSIATAFAMLSLGTK.A | HexNAc(4)Hex(5)Fuc(1)NeuAc(2) | N[+2351] | 1383.8739 | 4 | 5532.4738 | 5532.4654 | 0 | 1.5 | 64 | Specific | 497.16 | 485.27 | 485.27 | 10.11 | 35 | >sp|P01009|A1AT_HUMAN Alpha-1-antitrypsin OS=Homo sapiens OX=9606 GN=SERPINA1 PE=1 SV=3 | 642 | 20201115_10113_1ng_glyco_hcd203040_compound15.27263.27263.4 | controllerType=0 controllerNumber=1 scan=27263 | 55.7418 |
| 23849:3 | 18 | R.QLAHQSN[+2204.772]STNIFFSPVSIATAFAMLSLGTK.A | HexNAc(4)Hex(5)NeuAc(2) | N[+2205] | 1796.1491 | 3 | 5386.4328 | 5386.4075 | 0 | 4.7 | 64 | Specific | 535.35 | 520.15 | 520.15 | 10.12 | 35 | >sp|P01009|A1AT_HUMAN Alpha-1-antitrypsin OS=Homo sapiens OX=9606 GN=SERPINA1 PE=1 SV=3 | 642 | 20201115_10113_1ng_glyco_hcd203040_compound15.27340.27340.3 | controllerType=0 controllerNumber=1 scan=27340 | 55.9066 |
| 23851:3 | 18 | R.QLAHQSN[+2204.772]STNIFFSPVSIATAFAMLSLGTK.A | HexNAc(4)Hex(5)NeuAc(2) | N[+2205] | 1796.1491 | 3 | 5386.4328 | 5386.4075 | 0 | 4.7 | 64 | Specific | 567.01 | 507.63 | 507.63 | 10.43 | 35 | >sp|P01009|A1AT_HUMAN Alpha-1-antitrypsin OS=Homo sapiens OX=9606 GN=SERPINA1 PE=1 SV=3 | 642 | 20201115_10113_1ng_glyco_hcd203040_compound15.27342.27342.3 | controllerType=0 controllerNumber=1 scan=27342 | 55.9121 |
| 23778:4 | 18 | R.QLAHQSN[+2350.830]STNIFFSPVSIATAFAMLSLGTK.A | HexNAc(4)Hex(5)Fuc(1)NeuAc(2) | N[+2351] | 1383.8739 | 4 | 5532.4738 | 5532.4654 | 0 | 1.5 | 64 | Specific | 825.73 | 772.26 | 772.26 | 16.61 | 35 | >sp|P01009|A1AT_HUMAN Alpha-1-antitrypsin OS=Homo sapiens OX=9606 GN=SERPINA1 PE=1 SV=3 | 642 | 20201115_10113_1ng_glyco_hcd203040_compound15.27265.27265.4 | controllerType=0 controllerNumber=1 scan=27265 | 55.7456 |
| 23848:4 | 18 | R.QLAHQSN[+2204.772]STNIFFSPVSIATAFAMLSLGTK.A | HexNAc(4)Hex(5)NeuAc(2) | N[+2205] | 1347.3641 | 4 | 5386.4347 | 5386.4075 | 0 | 5.1 | 64 | Specific | 874.65 | 824.88 | 824.88 | 17.25 | 35 | >sp|P01009|A1AT_HUMAN Alpha-1-antitrypsin OS=Homo sapiens OX=9606 GN=SERPINA1 PE=1 SV=3 | 642 | 20201115_10113_1ng_glyco_hcd203040_compound15.27339.27339.4 | controllerType=0 controllerNumber=1 scan=27339 | 55.9048 |
| 23850:4 | 18 | R.QLAHQSN[+2204.772]STNIFFSPVSIATAFAMLSLGTK.A | HexNAc(4)Hex(5)NeuAc(2) | N[+2205] | 1347.3641 | 4 | 5386.4347 | 5386.4075 | 0 | 5.1 | 64 | Specific | 960.36 | 875.9 | 875.9 | 19.25 | 35 | >sp|P01009|A1AT_HUMAN Alpha-1-antitrypsin OS=Homo sapiens OX=9606 GN=SERPINA1 PE=1 SV=3 | 642 | 20201115_10113_1ng_glyco_hcd203040_compound15.27341.27341.4 | controllerType=0 controllerNumber=1 scan=27341 | 55.9102 |
| 20240:5 | 18 | K.YLGN[+2204.772]ATAIFFLPDEGKLQHLENELTHDIITK.F | HexNAc(4)Hex(5)NeuAc(2) | N[+2205] | 1149.9332 | 5 | 5745.637 | 5745.6098 | 0 | 4.7 | 268 | Specific | 781.2 | 600.97 | 600.97 | 14.95 | 35 | >sp|P01009|A1AT_HUMAN Alpha-1-antitrypsin OS=Homo sapiens OX=9606 GN=SERPINA1 PE=1 SV=3 | 642 | 20201115_10113_1ng_glyco_hcd203040_compound15.23556.23556.5 | controllerType=0 controllerNumber=1 scan=23556 | 47.5867 |
| 20241:4 | 18 | K.YLGN[+2204.772]ATAIFFLPDEGKLQHLENELTHDIITK.F | HexNAc(4)Hex(5)NeuAc(2) | N[+2205] | 1437.165 | 4 | 5745.6383 | 5745.6098 | 0 | 5 | 268 | Specific | 871.58 | 871.58 | 871.58 | 16.21 | 35 | >sp|P01009|A1AT_HUMAN Alpha-1-antitrypsin OS=Homo sapiens OX=9606 GN=SERPINA1 PE=1 SV=3 | 642 | 20201115_10113_1ng_glyco_hcd203040_compound15.23557.23557.4 | controllerType=0 controllerNumber=1 scan=23557 | 47.5918 |
| 20239:4 | 18 | K.YLGN[+2204.772]ATAIFFLPDEGKLQHLENELTHDIITK.F | HexNAc(4)Hex(5)NeuAc(2) | N[+2205] | 1437.165 | 4 | 5745.6383 | 5745.6098 | 0 | 5 | 268 | Specific | 1329.47 | 1185.07 | 1185.07 | 25.21 | 35 | >sp|P01009|A1AT_HUMAN Alpha-1-antitrypsin OS=Homo sapiens OX=9606 GN=SERPINA1 PE=1 SV=3 | 642 | 20201115_10113_1ng_glyco_hcd203040_compound15.23555.23555.4 | controllerType=0 controllerNumber=1 scan=23555 | 47.5855 |
| 24673:5 | 23 | K.DIDGYGGIALPELIC[+57.021]TMFHTSGYDTQAIVENN[+2350.830]ESTEYGLFQISNK. | HexNAc(4)Hex(5)Fuc(1)NeuAc(2) | C[+57], N[+2351] | 1473.238 | 5 | 7362.1607 | 7362.1592 | 0 | 0.2 | 33 | Specific | 294.08 | 294.08 | 294.08 | 7.34 | 25 | >sp|P00709|LALBA_HUMAN Alpha-lactalbumin OS=Homo sapiens OX=9606 GN=LALBA PE=1 SV=1 | 468 | 20201115_10113_1ng_glyco_hcd203040_compound15.28202.28202.2 | controllerType=0 controllerNumber=1 scan=28202 | 57.6464 |
| 09132:5 | 27 | K.SC[+57.021]QHN[+2204.772]GTMYQHGEIFSAHELFPSR.L | HexNAc(4)Hex(5)NeuAc(2) | C[+57], N[+2205] | 1008.4193 | 5 | 5038.0672 | 5038.0294 | 0 | 7.5 | 110 | Specific | 439.35 | 162 | 162 | 6.06 | 21 | >sp|Q6WN34|CRDL2_HUMAN Chordin-like protein 2 OS=Homo sapiens OX=9606 GN=CHRDL2 PE=1 SV=1 | 1189 | 20201115_10113_1ng_glyco_hcd203040_compound15.11993.11993.3 | controllerType=0 controllerNumber=1 scan=11993 | 25.5598 |
| 09106:5 | 27 | K.SC[+57.021]QHN[+2205.793]GTMYQHGEIFSAHELFPSR.L | HexNAc(4)Hex(5)Fuc(2)NeuAc(1) | C[+57], N[+2206] | 1008.6193 | 5 | 5039.0672 | 5039.0498 | 0 | 3.5 | 110 | Specific | 456.51 | 456.51 | 456.51 | 10.39 | 21 | >sp|Q6WN34|CRDL2_HUMAN Chordin-like protein 2 OS=Homo sapiens OX=9606 GN=CHRDL2 PE=1 SV=1 | 1189 | 20201115_10113_1ng_glyco_hcd203040_compound15.11966.11966.2 | controllerType=0 controllerNumber=1 scan=11966 | 25.5148 |
| 09130:5 | 27 | K.SC[+57.021]QHN[+2204.772]GTMYQHGEIFSAHELFPSR.L | HexNAc(4)Hex(5)NeuAc(2) | C[+57], N[+2205] | 1008.4193 | 5 | 5038.0672 | 5038.0294 | 0 | 7.5 | 110 | Specific | 462.65 | 119.03 | 119.03 | 5.88 | 21 | >sp|Q6WN34|CRDL2_HUMAN Chordin-like protein 2 OS=Homo sapiens OX=9606 GN=CHRDL2 PE=1 SV=1 | 1189 | 20201115_10113_1ng_glyco_hcd203040_compound15.11991.11991.3 | controllerType=0 controllerNumber=1 scan=11991 | 25.5554 |
| 09109:5 | 27 | K.SC[+57.021]QHN[+2205.793]GTMYQHGEIFSAHELFPSR.L | HexNAc(4)Hex(5)Fuc(2)NeuAc(1) | C[+57], N[+2206] | 1008.6193 | 5 | 5039.0672 | 5039.0498 | 0 | 3.5 | 110 | Specific | 597.61 | 597.61 | 597.61 | 11.99 | 21 | >sp|Q6WN34|CRDL2_HUMAN Chordin-like protein 2 OS=Homo sapiens OX=9606 GN=CHRDL2 PE=1 SV=1 | 1189 | 20201115_10113_1ng_glyco_hcd203040_compound15.11969.11969.2 | controllerType=0 controllerNumber=1 scan=11969 | 25.5203 |
| 02518:3 | 36 | R.LRN[+1216.423]VSWATGR.S | HexNAc(2)Hex(5) | N[+1216] | 792.6927 | 3 | 2376.0636 | 2376.0559 | 0 | 3.2 | 149 | Specific | 208.71 | 47.63 | 47.63 | 3.15 | 16 | >sp|P08571|CD14_HUMAN Monocyte differentiation antigen CD14 OS=Homo sapiens OX=9606 GN=CD14 PE=1 SV=2 | 1232 | 20201115_10113_1ng_glyco_hcd203040_compound15.5141.5141.3 | controllerType=0 controllerNumber=1 scan=5141 | 14.2971 |
| 13865:4 | 43 | R.IIVPLNNREN[+2204.772]ISDPTSPLR.T | HexNAc(4)Hex(5)NeuAc(2) | N[+2205] | 1088.995 | 4 | 4352.9583 | 4352.9496 | 0 | 2 | 62 | Specific | 273.82 | 207.87 | 207.87 | 5.95 | 17 | >sp|P01591|IGJ_HUMAN Immunoglobulin J chain OS=Homo sapiens OX=9606 GN=JCHAIN PE=1 SV=4 | 1392 | 20201115_10113_1ng_glyco_hcd203040_compound15.16913.16913.4 | controllerType=0 controllerNumber=1 scan=16913 | 34.0411 |
| 13807:4 | 43 | R.IIVPLNNREN[+2350.830]ISDPTSPLR.T | HexNAc(4)Hex(5)Fuc(1)NeuAc(2) | N[+2351] | 1125.5094 | 4 | 4499.0159 | 4499.0075 | 0 | 1.9 | 62 | Specific | 273.9 | 235.5 | 235.5 | 5.08 | 17 | >sp|P01591|IGJ_HUMAN Immunoglobulin J chain OS=Homo sapiens OX=9606 GN=JCHAIN PE=1 SV=4 | 1392 | 20201115_10113_1ng_glyco_hcd203040_compound15.16853.16853.4 | controllerType=0 controllerNumber=1 scan=16853 | 33.9443 |
| 12019:3 | 43 | R.IIVPLNNREN[+1694.603]ISDPTSPLR.T | HexNAc(3)Hex(4)Fuc(1)NeuAc(1) | N[+1695] | 1281.5898 | 3 | 3842.7548 | 3842.7799 | 0 | -6.5 | 62 | Specific | 277.77 | 277.77 | 277.77 | 3.41 | 17 | >sp|P01591|IGJ_HUMAN Immunoglobulin J chain OS=Homo sapiens OX=9606 GN=JCHAIN PE=1 SV=4 | 1392 | 20201115_10113_1ng_glyco_hcd203040_compound15.14993.14993.3 | controllerType=0 controllerNumber=1 scan=14993 | 30.7651 |
| 13868:4 | 43 | R.IIVPLNNREN[+2204.772]ISDPTSPLR.T | HexNAc(4)Hex(5)NeuAc(2) | N[+2205] | 1088.995 | 4 | 4352.9583 | 4352.9496 | 0 | 2 | 62 | Specific | 329.61 | 304.37 | 304.37 | 5.82 | 17 | >sp|P01591|IGJ_HUMAN Immunoglobulin J chain OS=Homo sapiens OX=9606 GN=JCHAIN PE=1 SV=4 | 1392 | 20201115_10113_1ng_glyco_hcd203040_compound15.16916.16916.4 | controllerType=0 controllerNumber=1 scan=16916 | 34.0484 |
| 12016:3 | 43 | R.IIVPLNNREN[+1694.603]ISDPTSPLR.T | HexNAc(3)Hex(4)Fuc(1)NeuAc(1) | N[+1695] | 1281.5898 | 3 | 3842.7548 | 3842.7799 | 0 | -6.5 | 62 | Specific | 330.55 | 315.27 | 315.27 | 5.1 | 17 | >sp|P01591|IGJ_HUMAN Immunoglobulin J chain OS=Homo sapiens OX=9606 GN=JCHAIN PE=1 SV=4 | 1392 | 20201115_10113_1ng_glyco_hcd203040_compound15.14990.14990.3 | controllerType=0 controllerNumber=1 scan=14990 | 30.7612 |
| 11817:3 | 43 | R.IIVPLNNREN[+2059.735]ISDPTSPLR.T | HexNAc(4)Hex(5)Fuc(1)NeuAc(1) | N[+2060] | 1403.3116 | 3 | 4207.9202 | 4207.9121 | 0 | 1.9 | 62 | Specific | 377.43 | 357.69 | 357.69 | 7.53 | 17 | >sp|P01591|IGJ_HUMAN Immunoglobulin J chain OS=Homo sapiens OX=9606 GN=JCHAIN PE=1 SV=4 | 1392 | 20201115_10113_1ng_glyco_hcd203040_compound15.14783.14783.3 | controllerType=0 controllerNumber=1 scan=14783 | 30.3538 |
| 11903:3 | 43 | R.IIVPLNNREN[+1856.656]ISDPTSPLR.T | HexNAc(3)Hex(5)Fuc(1)NeuAc(1) | N[+1857] | 1335.6185 | 3 | 4004.8408 | 4004.8327 | 0 | 2 | 62 | Specific | 469.79 | 463.73 | 463.73 | 9.65 | 17 | >sp|P01591|IGJ_HUMAN Immunoglobulin J chain OS=Homo sapiens OX=9606 GN=JCHAIN PE=1 SV=4 | 1392 | 20201115_10113_1ng_glyco_hcd203040_compound15.14873.14873.3 | controllerType=0 controllerNumber=1 scan=14873 | 30.5535 |
| 11819:3 | 43 | R.IIVPLNNREN[+2059.735]ISDPTSPLR.T | HexNAc(4)Hex(5)Fuc(1)NeuAc(1) | N[+2060] | 1403.3116 | 3 | 4207.9202 | 4207.9121 | 0 | 1.9 | 62 | Specific | 511.38 | 482.13 | 482.13 | 9.74 | 17 | >sp|P01591|IGJ_HUMAN Immunoglobulin J chain OS=Homo sapiens OX=9606 GN=JCHAIN PE=1 SV=4 | 1392 | 20201115_10113_1ng_glyco_hcd203040_compound15.14785.14785.3 | controllerType=0 controllerNumber=1 scan=14785 | 30.3607 |
| 11906:3 | 43 | R.IIVPLNNREN[+1856.656]ISDPTSPLR.T | HexNAc(3)Hex(5)Fuc(1)NeuAc(1) | N[+1857] | 1335.6185 | 3 | 4004.8408 | 4004.8327 | 0 | 2 | 62 | Specific | 538.62 | 538.62 | 538.62 | 10.06 | 17 | >sp|P01591|IGJ_HUMAN Immunoglobulin J chain OS=Homo sapiens OX=9606 GN=JCHAIN PE=1 SV=4 | 1392 | 20201115_10113_1ng_glyco_hcd203040_compound15.14876.14876.3 | controllerType=0 controllerNumber=1 scan=14876 | 30.5581 |
| 10226:3 | 43 | R.IIVPLNNREN[+1809.666]ISDPTSPLR.T | HexNAc(5)Hex(4)Fuc(1) | N[+1810] | 1319.955 | 3 | 3957.8505 | 3957.8432 | 0 | 1.8 | 62 | Specific | 544.93 | 468.24 | 468.24 | 10.06 | 17 | >sp|P01591|IGJ_HUMAN Immunoglobulin J chain OS=Homo sapiens OX=9606 GN=JCHAIN PE=1 SV=4 | 1392 | 20201115_10113_1ng_glyco_hcd203040_compound15.13135.13135.3 | controllerType=0 controllerNumber=1 scan=13135 | 27.4826 |
| 10224:3 | 43 | R.IIVPLNNREN[+1809.666]ISDPTSPLR.T | HexNAc(5)Hex(4)Fuc(1) | N[+1810] | 1319.955 | 3 | 3957.8505 | 3957.8432 | 0 | 1.8 | 62 | Specific | 552.29 | 437.71 | 437.71 | 10.06 | 17 | >sp|P01591|IGJ_HUMAN Immunoglobulin J chain OS=Homo sapiens OX=9606 GN=JCHAIN PE=1 SV=4 | 1392 | 20201115_10113_1ng_glyco_hcd203040_compound15.13133.13133.3 | controllerType=0 controllerNumber=1 scan=13133 | 27.4775 |
| 21653:3 | 67 | R.QFWIFDVQNPQEVMMN[+2350.830]SSNIQVK.Q | HexNAc(4)Hex(5)Fuc(1)NeuAc(2) | N[+2351] | 1711.7315 | 3 | 5133.18 | 5133.1631 | 0 | 3.3 | 64 | Specific | 221.37 | 216.1 | 216.1 | 6.86 | 11 | >sp|P16671|CD36_HUMAN Platelet glycoprotein 4 OS=Homo sapiens OX=9606 GN=CD36 PE=1 SV=2 | 812 | 20201115_10113_1ng_glyco_hcd203040_compound15.25035.25035.3 | controllerType=0 controllerNumber=1 scan=25035 | 50.8137 |
| 24398:3 | 103 | R.AVN[+2205.793]ITSENLIDDVVSLIR.G | HexNAc(4)Hex(5)Fuc(2)NeuAc(1) | N[+2206] | 1392.9624 | 3 | 4176.8726 | 4176.8685 | 0 | 1 | 413 | Specific | 480.7 | 438.91 | 438.91 | 9.71 | 9 | >sp|P02748|CO9_HUMAN Complement component C9 OS=Homo sapiens OX=9606 GN=C9 PE=1 SV=2 | 1547 | 20201115_10113_1ng_glyco_hcd203040_compound15.27915.27915.3 | controllerType=0 controllerNumber=1 scan=27915 | 57.0997 |
| 24400:3 | 103 | R.AVN[+2205.793]ITSENLIDDVVSLIR.G | HexNAc(4)Hex(5)Fuc(2)NeuAc(1) | N[+2206] | 1392.9624 | 3 | 4176.8726 | 4176.8685 | 0 | 1 | 413 | Specific | 640.54 | 368.7 | 368.7 | 10.82 | 9 | >sp|P02748|CO9_HUMAN Complement component C9 OS=Homo sapiens OX=9606 GN=C9 PE=1 SV=2 | 1547 | 20201115_10113_1ng_glyco_hcd203040_compound15.27917.27917.3 | controllerType=0 controllerNumber=1 scan=27917 | 57.1047 |
| 17420:4 | 1 | R.RTAGWNVPIGTLRPFLN[+2569.905]WTGPPEPIEAAVAR.F | HexNAc(5)Hex(6)NeuAc(2) | N[+2570] | 1489.9432 | 4 | 5956.7511 | 5956.7167 | 0 | 5.8 | 140 | Specific | 261.08 | 244.71 | 244.71 | 5.52 | 130 | >sp|P02788|TRFL_HUMAN Lactotransferrin OS=Homo sapiens OX=9606 GN=LTF PE=1 SV=6 | 1270 | 20201119_10132_1ng_glyco_hcd203040_compound15.21089.21089.4 | controllerType=0 controllerNumber=1 scan=21089 | 43.3832 |
| 17533:3 | 1 | R.RTAGWNVPIGTLRPFLN[+2059.735]WTGPPEPIEAAVAR.F | HexNAc(4)Hex(5)Fuc(1)NeuAc(1) | N[+2060] | 1816.1921 | 3 | 5446.5618 | 5446.547 | 0 | 2.7 | 140 | Specific | 360.86 | 360.86 | 360.86 | 7.64 | 130 | >sp|P02788|TRFL_HUMAN Lactotransferrin OS=Homo sapiens OX=9606 GN=LTF PE=1 SV=6 | 1270 | 20201119_10132_1ng_glyco_hcd203040_compound15.21206.21206.3 | controllerType=0 controllerNumber=1 scan=21206 | 43.615 |
| 17423:4 | 1 | R.RTAGWNVPIGTLRPFLN[+2569.905]WTGPPEPIEAAVAR.F | HexNAc(5)Hex(6)NeuAc(2) | N[+2570] | 1489.9432 | 4 | 5956.7511 | 5956.7167 | 0 | 5.8 | 140 | Specific | 423.07 | 364.3 | 364.3 | 8.19 | 130 | >sp|P02788|TRFL_HUMAN Lactotransferrin OS=Homo sapiens OX=9606 GN=LTF PE=1 SV=6 | 1270 | 20201119_10132_1ng_glyco_hcd203040_compound15.21092.21092.4 | controllerType=0 controllerNumber=1 scan=21092 | 43.3886 |

Table S4. Cont.

| Query #:z | Protein Rank | Glycopeptide | Glycan | Modification Type(s) | Observed m/z | z | Observed (M+H) | Calc. mass (M+H) | Off-by-x error | Mass error (ppm) | StartPosition | Cleavage | Score | Delta | Delta Mod | \|Log Prob\| | # of unique peptides | Protein Name | Protein DB number | Comment | Scan # | Scan Time |
|---|---|---|---|---|---|---|---|---|---|---|---|---|---|---|---|---|---|---|---|---|---|---|
| 17543:5 | 1 | R.RTAGWNVPIGTLRPFLN[+2059.735]WTGPPEPIEAAVAR.F | HexNAc(4)Hex(5)Fuc(1)NeuAc(1) | N[+2060] | 1090.1187 | 5 | 5446.5642 | 5446.547 | 0 | 3.2 | 140 | Specific | 502.23 | 502.23 | 502.23 | 10.22 | 130 | >sp\|P02788\|TRFL_HUMAN Lactotransferrin OS=Homo sapiens OX=9606 GN=LTF PE=1 SV=6 | 1270 | 20201119_10132_1ng_glyco_hcd203040_compound15.21216.21216.5 | controllerType=0 controllerNumber=1 scan=21216 | 43.6313 |
| 17519:4 | 1 | R.RTAGWNVPIGTLRPFLN[+2059.735]WTGPPEPIEAAVAR.F | HexNAc(4)Hex(5)Fuc(1)NeuAc(1) | N[+2060] | 1362.3954 | 4 | 5446.5597 | 5446.547 | 0 | 2.3 | 140 | Specific | 571.32 | 571.32 | 571.32 | 10.65 | 130 | >sp\|P02788\|TRFL_HUMAN Lactotransferrin OS=Homo sapiens OX=9606 GN=LTF PE=1 SV=6 | 1270 | 20201119_10132_1ng_glyco_hcd203040_compound15.21192.21192.4 | controllerType=0 controllerNumber=1 scan=21192 | 43.5853 |
| 17521:4 | 1 | R.RTAGWNVPIGTLRPFLN[+2059.735]WTGPPEPIEAAVAR.F | HexNAc(4)Hex(5)Fuc(1)NeuAc(1) | N[+2060] | 1362.3954 | 4 | 5446.5597 | 5446.547 | 0 | 2.3 | 140 | Specific | 623.78 | 535.78 | 535.78 | 12.8 | 130 | >sp\|P02788\|TRFL_HUMAN Lactotransferrin OS=Homo sapiens OX=9606 GN=LTF PE=1 SV=6 | 1270 | 20201119_10132_1ng_glyco_hcd203040_compound15.21194.21194.4 | controllerType=0 controllerNumber=1 scan=21194 | 43.5908 |
| 20118:3 | 1 | R.TAGWNVPIGTLRPFLN[+2350.830]WTGPPEPIEAAVAR.F | HexNAc(4)Hex(5)Fuc(1)NeuAc(2) | N[+2351] | 1861.1989 | 3 | 5581.5821 | 5581.5413 | 0 | 7.3 | 141 | Specific | 202.03 | 159.31 | 159.31 | 4.39 | 130 | >sp\|P02788\|TRFL_HUMAN Lactotransferrin OS=Homo sapiens OX=9606 GN=LTF PE=1 SV=6 | 1270 | 20201119_10132_1ng_glyco_hcd203040_compound15.23911.23911.3 | controllerType=0 controllerNumber=1 scan=23911 | 49.346 |
| 18663:3 | 1 | R.TAGWNVPIGTLRPFLN[+2570.925]WTGPPEPIEAAVAR.F | HexNAc(5)Hex(6)Fuc(2)NeuAc(1) | N[+2571] | 1934.5604 | 3 | 5801.6666 | 5801.636 | 0 | 5.3 | 141 | Specific | 236.43 | 214.44 | 214.44 | 6.3 | 130 | >sp\|P02788\|TRFL_HUMAN Lactotransferrin OS=Homo sapiens OX=9606 GN=LTF PE=1 SV=6 | 1270 | 20201119_10132_1ng_glyco_hcd203040_compound15.22387.22387.3 | controllerType=0 controllerNumber=1 scan=22387 | 46.1905 |
| 17625:4 | 1 | R.TAGWNVPIGTLRPFLN[+2791.020]WTGPPEPIEAAVAR.F | HexNAc(6)Hex(7)Fuc(3) | N[+2791] | 1506.1939 | 4 | 6021.7536 | 6021.7307 | 0 | 3.8 | 141 | Specific | 243.42 | 243.42 | 243.42 | 7.01 | 130 | >sp\|P02788\|TRFL_HUMAN Lactotransferrin OS=Homo sapiens OX=9606 GN=LTF PE=1 SV=6 | 1270 | 20201119_10132_1ng_glyco_hcd203040_compound15.21301.21301.4 | controllerType=0 controllerNumber=1 scan=21301 | 43.779 |
| 17593:4 | 1 | R.TAGWNVPIGTLRPFLN[+2644.962]WTGPPEPIEAAVAR.F | HexNAc(6)Hex(7)Fuc(2) | N[+2645] | 1469.6794 | 4 | 5875.696 | 5875.6727 | 0 | 4 | 141 | Specific | 268.3 | 268.3 | 268.3 | 7.27 | 130 | >sp\|P02788\|TRFL_HUMAN Lactotransferrin OS=Homo sapiens OX=9606 GN=LTF PE=1 SV=6 | 1270 | 20201119_10132_1ng_glyco_hcd203040_compound15.21268.21268.3 | controllerType=0 controllerNumber=1 scan=21268 | 43.7212 |
| 18649:4 | 1 | R.TAGWNVPIGTLRPFLN[+2569.905]WTGPPEPIEAAVAR.F | HexNAc(5)Hex(6)NeuAc(2) | N[+2570] | 1450.923 | 4 | 5800.6701 | 5800.6156 | 0 | 9.4 | 141 | Specific | 271.24 | 271.24 | 271.24 | 5.9 | 130 | >sp\|P02788\|TRFL_HUMAN Lactotransferrin OS=Homo sapiens OX=9606 GN=LTF PE=1 SV=6 | 1270 | 20201119_10132_1ng_glyco_hcd203040_compound15.22373.22373.4 | controllerType=0 controllerNumber=1 scan=22373 | 46.1594 |
| 17719:4 | 1 | R.[+42.011]TAGWNVPIGTLRPFLN[+1872.651]WTGPPEPIEAAVAR.F | HexNAc(3)Hex(6)NeuAc(1) | .[42], N[+1873] | 1287.1091 | 4 | 5145.4147 | 5145.372 | 0 | 8.3 | 141 | Specific | 281.54 | 261.48 | 261.48 | 6.25 | 130 | >sp\|P02788\|TRFL_HUMAN Lactotransferrin OS=Homo sapiens OX=9606 GN=LTF PE=1 SV=6 | 1270 | 20201119_10132_1ng_glyco_hcd203040_compound15.21400.21400.4 | controllerType=0 controllerNumber=1 scan=21400 | 44.0025 |
| 17720:3 | 1 | R.[+42.011]TAGWNVPIGTLRPFLN[+1872.651]WTGPPEPIEAAVAR.F | HexNAc(3)Hex(6)NeuAc(1) | .[42], N[+1873] | 1715.8092 | 3 | 5145.4129 | 5145.372 | 0 | 8 | 141 | Specific | 295.5 | 290.84 | 290.84 | 6.5 | 130 | >sp\|P02788\|TRFL_HUMAN Lactotransferrin OS=Homo sapiens OX=9606 GN=LTF PE=1 SV=6 | 1270 | 20201119_10132_1ng_glyco_hcd203040_compound15.21401.21401.3 | controllerType=0 controllerNumber=1 scan=21401 | 44.0066 |
| 20063:3 | 1 | R.TAGWNVPIGTLRPFLN[+2350.830]WTGPPEPIEAAVAR.F | HexNAc(4)Hex(5)Fuc(1)NeuAc(2) | N[+2351] | 1861.1949 | 3 | 5581.5702 | 5581.5413 | 0 | 5.2 | 141 | Specific | 304.95 | 171.37 | 171.37 | 5.87 | 130 | >sp\|P02788\|TRFL_HUMAN Lactotransferrin OS=Homo sapiens OX=9606 GN=LTF PE=1 SV=6 | 1270 | 20201119_10132_1ng_glyco_hcd203040_compound15.23854.23854.3 | controllerType=0 controllerNumber=1 scan=23854 | 49.2264 |
| 20121:3 | 1 | R.TAGWNVPIGTLRPFLN[+2350.830]WTGPPEPIEAAVAR.F | HexNAc(4)Hex(5)Fuc(1)NeuAc(2) | N[+2351] | 1861.1989 | 3 | 5581.5821 | 5581.5413 | 0 | 7.3 | 141 | Specific | 319.3 | 193.49 | 193.49 | 6.33 | 130 | >sp\|P02788\|TRFL_HUMAN Lactotransferrin OS=Homo sapiens OX=9606 GN=LTF PE=1 SV=6 | 1270 | 20201119_10132_1ng_glyco_hcd203040_compound15.23914.23914.3 | controllerType=0 controllerNumber=1 scan=23914 | 49.3537 |
| 17718:3 | 1 | R.[+42.011]TAGWNVPIGTLRPFLN[+1872.651]WTGPPEPIEAAVAR.F | HexNAc(3)Hex(6)NeuAc(1) | .[42], N[+1873] | 1715.8092 | 3 | 5145.4129 | 5145.372 | 0 | 8 | 141 | Specific | 345.32 | 265.14 | 265.14 | 6.83 | 130 | >sp\|P02788\|TRFL_HUMAN Lactotransferrin OS=Homo sapiens OX=9606 GN=LTF PE=1 SV=6 | 1270 | 20201119_10132_1ng_glyco_hcd203040_compound15.21399.21399.3 | controllerType=0 controllerNumber=1 scan=21399 | 44.0011 |
| 17686:3 | 1 | R.TAGWNVPIGTLRPFLN[+2425.887]WTGPPEPIEAAVAR.F | HexNAc(5)Hex(6)Fuc(3) | N[+2426] | 1886.2164 | 3 | 5656.6346 | 5656.5985 | 0 | 6.4 | 141 | Specific | 350.97 | 171.04 | 171.04 | 6.02 | 130 | >sp\|P02788\|TRFL_HUMAN Lactotransferrin OS=Homo sapiens OX=9606 GN=LTF PE=1 SV=6 | 1270 | 20201119_10132_1ng_glyco_hcd203040_compound15.21365.21365.3 | controllerType=0 controllerNumber=1 scan=21365 | 43.9313 |
| 17678:4 | 1 | R.TAGWNVPIGTLRPFLN[+2425.887]WTGPPEPIEAAVAR.F | HexNAc(5)Hex(6)Fuc(3) | N[+2426] | 1414.9152 | 4 | 5656.6388 | 5656.5985 | 0 | 7.1 | 141 | Specific | 374.35 | 223.03 | 223.03 | 6.76 | 130 | >sp\|P02788\|TRFL_HUMAN Lactotransferrin OS=Homo sapiens OX=9606 GN=LTF PE=1 SV=6 | 1270 | 20201119_10132_1ng_glyco_hcd203040_compound15.21357.21357.4 | controllerType=0 controllerNumber=1 scan=21357 | 43.9165 |
| 20198:4 | 1 | R.TAGWNVPIGTLRPFLN[+2350.830]WTGPPEPIEAAVAR.F | HexNAc(4)Hex(5)Fuc(1)NeuAc(2) | N[+2351] | 1396.1449 | 4 | 5581.5578 | 5581.5413 | 0 | 3 | 141 | Specific | 380.15 | 280.98 | 280.98 | 7.82 | 130 | >sp\|P02788\|TRFL_HUMAN Lactotransferrin OS=Homo sapiens OX=9606 GN=LTF PE=1 SV=6 | 1270 | 20201119_10132_1ng_glyco_hcd203040_compound15.23995.23995.4 | controllerType=0 controllerNumber=1 scan=23995 | 49.5417 |
| 18915:3 | 1 | R.TAGWNVPIGTLRPFLN[+2059.735]WTGPPEPIEAAVAR.F | HexNAc(4)Hex(5)Fuc(1)NeuAc(1) | N[+2060] | 1764.1594 | 3 | 5290.4636 | 5290.4459 | 0 | 3.4 | 141 | Specific | 454.06 | 328.05 | 328.05 | 8.37 | 130 | >sp\|P02788\|TRFL_HUMAN Lactotransferrin OS=Homo sapiens OX=9606 GN=LTF PE=1 SV=6 | 1270 | 20201119_10132_1ng_glyco_hcd203040_compound15.22655.22655.3 | controllerType=0 controllerNumber=1 scan=22655 | 46.7216 |
| 17708:3 | 1 | R.TAGWNVPIGTLRPFLN[+2279.830]WTGPPEPIEAAVAR.F | HexNAc(5)Hex(6)Fuc(2) | N[+2280] | 1837.5258 | 3 | 5510.563 | 5510.5406 | 0 | 4.1 | 141 | Specific | 457.3 | 369.59 | 369.59 | 8.84 | 130 | >sp\|P02788\|TRFL_HUMAN Lactotransferrin OS=Homo sapiens OX=9606 GN=LTF PE=1 SV=6 | 1270 | 20201119_10132_1ng_glyco_hcd203040_compound15.21388.21388.3 | controllerType=0 controllerNumber=1 scan=21388 | 43.9746 |
| 17706:3 | 1 | R.TAGWNVPIGTLRPFLN[+2279.830]WTGPPEPIEAAVAR.F | HexNAc(5)Hex(6)Fuc(2) | N[+2280] | 1837.5258 | 3 | 5510.563 | 5510.5406 | 0 | 4.1 | 141 | Specific | 467.43 | 396.79 | 396.79 | 10.21 | 130 | >sp\|P02788\|TRFL_HUMAN Lactotransferrin OS=Homo sapiens OX=9606 GN=LTF PE=1 SV=6 | 1270 | 20201119_10132_1ng_glyco_hcd203040_compound15.21386.21386.3 | controllerType=0 controllerNumber=1 scan=21386 | 43.9705 |
| 20200:4 | 1 | R.TAGWNVPIGTLRPFLN[+2350.830]WTGPPEPIEAAVAR.F | HexNAc(4)Hex(5)Fuc(1)NeuAc(2) | N[+2351] | 1396.1449 | 4 | 5581.5578 | 5581.5413 | 0 | 3 | 141 | Specific | 507.15 | 305.6 | 305.6 | 8.24 | 130 | >sp\|P02788\|TRFL_HUMAN Lactotransferrin OS=Homo sapiens OX=9606 GN=LTF PE=1 SV=6 | 1270 | 20201119_10132_1ng_glyco_hcd203040_compound15.23997.23997.4 | controllerType=0 controllerNumber=1 scan=23997 | 49.5455 |
| 18913:3 | 1 | R.TAGWNVPIGTLRPFLN[+2059.735]WTGPPEPIEAAVAR.F | HexNAc(4)Hex(5)Fuc(1)NeuAc(1) | N[+2060] | 1764.1594 | 3 | 5290.4636 | 5290.4459 | 0 | 3.4 | 141 | Specific | 521.42 | 471.99 | 471.99 | 10.27 | 130 | >sp\|P02788\|TRFL_HUMAN Lactotransferrin OS=Homo sapiens OX=9606 GN=LTF PE=1 SV=6 | 1270 | 20201119_10132_1ng_glyco_hcd203040_compound15.22653.22653.3 | controllerType=0 controllerNumber=1 scan=22653 | 46.7084 |
| 17628:4 | 1 | R.TAGWNVPIGTLRPFLN[+2791.020]WTGPPEPIEAAVAR.F | HexNAc(6)Hex(7)Fuc(3) | N[+2791] | 1506.1939 | 4 | 6021.7536 | 6021.7307 | 0 | 3.8 | 141 | Specific | 537.36 | 500.4 | 500.4 | 10.3 | 130 | >sp\|P02788\|TRFL_HUMAN Lactotransferrin OS=Homo sapiens OX=9606 GN=LTF PE=1 SV=6 | 1270 | 20201119_10132_1ng_glyco_hcd203040_compound15.21304.21304.4 | controllerType=0 controllerNumber=1 scan=21304 | 43.7836 |
| 17690:4 | 1 | R.TAGWNVPIGTLRPFLN[+2279.830]WTGPPEPIEAAVAR.F | HexNAc(5)Hex(6)Fuc(2) | N[+2280] | 1378.3989 | 4 | 5510.5739 | 5510.5406 | 0 | 6 | 141 | Specific | 574.91 | 528.8 | 528.8 | 11.07 | 130 | >sp\|P02788\|TRFL_HUMAN Lactotransferrin OS=Homo sapiens OX=9606 GN=LTF PE=1 SV=6 | 1270 | 20201119_10132_1ng_glyco_hcd203040_compound15.21369.21369.4 | controllerType=0 controllerNumber=1 scan=21369 | 43.9375 |
| 20065:3 | 1 | R.TAGWNVPIGTLRPFLN[+2350.830]WTGPPEPIEAAVAR.F | HexNAc(4)Hex(5)Fuc(1)NeuAc(2) | N[+2351] | 1861.1949 | 3 | 5581.5702 | 5581.5413 | 0 | 5.2 | 141 | Specific | 578.61 | 420.91 | 420.91 | 11.15 | 130 | >sp\|P02788\|TRFL_HUMAN Lactotransferrin OS=Homo sapiens OX=9606 GN=LTF PE=1 SV=6 | 1270 | 20201119_10132_1ng_glyco_hcd203040_compound15.23856.23856.3 | controllerType=0 controllerNumber=1 scan=23856 | 49.2344 |
| 18672:3 | 1 | R.TAGWNVPIGTLRPFLN[+2205.793]WTGPPEPIEAAVAR.F | HexNAc(4)Hex(5)Fuc(2)NeuAc(1) | N[+2206] | 1812.8482 | 3 | 5436.5301 | 5436.5038 | 0 | 4.8 | 141 | Specific | 585.84 | 496.5 | 496.5 | 11.73 | 130 | >sp\|P02788\|TRFL_HUMAN Lactotransferrin OS=Homo sapiens OX=9606 GN=LTF PE=1 SV=6 | 1270 | 20201119_10132_1ng_glyco_hcd203040_compound15.22397.22397.3 | controllerType=0 controllerNumber=1 scan=22397 | 46.224 |
| 17681:4 | 1 | R.TAGWNVPIGTLRPFLN[+2425.887]WTGPPEPIEAAVAR.F | HexNAc(5)Hex(6)Fuc(3) | N[+2426] | 1414.9152 | 4 | 5656.6388 | 5656.5985 | 0 | 7.1 | 141 | Specific | 594.09 | 360.01 | 360.01 | 9.46 | 130 | >sp\|P02788\|TRFL_HUMAN Lactotransferrin OS=Homo sapiens OX=9606 GN=LTF PE=1 SV=6 | 1270 | 20201119_10132_1ng_glyco_hcd203040_compound15.21360.21360.4 | controllerType=0 controllerNumber=1 scan=21360 | 43.9224 |
| 18711:3 | 1 | R.TAGWNVPIGTLRPFLN[+2059.735]WTGPPEPIEAAVAR.F | HexNAc(4)Hex(5)Fuc(1)NeuAc(1) | N[+2060] | 1764.1578 | 3 | 5290.4588 | 5290.4459 | 0 | 2.5 | 141 | Specific | 613.87 | 573.01 | 573.01 | 12.62 | 130 | >sp\|P02788\|TRFL_HUMAN Lactotransferrin OS=Homo sapiens OX=9606 GN=LTF PE=1 SV=6 | 1270 | 20201119_10132_1ng_glyco_hcd203040_compound15.22438.22438.2 | controllerType=0 controllerNumber=1 scan=22438 | 46.3407 |
| 18669:4 | 1 | R.TAGWNVPIGTLRPFLN[+2205.793]WTGPPEPIEAAVAR.F | HexNAc(4)Hex(5)Fuc(2)NeuAc(1) | N[+2206] | 1359.8889 | 4 | 5436.5338 | 5436.5038 | 0 | 5.5 | 141 | Specific | 652.52 | 592.67 | 592.67 | 13.07 | 130 | >sp\|P02788\|TRFL_HUMAN Lactotransferrin OS=Homo sapiens OX=9606 GN=LTF PE=1 SV=6 | 1270 | 20201119_10132_1ng_glyco_hcd203040_compound15.22394.22394.4 | controllerType=0 controllerNumber=1 scan=22394 | 46.2116 |
| 17767:4 | 1 | R.TAGWNVPIGTLRPFLN[+1768.640]WTGPPEPIEAAVAR.F | HexNAc(4)Hex(5)Fuc(1) | N[+1769] | 1250.6003 | 4 | 4999.3796 | 4999.3504 | 0 | 5.8 | 141 | Specific | 669.52 | 653.46 | 653.46 | 13.07 | 130 | >sp\|P02788\|TRFL_HUMAN Lactotransferrin OS=Homo sapiens OX=9606 GN=LTF PE=1 SV=6 | 1270 | 20201119_10132_1ng_glyco_hcd203040_compound15.21450.21450.4 | controllerType=0 controllerNumber=1 scan=21450 | 44.1104 |
| 18670:3 | 1 | R.TAGWNVPIGTLRPFLN[+2205.793]WTGPPEPIEAAVAR.F | HexNAc(4)Hex(5)Fuc(2)NeuAc(1) | N[+2206] | 1812.8482 | 3 | 5436.5301 | 5436.5038 | 0 | 4.8 | 141 | Specific | 674.76 | 590.65 | 590.65 | 13.73 | 130 | >sp\|P02788\|TRFL_HUMAN Lactotransferrin OS=Homo sapiens OX=9606 GN=LTF PE=1 SV=6 | 1270 | 20201119_10132_1ng_glyco_hcd203040_compound15.22395.22395.3 | controllerType=0 controllerNumber=1 scan=22395 | 46.2129 |
| 17747:3 | 1 | R.TAGWNVPIGTLRPFLN[+1768.640]WTGPPEPIEAAVAR.F | HexNAc(4)Hex(5)Fuc(1) | N[+1769] | 1667.1274 | 3 | 4999.3677 | 4999.3504 | 0 | 3.4 | 141 | Specific | 697.37 | 578.51 | 578.51 | 13.62 | 130 | >sp\|P02788\|TRFL_HUMAN Lactotransferrin OS=Homo sapiens OX=9606 GN=LTF PE=1 SV=6 | 1270 | 20201119_10132_1ng_glyco_hcd203040_compound15.21429.21429.3 | controllerType=0 controllerNumber=1 scan=21429 | 44.063 |
| 18708:3 | 1 | R.TAGWNVPIGTLRPFLN[+2059.735]WTGPPEPIEAAVAR.F | HexNAc(4)Hex(5)Fuc(1)NeuAc(1) | N[+2060] | 1764.1578 | 3 | 5290.4588 | 5290.4459 | 0 | 2.5 | 141 | Specific | 697.96 | 653.91 | 653.91 | 13.62 | 130 | >sp\|P02788\|TRFL_HUMAN Lactotransferrin OS=Homo sapiens OX=9606 GN=LTF PE=1 SV=6 | 1270 | 20201119_10132_1ng_glyco_hcd203040_compound15.22435.22435.2 | controllerType=0 controllerNumber=1 scan=22435 | 46.3357 |
| 17692:4 | 1 | R.TAGWNVPIGTLRPFLN[+2279.830]WTGPPEPIEAAVAR.F | HexNAc(5)Hex(6)Fuc(2) | N[+2280] | 1378.3989 | 4 | 5510.5739 | 5510.5406 | 0 | 6 | 141 | Specific | 704.74 | 617.86 | 617.86 | 14.07 | 130 | >sp\|P02788\|TRFL_HUMAN Lactotransferrin OS=Homo sapiens OX=9606 GN=LTF PE=1 SV=6 | 1270 | 20201119_10132_1ng_glyco_hcd203040_compound15.21371.21371.4 | controllerType=0 controllerNumber=1 scan=21371 | 43.9422 |
| 17745:3 | 1 | R.TAGWNVPIGTLRPFLN[+1768.640]WTGPPEPIEAAVAR.F | HexNAc(4)Hex(5)Fuc(1) | N[+1769] | 1667.1274 | 3 | 4999.3677 | 4999.3504 | 0 | 3.4 | 141 | Specific | 748.91 | 650.59 | 650.59 | 14.62 | 130 | >sp\|P02788\|TRFL_HUMAN Lactotransferrin OS=Homo sapiens OX=9606 GN=LTF PE=1 SV=6 | 1270 | 20201119_10132_1ng_glyco_hcd203040_compound15.21427.21427.3 | controllerType=0 controllerNumber=1 scan=21427 | 44.0566 |
| 18749:3 | 1 | R.TAGWNVPIGTLRPFLN[+2059.735]WTGPPEPIEAAVAR.F | HexNAc(4)Hex(5)Fuc(1)NeuAc(1) | N[+2060] | 1764.1587 | 3 | 5290.4614 | 5290.4459 | 0 | 2.9 | 141 | Specific | 753.72 | 705.65 | 705.65 | 15.62 | 130 | >sp\|P02788\|TRFL_HUMAN Lactotransferrin OS=Homo sapiens OX=9606 GN=LTF PE=1 SV=6 | 1270 | 20201119_10132_1ng_glyco_hcd203040_compound15.22477.22477.2 | controllerType=0 controllerNumber=1 scan=22477 | 46.3986 |
| 18651:4 | 1 | R.TAGWNVPIGTLRPFLN[+2569.905]WTGPPEPIEAAVAR.F | HexNAc(5)Hex(6)NeuAc(2) | N[+2570] | 1450.923 | 4 | 5800.6701 | 5800.6156 | 0 | 9.4 | 141 | Specific | 781.87 | 650.2 | 650.2 | 14.28 | 130 | >sp\|P02788\|TRFL_HUMAN Lactotransferrin OS=Homo sapiens OX=9606 GN=LTF PE=1 SV=6 | 1270 | 20201119_10132_1ng_glyco_hcd203040_compound15.22375.22375.4 | controllerType=0 controllerNumber=1 scan=22375 | 46.1633 |
| 20050:5 | 1 | R.TAGWNVPIGTLRPFLN[+2350.830]WTGPPEPIEAAVAR.F | HexNAc(4)Hex(5)Fuc(1)NeuAc(2) | N[+2351] | 1117.1185 | 5 | 5581.5636 | 5581.5413 | 0 | 4 | 141 | Specific | 798.06 | 772.93 | 772.93 | 15.8 | 130 | >sp\|P02788\|TRFL_HUMAN Lactotransferrin OS=Homo sapiens OX=9606 GN=LTF PE=1 SV=6 | 1270 | 20201119_10132_1ng_glyco_hcd203040_compound15.23840.23840.5 | controllerType=0 controllerNumber=1 scan=23840 | 49.191 |
| 18671:4 | 1 | R.TAGWNVPIGTLRPFLN[+2205.793]WTGPPEPIEAAVAR.F | HexNAc(4)Hex(5)Fuc(2)NeuAc(1) | N[+2206] | 1359.8889 | 4 | 5436.5338 | 5436.5038 | 0 | 5.5 | 141 | Specific | 806.87 | 680.61 | 680.61 | 16.07 | 130 | >sp\|P02788\|TRFL_HUMAN Lactotransferrin OS=Homo sapiens OX=9606 GN=LTF PE=1 SV=6 | 1270 | 20201119_10132_1ng_glyco_hcd203040_compound15.22396.22396.4 | controllerType=0 controllerNumber=1 scan=22396 | 46.2197 |
| 20052:5 | 1 | R.TAGWNVPIGTLRPFLN[+2350.830]WTGPPEPIEAAVAR.F | HexNAc(4)Hex(5)Fuc(1)NeuAc(2) | N[+2351] | 1117.1185 | 5 | 5581.5636 | 5581.5413 | 0 | 4 | 141 | Specific | 846.78 | 846.78 | 846.78 | 16.8 | 130 | >sp\|P02788\|TRFL_HUMAN Lactotransferrin OS=Homo sapiens OX=9606 GN=LTF PE=1 SV=6 | 1270 | 20201119_10132_1ng_glyco_hcd203040_compound15.23842.23842.5 | controllerType=0 controllerNumber=1 scan=23842 | 49.197 |
| 17769:4 | 1 | R.TAGWNVPIGTLRPFLN[+1768.640]WTGPPEPIEAAVAR.F | HexNAc(4)Hex(5)Fuc(1) | N[+1769] | 1250.6003 | 4 | 4999.3796 | 4999.3504 | 0 | 5.8 | 141 | Specific | 862.2 | 757.71 | 757.71 | 17.07 | 130 | >sp\|P02788\|TRFL_HUMAN Lactotransferrin OS=Homo sapiens OX=9606 GN=LTF PE=1 SV=6 | 1270 | 20201119_10132_1ng_glyco_hcd203040_compound15.21452.21452.4 | controllerType=0 controllerNumber=1 scan=21452 | 44.1155 |
| 16517:3 | 1 | R.TAGWNIPMGLLFN[+1460.529]QTGSC[+57.021]K.F | HexNAc(4)Hex(4) | C[+57], N[+1461] | 1185.8547 | 3 | 3555.5495 | 3555.5387 | 0 | 3 | 485 | Specific | 276.93 | 276.93 | 182.05 | 6.92 | 130 | >sp\|P02788\|TRFL_HUMAN Lactotransferrin OS=Homo sapiens OX=9606 GN=LTF PE=1 SV=6 | 1270 | 20201119_10132_1ng_glyco_hcd203040_compound15.20145.20145.3 | controllerType=0 controllerNumber=1 scan=20145 | 41.35 |
| 16447:3 | 1 | R.TAGWNIPMGLLFN[+1581.555]QTGSC[+57.021]K.F | HexNAc(3)Hex(6) | C[+57], N[+1582] | 1226.1979 | 3 | 3676.5792 | 3676.565 | 0 | 3.9 | 485 | Specific | 312.42 | 312.42 | 204.41 | 7.19 | 130 | >sp\|P02788\|TRFL_HUMAN Lactotransferrin OS=Homo sapiens OX=9606 GN=LTF PE=1 SV=6 | 1270 | 20201119_10132_1ng_glyco_hcd203040_compound15.20072.20072.3 | controllerType=0 controllerNumber=1 scan=20072 | 41.2042 |
| 15904:3 | 1 | R.TAGWNIPMGLLFN[+2279.830]QTGSC[+57.021]K.F | HexNAc(5)Hex(6)Fuc(2) | C[+57], N[+2280] | 1458.9514 | 3 | 4374.8395 | 4374.8396 | 0 | 0 | 485 | Specific | 334.16 | 300.61 | 227.56 | 6.87 | 130 | >sp\|P02788\|TRFL_HUMAN Lactotransferrin OS=Homo sapiens OX=9606 GN=LTF PE=1 SV=6 | 1270 | 20201119_10132_1ng_glyco_hcd203040_compound15.19507.19507.3 | controllerType=0 controllerNumber=1 scan=19507 | 40.1236 |
| 16602:3 | 1 | R.TAGWNIPMGLLFN[+1257.449]QTGSC[+57.021]K.F | HexNAc(3)Hex(4) | C[+57], N[+1257] | 1118.1649 | 3 | 3352.4801 | 3352.4594 | 0 | 6.2 | 485 | Specific | 416.19 | 416.19 | 212.36 | 9.06 | 130 | >sp\|P02788\|TRFL_HUMAN Lactotransferrin OS=Homo sapiens OX=9606 GN=LTF PE=1 SV=6 | 1270 | 20201119_10132_1ng_glyco_hcd203040_compound15.20234.20234.3 | controllerType=0 controllerNumber=1 scan=20234 | 41.5405 |
| 16438:3 | 1 | R.TAGWNIPMGLLFN[+1622.582]QTGSC[+57.021]K.F | HexNAc(4)Hex(5) | C[+57], N[+1623] | 1239.8737 | 3 | 3717.6066 | 3717.5916 | 0 | 4 | 485 | Specific | 433.94 | 365.74 | 278.68 | 8.29 | 130 | >sp\|P02788\|TRFL_HUMAN Lactotransferrin OS=Homo sapiens OX=9606 GN=LTF PE=1 SV=6 | 1270 | 20201119_10132_1ng_glyco_hcd203040_compound15.20063.20063.5 | controllerType=0 controllerNumber=1 scan=20063 | 41.1895 |
| 16519:3 | 1 | R.TAGWNIPMGLLFN[+1460.529]QTGSC[+57.021]K.F | HexNAc(4)Hex(4) | C[+57], N[+1461] | 1185.8547 | 3 | 3555.5495 | 3555.5387 | 0 | 3 | 485 | Specific | 511.38 | 511.38 | 315.48 | 9.8 | 130 | >sp\|P02788\|TRFL_HUMAN Lactotransferrin OS=Homo sapiens OX=9606 GN=LTF PE=1 SV=6 | 1270 | 20201119_10132_1ng_glyco_hcd203040_compound15.20147.20147.3 | controllerType=0 controllerNumber=1 scan=20147 | 41.3552 |
| 16182:2 | 1 | R.TAGWNIPMGLLFN[+1768.640]QTGSC[+57.021]K.F | HexNAc(4)Hex(5)Fuc(1) | C[+57], N[+1769] | 1932.3453 | 2 | 3863.6833 | 3863.6495 | 0 | 8.8 | 485 | Specific | 600.44 | 575.86 | 575.86 | 11.47 | 130 | >sp\|P02788\|TRFL_HUMAN Lactotransferrin OS=Homo sapiens OX=9606 GN=LTF PE=1 SV=6 | 1270 | 20201119_10132_1ng_glyco_hcd203040_compound15.19797.19797.2 | controllerType=0 controllerNumber=1 scan=19797 | 40.6677 |
| 16449:3 | 1 | R.TAGWNIPMGLLFN[+1581.555]QTGSC[+57.021]K.F | HexNAc(3)Hex(6) | C[+57], N[+1582] | 1226.1979 | 3 | 3676.5792 | 3676.565 | 0 | 3.9 | 485 | Specific | 611.49 | 601.6 | 353.66 | 12.41 | 130 | >sp\|P02788\|TRFL_HUMAN Lactotransferrin OS=Homo sapiens OX=9606 GN=LTF PE=1 SV=6 | 1270 | 20201119_10132_1ng_glyco_hcd203040_compound15.20074.20074.3 | controllerType=0 controllerNumber=1 scan=20074 | 41.2087 |
| 16604:3 | 1 | R.TAGWNIPMGLLFN[+1257.449]QTGSC[+57.021]K.F | HexNAc(3)Hex(4) | C[+57], N[+1257] | 1118.1649 | 3 | 3352.4801 | 3352.4594 | 0 | 6.2 | 485 | Specific | 660.56 | 533.08 | 406.29 | 12.83 | 130 | >sp\|P02788\|TRFL_HUMAN Lactotransferrin OS=Homo sapiens OX=9606 GN=LTF PE=1 SV=6 | 1270 | 20201119_10132_1ng_glyco_hcd203040_compound15.20236.20236.3 | controllerType=0 controllerNumber=1 scan=20236 | 41.5461 |
| 16185:2 | 1 | R.TAGWNIPMGLLFN[+1768.640]QTGSC[+57.021]K.F | HexNAc(4)Hex(5)Fuc(1) | C[+57], N[+1769] | 1932.3453 | 2 | 3863.6833 | 3863.6495 | 0 | 8.8 | 485 | Specific | 737.6 | 659.81 | 659.81 | 13.47 | 130 | >sp\|P02788\|TRFL_HUMAN Lactotransferrin OS=Homo sapiens OX=9606 GN=LTF PE=1 SV=6 | 1270 | 20201119_10132_1ng_glyco_hcd203040_compound15.19800.19800.2 | controllerType=0 controllerNumber=1 scan=19800 | 40.6723 |
| 16346:3 | 1 | R.TAGWNIPMGLLFN[+1622.582]QTGSC[+57.021]K.F | HexNAc(4)Hex(5) | C[+57], N[+1623] | 1239.872 | 3 | 3717.6015 | 3717.5916 | 0 | 2.7 | 485 | Specific | 749.75 | 749.75 | 297.08 | 14.3 | 130 | >sp\|P02788\|TRFL_HUMAN Lactotransferrin OS=Homo sapiens OX=9606 GN=LTF PE=1 SV=6 | 1270 | 20201119_10132_1ng_glyco_hcd203040_compound15.19968.19968.3 | controllerType=0 controllerNumber=1 scan=19968 | 41.0123 |
| 16440:3 | 1 | R.TAGWNIPMGLLFN[+1622.582]QTGSC[+57.021]K.F | HexNAc(4)Hex(5) | C[+57], N[+1623] | 1239.8737 | 3 | 3717.6066 | 3717.5916 | 0 | 4 | 485 | Specific | 828.12 | 803.53 | 390.66 | 16.41 | 130 | >sp\|P02788\|TRFL_HUMAN Lactotransferrin OS=Homo sapiens OX=9606 GN=LTF PE=1 SV=6 | 1270 | 20201119_10132_1ng_glyco_hcd203040_compound15.20065.20065.5 | controllerType=0 controllerNumber=1 scan=20065 | 41.1927 |
| 18274:3 | 1 | R.TAGWNIPMGLLFN[+1913.677]QTGSC[+57.021]K.F | HexNAc(4)Hex(5)NeuAc(1) | C[+57], N[+1914] | 1336.9053 | 3 | 4008.7014 | 4008.687 | 0 | 3.6 | 485 | Specific | 875.36 | 841.04 | 841.04 | 17.41 | 130 | >sp\|P02788\|TRFL_HUMAN Lactotransferrin OS=Homo sapiens OX=9606 GN=LTF PE=1 SV=6 | 1270 | 20201119_10132_1ng_glyco_hcd203040_compound15.21980.21980.3 | controllerType=0 controllerNumber=1 scan=21980 | 45.2465 |
| 16349:3 | 1 | R.TAGWNIPMGLLFN[+1622.582]QTGSC[+57.021]K.F | HexNAc(4)Hex(5) | C[+57], N[+1623] | 1239.872 | 3 | 3717.6015 | 3717.5916 | 0 | 2.7 | 485 | Specific | 878.06 | 870.9 | 424.88 | 17.3 | 130 | >sp\|P02788\|TRFL_HUMAN Lactotransferrin OS=Homo sapiens OX=9606 GN=LTF PE=1 SV=6 | 1270 | 20201119_10132_1ng_glyco_hcd203040_compound15.19971.19971.3 | controllerType=0 controllerNumber=1 scan=19971 | 41.0166 |
| 16280:3 | 1 | R.TAGWNIPMGLLFN[+1606.587]QTGSC[+57.021]K.F | HexNAc(4)Hex(4)Fuc(1) | C[+57], N[+1607] | 1234.5385 | 3 | 3701.601 | 3701.5966 | 0 | 1.2 | 485 | Specific | 889.09 | 757.05 | 757.05 | 17.09 | 130 | >sp\|P02788\|TRFL_HUMAN Lactotransferrin OS=Homo sapiens OX=9606 GN=LTF PE=1 SV=6 | 1270 | 20201119_10132_1ng_glyco_hcd203040_compound15.19899.19899.3 | controllerType=0 controllerNumber=1 scan=19899 | 40.8869 |
| 18276:3 | 1 | R.TAGWNIPMGLLFN[+1913.677]QTGSC[+57.021]K.F | HexNAc(4)Hex(5)NeuAc(1) | C[+57], N[+1914] | 1336.9053 | 3 | 4008.7014 | 4008.687 | 0 | 3.6 | 485 | Specific | 894.06 | 894.06 | 894.06 | 17.41 | 130 | >sp\|P02788\|TRFL_HUMAN Lactotransferrin OS=Homo sapiens OX=9606 GN=LTF PE=1 SV=6 | 1270 | 20201119_10132_1ng_glyco_hcd203040_compound15.21982.21982.3 | controllerType=0 controllerNumber=1 scan=21982 | 45.2514 |
| 16154:3 | 1 | R.TAGWNIPMGLLFN[+1768.640]QTGSC[+57.021]K.F | HexNAc(4)Hex(5)Fuc(1) | C[+57], N[+1769] | 1288.5562 | 3 | 3863.6541 | 3863.6495 | 0 | 1.2 | 485 | Specific | 901.7 | 901.7 | 901.7 | 18.09 | 130 | >sp\|P02788\|TRFL_HUMAN Lactotransferrin OS=Homo sapiens OX=9606 GN=LTF PE=1 SV=6 | 1270 | 20201119_10132_1ng_glyco_hcd203040_compound15.19768.19768.3 | controllerType=0 controllerNumber=1 scan=19768 | 40.5975 |
| 16567:3 | 1 | R.TAGWNIPMGLLFN[+1216.423]QTGSC[+57.021]K.F | HexNAc(2)Hex(5) | C[+57], N[+1216] | 1104.4845 | 3 | 3311.439 | 3311.4328 | 0 | 1.9 | 485 | Specific | 923 | 893.2 | 376.28 | 18.09 | 130 | >sp\|P02788\|TRFL_HUMAN Lactotransferrin OS=Homo sapiens OX=9606 GN=LTF PE=1 SV=6 | 1270 | 20201119_10132_1ng_glyco_hcd203040_compound15.20197.20197.3 | controllerType=0 controllerNumber=1 scan=20197 | 41.4575 |
| 18122:3 | 1 | R.TAGWNIPMGLLFN[+2059.735]QTGSC[+57.021]K.F | HexNAc(4)Hex(5)Fuc(1)NeuAc(1) | C[+57], N[+2060] | 1385.5866 | 3 | 4154.7453 | 4154.7449 | 0 | 0.1 | 485 | Specific | 924.5 | 774.53 | 774.53 | 17.79 | 130 | >sp\|P02788\|TRFL_HUMAN Lactotransferrin OS=Homo sapiens OX=9606 GN=LTF PE=1 SV=6 | 1270 | 20201119_10132_1ng_glyco_hcd203040_compound15.21821.21821.3 | controllerType=0 controllerNumber=1 scan=21821 | 44.8887 |
| 16278:3 | 1 | R.TAGWNIPMGLLFN[+1606.587]QTGSC[+57.021]K.F | HexNAc(4)Hex(4)Fuc(1) | C[+57], N[+1607] | 1234.5385 | 3 | 3701.601 | 3701.5966 | 0 | 1.2 | 485 | Specific | 929.66 | 836.07 | 836.07 | 18.09 | 130 | >sp\|P02788\|TRFL_HUMAN Lactotransferrin OS=Homo sapiens OX=9606 GN=LTF PE=1 SV=6 | 1270 | 20201119_10132_1ng_glyco_hcd203040_compound15.19897.19897.3 | controllerType=0 controllerNumber=1 scan=19897 | 40.8819 |
| 16119:3 | 1 | R.TAGWNIPMGLLFN[+1768.640]QTGSC[+57.021]K.F | HexNAc(4)Hex(5)Fuc(1) | C[+57], N[+1769] | 1288.5606 | 3 | 3863.6673 | 3863.6495 | 0 | 4.6 | 485 | Specific | 968.47 | 945 | 945 | 19.41 | 130 | >sp\|P02788\|TRFL_HUMAN Lactotransferrin OS=Homo sapiens OX=9606 GN=LTF PE=1 SV=6 | 1270 | 20201119_10132_1ng_glyco_hcd203040_compound15.19731.19731.3 | controllerType=0 controllerNumber=1 scan=19731 | 40.5221 |
| 16117:3 | 1 | R.TAGWNIPMGLLFN[+1768.640]QTGSC[+57.021]K.F | HexNAc(4)Hex(5)Fuc(1) | C[+57], N[+1769] | 1288.5606 | 3 | 3863.6673 | 3863.6495 | 0 | 4.6 | 485 | Specific | 971.71 | 971.71 | 971.71 | 19.41 | 130 | >sp\|P02788\|TRFL_HUMAN Lactotransferrin OS=Homo sapiens OX=9606 GN=LTF PE=1 SV=6 | 1270 | 20201119_10132_1ng_glyco_hcd203040_compound15.19729.19729.3 | controllerType=0 controllerNumber=1 scan=19729 | 40.5168 |
| 18120:3 | 1 | R.TAGWNIPMGLLFN[+2059.735]QTGSC[+57.021]K.F | HexNAc(4)Hex(5)Fuc(1)NeuAc(1) | C[+57], N[+2060] | 1385.5866 | 3 | 4154.7453 | 4154.7449 | 0 | 0.1 | 485 | Specific | 989.45 | 882.19 | 882.19 | 18.79 | 130 | >sp\|P02788\|TRFL_HUMAN Lactotransferrin OS=Homo sapiens OX=9606 GN=LTF PE=1 SV=6 | 1270 | 20201119_10132_1ng_glyco_hcd203040_compound15.21819.21819.3 | controllerType=0 controllerNumber=1 scan=21819 | 44.8775 |
| 16152:3 | 1 | R.TAGWNIPMGLLFN[+1768.640]QTGSC[+57.021]K.F | HexNAc(4)Hex(5)Fuc(1) | C[+57], N[+1769] | 1288.5562 | 3 | 3863.6541 | 3863.6495 | 0 | 1.2 | 485 | Specific | 995.5 | 995.5 | 995.5 | 19.09 | 130 | >sp\|P02788\|TRFL_HUMAN Lactotransferrin OS=Homo sapiens OX=9606 GN=LTF PE=1 SV=6 | 1270 | 20201119_10132_1ng_glyco_hcd203040_compound15.19766.19766.3 | controllerType=0 controllerNumber=1 scan=19766 | 40.5919 |
| 16569:3 | 1 | R.TAGWNIPMGLLFN[+1216.423]QTGSC[+57.021]K.F | HexNAc(2)Hex(5) | C[+57], N[+1216] | 1104.4845 | 3 | 3311.439 | 3311.4328 | 0 | 1.9 | 485 | Specific | 1041.1 | 926.83 | 484.94 | 20.09 | 130 | >sp\|P02788\|TRFL_HUMAN Lactotransferrin OS=Homo sapiens OX=9606 GN=LTF PE=1 SV=6 | 1270 | 20201119_10132_1ng_glyco_hcd203040_compound15.20199.20199.3 | controllerType=0 controllerNumber=1 scan=20199 | 41.4626 |
| 13413:3 | 1 | R.TAGWNIPM[+15.995]GLLFN[+1768.640]QTGSC[+57.021]K.F | HexNAc(4)Hex(5)Fuc(1) | C[+57], M[+16], N[+1769] | 1293.8915 | 3 | 3879.6601 | 3879.6444 | 0 | 4 | 485 | Specific | 365.01 | 365.01 | 365.01 | 8.02 | 130 | >sp\|P02788\|TRFL_HUMAN Lactotransferrin OS=Homo sapiens OX=9606 GN=LTF PE=1 SV=6 | 1270 | 20201119_10132_1ng_glyco_hcd203040_compound15.16879.16879.3 | controllerType=0 controllerNumber=1 scan=16879 | 35.4428 |
| 16464:3 | 1 | R.TAGWNIPM[+15.995]GLLFN[+2059.735]QTGSC[+57.021]K.F | HexNAc(4)Hex(5)Fuc(1)NeuAc(1) | C[+57], M[+16], N[+2060] | 1390.9286 | 3 | 4170.7714 | 4170.7398 | 0 | 7.6 | 485 | Specific | 460.78 | 416.1 | 416.1 | 9.06 | 130 | >sp\|P02788\|TRFL_HUMAN Lactotransferrin OS=Homo sapiens OX=9606 GN=LTF PE=1 SV=6 | 1270 | 20201119_10132_1ng_glyco_hcd203040_compound15.20090.20090.3 | controllerType=0 controllerNumber=1 scan=20090 | 41.2423 |
| 16406:3 | 1 | R.TAGWNIPM[+15.995]GLLFN[+2059.735]QTGSC[+57.021]K.F | HexNAc(4)Hex(5)Fuc(1)NeuAc(1) | C[+57], M[+16], N[+2060] | 1390.9189 | 3 | 4170.7421 | 4170.7398 | 0 | 0.6 | 485 | Specific | 527.78 | 394.48 | 394.48 | 9.44 | 130 | >sp\|P02788\|TRFL_HUMAN Lactotransferrin OS=Homo sapiens OX=9606 GN=LTF PE=1 SV=6 | 1270 | 20201119_10132_1ng_glyco_hcd203040_compound15.20030.20030.3 | controllerType=0 controllerNumber=1 scan=20030 | 41.1276 |
| 16466:3 | 1 | R.TAGWNIPM[+15.995]GLLFN[+2059.735]QTGSC[+57.021]K.F | HexNAc(4)Hex(5)Fuc(1)NeuAc(1) | C[+57], M[+16], N[+2060] | 1390.9286 | 3 | 4170.7714 | 4170.7398 | 0 | 7.6 | 485 | Specific | 677.9 | 510.58 | 510.58 | 12.58 | 130 | >sp\|P02788\|TRFL_HUMAN Lactotransferrin OS=Homo sapiens OX=9606 GN=LTF PE=1 SV=6 | 1270 | 20201119_10132_1ng_glyco_hcd203040_compound15.20092.20092.3 | controllerType=0 controllerNumber=1 scan=20092 | 41.2483 |
| 16408:3 | 1 | R.TAGWNIPM[+15.995]GLLFN[+2059.735]QTGSC[+57.021]K.F | HexNAc(4)Hex(5)Fuc(1)NeuAc(1) | C[+57], M[+16], N[+2060] | 1390.9189 | 3 | 4170.7421 | 4170.7398 | 0 | 0.6 | 485 | Specific | 766.52 | 632.24 | 632.24 | 14.79 | 130 | >sp\|P02788\|TRFL_HUMAN Lactotransferrin OS=Homo sapiens OX=9606 GN=LTF PE=1 SV=6 | 1270 | 20201119_10132_1ng_glyco_hcd203040_compound15.20032.20032.3 | controllerType=0 controllerNumber=1 scan=20032 | 41.1324 |
| 18362:4 | 1 | R.TAGWNIPMGLLFN[+2076.750]QTGSC[+57.021]KFDEYFSQSC[+57.021]APGSDPR.S | HexNAc(4)Hex(6)Fuc(2) | C[+57]*2, N[+2077] | 1504.6409 | 4 | 6015.5417 | 6015.5074 | 0 | 5.7 | 485 | Specific | 379.71 | 369.19 | 369.19 | 7.79 | 130 | >sp\|P02788\|TRFL_HUMAN Lactotransferrin OS=Homo sapiens OX=9606 GN=LTF PE=1 SV=6 | 1270 | 20201119_10132_1ng_glyco_hcd203040_compound15.22072.22072.4 | controllerType=0 controllerNumber=1 scan=22072 | 45.473 |
| 19729:4 | 1 | R.TAGWNIPMGLLFN[+2350.830]QTGSC[+57.021]KFDEYFSQSC[+57.021]APGSDPR.S | HexNAc(4)Hex(5)Fuc(1)NeuAc(2) | C[+57]*2, N[+2351] | 1573.1566 | 4 | 6289.6046 | 6289.5875 | 0 | 2.7 | 485 | Specific | 401.69 | 388.68 | 388.68 | 9.72 | 130 | >sp\|P02788\|TRFL_HUMAN Lactotransferrin OS=Homo sapiens OX=9606 GN=LTF PE=1 SV=6 | 1270 | 20201119_10132_1ng_glyco_hcd203040_compound15.23505.23505.4 | controllerType=0 controllerNumber=1 scan=23505 | 48.4558 |
| 17355:3 | 1 | R.TAGWNIPMGLLFN[+1768.640]QTGSC[+57.021]KFDEYFSQSC[+57.021]APGSDPR.S | HexNAc(4)Hex(5)Fuc(1) | C[+57]*2, N[+1769] | 1903.143 | 3 | 5707.4145 | 5707.3966 | 0 | 3.1 | 485 | Specific | 455.02 | 395.73 | 395.73 | 9.82 | 130 | >sp\|P02788\|TRFL_HUMAN Lactotransferrin OS=Homo sapiens OX=9606 GN=LTF PE=1 SV=6 | 1270 | 20201119_10132_1ng_glyco_hcd203040_compound15.21021.21021.3 | controllerType=0 controllerNumber=1 scan=21021 | 43.225 |
| 18382:4 | 1 | R.TAGWNIPMGLLFN[+2205.793]QTGSC[+57.021]KFDEYFSQSC[+57.021]APGSDPR.S | HexNAc(4)Hex(5)Fuc(2)NeuAc(1) | C[+57]*2, N[+2206] | 1536.8991 | 4 | 6144.5744 | 6144.5499 | 0 | 4 | 485 | Specific | 581.87 | 569.88 | 569.88 | 11.6 | 130 | >sp\|P02788\|TRFL_HUMAN Lactotransferrin OS=Homo sapiens OX=9606 GN=LTF PE=1 SV=6 | 1270 | 20201119_10132_1ng_glyco_hcd203040_compound15.22093.22093.4 | controllerType=0 controllerNumber=1 scan=22093 | 45.5164 |
| 17380:4 | 1 | R.TAGWNIPMGLLFN[+1606.587]QTGSC[+57.021]KFDEYFSQSC[+57.021]APGSDPR.S | HexNAc(4)Hex(4)Fuc(1) | C[+57]*2, N[+1607] | 1387.0957 | 4 | 5545.361 | 5545.3438 | 0 | 3.1 | 485 | Specific | 589.98 | 589.98 | 589.98 | 11.49 | 130 | >sp\|P02788\|TRFL_HUMAN Lactotransferrin OS=Homo sapiens OX=9606 GN=LTF PE=1 SV=6 | 1270 | 20201119_10132_1ng_glyco_hcd203040_compound15.21047.21047.4 | controllerType=0 controllerNumber=1 scan=21047 | 43.2759 |
| 17352:4 | 1 | R.TAGWNIPMGLLFN[+1768.640]QTGSC[+57.021]KFDEYFSQSC[+57.021]APGSDPR.S | HexNAc(4)Hex(5)Fuc(1) | C[+57]*2, N[+1769] | 1427.6101 | 4 | 5707.4186 | 5707.3966 | 0 | 3.9 | 485 | Specific | 651.49 | 597.2 | 597.2 | 13.6 | 130 | >sp\|P02788\|TRFL_HUMAN Lactotransferrin OS=Homo sapiens OX=9606 GN=LTF PE=1 SV=6 | 1270 | 20201119_10132_1ng_glyco_hcd203040_compound15.21018.21018.4 | controllerType=0 controllerNumber=1 scan=21018 | 43.2199 |
| 17399:4 | 1 | R.TAGWNIPMGLLFN[+1622.582]QTGSC[+57.021]KFDEYFSQSC[+57.021]APGSDPR.S | HexNAc(4)Hex(5) | C[+57]*2, N[+1623] | 1391.0988 | 4 | 5561.3732 | 5561.3387 | 0 | 6.2 | 485 | Specific | 768.45 | 768.45 | 221.01 | 15.02 | 130 | >sp\|P02788\|TRFL_HUMAN Lactotransferrin OS=Homo sapiens OX=9606 GN=LTF PE=1 SV=6 | 1270 | 20201119_10132_1ng_glyco_hcd203040_compound15.21067.21067.4 | controllerType=0 controllerNumber=1 scan=21067 | 43.338 |
| 17357:3 | 1 | R.TAGWNIPMGLLFN[+1768.640]QTGSC[+57.021]KFDEYFSQSC[+57.021]APGSDPR.S | HexNAc(4)Hex(5)Fuc(1) | C[+57]*2, N[+1769] | 1903.143 | 3 | 5707.4145 | 5707.3966 | 0 | 3.1 | 485 | Specific | 774.44 | 643 | 643 | 15.57 | 130 | >sp\|P02788\|TRFL_HUMAN Lactotransferrin OS=Homo sapiens OX=9606 GN=LTF PE=1 SV=6 | 1270 | 20201119_10132_1ng_glyco_hcd203040_compound15.21023.21023.3 | controllerType=0 controllerNumber=1 scan=21023 | 43.2281 |
| 17402:4 | 1 | R.TAGWNIPMGLLFN[+1622.582]QTGSC[+57.021]KFDEYFSQSC[+57.021]APGSDPR.S | HexNAc(4)Hex(5) | C[+57]*2, N[+1623] | 1391.0988 | 4 | 5561.3732 | 5561.3387 | 0 | 6.2 | 485 | Specific | 864.03 | 749.7 | 337.45 | 17.02 | 130 | >sp\|P02788\|TRFL_HUMAN Lactotransferrin OS=Homo sapiens OX=9606 GN=LTF PE=1 SV=6 | 1270 | 20201119_10132_1ng_glyco_hcd203040_compound15.21070.21070.4 | controllerType=0 controllerNumber=1 scan=21070 | 43.3427 |
| 19731:4 | 1 | R.TAGWNIPMGLLFN[+2350.830]QTGSC[+57.021]KFDEYFSQSC[+57.021]APGSDPR.S | HexNAc(4)Hex(5)Fuc(1)NeuAc(2) | C[+57]*2, N[+2351] | 1573.1566 | 4 | 6289.6046 | 6289.5875 | 0 | 2.7 | 485 | Specific | 869.42 | 843.48 | 843.48 | 17.49 | 130 | >sp\|P02788\|TRFL_HUMAN Lactotransferrin OS=Homo sapiens OX=9606 GN=LTF PE=1 SV=6 | 1270 | 20201119_10132_1ng_glyco_hcd203040_compound15.23507.23507.4 | controllerType=0 controllerNumber=1 scan=23507 | 48.4591 |
| 18356:4 | 1 | R.TAGWNIPMGLLFN[+2059.735]QTGSC[+57.021]KFDEYFSQSC[+57.021]APGSDPR.S | HexNAc(4)Hex(5)Fuc(1)NeuAc(1) | C[+57]*2, N[+2060] | 1500.3864 | 4 | 5998.5236 | 5998.492 | 0 | 5.3 | 485 | Specific | 890.9 | 860.92 | 860.92 | 17.02 | 130 | >sp\|P02788\|TRFL_HUMAN Lactotransferrin OS=Homo sapiens OX=9606 GN=LTF PE=1 SV=6 | 1270 | 20201119_10132_1ng_glyco_hcd203040_compound15.22066.22066.4 | controllerType=0 controllerNumber=1 scan=22066 | 45.4609 |
| 18358:4 | 1 | R.TAGWNIPMGLLFN[+2059.735]QTGSC[+57.021]KFDEYFSQSC[+57.021]APGSDPR.S | HexNAc(4)Hex(5)Fuc(1)NeuAc(1) | C[+57]*2, N[+2060] | 1500.3864 | 4 | 5998.5236 | 5998.492 | 0 | 5.3 | 485 | Specific | 892.77 | 847.81 | 847.81 | 17.02 | 130 | >sp\|P02788\|TRFL_HUMAN Lactotransferrin OS=Homo sapiens OX=9606 GN=LTF PE=1 SV=6 | 1270 | 20201119_10132_1ng_glyco_hcd203040_compound15.22068.22068.4 | controllerType=0 controllerNumber=1 scan=22068 | 45.4665 |
| 18384:4 | 1 | R.TAGWNIPMGLLFN[+2205.793]QTGSC[+57.021]KFDEYFSQSC[+57.021]APGSDPR.S | HexNAc(4)Hex(5)Fuc(2)NeuAc(1) | C[+57]*2, N[+2206] | 1536.8991 | 4 | 6144.5744 | 6144.5499 | 0 | 4 | 485 | Specific | 916.3 | 863 | 863 | 18.6 | 130 | >sp\|P02788\|TRFL_HUMAN Lactotransferrin OS=Homo sapiens OX=9606 GN=LTF PE=1 SV=6 | 1270 | 20201119_10132_1ng_glyco_hcd203040_compound15.22095.22095.4 | controllerType=0 controllerNumber=1 scan=22095 | 45.5204 |
| 17354:4 | 1 | R.TAGWNIPMGLLFN[+1768.640]QTGSC[+57.021]KFDEYFSQSC[+57.021]APGSDPR.S | HexNAc(4)Hex(5)Fuc(1) | C[+57]*2, N[+1769] | 1427.6101 | 4 | 5707.4186 | 5707.3966 | 0 | 3.9 | 485 | Specific | 974 | 922.33 | 922.33 | 19.6 | 130 | >sp\|P02788\|TRFL_HUMAN Lactotransferrin OS=Homo sapiens OX=9606 GN=LTF PE=1 SV=6 | 1270 | 20201119_10132_1ng_glyco_hcd203040_compound15.21020.21020.4 | controllerType=0 controllerNumber=1 scan=21020 | 43.2239 |
| 08599:3 | 1 | R.N[+2059.735]GSDC[+57.021]PDKFC[+57.021]LFQSETK.N | HexNAc(4)Hex(5)Fuc(1)NeuAc(1) | C[+57]*2, N[+2060] | 1364.8773 | 3 | 4092.6172 | 4092.6088 | 0 | 2 | 642 | Specific | 210.34 | 101.32 | 101.32 | 3.45 | 130 | >sp\|P02788\|TRFL_HUMAN Lactotransferrin OS=Homo sapiens OX=9606 GN=LTF PE=1 SV=6 | 1270 | 20201119_10132_1ng_glyco_hcd203040_compound15.11866.11866.3 | controllerType=0 controllerNumber=1 scan=11866 | 26.7793 |
| 08529:3 | 1 | R.N[+2059.735]GSDC[+57.021]PDKFC[+57.021]LFQSETK.N | HexNAc(4)Hex(5)Fuc(1)NeuAc(1) | C[+57]*2, N[+2060] | 1364.8819 | 3 | 4092.6311 | 4092.6088 | 0 | 5.4 | 642 | Specific | 315.73 | 242.56 | 242.56 | 6.1 | 130 | >sp\|P02788\|TRFL_HUMAN Lactotransferrin OS=Homo sapiens OX=9606 GN=LTF PE=1 SV=6 | 1270 | 20201119_10132_1ng_glyco_hcd203040_compound15.11793.11793.3 | controllerType=0 controllerNumber=1 scan=11793 | 26.6549 |
| 08527:3 | 1 | R.N[+2059.735]GSDC[+57.021]PDKFC[+57.021]LFQSETK.N | HexNAc(4)Hex(5)Fuc(1)NeuAc(1) | C[+57]*2, N[+2060] | 1364.8819 | 3 | 4092.6311 | 4092.6088 | 0 | 5.4 | 642 | Specific | 333.92 | 291.54 | 291.54 | 6.3 | 130 | >sp\|P02788\|TRFL_HUMAN Lactotransferrin OS=Homo sapiens OX=9606 GN=LTF PE=1 SV=6 | 1270 | 20201119_10132_1ng_glyco_hcd203040_compound15.11791.11791.3 | controllerType=0 controllerNumber=1 scan=11791 | 26.6507 |
| 08437:3 | 1 | R.N[+2205.793]GSDC[+57.021]PDKFC[+57.021]LFQSETK.N | HexNAc(4)Hex(5)Fuc(2)NeuAc(1) | C[+57]*2, N[+2206] | 1413.5673 | 3 | 4238.6874 | 4238.6667 | 0 | 4.9 | 642 | Specific | 517.13 | 402.19 | 402.19 | 8.87 | 130 | >sp\|P02788\|TRFL_HUMAN Lactotransferrin OS=Homo sapiens OX=9606 GN=LTF PE=1 SV=6 | 1270 | 20201119_10132_1ng_glyco_hcd203040_compound15.11698.11698.3 | controllerType=0 controllerNumber=1 scan=11698 | 26.4966 |
| 08435:3 | 1 | R.N[+2205.793]GSDC[+57.021]PDKFC[+57.021]LFQSETK.N | HexNAc(4)Hex(5)Fuc(2)NeuAc(1) | C[+57]*2, N[+2206] | 1413.5673 | 3 | 4238.6874 | 4238.6667 | 0 | 4.9 | 642 | Specific | 552.16 | 492.71 | 492.71 | 9.07 | 130 | >sp\|P02788\|TRFL_HUMAN Lactotransferrin OS=Homo sapiens OX=9606 GN=LTF PE=1 SV=6 | 1270 | 20201119_10132_1ng_glyco_hcd203040_compound15.11696.11696.3 | controllerType=0 controllerNumber=1 scan=11696 | 26.4924 |
| 08462:3 | 1 | R.N[+2205.793]GSDC[+57.021]PDKFC[+57.021]LFQSETK.N | HexNAc(4)Hex(5)Fuc(2)NeuAc(1) | C[+57]*2, N[+2206] | 1413.5644 | 3 | 4238.6786 | 4238.6667 | 0 | 2.8 | 642 | Specific | 556.56 | 415.11 | 415.11 | 9.54 | 130 | >sp\|P02788\|TRFL_HUMAN Lactotransferrin OS=Homo sapiens OX=9606 GN=LTF PE=1 SV=6 | 1270 | 20201119_10132_1ng_glyco_hcd203040_compound15.11724.11724.3 | controllerType=0 controllerNumber=1 scan=11724 | 26.5403 |
| 08464:3 | 1 | R.N[+2205.793]GSDC[+57.021]PDKFC[+57.021]LFQSETK.N | HexNAc(4)Hex(5)Fuc(2)NeuAc(1) | C[+57]*2, N[+2206] | 1413.5644 | 3 | 4238.6786 | 4238.6667 | 0 | 2.8 | 642 | Specific | 651.83 | 475.81 | 475.81 | 12.69 | 130 | >sp\|P02788\|TRFL_HUMAN Lactotransferrin OS=Homo sapiens OX=9606 GN=LTF PE=1 SV=6 | 1270 | 20201119_10132_1ng_glyco_hcd203040_compound15.11726.11726.3 | controllerType=0 controllerNumber=1 scan=11726 | 26.5451 |
| 12486:4 | 5 | K.KGN[+2204.772]TTLNSFVIPSESDVPTHC[+57.021]PSQWWPYAGHC[+57.021]YK.I | HexNAc(4)Hex(5)NeuAc(2) | C[+57]*2, N[+2205] | 1542.9124 | 4 | 6168.6276 | 6168.5942 | 0 | 5.4 | 342 | Specific | 241.23 | 241.23 | 241.23 | 5.91 | 72 | >sp\|P22897\|MRC1_HUMAN Macrophage mannose receptor 1 OS=Homo sapiens OX=9606 GN=MRC1 PE=1 SV=1 | 1049 | 20201119_10132_1ng_glyco_hcd203040_compound15.15917.15917.4 | controllerType=0 controllerNumber=1 scan=15917 | 33.7242 |
| 07724:3 | 5 | K.TAHC[+57.021]N[+2205.793]ESFYFLC[+57.021]K.R | HexNAc(4)Hex(5)Fuc(2)NeuAc(1) | C[+57]*2, N[+2206] | 1294.8466 | 3 | 3882.5253 | 3882.5124 | 0 | 3.3 | 1201 | Specific | 376.5 | 287.48 | 287.48 | 7.53 | 72 | >sp\|P22897\|MRC1_HUMAN Macrophage mannose receptor 1 OS=Homo sapiens OX=9606 GN=MRC1 PE=1 SV=1 | 1049 | 20201119_10132_1ng_glyco_hcd203040_compound15.10940.10940.3 | controllerType=0 controllerNumber=1 scan=10940 | 25.163 |
| 07738:3 | 5 | K.TAHC[+57.021]N[+2205.793]ESFYFLC[+57.021]K.R | HexNAc(4)Hex(5)Fuc(2)NeuAc(1) | C[+57]*2, N[+2206] | 1294.847 | 3 | 3882.5264 | 3882.5124 | 0 | 3.6 | 1201 | Specific | 462.2 | 383.71 | 383.71 | 9.77 | 72 | >sp\|P22897\|MRC1_HUMAN Macrophage mannose receptor 1 OS=Homo sapiens OX=9606 GN=MRC1 PE=1 SV=1 | 1049 | 20201119_10132_1ng_glyco_hcd203040_compound15.10955.10955.3 | controllerType=0 controllerNumber=1 scan=10955 | 25.1903 |
| 07740:3 | 5 | K.TAHC[+57.021]N[+2205.793]ESFYFLC[+57.021]K.R | HexNAc(4)Hex(5)Fuc(2)NeuAc(1) | C[+57]*2, N[+2206] | 1294.847 | 3 | 3882.5264 | 3882.5124 | 0 | 3.6 | 1201 | Specific | 462.52 | 391.48 | 391.48 | 9.77 | 72 | >sp\|P22897\|MRC1_HUMAN Macrophage mannose receptor 1 OS=Homo sapiens OX=9606 GN=MRC1 PE=1 SV=1 | 1049 | 20201119_10132_1ng_glyco_hcd203040_compound15.10957.10957.3 | controllerType=0 controllerNumber=1 scan=10957 | 25.1944 |
| 17166:3 | 6 | R.LSLHRPALEDLLLGSEAN[+1460.529]LTC[+57.021]TLTGLR.D | HexNAc(4)Hex(4) | C[+57], N[+1461] | 1475.3878 | 3 | 4424.1487 | 4424.1271 | 0 | 4.9 | 127 | Specific | 260.43 | 260.43 | 260.43 | 5.99 | 57 | >sp\|P01876\|IGHA1_HUMAN Immunoglobulin heavy constant alpha 1 OS=Homo sapiens OX=9606 GN=IGHA1 PE=1 SV=2 | 255 | 20201119_10132_1ng_glyco_hcd203040_compound15.20824.20824.3 | controllerType=0 controllerNumber=1 scan=20824 | 42.8017 |
| 17114:3 | 6 | R.LSLHRPALEDLLLGSEAN[+1702.581]LTC[+57.021]TLTGLR.D | HexNAc(2)Hex(8) | C[+57], N[+1703] | 1556.074 | 3 | 4666.2075 | 4666.1796 | 0 | 6 | 127 | Specific | 359.72 | 359.72 | 359.72 | 6.99 | 57 | >sp\|P01876\|IGHA1_HUMAN Immunoglobulin heavy constant alpha 1 OS=Homo sapiens OX=9606 GN=IGHA1 PE=1 SV=2 | 255 | 20201119_10132_1ng_glyco_hcd203040_compound15.20769.20769.3 | controllerType=0 controllerNumber=1 scan=20769 | 42.637 |
| 17231:4 | 6 | R.LSLHRPALEDLLLGSEAN[+1095.397]LTC[+57.021]TLTGLR.D | HexNAc(3)Hex(3) | C[+57], N[+1095] | 1015.5092 | 4 | 4059.015 | 4058.9949 | 0 | 5 | 127 | Specific | 366.67 | 366.67 | 366.67 | 7.76 | 57 | >sp\|P01876\|IGHA1_HUMAN Immunoglobulin heavy constant alpha 1 OS=Homo sapiens OX=9606 GN=IGHA1 PE=1 SV=2 | 255 | 20201119_10132_1ng_glyco_hcd203040_compound15.20892.20892.4 | controllerType=0 controllerNumber=1 scan=20892 | 42.9651 |
| 17168:3 | 6 | R.LSLHRPALEDLLLGSEAN[+1378.476]LTC[+57.021]TLTGLR.D | HexNAc(2)Hex(6) | C[+57], N[+1378] | 1448.0402 | 3 | 4342.1061 | 4342.074 | 0 | 7.4 | 127 | Specific | 439.65 | 410.83 | 410.83 | 7.08 | 57 | >sp\|P01876\|IGHA1_HUMAN Immunoglobulin heavy constant alpha 1 OS=Homo sapiens OX=9606 GN=IGHA1 PE=1 SV=2 | 255 | 20201119_10132_1ng_glyco_hcd203040_compound15.20826.20826.3 | controllerType=0 controllerNumber=1 scan=20826 | 42.8087 |
| 17137:3 | 6 | R.LSLHRPALEDLLLGSEAN[+1622.582]LTC[+57.021]TLTGLR.D | HexNAc(4)Hex(5) | C[+57], N[+1623] | 1529.4086 | 3 | 4586.2113 | 4586.1799 | 0 | 6.9 | 127 | Specific | 455.81 | 390.88 | 390.88 | 8.38 | 57 | >sp\|P01876\|IGHA1_HUMAN Immunoglobulin heavy constant alpha 1 OS=Homo sapiens OX=9606 GN=IGHA1 PE=1 SV=2 | 255 | 20201119_10132_1ng_glyco_hcd203040_compound15.20793.20793.3 | controllerType=0 controllerNumber=1 scan=20793 | 42.6958 |
| 17179:5 | 6 | R.LSLHRPALEDLLLGSEAN[+1501.555]LTC[+57.021]TLTGLR.D | HexNAc(5)Hex(3) | C[+57], N[+1502] | 893.8408 | 5 | 4465.1748 | 4465.1536 | 0 | 4.7 | 127 | Specific | 500.84 | 500.84 | 500.84 | 10.14 | 57 | >sp\|P01876\|IGHA1_HUMAN Immunoglobulin heavy constant alpha 1 OS=Homo sapiens OX=9606 GN=IGHA1 PE=1 SV=2 | 255 | 20201119_10132_1ng_glyco_hcd203040_compound15.20837.20837.5 | controllerType=0 controllerNumber=1 scan=20837 | 42.8326 |
| 17117:3 | 6 | R.LSLHRPALEDLLLGSEAN[+1702.581]LTC[+57.021]TLTGLR.D | HexNAc(2)Hex(8) | C[+57], N[+1703] | 1556.074 | 3 | 4666.2075 | 4666.1796 | 0 | 6 | 127 | Specific | 573.81 | 573.81 | 573.81 | 10.37 | 57 | >sp\|P01876\|IGHA1_HUMAN Immunoglobulin heavy constant alpha 1 OS=Homo sapiens OX=9606 GN=IGHA1 PE=1 SV=2 | 255 | 20201119_10132_1ng_glyco_hcd203040_compound15.20772.20772.3 | controllerType=0 controllerNumber=1 scan=20772 | 42.6417 |
| 18088:3 | 6 | R.LSLHRPALEDLLLGSEAN[+1913.677]LTC[+57.021]TLTGLR.D | HexNAc(4)Hex(5)NeuAc(1) | C[+57], N[+1914] | 1626.4357 | 3 | 4877.2926 | 4877.2753 | 0 | 3.6 | 127 | Specific | 599.4 | 560.7 | 560.7 | 10.85 | 57 | >sp\|P01876\|IGHA1_HUMAN Immunoglobulin heavy constant alpha 1 OS=Homo sapiens OX=9606 GN=IGHA1 PE=1 SV=2 | 255 | 20201119_10132_1ng_glyco_hcd203040_compound15.21785.21785.3 | controllerType=0 controllerNumber=1 scan=21785 | 44.8027 |
| 17161:3 | 6 | R.LSLHRPALEDLLLGSEAN[+1216.423]LTC[+57.021]TLTGLR.D | HexNAc(2)Hex(5) | C[+57], N[+1216] | 1394.0178 | 3 | 4180.0387 | 4180.0211 | 0 | 4.2 | 127 | Specific | 601.54 | 601.54 | 601.54 | 11.95 | 57 | >sp\|P01876\|IGHA1_HUMAN Immunoglobulin heavy constant alpha 1 OS=Homo sapiens OX=9606 GN=IGHA1 PE=1 SV=2 | 255 | 20201119_10132_1ng_glyco_hcd203040_compound15.20819.20819.3 | controllerType=0 controllerNumber=1 scan=20819 | 42.792 |
| 17118:3 | 6 | R.LSLHRPALEDLLLGSEAN[+1663.608]LTC[+57.021]TLTGLR.D | HexNAc(5)Hex(4) | C[+57], N[+1664] | 1543.0801 | 3 | 4627.2259 | 4627.2064 | 0 | 4.2 | 127 | Specific | 620.39 | 557.24 | 557.24 | 11.95 | 57 | >sp\|P01876\|IGHA1_HUMAN Immunoglobulin heavy constant alpha 1 OS=Homo sapiens OX=9606 GN=IGHA1 PE=1 SV=2 | 255 | 20201119_10132_1ng_glyco_hcd203040_compound15.20773.20773.3 | controllerType=0 controllerNumber=1 scan=20773 | 42.647 |
| 18146:3 | 6 | R.LSLHRPALEDLLLGSEAN[+1751.624]LTC[+57.021]TLTGLR.D | HexNAc(4)Hex(4)NeuAc(1) | C[+57], N[+1752] | 1572.4189 | 3 | 4715.2421 | 4715.2225 | 0 | 4.2 | 127 | Specific | 638.9 | 638.9 | 638.9 | 11.95 | 57 | >sp\|P01876\|IGHA1_HUMAN Immunoglobulin heavy constant alpha 1 OS=Homo sapiens OX=9606 GN=IGHA1 PE=1 SV=2 | 255 | 20201119_10132_1ng_glyco_hcd203040_compound15.21846.21846.3 | controllerType=0 controllerNumber=1 scan=21846 | 44.9456 |
| 17116:3 | 6 | R.LSLHRPALEDLLLGSEAN[+1663.608]LTC[+57.021]TLTGLR.D | HexNAc(5)Hex(4) | C[+57], N[+1664] | 1543.0801 | 3 | 4627.2259 | 4627.2064 | 0 | 4.2 | 127 | Specific | 642.44 | 621.74 | 621.74 | 11.95 | 57 | >sp\|P01876\|IGHA1_HUMAN Immunoglobulin heavy constant alpha 1 OS=Homo sapiens OX=9606 GN=IGHA1 PE=1 SV=2 | 255 | 20201119_10132_1ng_glyco_hcd203040_compound15.20771.20771.3 | controllerType=0 controllerNumber=1 scan=20771 | 42.6395 |
| 17176:5 | 6 | R.LSLHRPALEDLLLGSEAN[+1501.555]LTC[+57.021]TLTGLR.D | HexNAc(5)Hex(3) | C[+57], N[+1502] | 893.8408 | 5 | 4465.1748 | 4465.1536 | 0 | 4.7 | 127 | Specific | 642.8 | 642.8 | 642.8 | 12.65 | 57 | >sp\|P01876\|IGHA1_HUMAN Immunoglobulin heavy constant alpha 1 OS=Homo sapiens OX=9606 GN=IGHA1 PE=1 SV=2 | 255 | 20201119_10132_1ng_glyco_hcd203040_compound15.20834.20834.5 | controllerType=0 controllerNumber=1 scan=20834 | 42.8222 |
| 18090:3 | 6 | R.LSLHRPALEDLLLGSEAN[+1913.677]LTC[+57.021]TLTGLR.D | HexNAc(4)Hex(5)NeuAc(1) | C[+57], N[+1914] | 1626.4357 | 3 | 4877.2926 | 4877.2753 | 0 | 3.6 | 127 | Specific | 665.82 | 606.54 | 606.54 | 12.85 | 57 | >sp\|P01876\|IGHA1_HUMAN Immunoglobulin heavy constant alpha 1 OS=Homo sapiens OX=9606 GN=IGHA1 PE=1 SV=2 | 255 | 20201119_10132_1ng_glyco_hcd203040_compound15.21787.21787.3 | controllerType=0 controllerNumber=1 scan=21787 | 44.8096 |
| 17157:3 | 6 | R.LSLHRPALEDLLLGSEAN[+1501.555]LTC[+57.021]TLTGLR.D | HexNAc(5)Hex(3) | C[+57], N[+1502] | 1489.0606 | 3 | 4465.1673 | 4465.1536 | 0 | 3.1 | 127 | Specific | 666.6 | 637.61 | 637.61 | 12.85 | 57 | >sp\|P01876\|IGHA1_HUMAN Immunoglobulin heavy constant alpha 1 OS=Homo sapiens OX=9606 GN=IGHA1 PE=1 SV=2 | 255 | 20201119_10132_1ng_glyco_hcd203040_compound15.20814.20814.3 | controllerType=0 controllerNumber=1 scan=20814 | 42.7776 |
| 17111:4 | 6 | R.LSLHRPALEDLLLGSEAN[+1663.608]LTC[+57.021]TLTGLR.D | HexNAc(5)Hex(4) | C[+57], N[+1664] | 1157.5618 | 4 | 4627.2254 | 4627.2064 | 0 | 4.1 | 127 | Specific | 667.34 | 667.34 | 667.34 | 13.73 | 57 | >sp\|P01876\|IGHA1_HUMAN Immunoglobulin heavy constant alpha 1 OS=Homo sapiens OX=9606 GN=IGHA1 PE=1 SV=2 | 255 | 20201119_10132_1ng_glyco_hcd203040_compound15.20766.20766.4 | controllerType=0 controllerNumber=1 scan=20766 | 42.6275 |
| 17154:3 | 6 | R.LSLHRPALEDLLLGSEAN[+1501.555]LTC[+57.021]TLTGLR.D | HexNAc(5)Hex(3) | C[+57], N[+1502] | 1489.0606 | 3 | 4465.1673 | 4465.1536 | 0 | 3.1 | 127 | Specific | 672.22 | 672.22 | 672.22 | 12.85 | 57 | >sp\|P01876\|IGHA1_HUMAN Immunoglobulin heavy constant alpha 1 OS=Homo sapiens OX=9606 GN=IGHA1 PE=1 SV=2 | 255 | 20201119_10132_1ng_glyco_hcd203040_compound15.20811.20811.3 | controllerType=0 controllerNumber=1 scan=20811 | 42.7644 |
| 17169:3 | 6 | R.LSLHRPALEDLLLGSEAN[+1460.529]LTC[+57.021]TLTGLR.D | HexNAc(4)Hex(4) | C[+57], N[+1461] | 1475.3878 | 3 | 4424.1487 | 4424.1271 | 0 | 4.9 | 127 | Specific | 686.49 | 672.91 | 672.91 | 12.37 | 57 | >sp\|P01876\|IGHA1_HUMAN Immunoglobulin heavy constant alpha 1 OS=Homo sapiens OX=9606 GN=IGHA1 PE=1 SV=2 | 255 | 20201119_10132_1ng_glyco_hcd203040_compound15.20827.20827.3 | controllerType=0 controllerNumber=1 scan=20827 | 42.8106 |
| 17139:3 | 6 | R.LSLHRPALEDLLLGSEAN[+1622.582]LTC[+57.021]TLTGLR.D | HexNAc(4)Hex(5) | C[+57], N[+1623] | 1529.4086 | 3 | 4586.2113 | 4586.1799 | 0 | 6.9 | 127 | Specific | 690.74 | 568.7 | 568.7 | 12.13 | 57 | >sp\|P01876\|IGHA1_HUMAN Immunoglobulin heavy constant alpha 1 OS=Homo sapiens OX=9606 GN=IGHA1 PE=1 SV=2 | 255 | 20201119_10132_1ng_glyco_hcd203040_compound15.20795.20795.3 | controllerType=0 controllerNumber=1 scan=20795 | 42.7014 |
| 17133:4 | 6 | R.LSLHRPALEDLLLGSEAN[+1581.555]LTC[+57.021]TLTGLR.D | HexNAc(3)Hex(6) | C[+57], N[+1582] | 1137.0486 | 4 | 4545.1727 | 4545.1533 | 0 | 4.3 | 127 | Specific | 691.67 | 666.36 | 666.36 | 13.73 | 57 | >sp\|P01876\|IGHA1_HUMAN Immunoglobulin heavy constant alpha 1 OS=Homo sapiens OX=9606 GN=IGHA1 PE=1 SV=2 | 255 | 20201119_10132_1ng_glyco_hcd203040_compound15.20789.20789.4 | controllerType=0 controllerNumber=1 scan=20789 | 42.6877 |
| 17132:4 | 6 | R.LSLHRPALEDLLLGSEAN[+1622.582]LTC[+57.021]TLTGLR.D | HexNAc(4)Hex(5) | C[+57], N[+1623] | 1147.3083 | 4 | 4586.2113 | 4586.1799 | 0 | 6.8 | 127 | Specific | 698.52 | 698.52 | 698.52 | 12.91 | 57 | >sp\|P01876\|IGHA1_HUMAN Immunoglobulin heavy constant alpha 1 OS=Homo sapiens OX=9606 GN=IGHA1 PE=1 SV=2 | 255 | 20201119_10132_1ng_glyco_hcd203040_compound15.20788.20788.4 | controllerType=0 controllerNumber=1 scan=20788 | 42.6859 |
| 17150:4 | 6 | R.LSLHRPALEDLLLGSEAN[+1501.555]LTC[+57.021]TLTGLR.D | HexNAc(5)Hex(3) | C[+57], N[+1502] | 1117.0479 | 4 | 4465.1697 | 4465.1536 | 0 | 3.6 | 127 | Specific | 699.37 | 699.37 | 699.37 | 13.73 | 57 | >sp\|P01876\|IGHA1_HUMAN Immunoglobulin heavy constant alpha 1 OS=Homo sapiens OX=9606 GN=IGHA1 PE=1 SV=2 | 255 | 20201119_10132_1ng_glyco_hcd203040_compound15.20807.20807.4 | controllerType=0 controllerNumber=1 scan=20807 | 42.7347 |
| 17165:3 | 6 | R.LSLHRPALEDLLLGSEAN[+1378.476]LTC[+57.021]TLTGLR.D | HexNAc(2)Hex(6) | C[+57], N[+1378] | 1448.0402 | 3 | 4342.1061 | 4342.074 | 0 | 7.4 | 127 | Specific | 712.5 | 670.25 | 670.25 | 13.13 | 57 | >sp\|P01876\|IGHA1_HUMAN Immunoglobulin heavy constant alpha 1 OS=Homo sapiens OX=9606 GN=IGHA1 PE=1 SV=2 | 255 | 20201119_10132_1ng_glyco_hcd203040_compound15.20823.20823.3 | controllerType=0 controllerNumber=1 scan=20823 | 42.8004 |
| 17135:4 | 6 | R.LSLHRPALEDLLLGSEAN[+1581.555]LTC[+57.021]TLTGLR.D | HexNAc(3)Hex(6) | C[+57], N[+1582] | 1137.0486 | 4 | 4545.1727 | 4545.1533 | 0 | 4.3 | 127 | Specific | 735.36 | 680.59 | 680.59 | 14.73 | 57 | >sp\|P01876\|IGHA1_HUMAN Immunoglobulin heavy constant alpha 1 OS=Homo sapiens OX=9606 GN=IGHA1 PE=1 SV=2 | 255 | 20201119_10132_1ng_glyco_hcd203040_compound15.20791.20791.4 | controllerType=0 controllerNumber=1 scan=20791 | 42.6928 |
| 18077:4 | 6 | R.LSLHRPALEDLLLGSEAN[+1913.677]LTC[+57.021]TLTGLR.D | HexNAc(4)Hex(5)NeuAc(1) | C[+57], N[+1914] | 1220.0761 | 4 | 4877.2825 | 4877.2753 | 0 | 1.5 | 127 | Specific | 737.86 | 737.86 | 737.86 | 14.41 | 57 | >sp\|P01876\|IGHA1_HUMAN Immunoglobulin heavy constant alpha 1 OS=Homo sapiens OX=9606 GN=IGHA1 PE=1 SV=2 | 255 | 20201119_10132_1ng_glyco_hcd203040_compound15.21774.21774.4 | controllerType=0 controllerNumber=1 scan=21774 | 44.7715 |
| 17113:4 | 6 | R.LSLHRPALEDLLLGSEAN[+1663.608]LTC[+57.021]TLTGLR.D | HexNAc(5)Hex(4) | C[+57], N[+1664] | 1157.5618 | 4 | 4627.2254 | 4627.2064 | 0 | 4.1 | 127 | Specific | 755.72 | 755.72 | 755.72 | 15.73 | 57 | >sp\|P01876\|IGHA1_HUMAN Immunoglobulin heavy constant alpha 1 OS=Homo sapiens OX=9606 GN=IGHA1 PE=1 SV=2 | 255 | 20201119_10132_1ng_glyco_hcd203040_compound15.20768.20768.4 | controllerType=0 controllerNumber=1 scan=20768 | 42.6358 |
| 18079:4 | 6 | R.LSLHRPALEDLLLGSEAN[+1913.677]LTC[+57.021]TLTGLR.D | HexNAc(4)Hex(5)NeuAc(1) | C[+57], N[+1914] | 1220.0761 | 4 | 4877.2825 | 4877.2753 | 0 | 1.5 | 127 | Specific | 768.84 | 733.79 | 733.79 | 15.41 | 57 | >sp\|P01876\|IGHA1_HUMAN Immunoglobulin heavy constant alpha 1 OS=Homo sapiens OX=9606 GN=IGHA1 PE=1 SV=2 | 255 | 20201119_10132_1ng_glyco_hcd203040_compound15.21776.21776.4 | controllerType=0 controllerNumber=1 scan=21776 | 44.7783 |
| 17136:4 | 6 | R.LSLHRPALEDLLLGSEAN[+1419.502]LTC[+57.021]TLTGLR.D | HexNAc(3)Hex(5) | C[+57], N[+1420] | 1096.5384 | 4 | 4383.1317 | 4383.1005 | 0 | 7.1 | 127 | Specific | 785.92 | 785.92 | 785.92 | 14.91 | 57 | >sp\|P01876\|IGHA1_HUMAN Immunoglobulin heavy constant alpha 1 OS=Homo sapiens OX=9606 GN=IGHA1 PE=1 SV=2 | 255 | 20201119_10132_1ng_glyco_hcd203040_compound15.20792.20792.4 | controllerType=0 controllerNumber=1 scan=20792 | 42.6947 |
| 17164:4 | 6 | R.LSLHRPALEDLLLGSEAN[+1298.476]LTC[+57.021]TLTGLR.D | HexNAc(4)Hex(3) | C[+57], N[+1298] | 1066.2787 | 4 | 4262.0931 | 4262.0742 | 0 | 4.4 | 127 | Specific | 807.23 | 807.23 | 807.23 | 16.73 | 57 | >sp\|P01876\|IGHA1_HUMAN Immunoglobulin heavy constant alpha 1 OS=Homo sapiens OX=9606 GN=IGHA1 PE=1 SV=2 | 255 | 20201119_10132_1ng_glyco_hcd203040_compound15.20822.20822.4 | controllerType=0 controllerNumber=1 scan=20822 | 42.7994 |
| 17170:4 | 6 | R.LSLHRPALEDLLLGSEAN[+1257.449]LTC[+57.021]TLTGLR.D | HexNAc(3)Hex(4) | C[+57], N[+1257] | 1056.0216 | 4 | 4221.0648 | 4221.0477 | 0 | 4 | 127 | Specific | 812.17 | 812.17 | 812.17 | 16.73 | 57 | >sp\|P01876\|IGHA1_HUMAN Immunoglobulin heavy constant alpha 1 OS=Homo sapiens OX=9606 GN=IGHA1 PE=1 SV=2 | 255 | 20201119_10132_1ng_glyco_hcd203040_compound15.20828.20828.4 | controllerType=0 controllerNumber=1 scan=20828 | 42.8124 |
| 18137:4 | 6 | R.LSLHRPALEDLLLGSEAN[+1751.624]LTC[+57.021]TLTGLR.D | HexNAc(4)Hex(4)NeuAc(1) | C[+57], N[+1752] | 1179.5668 | 4 | 4715.2454 | 4715.2225 | 0 | 4.9 | 127 | Specific | 828.12 | 828.12 | 828.12 | 16.73 | 57 | >sp\|P01876\|IGHA1_HUMAN Immunoglobulin heavy constant alpha 1 OS=Homo sapiens OX=9606 GN=IGHA1 PE=1 SV=2 | 255 | 20201119_10132_1ng_glyco_hcd203040_compound15.21837.21837.4 | controllerType=0 controllerNumber=1 scan=21837 | 44.933 |
| 17134:4 | 6 | R.LSLHRPALEDLLLGSEAN[+1622.582]LTC[+57.021]TLTGLR.D | HexNAc(4)Hex(5) | C[+57], N[+1623] | 1147.3083 | 4 | 4586.2113 | 4586.1799 | 0 | 6.8 | 127 | Specific | 833.79 | 726.78 | 726.78 | 15.91 | 57 | >sp\|P01876\|IGHA1_HUMAN Immunoglobulin heavy constant alpha 1 OS=Homo sapiens OX=9606 GN=IGHA1 PE=1 SV=2 | 255 | 20201119_10132_1ng_glyco_hcd203040_compound15.20790.20790.4 | controllerType=0 controllerNumber=1 scan=20790 | 42.6909 |
| 17163:3 | 6 | R.LSLHRPALEDLLLGSEAN[+1216.423]LTC[+57.021]TLTGLR.D | HexNAc(2)Hex(5) | C[+57], N[+1216] | 1394.0178 | 3 | 4180.0387 | 4180.0211 | 0 | 4.2 | 127 | Specific | 848.75 | 703.95 | 703.95 | 15.95 | 57 | >sp\|P01876\|IGHA1_HUMAN Immunoglobulin heavy constant alpha 1 OS=Homo sapiens OX=9606 GN=IGHA1 PE=1 SV=2 | 255 | 20201119_10132_1ng_glyco_hcd203040_compound15.20821.20821.3 | controllerType=0 controllerNumber=1 scan=20821 | 42.7974 |
| 17110:4 | 6 | R.LSLHRPALEDLLLGSEAN[+1702.581]LTC[+57.021]TLTGLR.D | HexNAc(2)Hex(8) | C[+57], N[+1703] | 1167.304 | 4 | 4666.1942 | 4666.1796 | 0 | 3.1 | 127 | Specific | 878.59 | 878.59 | 878.59 | 17.62 | 57 | >sp\|P01876\|IGHA1_HUMAN Immunoglobulin heavy constant alpha 1 OS=Homo sapiens OX=9606 GN=IGHA1 PE=1 SV=2 | 255 | 20201119_10132_1ng_glyco_hcd203040_compound15.20765.20765.4 | controllerType=0 controllerNumber=1 scan=20765 | 42.626 |
| 17152:4 | 6 | R.LSLHRPALEDLLLGSEAN[+1501.555]LTC[+57.021]TLTGLR.D | HexNAc(5)Hex(3) | C[+57], N[+1502] | 1117.0479 | 4 | 4465.1697 | 4465.1536 | 0 | 3.6 | 127 | Specific | 879.45 | 879.45 | 879.45 | 17.73 | 57 | >sp\|P01876\|IGHA1_HUMAN Immunoglobulin heavy constant alpha 1 OS=Homo sapiens OX=9606 GN=IGHA1 PE=1 SV=2 | 255 | 20201119_10132_1ng_glyco_hcd203040_compound15.20809.20809.4 | controllerType=0 controllerNumber=1 scan=20809 | 42.7478 |
| 17138:4 | 6 | R.LSLHRPALEDLLLGSEAN[+1419.502]LTC[+57.021]TLTGLR.D | HexNAc(3)Hex(5) | C[+57], N[+1420] | 1096.5384 | 4 | 4383.1317 | 4383.1005 | 0 | 7.1 | 127 | Specific | 884.62 | 881.59 | 881.59 | 16.91 | 57 | >sp\|P01876\|IGHA1_HUMAN Immunoglobulin heavy constant alpha 1 OS=Homo sapiens OX=9606 GN=IGHA1 PE=1 SV=2 | 255 | 20201119_10132_1ng_glyco_hcd203040_compound15.20794.20794.4 | controllerType=0 controllerNumber=1 scan=20794 | 42.6995 |
| 17167:4 | 6 | R.LSLHRPALEDLLLGSEAN[+1298.476]LTC[+57.021]TLTGLR.D | HexNAc(4)Hex(3) | C[+57], N[+1298] | 1066.2787 | 4 | 4262.0931 | 4262.0742 | 0 | 4.4 | 127 | Specific | 885.38 | 885.17 | 885.17 | 17.73 | 57 | >sp\|P01876\|IGHA1_HUMAN Immunoglobulin heavy constant alpha 1 OS=Homo sapiens OX=9606 GN=IGHA1 PE=1 SV=2 | 255 | 20201119_10132_1ng_glyco_hcd203040_compound15.20825.20825.4 | controllerType=0 controllerNumber=1 scan=20825 | 42.8068 |
| 17162:4 | 6 | R.LSLHRPALEDLLLGSEAN[+1460.529]LTC[+57.021]TLTGLR.D | HexNAc(4)Hex(4) | C[+57], N[+1461] | 1106.797 | 4 | 4424.1663 | 4424.1271 | 0 | 8.9 | 127 | Specific | 900.7 | 849.84 | 849.84 | 17.71 | 57 | >sp\|P01876\|IGHA1_HUMAN Immunoglobulin heavy constant alpha 1 OS=Homo sapiens OX=9606 GN=IGHA1 PE=1 SV=2 | 255 | 20201119_10132_1ng_glyco_hcd203040_compound15.20820.20820.4 | controllerType=0 controllerNumber=1 scan=20820 | 42.7942 |
| 17234:4 | 6 | R.LSLHRPALEDLLLGSEAN[+1095.397]LTC[+57.021]TLTGLR.D | HexNAc(3)Hex(3) | C[+57], N[+1095] | 1015.5092 | 4 | 4059.015 | 4058.9949 | 0 | 5 | 127 | Specific | 906.09 | 634.44 | 634.44 | 18.15 | 57 | >sp\|P01876\|IGHA1_HUMAN Immunoglobulin heavy constant alpha 1 OS=Homo sapiens OX=9606 GN=IGHA1 PE=1 SV=2 | 255 | 20201119_10132_1ng_glyco_hcd203040_compound15.20895.20895.4 | controllerType=0 controllerNumber=1 scan=20895 | 42.9693 |
| 17156:4 | 6 | R.LSLHRPALEDLLLGSEAN[+1216.423]LTC[+57.021]TLTGLR.D | HexNAc(2)Hex(5) | C[+57], N[+1216] | 1045.7635 | 4 | 4180.0321 | 4180.0211 | 0 | 2.6 | 127 | Specific | 928.05 | 864.64 | 864.64 | 18.62 | 57 | >sp\|P01876\|IGHA1_HUMAN Immunoglobulin heavy constant alpha 1 OS=Homo sapiens OX=9606 GN=IGHA1 PE=1 SV=2 | 255 | 20201119_10132_1ng_glyco_hcd203040_compound15.20813.20813.4 | controllerType=0 controllerNumber=1 scan=20813 | 42.7679 |

Table S4. Cont.

| Query #:z | Protein Rank | Glycopeptide | Glycan | Modification Type(s) | Observed m/z | z | Observed (M+H) | Calc. mass (M+H) | Off-by-x error | Mass error (ppm) | StartPosition | Cleavage | Score | Delta | Delta Mod | \|Log Prob\| | # of unique peptides | Protein Name | Protein DB number | Comment | Scan # | Scan Time |
|---|---|---|---|---|---|---|---|---|---|---|---|---|---|---|---|---|---|---|---|---|---|---|
| 17153:4 | 6 | R.LSLHRPALEDLLLGSEAN[+1378.476]LTC[+57.021]TLTGLR.D | HexNAc(2)Hex(6) | C[+57], N[+1378] | 1086.2769 | 4 | 4342.0858 | 4342.074 | 0 | 2.7 | 127 | Specific | 929.77 | 929.77 | 929.77 | 18.62 | 57 | >sp\|P01876\|IGHA1_HUMAN Immunoglobulin heavy constant alpha 1 OS=Homo sapiens OX=9606 GN=IGHA1 PE=1 SV=2 | 255 | 20201119_10132_1ng_glyco_hcd203040_compound15.20810.20810.4 | controllerType=0 controllerNumber=1 scan=20810 | 42.7633 |
| 17159:4 | 6 | R.LSLHRPALEDLLLGSEAN[+1216.423]LTC[+57.021]TLTGLR.D | HexNAc(2)Hex(5) | C[+57], N[+1216] | 1045.7635 | 4 | 4180.0321 | 4180.0211 | 0 | 2.6 | 127 | Specific | 972.64 | 866.34 | 866.34 | 19.62 | 57 | >sp\|P01876\|IGHA1_HUMAN Immunoglobulin heavy constant alpha 1 OS=Homo sapiens OX=9606 GN=IGHA1 PE=1 SV=2 | 255 | 20201119_10132_1ng_glyco_hcd203040_compound15.20816.20816.4 | controllerType=0 controllerNumber=1 scan=20816 | 42.7869 |
| 17068:4 | 6 | R.LSLHRPALEDLLLGSEAN[+1540.529]LTC[+57.021]TLTGLR.D | HexNAc(2)Hex(7) | C[+57], N[+1541] | 1126.788 | 4 | 4504.1302 | 4504.1268 | 0 | 0.8 | 127 | Specific | 989.31 | 900.47 | 900.47 | 19.11 | 57 | >sp\|P01876\|IGHA1_HUMAN Immunoglobulin heavy constant alpha 1 OS=Homo sapiens OX=9606 GN=IGHA1 PE=1 SV=2 | 255 | 20201119_10132_1ng_glyco_hcd203040_compound15.20721.20721.4 | controllerType=0 controllerNumber=1 scan=20721 | 42.5212 |
| 17071:4 | 6 | R.LSLHRPALEDLLLGSEAN[+1540.529]LTC[+57.021]TLTGLR.D | HexNAc(2)Hex(7) | C[+57], N[+1541] | 1126.788 | 4 | 4504.1302 | 4504.1268 | 0 | 0.8 | 127 | Specific | 1001.25 | 935.63 | 935.63 | 20.11 | 57 | >sp\|P01876\|IGHA1_HUMAN Immunoglobulin heavy constant alpha 1 OS=Homo sapiens OX=9606 GN=IGHA1 PE=1 SV=2 | 255 | 20201119_10132_1ng_glyco_hcd203040_compound15.20724.20724.4 | controllerType=0 controllerNumber=1 scan=20724 | 42.5448 |
| 17151:4 | 6 | R.LSLHRPALEDLLLGSEAN[+1378.476]LTC[+57.021]TLTGLR.D | HexNAc(2)Hex(6) | C[+57], N[+1378] | 1086.2769 | 4 | 4342.0858 | 4342.074 | 0 | 2.7 | 127 | Specific | 1001.43 | 916.05 | 916.05 | 20.62 | 57 | >sp\|P01876\|IGHA1_HUMAN Immunoglobulin heavy constant alpha 1 OS=Homo sapiens OX=9606 GN=IGHA1 PE=1 SV=2 | 255 | 20201119_10132_1ng_glyco_hcd203040_compound15.20808.20808.4 | controllerType=0 controllerNumber=1 scan=20808 | 42.7362 |
| 17112:4 | 6 | R.LSLHRPALEDLLLGSEAN[+1702.581]LTC[+57.021]TLTGLR.D | HexNAc(2)Hex(8) | C[+57], N[+1703] | 1167.304 | 4 | 4666.1942 | 4666.1796 | 0 | 3.1 | 127 | Specific | 1063.62 | 1063.62 | 1063.62 | 21.62 | 57 | >sp\|P01876\|IGHA1_HUMAN Immunoglobulin heavy constant alpha 1 OS=Homo sapiens OX=9606 GN=IGHA1 PE=1 SV=2 | 255 | 20201119_10132_1ng_glyco_hcd203040_compound15.20767.20767.4 | controllerType=0 controllerNumber=1 scan=20767 | 42.6319 |
| 17173:4 | 6 | R.LSLHRPALEDLLLGSEAN[+1257.449]LTC[+57.021]TLTGLR.D | HexNAc(3)Hex(4) | C[+57], N[+1257] | 1056.0216 | 4 | 4221.0648 | 4221.0477 | 0 | 4 | 127 | Specific | 1095.85 | 1095.85 | 1095.85 | 21.73 | 57 | >sp\|P01876\|IGHA1_HUMAN Immunoglobulin heavy constant alpha 1 OS=Homo sapiens OX=9606 GN=IGHA1 PE=1 SV=2 | 255 | 20201119_10132_1ng_glyco_hcd203040_compound15.20831.20831.4 | controllerType=0 controllerNumber=1 scan=20831 | 42.8177 |
| 12467:3 | 6 | R.LAGKPTHVN[+2263.835]VSVVMAEVDGTC[+57.021].Y | HexNAc(5)Hex(5)Fuc(3) | C[+57], N[+2264] | 1483.3118 | 3 | 4447.9209 | 4447.9135 | 0 | 1.7 | 332 | CRagged | 276.3 | 276.3 | 276.3 | 4.85 | 57 | >sp\|P01876\|IGHA1_HUMAN Immunoglobulin heavy constant alpha 1 OS=Homo sapiens OX=9606 GN=IGHA1 PE=1 SV=2 | 255 | 20201119_10132_1ng_glyco_hcd203040_compound15.15898.15898.3 | controllerType=0 controllerNumber=1 scan=15898 | 33.6904 |
| 12465:3 | 6 | R.LAGKPTHVN[+2263.835]VSVVMAEVDGTC[+57.021].Y | HexNAc(5)Hex(5)Fuc(3) | C[+57], N[+2264] | 1483.3118 | 3 | 4447.9209 | 4447.9135 | 0 | 1.7 | 332 | CRagged | 290.38 | 283.39 | 283.39 | 6.23 | 57 | >sp\|P01876\|IGHA1_HUMAN Immunoglobulin heavy constant alpha 1 OS=Homo sapiens OX=9606 GN=IGHA1 PE=1 SV=2 | 255 | 20201119_10132_1ng_glyco_hcd203040_compound15.15896.15896.3 | controllerType=0 controllerNumber=1 scan=15896 | 33.6838 |
| 12553:3 | 6 | R.LAGKPTHVN[+2059.735]VSVVMAEVDGTC[+57.021]Y.- | HexNAc(4)Hex(5)Fuc(1)NeuAc(1) | C[+57], N[+2060] | 1469.6346 | 3 | 4406.8892 | 4406.877 | 0 | 2.8 | 332 | CRagged | 217.25 | 217.25 | 217.25 | 5.96 | 57 | >sp\|P01876\|IGHA1_HUMAN Immunoglobulin heavy constant alpha 1 OS=Homo sapiens OX=9606 GN=IGHA1 PE=1 SV=2 | 255 | 20201119_10132_1ng_glyco_hcd203040_compound15.15987.15987.3 | controllerType=0 controllerNumber=1 scan=15987 | 33.8512 |
| 10862:3 | 6 | R.LAGKPTHVN[+1257.449]VSVVMAEVDGTC[+57.021]Y.- | HexNAc(3)Hex(4) | C[+57], N[+1257] | 1202.209 | 3 | 3604.6126 | 3604.5915 | 0 | 5.8 | 332 | CRagged | 237.24 | 237.24 | 237.24 | 6.73 | 57 | >sp\|P01876\|IGHA1_HUMAN Immunoglobulin heavy constant alpha 1 OS=Homo sapiens OX=9606 GN=IGHA1 PE=1 SV=2 | 255 | 20201119_10132_1ng_glyco_hcd203040_compound15.14211.14211.3 | controllerType=0 controllerNumber=1 scan=14211 | 30.8182 |
| 10812:3 | 6 | R.LAGKPTHVN[+1501.555]VSVVMAEVDGTC[+57.021]Y.- | HexNAc(5)Hex(3) | C[+57], N[+1502] | 1283.574 | 3 | 3848.7075 | 3848.6974 | 0 | 2.6 | 332 | CRagged | 242.41 | 242.41 | 242.41 | 7.21 | 57 | >sp\|P01876\|IGHA1_HUMAN Immunoglobulin heavy constant alpha 1 OS=Homo sapiens OX=9606 GN=IGHA1 PE=1 SV=2 | 255 | 20201119_10132_1ng_glyco_hcd203040_compound15.14159.14159.3 | controllerType=0 controllerNumber=1 scan=14159 | 30.7199 |
| 14568:3 | 6 | R.LAGKPTHVN[+2204.772]VSVVMAEVDGTC[+57.021]Y.- | HexNAc(4)Hex(5)NeuAc(2) | C[+57], N[+2205] | 1517.9828 | 3 | 4551.934 | 4551.9145 | 0 | 4.3 | 332 | CRagged | 258.18 | 258.18 | 258.18 | 7.57 | 57 | >sp\|P01876\|IGHA1_HUMAN Immunoglobulin heavy constant alpha 1 OS=Homo sapiens OX=9606 GN=IGHA1 PE=1 SV=2 | 255 | 20201119_10132_1ng_glyco_hcd203040_compound15.18081.18081.3 | controllerType=0 controllerNumber=1 scan=18081 | 37.5936 |
| 12626:3 | 6 | R.LAGKPTHVN[+1710.598]VSVVMAEVDGTC[+57.021]Y.- | HexNAc(3)Hex(5)NeuAc(1) | C[+57], N[+1711] | 1353.2614 | 3 | 4057.7697 | 4057.7397 | 0 | 7.4 | 332 | CRagged | 285.13 | 285.13 | 234.93 | 6.75 | 57 | >sp\|P01876\|IGHA1_HUMAN Immunoglobulin heavy constant alpha 1 OS=Homo sapiens OX=9606 GN=IGHA1 PE=1 SV=2 | 255 | 20201119_10132_1ng_glyco_hcd203040_compound15.16062.16062.3 | controllerType=0 controllerNumber=1 scan=16062 | 33.9756 |
| 14458:3 | 6 | R.LAGKPTHVN[+2350.830]VSVVMAEVDGTC[+57.021]Y.- | HexNAc(4)Hex(5)Fuc(1)NeuAc(2) | C[+57], N[+2351] | 1566.6751 | 3 | 4698.0108 | 4697.9724 | 0 | 8.2 | 332 | CRagged | 288.86 | 263.63 | 263.63 | 6.66 | 57 | >sp\|P01876\|IGHA1_HUMAN Immunoglobulin heavy constant alpha 1 OS=Homo sapiens OX=9606 GN=IGHA1 PE=1 SV=2 | 255 | 20201119_10132_1ng_glyco_hcd203040_compound15.17967.17967.3 | controllerType=0 controllerNumber=1 scan=17967 | 37.3868 |
| 14570:3 | 6 | R.LAGKPTHVN[+2204.772]VSVVMAEVDGTC[+57.021]Y.- | HexNAc(4)Hex(5)NeuAc(2) | C[+57], N[+2205] | 1517.9828 | 3 | 4551.934 | 4551.9145 | 0 | 4.3 | 332 | CRagged | 297.42 | 297.42 | 297.42 | 7.74 | 57 | >sp\|P01876\|IGHA1_HUMAN Immunoglobulin heavy constant alpha 1 OS=Homo sapiens OX=9606 GN=IGHA1 PE=1 SV=2 | 255 | 20201119_10132_1ng_glyco_hcd203040_compound15.18083.18083.3 | controllerType=0 controllerNumber=1 scan=18083 | 37.5975 |
| 12628:3 | 6 | R.LAGKPTHVN[+1710.598]VSVVMAEVDGTC[+57.021]Y.- | HexNAc(3)Hex(5)NeuAc(1) | C[+57], N[+1711] | 1353.2614 | 3 | 4057.7697 | 4057.7397 | 0 | 7.4 | 332 | CRagged | 310.78 | 310.78 | 226.68 | 6.92 | 57 | >sp\|P01876\|IGHA1_HUMAN Immunoglobulin heavy constant alpha 1 OS=Homo sapiens OX=9606 GN=IGHA1 PE=1 SV=2 | 255 | 20201119_10132_1ng_glyco_hcd203040_compound15.16064.16064.3 | controllerType=0 controllerNumber=1 scan=16064 | 33.979 |
| 12642:3 | 6 | R.LAGKPTHVN[+1710.598]VSVVMAEVDGTC[+57.021]Y.- | HexNAc(3)Hex(5)NeuAc(1) | C[+57], N[+1711] | 1353.2563 | 3 | 4057.7543 | 4057.7397 | 0 | 3.6 | 332 | CRagged | 311.21 | 310.48 | 310.48 | 7.74 | 57 | >sp\|P01876\|IGHA1_HUMAN Immunoglobulin heavy constant alpha 1 OS=Homo sapiens OX=9606 GN=IGHA1 PE=1 SV=2 | 255 | 20201119_10132_1ng_glyco_hcd203040_compound15.16079.16079.3 | controllerType=0 controllerNumber=1 scan=16079 | 34.006 |
| 09636:3 | 6 | R.LAGKPTHVN[+1216.423]VSVVMAEVDGTC[+57.021]Y.- | HexNAc(2)Hex(5) | C[+57], N[+1216] | 1188.5296 | 3 | 3563.5742 | 3563.5649 | 0 | 2.6 | 332 | CRagged | 315.76 | 315.76 | 266.44 | 8.22 | 57 | >sp\|P01876\|IGHA1_HUMAN Immunoglobulin heavy constant alpha 1 OS=Homo sapiens OX=9606 GN=IGHA1 PE=1 SV=2 | 255 | 20201119_10132_1ng_glyco_hcd203040_compound15.12941.12941.3 | controllerType=0 controllerNumber=1 scan=12941 | 28.6041 |
| 12555:3 | 6 | R.LAGKPTHVN[+2059.735]VSVVMAEVDGTC[+57.021]Y.- | HexNAc(4)Hex(5)Fuc(1)NeuAc(1) | C[+57], N[+2060] | 1469.6346 | 3 | 4406.8892 | 4406.877 | 0 | 2.8 | 332 | CRagged | 319.48 | 316.2 | 316.2 | 8.22 | 57 | >sp\|P01876\|IGHA1_HUMAN Immunoglobulin heavy constant alpha 1 OS=Homo sapiens OX=9606 GN=IGHA1 PE=1 SV=2 | 255 | 20201119_10132_1ng_glyco_hcd203040_compound15.15989.15989.3 | controllerType=0 controllerNumber=1 scan=15989 | 33.8552 |
| 10864:3 | 6 | R.LAGKPTHVN[+1257.449]VSVVMAEVDGTC[+57.021]Y.- | HexNAc(3)Hex(4) | C[+57], N[+1257] | 1202.209 | 3 | 3604.6126 | 3604.5915 | 0 | 5.8 | 332 | CRagged | 372.7 | 372.7 | 251.09 | 9.51 | 57 | >sp\|P01876\|IGHA1_HUMAN Immunoglobulin heavy constant alpha 1 OS=Homo sapiens OX=9606 GN=IGHA1 PE=1 SV=2 | 255 | 20201119_10132_1ng_glyco_hcd203040_compound15.14213.14213.3 | controllerType=0 controllerNumber=1 scan=14213 | 30.8219 |
| 10815:3 | 6 | R.LAGKPTHVN[+1419.502]VSVVMAEVDGTC[+57.021]Y.- | HexNAc(3)Hex(5) | C[+57], N[+1420] | 1256.2217 | 3 | 3766.6507 | 3766.6443 | 0 | 1.7 | 332 | CRagged | 480.03 | 383.72 | 284.16 | 10.11 | 57 | >sp\|P01876\|IGHA1_HUMAN Immunoglobulin heavy constant alpha 1 OS=Homo sapiens OX=9606 GN=IGHA1 PE=1 SV=2 | 255 | 20201119_10132_1ng_glyco_hcd203040_compound15.14162.14162.3 | controllerType=0 controllerNumber=1 scan=14162 | 30.726 |
| 12639:3 | 6 | R.LAGKPTHVN[+1751.624]VSVVMAEVDGTC[+57.021]Y.- | HexNAc(4)Hex(4)NeuAc(1) | C[+57], N[+1752] | 1366.9312 | 3 | 4098.7791 | 4098.7663 | 0 | 3.1 | 332 | CRagged | 491.66 | 481.62 | 481.62 | 10.34 | 57 | >sp\|P01876\|IGHA1_HUMAN Immunoglobulin heavy constant alpha 1 OS=Homo sapiens OX=9606 GN=IGHA1 PE=1 SV=2 | 255 | 20201119_10132_1ng_glyco_hcd203040_compound15.16076.16076.3 | controllerType=0 controllerNumber=1 scan=16076 | 34.0003 |
| 12640:3 | 6 | R.LAGKPTHVN[+1710.598]VSVVMAEVDGTC[+57.021]Y.- | HexNAc(3)Hex(5)NeuAc(1) | C[+57], N[+1711] | 1353.2563 | 3 | 4057.7543 | 4057.7397 | 0 | 3.6 | 332 | CRagged | 500.1 | 500.1 | 352.81 | 10.45 | 57 | >sp\|P01876\|IGHA1_HUMAN Immunoglobulin heavy constant alpha 1 OS=Homo sapiens OX=9606 GN=IGHA1 PE=1 SV=2 | 255 | 20201119_10132_1ng_glyco_hcd203040_compound15.16077.16077.3 | controllerType=0 controllerNumber=1 scan=16077 | 34.0023 |
| 12610:3 | 6 | R.LAGKPTHVN[+1751.624]VSVVMAEVDGTC[+57.021]Y.- | HexNAc(4)Hex(4)NeuAc(1) | C[+57], N[+1752] | 1366.9368 | 3 | 4098.7959 | 4098.7663 | 0 | 7.2 | 332 | CRagged | 544.05 | 544.05 | 544.05 | 9.98 | 57 | >sp\|P01876\|IGHA1_HUMAN Immunoglobulin heavy constant alpha 1 OS=Homo sapiens OX=9606 GN=IGHA1 PE=1 SV=2 | 255 | 20201119_10132_1ng_glyco_hcd203040_compound15.16046.16046.3 | controllerType=0 controllerNumber=1 scan=16046 | 33.9515 |
| 10879:3 | 6 | R.LAGKPTHVN[+1444.534]VSVVMAEVDGTC[+57.021]Y.- | HexNAc(4)Hex(3)Fuc(1) | C[+57], N[+1445] | 1264.5702 | 3 | 3791.6962 | 3791.6759 | 0 | 5.3 | 332 | CRagged | 552.86 | 511.11 | 511.11 | 10.22 | 57 | >sp\|P01876\|IGHA1_HUMAN Immunoglobulin heavy constant alpha 1 OS=Homo sapiens OX=9606 GN=IGHA1 PE=1 SV=2 | 255 | 20201119_10132_1ng_glyco_hcd203040_compound15.14228.14228.3 | controllerType=0 controllerNumber=1 scan=14228 | 30.8579 |
| 12637:3 | 6 | R.LAGKPTHVN[+1751.624]VSVVMAEVDGTC[+57.021]Y.- | HexNAc(4)Hex(4)NeuAc(1) | C[+57], N[+1752] | 1366.9312 | 3 | 4098.7791 | 4098.7663 | 0 | 3.1 | 332 | CRagged | 563.36 | 563.36 | 563.36 | 10.69 | 57 | >sp\|P01876\|IGHA1_HUMAN Immunoglobulin heavy constant alpha 1 OS=Homo sapiens OX=9606 GN=IGHA1 PE=1 SV=2 | 255 | 20201119_10132_1ng_glyco_hcd203040_compound15.16074.16074.3 | controllerType=0 controllerNumber=1 scan=16074 | 33.9965 |
| 12547:3 | 6 | R.LAGKPTHVN[+2262.814]VSVVMAEVDGTC[+57.021]Y.- | HexNAc(5)Hex(5)Fuc(1)NeuAc(1) | C[+57], N[+2263] | 1537.3272 | 3 | 4609.9671 | 4609.9564 | 0 | 2.3 | 332 | CRagged | 582.4 | 508.11 | 508.11 | 11.85 | 57 | >sp\|P01876\|IGHA1_HUMAN Immunoglobulin heavy constant alpha 1 OS=Homo sapiens OX=9606 GN=IGHA1 PE=1 SV=2 | 255 | 20201119_10132_1ng_glyco_hcd203040_compound15.15981.15981.3 | controllerType=0 controllerNumber=1 scan=15981 | 33.8409 |
| 12545:3 | 6 | R.LAGKPTHVN[+2262.814]VSVVMAEVDGTC[+57.021]Y.- | HexNAc(5)Hex(5)Fuc(1)NeuAc(1) | C[+57], N[+2263] | 1537.3272 | 3 | 4609.9671 | 4609.9564 | 0 | 2.3 | 332 | CRagged | 587.91 | 466.01 | 466.01 | 11.85 | 57 | >sp\|P01876\|IGHA1_HUMAN Immunoglobulin heavy constant alpha 1 OS=Homo sapiens OX=9606 GN=IGHA1 PE=1 SV=2 | 255 | 20201119_10132_1ng_glyco_hcd203040_compound15.15979.15979.3 | controllerType=0 controllerNumber=1 scan=15979 | 33.8374 |
| 10877:3 | 6 | R.LAGKPTHVN[+1444.534]VSVVMAEVDGTC[+57.021]Y.- | HexNAc(4)Hex(3)Fuc(1) | C[+57], N[+1445] | 1264.5702 | 3 | 3791.6962 | 3791.6759 | 0 | 5.3 | 332 | CRagged | 590.04 | 537.79 | 537.79 | 11.37 | 57 | >sp\|P01876\|IGHA1_HUMAN Immunoglobulin heavy constant alpha 1 OS=Homo sapiens OX=9606 GN=IGHA1 PE=1 SV=2 | 255 | 20201119_10132_1ng_glyco_hcd203040_compound15.14226.14226.3 | controllerType=0 controllerNumber=1 scan=14226 | 30.8524 |
| 12607:3 | 6 | R.LAGKPTHVN[+1751.624]VSVVMAEVDGTC[+57.021]Y.- | HexNAc(4)Hex(4)NeuAc(1) | C[+57], N[+1752] | 1366.9368 | 3 | 4098.7959 | 4098.7663 | 0 | 7.2 | 332 | CRagged | 613.87 | 543.76 | 543.76 | 12.13 | 57 | >sp\|P01876\|IGHA1_HUMAN Immunoglobulin heavy constant alpha 1 OS=Homo sapiens OX=9606 GN=IGHA1 PE=1 SV=2 | 255 | 20201119_10132_1ng_glyco_hcd203040_compound15.16043.16043.3 | controllerType=0 controllerNumber=1 scan=16043 | 33.9473 |
| 10804:3 | 6 | R.LAGKPTHVN[+1809.666]VSVVMAEVDGTC[+57.021]Y.- | HexNAc(5)Hex(4)Fuc(1) | C[+57], N[+1810] | 1386.277 | 3 | 4156.8166 | 4156.8081 | 0 | 2 | 332 | CRagged | 686.47 | 686.47 | 462.56 | 13.64 | 57 | >sp\|P01876\|IGHA1_HUMAN Immunoglobulin heavy constant alpha 1 OS=Homo sapiens OX=9606 GN=IGHA1 PE=1 SV=2 | 255 | 20201119_10132_1ng_glyco_hcd203040_compound15.14151.14151.3 | controllerType=0 controllerNumber=1 scan=14151 | 30.7042 |
| 10806:3 | 6 | R.LAGKPTHVN[+1809.666]VSVVMAEVDGTC[+57.021]Y.- | HexNAc(5)Hex(4)Fuc(1) | C[+57], N[+1810] | 1386.277 | 3 | 4156.8166 | 4156.8081 | 0 | 2 | 332 | CRagged | 782.15 | 732.09 | 512.76 | 15.64 | 57 | >sp\|P01876\|IGHA1_HUMAN Immunoglobulin heavy constant alpha 1 OS=Homo sapiens OX=9606 GN=IGHA1 PE=1 SV=2 | 255 | 20201119_10132_1ng_glyco_hcd203040_compound15.14153.14153.3 | controllerType=0 controllerNumber=1 scan=14153 | 30.7114 |
| 10805:3 | 6 | R.LAGKPTHVN[+1216.423]VSVVMAEVDGTC[+57.021]Y.- | HexNAc(2)Hex(5) | C[+57], N[+1216] | 1188.529 | 3 | 3563.5724 | 3563.5649 | 0 | 2.1 | 332 | CRagged | 798.68 | 779.48 | 511.84 | 15.64 | 57 | >sp\|P01876\|IGHA1_HUMAN Immunoglobulin heavy constant alpha 1 OS=Homo sapiens OX=9606 GN=IGHA1 PE=1 SV=2 | 255 | 20201119_10132_1ng_glyco_hcd203040_compound15.14152.14152.3 | controllerType=0 controllerNumber=1 scan=14152 | 30.71 |
| 10667:3 | 6 | R.LAGKPTHVN[+1702.581]VSVVMAEVDGTC[+57.021]Y.- | HexNAc(2)Hex(8) | C[+57], N[+1703] | 1350.5833 | 3 | 4049.7354 | 4049.7234 | 0 | 3 | 332 | CRagged | 826.87 | 826.87 | 487.67 | 16.85 | 57 | >sp\|P01876\|IGHA1_HUMAN Immunoglobulin heavy constant alpha 1 OS=Homo sapiens OX=9606 GN=IGHA1 PE=1 SV=2 | 255 | 20201119_10132_1ng_glyco_hcd203040_compound15.14008.14008.3 | controllerType=0 controllerNumber=1 scan=14008 | 30.3902 |
| 10836:3 | 6 | R.LAGKPTHVN[+1647.613]VSVVMAEVDGTC[+57.021]Y.- | HexNAc(5)Hex(3)Fuc(1) | C[+57], N[+1648] | 1332.2618 | 3 | 3994.7708 | 3994.7553 | 0 | 3.9 | 332 | CRagged | 841.85 | 725.16 | 725.16 | 16.95 | 57 | >sp\|P01876\|IGHA1_HUMAN Immunoglobulin heavy constant alpha 1 OS=Homo sapiens OX=9606 GN=IGHA1 PE=1 SV=2 | 255 | 20201119_10132_1ng_glyco_hcd203040_compound15.14184.14184.3 | controllerType=0 controllerNumber=1 scan=14184 | 30.7712 |
| 10715:3 | 6 | R.LAGKPTHVN[+1540.529]VSVVMAEVDGTC[+57.021]Y.- | HexNAc(2)Hex(7) | C[+57], N[+1541] | 1296.5651 | 3 | 3887.6808 | 3887.6706 | 0 | 2.6 | 332 | CRagged | 843.24 | 838.66 | 547.01 | 16.85 | 57 | >sp\|P01876\|IGHA1_HUMAN Immunoglobulin heavy constant alpha 1 OS=Homo sapiens OX=9606 GN=IGHA1 PE=1 SV=2 | 255 | 20201119_10132_1ng_glyco_hcd203040_compound15.14057.14057.3 | controllerType=0 controllerNumber=1 scan=14057 | 30.4796 |
| 10814:3 | 6 | R.LAGKPTHVN[+1501.555]VSVVMAEVDGTC[+57.021]Y.- | HexNAc(5)Hex(3) | C[+57], N[+1502] | 1283.574 | 3 | 3848.7075 | 3848.6974 | 0 | 2.6 | 332 | CRagged | 851.42 | 831.26 | 831.26 | 17.85 | 57 | >sp\|P01876\|IGHA1_HUMAN Immunoglobulin heavy constant alpha 1 OS=Homo sapiens OX=9606 GN=IGHA1 PE=1 SV=2 | 255 | 20201119_10132_1ng_glyco_hcd203040_compound15.14161.14161.3 | controllerType=0 controllerNumber=1 scan=14161 | 30.7239 |
| 10756:3 | 6 | R.LAGKPTHVN[+1663.608]VSVVMAEVDGTC[+57.021]Y.- | HexNAc(5)Hex(4) | C[+57], N[+1664] | 1337.5912 | 3 | 4010.7592 | 4010.7502 | 0 | 2.2 | 332 | CRagged | 873.89 | 873.89 | 536.26 | 17.85 | 57 | >sp\|P01876\|IGHA1_HUMAN Immunoglobulin heavy constant alpha 1 OS=Homo sapiens OX=9606 GN=IGHA1 PE=1 SV=2 | 255 | 20201119_10132_1ng_glyco_hcd203040_compound15.14100.14100.3 | controllerType=0 controllerNumber=1 scan=14100 | 30.5647 |
| 10803:3 | 6 | R.LAGKPTHVN[+1216.423]VSVVMAEVDGTC[+57.021]Y.- | HexNAc(2)Hex(5) | C[+57], N[+1216] | 1188.529 | 3 | 3563.5724 | 3563.5649 | 0 | 2.1 | 332 | CRagged | 879.42 | 879.42 | 569.88 | 17.64 | 57 | >sp\|P01876\|IGHA1_HUMAN Immunoglobulin heavy constant alpha 1 OS=Homo sapiens OX=9606 GN=IGHA1 PE=1 SV=2 | 255 | 20201119_10132_1ng_glyco_hcd203040_compound15.14150.14150.3 | controllerType=0 controllerNumber=1 scan=14150 | 30.7028 |
| 10754:3 | 6 | R.LAGKPTHVN[+1663.608]VSVVMAEVDGTC[+57.021]Y.- | HexNAc(5)Hex(4) | C[+57], N[+1664] | 1337.5912 | 3 | 4010.7592 | 4010.7502 | 0 | 2.2 | 332 | CRagged | 894.38 | 757.87 | 577.77 | 17.85 | 57 | >sp\|P01876\|IGHA1_HUMAN Immunoglobulin heavy constant alpha 1 OS=Homo sapiens OX=9606 GN=IGHA1 PE=1 SV=2 | 255 | 20201119_10132_1ng_glyco_hcd203040_compound15.14098.14098.3 | controllerType=0 controllerNumber=1 scan=14098 | 30.5583 |
| 10713:3 | 6 | R.LAGKPTHVN[+1540.529]VSVVMAEVDGTC[+57.021]Y.- | HexNAc(2)Hex(7) | C[+57], N[+1541] | 1296.5651 | 3 | 3887.6808 | 3887.6706 | 0 | 2.6 | 332 | CRagged | 910.79 | 910.79 | 578.21 | 18.85 | 57 | >sp\|P01876\|IGHA1_HUMAN Immunoglobulin heavy constant alpha 1 OS=Homo sapiens OX=9606 GN=IGHA1 PE=1 SV=2 | 255 | 20201119_10132_1ng_glyco_hcd203040_compound15.14055.14055.3 | controllerType=0 controllerNumber=1 scan=14055 | 30.4722 |
| 10751:3 | 6 | R.LAGKPTHVN[+1378.476]VSVVMAEVDGTC[+57.021]Y.- | HexNAc(2)Hex(6) | C[+57], N[+1378] | 1242.5466 | 3 | 3725.6251 | 3725.6178 | 0 | 2 | 332 | CRagged | 916.18 | 875.1 | 531.28 | 18.64 | 57 | >sp\|P01876\|IGHA1_HUMAN Immunoglobulin heavy constant alpha 1 OS=Homo sapiens OX=9606 GN=IGHA1 PE=1 SV=2 | 255 | 20201119_10132_1ng_glyco_hcd203040_compound15.14095.14095.3 | controllerType=0 controllerNumber=1 scan=14095 | 30.5544 |
| 10834:3 | 6 | R.LAGKPTHVN[+1647.613]VSVVMAEVDGTC[+57.021]Y.- | HexNAc(5)Hex(3)Fuc(1) | C[+57], N[+1648] | 1332.2618 | 3 | 3994.7708 | 3994.7553 | 0 | 3.9 | 332 | CRagged | 925.78 | 925.78 | 925.78 | 18.95 | 57 | >sp\|P01876\|IGHA1_HUMAN Immunoglobulin heavy constant alpha 1 OS=Homo sapiens OX=9606 GN=IGHA1 PE=1 SV=2 | 255 | 20201119_10132_1ng_glyco_hcd203040_compound15.14182.14182.3 | controllerType=0 controllerNumber=1 scan=14182 | 30.7619 |
| 10748:3 | 6 | R.LAGKPTHVN[+1378.476]VSVVMAEVDGTC[+57.021]Y.- | HexNAc(2)Hex(6) | C[+57], N[+1378] | 1242.5466 | 3 | 3725.6251 | 3725.6178 | 0 | 2 | 332 | CRagged | 944.05 | 944.05 | 583.13 | 18.64 | 57 | >sp\|P01876\|IGHA1_HUMAN Immunoglobulin heavy constant alpha 1 OS=Homo sapiens OX=9606 GN=IGHA1 PE=1 SV=2 | 255 | 20201119_10132_1ng_glyco_hcd203040_compound15.14092.14092.3 | controllerType=0 controllerNumber=1 scan=14092 | 30.5453 |
| 06739:3 | 7 | R.FSDGLESN[+1864.634]SSTQFEVK.K | HexNAc(2)Hex(9) | N[+1865] | 1213.8234 | 3 | 3639.4557 | 3639.4472 | 0 | 2.4 | 219 | Specific | 397.19 | 397.19 | 397.19 | 9.25 | 65 | >sp\|P0C0L5\|CO4B_HUMAN Complement C4-B OS=Homo sapiens OX=9606 GN=C4B PE=1 SV=2 | 1424 | 20201119_10132_1ng_glyco_hcd203040_compound15.9914.9914.3 | controllerType=0 controllerNumber=1 scan=9914 | 23.4518 |
| 04266:3 | 7 | R.FSDGLESN[+1864.634]SSTQFEVKK.Y | HexNAc(2)Hex(9) | N[+1865] | 1256.5222 | 3 | 3767.5519 | 3767.5421 | 0 | 2.6 | 219 | Specific | 315.09 | 305.66 | 305.66 | 6.78 | 65 | >sp\|P0C0L5\|CO4B_HUMAN Complement C4-B OS=Homo sapiens OX=9606 GN=C4B PE=1 SV=2 | 1424 | 20201119_10132_1ng_glyco_hcd203040_compound15.7351.7351.3 | controllerType=0 controllerNumber=1 scan=7351 | 19.1412 |
| 09387:3 | 8 | K.TVLTPATNHMGN[+1540.529]VTFTIPANR.E | HexNAc(2)Hex(7) | N[+1541] | 1265.9048 | 3 | 3795.7 | 3795.6886 | 0 | 3 | 74 | Specific | 204.59 | 200.39 | 100.04 | 5.25 | 75 | >sp\|P01024\|CO3_HUMAN Complement C3 OS=Homo sapiens OX=9606 GN=C3 PE=1 SV=2 | 492 | 20201119_10132_1ng_glyco_hcd203040_compound15.12683.12683.3 | controllerType=0 controllerNumber=1 scan=12683 | 28.1502 |
| 09389:3 | 8 | K.TVLTPATNHMGN[+1540.529]VTFTIPANR.E | HexNAc(2)Hex(7) | N[+1541] | 1265.9048 | 3 | 3795.7 | 3795.6886 | 0 | 3 | 74 | Specific | 359.89 | 336.1 | 163.98 | 5.95 | 75 | >sp\|P01024\|CO3_HUMAN Complement C3 OS=Homo sapiens OX=9606 GN=C3 PE=1 SV=2 | 492 | 20201119_10132_1ng_glyco_hcd203040_compound15.12685.12685.3 | controllerType=0 controllerNumber=1 scan=12685 | 28.1539 |
| 18361:3 | 9 | K.QIGLYPVLVIDSSGYVNPN[+1913.677]YTGR.I | HexNAc(4)Hex(5)NeuAc(1) | N[+1914] | 1480.3397 | 3 | 4439.0044 | 4438.9805 | 0 | 5.4 | 168 | Specific | 395.58 | 368.79 | 368.79 | 8.06 | 55 | >sp\|P01833\|PIGR_HUMAN Polymeric immunoglobulin receptor OS=Homo sapiens OX=9606 GN=PIGR PE=1 SV=4 | 902 | 20201119_10132_1ng_glyco_hcd203040_compound15.22071.22071.3 | controllerType=0 controllerNumber=1 scan=22071 | 45.4705 |
| 18363:3 | 9 | K.QIGLYPVLVIDSSGYVNPN[+1913.677]YTGR.I | HexNAc(4)Hex(5)NeuAc(1) | N[+1914] | 1480.3397 | 3 | 4439.0044 | 4438.9805 | 0 | 5.4 | 168 | Specific | 563.56 | 506.39 | 506.39 | 10.14 | 55 | >sp\|P01833\|PIGR_HUMAN Polymeric immunoglobulin receptor OS=Homo sapiens OX=9606 GN=PIGR PE=1 SV=4 | 902 | 20201119_10132_1ng_glyco_hcd203040_compound15.22073.22073.3 | controllerType=0 controllerNumber=1 scan=22073 | 45.4751 |
| 22450:3 | 9 | R.LSLLEEPGN[+2351.851]GTFTVILNQLTSR.D | HexNAc(4)Hex(5)Fuc(3)NeuAc(1) | N[+2352] | 1585.3887 | 3 | 4754.1515 | 4754.1433 | 0 | 1.7 | 413 | Specific | 291.69 | 174.32 | 174.32 | 6.69 | 55 | >sp\|P01833\|PIGR_HUMAN Polymeric immunoglobulin receptor OS=Homo sapiens OX=9606 GN=PIGR PE=1 SV=4 | 902 | 20201119_10132_1ng_glyco_hcd203040_compound15.26339.26339.3 | controllerType=0 controllerNumber=1 scan=26339 | 53.7057 |
| 19861:4 | 9 | R.LSLLEEPGN[+2205.793]GTFTVILNQLTSR.D | HexNAc(4)Hex(5)Fuc(2)NeuAc(1) | N[+2206] | 1152.7798 | 4 | 4608.0975 | 4608.0854 | 0 | 2.6 | 413 | Specific | 796.12 | 644.03 | 644.03 | 15.91 | 55 | >sp\|P01833\|PIGR_HUMAN Polymeric immunoglobulin receptor OS=Homo sapiens OX=9606 GN=PIGR PE=1 SV=4 | 902 | 20201119_10132_1ng_glyco_hcd203040_compound15.23642.23642.3 | controllerType=0 controllerNumber=1 scan=23642 | 48.738 |
| 19864:4 | 9 | R.LSLLEEPGN[+2205.793]GTFTVILNQLTSR.D | HexNAc(4)Hex(5)Fuc(2)NeuAc(1) | N[+2206] | 1152.7798 | 4 | 4608.0975 | 4608.0854 | 0 | 2.6 | 413 | Specific | 834.37 | 633.44 | 633.44 | 16.91 | 55 | >sp\|P01833\|PIGR_HUMAN Polymeric immunoglobulin receptor OS=Homo sapiens OX=9606 GN=PIGR PE=1 SV=4 | 902 | 20201119_10132_1ng_glyco_hcd203040_compound15.23645.23645.3 | controllerType=0 controllerNumber=1 scan=23645 | 48.7464 |
| 22261:4 | 9 | R.LSLLEEPGN[+2350.830]GTFTVILNQLTSR.D | HexNAc(4)Hex(5)Fuc(1)NeuAc(2) | N[+2351] | 1189.0426 | 4 | 4753.1488 | 4753.1229 | 0 | 5.4 | 413 | Specific | 879.65 | 682.14 | 682.14 | 17.44 | 55 | >sp\|P01833\|PIGR_HUMAN Polymeric immunoglobulin receptor OS=Homo sapiens OX=9606 GN=PIGR PE=1 SV=4 | 902 | 20201119_10132_1ng_glyco_hcd203040_compound15.26142.26142.4 | controllerType=0 controllerNumber=1 scan=26142 | 53.3145 |
| 22262:4 | 9 | R.LSLLEEPGN[+2350.830]GTFTVILNQLTSR.D | HexNAc(4)Hex(5)Fuc(1)NeuAc(2) | N[+2351] | 1189.0426 | 4 | 4753.1488 | 4753.1229 | 0 | 5.4 | 413 | Specific | 896.63 | 687.33 | 687.33 | 17.44 | 55 | >sp\|P01833\|PIGR_HUMAN Polymeric immunoglobulin receptor OS=Homo sapiens OX=9606 GN=PIGR PE=1 SV=4 | 902 | 20201119_10132_1ng_glyco_hcd203040_compound15.26143.26143.4 | controllerType=0 controllerNumber=1 scan=26143 | 53.3197 |
| 19884:4 | 9 | R.LSLLEEPGN[+2205.793]GTFTVILNQLTSR.D | HexNAc(4)Hex(5)Fuc(2)NeuAc(1) | N[+2206] | 1152.7812 | 4 | 4608.1029 | 4608.0854 | 0 | 3.8 | 413 | Specific | 902.34 | 718.98 | 718.98 | 19.02 | 55 | >sp\|P01833\|PIGR_HUMAN Polymeric immunoglobulin receptor OS=Homo sapiens OX=9606 GN=PIGR PE=1 SV=4 | 902 | 20201119_10132_1ng_glyco_hcd203040_compound15.23666.23666.4 | controllerType=0 controllerNumber=1 scan=23666 | 48.7877 |
| 19887:4 | 9 | R.LSLLEEPGN[+2205.793]GTFTVILNQLTSR.D | HexNAc(4)Hex(5)Fuc(2)NeuAc(1) | N[+2206] | 1152.7812 | 4 | 4608.1029 | 4608.0854 | 0 | 3.8 | 413 | Specific | 942.88 | 728.06 | 728.06 | 19.02 | 55 | >sp\|P01833\|PIGR_HUMAN Polymeric immunoglobulin receptor OS=Homo sapiens OX=9606 GN=PIGR PE=1 SV=4 | 902 | 20201119_10132_1ng_glyco_hcd203040_compound15.23669.23669.4 | controllerType=0 controllerNumber=1 scan=23669 | 48.7928 |
| 19881:3 | 9 | R.LSLLEEPGN[+2205.793]GTFTVILNQLTSR.D | HexNAc(4)Hex(5)Fuc(2)NeuAc(1) | N[+2206] | 1536.7061 | 3 | 4608.1038 | 4608.0854 | 0 | 4 | 413 | Specific | 965.03 | 693.63 | 693.63 | 19.87 | 55 | >sp\|P01833\|PIGR_HUMAN Polymeric immunoglobulin receptor OS=Homo sapiens OX=9606 GN=PIGR PE=1 SV=4 | 902 | 20201119_10132_1ng_glyco_hcd203040_compound15.23663.23663.3 | controllerType=0 controllerNumber=1 scan=23663 | 48.7841 |
| 19948:3 | 9 | R.LSLLEEPGN[+2059.735]GTFTVILNQLTSR.D | HexNAc(4)Hex(5)Fuc(1)NeuAc(1) | N[+2060] | 1488.02 | 3 | 4462.0453 | 4462.0275 | 0 | 4 | 413 | Specific | 992.53 | 740.89 | 740.89 | 19.87 | 55 | >sp\|P01833\|PIGR_HUMAN Polymeric immunoglobulin receptor OS=Homo sapiens OX=9606 GN=PIGR PE=1 SV=4 | 902 | 20201119_10132_1ng_glyco_hcd203040_compound15.23734.23734.3 | controllerType=0 controllerNumber=1 scan=23734 | 48.9512 |
| 19946:3 | 9 | R.LSLLEEPGN[+2059.735]GTFTVILNQLTSR.D | HexNAc(4)Hex(5)Fuc(1)NeuAc(1) | N[+2060] | 1488.02 | 3 | 4462.0453 | 4462.0275 | 0 | 4 | 413 | Specific | 1048.41 | 794.69 | 794.69 | 20.87 | 55 | >sp\|P01833\|PIGR_HUMAN Polymeric immunoglobulin receptor OS=Homo sapiens OX=9606 GN=PIGR PE=1 SV=4 | 902 | 20201119_10132_1ng_glyco_hcd203040_compound15.23732.23732.3 | controllerType=0 controllerNumber=1 scan=23732 | 48.9446 |
| 19879:3 | 9 | R.LSLLEEPGN[+2205.793]GTFTVILNQLTSR.D | HexNAc(4)Hex(5)Fuc(2)NeuAc(1) | N[+2206] | 1536.7061 | 3 | 4608.1038 | 4608.0854 | 0 | 4 | 413 | Specific | 1071.61 | 828.71 | 828.71 | 21.87 | 55 | >sp\|P01833\|PIGR_HUMAN Polymeric immunoglobulin receptor OS=Homo sapiens OX=9606 GN=PIGR PE=1 SV=4 | 902 | 20201119_10132_1ng_glyco_hcd203040_compound15.23661.23661.3 | controllerType=0 controllerNumber=1 scan=23661 | 48.7788 |
| 19921:4 | 9 | R.LSLLEEPGN[+2059.735]GTFTVILNQLTSR.D | HexNAc(4)Hex(5)Fuc(1)NeuAc(1) | N[+2060] | 1116.2642 | 4 | 4462.035 | 4462.0275 | 0 | 1.7 | 413 | Specific | 1090.13 | 857.25 | 857.25 | 21.7 | 55 | >sp\|P01833\|PIGR_HUMAN Polymeric immunoglobulin receptor OS=Homo sapiens OX=9606 GN=PIGR PE=1 SV=4 | 902 | 20201119_10132_1ng_glyco_hcd203040_compound15.23704.23704.4 | controllerType=0 controllerNumber=1 scan=23704 | 48.8697 |
| 15381:3 | 9 | K.IIEGEPNLKVPGN[+2204.772]VTAVLGETLK.V | HexNAc(4)Hex(5)NeuAc(2) | N[+2205] | 1532.7211 | 3 | 4596.1488 | 4596.1218 | 0 | 5.9 | 457 | Specific | 514.39 | 424.42 | 424.42 | 9.53 | 55 | >sp\|P01833\|PIGR_HUMAN Polymeric immunoglobulin receptor OS=Homo sapiens OX=9606 GN=PIGR PE=1 SV=4 | 902 | 20201119_10132_1ng_glyco_hcd203040_compound15.18959.18959.3 | controllerType=0 controllerNumber=1 scan=18959 | [illegible] |
| 15384:3 | 9 | K.IIEGEPNLKVPGN[+2204.772]VTAVLGETLK.V | HexNAc(4)Hex(5)NeuAc(2) | N[+2205] | 1532.7211 | 3 | 4596.1488 | 4596.1218 | 0 | 5.9 | 457 | Specific | 609.71 | 546.35 | 546.35 | 11.88 | 55 | >sp\|P01833\|PIGR_HUMAN Polymeric immunoglobulin receptor OS=Homo sapiens OX=9606 GN=PIGR PE=1 SV=4 | 902 | 20201119_10132_1ng_glyco_hcd203040_compound15.18962.18962.3 | controllerType=0 controllerNumber=1 scan=18962 | [illegible] |
| 16489:5 | 9 | K.IIEGEPNLKVPGN[+2350.830]VTAVLGETLKVPC[+57.021]HFPC[+57.021]K.F | HexNAc(4)Hex(5)Fuc(1)NeuAc(2) | C[+57]*2, N[+2351] | 1154.3379 | 5 | 5767.6604 | 5767.6372 | 0 | 4 | 457 | Specific | 314.56 | 253.7 | 253.7 | 7.19 | 55 | >sp\|P01833\|PIGR_HUMAN Polymeric immunoglobulin receptor OS=Homo sapiens OX=9606 GN=PIGR PE=1 SV=4 | 902 | 20201119_10132_1ng_glyco_hcd203040_compound15.20116.20116.5 | controllerType=0 controllerNumber=1 scan=20116 | [illegible] |
| 15012:5 | 9 | K.IIEGEPNLKVPGN[+2205.793]VTAVLGETLKVPC[+57.021]HFPC[+57.021]K.F | HexNAc(4)Hex(5)Fuc(2)NeuAc(1) | C[+57]*2, N[+2206] | 1125.337 | 5 | 5622.6559 | 5622.5997 | 0 | 10 | 457 | Specific | 380.31 | 380.31 | 380.31 | 8.08 | 55 | >sp\|P01833\|PIGR_HUMAN Polymeric immunoglobulin receptor OS=Homo sapiens OX=9606 GN=PIGR PE=1 SV=4 | 902 | 20201119_10132_1ng_glyco_hcd203040_compound15.18541.18541.5 | controllerType=0 controllerNumber=1 scan=18541 | 38.4093 |
| 13697:5 | 9 | K.IIEGEPNLKVPGN[+2059.735]VTAVLGETLKVPC[+57.021]HFPC[+57.021]K.F | HexNAc(4)Hex(5)Fuc(1)NeuAc(1) | C[+57]*2, N[+2060] | 1096.1207 | 5 | 5476.5745 | 5476.5418 | 0 | 6 | 457 | Specific | 424.38 | 424.38 | 424.38 | 7.66 | 55 | >sp\|P01833\|PIGR_HUMAN Polymeric immunoglobulin receptor OS=Homo sapiens OX=9606 GN=PIGR PE=1 SV=4 | 902 | 20201119_10132_1ng_glyco_hcd203040_compound15.17175.17175.5 | controllerType=0 controllerNumber=1 scan=17175 | 35.9966 |
| 15010:5 | 9 | K.IIEGEPNLKVPGN[+2205.793]VTAVLGETLKVPC[+57.021]HFPC[+57.021]K.F | HexNAc(4)Hex(5)Fuc(2)NeuAc(1) | C[+57]*2, N[+2206] | 1125.337 | 5 | 5622.6559 | 5622.5997 | 0 | 10 | 457 | Specific | 455.42 | 455.42 | 455.42 | 8.2 | 55 | >sp\|P01833\|PIGR_HUMAN Polymeric immunoglobulin receptor OS=Homo sapiens OX=9606 GN=PIGR PE=1 SV=4 | 902 | 20201119_10132_1ng_glyco_hcd203040_compound15.18539.18539.5 | controllerType=0 controllerNumber=1 scan=18539 | 38.4057 |
| 15013:4 | 9 | K.IIEGEPNLKVPGN[+2205.793]VTAVLGETLKVPC[+57.021]HFPC[+57.021]K.F | HexNAc(4)Hex(5)Fuc(2)NeuAc(1) | C[+57]*2, N[+2206] | 1406.4194 | 4 | 5622.6558 | 5622.5997 | 0 | 10 | 457 | Specific | 546.94 | 524.22 | 524.22 | 8.02 | 55 | >sp\|P01833\|PIGR_HUMAN Polymeric immunoglobulin receptor OS=Homo sapiens OX=9606 GN=PIGR PE=1 SV=4 | 902 | 20201119_10132_1ng_glyco_hcd203040_compound15.18542.18542.4 | controllerType=0 controllerNumber=1 scan=18542 | 38.4123 |
| 14990:5 | 9 | K.IIEGEPNLKVPGN[+2059.735]VTAVLGETLKVPC[+57.021]HFPC[+57.021]K.F | HexNAc(4)Hex(5)Fuc(1)NeuAc(1) | C[+57]*2, N[+2060] | 1096.124 | 5 | 5476.591 | 5476.5418 | 0 | 9 | 457 | Specific | 551.31 | 551.31 | 551.31 | 9.15 | 55 | >sp\|P01833\|PIGR_HUMAN Polymeric immunoglobulin receptor OS=Homo sapiens OX=9606 GN=PIGR PE=1 SV=4 | 902 | 20201119_10132_1ng_glyco_hcd203040_compound15.18518.18518.5 | controllerType=0 controllerNumber=1 scan=18518 | 38.3594 |
| 14988:5 | 9 | K.IIEGEPNLKVPGN[+2059.735]VTAVLGETLKVPC[+57.021]HFPC[+57.021]K.F | HexNAc(4)Hex(5)Fuc(1)NeuAc(1) | C[+57]*2, N[+2060] | 1096.124 | 5 | 5476.591 | 5476.5418 | 0 | 9 | 457 | Specific | 557.86 | 503.26 | 503.26 | 9.15 | 55 | >sp\|P01833\|PIGR_HUMAN Polymeric immunoglobulin receptor OS=Homo sapiens OX=9606 GN=PIGR PE=1 SV=4 | 902 | 20201119_10132_1ng_glyco_hcd203040_compound15.18516.18516.5 | controllerType=0 controllerNumber=1 scan=18516 | 38.3549 |
| 13695:5 | 9 | K.IIEGEPNLKVPGN[+2059.735]VTAVLGETLKVPC[+57.021]HFPC[+57.021]K.F | HexNAc(4)Hex(5)Fuc(1)NeuAc(1) | C[+57]*2, N[+2060] | 1096.1207 | 5 | 5476.5745 | 5476.5418 | 0 | 6 | 457 | Specific | 657.98 | 650.19 | 650.19 | 12.73 | 55 | >sp\|P01833\|PIGR_HUMAN Polymeric immunoglobulin receptor OS=Homo sapiens OX=9606 GN=PIGR PE=1 SV=4 | 902 | 20201119_10132_1ng_glyco_hcd203040_compound15.17173.17173.5 | controllerType=0 controllerNumber=1 scan=17173 | 35.9863 |
| 15011:4 | 9 | K.IIEGEPNLKVPGN[+2205.793]VTAVLGETLKVPC[+57.021]HFPC[+57.021]K.F | HexNAc(4)Hex(5)Fuc(2)NeuAc(1) | C[+57]*2, N[+2206] | 1406.4194 | 4 | 5622.6558 | 5622.5997 | 0 | 10 | 457 | Specific | 679.73 | 679.73 | 679.73 | 11.17 | 55 | >sp\|P01833\|PIGR_HUMAN Polymeric immunoglobulin receptor OS=Homo sapiens OX=9606 GN=PIGR PE=1 SV=4 | 902 | 20201119_10132_1ng_glyco_hcd203040_compound15.18540.18540.4 | controllerType=0 controllerNumber=1 scan=18540 | 38.4072 |
| 14998:4 | 9 | K.IIEGEPNLKVPGN[+2059.735]VTAVLGETLKVPC[+57.021]HFPC[+57.021]K.F | HexNAc(4)Hex(5)Fuc(1)NeuAc(1) | C[+57]*2, N[+2060] | 1369.8984 | 4 | 5476.5719 | 5476.5418 | 0 | 5.5 | 457 | Specific | 686.3 | 686.3 | 686.3 | 11.95 | 55 | >sp\|P01833\|PIGR_HUMAN Polymeric immunoglobulin receptor OS=Homo sapiens OX=9606 GN=PIGR PE=1 SV=4 | 902 | 20201119_10132_1ng_glyco_hcd203040_compound15.18526.18526.4 | controllerType=0 controllerNumber=1 scan=18526 | 38.3842 |
| 15000:4 | 9 | K.IIEGEPNLKVPGN[+2059.735]VTAVLGETLKVPC[+57.021]HFPC[+57.021]K.F | HexNAc(4)Hex(5)Fuc(1)NeuAc(1) | C[+57]*2, N[+2060] | 1369.8984 | 4 | 5476.5719 | 5476.5418 | 0 | 5.5 | 457 | Specific | 690.65 | 627.81 | 627.81 | 11.95 | 55 | >sp\|P01833\|PIGR_HUMAN Polymeric immunoglobulin receptor OS=Homo sapiens OX=9606 GN=PIGR PE=1 SV=4 | 902 | 20201119_10132_1ng_glyco_hcd203040_compound15.18528.18528.4 | controllerType=0 controllerNumber=1 scan=18528 | 38.3884 |
| 14967:4 | 9 | K.IIEGEPNLKVPGN[+2205.793]VTAVLGETLKVPC[+57.021]HFPC[+57.021]K.F | HexNAc(4)Hex(5)Fuc(2)NeuAc(1) | C[+57]*2, N[+2206] | 1406.4175 | 4 | 5622.6481 | 5622.5997 | 0 | 8.6 | 457 | Specific | 852.23 | 852.23 | 852.23 | 15.52 | 55 | >sp\|P01833\|PIGR_HUMAN Polymeric immunoglobulin receptor OS=Homo sapiens OX=9606 GN=PIGR PE=1 SV=4 | 902 | 20201119_10132_1ng_glyco_hcd203040_compound15.18494.18494.4 | controllerType=0 controllerNumber=1 scan=18494 | 38.3103 |
| 14965:4 | 9 | K.IIEGEPNLKVPGN[+2205.793]VTAVLGETLKVPC[+57.021]HFPC[+57.021]K.F | HexNAc(4)Hex(5)Fuc(2)NeuAc(1) | C[+57]*2, N[+2206] | 1406.4175 | 4 | 5622.6481 | 5622.5997 | 0 | 8.6 | 457 | Specific | 1002.49 | 981.47 | 981.47 | 18.52 | 55 | >sp\|P01833\|PIGR_HUMAN Polymeric immunoglobulin receptor OS=Homo sapiens OX=9606 GN=PIGR PE=1 SV=4 | 902 | 20201119_10132_1ng_glyco_hcd203040_compound15.18492.18492.4 | controllerType=0 controllerNumber=1 scan=18492 | 38.3054 |
| 13016:3 | 9 | K.VPGN[+2059.735]VTAVLGETLK.V | HexNAc(4)Hex(5)Fuc(1)NeuAc(1) | N[+2060] | 1153.1857 | 3 | 3457.5426 | 3457.5348 | 0 | 2.3 | 466 | Specific | 316.23 | 159.98 | 159.98 | 6.12 | 55 | >sp\|P01833\|PIGR_HUMAN Polymeric immunoglobulin receptor OS=Homo sapiens OX=9606 GN=PIGR PE=1 SV=4 | 902 | 20201119_10132_1ng_glyco_hcd203040_compound15.16467.16467.3 | controllerType=0 controllerNumber=1 scan=16467 | 34.7228 |
| 13014:3 | 9 | K.VPGN[+2059.735]VTAVLGETLK.V | HexNAc(4)Hex(5)Fuc(1)NeuAc(1) | N[+2060] | 1153.1857 | 3 | 3457.5426 | 3457.5348 | 0 | 2.3 | 466 | Specific | 478.84 | 464.46 | 464.46 | 9.58 | 55 | >sp\|P01833\|PIGR_HUMAN Polymeric immunoglobulin receptor OS=Homo sapiens OX=9606 GN=PIGR PE=1 SV=4 | 902 | 20201119_10132_1ng_glyco_hcd203040_compound15.16465.16465.3 | controllerType=0 controllerNumber=1 scan=16465 | 34.7191 |
| 12869:3 | 9 | K.VPGN[+2205.793]VTAVLGETLK.V | HexNAc(4)Hex(5)Fuc(2)NeuAc(1) | N[+2206] | 1201.8719 | 3 | 3603.6011 | 3603.5927 | 0 | 2.3 | 466 | Specific | 618.52 | 530.24 | 530.24 | 12.11 | 55 | >sp\|P01833\|PIGR_HUMAN Polymeric immunoglobulin receptor OS=Homo sapiens OX=9606 GN=PIGR PE=1 SV=4 | 902 | 20201119_10132_1ng_glyco_hcd203040_compound15.16314.16314.3 | controllerType=0 controllerNumber=1 scan=16314 | 34.4171 |
| 12867:3 | 9 | K.VPGN[+2205.793]VTAVLGETLK.V | HexNAc(4)Hex(5)Fuc(2)NeuAc(1) | N[+2206] | 1201.8719 | 3 | 3603.6011 | 3603.5927 | 0 | 2.3 | 466 | Specific | 631.78 | 602.75 | 602.75 | 12.11 | 55 | >sp\|P01833\|PIGR_HUMAN Polymeric immunoglobulin receptor OS=Homo sapiens OX=9606 GN=PIGR PE=1 SV=4 | 902 | 20201119_10132_1ng_glyco_hcd203040_compound15.16312.16312.3 | controllerType=0 controllerNumber=1 scan=16312 | 34.4129 |
| 12952:4 | 9 | K.VPGN[+2205.793]VTAVLGETLKVPC[+57.021]HFPC[+57.021]K.F | HexNAc(4)Hex(5)Fuc(2)NeuAc(1) | C[+57]*2, N[+2206] | 1158.0213 | 4 | 4629.0633 | 4629.0502 | 0 | 2.8 | 466 | Specific | 426.29 | 240.37 | 240.37 | 7.59 | 55 | >sp\|P01833\|PIGR_HUMAN Polymeric immunoglobulin receptor OS=Homo sapiens OX=9606 GN=PIGR PE=1 SV=4 | 902 | 20201119_10132_1ng_glyco_hcd203040_compound15.16400.16400.3 | controllerType=0 controllerNumber=1 scan=16400 | 34.5747 |
| 12960:4 | 9 | K.VPGN[+2205.793]VTAVLGETLKVPC[+57.021]HFPC[+57.021]K.F | HexNAc(4)Hex(5)Fuc(2)NeuAc(1) | C[+57]*2, N[+2206] | 1158.0213 | 4 | 4629.0633 | 4629.0502 | 0 | 2.8 | 466 | Specific | 553.26 | 376.25 | 376.25 | 10.2 | 55 | >sp\|P01833\|PIGR_HUMAN Polymeric immunoglobulin receptor OS=Homo sapiens OX=9606 GN=PIGR PE=1 SV=4 | 902 | 20201119_10132_1ng_glyco_hcd203040_compound15.16408.16408.3 | controllerType=0 controllerNumber=1 scan=16408 | 34.5901 |
| 12950:4 | 9 | K.VPGN[+2205.793]VTAVLGETLKVPC[+57.021]HFPC[+57.021]K.F | HexNAc(4)Hex(5)Fuc(2)NeuAc(1) | C[+57]*2, N[+2206] | 1158.0213 | 4 | 4629.0633 | 4629.0502 | 0 | 2.8 | 466 | Specific | 679.49 | 448.75 | 448.75 | 13.35 | 55 | >sp\|P01833\|PIGR_HUMAN Polymeric immunoglobulin receptor OS=Homo sapiens OX=9606 GN=PIGR PE=1 SV=4 | 902 | 20201119_10132_1ng_glyco_hcd203040_compound15.16398.16398.3 | controllerType=0 controllerNumber=1 scan=16398 | 34.566 |
| 12958:4 | 9 | K.VPGN[+2205.793]VTAVLGETLKVPC[+57.021]HFPC[+57.021]K.F | HexNAc(4)Hex(5)Fuc(2)NeuAc(1) | C[+57]*2, N[+2206] | 1158.0213 | 4 | 4629.0633 | 4629.0502 | 0 | 2.8 | 466 | Specific | 716.16 | 474.17 | 474.17 | 14.35 | 55 | >sp\|P01833\|PIGR_HUMAN Polymeric immunoglobulin receptor OS=Homo sapiens OX=9606 GN=PIGR PE=1 SV=4 | 902 | 20201119_10132_1ng_glyco_hcd203040_compound15.16406.16406.3 | controllerType=0 controllerNumber=1 scan=16406 | 34.5835 |
| 03509:3 | 9 | K.WN[+1768.640]NTGC[+57.021]QALPSQDEGPSK.A | HexNAc(4)Hex(5)Fuc(1) | C[+57], N[+1769] | 1253.18 | 3 | 3757.5254 | 3757.5162 | 0 | 2.5 | 498 | Specific | 262.93 | 247.24 | 247.24 | 6.78 | 55 | >sp\|P01833\|PIGR_HUMAN Polymeric immunoglobulin receptor OS=Homo sapiens OX=9606 GN=PIGR PE=1 SV=4 | 902 | 20201119_10132_1ng_glyco_hcd203040_compound15.6564.6564.3 | controllerType=0 controllerNumber=1 scan=6564 | 17.8223 |
| 05261:3 | 9 | K.WN[+1913.677]NTGC[+57.021]QALPSQDEGPSK.A | HexNAc(4)Hex(5)NeuAc(1) | C[+57], N[+1914] | 1301.527 | 3 | 3902.5666 | 3902.5537 | 0 | 3.3 | 498 | Specific | 266.59 | 237.12 | 237.12 | 6.78 | 55 | >sp\|P01833\|PIGR_HUMAN Polymeric immunoglobulin receptor OS=Homo sapiens OX=9606 GN=PIGR PE=1 SV=4 | 902 | 20201119_10132_1ng_glyco_hcd203040_compound15.8381.8381.3 | controllerType=0 controllerNumber=1 scan=8381 | 20.8676 |
| 05263:3 | 9 | K.WN[+1913.677]NTGC[+57.021]QALPSQDEGPSK.A | HexNAc(4)Hex(5)NeuAc(1) | C[+57], N[+1914] | 1301.527 | 3 | 3902.5666 | 3902.5537 | 0 | 3.3 | 498 | Specific | 334.27 | 290.06 | 290.06 | 7.38 | 55 | >sp\|P01833\|PIGR_HUMAN Polymeric immunoglobulin receptor OS=Homo sapiens OX=9606 GN=PIGR PE=1 SV=4 | 902 | 20201119_10132_1ng_glyco_hcd203040_compound15.8383.8383.3 | controllerType=0 controllerNumber=1 scan=8383 | 20.8716 |
| 05142:3 | 9 | K.WN[+2059.735]NTGC[+57.021]QALPSQDEGPSK.A | HexNAc(4)Hex(5)Fuc(1)NeuAc(1) | C[+57], N[+2060] | 1350.2127 | 3 | 4048.6236 | 4048.6116 | 0 | 3 | 498 | Specific | 538.95 | 482.23 | 482.23 | 9.95 | 55 | >sp\|P01833\|PIGR_HUMAN Polymeric immunoglobulin receptor OS=Homo sapiens OX=9606 GN=PIGR PE=1 SV=4 | 902 | 20201119_10132_1ng_glyco_hcd203040_compound15.8258.8258.3 | controllerType=0 controllerNumber=1 scan=8258 | 20.6618 |
| 05140:3 | 9 | K.WN[+2059.735]NTGC[+57.021]QALPSQDEGPSK.A | HexNAc(4)Hex(5)Fuc(1)NeuAc(1) | C[+57], N[+2060] | 1350.2127 | 3 | 4048.6236 | 4048.6116 | 0 | 3 | 498 | Specific | 552.99 | 490.21 | 490.21 | 10.15 | 55 | >sp\|P01833\|PIGR_HUMAN Polymeric immunoglobulin receptor OS=Homo sapiens OX=9606 GN=PIGR PE=1 SV=4 | 902 | 20201119_10132_1ng_glyco_hcd203040_compound15.8256.8256.3 | controllerType=0 controllerNumber=1 scan=8256 | 20.657 |
| 16593:4 | 10 | K.RNIAAFGGDPNN[+1913.677]ITLFGESAGGASVSLQTLSPYNK.G | HexNAc(4)Hex(5)NeuAc(1) | N[+1914] | 1370.8782 | 4 | 5480.4909 | 5480.4644 | 0 | 4.8 | 196 | Specific | 204.8 | 194.87 | 194.87 | 5.96 | 44 | >sp\|P19835\|CEL_HUMAN Bile salt-activated lipase OS=Homo sapiens OX=9606 GN=CEL PE=1 SV=3 | 476 | 20201119_10132_1ng_glyco_hcd203040_compound15.20224.20224.2 | controllerType=0 controllerNumber=1 scan=20224 | 41.5234 |
| 05085:3 | 11 | R.LSPN[+2351.851]ASAEHLELR.W | HexNAc(4)Hex(5)Fuc(3)NeuAc(1) | N[+2352] | 1263.5444 | 3 | 3788.6186 | 3788.6 | 0 | 4.9 | 52 | Specific | 256.77 | 256.77 | 256.77 | 6.07 | 42 | >sp\|Q13410\|BT1A1_HUMAN Butyrophilin subfamily 1 member A1 OS=Homo sapiens OX=9606 GN=BTN1A1 PE=1 SV=3 | 179 | 20201119_10132_1ng_glyco_hcd203040_compound15.8199.8199.3 | controllerType=0 controllerNumber=1 scan=8199 | 20.5621 |
| 07109:3 | 11 | R.LSPN[+2350.830]ASAEHLELR.W | HexNAc(4)Hex(5)Fuc(1)NeuAc(2) | N[+2351] | 1263.2021 | 3 | 3787.5917 | 3787.5796 | 0 | 3.2 | 52 | Specific | 468.46 | 404.11 | 404.11 | 9.66 | 42 | >sp\|Q13410\|BT1A1_HUMAN Butyrophilin subfamily 1 member A1 OS=Homo sapiens OX=9606 GN=BTN1A1 PE=1 SV=3 | 179 | 20201119_10132_1ng_glyco_hcd203040_compound15.10298.10298.3 | controllerType=0 controllerNumber=1 scan=10298 | 24.0973 |
| 07081:3 | 11 | R.LSPN[+2350.830]ASAEHLELR.W | HexNAc(4)Hex(5)Fuc(1)NeuAc(2) | N[+2351] | 1263.2009 | 3 | 3787.588 | 3787.5796 | 0 | 2.2 | 52 | Specific | 564.34 | 477.77 | 477.77 | 10.03 | 42 | >sp\|Q13410\|BT1A1_HUMAN Butyrophilin subfamily 1 member A1 OS=Homo sapiens OX=9606 GN=BTN1A1 PE=1 SV=3 | 179 | 20201119_10132_1ng_glyco_hcd203040_compound15.10269.10269.3 | controllerType=0 controllerNumber=1 scan=10269 | 24.0487 |
| 05082:3 | 11 | R.LSPN[+2205.793]ASAEHLELR.W | HexNAc(4)Hex(5)Fuc(2)NeuAc(1) | N[+2206] | 1214.8574 | 3 | 3642.5575 | 3642.5421 | 0 | 4.2 | 52 | Specific | 565.78 | 487.59 | 487.59 | 10.14 | 42 | >sp\|Q13410\|BT1A1_HUMAN Butyrophilin subfamily 1 member A1 OS=Homo sapiens OX=9606 GN=BTN1A1 PE=1 SV=3 | 179 | 20201119_10132_1ng_glyco_hcd203040_compound15.8196.8196.3 | controllerType=0 controllerNumber=1 scan=8196 | 20.5548 |
| 07107:3 | 11 | R.LSPN[+2350.830]ASAEHLELR.W | HexNAc(4)Hex(5)Fuc(1)NeuAc(2) | N[+2351] | 1263.2021 | 3 | 3787.5917 | 3787.5796 | 0 | 3.2 | 52 | Specific | 567.77 | 517.29 | 517.29 | 10.03 | 42 | >sp\|Q13410\|BT1A1_HUMAN Butyrophilin subfamily 1 member A1 OS=Homo sapiens OX=9606 GN=BTN1A1 PE=1 SV=3 | 179 | 20201119_10132_1ng_glyco_hcd203040_compound15.10296.10296.3 | controllerType=0 controllerNumber=1 scan=10296 | 24.0933 |
| 05084:3 | 11 | R.LSPN[+2205.793]ASAEHLELR.W | HexNAc(4)Hex(5)Fuc(2)NeuAc(1) | N[+2206] | 1214.8574 | 3 | 3642.5575 | 3642.5421 | 0 | 4.2 | 52 | Specific | 600.71 | 447.32 | 447.32 | 12.29 | 42 | >sp\|Q13410\|BT1A1_HUMAN Butyrophilin subfamily 1 member A1 OS=Homo sapiens OX=9606 GN=BTN1A1 PE=1 SV=3 | 179 | 20201119_10132_1ng_glyco_hcd203040_compound15.8198.8198.3 | controllerType=0 controllerNumber=1 scan=8198 | 20.5602 |
| 07079:3 | 11 | R.LSPN[+2350.830]ASAEHLELR.W | HexNAc(4)Hex(5)Fuc(1)NeuAc(2) | N[+2351] | 1263.2009 | 3 | 3787.588 | 3787.5796 | 0 | 2.2 | 52 | Specific | 613.74 | 450.14 | 450.14 | 12.19 | 42 | >sp\|Q13410\|BT1A1_HUMAN Butyrophilin subfamily 1 member A1 OS=Homo sapiens OX=9606 GN=BTN1A1 PE=1 SV=3 | 179 | 20201119_10132_1ng_glyco_hcd203040_compound15.10267.10267.3 | controllerType=0 controllerNumber=1 scan=10267 | 24.0443 |
| 17250:3 | 11 | R.DTSAKN[+2059.735]VSC[+57.021]YIQNLLLGQEK.K | HexNAc(4)Hex(5)Fuc(1)NeuAc(1) | C[+57], N[+2060] | 1447.6395 | 3 | 4340.9038 | 4340.8842 | 0 | 4.5 | 210 | Specific | 293.11 | 114.67 | 114.67 | 4.96 | 42 | >sp\|Q13410\|BT1A1_HUMAN Butyrophilin subfamily 1 member A1 OS=Homo sapiens OX=9606 GN=BTN1A1 PE=1 SV=3 | 179 | 20201119_10132_1ng_glyco_hcd203040_compound15.20911.20911.3 | controllerType=0 controllerNumber=1 scan=20911 | 42.9946 |
| 17247:3 | 11 | R.DTSAKN[+2059.735]VSC[+57.021]YIQNLLLGQEK.K | HexNAc(4)Hex(5)Fuc(1)NeuAc(1) | C[+57], N[+2060] | 1447.6395 | 3 | 4340.9038 | 4340.8842 | 0 | 4.5 | 210 | Specific | 431.61 | 399.01 | 399.01 | 9.25 | 42 | >sp\|Q13410\|BT1A1_HUMAN Butyrophilin subfamily 1 member A1 OS=Homo sapiens OX=9606 GN=BTN1A1 PE=1 SV=3 | 179 | 20201119_10132_1ng_glyco_hcd203040_compound15.20908.20908.3 | controllerType=0 controllerNumber=1 scan=20908 | 42.9885 |
| 17089:3 | 11 | K.N[+1913.677]VSC[+57.021]YIQNLLLGQEK.K | HexNAc(4)Hex(5)NeuAc(1) | C[+57], N[+1914] | 1231.5455 | 3 | 3692.6218 | 3692.5876 | 0 | 9.3 | 215 | Specific | 387.31 | 379.83 | 379.83 | 7.99 | 42 | >sp\|Q13410\|BT1A1_HUMAN Butyrophilin subfamily 1 member A1 OS=Homo sapiens OX=9606 GN=BTN1A1 PE=1 SV=3 | 179 | 20201119_10132_1ng_glyco_hcd203040_compound15.20743.20743.3 | controllerType=0 controllerNumber=1 scan=20743 | 42.584 |
| 17092:3 | 11 | K.N[+1913.677]VSC[+57.021]YIQNLLLGQEK.K | HexNAc(4)Hex(5)NeuAc(1) | C[+57], N[+1914] | 1231.5455 | 3 | 3692.6218 | 3692.5876 | 0 | 9.3 | 215 | Specific | 489.59 | 409.08 | 409.08 | 8.34 | 42 | >sp\|Q13410\|BT1A1_HUMAN Butyrophilin subfamily 1 member A1 OS=Homo sapiens OX=9606 GN=BTN1A1 PE=1 SV=3 | 179 | 20201119_10132_1ng_glyco_hcd203040_compound15.20746.20746.3 | controllerType=0 controllerNumber=1 scan=20746 | 42.589 |
| 17002:3 | 11 | K.N[+2059.735]VSC[+57.021]YIQNLLLGQEK.K | HexNAc(4)Hex(5)Fuc(1)NeuAc(1) | C[+57], N[+2060] | 1280.2278 | 3 | 3838.669 | 3838.6455 | 0 | 6.1 | 215 | Specific | 521.62 | 427.54 | 427.54 | 9.28 | 42 | >sp\|Q13410\|BT1A1_HUMAN Butyrophilin subfamily 1 member A1 OS=Homo sapiens OX=9606 GN=BTN1A1 PE=1 SV=3 | 179 | 20201119_10132_1ng_glyco_hcd203040_compound15.20652.20652.3 | controllerType=0 controllerNumber=1 scan=20652 | 42.3721 |
| 17004:3 | 11 | K.N[+2059.735]VSC[+57.021]YIQNLLLGQEK.K | HexNAc(4)Hex(5)Fuc(1)NeuAc(1) | C[+57], N[+2060] | 1280.2278 | 3 | 3838.669 | 3838.6455 | 0 | 6.1 | 215 | Specific | 632.4 | 427.08 | 427.08 | 11.63 | 42 | >sp\|Q13410\|BT1A1_HUMAN Butyrophilin subfamily 1 member A1 OS=Homo sapiens OX=9606 GN=BTN1A1 PE=1 SV=3 | 179 | 20201119_10132_1ng_glyco_hcd203040_compound15.20654.20654.3 | controllerType=0 controllerNumber=1 scan=20654 | 42.3764 |
| 21440:4 | 13 | K.VFPLSLDSTPQDGNVVVAC[+57.021]LVQGFFPQEPLSVTWSESGQN[+1809.666]VTAI | HexNAc(5)Hex(4)Fuc(1) | C[+57], N[+1810] | 1647.2697 | 4 | 6586.0568 | 6586.0275 | 0 | 4.5 | 8 | Specific | 261.26 | 261.26 | 261.26 | 7.85 | 36 | >sp\|P01877\|IGHA2_HUMAN Immunoglobulin heavy constant alpha 2 OS=Homo sapiens OX=9606 GN=IGHA2 PE=1 SV=4 | 576 | 20201119_10132_1ng_glyco_hcd203040_compound15.25285.25285.3 | controllerType=0 controllerNumber=1 scan=25285 | 51.8073 |
| 23259:5 | 13 | K.VFPLSLDSTPQDGNVVVAC[+57.021]LVQGFFPQEPLSVTWSESGQN[+2262.814]VTAI | HexNAc(5)Hex(5)Fuc(1)NeuAc(1) | C[+57], N[+2263] | 1408.6473 | 5 | 7039.2072 | 7039.1757 | 0 | 4.5 | 8 | Specific | 268.05 | 268.05 | 268.05 | 7.85 | 36 | >sp\|P01877\|IGHA2_HUMAN Immunoglobulin heavy constant alpha 2 OS=Homo sapiens OX=9606 GN=IGHA2 PE=1 SV=4 | 576 | 20201119_10132_1ng_glyco_hcd203040_compound15.27180.27180.5 | controllerType=0 controllerNumber=1 scan=27180 | 55.2295 |
| 21463:4 | 13 | K.[+42.011]VFPLSLDSTPQDGNVVVAC[+57.021]LVQGFFPQEPLSVTWSESGQN[+1768 | HexNAc(4)Hex(5)Fuc(1) | .[42], C[+57], N[+1769] | 1647.5166 | 4 | 6587.0446 | 6587.0115 | 0 | 5 | 8 | Specific | 344.13 | 295.37 | 295.37 | 6.21 | 36 | >sp\|P01877\|IGHA2_HUMAN Immunoglobulin heavy constant alpha 2 OS=Homo sapiens OX=9606 GN=IGHA2 PE=1 SV=4 | 576 | 20201119_10132_1ng_glyco_hcd203040_compound15.25309.25309.4 | controllerType=0 controllerNumber=1 scan=25309 | 51.8636 |
| 21780:4 | 13 | K.VFPLSLDSTPQDGNVVVAC[+57.021]LVQGFFPQEPLSVTWSESGQN[+1647.613]VTAI | HexNAc(5)Hex(3)Fuc(1) | C[+57], N[+1648] | 1606.7576 | 4 | 6424.0085 | 6423.9746 | 0 | 5.3 | 8 | Specific | 394.85 | 394.85 | 394.85 | 9.79 | 36 | >sp\|P01877\|IGHA2_HUMAN Immunoglobulin heavy constant alpha 2 OS=Homo sapiens OX=9606 GN=IGHA2 PE=1 SV=4 | 576 | 20201119_10132_1ng_glyco_hcd203040_compound15.25644.25644.4 | controllerType=0 controllerNumber=1 scan=25644 | 52.4417 |
| 21461:4 | 13 | K.[+42.011]VFPLSLDSTPQDGNVVVAC[+57.021]LVQGFFPQEPLSVTWSESGQN[+1768 | HexNAc(4)Hex(5)Fuc(1) | .[42], C[+57], N[+1769] | 1647.5166 | 4 | 6587.0446 | 6587.0115 | 0 | 5 | 8 | Specific | 425.08 | 385.65 | 385.65 | 8.59 | 36 | >sp\|P01877\|IGHA2_HUMAN Immunoglobulin heavy constant alpha 2 OS=Homo sapiens OX=9606 GN=IGHA2 PE=1 SV=4 | 576 | 20201119_10132_1ng_glyco_hcd203040_compound15.25307.25307.4 | controllerType=0 controllerNumber=1 scan=25307 | 51.8532 |
| 21442:4 | 13 | K.VFPLSLDSTPQDGNVVVAC[+57.021]LVQGFFPQEPLSVTWSESGQN[+1809.666]VTAI | HexNAc(5)Hex(4)Fuc(1) | C[+57], N[+1810] | 1647.2697 | 4 | 6586.0568 | 6586.0275 | 0 | 4.5 | 8 | Specific | 475.13 | 473.77 | 473.77 | 10.71 | 36 | >sp\|P01877\|IGHA2_HUMAN Immunoglobulin heavy constant alpha 2 OS=Homo sapiens OX=9606 GN=IGHA2 PE=1 SV=4 | 576 | 20201119_10132_1ng_glyco_hcd203040_compound15.25287.25287.3 | controllerType=0 controllerNumber=1 scan=25287 | 51.8135 |
| 21782:4 | 13 | K.VFPLSLDSTPQDGNVVVAC[+57.021]LVQGFFPQEPLSVTWSESGQN[+1647.613]VTAI | HexNAc(5)Hex(3)Fuc(1) | C[+57], N[+1648] | 1606.7576 | 4 | 6424.0085 | 6423.9746 | 0 | 5.3 | 8 | Specific | 738.56 | 676.55 | 676.55 | 14.65 | 36 | >sp\|P01877\|IGHA2_HUMAN Immunoglobulin heavy constant alpha 2 OS=Homo sapiens OX=9606 GN=IGHA2 PE=1 SV=4 | 576 | 20201119_10132_1ng_glyco_hcd203040_compound15.25646.25646.4 | controllerType=0 controllerNumber=1 scan=25646 | 52.447 |
| 00891:3 | 13 | K.HYTN[+1444.534]SSQDVTVPC[+57.021]R.V | HexNAc(4)Hex(3)Fuc(1) | C[+57], N[+1445] | 1036.7688 | 3 | 3108.2918 | 3108.2832 | 0 | 2.8 | 89 | Specific | 274.27 | 238.72 | 238.72 | 6.67 | 36 | >sp\|P01877\|IGHA2_HUMAN Immunoglobulin heavy constant alpha 2 OS=Homo sapiens OX=9606 GN=IGHA2 PE=1 SV=4 | 576 | 20201119_10132_1ng_glyco_hcd203040_compound15.3838.3838.3 | controllerType=0 controllerNumber=1 scan=3838 | 12.8867 |
| 01683:3 | 13 | K.HYTN[+2262.814]SSQDVTVPC[+57.021]R.V | HexNAc(5)Hex(5)Fuc(1)NeuAc(1) | C[+57], N[+2263] | 1309.5285 | 3 | 3926.5709 | 3926.5636 | 0 | 1.9 | 89 | Specific | 286.78 | 235.04 | 235.04 | 6.56 | 36 | >sp\|P01877\|IGHA2_HUMAN Immunoglobulin heavy constant alpha 2 OS=Homo sapiens OX=9606 GN=IGHA2 PE=1 SV=4 | 576 | 20201119_10132_1ng_glyco_hcd203040_compound15.4668.4668.3 | controllerType=0 controllerNumber=1 scan=4668 | 14.4243 |
| 01681:3 | 13 | K.HYTN[+2262.814]SSQDVTVPC[+57.021]R.V | HexNAc(5)Hex(5)Fuc(1)NeuAc(1) | C[+57], N[+2263] | 1309.5285 | 3 | 3926.5709 | 3926.5636 | 0 | 1.9 | 89 | Specific | 315.79 | 267.65 | 267.65 | 6.99 | 36 | >sp\|P01877\|IGHA2_HUMAN Immunoglobulin heavy constant alpha 2 OS=Homo sapiens OX=9606 GN=IGHA2 PE=1 SV=4 | 576 | 20201119_10132_1ng_glyco_hcd203040_compound15.4666.4666.3 | controllerType=0 controllerNumber=1 scan=4666 | 14.4197 |
| 00893:3 | 13 | K.HYTN[+1444.534]SSQDVTVPC[+57.021]R.V | HexNAc(4)Hex(3)Fuc(1) | C[+57], N[+1445] | 1036.7688 | 3 | 3108.2918 | 3108.2832 | 0 | 2.8 | 89 | Specific | 332.32 | 260.41 | 260.41 | 7.2 | 36 | >sp\|P01877\|IGHA2_HUMAN Immunoglobulin heavy constant alpha 2 OS=Homo sapiens OX=9606 GN=IGHA2 PE=1 SV=4 | 576 | 20201119_10132_1ng_glyco_hcd203040_compound15.3840.3840.3 | controllerType=0 controllerNumber=1 scan=3840 | 12.8906 |
| 00855:3 | 13 | K.HYTN[+1647.613]SSQDVTVPC[+57.021]R.V | HexNAc(5)Hex(3)Fuc(1) | C[+57], N[+1648] | 1104.4627 | 3 | 3311.3735 | 3311.3625 | 0 | 3.3 | 89 | Specific | 434.76 | 316.24 | 316.24 | 7.93 | 36 | >sp\|P01877\|IGHA2_HUMAN Immunoglobulin heavy constant alpha 2 OS=Homo sapiens OX=9606 GN=IGHA2 PE=1 SV=4 | 576 | 20201119_10132_1ng_glyco_hcd203040_compound15.3801.3801.3 | controllerType=0 controllerNumber=1 scan=3801 | 12.8221 |
| 00851:3 | 13 | K.HYTN[+1809.666]SSQDVTVPC[+57.021]R.V | HexNAc(5)Hex(4)Fuc(1) | C[+57], N[+1810] | 1158.4803 | 3 | 3473.4263 | 3473.4154 | 0 | 3.1 | 89 | Specific | 474.86 | 294.81 | 294.81 | 7.87 | 36 | >sp\|P01877\|IGHA2_HUMAN Immunoglobulin heavy constant alpha 2 OS=Homo sapiens OX=9606 GN=IGHA2 PE=1 SV=4 | 576 | 20201119_10132_1ng_glyco_hcd203040_compound15.3797.3797.3 | controllerType=0 controllerNumber=1 scan=3797 | 12.8158 |
| 00849:3 | 13 | K.HYTN[+1809.666]SSQDVTVPC[+57.021]R.V | HexNAc(5)Hex(4)Fuc(1) | C[+57], N[+1810] | 1158.4803 | 3 | 3473.4263 | 3473.4154 | 0 | 3.1 | 89 | Specific | 497.15 | 387.03 | 387.03 | 9.68 | 36 | >sp\|P01877\|IGHA2_HUMAN Immunoglobulin heavy constant alpha 2 OS=Homo sapiens OX=9606 GN=IGHA2 PE=1 SV=4 | 576 | 20201119_10132_1ng_glyco_hcd203040_compound15.3795.3795.3 | controllerType=0 controllerNumber=1 scan=3795 | 12.8122 |
| 00857:3 | 13 | K.HYTN[+1647.613]SSQDVTVPC[+57.021]R.V | HexNAc(5)Hex(3)Fuc(1) | C[+57], N[+1648] | 1104.4627 | 3 | 3311.3735 | 3311.3625 | 0 | 3.3 | 89 | Specific | 530.48 | 374.43 | 374.43 | 9.83 | 36 | >sp\|P01877\|IGHA2_HUMAN Immunoglobulin heavy constant alpha 2 OS=Homo sapiens OX=9606 GN=IGHA2 PE=1 SV=4 | 576 | 20201119_10132_1ng_glyco_hcd203040_compound15.3803.3803.3 | controllerType=0 controllerNumber=1 scan=3803 | 12.8256 |
| 02817:4 | 13 | K.HYTN[+1647.613]SSQDVTVPC[+57.021]RVPPPPPC[+57.021]C[+57.021]HPR.L | HexNAc(5)Hex(3)Fuc(1) | C[+57]*3, N[+1648] | 1152.2518 | 4 | 4605.9852 | 4605.9689 | 0 | 3.5 | 89 | Specific | 404.88 | 246.82 | 246.82 | 6.91 | 36 | >sp\|P01877\|IGHA2_HUMAN Immunoglobulin heavy constant alpha 2 OS=Homo sapiens OX=9606 GN=IGHA2 PE=1 SV=4 | 576 | 20201119_10132_1ng_glyco_hcd203040_compound15.5845.5845.4 | controllerType=0 controllerNumber=1 scan=5845 | 16.5562 |
| 02815:4 | 13 | K.HYTN[+1647.613]SSQDVTVPC[+57.021]RVPPPPPC[+57.021]C[+57.021]HPR.L | HexNAc(5)Hex(3)Fuc(1) | C[+57]*3, N[+1648] | 1152.2518 | 4 | 4605.9852 | 4605.9689 | 0 | 3.5 | 89 | Specific | 429.98 | 426.49 | 426.49 | 8.93 | 36 | >sp\|P01877\|IGHA2_HUMAN Immunoglobulin heavy constant alpha 2 OS=Homo sapiens OX=9606 GN=IGHA2 PE=1 SV=4 | 576 | 20201119_10132_1ng_glyco_hcd203040_compound15.5843.5843.4 | controllerType=0 controllerNumber=1 scan=5843 | 16.5526 |
| 02810:5 | 13 | K.HYTN[+1647.613]SSQDVTVPC[+57.021]RVPPPPPC[+57.021]C[+57.021]HPR.L | HexNAc(5)Hex(3)Fuc(1) | C[+57]*3, N[+1648] | 922.002 | 5 | 4605.9808 | 4605.9689 | 0 | 2.6 | 89 | Specific | 462.55 | 301.67 | 301.67 | 8.03 | 36 | >sp\|P01877\|IGHA2_HUMAN Immunoglobulin heavy constant alpha 2 OS=Homo sapiens OX=9606 GN=IGHA2 PE=1 SV=4 | 576 | 20201119_10132_1ng_glyco_hcd203040_compound15.5838.5838.5 | controllerType=0 controllerNumber=1 scan=5838 | 16.5441 |
| 02812:5 | 13 | K.HYTN[+1647.613]SSQDVTVPC[+57.021]RVPPPPPC[+57.021]C[+57.021]HPR.L | HexNAc(5)Hex(3)Fuc(1) | C[+57]*3, N[+1648] | 922.002 | 5 | 4605.9808 | 4605.9689 | 0 | 2.6 | 89 | Specific | 620.56 | 390.3 | 390.3 | 12.35 | 36 | >sp\|P01877\|IGHA2_HUMAN Immunoglobulin heavy constant alpha 2 OS=Homo sapiens OX=9606 GN=IGHA2 PE=1 SV=4 | 576 | 20201119_10132_1ng_glyco_hcd203040_compound15.5840.5840.5 | controllerType=0 controllerNumber=1 scan=5840 | 16.5481 |
| 04241:3 | 13 | K.TPLTAN[+2262.814]ITK.S | HexNAc(5)Hex(5)Fuc(1)NeuAc(1) | N[+2263] | 1074.4644 | 3 | 3221.3786 | 3221.3711 | 0 | 2.3 | 200 | Specific | 216.9 | 112.1 | 112.1 | 3.59 | 36 | >sp\|P01877\|IGHA2_HUMAN Immunoglobulin heavy constant alpha 2 OS=Homo sapiens OX=9606 GN=IGHA2 PE=1 SV=4 | 576 | 20201119_10132_1ng_glyco_hcd203040_compound15.7325.7325.3 | controllerType=0 controllerNumber=1 scan=7325 | 19.0974 |
| 02411:3 | 13 | K.TPLTAN[+1971.719]ITK.S | HexNAc(5)Hex(5)Fuc(1) | N[+1972] | 977.4325 | 3 | 2930.283 | 2930.2757 | 0 | 2.5 | 200 | Specific | 281.06 | 193.08 | 193.08 | 5.74 | 36 | >sp\|P01877\|IGHA2_HUMAN Immunoglobulin heavy constant alpha 2 OS=Homo sapiens OX=9606 GN=IGHA2 PE=1 SV=4 | 576 | 20201119_10132_1ng_glyco_hcd203040_compound15.5424.5424.3 | controllerType=0 controllerNumber=1 scan=5424 | 15.789 |
| 02522:2 | 13 | K.TPLTAN[+1501.555]ITK.S | HexNAc(5)Hex(3) | N[+1502] | 1230.5632 | 2 | 2460.119 | 2460.1121 | 0 | 2.8 | 200 | Specific | 441.11 | 282.33 | 282.33 | 7.18 | 36 | >sp\|P01877\|IGHA2_HUMAN Immunoglobulin heavy constant alpha 2 OS=Homo sapiens OX=9606 GN=IGHA2 PE=1 SV=4 | 576 | 20201119_10132_1ng_glyco_hcd203040_compound15.5539.5539.2 | controllerType=0 controllerNumber=1 scan=5539 | 16.003 |
| 02533:3 | 13 | K.TPLTAN[+1647.613]ITK.S | HexNAc(5)Hex(3)Fuc(1) | N[+1648] | 869.4015 | 3 | 2606.19 | 2606.17 | 0 | 7.7 | 200 | Specific | 443.43 | 177.43 | 177.43 | 5.6 | 36 | >sp\|P01877\|IGHA2_HUMAN Immunoglobulin heavy constant alpha 2 OS=Homo sapiens OX=9606 GN=IGHA2 PE=1 SV=4 | 576 | 20201119_10132_1ng_glyco_hcd203040_compound15.5551.5551.3 | controllerType=0 controllerNumber=1 scan=5551 | 16.0266 |
| 02524:2 | 13 | K.TPLTAN[+1501.555]ITK.S | HexNAc(5)Hex(3) | N[+1502] | 1230.5632 | 2 | 2460.119 | 2460.1121 | 0 | 2.8 | 200 | Specific | 456.44 | 410.54 | 410.54 | 9.05 | 36 | >sp\|P01877\|IGHA2_HUMAN Immunoglobulin heavy constant alpha 2 OS=Homo sapiens OX=9606 GN=IGHA2 PE=1 SV=4 | 576 | 20201119_10132_1ng_glyco_hcd203040_compound15.5541.5541.2 | controllerType=0 controllerNumber=1 scan=5541 | 16.0073 |
| 02463:2 | 13 | K.TPLTAN[+1444.534]ITK.S | HexNAc(4)Hex(3)Fuc(1) | N[+1445] | 1202.0541 | 2 | 2403.101 | 2403.0906 | 0 | 4.3 | 200 | Specific | 461.81 | 287.63 | 287.63 | 7.58 | 36 | >sp\|P01877\|IGHA2_HUMAN Immunoglobulin heavy constant alpha 2 OS=Homo sapiens OX=9606 GN=IGHA2 PE=1 SV=4 | 576 | 20201119_10132_1ng_glyco_hcd203040_compound15.5478.5478.2 | controllerType=0 controllerNumber=1 scan=5478 | 15.8896 |
| 02465:2 | 13 | K.TPLTAN[+1444.534]ITK.S | HexNAc(4)Hex(3)Fuc(1) | N[+1445] | 1202.0541 | 2 | 2403.101 | 2403.0906 | 0 | 4.3 | 200 | Specific | 481.04 | 250.92 | 250.92 | 7.38 | 36 | >sp\|P01877\|IGHA2_HUMAN Immunoglobulin heavy constant alpha 2 OS=Homo sapiens OX=9606 GN=IGHA2 PE=1 SV=4 | 576 | 20201119_10132_1ng_glyco_hcd203040_compound15.5480.5480.2 | controllerType=0 controllerNumber=1 scan=5480 | 15.8933 |
| 02535:3 | 13 | K.TPLTAN[+1647.613]ITK.S | HexNAc(5)Hex(3)Fuc(1) | N[+1648] | 869.4015 | 3 | 2606.19 | 2606.17 | 0 | 7.7 | 200 | Specific | 545.83 | 273.07 | 273.07 | 6.79 | 36 | >sp\|P01877\|IGHA2_HUMAN Immunoglobulin heavy constant alpha 2 OS=Homo sapiens OX=9606 GN=IGHA2 PE=1 SV=4 | 576 | 20201119_10132_1ng_glyco_hcd203040_compound15.5553.5553.3 | controllerType=0 controllerNumber=1 scan=5553 | 16.0317 |
| 02514:2 | 13 | K.TPLTAN[+1647.613]ITK.S | HexNAc(5)Hex(3)Fuc(1) | N[+1648] | 1303.5918 | 2 | 2606.1764 | 2606.17 | 0 | 2.5 | 200 | Specific | 594.42 | 297.65 | 297.65 | 9 | 36 | >sp\|P01877\|IGHA2_HUMAN Immunoglobulin heavy constant alpha 2 OS=Homo sapiens OX=9606 GN=IGHA2 PE=1 SV=4 | 576 | 20201119_10132_1ng_glyco_hcd203040_compound15.5531.5531.2 | controllerType=0 controllerNumber=1 scan=5531 | 15.9894 |
| 02512:2 | 13 | K.TPLTAN[+1647.613]ITK.S | HexNAc(5)Hex(3)Fuc(1) | N[+1648] | 1303.5918 | 2 | 2606.1764 | 2606.17 | 0 | 2.5 | 200 | Specific | 634.47 | 340.79 | 340.79 | 10.29 | 36 | >sp\|P01877\|IGHA2_HUMAN Immunoglobulin heavy constant alpha 2 OS=Homo sapiens OX=9606 GN=IGHA2 PE=1 SV=4 | 576 | 20201119_10132_1ng_glyco_hcd203040_compound15.5529.5529.2 | controllerType=0 controllerNumber=1 scan=5529 | 15.9845 |
| 09748:3 | 13 | R.MAGKPTHIN[+1809.666]VSVVMAEADGTC[+57.021]Y.- | HexNAc(5)Hex(4)Fuc(1) | C[+57], N[+1810] | 1387.5894 | 3 | 4160.7537 | 4160.7489 | 0 | 1.1 | 319 | CRagged | 245.54 | 209.74 | 209.74 | 6.79 | 36 | >sp\|P01877\|IGHA2_HUMAN Immunoglobulin heavy constant alpha 2 OS=Homo sapiens OX=9606 GN=IGHA2 PE=1 SV=4 | 576 | 20201119_10132_1ng_glyco_hcd203040_compound15.13057.13057.3 | controllerType=0 controllerNumber=1 scan=13057 | 28.8068 |

Table S4. Cont.

| Query #:z | Protein Rank | Glycopeptide | Glycan | Modification Type(s) | Observed m/z | z | Observed (M+H) | Calc. mass (M+H) | Off-by-x error | Mass error (ppm) | StartPosition | Cleavage | Score | Delta | Delta Mod | \|Log Prob\| | # of unique peptides | Protein Name | Protein DB number | Comment | Scan # | Scan Time |
|---|---|---|---|---|---|---|---|---|---|---|---|---|---|---|---|---|---|---|---|---|---|---|
| 09846:3 | 13 | R.MAGKPTHIN[+1501.555]VSVVMAEADGTC[+57.021]Y.- | HexNAc(5)Hex(3) | C[+57], N[+1502] | 1284.8893 | 3 | 3852.6535 | 3852.6382 | 0 | 4 | 319 | CRagged | 287.43 | 287.43 | 287.43 | 7.74 | 36 | >sp\|P01877\|IGHA2_HUMAN Immunoglobulin heavy constant alpha 2 OS=Homo sapiens OX=9606 GN=IGHA2 PE=1 SV=4 | 576 | 20201119_10132_1ng_glyco_hcd203040_compound15.13159.13159.3 | controllerType=0 controllerNumber=1 scan=13159 | 28.9809 |
| 09848:3 | 13 | R.MAGKPTHIN[+1501.555]VSVVMAEADGTC[+57.021]Y.- | HexNAc(5)Hex(3) | C[+57], N[+1502] | 1284.8893 | 3 | 3852.6535 | 3852.6382 | 0 | 4 | 319 | CRagged | 345.77 | 340.93 | 340.93 | 8.43 | 36 | >sp\|P01877\|IGHA2_HUMAN Immunoglobulin heavy constant alpha 2 OS=Homo sapiens OX=9606 GN=IGHA2 PE=1 SV=4 | 576 | 20201119_10132_1ng_glyco_hcd203040_compound15.13161.13161.3 | controllerType=0 controllerNumber=1 scan=13161 | 28.9848 |
| 09851:3 | 13 | R.MAGKPTHIN[+1647.613]VSVVMAEADGTC[+57.021]Y.- | HexNAc(5)Hex(3)Fuc(1) | C[+57], N[+1648] | 1333.5748 | 3 | 3998.7097 | 3998.6961 | 0 | 3.4 | 319 | CRagged | 370.22 | 370.22 | 370.22 | 8.46 | 36 | >sp\|P01877\|IGHA2_HUMAN Immunoglobulin heavy constant alpha 2 OS=Homo sapiens OX=9606 GN=IGHA2 PE=1 SV=4 | 576 | 20201119_10132_1ng_glyco_hcd203040_compound15.13164.13164.3 | controllerType=0 controllerNumber=1 scan=13164 | 28.9906 |
| 09625:3 | 13 | R.MAGKPTHIN[+1702.581]VSVVMAEADGTC[+57.021]Y.- | HexNAc(2)Hex(8) | C[+57], N[+1703] | 1351.8975 | 3 | 4053.678 | 4053.6642 | 0 | 3.4 | 319 | CRagged | 751.26 | 751.26 | 409.32 | 15.85 | 36 | >sp\|P01877\|IGHA2_HUMAN Immunoglobulin heavy constant alpha 2 OS=Homo sapiens OX=9606 GN=IGHA2 PE=1 SV=4 | 576 | 20201119_10132_1ng_glyco_hcd203040_compound15.12930.12930.3 | controllerType=0 controllerNumber=1 scan=12930 | 28.582 |
| 09764:3 | 13 | R.MAGKPTHIN[+1216.423]VSVVMAEADGTC[+57.021]Y.- | HexNAc(2)Hex(5) | C[+57], N[+1216] | 1189.8422 | 3 | 3567.5121 | 3567.5057 | 0 | 1.8 | 319 | CRagged | 787.95 | 787.95 | 439.55 | 15.64 | 36 | >sp\|P01877\|IGHA2_HUMAN Immunoglobulin heavy constant alpha 2 OS=Homo sapiens OX=9606 GN=IGHA2 PE=1 SV=4 | 576 | 20201119_10132_1ng_glyco_hcd203040_compound15.13074.13074.3 | controllerType=0 controllerNumber=1 scan=13074 | 28.8354 |
| 09680:3 | 13 | R.MAGKPTHIN[+1378.476]VSVVMAEADGTC[+57.021]Y.- | HexNAc(2)Hex(6) | C[+57], N[+1378] | 1243.8608 | 3 | 3729.5678 | 3729.5585 | 0 | 2.5 | 319 | CRagged | 815.18 | 815.18 | 443.35 | 16.85 | 36 | >sp\|P01877\|IGHA2_HUMAN Immunoglobulin heavy constant alpha 2 OS=Homo sapiens OX=9606 GN=IGHA2 PE=1 SV=4 | 576 | 20201119_10132_1ng_glyco_hcd203040_compound15.12987.12987.3 | controllerType=0 controllerNumber=1 scan=12987 | 28.6838 |
| 09623:3 | 13 | R.MAGKPTHIN[+1702.581]VSVVMAEADGTC[+57.021]Y.- | HexNAc(2)Hex(8) | C[+57], N[+1703] | 1351.8975 | 3 | 4053.678 | 4053.6642 | 0 | 3.4 | 319 | CRagged | 819.48 | 819.48 | 492.91 | 16.85 | 36 | >sp\|P01877\|IGHA2_HUMAN Immunoglobulin heavy constant alpha 2 OS=Homo sapiens OX=9606 GN=IGHA2 PE=1 SV=4 | 576 | 20201119_10132_1ng_glyco_hcd203040_compound15.12928.12928.3 | controllerType=0 controllerNumber=1 scan=12928 | 28.5748 |
| 09678:3 | 13 | R.MAGKPTHIN[+1378.476]VSVVMAEADGTC[+57.021]Y.- | HexNAc(2)Hex(6) | C[+57], N[+1378] | 1243.8608 | 3 | 3729.5678 | 3729.5585 | 0 | 2.5 | 319 | CRagged | 832.18 | 669.13 | 501.3 | 16.85 | 36 | >sp\|P01877\|IGHA2_HUMAN Immunoglobulin heavy constant alpha 2 OS=Homo sapiens OX=9606 GN=IGHA2 PE=1 SV=4 | 576 | 20201119_10132_1ng_glyco_hcd203040_compound15.12985.12985.3 | controllerType=0 controllerNumber=1 scan=12985 | 28.678 |
| 09766:3 | 13 | R.MAGKPTHIN[+1216.423]VSVVMAEADGTC[+57.021]Y.- | HexNAc(2)Hex(5) | C[+57], N[+1216] | 1189.8422 | 3 | 3567.5121 | 3567.5057 | 0 | 1.8 | 319 | CRagged | 850.28 | 783.41 | 475.76 | 17.64 | 36 | >sp\|P01877\|IGHA2_HUMAN Immunoglobulin heavy constant alpha 2 OS=Homo sapiens OX=9606 GN=IGHA2 PE=1 SV=4 | 576 | 20201119_10132_1ng_glyco_hcd203040_compound15.13076.13076.3 | controllerType=0 controllerNumber=1 scan=13076 | 28.84 |
| 13388:4 | 17 | K.LKELPGVC[+57.021]N[+2059.735]ETMMALWEEC[+57.021]KPC[+57.021]LK.Q | HexNAc(4)Hex(5)Fuc(1)NeuAc(1) | C[+57]*3, N[+2060] | 1249.7887 | 4 | 4996.133 | 4996.1196 | 0 | 2.7 | 95 | Specific | 779.01 | 667.28 | 667.28 | 15.35 | 26 | >sp\|P10909\|CLUS_HUMAN Clusterin OS=Homo sapiens OX=9606 GN=CLU PE=1 SV=1 | 1467 | 20201119_10132_1ng_glyco_hcd203040_compound15.16853.16853.4 | controllerType=0 controllerNumber=1 scan=16853 | 35.3885 |
| 13341:4 | 17 | K.LKELPGVC[+57.021]N[+2205.793]ETMMALWEEC[+57.021]KPC[+57.021]LK.Q | HexNAc(4)Hex(5)Fuc(2)NeuAc(1) | C[+57]*3, N[+2206] | 1286.3052 | 4 | 5142.1989 | 5142.1776 | 0 | 4.1 | 95 | Specific | 840.64 | 840.64 | 840.64 | 16.46 | 26 | >sp\|P10909\|CLUS_HUMAN Clusterin OS=Homo sapiens OX=9606 GN=CLU PE=1 SV=1 | 1467 | 20201119_10132_1ng_glyco_hcd203040_compound15.16804.16804.4 | controllerType=0 controllerNumber=1 scan=16804 | 35.2974 |
| 13339:4 | 17 | K.LKELPGVC[+57.021]N[+2205.793]ETMMALWEEC[+57.021]KPC[+57.021]LK.Q | HexNAc(4)Hex(5)Fuc(2)NeuAc(1) | C[+57]*3, N[+2206] | 1286.3052 | 4 | 5142.1989 | 5142.1776 | 0 | 4.1 | 95 | Specific | 900.2 | 900.2 | 900.2 | 18.46 | 26 | >sp\|P10909\|CLUS_HUMAN Clusterin OS=Homo sapiens OX=9606 GN=CLU PE=1 SV=1 | 1467 | 20201119_10132_1ng_glyco_hcd203040_compound15.16802.16802.4 | controllerType=0 controllerNumber=1 scan=16802 | 35.2927 |
| 13386:4 | 17 | K.LKELPGVC[+57.021]N[+2059.735]ETMMALWEEC[+57.021]KPC[+57.021]LK.Q | HexNAc(4)Hex(5)Fuc(1)NeuAc(1) | C[+57]*3, N[+2060] | 1249.7887 | 4 | 4996.133 | 4996.1196 | 0 | 2.7 | 95 | Specific | 952.24 | 952.24 | 952.24 | 19.35 | 26 | >sp\|P10909\|CLUS_HUMAN Clusterin OS=Homo sapiens OX=9606 GN=CLU PE=1 SV=1 | 1467 | 20201119_10132_1ng_glyco_hcd203040_compound15.16851.16851.4 | controllerType=0 controllerNumber=1 scan=16851 | 35.3836 |
| 13368:4 | 17 | K.LKELPGVC[+57.021]N[+2190.793]ETM[+15.995]MALWEEC[+57.021]KPC[+57.021 | HexNAc(6)Hex(6) | C[+57]*3, M[+16], N[+2191] | 1286.5562 | 4 | 5143.2028 | 5143.1728 | 0 | 5.8 | 95 | Specific | 230.02 | 197.52 | 13.09 | 3.73 | 26 | >sp\|P10909\|CLUS_HUMAN Clusterin OS=Homo sapiens OX=9606 GN=CLU PE=1 SV=1 | 1467 | 20201119_10132_1ng_glyco_hcd203040_compound15.16832.16832.2 | controllerType=0 controllerNumber=1 scan=16832 | 35.3487 |
| 13366:4 | 17 | K.LKELPGVC[+57.021]N[+2190.793]ETM[+15.995]MALWEEC[+57.021]KPC[+57.021 | HexNAc(6)Hex(6) | C[+57]*3, M[+16], N[+2191] | 1286.5562 | 4 | 5143.2028 | 5143.1728 | 0 | 5.8 | 95 | Specific | 303.2 | 271.92 | 14.56 | 6.6 | 26 | >sp\|P10909\|CLUS_HUMAN Clusterin OS=Homo sapiens OX=9606 GN=CLU PE=1 SV=1 | 1467 | 20201119_10132_1ng_glyco_hcd203040_compound15.16830.16830.2 | controllerType=0 controllerNumber=1 scan=16830 | 35.3436 |
| 22842:4 | 17 | K.MLN[+2350.830]TSSLLEQLNEQFNWVSR.L | HexNAc(4)Hex(5)Fuc(1)NeuAc(2) | N[+2351] | 1190.7681 | 4 | 4760.0506 | 4760.0171 | 0 | 7 | 352 | Specific | 319.32 | 215.39 | 215.39 | 6.41 | 26 | >sp\|P10909\|CLUS_HUMAN Clusterin OS=Homo sapiens OX=9606 GN=CLU PE=1 SV=1 | 1467 | 20201119_10132_1ng_glyco_hcd203040_compound15.26747.26747.4 | controllerType=0 controllerNumber=1 scan=26747 | 54.4551 |
| 20347:4 | 17 | K.MLN[+2059.735]TSSLLEQLNEQFNWVSR.L | HexNAc(4)Hex(5)Fuc(1)NeuAc(1) | N[+2060] | 1117.9924 | 4 | 4468.9476 | 4468.9217 | 0 | 5.8 | 352 | Specific | 360.59 | 325.2 | 325.2 | 7.45 | 26 | >sp\|P10909\|CLUS_HUMAN Clusterin OS=Homo sapiens OX=9606 GN=CLU PE=1 SV=1 | 1467 | 20201119_10132_1ng_glyco_hcd203040_compound15.24149.24149.4 | controllerType=0 controllerNumber=1 scan=24149 | 49.8364 |
| 20349:4 | 17 | K.MLN[+2059.735]TSSLLEQLNEQFNWVSR.L | HexNAc(4)Hex(5)Fuc(1)NeuAc(1) | N[+2060] | 1117.9924 | 4 | 4468.9476 | 4468.9217 | 0 | 5.8 | 352 | Specific | 461.5 | 461.5 | 461.5 | 9.45 | 26 | >sp\|P10909\|CLUS_HUMAN Clusterin OS=Homo sapiens OX=9606 GN=CLU PE=1 SV=1 | 1467 | 20201119_10132_1ng_glyco_hcd203040_compound15.24151.24151.4 | controllerType=0 controllerNumber=1 scan=24151 | 49.8407 |
| 22844:4 | 17 | K.MLN[+2350.830]TSSLLEQLNEQFNWVSR.L | HexNAc(4)Hex(5)Fuc(1)NeuAc(2) | N[+2351] | 1190.7681 | 4 | 4760.0506 | 4760.0171 | 0 | 7 | 352 | Specific | 521.19 | 379.53 | 379.53 | 9.38 | 26 | >sp\|P10909\|CLUS_HUMAN Clusterin OS=Homo sapiens OX=9606 GN=CLU PE=1 SV=1 | 1467 | 20201119_10132_1ng_glyco_hcd203040_compound15.26749.26749.4 | controllerType=0 controllerNumber=1 scan=26749 | 54.4596 |
| 12405:3 | 17 | R.LAN[+2350.830]LTQGEDQYYLR.V | HexNAc(4)Hex(5)Fuc(1)NeuAc(2) | N[+2351] | 1345.5638 | 3 | 4034.6768 | 4034.664 | 0 | 3.2 | 372 | Specific | 261.02 | 243.72 | 243.72 | 6.59 | 26 | >sp\|P10909\|CLUS_HUMAN Clusterin OS=Homo sapiens OX=9606 GN=CLU PE=1 SV=1 | 1467 | 20201119_10132_1ng_glyco_hcd203040_compound15.15833.15833.3 | controllerType=0 controllerNumber=1 scan=15833 | 33.5767 |
| 09733:3 | 17 | R.LAN[+2205.793]LTQGEDQYYLR.V | HexNAc(4)Hex(5)Fuc(2)NeuAc(1) | N[+2206] | 1297.2138 | 3 | 3889.6269 | 3889.6265 | 0 | 0.1 | 372 | Specific | 284.72 | 213.47 | 213.47 | 5.87 | 26 | >sp\|P10909\|CLUS_HUMAN Clusterin OS=Homo sapiens OX=9606 GN=CLU PE=1 SV=1 | 1467 | 20201119_10132_1ng_glyco_hcd203040_compound15.13042.13042.3 | controllerType=0 controllerNumber=1 scan=13042 | 28.781 |
| 07951:5 | 24 | K.QTDEIKDTRN[+1768.640]ESTQNC[+57.021]VVAEPEKMESSISSSSEEMSLSK.C | HexNAc(4)Hex(5)Fuc(1) | C[+57], N[+1769] | 1232.7332 | 5 | 6159.6367 | 6159.6202 | 0 | 2.7 | 60 | Specific | 336.97 | 336.97 | 336.97 | 6.8 | 26 | >sp\|P47710\|CASA1_HUMAN Alpha-S1-casein OS=Homo sapiens OX=9606 GN=CSN1S1 PE=1 SV=1 | 989 | 20201119_10132_1ng_glyco_hcd203040_compound15.11175.11175.5 | controllerType=0 controllerNumber=1 scan=11175 | 25.5692 |
| 07949:5 | 24 | K.QTDEIKDTRN[+1768.640]ESTQNC[+57.021]VVAEPEKMESSISSSSEEMSLSK.C | HexNAc(4)Hex(5)Fuc(1) | C[+57], N[+1769] | 1232.7332 | 5 | 6159.6367 | 6159.6202 | 0 | 2.7 | 60 | Specific | 407.85 | 323.06 | 323.06 | 7.15 | 26 | >sp\|P47710\|CASA1_HUMAN Alpha-S1-casein OS=Homo sapiens OX=9606 GN=CSN1S1 PE=1 SV=1 | 989 | 20201119_10132_1ng_glyco_hcd203040_compound15.11173.11173.5 | controllerType=0 controllerNumber=1 scan=11173 | 25.5536 |
| 13672:3 | 25 | K.VVLHPN[+2205.793]YSQVDIGLIK.L | HexNAc(4)Hex(5)Fuc(2)NeuAc(1) | N[+2206] | 1334.2731 | 3 | 4000.8047 | 4000.8041 | 0 | 0.2 | 236 | Specific | 620.67 | 493.52 | 493.52 | 11.67 | 24 | >sp\|P00738\|HPT_HUMAN Haptoglobin OS=Homo sapiens OX=9606 GN=HP PE=1 SV=1 | 407 | 20201119_10132_1ng_glyco_hcd203040_compound15.17147.17147.3 | controllerType=0 controllerNumber=1 scan=17147 | 35.9202 |
| 13674:3 | 25 | K.VVLHPN[+2205.793]YSQVDIGLIK.L | HexNAc(4)Hex(5)Fuc(2)NeuAc(1) | N[+2206] | 1334.2731 | 3 | 4000.8047 | 4000.8041 | 0 | 0.2 | 236 | Specific | 723.56 | 641.81 | 641.81 | 13.67 | 24 | >sp\|P00738\|HPT_HUMAN Haptoglobin OS=Homo sapiens OX=9606 GN=HP PE=1 SV=1 | 407 | 20201119_10132_1ng_glyco_hcd203040_compound15.17149.17149.3 | controllerType=0 controllerNumber=1 scan=17149 | 35.9241 |
| 09422:4 | 30 | K.SC[+57.021]QHN[+2350.830]GTMYQHGEIFSAHELFPSR.L | HexNAc(4)Hex(5)Fuc(1)NeuAc(2) | C[+57], N[+2351] | 1296.7788 | 4 | 5184.0934 | 5184.0873 | 0 | 1.2 | 110 | Specific | 206.9 | 206.9 | 206.9 | 4.98 | 21 | >sp\|Q6WN34\|CRDL2_HUMAN Chordin-like protein 2 OS=Homo sapiens OX=9606 GN=CHRDL2 PE=1 SV=1 | 1189 | 20201119_10132_1ng_glyco_hcd203040_compound15.12719.12719.2 | controllerType=0 controllerNumber=1 scan=12719 | 28.2108 |
| 07992:5 | 30 | K.SC[+57.021]QHN[+2204.772]GTMYQHGEIFSAHELFPSR.L | HexNAc(4)Hex(5)NeuAc(2) | C[+57], N[+2205] | 1008.4193 | 5 | 5038.0675 | 5038.0294 | 0 | 7.6 | 110 | Specific | 391.54 | 391.54 | 391.54 | 9.27 | 21 | >sp\|Q6WN34\|CRDL2_HUMAN Chordin-like protein 2 OS=Homo sapiens OX=9606 GN=CHRDL2 PE=1 SV=1 | 1189 | 20201119_10132_1ng_glyco_hcd203040_compound15.11218.11218.5 | controllerType=0 controllerNumber=1 scan=11218 | 25.6595 |
| 07964:4 | 30 | K.SC[+57.021]QHN[+2205.793]GTMYQHGEIFSAHELFPSR.L | HexNAc(4)Hex(5)Fuc(2)NeuAc(1) | C[+57], N[+2206] | 1260.5259 | 4 | 5039.0817 | 5039.0498 | 0 | 6.3 | 110 | Specific | 478.2 | 478.2 | 478.2 | 8.83 | 21 | >sp\|Q6WN34\|CRDL2_HUMAN Chordin-like protein 2 OS=Homo sapiens OX=9606 GN=CHRDL2 PE=1 SV=1 | 1189 | 20201119_10132_1ng_glyco_hcd203040_compound15.11189.11189.4 | controllerType=0 controllerNumber=1 scan=11189 | 25.593 |
| 07994:5 | 30 | K.SC[+57.021]QHN[+2204.772]GTMYQHGEIFSAHELFPSR.L | HexNAc(4)Hex(5)NeuAc(2) | C[+57], N[+2205] | 1008.4193 | 5 | 5038.0675 | 5038.0294 | 0 | 7.6 | 110 | Specific | 534.2 | 534.2 | 534.2 | 9.78 | 21 | >sp\|Q6WN34\|CRDL2_HUMAN Chordin-like protein 2 OS=Homo sapiens OX=9606 GN=CHRDL2 PE=1 SV=1 | 1189 | 20201119_10132_1ng_glyco_hcd203040_compound15.11220.11220.5 | controllerType=0 controllerNumber=1 scan=11220 | 25.6635 |
| 07966:4 | 30 | K.SC[+57.021]QHN[+2205.793]GTMYQHGEIFSAHELFPSR.L | HexNAc(4)Hex(5)Fuc(2)NeuAc(1) | C[+57], N[+2206] | 1260.5259 | 4 | 5039.0817 | 5039.0498 | 0 | 6.3 | 110 | Specific | 559.62 | 559.62 | 559.62 | 9.2 | 21 | >sp\|Q6WN34\|CRDL2_HUMAN Chordin-like protein 2 OS=Homo sapiens OX=9606 GN=CHRDL2 PE=1 SV=1 | 1189 | 20201119_10132_1ng_glyco_hcd203040_compound15.11191.11191.4 | controllerType=0 controllerNumber=1 scan=11191 | 25.5981 |
| 23610:3 | 35 | R.QLAHQSN[+2204.772]STNIFFSPVSIATAFAMLSLGTK.A | HexNAc(4)Hex(5)NeuAc(2) | N[+2205] | 1796.1478 | 3 | 5386.4288 | 5386.4075 | 0 | 4 | 64 | Specific | 319.21 | 252.24 | 252.24 | 7.61 | 20 | >sp\|P01009\|A1AT_HUMAN Alpha-1-antitrypsin OS=Homo sapiens OX=9606 GN=SERPINA1 PE=1 SV=3 | 642 | 20201119_10132_1ng_glyco_hcd203040_compound15.27547.27547.3 | controllerType=0 controllerNumber=1 scan=27547 | 55.9164 |
| 08959:3 | 40 | R.IIVPLNNREN[+1606.587]ISDPTSPLR.T | HexNAc(4)Hex(4)Fuc(1) | N[+1607] | 1252.2624 | 3 | 3754.7726 | 3754.7638 | 0 | 2.3 | 62 | Specific | 260.39 | 208.62 | 208.62 | 6.16 | 23 | >sp\|P01591\|IGJ_HUMAN Immunoglobulin J chain OS=Homo sapiens OX=9606 GN=JCHAIN PE=1 SV=4 | 1392 | 20201119_10132_1ng_glyco_hcd203040_compound15.12240.12240.3 | controllerType=0 controllerNumber=1 scan=12240 | 27.4045 |
| 08993:3 | 40 | R.IIVPLNNREN[+1460.529]ISDPTSPLR.T | HexNAc(4)Hex(4) | N[+1461] | 1203.5776 | 3 | 3608.7182 | 3608.7059 | 0 | 3.4 | 62 | Specific | 266.35 | 258.63 | 258.63 | 6.51 | 23 | >sp\|P01591\|IGJ_HUMAN Immunoglobulin J chain OS=Homo sapiens OX=9606 GN=JCHAIN PE=1 SV=4 | 1392 | 20201119_10132_1ng_glyco_hcd203040_compound15.12275.12275.3 | controllerType=0 controllerNumber=1 scan=12275 | 27.4628 |
| 08961:3 | 40 | R.IIVPLNNREN[+1606.587]ISDPTSPLR.T | HexNAc(4)Hex(4)Fuc(1) | N[+1607] | 1252.2624 | 3 | 3754.7726 | 3754.7638 | 0 | 2.3 | 62 | Specific | 293.03 | 187.85 | 187.85 | 6.02 | 23 | >sp\|P01591\|IGJ_HUMAN Immunoglobulin J chain OS=Homo sapiens OX=9606 GN=JCHAIN PE=1 SV=4 | 1392 | 20201119_10132_1ng_glyco_hcd203040_compound15.12242.12242.3 | controllerType=0 controllerNumber=1 scan=12242 | 27.4081 |
| 08995:3 | 40 | R.IIVPLNNREN[+1460.529]ISDPTSPLR.T | HexNAc(4)Hex(4) | N[+1461] | 1203.5776 | 3 | 3608.7182 | 3608.7059 | 0 | 3.4 | 62 | Specific | 310.7 | 279.14 | 279.14 | 6.61 | 23 | >sp\|P01591\|IGJ_HUMAN Immunoglobulin J chain OS=Homo sapiens OX=9606 GN=JCHAIN PE=1 SV=4 | 1392 | 20201119_10132_1ng_glyco_hcd203040_compound15.12277.12277.3 | controllerType=0 controllerNumber=1 scan=12277 | 27.4664 |
| 12668:3 | 40 | R.IIVPLNNREN[+2204.772]ISDPTSPLR.T | HexNAc(4)Hex(5)NeuAc(2) | N[+2205] | 1451.6633 | 3 | 4352.9752 | 4352.9496 | 0 | 5.9 | 62 | Specific | 354.97 | 341.05 | 341.05 | 6.89 | 23 | >sp\|P01591\|IGJ_HUMAN Immunoglobulin J chain OS=Homo sapiens OX=9606 GN=JCHAIN PE=1 SV=4 | 1392 | 20201119_10132_1ng_glyco_hcd203040_compound15.16106.16106.3 | controllerType=0 controllerNumber=1 scan=16106 | 34.0606 |
| 10605:3 | 40 | R.IIVPLNNREN[+2059.735]ISDPTSPLR.T | HexNAc(4)Hex(5)Fuc(1)NeuAc(1) | N[+2060] | 1403.3112 | 3 | 4207.9191 | 4207.9121 | 0 | 1.7 | 62 | Specific | 382.49 | 347.06 | 347.06 | 7.45 | 23 | >sp\|P01591\|IGJ_HUMAN Immunoglobulin J chain OS=Homo sapiens OX=9606 GN=JCHAIN PE=1 SV=4 | 1392 | 20201119_10132_1ng_glyco_hcd203040_compound15.13943.13943.3 | controllerType=0 controllerNumber=1 scan=13943 | 30.2742 |
| 10773:3 | 40 | R.IIVPLNNREN[+1856.656]ISDPTSPLR.T | HexNAc(3)Hex(5)Fuc(1)NeuAc(1) | N[+1857] | 1335.6212 | 3 | 4004.8489 | 4004.8327 | 0 | 4 | 62 | Specific | 403 | 336.97 | 336.97 | 7.72 | 23 | >sp\|P01591\|IGJ_HUMAN Immunoglobulin J chain OS=Homo sapiens OX=9606 GN=JCHAIN PE=1 SV=4 | 1392 | 20201119_10132_1ng_glyco_hcd203040_compound15.14118.14118.3 | controllerType=0 controllerNumber=1 scan=14118 | 30.6036 |
| 08963:3 | 40 | R.IIVPLNNREN[+1809.666]ISDPTSPLR.T | HexNAc(5)Hex(4)Fuc(1) | N[+1810] | 1319.9564 | 3 | 3957.8545 | 3957.8432 | 0 | 2.9 | 62 | Specific | 404.17 | 404.17 | 404.17 | 9.12 | 23 | >sp\|P01591\|IGJ_HUMAN Immunoglobulin J chain OS=Homo sapiens OX=9606 GN=JCHAIN PE=1 SV=4 | 1392 | 20201119_10132_1ng_glyco_hcd203040_compound15.12244.12244.3 | controllerType=0 controllerNumber=1 scan=12244 | 27.4115 |
| 10770:3 | 40 | R.IIVPLNNREN[+1856.656]ISDPTSPLR.T | HexNAc(3)Hex(5)Fuc(1)NeuAc(1) | N[+1857] | 1335.6212 | 3 | 4004.8489 | 4004.8327 | 0 | 4 | 62 | Specific | 411.94 | 330.04 | 330.04 | 7.72 | 23 | >sp\|P01591\|IGJ_HUMAN Immunoglobulin J chain OS=Homo sapiens OX=9606 GN=JCHAIN PE=1 SV=4 | 1392 | 20201119_10132_1ng_glyco_hcd203040_compound15.14115.14115.3 | controllerType=0 controllerNumber=1 scan=14115 | 30.5964 |
| 12573:3 | 40 | R.IIVPLNNREN[+2350.830]ISDPTSPLR.T | HexNAc(4)Hex(5)Fuc(1)NeuAc(2) | N[+2351] | 1500.3427 | 3 | 4499.0136 | 4499.0075 | 0 | 1.4 | 62 | Specific | 416.9 | 340.4 | 340.4 | 7.4 | 23 | >sp\|P01591\|IGJ_HUMAN Immunoglobulin J chain OS=Homo sapiens OX=9606 GN=JCHAIN PE=1 SV=4 | 1392 | 20201119_10132_1ng_glyco_hcd203040_compound15.16008.16008.3 | controllerType=0 controllerNumber=1 scan=16008 | 33.8911 |
| 12571:3 | 40 | R.IIVPLNNREN[+2350.830]ISDPTSPLR.T | HexNAc(4)Hex(5)Fuc(1)NeuAc(2) | N[+2351] | 1500.3427 | 3 | 4499.0136 | 4499.0075 | 0 | 1.4 | 62 | Specific | 428.31 | 405.11 | 405.11 | 8.91 | 23 | >sp\|P01591\|IGJ_HUMAN Immunoglobulin J chain OS=Homo sapiens OX=9606 GN=JCHAIN PE=1 SV=4 | 1392 | 20201119_10132_1ng_glyco_hcd203040_compound15.16006.16006.3 | controllerType=0 controllerNumber=1 scan=16006 | 33.8865 |
| 12670:3 | 40 | R.IIVPLNNREN[+2204.772]ISDPTSPLR.T | HexNAc(4)Hex(5)NeuAc(2) | N[+2205] | 1451.6633 | 3 | 4352.9752 | 4352.9496 | 0 | 5.9 | 62 | Specific | 430.5 | 430.5 | 430.5 | 8.66 | 23 | >sp\|P01591\|IGJ_HUMAN Immunoglobulin J chain OS=Homo sapiens OX=9606 GN=JCHAIN PE=1 SV=4 | 1392 | 20201119_10132_1ng_glyco_hcd203040_compound15.16108.16108.3 | controllerType=0 controllerNumber=1 scan=16108 | 34.0645 |
| 10603:3 | 40 | R.IIVPLNNREN[+2059.735]ISDPTSPLR.T | HexNAc(4)Hex(5)Fuc(1)NeuAc(1) | N[+2060] | 1403.3112 | 3 | 4207.9191 | 4207.9121 | 0 | 1.7 | 62 | Specific | 438.41 | 421.66 | 421.66 | 8.93 | 23 | >sp\|P01591\|IGJ_HUMAN Immunoglobulin J chain OS=Homo sapiens OX=9606 GN=JCHAIN PE=1 SV=4 | 1392 | 20201119_10132_1ng_glyco_hcd203040_compound15.13941.13941.3 | controllerType=0 controllerNumber=1 scan=13941 | 30.27 |
| 09007:3 | 40 | R.IIVPLNNREN[+1663.608]ISDPTSPLR.T | HexNAc(5)Hex(4) | N[+1664] | 1271.2692 | 3 | 3811.7931 | 3811.7853 | 0 | 2 | 62 | Specific | 466.39 | 466.39 | 466.39 | 9.16 | 23 | >sp\|P01591\|IGJ_HUMAN Immunoglobulin J chain OS=Homo sapiens OX=9606 GN=JCHAIN PE=1 SV=4 | 1392 | 20201119_10132_1ng_glyco_hcd203040_compound15.12290.12290.3 | controllerType=0 controllerNumber=1 scan=12290 | 27.4888 |
| 08965:3 | 40 | R.IIVPLNNREN[+1809.666]ISDPTSPLR.T | HexNAc(5)Hex(4)Fuc(1) | N[+1810] | 1319.9564 | 3 | 3957.8545 | 3957.8432 | 0 | 2.9 | 62 | Specific | 501.14 | 375.33 | 375.33 | 9.38 | 23 | >sp\|P01591\|IGJ_HUMAN Immunoglobulin J chain OS=Homo sapiens OX=9606 GN=JCHAIN PE=1 SV=4 | 1392 | 20201119_10132_1ng_glyco_hcd203040_compound15.12246.12246.3 | controllerType=0 controllerNumber=1 scan=12246 | 27.4156 |
| 09009:3 | 40 | R.IIVPLNNREN[+1663.608]ISDPTSPLR.T | HexNAc(5)Hex(4) | N[+1664] | 1271.2692 | 3 | 3811.7931 | 3811.7853 | 0 | 2 | 62 | Specific | 518.9 | 489.6 | 489.6 | 9.32 | 23 | >sp\|P01591\|IGJ_HUMAN Immunoglobulin J chain OS=Homo sapiens OX=9606 GN=JCHAIN PE=1 SV=4 | 1392 | 20201119_10132_1ng_glyco_hcd203040_compound15.12292.12292.3 | controllerType=0 controllerNumber=1 scan=12292 | 27.4937 |
| 17209:3 | 52 | R.NYIVPILWLN[+1864.634]ETGTIGDEK.A | HexNAc(2)Hex(9) | N[+1865] | 1347.2634 | 3 | 4039.7756 | 4039.7674 | 0 | 2 | 408 | Specific | 247.18 | 247.18 | 247.18 | 6.45 | 13 | >sp\|P16671\|CD36_HUMAN Platelet glycoprotein 4 OS=Homo sapiens OX=9606 GN=CD36 PE=1 SV=2 | 812 | 20201119_10132_1ng_glyco_hcd203040_compound15.20869.20869.3 | controllerType=0 controllerNumber=1 scan=20869 | 42.9166 |
| 17211:3 | 52 | R.NYIVPILWLN[+1864.634]ETGTIGDEK.A | HexNAc(2)Hex(9) | N[+1865] | 1347.2634 | 3 | 4039.7756 | 4039.7674 | 0 | 2 | 408 | Specific | 396.79 | 381.89 | 381.89 | 9.23 | 13 | >sp\|P16671\|CD36_HUMAN Platelet glycoprotein 4 OS=Homo sapiens OX=9606 GN=CD36 PE=1 SV=2 | 812 | 20201119_10132_1ng_glyco_hcd203040_compound15.20871.20871.3 | controllerType=0 controllerNumber=1 scan=20871 | 42.9201 |
| 06091:3 | 459 | R.[+42.011]VEDAYILTC[+57.021]N[+730.264]VSLEYEK.T | HexNAc(2)Hex(2) | .[42], C[+57], N[+730] | 940.0957 | 3 | 2818.2726 | 2818.2486 | 0 | 8.5 | 225 | Specific | 281.36 | 224.39 | 224.39 | 3.14 | 2 | >sp\|P40227\|TCPZ_HUMAN T-complex protein 1 subunit zeta OS=Homo sapiens OX=9606 GN=CCT6A PE=1 SV=3 | 1115 | 20201119_10132_1ng_glyco_hcd203040_compound15.9239.9239.3 | controllerType=0 controllerNumber=1 scan=9239 | 22.3195 |
| 06089:3 | 459 | R.[+42.011]VEDAYILTC[+57.021]N[+730.264]VSLEYEK.T | HexNAc(2)Hex(2) | .[42], C[+57], N[+730] | 940.0957 | 3 | 2818.2726 | 2818.2486 | 0 | 8.5 | 225 | Specific | 285.3 | 176.58 | 176.58 | 3.12 | 2 | >sp\|P40227\|TCPZ_HUMAN T-complex protein 1 subunit zeta OS=Homo sapiens OX=9606 GN=CCT6A PE=1 SV=3 | 1115 | 20201119_10132_1ng_glyco_hcd203040_compound15.9237.9237.3 | controllerType=0 controllerNumber=1 scan=9237 | 22.3155 |
| 18436:4 | 1 | R.RTAGWNVPIGTLRPFLN[+2059.735]WTGPPEPIEAAVAR.F | HexNAc(4)Hex(5)Fuc(1)NeuAc(1) | N[+2060] | 1362.3976 | 4 | 5446.5685 | 5446.547 | 0 | 4 | 140 | Specific | 286.99 | 286.99 | 286.99 | 6.36 | 162 | >sp\|P02788\|TRFL_HUMAN Lactotransferrin OS=Homo sapiens OX=9606 GN=LTF PE=1 SV=6 | 1270 | 20201112_10613_1ng_glyco_hcd203040_compound15.22064.22064.4 | controllerType=0 controllerNumber=1 scan=22064 | 44.1237 |
| 18083:4 | 1 | R.RTAGWNVPIGTLRPFLN[+2716.983]WTGPPEPIEAAVAR.F | HexNAc(5)Hex(6)Fuc(3)NeuAc(1) | N[+2717] | 1526.7081 | 4 | 6103.8107 | 6103.795 | 0 | 2.6 | 140 | Specific | 394.78 | 394.78 | 394.78 | 9.27 | 162 | >sp\|P02788\|TRFL_HUMAN Lactotransferrin OS=Homo sapiens OX=9606 GN=LTF PE=1 SV=6 | 1270 | 20201112_10613_1ng_glyco_hcd203040_compound15.21697.21697.4 | controllerType=0 controllerNumber=1 scan=21697 | 43.4226 |
| 18364:4 | 1 | R.RTAGWNVPIGTLRPFLN[+2059.735]WTGPPEPIEAAVAR.F | HexNAc(4)Hex(5)Fuc(1)NeuAc(1) | N[+2060] | 1362.3927 | 4 | 5446.549 | 5446.547 | 0 | 0.4 | 140 | Specific | 457.43 | 457.43 | 457.43 | 9.81 | 162 | >sp\|P02788\|TRFL_HUMAN Lactotransferrin OS=Homo sapiens OX=9606 GN=LTF PE=1 SV=6 | 1270 | 20201112_10613_1ng_glyco_hcd203040_compound15.21989.21989.2 | controllerType=0 controllerNumber=1 scan=21989 | 43.9686 |
| 18438:4 | 1 | R.RTAGWNVPIGTLRPFLN[+2059.735]WTGPPEPIEAAVAR.F | HexNAc(4)Hex(5)Fuc(1)NeuAc(1) | N[+2060] | 1362.3976 | 4 | 5446.5685 | 5446.547 | 0 | 4 | 140 | Specific | 489.13 | 489.13 | 489.13 | 9.26 | 162 | >sp\|P02788\|TRFL_HUMAN Lactotransferrin OS=Homo sapiens OX=9606 GN=LTF PE=1 SV=6 | 1270 | 20201112_10613_1ng_glyco_hcd203040_compound15.22066.22066.4 | controllerType=0 controllerNumber=1 scan=22066 | 44.1285 |
| 18084:4 | 1 | R.RTAGWNVPIGTLRPFLN[+2716.983]WTGPPEPIEAAVAR.F | HexNAc(5)Hex(6)Fuc(3)NeuAc(1) | N[+2717] | 1526.7081 | 4 | 6103.8107 | 6103.795 | 0 | 2.6 | 140 | Specific | 525.01 | 394.99 | 394.99 | 10.03 | 162 | >sp\|P02788\|TRFL_HUMAN Lactotransferrin OS=Homo sapiens OX=9606 GN=LTF PE=1 SV=6 | 1270 | 20201112_10613_1ng_glyco_hcd203040_compound15.21698.21698.4 | controllerType=0 controllerNumber=1 scan=21698 | 43.4273 |
| 17290:4 | 1 | R.RTAGWNVPIGTLRPFLN[+1768.640]WTGPPEPIEAAVAR.F | HexNAc(4)Hex(5)Fuc(1) | N[+1769] | 1289.6185 | 4 | 5155.4523 | 5155.4516 | 0 | 0.1 | 140 | Specific | 563.72 | 474.45 | 474.45 | 10.42 | 162 | >sp\|P02788\|TRFL_HUMAN Lactotransferrin OS=Homo sapiens OX=9606 GN=LTF PE=1 SV=6 | 1270 | 20201112_10613_1ng_glyco_hcd203040_compound15.20870.20870.4 | controllerType=0 controllerNumber=1 scan=20870 | 41.6494 |
| 17293:4 | 1 | R.RTAGWNVPIGTLRPFLN[+1768.640]WTGPPEPIEAAVAR.F | HexNAc(4)Hex(5)Fuc(1) | N[+1769] | 1289.6185 | 4 | 5155.4523 | 5155.4516 | 0 | 0.1 | 140 | Specific | 619.44 | 499.82 | 499.82 | 12.65 | 162 | >sp\|P02788\|TRFL_HUMAN Lactotransferrin OS=Homo sapiens OX=9606 GN=LTF PE=1 SV=6 | 1270 | 20201112_10613_1ng_glyco_hcd203040_compound15.20873.20873.4 | controllerType=0 controllerNumber=1 scan=20873 | 41.6558 |
| 18367:4 | 1 | R.RTAGWNVPIGTLRPFLN[+2059.735]WTGPPEPIEAAVAR.F | HexNAc(4)Hex(5)Fuc(1)NeuAc(1) | N[+2060] | 1362.3927 | 4 | 5446.549 | 5446.547 | 0 | 0.4 | 140 | Specific | 693.38 | 568.39 | 568.39 | 13.65 | 162 | >sp\|P02788\|TRFL_HUMAN Lactotransferrin OS=Homo sapiens OX=9606 GN=LTF PE=1 SV=6 | 1270 | 20201112_10613_1ng_glyco_hcd203040_compound15.21992.21992.2 | controllerType=0 controllerNumber=1 scan=21992 | 43.973 |
| 20584:4 | 1 | R.TAGWNVPIGTLRPFLN[+2863.041]WTGPPEPIEAAVAR.F | HexNAc(5)Hex(6)Fuc(4)NeuAc(1) | N[+2863] | 1524.1929 | 4 | 6093.7497 | 6093.7518 | 0 | -0.3 | 141 | Specific | 214.62 | 214.62 | 214.62 | 5.33 | 162 | >sp\|P02788\|TRFL_HUMAN Lactotransferrin OS=Homo sapiens OX=9606 GN=LTF PE=1 SV=6 | 1270 | 20201112_10613_1ng_glyco_hcd203040_compound15.24305.24305.4 | controllerType=0 controllerNumber=1 scan=24305 | 48.8163 |
| 18460:3 | 1 | R.TAGWNVPIGTLRPFLN[+2279.830]WTGPPEPIEAAVAR.F | HexNAc(5)Hex(6)Fuc(2) | N[+2280] | 1837.5214 | 3 | 5510.5498 | 5510.5406 | 0 | 1.7 | 141 | Specific | 226.35 | 226.35 | 226.35 | 6.82 | 162 | >sp\|P02788\|TRFL_HUMAN Lactotransferrin OS=Homo sapiens OX=9606 GN=LTF PE=1 SV=6 | 1270 | 20201112_10613_1ng_glyco_hcd203040_compound15.22089.22089.3 | controllerType=0 controllerNumber=1 scan=22089 | 44.1706 |
| 19369:4 | 1 | R.TAGWNVPIGTLRPFLN[+2716.983]WTGPPEPIEAAVAR.F | HexNAc(5)Hex(6)Fuc(3)NeuAc(1) | N[+2717] | 1487.6819 | 4 | 5947.7057 | 5947.6939 | 0 | 2 | 141 | Specific | 273.24 | 273.24 | 273.24 | 6.87 | 162 | >sp\|P02788\|TRFL_HUMAN Lactotransferrin OS=Homo sapiens OX=9606 GN=LTF PE=1 SV=6 | 1270 | 20201112_10613_1ng_glyco_hcd203040_compound15.23037.23037.4 | controllerType=0 controllerNumber=1 scan=23037 | 46.2084 |
| 19274:4 | 1 | R.TAGWNVPIGTLRPFLN[+2791.020]WTGPPEPIEAAVAR.F | HexNAc(6)Hex(7)Fuc(3) | N[+2791] | 1506.1875 | 4 | 6021.7282 | 6021.7307 | 0 | -0.4 | 141 | Specific | 282.83 | 282.83 | 282.83 | 5.11 | 162 | >sp\|P02788\|TRFL_HUMAN Lactotransferrin OS=Homo sapiens OX=9606 GN=LTF PE=1 SV=6 | 1270 | 20201112_10613_1ng_glyco_hcd203040_compound15.22938.22938.4 | controllerType=0 controllerNumber=1 scan=22938 | 46.0137 |
| 19422:3 | 1 | R.TAGWNVPIGTLRPFLN[+2424.867]WTGPPEPIEAAVAR.F | HexNAc(5)Hex(6)Fuc(1)NeuAc(1) | N[+2425] | 1885.8668 | 3 | 5655.5858 | 5655.5781 | 0 | 1.4 | 141 | Specific | 298.45 | 254.49 | 254.49 | 7.09 | 162 | >sp\|P02788\|TRFL_HUMAN Lactotransferrin OS=Homo sapiens OX=9606 GN=LTF PE=1 SV=6 | 1270 | 20201112_10613_1ng_glyco_hcd203040_compound15.23092.23092.3 | controllerType=0 controllerNumber=1 scan=23092 | 46.[illegible]241 |
| 20586:4 | 1 | R.TAGWNVPIGTLRPFLN[+2863.041]WTGPPEPIEAAVAR.F | HexNAc(5)Hex(6)Fuc(4)NeuAc(1) | N[+2863] | 1524.1929 | 4 | 6093.7497 | 6093.7518 | 0 | -0.3 | 141 | Specific | 305.28 | 305.28 | 305.28 | 7.36 | 162 | >sp\|P02788\|TRFL_HUMAN Lactotransferrin OS=Homo sapiens OX=9606 GN=LTF PE=1 SV=6 | 1270 | 20201112_10613_1ng_glyco_hcd203040_compound15.24307.24307.4 | controllerType=0 controllerNumber=1 scan=24307 | 4[illegible]09 |
| 20820:4 | 1 | R.TAGWNVPIGTLRPFLN[+2350.830]WTGPPEPIEAAVAR.F | HexNAc(4)Hex(5)Fuc(1)NeuAc(2) | N[+2351] | 1396.1417 | 4 | 5581.5451 | 5581.5413 | 0 | 0.7 | 141 | Specific | 320.27 | 199.84 | 199.84 | 6.41 | 162 | >sp\|P02788\|TRFL_HUMAN Lactotransferrin OS=Homo sapiens OX=9606 GN=LTF PE=1 SV=6 | 1270 | 20201112_10613_1ng_glyco_hcd203040_compound15.24552.24552.4 | controllerType=0 controllerNumber=1 scan=24552 | 49.[illegible]29 |
| 19485:3 | 1 | R.TAGWNVPIGTLRPFLN[+2205.793]WTGPPEPIEAAVAR.F | HexNAc(4)Hex(5)Fuc(2)NeuAc(1) | N[+2206] | 1812.8458 | 3 | 5436.5228 | 5436.5038 | 0 | 3.5 | 141 | Specific | 336.71 | 336.71 | 336.71 | 7.54 | 162 | >sp\|P02788\|TRFL_HUMAN Lactotransferrin OS=Homo sapiens OX=9606 GN=LTF PE=1 SV=6 | 1270 | 20201112_10613_1ng_glyco_hcd203040_compound15.23158.23158.3 | controllerType=0 controllerNumber=1 scan=23158 | 46.4785 |
| 20848:4 | 1 | R.[+42.011]TAGWNVPIGTLRPFLN[+2336.851]WTGPPEPIEAAVAR.F | HexNAc(6)Hex(6)Fuc(1) | .[42], N[+2337] | 1403.1414 | 4 | 5609.5436 | 5609.5726 | 0 | -5.2 | 141 | Specific | 338.38 | 338.38 | 338.38 | 6.22 | 162 | >sp\|P02788\|TRFL_HUMAN Lactotransferrin OS=Homo sapiens OX=9606 GN=LTF PE=1 SV=6 | 1270 | 20201112_10613_1ng_glyco_hcd203040_compound15.24581.24581.4 | controllerType=0 controllerNumber=1 scan=24581 | 49.4295 |
| 19647:4 | 1 | R.[+42.011]TAGWNVPIGTLRPFLN[+2018.708]WTGPPEPIEAAVAR.F | HexNAc(3)Hex(6)Fuc(1)NeuAc(1) | .[42], N[+2019] | 1323.6187 | 4 | 5291.4528 | 5291.4299 | 0 | 4.3 | 141 | Specific | 383.03 | 383.03 | 383.03 | 8.66 | 162 | >sp\|P02788\|TRFL_HUMAN Lactotransferrin OS=Homo sapiens OX=9606 GN=LTF PE=1 SV=6 | 1270 | 20201112_10613_1ng_glyco_hcd203040_compound15.23330.23330.2 | controllerType=0 controllerNumber=1 scan=23330 | 46.7982 |
| 18458:4 | 1 | R.TAGWNVPIGTLRPFLN[+2279.830]WTGPPEPIEAAVAR.F | HexNAc(5)Hex(6)Fuc(2) | N[+2280] | 1378.3933 | 4 | 5510.5514 | 5510.5406 | 0 | 2 | 141 | Specific | 386.96 | 386.96 | 386.96 | 9.5 | 162 | >sp\|P02788\|TRFL_HUMAN Lactotransferrin OS=Homo sapiens OX=9606 GN=LTF PE=1 SV=6 | 1270 | 20201112_10613_1ng_glyco_hcd203040_compound15.22087.22087.4 | controllerType=0 controllerNumber=1 scan=22087 | 44.1683 |
| 19977:4 | 1 | R.TAGWNVPIGTLRPFLN[+2059.735]WTGPPEPIEAAVAR.F | HexNAc(4)Hex(5)Fuc(1)NeuAc(1) | N[+2060] | 1323.3726 | 4 | 5290.4684 | 5290.4459 | 0 | 4.3 | 141 | Specific | 389.58 | 389.58 | 389.58 | 8.66 | 162 | >sp\|P02788\|TRFL_HUMAN Lactotransferrin OS=Homo sapiens OX=9606 GN=LTF PE=1 SV=6 | 1270 | 20201112_10613_1ng_glyco_hcd203040_compound15.23673.23673.4 | controllerType=0 controllerNumber=1 scan=23673 | 47.472 |
| 19666:3 | 1 | R.[+42.011]TAGWNVPIGTLRPFLN[+2018.708]WTGPPEPIEAAVAR.F | HexNAc(3)Hex(6)Fuc(1)NeuAc(1) | .[42], N[+2019] | 1764.4882 | 3 | 5291.4502 | 5291.4299 | 0 | 3.8 | 141 | Specific | 409.22 | 310.1 | 310.1 | 7.1 | 162 | >sp\|P02788\|TRFL_HUMAN Lactotransferrin OS=Homo sapiens OX=9606 GN=LTF PE=1 SV=6 | 1270 | 20201112_10613_1ng_glyco_hcd203040_compound15.23350.23350.3 | controllerType=0 controllerNumber=1 scan=23350 | 46.8372 |
| 19650:4 | 1 | R.[+42.011]TAGWNVPIGTLRPFLN[+2018.708]WTGPPEPIEAAVAR.F | HexNAc(3)Hex(6)Fuc(1)NeuAc(1) | .[42], N[+2019] | 1323.6187 | 4 | 5291.4528 | 5291.4299 | 0 | 4.3 | 141 | Specific | 431.71 | 419.33 | 419.33 | 8.91 | 162 | >sp\|P02788\|TRFL_HUMAN Lactotransferrin OS=Homo sapiens OX=9606 GN=LTF PE=1 SV=6 | 1270 | 20201112_10613_1ng_glyco_hcd203040_compound15.23333.23333.2 | controllerType=0 controllerNumber=1 scan=23333 | 46.8038 |
| 18462:3 | 1 | R.TAGWNVPIGTLRPFLN[+2279.830]WTGPPEPIEAAVAR.F | HexNAc(5)Hex(6)Fuc(2) | N[+2280] | 1837.5214 | 3 | 5510.5498 | 5510.5406 | 0 | 1.7 | 141 | Specific | 432.89 | 344.89 | 344.89 | 8.17 | 162 | >sp\|P02788\|TRFL_HUMAN Lactotransferrin OS=Homo sapiens OX=9606 GN=LTF PE=1 SV=6 | 1270 | 20201112_10613_1ng_glyco_hcd203040_compound15.22091.22091.3 | controllerType=0 controllerNumber=1 scan=22091 | 44.1748 |
| 18416:4 | 1 | R.TAGWNVPIGTLRPFLN[+2498.904]WTGPPEPIEAAVAR.F | HexNAc(6)Hex(7)Fuc(1) | N[+2499] | 1433.1626 | 4 | 5729.6286 | 5729.6148 | 0 | 2.4 | 141 | Specific | 435.32 | 398.83 | 398.83 | 9.44 | 162 | >sp\|P02788\|TRFL_HUMAN Lactotransferrin OS=Homo sapiens OX=9606 GN=LTF PE=1 SV=6 | 1270 | 20201112_10613_1ng_glyco_hcd203040_compound15.22043.22043.4 | controllerType=0 controllerNumber=1 scan=22043 | 44.066 |
| 19359:3 | 1 | R.TAGWNVPIGTLRPFLN[+2424.867]WTGPPEPIEAAVAR.F | HexNAc(5)Hex(6)Fuc(1)NeuAc(1) | N[+2425] | 1885.8674 | 3 | 5655.5876 | 5655.5781 | 0 | 1.7 | 141 | Specific | 437.61 | 302.17 | 302.17 | 8.07 | 162 | >sp\|P02788\|TRFL_HUMAN Lactotransferrin OS=Homo sapiens OX=9606 GN=LTF PE=1 SV=6 | 1270 | 20201112_10613_1ng_glyco_hcd203040_compound15.23026.23026.3 | controllerType=0 controllerNumber=1 scan=23026 | 46.1881 |
| 19648:3 | 1 | R.TAGWNVPIGTLRPFLN[+2059.735]WTGPPEPIEAAVAR.F | HexNAc(4)Hex(5)Fuc(1)NeuAc(1) | N[+2060] | 1764.1561 | 3 | 5290.4537 | 5290.4459 | 0 | 1.5 | 141 | Specific | 490.82 | 479.67 | 479.67 | 10.1 | 162 | >sp\|P02788\|TRFL_HUMAN Lactotransferrin OS=Homo sapiens OX=9606 GN=LTF PE=1 SV=6 | 1270 | 20201112_10613_1ng_glyco_hcd203040_compound15.23331.23331.3 | controllerType=0 controllerNumber=1 scan=23331 | 46.7993 |
| 19665:4 | 1 | R.TAGWNVPIGTLRPFLN[+2059.735]WTGPPEPIEAAVAR.F | HexNAc(4)Hex(5)Fuc(1)NeuAc(1) | N[+2060] | 1323.3781 | 4 | 5290.4904 | 5290.4459 | 0 | 8.4 | 141 | Specific | 490.82 | 490.82 | 490.82 | 8.47 | 162 | >sp\|P02788\|TRFL_HUMAN Lactotransferrin OS=Homo sapiens OX=9606 GN=LTF PE=1 SV=6 | 1270 | 20201112_10613_1ng_glyco_hcd203040_compound15.23349.23349.2 | controllerType=0 controllerNumber=1 scan=23349 | 46.8353 |
| 20810:5 | 1 | R.TAGWNVPIGTLRPFLN[+2350.830]WTGPPEPIEAAVAR.F | HexNAc(4)Hex(5)Fuc(1)NeuAc(2) | N[+2351] | 1117.1165 | 5 | 5581.5532 | 5581.5413 | 0 | 2.1 | 141 | Specific | 513.64 | 502.66 | 502.66 | 10.08 | 162 | >sp\|P02788\|TRFL_HUMAN Lactotransferrin OS=Homo sapiens OX=9606 GN=LTF PE=1 SV=6 | 1270 | 20201112_10613_1ng_glyco_hcd203040_compound15.24541.24541.5 | controllerType=0 controllerNumber=1 scan=24541 | 49.3413 |
| 19979:4 | 1 | R.TAGWNVPIGTLRPFLN[+2059.735]WTGPPEPIEAAVAR.F | HexNAc(4)Hex(5)Fuc(1)NeuAc(1) | N[+2060] | 1323.3726 | 4 | 5290.4684 | 5290.4459 | 0 | 4.3 | 141 | Specific | 514.01 | 457.67 | 457.67 | 9.14 | 162 | >sp\|P02788\|TRFL_HUMAN Lactotransferrin OS=Homo sapiens OX=9606 GN=LTF PE=1 SV=6 | 1270 | 20201112_10613_1ng_glyco_hcd203040_compound15.23675.23675.4 | controllerType=0 controllerNumber=1 scan=23675 | 47.4758 |
| 19480:3 | 1 | R.TAGWNVPIGTLRPFLN[+2059.735]WTGPPEPIEAAVAR.F | HexNAc(4)Hex(5)Fuc(1)NeuAc(1) | N[+2060] | 1764.1612 | 3 | 5290.4691 | 5290.4459 | 0 | 4.4 | 141 | Specific | 543.42 | 486.79 | 486.79 | 9.5 | 162 | >sp\|P02788\|TRFL_HUMAN Lactotransferrin OS=Homo sapiens OX=9606 GN=LTF PE=1 SV=6 | 1270 | 20201112_10613_1ng_glyco_hcd203040_compound15.23153.23153.3 | controllerType=0 controllerNumber=1 scan=23153 | 46.4666 |
| 19418:4 | 1 | R.TAGWNVPIGTLRPFLN[+2424.867]WTGPPEPIEAAVAR.F | HexNAc(5)Hex(6)Fuc(1)NeuAc(1) | N[+2425] | 1414.6522 | 4 | 5655.5871 | 5655.5781 | 0 | 1.6 | 141 | Specific | 543.7 | 318.39 | 318.39 | 8.5 | 162 | >sp\|P02788\|TRFL_HUMAN Lactotransferrin OS=Homo sapiens OX=9606 GN=LTF PE=1 SV=6 | 1270 | 20201112_10613_1ng_glyco_hcd203040_compound15.23088.23088.4 | controllerType=0 controllerNumber=1 scan=23088 | 46.3153 |
| 18418:4 | 1 | R.TAGWNVPIGTLRPFLN[+2498.904]WTGPPEPIEAAVAR.F | HexNAc(6)Hex(7)Fuc(1) | N[+2499] | 1433.1626 | 4 | 5729.6286 | 5729.6148 | 0 | 2.4 | 141 | Specific | 559.79 | 515.31 | 515.31 | 10.1 | 162 | >sp\|P02788\|TRFL_HUMAN Lactotransferrin OS=Homo sapiens OX=9606 GN=LTF PE=1 SV=6 | 1270 | 20201112_10613_1ng_glyco_hcd203040_compound15.22045.22045.4 | controllerType=0 controllerNumber=1 scan=22045 | 44.0701 |
| 19362:3 | 1 | R.TAGWNVPIGTLRPFLN[+2424.867]WTGPPEPIEAAVAR.F | HexNAc(5)Hex(6)Fuc(1)NeuAc(1) | N[+2425] | 1885.8674 | 3 | 5655.5876 | 5655.5781 | 0 | 1.7 | 141 | Specific | 572.41 | 428.81 | 428.81 | 10.49 | 162 | >sp\|P02788\|TRFL_HUMAN Lactotransferrin OS=Homo sapiens OX=9606 GN=LTF PE=1 SV=6 | 1270 | 20201112_10613_1ng_glyco_hcd203040_compound15.23029.23029.3 | controllerType=0 controllerNumber=1 scan=23029 | 46.1934 |
| 19348:4 | 1 | R.TAGWNVPIGTLRPFLN[+2424.867]WTGPPEPIEAAVAR.F | HexNAc(5)Hex(6)Fuc(1)NeuAc(1) | N[+2425] | 1414.6522 | 4 | 5655.5871 | 5655.5781 | 0 | 1.6 | 141 | Specific | 573.56 | 466.04 | 466.04 | 10.41 | 162 | >sp\|P02788\|TRFL_HUMAN Lactotransferrin OS=Homo sapiens OX=9606 GN=LTF PE=1 SV=6 | 1270 | 20201112_10613_1ng_glyco_hcd203040_compound15.23015.23015.4 | controllerType=0 controllerNumber=1 scan=23015 | 46.168 |
| 19461:3 | 1 | R.TAGWNVPIGTLRPFLN[+2059.735]WTGPPEPIEAAVAR.F | HexNAc(4)Hex(5)Fuc(1)NeuAc(1) | N[+2060] | 1764.1579 | 3 | 5290.4592 | 5290.4459 | 0 | 2.5 | 141 | Specific | 573.57 | 517.27 | 517.27 | 10.18 | 162 | >sp\|P02788\|TRFL_HUMAN Lactotransferrin OS=Homo sapiens OX=9606 GN=LTF PE=1 SV=6 | 1270 | 20201112_10613_1ng_glyco_hcd203040_compound15.23133.23133.3 | controllerType=0 controllerNumber=1 scan=23133 | 46.4245 |
| 19416:4 | 1 | R.TAGWNVPIGTLRPFLN[+2424.867]WTGPPEPIEAAVAR.F | HexNAc(5)Hex(6)Fuc(1)NeuAc(1) | N[+2425] | 1414.6522 | 4 | 5655.5871 | 5655.5781 | 0 | 1.6 | 141 | Specific | 585.77 | 464.75 | 464.75 | 11.64 | 162 | >sp\|P02788\|TRFL_HUMAN Lactotransferrin OS=Homo sapiens OX=9606 GN=LTF PE=1 SV=6 | 1270 | 20201112_10613_1ng_glyco_hcd203040_compound15.23086.23086.4 | controllerType=0 controllerNumber=1 scan=23086 | 46.3111 |
| 19443:3 | 1 | R.TAGWNVPIGTLRPFLN[+2205.793]WTGPPEPIEAAVAR.F | HexNAc(4)Hex(5)Fuc(2)NeuAc(1) | N[+2206] | 1812.8387 | 3 | 5436.5016 | 5436.5038 | 0 | -0.4 | 141 | Specific | 587.31 | 542.72 | 542.72 | 11.65 | 162 | >sp\|P02788\|TRFL_HUMAN Lactotransferrin OS=Homo sapiens OX=9606 GN=LTF PE=1 SV=6 | 1270 | 20201112_10613_1ng_glyco_hcd203040_compound15.23114.23114.3 | controllerType=0 controllerNumber=1 scan=23114 | 46.3737 |
| 19440:3 | 1 | R.TAGWNVPIGTLRPFLN[+2205.793]WTGPPEPIEAAVAR.F | HexNAc(4)Hex(5)Fuc(2)NeuAc(1) | N[+2206] | 1812.8387 | 3 | 5436.5016 | 5436.5038 | 0 | -0.4 | 141 | Specific | 589.57 | 486.97 | 486.97 | 11.65 | 162 | >sp\|P02788\|TRFL_HUMAN Lactotransferrin OS=Homo sapiens OX=9606 GN=LTF PE=1 SV=6 | 1270 | 20201112_10613_1ng_glyco_hcd203040_compound15.23111.23111.3 | controllerType=0 controllerNumber=1 scan=23111 | 46.3679 |
| 19356:5 | 1 | R.TAGWNVPIGTLRPFLN[+2570.925]WTGPPEPIEAAVAR.F | HexNAc(5)Hex(6)Fuc(2)NeuAc(1) | N[+2571] | 1161.1318 | 5 | 5801.6301 | 5801.636 | 0 | -1 | 141 | Specific | 626.88 | 602.55 | 602.55 | 12.36 | 162 | >sp\|P02788\|TRFL_HUMAN Lactotransferrin OS=Homo sapiens OX=9606 GN=LTF PE=1 SV=6 | 1270 | 20201112_10613_1ng_glyco_hcd203040_compound15.23023.23023.5 | controllerType=0 controllerNumber=1 scan=23023 | 46.1821 |
| 19488:3 | 1 | R.TAGWNVPIGTLRPFLN[+2205.793]WTGPPEPIEAAVAR.F | HexNAc(4)Hex(5)Fuc(2)NeuAc(1) | N[+2206] | 1812.8458 | 3 | 5436.5228 | 5436.5038 | 0 | 3.5 | 141 | Specific | 634.16 | 592.66 | 592.66 | 12.41 | 162 | >sp\|P02788\|TRFL_HUMAN Lactotransferrin OS=Homo sapiens OX=9606 GN=LTF PE=1 SV=6 | 1270 | 20201112_10613_1ng_glyco_hcd203040_compound15.23161.23161.3 | controllerType=0 controllerNumber=1 scan=23161 | 46.4876 |
| 19651:3 | 1 | R.TAGWNVPIGTLRPFLN[+2059.735]WTGPPEPIEAAVAR.F | HexNAc(4)Hex(5)Fuc(1)NeuAc(1) | N[+2060] | 1764.1561 | 3 | 5290.4537 | 5290.4459 | 0 | 1.5 | 141 | Specific | 643.37 | 526.98 | 526.98 | 12.72 | 162 | >sp\|P02788\|TRFL_HUMAN Lactotransferrin OS=Homo sapiens OX=9606 GN=LTF PE=1 SV=6 | 1270 | 20201112_10613_1ng_glyco_hcd203040_compound15.23334.23334.3 | controllerType=0 controllerNumber=1 scan=23334 | 46.8054 |
| 19437:4 | 1 | R.TAGWNVPIGTLRPFLN[+2205.793]WTGPPEPIEAAVAR.F | HexNAc(4)Hex(5)Fuc(2)NeuAc(1) | N[+2206] | 1359.8809 | 4 | 5436.5016 | 5436.5038 | 0 | -0.4 | 141 | Specific | 648.22 | 567.23 | 567.23 | 12.57 | 162 | >sp\|P02788\|TRFL_HUMAN Lactotransferrin OS=Homo sapiens OX=9606 GN=LTF PE=1 SV=6 | 1270 | 20201112_10613_1ng_glyco_hcd203040_compound15.23108.23108.4 | controllerType=0 controllerNumber=1 scan=23108 | 46.3602 |
| 18461:4 | 1 | R.TAGWNVPIGTLRPFLN[+2279.830]WTGPPEPIEAAVAR.F | HexNAc(5)Hex(6)Fuc(2) | N[+2280] | 1378.3933 | 4 | 5510.5514 | 5510.5406 | 0 | 2 | 141 | Specific | 657.98 | 557.27 | 557.27 | 13.64 | 162 | >sp\|P02788\|TRFL_HUMAN Lactotransferrin OS=Homo sapiens OX=9606 GN=LTF PE=1 SV=6 | 1270 | 20201112_10613_1ng_glyco_hcd203040_compound15.22090.22090.4 | controllerType=0 controllerNumber=1 scan=22090 | 44.1727 |
| 19371:4 | 1 | R.TAGWNVPIGTLRPFLN[+2716.983]WTGPPEPIEAAVAR.F | HexNAc(5)Hex(6)Fuc(3)NeuAc(1) | N[+2717] | 1487.6819 | 4 | 5947.7057 | 5947.6939 | 0 | 2 | 141 | Specific | 673.53 | 555.07 | 555.07 | 13.64 | 162 | >sp\|P02788\|TRFL_HUMAN Lactotransferrin OS=Homo sapiens OX=9606 GN=LTF PE=1 SV=6 | 1270 | 20201112_10613_1ng_glyco_hcd203040_compound15.23039.23039.4 | controllerType=0 controllerNumber=1 scan=23039 | 46.2119 |
| 19482:3 | 1 | R.TAGWNVPIGTLRPFLN[+2059.735]WTGPPEPIEAAVAR.F | HexNAc(4)Hex(5)Fuc(1)NeuAc(1) | N[+2060] | 1764.1612 | 3 | 5290.4691 | 5290.4459 | 0 | 4.4 | 141 | Specific | 675.1 | 547.69 | 547.69 | 12.88 | 162 | >sp\|P02788\|TRFL_HUMAN Lactotransferrin OS=Homo sapiens OX=9606 GN=LTF PE=1 SV=6 | 1270 | 20201112_10613_1ng_glyco_hcd203040_compound15.23155.23155.3 | controllerType=0 controllerNumber=1 scan=23155 | 46.4715 |
| 19351:4 | 1 | R.TAGWNVPIGTLRPFLN[+2424.867]WTGPPEPIEAAVAR.F | HexNAc(5)Hex(6)Fuc(1)NeuAc(1) | N[+2425] | 1414.6522 | 4 | 5655.5871 | 5655.5781 | 0 | 1.6 | 141 | Specific | 687.83 | 467.17 | 467.17 | 13.64 | 162 | >sp\|P02788\|TRFL_HUMAN Lactotransferrin OS=Homo sapiens OX=9606 GN=LTF PE=1 SV=6 | 1270 | 20201112_10613_1ng_glyco_hcd203040_compound15.23018.23018.4 | controllerType=0 controllerNumber=1 scan=23018 | 46.1731 |
| 18500:4 | 1 | R.TAGWNVPIGTLRPFLN[+1768.640]WTGPPEPIEAAVAR.F | HexNAc(4)Hex(5)Fuc(1) | N[+1769] | 1250.5936 | 4 | 4999.3527 | 4999.3504 | 0 | 0.4 | 141 | Specific | 708.52 | 665.71 | 665.71 | 14.57 | 162 | >sp\|P02788\|TRFL_HUMAN Lactotransferrin OS=Homo sapiens OX=9606 GN=LTF PE=1 SV=6 | 1270 | 20201112_10613_1ng_glyco_hcd203040_compound15.22131.22131.4 | controllerType=0 controllerNumber=1 scan=22131 | 44.2625 |
| 18501:3 | 1 | R.TAGWNVPIGTLRPFLN[+1768.640]WTGPPEPIEAAVAR.F | HexNAc(4)Hex(5)Fuc(1) | N[+1769] | 1667.1239 | 3 | 4999.357 | 4999.3504 | 0 | 1.3 | 141 | Specific | 709.12 | 597.31 | 597.31 | 14.72 | 162 | >sp\|P02788\|TRFL_HUMAN Lactotransferrin OS=Homo sapiens OX=9606 GN=LTF PE=1 SV=6 | 1270 | 20201112_10613_1ng_glyco_hcd203040_compound15.22132.22132.3 | controllerType=0 controllerNumber=1 scan=22132 | 44.264 |
| 18503:3 | 1 | R.TAGWNVPIGTLRPFLN[+1768.640]WTGPPEPIEAAVAR.F | HexNAc(4)Hex(5)Fuc(1) | N[+1769] | 1667.1239 | 3 | 4999.357 | 4999.3504 | 0 | 1.3 | 141 | Specific | 716.34 | 598.71 | 598.71 | 14.72 | 162 | >sp\|P02788\|TRFL_HUMAN Lactotransferrin OS=Homo sapiens OX=9606 GN=LTF PE=1 SV=6 | 1270 | 20201112_10613_1ng_glyco_hcd203040_compound15.22134.22134.3 | controllerType=0 controllerNumber=1 scan=22134 | 44.2715 |
| 19458:4 | 1 | R.TAGWNVPIGTLRPFLN[+2059.735]WTGPPEPIEAAVAR.F | HexNAc(4)Hex(5)Fuc(1)NeuAc(1) | N[+2060] | 1323.3709 | 4 | 5290.4616 | 5290.4459 | 0 | 3 | 141 | Specific | 717.49 | 685.48 | 685.48 | 14.33 | 162 | >sp\|P02788\|TRFL_HUMAN Lactotransferrin OS=Homo sapiens OX=9606 GN=LTF PE=1 SV=6 | 1270 | 20201112_10613_1ng_glyco_hcd203040_compound15.23130.23130.4 | controllerType=0 controllerNumber=1 scan=23130 | 46.4122 |
| 19434:4 | 1 | R.TAGWNVPIGTLRPFLN[+2351.851]WTGPPEPIEAAVAR.F | HexNAc(4)Hex(5)Fuc(3)NeuAc(1) | N[+2352] | 1396.3972 | 4 | 5582.5671 | 5582.5617 | 0 | 1 | 141 | Specific | 726.23 | 647 | 647 | 14.64 | 162 | >sp\|P02788\|TRFL_HUMAN Lactotransferrin OS=Homo sapiens OX=9606 GN=LTF PE=1 SV=6 | 1270 | 20201112_10613_1ng_glyco_hcd203040_compound15.23104.23104.4 | controllerType=0 controllerNumber=1 scan=23104 | 46.3487 |
| 19459:3 | 1 | R.TAGWNVPIGTLRPFLN[+2059.735]WTGPPEPIEAAVAR.F | HexNAc(4)Hex(5)Fuc(1)NeuAc(1) | N[+2060] | 1764.1579 | 3 | 5290.4592 | 5290.4459 | 0 | 2.5 | 141 | Specific | 733.26 | 591.72 | 591.72 | 14.41 | 162 | >sp\|P02788\|TRFL_HUMAN Lactotransferrin OS=Homo sapiens OX=9606 GN=LTF PE=1 SV=6 | 1270 | 20201112_10613_1ng_glyco_hcd203040_compound15.23131.23131.3 | controllerType=0 controllerNumber=1 scan=23131 | 46.4137 |
| 20822:4 | 1 | R.TAGWNVPIGTLRPFLN[+2350.830]WTGPPEPIEAAVAR.F | HexNAc(4)Hex(5)Fuc(1)NeuAc(2) | N[+2351] | 1396.1417 | 4 | 5581.5451 | 5581.5413 | 0 | 0.7 | 141 | Specific | 769.25 | 639.56 | 639.56 | 15.57 | 162 | >sp\|P02788\|TRFL_HUMAN Lactotransferrin OS=Homo sapiens OX=9606 GN=LTF PE=1 SV=6 | 1270 | 20201112_10613_1ng_glyco_hcd203040_compound15.24554.24554.4 | controllerType=0 controllerNumber=1 scan=24554 | 49.3668 |
| 19436:4 | 1 | R.TAGWNVPIGTLRPFLN[+2351.851]WTGPPEPIEAAVAR.F | HexNAc(4)Hex(5)Fuc(3)NeuAc(1) | N[+2352] | 1396.3972 | 4 | 5582.5671 | 5582.5617 | 0 | 1 | 141 | Specific | 824.42 | 695.39 | 695.39 | 16.64 | 162 | >sp\|P02788\|TRFL_HUMAN Lactotransferrin OS=Homo sapiens OX=9606 GN=LTF PE=1 SV=6 | 1270 | 20201112_10613_1ng_glyco_hcd203040_compound15.23106.23106.4 | controllerType=0 controllerNumber=1 scan=23106 | 46.3557 |
| 19439:4 | 1 | R.TAGWNVPIGTLRPFLN[+2205.793]WTGPPEPIEAAVAR.F | HexNAc(4)Hex(5)Fuc(2)NeuAc(1) | N[+2206] | 1359.8809 | 4 | 5436.5016 | 5436.5038 | 0 | -0.4 | 141 | Specific | 839.08 | 688.2 | 688.2 | 16.57 | 162 | >sp\|P02788\|TRFL_HUMAN Lactotransferrin OS=Homo sapiens OX=9606 GN=LTF PE=1 SV=6 | 1270 | 20201112_10613_1ng_glyco_hcd203040_compound15.23110.23110.4 | controllerType=0 controllerNumber=1 scan=23110 | 46.3671 |
| 19460:4 | 1 | R.TAGWNVPIGTLRPFLN[+2059.735]WTGPPEPIEAAVAR.F | HexNAc(4)Hex(5)Fuc(1)NeuAc(1) | N[+2060] | 1323.3709 | 4 | 5290.4616 | 5290.4459 | 0 | 3 | 141 | Specific | 860.07 | 707.02 | 707.02 | 17.33 | 162 | >sp\|P02788\|TRFL_HUMAN Lactotransferrin OS=Homo sapiens OX=9606 GN=LTF PE=1 SV=6 | 1270 | 20201112_10613_1ng_glyco_hcd203040_compound15.23132.23132.4 | controllerType=0 controllerNumber=1 scan=23132 | 46.4208 |
| 19500:5 | 1 | R.TAGWNVPIGTLRPFLN[+2059.735]WTGPPEPIEAAVAR.F | HexNAc(4)Hex(5)Fuc(1)NeuAc(1) | N[+2060] | 1058.8955 | 5 | 5290.4484 | 5290.4459 | 0 | 0.5 | 141 | Specific | 874.95 | 837.44 | 837.44 | 17.98 | 162 | >sp\|P02788\|TRFL_HUMAN Lactotransferrin OS=Homo sapiens OX=9606 GN=LTF PE=1 SV=6 | 1270 | 20201112_10613_1ng_glyco_hcd203040_compound15.23174.23174.5 | controllerType=0 controllerNumber=1 scan=23174 | 46.5165 |
| 18502:4 | 1 | R.TAGWNVPIGTLRPFLN[+1768.640]WTGPPEPIEAAVAR.F | HexNAc(4)Hex(5)Fuc(1) | N[+1769] | 1250.5936 | 4 | 4999.3527 | 4999.3504 | 0 | 0.4 | 141 | Specific | 875.26 | 733.11 | 733.11 | 17.57 | 162 | >sp\|P02788\|TRFL_HUMAN Lactotransferrin OS=Homo sapiens OX=9606 GN=LTF PE=1 SV=6 | 1270 | 20201112_10613_1ng_glyco_hcd203040_compound15.22133.22133.4 | controllerType=0 controllerNumber=1 scan=22133 | 44.2687 |
| 19667:4 | 1 | R.TAGWNVPIGTLRPFLN[+2059.735]WTGPPEPIEAAVAR.F | HexNAc(4)Hex(5)Fuc(1)NeuAc(1) | N[+2060] | 1323.3781 | 4 | 5290.4904 | 5290.4459 | 0 | 8.4 | 141 | Specific | 875.92 | 863.99 | 863.99 | 16.09 | 162 | >sp\|P02788\|TRFL_HUMAN Lactotransferrin OS=Homo sapiens OX=9606 GN=LTF PE=1 SV=6 | 1270 | 20201112_10613_1ng_glyco_hcd203040_compound15.23351.23351.2 | controllerType=0 controllerNumber=1 scan=23351 | 46.8404 |
| 19502:5 | 1 | R.TAGWNVPIGTLRPFLN[+2059.735]WTGPPEPIEAAVAR.F | HexNAc(4)Hex(5)Fuc(1)NeuAc(1) | N[+2060] | 1058.8955 | 5 | 5290.4484 | 5290.4459 | 0 | 0.5 | 141 | Specific | 907.28 | 863.89 | 863.89 | 18.98 | 162 | >sp\|P02788\|TRFL_HUMAN Lactotransferrin OS=Homo sapiens OX=9606 GN=LTF PE=1 SV=6 | 1270 | 20201112_10613_1ng_glyco_hcd203040_compound15.23176.23176.5 | controllerType=0 controllerNumber=1 scan=23176 | 46.5234 |
| 20812:5 | 1 | R.TAGWNVPIGTLRPFLN[+2350.830]WTGPPEPIEAAVAR.F | HexNAc(4)Hex(5)Fuc(1)NeuAc(2) | N[+2351] | 1117.1165 | 5 | 5581.5532 | 5581.5413 | 0 | 2.1 | 141 | Specific | 910.22 | 741.09 | 741.09 | 18.74 | 162 | >sp\|P02788\|TRFL_HUMAN Lactotransferrin OS=Homo sapiens OX=9606 GN=LTF PE=1 SV=6 | 1270 | 20201112_10613_1ng_glyco_hcd203040_compound15.24543.24543.5 | controllerType=0 controllerNumber=1 scan=24543 | 49.3448 |
| 16890:3 | 1 | R.[+42.011]TAGWNIPMGLLFN[+1872.651]QTGSC[+57.021]K.F | HexNAc(3)Hex(6)NeuAc(1) | .[42], C[+57], N[+1873] | 1337.2397 | 3 | 4009.7045 | 4009.671 | 0 | 8.4 | 485 | Specific | 218.52 | 187.9 | 187.9 | 2.08 | 162 | >sp\|P02788\|TRFL_HUMAN Lactotransferrin OS=Homo sapiens OX=9606 GN=LTF PE=1 SV=6 | 1270 | 20201112_10613_1ng_glyco_hcd203040_compound15.20453.20453.3 | controllerType=0 controllerNumber=1 scan=20453 | 40.7864 |
| 16983:2 | 1 | R.TAGWNIPMGLLFN[+1768.640]QTGSC[+57.021]K.F | HexNAc(4)Hex(5)Fuc(1) | C[+57], N[+1769] | 1932.3325 | 2 | 3863.6577 | 3863.6495 | 0 | 2.1 | 485 | Specific | 429.74 | 423.61 | 423.61 | 9.33 | 162 | >sp\|P02788\|TRFL_HUMAN Lactotransferrin OS=Homo sapiens OX=9606 GN=LTF PE=1 SV=6 | 1270 | 20201112_10613_1ng_glyco_hcd203040_compound15.20550.20550.2 | controllerType=0 controllerNumber=1 scan=20550 | 40.9968 |
| 16985:2 | 1 | R.TAGWNIPMGLLFN[+1768.640]QTGSC[+57.021]K.F | HexNAc(4)Hex(5)Fuc(1) | C[+57], N[+1769] | 1932.3325 | 2 | 3863.6577 | 3863.6495 | 0 | 2.1 | 485 | Specific | 466.67 | 450.36 | 450.36 | 9.38 | 162 | >sp\|P02788\|TRFL_HUMAN Lactotransferrin OS=Homo sapiens OX=9606 GN=LTF PE=1 SV=6 | 1270 | 20201112_10613_1ng_glyco_hcd203040_compound15.20552.20552.2 | controllerType=0 controllerNumber=1 scan=20552 | 41.0001 |
| 17287:3 | 1 | R.TAGWNIPMGLLFN[+1460.529]QTGSC[+57.021]K.F | HexNAc(4)Hex(4) | C[+57], N[+1461] | 1185.8509 | 3 | 3555.5381 | 3555.5387 | 0 | -0.2 | 485 | Specific | 477.4 | 459.04 | 180.35 | 9.77 | 162 | >sp\|P02788\|TRFL_HUMAN Lactotransferrin OS=Homo sapiens OX=9606 GN=LTF PE=1 SV=6 | 1270 | 20201112_10613_1ng_glyco_hcd203040_compound15.20867.20867.3 | controllerType=0 controllerNumber=1 scan=20867 | 41.6408 |
| 18914:3 | 1 | R.TAGWNIPMGLLFN[+2059.735]QTGSC[+57.021]K.F | HexNAc(4)Hex(5)Fuc(1)NeuAc(1) | C[+57], N[+2060] | 1385.5859 | 3 | 4154.7431 | 4154.7449 | 0 | -0.4 | 485 | Specific | 481.25 | 481.25 | 481.25 | 9.77 | 162 | >sp\|P02788\|TRFL_HUMAN Lactotransferrin OS=Homo sapiens OX=9606 GN=LTF PE=1 SV=6 | 1270 | 20201112_10613_1ng_glyco_hcd203040_compound15.22563.22563.4 | controllerType=0 controllerNumber=1 scan=22563 | 45.1893 |
| 20737:3 | 1 | R.TAGWNIPMGLLFN[+2350.830]QTGSC[+57.021]K.F | HexNAc(4)Hex(5)Fuc(1)NeuAc(2) | C[+57], N[+2351] | 1482.6243 | 3 | 4445.8584 | 4445.8403 | 0 | 4.1 | 485 | Specific | 494.6 | 369.96 | 369.96 | 7.59 | 162 | >sp\|P02788\|TRFL_HUMAN Lactotransferrin OS=Homo sapiens OX=9606 GN=LTF PE=1 SV=6 | 1270 | 20201112_10613_1ng_glyco_hcd203040_compound15.24465.24465.3 | controllerType=0 controllerNumber=1 scan=24465 | 49.1712 |
| 19065:3 | 1 | R.TAGWNIPMGLLFN[+1751.624]QTGSC[+57.021]K.F | HexNAc(4)Hex(4)NeuAc(1) | C[+57], N[+1752] | 1282.8847 | 3 | 3846.6395 | 3846.6341 | 0 | 1.4 | 485 | Specific | 545.19 | 545.19 | 545.19 | 10.07 | 162 | >sp\|P02788\|TRFL_HUMAN Lactotransferrin OS=Homo sapiens OX=9606 GN=LTF PE=1 SV=6 | 1270 | 20201112_10613_1ng_glyco_hcd203040_compound15.22721.22721.3 | controllerType=0 controllerNumber=1 scan=22721 | 45.5684 |
| 20824:3 | 1 | R.TAGWNIPMGLLFN[+2350.830]QTGSC[+57.021]K.F | HexNAc(4)Hex(5)Fuc(1)NeuAc(2) | C[+57], N[+2351] | 1482.6194 | 3 | 4445.8438 | 4445.8403 | 0 | 0.8 | 485 | Specific | 574.32 | 548.38 | 548.38 | 10.22 | 162 | >sp\|P02788\|TRFL_HUMAN Lactotransferrin OS=Homo sapiens OX=9606 GN=LTF PE=1 SV=6 | 1270 | 20201112_10613_1ng_glyco_hcd203040_compound15.24556.24556.3 | controllerType=0 controllerNumber=1 scan=24556 | 49.3697 |
| 20739:3 | 1 | R.TAGWNIPMGLLFN[+2350.830]QTGSC[+57.021]K.F | HexNAc(4)Hex(5)Fuc(1)NeuAc(2) | C[+57], N[+2351] | 1482.6243 | 3 | 4445.8584 | 4445.8403 | 0 | 4.1 | 485 | Specific | 747.72 | 674.2 | 674.2 | 13.61 | 162 | >sp\|P02788\|TRFL_HUMAN Lactotransferrin OS=Homo sapiens OX=9606 GN=LTF PE=1 SV=6 | 1270 | 20201112_10613_1ng_glyco_hcd203040_compound15.24467.24467.3 | controllerType=0 controllerNumber=1 scan=24467 | 49.1755 |
| 17347:2 | 1 | R.TAGWNIPMGLLFN[+1216.423]QTGSC[+57.021]K.F | HexNAc(2)Hex(5) | C[+57], N[+1216] | 1656.2196 | 2 | 3311.4319 | 3311.4328 | 0 | -0.3 | 485 | Specific | 764.02 | 755.12 | 245.78 | 15.46 | 162 | >sp\|P02788\|TRFL_HUMAN Lactotransferrin OS=Homo sapiens OX=9606 GN=LTF PE=1 SV=6 | 1270 | 20201112_10613_1ng_glyco_hcd203040_compound15.20929.20929.2 | controllerType=0 controllerNumber=1 scan=20929 | 41.77 |
| 17289:3 | 1 | R.TAGWNIPMGLLFN[+1460.529]QTGSC[+57.021]K.F | HexNAc(4)Hex(4) | C[+57], N[+1461] | 1185.8509 | 3 | 3555.5381 | 3555.5387 | 0 | -0.2 | 485 | Specific | 771.98 | 744.72 | 426.81 | 15.38 | 162 | >sp\|P02788\|TRFL_HUMAN Lactotransferrin OS=Homo sapiens OX=9606 GN=LTF PE=1 SV=6 | 1270 | 20201112_10613_1ng_glyco_hcd203040_compound15.20869.20869.3 | controllerType=0 controllerNumber=1 scan=20869 | 41.6475 |
| 17072:3 | 1 | R.TAGWNIPMGLLFN[+1606.587]QTGSC[+57.021]K.F | HexNAc(4)Hex(4)Fuc(1) | C[+57], N[+1607] | 1234.539 | 3 | 3701.6024 | 3701.5966 | 0 | 1.6 | 485 | Specific | 788.29 | 788.29 | 788.29 | 15.45 | 162 | >sp\|P02788\|TRFL_HUMAN Lactotransferrin OS=Homo sapiens OX=9606 GN=LTF PE=1 SV=6 | 1270 | 20201112_10613_1ng_glyco_hcd203040_compound15.20643.20643.3 | controllerType=0 controllerNumber=1 scan=20643 | 41.1785 |
| 17384:3 | 1 | R.TAGWNIPMGLLFN[+1216.423]QTGSC[+57.021]K.F | HexNAc(2)Hex(5) | C[+57], N[+1216] | 1104.4846 | 3 | 3311.4394 | 3311.4328 | 0 | 2 | 485 | Specific | 804.53 | 804.53 | 302.84 | 16.45 | 162 | >sp\|P02788\|TRFL_HUMAN Lactotransferrin OS=Homo sapiens OX=9606 GN=LTF PE=1 SV=6 | 1270 | 20201112_10613_1ng_glyco_hcd203040_compound15.20968.20968.3 | controllerType=0 controllerNumber=1 scan=20968 | 41.8538 |
| 18972:4 | 1 | R.TAGWNIPMGLLFN[+1913.677]QTGSC[+57.021]K.F | HexNAc(4)Hex(5)NeuAc(1) | C[+57], N[+1914] | 1002.9275 | 4 | 4008.6881 | 4008.687 | 0 | 0.3 | 485 | Specific | 868.67 | 657.97 | 657.97 | 17.79 | 162 | >sp\|P02788\|TRFL_HUMAN Lactotransferrin OS=Homo sapiens OX=9606 GN=LTF PE=1 SV=6 | 1270 | 20201112_10613_1ng_glyco_hcd203040_compound15.22623.22623.4 | controllerType=0 controllerNumber=1 scan=22623 | 45.3191 |
| 17319:3 | 1 | R.TAGWNIPMGLLFN[+1216.423]QTGSC[+57.021]K.F | HexNAc(2)Hex(5) | C[+57], N[+1216] | 1104.4821 | 3 | 3311.4317 | 3311.4328 | 0 | -0.3 | 485 | Specific | 876.16 | 876.16 | 366.19 | 17.38 | 162 | >sp\|P02788\|TRFL_HUMAN Lactotransferrin OS=Homo sapiens OX=9606 GN=LTF PE=1 SV=6 | 1270 | 20201112_10613_1ng_glyco_hcd203040_compound15.20900.20900.3 | controllerType=0 controllerNumber=1 scan=20900 | 41.7054 |
| 17164:3 | 1 | R.TAGWNIPMGLLFN[+1622.582]QTGSC[+57.021]K.F | HexNAc(4)Hex(5) | C[+57], N[+1623] | 1239.8704 | 3 | 3717.5967 | 3717.5916 | 0 | 1.4 | 485 | Specific | 876.96 | 863.12 | 433.82 | 17.45 | 162 | >sp\|P02788\|TRFL_HUMAN Lactotransferrin OS=Homo sapiens OX=9606 GN=LTF PE=1 SV=6 | 1270 | 20201112_10613_1ng_glyco_hcd203040_compound15.20739.20739.3 | controllerType=0 controllerNumber=1 scan=20739 | 41.378 |
| 17074:3 | 1 | R.TAGWNIPMGLLFN[+1606.587]QTGSC[+57.021]K.F | HexNAc(4)Hex(4)Fuc(1) | C[+57], N[+1607] | 1234.539 | 3 | 3701.6024 | 3701.5966 | 0 | 1.6 | 485 | Specific | 883.41 | 738.22 | 738.22 | 17.45 | 162 | >sp\|P02788\|TRFL_HUMAN Lactotransferrin OS=Homo sapiens OX=9606 GN=LTF PE=1 SV=6 | 1270 | 20201112_10613_1ng_glyco_hcd203040_compound15.20645.20645.3 | controllerType=0 controllerNumber=1 scan=20645 | 41.184 |
| 18815:4 | 1 | R.TAGWNIPMGLLFN[+2059.735]QTGSC[+57.021]K.F | HexNAc(4)Hex(5)Fuc(1)NeuAc(1) | C[+57], N[+2060] | 1039.4424 | 4 | 4154.7479 | 4154.7449 | 0 | 0.7 | 485 | Specific | 884.73 | 671.55 | 671.55 | 17.86 | 162 | >sp\|P02788\|TRFL_HUMAN Lactotransferrin OS=Homo sapiens OX=9606 GN=LTF PE=1 SV=6 | 1270 | 20201112_10613_1ng_glyco_hcd203040_compound15.22459.22459.4 | controllerType=0 controllerNumber=1 scan=22459 | 44.9554 |
| 17162:3 | 1 | R.TAGWNIPMGLLFN[+1622.582]QTGSC[+57.021]K.F | HexNAc(4)Hex(5) | C[+57], N[+1623] | 1239.8704 | 3 | 3717.5967 | 3717.5916 | 0 | 1.4 | 485 | Specific | 892.01 | 808.89 | 399.28 | 17.45 | 162 | >sp\|P02788\|TRFL_HUMAN Lactotransferrin OS=Homo sapiens OX=9606 GN=LTF PE=1 SV=6 | 1270 | 20201112_10613_1ng_glyco_hcd203040_compound15.20737.20737.3 | controllerType=0 controllerNumber=1 scan=20737 | 41.3729 |
| 18789:3 | 1 | R.TAGWNIPMGLLFN[+2059.735]QTGSC[+57.021]K.F | HexNAc(4)Hex(5)Fuc(1)NeuAc(1) | C[+57], N[+2060] | 1385.5895 | 3 | 4154.7541 | 4154.7449 | 0 | 2.2 | 485 | Specific | 892.07 | 847.31 | 847.31 | 17.14 | 162 | >sp\|P02788\|TRFL_HUMAN Lactotransferrin OS=Homo sapiens OX=9606 GN=LTF PE=1 SV=6 | 1270 | 20201112_10613_1ng_glyco_hcd203040_compound15.22432.22432.3 | controllerType=0 controllerNumber=1 scan=22432 | 44.8975 |
| 16930:3 | 1 | R.TAGWNIPMGLLFN[+1768.640]QTGSC[+57.021]K.F | HexNAc(4)Hex(5)Fuc(1) | C[+57], N[+1769] | 1288.5574 | 3 | 3863.6577 | 3863.6495 | 0 | 2.1 | 485 | Specific | 894.75 | 731.1 | 731.1 | 17.14 | 162 | >sp\|P02788\|TRFL_HUMAN Lactotransferrin OS=Homo sapiens OX=9606 GN=LTF PE=1 SV=6 | 1270 | 20201112_10613_1ng_glyco_hcd203040_compound15.20495.20495.3 | controllerType=0 controllerNumber=1 scan=20495 | 40.8705 |
| 18778:3 | 1 | R.TAGWNIPMGLLFN[+2205.793]QTGSC[+57.021]K.F | HexNAc(4)Hex(5)Fuc(2)NeuAc(1) | C[+57], N[+2206] | 1434.2754 | 3 | 4300.8118 | 4300.8028 | 0 | 2.1 | 485 | Specific | 899.75 | 889.16 | 889.16 | 17.14 | 162 | >sp\|P02788\|TRFL_HUMAN Lactotransferrin OS=Homo sapiens OX=9606 GN=LTF PE=1 SV=6 | 1270 | 20201112_10613_1ng_glyco_hcd203040_compound15.22421.22421.3 | controllerType=0 controllerNumber=1 scan=22421 | 44.8712 |
| 17345:2 | 1 | R.TAGWNIPMGLLFN[+1216.423]QTGSC[+57.021]K.F | HexNAc(2)Hex(5) | C[+57], N[+1216] | 1656.2196 | 2 | 3311.4319 | 3311.4328 | 0 | -0.3 | 485 | Specific | 900.39 | 900.39 | 325.56 | 18.46 | 162 | >sp\|P02788\|TRFL_HUMAN Lactotransferrin OS=Homo sapiens OX=9606 GN=LTF PE=1 SV=6 | 1270 | 20201112_10613_1ng_glyco_hcd203040_compound15.20927.20927.2 | controllerType=0 controllerNumber=1 scan=20927 | 41.7632 |

Table S4. Cont.

| Query #:z | Protein Rank | Glycopeptide | Glycan | Modification Type(s) | Observed m/z | z | Observed (M+H) | Calc. mass (M+H) | Off-by-x error | Mass error (ppm) | StartPosition | Cleavage | Score | Delta | Delta Mod | \|Log Prob\| | # of unique peptides | Protein Name | Protein DB number | Comment | Scan # | Scan Time |
|---|---|---|---|---|---|---|---|---|---|---|---|---|---|---|---|---|---|---|---|---|---|---|
| 20826:3 | 1 | R.TAGWNIPMGLLFN[+2350.830]QTGSC[+57.021]K.F | HexNAc(4)Hex(5)Fuc(1)NeuAc(2) | C[+57], N[+2351] | 1482.6194 | 3 | 4445.8438 | 4445.8403 | 0 | 0.8 | 485 | Specific | 906.16 | 765.55 | 765.55 | 18.45 | 162 | >sp\|P02788\|TRFL_HUMAN Lactotransferrin OS=Homo sapiens OX=9606 GN=LTF PE=1 SV=6 | 1270 | 20201112_10613_1ng_glyco_hcd203040_compound15.24558.24558.3 | controllerType=0 controllerNumber=1 scan=24558 | 49.3733 |
| 18969:4 | 1 | R.TAGWNIPMGLLFN[+1913.677]QTGSC[+57.021]K.F | HexNAc(4)Hex(5)NeuAc(1) | C[+57], N[+1914] | 1002.9275 | 4 | 4008.6881 | 4008.687 | 0 | 0.3 | 485 | Specific | 907.81 | 759.66 | 759.66 | 18.79 | 162 | >sp\|P02788\|TRFL_HUMAN Lactotransferrin OS=Homo sapiens OX=9606 GN=LTF PE=1 SV=6 | 1270 | 20201112_10613_1ng_glyco_hcd203040_compound15.22620.22620.4 | controllerType=0 controllerNumber=1 scan=22620 | 45.307 |
| 18962:3 | 1 | R.TAGWNIPMGLLFN[+1913.677]QTGSC[+57.021]K.F | HexNAc(4)Hex(5)NeuAc(1) | C[+57], N[+1914] | 1336.903 | 3 | 4008.6945 | 4008.687 | 0 | 1.9 | 485 | Specific | 909.77 | 872.82 | 872.82 | 18.45 | 162 | >sp\|P02788\|TRFL_HUMAN Lactotransferrin OS=Homo sapiens OX=9606 GN=LTF PE=1 SV=6 | 1270 | 20201112_10613_1ng_glyco_hcd203040_compound15.22613.22613.3 | controllerType=0 controllerNumber=1 scan=22613 | 45.2917 |
| 19067:3 | 1 | R.TAGWNIPMGLLFN[+1751.624]QTGSC[+57.021]K.F | HexNAc(4)Hex(4)NeuAc(1) | C[+57], N[+1752] | 1282.8847 | 3 | 3846.6395 | 3846.6341 | 0 | 1.4 | 485 | Specific | 920.87 | 823.55 | 823.55 | 18.45 | 162 | >sp\|P02788\|TRFL_HUMAN Lactotransferrin OS=Homo sapiens OX=9606 GN=LTF PE=1 SV=6 | 1270 | 20201112_10613_1ng_glyco_hcd203040_compound15.22723.22723.3 | controllerType=0 controllerNumber=1 scan=22723 | 45.5721 |
| 16927:3 | 1 | R.TAGWNIPMGLLFN[+1768.640]QTGSC[+57.021]K.F | HexNAc(4)Hex(5)Fuc(1) | C[+57], N[+1769] | 1288.5574 | 3 | 3863.6577 | 3863.6495 | 0 | 2.1 | 485 | Specific | 928.58 | 826.51 | 826.51 | 18.14 | 162 | >sp\|P02788\|TRFL_HUMAN Lactotransferrin OS=Homo sapiens OX=9606 GN=LTF PE=1 SV=6 | 1270 | 20201112_10613_1ng_glyco_hcd203040_compound15.20492.20492.3 | controllerType=0 controllerNumber=1 scan=20492 | 40.8625 |
| 18792:3 | 1 | R.TAGWNIPMGLLFN[+2059.735]QTGSC[+57.021]K.F | HexNAc(4)Hex(5)Fuc(1)NeuAc(1) | C[+57], N[+2060] | 1385.5895 | 3 | 4154.7541 | 4154.7449 | 0 | 2.2 | 485 | Specific | 933.05 | 929.89 | 929.89 | 18.14 | 162 | >sp\|P02788\|TRFL_HUMAN Lactotransferrin OS=Homo sapiens OX=9606 GN=LTF PE=1 SV=6 | 1270 | 20201112_10613_1ng_glyco_hcd203040_compound15.22435.22435.3 | controllerType=0 controllerNumber=1 scan=22435 | 44.9084 |
| 18812:4 | 1 | R.TAGWNIPMGLLFN[+2059.735]QTGSC[+57.021]K.F | HexNAc(4)Hex(5)Fuc(1)NeuAc(1) | C[+57], N[+2060] | 1039.4424 | 4 | 4154.7479 | 4154.7449 | 0 | 0.7 | 485 | Specific | 947.09 | 830.01 | 830.01 | 18.86 | 162 | >sp\|P02788\|TRFL_HUMAN Lactotransferrin OS=Homo sapiens OX=9606 GN=LTF PE=1 SV=6 | 1270 | 20201112_10613_1ng_glyco_hcd203040_compound15.22456.22456.4 | controllerType=0 controllerNumber=1 scan=22456 | 44.9484 |
| 17321:3 | 1 | R.TAGWNIPMGLLFN[+1216.423]QTGSC[+57.021]K.F | HexNAc(2)Hex(5) | C[+57], N[+1216] | 1104.4821 | 3 | 3311.4317 | 3311.4328 | 0 | -0.3 | 485 | Specific | 952.41 | 859.39 | 433.44 | 19.38 | 162 | >sp\|P02788\|TRFL_HUMAN Lactotransferrin OS=Homo sapiens OX=9606 GN=LTF PE=1 SV=6 | 1270 | 20201112_10613_1ng_glyco_hcd203040_compound15.20902.20902.3 | controllerType=0 controllerNumber=1 scan=20902 | 41.7112 |
| 18960:3 | 1 | R.TAGWNIPMGLLFN[+1913.677]QTGSC[+57.021]K.F | HexNAc(4)Hex(5)NeuAc(1) | C[+57], N[+1914] | 1336.903 | 3 | 4008.6945 | 4008.687 | 0 | 1.9 | 485 | Specific | 955.23 | 942.38 | 942.38 | 19.45 | 162 | >sp\|P02788\|TRFL_HUMAN Lactotransferrin OS=Homo sapiens OX=9606 GN=LTF PE=1 SV=6 | 1270 | 20201112_10613_1ng_glyco_hcd203040_compound15.22611.22611.3 | controllerType=0 controllerNumber=1 scan=22611 | 45.2858 |
| 18776:3 | 1 | R.TAGWNIPMGLLFN[+2205.793]QTGSC[+57.021]K.F | HexNAc(4)Hex(5)Fuc(2)NeuAc(1) | C[+57], N[+2206] | 1434.2754 | 3 | 4300.8118 | 4300.8028 | 0 | 2.1 | 485 | Specific | 968.37 | 813.31 | 813.31 | 19.14 | 162 | >sp\|P02788\|TRFL_HUMAN Lactotransferrin OS=Homo sapiens OX=9606 GN=LTF PE=1 SV=6 | 1270 | 20201112_10613_1ng_glyco_hcd203040_compound15.22419.22419.3 | controllerType=0 controllerNumber=1 scan=22419 | 44.8659 |
| 17387:3 | 1 | R.TAGWNIPMGLLFN[+1216.423]QTGSC[+57.021]K.F | HexNAc(2)Hex(5) | C[+57], N[+1216] | 1104.4846 | 3 | 3311.4394 | 3311.4328 | 0 | 2 | 485 | Specific | 1034.71 | 827.44 | 456.21 | 20.45 | 162 | >sp\|P02788\|TRFL_HUMAN Lactotransferrin OS=Homo sapiens OX=9606 GN=LTF PE=1 SV=6 | 1270 | 20201112_10613_1ng_glyco_hcd203040_compound15.20971.20971.3 | controllerType=0 controllerNumber=1 scan=20971 | 41.8606 |
| 17512:3 | 1 | R.TAGWNIPM[+15.995]GLLFN[+2059.735]QTGSC[+57.021]K.F | HexNAc(4)Hex(5)Fuc(1)NeuAc(1) | C[+57], M[+16], N[+2060] | 1390.9217 | 3 | 4170.7505 | 4170.7398 | 0 | 2.6 | 485 | Specific | 299.81 | 285.21 | 285.21 | 6.63 | 162 | >sp\|P02788\|TRFL_HUMAN Lactotransferrin OS=Homo sapiens OX=9606 GN=LTF PE=1 SV=6 | 1270 | 20201112_10613_1ng_glyco_hcd203040_compound15.21101.21101.3 | controllerType=0 controllerNumber=1 scan=21101 | 42.1205 |
| 16991:3 | 1 | R.TAGWNIPM[+15.995]GLLFN[+2059.735]QTGSC[+57.021]K.F | HexNAc(4)Hex(5)Fuc(1)NeuAc(1) | C[+57], M[+16], N[+2060] | 1390.9228 | 3 | 4170.7538 | 4170.7398 | 0 | 3.4 | 485 | Specific | 352.81 | 310.21 | 310.21 | 7.15 | 162 | >sp\|P02788\|TRFL_HUMAN Lactotransferrin OS=Homo sapiens OX=9606 GN=LTF PE=1 SV=6 | 1270 | 20201112_10613_1ng_glyco_hcd203040_compound15.20559.20559.3 | controllerType=0 controllerNumber=1 scan=20559 | 41.017 |
| 16994:3 | 1 | R.TAGWNIPM[+15.995]GLLFN[+2059.735]QTGSC[+57.021]K.F | HexNAc(4)Hex(5)Fuc(1)NeuAc(1) | C[+57], M[+16], N[+2060] | 1390.9228 | 3 | 4170.7538 | 4170.7398 | 0 | 3.4 | 485 | Specific | 565.06 | 475.49 | 475.49 | 9.92 | 162 | >sp\|P02788\|TRFL_HUMAN Lactotransferrin OS=Homo sapiens OX=9606 GN=LTF PE=1 SV=6 | 1270 | 20201112_10613_1ng_glyco_hcd203040_compound15.20562.20562.3 | controllerType=0 controllerNumber=1 scan=20562 | 41.0222 |
| 17515:3 | 1 | R.TAGWNIPM[+15.995]GLLFN[+2059.735]QTGSC[+57.021]K.F | HexNAc(4)Hex(5)Fuc(1)NeuAc(1) | C[+57], M[+16], N[+2060] | 1390.9217 | 3 | 4170.7505 | 4170.7398 | 0 | 2.6 | 485 | Specific | 571.69 | 477.95 | 477.95 | 9.92 | 162 | >sp\|P02788\|TRFL_HUMAN Lactotransferrin OS=Homo sapiens OX=9606 GN=LTF PE=1 SV=6 | 1270 | 20201112_10613_1ng_glyco_hcd203040_compound15.21104.21104.3 | controllerType=0 controllerNumber=1 scan=21104 | 42.126 |
| 20506:4 | 1 | R.TAGWNIPMGLLFN[+2350.830]QTGSC[+57.021]KFDEYFSQSC[+57.021]APGSDPR.S | HexNAc(4)Hex(5)Fuc(1)NeuAc(2) | C[+57]*2, N[+2351] | 1573.1585 | 4 | 6289.612 | 6289.5875 | 0 | 3.9 | 485 | Specific | 264.16 | 264.16 | 264.16 | 5.76 | 162 | >sp\|P02788\|TRFL_HUMAN Lactotransferrin OS=Homo sapiens OX=9606 GN=LTF PE=1 SV=6 | 1270 | 20201112_10613_1ng_glyco_hcd203040_compound15.24224.24224.4 | controllerType=0 controllerNumber=1 scan=24224 | 48.6457 |
| 15583:4 | 1 | R.TAGWNIPMGLLFN[+2116.756]QTGSC[+57.021]KFDEYFSQSC[+57.021]APGSDPR.S | HexNAc(5)Hex(5)NeuAc(1) | C[+57]*2, N[+2117] | 1514.637 | 4 | 6055.526 | 6055.5135 | 0 | 2.1 | 485 | Specific | 287.03 | 264.25 | 264.25 | 6.74 | 162 | >sp\|P02788\|TRFL_HUMAN Lactotransferrin OS=Homo sapiens OX=9606 GN=LTF PE=1 SV=6 | 1270 | 20201112_10613_1ng_glyco_hcd203040_compound15.19062.19062.4 | controllerType=0 controllerNumber=1 scan=19062 | 38.202 |
| 19189:4 | 1 | R.TAGWNIPMGLLFN[+2059.735]QTGSC[+57.021]KFDEYFSQSC[+57.021]APGSDPR.S | HexNAc(4)Hex(5)Fuc(1)NeuAc(1) | C[+57]*2, N[+2060] | 1500.3771 | 4 | 5998.4865 | 5998.492 | 0 | -0.9 | 485 | Specific | 300.11 | 295.93 | 295.93 | 6.26 | 162 | >sp\|P02788\|TRFL_HUMAN Lactotransferrin OS=Homo sapiens OX=9606 GN=LTF PE=1 SV=6 | 1270 | 20201112_10613_1ng_glyco_hcd203040_compound15.22850.22850.4 | controllerType=0 controllerNumber=1 scan=22850 | 45.8341 |
| 19062:4 | 1 | R.TAGWNIPMGLLFN[+2076.750]QTGSC[+57.021]KFDEYFSQSC[+57.021]APGSDPR.S | HexNAc(4)Hex(6)Fuc(2) | C[+57]*2, N[+2077] | 1504.6361 | 4 | 6015.5226 | 6015.5074 | 0 | 2.5 | 485 | Specific | 320.84 | 320.84 | 320.84 | 6.84 | 162 | >sp\|P02788\|TRFL_HUMAN Lactotransferrin OS=Homo sapiens OX=9606 GN=LTF PE=1 SV=6 | 1270 | 20201112_10613_1ng_glyco_hcd203040_compound15.22718.22718.4 | controllerType=0 controllerNumber=1 scan=22718 | 45.5613 |
| 15585:4 | 1 | R.TAGWNIPMGLLFN[+2116.756]QTGSC[+57.021]KFDEYFSQSC[+57.021]APGSDPR.S | HexNAc(5)Hex(5)NeuAc(1) | C[+57]*2, N[+2117] | 1514.637 | 4 | 6055.526 | 6055.5135 | 0 | 2.1 | 485 | Specific | 331.47 | 316.37 | 316.37 | 7.39 | 162 | >sp\|P02788\|TRFL_HUMAN Lactotransferrin OS=Homo sapiens OX=9606 GN=LTF PE=1 SV=6 | 1270 | 20201112_10613_1ng_glyco_hcd203040_compound15.19064.19064.4 | controllerType=0 controllerNumber=1 scan=19064 | 38.2078 |
| 18234:3 | 1 | R.TAGWNIPMGLLFN[+1216.423]QTGSC[+57.021]KFDEYFSQSC[+57.021]APGSDPR.S | HexNAc(2)Hex(5) | C[+57]*2, N[+1216] | 1719.0694 | 3 | 5155.1937 | 5155.18 | 0 | 2.7 | 485 | Specific | 343.92 | 342.18 | 95.06 | 6.22 | 162 | >sp\|P02788\|TRFL_HUMAN Lactotransferrin OS=Homo sapiens OX=9606 GN=LTF PE=1 SV=6 | 1270 | 20201112_10613_1ng_glyco_hcd203040_compound15.21854.21854.3 | controllerType=0 controllerNumber=1 scan=21854 | 43.7215 |
| 19243:4 | 1 | R.TAGWNIPMGLLFN[+1913.677]QTGSC[+57.021]KFDEYFSQSC[+57.021]APGSDPR.S | HexNAc(4)Hex(5)NeuAc(1) | C[+57]*2, N[+1914] | 1463.8643 | 4 | 5852.4352 | 5852.4341 | 0 | 0.2 | 485 | Specific | 382.54 | 336.99 | 336.99 | 7.5 | 162 | >sp\|P02788\|TRFL_HUMAN Lactotransferrin OS=Homo sapiens OX=9606 GN=LTF PE=1 SV=6 | 1270 | 20201112_10613_1ng_glyco_hcd203040_compound15.22906.22906.4 | controllerType=0 controllerNumber=1 scan=22906 | 45.9446 |
| 19109:5 | 1 | R.TAGWNIPMGLLFN[+2059.735]QTGSC[+57.021]KFDEYFSQSC[+57.021]APGSDPR.S | HexNAc(4)Hex(5)Fuc(1)NeuAc(1) | C[+57]*2, N[+2060] | 1200.5073 | 5 | 5998.5075 | 5998.492 | 0 | 2.6 | 485 | Specific | 389.74 | 389.74 | 389.74 | 9.33 | 162 | >sp\|P02788\|TRFL_HUMAN Lactotransferrin OS=Homo sapiens OX=9606 GN=LTF PE=1 SV=6 | 1270 | 20201112_10613_1ng_glyco_hcd203040_compound15.22767.22767.5 | controllerType=0 controllerNumber=1 scan=22767 | 45.6711 |
| 19085:4 | 1 | R.TAGWNIPMGLLFN[+2205.793]QTGSC[+57.021]KFDEYFSQSC[+57.021]APGSDPR.S | HexNAc(4)Hex(5)Fuc(2)NeuAc(1) | C[+57]*2, N[+2206] | 1536.8944 | 4 | 6144.5558 | 6144.5499 | 0 | 1 | 485 | Specific | 393.87 | 368.39 | 368.39 | 7.82 | 162 | >sp\|P02788\|TRFL_HUMAN Lactotransferrin OS=Homo sapiens OX=9606 GN=LTF PE=1 SV=6 | 1270 | 20201112_10613_1ng_glyco_hcd203040_compound15.22742.22742.4 | controllerType=0 controllerNumber=1 scan=22742 | 45.6145 |
| 18046:3 | 1 | R.TAGWNIPMGLLFN[+1768.640]QTGSC[+57.021]KFDEYFSQSC[+57.021]APGSDPR.S | HexNAc(4)Hex(5)Fuc(1) | C[+57]*2, N[+1769] | 1903.1408 | 3 | 5707.4079 | 5707.3966 | 0 | 2 | 485 | Specific | 398.03 | 333.5 | 333.5 | 7.65 | 162 | >sp\|P02788\|TRFL_HUMAN Lactotransferrin OS=Homo sapiens OX=9606 GN=LTF PE=1 SV=6 | 1270 | 20201112_10613_1ng_glyco_hcd203040_compound15.21659.21659.3 | controllerType=0 controllerNumber=1 scan=21659 | 43.3304 |
| 18116:3 | 1 | R.TAGWNIPMGLLFN[+1622.582]QTGSC[+57.021]KFDEYFSQSC[+57.021]APGSDPR.S | HexNAc(4)Hex(5) | C[+57]*2, N[+1623] | 1854.45 | 3 | 5561.3356 | 5561.3387 | 0 | -0.6 | 485 | Specific | 434.53 | 335.84 | 99.35 | 7.82 | 162 | >sp\|P02788\|TRFL_HUMAN Lactotransferrin OS=Homo sapiens OX=9606 GN=LTF PE=1 SV=6 | 1270 | 20201112_10613_1ng_glyco_hcd203040_compound15.21732.21732.3 | controllerType=0 controllerNumber=1 scan=21732 | 43.491 |
| 18119:3 | 1 | R.TAGWNIPMGLLFN[+1622.582]QTGSC[+57.021]KFDEYFSQSC[+57.021]APGSDPR.S | HexNAc(4)Hex(5) | C[+57]*2, N[+1623] | 1854.45 | 3 | 5561.3356 | 5561.3387 | 0 | -0.6 | 485 | Specific | 440.85 | 350.42 | 178.45 | 8.07 | 162 | >sp\|P02788\|TRFL_HUMAN Lactotransferrin OS=Homo sapiens OX=9606 GN=LTF PE=1 SV=6 | 1270 | 20201112_10613_1ng_glyco_hcd203040_compound15.21735.21735.3 | controllerType=0 controllerNumber=1 scan=21735 | 43.5037 |
| 18237:3 | 1 | R.TAGWNIPMGLLFN[+1216.423]QTGSC[+57.021]KFDEYFSQSC[+57.021]APGSDPR.S | HexNAc(2)Hex(5) | C[+57]*2, N[+1216] | 1719.0694 | 3 | 5155.1937 | 5155.18 | 0 | 2.7 | 485 | Specific | 546.23 | 494.88 | 117.93 | 9.75 | 162 | >sp\|P02788\|TRFL_HUMAN Lactotransferrin OS=Homo sapiens OX=9606 GN=LTF PE=1 SV=6 | 1270 | 20201112_10613_1ng_glyco_hcd203040_compound15.21857.21857.3 | controllerType=0 controllerNumber=1 scan=21857 | 43.7265 |
| 19141:3 | 1 | R.TAGWNIPMGLLFN[+1913.677]QTGSC[+57.021]KFDEYFSQSC[+57.021]APGSDPR.S | HexNAc(4)Hex(5)NeuAc(1) | C[+57]*2, N[+1914] | 1951.4834 | 3 | 5852.4355 | 5852.4341 | 0 | 0.2 | 485 | Specific | 607.29 | 493.81 | 493.81 | 12.37 | 162 | >sp\|P02788\|TRFL_HUMAN Lactotransferrin OS=Homo sapiens OX=9606 GN=LTF PE=1 SV=6 | 1270 | 20201112_10613_1ng_glyco_hcd203040_compound15.22800.22800.3 | controllerType=0 controllerNumber=1 scan=22800 | 45.735 |
| 19245:4 | 1 | R.TAGWNIPMGLLFN[+1913.677]QTGSC[+57.021]KFDEYFSQSC[+57.021]APGSDPR.S | HexNAc(4)Hex(5)NeuAc(1) | C[+57]*2, N[+1914] | 1463.8643 | 4 | 5852.4352 | 5852.4341 | 0 | 0.2 | 485 | Specific | 609.81 | 570.64 | 570.64 | 12.29 | 162 | >sp\|P02788\|TRFL_HUMAN Lactotransferrin OS=Homo sapiens OX=9606 GN=LTF PE=1 SV=6 | 1270 | 20201112_10613_1ng_glyco_hcd203040_compound15.22908.22908.4 | controllerType=0 controllerNumber=1 scan=22908 | 45.9517 |
| 19152:5 | 1 | R.TAGWNIPMGLLFN[+1913.677]QTGSC[+57.021]KFDEYFSQSC[+57.021]APGSDPR.S | HexNAc(4)Hex(5)NeuAc(1) | C[+57]*2, N[+1914] | 1171.2941 | 5 | 5852.4412 | 5852.4341 | 0 | 1.2 | 485 | Specific | 644.98 | 609.43 | 609.43 | 12.77 | 162 | >sp\|P02788\|TRFL_HUMAN Lactotransferrin OS=Homo sapiens OX=9606 GN=LTF PE=1 SV=6 | 1270 | 20201112_10613_1ng_glyco_hcd203040_compound15.22811.22811.5 | controllerType=0 controllerNumber=1 scan=22811 | 45.7542 |
| 18085:4 | 1 | R.TAGWNIPMGLLFN[+1622.582]QTGSC[+57.021]KFDEYFSQSC[+57.021]APGSDPR.S | HexNAc(4)Hex(5) | C[+57]*2, N[+1623] | 1391.0945 | 4 | 5561.3561 | 5561.3387 | 0 | 3.1 | 485 | Specific | 668.19 | 636.82 | 186.43 | 13.05 | 162 | >sp\|P02788\|TRFL_HUMAN Lactotransferrin OS=Homo sapiens OX=9606 GN=LTF PE=1 SV=6 | 1270 | 20201112_10613_1ng_glyco_hcd203040_compound15.21700.21700.4 | controllerType=0 controllerNumber=1 scan=21700 | 43.4319 |
| 18040:4 | 1 | R.TAGWNIPMGLLFN[+1768.640]QTGSC[+57.021]KFDEYFSQSC[+57.021]APGSDPR.S | HexNAc(4)Hex(5)Fuc(1) | C[+57]*2, N[+1769] | 1427.6074 | 4 | 5707.4079 | 5707.3966 | 0 | 2 | 485 | Specific | 709.22 | 657.86 | 657.86 | 14.36 | 162 | >sp\|P02788\|TRFL_HUMAN Lactotransferrin OS=Homo sapiens OX=9606 GN=LTF PE=1 SV=6 | 1270 | 20201112_10613_1ng_glyco_hcd203040_compound15.21653.21653.4 | controllerType=0 controllerNumber=1 scan=21653 | 43.3213 |
| 19134:4 | 1 | R.TAGWNIPMGLLFN[+1913.677]QTGSC[+57.021]KFDEYFSQSC[+57.021]APGSDPR.S | HexNAc(4)Hex(5)NeuAc(1) | C[+57]*2, N[+1914] | 1463.8673 | 4 | 5852.4474 | 5852.4341 | 0 | 2.3 | 485 | Specific | 715.1 | 654.86 | 654.86 | 14.05 | 162 | >sp\|P02788\|TRFL_HUMAN Lactotransferrin OS=Homo sapiens OX=9606 GN=LTF PE=1 SV=6 | 1270 | 20201112_10613_1ng_glyco_hcd203040_compound15.22793.22793.4 | controllerType=0 controllerNumber=1 scan=22793 | 45.7247 |
| 20477:4 | 1 | R.TAGWNIPMGLLFN[+2204.772]QTGSC[+57.021]KFDEYFSQSC[+57.021]APGSDPR.S | HexNAc(4)Hex(5)NeuAc(2) | C[+57]*2, N[+2205] | 1536.6398 | 4 | 6143.5373 | 6143.5295 | 0 | 1.3 | 485 | Specific | 730.15 | 678.7 | 296.96 | 14.36 | 162 | >sp\|P02788\|TRFL_HUMAN Lactotransferrin OS=Homo sapiens OX=9606 GN=LTF PE=1 SV=6 | 1270 | 20201112_10613_1ng_glyco_hcd203040_compound15.24194.24194.4 | controllerType=0 controllerNumber=1 scan=24194 | 48.5858 |
| 18049:3 | 1 | R.TAGWNIPMGLLFN[+1768.640]QTGSC[+57.021]KFDEYFSQSC[+57.021]APGSDPR.S | HexNAc(4)Hex(5)Fuc(1) | C[+57]*2, N[+1769] | 1903.1408 | 3 | 5707.4079 | 5707.3966 | 0 | 2 | 485 | Specific | 733.59 | 620.99 | 620.99 | 14.44 | 162 | >sp\|P02788\|TRFL_HUMAN Lactotransferrin OS=Homo sapiens OX=9606 GN=LTF PE=1 SV=6 | 1270 | 20201112_10613_1ng_glyco_hcd203040_compound15.21662.21662.3 | controllerType=0 controllerNumber=1 scan=21662 | 43.3354 |
| 19144:3 | 1 | R.TAGWNIPMGLLFN[+1913.677]QTGSC[+57.021]KFDEYFSQSC[+57.021]APGSDPR.S | HexNAc(4)Hex(5)NeuAc(1) | C[+57]*2, N[+1914] | 1951.4834 | 3 | 5852.4355 | 5852.4341 | 0 | 0.2 | 485 | Specific | 755.68 | 618.41 | 618.41 | 15.37 | 162 | >sp\|P02788\|TRFL_HUMAN Lactotransferrin OS=Homo sapiens OX=9606 GN=LTF PE=1 SV=6 | 1270 | 20201112_10613_1ng_glyco_hcd203040_compound15.22803.22803.3 | controllerType=0 controllerNumber=1 scan=22803 | 45.7419 |
| 19192:4 | 1 | R.TAGWNIPMGLLFN[+2059.735]QTGSC[+57.021]KFDEYFSQSC[+57.021]APGSDPR.S | HexNAc(4)Hex(5)Fuc(1)NeuAc(1) | C[+57]*2, N[+2060] | 1500.3771 | 4 | 5998.4865 | 5998.492 | 0 | -0.9 | 485 | Specific | 762.3 | 718.8 | 718.8 | 14.67 | 162 | >sp\|P02788\|TRFL_HUMAN Lactotransferrin OS=Homo sapiens OX=9606 GN=LTF PE=1 SV=6 | 1270 | 20201112_10613_1ng_glyco_hcd203040_compound15.22853.22853.4 | controllerType=0 controllerNumber=1 scan=22853 | 45.8393 |
| 19087:4 | 1 | R.TAGWNIPMGLLFN[+2205.793]QTGSC[+57.021]KFDEYFSQSC[+57.021]APGSDPR.S | HexNAc(4)Hex(5)Fuc(2)NeuAc(1) | C[+57]*2, N[+2206] | 1536.8944 | 4 | 6144.5558 | 6144.5499 | 0 | 1 | 485 | Specific | 782.84 | 757.18 | 757.18 | 15.36 | 162 | >sp\|P02788\|TRFL_HUMAN Lactotransferrin OS=Homo sapiens OX=9606 GN=LTF PE=1 SV=6 | 1270 | 20201112_10613_1ng_glyco_hcd203040_compound15.22744.22744.4 | controllerType=0 controllerNumber=1 scan=22744 | 45.6209 |
| 20480:4 | 1 | R.TAGWNIPMGLLFN[+2204.772]QTGSC[+57.021]KFDEYFSQSC[+57.021]APGSDPR.S | HexNAc(4)Hex(5)NeuAc(2) | C[+57]*2, N[+2205] | 1536.6398 | 4 | 6143.5373 | 6143.5295 | 0 | 1.3 | 485 | Specific | 788.2 | 781.57 | 413.96 | 15.36 | 162 | >sp\|P02788\|TRFL_HUMAN Lactotransferrin OS=Homo sapiens OX=9606 GN=LTF PE=1 SV=6 | 1270 | 20201112_10613_1ng_glyco_hcd203040_compound15.24197.24197.4 | controllerType=0 controllerNumber=1 scan=24197 | 48.5911 |
| 20365:4 | 1 | R.TAGWNIPMGLLFN[+2350.830]QTGSC[+57.021]KFDEYFSQSC[+57.021]APGSDPR.S | HexNAc(4)Hex(5)Fuc(1)NeuAc(2) | C[+57]*2, N[+2351] | 1573.1552 | 4 | 6289.5988 | 6289.5875 | 0 | 1.8 | 485 | Specific | 828.73 | 795.4 | 795.4 | 16.36 | 162 | >sp\|P02788\|TRFL_HUMAN Lactotransferrin OS=Homo sapiens OX=9606 GN=LTF PE=1 SV=6 | 1270 | 20201112_10613_1ng_glyco_hcd203040_compound15.24077.24077.4 | controllerType=0 controllerNumber=1 scan=24077 | 48.3223 |
| 20367:4 | 1 | R.TAGWNIPMGLLFN[+2350.830]QTGSC[+57.021]KFDEYFSQSC[+57.021]APGSDPR.S | HexNAc(4)Hex(5)Fuc(1)NeuAc(2) | C[+57]*2, N[+2351] | 1573.1552 | 4 | 6289.5988 | 6289.5875 | 0 | 1.8 | 485 | Specific | 842.66 | 810.67 | 810.67 | 16.36 | 162 | >sp\|P02788\|TRFL_HUMAN Lactotransferrin OS=Homo sapiens OX=9606 GN=LTF PE=1 SV=6 | 1270 | 20201112_10613_1ng_glyco_hcd203040_compound15.24079.24079.4 | controllerType=0 controllerNumber=1 scan=24079 | 48.3295 |
| 19136:4 | 1 | R.TAGWNIPMGLLFN[+1913.677]QTGSC[+57.021]KFDEYFSQSC[+57.021]APGSDPR.S | HexNAc(4)Hex(5)NeuAc(1) | C[+57]*2, N[+1914] | 1463.8673 | 4 | 5852.4474 | 5852.4341 | 0 | 2.3 | 485 | Specific | 891.09 | 841.57 | 841.57 | 17.05 | 162 | >sp\|P02788\|TRFL_HUMAN Lactotransferrin OS=Homo sapiens OX=9606 GN=LTF PE=1 SV=6 | 1270 | 20201112_10613_1ng_glyco_hcd203040_compound15.22795.22795.4 | controllerType=0 controllerNumber=1 scan=22795 | 45.7296 |
| 18087:4 | 1 | R.TAGWNIPMGLLFN[+1622.582]QTGSC[+57.021]KFDEYFSQSC[+57.021]APGSDPR.S | HexNAc(4)Hex(5) | C[+57]*2, N[+1623] | 1391.0945 | 4 | 5561.3561 | 5561.3387 | 0 | 3.1 | 485 | Specific | 931.88 | 895.66 | 388.99 | 18.05 | 162 | >sp\|P02788\|TRFL_HUMAN Lactotransferrin OS=Homo sapiens OX=9606 GN=LTF PE=1 SV=6 | 1270 | 20201112_10613_1ng_glyco_hcd203040_compound15.21702.21702.4 | controllerType=0 controllerNumber=1 scan=21702 | 43.4364 |
| 19061:4 | 1 | R.TAGWNIPMGLLFN[+2059.735]QTGSC[+57.021]KFDEYFSQSC[+57.021]APGSDPR.S | HexNAc(4)Hex(5)Fuc(1)NeuAc(1) | C[+57]*2, N[+2060] | 1500.3862 | 4 | 5998.5231 | 5998.492 | 0 | 5.2 | 485 | Specific | 934.76 | 909.79 | 909.79 | 17.19 | 162 | >sp\|P02788\|TRFL_HUMAN Lactotransferrin OS=Homo sapiens OX=9606 GN=LTF PE=1 SV=6 | 1270 | 20201112_10613_1ng_glyco_hcd203040_compound15.22717.22717.4 | controllerType=0 controllerNumber=1 scan=22717 | 45.5599 |
| 19155:5 | 1 | R.TAGWNIPMGLLFN[+1913.677]QTGSC[+57.021]KFDEYFSQSC[+57.021]APGSDPR.S | HexNAc(4)Hex(5)NeuAc(1) | C[+57]*2, N[+1914] | 1171.2941 | 5 | 5852.4412 | 5852.4341 | 0 | 1.2 | 485 | Specific | 941.13 | 888.85 | 888.85 | 18.77 | 162 | >sp\|P02788\|TRFL_HUMAN Lactotransferrin OS=Homo sapiens OX=9606 GN=LTF PE=1 SV=6 | 1270 | 20201112_10613_1ng_glyco_hcd203040_compound15.22814.22814.5 | controllerType=0 controllerNumber=1 scan=22814 | 45.7622 |
| 19063:4 | 1 | R.TAGWNIPMGLLFN[+2059.735]QTGSC[+57.021]KFDEYFSQSC[+57.021]APGSDPR.S | HexNAc(4)Hex(5)Fuc(1)NeuAc(1) | C[+57]*2, N[+2060] | 1500.3862 | 4 | 5998.5231 | 5998.492 | 0 | 5.2 | 485 | Specific | 963.65 | 925.32 | 925.32 | 18.19 | 162 | >sp\|P02788\|TRFL_HUMAN Lactotransferrin OS=Homo sapiens OX=9606 GN=LTF PE=1 SV=6 | 1270 | 20201112_10613_1ng_glyco_hcd203040_compound15.22719.22719.4 | controllerType=0 controllerNumber=1 scan=22719 | 45.5666 |
| 19111:5 | 1 | R.TAGWNIPMGLLFN[+2059.735]QTGSC[+57.021]KFDEYFSQSC[+57.021]APGSDPR.S | HexNAc(4)Hex(5)Fuc(1)NeuAc(1) | C[+57]*2, N[+2060] | 1200.5073 | 5 | 5998.5075 | 5998.492 | 0 | 2.6 | 485 | Specific | 970.16 | 945.02 | 945.02 | 19.46 | 162 | >sp\|P02788\|TRFL_HUMAN Lactotransferrin OS=Homo sapiens OX=9606 GN=LTF PE=1 SV=6 | 1270 | 20201112_10613_1ng_glyco_hcd203040_compound15.22769.22769.5 | controllerType=0 controllerNumber=1 scan=22769 | 45.6749 |
| 18042:4 | 1 | R.TAGWNIPMGLLFN[+1768.640]QTGSC[+57.021]KFDEYFSQSC[+57.021]APGSDPR.S | HexNAc(4)Hex(5)Fuc(1) | C[+57]*2, N[+1769] | 1427.6074 | 4 | 5707.4079 | 5707.3966 | 0 | 2 | 485 | Specific | 978.55 | 917.73 | 917.73 | 19.36 | 162 | >sp\|P02788\|TRFL_HUMAN Lactotransferrin OS=Homo sapiens OX=9606 GN=LTF PE=1 SV=6 | 1270 | 20201112_10613_1ng_glyco_hcd203040_compound15.21655.21655.4 | controllerType=0 controllerNumber=1 scan=21655 | 43.326 |
| 18032:4 | 1 | R.TAGWNIPM[+15.995]GLLFN[+2059.735]QTGSC[+57.021]KFDEYFSQSC[+57.021]A | HexNAc(4)Hex(5)Fuc(1)NeuAc(1) | C[+57]*2, M[+16], N[+2060] | 1504.3824 | 4 | 6014.508 | 6014.487 | 0 | 3.5 | 485 | Specific | 472.47 | 408.77 | 408.77 | 9.21 | 162 | >sp\|P02788\|TRFL_HUMAN Lactotransferrin OS=Homo sapiens OX=9606 GN=LTF PE=1 SV=6 | 1270 | 20201112_10613_1ng_glyco_hcd203040_compound15.21644.21644.4 | controllerType=0 controllerNumber=1 scan=21644 | 43.2995 |
| 18034:4 | 1 | R.TAGWNIPM[+15.995]GLLFN[+2059.735]QTGSC[+57.021]KFDEYFSQSC[+57.021]A | HexNAc(4)Hex(5)Fuc(1)NeuAc(1) | C[+57]*2, M[+16], N[+2060] | 1504.3824 | 4 | 6014.508 | 6014.487 | 0 | 3.5 | 485 | Specific | 637.53 | 572.94 | 572.94 | 12.05 | 162 | >sp\|P02788\|TRFL_HUMAN Lactotransferrin OS=Homo sapiens OX=9606 GN=LTF PE=1 SV=6 | 1270 | 20201112_10613_1ng_glyco_hcd203040_compound15.21646.21646.4 | controllerType=0 controllerNumber=1 scan=21646 | 43.3034 |
| 17830:4 | 1 | R.TAGWNIPM[+15.995]GLLFN[+2059.735]QTGSC[+57.021]KFDEYFSQSC[+57.021]A | HexNAc(4)Hex(5)Fuc(1)NeuAc(1) | C[+57]*2, M[+16], N[+2060] | 1504.3795 | 4 | 6014.4962 | 6014.487 | 0 | 1.5 | 485 | Specific | 772.83 | 659.76 | 659.76 | 15.36 | 162 | >sp\|P02788\|TRFL_HUMAN Lactotransferrin OS=Homo sapiens OX=9606 GN=LTF PE=1 SV=6 | 1270 | 20201112_10613_1ng_glyco_hcd203040_compound15.21434.21434.4 | controllerType=0 controllerNumber=1 scan=21434 | 42.8786 |
| 17833:4 | 1 | R.TAGWNIPM[+15.995]GLLFN[+2059.735]QTGSC[+57.021]KFDEYFSQSC[+57.021]A | HexNAc(4)Hex(5)Fuc(1)NeuAc(1) | C[+57]*2, M[+16], N[+2060] | 1504.3795 | 4 | 6014.4962 | 6014.487 | 0 | 1.5 | 485 | Specific | 923.17 | 645.81 | 645.81 | 18.36 | 162 | >sp\|P02788\|TRFL_HUMAN Lactotransferrin OS=Homo sapiens OX=9606 GN=LTF PE=1 SV=6 | 1270 | 20201112_10613_1ng_glyco_hcd203040_compound15.21437.21437.4 | controllerType=0 controllerNumber=1 scan=21437 | 42.8839 |
| 09166:3 | 1 | R.N[+2205.793]GSDC[+57.021]PDKFC[+57.021]LFQSETK.N | HexNAc(4)Hex(5)Fuc(2)NeuAc(1) | C[+57]*2, N[+2206] | 1413.5556 | 3 | 4238.6522 | 4238.6667 | 0 | -3.4 | 642 | Specific | 412.85 | 249.56 | 249.56 | 5.78 | 162 | >sp\|P02788\|TRFL_HUMAN Lactotransferrin OS=Homo sapiens OX=9606 GN=LTF PE=1 SV=6 | 1270 | 20201112_10613_1ng_glyco_hcd203040_compound15.12392.12392.3 | controllerType=0 controllerNumber=1 scan=12392 | 26.683 |
| 09233:3 | 1 | R.N[+2059.735]GSDC[+57.021]PDKFC[+57.021]LFQSETK.N | HexNAc(4)Hex(5)Fuc(1)NeuAc(1) | C[+57]*2, N[+2060] | 1364.8752 | 3 | 4092.611 | 4092.6088 | 0 | 0.5 | 642 | Specific | 467.95 | 294.94 | 294.94 | 7.33 | 162 | >sp\|P02788\|TRFL_HUMAN Lactotransferrin OS=Homo sapiens OX=9606 GN=LTF PE=1 SV=6 | 1270 | 20201112_10613_1ng_glyco_hcd203040_compound15.12463.12463.3 | controllerType=0 controllerNumber=1 scan=12463 | 26.8147 |
| 09178:3 | 1 | R.N[+2205.793]GSDC[+57.021]PDKFC[+57.021]LFQSETK.N | HexNAc(4)Hex(5)Fuc(2)NeuAc(1) | C[+57]*2, N[+2206] | 1413.5574 | 3 | 4238.6577 | 4238.6667 | 0 | -2.1 | 642 | Specific | 472.99 | 293.45 | 293.45 | 6.71 | 162 | >sp\|P02788\|TRFL_HUMAN Lactotransferrin OS=Homo sapiens OX=9606 GN=LTF PE=1 SV=6 | 1270 | 20201112_10613_1ng_glyco_hcd203040_compound15.12405.12405.3 | controllerType=0 controllerNumber=1 scan=12405 | 26.7085 |
| 09168:3 | 1 | R.N[+2205.793]GSDC[+57.021]PDKFC[+57.021]LFQSETK.N | HexNAc(4)Hex(5)Fuc(2)NeuAc(1) | C[+57]*2, N[+2206] | 1413.5556 | 3 | 4238.6522 | 4238.6667 | 0 | -3.4 | 642 | Specific | 476.51 | 283.66 | 283.66 | 6.41 | 162 | >sp\|P02788\|TRFL_HUMAN Lactotransferrin OS=Homo sapiens OX=9606 GN=LTF PE=1 SV=6 | 1270 | 20201112_10613_1ng_glyco_hcd203040_compound15.12394.12394.3 | controllerType=0 controllerNumber=1 scan=12394 | 26.6871 |
| 09176:3 | 1 | R.N[+2205.793]GSDC[+57.021]PDKFC[+57.021]LFQSETK.N | HexNAc(4)Hex(5)Fuc(2)NeuAc(1) | C[+57]*2, N[+2206] | 1413.5574 | 3 | 4238.6577 | 4238.6667 | 0 | -2.1 | 642 | Specific | 521.92 | 429.8 | 429.8 | 8.84 | 162 | >sp\|P02788\|TRFL_HUMAN Lactotransferrin OS=Homo sapiens OX=9606 GN=LTF PE=1 SV=6 | 1270 | 20201112_10613_1ng_glyco_hcd203040_compound15.12403.12403.3 | controllerType=0 controllerNumber=1 scan=12403 | 26.7042 |
| 09231:3 | 1 | R.N[+2059.735]GSDC[+57.021]PDKFC[+57.021]LFQSETK.N | HexNAc(4)Hex(5)Fuc(1)NeuAc(1) | C[+57]*2, N[+2060] | 1364.8752 | 3 | 4092.611 | 4092.6088 | 0 | 0.5 | 642 | Specific | 585.39 | 416.33 | 416.33 | 11.12 | 162 | >sp\|P02788\|TRFL_HUMAN Lactotransferrin OS=Homo sapiens OX=9606 GN=LTF PE=1 SV=6 | 1270 | 20201112_10613_1ng_glyco_hcd203040_compound15.12461.12461.3 | controllerType=0 controllerNumber=1 scan=12461 | 26.8094 |
| 12092:3 | 2 | R.QSGVN[+2059.735]ATLPEENQPVVFNHVYNIK.L | HexNAc(4)Hex(5)Fuc(1)NeuAc(1) | N[+2060] | 1586.3742 | 3 | 4757.1081 | 4757.098 | 0 | 2.1 | 34 | Specific | 438.74 | 424.48 | 424.48 | 10.28 | 125 | >sp\|P24821\|TENA_HUMAN Tenascin OS=Homo sapiens OX=9606 GN=TNC PE=1 SV=3 | 486 | 20201112_10613_1ng_glyco_hcd203040_compound15.15423.15423.3 | controllerType=0 controllerNumber=1 scan=15423 | 31.8241 |
| 12094:3 | 2 | R.QSGVN[+2059.735]ATLPEENQPVVFNHVYNIK.L | HexNAc(4)Hex(5)Fuc(1)NeuAc(1) | N[+2060] | 1586.3742 | 3 | 4757.1081 | 4757.098 | 0 | 2.1 | 34 | Specific | 491.13 | 426.83 | 426.83 | 10.55 | 125 | >sp\|P24821\|TENA_HUMAN Tenascin OS=Homo sapiens OX=9606 GN=TNC PE=1 SV=3 | 486 | 20201112_10613_1ng_glyco_hcd203040_compound15.15425.15425.3 | controllerType=0 controllerNumber=1 scan=15425 | 31.8277 |
| 01776:3 | 2 | K.GPN[+2059.735]C[+57.021]SEPEC[+57.021]PGNC[+57.021]HLR.G | HexNAc(4)Hex(5)Fuc(1)NeuAc(1) | C[+57]*3, N[+2060] | 1315.1692 | 3 | 3943.4932 | 3943.4931 | 0 | 0 | 182 | Specific | 294.66 | 224.95 | 224.95 | 6.58 | 125 | >sp\|P24821\|TENA_HUMAN Tenascin OS=Homo sapiens OX=9606 GN=TNC PE=1 SV=3 | 486 | 20201112_10613_1ng_glyco_hcd203040_compound15.4724.4724.3 | controllerType=0 controllerNumber=1 scan=4724 | 13.9029 |
| 01774:3 | 2 | K.GPN[+2059.735]C[+57.021]SEPEC[+57.021]PGNC[+57.021]HLR.G | HexNAc(4)Hex(5)Fuc(1)NeuAc(1) | C[+57]*3, N[+2060] | 1315.1692 | 3 | 3943.4932 | 3943.4931 | 0 | 0 | 182 | Specific | 501.92 | 458.18 | 458.18 | 9.8 | 125 | >sp\|P24821\|TENA_HUMAN Tenascin OS=Homo sapiens OX=9606 GN=TNC PE=1 SV=3 | 486 | 20201112_10613_1ng_glyco_hcd203040_compound15.4722.4722.3 | controllerType=0 controllerNumber=1 scan=4722 | 13.8988 |
| 03999:4 | 2 | R.C[+57.021]IN[+2059.735]GTC[+57.021]YC[+57.021]EEGFTGEDC[+57.021]GKPTC[ | HexNAc(4)Hex(5)Fuc(1)NeuAc(1) | C[+57]*6, N[+2060] | 1430.5475 | 4 | 5719.1681 | 5719.1633 | 0 | 0.8 | 325 | Specific | 296.34 | 296.34 | 296.34 | 6.73 | 125 | >sp\|P24821\|TENA_HUMAN Tenascin OS=Homo sapiens OX=9606 GN=TNC PE=1 SV=3 | 486 | 20201112_10613_1ng_glyco_hcd203040_compound15.7037.7037.4 | controllerType=0 controllerNumber=1 scan=7037 | 17.8703 |
| 03997:4 | 2 | R.C[+57.021]IN[+2059.735]GTC[+57.021]YC[+57.021]EEGFTGEDC[+57.021]GKPTC[ | HexNAc(4)Hex(5)Fuc(1)NeuAc(1) | C[+57]*6, N[+2060] | 1430.5475 | 4 | 5719.1681 | 5719.1633 | 0 | 0.8 | 325 | Specific | 408.22 | 408.22 | 408.22 | 9.23 | 125 | >sp\|P24821\|TENA_HUMAN Tenascin OS=Homo sapiens OX=9606 GN=TNC PE=1 SV=3 | 486 | 20201112_10613_1ng_glyco_hcd203040_compound15.7035.7035.4 | controllerType=0 controllerNumber=1 scan=7035 | 17.8663 |
| 09157:4 | 2 | R.QTGLAPGQEYEISLHIVKN[+2059.735]NTR.G | HexNAc(4)Hex(5)Fuc(1)NeuAc(1) | N[+2060] | 1132.7627 | 4 | 4528.0291 | 4528.0241 | 0 | 1.1 | 770 | Specific | 403.28 | 268.47 | 268.47 | 7.52 | 125 | >sp\|P24821\|TENA_HUMAN Tenascin OS=Homo sapiens OX=9606 GN=TNC PE=1 SV=3 | 486 | 20201112_10613_1ng_glyco_hcd203040_compound15.12382.12382.2 | controllerType=0 controllerNumber=1 scan=12382 | 26.6647 |
| 09155:4 | 2 | R.QTGLAPGQEYEISLHIVKN[+2059.735]NTR.G | HexNAc(4)Hex(5)Fuc(1)NeuAc(1) | N[+2060] | 1132.7627 | 4 | 4528.0291 | 4528.0241 | 0 | 1.1 | 770 | Specific | 475.99 | 314.76 | 314.76 | 8.23 | 125 | >sp\|P24821\|TENA_HUMAN Tenascin OS=Homo sapiens OX=9606 GN=TNC PE=1 SV=3 | 486 | 20201112_10613_1ng_glyco_hcd203040_compound15.12380.12380.2 | controllerType=0 controllerNumber=1 scan=12380 | 26.6598 |
| 09185:3 | 2 | R.QTGLAPGQEYEISLHIVKN[+2059.735]NTR.G | HexNAc(4)Hex(5)Fuc(1)NeuAc(1) | N[+2060] | 1510.0143 | 3 | 4528.0285 | 4528.0241 | 0 | 1 | 770 | Specific | 486.14 | 279.67 | 279.67 | 7.57 | 125 | >sp\|P24821\|TENA_HUMAN Tenascin OS=Homo sapiens OX=9606 GN=TNC PE=1 SV=3 | 486 | 20201112_10613_1ng_glyco_hcd203040_compound15.12413.12413.3 | controllerType=0 controllerNumber=1 scan=12413 | 26.7258 |
| 09171:3 | 2 | R.QTGLAPGQEYEISLHIVKN[+2059.735]NTR.G | HexNAc(4)Hex(5)Fuc(1)NeuAc(1) | N[+2060] | 1510.0115 | 3 | 4528.02 | 4528.0241 | 0 | -0.9 | 770 | Specific | 487.77 | 302.83 | 302.83 | 7.07 | 125 | >sp\|P24821\|TENA_HUMAN Tenascin OS=Homo sapiens OX=9606 GN=TNC PE=1 SV=3 | 486 | 20201112_10613_1ng_glyco_hcd203040_compound15.12398.12398.3 | controllerType=0 controllerNumber=1 scan=12398 | 26.6971 |
| 09183:3 | 2 | R.QTGLAPGQEYEISLHIVKN[+2059.735]NTR.G | HexNAc(4)Hex(5)Fuc(1)NeuAc(1) | N[+2060] | 1510.0143 | 3 | 4528.0285 | 4528.0241 | 0 | 1 | 770 | Specific | 490.25 | 344.4 | 344.4 | 7.86 | 125 | >sp\|P24821\|TENA_HUMAN Tenascin OS=Homo sapiens OX=9606 GN=TNC PE=1 SV=3 | 486 | 20201112_10613_1ng_glyco_hcd203040_compound15.12411.12411.3 | controllerType=0 controllerNumber=1 scan=12411 | 26.7209 |
| 09169:3 | 2 | R.QTGLAPGQEYEISLHIVKN[+2059.735]NTR.G | HexNAc(4)Hex(5)Fuc(1)NeuAc(1) | N[+2060] | 1510.0115 | 3 | 4528.02 | 4528.0241 | 0 | -0.9 | 770 | Specific | 644.74 | 366.04 | 366.04 | 10.04 | 125 | >sp\|P24821\|TENA_HUMAN Tenascin OS=Homo sapiens OX=9606 GN=TNC PE=1 SV=3 | 486 | 20201112_10613_1ng_glyco_hcd203040_compound15.12396.12396.3 | controllerType=0 controllerNumber=1 scan=12396 | 26.6917 |
| 15620:2 | 2 | R.LN[+1378.476]YSLPTGQWVGVQLPR.N | HexNAc(2)Hex(6) | N[+1378] | 1653.7669 | 2 | 3306.5265 | 3306.5145 | 0 | 3.6 | 1017 | Specific | 426.79 | 385.23 | 385.23 | 8.8 | 125 | >sp\|P24821\|TENA_HUMAN Tenascin OS=Homo sapiens OX=9606 GN=TNC PE=1 SV=3 | 486 | 20201112_10613_1ng_glyco_hcd203040_compound15.19100.19100.2 | controllerType=0 controllerNumber=1 scan=19100 | 38.2876 |
| 15618:2 | 2 | R.LN[+1378.476]YSLPTGQWVGVQLPR.N | HexNAc(2)Hex(6) | N[+1378] | 1653.7669 | 2 | 3306.5265 | 3306.5145 | 0 | 3.6 | 1017 | Specific | 521.46 | 516.63 | 516.63 | 9.03 | 125 | >sp\|P24821\|TENA_HUMAN Tenascin OS=Homo sapiens OX=9606 GN=TNC PE=1 SV=3 | 486 | 20201112_10613_1ng_glyco_hcd203040_compound15.19098.19098.2 | controllerType=0 controllerNumber=1 scan=19098 | 38.2752 |
| 15541:2 | 2 | R.LN[+1540.529]YSLPTGQWVGVQLPR.N | HexNAc(2)Hex(7) | N[+1541] | 1734.7819 | 2 | 3468.5566 | 3468.5674 | 0 | -3.1 | 1017 | Specific | 606.03 | 485.96 | 485.96 | 11.31 | 125 | >sp\|P24821\|TENA_HUMAN Tenascin OS=Homo sapiens OX=9606 GN=TNC PE=1 SV=3 | 486 | 20201112_10613_1ng_glyco_hcd203040_compound15.19018.19018.2 | controllerType=0 controllerNumber=1 scan=19018 | 38.1125 |
| 15539:2 | 2 | R.LN[+1540.529]YSLPTGQWVGVQLPR.N | HexNAc(2)Hex(7) | N[+1541] | 1734.7819 | 2 | 3468.5566 | 3468.5674 | 0 | -3.1 | 1017 | Specific | 632.13 | 536.85 | 536.85 | 11.31 | 125 | >sp\|P24821\|TENA_HUMAN Tenascin OS=Homo sapiens OX=9606 GN=TNC PE=1 SV=3 | 486 | 20201112_10613_1ng_glyco_hcd203040_compound15.19016.19016.2 | controllerType=0 controllerNumber=1 scan=19016 | 38.1091 |
| 15466:2 | 2 | R.LN[+1702.581]YSLPTGQWVGVQLPR.N | HexNAc(2)Hex(8) | N[+1703] | 1815.8142 | 2 | 3630.6211 | 3630.6202 | 0 | 0.2 | 1017 | Specific | 681.59 | 581.51 | 581.51 | 13.46 | 125 | >sp\|P24821\|TENA_HUMAN Tenascin OS=Homo sapiens OX=9606 GN=TNC PE=1 SV=3 | 486 | 20201112_10613_1ng_glyco_hcd203040_compound15.18940.18940.2 | controllerType=0 controllerNumber=1 scan=18940 | 37.9699 |
| 15468:2 | 2 | R.LN[+1702.581]YSLPTGQWVGVQLPR.N | HexNAc(2)Hex(8) | N[+1703] | 1815.8142 | 2 | 3630.6211 | 3630.6202 | 0 | 0.2 | 1017 | Specific | 738.26 | 600.3 | 600.3 | 14.46 | 125 | >sp\|P24821\|TENA_HUMAN Tenascin OS=Homo sapiens OX=9606 GN=TNC PE=1 SV=3 | 486 | 20201112_10613_1ng_glyco_hcd203040_compound15.18942.18942.2 | controllerType=0 controllerNumber=1 scan=18942 | 37.9736 |
| 15626:3 | 2 | R.LN[+1378.476]YSLPTGQWVGVQLPR.N | HexNAc(2)Hex(6) | N[+1378] | 1102.844 | 3 | 3306.5175 | 3306.5145 | 0 | 0.9 | 1017 | Specific | 749.84 | 725.51 | 725.51 | 14.45 | 125 | >sp\|P24821\|TENA_HUMAN Tenascin OS=Homo sapiens OX=9606 GN=TNC PE=1 SV=3 | 486 | 20201112_10613_1ng_glyco_hcd203040_compound15.19107.19107.3 | controllerType=0 controllerNumber=1 scan=19107 | 38.2998 |
| 15662:3 | 2 | R.LN[+1216.423]YSLPTGQWVGVQLPR.N | HexNAc(2)Hex(5) | N[+1216] | 1048.8274 | 3 | 3144.4677 | 3144.4617 | 0 | 1.9 | 1017 | Specific | 800.45 | 790 | 790 | 16.45 | 125 | >sp\|P24821\|TENA_HUMAN Tenascin OS=Homo sapiens OX=9606 GN=TNC PE=1 SV=3 | 486 | 20201112_10613_1ng_glyco_hcd203040_compound15.19144.19144.3 | controllerType=0 controllerNumber=1 scan=19144 | 38.3688 |
| 15629:3 | 2 | R.LN[+1378.476]YSLPTGQWVGVQLPR.N | HexNAc(2)Hex(6) | N[+1378] | 1102.844 | 3 | 3306.5175 | 3306.5145 | 0 | 0.9 | 1017 | Specific | 868.09 | 712.42 | 712.42 | 17.45 | 125 | >sp\|P24821\|TENA_HUMAN Tenascin OS=Homo sapiens OX=9606 GN=TNC PE=1 SV=3 | 486 | 20201112_10613_1ng_glyco_hcd203040_compound15.19110.19110.3 | controllerType=0 controllerNumber=1 scan=19110 | 38.3086 |
| 15664:3 | 2 | R.LN[+1216.423]YSLPTGQWVGVQLPR.N | HexNAc(2)Hex(5) | N[+1216] | 1048.8274 | 3 | 3144.4677 | 3144.4617 | 0 | 1.9 | 1017 | Specific | 939.44 | 928.09 | 928.09 | 18.45 | 125 | >sp\|P24821\|TENA_HUMAN Tenascin OS=Homo sapiens OX=9606 GN=TNC PE=1 SV=3 | 486 | 20201112_10613_1ng_glyco_hcd203040_compound15.19146.19146.3 | controllerType=0 controllerNumber=1 scan=19146 | 38.3736 |
| 15525:3 | 2 | R.LN[+1540.529]YSLPTGQWVGVQLPR.N | HexNAc(2)Hex(7) | N[+1541] | 1156.8594 | 3 | 3468.5637 | 3468.5674 | 0 | -1 | 1017 | Specific | 985.35 | 883.91 | 883.91 | 18.76 | 125 | >sp\|P24821\|TENA_HUMAN Tenascin OS=Homo sapiens OX=9606 GN=TNC PE=1 SV=3 | 486 | 20201112_10613_1ng_glyco_hcd203040_compound15.19002.19002.3 | controllerType=0 controllerNumber=1 scan=19002 | 38.0824 |
| 15433:3 | 2 | R.LN[+1702.581]YSLPTGQWVGVQLPR.N | HexNAc(2)Hex(8) | N[+1703] | 1210.8777 | 3 | 3630.6187 | 3630.6202 | 0 | -0.4 | 1017 | Specific | 1026.08 | 923.63 | 923.63 | 20.38 | 125 | >sp\|P24821\|TENA_HUMAN Tenascin OS=Homo sapiens OX=9606 GN=TNC PE=1 SV=3 | 486 | 20201112_10613_1ng_glyco_hcd203040_compound15.18906.18906.3 | controllerType=0 controllerNumber=1 scan=18906 | 37.9067 |
| 15435:3 | 2 | R.LN[+1702.581]YSLPTGQWVGVQLPR.N | HexNAc(2)Hex(8) | N[+1703] | 1210.8777 | 3 | 3630.6187 | 3630.6202 | 0 | -0.4 | 1017 | Specific | 1047.42 | 841.01 | 841.01 | 20.38 | 125 | >sp\|P24821\|TENA_HUMAN Tenascin OS=Homo sapiens OX=9606 GN=TNC PE=1 SV=3 | 486 | 20201112_10613_1ng_glyco_hcd203040_compound15.18908.18908.3 | controllerType=0 controllerNumber=1 scan=18908 | 37.9121 |
| 15527:3 | 2 | R.LN[+1540.529]YSLPTGQWVGVQLPR.N | HexNAc(2)Hex(7) | N[+1541] | 1156.8594 | 3 | 3468.5637 | 3468.5674 | 0 | -1 | 1017 | Specific | 1056.51 | 884.24 | 884.24 | 20.76 | 125 | >sp\|P24821\|TENA_HUMAN Tenascin OS=Homo sapiens OX=9606 GN=TNC PE=1 SV=3 | 486 | 20201112_10613_1ng_glyco_hcd203040_compound15.19004.19004.3 | controllerType=0 controllerNumber=1 scan=19004 | 38.0879 |
| 03988:3 | 2 | R.N[+2059.735]TTSYVLR.G | HexNAc(4)Hex(5)Fuc(1)NeuAc(1) | N[+2060] | 1005.0858 | 3 | 3013.2428 | 3013.24 | 0 | 0.9 | 1034 | Specific | 207.89 | 119.78 | 119.78 | 3.65 | 125 | >sp\|P24821\|TENA_HUMAN Tenascin OS=Homo sapiens OX=9606 GN=TNC PE=1 SV=3 | 486 | 20201112_10613_1ng_glyco_hcd203040_compound15.7025.7025.3 | controllerType=0 controllerNumber=1 scan=7025 | 17.8475 |
| 04008:2 | 2 | R.N[+2059.735]TTSYVLR.G | HexNAc(4)Hex(5)Fuc(1)NeuAc(1) | N[+2060] | 1507.126 | 2 | 3013.2448 | 3013.24 | 0 | 1.6 | 1034 | Specific | 213.16 | 62.04 | 62.04 | 3.41 | 125 | >sp\|P24821\|TENA_HUMAN Tenascin OS=Homo sapiens OX=9606 GN=TNC PE=1 SV=3 | 486 | 20201112_10613_1ng_glyco_hcd203040_compound15.7046.7046.2 | controllerType=0 controllerNumber=1 scan=7046 | 17.8856 |
| 03990:3 | 2 | R.N[+2059.735]TTSYVLR.G | HexNAc(4)Hex(5)Fuc(1)NeuAc(1) | N[+2060] | 1005.0858 | 3 | 3013.2428 | 3013.24 | 0 | 0.9 | 1034 | Specific | 292.37 | 180.66 | 180.66 | 5.8 | 125 | >sp\|P24821\|TENA_HUMAN Tenascin OS=Homo sapiens OX=9606 GN=TNC PE=1 SV=3 | 486 | 20201112_10613_1ng_glyco_hcd203040_compound15.7027.7027.3 | controllerType=0 controllerNumber=1 scan=7027 | 17.8516 |
| 17804:4 | 2 | R.LN[+1913.677]WTAADQAYEHFIIQVQEANKVEAAR.N | HexNAc(4)Hex(5)NeuAc(1) | N[+1914] | 1258.0656 | 4 | 5029.2404 | 5029.2366 | 0 | 0.8 | 1092 | Specific | 927.25 | 750.4 | 750.4 | 18.83 | 125 | >sp\|P24821\|TENA_HUMAN Tenascin OS=Homo sapiens OX=9606 GN=TNC PE=1 SV=3 | 486 | 20201112_10613_1ng_glyco_hcd203040_compound15.21406.21406.4 | controllerType=0 controllerNumber=1 scan=21406 | 42.7998 |
| 17802:4 | 2 | R.LN[+1913.677]WTAADQAYEHFIIQVQEANKVEAAR.N | HexNAc(4)Hex(5)NeuAc(1) | N[+1914] | 1258.0656 | 4 | 5029.2404 | 5029.2366 | 0 | 0.8 | 1092 | Specific | 1005.81 | 921.86 | 921.86 | 20.83 | 125 | >sp\|P24821\|TENA_HUMAN Tenascin OS=Homo sapiens OX=9606 GN=TNC PE=1 SV=3 | 486 | 20201112_10613_1ng_glyco_hcd203040_compound15.21404.21404.4 | controllerType=0 controllerNumber=1 scan=21404 | 42.7954 |
| 07892:3 | 2 | R.N[+2059.735]LTVPGSLR.A | HexNAc(4)Hex(5)Fuc(1)NeuAc(1) | N[+2060] | 1006.1018 | 3 | 3016.2908 | 3016.2873 | 0 | 1.2 | 1119 | Specific | 245.13 | 161.22 | 161.22 | 5.45 | 125 | >sp\|P24821\|TENA_HUMAN Tenascin OS=Homo sapiens OX=9606 GN=TNC PE=1 SV=3 | 486 | 20201112_10613_1ng_glyco_hcd203040_compound15.11063.11063.3 | controllerType=0 controllerNumber=1 scan=11063 | 24.4748 |
| 07909:2 | 2 | R.N[+2059.735]LTVPGSLR.A | HexNAc(4)Hex(5)Fuc(1)NeuAc(1) | N[+2060] | 1508.6477 | 2 | 3016.2882 | 3016.2873 | 0 | 0.3 | 1119 | Specific | 263.1 | 122.17 | 122.17 | 4.71 | 125 | >sp\|P24821\|TENA_HUMAN Tenascin OS=Homo sapiens OX=9606 GN=TNC PE=1 SV=3 | 486 | 20201112_10613_1ng_glyco_hcd203040_compound15.11080.11080.2 | controllerType=0 controllerNumber=1 scan=11080 | 24.5031 |
| 07894:3 | 2 | R.N[+2059.735]LTVPGSLR.A | HexNAc(4)Hex(5)Fuc(1)NeuAc(1) | N[+2060] | 1006.1018 | 3 | 3016.2908 | 3016.2873 | 0 | 1.2 | 1119 | Specific | 328.9 | 235.58 | 235.58 | 6.68 | 125 | >sp\|P24821\|TENA_HUMAN Tenascin OS=Homo sapiens OX=9606 GN=TNC PE=1 SV=3 | 486 | 20201112_10613_1ng_glyco_hcd203040_compound15.11065.11065.3 | controllerType=0 controllerNumber=1 scan=11065 | 24.4788 |
| 11185:3 | 2 | R.LLETVEYN[+2059.735]ISGAER.T | HexNAc(4)Hex(5)Fuc(1)NeuAc(1) | N[+2060] | 1218.5231 | 3 | 3653.5548 | 3653.5468 | 0 | 2.2 | 1478 | Specific | 362.76 | 351.25 | 351.25 | 7.35 | 125 | >sp\|P24821\|TENA_HUMAN Tenascin OS=Homo sapiens OX=9606 GN=TNC PE=1 SV=3 | 486 | 20201112_10613_1ng_glyco_hcd203040_compound15.14483.14483.3 | controllerType=0 controllerNumber=1 scan=14483 | 30.1702 |
| 11188:3 | 2 | R.LLETVEYN[+2059.735]ISGAER.T | HexNAc(4)Hex(5)Fuc(1)NeuAc(1) | N[+2060] | 1218.5231 | 3 | 3653.5548 | 3653.5468 | 0 | 2.2 | 1478 | Specific | 581.05 | 404.26 | 404.26 | 10.99 | 125 | >sp\|P24821\|TENA_HUMAN Tenascin OS=Homo sapiens OX=9606 GN=TNC PE=1 SV=3 | 486 | 20201112_10613_1ng_glyco_hcd203040_compound15.14486.14486.3 | controllerType=0 controllerNumber=1 scan=14486 | 30.175 |
| 19360:5 | 2 | R.LLETVEYN[+2336.851]ISGAERTAHISGLPPSTDFIVYLSGLAPSIR.T | HexNAc(6)Hex(6)Fuc(1) | N[+2337] | 1305.6158 | 5 | 6524.0501 | 6524.0534 | 0 | -0.5 | 1478 | Specific | 314.19 | 314.19 | 314.19 | 7.73 | 125 | >sp\|P24821\|TENA_HUMAN Tenascin OS=Homo sapiens OX=9606 GN=TNC PE=1 SV=3 | 486 | 20201112_10613_1ng_glyco_hcd203040_compound15.23027.23027.2 | controllerType=0 controllerNumber=1 scan=23027 | 46.1896 |
| 19363:5 | 2 | R.LLETVEYN[+2336.851]ISGAERTAHISGLPPSTDFIVYLSGLAPSIR.T | HexNAc(6)Hex(6)Fuc(1) | N[+2337] | 1305.6158 | 5 | 6524.0501 | 6524.0534 | 0 | -0.5 | 1478 | Specific | 412.89 | 412.89 | 412.89 | 9.92 | 125 | >sp\|P24821\|TENA_HUMAN Tenascin OS=Homo sapiens OX=9606 GN=TNC PE=1 SV=3 | 486 | 20201112_10613_1ng_glyco_hcd203040_compound15.23030.23030.2 | controllerType=0 controllerNumber=1 scan=23030 | 46.1956 |
| 10315:3 | 4 | K.TVLTPATNHMGN[+1378.476]VTFTIPANR.E | HexNAc(2)Hex(6) | N[+1378] | 1211.8878 | 3 | 3633.6487 | 3633.6358 | 0 | 3.6 | 74 | Specific | 296.89 | 296.89 | 226.1 | 6.59 | 100 | >sp\|P01024\|CO3_HUMAN Complement C3 OS=Homo sapiens OX=9606 GN=C3 PE=1 SV=2 | 492 | 20201112_10613_1ng_glyco_hcd203040_compound15.13582.13582.3 | controllerType=0 controllerNumber=1 scan=13582 | 28.6687 |
| 10313:3 | 4 | K.TVLTPATNHMGN[+1378.476]VTFTIPANR.E | HexNAc(2)Hex(6) | N[+1378] | 1211.8878 | 3 | 3633.6487 | 3633.6358 | 0 | 3.6 | 74 | Specific | 350.05 | 350.05 | 214.58 | 7.21 | 100 | >sp\|P01024\|CO3_HUMAN Complement C3 OS=Homo sapiens OX=9606 GN=C3 PE=1 SV=2 | 492 | 20201112_10613_1ng_glyco_hcd203040_compound15.13580.13580.3 | controllerType=0 controllerNumber=1 scan=13580 | 28.6646 |
| 10216:3 | 4 | K.TVLTPATNHMGN[+1540.529]VTFTIPANR.E | HexNAc(2)Hex(7) | N[+1541] | 1265.9053 | 3 | 3795.7014 | 3795.6886 | 0 | 3.4 | 74 | Specific | 717.59 | 686.63 | 319.97 | 14.63 | 100 | >sp\|P01024\|CO3_HUMAN Complement C3 OS=Homo sapiens OX=9606 GN=C3 PE=1 SV=2 | 492 | 20201112_10613_1ng_glyco_hcd203040_compound15.13479.13479.3 | controllerType=0 controllerNumber=1 scan=13479 | 28.4697 |
| 10214:3 | 4 | K.TVLTPATNHMGN[+1540.529]VTFTIPANR.E | HexNAc(2)Hex(7) | N[+1541] | 1265.9053 | 3 | 3795.7014 | 3795.6886 | 0 | 3.4 | 74 | Specific | 790.47 | 790.47 | 440.35 | 15.63 | 100 | >sp\|P01024\|CO3_HUMAN Complement C3 OS=Homo sapiens OX=9606 GN=C3 PE=1 SV=2 | 492 | 20201112_10613_1ng_glyco_hcd203040_compound15.13477.13477.3 | controllerType=0 controllerNumber=1 scan=13477 | 28.4655 |
| 17564:3 | 6 | R.GLSFDVSLEVSQGPGLLN[+2570.925]DTK.V | HexNAc(5)Hex(6)Fuc(2)NeuAc(1) | N[+2571] | 1583.025 | 3 | 4747.0603 | 4747.0382 | 0 | 4.7 | 118 | Specific | 658.23 | 619.09 | 619.09 | 13.02 | 79 | >sp\|P01833\|PIGR_HUMAN Polymeric immunoglobulin receptor OS=Homo sapiens OX=9606 GN=PIGR PE=1 SV=4 | 902 | 20201112_10613_1ng_glyco_hcd203040_compound15.21156.21156.3 | controllerType=0 controllerNumber=1 scan=21156 | 42.2382 |
| 17566:3 | 6 | R.GLSFDVSLEVSQGPGLLN[+2570.925]DTK.V | HexNAc(5)Hex(6)Fuc(2)NeuAc(1) | N[+2571] | 1583.025 | 3 | 4747.0603 | 4747.0382 | 0 | 4.7 | 118 | Specific | 740.93 | 635.51 | 635.51 | 14.02 | 79 | >sp\|P01833\|PIGR_HUMAN Polymeric immunoglobulin receptor OS=Homo sapiens OX=9606 GN=PIGR PE=1 SV=4 | 902 | 20201112_10613_1ng_glyco_hcd203040_compound15.21158.21158.3 | controllerType=0 controllerNumber=1 scan=21158 | 42.2426 |
| 17777:3 | 6 | R.GLSFDVSLEVSQGPGLLN[+2059.735]DTK.V | HexNAc(4)Hex(5)Fuc(1)NeuAc(1) | N[+2060] | 1412.6243 | 3 | 4235.8584 | 4235.8481 | 0 | 2.4 | 118 | Specific | 850.44 | 791.69 | 791.69 | 17.55 | 79 | >sp\|P01833\|PIGR_HUMAN Polymeric immunoglobulin receptor OS=Homo sapiens OX=9606 GN=PIGR PE=1 SV=4 | 902 | 20201112_10613_1ng_glyco_hcd203040_compound15.21378.21378.3 | controllerType=0 controllerNumber=1 scan=21378 | 42.7372 |
| 17779:3 | 6 | R.GLSFDVSLEVSQGPGLLN[+2059.735]DTK.V | HexNAc(4)Hex(5)Fuc(1)NeuAc(1) | N[+2060] | 1412.6243 | 3 | 4235.8584 | 4235.8481 | 0 | 2.4 | 118 | Specific | 962.07 | 881.83 | 881.83 | 19.55 | 79 | >sp\|P01833\|PIGR_HUMAN Polymeric immunoglobulin receptor OS=Homo sapiens OX=9606 GN=PIGR PE=1 SV=4 | 902 | 20201112_10613_1ng_glyco_hcd203040_compound15.21380.21380.3 | controllerType=0 controllerNumber=1 scan=21380 | 42.7445 |
| 19047:3 | 6 | K.QIGLYPVLVIDSSGYVNPN[+1913.677]YTGR.I | HexNAc(4)Hex(5)NeuAc(1) | N[+1914] | 1480.3361 | 3 | 4438.9938 | 4438.9805 | 0 | 3 | 168 | Specific | 311.84 | 311.84 | 311.84 | 7.34 | 79 | >sp\|P01833\|PIGR_HUMAN Polymeric immunoglobulin receptor OS=Homo sapiens OX=9606 GN=PIGR PE=1 SV=4 | 902 | 20201112_10613_1ng_glyco_hcd203040_compound15.22702.22702.3 | controllerType=0 controllerNumber=1 scan=22702 | 45.5147 |
| 18926:3 | 6 | K.QIGLYPVLVIDSSGYVNPN[+2205.793]YTGR.I | HexNAc(4)Hex(5)Fuc(2)NeuAc(1) | N[+2206] | 1577.7055 | 3 | 4731.102 | 4731.0963 | 0 | 1.2 | 168 | Specific | 635.48 | 635.48 | 635.48 | 12.86 | 79 | >sp\|P01833\|PIGR_HUMAN Polymeric immunoglobulin receptor OS=Homo sapiens OX=9606 GN=PIGR PE=1 SV=4 | 902 | 20201112_10613_1ng_glyco_hcd203040_compound15.22575.22575.3 | controllerType=0 controllerNumber=1 scan=22575 | 45.2136 |
| 19049:3 | 6 | K.QIGLYPVLVIDSSGYVNPN[+1913.677]YTGR.I | HexNAc(4)Hex(5)NeuAc(1) | N[+1914] | 1480.3361 | 3 | 4438.9938 | 4438.9805 | 0 | 3 | 168 | Specific | 681.7 | 548.59 | 548.59 | 13.55 | 79 | >sp\|P01833\|PIGR_HUMAN Polymeric immunoglobulin receptor OS=Homo sapiens OX=9606 GN=PIGR PE=1 SV=4 | 902 | 20201112_10613_1ng_glyco_hcd203040_compound15.22704.22704.3 | controllerType=0 controllerNumber=1 scan=22704 | 45.5201 |
| 20986:3 | 6 | K.QIGLYPVLVIDSSGYVNPN[+2204.772]YTGR.I | HexNAc(4)Hex(5)NeuAc(2) | N[+2205] | 1577.3622 | 3 | 4730.0722 | 4730.0759 | 0 | -0.8 | 168 | Specific | 711.27 | 685.21 | 685.21 | 14.79 | 79 | >sp\|P01833\|PIGR_HUMAN Polymeric immunoglobulin receptor OS=Homo sapiens OX=9606 GN=PIGR PE=1 SV=4 | 902 | 20201112_10613_1ng_glyco_hcd203040_compound15.24725.24725.4 | controllerType=0 controllerNumber=1 scan=24725 | 49.7558 |
| 20984:3 | 6 | K.QIGLYPVLVIDSSGYVNPN[+2204.772]YTGR.I | HexNAc(4)Hex(5)NeuAc(2) | N[+2205] | 1577.3622 | 3 | 4730.0722 | 4730.0759 | 0 | -0.8 | 168 | Specific | 747.42 | 546.75 | 546.75 | 14.79 | 79 | >sp\|P01833\|PIGR_HUMAN Polymeric immunoglobulin receptor OS=Homo sapiens OX=9606 GN=PIGR PE=1 SV=4 | 902 | 20201112_10613_1ng_glyco_hcd203040_compound15.24723.24723.4 | controllerType=0 controllerNumber=1 scan=24723 | 49.7514 |
| 18928:3 | 6 | K.QIGLYPVLVIDSSGYVNPN[+2205.793]YTGR.I | HexNAc(4)Hex(5)Fuc(2)NeuAc(1) | N[+2206] | 1577.7055 | 3 | 4731.102 | 4731.0963 | 0 | 1.2 | 168 | Specific | 756.97 | 694.38 | 694.38 | 15.86 | 79 | >sp\|P01833\|PIGR_HUMAN Polymeric immunoglobulin receptor OS=Homo sapiens OX=9606 GN=PIGR PE=1 SV=4 | 902 | 20201112_10613_1ng_glyco_hcd203040_compound15.22577.22577.3 | controllerType=0 controllerNumber=1 scan=22577 | 45.2188 |
| 28788:3 | 6 | R.[+42.011]LSLLEEPGN[+2351.851]GTFTVILNQLTSR.D | HexNAc(4)Hex(5)Fuc(3)NeuAc(1) | .[42], N[+2352] | 1599.3823 | 3 | 4796.1322 | 4796.1539 | 0 | -4.5 | 413 | Specific | 200.71 | 174.9 | 174.9 | 3.62 | 79 | >sp\|P01833\|PIGR_HUMAN Polymeric immunoglobulin receptor OS=Homo sapiens OX=9606 GN=PIGR PE=1 SV=4 | 902 | 20201112_10613_1ng_glyco_hcd203040_compound15.32855.32855.3 | controllerType=0 controllerNumber=1 scan=32855 | 64.7448 |
| 28781:3 | 6 | R.[+42.011]LSLLEEPGN[+2351.851]GTFTVILNQLTSR.D | HexNAc(4)Hex(5)Fuc(3)NeuAc(1) | .[42], N[+2352] | 1599.3869 | 3 | 4796.1461 | 4796.1539 | 0 | -1.6 | 413 | Specific | 206.25 | 206.25 | 206.25 | 4.93 | 79 | >sp\|P01833\|PIGR_HUMAN Polymeric immunoglobulin receptor OS=Homo sapiens OX=9606 GN=PIGR PE=1 SV=4 | 902 | 20201112_10613_1ng_glyco_hcd203040_compound15.32847.32847.3 | controllerType=0 controllerNumber=1 scan=32847 | 64.7294 |
| 24674:3 | 6 | R.[+42.011]LSLLEEPGN[+2059.735]GTFTVILNQLTSR.D | HexNAc(4)Hex(5)Fuc(1)NeuAc(1) | .[42], N[+2060] | 1502.0168 | 3 | 4504.0358 | 4504.0381 | 0 | -0.5 | 413 | Specific | 255.45 | 242.62 | 242.62 | 6.91 | 79 | >sp\|P01833\|PIGR_HUMAN Polymeric immunoglobulin receptor OS=Homo sapiens OX=9606 GN=PIGR PE=1 SV=4 | 902 | 20201112_10613_1ng_glyco_hcd203040_compound15.28564.28564.3 | controllerType=0 controllerNumber=1 scan=28564 | 57.0764 |
| 22992:3 | 6 | R.LSLLEEPGN[+2350.830]GTFTVILNQLTSR.D | HexNAc(4)Hex(5)Fuc(1)NeuAc(2) | N[+2351] | 1585.0457 | 3 | 4753.1226 | 4753.1229 | 0 | -0.1 | 413 | Specific | 261.25 | 261.25 | 261.25 | 7.03 | 79 | >sp\|P01833\|PIGR_HUMAN Polymeric immunoglobulin receptor OS=Homo sapiens OX=9606 GN=PIGR PE=1 SV=4 | 902 | 20201112_10613_1ng_glyco_hcd203040_compound15.26818.26818.3 | controllerType=0 controllerNumber=1 scan=26818 | 53.7822 |
| 22873:3 | 6 | R.[+42.011]LSLLEEPGN[+2336.851]GTFTVILNQLTSR.D | HexNAc(6)Hex(6)Fuc(1) | .[42], N[+2337] | 1594.3884 | 3 | 4781.1505 | 4781.1542 | 0 | -0.8 | 413 | Specific | 306.35 | 306.35 | 306.35 | 7.58 | 79 | >sp\|P01833\|PIGR_HUMAN Polymeric immunoglobulin receptor OS=Homo sapiens OX=9606 GN=PIGR PE=1 SV=4 | 902 | 20201112_10613_1ng_glyco_hcd203040_compound15.26694.26694.3 | controllerType=0 controllerNumber=1 scan=26694 | 53.5202 |
| 22871:3 | 6 | R.[+42.011]LSLLEEPGN[+2336.851]GTFTVILNQLTSR.D | HexNAc(6)Hex(6)Fuc(1) | .[42], N[+2337] | 1594.3884 | 3 | 4781.1505 | 4781.1542 | 0 | -0.8 | 413 | Specific | 329.35 | 197.77 | 197.77 | 6.87 | 79 | >sp\|P01833\|PIGR_HUMAN Polymeric immunoglobulin receptor OS=Homo sapiens OX=9606 GN=PIGR PE=1 SV=4 | 902 | 20201112_10613_1ng_glyco_hcd203040_compound15.26692.26692.3 | controllerType=0 controllerNumber=1 scan=26692 | 53.5151 |
| 24676:3 | 6 | R.[+42.011]LSLLEEPGN[+2059.735]GTFTVILNQLTSR.D | HexNAc(4)Hex(5)Fuc(1)NeuAc(1) | .[42], N[+2060] | 1502.0168 | 3 | 4504.0358 | 4504.0381 | 0 | -0.5 | 413 | Specific | 340.94 | 340.94 | 340.94 | 7.93 | 79 | >sp\|P01833\|PIGR_HUMAN Polymeric immunoglobulin receptor OS=Homo sapiens OX=9606 GN=PIGR PE=1 SV=4 | 902 | 20201112_10613_1ng_glyco_hcd203040_compound15.28566.28566.3 | controllerType=0 controllerNumber=1 scan=28566 | 57.0799 |
| 22994:3 | 6 | R.LSLLEEPGN[+2350.830]GTFTVILNQLTSR.D | HexNAc(4)Hex(5)Fuc(1)NeuAc(2) | N[+2351] | 1585.0457 | 3 | 4753.1226 | 4753.1229 | 0 | -0.1 | 413 | Specific | 456.2 | 218.59 | 218.59 | 7.63 | 79 | >sp\|P01833\|PIGR_HUMAN Polymeric immunoglobulin receptor OS=Homo sapiens OX=9606 GN=PIGR PE=1 SV=4 | 902 | 20201112_10613_1ng_glyco_hcd203040_compound15.26820.26820.3 | controllerType=0 controllerNumber=1 scan=26820 | 53.7884 |
| 20657:4 | 6 | R.LSLLEEPGN[+2076.750]GTFTVILNQLTSR.D | HexNAc(4)Hex(6)Fuc(2) | N[+2077] | 1120.5203 | 4 | 4479.0594 | 4479.0428 | 0 | 3.7 | 413 | Specific | 501.71 | 455.56 | 455.56 | 9.77 | 79 | >sp\|P01833\|PIGR_HUMAN Polymeric immunoglobulin receptor OS=Homo sapiens OX=9606 GN=PIGR PE=1 SV=4 | 902 | 20201112_10613_1ng_glyco_hcd203040_compound15.24381.24381.4 | controllerType=0 controllerNumber=1 scan=24381 | 48.9875 |
| 20655:4 | 6 | R.LSLLEEPGN[+2076.750]GTFTVILNQLTSR.D | HexNAc(4)Hex(6)Fuc(2) | N[+2077] | 1120.5203 | 4 | 4479.0594 | 4479.0428 | 0 | 3.7 | 413 | Specific | 602.94 | 493.24 | 493.24 | 12.43 | 79 | >sp\|P01833\|PIGR_HUMAN Polymeric immunoglobulin receptor OS=Homo sapiens OX=9606 GN=PIGR PE=1 SV=4 | 902 | 20201112_10613_1ng_glyco_hcd203040_compound15.24379.24379.4 | controllerType=0 controllerNumber=1 scan=24379 | 48.9744 |
| 20578:3 | 6 | R.LSLLEEPGN[+2350.830]GTFTVILNQLTSR.D | HexNAc(4)Hex(5)Fuc(1)NeuAc(2) | N[+2351] | 1585.06 | 3 | 4753.1654 | 4753.1229 | 0 | 8.9 | 413 | Specific | 625.81 | 494.33 | 494.33 | 11.31 | 79 | >sp\|P01833\|PIGR_HUMAN Polymeric immunoglobulin receptor OS=Homo sapiens OX=9606 GN=PIGR PE=1 SV=4 | 902 | 20201112_10613_1ng_glyco_hcd203040_compound15.24299.24299.3 | controllerType=0 controllerNumber=1 scan=24299 | 48.8036 |
| 20613:4 | 6 | R.LSLLEEPGN[+2205.793]GTFTVILNQLTSR.D | HexNAc(4)Hex(5)Fuc(2)NeuAc(1) | N[+2206] | 1152.781 | 4 | 4608.1024 | 4608.0854 | 0 | 3.7 | 413 | Specific | 782.21 | 590.22 | 590.22 | 15.43 | 79 | >sp\|P01833\|PIGR_HUMAN Polymeric immunoglobulin receptor OS=Homo sapiens OX=9606 GN=PIGR PE=1 SV=4 | 902 | 20201112_10613_1ng_glyco_hcd203040_compound15.24336.24336.4 | controllerType=0 controllerNumber=1 scan=24336 | 48.8786 |
| 20612:3 | 6 | R.LSLLEEPGN[+2205.793]GTFTVILNQLTSR.D | HexNAc(4)Hex(5)Fuc(2)NeuAc(1) | N[+2206] | 1536.7044 | 3 | 4608.0987 | 4608.0854 | 0 | 2.9 | 413 | Specific | 798.83 | 608.44 | 608.44 | 15.55 | 79 | >sp\|P01833\|PIGR_HUMAN Polymeric immunoglobulin receptor OS=Homo sapiens OX=9606 GN=PIGR PE=1 SV=4 | 902 | 20201112_10613_1ng_glyco_hcd203040_compound15.24335.24335.3 | controllerType=0 controllerNumber=1 scan=24335 | 48.8772 |
| 20581:3 | 6 | R.LSLLEEPGN[+2350.830]GTFTVILNQLTSR.D | HexNAc(4)Hex(5)Fuc(1)NeuAc(2) | N[+2351] | 1585.06 | 3 | 4753.1654 | 4753.1229 | 0 | 8.9 | 413 | Specific | 843.28 | 617.62 | 617.62 | 15.31 | 79 | >sp\|P01833\|PIGR_HUMAN Polymeric immunoglobulin receptor OS=Homo sapiens OX=9606 GN=PIGR PE=1 SV=4 | 902 | 20201112_10613_1ng_glyco_hcd203040_compound15.24302.24302.3 | controllerType=0 controllerNumber=1 scan=24302 | 48.8091 |
| 20615:4 | 6 | R.LSLLEEPGN[+2205.793]GTFTVILNQLTSR.D | HexNAc(4)Hex(5)Fuc(2)NeuAc(1) | N[+2206] | 1152.781 | 4 | 4608.1024 | 4608.0854 | 0 | 3.7 | 413 | Specific | 917.95 | 917.95 | 917.95 | 18.43 | 79 | >sp\|P01833\|PIGR_HUMAN Polymeric immunoglobulin receptor OS=Homo sapiens OX=9606 GN=PIGR PE=1 SV=4 | 902 | 20201112_10613_1ng_glyco_hcd203040_compound15.24338.24338.4 | controllerType=0 controllerNumber=1 scan=24338 | 48.8833 |
| 20614:3 | 6 | R.LSLLEEPGN[+2205.793]GTFTVILNQLTSR.D | HexNAc(4)Hex(5)Fuc(2)NeuAc(1) | N[+2206] | 1536.7044 | 3 | 4608.0987 | 4608.0854 | 0 | 2.9 | 413 | Specific | 936.2 | 666.49 | 666.49 | 18.55 | 79 | >sp\|P01833\|PIGR_HUMAN Polymeric immunoglobulin receptor OS=Homo sapiens OX=9606 GN=PIGR PE=1 SV=4 | 902 | 20201112_10613_1ng_glyco_hcd203040_compound15.24337.24337.3 | controllerType=0 controllerNumber=1 scan=24337 | 48.8814 |
| 20618:3 | 6 | R.LSLLEEPGN[+2059.735]GTFTVILNQLTSR.D | HexNAc(4)Hex(5)Fuc(1)NeuAc(1) | N[+2060] | 1488.0184 | 3 | 4462.0405 | 4462.0275 | 0 | 2.9 | 413 | Specific | 942.76 | 690.42 | 690.42 | 18.55 | 79 | >sp\|P01833\|PIGR_HUMAN Polymeric immunoglobulin receptor OS=Homo sapiens OX=9606 GN=PIGR PE=1 SV=4 | 902 | 20201112_10613_1ng_glyco_hcd203040_compound15.24341.24341.3 | controllerType=0 controllerNumber=1 scan=24341 | 48.8918 |
| 20616:3 | 6 | R.LSLLEEPGN[+2059.735]GTFTVILNQLTSR.D | HexNAc(4)Hex(5)Fuc(1)NeuAc(1) | N[+2060] | 1488.0184 | 3 | 4462.0405 | 4462.0275 | 0 | 2.9 | 413 | Specific | 1069.81 | 837.36 | 837.36 | 21.55 | 79 | >sp\|P01833\|PIGR_HUMAN Polymeric immunoglobulin receptor OS=Homo sapiens OX=9606 GN=PIGR PE=1 SV=4 | 902 | 20201112_10613_1ng_glyco_hcd203040_compound15.24339.24339.3 | controllerType=0 controllerNumber=1 scan=24339 | 48.8852 |
| 16220:3 | 6 | K.IIEGEPNLKVPGN[+2059.735]VTAVLGETLK.V | HexNAc(4)Hex(5)Fuc(1)NeuAc(1) | N[+2060] | 1484.3671 | 3 | 4451.0868 | 4451.0843 | 0 | 0.6 | 457 | Specific | 597.62 | 461.19 | 461.19 | 11.53 | 79 | >sp\|P01833\|PIGR_HUMAN Polymeric immunoglobulin receptor OS=Homo sapiens OX=9606 GN=PIGR PE=1 SV=4 | 902 | 20201112_10613_1ng_glyco_hcd203040_compound15.19755.19755.3 | controllerType=0 controllerNumber=1 scan=19755 | 39.5017 |
| 18000:3 | 6 | K.IIEGEPNLKVPGN[+2350.830]VTAVLGETLK.V | HexNAc(4)Hex(5)Fuc(1)NeuAc(2) | N[+2351] | 1581.4025 | 3 | 4742.193 | 4742.1797 | 0 | 2.8 | 457 | Specific | 746.61 | 568.67 | 568.67 | 14.29 | 79 | >sp\|P01833\|PIGR_HUMAN Polymeric immunoglobulin receptor OS=Homo sapiens OX=9606 GN=PIGR PE=1 SV=4 | 902 | 20201112_10613_1ng_glyco_hcd203040_compound15.21611.21611.3 | controllerType=0 controllerNumber=1 scan=21611 | 43.2423 |
| 18006:4 | 6 | K.IIEGEPNLKVPGN[+2350.830]VTAVLGETLK.V | HexNAc(4)Hex(5)Fuc(1)NeuAc(2) | N[+2351] | 1186.3016 | 4 | 4742.1844 | 4742.1797 | 0 | 1 | 457 | Specific | 761.16 | 677.08 | 677.08 | 15.52 | 79 | >sp\|P01833\|PIGR_HUMAN Polymeric immunoglobulin receptor OS=Homo sapiens OX=9606 GN=PIGR PE=1 SV=4 | 902 | 20201112_10613_1ng_glyco_hcd203040_compound15.21617.21617.2 | controllerType=0 controllerNumber=1 scan=21617 | 43.2531 |
| 16192:3 | 6 | K.IIEGEPNLKVPGN[+2205.793]VTAVLGETLK.V | HexNAc(4)Hex(5)Fuc(2)NeuAc(1) | N[+2206] | 1533.0514 | 3 | 4597.1398 | 4597.1422 | 0 | -0.5 | 457 | Specific | 769.34 | 769.34 | 769.34 | 15.53 | 79 | >sp\|P01833\|PIGR_HUMAN Polymeric immunoglobulin receptor OS=Homo sapiens OX=9606 GN=PIGR PE=1 SV=4 | 902 | 20201112_10613_1ng_glyco_hcd203040_compound15.19724.19724.3 | controllerType=0 controllerNumber=1 scan=19724 | 39.4494 |
| 16217:3 | 6 | K.IIEGEPNLKVPGN[+2059.735]VTAVLGETLK.V | HexNAc(4)Hex(5)Fuc(1)NeuAc(1) | N[+2060] | 1484.3671 | 3 | 4451.0868 | 4451.0843 | 0 | 0.6 | 457 | Specific | 781.61 | 611.06 | 611.06 | 15.53 | 79 | >sp\|P01833\|PIGR_HUMAN Polymeric immunoglobulin receptor OS=Homo sapiens OX=9606 GN=PIGR PE=1 SV=4 | 902 | 20201112_10613_1ng_glyco_hcd203040_compound15.19752.19752.3 | controllerType=0 controllerNumber=1 scan=19752 | 39.4969 |
| 16195:3 | 6 | K.IIEGEPNLKVPGN[+2205.793]VTAVLGETLK.V | HexNAc(4)Hex(5)Fuc(2)NeuAc(1) | N[+2206] | 1533.0514 | 3 | 4597.1398 | 4597.1422 | 0 | -0.5 | 457 | Specific | 790.39 | 723.53 | 723.53 | 15.53 | 79 | >sp\|P01833\|PIGR_HUMAN Polymeric immunoglobulin receptor OS=Homo sapiens OX=9606 GN=PIGR PE=1 SV=4 | 902 | 20201112_10613_1ng_glyco_hcd203040_compound15.19727.19727.3 | controllerType=0 controllerNumber=1 scan=19727 | 39.4561 |

Table S4. Cont.

| Query #:z | Protein Rank | Glycopeptide | Glycan | Modification Type(s) | Observed m/z | z | Observed (M+H) | Calc. mass (M+H) | Off-by-x error | Mass error (ppm) | StartPosition | Cleavage | Score | Delta | Delta Mod | \|Log Prob\| | # of unique peptides | Protein Name | Protein DB number | Comment | Scan # | Scan Time |
|---|---|---|---|---|---|---|---|---|---|---|---|---|---|---|---|---|---|---|---|---|---|---|
| 17997:3 | 6 | K.IIEGEPNLKVPGN[+2350.830]VTAVLGETLK.V | HexNAc(4)Hex(5)Fuc(1)NeuAc(2) | N[+2351] | 1581.4025 | 3 | 4742.193 | 4742.1797 | 0 | 2.8 | 457 | Specific | 921.24 | 825.91 | 825.91 | 18.29 | 79 | >sp\|P01833\|PIGR_HUMAN Polymeric immunoglobulin receptor OS=Homo sapiens OX=9606 GN=PIGR PE=1 SV=4 | 902 | 20201112_10613_1ng_glyco_hcd203040_compound15.21608.21608.3 | controllerType=0 controllerNumber=1 scan=21608 | 43.232 |
| 18004:4 | 6 | K.IIEGEPNLKVPGN[+2350.830]VTAVLGETLK.V | HexNAc(4)Hex(5)Fuc(1)NeuAc(2) | N[+2351] | 1186.3016 | 4 | 4742.1844 | 4742.1797 | 0 | 1 | 457 | Specific | 989.95 | 896.76 | 896.76 | 19.52 | 79 | >sp\|P01833\|PIGR_HUMAN Polymeric immunoglobulin receptor OS=Homo sapiens OX=9606 GN=PIGR PE=1 SV=4 | 902 | 20201112_10613_1ng_glyco_hcd203040_compound15.21615.21615.2 | controllerType=0 controllerNumber=1 scan=21615 | 43.2475 |
| 15779:4 | 6 | K.[+42.011]IIEGEPNLKVPGN[+2018.708]VTAVLGETLKVPC[+57.021]HFPC[+57.021] | HexNAc(3)Hex(6)Fuc(1)NeuAc(1) | .[42], C[+57]*2, N[+2019] | 1370.1421 | 4 | 5477.5465 | 5477.5258 | 0 | 3.8 | 457 | Specific | 239.01 | 139.36 | 139.36 | 3.45 | 79 | >sp\|P01833\|PIGR_HUMAN Polymeric immunoglobulin receptor OS=Homo sapiens OX=9606 GN=PIGR PE=1 SV=4 | 902 | 20201112_10613_1ng_glyco_hcd203040_compound15.19265.19265.2 | controllerType=0 controllerNumber=1 scan=19265 | 38.5796 |
| 15925:5 | 6 | K.IIEGEPNLKVPGN[+2059.735]VTAVLGETLKVPC[+57.021]HFPC[+57.021]K.F | HexNAc(4)Hex(5)Fuc(1)NeuAc(1) | C[+57]*2, N[+2060] | 1096.1151 | 5 | 5476.5465 | 5476.5418 | 0 | 0.8 | 457 | Specific | 320.07 | 247.39 | 247.39 | 4.73 | 79 | >sp\|P01833\|PIGR_HUMAN Polymeric immunoglobulin receptor OS=Homo sapiens OX=9606 GN=PIGR PE=1 SV=4 | 902 | 20201112_10613_1ng_glyco_hcd203040_compound15.19417.19417.3 | controllerType=0 controllerNumber=1 scan=19417 | 38.8497 |
| 15868:5 | 6 | K.IIEGEPNLKVPGN[+2059.735]VTAVLGETLKVPC[+57.021]HFPC[+57.021]K.F | HexNAc(4)Hex(5)Fuc(1)NeuAc(1) | C[+57]*2, N[+2060] | 1096.115 | 5 | 5476.5459 | 5476.5418 | 0 | 0.7 | 457 | Specific | 362.86 | 362.86 | 362.86 | 7.67 | 79 | >sp\|P01833\|PIGR_HUMAN Polymeric immunoglobulin receptor OS=Homo sapiens OX=9606 GN=PIGR PE=1 SV=4 | 902 | 20201112_10613_1ng_glyco_hcd203040_compound15.19357.19357.2 | controllerType=0 controllerNumber=1 scan=19357 | 38.7469 |
| 15776:4 | 6 | K.[+42.011]IIEGEPNLKVPGN[+2018.708]VTAVLGETLKVPC[+57.021]HFPC[+57.021] | HexNAc(3)Hex(6)Fuc(1)NeuAc(1) | .[42], C[+57]*2, N[+2019] | 1370.1421 | 4 | 5477.5465 | 5477.5258 | 0 | 3.8 | 457 | Specific | 448.68 | 372.08 | 372.08 | 6.7 | 79 | >sp\|P01833\|PIGR_HUMAN Polymeric immunoglobulin receptor OS=Homo sapiens OX=9606 GN=PIGR PE=1 SV=4 | 902 | 20201112_10613_1ng_glyco_hcd203040_compound15.19262.19262.2 | controllerType=0 controllerNumber=1 scan=19262 | 38.5737 |
| 17193:4 | 6 | K.IIEGEPNLKVPGN[+2350.830]VTAVLGETLKVPC[+57.021]HFPC[+57.021]K.F | HexNAc(4)Hex(5)Fuc(1)NeuAc(2) | C[+57]*2, N[+2351] | 1442.6677 | 4 | 5767.6491 | 5767.6372 | 0 | 2.1 | 457 | Specific | 730.34 | 730.34 | 730.34 | 13.84 | 79 | >sp\|P01833\|PIGR_HUMAN Polymeric immunoglobulin receptor OS=Homo sapiens OX=9606 GN=PIGR PE=1 SV=4 | 902 | 20201112_10613_1ng_glyco_hcd203040_compound15.20769.20769.4 | controllerType=0 controllerNumber=1 scan=20769 | 41.4411 |
| 17159:4 | 6 | K.IIEGEPNLKVPGN[+2350.830]VTAVLGETLKVPC[+57.021]HFPC[+57.021]K.F | HexNAc(4)Hex(5)Fuc(1)NeuAc(2) | C[+57]*2, N[+2351] | 1442.6692 | 4 | 5767.6549 | 5767.6372 | 0 | 3.1 | 457 | Specific | 749.95 | 660.93 | 660.93 | 13.53 | 79 | >sp\|P01833\|PIGR_HUMAN Polymeric immunoglobulin receptor OS=Homo sapiens OX=9606 GN=PIGR PE=1 SV=4 | 902 | 20201112_10613_1ng_glyco_hcd203040_compound15.20733.20733.2 | controllerType=0 controllerNumber=1 scan=20733 | 41.3651 |
| 15870:5 | 6 | K.IIEGEPNLKVPGN[+2059.735]VTAVLGETLKVPC[+57.021]HFPC[+57.021]K.F | HexNAc(4)Hex(5)Fuc(1)NeuAc(1) | C[+57]*2, N[+2060] | 1096.115 | 5 | 5476.5459 | 5476.5418 | 0 | 0.7 | 457 | Specific | 788.68 | 693.49 | 693.49 | 15.3 | 79 | >sp\|P01833\|PIGR_HUMAN Polymeric immunoglobulin receptor OS=Homo sapiens OX=9606 GN=PIGR PE=1 SV=4 | 902 | 20201112_10613_1ng_glyco_hcd203040_compound15.19359.19359.2 | controllerType=0 controllerNumber=1 scan=19359 | 38.7502 |
| 17189:5 | 6 | K.IIEGEPNLKVPGN[+2350.830]VTAVLGETLKVPC[+57.021]HFPC[+57.021]K.F | HexNAc(4)Hex(5)Fuc(1)NeuAc(2) | C[+57]*2, N[+2351] | 1154.3346 | 5 | 5767.6439 | 5767.6372 | 0 | 1.2 | 457 | Specific | 826.26 | 647.93 | 647.93 | 16.3 | 79 | >sp\|P01833\|PIGR_HUMAN Polymeric immunoglobulin receptor OS=Homo sapiens OX=9606 GN=PIGR PE=1 SV=4 | 902 | 20201112_10613_1ng_glyco_hcd203040_compound15.20765.20765.5 | controllerType=0 controllerNumber=1 scan=20765 | 41.4328 |
| 17187:5 | 6 | K.IIEGEPNLKVPGN[+2350.830]VTAVLGETLKVPC[+57.021]HFPC[+57.021]K.F | HexNAc(4)Hex(5)Fuc(1)NeuAc(2) | C[+57]*2, N[+2351] | 1154.3346 | 5 | 5767.6439 | 5767.6372 | 0 | 1.2 | 457 | Specific | 835.64 | 687.07 | 687.07 | 16.3 | 79 | >sp\|P01833\|PIGR_HUMAN Polymeric immunoglobulin receptor OS=Homo sapiens OX=9606 GN=PIGR PE=1 SV=4 | 902 | 20201112_10613_1ng_glyco_hcd203040_compound15.20763.20763.5 | controllerType=0 controllerNumber=1 scan=20763 | 41.4279 |
| 17190:4 | 6 | K.IIEGEPNLKVPGN[+2350.830]VTAVLGETLKVPC[+57.021]HFPC[+57.021]K.F | HexNAc(4)Hex(5)Fuc(1)NeuAc(2) | C[+57]*2, N[+2351] | 1442.6677 | 4 | 5767.6491 | 5767.6372 | 0 | 2.1 | 457 | Specific | 966.58 | 815.97 | 815.97 | 18.84 | 79 | >sp\|P01833\|PIGR_HUMAN Polymeric immunoglobulin receptor OS=Homo sapiens OX=9606 GN=PIGR PE=1 SV=4 | 902 | 20201112_10613_1ng_glyco_hcd203040_compound15.20766.20766.4 | controllerType=0 controllerNumber=1 scan=20766 | 41.4344 |
| 17156:4 | 6 | K.IIEGEPNLKVPGN[+2350.830]VTAVLGETLKVPC[+57.021]HFPC[+57.021]K.F | HexNAc(4)Hex(5)Fuc(1)NeuAc(2) | C[+57]*2, N[+2351] | 1442.6692 | 4 | 5767.6549 | 5767.6372 | 0 | 3.1 | 457 | Specific | 1042.85 | 945.42 | 945.42 | 19.53 | 79 | >sp\|P01833\|PIGR_HUMAN Polymeric immunoglobulin receptor OS=Homo sapiens OX=9606 GN=PIGR PE=1 SV=4 | 902 | 20201112_10613_1ng_glyco_hcd203040_compound15.20730.20730.2 | controllerType=0 controllerNumber=1 scan=20730 | 41.3512 |
| 13655:3 | 6 | K.VPGN[+2205.793]VTAVLGETLK.V | HexNAc(4)Hex(5)Fuc(2)NeuAc(1) | N[+2206] | 1201.872 | 3 | 3603.6015 | 3603.5927 | 0 | 2.4 | 466 | Specific | 474.8 | 443.67 | 443.67 | 9.15 | 79 | >sp\|P01833\|PIGR_HUMAN Polymeric immunoglobulin receptor OS=Homo sapiens OX=9606 GN=PIGR PE=1 SV=4 | 902 | 20201112_10613_1ng_glyco_hcd203040_compound15.17063.17063.3 | controllerType=0 controllerNumber=1 scan=17063 | 34.5913 |
| 13657:3 | 6 | K.VPGN[+2205.793]VTAVLGETLK.V | HexNAc(4)Hex(5)Fuc(2)NeuAc(1) | N[+2206] | 1201.872 | 3 | 3603.6015 | 3603.5927 | 0 | 2.4 | 466 | Specific | 594.14 | 594.14 | 594.14 | 10.99 | 79 | >sp\|P01833\|PIGR_HUMAN Polymeric immunoglobulin receptor OS=Homo sapiens OX=9606 GN=PIGR PE=1 SV=4 | 902 | 20201112_10613_1ng_glyco_hcd203040_compound15.17065.17065.3 | controllerType=0 controllerNumber=1 scan=17065 | 34.5953 |
| 16705:3 | 6 | K.VPGN[+2350.830]VTAVLGETLK.V | HexNAc(4)Hex(5)Fuc(1)NeuAc(2) | N[+2351] | 1250.2171 | 3 | 3748.6367 | 3748.6302 | 0 | 1.7 | 466 | Specific | 635.69 | 472.45 | 472.45 | 12.3 | 79 | >sp\|P01833\|PIGR_HUMAN Polymeric immunoglobulin receptor OS=Homo sapiens OX=9606 GN=PIGR PE=1 SV=4 | 902 | 20201112_10613_1ng_glyco_hcd203040_compound15.20261.20261.3 | controllerType=0 controllerNumber=1 scan=20261 | 40.4312 |
| 16707:3 | 6 | K.VPGN[+2350.830]VTAVLGETLK.V | HexNAc(4)Hex(5)Fuc(1)NeuAc(2) | N[+2351] | 1250.2171 | 3 | 3748.6367 | 3748.6302 | 0 | 1.7 | 466 | Specific | 690.3 | 474.77 | 474.77 | 13.3 | 79 | >sp\|P01833\|PIGR_HUMAN Polymeric immunoglobulin receptor OS=Homo sapiens OX=9606 GN=PIGR PE=1 SV=4 | 902 | 20201112_10613_1ng_glyco_hcd203040_compound15.20263.20263.3 | controllerType=0 controllerNumber=1 scan=20263 | 40.4369 |
| 13760:4 | 6 | K.VPGN[+2205.793]VTAVLGETLKVPC[+57.021]HFPC[+57.021]K.F | HexNAc(4)Hex(5)Fuc(2)NeuAc(1) | C[+57]*2, N[+2206] | 1158.0196 | 4 | 4629.0565 | 4629.0502 | 0 | 1.3 | 466 | Specific | 554.52 | 550.83 | 550.83 | 10.38 | 79 | >sp\|P01833\|PIGR_HUMAN Polymeric immunoglobulin receptor OS=Homo sapiens OX=9606 GN=PIGR PE=1 SV=4 | 902 | 20201112_10613_1ng_glyco_hcd203040_compound15.17172.17172.4 | controllerType=0 controllerNumber=1 scan=17172 | 34.7871 |
| 15657:4 | 6 | K.VPGN[+2351.851]VTAVLGETLKVPC[+57.021]HFPC[+57.021]K.F | HexNAc(4)Hex(5)Fuc(3)NeuAc(1) | C[+57]*2, N[+2352] | 1194.5289 | 4 | 4775.0936 | 4775.1081 | 0 | -3 | 466 | Specific | 568.06 | 517.03 | 517.03 | 9.16 | 79 | >sp\|P01833\|PIGR_HUMAN Polymeric immunoglobulin receptor OS=Homo sapiens OX=9606 GN=PIGR PE=1 SV=4 | 902 | 20201112_10613_1ng_glyco_hcd203040_compound15.19139.19139.4 | controllerType=0 controllerNumber=1 scan=19139 | 38.3578 |
| 13736:4 | 6 | K.VPGN[+2205.793]VTAVLGETLKVPC[+57.021]HFPC[+57.021]K.F | HexNAc(4)Hex(5)Fuc(2)NeuAc(1) | C[+57]*2, N[+2206] | 1158.0193 | 4 | 4629.0555 | 4629.0502 | 0 | 1.1 | 466 | Specific | 615.84 | 494.17 | 494.17 | 12.6 | 79 | >sp\|P01833\|PIGR_HUMAN Polymeric immunoglobulin receptor OS=Homo sapiens OX=9606 GN=PIGR PE=1 SV=4 | 902 | 20201112_10613_1ng_glyco_hcd203040_compound15.17147.17147.3 | controllerType=0 controllerNumber=1 scan=17147 | 34.7448 |
| 13734:4 | 6 | K.VPGN[+2205.793]VTAVLGETLKVPC[+57.021]HFPC[+57.021]K.F | HexNAc(4)Hex(5)Fuc(2)NeuAc(1) | C[+57]*2, N[+2206] | 1158.0193 | 4 | 4629.0555 | 4629.0502 | 0 | 1.1 | 466 | Specific | 641.43 | 476.59 | 476.59 | 12.6 | 79 | >sp\|P01833\|PIGR_HUMAN Polymeric immunoglobulin receptor OS=Homo sapiens OX=9606 GN=PIGR PE=1 SV=4 | 902 | 20201112_10613_1ng_glyco_hcd203040_compound15.17145.17145.3 | controllerType=0 controllerNumber=1 scan=17145 | 34.7393 |
| 13763:4 | 6 | K.VPGN[+2205.793]VTAVLGETLKVPC[+57.021]HFPC[+57.021]K.F | HexNAc(4)Hex(5)Fuc(2)NeuAc(1) | C[+57]*2, N[+2206] | 1158.0196 | 4 | 4629.0565 | 4629.0502 | 0 | 1.3 | 466 | Specific | 655.07 | 528.87 | 528.87 | 13.6 | 79 | >sp\|P01833\|PIGR_HUMAN Polymeric immunoglobulin receptor OS=Homo sapiens OX=9606 GN=PIGR PE=1 SV=4 | 902 | 20201112_10613_1ng_glyco_hcd203040_compound15.17175.17175.4 | controllerType=0 controllerNumber=1 scan=17175 | 34.7952 |
| 15660:4 | 6 | K.VPGN[+2351.851]VTAVLGETLKVPC[+57.021]HFPC[+57.021]K.F | HexNAc(4)Hex(5)Fuc(3)NeuAc(1) | C[+57]*2, N[+2352] | 1194.5289 | 4 | 4775.0936 | 4775.1081 | 0 | -3 | 466 | Specific | 757.04 | 513.53 | 513.53 | 14.38 | 79 | >sp\|P01833\|PIGR_HUMAN Polymeric immunoglobulin receptor OS=Homo sapiens OX=9606 GN=PIGR PE=1 SV=4 | 902 | 20201112_10613_1ng_glyco_hcd203040_compound15.19142.19142.4 | controllerType=0 controllerNumber=1 scan=19142 | 38.365 |
| 13816:4 | 6 | K.VPGN[+2059.735]VTAVLGETLKVPC[+57.021]HFPC[+57.021]K.F | HexNAc(4)Hex(5)Fuc(1)NeuAc(1) | C[+57]*2, N[+2060] | 1121.5042 | 4 | 4482.9949 | 4482.9923 | 0 | 0.6 | 466 | Specific | 804.73 | 559.66 | 559.66 | 16.6 | 79 | >sp\|P01833\|PIGR_HUMAN Polymeric immunoglobulin receptor OS=Homo sapiens OX=9606 GN=PIGR PE=1 SV=4 | 902 | 20201112_10613_1ng_glyco_hcd203040_compound15.17230.17230.4 | controllerType=0 controllerNumber=1 scan=17230 | 34.9175 |
| 13814:4 | 6 | K.VPGN[+2059.735]VTAVLGETLKVPC[+57.021]HFPC[+57.021]K.F | HexNAc(4)Hex(5)Fuc(1)NeuAc(1) | C[+57]*2, N[+2060] | 1121.5042 | 4 | 4482.9949 | 4482.9923 | 0 | 0.6 | 466 | Specific | 929.01 | 780.22 | 780.22 | 18.6 | 79 | >sp\|P01833\|PIGR_HUMAN Polymeric immunoglobulin receptor OS=Homo sapiens OX=9606 GN=PIGR PE=1 SV=4 | 902 | 20201112_10613_1ng_glyco_hcd203040_compound15.17228.17228.4 | controllerType=0 controllerNumber=1 scan=17228 | 34.9064 |
| 05810:3 | 6 | K.WN[+1872.651]NTGC[+57.021]QALPSQDEGPSK.A | HexNAc(3)Hex(6)NeuAc(1) | C[+57], N[+1873] | 1287.8476 | 3 | 3861.5282 | 3861.5272 | 0 | 0.3 | 498 | Specific | 318.23 | 286.76 | 286.76 | 6.98 | 79 | >sp\|P01833\|PIGR_HUMAN Polymeric immunoglobulin receptor OS=Homo sapiens OX=9606 GN=PIGR PE=1 SV=4 | 902 | 20201112_10613_1ng_glyco_hcd203040_compound15.8910.8910.3 | controllerType=0 controllerNumber=1 scan=8910 | 20.9422 |
| 05963:3 | 6 | K.WN[+1548.545]NTGC[+57.021]QALPSQDEGPSK.A | HexNAc(3)Hex(4)NeuAc(1) | C[+57], N[+1549] | 1179.8157 | 3 | 3537.4326 | 3537.4215 | 0 | 3.1 | 498 | Specific | 368.47 | 327.85 | 327.85 | 7.25 | 79 | >sp\|P01833\|PIGR_HUMAN Polymeric immunoglobulin receptor OS=Homo sapiens OX=9606 GN=PIGR PE=1 SV=4 | 902 | 20201112_10613_1ng_glyco_hcd203040_compound15.9068.9068.3 | controllerType=0 controllerNumber=1 scan=9068 | 21.1995 |
| 05965:3 | 6 | K.WN[+1548.545]NTGC[+57.021]QALPSQDEGPSK.A | HexNAc(3)Hex(4)NeuAc(1) | C[+57], N[+1549] | 1179.8157 | 3 | 3537.4326 | 3537.4215 | 0 | 3.1 | 498 | Specific | 376.43 | 323.96 | 323.96 | 7.25 | 79 | >sp\|P01833\|PIGR_HUMAN Polymeric immunoglobulin receptor OS=Homo sapiens OX=9606 GN=PIGR PE=1 SV=4 | 902 | 20201112_10613_1ng_glyco_hcd203040_compound15.9070.9070.3 | controllerType=0 controllerNumber=1 scan=9070 | 21.2036 |
| 05716:3 | 6 | K.WN[+2059.735]NTGC[+57.021]QALPSQDEGPSK.A | HexNAc(4)Hex(5)Fuc(1)NeuAc(1) | C[+57], N[+2060] | 1350.2112 | 3 | 4048.6192 | 4048.6116 | 0 | 1.9 | 498 | Specific | 441.38 | 386.27 | 386.27 | 9.56 | 79 | >sp\|P01833\|PIGR_HUMAN Polymeric immunoglobulin receptor OS=Homo sapiens OX=9606 GN=PIGR PE=1 SV=4 | 902 | 20201112_10613_1ng_glyco_hcd203040_compound15.8813.8813.3 | controllerType=0 controllerNumber=1 scan=8813 | 20.7795 |
| 05864:3 | 6 | K.WN[+1710.598]NTGC[+57.021]QALPSQDEGPSK.A | HexNAc(3)Hex(5)NeuAc(1) | C[+57], N[+1711] | 1233.831 | 3 | 3699.4784 | 3699.4743 | 0 | 1.1 | 498 | Specific | 496.95 | 422.88 | 422.88 | 9.84 | 79 | >sp\|P01833\|PIGR_HUMAN Polymeric immunoglobulin receptor OS=Homo sapiens OX=9606 GN=PIGR PE=1 SV=4 | 902 | 20201112_10613_1ng_glyco_hcd203040_compound15.8966.8966.3 | controllerType=0 controllerNumber=1 scan=8966 | 21.0354 |
| 05718:3 | 6 | K.WN[+2059.735]NTGC[+57.021]QALPSQDEGPSK.A | HexNAc(4)Hex(5)Fuc(1)NeuAc(1) | C[+57], N[+2060] | 1350.2112 | 3 | 4048.6192 | 4048.6116 | 0 | 1.9 | 498 | Specific | 527.7 | 468.62 | 468.62 | 10.07 | 79 | >sp\|P01833\|PIGR_HUMAN Polymeric immunoglobulin receptor OS=Homo sapiens OX=9606 GN=PIGR PE=1 SV=4 | 902 | 20201112_10613_1ng_glyco_hcd203040_compound15.8815.8815.3 | controllerType=0 controllerNumber=1 scan=8815 | 20.783 |
| 05866:3 | 6 | K.WN[+1710.598]NTGC[+57.021]QALPSQDEGPSK.A | HexNAc(3)Hex(5)NeuAc(1) | C[+57], N[+1711] | 1233.831 | 3 | 3699.4784 | 3699.4743 | 0 | 1.1 | 498 | Specific | 584.9 | 506.93 | 506.93 | 11.45 | 79 | >sp\|P01833\|PIGR_HUMAN Polymeric immunoglobulin receptor OS=Homo sapiens OX=9606 GN=PIGR PE=1 SV=4 | 902 | 20201112_10613_1ng_glyco_hcd203040_compound15.8968.8968.3 | controllerType=0 controllerNumber=1 scan=8968 | 21.039 |
| 05805:3 | 6 | K.WN[+1913.677]NTGC[+57.021]QALPSQDEGPSK.A | HexNAc(4)Hex(5)NeuAc(1) | C[+57], N[+1914] | 1301.5234 | 3 | 3902.5556 | 3902.5537 | 0 | 0.5 | 498 | Specific | 623.95 | 544.22 | 544.22 | 12.38 | 79 | >sp\|P01833\|PIGR_HUMAN Polymeric immunoglobulin receptor OS=Homo sapiens OX=9606 GN=PIGR PE=1 SV=4 | 902 | 20201112_10613_1ng_glyco_hcd203040_compound15.8905.8905.3 | controllerType=0 controllerNumber=1 scan=8905 | 20.9333 |
| 05803:3 | 6 | K.WN[+1913.677]NTGC[+57.021]QALPSQDEGPSK.A | HexNAc(4)Hex(5)NeuAc(1) | C[+57], N[+1914] | 1301.5234 | 3 | 3902.5556 | 3902.5537 | 0 | 0.5 | 498 | Specific | 676.01 | 580.9 | 580.9 | 13.38 | 79 | >sp\|P01833\|PIGR_HUMAN Polymeric immunoglobulin receptor OS=Homo sapiens OX=9606 GN=PIGR PE=1 SV=4 | 902 | 20201112_10613_1ng_glyco_hcd203040_compound15.8903.8903.3 | controllerType=0 controllerNumber=1 scan=8903 | 20.9291 |
| 17260:3 | 10 | K.RNIAAFGGDPNN[+1913.677]ITLFGESAGGASVSLQTLSPYNK.G | HexNAc(4)Hex(5)NeuAc(1) | N[+1914] | 1827.4983 | 3 | 5480.4802 | 5480.4644 | 0 | 2.9 | 196 | Specific | 235.2 | 72.96 | 72.96 | 4.59 | 44 | >sp\|P19835\|CEL_HUMAN Bile salt-activated lipase OS=Homo sapiens OX=9606 GN=CEL PE=1 SV=3 | 476 | 20201112_10613_1ng_glyco_hcd203040_compound15.20839.20839.3 | controllerType=0 controllerNumber=1 scan=20839 | 41.5893 |
| 17258:3 | 10 | K.RNIAAFGGDPNN[+1913.677]ITLFGESAGGASVSLQTLSPYNK.G | HexNAc(4)Hex(5)NeuAc(1) | N[+1914] | 1827.4983 | 3 | 5480.4802 | 5480.4644 | 0 | 2.9 | 196 | Specific | 353.02 | 268.86 | 268.86 | 7.17 | 44 | >sp\|P19835\|CEL_HUMAN Bile salt-activated lipase OS=Homo sapiens OX=9606 GN=CEL PE=1 SV=3 | 476 | 20201112_10613_1ng_glyco_hcd203040_compound15.20837.20837.3 | controllerType=0 controllerNumber=1 scan=20837 | 41.5852 |
| 17789:5 | 12 | R.LSLHRPALEDLLLGSEAN[+1524.534]LTC[+57.021]TLTGLR.D | HexNAc(2)Hex(6)Fuc(1) | C[+57], N[+1525] | 898.4287 | 5 | 4488.1146 | 4488.1319 | 0 | -3.9 | 127 | Specific | 262.69 | 205.04 | 205.04 | 4.43 | 48 | >sp\|P01876\|IGHA1_HUMAN Immunoglobulin heavy constant alpha 1 OS=Homo sapiens OX=9606 GN=IGHA1 PE=1 SV=2 | 255 | 20201112_10613_1ng_glyco_hcd203040_compound15.21391.21391.5 | controllerType=0 controllerNumber=1 scan=21391 | 42.7682 |
| 17786:5 | 12 | R.LSLHRPALEDLLLGSEAN[+1524.534]LTC[+57.021]TLTGLR.D | HexNAc(2)Hex(6)Fuc(1) | C[+57], N[+1525] | 898.4287 | 5 | 4488.1146 | 4488.1319 | 0 | -3.9 | 127 | Specific | 298.37 | 277.78 | 277.78 | 4.92 | 48 | >sp\|P01876\|IGHA1_HUMAN Immunoglobulin heavy constant alpha 1 OS=Homo sapiens OX=9606 GN=IGHA1 PE=1 SV=2 | 255 | 20201112_10613_1ng_glyco_hcd203040_compound15.21388.21388.5 | controllerType=0 controllerNumber=1 scan=21388 | 42.7616 |
| 18737:3 | 12 | R.LSLHRPALEDLLLGSEAN[+2116.756]LTC[+57.021]TLTGLR.D | HexNAc(5)Hex(5)NeuAc(1) | C[+57], N[+2117] | 1694.1264 | 3 | 5080.3647 | 5080.3547 | 0 | 2 | 127 | Specific | 313.1 | 215.88 | 215.88 | 6.93 | 48 | >sp\|P01876\|IGHA1_HUMAN Immunoglobulin heavy constant alpha 1 OS=Homo sapiens OX=9606 GN=IGHA1 PE=1 SV=2 | 255 | 20201112_10613_1ng_glyco_hcd203040_compound15.22377.22377.3 | controllerType=0 controllerNumber=1 scan=22377 | 44.7394 |
| 17992:4 | 12 | R.LSLHRPALEDLLLGSEAN[+1501.555]LTC[+57.021]TLTGLR.D | HexNAc(5)Hex(3) | C[+57], N[+1502] | 1117.0409 | 4 | 4465.1419 | 4465.1536 | 0 | -2.6 | 127 | Specific | 353.84 | 353.84 | 353.84 | 7.31 | 48 | >sp\|P01876\|IGHA1_HUMAN Immunoglobulin heavy constant alpha 1 OS=Homo sapiens OX=9606 GN=IGHA1 PE=1 SV=2 | 255 | 20201112_10613_1ng_glyco_hcd203040_compound15.21603.21603.4 | controllerType=0 controllerNumber=1 scan=21603 | 43.2228 |
| 18739:3 | 12 | R.LSLHRPALEDLLLGSEAN[+2116.756]LTC[+57.021]TLTGLR.D | HexNAc(5)Hex(5)NeuAc(1) | C[+57], N[+2117] | 1694.1264 | 3 | 5080.3647 | 5080.3547 | 0 | 2 | 127 | Specific | 461.43 | 324.5 | 324.5 | 8.1 | 48 | >sp\|P01876\|IGHA1_HUMAN Immunoglobulin heavy constant alpha 1 OS=Homo sapiens OX=9606 GN=IGHA1 PE=1 SV=2 | 255 | 20201112_10613_1ng_glyco_hcd203040_compound15.22379.22379.3 | controllerType=0 controllerNumber=1 scan=22379 | 44.7435 |
| 18793:3 | 12 | R.LSLHRPALEDLLLGSEAN[+1751.624]LTC[+57.021]TLTGLR.D | HexNAc(4)Hex(4)NeuAc(1) | C[+57], N[+1752] | 1572.4162 | 3 | 4715.234 | 4715.2225 | 0 | 2.5 | 127 | Specific | 466.87 | 466.87 | 466.87 | 9.55 | 48 | >sp\|P01876\|IGHA1_HUMAN Immunoglobulin heavy constant alpha 1 OS=Homo sapiens OX=9606 GN=IGHA1 PE=1 SV=2 | 255 | 20201112_10613_1ng_glyco_hcd203040_compound15.22436.22436.3 | controllerType=0 controllerNumber=1 scan=22436 | 44.9094 |
| 18795:3 | 12 | R.LSLHRPALEDLLLGSEAN[+1751.624]LTC[+57.021]TLTGLR.D | HexNAc(4)Hex(4)NeuAc(1) | C[+57], N[+1752] | 1572.4162 | 3 | 4715.234 | 4715.2225 | 0 | 2.5 | 127 | Specific | 496.36 | 496.36 | 496.36 | 9.78 | 48 | >sp\|P01876\|IGHA1_HUMAN Immunoglobulin heavy constant alpha 1 OS=Homo sapiens OX=9606 GN=IGHA1 PE=1 SV=2 | 255 | 20201112_10613_1ng_glyco_hcd203040_compound15.22438.22438.3 | controllerType=0 controllerNumber=1 scan=22438 | 44.9154 |
| 17746:3 | 12 | R.LSLHRPALEDLLLGSEAN[+1622.582]LTC[+57.021]TLTGLR.D | HexNAc(4)Hex(5) | C[+57], N[+1623] | 1529.399 | 3 | 4586.1824 | 4586.1799 | 0 | 0.5 | 127 | Specific | 502.84 | 444.99 | 444.99 | 10.04 | 48 | >sp\|P01876\|IGHA1_HUMAN Immunoglobulin heavy constant alpha 1 OS=Homo sapiens OX=9606 GN=IGHA1 PE=1 SV=2 | 255 | 20201112_10613_1ng_glyco_hcd203040_compound15.21346.21346.3 | controllerType=0 controllerNumber=1 scan=21346 | 42.6537 |
| 17726:3 | 12 | R.LSLHRPALEDLLLGSEAN[+1540.529]LTC[+57.021]TLTGLR.D | HexNAc(2)Hex(7) | C[+57], N[+1541] | 1502.0478 | 3 | 4504.1288 | 4504.1268 | 0 | 0.5 | 127 | Specific | 509.24 | 509.24 | 509.24 | 9.97 | 48 | >sp\|P01876\|IGHA1_HUMAN Immunoglobulin heavy constant alpha 1 OS=Homo sapiens OX=9606 GN=IGHA1 PE=1 SV=2 | 255 | 20201112_10613_1ng_glyco_hcd203040_compound15.21325.21325.3 | controllerType=0 controllerNumber=1 scan=21325 | 42.604 |
| 17793:3 | 12 | R.LSLHRPALEDLLLGSEAN[+1298.476]LTC[+57.021]TLTGLR.D | HexNAc(4)Hex(3) | C[+57], N[+1298] | 1421.3621 | 3 | 4262.0718 | 4262.0742 | 0 | -0.6 | 127 | Specific | 552.95 | 472.5 | 472.5 | 10.41 | 48 | >sp\|P01876\|IGHA1_HUMAN Immunoglobulin heavy constant alpha 1 OS=Homo sapiens OX=9606 GN=IGHA1 PE=1 SV=2 | 255 | 20201112_10613_1ng_glyco_hcd203040_compound15.21395.21395.3 | controllerType=0 controllerNumber=1 scan=21395 | 42.7779 |
| 17791:3 | 12 | R.LSLHRPALEDLLLGSEAN[+1298.476]LTC[+57.021]TLTGLR.D | HexNAc(4)Hex(3) | C[+57], N[+1298] | 1421.3621 | 3 | 4262.0718 | 4262.0742 | 0 | -0.6 | 127 | Specific | 560.7 | 560.7 | 560.7 | 10.41 | 48 | >sp\|P01876\|IGHA1_HUMAN Immunoglobulin heavy constant alpha 1 OS=Homo sapiens OX=9606 GN=IGHA1 PE=1 SV=2 | 255 | 20201112_10613_1ng_glyco_hcd203040_compound15.21393.21393.3 | controllerType=0 controllerNumber=1 scan=21393 | 42.7729 |
| 17766:4 | 12 | R.LSLHRPALEDLLLGSEAN[+1622.582]LTC[+57.021]TLTGLR.D | HexNAc(4)Hex(5) | C[+57], N[+1623] | 1147.3013 | 4 | 4586.1834 | 4586.1799 | 0 | 0.8 | 127 | Specific | 567.98 | 567.98 | 567.98 | 10.94 | 48 | >sp\|P01876\|IGHA1_HUMAN Immunoglobulin heavy constant alpha 1 OS=Homo sapiens OX=9606 GN=IGHA1 PE=1 SV=2 | 255 | 20201112_10613_1ng_glyco_hcd203040_compound15.21367.21367.4 | controllerType=0 controllerNumber=1 scan=21367 | 42.7146 |
| 17728:3 | 12 | R.LSLHRPALEDLLLGSEAN[+1540.529]LTC[+57.021]TLTGLR.D | HexNAc(2)Hex(7) | C[+57], N[+1541] | 1502.0478 | 3 | 4504.1288 | 4504.1268 | 0 | 0.5 | 127 | Specific | 577.04 | 544.51 | 544.51 | 11.63 | 48 | >sp\|P01876\|IGHA1_HUMAN Immunoglobulin heavy constant alpha 1 OS=Homo sapiens OX=9606 GN=IGHA1 PE=1 SV=2 | 255 | 20201112_10613_1ng_glyco_hcd203040_compound15.21327.21327.3 | controllerType=0 controllerNumber=1 scan=21327 | 42.6081 |
| 17711:3 | 12 | R.LSLHRPALEDLLLGSEAN[+1622.582]LTC[+57.021]TLTGLR.D | HexNAc(4)Hex(5) | C[+57], N[+1623] | 1529.3976 | 3 | 4586.1784 | 4586.1799 | 0 | -0.3 | 127 | Specific | 589.25 | 551 | 551 | 11.63 | 48 | >sp\|P01876\|IGHA1_HUMAN Immunoglobulin heavy constant alpha 1 OS=Homo sapiens OX=9606 GN=IGHA1 PE=1 SV=2 | 255 | 20201112_10613_1ng_glyco_hcd203040_compound15.21309.21309.3 | controllerType=0 controllerNumber=1 scan=21309 | 42.5713 |
| 17775:3 | 12 | R.LSLHRPALEDLLLGSEAN[+1460.529]LTC[+57.021]TLTGLR.D | HexNAc(4)Hex(4) | C[+57], N[+1461] | 1475.3842 | 3 | 4424.1381 | 4424.1271 | 0 | 2.5 | 127 | Specific | 595.87 | 528.36 | 528.36 | 11.39 | 48 | >sp\|P01876\|IGHA1_HUMAN Immunoglobulin heavy constant alpha 1 OS=Homo sapiens OX=9606 GN=IGHA1 PE=1 SV=2 | 255 | 20201112_10613_1ng_glyco_hcd203040_compound15.21376.21376.3 | controllerType=0 controllerNumber=1 scan=21376 | 42.7343 |
| 17783:5 | 12 | R.LSLHRPALEDLLLGSEAN[+1501.555]LTC[+57.021]TLTGLR.D | HexNAc(5)Hex(3) | C[+57], N[+1502] | 893.8366 | 5 | 4465.1537 | 4465.1536 | 0 | 0 | 127 | Specific | 597.73 | 597.73 | 597.73 | 12.02 | 48 | >sp\|P01876\|IGHA1_HUMAN Immunoglobulin heavy constant alpha 1 OS=Homo sapiens OX=9606 GN=IGHA1 PE=1 SV=2 | 255 | 20201112_10613_1ng_glyco_hcd203040_compound15.21385.21385.5 | controllerType=0 controllerNumber=1 scan=21385 | 42.7552 |
| 17729:3 | 12 | R.LSLHRPALEDLLLGSEAN[+1663.608]LTC[+57.021]TLTGLR.D | HexNAc(5)Hex(4) | C[+57], N[+1664] | 1543.0753 | 3 | 4627.2112 | 4627.2064 | 0 | 1 | 127 | Specific | 610.68 | 591.21 | 591.21 | 12.7 | 48 | >sp\|P01876\|IGHA1_HUMAN Immunoglobulin heavy constant alpha 1 OS=Homo sapiens OX=9606 GN=IGHA1 PE=1 SV=2 | 255 | 20201112_10613_1ng_glyco_hcd203040_compound15.21328.21328.3 | controllerType=0 controllerNumber=1 scan=21328 | 42.6097 |
| 17772:3 | 12 | R.LSLHRPALEDLLLGSEAN[+1460.529]LTC[+57.021]TLTGLR.D | HexNAc(4)Hex(4) | C[+57], N[+1461] | 1475.3842 | 3 | 4424.1381 | 4424.1271 | 0 | 2.5 | 127 | Specific | 612.38 | 547.12 | 547.12 | 12.39 | 48 | >sp\|P01876\|IGHA1_HUMAN Immunoglobulin heavy constant alpha 1 OS=Homo sapiens OX=9606 GN=IGHA1 PE=1 SV=2 | 255 | 20201112_10613_1ng_glyco_hcd203040_compound15.21373.21373.3 | controllerType=0 controllerNumber=1 scan=21373 | 42.7264 |
| 17732:3 | 12 | R.LSLHRPALEDLLLGSEAN[+1663.608]LTC[+57.021]TLTGLR.D | HexNAc(5)Hex(4) | C[+57], N[+1664] | 1543.0753 | 3 | 4627.2112 | 4627.2064 | 0 | 1 | 127 | Specific | 620.96 | 550.54 | 550.54 | 12.7 | 48 | >sp\|P01876\|IGHA1_HUMAN Immunoglobulin heavy constant alpha 1 OS=Homo sapiens OX=9606 GN=IGHA1 PE=1 SV=2 | 255 | 20201112_10613_1ng_glyco_hcd203040_compound15.21331.21331.3 | controllerType=0 controllerNumber=1 scan=21331 | 42.6148 |
| 17785:5 | 12 | R.LSLHRPALEDLLLGSEAN[+1501.555]LTC[+57.021]TLTGLR.D | HexNAc(5)Hex(3) | C[+57], N[+1502] | 893.8366 | 5 | 4465.1537 | 4465.1536 | 0 | 0 | 127 | Specific | 623.82 | 623.82 | 623.82 | 13.02 | 48 | >sp\|P01876\|IGHA1_HUMAN Immunoglobulin heavy constant alpha 1 OS=Homo sapiens OX=9606 GN=IGHA1 PE=1 SV=2 | 255 | 20201112_10613_1ng_glyco_hcd203040_compound15.21387.21387.5 | controllerType=0 controllerNumber=1 scan=21387 | 42.7603 |
| 17768:3 | 12 | R.LSLHRPALEDLLLGSEAN[+1501.555]LTC[+57.021]TLTGLR.D | HexNAc(5)Hex(3) | C[+57], N[+1502] | 1489.0589 | 3 | 4465.1621 | 4465.1536 | 0 | 1.9 | 127 | Specific | 624.85 | 576.46 | 576.46 | 12.7 | 48 | >sp\|P01876\|IGHA1_HUMAN Immunoglobulin heavy constant alpha 1 OS=Homo sapiens OX=9606 GN=IGHA1 PE=1 SV=2 | 255 | 20201112_10613_1ng_glyco_hcd203040_compound15.21369.21369.3 | controllerType=0 controllerNumber=1 scan=21369 | 42.7202 |
| 18728:4 | 12 | R.LSLHRPALEDLLLGSEAN[+1913.677]LTC[+57.021]TLTGLR.D | HexNAc(4)Hex(5)NeuAc(1) | C[+57], N[+1914] | 1220.0776 | 4 | 4877.2884 | 4877.2753 | 0 | 2.7 | 127 | Specific | 634.33 | 634.33 | 634.33 | 12.86 | 48 | >sp\|P01876\|IGHA1_HUMAN Immunoglobulin heavy constant alpha 1 OS=Homo sapiens OX=9606 GN=IGHA1 PE=1 SV=2 | 255 | 20201112_10613_1ng_glyco_hcd203040_compound15.22368.22368.4 | controllerType=0 controllerNumber=1 scan=22368 | 44.7028 |
| 17713:3 | 12 | R.LSLHRPALEDLLLGSEAN[+1622.582]LTC[+57.021]TLTGLR.D | HexNAc(4)Hex(5) | C[+57], N[+1623] | 1529.3976 | 3 | 4586.1784 | 4586.1799 | 0 | -0.3 | 127 | Specific | 638.33 | 549.21 | 549.21 | 12.63 | 48 | >sp\|P01876\|IGHA1_HUMAN Immunoglobulin heavy constant alpha 1 OS=Homo sapiens OX=9606 GN=IGHA1 PE=1 SV=2 | 255 | 20201112_10613_1ng_glyco_hcd203040_compound15.21311.21311.3 | controllerType=0 controllerNumber=1 scan=21311 | 42.5761 |
| 17994:4 | 12 | R.LSLHRPALEDLLLGSEAN[+1501.555]LTC[+57.021]TLTGLR.D | HexNAc(5)Hex(3) | C[+57], N[+1502] | 1117.0409 | 4 | 4465.1419 | 4465.1536 | 0 | -2.6 | 127 | Specific | 660.15 | 520.97 | 520.97 | 12.95 | 48 | >sp\|P01876\|IGHA1_HUMAN Immunoglobulin heavy constant alpha 1 OS=Homo sapiens OX=9606 GN=IGHA1 PE=1 SV=2 | 255 | 20201112_10613_1ng_glyco_hcd203040_compound15.21605.21605.4 | controllerType=0 controllerNumber=1 scan=21605 | 43.2275 |
| 17688:4 | 12 | R.LSLHRPALEDLLLGSEAN[+1663.608]LTC[+57.021]TLTGLR.D | HexNAc(5)Hex(4) | C[+57], N[+1664] | 1157.5586 | 4 | 4627.2127 | 4627.2064 | 0 | 1.4 | 127 | Specific | 661.91 | 658.56 | 658.56 | 14.17 | 48 | >sp\|P01876\|IGHA1_HUMAN Immunoglobulin heavy constant alpha 1 OS=Homo sapiens OX=9606 GN=IGHA1 PE=1 SV=2 | 255 | 20201112_10613_1ng_glyco_hcd203040_compound15.21285.21285.4 | controllerType=0 controllerNumber=1 scan=21285 | 42.5154 |
| 17774:3 | 12 | R.LSLHRPALEDLLLGSEAN[+1216.423]LTC[+57.021]TLTGLR.D | HexNAc(2)Hex(5) | C[+57], N[+1216] | 1394.0136 | 3 | 4180.0263 | 4180.0211 | 0 | 1.2 | 127 | Specific | 670.82 | 588.69 | 588.69 | 13.7 | 48 | >sp\|P01876\|IGHA1_HUMAN Immunoglobulin heavy constant alpha 1 OS=Homo sapiens OX=9606 GN=IGHA1 PE=1 SV=2 | 255 | 20201112_10613_1ng_glyco_hcd203040_compound15.21375.21375.3 | controllerType=0 controllerNumber=1 scan=21375 | 42.7318 |
| 17748:3 | 12 | R.LSLHRPALEDLLLGSEAN[+1622.582]LTC[+57.021]TLTGLR.D | HexNAc(4)Hex(5) | C[+57], N[+1623] | 1529.399 | 3 | 4586.1824 | 4586.1799 | 0 | 0.5 | 127 | Specific | 672.9 | 553.59 | 553.59 | 13.7 | 48 | >sp\|P01876\|IGHA1_HUMAN Immunoglobulin heavy constant alpha 1 OS=Homo sapiens OX=9606 GN=IGHA1 PE=1 SV=2 | 255 | 20201112_10613_1ng_glyco_hcd203040_compound15.21348.21348.3 | controllerType=0 controllerNumber=1 scan=21348 | 42.6576 |
| 17765:3 | 12 | R.LSLHRPALEDLLLGSEAN[+1501.555]LTC[+57.021]TLTGLR.D | HexNAc(5)Hex(3) | C[+57], N[+1502] | 1489.0589 | 3 | 4465.1621 | 4465.1536 | 0 | 1.9 | 127 | Specific | 678.72 | 678.72 | 678.72 | 13.7 | 48 | >sp\|P01876\|IGHA1_HUMAN Immunoglobulin heavy constant alpha 1 OS=Homo sapiens OX=9606 GN=IGHA1 PE=1 SV=2 | 255 | 20201112_10613_1ng_glyco_hcd203040_compound15.21366.21366.3 | controllerType=0 controllerNumber=1 scan=21366 | 42.7135 |
| 17761:4 | 12 | R.LSLHRPALEDLLLGSEAN[+1501.555]LTC[+57.021]TLTGLR.D | HexNAc(5)Hex(3) | C[+57], N[+1502] | 1117.0466 | 4 | 4465.1644 | 4465.1536 | 0 | 2.4 | 127 | Specific | 688.45 | 688.45 | 688.45 | 13.86 | 48 | >sp\|P01876\|IGHA1_HUMAN Immunoglobulin heavy constant alpha 1 OS=Homo sapiens OX=9606 GN=IGHA1 PE=1 SV=2 | 255 | 20201112_10613_1ng_glyco_hcd203040_compound15.21362.21362.4 | controllerType=0 controllerNumber=1 scan=21362 | 42.702 |
| 17734:3 | 12 | R.LSLHRPALEDLLLGSEAN[+1378.476]LTC[+57.021]TLTGLR.D | HexNAc(2)Hex(6) | C[+57], N[+1378] | 1448.0311 | 3 | 4342.0786 | 4342.074 | 0 | 1.1 | 127 | Specific | 688.57 | 659.92 | 659.92 | 13.7 | 48 | >sp\|P01876\|IGHA1_HUMAN Immunoglobulin heavy constant alpha 1 OS=Homo sapiens OX=9606 GN=IGHA1 PE=1 SV=2 | 255 | 20201112_10613_1ng_glyco_hcd203040_compound15.21333.21333.3 | controllerType=0 controllerNumber=1 scan=21333 | 42.623 |
| 17725:4 | 12 | R.LSLHRPALEDLLLGSEAN[+1540.529]LTC[+57.021]TLTGLR.D | HexNAc(2)Hex(7) | C[+57], N[+1541] | 1126.7878 | 4 | 4504.1292 | 4504.1268 | 0 | 0.5 | 127 | Specific | 696.85 | 641.48 | 641.48 | 14.1 | 48 | >sp\|P01876\|IGHA1_HUMAN Immunoglobulin heavy constant alpha 1 OS=Homo sapiens OX=9606 GN=IGHA1 PE=1 SV=2 | 255 | 20201112_10613_1ng_glyco_hcd203040_compound15.21324.21324.4 | controllerType=0 controllerNumber=1 scan=21324 | 42.6026 |
| 17747:4 | 12 | R.LSLHRPALEDLLLGSEAN[+1419.502]LTC[+57.021]TLTGLR.D | HexNAc(3)Hex(5) | C[+57], N[+1420] | 1096.5313 | 4 | 4383.1033 | 4383.1005 | 0 | 0.6 | 127 | Specific | 702.1 | 702.1 | 702.1 | 15.17 | 48 | >sp\|P01876\|IGHA1_HUMAN Immunoglobulin heavy constant alpha 1 OS=Homo sapiens OX=9606 GN=IGHA1 PE=1 SV=2 | 255 | 20201112_10613_1ng_glyco_hcd203040_compound15.21347.21347.4 | controllerType=0 controllerNumber=1 scan=21347 | 42.6555 |
| 17771:3 | 12 | R.LSLHRPALEDLLLGSEAN[+1216.423]LTC[+57.021]TLTGLR.D | HexNAc(2)Hex(5) | C[+57], N[+1216] | 1394.0136 | 3 | 4180.0263 | 4180.0211 | 0 | 1.2 | 127 | Specific | 719.62 | 685.82 | 685.82 | 14.7 | 48 | >sp\|P01876\|IGHA1_HUMAN Immunoglobulin heavy constant alpha 1 OS=Homo sapiens OX=9606 GN=IGHA1 PE=1 SV=2 | 255 | 20201112_10613_1ng_glyco_hcd203040_compound15.21372.21372.3 | controllerType=0 controllerNumber=1 scan=21372 | 42.7253 |
| 17757:4 | 12 | R.LSLHRPALEDLLLGSEAN[+1460.529]LTC[+57.021]TLTGLR.D | HexNAc(4)Hex(4) | C[+57], N[+1461] | 1106.7886 | 4 | 4424.1326 | 4424.1271 | 0 | 1.3 | 127 | Specific | 720.7 | 718.06 | 718.06 | 15.17 | 48 | >sp\|P01876\|IGHA1_HUMAN Immunoglobulin heavy constant alpha 1 OS=Homo sapiens OX=9606 GN=IGHA1 PE=1 SV=2 | 255 | 20201112_10613_1ng_glyco_hcd203040_compound15.21357.21357.4 | controllerType=0 controllerNumber=1 scan=21357 | 42.6889 |
| 17763:4 | 12 | R.LSLHRPALEDLLLGSEAN[+1501.555]LTC[+57.021]TLTGLR.D | HexNAc(5)Hex(3) | C[+57], N[+1502] | 1117.0466 | 4 | 4465.1644 | 4465.1536 | 0 | 2.4 | 127 | Specific | 738.92 | 684.52 | 684.52 | 14.86 | 48 | >sp\|P01876\|IGHA1_HUMAN Immunoglobulin heavy constant alpha 1 OS=Homo sapiens OX=9606 GN=IGHA1 PE=1 SV=2 | 255 | 20201112_10613_1ng_glyco_hcd203040_compound15.21364.21364.4 | controllerType=0 controllerNumber=1 scan=21364 | 42.7086 |
| 17731:3 | 12 | R.LSLHRPALEDLLLGSEAN[+1378.476]LTC[+57.021]TLTGLR.D | HexNAc(2)Hex(6) | C[+57], N[+1378] | 1448.0311 | 3 | 4342.0786 | 4342.074 | 0 | 1.1 | 127 | Specific | 748.79 | 667.16 | 667.16 | 14.7 | 48 | >sp\|P01876\|IGHA1_HUMAN Immunoglobulin heavy constant alpha 1 OS=Homo sapiens OX=9606 GN=IGHA1 PE=1 SV=2 | 255 | 20201112_10613_1ng_glyco_hcd203040_compound15.21330.21330.3 | controllerType=0 controllerNumber=1 scan=21330 | 42.612 |
| 17740:4 | 12 | R.LSLHRPALEDLLLGSEAN[+1419.502]LTC[+57.021]TLTGLR.D | HexNAc(3)Hex(5) | C[+57], N[+1420] | 1096.5325 | 4 | 4383.1082 | 4383.1005 | 0 | 1.8 | 127 | Specific | 749.38 | 749.38 | 749.38 | 15.17 | 48 | >sp\|P01876\|IGHA1_HUMAN Immunoglobulin heavy constant alpha 1 OS=Homo sapiens OX=9606 GN=IGHA1 PE=1 SV=2 | 255 | 20201112_10613_1ng_glyco_hcd203040_compound15.21339.21339.3 | controllerType=0 controllerNumber=1 scan=21339 | 42.6358 |
| 17856:4 | 12 | R.LSLHRPALEDLLLGSEAN[+1460.529]LTC[+57.021]TLTGLR.D | HexNAc(4)Hex(4) | C[+57], N[+1461] | 1106.7875 | 4 | 4424.1282 | 4424.1271 | 0 | 0.3 | 127 | Specific | 764.69 | 631.56 | 631.56 | 16.1 | 48 | >sp\|P01876\|IGHA1_HUMAN Immunoglobulin heavy constant alpha 1 OS=Homo sapiens OX=9606 GN=IGHA1 PE=1 SV=2 | 255 | 20201112_10613_1ng_glyco_hcd203040_compound15.21461.21461.4 | controllerType=0 controllerNumber=1 scan=21461 | 42.9338 |
| 17759:4 | 12 | R.LSLHRPALEDLLLGSEAN[+1460.529]LTC[+57.021]TLTGLR.D | HexNAc(4)Hex(4) | C[+57], N[+1461] | 1106.7886 | 4 | 4424.1326 | 4424.1271 | 0 | 1.3 | 127 | Specific | 770.28 | 702.29 | 702.29 | 16.17 | 48 | >sp\|P01876\|IGHA1_HUMAN Immunoglobulin heavy constant alpha 1 OS=Homo sapiens OX=9606 GN=IGHA1 PE=1 SV=2 | 255 | 20201112_10613_1ng_glyco_hcd203040_compound15.21359.21359.4 | controllerType=0 controllerNumber=1 scan=21359 | 42.6946 |
| 17769:4 | 12 | R.LSLHRPALEDLLLGSEAN[+1622.582]LTC[+57.021]TLTGLR.D | HexNAc(4)Hex(5) | C[+57], N[+1623] | 1147.3013 | 4 | 4586.1834 | 4586.1799 | 0 | 0.8 | 127 | Specific | 773.14 | 642.72 | 642.72 | 16.17 | 48 | >sp\|P01876\|IGHA1_HUMAN Immunoglobulin heavy constant alpha 1 OS=Homo sapiens OX=9606 GN=IGHA1 PE=1 SV=2 | 255 | 20201112_10613_1ng_glyco_hcd203040_compound15.21370.21370.4 | controllerType=0 controllerNumber=1 scan=21370 | 42.7221 |
| 18730:4 | 12 | R.LSLHRPALEDLLLGSEAN[+1913.677]LTC[+57.021]TLTGLR.D | HexNAc(4)Hex(5)NeuAc(1) | C[+57], N[+1914] | 1220.0776 | 4 | 4877.2884 | 4877.2753 | 0 | 2.7 | 127 | Specific | 775.34 | 745.47 | 745.47 | 15.86 | 48 | >sp\|P01876\|IGHA1_HUMAN Immunoglobulin heavy constant alpha 1 OS=Homo sapiens OX=9606 GN=IGHA1 PE=1 SV=2 | 255 | 20201112_10613_1ng_glyco_hcd203040_compound15.22370.22370.4 | controllerType=0 controllerNumber=1 scan=22370 | 44.7209 |
| 17690:4 | 12 | R.LSLHRPALEDLLLGSEAN[+1663.608]LTC[+57.021]TLTGLR.D | HexNAc(5)Hex(4) | C[+57], N[+1664] | 1157.5586 | 4 | 4627.2127 | 4627.2064 | 0 | 1.4 | 127 | Specific | 778.52 | 700.12 | 700.12 | 16.17 | 48 | >sp\|P01876\|IGHA1_HUMAN Immunoglobulin heavy constant alpha 1 OS=Homo sapiens OX=9606 GN=IGHA1 PE=1 SV=2 | 255 | 20201112_10613_1ng_glyco_hcd203040_compound15.21287.21287.4 | controllerType=0 controllerNumber=1 scan=21287 | 42.5209 |
| 17787:4 | 12 | R.LSLHRPALEDLLLGSEAN[+1298.476]LTC[+57.021]TLTGLR.D | HexNAc(4)Hex(3) | C[+57], N[+1298] | 1066.2741 | 4 | 4262.0745 | 4262.0742 | 0 | 0.1 | 127 | Specific | 785.56 | 785.56 | 785.56 | 16.1 | 48 | >sp\|P01876\|IGHA1_HUMAN Immunoglobulin heavy constant alpha 1 OS=Homo sapiens OX=9606 GN=IGHA1 PE=1 SV=2 | 255 | 20201112_10613_1ng_glyco_hcd203040_compound15.21389.21389.4 | controllerType=0 controllerNumber=1 scan=21389 | 42.7627 |
| 17835:4 | 12 | R.LSLHRPALEDLLLGSEAN[+1095.397]LTC[+57.021]TLTGLR.D | HexNAc(3)Hex(3) | C[+57], N[+1095] | 1015.5046 | 4 | 4058.9964 | 4058.9949 | 0 | 0.4 | 127 | Specific | 819.55 | 819.55 | 819.55 | 17.1 | 48 | >sp\|P01876\|IGHA1_HUMAN Immunoglobulin heavy constant alpha 1 OS=Homo sapiens OX=9606 GN=IGHA1 PE=1 SV=2 | 255 | 20201112_10613_1ng_glyco_hcd203040_compound15.21439.21439.4 | controllerType=0 controllerNumber=1 scan=21439 | 42.887 |
| 17730:4 | 12 | R.LSLHRPALEDLLLGSEAN[+1378.476]LTC[+57.021]TLTGLR.D | HexNAc(2)Hex(6) | C[+57], N[+1378] | 1086.2749 | 4 | 4342.078 | 4342.074 | 0 | 0.9 | 127 | Specific | 862.79 | 747.82 | 747.82 | 18.17 | 48 | >sp\|P01876\|IGHA1_HUMAN Immunoglobulin heavy constant alpha 1 OS=Homo sapiens OX=9606 GN=IGHA1 PE=1 SV=2 | 255 | 20201112_10613_1ng_glyco_hcd203040_compound15.21329.21329.4 | controllerType=0 controllerNumber=1 scan=21329 | 42.6108 |
| 17790:4 | 12 | R.LSLHRPALEDLLLGSEAN[+1298.476]LTC[+57.021]TLTGLR.D | HexNAc(4)Hex(3) | C[+57], N[+1298] | 1066.2741 | 4 | 4262.0745 | 4262.0742 | 0 | 0.1 | 127 | Specific | 871.63 | 800.25 | 800.25 | 18.1 | 48 | >sp\|P01876\|IGHA1_HUMAN Immunoglobulin heavy constant alpha 1 OS=Homo sapiens OX=9606 GN=IGHA1 PE=1 SV=2 | 255 | 20201112_10613_1ng_glyco_hcd203040_compound15.21392.21392.4 | controllerType=0 controllerNumber=1 scan=21392 | 42.7714 |
| 17837:4 | 12 | R.LSLHRPALEDLLLGSEAN[+1095.397]LTC[+57.021]TLTGLR.D | HexNAc(3)Hex(3) | C[+57], N[+1095] | 1015.5046 | 4 | 4058.9964 | 4058.9949 | 0 | 0.4 | 127 | Specific | 899.69 | 847.57 | 847.57 | 18.1 | 48 | >sp\|P01876\|IGHA1_HUMAN Immunoglobulin heavy constant alpha 1 OS=Homo sapiens OX=9606 GN=IGHA1 PE=1 SV=2 | 255 | 20201112_10613_1ng_glyco_hcd203040_compound15.21441.21441.4 | controllerType=0 controllerNumber=1 scan=21441 | 42.8934 |
| 17753:4 | 12 | R.LSLHRPALEDLLLGSEAN[+1216.423]LTC[+57.021]TLTGLR.D | HexNAc(2)Hex(5) | C[+57], N[+1216] | 1045.7614 | 4 | 4180.0238 | 4180.0211 | 0 | 0.6 | 127 | Specific | 900.96 | 808.16 | 808.16 | 19.17 | 48 | >sp\|P01876\|IGHA1_HUMAN Immunoglobulin heavy constant alpha 1 OS=Homo sapiens OX=9606 GN=IGHA1 PE=1 SV=2 | 255 | 20201112_10613_1ng_glyco_hcd203040_compound15.21353.21353.4 | controllerType=0 controllerNumber=1 scan=21353 | 42.6653 |
| 17743:4 | 12 | R.LSLHRPALEDLLLGSEAN[+1419.502]LTC[+57.021]TLTGLR.D | HexNAc(3)Hex(5) | C[+57], N[+1420] | 1096.5325 | 4 | 4383.1082 | 4383.1005 | 0 | 1.8 | 127 | Specific | 913.3 | 817.08 | 817.08 | 19.17 | 48 | >sp\|P01876\|IGHA1_HUMAN Immunoglobulin heavy constant alpha 1 OS=Homo sapiens OX=9606 GN=IGHA1 PE=1 SV=2 | 255 | 20201112_10613_1ng_glyco_hcd203040_compound15.21342.21342.3 | controllerType=0 controllerNumber=1 scan=21342 | 42.6458 |
| 17749:4 | 12 | R.LSLHRPALEDLLLGSEAN[+1419.502]LTC[+57.021]TLTGLR.D | HexNAc(3)Hex(5) | C[+57], N[+1420] | 1096.5313 | 4 | 4383.1033 | 4383.1005 | 0 | 0.6 | 127 | Specific | 921.36 | 686.57 | 686.57 | 19.17 | 48 | >sp\|P01876\|IGHA1_HUMAN Immunoglobulin heavy constant alpha 1 OS=Homo sapiens OX=9606 GN=IGHA1 PE=1 SV=2 | 255 | 20201112_10613_1ng_glyco_hcd203040_compound15.21349.21349.4 | controllerType=0 controllerNumber=1 scan=21349 | 42.6598 |
| 17727:4 | 12 | R.LSLHRPALEDLLLGSEAN[+1540.529]LTC[+57.021]TLTGLR.D | HexNAc(2)Hex(7) | C[+57], N[+1541] | 1126.7878 | 4 | 4504.1292 | 4504.1268 | 0 | 0.5 | 127 | Specific | 972.15 | 939.57 | 939.57 | 20.1 | 48 | >sp\|P01876\|IGHA1_HUMAN Immunoglobulin heavy constant alpha 1 OS=Homo sapiens OX=9606 GN=IGHA1 PE=1 SV=2 | 255 | 20201112_10613_1ng_glyco_hcd203040_compound15.21326.21326.4 | controllerType=0 controllerNumber=1 scan=21326 | 42.6069 |
| 17756:4 | 12 | R.LSLHRPALEDLLLGSEAN[+1216.423]LTC[+57.021]TLTGLR.D | HexNAc(2)Hex(5) | C[+57], N[+1216] | 1045.7614 | 4 | 4180.0238 | 4180.0211 | 0 | 0.6 | 127 | Specific | 993.95 | 993.95 | 993.95 | 20.17 | 48 | >sp\|P01876\|IGHA1_HUMAN Immunoglobulin heavy constant alpha 1 OS=Homo sapiens OX=9606 GN=IGHA1 PE=1 SV=2 | 255 | 20201112_10613_1ng_glyco_hcd203040_compound15.21356.21356.4 | controllerType=0 controllerNumber=1 scan=21356 | 42.6872 |
| 17733:4 | 12 | R.LSLHRPALEDLLLGSEAN[+1378.476]LTC[+57.021]TLTGLR.D | HexNAc(2)Hex(6) | C[+57], N[+1378] | 1086.2749 | 4 | 4342.078 | 4342.074 | 0 | 0.9 | 127 | Specific | 1001.61 | 994.67 | 994.67 | 21.17 | 48 | >sp\|P01876\|IGHA1_HUMAN Immunoglobulin heavy constant alpha 1 OS=Homo sapiens OX=9606 GN=IGHA1 PE=1 SV=2 | 255 | 20201112_10613_1ng_glyco_hcd203040_compound15.21332.21332.4 | controllerType=0 controllerNumber=1 scan=21332 | 42.622 |
| 13365:3 | 12 | R.LAGKPTHVN[+1548.545]VSVVMAEVDGTC[+57.021]Y.- | HexNAc(3)Hex(4)NeuAc(1) | C[+57], N[+1549] | 1299.2357 | 3 | 3895.6924 | 3895.6869 | 0 | 1.4 | 332 | CRagged | 226.35 | 220.49 | 220.49 | 6.9 | 48 | >sp\|P01876\|IGHA1_HUMAN Immunoglobulin heavy constant alpha 1 OS=Homo sapiens OX=9606 GN=IGHA1 PE=1 SV=2 | 255 | 20201112_10613_1ng_glyco_hcd203040_compound15.16763.16763.3 | controllerType=0 controllerNumber=1 scan=16763 | 34.0903 |
| 11454:4 | 12 | R.LAGKPTHVN[+1809.666]VSVVMAEVDGTC[+57.021]Y.- | HexNAc(5)Hex(4)Fuc(1) | C[+57], N[+1810] | 1039.9576 | 4 | 4156.8084 | 4156.8081 | 0 | 0.1 | 332 | CRagged | 256.47 | 175.5 | 175.5 | 4.45 | 48 | >sp\|P01876\|IGHA1_HUMAN Immunoglobulin heavy constant alpha 1 OS=Homo sapiens OX=9606 GN=IGHA1 PE=1 SV=2 | 255 | 20201112_10613_1ng_glyco_hcd203040_compound15.14762.14762.4 | controllerType=0 controllerNumber=1 scan=14762 | 30.6963 |
| 15212:3 | 12 | R.LAGKPTHVN[+2204.772]VSVVMAEVDGTC[+57.021]Y.- | HexNAc(4)Hex(5)NeuAc(2) | C[+57], N[+2205] | 1517.9776 | 3 | 4551.9182 | 4551.9145 | 0 | 0.8 | 332 | CRagged | 304.32 | 304.32 | 304.32 | 7.73 | 48 | >sp\|P01876\|IGHA1_HUMAN Immunoglobulin heavy constant alpha 1 OS=Homo sapiens OX=9606 GN=IGHA1 PE=1 SV=2 | 255 | 20201112_10613_1ng_glyco_hcd203040_compound15.18677.18677.3 | controllerType=0 controllerNumber=1 scan=18677 | 37.4915 |
| 11578:4 | 12 | R.LAGKPTHVN[+1647.613]VSVVMAEVDGTC[+57.021]Y.- | HexNAc(5)Hex(3)Fuc(1) | C[+57], N[+1648] | 999.4445 | 4 | 3994.7562 | 3994.7553 | 0 | 0.2 | 332 | CRagged | 310.36 | 310.36 | 310.36 | 7.58 | 48 | >sp\|P01876\|IGHA1_HUMAN Immunoglobulin heavy constant alpha 1 OS=Homo sapiens OX=9606 GN=IGHA1 PE=1 SV=2 | 255 | 20201112_10613_1ng_glyco_hcd203040_compound15.14891.14891.4 | controllerType=0 controllerNumber=1 scan=14891 | 30.9243 |
| 11612:3 | 12 | R.LAGKPTHVN[+1216.423]VSVVMAEVDGTC[+57.021]Y.- | HexNAc(2)Hex(5) | C[+57], N[+1216] | 1188.5171 | 3 | 3563.5369 | 3563.5649 | 0 | -7.9 | 332 | CRagged | 318.59 | 276.1 | 225.04 | 3.77 | 48 | >sp\|P01876\|IGHA1_HUMAN Immunoglobulin heavy constant alpha 1 OS=Homo sapiens OX=9606 GN=IGHA1 PE=1 SV=2 | 255 | 20201112_10613_1ng_glyco_hcd203040_compound15.14926.14926.3 | controllerType=0 controllerNumber=1 scan=14926 | 30.9908 |
| 11580:4 | 12 | R.LAGKPTHVN[+1647.613]VSVVMAEVDGTC[+57.021]Y.- | HexNAc(5)Hex(3)Fuc(1) | C[+57], N[+1648] | 999.4445 | 4 | 3994.7562 | 3994.7553 | 0 | 0.2 | 332 | CRagged | 354.72 | 354.72 | 354.72 | 6.75 | 48 | >sp\|P01876\|IGHA1_HUMAN Immunoglobulin heavy constant alpha 1 OS=Homo sapiens OX=9606 GN=IGHA1 PE=1 SV=2 | 255 | 20201112_10613_1ng_glyco_hcd203040_compound15.14893.14893.4 | controllerType=0 controllerNumber=1 scan=14893 | 30.9281 |
| 11451:4 | 12 | R.LAGKPTHVN[+1809.666]VSVVMAEVDGTC[+57.021]Y.- | HexNAc(5)Hex(4)Fuc(1) | C[+57], N[+1810] | 1039.9576 | 4 | 4156.8084 | 4156.8081 | 0 | 0.1 | 332 | CRagged | 366.02 | 352.34 | 325.97 | 7.1 | 48 | >sp\|P01876\|IGHA1_HUMAN Immunoglobulin heavy constant alpha 1 OS=Homo sapiens OX=9606 GN=IGHA1 PE=1 SV=2 | 255 | 20201112_10613_1ng_glyco_hcd203040_compound15.14759.14759.4 | controllerType=0 controllerNumber=1 scan=14759 | 30.6915 |
| 11553:3 | 12 | R.LAGKPTHVN[+1809.666]VSVVMAEVDGTC[+57.021]Y.- | HexNAc(5)Hex(4)Fuc(1) | C[+57], N[+1810] | 1386.2803 | 3 | 4156.8264 | 4156.8081 | 0 | 4.4 | 332 | CRagged | 379.55 | 333.23 | 285.86 | 5.9 | 48 | >sp\|P01876\|IGHA1_HUMAN Immunoglobulin heavy constant alpha 1 OS=Homo sapiens OX=9606 GN=IGHA1 PE=1 SV=2 | 255 | 20201112_10613_1ng_glyco_hcd203040_compound15.14865.14865.3 | controllerType=0 controllerNumber=1 scan=14865 | 30.8773 |
| 11550:3 | 12 | R.LAGKPTHVN[+1809.666]VSVVMAEVDGTC[+57.021]Y.- | HexNAc(5)Hex(4)Fuc(1) | C[+57], N[+1810] | 1386.2803 | 3 | 4156.8264 | 4156.8081 | 0 | 4.4 | 332 | CRagged | 383.1 | 294.05 | 281.99 | 7.01 | 48 | >sp\|P01876\|IGHA1_HUMAN Immunoglobulin heavy constant alpha 1 OS=Homo sapiens OX=9606 GN=IGHA1 PE=1 SV=2 | 255 | 20201112_10613_1ng_glyco_hcd203040_compound15.14862.14862.3 | controllerType=0 controllerNumber=1 scan=14862 | 30.8727 |
| 11437:3 | 12 | R.LAGKPTHVN[+1971.719]VSVVMAEVDGTC[+57.021]Y.- | HexNAc(5)Hex(5)Fuc(1) | C[+57], N[+1972] | 1440.2922 | 3 | 4318.862 | 4318.861 | 0 | 0.2 | 332 | CRagged | 437.62 | 431.61 | 335.06 | 9.98 | 48 | >sp\|P01876\|IGHA1_HUMAN Immunoglobulin heavy constant alpha 1 OS=Homo sapiens OX=9606 GN=IGHA1 PE=1 SV=2 | 255 | 20201112_10613_1ng_glyco_hcd203040_compound15.14744.14744.3 | controllerType=0 controllerNumber=1 scan=14744 | 30.6634 |
| 11435:3 | 12 | R.LAGKPTHVN[+1971.719]VSVVMAEVDGTC[+57.021]Y.- | HexNAc(5)Hex(5)Fuc(1) | C[+57], N[+1972] | 1440.2922 | 3 | 4318.862 | 4318.861 | 0 | 0.2 | 332 | CRagged | 542.31 | 542.31 | 366.35 | 10.49 | 48 | >sp\|P01876\|IGHA1_HUMAN Immunoglobulin heavy constant alpha 1 OS=Homo sapiens OX=9606 GN=IGHA1 PE=1 SV=2 | 255 | 20201112_10613_1ng_glyco_hcd203040_compound15.14742.14742.3 | controllerType=0 controllerNumber=1 scan=14742 | 30.6579 |
| 11577:3 | 12 | R.LAGKPTHVN[+1501.555]VSVVMAEVDGTC[+57.021]Y.- | HexNAc(5)Hex(3) | C[+57], N[+1502] | 1283.572 | 3 | 3848.7013 | 3848.6974 | 0 | 1 | 332 | CRagged | 597.71 | 554.82 | 554.82 | 11.94 | 48 | >sp\|P01876\|IGHA1_HUMAN Immunoglobulin heavy constant alpha 1 OS=Homo sapiens OX=9606 GN=IGHA1 PE=1 SV=2 | 255 | 20201112_10613_1ng_glyco_hcd203040_compound15.14890.14890.3 | controllerType=0 controllerNumber=1 scan=14890 | 30.9224 |
| 11383:3 | 12 | R.LAGKPTHVN[+1702.581]VSVVMAEVDGTC[+57.021]Y.- | HexNAc(2)Hex(8) | C[+57], N[+1703] | 1350.5793 | 3 | 4049.7233 | 4049.7234 | 0 | 0 | 332 | CRagged | 621.21 | 621.21 | 383.82 | 12.87 | 48 | >sp\|P01876\|IGHA1_HUMAN Immunoglobulin heavy constant alpha 1 OS=Homo sapiens OX=9606 GN=IGHA1 PE=1 SV=2 | 255 | 20201112_10613_1ng_glyco_hcd203040_compound15.14688.14688.3 | controllerType=0 controllerNumber=1 scan=14688 | 30.5147 |
| 11410:3 | 12 | R.LAGKPTHVN[+1540.529]VSVVMAEVDGTC[+57.021]Y.- | HexNAc(2)Hex(7) | C[+57], N[+1541] | 1296.5627 | 3 | 3887.6735 | 3887.6706 | 0 | 0.7 | 332 | CRagged | 651.26 | 651.26 | 386.89 | 13.94 | 48 | >sp\|P01876\|IGHA1_HUMAN Immunoglobulin heavy constant alpha 1 OS=Homo sapiens OX=9606 GN=IGHA1 PE=1 SV=2 | 255 | 20201112_10613_1ng_glyco_hcd203040_compound15.14716.14716.3 | controllerType=0 controllerNumber=1 scan=14716 | 30.5698 |
| 11438:3 | 12 | R.LAGKPTHVN[+1809.666]VSVVMAEVDGTC[+57.021]Y.- | HexNAc(5)Hex(4)Fuc(1) | C[+57], N[+1810] | 1386.2752 | 3 | 4156.8111 | 4156.8081 | 0 | 0.7 | 332 | CRagged | 690.12 | 690.12 | 455.09 | 13.94 | 48 | >sp\|P01876\|IGHA1_HUMAN Immunoglobulin heavy constant alpha 1 OS=Homo sapiens OX=9606 GN=IGHA1 PE=1 SV=2 | 255 | 20201112_10613_1ng_glyco_hcd203040_compound15.14746.14746.3 | controllerType=0 controllerNumber=1 scan=14746 | 30.6679 |
| 11429:3 | 12 | R.LAGKPTHVN[+1663.608]VSVVMAEVDGTC[+57.021]Y.- | HexNAc(5)Hex(4) | C[+57], N[+1664] | 1337.5884 | 3 | 4010.7508 | 4010.7502 | 0 | 0.1 | 332 | CRagged | 718.86 | 704.23 | 429.82 | 14.87 | 48 | >sp\|P01876\|IGHA1_HUMAN Immunoglobulin heavy constant alpha 1 OS=Homo sapiens OX=9606 GN=IGHA1 PE=1 SV=2 | 255 | 20201112_10613_1ng_glyco_hcd203040_compound15.14736.14736.3 | controllerType=0 controllerNumber=1 scan=14736 | 30.6463 |
| 11540:3 | 12 | R.LAGKPTHVN[+1647.613]VSVVMAEVDGTC[+57.021]Y.- | HexNAc(5)Hex(3)Fuc(1) | C[+57], N[+1648] | 1332.2598 | 3 | 3994.7649 | 3994.7553 | 0 | 2.4 | 332 | CRagged | 759.39 | 612.83 | 612.83 | 15.63 | 48 | >sp\|P01876\|IGHA1_HUMAN Immunoglobulin heavy constant alpha 1 OS=Homo sapiens OX=9606 GN=IGHA1 PE=1 SV=2 | 255 | 20201112_10613_1ng_glyco_hcd203040_compound15.14852.14852.3 | controllerType=0 controllerNumber=1 scan=14852 | 30.8594 |
| 11538:3 | 12 | R.LAGKPTHVN[+1647.613]VSVVMAEVDGTC[+57.021]Y.- | HexNAc(5)Hex(3)Fuc(1) | C[+57], N[+1648] | 1332.2598 | 3 | 3994.7649 | 3994.7553 | 0 | 2.4 | 332 | CRagged | 792 | 792 | 792 | 15.63 | 48 | >sp\|P01876\|IGHA1_HUMAN Immunoglobulin heavy constant alpha 1 OS=Homo sapiens OX=9606 GN=IGHA1 PE=1 SV=2 | 255 | 20201112_10613_1ng_glyco_hcd203040_compound15.14850.14850.3 | controllerType=0 controllerNumber=1 scan=14850 | 30.8548 |
| 11440:3 | 12 | R.LAGKPTHVN[+1809.666]VSVVMAEVDGTC[+57.021]Y.- | HexNAc(5)Hex(4)Fuc(1) | C[+57], N[+1810] | 1386.2752 | 3 | 4156.8111 | 4156.8081 | 0 | 0.7 | 332 | CRagged | 793.53 | 773.3 | 488.7 | 15.94 | 48 | >sp\|P01876\|IGHA1_HUMAN Immunoglobulin heavy constant alpha 1 OS=Homo sapiens OX=9606 GN=IGHA1 PE=1 SV=2 | 255 | 20201112_10613_1ng_glyco_hcd203040_compound15.14748.14748.3 | controllerType=0 controllerNumber=1 scan=14748 | 30.673 |
| 11422:3 | 12 | R.LAGKPTHVN[+1378.476]VSVVMAEVDGTC[+57.021]Y.- | HexNAc(2)Hex(6) | C[+57], N[+1378] | 1242.5449 | 3 | 3725.62 | 3725.6178 | 0 | 0.6 | 332 | CRagged | 800.53 | 773.17 | 485.54 | 16.94 | 48 | >sp\|P01876\|IGHA1_HUMAN Immunoglobulin heavy constant alpha 1 OS=Homo sapiens OX=9606 GN=IGHA1 PE=1 SV=2 | 255 | 20201112_10613_1ng_glyco_hcd203040_compound15.14729.14729.3 | controllerType=0 controllerNumber=1 scan=14729 | 30.6276 |
| 11385:3 | 12 | R.LAGKPTHVN[+1702.581]VSVVMAEVDGTC[+57.021]Y.- | HexNAc(2)Hex(8) | C[+57], N[+1703] | 1350.5793 | 3 | 4049.7233 | 4049.7234 | 0 | 0 | 332 | CRagged | 807.7 | 781.16 | 478.21 | 16.87 | 48 | >sp\|P01876\|IGHA1_HUMAN Immunoglobulin heavy constant alpha 1 OS=Homo sapiens OX=9606 GN=IGHA1 PE=1 SV=2 | 255 | 20201112_10613_1ng_glyco_hcd203040_compound15.14690.14690.3 | controllerType=0 controllerNumber=1 scan=14690 | 30.519 |
| 11427:3 | 12 | R.LAGKPTHVN[+1663.608]VSVVMAEVDGTC[+57.021]Y.- | HexNAc(5)Hex(4) | C[+57], N[+1664] | 1337.5884 | 3 | 4010.7508 | 4010.7502 | 0 | 0.1 | 332 | CRagged | 818.33 | 713.93 | 541.96 | 16.87 | 48 | >sp\|P01876\|IGHA1_HUMAN Immunoglobulin heavy constant alpha 1 OS=Homo sapiens OX=9606 GN=IGHA1 PE=1 SV=2 | 255 | 20201112_10613_1ng_glyco_hcd203040_compound15.14734.14734.3 | controllerType=0 controllerNumber=1 scan=14734 | 30.6398 |
| 11413:3 | 12 | R.LAGKPTHVN[+1540.529]VSVVMAEVDGTC[+57.021]Y.- | HexNAc(2)Hex(7) | C[+57], N[+1541] | 1296.5627 | 3 | 3887.6735 | 3887.6706 | 0 | 0.7 | 332 | CRagged | 823.86 | 779.15 | 488.7 | 16.94 | 48 | >sp\|P01876\|IGHA1_HUMAN Immunoglobulin heavy constant alpha 1 OS=Homo sapiens OX=9606 GN=IGHA1 PE=1 SV=2 | 255 | 20201112_10613_1ng_glyco_hcd203040_compound15.14719.14719.3 | controllerType=0 controllerNumber=1 scan=14719 | 30.576 |
| 11449:3 | 12 | R.LAGKPTHVN[+1216.423]VSVVMAEVDGTC[+57.021]Y.- | HexNAc(2)Hex(5) | C[+57], N[+1216] | 1188.5275 | 3 | 3563.568 | 3563.5649 | 0 | 0.9 | 332 | CRagged | 841.33 | 819.56 | 499.27 | 16.94 | 48 | >sp\|P01876\|IGHA1_HUMAN Immunoglobulin heavy constant alpha 1 OS=Homo sapiens OX=9606 GN=IGHA1 PE=1 SV=2 | 255 | 20201112_10613_1ng_glyco_hcd203040_compound15.14757.14757.3 | controllerType=0 controllerNumber=1 scan=14757 | 30.6889 |
| 11420:3 | 12 | R.LAGKPTHVN[+1378.476]VSVVMAEVDGTC[+57.021]Y.- | HexNAc(2)Hex(6) | C[+57], N[+1378] | 1242.5449 | 3 | 3725.62 | 3725.6178 | 0 | 0.6 | 332 | CRagged | 867.71 | 867.71 | 545.04 | 17.94 | 48 | >sp\|P01876\|IGHA1_HUMAN Immunoglobulin heavy constant alpha 1 OS=Homo sapiens OX=9606 GN=IGHA1 PE=1 SV=2 | 255 | 20201112_10613_1ng_glyco_hcd203040_compound15.14727.14727.3 | controllerType=0 controllerNumber=1 scan=14727 | 30.6172 |
| 11446:3 | 12 | R.LAGKPTHVN[+1216.423]VSVVMAEVDGTC[+57.021]Y.- | HexNAc(2)Hex(5) | C[+57], N[+1216] | 1188.5275 | 3 | 3563.568 | 3563.5649 | 0 | 0.9 | 332 | CRagged | 917.91 | 917.91 | 577.46 | 18.94 | 48 | >sp\|P01876\|IGHA1_HUMAN Immunoglobulin heavy constant alpha 1 OS=Homo sapiens OX=9606 GN=IGHA1 PE=1 SV=2 | 255 | 20201112_10613_1ng_glyco_hcd203040_compound15.14754.14754.3 | controllerType=0 controllerNumber=1 scan=14754 | 30.6797 |
| 07698:3 | 13 | R.LSPN[+2350.830]ASAEHLELR.W | HexNAc(4)Hex(5)Fuc(1)NeuAc(2) | N[+2351] | 1263.1998 | 3 | 3787.5847 | 3787.5796 | 0 | 1.4 | 52 | Specific | 565.41 | 465 | 465 | 10.15 | 37 | >sp\|Q13410\|BT1A1_HUMAN Butyrophilin subfamily 1 member A1 OS=Homo sapiens OX=9606 GN=BTN1A1 PE=1 SV=3 | 179 | 20201112_10613_1ng_glyco_hcd203040_compound15.10862.10862.3 | controllerType=0 controllerNumber=1 scan=10862 | 24.1396 |
| 07696:3 | 13 | R.LSPN[+2350.830]ASAEHLELR.W | HexNAc(4)Hex(5)Fuc(1)NeuAc(2) | N[+2351] | 1263.1998 | 3 | 3787.5847 | 3787.5796 | 0 | 1.4 | 52 | Specific | 589.52 | 513.43 | 513.43 | 11.38 | 37 | >sp\|Q13410\|BT1A1_HUMAN Butyrophilin subfamily 1 member A1 OS=Homo sapiens OX=9606 GN=BTN1A1 PE=1 SV=3 | 179 | 20201112_10613_1ng_glyco_hcd203040_compound15.10860.10860.3 | controllerType=0 controllerNumber=1 scan=10860 | 24.1353 |
| 05705:3 | 13 | R.LSPN[+2205.793]ASAEHLELR.W | HexNAc(4)Hex(5)Fuc(2)NeuAc(1) | N[+2206] | 1214.855 | 3 | 3642.5506 | 3642.5421 | 0 | 2.3 | 52 | Specific | 641.05 | 450.34 | 450.34 | 12.07 | 37 | >sp\|Q13410\|BT1A1_HUMAN Butyrophilin subfamily 1 member A1 OS=Homo sapiens OX=9606 GN=BTN1A1 PE=1 SV=3 | 179 | 20201112_10613_1ng_glyco_hcd203040_compound15.8801.8801.3 | controllerType=0 controllerNumber=1 scan=8801 | 20.7568 |
| 05703:3 | 13 | R.LSPN[+2205.793]ASAEHLELR.W | HexNAc(4)Hex(5)Fuc(2)NeuAc(1) | N[+2206] | 1214.855 | 3 | 3642.5506 | 3642.5421 | 0 | 2.3 | 52 | Specific | 657.88 | 587.16 | 587.16 | 13.07 | 37 | >sp\|Q13410\|BT1A1_HUMAN Butyrophilin subfamily 1 member A1 OS=Homo sapiens OX=9606 GN=BTN1A1 PE=1 SV=3 | 179 | 20201112_10613_1ng_glyco_hcd203040_compound15.8799.8799.3 | controllerType=0 controllerNumber=1 scan=8799 | 20.7525 |
| 17704:3 | 13 | K.N[+1913.677]VSC[+57.021]YIQNLLLGQEK.K | HexNAc(4)Hex(5)NeuAc(1) | C[+57], N[+1914] | 1231.535 | 3 | 3692.5904 | 3692.5876 | 0 | 0.8 | 215 | Specific | 563.38 | 519.83 | 519.83 | 10.07 | 37 | >sp\|Q13410\|BT1A1_HUMAN Butyrophilin subfamily 1 member A1 OS=Homo sapiens OX=9606 GN=BTN1A1 PE=1 SV=3 | 179 | 20201112_10613_1ng_glyco_hcd203040_compound15.21302.21302.3 | controllerType=0 controllerNumber=1 scan=21302 | 42.5548 |
| 17706:3 | 13 | K.N[+1913.677]VSC[+57.021]YIQNLLLGQEK.K | HexNAc(4)Hex(5)NeuAc(1) | C[+57], N[+1914] | 1231.535 | 3 | 3692.5904 | 3692.5876 | 0 | 0.8 | 215 | Specific | 613.42 | 453.83 | 453.83 | 12.3 | 37 | >sp\|Q13410\|BT1A1_HUMAN Butyrophilin subfamily 1 member A1 OS=Homo sapiens OX=9606 GN=BTN1A1 PE=1 SV=3 | 179 | 20201112_10613_1ng_glyco_hcd203040_compound15.21304.21304.3 | controllerType=0 controllerNumber=1 scan=21304 | 42.562 |
| 24149:4 | 14 | R.QLAHQSN[+2351.851]STNIFFSPVSIATAFAMLSLGTK.A | HexNAc(4)Hex(5)Fuc(3)NeuAc(1) | N[+2352] | 1384.1261 | 4 | 5533.4826 | 5533.4858 | 0 | -0.6 | 64 | Specific | 209.83 | 209.83 | 209.83 | 5.33 | 41 | >sp\|P01009\|A1AT_HUMAN Alpha-1-antitrypsin OS=Homo sapiens OX=9606 GN=SERPINA1 PE=1 SV=3 | 642 | 20201112_10613_1ng_glyco_hcd203040_compound15.28019.28019.4 | controllerType=0 controllerNumber=1 scan=28019 | 56.0697 |
| 26870:4 | 14 | R.QLAHQSN[+2861.000]STNIFFSPVSIATAFAMLSLGTK.A | HexNAc(5)Hex(6)NeuAc(3) | N[+2861] | 1511.4169 | 4 | 6042.6457 | 6042.6351 | 0 | 1.7 | 64 | Specific | 225.51 | 225.51 | 225.51 | 6.74 | 41 | >sp\|P01009\|A1AT_HUMAN Alpha-1-antitrypsin OS=Homo sapiens OX=9606 GN=SERPINA1 PE=1 SV=3 | 642 | 20201112_10613_1ng_glyco_hcd203040_compound15.30842.30842.5 | controllerType=0 controllerNumber=1 scan=30842 | 61.1387 |
| 24066:3 | 14 | R.QLAHQSN[+2350.830]STNIFFSPVSIATAFAMLSLGTK.A | HexNAc(4)Hex(5)Fuc(1)NeuAc(2) | N[+2351] | 1844.8413 | 3 | 5532.5093 | 5532.4654 | 0 | 7.9 | 64 | Specific | 277.11 | 254.13 | 254.13 | 5.55 | 41 | >sp\|P01009\|A1AT_HUMAN Alpha-1-antitrypsin OS=Homo sapiens OX=9606 GN=SERPINA1 PE=1 SV=3 | 642 | 20201112_10613_1ng_glyco_hcd203040_compound15.27933.27933.3 | controllerType=0 controllerNumber=1 scan=27933 | 55.8971 |
| 24068:3 | 14 | R.QLAHQSN[+2350.830]STNIFFSPVSIATAFAMLSLGTK.A | HexNAc(4)Hex(5)Fuc(1)NeuAc(2) | N[+2351] | 1844.8413 | 3 | 5532.5093 | 5532.4654 | 0 | 7.9 | 64 | Specific | 333.17 | 333.17 | 333.17 | 6.3 | 41 | >sp\|P01009\|A1AT_HUMAN Alpha-1-antitrypsin OS=Homo sapiens OX=9606 GN=SERPINA1 PE=1 SV=3 | 642 | 20201112_10613_1ng_glyco_hcd203040_compound15.27935.27935.3 | controllerType=0 controllerNumber=1 scan=27935 | 55.9011 |
| 24163:3 | 14 | R.QLAHQSN[+2204.772]STNIFFSPVSIATAFAMLSLGTK.A | HexNAc(4)Hex(5)NeuAc(2) | N[+2205] | 1796.1438 | 3 | 5386.4167 | 5386.4075 | 0 | 1.7 | 64 | Specific | 341.3 | 312.75 | 312.75 | 7.75 | 41 | >sp\|P01009\|A1AT_HUMAN Alpha-1-antitrypsin OS=Homo sapiens OX=9606 GN=SERPINA1 PE=1 SV=3 | 642 | 20201112_10613_1ng_glyco_hcd203040_compound15.28034.28034.3 | controllerType=0 controllerNumber=1 scan=28034 | 56.0983 |
| 24165:3 | 14 | R.QLAHQSN[+2204.772]STNIFFSPVSIATAFAMLSLGTK.A | HexNAc(4)Hex(5)NeuAc(2) | N[+2205] | 1796.1438 | 3 | 5386.4167 | 5386.4075 | 0 | 1.7 | 64 | Specific | 378.8 | 351.52 | 316.61 | 8.18 | 41 | >sp\|P01009\|A1AT_HUMAN Alpha-1-antitrypsin OS=Homo sapiens OX=9606 GN=SERPINA1 PE=1 SV=3 | 642 | 20201112_10613_1ng_glyco_hcd203040_compound15.28036.28036.3 | controllerType=0 controllerNumber=1 scan=28036 | 56.1018 |
| 26872:4 | 14 | R.QLAHQSN[+2861.000]STNIFFSPVSIATAFAMLSLGTK.A | HexNAc(5)Hex(6)NeuAc(3) | N[+2861] | 1511.4169 | 4 | 6042.6457 | 6042.6351 | 0 | 1.7 | 64 | Specific | 406.21 | 348.58 | 348.58 | 7.96 | 41 | >sp\|P01009\|A1AT_HUMAN Alpha-1-antitrypsin OS=Homo sapiens OX=9606 GN=SERPINA1 PE=1 SV=3 | 642 | 20201112_10613_1ng_glyco_hcd203040_compound15.30844.30844.5 | controllerType=0 controllerNumber=1 scan=30844 | 61.1428 |
| 24160:4 | 14 | R.QLAHQSN[+2204.772]STNIFFSPVSIATAFAMLSLGTK.A | HexNAc(4)Hex(5)NeuAc(2) | N[+2205] | 1347.3579 | 4 | 5386.4098 | 5386.4075 | 0 | 0.4 | 64 | Specific | 439.31 | 433.82 | 421.83 | 9.68 | 41 | >sp\|P01009\|A1AT_HUMAN Alpha-1-antitrypsin OS=Homo sapiens OX=9606 GN=SERPINA1 PE=1 SV=3 | 642 | 20201112_10613_1ng_glyco_hcd203040_compound15.28031.28031.4 | controllerType=0 controllerNumber=1 scan=28031 | 56.092 |
| 24162:4 | 14 | R.QLAHQSN[+2204.772]STNIFFSPVSIATAFAMLSLGTK.A | HexNAc(4)Hex(5)NeuAc(2) | N[+2205] | 1347.3579 | 4 | 5386.4098 | 5386.4075 | 0 | 0.4 | 64 | Specific | 662.77 | 595.57 | 436.48 | 13.57 | 41 | >sp\|P01009\|A1AT_HUMAN Alpha-1-antitrypsin OS=Homo sapiens OX=9606 GN=SERPINA1 PE=1 SV=3 | 642 | 20201112_10613_1ng_glyco_hcd203040_compound15.28033.28033.4 | controllerType=0 controllerNumber=1 scan=28033 | 56.0965 |
| 22744:5 | 14 | K.ADTHDEILEGLNFN[+2862.021]LTEIPEAQIHEGFQELLR.T | HexNAc(5)Hex(6)Fuc(2)NeuAc(2) | N[+2862] | 1311.5752 | 5 | 6553.8469 | 6553.8443 | 0 | 0.4 | 94 | Specific | 407.52 | 356.42 | 356.42 | 8.2 | 41 | >sp\|P01009\|A1AT_HUMAN Alpha-1-antitrypsin OS=Homo sapiens OX=9606 GN=SERPINA1 PE=1 SV=3 | 642 | 20201112_10613_1ng_glyco_hcd203040_compound15.26559.26559.5 | controllerType=0 controllerNumber=1 scan=26559 | 53.2549 |
| 22745:5 | 14 | K.ADTHDEILEGLNFN[+2862.021]LTEIPEAQIHEGFQELLR.T | HexNAc(5)Hex(6)Fuc(2)NeuAc(2) | N[+2862] | 1311.5752 | 5 | 6553.8469 | 6553.8443 | 0 | 0.4 | 94 | Specific | 416.97 | 381.75 | 381.75 | 9.6 | 41 | >sp\|P01009\|A1AT_HUMAN Alpha-1-antitrypsin OS=Homo sapiens OX=9606 GN=SERPINA1 PE=1 SV=3 | 642 | 20201112_10613_1ng_glyco_hcd203040_compound15.26560.26560.5 | controllerType=0 controllerNumber=1 scan=26560 | 53.2582 |
| 22722:5 | 14 | K.ADTHDEILEGLNFN[+2861.000]LTEIPEAQIHEGFQELLR.T | HexNAc(5)Hex(6)NeuAc(3) | N[+2861] | 1311.3765 | 5 | 6552.8532 | 6552.8239 | 0 | 4.5 | 94 | Specific | 532.31 | 464.12 | 464.12 | 9.48 | 41 | >sp\|P01009\|A1AT_HUMAN Alpha-1-antitrypsin OS=Homo sapiens OX=9606 GN=SERPINA1 PE=1 SV=3 | 642 | 20201112_10613_1ng_glyco_hcd203040_compound15.26536.26536.4 | controllerType=0 controllerNumber=1 scan=26536 | 53.1943 |
| 22724:5 | 14 | K.ADTHDEILEGLNFN[+2861.000]LTEIPEAQIHEGFQELLR.T | HexNAc(5)Hex(6)NeuAc(3) | N[+2861] | 1311.3765 | 5 | 6552.8532 | 6552.8239 | 0 | 4.5 | 94 | Specific | 587.56 | 587.56 | 587.56 | 10.85 | 41 | >sp\|P01009\|A1AT_HUMAN Alpha-1-antitrypsin OS=Homo sapiens OX=9606 GN=SERPINA1 PE=1 SV=3 | 642 | 20201112_10613_1ng_glyco_hcd203040_compound15.26538.26538.4 | controllerType=0 controllerNumber=1 scan=26538 | 53.1995 |
| 20259:3 | 14 | K.YLGN[+2204.772]ATAIFFLPDEGK.L | HexNAc(4)Hex(5)NeuAc(2) | N[+2205] | 1320.8946 | 3 | 3960.6692 | 3960.6677 | 0 | 0.4 | 268 | Specific | 559.25 | 480.62 | 480.62 | 10.15 | 41 | >sp\|P01009\|A1AT_HUMAN Alpha-1-antitrypsin OS=Homo sapiens OX=9606 GN=SERPINA1 PE=1 SV=3 | 642 | 20201112_10613_1ng_glyco_hcd203040_compound15.23967.23967.3 | controllerType=0 controllerNumber=1 scan=23967 | 48.0959 |

Table S4. Cont.

| Query #:z | Protein Rank | Glycopeptide | Glycan | Modification Type(s) | Observed m/z | z | Observed (M+H) | Calc. mass (M+H) | Off-by-x error | Mass error (ppm) | StartPosition | Cleavage | Score | Delta | Delta Mod | \|Log Prob\| | # of unique peptides | Protein Name | Protein DB number | Comment | Scan # | Scan Time |
|---|---|---|---|---|---|---|---|---|---|---|---|---|---|---|---|---|---|---|---|---|---|---|
| 20262:3 | 14 | K.YLGN[+2204.772]ATAIFFLPDEGK.L | HexNAc(4)Hex(5)NeuAc(2) | N[+2205] | 1320.8946 | 3 | 3960.6692 | 3960.6677 | 0 | 0.4 | 268 | Specific | 696.69 | 574.76 | 574.76 | 13.38 | 41 | >sp\|P01009\|A1AT_HUMAN Alpha-1-antitrypsin OS=Homo sapiens OX=9606 GN=SERPINA1 PE=1 SV=3 | 642 | 20201112_10613_1ng_glyco_hcd203040_compound15.23970.23970.3 | controllerType=0 controllerNumber=1 scan=23970 | 48.1007 |
| 00348:3 | 15 | K.KKEDALN[+2059.735]ETRESETK.L | HexNAc(4)Hex(5)Fuc(1)NeuAc(1) | N[+2060] | 1279.8831 | 3 | 3837.6348 | 3837.6276 | 0 | 1.9 | 80 | Specific | 285.13 | 219.68 | 219.68 | 5.26 | 32 | >sp\|P10909\|CLUS_HUMAN Clusterin OS=Homo sapiens OX=9606 GN=CLU PE=1 SV=1 | 1467 | 20201112_10613_1ng_glyco_hcd203040_compound15.3237.3237.3 | controllerType=0 controllerNumber=1 scan=3237 | 11.2932 |
| 14128:4 | 15 | K.LKELPGVC[+57.021]N[+2059.735]ETMMALWEEC[+57.021]KPC[+57.021]LK.Q | HexNAc(4)Hex(5)Fuc(1)NeuAc(1) | C[+57]*3, N[+2060] | 1249.7852 | 4 | 4996.1188 | 4996.1196 | 0 | -0.2 | 95 | Specific | 239.62 | 239.62 | 239.62 | 4.65 | 32 | >sp\|P10909\|CLUS_HUMAN Clusterin OS=Homo sapiens OX=9606 GN=CLU PE=1 SV=1 | 1467 | 20201112_10613_1ng_glyco_hcd203040_compound15.17554.17554.4 | controllerType=0 controllerNumber=1 scan=17554 | 35.5092 |
| 14189:3 | 15 | K.LKELPGVC[+57.021]N[+2059.735]ETMMALWEEC[+57.021]KPC[+57.021]LK.Q | HexNAc(4)Hex(5)Fuc(1)NeuAc(1) | C[+57]*3, N[+2060] | 1666.0438 | 3 | 4996.1168 | 4996.1196 | 0 | -0.6 | 95 | Specific | 315.88 | 286.69 | 286.69 | 6.89 | 32 | >sp\|P10909\|CLUS_HUMAN Clusterin OS=Homo sapiens OX=9606 GN=CLU PE=1 SV=1 | 1467 | 20201112_10613_1ng_glyco_hcd203040_compound15.17617.17617.3 | controllerType=0 controllerNumber=1 scan=17617 | 35.6168 |
| 14137:4 | 15 | K.LKELPGVC[+57.021]N[+2205.793]ETMMALWEEC[+57.021]KPC[+57.021]LK.Q | HexNAc(4)Hex(5)Fuc(2)NeuAc(1) | C[+57]*3, N[+2206] | 1286.3021 | 4 | 5142.1867 | 5142.1776 | 0 | 1.8 | 95 | Specific | 325.49 | 325.49 | 325.49 | 7.97 | 32 | >sp\|P10909\|CLUS_HUMAN Clusterin OS=Homo sapiens OX=9606 GN=CLU PE=1 SV=1 | 1467 | 20201112_10613_1ng_glyco_hcd203040_compound15.17563.17563.4 | controllerType=0 controllerNumber=1 scan=17563 | 35.5222 |
| 14186:3 | 15 | K.LKELPGVC[+57.021]N[+2059.735]ETMMALWEEC[+57.021]KPC[+57.021]LK.Q | HexNAc(4)Hex(5)Fuc(1)NeuAc(1) | C[+57]*3, N[+2060] | 1666.0438 | 3 | 4996.1168 | 4996.1196 | 0 | -0.6 | 95 | Specific | 327.62 | 327.62 | 327.62 | 7.43 | 32 | >sp\|P10909\|CLUS_HUMAN Clusterin OS=Homo sapiens OX=9606 GN=CLU PE=1 SV=1 | 1467 | 20201112_10613_1ng_glyco_hcd203040_compound15.17614.17614.3 | controllerType=0 controllerNumber=1 scan=17614 | 35.6123 |
| 14139:4 | 15 | K.LKELPGVC[+57.021]N[+2205.793]ETMMALWEEC[+57.021]KPC[+57.021]LK.Q | HexNAc(4)Hex(5)Fuc(2)NeuAc(1) | C[+57]*3, N[+2206] | 1286.3021 | 4 | 5142.1867 | 5142.1776 | 0 | 1.8 | 95 | Specific | 527.71 | 479.05 | 479.05 | 10.45 | 32 | >sp\|P10909\|CLUS_HUMAN Clusterin OS=Homo sapiens OX=9606 GN=CLU PE=1 SV=1 | 1467 | 20201112_10613_1ng_glyco_hcd203040_compound15.17565.17565.4 | controllerType=0 controllerNumber=1 scan=17565 | 35.5257 |
| 14182:4 | 15 | K.LKELPGVC[+57.021]N[+2059.735]ETMMALWEEC[+57.021]KPC[+57.021]LK.Q | HexNAc(4)Hex(5)Fuc(1)NeuAc(1) | C[+57]*3, N[+2060] | 1249.7863 | 4 | 4996.1232 | 4996.1196 | 0 | 0.7 | 95 | Specific | 530.3 | 476.84 | 476.84 | 10.45 | 32 | >sp\|P10909\|CLUS_HUMAN Clusterin OS=Homo sapiens OX=9606 GN=CLU PE=1 SV=1 | 1467 | 20201112_10613_1ng_glyco_hcd203040_compound15.17610.17610.4 | controllerType=0 controllerNumber=1 scan=17610 | 35.6051 |
| 14184:4 | 15 | K.LKELPGVC[+57.021]N[+2059.735]ETMMALWEEC[+57.021]KPC[+57.021]LK.Q | HexNAc(4)Hex(5)Fuc(1)NeuAc(1) | C[+57]*3, N[+2060] | 1249.7863 | 4 | 4996.1232 | 4996.1196 | 0 | 0.7 | 95 | Specific | 820.56 | 791.89 | 791.89 | 16.83 | 32 | >sp\|P10909\|CLUS_HUMAN Clusterin OS=Homo sapiens OX=9606 GN=CLU PE=1 SV=1 | 1467 | 20201112_10613_1ng_glyco_hcd203040_compound15.17612.17612.4 | controllerType=0 controllerNumber=1 scan=17612 | 35.609 |
| 23345:3 | 15 | K.[+42.011]MLN[+2336.851]TSSLLEQLNEQFNWVSR.L | HexNAc(6)Hex(6)Fuc(1) | .[42], N[+2337] | 1596.678 | 3 | 4788.0196 | 4788.0484 | 0 | -6 | 352 | Specific | 206.2 | 134.51 | 134.51 | 3.53 | 32 | >sp\|P10909\|CLUS_HUMAN Clusterin OS=Homo sapiens OX=9606 GN=CLU PE=1 SV=1 | 1467 | 20201112_10613_1ng_glyco_hcd203040_compound15.27184.27184.3 | controllerType=0 controllerNumber=1 scan=27184 | 54.4931 |
| 23343:3 | 15 | K.[+42.011]MLN[+2336.851]TSSLLEQLNEQFNWVSR.L | HexNAc(6)Hex(6)Fuc(1) | .[42], N[+2337] | 1596.678 | 3 | 4788.0196 | 4788.0484 | 0 | -6 | 352 | Specific | 214.4 | 214.4 | 214.4 | 4.11 | 32 | >sp\|P10909\|CLUS_HUMAN Clusterin OS=Homo sapiens OX=9606 GN=CLU PE=1 SV=1 | 1467 | 20201112_10613_1ng_glyco_hcd203040_compound15.27182.27182.3 | controllerType=0 controllerNumber=1 scan=27182 | 54.4891 |
| 23234:4 | 15 | K.MLN[+2350.830]TSSLLEQLNEQFNWVSR.L | HexNAc(4)Hex(5)Fuc(1)NeuAc(2) | N[+2351] | 1190.7622 | 4 | 4760.0272 | 4760.0171 | 0 | 2.1 | 352 | Specific | 482.7 | 359.34 | 359.34 | 8.94 | 32 | >sp\|P10909\|CLUS_HUMAN Clusterin OS=Homo sapiens OX=9606 GN=CLU PE=1 SV=1 | 1467 | 20201112_10613_1ng_glyco_hcd203040_compound15.27069.27069.4 | controllerType=0 controllerNumber=1 scan=27069 | 54.2663 |
| 20996:3 | 15 | K.MLN[+2059.735]TSSLLEQLNEQFNWVSR.L | HexNAc(4)Hex(5)Fuc(1)NeuAc(1) | N[+2060] | 1490.3171 | 3 | 4468.9367 | 4468.9217 | 0 | 3.4 | 352 | Specific | 736.87 | 539.19 | 539.19 | 14.55 | 32 | >sp\|P10909\|CLUS_HUMAN Clusterin OS=Homo sapiens OX=9606 GN=CLU PE=1 SV=1 | 1467 | 20201112_10613_1ng_glyco_hcd203040_compound15.24735.24735.5 | controllerType=0 controllerNumber=1 scan=24735 | 49.7825 |
| 23232:4 | 15 | K.MLN[+2350.830]TSSLLEQLNEQFNWVSR.L | HexNAc(4)Hex(5)Fuc(1)NeuAc(2) | N[+2351] | 1190.7622 | 4 | 4760.0272 | 4760.0171 | 0 | 2.1 | 352 | Specific | 770.5 | 579.89 | 579.89 | 15.96 | 32 | >sp\|P10909\|CLUS_HUMAN Clusterin OS=Homo sapiens OX=9606 GN=CLU PE=1 SV=1 | 1467 | 20201112_10613_1ng_glyco_hcd203040_compound15.27067.27067.4 | controllerType=0 controllerNumber=1 scan=27067 | 54.2618 |
| 20994:3 | 15 | K.MLN[+2059.735]TSSLLEQLNEQFNWVSR.L | HexNAc(4)Hex(5)Fuc(1)NeuAc(1) | N[+2060] | 1490.3171 | 3 | 4468.9367 | 4468.9217 | 0 | 3.4 | 352 | Specific | 795.73 | 548.11 | 548.11 | 15.55 | 32 | >sp\|P10909\|CLUS_HUMAN Clusterin OS=Homo sapiens OX=9606 GN=CLU PE=1 SV=1 | 1467 | 20201112_10613_1ng_glyco_hcd203040_compound15.24733.24733.5 | controllerType=0 controllerNumber=1 scan=24733 | 49.7765 |
| 21010:4 | 15 | K.MLN[+2059.735]TSSLLEQLNEQFNWVSR.L | HexNAc(4)Hex(5)Fuc(1)NeuAc(1) | N[+2060] | 1117.9859 | 4 | 4468.9217 | 4468.9217 | 0 | 0 | 352 | Specific | 800.99 | 618.44 | 618.44 | 17.2 | 32 | >sp\|P10909\|CLUS_HUMAN Clusterin OS=Homo sapiens OX=9606 GN=CLU PE=1 SV=1 | 1467 | 20201112_10613_1ng_glyco_hcd203040_compound15.24750.24750.4 | controllerType=0 controllerNumber=1 scan=24750 | 49.8102 |
| 21013:4 | 15 | K.MLN[+2059.735]TSSLLEQLNEQFNWVSR.L | HexNAc(4)Hex(5)Fuc(1)NeuAc(1) | N[+2060] | 1117.9859 | 4 | 4468.9217 | 4468.9217 | 0 | 0 | 352 | Specific | 841.78 | 510.5 | 510.5 | 17.2 | 32 | >sp\|P10909\|CLUS_HUMAN Clusterin OS=Homo sapiens OX=9606 GN=CLU PE=1 SV=1 | 1467 | 20201112_10613_1ng_glyco_hcd203040_compound15.24753.24753.4 | controllerType=0 controllerNumber=1 scan=24753 | 49.8208 |
| 23227:3 | 15 | K.MLN[+2350.830]TSSLLEQLNEQFNWVSR.L | HexNAc(4)Hex(5)Fuc(1)NeuAc(2) | N[+2351] | 1587.348 | 3 | 4760.0293 | 4760.0171 | 0 | 2.6 | 352 | Specific | 847.12 | 649.69 | 649.69 | 16.55 | 32 | >sp\|P10909\|CLUS_HUMAN Clusterin OS=Homo sapiens OX=9606 GN=CLU PE=1 SV=1 | 1467 | 20201112_10613_1ng_glyco_hcd203040_compound15.27062.27062.3 | controllerType=0 controllerNumber=1 scan=27062 | 54.2518 |
| 23229:3 | 15 | K.MLN[+2350.830]TSSLLEQLNEQFNWVSR.L | HexNAc(4)Hex(5)Fuc(1)NeuAc(2) | N[+2351] | 1587.348 | 3 | 4760.0293 | 4760.0171 | 0 | 2.6 | 352 | Specific | 917.47 | 607.88 | 607.88 | 18.55 | 32 | >sp\|P10909\|CLUS_HUMAN Clusterin OS=Homo sapiens OX=9606 GN=CLU PE=1 SV=1 | 1467 | 20201112_10613_1ng_glyco_hcd203040_compound15.27064.27064.3 | controllerType=0 controllerNumber=1 scan=27064 | 54.2564 |
| 21007:3 | 15 | K.MLN[+2059.735]TSSLLEQLNEQFNWVSR.L | HexNAc(4)Hex(5)Fuc(1)NeuAc(1) | N[+2060] | 1490.3148 | 3 | 4468.9297 | 4468.9217 | 0 | 1.8 | 352 | Specific | 973.08 | 710.75 | 710.75 | 19.86 | 32 | >sp\|P10909\|CLUS_HUMAN Clusterin OS=Homo sapiens OX=9606 GN=CLU PE=1 SV=1 | 1467 | 20201112_10613_1ng_glyco_hcd203040_compound15.24747.24747.3 | controllerType=0 controllerNumber=1 scan=24747 | 49.806 |
| 21005:3 | 15 | K.MLN[+2059.735]TSSLLEQLNEQFNWVSR.L | HexNAc(4)Hex(5)Fuc(1)NeuAc(1) | N[+2060] | 1490.3148 | 3 | 4468.9297 | 4468.9217 | 0 | 1.8 | 352 | Specific | 1016.02 | 759.72 | 759.72 | 20.86 | 32 | >sp\|P10909\|CLUS_HUMAN Clusterin OS=Homo sapiens OX=9606 GN=CLU PE=1 SV=1 | 1467 | 20201112_10613_1ng_glyco_hcd203040_compound15.24745.24745.3 | controllerType=0 controllerNumber=1 scan=24745 | 49.801 |
| 20523:3 | 15 | K.M[+15.995]LN[+2059.735]TSSLLEQLNEQFNWVSR.L | HexNAc(4)Hex(5)Fuc(1)NeuAc(1) | M[+16], N[+2060] | 1495.6423 | 3 | 4484.9123 | 4484.9166 | 0 | -1 | 352 | Specific | 750.28 | 532.16 | 532.16 | 15.17 | 32 | >sp\|P10909\|CLUS_HUMAN Clusterin OS=Homo sapiens OX=9606 GN=CLU PE=1 SV=1 | 1467 | 20201112_10613_1ng_glyco_hcd203040_compound15.24242.24242.3 | controllerType=0 controllerNumber=1 scan=24242 | 48.6834 |
| 20526:3 | 15 | K.M[+15.995]LN[+2059.735]TSSLLEQLNEQFNWVSR.L | HexNAc(4)Hex(5)Fuc(1)NeuAc(1) | M[+16], N[+2060] | 1495.6423 | 3 | 4484.9123 | 4484.9166 | 0 | -1 | 352 | Specific | 894.33 | 588.38 | 588.38 | 17.17 | 32 | >sp\|P10909\|CLUS_HUMAN Clusterin OS=Homo sapiens OX=9606 GN=CLU PE=1 SV=1 | 1467 | 20201112_10613_1ng_glyco_hcd203040_compound15.24245.24245.3 | controllerType=0 controllerNumber=1 scan=24245 | 48.6915 |
| 10531:3 | 15 | R.LAN[+2059.735]LTQGEDQYYLR.V | HexNAc(4)Hex(5)Fuc(1)NeuAc(1) | N[+2060] | 1248.5275 | 3 | 3743.568 | 3743.5686 | 0 | -0.2 | 372 | Specific | 207.73 | 207.73 | 207.73 | 4.99 | 32 | >sp\|P10909\|CLUS_HUMAN Clusterin OS=Homo sapiens OX=9606 GN=CLU PE=1 SV=1 | 1467 | 20201112_10613_1ng_glyco_hcd203040_compound15.13805.13805.3 | controllerType=0 controllerNumber=1 scan=13805 | 29.0441 |
| 10533:3 | 15 | R.LAN[+2059.735]LTQGEDQYYLR.V | HexNAc(4)Hex(5)Fuc(1)NeuAc(1) | N[+2060] | 1248.5275 | 3 | 3743.568 | 3743.5686 | 0 | -0.2 | 372 | Specific | 224.5 | 181.24 | 181.24 | 3.7 | 32 | >sp\|P10909\|CLUS_HUMAN Clusterin OS=Homo sapiens OX=9606 GN=CLU PE=1 SV=1 | 1467 | 20201112_10613_1ng_glyco_hcd203040_compound15.13807.13807.3 | controllerType=0 controllerNumber=1 scan=13807 | 29.0476 |
| 13020:3 | 15 | R.LAN[+2350.830]LTQGEDQYYLR.V | HexNAc(4)Hex(5)Fuc(1)NeuAc(2) | N[+2351] | 1345.5635 | 3 | 4034.676 | 4034.664 | 0 | 3 | 372 | Specific | 574.54 | 512.2 | 512.2 | 9.76 | 32 | >sp\|P10909\|CLUS_HUMAN Clusterin OS=Homo sapiens OX=9606 GN=CLU PE=1 SV=1 | 1467 | 20201112_10613_1ng_glyco_hcd203040_compound15.16384.16384.3 | controllerType=0 controllerNumber=1 scan=16384 | 33.3897 |
| 13022:3 | 15 | R.LAN[+2350.830]LTQGEDQYYLR.V | HexNAc(4)Hex(5)Fuc(1)NeuAc(2) | N[+2351] | 1345.5635 | 3 | 4034.676 | 4034.664 | 0 | 3 | 372 | Specific | 623.3 | 483.35 | 483.35 | 11.99 | 32 | >sp\|P10909\|CLUS_HUMAN Clusterin OS=Homo sapiens OX=9606 GN=CLU PE=1 SV=1 | 1467 | 20201112_10613_1ng_glyco_hcd203040_compound15.16386.16386.3 | controllerType=0 controllerNumber=1 scan=16386 | 33.3936 |
| 02118:4 | 17 | R.EKQTDEIKDTRN[+2059.735]ESTQNC[+57.021]VVAEPEK.M | HexNAc(4)Hex(5)Fuc(1)NeuAc(1) | C[+57], N[+2060] | 1252.7889 | 4 | 5008.1339 | 5008.1251 | 0 | 1.8 | 58 | Specific | 351.17 | 343.04 | 343.04 | 7.06 | 34 | >sp\|P47710\|CASA1_HUMAN Alpha-S1-casein OS=Homo sapiens OX=9606 GN=CSN1S1 PE=1 SV=1 | 989 | 20201112_10613_1ng_glyco_hcd203040_compound15.5080.5080.4 | controllerType=0 controllerNumber=1 scan=5080 | 14.5095 |
| 02116:4 | 17 | R.EKQTDEIKDTRN[+2059.735]ESTQNC[+57.021]VVAEPEK.M | HexNAc(4)Hex(5)Fuc(1)NeuAc(1) | C[+57], N[+2060] | 1252.7889 | 4 | 5008.1339 | 5008.1251 | 0 | 1.8 | 58 | Specific | 421.29 | 421.29 | 421.29 | 8.93 | 34 | >sp\|P47710\|CASA1_HUMAN Alpha-S1-casein OS=Homo sapiens OX=9606 GN=CSN1S1 PE=1 SV=1 | 989 | 20201112_10613_1ng_glyco_hcd203040_compound15.5078.5078.4 | controllerType=0 controllerNumber=1 scan=5078 | 14.505 |
| 09510:5 | 17 | K.QTDEIKDTRN[+2059.735]ESTQNC[+57.021]VVAEPEKMESSISSSSEEMSLSK.C | HexNAc(4)Hex(5)Fuc(1)NeuAc(1) | C[+57], N[+2060] | 1290.9487 | 5 | 6450.7146 | 6450.7156 | 0 | -0.2 | 60 | Specific | 595.03 | 475.23 | 475.23 | 11.61 | 34 | >sp\|P47710\|CASA1_HUMAN Alpha-S1-casein OS=Homo sapiens OX=9606 GN=CSN1S1 PE=1 SV=1 | 989 | 20201112_10613_1ng_glyco_hcd203040_compound15.12749.12749.5 | controllerType=0 controllerNumber=1 scan=12749 | 27.2959 |
| 09512:5 | 17 | K.QTDEIKDTRN[+2059.735]ESTQNC[+57.021]VVAEPEKMESSISSSSEEMSLSK.C | HexNAc(4)Hex(5)Fuc(1)NeuAc(1) | C[+57], N[+2060] | 1290.9487 | 5 | 6450.7146 | 6450.7156 | 0 | -0.2 | 60 | Specific | 763.33 | 582.59 | 582.59 | 15.61 | 34 | >sp\|P47710\|CASA1_HUMAN Alpha-S1-casein OS=Homo sapiens OX=9606 GN=CSN1S1 PE=1 SV=1 | 989 | 20201112_10613_1ng_glyco_hcd203040_compound15.12751.12751.5 | controllerType=0 controllerNumber=1 scan=12751 | 27.3011 |
| 10105:4 | 21 | K.SC[+57.021]QHN[+2350.830]GTMYQHGEIFSAHELFPSR.L | HexNAc(4)Hex(5)Fuc(1)NeuAc(2) | C[+57], N[+2351] | 1296.7792 | 4 | 5184.0949 | 5184.0873 | 0 | 1.5 | 110 | Specific | 545.28 | 545.28 | 545.28 | 10.32 | 24 | >sp\|Q6WN34\|CRDL2_HUMAN Chordin-like protein 2 OS=Homo sapiens OX=9606 GN=CHRDL2 PE=1 SV=1 | 1189 | 20201112_10613_1ng_glyco_hcd203040_compound15.13364.13364.4 | controllerType=0 controllerNumber=1 scan=13364 | 28.2873 |
| 10107:4 | 21 | K.SC[+57.021]QHN[+2350.830]GTMYQHGEIFSAHELFPSR.L | HexNAc(4)Hex(5)Fuc(1)NeuAc(2) | C[+57], N[+2351] | 1296.7792 | 4 | 5184.0949 | 5184.0873 | 0 | 1.5 | 110 | Specific | 550.62 | 550.62 | 550.62 | 10.48 | 24 | >sp\|Q6WN34\|CRDL2_HUMAN Chordin-like protein 2 OS=Homo sapiens OX=9606 GN=CHRDL2 PE=1 SV=1 | 1189 | 20201112_10613_1ng_glyco_hcd203040_compound15.13366.13366.4 | controllerType=0 controllerNumber=1 scan=13366 | 28.2912 |
| 14364:2 | 22 | K.LC[+57.021]DN[+1866.688]LSTK.N | HexNAc(6)Hex(4) | C[+57], N[+1867] | 1409.0695 | 2 | 2817.1317 | 2817.1487 | 0 | -6 | 285 | Specific | 232.68 | 192.13 | 192.13 | 2.16 | 25 | >sp\|P02774\|VTDB_HUMAN Vitamin D-binding protein OS=Homo sapiens OX=9606 GN=GC PE=1 SV=2 | 1103 | 20201112_10613_1ng_glyco_hcd203040_compound15.17798.17798.2 | controllerType=0 controllerNumber=1 scan=17798 | 35.918 |
| 14366:2 | 22 | K.LC[+57.021]DN[+1866.688]LSTK.N | HexNAc(6)Hex(4) | C[+57], N[+1867] | 1409.0695 | 2 | 2817.1317 | 2817.1487 | 0 | -6 | 285 | Specific | 255.59 | 162.75 | 162.75 | 2.1 | 25 | >sp\|P02774\|VTDB_HUMAN Vitamin D-binding protein OS=Homo sapiens OX=9606 GN=GC PE=1 SV=2 | 1103 | 20201112_10613_1ng_glyco_hcd203040_compound15.17800.17800.2 | controllerType=0 controllerNumber=1 scan=17800 | 35.9253 |
| 17923:3 | 41 | R.NYIVPILWLN[+1864.634]ETGTIGDEK.A | HexNAc(2)Hex(9) | N[+1865] | 1347.2623 | 3 | 4039.7722 | 4039.7674 | 0 | 1.2 | 408 | Specific | 480.81 | 478.68 | 478.68 | 9.84 | 14 | >sp\|P16671\|CD36_HUMAN Platelet glycoprotein 4 OS=Homo sapiens OX=9606 GN=CD36 PE=1 SV=2 | 812 | 20201112_10613_1ng_glyco_hcd203040_compound15.21531.21531.3 | controllerType=0 controllerNumber=1 scan=21531 | 43.0805 |
| 17925:3 | 41 | R.NYIVPILWLN[+1864.634]ETGTIGDEK.A | HexNAc(2)Hex(9) | N[+1865] | 1347.2623 | 3 | 4039.7722 | 4039.7674 | 0 | 1.2 | 408 | Specific | 708.17 | 697.89 | 697.89 | 14.45 | 14 | >sp\|P16671\|CD36_HUMAN Platelet glycoprotein 4 OS=Homo sapiens OX=9606 GN=CD36 PE=1 SV=2 | 812 | 20201112_10613_1ng_glyco_hcd203040_compound15.21533.21533.3 | controllerType=0 controllerNumber=1 scan=21533 | 43.0844 |
| 16700:3 | 41 | R.NYIVPILWLN[+1864.634]ETGTIGDEKANM[+15.995]FR.S | HexNAc(2)Hex(9) | M[+16], N[+1865] | 1559.0254 | 3 | 4675.0618 | 4675.0523 | 0 | 2 | 408 | Specific | 763.47 | 763.47 | 763.47 | 15.52 | 14 | >sp\|P16671\|CD36_HUMAN Platelet glycoprotein 4 OS=Homo sapiens OX=9606 GN=CD36 PE=1 SV=2 | 812 | 20201112_10613_1ng_glyco_hcd203040_compound15.20256.20256.3 | controllerType=0 controllerNumber=1 scan=20256 | 40.4228 |
| 16698:3 | 41 | R.NYIVPILWLN[+1864.634]ETGTIGDEKANM[+15.995]FR.S | HexNAc(2)Hex(9) | M[+16], N[+1865] | 1559.0254 | 3 | 4675.0618 | 4675.0523 | 0 | 2 | 408 | Specific | 770.6 | 741.3 | 741.3 | 15.52 | 14 | >sp\|P16671\|CD36_HUMAN Platelet glycoprotein 4 OS=Homo sapiens OX=9606 GN=CD36 PE=1 SV=2 | 812 | 20201112_10613_1ng_glyco_hcd203040_compound15.20254.20254.3 | controllerType=0 controllerNumber=1 scan=20254 | 40.3985 |
| 23698:4 | 42 | K.[+42.011]VFPLSLDSTPQDGNVVVAC[+57.021]LVQGFFPQEPLSVTWSESGQN[+2059 | HexNAc(4)Hex(5)Fuc(1)NeuAc(1) | .[42], C[+57], N[+2060] | 1720.2873 | 4 | 6878.1274 | 6878.1069 | 0 | 3 | 123 | Specific | 237.04 | 237.04 | 237.04 | 7.55 | 21 | >sp\|P0DOX2\|IGA2_HUMAN Immunoglobulin alpha-2 heavy chain OS=Homo sapiens OX=9606 PE=1 SV=2 | 631 | 20201112_10613_1ng_glyco_hcd203040_compound15.27551.27551.4 | controllerType=0 controllerNumber=1 scan=27551 | 55.1874 |
| 23618:4 | 42 | K.VFPLSLDSTPQDGNVVVAC[+57.021]LVQGFFPQEPLSVTWSESGQN[+2262.814]VTAI | HexNAc(5)Hex(5)Fuc(1)NeuAc(1) | C[+57], N[+2263] | 1760.5507 | 4 | 7039.1811 | 7039.1757 | 0 | 0.8 | 123 | Specific | 240.7 | 218.71 | 218.71 | 7.72 | 21 | >sp\|P0DOX2\|IGA2_HUMAN Immunoglobulin alpha-2 heavy chain OS=Homo sapiens OX=9606 PE=1 SV=2 | 631 | 20201112_10613_1ng_glyco_hcd203040_compound15.27468.27468.4 | controllerType=0 controllerNumber=1 scan=27468 | 55.0274 |
| 23620:4 | 42 | K.VFPLSLDSTPQDGNVVVAC[+57.021]LVQGFFPQEPLSVTWSESGQN[+2262.814]VTAI | HexNAc(5)Hex(5)Fuc(1)NeuAc(1) | C[+57], N[+2263] | 1760.5507 | 4 | 7039.1811 | 7039.1757 | 0 | 0.8 | 123 | Specific | 320.82 | 309.64 | 309.64 | 8.55 | 21 | >sp\|P0DOX2\|IGA2_HUMAN Immunoglobulin alpha-2 heavy chain OS=Homo sapiens OX=9606 PE=1 SV=2 | 631 | 20201112_10613_1ng_glyco_hcd203040_compound15.27470.27470.4 | controllerType=0 controllerNumber=1 scan=27470 | 55.0323 |
| 22094:4 | 42 | K.VFPLSLDSTPQDGNVVVAC[+57.021]LVQGFFPQEPLSVTWSESGQN[+1809.666]VTAI | HexNAc(5)Hex(4)Fuc(1) | C[+57], N[+1810] | 1647.2764 | 4 | 6586.0837 | 6586.0275 | 0 | 8.5 | 123 | Specific | 440.03 | 433.37 | 433.37 | 8.28 | 21 | >sp\|P0DOX2\|IGA2_HUMAN Immunoglobulin alpha-2 heavy chain OS=Homo sapiens OX=9606 PE=1 SV=2 | 631 | 20201112_10613_1ng_glyco_hcd203040_compound15.25876.25876.2 | controllerType=0 controllerNumber=1 scan=25876 | 51.9662 |
| 22159:4 | 42 | K.VFPLSLDSTPQDGNVVVAC[+57.021]LVQGFFPQEPLSVTWSESGQN[+1809.666]VTAI | HexNAc(5)Hex(4)Fuc(1) | C[+57], N[+1810] | 1647.2711 | 4 | 6586.0627 | 6586.0275 | 0 | 5.3 | 123 | Specific | 616.29 | 604.76 | 604.76 | 12.59 | 21 | >sp\|P0DOX2\|IGA2_HUMAN Immunoglobulin alpha-2 heavy chain OS=Homo sapiens OX=9606 PE=1 SV=2 | 631 | 20201112_10613_1ng_glyco_hcd203040_compound15.25944.25944.4 | controllerType=0 controllerNumber=1 scan=25944 | 52.0948 |
| 22162:4 | 42 | K.VFPLSLDSTPQDGNVVVAC[+57.021]LVQGFFPQEPLSVTWSESGQN[+1809.666]VTAI | HexNAc(5)Hex(4)Fuc(1) | C[+57], N[+1810] | 1647.2711 | 4 | 6586.0627 | 6586.0275 | 0 | 5.3 | 123 | Specific | 672.3 | 661.44 | 661.44 | 13.59 | 21 | >sp\|P0DOX2\|IGA2_HUMAN Immunoglobulin alpha-2 heavy chain OS=Homo sapiens OX=9606 PE=1 SV=2 | 631 | 20201112_10613_1ng_glyco_hcd203040_compound15.25947.25947.4 | controllerType=0 controllerNumber=1 scan=25947 | 52.1022 |
| 03136:2 | 42 | K.TPLTAN[+1501.555]ITK.S | HexNAc(5)Hex(3) | N[+1502] | 1230.5627 | 2 | 2460.1181 | 2460.1121 | 0 | 2.4 | 315 | Specific | 215.87 | 86.26 | 86.26 | 3.41 | 21 | >sp\|P0DOX2\|IGA2_HUMAN Immunoglobulin alpha-2 heavy chain OS=Homo sapiens OX=9606 PE=1 SV=2 | 631 | 20201112_10613_1ng_glyco_hcd203040_compound15.6137.6137.2 | controllerType=0 controllerNumber=1 scan=6137 | 16.331 |
| 04830:3 | 42 | K.TPLTAN[+2262.814]ITK.S | HexNAc(5)Hex(5)Fuc(1)NeuAc(1) | N[+2263] | 1074.4643 | 3 | 3221.3782 | 3221.3711 | 0 | 2.2 | 315 | Specific | 242.04 | 106.9 | 106.9 | 4.39 | 21 | >sp\|P0DOX2\|IGA2_HUMAN Immunoglobulin alpha-2 heavy chain OS=Homo sapiens OX=9606 PE=1 SV=2 | 631 | 20201112_10613_1ng_glyco_hcd203040_compound15.7897.7897.3 | controllerType=0 controllerNumber=1 scan=7897 | 19.2726 |
| 03138:2 | 42 | K.TPLTAN[+1501.555]ITK.S | HexNAc(5)Hex(3) | N[+1502] | 1230.5627 | 2 | 2460.1181 | 2460.1121 | 0 | 2.4 | 315 | Specific | 281.81 | 156.67 | 156.67 | 5.38 | 21 | >sp\|P0DOX2\|IGA2_HUMAN Immunoglobulin alpha-2 heavy chain OS=Homo sapiens OX=9606 PE=1 SV=2 | 631 | 20201112_10613_1ng_glyco_hcd203040_compound15.6139.6139.2 | controllerType=0 controllerNumber=1 scan=6139 | 16.335 |
| 03063:3 | 42 | K.TPLTAN[+1809.666]ITK.S | HexNAc(5)Hex(4)Fuc(1) | N[+1810] | 923.4138 | 3 | 2768.227 | 2768.2228 | 0 | 1.5 | 315 | Specific | 414.67 | 272.44 | 272.44 | 6.98 | 21 | >sp\|P0DOX2\|IGA2_HUMAN Immunoglobulin alpha-2 heavy chain OS=Homo sapiens OX=9606 PE=1 SV=2 | 631 | 20201112_10613_1ng_glyco_hcd203040_compound15.6062.6062.3 | controllerType=0 controllerNumber=1 scan=6062 | 16.1963 |
| 03126:3 | 42 | K.TPLTAN[+1647.613]ITK.S | HexNAc(5)Hex(3)Fuc(1) | N[+1648] | 869.3952 | 3 | 2606.1711 | 2606.17 | 0 | 0.4 | 315 | Specific | 514.89 | 198.21 | 198.21 | 6.63 | 21 | >sp\|P0DOX2\|IGA2_HUMAN Immunoglobulin alpha-2 heavy chain OS=Homo sapiens OX=9606 PE=1 SV=2 | 631 | 20201112_10613_1ng_glyco_hcd203040_compound15.6127.6127.3 | controllerType=0 controllerNumber=1 scan=6127 | 16.3128 |
| 03128:3 | 42 | K.TPLTAN[+1647.613]ITK.S | HexNAc(5)Hex(3)Fuc(1) | N[+1648] | 869.3952 | 3 | 2606.1711 | 2606.17 | 0 | 0.4 | 315 | Specific | 538.75 | 254.97 | 254.97 | 7.56 | 21 | >sp\|P0DOX2\|IGA2_HUMAN Immunoglobulin alpha-2 heavy chain OS=Homo sapiens OX=9606 PE=1 SV=2 | 631 | 20201112_10613_1ng_glyco_hcd203040_compound15.6129.6129.3 | controllerType=0 controllerNumber=1 scan=6129 | 16.3172 |
| 03067:2 | 42 | K.TPLTAN[+1809.666]ITK.S | HexNAc(5)Hex(4)Fuc(1) | N[+1810] | 1384.6171 | 2 | 2768.2269 | 2768.2228 | 0 | 1.5 | 315 | Specific | 544.96 | 283.82 | 283.82 | 7.82 | 21 | >sp\|P0DOX2\|IGA2_HUMAN Immunoglobulin alpha-2 heavy chain OS=Homo sapiens OX=9606 PE=1 SV=2 | 631 | 20201112_10613_1ng_glyco_hcd203040_compound15.6066.6066.2 | controllerType=0 controllerNumber=1 scan=6066 | 16.2046 |
| 03065:3 | 42 | K.TPLTAN[+1809.666]ITK.S | HexNAc(5)Hex(4)Fuc(1) | N[+1810] | 923.4138 | 3 | 2768.227 | 2768.2228 | 0 | 1.5 | 315 | Specific | 551.51 | 272.9 | 272.9 | 7.78 | 21 | >sp\|P0DOX2\|IGA2_HUMAN Immunoglobulin alpha-2 heavy chain OS=Homo sapiens OX=9606 PE=1 SV=2 | 631 | 20201112_10613_1ng_glyco_hcd203040_compound15.6064.6064.3 | controllerType=0 controllerNumber=1 scan=6064 | 16.2013 |
| 03125:2 | 42 | K.TPLTAN[+1647.613]ITK.S | HexNAc(5)Hex(3)Fuc(1) | N[+1648] | 1303.5896 | 2 | 2606.172 | 2606.17 | 0 | 0.8 | 315 | Specific | 573.3 | 286.1 | 286.1 | 7.97 | 21 | >sp\|P0DOX2\|IGA2_HUMAN Immunoglobulin alpha-2 heavy chain OS=Homo sapiens OX=9606 PE=1 SV=2 | 631 | 20201112_10613_1ng_glyco_hcd203040_compound15.6126.6126.2 | controllerType=0 controllerNumber=1 scan=6126 | 16.3109 |
| 03123:2 | 42 | K.TPLTAN[+1647.613]ITK.S | HexNAc(5)Hex(3)Fuc(1) | N[+1648] | 1303.5896 | 2 | 2606.172 | 2606.17 | 0 | 0.8 | 315 | Specific | 590.64 | 320.29 | 320.29 | 9.39 | 21 | >sp\|P0DOX2\|IGA2_HUMAN Immunoglobulin alpha-2 heavy chain OS=Homo sapiens OX=9606 PE=1 SV=2 | 631 | 20201112_10613_1ng_glyco_hcd203040_compound15.6124.6124.2 | controllerType=0 controllerNumber=1 scan=6124 | 16.3056 |
| 03069:2 | 42 | K.TPLTAN[+1809.666]ITK.S | HexNAc(5)Hex(4)Fuc(1) | N[+1810] | 1384.6171 | 2 | 2768.2269 | 2768.2228 | 0 | 1.5 | 315 | Specific | 619.84 | 352.83 | 352.83 | 10.74 | 21 | >sp\|P0DOX2\|IGA2_HUMAN Immunoglobulin alpha-2 heavy chain OS=Homo sapiens OX=9606 PE=1 SV=2 | 631 | 20201112_10613_1ng_glyco_hcd203040_compound15.6068.6068.2 | controllerType=0 controllerNumber=1 scan=6068 | 16.2089 |
| 11585:3 | 46 | R.IIVPLNNREN[+1694.603]ISDPTSPLR.T | HexNAc(3)Hex(4)Fuc(1)NeuAc(1) | N[+1695] | 1281.5989 | 3 | 3842.7822 | 3842.7799 | 0 | 0.6 | 62 | Specific | 304.75 | 241.56 | 241.56 | 6.56 | 16 | >sp\|P01591\|IGJ_HUMAN Immunoglobulin J chain OS=Homo sapiens OX=9606 GN=JCHAIN PE=1 SV=4 | 1392 | 20201112_10613_1ng_glyco_hcd203040_compound15.14898.14898.3 | controllerType=0 controllerNumber=1 scan=14898 | 30.9357 |
| 11587:3 | 46 | R.IIVPLNNREN[+1694.603]ISDPTSPLR.T | HexNAc(3)Hex(4)Fuc(1)NeuAc(1) | N[+1695] | 1281.5989 | 3 | 3842.7822 | 3842.7799 | 0 | 0.6 | 62 | Specific | 331.76 | 260.39 | 260.39 | 5.52 | 16 | >sp\|P01591\|IGJ_HUMAN Immunoglobulin J chain OS=Homo sapiens OX=9606 GN=JCHAIN PE=1 SV=4 | 1392 | 20201112_10613_1ng_glyco_hcd203040_compound15.14900.14900.3 | controllerType=0 controllerNumber=1 scan=14900 | 30.9396 |
| 11418:3 | 46 | R.IIVPLNNREN[+2059.735]ISDPTSPLR.T | HexNAc(4)Hex(5)Fuc(1)NeuAc(1) | N[+2060] | 1403.3099 | 3 | 4207.9151 | 4207.9121 | 0 | 0.7 | 62 | Specific | 361.25 | 329.5 | 329.5 | 7.3 | 16 | >sp\|P01591\|IGJ_HUMAN Immunoglobulin J chain OS=Homo sapiens OX=9606 GN=JCHAIN PE=1 SV=4 | 1392 | 20201112_10613_1ng_glyco_hcd203040_compound15.14724.14724.3 | controllerType=0 controllerNumber=1 scan=14724 | 30.6016 |
| 11415:3 | 46 | R.IIVPLNNREN[+2059.735]ISDPTSPLR.T | HexNAc(4)Hex(5)Fuc(1)NeuAc(1) | N[+2060] | 1403.3099 | 3 | 4207.9151 | 4207.9121 | 0 | 0.7 | 62 | Specific | 364.41 | 364.41 | 364.41 | 7.55 | 16 | >sp\|P01591\|IGJ_HUMAN Immunoglobulin J chain OS=Homo sapiens OX=9606 GN=JCHAIN PE=1 SV=4 | 1392 | 20201112_10613_1ng_glyco_hcd203040_compound15.14721.14721.3 | controllerType=0 controllerNumber=1 scan=14721 | 30.5951 |
| 09834:3 | 46 | R.IIVPLNNREN[+1663.608]ISDPTSPLR.T | HexNAc(5)Hex(4) | N[+1664] | 1271.2657 | 3 | 3811.7825 | 3811.7853 | 0 | -0.7 | 62 | Specific | 377.97 | 377.97 | 377.97 | 8.99 | 16 | >sp\|P01591\|IGJ_HUMAN Immunoglobulin J chain OS=Homo sapiens OX=9606 GN=JCHAIN PE=1 SV=4 | 1392 | 20201112_10613_1ng_glyco_hcd203040_compound15.13084.13084.3 | controllerType=0 controllerNumber=1 scan=13084 | 27.8266 |
| 09837:3 | 46 | R.IIVPLNNREN[+1663.608]ISDPTSPLR.T | HexNAc(5)Hex(4) | N[+1664] | 1271.2657 | 3 | 3811.7825 | 3811.7853 | 0 | -0.7 | 62 | Specific | 398.84 | 398.84 | 398.84 | 8.99 | 16 | >sp\|P01591\|IGJ_HUMAN Immunoglobulin J chain OS=Homo sapiens OX=9606 GN=JCHAIN PE=1 SV=4 | 1392 | 20201112_10613_1ng_glyco_hcd203040_compound15.13087.13087.3 | controllerType=0 controllerNumber=1 scan=13087 | 27.8318 |
| 11499:3 | 46 | R.IIVPLNNREN[+1856.656]ISDPTSPLR.T | HexNAc(3)Hex(5)Fuc(1)NeuAc(1) | N[+1857] | 1335.6169 | 3 | 4004.8361 | 4004.8327 | 0 | 0.8 | 62 | Specific | 415.14 | 415.14 | 415.14 | 9.17 | 16 | >sp\|P01591\|IGJ_HUMAN Immunoglobulin J chain OS=Homo sapiens OX=9606 GN=JCHAIN PE=1 SV=4 | 1392 | 20201112_10613_1ng_glyco_hcd203040_compound15.14809.14809.3 | controllerType=0 controllerNumber=1 scan=14809 | 30.7798 |
| 11502:3 | 46 | R.IIVPLNNREN[+1856.656]ISDPTSPLR.T | HexNAc(3)Hex(5)Fuc(1)NeuAc(1) | N[+1857] | 1335.6169 | 3 | 4004.8361 | 4004.8327 | 0 | 0.8 | 62 | Specific | 536.73 | 526.48 | 526.48 | 9.81 | 16 | >sp\|P01591\|IGJ_HUMAN Immunoglobulin J chain OS=Homo sapiens OX=9606 GN=JCHAIN PE=1 SV=4 | 1392 | 20201112_10613_1ng_glyco_hcd203040_compound15.14812.14812.3 | controllerType=0 controllerNumber=1 scan=14812 | 30.7849 |
| 24649:3 | 96 | R.AVN[+2204.772]ITSENLIDDVVSLIR.G | HexNAc(4)Hex(5)NeuAc(2) | N[+2205] | 1392.6218 | 3 | 4175.8507 | 4175.8481 | 0 | 0.6 | 413 | Specific | 477.9 | 431.31 | 431.31 | 9.77 | 8 | >sp\|P02748\|CO9_HUMAN Complement component C9 OS=Homo sapiens OX=9606 GN=C9 PE=1 SV=2 | 1547 | 20201112_10613_1ng_glyco_hcd203040_compound15.28538.28538.3 | controllerType=0 controllerNumber=1 scan=28538 | 57.0262 |
| 24651:3 | 96 | R.AVN[+2204.772]ITSENLIDDVVSLIR.G | HexNAc(4)Hex(5)NeuAc(2) | N[+2205] | 1392.6218 | 3 | 4175.8507 | 4175.8481 | 0 | 0.6 | 413 | Specific | 501.18 | 252.72 | 252.72 | 7.66 | 8 | >sp\|P02748\|CO9_HUMAN Complement component C9 OS=Homo sapiens OX=9606 GN=C9 PE=1 SV=2 | 1547 | 20201112_10613_1ng_glyco_hcd203040_compound15.28540.28540.3 | controllerType=0 controllerNumber=1 scan=28540 | 57.0298 |
| 18480:5 | 1 | R.RTAGWNVPIGTLRPFLN[+2059.735]WTGPPEPIEAAVAR.F | HexNAc(4)Hex(5)Fuc(1)NeuAc(1) | N[+2060] | 1090.1143 | 5 | 5446.5422 | 5446.547 | 0 | -0.9 | 140 | Specific | 213.96 | 201.53 | 201.53 | 4.51 | 139 | >sp\|P02788\|TRFL_HUMAN Lactotransferrin OS=Homo sapiens OX=9606 GN=LTF PE=1 SV=6 | 1270 | 20201113_10633_1ng_glyco_hcd203040_compound15.21972.21972.5 | controllerType=0 controllerNumber=1 scan=21972 | 43.7392 |
| 19851:3 | 1 | R.[+42.011]RTAGWNVPIGTLRPFLN[+1864.634]WTGPPEPIEAAVAR.F | HexNAc(2)Hex(9) | .[42], N[+1865] | 1765.1597 | 3 | 5293.4645 | 5293.4568 | 0 | 1.5 | 140 | Specific | 227.94 | 227.94 | 227.94 | 6.28 | 139 | >sp\|P02788\|TRFL_HUMAN Lactotransferrin OS=Homo sapiens OX=9606 GN=LTF PE=1 SV=6 | 1270 | 20201113_10633_1ng_glyco_hcd203040_compound15.23397.23397.3 | controllerType=0 controllerNumber=1 scan=23397 | 46.5828 |
| 18465:3 | 1 | R.RTAGWNVPIGTLRPFLN[+2059.735]WTGPPEPIEAAVAR.F | HexNAc(4)Hex(5)Fuc(1)NeuAc(1) | N[+2060] | 1816.198 | 3 | 5446.5793 | 5446.547 | 0 | 5.9 | 140 | Specific | 280.78 | 280.78 | 280.78 | 4.65 | 139 | >sp\|P02788\|TRFL_HUMAN Lactotransferrin OS=Homo sapiens OX=9606 GN=LTF PE=1 SV=6 | 1270 | 20201113_10633_1ng_glyco_hcd203040_compound15.21956.21956.3 | controllerType=0 controllerNumber=1 scan=21956 | 43.7011 |
| 18614:4 | 1 | R.RTAGWNVPIGTLRPFLN[+2059.735]WTGPPEPIEAAVAR.F | HexNAc(4)Hex(5)Fuc(1)NeuAc(1) | N[+2060] | 1362.4043 | 4 | 5446.5954 | 5446.547 | 0 | 8.9 | 140 | Specific | 314.97 | 314.97 | 314.97 | 6.67 | 139 | >sp\|P02788\|TRFL_HUMAN Lactotransferrin OS=Homo sapiens OX=9606 GN=LTF PE=1 SV=6 | 1270 | 20201113_10633_1ng_glyco_hcd203040_compound15.22110.22110.2 | controllerType=0 controllerNumber=1 scan=22110 | 43.9721 |
| 18616:4 | 1 | R.RTAGWNVPIGTLRPFLN[+2059.735]WTGPPEPIEAAVAR.F | HexNAc(4)Hex(5)Fuc(1)NeuAc(1) | N[+2060] | 1362.4043 | 4 | 5446.5954 | 5446.547 | 0 | 8.9 | 140 | Specific | 377.62 | 377.62 | 377.62 | 8.6 | 139 | >sp\|P02788\|TRFL_HUMAN Lactotransferrin OS=Homo sapiens OX=9606 GN=LTF PE=1 SV=6 | 1270 | 20201113_10633_1ng_glyco_hcd203040_compound15.22112.22112.2 | controllerType=0 controllerNumber=1 scan=22112 | 43.9761 |
| 19469:4 | 1 | R.RTAGWNVPIGTLRPFLN[+2350.830]WTGPPEPIEAAVAR.F | HexNAc(4)Hex(5)Fuc(1)NeuAc(2) | N[+2351] | 1435.1659 | 4 | 5737.6418 | 5737.6424 | 0 | -0.1 | 140 | Specific | 438.4 | 372.3 | 372.3 | 8.4 | 139 | >sp\|P02788\|TRFL_HUMAN Lactotransferrin OS=Homo sapiens OX=9606 GN=LTF PE=1 SV=6 | 1270 | 20201113_10633_1ng_glyco_hcd203040_compound15.23001.23001.4 | controllerType=0 controllerNumber=1 scan=23001 | 45.8532 |
| 18451:4 | 1 | R.RTAGWNVPIGTLRPFLN[+2059.735]WTGPPEPIEAAVAR.F | HexNAc(4)Hex(5)Fuc(1)NeuAc(1) | N[+2060] | 1362.3973 | 4 | 5446.5675 | 5446.547 | 0 | 3.8 | 140 | Specific | 487.44 | 487.44 | 487.44 | 9.98 | 139 | >sp\|P02788\|TRFL_HUMAN Lactotransferrin OS=Homo sapiens OX=9606 GN=LTF PE=1 SV=6 | 1270 | 20201113_10633_1ng_glyco_hcd203040_compound15.21942.21942.4 | controllerType=0 controllerNumber=1 scan=21942 | 43.6784 |
| 19472:4 | 1 | R.RTAGWNVPIGTLRPFLN[+2350.830]WTGPPEPIEAAVAR.F | HexNAc(4)Hex(5)Fuc(1)NeuAc(2) | N[+2351] | 1435.1659 | 4 | 5737.6418 | 5737.6424 | 0 | -0.1 | 140 | Specific | 504.86 | 426.26 | 426.26 | 10.17 | 139 | >sp\|P02788\|TRFL_HUMAN Lactotransferrin OS=Homo sapiens OX=9606 GN=LTF PE=1 SV=6 | 1270 | 20201113_10633_1ng_glyco_hcd203040_compound15.23004.23004.4 | controllerType=0 controllerNumber=1 scan=23004 | 45.8583 |
| 18463:5 | 1 | R.RTAGWNVPIGTLRPFLN[+2059.735]WTGPPEPIEAAVAR.F | HexNAc(4)Hex(5)Fuc(1)NeuAc(1) | N[+2060] | 1090.1188 | 5 | 5446.5648 | 5446.547 | 0 | 3.3 | 140 | Specific | 513.31 | 502.13 | 502.13 | 9.91 | 139 | >sp\|P02788\|TRFL_HUMAN Lactotransferrin OS=Homo sapiens OX=9606 GN=LTF PE=1 SV=6 | 1270 | 20201113_10633_1ng_glyco_hcd203040_compound15.21954.21954.5 | controllerType=0 controllerNumber=1 scan=21954 | 43.6976 |
| 18482:5 | 1 | R.RTAGWNVPIGTLRPFLN[+2059.735]WTGPPEPIEAAVAR.F | HexNAc(4)Hex(5)Fuc(1)NeuAc(1) | N[+2060] | 1090.1143 | 5 | 5446.5422 | 5446.547 | 0 | -0.9 | 140 | Specific | 561.82 | 525.51 | 525.51 | 9.75 | 139 | >sp\|P02788\|TRFL_HUMAN Lactotransferrin OS=Homo sapiens OX=9606 GN=LTF PE=1 SV=6 | 1270 | 20201113_10633_1ng_glyco_hcd203040_compound15.21974.21974.5 | controllerType=0 controllerNumber=1 scan=21974 | 43.7429 |
| 17488:4 | 1 | R.RTAGWNVPIGTLRPFLN[+1768.640]WTGPPEPIEAAVAR.F | HexNAc(4)Hex(5)Fuc(1) | N[+1769] | 1289.619 | 4 | 5155.4543 | 5155.4516 | 0 | 0.5 | 140 | Specific | 567.12 | 402.75 | 402.75 | 10.49 | 139 | >sp\|P02788\|TRFL_HUMAN Lactotransferrin OS=Homo sapiens OX=9606 GN=LTF PE=1 SV=6 | 1270 | 20201113_10633_1ng_glyco_hcd203040_compound15.20939.20939.4 | controllerType=0 controllerNumber=1 scan=20939 | 41.5906 |
| 18414:4 | 1 | R.RTAGWNVPIGTLRPFLN[+2205.793]WTGPPEPIEAAVAR.F | HexNAc(4)Hex(5)Fuc(2)NeuAc(1) | N[+2206] | 1398.9076 | 4 | 5592.6086 | 5592.6049 | 0 | 0.7 | 140 | Specific | 575.78 | 436.9 | 436.9 | 11.95 | 139 | >sp\|P02788\|TRFL_HUMAN Lactotransferrin OS=Homo sapiens OX=9606 GN=LTF PE=1 SV=6 | 1270 | 20201113_10633_1ng_glyco_hcd203040_compound15.21903.21903.4 | controllerType=0 controllerNumber=1 scan=21903 | 43.5859 |
| 18466:5 | 1 | R.RTAGWNVPIGTLRPFLN[+2059.735]WTGPPEPIEAAVAR.F | HexNAc(4)Hex(5)Fuc(1)NeuAc(1) | N[+2060] | 1090.1188 | 5 | 5446.5648 | 5446.547 | 0 | 3.3 | 140 | Specific | 579.27 | 563.71 | 563.71 | 11.47 | 139 | >sp\|P02788\|TRFL_HUMAN Lactotransferrin OS=Homo sapiens OX=9606 GN=LTF PE=1 SV=6 | 1270 | 20201113_10633_1ng_glyco_hcd203040_compound15.21957.21957.5 | controllerType=0 controllerNumber=1 scan=21957 | 43.7031 |
| 18435:4 | 1 | R.RTAGWNVPIGTLRPFLN[+2059.735]WTGPPEPIEAAVAR.F | HexNAc(4)Hex(5)Fuc(1)NeuAc(1) | N[+2060] | 1362.3936 | 4 | 5446.5524 | 5446.547 | 0 | 1 | 140 | Specific | 612.1 | 488.45 | 488.45 | 12.95 | 139 | >sp\|P02788\|TRFL_HUMAN Lactotransferrin OS=Homo sapiens OX=9606 GN=LTF PE=1 SV=6 | 1270 | 20201113_10633_1ng_glyco_hcd203040_compound15.21925.21925.4 | controllerType=0 controllerNumber=1 scan=21925 | 43.6404 |
| 18453:4 | 1 | R.RTAGWNVPIGTLRPFLN[+2059.735]WTGPPEPIEAAVAR.F | HexNAc(4)Hex(5)Fuc(1)NeuAc(1) | N[+2060] | 1362.3973 | 4 | 5446.5675 | 5446.547 | 0 | 3.8 | 140 | Specific | 613.29 | 483.29 | 483.29 | 12.63 | 139 | >sp\|P02788\|TRFL_HUMAN Lactotransferrin OS=Homo sapiens OX=9606 GN=LTF PE=1 SV=6 | 1270 | 20201113_10633_1ng_glyco_hcd203040_compound15.21944.21944.4 | controllerType=0 controllerNumber=1 scan=21944 | 43.683 |
| 18412:4 | 1 | R.RTAGWNVPIGTLRPFLN[+2205.793]WTGPPEPIEAAVAR.F | HexNAc(4)Hex(5)Fuc(2)NeuAc(1) | N[+2206] | 1398.9076 | 4 | 5592.6086 | 5592.6049 | 0 | 0.7 | 140 | Specific | 630.29 | 570.87 | 570.87 | 12.95 | 139 | >sp\|P02788\|TRFL_HUMAN Lactotransferrin OS=Homo sapiens OX=9606 GN=LTF PE=1 SV=6 | 1270 | 20201113_10633_1ng_glyco_hcd203040_compound15.21901.21901.4 | controllerType=0 controllerNumber=1 scan=21901 | 43.5783 |
| 17490:4 | 1 | R.RTAGWNVPIGTLRPFLN[+1768.640]WTGPPEPIEAAVAR.F | HexNAc(4)Hex(5)Fuc(1) | N[+1769] | 1289.619 | 4 | 5155.4543 | 5155.4516 | 0 | 0.5 | 140 | Specific | 654.31 | 610.84 | 610.84 | 13.73 | 139 | >sp\|P02788\|TRFL_HUMAN Lactotransferrin OS=Homo sapiens OX=9606 GN=LTF PE=1 SV=6 | 1270 | 20201113_10633_1ng_glyco_hcd203040_compound15.20941.20941.4 | controllerType=0 controllerNumber=1 scan=20941 | 41.5961 |
| 18438:4 | 1 | R.RTAGWNVPIGTLRPFLN[+2059.735]WTGPPEPIEAAVAR.F | HexNAc(4)Hex(5)Fuc(1)NeuAc(1) | N[+2060] | 1362.3936 | 4 | 5446.5524 | 5446.547 | 0 | 1 | 140 | Specific | 664.35 | 530.27 | 530.27 | 13.95 | 139 | >sp\|P02788\|TRFL_HUMAN Lactotransferrin OS=Homo sapiens OX=9606 GN=LTF PE=1 SV=6 | 1270 | 20201113_10633_1ng_glyco_hcd203040_compound15.21928.21928.4 | controllerType=0 controllerNumber=1 scan=21928 | 43.6493 |
| 19784:4 | 1 | R.TAGWNVPIGTLRPFLN[+2059.735]WTGPPEPIEAAVAR.F | HexNAc(4)Hex(5)Fuc(1)NeuAc(1) | N[+2060] | 1323.3684 | 4 | 5290.4518 | 5290.4459 | 0 | 1.1 | 141 | Specific | 227.68 | 224.67 | 224.67 | 6.36 | 139 | >sp\|P02788\|TRFL_HUMAN Lactotransferrin OS=Homo sapiens OX=9606 GN=LTF PE=1 SV=6 | 1270 | 20201113_10633_1ng_glyco_hcd203040_compound15.23328.23328.4 | controllerType=0 controllerNumber=1 scan=23328 | 46.4787 |
| 18716:3 | 1 | R.[+42.011]TAGWNVPIGTLRPFLN[+1872.651]WTGPPEPIEAAVAR.F | HexNAc(3)Hex(6)NeuAc(1) | .[42], N[+1873] | 1715.8103 | 3 | 5145.4162 | 5145.372 | 0 | 8.6 | 141 | Specific | 242.38 | 121.51 | 121.51 | 3.35 | 139 | >sp\|P02788\|TRFL_HUMAN Lactotransferrin OS=Homo sapiens OX=9606 GN=LTF PE=1 SV=6 | 1270 | 20201113_10633_1ng_glyco_hcd203040_compound15.22217.22217.3 | controllerType=0 controllerNumber=1 scan=22217 | 44.2091 |
| 17485:4 | 1 | R.[+42.011]TAGWNVPIGTLRPFLN[+2028.740]WTGPPEPIEAAVAR.F | HexNAc(6)Hex(5) | .[42], N[+2029] | 1326.1337 | 4 | 5301.5129 | 5301.4618 | 0 | 9.6 | 141 | Specific | 243.06 | 119.18 | 119.18 | 2.1 | 139 | >sp\|P02788\|TRFL_HUMAN Lactotransferrin OS=Homo sapiens OX=9606 GN=LTF PE=1 SV=6 | 1270 | 20201113_10633_1ng_glyco_hcd203040_compound15.20936.20936.4 | controllerType=0 controllerNumber=1 scan=20936 | 41.5845 |
| 18715:4 | 1 | R.[+42.011]TAGWNVPIGTLRPFLN[+1872.651]WTGPPEPIEAAVAR.F | HexNAc(3)Hex(6)NeuAc(1) | .[42], N[+1873] | 1287.1099 | 4 | 5145.4176 | 5145.372 | 0 | 8.9 | 141 | Specific | 326.94 | 206.18 | 206.18 | 4.92 | 139 | >sp\|P02788\|TRFL_HUMAN Lactotransferrin OS=Homo sapiens OX=9606 GN=LTF PE=1 SV=6 | 1270 | 20201113_10633_1ng_glyco_hcd203040_compound15.22216.22216.4 | controllerType=0 controllerNumber=1 scan=22216 | 44.2078 |
| 19762:5 | 1 | R.TAGWNVPIGTLRPFLN[+2059.735]WTGPPEPIEAAVAR.F | HexNAc(4)Hex(5)Fuc(1)NeuAc(1) | N[+2060] | 1058.896 | 5 | 5290.4509 | 5290.4459 | 0 | 0.9 | 141 | Specific | 503.76 | 398.08 | 398.08 | 10.54 | 139 | >sp\|P02788\|TRFL_HUMAN Lactotransferrin OS=Homo sapiens OX=9606 GN=LTF PE=1 SV=6 | 1270 | 20201113_10633_1ng_glyco_hcd203040_compound15.23305.23305.5 | controllerType=0 controllerNumber=1 scan=23305 | 46.4343 |
| 19787:3 | 1 | R.TAGWNVPIGTLRPFLN[+2059.735]WTGPPEPIEAAVAR.F | HexNAc(4)Hex(5)Fuc(1)NeuAc(1) | N[+2060] | 1764.1546 | 3 | 5290.4493 | 5290.4459 | 0 | 0.7 | 141 | Specific | 509.89 | 456.52 | 456.52 | 10.15 | 139 | >sp\|P02788\|TRFL_HUMAN Lactotransferrin OS=Homo sapiens OX=9606 GN=LTF PE=1 SV=6 | 1270 | 20201113_10633_1ng_glyco_hcd203040_compound15.23331.23331.3 | controllerType=0 controllerNumber=1 scan=23331 | 46.4834 |
| 21163:3 | 1 | R.TAGWNVPIGTLRPFLN[+2350.830]WTGPPEPIEAAVAR.F | HexNAc(4)Hex(5)Fuc(1)NeuAc(2) | N[+2351] | 1861.1904 | 3 | 5581.5566 | 5581.5413 | 0 | 2.7 | 141 | Specific | 517.12 | 344.12 | 344.12 | 8.57 | 139 | >sp\|P02788\|TRFL_HUMAN Lactotransferrin OS=Homo sapiens OX=9606 GN=LTF PE=1 SV=6 | 1270 | 20201113_10633_1ng_glyco_hcd203040_compound15.24767.24767.3 | controllerType=0 controllerNumber=1 scan=24767 | 49.2377 |
| 21166:3 | 1 | R.TAGWNVPIGTLRPFLN[+2350.830]WTGPPEPIEAAVAR.F | HexNAc(4)Hex(5)Fuc(1)NeuAc(2) | N[+2351] | 1861.1904 | 3 | 5581.5566 | 5581.5413 | 0 | 2.7 | 141 | Specific | 560.47 | 527.18 | 527.18 | 10.5 | 139 | >sp\|P02788\|TRFL_HUMAN Lactotransferrin OS=Homo sapiens OX=9606 GN=LTF PE=1 SV=6 | 1270 | 20201113_10633_1ng_glyco_hcd203040_compound15.24770.24770.3 | controllerType=0 controllerNumber=1 scan=24770 | 49.242 |
| 21201:4 | 1 | R.TAGWNVPIGTLRPFLN[+2350.830]WTGPPEPIEAAVAR.F | HexNAc(4)Hex(5)Fuc(1)NeuAc(2) | N[+2351] | 1396.1422 | 4 | 5581.547 | 5581.5413 | 0 | 1 | 141 | Specific | 569.55 | 435.64 | 435.64 | 10.32 | 139 | >sp\|P02788\|TRFL_HUMAN Lactotransferrin OS=Homo sapiens OX=9606 GN=LTF PE=1 SV=6 | 1270 | 20201113_10633_1ng_glyco_hcd203040_compound15.24806.24806.6 | controllerType=0 controllerNumber=1 scan=24806 | 49.3157 |
| 19714:4 | 1 | R.TAGWNVPIGTLRPFLN[+2205.793]WTGPPEPIEAAVAR.F | HexNAc(4)Hex(5)Fuc(2)NeuAc(1) | N[+2206] | 1359.8854 | 4 | 5436.5197 | 5436.5038 | 0 | 2.9 | 141 | Specific | 661.71 | 610.2 | 610.2 | 13.58 | 139 | >sp\|P02788\|TRFL_HUMAN Lactotransferrin OS=Homo sapiens OX=9606 GN=LTF PE=1 SV=6 | 1270 | 20201113_10633_1ng_glyco_hcd203040_compound15.23255.23255.4 | controllerType=0 controllerNumber=1 scan=23255 | 46.3206 |
| 19791:3 | 1 | R.TAGWNVPIGTLRPFLN[+2205.793]WTGPPEPIEAAVAR.F | HexNAc(4)Hex(5)Fuc(2)NeuAc(1) | N[+2206] | 1812.8441 | 3 | 5436.5177 | 5436.5038 | 0 | 2.6 | 141 | Specific | 662.55 | 572.85 | 572.85 | 13.74 | 139 | >sp\|P02788\|TRFL_HUMAN Lactotransferrin OS=Homo sapiens OX=9606 GN=LTF PE=1 SV=6 | 1270 | 20201113_10633_1ng_glyco_hcd203040_compound15.23335.23335.3 | controllerType=0 controllerNumber=1 scan=23335 | 46.4877 |
| 21160:4 | 1 | R.TAGWNVPIGTLRPFLN[+2350.830]WTGPPEPIEAAVAR.F | HexNAc(4)Hex(5)Fuc(1)NeuAc(2) | N[+2351] | 1396.1449 | 4 | 5581.5578 | 5581.5413 | 0 | 3 | 141 | Specific | 669.34 | 620.55 | 620.55 | 13.58 | 139 | >sp\|P02788\|TRFL_HUMAN Lactotransferrin OS=Homo sapiens OX=9606 GN=LTF PE=1 SV=6 | 1270 | 20201113_10633_1ng_glyco_hcd203040_compound15.24764.24764.4 | controllerType=0 controllerNumber=1 scan=24764 | 49.232 |
| 18717:4 | 1 | R.TAGWNVPIGTLRPFLN[+1768.640]WTGPPEPIEAAVAR.F | HexNAc(4)Hex(5)Fuc(1) | N[+1769] | 1250.5953 | 4 | 4999.3595 | 4999.3504 | 0 | 1.8 | 141 | Specific | 684.08 | 659.44 | 659.44 | 13.56 | 139 | >sp\|P02788\|TRFL_HUMAN Lactotransferrin OS=Homo sapiens OX=9606 GN=LTF PE=1 SV=6 | 1270 | 20201113_10633_1ng_glyco_hcd203040_compound15.22218.22218.4 | controllerType=0 controllerNumber=1 scan=22218 | 44.2104 |
| 18725:3 | 1 | R.TAGWNVPIGTLRPFLN[+1768.640]WTGPPEPIEAAVAR.F | HexNAc(4)Hex(5)Fuc(1) | N[+1769] | 1667.1248 | 3 | 4999.36 | 4999.3504 | 0 | 1.9 | 141 | Specific | 687.34 | 540.81 | 540.81 | 13.74 | 139 | >sp\|P02788\|TRFL_HUMAN Lactotransferrin OS=Homo sapiens OX=9606 GN=LTF PE=1 SV=6 | 1270 | 20201113_10633_1ng_glyco_hcd203040_compound15.22226.22226.3 | controllerType=0 controllerNumber=1 scan=22226 | 44.225 |
| 18722:3 | 1 | R.TAGWNVPIGTLRPFLN[+1768.640]WTGPPEPIEAAVAR.F | HexNAc(4)Hex(5)Fuc(1) | N[+1769] | 1667.1248 | 3 | 4999.36 | 4999.3504 | 0 | 1.9 | 141 | Specific | 689.29 | 598.42 | 598.42 | 13.74 | 139 | >sp\|P02788\|TRFL_HUMAN Lactotransferrin OS=Homo sapiens OX=9606 GN=LTF PE=1 SV=6 | 1270 | 20201113_10633_1ng_glyco_hcd203040_compound15.22223.22223.3 | controllerType=0 controllerNumber=1 scan=22223 | 44.2201 |
| 21162:4 | 1 | R.TAGWNVPIGTLRPFLN[+2350.830]WTGPPEPIEAAVAR.F | HexNAc(4)Hex(5)Fuc(1)NeuAc(2) | N[+2351] | 1396.1449 | 4 | 5581.5578 | 5581.5413 | 0 | 3 | 141 | Specific | 785.94 | 611.87 | 611.87 | 15.58 | 139 | >sp\|P02788\|TRFL_HUMAN Lactotransferrin OS=Homo sapiens OX=9606 GN=LTF PE=1 SV=6 | 1270 | 20201113_10633_1ng_glyco_hcd203040_compound15.24766.24766.4 | controllerType=0 controllerNumber=1 scan=24766 | 49.2369 |
| 19759:5 | 1 | R.TAGWNVPIGTLRPFLN[+2059.735]WTGPPEPIEAAVAR.F | HexNAc(4)Hex(5)Fuc(1)NeuAc(1) | N[+2060] | 1058.896 | 5 | 5290.4509 | 5290.4459 | 0 | 0.9 | 141 | Specific | 833.36 | 709.95 | 709.95 | 17.1 | 139 | >sp\|P02788\|TRFL_HUMAN Lactotransferrin OS=Homo sapiens OX=9606 GN=LTF PE=1 SV=6 | 1270 | 20201113_10633_1ng_glyco_hcd203040_compound15.23302.23302.5 | controllerType=0 controllerNumber=1 scan=23302 | 46.4282 |
| 19786:4 | 1 | R.TAGWNVPIGTLRPFLN[+2059.735]WTGPPEPIEAAVAR.F | HexNAc(4)Hex(5)Fuc(1)NeuAc(1) | N[+2060] | 1323.3684 | 4 | 5290.4518 | 5290.4459 | 0 | 1.1 | 141 | Specific | 833.72 | 697.33 | 697.33 | 16.56 | 139 | >sp\|P02788\|TRFL_HUMAN Lactotransferrin OS=Homo sapiens OX=9606 GN=LTF PE=1 SV=6 | 1270 | 20201113_10633_1ng_glyco_hcd203040_compound15.23330.23330.4 | controllerType=0 controllerNumber=1 scan=23330 | 46.4821 |
| 19717:4 | 1 | R.TAGWNVPIGTLRPFLN[+2205.793]WTGPPEPIEAAVAR.F | HexNAc(4)Hex(5)Fuc(2)NeuAc(1) | N[+2206] | 1359.8854 | 4 | 5436.5197 | 5436.5038 | 0 | 2.9 | 141 | Specific | 836.58 | 695.56 | 695.56 | 16.58 | 139 | >sp\|P02788\|TRFL_HUMAN Lactotransferrin OS=Homo sapiens OX=9606 GN=LTF PE=1 SV=6 | 1270 | 20201113_10633_1ng_glyco_hcd203040_compound15.23258.23258.4 | controllerType=0 controllerNumber=1 scan=23258 | 46.3385 |
| 18720:4 | 1 | R.TAGWNVPIGTLRPFLN[+1768.640]WTGPPEPIEAAVAR.F | HexNAc(4)Hex(5)Fuc(1) | N[+1769] | 1250.5953 | 4 | 4999.3595 | 4999.3504 | 0 | 1.8 | 141 | Specific | 928.12 | 828.2 | 828.2 | 18.56 | 139 | >sp\|P02788\|TRFL_HUMAN Lactotransferrin OS=Homo sapiens OX=9606 GN=LTF PE=1 SV=6 | 1270 | 20201113_10633_1ng_glyco_hcd203040_compound15.22221.22221.4 | controllerType=0 controllerNumber=1 scan=22221 | 44.2153 |
| 17092:3 | 1 | R.[+42.011]TAGWNIPMGLLFN[+1872.651]QTGSC[+57.021]K.F | HexNAc(3)Hex(6)NeuAc(1) | .[42], C[+57], N[+1873] | 1337.2416 | 3 | 4009.7103 | 4009.671 | 0 | 9.8 | 485 | Specific | 235.54 | 176.71 | 176.71 | 3.28 | 139 | >sp\|P02788\|TRFL_HUMAN Lactotransferrin OS=Homo sapiens OX=9606 GN=LTF PE=1 SV=6 | 1270 | 20201113_10633_1ng_glyco_hcd203040_compound15.20527.20527.3 | controllerType=0 controllerNumber=1 scan=20527 | 40.7737 |
| 17450:3 | 1 | R.TAGWNIPMGLLFN[+1622.582]QTGSC[+57.021]K.F | HexNAc(4)Hex(5) | C[+57], N[+1623] | 1239.8713 | 3 | 3717.5993 | 3717.5916 | 0 | 2.1 | 485 | Specific | 328.8 | 318.55 | 245.16 | 7.48 | 139 | >sp\|P02788\|TRFL_HUMAN Lactotransferrin OS=Homo sapiens OX=9606 GN=LTF PE=1 SV=6 | 1270 | 20201113_10633_1ng_glyco_hcd203040_compound15.20899.20899.7 | controllerType=0 controllerNumber=1 scan=20899 | 41.5033 |
| 17189:2 | 1 | R.TAGWNIPMGLLFN[+1768.640]QTGSC[+57.021]K.F | HexNAc(4)Hex(5)Fuc(1) | C[+57], N[+1769] | 1932.3339 | 2 | 3863.6606 | 3863.6495 | 0 | 2.9 | 485 | Specific | 393.32 | 364.53 | 364.53 | 7.96 | 139 | >sp\|P02788\|TRFL_HUMAN Lactotransferrin OS=Homo sapiens OX=9606 GN=LTF PE=1 SV=6 | 1270 | 20201113_10633_1ng_glyco_hcd203040_compound15.20628.20628.2 | controllerType=0 controllerNumber=1 scan=20628 | 40.9755 |
| 17187:2 | 1 | R.TAGWNIPMGLLFN[+1768.640]QTGSC[+57.021]K.F | HexNAc(4)Hex(5)Fuc(1) | C[+57], N[+1769] | 1932.3339 | 2 | 3863.6606 | 3863.6495 | 0 | 2.9 | 485 | Specific | 404.1 | 359.37 | 359.37 | 8.03 | 139 | >sp\|P02788\|TRFL_HUMAN Lactotransferrin OS=Homo sapiens OX=9606 GN=LTF PE=1 SV=6 | 1270 | 20201113_10633_1ng_glyco_hcd203040_compound15.20626.20626.2 | controllerType=0 controllerNumber=1 scan=20626 | 40.9709 |
| 17193:4 | 1 | R.TAGWNIPMGLLFN[+1768.640]QTGSC[+57.021]K.F | HexNAc(4)Hex(5)Fuc(1) | C[+57], N[+1769] | 966.6733 | 4 | 3863.6715 | 3863.6495 | 0 | 5.7 | 485 | Specific | 422.97 | 300.14 | 300.14 | 6.22 | 139 | >sp\|P02788\|TRFL_HUMAN Lactotransferrin OS=Homo sapiens OX=9606 GN=LTF PE=1 SV=6 | 1270 | 20201113_10633_1ng_glyco_hcd203040_compound15.20632.20632.4 | controllerType=0 controllerNumber=1 scan=20632 | 40.99 |
| 17471:3 | 1 | R.TAGWNIPMGLLFN[+1460.529]QTGSC[+57.021]K.F | HexNAc(4)Hex(4) | C[+57], N[+1461] | 1185.8528 | 3 | 3555.544 | 3555.5387 | 0 | 1.5 | 485 | Specific | 481.48 | 481.48 | 224.96 | 9.7 | 139 | >sp\|P02788\|TRFL_HUMAN Lactotransferrin OS=Homo sapiens OX=9606 GN=LTF PE=1 SV=6 | 1270 | 20201113_10633_1ng_glyco_hcd203040_compound15.20921.20921.3 | controllerType=0 controllerNumber=1 scan=20921 | 41.5533 |
| 17191:4 | 1 | R.TAGWNIPMGLLFN[+1768.640]QTGSC[+57.021]K.F | HexNAc(4)Hex(5)Fuc(1) | C[+57], N[+1769] | 966.6733 | 4 | 3863.6715 | 3863.6495 | 0 | 5.7 | 485 | Specific | 490.08 | 464.54 | 464.54 | 9.47 | 139 | >sp\|P02788\|TRFL_HUMAN Lactotransferrin OS=Homo sapiens OX=9606 GN=LTF PE=1 SV=6 | 1270 | 20201113_10633_1ng_glyco_hcd203040_compound15.20630.20630.4 | controllerType=0 controllerNumber=1 scan=20630 | 40.9789 |
| 17314:3 | 1 | R.TAGWNIPMGLLFN[+1622.582]QTGSC[+57.021]K.F | HexNAc(4)Hex(5) | C[+57], N[+1623] | 1239.8736 | 3 | 3717.6062 | 3717.5916 | 0 | 3.9 | 485 | Specific | 495.5 | 495.5 | 164.95 | 9.38 | 139 | >sp\|P02788\|TRFL_HUMAN Lactotransferrin OS=Homo sapiens OX=9606 GN=LTF PE=1 SV=6 | 1270 | 20201113_10633_1ng_glyco_hcd203040_compound15.20758.20758.3 | controllerType=0 controllerNumber=1 scan=20758 | 41.227 |
| 19201:3 | 1 | R.TAGWNIPMGLLFN[+2059.735]QTGSC[+57.021]K.F | HexNAc(4)Hex(5)Fuc(1)NeuAc(1) | C[+57], N[+2060] | 1385.5892 | 3 | 4154.753 | 4154.7449 | 0 | 1.9 | 485 | Specific | 523.89 | 510.75 | 510.75 | 9.8 | 139 | >sp\|P02788\|TRFL_HUMAN Lactotransferrin OS=Homo sapiens OX=9606 GN=LTF PE=1 SV=6 | 1270 | 20201113_10633_1ng_glyco_hcd203040_compound15.22722.22722.3 | controllerType=0 controllerNumber=1 scan=22722 | 45.2684 |
| 21226:3 | 1 | R.TAGWNIPMGLLFN[+2351.851]QTGSC[+57.021]K.F | HexNAc(4)Hex(5)Fuc(3)NeuAc(1) | C[+57], N[+2352] | 1482.9645 | 3 | 4446.8789 | 4446.8607 | 0 | 4.1 | 485 | Specific | 631.95 | 600.62 | 600.62 | 12.03 | 139 | >sp\|P02788\|TRFL_HUMAN Lactotransferrin OS=Homo sapiens OX=9606 GN=LTF PE=1 SV=6 | 1270 | 20201113_10633_1ng_glyco_hcd203040_compound15.24832.24832.3 | controllerType=0 controllerNumber=1 scan=24832 | 49.3651 |
| 17473:3 | 1 | R.TAGWNIPMGLLFN[+1460.529]QTGSC[+57.021]K.F | HexNAc(4)Hex(4) | C[+57], N[+1461] | 1185.8528 | 3 | 3555.544 | 3555.5387 | 0 | 1.5 | 485 | Specific | 717.34 | 681.03 | 355.68 | 14.35 | 139 | >sp\|P02788\|TRFL_HUMAN Lactotransferrin OS=Homo sapiens OX=9606 GN=LTF PE=1 SV=6 | 1270 | 20201113_10633_1ng_glyco_hcd203040_compound15.20923.20923.3 | controllerType=0 controllerNumber=1 scan=20923 | 41.5593 |
| 17363:3 | 1 | R.TAGWNIPMGLLFN[+1622.582]QTGSC[+57.021]K.F | HexNAc(4)Hex(5) | C[+57], N[+1623] | 1239.8713 | 3 | 3717.5993 | 3717.5916 | 0 | 2.1 | 485 | Specific | 718.59 | 718.59 | 309.01 | 14.36 | 139 | >sp\|P02788\|TRFL_HUMAN Lactotransferrin OS=Homo sapiens OX=9606 GN=LTF PE=1 SV=6 | 1270 | 20201113_10633_1ng_glyco_hcd203040_compound15.20809.20809.3 | controllerType=0 controllerNumber=1 scan=20809 | 41.3225 |
| 17604:3 | 1 | R.TAGWNIPMGLLFN[+1216.423]QTGSC[+57.021]K.F | HexNAc(2)Hex(5) | C[+57], N[+1216] | 1104.4883 | 3 | 3311.4504 | 3311.4328 | 0 | 5.3 | 485 | Specific | 726.27 | 726.27 | 240.91 | 13.54 | 139 | >sp\|P02788\|TRFL_HUMAN Lactotransferrin OS=Homo sapiens OX=9606 GN=LTF PE=1 SV=6 | 1270 | 20201113_10633_1ng_glyco_hcd203040_compound15.21059.21059.3 | controllerType=0 controllerNumber=1 scan=21059 | 41.8166 |
| 19016:3 | 1 | R.TAGWNIPMGLLFN[+2204.772]QTGSC[+57.021]K.F | HexNAc(4)Hex(5)NeuAc(2) | C[+57], N[+2205] | 1433.9398 | 3 | 4299.8047 | 4299.7824 | 0 | 5.2 | 485 | Specific | 739.6 | 719.57 | 719.57 | 13.54 | 139 | >sp\|P02788\|TRFL_HUMAN Lactotransferrin OS=Homo sapiens OX=9606 GN=LTF PE=1 SV=6 | 1270 | 20201113_10633_1ng_glyco_hcd203040_compound15.22529.22529.3 | controllerType=0 controllerNumber=1 scan=22529 | 44.8536 |
| 19203:3 | 1 | R.TAGWNIPMGLLFN[+2059.735]QTGSC[+57.021]K.F | HexNAc(4)Hex(5)Fuc(1)NeuAc(1) | C[+57], N[+2060] | 1385.5892 | 3 | 4154.753 | 4154.7449 | 0 | 1.9 | 485 | Specific | 748.69 | 732.97 | 732.97 | 14.36 | 139 | >sp\|P02788\|TRFL_HUMAN Lactotransferrin OS=Homo sapiens OX=9606 GN=LTF PE=1 SV=6 | 1270 | 20201113_10633_1ng_glyco_hcd203040_compound15.22724.22724.3 | controllerType=0 controllerNumber=1 scan=22724 | 45.2732 |
| 19058:3 | 1 | R.TAGWNIPMGLLFN[+2059.735]QTGSC[+57.021]K.F | HexNAc(4)Hex(5)Fuc(1)NeuAc(1) | C[+57], N[+2060] | 1385.5899 | 3 | 4154.7551 | 4154.7449 | 0 | 2.5 | 485 | Specific | 759.17 | 671.64 | 671.64 | 15.36 | 139 | >sp\|P02788\|TRFL_HUMAN Lactotransferrin OS=Homo sapiens OX=9606 GN=LTF PE=1 SV=6 | 1270 | 20201113_10633_1ng_glyco_hcd203040_compound15.22573.22573.3 | controllerType=0 controllerNumber=1 scan=22573 | 44.942 |
| 17237:3 | 1 | R.TAGWNIPMGLLFN[+1606.587]QTGSC[+57.021]K.F | HexNAc(4)Hex(4)Fuc(1) | C[+57], N[+1607] | 1234.5372 | 3 | 3701.5969 | 3701.5966 | 0 | 0.1 | 485 | Specific | 864.07 | 792.03 | 792.03 | 17.13 | 139 | >sp\|P02788\|TRFL_HUMAN Lactotransferrin OS=Homo sapiens OX=9606 GN=LTF PE=1 SV=6 | 1270 | 20201113_10633_1ng_glyco_hcd203040_compound15.20678.20678.3 | controllerType=0 controllerNumber=1 scan=20678 | 41.08 |
| 17317:3 | 1 | R.TAGWNIPMGLLFN[+1622.582]QTGSC[+57.021]K.F | HexNAc(4)Hex(5) | C[+57], N[+1623] | 1239.8736 | 3 | 3717.6062 | 3717.5916 | 0 | 3.9 | 485 | Specific | 867.08 | 810.93 | 404.99 | 17.03 | 139 | >sp\|P02788\|TRFL_HUMAN Lactotransferrin OS=Homo sapiens OX=9606 GN=LTF PE=1 SV=6 | 1270 | 20201113_10633_1ng_glyco_hcd203040_compound15.20761.20761.3 | controllerType=0 controllerNumber=1 scan=20761 | 41.2315 |
| 17103:3 | 1 | R.TAGWNIPMGLLFN[+1768.640]QTGSC[+57.021]K.F | HexNAc(4)Hex(5)Fuc(1) | C[+57], N[+1769] | 1288.5532 | 3 | 3863.6449 | 3863.6495 | 0 | -1.2 | 485 | Specific | 908.29 | 908.29 | 908.29 | 17.55 | 139 | >sp\|P02788\|TRFL_HUMAN Lactotransferrin OS=Homo sapiens OX=9606 GN=LTF PE=1 SV=6 | 1270 | 20201113_10633_1ng_glyco_hcd203040_compound15.20539.20539.3 | controllerType=0 controllerNumber=1 scan=20539 | 40.7962 |
| 17365:3 | 1 | R.TAGWNIPMGLLFN[+1622.582]QTGSC[+57.021]K.F | HexNAc(4)Hex(5) | C[+57], N[+1623] | 1239.8713 | 3 | 3717.5993 | 3717.5916 | 0 | 2.1 | 485 | Specific | 916.71 | 879.03 | 427.91 | 18.36 | 139 | >sp\|P02788\|TRFL_HUMAN Lactotransferrin OS=Homo sapiens OX=9606 GN=LTF PE=1 SV=6 | 1270 | 20201113_10633_1ng_glyco_hcd203040_compound15.20811.20811.3 | controllerType=0 controllerNumber=1 scan=20811 | 41.3265 |
| 17239:3 | 1 | R.TAGWNIPMGLLFN[+1606.587]QTGSC[+57.021]K.F | HexNAc(4)Hex(4)Fuc(1) | C[+57], N[+1607] | 1234.5372 | 3 | 3701.5969 | 3701.5966 | 0 | 0.1 | 485 | Specific | 921.39 | 919.16 | 919.16 | 18.13 | 139 | >sp\|P02788\|TRFL_HUMAN Lactotransferrin OS=Homo sapiens OX=9606 GN=LTF PE=1 SV=6 | 1270 | 20201113_10633_1ng_glyco_hcd203040_compound15.20680.20680.3 | controllerType=0 controllerNumber=1 scan=20680 | 41.0846 |
| 17105:3 | 1 | R.TAGWNIPMGLLFN[+1768.640]QTGSC[+57.021]K.F | HexNAc(4)Hex(5)Fuc(1) | C[+57], N[+1769] | 1288.5532 | 3 | 3863.6449 | 3863.6495 | 0 | -1.2 | 485 | Specific | 924.17 | 924.17 | 924.17 | 17.55 | 139 | >sp\|P02788\|TRFL_HUMAN Lactotransferrin OS=Homo sapiens OX=9606 GN=LTF PE=1 SV=6 | 1270 | 20201113_10633_1ng_glyco_hcd203040_compound15.20541.20541.3 | controllerType=0 controllerNumber=1 scan=20541 | 40.802 |
| 17605:3 | 1 | R.TAGWNIPMGLLFN[+1216.423]QTGSC[+57.021]K.F | HexNAc(2)Hex(5) | C[+57], N[+1216] | 1104.4883 | 3 | 3311.4504 | 3311.4328 | 0 | 5.3 | 485 | Specific | 925.59 | 898.77 | 439.53 | 17.54 | 139 | >sp\|P02788\|TRFL_HUMAN Lactotransferrin OS=Homo sapiens OX=9606 GN=LTF PE=1 SV=6 | 1270 | 20201113_10633_1ng_glyco_hcd203040_compound15.21060.21060.3 | controllerType=0 controllerNumber=1 scan=21060 | 41.821 |
| 19014:3 | 1 | R.TAGWNIPMGLLFN[+2204.772]QTGSC[+57.021]K.F | HexNAc(4)Hex(5)NeuAc(2) | C[+57], N[+2205] | 1433.9398 | 3 | 4299.8047 | 4299.7824 | 0 | 5.2 | 485 | Specific | 935.97 | 935.97 | 935.97 | 17.54 | 139 | >sp\|P02788\|TRFL_HUMAN Lactotransferrin OS=Homo sapiens OX=9606 GN=LTF PE=1 SV=6 | 1270 | 20201113_10633_1ng_glyco_hcd203040_compound15.22527.22527.3 | controllerType=0 controllerNumber=1 scan=22527 | 44.8486 |
| 19060:3 | 1 | R.TAGWNIPMGLLFN[+2059.735]QTGSC[+57.021]K.F | HexNAc(4)Hex(5)Fuc(1)NeuAc(1) | C[+57], N[+2060] | 1385.5899 | 3 | 4154.7551 | 4154.7449 | 0 | 2.5 | 485 | Specific | 939.91 | 916.92 | 916.92 | 18.36 | 139 | >sp\|P02788\|TRFL_HUMAN Lactotransferrin OS=Homo sapiens OX=9606 GN=LTF PE=1 SV=6 | 1270 | 20201113_10633_1ng_glyco_hcd203040_compound15.22575.22575.3 | controllerType=0 controllerNumber=1 scan=22575 | 44.9465 |
| 19357:5 | 1 | R.TAGWNIPMGLLFN[+2076.750]QTGSC[+57.021]KFDEYFSQSC[+57.021]APGSDPR.S | HexNAc(4)Hex(6)Fuc(2) | C[+57]*2, N[+2077] | 1203.917 | 5 | 6015.5559 | 6015.5074 | 0 | 8.1 | 485 | Specific | 229.09 | 203.14 | 203.14 | 4.14 | 139 | >sp\|P02788\|TRFL_HUMAN Lactotransferrin OS=Homo sapiens OX=9606 GN=LTF PE=1 SV=6 | 1270 | 20201113_10633_1ng_glyco_hcd203040_compound15.22883.22883.3 | controllerType=0 controllerNumber=1 scan=22883 | 45.585 |
| 18264:4 | 1 | R.[+42.011]TAGWNIPMGLLFN[+1872.651]QTGSC[+57.021]KFDEYFSQSC[+57.021]A | HexNAc(3)Hex(6)NeuAc(1) | .[42], C[+57]*2, N[+1873] | 1464.1232 | 4 | 5853.4709 | 5853.4181 | 0 | 9 | 485 | Specific | 242.57 | 219.87 | 219.87 | 3.71 | 139 | >sp\|P02788\|TRFL_HUMAN Lactotransferrin OS=Homo sapiens OX=9606 GN=LTF PE=1 SV=6 | 1270 | 20201113_10633_1ng_glyco_hcd203040_compound15.21747.21747.4 | controllerType=0 controllerNumber=1 scan=21747 | 43.2717 |
| 18352:3 | 1 | R.TAGWNIPMGLLFN[+1606.587]QTGSC[+57.021]KFDEYFSQSC[+57.021]APGSDPR.S | HexNAc(4)Hex(4)Fuc(1) | C[+57]*2, N[+1607] | 1849.1229 | 3 | 5545.3541 | 5545.3438 | 0 | 1.9 | 485 | Specific | 332.9 | 302.5 | 302.5 | 7.76 | 139 | >sp\|P02788\|TRFL_HUMAN Lactotransferrin OS=Homo sapiens OX=9606 GN=LTF PE=1 SV=6 | 1270 | 20201113_10633_1ng_glyco_hcd203040_compound15.21838.21838.3 | controllerType=0 controllerNumber=1 scan=21838 | 43.4474 |
| 18261:4 | 1 | R.[+42.011]TAGWNIPMGLLFN[+1872.651]QTGSC[+57.021]KFDEYFSQSC[+57.021]A | HexNAc(3)Hex(6)NeuAc(1) | .[42], C[+57]*2, N[+1873] | 1464.1232 | 4 | 5853.4709 | 5853.4181 | 0 | 9 | 485 | Specific | 389.23 | 347.74 | 347.74 | 6.74 | 139 | >sp\|P02788\|TRFL_HUMAN Lactotransferrin OS=Homo sapiens OX=9606 GN=LTF PE=1 SV=6 | 1270 | 20201113_10633_1ng_glyco_hcd203040_compound15.21744.21744.4 | controllerType=0 controllerNumber=1 scan=21744 | 43.2661 |
| 18442:3 | 1 | R.TAGWNIPMGLLFN[+1216.423]QTGSC[+57.021]KFDEYFSQSC[+57.021]APGSDPR.S | HexNAc(2)Hex(5) | C[+57]*2, N[+1216] | 1719.0705 | 3 | 5155.197 | 5155.18 | 0 | 3.3 | 485 | Specific | 397.31 | 393.66 | 117.66 | 9.41 | 139 | >sp\|P02788\|TRFL_HUMAN Lactotransferrin OS=Homo sapiens OX=9606 GN=LTF PE=1 SV=6 | 1270 | 20201113_10633_1ng_glyco_hcd203040_compound15.21932.21932.3 | controllerType=0 controllerNumber=1 scan=21932 | 43.6548 |
| 18440:3 | 1 | R.TAGWNIPMGLLFN[+1216.423]QTGSC[+57.021]KFDEYFSQSC[+57.021]APGSDPR.S | HexNAc(2)Hex(5) | C[+57]*2, N[+1216] | 1719.0705 | 3 | 5155.197 | 5155.18 | 0 | 3.3 | 485 | Specific | 416.5 | 397.88 | 126.47 | 9.48 | 139 | >sp\|P02788\|TRFL_HUMAN Lactotransferrin OS=Homo sapiens OX=9606 GN=LTF PE=1 SV=6 | 1270 | 20201113_10633_1ng_glyco_hcd203040_compound15.21930.21930.3 | controllerType=0 controllerNumber=1 scan=21930 | 43.6513 |

Table S4. Cont.

| Query #:z | Protein Rank | Glycopeptide | Glycan | Modification Type(s) | Observed m/z | z | Observed (M+H) | Calc. mass (M+H) | Off-by-x error | Mass error (ppm) | StartPosition | Cleavage | Score | Delta | Delta Mod | \|Log Prob\| | # of unique peptides | Protein Name | Protein DB number | Comment | Scan # | Scan Time |
|---|---|---|---|---|---|---|---|---|---|---|---|---|---|---|---|---|---|---|---|---|---|---|
| 18350:3 | 1 | R.TAGWNIPMGLLFN[+1606.587]QTGSC[+57.021]KFDEYFSQSC[+57.021]APGSDPR.S | HexNAc(4)Hex(4)Fuc(1) | C[+57]*2, N[+1607] | 1849.1229 | 3 | 5545.3541 | 5545.3438 | 0 | 1.9 | 485 | Specific | 497.58 | 471.05 | 471.05 | 10.21 | 139 | >sp\|P02788\|TRFL_HUMAN Lactotransferrin OS=Homo sapiens OX=9606 GN=LTF PE=1 SV=6 | 1270 | 20201113_10633_1ng_glyco_hcd203040_compound15.21836.21836.3 | controllerType=0 controllerNumber=1 scan=21836 | 43.4439 |
| 19336:4 | 1 | R.TAGWNIPMGLLFN[+2205.793]QTGSC[+57.021]KFDEYFSQSC[+57.021]APGSDPR.S | HexNAc(4)Hex(5)Fuc(2)NeuAc(1) | C[+57]*2, N[+2206] | 1536.8978 | 4 | 6144.5695 | 6144.5499 | 0 | 3.2 | 485 | Specific | 535.9 | 526.3 | 526.3 | 9.96 | 139 | >sp\|P02788\|TRFL_HUMAN Lactotransferrin OS=Homo sapiens OX=9606 GN=LTF PE=1 SV=6 | 1270 | 20201113_10633_1ng_glyco_hcd203040_compound15.22862.22862.4 | controllerType=0 controllerNumber=1 scan=22862 | 45.5471 |
| 18306:5 | 1 | R.TAGWNIPMGLLFN[+1768.640]QTGSC[+57.021]KFDEYFSQSC[+57.021]APGSDPR.S | HexNAc(4)Hex(5)Fuc(1) | C[+57]*2, N[+1769] | 1142.2898 | 5 | 5707.4199 | 5707.3966 | 0 | 4.1 | 485 | Specific | 538.68 | 538.68 | 538.68 | 10.5 | 139 | >sp\|P02788\|TRFL_HUMAN Lactotransferrin OS=Homo sapiens OX=9606 GN=LTF PE=1 SV=6 | 1270 | 20201113_10633_1ng_glyco_hcd203040_compound15.21790.21790.5 | controllerType=0 controllerNumber=1 scan=21790 | 43.3532 |
| 18342:3 | 1 | R.TAGWNIPMGLLFN[+1622.582]QTGSC[+57.021]KFDEYFSQSC[+57.021]APGSDPR.S | HexNAc(4)Hex(5) | C[+57]*2, N[+1623] | 1854.455 | 3 | 5561.3506 | 5561.3387 | 0 | 2.1 | 485 | Specific | 547.19 | 464.58 | 168.85 | 10.45 | 139 | >sp\|P02788\|TRFL_HUMAN Lactotransferrin OS=Homo sapiens OX=9606 GN=LTF PE=1 SV=6 | 1270 | 20201113_10633_1ng_glyco_hcd203040_compound15.21828.21828.3 | controllerType=0 controllerNumber=1 scan=21828 | 43.4302 |
| 18345:3 | 1 | R.TAGWNIPMGLLFN[+1622.582]QTGSC[+57.021]KFDEYFSQSC[+57.021]APGSDPR.S | HexNAc(4)Hex(5) | C[+57]*2, N[+1623] | 1854.455 | 3 | 5561.3506 | 5561.3387 | 0 | 2.1 | 485 | Specific | 554.15 | 427.08 | 171.58 | 10.62 | 139 | >sp\|P02788\|TRFL_HUMAN Lactotransferrin OS=Homo sapiens OX=9606 GN=LTF PE=1 SV=6 | 1270 | 20201113_10633_1ng_glyco_hcd203040_compound15.21831.21831.3 | controllerType=0 controllerNumber=1 scan=21831 | 43.4343 |
| 18304:5 | 1 | R.TAGWNIPMGLLFN[+1768.640]QTGSC[+57.021]KFDEYFSQSC[+57.021]APGSDPR.S | HexNAc(4)Hex(5)Fuc(1) | C[+57]*2, N[+1769] | 1142.2898 | 5 | 5707.4199 | 5707.3966 | 0 | 4.1 | 485 | Specific | 556.27 | 556.27 | 556.27 | 10.67 | 139 | >sp\|P02788\|TRFL_HUMAN Lactotransferrin OS=Homo sapiens OX=9606 GN=LTF PE=1 SV=6 | 1270 | 20201113_10633_1ng_glyco_hcd203040_compound15.21788.21788.5 | controllerType=0 controllerNumber=1 scan=21788 | 43.3494 |
| 18334:4 | 1 | R.TAGWNIPMGLLFN[+1622.582]QTGSC[+57.021]KFDEYFSQSC[+57.021]APGSDPR.S | HexNAc(4)Hex(5) | C[+57]*2, N[+1623] | 1391.0945 | 4 | 5561.3561 | 5561.3387 | 0 | 3.1 | 485 | Specific | 612.16 | 555.02 | 164.33 | 12.37 | 139 | >sp\|P02788\|TRFL_HUMAN Lactotransferrin OS=Homo sapiens OX=9606 GN=LTF PE=1 SV=6 | 1270 | 20201113_10633_1ng_glyco_hcd203040_compound15.21820.21820.4 | controllerType=0 controllerNumber=1 scan=21820 | 43.4138 |
| 20786:4 | 1 | R.TAGWNIPMGLLFN[+2350.830]QTGSC[+57.021]KFDEYFSQSC[+57.021]APGSDPR.S | HexNAc(4)Hex(5)Fuc(1)NeuAc(2) | C[+57]*2, N[+2351] | 1573.1498 | 4 | 6289.5773 | 6289.5875 | 0 | -1.6 | 485 | Specific | 739.93 | 654.53 | 654.53 | 13.89 | 139 | >sp\|P02788\|TRFL_HUMAN Lactotransferrin OS=Homo sapiens OX=9606 GN=LTF PE=1 SV=6 | 1270 | 20201113_10633_1ng_glyco_hcd203040_compound15.24374.24374.4 | controllerType=0 controllerNumber=1 scan=24374 | 48.4015 |
| 18317:4 | 1 | R.TAGWNIPMGLLFN[+1606.587]QTGSC[+57.021]KFDEYFSQSC[+57.021]APGSDPR.S | HexNAc(4)Hex(4)Fuc(1) | C[+57]*2, N[+1607] | 1387.1024 | 4 | 5545.3879 | 5545.3438 | 0 | 7.9 | 485 | Specific | 750.26 | 750.26 | 750.26 | 14.54 | 139 | >sp\|P02788\|TRFL_HUMAN Lactotransferrin OS=Homo sapiens OX=9606 GN=LTF PE=1 SV=6 | 1270 | 20201113_10633_1ng_glyco_hcd203040_compound15.21802.21802.4 | controllerType=0 controllerNumber=1 scan=21802 | 43.3773 |
| 18270:3 | 1 | R.TAGWNIPMGLLFN[+1768.640]QTGSC[+57.021]KFDEYFSQSC[+57.021]APGSDPR.S | HexNAc(4)Hex(5)Fuc(1) | C[+57]*2, N[+1769] | 1903.1417 | 3 | 5707.4105 | 5707.3966 | 0 | 2.4 | 485 | Specific | 775.24 | 632.89 | 632.89 | 15.86 | 139 | >sp\|P02788\|TRFL_HUMAN Lactotransferrin OS=Homo sapiens OX=9606 GN=LTF PE=1 SV=6 | 1270 | 20201113_10633_1ng_glyco_hcd203040_compound15.21753.21753.3 | controllerType=0 controllerNumber=1 scan=21753 | 43.2819 |
| 18336:4 | 1 | R.TAGWNIPMGLLFN[+1622.582]QTGSC[+57.021]KFDEYFSQSC[+57.021]APGSDPR.S | HexNAc(4)Hex(5) | C[+57]*2, N[+1623] | 1391.0945 | 4 | 5561.3561 | 5561.3387 | 0 | 3.1 | 485 | Specific | 864.26 | 740.05 | 295.04 | 17.37 | 139 | >sp\|P02788\|TRFL_HUMAN Lactotransferrin OS=Homo sapiens OX=9606 GN=LTF PE=1 SV=6 | 1270 | 20201113_10633_1ng_glyco_hcd203040_compound15.21822.21822.4 | controllerType=0 controllerNumber=1 scan=21822 | 43.4183 |
| 20788:4 | 1 | R.TAGWNIPMGLLFN[+2350.830]QTGSC[+57.021]KFDEYFSQSC[+57.021]APGSDPR.S | HexNAc(4)Hex(5)Fuc(1)NeuAc(2) | C[+57]*2, N[+2351] | 1573.1498 | 4 | 6289.5773 | 6289.5875 | 0 | -1.6 | 485 | Specific | 876.61 | 815.71 | 815.71 | 16.89 | 139 | >sp\|P02788\|TRFL_HUMAN Lactotransferrin OS=Homo sapiens OX=9606 GN=LTF PE=1 SV=6 | 1270 | 20201113_10633_1ng_glyco_hcd203040_compound15.24376.24376.4 | controllerType=0 controllerNumber=1 scan=24376 | 48.4063 |
| 18268:3 | 1 | R.TAGWNIPMGLLFN[+1768.640]QTGSC[+57.021]KFDEYFSQSC[+57.021]APGSDPR.S | HexNAc(4)Hex(5)Fuc(1) | C[+57]*2, N[+1769] | 1903.1417 | 3 | 5707.4105 | 5707.3966 | 0 | 2.4 | 485 | Specific | 876.64 | 774.94 | 774.94 | 17.86 | 139 | >sp\|P02788\|TRFL_HUMAN Lactotransferrin OS=Homo sapiens OX=9606 GN=LTF PE=1 SV=6 | 1270 | 20201113_10633_1ng_glyco_hcd203040_compound15.21751.21751.3 | controllerType=0 controllerNumber=1 scan=21751 | 43.2775 |
| 19339:4 | 1 | R.TAGWNIPMGLLFN[+2205.793]QTGSC[+57.021]KFDEYFSQSC[+57.021]APGSDPR.S | HexNAc(4)Hex(5)Fuc(2)NeuAc(1) | C[+57]*2, N[+2206] | 1536.8978 | 4 | 6144.5695 | 6144.5499 | 0 | 3.2 | 485 | Specific | 898.89 | 852.29 | 852.29 | 17.37 | 139 | >sp\|P02788\|TRFL_HUMAN Lactotransferrin OS=Homo sapiens OX=9606 GN=LTF PE=1 SV=6 | 1270 | 20201113_10633_1ng_glyco_hcd203040_compound15.22865.22865.4 | controllerType=0 controllerNumber=1 scan=22865 | 45.5513 |
| 18320:4 | 1 | R.TAGWNIPMGLLFN[+1606.587]QTGSC[+57.021]KFDEYFSQSC[+57.021]APGSDPR.S | HexNAc(4)Hex(4)Fuc(1) | C[+57]*2, N[+1607] | 1387.1024 | 4 | 5545.3879 | 5545.3438 | 0 | 7.9 | 485 | Specific | 926 | 869.48 | 869.48 | 17.54 | 139 | >sp\|P02788\|TRFL_HUMAN Lactotransferrin OS=Homo sapiens OX=9606 GN=LTF PE=1 SV=6 | 1270 | 20201113_10633_1ng_glyco_hcd203040_compound15.21805.21805.4 | controllerType=0 controllerNumber=1 scan=21805 | 43.3827 |
| 19403:4 | 1 | R.TAGWNIPMGLLFN[+1913.677]QTGSC[+57.021]KFDEYFSQSC[+57.021]APGSDPR.S | HexNAc(4)Hex(5)NeuAc(1) | C[+57]*2, N[+1914] | 1463.8699 | 4 | 5852.4577 | 5852.4341 | 0 | 4 | 485 | Specific | 936.74 | 891.16 | 891.16 | 18.37 | 139 | >sp\|P02788\|TRFL_HUMAN Lactotransferrin OS=Homo sapiens OX=9606 GN=LTF PE=1 SV=6 | 1270 | 20201113_10633_1ng_glyco_hcd203040_compound15.22932.22932.4 | controllerType=0 controllerNumber=1 scan=22932 | 45.7035 |
| 19334:4 | 1 | R.TAGWNIPMGLLFN[+2059.735]QTGSC[+57.021]KFDEYFSQSC[+57.021]APGSDPR.S | HexNAc(4)Hex(5)Fuc(1)NeuAc(1) | C[+57]*2, N[+2060] | 1500.3862 | 4 | 5998.5231 | 5998.492 | 0 | 5.2 | 485 | Specific | 948.64 | 922.82 | 922.82 | 17.88 | 139 | >sp\|P02788\|TRFL_HUMAN Lactotransferrin OS=Homo sapiens OX=9606 GN=LTF PE=1 SV=6 | 1270 | 20201113_10633_1ng_glyco_hcd203040_compound15.22860.22860.4 | controllerType=0 controllerNumber=1 scan=22860 | 45.5445 |
| 19401:4 | 1 | R.TAGWNIPMGLLFN[+1913.677]QTGSC[+57.021]KFDEYFSQSC[+57.021]APGSDPR.S | HexNAc(4)Hex(5)NeuAc(1) | C[+57]*2, N[+1914] | 1463.8699 | 4 | 5852.4577 | 5852.4341 | 0 | 4 | 485 | Specific | 957.45 | 864.27 | 864.27 | 19.37 | 139 | >sp\|P02788\|TRFL_HUMAN Lactotransferrin OS=Homo sapiens OX=9606 GN=LTF PE=1 SV=6 | 1270 | 20201113_10633_1ng_glyco_hcd203040_compound15.22930.22930.4 | controllerType=0 controllerNumber=1 scan=22930 | 45.6962 |
| 18260:4 | 1 | R.TAGWNIPMGLLFN[+1768.640]QTGSC[+57.021]KFDEYFSQSC[+57.021]APGSDPR.S | HexNAc(4)Hex(5)Fuc(1) | C[+57]*2, N[+1769] | 1427.6093 | 4 | 5707.4152 | 5707.3966 | 0 | 3.3 | 485 | Specific | 998.37 | 952.69 | 952.69 | 19.37 | 139 | >sp\|P02788\|TRFL_HUMAN Lactotransferrin OS=Homo sapiens OX=9606 GN=LTF PE=1 SV=6 | 1270 | 20201113_10633_1ng_glyco_hcd203040_compound15.21743.21743.4 | controllerType=0 controllerNumber=1 scan=21743 | 43.265 |
| 19332:4 | 1 | R.TAGWNIPMGLLFN[+2059.735]QTGSC[+57.021]KFDEYFSQSC[+57.021]APGSDPR.S | HexNAc(4)Hex(5)Fuc(1)NeuAc(1) | C[+57]*2, N[+2060] | 1500.3862 | 4 | 5998.5231 | 5998.492 | 0 | 5.2 | 485 | Specific | 1004.99 | 956.58 | 956.58 | 19.88 | 139 | >sp\|P02788\|TRFL_HUMAN Lactotransferrin OS=Homo sapiens OX=9606 GN=LTF PE=1 SV=6 | 1270 | 20201113_10633_1ng_glyco_hcd203040_compound15.22858.22858.4 | controllerType=0 controllerNumber=1 scan=22858 | 45.5386 |
| 18258:4 | 1 | R.TAGWNIPMGLLFN[+1768.640]QTGSC[+57.021]KFDEYFSQSC[+57.021]APGSDPR.S | HexNAc(4)Hex(5)Fuc(1) | C[+57]*2, N[+1769] | 1427.6093 | 4 | 5707.4152 | 5707.3966 | 0 | 3.3 | 485 | Specific | 1026.42 | 973.82 | 973.82 | 20.37 | 139 | >sp\|P02788\|TRFL_HUMAN Lactotransferrin OS=Homo sapiens OX=9606 GN=LTF PE=1 SV=6 | 1270 | 20201113_10633_1ng_glyco_hcd203040_compound15.21741.21741.4 | controllerType=0 controllerNumber=1 scan=21741 | 43.2582 |
| 09372:3 | 1 | R.N[+2059.735]GSDC[+57.021]PDKFC[+57.021]LFQSETK.N | HexNAc(4)Hex(5)Fuc(1)NeuAc(1) | C[+57]*2, N[+2060] | 1364.8785 | 3 | 4092.6209 | 4092.6088 | 0 | 2.9 | 642 | Specific | 210.81 | 100.61 | 100.61 | 3.62 | 139 | >sp\|P02788\|TRFL_HUMAN Lactotransferrin OS=Homo sapiens OX=9606 GN=LTF PE=1 SV=6 | 1270 | 20201113_10633_1ng_glyco_hcd203040_compound15.12437.12437.3 | controllerType=0 controllerNumber=1 scan=12437 | 26.9036 |
| 09370:3 | 1 | R.N[+2059.735]GSDC[+57.021]PDKFC[+57.021]LFQSETK.N | HexNAc(4)Hex(5)Fuc(1)NeuAc(1) | C[+57]*2, N[+2060] | 1364.8785 | 3 | 4092.6209 | 4092.6088 | 0 | 2.9 | 642 | Specific | 326.53 | 210.23 | 210.23 | 6.67 | 139 | >sp\|P02788\|TRFL_HUMAN Lactotransferrin OS=Homo sapiens OX=9606 GN=LTF PE=1 SV=6 | 1270 | 20201113_10633_1ng_glyco_hcd203040_compound15.12435.12435.3 | controllerType=0 controllerNumber=1 scan=12435 | 26.8995 |
| 09216:3 | 1 | R.N[+2205.793]GSDC[+57.021]PDKFC[+57.021]LFQSETK.N | HexNAc(4)Hex(5)Fuc(2)NeuAc(1) | C[+57]*2, N[+2206] | 1413.5613 | 3 | 4238.6695 | 4238.6667 | 0 | 0.6 | 642 | Specific | 396.63 | 242.32 | 242.32 | 6.75 | 139 | >sp\|P02788\|TRFL_HUMAN Lactotransferrin OS=Homo sapiens OX=9606 GN=LTF PE=1 SV=6 | 1270 | 20201113_10633_1ng_glyco_hcd203040_compound15.12274.12274.3 | controllerType=0 controllerNumber=1 scan=12274 | 26.6109 |
| 09213:3 | 1 | R.N[+2205.793]GSDC[+57.021]PDKFC[+57.021]LFQSETK.N | HexNAc(4)Hex(5)Fuc(2)NeuAc(1) | C[+57]*2, N[+2206] | 1413.5613 | 3 | 4238.6695 | 4238.6667 | 0 | 0.6 | 642 | Specific | 448.39 | 382.48 | 382.48 | 9.28 | 139 | >sp\|P02788\|TRFL_HUMAN Lactotransferrin OS=Homo sapiens OX=9606 GN=LTF PE=1 SV=6 | 1270 | 20201113_10633_1ng_glyco_hcd203040_compound15.12271.12271.3 | controllerType=0 controllerNumber=1 scan=12271 | 26.6061 |
| 13255:4 | 6 | K.KGN[+2205.793]TTLNSFVIPSESDVPTHC[+57.021]PSQWWPYAGHC[+57.021]YK.I | HexNAc(4)Hex(5)Fuc(2)NeuAc(1) | C[+57]*2, N[+2206] | 1543.1597 | 4 | 6169.6169 | 6169.6146 | 0 | 0.4 | 342 | Specific | 369.64 | 369.64 | 369.64 | 7.76 | 81 | >sp\|P22897\|MRC1_HUMAN Macrophage mannose receptor 1 OS=Homo sapiens OX=9606 GN=MRC1 PE=1 SV=1 | 1049 | 20201113_10633_1ng_glyco_hcd203040_compound15.16498.16498.2 | controllerType=0 controllerNumber=1 scan=16498 | 33.7992 |
| 13253:4 | 6 | K.KGN[+2205.793]TTLNSFVIPSESDVPTHC[+57.021]PSQWWPYAGHC[+57.021]YK.I | HexNAc(4)Hex(5)Fuc(2)NeuAc(1) | C[+57]*2, N[+2206] | 1543.1597 | 4 | 6169.6169 | 6169.6146 | 0 | 0.4 | 342 | Specific | 389.33 | 389.33 | 389.33 | 9.15 | 81 | >sp\|P22897\|MRC1_HUMAN Macrophage mannose receptor 1 OS=Homo sapiens OX=9606 GN=MRC1 PE=1 SV=1 | 1049 | 20201113_10633_1ng_glyco_hcd203040_compound15.16496.16496.2 | controllerType=0 controllerNumber=1 scan=16496 | 33.7955 |
| 17470:3 | 8 | K.RNIAAFGGDPNN[+1913.677]ITLFGESAGGASVSLQTLSPYNK.G | HexNAc(4)Hex(5)NeuAc(1) | N[+1914] | 1827.5001 | 3 | 5480.4857 | 5480.4644 | 0 | 3.9 | 196 | Specific | 215.7 | 143.79 | 143.79 | 4.83 | 61 | >sp\|P19835\|CEL_HUMAN Bile salt-activated lipase OS=Homo sapiens OX=9606 GN=CEL PE=1 SV=3 | 476 | 20201113_10633_1ng_glyco_hcd203040_compound15.20920.20920.3 | controllerType=0 controllerNumber=1 scan=20920 | 41.5518 |
| 17472:3 | 8 | K.RNIAAFGGDPNN[+1913.677]ITLFGESAGGASVSLQTLSPYNK.G | HexNAc(4)Hex(5)NeuAc(1) | N[+1914] | 1827.5001 | 3 | 5480.4857 | 5480.4644 | 0 | 3.9 | 196 | Specific | 244.01 | 68.72 | 68.72 | 5.16 | 61 | >sp\|P19835\|CEL_HUMAN Bile salt-activated lipase OS=Homo sapiens OX=9606 GN=CEL PE=1 SV=3 | 476 | 20201113_10633_1ng_glyco_hcd203040_compound15.20922.20922.3 | controllerType=0 controllerNumber=1 scan=20922 | 41.5559 |
| 19309:4 | 8 | R.NIAAFGGDPNN[+1913.677]ITLFGESAGGASVSLQTLSPYNK.G | HexNAc(4)Hex(5)NeuAc(1) | N[+1914] | 1331.8505 | 4 | 5324.38 | 5324.3633 | 0 | 3.1 | 197 | Specific | 201.04 | 201.04 | 201.04 | 5.52 | 61 | >sp\|P19835\|CEL_HUMAN Bile salt-activated lipase OS=Homo sapiens OX=9606 GN=CEL PE=1 SV=3 | 476 | 20201113_10633_1ng_glyco_hcd203040_compound15.22834.22834.4 | controllerType=0 controllerNumber=1 scan=22834 | 45.4929 |
| 19312:4 | 8 | R.NIAAFGGDPNN[+1913.677]ITLFGESAGGASVSLQTLSPYNK.G | HexNAc(4)Hex(5)NeuAc(1) | N[+1914] | 1331.8505 | 4 | 5324.38 | 5324.3633 | 0 | 3.1 | 197 | Specific | 841.12 | 739.46 | 739.46 | 17.01 | 61 | >sp\|P19835\|CEL_HUMAN Bile salt-activated lipase OS=Homo sapiens OX=9606 GN=CEL PE=1 SV=3 | 476 | 20201113_10633_1ng_glyco_hcd203040_compound15.22837.22837.4 | controllerType=0 controllerNumber=1 scan=22837 | 45.4963 |
| 18011:3 | 9 | R.GLSFDVSLEVSQGPGLLN[+2059.735]DTK.V | HexNAc(4)Hex(5)Fuc(1)NeuAc(1) | N[+2060] | 1412.6236 | 3 | 4235.8562 | 4235.8481 | 0 | 1.9 | 118 | Specific | 871.07 | 810.57 | 810.57 | 17.67 | 60 | >sp\|P01833\|PIGR_HUMAN Polymeric immunoglobulin receptor OS=Homo sapiens OX=9606 GN=PIGR PE=1 SV=4 | 902 | 20201113_10633_1ng_glyco_hcd203040_compound15.21483.21483.3 | controllerType=0 controllerNumber=1 scan=21483 | 42.7216 |
| 18013:3 | 9 | R.GLSFDVSLEVSQGPGLLN[+2059.735]DTK.V | HexNAc(4)Hex(5)Fuc(1)NeuAc(1) | N[+2060] | 1412.6236 | 3 | 4235.8562 | 4235.8481 | 0 | 1.9 | 118 | Specific | 1000.56 | 882.2 | 882.2 | 20.67 | 60 | >sp\|P01833\|PIGR_HUMAN Polymeric immunoglobulin receptor OS=Homo sapiens OX=9606 GN=PIGR PE=1 SV=4 | 902 | 20201113_10633_1ng_glyco_hcd203040_compound15.21485.21485.3 | controllerType=0 controllerNumber=1 scan=21485 | 42.7265 |
| 19304:3 | 9 | K.QIGLYPVLVIDSSGYVNPN[+1913.677]YTGR.I | HexNAc(4)Hex(5)NeuAc(1) | N[+1914] | 1480.3343 | 3 | 4438.9883 | 4438.9805 | 0 | 1.8 | 168 | Specific | 502.98 | 502.98 | 502.98 | 10.09 | 60 | >sp\|P01833\|PIGR_HUMAN Polymeric immunoglobulin receptor OS=Homo sapiens OX=9606 GN=PIGR PE=1 SV=4 | 902 | 20201113_10633_1ng_glyco_hcd203040_compound15.22828.22828.3 | controllerType=0 controllerNumber=1 scan=22828 | 45.4799 |
| 19305:3 | 9 | K.QIGLYPVLVIDSSGYVNPN[+1913.677]YTGR.I | HexNAc(4)Hex(5)NeuAc(1) | N[+1914] | 1480.3343 | 3 | 4438.9883 | 4438.9805 | 0 | 1.8 | 168 | Specific | 577.27 | 415.14 | 415.14 | 11.66 | 60 | >sp\|P01833\|PIGR_HUMAN Polymeric immunoglobulin receptor OS=Homo sapiens OX=9606 GN=PIGR PE=1 SV=4 | 902 | 20201113_10633_1ng_glyco_hcd203040_compound15.22829.22829.3 | controllerType=0 controllerNumber=1 scan=22829 | 45.4836 |
| 19235:3 | 9 | K.QIGLYPVLVIDSSGYVNPN[+2059.735]YTGR.I | HexNAc(4)Hex(5)Fuc(1)NeuAc(1) | N[+2060] | 1529.0208 | 3 | 4585.0479 | 4585.0384 | 0 | 2.1 | 168 | Specific | 697.36 | 593.93 | 593.93 | 13.67 | 60 | >sp\|P01833\|PIGR_HUMAN Polymeric immunoglobulin receptor OS=Homo sapiens OX=9606 GN=PIGR PE=1 SV=4 | 902 | 20201113_10633_1ng_glyco_hcd203040_compound15.22757.22757.3 | controllerType=0 controllerNumber=1 scan=22757 | 45.3446 |
| 19238:3 | 9 | K.QIGLYPVLVIDSSGYVNPN[+2059.735]YTGR.I | HexNAc(4)Hex(5)Fuc(1)NeuAc(1) | N[+2060] | 1529.0208 | 3 | 4585.0479 | 4585.0384 | 0 | 2.1 | 168 | Specific | 801.09 | 642.02 | 642.02 | 16.67 | 60 | >sp\|P01833\|PIGR_HUMAN Polymeric immunoglobulin receptor OS=Homo sapiens OX=9606 GN=PIGR PE=1 SV=4 | 902 | 20201113_10633_1ng_glyco_hcd203040_compound15.22760.22760.3 | controllerType=0 controllerNumber=1 scan=22760 | 45.3489 |
| 21034:4 | 9 | R.LSLLEEPGN[+2076.750]GTFTVILNQLTSR.D | HexNAc(4)Hex(6)Fuc(2) | N[+2077] | 1120.5262 | 4 | 4479.0828 | 4479.0428 | 0 | 8.9 | 413 | Specific | 293.44 | 237.3 | 237.3 | 4.53 | 60 | >sp\|P01833\|PIGR_HUMAN Polymeric immunoglobulin receptor OS=Homo sapiens OX=9606 GN=PIGR PE=1 SV=4 | 902 | 20201113_10633_1ng_glyco_hcd203040_compound15.24632.24632.4 | controllerType=0 controllerNumber=1 scan=24632 | 48.9376 |
| 21020:3 | 9 | R.LSLLEEPGN[+2059.735]GTFTVILNQLTSR.D | HexNAc(4)Hex(5)Fuc(1)NeuAc(1) | N[+2060] | 1488.025 | 3 | 4462.0603 | 4462.0275 | 0 | 7.4 | 413 | Specific | 955.38 | 745.67 | 745.67 | 18.51 | 60 | >sp\|P01833\|PIGR_HUMAN Polymeric immunoglobulin receptor OS=Homo sapiens OX=9606 GN=PIGR PE=1 SV=4 | 902 | 20201113_10633_1ng_glyco_hcd203040_compound15.24618.24618.3 | controllerType=0 controllerNumber=1 scan=24618 | 48.9105 |
| 21026:4 | 9 | R.LSLLEEPGN[+2059.735]GTFTVILNQLTSR.D | HexNAc(4)Hex(5)Fuc(1)NeuAc(1) | N[+2060] | 1116.2696 | 4 | 4462.0565 | 4462.0275 | 0 | 6.5 | 413 | Specific | 963.53 | 794.73 | 794.73 | 19.43 | 60 | >sp\|P01833\|PIGR_HUMAN Polymeric immunoglobulin receptor OS=Homo sapiens OX=9606 GN=PIGR PE=1 SV=4 | 902 | 20201113_10633_1ng_glyco_hcd203040_compound15.24624.24624.4 | controllerType=0 controllerNumber=1 scan=24624 | 48.9201 |
| 21029:4 | 9 | R.LSLLEEPGN[+2059.735]GTFTVILNQLTSR.D | HexNAc(4)Hex(5)Fuc(1)NeuAc(1) | N[+2060] | 1116.2696 | 4 | 4462.0565 | 4462.0275 | 0 | 6.5 | 413 | Specific | 978.62 | 712.83 | 712.83 | 19.43 | 60 | >sp\|P01833\|PIGR_HUMAN Polymeric immunoglobulin receptor OS=Homo sapiens OX=9606 GN=PIGR PE=1 SV=4 | 902 | 20201113_10633_1ng_glyco_hcd203040_compound15.24627.24627.4 | controllerType=0 controllerNumber=1 scan=24627 | 48.9271 |
| 23386:4 | 9 | R.LSLLEEPGN[+2350.830]GTFTVILNQLTSR.D | HexNAc(4)Hex(5)Fuc(1)NeuAc(2) | N[+2351] | 1189.04 | 4 | 4753.138 | 4753.1229 | 0 | 3.2 | 413 | Specific | 985.42 | 763.61 | 763.61 | 19.88 | 60 | >sp\|P01833\|PIGR_HUMAN Polymeric immunoglobulin receptor OS=Homo sapiens OX=9606 GN=PIGR PE=1 SV=4 | 902 | 20201113_10633_1ng_glyco_hcd203040_compound15.27087.27087.4 | controllerType=0 controllerNumber=1 scan=27087 | 53.1982 |
| 21022:3 | 9 | R.LSLLEEPGN[+2059.735]GTFTVILNQLTSR.D | HexNAc(4)Hex(5)Fuc(1)NeuAc(1) | N[+2060] | 1488.025 | 3 | 4462.0603 | 4462.0275 | 0 | 7.4 | 413 | Specific | 988.71 | 718.41 | 718.41 | 18.51 | 60 | >sp\|P01833\|PIGR_HUMAN Polymeric immunoglobulin receptor OS=Homo sapiens OX=9606 GN=PIGR PE=1 SV=4 | 902 | 20201113_10633_1ng_glyco_hcd203040_compound15.24620.24620.3 | controllerType=0 controllerNumber=1 scan=24620 | 48.9154 |
| 23384:4 | 9 | R.LSLLEEPGN[+2350.830]GTFTVILNQLTSR.D | HexNAc(4)Hex(5)Fuc(1)NeuAc(2) | N[+2351] | 1189.04 | 4 | 4753.138 | 4753.1229 | 0 | 3.2 | 413 | Specific | 1030.1 | 810.39 | 810.39 | 20.88 | 60 | >sp\|P01833\|PIGR_HUMAN Polymeric immunoglobulin receptor OS=Homo sapiens OX=9606 GN=PIGR PE=1 SV=4 | 902 | 20201113_10633_1ng_glyco_hcd203040_compound15.27085.27085.4 | controllerType=0 controllerNumber=1 scan=27085 | 53.1932 |
| 18307:3 | 9 | K.IIEGEPNLKVPGN[+2350.830]VTAVLGETLK.V | HexNAc(4)Hex(5)Fuc(1)NeuAc(2) | N[+2351] | 1581.4039 | 3 | 4742.197 | 4742.1797 | 0 | 3.7 | 457 | Specific | 250.5 | 250.5 | 250.5 | 6.09 | 60 | >sp\|P01833\|PIGR_HUMAN Polymeric immunoglobulin receptor OS=Homo sapiens OX=9606 GN=PIGR PE=1 SV=4 | 902 | 20201113_10633_1ng_glyco_hcd203040_compound15.21791.21791.2 | controllerType=0 controllerNumber=1 scan=21791 | 43.3559 |
| 18305:3 | 9 | K.IIEGEPNLKVPGN[+2350.830]VTAVLGETLK.V | HexNAc(4)Hex(5)Fuc(1)NeuAc(2) | N[+2351] | 1581.4039 | 3 | 4742.197 | 4742.1797 | 0 | 3.7 | 457 | Specific | 379.06 | 301.23 | 301.23 | 7.09 | 60 | >sp\|P01833\|PIGR_HUMAN Polymeric immunoglobulin receptor OS=Homo sapiens OX=9606 GN=PIGR PE=1 SV=4 | 902 | 20201113_10633_1ng_glyco_hcd203040_compound15.21789.21789.2 | controllerType=0 controllerNumber=1 scan=21789 | 43.3509 |
| 16385:3 | 9 | K.IIEGEPNLKVPGN[+2059.735]VTAVLGETLK.V | HexNAc(4)Hex(5)Fuc(1)NeuAc(1) | N[+2060] | 1484.3701 | 3 | 4451.0956 | 4451.0843 | 0 | 2.5 | 457 | Specific | 800.59 | 702.28 | 702.28 | 16.49 | 60 | >sp\|P01833\|PIGR_HUMAN Polymeric immunoglobulin receptor OS=Homo sapiens OX=9606 GN=PIGR PE=1 SV=4 | 902 | 20201113_10633_1ng_glyco_hcd203040_compound15.19789.19789.3 | controllerType=0 controllerNumber=1 scan=19789 | 39.5003 |
| 16382:3 | 9 | K.IIEGEPNLKVPGN[+2059.735]VTAVLGETLK.V | HexNAc(4)Hex(5)Fuc(1)NeuAc(1) | N[+2060] | 1484.3701 | 3 | 4451.0956 | 4451.0843 | 0 | 2.5 | 457 | Specific | 811.33 | 759.71 | 759.71 | 16.49 | 60 | >sp\|P01833\|PIGR_HUMAN Polymeric immunoglobulin receptor OS=Homo sapiens OX=9606 GN=PIGR PE=1 SV=4 | 902 | 20201113_10633_1ng_glyco_hcd203040_compound15.19786.19786.3 | controllerType=0 controllerNumber=1 scan=19786 | 39.4936 |
| 16314:3 | 9 | K.IIEGEPNLKVPGN[+2205.793]VTAVLGETLK.V | HexNAc(4)Hex(5)Fuc(2)NeuAc(1) | N[+2206] | 1533.0544 | 3 | 4597.1486 | 4597.1422 | 0 | 1.4 | 457 | Specific | 832.47 | 696.48 | 696.48 | 16.47 | 60 | >sp\|P01833\|PIGR_HUMAN Polymeric immunoglobulin receptor OS=Homo sapiens OX=9606 GN=PIGR PE=1 SV=4 | 902 | 20201113_10633_1ng_glyco_hcd203040_compound15.19712.19712.3 | controllerType=0 controllerNumber=1 scan=19712 | 39.3636 |
| 16312:3 | 9 | K.IIEGEPNLKVPGN[+2205.793]VTAVLGETLK.V | HexNAc(4)Hex(5)Fuc(2)NeuAc(1) | N[+2206] | 1533.0544 | 3 | 4597.1486 | 4597.1422 | 0 | 1.4 | 457 | Specific | 843.52 | 739.67 | 739.67 | 16.47 | 60 | >sp\|P01833\|PIGR_HUMAN Polymeric immunoglobulin receptor OS=Homo sapiens OX=9606 GN=PIGR PE=1 SV=4 | 902 | 20201113_10633_1ng_glyco_hcd203040_compound15.19710.19710.3 | controllerType=0 controllerNumber=1 scan=19710 | 39.3585 |
| 15966:4 | 9 | K.IIEGEPNLKVPGN[+2205.793]VTAVLGETLKVPC[+57.021]HFPC[+57.021]K.F | HexNAc(4)Hex(5)Fuc(2)NeuAc(1) | C[+57]*2, N[+2206] | 1406.4082 | 4 | 5622.611 | 5622.5997 | 0 | 2 | 457 | Specific | 265.01 | 265.01 | 265.01 | 6.26 | 60 | >sp\|P01833\|PIGR_HUMAN Polymeric immunoglobulin receptor OS=Homo sapiens OX=9606 GN=PIGR PE=1 SV=4 | 902 | 20201113_10633_1ng_glyco_hcd203040_compound15.19305.19305.2 | controllerType=0 controllerNumber=1 scan=19305 | 38.5913 |
| 15964:4 | 9 | K.IIEGEPNLKVPGN[+2205.793]VTAVLGETLKVPC[+57.021]HFPC[+57.021]K.F | HexNAc(4)Hex(5)Fuc(2)NeuAc(1) | C[+57]*2, N[+2206] | 1406.4082 | 4 | 5622.611 | 5622.5997 | 0 | 2 | 457 | Specific | 555.23 | 555.23 | 555.23 | 9.79 | 60 | >sp\|P01833\|PIGR_HUMAN Polymeric immunoglobulin receptor OS=Homo sapiens OX=9606 GN=PIGR PE=1 SV=4 | 902 | 20201113_10633_1ng_glyco_hcd203040_compound15.19303.19303.2 | controllerType=0 controllerNumber=1 scan=19303 | 38.5858 |
| 17444:5 | 9 | K.IIEGEPNLKVPGN[+2350.830]VTAVLGETLKVPC[+57.021]HFPC[+57.021]K.F | HexNAc(4)Hex(5)Fuc(1)NeuAc(2) | C[+57]*2, N[+2351] | 1154.3373 | 5 | 5767.6573 | 5767.6372 | 0 | 3.5 | 457 | Specific | 592.12 | 490.57 | 490.57 | 11.17 | 60 | >sp\|P01833\|PIGR_HUMAN Polymeric immunoglobulin receptor OS=Homo sapiens OX=9606 GN=PIGR PE=1 SV=4 | 902 | 20201113_10633_1ng_glyco_hcd203040_compound15.20893.20893.5 | controllerType=0 controllerNumber=1 scan=20893 | 41.4906 |
| 15965:4 | 9 | K.IIEGEPNLKVPGN[+2205.793]VTAVLGETLKVPC[+57.021]HFPC[+57.021]K.F | HexNAc(4)Hex(5)Fuc(2)NeuAc(1) | C[+57]*2, N[+2206] | 1406.4082 | 4 | 5622.611 | 5622.5997 | 0 | 2 | 457 | Specific | 681.8 | 587.97 | 587.97 | 13.03 | 60 | >sp\|P01833\|PIGR_HUMAN Polymeric immunoglobulin receptor OS=Homo sapiens OX=9606 GN=PIGR PE=1 SV=4 | 902 | 20201113_10633_1ng_glyco_hcd203040_compound15.19304.19304.3 | controllerType=0 controllerNumber=1 scan=19304 | 38.5884 |
| 15951:4 | 9 | K.IIEGEPNLKVPGN[+2205.793]VTAVLGETLKVPC[+57.021]HFPC[+57.021]K.F | HexNAc(4)Hex(5)Fuc(2)NeuAc(1) | C[+57]*2, N[+2206] | 1406.4085 | 4 | 5622.612 | 5622.5997 | 0 | 2.2 | 457 | Specific | 731.18 | 731.18 | 731.18 | 14.03 | 60 | >sp\|P01833\|PIGR_HUMAN Polymeric immunoglobulin receptor OS=Homo sapiens OX=9606 GN=PIGR PE=1 SV=4 | 902 | 20201113_10633_1ng_glyco_hcd203040_compound15.19289.19289.2 | controllerType=0 controllerNumber=1 scan=19289 | 38.5612 |
| 15962:4 | 9 | K.IIEGEPNLKVPGN[+2205.793]VTAVLGETLKVPC[+57.021]HFPC[+57.021]K.F | HexNAc(4)Hex(5)Fuc(2)NeuAc(1) | C[+57]*2, N[+2206] | 1406.4082 | 4 | 5622.611 | 5622.5997 | 0 | 2 | 457 | Specific | 737.53 | 674.47 | 674.47 | 14.03 | 60 | >sp\|P01833\|PIGR_HUMAN Polymeric immunoglobulin receptor OS=Homo sapiens OX=9606 GN=PIGR PE=1 SV=4 | 902 | 20201113_10633_1ng_glyco_hcd203040_compound15.19301.19301.3 | controllerType=0 controllerNumber=1 scan=19301 | 38.5835 |
| 17383:5 | 9 | K.IIEGEPNLKVPGN[+2350.830]VTAVLGETLKVPC[+57.021]HFPC[+57.021]K.F | HexNAc(4)Hex(5)Fuc(1)NeuAc(2) | C[+57]*2, N[+2351] | 1154.3346 | 5 | 5767.6439 | 5767.6372 | 0 | 1.2 | 457 | Specific | 780.27 | 557.61 | 557.61 | 15.49 | 60 | >sp\|P01833\|PIGR_HUMAN Polymeric immunoglobulin receptor OS=Homo sapiens OX=9606 GN=PIGR PE=1 SV=4 | 902 | 20201113_10633_1ng_glyco_hcd203040_compound15.20829.20829.5 | controllerType=0 controllerNumber=1 scan=20829 | 41.3624 |
| 15953:4 | 9 | K.IIEGEPNLKVPGN[+2205.793]VTAVLGETLKVPC[+57.021]HFPC[+57.021]K.F | HexNAc(4)Hex(5)Fuc(2)NeuAc(1) | C[+57]*2, N[+2206] | 1406.4085 | 4 | 5622.612 | 5622.5997 | 0 | 2.2 | 457 | Specific | 785.41 | 658.38 | 658.38 | 15.03 | 60 | >sp\|P01833\|PIGR_HUMAN Polymeric immunoglobulin receptor OS=Homo sapiens OX=9606 GN=PIGR PE=1 SV=4 | 902 | 20201113_10633_1ng_glyco_hcd203040_compound15.19291.19291.2 | controllerType=0 controllerNumber=1 scan=19291 | 38.5669 |
| 17385:5 | 9 | K.IIEGEPNLKVPGN[+2350.830]VTAVLGETLKVPC[+57.021]HFPC[+57.021]K.F | HexNAc(4)Hex(5)Fuc(1)NeuAc(2) | C[+57]*2, N[+2351] | 1154.3346 | 5 | 5767.6439 | 5767.6372 | 0 | 1.2 | 457 | Specific | 814.33 | 767.44 | 767.44 | 16.49 | 60 | >sp\|P01833\|PIGR_HUMAN Polymeric immunoglobulin receptor OS=Homo sapiens OX=9606 GN=PIGR PE=1 SV=4 | 902 | 20201113_10633_1ng_glyco_hcd203040_compound15.20831.20831.5 | controllerType=0 controllerNumber=1 scan=20831 | 41.367 |
| 16927:3 | 9 | K.VPGN[+2350.830]VTAVLGETLK.V | HexNAc(4)Hex(5)Fuc(1)NeuAc(2) | N[+2351] | 1250.214 | 3 | 3748.6276 | 3748.6302 | 0 | -0.7 | 466 | Specific | 235.34 | 146.34 | 146.34 | 3.88 | 60 | >sp\|P01833\|PIGR_HUMAN Polymeric immunoglobulin receptor OS=Homo sapiens OX=9606 GN=PIGR PE=1 SV=4 | 902 | 20201113_10633_1ng_glyco_hcd203040_compound15.20354.20354.3 | controllerType=0 controllerNumber=1 scan=20354 | 40.485 |
| 13729:3 | 9 | K.VPGN[+2205.793]VTAVLGETLK.V | HexNAc(4)Hex(5)Fuc(2)NeuAc(1) | N[+2206] | 1201.8715 | 3 | 3603.6 | 3603.5927 | 0 | 2 | 466 | Specific | 390.25 | 371.4 | 371.4 | 7.69 | 60 | >sp\|P01833\|PIGR_HUMAN Polymeric immunoglobulin receptor OS=Homo sapiens OX=9606 GN=PIGR PE=1 SV=4 | 902 | 20201113_10633_1ng_glyco_hcd203040_compound15.16988.16988.3 | controllerType=0 controllerNumber=1 scan=16988 | 34.6002 |
| 13727:3 | 9 | K.VPGN[+2205.793]VTAVLGETLK.V | HexNAc(4)Hex(5)Fuc(2)NeuAc(1) | N[+2206] | 1201.8715 | 3 | 3603.6 | 3603.5927 | 0 | 2 | 466 | Specific | 441.89 | 382.58 | 382.58 | 9.37 | 60 | >sp\|P01833\|PIGR_HUMAN Polymeric immunoglobulin receptor OS=Homo sapiens OX=9606 GN=PIGR PE=1 SV=4 | 902 | 20201113_10633_1ng_glyco_hcd203040_compound15.16986.16986.3 | controllerType=0 controllerNumber=1 scan=16986 | 34.5936 |
| 13969:4 | 9 | K.VPGN[+2059.735]VTAVLGETLKVPC[+57.021]HFPC[+57.021]K.F | HexNAc(4)Hex(5)Fuc(1)NeuAc(1) | C[+57]*2, N[+2060] | 1121.5044 | 4 | 4482.9959 | 4482.9923 | 0 | 0.8 | 466 | Specific | 324.63 | 305.86 | 305.86 | 7.43 | 60 | >sp\|P01833\|PIGR_HUMAN Polymeric immunoglobulin receptor OS=Homo sapiens OX=9606 GN=PIGR PE=1 SV=4 | 902 | 20201113_10633_1ng_glyco_hcd203040_compound15.17236.17236.3 | controllerType=0 controllerNumber=1 scan=17236 | 35.046 |
| 13828:3 | 9 | K.VPGN[+2205.793]VTAVLGETLKVPC[+57.021]HFPC[+57.021]K.F | HexNAc(4)Hex(5)Fuc(2)NeuAc(1) | C[+57]*2, N[+2206] | 1543.6904 | 3 | 4629.0565 | 4629.0502 | 0 | 1.4 | 466 | Specific | 397.4 | 337.06 | 337.06 | 7.28 | 60 | >sp\|P01833\|PIGR_HUMAN Polymeric immunoglobulin receptor OS=Homo sapiens OX=9606 GN=PIGR PE=1 SV=4 | 902 | 20201113_10633_1ng_glyco_hcd203040_compound15.17090.17090.3 | controllerType=0 controllerNumber=1 scan=17090 | 34.798 |
| 13971:4 | 9 | K.VPGN[+2059.735]VTAVLGETLKVPC[+57.021]HFPC[+57.021]K.F | HexNAc(4)Hex(5)Fuc(1)NeuAc(1) | C[+57]*2, N[+2060] | 1121.5044 | 4 | 4482.9959 | 4482.9923 | 0 | 0.8 | 466 | Specific | 403.59 | 291.94 | 291.94 | 7.48 | 60 | >sp\|P01833\|PIGR_HUMAN Polymeric immunoglobulin receptor OS=Homo sapiens OX=9606 GN=PIGR PE=1 SV=4 | 902 | 20201113_10633_1ng_glyco_hcd203040_compound15.17238.17238.3 | controllerType=0 controllerNumber=1 scan=17238 | 35.0497 |
| 13826:3 | 9 | K.VPGN[+2205.793]VTAVLGETLKVPC[+57.021]HFPC[+57.021]K.F | HexNAc(4)Hex(5)Fuc(2)NeuAc(1) | C[+57]*2, N[+2206] | 1543.6904 | 3 | 4629.0565 | 4629.0502 | 0 | 1.4 | 466 | Specific | 497.86 | 437.37 | 437.37 | 9.35 | 60 | >sp\|P01833\|PIGR_HUMAN Polymeric immunoglobulin receptor OS=Homo sapiens OX=9606 GN=PIGR PE=1 SV=4 | 902 | 20201113_10633_1ng_glyco_hcd203040_compound15.17088.17088.3 | controllerType=0 controllerNumber=1 scan=17088 | [illegible] |
| 05937:3 | 9 | K.WN[+1913.677]NTGC[+57.021]QALPSQDEGPSK.A | HexNAc(4)Hex(5)NeuAc(1) | C[+57], N[+1914] | 1301.5251 | 3 | 3902.5607 | 3902.5537 | 0 | 1.8 | 498 | Specific | 241.18 | 193.85 | 193.85 | 5.93 | 60 | >sp\|P01833\|PIGR_HUMAN Polymeric immunoglobulin receptor OS=Homo sapiens OX=9606 GN=PIGR PE=1 SV=4 | 902 | 20201113_10633_1ng_glyco_hcd203040_compound15.8855.8855.3 | controllerType=0 controllerNumber=1 scan=8855 | 20.9487 |
| 05939:3 | 9 | K.WN[+1913.677]NTGC[+57.021]QALPSQDEGPSK.A | HexNAc(4)Hex(5)NeuAc(1) | C[+57], N[+1914] | 1301.5251 | 3 | 3902.5607 | 3902.5537 | 0 | 1.8 | 498 | Specific | 330.37 | 270.8 | 270.8 | 7.2 | 60 | >sp\|P01833\|PIGR_HUMAN Polymeric immunoglobulin receptor OS=Homo sapiens OX=9606 GN=PIGR PE=1 SV=4 | 902 | 20201113_10633_1ng_glyco_hcd203040_compound15.8857.8857.3 | controllerType=0 controllerNumber=1 scan=8857 | 20.9526 |
| 05999:3 | 9 | K.WN[+1710.598]NTGC[+57.021]QALPSQDEGPSK.A | HexNAc(3)Hex(5)NeuAc(1) | C[+57], N[+1711] | 1233.8358 | 3 | 3699.4927 | 3699.4743 | 0 | 5 | 498 | Specific | 374.68 | 326.71 | 326.71 | 6.77 | 60 | >sp\|P01833\|PIGR_HUMAN Polymeric immunoglobulin receptor OS=Homo sapiens OX=9606 GN=PIGR PE=1 SV=4 | 902 | 20201113_10633_1ng_glyco_hcd203040_compound15.8919.8919.3 | controllerType=0 controllerNumber=1 scan=8919 | 21.0558 |
| 05871:3 | 9 | K.WN[+1872.651]NTGC[+57.021]QALPSQDEGPSK.A | HexNAc(3)Hex(6)NeuAc(1) | C[+57], N[+1873] | 1287.8461 | 3 | 3861.5238 | 3861.5272 | 0 | -0.9 | 498 | Specific | 394.26 | 367.83 | 367.83 | 6.98 | 60 | >sp\|P01833\|PIGR_HUMAN Polymeric immunoglobulin receptor OS=Homo sapiens OX=9606 GN=PIGR PE=1 SV=4 | 902 | 20201113_10633_1ng_glyco_hcd203040_compound15.8786.8786.3 | controllerType=0 controllerNumber=1 scan=8786 | 20.8316 |
| 05873:3 | 9 | K.WN[+1872.651]NTGC[+57.021]QALPSQDEGPSK.A | HexNAc(3)Hex(6)NeuAc(1) | C[+57], N[+1873] | 1287.8461 | 3 | 3861.5238 | 3861.5272 | 0 | -0.9 | 498 | Specific | 444.4 | 412.48 | 412.48 | 8.67 | 60 | >sp\|P01833\|PIGR_HUMAN Polymeric immunoglobulin receptor OS=Homo sapiens OX=9606 GN=PIGR PE=1 SV=4 | 902 | 20201113_10633_1ng_glyco_hcd203040_compound15.8788.8788.3 | controllerType=0 controllerNumber=1 scan=8788 | 20.8357 |
| 05779:3 | 9 | K.WN[+2059.735]NTGC[+57.021]QALPSQDEGPSK.A | HexNAc(4)Hex(5)Fuc(1)NeuAc(1) | C[+57], N[+2060] | 1350.2111 | 3 | 4048.6188 | 4048.6116 | 0 | 1.8 | 498 | Specific | 660.43 | 592.76 | 592.76 | 13.36 | 60 | >sp\|P01833\|PIGR_HUMAN Polymeric immunoglobulin receptor OS=Homo sapiens OX=9606 GN=PIGR PE=1 SV=4 | 902 | 20201113_10633_1ng_glyco_hcd203040_compound15.8691.8691.3 | controllerType=0 controllerNumber=1 scan=8691 | 20.6643 |
| 05781:3 | 9 | K.WN[+2059.735]NTGC[+57.021]QALPSQDEGPSK.A | HexNAc(4)Hex(5)Fuc(1)NeuAc(1) | C[+57], N[+2060] | 1350.2111 | 3 | 4048.6188 | 4048.6116 | 0 | 1.8 | 498 | Specific | 697.33 | 620.46 | 620.46 | 13.36 | 60 | >sp\|P01833\|PIGR_HUMAN Polymeric immunoglobulin receptor OS=Homo sapiens OX=9606 GN=PIGR PE=1 SV=4 | 902 | 20201113_10633_1ng_glyco_hcd203040_compound15.8693.8693.3 | controllerType=0 controllerNumber=1 scan=8693 | 20.6698 |
| 10305:3 | 10 | K.TVLTPATNHMGN[+1378.476]VTFTIPANR.E | HexNAc(2)Hex(6) | N[+1378] | 1211.8845 | 3 | 3633.6388 | 3633.6358 | 0 | 0.8 | 74 | Specific | 328.8 | 328.8 | 228.93 | 8.05 | 78 | >sp\|P01024\|CO3_HUMAN Complement C3 OS=Homo sapiens OX=9606 GN=C3 PE=1 SV=2 | 492 | 20201113_10633_1ng_glyco_hcd203040_compound15.13403.13403.4 | controllerType=0 controllerNumber=1 scan=13403 | 28.5373 |
| 18050:3 | 11 | R.LSLHRPALEDLLLGSEAN[+1257.449]LTC[+57.021]TLTGLR.D | HexNAc(3)Hex(4) | C[+57], N[+1257] | 1407.6921 | 3 | 4221.0617 | 4221.0477 | 0 | 3.3 | 127 | Specific | 235.4 | 235.4 | 235.4 | 5.79 | 40 | >sp\|P01876\|IGHA1_HUMAN Immunoglobulin heavy constant alpha 1 OS=Homo sapiens OX=9606 GN=IGHA1 PE=1 SV=2 | 255 | 20201113_10633_1ng_glyco_hcd203040_compound15.21524.21524.3 | controllerType=0 controllerNumber=1 scan=21524 | 42.8114 |
| 18171:3 | 11 | R.LSLHRPALEDLLLGSEAN[+1501.555]LTC[+57.021]TLTGLR.D | HexNAc(5)Hex(3) | C[+57], N[+1502] | 1489.0534 | 3 | 4465.1457 | 4465.1536 | 0 | -1.8 | 127 | Specific | 251.43 | 238.25 | 238.25 | 5.39 | 40 | >sp\|P01876\|IGHA1_HUMAN Immunoglobulin heavy constant alpha 1 OS=Homo sapiens OX=9606 GN=IGHA1 PE=1 SV=2 | 255 | 20201113_10633_1ng_glyco_hcd203040_compound15.21650.21650.3 | controllerType=0 controllerNumber=1 scan=21650 | 43.0737 |
| 18174:3 | 11 | R.LSLHRPALEDLLLGSEAN[+1501.555]LTC[+57.021]TLTGLR.D | HexNAc(5)Hex(3) | C[+57], N[+1502] | 1489.0534 | 3 | 4465.1457 | 4465.1536 | 0 | -1.8 | 127 | Specific | 282.56 | 253.32 | 253.32 | 5.59 | 40 | >sp\|P01876\|IGHA1_HUMAN Immunoglobulin heavy constant alpha 1 OS=Homo sapiens OX=9606 GN=IGHA1 PE=1 SV=2 | 255 | 20201113_10633_1ng_glyco_hcd203040_compound15.21653.21653.3 | controllerType=0 controllerNumber=1 scan=21653 | 43.0802 |
| 18052:3 | 11 | R.LSLHRPALEDLLLGSEAN[+1257.449]LTC[+57.021]TLTGLR.D | HexNAc(3)Hex(4) | C[+57], N[+1257] | 1407.6921 | 3 | 4221.0617 | 4221.0477 | 0 | 3.3 | 127 | Specific | 296.55 | 293.25 | 293.25 | 5.19 | 40 | >sp\|P01876\|IGHA1_HUMAN Immunoglobulin heavy constant alpha 1 OS=Homo sapiens OX=9606 GN=IGHA1 PE=1 SV=2 | 255 | 20201113_10633_1ng_glyco_hcd203040_compound15.21526.21526.3 | controllerType=0 controllerNumber=1 scan=21526 | 42.8148 |
| 19041:4 | 11 | R.LSLHRPALEDLLLGSEAN[+1751.624]LTC[+57.021]TLTGLR.D | HexNAc(4)Hex(4)NeuAc(1) | C[+57], N[+1752] | 1179.5617 | 4 | 4715.2249 | 4715.2225 | 0 | 0.5 | 127 | Specific | 306.57 | 306.57 | 306.57 | 7.46 | 40 | >sp\|P01876\|IGHA1_HUMAN Immunoglobulin heavy constant alpha 1 OS=Homo sapiens OX=9606 GN=IGHA1 PE=1 SV=2 | 255 | 20201113_10633_1ng_glyco_hcd203040_compound15.22555.22555.4 | controllerType=0 controllerNumber=1 scan=22555 | 44.9049 |
| 18237:4 | 11 | R.LSLHRPALEDLLLGSEAN[+1501.555]LTC[+57.021]TLTGLR.D | HexNAc(5)Hex(3) | C[+57], N[+1502] | 1117.0446 | 4 | 4465.1566 | 4465.1536 | 0 | 0.7 | 127 | Specific | 419.02 | 419.02 | 419.02 | 9.67 | 40 | >sp\|P01876\|IGHA1_HUMAN Immunoglobulin heavy constant alpha 1 OS=Homo sapiens OX=9606 GN=IGHA1 PE=1 SV=2 | 255 | 20201113_10633_1ng_glyco_hcd203040_compound15.21719.21719.4 | controllerType=0 controllerNumber=1 scan=21719 | 43.2129 |
| 17980:3 | 11 | R.LSLHRPALEDLLLGSEAN[+1540.529]LTC[+57.021]TLTGLR.D | HexNAc(2)Hex(7) | C[+57], N[+1541] | 1502.0514 | 3 | 4504.1398 | 4504.1268 | 0 | 2.9 | 127 | Specific | 430.3 | 430.3 | 430.3 | 9.38 | 40 | >sp\|P01876\|IGHA1_HUMAN Immunoglobulin heavy constant alpha 1 OS=Homo sapiens OX=9606 GN=IGHA1 PE=1 SV=2 | 255 | 20201113_10633_1ng_glyco_hcd203040_compound15.21451.21451.3 | controllerType=0 controllerNumber=1 scan=21451 | 42.6354 |
| 18080:3 | 11 | R.LSLHRPALEDLLLGSEAN[+1095.397]LTC[+57.021]TLTGLR.D | HexNAc(3)Hex(3) | C[+57], N[+1095] | 1353.6741 | 3 | 4059.0078 | 4058.9949 | 0 | 3.2 | 127 | Specific | 483.78 | 417.31 | 417.31 | 9.28 | 40 | >sp\|P01876\|IGHA1_HUMAN Immunoglobulin heavy constant alpha 1 OS=Homo sapiens OX=9606 GN=IGHA1 PE=1 SV=2 | 255 | 20201113_10633_1ng_glyco_hcd203040_compound15.21555.21555.3 | controllerType=0 controllerNumber=1 scan=21555 | 42.8895 |
| 18000:3 | 11 | R.LSLHRPALEDLLLGSEAN[+1622.582]LTC[+57.021]TLTGLR.D | HexNAc(4)Hex(5) | C[+57], N[+1623] | 1529.3998 | 3 | 4586.185 | 4586.1799 | 0 | 1.1 | 127 | Specific | 498.07 | 443.94 | 443.94 | 9.6 | 40 | >sp\|P01876\|IGHA1_HUMAN Immunoglobulin heavy constant alpha 1 OS=Homo sapiens OX=9606 GN=IGHA1 PE=1 SV=2 | 255 | 20201113_10633_1ng_glyco_hcd203040_compound15.21472.21472.3 | controllerType=0 controllerNumber=1 scan=21472 | 42.6908 |
| 18015:3 | 11 | R.LSLHRPALEDLLLGSEAN[+1460.529]LTC[+57.021]TLTGLR.D | HexNAc(4)Hex(4) | C[+57], N[+1461] | 1475.386 | 3 | 4424.1436 | 4424.1271 | 0 | 3.7 | 127 | Specific | 498.59 | 483.55 | 483.55 | 9.28 | 40 | >sp\|P01876\|IGHA1_HUMAN Immunoglobulin heavy constant alpha 1 OS=Homo sapiens OX=9606 GN=IGHA1 PE=1 SV=2 | 255 | 20201113_10633_1ng_glyco_hcd203040_compound15.21488.21488.3 | controllerType=0 controllerNumber=1 scan=21488 | 42.7325 |
| 18219:4 | 11 | R.LSLHRPALEDLLLGSEAN[+1501.555]LTC[+57.021]TLTGLR.D | HexNAc(5)Hex(3) | C[+57], N[+1502] | 1117.0401 | 4 | 4465.1385 | 4465.1536 | 0 | -3.4 | 127 | Specific | 514.01 | 514.01 | 514.01 | 8.78 | 40 | >sp\|P01876\|IGHA1_HUMAN Immunoglobulin heavy constant alpha 1 OS=Homo sapiens OX=9606 GN=IGHA1 PE=1 SV=2 | 255 | 20201113_10633_1ng_glyco_hcd203040_compound15.21700.21700.4 | controllerType=0 controllerNumber=1 scan=21700 | 43.1733 |
| 17967:3 | 11 | R.LSLHRPALEDLLLGSEAN[+1663.608]LTC[+57.021]TLTGLR.D | HexNAc(5)Hex(4) | C[+57], N[+1664] | 1543.0748 | 3 | 4627.2097 | 4627.2064 | 0 | 0.7 | 127 | Specific | 544.32 | 509.18 | 509.18 | 9.84 | 40 | >sp\|P01876\|IGHA1_HUMAN Immunoglobulin heavy constant alpha 1 OS=Homo sapiens OX=9606 GN=IGHA1 PE=1 SV=2 | 255 | 20201113_10633_1ng_glyco_hcd203040_compound15.21437.21437.3 | controllerType=0 controllerNumber=1 scan=21437 | 42.5944 |
| 18002:3 | 11 | R.LSLHRPALEDLLLGSEAN[+1622.582]LTC[+57.021]TLTGLR.D | HexNAc(4)Hex(5) | C[+57], N[+1623] | 1529.3998 | 3 | 4586.185 | 4586.1799 | 0 | 1.1 | 127 | Specific | 562.56 | 482.76 | 482.76 | 10.01 | 40 | >sp\|P01876\|IGHA1_HUMAN Immunoglobulin heavy constant alpha 1 OS=Homo sapiens OX=9606 GN=IGHA1 PE=1 SV=2 | 255 | 20201113_10633_1ng_glyco_hcd203040_compound15.21474.21474.3 | controllerType=0 controllerNumber=1 scan=21474 | 42.6977 |
| 17981:3 | 11 | R.LSLHRPALEDLLLGSEAN[+1501.555]LTC[+57.021]TLTGLR.D | HexNAc(5)Hex(3) | C[+57], N[+1502] | 1489.0605 | 3 | 4465.1669 | 4465.1536 | 0 | 3 | 127 | Specific | 601.43 | 601.43 | 601.43 | 12.26 | 40 | >sp\|P01876\|IGHA1_HUMAN Immunoglobulin heavy constant alpha 1 OS=Homo sapiens OX=9606 GN=IGHA1 PE=1 SV=2 | 255 | 20201113_10633_1ng_glyco_hcd203040_compound15.21452.21452.3 | controllerType=0 controllerNumber=1 scan=21452 | 42.6365 |
| 17953:3 | 11 | R.LSLHRPALEDLLLGSEAN[+1702.581]LTC[+57.021]TLTGLR.D | HexNAc(2)Hex(8) | C[+57], N[+1703] | 1556.066 | 3 | 4666.1834 | 4666.1796 | 0 | 0.8 | 127 | Specific | 617.06 | 570.83 | 570.83 | 12.25 | 40 | >sp\|P01876\|IGHA1_HUMAN Immunoglobulin heavy constant alpha 1 OS=Homo sapiens OX=9606 GN=IGHA1 PE=1 SV=2 | 255 | 20201113_10633_1ng_glyco_hcd203040_compound15.21422.21422.3 | controllerType=0 controllerNumber=1 scan=21422 | 42.5534 |
| 17969:3 | 11 | R.LSLHRPALEDLLLGSEAN[+1663.608]LTC[+57.021]TLTGLR.D | HexNAc(5)Hex(4) | C[+57], N[+1664] | 1543.0748 | 3 | 4627.2097 | 4627.2064 | 0 | 0.7 | 127 | Specific | 618.7 | 509.25 | 509.25 | 12.25 | 40 | >sp\|P01876\|IGHA1_HUMAN Immunoglobulin heavy constant alpha 1 OS=Homo sapiens OX=9606 GN=IGHA1 PE=1 SV=2 | 255 | 20201113_10633_1ng_glyco_hcd203040_compound15.21439.21439.3 | controllerType=0 controllerNumber=1 scan=21439 | 42.599 |
| 18017:3 | 11 | R.LSLHRPALEDLLLGSEAN[+1460.529]LTC[+57.021]TLTGLR.D | HexNAc(4)Hex(4) | C[+57], N[+1461] | 1475.386 | 3 | 4424.1436 | 4424.1271 | 0 | 3.7 | 127 | Specific | 621.67 | 546.37 | 546.37 | 11.93 | 40 | >sp\|P01876\|IGHA1_HUMAN Immunoglobulin heavy constant alpha 1 OS=Homo sapiens OX=9606 GN=IGHA1 PE=1 SV=2 | 255 | 20201113_10633_1ng_glyco_hcd203040_compound15.21490.21490.3 | controllerType=0 controllerNumber=1 scan=21490 | 42.7368 |
| 18079:3 | 11 | R.LSLHRPALEDLLLGSEAN[+1095.397]LTC[+57.021]TLTGLR.D | HexNAc(3)Hex(3) | C[+57], N[+1095] | 1353.6741 | 3 | 4059.0078 | 4058.9949 | 0 | 3.2 | 127 | Specific | 622.88 | 615.94 | 615.94 | 11.93 | 40 | >sp\|P01876\|IGHA1_HUMAN Immunoglobulin heavy constant alpha 1 OS=Homo sapiens OX=9606 GN=IGHA1 PE=1 SV=2 | 255 | 20201113_10633_1ng_glyco_hcd203040_compound15.21554.21554.3 | controllerType=0 controllerNumber=1 scan=21554 | 42.8852 |
| 17951:3 | 11 | R.LSLHRPALEDLLLGSEAN[+1702.581]LTC[+57.021]TLTGLR.D | HexNAc(2)Hex(8) | C[+57], N[+1703] | 1556.066 | 3 | 4666.1834 | 4666.1796 | 0 | 0.8 | 127 | Specific | 626.6 | 617.49 | 617.49 | 12.25 | 40 | >sp\|P01876\|IGHA1_HUMAN Immunoglobulin heavy constant alpha 1 OS=Homo sapiens OX=9606 GN=IGHA1 PE=1 SV=2 | 255 | 20201113_10633_1ng_glyco_hcd203040_compound15.21420.21420.3 | controllerType=0 controllerNumber=1 scan=21420 | 42.5487 |
| 17982:3 | 11 | R.LSLHRPALEDLLLGSEAN[+1540.529]LTC[+57.021]TLTGLR.D | HexNAc(2)Hex(7) | C[+57], N[+1541] | 1502.0514 | 3 | 4504.1398 | 4504.1268 | 0 | 2.9 | 127 | Specific | 630.09 | 570.9 | 570.9 | 12.26 | 40 | >sp\|P01876\|IGHA1_HUMAN Immunoglobulin heavy constant alpha 1 OS=Homo sapiens OX=9606 GN=IGHA1 PE=1 SV=2 | 255 | 20201113_10633_1ng_glyco_hcd203040_compound15.21453.21453.3 | controllerType=0 controllerNumber=1 scan=21453 | 42.6388 |
| 18221:4 | 11 | R.LSLHRPALEDLLLGSEAN[+1501.555]LTC[+57.021]TLTGLR.D | HexNAc(5)Hex(3) | C[+57], N[+1502] | 1117.0401 | 4 | 4465.1385 | 4465.1536 | 0 | -3.4 | 127 | Specific | 639.82 | 381.68 | 381.68 | 11.34 | 40 | >sp\|P01876\|IGHA1_HUMAN Immunoglobulin heavy constant alpha 1 OS=Homo sapiens OX=9606 GN=IGHA1 PE=1 SV=2 | 255 | 20201113_10633_1ng_glyco_hcd203040_compound15.21702.21702.4 | controllerType=0 controllerNumber=1 scan=21702 | 43.178 |
| 17983:3 | 11 | R.LSLHRPALEDLLLGSEAN[+1501.555]LTC[+57.021]TLTGLR.D | HexNAc(5)Hex(3) | C[+57], N[+1502] | 1489.0605 | 3 | 4465.1669 | 4465.1536 | 0 | 3 | 127 | Specific | 642.36 | 596.65 | 596.65 | 12.26 | 40 | >sp\|P01876\|IGHA1_HUMAN Immunoglobulin heavy constant alpha 1 OS=Homo sapiens OX=9606 GN=IGHA1 PE=1 SV=2 | 255 | 20201113_10633_1ng_glyco_hcd203040_compound15.21454.21454.3 | controllerType=0 controllerNumber=1 scan=21454 | 42.6449 |
| 18014:4 | 11 | R.LSLHRPALEDLLLGSEAN[+1460.529]LTC[+57.021]TLTGLR.D | HexNAc(4)Hex(4) | C[+57], N[+1461] | 1106.7923 | 4 | 4424.1473 | 4424.1271 | 0 | 4.6 | 127 | Specific | 647.2 | 647.2 | 647.2 | 11.91 | 40 | >sp\|P01876\|IGHA1_HUMAN Immunoglobulin heavy constant alpha 1 OS=Homo sapiens OX=9606 GN=IGHA1 PE=1 SV=2 | 255 | 20201113_10633_1ng_glyco_hcd203040_compound15.21487.21487.4 | controllerType=0 controllerNumber=1 scan=21487 | 42.7311 |
| 18240:4 | 11 | R.LSLHRPALEDLLLGSEAN[+1501.555]LTC[+57.021]TLTGLR.D | HexNAc(5)Hex(3) | C[+57], N[+1502] | 1117.0446 | 4 | 4465.1566 | 4465.1536 | 0 | 0.7 | 127 | Specific | 648.2 | 581.6 | 581.6 | 12.72 | 40 | >sp\|P01876\|IGHA1_HUMAN Immunoglobulin heavy constant alpha 1 OS=Homo sapiens OX=9606 GN=IGHA1 PE=1 SV=2 | 255 | 20201113_10633_1ng_glyco_hcd203040_compound15.21722.21722.4 | controllerType=0 controllerNumber=1 scan=21722 | 43.2186 |
| 18044:4 | 11 | R.LSLHRPALEDLLLGSEAN[+1298.476]LTC[+57.021]TLTGLR.D | HexNAc(4)Hex(3) | C[+57], N[+1298] | 1066.2771 | 4 | 4262.0867 | 4262.0742 | 0 | 2.9 | 127 | Specific | 671.09 | 671.09 | 671.09 | 13.74 | 40 | >sp\|P01876\|IGHA1_HUMAN Immunoglobulin heavy constant alpha 1 OS=Homo sapiens OX=9606 GN=IGHA1 PE=1 SV=2 | 255 | 20201113_10633_1ng_glyco_hcd203040_compound15.21518.21518.4 | controllerType=0 controllerNumber=1 scan=21518 | 42.7987 |
| 19044:4 | 11 | R.LSLHRPALEDLLLGSEAN[+1751.624]LTC[+57.021]TLTGLR.D | HexNAc(4)Hex(4)NeuAc(1) | C[+57], N[+1752] | 1179.5617 | 4 | 4715.2249 | 4715.2225 | 0 | 0.5 | 127 | Specific | 677.85 | 676.64 | 676.64 | 13.5 | 40 | >sp\|P01876\|IGHA1_HUMAN Immunoglobulin heavy constant alpha 1 OS=Homo sapiens OX=9606 GN=IGHA1 PE=1 SV=2 | 255 | 20201113_10633_1ng_glyco_hcd203040_compound15.22558.22558.4 | controllerType=0 controllerNumber=1 scan=22558 | 44.9104 |
| 18008:3 | 11 | R.LSLHRPALEDLLLGSEAN[+1216.423]LTC[+57.021]TLTGLR.D | HexNAc(2)Hex(5) | C[+57], N[+1216] | 1394.0134 | 3 | 4180.0255 | 4180.0211 | 0 | 1.1 | 127 | Specific | 685.06 | 685.06 | 685.06 | 13.25 | 40 | >sp\|P01876\|IGHA1_HUMAN Immunoglobulin heavy constant alpha 1 OS=Homo sapiens OX=9606 GN=IGHA1 PE=1 SV=2 | 255 | 20201113_10633_1ng_glyco_hcd203040_compound15.21480.21480.3 | controllerType=0 controllerNumber=1 scan=21480 | 42.712 |
| 17958:4 | 11 | R.LSLHRPALEDLLLGSEAN[+1663.608]LTC[+57.021]TLTGLR.D | HexNAc(5)Hex(4) | C[+57], N[+1664] | 1157.559 | 4 | 4627.2142 | 4627.2064 | 0 | 1.7 | 127 | Specific | 694.7 | 681.68 | 681.68 | 13.72 | 40 | >sp\|P01876\|IGHA1_HUMAN Immunoglobulin heavy constant alpha 1 OS=Homo sapiens OX=9606 GN=IGHA1 PE=1 SV=2 | 255 | 20201113_10633_1ng_glyco_hcd203040_compound15.21428.21428.4 | controllerType=0 controllerNumber=1 scan=21428 | 42.5711 |
| 18073:4 | 11 | R.LSLHRPALEDLLLGSEAN[+1095.397]LTC[+57.021]TLTGLR.D | HexNAc(3)Hex(3) | C[+57], N[+1095] | 1015.507 | 4 | 4059.0062 | 4058.9949 | 0 | 2.8 | 127 | Specific | 703.15 | 703.15 | 703.15 | 14.74 | 40 | >sp\|P01876\|IGHA1_HUMAN Immunoglobulin heavy constant alpha 1 OS=Homo sapiens OX=9606 GN=IGHA1 PE=1 SV=2 | 255 | 20201113_10633_1ng_glyco_hcd203040_compound15.21548.21548.4 | controllerType=0 controllerNumber=1 scan=21548 | 42.874 |
| 17976:4 | 11 | R.LSLHRPALEDLLLGSEAN[+1501.555]LTC[+57.021]TLTGLR.D | HexNAc(5)Hex(3) | C[+57], N[+1502] | 1117.0474 | 4 | 4465.1678 | 4465.1536 | 0 | 3.2 | 127 | Specific | 704.82 | 704.82 | 704.82 | 14.4 | 40 | >sp\|P01876\|IGHA1_HUMAN Immunoglobulin heavy constant alpha 1 OS=Homo sapiens OX=9606 GN=IGHA1 PE=1 SV=2 | 255 | 20201113_10633_1ng_glyco_hcd203040_compound15.21447.21447.4 | controllerType=0 controllerNumber=1 scan=21447 | 42.6218 |
| 18006:3 | 11 | R.LSLHRPALEDLLLGSEAN[+1216.423]LTC[+57.021]TLTGLR.D | HexNAc(2)Hex(5) | C[+57], N[+1216] | 1394.0134 | 3 | 4180.0255 | 4180.0211 | 0 | 1.1 | 127 | Specific | 750.44 | 675.27 | 675.27 | 15.25 | 40 | >sp\|P01876\|IGHA1_HUMAN Immunoglobulin heavy constant alpha 1 OS=Homo sapiens OX=9606 GN=IGHA1 PE=1 SV=2 | 255 | 20201113_10633_1ng_glyco_hcd203040_compound15.21478.21478.3 | controllerType=0 controllerNumber=1 scan=21478 | 42.7057 |
| 17919:4 | 11 | R.LSLHRPALEDLLLGSEAN[+1864.634]LTC[+57.021]TLTGLR.D | HexNAc(2)Hex(9) | C[+57], N[+1865] | 1207.8232 | 4 | 4828.2708 | 4828.2324 | 0 | 8 | 127 | Specific | 762.77 | 762.77 | 762.77 | 14.57 | 40 | >sp\|P01876\|IGHA1_HUMAN Immunoglobulin heavy constant alpha 1 OS=Homo sapiens OX=9606 GN=IGHA1 PE=1 SV=2 | 255 | 20201113_10633_1ng_glyco_hcd203040_compound15.21387.21387.4 | controllerType=0 controllerNumber=1 scan=21387 | 42.4737 |
| 17978:4 | 11 | R.LSLHRPALEDLLLGSEAN[+1501.555]LTC[+57.021]TLTGLR.D | HexNAc(5)Hex(3) | C[+57], N[+1502] | 1117.0474 | 4 | 4465.1678 | 4465.1536 | 0 | 3.2 | 127 | Specific | 777.36 | 777.36 | 777.36 | 15.4 | 40 | >sp\|P01876\|IGHA1_HUMAN Immunoglobulin heavy constant alpha 1 OS=Homo sapiens OX=9606 GN=IGHA1 PE=1 SV=2 | 255 | 20201113_10633_1ng_glyco_hcd203040_compound15.21449.21449.4 | controllerType=0 controllerNumber=1 scan=21449 | 42.6277 |
| 17960:4 | 11 | R.LSLHRPALEDLLLGSEAN[+1663.608]LTC[+57.021]TLTGLR.D | HexNAc(5)Hex(4) | C[+57], N[+1664] | 1157.559 | 4 | 4627.2142 | 4627.2064 | 0 | 1.7 | 127 | Specific | 787.31 | 714.77 | 714.77 | 15.72 | 40 | >sp\|P01876\|IGHA1_HUMAN Immunoglobulin heavy constant alpha 1 OS=Homo sapiens OX=9606 GN=IGHA1 PE=1 SV=2 | 255 | 20201113_10633_1ng_glyco_hcd203040_compound15.21430.21430.4 | controllerType=0 controllerNumber=1 scan=21430 | 42.5778 |
| 18016:4 | 11 | R.LSLHRPALEDLLLGSEAN[+1460.529]LTC[+57.021]TLTGLR.D | HexNAc(4)Hex(4) | C[+57], N[+1461] | 1106.7923 | 4 | 4424.1473 | 4424.1271 | 0 | 4.6 | 127 | Specific | 808.86 | 647.78 | 647.78 | 15.91 | 40 | >sp\|P01876\|IGHA1_HUMAN Immunoglobulin heavy constant alpha 1 OS=Homo sapiens OX=9606 GN=IGHA1 PE=1 SV=2 | 255 | 20201113_10633_1ng_glyco_hcd203040_compound15.21489.21489.4 | controllerType=0 controllerNumber=1 scan=21489 | 42.7352 |
| 17995:4 | 11 | R.LSLHRPALEDLLLGSEAN[+1216.423]LTC[+57.021]TLTGLR.D | HexNAc(2)Hex(5) | C[+57], N[+1216] | 1045.762 | 4 | 4180.0262 | 4180.0211 | 0 | 1.2 | 127 | Specific | 841.95 | 794.49 | 794.49 | 16.72 | 40 | >sp\|P01876\|IGHA1_HUMAN Immunoglobulin heavy constant alpha 1 OS=Homo sapiens OX=9606 GN=IGHA1 PE=1 SV=2 | 255 | 20201113_10633_1ng_glyco_hcd203040_compound15.21467.21467.4 | controllerType=0 controllerNumber=1 scan=21467 | 42.6789 |
| 17943:4 | 11 | R.LSLHRPALEDLLLGSEAN[+1702.581]LTC[+57.021]TLTGLR.D | HexNAc(2)Hex(8) | C[+57], N[+1703] | 1167.3025 | 4 | 4666.1883 | 4666.1796 | 0 | 1.9 | 127 | Specific | 868.04 | 868.04 | 868.04 | 17.74 | 40 | >sp\|P01876\|IGHA1_HUMAN Immunoglobulin heavy constant alpha 1 OS=Homo sapiens OX=9606 GN=IGHA1 PE=1 SV=2 | 255 | 20201113_10633_1ng_glyco_hcd203040_compound15.21412.21412.4 | controllerType=0 controllerNumber=1 scan=21412 | 42.5263 |
| 17947:4 | 11 | R.LSLHRPALEDLLLGSEAN[+1540.529]LTC[+57.021]TLTGLR.D | HexNAc(2)Hex(7) | C[+57], N[+1541] | 1126.7889 | 4 | 4504.1336 | 4504.1268 | 0 | 1.5 | 127 | Specific | 885.99 | 830.57 | 830.57 | 17.72 | 40 | >sp\|P01876\|IGHA1_HUMAN Immunoglobulin heavy constant alpha 1 OS=Homo sapiens OX=9606 GN=IGHA1 PE=1 SV=2 | 255 | 20201113_10633_1ng_glyco_hcd203040_compound15.21416.21416.4 | controllerType=0 controllerNumber=1 scan=21416 | 42.5369 |
| 18047:4 | 11 | R.LSLHRPALEDLLLGSEAN[+1298.476]LTC[+57.021]TLTGLR.D | HexNAc(4)Hex(3) | C[+57], N[+1298] | 1066.2771 | 4 | 4262.0867 | 4262.0742 | 0 | 2.9 | 127 | Specific | 886.77 | 820.99 | 820.99 | 17.74 | 40 | >sp\|P01876\|IGHA1_HUMAN Immunoglobulin heavy constant alpha 1 OS=Homo sapiens OX=9606 GN=IGHA1 PE=1 SV=2 | 255 | 20201113_10633_1ng_glyco_hcd203040_compound15.21521.21521.4 | controllerType=0 controllerNumber=1 scan=21521 | 42.8036 |
| 17921:4 | 11 | R.LSLHRPALEDLLLGSEAN[+1864.634]LTC[+57.021]TLTGLR.D | HexNAc(2)Hex(9) | C[+57], N[+1865] | 1207.8232 | 4 | 4828.2708 | 4828.2324 | 0 | 8 | 127 | Specific | 889.54 | 889.54 | 889.54 | 16.57 | 40 | >sp\|P01876\|IGHA1_HUMAN Immunoglobulin heavy constant alpha 1 OS=Homo sapiens OX=9606 GN=IGHA1 PE=1 SV=2 | 255 | 20201113_10633_1ng_glyco_hcd203040_compound15.21389.21389.4 | controllerType=0 controllerNumber=1 scan=21389 | 42.4778 |
| 18076:4 | 11 | R.LSLHRPALEDLLLGSEAN[+1095.397]LTC[+57.021]TLTGLR.D | HexNAc(3)Hex(3) | C[+57], N[+1095] | 1015.507 | 4 | 4059.0062 | 4058.9949 | 0 | 2.8 | 127 | Specific | 917.53 | 865.12 | 865.12 | 18.74 | 40 | >sp\|P01876\|IGHA1_HUMAN Immunoglobulin heavy constant alpha 1 OS=Homo sapiens OX=9606 GN=IGHA1 PE=1 SV=2 | 255 | 20201113_10633_1ng_glyco_hcd203040_compound15.21551.21551.4 | controllerType=0 controllerNumber=1 scan=21551 | 42.88 |
| 18020:4 | 11 | R.LSLHRPALEDLLLGSEAN[+1257.449]LTC[+57.021]TLTGLR.D | HexNAc(3)Hex(4) | C[+57], N[+1257] | 1056.0191 | 4 | 4221.0545 | 4221.0477 | 0 | 1.6 | 127 | Specific | 944.54 | 878.79 | 878.79 | 18.72 | 40 | >sp\|P01876\|IGHA1_HUMAN Immunoglobulin heavy constant alpha 1 OS=Homo sapiens OX=9606 GN=IGHA1 PE=1 SV=2 | 255 | 20201113_10633_1ng_glyco_hcd203040_compound15.21493.21493.4 | controllerType=0 controllerNumber=1 scan=21493 | 42.741 |
| 17997:4 | 11 | R.LSLHRPALEDLLLGSEAN[+1216.423]LTC[+57.021]TLTGLR.D | HexNAc(2)Hex(5) | C[+57], N[+1216] | 1045.762 | 4 | 4180.0262 | 4180.0211 | 0 | 1.2 | 127 | Specific | 952.89 | 952.89 | 952.89 | 19.72 | 40 | >sp\|P01876\|IGHA1_HUMAN Immunoglobulin heavy constant alpha 1 OS=Homo sapiens OX=9606 GN=IGHA1 PE=1 SV=2 | 255 | 20201113_10633_1ng_glyco_hcd203040_compound15.21469.21469.4 | controllerType=0 controllerNumber=1 scan=21469 | 42.6866 |
| 17914:4 | 11 | R.LSLHRPALEDLLLGSEAN[+1540.529]LTC[+57.021]TLTGLR.D | HexNAc(2)Hex(7) | C[+57], N[+1541] | 1126.7898 | 4 | 4504.1375 | 4504.1268 | 0 | 2.4 | 127 | Specific | 963.74 | 894.25 | 894.25 | 19.74 | 40 | >sp\|P01876\|IGHA1_HUMAN Immunoglobulin heavy constant alpha 1 OS=Homo sapiens OX=9606 GN=IGHA1 PE=1 SV=2 | 255 | 20201113_10633_1ng_glyco_hcd203040_compound15.21381.21381.2 | controllerType=0 controllerNumber=1 scan=21381 | 42.4542 |
| 17945:4 | 11 | R.LSLHRPALEDLLLGSEAN[+1702.581]LTC[+57.021]TLTGLR.D | HexNAc(2)Hex(8) | C[+57], N[+1703] | 1167.3025 | 4 | 4666.1883 | 4666.1796 | 0 | 1.9 | 127 | Specific | 984.68 | 938.28 | 938.28 | 19.74 | 40 | >sp\|P01876\|IGHA1_HUMAN Immunoglobulin heavy constant alpha 1 OS=Homo sapiens OX=9606 GN=IGHA1 PE=1 SV=2 | 255 | 20201113_10633_1ng_glyco_hcd203040_compound15.21414.21414.4 | controllerType=0 controllerNumber=1 scan=21414 | 42.5336 |
| 17949:4 | 11 | R.LSLHRPALEDLLLGSEAN[+1540.529]LTC[+57.021]TLTGLR.D | HexNAc(2)Hex(7) | C[+57], N[+1541] | 1126.7889 | 4 | 4504.1336 | 4504.1268 | 0 | 1.5 | 127 | Specific | 999.92 | 925.44 | 925.44 | 19.72 | 40 | >sp\|P01876\|IGHA1_HUMAN Immunoglobulin heavy constant alpha 1 OS=Homo sapiens OX=9606 GN=IGHA1 PE=1 SV=2 | 255 | 20201113_10633_1ng_glyco_hcd203040_compound15.21418.21418.4 | controllerType=0 controllerNumber=1 scan=21418 | 42.545 |
| 18022:4 | 11 | R.LSLHRPALEDLLLGSEAN[+1257.449]LTC[+57.021]TLTGLR.D | HexNAc(3)Hex(4) | C[+57], N[+1257] | 1056.0191 | 4 | 4221.0545 | 4221.0477 | 0 | 1.6 | 127 | Specific | 1067.37 | 1006.54 | 1006.54 | 21.72 | 40 | >sp\|P01876\|IGHA1_HUMAN Immunoglobulin heavy constant alpha 1 OS=Homo sapiens OX=9606 GN=IGHA1 PE=1 SV=2 | 255 | 20201113_10633_1ng_glyco_hcd203040_compound15.21495.21495.4 | controllerType=0 controllerNumber=1 scan=21495 | 42.7478 |
| 13266:3 | 11 | R.LAGKPTHVN[+2262.814]VSVVMAEVDGTC[+57.021]Y.- | HexNAc(5)Hex(5)Fuc(1)NeuAc(1) | C[+57], N[+2263] | 1537.3278 | 3 | 4609.9689 | 4609.9564 | 0 | 2.7 | 332 | CRagged | 207.28 | 207.28 | 207.28 | 5.57 | 40 | >sp\|P01876\|IGHA1_HUMAN Immunoglobulin heavy constant alpha 1 OS=Homo sapiens OX=9606 GN=IGHA1 PE=1 SV=2 | 255 | 20201113_10633_1ng_glyco_hcd203040_compound15.16509.16509.3 | controllerType=0 controllerNumber=1 scan=16509 | 33.818 |
| 11540:3 | 11 | R.LAGKPTHVN[+1540.529]VSVVMAEVDGTC[+57.021]Y.- | HexNAc(2)Hex(7) | C[+57], N[+1541] | 1296.5636 | 3 | 3887.6762 | 3887.6706 | 0 | 1.5 | 332 | CRagged | 221.54 | 76.85 | 76.85 | 4.08 | 40 | >sp\|P01876\|IGHA1_HUMAN Immunoglobulin heavy constant alpha 1 OS=Homo sapiens OX=9606 GN=IGHA1 PE=1 SV=2 | 255 | 20201113_10633_1ng_glyco_hcd203040_compound15.14680.14680.3 | controllerType=0 controllerNumber=1 scan=14680 | 30.6888 |
| 11527:3 | 11 | R.LAGKPTHVN[+1378.476]VSVVMAEVDGTC[+57.021]Y.- | HexNAc(2)Hex(6) | C[+57], N[+1378] | 1242.5455 | 3 | 3725.6218 | 3725.6178 | 0 | 1.1 | 332 | CRagged | 491.87 | 491.87 | 345.85 | 10.16 | 40 | >sp\|P01876\|IGHA1_HUMAN Immunoglobulin heavy constant alpha 1 OS=Homo sapiens OX=9606 GN=IGHA1 PE=1 SV=2 | 255 | 20201113_10633_1ng_glyco_hcd203040_compound15.14667.14667.3 | controllerType=0 controllerNumber=1 scan=14667 | 30.6669 |
| 11535:3 | 11 | R.LAGKPTHVN[+1663.608]VSVVMAEVDGTC[+57.021]Y.- | HexNAc(5)Hex(4) | C[+57], N[+1664] | 1337.5858 | 3 | 4010.7427 | 4010.7502 | 0 | -1.9 | 332 | CRagged | 508.97 | 508.97 | 378.16 | 9.45 | 40 | >sp\|P01876\|IGHA1_HUMAN Immunoglobulin heavy constant alpha 1 OS=Homo sapiens OX=9606 GN=IGHA1 PE=1 SV=2 | 255 | 20201113_10633_1ng_glyco_hcd203040_compound15.14675.14675.3 | controllerType=0 controllerNumber=1 scan=14675 | 30.6778 |
| 11528:3 | 11 | R.LAGKPTHVN[+1809.666]VSVVMAEVDGTC[+57.021]Y.- | HexNAc(5)Hex(4)Fuc(1) | C[+57], N[+1810] | 1386.2743 | 3 | 4156.8085 | 4156.8081 | 0 | 0.1 | 332 | CRagged | 571.91 | 571.91 | 398.49 | 10.36 | 40 | >sp\|P01876\|IGHA1_HUMAN Immunoglobulin heavy constant alpha 1 OS=Homo sapiens OX=9606 GN=IGHA1 PE=1 SV=2 | 255 | 20201113_10633_1ng_glyco_hcd203040_compound15.14668.14668.3 | controllerType=0 controllerNumber=1 scan=14668 | 30.6677 |
| 11415:3 | 11 | R.LAGKPTHVN[+1702.581]VSVVMAEVDGTC[+57.021]Y.- | HexNAc(2)Hex(8) | C[+57], N[+1703] | 1350.5809 | 3 | 4049.728 | 4049.7234 | 0 | 1.1 | 332 | CRagged | 659.58 | 659.58 | 425.59 | 13.81 | 40 | >sp\|P01876\|IGHA1_HUMAN Immunoglobulin heavy constant alpha 1 OS=Homo sapiens OX=9606 GN=IGHA1 PE=1 SV=2 | 255 | 20201113_10633_1ng_glyco_hcd203040_compound15.14551.14551.3 | controllerType=0 controllerNumber=1 scan=14551 | 30.4691 |
| 11577:3 | 11 | R.LAGKPTHVN[+1501.555]VSVVMAEVDGTC[+57.021]Y.- | HexNAc(5)Hex(3) | C[+57], N[+1502] | 1283.5757 | 3 | 3848.7127 | 3848.6974 | 0 | 4 | 332 | CRagged | 700.15 | 700.15 | 700.15 | 14.49 | 40 | >sp\|P01876\|IGHA1_HUMAN Immunoglobulin heavy constant alpha 1 OS=Homo sapiens OX=9606 GN=IGHA1 PE=1 SV=2 | 255 | 20201113_10633_1ng_glyco_hcd203040_compound15.14719.14719.3 | controllerType=0 controllerNumber=1 scan=14719 | 30.7582 |
| 11579:3 | 11 | R.LAGKPTHVN[+1501.555]VSVVMAEVDGTC[+57.021]Y.- | HexNAc(5)Hex(3) | C[+57], N[+1502] | 1283.5757 | 3 | 3848.7127 | 3848.6974 | 0 | 4 | 332 | CRagged | 733.79 | 733.79 | 733.79 | 14.49 | 40 | >sp\|P01876\|IGHA1_HUMAN Immunoglobulin heavy constant alpha 1 OS=Homo sapiens OX=9606 GN=IGHA1 PE=1 SV=2 | 255 | 20201113_10633_1ng_glyco_hcd203040_compound15.14721.14721.3 | controllerType=0 controllerNumber=1 scan=14721 | 30.7642 |
| 11566:3 | 11 | R.LAGKPTHVN[+1501.555]VSVVMAEVDGTC[+57.021]Y.- | HexNAc(5)Hex(3) | C[+57], N[+1502] | 1283.5717 | 3 | 3848.7006 | 3848.6974 | 0 | 0.8 | 332 | CRagged | 734.28 | 734.28 | 734.28 | 14.81 | 40 | >sp\|P01876\|IGHA1_HUMAN Immunoglobulin heavy constant alpha 1 OS=Homo sapiens OX=9606 GN=IGHA1 PE=1 SV=2 | 255 | 20201113_10633_1ng_glyco_hcd203040_compound15.14707.14707.3 | controllerType=0 controllerNumber=1 scan=14707 | 30.7354 |
| 11568:3 | 11 | R.LAGKPTHVN[+1501.555]VSVVMAEVDGTC[+57.021]Y.- | HexNAc(5)Hex(3) | C[+57], N[+1502] | 1283.5717 | 3 | 3848.7006 | 3848.6974 | 0 | 0.8 | 332 | CRagged | 756.85 | 599.7 | 599.7 | 15.81 | 40 | >sp\|P01876\|IGHA1_HUMAN Immunoglobulin heavy constant alpha 1 OS=Homo sapiens OX=9606 GN=IGHA1 PE=1 SV=2 | 255 | 20201113_10633_1ng_glyco_hcd203040_compound15.14709.14709.3 | controllerType=0 controllerNumber=1 scan=14709 | 30.7403 |
| 11580:3 | 11 | R.LAGKPTHVN[+1647.613]VSVVMAEVDGTC[+57.021]Y.- | HexNAc(5)Hex(3)Fuc(1) | C[+57], N[+1648] | 1332.261 | 3 | 3994.7686 | 3994.7553 | 0 | 3.3 | 332 | CRagged | 785.77 | 669.37 | 669.37 | 15.49 | 40 | >sp\|P01876\|IGHA1_HUMAN Immunoglobulin heavy constant alpha 1 OS=Homo sapiens OX=9606 GN=IGHA1 PE=1 SV=2 | 255 | 20201113_10633_1ng_glyco_hcd203040_compound15.14722.14722.3 | controllerType=0 controllerNumber=1 scan=14722 | 30.7675 |
| 11530:3 | 11 | R.LAGKPTHVN[+1378.476]VSVVMAEVDGTC[+57.021]Y.- | HexNAc(2)Hex(6) | C[+57], N[+1378] | 1242.5455 | 3 | 3725.6218 | 3725.6178 | 0 | 1.1 | 332 | CRagged | 806.96 | 724.48 | 488.79 | 16.81 | 40 | >sp\|P01876\|IGHA1_HUMAN Immunoglobulin heavy constant alpha 1 OS=Homo sapiens OX=9606 GN=IGHA1 PE=1 SV=2 | 255 | 20201113_10633_1ng_glyco_hcd203040_compound15.14670.14670.3 | controllerType=0 controllerNumber=1 scan=14670 | 30.6711 |
| 11418:3 | 11 | R.LAGKPTHVN[+1702.581]VSVVMAEVDGTC[+57.021]Y.- | HexNAc(2)Hex(8) | C[+57], N[+1703] | 1350.5809 | 3 | 4049.728 | 4049.7234 | 0 | 1.1 | 332 | CRagged | 808.7 | 737.77 | 457.11 | 16.81 | 40 | >sp\|P01876\|IGHA1_HUMAN Immunoglobulin heavy constant alpha 1 OS=Homo sapiens OX=9606 GN=IGHA1 PE=1 SV=2 | 255 | 20201113_10633_1ng_glyco_hcd203040_compound15.14554.14554.3 | controllerType=0 controllerNumber=1 scan=14554 | 30.474 |
| 11534:3 | 11 | R.LAGKPTHVN[+1216.423]VSVVMAEVDGTC[+57.021]Y.- | HexNAc(2)Hex(5) | C[+57], N[+1216] | 1188.5287 | 3 | 3563.5717 | 3563.5649 | 0 | 1.9 | 332 | CRagged | 812 | 812 | 463.76 | 16.83 | 40 | >sp\|P01876\|IGHA1_HUMAN Immunoglobulin heavy constant alpha 1 OS=Homo sapiens OX=9606 GN=IGHA1 PE=1 SV=2 | 255 | 20201113_10633_1ng_glyco_hcd203040_compound15.14674.14674.3 | controllerType=0 controllerNumber=1 scan=14674 | 30.6763 |

Table S4. Cont.

| Query #:z | Protein Rank | Glycopeptide | Glycan | Modification Type(s) | Observed m/z | z | Observed (M+H) | Calc. mass (M+H) | Off-by-x error | Mass error (ppm) | StartPosition | Cleavage | Score | Delta | Delta Mod | \|Log Prob\| | # of unique peptides | Protein Name | Protein DB number | Comment | Scan # | Scan Time |
|---|---|---|---|---|---|---|---|---|---|---|---|---|---|---|---|---|---|---|---|---|---|---|
| 11531:3 | 11 | R.LAGKPTHVN[+1809.666]VSVVMAEVDGTC[+57.021]Y.- | HexNAc(5)Hex(4)Fuc(1) | C[+57], N[+1810] | 1386.2743 | 3 | 4156.8085 | 4156.8081 | 0 | 0.1 | 332 | CRagged | 829.41 | 804.24 | 496.06 | 16.6 | 40 | >sp\|P01876\|IGHA1_HUMAN Immunoglobulin heavy constant alpha 1 OS=Homo sapiens OX=9606 GN=IGHA1 PE=1 SV=2 | 255 | 20201113_10633_1ng_glyco_hcd203040_compound15.14671.14671.3 | controllerType=0 controllerNumber=1 scan=14671 | 30.6724 |
| 11536:3 | 11 | R.LAGKPTHVN[+1216.423]VSVVMAEVDGTC[+57.021]Y.- | HexNAc(2)Hex(5) | C[+57], N[+1216] | 1188.5287 | 3 | 3563.5717 | 3563.5649 | 0 | 1.9 | 332 | CRagged | 850.76 | 829.84 | 504.05 | 17.83 | 40 | >sp\|P01876\|IGHA1_HUMAN Immunoglobulin heavy constant alpha 1 OS=Homo sapiens OX=9606 GN=IGHA1 PE=1 SV=2 | 255 | 20201113_10633_1ng_glyco_hcd203040_compound15.14676.14676.3 | controllerType=0 controllerNumber=1 scan=14676 | 30.6828 |
| 11578:3 | 11 | R.LAGKPTHVN[+1647.613]VSVVMAEVDGTC[+57.021]Y.- | HexNAc(5)Hex(3)Fuc(1) | C[+57], N[+1648] | 1332.261 | 3 | 3994.7686 | 3994.7553 | 0 | 3.3 | 332 | CRagged | 852.74 | 711.51 | 711.51 | 17.49 | 40 | >sp\|P01876\|IGHA1_HUMAN Immunoglobulin heavy constant alpha 1 OS=Homo sapiens OX=9606 GN=IGHA1 PE=1 SV=2 | 255 | 20201113_10633_1ng_glyco_hcd203040_compound15.14720.14720.3 | controllerType=0 controllerNumber=1 scan=14720 | 30.7603 |
| 05819:3 | 12 | R.LSPN[+2205.793]ASAEHLELR.W | HexNAc(4)Hex(5)Fuc(2)NeuAc(1) | N[+2206] | 1214.8549 | 3 | 3642.5502 | 3642.5421 | 0 | 2.2 | 52 | Specific | 589.68 | 382.43 | 382.43 | 11.41 | 39 | >sp\|Q13410\|BT1A1_HUMAN Butyrophilin subfamily 1 member A1 OS=Homo sapiens OX=9606 GN=BTN1A1 PE=1 SV=3 | 179 | 20201113_10633_1ng_glyco_hcd203040_compound15.8733.8733.3 | controllerType=0 controllerNumber=1 scan=8733 | 20.7369 |
| 05821:3 | 12 | R.LSPN[+2205.793]ASAEHLELR.W | HexNAc(4)Hex(5)Fuc(2)NeuAc(1) | N[+2206] | 1214.8549 | 3 | 3642.5502 | 3642.5421 | 0 | 2.2 | 52 | Specific | 671.9 | 519.85 | 519.85 | 13.41 | 39 | >sp\|Q13410\|BT1A1_HUMAN Butyrophilin subfamily 1 member A1 OS=Homo sapiens OX=9606 GN=BTN1A1 PE=1 SV=3 | 179 | 20201113_10633_1ng_glyco_hcd203040_compound15.8735.8735.3 | controllerType=0 controllerNumber=1 scan=8735 | 20.7458 |
| 17875:3 | 12 | K.N[+2059.735]VSC[+57.021]YIQNLLLGQEK.K | HexNAc(4)Hex(5)Fuc(1)NeuAc(1) | C[+57], N[+2060] | 1280.2199 | 3 | 3838.6452 | 3838.6455 | 0 | -0.1 | 215 | Specific | 255.05 | 240.3 | 240.3 | 5.95 | 39 | >sp\|Q13410\|BT1A1_HUMAN Butyrophilin subfamily 1 member A1 OS=Homo sapiens OX=9606 GN=BTN1A1 PE=1 SV=3 | 179 | 20201113_10633_1ng_glyco_hcd203040_compound15.21341.21341.3 | controllerType=0 controllerNumber=1 scan=21341 | 42.3754 |
| 17878:3 | 12 | K.N[+2059.735]VSC[+57.021]YIQNLLLGQEK.K | HexNAc(4)Hex(5)Fuc(1)NeuAc(1) | C[+57], N[+2060] | 1280.2199 | 3 | 3838.6452 | 3838.6455 | 0 | -0.1 | 215 | Specific | 393.44 | 229.82 | 229.82 | 5.38 | 39 | >sp\|Q13410\|BT1A1_HUMAN Butyrophilin subfamily 1 member A1 OS=Homo sapiens OX=9606 GN=BTN1A1 PE=1 SV=3 | 179 | 20201113_10633_1ng_glyco_hcd203040_compound15.21344.21344.3 | controllerType=0 controllerNumber=1 scan=21344 | 42.3835 |
| 24825:3 | 15 | R.QLAHQSN[+2204.772]STNIFFSPVSIATAFAMLSLGTK.A | HexNAc(4)Hex(5)NeuAc(2) | N[+2205] | 1796.1462 | 3 | 5386.4241 | 5386.4075 | 0 | 3.1 | 64 | Specific | 603.62 | 603.62 | 603.62 | 12.74 | 33 | >sp\|P01009\|A1AT_HUMAN Alpha-1-antitrypsin OS=Homo sapiens OX=9606 GN=SERPINA1 PE=1 SV=3 | 642 | 20201113_10633_1ng_glyco_hcd203040_compound15.28582.28582.3 | controllerType=0 controllerNumber=1 scan=28582 | 55.9139 |
| 24827:3 | 15 | R.QLAHQSN[+2204.772]STNIFFSPVSIATAFAMLSLGTK.A | HexNAc(4)Hex(5)NeuAc(2) | N[+2205] | 1796.1462 | 3 | 5386.4241 | 5386.4075 | 0 | 3.1 | 64 | Specific | 697.1 | 618.52 | 618.52 | 13.74 | 33 | >sp\|P01009\|A1AT_HUMAN Alpha-1-antitrypsin OS=Homo sapiens OX=9606 GN=SERPINA1 PE=1 SV=3 | 642 | 20201113_10633_1ng_glyco_hcd203040_compound15.28584.28584.3 | controllerType=0 controllerNumber=1 scan=28584 | 55.9178 |
| 24822:4 | 15 | R.QLAHQSN[+2204.772]STNIFFSPVSIATAFAMLSLGTK.A | HexNAc(4)Hex(5)NeuAc(2) | N[+2205] | 1347.3589 | 4 | 5386.4137 | 5386.4075 | 0 | 1.2 | 64 | Specific | 806.46 | 710.43 | 517.37 | 16.56 | 33 | >sp\|P01009\|A1AT_HUMAN Alpha-1-antitrypsin OS=Homo sapiens OX=9606 GN=SERPINA1 PE=1 SV=3 | 642 | 20201113_10633_1ng_glyco_hcd203040_compound15.28579.28579.4 | controllerType=0 controllerNumber=1 scan=28579 | 55.906 |
| 24824:4 | 15 | R.QLAHQSN[+2204.772]STNIFFSPVSIATAFAMLSLGTK.A | HexNAc(4)Hex(5)NeuAc(2) | N[+2205] | 1347.3589 | 4 | 5386.4137 | 5386.4075 | 0 | 1.2 | 64 | Specific | 840.7 | 784.68 | 549.55 | 16.56 | 33 | >sp\|P01009\|A1AT_HUMAN Alpha-1-antitrypsin OS=Homo sapiens OX=9606 GN=SERPINA1 PE=1 SV=3 | 642 | 20201113_10633_1ng_glyco_hcd203040_compound15.28581.28581.4 | controllerType=0 controllerNumber=1 scan=28581 | 55.912 |
| 23366:4 | 15 | K.ADTHDEILEGLNFN[+2861.000]LTEIPEAQIHEGFQELLR.T | HexNAc(5)Hex(6)NeuAc(3) | N[+2861] | 1638.9667 | 4 | 6552.8449 | 6552.8239 | 0 | 3.2 | 94 | Specific | 625.82 | 532.16 | 532.16 | 12.63 | 33 | >sp\|P01009\|A1AT_HUMAN Alpha-1-antitrypsin OS=Homo sapiens OX=9606 GN=SERPINA1 PE=1 SV=3 | 642 | 20201113_10633_1ng_glyco_hcd203040_compound15.27066.27066.4 | controllerType=0 controllerNumber=1 scan=27066 | 53.153 |
| 23369:4 | 15 | K.ADTHDEILEGLNFN[+2861.000]LTEIPEAQIHEGFQELLR.T | HexNAc(5)Hex(6)NeuAc(3) | N[+2861] | 1638.9667 | 4 | 6552.8449 | 6552.8239 | 0 | 3.2 | 94 | Specific | 703.5 | 692.64 | 692.64 | 14.63 | 33 | >sp\|P01009\|A1AT_HUMAN Alpha-1-antitrypsin OS=Homo sapiens OX=9606 GN=SERPINA1 PE=1 SV=3 | 642 | 20201113_10633_1ng_glyco_hcd203040_compound15.27069.27069.4 | controllerType=0 controllerNumber=1 scan=27069 | 53.16 |
| 23383:4 | 15 | K.ADTHDEILEGLNFN[+2861.000]LTEIPEAQIHEGFQELLR.T | HexNAc(5)Hex(6)NeuAc(3) | N[+2861] | 1638.9716 | 4 | 6552.8644 | 6552.8239 | 0 | 6.2 | 94 | Specific | 735.31 | 735.31 | 735.31 | 14.18 | 33 | >sp\|P01009\|A1AT_HUMAN Alpha-1-antitrypsin OS=Homo sapiens OX=9606 GN=SERPINA1 PE=1 SV=3 | 642 | 20201113_10633_1ng_glyco_hcd203040_compound15.27084.27084.4 | controllerType=0 controllerNumber=1 scan=27084 | 53.1917 |
| 23385:4 | 15 | K.ADTHDEILEGLNFN[+2861.000]LTEIPEAQIHEGFQELLR.T | HexNAc(5)Hex(6)NeuAc(3) | N[+2861] | 1638.9716 | 4 | 6552.8644 | 6552.8239 | 0 | 6.2 | 94 | Specific | 773.05 | 773.05 | 773.05 | 15.18 | 33 | >sp\|P01009\|A1AT_HUMAN Alpha-1-antitrypsin OS=Homo sapiens OX=9606 GN=SERPINA1 PE=1 SV=3 | 642 | 20201113_10633_1ng_glyco_hcd203040_compound15.27086.27086.4 | controllerType=0 controllerNumber=1 scan=27086 | 53.1958 |
| 20467:4 | 15 | K.YLGN[+2204.772]ATAIFFLPDEGKLQHLENELTHDIITK.F | HexNAc(4)Hex(5)NeuAc(2) | N[+2205] | 1437.1646 | 4 | 5745.6364 | 5745.6098 | 0 | 4.6 | 268 | Specific | 536.27 | 397.28 | 397.28 | 8.92 | 33 | >sp\|P01009\|A1AT_HUMAN Alpha-1-antitrypsin OS=Homo sapiens OX=9606 GN=SERPINA1 PE=1 SV=3 | 642 | 20201113_10633_1ng_glyco_hcd203040_compound15.24043.24043.4 | controllerType=0 controllerNumber=1 scan=24043 | 47.7665 |
| 20468:5 | 15 | K.YLGN[+2204.772]ATAIFFLPDEGKLQHLENELTHDIITK.F | HexNAc(4)Hex(5)NeuAc(2) | N[+2205] | 1149.9312 | 5 | 5745.6267 | 5745.6098 | 0 | 2.9 | 268 | Specific | 686.3 | 588.31 | 588.31 | 13.62 | 33 | >sp\|P01009\|A1AT_HUMAN Alpha-1-antitrypsin OS=Homo sapiens OX=9606 GN=SERPINA1 PE=1 SV=3 | 642 | 20201113_10633_1ng_glyco_hcd203040_compound15.24044.24044.5 | controllerType=0 controllerNumber=1 scan=24044 | 47.7685 |
| 14309:4 | 23 | K.LKELPGVC[+57.021]N[+2059.735]ETMMALWEEC[+57.021]KPC[+57.021]LK.Q | HexNAc(4)Hex(5)Fuc(1)NeuAc(1) | C[+57]*3, N[+2060] | 1249.7899 | 4 | 4996.1379 | 4996.1196 | 0 | 3.6 | 95 | Specific | 293.74 | 293.74 | 293.74 | 6.52 | 24 | >sp\|P10909\|CLUS_HUMAN Clusterin OS=Homo sapiens OX=9606 GN=CLU PE=1 SV=1 | 1467 | 20201113_10633_1ng_glyco_hcd203040_compound15.17588.17588.4 | controllerType=0 controllerNumber=1 scan=17588 | 35.6342 |
| 00510:3 | 23 | R.EIRHN[+2059.735]STGC[+57.021]LR.M | HexNAc(4)Hex(5)Fuc(1)NeuAc(1) | C[+57], N[+2060] | 1134.8054 | 3 | 3402.4018 | 3402.3994 | 0 | 0.7 | 287 | Specific | 294.18 | 184.52 | 184.52 | 5.11 | 24 | >sp\|P10909\|CLUS_HUMAN Clusterin OS=Homo sapiens OX=9606 GN=CLU PE=1 SV=1 | 1467 | 20201113_10633_1ng_glyco_hcd203040_compound15.3229.3229.3 | controllerType=0 controllerNumber=1 scan=3229 | 11.2875 |
| 00512:3 | 23 | R.EIRHN[+2059.735]STGC[+57.021]LR.M | HexNAc(4)Hex(5)Fuc(1)NeuAc(1) | C[+57], N[+2060] | 1134.8054 | 3 | 3402.4018 | 3402.3994 | 0 | 0.7 | 287 | Specific | 311.01 | 206.06 | 206.06 | 5.58 | 24 | >sp\|P10909\|CLUS_HUMAN Clusterin OS=Homo sapiens OX=9606 GN=CLU PE=1 SV=1 | 1467 | 20201113_10633_1ng_glyco_hcd203040_compound15.3231.3231.3 | controllerType=0 controllerNumber=1 scan=3231 | 11.292 |
| 24007:4 | 23 | K.MLN[+2350.830]TSSLLEQLNEQFNWVSR.L | HexNAc(4)Hex(5)Fuc(1)NeuAc(2) | N[+2351] | 1190.7621 | 4 | 4760.0267 | 4760.0171 | 0 | 2 | 352 | Specific | 306.5 | 298.8 | 298.8 | 7.95 | 24 | >sp\|P10909\|CLUS_HUMAN Clusterin OS=Homo sapiens OX=9606 GN=CLU PE=1 SV=1 | 1467 | 20201113_10633_1ng_glyco_hcd203040_compound15.27732.27732.4 | controllerType=0 controllerNumber=1 scan=27732 | 54.3926 |
| 24009:4 | 23 | K.MLN[+2350.830]TSSLLEQLNEQFNWVSR.L | HexNAc(4)Hex(5)Fuc(1)NeuAc(2) | N[+2351] | 1190.7621 | 4 | 4760.0267 | 4760.0171 | 0 | 2 | 352 | Specific | 321.25 | 271.13 | 271.13 | 6.99 | 24 | >sp\|P10909\|CLUS_HUMAN Clusterin OS=Homo sapiens OX=9606 GN=CLU PE=1 SV=1 | 1467 | 20201113_10633_1ng_glyco_hcd203040_compound15.27734.27734.4 | controllerType=0 controllerNumber=1 scan=27734 | 54.3973 |
| 23994:3 | 23 | K.MLN[+2350.830]TSSLLEQLNEQFNWVSR.L | HexNAc(4)Hex(5)Fuc(1)NeuAc(2) | N[+2351] | 1587.3478 | 3 | 4760.029 | 4760.0171 | 0 | 2.5 | 352 | Specific | 562.83 | 483.56 | 483.56 | 10.43 | 24 | >sp\|P10909\|CLUS_HUMAN Clusterin OS=Homo sapiens OX=9606 GN=CLU PE=1 SV=1 | 1467 | 20201113_10633_1ng_glyco_hcd203040_compound15.27719.27719.3 | controllerType=0 controllerNumber=1 scan=27719 | 54.3696 |
| 23997:3 | 23 | K.MLN[+2350.830]TSSLLEQLNEQFNWVSR.L | HexNAc(4)Hex(5)Fuc(1)NeuAc(2) | N[+2351] | 1587.3478 | 3 | 4760.029 | 4760.0171 | 0 | 2.5 | 352 | Specific | 715.75 | 519.23 | 519.23 | 14.67 | 24 | >sp\|P10909\|CLUS_HUMAN Clusterin OS=Homo sapiens OX=9606 GN=CLU PE=1 SV=1 | 1467 | 20201113_10633_1ng_glyco_hcd203040_compound15.27722.27722.3 | controllerType=0 controllerNumber=1 scan=27722 | 54.3744 |
| 10472:3 | 23 | R.LAN[+2205.793]LTQGEDQYYLR.V | HexNAc(4)Hex(5)Fuc(2)NeuAc(1) | N[+2206] | 1297.2166 | 3 | 3889.6353 | 3889.6265 | 0 | 2.3 | 372 | Specific | 234.37 | 234.37 | 234.37 | 6.11 | 24 | >sp\|P10909\|CLUS_HUMAN Clusterin OS=Homo sapiens OX=9606 GN=CLU PE=1 SV=1 | 1467 | 20201113_10633_1ng_glyco_hcd203040_compound15.13576.13576.3 | controllerType=0 controllerNumber=1 scan=13576 | 28.8318 |
| 10474:3 | 23 | R.LAN[+2205.793]LTQGEDQYYLR.V | HexNAc(4)Hex(5)Fuc(2)NeuAc(1) | N[+2206] | 1297.2166 | 3 | 3889.6353 | 3889.6265 | 0 | 2.3 | 372 | Specific | 358.91 | 346.34 | 346.34 | 7.59 | 24 | >sp\|P10909\|CLUS_HUMAN Clusterin OS=Homo sapiens OX=9606 GN=CLU PE=1 SV=1 | 1467 | 20201113_10633_1ng_glyco_hcd203040_compound15.13578.13578.3 | controllerType=0 controllerNumber=1 scan=13578 | 28.8359 |
| 14475:2 | 25 | K.LC[+57.021]DN[+1866.688]LSTK.N | HexNAc(6)Hex(4) | C[+57], N[+1867] | 1409.0729 | 2 | 2817.1386 | 2817.1487 | 0 | -3.6 | 285 | Specific | 233.67 | 233.67 | 233.67 | 2.09 | 23 | >sp\|P02774\|VTDB_HUMAN Vitamin D-binding protein OS=Homo sapiens OX=9606 GN=GC PE=1 SV=2 | 1103 | 20201113_10633_1ng_glyco_hcd203040_compound15.17760.17760.2 | controllerType=0 controllerNumber=1 scan=17760 | 35.9094 |
| 22717:4 | 41 | K.VFPLSLDSTPQDGNVVVAC[+57.021]LVQGFFPQEPLSVTWSESGQN[+1647.613]VTAI | HexNAc(5)Hex(3)Fuc(1) | C[+57], N[+1648] | 1606.7611 | 4 | 6424.0226 | 6423.9746 | 0 | 7.5 | 123 | Specific | 735.57 | 675.6 | 675.6 | 14.98 | 16 | >sp\|P0DOX2\|IGA2_HUMAN Immunoglobulin alpha-2 heavy chain OS=Homo sapiens OX=9606 PE=1 SV=2 | 631 | 20201113_10633_1ng_glyco_hcd203040_compound15.26379.26379.4 | controllerType=0 controllerNumber=1 scan=26379 | 52.023 |
| 22702:4 | 41 | K.VFPLSLDSTPQDGNVVVAC[+57.021]LVQGFFPQEPLSVTWSESGQN[+1647.613]VTAI | HexNAc(5)Hex(3)Fuc(1) | C[+57], N[+1648] | 1606.7551 | 4 | 6423.9987 | 6423.9746 | 0 | 3.7 | 123 | Specific | 765.19 | 674.26 | 674.26 | 16.8 | 16 | >sp\|P0DOX2\|IGA2_HUMAN Immunoglobulin alpha-2 heavy chain OS=Homo sapiens OX=9606 PE=1 SV=2 | 631 | 20201113_10633_1ng_glyco_hcd203040_compound15.26363.26363.4 | controllerType=0 controllerNumber=1 scan=26363 | 51.9876 |
| 22719:4 | 41 | K.VFPLSLDSTPQDGNVVVAC[+57.021]LVQGFFPQEPLSVTWSESGQN[+1647.613]VTAI | HexNAc(5)Hex(3)Fuc(1) | C[+57], N[+1648] | 1606.7611 | 4 | 6424.0226 | 6423.9746 | 0 | 7.5 | 123 | Specific | 905.09 | 905.09 | 905.09 | 18.98 | 16 | >sp\|P0DOX2\|IGA2_HUMAN Immunoglobulin alpha-2 heavy chain OS=Homo sapiens OX=9606 PE=1 SV=2 | 631 | 20201113_10633_1ng_glyco_hcd203040_compound15.26381.26381.4 | controllerType=0 controllerNumber=1 scan=26381 | 52.0313 |
| 22705:4 | 41 | K.VFPLSLDSTPQDGNVVVAC[+57.021]LVQGFFPQEPLSVTWSESGQN[+1647.613]VTAI | HexNAc(5)Hex(3)Fuc(1) | C[+57], N[+1648] | 1606.7551 | 4 | 6423.9987 | 6423.9746 | 0 | 3.7 | 123 | Specific | 919.82 | 777.84 | 777.84 | 19.8 | 16 | >sp\|P0DOX2\|IGA2_HUMAN Immunoglobulin alpha-2 heavy chain OS=Homo sapiens OX=9606 PE=1 SV=2 | 631 | 20201113_10633_1ng_glyco_hcd203040_compound15.26366.26366.4 | controllerType=0 controllerNumber=1 scan=26366 | 52.0037 |
| 03211:2 | 41 | K.TPLTAN[+1501.555]ITK.S | HexNAc(5)Hex(3) | N[+1502] | 1230.5629 | 2 | 2460.1185 | 2460.1121 | 0 | 2.6 | 315 | Specific | 228.13 | 84.63 | 84.63 | 4.45 | 16 | >sp\|P0DOX2\|IGA2_HUMAN Immunoglobulin alpha-2 heavy chain OS=Homo sapiens OX=9606 PE=1 SV=2 | 631 | 20201113_10633_1ng_glyco_hcd203040_compound15.6033.6033.2 | controllerType=0 controllerNumber=1 scan=6033 | 16.2722 |
| 03133:3 | 41 | K.TPLTAN[+1809.666]ITK.S | HexNAc(5)Hex(4)Fuc(1) | N[+1810] | 923.4138 | 3 | 2768.2268 | 2768.2228 | 0 | 1.4 | 315 | Specific | 282.1 | 155.86 | 155.86 | 5.21 | 16 | >sp\|P0DOX2\|IGA2_HUMAN Immunoglobulin alpha-2 heavy chain OS=Homo sapiens OX=9606 PE=1 SV=2 | 631 | 20201113_10633_1ng_glyco_hcd203040_compound15.5952.5952.3 | controllerType=0 controllerNumber=1 scan=5952 | 16.1254 |
| 03214:2 | 41 | K.TPLTAN[+1501.555]ITK.S | HexNAc(5)Hex(3) | N[+1502] | 1230.5629 | 2 | 2460.1185 | 2460.1121 | 0 | 2.6 | 315 | Specific | 331.03 | 307.02 | 307.02 | 7.21 | 16 | >sp\|P0DOX2\|IGA2_HUMAN Immunoglobulin alpha-2 heavy chain OS=Homo sapiens OX=9606 PE=1 SV=2 | 631 | 20201113_10633_1ng_glyco_hcd203040_compound15.6036.6036.2 | controllerType=0 controllerNumber=1 scan=6036 | 16.2766 |
| 03173:2 | 41 | K.TPLTAN[+1444.534]ITK.S | HexNAc(4)Hex(3)Fuc(1) | N[+1445] | 1202.048 | 2 | 2403.0888 | 2403.0906 | 0 | -0.8 | 315 | Specific | 419.92 | 223.27 | 223.27 | 6.46 | 16 | >sp\|P0DOX2\|IGA2_HUMAN Immunoglobulin alpha-2 heavy chain OS=Homo sapiens OX=9606 PE=1 SV=2 | 631 | 20201113_10633_1ng_glyco_hcd203040_compound15.5993.5993.2 | controllerType=0 controllerNumber=1 scan=5993 | 16.1985 |
| 03204:3 | 41 | K.TPLTAN[+1647.613]ITK.S | HexNAc(5)Hex(3)Fuc(1) | N[+1648] | 869.3961 | 3 | 2606.1739 | 2606.17 | 0 | 1.5 | 315 | Specific | 462.54 | 198.37 | 198.37 | 6.48 | 16 | >sp\|P0DOX2\|IGA2_HUMAN Immunoglobulin alpha-2 heavy chain OS=Homo sapiens OX=9606 PE=1 SV=2 | 631 | 20201113_10633_1ng_glyco_hcd203040_compound15.6026.6026.3 | controllerType=0 controllerNumber=1 scan=6026 | 16.2584 |
| 03135:3 | 41 | K.TPLTAN[+1809.666]ITK.S | HexNAc(5)Hex(4)Fuc(1) | N[+1810] | 923.4138 | 3 | 2768.2268 | 2768.2228 | 0 | 1.4 | 315 | Specific | 500.59 | 256.15 | 256.15 | 7.35 | 16 | >sp\|P0DOX2\|IGA2_HUMAN Immunoglobulin alpha-2 heavy chain OS=Homo sapiens OX=9606 PE=1 SV=2 | 631 | 20201113_10633_1ng_glyco_hcd203040_compound15.5954.5954.3 | controllerType=0 controllerNumber=1 scan=5954 | 16.1294 |
| 03175:2 | 41 | K.TPLTAN[+1444.534]ITK.S | HexNAc(4)Hex(3)Fuc(1) | N[+1445] | 1202.048 | 2 | 2403.0888 | 2403.0906 | 0 | -0.8 | 315 | Specific | 503.57 | 300.18 | 300.18 | 7.35 | 16 | >sp\|P0DOX2\|IGA2_HUMAN Immunoglobulin alpha-2 heavy chain OS=Homo sapiens OX=9606 PE=1 SV=2 | 631 | 20201113_10633_1ng_glyco_hcd203040_compound15.5995.5995.2 | controllerType=0 controllerNumber=1 scan=5995 | 16.2031 |
| 03141:2 | 41 | K.TPLTAN[+1809.666]ITK.S | HexNAc(5)Hex(4)Fuc(1) | N[+1810] | 1384.6166 | 2 | 2768.226 | 2768.2228 | 0 | 1.1 | 315 | Specific | 514.46 | 298.32 | 298.32 | 7.56 | 16 | >sp\|P0DOX2\|IGA2_HUMAN Immunoglobulin alpha-2 heavy chain OS=Homo sapiens OX=9606 PE=1 SV=2 | 631 | 20201113_10633_1ng_glyco_hcd203040_compound15.5960.5960.2 | controllerType=0 controllerNumber=1 scan=5960 | 16.1389 |
| 03206:3 | 41 | K.TPLTAN[+1647.613]ITK.S | HexNAc(5)Hex(3)Fuc(1) | N[+1648] | 869.3961 | 3 | 2606.1739 | 2606.17 | 0 | 1.5 | 315 | Specific | 548.62 | 267.42 | 267.42 | 7.51 | 16 | >sp\|P0DOX2\|IGA2_HUMAN Immunoglobulin alpha-2 heavy chain OS=Homo sapiens OX=9606 PE=1 SV=2 | 631 | 20201113_10633_1ng_glyco_hcd203040_compound15.6028.6028.3 | controllerType=0 controllerNumber=1 scan=6028 | 16.2639 |
| 03143:2 | 41 | K.TPLTAN[+1809.666]ITK.S | HexNAc(5)Hex(4)Fuc(1) | N[+1810] | 1384.6166 | 2 | 2768.226 | 2768.2228 | 0 | 1.1 | 315 | Specific | 594.57 | 287.9 | 287.9 | 9.13 | 16 | >sp\|P0DOX2\|IGA2_HUMAN Immunoglobulin alpha-2 heavy chain OS=Homo sapiens OX=9606 PE=1 SV=2 | 631 | 20201113_10633_1ng_glyco_hcd203040_compound15.5962.5962.2 | controllerType=0 controllerNumber=1 scan=5962 | 16.143 |
| 03186:2 | 41 | K.TPLTAN[+1647.613]ITK.S | HexNAc(5)Hex(3)Fuc(1) | N[+1648] | 1303.5911 | 2 | 2606.1749 | 2606.17 | 0 | 1.9 | 315 | Specific | 670.6 | 282.82 | 282.82 | 11.14 | 16 | >sp\|P0DOX2\|IGA2_HUMAN Immunoglobulin alpha-2 heavy chain OS=Homo sapiens OX=9606 PE=1 SV=2 | 631 | 20201113_10633_1ng_glyco_hcd203040_compound15.6007.6007.2 | controllerType=0 controllerNumber=1 scan=6007 | 16.2238 |
| 03188:2 | 41 | K.TPLTAN[+1647.613]ITK.S | HexNAc(5)Hex(3)Fuc(1) | N[+1648] | 1303.5911 | 2 | 2606.1749 | 2606.17 | 0 | 1.9 | 315 | Specific | 678.15 | 301.55 | 301.55 | 11.14 | 16 | >sp\|P0DOX2\|IGA2_HUMAN Immunoglobulin alpha-2 heavy chain OS=Homo sapiens OX=9606 PE=1 SV=2 | 631 | 20201113_10633_1ng_glyco_hcd203040_compound15.6009.6009.2 | controllerType=0 controllerNumber=1 scan=6009 | 16.2292 |
| 13441:3 | 56 | R.IIVPLNNREN[+2204.772]ISDPTSPLR.T | HexNAc(4)Hex(5)NeuAc(2) | N[+2205] | 1451.6591 | 3 | 4352.9628 | 4352.9496 | 0 | 3 | 62 | Specific | 219.04 | 219.04 | 219.04 | 4.92 | 15 | >sp\|P01591\|IGJ_HUMAN Immunoglobulin J chain OS=Homo sapiens OX=9606 GN=JCHAIN PE=1 SV=4 | 1392 | 20201113_10633_1ng_glyco_hcd203040_compound15.16690.16690.3 | controllerType=0 controllerNumber=1 scan=16690 | 34.1172 |
| 11633:3 | 56 | R.IIVPLNNREN[+1694.603]ISDPTSPLR.T | HexNAc(3)Hex(4)Fuc(1)NeuAc(1) | N[+1695] | 1281.6022 | 3 | 3842.7921 | 3842.7799 | 0 | 3.2 | 62 | Specific | 245.54 | 190.14 | 190.14 | 3.92 | 15 | >sp\|P01591\|IGJ_HUMAN Immunoglobulin J chain OS=Homo sapiens OX=9606 GN=JCHAIN PE=1 SV=4 | 1392 | 20201113_10633_1ng_glyco_hcd203040_compound15.14777.14777.3 | controllerType=0 controllerNumber=1 scan=14777 | 30.8963 |
| 13443:3 | 56 | R.IIVPLNNREN[+2204.772]ISDPTSPLR.T | HexNAc(4)Hex(5)NeuAc(2) | N[+2205] | 1451.6591 | 3 | 4352.9628 | 4352.9496 | 0 | 3 | 62 | Specific | 312.49 | 296.8 | 296.8 | 6.91 | 15 | >sp\|P01591\|IGJ_HUMAN Immunoglobulin J chain OS=Homo sapiens OX=9606 GN=JCHAIN PE=1 SV=4 | 1392 | 20201113_10633_1ng_glyco_hcd203040_compound15.16692.16692.3 | controllerType=0 controllerNumber=1 scan=16692 | 34.1207 |
| 11631:3 | 56 | R.IIVPLNNREN[+1694.603]ISDPTSPLR.T | HexNAc(3)Hex(4)Fuc(1)NeuAc(1) | N[+1695] | 1281.6022 | 3 | 3842.7921 | 3842.7799 | 0 | 3.2 | 62 | Specific | 315.52 | 315.52 | 315.52 | 6.8 | 15 | >sp\|P01591\|IGJ_HUMAN Immunoglobulin J chain OS=Homo sapiens OX=9606 GN=JCHAIN PE=1 SV=4 | 1392 | 20201113_10633_1ng_glyco_hcd203040_compound15.14775.14775.3 | controllerType=0 controllerNumber=1 scan=14775 | 30.8924 |
| 11452:3 | 56 | R.IIVPLNNREN[+1913.677]ISDPTSPLR.T | HexNAc(4)Hex(5)NeuAc(1) | N[+1914] | 1354.624 | 3 | 4061.8573 | 4061.8542 | 0 | 0.8 | 62 | Specific | 322.04 | 315.51 | 315.51 | 7.12 | 15 | >sp\|P01591\|IGJ_HUMAN Immunoglobulin J chain OS=Homo sapiens OX=9606 GN=JCHAIN PE=1 SV=4 | 1392 | 20201113_10633_1ng_glyco_hcd203040_compound15.14589.14589.3 | controllerType=0 controllerNumber=1 scan=14589 | 30.531 |
| 11454:3 | 56 | R.IIVPLNNREN[+1913.677]ISDPTSPLR.T | HexNAc(4)Hex(5)NeuAc(1) | N[+1914] | 1354.624 | 3 | 4061.8573 | 4061.8542 | 0 | 0.8 | 62 | Specific | 413.45 | 342.48 | 342.48 | 7.51 | 15 | >sp\|P01591\|IGJ_HUMAN Immunoglobulin J chain OS=Homo sapiens OX=9606 GN=JCHAIN PE=1 SV=4 | 1392 | 20201113_10633_1ng_glyco_hcd203040_compound15.14591.14591.3 | controllerType=0 controllerNumber=1 scan=14591 | 30.5361 |
| 06776:3 | 364 | R.[+42.011]VEDAYILTC[+57.021]N[+730.264]VSLEYEK.T | HexNAc(2)Hex(2) | .[42], C[+57], N[+730] | 940.0938 | 3 | 2818.2668 | 2818.2486 | 0 | 6.5 | 225 | Specific | 258.59 | 193.87 | 193.87 | 4.34 | 3 | >sp\|P40227\|TCPZ_HUMAN T-complex protein 1 subunit zeta OS=Homo sapiens OX=9606 GN=CCT6A PE=1 SV=3 | 1115 | 20201113_10633_1ng_glyco_hcd203040_compound15.9721.9721.3 | controllerType=0 controllerNumber=1 scan=9721 | 22.3476 |
| 06778:3 | 364 | R.[+42.011]VEDAYILTC[+57.021]N[+730.264]VSLEYEK.T | HexNAc(2)Hex(2) | .[42], C[+57], N[+730] | 940.0938 | 3 | 2818.2668 | 2818.2486 | 0 | 6.5 | 225 | Specific | 263.23 | 122.62 | 122.62 | 2.77 | 3 | >sp\|P40227\|TCPZ_HUMAN T-complex protein 1 subunit zeta OS=Homo sapiens OX=9606 GN=CCT6A PE=1 SV=3 | 1115 | 20201113_10633_1ng_glyco_hcd203040_compound15.9723.9723.3 | controllerType=0 controllerNumber=1 scan=9723 | 22.3518 |
| 13814:3 | 395 | R.[+42.011]TVLISKN[+2059.735]LTNPR.G | HexNAc(4)Hex(5)Fuc(1)NeuAc(1) | .[42], N[+2060] | 1153.1949 | 3 | 3457.5701 | 3457.546 | 0 | 7 | 1544 | Specific | 237.98 | 170.55 | 170.55 | 3.37 | 4 | >sp\|P98164\|LRP2_HUMAN Low-density lipoprotein receptor-related protein 2 OS=Homo sapiens OX=9606 GN=LRP2 PE=1 SV=3 | 656 | 20201113_10633_1ng_glyco_hcd203040_compound15.17076.17076.3 | controllerType=0 controllerNumber=1 scan=17076 | 34.775 |
| 13817:3 | 395 | R.[+42.011]TVLISKN[+2059.735]LTNPR.G | HexNAc(4)Hex(5)Fuc(1)NeuAc(1) | .[42], N[+2060] | 1153.1949 | 3 | 3457.5701 | 3457.546 | 0 | 7 | 1544 | Specific | 308.24 | 124.71 | 124.71 | 3.21 | 4 | >sp\|P98164\|LRP2_HUMAN Low-density lipoprotein receptor-related protein 2 OS=Homo sapiens OX=9606 GN=LRP2 PE=1 SV=3 | 656 | 20201113_10633_1ng_glyco_hcd203040_compound15.17079.17079.3 | controllerType=0 controllerNumber=1 scan=17079 | 34.7797 |
| 17867:4 | 1 | R.RTAGWNVPIGTLRPFLN[+2351.851]WTGPPEPIEAAVAR.F | HexNAc(4)Hex(5)Fuc(3)NeuAc(1) | N[+2352] | 1435.4252 | 4 | 5738.6789 | 5738.6628 | 0 | 2.8 | 140 | Specific | 392.06 | 310.16 | 310.16 | 7.2 | 149 | >sp\|P02788\|TRFL_HUMAN Lactotransferrin OS=Homo sapiens OX=9606 GN=LTF PE=1 SV=6 | 1270 | 20201113_12413_1ng_glyco_hcd203040_compound15.21394.21394.4 | controllerType=0 controllerNumber=1 scan=21394 | 43.5499 |
| 17749:4 | 1 | R.RTAGWNVPIGTLRPFLN[+2716.983]WTGPPEPIEAAVAR.F | HexNAc(5)Hex(6)Fuc(3)NeuAc(1) | N[+2717] | 1526.7095 | 4 | 6103.8161 | 6103.795 | 0 | 3.5 | 140 | Specific | 440.99 | 431.57 | 431.57 | 9.1 | 149 | >sp\|P02788\|TRFL_HUMAN Lactotransferrin OS=Homo sapiens OX=9606 GN=LTF PE=1 SV=6 | 1270 | 20201113_12413_1ng_glyco_hcd203040_compound15.21271.21271.4 | controllerType=0 controllerNumber=1 scan=21271 | 43.2982 |
| 16962:4 | 1 | R.RTAGWNVPIGTLRPFLN[+1768.640]WTGPPEPIEAAVAR.F | HexNAc(4)Hex(5)Fuc(1) | N[+1769] | 1289.6209 | 4 | 5155.4616 | 5155.4516 | 0 | 1.9 | 140 | Specific | 444.21 | 392.67 | 392.67 | 9.01 | 149 | >sp\|P02788\|TRFL_HUMAN Lactotransferrin OS=Homo sapiens OX=9606 GN=LTF PE=1 SV=6 | 1270 | 20201113_12413_1ng_glyco_hcd203040_compound15.20448.20448.4 | controllerType=0 controllerNumber=1 scan=20448 | 41.5044 |
| 17752:4 | 1 | R.RTAGWNVPIGTLRPFLN[+2716.983]WTGPPEPIEAAVAR.F | HexNAc(5)Hex(6)Fuc(3)NeuAc(1) | N[+2717] | 1526.7095 | 4 | 6103.8161 | 6103.795 | 0 | 3.5 | 140 | Specific | 463.56 | 374.27 | 374.27 | 9.1 | 149 | >sp\|P02788\|TRFL_HUMAN Lactotransferrin OS=Homo sapiens OX=9606 GN=LTF PE=1 SV=6 | 1270 | 20201113_12413_1ng_glyco_hcd203040_compound15.21274.21274.4 | controllerType=0 controllerNumber=1 scan=21274 | 43.3105 |
| 17869:4 | 1 | R.RTAGWNVPIGTLRPFLN[+2351.851]WTGPPEPIEAAVAR.F | HexNAc(4)Hex(5)Fuc(3)NeuAc(1) | N[+2352] | 1435.4252 | 4 | 5738.6789 | 5738.6628 | 0 | 2.8 | 140 | Specific | 522.5 | 346.13 | 346.13 | 7.97 | 149 | >sp\|P02788\|TRFL_HUMAN Lactotransferrin OS=Homo sapiens OX=9606 GN=LTF PE=1 SV=6 | 1270 | 20201113_12413_1ng_glyco_hcd203040_compound15.21396.21396.4 | controllerType=0 controllerNumber=1 scan=21396 | 43.5589 |
| 18879:4 | 1 | R.RTAGWNVPIGTLRPFLN[+2350.830]WTGPPEPIEAAVAR.F | HexNAc(4)Hex(5)Fuc(1)NeuAc(2) | N[+2351] | 1435.1801 | 4 | 5737.6984 | 5737.6424 | 0 | 9.8 | 140 | Specific | 524.69 | 516.02 | 516.02 | 8.55 | 149 | >sp\|P02788\|TRFL_HUMAN Lactotransferrin OS=Homo sapiens OX=9606 GN=LTF PE=1 SV=6 | 1270 | 20201113_12413_1ng_glyco_hcd203040_compound15.22454.22454.4 | controllerType=0 controllerNumber=1 scan=22454 | 45.8547 |
| 16964:4 | 1 | R.RTAGWNVPIGTLRPFLN[+1768.640]WTGPPEPIEAAVAR.F | HexNAc(4)Hex(5)Fuc(1) | N[+1769] | 1289.6209 | 4 | 5155.4616 | 5155.4516 | 0 | 1.9 | 140 | Specific | 572.04 | 501.52 | 501.52 | 9.61 | 149 | >sp\|P02788\|TRFL_HUMAN Lactotransferrin OS=Homo sapiens OX=9606 GN=LTF PE=1 SV=6 | 1270 | 20201113_12413_1ng_glyco_hcd203040_compound15.20450.20450.4 | controllerType=0 controllerNumber=1 scan=20450 | 41.5081 |
| 18961:3 | 1 | R.TAGWNVPIGTLRPFLN[+2716.983]WTGPPEPIEAAVAR.F | HexNAc(5)Hex(6)Fuc(3)NeuAc(1) | N[+2717] | 1983.2417 | 3 | 5947.7104 | 5947.6939 | 0 | 2.8 | 141 | Specific | 206.91 | 206.91 | 206.91 | 4.64 | 149 | >sp\|P02788\|TRFL_HUMAN Lactotransferrin OS=Homo sapiens OX=9606 GN=LTF PE=1 SV=6 | 1270 | 20201113_12413_1ng_glyco_hcd203040_compound15.22540.22540.3 | controllerType=0 controllerNumber=1 scan=22540 | 46.0444 |
| 18083:3 | 1 | R.TAGWNVPIGTLRPFLN[+2279.830]WTGPPEPIEAAVAR.F | HexNAc(5)Hex(6)Fuc(2) | N[+2280] | 1837.5184 | 3 | 5510.5406 | 5510.5406 | 0 | 0 | 141 | Specific | 290.54 | 276.45 | 276.45 | 6.28 | 149 | >sp\|P02788\|TRFL_HUMAN Lactotransferrin OS=Homo sapiens OX=9606 GN=LTF PE=1 SV=6 | 1270 | 20201113_12413_1ng_glyco_hcd203040_compound15.21619.21619.3 | controllerType=0 controllerNumber=1 scan=21619 | 44.0002 |
| 18127:3 | 1 | R.TAGWNVPIGTLRPFLN[+1768.640]WTGPPEPIEAAVAR.F | HexNAc(4)Hex(5)Fuc(1) | N[+1769] | 1667.1258 | 3 | 4999.3629 | 4999.3504 | 0 | 2.5 | 141 | Specific | 296.61 | 244.39 | 244.39 | 6.54 | 149 | >sp\|P02788\|TRFL_HUMAN Lactotransferrin OS=Homo sapiens OX=9606 GN=LTF PE=1 SV=6 | 1270 | 20201113_12413_1ng_glyco_hcd203040_compound15.21666.21666.3 | controllerType=0 controllerNumber=1 scan=21666 | 44.1134 |
| 18963:3 | 1 | R.TAGWNVPIGTLRPFLN[+2716.983]WTGPPEPIEAAVAR.F | HexNAc(5)Hex(6)Fuc(3)NeuAc(1) | N[+2717] | 1983.2417 | 3 | 5947.7104 | 5947.6939 | 0 | 2.8 | 141 | Specific | 301.41 | 297.21 | 297.21 | 6.72 | 149 | >sp\|P02788\|TRFL_HUMAN Lactotransferrin OS=Homo sapiens OX=9606 GN=LTF PE=1 SV=6 | 1270 | 20201113_12413_1ng_glyco_hcd203040_compound15.22542.22542.3 | controllerType=0 controllerNumber=1 scan=22542 | 46.0488 |
| 19135:3 | 1 | R.TAGWNVPIGTLRPFLN[+2059.735]WTGPPEPIEAAVAR.F | HexNAc(4)Hex(5)Fuc(1)NeuAc(1) | N[+2060] | 1764.1646 | 3 | 5290.4794 | 5290.4459 | 0 | 6.3 | 141 | Specific | 312.91 | 312.91 | 312.91 | 6.13 | 149 | >sp\|P02788\|TRFL_HUMAN Lactotransferrin OS=Homo sapiens OX=9606 GN=LTF PE=1 SV=6 | 1270 | 20201113_12413_1ng_glyco_hcd203040_compound15.22721.22721.3 | controllerType=0 controllerNumber=1 scan=22721 | 46.4369 |
| 19076:3 | 1 | R.[+42.011]TAGWNVPIGTLRPFLN[+2190.793]WTGPPEPIEAAVAR.F | HexNAc(6)Hex(6) | .[42], N[+2191] | 1821.8508 | 3 | 5463.5378 | 5463.5147 | 0 | 4.2 | 141 | Specific | 355.85 | 88.01 | 88.01 | 4.93 | 149 | >sp\|P02788\|TRFL_HUMAN Lactotransferrin OS=Homo sapiens OX=9606 GN=LTF PE=1 SV=6 | 1270 | 20201113_12413_1ng_glyco_hcd203040_compound15.22660.22660.3 | controllerType=0 controllerNumber=1 scan=22660 | 46.3114 |
| 18075:4 | 1 | R.TAGWNVPIGTLRPFLN[+2278.809]WTGPPEPIEAAVAR.F | HexNAc(5)Hex(6)NeuAc(1) | N[+2279] | 1378.1377 | 4 | 5509.529 | 5509.5201 | 0 | 1.6 | 141 | Specific | 360.13 | 291.86 | 291.86 | 6.96 | 149 | >sp\|P02788\|TRFL_HUMAN Lactotransferrin OS=Homo sapiens OX=9606 GN=LTF PE=1 SV=6 | 1270 | 20201113_12413_1ng_glyco_hcd203040_compound15.21611.21611.4 | controllerType=0 controllerNumber=1 scan=21611 | 43.9816 |
| 18077:4 | 1 | R.TAGWNVPIGTLRPFLN[+2278.809]WTGPPEPIEAAVAR.F | HexNAc(5)Hex(6)NeuAc(1) | N[+2279] | 1378.1377 | 4 | 5509.529 | 5509.5201 | 0 | 1.6 | 141 | Specific | 363.96 | 159.37 | 159.37 | 5.9 | 149 | >sp\|P02788\|TRFL_HUMAN Lactotransferrin OS=Homo sapiens OX=9606 GN=LTF PE=1 SV=6 | 1270 | 20201113_12413_1ng_glyco_hcd203040_compound15.21613.21613.4 | controllerType=0 controllerNumber=1 scan=21613 | 43.9862 |
| 19078:3 | 1 | R.[+42.011]TAGWNVPIGTLRPFLN[+2190.793]WTGPPEPIEAAVAR.F | HexNAc(6)Hex(6) | .[42], N[+2191] | 1821.8508 | 3 | 5463.5378 | 5463.5147 | 0 | 4.2 | 141 | Specific | 366.9 | 56.06 | 56.06 | 3.73 | 149 | >sp\|P02788\|TRFL_HUMAN Lactotransferrin OS=Homo sapiens OX=9606 GN=LTF PE=1 SV=6 | 1270 | 20201113_12413_1ng_glyco_hcd203040_compound15.22662.22662.3 | controllerType=0 controllerNumber=1 scan=22662 | 46.3172 |
| 18039:3 | 1 | R.TAGWNVPIGTLRPFLN[+2279.830]WTGPPEPIEAAVAR.F | HexNAc(5)Hex(6)Fuc(2) | N[+2280] | 1837.5219 | 3 | 5510.5513 | 5510.5406 | 0 | 1.9 | 141 | Specific | 369.33 | 369.33 | 369.33 | 7.38 | 149 | >sp\|P02788\|TRFL_HUMAN Lactotransferrin OS=Homo sapiens OX=9606 GN=LTF PE=1 SV=6 | 1270 | 20201113_12413_1ng_glyco_hcd203040_compound15.21573.21573.2 | controllerType=0 controllerNumber=1 scan=21573 | 43.9015 |
| 20436:3 | 1 | R.TAGWNVPIGTLRPFLN[+2350.830]WTGPPEPIEAAVAR.F | HexNAc(4)Hex(5)Fuc(1)NeuAc(2) | N[+2351] | 1861.1893 | 3 | 5581.5533 | 5581.5413 | 0 | 2.2 | 141 | Specific | 387.64 | 226.94 | 226.94 | 6.72 | 149 | >sp\|P02788\|TRFL_HUMAN Lactotransferrin OS=Homo sapiens OX=9606 GN=LTF PE=1 SV=6 | 1270 | 20201113_12413_1ng_glyco_hcd203040_compound15.24081.24081.3 | controllerType=0 controllerNumber=1 scan=24081 | 49.4188 |
| 18085:3 | 1 | R.TAGWNVPIGTLRPFLN[+2279.830]WTGPPEPIEAAVAR.F | HexNAc(5)Hex(6)Fuc(2) | N[+2280] | 1837.5184 | 3 | 5510.5406 | 5510.5406 | 0 | 0 | 141 | Specific | 397.19 | 292.61 | 292.61 | 6.76 | 149 | >sp\|P02788\|TRFL_HUMAN Lactotransferrin OS=Homo sapiens OX=9606 GN=LTF PE=1 SV=6 | 1270 | 20201113_12413_1ng_glyco_hcd203040_compound15.21621.21621.3 | controllerType=0 controllerNumber=1 scan=21621 | 44.0065 |
| 19574:4 | 1 | R.TAGWNVPIGTLRPFLN[+2059.735]WTGPPEPIEAAVAR.F | HexNAc(4)Hex(5)Fuc(1)NeuAc(1) | N[+2060] | 1323.372 | 4 | 5290.466 | 5290.4459 | 0 | 3.8 | 141 | Specific | 398.42 | 398.42 | 398.42 | 8.82 | 149 | >sp\|P02788\|TRFL_HUMAN Lactotransferrin OS=Homo sapiens OX=9606 GN=LTF PE=1 SV=6 | 1270 | 20201113_12413_1ng_glyco_hcd203040_compound15.23180.23180.4 | controllerType=0 controllerNumber=1 scan=23180 | 47.3879 |
| 19573:3 | 1 | R.TAGWNVPIGTLRPFLN[+2059.735]WTGPPEPIEAAVAR.F | HexNAc(4)Hex(5)Fuc(1)NeuAc(1) | N[+2060] | 1764.1609 | 3 | 5290.468 | 5290.4459 | 0 | 4.2 | 141 | Specific | 431.84 | 397.56 | 397.56 | 8.96 | 149 | >sp\|P02788\|TRFL_HUMAN Lactotransferrin OS=Homo sapiens OX=9606 GN=LTF PE=1 SV=6 | 1270 | 20201113_12413_1ng_glyco_hcd203040_compound15.23179.23179.3 | controllerType=0 controllerNumber=1 scan=23179 | 47.3865 |
| 19116:4 | 1 | R.TAGWNVPIGTLRPFLN[+2076.750]WTGPPEPIEAAVAR.F | HexNAc(4)Hex(6)Fuc(2) | N[+2077] | 1327.6265 | 4 | 5307.484 | 5307.4612 | 0 | 4.3 | 141 | Specific | 485.92 | 485.92 | 485.92 | 9.19 | 149 | >sp\|P02788\|TRFL_HUMAN Lactotransferrin OS=Homo sapiens OX=9606 GN=LTF PE=1 SV=6 | 1270 | 20201113_12413_1ng_glyco_hcd203040_compound15.22702.22702.4 | controllerType=0 controllerNumber=1 scan=22702 | 46.3979 |
| 19072:4 | 1 | R.TAGWNVPIGTLRPFLN[+2076.750]WTGPPEPIEAAVAR.F | HexNAc(4)Hex(6)Fuc(2) | N[+2077] | 1327.6274 | 4 | 5307.4879 | 5307.4612 | 0 | 5 | 141 | Specific | 530.04 | 530.04 | 530.04 | 8.79 | 149 | >sp\|P02788\|TRFL_HUMAN Lactotransferrin OS=Homo sapiens OX=9606 GN=LTF PE=1 SV=6 | 1270 | 20201113_12413_1ng_glyco_hcd203040_compound15.22656.22656.4 | controllerType=0 controllerNumber=1 scan=22656 | 46.3047 |
| 19143:3 | 1 | R.TAGWNVPIGTLRPFLN[+2059.735]WTGPPEPIEAAVAR.F | HexNAc(4)Hex(5)Fuc(1)NeuAc(1) | N[+2060] | 1764.1591 | 3 | 5290.4629 | 5290.4459 | 0 | 3.2 | 141 | Specific | 531.11 | 474.09 | 474.09 | 9.53 | 149 | >sp\|P02788\|TRFL_HUMAN Lactotransferrin OS=Homo sapiens OX=9606 GN=LTF PE=1 SV=6 | 1270 | 20201113_12413_1ng_glyco_hcd203040_compound15.22730.22730.2 | controllerType=0 controllerNumber=1 scan=22730 | 46.4539 |
| 19054:3 | 1 | R.TAGWNVPIGTLRPFLN[+2351.851]WTGPPEPIEAAVAR.F | HexNAc(4)Hex(5)Fuc(3)NeuAc(1) | N[+2352] | 1861.5296 | 3 | 5582.5743 | 5582.5617 | 0 | 2.3 | 141 | Specific | 533.75 | 449.65 | 449.65 | 9.53 | 149 | >sp\|P02788\|TRFL_HUMAN Lactotransferrin OS=Homo sapiens OX=9606 GN=LTF PE=1 SV=6 | 1270 | 20201113_12413_1ng_glyco_hcd203040_compound15.22637.22637.3 | controllerType=0 controllerNumber=1 scan=22637 | 46.2585 |
| 19068:3 | 1 | R.TAGWNVPIGTLRPFLN[+2059.735]WTGPPEPIEAAVAR.F | HexNAc(4)Hex(5)Fuc(1)NeuAc(1) | N[+2060] | 1764.1688 | 3 | 5290.4918 | 5290.4459 | 0 | 8.7 | 141 | Specific | 579.31 | 502.14 | 502.14 | 10.06 | 149 | >sp\|P02788\|TRFL_HUMAN Lactotransferrin OS=Homo sapiens OX=9606 GN=LTF PE=1 SV=6 | 1270 | 20201113_12413_1ng_glyco_hcd203040_compound15.22652.22652.3 | controllerType=0 controllerNumber=1 scan=22652 | 46.2949 |
| 19056:3 | 1 | R.TAGWNVPIGTLRPFLN[+2351.851]WTGPPEPIEAAVAR.F | HexNAc(4)Hex(5)Fuc(3)NeuAc(1) | N[+2352] | 1861.5296 | 3 | 5582.5743 | 5582.5617 | 0 | 2.3 | 141 | Specific | 595.29 | 572.12 | 572.12 | 10.92 | 149 | >sp\|P02788\|TRFL_HUMAN Lactotransferrin OS=Homo sapiens OX=9606 GN=LTF PE=1 SV=6 | 1270 | 20201113_12413_1ng_glyco_hcd203040_compound15.22639.22639.3 | controllerType=0 controllerNumber=1 scan=22639 | 46.2628 |
| 19575:3 | 1 | R.TAGWNVPIGTLRPFLN[+2059.735]WTGPPEPIEAAVAR.F | HexNAc(4)Hex(5)Fuc(1)NeuAc(1) | N[+2060] | 1764.1609 | 3 | 5290.468 | 5290.4459 | 0 | 4.2 | 141 | Specific | 675.94 | 607.93 | 607.93 | 12.79 | 149 | >sp\|P02788\|TRFL_HUMAN Lactotransferrin OS=Homo sapiens OX=9606 GN=LTF PE=1 SV=6 | 1270 | 20201113_12413_1ng_glyco_hcd203040_compound15.23181.23181.3 | controllerType=0 controllerNumber=1 scan=23181 | 47.3904 |
| 19138:3 | 1 | R.TAGWNVPIGTLRPFLN[+2059.735]WTGPPEPIEAAVAR.F | HexNAc(4)Hex(5)Fuc(1)NeuAc(1) | N[+2060] | 1764.1646 | 3 | 5290.4794 | 5290.4459 | 0 | 6.3 | 141 | Specific | 678.79 | 632.04 | 632.04 | 12.13 | 149 | >sp\|P02788\|TRFL_HUMAN Lactotransferrin OS=Homo sapiens OX=9606 GN=LTF PE=1 SV=6 | 1270 | 20201113_12413_1ng_glyco_hcd203040_compound15.22724.22724.3 | controllerType=0 controllerNumber=1 scan=22724 | 46.4441 |
| 19093:3 | 1 | R.TAGWNVPIGTLRPFLN[+2059.735]WTGPPEPIEAAVAR.F | HexNAc(4)Hex(5)Fuc(1)NeuAc(1) | N[+2060] | 1764.16 | 3 | 5290.4654 | 5290.4459 | 0 | 3.7 | 141 | Specific | 693.31 | 663.03 | 663.03 | 12.79 | 149 | >sp\|P02788\|TRFL_HUMAN Lactotransferrin OS=Homo sapiens OX=9606 GN=LTF PE=1 SV=6 | 1270 | 20201113_12413_1ng_glyco_hcd203040_compound15.22678.22678.2 | controllerType=0 controllerNumber=1 scan=22678 | 46.348 |
| 19090:3 | 1 | R.TAGWNVPIGTLRPFLN[+2059.735]WTGPPEPIEAAVAR.F | HexNAc(4)Hex(5)Fuc(1)NeuAc(1) | N[+2060] | 1764.16 | 3 | 5290.4654 | 5290.4459 | 0 | 3.7 | 141 | Specific | 706.06 | 608.77 | 608.77 | 13.79 | 149 | >sp\|P02788\|TRFL_HUMAN Lactotransferrin OS=Homo sapiens OX=9606 GN=LTF PE=1 SV=6 | 1270 | 20201113_12413_1ng_glyco_hcd203040_compound15.22675.22675.2 | controllerType=0 controllerNumber=1 scan=22675 | 46.3438 |
| 19066:3 | 1 | R.TAGWNVPIGTLRPFLN[+2059.735]WTGPPEPIEAAVAR.F | HexNAc(4)Hex(5)Fuc(1)NeuAc(1) | N[+2060] | 1764.1688 | 3 | 5290.4918 | 5290.4459 | 0 | 8.7 | 141 | Specific | 706.94 | 663.97 | 663.97 | 13.06 | 149 | >sp\|P02788\|TRFL_HUMAN Lactotransferrin OS=Homo sapiens OX=9606 GN=LTF PE=1 SV=6 | 1270 | 20201113_12413_1ng_glyco_hcd203040_compound15.22650.22650.3 | controllerType=0 controllerNumber=1 scan=22650 | 46.2868 |
| 18133:4 | 1 | R.TAGWNVPIGTLRPFLN[+1768.640]WTGPPEPIEAAVAR.F | HexNAc(4)Hex(5)Fuc(1) | N[+1769] | 1250.5964 | 4 | 4999.3639 | 4999.3504 | 0 | 2.7 | 141 | Specific | 724.67 | 678.79 | 678.79 | 13.94 | 149 | >sp\|P02788\|TRFL_HUMAN Lactotransferrin OS=Homo sapiens OX=9606 GN=LTF PE=1 SV=6 | 1270 | 20201113_12413_1ng_glyco_hcd203040_compound15.21672.21672.4 | controllerType=0 controllerNumber=1 scan=21672 | 44.1244 |
| 18129:3 | 1 | R.TAGWNVPIGTLRPFLN[+1768.640]WTGPPEPIEAAVAR.F | HexNAc(4)Hex(5)Fuc(1) | N[+1769] | 1667.1258 | 3 | 4999.3629 | 4999.3504 | 0 | 2.5 | 141 | Specific | 747.82 | 641.33 | 641.33 | 13.92 | 149 | >sp\|P02788\|TRFL_HUMAN Lactotransferrin OS=Homo sapiens OX=9606 GN=LTF PE=1 SV=6 | 1270 | 20201113_12413_1ng_glyco_hcd203040_compound15.21668.21668.3 | controllerType=0 controllerNumber=1 scan=21668 | 44.1182 |
| 19146:3 | 1 | R.TAGWNVPIGTLRPFLN[+2059.735]WTGPPEPIEAAVAR.F | HexNAc(4)Hex(5)Fuc(1)NeuAc(1) | N[+2060] | 1764.1591 | 3 | 5290.4629 | 5290.4459 | 0 | 3.2 | 141 | Specific | 760.29 | 641.07 | 641.07 | 14.92 | 149 | >sp\|P02788\|TRFL_HUMAN Lactotransferrin OS=Homo sapiens OX=9606 GN=LTF PE=1 SV=6 | 1270 | 20201113_12413_1ng_glyco_hcd203040_compound15.22733.22733.2 | controllerType=0 controllerNumber=1 scan=22733 | 46.4581 |
| 19267:4 | 1 | R.TAGWNVPIGTLRPFLN[+2059.735]WTGPPEPIEAAVAR.F | HexNAc(4)Hex(5)Fuc(1)NeuAc(1) | N[+2060] | 1323.3737 | 4 | 5290.4728 | 5290.4459 | 0 | 5.1 | 141 | Specific | 771 | 611.27 | 611.27 | 14.18 | 149 | >sp\|P02788\|TRFL_HUMAN Lactotransferrin OS=Homo sapiens OX=9606 GN=LTF PE=1 SV=6 | 1270 | 20201113_12413_1ng_glyco_hcd203040_compound15.22860.22860.4 | controllerType=0 controllerNumber=1 scan=22860 | 46.7074 |
| 19576:4 | 1 | R.TAGWNVPIGTLRPFLN[+2059.735]WTGPPEPIEAAVAR.F | HexNAc(4)Hex(5)Fuc(1)NeuAc(1) | N[+2060] | 1323.372 | 4 | 5290.466 | 5290.4459 | 0 | 3.8 | 141 | Specific | 798.55 | 664.84 | 664.84 | 14.8 | 149 | >sp\|P02788\|TRFL_HUMAN Lactotransferrin OS=Homo sapiens OX=9606 GN=LTF PE=1 SV=6 | 1270 | 20201113_12413_1ng_glyco_hcd203040_compound15.23182.23182.4 | controllerType=0 controllerNumber=1 scan=23182 | 47.3931 |
| 18136:4 | 1 | R.TAGWNVPIGTLRPFLN[+1768.640]WTGPPEPIEAAVAR.F | HexNAc(4)Hex(5)Fuc(1) | N[+1769] | 1250.5964 | 4 | 4999.3639 | 4999.3504 | 0 | 2.7 | 141 | Specific | 873.09 | 764.08 | 764.08 | 16.94 | 149 | >sp\|P02788\|TRFL_HUMAN Lactotransferrin OS=Homo sapiens OX=9606 GN=LTF PE=1 SV=6 | 1270 | 20201113_12413_1ng_glyco_hcd203040_compound15.21675.21675.4 | controllerType=0 controllerNumber=1 scan=21675 | 44.1313 |
| 20493:3 | 1 | R.[+42.011]TAGWNIPMGLLFN[+2336.851]QTGSC[+57.021]K.F | HexNAc(6)Hex(6)Fuc(1) | .[42], C[+57], N[+2337] | 1491.95 | 3 | 4473.8355 | 4473.8716 | 0 | -8.1 | 485 | Specific | 283.01 | 239.39 | 239.39 | 3.79 | 149 | >sp\|P02788\|TRFL_HUMAN Lactotransferrin OS=Homo sapiens OX=9606 GN=LTF PE=1 SV=6 | 1270 | 20201113_12413_1ng_glyco_hcd203040_compound15.24141.24141.5 | controllerType=0 controllerNumber=1 scan=24141 | 49.5397 |
| 16334:3 | 1 | R.TAGWNIPMGLLFN[+2279.830]QTGSC[+57.021]K.F | HexNAc(5)Hex(6)Fuc(2) | C[+57], N[+2280] | 1458.9527 | 3 | 4374.8435 | 4374.8396 | 0 | 0.9 | 485 | Specific | 316.9 | 316.9 | 234.99 | 7.02 | 149 | >sp\|P02788\|TRFL_HUMAN Lactotransferrin OS=Homo sapiens OX=9606 GN=LTF PE=1 SV=6 | 1270 | 20201113_12413_1ng_glyco_hcd203040_compound15.19794.19794.3 | controllerType=0 controllerNumber=1 scan=19794 | 40.2413 |
| 18779:3 | 1 | R.TAGWNIPMGLLFN[+1548.545]QTGSC[+57.021]K.F | HexNAc(3)Hex(4)NeuAc(1) | C[+57], N[+1549] | 1215.1918 | 3 | 3643.5609 | 3643.5548 | 0 | 1.7 | 485 | Specific | 362.64 | 362.64 | 362.64 | 7.58 | 149 | >sp\|P02788\|TRFL_HUMAN Lactotransferrin OS=Homo sapiens OX=9606 GN=LTF PE=1 SV=6 | 1270 | 20201113_12413_1ng_glyco_hcd203040_compound15.22350.22350.3 | controllerType=0 controllerNumber=1 scan=22350 | 45.6381 |
| 16717:3 | 1 | R.TAGWNIPMGLLFN[+1606.587]QTGSC[+57.021]K.F | HexNAc(4)Hex(4)Fuc(1) | C[+57], N[+1607] | 1234.538 | 3 | 3701.5995 | 3701.5966 | 0 | 0.8 | 485 | Specific | 373.17 | 373.17 | 373.17 | 7.38 | 149 | >sp\|P02788\|TRFL_HUMAN Lactotransferrin OS=Homo sapiens OX=9606 GN=LTF PE=1 SV=6 | 1270 | 20201113_12413_1ng_glyco_hcd203040_compound15.20193.20193.3 | controllerType=0 controllerNumber=1 scan=20193 | 40.9833 |
| 16336:3 | 1 | R.TAGWNIPMGLLFN[+2279.830]QTGSC[+57.021]K.F | HexNAc(5)Hex(6)Fuc(2) | C[+57], N[+2280] | 1458.9527 | 3 | 4374.8435 | 4374.8396 | 0 | 0.9 | 485 | Specific | 434.59 | 424.77 | 355.63 | 9.2 | 149 | >sp\|P02788\|TRFL_HUMAN Lactotransferrin OS=Homo sapiens OX=9606 GN=LTF PE=1 SV=6 | 1270 | 20201113_12413_1ng_glyco_hcd203040_compound15.19796.19796.3 | controllerType=0 controllerNumber=1 scan=19796 | 40.2454 |
| 16627:2 | 1 | R.TAGWNIPMGLLFN[+1768.640]QTGSC[+57.021]K.F | HexNAc(4)Hex(5)Fuc(1) | C[+57], N[+1769] | 1932.3275 | 2 | 3863.6477 | 3863.6495 | 0 | -0.5 | 485 | Specific | 450.26 | 376.24 | 376.24 | 8.36 | 149 | >sp\|P02788\|TRFL_HUMAN Lactotransferrin OS=Homo sapiens OX=9606 GN=LTF PE=1 SV=6 | 1270 | 20201113_12413_1ng_glyco_hcd203040_compound15.20099.20099.2 | controllerType=0 controllerNumber=1 scan=20099 | 40.8054 |
| 16435:3 | 1 | R.TAGWNIPMGLLFN[+2133.772]QTGSC[+57.021]K.F | HexNAc(5)Hex(6)Fuc(1) | C[+57], N[+2134] | 1410.2645 | 3 | 4228.7788 | 4228.7817 | 0 | -0.7 | 485 | Specific | 468.41 | 468.41 | 238.64 | 8.58 | 149 | >sp\|P02788\|TRFL_HUMAN Lactotransferrin OS=Homo sapiens OX=9606 GN=LTF PE=1 SV=6 | 1270 | 20201113_12413_1ng_glyco_hcd203040_compound15.19899.19899.3 | controllerType=0 controllerNumber=1 scan=19899 | 40.4175 |
| 18578:3 | 1 | R.TAGWNIPMGLLFN[+1856.656]QTGSC[+57.021]K.F | HexNAc(3)Hex(5)Fuc(1)NeuAc(1) | C[+57], N[+1857] | 1317.8995 | 3 | 3951.6839 | 3951.6655 | 0 | 4.6 | 485 | Specific | 500.63 | 466.23 | 228.67 | 9.46 | 149 | >sp\|P02788\|TRFL_HUMAN Lactotransferrin OS=Homo sapiens OX=9606 GN=LTF PE=1 SV=6 | 1270 | 20201113_12413_1ng_glyco_hcd203040_compound15.22140.22140.3 | controllerType=0 controllerNumber=1 scan=22140 | 45.1779 |
| 18420:3 | 1 | R.TAGWNIPMGLLFN[+2351.851]QTGSC[+57.021]K.F | HexNAc(4)Hex(5)Fuc(3)NeuAc(1) | C[+57], N[+2352] | 1482.9583 | 3 | 4446.8604 | 4446.8607 | 0 | -0.1 | 485 | Specific | 511.68 | 439.72 | 439.72 | 9.31 | 149 | >sp\|P02788\|TRFL_HUMAN Lactotransferrin OS=Homo sapiens OX=9606 GN=LTF PE=1 SV=6 | 1270 | 20201113_12413_1ng_glyco_hcd203040_compound15.21974.21974.3 | controllerType=0 controllerNumber=1 scan=21974 | 44.7862 |
| 17078:3 | 1 | R.TAGWNIPMGLLFN[+1257.449]QTGSC[+57.021]K.F | HexNAc(3)Hex(4) | C[+57], N[+1257] | 1118.162 | 3 | 3352.4715 | 3352.4594 | 0 | 3.6 | 485 | Specific | 515.3 | 365.54 | 223.51 | 8.01 | 149 | >sp\|P02788\|TRFL_HUMAN Lactotransferrin OS=Homo sapiens OX=9606 GN=LTF PE=1 SV=6 | 1270 | 20201113_12413_1ng_glyco_hcd203040_compound15.20569.20569.3 | controllerType=0 controllerNumber=1 scan=20569 | 41.7396 |
| 18706:3 | 1 | R.TAGWNIPMGLLFN[+1710.598]QTGSC[+57.021]K.F | HexNAc(3)Hex(5)NeuAc(1) | C[+57], N[+1711] | 1269.2077 | 3 | 3805.6085 | 3805.6076 | 0 | 0.2 | 485 | Specific | 587.33 | 535.38 | 222.39 | 10.83 | 149 | >sp\|P02788\|TRFL_HUMAN Lactotransferrin OS=Homo sapiens OX=9606 GN=LTF PE=1 SV=6 | 1270 | 20201113_12413_1ng_glyco_hcd203040_compound15.22273.22273.3 | controllerType=0 controllerNumber=1 scan=22273 | 45.4673 |
| 16912:3 | 1 | R.TAGWNIPMGLLFN[+1378.476]QTGSC[+57.021]K.F | HexNAc(2)Hex(6) | C[+57], N[+1378] | 1158.5014 | 3 | 3473.4896 | 3473.4856 | 0 | 1.2 | 485 | Specific | 597.23 | 597.23 | 226.2 | 11.03 | 149 | >sp\|P02788\|TRFL_HUMAN Lactotransferrin OS=Homo sapiens OX=9606 GN=LTF PE=1 SV=6 | 1270 | 20201113_12413_1ng_glyco_hcd203040_compound15.20396.20396.3 | controllerType=0 controllerNumber=1 scan=20396 | 41.3997 |
| 16437:3 | 1 | R.TAGWNIPMGLLFN[+2133.772]QTGSC[+57.021]K.F | HexNAc(5)Hex(6)Fuc(1) | C[+57], N[+2134] | 1410.2645 | 3 | 4228.7788 | 4228.7817 | 0 | -0.7 | 485 | Specific | 605.97 | 605.97 | 376.3 | 11.19 | 149 | >sp\|P02788\|TRFL_HUMAN Lactotransferrin OS=Homo sapiens OX=9606 GN=LTF PE=1 SV=6 | 1270 | 20201113_12413_1ng_glyco_hcd203040_compound15.19901.19901.3 | controllerType=0 controllerNumber=1 scan=19901 | 40.4211 |
| 16810:3 | 1 | R.TAGWNIPMGLLFN[+1622.582]QTGSC[+57.021]K.F | HexNAc(4)Hex(5) | C[+57], N[+1623] | 1239.8773 | 3 | 3717.6172 | 3717.5916 | 0 | 6.9 | 485 | Specific | 638.71 | 638.71 | 268.55 | 11.33 | 149 | >sp\|P02788\|TRFL_HUMAN Lactotransferrin OS=Homo sapiens OX=9606 GN=LTF PE=1 SV=6 | 1270 | 20201113_12413_1ng_glyco_hcd203040_compound15.20290.20290.3 | controllerType=0 controllerNumber=1 scan=20290 | 41.1904 |
| 17080:3 | 1 | R.TAGWNIPMGLLFN[+1257.449]QTGSC[+57.021]K.F | HexNAc(3)Hex(4) | C[+57], N[+1257] | 1118.162 | 3 | 3352.4715 | 3352.4594 | 0 | 3.6 | 485 | Specific | 663.75 | 538.12 | 333.68 | 12.99 | 149 | >sp\|P02788\|TRFL_HUMAN Lactotransferrin OS=Homo sapiens OX=9606 GN=LTF PE=1 SV=6 | 1270 | 20201113_12413_1ng_glyco_hcd203040_compound15.20571.20571.3 | controllerType=0 controllerNumber=1 scan=20571 | 41.7451 |
| 18734:3 | 1 | R.TAGWNIPMGLLFN[+2059.735]QTGSC[+57.021]K.F | HexNAc(4)Hex(5)Fuc(1)NeuAc(1) | C[+57], N[+2060] | 1385.5938 | 3 | 4154.7669 | 4154.7449 | 0 | 5.3 | 485 | Specific | 666.43 | 525.82 | 525.82 | 12.36 | 149 | >sp\|P02788\|TRFL_HUMAN Lactotransferrin OS=Homo sapiens OX=9606 GN=LTF PE=1 SV=6 | 1270 | 20201113_12413_1ng_glyco_hcd203040_compound15.22302.22302.3 | controllerType=0 controllerNumber=1 scan=22302 | 45.5232 |
| 16612:3 | 1 | R.TAGWNIPMGLLFN[+1768.640]QTGSC[+57.021]K.F | HexNAc(4)Hex(5)Fuc(1) | C[+57], N[+1769] | 1288.5539 | 3 | 3863.6471 | 3863.6495 | 0 | -0.6 | 485 | Specific | 680.72 | 680.72 | 680.72 | 12.19 | 149 | >sp\|P02788\|TRFL_HUMAN Lactotransferrin OS=Homo sapiens OX=9606 GN=LTF PE=1 SV=6 | 1270 | 20201113_12413_1ng_glyco_hcd203040_compound15.20084.20084.3 | controllerType=0 controllerNumber=1 scan=20084 | 40.7798 |
| 18623:4 | 1 | R.TAGWNIPMGLLFN[+1913.677]QTGSC[+57.021]K.F | HexNAc(4)Hex(5)NeuAc(1) | C[+57], N[+1914] | 1002.9288 | 4 | 4008.6932 | 4008.687 | 0 | 1.6 | 485 | Specific | 699.77 | 517.53 | 517.53 | 12.86 | 149 | >sp\|P02788\|TRFL_HUMAN Lactotransferrin OS=Homo sapiens OX=9606 GN=LTF PE=1 SV=6 | 1270 | 20201113_12413_1ng_glyco_hcd203040_compound15.22187.22187.4 | controllerType=0 controllerNumber=1 scan=22187 | 45.2815 |
| 18626:4 | 1 | R.TAGWNIPMGLLFN[+1913.677]QTGSC[+57.021]K.F | HexNAc(4)Hex(5)NeuAc(1) | C[+57], N[+1914] | 1002.9288 | 4 | 4008.6932 | 4008.687 | 0 | 1.6 | 485 | Specific | 710.9 | 633.93 | 633.93 | 13.86 | 149 | >sp\|P02788\|TRFL_HUMAN Lactotransferrin OS=Homo sapiens OX=9606 GN=LTF PE=1 SV=6 | 1270 | 20201113_12413_1ng_glyco_hcd203040_compound15.22190.22190.4 | controllerType=0 controllerNumber=1 scan=22190 | 45.2902 |
| 16939:3 | 1 | R.TAGWNIPMGLLFN[+1460.529]QTGSC[+57.021]K.F | HexNAc(4)Hex(4) | C[+57], N[+1461] | 1185.8559 | 3 | 3555.5531 | 3555.5387 | 0 | 4 | 485 | Specific | 723.09 | 705.81 | 339.33 | 13.99 | 149 | >sp\|P02788\|TRFL_HUMAN Lactotransferrin OS=Homo sapiens OX=9606 GN=LTF PE=1 SV=6 | 1270 | 20201113_12413_1ng_glyco_hcd203040_compound15.20424.20424.3 | controllerType=0 controllerNumber=1 scan=20424 | 41.4549 |
| 18581:3 | 1 | R.TAGWNIPMGLLFN[+1856.656]QTGSC[+57.021]K.F | HexNAc(3)Hex(5)Fuc(1)NeuAc(1) | C[+57], N[+1857] | 1317.8995 | 3 | 3951.6839 | 3951.6655 | 0 | 4.6 | 485 | Specific | 733.45 | 563.31 | 325.44 | 13.99 | 149 | >sp\|P02788\|TRFL_HUMAN Lactotransferrin OS=Homo sapiens OX=9606 GN=LTF PE=1 SV=6 | 1270 | 20201113_12413_1ng_glyco_hcd203040_compound15.22143.22143.3 | controllerType=0 controllerNumber=1 scan=22143 | 45.1821 |
| 16941:3 | 1 | R.TAGWNIPMGLLFN[+1460.529]QTGSC[+57.021]K.F | HexNAc(4)Hex(4) | C[+57], N[+1461] | 1185.8559 | 3 | 3555.5531 | 3555.5387 | 0 | 4 | 485 | Specific | 744.26 | 738.84 | 374.2 | 13.99 | 149 | >sp\|P02788\|TRFL_HUMAN Lactotransferrin OS=Homo sapiens OX=9606 GN=LTF PE=1 SV=6 | 1270 | 20201113_12413_1ng_glyco_hcd203040_compound15.20426.20426.3 | controllerType=0 controllerNumber=1 scan=20426 | 41.4602 |
| 17146:3 | 1 | R.TAGWNIPMGLLFN[+1095.397]QTGSC[+57.021]K.F | HexNAc(3)Hex(3) | C[+57], N[+1095] | 1064.1438 | 3 | 3190.4169 | 3190.4065 | 0 | 3.3 | 485 | Specific | 760.54 | 754.54 | 754.54 | 15.12 | 149 | >sp\|P02788\|TRFL_HUMAN Lactotransferrin OS=Homo sapiens OX=9606 GN=LTF PE=1 SV=6 | 1270 | 20201113_12413_1ng_glyco_hcd203040_compound15.20640.20640.3 | controllerType=0 controllerNumber=1 scan=20640 | 41.8818 |
| 18725:3 | 1 | R.TAGWNIPMGLLFN[+1710.598]QTGSC[+57.021]K.F | HexNAc(3)Hex(5)NeuAc(1) | C[+57], N[+1711] | 1269.2136 | 3 | 3805.6261 | 3805.6076 | 0 | 4.9 | 485 | Specific | 794.37 | 741.23 | 325.22 | 14.36 | 149 | >sp\|P02788\|TRFL_HUMAN Lactotransferrin OS=Homo sapiens OX=9606 GN=LTF PE=1 SV=6 | 1270 | 20201113_12413_1ng_glyco_hcd203040_compound15.22293.22293.3 | controllerType=0 controllerNumber=1 scan=22293 | 45.5059 |
| 16780:3 | 1 | R.TAGWNIPMGLLFN[+1622.582]QTGSC[+57.021]K.F | HexNAc(4)Hex(5) | C[+57], N[+1623] | 1239.8718 | 3 | 3717.6007 | 3717.5916 | 0 | 2.5 | 485 | Specific | 796.03 | 786.81 | 346.25 | 15.12 | 149 | >sp\|P02788\|TRFL_HUMAN Lactotransferrin OS=Homo sapiens OX=9606 GN=LTF PE=1 SV=6 | 1270 | 20201113_12413_1ng_glyco_hcd203040_compound15.20258.20258.3 | controllerType=0 controllerNumber=1 scan=20258 | 41.1016 |
| 16940:3 | 1 | R.TAGWNIPMGLLFN[+1419.502]QTGSC[+57.021]K.F | HexNAc(3)Hex(5) | C[+57], N[+1420] | 1172.1807 | 3 | 3514.5276 | 3514.5122 | 0 | 4.4 | 485 | Specific | 817.41 | 775.18 | 367.16 | 15.99 | 149 | >sp\|P02788\|TRFL_HUMAN Lactotransferrin OS=Homo sapiens OX=9606 GN=LTF PE=1 SV=6 | 1270 | 20201113_12413_1ng_glyco_hcd203040_compound15.20425.20425.3 | controllerType=0 controllerNumber=1 scan=20425 | 41.4562 |
| 18782:3 | 1 | R.TAGWNIPMGLLFN[+1548.545]QTGSC[+57.021]K.F | HexNAc(3)Hex(4)NeuAc(1) | C[+57], N[+1549] | 1215.1918 | 3 | 3643.5609 | 3643.5548 | 0 | 1.7 | 485 | Specific | 826.78 | 798.62 | 798.62 | 16.03 | 149 | >sp\|P02788\|TRFL_HUMAN Lactotransferrin OS=Homo sapiens OX=9606 GN=LTF PE=1 SV=6 | 1270 | 20201113_12413_1ng_glyco_hcd203040_compound15.22353.22353.3 | controllerType=0 controllerNumber=1 scan=22353 | 45.6464 |
| 16856:3 | 1 | R.TAGWNIPMGLLFN[+1581.555]QTGSC[+57.021]K.F | HexNAc(3)Hex(6) | C[+57], N[+1582] | 1226.1959 | 3 | 3676.573 | 3676.565 | 0 | 2.2 | 485 | Specific | 838.82 | 820.89 | 393.85 | 16.03 | 149 | >sp\|P02788\|TRFL_HUMAN Lactotransferrin OS=Homo sapiens OX=9606 GN=LTF PE=1 SV=6 | 1270 | 20201113_12413_1ng_glyco_hcd203040_compound15.20337.20337.3 | controllerType=0 controllerNumber=1 scan=20337 | 41.2837 |
| 20557:3 | 1 | R.TAGWNIPMGLLFN[+2204.772]QTGSC[+57.021]K.F | HexNAc(4)Hex(5)NeuAc(2) | C[+57], N[+2205] | 1433.9399 | 3 | 4299.8051 | 4299.7824 | 0 | 5.3 | 485 | Specific | 859.09 | 774.79 | 774.79 | 16.36 | 149 | >sp\|P02788\|TRFL_HUMAN Lactotransferrin OS=Homo sapiens OX=9606 GN=LTF PE=1 SV=6 | 1270 | 20201113_12413_1ng_glyco_hcd203040_compound15.24208.24208.3 | controllerType=0 controllerNumber=1 scan=24208 | 49.6968 |
| 16990:2 | 1 | R.TAGWNIPMGLLFN[+1216.423]QTGSC[+57.021]K.F | HexNAc(2)Hex(5) | C[+57], N[+1216] | 1656.2231 | 2 | 3311.4389 | 3311.4328 | 0 | 1.8 | 485 | Specific | 866.72 | 829.44 | 295.44 | 17.01 | 149 | >sp\|P02788\|TRFL_HUMAN Lactotransferrin OS=Homo sapiens OX=9606 GN=LTF PE=1 SV=6 | 1270 | 20201113_12413_1ng_glyco_hcd203040_compound15.20477.20477.2 | controllerType=0 controllerNumber=1 scan=20477 | 41.5676 |
| 16782:3 | 1 | R.TAGWNIPMGLLFN[+1622.582]QTGSC[+57.021]K.F | HexNAc(4)Hex(5) | C[+57], N[+1623] | 1239.8718 | 3 | 3717.6007 | 3717.5916 | 0 | 2.5 | 485 | Specific | 866.91 | 815.65 | 392.7 | 17.12 | 149 | >sp\|P02788\|TRFL_HUMAN Lactotransferrin OS=Homo sapiens OX=9606 GN=LTF PE=1 SV=6 | 1270 | 20201113_12413_1ng_glyco_hcd203040_compound15.20260.20260.3 | controllerType=0 controllerNumber=1 scan=20260 | 41.1065 |
| 16942:3 | 1 | R.TAGWNIPMGLLFN[+1419.502]QTGSC[+57.021]K.F | HexNAc(3)Hex(5) | C[+57], N[+1420] | 1172.1807 | 3 | 3514.5276 | 3514.5122 | 0 | 4.4 | 485 | Specific | 875.76 | 853.89 | 372.56 | 16.99 | 149 | >sp\|P02788\|TRFL_HUMAN Lactotransferrin OS=Homo sapiens OX=9606 GN=LTF PE=1 SV=6 | 1270 | 20201113_12413_1ng_glyco_hcd203040_compound15.20427.20427.3 | controllerType=0 controllerNumber=1 scan=20427 | 41.4621 |
| 16832:3 | 1 | R.TAGWNIPMGLLFN[+1581.555]QTGSC[+57.021]K.F | HexNAc(3)Hex(6) | C[+57], N[+1582] | 1226.2034 | 3 | 3676.5957 | 3676.565 | 0 | 8.4 | 485 | Specific | 879.02 | 864.55 | 394.79 | 16.26 | 149 | >sp\|P02788\|TRFL_HUMAN Lactotransferrin OS=Homo sapiens OX=9606 GN=LTF PE=1 SV=6 | 1270 | 20201113_12413_1ng_glyco_hcd203040_compound15.20312.20312.3 | controllerType=0 controllerNumber=1 scan=20312 | 41.2274 |
| 16614:3 | 1 | R.TAGWNIPMGLLFN[+1768.640]QTGSC[+57.021]K.F | HexNAc(4)Hex(5)Fuc(1) | C[+57], N[+1769] | 1288.5539 | 3 | 3863.6471 | 3863.6495 | 0 | -0.6 | 485 | Specific | 880.95 | 871.65 | 871.65 | 16.19 | 149 | >sp\|P02788\|TRFL_HUMAN Lactotransferrin OS=Homo sapiens OX=9606 GN=LTF PE=1 SV=6 | 1270 | 20201113_12413_1ng_glyco_hcd203040_compound15.20086.20086.3 | controllerType=0 controllerNumber=1 scan=20086 | 40.7846 |
| 18687:3 | 1 | R.TAGWNIPMGLLFN[+1751.624]QTGSC[+57.021]K.F | HexNAc(4)Hex(4)NeuAc(1) | C[+57], N[+1752] | 1282.8881 | 3 | 3846.6498 | 3846.6341 | 0 | 4.1 | 485 | Specific | 885.47 | 885.47 | 885.47 | 16.99 | 149 | >sp\|P02788\|TRFL_HUMAN Lactotransferrin OS=Homo sapiens OX=9606 GN=LTF PE=1 SV=6 | 1270 | 20201113_12413_1ng_glyco_hcd203040_compound15.22253.22253.3 | controllerType=0 controllerNumber=1 scan=22253 | 45.4293 |
| 16835:3 | 1 | R.TAGWNIPMGLLFN[+1581.555]QTGSC[+57.021]K.F | HexNAc(3)Hex(6) | C[+57], N[+1582] | 1226.2034 | 3 | 3676.5957 | 3676.565 | 0 | 8.4 | 485 | Specific | 886.16 | 824.53 | 426.12 | 16.26 | 149 | >sp\|P02788\|TRFL_HUMAN Lactotransferrin OS=Homo sapiens OX=9606 GN=LTF PE=1 SV=6 | 1270 | 20201113_12413_1ng_glyco_hcd203040_compound15.20315.20315.3 | controllerType=0 controllerNumber=1 scan=20315 | 41.2331 |
| 16914:3 | 1 | R.TAGWNIPMGLLFN[+1378.476]QTGSC[+57.021]K.F | HexNAc(2)Hex(6) | C[+57], N[+1378] | 1158.5014 | 3 | 3473.4896 | 3473.4856 | 0 | 1.2 | 485 | Specific | 889.05 | 889.05 | 380.11 | 17.03 | 149 | >sp\|P02788\|TRFL_HUMAN Lactotransferrin OS=Homo sapiens OX=9606 GN=LTF PE=1 SV=6 | 1270 | 20201113_12413_1ng_glyco_hcd203040_compound15.20398.20398.3 | controllerType=0 controllerNumber=1 scan=20398 | 41.4038 |
| 18697:3 | 1 | R.TAGWNIPMGLLFN[+1751.624]QTGSC[+57.021]K.F | HexNAc(4)Hex(4)NeuAc(1) | C[+57], N[+1752] | 1282.8832 | 3 | 3846.6352 | 3846.6341 | 0 | 0.3 | 485 | Specific | 889.29 | 785.04 | 785.04 | 16.83 | 149 | >sp\|P02788\|TRFL_HUMAN Lactotransferrin OS=Homo sapiens OX=9606 GN=LTF PE=1 SV=6 | 1270 | 20201113_12413_1ng_glyco_hcd203040_compound15.22264.22264.3 | controllerType=0 controllerNumber=1 scan=22264 | 45.453 |
| 16858:3 | 1 | R.TAGWNIPMGLLFN[+1581.555]QTGSC[+57.021]K.F | HexNAc(3)Hex(6) | C[+57], N[+1582] | 1226.1959 | 3 | 3676.573 | 3676.565 | 0 | 2.2 | 485 | Specific | 890.79 | 858.22 | 392.97 | 17.03 | 149 | >sp\|P02788\|TRFL_HUMAN Lactotransferrin OS=Homo sapiens OX=9606 GN=LTF PE=1 SV=6 | 1270 | 20201113_12413_1ng_glyco_hcd203040_compound15.20339.20339.3 | controllerType=0 controllerNumber=1 scan=20339 | 41.2895 |
| 18426:3 | 1 | R.TAGWNIPMGLLFN[+2059.735]QTGSC[+57.021]K.F | HexNAc(4)Hex(5)Fuc(1)NeuAc(1) | C[+57], N[+2060] | 1385.5933 | 3 | 4154.7654 | 4154.7449 | 0 | 4.9 | 485 | Specific | 891.26 | 880.47 | 880.47 | 16.36 | 149 | >sp\|P02788\|TRFL_HUMAN Lactotransferrin OS=Homo sapiens OX=9606 GN=LTF PE=1 SV=6 | 1270 | 20201113_12413_1ng_glyco_hcd203040_compound15.21981.21981.3 | controllerType=0 controllerNumber=1 scan=21981 | 44.8058 |
| 16719:3 | 1 | R.TAGWNIPMGLLFN[+1606.587]QTGSC[+57.021]K.F | HexNAc(4)Hex(4)Fuc(1) | C[+57], N[+1607] | 1234.538 | 3 | 3701.5995 | 3701.5966 | 0 | 0.8 | 485 | Specific | 892.69 | 747.05 | 747.05 | 16.83 | 149 | >sp\|P02788\|TRFL_HUMAN Lactotransferrin OS=Homo sapiens OX=9606 GN=LTF PE=1 SV=6 | 1270 | 20201113_12413_1ng_glyco_hcd203040_compound15.20195.20195.3 | controllerType=0 controllerNumber=1 scan=20195 | 40.9867 |
| 18576:3 | 1 | R.TAGWNIPMGLLFN[+1913.677]QTGSC[+57.021]K.F | HexNAc(4)Hex(5)NeuAc(1) | C[+57], N[+1914] | 1336.9046 | 3 | 4008.6992 | 4008.687 | 0 | 3.1 | 485 | Specific | 897.69 | 897.69 | 897.69 | 17.12 | 149 | >sp\|P02788\|TRFL_HUMAN Lactotransferrin OS=Homo sapiens OX=9606 GN=LTF PE=1 SV=6 | 1270 | 20201113_12413_1ng_glyco_hcd203040_compound15.22138.22138.3 | controllerType=0 controllerNumber=1 scan=22138 | 45.1756 |
| 18690:3 | 1 | R.TAGWNIPMGLLFN[+1751.624]QTGSC[+57.021]K.F | HexNAc(4)Hex(4)NeuAc(1) | C[+57], N[+1752] | 1282.8881 | 3 | 3846.6498 | 3846.6341 | 0 | 4.1 | 485 | Specific | 911.77 | 911.77 | 911.77 | 17.99 | 149 | >sp\|P02788\|TRFL_HUMAN Lactotransferrin OS=Homo sapiens OX=9606 GN=LTF PE=1 SV=6 | 1270 | 20201113_12413_1ng_glyco_hcd203040_compound15.22256.22256.3 | controllerType=0 controllerNumber=1 scan=22256 | 45.4374 |
| 16812:3 | 1 | R.TAGWNIPMGLLFN[+1622.582]QTGSC[+57.021]K.F | HexNAc(4)Hex(5) | C[+57], N[+1623] | 1239.8773 | 3 | 3717.6172 | 3717.5916 | 0 | 6.9 | 485 | Specific | 916.82 | 908.04 | 417.06 | 17.33 | 149 | >sp\|P02788\|TRFL_HUMAN Lactotransferrin OS=Homo sapiens OX=9606 GN=LTF PE=1 SV=6 | 1270 | 20201113_12413_1ng_glyco_hcd203040_compound15.20292.20292.3 | controllerType=0 controllerNumber=1 scan=20292 | 41.1946 |

Table S4. Cont.

| Query #:z | Protein Rank | Glycopeptide | Glycan | Modification Type(s) | Observed m/z | z | Observed (M+H) | Calc. mass (M+H) | Off-by-x error | Mass error (ppm) | StartPosition | Cleavage | Score | Delta | Delta Mod | \|Log Prob\| | # of unique peptides | Protein Name | Protein DB number | Comment | Scan # | Scan Time |
|---|---|---|---|---|---|---|---|---|---|---|---|---|---|---|---|---|---|---|---|---|---|---|
| 18708:3 | 1 | R.TAGWNIPMGLLFN[+1710.598]QTGSC[+57.021]K.F | HexNAc(3)Hex(5)NeuAc(1) | C[+57], N[+1711] | 1269.2077 | 3 | 3805.6085 | 3805.6076 | 0 | 0.2 | 485 | Specific | 919.28 | 696.42 | 403.31 | 17.83 | 149 | >sp\|P02788\|TRFL_HUMAN Lactotransferrin OS=Homo sapiens OX=9606 GN=LTF PE=1 SV=6 | 1270 | 20201113_12413_1ng_glyco_hcd203040_compound15.22275.22275.3 | controllerType=0 controllerNumber=1 scan=22275 | 45.4739 |
| 18422:3 | 1 | R.TAGWNIPMGLLFN[+2059.735]QTGSC[+57.021]K.F | HexNAc(4)Hex(5)Fuc(1)NeuAc(1) | C[+57], N[+2060] | 1385.5889 | 3 | 4154.7522 | 4154.7449 | 0 | 1.8 | 485 | Specific | 931.58 | 690.43 | 690.43 | 18.03 | 149 | >sp\|P02788\|TRFL_HUMAN Lactotransferrin OS=Homo sapiens OX=9606 GN=LTF PE=1 SV=6 | 1270 | 20201113_12413_1ng_glyco_hcd203040_compound15.21976.21976.3 | controllerType=0 controllerNumber=1 scan=21976 | 44.7909 |
| 16988:2 | 1 | R.TAGWNIPMGLLFN[+1216.423]QTGSC[+57.021]K.F | HexNAc(2)Hex(5) | C[+57], N[+1216] | 1656.2231 | 2 | 3311.4389 | 3311.4328 | 0 | 1.8 | 485 | Specific | 935.1 | 896.04 | 361.02 | 18.01 | 149 | >sp\|P02788\|TRFL_HUMAN Lactotransferrin OS=Homo sapiens OX=9606 GN=LTF PE=1 SV=6 | 1270 | 20201113_12413_1ng_glyco_hcd203040_compound15.20475.20475.2 | controllerType=0 controllerNumber=1 scan=20475 | 41.5596 |
| 20555:3 | 1 | R.TAGWNIPMGLLFN[+2204.772]QTGSC[+57.021]K.F | HexNAc(4)Hex(5)NeuAc(2) | C[+57], N[+2205] | 1433.9399 | 3 | 4299.8051 | 4299.7824 | 0 | 5.3 | 485 | Specific | 937.75 | 916.55 | 916.55 | 17.36 | 149 | >sp\|P02788\|TRFL_HUMAN Lactotransferrin OS=Homo sapiens OX=9606 GN=LTF PE=1 SV=6 | 1270 | 20201113_12413_1ng_glyco_hcd203040_compound15.24206.24206.3 | controllerType=0 controllerNumber=1 scan=24206 | 49.6884 |
| 18728:3 | 1 | R.TAGWNIPMGLLFN[+1710.598]QTGSC[+57.021]K.F | HexNAc(3)Hex(5)NeuAc(1) | C[+57], N[+1711] | 1269.2136 | 3 | 3805.6261 | 3805.6076 | 0 | 4.9 | 485 | Specific | 961.13 | 851.62 | 445.07 | 18.36 | 149 | >sp\|P02788\|TRFL_HUMAN Lactotransferrin OS=Homo sapiens OX=9606 GN=LTF PE=1 SV=6 | 1270 | 20201113_12413_1ng_glyco_hcd203040_compound15.22296.22296.3 | controllerType=0 controllerNumber=1 scan=22296 | 45.5123 |
| 18574:3 | 1 | R.TAGWNIPMGLLFN[+1913.677]QTGSC[+57.021]K.F | HexNAc(4)Hex(5)NeuAc(1) | C[+57], N[+1914] | 1336.9046 | 3 | 4008.6992 | 4008.687 | 0 | 3.1 | 485 | Specific | 969.78 | 969.78 | 969.78 | 19.12 | 149 | >sp\|P02788\|TRFL_HUMAN Lactotransferrin OS=Homo sapiens OX=9606 GN=LTF PE=1 SV=6 | 1270 | 20201113_12413_1ng_glyco_hcd203040_compound15.22136.22136.3 | controllerType=0 controllerNumber=1 scan=22136 | 45.1684 |
| 18700:3 | 1 | R.TAGWNIPMGLLFN[+1751.624]QTGSC[+57.021]K.F | HexNAc(4)Hex(4)NeuAc(1) | C[+57], N[+1752] | 1282.8832 | 3 | 3846.6352 | 3846.6341 | 0 | 0.3 | 485 | Specific | 973.02 | 864.41 | 864.41 | 18.83 | 149 | >sp\|P02788\|TRFL_HUMAN Lactotransferrin OS=Homo sapiens OX=9606 GN=LTF PE=1 SV=6 | 1270 | 20201113_12413_1ng_glyco_hcd203040_compound15.22267.22267.3 | controllerType=0 controllerNumber=1 scan=22267 | 45.4581 |
| 16970:3 | 1 | R.TAGWNIPMGLLFN[+1216.423]QTGSC[+57.021]K.F | HexNAc(2)Hex(5) | C[+57], N[+1216] | 1104.4855 | 3 | 3311.442 | 3311.4328 | 0 | 2.8 | 485 | Specific | 978.24 | 978.24 | 442.09 | 19.12 | 149 | >sp\|P02788\|TRFL_HUMAN Lactotransferrin OS=Homo sapiens OX=9606 GN=LTF PE=1 SV=6 | 1270 | 20201113_12413_1ng_glyco_hcd203040_compound15.20456.20456.3 | controllerType=0 controllerNumber=1 scan=20456 | 41.5173 |
| 18403:3 | 1 | R.TAGWNIPMGLLFN[+2205.793]QTGSC[+57.021]K.F | HexNAc(4)Hex(5)Fuc(2)NeuAc(1) | C[+57], N[+2206] | 1434.2679 | 3 | 4300.7891 | 4300.8028 | 0 | -3.2 | 485 | Specific | 978.94 | 862.12 | 862.12 | 17.65 | 149 | >sp\|P02788\|TRFL_HUMAN Lactotransferrin OS=Homo sapiens OX=9606 GN=LTF PE=1 SV=6 | 1270 | 20201113_12413_1ng_glyco_hcd203040_compound15.21956.21956.3 | controllerType=0 controllerNumber=1 scan=21956 | 44.7533 |
| 18424:3 | 1 | R.TAGWNIPMGLLFN[+2059.735]QTGSC[+57.021]K.F | HexNAc(4)Hex(5)Fuc(1)NeuAc(1) | C[+57], N[+2060] | 1385.5933 | 3 | 4154.7654 | 4154.7449 | 0 | 4.9 | 485 | Specific | 983.45 | 924.53 | 924.53 | 18.36 | 149 | >sp\|P02788\|TRFL_HUMAN Lactotransferrin OS=Homo sapiens OX=9606 GN=LTF PE=1 SV=6 | 1270 | 20201113_12413_1ng_glyco_hcd203040_compound15.21979.21979.3 | controllerType=0 controllerNumber=1 scan=21979 | 44.7978 |
| 18419:3 | 1 | R.TAGWNIPMGLLFN[+2059.735]QTGSC[+57.021]K.F | HexNAc(4)Hex(5)Fuc(1)NeuAc(1) | C[+57], N[+2060] | 1385.5889 | 3 | 4154.7522 | 4154.7449 | 0 | 1.8 | 485 | Specific | 999.92 | 885.02 | 885.02 | 19.03 | 149 | >sp\|P02788\|TRFL_HUMAN Lactotransferrin OS=Homo sapiens OX=9606 GN=LTF PE=1 SV=6 | 1270 | 20201113_12413_1ng_glyco_hcd203040_compound15.21973.21973.3 | controllerType=0 controllerNumber=1 scan=21973 | 44.785 |
| 18400:3 | 1 | R.TAGWNIPMGLLFN[+2205.793]QTGSC[+57.021]K.F | HexNAc(4)Hex(5)Fuc(2)NeuAc(1) | C[+57], N[+2206] | 1434.2679 | 3 | 4300.7891 | 4300.8028 | 0 | -3.2 | 485 | Specific | 1020.43 | 890.23 | 890.23 | 18.65 | 149 | >sp\|P02788\|TRFL_HUMAN Lactotransferrin OS=Homo sapiens OX=9606 GN=LTF PE=1 SV=6 | 1270 | 20201113_12413_1ng_glyco_hcd203040_compound15.21953.21953.3 | controllerType=0 controllerNumber=1 scan=21953 | 44.7421 |
| 16973:3 | 1 | R.TAGWNIPMGLLFN[+1216.423]QTGSC[+57.021]K.F | HexNAc(2)Hex(5) | C[+57], N[+1216] | 1104.4855 | 3 | 3311.442 | 3311.4328 | 0 | 2.8 | 485 | Specific | 1030.24 | 1030.24 | 512.86 | 20.12 | 149 | >sp\|P02788\|TRFL_HUMAN Lactotransferrin OS=Homo sapiens OX=9606 GN=LTF PE=1 SV=6 | 1270 | 20201113_12413_1ng_glyco_hcd203040_compound15.20459.20459.3 | controllerType=0 controllerNumber=1 scan=20459 | 41.5256 |
| 16444:3 | 1 | R.TAGWNIPM[+15.995]GLLFN[+1913.677]QTGSC[+57.021]K.F | HexNAc(4)Hex(5)NeuAc(1) | C[+57], M[+16], N[+1914] | 1342.24 | 3 | 4024.7054 | 4024.6819 | 0 | 5.9 | 485 | Specific | 603.06 | 576.7 | 576.7 | 11.36 | 149 | >sp\|P02788\|TRFL_HUMAN Lactotransferrin OS=Homo sapiens OX=9606 GN=LTF PE=1 SV=6 | 1270 | 20201113_12413_1ng_glyco_hcd203040_compound15.19908.19908.3 | controllerType=0 controllerNumber=1 scan=19908 | 40.4376 |
| 16865:3 | 1 | R.TAGWNIPM[+15.995]GLLFN[+2059.735]QTGSC[+57.021]K.F | HexNAc(4)Hex(5)Fuc(1)NeuAc(1) | C[+57], M[+16], N[+2060] | 1390.9207 | 3 | 4170.7476 | 4170.7398 | 0 | 1.9 | 485 | Specific | 686.85 | 623.92 | 623.92 | 13.03 | 149 | >sp\|P02788\|TRFL_HUMAN Lactotransferrin OS=Homo sapiens OX=9606 GN=LTF PE=1 SV=6 | 1270 | 20201113_12413_1ng_glyco_hcd203040_compound15.20347.20347.3 | controllerType=0 controllerNumber=1 scan=20347 | 41.3064 |
| 16867:3 | 1 | R.TAGWNIPM[+15.995]GLLFN[+2059.735]QTGSC[+57.021]K.F | HexNAc(4)Hex(5)Fuc(1)NeuAc(1) | C[+57], M[+16], N[+2060] | 1390.9207 | 3 | 4170.7476 | 4170.7398 | 0 | 1.9 | 485 | Specific | 769.92 | 701.77 | 701.77 | 15.03 | 149 | >sp\|P02788\|TRFL_HUMAN Lactotransferrin OS=Homo sapiens OX=9606 GN=LTF PE=1 SV=6 | 1270 | 20201113_12413_1ng_glyco_hcd203040_compound15.20349.20349.3 | controllerType=0 controllerNumber=1 scan=20349 | 41.311 |
| 17892:4 | 1 | R.TAGWNIPMGLLFN[+1216.423]QTGSC[+57.021]KFDEYFSQSC[+57.021]APGSDPR.S | HexNAc(2)Hex(5) | C[+57]*2, N[+1216] | 1289.5565 | 4 | 5155.2043 | 5155.18 | 0 | 4.7 | 485 | Specific | 206.26 | 206.26 | 122.48 | 3.52 | 149 | >sp\|P02788\|TRFL_HUMAN Lactotransferrin OS=Homo sapiens OX=9606 GN=LTF PE=1 SV=6 | 1270 | 20201113_12413_1ng_glyco_hcd203040_compound15.21420.21420.4 | controllerType=0 controllerNumber=1 scan=21420 | 43.6137 |
| 20101:4 | 1 | R.TAGWNIPMGLLFN[+2204.772]QTGSC[+57.021]KFDEYFSQSC[+57.021]APGSDPR.S | HexNAc(4)Hex(5)NeuAc(2) | C[+57]*2, N[+2205] | 1536.6482 | 4 | 6143.571 | 6143.5295 | 0 | 6.7 | 485 | Specific | 313.58 | 302.3 | 302.3 | 5.77 | 149 | >sp\|P02788\|TRFL_HUMAN Lactotransferrin OS=Homo sapiens OX=9606 GN=LTF PE=1 SV=6 | 1270 | 20201113_12413_1ng_glyco_hcd203040_compound15.23732.23732.4 | controllerType=0 controllerNumber=1 scan=23732 | 48.6249 |
| 20104:4 | 1 | R.TAGWNIPMGLLFN[+2204.772]QTGSC[+57.021]KFDEYFSQSC[+57.021]APGSDPR.S | HexNAc(4)Hex(5)NeuAc(2) | C[+57]*2, N[+2205] | 1536.6482 | 4 | 6143.571 | 6143.5295 | 0 | 6.7 | 485 | Specific | 333.33 | 333.33 | 333.33 | 5.95 | 149 | >sp\|P02788\|TRFL_HUMAN Lactotransferrin OS=Homo sapiens OX=9606 GN=LTF PE=1 SV=6 | 1270 | 20201113_12413_1ng_glyco_hcd203040_compound15.23735.23735.4 | controllerType=0 controllerNumber=1 scan=23735 | 48.629 |
| 18807:3 | 1 | R.TAGWNIPMGLLFN[+1913.677]QTGSC[+57.021]KFDEYFSQSC[+57.021]APGSDPR.S | HexNAc(4)Hex(5)NeuAc(1) | C[+57]*2, N[+1914] | 1951.4834 | 3 | 5852.4355 | 5852.4341 | 0 | 0.2 | 485 | Specific | 557.27 | 453.87 | 453.87 | 9.04 | 149 | >sp\|P02788\|TRFL_HUMAN Lactotransferrin OS=Homo sapiens OX=9606 GN=LTF PE=1 SV=6 | 1270 | 20201113_12413_1ng_glyco_hcd203040_compound15.22379.22379.3 | controllerType=0 controllerNumber=1 scan=22379 | 45.6925 |
| 18810:3 | 1 | R.TAGWNIPMGLLFN[+1913.677]QTGSC[+57.021]KFDEYFSQSC[+57.021]APGSDPR.S | HexNAc(4)Hex(5)NeuAc(1) | C[+57]*2, N[+1914] | 1951.4834 | 3 | 5852.4355 | 5852.4341 | 0 | 0.2 | 485 | Specific | 612.94 | 500.98 | 500.98 | 11.26 | 149 | >sp\|P02788\|TRFL_HUMAN Lactotransferrin OS=Homo sapiens OX=9606 GN=LTF PE=1 SV=6 | 1270 | 20201113_12413_1ng_glyco_hcd203040_compound15.22382.22382.3 | controllerType=0 controllerNumber=1 scan=22382 | 45.6975 |
| 18802:4 | 1 | R.TAGWNIPMGLLFN[+1913.677]QTGSC[+57.021]KFDEYFSQSC[+57.021]APGSDPR.S | HexNAc(4)Hex(5)NeuAc(1) | C[+57]*2, N[+1914] | 1463.8665 | 4 | 5852.444 | 5852.4341 | 0 | 1.7 | 485 | Specific | 621.61 | 559.68 | 559.68 | 11.47 | 149 | >sp\|P02788\|TRFL_HUMAN Lactotransferrin OS=Homo sapiens OX=9606 GN=LTF PE=1 SV=6 | 1270 | 20201113_12413_1ng_glyco_hcd203040_compound15.22374.22374.4 | controllerType=0 controllerNumber=1 scan=22374 | 45.6849 |
| 18721:4 | 1 | R.TAGWNIPMGLLFN[+2059.735]QTGSC[+57.021]KFDEYFSQSC[+57.021]APGSDPR.S | HexNAc(4)Hex(5)Fuc(1)NeuAc(1) | C[+57]*2, N[+2060] | 1500.3894 | 4 | 5998.5358 | 5998.492 | 0 | 7.3 | 485 | Specific | 639.87 | 629.56 | 629.56 | 10.77 | 149 | >sp\|P02788\|TRFL_HUMAN Lactotransferrin OS=Homo sapiens OX=9606 GN=LTF PE=1 SV=6 | 1270 | 20201113_12413_1ng_glyco_hcd203040_compound15.22289.22289.4 | controllerType=0 controllerNumber=1 scan=22289 | 45.4971 |
| 18733:4 | 1 | R.TAGWNIPMGLLFN[+2205.793]QTGSC[+57.021]KFDEYFSQSC[+57.021]APGSDPR.S | HexNAc(4)Hex(5)Fuc(2)NeuAc(1) | C[+57]*2, N[+2206] | 1536.9008 | 4 | 6144.5812 | 6144.5499 | 0 | 5.1 | 485 | Specific | 758.62 | 684.85 | 684.85 | 13.8 | 149 | >sp\|P02788\|TRFL_HUMAN Lactotransferrin OS=Homo sapiens OX=9606 GN=LTF PE=1 SV=6 | 1270 | 20201113_12413_1ng_glyco_hcd203040_compound15.22301.22301.4 | controllerType=0 controllerNumber=1 scan=22301 | 45.5213 |
| 19988:4 | 1 | R.TAGWNIPMGLLFN[+2350.830]QTGSC[+57.021]KFDEYFSQSC[+57.021]APGSDPR.S | HexNAc(4)Hex(5)Fuc(1)NeuAc(2) | C[+57]*2, N[+2351] | 1573.1534 | 4 | 6289.592 | 6289.5875 | 0 | 0.7 | 485 | Specific | 883.55 | 828.18 | 828.18 | 16.27 | 149 | >sp\|P02788\|TRFL_HUMAN Lactotransferrin OS=Homo sapiens OX=9606 GN=LTF PE=1 SV=6 | 1270 | 20201113_12413_1ng_glyco_hcd203040_compound15.23614.23614.4 | controllerType=0 controllerNumber=1 scan=23614 | 48.3878 |
| 18723:4 | 1 | R.TAGWNIPMGLLFN[+2059.735]QTGSC[+57.021]KFDEYFSQSC[+57.021]APGSDPR.S | HexNAc(4)Hex(5)Fuc(1)NeuAc(1) | C[+57]*2, N[+2060] | 1500.3894 | 4 | 5998.5358 | 5998.492 | 0 | 7.3 | 485 | Specific | 940.44 | 906.44 | 906.44 | 16.77 | 149 | >sp\|P02788\|TRFL_HUMAN Lactotransferrin OS=Homo sapiens OX=9606 GN=LTF PE=1 SV=6 | 1270 | 20201113_12413_1ng_glyco_hcd203040_compound15.22291.22291.4 | controllerType=0 controllerNumber=1 scan=22291 | 45.5015 |
| 18804:4 | 1 | R.TAGWNIPMGLLFN[+1913.677]QTGSC[+57.021]KFDEYFSQSC[+57.021]APGSDPR.S | HexNAc(4)Hex(5)NeuAc(1) | C[+57]*2, N[+1914] | 1463.8665 | 4 | 5852.444 | 5852.4341 | 0 | 1.7 | 485 | Specific | 970.45 | 896.4 | 896.4 | 18.47 | 149 | >sp\|P02788\|TRFL_HUMAN Lactotransferrin OS=Homo sapiens OX=9606 GN=LTF PE=1 SV=6 | 1270 | 20201113_12413_1ng_glyco_hcd203040_compound15.22376.22376.4 | controllerType=0 controllerNumber=1 scan=22376 | 45.6887 |
| 19986:4 | 1 | R.TAGWNIPMGLLFN[+2350.830]QTGSC[+57.021]KFDEYFSQSC[+57.021]APGSDPR.S | HexNAc(4)Hex(5)Fuc(1)NeuAc(2) | C[+57]*2, N[+2351] | 1573.1534 | 4 | 6289.592 | 6289.5875 | 0 | 0.7 | 485 | Specific | 981.51 | 935.26 | 935.26 | 18.27 | 149 | >sp\|P02788\|TRFL_HUMAN Lactotransferrin OS=Homo sapiens OX=9606 GN=LTF PE=1 SV=6 | 1270 | 20201113_12413_1ng_glyco_hcd203040_compound15.23612.23612.4 | controllerType=0 controllerNumber=1 scan=23612 | 48.3822 |
| 08778:3 | 1 | R.N[+2204.772]GSDC[+57.021]PDKFC[+57.021]LFQSETK.N | HexNAc(4)Hex(5)NeuAc(2) | C[+57]*2, N[+2205] | 1413.2267 | 3 | 4237.6657 | 4237.6463 | 0 | 4.6 | 642 | Specific | 225.48 | 151.24 | 151.24 | 3.77 | 149 | >sp\|P02788\|TRFL_HUMAN Lactotransferrin OS=Homo sapiens OX=9606 GN=LTF PE=1 SV=6 | 1270 | 20201113_12413_1ng_glyco_hcd203040_compound15.11915.11915.2 | controllerType=0 controllerNumber=1 scan=11915 | 26.4744 |
| 08776:3 | 1 | R.N[+2204.772]GSDC[+57.021]PDKFC[+57.021]LFQSETK.N | HexNAc(4)Hex(5)NeuAc(2) | C[+57]*2, N[+2205] | 1413.2267 | 3 | 4237.6657 | 4237.6463 | 0 | 4.6 | 642 | Specific | 273.57 | 168.4 | 168.4 | 5.33 | 149 | >sp\|P02788\|TRFL_HUMAN Lactotransferrin OS=Homo sapiens OX=9606 GN=LTF PE=1 SV=6 | 1270 | 20201113_12413_1ng_glyco_hcd203040_compound15.11913.11913.2 | controllerType=0 controllerNumber=1 scan=11913 | 26.4702 |
| 08648:3 | 1 | R.N[+2570.925]GSDC[+57.021]PDKFC[+57.021]LFQSETK.N | HexNAc(5)Hex(6)Fuc(2)NeuAc(1) | C[+57]*2, N[+2571] | 1535.2726 | 3 | 4603.8034 | 4603.7989 | 0 | 1 | 642 | Specific | 341.7 | 325.07 | 325.07 | 6.78 | 149 | >sp\|P02788\|TRFL_HUMAN Lactotransferrin OS=Homo sapiens OX=9606 GN=LTF PE=1 SV=6 | 1270 | 20201113_12413_1ng_glyco_hcd203040_compound15.11777.11777.3 | controllerType=0 controllerNumber=1 scan=11777 | 26.241 |
| 08646:3 | 1 | R.N[+2570.925]GSDC[+57.021]PDKFC[+57.021]LFQSETK.N | HexNAc(5)Hex(6)Fuc(2)NeuAc(1) | C[+57]*2, N[+2571] | 1535.2726 | 3 | 4603.8034 | 4603.7989 | 0 | 1 | 642 | Specific | 380.69 | 322.02 | 322.02 | 6.94 | 149 | >sp\|P02788\|TRFL_HUMAN Lactotransferrin OS=Homo sapiens OX=9606 GN=LTF PE=1 SV=6 | 1270 | 20201113_12413_1ng_glyco_hcd203040_compound15.11775.11775.3 | controllerType=0 controllerNumber=1 scan=11775 | 26.2372 |
| 11263:3 | 1 | R.N[+2350.830]GSDC[+57.021]PDKFC[+57.021]LFQSETK.N | HexNAc(4)Hex(5)Fuc(1)NeuAc(2) | C[+57]*2, N[+2351] | 1461.9109 | 3 | 4383.7183 | 4383.7043 | 0 | 3.2 | 642 | Specific | 438.53 | 390.97 | 390.97 | 8.76 | 149 | >sp\|P02788\|TRFL_HUMAN Lactotransferrin OS=Homo sapiens OX=9606 GN=LTF PE=1 SV=6 | 1270 | 20201113_12413_1ng_glyco_hcd203040_compound15.14490.14490.3 | controllerType=0 controllerNumber=1 scan=14490 | 30.8768 |
| 08862:3 | 1 | R.N[+2205.793]GSDC[+57.021]PDKFC[+57.021]LFQSETK.N | HexNAc(4)Hex(5)Fuc(2)NeuAc(1) | C[+57]*2, N[+2206] | 1413.5639 | 3 | 4238.6771 | 4238.6667 | 0 | 2.5 | 642 | Specific | 456.56 | 312.06 | 312.06 | 7.04 | 149 | >sp\|P02788\|TRFL_HUMAN Lactotransferrin OS=Homo sapiens OX=9606 GN=LTF PE=1 SV=6 | 1270 | 20201113_12413_1ng_glyco_hcd203040_compound15.12003.12003.3 | controllerType=0 controllerNumber=1 scan=12003 | 26.6281 |
| 08810:3 | 1 | R.N[+2204.772]GSDC[+57.021]PDKFC[+57.021]LFQSETK.N | HexNAc(4)Hex(5)NeuAc(2) | C[+57]*2, N[+2205] | 1413.2274 | 3 | 4237.6675 | 4237.6463 | 0 | 5 | 642 | Specific | 477.07 | 381.44 | 381.44 | 8.21 | 149 | >sp\|P02788\|TRFL_HUMAN Lactotransferrin OS=Homo sapiens OX=9606 GN=LTF PE=1 SV=6 | 1270 | 20201113_12413_1ng_glyco_hcd203040_compound15.11948.11948.3 | controllerType=0 controllerNumber=1 scan=11948 | 26.5309 |
| 08888:3 | 1 | R.N[+2059.735]GSDC[+57.021]PDKFC[+57.021]LFQSETK.N | HexNAc(4)Hex(5)Fuc(1)NeuAc(1) | C[+57]*2, N[+2060] | 1364.8771 | 3 | 4092.6168 | 4092.6088 | 0 | 2 | 642 | Specific | 535.35 | 389.97 | 389.97 | 9.1 | 149 | >sp\|P02788\|TRFL_HUMAN Lactotransferrin OS=Homo sapiens OX=9606 GN=LTF PE=1 SV=6 | 1270 | 20201113_12413_1ng_glyco_hcd203040_compound15.12030.12030.3 | controllerType=0 controllerNumber=1 scan=12030 | 26.6759 |
| 08886:3 | 1 | R.N[+2059.735]GSDC[+57.021]PDKFC[+57.021]LFQSETK.N | HexNAc(4)Hex(5)Fuc(1)NeuAc(1) | C[+57]*2, N[+2060] | 1364.8771 | 3 | 4092.6168 | 4092.6088 | 0 | 2 | 642 | Specific | 538.28 | 391.85 | 391.85 | 9.1 | 149 | >sp\|P02788\|TRFL_HUMAN Lactotransferrin OS=Homo sapiens OX=9606 GN=LTF PE=1 SV=6 | 1270 | 20201113_12413_1ng_glyco_hcd203040_compound15.12028.12028.3 | controllerType=0 controllerNumber=1 scan=12028 | 26.6714 |
| 08809:3 | 1 | R.N[+2204.772]GSDC[+57.021]PDKFC[+57.021]LFQSETK.N | HexNAc(4)Hex(5)NeuAc(2) | C[+57]*2, N[+2205] | 1413.2274 | 3 | 4237.6675 | 4237.6463 | 0 | 5 | 642 | Specific | 550.46 | 493.33 | 493.33 | 8.6 | 149 | >sp\|P02788\|TRFL_HUMAN Lactotransferrin OS=Homo sapiens OX=9606 GN=LTF PE=1 SV=6 | 1270 | 20201113_12413_1ng_glyco_hcd203040_compound15.11947.11947.3 | controllerType=0 controllerNumber=1 scan=11947 | 26.5272 |
| 11261:3 | 1 | R.N[+2350.830]GSDC[+57.021]PDKFC[+57.021]LFQSETK.N | HexNAc(4)Hex(5)Fuc(1)NeuAc(2) | C[+57]*2, N[+2351] | 1461.9109 | 3 | 4383.7183 | 4383.7043 | 0 | 3.2 | 642 | Specific | 573.44 | 440.77 | 440.77 | 10.59 | 149 | >sp\|P02788\|TRFL_HUMAN Lactotransferrin OS=Homo sapiens OX=9606 GN=LTF PE=1 SV=6 | 1270 | 20201113_12413_1ng_glyco_hcd203040_compound15.14488.14488.3 | controllerType=0 controllerNumber=1 scan=14488 | 30.8713 |
| 08860:3 | 1 | R.N[+2205.793]GSDC[+57.021]PDKFC[+57.021]LFQSETK.N | HexNAc(4)Hex(5)Fuc(2)NeuAc(1) | C[+57]*2, N[+2206] | 1413.5639 | 3 | 4238.6771 | 4238.6667 | 0 | 2.5 | 642 | Specific | 622.03 | 517.69 | 517.69 | 11.59 | 149 | >sp\|P02788\|TRFL_HUMAN Lactotransferrin OS=Homo sapiens OX=9606 GN=LTF PE=1 SV=6 | 1270 | 20201113_12413_1ng_glyco_hcd203040_compound15.12001.12001.3 | controllerType=0 controllerNumber=1 scan=12001 | 26.6237 |
| 09943:3 | 3 | K.TVLTPATNHMGN[+1378.476]VTFTIPANR.E | HexNAc(2)Hex(6) | N[+1378] | 1211.8862 | 3 | 3633.6439 | 3633.6358 | 0 | 2.2 | 74 | Specific | 206.68 | 148.7 | 147.23 | 4.35 | 102 | >sp\|P01024\|CO3_HUMAN Complement C3 OS=Homo sapiens OX=9606 GN=C3 PE=1 SV=2 | 492 | 20201113_12413_1ng_glyco_hcd203040_compound15.13121.13121.2 | controllerType=0 controllerNumber=1 scan=13121 | 28.5084 |
| 09843:3 | 3 | K.TVLTPATNHMGN[+1540.529]VTFTIPANR.E | HexNAc(2)Hex(7) | N[+1541] | 1265.9039 | 3 | 3795.697 | 3795.6886 | 0 | 2.2 | 74 | Specific | 774.18 | 705.23 | 391.29 | 15.01 | 102 | >sp\|P01024\|CO3_HUMAN Complement C3 OS=Homo sapiens OX=9606 GN=C3 PE=1 SV=2 | 492 | 20201113_12413_1ng_glyco_hcd203040_compound15.13018.13018.3 | controllerType=0 controllerNumber=1 scan=13018 | 28.3191 |
| 09841:3 | 3 | K.TVLTPATNHMGN[+1540.529]VTFTIPANR.E | HexNAc(2)Hex(7) | N[+1541] | 1265.9039 | 3 | 3795.697 | 3795.6886 | 0 | 2.2 | 74 | Specific | 804.33 | 804.33 | 470.8 | 16.01 | 102 | >sp\|P01024\|CO3_HUMAN Complement C3 OS=Homo sapiens OX=9606 GN=C3 PE=1 SV=2 | 492 | 20201113_12413_1ng_glyco_hcd203040_compound15.13016.13016.3 | controllerType=0 controllerNumber=1 scan=13016 | 28.3153 |
| 15114:3 | 4 | R.LN[+1702.581]YSLPTGQWVGVQLPR.N | HexNAc(2)Hex(8) | N[+1703] | 1210.8791 | 3 | 3630.6227 | 3630.6202 | 0 | 0.7 | 1017 | Specific | 502.34 | 502.34 | 502.34 | 9.23 | 94 | >sp\|P24821\|TENA_HUMAN Tenascin OS=Homo sapiens OX=9606 GN=TNC PE=1 SV=3 | 486 | 20201113_12413_1ng_glyco_hcd203040_compound15.18498.18498.3 | controllerType=0 controllerNumber=1 scan=18498 | 37.8253 |
| 15116:3 | 4 | R.LN[+1702.581]YSLPTGQWVGVQLPR.N | HexNAc(2)Hex(8) | N[+1703] | 1210.8791 | 3 | 3630.6227 | 3630.6202 | 0 | 0.7 | 1017 | Specific | 751.2 | 606.26 | 606.26 | 14.75 | 94 | >sp\|P24821\|TENA_HUMAN Tenascin OS=Homo sapiens OX=9606 GN=TNC PE=1 SV=3 | 486 | 20201113_12413_1ng_glyco_hcd203040_compound15.18500.18500.3 | controllerType=0 controllerNumber=1 scan=18500 | 37.8301 |
| 17511:3 | 6 | R.GLSFDVSLEVSQGPGLLN[+2059.735]DTK.V | HexNAc(4)Hex(5)Fuc(1)NeuAc(1) | N[+2060] | 1412.6275 | 3 | 4235.8679 | 4235.8481 | 0 | 4.7 | 118 | Specific | 324.75 | 324.75 | 324.75 | 7.16 | 74 | >sp\|P01833\|PIGR_HUMAN Polymeric immunoglobulin receptor OS=Homo sapiens OX=9606 GN=PIGR PE=1 SV=4 | 902 | 20201113_12413_1ng_glyco_hcd203040_compound15.21022.21022.3 | controllerType=0 controllerNumber=1 scan=21022 | 42.7373 |
| 17508:3 | 6 | R.GLSFDVSLEVSQGPGLLN[+2059.735]DTK.V | HexNAc(4)Hex(5)Fuc(1)NeuAc(1) | N[+2060] | 1412.6275 | 3 | 4235.8679 | 4235.8481 | 0 | 4.7 | 118 | Specific | 427.2 | 427.2 | 427.2 | 9.16 | 74 | >sp\|P01833\|PIGR_HUMAN Polymeric immunoglobulin receptor OS=Homo sapiens OX=9606 GN=PIGR PE=1 SV=4 | 902 | 20201113_12413_1ng_glyco_hcd203040_compound15.21019.21019.3 | controllerType=0 controllerNumber=1 scan=21019 | 42.7328 |
| 20604:4 | 6 | K.QIGLYPVLVIDSSGYVNPN[+2204.772]YTGR.I | HexNAc(4)Hex(5)NeuAc(2) | N[+2205] | 1183.283 | 4 | 4730.1102 | 4730.0759 | 0 | 7.2 | 168 | Specific | 344.91 | 344.91 | 344.91 | 6.72 | 74 | >sp\|P01833\|PIGR_HUMAN Polymeric immunoglobulin receptor OS=Homo sapiens OX=9606 GN=PIGR PE=1 SV=4 | 902 | 20201113_12413_1ng_glyco_hcd203040_compound15.24257.24257.4 | controllerType=0 controllerNumber=1 scan=24257 | 49.8104 |
| 20606:4 | 6 | K.QIGLYPVLVIDSSGYVNPN[+2204.772]YTGR.I | HexNAc(4)Hex(5)NeuAc(2) | N[+2205] | 1183.283 | 4 | 4730.1102 | 4730.0759 | 0 | 7.2 | 168 | Specific | 644.62 | 513.44 | 513.44 | 11.43 | 74 | >sp\|P01833\|PIGR_HUMAN Polymeric immunoglobulin receptor OS=Homo sapiens OX=9606 GN=PIGR PE=1 SV=4 | 902 | 20201113_12413_1ng_glyco_hcd203040_compound15.24259.24259.4 | controllerType=0 controllerNumber=1 scan=24259 | 49.8153 |
| 18672:3 | 6 | K.QIGLYPVLVIDSSGYVNPN[+1913.677]YTGR.I | HexNAc(4)Hex(5)NeuAc(1) | N[+1914] | 1480.3364 | 3 | 4438.9945 | 4438.9805 | 0 | 3.2 | 168 | Specific | 846.69 | 735 | 735 | 16.38 | 74 | >sp\|P01833\|PIGR_HUMAN Polymeric immunoglobulin receptor OS=Homo sapiens OX=9606 GN=PIGR PE=1 SV=4 | 902 | 20201113_12413_1ng_glyco_hcd203040_compound15.22238.22238.3 | controllerType=0 controllerNumber=1 scan=22238 | 45.4002 |
| 18674:3 | 6 | K.QIGLYPVLVIDSSGYVNPN[+1913.677]YTGR.I | HexNAc(4)Hex(5)NeuAc(1) | N[+1914] | 1480.3364 | 3 | 4438.9945 | 4438.9805 | 0 | 3.2 | 168 | Specific | 896.52 | 641.03 | 641.03 | 17.38 | 74 | >sp\|P01833\|PIGR_HUMAN Polymeric immunoglobulin receptor OS=Homo sapiens OX=9606 GN=PIGR PE=1 SV=4 | 902 | 20201113_12413_1ng_glyco_hcd203040_compound15.22240.22240.3 | controllerType=0 controllerNumber=1 scan=22240 | 45.4061 |
| 22507:3 | 6 | R.[+42.011]LSLLEEPGN[+2336.851]GTFTVILNQLTSR.D | HexNAc(6)Hex(6)Fuc(1) | .[42], N[+2337] | 1594.3967 | 3 | 4781.1754 | 4781.1542 | 0 | 4.4 | 413 | Specific | 367.13 | 331.13 | 331.13 | 7.43 | 74 | >sp\|P01833\|PIGR_HUMAN Polymeric immunoglobulin receptor OS=Homo sapiens OX=9606 GN=PIGR PE=1 SV=4 | 902 | 20201113_12413_1ng_glyco_hcd203040_compound15.26240.26240.3 | controllerType=0 controllerNumber=1 scan=26240 | 53.4724 |
| 22505:3 | 6 | R.[+42.011]LSLLEEPGN[+2336.851]GTFTVILNQLTSR.D | HexNAc(6)Hex(6)Fuc(1) | .[42], N[+2337] | 1594.3967 | 3 | 4781.1754 | 4781.1542 | 0 | 4.4 | 413 | Specific | 387.01 | 352.7 | 352.7 | 7.55 | 74 | >sp\|P01833\|PIGR_HUMAN Polymeric immunoglobulin receptor OS=Homo sapiens OX=9606 GN=PIGR PE=1 SV=4 | 902 | 20201113_12413_1ng_glyco_hcd203040_compound15.26238.26238.3 | controllerType=0 controllerNumber=1 scan=26238 | 53.463 |
| 20283:3 | 6 | R.LSLLEEPGN[+2059.735]GTFTVILNQLTSR.D | HexNAc(4)Hex(5)Fuc(1)NeuAc(1) | N[+2060] | 1488.0239 | 3 | 4462.057 | 4462.0275 | 0 | 6.6 | 413 | Specific | 450.59 | 439.26 | 439.26 | 8.51 | 74 | >sp\|P01833\|PIGR_HUMAN Polymeric immunoglobulin receptor OS=Homo sapiens OX=9606 GN=PIGR PE=1 SV=4 | 902 | 20201113_12413_1ng_glyco_hcd203040_compound15.23922.23922.3 | controllerType=0 controllerNumber=1 scan=23922 | 49.0445 |
| 20249:3 | 6 | R.LSLLEEPGN[+2205.793]GTFTVILNQLTSR.D | HexNAc(4)Hex(5)Fuc(2)NeuAc(1) | N[+2206] | 1536.6983 | 3 | 4608.0803 | 4608.0854 | 0 | -1.1 | 413 | Specific | 559.51 | 421.34 | 421.34 | 8.97 | 74 | >sp\|P01833\|PIGR_HUMAN Polymeric immunoglobulin receptor OS=Homo sapiens OX=9606 GN=PIGR PE=1 SV=4 | 902 | 20201113_12413_1ng_glyco_hcd203040_compound15.23886.23886.3 | controllerType=0 controllerNumber=1 scan=23886 | 48.9732 |
| 22619:3 | 6 | R.LSLLEEPGN[+2350.830]GTFTVILNQLTSR.D | HexNAc(4)Hex(5)Fuc(1)NeuAc(2) | N[+2351] | 1585.0558 | 3 | 4753.153 | 4753.1229 | 0 | 6.3 | 413 | Specific | 572.13 | 501.53 | 501.53 | 9.11 | 74 | >sp\|P01833\|PIGR_HUMAN Polymeric immunoglobulin receptor OS=Homo sapiens OX=9606 GN=PIGR PE=1 SV=4 | 902 | 20201113_12413_1ng_glyco_hcd203040_compound15.26356.26356.3 | controllerType=0 controllerNumber=1 scan=26356 | 53.6868 |
| 20251:3 | 6 | R.LSLLEEPGN[+2205.793]GTFTVILNQLTSR.D | HexNAc(4)Hex(5)Fuc(2)NeuAc(1) | N[+2206] | 1536.6983 | 3 | 4608.0803 | 4608.0854 | 0 | -1.1 | 413 | Specific | 620.68 | 380.22 | 380.22 | 11.19 | 74 | >sp\|P01833\|PIGR_HUMAN Polymeric immunoglobulin receptor OS=Homo sapiens OX=9606 GN=PIGR PE=1 SV=4 | 902 | 20201113_12413_1ng_glyco_hcd203040_compound15.23888.23888.3 | controllerType=0 controllerNumber=1 scan=23888 | 48.9769 |
| 22621:3 | 6 | R.LSLLEEPGN[+2350.830]GTFTVILNQLTSR.D | HexNAc(4)Hex(5)Fuc(1)NeuAc(2) | N[+2351] | 1585.0558 | 3 | 4753.153 | 4753.1229 | 0 | 6.3 | 413 | Specific | 697.36 | 464.03 | 464.03 | 12.33 | 74 | >sp\|P01833\|PIGR_HUMAN Polymeric immunoglobulin receptor OS=Homo sapiens OX=9606 GN=PIGR PE=1 SV=4 | 902 | 20201113_12413_1ng_glyco_hcd203040_compound15.26358.26358.3 | controllerType=0 controllerNumber=1 scan=26358 | 53.6903 |
| 20036:3 | 6 | R.LSLLEEPGN[+2570.925]GTFTVILNQLTSR.D | HexNAc(5)Hex(6)Fuc(2)NeuAc(1) | N[+2571] | 1658.4166 | 3 | 4973.2352 | 4973.2176 | 0 | 3.5 | 413 | Specific | 721.22 | 388.88 | 388.88 | 13.99 | 74 | >sp\|P01833\|PIGR_HUMAN Polymeric immunoglobulin receptor OS=Homo sapiens OX=9606 GN=PIGR PE=1 SV=4 | 902 | 20201113_12413_1ng_glyco_hcd203040_compound15.23664.23664.3 | controllerType=0 controllerNumber=1 scan=23664 | 48.4814 |
| 20286:3 | 6 | R.LSLLEEPGN[+2059.735]GTFTVILNQLTSR.D | HexNAc(4)Hex(5)Fuc(1)NeuAc(1) | N[+2060] | 1488.0239 | 3 | 4462.057 | 4462.0275 | 0 | 6.6 | 413 | Specific | 771.15 | 574.4 | 574.4 | 14.33 | 74 | >sp\|P01833\|PIGR_HUMAN Polymeric immunoglobulin receptor OS=Homo sapiens OX=9606 GN=PIGR PE=1 SV=4 | 902 | 20201113_12413_1ng_glyco_hcd203040_compound15.23925.23925.3 | controllerType=0 controllerNumber=1 scan=23925 | 49.0541 |
| 15921:3 | 6 | K.IIEGEPNLKVPGN[+1913.677]VTAVLGETLK.V | HexNAc(4)Hex(5)NeuAc(1) | N[+1914] | 1435.6898 | 3 | 4305.0548 | 4305.0264 | 0 | 6.6 | 457 | Specific | 275.78 | 275.78 | 275.78 | 5.78 | 74 | >sp\|P01833\|PIGR_HUMAN Polymeric immunoglobulin receptor OS=Homo sapiens OX=9606 GN=PIGR PE=1 SV=4 | 902 | 20201113_12413_1ng_glyco_hcd203040_compound15.19358.19358.3 | controllerType=0 controllerNumber=1 scan=19358 | 39.4532 |
| 15884:3 | 6 | K.IIEGEPNLKVPGN[+2059.735]VTAVLGETLK.V | HexNAc(4)Hex(5)Fuc(1)NeuAc(1) | N[+2060] | 1484.3776 | 3 | 4451.1183 | 4451.0843 | 0 | 7.6 | 457 | Specific | 376.7 | 376.7 | 376.7 | 8.07 | 74 | >sp\|P01833\|PIGR_HUMAN Polymeric immunoglobulin receptor OS=Homo sapiens OX=9606 GN=PIGR PE=1 SV=4 | 902 | 20201113_12413_1ng_glyco_hcd203040_compound15.19320.19320.3 | controllerType=0 controllerNumber=1 scan=19320 | 39.369 |
| 14133:3 | 6 | K.IIEGEPNLKVPGN[+1913.677]VTAVLGETLK.V | HexNAc(4)Hex(5)NeuAc(1) | N[+1914] | 1435.6899 | 3 | 4305.0551 | 4305.0264 | 0 | 6.7 | 457 | Specific | 455.92 | 426.52 | 426.52 | 8.32 | 74 | >sp\|P01833\|PIGR_HUMAN Polymeric immunoglobulin receptor OS=Homo sapiens OX=9606 GN=PIGR PE=1 SV=4 | 902 | 20201113_12413_1ng_glyco_hcd203040_compound15.17480.17480.3 | controllerType=0 controllerNumber=1 scan=17480 | 35.9931 |
| 15819:3 | 6 | K.IIEGEPNLKVPGN[+2205.793]VTAVLGETLK.V | HexNAc(4)Hex(5)Fuc(2)NeuAc(1) | N[+2206] | 1533.0591 | 3 | 4597.1629 | 4597.1422 | 0 | 4.5 | 457 | Specific | 462.14 | 462.14 | 462.14 | 8.98 | 74 | >sp\|P01833\|PIGR_HUMAN Polymeric immunoglobulin receptor OS=Homo sapiens OX=9606 GN=PIGR PE=1 SV=4 | 902 | 20201113_12413_1ng_glyco_hcd203040_compound15.19250.19250.2 | controllerType=0 controllerNumber=1 scan=19250 | 39.2363 |
| 15817:3 | 6 | K.IIEGEPNLKVPGN[+2205.793]VTAVLGETLK.V | HexNAc(4)Hex(5)Fuc(2)NeuAc(1) | N[+2206] | 1533.0591 | 3 | 4597.1629 | 4597.1422 | 0 | 4.5 | 457 | Specific | 493.8 | 463.5 | 463.5 | 9.27 | 74 | >sp\|P01833\|PIGR_HUMAN Polymeric immunoglobulin receptor OS=Homo sapiens OX=9606 GN=PIGR PE=1 SV=4 | 902 | 20201113_12413_1ng_glyco_hcd203040_compound15.19248.19248.2 | controllerType=0 controllerNumber=1 scan=19248 | 39.[illegible]308 |
| 15814:3 | 6 | K.IIEGEPNLKVPGN[+2205.793]VTAVLGETLK.V | HexNAc(4)Hex(5)Fuc(2)NeuAc(1) | N[+2206] | 1533.0529 | 3 | 4597.1442 | 4597.1422 | 0 | 0.4 | 457 | Specific | 609.86 | 539.66 | 539.66 | 11.64 | 74 | >sp\|P01833\|PIGR_HUMAN Polymeric immunoglobulin receptor OS=Homo sapiens OX=9606 GN=PIGR PE=1 SV=4 | 902 | 20201113_12413_1ng_glyco_hcd203040_compound15.19244.19244.3 | controllerType=0 controllerNumber=1 scan=19244 | 39.2217 |
| 15919:3 | 6 | K.IIEGEPNLKVPGN[+1913.677]VTAVLGETLK.V | HexNAc(4)Hex(5)NeuAc(1) | N[+1914] | 1435.6898 | 3 | 4305.0548 | 4305.0264 | 0 | 6.6 | 457 | Specific | 609.94 | 609.4 | 609.4 | 11.14 | 74 | >sp\|P01833\|PIGR_HUMAN Polymeric immunoglobulin receptor OS=Homo sapiens OX=9606 GN=PIGR PE=1 SV=4 | 902 | 20201113_12413_1ng_glyco_hcd203040_compound15.19356.19356.3 | controllerType=0 controllerNumber=1 scan=19356 | 39.4495 |
| 15816:3 | 6 | K.IIEGEPNLKVPGN[+2205.793]VTAVLGETLK.V | HexNAc(4)Hex(5)Fuc(2)NeuAc(1) | N[+2206] | 1533.0529 | 3 | 4597.1442 | 4597.1422 | 0 | 0.4 | 457 | Specific | 611.38 | 485.42 | 485.42 | 11.64 | 74 | >sp\|P01833\|PIGR_HUMAN Polymeric immunoglobulin receptor OS=Homo sapiens OX=9606 GN=PIGR PE=1 SV=4 | 902 | 20201113_12413_1ng_glyco_hcd203040_compound15.19246.19246.3 | controllerType=0 controllerNumber=1 scan=19246 | 39.2263 |
| 15868:3 | 6 | K.IIEGEPNLKVPGN[+2059.735]VTAVLGETLK.V | HexNAc(4)Hex(5)Fuc(1)NeuAc(1) | N[+2060] | 1484.3705 | 3 | 4451.0971 | 4451.0843 | 0 | 2.9 | 457 | Specific | 697.93 | 697.93 | 697.93 | 12.93 | 74 | >sp\|P01833\|PIGR_HUMAN Polymeric immunoglobulin receptor OS=Homo sapiens OX=9606 GN=PIGR PE=1 SV=4 | 902 | 20201113_12413_1ng_glyco_hcd203040_compound15.19303.19303.3 | controllerType=0 controllerNumber=1 scan=19303 | 39.3397 |
| 15886:3 | 6 | K.IIEGEPNLKVPGN[+2059.735]VTAVLGETLK.V | HexNAc(4)Hex(5)Fuc(1)NeuAc(1) | N[+2060] | 1484.3776 | 3 | 4451.1183 | 4451.0843 | 0 | 7.6 | 457 | Specific | 790.1 | 703.58 | 703.58 | 14.07 | 74 | >sp\|P01833\|PIGR_HUMAN Polymeric immunoglobulin receptor OS=Homo sapiens OX=9606 GN=PIGR PE=1 SV=4 | 902 | 20201113_12413_1ng_glyco_hcd203040_compound15.19322.19322.3 | controllerType=0 controllerNumber=1 scan=19322 | 39.3756 |
| 15862:3 | 6 | K.IIEGEPNLKVPGN[+2059.735]VTAVLGETLK.V | HexNAc(4)Hex(5)Fuc(1)NeuAc(1) | N[+2060] | 1484.3712 | 3 | 4451.0989 | 4451.0843 | 0 | 3.3 | 457 | Specific | 890.52 | 872.59 | 872.59 | 16.93 | 74 | >sp\|P01833\|PIGR_HUMAN Polymeric immunoglobulin receptor OS=Homo sapiens OX=9606 GN=PIGR PE=1 SV=4 | 902 | 20201113_12413_1ng_glyco_hcd203040_compound15.19296.19296.3 | controllerType=0 controllerNumber=1 scan=19296 | 39.3241 |
| 15866:3 | 6 | K.IIEGEPNLKVPGN[+2059.735]VTAVLGETLK.V | HexNAc(4)Hex(5)Fuc(1)NeuAc(1) | N[+2060] | 1484.3705 | 3 | 4451.0971 | 4451.0843 | 0 | 2.9 | 457 | Specific | 973.77 | 897.4 | 897.4 | 18.93 | 74 | >sp\|P01833\|PIGR_HUMAN Polymeric immunoglobulin receptor OS=Homo sapiens OX=9606 GN=PIGR PE=1 SV=4 | 902 | 20201113_12413_1ng_glyco_hcd203040_compound15.19301.19301.3 | controllerType=0 controllerNumber=1 scan=19301 | 39.335 |
| 15864:3 | 6 | K.IIEGEPNLKVPGN[+2059.735]VTAVLGETLK.V | HexNAc(4)Hex(5)Fuc(1)NeuAc(1) | N[+2060] | 1484.3712 | 3 | 4451.0989 | 4451.0843 | 0 | 3.3 | 457 | Specific | 1001.29 | 913.32 | 913.32 | 19.93 | 74 | >sp\|P01833\|PIGR_HUMAN Polymeric immunoglobulin receptor OS=Homo sapiens OX=9606 GN=PIGR PE=1 SV=4 | 902 | 20201113_12413_1ng_glyco_hcd203040_compound15.19298.19298.3 | controllerType=0 controllerNumber=1 scan=19298 | 39.329 |
| 16800:4 | 6 | K.IIEGEPNLKVPGN[+2350.830]VTAVLGETLKVPC[+57.021]HFPC[+57.021]K.F | HexNAc(4)Hex(5)Fuc(1)NeuAc(2) | C[+57]*2, N[+2351] | 1442.6699 | 4 | 5767.6579 | 5767.6372 | 0 | 3.6 | 457 | Specific | 254.34 | 254.34 | 254.34 | 5.06 | 74 | >sp\|P01833\|PIGR_HUMAN Polymeric immunoglobulin receptor OS=Homo sapiens OX=9606 GN=PIGR PE=1 SV=4 | 902 | 20201113_12413_1ng_glyco_hcd203040_compound15.20279.20279.4 | controllerType=0 controllerNumber=1 scan=20279 | 41.165 |
| 16798:4 | 6 | K.IIEGEPNLKVPGN[+2350.830]VTAVLGETLKVPC[+57.021]HFPC[+57.021]K.F | HexNAc(4)Hex(5)Fuc(1)NeuAc(2) | C[+57]*2, N[+2351] | 1442.6699 | 4 | 5767.6579 | 5767.6372 | 0 | 3.6 | 457 | Specific | 256.87 | 256.87 | 256.87 | 5.06 | 74 | >sp\|P01833\|PIGR_HUMAN Polymeric immunoglobulin receptor OS=Homo sapiens OX=9606 GN=PIGR PE=1 SV=4 | 902 | 20201113_12413_1ng_glyco_hcd203040_compound15.20277.20277.4 | controllerType=0 controllerNumber=1 scan=20277 | 41.1574 |
| 15373:4 | 6 | K.IIEGEPNLKVPGN[+2205.793]VTAVLGETLKVPC[+57.021]HFPC[+57.021]K.F | HexNAc(4)Hex(5)Fuc(2)NeuAc(1) | C[+57]*2, N[+2206] | 1406.4033 | 4 | 5622.5915 | 5622.5997 | 0 | -1.5 | 457 | Specific | 270.16 | 260 | 260 | 4.29 | 74 | >sp\|P01833\|PIGR_HUMAN Polymeric immunoglobulin receptor OS=Homo sapiens OX=9606 GN=PIGR PE=1 SV=4 | 902 | 20201113_12413_1ng_glyco_hcd203040_compound15.18767.18767.4 | controllerType=0 controllerNumber=1 scan=18767 | 38.3423 |
| 15494:4 | 6 | K.IIEGEPNLKVPGN[+2059.735]VTAVLGETLKVPC[+57.021]HFPC[+57.021]K.F | HexNAc(4)Hex(5)Fuc(1)NeuAc(1) | C[+57]*2, N[+2060] | 1369.8938 | 4 | 5476.5534 | 5476.5418 | 0 | 2.1 | 457 | Specific | 453.31 | 361.52 | 361.52 | 6.22 | 74 | >sp\|P01833\|PIGR_HUMAN Polymeric immunoglobulin receptor OS=Homo sapiens OX=9606 GN=PIGR PE=1 SV=4 | 902 | 20201113_12413_1ng_glyco_hcd203040_compound15.18892.18892.4 | controllerType=0 controllerNumber=1 scan=18892 | 38.5725 |
| 15375:4 | 6 | K.IIEGEPNLKVPGN[+2205.793]VTAVLGETLKVPC[+57.021]HFPC[+57.021]K.F | HexNAc(4)Hex(5)Fuc(2)NeuAc(1) | C[+57]*2, N[+2206] | 1406.4033 | 4 | 5622.5915 | 5622.5997 | 0 | -1.5 | 457 | Specific | 699.81 | 674.53 | 674.53 | 10.65 | 74 | >sp\|P01833\|PIGR_HUMAN Polymeric immunoglobulin receptor OS=Homo sapiens OX=9606 GN=PIGR PE=1 SV=4 | 902 | 20201113_12413_1ng_glyco_hcd203040_compound15.18769.18769.4 | controllerType=0 controllerNumber=1 scan=18769 | 38.3464 |
| 16848:5 | 6 | K.IIEGEPNLKVPGN[+2350.830]VTAVLGETLKVPC[+57.021]HFPC[+57.021]K.F | HexNAc(4)Hex(5)Fuc(1)NeuAc(2) | C[+57]*2, N[+2351] | 1154.3352 | 5 | 5767.6469 | 5767.6372 | 0 | 1.7 | 457 | Specific | 775.79 | 586.53 | 586.53 | 14.06 | 74 | >sp\|P01833\|PIGR_HUMAN Polymeric immunoglobulin receptor OS=Homo sapiens OX=9606 GN=PIGR PE=1 SV=4 | 902 | 20201113_12413_1ng_glyco_hcd203040_compound15.20329.20329.5 | controllerType=0 controllerNumber=1 scan=20329 | 41.2672 |
| 15423:4 | 6 | K.IIEGEPNLKVPGN[+2205.793]VTAVLGETLKVPC[+57.021]HFPC[+57.021]K.F | HexNAc(4)Hex(5)Fuc(2)NeuAc(1) | C[+57]*2, N[+2206] | 1406.4166 | 4 | 5622.6447 | 5622.5997 | 0 | 8 | 457 | Specific | 785.02 | 785.02 | 785.02 | 12.72 | 74 | >sp\|P01833\|PIGR_HUMAN Polymeric immunoglobulin receptor OS=Homo sapiens OX=9606 GN=PIGR PE=1 SV=4 | 902 | 20201113_12413_1ng_glyco_hcd203040_compound15.18819.18819.4 | controllerType=0 controllerNumber=1 scan=18819 | 38.4376 |
| 15486:5 | 6 | K.IIEGEPNLKVPGN[+2059.735]VTAVLGETLKVPC[+57.021]HFPC[+57.021]K.F | HexNAc(4)Hex(5)Fuc(1)NeuAc(1) | C[+57]*2, N[+2060] | 1096.1174 | 5 | 5476.5581 | 5476.5418 | 0 | 3 | 457 | Specific | 803.02 | 634.36 | 634.36 | 15.15 | 74 | >sp\|P01833\|PIGR_HUMAN Polymeric immunoglobulin receptor OS=Homo sapiens OX=9606 GN=PIGR PE=1 SV=4 | 902 | 20201113_12413_1ng_glyco_hcd203040_compound15.18884.18884.5 | controllerType=0 controllerNumber=1 scan=18884 | 38.5568 |
| 15440:5 | 6 | K.IIEGEPNLKVPGN[+2059.735]VTAVLGETLKVPC[+57.021]HFPC[+57.021]K.F | HexNAc(4)Hex(5)Fuc(1)NeuAc(1) | C[+57]*2, N[+2060] | 1096.122 | 5 | 5476.5806 | 5476.5418 | 0 | 7.1 | 457 | Specific | 847.71 | 833.33 | 833.33 | 14.36 | 74 | >sp\|P01833\|PIGR_HUMAN Polymeric immunoglobulin receptor OS=Homo sapiens OX=9606 GN=PIGR PE=1 SV=4 | 902 | 20201113_12413_1ng_glyco_hcd203040_compound15.18836.18836.3 | controllerType=0 controllerNumber=1 scan=18836 | 38.4701 |
| 15421:4 | 6 | K.IIEGEPNLKVPGN[+2205.793]VTAVLGETLKVPC[+57.021]HFPC[+57.021]K.F | HexNAc(4)Hex(5)Fuc(2)NeuAc(1) | C[+57]*2, N[+2206] | 1406.4166 | 4 | 5622.6447 | 5622.5997 | 0 | 8 | 457 | Specific | 848.08 | 848.08 | 848.08 | 13.72 | 74 | >sp\|P01833\|PIGR_HUMAN Polymeric immunoglobulin receptor OS=Homo sapiens OX=9606 GN=PIGR PE=1 SV=4 | 902 | 20201113_12413_1ng_glyco_hcd203040_compound15.18817.18817.4 | controllerType=0 controllerNumber=1 scan=18817 | 38.4332 |
| 15438:5 | 6 | K.IIEGEPNLKVPGN[+2059.735]VTAVLGETLKVPC[+57.021]HFPC[+57.021]K.F | HexNAc(4)Hex(5)Fuc(1)NeuAc(1) | C[+57]*2, N[+2060] | 1096.122 | 5 | 5476.5806 | 5476.5418 | 0 | 7.1 | 457 | Specific | 868.65 | 815.46 | 815.46 | 15.36 | 74 | >sp\|P01833\|PIGR_HUMAN Polymeric immunoglobulin receptor OS=Homo sapiens OX=9606 GN=PIGR PE=1 SV=4 | 902 | 20201113_12413_1ng_glyco_hcd203040_compound15.18834.18834.3 | controllerType=0 controllerNumber=1 scan=18834 | 38.4662 |
| 15484:5 | 6 | K.IIEGEPNLKVPGN[+2059.735]VTAVLGETLKVPC[+57.021]HFPC[+57.021]K.F | HexNAc(4)Hex(5)Fuc(1)NeuAc(1) | C[+57]*2, N[+2060] | 1096.1174 | 5 | 5476.5581 | 5476.5418 | 0 | 3 | 457 | Specific | 967.15 | 832.67 | 832.67 | 18.15 | 74 | >sp\|P01833\|PIGR_HUMAN Polymeric immunoglobulin receptor OS=Homo sapiens OX=9606 GN=PIGR PE=1 SV=4 | 902 | 20201113_12413_1ng_glyco_hcd203040_compound15.18882.18882.5 | controllerType=0 controllerNumber=1 scan=18882 | 38.5504 |
| 16850:5 | 6 | K.IIEGEPNLKVPGN[+2350.830]VTAVLGETLKVPC[+57.021]HFPC[+57.021]K.F | HexNAc(4)Hex(5)Fuc(1)NeuAc(2) | C[+57]*2, N[+2351] | 1154.3352 | 5 | 5767.6469 | 5767.6372 | 0 | 1.7 | 457 | Specific | 967.33 | 726.76 | 726.76 | 18.06 | 74 | >sp\|P01833\|PIGR_HUMAN Polymeric immunoglobulin receptor OS=Homo sapiens OX=9606 GN=PIGR PE=1 SV=4 | 902 | 20201113_12413_1ng_glyco_hcd203040_compound15.20331.20331.5 | controllerType=0 controllerNumber=1 scan=20331 | 41.2731 |
| 15417:5 | 6 | K.IIEGEPNLKVPGN[+2059.735]VTAVLGETLKVPC[+57.021]HFPC[+57.021]K.F | HexNAc(4)Hex(5)Fuc(1)NeuAc(1) | C[+57]*2, N[+2060] | 1096.114 | 5 | 5476.541 | 5476.5418 | 0 | -0.2 | 457 | Specific | 978.78 | 843.57 | 843.57 | 17.86 | 74 | >sp\|P01833\|PIGR_HUMAN Polymeric immunoglobulin receptor OS=Homo sapiens OX=9606 GN=PIGR PE=1 SV=4 | 902 | 20201113_12413_1ng_glyco_hcd203040_compound15.18812.18812.5 | controllerType=0 controllerNumber=1 scan=18812 | 38.4209 |
| 13365:3 | 6 | K.VPGN[+2205.793]VTAVLGETLK.V | HexNAc(4)Hex(5)Fuc(2)NeuAc(1) | N[+2206] | 1201.8796 | 3 | 3603.6242 | 3603.5927 | 0 | 8.7 | 466 | Specific | 345.57 | 345.57 | 345.57 | 6.35 | 74 | >sp\|P01833\|PIGR_HUMAN Polymeric immunoglobulin receptor OS=Homo sapiens OX=9606 GN=PIGR PE=1 SV=4 | 902 | 20201113_12413_1ng_glyco_hcd203040_compound15.16685.16685.3 | controllerType=0 controllerNumber=1 scan=16685 | 34.5705 |
| 13367:3 | 6 | K.VPGN[+2205.793]VTAVLGETLK.V | HexNAc(4)Hex(5)Fuc(2)NeuAc(1) | N[+2206] | 1201.8796 | 3 | 3603.6242 | 3603.5927 | 0 | 8.7 | 466 | Specific | 388.75 | 384.96 | 384.96 | 8.07 | 74 | >sp\|P01833\|PIGR_HUMAN Polymeric immunoglobulin receptor OS=Homo sapiens OX=9606 GN=PIGR PE=1 SV=4 | 902 | 20201113_12413_1ng_glyco_hcd203040_compound15.16687.16687.3 | controllerType=0 controllerNumber=1 scan=16687 | 34.5742 |
| 13443:3 | 6 | K.VPGN[+2059.735]VTAVLGETLK.V | HexNAc(4)Hex(5)Fuc(1)NeuAc(1) | N[+2060] | 1153.1891 | 3 | 3457.5529 | 3457.5348 | 0 | 5.2 | 466 | Specific | 573.57 | 541.34 | 541.34 | 10.28 | 74 | >sp\|P01833\|PIGR_HUMAN Polymeric immunoglobulin receptor OS=Homo sapiens OX=9606 GN=PIGR PE=1 SV=4 | 902 | 20201113_12413_1ng_glyco_hcd203040_compound15.16766.16766.3 | controllerType=0 controllerNumber=1 scan=16766 | 34.736 |
| 13445:3 | 6 | K.VPGN[+2059.735]VTAVLGETLK.V | HexNAc(4)Hex(5)Fuc(1)NeuAc(1) | N[+2060] | 1153.1891 | 3 | 3457.5529 | 3457.5348 | 0 | 5.2 | 466 | Specific | 639.29 | 403.88 | 403.88 | 11.28 | 74 | >sp\|P01833\|PIGR_HUMAN Polymeric immunoglobulin receptor OS=Homo sapiens OX=9606 GN=PIGR PE=1 SV=4 | 902 | 20201113_12413_1ng_glyco_hcd203040_compound15.16768.16768.3 | controllerType=0 controllerNumber=1 scan=16768 | 34.7409 |
| 16433:3 | 6 | K.VPGN[+2350.830]VTAVLGETLK.V | HexNAc(4)Hex(5)Fuc(1)NeuAc(2) | N[+2351] | 1250.212 | 3 | 3748.6214 | 3748.6302 | 0 | -2.4 | 466 | Specific | 733.22 | 509.28 | 509.28 | 12.89 | 74 | >sp\|P01833\|PIGR_HUMAN Polymeric immunoglobulin receptor OS=Homo sapiens OX=9606 GN=PIGR PE=1 SV=4 | 902 | 20201113_12413_1ng_glyco_hcd203040_compound15.19897.19897.3 | controllerType=0 controllerNumber=1 scan=19897 | 40.4144 |
| 15436:4 | 6 | K.VPGN[+2350.830]VTAVLGETLKVPC[+57.021]HFPC[+57.021]K.F | HexNAc(4)Hex(5)Fuc(1)NeuAc(2) | C[+57]*2, N[+2351] | 1194.2853 | 4 | 4774.1195 | 4774.0877 | 0 | 6.6 | 466 | Specific | 356.77 | 274.37 | 274.37 | 4.01 | 74 | >sp\|P01833\|PIGR_HUMAN Polymeric immunoglobulin receptor OS=Homo sapiens OX=9606 GN=PIGR PE=1 SV=4 | 902 | 20201113_12413_1ng_glyco_hcd203040_compound15.18832.18832.4 | controllerType=0 controllerNumber=1 scan=18832 | 38.4619 |
| 15357:3 | 6 | K.VPGN[+2350.830]VTAVLGETLKVPC[+57.021]HFPC[+57.021]K.F | HexNAc(4)Hex(5)Fuc(1)NeuAc(2) | C[+57]*2, N[+2351] | 1592.0414 | 3 | 4774.1098 | 4774.0877 | 0 | 4.6 | 466 | Specific | 408.52 | 250.93 | 250.93 | 6.06 | 74 | >sp\|P01833\|PIGR_HUMAN Polymeric immunoglobulin receptor OS=Homo sapiens OX=9606 GN=PIGR PE=1 SV=4 | 902 | 20201113_12413_1ng_glyco_hcd203040_compound15.18750.18750.3 | controllerType=0 controllerNumber=1 scan=18750 | 38.3119 |
| 13503:3 | 6 | K.VPGN[+2059.735]VTAVLGETLKVPC[+57.021]HFPC[+57.021]K.F | HexNAc(4)Hex(5)Fuc(1)NeuAc(1) | C[+57]*2, N[+2060] | 1495.0034 | 3 | 4482.9955 | 4482.9923 | 0 | 0.7 | 466 | Specific | 476.15 | 476.15 | 476.15 | 8.19 | 74 | >sp\|P01833\|PIGR_HUMAN Polymeric immunoglobulin receptor OS=Homo sapiens OX=9606 GN=PIGR PE=1 SV=4 | 902 | 20201113_12413_1ng_glyco_hcd203040_compound15.16828.16828.3 | controllerType=0 controllerNumber=1 scan=16828 | 34.8665 |
| 13505:3 | 6 | K.VPGN[+2059.735]VTAVLGETLKVPC[+57.021]HFPC[+57.021]K.F | HexNAc(4)Hex(5)Fuc(1)NeuAc(1) | C[+57]*2, N[+2060] | 1495.0034 | 3 | 4482.9955 | 4482.9923 | 0 | 0.7 | 466 | Specific | 668.76 | 638.73 | 638.73 | 11.8 | 74 | >sp\|P01833\|PIGR_HUMAN Polymeric immunoglobulin receptor OS=Homo sapiens OX=9606 GN=PIGR PE=1 SV=4 | 902 | 20201113_12413_1ng_glyco_hcd203040_compound15.16830.16830.3 | controllerType=0 controllerNumber=1 scan=16830 | 34.8701 |
| 13397:4 | 6 | K.VPGN[+2205.793]VTAVLGETLKVPC[+57.021]HFPC[+57.021]K.F | HexNAc(4)Hex(5)Fuc(2)NeuAc(1) | C[+57]*2, N[+2206] | 1158.0201 | 4 | 4629.0584 | 4629.0502 | 0 | 1.8 | 466 | Specific | 793.05 | 708.71 | 708.71 | 14.57 | 74 | >sp\|P01833\|PIGR_HUMAN Polymeric immunoglobulin receptor OS=Homo sapiens OX=9606 GN=PIGR PE=1 SV=4 | 902 | 20201113_12413_1ng_glyco_hcd203040_compound15.16718.16718.3 | controllerType=0 controllerNumber=1 scan=16718 | 34.6291 |
| 13395:4 | 6 | K.VPGN[+2205.793]VTAVLGETLKVPC[+57.021]HFPC[+57.021]K.F | HexNAc(4)Hex(5)Fuc(2)NeuAc(1) | C[+57]*2, N[+2206] | 1158.0201 | 4 | 4629.0584 | 4629.0502 | 0 | 1.8 | 466 | Specific | 864.39 | 579.55 | 579.55 | 16.57 | 74 | >sp\|P01833\|PIGR_HUMAN Polymeric immunoglobulin receptor OS=Homo sapiens OX=9606 GN=PIGR PE=1 SV=4 | 902 | 20201113_12413_1ng_glyco_hcd203040_compound15.16716.16716.3 | controllerType=0 controllerNumber=1 scan=16716 | 34.6236 |
| 05676:3 | 6 | K.WN[+1548.545]NTGC[+57.021]QALPSQDEGPSK.A | HexNAc(3)Hex(4)NeuAc(1) | C[+57], N[+1549] | 1179.8139 | 3 | 3537.4271 | 3537.4215 | 0 | 1.6 | 498 | Specific | 248.75 | 194.98 | 194.98 | 5.74 | 74 | >sp\|P01833\|PIGR_HUMAN Polymeric immunoglobulin receptor OS=Homo sapiens OX=9606 GN=PIGR PE=1 SV=4 | 902 | 20201113_12413_1ng_glyco_hcd203040_compound15.8692.8692.3 | controllerType=0 controllerNumber=1 scan=8692 | 21.0913 |
| 07548:3 | 6 | K.WN[+2204.772]NTGC[+57.021]QALPSQDEGPSK.A | HexNAc(4)Hex(5)NeuAc(2) | C[+57], N[+2205] | 1398.5561 | 3 | 4193.6537 | 4193.6491 | 0 | 1.1 | 498 | Specific | 329.64 | 283.42 | 283.42 | 6.79 | 74 | >sp\|P01833\|PIGR_HUMAN Polymeric immunoglobulin receptor OS=Homo sapiens OX=9606 GN=PIGR PE=1 SV=4 | 902 | 20201113_12413_1ng_glyco_hcd203040_compound15.10632.10632.3 | controllerType=0 controllerNumber=1 scan=10632 | 24.3173 |
| 07551:3 | 6 | K.WN[+2204.772]NTGC[+57.021]QALPSQDEGPSK.A | HexNAc(4)Hex(5)NeuAc(2) | C[+57], N[+2205] | 1398.5561 | 3 | 4193.6537 | 4193.6491 | 0 | 1.1 | 498 | Specific | 375.26 | 304.65 | 304.65 | 7.21 | 74 | >sp\|P01833\|PIGR_HUMAN Polymeric immunoglobulin receptor OS=Homo sapiens OX=9606 GN=PIGR PE=1 SV=4 | 902 | 20201113_12413_1ng_glyco_hcd203040_compound15.10635.10635.3 | controllerType=0 controllerNumber=1 scan=10635 | 24.3217 |
| 05512:3 | 6 | K.WN[+1872.651]NTGC[+57.021]QALPSQDEGPSK.A | HexNAc(3)Hex(6)NeuAc(1) | C[+57], N[+1873] | 1287.8492 | 3 | 3861.533 | 3861.5272 | 0 | 1.5 | 498 | Specific | 399.58 | 380.2 | 380.2 | 8.97 | 74 | >sp\|P01833\|PIGR_HUMAN Polymeric immunoglobulin receptor OS=Homo sapiens OX=9606 GN=PIGR PE=1 SV=4 | 902 | 20201113_12413_1ng_glyco_hcd203040_compound15.8523.8523.3 | controllerType=0 controllerNumber=1 scan=8523 | 20.813 |
| 05678:3 | 6 | K.WN[+1548.545]NTGC[+57.021]QALPSQDEGPSK.A | HexNAc(3)Hex(4)NeuAc(1) | C[+57], N[+1549] | 1179.8139 | 3 | 3537.4271 | 3537.4215 | 0 | 1.6 | 498 | Specific | 430.27 | 343.5 | 343.5 | 7.57 | 74 | >sp\|P01833\|PIGR_HUMAN Polymeric immunoglobulin receptor OS=Homo sapiens OX=9606 GN=PIGR PE=1 SV=4 | 902 | 20201113_12413_1ng_glyco_hcd203040_compound15.8694.8694.3 | controllerType=0 controllerNumber=1 scan=8694 | 21.0952 |
| 05420:3 | 6 | K.WN[+2059.735]NTGC[+57.021]QALPSQDEGPSK.A | HexNAc(4)Hex(5)Fuc(1)NeuAc(1) | C[+57], N[+2060] | 1350.2104 | 3 | 4048.6166 | 4048.6116 | 0 | 1.2 | 498 | Specific | 532.02 | 476.77 | 476.77 | 9.56 | 74 | >sp\|P01833\|PIGR_HUMAN Polymeric immunoglobulin receptor OS=Homo sapiens OX=9606 GN=PIGR PE=1 SV=4 | 902 | 20201113_12413_1ng_glyco_hcd203040_compound15.8428.8428.3 | controllerType=0 controllerNumber=1 scan=8428 | 20.6507 |
| 05514:3 | 6 | K.WN[+1872.651]NTGC[+57.021]QALPSQDEGPSK.A | HexNAc(3)Hex(6)NeuAc(1) | C[+57], N[+1873] | 1287.8492 | 3 | 3861.533 | 3861.5272 | 0 | 1.5 | 498 | Specific | 535.01 | 502.75 | 502.75 | 9.56 | 74 | >sp\|P01833\|PIGR_HUMAN Polymeric immunoglobulin receptor OS=Homo sapiens OX=9606 GN=PIGR PE=1 SV=4 | 902 | 20201113_12413_1ng_glyco_hcd203040_compound15.8525.8525.3 | controllerType=0 controllerNumber=1 scan=8525 | 20.8166 |
| 07497:3 | 6 | K.WN[+2204.772]NTGC[+57.021]QALPSQDEGPSK.A | HexNAc(4)Hex(5)NeuAc(2) | C[+57], N[+2205] | 1398.5574 | 3 | 4193.6577 | 4193.6491 | 0 | 2.1 | 498 | Specific | 583.93 | 528.47 | 528.47 | 10.95 | 74 | >sp\|P01833\|PIGR_HUMAN Polymeric immunoglobulin receptor OS=Homo sapiens OX=9606 GN=PIGR PE=1 SV=4 | 902 | 20201113_12413_1ng_glyco_hcd203040_compound15.10579.10579.3 | controllerType=0 controllerNumber=1 scan=10579 | 24.2305 |
| 07499:3 | 6 | K.WN[+2204.772]NTGC[+57.021]QALPSQDEGPSK.A | HexNAc(4)Hex(5)NeuAc(2) | C[+57], N[+2205] | 1398.5574 | 3 | 4193.6577 | 4193.6491 | 0 | 2.1 | 498 | Specific | 589.38 | 514.39 | 514.39 | 10.95 | 74 | >sp\|P01833\|PIGR_HUMAN Polymeric immunoglobulin receptor OS=Homo sapiens OX=9606 GN=PIGR PE=1 SV=4 | 902 | 20201113_12413_1ng_glyco_hcd203040_compound15.10581.10581.3 | controllerType=0 controllerNumber=1 scan=10581 | 24.2343 |
| 05422:3 | 6 | K.WN[+2059.735]NTGC[+57.021]QALPSQDEGPSK.A | HexNAc(4)Hex(5)Fuc(1)NeuAc(1) | C[+57], N[+2060] | 1350.2104 | 3 | 4048.6166 | 4048.6116 | 0 | 1.2 | 498 | Specific | 632.78 | 573.12 | 573.12 | 11.95 | 74 | >sp\|P01833\|PIGR_HUMAN Polymeric immunoglobulin receptor OS=Homo sapiens OX=9606 GN=PIGR PE=1 SV=4 | 902 | 20201113_12413_1ng_glyco_hcd203040_compound15.8430.8430.3 | controllerType=0 controllerNumber=1 scan=8430 | 20.6548 |
| 05567:3 | 6 | K.WN[+1710.598]NTGC[+57.021]QALPSQDEGPSK.A | HexNAc(3)Hex(5)NeuAc(1) | C[+57], N[+1711] | 1233.8337 | 3 | 3699.4865 | 3699.4743 | 0 | 3.3 | 498 | Specific | 660.91 | 593.16 | 593.16 | 13.04 | 74 | >sp\|P01833\|PIGR_HUMAN Polymeric immunoglobulin receptor OS=Homo sapiens OX=9606 GN=PIGR PE=1 SV=4 | 902 | 20201113_12413_1ng_glyco_hcd203040_compound15.8580.8580.3 | controllerType=0 controllerNumber=1 scan=8580 | 20.9084 |
| 05569:3 | 6 | K.WN[+1710.598]NTGC[+57.021]QALPSQDEGPSK.A | HexNAc(3)Hex(5)NeuAc(1) | C[+57], N[+1711] | 1233.8337 | 3 | 3699.4865 | 3699.4743 | 0 | 3.3 | 498 | Specific | 742.2 | 662 | 662 | 14.04 | 74 | >sp\|P01833\|PIGR_HUMAN Polymeric immunoglobulin receptor OS=Homo sapiens OX=9606 GN=PIGR PE=1 SV=4 | 902 | 20201113_12413_1ng_glyco_hcd203040_compound15.8582.8582.3 | controllerType=0 controllerNumber=1 scan=8582 | 20.9128 |
| 05481:3 | 6 | K.WN[+1913.677]NTGC[+57.021]QALPSQDEGPSK.A | HexNAc(4)Hex(5)NeuAc(1) | C[+57], N[+1914] | 1301.5243 | 3 | 3902.5585 | 3902.5537 | 0 | 1.2 | 498 | Specific | 758.32 | 658.38 | 658.38 | 14.95 | 74 | >sp\|P01833\|PIGR_HUMAN Polymeric immunoglobulin receptor OS=Homo sapiens OX=9606 GN=PIGR PE=1 SV=4 | 902 | 20201113_12413_1ng_glyco_hcd203040_compound15.8491.8491.3 | controllerType=0 controllerNumber=1 scan=8491 | 20.7601 |
| 05479:3 | 6 | K.WN[+1913.677]NTGC[+57.021]QALPSQDEGPSK.A | HexNAc(4)Hex(5)NeuAc(1) | C[+57], N[+1914] | 1301.5243 | 3 | 3902.5585 | 3902.5537 | 0 | 1.2 | 498 | Specific | 772.13 | 699.44 | 699.44 | 14.95 | 74 | >sp\|P01833\|PIGR_HUMAN Polymeric immunoglobulin receptor OS=Homo sapiens OX=9606 GN=PIGR PE=1 SV=4 | 902 | 20201113_12413_1ng_glyco_hcd203040_compound15.8489.8489.3 | controllerType=0 controllerNumber=1 scan=8489 | 20.756 |
| 04615:3 | 7 | R.FSDGLESN[+1864.634]SSTQFEVKK.Y | HexNAc(2)Hex(9) | N[+1865] | 1256.5208 | 3 | 3767.5479 | 3767.5421 | 0 | 1.5 | 219 | Specific | 256.04 | 256.04 | 256.04 | 6.11 | 72 | >sp\|P0C0L4\|CO4A_HUMAN Complement C4-A OS=Homo sapiens OX=9606 GN=C4A PE=1 SV=2 | 1235 | 20201113_12413_1ng_glyco_hcd203040_compound15.7595.7595.3 | controllerType=0 controllerNumber=1 scan=7595 | 19.2581 |
| 18699:3 | 10 | R.NIAAFGGDPNN[+1913.677]ITLFGESAGGASVSLQTLSPYNK.G | HexNAc(4)Hex(5)NeuAc(1) | N[+1914] | 1775.4728 | 3 | 5324.4038 | 5324.3633 | 0 | 7.6 | 197 | Specific | 425.79 | 425.79 | 425.79 | 8.32 | 45 | >sp\|P19835\|CEL_HUMAN Bile salt-activated lipase OS=Homo sapiens OX=9606 GN=CEL PE=1 SV=3 | 476 | 20201113_12413_1ng_glyco_hcd203040_compound15.22266.22266.3 | controllerType=0 controllerNumber=1 scan=22266 | 45.4552 |
| 18701:3 | 10 | R.NIAAFGGDPNN[+1913.677]ITLFGESAGGASVSLQTLSPYNK.G | HexNAc(4)Hex(5)NeuAc(1) | N[+1914] | 1775.4728 | 3 | 5324.4038 | 5324.3633 | 0 | 7.6 | 197 | Specific | 571.54 | 550.84 | 550.84 | 8.92 | 45 | >sp\|P19835\|CEL_HUMAN Bile salt-activated lipase OS=Homo sapiens OX=9606 GN=CEL PE=1 SV=3 | 476 | 20201113_12413_1ng_glyco_hcd203040_compound15.22268.22268.3 | controllerType=0 controllerNumber=1 scan=22268 | 45.4605 |
| 18682:3 | 10 | R.NIAAFGGDPNN[+1913.677]ITLFGESAGGASVSLQTLSPYNK.G | HexNAc(4)Hex(5)NeuAc(1) | N[+1914] | 1775.4653 | 3 | 5324.3813 | 5324.3633 | 0 | 3.4 | 197 | Specific | 631.58 | 631.58 | 631.58 | 12 | 45 | >sp\|P19835\|CEL_HUMAN Bile salt-activated lipase OS=Homo sapiens OX=9606 GN=CEL PE=1 SV=3 | 476 | 20201113_12413_1ng_glyco_hcd203040_compound15.22248.22248.3 | controllerType=0 controllerNumber=1 scan=22248 | 45.4194 |
| 18684:3 | 10 | R.NIAAFGGDPNN[+1913.677]ITLFGESAGGASVSLQTLSPYNK.G | HexNAc(4)Hex(5)NeuAc(1) | N[+1914] | 1775.4653 | 3 | 5324.3813 | 5324.3633 | 0 | 3.4 | 197 | Specific | 847.74 | 810.23 | 810.23 | 16 | 45 | >sp\|P19835\|CEL_HUMAN Bile salt-activated lipase OS=Homo sapiens OX=9606 GN=CEL PE=1 SV=3 | 476 | 20201113_12413_1ng_glyco_hcd203040_compound15.22250.22250.3 | controllerType=0 controllerNumber=1 scan=22250 | 45.4228 |
| 17484:5 | 11 | R.LSLHRPALEDLLLGSEAN[+1524.534]LTC[+57.021]TLTGLR.D | HexNAc(2)Hex(6)Fuc(1) | C[+57], N[+1525] | 898.4309 | 5 | 4488.1252 | 4488.1319 | 0 | -1.5 | 127 | Specific | 231.09 | 152.89 | 152.89 | 3.92 | 51 | >sp\|P01876\|IGHA1_HUMAN Immunoglobulin heavy constant alpha 1 OS=Homo sapiens OX=9606 GN=IGHA1 PE=1 SV=2 | 255 | 20201113_12413_1ng_glyco_hcd203040_compound15.20994.20994.5 | controllerType=0 controllerNumber=1 scan=20994 | 42.6809 |
| 18410:4 | 11 | R.LSLHRPALEDLLLGSEAN[+1710.598]LTC[+57.021]TLTGLR.D | HexNAc(3)Hex(5)NeuAc(1) | C[+57], N[+1711] | 1169.309 | 4 | 4674.2142 | 4674.1959 | 0 | 3.9 | 127 | Specific | 331.54 | 331.54 | 331.54 | 7.41 | 51 | >sp\|P01876\|IGHA1_HUMAN Immunoglobulin heavy constant alpha 1 OS=Homo sapiens OX=9606 GN=IGHA1 PE=1 SV=2 | 255 | 20201113_12413_1ng_glyco_hcd203040_compound15.21963.21963.4 | controllerType=0 controllerNumber=1 scan=21963 | 44.7692 |
| 18371:3 | 11 | R.LSLHRPALEDLLLGSEAN[+2116.756]LTC[+57.021]TLTGLR.D | HexNAc(5)Hex(5)NeuAc(1) | C[+57], N[+2117] | 1694.1306 | 3 | 5080.3772 | 5080.3547 | 0 | 4.4 | 127 | Specific | 470.68 | 470.68 | 470.68 | 9.05 | 51 | >sp\|P01876\|IGHA1_HUMAN Immunoglobulin heavy constant alpha 1 OS=Homo sapiens OX=9606 GN=IGHA1 PE=1 SV=2 | 255 | 20201113_12413_1ng_glyco_hcd203040_compound15.21923.21923.3 | controllerType=0 controllerNumber=1 scan=21923 | 44.6736 |
| 19445:4 | 11 | R.LSLHRPALEDLLLGSEAN[+2204.772]LTC[+57.021]TLTGLR.D | HexNAc(4)Hex(5)NeuAc(2) | C[+57], N[+2205] | 1292.8527 | 4 | 5168.3888 | 5168.3707 | 0 | 3.5 | 127 | Specific | 485.73 | 474.17 | 474.17 | 9.75 | 51 | >sp\|P01876\|IGHA1_HUMAN Immunoglobulin heavy constant alpha 1 OS=Homo sapiens OX=9606 GN=IGHA1 PE=1 SV=2 | 255 | 20201113_12413_1ng_glyco_hcd203040_compound15.23046.23046.2 | controllerType=0 controllerNumber=1 scan=23046 | 47.1158 |
| 18452:3 | 11 | R.LSLHRPALEDLLLGSEAN[+1751.624]LTC[+57.021]TLTGLR.D | HexNAc(4)Hex(4)NeuAc(1) | C[+57], N[+1752] | 1572.418 | 3 | 4715.2395 | 4715.2225 | 0 | 3.6 | 127 | Specific | 575.19 | 503.46 | 503.46 | 10.66 | 51 | >sp\|P01876\|IGHA1_HUMAN Immunoglobulin heavy constant alpha 1 OS=Homo sapiens OX=9606 GN=IGHA1 PE=1 SV=2 | 255 | 20201113_12413_1ng_glyco_hcd203040_compound15.22008.22008.3 | controllerType=0 controllerNumber=1 scan=22008 | 44.8765 |
| 17449:4 | 11 | R.LSLHRPALEDLLLGSEAN[+1378.476]LTC[+57.021]TLTGLR.D | HexNAc(2)Hex(6) | C[+57], N[+1378] | 1086.2741 | 4 | 4342.0745 | 4342.074 | 0 | 0.1 | 127 | Specific | 590 | 590 | 590 | 11.08 | 51 | >sp\|P01876\|IGHA1_HUMAN Immunoglobulin heavy constant alpha 1 OS=Homo sapiens OX=9606 GN=IGHA1 PE=1 SV=2 | 255 | 20201113_12413_1ng_glyco_hcd203040_compound15.20958.20958.4 | controllerType=0 controllerNumber=1 scan=20958 | 42.6074 |
| 18376:3 | 11 | R.LSLHRPALEDLLLGSEAN[+1872.651]LTC[+57.021]TLTGLR.D | HexNAc(3)Hex(6)NeuAc(1) | C[+57], N[+1873] | 1612.7613 | 3 | 4836.2693 | 4836.2487 | 0 | 4.3 | 127 | Specific | 594.33 | 549.76 | 549.76 | 10.66 | 51 | >sp\|P01876\|IGHA1_HUMAN Immunoglobulin heavy constant alpha 1 OS=Homo sapiens OX=9606 GN=IGHA1 PE=1 SV=2 | 255 | 20201113_12413_1ng_glyco_hcd203040_compound15.21928.21928.3 | controllerType=0 controllerNumber=1 scan=21928 | 44.6883 |
| 17443:4 | 11 | R.LSLHRPALEDLLLGSEAN[+1419.502]LTC[+57.021]TLTGLR.D | HexNAc(3)Hex(5) | C[+57], N[+1420] | 1096.5357 | 4 | 4383.1209 | 4383.1005 | 0 | 4.7 | 127 | Specific | 608.24 | 608.24 | 608.24 | 12.23 | 51 | >sp\|P01876\|IGHA1_HUMAN Immunoglobulin heavy constant alpha 1 OS=Homo sapiens OX=9606 GN=IGHA1 PE=1 SV=2 | 255 | 20201113_12413_1ng_glyco_hcd203040_compound15.20951.20951.4 | controllerType=0 controllerNumber=1 scan=20951 | 42.5847 |
| 17503:3 | 11 | R.LSLHRPALEDLLLGSEAN[+1501.555]LTC[+57.021]TLTGLR.D | HexNAc(5)Hex(3) | C[+57], N[+1502] | 1489.0602 | 3 | 4465.1662 | 4465.1536 | 0 | 2.8 | 127 | Specific | 617.85 | 573.52 | 573.52 | 11.79 | 51 | >sp\|P01876\|IGHA1_HUMAN Immunoglobulin heavy constant alpha 1 OS=Homo sapiens OX=9606 GN=IGHA1 PE=1 SV=2 | 255 | 20201113_12413_1ng_glyco_hcd203040_compound15.21014.21014.3 | controllerType=0 controllerNumber=1 scan=21014 | 42.7217 |
| 17453:3 | 11 | R.LSLHRPALEDLLLGSEAN[+1216.423]LTC[+57.021]TLTGLR.D | HexNAc(2)Hex(5) | C[+57], N[+1216] | 1394.0108 | 3 | 4180.0178 | 4180.0211 | 0 | -0.8 | 127 | Specific | 621.81 | 621.81 | 621.81 | 10.87 | 51 | >sp\|P01876\|IGHA1_HUMAN Immunoglobulin heavy constant alpha 1 OS=Homo sapiens OX=9606 GN=IGHA1 PE=1 SV=2 | 255 | 20201113_12413_1ng_glyco_hcd203040_compound15.20962.20962.3 | controllerType=0 controllerNumber=1 scan=20962 | 42.6134 |
| 18360:4 | 11 | R.LSLHRPALEDLLLGSEAN[+1913.677]LTC[+57.021]TLTGLR.D | HexNAc(4)Hex(5)NeuAc(1) | C[+57], N[+1914] | 1220.0834 | 4 | 4877.3118 | 4877.2753 | 0 | 7.5 | 127 | Specific | 632.82 | 632.82 | 632.82 | 11.51 | 51 | >sp\|P01876\|IGHA1_HUMAN Immunoglobulin heavy constant alpha 1 OS=Homo sapiens OX=9606 GN=IGHA1 PE=1 SV=2 | 255 | 20201113_12413_1ng_glyco_hcd203040_compound15.21911.21911.4 | controllerType=0 controllerNumber=1 scan=21911 | 44.6348 |
| 18363:4 | 11 | R.LSLHRPALEDLLLGSEAN[+1913.677]LTC[+57.021]TLTGLR.D | HexNAc(4)Hex(5)NeuAc(1) | C[+57], N[+1914] | 1220.0834 | 4 | 4877.3118 | 4877.2753 | 0 | 7.5 | 127 | Specific | 634.19 | 622.66 | 622.66 | 11.51 | 51 | >sp\|P01876\|IGHA1_HUMAN Immunoglobulin heavy constant alpha 1 OS=Homo sapiens OX=9606 GN=IGHA1 PE=1 SV=2 | 255 | 20201113_12413_1ng_glyco_hcd203040_compound15.21914.21914.4 | controllerType=0 controllerNumber=1 scan=21914 | 44.6432 |
| 17500:4 | 11 | R.LSLHRPALEDLLLGSEAN[+1647.613]LTC[+57.021]TLTGLR.D | HexNAc(5)Hex(3)Fuc(1) | C[+57], N[+1648] | 1153.5591 | 4 | 4611.2147 | 4611.2115 | 0 | 0.7 | 127 | Specific | 643.05 | 643.05 | 643.05 | 12.08 | 51 | >sp\|P01876\|IGHA1_HUMAN Immunoglobulin heavy constant alpha 1 OS=Homo sapiens OX=9606 GN=IGHA1 PE=1 SV=2 | 255 | 20201113_12413_1ng_glyco_hcd203040_compound15.21010.21010.4 | controllerType=0 controllerNumber=1 scan=21010 | 42.7104 |
| 17458:3 | 11 | R.LSLHRPALEDLLLGSEAN[+1460.529]LTC[+57.021]TLTGLR.D | HexNAc(4)Hex(4) | C[+57], N[+1461] | 1475.3782 | 3 | 4424.1201 | 4424.1271 | 0 | -1.6 | 127 | Specific | 646.12 | 581.66 | 581.66 | 10.87 | 51 | >sp\|P01876\|IGHA1_HUMAN Immunoglobulin heavy constant alpha 1 OS=Homo sapiens OX=9606 GN=IGHA1 PE=1 SV=2 | 255 | 20201113_12413_1ng_glyco_hcd203040_compound15.20967.20967.3 | controllerType=0 controllerNumber=1 scan=20967 | 42.6235 |
| 18401:3 | 11 | R.LSLHRPALEDLLLGSEAN[+1710.598]LTC[+57.021]TLTGLR.D | HexNAc(3)Hex(5)NeuAc(1) | C[+57], N[+1711] | 1558.7429 | 3 | 4674.2141 | 4674.1959 | 0 | 3.9 | 127 | Specific | 646.77 | 643.91 | 643.91 | 11.66 | 51 | >sp\|P01876\|IGHA1_HUMAN Immunoglobulin heavy constant alpha 1 OS=Homo sapiens OX=9606 GN=IGHA1 PE=1 SV=2 | 255 | 20201113_12413_1ng_glyco_hcd203040_compound15.21954.21954.3 | controllerType=0 controllerNumber=1 scan=21954 | 44.7433 |
| 18374:3 | 11 | R.LSLHRPALEDLLLGSEAN[+1872.651]LTC[+57.021]TLTGLR.D | HexNAc(3)Hex(6)NeuAc(1) | C[+57], N[+1873] | 1612.7613 | 3 | 4836.2693 | 4836.2487 | 0 | 4.3 | 127 | Specific | 654.13 | 582.58 | 582.58 | 12.66 | 51 | >sp\|P01876\|IGHA1_HUMAN Immunoglobulin heavy constant alpha 1 OS=Homo sapiens OX=9606 GN=IGHA1 PE=1 SV=2 | 255 | 20201113_12413_1ng_glyco_hcd203040_compound15.21926.21926.3 | controllerType=0 controllerNumber=1 scan=21926 | 44.6802 |
| 17501:3 | 11 | R.LSLHRPALEDLLLGSEAN[+1501.555]LTC[+57.021]TLTGLR.D | HexNAc(5)Hex(3) | C[+57], N[+1502] | 1489.0602 | 3 | 4465.1662 | 4465.1536 | 0 | 2.8 | 127 | Specific | 658.93 | 658.93 | 658.93 | 12.79 | 51 | >sp\|P01876\|IGHA1_HUMAN Immunoglobulin heavy constant alpha 1 OS=Homo sapiens OX=9606 GN=IGHA1 PE=1 SV=2 | 255 | 20201113_12413_1ng_glyco_hcd203040_compound15.21012.21012.3 | controllerType=0 controllerNumber=1 scan=21012 | 42.715 |
| 18447:4 | 11 | R.LSLHRPALEDLLLGSEAN[+1751.624]LTC[+57.021]TLTGLR.D | HexNAc(4)Hex(4)NeuAc(1) | C[+57], N[+1752] | 1179.5651 | 4 | 4715.2386 | 4715.2225 | 0 | 3.4 | 127 | Specific | 660.7 | 618.48 | 618.48 | 13.37 | 51 | >sp\|P01876\|IGHA1_HUMAN Immunoglobulin heavy constant alpha 1 OS=Homo sapiens OX=9606 GN=IGHA1 PE=1 SV=2 | 255 | 20201113_12413_1ng_glyco_hcd203040_compound15.22003.22003.4 | controllerType=0 controllerNumber=1 scan=22003 | 44.8663 |

Table S4. Cont.

| Query #:z | Protein Rank | Glycopeptide | Glycan | Modification Type(s) | Observed m/z | z | Observed (M+H) | Calc. mass (M+H) | Off-by-x error | Mass error (ppm) | StartPosition | Cleavage | Score | Delta | Delta Mod | \|Log Prob\| | # of unique peptides | Protein Name | Protein DB number | Comment | Scan # | Scan Time |
|---|---|---|---|---|---|---|---|---|---|---|---|---|---|---|---|---|---|---|---|---|---|---|
| 17690:4 | 11 | R.LSLHRPALEDLLLGSEAN[+1501.555]LTC[+57.021]TLTGLR.D | HexNAc(5)Hex(3) | C[+57], N[+1502] | 1117.0492 | 4 | 4465.1751 | 4465.1536 | 0 | 4.8 | 127 | Specific | 665.81 | 648.32 | 648.32 | 13.23 | 51 | >sp\|P01876\|IGHA1_HUMAN Immunoglobulin heavy constant alpha 1 OS=Homo sapiens OX=9606 GN=IGHA1 PE=1 SV=2 | 255 | 20201113_12413_1ng_glyco_hcd203040_compound15.21209.21209.2 | controllerType=0 controllerNumber=1 scan=21209 | 43.1806 |
| 17455:3 | 11 | R.LSLHRPALEDLLLGSEAN[+1460.529]LTC[+57.021]TLTGLR.D | HexNAc(4)Hex(4) | C[+57], N[+1461] | 1475.3782 | 3 | 4424.1201 | 4424.1271 | 0 | -1.6 | 127 | Specific | 668.51 | 601.24 | 601.24 | 11.87 | 51 | >sp\|P01876\|IGHA1_HUMAN Immunoglobulin heavy constant alpha 1 OS=Homo sapiens OX=9606 GN=IGHA1 PE=1 SV=2 | 255 | 20201113_12413_1ng_glyco_hcd203040_compound15.20964.20964.3 | controllerType=0 controllerNumber=1 scan=20964 | 42.616 |
| 17498:4 | 11 | R.LSLHRPALEDLLLGSEAN[+1647.613]LTC[+57.021]TLTGLR.D | HexNAc(5)Hex(3)Fuc(1) | C[+57], N[+1648] | 1153.5591 | 4 | 4611.2147 | 4611.2115 | 0 | 0.7 | 127 | Specific | 686.52 | 686.52 | 686.52 | 13.08 | 51 | >sp\|P01876\|IGHA1_HUMAN Immunoglobulin heavy constant alpha 1 OS=Homo sapiens OX=9606 GN=IGHA1 PE=1 SV=2 | 255 | 20201113_12413_1ng_glyco_hcd203040_compound15.21008.21008.4 | controllerType=0 controllerNumber=1 scan=21008 | 42.706 |
| 19448:4 | 11 | R.LSLHRPALEDLLLGSEAN[+2204.772]LTC[+57.021]TLTGLR.D | HexNAc(4)Hex(5)NeuAc(2) | C[+57], N[+2205] | 1292.8527 | 4 | 5168.3888 | 5168.3707 | 0 | 3.5 | 127 | Specific | 693.24 | 644.86 | 644.86 | 13.37 | 51 | >sp\|P01876\|IGHA1_HUMAN Immunoglobulin heavy constant alpha 1 OS=Homo sapiens OX=9606 GN=IGHA1 PE=1 SV=2 | 255 | 20201113_12413_1ng_glyco_hcd203040_compound15.23049.23049.2 | controllerType=0 controllerNumber=1 scan=23049 | 47.1333 |
| 17435:4 | 11 | R.LSLHRPALEDLLLGSEAN[+1501.555]LTC[+57.021]TLTGLR.D | HexNAc(5)Hex(3) | C[+57], N[+1502] | 1117.0503 | 4 | 4465.1795 | 4465.1536 | 0 | 5.8 | 127 | Specific | 703.11 | 703.11 | 703.11 | 13.61 | 51 | >sp\|P01876\|IGHA1_HUMAN Immunoglobulin heavy constant alpha 1 OS=Homo sapiens OX=9606 GN=IGHA1 PE=1 SV=2 | 255 | 20201113_12413_1ng_glyco_hcd203040_compound15.20943.20943.4 | controllerType=0 controllerNumber=1 scan=20943 | 42.5564 |
| 17688:4 | 11 | R.LSLHRPALEDLLLGSEAN[+1501.555]LTC[+57.021]TLTGLR.D | HexNAc(5)Hex(3) | C[+57], N[+1502] | 1117.0492 | 4 | 4465.1751 | 4465.1536 | 0 | 4.8 | 127 | Specific | 729.73 | 729.73 | 729.73 | 14.23 | 51 | >sp\|P01876\|IGHA1_HUMAN Immunoglobulin heavy constant alpha 1 OS=Homo sapiens OX=9606 GN=IGHA1 PE=1 SV=2 | 255 | 20201113_12413_1ng_glyco_hcd203040_compound15.21207.21207.2 | controllerType=0 controllerNumber=1 scan=21207 | 43.1754 |
| 17456:3 | 11 | R.LSLHRPALEDLLLGSEAN[+1216.423]LTC[+57.021]TLTGLR.D | HexNAc(2)Hex(5) | C[+57], N[+1216] | 1394.0108 | 3 | 4180.0178 | 4180.0211 | 0 | -0.8 | 127 | Specific | 732.68 | 732.68 | 732.68 | 12.87 | 51 | >sp\|P01876\|IGHA1_HUMAN Immunoglobulin heavy constant alpha 1 OS=Homo sapiens OX=9606 GN=IGHA1 PE=1 SV=2 | 255 | 20201113_12413_1ng_glyco_hcd203040_compound15.20965.20965.3 | controllerType=0 controllerNumber=1 scan=20965 | 42.6193 |
| 17439:4 | 11 | R.LSLHRPALEDLLLGSEAN[+1501.555]LTC[+57.021]TLTGLR.D | HexNAc(5)Hex(3) | C[+57], N[+1502] | 1117.0503 | 4 | 4465.1795 | 4465.1536 | 0 | 5.8 | 127 | Specific | 740.58 | 721.57 | 721.57 | 13.61 | 51 | >sp\|P01876\|IGHA1_HUMAN Immunoglobulin heavy constant alpha 1 OS=Homo sapiens OX=9606 GN=IGHA1 PE=1 SV=2 | 255 | 20201113_12413_1ng_glyco_hcd203040_compound15.20947.20947.4 | controllerType=0 controllerNumber=1 scan=20947 | 42.5709 |
| 17454:4 | 11 | R.LSLHRPALEDLLLGSEAN[+1216.423]LTC[+57.021]TLTGLR.D | HexNAc(2)Hex(5) | C[+57], N[+1216] | 1045.7605 | 4 | 4180.0203 | 4180.0211 | 0 | -0.2 | 127 | Specific | 757.5 | 757.5 | 757.5 | 15.08 | 51 | >sp\|P01876\|IGHA1_HUMAN Immunoglobulin heavy constant alpha 1 OS=Homo sapiens OX=9606 GN=IGHA1 PE=1 SV=2 | 255 | 20201113_12413_1ng_glyco_hcd203040_compound15.20963.20963.4 | controllerType=0 controllerNumber=1 scan=20963 | 42.6149 |
| 18372:4 | 11 | R.LSLHRPALEDLLLGSEAN[+2116.756]LTC[+57.021]TLTGLR.D | HexNAc(5)Hex(5)NeuAc(1) | C[+57], N[+2117] | 1270.8484 | 4 | 5080.3717 | 5080.3547 | 0 | 3.4 | 127 | Specific | 764.54 | 618.13 | 618.13 | 15.37 | 51 | >sp\|P01876\|IGHA1_HUMAN Immunoglobulin heavy constant alpha 1 OS=Homo sapiens OX=9606 GN=IGHA1 PE=1 SV=2 | 255 | 20201113_12413_1ng_glyco_hcd203040_compound15.21924.21924.4 | controllerType=0 controllerNumber=1 scan=21924 | 44.6764 |
| 18408:4 | 11 | R.LSLHRPALEDLLLGSEAN[+1710.598]LTC[+57.021]TLTGLR.D | HexNAc(3)Hex(5)NeuAc(1) | C[+57], N[+1711] | 1169.309 | 4 | 4674.2142 | 4674.1959 | 0 | 3.9 | 127 | Specific | 770.72 | 770.72 | 770.72 | 15.23 | 51 | >sp\|P01876\|IGHA1_HUMAN Immunoglobulin heavy constant alpha 1 OS=Homo sapiens OX=9606 GN=IGHA1 PE=1 SV=2 | 255 | 20201113_12413_1ng_glyco_hcd203040_compound15.21961.21961.4 | controllerType=0 controllerNumber=1 scan=21961 | 44.7597 |
| 18449:4 | 11 | R.LSLHRPALEDLLLGSEAN[+1751.624]LTC[+57.021]TLTGLR.D | HexNAc(4)Hex(4)NeuAc(1) | C[+57], N[+1752] | 1179.5651 | 4 | 4715.2386 | 4715.2225 | 0 | 3.4 | 127 | Specific | 773.92 | 773.92 | 773.92 | 15.37 | 51 | >sp\|P01876\|IGHA1_HUMAN Immunoglobulin heavy constant alpha 1 OS=Homo sapiens OX=9606 GN=IGHA1 PE=1 SV=2 | 255 | 20201113_12413_1ng_glyco_hcd203040_compound15.22005.22005.4 | controllerType=0 controllerNumber=1 scan=22005 | 44.8712 |
| 17462:4 | 11 | R.LSLHRPALEDLLLGSEAN[+1460.529]LTC[+57.021]TLTGLR.D | HexNAc(4)Hex(4) | C[+57], N[+1461] | 1106.7875 | 4 | 4424.1282 | 4424.1271 | 0 | 0.3 | 127 | Specific | 782.32 | 609.18 | 609.18 | 15.08 | 51 | >sp\|P01876\|IGHA1_HUMAN Immunoglobulin heavy constant alpha 1 OS=Homo sapiens OX=9606 GN=IGHA1 PE=1 SV=2 | 255 | 20201113_12413_1ng_glyco_hcd203040_compound15.20971.20971.4 | controllerType=0 controllerNumber=1 scan=20971 | 42.633 |
| 17441:4 | 11 | R.LSLHRPALEDLLLGSEAN[+1460.529]LTC[+57.021]TLTGLR.D | HexNAc(4)Hex(4) | C[+57], N[+1461] | 1106.7935 | 4 | 4424.1522 | 4424.1271 | 0 | 5.7 | 127 | Specific | 803.51 | 803.51 | 803.51 | 15.61 | 51 | >sp\|P01876\|IGHA1_HUMAN Immunoglobulin heavy constant alpha 1 OS=Homo sapiens OX=9606 GN=IGHA1 PE=1 SV=2 | 255 | 20201113_12413_1ng_glyco_hcd203040_compound15.20949.20949.2 | controllerType=0 controllerNumber=1 scan=20949 | 42.5822 |
| 17459:4 | 11 | R.LSLHRPALEDLLLGSEAN[+1460.529]LTC[+57.021]TLTGLR.D | HexNAc(4)Hex(4) | C[+57], N[+1461] | 1106.7875 | 4 | 4424.1282 | 4424.1271 | 0 | 0.3 | 127 | Specific | 834.33 | 834.33 | 834.33 | 16.08 | 51 | >sp\|P01876\|IGHA1_HUMAN Immunoglobulin heavy constant alpha 1 OS=Homo sapiens OX=9606 GN=IGHA1 PE=1 SV=2 | 255 | 20201113_12413_1ng_glyco_hcd203040_compound15.20968.20968.4 | controllerType=0 controllerNumber=1 scan=20968 | 42.6246 |
| 17446:4 | 11 | R.LSLHRPALEDLLLGSEAN[+1419.502]LTC[+57.021]TLTGLR.D | HexNAc(3)Hex(5) | C[+57], N[+1420] | 1096.5357 | 4 | 4383.1209 | 4383.1005 | 0 | 4.7 | 127 | Specific | 888.32 | 817.18 | 817.18 | 17.23 | 51 | >sp\|P01876\|IGHA1_HUMAN Immunoglobulin heavy constant alpha 1 OS=Homo sapiens OX=9606 GN=IGHA1 PE=1 SV=2 | 255 | 20201113_12413_1ng_glyco_hcd203040_compound15.20954.20954.4 | controllerType=0 controllerNumber=1 scan=20954 | 42.5955 |
| 17451:4 | 11 | R.LSLHRPALEDLLLGSEAN[+1378.476]LTC[+57.021]TLTGLR.D | HexNAc(2)Hex(6) | C[+57], N[+1378] | 1086.2741 | 4 | 4342.0745 | 4342.074 | 0 | 0.1 | 127 | Specific | 901.88 | 889.73 | 889.73 | 18.08 | 51 | >sp\|P01876\|IGHA1_HUMAN Immunoglobulin heavy constant alpha 1 OS=Homo sapiens OX=9606 GN=IGHA1 PE=1 SV=2 | 255 | 20201113_12413_1ng_glyco_hcd203040_compound15.20960.20960.4 | controllerType=0 controllerNumber=1 scan=20960 | 42.6114 |
| 17376:4 | 11 | R.LSLHRPALEDLLLGSEAN[+1702.581]LTC[+57.021]TLTGLR.D | HexNAc(2)Hex(8) | C[+57], N[+1703] | 1167.3036 | 4 | 4666.1927 | 4666.1796 | 0 | 2.8 | 127 | Specific | 916.49 | 916.49 | 916.49 | 18.37 | 51 | >sp\|P01876\|IGHA1_HUMAN Immunoglobulin heavy constant alpha 1 OS=Homo sapiens OX=9606 GN=IGHA1 PE=1 SV=2 | 255 | 20201113_12413_1ng_glyco_hcd203040_compound15.20880.20880.2 | controllerType=0 controllerNumber=1 scan=20880 | 42.3848 |
| 17370:4 | 11 | R.LSLHRPALEDLLLGSEAN[+1540.529]LTC[+57.021]TLTGLR.D | HexNAc(2)Hex(7) | C[+57], N[+1541] | 1126.7889 | 4 | 4504.1336 | 4504.1268 | 0 | 1.5 | 127 | Specific | 936 | 881.82 | 881.82 | 18.27 | 51 | >sp\|P01876\|IGHA1_HUMAN Immunoglobulin heavy constant alpha 1 OS=Homo sapiens OX=9606 GN=IGHA1 PE=1 SV=2 | 255 | 20201113_12413_1ng_glyco_hcd203040_compound15.20874.20874.2 | controllerType=0 controllerNumber=1 scan=20874 | 42.3688 |
| 17378:4 | 11 | R.LSLHRPALEDLLLGSEAN[+1702.581]LTC[+57.021]TLTGLR.D | HexNAc(2)Hex(8) | C[+57], N[+1703] | 1167.3036 | 4 | 4666.1927 | 4666.1796 | 0 | 2.8 | 127 | Specific | 937.38 | 776.06 | 776.06 | 18.37 | 51 | >sp\|P01876\|IGHA1_HUMAN Immunoglobulin heavy constant alpha 1 OS=Homo sapiens OX=9606 GN=IGHA1 PE=1 SV=2 | 255 | 20201113_12413_1ng_glyco_hcd203040_compound15.20882.20882.2 | controllerType=0 controllerNumber=1 scan=20882 | 42.3942 |
| 17444:4 | 11 | R.LSLHRPALEDLLLGSEAN[+1460.529]LTC[+57.021]TLTGLR.D | HexNAc(4)Hex(4) | C[+57], N[+1461] | 1106.7935 | 4 | 4424.1522 | 4424.1271 | 0 | 5.7 | 127 | Specific | 948.45 | 899.93 | 899.93 | 17.61 | 51 | >sp\|P01876\|IGHA1_HUMAN Immunoglobulin heavy constant alpha 1 OS=Homo sapiens OX=9606 GN=IGHA1 PE=1 SV=2 | 255 | 20201113_12413_1ng_glyco_hcd203040_compound15.20952.20952.2 | controllerType=0 controllerNumber=1 scan=20952 | 42.5903 |
| 17457:4 | 11 | R.LSLHRPALEDLLLGSEAN[+1216.423]LTC[+57.021]TLTGLR.D | HexNAc(2)Hex(5) | C[+57], N[+1216] | 1045.7605 | 4 | 4180.0203 | 4180.0211 | 0 | -0.2 | 127 | Specific | 951.54 | 951.54 | 951.54 | 19.08 | 51 | >sp\|P01876\|IGHA1_HUMAN Immunoglobulin heavy constant alpha 1 OS=Homo sapiens OX=9606 GN=IGHA1 PE=1 SV=2 | 255 | 20201113_12413_1ng_glyco_hcd203040_compound15.20966.20966.4 | controllerType=0 controllerNumber=1 scan=20966 | 42.6227 |
| 17373:4 | 11 | R.LSLHRPALEDLLLGSEAN[+1540.529]LTC[+57.021]TLTGLR.D | HexNAc(2)Hex(7) | C[+57], N[+1541] | 1126.7889 | 4 | 4504.1336 | 4504.1268 | 0 | 1.5 | 127 | Specific | 1095.55 | 1085.64 | 1085.64 | 21.27 | 51 | >sp\|P01876\|IGHA1_HUMAN Immunoglobulin heavy constant alpha 1 OS=Homo sapiens OX=9606 GN=IGHA1 PE=1 SV=2 | 255 | 20201113_12413_1ng_glyco_hcd203040_compound15.20877.20877.2 | controllerType=0 controllerNumber=1 scan=20877 | 42.3789 |
| 14899:3 | 11 | R.LAGKPTHVN[+2553.910]VSVVMAEVDGTC[+57.021]Y.- | HexNAc(5)Hex(5)Fuc(1)NeuAc(2) | C[+57], N[+2554] | 1634.3599 | 3 | 4901.0651 | 4901.0518 | 0 | 2.7 | 332 | CRagged | 204.11 | 196.06 | 196.06 | 4.81 | 51 | >sp\|P01876\|IGHA1_HUMAN Immunoglobulin heavy constant alpha 1 OS=Homo sapiens OX=9606 GN=IGHA1 PE=1 SV=2 | 255 | 20201113_12413_1ng_glyco_hcd203040_compound15.18275.18275.3 | controllerType=0 controllerNumber=1 scan=18275 | 37.4044 |
| 11094:3 | 11 | R.LAGKPTHVN[+1581.555]VSVVMAEVDGTC[+57.021]Y.- | HexNAc(3)Hex(6) | C[+57], N[+1582] | 1310.2393 | 3 | 3928.7034 | 3928.6971 | 0 | 1.6 | 332 | CRagged | 224.24 | 151 | 151 | 4.26 | 51 | >sp\|P01876\|IGHA1_HUMAN Immunoglobulin heavy constant alpha 1 OS=Homo sapiens OX=9606 GN=IGHA1 PE=1 SV=2 | 255 | 20201113_12413_1ng_glyco_hcd203040_compound15.14312.14312.3 | controllerType=0 controllerNumber=1 scan=14312 | 30.525 |
| 11023:3 | 11 | R.LAGKPTHVN[+1702.581]VSVVMAEVDGTC[+57.021]Y.- | HexNAc(2)Hex(8) | C[+57], N[+1703] | 1350.5823 | 3 | 4049.7324 | 4049.7234 | 0 | 2.2 | 332 | CRagged | 241.42 | 241.42 | 209.07 | 6.23 | 51 | >sp\|P01876\|IGHA1_HUMAN Immunoglobulin heavy constant alpha 1 OS=Homo sapiens OX=9606 GN=IGHA1 PE=1 SV=2 | 255 | 20201113_12413_1ng_glyco_hcd203040_compound15.14239.14239.3 | controllerType=0 controllerNumber=1 scan=14239 | 30.3856 |
| 11112:3 | 11 | R.LAGKPTHVN[+1971.719]VSVVMAEVDGTC[+57.021]Y.- | HexNAc(5)Hex(5)Fuc(1) | C[+57], N[+1972] | 1440.2954 | 3 | 4318.8716 | 4318.861 | 0 | 2.5 | 332 | CRagged | 245.82 | 245.82 | 245.82 | 6.44 | 51 | >sp\|P01876\|IGHA1_HUMAN Immunoglobulin heavy constant alpha 1 OS=Homo sapiens OX=9606 GN=IGHA1 PE=1 SV=2 | 255 | 20201113_12413_1ng_glyco_hcd203040_compound15.14331.14331.3 | controllerType=0 controllerNumber=1 scan=14331 | 30.5694 |
| 12851:3 | 11 | R.LAGKPTHVN[+2262.814]VSVVMAEVDGTC[+57.021]Y.- | HexNAc(5)Hex(5)Fuc(1)NeuAc(1) | C[+57], N[+2263] | 1537.3287 | 3 | 4609.9715 | 4609.9564 | 0 | 3.3 | 332 | CRagged | 325.67 | 325.67 | 325.67 | 7.28 | 51 | >sp\|P01876\|IGHA1_HUMAN Immunoglobulin heavy constant alpha 1 OS=Homo sapiens OX=9606 GN=IGHA1 PE=1 SV=2 | 255 | 20201113_12413_1ng_glyco_hcd203040_compound15.16153.16153.3 | controllerType=0 controllerNumber=1 scan=16153 | 33.6648 |
| 11114:3 | 11 | R.LAGKPTHVN[+1971.719]VSVVMAEVDGTC[+57.021]Y.- | HexNAc(5)Hex(5)Fuc(1) | C[+57], N[+1972] | 1440.2954 | 3 | 4318.8716 | 4318.861 | 0 | 2.5 | 332 | CRagged | 333.58 | 333.58 | 304.3 | 7.39 | 51 | >sp\|P01876\|IGHA1_HUMAN Immunoglobulin heavy constant alpha 1 OS=Homo sapiens OX=9606 GN=IGHA1 PE=1 SV=2 | 255 | 20201113_12413_1ng_glyco_hcd203040_compound15.14333.14333.3 | controllerType=0 controllerNumber=1 scan=14333 | 30.575 |
| 14808:4 | 11 | R.LAGKPTHVN[+2553.910]VSVVMAEVDGTC[+57.021]Y.- | HexNAc(5)Hex(5)Fuc(1)NeuAc(2) | C[+57], N[+2554] | 1226.0192 | 4 | 4901.055 | 4901.0518 | 0 | 0.7 | 332 | CRagged | 343.97 | 289.85 | 289.85 | 6.79 | 51 | >sp\|P01876\|IGHA1_HUMAN Immunoglobulin heavy constant alpha 1 OS=Homo sapiens OX=9606 GN=IGHA1 PE=1 SV=2 | 255 | 20201113_12413_1ng_glyco_hcd203040_compound15.18180.18180.2 | controllerType=0 controllerNumber=1 scan=18180 | 37.2369 |
| 11092:3 | 11 | R.LAGKPTHVN[+1581.555]VSVVMAEVDGTC[+57.021]Y.- | HexNAc(3)Hex(6) | C[+57], N[+1582] | 1310.2393 | 3 | 3928.7034 | 3928.6971 | 0 | 1.6 | 332 | CRagged | 395.26 | 395.26 | 317.73 | 9.03 | 51 | >sp\|P01876\|IGHA1_HUMAN Immunoglobulin heavy constant alpha 1 OS=Homo sapiens OX=9606 GN=IGHA1 PE=1 SV=2 | 255 | 20201113_12413_1ng_glyco_hcd203040_compound15.14310.14310.3 | controllerType=0 controllerNumber=1 scan=14310 | 30.5214 |
| 09881:3 | 11 | R.LAGKPTHVN[+1378.476]VSVVMAEVDGTC[+57.021]Y.- | HexNAc(2)Hex(6) | C[+57], N[+1378] | 1242.545 | 3 | 3725.6204 | 3725.6178 | 0 | 0.7 | 332 | CRagged | 436.19 | 427.34 | 231 | 8.99 | 51 | >sp\|P01876\|IGHA1_HUMAN Immunoglobulin heavy constant alpha 1 OS=Homo sapiens OX=9606 GN=IGHA1 PE=1 SV=2 | 255 | 20201113_12413_1ng_glyco_hcd203040_compound15.13057.13057.2 | controllerType=0 controllerNumber=1 scan=13057 | 28.3841 |
| 09883:3 | 11 | R.LAGKPTHVN[+1378.476]VSVVMAEVDGTC[+57.021]Y.- | HexNAc(2)Hex(6) | C[+57], N[+1378] | 1242.545 | 3 | 3725.6204 | 3725.6178 | 0 | 0.7 | 332 | CRagged | 443.13 | 443.13 | 404.97 | 9 | 51 | >sp\|P01876\|IGHA1_HUMAN Immunoglobulin heavy constant alpha 1 OS=Homo sapiens OX=9606 GN=IGHA1 PE=1 SV=2 | 255 | 20201113_12413_1ng_glyco_hcd203040_compound15.13059.13059.2 | controllerType=0 controllerNumber=1 scan=13059 | 28.3881 |
| 14825:3 | 11 | R.LAGKPTHVN[+2351.851]VSVVMAEVDGTC[+57.021]Y.- | HexNAc(4)Hex(5)Fuc(3)NeuAc(1) | C[+57], N[+2352] | 1566.9983 | 3 | 4698.9803 | 4698.9928 | 0 | -2.7 | 332 | CRagged | 495.13 | 490.31 | 490.31 | 8.42 | 51 | >sp\|P01876\|IGHA1_HUMAN Immunoglobulin heavy constant alpha 1 OS=Homo sapiens OX=9606 GN=IGHA1 PE=1 SV=2 | 255 | 20201113_12413_1ng_glyco_hcd203040_compound15.18198.18198.3 | controllerType=0 controllerNumber=1 scan=18198 | 37.2725 |
| 14810:4 | 11 | R.LAGKPTHVN[+2553.910]VSVVMAEVDGTC[+57.021]Y.- | HexNAc(5)Hex(5)Fuc(1)NeuAc(2) | C[+57], N[+2554] | 1226.0192 | 4 | 4901.055 | 4901.0518 | 0 | 0.7 | 332 | CRagged | 520.54 | 426.22 | 426.22 | 9.44 | 51 | >sp\|P01876\|IGHA1_HUMAN Immunoglobulin heavy constant alpha 1 OS=Homo sapiens OX=9606 GN=IGHA1 PE=1 SV=2 | 255 | 20201113_12413_1ng_glyco_hcd203040_compound15.18182.18182.2 | controllerType=0 controllerNumber=1 scan=18182 | 37.2407 |
| 12849:3 | 11 | R.LAGKPTHVN[+2262.814]VSVVMAEVDGTC[+57.021]Y.- | HexNAc(5)Hex(5)Fuc(1)NeuAc(1) | C[+57], N[+2263] | 1537.3287 | 3 | 4609.9715 | 4609.9564 | 0 | 3.3 | 332 | CRagged | 531.55 | 299.52 | 299.52 | 7.98 | 51 | >sp\|P01876\|IGHA1_HUMAN Immunoglobulin heavy constant alpha 1 OS=Homo sapiens OX=9606 GN=IGHA1 PE=1 SV=2 | 255 | 20201113_12413_1ng_glyco_hcd203040_compound15.16151.16151.3 | controllerType=0 controllerNumber=1 scan=16151 | 33.6603 |
| 12854:3 | 11 | R.LAGKPTHVN[+2059.735]VSVVMAEVDGTC[+57.021]Y.- | HexNAc(4)Hex(5)Fuc(1)NeuAc(1) | C[+57], N[+2060] | 1469.6358 | 3 | 4406.8928 | 4406.877 | 0 | 3.6 | 332 | CRagged | 569.74 | 301.27 | 301.27 | 8.02 | 51 | >sp\|P01876\|IGHA1_HUMAN Immunoglobulin heavy constant alpha 1 OS=Homo sapiens OX=9606 GN=IGHA1 PE=1 SV=2 | 255 | 20201113_12413_1ng_glyco_hcd203040_compound15.16156.16156.3 | controllerType=0 controllerNumber=1 scan=16156 | 33.669 |
| 11081:3 | 11 | R.LAGKPTHVN[+1809.666]VSVVMAEVDGTC[+57.021]Y.- | HexNAc(5)Hex(4)Fuc(1) | C[+57], N[+1810] | 1386.2762 | 3 | 4156.814 | 4156.8081 | 0 | 1.4 | 332 | CRagged | 578.02 | 505.78 | 433.9 | 11.01 | 51 | >sp\|P01876\|IGHA1_HUMAN Immunoglobulin heavy constant alpha 1 OS=Homo sapiens OX=9606 GN=IGHA1 PE=1 SV=2 | 255 | 20201113_12413_1ng_glyco_hcd203040_compound15.14299.14299.3 | controllerType=0 controllerNumber=1 scan=14299 | 30.5019 |
| 11160:3 | 11 | R.LAGKPTHVN[+1501.555]VSVVMAEVDGTC[+57.021]Y.- | HexNAc(5)Hex(3) | C[+57], N[+1502] | 1283.5742 | 3 | 3848.7079 | 3848.6974 | 0 | 2.7 | 332 | CRagged | 602.45 | 473.02 | 473.02 | 12.1 | 51 | >sp\|P01876\|IGHA1_HUMAN Immunoglobulin heavy constant alpha 1 OS=Homo sapiens OX=9606 GN=IGHA1 PE=1 SV=2 | 255 | 20201113_12413_1ng_glyco_hcd203040_compound15.14383.14383.3 | controllerType=0 controllerNumber=1 scan=14383 | 30.6811 |
| 11157:3 | 11 | R.LAGKPTHVN[+1501.555]VSVVMAEVDGTC[+57.021]Y.- | HexNAc(5)Hex(3) | C[+57], N[+1502] | 1283.5742 | 3 | 3848.7079 | 3848.6974 | 0 | 2.7 | 332 | CRagged | 652.33 | 652.33 | 652.33 | 13.1 | 51 | >sp\|P01876\|IGHA1_HUMAN Immunoglobulin heavy constant alpha 1 OS=Homo sapiens OX=9606 GN=IGHA1 PE=1 SV=2 | 255 | 20201113_12413_1ng_glyco_hcd203040_compound15.14380.14380.3 | controllerType=0 controllerNumber=1 scan=14380 | 30.6763 |
| 11068:3 | 11 | R.LAGKPTHVN[+1663.608]VSVVMAEVDGTC[+57.021]Y.- | HexNAc(5)Hex(4) | C[+57], N[+1664] | 1337.5908 | 3 | 4010.7577 | 4010.7502 | 0 | 1.9 | 332 | CRagged | 762.29 | 762.29 | 480.71 | 15.01 | 51 | >sp\|P01876\|IGHA1_HUMAN Immunoglobulin heavy constant alpha 1 OS=Homo sapiens OX=9606 GN=IGHA1 PE=1 SV=2 | 255 | 20201113_12413_1ng_glyco_hcd203040_compound15.14285.14285.3 | controllerType=0 controllerNumber=1 scan=14285 | 30.4725 |
| 11078:3 | 11 | R.LAGKPTHVN[+1809.666]VSVVMAEVDGTC[+57.021]Y.- | HexNAc(5)Hex(4)Fuc(1) | C[+57], N[+1810] | 1386.2762 | 3 | 4156.814 | 4156.8081 | 0 | 1.4 | 332 | CRagged | 773.75 | 773.75 | 500.77 | 15.01 | 51 | >sp\|P01876\|IGHA1_HUMAN Immunoglobulin heavy constant alpha 1 OS=Homo sapiens OX=9606 GN=IGHA1 PE=1 SV=2 | 255 | 20201113_12413_1ng_glyco_hcd203040_compound15.14296.14296.3 | controllerType=0 controllerNumber=1 scan=14296 | 30.4958 |
| 11155:3 | 11 | R.LAGKPTHVN[+1647.613]VSVVMAEVDGTC[+57.021]Y.- | HexNAc(5)Hex(3)Fuc(1) | C[+57], N[+1648] | 1332.2593 | 3 | 3994.7635 | 3994.7553 | 0 | 2 | 332 | CRagged | 782.4 | 782.4 | 782.4 | 15.01 | 51 | >sp\|P01876\|IGHA1_HUMAN Immunoglobulin heavy constant alpha 1 OS=Homo sapiens OX=9606 GN=IGHA1 PE=1 SV=2 | 255 | 20201113_12413_1ng_glyco_hcd203040_compound15.14378.14378.3 | controllerType=0 controllerNumber=1 scan=14378 | 30.6738 |
| 12856:3 | 11 | R.LAGKPTHVN[+2059.735]VSVVMAEVDGTC[+57.021]Y.- | HexNAc(4)Hex(5)Fuc(1)NeuAc(1) | C[+57], N[+2060] | 1469.6358 | 3 | 4406.8928 | 4406.877 | 0 | 3.6 | 332 | CRagged | 804.86 | 758.93 | 758.93 | 15.97 | 51 | >sp\|P01876\|IGHA1_HUMAN Immunoglobulin heavy constant alpha 1 OS=Homo sapiens OX=9606 GN=IGHA1 PE=1 SV=2 | 255 | 20201113_12413_1ng_glyco_hcd203040_compound15.16158.16158.3 | controllerType=0 controllerNumber=1 scan=16158 | 33.673 |
| 11153:3 | 11 | R.LAGKPTHVN[+1647.613]VSVVMAEVDGTC[+57.021]Y.- | HexNAc(5)Hex(3)Fuc(1) | C[+57], N[+1648] | 1332.2593 | 3 | 3994.7635 | 3994.7553 | 0 | 2 | 332 | CRagged | 837.48 | 837.48 | 837.48 | 16.01 | 51 | >sp\|P01876\|IGHA1_HUMAN Immunoglobulin heavy constant alpha 1 OS=Homo sapiens OX=9606 GN=IGHA1 PE=1 SV=2 | 255 | 20201113_12413_1ng_glyco_hcd203040_compound15.14376.14376.3 | controllerType=0 controllerNumber=1 scan=14376 | 30.663 |
| 11066:3 | 11 | R.LAGKPTHVN[+1663.608]VSVVMAEVDGTC[+57.021]Y.- | HexNAc(5)Hex(4) | C[+57], N[+1664] | 1337.5908 | 3 | 4010.7577 | 4010.7502 | 0 | 1.9 | 332 | CRagged | 839.28 | 839.28 | 567.71 | 16.01 | 51 | >sp\|P01876\|IGHA1_HUMAN Immunoglobulin heavy constant alpha 1 OS=Homo sapiens OX=9606 GN=IGHA1 PE=1 SV=2 | 255 | 20201113_12413_1ng_glyco_hcd203040_compound15.14283.14283.3 | controllerType=0 controllerNumber=1 scan=14283 | 30.4662 |
| 11108:3 | 11 | R.LAGKPTHVN[+1216.423]VSVVMAEVDGTC[+57.021]Y.- | HexNAc(2)Hex(5) | C[+57], N[+1216] | 1188.529 | 3 | 3563.5724 | 3563.5649 | 0 | 2.1 | 332 | CRagged | 858.48 | 812.72 | 522.56 | 17.01 | 51 | >sp\|P01876\|IGHA1_HUMAN Immunoglobulin heavy constant alpha 1 OS=Homo sapiens OX=9606 GN=IGHA1 PE=1 SV=2 | 255 | 20201113_12413_1ng_glyco_hcd203040_compound15.14327.14327.3 | controllerType=0 controllerNumber=1 scan=14327 | 30.5583 |
| 11025:3 | 11 | R.LAGKPTHVN[+1702.581]VSVVMAEVDGTC[+57.021]Y.- | HexNAc(2)Hex(8) | C[+57], N[+1703] | 1350.5823 | 3 | 4049.7324 | 4049.7234 | 0 | 2.2 | 332 | CRagged | 900.79 | 755.19 | 533.92 | 18.1 | 51 | >sp\|P01876\|IGHA1_HUMAN Immunoglobulin heavy constant alpha 1 OS=Homo sapiens OX=9606 GN=IGHA1 PE=1 SV=2 | 255 | 20201113_12413_1ng_glyco_hcd203040_compound15.14241.14241.3 | controllerType=0 controllerNumber=1 scan=14241 | 30.3884 |
| 11098:3 | 11 | R.LAGKPTHVN[+1216.423]VSVVMAEVDGTC[+57.021]Y.- | HexNAc(2)Hex(5) | C[+57], N[+1216] | 1188.5287 | 3 | 3563.5717 | 3563.5649 | 0 | 1.9 | 332 | CRagged | 909.18 | 895.84 | 563.35 | 18.01 | 51 | >sp\|P01876\|IGHA1_HUMAN Immunoglobulin heavy constant alpha 1 OS=Homo sapiens OX=9606 GN=IGHA1 PE=1 SV=2 | 255 | 20201113_12413_1ng_glyco_hcd203040_compound15.14316.14316.3 | controllerType=0 controllerNumber=1 scan=14316 | 30.5388 |
| 11096:3 | 11 | R.LAGKPTHVN[+1216.423]VSVVMAEVDGTC[+57.021]Y.- | HexNAc(2)Hex(5) | C[+57], N[+1216] | 1188.5287 | 3 | 3563.5717 | 3563.5649 | 0 | 1.9 | 332 | CRagged | 910.91 | 910.91 | 556.15 | 18.01 | 51 | >sp\|P01876\|IGHA1_HUMAN Immunoglobulin heavy constant alpha 1 OS=Homo sapiens OX=9606 GN=IGHA1 PE=1 SV=2 | 255 | 20201113_12413_1ng_glyco_hcd203040_compound15.14314.14314.3 | controllerType=0 controllerNumber=1 scan=14314 | 30.5284 |
| 11076:3 | 11 | R.LAGKPTHVN[+1378.476]VSVVMAEVDGTC[+57.021]Y.- | HexNAc(2)Hex(6) | C[+57], N[+1378] | 1242.5466 | 3 | 3725.6251 | 3725.6178 | 0 | 2 | 332 | CRagged | 916.66 | 911.55 | 570.22 | 18.01 | 51 | >sp\|P01876\|IGHA1_HUMAN Immunoglobulin heavy constant alpha 1 OS=Homo sapiens OX=9606 GN=IGHA1 PE=1 SV=2 | 255 | 20201113_12413_1ng_glyco_hcd203040_compound15.14294.14294.3 | controllerType=0 controllerNumber=1 scan=14294 | 30.4935 |
| 11074:3 | 11 | R.LAGKPTHVN[+1378.476]VSVVMAEVDGTC[+57.021]Y.- | HexNAc(2)Hex(6) | C[+57], N[+1378] | 1242.5466 | 3 | 3725.6251 | 3725.6178 | 0 | 2 | 332 | CRagged | 935.65 | 935.65 | 581.52 | 18.01 | 51 | >sp\|P01876\|IGHA1_HUMAN Immunoglobulin heavy constant alpha 1 OS=Homo sapiens OX=9606 GN=IGHA1 PE=1 SV=2 | 255 | 20201113_12413_1ng_glyco_hcd203040_compound15.14292.14292.3 | controllerType=0 controllerNumber=1 scan=14292 | 30.487 |
| 11105:3 | 11 | R.LAGKPTHVN[+1216.423]VSVVMAEVDGTC[+57.021]Y.- | HexNAc(2)Hex(5) | C[+57], N[+1216] | 1188.529 | 3 | 3563.5724 | 3563.5649 | 0 | 2.1 | 332 | CRagged | 937.32 | 937.32 | 583.13 | 18.01 | 51 | >sp\|P01876\|IGHA1_HUMAN Immunoglobulin heavy constant alpha 1 OS=Homo sapiens OX=9606 GN=IGHA1 PE=1 SV=2 | 255 | 20201113_12413_1ng_glyco_hcd203040_compound15.14324.14324.3 | controllerType=0 controllerNumber=1 scan=14324 | 30.5512 |
| 05448:3 | 12 | R.LSPN[+2205.793]ASAEHLELR.W | HexNAc(4)Hex(5)Fuc(2)NeuAc(1) | N[+2206] | 1214.8549 | 3 | 3642.5502 | 3642.5421 | 0 | 2.2 | 52 | Specific | 299.96 | 139.22 | 139.22 | 4.83 | 40 | >sp\|Q13410\|BT1A1_HUMAN Butyrophilin subfamily 1 member A1 OS=Homo sapiens OX=9606 GN=BTN1A1 PE=1 SV=3 | 179 | 20201113_12413_1ng_glyco_hcd203040_compound15.8457.8457.3 | controllerType=0 controllerNumber=1 scan=8457 | 20.7013 |
| 07428:3 | 12 | R.LSPN[+2350.830]ASAEHLELR.W | HexNAc(4)Hex(5)Fuc(1)NeuAc(2) | N[+2351] | 1263.2 | 3 | 3787.5855 | 3787.5796 | 0 | 1.6 | 52 | Specific | 578.27 | 384.72 | 384.72 | 10.6 | 40 | >sp\|Q13410\|BT1A1_HUMAN Butyrophilin subfamily 1 member A1 OS=Homo sapiens OX=9606 GN=BTN1A1 PE=1 SV=3 | 179 | 20201113_12413_1ng_glyco_hcd203040_compound15.10508.10508.3 | controllerType=0 controllerNumber=1 scan=10508 | 24.1167 |
| 07430:3 | 12 | R.LSPN[+2350.830]ASAEHLELR.W | HexNAc(4)Hex(5)Fuc(1)NeuAc(2) | N[+2351] | 1263.2 | 3 | 3787.5855 | 3787.5796 | 0 | 1.6 | 52 | Specific | 638.9 | 541.02 | 541.02 | 11.6 | 40 | >sp\|Q13410\|BT1A1_HUMAN Butyrophilin subfamily 1 member A1 OS=Homo sapiens OX=9606 GN=BTN1A1 PE=1 SV=3 | 179 | 20201113_12413_1ng_glyco_hcd203040_compound15.10510.10510.3 | controllerType=0 controllerNumber=1 scan=10510 | 24.1214 |
| 17346:3 | 12 | K.N[+2059.735]VSC[+57.021]YIQNLLLGQEK.K | HexNAc(4)Hex(5)Fuc(1)NeuAc(1) | C[+57], N[+2060] | 1280.223 | 3 | 3838.6543 | 3838.6455 | 0 | 2.3 | 215 | Specific | 496.02 | 449.24 | 449.24 | 9.52 | 40 | >sp\|Q13410\|BT1A1_HUMAN Butyrophilin subfamily 1 member A1 OS=Homo sapiens OX=9606 GN=BTN1A1 PE=1 SV=3 | 179 | 20201113_12413_1ng_glyco_hcd203040_compound15.20849.20849.3 | controllerType=0 controllerNumber=1 scan=20849 | 42.3142 |
| 17419:3 | 12 | K.N[+1913.677]VSC[+57.021]YIQNLLLGQEK.K | HexNAc(4)Hex(5)NeuAc(1) | C[+57], N[+1914] | 1231.5368 | 3 | 3692.5958 | 3692.5876 | 0 | 2.2 | 215 | Specific | 697.21 | 508.58 | 508.58 | 13.04 | 40 | >sp\|Q13410\|BT1A1_HUMAN Butyrophilin subfamily 1 member A1 OS=Homo sapiens OX=9606 GN=BTN1A1 PE=1 SV=3 | 179 | 20201113_12413_1ng_glyco_hcd203040_compound15.20926.20926.3 | controllerType=0 controllerNumber=1 scan=20926 | 42.5204 |
| 23804:3 | 14 | R.QLAHQSN[+2204.772]STNIFFSPVSIATAFAMLSLGTK.A | HexNAc(4)Hex(5)NeuAc(2) | N[+2205] | 1796.1551 | 3 | 5386.4508 | 5386.4075 | 0 | 8 | 64 | Specific | 271.57 | 214.69 | 214.69 | 5.13 | 36 | >sp\|P01009\|A1AT_HUMAN Alpha-1-antitrypsin OS=Homo sapiens OX=9606 GN=SERPINA1 PE=1 SV=3 | 642 | 20201113_12413_1ng_glyco_hcd203040_compound15.27594.27594.3 | controllerType=0 controllerNumber=1 scan=27594 | 56.0614 |
| 23721:4 | 14 | R.QLAHQSN[+2204.772]STNIFFSPVSIATAFAMLSLGTK.A | HexNAc(4)Hex(5)NeuAc(2) | N[+2205] | 1347.3577 | 4 | 5386.4088 | 5386.4075 | 0 | 0.3 | 64 | Specific | 451.13 | 451.13 | 443.65 | 8.83 | 36 | >sp\|P01009\|A1AT_HUMAN Alpha-1-antitrypsin OS=Homo sapiens OX=9606 GN=SERPINA1 PE=1 SV=3 | 642 | 20201113_12413_1ng_glyco_hcd203040_compound15.27508.27508.4 | controllerType=0 controllerNumber=1 scan=27508 | 55.8967 |
| 23810:4 | 14 | R.QLAHQSN[+2204.772]STNIFFSPVSIATAFAMLSLGTK.A | HexNAc(4)Hex(5)NeuAc(2) | N[+2205] | 1347.3663 | 4 | 5386.4435 | 5386.4075 | 0 | 6.7 | 64 | Specific | 647.73 | 622.37 | 622.37 | 11.15 | 36 | >sp\|P01009\|A1AT_HUMAN Alpha-1-antitrypsin OS=Homo sapiens OX=9606 GN=SERPINA1 PE=1 SV=3 | 642 | 20201113_12413_1ng_glyco_hcd203040_compound15.27600.27600.4 | controllerType=0 controllerNumber=1 scan=27600 | 56.0722 |
| 23724:3 | 14 | R.QLAHQSN[+2204.772]STNIFFSPVSIATAFAMLSLGTK.A | HexNAc(4)Hex(5)NeuAc(2) | N[+2205] | 1796.1435 | 3 | 5386.416 | 5386.4075 | 0 | 1.6 | 64 | Specific | 841.09 | 823.1 | 546.39 | 15.83 | 36 | >sp\|P01009\|A1AT_HUMAN Alpha-1-antitrypsin OS=Homo sapiens OX=9606 GN=SERPINA1 PE=1 SV=3 | 642 | 20201113_12413_1ng_glyco_hcd203040_compound15.27511.27511.3 | controllerType=0 controllerNumber=1 scan=27511 | 55.9026 |
| 23723:4 | 14 | R.QLAHQSN[+2204.772]STNIFFSPVSIATAFAMLSLGTK.A | HexNAc(4)Hex(5)NeuAc(2) | N[+2205] | 1347.3577 | 4 | 5386.4088 | 5386.4075 | 0 | 0.3 | 64 | Specific | 888 | 813.39 | 559.74 | 16.65 | 36 | >sp\|P01009\|A1AT_HUMAN Alpha-1-antitrypsin OS=Homo sapiens OX=9606 GN=SERPINA1 PE=1 SV=3 | 642 | 20201113_12413_1ng_glyco_hcd203040_compound15.27510.27510.4 | controllerType=0 controllerNumber=1 scan=27510 | 55.901 |
| 22287:4 | 14 | K.ADTHDEILEGLNFN[+2861.000]LTEIPEAQIHEGFQELLR.T | HexNAc(5)Hex(6)NeuAc(3) | N[+2861] | 1638.9733 | 4 | 6552.8712 | 6552.8239 | 0 | 7.2 | 94 | Specific | 780.31 | 549.5 | 549.5 | 14.13 | 36 | >sp\|P01009\|A1AT_HUMAN Alpha-1-antitrypsin OS=Homo sapiens OX=9606 GN=SERPINA1 PE=1 SV=3 | 642 | 20201113_12413_1ng_glyco_hcd203040_compound15.26011.26011.5 | controllerType=0 controllerNumber=1 scan=26011 | 53.0221 |
| 01998:3 | 15 | K.N[+1216.423]NSIPDKQITASSSYK.T | HexNAc(2)Hex(5) | N[+1216] | 990.4407 | 3 | 2969.3076 | 2969.2991 | 0 | 2.8 | 238 | Specific | 265 | 166.5 | 166.5 | 5.46 | 30 | >sp\|Q08431\|MFGM_HUMAN Lactadherin OS=Homo sapiens OX=9606 GN=MFGE8 PE=1 SV=3 | 1064 | 20201113_12413_1ng_glyco_hcd203040_compound15.4880.4880.3 | controllerType=0 controllerNumber=1 scan=4880 | [illegible] |
| 01866:3 | 15 | K.N[+1540.529]NSIPDKQITASSSYK.T | HexNAc(2)Hex(7) | N[+1541] | 1098.4743 | 3 | 3293.4083 | 3293.4048 | 0 | 1.1 | 238 | Specific | 312.13 | 312.13 | 312.13 | 6.49 | 30 | >sp\|Q08431\|MFGM_HUMAN Lactadherin OS=Homo sapiens OX=9606 GN=MFGE8 PE=1 SV=3 | 1064 | 20201113_12413_1ng_glyco_hcd203040_compound15.4743.4743.3 | controllerType=0 controllerNumber=1 scan=4743 | [illegible] |
| 01864:3 | 15 | K.N[+1540.529]NSIPDKQITASSSYK.T | HexNAc(2)Hex(7) | N[+1541] | 1098.4743 | 3 | 3293.4083 | 3293.4048 | 0 | 1.1 | 238 | Specific | 372.88 | 135.76 | 135.76 | 5.08 | 30 | >sp\|Q08431\|MFGM_HUMAN Lactadherin OS=Homo sapiens OX=9606 GN=MFGE8 PE=1 SV=3 | 1064 | 20201113_12413_1ng_glyco_hcd203040_compound15.4741.4741.3 | controllerType=0 controllerNumber=1 scan=4741 | [illegible] |
| 01984:3 | 15 | K.N[+1216.423]NSIPDKQITASSSYK.T | HexNAc(2)Hex(5) | N[+1216] | 990.4398 | 3 | 2969.305 | 2969.2991 | 0 | 2 | 238 | Specific | 388.39 | 129.32 | 129.32 | 5.1 | 30 | >sp\|Q08431\|MFGM_HUMAN Lactadherin OS=Homo sapiens OX=9606 GN=MFGE8 PE=1 SV=3 | 1064 | 20201113_12413_1ng_glyco_hcd203040_compound15.4865.4865.3 | controllerType=0 controllerNumber=1 scan=4865 | 14.5461 |
| 01986:3 | 15 | K.N[+1216.423]NSIPDKQITASSSYK.T | HexNAc(2)Hex(5) | N[+1216] | 990.4398 | 3 | 2969.305 | 2969.2991 | 0 | 2 | 238 | Specific | 431.74 | 381.53 | 381.53 | 8.67 | 30 | >sp\|Q08431\|MFGM_HUMAN Lactadherin OS=Homo sapiens OX=9606 GN=MFGE8 PE=1 SV=3 | 1064 | 20201113_12413_1ng_glyco_hcd203040_compound15.4867.4867.3 | controllerType=0 controllerNumber=1 scan=4867 | 14.5497 |
| 13871:3 | 16 | K.LKELPGVC[+57.021]N[+2059.735]ETMMALWEEC[+57.021]KPC[+57.021]LK.Q | HexNAc(4)Hex(5)Fuc(1)NeuAc(1) | C[+57]*3, N[+2060] | 1666.0444 | 3 | 4996.1186 | 4996.1196 | 0 | -0.2 | 95 | Specific | 220.23 | 187.69 | 187.69 | 3.21 | 32 | >sp\|P10909\|CLUS_HUMAN Clusterin OS=Homo sapiens OX=9606 GN=CLU PE=1 SV=1 | 1467 | 20201113_12413_1ng_glyco_hcd203040_compound15.17209.17209.3 | controllerType=0 controllerNumber=1 scan=17209 | 35.5203 |
| 13703:4 | 16 | K.LKELPGVC[+57.021]N[+2424.867]ETMMALWEEC[+57.021]KPC[+57.021]LK.Q | HexNAc(5)Hex(6)Fuc(1)NeuAc(1) | C[+57]*3, N[+2425] | 1341.0731 | 4 | 5361.2706 | 5361.2518 | 0 | 3.5 | 95 | Specific | 289.3 | 289.3 | 272.47 | 6.58 | 32 | >sp\|P10909\|CLUS_HUMAN Clusterin OS=Homo sapiens OX=9606 GN=CLU PE=1 SV=1 | 1467 | 20201113_12413_1ng_glyco_hcd203040_compound15.17035.17035.4 | controllerType=0 controllerNumber=1 scan=17035 | 35.2255 |
| 15510:4 | 16 | K.LKELPGVC[+57.021]N[+2351.851]ETMMALWEEC[+57.021]KPC[+57.021]LK.Q | HexNAc(4)Hex(5)Fuc(3)NeuAc(1) | C[+57]*3, N[+2352] | 1322.811 | 4 | 5288.2222 | 5288.2355 | 0 | -2.5 | 95 | Specific | 335.33 | 335.33 | 335.33 | 6.06 | 32 | >sp\|P10909\|CLUS_HUMAN Clusterin OS=Homo sapiens OX=9606 GN=CLU PE=1 SV=1 | 1467 | 20201113_12413_1ng_glyco_hcd203040_compound15.18909.18909.4 | controllerType=0 controllerNumber=1 scan=18909 | 38.6067 |
| 13869:3 | 16 | K.LKELPGVC[+57.021]N[+2059.735]ETMMALWEEC[+57.021]KPC[+57.021]LK.Q | HexNAc(4)Hex(5)Fuc(1)NeuAc(1) | C[+57]*3, N[+2060] | 1666.0444 | 3 | 4996.1186 | 4996.1196 | 0 | -0.2 | 95 | Specific | 375.17 | 375.17 | 375.17 | 8.06 | 32 | >sp\|P10909\|CLUS_HUMAN Clusterin OS=Homo sapiens OX=9606 GN=CLU PE=1 SV=1 | 1467 | 20201113_12413_1ng_glyco_hcd203040_compound15.17207.17207.3 | controllerType=0 controllerNumber=1 scan=17207 | 35.5168 |
| 13773:4 | 16 | K.LKELPGVC[+57.021]N[+2204.772]ETMMALWEEC[+57.021]KPC[+57.021]LK.Q | HexNAc(4)Hex(5)NeuAc(2) | C[+57]*3, N[+2205] | 1286.0529 | 4 | 5141.1896 | 5141.1572 | 0 | 6.3 | 95 | Specific | 688.34 | 502.64 | 502.64 | 12.14 | 32 | >sp\|P10909\|CLUS_HUMAN Clusterin OS=Homo sapiens OX=9606 GN=CLU PE=1 SV=1 | 1467 | 20201113_12413_1ng_glyco_hcd203040_compound15.17108.17108.4 | controllerType=0 controllerNumber=1 scan=17108 | 35.3466 |
| 13771:4 | 16 | K.LKELPGVC[+57.021]N[+2204.772]ETMMALWEEC[+57.021]KPC[+57.021]LK.Q | HexNAc(4)Hex(5)NeuAc(2) | C[+57]*3, N[+2205] | 1286.0529 | 4 | 5141.1896 | 5141.1572 | 0 | 6.3 | 95 | Specific | 728.2 | 691.73 | 691.73 | 13.14 | 32 | >sp\|P10909\|CLUS_HUMAN Clusterin OS=Homo sapiens OX=9606 GN=CLU PE=1 SV=1 | 1467 | 20201113_12413_1ng_glyco_hcd203040_compound15.17106.17106.4 | controllerType=0 controllerNumber=1 scan=17106 | 35.342 |
| 13855:4 | 16 | K.LKELPGVC[+57.021]N[+2059.735]ETMMALWEEC[+57.021]KPC[+57.021]LK.Q | HexNAc(4)Hex(5)Fuc(1)NeuAc(1) | C[+57]*3, N[+2060] | 1249.7876 | 4 | 4996.1286 | 4996.1196 | 0 | 1.8 | 95 | Specific | 846.47 | 764.24 | 764.24 | 15.83 | 32 | >sp\|P10909\|CLUS_HUMAN Clusterin OS=Homo sapiens OX=9606 GN=CLU PE=1 SV=1 | 1467 | 20201113_12413_1ng_glyco_hcd203040_compound15.17193.17193.4 | controllerType=0 controllerNumber=1 scan=17193 | 35.4963 |
| 13853:4 | 16 | K.LKELPGVC[+57.021]N[+2059.735]ETMMALWEEC[+57.021]KPC[+57.021]LK.Q | HexNAc(4)Hex(5)Fuc(1)NeuAc(1) | C[+57]*3, N[+2060] | 1249.7876 | 4 | 4996.1286 | 4996.1196 | 0 | 1.8 | 95 | Specific | 864.74 | 829.49 | 829.49 | 16.83 | 32 | >sp\|P10909\|CLUS_HUMAN Clusterin OS=Homo sapiens OX=9606 GN=CLU PE=1 SV=1 | 1467 | 20201113_12413_1ng_glyco_hcd203040_compound15.17191.17191.4 | controllerType=0 controllerNumber=1 scan=17191 | 35.4921 |
| 17512:3 | 16 | K.ELPGVC[+57.021]N[+2350.830]ETMMALWEEC[+57.021]KPC[+57.021]LK.Q | HexNAc(4)Hex(5)Fuc(1)NeuAc(2) | C[+57]*3, N[+2351] | 1682.6886 | 3 | 5046.0511 | 5046.036 | 0 | 3 | 97 | Specific | 236.48 | 220.48 | 220.48 | 5.13 | 32 | >sp\|P10909\|CLUS_HUMAN Clusterin OS=Homo sapiens OX=9606 GN=CLU PE=1 SV=1 | 1467 | 20201113_12413_1ng_glyco_hcd203040_compound15.21023.21023.3 | controllerType=0 controllerNumber=1 scan=21023 | 42.7392 |
| 20878:3 | 16 | K.MLN[+1913.677]TSSLLEQLNEQFNWVSR.L | HexNAc(4)Hex(5)NeuAc(1) | N[+1914] | 1441.6314 | 3 | 4322.8797 | 4322.8637 | 0 | 3.7 | 352 | Specific | 266.75 | 266.75 | 266.75 | 6.6 | 32 | >sp\|P10909\|CLUS_HUMAN Clusterin OS=Homo sapiens OX=9606 GN=CLU PE=1 SV=1 | 1467 | 20201113_12413_1ng_glyco_hcd203040_compound15.24542.24542.3 | controllerType=0 controllerNumber=1 scan=24542 | 50.3745 |
| 20571:3 | 16 | K.MLN[+2059.735]TSSLLEQLNEQFNWVSR.L | HexNAc(4)Hex(5)Fuc(1)NeuAc(1) | N[+2060] | 1490.3207 | 3 | 4468.9477 | 4468.9217 | 0 | 5.8 | 352 | Specific | 560.99 | 551.07 | 551.07 | 9.14 | 32 | >sp\|P10909\|CLUS_HUMAN Clusterin OS=Homo sapiens OX=9606 GN=CLU PE=1 SV=1 | 1467 | 20201113_12413_1ng_glyco_hcd203040_compound15.24222.24222.3 | controllerType=0 controllerNumber=1 scan=24222 | 49.7275 |
| 20881:3 | 16 | K.MLN[+1913.677]TSSLLEQLNEQFNWVSR.L | HexNAc(4)Hex(5)NeuAc(1) | N[+1914] | 1441.6314 | 3 | 4322.8797 | 4322.8637 | 0 | 3.7 | 352 | Specific | 563.15 | 521.78 | 521.78 | 9.77 | 32 | >sp\|P10909\|CLUS_HUMAN Clusterin OS=Homo sapiens OX=9606 GN=CLU PE=1 SV=1 | 1467 | 20201113_12413_1ng_glyco_hcd203040_compound15.24545.24545.3 | controllerType=0 controllerNumber=1 scan=24545 | 50.3826 |
| 20573:3 | 16 | K.MLN[+2059.735]TSSLLEQLNEQFNWVSR.L | HexNAc(4)Hex(5)Fuc(1)NeuAc(1) | N[+2060] | 1490.3207 | 3 | 4468.9477 | 4468.9217 | 0 | 5.8 | 352 | Specific | 580.63 | 567.96 | 567.96 | 10.36 | 32 | >sp\|P10909\|CLUS_HUMAN Clusterin OS=Homo sapiens OX=9606 GN=CLU PE=1 SV=1 | 1467 | 20201113_12413_1ng_glyco_hcd203040_compound15.24224.24224.3 | controllerType=0 controllerNumber=1 scan=24224 | 49.733 |
| 20602:3 | 16 | K.MLN[+2059.735]TSSLLEQLNEQFNWVSR.L | HexNAc(4)Hex(5)Fuc(1)NeuAc(1) | N[+2060] | 1490.3234 | 3 | 4468.9557 | 4468.9217 | 0 | 7.6 | 352 | Specific | 629.61 | 419.29 | 419.29 | 11.26 | 32 | >sp\|P10909\|CLUS_HUMAN Clusterin OS=Homo sapiens OX=9606 GN=CLU PE=1 SV=1 | 1467 | 20201113_12413_1ng_glyco_hcd203040_compound15.24255.24255.3 | controllerType=0 controllerNumber=1 scan=24255 | 49.8081 |
| 22963:3 | 16 | K.MLN[+2350.830]TSSLLEQLNEQFNWVSR.L | HexNAc(4)Hex(5)Fuc(1)NeuAc(2) | N[+2351] | 1587.3484 | 3 | 4760.0308 | 4760.0171 | 0 | 2.9 | 352 | Specific | 717.73 | 475.46 | 475.46 | 14.12 | 32 | >sp\|P10909\|CLUS_HUMAN Clusterin OS=Homo sapiens OX=9606 GN=CLU PE=1 SV=1 | 1467 | 20201113_12413_1ng_glyco_hcd203040_compound15.26715.26715.3 | controllerType=0 controllerNumber=1 scan=26715 | 54.3996 |
| 22928:3 | 16 | K.MLN[+2350.830]TSSLLEQLNEQFNWVSR.L | HexNAc(4)Hex(5)Fuc(1)NeuAc(2) | N[+2351] | 1587.351 | 3 | 4760.0385 | 4760.0171 | 0 | 4.5 | 352 | Specific | 755.85 | 543.24 | 543.24 | 14.99 | 32 | >sp\|P10909\|CLUS_HUMAN Clusterin OS=Homo sapiens OX=9606 GN=CLU PE=1 SV=1 | 1467 | 20201113_12413_1ng_glyco_hcd203040_compound15.26677.26677.3 | controllerType=0 controllerNumber=1 scan=26677 | 54.3262 |
| 22965:3 | 16 | K.MLN[+2350.830]TSSLLEQLNEQFNWVSR.L | HexNAc(4)Hex(5)Fuc(1)NeuAc(2) | N[+2351] | 1587.3484 | 3 | 4760.0308 | 4760.0171 | 0 | 2.9 | 352 | Specific | 818.26 | 555.66 | 555.66 | 16.12 | 32 | >sp\|P10909\|CLUS_HUMAN Clusterin OS=Homo sapiens OX=9606 GN=CLU PE=1 SV=1 | 1467 | 20201113_12413_1ng_glyco_hcd203040_compound15.26717.26717.3 | controllerType=0 controllerNumber=1 scan=26717 | 54.4034 |
| 22930:3 | 16 | K.MLN[+2350.830]TSSLLEQLNEQFNWVSR.L | HexNAc(4)Hex(5)Fuc(1)NeuAc(2) | N[+2351] | 1587.351 | 3 | 4760.0385 | 4760.0171 | 0 | 4.5 | 352 | Specific | 895.26 | 642.9 | 642.9 | 16.99 | 32 | >sp\|P10909\|CLUS_HUMAN Clusterin OS=Homo sapiens OX=9606 GN=CLU PE=1 SV=1 | 1467 | 20201113_12413_1ng_glyco_hcd203040_compound15.26679.26679.3 | controllerType=0 controllerNumber=1 scan=26679 | 54.3302 |
| 20605:3 | 16 | K.MLN[+2059.735]TSSLLEQLNEQFNWVSR.L | HexNAc(4)Hex(5)Fuc(1)NeuAc(1) | N[+2060] | 1490.3234 | 3 | 4468.9557 | 4468.9217 | 0 | 7.6 | 352 | Specific | 1003.97 | 746.49 | 746.49 | 19.26 | 32 | >sp\|P10909\|CLUS_HUMAN Clusterin OS=Homo sapiens OX=9606 GN=CLU PE=1 SV=1 | 1467 | 20201113_12413_1ng_glyco_hcd203040_compound15.24258.24258.3 | controllerType=0 controllerNumber=1 scan=24258 | 49.8122 |
| 10061:3 | 16 | R.LAN[+2205.793]LTQGEDQYYLR.V | HexNAc(4)Hex(5)Fuc(2)NeuAc(1) | N[+2206] | 1297.2164 | 3 | 3889.6345 | 3889.6265 | 0 | 2.1 | 372 | Specific | 251.34 | 251.34 | 251.34 | 6.29 | 32 | >sp\|P10909\|CLUS_HUMAN Clusterin OS=Homo sapiens OX=9606 GN=CLU PE=1 SV=1 | 1467 | 20201113_12413_1ng_glyco_hcd203040_compound15.13244.13244.3 | controllerType=0 controllerNumber=1 scan=13244 | 28.7369 |
| 10063:3 | 16 | R.LAN[+2205.793]LTQGEDQYYLR.V | HexNAc(4)Hex(5)Fuc(2)NeuAc(1) | N[+2206] | 1297.2164 | 3 | 3889.6345 | 3889.6265 | 0 | 2.1 | 372 | Specific | 316.96 | 316.96 | 316.96 | 6.95 | 32 | >sp\|P10909\|CLUS_HUMAN Clusterin OS=Homo sapiens OX=9606 GN=CLU PE=1 SV=1 | 1467 | 20201113_12413_1ng_glyco_hcd203040_compound15.13246.13246.3 | controllerType=0 controllerNumber=1 scan=13246 | 28.7424 |
| 15519:3 | 18 | R.[+42.011]TLN[+2018.708]QSSDELQLSMGNAMFVK.E | HexNAc(3)Hex(6)Fuc(1)NeuAc(1) | .[42], N[+2019] | 1425.275 | 3 | 4273.8104 | 4273.7766 | 0 | 7.9 | 125 | Specific | 375.61 | 251.17 | 251.17 | 6.35 | 31 | >sp\|P01011\|AACT_HUMAN Alpha-1-antichymotrypsin OS=Homo sapiens OX=9606 GN=SERPINA3 PE=1 SV=2 | 1314 | 20201113_12413_1ng_glyco_hcd203040_compound15.18918.18918.3 | controllerType=0 controllerNumber=1 scan=18918 | 38.6224 |
| 15517:3 | 18 | R.[+42.011]TLN[+2018.708]QSSDELQLSMGNAMFVK.E | HexNAc(3)Hex(6)Fuc(1)NeuAc(1) | .[42], N[+2019] | 1425.275 | 3 | 4273.8104 | 4273.7766 | 0 | 7.9 | 125 | Specific | 419.6 | 369.2 | 369.2 | 6.99 | 31 | >sp\|P01011\|AACT_HUMAN Alpha-1-antichymotrypsin OS=Homo sapiens OX=9606 GN=SERPINA3 PE=1 SV=2 | 1314 | 20201113_12413_1ng_glyco_hcd203040_compound15.18916.18916.3 | controllerType=0 controllerNumber=1 scan=18916 | 38.6188 |
| 01969:4 | 19 | R.EKQTDEIKDTRN[+2059.735]ESTQNC[+57.021]VVAEPEK.M | HexNAc(4)Hex(5)Fuc(1)NeuAc(1) | C[+57], N[+2060] | 1252.79 | 4 | 5008.1383 | 5008.1251 | 0 | 2.6 | 58 | Specific | 319.98 | 248.77 | 248.77 | 5.49 | 30 | >sp\|P47710\|CASA1_HUMAN Alpha-S1-casein OS=Homo sapiens OX=9606 GN=CSN1S1 PE=1 SV=1 | 989 | 20201113_12413_1ng_glyco_hcd203040_compound15.4850.4850.4 | controllerType=0 controllerNumber=1 scan=4850 | 14.5211 |
| 01967:4 | 19 | R.EKQTDEIKDTRN[+2059.735]ESTQNC[+57.021]VVAEPEK.M | HexNAc(4)Hex(5)Fuc(1)NeuAc(1) | C[+57], N[+2060] | 1252.79 | 4 | 5008.1383 | 5008.1251 | 0 | 2.6 | 58 | Specific | 495.84 | 377.54 | 377.54 | 8.15 | 30 | >sp\|P47710\|CASA1_HUMAN Alpha-S1-casein OS=Homo sapiens OX=9606 GN=CSN1S1 PE=1 SV=1 | 989 | 20201113_12413_1ng_glyco_hcd203040_compound15.4848.4848.4 | controllerType=0 controllerNumber=1 scan=4848 | 14.5169 |
| 23264:4 | 20 | K.VFPLSLDSTPQDGNVVVAC[+57.021]LVQGFFPQEPLSVTWSESGQN[+2262.814]VTAI | HexNAc(5)Hex(5)Fuc(1)NeuAc(1) | C[+57], N[+2263] | 1760.5655 | 4 | 7039.2402 | 7039.1757 | 0 | 9.2 | 8 | Specific | 327.39 | 310.99 | 310.99 | 6.44 | 30 | >sp\|P01877\|IGHA2_HUMAN Immunoglobulin heavy constant alpha 2 OS=Homo sapiens OX=9606 GN=IGHA2 PE=1 SV=4 | 576 | 20201113_12413_1ng_glyco_hcd203040_compound15.27029.27029.2 | controllerType=0 controllerNumber=1 scan=27029 | 54.9774 |
| 23406:4 | 20 | K.VFPLSLDSTPQDGNVVVAC[+57.021]LVQGFFPQEPLSVTWSESGQN[+2262.814]VTAI | HexNAc(5)Hex(5)Fuc(1)NeuAc(1) | C[+57], N[+2263] | 1760.5515 | 4 | 7039.1841 | 7039.1757 | 0 | 1.2 | 8 | Specific | 475.36 | 422.92 | 422.92 | 9.72 | 30 | >sp\|P01877\|IGHA2_HUMAN Immunoglobulin heavy constant alpha 2 OS=Homo sapiens OX=9606 GN=IGHA2 PE=1 SV=4 | 576 | 20201113_12413_1ng_glyco_hcd203040_compound15.27177.27177.4 | controllerType=0 controllerNumber=1 scan=27177 | 55.2741 |
| 23330:4 | 20 | K.VFPLSLDSTPQDGNVVVAC[+57.021]LVQGFFPQEPLSVTWSESGQN[+2262.814]VTAI | HexNAc(5)Hex(5)Fuc(1)NeuAc(1) | C[+57], N[+2263] | 1760.5557 | 4 | 7039.2012 | 7039.1757 | 0 | 3.6 | 8 | Specific | 624.48 | 598.02 | 598.02 | 12.3 | 30 | >sp\|P01877\|IGHA2_HUMAN Immunoglobulin heavy constant alpha 2 OS=Homo sapiens OX=9606 GN=IGHA2 PE=1 SV=4 | 576 | 20201113_12413_1ng_glyco_hcd203040_compound15.27098.27098.4 | controllerType=0 controllerNumber=1 scan=27098 | 55.116 |
| 23404:4 | 20 | K.VFPLSLDSTPQDGNVVVAC[+57.021]LVQGFFPQEPLSVTWSESGQN[+2262.814]VTAI | HexNAc(5)Hex(5)Fuc(1)NeuAc(1) | C[+57], N[+2263] | 1760.5515 | 4 | 7039.1841 | 7039.1757 | 0 | 1.2 | 8 | Specific | 657.44 | 612.82 | 612.82 | 13.34 | 30 | >sp\|P01877\|IGHA2_HUMAN Immunoglobulin heavy constant alpha 2 OS=Homo sapiens OX=9606 GN=IGHA2 PE=1 SV=4 | 576 | 20201113_12413_1ng_glyco_hcd203040_compound15.27175.27175.4 | controllerType=0 controllerNumber=1 scan=27175 | 55.2687 |
| 01098:3 | 20 | K.HYTN[+1971.719]SSQDVTVPC[+57.021]R.V | HexNAc(5)Hex(5)Fuc(1) | C[+57], N[+1972] | 1212.494 | 3 | 3635.4673 | 3635.4682 | 0 | -0.2 | 89 | Specific | 324.31 | 161.78 | 161.78 | 5.52 | 30 | >sp\|P01877\|IGHA2_HUMAN Immunoglobulin heavy constant alpha 2 OS=Homo sapiens OX=9606 GN=IGHA2 PE=1 SV=4 | 576 | 20201113_12413_1ng_glyco_hcd203040_compound15.3943.3943.3 | controllerType=0 controllerNumber=1 scan=3943 | 12.8791 |
| 01096:3 | 20 | K.HYTN[+1971.719]SSQDVTVPC[+57.021]R.V | HexNAc(5)Hex(5)Fuc(1) | C[+57], N[+1972] | 1212.494 | 3 | 3635.4673 | 3635.4682 | 0 | -0.2 | 89 | Specific | 325.38 | 212.77 | 212.77 | 6 | 30 | >sp\|P01877\|IGHA2_HUMAN Immunoglobulin heavy constant alpha 2 OS=Homo sapiens OX=9606 GN=IGHA2 PE=1 SV=4 | 576 | 20201113_12413_1ng_glyco_hcd203040_compound15.3941.3941.3 | controllerType=0 controllerNumber=1 scan=3941 | 12.8746 |
| 01073:3 | 20 | K.HYTN[+1809.666]SSQDVTVPC[+57.021]R.V | HexNAc(5)Hex(4)Fuc(1) | C[+57], N[+1810] | 1158.48 | 3 | 3473.4256 | 3473.4154 | 0 | 2.9 | 89 | Specific | 433.17 | 344.23 | 344.23 | 7.65 | 30 | >sp\|P01877\|IGHA2_HUMAN Immunoglobulin heavy constant alpha 2 OS=Homo sapiens OX=9606 GN=IGHA2 PE=1 SV=4 | 576 | 20201113_12413_1ng_glyco_hcd203040_compound15.3917.3917.3 | controllerType=0 controllerNumber=1 scan=3917 | 12.8317 |
| 01101:3 | 20 | K.HYTN[+1647.613]SSQDVTVPC[+57.021]R.V | HexNAc(5)Hex(3)Fuc(1) | C[+57], N[+1648] | 1104.4597 | 3 | 3311.3647 | 3311.3625 | 0 | 0.6 | 89 | Specific | 519.19 | 519.19 | 519.19 | 9.21 | 30 | >sp\|P01877\|IGHA2_HUMAN Immunoglobulin heavy constant alpha 2 OS=Homo sapiens OX=9606 GN=IGHA2 PE=1 SV=4 | 576 | 20201113_12413_1ng_glyco_hcd203040_compound15.3946.3946.3 | controllerType=0 controllerNumber=1 scan=3946 | 12.884 |
| 01103:3 | 20 | K.HYTN[+1647.613]SSQDVTVPC[+57.021]R.V | HexNAc(5)Hex(3)Fuc(1) | C[+57], N[+1648] | 1104.4597 | 3 | 3311.3647 | 3311.3625 | 0 | 0.6 | 89 | Specific | 580.49 | 467.66 | 467.66 | 10.74 | 30 | >sp\|P01877\|IGHA2_HUMAN Immunoglobulin heavy constant alpha 2 OS=Homo sapiens OX=9606 GN=IGHA2 PE=1 SV=4 | 576 | 20201113_12413_1ng_glyco_hcd203040_compound15.3948.3948.3 | controllerType=0 controllerNumber=1 scan=3948 | 12.8899 |
| 01075:3 | 20 | K.HYTN[+1809.666]SSQDVTVPC[+57.021]R.V | HexNAc(5)Hex(4)Fuc(1) | C[+57], N[+1810] | 1158.48 | 3 | 3473.4256 | 3473.4154 | 0 | 2.9 | 89 | Specific | 606.24 | 428.97 | 428.97 | 12.02 | 30 | >sp\|P01877\|IGHA2_HUMAN Immunoglobulin heavy constant alpha 2 OS=Homo sapiens OX=9606 GN=IGHA2 PE=1 SV=4 | 576 | 20201113_12413_1ng_glyco_hcd203040_compound15.3919.3919.3 | controllerType=0 controllerNumber=1 scan=3919 | 12.8367 |
| 03131:4 | 20 | K.HYTN[+1647.613]SSQDVTVPC[+57.021]RVPPPPPC[+57.021]C[+57.021]HPR.L | HexNAc(5)Hex(3)Fuc(1) | C[+57]*3, N[+1648] | 1152.2499 | 4 | 4605.9779 | 4605.9689 | 0 | 2 | 89 | Specific | 218.85 | 196.22 | 196.22 | 4.02 | 30 | >sp\|P01877\|IGHA2_HUMAN Immunoglobulin heavy constant alpha 2 OS=Homo sapiens OX=9606 GN=IGHA2 PE=1 SV=4 | 576 | 20201113_12413_1ng_glyco_hcd203040_compound15.6053.6053.4 | controllerType=0 controllerNumber=1 scan=6053 | 16.6516 |
| 03055:4 | 20 | K.HYTN[+1647.613]SSQDVTVPC[+57.021]RVPPPPPC[+57.021]C[+57.021]HPR.L | HexNAc(5)Hex(3)Fuc(1) | C[+57]*3, N[+1648] | 1152.2504 | 4 | 4605.9798 | 4605.9689 | 0 | 2.4 | 89 | Specific | 314.93 | 293.47 | 293.47 | 6.2 | 30 | >sp\|P01877\|IGHA2_HUMAN Immunoglobulin heavy constant alpha 2 OS=Homo sapiens OX=9606 GN=IGHA2 PE=1 SV=4 | 576 | 20201113_12413_1ng_glyco_hcd203040_compound15.5974.5974.4 | controllerType=0 controllerNumber=1 scan=5974 | 16.515 |
| 03053:4 | 20 | K.HYTN[+1647.613]SSQDVTVPC[+57.021]RVPPPPPC[+57.021]C[+57.021]HPR.L | HexNAc(5)Hex(3)Fuc(1) | C[+57]*3, N[+1648] | 1152.2504 | 4 | 4605.9798 | 4605.9689 | 0 | 2.4 | 89 | Specific | 323.55 | 323.55 | 323.55 | 6.53 | 30 | >sp\|P01877\|IGHA2_HUMAN Immunoglobulin heavy constant alpha 2 OS=Homo sapiens OX=9606 GN=IGHA2 PE=1 SV=4 | 576 | 20201113_12413_1ng_glyco_hcd203040_compound15.5972.5972.4 | controllerType=0 controllerNumber=1 scan=5972 | 16.5109 |
| 03128:4 | 20 | K.HYTN[+1647.613]SSQDVTVPC[+57.021]RVPPPPPC[+57.021]C[+57.021]HPR.L | HexNAc(5)Hex(3)Fuc(1) | C[+57]*3, N[+1648] | 1152.2499 | 4 | 4605.9779 | 4605.9689 | 0 | 2 | 89 | Specific | 381.64 | 334.53 | 334.53 | 6.7 | 30 | >sp\|P01877\|IGHA2_HUMAN Immunoglobulin heavy constant alpha 2 OS=Homo sapiens OX=9606 GN=IGHA2 PE=1 SV=4 | 576 | 20201113_12413_1ng_glyco_hcd203040_compound15.6050.6050.4 | controllerType=0 controllerNumber=1 scan=6050 | 16.6468 |
| 03124:5 | 20 | K.HYTN[+1647.613]SSQDVTVPC[+57.021]RVPPPPPC[+57.021]C[+57.021]HPR.L | HexNAc(5)Hex(3)Fuc(1) | C[+57]*3, N[+1648] | 922.0021 | 5 | 4605.9814 | 4605.9689 | 0 | 2.7 | 89 | Specific | 408.36 | 307.2 | 307.2 | 7.21 | 30 | >sp\|P01877\|IGHA2_HUMAN Immunoglobulin heavy constant alpha 2 OS=Homo sapiens OX=9606 GN=IGHA2 PE=1 SV=4 | 576 | 20201113_12413_1ng_glyco_hcd203040_compound15.6046.6046.5 | controllerType=0 controllerNumber=1 scan=6046 | 16.6407 |
| 03122:5 | 20 | K.HYTN[+1647.613]SSQDVTVPC[+57.021]RVPPPPPC[+57.021]C[+57.021]HPR.L | HexNAc(5)Hex(3)Fuc(1) | C[+57]*3, N[+1648] | 922.0021 | 5 | 4605.9814 | 4605.9689 | 0 | 2.7 | 89 | Specific | 421.77 | 310.37 | 310.37 | 7.37 | 30 | >sp\|P01877\|IGHA2_HUMAN Immunoglobulin heavy constant alpha 2 OS=Homo sapiens OX=9606 GN=IGHA2 PE=1 SV=4 | 576 | 20201113_12413_1ng_glyco_hcd203040_compound15.6044.6044.5 | controllerType=0 controllerNumber=1 scan=6044 | 16.637 |
| 04531:2 | 20 | K.TPLTAN[+2059.735]ITK.S | HexNAc(4)Hex(5)Fuc(1)NeuAc(1) | N[+2060] | 1509.653 | 2 | 3018.2987 | 3018.2917 | 0 | 2.3 | 200 | Specific | 224.8 | 154.18 | 154.18 | 3.97 | 30 | >sp\|P01877\|IGHA2_HUMAN Immunoglobulin heavy constant alpha 2 OS=Homo sapiens OX=9606 GN=IGHA2 PE=1 SV=4 | 576 | 20201113_12413_1ng_glyco_hcd203040_compound15.7508.7508.2 | controllerType=0 controllerNumber=1 scan=7508 | 19.1097 |
| 02785:3 | 20 | K.TPLTAN[+1809.666]ITK.S | HexNAc(5)Hex(4)Fuc(1) | N[+1810] | 923.4155 | 3 | 2768.2319 | 2768.2228 | 0 | 3.3 | 200 | Specific | 227.34 | 60.88 | 60.88 | 3.72 | 30 | >sp\|P01877\|IGHA2_HUMAN Immunoglobulin heavy constant alpha 2 OS=Homo sapiens OX=9606 GN=IGHA2 PE=1 SV=4 | 576 | 20201113_12413_1ng_glyco_hcd203040_compound15.5695.5695.3 | controllerType=0 controllerNumber=1 scan=5695 | 16.0227 |
| 02727:3 | 20 | K.TPLTAN[+1971.719]ITK.S | HexNAc(5)Hex(5)Fuc(1) | N[+1972] | 977.4309 | 3 | 2930.2781 | 2930.2757 | 0 | 0.8 | 200 | Specific | 243.85 | 66.89 | 66.89 | 3.93 | 30 | >sp\|P01877\|IGHA2_HUMAN Immunoglobulin heavy constant alpha 2 OS=Homo sapiens OX=9606 GN=IGHA2 PE=1 SV=4 | 576 | 20201113_12413_1ng_glyco_hcd203040_compound15.5635.5635.3 | controllerType=0 controllerNumber=1 scan=5635 | 15.9137 |
| 02730:2 | 20 | K.TPLTAN[+1971.719]ITK.S | HexNAc(5)Hex(5)Fuc(1) | N[+1972] | 1465.6416 | 2 | 2930.276 | 2930.2757 | 0 | 0.1 | 200 | Specific | 268.09 | 95.95 | 95.95 | 4.25 | 30 | >sp\|P01877\|IGHA2_HUMAN Immunoglobulin heavy constant alpha 2 OS=Homo sapiens OX=9606 GN=IGHA2 PE=1 SV=4 | 576 | 20201113_12413_1ng_glyco_hcd203040_compound15.5638.5638.2 | controllerType=0 controllerNumber=1 scan=5638 | 15.9185 |
| 02732:2 | 20 | K.TPLTAN[+1971.719]ITK.S | HexNAc(5)Hex(5)Fuc(1) | N[+1972] | 1465.6416 | 2 | 2930.276 | 2930.2757 | 0 | 0.1 | 200 | Specific | 312.16 | 239.92 | 239.92 | 6.01 | 30 | >sp\|P01877\|IGHA2_HUMAN Immunoglobulin heavy constant alpha 2 OS=Homo sapiens OX=9606 GN=IGHA2 PE=1 SV=4 | 576 | 20201113_12413_1ng_glyco_hcd203040_compound15.5640.5640.2 | controllerType=0 controllerNumber=1 scan=5640 | 15.9224 |
| 02803:2 | 20 | K.TPLTAN[+1444.534]ITK.S | HexNAc(4)Hex(3)Fuc(1) | N[+1445] | 1202.0525 | 2 | 2403.0978 | 2403.0906 | 0 | 3 | 200 | Specific | 349.53 | 160.46 | 160.46 | 5.6 | 30 | >sp\|P01877\|IGHA2_HUMAN Immunoglobulin heavy constant alpha 2 OS=Homo sapiens OX=9606 GN=IGHA2 PE=1 SV=4 | 576 | 20201113_12413_1ng_glyco_hcd203040_compound15.5713.5713.2 | controllerType=0 controllerNumber=1 scan=5713 | 16.0553 |
| 02864:3 | 20 | K.TPLTAN[+1647.613]ITK.S | HexNAc(5)Hex(3)Fuc(1) | N[+1648] | 869.3962 | 3 | 2606.1741 | 2606.17 | 0 | 1.6 | 200 | Specific | 371.79 | 173.2 | 173.2 | 5.97 | 30 | >sp\|P01877\|IGHA2_HUMAN Immunoglobulin heavy constant alpha 2 OS=Homo sapiens OX=9606 GN=IGHA2 PE=1 SV=4 | 576 | 20201113_12413_1ng_glyco_hcd203040_compound15.5777.5777.3 | controllerType=0 controllerNumber=1 scan=5777 | 16.1728 |
| 02804:2 | 20 | K.TPLTAN[+1444.534]ITK.S | HexNAc(4)Hex(3)Fuc(1) | N[+1445] | 1202.0525 | 2 | 2403.0978 | 2403.0906 | 0 | 3 | 200 | Specific | 435.51 | 232.55 | 232.55 | 6.74 | 30 | >sp\|P01877\|IGHA2_HUMAN Immunoglobulin heavy constant alpha 2 OS=Homo sapiens OX=9606 GN=IGHA2 PE=1 SV=4 | 576 | 20201113_12413_1ng_glyco_hcd203040_compound15.5714.5714.2 | controllerType=0 controllerNumber=1 scan=5714 | 16.0591 |
| 02787:3 | 20 | K.TPLTAN[+1809.666]ITK.S | HexNAc(5)Hex(4)Fuc(1) | N[+1810] | 923.4155 | 3 | 2768.2319 | 2768.2228 | 0 | 3.3 | 200 | Specific | 469.53 | 213.96 | 213.96 | 6.64 | 30 | >sp\|P01877\|IGHA2_HUMAN Immunoglobulin heavy constant alpha 2 OS=Homo sapiens OX=9606 GN=IGHA2 PE=1 SV=4 | 576 | 20201113_12413_1ng_glyco_hcd203040_compound15.5697.5697.3 | controllerType=0 controllerNumber=1 scan=5697 | 16.0265 |
| 02866:3 | 20 | K.TPLTAN[+1647.613]ITK.S | HexNAc(5)Hex(3)Fuc(1) | N[+1648] | 869.3962 | 3 | 2606.1741 | 2606.17 | 0 | 1.6 | 200 | Specific | 476.37 | 241.4 | 241.4 | 6.88 | 30 | >sp\|P01877\|IGHA2_HUMAN Immunoglobulin heavy constant alpha 2 OS=Homo sapiens OX=9606 GN=IGHA2 PE=1 SV=4 | 576 | 20201113_12413_1ng_glyco_hcd203040_compound15.5779.5779.3 | controllerType=0 controllerNumber=1 scan=5779 | 16.1767 |
| 02839:2 | 20 | K.TPLTAN[+1647.613]ITK.S | HexNAc(5)Hex(3)Fuc(1) | N[+1648] | 1303.5928 | 2 | 2606.1784 | 2606.17 | 0 | 3.2 | 200 | Specific | 597.19 | 318.63 | 318.63 | 8.96 | 30 | >sp\|P01877\|IGHA2_HUMAN Immunoglobulin heavy constant alpha 2 OS=Homo sapiens OX=9606 GN=IGHA2 PE=1 SV=4 | 576 | 20201113_12413_1ng_glyco_hcd203040_compound15.5751.5751.2 | controllerType=0 controllerNumber=1 scan=5751 | 16.1277 |
| 02784:2 | 20 | K.TPLTAN[+1809.666]ITK.S | HexNAc(5)Hex(4)Fuc(1) | N[+1810] | 1384.6185 | 2 | 2768.2296 | 2768.2228 | 0 | 2.5 | 200 | Specific | 605.71 | 335.09 | 335.09 | 10.14 | 30 | >sp\|P01877\|IGHA2_HUMAN Immunoglobulin heavy constant alpha 2 OS=Homo sapiens OX=9606 GN=IGHA2 PE=1 SV=4 | 576 | 20201113_12413_1ng_glyco_hcd203040_compound15.5694.5694.2 | controllerType=0 controllerNumber=1 scan=5694 | 16.0208 |
| 02837:2 | 20 | K.TPLTAN[+1647.613]ITK.S | HexNAc(5)Hex(3)Fuc(1) | N[+1648] | 1303.5928 | 2 | 2606.1784 | 2606.17 | 0 | 3.2 | 200 | Specific | 618.42 | 360.1 | 360.1 | 10.25 | 30 | >sp\|P01877\|IGHA2_HUMAN Immunoglobulin heavy constant alpha 2 OS=Homo sapiens OX=9606 GN=IGHA2 PE=1 SV=4 | 576 | 20201113_12413_1ng_glyco_hcd203040_compound15.5749.5749.2 | controllerType=0 controllerNumber=1 scan=5749 | 16.1227 |
| 02782:2 | 20 | K.TPLTAN[+1809.666]ITK.S | HexNAc(5)Hex(4)Fuc(1) | N[+1810] | 1384.6185 | 2 | 2768.2296 | 2768.2228 | 0 | 2.5 | 200 | Specific | 629.56 | 363.7 | 363.7 | 10.25 | 30 | >sp\|P01877\|IGHA2_HUMAN Immunoglobulin heavy constant alpha 2 OS=Homo sapiens OX=9606 GN=IGHA2 PE=1 SV=4 | 576 | 20201113_12413_1ng_glyco_hcd203040_compound15.5692.5692.2 | controllerType=0 controllerNumber=1 scan=5692 | 16.0165 |
| 11698:3 | 20 | R.MAGKPTHIN[+1913.677]VSVVMAEADGTC[+57.021]Y.- | HexNAc(4)Hex(5)NeuAc(1) | C[+57], N[+1914] | 1422.2618 | 3 | 4264.7708 | 4264.7599 | 0 | 2.6 | 319 | CRagged | 216.64 | 216.64 | 216.64 | 4.81 | 30 | >sp\|P01877\|IGHA2_HUMAN Immunoglobulin heavy constant alpha 2 OS=Homo sapiens OX=9606 GN=IGHA2 PE=1 SV=4 | 576 | 20201113_12413_1ng_glyco_hcd203040_compound15.14942.14942.3 | controllerType=0 controllerNumber=1 scan=14942 | 31.6464 |
| 10009:3 | 20 | R.MAGKPTHIN[+1540.529]VSVVMAEADGTC[+57.021]Y.- | HexNAc(2)Hex(7) | C[+57], N[+1541] | 1297.8782 | 3 | 3891.6201 | 3891.6114 | 0 | 2.3 | 319 | CRagged | 599.02 | 573.71 | 375.62 | 11.1 | 30 | >sp\|P01877\|IGHA2_HUMAN Immunoglobulin heavy constant alpha 2 OS=Homo sapiens OX=9606 GN=IGHA2 PE=1 SV=4 | 576 | 20201113_12413_1ng_glyco_hcd203040_compound15.13190.13190.3 | controllerType=0 controllerNumber=1 scan=13190 | 28.6416 |
| 10006:3 | 20 | R.MAGKPTHIN[+1540.529]VSVVMAEADGTC[+57.021]Y.- | HexNAc(2)Hex(7) | C[+57], N[+1541] | 1297.8782 | 3 | 3891.6201 | 3891.6114 | 0 | 2.3 | 319 | CRagged | 668.17 | 668.17 | 395.97 | 13.1 | 30 | >sp\|P01877\|IGHA2_HUMAN Immunoglobulin heavy constant alpha 2 OS=Homo sapiens OX=9606 GN=IGHA2 PE=1 SV=4 | 576 | 20201113_12413_1ng_glyco_hcd203040_compound15.13187.13187.3 | controllerType=0 controllerNumber=1 scan=13187 | 28.6343 |
| 10060:3 | 20 | R.MAGKPTHIN[+1216.423]VSVVMAEADGTC[+57.021]Y.- | HexNAc(2)Hex(5) | C[+57], N[+1216] | 1189.8425 | 3 | 3567.5128 | 3567.5057 | 0 | 2 | 319 | CRagged | 768.13 | 768.13 | 434.4 | 15.01 | 30 | >sp\|P01877\|IGHA2_HUMAN Immunoglobulin heavy constant alpha 2 OS=Homo sapiens OX=9606 GN=IGHA2 PE=1 SV=4 | 576 | 20201113_12413_1ng_glyco_hcd203040_compound15.13243.13243.3 | controllerType=0 controllerNumber=1 scan=13243 | 28.7349 |
| 09936:3 | 20 | R.MAGKPTHIN[+1702.581]VSVVMAEADGTC[+57.021]Y.- | HexNAc(2)Hex(8) | C[+57], N[+1703] | 1351.8979 | 3 | 4053.6791 | 4053.6642 | 0 | 3.7 | 319 | CRagged | 805.36 | 654.18 | 464.8 | 15.97 | 30 | >sp\|P01877\|IGHA2_HUMAN Immunoglobulin heavy constant alpha 2 OS=Homo sapiens OX=9606 GN=IGHA2 PE=1 SV=4 | 576 | 20201113_12413_1ng_glyco_hcd203040_compound15.13114.13114.3 | controllerType=0 controllerNumber=1 scan=13114 | 28.4974 |
| 09934:3 | 20 | R.MAGKPTHIN[+1702.581]VSVVMAEADGTC[+57.021]Y.- | HexNAc(2)Hex(8) | C[+57], N[+1703] | 1351.8979 | 3 | 4053.6791 | 4053.6642 | 0 | 3.7 | 319 | CRagged | 812.81 | 812.81 | 526 | 15.97 | 30 | >sp\|P01877\|IGHA2_HUMAN Immunoglobulin heavy constant alpha 2 OS=Homo sapiens OX=9606 GN=IGHA2 PE=1 SV=4 | 576 | 20201113_12413_1ng_glyco_hcd203040_compound15.13112.13112.3 | controllerType=0 controllerNumber=1 scan=13112 | 28.4914 |
| 10008:3 | 20 | R.MAGKPTHIN[+1378.476]VSVVMAEADGTC[+57.021]Y.- | HexNAc(2)Hex(6) | C[+57], N[+1378] | 1243.8607 | 3 | 3729.5674 | 3729.5585 | 0 | 2.4 | 319 | CRagged | 841.39 | 714.3 | 491.43 | 16.1 | 30 | >sp\|P01877\|IGHA2_HUMAN Immunoglobulin heavy constant alpha 2 OS=Homo sapiens OX=9606 GN=IGHA2 PE=1 SV=4 | 576 | 20201113_12413_1ng_glyco_hcd203040_compound15.13189.13189.3 | controllerType=0 controllerNumber=1 scan=13189 | 28.6396 |
| 10062:3 | 20 | R.MAGKPTHIN[+1216.423]VSVVMAEADGTC[+57.021]Y.- | HexNAc(2)Hex(5) | C[+57], N[+1216] | 1189.8425 | 3 | 3567.5128 | 3567.5057 | 0 | 2 | 319 | CRagged | 865.01 | 854.05 | 466.14 | 17.01 | 30 | >sp\|P01877\|IGHA2_HUMAN Immunoglobulin heavy constant alpha 2 OS=Homo sapiens OX=9606 GN=IGHA2 PE=1 SV=4 | 576 | 20201113_12413_1ng_glyco_hcd203040_compound15.13245.13245.3 | controllerType=0 controllerNumber=1 scan=13245 | 28.7405 |
| 10005:3 | 20 | R.MAGKPTHIN[+1378.476]VSVVMAEADGTC[+57.021]Y.- | HexNAc(2)Hex(6) | C[+57], N[+1378] | 1243.8607 | 3 | 3729.5674 | 3729.5585 | 0 | 2.4 | 319 | CRagged | 920.42 | 771.01 | 525.83 | 18.1 | 30 | >sp\|P01877\|IGHA2_HUMAN Immunoglobulin heavy constant alpha 2 OS=Homo sapiens OX=9606 GN=IGHA2 PE=1 SV=4 | 576 | 20201113_12413_1ng_glyco_hcd203040_compound15.13186.13186.3 | controllerType=0 controllerNumber=1 scan=13186 | 28.6332 |
| 08237:5 | 23 | K.SC[+57.021]QHN[+2059.735]GTMYQHGEIFSAHELFPSR.L | HexNAc(4)Hex(5)Fuc(1)NeuAc(1) | C[+57], N[+2060] | 979.4094 | 5 | 4893.0178 | 4892.9919 | 0 | 5.3 | 110 | Specific | 460.38 | 434.45 | 434.45 | 8.79 | 24 | >sp\|Q6WN34\|CRDL2_HUMAN Chordin-like protein 2 OS=Homo sapiens OX=9606 GN=CHRDL2 PE=1 SV=1 | 1189 | 20201113_12413_1ng_glyco_hcd203040_compound15.11349.11349.3 | controllerType=0 controllerNumber=1 scan=11349 | 25.4944 |
| 08239:5 | 23 | K.SC[+57.021]QHN[+2059.735]GTMYQHGEIFSAHELFPSR.L | HexNAc(4)Hex(5)Fuc(1)NeuAc(1) | C[+57], N[+2060] | 979.4094 | 5 | 4893.0178 | 4892.9919 | 0 | 5.3 | 110 | Specific | 515.77 | 515.77 | 515.77 | 9.08 | 24 | >sp\|Q6WN34\|CRDL2_HUMAN Chordin-like protein 2 OS=Homo sapiens OX=9606 GN=CHRDL2 PE=1 SV=1 | 1189 | 20201113_12413_1ng_glyco_hcd203040_compound15.11351.11351.3 | controllerType=0 controllerNumber=1 scan=11351 | 25.4984 |

Table S4. Cont.

| Query #:z | Protein Rank | Glycopeptide | Glycan | Modification Type(s) | Observed m/z | z | Observed (M+H) | Calc. mass (M+H) | Off-by-x error | Mass error (ppm) | StartPosition | Cleavage | Score | Delta | Delta Mod | \|Log Prob\| | # of unique peptides | Protein Name | Protein DB number | Comment | Scan # | Scan Time |
|---|---|---|---|---|---|---|---|---|---|---|---|---|---|---|---|---|---|---|---|---|---|---|
| 08363:4 | 23 | K.SC[+57.021]QHN[+2059.735]GTMYQHGEIFSAHELFPSR.L | HexNAc(4)Hex(5)Fuc(1)NeuAc(1) | C[+57], N[+2060] | 1224.0057 | 4 | 4893.0008 | 4892.9919 | 0 | 1.8 | 110 | Specific | 518.17 | 518.17 | 518.17 | 9.17 | 24 | >sp\|Q6WN34\|CRDL2_HUMAN Chordin-like protein 2 OS=Homo sapiens OX=9606 GN=CHRDL2 PE=1 SV=1 | 1189 | 20201113_12413_1ng_glyco_hcd203040_compound15.11480.11480.4 | controllerType=0 controllerNumber=1 scan=11480 | 25.7162 |
| 08361:4 | 23 | K.SC[+57.021]QHN[+2059.735]GTMYQHGEIFSAHELFPSR.L | HexNAc(4)Hex(5)Fuc(1)NeuAc(1) | C[+57], N[+2060] | 1224.0057 | 4 | 4893.0008 | 4892.9919 | 0 | 1.8 | 110 | Specific | 604.69 | 604.69 | 604.69 | 11.7 | 24 | >sp\|Q6WN34\|CRDL2_HUMAN Chordin-like protein 2 OS=Homo sapiens OX=9606 GN=CHRDL2 PE=1 SV=1 | 1189 | 20201113_12413_1ng_glyco_hcd203040_compound15.11478.11478.4 | controllerType=0 controllerNumber=1 scan=11478 | 25.7124 |
| 08304:4 | 23 | K.SC[+57.021]QHN[+2059.735]GTMYQHGEIFSAHELFPSR.L | HexNAc(4)Hex(5)Fuc(1)NeuAc(1) | C[+57], N[+2060] | 1224.0054 | 4 | 4892.9998 | 4892.9919 | 0 | 1.6 | 110 | Specific | 754.11 | 754.11 | 754.11 | 14.7 | 24 | >sp\|Q6WN34\|CRDL2_HUMAN Chordin-like protein 2 OS=Homo sapiens OX=9606 GN=CHRDL2 PE=1 SV=1 | 1189 | 20201113_12413_1ng_glyco_hcd203040_compound15.11419.11419.4 | controllerType=0 controllerNumber=1 scan=11419 | 25.6138 |
| 08306:4 | 23 | K.SC[+57.021]QHN[+2059.735]GTMYQHGEIFSAHELFPSR.L | HexNAc(4)Hex(5)Fuc(1)NeuAc(1) | C[+57], N[+2060] | 1224.0054 | 4 | 4892.9998 | 4892.9919 | 0 | 1.6 | 110 | Specific | 833.06 | 833.06 | 833.06 | 15.7 | 24 | >sp\|Q6WN34\|CRDL2_HUMAN Chordin-like protein 2 OS=Homo sapiens OX=9606 GN=CHRDL2 PE=1 SV=1 | 1189 | 20201113_12413_1ng_glyco_hcd203040_compound15.11421.11421.4 | controllerType=0 controllerNumber=1 scan=11421 | 25.6179 |
| 23507:5 | 24 | R.FFVPLFLVGILFPAILAKQFTKC[+57.021]ELSQLLKDIDGYGGIALPELIC[+57.021]TMFH | HexNAc(2)Hex(3) | C[+57]*2, N[+892] | 1873.9386 | 5 | 9365.6639 | 9365.6377 | 0 | 2.8 | 3 | Specific | 506.53 | 506.53 | 269.54 | 7.01 | 24 | >sp\|P00709\|LALBA_HUMAN Alpha-lactalbumin OS=Homo sapiens OX=9606 GN=LALBA PE=1 SV=1 | 468 | 20201113_12413_1ng_glyco_hcd203040_compound15.27282.27282.5 | controllerType=0 controllerNumber=1 scan=27282 | 55.474 |
| 23509:5 | 24 | R.FFVPLFLVGILFPAILAKQFTKC[+57.021]ELSQLLKDIDGYGGIALPELIC[+57.021]TMFH | HexNAc(2)Hex(3) | C[+57]*2, N[+892] | 1873.9386 | 5 | 9365.6639 | 9365.6377 | 0 | 2.8 | 3 | Specific | 521.82 | 521.82 | 328.22 | 7.14 | 24 | >sp\|P00709\|LALBA_HUMAN Alpha-lactalbumin OS=Homo sapiens OX=9606 GN=LALBA PE=1 SV=1 | 468 | 20201113_12413_1ng_glyco_hcd203040_compound15.27284.27284.5 | controllerType=0 controllerNumber=1 scan=27284 | 55.4797 |
| 13089:3 | 46 | R.IIVPLNNREN[+2204.772]ISDPTSPLR.T | HexNAc(4)Hex(5)NeuAc(2) | N[+2205] | 1451.6591 | 3 | 4352.9628 | 4352.9496 | 0 | 3 | 62 | Specific | 238.23 | 238.23 | 238.23 | 5.79 | 15 | >sp\|P01591\|IGJ_HUMAN Immunoglobulin J chain OS=Homo sapiens OX=9606 GN=JCHAIN PE=1 SV=4 | 1392 | 20201113_12413_1ng_glyco_hcd203040_compound15.16399.16399.3 | controllerType=0 controllerNumber=1 scan=16399 | 34.0804 |
| 11059:3 | 46 | R.IIVPLNNREN[+2059.735]ISDPTSPLR.T | HexNAc(4)Hex(5)Fuc(1)NeuAc(1) | N[+2060] | 1403.3115 | 3 | 4207.9198 | 4207.9121 | 0 | 1.8 | 62 | Specific | 295.02 | 274 | 274 | 6.22 | 15 | >sp\|P01591\|IGJ_HUMAN Immunoglobulin J chain OS=Homo sapiens OX=9606 GN=JCHAIN PE=1 SV=4 | 1392 | 20201113_12413_1ng_glyco_hcd203040_compound15.14276.14276.3 | controllerType=0 controllerNumber=1 scan=14276 | 30.453 |
| 12971:3 | 46 | R.IIVPLNNREN[+2350.830]ISDPTSPLR.T | HexNAc(4)Hex(5)Fuc(1)NeuAc(2) | N[+2351] | 1500.3519 | 3 | 4499.041 | 4499.0075 | 0 | 7.5 | 62 | Specific | 307.64 | 307.64 | 307.64 | 5.8 | 15 | >sp\|P01591\|IGJ_HUMAN Immunoglobulin J chain OS=Homo sapiens OX=9606 GN=JCHAIN PE=1 SV=4 | 1392 | 20201113_12413_1ng_glyco_hcd203040_compound15.16277.16277.3 | controllerType=0 controllerNumber=1 scan=16277 | 33.8764 |
| 11061:3 | 46 | R.IIVPLNNREN[+2059.735]ISDPTSPLR.T | HexNAc(4)Hex(5)Fuc(1)NeuAc(1) | N[+2060] | 1403.3115 | 3 | 4207.9198 | 4207.9121 | 0 | 1.8 | 62 | Specific | 309.79 | 309.79 | 309.79 | 6.57 | 15 | >sp\|P01591\|IGJ_HUMAN Immunoglobulin J chain OS=Homo sapiens OX=9606 GN=JCHAIN PE=1 SV=4 | 1392 | 20201113_12413_1ng_glyco_hcd203040_compound15.14278.14278.3 | controllerType=0 controllerNumber=1 scan=14278 | 30.4569 |
| 11193:3 | 46 | R.IIVPLNNREN[+1694.603]ISDPTSPLR.T | HexNAc(3)Hex(4)Fuc(1)NeuAc(1) | N[+1695] | 1281.5981 | 3 | 3842.7799 | 3842.7799 | 0 | 0 | 62 | Specific | 385.06 | 359.08 | 359.08 | 6.92 | 15 | >sp\|P01591\|IGJ_HUMAN Immunoglobulin J chain OS=Homo sapiens OX=9606 GN=JCHAIN PE=1 SV=4 | 1392 | 20201113_12413_1ng_glyco_hcd203040_compound15.14417.14417.3 | controllerType=0 controllerNumber=1 scan=14417 | 30.7471 |
| 13056:3 | 46 | R.IIVPLNNREN[+2204.772]ISDPTSPLR.T | HexNAc(4)Hex(5)NeuAc(2) | N[+2205] | 1451.6583 | 3 | 4352.9602 | 4352.9496 | 0 | 2.4 | 62 | Specific | 416.7 | 391.67 | 391.67 | 8.84 | 15 | >sp\|P01591\|IGJ_HUMAN Immunoglobulin J chain OS=Homo sapiens OX=9606 GN=JCHAIN PE=1 SV=4 | 1392 | 20201113_12413_1ng_glyco_hcd203040_compound15.16365.16365.3 | controllerType=0 controllerNumber=1 scan=16365 | 34.0257 |
| 11195:3 | 46 | R.IIVPLNNREN[+1694.603]ISDPTSPLR.T | HexNAc(3)Hex(4)Fuc(1)NeuAc(1) | N[+1695] | 1281.5981 | 3 | 3842.7799 | 3842.7799 | 0 | 0 | 62 | Specific | 422.29 | 340.45 | 340.45 | 7 | 15 | >sp\|P01591\|IGJ_HUMAN Immunoglobulin J chain OS=Homo sapiens OX=9606 GN=JCHAIN PE=1 SV=4 | 1392 | 20201113_12413_1ng_glyco_hcd203040_compound15.14419.14419.3 | controllerType=0 controllerNumber=1 scan=14419 | 30.7512 |
| 13053:3 | 46 | R.IIVPLNNREN[+2204.772]ISDPTSPLR.T | HexNAc(4)Hex(5)NeuAc(2) | N[+2205] | 1451.6583 | 3 | 4352.9602 | 4352.9496 | 0 | 2.4 | 62 | Specific | 430.78 | 395.77 | 395.77 | 8.84 | 15 | >sp\|P01591\|IGJ_HUMAN Immunoglobulin J chain OS=Homo sapiens OX=9606 GN=JCHAIN PE=1 SV=4 | 1392 | 20201113_12413_1ng_glyco_hcd203040_compound15.16362.16362.3 | controllerType=0 controllerNumber=1 scan=16362 | 34.0208 |
| 12973:3 | 46 | R.IIVPLNNREN[+2350.830]ISDPTSPLR.T | HexNAc(4)Hex(5)Fuc(1)NeuAc(2) | N[+2351] | 1500.3519 | 3 | 4499.041 | 4499.0075 | 0 | 7.5 | 62 | Specific | 440.07 | 388.17 | 388.17 | 7.99 | 15 | >sp\|P01591\|IGJ_HUMAN Immunoglobulin J chain OS=Homo sapiens OX=9606 GN=JCHAIN PE=1 SV=4 | 1392 | 20201113_12413_1ng_glyco_hcd203040_compound15.16279.16279.3 | controllerType=0 controllerNumber=1 scan=16279 | 33.8802 |
| 13092:3 | 46 | R.IIVPLNNREN[+2204.772]ISDPTSPLR.T | HexNAc(4)Hex(5)NeuAc(2) | N[+2205] | 1451.6591 | 3 | 4352.9628 | 4352.9496 | 0 | 3 | 62 | Specific | 475.7 | 460.78 | 460.78 | 9.05 | 15 | >sp\|P01591\|IGJ_HUMAN Immunoglobulin J chain OS=Homo sapiens OX=9606 GN=JCHAIN PE=1 SV=4 | 1392 | 20201113_12413_1ng_glyco_hcd203040_compound15.16402.16402.3 | controllerType=0 controllerNumber=1 scan=16402 | 34.0858 |
| 11058:3 | 46 | R.IIVPLNNREN[+1913.677]ISDPTSPLR.T | HexNAc(4)Hex(5)NeuAc(1) | N[+1914] | 1354.6254 | 3 | 4061.8617 | 4061.8542 | 0 | 1.9 | 62 | Specific | 504.94 | 465.77 | 465.77 | 9.05 | 15 | >sp\|P01591\|IGJ_HUMAN Immunoglobulin J chain OS=Homo sapiens OX=9606 GN=JCHAIN PE=1 SV=4 | 1392 | 20201113_12413_1ng_glyco_hcd203040_compound15.14275.14275.3 | controllerType=0 controllerNumber=1 scan=14275 | 30.4512 |
| 11056:3 | 46 | R.IIVPLNNREN[+1913.677]ISDPTSPLR.T | HexNAc(4)Hex(5)NeuAc(1) | N[+1914] | 1354.6254 | 3 | 4061.8617 | 4061.8542 | 0 | 1.9 | 62 | Specific | 511.19 | 491.54 | 491.54 | 9.05 | 15 | >sp\|P01591\|IGJ_HUMAN Immunoglobulin J chain OS=Homo sapiens OX=9606 GN=JCHAIN PE=1 SV=4 | 1392 | 20201113_12413_1ng_glyco_hcd203040_compound15.14273.14273.3 | controllerType=0 controllerNumber=1 scan=14273 | 30.4468 |
| 11000:3 | 46 | R.IIVPLNNREN[+2059.735]ISDPTSPLR.T | HexNAc(4)Hex(5)Fuc(1)NeuAc(1) | N[+2060] | 1403.3097 | 3 | 4207.9147 | 4207.9121 | 0 | 0.6 | 62 | Specific | 515.99 | 485.79 | 485.79 | 8.85 | 15 | >sp\|P01591\|IGJ_HUMAN Immunoglobulin J chain OS=Homo sapiens OX=9606 GN=JCHAIN PE=1 SV=4 | 1392 | 20201113_12413_1ng_glyco_hcd203040_compound15.14215.14215.3 | controllerType=0 controllerNumber=1 scan=14215 | 30.3388 |
| 11002:3 | 46 | R.IIVPLNNREN[+2059.735]ISDPTSPLR.T | HexNAc(4)Hex(5)Fuc(1)NeuAc(1) | N[+2060] | 1403.3097 | 3 | 4207.9147 | 4207.9121 | 0 | 0.6 | 62 | Specific | 517.93 | 474.91 | 474.91 | 8.85 | 15 | >sp\|P01591\|IGJ_HUMAN Immunoglobulin J chain OS=Homo sapiens OX=9606 GN=JCHAIN PE=1 SV=4 | 1392 | 20201113_12413_1ng_glyco_hcd203040_compound15.14217.14217.3 | controllerType=0 controllerNumber=1 scan=14217 | 30.3434 |
| 16352:3 | 51 | R.NYIVPILWLN[+1864.634]ETGTIGDEKANM[+15.995]FR.S | HexNAc(2)Hex(9) | M[+16], N[+1865] | 1559.0242 | 3 | 4675.0581 | 4675.0523 | 0 | 1.2 | 408 | Specific | 343.54 | 343.54 | 343.54 | 7.12 | 13 | >sp\|P16671\|CD36_HUMAN Platelet glycoprotein 4 OS=Homo sapiens OX=9606 GN=CD36 PE=1 SV=2 | 812 | 20201113_12413_1ng_glyco_hcd203040_compound15.19813.19813.3 | controllerType=0 controllerNumber=1 scan=19813 | 40.2774 |
| 16350:3 | 51 | R.NYIVPILWLN[+1864.634]ETGTIGDEKANM[+15.995]FR.S | HexNAc(2)Hex(9) | M[+16], N[+1865] | 1559.0242 | 3 | 4675.0581 | 4675.0523 | 0 | 1.2 | 408 | Specific | 442.39 | 442.39 | 442.39 | 9.02 | 13 | >sp\|P16671\|CD36_HUMAN Platelet glycoprotein 4 OS=Homo sapiens OX=9606 GN=CD36 PE=1 SV=2 | 812 | 20201113_12413_1ng_glyco_hcd203040_compound15.19811.19811.3 | controllerType=0 controllerNumber=1 scan=19811 | 40.2737 |
| 19699:4 | 1 | R.RTAGWNVPIGTLRPFLN[+2350.830]WTGPPEPIEAAVAR.F | HexNAc(4)Hex(5)Fuc(1)NeuAc(2) | N[+2351] | 1435.1722 | 4 | 5737.6671 | 5737.6424 | 0 | 4.3 | 140 | Specific | 253.97 | 253.97 | 253.97 | 5.48 | 142 | >sp\|P02788\|TRFL_HUMAN Lactotransferrin OS=Homo sapiens OX=9606 GN=LTF PE=1 SV=6 | 1270 | 20201113_12433_1ng_glyco_hcd203040_compound15.23144.23144.4 | controllerType=0 controllerNumber=1 scan=23144 | 46.0111 |
| 19890:4 | 1 | R.RTAGWNVPIGTLRPFLN[+1930.692]WTGPPEPIEAAVAR.F | HexNAc(4)Hex(6)Fuc(1) | N[+1931] | 1330.1185 | 4 | 5317.4523 | 5317.5044 | 0 | -9.8 | 140 | Specific | 284.01 | 266.79 | 266.79 | 4.07 | 142 | >sp\|P02788\|TRFL_HUMAN Lactotransferrin OS=Homo sapiens OX=9606 GN=LTF PE=1 SV=6 | 1270 | 20201113_12433_1ng_glyco_hcd203040_compound15.23343.23343.2 | controllerType=0 controllerNumber=1 scan=23343 | 46.4317 |
| 19892:4 | 1 | R.RTAGWNVPIGTLRPFLN[+1930.692]WTGPPEPIEAAVAR.F | HexNAc(4)Hex(6)Fuc(1) | N[+1931] | 1330.1185 | 4 | 5317.4523 | 5317.5044 | 0 | -9.8 | 140 | Specific | 313.33 | 313.33 | 313.33 | 4.33 | 142 | >sp\|P02788\|TRFL_HUMAN Lactotransferrin OS=Homo sapiens OX=9606 GN=LTF PE=1 SV=6 | 1270 | 20201113_12433_1ng_glyco_hcd203040_compound15.23345.23345.2 | controllerType=0 controllerNumber=1 scan=23345 | 46.4378 |
| 18499:4 | 1 | R.RTAGWNVPIGTLRPFLN[+2716.983]WTGPPEPIEAAVAR.F | HexNAc(5)Hex(6)Fuc(3)NeuAc(1) | N[+2717] | 1526.7075 | 4 | 6103.8083 | 6103.795 | 0 | 2.2 | 140 | Specific | 403.12 | 403.12 | 403.12 | 9.81 | 142 | >sp\|P02788\|TRFL_HUMAN Lactotransferrin OS=Homo sapiens OX=9606 GN=LTF PE=1 SV=6 | 1270 | 20201113_12433_1ng_glyco_hcd203040_compound15.21893.21893.4 | controllerType=0 controllerNumber=1 scan=21893 | 43.3219 |
| 18502:4 | 1 | R.RTAGWNVPIGTLRPFLN[+2716.983]WTGPPEPIEAAVAR.F | HexNAc(5)Hex(6)Fuc(3)NeuAc(1) | N[+2717] | 1526.7075 | 4 | 6103.8083 | 6103.795 | 0 | 2.2 | 140 | Specific | 481.75 | 410.04 | 410.04 | 10.08 | 142 | >sp\|P02788\|TRFL_HUMAN Lactotransferrin OS=Homo sapiens OX=9606 GN=LTF PE=1 SV=6 | 1270 | 20201113_12433_1ng_glyco_hcd203040_compound15.21896.21896.4 | controllerType=0 controllerNumber=1 scan=21896 | 43.3297 |
| 18538:4 | 1 | R.RTAGWNVPIGTLRPFLN[+2570.925]WTGPPEPIEAAVAR.F | HexNAc(5)Hex(6)Fuc(2)NeuAc(1) | N[+2571] | 1490.1957 | 4 | 5957.7609 | 5957.7371 | 0 | 4 | 140 | Specific | 519.59 | 519.59 | 519.59 | 9.91 | 142 | >sp\|P02788\|TRFL_HUMAN Lactotransferrin OS=Homo sapiens OX=9606 GN=LTF PE=1 SV=6 | 1270 | 20201113_12433_1ng_glyco_hcd203040_compound15.21934.21934.4 | controllerType=0 controllerNumber=1 scan=21934 | 43.4076 |
| 18540:4 | 1 | R.RTAGWNVPIGTLRPFLN[+2570.925]WTGPPEPIEAAVAR.F | HexNAc(5)Hex(6)Fuc(2)NeuAc(1) | N[+2571] | 1490.1957 | 4 | 5957.7609 | 5957.7371 | 0 | 4 | 140 | Specific | 536.35 | 536.35 | 536.35 | 10.03 | 142 | >sp\|P02788\|TRFL_HUMAN Lactotransferrin OS=Homo sapiens OX=9606 GN=LTF PE=1 SV=6 | 1270 | 20201113_12433_1ng_glyco_hcd203040_compound15.21936.21936.4 | controllerType=0 controllerNumber=1 scan=21936 | 43.4133 |
| 17690:4 | 1 | R.RTAGWNVPIGTLRPFLN[+1768.640]WTGPPEPIEAAVAR.F | HexNAc(4)Hex(5)Fuc(1) | N[+1769] | 1289.6213 | 4 | 5155.4635 | 5155.4516 | 0 | 2.3 | 140 | Specific | 544.21 | 467.72 | 467.72 | 10.38 | 142 | >sp\|P02788\|TRFL_HUMAN Lactotransferrin OS=Homo sapiens OX=9606 GN=LTF PE=1 SV=6 | 1270 | 20201113_12433_1ng_glyco_hcd203040_compound15.21050.21050.4 | controllerType=0 controllerNumber=1 scan=21050 | 41.5411 |
| 17692:4 | 1 | R.RTAGWNVPIGTLRPFLN[+1768.640]WTGPPEPIEAAVAR.F | HexNAc(4)Hex(5)Fuc(1) | N[+1769] | 1289.6213 | 4 | 5155.4635 | 5155.4516 | 0 | 2.3 | 140 | Specific | 721.02 | 556.8 | 556.8 | 14.85 | 142 | >sp\|P02788\|TRFL_HUMAN Lactotransferrin OS=Homo sapiens OX=9606 GN=LTF PE=1 SV=6 | 1270 | 20201113_12433_1ng_glyco_hcd203040_compound15.21052.21052.4 | controllerType=0 controllerNumber=1 scan=21052 | 41.5463 |
| 19863:3 | 1 | R.TAGWNVPIGTLRPFLN[+2205.793]WTGPPEPIEAAVAR.F | HexNAc(4)Hex(5)Fuc(2)NeuAc(1) | N[+2206] | 1812.8432 | 3 | 5436.5151 | 5436.5038 | 0 | 2.1 | 141 | Specific | 249.98 | 249.98 | 249.98 | 7.12 | 142 | >sp\|P02788\|TRFL_HUMAN Lactotransferrin OS=Homo sapiens OX=9606 GN=LTF PE=1 SV=6 | 1270 | 20201113_12433_1ng_glyco_hcd203040_compound15.23315.23315.3 | controllerType=0 controllerNumber=1 scan=23315 | 46.3776 |
| 19840:3 | 1 | R.TAGWNVPIGTLRPFLN[+2351.851]WTGPPEPIEAAVAR.F | HexNAc(4)Hex(5)Fuc(3)NeuAc(1) | N[+2352] | 1861.5311 | 3 | 5582.5787 | 5582.5617 | 0 | 3.1 | 141 | Specific | 303.11 | 297.36 | 297.36 | 7.29 | 142 | >sp\|P02788\|TRFL_HUMAN Lactotransferrin OS=Homo sapiens OX=9606 GN=LTF PE=1 SV=6 | 1270 | 20201113_12433_1ng_glyco_hcd203040_compound15.23291.23291.3 | controllerType=0 controllerNumber=1 scan=23291 | 46.3252 |
| 21269:3 | 1 | R.TAGWNVPIGTLRPFLN[+2350.830]WTGPPEPIEAAVAR.F | HexNAc(4)Hex(5)Fuc(1)NeuAc(2) | N[+2351] | 1861.1849 | 3 | 5581.5401 | 5581.5413 | 0 | -0.2 | 141 | Specific | 341.83 | 168.17 | 168.17 | 6.18 | 142 | >sp\|P02788\|TRFL_HUMAN Lactotransferrin OS=Homo sapiens OX=9606 GN=LTF PE=1 SV=6 | 1270 | 20201113_12433_1ng_glyco_hcd203040_compound15.24782.24782.2 | controllerType=0 controllerNumber=1 scan=24782 | 49.2441 |
| 19889:3 | 1 | R.TAGWNVPIGTLRPFLN[+2059.735]WTGPPEPIEAAVAR.F | HexNAc(4)Hex(5)Fuc(1)NeuAc(1) | N[+2060] | 1764.16 | 3 | 5290.4654 | 5290.4459 | 0 | 3.7 | 141 | Specific | 357.36 | 323.67 | 323.67 | 7.41 | 142 | >sp\|P02788\|TRFL_HUMAN Lactotransferrin OS=Homo sapiens OX=9606 GN=LTF PE=1 SV=6 | 1270 | 20201113_12433_1ng_glyco_hcd203040_compound15.23342.23342.2 | controllerType=0 controllerNumber=1 scan=23342 | 46.4303 |
| 19796:4 | 1 | R.TAGWNVPIGTLRPFLN[+2424.867]WTGPPEPIEAAVAR.F | HexNAc(5)Hex(6)Fuc(1)NeuAc(1) | N[+2425] | 1414.6545 | 4 | 5655.5963 | 5655.5781 | 0 | 3.2 | 141 | Specific | 364.46 | 265.53 | 265.53 | 7.52 | 142 | >sp\|P02788\|TRFL_HUMAN Lactotransferrin OS=Homo sapiens OX=9606 GN=LTF PE=1 SV=6 | 1270 | 20201113_12433_1ng_glyco_hcd203040_compound15.23245.23245.4 | controllerType=0 controllerNumber=1 scan=23245 | 46.2365 |
| 19862:4 | 1 | R.TAGWNVPIGTLRPFLN[+2205.793]WTGPPEPIEAAVAR.F | HexNAc(4)Hex(5)Fuc(2)NeuAc(1) | N[+2206] | 1359.8842 | 4 | 5436.5148 | 5436.5038 | 0 | 2 | 141 | Specific | 373.73 | 355.88 | 355.88 | 8.32 | 142 | >sp\|P02788\|TRFL_HUMAN Lactotransferrin OS=Homo sapiens OX=9606 GN=LTF PE=1 SV=6 | 1270 | 20201113_12433_1ng_glyco_hcd203040_compound15.23314.23314.4 | controllerType=0 controllerNumber=1 scan=23314 | 46.3762 |
| 19829:4 | 1 | R.TAGWNVPIGTLRPFLN[+2424.867]WTGPPEPIEAAVAR.F | HexNAc(5)Hex(6)Fuc(1)NeuAc(1) | N[+2425] | 1414.6565 | 4 | 5655.6042 | 5655.5781 | 0 | 4.6 | 141 | Specific | 405.36 | 260.79 | 260.79 | 7.42 | 142 | >sp\|P02788\|TRFL_HUMAN Lactotransferrin OS=Homo sapiens OX=9606 GN=LTF PE=1 SV=6 | 1270 | 20201113_12433_1ng_glyco_hcd203040_compound15.23279.23279.4 | controllerType=0 controllerNumber=1 scan=23279 | 46.2984 |
| 19706:3 | 1 | R.TAGWNVPIGTLRPFLN[+2716.983]WTGPPEPIEAAVAR.F | HexNAc(5)Hex(6)Fuc(3)NeuAc(1) | N[+2717] | 1983.2408 | 3 | 5947.7079 | 5947.6939 | 0 | 2.4 | 141 | Specific | 439.37 | 386.97 | 386.97 | 9.91 | 142 | >sp\|P02788\|TRFL_HUMAN Lactotransferrin OS=Homo sapiens OX=9606 GN=LTF PE=1 SV=6 | 1270 | 20201113_12433_1ng_glyco_hcd203040_compound15.23151.23151.3 | controllerType=0 controllerNumber=1 scan=23151 | 46.0221 |
| 20076:4 | 1 | R.TAGWNVPIGTLRPFLN[+2205.793]WTGPPEPIEAAVAR.F | HexNAc(4)Hex(5)Fuc(2)NeuAc(1) | N[+2206] | 1359.8822 | 4 | 5436.507 | 5436.5038 | 0 | 0.6 | 141 | Specific | 472.51 | 315.58 | 315.58 | 7.87 | 142 | >sp\|P02788\|TRFL_HUMAN Lactotransferrin OS=Homo sapiens OX=9606 GN=LTF PE=1 SV=6 | 1270 | 20201113_12433_1ng_glyco_hcd203040_compound15.23541.23541.3 | controllerType=0 controllerNumber=1 scan=23541 | 46.7918 |
| 21280:4 | 1 | R.TAGWNVPIGTLRPFLN[+2350.830]WTGPPEPIEAAVAR.F | HexNAc(4)Hex(5)Fuc(1)NeuAc(2) | N[+2351] | 1396.1397 | 4 | 5581.5368 | 5581.5413 | 0 | -0.8 | 141 | Specific | 481.47 | 481.47 | 481.47 | 9.41 | 142 | >sp\|P02788\|TRFL_HUMAN Lactotransferrin OS=Homo sapiens OX=9606 GN=LTF PE=1 SV=6 | 1270 | 20201113_12433_1ng_glyco_hcd203040_compound15.24793.24793.6 | controllerType=0 controllerNumber=1 scan=24793 | 49.2661 |
| 19754:4 | 1 | R.TAGWNVPIGTLRPFLN[+2716.983]WTGPPEPIEAAVAR.F | HexNAc(5)Hex(6)Fuc(3)NeuAc(1) | N[+2717] | 1487.6818 | 4 | 5947.7052 | 5947.6939 | 0 | 1.9 | 141 | Specific | 522.88 | 514.74 | 514.74 | 10.24 | 142 | >sp\|P02788\|TRFL_HUMAN Lactotransferrin OS=Homo sapiens OX=9606 GN=LTF PE=1 SV=6 | 1270 | 20201113_12433_1ng_glyco_hcd203040_compound15.23201.23201.4 | controllerType=0 controllerNumber=1 scan=23201 | 46.1504 |
| 19709:3 | 1 | R.TAGWNVPIGTLRPFLN[+2716.983]WTGPPEPIEAAVAR.F | HexNAc(5)Hex(6)Fuc(3)NeuAc(1) | N[+2717] | 1983.2408 | 3 | 5947.7079 | 5947.6939 | 0 | 2.4 | 141 | Specific | 523.96 | 483.71 | 483.71 | 10.25 | 142 | >sp\|P02788\|TRFL_HUMAN Lactotransferrin OS=Homo sapiens OX=9606 GN=LTF PE=1 SV=6 | 1270 | 20201113_12433_1ng_glyco_hcd203040_compound15.23154.23154.3 | controllerType=0 controllerNumber=1 scan=23154 | 46.0272 |
| 19842:3 | 1 | R.TAGWNVPIGTLRPFLN[+2351.851]WTGPPEPIEAAVAR.F | HexNAc(4)Hex(5)Fuc(3)NeuAc(1) | N[+2352] | 1861.5311 | 3 | 5582.5787 | 5582.5617 | 0 | 3.1 | 141 | Specific | 531.22 | 476.75 | 476.75 | 10.37 | 142 | >sp\|P02788\|TRFL_HUMAN Lactotransferrin OS=Homo sapiens OX=9606 GN=LTF PE=1 SV=6 | 1270 | 20201113_12433_1ng_glyco_hcd203040_compound15.23293.23293.3 | controllerType=0 controllerNumber=1 scan=23293 | 46.3293 |
| 19766:4 | 1 | R.TAGWNVPIGTLRPFLN[+2716.983]WTGPPEPIEAAVAR.F | HexNAc(5)Hex(6)Fuc(3)NeuAc(1) | N[+2717] | 1487.6847 | 4 | 5947.717 | 5947.6939 | 0 | 3.9 | 141 | Specific | 548.26 | 515.32 | 515.32 | 10.01 | 142 | >sp\|P02788\|TRFL_HUMAN Lactotransferrin OS=Homo sapiens OX=9606 GN=LTF PE=1 SV=6 | 1270 | 20201113_12433_1ng_glyco_hcd203040_compound15.23214.23214.4 | controllerType=0 controllerNumber=1 scan=23214 | 46.1731 |
| 19799:4 | 1 | R.TAGWNVPIGTLRPFLN[+2424.867]WTGPPEPIEAAVAR.F | HexNAc(5)Hex(6)Fuc(1)NeuAc(1) | N[+2425] | 1414.6545 | 4 | 5655.5963 | 5655.5781 | 0 | 3.2 | 141 | Specific | 559.05 | 404.14 | 404.14 | 10.56 | 142 | >sp\|P02788\|TRFL_HUMAN Lactotransferrin OS=Homo sapiens OX=9606 GN=LTF PE=1 SV=6 | 1270 | 20201113_12433_1ng_glyco_hcd203040_compound15.23248.23248.4 | controllerType=0 controllerNumber=1 scan=23248 | 46.2435 |
| 19831:4 | 1 | R.TAGWNVPIGTLRPFLN[+2424.867]WTGPPEPIEAAVAR.F | HexNAc(5)Hex(6)Fuc(1)NeuAc(1) | N[+2425] | 1414.6565 | 4 | 5655.6042 | 5655.5781 | 0 | 4.6 | 141 | Specific | 591.74 | 431.54 | 431.54 | 11.48 | 142 | >sp\|P02788\|TRFL_HUMAN Lactotransferrin OS=Homo sapiens OX=9606 GN=LTF PE=1 SV=6 | 1270 | 20201113_12433_1ng_glyco_hcd203040_compound15.23281.23281.4 | controllerType=0 controllerNumber=1 scan=23281 | 46.306 |
| 21272:3 | 1 | R.TAGWNVPIGTLRPFLN[+2350.830]WTGPPEPIEAAVAR.F | HexNAc(4)Hex(5)Fuc(1)NeuAc(2) | N[+2351] | 1861.1849 | 3 | 5581.5401 | 5581.5413 | 0 | -0.2 | 141 | Specific | 632.22 | 457.98 | 457.98 | 12.2 | 142 | >sp\|P02788\|TRFL_HUMAN Lactotransferrin OS=Homo sapiens OX=9606 GN=LTF PE=1 SV=6 | 1270 | 20201113_12433_1ng_glyco_hcd203040_compound15.24785.24785.2 | controllerType=0 controllerNumber=1 scan=24785 | 49.2484 |
| 19865:3 | 1 | R.TAGWNVPIGTLRPFLN[+2205.793]WTGPPEPIEAAVAR.F | HexNAc(4)Hex(5)Fuc(2)NeuAc(1) | N[+2206] | 1812.8432 | 3 | 5436.5151 | 5436.5038 | 0 | 2.1 | 141 | Specific | 635.65 | 572.21 | 572.21 | 12.85 | 142 | >sp\|P02788\|TRFL_HUMAN Lactotransferrin OS=Homo sapiens OX=9606 GN=LTF PE=1 SV=6 | 1270 | 20201113_12433_1ng_glyco_hcd203040_compound15.23317.23317.3 | controllerType=0 controllerNumber=1 scan=23317 | 46.381 |
| 19757:4 | 1 | R.TAGWNVPIGTLRPFLN[+2716.983]WTGPPEPIEAAVAR.F | HexNAc(5)Hex(6)Fuc(3)NeuAc(1) | N[+2717] | 1487.6818 | 4 | 5947.7052 | 5947.6939 | 0 | 1.9 | 141 | Specific | 644.57 | 559.63 | 559.63 | 12.83 | 142 | >sp\|P02788\|TRFL_HUMAN Lactotransferrin OS=Homo sapiens OX=9606 GN=LTF PE=1 SV=6 | 1270 | 20201113_12433_1ng_glyco_hcd203040_compound15.23204.23204.4 | controllerType=0 controllerNumber=1 scan=23204 | 46.156 |
| 18893:3 | 1 | R.TAGWNVPIGTLRPFLN[+1768.640]WTGPPEPIEAAVAR.F | HexNAc(4)Hex(5)Fuc(1) | N[+1769] | 1667.1204 | 3 | 4999.3468 | 4999.3504 | 0 | -0.7 | 141 | Specific | 647.04 | 513.37 | 513.37 | 12.2 | 142 | >sp\|P02788\|TRFL_HUMAN Lactotransferrin OS=Homo sapiens OX=9606 GN=LTF PE=1 SV=6 | 1270 | 20201113_12433_1ng_glyco_hcd203040_compound15.22304.22304.3 | controllerType=0 controllerNumber=1 scan=22304 | 44.1855 |
| 21372:4 | 1 | R.TAGWNVPIGTLRPFLN[+2350.830]WTGPPEPIEAAVAR.F | HexNAc(4)Hex(5)Fuc(1)NeuAc(2) | N[+2351] | 1396.1449 | 4 | 5581.5578 | 5581.5413 | 0 | 3 | 141 | Specific | 653.93 | 523.26 | 523.26 | 13.82 | 142 | >sp\|P02788\|TRFL_HUMAN Lactotransferrin OS=Homo sapiens OX=9606 GN=LTF PE=1 SV=6 | 1270 | 20201113_12433_1ng_glyco_hcd203040_compound15.24889.24889.4 | controllerType=0 controllerNumber=1 scan=24889 | 49.4702 |
| 19867:5 | 1 | R.TAGWNVPIGTLRPFLN[+2205.793]WTGPPEPIEAAVAR.F | HexNAc(4)Hex(5)Fuc(2)NeuAc(1) | N[+2206] | 1088.1095 | 5 | 5436.5184 | 5436.5038 | 0 | 2.7 | 141 | Specific | 665.39 | 545.14 | 545.14 | 13.93 | 142 | >sp\|P02788\|TRFL_HUMAN Lactotransferrin OS=Homo sapiens OX=9606 GN=LTF PE=1 SV=6 | 1270 | 20201113_12433_1ng_glyco_hcd203040_compound15.23319.23319.5 | controllerType=0 controllerNumber=1 scan=23319 | 46.3833 |
| 18890:3 | 1 | R.TAGWNVPIGTLRPFLN[+1768.640]WTGPPEPIEAAVAR.F | HexNAc(4)Hex(5)Fuc(1) | N[+1769] | 1667.1204 | 3 | 4999.3468 | 4999.3504 | 0 | -0.7 | 141 | Specific | 690.06 | 572.17 | 572.17 | 13.2 | 142 | >sp\|P02788\|TRFL_HUMAN Lactotransferrin OS=Homo sapiens OX=9606 GN=LTF PE=1 SV=6 | 1270 | 20201113_12433_1ng_glyco_hcd203040_compound15.22301.22301.3 | controllerType=0 controllerNumber=1 scan=22301 | 44.1807 |
| 18892:4 | 1 | R.TAGWNVPIGTLRPFLN[+1768.640]WTGPPEPIEAAVAR.F | HexNAc(4)Hex(5)Fuc(1) | N[+1769] | 1250.5918 | 4 | 4999.3454 | 4999.3504 | 0 | -1 | 141 | Specific | 726.09 | 674.79 | 674.79 | 14.18 | 142 | >sp\|P02788\|TRFL_HUMAN Lactotransferrin OS=Homo sapiens OX=9606 GN=LTF PE=1 SV=6 | 1270 | 20201113_12433_1ng_glyco_hcd203040_compound15.22303.22303.4 | controllerType=0 controllerNumber=1 scan=22303 | 44.1828 |
| 19850:4 | 1 | R.TAGWNVPIGTLRPFLN[+2059.735]WTGPPEPIEAAVAR.F | HexNAc(4)Hex(5)Fuc(1)NeuAc(1) | N[+2060] | 1323.3718 | 4 | 5290.4655 | 5290.4459 | 0 | 3.7 | 141 | Specific | 726.11 | 654.48 | 654.48 | 14.48 | 142 | >sp\|P02788\|TRFL_HUMAN Lactotransferrin OS=Homo sapiens OX=9606 GN=LTF PE=1 SV=6 | 1270 | 20201113_12433_1ng_glyco_hcd203040_compound15.23301.23301.4 | controllerType=0 controllerNumber=1 scan=23301 | 46.3433 |
| 19717:4 | 1 | R.TAGWNVPIGTLRPFLN[+2570.925]WTGPPEPIEAAVAR.F | HexNAc(5)Hex(6)Fuc(2)NeuAc(1) | N[+2571] | 1451.1696 | 4 | 5801.6564 | 5801.636 | 0 | 3.5 | 141 | Specific | 728.72 | 629.28 | 629.28 | 14.82 | 142 | >sp\|P02788\|TRFL_HUMAN Lactotransferrin OS=Homo sapiens OX=9606 GN=LTF PE=1 SV=6 | 1270 | 20201113_12433_1ng_glyco_hcd203040_compound15.23162.23162.2 | controllerType=0 controllerNumber=1 scan=23162 | 46.0434 |
| 21282:4 | 1 | R.TAGWNVPIGTLRPFLN[+2350.830]WTGPPEPIEAAVAR.F | HexNAc(4)Hex(5)Fuc(1)NeuAc(2) | N[+2351] | 1396.1397 | 4 | 5581.5368 | 5581.5413 | 0 | -0.8 | 141 | Specific | 731.02 | 554.06 | 554.06 | 14.18 | 142 | >sp\|P02788\|TRFL_HUMAN Lactotransferrin OS=Homo sapiens OX=9606 GN=LTF PE=1 SV=6 | 1270 | 20201113_12433_1ng_glyco_hcd203040_compound15.24795.24795.6 | controllerType=0 controllerNumber=1 scan=24795 | 49.2699 |
| 19901:3 | 1 | R.TAGWNVPIGTLRPFLN[+2205.793]WTGPPEPIEAAVAR.F | HexNAc(4)Hex(5)Fuc(2)NeuAc(1) | N[+2206] | 1812.8454 | 3 | 5436.5217 | 5436.5038 | 0 | 3.3 | 141 | Specific | 743.38 | 644.21 | 644.21 | 14.84 | 142 | >sp\|P02788\|TRFL_HUMAN Lactotransferrin OS=Homo sapiens OX=9606 GN=LTF PE=1 SV=6 | 1270 | 20201113_12433_1ng_glyco_hcd203040_compound15.23354.23354.2 | controllerType=0 controllerNumber=1 scan=23354 | 46.4541 |
| 19891:3 | 1 | R.TAGWNVPIGTLRPFLN[+2059.735]WTGPPEPIEAAVAR.F | HexNAc(4)Hex(5)Fuc(1)NeuAc(1) | N[+2060] | 1764.16 | 3 | 5290.4654 | 5290.4459 | 0 | 3.7 | 141 | Specific | 746.72 | 670.62 | 670.62 | 14.5 | 142 | >sp\|P02788\|TRFL_HUMAN Lactotransferrin OS=Homo sapiens OX=9606 GN=LTF PE=1 SV=6 | 1270 | 20201113_12433_1ng_glyco_hcd203040_compound15.23344.23344.2 | controllerType=0 controllerNumber=1 scan=23344 | 46.4341 |
| 19843:5 | 1 | R.TAGWNVPIGTLRPFLN[+2205.793]WTGPPEPIEAAVAR.F | HexNAc(4)Hex(5)Fuc(2)NeuAc(1) | N[+2206] | 1088.1097 | 5 | 5436.5196 | 5436.5038 | 0 | 2.9 | 141 | Specific | 747.7 | 641.12 | 641.12 | 14.93 | 142 | >sp\|P02788\|TRFL_HUMAN Lactotransferrin OS=Homo sapiens OX=9606 GN=LTF PE=1 SV=6 | 1270 | 20201113_12433_1ng_glyco_hcd203040_compound15.23294.23294.4 | controllerType=0 controllerNumber=1 scan=23294 | 46.3312 |
| 19719:4 | 1 | R.TAGWNVPIGTLRPFLN[+2570.925]WTGPPEPIEAAVAR.F | HexNAc(5)Hex(6)Fuc(2)NeuAc(1) | N[+2571] | 1451.1696 | 4 | 5801.6564 | 5801.636 | 0 | 3.5 | 141 | Specific | 768.31 | 624.81 | 624.81 | 15.82 | 142 | >sp\|P02788\|TRFL_HUMAN Lactotransferrin OS=Homo sapiens OX=9606 GN=LTF PE=1 SV=6 | 1270 | 20201113_12433_1ng_glyco_hcd203040_compound15.23164.23164.2 | controllerType=0 controllerNumber=1 scan=23164 | 46.0507 |
| 21374:4 | 1 | R.TAGWNVPIGTLRPFLN[+2350.830]WTGPPEPIEAAVAR.F | HexNAc(4)Hex(5)Fuc(1)NeuAc(2) | N[+2351] | 1396.1449 | 4 | 5581.5578 | 5581.5413 | 0 | 3 | 141 | Specific | 774.6 | 643.71 | 643.71 | 15.82 | 142 | >sp\|P02788\|TRFL_HUMAN Lactotransferrin OS=Homo sapiens OX=9606 GN=LTF PE=1 SV=6 | 1270 | 20201113_12433_1ng_glyco_hcd203040_compound15.24891.24891.4 | controllerType=0 controllerNumber=1 scan=24891 | 49.4746 |
| 19864:4 | 1 | R.TAGWNVPIGTLRPFLN[+2205.793]WTGPPEPIEAAVAR.F | HexNAc(4)Hex(5)Fuc(2)NeuAc(1) | N[+2206] | 1359.8842 | 4 | 5436.5148 | 5436.5038 | 0 | 2 | 141 | Specific | 817.36 | 704.19 | 704.19 | 16.83 | 142 | >sp\|P02788\|TRFL_HUMAN Lactotransferrin OS=Homo sapiens OX=9606 GN=LTF PE=1 SV=6 | 1270 | 20201113_12433_1ng_glyco_hcd203040_compound15.23316.23316.4 | controllerType=0 controllerNumber=1 scan=23316 | 46.3801 |
| 19853:4 | 1 | R.TAGWNVPIGTLRPFLN[+2059.735]WTGPPEPIEAAVAR.F | HexNAc(4)Hex(5)Fuc(1)NeuAc(1) | N[+2060] | 1323.3718 | 4 | 5290.4655 | 5290.4459 | 0 | 3.7 | 141 | Specific | 868.44 | 733.89 | 733.89 | 17.48 | 142 | >sp\|P02788\|TRFL_HUMAN Lactotransferrin OS=Homo sapiens OX=9606 GN=LTF PE=1 SV=6 | 1270 | 20201113_12433_1ng_glyco_hcd203040_compound15.23304.23304.4 | controllerType=0 controllerNumber=1 scan=23304 | 46.3538 |
| 18895:4 | 1 | R.TAGWNVPIGTLRPFLN[+1768.640]WTGPPEPIEAAVAR.F | HexNAc(4)Hex(5)Fuc(1) | N[+1769] | 1250.5918 | 4 | 4999.3454 | 4999.3504 | 0 | -1 | 141 | Specific | 881.53 | 724.02 | 724.02 | 17.18 | 142 | >sp\|P02788\|TRFL_HUMAN Lactotransferrin OS=Homo sapiens OX=9606 GN=LTF PE=1 SV=6 | 1270 | 20201113_12433_1ng_glyco_hcd203040_compound15.22306.22306.4 | controllerType=0 controllerNumber=1 scan=22306 | 44.1918 |
| 19870:5 | 1 | R.TAGWNVPIGTLRPFLN[+2205.793]WTGPPEPIEAAVAR.F | HexNAc(4)Hex(5)Fuc(2)NeuAc(1) | N[+2206] | 1088.1095 | 5 | 5436.5184 | 5436.5038 | 0 | 2.7 | 141 | Specific | 937.8 | 759.82 | 759.82 | 18.93 | 142 | >sp\|P02788\|TRFL_HUMAN Lactotransferrin OS=Homo sapiens OX=9606 GN=LTF PE=1 SV=6 | 1270 | 20201113_12433_1ng_glyco_hcd203040_compound15.23322.23322.5 | controllerType=0 controllerNumber=1 scan=23322 | 46.3881 |
| 19845:5 | 1 | R.TAGWNVPIGTLRPFLN[+2205.793]WTGPPEPIEAAVAR.F | HexNAc(4)Hex(5)Fuc(2)NeuAc(1) | N[+2206] | 1088.1097 | 5 | 5436.5196 | 5436.5038 | 0 | 2.9 | 141 | Specific | 991.26 | 783.09 | 783.09 | 19.93 | 142 | >sp\|P02788\|TRFL_HUMAN Lactotransferrin OS=Homo sapiens OX=9606 GN=LTF PE=1 SV=6 | 1270 | 20201113_12433_1ng_glyco_hcd203040_compound15.23296.23296.4 | controllerType=0 controllerNumber=1 scan=23296 | 46.3353 |
| 17695:3 | 1 | R.TAGWNIPMGLLFN[+1460.529]QTGSC[+57.021]K.F | HexNAc(4)Hex(4) | C[+57], N[+1461] | 1185.8517 | 3 | 3555.5407 | 3555.5387 | 0 | 0.5 | 485 | Specific | 300.02 | 300.02 | 300.02 | 6.56 | 142 | >sp\|P02788\|TRFL_HUMAN Lactotransferrin OS=Homo sapiens OX=9606 GN=LTF PE=1 SV=6 | 1270 | 20201113_12433_1ng_glyco_hcd203040_compound15.21055.21055.3 | controllerType=0 controllerNumber=1 scan=21055 | 41.5511 |
| 19424:3 | 1 | R.TAGWNIPMGLLFN[+1872.651]QTGSC[+57.021]K.F | HexNAc(3)Hex(6)NeuAc(1) | C[+57], N[+1873] | 1323.2254 | 3 | 3967.6617 | 3967.6604 | 0 | 0.3 | 485 | Specific | 319.97 | 318.19 | 222.05 | 6.81 | 142 | >sp\|P02788\|TRFL_HUMAN Lactotransferrin OS=Homo sapiens OX=9606 GN=LTF PE=1 SV=6 | 1270 | 20201113_12433_1ng_glyco_hcd203040_compound15.22858.22858.3 | controllerType=0 controllerNumber=1 scan=22858 | 45.4346 |
| 17486:3 | 1 | R.TAGWNIPMGLLFN[+1606.587]QTGSC[+57.021]K.F | HexNAc(4)Hex(4)Fuc(1) | C[+57], N[+1607] | 1234.5375 | 3 | 3701.598 | 3701.5966 | 0 | 0.4 | 485 | Specific | 404.34 | 404.34 | 404.34 | 9.07 | 142 | >sp\|P02788\|TRFL_HUMAN Lactotransferrin OS=Homo sapiens OX=9606 GN=LTF PE=1 SV=6 | 1270 | 20201113_12433_1ng_glyco_hcd203040_compound15.20838.20838.3 | controllerType=0 controllerNumber=1 scan=20838 | 41.1014 |
| 21509:3 | 1 | R.TAGWNIPMGLLFN[+2204.772]QTGSC[+57.021]K.F | HexNAc(4)Hex(5)NeuAc(2) | C[+57], N[+2205] | 1433.9372 | 3 | 4299.797 | 4299.7824 | 0 | 3.4 | 485 | Specific | 425.21 | 425.21 | 425.21 | 9.46 | 142 | >sp\|P02788\|TRFL_HUMAN Lactotransferrin OS=Homo sapiens OX=9606 GN=LTF PE=1 SV=6 | 1270 | 20201113_12433_1ng_glyco_hcd203040_compound15.25032.25032.3 | controllerType=0 controllerNumber=1 scan=25032 | 49.7565 |
| 17386:2 | 1 | R.TAGWNIPMGLLFN[+1768.640]QTGSC[+57.021]K.F | HexNAc(4)Hex(5)Fuc(1) | C[+57], N[+1769] | 1932.3361 | 2 | 3863.665 | 3863.6495 | 0 | 4 | 485 | Specific | 431.93 | 406.64 | 406.64 | 9.13 | 142 | >sp\|P02788\|TRFL_HUMAN Lactotransferrin OS=Homo sapiens OX=9606 GN=LTF PE=1 SV=6 | 1270 | 20201113_12433_1ng_glyco_hcd203040_compound15.20734.20734.2 | controllerType=0 controllerNumber=1 scan=20734 | 40.8997 |
| 17388:2 | 1 | R.TAGWNIPMGLLFN[+1768.640]QTGSC[+57.021]K.F | HexNAc(4)Hex(5)Fuc(1) | C[+57], N[+1769] | 1932.3361 | 2 | 3863.665 | 3863.6495 | 0 | 4 | 485 | Specific | 447.19 | 428.12 | 428.12 | 9.22 | 142 | >sp\|P02788\|TRFL_HUMAN Lactotransferrin OS=Homo sapiens OX=9606 GN=LTF PE=1 SV=6 | 1270 | 20201113_12433_1ng_glyco_hcd203040_compound15.20736.20736.2 | controllerType=0 controllerNumber=1 scan=20736 | 40.9036 |
| 19426:3 | 1 | R.TAGWNIPMGLLFN[+1872.651]QTGSC[+57.021]K.F | HexNAc(3)Hex(6)NeuAc(1) | C[+57], N[+1873] | 1323.2254 | 3 | 3967.6617 | 3967.6604 | 0 | 0.3 | 485 | Specific | 491.25 | 287.15 | 287.15 | 7.29 | 142 | >sp\|P02788\|TRFL_HUMAN Lactotransferrin OS=Homo sapiens OX=9606 GN=LTF PE=1 SV=6 | 1270 | 20201113_12433_1ng_glyco_hcd203040_compound15.22860.22860.3 | controllerType=0 controllerNumber=1 scan=22860 | 45.4408 |
| 17399:4 | 1 | R.TAGWNIPMGLLFN[+1768.640]QTGSC[+57.021]K.F | HexNAc(4)Hex(5)Fuc(1) | C[+57], N[+1769] | 966.6718 | 4 | 3863.6654 | 3863.6495 | 0 | 4.1 | 485 | Specific | 553.26 | 518.18 | 518.18 | 9.99 | 142 | >sp\|P02788\|TRFL_HUMAN Lactotransferrin OS=Homo sapiens OX=9606 GN=LTF PE=1 SV=6 | 1270 | 20201113_12433_1ng_glyco_hcd203040_compound15.20747.20747.4 | controllerType=0 controllerNumber=1 scan=20747 | 40.9254 |
| 19314:3 | 1 | R.TAGWNIPMGLLFN[+2059.735]QTGSC[+57.021]K.F | HexNAc(4)Hex(5)Fuc(1)NeuAc(1) | C[+57], N[+2060] | 1385.5969 | 3 | 4154.776 | 4154.7449 | 0 | 7.5 | 485 | Specific | 557.12 | 534.92 | 534.92 | 9.11 | 142 | >sp\|P02788\|TRFL_HUMAN Lactotransferrin OS=Homo sapiens OX=9606 GN=LTF PE=1 SV=6 | 1270 | 20201113_12433_1ng_glyco_hcd203040_compound15.22743.22743.4 | controllerType=0 controllerNumber=1 scan=22743 | 45.1748 |
| 17396:4 | 1 | R.TAGWNIPMGLLFN[+1768.640]QTGSC[+57.021]K.F | HexNAc(4)Hex(5)Fuc(1) | C[+57], N[+1769] | 966.6718 | 4 | 3863.6654 | 3863.6495 | 0 | 4.1 | 485 | Specific | 564.65 | 528.55 | 528.55 | 9.99 | 142 | >sp\|P02788\|TRFL_HUMAN Lactotransferrin OS=Homo sapiens OX=9606 GN=LTF PE=1 SV=6 | 1270 | 20201113_12433_1ng_glyco_hcd203040_compound15.20744.20744.4 | controllerType=0 controllerNumber=1 scan=20744 | 40.9188 |
| 17790:2 | 1 | R.TAGWNIPMGLLFN[+1216.423]QTGSC[+57.021]K.F | HexNAc(2)Hex(5) | C[+57], N[+1216] | 1656.2299 | 2 | 3311.4525 | 3311.4328 | 0 | 6 | 485 | Specific | 598.05 | 579.95 | 216.13 | 10.87 | 142 | >sp\|P02788\|TRFL_HUMAN Lactotransferrin OS=Homo sapiens OX=9606 GN=LTF PE=1 SV=6 | 1270 | 20201113_12433_1ng_glyco_hcd203040_compound15.21154.21154.2 | controllerType=0 controllerNumber=1 scan=21154 | 41.7494 |
| 19458:3 | 1 | R.TAGWNIPMGLLFN[+1751.624]QTGSC[+57.021]K.F | HexNAc(4)Hex(4)NeuAc(1) | C[+57], N[+1752] | 1282.8881 | 3 | 3846.6498 | 3846.6341 | 0 | 4.1 | 485 | Specific | 696.27 | 696.27 | 696.27 | 13.13 | 142 | >sp\|P02788\|TRFL_HUMAN Lactotransferrin OS=Homo sapiens OX=9606 GN=LTF PE=1 SV=6 | 1270 | 20201113_12433_1ng_glyco_hcd203040_compound15.22894.22894.3 | controllerType=0 controllerNumber=1 scan=22894 | 45.5264 |
| 19370:4 | 1 | R.TAGWNIPMGLLFN[+1913.677]QTGSC[+57.021]K.F | HexNAc(4)Hex(5)NeuAc(1) | C[+57], N[+1914] | 1002.9283 | 4 | 4008.6912 | 4008.687 | 0 | 1.1 | 485 | Specific | 737.58 | 685.44 | 685.44 | 14.22 | 142 | >sp\|P02788\|TRFL_HUMAN Lactotransferrin OS=Homo sapiens OX=9606 GN=LTF PE=1 SV=6 | 1270 | 20201113_12433_1ng_glyco_hcd203040_compound15.22802.22802.4 | controllerType=0 controllerNumber=1 scan=22802 | 45.3321 |
| 17698:3 | 1 | R.TAGWNIPMGLLFN[+1460.529]QTGSC[+57.021]K.F | HexNAc(4)Hex(4) | C[+57], N[+1461] | 1185.8517 | 3 | 3555.5407 | 3555.5387 | 0 | 0.5 | 485 | Specific | 741.66 | 588.24 | 348.73 | 14.11 | 142 | >sp\|P02788\|TRFL_HUMAN Lactotransferrin OS=Homo sapiens OX=9606 GN=LTF PE=1 SV=6 | 1270 | 20201113_12433_1ng_glyco_hcd203040_compound15.21058.21058.3 | controllerType=0 controllerNumber=1 scan=21058 | 41.5551 |
| 17788:2 | 1 | R.TAGWNIPMGLLFN[+1216.423]QTGSC[+57.021]K.F | HexNAc(2)Hex(5) | C[+57], N[+1216] | 1656.2299 | 2 | 3311.4525 | 3311.4328 | 0 | 6 | 485 | Specific | 792.91 | 792.91 | 252.59 | 14.87 | 142 | >sp\|P02788\|TRFL_HUMAN Lactotransferrin OS=Homo sapiens OX=9606 GN=LTF PE=1 SV=6 | 1270 | 20201113_12433_1ng_glyco_hcd203040_compound15.21152.21152.2 | controllerType=0 controllerNumber=1 scan=21152 | 41.7454 |
| 19373:4 | 1 | R.TAGWNIPMGLLFN[+1913.677]QTGSC[+57.021]K.F | HexNAc(4)Hex(5)NeuAc(1) | C[+57], N[+1914] | 1002.9283 | 4 | 4008.6912 | 4008.687 | 0 | 1.1 | 485 | Specific | 842.59 | 660.88 | 660.88 | 16.22 | 142 | >sp\|P02788\|TRFL_HUMAN Lactotransferrin OS=Homo sapiens OX=9606 GN=LTF PE=1 SV=6 | 1270 | 20201113_12433_1ng_glyco_hcd203040_compound15.22805.22805.4 | controllerType=0 controllerNumber=1 scan=22805 | 45.3375 |
| 17750:3 | 1 | R.TAGWNIPMGLLFN[+1216.423]QTGSC[+57.021]K.F | HexNAc(2)Hex(5) | C[+57], N[+1216] | 1104.4857 | 3 | 3311.4427 | 3311.4328 | 0 | 3 | 485 | Specific | 850.35 | 850.35 | 353.46 | 17.48 | 142 | >sp\|P02788\|TRFL_HUMAN Lactotransferrin OS=Homo sapiens OX=9606 GN=LTF PE=1 SV=6 | 1270 | 20201113_12433_1ng_glyco_hcd203040_compound15.21112.21112.3 | controllerType=0 controllerNumber=1 scan=21112 | 41.6607 |
| 19188:3 | 1 | R.TAGWNIPMGLLFN[+2059.735]QTGSC[+57.021]K.F | HexNAc(4)Hex(5)Fuc(1)NeuAc(1) | C[+57], N[+2060] | 1385.5889 | 3 | 4154.7522 | 4154.7449 | 0 | 1.8 | 485 | Specific | 865.87 | 779.99 | 779.99 | 17.48 | 142 | >sp\|P02788\|TRFL_HUMAN Lactotransferrin OS=Homo sapiens OX=9606 GN=LTF PE=1 SV=6 | 1270 | 20201113_12433_1ng_glyco_hcd203040_compound15.22612.22612.3 | controllerType=0 controllerNumber=1 scan=22612 | 44.9068 |
| 21511:3 | 1 | R.TAGWNIPMGLLFN[+2204.772]QTGSC[+57.021]K.F | HexNAc(4)Hex(5)NeuAc(2) | C[+57], N[+2205] | 1433.9372 | 3 | 4299.797 | 4299.7824 | 0 | 3.4 | 485 | Specific | 870.01 | 742.06 | 742.06 | 17.48 | 142 | >sp\|P02788\|TRFL_HUMAN Lactotransferrin OS=Homo sapiens OX=9606 GN=LTF PE=1 SV=6 | 1270 | 20201113_12433_1ng_glyco_hcd203040_compound15.25034.25034.3 | controllerType=0 controllerNumber=1 scan=25034 | 49.7604 |
| 19179:3 | 1 | R.TAGWNIPMGLLFN[+2205.793]QTGSC[+57.021]K.F | HexNAc(4)Hex(5)Fuc(2)NeuAc(1) | C[+57], N[+2206] | 1434.2761 | 3 | 4300.8136 | 4300.8028 | 0 | 2.5 | 485 | Specific | 870.29 | 831.35 | 831.35 | 17.48 | 142 | >sp\|P02788\|TRFL_HUMAN Lactotransferrin OS=Homo sapiens OX=9606 GN=LTF PE=1 SV=6 | 1270 | 20201113_12433_1ng_glyco_hcd203040_compound15.22603.22603.3 | controllerType=0 controllerNumber=1 scan=22603 | 44.8811 |
| 17563:3 | 1 | R.TAGWNIPMGLLFN[+1622.582]QTGSC[+57.021]K.F | HexNAc(4)Hex(5) | C[+57], N[+1623] | 1239.874 | 3 | 3717.6073 | 3717.5916 | 0 | 4.2 | 485 | Specific | 872.75 | 870.6 | 405.63 | 17.13 | 142 | >sp\|P02788\|TRFL_HUMAN Lactotransferrin OS=Homo sapiens OX=9606 GN=LTF PE=1 SV=6 | 1270 | 20201113_12433_1ng_glyco_hcd203040_compound15.20918.20918.3 | controllerType=0 controllerNumber=1 scan=20918 | 41.2531 |
| 17565:3 | 1 | R.TAGWNIPMGLLFN[+1622.582]QTGSC[+57.021]K.F | HexNAc(4)Hex(5) | C[+57], N[+1623] | 1239.874 | 3 | 3717.6073 | 3717.5916 | 0 | 4.2 | 485 | Specific | 881.25 | 850.76 | 424.92 | 17.13 | 142 | >sp\|P02788\|TRFL_HUMAN Lactotransferrin OS=Homo sapiens OX=9606 GN=LTF PE=1 SV=6 | 1270 | 20201113_12433_1ng_glyco_hcd203040_compound15.20920.20920.3 | controllerType=0 controllerNumber=1 scan=20920 | 41.2581 |
| 19461:3 | 1 | R.TAGWNIPMGLLFN[+1751.624]QTGSC[+57.021]K.F | HexNAc(4)Hex(4)NeuAc(1) | C[+57], N[+1752] | 1282.8881 | 3 | 3846.6498 | 3846.6341 | 0 | 4.1 | 485 | Specific | 889.82 | 810.83 | 810.83 | 17.13 | 142 | >sp\|P02788\|TRFL_HUMAN Lactotransferrin OS=Homo sapiens OX=9606 GN=LTF PE=1 SV=6 | 1270 | 20201113_12433_1ng_glyco_hcd203040_compound15.22897.22897.3 | controllerType=0 controllerNumber=1 scan=22897 | 45.5307 |
| 19191:3 | 1 | R.TAGWNIPMGLLFN[+2059.735]QTGSC[+57.021]K.F | HexNAc(4)Hex(5)Fuc(1)NeuAc(1) | C[+57], N[+2060] | 1385.5889 | 3 | 4154.7522 | 4154.7449 | 0 | 1.8 | 485 | Specific | 894.71 | 694.93 | 694.93 | 17.48 | 142 | >sp\|P02788\|TRFL_HUMAN Lactotransferrin OS=Homo sapiens OX=9606 GN=LTF PE=1 SV=6 | 1270 | 20201113_12433_1ng_glyco_hcd203040_compound15.22615.22615.3 | controllerType=0 controllerNumber=1 scan=22615 | 44.9199 |
| 17753:3 | 1 | R.TAGWNIPMGLLFN[+1216.423]QTGSC[+57.021]K.F | HexNAc(2)Hex(5) | C[+57], N[+1216] | 1104.4857 | 3 | 3311.4427 | 3311.4328 | 0 | 3 | 485 | Specific | 899.66 | 898.45 | 395.48 | 17.48 | 142 | >sp\|P02788\|TRFL_HUMAN Lactotransferrin OS=Homo sapiens OX=9606 GN=LTF PE=1 SV=6 | 1270 | 20201113_12433_1ng_glyco_hcd203040_compound15.21115.21115.3 | controllerType=0 controllerNumber=1 scan=21115 | 41.6664 |
| 21291:3 | 1 | R.TAGWNIPMGLLFN[+2350.830]QTGSC[+57.021]K.F | HexNAc(4)Hex(5)Fuc(1)NeuAc(2) | C[+57], N[+2351] | 1482.6197 | 3 | 4445.8445 | 4445.8403 | 0 | 0.9 | 485 | Specific | 908.96 | 774.01 | 774.01 | 18.11 | 142 | >sp\|P02788\|TRFL_HUMAN Lactotransferrin OS=Homo sapiens OX=9606 GN=LTF PE=1 SV=6 | 1270 | 20201113_12433_1ng_glyco_hcd203040_compound15.24805.24805.3 | controllerType=0 controllerNumber=1 scan=24805 | 49.2888 |
| 17336:3 | 1 | R.TAGWNIPMGLLFN[+1768.640]QTGSC[+57.021]K.F | HexNAc(4)Hex(5)Fuc(1) | C[+57], N[+1769] | 1288.558 | 3 | 3863.6596 | 3863.6495 | 0 | 2.6 | 485 | Specific | 917.39 | 917.39 | 917.39 | 18.48 | 142 | >sp\|P02788\|TRFL_HUMAN Lactotransferrin OS=Homo sapiens OX=9606 GN=LTF PE=1 SV=6 | 1270 | 20201113_12433_1ng_glyco_hcd203040_compound15.20682.20682.3 | controllerType=0 controllerNumber=1 scan=20682 | 40.8002 |
| 19330:3 | 1 | R.TAGWNIPMGLLFN[+1913.677]QTGSC[+57.021]K.F | HexNAc(4)Hex(5)NeuAc(1) | C[+57], N[+1914] | 1336.9055 | 3 | 4008.7018 | 4008.687 | 0 | 3.7 | 485 | Specific | 919.92 | 905.95 | 905.95 | 18.13 | 142 | >sp\|P02788\|TRFL_HUMAN Lactotransferrin OS=Homo sapiens OX=9606 GN=LTF PE=1 SV=6 | 1270 | 20201113_12433_1ng_glyco_hcd203040_compound15.22760.22760.3 | controllerType=0 controllerNumber=1 scan=22760 | 45.2202 |
| 17333:3 | 1 | R.TAGWNIPMGLLFN[+1768.640]QTGSC[+57.021]K.F | HexNAc(4)Hex(5)Fuc(1) | C[+57], N[+1769] | 1288.558 | 3 | 3863.6596 | 3863.6495 | 0 | 2.6 | 485 | Specific | 929.32 | 929.32 | 929.32 | 18.48 | 142 | >sp\|P02788\|TRFL_HUMAN Lactotransferrin OS=Homo sapiens OX=9606 GN=LTF PE=1 SV=6 | 1270 | 20201113_12433_1ng_glyco_hcd203040_compound15.20679.20679.3 | controllerType=0 controllerNumber=1 scan=20679 | 40.7938 |
| 21288:3 | 1 | R.TAGWNIPMGLLFN[+2350.830]QTGSC[+57.021]K.F | HexNAc(4)Hex(5)Fuc(1)NeuAc(2) | C[+57], N[+2351] | 1482.6197 | 3 | 4445.8445 | 4445.8403 | 0 | 0.9 | 485 | Specific | 934.33 | 805.24 | 805.24 | 18.11 | 142 | >sp\|P02788\|TRFL_HUMAN Lactotransferrin OS=Homo sapiens OX=9606 GN=LTF PE=1 SV=6 | 1270 | 20201113_12433_1ng_glyco_hcd203040_compound15.24802.24802.3 | controllerType=0 controllerNumber=1 scan=24802 | 49.2829 |
| 19328:3 | 1 | R.TAGWNIPMGLLFN[+1913.677]QTGSC[+57.021]K.F | HexNAc(4)Hex(5)NeuAc(1) | C[+57], N[+1914] | 1336.9055 | 3 | 4008.7018 | 4008.687 | 0 | 3.7 | 485 | Specific | 981.67 | 967.27 | 967.27 | 19.13 | 142 | >sp\|P02788\|TRFL_HUMAN Lactotransferrin OS=Homo sapiens OX=9606 GN=LTF PE=1 SV=6 | 1270 | 20201113_12433_1ng_glyco_hcd203040_compound15.22758.22758.3 | controllerType=0 controllerNumber=1 scan=22758 | 45.2093 |
| 17488:3 | 1 | R.TAGWNIPMGLLFN[+1606.587]QTGSC[+57.021]K.F | HexNAc(4)Hex(4)Fuc(1) | C[+57], N[+1607] | 1234.5375 | 3 | 3701.598 | 3701.5966 | 0 | 0.4 | 485 | Specific | 981.94 | 747.26 | 747.26 | 19.11 | 142 | >sp\|P02788\|TRFL_HUMAN Lactotransferrin OS=Homo sapiens OX=9606 GN=LTF PE=1 SV=6 | 1270 | 20201113_12433_1ng_glyco_hcd203040_compound15.20840.20840.3 | controllerType=0 controllerNumber=1 scan=20840 | 41.1049 |
| 19177:3 | 1 | R.TAGWNIPMGLLFN[+2205.793]QTGSC[+57.021]K.F | HexNAc(4)Hex(5)Fuc(2)NeuAc(1) | C[+57], N[+2206] | 1434.2761 | 3 | 4300.8136 | 4300.8028 | 0 | 2.5 | 485 | Specific | 991.02 | 857.03 | 857.03 | 19.48 | 142 | >sp\|P02788\|TRFL_HUMAN Lactotransferrin OS=Homo sapiens OX=9606 GN=LTF PE=1 SV=6 | 1270 | 20201113_12433_1ng_glyco_hcd203040_compound15.22601.22601.3 | controllerType=0 controllerNumber=1 scan=22601 | 44.8661 |
| 18445:3 | 1 | R.TAGWNIPM[+15.995]GLLFN[+2059.735]QTGSC[+57.021]K.F | HexNAc(4)Hex(5)Fuc(1)NeuAc(1) | C[+57], M[+16], N[+2060] | 1390.9268 | 3 | 4170.7659 | 4170.7398 | 0 | 6.3 | 485 | Specific | 256.06 | 151.85 | 151.85 | 3.76 | 142 | >sp\|P02788\|TRFL_HUMAN Lactotransferrin OS=Homo sapiens OX=9606 GN=LTF PE=1 SV=6 | 1270 | 20201113_12433_1ng_glyco_hcd203040_compound15.21837.21837.3 | controllerType=0 controllerNumber=1 scan=21837 | 43.2155 |
| 18560:4 | 1 | R.TAGWNIPMGLLFN[+1606.587]QTGSC[+57.021]KFDEYFSQSC[+57.021]APGSDPR.S | HexNAc(4)Hex(4)Fuc(1) | C[+57]*2, N[+1607] | 1387.0978 | 4 | 5545.3693 | 5545.3438 | 0 | 4.6 | 485 | Specific | 362.06 | 362.06 | 362.06 | 6.99 | 142 | >sp\|P02788\|TRFL_HUMAN Lactotransferrin OS=Homo sapiens OX=9606 GN=LTF PE=1 SV=6 | 1270 | 20201113_12433_1ng_glyco_hcd203040_compound15.21957.21957.4 | controllerType=0 controllerNumber=1 scan=21957 | 43.4535 |
| 18486:3 | 1 | R.TAGWNIPMGLLFN[+1768.640]QTGSC[+57.021]KFDEYFSQSC[+57.021]APGSDPR.S | HexNAc(4)Hex(5)Fuc(1) | C[+57]*2, N[+1769] | 1903.1417 | 3 | 5707.4105 | 5707.3966 | 0 | 2.4 | 485 | Specific | 388.28 | 348.87 | 348.87 | 7.38 | 142 | >sp\|P02788\|TRFL_HUMAN Lactotransferrin OS=Homo sapiens OX=9606 GN=LTF PE=1 SV=6 | 1270 | 20201113_12433_1ng_glyco_hcd203040_compound15.21880.21880.3 | controllerType=0 controllerNumber=1 scan=21880 | 43.3009 |
| 18544:3 | 1 | R.TAGWNIPMGLLFN[+1622.582]QTGSC[+57.021]KFDEYFSQSC[+57.021]APGSDPR.S | HexNAc(4)Hex(5) | C[+57]*2, N[+1623] | 1854.4594 | 3 | 5561.3638 | 5561.3387 | 0 | 4.5 | 485 | Specific | 411.6 | 340.54 | 177.44 | 7.04 | 142 | >sp\|P02788\|TRFL_HUMAN Lactotransferrin OS=Homo sapiens OX=9606 GN=LTF PE=1 SV=6 | 1270 | 20201113_12433_1ng_glyco_hcd203040_compound15.21940.21940.3 | controllerType=0 controllerNumber=1 scan=21940 | 43.4256 |
| 19573:3 | 1 | R.TAGWNIPMGLLFN[+1913.677]QTGSC[+57.021]KFDEYFSQSC[+57.021]APGSDPR.S | HexNAc(4)Hex(5)NeuAc(1) | C[+57]*2, N[+1914] | 1951.4964 | 3 | 5852.4747 | 5852.4341 | 0 | 6.9 | 485 | Specific | 420.4 | 411.68 | 411.68 | 8.15 | 142 | >sp\|P02788\|TRFL_HUMAN Lactotransferrin OS=Homo sapiens OX=9606 GN=LTF PE=1 SV=6 | 1270 | 20201113_12433_1ng_glyco_hcd203040_compound15.23013.23013.3 | controllerType=0 controllerNumber=1 scan=23013 | 45.7583 |
| 21281:4 | 1 | R.[+42.011]TAGWNIPMGLLFN[+2059.735]QTGSC[+57.021]KFDEYFSQSC[+57.021]A | HexNAc(4)Hex(5)Fuc(1)NeuAc(1) | .[42], C[+57]*2, N[+2060] | 1510.8777 | 4 | 6040.4889 | 6040.5026 | 0 | -2.3 | 485 | Specific | 459.64 | 407.06 | 407.06 | 8 | 142 | >sp\|P02788\|TRFL_HUMAN Lactotransferrin OS=Homo sapiens OX=9606 GN=LTF PE=1 SV=6 | 1270 | 20201113_12433_1ng_glyco_hcd203040_compound15.24794.24794.4 | controllerType=0 controllerNumber=1 scan=24794 | 49.2672 |
| 18562:4 | 1 | R.TAGWNIPMGLLFN[+1606.587]QTGSC[+57.021]KFDEYFSQSC[+57.021]APGSDPR.S | HexNAc(4)Hex(4)Fuc(1) | C[+57]*2, N[+1607] | 1387.0978 | 4 | 5545.3693 | 5545.3438 | 0 | 4.6 | 485 | Specific | 480.56 | 474.73 | 474.73 | 8.92 | 142 | >sp\|P02788\|TRFL_HUMAN Lactotransferrin OS=Homo sapiens OX=9606 GN=LTF PE=1 SV=6 | 1270 | 20201113_12433_1ng_glyco_hcd203040_compound15.21959.21959.4 | controllerType=0 controllerNumber=1 scan=21959 | 43.4685 |
| 21283:4 | 1 | R.[+42.011]TAGWNIPMGLLFN[+2059.735]QTGSC[+57.021]KFDEYFSQSC[+57.021]A | HexNAc(4)Hex(5)Fuc(1)NeuAc(1) | .[42], C[+57]*2, N[+2060] | 1510.8777 | 4 | 6040.4889 | 6040.5026 | 0 | -2.3 | 485 | Specific | 492.89 | 457.39 | 457.39 | 8 | 142 | >sp\|P02788\|TRFL_HUMAN Lactotransferrin OS=Homo sapiens OX=9606 GN=LTF PE=1 SV=6 | 1270 | 20201113_12433_1ng_glyco_hcd203040_compound15.24796.24796.4 | controllerType=0 controllerNumber=1 scan=24796 | 49.2716 |
| 20910:4 | 1 | R.TAGWNIPMGLLFN[+2350.830]QTGSC[+57.021]KFDEYFSQSC[+57.021]APGSDPR.S | HexNAc(4)Hex(5)Fuc(1)NeuAc(2) | C[+57]*2, N[+2351] | 1573.156 | 4 | 6289.6022 | 6289.5875 | 0 | 2.3 | 485 | Specific | 524.69 | 474.29 | 474.29 | 9.46 | 142 | >sp\|P02788\|TRFL_HUMAN Lactotransferrin OS=Homo sapiens OX=9606 GN=LTF PE=1 SV=6 | 1270 | 20201113_12433_1ng_glyco_hcd203040_compound15.24408.24408.4 | controllerType=0 controllerNumber=1 scan=24408 | 48.4595 |
| 20866:4 | 1 | R.TAGWNIPMGLLFN[+2350.830]QTGSC[+57.021]KFDEYFSQSC[+57.021]APGSDPR.S | HexNAc(4)Hex(5)Fuc(1)NeuAc(2) | C[+57]*2, N[+2351] | 1573.1583 | 4 | 6289.6115 | 6289.5875 | 0 | 3.8 | 485 | Specific | 526.84 | 526.84 | 526.84 | 9.11 | 142 | >sp\|P02788\|TRFL_HUMAN Lactotransferrin OS=Homo sapiens OX=9606 GN=LTF PE=1 SV=6 | 1270 | 20201113_12433_1ng_glyco_hcd203040_compound15.24362.24362.4 | controllerType=0 controllerNumber=1 scan=24362 | 48.371 |
| 19568:4 | 1 | R.TAGWNIPMGLLFN[+1913.677]QTGSC[+57.021]KFDEYFSQSC[+57.021]APGSDPR.S | HexNAc(4)Hex(5)NeuAc(1) | C[+57]*2, N[+1914] | 1463.876 | 4 | 5852.4821 | 5852.4341 | 0 | 8.2 | 485 | Specific | 532.55 | 532.55 | 532.55 | 8.46 | 142 | >sp\|P02788\|TRFL_HUMAN Lactotransferrin OS=Homo sapiens OX=9606 GN=LTF PE=1 SV=6 | 1270 | 20201113_12433_1ng_glyco_hcd203040_compound15.23008.23008.4 | controllerType=0 controllerNumber=1 scan=23008 | 45.7505 |
| 19576:3 | 1 | R.TAGWNIPMGLLFN[+1913.677]QTGSC[+57.021]KFDEYFSQSC[+57.021]APGSDPR.S | HexNAc(4)Hex(5)NeuAc(1) | C[+57]*2, N[+1914] | 1951.4964 | 3 | 5852.4747 | 5852.4341 | 0 | 6.9 | 485 | Specific | 538.76 | 498.89 | 498.89 | 8.7 | 142 | >sp\|P02788\|TRFL_HUMAN Lactotransferrin OS=Homo sapiens OX=9606 GN=LTF PE=1 SV=6 | 1270 | 20201113_12433_1ng_glyco_hcd203040_compound15.23016.23016.3 | controllerType=0 controllerNumber=1 scan=23016 | 45.7632 |
| 18542:3 | 1 | R.TAGWNIPMGLLFN[+1622.582]QTGSC[+57.021]KFDEYFSQSC[+57.021]APGSDPR.S | HexNAc(4)Hex(5) | C[+57]*2, N[+1623] | 1854.4594 | 3 | 5561.3638 | 5561.3387 | 0 | 4.5 | 485 | Specific | 598.2 | 542.58 | 185.68 | 10.71 | 142 | >sp\|P02788\|TRFL_HUMAN Lactotransferrin OS=Homo sapiens OX=9606 GN=LTF PE=1 SV=6 | 1270 | 20201113_12433_1ng_glyco_hcd203040_compound15.21938.21938.3 | controllerType=0 controllerNumber=1 scan=21938 | 43.4166 |
| 19604:4 | 1 | R.TAGWNIPMGLLFN[+2059.735]QTGSC[+57.021]KFDEYFSQSC[+57.021]APGSDPR.S | HexNAc(4)Hex(5)Fuc(1)NeuAc(1) | C[+57]*2, N[+2060] | 1500.3768 | 4 | 5998.4855 | 5998.492 | 0 | -1.1 | 485 | Specific | 599.14 | 544.68 | 544.68 | 10.39 | 142 | >sp\|P02788\|TRFL_HUMAN Lactotransferrin OS=Homo sapiens OX=9606 GN=LTF PE=1 SV=6 | 1270 | 20201113_12433_1ng_glyco_hcd203040_compound15.23045.23045.4 | controllerType=0 controllerNumber=1 scan=23045 | 45.8277 |
| 20912:4 | 1 | R.TAGWNIPMGLLFN[+2350.830]QTGSC[+57.021]KFDEYFSQSC[+57.021]APGSDPR.S | HexNAc(4)Hex(5)Fuc(1)NeuAc(2) | C[+57]*2, N[+2351] | 1573.156 | 4 | 6289.6022 | 6289.5875 | 0 | 2.3 | 485 | Specific | 662.36 | 545.15 | 545.15 | 13.04 | 142 | >sp\|P02788\|TRFL_HUMAN Lactotransferrin OS=Homo sapiens OX=9606 GN=LTF PE=1 SV=6 | 1270 | 20201113_12433_1ng_glyco_hcd203040_compound15.24410.24410.4 | controllerType=0 controllerNumber=1 scan=24410 | 48.4635 |
| 18471:3 | 1 | R.TAGWNIPMGLLFN[+1768.640]QTGSC[+57.021]KFDEYFSQSC[+57.021]APGSDPR.S | HexNAc(4)Hex(5)Fuc(1) | C[+57]*2, N[+1769] | 1903.1412 | 3 | 5707.409 | 5707.3966 | 0 | 2.2 | 485 | Specific | 684.97 | 605.57 | 605.57 | 13.07 | 142 | >sp\|P02788\|TRFL_HUMAN Lactotransferrin OS=Homo sapiens OX=9606 GN=LTF PE=1 SV=6 | 1270 | 20201113_12433_1ng_glyco_hcd203040_compound15.21864.21864.3 | controllerType=0 controllerNumber=1 scan=21864 | 43.2689 |
| 18489:3 | 1 | R.TAGWNIPMGLLFN[+1768.640]QTGSC[+57.021]KFDEYFSQSC[+57.021]APGSDPR.S | HexNAc(4)Hex(5)Fuc(1) | C[+57]*2, N[+1769] | 1903.1417 | 3 | 5707.4105 | 5707.3966 | 0 | 2.4 | 485 | Specific | 687.19 | 537.84 | 537.84 | 13.06 | 142 | >sp\|P02788\|TRFL_HUMAN Lactotransferrin OS=Homo sapiens OX=9606 GN=LTF PE=1 SV=6 | 1270 | 20201113_12433_1ng_glyco_hcd203040_compound15.21883.21883.3 | controllerType=0 controllerNumber=1 scan=21883 | 43.3052 |
| 19455:4 | 1 | R.TAGWNIPMGLLFN[+2205.793]QTGSC[+57.021]KFDEYFSQSC[+57.021]APGSDPR.S | HexNAc(4)Hex(5)Fuc(2)NeuAc(1) | C[+57]*2, N[+2206] | 1536.8994 | 4 | 6144.5758 | 6144.5499 | 0 | 4.2 | 485 | Specific | 692.77 | 663.9 | 663.9 | 12.69 | 142 | >sp\|P02788\|TRFL_HUMAN Lactotransferrin OS=Homo sapiens OX=9606 GN=LTF PE=1 SV=6 | 1270 | 20201113_12433_1ng_glyco_hcd203040_compound15.22891.22891.4 | controllerType=0 controllerNumber=1 scan=22891 | 45.5208 |
| 18482:4 | 1 | R.TAGWNIPMGLLFN[+1768.640]QTGSC[+57.021]KFDEYFSQSC[+57.021]APGSDPR.S | HexNAc(4)Hex(5)Fuc(1) | C[+57]*2, N[+1769] | 1427.6095 | 4 | 5707.4162 | 5707.3966 | 0 | 3.4 | 485 | Specific | 728.98 | 689 | 689 | 14.04 | 142 | >sp\|P02788\|TRFL_HUMAN Lactotransferrin OS=Homo sapiens OX=9606 GN=LTF PE=1 SV=6 | 1270 | 20201113_12433_1ng_glyco_hcd203040_compound15.21876.21876.4 | controllerType=0 controllerNumber=1 scan=21876 | 43.2942 |
| 19606:4 | 1 | R.TAGWNIPMGLLFN[+2059.735]QTGSC[+57.021]KFDEYFSQSC[+57.021]APGSDPR.S | HexNAc(4)Hex(5)Fuc(1)NeuAc(1) | C[+57]*2, N[+2060] | 1500.3768 | 4 | 5998.4855 | 5998.492 | 0 | -1.1 | 485 | Specific | 730.83 | 681.43 | 681.43 | 13.39 | 142 | >sp\|P02788\|TRFL_HUMAN Lactotransferrin OS=Homo sapiens OX=9606 GN=LTF PE=1 SV=6 | 1270 | 20201113_12433_1ng_glyco_hcd203040_compound15.23047.23047.4 | controllerType=0 controllerNumber=1 scan=23047 | 45.8318 |
| 19483:4 | 1 | R.TAGWNIPMGLLFN[+2059.735]QTGSC[+57.021]KFDEYFSQSC[+57.021]APGSDPR.S | HexNAc(4)Hex(5)Fuc(1)NeuAc(1) | C[+57]*2, N[+2060] | 1500.381 | 4 | 5998.5021 | 5998.492 | 0 | 1.7 | 485 | Specific | 757.67 | 674.87 | 674.87 | 15.04 | 142 | >sp\|P02788\|TRFL_HUMAN Lactotransferrin OS=Homo sapiens OX=9606 GN=LTF PE=1 SV=6 | 1270 | 20201113_12433_1ng_glyco_hcd203040_compound15.22920.22920.4 | controllerType=0 controllerNumber=1 scan=22920 | 45.5714 |
| 18473:3 | 1 | R.TAGWNIPMGLLFN[+1768.640]QTGSC[+57.021]KFDEYFSQSC[+57.021]APGSDPR.S | HexNAc(4)Hex(5)Fuc(1) | C[+57]*2, N[+1769] | 1903.1412 | 3 | 5707.409 | 5707.3966 | 0 | 2.2 | 485 | Specific | 786.04 | 657.47 | 657.47 | 15.07 | 142 | >sp\|P02788\|TRFL_HUMAN Lactotransferrin OS=Homo sapiens OX=9606 GN=LTF PE=1 SV=6 | 1270 | 20201113_12433_1ng_glyco_hcd203040_compound15.21866.21866.3 | controllerType=0 controllerNumber=1 scan=21866 | 43.2731 |
| 18533:4 | 1 | R.TAGWNIPMGLLFN[+1622.582]QTGSC[+57.021]KFDEYFSQSC[+57.021]APGSDPR.S | HexNAc(4)Hex(5) | C[+57]*2, N[+1623] | 1391.0967 | 4 | 5561.3649 | 5561.3387 | 0 | 4.7 | 485 | Specific | 841.14 | 790.41 | 243.92 | 15.69 | 142 | >sp\|P02788\|TRFL_HUMAN Lactotransferrin OS=Homo sapiens OX=9606 GN=LTF PE=1 SV=6 | 1270 | 20201113_12433_1ng_glyco_hcd203040_compound15.21929.21929.4 | controllerType=0 controllerNumber=1 scan=21929 | 43.3974 |
| 19457:4 | 1 | R.TAGWNIPMGLLFN[+2205.793]QTGSC[+57.021]KFDEYFSQSC[+57.021]APGSDPR.S | HexNAc(4)Hex(5)Fuc(2)NeuAc(1) | C[+57]*2, N[+2206] | 1536.8994 | 4 | 6144.5758 | 6144.5499 | 0 | 4.2 | 485 | Specific | 884 | 853.26 | 853.26 | 16.69 | 142 | >sp\|P02788\|TRFL_HUMAN Lactotransferrin OS=Homo sapiens OX=9606 GN=LTF PE=1 SV=6 | 1270 | 20201113_12433_1ng_glyco_hcd203040_compound15.22893.22893.4 | controllerType=0 controllerNumber=1 scan=22893 | 45.5252 |
| 19571:4 | 1 | R.TAGWNIPMGLLFN[+1913.677]QTGSC[+57.021]KFDEYFSQSC[+57.021]APGSDPR.S | HexNAc(4)Hex(5)NeuAc(1) | C[+57]*2, N[+1914] | 1463.876 | 4 | 5852.4821 | 5852.4341 | 0 | 8.2 | 485 | Specific | 886.95 | 886.95 | 886.95 | 15.93 | 142 | >sp\|P02788\|TRFL_HUMAN Lactotransferrin OS=Homo sapiens OX=9606 GN=LTF PE=1 SV=6 | 1270 | 20201113_12433_1ng_glyco_hcd203040_compound15.23011.23011.4 | controllerType=0 controllerNumber=1 scan=23011 | 45.755 |
| 18535:4 | 1 | R.TAGWNIPMGLLFN[+1622.582]QTGSC[+57.021]KFDEYFSQSC[+57.021]APGSDPR.S | HexNAc(4)Hex(5) | C[+57]*2, N[+1623] | 1391.0967 | 4 | 5561.3649 | 5561.3387 | 0 | 4.7 | 485 | Specific | 899.57 | 734.18 | 388.08 | 16.69 | 142 | >sp\|P02788\|TRFL_HUMAN Lactotransferrin OS=Homo sapiens OX=9606 GN=LTF PE=1 SV=6 | 1270 | 20201113_12433_1ng_glyco_hcd203040_compound15.21931.21931.4 | controllerType=0 controllerNumber=1 scan=21931 | 43.4032 |
| 18484:4 | 1 | R.TAGWNIPMGLLFN[+1768.640]QTGSC[+57.021]KFDEYFSQSC[+57.021]APGSDPR.S | HexNAc(4)Hex(5)Fuc(1) | C[+57]*2, N[+1769] | 1427.6095 | 4 | 5707.4162 | 5707.3966 | 0 | 3.4 | 485 | Specific | 904.97 | 865.26 | 865.26 | 18.04 | 142 | >sp\|P02788\|TRFL_HUMAN Lactotransferrin OS=Homo sapiens OX=9606 GN=LTF PE=1 SV=6 | 1270 | 20201113_12433_1ng_glyco_hcd203040_compound15.21878.21878.4 | controllerType=0 controllerNumber=1 scan=21878 | 43.2987 |
| 20864:4 | 1 | R.TAGWNIPMGLLFN[+2350.830]QTGSC[+57.021]KFDEYFSQSC[+57.021]APGSDPR.S | HexNAc(4)Hex(5)Fuc(1)NeuAc(2) | C[+57]*2, N[+2351] | 1573.1583 | 4 | 6289.6115 | 6289.5875 | 0 | 3.8 | 485 | Specific | 952.98 | 920.21 | 920.21 | 18.69 | 142 | >sp\|P02788\|TRFL_HUMAN Lactotransferrin OS=Homo sapiens OX=9606 GN=LTF PE=1 SV=6 | 1270 | 20201113_12433_1ng_glyco_hcd203040_compound15.24360.24360.4 | controllerType=0 controllerNumber=1 scan=24360 | 48.3649 |
| 19485:4 | 1 | R.TAGWNIPMGLLFN[+2059.735]QTGSC[+57.021]KFDEYFSQSC[+57.021]APGSDPR.S | HexNAc(4)Hex(5)Fuc(1)NeuAc(1) | C[+57]*2, N[+2060] | 1500.381 | 4 | 5998.5021 | 5998.492 | 0 | 1.7 | 485 | Specific | 1001.07 | 944.41 | 944.41 | 20.04 | 142 | >sp\|P02788\|TRFL_HUMAN Lactotransferrin OS=Homo sapiens OX=9606 GN=LTF PE=1 SV=6 | 1270 | 20201113_12433_1ng_glyco_hcd203040_compound15.22922.22922.4 | controllerType=0 controllerNumber=1 scan=22922 | 45.5755 |
| 08522:3 | 1 | K.FGRN[+1501.555]GSDC[+57.021]PDKFC[+57.021]LFQSETK.N | HexNAc(5)Hex(3) | C[+57]*2, N[+1502] | 1298.8847 | 3 | 3894.6395 | 3894.6202 | 0 | 5 | 639 | Specific | 201.94 | 201.94 | 201.94 | 3.39 | 142 | >sp\|P02788\|TRFL_HUMAN Lactotransferrin OS=Homo sapiens OX=9606 GN=LTF PE=1 SV=6 | 1270 | 20201113_12433_1ng_glyco_hcd203040_compound15.11503.11503.3 | controllerType=0 controllerNumber=1 scan=11503 | 25.2122 |
| 09436:3 | 1 | R.N[+2059.735]GSDC[+57.021]PDKFC[+57.021]LFQSETK.N | HexNAc(4)Hex(5)Fuc(1)NeuAc(1) | C[+57]*2, N[+2060] | 1364.8797 | 3 | 4092.6245 | 4092.6088 | 0 | 3.8 | 642 | Specific | 365.29 | 303.88 | 303.88 | 6.38 | 142 | >sp\|P02788\|TRFL_HUMAN Lactotransferrin OS=Homo sapiens OX=9606 GN=LTF PE=1 SV=6 | 1270 | 20201113_12433_1ng_glyco_hcd203040_compound15.12454.12454.3 | controllerType=0 controllerNumber=1 scan=12454 | 26.7997 |
| 09438:3 | 1 | R.N[+2059.735]GSDC[+57.021]PDKFC[+57.021]LFQSETK.N | HexNAc(4)Hex(5)Fuc(1)NeuAc(1) | C[+57]*2, N[+2060] | 1364.8797 | 3 | 4092.6245 | 4092.6088 | 0 | 3.8 | 642 | Specific | 400.67 | 267.88 | 267.88 | 6.46 | 142 | >sp\|P02788\|TRFL_HUMAN Lactotransferrin OS=Homo sapiens OX=9606 GN=LTF PE=1 SV=6 | 1270 | 20201113_12433_1ng_glyco_hcd203040_compound15.12456.12456.3 | controllerType=0 controllerNumber=1 scan=12456 | 26.8036 |
| 09358:3 | 1 | R.N[+2205.793]GSDC[+57.021]PDKFC[+57.021]LFQSETK.N | HexNAc(4)Hex(5)Fuc(2)NeuAc(1) | C[+57]*2, N[+2206] | 1413.5629 | 3 | 4238.6742 | 4238.6667 | 0 | 1.8 | 642 | Specific | 402.84 | 255.6 | 255.6 | 6.78 | 142 | >sp\|P02788\|TRFL_HUMAN Lactotransferrin OS=Homo sapiens OX=9606 GN=LTF PE=1 SV=6 | 1270 | 20201113_12433_1ng_glyco_hcd203040_compound15.12374.12374.3 | controllerType=0 controllerNumber=1 scan=12374 | 26.6712 |
| 09360:3 | 1 | R.N[+2205.793]GSDC[+57.021]PDKFC[+57.021]LFQSETK.N | HexNAc(4)Hex(5)Fuc(2)NeuAc(1) | C[+57]*2, N[+2206] | 1413.5629 | 3 | 4238.6742 | 4238.6667 | 0 | 1.8 | 642 | Specific | 594.37 | 423.01 | 423.01 | 10.87 | 142 | >sp\|P02788\|TRFL_HUMAN Lactotransferrin OS=Homo sapiens OX=9606 GN=LTF PE=1 SV=6 | 1270 | 20201113_12433_1ng_glyco_hcd203040_compound15.12376.12376.3 | controllerType=0 controllerNumber=1 scan=12376 | 26.675 |

Table S4. Cont.

| Query #:z | Protein Rank | Glycopeptide | Glycan | Modification Type(s) | Observed m/z | z | Observed (M+H) | Calc. mass (M+H) | Off-by-x error | Mass error (ppm) | StartPosition | Cleavage | Score | Delta | Delta Mod | \|Log Prob\| | # of unique peptides | Protein Name | Protein DB number | Comment | Scan # | Scan Time |
|---|---|---|---|---|---|---|---|---|---|---|---|---|---|---|---|---|---|---|---|---|---|---|
| 07589:3 | 4 | R.FSDGLESN[+1864.634]SSTQFEVK.K | HexNAc(2)Hex(9) | N[+1865] | 1213.8223 | 3 | 3639.4524 | 3639.4472 | 0 | 1.4 | 219 | Specific | 618.33 | 618.33 | 618.33 | 12.28 | 89 | >sp\|P0C0L5\|CO4B_HUMAN Complement C4-B OS=Homo sapiens OX=9606 GN=C4B PE=1 SV=2 | 1424 | 20201113_12433_1ng_glyco_hcd203040_compound15.10531.10531.3 | controllerType=0 controllerNumber=1 scan=10531 | 23.6216 |
| 07591:3 | 4 | R.FSDGLESN[+1864.634]SSTQFEVK.K | HexNAc(2)Hex(9) | N[+1865] | 1213.8223 | 3 | 3639.4524 | 3639.4472 | 0 | 1.4 | 219 | Specific | 648.1 | 623.9 | 623.9 | 12.28 | 89 | >sp\|P0C0L5\|CO4B_HUMAN Complement C4-B OS=Homo sapiens OX=9606 GN=C4B PE=1 SV=2 | 1424 | 20201113_12433_1ng_glyco_hcd203040_compound15.10533.10533.3 | controllerType=0 controllerNumber=1 scan=10533 | 23.6252 |
| 05058:3 | 4 | R.FSDGLESN[+1864.634]SSTQFEVKK.Y | HexNAc(2)Hex(9) | N[+1865] | 1256.5217 | 3 | 3767.5504 | 3767.5421 | 0 | 2.2 | 219 | Specific | 460.04 | 416.68 | 416.68 | 9.1 | 89 | >sp\|P0C0L5\|CO4B_HUMAN Complement C4-B OS=Homo sapiens OX=9606 GN=C4B PE=1 SV=2 | 1424 | 20201113_12433_1ng_glyco_hcd203040_compound15.7912.7912.3 | controllerType=0 controllerNumber=1 scan=7912 | 19.3423 |
| 05055:3 | 4 | R.FSDGLESN[+1864.634]SSTQFEVKK.Y | HexNAc(2)Hex(9) | N[+1865] | 1256.5217 | 3 | 3767.5504 | 3767.5421 | 0 | 2.2 | 219 | Specific | 641.13 | 487.94 | 487.94 | 11.87 | 89 | >sp\|P0C0L5\|CO4B_HUMAN Complement C4-B OS=Homo sapiens OX=9606 GN=C4B PE=1 SV=2 | 1424 | 20201113_12433_1ng_glyco_hcd203040_compound15.7909.7909.3 | controllerType=0 controllerNumber=1 scan=7909 | 19.3367 |
| 10473:3 | 5 | K.TVLTPATNHMGN[+1378.476]VTFTIPANR.E | HexNAc(2)Hex(6) | N[+1378] | 1211.8914 | 3 | 3633.6597 | 3633.6358 | 0 | 6.6 | 74 | Specific | 223.18 | 223.18 | 185.12 | 4.12 | 102 | >sp\|P01024\|CO3_HUMAN Complement C3 OS=Homo sapiens OX=9606 GN=C3 PE=1 SV=2 | 492 | 20201113_12433_1ng_glyco_hcd203040_compound15.13526.13526.3 | controllerType=0 controllerNumber=1 scan=13526 | 28.5501 |
| 10475:3 | 5 | K.TVLTPATNHMGN[+1378.476]VTFTIPANR.E | HexNAc(2)Hex(6) | N[+1378] | 1211.8914 | 3 | 3633.6597 | 3633.6358 | 0 | 6.6 | 74 | Specific | 237.26 | 138.72 | 138.72 | 4.34 | 102 | >sp\|P01024\|CO3_HUMAN Complement C3 OS=Homo sapiens OX=9606 GN=C3 PE=1 SV=2 | 492 | 20201113_12433_1ng_glyco_hcd203040_compound15.13528.13528.3 | controllerType=0 controllerNumber=1 scan=13528 | 28.5542 |
| 07970:3 | 6 | K.WEC[+57.021]KN[+2059.735]DTLLGIK.G | HexNAc(4)Hex(5)Fuc(1)NeuAc(1) | C[+57], N[+2060] | 1179.5037 | 3 | 3536.4966 | 3536.4865 | 0 | 2.9 | 100 | Specific | 313.49 | 275.19 | 275.19 | 6.34 | 86 | >sp\|P22897\|MRC1_HUMAN Macrophage mannose receptor 1 OS=Homo sapiens OX=9606 GN=MRC1 PE=1 SV=1 | 1049 | 20201113_12433_1ng_glyco_hcd203040_compound15.10924.10924.3 | controllerType=0 controllerNumber=1 scan=10924 | 24.2559 |
| 07968:3 | 6 | K.WEC[+57.021]KN[+2059.735]DTLLGIK.G | HexNAc(4)Hex(5)Fuc(1)NeuAc(1) | C[+57], N[+2060] | 1179.5037 | 3 | 3536.4966 | 3536.4865 | 0 | 2.9 | 100 | Specific | 411.14 | 191.5 | 191.5 | 6.3 | 86 | >sp\|P22897\|MRC1_HUMAN Macrophage mannose receptor 1 OS=Homo sapiens OX=9606 GN=MRC1 PE=1 SV=1 | 1049 | 20201113_12433_1ng_glyco_hcd203040_compound15.10922.10922.3 | controllerType=0 controllerNumber=1 scan=10922 | 24.2524 |
| 14809:4 | 6 | K.KGN[+2351.851]TTLNSFVIPSESDVPTHC[+57.021]PSQWWPYAGHC[+57.021]YK.I | HexNAc(4)Hex(5)Fuc(3)NeuAc(1) | C[+57]*2, N[+2352] | 1579.6753 | 4 | 6315.6794 | 6315.6725 | 0 | 1.1 | 342 | Specific | 343.74 | 276.42 | 276.42 | 6.27 | 86 | >sp\|P22897\|MRC1_HUMAN Macrophage mannose receptor 1 OS=Homo sapiens OX=9606 GN=MRC1 PE=1 SV=1 | 1049 | 20201113_12433_1ng_glyco_hcd203040_compound15.18034.18034.4 | controllerType=0 controllerNumber=1 scan=18034 | 36.1262 |
| 13534:5 | 6 | K.KGN[+2205.793]TTLNSFVIPSESDVPTHC[+57.021]PSQWWPYAGHC[+57.021]YK.I | HexNAc(4)Hex(5)Fuc(2)NeuAc(1) | C[+57]*2, N[+2206] | 1234.7327 | 5 | 6169.6342 | 6169.6146 | 0 | 3.2 | 342 | Specific | 347.19 | 347.19 | 347.19 | 7.57 | 86 | >sp\|P22897\|MRC1_HUMAN Macrophage mannose receptor 1 OS=Homo sapiens OX=9606 GN=MRC1 PE=1 SV=1 | 1049 | 20201113_12433_1ng_glyco_hcd203040_compound15.16715.16715.5 | controllerType=0 controllerNumber=1 scan=16715 | 33.8637 |
| 14811:4 | 6 | K.KGN[+2351.851]TTLNSFVIPSESDVPTHC[+57.021]PSQWWPYAGHC[+57.021]YK.I | HexNAc(4)Hex(5)Fuc(3)NeuAc(1) | C[+57]*2, N[+2352] | 1579.6753 | 4 | 6315.6794 | 6315.6725 | 0 | 1.1 | 342 | Specific | 417.84 | 417.84 | 417.84 | 8.56 | 86 | >sp\|P22897\|MRC1_HUMAN Macrophage mannose receptor 1 OS=Homo sapiens OX=9606 GN=MRC1 PE=1 SV=1 | 1049 | 20201113_12433_1ng_glyco_hcd203040_compound15.18036.18036.4 | controllerType=0 controllerNumber=1 scan=18036 | 36.1303 |
| 08729:3 | 6 | K.TAHC[+57.021]N[+2059.735]ESFYFLC[+57.021]K.R | HexNAc(4)Hex(5)Fuc(1)NeuAc(1) | C[+57]*2, N[+2060] | 1246.1569 | 3 | 3736.4562 | 3736.4545 | 0 | 0.5 | 1201 | Specific | 558.62 | 418.66 | 418.66 | 9.66 | 86 | >sp\|P22897\|MRC1_HUMAN Macrophage mannose receptor 1 OS=Homo sapiens OX=9606 GN=MRC1 PE=1 SV=1 | 1049 | 20201113_12433_1ng_glyco_hcd203040_compound15.11717.11717.3 | controllerType=0 controllerNumber=1 scan=11717 | 25.5543 |
| 08731:3 | 6 | K.TAHC[+57.021]N[+2059.735]ESFYFLC[+57.021]K.R | HexNAc(4)Hex(5)Fuc(1)NeuAc(1) | C[+57]*2, N[+2060] | 1246.1569 | 3 | 3736.4562 | 3736.4545 | 0 | 0.5 | 1201 | Specific | 654.53 | 551.8 | 551.8 | 12.92 | 86 | >sp\|P22897\|MRC1_HUMAN Macrophage mannose receptor 1 OS=Homo sapiens OX=9606 GN=MRC1 PE=1 SV=1 | 1049 | 20201113_12433_1ng_glyco_hcd203040_compound15.11719.11719.3 | controllerType=0 controllerNumber=1 scan=11719 | 25.5583 |
| 18018:3 | 8 | R.GLSFDVSLEVSQGPGLLN[+2716.983]DTK.V | HexNAc(5)Hex(6)Fuc(3)NeuAc(1) | N[+2717] | 1631.7138 | 3 | 4893.1269 | 4893.0962 | 0 | 6.3 | 118 | Specific | 476.04 | 476.04 | 476.04 | 8.82 | 61 | >sp\|P01833\|PIGR_HUMAN Polymeric immunoglobulin receptor OS=Homo sapiens OX=9606 GN=PIGR PE=1 SV=4 | 902 | 20201113_12433_1ng_glyco_hcd203040_compound15.21391.21391.3 | controllerType=0 controllerNumber=1 scan=21391 | 42.2222 |
| 18280:3 | 8 | R.GLSFDVSLEVSQGPGLLN[+2059.735]DTK.V | HexNAc(4)Hex(5)Fuc(1)NeuAc(1) | N[+2060] | 1412.6287 | 3 | 4235.8716 | 4235.8481 | 0 | 5.5 | 118 | Specific | 481.78 | 481.78 | 481.78 | 9.08 | 61 | >sp\|P01833\|PIGR_HUMAN Polymeric immunoglobulin receptor OS=Homo sapiens OX=9606 GN=PIGR PE=1 SV=4 | 902 | 20201113_12433_1ng_glyco_hcd203040_compound15.21665.21665.3 | controllerType=0 controllerNumber=1 scan=21665 | 42.8429 |
| 19381:3 | 8 | K.QIGLYPVLVIDSSGYVNPN[+2059.735]YTGR.I | HexNAc(4)Hex(5)Fuc(1)NeuAc(1) | N[+2060] | 1529.0203 | 3 | 4585.0464 | 4585.0384 | 0 | 1.7 | 168 | Specific | 324.95 | 252.64 | 252.64 | 7.61 | 61 | >sp\|P01833\|PIGR_HUMAN Polymeric immunoglobulin receptor OS=Homo sapiens OX=9606 GN=PIGR PE=1 SV=4 | 902 | 20201113_12433_1ng_glyco_hcd203040_compound15.22814.22814.3 | controllerType=0 controllerNumber=1 scan=22814 | 45.3584 |
| 19383:3 | 8 | K.QIGLYPVLVIDSSGYVNPN[+2059.735]YTGR.I | HexNAc(4)Hex(5)Fuc(1)NeuAc(1) | N[+2060] | 1529.0203 | 3 | 4585.0464 | 4585.0384 | 0 | 1.7 | 168 | Specific | 886.93 | 807.49 | 807.49 | 18.16 | 61 | >sp\|P01833\|PIGR_HUMAN Polymeric immunoglobulin receptor OS=Homo sapiens OX=9606 GN=PIGR PE=1 SV=4 | 902 | 20201113_12433_1ng_glyco_hcd203040_compound15.22816.22816.3 | controllerType=0 controllerNumber=1 scan=22816 | 45.362 |
| 29650:3 | 8 | R.[+42.011]LSLLEEPGN[+2351.851]GTFTVILNQLTSR.D | HexNAc(4)Hex(5)Fuc(3)NeuAc(1) | .[42], N[+2352] | 1599.3843 | 3 | 4796.1385 | 4796.1539 | 0 | -3.2 | 413 | Specific | 308.53 | 263.18 | 263.18 | 6.02 | 61 | >sp\|P01833\|PIGR_HUMAN Polymeric immunoglobulin receptor OS=Homo sapiens OX=9606 GN=PIGR PE=1 SV=4 | 902 | 20201113_12433_1ng_glyco_hcd203040_compound15.33514.33514.3 | controllerType=0 controllerNumber=1 scan=33514 | 64.9619 |
| 23536:3 | 8 | R.[+42.011]LSLLEEPGN[+2336.851]GTFTVILNQLTSR.D | HexNAc(6)Hex(6)Fuc(1) | .[42], N[+2337] | 1594.3842 | 3 | 4781.1381 | 4781.1542 | 0 | -3.4 | 413 | Specific | 326.56 | 268.81 | 268.81 | 6.02 | 61 | >sp\|P01833\|PIGR_HUMAN Polymeric immunoglobulin receptor OS=Homo sapiens OX=9606 GN=PIGR PE=1 SV=4 | 902 | 20201113_12433_1ng_glyco_hcd203040_compound15.27143.27143.3 | controllerType=0 controllerNumber=1 scan=27143 | 53.4603 |
| 23534:3 | 8 | R.[+42.011]LSLLEEPGN[+2336.851]GTFTVILNQLTSR.D | HexNAc(6)Hex(6)Fuc(1) | .[42], N[+2337] | 1594.3842 | 3 | 4781.1381 | 4781.1542 | 0 | -3.4 | 413 | Specific | 354.41 | 344.53 | 344.53 | 6.61 | 61 | >sp\|P01833\|PIGR_HUMAN Polymeric immunoglobulin receptor OS=Homo sapiens OX=9606 GN=PIGR PE=1 SV=4 | 902 | 20201113_12433_1ng_glyco_hcd203040_compound15.27141.27141.3 | controllerType=0 controllerNumber=1 scan=27141 | 53.4562 |
| 23550:4 | 8 | R.LSLLEEPGN[+2351.851]GTFTVILNQLTSR.D | HexNAc(4)Hex(5)Fuc(3)NeuAc(1) | N[+2352] | 1189.2991 | 4 | 4754.1746 | 4754.1433 | 0 | 6.6 | 413 | Specific | 360.4 | 360.13 | 360.13 | 7.68 | 61 | >sp\|P01833\|PIGR_HUMAN Polymeric immunoglobulin receptor OS=Homo sapiens OX=9606 GN=PIGR PE=1 SV=4 | 902 | 20201113_12433_1ng_glyco_hcd203040_compound15.27157.27157.4 | controllerType=0 controllerNumber=1 scan=27157 | 53.483 |
| 23508:4 | 8 | R.LSLLEEPGN[+2351.851]GTFTVILNQLTSR.D | HexNAc(4)Hex(5)Fuc(3)NeuAc(1) | N[+2352] | 1189.3007 | 4 | 4754.181 | 4754.1433 | 0 | 7.9 | 413 | Specific | 486.82 | 466.53 | 466.53 | 9.4 | 61 | >sp\|P01833\|PIGR_HUMAN Polymeric immunoglobulin receptor OS=Homo sapiens OX=9606 GN=PIGR PE=1 SV=4 | 902 | 20201113_12433_1ng_glyco_hcd203040_compound15.27114.27114.4 | controllerType=0 controllerNumber=1 scan=27114 | 53.3999 |
| 23506:4 | 8 | R.LSLLEEPGN[+2351.851]GTFTVILNQLTSR.D | HexNAc(4)Hex(5)Fuc(3)NeuAc(1) | N[+2352] | 1189.3007 | 4 | 4754.181 | 4754.1433 | 0 | 7.9 | 413 | Specific | 547.74 | 446.19 | 446.19 | 9.7 | 61 | >sp\|P01833\|PIGR_HUMAN Polymeric immunoglobulin receptor OS=Homo sapiens OX=9606 GN=PIGR PE=1 SV=4 | 902 | 20201113_12433_1ng_glyco_hcd203040_compound15.27112.27112.4 | controllerType=0 controllerNumber=1 scan=27112 | 53.3952 |
| 18501:4 | 8 | K.IIEGEPNLKVPGN[+2350.830]VTAVLGETLK.V | HexNAc(4)Hex(5)Fuc(1)NeuAc(2) | N[+2351] | 1186.3034 | 4 | 4742.1917 | 4742.1797 | 0 | 2.5 | 457 | Specific | 212.25 | 184.56 | 184.56 | 3.65 | 61 | >sp\|P01833\|PIGR_HUMAN Polymeric immunoglobulin receptor OS=Homo sapiens OX=9606 GN=PIGR PE=1 SV=4 | 902 | 20201113_12433_1ng_glyco_hcd203040_compound15.21895.21895.4 | controllerType=0 controllerNumber=1 scan=21895 | 43.3278 |
| 18490:3 | 8 | K.IIEGEPNLKVPGN[+2350.830]VTAVLGETLK.V | HexNAc(4)Hex(5)Fuc(1)NeuAc(2) | N[+2351] | 1581.4037 | 3 | 4742.1967 | 4742.1797 | 0 | 3.6 | 457 | Specific | 665.69 | 603.9 | 603.9 | 13.4 | 61 | >sp\|P01833\|PIGR_HUMAN Polymeric immunoglobulin receptor OS=Homo sapiens OX=9606 GN=PIGR PE=1 SV=4 | 902 | 20201113_12433_1ng_glyco_hcd203040_compound15.21884.21884.3 | controllerType=0 controllerNumber=1 scan=21884 | 43.3092 |
| 18487:3 | 8 | K.IIEGEPNLKVPGN[+2350.830]VTAVLGETLK.V | HexNAc(4)Hex(5)Fuc(1)NeuAc(2) | N[+2351] | 1581.4037 | 3 | 4742.1967 | 4742.1797 | 0 | 3.6 | 457 | Specific | 693.35 | 670.86 | 670.86 | 13.4 | 61 | >sp\|P01833\|PIGR_HUMAN Polymeric immunoglobulin receptor OS=Homo sapiens OX=9606 GN=PIGR PE=1 SV=4 | 902 | 20201113_12433_1ng_glyco_hcd203040_compound15.21881.21881.3 | controllerType=0 controllerNumber=1 scan=21881 | 43.3021 |
| 16561:3 | 8 | K.IIEGEPNLKVPGN[+2205.793]VTAVLGETLK.V | HexNAc(4)Hex(5)Fuc(2)NeuAc(1) | N[+2206] | 1533.0571 | 3 | 4597.1566 | 4597.1422 | 0 | 3.1 | 457 | Specific | 783.2 | 684.93 | 684.93 | 15.75 | 61 | >sp\|P01833\|PIGR_HUMAN Polymeric immunoglobulin receptor OS=Homo sapiens OX=9606 GN=PIGR PE=1 SV=4 | 902 | 20201113_12433_1ng_glyco_hcd203040_compound15.19876.19876.3 | controllerType=0 controllerNumber=1 scan=19876 | 39.3897 |
| 16559:3 | 8 | K.IIEGEPNLKVPGN[+2205.793]VTAVLGETLK.V | HexNAc(4)Hex(5)Fuc(2)NeuAc(1) | N[+2206] | 1533.0571 | 3 | 4597.1566 | 4597.1422 | 0 | 3.1 | 457 | Specific | 844.84 | 788.87 | 788.87 | 16.75 | 61 | >sp\|P01833\|PIGR_HUMAN Polymeric immunoglobulin receptor OS=Homo sapiens OX=9606 GN=PIGR PE=1 SV=4 | 902 | 20201113_12433_1ng_glyco_hcd203040_compound15.19874.19874.3 | controllerType=0 controllerNumber=1 scan=19874 | 39.385 |
| 16041:5 | 8 | K.IIEGEPNLKVPGN[+2059.735]VTAVLGETLKVPC[+57.021]HFPC[+57.021]K.F | HexNAc(4)Hex(5)Fuc(1)NeuAc(1) | C[+57]*2, N[+2060] | 1096.1166 | 5 | 5476.5538 | 5476.5418 | 0 | 2.2 | 457 | Specific | 440.93 | 387.16 | 387.16 | 9.32 | 61 | >sp\|P01833\|PIGR_HUMAN Polymeric immunoglobulin receptor OS=Homo sapiens OX=9606 GN=PIGR PE=1 SV=4 | 902 | 20201113_12433_1ng_glyco_hcd203040_compound15.19310.19310.5 | controllerType=0 controllerNumber=1 scan=19310 | 38.3625 |
| 16038:5 | 8 | K.IIEGEPNLKVPGN[+2059.735]VTAVLGETLKVPC[+57.021]HFPC[+57.021]K.F | HexNAc(4)Hex(5)Fuc(1)NeuAc(1) | C[+57]*2, N[+2060] | 1096.1166 | 5 | 5476.5538 | 5476.5418 | 0 | 2.2 | 457 | Specific | 579.12 | 481.44 | 481.44 | 11.25 | 61 | >sp\|P01833\|PIGR_HUMAN Polymeric immunoglobulin receptor OS=Homo sapiens OX=9606 GN=PIGR PE=1 SV=4 | 902 | 20201113_12433_1ng_glyco_hcd203040_compound15.19307.19307.5 | controllerType=0 controllerNumber=1 scan=19307 | 38.3539 |
| 16053:4 | 8 | K.IIEGEPNLKVPGN[+2205.793]VTAVLGETLKVPC[+57.021]HFPC[+57.021]K.F | HexNAc(4)Hex(5)Fuc(2)NeuAc(1) | C[+57]*2, N[+2206] | 1406.4069 | 4 | 5622.6056 | 5622.5997 | 0 | 1 | 457 | Specific | 762.2 | 698.78 | 698.78 | 14.33 | 61 | >sp\|P01833\|PIGR_HUMAN Polymeric immunoglobulin receptor OS=Homo sapiens OX=9606 GN=PIGR PE=1 SV=4 | 902 | 20201113_12433_1ng_glyco_hcd203040_compound15.19323.19323.4 | controllerType=0 controllerNumber=1 scan=19323 | 38.3867 |
| 16052:5 | 8 | K.IIEGEPNLKVPGN[+2204.772]VTAVLGETLKVPC[+57.021]HFPC[+57.021]K.F | HexNAc(4)Hex(5)NeuAc(2) | C[+57]*2, N[+2205] | 1125.1271 | 5 | 5621.6063 | 5621.5793 | 0 | 4.8 | 457 | Specific | 769.49 | 561.51 | 561.51 | 14.61 | 61 | >sp\|P01833\|PIGR_HUMAN Polymeric immunoglobulin receptor OS=Homo sapiens OX=9606 GN=PIGR PE=1 SV=4 | 902 | 20201113_12433_1ng_glyco_hcd203040_compound15.19322.19322.5 | controllerType=0 controllerNumber=1 scan=19322 | 38.3847 |
| 17606:5 | 8 | K.IIEGEPNLKVPGN[+2350.830]VTAVLGETLKVPC[+57.021]HFPC[+57.021]K.F | HexNAc(4)Hex(5)Fuc(1)NeuAc(2) | C[+57]*2, N[+2351] | 1154.3356 | 5 | 5767.6488 | 5767.6372 | 0 | 2 | 457 | Specific | 806.32 | 562.41 | 562.41 | 16.25 | 61 | >sp\|P01833\|PIGR_HUMAN Polymeric immunoglobulin receptor OS=Homo sapiens OX=9606 GN=PIGR PE=1 SV=4 | 902 | 20201113_12433_1ng_glyco_hcd203040_compound15.20963.20963.5 | controllerType=0 controllerNumber=1 scan=20963 | 41.3619 |
| 17608:5 | 8 | K.IIEGEPNLKVPGN[+2350.830]VTAVLGETLKVPC[+57.021]HFPC[+57.021]K.F | HexNAc(4)Hex(5)Fuc(1)NeuAc(2) | C[+57]*2, N[+2351] | 1154.3356 | 5 | 5767.6488 | 5767.6372 | 0 | 2 | 457 | Specific | 862.47 | 834.35 | 834.35 | 17.25 | 61 | >sp\|P01833\|PIGR_HUMAN Polymeric immunoglobulin receptor OS=Homo sapiens OX=9606 GN=PIGR PE=1 SV=4 | 902 | 20201113_12433_1ng_glyco_hcd203040_compound15.20965.20965.5 | controllerType=0 controllerNumber=1 scan=20965 | 41.3664 |
| 16050:4 | 8 | K.IIEGEPNLKVPGN[+2205.793]VTAVLGETLKVPC[+57.021]HFPC[+57.021]K.F | HexNAc(4)Hex(5)Fuc(2)NeuAc(1) | C[+57]*2, N[+2206] | 1406.4069 | 4 | 5622.6056 | 5622.5997 | 0 | 1 | 457 | Specific | 865.33 | 776.13 | 776.13 | 16.33 | 61 | >sp\|P01833\|PIGR_HUMAN Polymeric immunoglobulin receptor OS=Homo sapiens OX=9606 GN=PIGR PE=1 SV=4 | 902 | 20201113_12433_1ng_glyco_hcd203040_compound15.19320.19320.4 | controllerType=0 controllerNumber=1 scan=19320 | 38.3799 |
| 16049:5 | 8 | K.IIEGEPNLKVPGN[+2204.772]VTAVLGETLKVPC[+57.021]HFPC[+57.021]K.F | HexNAc(4)Hex(5)NeuAc(2) | C[+57]*2, N[+2205] | 1125.1271 | 5 | 5621.6063 | 5621.5793 | 0 | 4.8 | 457 | Specific | 908.3 | 760.51 | 760.51 | 17.61 | 61 | >sp\|P01833\|PIGR_HUMAN Polymeric immunoglobulin receptor OS=Homo sapiens OX=9606 GN=PIGR PE=1 SV=4 | 902 | 20201113_12433_1ng_glyco_hcd203040_compound15.19319.19319.5 | controllerType=0 controllerNumber=1 scan=19319 | 38.379 |
| 17572:5 | 8 | K.IIEGEPNLKVPGN[+2350.830]VTAVLGETLKVPC[+57.021]HFPC[+57.021]K.F | HexNAc(4)Hex(5)Fuc(1)NeuAc(2) | C[+57]*2, N[+2351] | 1154.3385 | 5 | 5767.6634 | 5767.6372 | 0 | 4.5 | 457 | Specific | 944.67 | 758.43 | 758.43 | 17.9 | 61 | >sp\|P01833\|PIGR_HUMAN Polymeric immunoglobulin receptor OS=Homo sapiens OX=9606 GN=PIGR PE=1 SV=4 | 902 | 20201113_12433_1ng_glyco_hcd203040_compound15.20927.20927.5 | controllerType=0 controllerNumber=1 scan=20927 | 41.2679 |
| 17168:3 | 8 | K.VPGN[+2350.830]VTAVLGETLK.V | HexNAc(4)Hex(5)Fuc(1)NeuAc(2) | N[+2351] | 1250.2204 | 3 | 3748.6466 | 3748.6302 | 0 | 4.4 | 466 | Specific | 264.36 | 124.28 | 124.28 | 4.06 | 61 | >sp\|P01833\|PIGR_HUMAN Polymeric immunoglobulin receptor OS=Homo sapiens OX=9606 GN=PIGR PE=1 SV=4 | 902 | 20201113_12433_1ng_glyco_hcd203040_compound15.20508.20508.3 | controllerType=0 controllerNumber=1 scan=20508 | 40.4948 |
| 13969:3 | 8 | K.VPGN[+2205.793]VTAVLGETLK.V | HexNAc(4)Hex(5)Fuc(2)NeuAc(1) | N[+2206] | 1201.8777 | 3 | 3603.6187 | 3603.5927 | 0 | 7.2 | 466 | Specific | 347.12 | 347.12 | 347.12 | 6.45 | 61 | >sp\|P01833\|PIGR_HUMAN Polymeric immunoglobulin receptor OS=Homo sapiens OX=9606 GN=PIGR PE=1 SV=4 | 902 | 20201113_12433_1ng_glyco_hcd203040_compound15.17165.17165.3 | controllerType=0 controllerNumber=1 scan=17165 | 34.6061 |
| 13971:3 | 8 | K.VPGN[+2205.793]VTAVLGETLK.V | HexNAc(4)Hex(5)Fuc(2)NeuAc(1) | N[+2206] | 1201.8777 | 3 | 3603.6187 | 3603.5927 | 0 | 7.2 | 466 | Specific | 449.72 | 382.85 | 382.85 | 8.44 | 61 | >sp\|P01833\|PIGR_HUMAN Polymeric immunoglobulin receptor OS=Homo sapiens OX=9606 GN=PIGR PE=1 SV=4 | 902 | 20201113_12433_1ng_glyco_hcd203040_compound15.17167.17167.3 | controllerType=0 controllerNumber=1 scan=17167 | 34.6098 |
| 14133:4 | 8 | K.VPGN[+2059.735]VTAVLGETLKVPC[+57.021]HFPC[+57.021]K.F | HexNAc(4)Hex(5)Fuc(1)NeuAc(1) | C[+57]*2, N[+2060] | 1121.505 | 4 | 4482.9984 | 4482.9923 | 0 | 1.3 | 466 | Specific | 329.83 | 284.1 | 284.1 | 7.2 | 61 | >sp\|P01833\|PIGR_HUMAN Polymeric immunoglobulin receptor OS=Homo sapiens OX=9606 GN=PIGR PE=1 SV=4 | 902 | 20201113_12433_1ng_glyco_hcd203040_compound15.17335.17335.4 | controllerType=0 controllerNumber=1 scan=17335 | 34.9575 |
| 14135:4 | 8 | K.VPGN[+2059.735]VTAVLGETLKVPC[+57.021]HFPC[+57.021]K.F | HexNAc(4)Hex(5)Fuc(1)NeuAc(1) | C[+57]*2, N[+2060] | 1121.505 | 4 | 4482.9984 | 4482.9923 | 0 | 1.3 | 466 | Specific | 357.05 | 262.83 | 262.83 | 7.45 | 61 | >sp\|P01833\|PIGR_HUMAN Polymeric immunoglobulin receptor OS=Homo sapiens OX=9606 GN=PIGR PE=1 SV=4 | 902 | 20201113_12433_1ng_glyco_hcd203040_compound15.17337.17337.4 | controllerType=0 controllerNumber=1 scan=17337 | 34.9617 |
| 05986:3 | 8 | K.WN[+1872.651]NTGC[+57.021]QALPSQDEGPSK.A | HexNAc(3)Hex(6)NeuAc(1) | C[+57], N[+1873] | 1287.86 | 3 | 3861.5656 | 3861.5272 | 0 | 9.9 | 498 | Specific | 210.21 | 210.21 | 210.21 | 3.35 | 61 | >sp\|P01833\|PIGR_HUMAN Polymeric immunoglobulin receptor OS=Homo sapiens OX=9606 GN=PIGR PE=1 SV=4 | 902 | 20201113_12433_1ng_glyco_hcd203040_compound15.8872.8872.3 | controllerType=0 controllerNumber=1 scan=8872 | 20.9178 |
| 05988:3 | 8 | K.WN[+1872.651]NTGC[+57.021]QALPSQDEGPSK.A | HexNAc(3)Hex(6)NeuAc(1) | C[+57], N[+1873] | 1287.86 | 3 | 3861.5656 | 3861.5272 | 0 | 9.9 | 498 | Specific | 289.25 | 289.25 | 289.25 | 4.96 | 61 | >sp\|P01833\|PIGR_HUMAN Polymeric immunoglobulin receptor OS=Homo sapiens OX=9606 GN=PIGR PE=1 SV=4 | 902 | 20201113_12433_1ng_glyco_hcd203040_compound15.8874.8874.3 | controllerType=0 controllerNumber=1 scan=8874 | 20.9209 |
| 06142:3 | 8 | K.WN[+1548.545]NTGC[+57.021]QALPSQDEGPSK.A | HexNAc(3)Hex(4)NeuAc(1) | C[+57], N[+1549] | 1179.8138 | 3 | 3537.4268 | 3537.4215 | 0 | 1.5 | 498 | Specific | 303.93 | 273.28 | 273.28 | 6.77 | 61 | >sp\|P01833\|PIGR_HUMAN Polymeric immunoglobulin receptor OS=Homo sapiens OX=9606 GN=PIGR PE=1 SV=4 | 902 | 20201113_12433_1ng_glyco_hcd203040_compound15.9034.9034.3 | controllerType=0 controllerNumber=1 scan=9034 | 21.1882 |
| 06144:3 | 8 | K.WN[+1548.545]NTGC[+57.021]QALPSQDEGPSK.A | HexNAc(3)Hex(4)NeuAc(1) | C[+57], N[+1549] | 1179.8138 | 3 | 3537.4268 | 3537.4215 | 0 | 1.5 | 498 | Specific | 411.04 | 327.26 | 327.26 | 7.44 | 61 | >sp\|P01833\|PIGR_HUMAN Polymeric immunoglobulin receptor OS=Homo sapiens OX=9606 GN=PIGR PE=1 SV=4 | 902 | 20201113_12433_1ng_glyco_hcd203040_compound15.9036.9036.3 | controllerType=0 controllerNumber=1 scan=9036 | 21.1922 |
| 06070:3 | 8 | K.WN[+1710.598]NTGC[+57.021]QALPSQDEGPSK.A | HexNAc(3)Hex(5)NeuAc(1) | C[+57], N[+1711] | 1233.8338 | 3 | 3699.4868 | 3699.4743 | 0 | 3.4 | 498 | Specific | 484.24 | 428.02 | 428.02 | 9.51 | 61 | >sp\|P01833\|PIGR_HUMAN Polymeric immunoglobulin receptor OS=Homo sapiens OX=9606 GN=PIGR PE=1 SV=4 | 902 | 20201113_12433_1ng_glyco_hcd203040_compound15.8960.8960.3 | controllerType=0 controllerNumber=1 scan=8960 | 21.0671 |
| 06072:3 | 8 | K.WN[+1710.598]NTGC[+57.021]QALPSQDEGPSK.A | HexNAc(3)Hex(5)NeuAc(1) | C[+57], N[+1711] | 1233.8338 | 3 | 3699.4868 | 3699.4743 | 0 | 3.4 | 498 | Specific | 512.64 | 448.33 | 448.33 | 9.69 | 61 | >sp\|P01833\|PIGR_HUMAN Polymeric immunoglobulin receptor OS=Homo sapiens OX=9606 GN=PIGR PE=1 SV=4 | 902 | 20201113_12433_1ng_glyco_hcd203040_compound15.8962.8962.3 | controllerType=0 controllerNumber=1 scan=8962 | 21.0712 |
| 08016:3 | 8 | K.WN[+2204.772]NTGC[+57.021]QALPSQDEGPSK.A | HexNAc(4)Hex(5)NeuAc(2) | C[+57], N[+2205] | 1398.5543 | 3 | 4193.6482 | 4193.6491 | 0 | -0.2 | 498 | Specific | 583.76 | 479.66 | 479.66 | 10.63 | 61 | >sp\|P01833\|PIGR_HUMAN Polymeric immunoglobulin receptor OS=Homo sapiens OX=9606 GN=PIGR PE=1 SV=4 | 902 | 20201113_12433_1ng_glyco_hcd203040_compound15.10972.10972.3 | controllerType=0 controllerNumber=1 scan=10972 | 24.3312 |
| 08019:3 | 8 | K.WN[+2204.772]NTGC[+57.021]QALPSQDEGPSK.A | HexNAc(4)Hex(5)NeuAc(2) | C[+57], N[+2205] | 1398.5543 | 3 | 4193.6482 | 4193.6491 | 0 | -0.2 | 498 | Specific | 589.42 | 501.62 | 501.62 | 10.63 | 61 | >sp\|P01833\|PIGR_HUMAN Polymeric immunoglobulin receptor OS=Homo sapiens OX=9606 GN=PIGR PE=1 SV=4 | 902 | 20201113_12433_1ng_glyco_hcd203040_compound15.10975.10975.3 | controllerType=0 controllerNumber=1 scan=10975 | 24.3367 |
| 05941:3 | 8 | K.WN[+1913.677]NTGC[+57.021]QALPSQDEGPSK.A | HexNAc(4)Hex(5)NeuAc(1) | C[+57], N[+1914] | 1301.5263 | 3 | 3902.5644 | 3902.5537 | 0 | 2.7 | 498 | Specific | 731.27 | 665.68 | 665.68 | 14.28 | 61 | >sp\|P01833\|PIGR_HUMAN Polymeric immunoglobulin receptor OS=Homo sapiens OX=9606 GN=PIGR PE=1 SV=4 | 902 | 20201113_12433_1ng_glyco_hcd203040_compound15.8825.8825.3 | controllerType=0 controllerNumber=1 scan=8825 | 20.8394 |
| 05843:3 | 8 | K.WN[+2059.735]NTGC[+57.021]QALPSQDEGPSK.A | HexNAc(4)Hex(5)Fuc(1)NeuAc(1) | C[+57], N[+2060] | 1350.212 | 3 | 4048.6214 | 4048.6116 | 0 | 2.4 | 498 | Specific | 804.03 | 701.89 | 701.89 | 16.28 | 61 | >sp\|P01833\|PIGR_HUMAN Polymeric immunoglobulin receptor OS=Homo sapiens OX=9606 GN=PIGR PE=1 SV=4 | 902 | 20201113_12433_1ng_glyco_hcd203040_compound15.8723.8723.3 | controllerType=0 controllerNumber=1 scan=8723 | 20.668 |
| 05845:3 | 8 | K.WN[+2059.735]NTGC[+57.021]QALPSQDEGPSK.A | HexNAc(4)Hex(5)Fuc(1)NeuAc(1) | C[+57], N[+2060] | 1350.212 | 3 | 4048.6214 | 4048.6116 | 0 | 2.4 | 498 | Specific | 812.03 | 722.18 | 722.18 | 16.28 | 61 | >sp\|P01833\|PIGR_HUMAN Polymeric immunoglobulin receptor OS=Homo sapiens OX=9606 GN=PIGR PE=1 SV=4 | 902 | 20201113_12433_1ng_glyco_hcd203040_compound15.8725.8725.3 | controllerType=0 controllerNumber=1 scan=8725 | 20.6731 |
| 05943:3 | 8 | K.WN[+1913.677]NTGC[+57.021]QALPSQDEGPSK.A | HexNAc(4)Hex(5)NeuAc(1) | C[+57], N[+1914] | 1301.5263 | 3 | 3902.5644 | 3902.5537 | 0 | 2.7 | 498 | Specific | 844.22 | 748.56 | 748.56 | 16.28 | 61 | >sp\|P01833\|PIGR_HUMAN Polymeric immunoglobulin receptor OS=Homo sapiens OX=9606 GN=PIGR PE=1 SV=4 | 902 | 20201113_12433_1ng_glyco_hcd203040_compound15.8827.8827.3 | controllerType=0 controllerNumber=1 scan=8827 | 20.8436 |
| 17714:4 | 10 | K.RNIAAFGGDPNN[+1913.677]ITLFGESAGGASVSLQTLSPYNK.G | HexNAc(4)Hex(5)NeuAc(1) | N[+1914] | 1370.8744 | 4 | 5480.4757 | 5480.4644 | 0 | 2.1 | 196 | Specific | 204.65 | 204.65 | 204.65 | 4.91 | 47 | >sp\|P19835\|CEL_HUMAN Bile salt-activated lipase OS=Homo sapiens OX=9606 GN=CEL PE=1 SV=3 | 476 | 20201113_12433_1ng_glyco_hcd203040_compound15.21075.21075.4 | controllerType=0 controllerNumber=1 scan=21075 | 41.5937 |
| 17716:4 | 10 | K.RNIAAFGGDPNN[+1913.677]ITLFGESAGGASVSLQTLSPYNK.G | HexNAc(4)Hex(5)NeuAc(1) | N[+1914] | 1370.8744 | 4 | 5480.4757 | 5480.4644 | 0 | 2.1 | 196 | Specific | 536.84 | 426.88 | 426.88 | 10.01 | 47 | >sp\|P19835\|CEL_HUMAN Bile salt-activated lipase OS=Homo sapiens OX=9606 GN=CEL PE=1 SV=3 | 476 | 20201113_12433_1ng_glyco_hcd203040_compound15.21077.21077.4 | controllerType=0 controllerNumber=1 scan=21077 | 41.5977 |
| 18289:3 | 11 | R.LSLHRPALEDLLLGSEAN[+1257.449]LTC[+57.021]TLTGLR.D | HexNAc(3)Hex(4) | C[+57], N[+1257] | 1407.694 | 3 | 4221.0675 | 4221.0477 | 0 | 4.7 | 127 | Specific | 231.41 | 231.41 | 231.41 | 5.34 | 47 | >sp\|P01876\|IGHA1_HUMAN Immunoglobulin heavy constant alpha 1 OS=Homo sapiens OX=9606 GN=IGHA1 PE=1 SV=2 | 255 | 20201113_12433_1ng_glyco_hcd203040_compound15.21674.21674.3 | controllerType=0 controllerNumber=1 scan=21674 | 42.8613 |
| 18291:3 | 11 | R.LSLHRPALEDLLLGSEAN[+1257.449]LTC[+57.021]TLTGLR.D | HexNAc(3)Hex(4) | C[+57], N[+1257] | 1407.694 | 3 | 4221.0675 | 4221.0477 | 0 | 4.7 | 127 | Specific | 270.97 | 270.97 | 270.97 | 5.96 | 47 | >sp\|P01876\|IGHA1_HUMAN Immunoglobulin heavy constant alpha 1 OS=Homo sapiens OX=9606 GN=IGHA1 PE=1 SV=2 | 255 | 20201113_12433_1ng_glyco_hcd203040_compound15.21676.21676.3 | controllerType=0 controllerNumber=1 scan=21676 | 42.8647 |
| 18146:4 | 11 | R.LSLHRPALEDLLLGSEAN[+1864.634]LTC[+57.021]TLTGLR.D | HexNAc(2)Hex(9) | C[+57], N[+1865] | 1207.8207 | 4 | 4828.2611 | 4828.2324 | 0 | 5.9 | 127 | Specific | 314.26 | 314.26 | 314.26 | 6.59 | 47 | >sp\|P01876\|IGHA1_HUMAN Immunoglobulin heavy constant alpha 1 OS=Homo sapiens OX=9606 GN=IGHA1 PE=1 SV=2 | 255 | 20201113_12433_1ng_glyco_hcd203040_compound15.21525.21525.4 | controllerType=0 controllerNumber=1 scan=21525 | 42.4983 |
| 18294:4 | 11 | R.LSLHRPALEDLLLGSEAN[+1095.397]LTC[+57.021]TLTGLR.D | HexNAc(3)Hex(3) | C[+57], N[+1095] | 1015.5083 | 4 | 4059.0115 | 4058.9949 | 0 | 4.1 | 127 | Specific | 384 | 384 | 384 | 9.1 | 47 | >sp\|P01876\|IGHA1_HUMAN Immunoglobulin heavy constant alpha 1 OS=Homo sapiens OX=9606 GN=IGHA1 PE=1 SV=2 | 255 | 20201113_12433_1ng_glyco_hcd203040_compound15.21680.21680.4 | controllerType=0 controllerNumber=1 scan=21680 | 42.8734 |
| 18150:3 | 11 | R.LSLHRPALEDLLLGSEAN[+1864.634]LTC[+57.021]TLTGLR.D | HexNAc(2)Hex(9) | C[+57], N[+1865] | 1610.093 | 3 | 4828.2643 | 4828.2324 | 0 | 6.6 | 127 | Specific | 417.34 | 417.34 | 417.34 | 8.06 | 47 | >sp\|P01876\|IGHA1_HUMAN Immunoglobulin heavy constant alpha 1 OS=Homo sapiens OX=9606 GN=IGHA1 PE=1 SV=2 | 255 | 20201113_12433_1ng_glyco_hcd203040_compound15.21529.21529.3 | controllerType=0 controllerNumber=1 scan=21529 | 42.5062 |
| 18245:3 | 11 | R.LSLHRPALEDLLLGSEAN[+1501.555]LTC[+57.021]TLTGLR.D | HexNAc(5)Hex(3) | C[+57], N[+1502] | 1489.0607 | 3 | 4465.1676 | 4465.1536 | 0 | 3.1 | 127 | Specific | 443.36 | 443.36 | 443.36 | 9.05 | 47 | >sp\|P01876\|IGHA1_HUMAN Immunoglobulin heavy constant alpha 1 OS=Homo sapiens OX=9606 GN=IGHA1 PE=1 SV=2 | 255 | 20201113_12433_1ng_glyco_hcd203040_compound15.21629.21629.3 | controllerType=0 controllerNumber=1 scan=21629 | 42.77 |
| 19150:3 | 11 | R.LSLHRPALEDLLLGSEAN[+1913.677]LTC[+57.021]TLTGLR.D | HexNAc(4)Hex(5)NeuAc(1) | C[+57], N[+1914] | 1626.4295 | 3 | 4877.2739 | 4877.2753 | 0 | -0.3 | 127 | Specific | 454.89 | 448.62 | 448.62 | 8.4 | 47 | >sp\|P01876\|IGHA1_HUMAN Immunoglobulin heavy constant alpha 1 OS=Homo sapiens OX=9606 GN=IGHA1 PE=1 SV=2 | 255 | 20201113_12433_1ng_glyco_hcd203040_compound15.22572.22572.3 | controllerType=0 controllerNumber=1 scan=22572 | 44.7934 |
| 18182:4 | 11 | R.LSLHRPALEDLLLGSEAN[+1663.608]LTC[+57.021]TLTGLR.D | HexNAc(5)Hex(4) | C[+57], N[+1664] | 1157.5614 | 4 | 4627.2239 | 4627.2064 | 0 | 3.8 | 127 | Specific | 491.58 | 441.66 | 441.66 | 9.41 | 47 | >sp\|P01876\|IGHA1_HUMAN Immunoglobulin heavy constant alpha 1 OS=Homo sapiens OX=9606 GN=IGHA1 PE=1 SV=2 | 255 | 20201113_12433_1ng_glyco_hcd203040_compound15.21563.21563.4 | controllerType=0 controllerNumber=1 scan=21563 | 42.6091 |
| 18225:3 | 11 | R.LSLHRPALEDLLLGSEAN[+1622.582]LTC[+57.021]TLTGLR.D | HexNAc(4)Hex(5) | C[+57], N[+1623] | 1529.3915 | 3 | 4586.1601 | 4586.1799 | 0 | -4.3 | 127 | Specific | 550.76 | 495.97 | 495.97 | 7.94 | 47 | >sp\|P01876\|IGHA1_HUMAN Immunoglobulin heavy constant alpha 1 OS=Homo sapiens OX=9606 GN=IGHA1 PE=1 SV=2 | 255 | 20201113_12433_1ng_glyco_hcd203040_compound15.21608.21608.3 | controllerType=0 controllerNumber=1 scan=21608 | [illegible] |
| 18329:4 | 11 | R.LSLHRPALEDLLLGSEAN[+1216.423]LTC[+57.021]TLTGLR.D | HexNAc(2)Hex(5) | C[+57], N[+1216] | 1045.7666 | 4 | 4180.0447 | 4180.0211 | 0 | 5.7 | 127 | Specific | 556.69 | 556.69 | 556.69 | 9.63 | 47 | >sp\|P01876\|IGHA1_HUMAN Immunoglobulin heavy constant alpha 1 OS=Homo sapiens OX=9606 GN=IGHA1 PE=1 SV=2 | 255 | 20201113_12433_1ng_glyco_hcd203040_compound15.21716.21716.3 | controllerType=0 controllerNumber=1 scan=21716 | 42.9696 |
| 18284:4 | 11 | R.LSLHRPALEDLLLGSEAN[+1257.449]LTC[+57.021]TLTGLR.D | HexNAc(3)Hex(4) | C[+57], N[+1257] | 1056.0195 | 4 | 4221.056 | 4221.0477 | 0 | 2 | 127 | Specific | 560.22 | 507.3 | 507.3 | 10.27 | 47 | >sp\|P01876\|IGHA1_HUMAN Immunoglobulin heavy constant alpha 1 OS=Homo sapiens OX=9606 GN=IGHA1 PE=1 SV=2 | 255 | 20201113_12433_1ng_glyco_hcd203040_compound15.21669.21669.4 | controllerType=0 controllerNumber=1 scan=21669 | 42.8495 |
| 18300:3 | 11 | R.LSLHRPALEDLLLGSEAN[+1095.397]LTC[+57.021]TLTGLR.D | HexNAc(3)Hex(3) | C[+57], N[+1095] | 1353.6789 | 3 | 4059.0221 | 4058.9949 | 0 | 6.7 | 127 | Specific | 564.43 | 548.67 | 548.67 | 8.83 | 47 | >sp\|P01876\|IGHA1_HUMAN Immunoglobulin heavy constant alpha 1 OS=Homo sapiens OX=9606 GN=IGHA1 PE=1 SV=2 | 255 | 20201113_12433_1ng_glyco_hcd203040_compound15.21686.21686.3 | controllerType=0 controllerNumber=1 scan=21686 | 42.8865 |
| 18231:3 | 11 | R.LSLHRPALEDLLLGSEAN[+1298.476]LTC[+57.021]TLTGLR.D | HexNAc(4)Hex(3) | C[+57], N[+1298] | 1421.366 | 3 | 4262.0835 | 4262.0742 | 0 | 2.2 | 127 | Specific | 576.12 | 576.12 | 576.12 | 9.72 | 47 | >sp\|P01876\|IGHA1_HUMAN Immunoglobulin heavy constant alpha 1 OS=Homo sapiens OX=9606 GN=IGHA1 PE=1 SV=2 | 255 | 20201113_12433_1ng_glyco_hcd203040_compound15.21614.21614.3 | controllerType=0 controllerNumber=1 scan=21614 | 42.7281 |
| 18287:4 | 11 | R.LSLHRPALEDLLLGSEAN[+1647.613]LTC[+57.021]TLTGLR.D | HexNAc(5)Hex(3)Fuc(1) | C[+57], N[+1648] | 1153.5617 | 4 | 4611.2249 | 4611.2115 | 0 | 2.9 | 127 | Specific | 596.04 | 575.71 | 575.71 | 11.52 | 47 | >sp\|P01876\|IGHA1_HUMAN Immunoglobulin heavy constant alpha 1 OS=Homo sapiens OX=9606 GN=IGHA1 PE=1 SV=2 | 255 | 20201113_12433_1ng_glyco_hcd203040_compound15.21672.21672.4 | controllerType=0 controllerNumber=1 scan=21672 | 42.8553 |
| 18227:3 | 11 | R.LSLHRPALEDLLLGSEAN[+1622.582]LTC[+57.021]TLTGLR.D | HexNAc(4)Hex(5) | C[+57], N[+1623] | 1529.3915 | 3 | 4586.1601 | 4586.1799 | 0 | -4.3 | 127 | Specific | 597.22 | 522.15 | 522.15 | 9.41 | 47 | >sp\|P01876\|IGHA1_HUMAN Immunoglobulin heavy constant alpha 1 OS=Homo sapiens OX=9606 GN=IGHA1 PE=1 SV=2 | 255 | 20201113_12433_1ng_glyco_hcd203040_compound15.21610.21610.3 | controllerType=0 controllerNumber=1 scan=21610 | 42.7221 |
| 18202:4 | 11 | R.LSLHRPALEDLLLGSEAN[+1419.502]LTC[+57.021]TLTGLR.D | HexNAc(3)Hex(5) | C[+57], N[+1420] | 1096.5353 | 4 | 4383.1195 | 4383.1005 | 0 | 4.3 | 127 | Specific | 602.6 | 602.6 | 602.6 | 12.18 | 47 | >sp\|P01876\|IGHA1_HUMAN Immunoglobulin heavy constant alpha 1 OS=Homo sapiens OX=9606 GN=IGHA1 PE=1 SV=2 | 255 | 20201113_12433_1ng_glyco_hcd203040_compound15.21584.21584.4 | controllerType=0 controllerNumber=1 scan=21584 | 42.6663 |
| 18184:3 | 11 | R.LSLHRPALEDLLLGSEAN[+1663.608]LTC[+57.021]TLTGLR.D | HexNAc(5)Hex(4) | C[+57], N[+1664] | 1543.0796 | 3 | 4627.2244 | 4627.2064 | 0 | 3.9 | 127 | Specific | 616.84 | 574.94 | 574.94 | 11.63 | 47 | >sp\|P01876\|IGHA1_HUMAN Immunoglobulin heavy constant alpha 1 OS=Homo sapiens OX=9606 GN=IGHA1 PE=1 SV=2 | 255 | 20201113_12433_1ng_glyco_hcd203040_compound15.21565.21565.3 | controllerType=0 controllerNumber=1 scan=21565 | 42.6179 |
| 18153:3 | 11 | R.LSLHRPALEDLLLGSEAN[+1864.634]LTC[+57.021]TLTGLR.D | HexNAc(2)Hex(9) | C[+57], N[+1865] | 1610.093 | 3 | 4828.2643 | 4828.2324 | 0 | 6.6 | 127 | Specific | 625.21 | 605.66 | 605.66 | 11.08 | 47 | >sp\|P01876\|IGHA1_HUMAN Immunoglobulin heavy constant alpha 1 OS=Homo sapiens OX=9606 GN=IGHA1 PE=1 SV=2 | 255 | 20201113_12433_1ng_glyco_hcd203040_compound15.21532.21532.3 | controllerType=0 controllerNumber=1 scan=21532 | 42.5104 |
| 18171:4 | 11 | R.LSLHRPALEDLLLGSEAN[+1622.582]LTC[+57.021]TLTGLR.D | HexNAc(4)Hex(5) | C[+57], N[+1623] | 1147.3075 | 4 | 4586.2083 | 4586.1799 | 0 | 6.2 | 127 | Specific | 630.62 | 387.66 | 387.66 | 11.63 | 47 | >sp\|P01876\|IGHA1_HUMAN Immunoglobulin heavy constant alpha 1 OS=Homo sapiens OX=9606 GN=IGHA1 PE=1 SV=2 | 255 | 20201113_12433_1ng_glyco_hcd203040_compound15.21551.21551.4 | controllerType=0 controllerNumber=1 scan=21551 | 42.5768 |
| 18169:4 | 11 | R.LSLHRPALEDLLLGSEAN[+1622.582]LTC[+57.021]TLTGLR.D | HexNAc(4)Hex(5) | C[+57], N[+1623] | 1147.3075 | 4 | 4586.2083 | 4586.1799 | 0 | 6.2 | 127 | Specific | 635.05 | 565.37 | 565.37 | 11.63 | 47 | >sp\|P01876\|IGHA1_HUMAN Immunoglobulin heavy constant alpha 1 OS=Homo sapiens OX=9606 GN=IGHA1 PE=1 SV=2 | 255 | 20201113_12433_1ng_glyco_hcd203040_compound15.21549.21549.4 | controllerType=0 controllerNumber=1 scan=21549 | 42.5674 |
| 18233:3 | 11 | R.LSLHRPALEDLLLGSEAN[+1298.476]LTC[+57.021]TLTGLR.D | HexNAc(4)Hex(3) | C[+57], N[+1298] | 1421.366 | 3 | 4262.0835 | 4262.0742 | 0 | 2.2 | 127 | Specific | 636.7 | 580.42 | 580.42 | 11.98 | 47 | >sp\|P01876\|IGHA1_HUMAN Immunoglobulin heavy constant alpha 1 OS=Homo sapiens OX=9606 GN=IGHA1 PE=1 SV=2 | 255 | 20201113_12433_1ng_glyco_hcd203040_compound15.21616.21616.3 | controllerType=0 controllerNumber=1 scan=21616 | 42.7403 |
| 18303:3 | 11 | R.LSLHRPALEDLLLGSEAN[+1095.397]LTC[+57.021]TLTGLR.D | HexNAc(3)Hex(3) | C[+57], N[+1095] | 1353.6789 | 3 | 4059.0221 | 4058.9949 | 0 | 6.7 | 127 | Specific | 645.5 | 598.8 | 598.8 | 11.08 | 47 | >sp\|P01876\|IGHA1_HUMAN Immunoglobulin heavy constant alpha 1 OS=Homo sapiens OX=9606 GN=IGHA1 PE=1 SV=2 | 255 | 20201113_12433_1ng_glyco_hcd203040_compound15.21689.21689.3 | controllerType=0 controllerNumber=1 scan=21689 | 42.8981 |
| 18247:3 | 11 | R.LSLHRPALEDLLLGSEAN[+1501.555]LTC[+57.021]TLTGLR.D | HexNAc(5)Hex(3) | C[+57], N[+1502] | 1489.0607 | 3 | 4465.1676 | 4465.1536 | 0 | 3.1 | 127 | Specific | 652.6 | 627.25 | 627.25 | 12.98 | 47 | >sp\|P01876\|IGHA1_HUMAN Immunoglobulin heavy constant alpha 1 OS=Homo sapiens OX=9606 GN=IGHA1 PE=1 SV=2 | 255 | 20201113_12433_1ng_glyco_hcd203040_compound15.21631.21631.3 | controllerType=0 controllerNumber=1 scan=21631 | 42.7741 |
| 18331:4 | 11 | R.LSLHRPALEDLLLGSEAN[+1216.423]LTC[+57.021]TLTGLR.D | HexNAc(2)Hex(5) | C[+57], N[+1216] | 1045.7666 | 4 | 4180.0447 | 4180.0211 | 0 | 5.7 | 127 | Specific | 653.05 | 471.59 | 471.59 | 12.89 | 47 | >sp\|P01876\|IGHA1_HUMAN Immunoglobulin heavy constant alpha 1 OS=Homo sapiens OX=9606 GN=IGHA1 PE=1 SV=2 | 255 | 20201113_12433_1ng_glyco_hcd203040_compound15.21718.21718.3 | controllerType=0 controllerNumber=1 scan=21718 | 42.9751 |
| 18186:3 | 11 | R.LSLHRPALEDLLLGSEAN[+1663.608]LTC[+57.021]TLTGLR.D | HexNAc(5)Hex(4) | C[+57], N[+1664] | 1543.0796 | 3 | 4627.2244 | 4627.2064 | 0 | 3.9 | 127 | Specific | 657.77 | 605.37 | 605.37 | 12.63 | 47 | >sp\|P01876\|IGHA1_HUMAN Immunoglobulin heavy constant alpha 1 OS=Homo sapiens OX=9606 GN=IGHA1 PE=1 SV=2 | 255 | 20201113_12433_1ng_glyco_hcd203040_compound15.21567.21567.3 | controllerType=0 controllerNumber=1 scan=21567 | 42.6232 |
| 18209:3 | 11 | R.LSLHRPALEDLLLGSEAN[+1460.529]LTC[+57.021]TLTGLR.D | HexNAc(4)Hex(4) | C[+57], N[+1461] | 1475.3858 | 3 | 4424.1428 | 4424.1271 | 0 | 3.6 | 127 | Specific | 666.02 | 582.58 | 582.58 | 12.63 | 47 | >sp\|P01876\|IGHA1_HUMAN Immunoglobulin heavy constant alpha 1 OS=Homo sapiens OX=9606 GN=IGHA1 PE=1 SV=2 | 255 | 20201113_12433_1ng_glyco_hcd203040_compound15.21591.21591.3 | controllerType=0 controllerNumber=1 scan=21591 | 42.6824 |
| 19189:4 | 11 | R.LSLHRPALEDLLLGSEAN[+1751.624]LTC[+57.021]TLTGLR.D | HexNAc(4)Hex(4)NeuAc(1) | C[+57], N[+1752] | 1179.5634 | 4 | 4715.2318 | 4715.2225 | 0 | 2 | 127 | Specific | 672.03 | 626.87 | 626.87 | 13.53 | 47 | >sp\|P01876\|IGHA1_HUMAN Immunoglobulin heavy constant alpha 1 OS=Homo sapiens OX=9606 GN=IGHA1 PE=1 SV=2 | 255 | 20201113_12433_1ng_glyco_hcd203040_compound15.22613.22613.4 | controllerType=0 controllerNumber=1 scan=22613 | 44.9082 |
| 18246:4 | 11 | R.LSLHRPALEDLLLGSEAN[+1298.476]LTC[+57.021]TLTGLR.D | HexNAc(4)Hex(3) | C[+57], N[+1298] | 1066.276 | 4 | 4262.0823 | 4262.0742 | 0 | 1.9 | 127 | Specific | 674.24 | 674.24 | 674.24 | 13.53 | 47 | >sp\|P01876\|IGHA1_HUMAN Immunoglobulin heavy constant alpha 1 OS=Homo sapiens OX=9606 GN=IGHA1 PE=1 SV=2 | 255 | 20201113_12433_1ng_glyco_hcd203040_compound15.21630.21630.4 | controllerType=0 controllerNumber=1 scan=21630 | 42.7714 |
| 18180:4 | 11 | R.LSLHRPALEDLLLGSEAN[+1663.608]LTC[+57.021]TLTGLR.D | HexNAc(5)Hex(4) | C[+57], N[+1664] | 1157.5614 | 4 | 4627.2239 | 4627.2064 | 0 | 3.8 | 127 | Specific | 683.46 | 683.46 | 683.46 | 13.18 | 47 | >sp\|P01876\|IGHA1_HUMAN Immunoglobulin heavy constant alpha 1 OS=Homo sapiens OX=9606 GN=IGHA1 PE=1 SV=2 | 255 | 20201113_12433_1ng_glyco_hcd203040_compound15.21561.21561.4 | controllerType=0 controllerNumber=1 scan=21561 | 42.6023 |
| 18207:3 | 11 | R.LSLHRPALEDLLLGSEAN[+1460.529]LTC[+57.021]TLTGLR.D | HexNAc(4)Hex(4) | C[+57], N[+1461] | 1475.3858 | 3 | 4424.1428 | 4424.1271 | 0 | 3.6 | 127 | Specific | 689.08 | 624.44 | 624.44 | 12.63 | 47 | >sp\|P01876\|IGHA1_HUMAN Immunoglobulin heavy constant alpha 1 OS=Homo sapiens OX=9606 GN=IGHA1 PE=1 SV=2 | 255 | 20201113_12433_1ng_glyco_hcd203040_compound15.21589.21589.3 | controllerType=0 controllerNumber=1 scan=21589 | 42.676 |
| 18189:4 | 11 | R.LSLHRPALEDLLLGSEAN[+1460.529]LTC[+57.021]TLTGLR.D | HexNAc(4)Hex(4) | C[+57], N[+1461] | 1106.7912 | 4 | 4424.1429 | 4424.1271 | 0 | 3.6 | 127 | Specific | 701.33 | 693.34 | 693.34 | 14.18 | 47 | >sp\|P01876\|IGHA1_HUMAN Immunoglobulin heavy constant alpha 1 OS=Homo sapiens OX=9606 GN=IGHA1 PE=1 SV=2 | 255 | 20201113_12433_1ng_glyco_hcd203040_compound15.21570.21570.4 | controllerType=0 controllerNumber=1 scan=21570 | 42.6284 |
| 18173:4 | 11 | R.LSLHRPALEDLLLGSEAN[+1501.555]LTC[+57.021]TLTGLR.D | HexNAc(5)Hex(3) | C[+57], N[+1502] | 1117.0506 | 4 | 4465.1805 | 4465.1536 | 0 | 6 | 127 | Specific | 706.58 | 706.58 | 706.58 | 13.89 | 47 | >sp\|P01876\|IGHA1_HUMAN Immunoglobulin heavy constant alpha 1 OS=Homo sapiens OX=9606 GN=IGHA1 PE=1 SV=2 | 255 | 20201113_12433_1ng_glyco_hcd203040_compound15.21553.21553.4 | controllerType=0 controllerNumber=1 scan=21553 | 42.58 |
| 18154:3 | 11 | R.LSLHRPALEDLLLGSEAN[+1540.529]LTC[+57.021]TLTGLR.D | HexNAc(2)Hex(7) | C[+57], N[+1541] | 1502.0567 | 3 | 4504.1555 | 4504.1268 | 0 | 6.4 | 127 | Specific | 725.88 | 725.88 | 725.88 | 13.08 | 47 | >sp\|P01876\|IGHA1_HUMAN Immunoglobulin heavy constant alpha 1 OS=Homo sapiens OX=9606 GN=IGHA1 PE=1 SV=2 | 255 | 20201113_12433_1ng_glyco_hcd203040_compound15.21533.21533.3 | controllerType=0 controllerNumber=1 scan=21533 | 42.5162 |
| 19192:4 | 11 | R.LSLHRPALEDLLLGSEAN[+1751.624]LTC[+57.021]TLTGLR.D | HexNAc(4)Hex(4)NeuAc(1) | C[+57], N[+1752] | 1179.5634 | 4 | 4715.2318 | 4715.2225 | 0 | 2 | 127 | Specific | 725.92 | 663.8 | 663.8 | 14.53 | 47 | >sp\|P01876\|IGHA1_HUMAN Immunoglobulin heavy constant alpha 1 OS=Homo sapiens OX=9606 GN=IGHA1 PE=1 SV=2 | 255 | 20201113_12433_1ng_glyco_hcd203040_compound15.22616.22616.4 | controllerType=0 controllerNumber=1 scan=22616 | 44.9207 |
| 18152:3 | 11 | R.LSLHRPALEDLLLGSEAN[+1540.529]LTC[+57.021]TLTGLR.D | HexNAc(2)Hex(7) | C[+57], N[+1541] | 1502.0567 | 3 | 4504.1555 | 4504.1268 | 0 | 6.4 | 127 | Specific | 729.54 | 669.89 | 669.89 | 13.08 | 47 | >sp\|P01876\|IGHA1_HUMAN Immunoglobulin heavy constant alpha 1 OS=Homo sapiens OX=9606 GN=IGHA1 PE=1 SV=2 | 255 | 20201113_12433_1ng_glyco_hcd203040_compound15.21531.21531.3 | controllerType=0 controllerNumber=1 scan=21531 | 42.5085 |
| 19144:4 | 11 | R.LSLHRPALEDLLLGSEAN[+1913.677]LTC[+57.021]TLTGLR.D | HexNAc(4)Hex(5)NeuAc(1) | C[+57], N[+1914] | 1220.0798 | 4 | 4877.2972 | 4877.2753 | 0 | 4.5 | 127 | Specific | 751.39 | 738.44 | 738.44 | 15.18 | 47 | >sp\|P01876\|IGHA1_HUMAN Immunoglobulin heavy constant alpha 1 OS=Homo sapiens OX=9606 GN=IGHA1 PE=1 SV=2 | 255 | 20201113_12433_1ng_glyco_hcd203040_compound15.22566.22566.4 | controllerType=0 controllerNumber=1 scan=22566 | 44.7726 |
| 18192:4 | 11 | R.LSLHRPALEDLLLGSEAN[+1419.502]LTC[+57.021]TLTGLR.D | HexNAc(3)Hex(5) | C[+57], N[+1420] | 1096.5346 | 4 | 4383.1165 | 4383.1005 | 0 | 3.7 | 127 | Specific | 760.35 | 760.35 | 760.35 | 15.18 | 47 | >sp\|P01876\|IGHA1_HUMAN Immunoglobulin heavy constant alpha 1 OS=Homo sapiens OX=9606 GN=IGHA1 PE=1 SV=2 | 255 | 20201113_12433_1ng_glyco_hcd203040_compound15.21573.21573.4 | controllerType=0 controllerNumber=1 scan=21573 | 42.6372 |
| 18175:4 | 11 | R.LSLHRPALEDLLLGSEAN[+1501.555]LTC[+57.021]TLTGLR.D | HexNAc(5)Hex(3) | C[+57], N[+1502] | 1117.0506 | 4 | 4465.1805 | 4465.1536 | 0 | 6 | 127 | Specific | 801.1 | 796.57 | 796.57 | 15.89 | 47 | >sp\|P01876\|IGHA1_HUMAN Immunoglobulin heavy constant alpha 1 OS=Homo sapiens OX=9606 GN=IGHA1 PE=1 SV=2 | 255 | 20201113_12433_1ng_glyco_hcd203040_compound15.21555.21555.4 | controllerType=0 controllerNumber=1 scan=21555 | 42.5877 |
| 18185:4 | 11 | R.LSLHRPALEDLLLGSEAN[+1702.581]LTC[+57.021]TLTGLR.D | HexNAc(2)Hex(8) | C[+57], N[+1703] | 1167.3042 | 4 | 4666.1951 | 4666.1796 | 0 | 3.3 | 127 | Specific | 814.18 | 814.18 | 814.18 | 16.52 | 47 | >sp\|P01876\|IGHA1_HUMAN Immunoglobulin heavy constant alpha 1 OS=Homo sapiens OX=9606 GN=IGHA1 PE=1 SV=2 | 255 | 20201113_12433_1ng_glyco_hcd203040_compound15.21566.21566.4 | controllerType=0 controllerNumber=1 scan=21566 | 42.6194 |
| 18147:4 | 11 | R.LSLHRPALEDLLLGSEAN[+1540.529]LTC[+57.021]TLTGLR.D | HexNAc(2)Hex(7) | C[+57], N[+1541] | 1126.7939 | 4 | 4504.1536 | 4504.1268 | 0 | 6 | 127 | Specific | 817.75 | 817.75 | 817.75 | 15.89 | 47 | >sp\|P01876\|IGHA1_HUMAN Immunoglobulin heavy constant alpha 1 OS=Homo sapiens OX=9606 GN=IGHA1 PE=1 SV=2 | 255 | 20201113_12433_1ng_glyco_hcd203040_compound15.21526.21526.4 | controllerType=0 controllerNumber=1 scan=21526 | 42.5001 |
| 18281:4 | 11 | R.LSLHRPALEDLLLGSEAN[+1257.449]LTC[+57.021]TLTGLR.D | HexNAc(3)Hex(4) | C[+57], N[+1257] | 1056.0195 | 4 | 4221.056 | 4221.0477 | 0 | 2 | 127 | Specific | 824.88 | 824.44 | 824.44 | 16.53 | 47 | >sp\|P01876\|IGHA1_HUMAN Immunoglobulin heavy constant alpha 1 OS=Homo sapiens OX=9606 GN=IGHA1 PE=1 SV=2 | 255 | 20201113_12433_1ng_glyco_hcd203040_compound15.21666.21666.4 | controllerType=0 controllerNumber=1 scan=21666 | 42.8444 |
| 18248:4 | 11 | R.LSLHRPALEDLLLGSEAN[+1298.476]LTC[+57.021]TLTGLR.D | HexNAc(4)Hex(3) | C[+57], N[+1298] | 1066.276 | 4 | 4262.0823 | 4262.0742 | 0 | 1.9 | 127 | Specific | 865.22 | 795.43 | 795.43 | 17.53 | 47 | >sp\|P01876\|IGHA1_HUMAN Immunoglobulin heavy constant alpha 1 OS=Homo sapiens OX=9606 GN=IGHA1 PE=1 SV=2 | 255 | 20201113_12433_1ng_glyco_hcd203040_compound15.21632.21632.4 | controllerType=0 controllerNumber=1 scan=21632 | 42.7757 |
| 18204:4 | 11 | R.LSLHRPALEDLLLGSEAN[+1419.502]LTC[+57.021]TLTGLR.D | HexNAc(3)Hex(5) | C[+57], N[+1420] | 1096.5353 | 4 | 4383.1195 | 4383.1005 | 0 | 4.3 | 127 | Specific | 872.69 | 785.88 | 785.88 | 17.18 | 47 | >sp\|P01876\|IGHA1_HUMAN Immunoglobulin heavy constant alpha 1 OS=Homo sapiens OX=9606 GN=IGHA1 PE=1 SV=2 | 255 | 20201113_12433_1ng_glyco_hcd203040_compound15.21586.21586.4 | controllerType=0 controllerNumber=1 scan=21586 | 42.6723 |
| 18191:4 | 11 | R.LSLHRPALEDLLLGSEAN[+1460.529]LTC[+57.021]TLTGLR.D | HexNAc(4)Hex(4) | C[+57], N[+1461] | 1106.7912 | 4 | 4424.1429 | 4424.1271 | 0 | 3.6 | 127 | Specific | 884.5 | 773.62 | 773.62 | 17.18 | 47 | >sp\|P01876\|IGHA1_HUMAN Immunoglobulin heavy constant alpha 1 OS=Homo sapiens OX=9606 GN=IGHA1 PE=1 SV=2 | 255 | 20201113_12433_1ng_glyco_hcd203040_compound15.21572.21572.4 | controllerType=0 controllerNumber=1 scan=21572 | 42.6355 |
| 18194:4 | 11 | R.LSLHRPALEDLLLGSEAN[+1419.502]LTC[+57.021]TLTGLR.D | HexNAc(3)Hex(5) | C[+57], N[+1420] | 1096.5346 | 4 | 4383.1165 | 4383.1005 | 0 | 3.7 | 127 | Specific | 904.01 | 832.89 | 832.89 | 18.18 | 47 | >sp\|P01876\|IGHA1_HUMAN Immunoglobulin heavy constant alpha 1 OS=Homo sapiens OX=9606 GN=IGHA1 PE=1 SV=2 | 255 | 20201113_12433_1ng_glyco_hcd203040_compound15.21575.21575.4 | controllerType=0 controllerNumber=1 scan=21575 | 42.6441 |
| 18201:4 | 11 | R.LSLHRPALEDLLLGSEAN[+1216.423]LTC[+57.021]TLTGLR.D | HexNAc(2)Hex(5) | C[+57], N[+1216] | 1045.7635 | 4 | 4180.0321 | 4180.0211 | 0 | 2.6 | 127 | Specific | 914.3 | 845.76 | 845.76 | 18.52 | 47 | >sp\|P01876\|IGHA1_HUMAN Immunoglobulin heavy constant alpha 1 OS=Homo sapiens OX=9606 GN=IGHA1 PE=1 SV=2 | 255 | 20201113_12433_1ng_glyco_hcd203040_compound15.21583.21583.4 | controllerType=0 controllerNumber=1 scan=21583 | 42.6646 |
| 18187:4 | 11 | R.LSLHRPALEDLLLGSEAN[+1702.581]LTC[+57.021]TLTGLR.D | HexNAc(2)Hex(8) | C[+57], N[+1703] | 1167.3042 | 4 | 4666.1951 | 4666.1796 | 0 | 3.3 | 127 | Specific | 919.5 | 895.43 | 895.43 | 18.52 | 47 | >sp\|P01876\|IGHA1_HUMAN Immunoglobulin heavy constant alpha 1 OS=Homo sapiens OX=9606 GN=IGHA1 PE=1 SV=2 | 255 | 20201113_12433_1ng_glyco_hcd203040_compound15.21568.21568.4 | controllerType=0 controllerNumber=1 scan=21568 | 42.625 |
| 18296:4 | 11 | R.LSLHRPALEDLLLGSEAN[+1095.397]LTC[+57.021]TLTGLR.D | HexNAc(3)Hex(3) | C[+57], N[+1095] | 1015.5083 | 4 | 4059.0115 | 4058.9949 | 0 | 4.1 | 127 | Specific | 945.49 | 945.49 | 945.49 | 18.18 | 47 | >sp\|P01876\|IGHA1_HUMAN Immunoglobulin heavy constant alpha 1 OS=Homo sapiens OX=9606 GN=IGHA1 PE=1 SV=2 | 255 | 20201113_12433_1ng_glyco_hcd203040_compound15.21682.21682.4 | controllerType=0 controllerNumber=1 scan=21682 | 42.8771 |
| 18203:4 | 11 | R.LSLHRPALEDLLLGSEAN[+1216.423]LTC[+57.021]TLTGLR.D | HexNAc(2)Hex(5) | C[+57], N[+1216] | 1045.7635 | 4 | 4180.0321 | 4180.0211 | 0 | 2.6 | 127 | Specific | 954.58 | 916.72 | 916.72 | 19.52 | 47 | >sp\|P01876\|IGHA1_HUMAN Immunoglobulin heavy constant alpha 1 OS=Homo sapiens OX=9606 GN=IGHA1 PE=1 SV=2 | 255 | 20201113_12433_1ng_glyco_hcd203040_compound15.21585.21585.4 | controllerType=0 controllerNumber=1 scan=21585 | 42.6709 |
| 18148:4 | 11 | R.LSLHRPALEDLLLGSEAN[+1864.634]LTC[+57.021]TLTGLR.D | HexNAc(2)Hex(9) | C[+57], N[+1865] | 1207.8207 | 4 | 4828.2611 | 4828.2324 | 0 | 5.9 | 127 | Specific | 994.73 | 921.47 | 921.47 | 18.89 | 47 | >sp\|P01876\|IGHA1_HUMAN Immunoglobulin heavy constant alpha 1 OS=Homo sapiens OX=9606 GN=IGHA1 PE=1 SV=2 | 255 | 20201113_12433_1ng_glyco_hcd203040_compound15.21527.21527.4 | controllerType=0 controllerNumber=1 scan=21527 | 42.5015 |
| 18149:4 | 11 | R.LSLHRPALEDLLLGSEAN[+1540.529]LTC[+57.021]TLTGLR.D | HexNAc(2)Hex(7) | C[+57], N[+1541] | 1126.7939 | 4 | 4504.1536 | 4504.1268 | 0 | 6 | 127 | Specific | 1016.99 | 975.37 | 975.37 | 19.89 | 47 | >sp\|P01876\|IGHA1_HUMAN Immunoglobulin heavy constant alpha 1 OS=Homo sapiens OX=9606 GN=IGHA1 PE=1 SV=2 | 255 | 20201113_12433_1ng_glyco_hcd203040_compound15.21528.21528.4 | controllerType=0 controllerNumber=1 scan=21528 | 42.5052 |
| 15491:4 | 11 | R.LAGKPTHVN[+2553.910]VSVVMAEVDGTC[+57.021]Y.- | HexNAc(5)Hex(5)Fuc(1)NeuAc(2) | C[+57], N[+2554] | 1226.0295 | 4 | 4901.0962 | 4901.0518 | 0 | 9.1 | 332 | CRagged | 344.92 | 273.18 | 273.18 | 6.49 | 47 | >sp\|P01876\|IGHA1_HUMAN Immunoglobulin heavy constant alpha 1 OS=Homo sapiens OX=9606 GN=IGHA1 PE=1 SV=2 | 255 | 20201113_12433_1ng_glyco_hcd203040_compound15.18739.18739.4 | controllerType=0 controllerNumber=1 scan=18739 | 37.3407 |
| 15481:3 | 11 | R.LAGKPTHVN[+2553.910]VSVVMAEVDGTC[+57.021]Y.- | HexNAc(5)Hex(5)Fuc(1)NeuAc(2) | C[+57], N[+2554] | 1634.3704 | 3 | 4901.0966 | 4901.0518 | 0 | 9.2 | 332 | CRagged | 351.29 | 248.57 | 248.57 | 6.47 | 47 | >sp\|P01876\|IGHA1_HUMAN Immunoglobulin heavy constant alpha 1 OS=Homo sapiens OX=9606 GN=IGHA1 PE=1 SV=2 | 255 | 20201113_12433_1ng_glyco_hcd203040_compound15.18729.18729.3 | controllerType=0 controllerNumber=1 scan=18729 | 37.3209 |
| 13479:3 | 11 | R.LAGKPTHVN[+2059.735]VSVVMAEVDGTC[+57.021]Y.- | HexNAc(4)Hex(5)Fuc(1)NeuAc(1) | C[+57], N[+2060] | 1469.6306 | 3 | 4406.8771 | 4406.877 | 0 | 0 | 332 | CRagged | 371.88 | 371.88 | 371.88 | 8.31 | 47 | >sp\|P01876\|IGHA1_HUMAN Immunoglobulin heavy constant alpha 1 OS=Homo sapiens OX=9606 GN=IGHA1 PE=1 SV=2 | 255 | 20201113_12433_1ng_glyco_hcd203040_compound15.16658.16658.3 | controllerType=0 controllerNumber=1 scan=16658 | 33.7647 |
| 13481:3 | 11 | R.LAGKPTHVN[+2059.735]VSVVMAEVDGTC[+57.021]Y.- | HexNAc(4)Hex(5)Fuc(1)NeuAc(1) | C[+57], N[+2060] | 1469.6306 | 3 | 4406.8771 | 4406.877 | 0 | 0 | 332 | CRagged | 389.64 | 270.21 | 270.21 | 7.75 | 47 | >sp\|P01876\|IGHA1_HUMAN Immunoglobulin heavy constant alpha 1 OS=Homo sapiens OX=9606 GN=IGHA1 PE=1 SV=2 | 255 | 20201113_12433_1ng_glyco_hcd203040_compound15.16660.16660.3 | controllerType=0 controllerNumber=1 scan=16660 | 33.768 |
| 13440:4 | 11 | R.LAGKPTHVN[+2262.814]VSVVMAEVDGTC[+57.021]Y.- | HexNAc(5)Hex(5)Fuc(1)NeuAc(1) | C[+57], N[+2263] | 1153.2466 | 4 | 4609.9647 | 4609.9564 | 0 | 1.8 | 332 | CRagged | 390.57 | 338.64 | 338.64 | 8.48 | 47 | >sp\|P01876\|IGHA1_HUMAN Immunoglobulin heavy constant alpha 1 OS=Homo sapiens OX=9606 GN=IGHA1 PE=1 SV=2 | 255 | 20201113_12433_1ng_glyco_hcd203040_compound15.16618.16618.2 | controllerType=0 controllerNumber=1 scan=16618 | 33.6925 |
| 15490:4 | 11 | R.LAGKPTHVN[+2553.910]VSVVMAEVDGTC[+57.021]Y.- | HexNAc(5)Hex(5)Fuc(1)NeuAc(2) | C[+57], N[+2554] | 1226.0295 | 4 | 4901.0962 | 4901.0518 | 0 | 9.1 | 332 | CRagged | 421.08 | 327.34 | 327.34 | 6.96 | 47 | >sp\|P01876\|IGHA1_HUMAN Immunoglobulin heavy constant alpha 1 OS=Homo sapiens OX=9606 GN=IGHA1 PE=1 SV=2 | 255 | 20201113_12433_1ng_glyco_hcd203040_compound15.18738.18738.4 | controllerType=0 controllerNumber=1 scan=18738 | 37.3371 |
| 13442:4 | 11 | R.LAGKPTHVN[+2262.814]VSVVMAEVDGTC[+57.021]Y.- | HexNAc(5)Hex(5)Fuc(1)NeuAc(1) | C[+57], N[+2263] | 1153.2466 | 4 | 4609.9647 | 4609.9564 | 0 | 1.8 | 332 | CRagged | 428.74 | 420 | 420 | 9.37 | 47 | >sp\|P01876\|IGHA1_HUMAN Immunoglobulin heavy constant alpha 1 OS=Homo sapiens OX=9606 GN=IGHA1 PE=1 SV=2 | 255 | 20201113_12433_1ng_glyco_hcd203040_compound15.16620.16620.2 | controllerType=0 controllerNumber=1 scan=16620 | 33.6984 |
| 15479:3 | 11 | R.LAGKPTHVN[+2553.910]VSVVMAEVDGTC[+57.021]Y.- | HexNAc(5)Hex(5)Fuc(1)NeuAc(2) | C[+57], N[+2554] | 1634.3704 | 3 | 4901.0966 | 4901.0518 | 0 | 9.2 | 332 | CRagged | 451.42 | 312.85 | 312.85 | 7.07 | 47 | >sp\|P01876\|IGHA1_HUMAN Immunoglobulin heavy constant alpha 1 OS=Homo sapiens OX=9606 GN=IGHA1 PE=1 SV=2 | 255 | 20201113_12433_1ng_glyco_hcd203040_compound15.18727.18727.3 | controllerType=0 controllerNumber=1 scan=18727 | 37.3172 |
| 13476:3 | 11 | R.LAGKPTHVN[+1913.677]VSVVMAEVDGTC[+57.021]Y.- | HexNAc(4)Hex(5)NeuAc(1) | C[+57], N[+1914] | 1420.9474 | 3 | 4260.8278 | 4260.8191 | 0 | 2 | 332 | CRagged | 519.25 | 404.43 | 404.43 | 10.6 | 47 | >sp\|P01876\|IGHA1_HUMAN Immunoglobulin heavy constant alpha 1 OS=Homo sapiens OX=9606 GN=IGHA1 PE=1 SV=2 | 255 | 20201113_12433_1ng_glyco_hcd203040_compound15.16655.16655.3 | controllerType=0 controllerNumber=1 scan=16655 | 33.7582 |
| 13478:3 | 11 | R.LAGKPTHVN[+1913.677]VSVVMAEVDGTC[+57.021]Y.- | HexNAc(4)Hex(5)NeuAc(1) | C[+57], N[+1914] | 1420.9474 | 3 | 4260.8278 | 4260.8191 | 0 | 2 | 332 | CRagged | 542.48 | 538.67 | 538.67 | 10.72 | 47 | >sp\|P01876\|IGHA1_HUMAN Immunoglobulin heavy constant alpha 1 OS=Homo sapiens OX=9606 GN=IGHA1 PE=1 SV=2 | 255 | 20201113_12433_1ng_glyco_hcd203040_compound15.16657.16657.3 | controllerType=0 controllerNumber=1 scan=16657 | 33.7628 |
| 11745:3 | 11 | R.LAGKPTHVN[+1501.555]VSVVMAEVDGTC[+57.021]Y.- | HexNAc(5)Hex(3) | C[+57], N[+1502] | 1283.5733 | 3 | 3848.7053 | 3848.6974 | 0 | 2.1 | 332 | CRagged | 725.92 | 713.88 | 713.88 | 15.18 | 47 | >sp\|P01876\|IGHA1_HUMAN Immunoglobulin heavy constant alpha 1 OS=Homo sapiens OX=9606 GN=IGHA1 PE=1 SV=2 | 255 | 20201113_12433_1ng_glyco_hcd203040_compound15.14842.14842.3 | controllerType=0 controllerNumber=1 scan=14842 | 30.8079 |
| 11736:3 | 11 | R.LAGKPTHVN[+1647.613]VSVVMAEVDGTC[+57.021]Y.- | HexNAc(5)Hex(3)Fuc(1) | C[+57], N[+1648] | 1332.2598 | 3 | 3994.7649 | 3994.7553 | 0 | 2.4 | 332 | CRagged | 734.59 | 569.48 | 569.48 | 15.18 | 47 | >sp\|P01876\|IGHA1_HUMAN Immunoglobulin heavy constant alpha 1 OS=Homo sapiens OX=9606 GN=IGHA1 PE=1 SV=2 | 255 | 20201113_12433_1ng_glyco_hcd203040_compound15.14832.14832.3 | controllerType=0 controllerNumber=1 scan=14832 | 30.7886 |
| 11769:3 | 11 | R.LAGKPTHVN[+1647.613]VSVVMAEVDGTC[+57.021]Y.- | HexNAc(5)Hex(3)Fuc(1) | C[+57], N[+1648] | 1332.2601 | 3 | 3994.7656 | 3994.7553 | 0 | 2.6 | 332 | CRagged | 737.21 | 657.57 | 657.57 | 15.18 | 47 | >sp\|P01876\|IGHA1_HUMAN Immunoglobulin heavy constant alpha 1 OS=Homo sapiens OX=9606 GN=IGHA1 PE=1 SV=2 | 255 | 20201113_12433_1ng_glyco_hcd203040_compound15.14867.14867.3 | controllerType=0 controllerNumber=1 scan=14867 | 30.8527 |
| 11743:3 | 11 | R.LAGKPTHVN[+1501.555]VSVVMAEVDGTC[+57.021]Y.- | HexNAc(5)Hex(3) | C[+57], N[+1502] | 1283.5733 | 3 | 3848.7053 | 3848.6974 | 0 | 2.1 | 332 | CRagged | 767.64 | 612.71 | 612.71 | 16.18 | 47 | >sp\|P01876\|IGHA1_HUMAN Immunoglobulin heavy constant alpha 1 OS=Homo sapiens OX=9606 GN=IGHA1 PE=1 SV=2 | 255 | 20201113_12433_1ng_glyco_hcd203040_compound15.14840.14840.3 | controllerType=0 controllerNumber=1 scan=14840 | 30.8035 |
| 11671:3 | 11 | R.LAGKPTHVN[+1216.423]VSVVMAEVDGTC[+57.021]Y.- | HexNAc(2)Hex(5) | C[+57], N[+1216] | 1188.5284 | 3 | 3563.5706 | 3563.5649 | 0 | 1.6 | 332 | CRagged | 786.07 | 719.4 | 466.4 | 16.18 | 47 | >sp\|P01876\|IGHA1_HUMAN Immunoglobulin heavy constant alpha 1 OS=Homo sapiens OX=9606 GN=IGHA1 PE=1 SV=2 | 255 | 20201113_12433_1ng_glyco_hcd203040_compound15.14765.14765.3 | controllerType=0 controllerNumber=1 scan=14765 | 30.6551 |
| 11771:3 | 11 | R.LAGKPTHVN[+1647.613]VSVVMAEVDGTC[+57.021]Y.- | HexNAc(5)Hex(3)Fuc(1) | C[+57], N[+1648] | 1332.2601 | 3 | 3994.7656 | 3994.7553 | 0 | 2.6 | 332 | CRagged | 829.09 | 685.39 | 685.39 | 17.18 | 47 | >sp\|P01876\|IGHA1_HUMAN Immunoglobulin heavy constant alpha 1 OS=Homo sapiens OX=9606 GN=IGHA1 PE=1 SV=2 | 255 | 20201113_12433_1ng_glyco_hcd203040_compound15.14869.14869.3 | controllerType=0 controllerNumber=1 scan=14869 | 30.8572 |
| 11616:3 | 11 | R.LAGKPTHVN[+1540.529]VSVVMAEVDGTC[+57.021]Y.- | HexNAc(2)Hex(7) | C[+57], N[+1541] | 1296.5652 | 3 | 3887.6812 | 3887.6706 | 0 | 2.7 | 332 | CRagged | 833.42 | 777.88 | 492.26 | 17.18 | 47 | >sp\|P01876\|IGHA1_HUMAN Immunoglobulin heavy constant alpha 1 OS=Homo sapiens OX=9606 GN=IGHA1 PE=1 SV=2 | 255 | 20201113_12433_1ng_glyco_hcd203040_compound15.14708.14708.3 | controllerType=0 controllerNumber=1 scan=14708 | 30.5542 |
| 11614:3 | 11 | R.LAGKPTHVN[+1540.529]VSVVMAEVDGTC[+57.021]Y.- | HexNAc(2)Hex(7) | C[+57], N[+1541] | 1296.5652 | 3 | 3887.6812 | 3887.6706 | 0 | 2.7 | 332 | CRagged | 838.56 | 838.56 | 531.89 | 17.18 | 47 | >sp\|P01876\|IGHA1_HUMAN Immunoglobulin heavy constant alpha 1 OS=Homo sapiens OX=9606 GN=IGHA1 PE=1 SV=2 | 255 | 20201113_12433_1ng_glyco_hcd203040_compound15.14706.14706.3 | controllerType=0 controllerNumber=1 scan=14706 | 30.5483 |
| 11733:3 | 11 | R.LAGKPTHVN[+1647.613]VSVVMAEVDGTC[+57.021]Y.- | HexNAc(5)Hex(3)Fuc(1) | C[+57], N[+1648] | 1332.2598 | 3 | 3994.7649 | 3994.7553 | 0 | 2.4 | 332 | CRagged | 868.73 | 868.73 | 868.73 | 18.18 | 47 | >sp\|P01876\|IGHA1_HUMAN Immunoglobulin heavy constant alpha 1 OS=Homo sapiens OX=9606 GN=IGHA1 PE=1 SV=2 | 255 | 20201113_12433_1ng_glyco_hcd203040_compound15.14829.14829.3 | controllerType=0 controllerNumber=1 scan=14829 | 30.7798 |
| 11636:3 | 11 | R.LAGKPTHVN[+1378.476]VSVVMAEVDGTC[+57.021]Y.- | HexNAc(2)Hex(6) | C[+57], N[+1378] | 1242.5471 | 3 | 3725.6266 | 3725.6178 | 0 | 2.4 | 332 | CRagged | 874.87 | 874.87 | 514.91 | 18.18 | 47 | >sp\|P01876\|IGHA1_HUMAN Immunoglobulin heavy constant alpha 1 OS=Homo sapiens OX=9606 GN=IGHA1 PE=1 SV=2 | 255 | 20201113_12433_1ng_glyco_hcd203040_compound15.14729.14729.3 | controllerType=0 controllerNumber=1 scan=14729 | 30.5946 |
| 11638:3 | 11 | R.LAGKPTHVN[+1378.476]VSVVMAEVDGTC[+57.021]Y.- | HexNAc(2)Hex(6) | C[+57], N[+1378] | 1242.5471 | 3 | 3725.6266 | 3725.6178 | 0 | 2.4 | 332 | CRagged | 894.75 | 847.56 | 531.92 | 18.18 | 47 | >sp\|P01876\|IGHA1_HUMAN Immunoglobulin heavy constant alpha 1 OS=Homo sapiens OX=9606 GN=IGHA1 PE=1 SV=2 | 255 | 20201113_12433_1ng_glyco_hcd203040_compound15.14731.14731.3 | controllerType=0 controllerNumber=1 scan=14731 | 30.6005 |
| 11674:3 | 11 | R.LAGKPTHVN[+1216.423]VSVVMAEVDGTC[+57.021]Y.- | HexNAc(2)Hex(5) | C[+57], N[+1216] | 1188.5284 | 3 | 3563.5706 | 3563.5649 | 0 | 1.6 | 332 | CRagged | 909.77 | 892.92 | 568.54 | 19.18 | 47 | >sp\|P01876\|IGHA1_HUMAN Immunoglobulin heavy constant alpha 1 OS=Homo sapiens OX=9606 GN=IGHA1 PE=1 SV=2 | 255 | 20201113_12433_1ng_glyco_hcd203040_compound15.14768.14768.3 | controllerType=0 controllerNumber=1 scan=14768 | 30.6602 |
| 07951:3 | 12 | R.LSPN[+2350.830]ASAEHLELR.W | HexNAc(4)Hex(5)Fuc(1)NeuAc(2) | N[+2351] | 1263.2012 | 3 | 3787.5891 | 3787.5796 | 0 | 2.5 | 52 | Specific | 384.13 | 380.24 | 380.24 | 9.21 | 39 | >sp\|Q13410\|BT1A1_HUMAN Butyrophilin subfamily 1 member A1 OS=Homo sapiens OX=9606 GN=BTN1A1 PE=1 SV=3 | 179 | 20201113_12433_1ng_glyco_hcd203040_compound15.10905.10905.3 | controllerType=0 controllerNumber=1 scan=10905 | 24.2269 |
| 07953:3 | 12 | R.LSPN[+2350.830]ASAEHLELR.W | HexNAc(4)Hex(5)Fuc(1)NeuAc(2) | N[+2351] | 1263.2012 | 3 | 3787.5891 | 3787.5796 | 0 | 2.5 | 52 | Specific | 509.5 | 405.84 | 405.84 | 9.7 | 39 | >sp\|Q13410\|BT1A1_HUMAN Butyrophilin subfamily 1 member A1 OS=Homo sapiens OX=9606 GN=BTN1A1 PE=1 SV=3 | 179 | 20201113_12433_1ng_glyco_hcd203040_compound15.10907.10907.3 | controllerType=0 controllerNumber=1 scan=10907 | 24.2304 |
| 05844:3 | 12 | R.LSPN[+2205.793]ASAEHLELR.W | HexNAc(4)Hex(5)Fuc(2)NeuAc(1) | N[+2206] | 1214.8554 | 3 | 3642.5517 | 3642.5421 | 0 | 2.6 | 52 | Specific | 579.63 | 426.82 | 426.82 | 11.29 | 39 | >sp\|Q13410\|BT1A1_HUMAN Butyrophilin subfamily 1 member A1 OS=Homo sapiens OX=9606 GN=BTN1A1 PE=1 SV=3 | 179 | 20201113_12433_1ng_glyco_hcd203040_compound15.8724.8724.3 | controllerType=0 controllerNumber=1 scan=8724 | 20.6714 |
| 05842:3 | 12 | R.LSPN[+2205.793]ASAEHLELR.W | HexNAc(4)Hex(5)Fuc(2)NeuAc(1) | N[+2206] | 1214.8554 | 3 | 3642.5517 | 3642.5421 | 0 | 2.6 | 52 | Specific | 681.21 | 577.72 | 577.72 | 13.29 | 39 | >sp\|Q13410\|BT1A1_HUMAN Butyrophilin subfamily 1 member A1 OS=Homo sapiens OX=9606 GN=BTN1A1 PE=1 SV=3 | 179 | 20201113_12433_1ng_glyco_hcd203040_compound15.8722.8722.3 | controllerType=0 controllerNumber=1 scan=8722 | 20.6661 |
| 18101:3 | 12 | K.N[+2059.735]VSC[+57.021]YIQNLLLGQEK.K | HexNAc(4)Hex(5)Fuc(1)NeuAc(1) | C[+57], N[+2060] | 1280.2212 | 3 | 3838.6492 | 3838.6455 | 0 | 1 | 215 | Specific | 391.21 | 312 | 312 | 7.07 | 39 | >sp\|Q13410\|BT1A1_HUMAN Butyrophilin subfamily 1 member A1 OS=Homo sapiens OX=9606 GN=BTN1A1 PE=1 SV=3 | 179 | 20201113_12433_1ng_glyco_hcd203040_compound15.21478.21478.3 | controllerType=0 controllerNumber=1 scan=21478 | 42.3994 |
| 18104:3 | 12 | K.N[+2059.735]VSC[+57.021]YIQNLLLGQEK.K | HexNAc(4)Hex(5)Fuc(1)NeuAc(1) | C[+57], N[+2060] | 1280.2212 | 3 | 3838.6492 | 3838.6455 | 0 | 1 | 215 | Specific | 486.33 | 339.06 | 339.06 | 7.5 | 39 | >sp\|Q13410\|BT1A1_HUMAN Butyrophilin subfamily 1 member A1 OS=Homo sapiens OX=9606 GN=BTN1A1 PE=1 SV=3 | 179 | 20201113_12433_1ng_glyco_hcd203040_compound15.21481.21481.3 | controllerType=0 controllerNumber=1 scan=21481 | 42.404 |

Table S4. Cont.

| Query #:z | Protein Rank | Glycopeptide | Glycan | Modification Type(s) | Observed m/z | z | Observed (M+H) | Calc. mass (M+H) | Off-by-x error | Mass error (ppm) | StartPosition | Cleavage | Score | Delta | Delta Mod | \|Log Prob\| | # of unique peptides | Protein Name | Protein DB number | Comment | Scan # | Scan Time |
|---|---|---|---|---|---|---|---|---|---|---|---|---|---|---|---|---|---|---|---|---|---|---|
| 24535:4 | 14 | K.VFPLSLDSTPQDGNVVVAC[+57.021]LVQGFFPQEPLSVTWSESGQN[+2117.777]VTAI | HexNAc(5)Hex(5)Fuc(2) | C[+57], N[+2118] | 1724.2974 | 4 | 6894.168 | 6894.1382 | 0 | 4.3 | 8 | Specific | 240.72 | 240.72 | 240.72 | 7.15 | 38 | >sp\|P01877\|IGHA2_HUMAN Immunoglobulin heavy constant alpha 2 OS=Homo sapiens OX=9606 GN=IGHA2 PE=1 SV=4 | 576 | 20201113_12433_1ng_glyco_hcd203040_compound15.28184.28184.4 | controllerType=0 controllerNumber=1 scan=28184 | 55.3774 |
| 24429:4 | 14 | K.VFPLSLDSTPQDGNVVVAC[+57.021]LVQGFFPQEPLSVTWSESGQN[+2059.735]VTAI | HexNAc(4)Hex(5)Fuc(1)NeuAc(1) | C[+57], N[+2060] | 1709.7888 | 4 | 6836.1333 | 6836.0963 | 0 | 5.4 | 8 | Specific | 271.46 | 271.46 | 271.46 | 7.07 | 38 | >sp\|P01877\|IGHA2_HUMAN Immunoglobulin heavy constant alpha 2 OS=Homo sapiens OX=9606 GN=IGHA2 PE=1 SV=4 | 576 | 20201113_12433_1ng_glyco_hcd203040_compound15.28073.28073.4 | controllerType=0 controllerNumber=1 scan=28073 | 55.1635 |
| 24467:4 | 14 | K.VFPLSLDSTPQDGNVVVAC[+57.021]LVQGFFPQEPLSVTWSESGQN[+2262.814]VTAI | HexNAc(5)Hex(5)Fuc(1)NeuAc(1) | C[+57], N[+2263] | 1760.5598 | 4 | 7039.2173 | 7039.1757 | 0 | 5.9 | 8 | Specific | 393.45 | 391.84 | 391.84 | 9.71 | 38 | >sp\|P01877\|IGHA2_HUMAN Immunoglobulin heavy constant alpha 2 OS=Homo sapiens OX=9606 GN=IGHA2 PE=1 SV=4 | 576 | 20201113_12433_1ng_glyco_hcd203040_compound15.28113.28113.4 | controllerType=0 controllerNumber=1 scan=28113 | 55.2418 |
| 22666:4 | 14 | K.VFPLSLDSTPQDGNVVVAC[+57.021]LVQGFFPQEPLSVTWSESGQN[+1809.666]VTAI | HexNAc(5)Hex(4)Fuc(1) | C[+57], N[+1810] | 1647.2725 | 4 | 6586.068 | 6586.0275 | 0 | 6.2 | 8 | Specific | 597.33 | 575.58 | 575.58 | 11.48 | 38 | >sp\|P01877\|IGHA2_HUMAN Immunoglobulin heavy constant alpha 2 OS=Homo sapiens OX=9606 GN=IGHA2 PE=1 SV=4 | 576 | 20201113_12433_1ng_glyco_hcd203040_compound15.26231.26231.4 | controllerType=0 controllerNumber=1 scan=26231 | 51.8691 |
| 22668:4 | 14 | K.VFPLSLDSTPQDGNVVVAC[+57.021]LVQGFFPQEPLSVTWSESGQN[+1809.666]VTAI | HexNAc(5)Hex(4)Fuc(1) | C[+57], N[+1810] | 1647.2725 | 4 | 6586.068 | 6586.0275 | 0 | 6.2 | 8 | Specific | 693.91 | 683.18 | 683.18 | 13.48 | 38 | >sp\|P01877\|IGHA2_HUMAN Immunoglobulin heavy constant alpha 2 OS=Homo sapiens OX=9606 GN=IGHA2 PE=1 SV=4 | 576 | 20201113_12433_1ng_glyco_hcd203040_compound15.26233.26233.4 | controllerType=0 controllerNumber=1 scan=26233 | 51.8749 |
| 22708:4 | 14 | K.VFPLSLDSTPQDGNVVVAC[+57.021]LVQGFFPQEPLSVTWSESGQN[+1647.613]VTAI | HexNAc(5)Hex(3)Fuc(1) | C[+57], N[+1648] | 1606.7551 | 4 | 6423.9987 | 6423.9746 | 0 | 3.7 | 8 | Specific | 742.15 | 720.92 | 720.92 | 15.03 | 38 | >sp\|P01877\|IGHA2_HUMAN Immunoglobulin heavy constant alpha 2 OS=Homo sapiens OX=9606 GN=IGHA2 PE=1 SV=4 | 576 | 20201113_12433_1ng_glyco_hcd203040_compound15.26275.26275.2 | controllerType=0 controllerNumber=1 scan=26275 | 51.9546 |
| 22710:4 | 14 | K.VFPLSLDSTPQDGNVVVAC[+57.021]LVQGFFPQEPLSVTWSESGQN[+1647.613]VTAI | HexNAc(5)Hex(3)Fuc(1) | C[+57], N[+1648] | 1606.7551 | 4 | 6423.9987 | 6423.9746 | 0 | 3.7 | 8 | Specific | 866.47 | 723.95 | 723.95 | 18.03 | 38 | >sp\|P01877\|IGHA2_HUMAN Immunoglobulin heavy constant alpha 2 OS=Homo sapiens OX=9606 GN=IGHA2 PE=1 SV=4 | 576 | 20201113_12433_1ng_glyco_hcd203040_compound15.26277.26277.2 | controllerType=0 controllerNumber=1 scan=26277 | 51.9622 |
| 02263:3 | 14 | K.HYTN[+2059.735]SSQDVTVPC[+57.021]R.V | HexNAc(4)Hex(5)Fuc(1)NeuAc(1) | C[+57], N[+2060] | 1241.8337 | 3 | 3723.4865 | 3723.4842 | 0 | 0.6 | 89 | Specific | 236.81 | 180.02 | 180.02 | 5.36 | 38 | >sp\|P01877\|IGHA2_HUMAN Immunoglobulin heavy constant alpha 2 OS=Homo sapiens OX=9606 GN=IGHA2 PE=1 SV=4 | 576 | 20201113_12433_1ng_glyco_hcd203040_compound15.5016.5016.3 | controllerType=0 controllerNumber=1 scan=5016 | 14.4192 |
| 01432:3 | 14 | K.HYTN[+1971.719]SSQDVTVPC[+57.021]R.V | HexNAc(5)Hex(5)Fuc(1) | C[+57], N[+1972] | 1212.496 | 3 | 3635.4735 | 3635.4682 | 0 | 1.5 | 89 | Specific | 311 | 165.5 | 165.5 | 6.07 | 38 | >sp\|P01877\|IGHA2_HUMAN Immunoglobulin heavy constant alpha 2 OS=Homo sapiens OX=9606 GN=IGHA2 PE=1 SV=4 | 576 | 20201113_12433_1ng_glyco_hcd203040_compound15.4153.4153.3 | controllerType=0 controllerNumber=1 scan=4153 | 12.9409 |
| 01434:3 | 14 | K.HYTN[+1971.719]SSQDVTVPC[+57.021]R.V | HexNAc(5)Hex(5)Fuc(1) | C[+57], N[+1972] | 1212.496 | 3 | 3635.4735 | 3635.4682 | 0 | 1.5 | 89 | Specific | 312.94 | 242.47 | 242.47 | 6.74 | 38 | >sp\|P01877\|IGHA2_HUMAN Immunoglobulin heavy constant alpha 2 OS=Homo sapiens OX=9606 GN=IGHA2 PE=1 SV=4 | 576 | 20201113_12433_1ng_glyco_hcd203040_compound15.4155.4155.3 | controllerType=0 controllerNumber=1 scan=4155 | 12.9449 |
| 02261:3 | 14 | K.HYTN[+2059.735]SSQDVTVPC[+57.021]R.V | HexNAc(4)Hex(5)Fuc(1)NeuAc(1) | C[+57], N[+2060] | 1241.8337 | 3 | 3723.4865 | 3723.4842 | 0 | 0.6 | 89 | Specific | 321.32 | 212.05 | 212.05 | 5.9 | 38 | >sp\|P01877\|IGHA2_HUMAN Immunoglobulin heavy constant alpha 2 OS=Homo sapiens OX=9606 GN=IGHA2 PE=1 SV=4 | 576 | 20201113_12433_1ng_glyco_hcd203040_compound15.5014.5014.3 | controllerType=0 controllerNumber=1 scan=5014 | 14.4153 |
| 01409:3 | 14 | K.HYTN[+1809.666]SSQDVTVPC[+57.021]R.V | HexNAc(5)Hex(4)Fuc(1) | C[+57], N[+1810] | 1158.4794 | 3 | 3473.4237 | 3473.4154 | 0 | 2.4 | 89 | Specific | 584.01 | 393.94 | 393.94 | 11.29 | 38 | >sp\|P01877\|IGHA2_HUMAN Immunoglobulin heavy constant alpha 2 OS=Homo sapiens OX=9606 GN=IGHA2 PE=1 SV=4 | 576 | 20201113_12433_1ng_glyco_hcd203040_compound15.4129.4129.3 | controllerType=0 controllerNumber=1 scan=4129 | 12.8975 |
| 01411:3 | 14 | K.HYTN[+1809.666]SSQDVTVPC[+57.021]R.V | HexNAc(5)Hex(4)Fuc(1) | C[+57], N[+1810] | 1158.4794 | 3 | 3473.4237 | 3473.4154 | 0 | 2.4 | 89 | Specific | 593.31 | 462.62 | 462.62 | 11.29 | 38 | >sp\|P01877\|IGHA2_HUMAN Immunoglobulin heavy constant alpha 2 OS=Homo sapiens OX=9606 GN=IGHA2 PE=1 SV=4 | 576 | 20201113_12433_1ng_glyco_hcd203040_compound15.4131.4131.3 | controllerType=0 controllerNumber=1 scan=4131 | 12.9015 |
| 01482:3 | 14 | K.HYTN[+1647.613]SSQDVTVPC[+57.021]R.V | HexNAc(5)Hex(3)Fuc(1) | C[+57], N[+1648] | 1104.4605 | 3 | 3311.3669 | 3311.3625 | 0 | 1.3 | 89 | Specific | 628.34 | 529.21 | 529.21 | 12.29 | 38 | >sp\|P01877\|IGHA2_HUMAN Immunoglobulin heavy constant alpha 2 OS=Homo sapiens OX=9606 GN=IGHA2 PE=1 SV=4 | 576 | 20201113_12433_1ng_glyco_hcd203040_compound15.4205.4205.3 | controllerType=0 controllerNumber=1 scan=4205 | 13.0289 |
| 01484:3 | 14 | K.HYTN[+1647.613]SSQDVTVPC[+57.021]R.V | HexNAc(5)Hex(3)Fuc(1) | C[+57], N[+1648] | 1104.4605 | 3 | 3311.3669 | 3311.3625 | 0 | 1.3 | 89 | Specific | 719.55 | 609.16 | 609.16 | 14.29 | 38 | >sp\|P01877\|IGHA2_HUMAN Immunoglobulin heavy constant alpha 2 OS=Homo sapiens OX=9606 GN=IGHA2 PE=1 SV=4 | 576 | 20201113_12433_1ng_glyco_hcd203040_compound15.4207.4207.3 | controllerType=0 controllerNumber=1 scan=4207 | 13.0334 |
| 03555:5 | 14 | K.HYTN[+1647.613]SSQDVTVPC[+57.021]RVPPPPPC[+57.021]C[+57.021]HPR.L | HexNAc(5)Hex(3)Fuc(1) | C[+57]*3, N[+1648] | 921.9995 | 5 | 4605.9686 | 4605.9689 | 0 | -0.1 | 89 | Specific | 473.35 | 244.09 | 244.09 | 7.55 | 38 | >sp\|P01877\|IGHA2_HUMAN Immunoglobulin heavy constant alpha 2 OS=Homo sapiens OX=9606 GN=IGHA2 PE=1 SV=4 | 576 | 20201113_12433_1ng_glyco_hcd203040_compound15.6355.6355.5 | controllerType=0 controllerNumber=1 scan=6355 | 16.7817 |
| 03557:5 | 14 | K.HYTN[+1647.613]SSQDVTVPC[+57.021]RVPPPPPC[+57.021]C[+57.021]HPR.L | HexNAc(5)Hex(3)Fuc(1) | C[+57]*3, N[+1648] | 921.9995 | 5 | 4605.9686 | 4605.9689 | 0 | -0.1 | 89 | Specific | 572.45 | 427.72 | 427.72 | 10.12 | 38 | >sp\|P01877\|IGHA2_HUMAN Immunoglobulin heavy constant alpha 2 OS=Homo sapiens OX=9606 GN=IGHA2 PE=1 SV=4 | 576 | 20201113_12433_1ng_glyco_hcd203040_compound15.6357.6357.5 | controllerType=0 controllerNumber=1 scan=6357 | 16.7856 |
| 03149:2 | 14 | K.TPLTAN[+1971.719]ITK.S | HexNAc(5)Hex(5)Fuc(1) | N[+1972] | 1465.6454 | 2 | 2930.2836 | 2930.2757 | 0 | 2.7 | 200 | Specific | 233.6 | 179.18 | 179.18 | 5.49 | 38 | >sp\|P01877\|IGHA2_HUMAN Immunoglobulin heavy constant alpha 2 OS=Homo sapiens OX=9606 GN=IGHA2 PE=1 SV=4 | 576 | 20201113_12433_1ng_glyco_hcd203040_compound15.5934.5934.2 | controllerType=0 controllerNumber=1 scan=5934 | 16.026 |
| 03128:2 | 14 | K.TPLTAN[+1606.587]ITK.S | HexNAc(4)Hex(4)Fuc(1) | N[+1607] | 1283.0802 | 2 | 2565.1532 | 2565.1435 | 0 | 3.8 | 200 | Specific | 246.05 | 106.81 | 106.81 | 4.16 | 38 | >sp\|P01877\|IGHA2_HUMAN Immunoglobulin heavy constant alpha 2 OS=Homo sapiens OX=9606 GN=IGHA2 PE=1 SV=4 | 576 | 20201113_12433_1ng_glyco_hcd203040_compound15.5912.5912.2 | controllerType=0 controllerNumber=1 scan=5912 | 15.9791 |
| 04959:2 | 14 | K.TPLTAN[+2059.735]ITK.S | HexNAc(4)Hex(5)Fuc(1)NeuAc(1) | N[+2060] | 1509.651 | 2 | 3018.2948 | 3018.2917 | 0 | 1 | 200 | Specific | 255.72 | 107.88 | 107.88 | 4.29 | 38 | >sp\|P01877\|IGHA2_HUMAN Immunoglobulin heavy constant alpha 2 OS=Homo sapiens OX=9606 GN=IGHA2 PE=1 SV=4 | 576 | 20201113_12433_1ng_glyco_hcd203040_compound15.7809.7809.2 | controllerType=0 controllerNumber=1 scan=7809 | 19.1586 |
| 03268:2 | 14 | K.TPLTAN[+1501.555]ITK.S | HexNAc(5)Hex(3) | N[+1502] | 1230.5641 | 2 | 2460.121 | 2460.1121 | 0 | 3.6 | 200 | Specific | 256.57 | 99.76 | 99.76 | 4.31 | 38 | >sp\|P01877\|IGHA2_HUMAN Immunoglobulin heavy constant alpha 2 OS=Homo sapiens OX=9606 GN=IGHA2 PE=1 SV=4 | 576 | 20201113_12433_1ng_glyco_hcd203040_compound15.6058.6058.2 | controllerType=0 controllerNumber=1 scan=6058 | 16.269 |
| 04971:3 | 14 | K.TPLTAN[+2262.814]ITK.S | HexNAc(5)Hex(5)Fuc(1)NeuAc(1) | N[+2263] | 1074.4644 | 3 | 3221.3786 | 3221.3711 | 0 | 2.3 | 200 | Specific | 264.59 | 231.05 | 231.05 | 5.9 | 38 | >sp\|P01877\|IGHA2_HUMAN Immunoglobulin heavy constant alpha 2 OS=Homo sapiens OX=9606 GN=IGHA2 PE=1 SV=4 | 576 | 20201113_12433_1ng_glyco_hcd203040_compound15.7822.7822.3 | controllerType=0 controllerNumber=1 scan=7822 | 19.182 |
| 04961:2 | 14 | K.TPLTAN[+2059.735]ITK.S | HexNAc(4)Hex(5)Fuc(1)NeuAc(1) | N[+2060] | 1509.651 | 2 | 3018.2948 | 3018.2917 | 0 | 1 | 200 | Specific | 294.55 | 131.95 | 131.95 | 4.73 | 38 | >sp\|P01877\|IGHA2_HUMAN Immunoglobulin heavy constant alpha 2 OS=Homo sapiens OX=9606 GN=IGHA2 PE=1 SV=4 | 576 | 20201113_12433_1ng_glyco_hcd203040_compound15.7811.7811.2 | controllerType=0 controllerNumber=1 scan=7811 | 19.1627 |
| 03270:2 | 14 | K.TPLTAN[+1501.555]ITK.S | HexNAc(5)Hex(3) | N[+1502] | 1230.5641 | 2 | 2460.121 | 2460.1121 | 0 | 3.6 | 200 | Specific | 316.63 | 316.63 | 316.63 | 6.4 | 38 | >sp\|P01877\|IGHA2_HUMAN Immunoglobulin heavy constant alpha 2 OS=Homo sapiens OX=9606 GN=IGHA2 PE=1 SV=4 | 576 | 20201113_12433_1ng_glyco_hcd203040_compound15.6060.6060.2 | controllerType=0 controllerNumber=1 scan=6060 | 16.2731 |
| 04974:3 | 14 | K.TPLTAN[+2262.814]ITK.S | HexNAc(5)Hex(5)Fuc(1)NeuAc(1) | N[+2263] | 1074.4644 | 3 | 3221.3786 | 3221.3711 | 0 | 2.3 | 200 | Specific | 366.04 | 224.1 | 224.1 | 6.48 | 38 | >sp\|P01877\|IGHA2_HUMAN Immunoglobulin heavy constant alpha 2 OS=Homo sapiens OX=9606 GN=IGHA2 PE=1 SV=4 | 576 | 20201113_12433_1ng_glyco_hcd203040_compound15.7825.7825.3 | controllerType=0 controllerNumber=1 scan=7825 | 19.1863 |
| 03134:3 | 14 | K.TPLTAN[+1971.719]ITK.S | HexNAc(5)Hex(5)Fuc(1) | N[+1972] | 977.4344 | 3 | 2930.2887 | 2930.2757 | 0 | 4.4 | 200 | Specific | 400.15 | 257.94 | 257.94 | 6.59 | 38 | >sp\|P01877\|IGHA2_HUMAN Immunoglobulin heavy constant alpha 2 OS=Homo sapiens OX=9606 GN=IGHA2 PE=1 SV=4 | 576 | 20201113_12433_1ng_glyco_hcd203040_compound15.5918.5918.3 | controllerType=0 controllerNumber=1 scan=5918 | 15.9908 |
| 03254:3 | 14 | K.TPLTAN[+1647.613]ITK.S | HexNAc(5)Hex(3)Fuc(1) | N[+1648] | 869.3971 | 3 | 2606.1768 | 2606.17 | 0 | 2.6 | 200 | Specific | 467.15 | 193.6 | 193.6 | 6.74 | 38 | >sp\|P01877\|IGHA2_HUMAN Immunoglobulin heavy constant alpha 2 OS=Homo sapiens OX=9606 GN=IGHA2 PE=1 SV=4 | 576 | 20201113_12433_1ng_glyco_hcd203040_compound15.6043.6043.3 | controllerType=0 controllerNumber=1 scan=6043 | 16.2343 |
| 03186:3 | 14 | K.TPLTAN[+1809.666]ITK.S | HexNAc(5)Hex(4)Fuc(1) | N[+1810] | 923.4151 | 3 | 2768.2308 | 2768.2228 | 0 | 2.9 | 200 | Specific | 484.27 | 186.56 | 186.56 | 6.53 | 38 | >sp\|P01877\|IGHA2_HUMAN Immunoglobulin heavy constant alpha 2 OS=Homo sapiens OX=9606 GN=IGHA2 PE=1 SV=4 | 576 | 20201113_12433_1ng_glyco_hcd203040_compound15.5972.5972.3 | controllerType=0 controllerNumber=1 scan=5972 | 16.0919 |
| 03200:2 | 14 | K.TPLTAN[+1444.534]ITK.S | HexNAc(4)Hex(3)Fuc(1) | N[+1445] | 1202.0506 | 2 | 2403.0939 | 2403.0906 | 0 | 1.3 | 200 | Specific | 525.18 | 258.29 | 258.29 | 7.4 | 38 | >sp\|P01877\|IGHA2_HUMAN Immunoglobulin heavy constant alpha 2 OS=Homo sapiens OX=9606 GN=IGHA2 PE=1 SV=4 | 576 | 20201113_12433_1ng_glyco_hcd203040_compound15.5987.5987.2 | controllerType=0 controllerNumber=1 scan=5987 | 16.1202 |
| 03256:3 | 14 | K.TPLTAN[+1647.613]ITK.S | HexNAc(5)Hex(3)Fuc(1) | N[+1648] | 869.3971 | 3 | 2606.1768 | 2606.17 | 0 | 2.6 | 200 | Specific | 553.36 | 276.44 | 276.44 | 7.75 | 38 | >sp\|P01877\|IGHA2_HUMAN Immunoglobulin heavy constant alpha 2 OS=Homo sapiens OX=9606 GN=IGHA2 PE=1 SV=4 | 576 | 20201113_12433_1ng_glyco_hcd203040_compound15.6045.6045.3 | controllerType=0 controllerNumber=1 scan=6045 | 16.244 |
| 03188:3 | 14 | K.TPLTAN[+1809.666]ITK.S | HexNAc(5)Hex(4)Fuc(1) | N[+1810] | 923.4151 | 3 | 2768.2308 | 2768.2228 | 0 | 2.9 | 200 | Specific | 558.72 | 274.48 | 274.48 | 7.75 | 38 | >sp\|P01877\|IGHA2_HUMAN Immunoglobulin heavy constant alpha 2 OS=Homo sapiens OX=9606 GN=IGHA2 PE=1 SV=4 | 576 | 20201113_12433_1ng_glyco_hcd203040_compound15.5974.5974.3 | controllerType=0 controllerNumber=1 scan=5974 | 16.0964 |
| 03168:2 | 14 | K.TPLTAN[+1809.666]ITK.S | HexNAc(5)Hex(4)Fuc(1) | N[+1810] | 1384.6186 | 2 | 2768.2299 | 2768.2228 | 0 | 2.5 | 200 | Specific | 559.21 | 294.21 | 294.21 | 7.73 | 38 | >sp\|P01877\|IGHA2_HUMAN Immunoglobulin heavy constant alpha 2 OS=Homo sapiens OX=9606 GN=IGHA2 PE=1 SV=4 | 576 | 20201113_12433_1ng_glyco_hcd203040_compound15.5954.5954.2 | controllerType=0 controllerNumber=1 scan=5954 | 16.0613 |
| 03198:2 | 14 | K.TPLTAN[+1444.534]ITK.S | HexNAc(4)Hex(3)Fuc(1) | N[+1445] | 1202.0506 | 2 | 2403.0939 | 2403.0906 | 0 | 1.3 | 200 | Specific | 568.78 | 370.7 | 370.7 | 8.37 | 38 | >sp\|P01877\|IGHA2_HUMAN Immunoglobulin heavy constant alpha 2 OS=Homo sapiens OX=9606 GN=IGHA2 PE=1 SV=4 | 576 | 20201113_12433_1ng_glyco_hcd203040_compound15.5985.5985.2 | controllerType=0 controllerNumber=1 scan=5985 | 16.1163 |
| 03247:2 | 14 | K.TPLTAN[+1647.613]ITK.S | HexNAc(5)Hex(3)Fuc(1) | N[+1648] | 1303.5918 | 2 | 2606.1764 | 2606.17 | 0 | 2.5 | 200 | Specific | 598.08 | 324.02 | 324.02 | 9.24 | 38 | >sp\|P01877\|IGHA2_HUMAN Immunoglobulin heavy constant alpha 2 OS=Homo sapiens OX=9606 GN=IGHA2 PE=1 SV=4 | 576 | 20201113_12433_1ng_glyco_hcd203040_compound15.6036.6036.2 | controllerType=0 controllerNumber=1 scan=6036 | 16.207 |
| 03170:2 | 14 | K.TPLTAN[+1809.666]ITK.S | HexNAc(5)Hex(4)Fuc(1) | N[+1810] | 1384.6186 | 2 | 2768.2299 | 2768.2228 | 0 | 2.5 | 200 | Specific | 611.2 | 300.1 | 300.1 | 9.99 | 38 | >sp\|P01877\|IGHA2_HUMAN Immunoglobulin heavy constant alpha 2 OS=Homo sapiens OX=9606 GN=IGHA2 PE=1 SV=4 | 576 | 20201113_12433_1ng_glyco_hcd203040_compound15.5956.5956.2 | controllerType=0 controllerNumber=1 scan=5956 | 16.0657 |
| 03249:2 | 14 | K.TPLTAN[+1647.613]ITK.S | HexNAc(5)Hex(3)Fuc(1) | N[+1648] | 1303.5918 | 2 | 2606.1764 | 2606.17 | 0 | 2.5 | 200 | Specific | 615.01 | 322.58 | 322.58 | 10.24 | 38 | >sp\|P01877\|IGHA2_HUMAN Immunoglobulin heavy constant alpha 2 OS=Homo sapiens OX=9606 GN=IGHA2 PE=1 SV=4 | 576 | 20201113_12433_1ng_glyco_hcd203040_compound15.6038.6038.2 | controllerType=0 controllerNumber=1 scan=6038 | 16.2268 |
| 10527:3 | 14 | R.MAGKPTHIN[+1540.529]VSVVMAEADGTC[+57.021]Y.- | HexNAc(2)Hex(7) | C[+57], N[+1541] | 1297.8771 | 3 | 3891.6168 | 3891.6114 | 0 | 1.4 | 319 | CRagged | 726.75 | 602.56 | 419.55 | 15.18 | 38 | >sp\|P01877\|IGHA2_HUMAN Immunoglobulin heavy constant alpha 2 OS=Homo sapiens OX=9606 GN=IGHA2 PE=1 SV=4 | 576 | 20201113_12433_1ng_glyco_hcd203040_compound15.13582.13582.3 | controllerType=0 controllerNumber=1 scan=13582 | 28.6451 |
| 10530:3 | 14 | R.MAGKPTHIN[+1540.529]VSVVMAEADGTC[+57.021]Y.- | HexNAc(2)Hex(7) | C[+57], N[+1541] | 1297.8771 | 3 | 3891.6168 | 3891.6114 | 0 | 1.4 | 319 | CRagged | 746.43 | 746.43 | 401.82 | 15.18 | 38 | >sp\|P01877\|IGHA2_HUMAN Immunoglobulin heavy constant alpha 2 OS=Homo sapiens OX=9606 GN=IGHA2 PE=1 SV=4 | 576 | 20201113_12433_1ng_glyco_hcd203040_compound15.13585.13585.3 | controllerType=0 controllerNumber=1 scan=13585 | 28.6499 |
| 10499:3 | 14 | R.MAGKPTHIN[+1702.581]VSVVMAEADGTC[+57.021]Y.- | HexNAc(2)Hex(8) | C[+57], N[+1703] | 1351.8964 | 3 | 4053.6747 | 4053.6642 | 0 | 2.6 | 319 | CRagged | 807.24 | 807.24 | 487.89 | 17.18 | 38 | >sp\|P01877\|IGHA2_HUMAN Immunoglobulin heavy constant alpha 2 OS=Homo sapiens OX=9606 GN=IGHA2 PE=1 SV=4 | 576 | 20201113_12433_1ng_glyco_hcd203040_compound15.13553.13553.3 | controllerType=0 controllerNumber=1 scan=13553 | 28.5961 |
| 10501:3 | 14 | R.MAGKPTHIN[+1702.581]VSVVMAEADGTC[+57.021]Y.- | HexNAc(2)Hex(8) | C[+57], N[+1703] | 1351.8964 | 3 | 4053.6747 | 4053.6642 | 0 | 2.6 | 319 | CRagged | 818.89 | 656.89 | 459.47 | 17.18 | 38 | >sp\|P01877\|IGHA2_HUMAN Immunoglobulin heavy constant alpha 2 OS=Homo sapiens OX=9606 GN=IGHA2 PE=1 SV=4 | 576 | 20201113_12433_1ng_glyco_hcd203040_compound15.13555.13555.3 | controllerType=0 controllerNumber=1 scan=13555 | 28.6008 |
| 10613:3 | 14 | R.MAGKPTHIN[+1216.423]VSVVMAEADGTC[+57.021]Y.- | HexNAc(2)Hex(5) | C[+57], N[+1216] | 1189.8431 | 3 | 3567.5147 | 3567.5057 | 0 | 2.5 | 319 | CRagged | 833.39 | 833.39 | 454.15 | 17.18 | 38 | >sp\|P01877\|IGHA2_HUMAN Immunoglobulin heavy constant alpha 2 OS=Homo sapiens OX=9606 GN=IGHA2 PE=1 SV=4 | 576 | 20201113_12433_1ng_glyco_hcd203040_compound15.13671.13671.3 | controllerType=0 controllerNumber=1 scan=13671 | 28.8275 |
| 10611:3 | 14 | R.MAGKPTHIN[+1216.423]VSVVMAEADGTC[+57.021]Y.- | HexNAc(2)Hex(5) | C[+57], N[+1216] | 1189.8431 | 3 | 3567.5147 | 3567.5057 | 0 | 2.5 | 319 | CRagged | 838.25 | 800.07 | 462.6 | 17.18 | 38 | >sp\|P01877\|IGHA2_HUMAN Immunoglobulin heavy constant alpha 2 OS=Homo sapiens OX=9606 GN=IGHA2 PE=1 SV=4 | 576 | 20201113_12433_1ng_glyco_hcd203040_compound15.13669.13669.3 | controllerType=0 controllerNumber=1 scan=13669 | 28.8122 |
| 10563:3 | 14 | R.MAGKPTHIN[+1378.476]VSVVMAEADGTC[+57.021]Y.- | HexNAc(2)Hex(6) | C[+57], N[+1378] | 1243.8626 | 3 | 3729.5733 | 3729.5585 | 0 | 3.9 | 319 | CRagged | 890.21 | 828.41 | 504.17 | 17.83 | 38 | >sp\|P01877\|IGHA2_HUMAN Immunoglobulin heavy constant alpha 2 OS=Homo sapiens OX=9606 GN=IGHA2 PE=1 SV=4 | 576 | 20201113_12433_1ng_glyco_hcd203040_compound15.13619.13619.3 | controllerType=0 controllerNumber=1 scan=13619 | 28.7154 |
| 10561:3 | 14 | R.MAGKPTHIN[+1378.476]VSVVMAEADGTC[+57.021]Y.- | HexNAc(2)Hex(6) | C[+57], N[+1378] | 1243.8626 | 3 | 3729.5733 | 3729.5585 | 0 | 3.9 | 319 | CRagged | 978.8 | 952.31 | 562.61 | 19.83 | 38 | >sp\|P01877\|IGHA2_HUMAN Immunoglobulin heavy constant alpha 2 OS=Homo sapiens OX=9606 GN=IGHA2 PE=1 SV=4 | 576 | 20201113_12433_1ng_glyco_hcd203040_compound15.13617.13617.3 | controllerType=0 controllerNumber=1 scan=13617 | 28.7091 |
| 02405:3 | 15 | K.N[+1216.423]NSIPDKQITASSSYK.T | HexNAc(2)Hex(5) | N[+1216] | 990.4396 | 3 | 2969.3043 | 2969.2991 | 0 | 1.7 | 238 | Specific | 295.57 | 157.1 | 157.1 | 5.44 | 31 | >sp\|Q08431\|MFGM_HUMAN Lactadherin OS=Homo sapiens OX=9606 GN=MFGE8 PE=1 SV=3 | 1064 | 20201113_12433_1ng_glyco_hcd203040_compound15.5163.5163.3 | controllerType=0 controllerNumber=1 scan=5163 | 14.6772 |
| 02287:3 | 15 | K.N[+1540.529]NSIPDKQITASSSYK.T | HexNAc(2)Hex(7) | N[+1541] | 1098.4771 | 3 | 3293.4167 | 3293.4048 | 0 | 3.6 | 238 | Specific | 298.58 | 298.58 | 298.58 | 5.97 | 31 | >sp\|Q08431\|MFGM_HUMAN Lactadherin OS=Homo sapiens OX=9606 GN=MFGE8 PE=1 SV=3 | 1064 | 20201113_12433_1ng_glyco_hcd203040_compound15.5041.5041.3 | controllerType=0 controllerNumber=1 scan=5041 | 14.4612 |
| 02403:3 | 15 | K.N[+1216.423]NSIPDKQITASSSYK.T | HexNAc(2)Hex(5) | N[+1216] | 990.4396 | 3 | 2969.3043 | 2969.2991 | 0 | 1.7 | 238 | Specific | 303.85 | 263.44 | 263.44 | 6.36 | 31 | >sp\|Q08431\|MFGM_HUMAN Lactadherin OS=Homo sapiens OX=9606 GN=MFGE8 PE=1 SV=3 | 1064 | 20201113_12433_1ng_glyco_hcd203040_compound15.5161.5161.3 | controllerType=0 controllerNumber=1 scan=5161 | 14.6724 |
| 08691:3 | 15 | K.VAYSN[+2205.793]DSANWTEYQDPR.T | HexNAc(4)Hex(5)Fuc(2)NeuAc(1) | N[+2206] | 1407.8986 | 3 | 4221.6813 | 4221.6658 | 0 | 3.7 | 321 | Specific | 248.04 | 187.62 | 4.16 | 5.54 | 31 | >sp\|Q08431\|MFGM_HUMAN Lactadherin OS=Homo sapiens OX=9606 GN=MFGE8 PE=1 SV=3 | 1064 | 20201113_12433_1ng_glyco_hcd203040_compound15.11677.11677.3 | controllerType=0 controllerNumber=1 scan=11677 | 25.4881 |
| 08693:3 | 15 | K.VAYSN[+2205.793]DSANWTEYQDPR.T | HexNAc(4)Hex(5)Fuc(2)NeuAc(1) | N[+2206] | 1407.8986 | 3 | 4221.6813 | 4221.6658 | 0 | 3.7 | 321 | Specific | 313.14 | 256.42 | 0 | 6.38 | 31 | >sp\|Q08431\|MFGM_HUMAN Lactadherin OS=Homo sapiens OX=9606 GN=MFGE8 PE=1 SV=3 | 1064 | 20201113_12433_1ng_glyco_hcd203040_compound15.11679.11679.3 | controllerType=0 controllerNumber=1 scan=11679 | 25.4921 |
| 27639:4 | 17 | R.QLAHQSN[+2861.000]STNIFFSPVSIATAFAMLSLGTK.A | HexNAc(5)Hex(6)NeuAc(3) | N[+2861] | 1511.4171 | 4 | 6042.6466 | 6042.6351 | 0 | 1.9 | 64 | Specific | 263.72 | 253.9 | 253.9 | 7.1 | 32 | >sp\|P01009\|A1AT_HUMAN Alpha-1-antitrypsin OS=Homo sapiens OX=9606 GN=SERPINA1 PE=1 SV=3 | 642 | 20201113_12433_1ng_glyco_hcd203040_compound15.31411.31411.4 | controllerType=0 controllerNumber=1 scan=31411 | 61.1735 |
| 24841:5 | 17 | R.QLAHQSN[+2204.772]STNIFFSPVSIATAFAMLSLGTK.A | HexNAc(4)Hex(5)NeuAc(2) | N[+2205] | 1078.0906 | 5 | 5386.4238 | 5386.4075 | 0 | 3 | 64 | Specific | 299.31 | 299.31 | 299.31 | 7.38 | 32 | >sp\|P01009\|A1AT_HUMAN Alpha-1-antitrypsin OS=Homo sapiens OX=9606 GN=SERPINA1 PE=1 SV=3 | 642 | 20201113_12433_1ng_glyco_hcd203040_compound15.28501.28501.5 | controllerType=0 controllerNumber=1 scan=28501 | 55.9511 |
| 24839:5 | 17 | R.QLAHQSN[+2204.772]STNIFFSPVSIATAFAMLSLGTK.A | HexNAc(4)Hex(5)NeuAc(2) | N[+2205] | 1078.0906 | 5 | 5386.4238 | 5386.4075 | 0 | 3 | 64 | Specific | 373.13 | 373.13 | 373.13 | 8.43 | 32 | >sp\|P01009\|A1AT_HUMAN Alpha-1-antitrypsin OS=Homo sapiens OX=9606 GN=SERPINA1 PE=1 SV=3 | 642 | 20201113_12433_1ng_glyco_hcd203040_compound15.28499.28499.5 | controllerType=0 controllerNumber=1 scan=28499 | 55.9473 |
| 27641:4 | 17 | R.QLAHQSN[+2861.000]STNIFFSPVSIATAFAMLSLGTK.A | HexNAc(5)Hex(6)NeuAc(3) | N[+2861] | 1511.4171 | 4 | 6042.6466 | 6042.6351 | 0 | 1.9 | 64 | Specific | 391.77 | 391.77 | 391.77 | 9.79 | 32 | >sp\|P01009\|A1AT_HUMAN Alpha-1-antitrypsin OS=Homo sapiens OX=9606 GN=SERPINA1 PE=1 SV=3 | 642 | 20201113_12433_1ng_glyco_hcd203040_compound15.31413.31413.4 | controllerType=0 controllerNumber=1 scan=31413 | 61.1776 |
| 24827:3 | 17 | R.QLAHQSN[+2204.772]STNIFFSPVSIATAFAMLSLGTK.A | HexNAc(4)Hex(5)NeuAc(2) | N[+2205] | 1796.1447 | 3 | 5386.4197 | 5386.4075 | 0 | 2.3 | 64 | Specific | 606.22 | 581.63 | 392.79 | 12.85 | 32 | >sp\|P01009\|A1AT_HUMAN Alpha-1-antitrypsin OS=Homo sapiens OX=9606 GN=SERPINA1 PE=1 SV=3 | 642 | 20201113_12433_1ng_glyco_hcd203040_compound15.28487.28487.3 | controllerType=0 controllerNumber=1 scan=28487 | 55.9258 |
| 24825:3 | 17 | R.QLAHQSN[+2204.772]STNIFFSPVSIATAFAMLSLGTK.A | HexNAc(4)Hex(5)NeuAc(2) | N[+2205] | 1796.1447 | 3 | 5386.4197 | 5386.4075 | 0 | 2.3 | 64 | Specific | 734.64 | 678.99 | 563.38 | 14.85 | 32 | >sp\|P01009\|A1AT_HUMAN Alpha-1-antitrypsin OS=Homo sapiens OX=9606 GN=SERPINA1 PE=1 SV=3 | 642 | 20201113_12433_1ng_glyco_hcd203040_compound15.28485.28485.3 | controllerType=0 controllerNumber=1 scan=28485 | 55.9215 |
| 24824:4 | 17 | R.QLAHQSN[+2204.772]STNIFFSPVSIATAFAMLSLGTK.A | HexNAc(4)Hex(5)NeuAc(2) | N[+2205] | 1347.3596 | 4 | 5386.4167 | 5386.4075 | 0 | 1.7 | 64 | Specific | 857.64 | 777.2 | 551.35 | 17.83 | 32 | >sp\|P01009\|A1AT_HUMAN Alpha-1-antitrypsin OS=Homo sapiens OX=9606 GN=SERPINA1 PE=1 SV=3 | 642 | 20201113_12433_1ng_glyco_hcd203040_compound15.28484.28484.4 | controllerType=0 controllerNumber=1 scan=28484 | 55.9198 |
| 24822:4 | 17 | R.QLAHQSN[+2204.772]STNIFFSPVSIATAFAMLSLGTK.A | HexNAc(4)Hex(5)NeuAc(2) | N[+2205] | 1347.3596 | 4 | 5386.4167 | 5386.4075 | 0 | 1.7 | 64 | Specific | 878.87 | 821.8 | 545.5 | 17.83 | 32 | >sp\|P01009\|A1AT_HUMAN Alpha-1-antitrypsin OS=Homo sapiens OX=9606 GN=SERPINA1 PE=1 SV=3 | 642 | 20201113_12433_1ng_glyco_hcd203040_compound15.28482.28482.4 | controllerType=0 controllerNumber=1 scan=28482 | 55.9133 |
| 14478:4 | 21 | K.LKELPGVC[+57.021]N[+2059.735]ETMMALWEEC[+57.021]KPC[+57.021]LK.Q | HexNAc(4)Hex(5)Fuc(1)NeuAc(1) | C[+57]*3, N[+2060] | 1249.7883 | 4 | 4996.1315 | 4996.1196 | 0 | 2.4 | 95 | Specific | 717.8 | 704.85 | 704.85 | 14.75 | 28 | >sp\|P10909\|CLUS_HUMAN Clusterin OS=Homo sapiens OX=9606 GN=CLU PE=1 SV=1 | 1467 | 20201113_12433_1ng_glyco_hcd203040_compound15.17692.17692.4 | controllerType=0 controllerNumber=1 scan=17692 | 35.5561 |
| 14395:4 | 21 | K.LKELPGVC[+57.021]N[+2205.793]ETMMALWEEC[+57.021]KPC[+57.021]LK.Q | HexNAc(4)Hex(5)Fuc(2)NeuAc(1) | C[+57]*3, N[+2206] | 1286.3029 | 4 | 5142.1896 | 5142.1776 | 0 | 2.3 | 95 | Specific | 765.16 | 744.89 | 744.89 | 15.75 | 28 | >sp\|P10909\|CLUS_HUMAN Clusterin OS=Homo sapiens OX=9606 GN=CLU PE=1 SV=1 | 1467 | 20201113_12433_1ng_glyco_hcd203040_compound15.17606.17606.4 | controllerType=0 controllerNumber=1 scan=17606 | 35.4043 |
| 14480:4 | 21 | K.LKELPGVC[+57.021]N[+2059.735]ETMMALWEEC[+57.021]KPC[+57.021]LK.Q | HexNAc(4)Hex(5)Fuc(1)NeuAc(1) | C[+57]*3, N[+2060] | 1249.7883 | 4 | 4996.1315 | 4996.1196 | 0 | 2.4 | 95 | Specific | 835.83 | 784.77 | 784.77 | 16.75 | 28 | >sp\|P10909\|CLUS_HUMAN Clusterin OS=Homo sapiens OX=9606 GN=CLU PE=1 SV=1 | 1467 | 20201113_12433_1ng_glyco_hcd203040_compound15.17694.17694.4 | controllerType=0 controllerNumber=1 scan=17694 | 35.5602 |
| 14393:4 | 21 | K.LKELPGVC[+57.021]N[+2205.793]ETMMALWEEC[+57.021]KPC[+57.021]LK.Q | HexNAc(4)Hex(5)Fuc(2)NeuAc(1) | C[+57]*3, N[+2206] | 1286.3029 | 4 | 5142.1896 | 5142.1776 | 0 | 2.3 | 95 | Specific | 968.97 | 968.97 | 968.97 | 19.75 | 28 | >sp\|P10909\|CLUS_HUMAN Clusterin OS=Homo sapiens OX=9606 GN=CLU PE=1 SV=1 | 1467 | 20201113_12433_1ng_glyco_hcd203040_compound15.17604.17604.4 | controllerType=0 controllerNumber=1 scan=17604 | 35.3994 |
| 21515:3 | 21 | K.MLN[+2205.793]TSSLLEQLNEQFNWVSR.L | HexNAc(4)Hex(5)Fuc(2)NeuAc(1) | N[+2206] | 1539.0045 | 3 | 4614.9988 | 4614.9796 | 0 | 4.2 | 352 | Specific | 512.75 | 434.48 | 434.48 | 9.54 | 28 | >sp\|P10909\|CLUS_HUMAN Clusterin OS=Homo sapiens OX=9606 GN=CLU PE=1 SV=1 | 1467 | 20201113_12433_1ng_glyco_hcd203040_compound15.25038.25038.3 | controllerType=0 controllerNumber=1 scan=25038 | 49.7663 |
| 21517:3 | 21 | K.MLN[+2205.793]TSSLLEQLNEQFNWVSR.L | HexNAc(4)Hex(5)Fuc(2)NeuAc(1) | N[+2206] | 1539.0045 | 3 | 4614.9988 | 4614.9796 | 0 | 4.2 | 352 | Specific | 568.55 | 455.29 | 455.29 | 8.48 | 28 | >sp\|P10909\|CLUS_HUMAN Clusterin OS=Homo sapiens OX=9606 GN=CLU PE=1 SV=1 | 1467 | 20201113_12433_1ng_glyco_hcd203040_compound15.25040.25040.3 | controllerType=0 controllerNumber=1 scan=25040 | 49.7721 |
| 24048:3 | 21 | K.MLN[+2350.830]TSSLLEQLNEQFNWVSR.L | HexNAc(4)Hex(5)Fuc(1)NeuAc(2) | N[+2351] | 1587.3499 | 3 | 4760.0351 | 4760.0171 | 0 | 3.8 | 352 | Specific | 688.26 | 368.02 | 368.02 | 11.71 | 28 | >sp\|P10909\|CLUS_HUMAN Clusterin OS=Homo sapiens OX=9606 GN=CLU PE=1 SV=1 | 1467 | 20201113_12433_1ng_glyco_hcd203040_compound15.27677.27677.3 | controllerType=0 controllerNumber=1 scan=27677 | 54.4431 |
| 11737:3 | 50 | R.IIVPLNNREN[+1856.656]ISDPTSPLR.T | HexNAc(3)Hex(5)Fuc(1)NeuAc(1) | N[+1857] | 1335.6185 | 3 | 4004.8408 | 4004.8327 | 0 | 2 | 62 | Specific | 212.85 | 212.85 | 212.85 | 3.85 | 15 | >sp\|P01591\|IGJ_HUMAN Immunoglobulin J chain OS=Homo sapiens OX=9606 GN=JCHAIN PE=1 SV=4 | 1392 | 20201113_12433_1ng_glyco_hcd203040_compound15.14833.14833.3 | controllerType=0 controllerNumber=1 scan=14833 | 30.7902 |
| 11758:3 | 50 | R.IIVPLNNREN[+1694.603]ISDPTSPLR.T | HexNAc(3)Hex(4)Fuc(1)NeuAc(1) | N[+1695] | 1281.5995 | 3 | 3842.7841 | 3842.7799 | 0 | 1.1 | 62 | Specific | 329.38 | 248.66 | 248.66 | 6.15 | 15 | >sp\|P01591\|IGJ_HUMAN Immunoglobulin J chain OS=Homo sapiens OX=9606 GN=JCHAIN PE=1 SV=4 | 1392 | 20201113_12433_1ng_glyco_hcd203040_compound15.14855.14855.3 | controllerType=0 controllerNumber=1 scan=14855 | 30.8316 |
| 11761:3 | 50 | R.IIVPLNNREN[+1694.603]ISDPTSPLR.T | HexNAc(3)Hex(4)Fuc(1)NeuAc(1) | N[+1695] | 1281.5995 | 3 | 3842.7841 | 3842.7799 | 0 | 1.1 | 62 | Specific | 349.65 | 301.48 | 301.48 | 6.36 | 15 | >sp\|P01591\|IGJ_HUMAN Immunoglobulin J chain OS=Homo sapiens OX=9606 GN=JCHAIN PE=1 SV=4 | 1392 | 20201113_12433_1ng_glyco_hcd203040_compound15.14858.14858.3 | controllerType=0 controllerNumber=1 scan=14858 | 30.8378 |
| 11552:3 | 50 | R.IIVPLNNREN[+2059.735]ISDPTSPLR.T | HexNAc(4)Hex(5)Fuc(1)NeuAc(1) | N[+2060] | 1403.3132 | 3 | 4207.9249 | 4207.9121 | 0 | 3.1 | 62 | Specific | 381.71 | 357.68 | 357.68 | 7.57 | 15 | >sp\|P01591\|IGJ_HUMAN Immunoglobulin J chain OS=Homo sapiens OX=9606 GN=JCHAIN PE=1 SV=4 | 1392 | 20201113_12433_1ng_glyco_hcd203040_compound15.14641.14641.3 | controllerType=0 controllerNumber=1 scan=14641 | 30.4167 |
| 11717:3 | 50 | R.IIVPLNNREN[+1856.656]ISDPTSPLR.T | HexNAc(3)Hex(5)Fuc(1)NeuAc(1) | N[+1857] | 1335.6182 | 3 | 4004.8401 | 4004.8327 | 0 | 1.8 | 62 | Specific | 397.84 | 397.84 | 397.84 | 9.04 | 15 | >sp\|P01591\|IGJ_HUMAN Immunoglobulin J chain OS=Homo sapiens OX=9606 GN=JCHAIN PE=1 SV=4 | 1392 | 20201113_12433_1ng_glyco_hcd203040_compound15.14812.14812.3 | controllerType=0 controllerNumber=1 scan=14812 | 30.7469 |
| 11554:3 | 50 | R.IIVPLNNREN[+2059.735]ISDPTSPLR.T | HexNAc(4)Hex(5)Fuc(1)NeuAc(1) | N[+2060] | 1403.3132 | 3 | 4207.9249 | 4207.9121 | 0 | 3.1 | 62 | Specific | 456.52 | 413.42 | 413.42 | 9.14 | 15 | >sp\|P01591\|IGJ_HUMAN Immunoglobulin J chain OS=Homo sapiens OX=9606 GN=JCHAIN PE=1 SV=4 | 1392 | 20201113_12433_1ng_glyco_hcd203040_compound15.14643.14643.3 | controllerType=0 controllerNumber=1 scan=14643 | 30.4224 |
| 11564:3 | 50 | R.IIVPLNNREN[+2059.735]ISDPTSPLR.T | HexNAc(4)Hex(5)Fuc(1)NeuAc(1) | N[+2060] | 1403.3119 | 3 | 4207.9213 | 4207.9121 | 0 | 2.2 | 62 | Specific | 486.81 | 469.34 | 469.34 | 9.31 | 15 | >sp\|P01591\|IGJ_HUMAN Immunoglobulin J chain OS=Homo sapiens OX=9606 GN=JCHAIN PE=1 SV=4 | 1392 | 20201113_12433_1ng_glyco_hcd203040_compound15.14654.14654.3 | controllerType=0 controllerNumber=1 scan=14654 | 30.4457 |
| 11566:3 | 50 | R.IIVPLNNREN[+2059.735]ISDPTSPLR.T | HexNAc(4)Hex(5)Fuc(1)NeuAc(1) | N[+2060] | 1403.3119 | 3 | 4207.9213 | 4207.9121 | 0 | 2.2 | 62 | Specific | 526.47 | 522.39 | 522.39 | 9.48 | 15 | >sp\|P01591\|IGJ_HUMAN Immunoglobulin J chain OS=Homo sapiens OX=9606 GN=JCHAIN PE=1 SV=4 | 1392 | 20201113_12433_1ng_glyco_hcd203040_compound15.14656.14656.3 | controllerType=0 controllerNumber=1 scan=14656 | 30.45 |
| 17825:4 | 1 | R.RTAGWNVPIGTLRPFLN[+2570.925]WTGPPEPIEAAVAR.F | HexNAc(5)Hex(6)Fuc(2)NeuAc(1) | N[+2571] | 1490.1952 | 4 | 5957.7589 | 5957.7371 | 0 | 3.7 | 140 | Specific | 476.01 | 378.65 | 378.65 | 9.78 | 155 | >sp\|P02788\|TRFL_HUMAN Lactotransferrin OS=Homo sapiens OX=9606 GN=LTF PE=1 SV=6 | 1270 | 20201118_12812_1ng_glyco_hcd203040_compound15.21412.21412.4 | controllerType=0 controllerNumber=1 scan=21412 | 43.3123 |
| 17808:4 | 1 | R.RTAGWNVPIGTLRPFLN[+2716.983]WTGPPEPIEAAVAR.F | HexNAc(5)Hex(6)Fuc(3)NeuAc(1) | N[+2717] | 1526.7094 | 4 | 6103.8156 | 6103.795 | 0 | 3.4 | 140 | Specific | 490.84 | 376.06 | 376.06 | 8.41 | 155 | >sp\|P02788\|TRFL_HUMAN Lactotransferrin OS=Homo sapiens OX=9606 GN=LTF PE=1 SV=6 | 1270 | 20201118_12812_1ng_glyco_hcd203040_compound15.21394.21394.4 | controllerType=0 controllerNumber=1 scan=21394 | 43.2673 |
| 17827:4 | 1 | R.RTAGWNVPIGTLRPFLN[+2570.925]WTGPPEPIEAAVAR.F | HexNAc(5)Hex(6)Fuc(2)NeuAc(1) | N[+2571] | 1490.1952 | 4 | 5957.7589 | 5957.7371 | 0 | 3.7 | 140 | Specific | 515.87 | 437.72 | 437.72 | 10.11 | 155 | >sp\|P02788\|TRFL_HUMAN Lactotransferrin OS=Homo sapiens OX=9606 GN=LTF PE=1 SV=6 | 1270 | 20201118_12812_1ng_glyco_hcd203040_compound15.21414.21414.4 | controllerType=0 controllerNumber=1 scan=21414 | 43.3182 |
| 17810:4 | 1 | R.RTAGWNVPIGTLRPFLN[+2716.983]WTGPPEPIEAAVAR.F | HexNAc(5)Hex(6)Fuc(3)NeuAc(1) | N[+2717] | 1526.7094 | 4 | 6103.8156 | 6103.795 | 0 | 3.4 | 140 | Specific | 522.27 | 416.9 | 416.9 | 10.11 | 155 | >sp\|P02788\|TRFL_HUMAN Lactotransferrin OS=Homo sapiens OX=9606 GN=LTF PE=1 SV=6 | 1270 | 20201118_12812_1ng_glyco_hcd203040_compound15.21396.21396.4 | controllerType=0 controllerNumber=1 scan=21396 | 43.2727 |
| 18933:4 | 1 | R.RTAGWNVPIGTLRPFLN[+2350.830]WTGPPEPIEAAVAR.F | HexNAc(4)Hex(5)Fuc(1)NeuAc(2) | N[+2351] | 1435.1685 | 4 | 5737.652 | 5737.6424 | 0 | 1.7 | 140 | Specific | 532.39 | 386.06 | 386.06 | 10.15 | 155 | >sp\|P02788\|TRFL_HUMAN Lactotransferrin OS=Homo sapiens OX=9606 GN=LTF PE=1 SV=6 | 1270 | 20201118_12812_1ng_glyco_hcd203040_compound15.22567.22567.4 | controllerType=0 controllerNumber=1 scan=22567 | 45.8134 |
| 18936:4 | 1 | R.RTAGWNVPIGTLRPFLN[+2350.830]WTGPPEPIEAAVAR.F | HexNAc(4)Hex(5)Fuc(1)NeuAc(2) | N[+2351] | 1435.1685 | 4 | 5737.652 | 5737.6424 | 0 | 1.7 | 140 | Specific | 559.36 | 493.04 | 493.04 | 10.34 | 155 | >sp\|P02788\|TRFL_HUMAN Lactotransferrin OS=Homo sapiens OX=9606 GN=LTF PE=1 SV=6 | 1270 | 20201118_12812_1ng_glyco_hcd203040_compound15.22570.22570.4 | controllerType=0 controllerNumber=1 scan=22570 | 45.8227 |
| 17003:4 | 1 | R.RTAGWNVPIGTLRPFLN[+1768.640]WTGPPEPIEAAVAR.F | HexNAc(4)Hex(5)Fuc(1) | N[+1769] | 1289.6207 | 4 | 5155.4611 | 5155.4516 | 0 | 1.8 | 140 | Specific | 651.73 | 477.01 | 477.01 | 13.56 | 155 | >sp\|P02788\|TRFL_HUMAN Lactotransferrin OS=Homo sapiens OX=9606 GN=LTF PE=1 SV=6 | 1270 | 20201118_12812_1ng_glyco_hcd203040_compound15.20552.20552.4 | controllerType=0 controllerNumber=1 scan=20552 | 41.4259 |
| 17005:4 | 1 | R.RTAGWNVPIGTLRPFLN[+1768.640]WTGPPEPIEAAVAR.F | HexNAc(4)Hex(5)Fuc(1) | N[+1769] | 1289.6207 | 4 | 5155.4611 | 5155.4516 | 0 | 1.8 | 140 | Specific | 661.23 | 529.13 | 529.13 | 13.56 | 155 | >sp\|P02788\|TRFL_HUMAN Lactotransferrin OS=Homo sapiens OX=9606 GN=LTF PE=1 SV=6 | 1270 | 20201118_12812_1ng_glyco_hcd203040_compound15.20554.20554.4 | controllerType=0 controllerNumber=1 scan=20554 | 41.431 |
| 19137:4 | 1 | R.TAGWNVPIGTLRPFLN[+2076.750]WTGPPEPIEAAVAR.F | HexNAc(4)Hex(6)Fuc(2) | N[+2077] | 1327.6248 | 4 | 5307.4772 | 5307.4612 | 0 | 3 | 141 | Specific | 224.28 | 224.28 | 224.28 | 6.52 | 155 | >sp\|P02788\|TRFL_HUMAN Lactotransferrin OS=Homo sapiens OX=9606 GN=LTF PE=1 SV=6 | 1270 | 20201118_12812_1ng_glyco_hcd203040_compound15.22780.22780.4 | controllerType=0 controllerNumber=1 scan=22780 | 46.2978 |
| 19135:4 | 1 | R.TAGWNVPIGTLRPFLN[+2076.750]WTGPPEPIEAAVAR.F | HexNAc(4)Hex(6)Fuc(2) | N[+2077] | 1327.6248 | 4 | 5307.4772 | 5307.4612 | 0 | 3 | 141 | Specific | 230.01 | 230.01 | 230.01 | 6.52 | 155 | >sp\|P02788\|TRFL_HUMAN Lactotransferrin OS=Homo sapiens OX=9606 GN=LTF PE=1 SV=6 | 1270 | 20201118_12812_1ng_glyco_hcd203040_compound15.22778.22778.4 | controllerType=0 controllerNumber=1 scan=22778 | 46.2915 |
| 19169:5 | 1 | R.TAGWNVPIGTLRPFLN[+2076.750]WTGPPEPIEAAVAR.F | HexNAc(4)Hex(6)Fuc(2) | N[+2077] | 1062.3032 | 5 | 5307.487 | 5307.4612 | 0 | 4.9 | 141 | Specific | 233.36 | 233.36 | 233.36 | 5.05 | 155 | >sp\|P02788\|TRFL_HUMAN Lactotransferrin OS=Homo sapiens OX=9606 GN=LTF PE=1 SV=6 | 1270 | 20201118_12812_1ng_glyco_hcd203040_compound15.22813.22813.5 | controllerType=0 controllerNumber=1 scan=22813 | 46.3755 |
| 18158:4 | 1 | R.[+42.011]TAGWNVPIGTLRPFLN[+1872.651]WTGPPEPIEAAVAR.F | HexNAc(3)Hex(6)NeuAc(1) | .[42], N[+1873] | 1287.1108 | 4 | 5145.4215 | 5145.372 | 0 | 9.6 | 141 | Specific | 244.84 | 151.82 | 151.82 | 2.29 | 155 | >sp\|P02788\|TRFL_HUMAN Lactotransferrin OS=Homo sapiens OX=9606 GN=LTF PE=1 SV=6 | 1270 | 20201118_12812_1ng_glyco_hcd203040_compound15.21759.21759.4 | controllerType=0 controllerNumber=1 scan=21759 | 44.0384 |
| 18155:4 | 1 | R.[+42.011]TAGWNVPIGTLRPFLN[+1872.651]WTGPPEPIEAAVAR.F | HexNAc(3)Hex(6)NeuAc(1) | .[42], N[+1873] | 1287.1108 | 4 | 5145.4215 | 5145.372 | 0 | 9.6 | 141 | Specific | 245.82 | 201.97 | 201.97 | 4.59 | 155 | >sp\|P02788\|TRFL_HUMAN Lactotransferrin OS=Homo sapiens OX=9606 GN=LTF PE=1 SV=6 | 1270 | 20201118_12812_1ng_glyco_hcd203040_compound15.21756.21756.4 | controllerType=0 controllerNumber=1 scan=21756 | 44.0334 |
| 19582:3 | 1 | R.TAGWNVPIGTLRPFLN[+2059.735]WTGPPEPIEAAVAR.F | HexNAc(4)Hex(5)Fuc(1)NeuAc(1) | N[+2060] | 1764.158 | 3 | 5290.4596 | 5290.4459 | 0 | 2.6 | 141 | Specific | 256.93 | 247.26 | 247.26 | 7.07 | 155 | >sp\|P02788\|TRFL_HUMAN Lactotransferrin OS=Homo sapiens OX=9606 GN=LTF PE=1 SV=6 | 1270 | 20201118_12812_1ng_glyco_hcd203040_compound15.23245.23245.3 | controllerType=0 controllerNumber=1 scan=23245 | 47.1986 |
| 19109:4 | 1 | R.TAGWNVPIGTLRPFLN[+2716.983]WTGPPEPIEAAVAR.F | HexNAc(5)Hex(6)Fuc(3)NeuAc(1) | N[+2717] | 1487.6915 | 4 | 5947.7443 | 5947.6939 | 0 | 8.5 | 141 | Specific | 258.78 | 258.78 | 258.78 | 5.4 | 155 | >sp\|P02788\|TRFL_HUMAN Lactotransferrin OS=Homo sapiens OX=9606 GN=LTF PE=1 SV=6 | 1270 | 20201118_12812_1ng_glyco_hcd203040_compound15.22750.22750.4 | controllerType=0 controllerNumber=1 scan=22750 | 46.2131 |
| 19612:3 | 1 | R.TAGWNVPIGTLRPFLN[+2059.735]WTGPPEPIEAAVAR.F | HexNAc(4)Hex(5)Fuc(1)NeuAc(1) | N[+2060] | 1764.1596 | 3 | 5290.4643 | 5290.4459 | 0 | 3.5 | 141 | Specific | 277.52 | 277.52 | 277.52 | 7.33 | 155 | >sp\|P02788\|TRFL_HUMAN Lactotransferrin OS=Homo sapiens OX=9606 GN=LTF PE=1 SV=6 | 1270 | 20201118_12812_1ng_glyco_hcd203040_compound15.23276.23276.3 | controllerType=0 controllerNumber=1 scan=23276 | 47.2488 |
| 20468:3 | 1 | R.TAGWNVPIGTLRPFLN[+2350.830]WTGPPEPIEAAVAR.F | HexNAc(4)Hex(5)Fuc(1)NeuAc(2) | N[+2351] | 1861.1927 | 3 | 5581.5636 | 5581.5413 | 0 | 4 | 141 | Specific | 282.05 | 165.42 | 165.42 | 6.43 | 155 | >sp\|P02788\|TRFL_HUMAN Lactotransferrin OS=Homo sapiens OX=9606 GN=LTF PE=1 SV=6 | 1270 | 20201118_12812_1ng_glyco_hcd203040_compound15.24171.24171.3 | controllerType=0 controllerNumber=1 scan=24171 | 49.2473 |
| 19580:3 | 1 | R.TAGWNVPIGTLRPFLN[+2059.735]WTGPPEPIEAAVAR.F | HexNAc(4)Hex(5)Fuc(1)NeuAc(1) | N[+2060] | 1764.158 | 3 | 5290.4596 | 5290.4459 | 0 | 2.6 | 141 | Specific | 283.5 | 283.5 | 283.5 | 7.46 | 155 | >sp\|P02788\|TRFL_HUMAN Lactotransferrin OS=Homo sapiens OX=9606 GN=LTF PE=1 SV=6 | 1270 | 20201118_12812_1ng_glyco_hcd203040_compound15.23243.23243.3 | controllerType=0 controllerNumber=1 scan=23243 | 47.195 |
| 19615:3 | 1 | R.TAGWNVPIGTLRPFLN[+2059.735]WTGPPEPIEAAVAR.F | HexNAc(4)Hex(5)Fuc(1)NeuAc(1) | N[+2060] | 1764.1596 | 3 | 5290.4643 | 5290.4459 | 0 | 3.5 | 141 | Specific | 290.64 | 252.88 | 252.88 | 7.26 | 155 | >sp\|P02788\|TRFL_HUMAN Lactotransferrin OS=Homo sapiens OX=9606 GN=LTF PE=1 SV=6 | 1270 | 20201118_12812_1ng_glyco_hcd203040_compound15.23279.23279.3 | controllerType=0 controllerNumber=1 scan=23279 | 47.2532 |
| 20539:4 | 1 | R.[+42.011]TAGWNVPIGTLRPFLN[+2336.851]WTGPPEPIEAAVAR.F | HexNAc(6)Hex(6)Fuc(1) | .[42], N[+2337] | 1403.1426 | 4 | 5609.5485 | 5609.5726 | 0 | -4.3 | 141 | Specific | 322.25 | 322.25 | 322.25 | 6.31 | 155 | >sp\|P02788\|TRFL_HUMAN Lactotransferrin OS=Homo sapiens OX=9606 GN=LTF PE=1 SV=6 | 1270 | 20201118_12812_1ng_glyco_hcd203040_compound15.24245.24245.4 | controllerType=0 controllerNumber=1 scan=24245 | 49.4211 |
| 18133:4 | 1 | R.[+42.011]TAGWNVPIGTLRPFLN[+1872.651]WTGPPEPIEAAVAR.F | HexNAc(3)Hex(6)NeuAc(1) | .[42], N[+1873] | 1287.1106 | 4 | 5145.4206 | 5145.372 | 0 | 9.4 | 141 | Specific | 329.17 | 319.37 | 319.37 | 5.87 | 155 | >sp\|P02788\|TRFL_HUMAN Lactotransferrin OS=Homo sapiens OX=9606 GN=LTF PE=1 SV=6 | 1270 | 20201118_12812_1ng_glyco_hcd203040_compound15.21733.21733.2 | controllerType=0 controllerNumber=1 scan=21733 | 43.976 |
| 18135:4 | 1 | R.[+42.011]TAGWNVPIGTLRPFLN[+1872.651]WTGPPEPIEAAVAR.F | HexNAc(3)Hex(6)NeuAc(1) | .[42], N[+1873] | 1287.1106 | 4 | 5145.4206 | 5145.372 | 0 | 9.4 | 141 | Specific | 343.07 | 244.33 | 244.33 | 5.65 | 155 | >sp\|P02788\|TRFL_HUMAN Lactotransferrin OS=Homo sapiens OX=9606 GN=LTF PE=1 SV=6 | 1270 | 20201118_12812_1ng_glyco_hcd203040_compound15.21735.21735.2 | controllerType=0 controllerNumber=1 scan=21735 | 43.984 |
| 19060:3 | 1 | R.TAGWNVPIGTLRPFLN[+2424.867]WTGPPEPIEAAVAR.F | HexNAc(5)Hex(6)Fuc(1)NeuAc(1) | N[+2425] | 1885.8718 | 3 | 5655.6008 | 5655.5781 | 0 | 4 | 141 | Specific | 372.3 | 261.55 | 261.55 | 7.38 | 155 | >sp\|P02788\|TRFL_HUMAN Lactotransferrin OS=Homo sapiens OX=9606 GN=LTF PE=1 SV=6 | 1270 | 20201118_12812_1ng_glyco_hcd203040_compound15.22699.22699.3 | controllerType=0 controllerNumber=1 scan=22699 | 46.0968 |
| 19038:3 | 1 | R.TAGWNVPIGTLRPFLN[+2716.983]WTGPPEPIEAAVAR.F | HexNAc(5)Hex(6)Fuc(3)NeuAc(1) | N[+2717] | 1983.2426 | 3 | 5947.7134 | 5947.6939 | 0 | 3.3 | 141 | Specific | 385.56 | 378.87 | 378.87 | 9.73 | 155 | >sp\|P02788\|TRFL_HUMAN Lactotransferrin OS=Homo sapiens OX=9606 GN=LTF PE=1 SV=6 | 1270 | 20201118_12812_1ng_glyco_hcd203040_compound15.22676.22676.3 | controllerType=0 controllerNumber=1 scan=22676 | 46.0406 |
| 20511:3 | 1 | R.TAGWNVPIGTLRPFLN[+2350.830]WTGPPEPIEAAVAR.F | HexNAc(4)Hex(5)Fuc(1)NeuAc(2) | N[+2351] | 1861.1925 | 3 | 5581.5629 | 5581.5413 | 0 | 3.9 | 141 | Specific | 402.36 | 237.31 | 237.31 | 7.42 | 155 | >sp\|P02788\|TRFL_HUMAN Lactotransferrin OS=Homo sapiens OX=9606 GN=LTF PE=1 SV=6 | 1270 | 20201118_12812_1ng_glyco_hcd203040_compound15.24215.24215.2 | controllerType=0 controllerNumber=1 scan=24215 | 49.3571 |
| 19062:3 | 1 | R.TAGWNVPIGTLRPFLN[+2424.867]WTGPPEPIEAAVAR.F | HexNAc(5)Hex(6)Fuc(1)NeuAc(1) | N[+2425] | 1885.8718 | 3 | 5655.6008 | 5655.5781 | 0 | 4 | 141 | Specific | 432.97 | 294.8 | 294.8 | 7.75 | 155 | >sp\|P02788\|TRFL_HUMAN Lactotransferrin OS=Homo sapiens OX=9606 GN=LTF PE=1 SV=6 | 1270 | 20201118_12812_1ng_glyco_hcd203040_compound15.22701.22701.3 | controllerType=0 controllerNumber=1 scan=22701 | 46.1028 |
| 20462:3 | 1 | R.TAGWNVPIGTLRPFLN[+2350.830]WTGPPEPIEAAVAR.F | HexNAc(4)Hex(5)Fuc(1)NeuAc(2) | N[+2351] | 1861.1886 | 3 | 5581.5511 | 5581.5413 | 0 | 1.8 | 141 | Specific | 435.27 | 267.85 | 267.85 | 7.78 | 155 | >sp\|P02788\|TRFL_HUMAN Lactotransferrin OS=Homo sapiens OX=9606 GN=LTF PE=1 SV=6 | 1270 | 20201118_12812_1ng_glyco_hcd203040_compound15.24164.24164.2 | controllerType=0 controllerNumber=1 scan=24164 | 49.2289 |
| 18193:3 | 1 | R.TAGWNVPIGTLRPFLN[+1768.640]WTGPPEPIEAAVAR.F | HexNAc(4)Hex(5)Fuc(1) | N[+1769] | 1667.1261 | 3 | 4999.3636 | 4999.3504 | 0 | 2.6 | 141 | Specific | 510.76 | 426.8 | 426.8 | 10.4 | 155 | >sp\|P02788\|TRFL_HUMAN Lactotransferrin OS=Homo sapiens OX=9606 GN=LTF PE=1 SV=6 | 1270 | 20201118_12812_1ng_glyco_hcd203040_compound15.21796.21796.3 | controllerType=0 controllerNumber=1 scan=21796 | 44.1335 |
| 19112:4 | 1 | R.TAGWNVPIGTLRPFLN[+2716.983]WTGPPEPIEAAVAR.F | HexNAc(5)Hex(6)Fuc(3)NeuAc(1) | N[+2717] | 1487.6915 | 4 | 5947.7443 | 5947.6939 | 0 | 8.5 | 141 | Specific | 522.14 | 447.39 | 447.39 | 8.73 | 155 | >sp\|P02788\|TRFL_HUMAN Lactotransferrin OS=Homo sapiens OX=9606 GN=LTF PE=1 SV=6 | 1270 | 20201118_12812_1ng_glyco_hcd203040_compound15.22753.22753.4 | controllerType=0 controllerNumber=1 scan=22753 | 46.2179 |
| 20470:3 | 1 | R.TAGWNVPIGTLRPFLN[+2350.830]WTGPPEPIEAAVAR.F | HexNAc(4)Hex(5)Fuc(1)NeuAc(2) | N[+2351] | 1861.1927 | 3 | 5581.5636 | 5581.5413 | 0 | 4 | 141 | Specific | 544.45 | 358.46 | 358.46 | 8.65 | 155 | >sp\|P02788\|TRFL_HUMAN Lactotransferrin OS=Homo sapiens OX=9606 GN=LTF PE=1 SV=6 | 1270 | 20201118_12812_1ng_glyco_hcd203040_compound15.24173.24173.3 | controllerType=0 controllerNumber=1 scan=24173 | 49.2553 |
| 19563:4 | 1 | R.TAGWNVPIGTLRPFLN[+2059.735]WTGPPEPIEAAVAR.F | HexNAc(4)Hex(5)Fuc(1)NeuAc(1) | N[+2060] | 1323.37 | 4 | 5290.4582 | 5290.4459 | 0 | 2.3 | 141 | Specific | 548.66 | 479.03 | 479.03 | 10.38 | 155 | >sp\|P02788\|TRFL_HUMAN Lactotransferrin OS=Homo sapiens OX=9606 GN=LTF PE=1 SV=6 | 1270 | 20201118_12812_1ng_glyco_hcd203040_compound15.23226.23226.4 | controllerType=0 controllerNumber=1 scan=23226 | 47.1597 |
| 19046:5 | 1 | R.TAGWNVPIGTLRPFLN[+2570.925]WTGPPEPIEAAVAR.F | HexNAc(5)Hex(6)Fuc(2)NeuAc(1) | N[+2571] | 1161.1365 | 5 | 5801.6533 | 5801.636 | 0 | 3 | 141 | Specific | 557.18 | 470.39 | 470.39 | 10.69 | 155 | >sp\|P02788\|TRFL_HUMAN Lactotransferrin OS=Homo sapiens OX=9606 GN=LTF PE=1 SV=6 | 1270 | 20201118_12812_1ng_glyco_hcd203040_compound15.22684.22684.5 | controllerType=0 controllerNumber=1 scan=22684 | 46.0649 |
| 19119:3 | 1 | R.TAGWNVPIGTLRPFLN[+2205.793]WTGPPEPIEAAVAR.F | HexNAc(4)Hex(5)Fuc(2)NeuAc(1) | N[+2206] | 1812.848 | 3 | 5436.5294 | 5436.5038 | 0 | 4.7 | 141 | Specific | 570.45 | 498.31 | 498.31 | 10.47 | 155 | >sp\|P02788\|TRFL_HUMAN Lactotransferrin OS=Homo sapiens OX=9606 GN=LTF PE=1 SV=6 | 1270 | 20201118_12812_1ng_glyco_hcd203040_compound15.22761.22761.3 | controllerType=0 controllerNumber=1 scan=22761 | 46.2515 |
| 19042:5 | 1 | R.TAGWNVPIGTLRPFLN[+2570.925]WTGPPEPIEAAVAR.F | HexNAc(5)Hex(6)Fuc(2)NeuAc(1) | N[+2571] | 1161.1365 | 5 | 5801.6533 | 5801.636 | 0 | 3 | 141 | Specific | 578.65 | 511.93 | 511.93 | 11.91 | 155 | >sp\|P02788\|TRFL_HUMAN Lactotransferrin OS=Homo sapiens OX=9606 GN=LTF PE=1 SV=6 | 1270 | 20201118_12812_1ng_glyco_hcd203040_compound15.22680.22680.5 | controllerType=0 controllerNumber=1 scan=22680 | 46.0459 |
| 20467:4 | 1 | R.TAGWNVPIGTLRPFLN[+2350.830]WTGPPEPIEAAVAR.F | HexNAc(4)Hex(5)Fuc(1)NeuAc(2) | N[+2351] | 1396.1458 | 4 | 5581.5612 | 5581.5413 | 0 | 3.6 | 141 | Specific | 589.36 | 515.96 | 515.96 | 11.89 | 155 | >sp\|P02788\|TRFL_HUMAN Lactotransferrin OS=Homo sapiens OX=9606 GN=LTF PE=1 SV=6 | 1270 | 20201118_12812_1ng_glyco_hcd203040_compound15.24170.24170.4 | controllerType=0 controllerNumber=1 scan=24170 | 49.2459 |
| 20465:3 | 1 | R.TAGWNVPIGTLRPFLN[+2350.830]WTGPPEPIEAAVAR.F | HexNAc(4)Hex(5)Fuc(1)NeuAc(2) | N[+2351] | 1861.1886 | 3 | 5581.5511 | 5581.5413 | 0 | 1.8 | 141 | Specific | 592.17 | 416.19 | 416.19 | 11.86 | 155 | >sp\|P02788\|TRFL_HUMAN Lactotransferrin OS=Homo sapiens OX=9606 GN=LTF PE=1 SV=6 | 1270 | 20201118_12812_1ng_glyco_hcd203040_compound15.24167.24167.2 | controllerType=0 controllerNumber=1 scan=24167 | 49.2358 |
| 19053:4 | 1 | R.TAGWNVPIGTLRPFLN[+2424.867]WTGPPEPIEAAVAR.F | HexNAc(5)Hex(6)Fuc(1)NeuAc(1) | N[+2425] | 1414.6564 | 4 | 5655.6037 | 5655.5781 | 0 | 4.5 | 141 | Specific | 643.01 | 549.38 | 549.38 | 12.62 | 155 | >sp\|P02788\|TRFL_HUMAN Lactotransferrin OS=Homo sapiens OX=9606 GN=LTF PE=1 SV=6 | 1270 | 20201118_12812_1ng_glyco_hcd203040_compound15.22692.22692.4 | controllerType=0 controllerNumber=1 scan=22692 | 46.0847 |
| 19122:3 | 1 | R.TAGWNVPIGTLRPFLN[+2205.793]WTGPPEPIEAAVAR.F | HexNAc(4)Hex(5)Fuc(2)NeuAc(1) | N[+2206] | 1812.848 | 3 | 5436.5294 | 5436.5038 | 0 | 4.7 | 141 | Specific | 648.93 | 579.12 | 579.12 | 12.69 | 155 | >sp\|P02788\|TRFL_HUMAN Lactotransferrin OS=Homo sapiens OX=9606 GN=LTF PE=1 SV=6 | 1270 | 20201118_12812_1ng_glyco_hcd203040_compound15.22764.22764.3 | controllerType=0 controllerNumber=1 scan=22764 | 46.2566 |
| 19141:3 | 1 | R.TAGWNVPIGTLRPFLN[+2059.735]WTGPPEPIEAAVAR.F | HexNAc(4)Hex(5)Fuc(1)NeuAc(1) | N[+2060] | 1764.1599 | 3 | 5290.4651 | 5290.4459 | 0 | 3.6 | 141 | Specific | 660.19 | 569.17 | 569.17 | 13.96 | 155 | >sp\|P02788\|TRFL_HUMAN Lactotransferrin OS=Homo sapiens OX=9606 GN=LTF PE=1 SV=6 | 1270 | 20201118_12812_1ng_glyco_hcd203040_compound15.22784.22784.3 | controllerType=0 controllerNumber=1 scan=22784 | 46.3052 |
| 18192:4 | 1 | R.TAGWNVPIGTLRPFLN[+1768.640]WTGPPEPIEAAVAR.F | HexNAc(4)Hex(5)Fuc(1) | N[+1769] | 1250.597 | 4 | 4999.3664 | 4999.3504 | 0 | 3.2 | 141 | Specific | 665.02 | 637.36 | 637.36 | 13.89 | 155 | >sp\|P02788\|TRFL_HUMAN Lactotransferrin OS=Homo sapiens OX=9606 GN=LTF PE=1 SV=6 | 1270 | 20201118_12812_1ng_glyco_hcd203040_compound15.21795.21795.4 | controllerType=0 controllerNumber=1 scan=21795 | 44.132 |
| 19116:4 | 1 | R.TAGWNVPIGTLRPFLN[+2205.793]WTGPPEPIEAAVAR.F | HexNAc(4)Hex(5)Fuc(2)NeuAc(1) | N[+2206] | 1359.8881 | 4 | 5436.5304 | 5436.5038 | 0 | 4.9 | 141 | Specific | 666.94 | 639.73 | 639.73 | 13.62 | 155 | >sp\|P02788\|TRFL_HUMAN Lactotransferrin OS=Homo sapiens OX=9606 GN=LTF PE=1 SV=6 | 1270 | 20201118_12812_1ng_glyco_hcd203040_compound15.22758.22758.4 | controllerType=0 controllerNumber=1 scan=22758 | 46.2387 |
| 19134:4 | 1 | R.TAGWNVPIGTLRPFLN[+2059.735]WTGPPEPIEAAVAR.F | HexNAc(4)Hex(5)Fuc(1)NeuAc(1) | N[+2060] | 1323.3717 | 4 | 5290.465 | 5290.4459 | 0 | 3.6 | 141 | Specific | 672.73 | 645.28 | 645.28 | 13.89 | 155 | >sp\|P02788\|TRFL_HUMAN Lactotransferrin OS=Homo sapiens OX=9606 GN=LTF PE=1 SV=6 | 1270 | 20201118_12812_1ng_glyco_hcd203040_compound15.22777.22777.4 | controllerType=0 controllerNumber=1 scan=22777 | 46.29 |
| 20473:5 | 1 | R.TAGWNVPIGTLRPFLN[+2350.830]WTGPPEPIEAAVAR.F | HexNAc(4)Hex(5)Fuc(1)NeuAc(2) | N[+2351] | 1117.1184 | 5 | 5581.5629 | 5581.5413 | 0 | 3.9 | 141 | Specific | 690.86 | 632.75 | 632.75 | 13.65 | 155 | >sp\|P02788\|TRFL_HUMAN Lactotransferrin OS=Homo sapiens OX=9606 GN=LTF PE=1 SV=6 | 1270 | 20201118_12812_1ng_glyco_hcd203040_compound15.24176.24176.5 | controllerType=0 controllerNumber=1 scan=24176 | 49.2587 |
| 18195:3 | 1 | R.TAGWNVPIGTLRPFLN[+1768.640]WTGPPEPIEAAVAR.F | HexNAc(4)Hex(5)Fuc(1) | N[+1769] | 1667.1261 | 3 | 4999.3636 | 4999.3504 | 0 | 2.6 | 141 | Specific | 720.45 | 564.1 | 564.1 | 14.96 | 155 | >sp\|P02788\|TRFL_HUMAN Lactotransferrin OS=Homo sapiens OX=9606 GN=LTF PE=1 SV=6 | 1270 | 20201118_12812_1ng_glyco_hcd203040_compound15.21798.21798.3 | controllerType=0 controllerNumber=1 scan=21798 | 44.1378 |
| 19138:3 | 1 | R.TAGWNVPIGTLRPFLN[+2059.735]WTGPPEPIEAAVAR.F | HexNAc(4)Hex(5)Fuc(1)NeuAc(1) | N[+2060] | 1764.1599 | 3 | 5290.4651 | 5290.4459 | 0 | 3.6 | 141 | Specific | 724.45 | 650.61 | 650.61 | 14.96 | 155 | >sp\|P02788\|TRFL_HUMAN Lactotransferrin OS=Homo sapiens OX=9606 GN=LTF PE=1 SV=6 | 1270 | 20201118_12812_1ng_glyco_hcd203040_compound15.22781.22781.3 | controllerType=0 controllerNumber=1 scan=22781 | 46.299 |
| 19056:4 | 1 | R.TAGWNVPIGTLRPFLN[+2424.867]WTGPPEPIEAAVAR.F | HexNAc(5)Hex(6)Fuc(1)NeuAc(1) | N[+2425] | 1414.6564 | 4 | 5655.6037 | 5655.5781 | 0 | 4.5 | 141 | Specific | 749.16 | 627.77 | 627.77 | 14.62 | 155 | >sp\|P02788\|TRFL_HUMAN Lactotransferrin OS=Homo sapiens OX=9606 GN=LTF PE=1 SV=6 | 1270 | 20201118_12812_1ng_glyco_hcd203040_compound15.22695.22695.4 | controllerType=0 controllerNumber=1 scan=22695 | 46.0897 |
| 19136:4 | 1 | R.TAGWNVPIGTLRPFLN[+2059.735]WTGPPEPIEAAVAR.F | HexNAc(4)Hex(5)Fuc(1)NeuAc(1) | N[+2060] | 1323.3717 | 4 | 5290.465 | 5290.4459 | 0 | 3.6 | 141 | Specific | 823.19 | 723.27 | 723.27 | 16.89 | 155 | >sp\|P02788\|TRFL_HUMAN Lactotransferrin OS=Homo sapiens OX=9606 GN=LTF PE=1 SV=6 | 1270 | 20201118_12812_1ng_glyco_hcd203040_compound15.22779.22779.4 | controllerType=0 controllerNumber=1 scan=22779 | 46.2971 |
| 20469:4 | 1 | R.TAGWNVPIGTLRPFLN[+2350.830]WTGPPEPIEAAVAR.F | HexNAc(4)Hex(5)Fuc(1)NeuAc(2) | N[+2351] | 1396.1458 | 4 | 5581.5612 | 5581.5413 | 0 | 3.6 | 141 | Specific | 825.39 | 660.86 | 660.86 | 16.89 | 155 | >sp\|P02788\|TRFL_HUMAN Lactotransferrin OS=Homo sapiens OX=9606 GN=LTF PE=1 SV=6 | 1270 | 20201118_12812_1ng_glyco_hcd203040_compound15.24172.24172.4 | controllerType=0 controllerNumber=1 scan=24172 | 49.2528 |
| 19139:5 | 1 | R.TAGWNVPIGTLRPFLN[+2205.793]WTGPPEPIEAAVAR.F | HexNAc(4)Hex(5)Fuc(2)NeuAc(1) | N[+2206] | 1088.1089 | 5 | 5436.5153 | 5436.5038 | 0 | 2.1 | 141 | Specific | 838.56 | 710.1 | 710.1 | 16.81 | 155 | >sp\|P02788\|TRFL_HUMAN Lactotransferrin OS=Homo sapiens OX=9606 GN=LTF PE=1 SV=6 | 1270 | 20201118_12812_1ng_glyco_hcd203040_compound15.22782.22782.5 | controllerType=0 controllerNumber=1 scan=22782 | 46.3001 |
| 19118:4 | 1 | R.TAGWNVPIGTLRPFLN[+2205.793]WTGPPEPIEAAVAR.F | HexNAc(4)Hex(5)Fuc(2)NeuAc(1) | N[+2206] | 1359.8881 | 4 | 5436.5304 | 5436.5038 | 0 | 4.9 | 141 | Specific | 860.56 | 712.74 | 712.74 | 17.62 | 155 | >sp\|P02788\|TRFL_HUMAN Lactotransferrin OS=Homo sapiens OX=9606 GN=LTF PE=1 SV=6 | 1270 | 20201118_12812_1ng_glyco_hcd203040_compound15.22760.22760.4 | controllerType=0 controllerNumber=1 scan=22760 | 46.2508 |
| 18194:4 | 1 | R.TAGWNVPIGTLRPFLN[+1768.640]WTGPPEPIEAAVAR.F | HexNAc(4)Hex(5)Fuc(1) | N[+1769] | 1250.597 | 4 | 4999.3664 | 4999.3504 | 0 | 3.2 | 141 | Specific | 862.13 | 787.23 | 787.23 | 17.89 | 155 | >sp\|P02788\|TRFL_HUMAN Lactotransferrin OS=Homo sapiens OX=9606 GN=LTF PE=1 SV=6 | 1270 | 20201118_12812_1ng_glyco_hcd203040_compound15.21797.21797.4 | controllerType=0 controllerNumber=1 scan=21797 | 44.1366 |
| 19149:5 | 1 | R.TAGWNVPIGTLRPFLN[+2059.735]WTGPPEPIEAAVAR.F | HexNAc(4)Hex(5)Fuc(1)NeuAc(1) | N[+2060] | 1058.8978 | 5 | 5290.46 | 5290.4459 | 0 | 2.7 | 141 | Specific | 867.57 | 841.06 | 841.06 | 17.91 | 155 | >sp\|P02788\|TRFL_HUMAN Lactotransferrin OS=Homo sapiens OX=9606 GN=LTF PE=1 SV=6 | 1270 | 20201118_12812_1ng_glyco_hcd203040_compound15.22792.22792.5 | controllerType=0 controllerNumber=1 scan=22792 | 46.3216 |
| 20476:5 | 1 | R.TAGWNVPIGTLRPFLN[+2350.830]WTGPPEPIEAAVAR.F | HexNAc(4)Hex(5)Fuc(1)NeuAc(2) | N[+2351] | 1117.1184 | 5 | 5581.5629 | 5581.5413 | 0 | 3.9 | 141 | Specific | 873.52 | 694.37 | 694.37 | 17.65 | 155 | >sp\|P02788\|TRFL_HUMAN Lactotransferrin OS=Homo sapiens OX=9606 GN=LTF PE=1 SV=6 | 1270 | 20201118_12812_1ng_glyco_hcd203040_compound15.24179.24179.5 | controllerType=0 controllerNumber=1 scan=24179 | 49.269 |
| 19565:4 | 1 | R.TAGWNVPIGTLRPFLN[+2059.735]WTGPPEPIEAAVAR.F | HexNAc(4)Hex(5)Fuc(1)NeuAc(1) | N[+2060] | 1323.37 | 4 | 5290.4582 | 5290.4459 | 0 | 2.3 | 141 | Specific | 895.05 | 817.37 | 817.37 | 17.79 | 155 | >sp\|P02788\|TRFL_HUMAN Lactotransferrin OS=Homo sapiens OX=9606 GN=LTF PE=1 SV=6 | 1270 | 20201118_12812_1ng_glyco_hcd203040_compound15.23228.23228.4 | controllerType=0 controllerNumber=1 scan=23228 | 47.1638 |
| 19142:5 | 1 | R.TAGWNVPIGTLRPFLN[+2205.793]WTGPPEPIEAAVAR.F | HexNAc(4)Hex(5)Fuc(2)NeuAc(1) | N[+2206] | 1088.1089 | 5 | 5436.5153 | 5436.5038 | 0 | 2.1 | 141 | Specific | 904.14 | 726.23 | 726.23 | 18.81 | 155 | >sp\|P02788\|TRFL_HUMAN Lactotransferrin OS=Homo sapiens OX=9606 GN=LTF PE=1 SV=6 | 1270 | 20201118_12812_1ng_glyco_hcd203040_compound15.22785.22785.5 | controllerType=0 controllerNumber=1 scan=22785 | 46.3078 |
| 19151:5 | 1 | R.TAGWNVPIGTLRPFLN[+2059.735]WTGPPEPIEAAVAR.F | HexNAc(4)Hex(5)Fuc(1)NeuAc(1) | N[+2060] | 1058.8978 | 5 | 5290.46 | 5290.4459 | 0 | 2.7 | 141 | Specific | 938.78 | 790.55 | 790.55 | 18.91 | 155 | >sp\|P02788\|TRFL_HUMAN Lactotransferrin OS=Homo sapiens OX=9606 GN=LTF PE=1 SV=6 | 1270 | 20201118_12812_1ng_glyco_hcd203040_compound15.22794.22794.5 | controllerType=0 controllerNumber=1 scan=22794 | 46.3293 |
| 18806:3 | 1 | R.TAGWNIPMGLLFN[+1751.624]QTGSC[+57.021]K.F | HexNAc(4)Hex(4)NeuAc(1) | C[+57], N[+1752] | 1282.8832 | 3 | 3846.6352 | 3846.6341 | 0 | 0.3 | 485 | Specific | 244.02 | 243.2 | 243.2 | 6.34 | 155 | >sp\|P02788\|TRFL_HUMAN Lactotransferrin OS=Homo sapiens OX=9606 GN=LTF PE=1 SV=6 | 1270 | 20201118_12812_1ng_glyco_hcd203040_compound15.22435.22435.3 | controllerType=0 controllerNumber=1 scan=22435 | 45.5547 |
| 20623:3 | 1 | R.[+42.011]TAGWNIPMGLLFN[+2336.851]QTGSC[+57.021]K.F | HexNAc(6)Hex(6)Fuc(1) | .[42], C[+57], N[+2337] | 1491.951 | 3 | 4473.8384 | 4473.8716 | 0 | -7.4 | 485 | Specific | 252.99 | 177.08 | 177.08 | 4.41 | 155 | >sp\|P02788\|TRFL_HUMAN Lactotransferrin OS=Homo sapiens OX=9606 GN=LTF PE=1 SV=6 | 1270 | 20201118_12812_1ng_glyco_hcd203040_compound15.24332.24332.3 | controllerType=0 controllerNumber=1 scan=24332 | 49.6037 |
| 16999:3 | 1 | R.TAGWNIPMGLLFN[+1460.529]QTGSC[+57.021]K.F | HexNAc(4)Hex(4) | C[+57], N[+1461] | 1185.8505 | 3 | 3555.537 | 3555.5387 | 0 | -0.5 | 485 | Specific | 292.78 | 244.77 | 244.77 | 4.86 | 155 | >sp\|P02788\|TRFL_HUMAN Lactotransferrin OS=Homo sapiens OX=9606 GN=LTF PE=1 SV=6 | 1270 | 20201118_12812_1ng_glyco_hcd203040_compound15.20547.20547.3 | controllerType=0 controllerNumber=1 scan=20547 | 41.4138 |
| 16446:3 | 1 | R.TAGWNIPMGLLFN[+2279.830]QTGSC[+57.021]K.F | HexNAc(5)Hex(6)Fuc(2) | C[+57], N[+2280] | 1458.9546 | 3 | 4374.8494 | 4374.8396 | 0 | 2.2 | 485 | Specific | 309.63 | 285.39 | 193.91 | 7.08 | 155 | >sp\|P02788\|TRFL_HUMAN Lactotransferrin OS=Homo sapiens OX=9606 GN=LTF PE=1 SV=6 | 1270 | 20201118_12812_1ng_glyco_hcd203040_compound15.19972.19972.3 | controllerType=0 controllerNumber=1 scan=19972 | 40.2628 |
| 17001:3 | 1 | R.TAGWNIPMGLLFN[+1460.529]QTGSC[+57.021]K.F | HexNAc(4)Hex(4) | C[+57], N[+1461] | 1185.8505 | 3 | 3555.537 | 3555.5387 | 0 | -0.5 | 485 | Specific | 366.18 | 352.12 | 297.46 | 5.53 | 155 | >sp\|P02788\|TRFL_HUMAN Lactotransferrin OS=Homo sapiens OX=9606 GN=LTF PE=1 SV=6 | 1270 | 20201118_12812_1ng_glyco_hcd203040_compound15.20549.20549.3 | controllerType=0 controllerNumber=1 scan=20549 | 41.4177 |
| 16448:3 | 1 | R.TAGWNIPMGLLFN[+2279.830]QTGSC[+57.021]K.F | HexNAc(5)Hex(6)Fuc(2) | C[+57], N[+2280] | 1458.9546 | 3 | 4374.8494 | 4374.8396 | 0 | 2.2 | 485 | Specific | 469.12 | 469.12 | 270.66 | 9.71 | 155 | >sp\|P02788\|TRFL_HUMAN Lactotransferrin OS=Homo sapiens OX=9606 GN=LTF PE=1 SV=6 | 1270 | 20201118_12812_1ng_glyco_hcd203040_compound15.19974.19974.3 | controllerType=0 controllerNumber=1 scan=19974 | 40.2663 |
| 18809:3 | 1 | R.TAGWNIPMGLLFN[+1751.624]QTGSC[+57.021]K.F | HexNAc(4)Hex(4)NeuAc(1) | C[+57], N[+1752] | 1282.8832 | 3 | 3846.6352 | 3846.6341 | 0 | 0.3 | 485 | Specific | 602.13 | 335 | 335 | 10.72 | 155 | >sp\|P02788\|TRFL_HUMAN Lactotransferrin OS=Homo sapiens OX=9606 GN=LTF PE=1 SV=6 | 1270 | 20201118_12812_1ng_glyco_hcd203040_compound15.22438.22438.3 | controllerType=0 controllerNumber=1 scan=22438 | 45.5617 |

Table S4. Cont.

| Query #:z | Protein Rank | Glycopeptide | Glycan | Modification Type(s) | Observed m/z | z | Observed (M+H) | Calc. mass (M+H) | Off-by-x error | Mass error (ppm) | StartPosition | Cleavage | Score | Delta | Delta Mod | \|Log Prob\| | # of unique peptides | Protein Name | Protein DB number | Comment | Scan # | Scan Time |
|---|---|---|---|---|---|---|---|---|---|---|---|---|---|---|---|---|---|---|---|---|---|---|
| 16863:3 | 1 | R.TAGWNIPMGLLFN[+1622.582]QTGSC[+57.021]K.F | HexNAc(4)Hex(5) | C[+57], N[+1623] | 1239.8715 | 3 | 3717.6 | 3717.5916 | 0 | 2.3 | 485 | Specific | 699.66 | 699.66 | 321.78 | 13.58 | 155 | >sp\|P02788\|TRFL_HUMAN Lactotransferrin OS=Homo sapiens OX=9606 GN=LTF PE=1 SV=6 | 1270 | 20201118_12812_1ng_glyco_hcd203040_compound15.20406.20406.3 | controllerType=0 controllerNumber=1 scan=20406 | 41.1106 |
| 16797:3 | 1 | R.TAGWNIPMGLLFN[+1622.582]QTGSC[+57.021]K.F | HexNAc(4)Hex(5) | C[+57], N[+1623] | 1239.8713 | 3 | 3717.5993 | 3717.5916 | 0 | 2.1 | 485 | Specific | 716.59 | 716.59 | 324.22 | 14.58 | 155 | >sp\|P02788\|TRFL_HUMAN Lactotransferrin OS=Homo sapiens OX=9606 GN=LTF PE=1 SV=6 | 1270 | 20201118_12812_1ng_glyco_hcd203040_compound15.20337.20337.3 | controllerType=0 controllerNumber=1 scan=20337 | 40.9766 |
| 18476:3 | 1 | R.TAGWNIPMGLLFN[+2205.793]QTGSC[+57.021]K.F | HexNAc(4)Hex(5)Fuc(2)NeuAc(1) | C[+57], N[+2206] | 1434.2796 | 3 | 4300.8242 | 4300.8028 | 0 | 5 | 485 | Specific | 822.52 | 822.52 | 822.52 | 15.77 | 155 | >sp\|P02788\|TRFL_HUMAN Lactotransferrin OS=Homo sapiens OX=9606 GN=LTF PE=1 SV=6 | 1270 | 20201118_12812_1ng_glyco_hcd203040_compound15.22090.22090.3 | controllerType=0 controllerNumber=1 scan=22090 | 44.7673 |
| 16625:3 | 1 | R.TAGWNIPMGLLFN[+1768.640]QTGSC[+57.021]K.F | HexNAc(4)Hex(5)Fuc(1) | C[+57], N[+1769] | 1288.5589 | 3 | 3863.6621 | 3863.6495 | 0 | 3.3 | 485 | Specific | 823.7 | 788.77 | 788.77 | 16.68 | 155 | >sp\|P02788\|TRFL_HUMAN Lactotransferrin OS=Homo sapiens OX=9606 GN=LTF PE=1 SV=6 | 1270 | 20201118_12812_1ng_glyco_hcd203040_compound15.20159.20159.3 | controllerType=0 controllerNumber=1 scan=20159 | 40.6386 |
| 16799:3 | 1 | R.TAGWNIPMGLLFN[+1622.582]QTGSC[+57.021]K.F | HexNAc(4)Hex(5) | C[+57], N[+1623] | 1239.8713 | 3 | 3717.5993 | 3717.5916 | 0 | 2.1 | 485 | Specific | 835.47 | 742.36 | 371.71 | 16.58 | 155 | >sp\|P02788\|TRFL_HUMAN Lactotransferrin OS=Homo sapiens OX=9606 GN=LTF PE=1 SV=6 | 1270 | 20201118_12812_1ng_glyco_hcd203040_compound15.20339.20339.3 | controllerType=0 controllerNumber=1 scan=20339 | 40.982 |
| 20519:3 | 1 | R.TAGWNIPMGLLFN[+2350.830]QTGSC[+57.021]K.F | HexNAc(4)Hex(5)Fuc(1)NeuAc(2) | C[+57], N[+2351] | 1482.6247 | 3 | 4445.8595 | 4445.8403 | 0 | 4.3 | 485 | Specific | 872.07 | 847.6 | 847.6 | 17.42 | 155 | >sp\|P02788\|TRFL_HUMAN Lactotransferrin OS=Homo sapiens OX=9606 GN=LTF PE=1 SV=6 | 1270 | 20201118_12812_1ng_glyco_hcd203040_compound15.24224.24224.3 | controllerType=0 controllerNumber=1 scan=24224 | 49.3762 |
| 20701:3 | 1 | R.TAGWNIPMGLLFN[+2204.772]QTGSC[+57.021]K.F | HexNAc(4)Hex(5)NeuAc(2) | C[+57], N[+2205] | 1433.9379 | 3 | 4299.7992 | 4299.7824 | 0 | 3.9 | 485 | Specific | 885.83 | 752.03 | 752.03 | 17.42 | 155 | >sp\|P02788\|TRFL_HUMAN Lactotransferrin OS=Homo sapiens OX=9606 GN=LTF PE=1 SV=6 | 1270 | 20201118_12812_1ng_glyco_hcd203040_compound15.24414.24414.3 | controllerType=0 controllerNumber=1 scan=24414 | 49.8132 |
| 20703:3 | 1 | R.TAGWNIPMGLLFN[+2204.772]QTGSC[+57.021]K.F | HexNAc(4)Hex(5)NeuAc(2) | C[+57], N[+2205] | 1433.9379 | 3 | 4299.7992 | 4299.7824 | 0 | 3.9 | 485 | Specific | 889.1 | 857.36 | 857.36 | 17.42 | 155 | >sp\|P02788\|TRFL_HUMAN Lactotransferrin OS=Homo sapiens OX=9606 GN=LTF PE=1 SV=6 | 1270 | 20201118_12812_1ng_glyco_hcd203040_compound15.24416.24416.3 | controllerType=0 controllerNumber=1 scan=24416 | 49.8181 |
| 18501:3 | 1 | R.TAGWNIPMGLLFN[+2059.735]QTGSC[+57.021]K.F | HexNAc(4)Hex(5)Fuc(1)NeuAc(1) | C[+57], N[+2060] | 1385.5905 | 3 | 4154.757 | 4154.7449 | 0 | 2.9 | 485 | Specific | 895.07 | 826.56 | 826.56 | 17.68 | 155 | >sp\|P02788\|TRFL_HUMAN Lactotransferrin OS=Homo sapiens OX=9606 GN=LTF PE=1 SV=6 | 1270 | 20201118_12812_1ng_glyco_hcd203040_compound15.22117.22117.3 | controllerType=0 controllerNumber=1 scan=22117 | 44.8337 |
| 17034:2 | 1 | R.TAGWNIPMGLLFN[+1216.423]QTGSC[+57.021]K.F | HexNAc(2)Hex(5) | C[+57], N[+1216] | 1656.2254 | 2 | 3311.4435 | 3311.4328 | 0 | 3.2 | 485 | Specific | 898.81 | 860.54 | 324.95 | 17.75 | 155 | >sp\|P02788\|TRFL_HUMAN Lactotransferrin OS=Homo sapiens OX=9606 GN=LTF PE=1 SV=6 | 1270 | 20201118_12812_1ng_glyco_hcd203040_compound15.20584.20584.2 | controllerType=0 controllerNumber=1 scan=20584 | 41.495 |
| 18661:3 | 1 | R.TAGWNIPMGLLFN[+1913.677]QTGSC[+57.021]K.F | HexNAc(4)Hex(5)NeuAc(1) | C[+57], N[+1914] | 1336.9042 | 3 | 4008.6981 | 4008.687 | 0 | 2.8 | 485 | Specific | 906.6 | 899.28 | 899.28 | 18.68 | 155 | >sp\|P02788\|TRFL_HUMAN Lactotransferrin OS=Homo sapiens OX=9606 GN=LTF PE=1 SV=6 | 1270 | 20201118_12812_1ng_glyco_hcd203040_compound15.22284.22284.3 | controllerType=0 controllerNumber=1 scan=22284 | 45.2181 |
| 17036:2 | 1 | R.TAGWNIPMGLLFN[+1216.423]QTGSC[+57.021]K.F | HexNAc(2)Hex(5) | C[+57], N[+1216] | 1656.2254 | 2 | 3311.4435 | 3311.4328 | 0 | 3.2 | 485 | Specific | 907.7 | 786.57 | 299.41 | 18.75 | 155 | >sp\|P02788\|TRFL_HUMAN Lactotransferrin OS=Homo sapiens OX=9606 GN=LTF PE=1 SV=6 | 1270 | 20201118_12812_1ng_glyco_hcd203040_compound15.20586.20586.2 | controllerType=0 controllerNumber=1 scan=20586 | 41.4994 |
| 18663:3 | 1 | R.TAGWNIPMGLLFN[+1913.677]QTGSC[+57.021]K.F | HexNAc(4)Hex(5)NeuAc(1) | C[+57], N[+1914] | 1336.9042 | 3 | 4008.6981 | 4008.687 | 0 | 2.8 | 485 | Specific | 914.32 | 907.87 | 907.87 | 18.68 | 155 | >sp\|P02788\|TRFL_HUMAN Lactotransferrin OS=Homo sapiens OX=9606 GN=LTF PE=1 SV=6 | 1270 | 20201118_12812_1ng_glyco_hcd203040_compound15.22286.22286.3 | controllerType=0 controllerNumber=1 scan=22286 | 45.2234 |
| 16865:3 | 1 | R.TAGWNIPMGLLFN[+1622.582]QTGSC[+57.021]K.F | HexNAc(4)Hex(5) | C[+57], N[+1623] | 1239.8715 | 3 | 3717.6 | 3717.5916 | 0 | 2.3 | 485 | Specific | 917.09 | 836.26 | 433.36 | 18.58 | 155 | >sp\|P02788\|TRFL_HUMAN Lactotransferrin OS=Homo sapiens OX=9606 GN=LTF PE=1 SV=6 | 1270 | 20201118_12812_1ng_glyco_hcd203040_compound15.20408.20408.3 | controllerType=0 controllerNumber=1 scan=20408 | 41.1151 |
| 18675:4 | 1 | R.TAGWNIPMGLLFN[+1913.677]QTGSC[+57.021]K.F | HexNAc(4)Hex(5)NeuAc(1) | C[+57], N[+1914] | 1002.9297 | 4 | 4008.6968 | 4008.687 | 0 | 2.5 | 485 | Specific | 922.36 | 917.02 | 917.02 | 18.61 | 155 | >sp\|P02788\|TRFL_HUMAN Lactotransferrin OS=Homo sapiens OX=9606 GN=LTF PE=1 SV=6 | 1270 | 20201118_12812_1ng_glyco_hcd203040_compound15.22298.22298.4 | controllerType=0 controllerNumber=1 scan=22298 | 45.2468 |
| 16628:3 | 1 | R.TAGWNIPMGLLFN[+1768.640]QTGSC[+57.021]K.F | HexNAc(4)Hex(5)Fuc(1) | C[+57], N[+1769] | 1288.5589 | 3 | 3863.6621 | 3863.6495 | 0 | 3.3 | 485 | Specific | 931.18 | 758.55 | 758.55 | 18.68 | 155 | >sp\|P02788\|TRFL_HUMAN Lactotransferrin OS=Homo sapiens OX=9606 GN=LTF PE=1 SV=6 | 1270 | 20201118_12812_1ng_glyco_hcd203040_compound15.20162.20162.3 | controllerType=0 controllerNumber=1 scan=20162 | 40.6436 |
| 20516:3 | 1 | R.TAGWNIPMGLLFN[+2350.830]QTGSC[+57.021]K.F | HexNAc(4)Hex(5)Fuc(1)NeuAc(2) | C[+57], N[+2351] | 1482.6247 | 3 | 4445.8595 | 4445.8403 | 0 | 4.3 | 485 | Specific | 950.18 | 908.39 | 908.39 | 19.42 | 155 | >sp\|P02788\|TRFL_HUMAN Lactotransferrin OS=Homo sapiens OX=9606 GN=LTF PE=1 SV=6 | 1270 | 20201118_12812_1ng_glyco_hcd203040_compound15.24221.24221.3 | controllerType=0 controllerNumber=1 scan=24221 | 49.3699 |
| 17007:3 | 1 | R.TAGWNIPMGLLFN[+1216.423]QTGSC[+57.021]K.F | HexNAc(2)Hex(5) | C[+57], N[+1216] | 1104.4856 | 3 | 3311.4423 | 3311.4328 | 0 | 2.9 | 485 | Specific | 964.65 | 964.65 | 394.71 | 19.68 | 155 | >sp\|P02788\|TRFL_HUMAN Lactotransferrin OS=Homo sapiens OX=9606 GN=LTF PE=1 SV=6 | 1270 | 20201118_12812_1ng_glyco_hcd203040_compound15.20556.20556.3 | controllerType=0 controllerNumber=1 scan=20556 | 41.434 |
| 18672:4 | 1 | R.TAGWNIPMGLLFN[+1913.677]QTGSC[+57.021]K.F | HexNAc(4)Hex(5)NeuAc(1) | C[+57], N[+1914] | 1002.9297 | 4 | 4008.6968 | 4008.687 | 0 | 2.5 | 485 | Specific | 976.03 | 945.66 | 945.66 | 19.61 | 155 | >sp\|P02788\|TRFL_HUMAN Lactotransferrin OS=Homo sapiens OX=9606 GN=LTF PE=1 SV=6 | 1270 | 20201118_12812_1ng_glyco_hcd203040_compound15.22295.22295.4 | controllerType=0 controllerNumber=1 scan=22295 | 45.2389 |
| 18499:3 | 1 | R.TAGWNIPMGLLFN[+2059.735]QTGSC[+57.021]K.F | HexNAc(4)Hex(5)Fuc(1)NeuAc(1) | C[+57], N[+2060] | 1385.5905 | 3 | 4154.757 | 4154.7449 | 0 | 2.9 | 485 | Specific | 980.38 | 942.8 | 942.8 | 19.68 | 155 | >sp\|P02788\|TRFL_HUMAN Lactotransferrin OS=Homo sapiens OX=9606 GN=LTF PE=1 SV=6 | 1270 | 20201118_12812_1ng_glyco_hcd203040_compound15.22115.22115.3 | controllerType=0 controllerNumber=1 scan=22115 | 44.8272 |
| 18473:3 | 1 | R.TAGWNIPMGLLFN[+2205.793]QTGSC[+57.021]K.F | HexNAc(4)Hex(5)Fuc(2)NeuAc(1) | C[+57], N[+2206] | 1434.2796 | 3 | 4300.8242 | 4300.8028 | 0 | 5 | 485 | Specific | 988.87 | 811.84 | 811.84 | 18.77 | 155 | >sp\|P02788\|TRFL_HUMAN Lactotransferrin OS=Homo sapiens OX=9606 GN=LTF PE=1 SV=6 | 1270 | 20201118_12812_1ng_glyco_hcd203040_compound15.22087.22087.3 | controllerType=0 controllerNumber=1 scan=22087 | 44.7554 |
| 17010:3 | 1 | R.TAGWNIPMGLLFN[+1216.423]QTGSC[+57.021]K.F | HexNAc(2)Hex(5) | C[+57], N[+1216] | 1104.4856 | 3 | 3311.4423 | 3311.4328 | 0 | 2.9 | 485 | Specific | 1081.82 | 901.38 | 517.15 | 21.68 | 155 | >sp\|P02788\|TRFL_HUMAN Lactotransferrin OS=Homo sapiens OX=9606 GN=LTF PE=1 SV=6 | 1270 | 20201118_12812_1ng_glyco_hcd203040_compound15.20559.20559.3 | controllerType=0 controllerNumber=1 scan=20559 | 41.441 |
| 17489:3 | 1 | R.TAGWNIPM[+15.995]GLLFN[+2059.735]QTGSC[+57.021]K.F | HexNAc(4)Hex(5)Fuc(1)NeuAc(1) | C[+57], M[+16], N[+2060] | 1390.9285 | 3 | 4170.771 | 4170.7398 | 0 | 7.5 | 485 | Specific | 299.22 | 190.75 | 190.75 | 5.05 | 155 | >sp\|P02788\|TRFL_HUMAN Lactotransferrin OS=Homo sapiens OX=9606 GN=LTF PE=1 SV=6 | 1270 | 20201118_12812_1ng_glyco_hcd203040_compound15.21059.21059.3 | controllerType=0 controllerNumber=1 scan=21059 | 42.4845 |
| 17231:3 | 1 | R.TAGWNIPM[+15.995]GLLFN[+1913.677]QTGSC[+57.021]K.F | HexNAc(4)Hex(5)NeuAc(1) | C[+57], M[+16], N[+1914] | 1342.235 | 3 | 4024.6906 | 4024.6819 | 0 | 2.2 | 485 | Specific | 381.48 | 343.98 | 343.98 | 7.71 | 155 | >sp\|P02788\|TRFL_HUMAN Lactotransferrin OS=Homo sapiens OX=9606 GN=LTF PE=1 SV=6 | 1270 | 20201118_12812_1ng_glyco_hcd203040_compound15.20790.20790.3 | controllerType=0 controllerNumber=1 scan=20790 | 41.9163 |
| 17491:3 | 1 | R.TAGWNIPM[+15.995]GLLFN[+2059.735]QTGSC[+57.021]K.F | HexNAc(4)Hex(5)Fuc(1)NeuAc(1) | C[+57], M[+16], N[+2060] | 1390.9285 | 3 | 4170.771 | 4170.7398 | 0 | 7.5 | 485 | Specific | 509.95 | 406.75 | 406.75 | 8.53 | 155 | >sp\|P02788\|TRFL_HUMAN Lactotransferrin OS=Homo sapiens OX=9606 GN=LTF PE=1 SV=6 | 1270 | 20201118_12812_1ng_glyco_hcd203040_compound15.21061.21061.3 | controllerType=0 controllerNumber=1 scan=21061 | 42.4892 |
| 17233:3 | 1 | R.TAGWNIPM[+15.995]GLLFN[+1913.677]QTGSC[+57.021]K.F | HexNAc(4)Hex(5)NeuAc(1) | C[+57], M[+16], N[+1914] | 1342.235 | 3 | 4024.6906 | 4024.6819 | 0 | 2.2 | 485 | Specific | 528.34 | 401.45 | 401.45 | 10.03 | 155 | >sp\|P02788\|TRFL_HUMAN Lactotransferrin OS=Homo sapiens OX=9606 GN=LTF PE=1 SV=6 | 1270 | 20201118_12812_1ng_glyco_hcd203040_compound15.20792.20792.3 | controllerType=0 controllerNumber=1 scan=20792 | 41.9202 |
| 18796:5 | 1 | R.TAGWNIPMGLLFN[+2059.735]QTGSC[+57.021]KFDEYFSQSC[+57.021]APGSDPR.S | HexNAc(4)Hex(5)Fuc(1)NeuAc(1) | C[+57]*2, N[+2060] | 1200.5064 | 5 | 5998.5026 | 5998.492 | 0 | 1.8 | 485 | Specific | 264.1 | 264.1 | 264.1 | 7 | 155 | >sp\|P02788\|TRFL_HUMAN Lactotransferrin OS=Homo sapiens OX=9606 GN=LTF PE=1 SV=6 | 1270 | 20201118_12812_1ng_glyco_hcd203040_compound15.22425.22425.5 | controllerType=0 controllerNumber=1 scan=22425 | 45.5426 |
| 17938:3 | 1 | R.TAGWNIPMGLLFN[+1216.423]QTGSC[+57.021]KFDEYFSQSC[+57.021]APGSDPR.S | HexNAc(2)Hex(5) | C[+57]*2, N[+1216] | 1719.0722 | 3 | 5155.2021 | 5155.18 | 0 | 4.3 | 485 | Specific | 287.85 | 285.85 | 47.66 | 5.98 | 155 | >sp\|P02788\|TRFL_HUMAN Lactotransferrin OS=Homo sapiens OX=9606 GN=LTF PE=1 SV=6 | 1270 | 20201118_12812_1ng_glyco_hcd203040_compound15.21529.21529.3 | controllerType=0 controllerNumber=1 scan=21529 | 43.5696 |
| 20164:4 | 1 | R.TAGWNIPMGLLFN[+2350.830]QTGSC[+57.021]KFDEYFSQSC[+57.021]APGSDPR.S | HexNAc(4)Hex(5)Fuc(1)NeuAc(2) | C[+57]*2, N[+2351] | 1573.1572 | 4 | 6289.6071 | 6289.5875 | 0 | 3.1 | 485 | Specific | 289.51 | 263.84 | 263.84 | 7.27 | 155 | >sp\|P02788\|TRFL_HUMAN Lactotransferrin OS=Homo sapiens OX=9606 GN=LTF PE=1 SV=6 | 1270 | 20201118_12812_1ng_glyco_hcd203040_compound15.23853.23853.4 | controllerType=0 controllerNumber=1 scan=23853 | 48.5383 |
| 17940:3 | 1 | R.TAGWNIPMGLLFN[+1216.423]QTGSC[+57.021]KFDEYFSQSC[+57.021]APGSDPR.S | HexNAc(2)Hex(5) | C[+57]*2, N[+1216] | 1719.0722 | 3 | 5155.2021 | 5155.18 | 0 | 4.3 | 485 | Specific | 317.46 | 317.46 | 143.79 | 6.14 | 155 | >sp\|P02788\|TRFL_HUMAN Lactotransferrin OS=Homo sapiens OX=9606 GN=LTF PE=1 SV=6 | 1270 | 20201118_12812_1ng_glyco_hcd203040_compound15.21531.21531.3 | controllerType=0 controllerNumber=1 scan=21531 | 43.5759 |
| 18879:3 | 1 | R.TAGWNIPMGLLFN[+1913.677]QTGSC[+57.021]KFDEYFSQSC[+57.021]APGSDPR.S | HexNAc(4)Hex(5)NeuAc(1) | C[+57]*2, N[+1914] | 1951.4843 | 3 | 5852.4385 | 5852.4341 | 0 | 0.7 | 485 | Specific | 394.18 | 307.14 | 307.14 | 7.76 | 155 | >sp\|P02788\|TRFL_HUMAN Lactotransferrin OS=Homo sapiens OX=9606 GN=LTF PE=1 SV=6 | 1270 | 20201118_12812_1ng_glyco_hcd203040_compound15.22511.22511.3 | controllerType=0 controllerNumber=1 scan=22511 | 45.7091 |
| 18877:3 | 1 | R.TAGWNIPMGLLFN[+1913.677]QTGSC[+57.021]KFDEYFSQSC[+57.021]APGSDPR.S | HexNAc(4)Hex(5)NeuAc(1) | C[+57]*2, N[+1914] | 1951.4843 | 3 | 5852.4385 | 5852.4341 | 0 | 0.7 | 485 | Specific | 395.1 | 291.44 | 291.44 | 7.57 | 155 | >sp\|P02788\|TRFL_HUMAN Lactotransferrin OS=Homo sapiens OX=9606 GN=LTF PE=1 SV=6 | 1270 | 20201118_12812_1ng_glyco_hcd203040_compound15.22509.22509.3 | controllerType=0 controllerNumber=1 scan=22509 | 45.7017 |
| 18785:4 | 1 | R.TAGWNIPMGLLFN[+2205.793]QTGSC[+57.021]KFDEYFSQSC[+57.021]APGSDPR.S | HexNAc(4)Hex(5)Fuc(2)NeuAc(1) | C[+57]*2, N[+2206] | 1536.898 | 4 | 6144.57 | 6144.5499 | 0 | 3.3 | 485 | Specific | 406.81 | 388.92 | 388.92 | 9.77 | 155 | >sp\|P02788\|TRFL_HUMAN Lactotransferrin OS=Homo sapiens OX=9606 GN=LTF PE=1 SV=6 | 1270 | 20201118_12812_1ng_glyco_hcd203040_compound15.22413.22413.3 | controllerType=0 controllerNumber=1 scan=22413 | 45.518 |
| 18907:5 | 1 | R.TAGWNIPMGLLFN[+1913.677]QTGSC[+57.021]KFDEYFSQSC[+57.021]APGSDPR.S | HexNAc(4)Hex(5)NeuAc(1) | C[+57]*2, N[+1914] | 1171.2941 | 5 | 5852.4414 | 5852.4341 | 0 | 1.2 | 485 | Specific | 450.42 | 406.83 | 406.83 | 9.96 | 155 | >sp\|P02788\|TRFL_HUMAN Lactotransferrin OS=Homo sapiens OX=9606 GN=LTF PE=1 SV=6 | 1270 | 20201118_12812_1ng_glyco_hcd203040_compound15.22540.22540.5 | controllerType=0 controllerNumber=1 scan=22540 | 45.7595 |
| 18850:3 | 1 | R.TAGWNIPMGLLFN[+1913.677]QTGSC[+57.021]KFDEYFSQSC[+57.021]APGSDPR.S | HexNAc(4)Hex(5)NeuAc(1) | C[+57]*2, N[+1914] | 1951.4871 | 3 | 5852.4469 | 5852.4341 | 0 | 2.2 | 485 | Specific | 462.95 | 355.62 | 355.62 | 8.36 | 155 | >sp\|P02788\|TRFL_HUMAN Lactotransferrin OS=Homo sapiens OX=9606 GN=LTF PE=1 SV=6 | 1270 | 20201118_12812_1ng_glyco_hcd203040_compound15.22481.22481.3 | controllerType=0 controllerNumber=1 scan=22481 | 45.6507 |
| 17804:4 | 1 | R.TAGWNIPMGLLFN[+1768.640]QTGSC[+57.021]KFDEYFSQSC[+57.021]APGSDPR.S | HexNAc(4)Hex(5)Fuc(1) | C[+57]*2, N[+1769] | 1427.6094 | 4 | 5707.4157 | 5707.3966 | 0 | 3.3 | 485 | Specific | 514.68 | 488.1 | 488.1 | 10.34 | 155 | >sp\|P02788\|TRFL_HUMAN Lactotransferrin OS=Homo sapiens OX=9606 GN=LTF PE=1 SV=6 | 1270 | 20201118_12812_1ng_glyco_hcd203040_compound15.21390.21390.4 | controllerType=0 controllerNumber=1 scan=21390 | 43.2593 |
| 18788:4 | 1 | R.TAGWNIPMGLLFN[+2205.793]QTGSC[+57.021]KFDEYFSQSC[+57.021]APGSDPR.S | HexNAc(4)Hex(5)Fuc(2)NeuAc(1) | C[+57]*2, N[+2206] | 1536.898 | 4 | 6144.57 | 6144.5499 | 0 | 3.3 | 485 | Specific | 545.29 | 523.63 | 523.63 | 10.48 | 155 | >sp\|P02788\|TRFL_HUMAN Lactotransferrin OS=Homo sapiens OX=9606 GN=LTF PE=1 SV=6 | 1270 | 20201118_12812_1ng_glyco_hcd203040_compound15.22416.22416.3 | controllerType=0 controllerNumber=1 scan=22416 | 45.5239 |
| 17833:4 | 1 | R.TAGWNIPMGLLFN[+1622.582]QTGSC[+57.021]KFDEYFSQSC[+57.021]APGSDPR.S | HexNAc(4)Hex(5) | C[+57]*2, N[+1623] | 1391.0956 | 4 | 5561.3605 | 5561.3387 | 0 | 3.9 | 485 | Specific | 583.58 | 548.87 | 160.41 | 11.63 | 155 | >sp\|P02788\|TRFL_HUMAN Lactotransferrin OS=Homo sapiens OX=9606 GN=LTF PE=1 SV=6 | 1270 | 20201118_12812_1ng_glyco_hcd203040_compound15.21420.21420.4 | controllerType=0 controllerNumber=1 scan=21420 | 43.3294 |
| 18870:4 | 1 | R.TAGWNIPMGLLFN[+1913.677]QTGSC[+57.021]KFDEYFSQSC[+57.021]APGSDPR.S | HexNAc(4)Hex(5)NeuAc(1) | C[+57]*2, N[+1914] | 1463.8655 | 4 | 5852.4401 | 5852.4341 | 0 | 1 | 485 | Specific | 642.27 | 581.31 | 581.31 | 12.57 | 155 | >sp\|P02788\|TRFL_HUMAN Lactotransferrin OS=Homo sapiens OX=9606 GN=LTF PE=1 SV=6 | 1270 | 20201118_12812_1ng_glyco_hcd203040_compound15.22502.22502.4 | controllerType=0 controllerNumber=1 scan=22502 | 45.6932 |
| 20199:4 | 1 | R.TAGWNIPMGLLFN[+2205.793]QTGSC[+57.021]KFDEYFSQSC[+57.021]APGSDPR.S | HexNAc(4)Hex(5)Fuc(2)NeuAc(1) | C[+57]*2, N[+2206] | 1536.89 | 4 | 6144.5382 | 6144.5499 | 0 | -1.9 | 485 | Specific | 713.48 | 641.59 | 641.59 | 13.71 | 155 | >sp\|P02788\|TRFL_HUMAN Lactotransferrin OS=Homo sapiens OX=9606 GN=LTF PE=1 SV=6 | 1270 | 20201118_12812_1ng_glyco_hcd203040_compound15.23889.23889.4 | controllerType=0 controllerNumber=1 scan=23889 | 48.612 |
| 18794:5 | 1 | R.TAGWNIPMGLLFN[+2059.735]QTGSC[+57.021]KFDEYFSQSC[+57.021]APGSDPR.S | HexNAc(4)Hex(5)Fuc(1)NeuAc(1) | C[+57]*2, N[+2060] | 1200.5064 | 5 | 5998.5026 | 5998.492 | 0 | 1.8 | 485 | Specific | 730.02 | 725.23 | 725.23 | 14.82 | 155 | >sp\|P02788\|TRFL_HUMAN Lactotransferrin OS=Homo sapiens OX=9606 GN=LTF PE=1 SV=6 | 1270 | 20201118_12812_1ng_glyco_hcd203040_compound15.22423.22423.5 | controllerType=0 controllerNumber=1 scan=22423 | 45.5379 |
| 18853:3 | 1 | R.TAGWNIPMGLLFN[+1913.677]QTGSC[+57.021]KFDEYFSQSC[+57.021]APGSDPR.S | HexNAc(4)Hex(5)NeuAc(1) | C[+57]*2, N[+1914] | 1951.4871 | 3 | 5852.4469 | 5852.4341 | 0 | 2.2 | 485 | Specific | 742.44 | 591.92 | 591.92 | 14.87 | 155 | >sp\|P02788\|TRFL_HUMAN Lactotransferrin OS=Homo sapiens OX=9606 GN=LTF PE=1 SV=6 | 1270 | 20201118_12812_1ng_glyco_hcd203040_compound15.22484.22484.3 | controllerType=0 controllerNumber=1 scan=22484 | 45.6586 |
| 18753:4 | 1 | R.TAGWNIPMGLLFN[+2205.793]QTGSC[+57.021]KFDEYFSQSC[+57.021]APGSDPR.S | HexNAc(4)Hex(5)Fuc(2)NeuAc(1) | C[+57]*2, N[+2206] | 1536.8986 | 4 | 6144.5724 | 6144.5499 | 0 | 3.7 | 485 | Specific | 751.18 | 704.85 | 704.85 | 15.9 | 155 | >sp\|P02788\|TRFL_HUMAN Lactotransferrin OS=Homo sapiens OX=9606 GN=LTF PE=1 SV=6 | 1270 | 20201118_12812_1ng_glyco_hcd203040_compound15.22380.22380.4 | controllerType=0 controllerNumber=1 scan=22380 | 45.4518 |
| 20201:4 | 1 | R.TAGWNIPMGLLFN[+2205.793]QTGSC[+57.021]KFDEYFSQSC[+57.021]APGSDPR.S | HexNAc(4)Hex(5)Fuc(2)NeuAc(1) | C[+57]*2, N[+2206] | 1536.89 | 4 | 6144.5382 | 6144.5499 | 0 | -1.9 | 485 | Specific | 757.92 | 745.5 | 745.5 | 14.71 | 155 | >sp\|P02788\|TRFL_HUMAN Lactotransferrin OS=Homo sapiens OX=9606 GN=LTF PE=1 SV=6 | 1270 | 20201118_12812_1ng_glyco_hcd203040_compound15.23891.23891.4 | controllerType=0 controllerNumber=1 scan=23891 | 48.6153 |
| 17836:4 | 1 | R.TAGWNIPMGLLFN[+1622.582]QTGSC[+57.021]KFDEYFSQSC[+57.021]APGSDPR.S | HexNAc(4)Hex(5) | C[+57]*2, N[+1623] | 1391.0956 | 4 | 5561.3605 | 5561.3387 | 0 | 3.9 | 485 | Specific | 795.53 | 746.13 | 307.5 | 15.63 | 155 | >sp\|P02788\|TRFL_HUMAN Lactotransferrin OS=Homo sapiens OX=9606 GN=LTF PE=1 SV=6 | 1270 | 20201118_12812_1ng_glyco_hcd203040_compound15.21423.21423.4 | controllerType=0 controllerNumber=1 scan=21423 | 43.335 |
| 20166:4 | 1 | R.TAGWNIPMGLLFN[+2350.830]QTGSC[+57.021]KFDEYFSQSC[+57.021]APGSDPR.S | HexNAc(4)Hex(5)Fuc(1)NeuAc(2) | C[+57]*2, N[+2351] | 1573.1572 | 4 | 6289.6071 | 6289.5875 | 0 | 3.1 | 485 | Specific | 820.96 | 627.28 | 627.28 | 16.9 | 155 | >sp\|P02788\|TRFL_HUMAN Lactotransferrin OS=Homo sapiens OX=9606 GN=LTF PE=1 SV=6 | 1270 | 20201118_12812_1ng_glyco_hcd203040_compound15.23855.23855.4 | controllerType=0 controllerNumber=1 scan=23855 | 48.542 |
| 18769:4 | 1 | R.TAGWNIPMGLLFN[+2059.735]QTGSC[+57.021]KFDEYFSQSC[+57.021]APGSDPR.S | HexNAc(4)Hex(5)Fuc(1)NeuAc(1) | C[+57]*2, N[+2060] | 1500.386 | 4 | 5998.5221 | 5998.492 | 0 | 5 | 485 | Specific | 865.62 | 784.99 | 784.99 | 17.63 | 155 | >sp\|P02788\|TRFL_HUMAN Lactotransferrin OS=Homo sapiens OX=9606 GN=LTF PE=1 SV=6 | 1270 | 20201118_12812_1ng_glyco_hcd203040_compound15.22397.22397.4 | controllerType=0 controllerNumber=1 scan=22397 | 45.4847 |
| 17806:4 | 1 | R.TAGWNIPMGLLFN[+1768.640]QTGSC[+57.021]KFDEYFSQSC[+57.021]APGSDPR.S | HexNAc(4)Hex(5)Fuc(1) | C[+57]*2, N[+1769] | 1427.6094 | 4 | 5707.4157 | 5707.3966 | 0 | 3.3 | 485 | Specific | 895.62 | 845.66 | 845.66 | 17.9 | 155 | >sp\|P02788\|TRFL_HUMAN Lactotransferrin OS=Homo sapiens OX=9606 GN=LTF PE=1 SV=6 | 1270 | 20201118_12812_1ng_glyco_hcd203040_compound15.21392.21392.4 | controllerType=0 controllerNumber=1 scan=21392 | 43.2632 |
| 18771:4 | 1 | R.TAGWNIPMGLLFN[+2059.735]QTGSC[+57.021]KFDEYFSQSC[+57.021]APGSDPR.S | HexNAc(4)Hex(5)Fuc(1)NeuAc(1) | C[+57]*2, N[+2060] | 1500.386 | 4 | 5998.5221 | 5998.492 | 0 | 5 | 485 | Specific | 951.1 | 930.29 | 930.29 | 19.63 | 155 | >sp\|P02788\|TRFL_HUMAN Lactotransferrin OS=Homo sapiens OX=9606 GN=LTF PE=1 SV=6 | 1270 | 20201118_12812_1ng_glyco_hcd203040_compound15.22399.22399.4 | controllerType=0 controllerNumber=1 scan=22399 | 45.4899 |
| 18756:4 | 1 | R.TAGWNIPMGLLFN[+2205.793]QTGSC[+57.021]KFDEYFSQSC[+57.021]APGSDPR.S | HexNAc(4)Hex(5)Fuc(2)NeuAc(1) | C[+57]*2, N[+2206] | 1536.8986 | 4 | 6144.5724 | 6144.5499 | 0 | 3.7 | 485 | Specific | 954.49 | 904.29 | 904.29 | 19.9 | 155 | >sp\|P02788\|TRFL_HUMAN Lactotransferrin OS=Homo sapiens OX=9606 GN=LTF PE=1 SV=6 | 1270 | 20201118_12812_1ng_glyco_hcd203040_compound15.22383.22383.4 | controllerType=0 controllerNumber=1 scan=22383 | 45.4571 |
| 18873:4 | 1 | R.TAGWNIPMGLLFN[+1913.677]QTGSC[+57.021]KFDEYFSQSC[+57.021]APGSDPR.S | HexNAc(4)Hex(5)NeuAc(1) | C[+57]*2, N[+1914] | 1463.8655 | 4 | 5852.4401 | 5852.4341 | 0 | 1 | 485 | Specific | 954.63 | 873.2 | 873.2 | 19.57 | 155 | >sp\|P02788\|TRFL_HUMAN Lactotransferrin OS=Homo sapiens OX=9606 GN=LTF PE=1 SV=6 | 1270 | 20201118_12812_1ng_glyco_hcd203040_compound15.22505.22505.4 | controllerType=0 controllerNumber=1 scan=22505 | 45.6967 |
| 17622:4 | 1 | R.TAGWNIPM[+15.995]GLLFN[+2059.735]QTGSC[+57.021]KFDEYFSQSC[+57.021]A | HexNAc(4)Hex(5)Fuc(1)NeuAc(1) | C[+57]*2, M[+16], N[+2060] | 1504.3816 | 4 | 6014.5046 | 6014.487 | 0 | 2.9 | 485 | Specific | 630.27 | 548.55 | 548.55 | 12.9 | 155 | >sp\|P02788\|TRFL_HUMAN Lactotransferrin OS=Homo sapiens OX=9606 GN=LTF PE=1 SV=6 | 1270 | 20201118_12812_1ng_glyco_hcd203040_compound15.21199.21199.4 | controllerType=0 controllerNumber=1 scan=21199 | 42.8275 |
| 17625:4 | 1 | R.TAGWNIPM[+15.995]GLLFN[+2059.735]QTGSC[+57.021]KFDEYFSQSC[+57.021]A | HexNAc(4)Hex(5)Fuc(1)NeuAc(1) | C[+57]*2, M[+16], N[+2060] | 1504.3816 | 4 | 6014.5046 | 6014.487 | 0 | 2.9 | 485 | Specific | 812.37 | 717.2 | 717.2 | 16.9 | 155 | >sp\|P02788\|TRFL_HUMAN Lactotransferrin OS=Homo sapiens OX=9606 GN=LTF PE=1 SV=6 | 1270 | 20201118_12812_1ng_glyco_hcd203040_compound15.21202.21202.4 | controllerType=0 controllerNumber=1 scan=21202 | 42.8357 |
| 08981:3 | 1 | R.N[+2059.735]GSDC[+57.021]PDKFC[+57.021]LFQSETK.N | HexNAc(4)Hex(5)Fuc(1)NeuAc(1) | C[+57]*2, N[+2060] | 1364.8784 | 3 | 4092.6205 | 4092.6088 | 0 | 2.9 | 642 | Specific | 593.36 | 463.07 | 463.07 | 11.28 | 155 | >sp\|P02788\|TRFL_HUMAN Lactotransferrin OS=Homo sapiens OX=9606 GN=LTF PE=1 SV=6 | 1270 | 20201118_12812_1ng_glyco_hcd203040_compound15.12193.12193.3 | controllerType=0 controllerNumber=1 scan=12193 | 26.6923 |
| 08979:3 | 1 | R.N[+2059.735]GSDC[+57.021]PDKFC[+57.021]LFQSETK.N | HexNAc(4)Hex(5)Fuc(1)NeuAc(1) | C[+57]*2, N[+2060] | 1364.8784 | 3 | 4092.6205 | 4092.6088 | 0 | 2.9 | 642 | Specific | 697.05 | 592.08 | 592.08 | 13.28 | 155 | >sp\|P02788\|TRFL_HUMAN Lactotransferrin OS=Homo sapiens OX=9606 GN=LTF PE=1 SV=6 | 1270 | 20201118_12812_1ng_glyco_hcd203040_compound15.12191.12191.3 | controllerType=0 controllerNumber=1 scan=12191 | 26.6876 |
| 07127:3 | 3 | R.FSDGLESN[+1864.634]SSTQFEVK.K | HexNAc(2)Hex(9) | N[+1865] | 1213.8267 | 3 | 3639.4656 | 3639.4472 | 0 | 5.1 | 219 | Specific | 448.5 | 448.5 | 448.5 | 8.91 | 83 | >sp\|P0C0L4\|CO4A_HUMAN Complement C4-A OS=Homo sapiens OX=9606 GN=C4A PE=1 SV=2 | 1235 | 20201118_12812_1ng_glyco_hcd203040_compound15.10256.10256.3 | controllerType=0 controllerNumber=1 scan=10256 | 23.4297 |
| 07125:3 | 3 | R.FSDGLESN[+1864.634]SSTQFEVK.K | HexNAc(2)Hex(9) | N[+1865] | 1213.8267 | 3 | 3639.4656 | 3639.4472 | 0 | 5.1 | 219 | Specific | 498.52 | 498.52 | 498.52 | 9.15 | 83 | >sp\|P0C0L4\|CO4A_HUMAN Complement C4-A OS=Homo sapiens OX=9606 GN=C4A PE=1 SV=2 | 1235 | 20201118_12812_1ng_glyco_hcd203040_compound15.10254.10254.3 | controllerType=0 controllerNumber=1 scan=10254 | 23.4262 |
| 04560:3 | 3 | R.FSDGLESN[+1864.634]SSTQFEVKK.Y | HexNAc(2)Hex(9) | N[+1865] | 1256.5234 | 3 | 3767.5556 | 3767.5421 | 0 | 3.6 | 219 | Specific | 376.07 | 362.42 | 362.42 | 7.42 | 83 | >sp\|P0C0L4\|CO4A_HUMAN Complement C4-A OS=Homo sapiens OX=9606 GN=C4A PE=1 SV=2 | 1235 | 20201118_12812_1ng_glyco_hcd203040_compound15.7601.7601.3 | controllerType=0 controllerNumber=1 scan=7601 | 19.1181 |
| 04558:3 | 3 | R.FSDGLESN[+1864.634]SSTQFEVKK.Y | HexNAc(2)Hex(9) | N[+1865] | 1256.5234 | 3 | 3767.5556 | 3767.5421 | 0 | 3.6 | 219 | Specific | 475.97 | 475.97 | 475.97 | 9.4 | 83 | >sp\|P0C0L4\|CO4A_HUMAN Complement C4-A OS=Homo sapiens OX=9606 GN=C4A PE=1 SV=2 | 1235 | 20201118_12812_1ng_glyco_hcd203040_compound15.7599.7599.3 | controllerType=0 controllerNumber=1 scan=7599 | 19.1145 |
| 09850:3 | 4 | K.TVLTPATNHMGN[+1540.529]VTFTIPANR.E | HexNAc(2)Hex(7) | N[+1541] | 1265.9029 | 3 | 3795.6941 | 3795.6886 | 0 | 1.4 | 74 | Specific | 471.01 | 299.42 | 167.01 | 6.58 | 91 | >sp\|P01024\|CO3_HUMAN Complement C3 OS=Homo sapiens OX=9606 GN=C3 PE=1 SV=2 | 492 | 20201118_12812_1ng_glyco_hcd203040_compound15.13092.13092.3 | controllerType=0 controllerNumber=1 scan=13092 | 28.1405 |
| 09848:3 | 4 | K.TVLTPATNHMGN[+1540.529]VTFTIPANR.E | HexNAc(2)Hex(7) | N[+1541] | 1265.9029 | 3 | 3795.6941 | 3795.6886 | 0 | 1.4 | 74 | Specific | 628.34 | 592.51 | 313.26 | 12.65 | 91 | >sp\|P01024\|CO3_HUMAN Complement C3 OS=Homo sapiens OX=9606 GN=C3 PE=1 SV=2 | 492 | 20201118_12812_1ng_glyco_hcd203040_compound15.13090.13090.3 | controllerType=0 controllerNumber=1 scan=13090 | 28.1359 |
| 15260:3 | 7 | R.LN[+1378.476]YSLPTGQWVGVQLPR.N | HexNAc(2)Hex(6) | N[+1378] | 1102.846 | 3 | 3306.5234 | 3306.5145 | 0 | 2.7 | 1017 | Specific | 801.97 | 752.97 | 752.97 | 16.68 | 78 | >sp\|P24821\|TENA_HUMAN Tenascin OS=Homo sapiens OX=9606 GN=TNC PE=1 SV=3 | 486 | 20201118_12812_1ng_glyco_hcd203040_compound15.18714.18714.3 | controllerType=0 controllerNumber=1 scan=18714 | 37.9677 |
| 15262:3 | 7 | R.LN[+1378.476]YSLPTGQWVGVQLPR.N | HexNAc(2)Hex(6) | N[+1378] | 1102.846 | 3 | 3306.5234 | 3306.5145 | 0 | 2.7 | 1017 | Specific | 872.88 | 757.37 | 757.37 | 17.68 | 78 | >sp\|P24821\|TENA_HUMAN Tenascin OS=Homo sapiens OX=9606 GN=TNC PE=1 SV=3 | 486 | 20201118_12812_1ng_glyco_hcd203040_compound15.18716.18716.3 | controllerType=0 controllerNumber=1 scan=18716 | 37.9724 |
| 17458:3 | 8 | R.GLSFDVSLEVSQGPGLLN[+2570.925]DTK.V | HexNAc(5)Hex(6)Fuc(2)NeuAc(1) | N[+2571] | 1583.0184 | 3 | 4747.0405 | 4747.0382 | 0 | 0.5 | 118 | Specific | 391.01 | 391.01 | 391.01 | 9.22 | 73 | >sp\|P01833\|PIGR_HUMAN Polymeric immunoglobulin receptor OS=Homo sapiens OX=9606 GN=PIGR PE=1 SV=4 | 902 | 20201118_12812_1ng_glyco_hcd203040_compound15.21026.21026.3 | controllerType=0 controllerNumber=1 scan=21026 | 42.4051 |
| 17516:3 | 8 | R.GLSFDVSLEVSQGPGLLN[+2351.851]DTK.V | HexNAc(4)Hex(5)Fuc(3)NeuAc(1) | N[+2352] | 1509.9981 | 3 | 4527.9798 | 4527.964 | 0 | 3.5 | 118 | Specific | 447.5 | 286.18 | 286.18 | 7.88 | 73 | >sp\|P01833\|PIGR_HUMAN Polymeric immunoglobulin receptor OS=Homo sapiens OX=9606 GN=PIGR PE=1 SV=4 | 902 | 20201118_12812_1ng_glyco_hcd203040_compound15.21087.21087.3 | controllerType=0 controllerNumber=1 scan=21087 | [illegible] |
| 19371:3 | 8 | R.GLSFDVSLEVSQGPGLLN[+2350.830]DTK.V | HexNAc(4)Hex(5)Fuc(1)NeuAc(2) | N[+2351] | 1509.6574 | 3 | 4526.9577 | 4526.9436 | 0 | 3.1 | 118 | Specific | 613.72 | 574.55 | 574.55 | 12.68 | 73 | >sp\|P01833\|PIGR_HUMAN Polymeric immunoglobulin receptor OS=Homo sapiens OX=9606 GN=PIGR PE=1 SV=4 | 902 | 20201118_12812_1ng_glyco_hcd203040_compound15.23026.23026.3 | controllerType=0 controllerNumber=1 scan=23026 | [illegible] |
| 17514:3 | 8 | R.GLSFDVSLEVSQGPGLLN[+2205.793]DTK.V | HexNAc(4)Hex(5)Fuc(2)NeuAc(1) | N[+2206] | 1461.3108 | 3 | 4381.918 | 4381.906 | 0 | 2.7 | 118 | Specific | 848.68 | 800.65 | 800.65 | 16.68 | 73 | >sp\|P01833\|PIGR_HUMAN Polymeric immunoglobulin receptor OS=Homo sapiens OX=9606 GN=PIGR PE=1 SV=4 | 902 | 20201118_12812_1ng_glyco_hcd203040_compound15.21085.21085.3 | controllerType=0 controllerNumber=1 scan=21085 | 42.5548 |
| 17511:3 | 8 | R.GLSFDVSLEVSQGPGLLN[+2205.793]DTK.V | HexNAc(4)Hex(5)Fuc(2)NeuAc(1) | N[+2206] | 1461.3108 | 3 | 4381.918 | 4381.906 | 0 | 2.7 | 118 | Specific | 880.31 | 676.04 | 676.04 | 17.68 | 73 | >sp\|P01833\|PIGR_HUMAN Polymeric immunoglobulin receptor OS=Homo sapiens OX=9606 GN=PIGR PE=1 SV=4 | 902 | 20201118_12812_1ng_glyco_hcd203040_compound15.21082.21082.3 | controllerType=0 controllerNumber=1 scan=21082 | 42.5457 |
| 17559:3 | 8 | R.GLSFDVSLEVSQGPGLLN[+2059.735]DTK.V | HexNAc(4)Hex(5)Fuc(1)NeuAc(1) | N[+2060] | 1412.6268 | 3 | 4235.8657 | 4235.8481 | 0 | 4.2 | 118 | Specific | 973.94 | 715.17 | 715.17 | 19.41 | 73 | >sp\|P01833\|PIGR_HUMAN Polymeric immunoglobulin receptor OS=Homo sapiens OX=9606 GN=PIGR PE=1 SV=4 | 902 | 20201118_12812_1ng_glyco_hcd203040_compound15.21133.21133.3 | controllerType=0 controllerNumber=1 scan=21133 | 42.6667 |
| 17561:3 | 8 | R.GLSFDVSLEVSQGPGLLN[+2059.735]DTK.V | HexNAc(4)Hex(5)Fuc(1)NeuAc(1) | N[+2060] | 1412.6268 | 3 | 4235.8657 | 4235.8481 | 0 | 4.2 | 118 | Specific | 999.34 | 915.05 | 915.05 | 19.41 | 73 | >sp\|P01833\|PIGR_HUMAN Polymeric immunoglobulin receptor OS=Homo sapiens OX=9606 GN=PIGR PE=1 SV=4 | 902 | 20201118_12812_1ng_glyco_hcd203040_compound15.21135.21135.3 | controllerType=0 controllerNumber=1 scan=21135 | 42.6752 |
| 19373:3 | 8 | R.GLSFDVSLEVSQGPGLLN[+2350.830]DTK.V | HexNAc(4)Hex(5)Fuc(1)NeuAc(2) | N[+2351] | 1509.6574 | 3 | 4526.9577 | 4526.9436 | 0 | 3.1 | 118 | Specific | 1014.01 | 954.24 | 954.24 | 20.68 | 73 | >sp\|P01833\|PIGR_HUMAN Polymeric immunoglobulin receptor OS=Homo sapiens OX=9606 GN=PIGR PE=1 SV=4 | 902 | 20201118_12812_1ng_glyco_hcd203040_compound15.23028.23028.3 | controllerType=0 controllerNumber=1 scan=23028 | 46.7347 |
| 18763:4 | 8 | K.QIGLYPVLVIDSSGYVNPN[+1913.677]YTGR.I | HexNAc(4)Hex(5)NeuAc(1) | N[+1914] | 1110.5038 | 4 | 4438.9935 | 4438.9805 | 0 | 2.9 | 168 | Specific | 311.98 | 311.98 | 311.98 | 7.09 | 73 | >sp\|P01833\|PIGR_HUMAN Polymeric immunoglobulin receptor OS=Homo sapiens OX=9606 GN=PIGR PE=1 SV=4 | 902 | 20201118_12812_1ng_glyco_hcd203040_compound15.22390.22390.4 | controllerType=0 controllerNumber=1 scan=22390 | 45.4698 |
| 18698:3 | 8 | K.QIGLYPVLVIDSSGYVNPN[+2205.793]YTGR.I | HexNAc(4)Hex(5)Fuc(2)NeuAc(1) | N[+2206] | 1577.7065 | 3 | 4731.1049 | 4731.0963 | 0 | 1.8 | 168 | Specific | 424.7 | 327.3 | 327.3 | 8.78 | 73 | >sp\|P01833\|PIGR_HUMAN Polymeric immunoglobulin receptor OS=Homo sapiens OX=9606 GN=PIGR PE=1 SV=4 | 902 | 20201118_12812_1ng_glyco_hcd203040_compound15.22322.22322.3 | controllerType=0 controllerNumber=1 scan=22322 | 45.3042 |
| 18696:3 | 8 | K.QIGLYPVLVIDSSGYVNPN[+2205.793]YTGR.I | HexNAc(4)Hex(5)Fuc(2)NeuAc(1) | N[+2206] | 1577.7065 | 3 | 4731.1049 | 4731.0963 | 0 | 1.8 | 168 | Specific | 488.8 | 488.8 | 488.8 | 10.91 | 73 | >sp\|P01833\|PIGR_HUMAN Polymeric immunoglobulin receptor OS=Homo sapiens OX=9606 GN=PIGR PE=1 SV=4 | 902 | 20201118_12812_1ng_glyco_hcd203040_compound15.22320.22320.3 | controllerType=0 controllerNumber=1 scan=22320 | 45.2978 |
| 18766:4 | 8 | K.QIGLYPVLVIDSSGYVNPN[+1913.677]YTGR.I | HexNAc(4)Hex(5)NeuAc(1) | N[+1914] | 1110.5038 | 4 | 4438.9935 | 4438.9805 | 0 | 2.9 | 168 | Specific | 495.67 | 381.83 | 381.83 | 11.04 | 73 | >sp\|P01833\|PIGR_HUMAN Polymeric immunoglobulin receptor OS=Homo sapiens OX=9606 GN=PIGR PE=1 SV=4 | 902 | 20201118_12812_1ng_glyco_hcd203040_compound15.22393.22393.4 | controllerType=0 controllerNumber=1 scan=22393 | 45.475 |
| 18700:4 | 8 | K.QIGLYPVLVIDSSGYVNPN[+2059.735]YTGR.I | HexNAc(4)Hex(5)Fuc(1)NeuAc(1) | N[+2060] | 1147.0168 | 4 | 4585.0452 | 4585.0384 | 0 | 1.5 | 168 | Specific | 661.02 | 640.85 | 640.85 | 14.55 | 73 | >sp\|P01833\|PIGR_HUMAN Polymeric immunoglobulin receptor OS=Homo sapiens OX=9606 GN=PIGR PE=1 SV=4 | 902 | 20201118_12812_1ng_glyco_hcd203040_compound15.22324.22324.4 | controllerType=0 controllerNumber=1 scan=22324 | 45.3074 |
| 18705:3 | 8 | K.QIGLYPVLVIDSSGYVNPN[+2059.735]YTGR.I | HexNAc(4)Hex(5)Fuc(1)NeuAc(1) | N[+2060] | 1529.0237 | 3 | 4585.0567 | 4585.0384 | 0 | 4 | 168 | Specific | 679.96 | 535.46 | 535.46 | 14.36 | 73 | >sp\|P01833\|PIGR_HUMAN Polymeric immunoglobulin receptor OS=Homo sapiens OX=9606 GN=PIGR PE=1 SV=4 | 902 | 20201118_12812_1ng_glyco_hcd203040_compound15.22330.22330.3 | controllerType=0 controllerNumber=1 scan=22330 | 45.3277 |
| 18744:3 | 8 | K.QIGLYPVLVIDSSGYVNPN[+1913.677]YTGR.I | HexNAc(4)Hex(5)NeuAc(1) | N[+1914] | 1480.3373 | 3 | 4438.9975 | 4438.9805 | 0 | 3.8 | 168 | Specific | 737.45 | 701.48 | 701.48 | 15.36 | 73 | >sp\|P01833\|PIGR_HUMAN Polymeric immunoglobulin receptor OS=Homo sapiens OX=9606 GN=PIGR PE=1 SV=4 | 902 | 20201118_12812_1ng_glyco_hcd203040_compound15.22371.22371.3 | controllerType=0 controllerNumber=1 scan=22371 | 45.4315 |
| 18746:3 | 8 | K.QIGLYPVLVIDSSGYVNPN[+1913.677]YTGR.I | HexNAc(4)Hex(5)NeuAc(1) | N[+1914] | 1480.3373 | 3 | 4438.9975 | 4438.9805 | 0 | 3.8 | 168 | Specific | 868.06 | 601.12 | 601.12 | 18.36 | 73 | >sp\|P01833\|PIGR_HUMAN Polymeric immunoglobulin receptor OS=Homo sapiens OX=9606 GN=PIGR PE=1 SV=4 | 902 | 20201118_12812_1ng_glyco_hcd203040_compound15.22373.22373.3 | controllerType=0 controllerNumber=1 scan=22373 | 45.4364 |
| 18703:4 | 8 | K.QIGLYPVLVIDSSGYVNPN[+2059.735]YTGR.I | HexNAc(4)Hex(5)Fuc(1)NeuAc(1) | N[+2060] | 1147.0168 | 4 | 4585.0452 | 4585.0384 | 0 | 1.5 | 168 | Specific | 894.36 | 804.19 | 804.19 | 18.55 | 73 | >sp\|P01833\|PIGR_HUMAN Polymeric immunoglobulin receptor OS=Homo sapiens OX=9606 GN=PIGR PE=1 SV=4 | 902 | 20201118_12812_1ng_glyco_hcd203040_compound15.22327.22327.4 | controllerType=0 controllerNumber=1 scan=22327 | 45.3214 |
| 18707:3 | 8 | K.QIGLYPVLVIDSSGYVNPN[+2059.735]YTGR.I | HexNAc(4)Hex(5)Fuc(1)NeuAc(1) | N[+2060] | 1529.0237 | 3 | 4585.0567 | 4585.0384 | 0 | 4 | 168 | Specific | 918.49 | 822.22 | 822.22 | 19.36 | 73 | >sp\|P01833\|PIGR_HUMAN Polymeric immunoglobulin receptor OS=Homo sapiens OX=9606 GN=PIGR PE=1 SV=4 | 902 | 20201118_12812_1ng_glyco_hcd203040_compound15.22332.22332.3 | controllerType=0 controllerNumber=1 scan=22332 | 45.3362 |
| 28734:4 | 8 | R.[+42.011]LSLLEEPGN[+2350.830]GTFTVILNQLTSR.D | HexNAc(4)Hex(5)Fuc(1)NeuAc(2) | .[42], N[+2351] | 1199.5422 | 4 | 4795.1468 | 4795.1335 | 0 | 2.8 | 413 | Specific | 205.46 | 205.46 | 205.46 | 5.26 | 73 | >sp\|P01833\|PIGR_HUMAN Polymeric immunoglobulin receptor OS=Homo sapiens OX=9606 GN=PIGR PE=1 SV=4 | 902 | 20201118_12812_1ng_glyco_hcd203040_compound15.32777.32777.4 | controllerType=0 controllerNumber=1 scan=32777 | 65.3975 |
| 28711:4 | 8 | R.[+42.011]LSLLEEPGN[+2351.851]GTFTVILNQLTSR.D | HexNAc(4)Hex(5)Fuc(3)NeuAc(1) | .[42], N[+2352] | 1199.7991 | 4 | 4796.1746 | 4796.1539 | 0 | 4.3 | 413 | Specific | 295.05 | 295.05 | 295.05 | 6.94 | 73 | >sp\|P01833\|PIGR_HUMAN Polymeric immunoglobulin receptor OS=Homo sapiens OX=9606 GN=PIGR PE=1 SV=4 | 902 | 20201118_12812_1ng_glyco_hcd203040_compound15.32753.32753.2 | controllerType=0 controllerNumber=1 scan=32753 | 65.3527 |
| 28780:3 | 8 | R.[+42.011]LSLLEEPGN[+2351.851]GTFTVILNQLTSR.D | HexNAc(4)Hex(5)Fuc(3)NeuAc(1) | .[42], N[+2352] | 1599.3867 | 3 | 4796.1454 | 4796.1539 | 0 | -1.8 | 413 | Specific | 337.76 | 280.47 | 280.47 | 6.1 | 73 | >sp\|P01833\|PIGR_HUMAN Polymeric immunoglobulin receptor OS=Homo sapiens OX=9606 GN=PIGR PE=1 SV=4 | 902 | 20201118_12812_1ng_glyco_hcd203040_compound15.32825.32825.3 | controllerType=0 controllerNumber=1 scan=32825 | 65.4837 |
| 20320:4 | 8 | R.LSLLEEPGN[+2076.750]GTFTVILNQLTSR.D | HexNAc(4)Hex(6)Fuc(2) | N[+2077] | 1120.5227 | 4 | 4479.0692 | 4479.0428 | 0 | 5.9 | 413 | Specific | 338.9 | 320.7 | 320.7 | 6.72 | 73 | >sp\|P01833\|PIGR_HUMAN Polymeric immunoglobulin receptor OS=Homo sapiens OX=9606 GN=PIGR PE=1 SV=4 | 902 | 20201118_12812_1ng_glyco_hcd203040_compound15.24017.24017.4 | controllerType=0 controllerNumber=1 scan=24017 | 48.9175 |
| 22571:3 | 8 | R.[+42.011]LSLLEEPGN[+2336.851]GTFTVILNQLTSR.D | HexNAc(6)Hex(6)Fuc(1) | .[42], N[+2337] | 1594.3843 | 3 | 4781.1385 | 4781.1542 | 0 | -3.3 | 413 | Specific | 339.47 | 310.59 | 310.59 | 6 | 73 | >sp\|P01833\|PIGR_HUMAN Polymeric immunoglobulin receptor OS=Homo sapiens OX=9606 GN=PIGR PE=1 SV=4 | 902 | 20201118_12812_1ng_glyco_hcd203040_compound15.26363.26363.3 | controllerType=0 controllerNumber=1 scan=26363 | 53.5313 |
| 28782:3 | 8 | R.[+42.011]LSLLEEPGN[+2351.851]GTFTVILNQLTSR.D | HexNAc(4)Hex(5)Fuc(3)NeuAc(1) | .[42], N[+2352] | 1599.3867 | 3 | 4796.1454 | 4796.1539 | 0 | -1.8 | 413 | Specific | 341.87 | 305.13 | 305.13 | 6.42 | 73 | >sp\|P01833\|PIGR_HUMAN Polymeric immunoglobulin receptor OS=Homo sapiens OX=9606 GN=PIGR PE=1 SV=4 | 902 | 20201118_12812_1ng_glyco_hcd203040_compound15.32827.32827.3 | controllerType=0 controllerNumber=1 scan=32827 | 65.4872 |
| 22569:3 | 8 | R.[+42.011]LSLLEEPGN[+2336.851]GTFTVILNQLTSR.D | HexNAc(6)Hex(6)Fuc(1) | .[42], N[+2337] | 1594.3843 | 3 | 4781.1385 | 4781.1542 | 0 | -3.3 | 413 | Specific | 413.4 | 386.21 | 386.21 | 8.06 | 73 | >sp\|P01833\|PIGR_HUMAN Polymeric immunoglobulin receptor OS=Homo sapiens OX=9606 GN=PIGR PE=1 SV=4 | 902 | 20201118_12812_1ng_glyco_hcd203040_compound15.26361.26361.3 | controllerType=0 controllerNumber=1 scan=26361 | 53.5256 |
| 20325:4 | 8 | R.LSLLEEPGN[+2076.750]GTFTVILNQLTSR.D | HexNAc(4)Hex(6)Fuc(2) | N[+2077] | 1120.5227 | 4 | 4479.0692 | 4479.0428 | 0 | 5.9 | 413 | Specific | 428.4 | 333.32 | 333.32 | 7.15 | 73 | >sp\|P01833\|PIGR_HUMAN Polymeric immunoglobulin receptor OS=Homo sapiens OX=9606 GN=PIGR PE=1 SV=4 | 902 | 20201118_12812_1ng_glyco_hcd203040_compound15.24022.24022.4 | controllerType=0 controllerNumber=1 scan=24022 | 48.9248 |
| 20328:4 | 8 | R.LSLLEEPGN[+2076.750]GTFTVILNQLTSR.D | HexNAc(4)Hex(6)Fuc(2) | N[+2077] | 1120.5227 | 4 | 4479.0692 | 4479.0428 | 0 | 5.9 | 413 | Specific | 442.67 | 350.67 | 350.67 | 7.29 | 73 | >sp\|P01833\|PIGR_HUMAN Polymeric immunoglobulin receptor OS=Homo sapiens OX=9606 GN=PIGR PE=1 SV=4 | 902 | 20201118_12812_1ng_glyco_hcd203040_compound15.24025.24025.4 | controllerType=0 controllerNumber=1 scan=24025 | 48.9347 |
| 20330:3 | 8 | R.LSLLEEPGN[+2076.750]GTFTVILNQLTSR.D | HexNAc(4)Hex(6)Fuc(2) | N[+2077] | 1493.6955 | 3 | 4479.0719 | 4479.0428 | 0 | 6.5 | 413 | Specific | 523.06 | 458.99 | 458.99 | 9 | 73 | >sp\|P01833\|PIGR_HUMAN Polymeric immunoglobulin receptor OS=Homo sapiens OX=9606 GN=PIGR PE=1 SV=4 | 902 | 20201118_12812_1ng_glyco_hcd203040_compound15.24027.24027.3 | controllerType=0 controllerNumber=1 scan=24027 | 48.9378 |
| 22721:3 | 8 | R.LSLLEEPGN[+2351.851]GTFTVILNQLTSR.D | HexNAc(4)Hex(5)Fuc(3)NeuAc(1) | N[+2352] | 1585.3846 | 3 | 4754.1392 | 4754.1433 | 0 | -0.9 | 413 | Specific | 547 | 519.4 | 519.4 | 9.23 | 73 | >sp\|P01833\|PIGR_HUMAN Polymeric immunoglobulin receptor OS=Homo sapiens OX=9606 GN=PIGR PE=1 SV=4 | 902 | 20201118_12812_1ng_glyco_hcd203040_compound15.26519.26519.3 | controllerType=0 controllerNumber=1 scan=26519 | 53.8569 |
| 22723:3 | 8 | R.LSLLEEPGN[+2351.851]GTFTVILNQLTSR.D | HexNAc(4)Hex(5)Fuc(3)NeuAc(1) | N[+2352] | 1585.3846 | 3 | 4754.1392 | 4754.1433 | 0 | -0.9 | 413 | Specific | 550.39 | 550.39 | 550.39 | 9.23 | 73 | >sp\|P01833\|PIGR_HUMAN Polymeric immunoglobulin receptor OS=Homo sapiens OX=9606 GN=PIGR PE=1 SV=4 | 902 | 20201118_12812_1ng_glyco_hcd203040_compound15.26521.26521.3 | controllerType=0 controllerNumber=1 scan=26521 | 53.8611 |
| 20158:3 | 8 | R.LSLLEEPGN[+2570.925]GTFTVILNQLTSR.D | HexNAc(5)Hex(6)Fuc(2)NeuAc(1) | N[+2571] | 1658.4111 | 3 | 4973.2187 | 4973.2176 | 0 | 0.2 | 413 | Specific | 627.34 | 340.33 | 340.33 | 10.71 | 73 | >sp\|P01833\|PIGR_HUMAN Polymeric immunoglobulin receptor OS=Homo sapiens OX=9606 GN=PIGR PE=1 SV=4 | 902 | 20201118_12812_1ng_glyco_hcd203040_compound15.23846.23846.3 | controllerType=0 controllerNumber=1 scan=23846 | 48.5241 |
| 20156:3 | 8 | R.LSLLEEPGN[+2570.925]GTFTVILNQLTSR.D | HexNAc(5)Hex(6)Fuc(2)NeuAc(1) | N[+2571] | 1658.4111 | 3 | 4973.2187 | 4973.2176 | 0 | 0.2 | 413 | Specific | 679.81 | 467.4 | 467.4 | 13.35 | 73 | >sp\|P01833\|PIGR_HUMAN Polymeric immunoglobulin receptor OS=Homo sapiens OX=9606 GN=PIGR PE=1 SV=4 | 902 | 20201118_12812_1ng_glyco_hcd203040_compound15.23844.23844.3 | controllerType=0 controllerNumber=1 scan=23844 | 48.5181 |
| 20322:4 | 8 | R.LSLLEEPGN[+2076.750]GTFTVILNQLTSR.D | HexNAc(4)Hex(6)Fuc(2) | N[+2077] | 1120.5227 | 4 | 4479.0692 | 4479.0428 | 0 | 5.9 | 413 | Specific | 712.33 | 570.71 | 570.71 | 13.79 | 73 | >sp\|P01833\|PIGR_HUMAN Polymeric immunoglobulin receptor OS=Homo sapiens OX=9606 GN=PIGR PE=1 SV=4 | 902 | 20201118_12812_1ng_glyco_hcd203040_compound15.24019.24019.4 | controllerType=0 controllerNumber=1 scan=24019 | 48.9205 |
| 20399:3 | 8 | R.LSLLEEPGN[+2059.735]GTFTVILNQLTSR.D | HexNAc(4)Hex(5)Fuc(1)NeuAc(1) | N[+2060] | 1488.0142 | 3 | 4462.0281 | 4462.0275 | 0 | 0.1 | 413 | Specific | 768.73 | 690.17 | 690.17 | 15.35 | 73 | >sp\|P01833\|PIGR_HUMAN Polymeric immunoglobulin receptor OS=Homo sapiens OX=9606 GN=PIGR PE=1 SV=4 | 902 | 20201118_12812_1ng_glyco_hcd203040_compound15.24099.24099.3 | controllerType=0 controllerNumber=1 scan=24099 | 49.0969 |
| 20401:3 | 8 | R.LSLLEEPGN[+2059.735]GTFTVILNQLTSR.D | HexNAc(4)Hex(5)Fuc(1)NeuAc(1) | N[+2060] | 1488.0142 | 3 | 4462.0281 | 4462.0275 | 0 | 0.1 | 413 | Specific | 856.66 | 588.52 | 588.52 | 17.35 | 73 | >sp\|P01833\|PIGR_HUMAN Polymeric immunoglobulin receptor OS=Homo sapiens OX=9606 GN=PIGR PE=1 SV=4 | 902 | 20201118_12812_1ng_glyco_hcd203040_compound15.24101.24101.3 | controllerType=0 controllerNumber=1 scan=24101 | 49.1007 |
| 22455:3 | 8 | R.LSLLEEPGN[+2350.830]GTFTVILNQLTSR.D | HexNAc(4)Hex(5)Fuc(1)NeuAc(2) | N[+2351] | 1585.0501 | 3 | 4753.1358 | 4753.1229 | 0 | 2.7 | 413 | Specific | 938.46 | 668.87 | 668.87 | 18.68 | 73 | >sp\|P01833\|PIGR_HUMAN Polymeric immunoglobulin receptor OS=Homo sapiens OX=9606 GN=PIGR PE=1 SV=4 | 902 | 20201118_12812_1ng_glyco_hcd203040_compound15.26242.26242.3 | controllerType=0 controllerNumber=1 scan=26242 | 53.2615 |
| 22458:4 | 8 | R.LSLLEEPGN[+2350.830]GTFTVILNQLTSR.D | HexNAc(4)Hex(5)Fuc(1)NeuAc(2) | N[+2351] | 1189.04 | 4 | 4753.138 | 4753.1229 | 0 | 3.2 | 413 | Specific | 960.49 | 746.35 | 746.35 | 19.7 | 73 | >sp\|P01833\|PIGR_HUMAN Polymeric immunoglobulin receptor OS=Homo sapiens OX=9606 GN=PIGR PE=1 SV=4 | 902 | 20201118_12812_1ng_glyco_hcd203040_compound15.26245.26245.4 | controllerType=0 controllerNumber=1 scan=26245 | 53.2673 |
| 22456:4 | 8 | R.LSLLEEPGN[+2350.830]GTFTVILNQLTSR.D | HexNAc(4)Hex(5)Fuc(1)NeuAc(2) | N[+2351] | 1189.04 | 4 | 4753.138 | 4753.1229 | 0 | 3.2 | 413 | Specific | 989.13 | 768.61 | 768.61 | 19.7 | 73 | >sp\|P01833\|PIGR_HUMAN Polymeric immunoglobulin receptor OS=Homo sapiens OX=9606 GN=PIGR PE=1 SV=4 | 902 | 20201118_12812_1ng_glyco_hcd203040_compound15.26243.26243.4 | controllerType=0 controllerNumber=1 scan=26243 | 53.2624 |
| 22453:3 | 8 | R.LSLLEEPGN[+2350.830]GTFTVILNQLTSR.D | HexNAc(4)Hex(5)Fuc(1)NeuAc(2) | N[+2351] | 1585.0501 | 3 | 4753.1358 | 4753.1229 | 0 | 2.7 | 413 | Specific | 1046.4 | 991.92 | 991.92 | 20.68 | 73 | >sp\|P01833\|PIGR_HUMAN Polymeric immunoglobulin receptor OS=Homo sapiens OX=9606 GN=PIGR PE=1 SV=4 | 902 | 20201118_12812_1ng_glyco_hcd203040_compound15.26240.26240.3 | controllerType=0 controllerNumber=1 scan=26240 | 53.2565 |
| 15947:3 | 8 | K.IIEGEPNLKVPGN[+1913.677]VTAVLGETLK.V | HexNAc(4)Hex(5)NeuAc(1) | N[+1914] | 1435.684 | 3 | 4305.0375 | 4305.0264 | 0 | 2.6 | 457 | Specific | 266.4 | 266.4 | 266.4 | 7.4 | 73 | >sp\|P01833\|PIGR_HUMAN Polymeric immunoglobulin receptor OS=Homo sapiens OX=9606 GN=PIGR PE=1 SV=4 | 902 | 20201118_12812_1ng_glyco_hcd203040_compound15.19449.19449.3 | controllerType=0 controllerNumber=1 scan=19449 | 39.3438 |
| 15945:3 | 8 | K.IIEGEPNLKVPGN[+1913.677]VTAVLGETLK.V | HexNAc(4)Hex(5)NeuAc(1) | N[+1914] | 1435.684 | 3 | 4305.0375 | 4305.0264 | 0 | 2.6 | 457 | Specific | 378.66 | 378.66 | 378.66 | 9.99 | 73 | >sp\|P01833\|PIGR_HUMAN Polymeric immunoglobulin receptor OS=Homo sapiens OX=9606 GN=PIGR PE=1 SV=4 | 902 | 20201118_12812_1ng_glyco_hcd203040_compound15.19447.19447.3 | controllerType=0 controllerNumber=1 scan=19447 | 39.3391 |
| 15887:4 | 8 | K.IIEGEPNLKVPGN[+2205.793]VTAVLGETLK.V | HexNAc(4)Hex(5)Fuc(2)NeuAc(1) | N[+2206] | 1150.0441 | 4 | 4597.1546 | 4597.1422 | 0 | 2.7 | 457 | Specific | 641.71 | 605.42 | 605.42 | 13.15 | 73 | >sp\|P01833\|PIGR_HUMAN Polymeric immunoglobulin receptor OS=Homo sapiens OX=9606 GN=PIGR PE=1 SV=4 | 902 | 20201118_12812_1ng_glyco_hcd203040_compound15.19387.19387.2 | controllerType=0 controllerNumber=1 scan=19387 | 39.206 |
| 15885:4 | 8 | K.IIEGEPNLKVPGN[+2205.793]VTAVLGETLK.V | HexNAc(4)Hex(5)Fuc(2)NeuAc(1) | N[+2206] | 1150.0441 | 4 | 4597.1546 | 4597.1422 | 0 | 2.7 | 457 | Specific | 673.76 | 629.95 | 629.95 | 14.15 | 73 | >sp\|P01833\|PIGR_HUMAN Polymeric immunoglobulin receptor OS=Homo sapiens OX=9606 GN=PIGR PE=1 SV=4 | 902 | 20201118_12812_1ng_glyco_hcd203040_compound15.19385.19385.2 | controllerType=0 controllerNumber=1 scan=19385 | 39.2013 |
| 17781:4 | 8 | K.IIEGEPNLKVPGN[+2350.830]VTAVLGETLK.V | HexNAc(4)Hex(5)Fuc(1)NeuAc(2) | N[+2351] | 1186.3031 | 4 | 4742.1907 | 4742.1797 | 0 | 2.3 | 457 | Specific | 758.64 | 708.8 | 708.8 | 16.05 | 73 | >sp\|P01833\|PIGR_HUMAN Polymeric immunoglobulin receptor OS=Homo sapiens OX=9606 GN=PIGR PE=1 SV=4 | 902 | 20201118_12812_1ng_glyco_hcd203040_compound15.21366.21366.4 | controllerType=0 controllerNumber=1 scan=21366 | 43.209 |
| 15908:3 | 8 | K.IIEGEPNLKVPGN[+2205.793]VTAVLGETLK.V | HexNAc(4)Hex(5)Fuc(2)NeuAc(1) | N[+2206] | 1533.0571 | 3 | 4597.1566 | 4597.1422 | 0 | 3.1 | 457 | Specific | 806.6 | 743.54 | 743.54 | 17.22 | 73 | >sp\|P01833\|PIGR_HUMAN Polymeric immunoglobulin receptor OS=Homo sapiens OX=9606 GN=PIGR PE=1 SV=4 | 902 | 20201118_12812_1ng_glyco_hcd203040_compound15.19409.19409.3 | controllerType=0 controllerNumber=1 scan=19409 | 39.2513 |
| 15910:3 | 8 | K.IIEGEPNLKVPGN[+2205.793]VTAVLGETLK.V | HexNAc(4)Hex(5)Fuc(2)NeuAc(1) | N[+2206] | 1533.0571 | 3 | 4597.1566 | 4597.1422 | 0 | 3.1 | 457 | Specific | 815.26 | 815.26 | 815.26 | 17.22 | 73 | >sp\|P01833\|PIGR_HUMAN Polymeric immunoglobulin receptor OS=Homo sapiens OX=9606 GN=PIGR PE=1 SV=4 | 902 | 20201118_12812_1ng_glyco_hcd203040_compound15.19411.19411.3 | controllerType=0 controllerNumber=1 scan=19411 | 39.2557 |
| 15888:3 | 8 | K.IIEGEPNLKVPGN[+2059.735]VTAVLGETLK.V | HexNAc(4)Hex(5)Fuc(1)NeuAc(1) | N[+2060] | 1484.3693 | 3 | 4451.0934 | 4451.0843 | 0 | 2 | 457 | Specific | 819.67 | 720.32 | 720.32 | 17.12 | 73 | >sp\|P01833\|PIGR_HUMAN Polymeric immunoglobulin receptor OS=Homo sapiens OX=9606 GN=PIGR PE=1 SV=4 | 902 | 20201118_12812_1ng_glyco_hcd203040_compound15.19388.19388.3 | controllerType=0 controllerNumber=1 scan=19388 | 39.2088 |
| 17784:4 | 8 | K.IIEGEPNLKVPGN[+2350.830]VTAVLGETLK.V | HexNAc(4)Hex(5)Fuc(1)NeuAc(2) | N[+2351] | 1186.3031 | 4 | 4742.1907 | 4742.1797 | 0 | 2.3 | 457 | Specific | 846.6 | 758.33 | 758.33 | 17.05 | 73 | >sp\|P01833\|PIGR_HUMAN Polymeric immunoglobulin receptor OS=Homo sapiens OX=9606 GN=PIGR PE=1 SV=4 | 902 | 20201118_12812_1ng_glyco_hcd203040_compound15.21369.21369.4 | controllerType=0 controllerNumber=1 scan=21369 | 43.2134 |
| 15914:3 | 8 | K.IIEGEPNLKVPGN[+2059.735]VTAVLGETLK.V | HexNAc(4)Hex(5)Fuc(1)NeuAc(1) | N[+2060] | 1484.3712 | 3 | 4451.0989 | 4451.0843 | 0 | 3.3 | 457 | Specific | 863.07 | 763.89 | 763.89 | 18.22 | 73 | >sp\|P01833\|PIGR_HUMAN Polymeric immunoglobulin receptor OS=Homo sapiens OX=9606 GN=PIGR PE=1 SV=4 | 902 | 20201118_12812_1ng_glyco_hcd203040_compound15.19415.19415.3 | controllerType=0 controllerNumber=1 scan=19415 | 39.2698 |
| 15886:3 | 8 | K.IIEGEPNLKVPGN[+2059.735]VTAVLGETLK.V | HexNAc(4)Hex(5)Fuc(1)NeuAc(1) | N[+2060] | 1484.3693 | 3 | 4451.0934 | 4451.0843 | 0 | 2 | 457 | Specific | 1018.3 | 911.76 | 911.76 | 21.12 | 73 | >sp\|P01833\|PIGR_HUMAN Polymeric immunoglobulin receptor OS=Homo sapiens OX=9606 GN=PIGR PE=1 SV=4 | 902 | 20201118_12812_1ng_glyco_hcd203040_compound15.19386.19386.3 | controllerType=0 controllerNumber=1 scan=19386 | 39.203 |
| 15912:3 | 8 | K.IIEGEPNLKVPGN[+2059.735]VTAVLGETLK.V | HexNAc(4)Hex(5)Fuc(1)NeuAc(1) | N[+2060] | 1484.3712 | 3 | 4451.0989 | 4451.0843 | 0 | 3.3 | 457 | Specific | 1068.36 | 991.21 | 991.21 | 22.22 | 73 | >sp\|P01833\|PIGR_HUMAN Polymeric immunoglobulin receptor OS=Homo sapiens OX=9606 GN=PIGR PE=1 SV=4 | 902 | 20201118_12812_1ng_glyco_hcd203040_compound15.19413.19413.3 | controllerType=0 controllerNumber=1 scan=19413 | 39.2586 |
| 16903:4 | 8 | K.IIEGEPNLKVPGN[+2350.830]VTAVLGETLKVPC[+57.021]HFPC[+57.021]K.F | HexNAc(4)Hex(5)Fuc(1)NeuAc(2) | C[+57]*2, N[+2351] | 1442.6685 | 4 | 5767.652 | 5767.6372 | 0 | 2.6 | 457 | Specific | 747.89 | 602.07 | 602.07 | 13.39 | 73 | >sp\|P01833\|PIGR_HUMAN Polymeric immunoglobulin receptor OS=Homo sapiens OX=9606 GN=PIGR PE=1 SV=4 | 902 | 20201118_12812_1ng_glyco_hcd203040_compound15.20448.20448.4 | controllerType=0 controllerNumber=1 scan=20448 | 41.2118 |
| 15423:4 | 8 | K.IIEGEPNLKVPGN[+2205.793]VTAVLGETLKVPC[+57.021]HFPC[+57.021]K.F | HexNAc(4)Hex(5)Fuc(2)NeuAc(1) | C[+57]*2, N[+2206] | 1406.4095 | 4 | 5622.6164 | 5622.5997 | 0 | 3 | 457 | Specific | 762.33 | 686.9 | 686.9 | 14.39 | 73 | >sp\|P01833\|PIGR_HUMAN Polymeric immunoglobulin receptor OS=Homo sapiens OX=9606 GN=PIGR PE=1 SV=4 | 902 | 20201118_12812_1ng_glyco_hcd203040_compound15.18883.18883.4 | controllerType=0 controllerNumber=1 scan=18883 | 38.2959 |
| 15492:4 | 8 | K.IIEGEPNLKVPGN[+2059.735]VTAVLGETLKVPC[+57.021]HFPC[+57.021]K.F | HexNAc(4)Hex(5)Fuc(1)NeuAc(1) | C[+57]*2, N[+2060] | 1369.897 | 4 | 5476.5661 | 5476.5418 | 0 | 4.4 | 457 | Specific | 783.4 | 758.49 | 758.49 | 14.12 | 73 | >sp\|P01833\|PIGR_HUMAN Polymeric immunoglobulin receptor OS=Homo sapiens OX=9606 GN=PIGR PE=1 SV=4 | 902 | 20201118_12812_1ng_glyco_hcd203040_compound15.18955.18955.4 | controllerType=0 controllerNumber=1 scan=18955 | 38.4347 |
| 15389:5 | 8 | K.IIEGEPNLKVPGN[+2205.793]VTAVLGETLKVPC[+57.021]HFPC[+57.021]K.F | HexNAc(4)Hex(5)Fuc(2)NeuAc(1) | C[+57]*2, N[+2206] | 1125.3348 | 5 | 5622.6451 | 5622.5997 | 0 | 8.1 | 457 | Specific | 848.13 | 848.13 | 848.13 | 14.41 | 73 | >sp\|P01833\|PIGR_HUMAN Polymeric immunoglobulin receptor OS=Homo sapiens OX=9606 GN=PIGR PE=1 SV=4 | 902 | 20201118_12812_1ng_glyco_hcd203040_compound15.18848.18848.5 | controllerType=0 controllerNumber=1 scan=18848 | 38.223 |
| 16901:4 | 8 | K.IIEGEPNLKVPGN[+2350.830]VTAVLGETLKVPC[+57.021]HFPC[+57.021]K.F | HexNAc(4)Hex(5)Fuc(1)NeuAc(2) | C[+57]*2, N[+2351] | 1442.6685 | 4 | 5767.652 | 5767.6372 | 0 | 2.6 | 457 | Specific | 873.05 | 666.02 | 666.02 | 16.39 | 73 | >sp\|P01833\|PIGR_HUMAN Polymeric immunoglobulin receptor OS=Homo sapiens OX=9606 GN=PIGR PE=1 SV=4 | 902 | 20201118_12812_1ng_glyco_hcd203040_compound15.20446.20446.4 | controllerType=0 controllerNumber=1 scan=20446 | 41.2061 |
| 15399:5 | 8 | K.IIEGEPNLKVPGN[+2059.735]VTAVLGETLKVPC[+57.021]HFPC[+57.021]K.F | HexNAc(4)Hex(5)Fuc(1)NeuAc(1) | C[+57]*2, N[+2060] | 1096.1239 | 5 | 5476.5904 | 5476.5418 | 0 | 8.9 | 457 | Specific | 876.67 | 793.47 | 793.47 | 15.06 | 73 | >sp\|P01833\|PIGR_HUMAN Polymeric immunoglobulin receptor OS=Homo sapiens OX=9606 GN=PIGR PE=1 SV=4 | 902 | 20201118_12812_1ng_glyco_hcd203040_compound15.18858.18858.5 | controllerType=0 controllerNumber=1 scan=18858 | 38.2389 |
| 15411:5 | 8 | K.IIEGEPNLKVPGN[+2059.735]VTAVLGETLKVPC[+57.021]HFPC[+57.021]K.F | HexNAc(4)Hex(5)Fuc(1)NeuAc(1) | C[+57]*2, N[+2060] | 1096.1168 | 5 | 5476.555 | 5476.5418 | 0 | 2.4 | 457 | Specific | 880.54 | 772.06 | 772.06 | 16.92 | 73 | >sp\|P01833\|PIGR_HUMAN Polymeric immunoglobulin receptor OS=Homo sapiens OX=9606 GN=PIGR PE=1 SV=4 | 902 | 20201118_12812_1ng_glyco_hcd203040_compound15.18871.18871.5 | controllerType=0 controllerNumber=1 scan=18871 | 38.269 |
| 16902:5 | 8 | K.IIEGEPNLKVPGN[+2350.830]VTAVLGETLKVPC[+57.021]HFPC[+57.021]K.F | HexNAc(4)Hex(5)Fuc(1)NeuAc(2) | C[+57]*2, N[+2351] | 1154.3357 | 5 | 5767.6494 | 5767.6372 | 0 | 2.1 | 457 | Specific | 884.52 | 810.1 | 810.1 | 16.92 | 73 | >sp\|P01833\|PIGR_HUMAN Polymeric immunoglobulin receptor OS=Homo sapiens OX=9606 GN=PIGR PE=1 SV=4 | 902 | 20201118_12812_1ng_glyco_hcd203040_compound15.20447.20447.5 | controllerType=0 controllerNumber=1 scan=20447 | 41.2098 |
| 15396:5 | 8 | K.IIEGEPNLKVPGN[+2059.735]VTAVLGETLKVPC[+57.021]HFPC[+57.021]K.F | HexNAc(4)Hex(5)Fuc(1)NeuAc(1) | C[+57]*2, N[+2060] | 1096.1239 | 5 | 5476.5904 | 5476.5418 | 0 | 8.9 | 457 | Specific | 941.52 | 819.58 | 819.58 | 16.06 | 73 | >sp\|P01833\|PIGR_HUMAN Polymeric immunoglobulin receptor OS=Homo sapiens OX=9606 GN=PIGR PE=1 SV=4 | 902 | 20201118_12812_1ng_glyco_hcd203040_compound15.18855.18855.5 | controllerType=0 controllerNumber=1 scan=18855 | 38.2329 |
| 15366:5 | 8 | K.IIEGEPNLKVPGN[+2205.793]VTAVLGETLKVPC[+57.021]HFPC[+57.021]K.F | HexNAc(4)Hex(5)Fuc(2)NeuAc(1) | C[+57]*2, N[+2206] | 1125.3285 | 5 | 5622.6134 | 5622.5997 | 0 | 2.4 | 457 | Specific | 950.1 | 892.04 | 892.04 | 18.92 | 73 | >sp\|P01833\|PIGR_HUMAN Polymeric immunoglobulin receptor OS=Homo sapiens OX=9606 GN=PIGR PE=1 SV=4 | 902 | 20201118_12812_1ng_glyco_hcd203040_compound15.18824.18824.5 | controllerType=0 controllerNumber=1 scan=18824 | 38.1749 |
| 15489:4 | 8 | K.IIEGEPNLKVPGN[+2059.735]VTAVLGETLKVPC[+57.021]HFPC[+57.021]K.F | HexNAc(4)Hex(5)Fuc(1)NeuAc(1) | C[+57]*2, N[+2060] | 1369.897 | 4 | 5476.5661 | 5476.5418 | 0 | 4.4 | 457 | Specific | 957.06 | 837.83 | 837.83 | 18.12 | 73 | >sp\|P01833\|PIGR_HUMAN Polymeric immunoglobulin receptor OS=Homo sapiens OX=9606 GN=PIGR PE=1 SV=4 | 902 | 20201118_12812_1ng_glyco_hcd203040_compound15.18952.18952.4 | controllerType=0 controllerNumber=1 scan=18952 | 38.4294 |
| 16900:5 | 8 | K.IIEGEPNLKVPGN[+2350.830]VTAVLGETLKVPC[+57.021]HFPC[+57.021]K.F | HexNAc(4)Hex(5)Fuc(1)NeuAc(2) | C[+57]*2, N[+2351] | 1154.3357 | 5 | 5767.6494 | 5767.6372 | 0 | 2.1 | 457 | Specific | 978.2 | 744.6 | 744.6 | 18.92 | 73 | >sp\|P01833\|PIGR_HUMAN Polymeric immunoglobulin receptor OS=Homo sapiens OX=9606 GN=PIGR PE=1 SV=4 | 902 | 20201118_12812_1ng_glyco_hcd203040_compound15.20445.20445.5 | controllerType=0 controllerNumber=1 scan=20445 | 41.2046 |
| 15363:5 | 8 | K.IIEGEPNLKVPGN[+2205.793]VTAVLGETLKVPC[+57.021]HFPC[+57.021]K.F | HexNAc(4)Hex(5)Fuc(2)NeuAc(1) | C[+57]*2, N[+2206] | 1125.3285 | 5 | 5622.6134 | 5622.5997 | 0 | 2.4 | 457 | Specific | 1052.6 | 928.71 | 928.71 | 20.92 | 73 | >sp\|P01833\|PIGR_HUMAN Polymeric immunoglobulin receptor OS=Homo sapiens OX=9606 GN=PIGR PE=1 SV=4 | 902 | 20201118_12812_1ng_glyco_hcd203040_compound15.18821.18821.5 | controllerType=0 controllerNumber=1 scan=18821 | 38.1674 |
| 15409:5 | 8 | K.IIEGEPNLKVPGN[+2059.735]VTAVLGETLKVPC[+57.021]HFPC[+57.021]K.F | HexNAc(4)Hex(5)Fuc(1)NeuAc(1) | C[+57]*2, N[+2060] | 1096.1168 | 5 | 5476.555 | 5476.5418 | 0 | 2.4 | 457 | Specific | 1054.7 | 916.31 | 916.31 | 20.92 | 73 | >sp\|P01833\|PIGR_HUMAN Polymeric immunoglobulin receptor OS=Homo sapiens OX=9606 GN=PIGR PE=1 SV=4 | 902 | 20201118_12812_1ng_glyco_hcd203040_compound15.18869.18869.5 | controllerType=0 controllerNumber=1 scan=18869 | 38.2613 |
| 15387:5 | 8 | K.IIEGEPNLKVPGN[+2205.793]VTAVLGETLKVPC[+57.021]HFPC[+57.021]K.F | HexNAc(4)Hex(5)Fuc(2)NeuAc(1) | C[+57]*2, N[+2206] | 1125.3348 | 5 | 5622.6451 | 5622.5997 | 0 | 8.1 | 457 | Specific | 1061.36 | 1061.36 | 1061.36 | 19.41 | 73 | >sp\|P01833\|PIGR_HUMAN Polymeric immunoglobulin receptor OS=Homo sapiens OX=9606 GN=PIGR PE=1 SV=4 | 902 | 20201118_12812_1ng_glyco_hcd203040_compound15.18846.18846.5 | controllerType=0 controllerNumber=1 scan=18846 | 38.2133 |
| 15421:4 | 8 | K.IIEGEPNLKVPGN[+2205.793]VTAVLGETLKVPC[+57.021]HFPC[+57.021]K.F | HexNAc(4)Hex(5)Fuc(2)NeuAc(1) | C[+57]*2, N[+2206] | 1406.4095 | 4 | 5622.6164 | 5622.5997 | 0 | 3 | 457 | Specific | 1074.58 | 970.49 | 970.49 | 20.39 | 73 | >sp\|P01833\|PIGR_HUMAN Polymeric immunoglobulin receptor OS=Homo sapiens OX=9606 GN=PIGR PE=1 SV=4 | 902 | 20201118_12812_1ng_glyco_hcd203040_compound15.18881.18881.4 | controllerType=0 controllerNumber=1 scan=18881 | 38.2908 |
| 16529:3 | 8 | K.VPGN[+2350.830]VTAVLGETLK.V | HexNAc(4)Hex(5)Fuc(1)NeuAc(2) | N[+2351] | 1250.2185 | 3 | 3748.6409 | 3748.6302 | 0 | 2.9 | 466 | Specific | 402.22 | 198.83 | 198.83 | 6.89 | 73 | >sp\|P01833\|PIGR_HUMAN Polymeric immunoglobulin receptor OS=Homo sapiens OX=9606 GN=PIGR PE=1 SV=4 | 902 | 20201118_12812_1ng_glyco_hcd203040_compound15.20059.20059.3 | controllerType=0 controllerNumber=1 scan=20059 | 40.4288 |
| 13558:3 | 8 | K.VPGN[+2059.735]VTAVLGETLK.V | HexNAc(4)Hex(5)Fuc(1)NeuAc(1) | N[+2060] | 1153.1868 | 3 | 3457.5459 | 3457.5348 | 0 | 3.2 | 466 | Specific | 439.82 | 277.79 | 277.79 | 7.57 | 73 | >sp\|P01833\|PIGR_HUMAN Polymeric immunoglobulin receptor OS=Homo sapiens OX=9606 GN=PIGR PE=1 SV=4 | 902 | 20201118_12812_1ng_glyco_hcd203040_compound15.16950.16950.3 | controllerType=0 controllerNumber=1 scan=16950 | 34.7738 |
| 16531:3 | 8 | K.VPGN[+2350.830]VTAVLGETLK.V | HexNAc(4)Hex(5)Fuc(1)NeuAc(2) | N[+2351] | 1250.2185 | 3 | 3748.6409 | 3748.6302 | 0 | 2.9 | 466 | Specific | 535.34 | 334.9 | 334.9 | 8.45 | 73 | >sp\|P01833\|PIGR_HUMAN Polymeric immunoglobulin receptor OS=Homo sapiens OX=9606 GN=PIGR PE=1 SV=4 | 902 | 20201118_12812_1ng_glyco_hcd203040_compound15.20061.20061.3 | controllerType=0 controllerNumber=1 scan=20061 | 40.4325 |
| 13374:3 | 8 | K.VPGN[+2205.793]VTAVLGETLK.V | HexNAc(4)Hex(5)Fuc(2)NeuAc(1) | N[+2206] | 1201.8751 | 3 | 3603.6106 | 3603.5927 | 0 | 5 | 466 | Specific | 659.62 | 636.66 | 636.66 | 13.24 | 73 | >sp\|P01833\|PIGR_HUMAN Polymeric immunoglobulin receptor OS=Homo sapiens OX=9606 GN=PIGR PE=1 SV=4 | 902 | 20201118_12812_1ng_glyco_hcd203040_compound15.16759.16759.3 | controllerType=0 controllerNumber=1 scan=16759 | 34.397 |
| 13376:3 | 8 | K.VPGN[+2205.793]VTAVLGETLK.V | HexNAc(4)Hex(5)Fuc(2)NeuAc(1) | N[+2206] | 1201.8751 | 3 | 3603.6106 | 3603.5927 | 0 | 5 | 466 | Specific | 772.63 | 485.85 | 485.85 | 15.24 | 73 | >sp\|P01833\|PIGR_HUMAN Polymeric immunoglobulin receptor OS=Homo sapiens OX=9606 GN=PIGR PE=1 SV=4 | 902 | 20201118_12812_1ng_glyco_hcd203040_compound15.16761.16761.3 | controllerType=0 controllerNumber=1 scan=16761 | 34.4011 |

Table S4. Cont.

| Query #:z | Protein Rank | Glycopeptide | Glycan | Modification Type(s) | Observed m/z | z | Observed (M+H) | Calc. mass (M+H) | Off-by-x error | Mass error (ppm) | StartPosition | Cleavage | Score | Delta | Delta Mod | \|Log Prob\| | # of unique peptides | Protein Name | Protein DB number | Comment | Scan # | Scan Time |
|---|---|---|---|---|---|---|---|---|---|---|---|---|---|---|---|---|---|---|---|---|---|---|
| 13545:3 | 8 | K.VPGN[+2059.735]VTAVLGETLKVPC[+57.021]HFPC[+57.021]K.F | HexNAc(4)Hex(5)Fuc(1)NeuAc(1) | C[+57]*2, N[+2060] | 1495.0098 | 3 | 4483.0149 | 4482.9923 | 0 | 5 | 466 | Specific | 524.91 | 495.72 | 495.72 | 8.17 | 73 | >sp\|P01833\|PIGR_HUMAN Polymeric immunoglobulin receptor OS=Homo sapiens OX=9606 GN=PIGR PE=1 SV=4 | 902 | 20201118_12812_1ng_glyco_hcd203040_compound15.16936.16936.3 | controllerType=0 controllerNumber=1 scan=16936 | 34.7487 |
| 15427:3 | 8 | K.VPGN[+2350.830]VTAVLGETLKVPC[+57.021]HFPC[+57.021]K.F | HexNAc(4)Hex(5)Fuc(1)NeuAc(2) | C[+57]*2, N[+2351] | 1592.0427 | 3 | 4774.1134 | 4774.0877 | 0 | 5.4 | 466 | Specific | 539.35 | 448.35 | 448.35 | 8.31 | 73 | >sp\|P01833\|PIGR_HUMAN Polymeric immunoglobulin receptor OS=Homo sapiens OX=9606 GN=PIGR PE=1 SV=4 | 902 | 20201118_12812_1ng_glyco_hcd203040_compound15.18887.18887.3 | controllerType=0 controllerNumber=1 scan=18887 | 38.3035 |
| 13547:3 | 8 | K.VPGN[+2059.735]VTAVLGETLKVPC[+57.021]HFPC[+57.021]K.F | HexNAc(4)Hex(5)Fuc(1)NeuAc(1) | C[+57]*2, N[+2060] | 1495.0098 | 3 | 4483.0149 | 4482.9923 | 0 | 5 | 466 | Specific | 604.19 | 587.73 | 587.73 | 10.73 | 73 | >sp\|P01833\|PIGR_HUMAN Polymeric immunoglobulin receptor OS=Homo sapiens OX=9606 GN=PIGR PE=1 SV=4 | 902 | 20201118_12812_1ng_glyco_hcd203040_compound15.16938.16938.3 | controllerType=0 controllerNumber=1 scan=16938 | 34.7543 |
| 15425:3 | 8 | K.VPGN[+2350.830]VTAVLGETLKVPC[+57.021]HFPC[+57.021]K.F | HexNAc(4)Hex(5)Fuc(1)NeuAc(2) | C[+57]*2, N[+2351] | 1592.0427 | 3 | 4774.1134 | 4774.0877 | 0 | 5.4 | 466 | Specific | 609.51 | 550.58 | 550.58 | 10.73 | 73 | >sp\|P01833\|PIGR_HUMAN Polymeric immunoglobulin receptor OS=Homo sapiens OX=9606 GN=PIGR PE=1 SV=4 | 902 | 20201118_12812_1ng_glyco_hcd203040_compound15.18885.18885.3 | controllerType=0 controllerNumber=1 scan=18885 | 38.2997 |
| 13500:4 | 8 | K.VPGN[+2205.793]VTAVLGETLKVPC[+57.021]HFPC[+57.021]K.F | HexNAc(4)Hex(5)Fuc(2)NeuAc(1) | C[+57]*2, N[+2206] | 1158.0212 | 4 | 4629.0628 | 4629.0502 | 0 | 2.7 | 466 | Specific | 659.96 | 603.79 | 603.79 | 13.27 | 73 | >sp\|P01833\|PIGR_HUMAN Polymeric immunoglobulin receptor OS=Homo sapiens OX=9606 GN=PIGR PE=1 SV=4 | 902 | 20201118_12812_1ng_glyco_hcd203040_compound15.16889.16889.3 | controllerType=0 controllerNumber=1 scan=16889 | 34.6483 |
| 13498:4 | 8 | K.VPGN[+2205.793]VTAVLGETLKVPC[+57.021]HFPC[+57.021]K.F | HexNAc(4)Hex(5)Fuc(2)NeuAc(1) | C[+57]*2, N[+2206] | 1158.0212 | 4 | 4629.0628 | 4629.0502 | 0 | 2.7 | 466 | Specific | 673.1 | 545.19 | 545.19 | 13.27 | 73 | >sp\|P01833\|PIGR_HUMAN Polymeric immunoglobulin receptor OS=Homo sapiens OX=9606 GN=PIGR PE=1 SV=4 | 902 | 20201118_12812_1ng_glyco_hcd203040_compound15.16887.16887.3 | controllerType=0 controllerNumber=1 scan=16887 | 34.6444 |
| 15416:4 | 8 | K.VPGN[+2350.830]VTAVLGETLKVPC[+57.021]HFPC[+57.021]K.F | HexNAc(4)Hex(5)Fuc(1)NeuAc(2) | C[+57]*2, N[+2351] | 1194.2826 | 4 | 4774.1087 | 4774.0877 | 0 | 4.4 | 466 | Specific | 792.49 | 694.59 | 694.59 | 15.01 | 73 | >sp\|P01833\|PIGR_HUMAN Polymeric immunoglobulin receptor OS=Homo sapiens OX=9606 GN=PIGR PE=1 SV=4 | 902 | 20201118_12812_1ng_glyco_hcd203040_compound15.18876.18876.4 | controllerType=0 controllerNumber=1 scan=18876 | 38.2817 |
| 13526:4 | 8 | K.VPGN[+2059.735]VTAVLGETLKVPC[+57.021]HFPC[+57.021]K.F | HexNAc(4)Hex(5)Fuc(1)NeuAc(1) | C[+57]*2, N[+2060] | 1121.5066 | 4 | 4483.0047 | 4482.9923 | 0 | 2.8 | 466 | Specific | 807.75 | 723.65 | 723.65 | 16.27 | 73 | >sp\|P01833\|PIGR_HUMAN Polymeric immunoglobulin receptor OS=Homo sapiens OX=9606 GN=PIGR PE=1 SV=4 | 902 | 20201118_12812_1ng_glyco_hcd203040_compound15.16917.16917.4 | controllerType=0 controllerNumber=1 scan=16917 | 34.7128 |
| 13529:4 | 8 | K.VPGN[+2059.735]VTAVLGETLKVPC[+57.021]HFPC[+57.021]K.F | HexNAc(4)Hex(5)Fuc(1)NeuAc(1) | C[+57]*2, N[+2060] | 1121.5066 | 4 | 4483.0047 | 4482.9923 | 0 | 2.8 | 466 | Specific | 814.66 | 725.48 | 725.48 | 16.27 | 73 | >sp\|P01833\|PIGR_HUMAN Polymeric immunoglobulin receptor OS=Homo sapiens OX=9606 GN=PIGR PE=1 SV=4 | 902 | 20201118_12812_1ng_glyco_hcd203040_compound15.16920.16920.4 | controllerType=0 controllerNumber=1 scan=16920 | 34.7182 |
| 15414:4 | 8 | K.VPGN[+2350.830]VTAVLGETLKVPC[+57.021]HFPC[+57.021]K.F | HexNAc(4)Hex(5)Fuc(1)NeuAc(2) | C[+57]*2, N[+2351] | 1194.2826 | 4 | 4774.1087 | 4774.0877 | 0 | 4.4 | 466 | Specific | 933.29 | 832.12 | 832.12 | 18.01 | 73 | >sp\|P01833\|PIGR_HUMAN Polymeric immunoglobulin receptor OS=Homo sapiens OX=9606 GN=PIGR PE=1 SV=4 | 902 | 20201118_12812_1ng_glyco_hcd203040_compound15.18874.18874.4 | controllerType=0 controllerNumber=1 scan=18874 | 38.2766 |
| 05724:3 | 8 | K.WN[+1548.545]NTGC[+57.021]QALPSQDEGPSK.A | HexNAc(3)Hex(4)NeuAc(1) | C[+57], N[+1549] | 1179.8132 | 3 | 3537.4249 | 3537.4215 | 0 | 1 | 498 | Specific | 238.74 | 191.08 | 191.08 | 6 | 73 | >sp\|P01833\|PIGR_HUMAN Polymeric immunoglobulin receptor OS=Homo sapiens OX=9606 GN=PIGR PE=1 SV=4 | 902 | 20201118_12812_1ng_glyco_hcd203040_compound15.8804.8804.3 | controllerType=0 controllerNumber=1 scan=8804 | 21.087 |
| 05726:3 | 8 | K.WN[+1548.545]NTGC[+57.021]QALPSQDEGPSK.A | HexNAc(3)Hex(4)NeuAc(1) | C[+57], N[+1549] | 1179.8132 | 3 | 3537.4249 | 3537.4215 | 0 | 1 | 498 | Specific | 414.27 | 336.66 | 336.66 | 7.72 | 73 | >sp\|P01833\|PIGR_HUMAN Polymeric immunoglobulin receptor OS=Homo sapiens OX=9606 GN=PIGR PE=1 SV=4 | 902 | 20201118_12812_1ng_glyco_hcd203040_compound15.8806.8806.3 | controllerType=0 controllerNumber=1 scan=8806 | 21.0909 |
| 05506:3 | 8 | K.WN[+1872.651]NTGC[+57.021]QALPSQDEGPSK.A | HexNAc(3)Hex(6)NeuAc(1) | C[+57], N[+1873] | 1287.8505 | 3 | 3861.537 | 3861.5272 | 0 | 2.6 | 498 | Specific | 582.06 | 545.13 | 545.13 | 11.68 | 73 | >sp\|P01833\|PIGR_HUMAN Polymeric immunoglobulin receptor OS=Homo sapiens OX=9606 GN=PIGR PE=1 SV=4 | 902 | 20201118_12812_1ng_glyco_hcd203040_compound15.8578.8578.3 | controllerType=0 controllerNumber=1 scan=8578 | 20.7079 |
| 05508:3 | 8 | K.WN[+1872.651]NTGC[+57.021]QALPSQDEGPSK.A | HexNAc(3)Hex(6)NeuAc(1) | C[+57], N[+1873] | 1287.8505 | 3 | 3861.537 | 3861.5272 | 0 | 2.6 | 498 | Specific | 586.74 | 525.68 | 525.68 | 11.68 | 73 | >sp\|P01833\|PIGR_HUMAN Polymeric immunoglobulin receptor OS=Homo sapiens OX=9606 GN=PIGR PE=1 SV=4 | 902 | 20201118_12812_1ng_glyco_hcd203040_compound15.8580.8580.3 | controllerType=0 controllerNumber=1 scan=8580 | 20.7116 |
| 05634:3 | 8 | K.WN[+1710.598]NTGC[+57.021]QALPSQDEGPSK.A | HexNAc(3)Hex(5)NeuAc(1) | C[+57], N[+1711] | 1233.8336 | 3 | 3699.4861 | 3699.4743 | 0 | 3.2 | 498 | Specific | 639.8 | 582.98 | 582.98 | 12.68 | 73 | >sp\|P01833\|PIGR_HUMAN Polymeric immunoglobulin receptor OS=Homo sapiens OX=9606 GN=PIGR PE=1 SV=4 | 902 | 20201118_12812_1ng_glyco_hcd203040_compound15.8711.8711.3 | controllerType=0 controllerNumber=1 scan=8711 | 20.9292 |
| 05632:3 | 8 | K.WN[+1710.598]NTGC[+57.021]QALPSQDEGPSK.A | HexNAc(3)Hex(5)NeuAc(1) | C[+57], N[+1711] | 1233.8336 | 3 | 3699.4861 | 3699.4743 | 0 | 3.2 | 498 | Specific | 645.25 | 576.34 | 576.34 | 12.68 | 73 | >sp\|P01833\|PIGR_HUMAN Polymeric immunoglobulin receptor OS=Homo sapiens OX=9606 GN=PIGR PE=1 SV=4 | 902 | 20201118_12812_1ng_glyco_hcd203040_compound15.8709.8709.3 | controllerType=0 controllerNumber=1 scan=8709 | 20.9257 |
| 05589:3 | 8 | K.WN[+1710.598]NTGC[+57.021]QALPSQDEGPSK.A | HexNAc(3)Hex(5)NeuAc(1) | C[+57], N[+1711] | 1233.8342 | 3 | 3699.4879 | 3699.4743 | 0 | 3.7 | 498 | Specific | 752.76 | 687.5 | 687.5 | 15.68 | 73 | >sp\|P01833\|PIGR_HUMAN Polymeric immunoglobulin receptor OS=Homo sapiens OX=9606 GN=PIGR PE=1 SV=4 | 902 | 20201118_12812_1ng_glyco_hcd203040_compound15.8664.8664.3 | controllerType=0 controllerNumber=1 scan=8664 | 20.8508 |
| 05591:3 | 8 | K.WN[+1710.598]NTGC[+57.021]QALPSQDEGPSK.A | HexNAc(3)Hex(5)NeuAc(1) | C[+57], N[+1711] | 1233.8342 | 3 | 3699.4879 | 3699.4743 | 0 | 3.7 | 498 | Specific | 781.23 | 685.69 | 685.69 | 15.68 | 73 | >sp\|P01833\|PIGR_HUMAN Polymeric immunoglobulin receptor OS=Homo sapiens OX=9606 GN=PIGR PE=1 SV=4 | 902 | 20201118_12812_1ng_glyco_hcd203040_compound15.8666.8666.3 | controllerType=0 controllerNumber=1 scan=8666 | 20.8548 |
| 07545:3 | 9 | K.WEC[+57.021]KN[+2059.735]DTLLGIK.G | HexNAc(4)Hex(5)Fuc(1)NeuAc(1) | C[+57], N[+2060] | 1179.5014 | 3 | 3536.4896 | 3536.4865 | 0 | 0.9 | 100 | Specific | 281.78 | 46.41 | 46.41 | 3.96 | 72 | >sp\|P22897\|MRC1_HUMAN Macrophage mannose receptor 1 OS=Homo sapiens OX=9606 GN=MRC1 PE=1 SV=1 | 1049 | 20201118_12812_1ng_glyco_hcd203040_compound15.10688.10688.3 | controllerType=0 controllerNumber=1 scan=10688 | 24.1672 |
| 16295:3 | 9 | R.VWIALNSN[+2204.772]LTDNQYTWTDKWR.V | HexNAc(4)Hex(5)NeuAc(2) | N[+2205] | 1610.3694 | 3 | 4829.0938 | 4829.0617 | 0 | 6.7 | 1153 | Specific | 497.56 | 328.02 | 328.02 | 6.78 | 72 | >sp\|P22897\|MRC1_HUMAN Macrophage mannose receptor 1 OS=Homo sapiens OX=9606 GN=MRC1 PE=1 SV=1 | 1049 | 20201118_12812_1ng_glyco_hcd203040_compound15.19816.19816.3 | controllerType=0 controllerNumber=1 scan=19816 | 39.9716 |
| 16297:3 | 9 | R.VWIALNSN[+2204.772]LTDNQYTWTDKWR.V | HexNAc(4)Hex(5)NeuAc(2) | N[+2205] | 1610.3694 | 3 | 4829.0938 | 4829.0617 | 0 | 6.7 | 1153 | Specific | 512.36 | 410.9 | 410.9 | 8.59 | 72 | >sp\|P22897\|MRC1_HUMAN Macrophage mannose receptor 1 OS=Homo sapiens OX=9606 GN=MRC1 PE=1 SV=1 | 1049 | 20201118_12812_1ng_glyco_hcd203040_compound15.19818.19818.3 | controllerType=0 controllerNumber=1 scan=19818 | 39.9774 |
| 08210:3 | 9 | K.[+42.011]TAHC[+57.021]N[+2018.708]ESFYFLC[+57.021]K.R | HexNAc(3)Hex(6)Fuc(1)NeuAc(1) | .[42], C[+57]*2, N[+2019] | 1246.4947 | 3 | 3737.4695 | 3737.4385 | 0 | 8.3 | 1201 | Specific | 249.27 | 122.34 | 122.34 | 4.01 | 72 | >sp\|P22897\|MRC1_HUMAN Macrophage mannose receptor 1 OS=Homo sapiens OX=9606 GN=MRC1 PE=1 SV=1 | 1049 | 20201118_12812_1ng_glyco_hcd203040_compound15.11384.11384.3 | controllerType=0 controllerNumber=1 scan=11384 | 25.3229 |
| 08244:3 | 9 | K.TAHC[+57.021]N[+2059.735]ESFYFLC[+57.021]K.R | HexNAc(4)Hex(5)Fuc(1)NeuAc(1) | C[+57]*2, N[+2060] | 1246.165 | 3 | 3736.4804 | 3736.4545 | 0 | 6.9 | 1201 | Specific | 482.55 | 439.54 | 439.54 | 8.83 | 72 | >sp\|P22897\|MRC1_HUMAN Macrophage mannose receptor 1 OS=Homo sapiens OX=9606 GN=MRC1 PE=1 SV=1 | 1049 | 20201118_12812_1ng_glyco_hcd203040_compound15.11419.11419.3 | controllerType=0 controllerNumber=1 scan=11419 | 25.3787 |
| 08246:3 | 9 | K.TAHC[+57.021]N[+2059.735]ESFYFLC[+57.021]K.R | HexNAc(4)Hex(5)Fuc(1)NeuAc(1) | C[+57]*2, N[+2060] | 1246.165 | 3 | 3736.4804 | 3736.4545 | 0 | 6.9 | 1201 | Specific | 482.85 | 433.66 | 433.66 | 8.83 | 72 | >sp\|P22897\|MRC1_HUMAN Macrophage mannose receptor 1 OS=Homo sapiens OX=9606 GN=MRC1 PE=1 SV=1 | 1049 | 20201118_12812_1ng_glyco_hcd203040_compound15.11421.11421.3 | controllerType=0 controllerNumber=1 scan=11421 | 25.3824 |
| 17591:4 | 10 | R.LSLHRPALEDLLLGSEAN[+1663.608]LTC[+57.021]TLTGLR.D | HexNAc(5)Hex(4) | C[+57], N[+1664] | 1157.5592 | 4 | 4627.2152 | 4627.2064 | 0 | 1.9 | 127 | Specific | 382.31 | 382.31 | 382.31 | 9.63 | 54 | >sp\|P01876\|IGHA1_HUMAN Immunoglobulin heavy constant alpha 1 OS=Homo sapiens OX=9606 GN=IGHA1 PE=1 SV=2 | 255 | 20201118_12812_1ng_glyco_hcd203040_compound15.21167.21167.4 | controllerType=0 controllerNumber=1 scan=21167 | 42.7521 |
| 18563:4 | 10 | R.LSLHRPALEDLLLGSEAN[+1751.624]LTC[+57.021]TLTGLR.D | HexNAc(4)Hex(4)NeuAc(1) | C[+57], N[+1752] | 1179.5625 | 4 | 4715.2283 | 4715.2225 | 0 | 1.2 | 127 | Specific | 417.54 | 417.54 | 417.54 | 9.86 | 54 | >sp\|P01876\|IGHA1_HUMAN Immunoglobulin heavy constant alpha 1 OS=Homo sapiens OX=9606 GN=IGHA1 PE=1 SV=2 | 255 | 20201118_12812_1ng_glyco_hcd203040_compound15.22182.22182.4 | controllerType=0 controllerNumber=1 scan=22182 | 44.9921 |
| 17592:3 | 10 | R.LSLHRPALEDLLLGSEAN[+1378.476]LTC[+57.021]TLTGLR.D | HexNAc(2)Hex(6) | C[+57], N[+1378] | 1448.0331 | 3 | 4342.0849 | 4342.074 | 0 | 2.5 | 127 | Specific | 516.35 | 516.35 | 516.35 | 9.78 | 54 | >sp\|P01876\|IGHA1_HUMAN Immunoglobulin heavy constant alpha 1 OS=Homo sapiens OX=9606 GN=IGHA1 PE=1 SV=2 | 255 | 20201118_12812_1ng_glyco_hcd203040_compound15.21168.21168.3 | controllerType=0 controllerNumber=1 scan=21168 | 42.7533 |
| 17567:3 | 10 | R.LSLHRPALEDLLLGSEAN[+1622.582]LTC[+57.021]TLTGLR.D | HexNAc(4)Hex(5) | C[+57], N[+1623] | 1529.3993 | 3 | 4586.1835 | 4586.1799 | 0 | 0.8 | 127 | Specific | 539.77 | 512.24 | 512.24 | 9.59 | 54 | >sp\|P01876\|IGHA1_HUMAN Immunoglobulin heavy constant alpha 1 OS=Homo sapiens OX=9606 GN=IGHA1 PE=1 SV=2 | 255 | 20201118_12812_1ng_glyco_hcd203040_compound15.21142.21142.3 | controllerType=0 controllerNumber=1 scan=21142 | 42.6901 |
| 17595:3 | 10 | R.LSLHRPALEDLLLGSEAN[+1378.476]LTC[+57.021]TLTGLR.D | HexNAc(2)Hex(6) | C[+57], N[+1378] | 1448.0331 | 3 | 4342.0849 | 4342.074 | 0 | 2.5 | 127 | Specific | 602.05 | 584.03 | 584.03 | 12.33 | 54 | >sp\|P01876\|IGHA1_HUMAN Immunoglobulin heavy constant alpha 1 OS=Homo sapiens OX=9606 GN=IGHA1 PE=1 SV=2 | 255 | 20201118_12812_1ng_glyco_hcd203040_compound15.21171.21171.3 | controllerType=0 controllerNumber=1 scan=21171 | 42.7684 |
| 17583:3 | 10 | R.LSLHRPALEDLLLGSEAN[+1460.529]LTC[+57.021]TLTGLR.D | HexNAc(4)Hex(4) | C[+57], N[+1461] | 1475.3846 | 3 | 4424.1392 | 4424.1271 | 0 | 2.7 | 127 | Specific | 608.2 | 581.42 | 581.42 | 12.33 | 54 | >sp\|P01876\|IGHA1_HUMAN Immunoglobulin heavy constant alpha 1 OS=Homo sapiens OX=9606 GN=IGHA1 PE=1 SV=2 | 255 | 20201118_12812_1ng_glyco_hcd203040_compound15.21158.21158.3 | controllerType=0 controllerNumber=1 scan=21158 | 42.725 |
| 17590:3 | 10 | R.LSLHRPALEDLLLGSEAN[+1501.555]LTC[+57.021]TLTGLR.D | HexNAc(5)Hex(3) | C[+57], N[+1502] | 1489.0595 | 3 | 4465.164 | 4465.1536 | 0 | 2.3 | 127 | Specific | 620.24 | 620.24 | 620.24 | 12.23 | 54 | >sp\|P01876\|IGHA1_HUMAN Immunoglobulin heavy constant alpha 1 OS=Homo sapiens OX=9606 GN=IGHA1 PE=1 SV=2 | 255 | 20201118_12812_1ng_glyco_hcd203040_compound15.21166.21166.3 | controllerType=0 controllerNumber=1 scan=21166 | 42.751 |
| 17570:3 | 10 | R.LSLHRPALEDLLLGSEAN[+1622.582]LTC[+57.021]TLTGLR.D | HexNAc(4)Hex(5) | C[+57], N[+1623] | 1529.3993 | 3 | 4586.1835 | 4586.1799 | 0 | 0.8 | 127 | Specific | 628.76 | 544.79 | 544.79 | 12 | 54 | >sp\|P01876\|IGHA1_HUMAN Immunoglobulin heavy constant alpha 1 OS=Homo sapiens OX=9606 GN=IGHA1 PE=1 SV=2 | 255 | 20201118_12812_1ng_glyco_hcd203040_compound15.21145.21145.3 | controllerType=0 controllerNumber=1 scan=21145 | 42.6967 |
| 18505:4 | 10 | R.LSLHRPALEDLLLGSEAN[+1872.651]LTC[+57.021]TLTGLR.D | HexNAc(3)Hex(6)NeuAc(1) | C[+57], N[+1873] | 1209.8223 | 4 | 4836.2674 | 4836.2487 | 0 | 3.9 | 127 | Specific | 629.07 | 629.07 | 629.07 | 12.69 | 54 | >sp\|P01876\|IGHA1_HUMAN Immunoglobulin heavy constant alpha 1 OS=Homo sapiens OX=9606 GN=IGHA1 PE=1 SV=2 | 255 | 20201118_12812_1ng_glyco_hcd203040_compound15.22121.22121.4 | controllerType=0 controllerNumber=1 scan=22121 | 44.839 |
| 17471:3 | 10 | R.LSLHRPALEDLLLGSEAN[+1864.634]LTC[+57.021]TLTGLR.D | HexNAc(2)Hex(9) | C[+57], N[+1865] | 1610.086 | 3 | 4828.2434 | 4828.2324 | 0 | 2.3 | 127 | Specific | 634.7 | 596.06 | 596.06 | 12.23 | 54 | >sp\|P01876\|IGHA1_HUMAN Immunoglobulin heavy constant alpha 1 OS=Homo sapiens OX=9606 GN=IGHA1 PE=1 SV=2 | 255 | 20201118_12812_1ng_glyco_hcd203040_compound15.21040.21040.3 | controllerType=0 controllerNumber=1 scan=21040 | 42.4397 |
| 17585:3 | 10 | R.LSLHRPALEDLLLGSEAN[+1460.529]LTC[+57.021]TLTGLR.D | HexNAc(4)Hex(4) | C[+57], N[+1461] | 1475.3846 | 3 | 4424.1392 | 4424.1271 | 0 | 2.7 | 127 | Specific | 642.88 | 525.62 | 525.62 | 12.33 | 54 | >sp\|P01876\|IGHA1_HUMAN Immunoglobulin heavy constant alpha 1 OS=Homo sapiens OX=9606 GN=IGHA1 PE=1 SV=2 | 255 | 20201118_12812_1ng_glyco_hcd203040_compound15.21160.21160.3 | controllerType=0 controllerNumber=1 scan=21160 | 42.7332 |
| 17593:3 | 10 | R.LSLHRPALEDLLLGSEAN[+1501.555]LTC[+57.021]TLTGLR.D | HexNAc(5)Hex(3) | C[+57], N[+1502] | 1489.0595 | 3 | 4465.164 | 4465.1536 | 0 | 2.3 | 127 | Specific | 663.2 | 613.05 | 613.05 | 13.23 | 54 | >sp\|P01876\|IGHA1_HUMAN Immunoglobulin heavy constant alpha 1 OS=Homo sapiens OX=9606 GN=IGHA1 PE=1 SV=2 | 255 | 20201118_12812_1ng_glyco_hcd203040_compound15.21169.21169.3 | controllerType=0 controllerNumber=1 scan=21169 | 42.7617 |
| 17513:3 | 10 | R.LSLHRPALEDLLLGSEAN[+1540.529]LTC[+57.021]TLTGLR.D | HexNAc(2)Hex(7) | C[+57], N[+1541] | 1502.0505 | 3 | 4504.1369 | 4504.1268 | 0 | 2.2 | 127 | Specific | 665.24 | 606.71 | 606.71 | 13.23 | 54 | >sp\|P01876\|IGHA1_HUMAN Immunoglobulin heavy constant alpha 1 OS=Homo sapiens OX=9606 GN=IGHA1 PE=1 SV=2 | 255 | 20201118_12812_1ng_glyco_hcd203040_compound15.21084.21084.3 | controllerType=0 controllerNumber=1 scan=21084 | 42.5534 |
| 17587:4 | 10 | R.LSLHRPALEDLLLGSEAN[+1378.476]LTC[+57.021]TLTGLR.D | HexNAc(2)Hex(6) | C[+57], N[+1378] | 1086.277 | 4 | 4342.0863 | 4342.074 | 0 | 2.8 | 127 | Specific | 666.83 | 666.83 | 666.83 | 13.96 | 54 | >sp\|P01876\|IGHA1_HUMAN Immunoglobulin heavy constant alpha 1 OS=Homo sapiens OX=9606 GN=IGHA1 PE=1 SV=2 | 255 | 20201118_12812_1ng_glyco_hcd203040_compound15.21163.21163.4 | controllerType=0 controllerNumber=1 scan=21163 | 42.7392 |
| 17468:3 | 10 | R.LSLHRPALEDLLLGSEAN[+1864.634]LTC[+57.021]TLTGLR.D | HexNAc(2)Hex(9) | C[+57], N[+1865] | 1610.086 | 3 | 4828.2434 | 4828.2324 | 0 | 2.3 | 127 | Specific | 670.8 | 670.8 | 670.8 | 13.23 | 54 | >sp\|P01876\|IGHA1_HUMAN Immunoglobulin heavy constant alpha 1 OS=Homo sapiens OX=9606 GN=IGHA1 PE=1 SV=2 | 255 | 20201118_12812_1ng_glyco_hcd203040_compound15.21037.21037.3 | controllerType=0 controllerNumber=1 scan=21037 | 42.4308 |
| 17612:5 | 10 | R.LSLHRPALEDLLLGSEAN[+1501.555]LTC[+57.021]TLTGLR.D | HexNAc(5)Hex(3) | C[+57], N[+1502] | 893.8392 | 5 | 4465.1669 | 4465.1536 | 0 | 3 | 127 | Specific | 673.97 | 673.97 | 673.97 | 13.89 | 54 | >sp\|P01876\|IGHA1_HUMAN Immunoglobulin heavy constant alpha 1 OS=Homo sapiens OX=9606 GN=IGHA1 PE=1 SV=2 | 255 | 20201118_12812_1ng_glyco_hcd203040_compound15.21189.21189.5 | controllerType=0 controllerNumber=1 scan=21189 | 42.808 |
| 17599:3 | 10 | R.LSLHRPALEDLLLGSEAN[+1216.423]LTC[+57.021]TLTGLR.D | HexNAc(2)Hex(5) | C[+57], N[+1216] | 1394.0149 | 3 | 4180.0303 | 4180.0211 | 0 | 2.2 | 127 | Specific | 691.88 | 672.86 | 672.86 | 13.23 | 54 | >sp\|P01876\|IGHA1_HUMAN Immunoglobulin heavy constant alpha 1 OS=Homo sapiens OX=9606 GN=IGHA1 PE=1 SV=2 | 255 | 20201118_12812_1ng_glyco_hcd203040_compound15.21175.21175.3 | controllerType=0 controllerNumber=1 scan=21175 | 42.7762 |
| 17586:4 | 10 | R.LSLHRPALEDLLLGSEAN[+1501.555]LTC[+57.021]TLTGLR.D | HexNAc(5)Hex(3) | C[+57], N[+1502] | 1117.0478 | 4 | 4465.1693 | 4465.1536 | 0 | 3.5 | 127 | Specific | 695.83 | 695.83 | 695.83 | 13.96 | 54 | >sp\|P01876\|IGHA1_HUMAN Immunoglobulin heavy constant alpha 1 OS=Homo sapiens OX=9606 GN=IGHA1 PE=1 SV=2 | 255 | 20201118_12812_1ng_glyco_hcd203040_compound15.21162.21162.4 | controllerType=0 controllerNumber=1 scan=21162 | 42.7377 |
| 18565:4 | 10 | R.LSLHRPALEDLLLGSEAN[+1751.624]LTC[+57.021]TLTGLR.D | HexNAc(4)Hex(4)NeuAc(1) | C[+57], N[+1752] | 1179.5625 | 4 | 4715.2283 | 4715.2225 | 0 | 1.2 | 127 | Specific | 698.14 | 698.14 | 698.14 | 13.86 | 54 | >sp\|P01876\|IGHA1_HUMAN Immunoglobulin heavy constant alpha 1 OS=Homo sapiens OX=9606 GN=IGHA1 PE=1 SV=2 | 255 | 20201118_12812_1ng_glyco_hcd203040_compound15.22184.22184.4 | controllerType=0 controllerNumber=1 scan=22184 | 44.9965 |
| 18467:4 | 10 | R.LSLHRPALEDLLLGSEAN[+1913.677]LTC[+57.021]TLTGLR.D | HexNAc(4)Hex(5)NeuAc(1) | C[+57], N[+1914] | 1220.0793 | 4 | 4877.2952 | 4877.2753 | 0 | 4.1 | 127 | Specific | 702.38 | 702.38 | 702.38 | 14.69 | 54 | >sp\|P01876\|IGHA1_HUMAN Immunoglobulin heavy constant alpha 1 OS=Homo sapiens OX=9606 GN=IGHA1 PE=1 SV=2 | 255 | 20201118_12812_1ng_glyco_hcd203040_compound15.22081.22081.4 | controllerType=0 controllerNumber=1 scan=22081 | 44.7429 |
| 17566:4 | 10 | R.LSLHRPALEDLLLGSEAN[+1419.502]LTC[+57.021]TLTGLR.D | HexNAc(3)Hex(5) | C[+57], N[+1420] | 1096.5328 | 4 | 4383.1092 | 4383.1005 | 0 | 2 | 127 | Specific | 706.91 | 660.33 | 660.33 | 14.86 | 54 | >sp\|P01876\|IGHA1_HUMAN Immunoglobulin heavy constant alpha 1 OS=Homo sapiens OX=9606 GN=IGHA1 PE=1 SV=2 | 255 | 20201118_12812_1ng_glyco_hcd203040_compound15.21141.21141.4 | controllerType=0 controllerNumber=1 scan=21141 | 42.6889 |
| 17544:4 | 10 | R.LSLHRPALEDLLLGSEAN[+1460.529]LTC[+57.021]TLTGLR.D | HexNAc(4)Hex(4) | C[+57], N[+1461] | 1106.7901 | 4 | 4424.1385 | 4424.1271 | 0 | 2.6 | 127 | Specific | 714.27 | 714.27 | 714.27 | 14.96 | 54 | >sp\|P01876\|IGHA1_HUMAN Immunoglobulin heavy constant alpha 1 OS=Homo sapiens OX=9606 GN=IGHA1 PE=1 SV=2 | 255 | 20201118_12812_1ng_glyco_hcd203040_compound15.21118.21118.4 | controllerType=0 controllerNumber=1 scan=21118 | 42.6244 |
| 17562:4 | 10 | R.LSLHRPALEDLLLGSEAN[+1622.582]LTC[+57.021]TLTGLR.D | HexNAc(4)Hex(5) | C[+57], N[+1623] | 1147.3017 | 4 | 4586.1849 | 4586.1799 | 0 | 1.1 | 127 | Specific | 714.89 | 714.89 | 714.89 | 14.63 | 54 | >sp\|P01876\|IGHA1_HUMAN Immunoglobulin heavy constant alpha 1 OS=Homo sapiens OX=9606 GN=IGHA1 PE=1 SV=2 | 255 | 20201118_12812_1ng_glyco_hcd203040_compound15.21137.21137.4 | controllerType=0 controllerNumber=1 scan=21137 | 42.6797 |
| 17609:5 | 10 | R.LSLHRPALEDLLLGSEAN[+1501.555]LTC[+57.021]TLTGLR.D | HexNAc(5)Hex(3) | C[+57], N[+1502] | 893.8392 | 5 | 4465.1669 | 4465.1536 | 0 | 3 | 127 | Specific | 719.64 | 719.64 | 719.64 | 14.89 | 54 | >sp\|P01876\|IGHA1_HUMAN Immunoglobulin heavy constant alpha 1 OS=Homo sapiens OX=9606 GN=IGHA1 PE=1 SV=2 | 255 | 20201118_12812_1ng_glyco_hcd203040_compound15.21186.21186.5 | controllerType=0 controllerNumber=1 scan=21186 | 42.8024 |
| 17510:3 | 10 | R.LSLHRPALEDLLLGSEAN[+1540.529]LTC[+57.021]TLTGLR.D | HexNAc(2)Hex(7) | C[+57], N[+1541] | 1502.0505 | 3 | 4504.1369 | 4504.1268 | 0 | 2.2 | 127 | Specific | 739.88 | 673.55 | 673.55 | 14.23 | 54 | >sp\|P01876\|IGHA1_HUMAN Immunoglobulin heavy constant alpha 1 OS=Homo sapiens OX=9606 GN=IGHA1 PE=1 SV=2 | 255 | 20201118_12812_1ng_glyco_hcd203040_compound15.21081.21081.3 | controllerType=0 controllerNumber=1 scan=21081 | 42.5446 |
| 17596:3 | 10 | R.LSLHRPALEDLLLGSEAN[+1216.423]LTC[+57.021]TLTGLR.D | HexNAc(2)Hex(5) | C[+57], N[+1216] | 1394.0149 | 3 | 4180.0303 | 4180.0211 | 0 | 2.2 | 127 | Specific | 744.23 | 678.13 | 678.13 | 14.23 | 54 | >sp\|P01876\|IGHA1_HUMAN Immunoglobulin heavy constant alpha 1 OS=Homo sapiens OX=9606 GN=IGHA1 PE=1 SV=2 | 255 | 20201118_12812_1ng_glyco_hcd203040_compound15.21172.21172.3 | controllerType=0 controllerNumber=1 scan=21172 | 42.7703 |
| 17606:4 | 10 | R.LSLHRPALEDLLLGSEAN[+1298.476]LTC[+57.021]TLTGLR.D | HexNAc(4)Hex(3) | C[+57], N[+1298] | 1066.2776 | 4 | 4262.0887 | 4262.0742 | 0 | 3.4 | 127 | Specific | 749.58 | 749.58 | 749.58 | 14.96 | 54 | >sp\|P01876\|IGHA1_HUMAN Immunoglobulin heavy constant alpha 1 OS=Homo sapiens OX=9606 GN=IGHA1 PE=1 SV=2 | 255 | 20201118_12812_1ng_glyco_hcd203040_compound15.21183.21183.4 | controllerType=0 controllerNumber=1 scan=21183 | 42.7958 |
| 17588:4 | 10 | R.LSLHRPALEDLLLGSEAN[+1501.555]LTC[+57.021]TLTGLR.D | HexNAc(5)Hex(3) | C[+57], N[+1502] | 1117.0478 | 4 | 4465.1693 | 4465.1536 | 0 | 3.5 | 127 | Specific | 761.27 | 761.27 | 761.27 | 15.96 | 54 | >sp\|P01876\|IGHA1_HUMAN Immunoglobulin heavy constant alpha 1 OS=Homo sapiens OX=9606 GN=IGHA1 PE=1 SV=2 | 255 | 20201118_12812_1ng_glyco_hcd203040_compound15.21164.21164.4 | controllerType=0 controllerNumber=1 scan=21164 | 42.7471 |
| 17546:4 | 10 | R.LSLHRPALEDLLLGSEAN[+1460.529]LTC[+57.021]TLTGLR.D | HexNAc(4)Hex(4) | C[+57], N[+1461] | 1106.7901 | 4 | 4424.1385 | 4424.1271 | 0 | 2.6 | 127 | Specific | 787.45 | 787.45 | 787.45 | 15.96 | 54 | >sp\|P01876\|IGHA1_HUMAN Immunoglobulin heavy constant alpha 1 OS=Homo sapiens OX=9606 GN=IGHA1 PE=1 SV=2 | 255 | 20201118_12812_1ng_glyco_hcd203040_compound15.21120.21120.4 | controllerType=0 controllerNumber=1 scan=21120 | 42.6294 |
| 17594:4 | 10 | R.LSLHRPALEDLLLGSEAN[+1663.608]LTC[+57.021]TLTGLR.D | HexNAc(5)Hex(4) | C[+57], N[+1664] | 1157.5592 | 4 | 4627.2152 | 4627.2064 | 0 | 1.9 | 127 | Specific | 800.24 | 730.74 | 730.74 | 16.86 | 54 | >sp\|P01876\|IGHA1_HUMAN Immunoglobulin heavy constant alpha 1 OS=Homo sapiens OX=9606 GN=IGHA1 PE=1 SV=2 | 255 | 20201118_12812_1ng_glyco_hcd203040_compound15.21170.21170.4 | controllerType=0 controllerNumber=1 scan=21170 | 42.7665 |
| 17564:4 | 10 | R.LSLHRPALEDLLLGSEAN[+1622.582]LTC[+57.021]TLTGLR.D | HexNAc(4)Hex(5) | C[+57], N[+1623] | 1147.3017 | 4 | 4586.1849 | 4586.1799 | 0 | 1.1 | 127 | Specific | 808.96 | 704.73 | 704.73 | 16.63 | 54 | >sp\|P01876\|IGHA1_HUMAN Immunoglobulin heavy constant alpha 1 OS=Homo sapiens OX=9606 GN=IGHA1 PE=1 SV=2 | 255 | 20201118_12812_1ng_glyco_hcd203040_compound15.21139.21139.4 | controllerType=0 controllerNumber=1 scan=21139 | 42.6847 |
| 18469:4 | 10 | R.LSLHRPALEDLLLGSEAN[+1913.677]LTC[+57.021]TLTGLR.D | HexNAc(4)Hex(5)NeuAc(1) | C[+57], N[+1914] | 1220.0793 | 4 | 4877.2952 | 4877.2753 | 0 | 4.1 | 127 | Specific | 823.63 | 784.1 | 784.1 | 16.69 | 54 | >sp\|P01876\|IGHA1_HUMAN Immunoglobulin heavy constant alpha 1 OS=Homo sapiens OX=9606 GN=IGHA1 PE=1 SV=2 | 255 | 20201118_12812_1ng_glyco_hcd203040_compound15.22083.22083.4 | controllerType=0 controllerNumber=1 scan=22083 | 44.7502 |
| 17545:4 | 10 | R.LSLHRPALEDLLLGSEAN[+1216.423]LTC[+57.021]TLTGLR.D | HexNAc(2)Hex(5) | C[+57], N[+1216] | 1045.7636 | 4 | 4180.0325 | 4180.0211 | 0 | 2.7 | 127 | Specific | 828.29 | 754.79 | 754.79 | 16.96 | 54 | >sp\|P01876\|IGHA1_HUMAN Immunoglobulin heavy constant alpha 1 OS=Homo sapiens OX=9606 GN=IGHA1 PE=1 SV=2 | 255 | 20201118_12812_1ng_glyco_hcd203040_compound15.21119.21119.4 | controllerType=0 controllerNumber=1 scan=21119 | 42.6259 |
| 17462:4 | 10 | R.LSLHRPALEDLLLGSEAN[+1864.634]LTC[+57.021]TLTGLR.D | HexNAc(2)Hex(9) | C[+57], N[+1865] | 1207.8169 | 4 | 4828.2459 | 4828.2324 | 0 | 2.8 | 127 | Specific | 831.36 | 831.36 | 831.36 | 16.96 | 54 | >sp\|P01876\|IGHA1_HUMAN Immunoglobulin heavy constant alpha 1 OS=Homo sapiens OX=9606 GN=IGHA1 PE=1 SV=2 | 255 | 20201118_12812_1ng_glyco_hcd203040_compound15.21031.21031.4 | controllerType=0 controllerNumber=1 scan=21031 | 42.4171 |
| 18508:4 | 10 | R.LSLHRPALEDLLLGSEAN[+1872.651]LTC[+57.021]TLTGLR.D | HexNAc(3)Hex(6)NeuAc(1) | C[+57], N[+1873] | 1209.8223 | 4 | 4836.2674 | 4836.2487 | 0 | 3.9 | 127 | Specific | 846.31 | 825.29 | 825.29 | 16.69 | 54 | >sp\|P01876\|IGHA1_HUMAN Immunoglobulin heavy constant alpha 1 OS=Homo sapiens OX=9606 GN=IGHA1 PE=1 SV=2 | 255 | 20201118_12812_1ng_glyco_hcd203040_compound15.22124.22124.4 | controllerType=0 controllerNumber=1 scan=22124 | 44.8459 |
| 17608:4 | 10 | R.LSLHRPALEDLLLGSEAN[+1298.476]LTC[+57.021]TLTGLR.D | HexNAc(4)Hex(3) | C[+57], N[+1298] | 1066.2776 | 4 | 4262.0887 | 4262.0742 | 0 | 3.4 | 127 | Specific | 861.51 | 836.27 | 836.27 | 17.96 | 54 | >sp\|P01876\|IGHA1_HUMAN Immunoglobulin heavy constant alpha 1 OS=Homo sapiens OX=9606 GN=IGHA1 PE=1 SV=2 | 255 | 20201118_12812_1ng_glyco_hcd203040_compound15.21185.21185.4 | controllerType=0 controllerNumber=1 scan=21185 | 42.8011 |
| 17569:4 | 10 | R.LSLHRPALEDLLLGSEAN[+1419.502]LTC[+57.021]TLTGLR.D | HexNAc(3)Hex(5) | C[+57], N[+1420] | 1096.5328 | 4 | 4383.1092 | 4383.1005 | 0 | 2 | 127 | Specific | 865.18 | 799.08 | 799.08 | 17.86 | 54 | >sp\|P01876\|IGHA1_HUMAN Immunoglobulin heavy constant alpha 1 OS=Homo sapiens OX=9606 GN=IGHA1 PE=1 SV=2 | 255 | 20201118_12812_1ng_glyco_hcd203040_compound15.21144.21144.4 | controllerType=0 controllerNumber=1 scan=21144 | 42.6953 |
| 17616:4 | 10 | R.LSLHRPALEDLLLGSEAN[+1257.449]LTC[+57.021]TLTGLR.D | HexNAc(3)Hex(4) | C[+57], N[+1257] | 1056.0196 | 4 | 4221.0565 | 4221.0477 | 0 | 2.1 | 127 | Specific | 887.99 | 812.59 | 812.59 | 17.86 | 54 | >sp\|P01876\|IGHA1_HUMAN Immunoglobulin heavy constant alpha 1 OS=Homo sapiens OX=9606 GN=IGHA1 PE=1 SV=2 | 255 | 20201118_12812_1ng_glyco_hcd203040_compound15.21193.21193.4 | controllerType=0 controllerNumber=1 scan=21193 | 42.8137 |
| 17473:4 | 10 | R.LSLHRPALEDLLLGSEAN[+1702.581]LTC[+57.021]TLTGLR.D | HexNAc(2)Hex(8) | C[+57], N[+1703] | 1167.3027 | 4 | 4666.1888 | 4666.1796 | 0 | 2 | 127 | Specific | 889.5 | 889.5 | 889.5 | 17.86 | 54 | >sp\|P01876\|IGHA1_HUMAN Immunoglobulin heavy constant alpha 1 OS=Homo sapiens OX=9606 GN=IGHA1 PE=1 SV=2 | 255 | 20201118_12812_1ng_glyco_hcd203040_compound15.21042.21042.4 | controllerType=0 controllerNumber=1 scan=21042 | 42.4435 |
| 17501:4 | 10 | R.LSLHRPALEDLLLGSEAN[+1540.529]LTC[+57.021]TLTGLR.D | HexNAc(2)Hex(7) | C[+57], N[+1541] | 1126.7897 | 4 | 4504.137 | 4504.1268 | 0 | 2.3 | 127 | Specific | 892.41 | 783.46 | 783.46 | 17.86 | 54 | >sp\|P01876\|IGHA1_HUMAN Immunoglobulin heavy constant alpha 1 OS=Homo sapiens OX=9606 GN=IGHA1 PE=1 SV=2 | 255 | 20201118_12812_1ng_glyco_hcd203040_compound15.21072.21072.4 | controllerType=0 controllerNumber=1 scan=21072 | 42.5139 |
| 17589:4 | 10 | R.LSLHRPALEDLLLGSEAN[+1378.476]LTC[+57.021]TLTGLR.D | HexNAc(2)Hex(6) | C[+57], N[+1378] | 1086.277 | 4 | 4342.0863 | 4342.074 | 0 | 2.8 | 127 | Specific | 980.88 | 940.45 | 940.45 | 19.96 | 54 | >sp\|P01876\|IGHA1_HUMAN Immunoglobulin heavy constant alpha 1 OS=Homo sapiens OX=9606 GN=IGHA1 PE=1 SV=2 | 255 | 20201118_12812_1ng_glyco_hcd203040_compound15.21165.21165.4 | controllerType=0 controllerNumber=1 scan=21165 | [illegible] |
| 17547:4 | 10 | R.LSLHRPALEDLLLGSEAN[+1216.423]LTC[+57.021]TLTGLR.D | HexNAc(2)Hex(5) | C[+57], N[+1216] | 1045.7636 | 4 | 4180.0325 | 4180.0211 | 0 | 2.7 | 127 | Specific | 1017.43 | 955.81 | 955.81 | 20.96 | 54 | >sp\|P01876\|IGHA1_HUMAN Immunoglobulin heavy constant alpha 1 OS=Homo sapiens OX=9606 GN=IGHA1 PE=1 SV=2 | 255 | 20201118_12812_1ng_glyco_hcd203040_compound15.21121.21121.4 | controllerType=0 controllerNumber=1 scan=21121 | [illegible] |
| 17465:4 | 10 | R.LSLHRPALEDLLLGSEAN[+1864.634]LTC[+57.021]TLTGLR.D | HexNAc(2)Hex(9) | C[+57], N[+1865] | 1207.8169 | 4 | 4828.2459 | 4828.2324 | 0 | 2.8 | 127 | Specific | 1027.52 | 1012.48 | 1012.48 | 20.96 | 54 | >sp\|P01876\|IGHA1_HUMAN Immunoglobulin heavy constant alpha 1 OS=Homo sapiens OX=9606 GN=IGHA1 PE=1 SV=2 | 255 | 20201118_12812_1ng_glyco_hcd203040_compound15.21034.21034.4 | controllerType=0 controllerNumber=1 scan=21034 | 42.4253 |
| 17475:4 | 10 | R.LSLHRPALEDLLLGSEAN[+1702.581]LTC[+57.021]TLTGLR.D | HexNAc(2)Hex(8) | C[+57], N[+1703] | 1167.3027 | 4 | 4666.1888 | 4666.1796 | 0 | 2 | 127 | Specific | 1034.57 | 1000.74 | 1000.74 | 20.86 | 54 | >sp\|P01876\|IGHA1_HUMAN Immunoglobulin heavy constant alpha 1 OS=Homo sapiens OX=9606 GN=IGHA1 PE=1 SV=2 | 255 | 20201118_12812_1ng_glyco_hcd203040_compound15.21044.21044.4 | controllerType=0 controllerNumber=1 scan=21044 | 42.452 |
| 17503:4 | 10 | R.LSLHRPALEDLLLGSEAN[+1540.529]LTC[+57.021]TLTGLR.D | HexNAc(2)Hex(7) | C[+57], N[+1541] | 1126.7897 | 4 | 4504.137 | 4504.1268 | 0 | 2.3 | 127 | Specific | 1044.7 | 1020.04 | 1020.04 | 20.86 | 54 | >sp\|P01876\|IGHA1_HUMAN Immunoglobulin heavy constant alpha 1 OS=Homo sapiens OX=9606 GN=IGHA1 PE=1 SV=2 | 255 | 20201118_12812_1ng_glyco_hcd203040_compound15.21074.21074.4 | controllerType=0 controllerNumber=1 scan=21074 | 42.5257 |
| 17619:4 | 10 | R.LSLHRPALEDLLLGSEAN[+1257.449]LTC[+57.021]TLTGLR.D | HexNAc(3)Hex(4) | C[+57], N[+1257] | 1056.0196 | 4 | 4221.0565 | 4221.0477 | 0 | 2.1 | 127 | Specific | 1101.44 | 1040.86 | 1040.86 | 22.86 | 54 | >sp\|P01876\|IGHA1_HUMAN Immunoglobulin heavy constant alpha 1 OS=Homo sapiens OX=9606 GN=IGHA1 PE=1 SV=2 | 255 | 20201118_12812_1ng_glyco_hcd203040_compound15.21196.21196.4 | controllerType=0 controllerNumber=1 scan=21196 | 42.823 |
| 11244:3 | 10 | R.LAGKPTHVN[+1825.661]VSVVMAEVDGTC[+57.021].Y | HexNAc(5)Hex(5) | C[+57], N[+1826] | 1337.2568 | 3 | 4009.7558 | 4009.7397 | 0 | 4 | 332 | CRagged | 275.62 | 135.57 | 37.58 | 4.15 | 54 | >sp\|P01876\|IGHA1_HUMAN Immunoglobulin heavy constant alpha 1 OS=Homo sapiens OX=9606 GN=IGHA1 PE=1 SV=2 | 255 | 20201118_12812_1ng_glyco_hcd203040_compound15.14535.14535.3 | controllerType=0 controllerNumber=1 scan=14535 | 30.5769 |
| 14947:4 | 10 | R.LAGKPTHVN[+2350.830]VSVVMAEVDGTC[+57.021]Y.- | HexNAc(4)Hex(5)Fuc(1)NeuAc(2) | C[+57], N[+2351] | 1175.252 | 4 | 4697.9862 | 4697.9724 | 0 | 2.9 | 332 | CRagged | 206.25 | 182.93 | 182.93 | 3.85 | 54 | >sp\|P01876\|IGHA1_HUMAN Immunoglobulin heavy constant alpha 1 OS=Homo sapiens OX=9606 GN=IGHA1 PE=1 SV=2 | 255 | 20201118_12812_1ng_glyco_hcd203040_compound15.18389.18389.4 | controllerType=0 controllerNumber=1 scan=18389 | 37.3505 |
| 14960:4 | 10 | R.LAGKPTHVN[+2553.910]VSVVMAEVDGTC[+57.021]Y.- | HexNAc(5)Hex(5)Fuc(1)NeuAc(2) | C[+57], N[+2554] | 1226.023 | 4 | 4901.0701 | 4901.0518 | 0 | 3.7 | 332 | CRagged | 215.14 | 215.14 | 215.14 | 4.96 | 54 | >sp\|P01876\|IGHA1_HUMAN Immunoglobulin heavy constant alpha 1 OS=Homo sapiens OX=9606 GN=IGHA1 PE=1 SV=2 | 255 | 20201118_12812_1ng_glyco_hcd203040_compound15.18403.18403.4 | controllerType=0 controllerNumber=1 scan=18403 | 37.3791 |
| 11268:3 | 10 | R.LAGKPTHVN[+1971.719]VSVVMAEVDGTC[+57.021]Y.- | HexNAc(5)Hex(5)Fuc(1) | C[+57], N[+1972] | 1440.2947 | 3 | 4318.8697 | 4318.861 | 0 | 2 | 332 | CRagged | 233.94 | 164.93 | 164.93 | 4.97 | 54 | >sp\|P01876\|IGHA1_HUMAN Immunoglobulin heavy constant alpha 1 OS=Homo sapiens OX=9606 GN=IGHA1 PE=1 SV=2 | 255 | 20201118_12812_1ng_glyco_hcd203040_compound15.14560.14560.3 | controllerType=0 controllerNumber=1 scan=14560 | 30.643 |
| 11307:4 | 10 | R.LAGKPTHVN[+1809.666]VSVVMAEVDGTC[+57.021]Y.- | HexNAc(5)Hex(4)Fuc(1) | C[+57], N[+1810] | 1039.9582 | 4 | 4156.8109 | 4156.8081 | 0 | 0.7 | 332 | CRagged | 245.56 | 126.87 | 126.87 | 3.37 | 54 | >sp\|P01876\|IGHA1_HUMAN Immunoglobulin heavy constant alpha 1 OS=Homo sapiens OX=9606 GN=IGHA1 PE=1 SV=2 | 255 | 20201118_12812_1ng_glyco_hcd203040_compound15.14601.14601.4 | controllerType=0 controllerNumber=1 scan=14601 | 30.725 |
| 14966:3 | 10 | R.LAGKPTHVN[+2553.910]VSVVMAEVDGTC[+57.021]Y.- | HexNAc(5)Hex(5)Fuc(1)NeuAc(2) | C[+57], N[+2554] | 1634.3598 | 3 | 4901.0648 | 4901.0518 | 0 | 2.7 | 332 | CRagged | 259.22 | 231.89 | 231.89 | 6.5 | 54 | >sp\|P01876\|IGHA1_HUMAN Immunoglobulin heavy constant alpha 1 OS=Homo sapiens OX=9606 GN=IGHA1 PE=1 SV=2 | 255 | 20201118_12812_1ng_glyco_hcd203040_compound15.18409.18409.3 | controllerType=0 controllerNumber=1 scan=18409 | 37.3933 |
| 13100:3 | 10 | R.LAGKPTHVN[+2262.814]VSVVMAEVDGTC[+57.021]Y.- | HexNAc(5)Hex(5)Fuc(1)NeuAc(1) | C[+57], N[+2263] | 1537.3282 | 3 | 4609.97 | 4609.9564 | 0 | 3 | 332 | CRagged | 276.84 | 276.84 | 276.84 | 7.12 | 54 | >sp\|P01876\|IGHA1_HUMAN Immunoglobulin heavy constant alpha 1 OS=Homo sapiens OX=9606 GN=IGHA1 PE=1 SV=2 | 255 | 20201118_12812_1ng_glyco_hcd203040_compound15.16474.16474.3 | controllerType=0 controllerNumber=1 scan=16474 | 33.8389 |
| 14939:3 | 10 | R.LAGKPTHVN[+2350.830]VSVVMAEVDGTC[+57.021]Y.- | HexNAc(4)Hex(5)Fuc(1)NeuAc(2) | C[+57], N[+2351] | 1566.6655 | 3 | 4697.9818 | 4697.9724 | 0 | 2 | 332 | CRagged | 283.93 | 215.5 | 215.5 | 6.6 | 54 | >sp\|P01876\|IGHA1_HUMAN Immunoglobulin heavy constant alpha 1 OS=Homo sapiens OX=9606 GN=IGHA1 PE=1 SV=2 | 255 | 20201118_12812_1ng_glyco_hcd203040_compound15.18381.18381.3 | controllerType=0 controllerNumber=1 scan=18381 | 37.3373 |
| 11261:4 | 10 | R.LAGKPTHVN[+1809.666]VSVVMAEVDGTC[+57.021]Y.- | HexNAc(5)Hex(4)Fuc(1) | C[+57], N[+1810] | 1039.9599 | 4 | 4156.8177 | 4156.8081 | 0 | 2.3 | 332 | CRagged | 304.74 | 238.09 | 238.09 | 4.97 | 54 | >sp\|P01876\|IGHA1_HUMAN Immunoglobulin heavy constant alpha 1 OS=Homo sapiens OX=9606 GN=IGHA1 PE=1 SV=2 | 255 | 20201118_12812_1ng_glyco_hcd203040_compound15.14553.14553.3 | controllerType=0 controllerNumber=1 scan=14553 | 30.6272 |
| 11259:4 | 10 | R.LAGKPTHVN[+1809.666]VSVVMAEVDGTC[+57.021]Y.- | HexNAc(5)Hex(4)Fuc(1) | C[+57], N[+1810] | 1039.9599 | 4 | 4156.8177 | 4156.8081 | 0 | 2.3 | 332 | CRagged | 345.32 | 188.31 | 188.31 | 6.96 | 54 | >sp\|P01876\|IGHA1_HUMAN Immunoglobulin heavy constant alpha 1 OS=Homo sapiens OX=9606 GN=IGHA1 PE=1 SV=2 | 255 | 20201118_12812_1ng_glyco_hcd203040_compound15.14551.14551.3 | controllerType=0 controllerNumber=1 scan=14551 | 30.6198 |
| 14964:3 | 10 | R.LAGKPTHVN[+2553.910]VSVVMAEVDGTC[+57.021]Y.- | HexNAc(5)Hex(5)Fuc(1)NeuAc(2) | C[+57], N[+2554] | 1634.3598 | 3 | 4901.0648 | 4901.0518 | 0 | 2.7 | 332 | CRagged | 354.82 | 274.96 | 274.96 | 7.54 | 54 | >sp\|P01876\|IGHA1_HUMAN Immunoglobulin heavy constant alpha 1 OS=Homo sapiens OX=9606 GN=IGHA1 PE=1 SV=2 | 255 | 20201118_12812_1ng_glyco_hcd203040_compound15.18407.18407.3 | controllerType=0 controllerNumber=1 scan=18407 | 37.3899 |
| 13102:3 | 10 | R.LAGKPTHVN[+2262.814]VSVVMAEVDGTC[+57.021]Y.- | HexNAc(5)Hex(5)Fuc(1)NeuAc(1) | C[+57], N[+2263] | 1537.3282 | 3 | 4609.97 | 4609.9564 | 0 | 3 | 332 | CRagged | 365.56 | 258.01 | 258.01 | 7.37 | 54 | >sp\|P01876\|IGHA1_HUMAN Immunoglobulin heavy constant alpha 1 OS=Homo sapiens OX=9606 GN=IGHA1 PE=1 SV=2 | 255 | 20201118_12812_1ng_glyco_hcd203040_compound15.16476.16476.3 | controllerType=0 controllerNumber=1 scan=16476 | 33.8441 |
| 14937:3 | 10 | R.LAGKPTHVN[+2350.830]VSVVMAEVDGTC[+57.021]Y.- | HexNAc(4)Hex(5)Fuc(1)NeuAc(2) | C[+57], N[+2351] | 1566.6655 | 3 | 4697.9818 | 4697.9724 | 0 | 2 | 332 | CRagged | 366.13 | 366.13 | 366.13 | 7.79 | 54 | >sp\|P01876\|IGHA1_HUMAN Immunoglobulin heavy constant alpha 1 OS=Homo sapiens OX=9606 GN=IGHA1 PE=1 SV=2 | 255 | 20201118_12812_1ng_glyco_hcd203040_compound15.18379.18379.3 | controllerType=0 controllerNumber=1 scan=18379 | 37.3337 |
| 11305:4 | 10 | R.LAGKPTHVN[+1809.666]VSVVMAEVDGTC[+57.021]Y.- | HexNAc(5)Hex(4)Fuc(1) | C[+57], N[+1810] | 1039.9582 | 4 | 4156.8109 | 4156.8081 | 0 | 0.7 | 332 | CRagged | 372.21 | 346.72 | 306.76 | 7.48 | 54 | >sp\|P01876\|IGHA1_HUMAN Immunoglobulin heavy constant alpha 1 OS=Homo sapiens OX=9606 GN=IGHA1 PE=1 SV=2 | 255 | 20201118_12812_1ng_glyco_hcd203040_compound15.14599.14599.4 | controllerType=0 controllerNumber=1 scan=14599 | 30.7207 |
| 14962:4 | 10 | R.LAGKPTHVN[+2553.910]VSVVMAEVDGTC[+57.021]Y.- | HexNAc(5)Hex(5)Fuc(1)NeuAc(2) | C[+57], N[+2554] | 1226.023 | 4 | 4901.0701 | 4901.0518 | 0 | 3.7 | 332 | CRagged | 376.78 | 271.61 | 271.61 | 7.11 | 54 | >sp\|P01876\|IGHA1_HUMAN Immunoglobulin heavy constant alpha 1 OS=Homo sapiens OX=9606 GN=IGHA1 PE=1 SV=2 | 255 | 20201118_12812_1ng_glyco_hcd203040_compound15.18405.18405.4 | controllerType=0 controllerNumber=1 scan=18405 | 37.3831 |
| 13211:3 | 10 | R.LAGKPTHVN[+1548.545]VSVVMAEVDGTC[+57.021]Y.- | HexNAc(3)Hex(4)NeuAc(1) | C[+57], N[+1549] | 1299.2374 | 3 | 3895.6975 | 3895.6869 | 0 | 2.7 | 332 | CRagged | 382.35 | 382.35 | 382.35 | 9.52 | 54 | >sp\|P01876\|IGHA1_HUMAN Immunoglobulin heavy constant alpha 1 OS=Homo sapiens OX=9606 GN=IGHA1 PE=1 SV=2 | 255 | 20201118_12812_1ng_glyco_hcd203040_compound15.16589.16589.3 | controllerType=0 controllerNumber=1 scan=16589 | 34.0707 |
| 13213:3 | 10 | R.LAGKPTHVN[+1548.545]VSVVMAEVDGTC[+57.021]Y.- | HexNAc(3)Hex(4)NeuAc(1) | C[+57], N[+1549] | 1299.2374 | 3 | 3895.6975 | 3895.6869 | 0 | 2.7 | 332 | CRagged | 396.45 | 338.29 | 338.29 | 6.37 | 54 | >sp\|P01876\|IGHA1_HUMAN Immunoglobulin heavy constant alpha 1 OS=Homo sapiens OX=9606 GN=IGHA1 PE=1 SV=2 | 255 | 20201118_12812_1ng_glyco_hcd203040_compound15.16591.16591.3 | controllerType=0 controllerNumber=1 scan=16591 | 34.0747 |
| 10073:3 | 10 | R.LAGKPTHVN[+1216.423]VSVVMAEVDGTC[+57.021]Y.- | HexNAc(2)Hex(5) | C[+57], N[+1216] | 1188.5289 | 3 | 3563.572 | 3563.5649 | 0 | 2 | 332 | CRagged | 442.54 | 442.54 | 265.73 | 9.79 | 54 | >sp\|P01876\|IGHA1_HUMAN Immunoglobulin heavy constant alpha 1 OS=Homo sapiens OX=9606 GN=IGHA1 PE=1 SV=2 | 255 | 20201118_12812_1ng_glyco_hcd203040_compound15.13323.13323.3 | controllerType=0 controllerNumber=1 scan=13323 | 28.5353 |
| 11325:3 | 10 | R.LAGKPTHVN[+1809.666]VSVVMAEVDGTC[+57.021]Y.- | HexNAc(5)Hex(4)Fuc(1) | C[+57], N[+1810] | 1386.2785 | 3 | 4156.8209 | 4156.8081 | 0 | 3.1 | 332 | CRagged | 474.44 | 474.44 | 359.52 | 9.87 | 54 | >sp\|P01876\|IGHA1_HUMAN Immunoglobulin heavy constant alpha 1 OS=Homo sapiens OX=9606 GN=IGHA1 PE=1 SV=2 | 255 | 20201118_12812_1ng_glyco_hcd203040_compound15.14620.14620.3 | controllerType=0 controllerNumber=1 scan=14620 | 30.7588 |
| 13158:3 | 10 | R.LAGKPTHVN[+1710.598]VSVVMAEVDGTC[+57.021]Y.- | HexNAc(3)Hex(5)NeuAc(1) | C[+57], N[+1711] | 1353.2566 | 3 | 4057.7554 | 4057.7397 | 0 | 3.9 | 332 | CRagged | 484.65 | 484.65 | 342.84 | 9.87 | 54 | >sp\|P01876\|IGHA1_HUMAN Immunoglobulin heavy constant alpha 1 OS=Homo sapiens OX=9606 GN=IGHA1 PE=1 SV=2 | 255 | 20201118_12812_1ng_glyco_hcd203040_compound15.16534.16534.3 | controllerType=0 controllerNumber=1 scan=16534 | 33.9525 |
| 11361:4 | 10 | R.LAGKPTHVN[+1647.613]VSVVMAEVDGTC[+57.021]Y.- | HexNAc(5)Hex(3)Fuc(1) | C[+57], N[+1648] | 999.4468 | 4 | 3994.7655 | 3994.7553 | 0 | 2.5 | 332 | CRagged | 487.33 | 385.79 | 385.79 | 10.06 | 54 | >sp\|P01876\|IGHA1_HUMAN Immunoglobulin heavy constant alpha 1 OS=Homo sapiens OX=9606 GN=IGHA1 PE=1 SV=2 | 255 | 20201118_12812_1ng_glyco_hcd203040_compound15.14657.14657.4 | controllerType=0 controllerNumber=1 scan=14657 | 30.8253 |
| 13160:3 | 10 | R.LAGKPTHVN[+1710.598]VSVVMAEVDGTC[+57.021]Y.- | HexNAc(3)Hex(5)NeuAc(1) | C[+57], N[+1711] | 1353.2566 | 3 | 4057.7554 | 4057.7397 | 0 | 3.9 | 332 | CRagged | 545.32 | 521.91 | 351.22 | 10.07 | 54 | >sp\|P01876\|IGHA1_HUMAN Immunoglobulin heavy constant alpha 1 OS=Homo sapiens OX=9606 GN=IGHA1 PE=1 SV=2 | 255 | 20201118_12812_1ng_glyco_hcd203040_compound15.16536.16536.3 | controllerType=0 controllerNumber=1 scan=16536 | 33.9573 |
| 10075:3 | 10 | R.LAGKPTHVN[+1216.423]VSVVMAEVDGTC[+57.021]Y.- | HexNAc(2)Hex(5) | C[+57], N[+1216] | 1188.5289 | 3 | 3563.572 | 3563.5649 | 0 | 2 | 332 | CRagged | 555.82 | 555.82 | 361.32 | 10.42 | 54 | >sp\|P01876\|IGHA1_HUMAN Immunoglobulin heavy constant alpha 1 OS=Homo sapiens OX=9606 GN=IGHA1 PE=1 SV=2 | 255 | 20201118_12812_1ng_glyco_hcd203040_compound15.13325.13325.3 | controllerType=0 controllerNumber=1 scan=13325 | 28.539 |
| 11330:3 | 10 | R.LAGKPTHVN[+1501.555]VSVVMAEVDGTC[+57.021]Y.- | HexNAc(5)Hex(3) | C[+57], N[+1502] | 1283.5745 | 3 | 3848.709 | 3848.6974 | 0 | 3 | 332 | CRagged | 570.18 | 570.18 | 570.18 | 10.52 | 54 | >sp\|P01876\|IGHA1_HUMAN Immunoglobulin heavy constant alpha 1 OS=Homo sapiens OX=9606 GN=IGHA1 PE=1 SV=2 | 255 | 20201118_12812_1ng_glyco_hcd203040_compound15.14625.14625.3 | controllerType=0 controllerNumber=1 scan=14625 | 30.7681 |
| 11359:4 | 10 | R.LAGKPTHVN[+1647.613]VSVVMAEVDGTC[+57.021]Y.- | HexNAc(5)Hex(3)Fuc(1) | C[+57], N[+1648] | 999.4468 | 4 | 3994.7655 | 3994.7553 | 0 | 2.5 | 332 | CRagged | 591.96 | 591.96 | 591.96 | 11.68 | 54 | >sp\|P01876\|IGHA1_HUMAN Immunoglobulin heavy constant alpha 1 OS=Homo sapiens OX=9606 GN=IGHA1 PE=1 SV=2 | 255 | 20201118_12812_1ng_glyco_hcd203040_compound15.14655.14655.4 | controllerType=0 controllerNumber=1 scan=14655 | 30.8193 |
| 13145:3 | 10 | R.LAGKPTHVN[+1751.624]VSVVMAEVDGTC[+57.021]Y.- | HexNAc(4)Hex(4)NeuAc(1) | C[+57], N[+1752] | 1366.93 | 3 | 4098.7754 | 4098.7663 | 0 | 2.2 | 332 | CRagged | 599.38 | 599.38 | 599.38 | 11.65 | 54 | >sp\|P01876\|IGHA1_HUMAN Immunoglobulin heavy constant alpha 1 OS=Homo sapiens OX=9606 GN=IGHA1 PE=1 SV=2 | 255 | 20201118_12812_1ng_glyco_hcd203040_compound15.16520.16520.3 | controllerType=0 controllerNumber=1 scan=16520 | 33.9198 |
| 13147:3 | 10 | R.LAGKPTHVN[+1751.624]VSVVMAEVDGTC[+57.021]Y.- | HexNAc(4)Hex(4)NeuAc(1) | C[+57], N[+1752] | 1366.93 | 3 | 4098.7754 | 4098.7663 | 0 | 2.2 | 332 | CRagged | 607.58 | 559.88 | 559.88 | 12.65 | 54 | >sp\|P01876\|IGHA1_HUMAN Immunoglobulin heavy constant alpha 1 OS=Homo sapiens OX=9606 GN=IGHA1 PE=1 SV=2 | 255 | 20201118_12812_1ng_glyco_hcd203040_compound15.16522.16522.3 | controllerType=0 controllerNumber=1 scan=16522 | 33.9244 |
| 11327:3 | 10 | R.LAGKPTHVN[+1809.666]VSVVMAEVDGTC[+57.021]Y.- | HexNAc(5)Hex(4)Fuc(1) | C[+57], N[+1810] | 1386.2785 | 3 | 4156.8209 | 4156.8081 | 0 | 3.1 | 332 | CRagged | 686.29 | 523.37 | 429.47 | 13.75 | 54 | >sp\|P01876\|IGHA1_HUMAN Immunoglobulin heavy constant alpha 1 OS=Homo sapiens OX=9606 GN=IGHA1 PE=1 SV=2 | 255 | 20201118_12812_1ng_glyco_hcd203040_compound15.14622.14622.3 | controllerType=0 controllerNumber=1 scan=14622 | 30.7626 |
| 11210:3 | 10 | R.LAGKPTHVN[+1663.608]VSVVMAEVDGTC[+57.021]Y.- | HexNAc(5)Hex(4) | C[+57], N[+1664] | 1337.5916 | 3 | 4010.7603 | 4010.7502 | 0 | 2.5 | 332 | CRagged | 729.29 | 707.9 | 463.37 | 14.75 | 54 | >sp\|P01876\|IGHA1_HUMAN Immunoglobulin heavy constant alpha 1 OS=Homo sapiens OX=9606 GN=IGHA1 PE=1 SV=2 | 255 | 20201118_12812_1ng_glyco_hcd203040_compound15.14500.14500.3 | controllerType=0 controllerNumber=1 scan=14500 | 30.513 |
| 11223:3 | 10 | R.LAGKPTHVN[+1378.476]VSVVMAEVDGTC[+57.021]Y.- | HexNAc(2)Hex(6) | C[+57], N[+1378] | 1242.5468 | 3 | 3725.6259 | 3725.6178 | 0 | 2.2 | 332 | CRagged | 738.95 | 738.95 | 414.89 | 14.65 | 54 | >sp\|P01876\|IGHA1_HUMAN Immunoglobulin heavy constant alpha 1 OS=Homo sapiens OX=9606 GN=IGHA1 PE=1 SV=2 | 255 | 20201118_12812_1ng_glyco_hcd203040_compound15.14514.14514.3 | controllerType=0 controllerNumber=1 scan=14514 | 30.541 |
| 11326:3 | 10 | R.LAGKPTHVN[+1647.613]VSVVMAEVDGTC[+57.021]Y.- | HexNAc(5)Hex(3)Fuc(1) | C[+57], N[+1648] | 1332.2599 | 3 | 3994.7653 | 3994.7553 | 0 | 2.5 | 332 | CRagged | 772.83 | 772.83 | 772.83 | 15.75 | 54 | >sp\|P01876\|IGHA1_HUMAN Immunoglobulin heavy constant alpha 1 OS=Homo sapiens OX=9606 GN=IGHA1 PE=1 SV=2 | 255 | 20201118_12812_1ng_glyco_hcd203040_compound15.14621.14621.3 | controllerType=0 controllerNumber=1 scan=14621 | 30.7602 |
| 11257:3 | 10 | R.LAGKPTHVN[+1809.666]VSVVMAEVDGTC[+57.021]Y.- | HexNAc(5)Hex(4)Fuc(1) | C[+57], N[+1810] | 1386.2768 | 3 | 4156.8158 | 4156.8081 | 0 | 1.8 | 332 | CRagged | 791.52 | 729.69 | 482.12 | 15.65 | 54 | >sp\|P01876\|IGHA1_HUMAN Immunoglobulin heavy constant alpha 1 OS=Homo sapiens OX=9606 GN=IGHA1 PE=1 SV=2 | 255 | 20201118_12812_1ng_glyco_hcd203040_compound15.14549.14549.3 | controllerType=0 controllerNumber=1 scan=14549 | 30.6029 |
| 11207:3 | 10 | R.LAGKPTHVN[+1663.608]VSVVMAEVDGTC[+57.021]Y.- | HexNAc(5)Hex(4) | C[+57], N[+1664] | 1337.5916 | 3 | 4010.7603 | 4010.7502 | 0 | 2.5 | 332 | CRagged | 792.79 | 690.38 | 493.35 | 15.75 | 54 | >sp\|P01876\|IGHA1_HUMAN Immunoglobulin heavy constant alpha 1 OS=Homo sapiens OX=9606 GN=IGHA1 PE=1 SV=2 | 255 | 20201118_12812_1ng_glyco_hcd203040_compound15.14497.14497.3 | controllerType=0 controllerNumber=1 scan=14497 | 30.5063 |
| 11333:3 | 10 | R.LAGKPTHVN[+1501.555]VSVVMAEVDGTC[+57.021]Y.- | HexNAc(5)Hex(3) | C[+57], N[+1502] | 1283.5745 | 3 | 3848.709 | 3848.6974 | 0 | 3 | 332 | CRagged | 819.38 | 784.52 | 784.52 | 16.75 | 54 | >sp\|P01876\|IGHA1_HUMAN Immunoglobulin heavy constant alpha 1 OS=Homo sapiens OX=9606 GN=IGHA1 PE=1 SV=2 | 255 | 20201118_12812_1ng_glyco_hcd203040_compound15.14628.14628.3 | controllerType=0 controllerNumber=1 scan=14628 | 30.7728 |
| 11254:3 | 10 | R.LAGKPTHVN[+1809.666]VSVVMAEVDGTC[+57.021]Y.- | HexNAc(5)Hex(4)Fuc(1) | C[+57], N[+1810] | 1386.2768 | 3 | 4156.8158 | 4156.8081 | 0 | 1.8 | 332 | CRagged | 839.32 | 839.32 | 538.57 | 16.65 | 54 | >sp\|P01876\|IGHA1_HUMAN Immunoglobulin heavy constant alpha 1 OS=Homo sapiens OX=9606 GN=IGHA1 PE=1 SV=2 | 255 | 20201118_12812_1ng_glyco_hcd203040_compound15.14546.14546.3 | controllerType=0 controllerNumber=1 scan=14546 | 30.5964 |
| 11328:3 | 10 | R.LAGKPTHVN[+1647.613]VSVVMAEVDGTC[+57.021]Y.- | HexNAc(5)Hex(3)Fuc(1) | C[+57], N[+1648] | 1332.2599 | 3 | 3994.7653 | 3994.7553 | 0 | 2.5 | 332 | CRagged | 847.48 | 762.27 | 762.27 | 16.75 | 54 | >sp\|P01876\|IGHA1_HUMAN Immunoglobulin heavy constant alpha 1 OS=Homo sapiens OX=9606 GN=IGHA1 PE=1 SV=2 | 255 | 20201118_12812_1ng_glyco_hcd203040_compound15.14623.14623.3 | controllerType=0 controllerNumber=1 scan=14623 | 30.7658 |
| 11269:3 | 10 | R.LAGKPTHVN[+1216.423]VSVVMAEVDGTC[+57.021]Y.- | HexNAc(2)Hex(5) | C[+57], N[+1216] | 1188.5283 | 3 | 3563.5702 | 3563.5649 | 0 | 1.5 | 332 | CRagged | 868.69 | 746.77 | 503.35 | 17.65 | 54 | >sp\|P01876\|IGHA1_HUMAN Immunoglobulin heavy constant alpha 1 OS=Homo sapiens OX=9606 GN=IGHA1 PE=1 SV=2 | 255 | 20201118_12812_1ng_glyco_hcd203040_compound15.14562.14562.3 | controllerType=0 controllerNumber=1 scan=14562 | 30.6479 |
| 11174:3 | 10 | R.LAGKPTHVN[+1540.529]VSVVMAEVDGTC[+57.021]Y.- | HexNAc(2)Hex(7) | C[+57], N[+1541] | 1296.5628 | 3 | 3887.6739 | 3887.6706 | 0 | 0.8 | 332 | CRagged | 868.71 | 775.63 | 528.63 | 17.42 | 54 | >sp\|P01876\|IGHA1_HUMAN Immunoglobulin heavy constant alpha 1 OS=Homo sapiens OX=9606 GN=IGHA1 PE=1 SV=2 | 255 | 20201118_12812_1ng_glyco_hcd203040_compound15.14463.14463.3 | controllerType=0 controllerNumber=1 scan=14463 | 30.4476 |
| 11128:3 | 10 | R.LAGKPTHVN[+1702.581]VSVVMAEVDGTC[+57.021]Y.- | HexNAc(2)Hex(8) | C[+57], N[+1703] | 1350.5831 | 3 | 4049.7346 | 4049.7234 | 0 | 2.8 | 332 | CRagged | 870.48 | 837.17 | 525.53 | 17.75 | 54 | >sp\|P01876\|IGHA1_HUMAN Immunoglobulin heavy constant alpha 1 OS=Homo sapiens OX=9606 GN=IGHA1 PE=1 SV=2 | 255 | 20201118_12812_1ng_glyco_hcd203040_compound15.14415.14415.3 | controllerType=0 controllerNumber=1 scan=14415 | 30.3564 |
| 11271:3 | 10 | R.LAGKPTHVN[+1216.423]VSVVMAEVDGTC[+57.021]Y.- | HexNAc(2)Hex(5) | C[+57], N[+1216] | 1188.5283 | 3 | 3563.5702 | 3563.5649 | 0 | 1.5 | 332 | CRagged | 875.44 | 818.65 | 518.99 | 17.65 | 54 | >sp\|P01876\|IGHA1_HUMAN Immunoglobulin heavy constant alpha 1 OS=Homo sapiens OX=9606 GN=IGHA1 PE=1 SV=2 | 255 | 20201118_12812_1ng_glyco_hcd203040_compound15.14564.14564.3 | controllerType=0 controllerNumber=1 scan=14564 | 30.6568 |
| 11226:3 | 10 | R.LAGKPTHVN[+1378.476]VSVVMAEVDGTC[+57.021]Y.- | HexNAc(2)Hex(6) | C[+57], N[+1378] | 1242.5468 | 3 | 3725.6259 | 3725.6178 | 0 | 2.2 | 332 | CRagged | 877.56 | 856.73 | 524.76 | 17.65 | 54 | >sp\|P01876\|IGHA1_HUMAN Immunoglobulin heavy constant alpha 1 OS=Homo sapiens OX=9606 GN=IGHA1 PE=1 SV=2 | 255 | 20201118_12812_1ng_glyco_hcd203040_compound15.14517.14517.3 | controllerType=0 controllerNumber=1 scan=14517 | 30.5468 |
| 11172:3 | 10 | R.LAGKPTHVN[+1540.529]VSVVMAEVDGTC[+57.021]Y.- | HexNAc(2)Hex(7) | C[+57], N[+1541] | 1296.5628 | 3 | 3887.6739 | 3887.6706 | 0 | 0.8 | 332 | CRagged | 912 | 912 | 582.98 | 18.42 | 54 | >sp\|P01876\|IGHA1_HUMAN Immunoglobulin heavy constant alpha 1 OS=Homo sapiens OX=9606 GN=IGHA1 PE=1 SV=2 | 255 | 20201118_12812_1ng_glyco_hcd203040_compound15.14461.14461.3 | controllerType=0 controllerNumber=1 scan=14461 | 30.4416 |
| 11125:3 | 10 | R.LAGKPTHVN[+1702.581]VSVVMAEVDGTC[+57.021]Y.- | HexNAc(2)Hex(8) | C[+57], N[+1703] | 1350.5831 | 3 | 4049.7346 | 4049.7234 | 0 | 2.8 | 332 | CRagged | 938.91 | 938.91 | 578 | 18.75 | 54 | >sp\|P01876\|IGHA1_HUMAN Immunoglobulin heavy constant alpha 1 OS=Homo sapiens OX=9606 GN=IGHA1 PE=1 SV=2 | 255 | 20201118_12812_1ng_glyco_hcd203040_compound15.14412.14412.3 | controllerType=0 controllerNumber=1 scan=14412 | 30.3491 |
| 18717:4 | 11 | R.NIAAFGGDPNN[+2059.735]ITLFGESAGGASVSLQTLSPYNK.G | HexNAc(4)Hex(5)Fuc(1)NeuAc(1) | N[+2060] | 1368.3639 | 4 | 5470.4337 | 5470.4212 | 0 | 2.3 | 197 | Specific | 415.58 | 415.58 | 415.58 | 9.52 | 50 | >sp\|P19835\|CEL_HUMAN Bile salt-activated lipase OS=Homo sapiens OX=9606 GN=CEL PE=1 SV=3 | 476 | 20201118_12812_1ng_glyco_hcd203040_compound15.22342.22342.4 | controllerType=0 controllerNumber=1 scan=22342 | 45.3691 |
| 18720:4 | 11 | R.NIAAFGGDPNN[+2059.735]ITLFGESAGGASVSLQTLSPYNK.G | HexNAc(4)Hex(5)Fuc(1)NeuAc(1) | N[+2060] | 1368.3639 | 4 | 5470.4337 | 5470.4212 | 0 | 2.3 | 197 | Specific | 557.84 | 492.18 | 492.18 | 10.29 | 50 | >sp\|P19835\|CEL_HUMAN Bile salt-activated lipase OS=Homo sapiens OX=9606 GN=CEL PE=1 SV=3 | 476 | 20201118_12812_1ng_glyco_hcd203040_compound15.22345.22345.4 | controllerType=0 controllerNumber=1 scan=22345 | 45.3761 |
| 18739:3 | 11 | R.NIAAFGGDPNN[+1913.677]ITLFGESAGGASVSLQTLSPYNK.G | HexNAc(4)Hex(5)NeuAc(1) | N[+1914] | 1775.4683 | 3 | 5324.3905 | 5324.3633 | 0 | 5.1 | 197 | Specific | 813.29 | 783.65 | 783.65 | 15.67 | 50 | >sp\|P19835\|CEL_HUMAN Bile salt-activated lipase OS=Homo sapiens OX=9606 GN=CEL PE=1 SV=3 | 476 | 20201118_12812_1ng_glyco_hcd203040_compound15.22365.22365.3 | controllerType=0 controllerNumber=1 scan=22365 | 45.4185 |
| 18721:4 | 11 | R.NIAAFGGDPNN[+1913.677]ITLFGESAGGASVSLQTLSPYNK.G | HexNAc(4)Hex(5)NeuAc(1) | N[+1914] | 1331.8505 | 4 | 5324.38 | 5324.3633 | 0 | 3.1 | 197 | Specific | 892.22 | 850.7 | 850.7 | 17.61 | 50 | >sp\|P19835\|CEL_HUMAN Bile salt-activated lipase OS=Homo sapiens OX=9606 GN=CEL PE=1 SV=3 | 476 | 20201118_12812_1ng_glyco_hcd203040_compound15.22347.22347.4 | controllerType=0 controllerNumber=1 scan=22347 | 45.3806 |
| 18723:4 | 11 | R.NIAAFGGDPNN[+1913.677]ITLFGESAGGASVSLQTLSPYNK.G | HexNAc(4)Hex(5)NeuAc(1) | N[+1914] | 1331.8505 | 4 | 5324.38 | 5324.3633 | 0 | 3.1 | 197 | Specific | 997.16 | 997.16 | 997.16 | 19.61 | 50 | >sp\|P19835\|CEL_HUMAN Bile salt-activated lipase OS=Homo sapiens OX=9606 GN=CEL PE=1 SV=3 | 476 | 20201118_12812_1ng_glyco_hcd203040_compound15.22349.22349.4 | controllerType=0 controllerNumber=1 scan=22349 | 45.3866 |
| 18742:3 | 11 | R.NIAAFGGDPNN[+1913.677]ITLFGESAGGASVSLQTLSPYNK.G | HexNAc(4)Hex(5)NeuAc(1) | N[+1914] | 1775.4683 | 3 | 5324.3905 | 5324.3633 | 0 | 5.1 | 197 | Specific | 1029.93 | 871.28 | 871.28 | 19.67 | 50 | >sp\|P19835\|CEL_HUMAN Bile salt-activated lipase OS=Homo sapiens OX=9606 GN=CEL PE=1 SV=3 | 476 | 20201118_12812_1ng_glyco_hcd203040_compound15.22368.22368.3 | controllerType=0 controllerNumber=1 scan=22368 | 45.4251 |
| 23683:3 | 14 | R.QLAHQSN[+2350.830]STNIFFSPVSIATAFAMLSLGTK.A | HexNAc(4)Hex(5)Fuc(1)NeuAc(2) | N[+2351] | 1844.8307 | 3 | 5532.4774 | 5532.4654 | 0 | 2.2 | 64 | Specific | 209.21 | 199.77 | 199.77 | 5.42 | 39 | >sp\|P01009\|A1AT_HUMAN Alpha-1-antitrypsin OS=Homo sapiens OX=9606 GN=SERPINA1 PE=1 SV=3 | 642 | 20201118_12812_1ng_glyco_hcd203040_compound15.27519.27519.3 | controllerType=0 controllerNumber=1 scan=27519 | 55.7913 |
| 26463:4 | 14 | R.QLAHQSN[+2862.021]STNIFFSPVSIATAFAMLSLGTK.A | HexNAc(5)Hex(6)Fuc(2)NeuAc(2) | N[+2862] | 1511.6755 | 4 | 6043.6803 | 6043.6555 | 0 | 4.1 | 64 | Specific | 238.05 | 238.05 | 238.05 | 6.6 | 39 | >sp\|P01009\|A1AT_HUMAN Alpha-1-antitrypsin OS=Homo sapiens OX=9606 GN=SERPINA1 PE=1 SV=3 | 642 | 20201118_12812_1ng_glyco_hcd203040_compound15.30406.30406.4 | controllerType=0 controllerNumber=1 scan=30406 | 61.0524 |
| 23821:4 | 14 | R.QLAHQSN[+2350.830]STNIFFSPVSIATAFAMLSLGTK.A | HexNAc(4)Hex(5)Fuc(1)NeuAc(2) | N[+2351] | 1383.8766 | 4 | 5532.4845 | 5532.4654 | 0 | 3.5 | 64 | Specific | 253.68 | 253.68 | 253.68 | 7 | 39 | >sp\|P01009\|A1AT_HUMAN Alpha-1-antitrypsin OS=Homo sapiens OX=9606 GN=SERPINA1 PE=1 SV=3 | 642 | 20201118_12812_1ng_glyco_hcd203040_compound15.27662.27662.4 | controllerType=0 controllerNumber=1 scan=27662 | 56.0596 |
| 23717:3 | 14 | R.QLAHQSN[+2350.830]STNIFFSPVSIATAFAMLSLGTK.A | HexNAc(4)Hex(5)Fuc(1)NeuAc(2) | N[+2351] | 1844.8335 | 3 | 5532.4858 | 5532.4654 | 0 | 3.7 | 64 | Specific | 257.01 | 257.01 | 257.01 | 7.07 | 39 | >sp\|P01009\|A1AT_HUMAN Alpha-1-antitrypsin OS=Homo sapiens OX=9606 GN=SERPINA1 PE=1 SV=3 | 642 | 20201118_12812_1ng_glyco_hcd203040_compound15.27554.27554.3 | controllerType=0 controllerNumber=1 scan=27554 | 55.8567 |
| 23790:5 | 14 | R.QLAHQSN[+2204.772]STNIFFSPVSIATAFAMLSLGTK.A | HexNAc(4)Hex(5)NeuAc(2) | N[+2205] | 1078.0895 | 5 | 5386.4183 | 5386.4075 | 0 | 2 | 64 | Specific | 269.41 | 269.41 | 269.41 | 6.99 | 39 | >sp\|P01009\|A1AT_HUMAN Alpha-1-antitrypsin OS=Homo sapiens OX=9606 GN=SERPINA1 PE=1 SV=3 | 642 | 20201118_12812_1ng_glyco_hcd203040_compound15.27630.27630.5 | controllerType=0 controllerNumber=1 scan=27630 | 55.9969 |
| 23850:4 | 14 | R.[+42.011]QLAHQSN[+2190.793]STNIFFSPVSIATAFAMLSLGTK.A | HexNAc(6)Hex(6) | .[42], N[+2191] | 1354.3605 | 4 | 5414.4201 | 5414.4388 | 0 | -3.5 | 64 | Specific | 280.53 | 280.53 | 280.53 | 5.64 | 39 | >sp\|P01009\|A1AT_HUMAN Alpha-1-antitrypsin OS=Homo sapiens OX=9606 GN=SERPINA1 PE=1 SV=3 | 642 | 20201118_12812_1ng_glyco_hcd203040_compound15.27693.27693.4 | controllerType=0 controllerNumber=1 scan=27693 | 56.1241 |
| 26465:4 | 14 | R.QLAHQSN[+2862.021]STNIFFSPVSIATAFAMLSLGTK.A | HexNAc(5)Hex(6)Fuc(2)NeuAc(2) | N[+2862] | 1511.6755 | 4 | 6043.6803 | 6043.6555 | 0 | 4.1 | 64 | Specific | 281.28 | 281.28 | 281.28 | 6.99 | 39 | >sp\|P01009\|A1AT_HUMAN Alpha-1-antitrypsin OS=Homo sapiens OX=9606 GN=SERPINA1 PE=1 SV=3 | 642 | 20201118_12812_1ng_glyco_hcd203040_compound15.30408.30408.4 | controllerType=0 controllerNumber=1 scan=30408 | 61.0571 |
| 23852:4 | 14 | R.[+42.011]QLAHQSN[+2190.793]STNIFFSPVSIATAFAMLSLGTK.A | HexNAc(6)Hex(6) | .[42], N[+2191] | 1354.3605 | 4 | 5414.4201 | 5414.4388 | 0 | -3.5 | 64 | Specific | 322.23 | 322.23 | 322.23 | 6.21 | 39 | >sp\|P01009\|A1AT_HUMAN Alpha-1-antitrypsin OS=Homo sapiens OX=9606 GN=SERPINA1 PE=1 SV=3 | 642 | 20201118_12812_1ng_glyco_hcd203040_compound15.27695.27695.4 | controllerType=0 controllerNumber=1 scan=27695 | 56.1299 |
| 26537:4 | 14 | R.QLAHQSN[+2861.000]STNIFFSPVSIATAFAMLSLGTK.A | HexNAc(5)Hex(6)NeuAc(3) | N[+2861] | 1511.4171 | 4 | 6042.6466 | 6042.6351 | 0 | 1.9 | 64 | Specific | 346.35 | 346.35 | 346.35 | 8.02 | 39 | >sp\|P01009\|A1AT_HUMAN Alpha-1-antitrypsin OS=Homo sapiens OX=9606 GN=SERPINA1 PE=1 SV=3 | 642 | 20201118_12812_1ng_glyco_hcd203040_compound15.30482.30482.2 | controllerType=0 controllerNumber=1 scan=30482 | 61.1924 |
| 23823:4 | 14 | R.QLAHQSN[+2350.830]STNIFFSPVSIATAFAMLSLGTK.A | HexNAc(4)Hex(5)Fuc(1)NeuAc(2) | N[+2351] | 1383.8766 | 4 | 5532.4845 | 5532.4654 | 0 | 3.5 | 64 | Specific | 348.48 | 348.48 | 348.48 | 8.12 | 39 | >sp\|P01009\|A1AT_HUMAN Alpha-1-antitrypsin OS=Homo sapiens OX=9606 GN=SERPINA1 PE=1 SV=3 | 642 | 20201118_12812_1ng_glyco_hcd203040_compound15.27664.27664.4 | controllerType=0 controllerNumber=1 scan=27664 | 56.0644 |
| 23674:4 | 14 | R.QLAHQSN[+2350.830]STNIFFSPVSIATAFAMLSLGTK.A | HexNAc(4)Hex(5)Fuc(1)NeuAc(2) | N[+2351] | 1383.875 | 4 | 5532.4782 | 5532.4654 | 0 | 2.3 | 64 | Specific | 590.98 | 554.78 | 554.78 | 11.79 | 39 | >sp\|P01009\|A1AT_HUMAN Alpha-1-antitrypsin OS=Homo sapiens OX=9606 GN=SERPINA1 PE=1 SV=3 | 642 | 20201118_12812_1ng_glyco_hcd203040_compound15.27510.27510.4 | controllerType=0 controllerNumber=1 scan=27510 | 55.7732 |
| 23854:4 | 14 | R.QLAHQSN[+2204.772]STNIFFSPVSIATAFAMLSLGTK.A | HexNAc(4)Hex(5)NeuAc(2) | N[+2205] | 1347.3573 | 4 | 5386.4074 | 5386.4075 | 0 | 0 | 64 | Specific | 699.38 | 633.47 | 510.5 | 12.85 | 39 | >sp\|P01009\|A1AT_HUMAN Alpha-1-antitrypsin OS=Homo sapiens OX=9606 GN=SERPINA1 PE=1 SV=3 | 642 | 20201118_12812_1ng_glyco_hcd203040_compound15.27697.27697.4 | controllerType=0 controllerNumber=1 scan=27697 | 56.1333 |
| 23700:4 | 14 | R.QLAHQSN[+2350.830]STNIFFSPVSIATAFAMLSLGTK.A | HexNAc(4)Hex(5)Fuc(1)NeuAc(2) | N[+2351] | 1383.8772 | 4 | 5532.487 | 5532.4654 | 0 | 3.9 | 64 | Specific | 749.04 | 711.21 | 711.21 | 14.62 | 39 | >sp\|P01009\|A1AT_HUMAN Alpha-1-antitrypsin OS=Homo sapiens OX=9606 GN=SERPINA1 PE=1 SV=3 | 642 | 20201118_12812_1ng_glyco_hcd203040_compound15.27537.27537.4 | controllerType=0 controllerNumber=1 scan=27537 | 55.8221 |
| 23702:4 | 14 | R.QLAHQSN[+2350.830]STNIFFSPVSIATAFAMLSLGTK.A | HexNAc(4)Hex(5)Fuc(1)NeuAc(2) | N[+2351] | 1383.8772 | 4 | 5532.487 | 5532.4654 | 0 | 3.9 | 64 | Specific | 749.96 | 705.32 | 705.32 | 14.62 | 39 | >sp\|P01009\|A1AT_HUMAN Alpha-1-antitrypsin OS=Homo sapiens OX=9606 GN=SERPINA1 PE=1 SV=3 | 642 | 20201118_12812_1ng_glyco_hcd203040_compound15.27539.27539.4 | controllerType=0 controllerNumber=1 scan=27539 | 55.8294 |
| 23676:4 | 14 | R.QLAHQSN[+2350.830]STNIFFSPVSIATAFAMLSLGTK.A | HexNAc(4)Hex(5)Fuc(1)NeuAc(2) | N[+2351] | 1383.875 | 4 | 5532.4782 | 5532.4654 | 0 | 2.3 | 64 | Specific | 768.88 | 672.83 | 672.83 | 15.79 | 39 | >sp\|P01009\|A1AT_HUMAN Alpha-1-antitrypsin OS=Homo sapiens OX=9606 GN=SERPINA1 PE=1 SV=3 | 642 | 20201118_12812_1ng_glyco_hcd203040_compound15.27512.27512.4 | controllerType=0 controllerNumber=1 scan=27512 | 55.7774 |
| 23856:4 | 14 | R.QLAHQSN[+2204.772]STNIFFSPVSIATAFAMLSLGTK.A | HexNAc(4)Hex(5)NeuAc(2) | N[+2205] | 1347.3573 | 4 | 5386.4074 | 5386.4075 | 0 | 0 | 64 | Specific | 781.47 | 712.38 | 538.65 | 14.85 | 39 | >sp\|P01009\|A1AT_HUMAN Alpha-1-antitrypsin OS=Homo sapiens OX=9606 GN=SERPINA1 PE=1 SV=3 | 642 | 20201118_12812_1ng_glyco_hcd203040_compound15.27699.27699.4 | controllerType=0 controllerNumber=1 scan=27699 | 56.1386 |

Table S4. Cont.

| Query #:z | Protein Rank | Glycopeptide | Glycan | Modification Type(s) | Observed m/z | z | Observed (M+H) | Calc. mass (M+H) | Off-by-x error | Mass error (ppm) | StartPosition | Cleavage | Score | Delta | Delta Mod | \|Log Prob\| | # of unique peptides | Protein Name | Protein DB number | Comment | Scan # | Scan Time |
|---|---|---|---|---|---|---|---|---|---|---|---|---|---|---|---|---|---|---|---|---|---|---|
| 23704:3 | 14 | R.QLAHQSN[+2204.772]STNIFFSPVSIATAFAMLSLGTK.A | HexNAc(4)Hex(5)NeuAc(2) | N[+2205] | 1796.1482 | 3 | 5386.4299 | 5386.4075 | 0 | 4.2 | 64 | Specific | 823.08 | 787.88 | 787.88 | 16.69 | 39 | >sp\|P01009\|A1AT_HUMAN Alpha-1-antitrypsin OS=Homo sapiens OX=9606 GN=SERPINA1 PE=1 SV=3 | 642 | 20201118_12812_1ng_glyco_hcd203040_compound15.27541.27541.3 | controllerType=0 controllerNumber=1 scan=27541 | 55.8327 |
| 23706:3 | 14 | R.QLAHQSN[+2204.772]STNIFFSPVSIATAFAMLSLGTK.A | HexNAc(4)Hex(5)NeuAc(2) | N[+2205] | 1796.1482 | 3 | 5386.4299 | 5386.4075 | 0 | 4.2 | 64 | Specific | 862.77 | 814.22 | 814.22 | 17.69 | 39 | >sp\|P01009\|A1AT_HUMAN Alpha-1-antitrypsin OS=Homo sapiens OX=9606 GN=SERPINA1 PE=1 SV=3 | 642 | 20201118_12812_1ng_glyco_hcd203040_compound15.27543.27543.3 | controllerType=0 controllerNumber=1 scan=27543 | 55.8378 |
| 23701:4 | 14 | R.QLAHQSN[+2204.772]STNIFFSPVSIATAFAMLSLGTK.A | HexNAc(4)Hex(5)NeuAc(2) | N[+2205] | 1347.3606 | 4 | 5386.4206 | 5386.4075 | 0 | 2.4 | 64 | Specific | 914.58 | 882.35 | 592.36 | 18.79 | 39 | >sp\|P01009\|A1AT_HUMAN Alpha-1-antitrypsin OS=Homo sapiens OX=9606 GN=SERPINA1 PE=1 SV=3 | 642 | 20201118_12812_1ng_glyco_hcd203040_compound15.27538.27538.4 | controllerType=0 controllerNumber=1 scan=27538 | 55.8275 |
| 23699:4 | 14 | R.QLAHQSN[+2204.772]STNIFFSPVSIATAFAMLSLGTK.A | HexNAc(4)Hex(5)NeuAc(2) | N[+2205] | 1347.3606 | 4 | 5386.4206 | 5386.4075 | 0 | 2.4 | 64 | Specific | 980.94 | 935.95 | 594.65 | 19.79 | 39 | >sp\|P01009\|A1AT_HUMAN Alpha-1-antitrypsin OS=Homo sapiens OX=9606 GN=SERPINA1 PE=1 SV=3 | 642 | 20201118_12812_1ng_glyco_hcd203040_compound15.27536.27536.4 | controllerType=0 controllerNumber=1 scan=27536 | 55.8206 |
| 20035:3 | 14 | K.YLGN[+2204.772]ATAIFFLPDEGK.L | HexNAc(4)Hex(5)NeuAc(2) | N[+2205] | 1320.8984 | 3 | 3960.6806 | 3960.6677 | 0 | 3.3 | 268 | Specific | 721.41 | 710.75 | 710.75 | 14.68 | 39 | >sp\|P01009\|A1AT_HUMAN Alpha-1-antitrypsin OS=Homo sapiens OX=9606 GN=SERPINA1 PE=1 SV=3 | 642 | 20201118_12812_1ng_glyco_hcd203040_compound15.23718.23718.3 | controllerType=0 controllerNumber=1 scan=23718 | 48.2296 |
| 20037:3 | 14 | K.YLGN[+2204.772]ATAIFFLPDEGK.L | HexNAc(4)Hex(5)NeuAc(2) | N[+2205] | 1320.8984 | 3 | 3960.6806 | 3960.6677 | 0 | 3.3 | 268 | Specific | 786.88 | 741.65 | 741.65 | 15.68 | 39 | >sp\|P01009\|A1AT_HUMAN Alpha-1-antitrypsin OS=Homo sapiens OX=9606 GN=SERPINA1 PE=1 SV=3 | 642 | 20201118_12812_1ng_glyco_hcd203040_compound15.23720.23720.3 | controllerType=0 controllerNumber=1 scan=23720 | 48.2341 |
| 19795:4 | 14 | K.YLGN[+2204.772]ATAIFFLPDEGKLQHLENELTHDIITK.F | HexNAc(4)Hex(5)NeuAc(2) | N[+2205] | 1437.1624 | 4 | 5745.6276 | 5745.6098 | 0 | 3.1 | 268 | Specific | 927 | 844.53 | 844.53 | 17.56 | 39 | >sp\|P01009\|A1AT_HUMAN Alpha-1-antitrypsin OS=Homo sapiens OX=9606 GN=SERPINA1 PE=1 SV=3 | 642 | 20201118_12812_1ng_glyco_hcd203040_compound15.23467.23467.4 | controllerType=0 controllerNumber=1 scan=23467 | 47.6793 |
| 19792:5 | 14 | K.YLGN[+2204.772]ATAIFFLPDEGKLQHLENELTHDIITK.F | HexNAc(4)Hex(5)NeuAc(2) | N[+2205] | 1149.9312 | 5 | 5745.6267 | 5745.6098 | 0 | 2.9 | 268 | Specific | 952.42 | 889.84 | 889.84 | 19.18 | 39 | >sp\|P01009\|A1AT_HUMAN Alpha-1-antitrypsin OS=Homo sapiens OX=9606 GN=SERPINA1 PE=1 SV=3 | 642 | 20201118_12812_1ng_glyco_hcd203040_compound15.23464.23464.5 | controllerType=0 controllerNumber=1 scan=23464 | 47.6694 |
| 19789:5 | 14 | K.YLGN[+2204.772]ATAIFFLPDEGKLQHLENELTHDIITK.F | HexNAc(4)Hex(5)NeuAc(2) | N[+2205] | 1149.9312 | 5 | 5745.6267 | 5745.6098 | 0 | 2.9 | 268 | Specific | 1149.64 | 1108.4 | 1108.4 | 22.18 | 39 | >sp\|P01009\|A1AT_HUMAN Alpha-1-antitrypsin OS=Homo sapiens OX=9606 GN=SERPINA1 PE=1 SV=3 | 642 | 20201118_12812_1ng_glyco_hcd203040_compound15.23461.23461.5 | controllerType=0 controllerNumber=1 scan=23461 | 47.653 |
| 19793:4 | 14 | K.YLGN[+2204.772]ATAIFFLPDEGKLQHLENELTHDIITK.F | HexNAc(4)Hex(5)NeuAc(2) | N[+2205] | 1437.1624 | 4 | 5745.6276 | 5745.6098 | 0 | 3.1 | 268 | Specific | 1287.18 | 1151.46 | 1151.46 | 24.56 | 39 | >sp\|P01009\|A1AT_HUMAN Alpha-1-antitrypsin OS=Homo sapiens OX=9606 GN=SERPINA1 PE=1 SV=3 | 642 | 20201118_12812_1ng_glyco_hcd203040_compound15.23465.23465.4 | controllerType=0 controllerNumber=1 scan=23465 | 47.6707 |
| 05452:3 | 15 | R.LSPN[+2205.793]ASAEHLELR.W | HexNAc(4)Hex(5)Fuc(2)NeuAc(1) | N[+2206] | 1214.8563 | 3 | 3642.5542 | 3642.5421 | 0 | 3.3 | 52 | Specific | 277.73 | 277.73 | 277.73 | 6.95 | 37 | >sp\|Q13410\|BT1A1_HUMAN Butyrophilin subfamily 1 member A1 OS=Homo sapiens OX=9606 GN=BTN1A1 PE=1 SV=3 | 179 | 20201118_12812_1ng_glyco_hcd203040_compound15.8523.8523.3 | controllerType=0 controllerNumber=1 scan=8523 | 20.6191 |
| 05454:3 | 15 | R.LSPN[+2205.793]ASAEHLELR.W | HexNAc(4)Hex(5)Fuc(2)NeuAc(1) | N[+2206] | 1214.8563 | 3 | 3642.5542 | 3642.5421 | 0 | 3.3 | 52 | Specific | 311.24 | 161 | 161 | 5.97 | 37 | >sp\|Q13410\|BT1A1_HUMAN Butyrophilin subfamily 1 member A1 OS=Homo sapiens OX=9606 GN=BTN1A1 PE=1 SV=3 | 179 | 20201118_12812_1ng_glyco_hcd203040_compound15.8525.8525.3 | controllerType=0 controllerNumber=1 scan=8525 | 20.6233 |
| 07471:3 | 15 | R.LSPN[+2350.830]ASAEHLELR.W | HexNAc(4)Hex(5)Fuc(1)NeuAc(2) | N[+2351] | 1263.2004 | 3 | 3787.5866 | 3787.5796 | 0 | 1.8 | 52 | Specific | 599.55 | 568.17 | 568.17 | 11.47 | 37 | >sp\|Q13410\|BT1A1_HUMAN Butyrophilin subfamily 1 member A1 OS=Homo sapiens OX=9606 GN=BTN1A1 PE=1 SV=3 | 179 | 20201118_12812_1ng_glyco_hcd203040_compound15.10612.10612.3 | controllerType=0 controllerNumber=1 scan=10612 | 24.0352 |
| 07469:3 | 15 | R.LSPN[+2350.830]ASAEHLELR.W | HexNAc(4)Hex(5)Fuc(1)NeuAc(2) | N[+2351] | 1263.2004 | 3 | 3787.5866 | 3787.5796 | 0 | 1.8 | 52 | Specific | 667.86 | 454.93 | 454.93 | 13.47 | 37 | >sp\|Q13410\|BT1A1_HUMAN Butyrophilin subfamily 1 member A1 OS=Homo sapiens OX=9606 GN=BTN1A1 PE=1 SV=3 | 179 | 20201118_12812_1ng_glyco_hcd203040_compound15.10610.10610.3 | controllerType=0 controllerNumber=1 scan=10610 | 24.0309 |
| 19537:3 | 15 | K.N[+2204.772]VSC[+57.021]YIQNLLLGQEK.K | HexNAc(4)Hex(5)NeuAc(2) | C[+57], N[+2205] | 1328.5724 | 3 | 3983.7028 | 3983.683 | 0 | 5 | 215 | Specific | 356.36 | 169.66 | 169.66 | 6.4 | 37 | >sp\|Q13410\|BT1A1_HUMAN Butyrophilin subfamily 1 member A1 OS=Homo sapiens OX=9606 GN=BTN1A1 PE=1 SV=3 | 179 | 20201118_12812_1ng_glyco_hcd203040_compound15.23198.23198.3 | controllerType=0 controllerNumber=1 scan=23198 | 47.0951 |
| 17492:3 | 15 | K.N[+1913.677]VSC[+57.021]YIQNLLLGQEK.K | HexNAc(4)Hex(5)NeuAc(1) | C[+57], N[+1914] | 1231.5384 | 3 | 3692.6006 | 3692.5876 | 0 | 3.5 | 215 | Specific | 606.81 | 525.17 | 525.17 | 12.5 | 37 | >sp\|Q13410\|BT1A1_HUMAN Butyrophilin subfamily 1 member A1 OS=Homo sapiens OX=9606 GN=BTN1A1 PE=1 SV=3 | 179 | 20201118_12812_1ng_glyco_hcd203040_compound15.21062.21062.3 | controllerType=0 controllerNumber=1 scan=21062 | 42.4911 |
| 17494:3 | 15 | K.N[+1913.677]VSC[+57.021]YIQNLLLGQEK.K | HexNAc(4)Hex(5)NeuAc(1) | C[+57], N[+1914] | 1231.5384 | 3 | 3692.6006 | 3692.5876 | 0 | 3.5 | 215 | Specific | 700.52 | 535.06 | 535.06 | 14.5 | 37 | >sp\|Q13410\|BT1A1_HUMAN Butyrophilin subfamily 1 member A1 OS=Homo sapiens OX=9606 GN=BTN1A1 PE=1 SV=3 | 179 | 20201118_12812_1ng_glyco_hcd203040_compound15.21064.21064.3 | controllerType=0 controllerNumber=1 scan=21064 | 42.4951 |
| 22572:5 | 17 | K.APDKNVIFSPLSISTALAFLSLGAHN[+2350.830]TTLTEILK.G | HexNAc(4)Hex(5)Fuc(1)NeuAc(2) | N[+2351] | 1187.5742 | 5 | 5933.842 | 5933.8085 | 0 | 5.6 | 68 | Specific | 393.72 | 393.72 | 393.72 | 8.07 | 34 | >sp\|P01011\|AACT_HUMAN Alpha-1-antichymotrypsin OS=Homo sapiens OX=9606 GN=SERPINA3 PE=1 SV=2 | 1314 | 20201118_12812_1ng_glyco_hcd203040_compound15.26364.26364.5 | controllerType=0 controllerNumber=1 scan=26364 | 53.5332 |
| 22574:5 | 17 | K.APDKNVIFSPLSISTALAFLSLGAHN[+2350.830]TTLTEILK.G | HexNAc(4)Hex(5)Fuc(1)NeuAc(2) | N[+2351] | 1187.5742 | 5 | 5933.842 | 5933.8085 | 0 | 5.6 | 68 | Specific | 503.65 | 503.65 | 503.65 | 8.59 | 34 | >sp\|P01011\|AACT_HUMAN Alpha-1-antichymotrypsin OS=Homo sapiens OX=9606 GN=SERPINA3 PE=1 SV=2 | 1314 | 20201118_12812_1ng_glyco_hcd203040_compound15.26366.26366.5 | controllerType=0 controllerNumber=1 scan=26366 | 53.5377 |
| 21176:4 | 17 | K.APDKNVIFSPLSISTALAFLSLGAHN[+2059.735]TTLTEILK.G | HexNAc(4)Hex(5)Fuc(1)NeuAc(1) | N[+2060] | 1411.4393 | 4 | 5642.7355 | 5642.7131 | 0 | 4 | 68 | Specific | 584.73 | 584.73 | 584.73 | 10.92 | 34 | >sp\|P01011\|AACT_HUMAN Alpha-1-antichymotrypsin OS=Homo sapiens OX=9606 GN=SERPINA3 PE=1 SV=2 | 1314 | 20201118_12812_1ng_glyco_hcd203040_compound15.24908.24908.4 | controllerType=0 controllerNumber=1 scan=24908 | 50.772 |
| 21173:4 | 17 | K.APDKNVIFSPLSISTALAFLSLGAHN[+2059.735]TTLTEILK.G | HexNAc(4)Hex(5)Fuc(1)NeuAc(1) | N[+2060] | 1411.4393 | 4 | 5642.7355 | 5642.7131 | 0 | 4 | 68 | Specific | 756.45 | 756.45 | 756.45 | 14.92 | 34 | >sp\|P01011\|AACT_HUMAN Alpha-1-antichymotrypsin OS=Homo sapiens OX=9606 GN=SERPINA3 PE=1 SV=2 | 1314 | 20201118_12812_1ng_glyco_hcd203040_compound15.24905.24905.4 | controllerType=0 controllerNumber=1 scan=24905 | 50.7675 |
| 24312:4 | 17 | K.NVIFSPLSISTALAFLSLGAHN[+2350.830]TTLTEILK.G | HexNAc(4)Hex(5)Fuc(1)NeuAc(2) | N[+2351] | 1381.4085 | 4 | 5522.612 | 5522.5967 | 0 | 2.8 | 72 | Specific | 245.75 | 245.75 | 245.75 | 6.87 | 34 | >sp\|P01011\|AACT_HUMAN Alpha-1-antichymotrypsin OS=Homo sapiens OX=9606 GN=SERPINA3 PE=1 SV=2 | 1314 | 20201118_12812_1ng_glyco_hcd203040_compound15.28173.28173.4 | controllerType=0 controllerNumber=1 scan=28173 | 57.0168 |
| 24305:4 | 17 | K.NVIFSPLSISTALAFLSLGAHN[+2350.830]TTLTEILK.G | HexNAc(4)Hex(5)Fuc(1)NeuAc(2) | N[+2351] | 1381.4098 | 4 | 5522.6173 | 5522.5967 | 0 | 3.7 | 72 | Specific | 256.41 | 256.41 | 256.41 | 7 | 34 | >sp\|P01011\|AACT_HUMAN Alpha-1-antichymotrypsin OS=Homo sapiens OX=9606 GN=SERPINA3 PE=1 SV=2 | 1314 | 20201118_12812_1ng_glyco_hcd203040_compound15.28165.28165.2 | controllerType=0 controllerNumber=1 scan=28165 | 57.0017 |
| 24307:4 | 17 | K.NVIFSPLSISTALAFLSLGAHN[+2350.830]TTLTEILK.G | HexNAc(4)Hex(5)Fuc(1)NeuAc(2) | N[+2351] | 1381.4098 | 4 | 5522.6173 | 5522.5967 | 0 | 3.7 | 72 | Specific | 263.07 | 263.07 | 263.07 | 7.07 | 34 | >sp\|P01011\|AACT_HUMAN Alpha-1-antichymotrypsin OS=Homo sapiens OX=9606 GN=SERPINA3 PE=1 SV=2 | 1314 | 20201118_12812_1ng_glyco_hcd203040_compound15.28167.28167.2 | controllerType=0 controllerNumber=1 scan=28167 | 57.0052 |
| 24310:4 | 17 | K.NVIFSPLSISTALAFLSLGAHN[+2350.830]TTLTEILK.G | HexNAc(4)Hex(5)Fuc(1)NeuAc(2) | N[+2351] | 1381.4085 | 4 | 5522.612 | 5522.5967 | 0 | 2.8 | 72 | Specific | 277.38 | 277.38 | 277.38 | 7.26 | 34 | >sp\|P01011\|AACT_HUMAN Alpha-1-antichymotrypsin OS=Homo sapiens OX=9606 GN=SERPINA3 PE=1 SV=2 | 1314 | 20201118_12812_1ng_glyco_hcd203040_compound15.28171.28171.4 | controllerType=0 controllerNumber=1 scan=28171 | 57.0133 |
| 20714:3 | 18 | K.MLN[+2059.735]TSSLLEQLNEQFNWVSR.L | HexNAc(4)Hex(5)Fuc(1)NeuAc(1) | N[+2060] | 1490.317 | 3 | 4468.9363 | 4468.9217 | 0 | 3.3 | 352 | Specific | 586.02 | 520.54 | 520.54 | 11.68 | 29 | >sp\|P10909\|CLUS_HUMAN Clusterin OS=Homo sapiens OX=9606 GN=CLU PE=1 SV=1 | 1467 | 20201118_12812_1ng_glyco_hcd203040_compound15.24428.24428.3 | controllerType=0 controllerNumber=1 scan=24428 | 49.8432 |
| 20726:4 | 18 | K.MLN[+2059.735]TSSLLEQLNEQFNWVSR.L | HexNAc(4)Hex(5)Fuc(1)NeuAc(1) | N[+2060] | 1117.9888 | 4 | 4468.9334 | 4468.9217 | 0 | 2.6 | 352 | Specific | 640.89 | 572.23 | 572.23 | 12.7 | 29 | >sp\|P10909\|CLUS_HUMAN Clusterin OS=Homo sapiens OX=9606 GN=CLU PE=1 SV=1 | 1467 | 20201118_12812_1ng_glyco_hcd203040_compound15.24440.24440.4 | controllerType=0 controllerNumber=1 scan=24440 | 49.8633 |
| 22994:4 | 18 | K.MLN[+2350.830]TSSLLEQLNEQFNWVSR.L | HexNAc(4)Hex(5)Fuc(1)NeuAc(2) | N[+2351] | 1190.7643 | 4 | 4760.0355 | 4760.0171 | 0 | 3.9 | 352 | Specific | 648.62 | 437.21 | 437.21 | 12.44 | 29 | >sp\|P10909\|CLUS_HUMAN Clusterin OS=Homo sapiens OX=9606 GN=CLU PE=1 SV=1 | 1467 | 20201118_12812_1ng_glyco_hcd203040_compound15.26803.26803.4 | controllerType=0 controllerNumber=1 scan=26803 | 54.4228 |
| 20729:4 | 18 | K.MLN[+2059.735]TSSLLEQLNEQFNWVSR.L | HexNAc(4)Hex(5)Fuc(1)NeuAc(1) | N[+2060] | 1117.9888 | 4 | 4468.9334 | 4468.9217 | 0 | 2.6 | 352 | Specific | 736.46 | 614.86 | 614.86 | 14.7 | 29 | >sp\|P10909\|CLUS_HUMAN Clusterin OS=Homo sapiens OX=9606 GN=CLU PE=1 SV=1 | 1467 | 20201118_12812_1ng_glyco_hcd203040_compound15.24443.24443.4 | controllerType=0 controllerNumber=1 scan=24443 | 49.8713 |
| 22996:4 | 18 | K.MLN[+2350.830]TSSLLEQLNEQFNWVSR.L | HexNAc(4)Hex(5)Fuc(1)NeuAc(2) | N[+2351] | 1190.7643 | 4 | 4760.0355 | 4760.0171 | 0 | 3.9 | 352 | Specific | 845.23 | 604.16 | 604.16 | 16.44 | 29 | >sp\|P10909\|CLUS_HUMAN Clusterin OS=Homo sapiens OX=9606 GN=CLU PE=1 SV=1 | 1467 | 20201118_12812_1ng_glyco_hcd203040_compound15.26805.26805.4 | controllerType=0 controllerNumber=1 scan=26805 | 54.4272 |
| 20717:3 | 18 | K.MLN[+2059.735]TSSLLEQLNEQFNWVSR.L | HexNAc(4)Hex(5)Fuc(1)NeuAc(1) | N[+2060] | 1490.317 | 3 | 4468.9363 | 4468.9217 | 0 | 3.3 | 352 | Specific | 874.61 | 526.49 | 526.49 | 17.68 | 29 | >sp\|P10909\|CLUS_HUMAN Clusterin OS=Homo sapiens OX=9606 GN=CLU PE=1 SV=1 | 1467 | 20201118_12812_1ng_glyco_hcd203040_compound15.24431.24431.3 | controllerType=0 controllerNumber=1 scan=24431 | 49.848 |
| 23001:3 | 18 | K.MLN[+2350.830]TSSLLEQLNEQFNWVSR.L | HexNAc(4)Hex(5)Fuc(1)NeuAc(2) | N[+2351] | 1587.351 | 3 | 4760.0385 | 4760.0171 | 0 | 4.5 | 352 | Specific | 882.04 | 608.91 | 608.91 | 17.41 | 29 | >sp\|P10909\|CLUS_HUMAN Clusterin OS=Homo sapiens OX=9606 GN=CLU PE=1 SV=1 | 1467 | 20201118_12812_1ng_glyco_hcd203040_compound15.26810.26810.3 | controllerType=0 controllerNumber=1 scan=26810 | 54.4385 |
| 22999:3 | 18 | K.MLN[+2350.830]TSSLLEQLNEQFNWVSR.L | HexNAc(4)Hex(5)Fuc(1)NeuAc(2) | N[+2351] | 1587.351 | 3 | 4760.0385 | 4760.0171 | 0 | 4.5 | 352 | Specific | 960.71 | 642.41 | 642.41 | 19.41 | 29 | >sp\|P10909\|CLUS_HUMAN Clusterin OS=Homo sapiens OX=9606 GN=CLU PE=1 SV=1 | 1467 | 20201118_12812_1ng_glyco_hcd203040_compound15.26808.26808.3 | controllerType=0 controllerNumber=1 scan=26808 | 54.4323 |
| 20245:3 | 18 | K.M[+15.995]LN[+2059.735]TSSLLEQLNEQFNWVSR.L | HexNAc(4)Hex(5)Fuc(1)NeuAc(1) | M[+16], N[+2060] | 1495.6481 | 3 | 4484.9298 | 4484.9166 | 0 | 3 | 352 | Specific | 465.52 | 366.31 | 366.31 | 8.17 | 29 | >sp\|P10909\|CLUS_HUMAN Clusterin OS=Homo sapiens OX=9606 GN=CLU PE=1 SV=1 | 1467 | 20201118_12812_1ng_glyco_hcd203040_compound15.23938.23938.3 | controllerType=0 controllerNumber=1 scan=23938 | 48.7234 |
| 20247:3 | 18 | K.M[+15.995]LN[+2059.735]TSSLLEQLNEQFNWVSR.L | HexNAc(4)Hex(5)Fuc(1)NeuAc(1) | M[+16], N[+2060] | 1495.6481 | 3 | 4484.9298 | 4484.9166 | 0 | 3 | 352 | Specific | 505.46 | 219.26 | 219.26 | 7.57 | 29 | >sp\|P10909\|CLUS_HUMAN Clusterin OS=Homo sapiens OX=9606 GN=CLU PE=1 SV=1 | 1467 | 20201118_12812_1ng_glyco_hcd203040_compound15.23940.23940.3 | controllerType=0 controllerNumber=1 scan=23940 | 48.7315 |
| 09975:3 | 18 | R.LAN[+2570.925]LTQGEDQYYLR.V | HexNAc(5)Hex(6)Fuc(2)NeuAc(1) | N[+2571] | 1418.9295 | 3 | 4254.7739 | 4254.7587 | 0 | 3.6 | 372 | Specific | 283.35 | 247.9 | 247.9 | 6.81 | 29 | >sp\|P10909\|CLUS_HUMAN Clusterin OS=Homo sapiens OX=9606 GN=CLU PE=1 SV=1 | 1467 | 20201118_12812_1ng_glyco_hcd203040_compound15.13221.13221.3 | controllerType=0 controllerNumber=1 scan=13221 | 28.3629 |
| 12945:3 | 18 | R.LAN[+2350.830]LTQGEDQYYLR.V | HexNAc(4)Hex(5)Fuc(1)NeuAc(2) | N[+2351] | 1345.5615 | 3 | 4034.6698 | 4034.664 | 0 | 1.4 | 372 | Specific | 340.03 | 320.84 | 320.84 | 7.34 | 29 | >sp\|P10909\|CLUS_HUMAN Clusterin OS=Homo sapiens OX=9606 GN=CLU PE=1 SV=1 | 1467 | 20201118_12812_1ng_glyco_hcd203040_compound15.16313.16313.3 | controllerType=0 controllerNumber=1 scan=16313 | 33.5628 |
| 12946:3 | 18 | R.LAN[+2350.830]LTQGEDQYYLR.V | HexNAc(4)Hex(5)Fuc(1)NeuAc(2) | N[+2351] | 1345.5615 | 3 | 4034.6698 | 4034.664 | 0 | 1.4 | 372 | Specific | 434.09 | 401.14 | 401.14 | 9.41 | 29 | >sp\|P10909\|CLUS_HUMAN Clusterin OS=Homo sapiens OX=9606 GN=CLU PE=1 SV=1 | 1467 | 20201118_12812_1ng_glyco_hcd203040_compound15.16314.16314.3 | controllerType=0 controllerNumber=1 scan=16314 | 33.5662 |
| 01887:5 | 20 | R.EKQTDEIKDTRN[+2059.735]ESTQNC[+57.021]VVAEPEK.M | HexNAc(4)Hex(5)Fuc(1)NeuAc(1) | C[+57], N[+2060] | 1002.4334 | 5 | 5008.1377 | 5008.1251 | 0 | 2.5 | 58 | Specific | 341.42 | 341.42 | 341.42 | 8.18 | 32 | >sp\|P47710\|CASA1_HUMAN Alpha-S1-casein OS=Homo sapiens OX=9606 GN=CSN1S1 PE=1 SV=1 | 989 | 20201118_12812_1ng_glyco_hcd203040_compound15.4830.4830.5 | controllerType=0 controllerNumber=1 scan=4830 | 14.4521 |
| 01889:5 | 20 | R.EKQTDEIKDTRN[+2059.735]ESTQNC[+57.021]VVAEPEK.M | HexNAc(4)Hex(5)Fuc(1)NeuAc(1) | C[+57], N[+2060] | 1002.4334 | 5 | 5008.1377 | 5008.1251 | 0 | 2.5 | 58 | Specific | 408.92 | 408.92 | 408.92 | 9.82 | 32 | >sp\|P47710\|CASA1_HUMAN Alpha-S1-casein OS=Homo sapiens OX=9606 GN=CSN1S1 PE=1 SV=1 | 989 | 20201118_12812_1ng_glyco_hcd203040_compound15.4832.4832.5 | controllerType=0 controllerNumber=1 scan=4832 | 14.4559 |
| 01875:4 | 20 | R.EKQTDEIKDTRN[+2059.735]ESTQNC[+57.021]VVAEPEK.M | HexNAc(4)Hex(5)Fuc(1)NeuAc(1) | C[+57], N[+2060] | 1252.7903 | 4 | 5008.1393 | 5008.1251 | 0 | 2.8 | 58 | Specific | 502.28 | 358.93 | 358.93 | 8.08 | 32 | >sp\|P47710\|CASA1_HUMAN Alpha-S1-casein OS=Homo sapiens OX=9606 GN=CSN1S1 PE=1 SV=1 | 989 | 20201118_12812_1ng_glyco_hcd203040_compound15.4818.4818.4 | controllerType=0 controllerNumber=1 scan=4818 | 14.4324 |
| 01877:4 | 20 | R.EKQTDEIKDTRN[+2059.735]ESTQNC[+57.021]VVAEPEK.M | HexNAc(4)Hex(5)Fuc(1)NeuAc(1) | C[+57], N[+2060] | 1252.7903 | 4 | 5008.1393 | 5008.1251 | 0 | 2.8 | 58 | Specific | 618.78 | 553.68 | 553.68 | 12.32 | 32 | >sp\|P47710\|CASA1_HUMAN Alpha-S1-casein OS=Homo sapiens OX=9606 GN=CSN1S1 PE=1 SV=1 | 989 | 20201118_12812_1ng_glyco_hcd203040_compound15.4820.4820.4 | controllerType=0 controllerNumber=1 scan=4820 | 14.4366 |
| 09237:4 | 20 | K.[+42.011]QTDEIKDTRN[+2018.708]ESTQNC[+57.021]VVAEPEKMESSISSSSEEMSLS | HexNAc(3)Hex(6)Fuc(1)NeuAc(1) | .[42], C[+57], N[+2019] | 1613.6879 | 4 | 6451.7296 | 6451.6996 | 0 | 4.7 | 60 | Specific | 204.88 | 198.21 | 198.21 | 3.67 | 32 | >sp\|P47710\|CASA1_HUMAN Alpha-S1-casein OS=Homo sapiens OX=9606 GN=CSN1S1 PE=1 SV=1 | 989 | 20201118_12812_1ng_glyco_hcd203040_compound15.12458.12458.2 | controllerType=0 controllerNumber=1 scan=12458 | 27.1337 |
| 09248:5 | 20 | K.[+42.011]QTDEIKDTRN[+2018.708]ESTQNC[+57.021]VVAEPEKMESSISSSSEEMSLS | HexNAc(3)Hex(6)Fuc(1)NeuAc(1) | .[42], C[+57], N[+2019] | 1291.1537 | 5 | 6451.7393 | 6451.6996 | 0 | 6.2 | 60 | Specific | 212.9 | 212.9 | 212.9 | 3.65 | 32 | >sp\|P47710\|CASA1_HUMAN Alpha-S1-casein OS=Homo sapiens OX=9606 GN=CSN1S1 PE=1 SV=1 | 989 | 20201118_12812_1ng_glyco_hcd203040_compound15.12469.12469.3 | controllerType=0 controllerNumber=1 scan=12469 | 27.152 |
| 09239:4 | 20 | K.[+42.011]QTDEIKDTRN[+2018.708]ESTQNC[+57.021]VVAEPEKMESSISSSSEEMSLS | HexNAc(3)Hex(6)Fuc(1)NeuAc(1) | .[42], C[+57], N[+2019] | 1613.6879 | 4 | 6451.7296 | 6451.6996 | 0 | 4.7 | 60 | Specific | 259.39 | 249.76 | 249.76 | 3.92 | 32 | >sp\|P47710\|CASA1_HUMAN Alpha-S1-casein OS=Homo sapiens OX=9606 GN=CSN1S1 PE=1 SV=1 | 989 | 20201118_12812_1ng_glyco_hcd203040_compound15.12460.12460.2 | controllerType=0 controllerNumber=1 scan=12460 | 27.1375 |
| 09325:5 | 20 | K.QTDEIKDTRN[+2059.735]ESTQNC[+57.021]VVAEPEKMESSISSSSEEMSLSK.C | HexNAc(4)Hex(5)Fuc(1)NeuAc(1) | C[+57], N[+2060] | 1290.9536 | 5 | 6450.739 | 6450.7156 | 0 | 3.6 | 60 | Specific | 643.91 | 510.73 | 510.73 | 11.86 | 32 | >sp\|P47710\|CASA1_HUMAN Alpha-S1-casein OS=Homo sapiens OX=9606 GN=CSN1S1 PE=1 SV=1 | 989 | 20201118_12812_1ng_glyco_hcd203040_compound15.12549.12549.5 | controllerType=0 controllerNumber=1 scan=12549 | 27.282 |
| 09323:5 | 20 | K.QTDEIKDTRN[+2059.735]ESTQNC[+57.021]VVAEPEKMESSISSSSEEMSLSK.C | HexNAc(4)Hex(5)Fuc(1)NeuAc(1) | C[+57], N[+2060] | 1290.9536 | 5 | 6450.739 | 6450.7156 | 0 | 3.6 | 60 | Specific | 819.83 | 545.94 | 545.94 | 15.86 | 32 | >sp\|P47710\|CASA1_HUMAN Alpha-S1-casein OS=Homo sapiens OX=9606 GN=CSN1S1 PE=1 SV=1 | 989 | 20201118_12812_1ng_glyco_hcd203040_compound15.12547.12547.5 | controllerType=0 controllerNumber=1 scan=12547 | 27.2774 |
| 01782:3 | 21 | K.N[+1540.529]NSIPDKQITASSSYK.T | HexNAc(2)Hex(7) | N[+1541] | 1098.4778 | 3 | 3293.4189 | 3293.4048 | 0 | 4.3 | 238 | Specific | 241.93 | 124.19 | 124.19 | 4.87 | 31 | >sp\|Q08431\|MFGM_HUMAN Lactadherin OS=Homo sapiens OX=9606 GN=MFGE8 PE=1 SV=3 | 1064 | 20201118_12812_1ng_glyco_hcd203040_compound15.4721.4721.2 | controllerType=0 controllerNumber=1 scan=4721 | 14.2636 |
| 12722:4 | 23 | K.MVSHHN[+1913.677]LTTGATLINEQWLLTTAK.N | HexNAc(4)Hex(5)NeuAc(1) | N[+1914] | 1149.0265 | 4 | 4593.0843 | 4593.0693 | 0 | 3.3 | 179 | Specific | 357.75 | 249.05 | 249.05 | 8.42 | 29 | >sp\|P00738\|HPT_HUMAN Haptoglobin OS=Homo sapiens OX=9606 GN=HP PE=1 SV=1 | 407 | 20201118_12812_1ng_glyco_hcd203040_compound15.16067.16067.4 | controllerType=0 controllerNumber=1 scan=16067 | 33.1586 |
| 12720:4 | 23 | K.MVSHHN[+1913.677]LTTGATLINEQWLLTTAK.N | HexNAc(4)Hex(5)NeuAc(1) | N[+1914] | 1149.0265 | 4 | 4593.0843 | 4593.0693 | 0 | 3.3 | 179 | Specific | 484.28 | 459.69 | 459.69 | 11.08 | 29 | >sp\|P00738\|HPT_HUMAN Haptoglobin OS=Homo sapiens OX=9606 GN=HP PE=1 SV=1 | 407 | 20201118_12812_1ng_glyco_hcd203040_compound15.16065.16065.4 | controllerType=0 controllerNumber=1 scan=16065 | 33.155 |
| 14439:4 | 23 | K.MVSHHN[+2204.772]LTTGATLINEQWLLTTAK.N | HexNAc(4)Hex(5)NeuAc(2) | N[+2205] | 1221.8022 | 4 | 4884.1868 | 4884.1647 | 0 | 4.5 | 179 | Specific | 561.27 | 449.14 | 449.14 | 11.21 | 29 | >sp\|P00738\|HPT_HUMAN Haptoglobin OS=Homo sapiens OX=9606 GN=HP PE=1 SV=1 | 407 | 20201118_12812_1ng_glyco_hcd203040_compound15.17863.17863.4 | controllerType=0 controllerNumber=1 scan=17863 | 36.4717 |
| 14437:4 | 23 | K.MVSHHN[+2204.772]LTTGATLINEQWLLTTAK.N | HexNAc(4)Hex(5)NeuAc(2) | N[+2205] | 1221.8022 | 4 | 4884.1868 | 4884.1647 | 0 | 4.5 | 179 | Specific | 628.62 | 628.62 | 628.62 | 13.43 | 29 | >sp\|P00738\|HPT_HUMAN Haptoglobin OS=Homo sapiens OX=9606 GN=HP PE=1 SV=1 | 407 | 20201118_12812_1ng_glyco_hcd203040_compound15.17861.17861.4 | controllerType=0 controllerNumber=1 scan=17861 | 36.4677 |
| 14254:3 | 23 | K.VVLHPN[+2205.793]YSQVDIGLIK.L | HexNAc(4)Hex(5)Fuc(2)NeuAc(1) | N[+2206] | 1334.2698 | 3 | 4000.7948 | 4000.8041 | 0 | -2.3 | 236 | Specific | 222.3 | 222.3 | 222.3 | 4.15 | 29 | >sp\|P00738\|HPT_HUMAN Haptoglobin OS=Homo sapiens OX=9606 GN=HP PE=1 SV=1 | 407 | 20201118_12812_1ng_glyco_hcd203040_compound15.17671.17671.3 | controllerType=0 controllerNumber=1 scan=17671 | 36.1007 |
| 14252:3 | 23 | K.VVLHPN[+2205.793]YSQVDIGLIK.L | HexNAc(4)Hex(5)Fuc(2)NeuAc(1) | N[+2206] | 1334.2698 | 3 | 4000.7948 | 4000.8041 | 0 | -2.3 | 236 | Specific | 261.1 | 261.1 | 261.1 | 5.75 | 29 | >sp\|P00738\|HPT_HUMAN Haptoglobin OS=Homo sapiens OX=9606 GN=HP PE=1 SV=1 | 407 | 20201118_12812_1ng_glyco_hcd203040_compound15.17669.17669.3 | controllerType=0 controllerNumber=1 scan=17669 | 36.0966 |
| 14217:3 | 23 | K.VVLHPN[+2204.772]YSQVDIGLIK.L | HexNAc(4)Hex(5)NeuAc(2) | N[+2205] | 1333.9428 | 3 | 3999.8139 | 3999.7837 | 0 | 7.5 | 236 | Specific | 332.52 | 319.39 | 319.39 | 6.09 | 29 | >sp\|P00738\|HPT_HUMAN Haptoglobin OS=Homo sapiens OX=9606 GN=HP PE=1 SV=1 | 407 | 20201118_12812_1ng_glyco_hcd203040_compound15.17633.17633.3 | controllerType=0 controllerNumber=1 scan=17633 | 36.0282 |
| 14215:3 | 23 | K.VVLHPN[+2204.772]YSQVDIGLIK.L | HexNAc(4)Hex(5)NeuAc(2) | N[+2205] | 1333.9428 | 3 | 3999.8139 | 3999.7837 | 0 | 7.5 | 236 | Specific | 455.46 | 455.04 | 455.04 | 8.29 | 29 | >sp\|P00738\|HPT_HUMAN Haptoglobin OS=Homo sapiens OX=9606 GN=HP PE=1 SV=1 | 407 | 20201118_12812_1ng_glyco_hcd203040_compound15.17631.17631.3 | controllerType=0 controllerNumber=1 scan=17631 | 36.0244 |
| 09907:5 | 27 | K.SC[+57.021]QHN[+2350.830]GTMYQHGEIFSAHELFPSR.L | HexNAc(4)Hex(5)Fuc(1)NeuAc(2) | C[+57], N[+2351] | 1037.6251 | 5 | 5184.0965 | 5184.0873 | 0 | 1.8 | 110 | Specific | 421.62 | 421.62 | 421.62 | 10.6 | 22 | >sp\|Q6WN34\|CRDL2_HUMAN Chordin-like protein 2 OS=Homo sapiens OX=9606 GN=CHRDL2 PE=1 SV=1 | 1189 | 20201118_12812_1ng_glyco_hcd203040_compound15.13151.13151.5 | controllerType=0 controllerNumber=1 scan=13151 | 28.245 |
| 09927:4 | 27 | K.SC[+57.021]QHN[+2350.830]GTMYQHGEIFSAHELFPSR.L | HexNAc(4)Hex(5)Fuc(1)NeuAc(2) | C[+57], N[+2351] | 1296.7831 | 4 | 5184.1105 | 5184.0873 | 0 | 4.5 | 110 | Specific | 545.52 | 545.52 | 545.52 | 10.39 | 22 | >sp\|Q6WN34\|CRDL2_HUMAN Chordin-like protein 2 OS=Homo sapiens OX=9606 GN=CHRDL2 PE=1 SV=1 | 1189 | 20201118_12812_1ng_glyco_hcd203040_compound15.13172.13172.4 | controllerType=0 controllerNumber=1 scan=13172 | 28.2814 |
| 09909:5 | 27 | K.SC[+57.021]QHN[+2350.830]GTMYQHGEIFSAHELFPSR.L | HexNAc(4)Hex(5)Fuc(1)NeuAc(2) | C[+57], N[+2351] | 1037.6251 | 5 | 5184.0965 | 5184.0873 | 0 | 1.8 | 110 | Specific | 564.03 | 533.16 | 533.16 | 11.37 | 22 | >sp\|Q6WN34\|CRDL2_HUMAN Chordin-like protein 2 OS=Homo sapiens OX=9606 GN=CHRDL2 PE=1 SV=1 | 1189 | 20201118_12812_1ng_glyco_hcd203040_compound15.13153.13153.5 | controllerType=0 controllerNumber=1 scan=13153 | 28.249 |
| 09929:4 | 27 | K.SC[+57.021]QHN[+2350.830]GTMYQHGEIFSAHELFPSR.L | HexNAc(4)Hex(5)Fuc(1)NeuAc(2) | C[+57], N[+2351] | 1296.7831 | 4 | 5184.1105 | 5184.0873 | 0 | 4.5 | 110 | Specific | 575.77 | 575.77 | 575.77 | 10.58 | 22 | >sp\|Q6WN34\|CRDL2_HUMAN Chordin-like protein 2 OS=Homo sapiens OX=9606 GN=CHRDL2 PE=1 SV=1 | 1189 | 20201118_12812_1ng_glyco_hcd203040_compound15.13174.13174.4 | controllerType=0 controllerNumber=1 scan=13174 | 28.2852 |
| 09883:4 | 27 | K.SC[+57.021]QHN[+2350.830]GTMYQHGEIFSAHELFPSR.L | HexNAc(4)Hex(5)Fuc(1)NeuAc(2) | C[+57], N[+2351] | 1296.7808 | 4 | 5184.1012 | 5184.0873 | 0 | 2.7 | 110 | Specific | 766.54 | 764.41 | 764.41 | 16.07 | 22 | >sp\|Q6WN34\|CRDL2_HUMAN Chordin-like protein 2 OS=Homo sapiens OX=9606 GN=CHRDL2 PE=1 SV=1 | 1189 | 20201118_12812_1ng_glyco_hcd203040_compound15.13126.13126.4 | controllerType=0 controllerNumber=1 scan=13126 | 28.1993 |
| 09881:4 | 27 | K.SC[+57.021]QHN[+2350.830]GTMYQHGEIFSAHELFPSR.L | HexNAc(4)Hex(5)Fuc(1)NeuAc(2) | C[+57], N[+2351] | 1296.7808 | 4 | 5184.1012 | 5184.0873 | 0 | 2.7 | 110 | Specific | 768.26 | 768.26 | 768.26 | 16.07 | 22 | >sp\|Q6WN34\|CRDL2_HUMAN Chordin-like protein 2 OS=Homo sapiens OX=9606 GN=CHRDL2 PE=1 SV=1 | 1189 | 20201118_12812_1ng_glyco_hcd203040_compound15.13124.13124.4 | controllerType=0 controllerNumber=1 scan=13124 | 28.1954 |
| 14110:3 | 30 | R.ALGFEN[+2350.830]ATQALGR.A | HexNAc(4)Hex(5)Fuc(1)NeuAc(2) | N[+2351] | 1233.5197 | 3 | 3698.5446 | 3698.5319 | 0 | 3.4 | 64 | Specific | 372.31 | 275.24 | 275.24 | 7.2 | 20 | >sp\|Q08380\|LG3BP_HUMAN Galectin-3-binding protein OS=Homo sapiens OX=9606 GN=LGALS3BP PE=1 SV=1 | 1562 | 20201118_12812_1ng_glyco_hcd203040_compound15.17522.17522.3 | controllerType=0 controllerNumber=1 scan=17522 | 35.8129 |
| 18483:3 | 36 | R.C[+57.021]MWSSALNSLN[+1864.634]LSFAGLEQVPK.G | HexNAc(2)Hex(9) | C[+57], N[+1865] | 1439.6183 | 3 | 4316.8405 | 4316.8341 | 0 | 1.5 | 272 | Specific | 634.33 | 606.06 | 606.06 | 12.58 | 17 | >sp\|P08571\|CD14_HUMAN Monocyte differentiation antigen CD14 OS=Homo sapiens OX=9606 GN=CD14 PE=1 SV=2 | 1232 | 20201118_12812_1ng_glyco_hcd203040_compound15.22098.22098.3 | controllerType=0 controllerNumber=1 scan=22098 | 44.7851 |
| 18486:3 | 36 | R.C[+57.021]MWSSALNSLN[+1864.634]LSFAGLEQVPK.G | HexNAc(2)Hex(9) | C[+57], N[+1865] | 1439.6183 | 3 | 4316.8405 | 4316.8341 | 0 | 1.5 | 272 | Specific | 884.82 | 773.5 | 773.5 | 17.58 | 17 | >sp\|P08571\|CD14_HUMAN Monocyte differentiation antigen CD14 OS=Homo sapiens OX=9606 GN=CD14 PE=1 SV=2 | 1232 | 20201118_12812_1ng_glyco_hcd203040_compound15.22101.22101.3 | controllerType=0 controllerNumber=1 scan=22101 | [illegible] |
| 22028:4 | 45 | K.[+42.011]VFPLSLDSTPQDGNVVVAC[+57.021]LVQGFFPQEPLSVTWSESGQN[+1606 | HexNAc(4)Hex(4)Fuc(1) | .[42], C[+57], N[+1607] | 1607.0043 | 4 | 6424.9953 | 6424.9587 | 0 | 5.7 | 123 | Specific | 202.92 | 202.92 | 202.92 | 4.35 | 19 | >sp\|P0DOX2\|IGA2_HUMAN Immunoglobulin alpha-2 heavy chain OS=Homo sapiens OX=9606 PE=1 SV=2 | 631 | 20201118_12812_1ng_glyco_hcd203040_compound15.25798.25798.4 | controllerType=0 controllerNumber=1 scan=25798 | [illegible] |
| 22031:4 | 45 | K.[+42.011]VFPLSLDSTPQDGNVVVAC[+57.021]LVQGFFPQEPLSVTWSESGQN[+1606 | HexNAc(4)Hex(4)Fuc(1) | .[42], C[+57], N[+1607] | 1607.0043 | 4 | 6424.9953 | 6424.9587 | 0 | 5.7 | 123 | Specific | 282.47 | 282.47 | 282.47 | 5.83 | 19 | >sp\|P0DOX2\|IGA2_HUMAN Immunoglobulin alpha-2 heavy chain OS=Homo sapiens OX=9606 PE=1 SV=2 | 631 | 20201118_12812_1ng_glyco_hcd203040_compound15.25801.25801.4 | controllerType=0 controllerNumber=1 scan=25801 | 52.4322 |
| 21742:4 | 45 | K.VFPLSLDSTPQDGNVVVAC[+57.021]LVQGFFPQEPLSVTWSESGQN[+1647.613]VTAI | HexNAc(5)Hex(3)Fuc(1) | C[+57], N[+1648] | 1606.7571 | 4 | 6424.0065 | 6423.9746 | 0 | 5 | 123 | Specific | 639.21 | 639.21 | 639.21 | 13.67 | 19 | >sp\|P0DOX2\|IGA2_HUMAN Immunoglobulin alpha-2 heavy chain OS=Homo sapiens OX=9606 PE=1 SV=2 | 631 | 20201118_12812_1ng_glyco_hcd203040_compound15.25495.25495.4 | controllerType=0 controllerNumber=1 scan=25495 | 51.8897 |
| 21748:4 | 45 | K.VFPLSLDSTPQDGNVVVAC[+57.021]LVQGFFPQEPLSVTWSESGQN[+1809.666]VTAI | HexNAc(5)Hex(4)Fuc(1) | C[+57], N[+1810] | 1647.2712 | 4 | 6586.0631 | 6586.0275 | 0 | 5.4 | 123 | Specific | 659.06 | 642.03 | 642.03 | 14.01 | 19 | >sp\|P0DOX2\|IGA2_HUMAN Immunoglobulin alpha-2 heavy chain OS=Homo sapiens OX=9606 PE=1 SV=2 | 631 | 20201118_12812_1ng_glyco_hcd203040_compound15.25502.25502.4 | controllerType=0 controllerNumber=1 scan=25502 | 51.9042 |
| 21744:4 | 45 | K.VFPLSLDSTPQDGNVVVAC[+57.021]LVQGFFPQEPLSVTWSESGQN[+1647.613]VTAI | HexNAc(5)Hex(3)Fuc(1) | C[+57], N[+1648] | 1606.7571 | 4 | 6424.0065 | 6423.9746 | 0 | 5 | 123 | Specific | 795.8 | 757.42 | 757.42 | 16.67 | 19 | >sp\|P0DOX2\|IGA2_HUMAN Immunoglobulin alpha-2 heavy chain OS=Homo sapiens OX=9606 PE=1 SV=2 | 631 | 20201118_12812_1ng_glyco_hcd203040_compound15.25497.25497.4 | controllerType=0 controllerNumber=1 scan=25497 | 51.8954 |
| 21750:4 | 45 | K.VFPLSLDSTPQDGNVVVAC[+57.021]LVQGFFPQEPLSVTWSESGQN[+1809.666]VTAI | HexNAc(5)Hex(4)Fuc(1) | C[+57], N[+1810] | 1647.2712 | 4 | 6586.0631 | 6586.0275 | 0 | 5.4 | 123 | Specific | 799.49 | 773.39 | 773.39 | 16.01 | 19 | >sp\|P0DOX2\|IGA2_HUMAN Immunoglobulin alpha-2 heavy chain OS=Homo sapiens OX=9606 PE=1 SV=2 | 631 | 20201118_12812_1ng_glyco_hcd203040_compound15.25504.25504.4 | controllerType=0 controllerNumber=1 scan=25504 | 51.9108 |
| 04533:3 | 45 | K.TPLTAN[+2263.835]ITK.S | HexNAc(5)Hex(5)Fuc(3) | N[+2264] | 1074.7985 | 3 | 3222.3809 | 3222.3915 | 0 | -3.3 | 315 | Specific | 238.46 | 174.95 | 174.95 | 4.11 | 19 | >sp\|P0DOX2\|IGA2_HUMAN Immunoglobulin alpha-2 heavy chain OS=Homo sapiens OX=9606 PE=1 SV=2 | 631 | 20201118_12812_1ng_glyco_hcd203040_compound15.7573.7573.3 | controllerType=0 controllerNumber=1 scan=7573 | 19.0692 |
| 02783:2 | 45 | K.TPLTAN[+1501.555]ITK.S | HexNAc(5)Hex(3) | N[+1502] | 1230.5641 | 2 | 2460.121 | 2460.1121 | 0 | 3.6 | 315 | Specific | 373.25 | 210.99 | 210.99 | 6.61 | 19 | >sp\|P0DOX2\|IGA2_HUMAN Immunoglobulin alpha-2 heavy chain OS=Homo sapiens OX=9606 PE=1 SV=2 | 631 | 20201118_12812_1ng_glyco_hcd203040_compound15.5758.5758.2 | controllerType=0 controllerNumber=1 scan=5758 | 16.0492 |
| 02697:3 | 45 | K.TPLTAN[+1809.666]ITK.S | HexNAc(5)Hex(4)Fuc(1) | N[+1810] | 923.4148 | 3 | 2768.2299 | 2768.2228 | 0 | 2.6 | 315 | Specific | 399.8 | 130.68 | 130.68 | 5.85 | 19 | >sp\|P0DOX2\|IGA2_HUMAN Immunoglobulin alpha-2 heavy chain OS=Homo sapiens OX=9606 PE=1 SV=2 | 631 | 20201118_12812_1ng_glyco_hcd203040_compound15.5669.5669.3 | controllerType=0 controllerNumber=1 scan=5669 | 15.8875 |
| 02785:2 | 45 | K.TPLTAN[+1501.555]ITK.S | HexNAc(5)Hex(3) | N[+1502] | 1230.5641 | 2 | 2460.121 | 2460.1121 | 0 | 3.6 | 315 | Specific | 404.97 | 233.77 | 233.77 | 6.7 | 19 | >sp\|P0DOX2\|IGA2_HUMAN Immunoglobulin alpha-2 heavy chain OS=Homo sapiens OX=9606 PE=1 SV=2 | 631 | 20201118_12812_1ng_glyco_hcd203040_compound15.5760.5760.2 | controllerType=0 controllerNumber=1 scan=5760 | 16.0534 |
| 02778:3 | 45 | K.TPLTAN[+1647.613]ITK.S | HexNAc(5)Hex(3)Fuc(1) | N[+1648] | 869.3977 | 3 | 2606.1785 | 2606.17 | 0 | 3.2 | 315 | Specific | 456.32 | 149.47 | 149.47 | 6.35 | 19 | >sp\|P0DOX2\|IGA2_HUMAN Immunoglobulin alpha-2 heavy chain OS=Homo sapiens OX=9606 PE=1 SV=2 | 631 | 20201118_12812_1ng_glyco_hcd203040_compound15.5753.5753.3 | controllerType=0 controllerNumber=1 scan=5753 | 16.0391 |
| 02699:3 | 45 | K.TPLTAN[+1809.666]ITK.S | HexNAc(5)Hex(4)Fuc(1) | N[+1810] | 923.4148 | 3 | 2768.2299 | 2768.2228 | 0 | 2.6 | 315 | Specific | 499.3 | 262.99 | 262.99 | 7.57 | 19 | >sp\|P0DOX2\|IGA2_HUMAN Immunoglobulin alpha-2 heavy chain OS=Homo sapiens OX=9606 PE=1 SV=2 | 631 | 20201118_12812_1ng_glyco_hcd203040_compound15.5671.5671.3 | controllerType=0 controllerNumber=1 scan=5671 | 15.8918 |
| 02745:2 | 45 | K.TPLTAN[+1444.534]ITK.S | HexNAc(4)Hex(3)Fuc(1) | N[+1445] | 1202.0513 | 2 | 2403.0953 | 2403.0906 | 0 | 2 | 315 | Specific | 536.14 | 302.78 | 302.78 | 7.87 | 19 | >sp\|P0DOX2\|IGA2_HUMAN Immunoglobulin alpha-2 heavy chain OS=Homo sapiens OX=9606 PE=1 SV=2 | 631 | 20201118_12812_1ng_glyco_hcd203040_compound15.5719.5719.2 | controllerType=0 controllerNumber=1 scan=5719 | 15.9754 |
| 02780:3 | 45 | K.TPLTAN[+1647.613]ITK.S | HexNAc(5)Hex(3)Fuc(1) | N[+1648] | 869.3977 | 3 | 2606.1785 | 2606.17 | 0 | 3.2 | 315 | Specific | 542.1 | 256.33 | 256.33 | 7.7 | 19 | >sp\|P0DOX2\|IGA2_HUMAN Immunoglobulin alpha-2 heavy chain OS=Homo sapiens OX=9606 PE=1 SV=2 | 631 | 20201118_12812_1ng_glyco_hcd203040_compound15.5755.5755.3 | controllerType=0 controllerNumber=1 scan=5755 | 16.0444 |
| 02747:2 | 45 | K.TPLTAN[+1444.534]ITK.S | HexNAc(4)Hex(3)Fuc(1) | N[+1445] | 1202.0513 | 2 | 2403.0953 | 2403.0906 | 0 | 2 | 315 | Specific | 563.99 | 275.83 | 275.83 | 7.93 | 19 | >sp\|P0DOX2\|IGA2_HUMAN Immunoglobulin alpha-2 heavy chain OS=Homo sapiens OX=9606 PE=1 SV=2 | 631 | 20201118_12812_1ng_glyco_hcd203040_compound15.5721.5721.2 | controllerType=0 controllerNumber=1 scan=5721 | 15.9794 |
| 02770:2 | 45 | K.TPLTAN[+1647.613]ITK.S | HexNAc(5)Hex(3)Fuc(1) | N[+1648] | 1303.5925 | 2 | 2606.1776 | 2606.17 | 0 | 2.9 | 315 | Specific | 596.86 | 332.35 | 332.35 | 9.68 | 19 | >sp\|P0DOX2\|IGA2_HUMAN Immunoglobulin alpha-2 heavy chain OS=Homo sapiens OX=9606 PE=1 SV=2 | 631 | 20201118_12812_1ng_glyco_hcd203040_compound15.5745.5745.2 | controllerType=0 controllerNumber=1 scan=5745 | 16.0257 |
| 02698:2 | 45 | K.TPLTAN[+1809.666]ITK.S | HexNAc(5)Hex(4)Fuc(1) | N[+1810] | 1384.6186 | 2 | 2768.2299 | 2768.2228 | 0 | 2.5 | 315 | Specific | 608.55 | 279.65 | 279.65 | 10.25 | 19 | >sp\|P0DOX2\|IGA2_HUMAN Immunoglobulin alpha-2 heavy chain OS=Homo sapiens OX=9606 PE=1 SV=2 | 631 | 20201118_12812_1ng_glyco_hcd203040_compound15.5670.5670.2 | controllerType=0 controllerNumber=1 scan=5670 | 15.889 |
| 02700:2 | 45 | K.TPLTAN[+1809.666]ITK.S | HexNAc(5)Hex(4)Fuc(1) | N[+1810] | 1384.6186 | 2 | 2768.2299 | 2768.2228 | 0 | 2.5 | 315 | Specific | 620.6 | 315.79 | 315.79 | 10.57 | 19 | >sp\|P0DOX2\|IGA2_HUMAN Immunoglobulin alpha-2 heavy chain OS=Homo sapiens OX=9606 PE=1 SV=2 | 631 | 20201118_12812_1ng_glyco_hcd203040_compound15.5672.5672.2 | controllerType=0 controllerNumber=1 scan=5672 | 15.8937 |
| 02768:2 | 45 | K.TPLTAN[+1647.613]ITK.S | HexNAc(5)Hex(3)Fuc(1) | N[+1648] | 1303.5925 | 2 | 2606.1776 | 2606.17 | 0 | 2.9 | 315 | Specific | 624.19 | 359.99 | 359.99 | 10.69 | 19 | >sp\|P0DOX2\|IGA2_HUMAN Immunoglobulin alpha-2 heavy chain OS=Homo sapiens OX=9606 PE=1 SV=2 | 631 | 20201118_12812_1ng_glyco_hcd203040_compound15.5743.5743.2 | controllerType=0 controllerNumber=1 scan=5743 | 16.0183 |
| 21246:3 | 49 | R.QFWIFDVQNPQEVMMN[+2351.851]SSNIQVK.Q | HexNAc(4)Hex(5)Fuc(3)NeuAc(1) | N[+2352] | 1712.065 | 3 | 5134.1805 | 5134.1835 | 0 | -0.6 | 64 | Specific | 344.18 | 344.18 | 344.18 | 7.82 | 14 | >sp\|P16671\|CD36_HUMAN Platelet glycoprotein 4 OS=Homo sapiens OX=9606 GN=CD36 PE=1 SV=2 | 812 | 20201118_12812_1ng_glyco_hcd203040_compound15.24981.24981.3 | controllerType=0 controllerNumber=1 scan=24981 | 50.9346 |
| 21248:3 | 49 | R.QFWIFDVQNPQEVMMN[+2351.851]SSNIQVK.Q | HexNAc(4)Hex(5)Fuc(3)NeuAc(1) | N[+2352] | 1712.065 | 3 | 5134.1805 | 5134.1835 | 0 | -0.6 | 64 | Specific | 559.57 | 538.33 | 538.33 | 10.37 | 14 | >sp\|P16671\|CD36_HUMAN Platelet glycoprotein 4 OS=Homo sapiens OX=9606 GN=CD36 PE=1 SV=2 | 812 | 20201118_12812_1ng_glyco_hcd203040_compound15.24983.24983.3 | controllerType=0 controllerNumber=1 scan=24983 | 50.9397 |
| 17641:3 | 49 | R.NYIVPILWLN[+1864.634]ETGTIGDEK.A | HexNAc(2)Hex(9) | N[+1865] | 1347.2646 | 3 | 4039.7792 | 4039.7674 | 0 | 2.9 | 408 | Specific | 374.09 | 374.09 | 374.09 | 7.82 | 14 | >sp\|P16671\|CD36_HUMAN Platelet glycoprotein 4 OS=Homo sapiens OX=9606 GN=CD36 PE=1 SV=2 | 812 | 20201118_12812_1ng_glyco_hcd203040_compound15.21219.21219.3 | controllerType=0 controllerNumber=1 scan=21219 | 42.8792 |
| 17644:3 | 49 | R.NYIVPILWLN[+1864.634]ETGTIGDEK.A | HexNAc(2)Hex(9) | N[+1865] | 1347.2646 | 3 | 4039.7792 | 4039.7674 | 0 | 2.9 | 408 | Specific | 834.82 | 834.82 | 834.82 | 16.68 | 14 | >sp\|P16671\|CD36_HUMAN Platelet glycoprotein 4 OS=Homo sapiens OX=9606 GN=CD36 PE=1 SV=2 | 812 | 20201118_12812_1ng_glyco_hcd203040_compound15.21222.21222.3 | controllerType=0 controllerNumber=1 scan=21222 | 42.8836 |
| 09461:3 | 54 | R.IIVPLNNREN[+1663.608]ISDPTSPLR.T | HexNAc(5)Hex(4) | N[+1664] | 1271.2687 | 3 | 3811.7916 | 3811.7853 | 0 | 1.7 | 62 | Specific | 369.99 | 364.47 | 364.47 | 7.32 | 14 | >sp\|P01591\|IGJ_HUMAN Immunoglobulin J chain OS=Homo sapiens OX=9606 GN=JCHAIN PE=1 SV=4 | 1392 | 20201118_12812_1ng_glyco_hcd203040_compound15.12690.12690.3 | controllerType=0 controllerNumber=1 scan=12690 | 27.5084 |
| 11100:3 | 54 | R.IIVPLNNREN[+2059.735]ISDPTSPLR.T | HexNAc(4)Hex(5)Fuc(1)NeuAc(1) | N[+2060] | 1403.3108 | 3 | 4207.918 | 4207.9121 | 0 | 1.4 | 62 | Specific | 462.17 | 418.53 | 418.53 | 9.3 | 14 | >sp\|P01591\|IGJ_HUMAN Immunoglobulin J chain OS=Homo sapiens OX=9606 GN=JCHAIN PE=1 SV=4 | 1392 | 20201118_12812_1ng_glyco_hcd203040_compound15.14386.14386.3 | controllerType=0 controllerNumber=1 scan=14386 | 30.3032 |
| 09464:3 | 54 | R.IIVPLNNREN[+1663.608]ISDPTSPLR.T | HexNAc(5)Hex(4) | N[+1664] | 1271.2687 | 3 | 3811.7916 | 3811.7853 | 0 | 1.7 | 62 | Specific | 474.63 | 472.04 | 472.04 | 9.3 | 14 | >sp\|P01591\|IGJ_HUMAN Immunoglobulin J chain OS=Homo sapiens OX=9606 GN=JCHAIN PE=1 SV=4 | 1392 | 20201118_12812_1ng_glyco_hcd203040_compound15.12693.12693.3 | controllerType=0 controllerNumber=1 scan=12693 | 27.5131 |
| 11103:3 | 54 | R.IIVPLNNREN[+2059.735]ISDPTSPLR.T | HexNAc(4)Hex(5)Fuc(1)NeuAc(1) | N[+2060] | 1403.3108 | 3 | 4207.918 | 4207.9121 | 0 | 1.4 | 62 | Specific | 477.34 | 440.83 | 440.83 | 9.3 | 14 | >sp\|P01591\|IGJ_HUMAN Immunoglobulin J chain OS=Homo sapiens OX=9606 GN=JCHAIN PE=1 SV=4 | 1392 | 20201118_12812_1ng_glyco_hcd203040_compound15.14389.14389.3 | controllerType=0 controllerNumber=1 scan=14389 | 30.3087 |
| 11129:3 | 54 | R.IIVPLNNREN[+1913.677]ISDPTSPLR.T | HexNAc(4)Hex(5)NeuAc(1) | N[+1914] | 1354.6265 | 3 | 4061.865 | 4061.8542 | 0 | 2.7 | 62 | Specific | 516.73 | 460.35 | 460.35 | 9.72 | 14 | >sp\|P01591\|IGJ_HUMAN Immunoglobulin J chain OS=Homo sapiens OX=9606 GN=JCHAIN PE=1 SV=4 | 1392 | 20201118_12812_1ng_glyco_hcd203040_compound15.14416.14416.3 | controllerType=0 controllerNumber=1 scan=14416 | 30.3584 |
| 11131:3 | 54 | R.IIVPLNNREN[+1913.677]ISDPTSPLR.T | HexNAc(4)Hex(5)NeuAc(1) | N[+1914] | 1354.6265 | 3 | 4061.865 | 4061.8542 | 0 | 2.7 | 62 | Specific | 629.04 | 629.04 | 629.04 | 12.28 | 14 | >sp\|P01591\|IGJ_HUMAN Immunoglobulin J chain OS=Homo sapiens OX=9606 GN=JCHAIN PE=1 SV=4 | 1392 | 20201118_12812_1ng_glyco_hcd203040_compound15.14418.14418.3 | controllerType=0 controllerNumber=1 scan=14418 | 30.3626 |
| 03358:2 | 62 | R.EEQFN[+1444.534]STFR.V | HexNAc(4)Hex(3)Fuc(1) | N[+1445] | 1301.5361 | 2 | 2602.0648 | 2602.056 | 0 | 3.4 | 172 | Specific | 237.98 | 104.1 | 104.1 | 4.57 | 10 | >sp\|P01859\|IGHG2_HUMAN Immunoglobulin heavy constant gamma 2 OS=Homo sapiens OX=9606 GN=IGHG2 PE=1 SV=2 | 387 | 20201118_12812_1ng_glyco_hcd203040_compound15.6356.6356.2 | controllerType=0 controllerNumber=1 scan=6356 | 17.0684 |
| 03355:2 | 62 | R.EEQFN[+1444.534]STFR.V | HexNAc(4)Hex(3)Fuc(1) | N[+1445] | 1301.5361 | 2 | 2602.0648 | 2602.056 | 0 | 3.4 | 172 | Specific | 240.36 | 175.65 | 175.65 | 5.74 | 10 | >sp\|P01859\|IGHG2_HUMAN Immunoglobulin heavy constant gamma 2 OS=Homo sapiens OX=9606 GN=IGHG2 PE=1 SV=2 | 387 | 20201118_12812_1ng_glyco_hcd203040_compound15.6353.6353.2 | controllerType=0 controllerNumber=1 scan=6353 | 17.064 |
| 24420:3 | 97 | R.AVN[+2204.772]ITSENLIDDVVSLIR.G | HexNAc(4)Hex(5)NeuAc(2) | N[+2205] | 1392.6268 | 3 | 4175.8657 | 4175.8481 | 0 | 4.2 | 413 | Specific | 515.85 | 462.04 | 462.04 | 9.86 | 12 | >sp\|P02748\|CO9_HUMAN Complement component C9 OS=Homo sapiens OX=9606 GN=C9 PE=1 SV=2 | 1547 | 20201118_12812_1ng_glyco_hcd203040_compound15.28285.28285.3 | controllerType=0 controllerNumber=1 scan=28285 | 57.2234 |
| 24422:3 | 97 | R.AVN[+2204.772]ITSENLIDDVVSLIR.G | HexNAc(4)Hex(5)NeuAc(2) | N[+2205] | 1392.6268 | 3 | 4175.8657 | 4175.8481 | 0 | 4.2 | 413 | Specific | 722.02 | 614.4 | 614.4 | 14.42 | 12 | >sp\|P02748\|CO9_HUMAN Complement component C9 OS=Homo sapiens OX=9606 GN=C9 PE=1 SV=2 | 1547 | 20201118_12812_1ng_glyco_hcd203040_compound15.28287.28287.3 | controllerType=0 controllerNumber=1 scan=28287 | 57.2286 |
| 16763:4 | 1 | R.RTAGWNVPIGTLRPFLN[+2278.809]WTGPPEPIEAAVAR.F | HexNAc(5)Hex(6)NeuAc(1) | N[+2279] | 1417.167 | 4 | 5665.6462 | 5665.6213 | 0 | 4.4 | 140 | Specific | 249.18 | 249.05 | 249.05 | 6.91 | 148 | >sp\|P02788\|TRFL_HUMAN Lactotransferrin OS=Homo sapiens OX=9606 GN=LTF PE=1 SV=6 | 1270 | 20201118_12833_1ng_glyco_hcd203040_compound15.20342.20342.4 | controllerType=0 controllerNumber=1 scan=20342 | 41.2063 |
| 16765:4 | 1 | R.RTAGWNVPIGTLRPFLN[+2278.809]WTGPPEPIEAAVAR.F | HexNAc(5)Hex(6)NeuAc(1) | N[+2279] | 1417.167 | 4 | 5665.6462 | 5665.6213 | 0 | 4.4 | 140 | Specific | 304.52 | 283.05 | 283.05 | 7.39 | 148 | >sp\|P02788\|TRFL_HUMAN Lactotransferrin OS=Homo sapiens OX=9606 GN=LTF PE=1 SV=6 | 1270 | 20201118_12833_1ng_glyco_hcd203040_compound15.20344.20344.4 | controllerType=0 controllerNumber=1 scan=20344 | 41.212 |
| 17911:4 | 1 | R.RTAGWNVPIGTLRPFLN[+2059.735]WTGPPEPIEAAVAR.F | HexNAc(4)Hex(5)Fuc(1)NeuAc(1) | N[+2060] | 1362.3947 | 4 | 5446.5568 | 5446.547 | 0 | 1.8 | 140 | Specific | 628.55 | 531.58 | 531.58 | 13.36 | 148 | >sp\|P02788\|TRFL_HUMAN Lactotransferrin OS=Homo sapiens OX=9606 GN=LTF PE=1 SV=6 | 1270 | 20201118_12833_1ng_glyco_hcd203040_compound15.21539.21539.4 | controllerType=0 controllerNumber=1 scan=21539 | 43.7692 |
| 18036:4 | 1 | R.[+42.011]TAGWNVPIGTLRPFLN[+1872.651]WTGPPEPIEAAVAR.F | HexNAc(3)Hex(6)NeuAc(1) | .[42], N[+1873] | 1287.1095 | 4 | 5145.4162 | 5145.372 | 0 | 8.6 | 141 | Specific | 214.25 | 187.9 | 187.9 | 3.42 | 148 | >sp\|P02788\|TRFL_HUMAN Lactotransferrin OS=Homo sapiens OX=9606 GN=LTF PE=1 SV=6 | 1270 | 20201118_12833_1ng_glyco_hcd203040_compound15.21669.21669.4 | controllerType=0 controllerNumber=1 scan=21669 | 44.0209 |
| 18030:3 | 1 | R.TAGWNVPIGTLRPFLN[+2279.830]WTGPPEPIEAAVAR.F | HexNAc(5)Hex(6)Fuc(2) | N[+2280] | 1837.5255 | 3 | 5510.5619 | 5510.5406 | 0 | 3.9 | 141 | Specific | 242.24 | 242.24 | 242.24 | 6.91 | 148 | >sp\|P02788\|TRFL_HUMAN Lactotransferrin OS=Homo sapiens OX=9606 GN=LTF PE=1 SV=6 | 1270 | 20201118_12833_1ng_glyco_hcd203040_compound15.21662.21662.3 | controllerType=0 controllerNumber=1 scan=21662 | 44.0073 |
| 19017:3 | 1 | R.TAGWNVPIGTLRPFLN[+2205.793]WTGPPEPIEAAVAR.F | HexNAc(4)Hex(5)Fuc(2)NeuAc(1) | N[+2206] | 1812.8443 | 3 | 5436.5184 | 5436.5038 | 0 | 2.7 | 141 | Specific | 257.5 | 257.5 | 257.5 | 7.66 | 148 | >sp\|P02788\|TRFL_HUMAN Lactotransferrin OS=Homo sapiens OX=9606 GN=LTF PE=1 SV=6 | 1270 | 20201118_12833_1ng_glyco_hcd203040_compound15.22695.22695.3 | controllerType=0 controllerNumber=1 scan=22695 | 46.3693 |
| 18042:3 | 1 | R.[+42.011]TAGWNVPIGTLRPFLN[+1872.651]WTGPPEPIEAAVAR.F | HexNAc(3)Hex(6)NeuAc(1) | .[42], N[+1873] | 1715.8116 | 3 | 5145.4203 | 5145.372 | 0 | 9.4 | 141 | Specific | 270.31 | 194.76 | 194.76 | 4.5 | 148 | >sp\|P02788\|TRFL_HUMAN Lactotransferrin OS=Homo sapiens OX=9606 GN=LTF PE=1 SV=6 | 1270 | 20201118_12833_1ng_glyco_hcd203040_compound15.21675.21675.3 | controllerType=0 controllerNumber=1 scan=21675 | 44.0316 |
| 18930:3 | 1 | R.TAGWNVPIGTLRPFLN[+2570.925]WTGPPEPIEAAVAR.F | HexNAc(5)Hex(6)Fuc(2)NeuAc(1) | N[+2571] | 1934.5573 | 3 | 5801.6573 | 5801.636 | 0 | 3.7 | 141 | Specific | 287.4 | 188.19 | 188.19 | 7.36 | 148 | >sp\|P02788\|TRFL_HUMAN Lactotransferrin OS=Homo sapiens OX=9606 GN=LTF PE=1 SV=6 | 1270 | 20201118_12833_1ng_glyco_hcd203040_compound15.22603.22603.3 | controllerType=0 controllerNumber=1 scan=22603 | 46.1404 |
| 18928:3 | 1 | R.TAGWNVPIGTLRPFLN[+2570.925]WTGPPEPIEAAVAR.F | HexNAc(5)Hex(6)Fuc(2)NeuAc(1) | N[+2571] | 1934.5573 | 3 | 5801.6573 | 5801.636 | 0 | 3.7 | 141 | Specific | 290.49 | 170.22 | 170.22 | 7.21 | 148 | >sp\|P02788\|TRFL_HUMAN Lactotransferrin OS=Homo sapiens OX=9606 GN=LTF PE=1 SV=6 | 1270 | 20201118_12833_1ng_glyco_hcd203040_compound15.22601.22601.3 | controllerType=0 controllerNumber=1 scan=22601 | 46.137 |
| 18039:3 | 1 | R.[+42.011]TAGWNVPIGTLRPFLN[+1872.651]WTGPPEPIEAAVAR.F | HexNAc(3)Hex(6)NeuAc(1) | .[42], N[+1873] | 1715.8116 | 3 | 5145.4203 | 5145.372 | 0 | 9.4 | 141 | Specific | 293.24 | 179.29 | 179.29 | 4.32 | 148 | >sp\|P02788\|TRFL_HUMAN Lactotransferrin OS=Homo sapiens OX=9606 GN=LTF PE=1 SV=6 | 1270 | 20201118_12833_1ng_glyco_hcd203040_compound15.21672.21672.3 | controllerType=0 controllerNumber=1 scan=21672 | 44.0266 |
| 18861:4 | 1 | R.TAGWNVPIGTLRPFLN[+2863.041]WTGPPEPIEAAVAR.F | HexNAc(5)Hex(6)Fuc(4)NeuAc(1) | N[+2863] | 1524.1982 | 4 | 6093.7712 | 6093.7518 | 0 | 3.2 | 141 | Specific | 298.72 | 298.72 | 298.72 | 7.89 | 148 | >sp\|P02788\|TRFL_HUMAN Lactotransferrin OS=Homo sapiens OX=9606 GN=LTF PE=1 SV=6 | 1270 | 20201118_12833_1ng_glyco_hcd203040_compound15.22531.22531.4 | controllerType=0 controllerNumber=1 scan=22531 | 45.9826 |
| 18038:4 | 1 | R.[+42.011]TAGWNVPIGTLRPFLN[+1872.651]WTGPPEPIEAAVAR.F | HexNAc(3)Hex(6)NeuAc(1) | .[42], N[+1873] | 1287.1095 | 4 | 5145.4162 | 5145.372 | 0 | 8.6 | 141 | Specific | 322.91 | 253.66 | 253.66 | 6.18 | 148 | >sp\|P02788\|TRFL_HUMAN Lactotransferrin OS=Homo sapiens OX=9606 GN=LTF PE=1 SV=6 | 1270 | 20201118_12833_1ng_glyco_hcd203040_compound15.21671.21671.4 | controllerType=0 controllerNumber=1 scan=21671 | 44.0253 |
| 18081:3 | 1 | R.TAGWNVPIGTLRPFLN[+1768.640]WTGPPEPIEAAVAR.F | HexNAc(4)Hex(5)Fuc(1) | N[+1769] | 1667.1269 | 3 | 4999.3662 | 4999.3504 | 0 | 3.2 | 141 | Specific | 335.62 | 321.42 | 321.42 | 8.62 | 148 | >sp\|P02788\|TRFL_HUMAN Lactotransferrin OS=Homo sapiens OX=9606 GN=LTF PE=1 SV=6 | 1270 | 20201118_12833_1ng_glyco_hcd203040_compound15.21716.21716.3 | controllerType=0 controllerNumber=1 scan=21716 | 44.1267 |
| 18091:4 | 1 | R.TAGWNVPIGTLRPFLN[+2059.735]WTGPPEPIEAAVAR.F | HexNAc(4)Hex(5)Fuc(1)NeuAc(1) | N[+2060] | 1323.3754 | 4 | 5290.4796 | 5290.4459 | 0 | 6.4 | 141 | Specific | 371.76 | 371.76 | 371.76 | 7.73 | 148 | >sp\|P02788\|TRFL_HUMAN Lactotransferrin OS=Homo sapiens OX=9606 GN=LTF PE=1 SV=6 | 1270 | 20201118_12833_1ng_glyco_hcd203040_compound15.21726.21726.4 | controllerType=0 controllerNumber=1 scan=21726 | 44.1488 |
| 18912:4 | 1 | R.TAGWNVPIGTLRPFLN[+2424.867]WTGPPEPIEAAVAR.F | HexNAc(5)Hex(6)Fuc(1)NeuAc(1) | N[+2425] | 1414.6542 | 4 | 5655.5949 | 5655.5781 | 0 | 3 | 141 | Specific | 440.06 | 312.82 | 312.82 | 8.7 | 148 | >sp\|P02788\|TRFL_HUMAN Lactotransferrin OS=Homo sapiens OX=9606 GN=LTF PE=1 SV=6 | 1270 | 20201118_12833_1ng_glyco_hcd203040_compound15.22584.22584.4 | controllerType=0 controllerNumber=1 scan=22584 | 46.0997 |
| 18933:4 | 1 | R.TAGWNVPIGTLRPFLN[+2425.887]WTGPPEPIEAAVAR.F | HexNAc(5)Hex(6)Fuc(3) | N[+2426] | 1414.9034 | 4 | 5656.5918 | 5656.5985 | 0 | -1.2 | 141 | Specific | 441.79 | 282.56 | 282.56 | 7.54 | 148 | >sp\|P02788\|TRFL_HUMAN Lactotransferrin OS=Homo sapiens OX=9606 GN=LTF PE=1 SV=6 | 1270 | 20201118_12833_1ng_glyco_hcd203040_compound15.22606.22606.4 | controllerType=0 controllerNumber=1 scan=22606 | 46.1459 |
| 18889:5 | 1 | R.TAGWNVPIGTLRPFLN[+2570.925]WTGPPEPIEAAVAR.F | HexNAc(5)Hex(6)Fuc(2)NeuAc(1) | N[+2571] | 1161.1366 | 5 | 5801.6539 | 5801.636 | 0 | 3.1 | 141 | Specific | 470.48 | 441.57 | 441.57 | 11.3 | 148 | >sp\|P02788\|TRFL_HUMAN Lactotransferrin OS=Homo sapiens OX=9606 GN=LTF PE=1 SV=6 | 1270 | 20201118_12833_1ng_glyco_hcd203040_compound15.22560.22560.5 | controllerType=0 controllerNumber=1 scan=22560 | 46.0461 |
| 18082:4 | 1 | R.TAGWNVPIGTLRPFLN[+1768.640]WTGPPEPIEAAVAR.F | HexNAc(4)Hex(5)Fuc(1) | N[+1769] | 1250.5978 | 4 | 4999.3693 | 4999.3504 | 0 | 3.8 | 141 | Specific | 522.44 | 488.58 | 488.58 | 10.86 | 148 | >sp\|P02788\|TRFL_HUMAN Lactotransferrin OS=Homo sapiens OX=9606 GN=LTF PE=1 SV=6 | 1270 | 20201118_12833_1ng_glyco_hcd203040_compound15.21717.21717.4 | controllerType=0 controllerNumber=1 scan=21717 | 44.1295 |
| 18008:4 | 1 | R.TAGWNVPIGTLRPFLN[+2279.830]WTGPPEPIEAAVAR.F | HexNAc(5)Hex(6)Fuc(2) | N[+2280] | 1378.3951 | 4 | 5510.5587 | 5510.5406 | 0 | 3.3 | 141 | Specific | 553.75 | 507.25 | 507.25 | 11.03 | 148 | >sp\|P02788\|TRFL_HUMAN Lactotransferrin OS=Homo sapiens OX=9606 GN=LTF PE=1 SV=6 | 1270 | 20201118_12833_1ng_glyco_hcd203040_compound15.21639.21639.4 | controllerType=0 controllerNumber=1 scan=21639 | 43.9591 |
| 18892:5 | 1 | R.TAGWNVPIGTLRPFLN[+2570.925]WTGPPEPIEAAVAR.F | HexNAc(5)Hex(6)Fuc(2)NeuAc(1) | N[+2571] | 1161.1366 | 5 | 5801.6539 | 5801.636 | 0 | 3.1 | 141 | Specific | 557.39 | 491.78 | 491.78 | 11.66 | 148 | >sp\|P02788\|TRFL_HUMAN Lactotransferrin OS=Homo sapiens OX=9606 GN=LTF PE=1 SV=6 | 1270 | 20201118_12833_1ng_glyco_hcd203040_compound15.22563.22563.5 | controllerType=0 controllerNumber=1 scan=22563 | 46.06 |
| 19020:3 | 1 | R.TAGWNVPIGTLRPFLN[+2205.793]WTGPPEPIEAAVAR.F | HexNAc(4)Hex(5)Fuc(2)NeuAc(1) | N[+2206] | 1812.8443 | 3 | 5436.5184 | 5436.5038 | 0 | 2.7 | 141 | Specific | 609.14 | 504.97 | 504.97 | 13.39 | 148 | >sp\|P02788\|TRFL_HUMAN Lactotransferrin OS=Homo sapiens OX=9606 GN=LTF PE=1 SV=6 | 1270 | 20201118_12833_1ng_glyco_hcd203040_compound15.22698.22698.3 | controllerType=0 controllerNumber=1 scan=22698 | 46.3751 |
| 18863:4 | 1 | R.TAGWNVPIGTLRPFLN[+2863.041]WTGPPEPIEAAVAR.F | HexNAc(5)Hex(6)Fuc(4)NeuAc(1) | N[+2863] | 1524.1982 | 4 | 6093.7712 | 6093.7518 | 0 | 3.2 | 141 | Specific | 622.67 | 503.57 | 503.57 | 13.22 | 148 | >sp\|P02788\|TRFL_HUMAN Lactotransferrin OS=Homo sapiens OX=9606 GN=LTF PE=1 SV=6 | 1270 | 20201118_12833_1ng_glyco_hcd203040_compound15.22533.22533.4 | controllerType=0 controllerNumber=1 scan=22533 | 45.987 |
| 19009:4 | 1 | R.TAGWNVPIGTLRPFLN[+2059.735]WTGPPEPIEAAVAR.F | HexNAc(4)Hex(5)Fuc(1)NeuAc(1) | N[+2060] | 1323.3723 | 4 | 5290.4674 | 5290.4459 | 0 | 4.1 | 141 | Specific | 647.93 | 619.2 | 619.2 | 12.67 | 148 | >sp\|P02788\|TRFL_HUMAN Lactotransferrin OS=Homo sapiens OX=9606 GN=LTF PE=1 SV=6 | 1270 | 20201118_12833_1ng_glyco_hcd203040_compound15.22687.22687.4 | controllerType=0 controllerNumber=1 scan=22687 | 46.3377 |
| 19012:3 | 1 | R.TAGWNVPIGTLRPFLN[+2059.735]WTGPPEPIEAAVAR.F | HexNAc(4)Hex(5)Fuc(1)NeuAc(1) | N[+2060] | 1764.1589 | 3 | 5290.4621 | 5290.4459 | 0 | 3.1 | 141 | Specific | 663 | 571.04 | 571.04 | 14.39 | 148 | >sp\|P02788\|TRFL_HUMAN Lactotransferrin OS=Homo sapiens OX=9606 GN=LTF PE=1 SV=6 | 1270 | 20201118_12833_1ng_glyco_hcd203040_compound15.22690.22690.3 | controllerType=0 controllerNumber=1 scan=22690 | 46.3491 |
| 18010:4 | 1 | R.TAGWNVPIGTLRPFLN[+2279.830]WTGPPEPIEAAVAR.F | HexNAc(5)Hex(6)Fuc(2) | N[+2280] | 1378.3951 | 4 | 5510.5587 | 5510.5406 | 0 | 3.3 | 141 | Specific | 671.1 | 603.18 | 603.18 | 14.22 | 148 | >sp\|P02788\|TRFL_HUMAN Lactotransferrin OS=Homo sapiens OX=9606 GN=LTF PE=1 SV=6 | 1270 | 20201118_12833_1ng_glyco_hcd203040_compound15.21641.21641.4 | controllerType=0 controllerNumber=1 scan=21641 | 43.9639 |
| 18083:3 | 1 | R.TAGWNVPIGTLRPFLN[+1768.640]WTGPPEPIEAAVAR.F | HexNAc(4)Hex(5)Fuc(1) | N[+1769] | 1667.1269 | 3 | 4999.3662 | 4999.3504 | 0 | 3.2 | 141 | Specific | 688.2 | 572.36 | 572.36 | 14.39 | 148 | >sp\|P02788\|TRFL_HUMAN Lactotransferrin OS=Homo sapiens OX=9606 GN=LTF PE=1 SV=6 | 1270 | 20201118_12833_1ng_glyco_hcd203040_compound15.21718.21718.3 | controllerType=0 controllerNumber=1 scan=21718 | 44.132 |
| 19010:3 | 1 | R.TAGWNVPIGTLRPFLN[+2059.735]WTGPPEPIEAAVAR.F | HexNAc(4)Hex(5)Fuc(1)NeuAc(1) | N[+2060] | 1764.1589 | 3 | 5290.4621 | 5290.4459 | 0 | 3.1 | 141 | Specific | 709.92 | 701.01 | 701.01 | 15.39 | 148 | >sp\|P02788\|TRFL_HUMAN Lactotransferrin OS=Homo sapiens OX=9606 GN=LTF PE=1 SV=6 | 1270 | 20201118_12833_1ng_glyco_hcd203040_compound15.22688.22688.3 | controllerType=0 controllerNumber=1 scan=22688 | 46.3392 |
| 18935:4 | 1 | R.TAGWNVPIGTLRPFLN[+2425.887]WTGPPEPIEAAVAR.F | HexNAc(5)Hex(6)Fuc(3) | N[+2426] | 1414.9034 | 4 | 5656.5918 | 5656.5985 | 0 | -1.2 | 141 | Specific | 717.31 | 424.04 | 424.04 | 14.18 | 148 | >sp\|P02788\|TRFL_HUMAN Lactotransferrin OS=Homo sapiens OX=9606 GN=LTF PE=1 SV=6 | 1270 | 20201118_12833_1ng_glyco_hcd203040_compound15.22608.22608.4 | controllerType=0 controllerNumber=1 scan=22608 | 46.1497 |
| 19170:4 | 1 | R.TAGWNVPIGTLRPFLN[+2059.735]WTGPPEPIEAAVAR.F | HexNAc(4)Hex(5)Fuc(1)NeuAc(1) | N[+2060] | 1323.3715 | 4 | 5290.464 | 5290.4459 | 0 | 3.4 | 141 | Specific | 746.29 | 655.48 | 655.48 | 15.22 | 148 | >sp\|P02788\|TRFL_HUMAN Lactotransferrin OS=Homo sapiens OX=9606 GN=LTF PE=1 SV=6 | 1270 | 20201118_12833_1ng_glyco_hcd203040_compound15.22858.22858.2 | controllerType=0 controllerNumber=1 scan=22858 | 46.6596 |
| 18915:4 | 1 | R.TAGWNVPIGTLRPFLN[+2424.867]WTGPPEPIEAAVAR.F | HexNAc(5)Hex(6)Fuc(1)NeuAc(1) | N[+2425] | 1414.6542 | 4 | 5655.5949 | 5655.5781 | 0 | 3 | 141 | Specific | 802.57 | 617.08 | 617.08 | 17.22 | 148 | >sp\|P02788\|TRFL_HUMAN Lactotransferrin OS=Homo sapiens OX=9606 GN=LTF PE=1 SV=6 | 1270 | 20201118_12833_1ng_glyco_hcd203040_compound15.22587.22587.4 | controllerType=0 controllerNumber=1 scan=22587 | 46.1042 |
| 19013:5 | 1 | R.TAGWNVPIGTLRPFLN[+2059.735]WTGPPEPIEAAVAR.F | HexNAc(4)Hex(5)Fuc(1)NeuAc(1) | N[+2060] | 1058.8982 | 5 | 5290.4619 | 5290.4459 | 0 | 3 | 141 | Specific | 807.6 | 739.65 | 739.65 | 17.85 | 148 | >sp\|P02788\|TRFL_HUMAN Lactotransferrin OS=Homo sapiens OX=9606 GN=LTF PE=1 SV=6 | 1270 | 20201118_12833_1ng_glyco_hcd203040_compound15.22691.22691.5 | controllerType=0 controllerNumber=1 scan=22691 | 46.3502 |
| 19015:5 | 1 | R.TAGWNVPIGTLRPFLN[+2059.735]WTGPPEPIEAAVAR.F | HexNAc(4)Hex(5)Fuc(1)NeuAc(1) | N[+2060] | 1058.8982 | 5 | 5290.4619 | 5290.4459 | 0 | 3 | 141 | Specific | 856.54 | 762.5 | 762.5 | 18.85 | 148 | >sp\|P02788\|TRFL_HUMAN Lactotransferrin OS=Homo sapiens OX=9606 GN=LTF PE=1 SV=6 | 1270 | 20201118_12833_1ng_glyco_hcd203040_compound15.22693.22693.5 | controllerType=0 controllerNumber=1 scan=22693 | 46.3577 |
| 19172:4 | 1 | R.TAGWNVPIGTLRPFLN[+2059.735]WTGPPEPIEAAVAR.F | HexNAc(4)Hex(5)Fuc(1)NeuAc(1) | N[+2060] | 1323.3715 | 4 | 5290.464 | 5290.4459 | 0 | 3.4 | 141 | Specific | 865.55 | 741.72 | 741.72 | 18.22 | 148 | >sp\|P02788\|TRFL_HUMAN Lactotransferrin OS=Homo sapiens OX=9606 GN=LTF PE=1 SV=6 | 1270 | 20201118_12833_1ng_glyco_hcd203040_compound15.22860.22860.2 | controllerType=0 controllerNumber=1 scan=22860 | 46.6646 |
| 18084:4 | 1 | R.TAGWNVPIGTLRPFLN[+1768.640]WTGPPEPIEAAVAR.F | HexNAc(4)Hex(5)Fuc(1) | N[+1769] | 1250.5978 | 4 | 4999.3693 | 4999.3504 | 0 | 3.8 | 141 | Specific | 890.41 | 809.97 | 809.97 | 18.22 | 148 | >sp\|P02788\|TRFL_HUMAN Lactotransferrin OS=Homo sapiens OX=9606 GN=LTF PE=1 SV=6 | 1270 | 20201118_12833_1ng_glyco_hcd203040_compound15.21719.21719.4 | controllerType=0 controllerNumber=1 scan=21719 | 44.1369 |
| 19011:4 | 1 | R.TAGWNVPIGTLRPFLN[+2059.735]WTGPPEPIEAAVAR.F | HexNAc(4)Hex(5)Fuc(1)NeuAc(1) | N[+2060] | 1323.3723 | 4 | 5290.4674 | 5290.4459 | 0 | 4.1 | 141 | Specific | 903.69 | 742.79 | 742.79 | 18.67 | 148 | >sp\|P02788\|TRFL_HUMAN Lactotransferrin OS=Homo sapiens OX=9606 GN=LTF PE=1 SV=6 | 1270 | 20201118_12833_1ng_glyco_hcd203040_compound15.22689.22689.4 | controllerType=0 controllerNumber=1 scan=22689 | 46.3429 |
| 20426:3 | 1 | R.TAGWNIPMGLLFN[+2350.830]QTGSC[+57.021]K.F | HexNAc(4)Hex(5)Fuc(1)NeuAc(2) | C[+57], N[+2351] | 1482.6197 | 3 | 4445.8445 | 4445.8403 | 0 | 0.9 | 485 | Specific | 641.14 | 555.25 | 555.25 | 12.18 | 148 | >sp\|P02788\|TRFL_HUMAN Lactotransferrin OS=Homo sapiens OX=9606 GN=LTF PE=1 SV=6 | 1270 | 20201118_12833_1ng_glyco_hcd203040_compound15.24168.24168.3 | controllerType=0 controllerNumber=1 scan=24168 | 49.4851 |
| 20428:3 | 1 | R.TAGWNIPMGLLFN[+2350.830]QTGSC[+57.021]K.F | HexNAc(4)Hex(5)Fuc(1)NeuAc(2) | C[+57], N[+2351] | 1482.6197 | 3 | 4445.8445 | 4445.8403 | 0 | 0.9 | 485 | Specific | 750.33 | 686.4 | 686.4 | 15.18 | 148 | >sp\|P02788\|TRFL_HUMAN Lactotransferrin OS=Homo sapiens OX=9606 GN=LTF PE=1 SV=6 | 1270 | 20201118_12833_1ng_glyco_hcd203040_compound15.24170.24170.3 | controllerType=0 controllerNumber=1 scan=24170 | 49.4895 |

Table S4. Cont.

| Query #:z | Protein Rank | Glycopeptide | Glycan | Modification Type(s) | Observed m/z | z | Observed (M+H) | Calc. mass (M+H) | Off-by-x error | Mass error (ppm) | StartPosition | Cleavage | Score | Delta | Delta Mod | \|Log Prob\| | # of unique peptides | Protein Name | Protein DB number | Comment | Scan # | Scan Time |
|---|---|---|---|---|---|---|---|---|---|---|---|---|---|---|---|---|---|---|---|---|---|---|
| 18531:3 | 1 | R.TAGWNIPMGLLFN[+1913.677]QTGSC[+57.021]K.F | HexNAc(4)Hex(5)NeuAc(1) | C[+57], N[+1914] | 1336.8996 | 3 | 4008.6842 | 4008.687 | 0 | -0.7 | 485 | Specific | 828.89 | 794.9 | 794.9 | 15.17 | 148 | >sp\|P02788\|TRFL_HUMAN Lactotransferrin OS=Homo sapiens OX=9606 GN=LTF PE=1 SV=6 | 1270 | 20201118_12833_1ng_glyco_hcd203040_compound15.22186.22186.3 | controllerType=0 controllerNumber=1 scan=22186 | 45.1882 |
| 16465:3 | 1 | R.TAGWNIPMGLLFN[+1768.640]QTGSC[+57.021]K.F | HexNAc(4)Hex(5)Fuc(1) | C[+57], N[+1769] | 1288.5576 | 3 | 3863.6581 | 3863.6495 | 0 | 2.2 | 485 | Specific | 882.89 | 691.87 | 691.87 | 17.18 | 148 | >sp\|P02788\|TRFL_HUMAN Lactotransferrin OS=Homo sapiens OX=9606 GN=LTF PE=1 SV=6 | 1270 | 20201118_12833_1ng_glyco_hcd203040_compound15.20033.20033.3 | controllerType=0 controllerNumber=1 scan=20033 | 40.629 |
| 18534:3 | 1 | R.TAGWNIPMGLLFN[+1913.677]QTGSC[+57.021]K.F | HexNAc(4)Hex(5)NeuAc(1) | C[+57], N[+1914] | 1336.8996 | 3 | 4008.6842 | 4008.687 | 0 | -0.7 | 485 | Specific | 899.28 | 873.56 | 873.56 | 16.17 | 148 | >sp\|P02788\|TRFL_HUMAN Lactotransferrin OS=Homo sapiens OX=9606 GN=LTF PE=1 SV=6 | 1270 | 20201118_12833_1ng_glyco_hcd203040_compound15.22189.22189.3 | controllerType=0 controllerNumber=1 scan=22189 | 45.1963 |
| 16463:3 | 1 | R.TAGWNIPMGLLFN[+1768.640]QTGSC[+57.021]K.F | HexNAc(4)Hex(5)Fuc(1) | C[+57], N[+1769] | 1288.5576 | 3 | 3863.6581 | 3863.6495 | 0 | 2.2 | 485 | Specific | 900.67 | 878.72 | 878.72 | 18.18 | 148 | >sp\|P02788\|TRFL_HUMAN Lactotransferrin OS=Homo sapiens OX=9606 GN=LTF PE=1 SV=6 | 1270 | 20201118_12833_1ng_glyco_hcd203040_compound15.20031.20031.3 | controllerType=0 controllerNumber=1 scan=20031 | 40.6233 |
| 16453:3 | 1 | R.TAGWNIPMGLLFN[+1768.640]QTGSC[+57.021]K.F | HexNAc(4)Hex(5)Fuc(1) | C[+57], N[+1769] | 1288.5579 | 3 | 3863.6592 | 3863.6495 | 0 | 2.5 | 485 | Specific | 901.72 | 901.72 | 901.72 | 18.21 | 148 | >sp\|P02788\|TRFL_HUMAN Lactotransferrin OS=Homo sapiens OX=9606 GN=LTF PE=1 SV=6 | 1270 | 20201118_12833_1ng_glyco_hcd203040_compound15.20020.20020.3 | controllerType=0 controllerNumber=1 scan=20020 | 40.6012 |
| 16451:3 | 1 | R.TAGWNIPMGLLFN[+1768.640]QTGSC[+57.021]K.F | HexNAc(4)Hex(5)Fuc(1) | C[+57], N[+1769] | 1288.5579 | 3 | 3863.6592 | 3863.6495 | 0 | 2.5 | 485 | Specific | 903.33 | 811.22 | 811.22 | 18.21 | 148 | >sp\|P02788\|TRFL_HUMAN Lactotransferrin OS=Homo sapiens OX=9606 GN=LTF PE=1 SV=6 | 1270 | 20201118_12833_1ng_glyco_hcd203040_compound15.20018.20018.3 | controllerType=0 controllerNumber=1 scan=20018 | 40.5967 |
| 18402:3 | 1 | R.TAGWNIPMGLLFN[+2059.735]QTGSC[+57.021]K.F | HexNAc(4)Hex(5)Fuc(1)NeuAc(1) | C[+57], N[+2060] | 1385.5908 | 3 | 4154.7577 | 4154.7449 | 0 | 3.1 | 485 | Specific | 933.82 | 807.82 | 807.82 | 18.21 | 148 | >sp\|P02788\|TRFL_HUMAN Lactotransferrin OS=Homo sapiens OX=9606 GN=LTF PE=1 SV=6 | 1270 | 20201118_12833_1ng_glyco_hcd203040_compound15.22051.22051.3 | controllerType=0 controllerNumber=1 scan=22051 | 44.8736 |
| 18404:3 | 1 | R.TAGWNIPMGLLFN[+2059.735]QTGSC[+57.021]K.F | HexNAc(4)Hex(5)Fuc(1)NeuAc(1) | C[+57], N[+2060] | 1385.5908 | 3 | 4154.7577 | 4154.7449 | 0 | 3.1 | 485 | Specific | 950.11 | 685.28 | 685.28 | 19.21 | 148 | >sp\|P02788\|TRFL_HUMAN Lactotransferrin OS=Homo sapiens OX=9606 GN=LTF PE=1 SV=6 | 1270 | 20201118_12833_1ng_glyco_hcd203040_compound15.22053.22053.3 | controllerType=0 controllerNumber=1 scan=22053 | 44.8791 |
| 18714:4 | 1 | R.TAGWNIPMGLLFN[+2076.750]QTGSC[+57.021]KFDEYFSQSC[+57.021]APGSDPR.S | HexNAc(4)Hex(6)Fuc(2) | C[+57]*2, N[+2077] | 1504.6401 | 4 | 6015.5387 | 6015.5074 | 0 | 5.2 | 485 | Specific | 264.77 | 254.01 | 254.01 | 6.51 | 148 | >sp\|P02788\|TRFL_HUMAN Lactotransferrin OS=Homo sapiens OX=9606 GN=LTF PE=1 SV=6 | 1270 | 20201118_12833_1ng_glyco_hcd203040_compound15.22378.22378.4 | controllerType=0 controllerNumber=1 scan=22378 | 45.6399 |
| 20068:4 | 1 | R.TAGWNIPMGLLFN[+2350.830]QTGSC[+57.021]KFDEYFSQSC[+57.021]APGSDPR.S | HexNAc(4)Hex(5)Fuc(1)NeuAc(2) | C[+57]*2, N[+2351] | 1573.1537 | 4 | 6289.5929 | 6289.5875 | 0 | 0.9 | 485 | Specific | 316.22 | 311 | 311 | 7.37 | 148 | >sp\|P02788\|TRFL_HUMAN Lactotransferrin OS=Homo sapiens OX=9606 GN=LTF PE=1 SV=6 | 1270 | 20201118_12833_1ng_glyco_hcd203040_compound15.23794.23794.4 | controllerType=0 controllerNumber=1 scan=23794 | 48.6481 |
| 18662:4 | 1 | R.TAGWNIPMGLLFN[+2350.830]QTGSC[+57.021]KFDEYFSQSC[+57.021]APGSDPR.S | HexNAc(4)Hex(5)Fuc(1)NeuAc(2) | C[+57]*2, N[+2351] | 1573.1666 | 4 | 6289.6447 | 6289.5875 | 0 | 9.1 | 485 | Specific | 476.5 | 460.9 | 460.9 | 8.58 | 148 | >sp\|P02788\|TRFL_HUMAN Lactotransferrin OS=Homo sapiens OX=9606 GN=LTF PE=1 SV=6 | 1270 | 20201118_12833_1ng_glyco_hcd203040_compound15.22324.22324.4 | controllerType=0 controllerNumber=1 scan=22324 | 45.5295 |
| 18683:5 | 1 | R.TAGWNIPMGLLFN[+2059.735]QTGSC[+57.021]KFDEYFSQSC[+57.021]APGSDPR.S | HexNAc(4)Hex(5)Fuc(1)NeuAc(1) | C[+57]*2, N[+2060] | 1200.5072 | 5 | 5998.5069 | 5998.492 | 0 | 2.5 | 485 | Specific | 497.07 | 497.07 | 497.07 | 10.93 | 148 | >sp\|P02788\|TRFL_HUMAN Lactotransferrin OS=Homo sapiens OX=9606 GN=LTF PE=1 SV=6 | 1270 | 20201118_12833_1ng_glyco_hcd203040_compound15.22346.22346.5 | controllerType=0 controllerNumber=1 scan=22346 | 45.577 |
| 20071:4 | 1 | R.TAGWNIPMGLLFN[+2350.830]QTGSC[+57.021]KFDEYFSQSC[+57.021]APGSDPR.S | HexNAc(4)Hex(5)Fuc(1)NeuAc(2) | C[+57]*2, N[+2351] | 1573.1537 | 4 | 6289.5929 | 6289.5875 | 0 | 0.9 | 485 | Specific | 520.36 | 497.72 | 497.72 | 9.94 | 148 | >sp\|P02788\|TRFL_HUMAN Lactotransferrin OS=Homo sapiens OX=9606 GN=LTF PE=1 SV=6 | 1270 | 20201118_12833_1ng_glyco_hcd203040_compound15.23797.23797.4 | controllerType=0 controllerNumber=1 scan=23797 | 48.6529 |
| 18644:5 | 1 | R.TAGWNIPMGLLFN[+2205.793]QTGSC[+57.021]KFDEYFSQSC[+57.021]APGSDPR.S | HexNAc(4)Hex(5)Fuc(2)NeuAc(1) | C[+57]*2, N[+2206] | 1229.7196 | 5 | 6144.5689 | 6144.5499 | 0 | 3.1 | 485 | Specific | 598.34 | 598.34 | 598.34 | 12.42 | 148 | >sp\|P02788\|TRFL_HUMAN Lactotransferrin OS=Homo sapiens OX=9606 GN=LTF PE=1 SV=6 | 1270 | 20201118_12833_1ng_glyco_hcd203040_compound15.22305.22305.5 | controllerType=0 controllerNumber=1 scan=22305 | 45.4894 |
| 18751:5 | 1 | R.TAGWNIPMGLLFN[+1913.677]QTGSC[+57.021]KFDEYFSQSC[+57.021]APGSDPR.S | HexNAc(4)Hex(5)NeuAc(1) | C[+57]*2, N[+1914] | 1171.2983 | 5 | 5852.4624 | 5852.4341 | 0 | 4.8 | 485 | Specific | 614.88 | 593.9 | 593.9 | 12.88 | 148 | >sp\|P02788\|TRFL_HUMAN Lactotransferrin OS=Homo sapiens OX=9606 GN=LTF PE=1 SV=6 | 1270 | 20201118_12833_1ng_glyco_hcd203040_compound15.22416.22416.5 | controllerType=0 controllerNumber=1 scan=22416 | 45.7181 |
| 17681:5 | 1 | R.TAGWNIPMGLLFN[+1768.640]QTGSC[+57.021]KFDEYFSQSC[+57.021]APGSDPR.S | HexNAc(4)Hex(5)Fuc(1) | C[+57]*2, N[+1769] | 1142.2881 | 5 | 5707.4113 | 5707.3966 | 0 | 2.6 | 485 | Specific | 642.08 | 561.43 | 561.43 | 13.42 | 148 | >sp\|P02788\|TRFL_HUMAN Lactotransferrin OS=Homo sapiens OX=9606 GN=LTF PE=1 SV=6 | 1270 | 20201118_12833_1ng_glyco_hcd203040_compound15.21299.21299.5 | controllerType=0 controllerNumber=1 scan=21299 | 43.2519 |
| 20060:4 | 1 | R.TAGWNIPMGLLFN[+2204.772]QTGSC[+57.021]KFDEYFSQSC[+57.021]APGSDPR.S | HexNAc(4)Hex(5)NeuAc(2) | C[+57]*2, N[+2205] | 1536.6436 | 4 | 6143.5524 | 6143.5295 | 0 | 3.7 | 485 | Specific | 661.45 | 640.76 | 244.21 | 13.79 | 148 | >sp\|P02788\|TRFL_HUMAN Lactotransferrin OS=Homo sapiens OX=9606 GN=LTF PE=1 SV=6 | 1270 | 20201118_12833_1ng_glyco_hcd203040_compound15.23786.23786.4 | controllerType=0 controllerNumber=1 scan=23786 | 48.6338 |
| 18636:4 | 1 | R.TAGWNIPMGLLFN[+2205.793]QTGSC[+57.021]KFDEYFSQSC[+57.021]APGSDPR.S | HexNAc(4)Hex(5)Fuc(2)NeuAc(1) | C[+57]*2, N[+2206] | 1536.8989 | 4 | 6144.5739 | 6144.5499 | 0 | 3.9 | 485 | Specific | 671.12 | 622.16 | 622.16 | 13.25 | 148 | >sp\|P02788\|TRFL_HUMAN Lactotransferrin OS=Homo sapiens OX=9606 GN=LTF PE=1 SV=6 | 1270 | 20201118_12833_1ng_glyco_hcd203040_compound15.22297.22297.4 | controllerType=0 controllerNumber=1 scan=22297 | 45.476 |
| 18664:4 | 1 | R.TAGWNIPMGLLFN[+2350.830]QTGSC[+57.021]KFDEYFSQSC[+57.021]APGSDPR.S | HexNAc(4)Hex(5)Fuc(1)NeuAc(2) | C[+57]*2, N[+2351] | 1573.1666 | 4 | 6289.6447 | 6289.5875 | 0 | 9.1 | 485 | Specific | 696.67 | 657.01 | 657.01 | 12.13 | 148 | >sp\|P02788\|TRFL_HUMAN Lactotransferrin OS=Homo sapiens OX=9606 GN=LTF PE=1 SV=6 | 1270 | 20201118_12833_1ng_glyco_hcd203040_compound15.22326.22326.4 | controllerType=0 controllerNumber=1 scan=22326 | 45.5374 |
| 18727:3 | 1 | R.TAGWNIPMGLLFN[+1913.677]QTGSC[+57.021]KFDEYFSQSC[+57.021]APGSDPR.S | HexNAc(4)Hex(5)NeuAc(1) | C[+57]*2, N[+1914] | 1951.4915 | 3 | 5852.4601 | 5852.4341 | 0 | 4.4 | 485 | Specific | 720.41 | 604.49 | 604.49 | 14.42 | 148 | >sp\|P02788\|TRFL_HUMAN Lactotransferrin OS=Homo sapiens OX=9606 GN=LTF PE=1 SV=6 | 1270 | 20201118_12833_1ng_glyco_hcd203040_compound15.22391.22391.3 | controllerType=0 controllerNumber=1 scan=22391 | 45.6623 |
| 18753:5 | 1 | R.TAGWNIPMGLLFN[+1913.677]QTGSC[+57.021]KFDEYFSQSC[+57.021]APGSDPR.S | HexNAc(4)Hex(5)NeuAc(1) | C[+57]*2, N[+1914] | 1171.2983 | 5 | 5852.4624 | 5852.4341 | 0 | 4.8 | 485 | Specific | 728.44 | 724.13 | 724.13 | 14.88 | 148 | >sp\|P02788\|TRFL_HUMAN Lactotransferrin OS=Homo sapiens OX=9606 GN=LTF PE=1 SV=6 | 1270 | 20201118_12833_1ng_glyco_hcd203040_compound15.22418.22418.5 | controllerType=0 controllerNumber=1 scan=22418 | 45.7225 |
| 18730:3 | 1 | R.TAGWNIPMGLLFN[+1913.677]QTGSC[+57.021]KFDEYFSQSC[+57.021]APGSDPR.S | HexNAc(4)Hex(5)NeuAc(1) | C[+57]*2, N[+1914] | 1951.4915 | 3 | 5852.4601 | 5852.4341 | 0 | 4.4 | 485 | Specific | 748.42 | 606.05 | 606.05 | 14.42 | 148 | >sp\|P02788\|TRFL_HUMAN Lactotransferrin OS=Homo sapiens OX=9606 GN=LTF PE=1 SV=6 | 1270 | 20201118_12833_1ng_glyco_hcd203040_compound15.22394.22394.3 | controllerType=0 controllerNumber=1 scan=22394 | 45.6664 |
| 19965:4 | 1 | R.TAGWNIPMGLLFN[+2350.830]QTGSC[+57.021]KFDEYFSQSC[+57.021]APGSDPR.S | HexNAc(4)Hex(5)Fuc(1)NeuAc(2) | C[+57]*2, N[+2351] | 1573.1585 | 4 | 6289.612 | 6289.5875 | 0 | 3.9 | 485 | Specific | 773.76 | 726.64 | 726.64 | 15.25 | 148 | >sp\|P02788\|TRFL_HUMAN Lactotransferrin OS=Homo sapiens OX=9606 GN=LTF PE=1 SV=6 | 1270 | 20201118_12833_1ng_glyco_hcd203040_compound15.23687.23687.4 | controllerType=0 controllerNumber=1 scan=23687 | 48.4162 |
| 20063:4 | 1 | R.TAGWNIPMGLLFN[+2204.772]QTGSC[+57.021]KFDEYFSQSC[+57.021]APGSDPR.S | HexNAc(4)Hex(5)NeuAc(2) | C[+57]*2, N[+2205] | 1536.6436 | 4 | 6143.5524 | 6143.5295 | 0 | 3.7 | 485 | Specific | 775.19 | 747.45 | 382.29 | 15.79 | 148 | >sp\|P02788\|TRFL_HUMAN Lactotransferrin OS=Homo sapiens OX=9606 GN=LTF PE=1 SV=6 | 1270 | 20201118_12833_1ng_glyco_hcd203040_compound15.23789.23789.4 | controllerType=0 controllerNumber=1 scan=23789 | 48.6401 |
| 17704:4 | 1 | R.TAGWNIPMGLLFN[+1622.582]QTGSC[+57.021]KFDEYFSQSC[+57.021]APGSDPR.S | HexNAc(4)Hex(5) | C[+57]*2, N[+1623] | 1391.0955 | 4 | 5561.36 | 5561.3387 | 0 | 3.8 | 485 | Specific | 775.73 | 730.5 | 211.07 | 15.79 | 148 | >sp\|P02788\|TRFL_HUMAN Lactotransferrin OS=Homo sapiens OX=9606 GN=LTF PE=1 SV=6 | 1270 | 20201118_12833_1ng_glyco_hcd203040_compound15.21323.21323.4 | controllerType=0 controllerNumber=1 scan=21323 | 43.3071 |
| 17640:4 | 1 | R.TAGWNIPMGLLFN[+1768.640]QTGSC[+57.021]KFDEYFSQSC[+57.021]APGSDPR.S | HexNAc(4)Hex(5)Fuc(1) | C[+57]*2, N[+1769] | 1427.6106 | 4 | 5707.4206 | 5707.3966 | 0 | 4.2 | 485 | Specific | 784.52 | 748.05 | 748.05 | 15.25 | 148 | >sp\|P02788\|TRFL_HUMAN Lactotransferrin OS=Homo sapiens OX=9606 GN=LTF PE=1 SV=6 | 1270 | 20201118_12833_1ng_glyco_hcd203040_compound15.21257.21257.4 | controllerType=0 controllerNumber=1 scan=21257 | 43.178 |
| 17641:3 | 1 | R.TAGWNIPMGLLFN[+1768.640]QTGSC[+57.021]KFDEYFSQSC[+57.021]APGSDPR.S | HexNAc(4)Hex(5)Fuc(1) | C[+57]*2, N[+1769] | 1903.1424 | 3 | 5707.4127 | 5707.3966 | 0 | 2.8 | 485 | Specific | 794.37 | 712 | 712 | 15.97 | 148 | >sp\|P02788\|TRFL_HUMAN Lactotransferrin OS=Homo sapiens OX=9606 GN=LTF PE=1 SV=6 | 1270 | 20201118_12833_1ng_glyco_hcd203040_compound15.21258.21258.3 | controllerType=0 controllerNumber=1 scan=21258 | 43.1794 |
| 17643:3 | 1 | R.TAGWNIPMGLLFN[+1768.640]QTGSC[+57.021]KFDEYFSQSC[+57.021]APGSDPR.S | HexNAc(4)Hex(5)Fuc(1) | C[+57]*2, N[+1769] | 1903.1424 | 3 | 5707.4127 | 5707.3966 | 0 | 2.8 | 485 | Specific | 814.89 | 653.64 | 653.64 | 16.97 | 148 | >sp\|P02788\|TRFL_HUMAN Lactotransferrin OS=Homo sapiens OX=9606 GN=LTF PE=1 SV=6 | 1270 | 20201118_12833_1ng_glyco_hcd203040_compound15.21260.21260.3 | controllerType=0 controllerNumber=1 scan=21260 | 43.1842 |
| 18685:5 | 1 | R.TAGWNIPMGLLFN[+2059.735]QTGSC[+57.021]KFDEYFSQSC[+57.021]APGSDPR.S | HexNAc(4)Hex(5)Fuc(1)NeuAc(1) | C[+57]*2, N[+2060] | 1200.5072 | 5 | 5998.5069 | 5998.492 | 0 | 2.5 | 485 | Specific | 851.38 | 828.65 | 828.65 | 18.42 | 148 | >sp\|P02788\|TRFL_HUMAN Lactotransferrin OS=Homo sapiens OX=9606 GN=LTF PE=1 SV=6 | 1270 | 20201118_12833_1ng_glyco_hcd203040_compound15.22348.22348.5 | controllerType=0 controllerNumber=1 scan=22348 | 45.5813 |
| 19967:4 | 1 | R.TAGWNIPMGLLFN[+2350.830]QTGSC[+57.021]KFDEYFSQSC[+57.021]APGSDPR.S | HexNAc(4)Hex(5)Fuc(1)NeuAc(2) | C[+57]*2, N[+2351] | 1573.1585 | 4 | 6289.612 | 6289.5875 | 0 | 3.9 | 485 | Specific | 851.48 | 828.52 | 828.52 | 17.25 | 148 | >sp\|P02788\|TRFL_HUMAN Lactotransferrin OS=Homo sapiens OX=9606 GN=LTF PE=1 SV=6 | 1270 | 20201118_12833_1ng_glyco_hcd203040_compound15.23689.23689.4 | controllerType=0 controllerNumber=1 scan=23689 | 48.4212 |
| 17706:4 | 1 | R.TAGWNIPMGLLFN[+1622.582]QTGSC[+57.021]KFDEYFSQSC[+57.021]APGSDPR.S | HexNAc(4)Hex(5) | C[+57]*2, N[+1623] | 1391.0955 | 4 | 5561.36 | 5561.3387 | 0 | 3.8 | 485 | Specific | 860.05 | 817.45 | 316.71 | 17.79 | 148 | >sp\|P02788\|TRFL_HUMAN Lactotransferrin OS=Homo sapiens OX=9606 GN=LTF PE=1 SV=6 | 1270 | 20201118_12833_1ng_glyco_hcd203040_compound15.21325.21325.4 | controllerType=0 controllerNumber=1 scan=21325 | 43.3111 |
| 18715:4 | 1 | R.TAGWNIPMGLLFN[+1913.677]QTGSC[+57.021]KFDEYFSQSC[+57.021]APGSDPR.S | HexNAc(4)Hex(5)NeuAc(1) | C[+57]*2, N[+1914] | 1463.8711 | 4 | 5852.4626 | 5852.4341 | 0 | 4.9 | 485 | Specific | 891.7 | 851.64 | 851.64 | 17.25 | 148 | >sp\|P02788\|TRFL_HUMAN Lactotransferrin OS=Homo sapiens OX=9606 GN=LTF PE=1 SV=6 | 1270 | 20201118_12833_1ng_glyco_hcd203040_compound15.22379.22379.4 | controllerType=0 controllerNumber=1 scan=22379 | 45.6412 |
| 18635:4 | 1 | R.TAGWNIPMGLLFN[+2059.735]QTGSC[+57.021]KFDEYFSQSC[+57.021]APGSDPR.S | HexNAc(4)Hex(5)Fuc(1)NeuAc(1) | C[+57]*2, N[+2060] | 1500.3882 | 4 | 5998.5309 | 5998.492 | 0 | 6.5 | 485 | Specific | 913.27 | 896.72 | 896.72 | 17.81 | 148 | >sp\|P02788\|TRFL_HUMAN Lactotransferrin OS=Homo sapiens OX=9606 GN=LTF PE=1 SV=6 | 1270 | 20201118_12833_1ng_glyco_hcd203040_compound15.22296.22296.4 | controllerType=0 controllerNumber=1 scan=22296 | 45.4746 |
| 18637:4 | 1 | R.TAGWNIPMGLLFN[+2059.735]QTGSC[+57.021]KFDEYFSQSC[+57.021]APGSDPR.S | HexNAc(4)Hex(5)Fuc(1)NeuAc(1) | C[+57]*2, N[+2060] | 1500.3882 | 4 | 5998.5309 | 5998.492 | 0 | 6.5 | 485 | Specific | 925.67 | 894.73 | 894.73 | 17.81 | 148 | >sp\|P02788\|TRFL_HUMAN Lactotransferrin OS=Homo sapiens OX=9606 GN=LTF PE=1 SV=6 | 1270 | 20201118_12833_1ng_glyco_hcd203040_compound15.22298.22298.4 | controllerType=0 controllerNumber=1 scan=22298 | 45.4807 |
| 18638:4 | 1 | R.TAGWNIPMGLLFN[+2205.793]QTGSC[+57.021]KFDEYFSQSC[+57.021]APGSDPR.S | HexNAc(4)Hex(5)Fuc(2)NeuAc(1) | C[+57]*2, N[+2206] | 1536.8989 | 4 | 6144.5739 | 6144.5499 | 0 | 3.9 | 485 | Specific | 927.03 | 864.6 | 864.6 | 18.25 | 148 | >sp\|P02788\|TRFL_HUMAN Lactotransferrin OS=Homo sapiens OX=9606 GN=LTF PE=1 SV=6 | 1270 | 20201118_12833_1ng_glyco_hcd203040_compound15.22299.22299.4 | controllerType=0 controllerNumber=1 scan=22299 | 45.482 |
| 18712:4 | 1 | R.TAGWNIPMGLLFN[+1913.677]QTGSC[+57.021]KFDEYFSQSC[+57.021]APGSDPR.S | HexNAc(4)Hex(5)NeuAc(1) | C[+57]*2, N[+1914] | 1463.8711 | 4 | 5852.4626 | 5852.4341 | 0 | 4.9 | 485 | Specific | 933.55 | 841.03 | 841.03 | 18.25 | 148 | >sp\|P02788\|TRFL_HUMAN Lactotransferrin OS=Homo sapiens OX=9606 GN=LTF PE=1 SV=6 | 1270 | 20201118_12833_1ng_glyco_hcd203040_compound15.22376.22376.4 | controllerType=0 controllerNumber=1 scan=22376 | 45.6344 |
| 18647:5 | 1 | R.TAGWNIPMGLLFN[+2205.793]QTGSC[+57.021]KFDEYFSQSC[+57.021]APGSDPR.S | HexNAc(4)Hex(5)Fuc(2)NeuAc(1) | C[+57]*2, N[+2206] | 1229.7196 | 5 | 6144.5689 | 6144.5499 | 0 | 3.1 | 485 | Specific | 946 | 831.58 | 831.58 | 19.42 | 148 | >sp\|P02788\|TRFL_HUMAN Lactotransferrin OS=Homo sapiens OX=9606 GN=LTF PE=1 SV=6 | 1270 | 20201118_12833_1ng_glyco_hcd203040_compound15.22308.22308.5 | controllerType=0 controllerNumber=1 scan=22308 | 45.4941 |
| 17642:4 | 1 | R.TAGWNIPMGLLFN[+1768.640]QTGSC[+57.021]KFDEYFSQSC[+57.021]APGSDPR.S | HexNAc(4)Hex(5)Fuc(1) | C[+57]*2, N[+1769] | 1427.6106 | 4 | 5707.4206 | 5707.3966 | 0 | 4.2 | 485 | Specific | 982.15 | 942.21 | 942.21 | 19.25 | 148 | >sp\|P02788\|TRFL_HUMAN Lactotransferrin OS=Homo sapiens OX=9606 GN=LTF PE=1 SV=6 | 1270 | 20201118_12833_1ng_glyco_hcd203040_compound15.21259.21259.4 | controllerType=0 controllerNumber=1 scan=21259 | 43.1827 |
| 19994:4 | 1 | R.TAGWNIPM[+15.995]GLLFN[+2351.851]QTGSC[+57.021]KFDEYFSQSC[+57.021]A | HexNAc(4)Hex(5)Fuc(3)NeuAc(1) | C[+57]*2, M[+16], N[+2352] | 1577.4133 | 4 | 6306.6315 | 6306.6028 | 0 | 4.6 | 485 | Specific | 233.87 | 210.05 | 210.05 | 6.05 | 148 | >sp\|P02788\|TRFL_HUMAN Lactotransferrin OS=Homo sapiens OX=9606 GN=LTF PE=1 SV=6 | 1270 | 20201118_12833_1ng_glyco_hcd203040_compound15.23717.23717.4 | controllerType=0 controllerNumber=1 scan=23717 | 48.4793 |
| 18542:4 | 1 | R.TAGWNIPM[+15.995]GLLFN[+2059.735]QTGSC[+57.021]KFDEYFSQSC[+57.021]A | HexNAc(4)Hex(5)Fuc(1)NeuAc(1) | C[+57]*2, M[+16], N[+2060] | 1504.3798 | 4 | 6014.4972 | 6014.487 | 0 | 1.7 | 485 | Specific | 320.42 | 243.5 | 243.5 | 7.39 | 148 | >sp\|P02788\|TRFL_HUMAN Lactotransferrin OS=Homo sapiens OX=9606 GN=LTF PE=1 SV=6 | 1270 | 20201118_12833_1ng_glyco_hcd203040_compound15.22198.22198.2 | controllerType=0 controllerNumber=1 scan=22198 | 45.2194 |
| 18545:4 | 1 | R.TAGWNIPM[+15.995]GLLFN[+2059.735]QTGSC[+57.021]KFDEYFSQSC[+57.021]A | HexNAc(4)Hex(5)Fuc(1)NeuAc(1) | C[+57]*2, M[+16], N[+2060] | 1504.3798 | 4 | 6014.4972 | 6014.487 | 0 | 1.7 | 485 | Specific | 483.59 | 365.48 | 365.48 | 8.64 | 148 | >sp\|P02788\|TRFL_HUMAN Lactotransferrin OS=Homo sapiens OX=9606 GN=LTF PE=1 SV=6 | 1270 | 20201118_12833_1ng_glyco_hcd203040_compound15.22201.22201.2 | controllerType=0 controllerNumber=1 scan=22201 | 45.2242 |
| 17635:4 | 1 | R.TAGWNIPM[+15.995]GLLFN[+2059.735]QTGSC[+57.021]KFDEYFSQSC[+57.021]A | HexNAc(4)Hex(5)Fuc(1)NeuAc(1) | C[+57]*2, M[+16], N[+2060] | 1504.381 | 4 | 6014.5021 | 6014.487 | 0 | 2.5 | 485 | Specific | 592.13 | 503.7 | 503.7 | 11.79 | 148 | >sp\|P02788\|TRFL_HUMAN Lactotransferrin OS=Homo sapiens OX=9606 GN=LTF PE=1 SV=6 | 1270 | 20201118_12833_1ng_glyco_hcd203040_compound15.21251.21251.4 | controllerType=0 controllerNumber=1 scan=21251 | 43.1624 |
| 17637:4 | 1 | R.TAGWNIPM[+15.995]GLLFN[+2059.735]QTGSC[+57.021]KFDEYFSQSC[+57.021]A | HexNAc(4)Hex(5)Fuc(1)NeuAc(1) | C[+57]*2, M[+16], N[+2060] | 1504.381 | 4 | 6014.5021 | 6014.487 | 0 | 2.5 | 485 | Specific | 778.35 | 680.32 | 680.32 | 15.79 | 148 | >sp\|P02788\|TRFL_HUMAN Lactotransferrin OS=Homo sapiens OX=9606 GN=LTF PE=1 SV=6 | 1270 | 20201118_12833_1ng_glyco_hcd203040_compound15.21253.21253.4 | controllerType=0 controllerNumber=1 scan=21253 | 43.1677 |
| 17679:4 | 1 | R.TAGWNIPM[+15.995]GLLFN[+2059.735]QTGSC[+57.021]KFDEYFSQSC[+57.021]A | HexNAc(4)Hex(5)Fuc(1)NeuAc(1) | C[+57]*2, M[+16], N[+2060] | 1504.3834 | 4 | 6014.5119 | 6014.487 | 0 | 4.1 | 485 | Specific | 794.59 | 699.89 | 699.89 | 15.25 | 148 | >sp\|P02788\|TRFL_HUMAN Lactotransferrin OS=Homo sapiens OX=9606 GN=LTF PE=1 SV=6 | 1270 | 20201118_12833_1ng_glyco_hcd203040_compound15.21297.21297.4 | controllerType=0 controllerNumber=1 scan=21297 | 43.2487 |
| 17682:4 | 1 | R.TAGWNIPM[+15.995]GLLFN[+2059.735]QTGSC[+57.021]KFDEYFSQSC[+57.021]A | HexNAc(4)Hex(5)Fuc(1)NeuAc(1) | C[+57]*2, M[+16], N[+2060] | 1504.3834 | 4 | 6014.5119 | 6014.487 | 0 | 4.1 | 485 | Specific | 891.62 | 821 | 821 | 17.25 | 148 | >sp\|P02788\|TRFL_HUMAN Lactotransferrin OS=Homo sapiens OX=9606 GN=LTF PE=1 SV=6 | 1270 | 20201118_12833_1ng_glyco_hcd203040_compound15.21300.21300.4 | controllerType=0 controllerNumber=1 scan=21300 | 43.2556 |
| 09577:3 | 1 | K.FGRN[+2350.830]GSDC[+57.021]PDKFC[+57.021]LFQSETK.N | HexNAc(4)Hex(5)Fuc(1)NeuAc(2) | C[+57]*2, N[+2351] | 1581.9759 | 3 | 4743.9131 | 4743.8952 | 0 | 3.8 | 639 | Specific | 216.6 | 168.4 | 168.4 | 4.42 | 148 | >sp\|P02788\|TRFL_HUMAN Lactotransferrin OS=Homo sapiens OX=9606 GN=LTF PE=1 SV=6 | 1270 | 20201118_12833_1ng_glyco_hcd203040_compound15.12829.12829.3 | controllerType=0 controllerNumber=1 scan=12829 | 28.0255 |
| 09537:3 | 1 | K.FGRN[+2350.830]GSDC[+57.021]PDKFC[+57.021]LFQSETK.N | HexNAc(4)Hex(5)Fuc(1)NeuAc(2) | C[+57]*2, N[+2351] | 1581.97 | 3 | 4743.8955 | 4743.8952 | 0 | 0.1 | 639 | Specific | 284.53 | 248.32 | 248.32 | 5.49 | 148 | >sp\|P02788\|TRFL_HUMAN Lactotransferrin OS=Homo sapiens OX=9606 GN=LTF PE=1 SV=6 | 1270 | 20201118_12833_1ng_glyco_hcd203040_compound15.12787.12787.3 | controllerType=0 controllerNumber=1 scan=12787 | 27.9551 |
| 09535:3 | 1 | K.FGRN[+2350.830]GSDC[+57.021]PDKFC[+57.021]LFQSETK.N | HexNAc(4)Hex(5)Fuc(1)NeuAc(2) | C[+57]*2, N[+2351] | 1581.97 | 3 | 4743.8955 | 4743.8952 | 0 | 0.1 | 639 | Specific | 347.53 | 347.53 | 347.53 | 6.5 | 148 | >sp\|P02788\|TRFL_HUMAN Lactotransferrin OS=Homo sapiens OX=9606 GN=LTF PE=1 SV=6 | 1270 | 20201118_12833_1ng_glyco_hcd203040_compound15.12785.12785.3 | controllerType=0 controllerNumber=1 scan=12785 | 27.9514 |
| 07797:3 | 1 | K.FGRN[+2059.735]GSDC[+57.021]PDKFC[+57.021]LFQSETK.N | HexNAc(4)Hex(5)Fuc(1)NeuAc(1) | C[+57]*2, N[+2060] | 1484.9434 | 3 | 4452.8157 | 4452.7998 | 0 | 3.6 | 639 | Specific | 355.22 | 355.22 | 355.22 | 7 | 148 | >sp\|P02788\|TRFL_HUMAN Lactotransferrin OS=Homo sapiens OX=9606 GN=LTF PE=1 SV=6 | 1270 | 20201118_12833_1ng_glyco_hcd203040_compound15.10970.10970.3 | controllerType=0 controllerNumber=1 scan=10970 | 24.8781 |
| 07811:3 | 1 | K.FGRN[+2059.735]GSDC[+57.021]PDKFC[+57.021]LFQSETK.N | HexNAc(4)Hex(5)Fuc(1)NeuAc(1) | C[+57]*2, N[+2060] | 1484.9415 | 3 | 4452.8098 | 4452.7998 | 0 | 2.2 | 639 | Specific | 370.98 | 370.98 | 370.98 | 7.24 | 148 | >sp\|P02788\|TRFL_HUMAN Lactotransferrin OS=Homo sapiens OX=9606 GN=LTF PE=1 SV=6 | 1270 | 20201118_12833_1ng_glyco_hcd203040_compound15.10985.10985.3 | controllerType=0 controllerNumber=1 scan=10985 | 24.9051 |
| 07813:3 | 1 | K.FGRN[+2059.735]GSDC[+57.021]PDKFC[+57.021]LFQSETK.N | HexNAc(4)Hex(5)Fuc(1)NeuAc(1) | C[+57]*2, N[+2060] | 1484.9415 | 3 | 4452.8098 | 4452.7998 | 0 | 2.2 | 639 | Specific | 420.85 | 380.66 | 380.66 | 8.97 | 148 | >sp\|P02788\|TRFL_HUMAN Lactotransferrin OS=Homo sapiens OX=9606 GN=LTF PE=1 SV=6 | 1270 | 20201118_12833_1ng_glyco_hcd203040_compound15.10987.10987.3 | controllerType=0 controllerNumber=1 scan=10987 | 24.9087 |
| 07799:3 | 1 | K.FGRN[+2059.735]GSDC[+57.021]PDKFC[+57.021]LFQSETK.N | HexNAc(4)Hex(5)Fuc(1)NeuAc(1) | C[+57]*2, N[+2060] | 1484.9434 | 3 | 4452.8157 | 4452.7998 | 0 | 3.6 | 639 | Specific | 450.05 | 375.94 | 375.94 | 9.01 | 148 | >sp\|P02788\|TRFL_HUMAN Lactotransferrin OS=Homo sapiens OX=9606 GN=LTF PE=1 SV=6 | 1270 | 20201118_12833_1ng_glyco_hcd203040_compound15.10972.10972.3 | controllerType=0 controllerNumber=1 scan=10972 | 24.8818 |
| 07750:4 | 1 | K.FGRN[+2205.793]GSDC[+57.021]PDKFC[+57.021]LFQSETK.N | HexNAc(4)Hex(5)Fuc(2)NeuAc(1) | C[+57]*2, N[+2206] | 1150.4715 | 4 | 4598.8641 | 4598.8577 | 0 | 1.4 | 639 | Specific | 486.42 | 486.42 | 486.42 | 9.66 | 148 | >sp\|P02788\|TRFL_HUMAN Lactotransferrin OS=Homo sapiens OX=9606 GN=LTF PE=1 SV=6 | 1270 | 20201118_12833_1ng_glyco_hcd203040_compound15.10922.10922.4 | controllerType=0 controllerNumber=1 scan=10922 | 24.7971 |
| 07720:4 | 1 | K.FGRN[+2205.793]GSDC[+57.021]PDKFC[+57.021]LFQSETK.N | HexNAc(4)Hex(5)Fuc(2)NeuAc(1) | C[+57]*2, N[+2206] | 1150.472 | 4 | 4598.866 | 4598.8577 | 0 | 1.8 | 639 | Specific | 556.14 | 556.14 | 556.14 | 10.02 | 148 | >sp\|P02788\|TRFL_HUMAN Lactotransferrin OS=Homo sapiens OX=9606 GN=LTF PE=1 SV=6 | 1270 | 20201118_12833_1ng_glyco_hcd203040_compound15.10891.10891.4 | controllerType=0 controllerNumber=1 scan=10891 | 24.7418 |
| 07722:4 | 1 | K.FGRN[+2205.793]GSDC[+57.021]PDKFC[+57.021]LFQSETK.N | HexNAc(4)Hex(5)Fuc(2)NeuAc(1) | C[+57]*2, N[+2206] | 1150.472 | 4 | 4598.866 | 4598.8577 | 0 | 1.8 | 639 | Specific | 574.99 | 574.99 | 574.99 | 11.21 | 148 | >sp\|P02788\|TRFL_HUMAN Lactotransferrin OS=Homo sapiens OX=9606 GN=LTF PE=1 SV=6 | 1270 | 20201118_12833_1ng_glyco_hcd203040_compound15.10893.10893.4 | controllerType=0 controllerNumber=1 scan=10893 | 24.7468 |
| 07748:4 | 1 | K.FGRN[+2205.793]GSDC[+57.021]PDKFC[+57.021]LFQSETK.N | HexNAc(4)Hex(5)Fuc(2)NeuAc(1) | C[+57]*2, N[+2206] | 1150.4715 | 4 | 4598.8641 | 4598.8577 | 0 | 1.4 | 639 | Specific | 644.2 | 644.2 | 644.2 | 12.21 | 148 | >sp\|P02788\|TRFL_HUMAN Lactotransferrin OS=Homo sapiens OX=9606 GN=LTF PE=1 SV=6 | 1270 | 20201118_12833_1ng_glyco_hcd203040_compound15.10920.10920.4 | controllerType=0 controllerNumber=1 scan=10920 | 24.7922 |
| 08754:3 | 1 | R.N[+2205.793]GSDC[+57.021]PDKFC[+57.021]LFQSETK.N | HexNAc(4)Hex(5)Fuc(2)NeuAc(1) | C[+57]*2, N[+2206] | 1413.5639 | 3 | 4238.6771 | 4238.6667 | 0 | 2.5 | 642 | Specific | 288.75 | 240.86 | 240.86 | 6.34 | 148 | >sp\|P02788\|TRFL_HUMAN Lactotransferrin OS=Homo sapiens OX=9606 GN=LTF PE=1 SV=6 | 1270 | 20201118_12833_1ng_glyco_hcd203040_compound15.11975.11975.3 | controllerType=0 controllerNumber=1 scan=11975 | 26.571 |
| 08705:3 | 1 | R.N[+2205.793]GSDC[+57.021]PDKFC[+57.021]LFQSETK.N | HexNAc(4)Hex(5)Fuc(2)NeuAc(1) | C[+57]*2, N[+2206] | 1413.56 | 3 | 4238.6654 | 4238.6667 | 0 | -0.3 | 642 | Specific | 311.09 | 205.65 | 205.65 | 6 | 148 | >sp\|P02788\|TRFL_HUMAN Lactotransferrin OS=Homo sapiens OX=9606 GN=LTF PE=1 SV=6 | 1270 | 20201118_12833_1ng_glyco_hcd203040_compound15.11924.11924.3 | controllerType=0 controllerNumber=1 scan=11924 | 26.4851 |
| 08840:3 | 1 | R.N[+2059.735]GSDC[+57.021]PDKFC[+57.021]LFQSETK.N | HexNAc(4)Hex(5)Fuc(1)NeuAc(1) | C[+57]*2, N[+2060] | 1364.879 | 3 | 4092.6223 | 4092.6088 | 0 | 3.3 | 642 | Specific | 326.8 | 242.72 | 242.72 | 6.75 | 148 | >sp\|P02788\|TRFL_HUMAN Lactotransferrin OS=Homo sapiens OX=9606 GN=LTF PE=1 SV=6 | 1270 | 20201118_12833_1ng_glyco_hcd203040_compound15.12064.12064.3 | controllerType=0 controllerNumber=1 scan=12064 | 26.7179 |
| 08842:3 | 1 | R.N[+2059.735]GSDC[+57.021]PDKFC[+57.021]LFQSETK.N | HexNAc(4)Hex(5)Fuc(1)NeuAc(1) | C[+57]*2, N[+2060] | 1364.879 | 3 | 4092.6223 | 4092.6088 | 0 | 3.3 | 642 | Specific | 340.59 | 216.62 | 216.62 | 6.66 | 148 | >sp\|P02788\|TRFL_HUMAN Lactotransferrin OS=Homo sapiens OX=9606 GN=LTF PE=1 SV=6 | 1270 | 20201118_12833_1ng_glyco_hcd203040_compound15.12066.12066.3 | controllerType=0 controllerNumber=1 scan=12066 | 26.7221 |
| 08707:3 | 1 | R.N[+2205.793]GSDC[+57.021]PDKFC[+57.021]LFQSETK.N | HexNAc(4)Hex(5)Fuc(2)NeuAc(1) | C[+57]*2, N[+2206] | 1413.56 | 3 | 4238.6654 | 4238.6667 | 0 | -0.3 | 642 | Specific | 403.94 | 308.52 | 308.52 | 6.9 | 148 | >sp\|P02788\|TRFL_HUMAN Lactotransferrin OS=Homo sapiens OX=9606 GN=LTF PE=1 SV=6 | 1270 | 20201118_12833_1ng_glyco_hcd203040_compound15.11926.11926.3 | controllerType=0 controllerNumber=1 scan=11926 | 26.4887 |
| 08756:3 | 1 | R.N[+2205.793]GSDC[+57.021]PDKFC[+57.021]LFQSETK.N | HexNAc(4)Hex(5)Fuc(2)NeuAc(1) | C[+57]*2, N[+2206] | 1413.5639 | 3 | 4238.6771 | 4238.6667 | 0 | 2.5 | 642 | Specific | 429.92 | 295.36 | 295.36 | 7.6 | 148 | >sp\|P02788\|TRFL_HUMAN Lactotransferrin OS=Homo sapiens OX=9606 GN=LTF PE=1 SV=6 | 1270 | 20201118_12833_1ng_glyco_hcd203040_compound15.11977.11977.3 | controllerType=0 controllerNumber=1 scan=11977 | 26.5747 |
| 06998:3 | 4 | R.FSDGLESN[+1864.634]SSTQFEVK.K | HexNAc(2)Hex(9) | N[+1865] | 1213.8232 | 3 | 3639.455 | 3639.4472 | 0 | 2.1 | 219 | Specific | 440.21 | 440.21 | 440.21 | 9.41 | 89 | >sp\|P0C0L4\|CO4A_HUMAN Complement C4-A OS=Homo sapiens OX=9606 GN=C4A PE=1 SV=2 | 1235 | 20201118_12833_1ng_glyco_hcd203040_compound15.10139.10139.3 | controllerType=0 controllerNumber=1 scan=10139 | 23.4649 |
| 06996:3 | 4 | R.FSDGLESN[+1864.634]SSTQFEVK.K | HexNAc(2)Hex(9) | N[+1865] | 1213.8232 | 3 | 3639.455 | 3639.4472 | 0 | 2.1 | 219 | Specific | 541.23 | 541.23 | 541.23 | 9.83 | 89 | >sp\|P0C0L4\|CO4A_HUMAN Complement C4-A OS=Homo sapiens OX=9606 GN=C4A PE=1 SV=2 | 1235 | 20201118_12833_1ng_glyco_hcd203040_compound15.10137.10137.3 | controllerType=0 controllerNumber=1 scan=10137 | 23.4603 |
| 04504:3 | 4 | R.FSDGLESN[+1864.634]SSTQFEVKK.Y | HexNAc(2)Hex(9) | N[+1865] | 1256.5215 | 3 | 3767.5501 | 3767.5421 | 0 | 2.1 | 219 | Specific | 375.27 | 271.46 | 271.46 | 7.28 | 89 | >sp\|P0C0L4\|CO4A_HUMAN Complement C4-A OS=Homo sapiens OX=9606 GN=C4A PE=1 SV=2 | 1235 | 20201118_12833_1ng_glyco_hcd203040_compound15.7557.7557.3 | controllerType=0 controllerNumber=1 scan=7557 | [illegible] |
| 04502:3 | 4 | R.FSDGLESN[+1864.634]SSTQFEVKK.Y | HexNAc(2)Hex(9) | N[+1865] | 1256.5215 | 3 | 3767.5501 | 3767.5421 | 0 | 2.1 | 219 | Specific | 548.68 | 355.37 | 355.37 | 8.34 | 89 | >sp\|P0C0L4\|CO4A_HUMAN Complement C4-A OS=Homo sapiens OX=9606 GN=C4A PE=1 SV=2 | 1235 | 20201118_12833_1ng_glyco_hcd203040_compound15.7555.7555.3 | controllerType=0 controllerNumber=1 scan=7555 | [illegible] |
| 16117:3 | 6 | R.VWIALNSN[+2205.793]LTDNQYTWTDKWR.V | HexNAc(4)Hex(5)Fuc(2)NeuAc(1) | N[+2206] | 1610.7056 | 3 | 4830.1023 | 4830.0821 | 0 | 4.2 | 1153 | Specific | 700.13 | 499.61 | 499.61 | 13.94 | 80 | >sp\|P22897\|MRC1_HUMAN Macrophage mannose receptor 1 OS=Homo sapiens OX=9606 GN=MRC1 PE=1 SV=1 | 1049 | 20201118_12833_1ng_glyco_hcd203040_compound15.19671.19671.3 | controllerType=0 controllerNumber=1 scan=19671 | 39.9328 |
| 16114:3 | 6 | R.VWIALNSN[+2205.793]LTDNQYTWTDKWR.V | HexNAc(4)Hex(5)Fuc(2)NeuAc(1) | N[+2206] | 1610.7056 | 3 | 4830.1023 | 4830.0821 | 0 | 4.2 | 1153 | Specific | 771.57 | 585.55 | 585.55 | 14.94 | 80 | >sp\|P22897\|MRC1_HUMAN Macrophage mannose receptor 1 OS=Homo sapiens OX=9606 GN=MRC1 PE=1 SV=1 | 1049 | 20201118_12833_1ng_glyco_hcd203040_compound15.19668.19668.3 | controllerType=0 controllerNumber=1 scan=19668 | 39.9276 |
| 08118:3 | 6 | K.TAHC[+57.021]N[+2059.735]ESFYFLC[+57.021]K.R | HexNAc(4)Hex(5)Fuc(1)NeuAc(1) | C[+57]*2, N[+2060] | 1246.16 | 3 | 3736.4654 | 3736.4545 | 0 | 2.9 | 1201 | Specific | 485.99 | 396.36 | 396.36 | 9.61 | 80 | >sp\|P22897\|MRC1_HUMAN Macrophage mannose receptor 1 OS=Homo sapiens OX=9606 GN=MRC1 PE=1 SV=1 | 1049 | 20201118_12833_1ng_glyco_hcd203040_compound15.11302.11302.3 | controllerType=0 controllerNumber=1 scan=11302 | 25.423 |
| 08091:3 | 6 | K.TAHC[+57.021]N[+2059.735]ESFYFLC[+57.021]K.R | HexNAc(4)Hex(5)Fuc(1)NeuAc(1) | C[+57]*2, N[+2060] | 1246.159 | 3 | 3736.4624 | 3736.4545 | 0 | 2.1 | 1201 | Specific | 510.93 | 488.89 | 488.89 | 9.63 | 80 | >sp\|P22897\|MRC1_HUMAN Macrophage mannose receptor 1 OS=Homo sapiens OX=9606 GN=MRC1 PE=1 SV=1 | 1049 | 20201118_12833_1ng_glyco_hcd203040_compound15.11274.11274.3 | controllerType=0 controllerNumber=1 scan=11274 | 25.3772 |
| 08116:3 | 6 | K.TAHC[+57.021]N[+2059.735]ESFYFLC[+57.021]K.R | HexNAc(4)Hex(5)Fuc(1)NeuAc(1) | C[+57]*2, N[+2060] | 1246.16 | 3 | 3736.4654 | 3736.4545 | 0 | 2.9 | 1201 | Specific | 540.69 | 515.2 | 515.2 | 9.8 | 80 | >sp\|P22897\|MRC1_HUMAN Macrophage mannose receptor 1 OS=Homo sapiens OX=9606 GN=MRC1 PE=1 SV=1 | 1049 | 20201118_12833_1ng_glyco_hcd203040_compound15.11300.11300.3 | controllerType=0 controllerNumber=1 scan=11300 | 25.4194 |
| 08093:3 | 6 | K.TAHC[+57.021]N[+2059.735]ESFYFLC[+57.021]K.R | HexNAc(4)Hex(5)Fuc(1)NeuAc(1) | C[+57]*2, N[+2060] | 1246.159 | 3 | 3736.4624 | 3736.4545 | 0 | 2.1 | 1201 | Specific | 606.8 | 503.47 | 503.47 | 12.12 | 80 | >sp\|P22897\|MRC1_HUMAN Macrophage mannose receptor 1 OS=Homo sapiens OX=9606 GN=MRC1 PE=1 SV=1 | 1049 | 20201118_12833_1ng_glyco_hcd203040_compound15.11276.11276.3 | controllerType=0 controllerNumber=1 scan=11276 | 25.381 |
| 19160:3 | 7 | R.GLSFDVSLEVSQGPGLLN[+2350.830]DTK.V | HexNAc(4)Hex(5)Fuc(1)NeuAc(2) | N[+2351] | 1509.6566 | 3 | 4526.9551 | 4526.9436 | 0 | 2.6 | 118 | Specific | 927.14 | 815.5 | 815.5 | 18.56 | 62 | >sp\|P01833\|PIGR_HUMAN Polymeric immunoglobulin receptor OS=Homo sapiens OX=9606 GN=PIGR PE=1 SV=4 | 902 | 20201118_12833_1ng_glyco_hcd203040_compound15.22848.22848.3 | controllerType=0 controllerNumber=1 scan=22848 | 46.6375 |
| 19163:3 | 7 | R.GLSFDVSLEVSQGPGLLN[+2350.830]DTK.V | HexNAc(4)Hex(5)Fuc(1)NeuAc(2) | N[+2351] | 1509.6566 | 3 | 4526.9551 | 4526.9436 | 0 | 2.6 | 118 | Specific | 965.65 | 915.51 | 915.51 | 19.56 | 62 | >sp\|P01833\|PIGR_HUMAN Polymeric immunoglobulin receptor OS=Homo sapiens OX=9606 GN=PIGR PE=1 SV=4 | 902 | 20201118_12833_1ng_glyco_hcd203040_compound15.22851.22851.3 | controllerType=0 controllerNumber=1 scan=22851 | 46.6456 |
| 19464:4 | 7 | R.GLSFDVSLEVSQGPGLLN[+2350.830]DTKVYTVDLGR.T | HexNAc(4)Hex(5)Fuc(1)NeuAc(2) | N[+2351] | 1358.3621 | 4 | 5430.4264 | 5430.425 | 0 | 0.3 | 118 | Specific | 565.75 | 466.29 | 466.29 | 10.27 | 62 | >sp\|P01833\|PIGR_HUMAN Polymeric immunoglobulin receptor OS=Homo sapiens OX=9606 GN=PIGR PE=1 SV=4 | 902 | 20201118_12833_1ng_glyco_hcd203040_compound15.23164.23164.4 | controllerType=0 controllerNumber=1 scan=23164 | 47.2716 |
| 19467:4 | 7 | R.GLSFDVSLEVSQGPGLLN[+2350.830]DTKVYTVDLGR.T | HexNAc(4)Hex(5)Fuc(1)NeuAc(2) | N[+2351] | 1358.3621 | 4 | 5430.4264 | 5430.425 | 0 | 0.3 | 118 | Specific | 619.16 | 512.33 | 512.33 | 12.45 | 62 | >sp\|P01833\|PIGR_HUMAN Polymeric immunoglobulin receptor OS=Homo sapiens OX=9606 GN=PIGR PE=1 SV=4 | 902 | 20201118_12833_1ng_glyco_hcd203040_compound15.23167.23167.4 | controllerType=0 controllerNumber=1 scan=23167 | 47.2771 |
| 20146:3 | 7 | R.LSLLEEPGN[+2205.793]GTFTVILNQLTSR.D | HexNAc(4)Hex(5)Fuc(2)NeuAc(1) | N[+2206] | 1536.7068 | 3 | 4608.106 | 4608.0854 | 0 | 4.5 | 413 | Specific | 259.85 | 259.85 | 259.85 | 6.28 | 62 | >sp\|P01833\|PIGR_HUMAN Polymeric immunoglobulin receptor OS=Homo sapiens OX=9606 GN=PIGR PE=1 SV=4 | 902 | 20201118_12833_1ng_glyco_hcd203040_compound15.23876.23876.3 | controllerType=0 controllerNumber=1 scan=23876 | 48.8468 |
| 20176:3 | 7 | R.LSLLEEPGN[+2076.750]GTFTVILNQLTSR.D | HexNAc(4)Hex(6)Fuc(2) | N[+2077] | 1493.6951 | 3 | 4479.0708 | 4479.0428 | 0 | 6.3 | 413 | Specific | 314.61 | 232.23 | 232.23 | 4.83 | 62 | >sp\|P01833\|PIGR_HUMAN Polymeric immunoglobulin receptor OS=Homo sapiens OX=9606 GN=PIGR PE=1 SV=4 | 902 | 20201118_12833_1ng_glyco_hcd203040_compound15.23907.23907.3 | controllerType=0 controllerNumber=1 scan=23907 | 48.9078 |
| 20183:4 | 7 | R.LSLLEEPGN[+2076.750]GTFTVILNQLTSR.D | HexNAc(4)Hex(6)Fuc(2) | N[+2077] | 1120.5213 | 4 | 4479.0633 | 4479.0428 | 0 | 4.6 | 413 | Specific | 335.85 | 330.63 | 330.63 | 6.52 | 62 | >sp\|P01833\|PIGR_HUMAN Polymeric immunoglobulin receptor OS=Homo sapiens OX=9606 GN=PIGR PE=1 SV=4 | 902 | 20201118_12833_1ng_glyco_hcd203040_compound15.23914.23914.4 | controllerType=0 controllerNumber=1 scan=23914 | 48.9284 |
| 22624:3 | 7 | R.LSLLEEPGN[+2350.830]GTFTVILNQLTSR.D | HexNAc(4)Hex(5)Fuc(1)NeuAc(2) | N[+2351] | 1585.049 | 3 | 4753.1325 | 4753.1229 | 0 | 2 | 413 | Specific | 341.61 | 313.09 | 313.09 | 7.58 | 62 | >sp\|P01833\|PIGR_HUMAN Polymeric immunoglobulin receptor OS=Homo sapiens OX=9606 GN=PIGR PE=1 SV=4 | 902 | 20201118_12833_1ng_glyco_hcd203040_compound15.26459.26459.3 | controllerType=0 controllerNumber=1 scan=26459 | 53.7124 |
| 20180:4 | 7 | R.LSLLEEPGN[+2076.750]GTFTVILNQLTSR.D | HexNAc(4)Hex(6)Fuc(2) | N[+2077] | 1120.5213 | 4 | 4479.0633 | 4479.0428 | 0 | 4.6 | 413 | Specific | 349.65 | 321.24 | 321.24 | 6.61 | 62 | >sp\|P01833\|PIGR_HUMAN Polymeric immunoglobulin receptor OS=Homo sapiens OX=9606 GN=PIGR PE=1 SV=4 | 902 | 20201118_12833_1ng_glyco_hcd203040_compound15.23911.23911.4 | controllerType=0 controllerNumber=1 scan=23911 | 48.9178 |
| 20178:3 | 7 | R.LSLLEEPGN[+2076.750]GTFTVILNQLTSR.D | HexNAc(4)Hex(6)Fuc(2) | N[+2077] | 1493.6951 | 3 | 4479.0708 | 4479.0428 | 0 | 6.3 | 413 | Specific | 378.93 | 297.34 | 297.34 | 5.64 | 62 | >sp\|P01833\|PIGR_HUMAN Polymeric immunoglobulin receptor OS=Homo sapiens OX=9606 GN=PIGR PE=1 SV=4 | 902 | 20201118_12833_1ng_glyco_hcd203040_compound15.23909.23909.3 | controllerType=0 controllerNumber=1 scan=23909 | 48.9145 |
| 22626:3 | 7 | R.LSLLEEPGN[+2350.830]GTFTVILNQLTSR.D | HexNAc(4)Hex(5)Fuc(1)NeuAc(2) | N[+2351] | 1585.049 | 3 | 4753.1325 | 4753.1229 | 0 | 2 | 413 | Specific | 535.19 | 535.19 | 535.19 | 10.18 | 62 | >sp\|P01833\|PIGR_HUMAN Polymeric immunoglobulin receptor OS=Homo sapiens OX=9606 GN=PIGR PE=1 SV=4 | 902 | 20201118_12833_1ng_glyco_hcd203040_compound15.26461.26461.3 | controllerType=0 controllerNumber=1 scan=26461 | 53.7165 |
| 20198:4 | 7 | R.LSLLEEPGN[+2059.735]GTFTVILNQLTSR.D | HexNAc(4)Hex(5)Fuc(1)NeuAc(1) | N[+2060] | 1116.2681 | 4 | 4462.0506 | 4462.0275 | 0 | 5.2 | 413 | Specific | 766.9 | 592.11 | 592.11 | 15.65 | 62 | >sp\|P01833\|PIGR_HUMAN Polymeric immunoglobulin receptor OS=Homo sapiens OX=9606 GN=PIGR PE=1 SV=4 | 902 | 20201118_12833_1ng_glyco_hcd203040_compound15.23930.23930.4 | controllerType=0 controllerNumber=1 scan=23930 | 48.9612 |
| 20194:3 | 7 | R.LSLLEEPGN[+2059.735]GTFTVILNQLTSR.D | HexNAc(4)Hex(5)Fuc(1)NeuAc(1) | N[+2060] | 1488.0206 | 3 | 4462.0471 | 4462.0275 | 0 | 4.4 | 413 | Specific | 777.52 | 570.9 | 570.9 | 15.02 | 62 | >sp\|P01833\|PIGR_HUMAN Polymeric immunoglobulin receptor OS=Homo sapiens OX=9606 GN=PIGR PE=1 SV=4 | 902 | 20201118_12833_1ng_glyco_hcd203040_compound15.23926.23926.3 | controllerType=0 controllerNumber=1 scan=23926 | 48.9543 |
| 20201:4 | 7 | R.LSLLEEPGN[+2059.735]GTFTVILNQLTSR.D | HexNAc(4)Hex(5)Fuc(1)NeuAc(1) | N[+2060] | 1116.2681 | 4 | 4462.0506 | 4462.0275 | 0 | 5.2 | 413 | Specific | 914.36 | 691.42 | 691.42 | 18.65 | 62 | >sp\|P01833\|PIGR_HUMAN Polymeric immunoglobulin receptor OS=Homo sapiens OX=9606 GN=PIGR PE=1 SV=4 | 902 | 20201118_12833_1ng_glyco_hcd203040_compound15.23933.23933.4 | controllerType=0 controllerNumber=1 scan=23933 | 48.9665 |
| 22415:3 | 7 | R.LSLLEEPGN[+2350.830]GTFTVILNQLTSR.D | HexNAc(4)Hex(5)Fuc(1)NeuAc(2) | N[+2351] | 1585.0521 | 3 | 4753.1416 | 4753.1229 | 0 | 3.9 | 413 | Specific | 943.39 | 657.89 | 657.89 | 18.02 | 62 | >sp\|P01833\|PIGR_HUMAN Polymeric immunoglobulin receptor OS=Homo sapiens OX=9606 GN=PIGR PE=1 SV=4 | 902 | 20201118_12833_1ng_glyco_hcd203040_compound15.26242.26242.3 | controllerType=0 controllerNumber=1 scan=26242 | 53.2799 |
| 22413:3 | 7 | R.LSLLEEPGN[+2350.830]GTFTVILNQLTSR.D | HexNAc(4)Hex(5)Fuc(1)NeuAc(2) | N[+2351] | 1585.0521 | 3 | 4753.1416 | 4753.1229 | 0 | 3.9 | 413 | Specific | 961.07 | 724.55 | 724.55 | 19.02 | 62 | >sp\|P01833\|PIGR_HUMAN Polymeric immunoglobulin receptor OS=Homo sapiens OX=9606 GN=PIGR PE=1 SV=4 | 902 | 20201118_12833_1ng_glyco_hcd203040_compound15.26240.26240.3 | controllerType=0 controllerNumber=1 scan=26240 | 53.2754 |
| 22421:4 | 7 | R.LSLLEEPGN[+2350.830]GTFTVILNQLTSR.D | HexNAc(4)Hex(5)Fuc(1)NeuAc(2) | N[+2351] | 1189.0419 | 4 | 4753.1458 | 4753.1229 | 0 | 4.8 | 413 | Specific | 971.43 | 760.22 | 760.22 | 19.65 | 62 | >sp\|P01833\|PIGR_HUMAN Polymeric immunoglobulin receptor OS=Homo sapiens OX=9606 GN=PIGR PE=1 SV=4 | 902 | 20201118_12833_1ng_glyco_hcd203040_compound15.26248.26248.4 | controllerType=0 controllerNumber=1 scan=26248 | 53.292 |
| 20196:3 | 7 | R.LSLLEEPGN[+2059.735]GTFTVILNQLTSR.D | HexNAc(4)Hex(5)Fuc(1)NeuAc(1) | N[+2060] | 1488.0206 | 3 | 4462.0471 | 4462.0275 | 0 | 4.4 | 413 | Specific | 1006.71 | 726.75 | 726.75 | 20.02 | 62 | >sp\|P01833\|PIGR_HUMAN Polymeric immunoglobulin receptor OS=Homo sapiens OX=9606 GN=PIGR PE=1 SV=4 | 902 | 20201118_12833_1ng_glyco_hcd203040_compound15.23928.23928.3 | controllerType=0 controllerNumber=1 scan=23928 | 48.9585 |
| 22418:4 | 7 | R.LSLLEEPGN[+2350.830]GTFTVILNQLTSR.D | HexNAc(4)Hex(5)Fuc(1)NeuAc(2) | N[+2351] | 1189.0419 | 4 | 4753.1458 | 4753.1229 | 0 | 4.8 | 413 | Specific | 1077.67 | 865.72 | 865.72 | 21.65 | 62 | >sp\|P01833\|PIGR_HUMAN Polymeric immunoglobulin receptor OS=Homo sapiens OX=9606 GN=PIGR PE=1 SV=4 | 902 | 20201118_12833_1ng_glyco_hcd203040_compound15.26245.26245.4 | controllerType=0 controllerNumber=1 scan=26245 | 53.2833 |
| 15722:3 | 7 | K.IIEGEPNLKVPGN[+2205.793]VTAVLGETLK.V | HexNAc(4)Hex(5)Fuc(2)NeuAc(1) | N[+2206] | 1533.0541 | 3 | 4597.1478 | 4597.1422 | 0 | 1.2 | 457 | Specific | 788.29 | 737.57 | 737.57 | 15.45 | 62 | >sp\|P01833\|PIGR_HUMAN Polymeric immunoglobulin receptor OS=Homo sapiens OX=9606 GN=PIGR PE=1 SV=4 | 902 | 20201118_12833_1ng_glyco_hcd203040_compound15.19256.19256.3 | controllerType=0 controllerNumber=1 scan=19256 | 39.1916 |
| 15720:3 | 7 | K.IIEGEPNLKVPGN[+2205.793]VTAVLGETLK.V | HexNAc(4)Hex(5)Fuc(2)NeuAc(1) | N[+2206] | 1533.0541 | 3 | 4597.1478 | 4597.1422 | 0 | 1.2 | 457 | Specific | 898.26 | 802.42 | 802.42 | 17.45 | 62 | >sp\|P01833\|PIGR_HUMAN Polymeric immunoglobulin receptor OS=Homo sapiens OX=9606 GN=PIGR PE=1 SV=4 | 902 | 20201118_12833_1ng_glyco_hcd203040_compound15.19254.19254.3 | controllerType=0 controllerNumber=1 scan=19254 | 39.187 |
| 14095:4 | 7 | K.IIEGEPNLKVPGN[+1913.677]VTAVLGETLKVPC[+57.021]HFPC[+57.021]K.F | HexNAc(4)Hex(5)NeuAc(1) | C[+57]*2, N[+1914] | 1333.3829 | 4 | 5330.5099 | 5330.4839 | 0 | 4.9 | 457 | Specific | 320.22 | 292.17 | 292.17 | 7.19 | 62 | >sp\|P01833\|PIGR_HUMAN Polymeric immunoglobulin receptor OS=Homo sapiens OX=9606 GN=PIGR PE=1 SV=4 | 902 | 20201118_12833_1ng_glyco_hcd203040_compound15.17531.17531.4 | controllerType=0 controllerNumber=1 scan=17531 | 36.1389 |
| 14093:4 | 7 | K.IIEGEPNLKVPGN[+1913.677]VTAVLGETLKVPC[+57.021]HFPC[+57.021]K.F | HexNAc(4)Hex(5)NeuAc(1) | C[+57]*2, N[+1914] | 1333.3829 | 4 | 5330.5099 | 5330.4839 | 0 | 4.9 | 457 | Specific | 453.4 | 444.1 | 444.1 | 9.47 | 62 | >sp\|P01833\|PIGR_HUMAN Polymeric immunoglobulin receptor OS=Homo sapiens OX=9606 GN=PIGR PE=1 SV=4 | 902 | 20201118_12833_1ng_glyco_hcd203040_compound15.17529.17529.4 | controllerType=0 controllerNumber=1 scan=17529 | 36.1353 |
| 15437:4 | 7 | K.IIEGEPNLKVPGN[+2059.735]VTAVLGETLKVPC[+57.021]HFPC[+57.021]K.F | HexNAc(4)Hex(5)Fuc(1)NeuAc(1) | C[+57]*2, N[+2060] | 1369.8918 | 4 | 5476.5456 | 5476.5418 | 0 | 0.7 | 457 | Specific | 497.35 | 497.35 | 497.35 | 9.79 | 62 | >sp\|P01833\|PIGR_HUMAN Polymeric immunoglobulin receptor OS=Homo sapiens OX=9606 GN=PIGR PE=1 SV=4 | 902 | 20201118_12833_1ng_glyco_hcd203040_compound15.18931.18931.2 | controllerType=0 controllerNumber=1 scan=18931 | 38.6025 |
| 15218:5 | 7 | K.IIEGEPNLKVPGN[+2059.735]VTAVLGETLKVPC[+57.021]HFPC[+57.021]K.F | HexNAc(4)Hex(5)Fuc(1)NeuAc(1) | C[+57]*2, N[+2060] | 1096.1157 | 5 | 5476.5494 | 5476.5418 | 0 | 1.4 | 457 | Specific | 533.25 | 413.62 | 413.62 | 10.86 | 62 | >sp\|P01833\|PIGR_HUMAN Polymeric immunoglobulin receptor OS=Homo sapiens OX=9606 GN=PIGR PE=1 SV=4 | 902 | 20201118_12833_1ng_glyco_hcd203040_compound15.18697.18697.5 | controllerType=0 controllerNumber=1 scan=18697 | 38.1893 |
| 15440:4 | 7 | K.IIEGEPNLKVPGN[+2059.735]VTAVLGETLKVPC[+57.021]HFPC[+57.021]K.F | HexNAc(4)Hex(5)Fuc(1)NeuAc(1) | C[+57]*2, N[+2060] | 1369.8918 | 4 | 5476.5456 | 5476.5418 | 0 | 0.7 | 457 | Specific | 566.41 | 566.41 | 566.41 | 10.1 | 62 | >sp\|P01833\|PIGR_HUMAN Polymeric immunoglobulin receptor OS=Homo sapiens OX=9606 GN=PIGR PE=1 SV=4 | 902 | 20201118_12833_1ng_glyco_hcd203040_compound15.18934.18934.2 | controllerType=0 controllerNumber=1 scan=18934 | 38.6069 |
| 15215:5 | 7 | K.IIEGEPNLKVPGN[+2059.735]VTAVLGETLKVPC[+57.021]HFPC[+57.021]K.F | HexNAc(4)Hex(5)Fuc(1)NeuAc(1) | C[+57]*2, N[+2060] | 1096.1157 | 5 | 5476.5494 | 5476.5418 | 0 | 1.4 | 457 | Specific | 663.43 | 571.3 | 571.3 | 14.21 | 62 | >sp\|P01833\|PIGR_HUMAN Polymeric immunoglobulin receptor OS=Homo sapiens OX=9606 GN=PIGR PE=1 SV=4 | 902 | 20201118_12833_1ng_glyco_hcd203040_compound15.18694.18694.5 | controllerType=0 controllerNumber=1 scan=18694 | 38.1839 |
| 15416:5 | 7 | K.IIEGEPNLKVPGN[+2205.793]VTAVLGETLKVPC[+57.021]HFPC[+57.021]K.F | HexNAc(4)Hex(5)Fuc(2)NeuAc(1) | C[+57]*2, N[+2206] | 1125.3293 | 5 | 5622.6176 | 5622.5997 | 0 | 3.2 | 457 | Specific | 751.87 | 751.87 | 751.87 | 16.24 | 62 | >sp\|P01833\|PIGR_HUMAN Polymeric immunoglobulin receptor OS=Homo sapiens OX=9606 GN=PIGR PE=1 SV=4 | 902 | 20201118_12833_1ng_glyco_hcd203040_compound15.18909.18909.5 | controllerType=0 controllerNumber=1 scan=18909 | 38.5624 |
| 15413:5 | 7 | K.IIEGEPNLKVPGN[+2205.793]VTAVLGETLKVPC[+57.021]HFPC[+57.021]K.F | HexNAc(4)Hex(5)Fuc(2)NeuAc(1) | C[+57]*2, N[+2206] | 1125.3293 | 5 | 5622.6176 | 5622.5997 | 0 | 3.2 | 457 | Specific | 760.15 | 760.15 | 760.15 | 16.24 | 62 | >sp\|P01833\|PIGR_HUMAN Polymeric immunoglobulin receptor OS=Homo sapiens OX=9606 GN=PIGR PE=1 SV=4 | 902 | 20201118_12833_1ng_glyco_hcd203040_compound15.18906.18906.5 | controllerType=0 controllerNumber=1 scan=18906 | 38.5569 |
| 15184:5 | 7 | K.IIEGEPNLKVPGN[+2205.793]VTAVLGETLKVPC[+57.021]HFPC[+57.021]K.F | HexNAc(4)Hex(5)Fuc(2)NeuAc(1) | C[+57]*2, N[+2206] | 1125.3304 | 5 | 5622.6231 | 5622.5997 | 0 | 4.2 | 457 | Specific | 809.12 | 787.99 | 787.99 | 16.7 | 62 | >sp\|P01833\|PIGR_HUMAN Polymeric immunoglobulin receptor OS=Homo sapiens OX=9606 GN=PIGR PE=1 SV=4 | 902 | 20201118_12833_1ng_glyco_hcd203040_compound15.18661.18661.5 | controllerType=0 controllerNumber=1 scan=18661 | 38.0902 |
| 15202:5 | 7 | K.IIEGEPNLKVPGN[+2205.793]VTAVLGETLKVPC[+57.021]HFPC[+57.021]K.F | HexNAc(4)Hex(5)Fuc(2)NeuAc(1) | C[+57]*2, N[+2206] | 1125.3281 | 5 | 5622.6115 | 5622.5997 | 0 | 2.1 | 457 | Specific | 814.63 | 814.63 | 814.63 | 17.21 | 62 | >sp\|P01833\|PIGR_HUMAN Polymeric immunoglobulin receptor OS=Homo sapiens OX=9606 GN=PIGR PE=1 SV=4 | 902 | 20201118_12833_1ng_glyco_hcd203040_compound15.18680.18680.5 | controllerType=0 controllerNumber=1 scan=18680 | 38.148 |
| 15203:4 | 7 | K.IIEGEPNLKVPGN[+2205.793]VTAVLGETLKVPC[+57.021]HFPC[+57.021]K.F | HexNAc(4)Hex(5)Fuc(2)NeuAc(1) | C[+57]*2, N[+2206] | 1406.4093 | 4 | 5622.6154 | 5622.5997 | 0 | 2.8 | 457 | Specific | 848.77 | 848.77 | 848.77 | 16.79 | 62 | >sp\|P01833\|PIGR_HUMAN Polymeric immunoglobulin receptor OS=Homo sapiens OX=9606 GN=PIGR PE=1 SV=4 | 902 | 20201118_12833_1ng_glyco_hcd203040_compound15.18681.18681.4 | controllerType=0 controllerNumber=1 scan=18681 | 38.1523 |
| 15182:5 | 7 | K.IIEGEPNLKVPGN[+2205.793]VTAVLGETLKVPC[+57.021]HFPC[+57.021]K.F | HexNAc(4)Hex(5)Fuc(2)NeuAc(1) | C[+57]*2, N[+2206] | 1125.3304 | 5 | 5622.6231 | 5622.5997 | 0 | 4.2 | 457 | Specific | 1045.65 | 945.83 | 945.83 | 20.7 | 62 | >sp\|P01833\|PIGR_HUMAN Polymeric immunoglobulin receptor OS=Homo sapiens OX=9606 GN=PIGR PE=1 SV=4 | 902 | 20201118_12833_1ng_glyco_hcd203040_compound15.18659.18659.5 | controllerType=0 controllerNumber=1 scan=18659 | 38.0831 |
| 15201:4 | 7 | K.IIEGEPNLKVPGN[+2205.793]VTAVLGETLKVPC[+57.021]HFPC[+57.021]K.F | HexNAc(4)Hex(5)Fuc(2)NeuAc(1) | C[+57]*2, N[+2206] | 1406.4093 | 4 | 5622.6154 | 5622.5997 | 0 | 2.8 | 457 | Specific | 1065.77 | 985.5 | 985.5 | 21.79 | 62 | >sp\|P01833\|PIGR_HUMAN Polymeric immunoglobulin receptor OS=Homo sapiens OX=9606 GN=PIGR PE=1 SV=4 | 902 | 20201118_12833_1ng_glyco_hcd203040_compound15.18679.18679.4 | controllerType=0 controllerNumber=1 scan=18679 | 38.1348 |
| 15200:5 | 7 | K.IIEGEPNLKVPGN[+2205.793]VTAVLGETLKVPC[+57.021]HFPC[+57.021]K.F | HexNAc(4)Hex(5)Fuc(2)NeuAc(1) | C[+57]*2, N[+2206] | 1125.3281 | 5 | 5622.6115 | 5622.5997 | 0 | 2.1 | 457 | Specific | 1155.94 | 1094.53 | 1094.53 | 24.21 | 62 | >sp\|P01833\|PIGR_HUMAN Polymeric immunoglobulin receptor OS=Homo sapiens OX=9606 GN=PIGR PE=1 SV=4 | 902 | 20201118_12833_1ng_glyco_hcd203040_compound15.18678.18678.5 | controllerType=0 controllerNumber=1 scan=18678 | 38.1333 |
| 13320:4 | 7 | K.VPGN[+2205.793]VTAVLGETLKVPC[+57.021]HFPC[+57.021]K.F | HexNAc(4)Hex(5)Fuc(2)NeuAc(1) | C[+57]*2, N[+2206] | 1158.0232 | 4 | 4629.0711 | 4629.0502 | 0 | 4.5 | 466 | Specific | 214.68 | 214.68 | 214.68 | 3.62 | 62 | >sp\|P01833\|PIGR_HUMAN Polymeric immunoglobulin receptor OS=Homo sapiens OX=9606 GN=PIGR PE=1 SV=4 | 902 | 20201118_12833_1ng_glyco_hcd203040_compound15.16727.16727.4 | controllerType=0 controllerNumber=1 scan=16727 | 34.7244 |
| 13318:4 | 7 | K.VPGN[+2205.793]VTAVLGETLKVPC[+57.021]HFPC[+57.021]K.F | HexNAc(4)Hex(5)Fuc(2)NeuAc(1) | C[+57]*2, N[+2206] | 1158.0232 | 4 | 4629.0711 | 4629.0502 | 0 | 4.5 | 466 | Specific | 322.65 | 322.65 | 322.65 | 7.19 | 62 | >sp\|P01833\|PIGR_HUMAN Polymeric immunoglobulin receptor OS=Homo sapiens OX=9606 GN=PIGR PE=1 SV=4 | 902 | 20201118_12833_1ng_glyco_hcd203040_compound15.16725.16725.4 | controllerType=0 controllerNumber=1 scan=16725 | 34.7206 |
| 07373:3 | 7 | K.YWC[+57.021]KWN[+1913.677]NTGC[+57.021]QALPSQDEGPSK.A | HexNAc(4)Hex(5)NeuAc(1) | C[+57]*2, N[+1914] | 1513.9482 | 3 | 4539.83 | 4539.822 | 0 | 1.8 | 494 | Specific | 441.25 | 408.21 | 408.21 | 9.69 | 62 | >sp\|P01833\|PIGR_HUMAN Polymeric immunoglobulin receptor OS=Homo sapiens OX=9606 GN=PIGR PE=1 SV=4 | 902 | 20201118_12833_1ng_glyco_hcd203040_compound15.10527.10527.3 | controllerType=0 controllerNumber=1 scan=10527 | 24.1138 |
| 07371:3 | 7 | K.YWC[+57.021]KWN[+1913.677]NTGC[+57.021]QALPSQDEGPSK.A | HexNAc(4)Hex(5)NeuAc(1) | C[+57]*2, N[+1914] | 1513.9482 | 3 | 4539.83 | 4539.822 | 0 | 1.8 | 494 | Specific | 465.07 | 455.32 | 455.32 | 9.69 | 62 | >sp\|P01833\|PIGR_HUMAN Polymeric immunoglobulin receptor OS=Homo sapiens OX=9606 GN=PIGR PE=1 SV=4 | 902 | 20201118_12833_1ng_glyco_hcd203040_compound15.10525.10525.3 | controllerType=0 controllerNumber=1 scan=10525 | 24.1102 |
| 05517:3 | 7 | K.WN[+1710.598]NTGC[+57.021]QALPSQDEGPSK.A | HexNAc(3)Hex(5)NeuAc(1) | C[+57], N[+1711] | 1233.8307 | 3 | 3699.4777 | 3699.4743 | 0 | 0.9 | 498 | Specific | 358.01 | 307.73 | 307.73 | 7.23 | 62 | >sp\|P01833\|PIGR_HUMAN Polymeric immunoglobulin receptor OS=Homo sapiens OX=9606 GN=PIGR PE=1 SV=4 | 902 | 20201118_12833_1ng_glyco_hcd203040_compound15.8604.8604.3 | controllerType=0 controllerNumber=1 scan=8604 | 20.9223 |
| 05515:3 | 7 | K.WN[+1710.598]NTGC[+57.021]QALPSQDEGPSK.A | HexNAc(3)Hex(5)NeuAc(1) | C[+57], N[+1711] | 1233.8307 | 3 | 3699.4777 | 3699.4743 | 0 | 0.9 | 498 | Specific | 387.61 | 325.71 | 325.71 | 7.63 | 62 | >sp\|P01833\|PIGR_HUMAN Polymeric immunoglobulin receptor OS=Homo sapiens OX=9606 GN=PIGR PE=1 SV=4 | 902 | 20201118_12833_1ng_glyco_hcd203040_compound15.8602.8602.3 | controllerType=0 controllerNumber=1 scan=8602 | 20.9183 |
| 05328:3 | 7 | K.WN[+2059.735]NTGC[+57.021]QALPSQDEGPSK.A | HexNAc(4)Hex(5)Fuc(1)NeuAc(1) | C[+57], N[+2060] | 1350.2109 | 3 | 4048.6181 | 4048.6116 | 0 | 1.6 | 498 | Specific | 537.94 | 464.45 | 464.45 | 9.83 | 62 | >sp\|P01833\|PIGR_HUMAN Polymeric immunoglobulin receptor OS=Homo sapiens OX=9606 GN=PIGR PE=1 SV=4 | 902 | 20201118_12833_1ng_glyco_hcd203040_compound15.8409.8409.3 | controllerType=0 controllerNumber=1 scan=8409 | 20.5955 |
| 05326:3 | 7 | K.WN[+2059.735]NTGC[+57.021]QALPSQDEGPSK.A | HexNAc(4)Hex(5)Fuc(1)NeuAc(1) | C[+57], N[+2060] | 1350.2109 | 3 | 4048.6181 | 4048.6116 | 0 | 1.6 | 498 | Specific | 568.9 | 491.4 | 491.4 | 10 | 62 | >sp\|P01833\|PIGR_HUMAN Polymeric immunoglobulin receptor OS=Homo sapiens OX=9606 GN=PIGR PE=1 SV=4 | 902 | 20201118_12833_1ng_glyco_hcd203040_compound15.8407.8407.3 | controllerType=0 controllerNumber=1 scan=8407 | 20.5919 |
| 05410:3 | 7 | K.WN[+1913.677]NTGC[+57.021]QALPSQDEGPSK.A | HexNAc(4)Hex(5)NeuAc(1) | C[+57], N[+1914] | 1301.5261 | 3 | 3902.5636 | 3902.5537 | 0 | 2.5 | 498 | Specific | 640.66 | 549.45 | 549.45 | 12.21 | 62 | >sp\|P01833\|PIGR_HUMAN Polymeric immunoglobulin receptor OS=Homo sapiens OX=9606 GN=PIGR PE=1 SV=4 | 902 | 20201118_12833_1ng_glyco_hcd203040_compound15.8494.8494.3 | controllerType=0 controllerNumber=1 scan=8494 | 20.7384 |
| 05412:3 | 7 | K.WN[+1913.677]NTGC[+57.021]QALPSQDEGPSK.A | HexNAc(4)Hex(5)NeuAc(1) | C[+57], N[+1914] | 1301.5261 | 3 | 3902.5636 | 3902.5537 | 0 | 2.5 | 498 | Specific | 695.81 | 621.11 | 621.11 | 13.21 | 62 | >sp\|P01833\|PIGR_HUMAN Polymeric immunoglobulin receptor OS=Homo sapiens OX=9606 GN=PIGR PE=1 SV=4 | 902 | 20201118_12833_1ng_glyco_hcd203040_compound15.8496.8496.3 | controllerType=0 controllerNumber=1 scan=8496 | 20.7427 |
| 09761:3 | 8 | K.TVLTPATNHMGN[+1378.476]VTFTIPANR.E | HexNAc(2)Hex(6) | N[+1378] | 1211.8858 | 3 | 3633.6428 | 3633.6358 | 0 | 1.9 | 74 | Specific | 328.22 | 328.22 | 176.05 | 7.97 | 74 | >sp\|P01024\|CO3_HUMAN Complement C3 OS=Homo sapiens OX=9606 GN=C3 PE=1 SV=2 | 492 | 20201118_12833_1ng_glyco_hcd203040_compound15.13019.13019.3 | controllerType=0 controllerNumber=1 scan=13019 | 28.3295 |
| 09763:3 | 8 | K.TVLTPATNHMGN[+1378.476]VTFTIPANR.E | HexNAc(2)Hex(6) | N[+1378] | 1211.8858 | 3 | 3633.6428 | 3633.6358 | 0 | 1.9 | 74 | Specific | 366.78 | 366.78 | 194.72 | 8.2 | 74 | >sp\|P01024\|CO3_HUMAN Complement C3 OS=Homo sapiens OX=9606 GN=C3 PE=1 SV=2 | 492 | 20201118_12833_1ng_glyco_hcd203040_compound15.13021.13021.3 | controllerType=0 controllerNumber=1 scan=13021 | 28.3329 |
| 18389:3 | 9 | R.LSLHRPALEDLLLGSEAN[+1913.677]LTC[+57.021]TLTGLR.D | HexNAc(4)Hex(5)NeuAc(1) | C[+57], N[+1914] | 1626.436 | 3 | 4877.2934 | 4877.2753 | 0 | 3.7 | 127 | Specific | 272.98 | 272.98 | 272.98 | 6.47 | 60 | >sp\|P01876\|IGHA1_HUMAN Immunoglobulin heavy constant alpha 1 OS=Homo sapiens OX=9606 GN=IGHA1 PE=1 SV=2 | 255 | 20201118_12833_1ng_glyco_hcd203040_compound15.22037.22037.3 | controllerType=0 controllerNumber=1 scan=22037 | 44.8428 |
| 17422:4 | 9 | R.LSLHRPALEDLLLGSEAN[+1825.661]LTC[+57.021]TLTGLR.D | HexNAc(5)Hex(5) | C[+57], N[+1826] | 1198.0728 | 4 | 4789.2694 | 4789.2592 | 0 | 2.1 | 127 | Specific | 304.77 | 304.77 | 304.77 | 6.98 | 60 | >sp\|P01876\|IGHA1_HUMAN Immunoglobulin heavy constant alpha 1 OS=Homo sapiens OX=9606 GN=IGHA1 PE=1 SV=2 | 255 | 20201118_12833_1ng_glyco_hcd203040_compound15.21028.21028.4 | controllerType=0 controllerNumber=1 scan=21028 | 42.6208 |
| 18378:4 | 9 | R.LSLHRPALEDLLLGSEAN[+1913.677]LTC[+57.021]TLTGLR.D | HexNAc(4)Hex(5)NeuAc(1) | C[+57], N[+1914] | 1220.0773 | 4 | 4877.2874 | 4877.2753 | 0 | 2.5 | 127 | Specific | 306.18 | 298.55 | 298.55 | 7.01 | 60 | >sp\|P01876\|IGHA1_HUMAN Immunoglobulin heavy constant alpha 1 OS=Homo sapiens OX=9606 GN=IGHA1 PE=1 SV=2 | 255 | 20201118_12833_1ng_glyco_hcd203040_compound15.22026.22026.4 | controllerType=0 controllerNumber=1 scan=22026 | 44.8222 |
| 17521:5 | 9 | R.LSLHRPALEDLLLGSEAN[+1524.534]LTC[+57.021]TLTGLR.D | HexNAc(2)Hex(6)Fuc(1) | C[+57], N[+1525] | 898.4291 | 5 | 4488.1164 | 4488.1319 | 0 | -3.4 | 127 | Specific | 314.19 | 223.85 | 223.85 | 3.65 | 60 | >sp\|P01876\|IGHA1_HUMAN Immunoglobulin heavy constant alpha 1 OS=Homo sapiens OX=9606 GN=IGHA1 PE=1 SV=2 | 255 | 20201118_12833_1ng_glyco_hcd203040_compound15.21132.21132.2 | controllerType=0 controllerNumber=1 scan=21132 | 42.9004 |
| 17734:4 | 9 | R.LSLHRPALEDLLLGSEAN[+1501.555]LTC[+57.021]TLTGLR.D | HexNAc(5)Hex(3) | C[+57], N[+1502] | 1117.0335 | 4 | 4465.1121 | 4465.1536 | 0 | -9.3 | 127 | Specific | 367.25 | 363.1 | 363.1 | 4.63 | 60 | >sp\|P01876\|IGHA1_HUMAN Immunoglobulin heavy constant alpha 1 OS=Homo sapiens OX=9606 GN=IGHA1 PE=1 SV=2 | 255 | 20201118_12833_1ng_glyco_hcd203040_compound15.21354.21354.4 | controllerType=0 controllerNumber=1 scan=21354 | 43.3698 |
| 17424:4 | 9 | R.LSLHRPALEDLLLGSEAN[+1825.661]LTC[+57.021]TLTGLR.D | HexNAc(5)Hex(5) | C[+57], N[+1826] | 1198.0728 | 4 | 4789.2694 | 4789.2592 | 0 | 2.1 | 127 | Specific | 397.7 | 355.21 | 355.21 | 7.76 | 60 | >sp\|P01876\|IGHA1_HUMAN Immunoglobulin heavy constant alpha 1 OS=Homo sapiens OX=9606 GN=IGHA1 PE=1 SV=2 | 255 | 20201118_12833_1ng_glyco_hcd203040_compound15.21030.21030.4 | controllerType=0 controllerNumber=1 scan=21030 | 42.6254 |
| 17389:4 | 9 | R.LSLHRPALEDLLLGSEAN[+1540.529]LTC[+57.021]TLTGLR.D | HexNAc(2)Hex(7) | C[+57], N[+1541] | 1126.7897 | 4 | 4504.137 | 4504.1268 | 0 | 2.3 | 127 | Specific | 429.35 | 429.35 | 429.35 | 9.53 | 60 | >sp\|P01876\|IGHA1_HUMAN Immunoglobulin heavy constant alpha 1 OS=Homo sapiens OX=9606 GN=IGHA1 PE=1 SV=2 | 255 | 20201118_12833_1ng_glyco_hcd203040_compound15.20994.20994.4 | controllerType=0 controllerNumber=1 scan=20994 | 42.5416 |
| 17493:3 | 9 | R.LSLHRPALEDLLLGSEAN[+1501.555]LTC[+57.021]TLTGLR.D | HexNAc(5)Hex(3) | C[+57], N[+1502] | 1489.0595 | 3 | 4465.164 | 4465.1536 | 0 | 2.3 | 127 | Specific | 444.54 | 444.54 | 444.54 | 9.09 | 60 | >sp\|P01876\|IGHA1_HUMAN Immunoglobulin heavy constant alpha 1 OS=Homo sapiens OX=9606 GN=IGHA1 PE=1 SV=2 | 255 | 20201118_12833_1ng_glyco_hcd203040_compound15.21103.21103.3 | controllerType=0 controllerNumber=1 scan=21103 | 42.8353 |
| 17488:3 | 9 | R.LSLHRPALEDLLLGSEAN[+1460.529]LTC[+57.021]TLTGLR.D | HexNAc(4)Hex(4) | C[+57], N[+1461] | 1475.3847 | 3 | 4424.1395 | 4424.1271 | 0 | 2.8 | 127 | Specific | 472.33 | 428.15 | 428.15 | 9.33 | 60 | >sp\|P01876\|IGHA1_HUMAN Immunoglobulin heavy constant alpha 1 OS=Homo sapiens OX=9606 GN=IGHA1 PE=1 SV=2 | 255 | 20201118_12833_1ng_glyco_hcd203040_compound15.21098.21098.3 | controllerType=0 controllerNumber=1 scan=21098 | 42.8188 |
| 18419:3 | 9 | R.LSLHRPALEDLLLGSEAN[+1751.624]LTC[+57.021]TLTGLR.D | HexNAc(4)Hex(4)NeuAc(1) | C[+57], N[+1752] | 1572.4175 | 3 | 4715.2381 | 4715.2225 | 0 | 3.3 | 127 | Specific | 501.57 | 501.57 | 501.57 | 9.39 | 60 | >sp\|P01876\|IGHA1_HUMAN Immunoglobulin heavy constant alpha 1 OS=Homo sapiens OX=9606 GN=IGHA1 PE=1 SV=2 | 255 | 20201118_12833_1ng_glyco_hcd203040_compound15.22068.22068.3 | controllerType=0 controllerNumber=1 scan=22068 | 44.9094 |
| 17736:4 | 9 | R.LSLHRPALEDLLLGSEAN[+1501.555]LTC[+57.021]TLTGLR.D | HexNAc(5)Hex(3) | C[+57], N[+1502] | 1117.0335 | 4 | 4465.1121 | 4465.1536 | 0 | -9.3 | 127 | Specific | 529.73 | 421.67 | 421.67 | 8.03 | 60 | >sp\|P01876\|IGHA1_HUMAN Immunoglobulin heavy constant alpha 1 OS=Homo sapiens OX=9606 GN=IGHA1 PE=1 SV=2 | 255 | 20201118_12833_1ng_glyco_hcd203040_compound15.21356.21356.4 | controllerType=0 controllerNumber=1 scan=21356 | 43.3754 |
| 17491:3 | 9 | R.LSLHRPALEDLLLGSEAN[+1460.529]LTC[+57.021]TLTGLR.D | HexNAc(4)Hex(4) | C[+57], N[+1461] | 1475.3847 | 3 | 4424.1395 | 4424.1271 | 0 | 2.8 | 127 | Specific | 569.04 | 540.53 | 540.53 | 9.7 | 60 | >sp\|P01876\|IGHA1_HUMAN Immunoglobulin heavy constant alpha 1 OS=Homo sapiens OX=9606 GN=IGHA1 PE=1 SV=2 | 255 | 20201118_12833_1ng_glyco_hcd203040_compound15.21101.21101.3 | controllerType=0 controllerNumber=1 scan=21101 | 42.8315 |
| 17484:3 | 9 | R.LSLHRPALEDLLLGSEAN[+1298.476]LTC[+57.021]TLTGLR.D | HexNAc(4)Hex(3) | C[+57], N[+1298] | 1421.3675 | 3 | 4262.0879 | 4262.0742 | 0 | 3.2 | 127 | Specific | 578.85 | 578.85 | 578.85 | 10.88 | 60 | >sp\|P01876\|IGHA1_HUMAN Immunoglobulin heavy constant alpha 1 OS=Homo sapiens OX=9606 GN=IGHA1 PE=1 SV=2 | 255 | 20201118_12833_1ng_glyco_hcd203040_compound15.21094.21094.3 | controllerType=0 controllerNumber=1 scan=21094 | 42.8114 |
| 17401:3 | 9 | R.LSLHRPALEDLLLGSEAN[+1540.529]LTC[+57.021]TLTGLR.D | HexNAc(2)Hex(7) | C[+57], N[+1541] | 1502.0508 | 3 | 4504.138 | 4504.1268 | 0 | 2.5 | 127 | Specific | 607.44 | 577.04 | 577.04 | 11.88 | 60 | >sp\|P01876\|IGHA1_HUMAN Immunoglobulin heavy constant alpha 1 OS=Homo sapiens OX=9606 GN=IGHA1 PE=1 SV=2 | 255 | 20201118_12833_1ng_glyco_hcd203040_compound15.21006.21006.3 | controllerType=0 controllerNumber=1 scan=21006 | 42.567 |
| 18422:3 | 9 | R.LSLHRPALEDLLLGSEAN[+1751.624]LTC[+57.021]TLTGLR.D | HexNAc(4)Hex(4)NeuAc(1) | C[+57], N[+1752] | 1572.4175 | 3 | 4715.2381 | 4715.2225 | 0 | 3.3 | 127 | Specific | 610.49 | 610.49 | 610.49 | 11.88 | 60 | >sp\|P01876\|IGHA1_HUMAN Immunoglobulin heavy constant alpha 1 OS=Homo sapiens OX=9606 GN=IGHA1 PE=1 SV=2 | 255 | 20201118_12833_1ng_glyco_hcd203040_compound15.22071.22071.3 | controllerType=0 controllerNumber=1 scan=22071 | 44.9139 |
| 17441:3 | 9 | R.LSLHRPALEDLLLGSEAN[+1622.582]LTC[+57.021]TLTGLR.D | HexNAc(4)Hex(5) | C[+57], N[+1623] | 1529.4024 | 3 | 4586.1926 | 4586.1799 | 0 | 2.8 | 127 | Specific | 619.03 | 516.36 | 516.36 | 11.88 | 60 | >sp\|P01876\|IGHA1_HUMAN Immunoglobulin heavy constant alpha 1 OS=Homo sapiens OX=9606 GN=IGHA1 PE=1 SV=2 | 255 | 20201118_12833_1ng_glyco_hcd203040_compound15.21048.21048.3 | controllerType=0 controllerNumber=1 scan=21048 | 42.6883 |

Table S4. Cont.

| Query #:z | Protein Rank | Glycopeptide | Glycan | Modification Type(s) | Observed m/z | z | Observed (M+H) | Calc. mass (M+H) | Off-by-x error | Mass error (ppm) | StartPosition | Cleavage | Score | Delta | Delta Mod | \|Log Prob\| | # of unique peptides | Protein Name | Protein DB number | Comment | Scan # | Scan Time |
|---|---|---|---|---|---|---|---|---|---|---|---|---|---|---|---|---|---|---|---|---|---|---|
| 17439:3 | 9 | R.LSLHRPALEDLLLGSEAN[+1622.582]LTC[+57.021]TLTGLR.D | HexNAc(4)Hex(5) | C[+57], N[+1623] | 1529.4024 | 3 | 4586.1926 | 4586.1799 | 0 | 2.8 | 127 | Specific | 619.24 | 523.1 | 523.1 | 11.88 | 60 | >sp\|P01876\|IGHA1_HUMAN Immunoglobulin heavy constant alpha 1 OS=Homo sapiens OX=9606 GN=IGHA1 PE=1 SV=2 | 255 | 20201118_12833_1ng_glyco_hcd203040_compound15.21046.21046.3 | controllerType=0 controllerNumber=1 scan=21046 | 42.6842 |
| 17496:3 | 9 | R.LSLHRPALEDLLLGSEAN[+1501.555]LTC[+57.021]TLTGLR.D | HexNAc(5)Hex(3) | C[+57], N[+1502] | 1489.0595 | 3 | 4465.164 | 4465.1536 | 0 | 2.3 | 127 | Specific | 623.58 | 580 | 580 | 11.85 | 60 | >sp\|P01876\|IGHA1_HUMAN Immunoglobulin heavy constant alpha 1 OS=Homo sapiens OX=9606 GN=IGHA1 PE=1 SV=2 | 255 | 20201118_12833_1ng_glyco_hcd203040_compound15.21106.21106.3 | controllerType=0 controllerNumber=1 scan=21106 | 42.8398 |
| 17414:3 | 9 | R.LSLHRPALEDLLLGSEAN[+1663.608]LTC[+57.021]TLTGLR.D | HexNAc(5)Hex(4) | C[+57], N[+1664] | 1543.0795 | 3 | 4627.224 | 4627.2064 | 0 | 3.8 | 127 | Specific | 624.66 | 544.32 | 544.32 | 11.88 | 60 | >sp\|P01876\|IGHA1_HUMAN Immunoglobulin heavy constant alpha 1 OS=Homo sapiens OX=9606 GN=IGHA1 PE=1 SV=2 | 255 | 20201118_12833_1ng_glyco_hcd203040_compound15.21020.21020.3 | controllerType=0 controllerNumber=1 scan=21020 | 42.6062 |
| 17412:3 | 9 | R.LSLHRPALEDLLLGSEAN[+1663.608]LTC[+57.021]TLTGLR.D | HexNAc(5)Hex(4) | C[+57], N[+1664] | 1543.0795 | 3 | 4627.224 | 4627.2064 | 0 | 3.8 | 127 | Specific | 625.77 | 567.21 | 567.21 | 11.88 | 60 | >sp\|P01876\|IGHA1_HUMAN Immunoglobulin heavy constant alpha 1 OS=Homo sapiens OX=9606 GN=IGHA1 PE=1 SV=2 | 255 | 20201118_12833_1ng_glyco_hcd203040_compound15.21018.21018.3 | controllerType=0 controllerNumber=1 scan=21018 | 42.5993 |
| 17447:3 | 9 | R.LSLHRPALEDLLLGSEAN[+1501.555]LTC[+57.021]TLTGLR.D | HexNAc(5)Hex(3) | C[+57], N[+1502] | 1489.0608 | 3 | 4465.168 | 4465.1536 | 0 | 3.2 | 127 | Specific | 640.49 | 640.49 | 640.49 | 11.88 | 60 | >sp\|P01876\|IGHA1_HUMAN Immunoglobulin heavy constant alpha 1 OS=Homo sapiens OX=9606 GN=IGHA1 PE=1 SV=2 | 255 | 20201118_12833_1ng_glyco_hcd203040_compound15.21055.21055.3 | controllerType=0 controllerNumber=1 scan=21055 | 42.7063 |
| 17535:3 | 9 | R.LSLHRPALEDLLLGSEAN[+1095.397]LTC[+57.021]TLTGLR.D | HexNAc(3)Hex(3) | C[+57], N[+1095] | 1353.6751 | 3 | 4059.0108 | 4058.9949 | 0 | 3.9 | 127 | Specific | 645.55 | 619.69 | 619.69 | 11.34 | 60 | >sp\|P01876\|IGHA1_HUMAN Immunoglobulin heavy constant alpha 1 OS=Homo sapiens OX=9606 GN=IGHA1 PE=1 SV=2 | 255 | 20201118_12833_1ng_glyco_hcd203040_compound15.21147.21147.3 | controllerType=0 controllerNumber=1 scan=21147 | 42.9297 |
| 17486:3 | 9 | R.LSLHRPALEDLLLGSEAN[+1298.476]LTC[+57.021]TLTGLR.D | HexNAc(4)Hex(3) | C[+57], N[+1298] | 1421.3675 | 3 | 4262.0879 | 4262.0742 | 0 | 3.2 | 127 | Specific | 652.55 | 588.93 | 588.93 | 12.88 | 60 | >sp\|P01876\|IGHA1_HUMAN Immunoglobulin heavy constant alpha 1 OS=Homo sapiens OX=9606 GN=IGHA1 PE=1 SV=2 | 255 | 20201118_12833_1ng_glyco_hcd203040_compound15.21096.21096.3 | controllerType=0 controllerNumber=1 scan=21096 | 42.8162 |
| 17443:4 | 9 | R.LSLHRPALEDLLLGSEAN[+1501.555]LTC[+57.021]TLTGLR.D | HexNAc(5)Hex(3) | C[+57], N[+1502] | 1117.0468 | 4 | 4465.1654 | 4465.1536 | 0 | 2.6 | 127 | Specific | 654.08 | 654.08 | 654.08 | 13.34 | 60 | >sp\|P01876\|IGHA1_HUMAN Immunoglobulin heavy constant alpha 1 OS=Homo sapiens OX=9606 GN=IGHA1 PE=1 SV=2 | 255 | 20201118_12833_1ng_glyco_hcd203040_compound15.21050.21050.4 | controllerType=0 controllerNumber=1 scan=21050 | 42.6916 |
| 17374:3 | 9 | R.LSLHRPALEDLLLGSEAN[+1702.581]LTC[+57.021]TLTGLR.D | HexNAc(2)Hex(8) | C[+57], N[+1703] | 1556.0689 | 3 | 4666.1922 | 4666.1796 | 0 | 2.7 | 127 | Specific | 655.55 | 606.42 | 606.42 | 12.88 | 60 | >sp\|P01876\|IGHA1_HUMAN Immunoglobulin heavy constant alpha 1 OS=Homo sapiens OX=9606 GN=IGHA1 PE=1 SV=2 | 255 | 20201118_12833_1ng_glyco_hcd203040_compound15.20978.20978.3 | controllerType=0 controllerNumber=1 scan=20978 | 42.4979 |
| 17538:3 | 9 | R.LSLHRPALEDLLLGSEAN[+1095.397]LTC[+57.021]TLTGLR.D | HexNAc(3)Hex(3) | C[+57], N[+1095] | 1353.6751 | 3 | 4059.0108 | 4058.9949 | 0 | 3.9 | 127 | Specific | 656.5 | 619.71 | 619.71 | 12.34 | 60 | >sp\|P01876\|IGHA1_HUMAN Immunoglobulin heavy constant alpha 1 OS=Homo sapiens OX=9606 GN=IGHA1 PE=1 SV=2 | 255 | 20201118_12833_1ng_glyco_hcd203040_compound15.21150.21150.3 | controllerType=0 controllerNumber=1 scan=21150 | 42.9371 |
| 17449:3 | 9 | R.LSLHRPALEDLLLGSEAN[+1501.555]LTC[+57.021]TLTGLR.D | HexNAc(5)Hex(3) | C[+57], N[+1502] | 1489.0608 | 3 | 4465.168 | 4465.1536 | 0 | 3.2 | 127 | Specific | 658.22 | 607.98 | 607.98 | 12.88 | 60 | >sp\|P01876\|IGHA1_HUMAN Immunoglobulin heavy constant alpha 1 OS=Homo sapiens OX=9606 GN=IGHA1 PE=1 SV=2 | 255 | 20201118_12833_1ng_glyco_hcd203040_compound15.21057.21057.3 | controllerType=0 controllerNumber=1 scan=21057 | 42.717 |
| 17431:3 | 9 | R.LSLHRPALEDLLLGSEAN[+1378.476]LTC[+57.021]TLTGLR.D | HexNAc(2)Hex(6) | C[+57], N[+1378] | 1448.0346 | 3 | 4342.0893 | 4342.074 | 0 | 3.5 | 127 | Specific | 665.97 | 610.72 | 610.72 | 12.88 | 60 | >sp\|P01876\|IGHA1_HUMAN Immunoglobulin heavy constant alpha 1 OS=Homo sapiens OX=9606 GN=IGHA1 PE=1 SV=2 | 255 | 20201118_12833_1ng_glyco_hcd203040_compound15.21038.21038.3 | controllerType=0 controllerNumber=1 scan=21038 | 42.6575 |
| 17372:3 | 9 | R.LSLHRPALEDLLLGSEAN[+1702.581]LTC[+57.021]TLTGLR.D | HexNAc(2)Hex(8) | C[+57], N[+1703] | 1556.0689 | 3 | 4666.1922 | 4666.1796 | 0 | 2.7 | 127 | Specific | 665.99 | 665.99 | 665.99 | 12.88 | 60 | >sp\|P01876\|IGHA1_HUMAN Immunoglobulin heavy constant alpha 1 OS=Homo sapiens OX=9606 GN=IGHA1 PE=1 SV=2 | 255 | 20201118_12833_1ng_glyco_hcd203040_compound15.20976.20976.3 | controllerType=0 controllerNumber=1 scan=20976 | 42.4925 |
| 17530:4 | 9 | R.LSLHRPALEDLLLGSEAN[+1095.397]LTC[+57.021]TLTGLR.D | HexNAc(3)Hex(3) | C[+57], N[+1095] | 1015.5073 | 4 | 4059.0074 | 4058.9949 | 0 | 3.1 | 127 | Specific | 667.01 | 667.01 | 667.01 | 13.34 | 60 | >sp\|P01876\|IGHA1_HUMAN Immunoglobulin heavy constant alpha 1 OS=Homo sapiens OX=9606 GN=IGHA1 PE=1 SV=2 | 255 | 20201118_12833_1ng_glyco_hcd203040_compound15.21142.21142.4 | controllerType=0 controllerNumber=1 scan=21142 | 42.9214 |
| 17398:3 | 9 | R.LSLHRPALEDLLLGSEAN[+1540.529]LTC[+57.021]TLTGLR.D | HexNAc(2)Hex(7) | C[+57], N[+1541] | 1502.0508 | 3 | 4504.138 | 4504.1268 | 0 | 2.5 | 127 | Specific | 670.66 | 611.82 | 611.82 | 12.88 | 60 | >sp\|P01876\|IGHA1_HUMAN Immunoglobulin heavy constant alpha 1 OS=Homo sapiens OX=9606 GN=IGHA1 PE=1 SV=2 | 255 | 20201118_12833_1ng_glyco_hcd203040_compound15.21003.21003.3 | controllerType=0 controllerNumber=1 scan=21003 | 42.5606 |
| 17429:3 | 9 | R.LSLHRPALEDLLLGSEAN[+1378.476]LTC[+57.021]TLTGLR.D | HexNAc(2)Hex(6) | C[+57], N[+1378] | 1448.0346 | 3 | 4342.0893 | 4342.074 | 0 | 3.5 | 127 | Specific | 674.56 | 648.26 | 648.26 | 12.88 | 60 | >sp\|P01876\|IGHA1_HUMAN Immunoglobulin heavy constant alpha 1 OS=Homo sapiens OX=9606 GN=IGHA1 PE=1 SV=2 | 255 | 20201118_12833_1ng_glyco_hcd203040_compound15.21036.21036.3 | controllerType=0 controllerNumber=1 scan=21036 | 42.6469 |
| 17457:3 | 9 | R.LSLHRPALEDLLLGSEAN[+1216.423]LTC[+57.021]TLTGLR.D | HexNAc(2)Hex(5) | C[+57], N[+1216] | 1394.0158 | 3 | 4180.0329 | 4180.0211 | 0 | 2.8 | 127 | Specific | 679.85 | 647.89 | 647.89 | 12.88 | 60 | >sp\|P01876\|IGHA1_HUMAN Immunoglobulin heavy constant alpha 1 OS=Homo sapiens OX=9606 GN=IGHA1 PE=1 SV=2 | 255 | 20201118_12833_1ng_glyco_hcd203040_compound15.21065.21065.3 | controllerType=0 controllerNumber=1 scan=21065 | 42.7357 |
| 17411:4 | 9 | R.LSLHRPALEDLLLGSEAN[+1622.582]LTC[+57.021]TLTGLR.D | HexNAc(4)Hex(5) | C[+57], N[+1623] | 1147.3044 | 4 | 4586.1956 | 4586.1799 | 0 | 3.4 | 127 | Specific | 703.76 | 703.76 | 703.76 | 14.34 | 60 | >sp\|P01876\|IGHA1_HUMAN Immunoglobulin heavy constant alpha 1 OS=Homo sapiens OX=9606 GN=IGHA1 PE=1 SV=2 | 255 | 20201118_12833_1ng_glyco_hcd203040_compound15.21017.21017.4 | controllerType=0 controllerNumber=1 scan=21017 | 42.5978 |
| 17403:4 | 9 | R.LSLHRPALEDLLLGSEAN[+1663.608]LTC[+57.021]TLTGLR.D | HexNAc(5)Hex(4) | C[+57], N[+1664] | 1157.5592 | 4 | 4627.2152 | 4627.2064 | 0 | 1.9 | 127 | Specific | 705.26 | 705.26 | 705.26 | 14.31 | 60 | >sp\|P01876\|IGHA1_HUMAN Immunoglobulin heavy constant alpha 1 OS=Homo sapiens OX=9606 GN=IGHA1 PE=1 SV=2 | 255 | 20201118_12833_1ng_glyco_hcd203040_compound15.21008.21008.4 | controllerType=0 controllerNumber=1 scan=21008 | 42.5719 |
| 17452:4 | 9 | R.LSLHRPALEDLLLGSEAN[+1419.502]LTC[+57.021]TLTGLR.D | HexNAc(3)Hex(5) | C[+57], N[+1420] | 1096.5336 | 4 | 4383.1126 | 4383.1005 | 0 | 2.8 | 127 | Specific | 716.25 | 716.25 | 716.25 | 14.34 | 60 | >sp\|P01876\|IGHA1_HUMAN Immunoglobulin heavy constant alpha 1 OS=Homo sapiens OX=9606 GN=IGHA1 PE=1 SV=2 | 255 | 20201118_12833_1ng_glyco_hcd203040_compound15.21060.21060.4 | controllerType=0 controllerNumber=1 scan=21060 | 42.7205 |
| 17483:4 | 9 | R.LSLHRPALEDLLLGSEAN[+1298.476]LTC[+57.021]TLTGLR.D | HexNAc(4)Hex(3) | C[+57], N[+1298] | 1066.276 | 4 | 4262.0823 | 4262.0742 | 0 | 1.9 | 127 | Specific | 720.14 | 720.14 | 720.14 | 14.31 | 60 | >sp\|P01876\|IGHA1_HUMAN Immunoglobulin heavy constant alpha 1 OS=Homo sapiens OX=9606 GN=IGHA1 PE=1 SV=2 | 255 | 20201118_12833_1ng_glyco_hcd203040_compound15.21093.21093.4 | controllerType=0 controllerNumber=1 scan=21093 | 42.81 |
| 18414:4 | 9 | R.LSLHRPALEDLLLGSEAN[+1710.598]LTC[+57.021]TLTGLR.D | HexNAc(3)Hex(5)NeuAc(1) | C[+57], N[+1711] | 1169.3091 | 4 | 4674.2147 | 4674.1959 | 0 | 4 | 127 | Specific | 733.58 | 733.58 | 733.58 | 13.79 | 60 | >sp\|P01876\|IGHA1_HUMAN Immunoglobulin heavy constant alpha 1 OS=Homo sapiens OX=9606 GN=IGHA1 PE=1 SV=2 | 255 | 20201118_12833_1ng_glyco_hcd203040_compound15.22063.22063.4 | controllerType=0 controllerNumber=1 scan=22063 | 44.9003 |
| 18408:4 | 9 | R.LSLHRPALEDLLLGSEAN[+1751.624]LTC[+57.021]TLTGLR.D | HexNAc(4)Hex(4)NeuAc(1) | C[+57], N[+1752] | 1179.5646 | 4 | 4715.2366 | 4715.2225 | 0 | 3 | 127 | Specific | 756.43 | 756.43 | 756.43 | 15.34 | 60 | >sp\|P01876\|IGHA1_HUMAN Immunoglobulin heavy constant alpha 1 OS=Homo sapiens OX=9606 GN=IGHA1 PE=1 SV=2 | 255 | 20201118_12833_1ng_glyco_hcd203040_compound15.22057.22057.4 | controllerType=0 controllerNumber=1 scan=22057 | 44.8834 |
| 17468:5 | 9 | R.LSLHRPALEDLLLGSEAN[+1501.555]LTC[+57.021]TLTGLR.D | HexNAc(5)Hex(3) | C[+57], N[+1502] | 893.8386 | 5 | 4465.1641 | 4465.1536 | 0 | 2.4 | 127 | Specific | 766.66 | 766.66 | 766.66 | 15.13 | 60 | >sp\|P01876\|IGHA1_HUMAN Immunoglobulin heavy constant alpha 1 OS=Homo sapiens OX=9606 GN=IGHA1 PE=1 SV=2 | 255 | 20201118_12833_1ng_glyco_hcd203040_compound15.21077.21077.5 | controllerType=0 controllerNumber=1 scan=21077 | 42.7682 |
| 18411:4 | 9 | R.LSLHRPALEDLLLGSEAN[+1751.624]LTC[+57.021]TLTGLR.D | HexNAc(4)Hex(4)NeuAc(1) | C[+57], N[+1752] | 1179.5646 | 4 | 4715.2366 | 4715.2225 | 0 | 3 | 127 | Specific | 773.21 | 773.21 | 773.21 | 15.34 | 60 | >sp\|P01876\|IGHA1_HUMAN Immunoglobulin heavy constant alpha 1 OS=Homo sapiens OX=9606 GN=IGHA1 PE=1 SV=2 | 255 | 20201118_12833_1ng_glyco_hcd203040_compound15.22060.22060.4 | controllerType=0 controllerNumber=1 scan=22060 | 44.8884 |
| 17471:5 | 9 | R.LSLHRPALEDLLLGSEAN[+1501.555]LTC[+57.021]TLTGLR.D | HexNAc(5)Hex(3) | C[+57], N[+1502] | 893.8386 | 5 | 4465.1641 | 4465.1536 | 0 | 2.4 | 127 | Specific | 773.52 | 773.52 | 773.52 | 15.13 | 60 | >sp\|P01876\|IGHA1_HUMAN Immunoglobulin heavy constant alpha 1 OS=Homo sapiens OX=9606 GN=IGHA1 PE=1 SV=2 | 255 | 20201118_12833_1ng_glyco_hcd203040_compound15.21080.21080.5 | controllerType=0 controllerNumber=1 scan=21080 | 42.7801 |
| 17445:4 | 9 | R.LSLHRPALEDLLLGSEAN[+1501.555]LTC[+57.021]TLTGLR.D | HexNAc(5)Hex(3) | C[+57], N[+1502] | 1117.0468 | 4 | 4465.1654 | 4465.1536 | 0 | 2.6 | 127 | Specific | 773.62 | 770.54 | 770.54 | 15.34 | 60 | >sp\|P01876\|IGHA1_HUMAN Immunoglobulin heavy constant alpha 1 OS=Homo sapiens OX=9606 GN=IGHA1 PE=1 SV=2 | 255 | 20201118_12833_1ng_glyco_hcd203040_compound15.21052.21052.4 | controllerType=0 controllerNumber=1 scan=21052 | 42.7012 |
| 17455:3 | 9 | R.LSLHRPALEDLLLGSEAN[+1216.423]LTC[+57.021]TLTGLR.D | HexNAc(2)Hex(5) | C[+57], N[+1216] | 1394.0158 | 3 | 4180.0329 | 4180.0211 | 0 | 2.8 | 127 | Specific | 775.38 | 712.84 | 712.84 | 14.88 | 60 | >sp\|P01876\|IGHA1_HUMAN Immunoglobulin heavy constant alpha 1 OS=Homo sapiens OX=9606 GN=IGHA1 PE=1 SV=2 | 255 | 20201118_12833_1ng_glyco_hcd203040_compound15.21063.21063.3 | controllerType=0 controllerNumber=1 scan=21063 | 42.7292 |
| 18380:4 | 9 | R.LSLHRPALEDLLLGSEAN[+1913.677]LTC[+57.021]TLTGLR.D | HexNAc(4)Hex(5)NeuAc(1) | C[+57], N[+1914] | 1220.0773 | 4 | 4877.2874 | 4877.2753 | 0 | 2.5 | 127 | Specific | 786.29 | 731.35 | 731.35 | 15.34 | 60 | >sp\|P01876\|IGHA1_HUMAN Immunoglobulin heavy constant alpha 1 OS=Homo sapiens OX=9606 GN=IGHA1 PE=1 SV=2 | 255 | 20201118_12833_1ng_glyco_hcd203040_compound15.22028.22028.4 | controllerType=0 controllerNumber=1 scan=22028 | 44.826 |
| 17405:4 | 9 | R.LSLHRPALEDLLLGSEAN[+1663.608]LTC[+57.021]TLTGLR.D | HexNAc(5)Hex(4) | C[+57], N[+1664] | 1157.5592 | 4 | 4627.2152 | 4627.2064 | 0 | 1.9 | 127 | Specific | 791.62 | 730.82 | 730.82 | 15.31 | 60 | >sp\|P01876\|IGHA1_HUMAN Immunoglobulin heavy constant alpha 1 OS=Homo sapiens OX=9606 GN=IGHA1 PE=1 SV=2 | 255 | 20201118_12833_1ng_glyco_hcd203040_compound15.21010.21010.4 | controllerType=0 controllerNumber=1 scan=21010 | 42.5789 |
| 17413:4 | 9 | R.LSLHRPALEDLLLGSEAN[+1622.582]LTC[+57.021]TLTGLR.D | HexNAc(4)Hex(5) | C[+57], N[+1623] | 1147.3044 | 4 | 4586.1956 | 4586.1799 | 0 | 3.4 | 127 | Specific | 833.1 | 738.14 | 738.14 | 16.34 | 60 | >sp\|P01876\|IGHA1_HUMAN Immunoglobulin heavy constant alpha 1 OS=Homo sapiens OX=9606 GN=IGHA1 PE=1 SV=2 | 255 | 20201118_12833_1ng_glyco_hcd203040_compound15.21019.21019.4 | controllerType=0 controllerNumber=1 scan=21019 | 42.6041 |
| 18417:4 | 9 | R.LSLHRPALEDLLLGSEAN[+1710.598]LTC[+57.021]TLTGLR.D | HexNAc(3)Hex(5)NeuAc(1) | C[+57], N[+1711] | 1169.3091 | 4 | 4674.2147 | 4674.1959 | 0 | 4 | 127 | Specific | 850.48 | 818.09 | 818.09 | 16.79 | 60 | >sp\|P01876\|IGHA1_HUMAN Immunoglobulin heavy constant alpha 1 OS=Homo sapiens OX=9606 GN=IGHA1 PE=1 SV=2 | 255 | 20201118_12833_1ng_glyco_hcd203040_compound15.22066.22066.4 | controllerType=0 controllerNumber=1 scan=22066 | 44.9055 |
| 17485:4 | 9 | R.LSLHRPALEDLLLGSEAN[+1298.476]LTC[+57.021]TLTGLR.D | HexNAc(4)Hex(3) | C[+57], N[+1298] | 1066.276 | 4 | 4262.0823 | 4262.0742 | 0 | 1.9 | 127 | Specific | 858.64 | 839.84 | 839.84 | 17.31 | 60 | >sp\|P01876\|IGHA1_HUMAN Immunoglobulin heavy constant alpha 1 OS=Homo sapiens OX=9606 GN=IGHA1 PE=1 SV=2 | 255 | 20201118_12833_1ng_glyco_hcd203040_compound15.21095.21095.4 | controllerType=0 controllerNumber=1 scan=21095 | 42.8148 |
| 17517:4 | 9 | R.LSLHRPALEDLLLGSEAN[+1257.449]LTC[+57.021]TLTGLR.D | HexNAc(3)Hex(4) | C[+57], N[+1257] | 1056.0186 | 4 | 4221.0526 | 4221.0477 | 0 | 1.2 | 127 | Specific | 864.31 | 838.52 | 838.52 | 17.31 | 60 | >sp\|P01876\|IGHA1_HUMAN Immunoglobulin heavy constant alpha 1 OS=Homo sapiens OX=9606 GN=IGHA1 PE=1 SV=2 | 255 | 20201118_12833_1ng_glyco_hcd203040_compound15.21128.21128.4 | controllerType=0 controllerNumber=1 scan=21128 | 42.8913 |
| 17446:4 | 9 | R.LSLHRPALEDLLLGSEAN[+1216.423]LTC[+57.021]TLTGLR.D | HexNAc(2)Hex(5) | C[+57], N[+1216] | 1045.7637 | 4 | 4180.033 | 4180.0211 | 0 | 2.8 | 127 | Specific | 873.19 | 768.44 | 768.44 | 17.34 | 60 | >sp\|P01876\|IGHA1_HUMAN Immunoglobulin heavy constant alpha 1 OS=Homo sapiens OX=9606 GN=IGHA1 PE=1 SV=2 | 255 | 20201118_12833_1ng_glyco_hcd203040_compound15.21054.21054.4 | controllerType=0 controllerNumber=1 scan=21054 | 42.7049 |
| 17454:4 | 9 | R.LSLHRPALEDLLLGSEAN[+1419.502]LTC[+57.021]TLTGLR.D | HexNAc(3)Hex(5) | C[+57], N[+1420] | 1096.5336 | 4 | 4383.1126 | 4383.1005 | 0 | 2.8 | 127 | Specific | 879.11 | 859.39 | 859.39 | 17.34 | 60 | >sp\|P01876\|IGHA1_HUMAN Immunoglobulin heavy constant alpha 1 OS=Homo sapiens OX=9606 GN=IGHA1 PE=1 SV=2 | 255 | 20201118_12833_1ng_glyco_hcd203040_compound15.21062.21062.4 | controllerType=0 controllerNumber=1 scan=21062 | 42.7273 |
| 17366:4 | 9 | R.LSLHRPALEDLLLGSEAN[+1702.581]LTC[+57.021]TLTGLR.D | HexNAc(2)Hex(8) | C[+57], N[+1703] | 1167.3036 | 4 | 4666.1927 | 4666.1796 | 0 | 2.8 | 127 | Specific | 893.62 | 893.62 | 893.62 | 17.34 | 60 | >sp\|P01876\|IGHA1_HUMAN Immunoglobulin heavy constant alpha 1 OS=Homo sapiens OX=9606 GN=IGHA1 PE=1 SV=2 | 255 | 20201118_12833_1ng_glyco_hcd203040_compound15.20970.20970.4 | controllerType=0 controllerNumber=1 scan=20970 | 42.4766 |
| 17532:4 | 9 | R.LSLHRPALEDLLLGSEAN[+1095.397]LTC[+57.021]TLTGLR.D | HexNAc(3)Hex(3) | C[+57], N[+1095] | 1015.5073 | 4 | 4059.0074 | 4058.9949 | 0 | 3.1 | 127 | Specific | 936.4 | 906.77 | 906.77 | 18.34 | 60 | >sp\|P01876\|IGHA1_HUMAN Immunoglobulin heavy constant alpha 1 OS=Homo sapiens OX=9606 GN=IGHA1 PE=1 SV=2 | 255 | 20201118_12833_1ng_glyco_hcd203040_compound15.21144.21144.4 | controllerType=0 controllerNumber=1 scan=21144 | 42.9256 |
| 17520:4 | 9 | R.LSLHRPALEDLLLGSEAN[+1257.449]LTC[+57.021]TLTGLR.D | HexNAc(3)Hex(4) | C[+57], N[+1257] | 1056.0186 | 4 | 4221.0526 | 4221.0477 | 0 | 1.2 | 127 | Specific | 937.99 | 883.81 | 883.81 | 18.31 | 60 | >sp\|P01876\|IGHA1_HUMAN Immunoglobulin heavy constant alpha 1 OS=Homo sapiens OX=9606 GN=IGHA1 PE=1 SV=2 | 255 | 20201118_12833_1ng_glyco_hcd203040_compound15.21131.21131.4 | controllerType=0 controllerNumber=1 scan=21131 | 42.8979 |
| 17387:4 | 9 | R.LSLHRPALEDLLLGSEAN[+1540.529]LTC[+57.021]TLTGLR.D | HexNAc(2)Hex(7) | C[+57], N[+1541] | 1126.7897 | 4 | 4504.137 | 4504.1268 | 0 | 2.3 | 127 | Specific | 938.23 | 823.44 | 823.44 | 18.31 | 60 | >sp\|P01876\|IGHA1_HUMAN Immunoglobulin heavy constant alpha 1 OS=Homo sapiens OX=9606 GN=IGHA1 PE=1 SV=2 | 255 | 20201118_12833_1ng_glyco_hcd203040_compound15.20992.20992.4 | controllerType=0 controllerNumber=1 scan=20992 | 42.5305 |
| 17407:4 | 9 | R.LSLHRPALEDLLLGSEAN[+1378.476]LTC[+57.021]TLTGLR.D | HexNAc(2)Hex(6) | C[+57], N[+1378] | 1086.2775 | 4 | 4342.0882 | 4342.074 | 0 | 3.3 | 127 | Specific | 940.37 | 870.21 | 870.21 | 18.34 | 60 | >sp\|P01876\|IGHA1_HUMAN Immunoglobulin heavy constant alpha 1 OS=Homo sapiens OX=9606 GN=IGHA1 PE=1 SV=2 | 255 | 20201118_12833_1ng_glyco_hcd203040_compound15.21013.21013.4 | controllerType=0 controllerNumber=1 scan=21013 | 42.5865 |
| 17448:4 | 9 | R.LSLHRPALEDLLLGSEAN[+1216.423]LTC[+57.021]TLTGLR.D | HexNAc(2)Hex(5) | C[+57], N[+1216] | 1045.7637 | 4 | 4180.033 | 4180.0211 | 0 | 2.8 | 127 | Specific | 982.96 | 982.96 | 982.96 | 19.34 | 60 | >sp\|P01876\|IGHA1_HUMAN Immunoglobulin heavy constant alpha 1 OS=Homo sapiens OX=9606 GN=IGHA1 PE=1 SV=2 | 255 | 20201118_12833_1ng_glyco_hcd203040_compound15.21056.21056.4 | controllerType=0 controllerNumber=1 scan=21056 | 42.7126 |
| 17409:4 | 9 | R.LSLHRPALEDLLLGSEAN[+1378.476]LTC[+57.021]TLTGLR.D | HexNAc(2)Hex(6) | C[+57], N[+1378] | 1086.2775 | 4 | 4342.0882 | 4342.074 | 0 | 3.3 | 127 | Specific | 1005.01 | 985.11 | 985.11 | 20.34 | 60 | >sp\|P01876\|IGHA1_HUMAN Immunoglobulin heavy constant alpha 1 OS=Homo sapiens OX=9606 GN=IGHA1 PE=1 SV=2 | 255 | 20201118_12833_1ng_glyco_hcd203040_compound15.21015.21015.4 | controllerType=0 controllerNumber=1 scan=21015 | 42.5949 |
| 17368:4 | 9 | R.LSLHRPALEDLLLGSEAN[+1702.581]LTC[+57.021]TLTGLR.D | HexNAc(2)Hex(8) | C[+57], N[+1703] | 1167.3036 | 4 | 4666.1927 | 4666.1796 | 0 | 2.8 | 127 | Specific | 1005.02 | 996.44 | 996.44 | 20.34 | 60 | >sp\|P01876\|IGHA1_HUMAN Immunoglobulin heavy constant alpha 1 OS=Homo sapiens OX=9606 GN=IGHA1 PE=1 SV=2 | 255 | 20201118_12833_1ng_glyco_hcd203040_compound15.20972.20972.4 | controllerType=0 controllerNumber=1 scan=20972 | 42.4833 |
| 11406:4 | 9 | K.TIDRLAGKPTHVN[+1216.423]VSVVMAEVDGTC[+57.021]Y.- | HexNAc(2)Hex(5) | C[+57], N[+1216] | 1012.9654 | 4 | 4048.8399 | 4048.8247 | 0 | 3.7 | 328 | CRagged | 268.4 | 268.4 | 268.4 | 5.21 | 60 | >sp\|P01876\|IGHA1_HUMAN Immunoglobulin heavy constant alpha 1 OS=Homo sapiens OX=9606 GN=IGHA1 PE=1 SV=2 | 255 | 20201118_12833_1ng_glyco_hcd203040_compound15.14725.14725.4 | controllerType=0 controllerNumber=1 scan=14725 | 31.3136 |
| 09948:3 | 9 | R.LAGKPTHVN[+1647.613]VSVVMAEVDGTC[+57.021]Y.- | HexNAc(5)Hex(3)Fuc(1) | C[+57], N[+1648] | 1332.2557 | 3 | 3994.7525 | 3994.7553 | 0 | -0.7 | 332 | CRagged | 204.28 | 180.02 | 180.02 | 3.43 | 60 | >sp\|P01876\|IGHA1_HUMAN Immunoglobulin heavy constant alpha 1 OS=Homo sapiens OX=9606 GN=IGHA1 PE=1 SV=2 | 255 | 20201118_12833_1ng_glyco_hcd203040_compound15.13212.13212.3 | controllerType=0 controllerNumber=1 scan=13212 | 28.6483 |
| 12989:3 | 9 | R.LAGKPTHVN[+1548.545]VSVVMAEVDGTC[+57.021]Y.- | HexNAc(3)Hex(4)NeuAc(1) | C[+57], N[+1549] | 1299.2353 | 3 | 3895.6913 | 3895.6869 | 0 | 1.1 | 332 | CRagged | 214.41 | 214.41 | 214.41 | 3.91 | 60 | >sp\|P01876\|IGHA1_HUMAN Immunoglobulin heavy constant alpha 1 OS=Homo sapiens OX=9606 GN=IGHA1 PE=1 SV=2 | 255 | 20201118_12833_1ng_glyco_hcd203040_compound15.16384.16384.3 | controllerType=0 controllerNumber=1 scan=16384 | 34.1321 |
| 09918:3 | 9 | R.LAGKPTHVN[+1216.423]VSVVMAEVDGTC[+57.021]Y.- | HexNAc(2)Hex(5) | C[+57], N[+1216] | 1188.5324 | 3 | 3563.5827 | 3563.5649 | 0 | 5 | 332 | CRagged | 233.83 | 233.83 | 233.83 | 6.19 | 60 | >sp\|P01876\|IGHA1_HUMAN Immunoglobulin heavy constant alpha 1 OS=Homo sapiens OX=9606 GN=IGHA1 PE=1 SV=2 | 255 | 20201118_12833_1ng_glyco_hcd203040_compound15.13181.13181.3 | controllerType=0 controllerNumber=1 scan=13181 | 28.5962 |
| 12830:3 | 9 | R.LAGKPTHVN[+2059.735]VSVVMAEVDGTC[+57.021]Y.- | HexNAc(4)Hex(5)Fuc(1)NeuAc(1) | C[+57], N[+2060] | 1469.6345 | 3 | 4406.8888 | 4406.877 | 0 | 2.7 | 332 | CRagged | 339.54 | 339.54 | 339.54 | 7.97 | 60 | >sp\|P01876\|IGHA1_HUMAN Immunoglobulin heavy constant alpha 1 OS=Homo sapiens OX=9606 GN=IGHA1 PE=1 SV=2 | 255 | 20201118_12833_1ng_glyco_hcd203040_compound15.16220.16220.3 | controllerType=0 controllerNumber=1 scan=16220 | 33.8413 |
| 12915:3 | 9 | R.LAGKPTHVN[+1751.624]VSVVMAEVDGTC[+57.021]Y.- | HexNAc(4)Hex(4)NeuAc(1) | C[+57], N[+1752] | 1366.934 | 3 | 4098.7875 | 4098.7663 | 0 | 5.2 | 332 | CRagged | 381.14 | 381.14 | 381.14 | 9.26 | 60 | >sp\|P01876\|IGHA1_HUMAN Immunoglobulin heavy constant alpha 1 OS=Homo sapiens OX=9606 GN=IGHA1 PE=1 SV=2 | 255 | 20201118_12833_1ng_glyco_hcd203040_compound15.16308.16308.3 | controllerType=0 controllerNumber=1 scan=16308 | 33.999 |
| 12832:3 | 9 | R.LAGKPTHVN[+2059.735]VSVVMAEVDGTC[+57.021]Y.- | HexNAc(4)Hex(5)Fuc(1)NeuAc(1) | C[+57], N[+2060] | 1469.6345 | 3 | 4406.8888 | 4406.877 | 0 | 2.7 | 332 | CRagged | 388.14 | 380.28 | 380.28 | 9.77 | 60 | >sp\|P01876\|IGHA1_HUMAN Immunoglobulin heavy constant alpha 1 OS=Homo sapiens OX=9606 GN=IGHA1 PE=1 SV=2 | 255 | 20201118_12833_1ng_glyco_hcd203040_compound15.16222.16222.3 | controllerType=0 controllerNumber=1 scan=16222 | 33.8453 |
| 11168:4 | 9 | R.LAGKPTHVN[+1647.613]VSVVMAEVDGTC[+57.021]Y.- | HexNAc(5)Hex(3)Fuc(1) | C[+57], N[+1648] | 999.4465 | 4 | 3994.764 | 3994.7553 | 0 | 2.2 | 332 | CRagged | 398.09 | 325.31 | 325.31 | 7.98 | 60 | >sp\|P01876\|IGHA1_HUMAN Immunoglobulin heavy constant alpha 1 OS=Homo sapiens OX=9606 GN=IGHA1 PE=1 SV=2 | 255 | 20201118_12833_1ng_glyco_hcd203040_compound15.14477.14477.4 | controllerType=0 controllerNumber=1 scan=14477 | 30.8648 |
| 12913:3 | 9 | R.LAGKPTHVN[+1751.624]VSVVMAEVDGTC[+57.021]Y.- | HexNAc(4)Hex(4)NeuAc(1) | C[+57], N[+1752] | 1366.934 | 3 | 4098.7875 | 4098.7663 | 0 | 5.2 | 332 | CRagged | 400.88 | 400.88 | 400.88 | 9.22 | 60 | >sp\|P01876\|IGHA1_HUMAN Immunoglobulin heavy constant alpha 1 OS=Homo sapiens OX=9606 GN=IGHA1 PE=1 SV=2 | 255 | 20201118_12833_1ng_glyco_hcd203040_compound15.16306.16306.3 | controllerType=0 controllerNumber=1 scan=16306 | 33.9954 |
| 12887:3 | 9 | R.LAGKPTHVN[+1710.598]VSVVMAEVDGTC[+57.021]Y.- | HexNAc(3)Hex(5)NeuAc(1) | C[+57], N[+1711] | 1353.2537 | 3 | 4057.7466 | 4057.7397 | 0 | 1.7 | 332 | CRagged | 416.32 | 315.89 | 296.19 | 8.19 | 60 | >sp\|P01876\|IGHA1_HUMAN Immunoglobulin heavy constant alpha 1 OS=Homo sapiens OX=9606 GN=IGHA1 PE=1 SV=2 | 255 | 20201118_12833_1ng_glyco_hcd203040_compound15.16279.16279.3 | controllerType=0 controllerNumber=1 scan=16279 | 33.9492 |
| 11055:3 | 9 | R.LAGKPTHVN[+1971.719]VSVVMAEVDGTC[+57.021]Y.- | HexNAc(5)Hex(5)Fuc(1) | C[+57], N[+1972] | 1440.2928 | 3 | 4318.8638 | 4318.861 | 0 | 0.7 | 332 | CRagged | 426.92 | 426.92 | 302.36 | 9.46 | 60 | >sp\|P01876\|IGHA1_HUMAN Immunoglobulin heavy constant alpha 1 OS=Homo sapiens OX=9606 GN=IGHA1 PE=1 SV=2 | 255 | 20201118_12833_1ng_glyco_hcd203040_compound15.14359.14359.3 | controllerType=0 controllerNumber=1 scan=14359 | 30.6162 |
| 12889:3 | 9 | R.LAGKPTHVN[+1710.598]VSVVMAEVDGTC[+57.021]Y.- | HexNAc(3)Hex(5)NeuAc(1) | C[+57], N[+1711] | 1353.2537 | 3 | 4057.7466 | 4057.7397 | 0 | 1.7 | 332 | CRagged | 438.6 | 438.6 | 298.48 | 9.94 | 60 | >sp\|P01876\|IGHA1_HUMAN Immunoglobulin heavy constant alpha 1 OS=Homo sapiens OX=9606 GN=IGHA1 PE=1 SV=2 | 255 | 20201118_12833_1ng_glyco_hcd203040_compound15.16281.16281.3 | controllerType=0 controllerNumber=1 scan=16281 | 33.9529 |
| 11057:3 | 9 | R.LAGKPTHVN[+1971.719]VSVVMAEVDGTC[+57.021]Y.- | HexNAc(5)Hex(5)Fuc(1) | C[+57], N[+1972] | 1440.2928 | 3 | 4318.8638 | 4318.861 | 0 | 0.7 | 332 | CRagged | 441.58 | 342.97 | 339.07 | 7.9 | 60 | >sp\|P01876\|IGHA1_HUMAN Immunoglobulin heavy constant alpha 1 OS=Homo sapiens OX=9606 GN=IGHA1 PE=1 SV=2 | 255 | 20201118_12833_1ng_glyco_hcd203040_compound15.14361.14361.3 | controllerType=0 controllerNumber=1 scan=14361 | 30.6197 |
| 12802:3 | 9 | R.LAGKPTHVN[+1913.677]VSVVMAEVDGTC[+57.021]Y.- | HexNAc(4)Hex(5)NeuAc(1) | C[+57], N[+1914] | 1420.9492 | 3 | 4260.8329 | 4260.8191 | 0 | 3.2 | 332 | CRagged | 444.49 | 333.22 | 333.22 | 8.4 | 60 | >sp\|P01876\|IGHA1_HUMAN Immunoglobulin heavy constant alpha 1 OS=Homo sapiens OX=9606 GN=IGHA1 PE=1 SV=2 | 255 | 20201118_12833_1ng_glyco_hcd203040_compound15.16191.16191.3 | controllerType=0 controllerNumber=1 scan=16191 | 33.79 |
| 11166:4 | 9 | R.LAGKPTHVN[+1647.613]VSVVMAEVDGTC[+57.021]Y.- | HexNAc(5)Hex(3)Fuc(1) | C[+57], N[+1648] | 999.4465 | 4 | 3994.764 | 3994.7553 | 0 | 2.2 | 332 | CRagged | 455.08 | 455.08 | 455.08 | 9.77 | 60 | >sp\|P01876\|IGHA1_HUMAN Immunoglobulin heavy constant alpha 1 OS=Homo sapiens OX=9606 GN=IGHA1 PE=1 SV=2 | 255 | 20201118_12833_1ng_glyco_hcd203040_compound15.14475.14475.4 | controllerType=0 controllerNumber=1 scan=14475 | 30.861 |
| 11186:3 | 9 | R.LAGKPTHVN[+1501.555]VSVVMAEVDGTC[+57.021]Y.- | HexNAc(5)Hex(3) | C[+57], N[+1502] | 1283.5705 | 3 | 3848.6969 | 3848.6974 | 0 | -0.1 | 332 | CRagged | 464.91 | 391.98 | 391.98 | 9.47 | 60 | >sp\|P01876\|IGHA1_HUMAN Immunoglobulin heavy constant alpha 1 OS=Homo sapiens OX=9606 GN=IGHA1 PE=1 SV=2 | 255 | 20201118_12833_1ng_glyco_hcd203040_compound15.14496.14496.3 | controllerType=0 controllerNumber=1 scan=14496 | 30.9 |
| 11188:3 | 9 | R.LAGKPTHVN[+1501.555]VSVVMAEVDGTC[+57.021]Y.- | HexNAc(5)Hex(3) | C[+57], N[+1502] | 1283.5705 | 3 | 3848.6969 | 3848.6974 | 0 | -0.1 | 332 | CRagged | 471.37 | 411.63 | 411.63 | 9.69 | 60 | >sp\|P01876\|IGHA1_HUMAN Immunoglobulin heavy constant alpha 1 OS=Homo sapiens OX=9606 GN=IGHA1 PE=1 SV=2 | 255 | 20201118_12833_1ng_glyco_hcd203040_compound15.14498.14498.3 | controllerType=0 controllerNumber=1 scan=14498 | 30.9042 |
| 12800:3 | 9 | R.LAGKPTHVN[+1913.677]VSVVMAEVDGTC[+57.021]Y.- | HexNAc(4)Hex(5)NeuAc(1) | C[+57], N[+1914] | 1420.9492 | 3 | 4260.8329 | 4260.8191 | 0 | 3.2 | 332 | CRagged | 511.97 | 422.61 | 422.61 | 10.25 | 60 | >sp\|P01876\|IGHA1_HUMAN Immunoglobulin heavy constant alpha 1 OS=Homo sapiens OX=9606 GN=IGHA1 PE=1 SV=2 | 255 | 20201118_12833_1ng_glyco_hcd203040_compound15.16189.16189.3 | controllerType=0 controllerNumber=1 scan=16189 | 33.786 |
| 11072:3 | 9 | R.LAGKPTHVN[+1663.608]VSVVMAEVDGTC[+57.021]Y.- | HexNAc(5)Hex(4) | C[+57], N[+1664] | 1337.592 | 3 | 4010.7614 | 4010.7502 | 0 | 2.8 | 332 | CRagged | 560.5 | 560.5 | 395.24 | 10.56 | 60 | >sp\|P01876\|IGHA1_HUMAN Immunoglobulin heavy constant alpha 1 OS=Homo sapiens OX=9606 GN=IGHA1 PE=1 SV=2 | 255 | 20201118_12833_1ng_glyco_hcd203040_compound15.14377.14377.3 | controllerType=0 controllerNumber=1 scan=14377 | 30.6507 |
| 11067:3 | 9 | R.LAGKPTHVN[+1809.666]VSVVMAEVDGTC[+57.021]Y.- | HexNAc(5)Hex(4)Fuc(1) | C[+57], N[+1810] | 1386.2781 | 3 | 4156.8198 | 4156.8081 | 0 | 2.8 | 332 | CRagged | 710.23 | 710.23 | 440.31 | 14.74 | 60 | >sp\|P01876\|IGHA1_HUMAN Immunoglobulin heavy constant alpha 1 OS=Homo sapiens OX=9606 GN=IGHA1 PE=1 SV=2 | 255 | 20201118_12833_1ng_glyco_hcd203040_compound15.14372.14372.3 | controllerType=0 controllerNumber=1 scan=14372 | 30.6419 |
| 11076:3 | 9 | R.LAGKPTHVN[+1663.608]VSVVMAEVDGTC[+57.021]Y.- | HexNAc(5)Hex(4) | C[+57], N[+1664] | 1337.592 | 3 | 4010.7614 | 4010.7502 | 0 | 2.8 | 332 | CRagged | 796.28 | 652.49 | 476.01 | 15.74 | 60 | >sp\|P01876\|IGHA1_HUMAN Immunoglobulin heavy constant alpha 1 OS=Homo sapiens OX=9606 GN=IGHA1 PE=1 SV=2 | 255 | 20201118_12833_1ng_glyco_hcd203040_compound15.14381.14381.3 | controllerType=0 controllerNumber=1 scan=14381 | 30.6574 |
| 11111:3 | 9 | R.LAGKPTHVN[+1647.613]VSVVMAEVDGTC[+57.021]Y.- | HexNAc(5)Hex(3)Fuc(1) | C[+57], N[+1648] | 1332.2608 | 3 | 3994.7678 | 3994.7553 | 0 | 3.1 | 332 | CRagged | 799.64 | 741.82 | 741.82 | 15.74 | 60 | >sp\|P01876\|IGHA1_HUMAN Immunoglobulin heavy constant alpha 1 OS=Homo sapiens OX=9606 GN=IGHA1 PE=1 SV=2 | 255 | 20201118_12833_1ng_glyco_hcd203040_compound15.14418.14418.3 | controllerType=0 controllerNumber=1 scan=14418 | [illegible] |
| 11033:3 | 9 | R.LAGKPTHVN[+1663.608]VSVVMAEVDGTC[+57.021]Y.- | HexNAc(5)Hex(4) | C[+57], N[+1664] | 1337.5909 | 3 | 4010.7581 | 4010.7502 | 0 | 2 | 332 | CRagged | 811.32 | 811.32 | 498.15 | 16.71 | 60 | >sp\|P01876\|IGHA1_HUMAN Immunoglobulin heavy constant alpha 1 OS=Homo sapiens OX=9606 GN=IGHA1 PE=1 SV=2 | 255 | 20201118_12833_1ng_glyco_hcd203040_compound15.14336.14336.3 | controllerType=0 controllerNumber=1 scan=14336 | 30.5735 |
| 11038:3 | 9 | R.LAGKPTHVN[+1378.476]VSVVMAEVDGTC[+57.021]Y.- | HexNAc(2)Hex(6) | C[+57], N[+1378] | 1242.5466 | 3 | 3725.6251 | 3725.6178 | 0 | 2 | 332 | CRagged | 821.44 | 821.44 | 492.22 | 16.71 | 60 | >sp\|P01876\|IGHA1_HUMAN Immunoglobulin heavy constant alpha 1 OS=Homo sapiens OX=9606 GN=IGHA1 PE=1 SV=2 | 255 | 20201118_12833_1ng_glyco_hcd203040_compound15.14342.14342.3 | controllerType=0 controllerNumber=1 scan=14342 | 30.5882 |
| 11069:3 | 9 | R.LAGKPTHVN[+1809.666]VSVVMAEVDGTC[+57.021]Y.- | HexNAc(5)Hex(4)Fuc(1) | C[+57], N[+1810] | 1386.2781 | 3 | 4156.8198 | 4156.8081 | 0 | 2.8 | 332 | CRagged | 822.74 | 799 | 531.13 | 16.74 | 60 | >sp\|P01876\|IGHA1_HUMAN Immunoglobulin heavy constant alpha 1 OS=Homo sapiens OX=9606 GN=IGHA1 PE=1 SV=2 | 255 | 20201118_12833_1ng_glyco_hcd203040_compound15.14374.14374.3 | controllerType=0 controllerNumber=1 scan=14374 | 30.647 |
| 10951:3 | 9 | R.LAGKPTHVN[+1702.581]VSVVMAEVDGTC[+57.021]Y.- | HexNAc(2)Hex(8) | C[+57], N[+1703] | 1350.5814 | 3 | 4049.7295 | 4049.7234 | 0 | 1.5 | 332 | CRagged | 837.02 | 803.58 | 496.96 | 16.71 | 60 | >sp\|P01876\|IGHA1_HUMAN Immunoglobulin heavy constant alpha 1 OS=Homo sapiens OX=9606 GN=IGHA1 PE=1 SV=2 | 255 | 20201118_12833_1ng_glyco_hcd203040_compound15.14251.14251.3 | controllerType=0 controllerNumber=1 scan=14251 | 30.4111 |
| 11040:3 | 9 | R.LAGKPTHVN[+1378.476]VSVVMAEVDGTC[+57.021]Y.- | HexNAc(2)Hex(6) | C[+57], N[+1378] | 1242.5466 | 3 | 3725.6251 | 3725.6178 | 0 | 2 | 332 | CRagged | 845.44 | 823.31 | 525.6 | 16.71 | 60 | >sp\|P01876\|IGHA1_HUMAN Immunoglobulin heavy constant alpha 1 OS=Homo sapiens OX=9606 GN=IGHA1 PE=1 SV=2 | 255 | 20201118_12833_1ng_glyco_hcd203040_compound15.14344.14344.3 | controllerType=0 controllerNumber=1 scan=14344 | 30.5941 |
| 11031:3 | 9 | R.LAGKPTHVN[+1663.608]VSVVMAEVDGTC[+57.021]Y.- | HexNAc(5)Hex(4) | C[+57], N[+1664] | 1337.5909 | 3 | 4010.7581 | 4010.7502 | 0 | 2 | 332 | CRagged | 859.61 | 853.88 | 556.11 | 17.71 | 60 | >sp\|P01876\|IGHA1_HUMAN Immunoglobulin heavy constant alpha 1 OS=Homo sapiens OX=9606 GN=IGHA1 PE=1 SV=2 | 255 | 20201118_12833_1ng_glyco_hcd203040_compound15.14334.14334.3 | controllerType=0 controllerNumber=1 scan=14334 | 30.5678 |
| 11075:3 | 9 | R.LAGKPTHVN[+1216.423]VSVVMAEVDGTC[+57.021]Y.- | HexNAc(2)Hex(5) | C[+57], N[+1216] | 1188.5296 | 3 | 3563.5742 | 3563.5649 | 0 | 2.6 | 332 | CRagged | 892.24 | 877.66 | 541.06 | 17.74 | 60 | >sp\|P01876\|IGHA1_HUMAN Immunoglobulin heavy constant alpha 1 OS=Homo sapiens OX=9606 GN=IGHA1 PE=1 SV=2 | 255 | 20201118_12833_1ng_glyco_hcd203040_compound15.14380.14380.3 | controllerType=0 controllerNumber=1 scan=14380 | 30.6663 |
| 10949:3 | 9 | R.LAGKPTHVN[+1702.581]VSVVMAEVDGTC[+57.021]Y.- | HexNAc(2)Hex(8) | C[+57], N[+1703] | 1350.5814 | 3 | 4049.7295 | 4049.7234 | 0 | 1.5 | 332 | CRagged | 905.7 | 905.7 | 596.8 | 18.71 | 60 | >sp\|P01876\|IGHA1_HUMAN Immunoglobulin heavy constant alpha 1 OS=Homo sapiens OX=9606 GN=IGHA1 PE=1 SV=2 | 255 | 20201118_12833_1ng_glyco_hcd203040_compound15.14249.14249.3 | controllerType=0 controllerNumber=1 scan=14249 | 30.401 |
| 11071:3 | 9 | R.LAGKPTHVN[+1216.423]VSVVMAEVDGTC[+57.021]Y.- | HexNAc(2)Hex(5) | C[+57], N[+1216] | 1188.5296 | 3 | 3563.5742 | 3563.5649 | 0 | 2.6 | 332 | CRagged | 911.05 | 911.05 | 553.18 | 18.74 | 60 | >sp\|P01876\|IGHA1_HUMAN Immunoglobulin heavy constant alpha 1 OS=Homo sapiens OX=9606 GN=IGHA1 PE=1 SV=2 | 255 | 20201118_12833_1ng_glyco_hcd203040_compound15.14376.14376.3 | controllerType=0 controllerNumber=1 scan=14376 | 30.6495 |
| 11108:3 | 9 | R.LAGKPTHVN[+1647.613]VSVVMAEVDGTC[+57.021]Y.- | HexNAc(5)Hex(3)Fuc(1) | C[+57], N[+1648] | 1332.2608 | 3 | 3994.7678 | 3994.7553 | 0 | 3.1 | 332 | CRagged | 951.37 | 951.37 | 951.37 | 19.74 | 60 | >sp\|P01876\|IGHA1_HUMAN Immunoglobulin heavy constant alpha 1 OS=Homo sapiens OX=9606 GN=IGHA1 PE=1 SV=2 | 255 | 20201118_12833_1ng_glyco_hcd203040_compound15.14415.14415.3 | controllerType=0 controllerNumber=1 scan=14415 | 30.7457 |
| 08396:3 | 9 | R.LAGKPTHVN[+1809.666]VSVVM[+15.995]AEVDGTC[+57.021]Y.- | HexNAc(5)Hex(4)Fuc(1) | C[+57], M[+16], N[+1810] | 1391.6069 | 3 | 4172.8061 | 4172.8031 | 0 | 0.7 | 332 | CRagged | 201.67 | 201.67 | 165.65 | 3.85 | 60 | >sp\|P01876\|IGHA1_HUMAN Immunoglobulin heavy constant alpha 1 OS=Homo sapiens OX=9606 GN=IGHA1 PE=1 SV=2 | 255 | 20201118_12833_1ng_glyco_hcd203040_compound15.11590.11590.3 | controllerType=0 controllerNumber=1 scan=11590 | 25.8952 |
| 08452:3 | 9 | R.LAGKPTHVN[+1216.423]VSVVM[+15.995]AEVDGTC[+57.021]Y.- | HexNAc(2)Hex(5) | C[+57], M[+16], N[+1216] | 1193.8597 | 3 | 3579.5645 | 3579.5599 | 0 | 1.3 | 332 | CRagged | 273.07 | 183.09 | 183.09 | 6.58 | 60 | >sp\|P01876\|IGHA1_HUMAN Immunoglobulin heavy constant alpha 1 OS=Homo sapiens OX=9606 GN=IGHA1 PE=1 SV=2 | 255 | 20201118_12833_1ng_glyco_hcd203040_compound15.11648.11648.3 | controllerType=0 controllerNumber=1 scan=11648 | 25.9974 |
| 08580:3 | 9 | R.LAGKPTHVN[+1216.423]VSVVM[+15.995]AEVDGTC[+57.021]Y.- | HexNAc(2)Hex(5) | C[+57], M[+16], N[+1216] | 1193.8615 | 3 | 3579.57 | 3579.5599 | 0 | 2.8 | 332 | CRagged | 362.91 | 362.91 | 362.91 | 8.23 | 60 | >sp\|P01876\|IGHA1_HUMAN Immunoglobulin heavy constant alpha 1 OS=Homo sapiens OX=9606 GN=IGHA1 PE=1 SV=2 | 255 | 20201118_12833_1ng_glyco_hcd203040_compound15.11782.11782.3 | controllerType=0 controllerNumber=1 scan=11782 | 26.2223 |
| 08582:3 | 9 | R.LAGKPTHVN[+1216.423]VSVVM[+15.995]AEVDGTC[+57.021]Y.- | HexNAc(2)Hex(5) | C[+57], M[+16], N[+1216] | 1193.8615 | 3 | 3579.57 | 3579.5599 | 0 | 2.8 | 332 | CRagged | 499.62 | 492.99 | 492.99 | 10.25 | 60 | >sp\|P01876\|IGHA1_HUMAN Immunoglobulin heavy constant alpha 1 OS=Homo sapiens OX=9606 GN=IGHA1 PE=1 SV=2 | 255 | 20201118_12833_1ng_glyco_hcd203040_compound15.11784.11784.3 | controllerType=0 controllerNumber=1 scan=11784 | 26.2261 |
| 18610:4 | 10 | R.NIAAFGGDPNN[+1913.677]ITLFGESAGGASVSLQTLSPYNK.G | HexNAc(4)Hex(5)NeuAc(1) | N[+1914] | 1331.8505 | 4 | 5324.38 | 5324.3633 | 0 | 3.1 | 197 | Specific | 938.06 | 938.06 | 938.06 | 19.68 | 54 | >sp\|P19835\|CEL_HUMAN Bile salt-activated lipase OS=Homo sapiens OX=9606 GN=CEL PE=1 SV=3 | 476 | 20201118_12833_1ng_glyco_hcd203040_compound15.22269.22269.4 | controllerType=0 controllerNumber=1 scan=22269 | 45.417 |
| 18608:4 | 10 | R.NIAAFGGDPNN[+1913.677]ITLFGESAGGASVSLQTLSPYNK.G | HexNAc(4)Hex(5)NeuAc(1) | N[+1914] | 1331.8505 | 4 | 5324.38 | 5324.3633 | 0 | 3.1 | 197 | Specific | 1013.98 | 962.84 | 962.84 | 21.68 | 54 | >sp\|P19835\|CEL_HUMAN Bile salt-activated lipase OS=Homo sapiens OX=9606 GN=CEL PE=1 SV=3 | 476 | 20201118_12833_1ng_glyco_hcd203040_compound15.22267.22267.4 | controllerType=0 controllerNumber=1 scan=22267 | 45.4063 |
| 05324:3 | 12 | R.LSPN[+2205.793]ASAEHLELR.W | HexNAc(4)Hex(5)Fuc(2)NeuAc(1) | N[+2206] | 1214.8554 | 3 | 3642.5517 | 3642.5421 | 0 | 2.6 | 52 | Specific | 570.89 | 420.36 | 420.36 | 9.97 | 40 | >sp\|Q13410\|BT1A1_HUMAN Butyrophilin subfamily 1 member A1 OS=Homo sapiens OX=9606 GN=BTN1A1 PE=1 SV=3 | 179 | 20201118_12833_1ng_glyco_hcd203040_compound15.8405.8405.3 | controllerType=0 controllerNumber=1 scan=8405 | 20.5886 |
| 05322:3 | 12 | R.LSPN[+2205.793]ASAEHLELR.W | HexNAc(4)Hex(5)Fuc(2)NeuAc(1) | N[+2206] | 1214.8554 | 3 | 3642.5517 | 3642.5421 | 0 | 2.6 | 52 | Specific | 584.92 | 543.7 | 543.7 | 11.15 | 40 | >sp\|Q13410\|BT1A1_HUMAN Butyrophilin subfamily 1 member A1 OS=Homo sapiens OX=9606 GN=BTN1A1 PE=1 SV=3 | 179 | 20201118_12833_1ng_glyco_hcd203040_compound15.8403.8403.3 | controllerType=0 controllerNumber=1 scan=8403 | 20.5837 |
| 07339:3 | 12 | R.LSPN[+2350.830]ASAEHLELR.W | HexNAc(4)Hex(5)Fuc(1)NeuAc(2) | N[+2351] | 1263.2006 | 3 | 3787.5873 | 3787.5796 | 0 | 2 | 52 | Specific | 602.1 | 545.43 | 545.43 | 12.12 | 40 | >sp\|Q13410\|BT1A1_HUMAN Butyrophilin subfamily 1 member A1 OS=Homo sapiens OX=9606 GN=BTN1A1 PE=1 SV=3 | 179 | 20201118_12833_1ng_glyco_hcd203040_compound15.10492.10492.3 | controllerType=0 controllerNumber=1 scan=10492 | 24.0523 |
| 07337:3 | 12 | R.LSPN[+2350.830]ASAEHLELR.W | HexNAc(4)Hex(5)Fuc(1)NeuAc(2) | N[+2351] | 1263.2006 | 3 | 3787.5873 | 3787.5796 | 0 | 2 | 52 | Specific | 666.63 | 627.69 | 627.69 | 13.12 | 40 | >sp\|Q13410\|BT1A1_HUMAN Butyrophilin subfamily 1 member A1 OS=Homo sapiens OX=9606 GN=BTN1A1 PE=1 SV=3 | 179 | 20201118_12833_1ng_glyco_hcd203040_compound15.10490.10490.3 | controllerType=0 controllerNumber=1 scan=10490 | 24.0478 |
| 26652:4 | 15 | R.QLAHQSN[+2861.000]STNIFFSPVSIATAFAMLSLGTK.A | HexNAc(5)Hex(6)NeuAc(3) | N[+2861] | 1511.4199 | 4 | 6042.6579 | 6042.6351 | 0 | 3.8 | 64 | Specific | 321.66 | 321.66 | 321.66 | 8.47 | 33 | >sp\|P01009\|A1AT_HUMAN Alpha-1-antitrypsin OS=Homo sapiens OX=9606 GN=SERPINA1 PE=1 SV=3 | 642 | 20201118_12833_1ng_glyco_hcd203040_compound15.30640.30640.2 | controllerType=0 controllerNumber=1 scan=30640 | 61.3058 |
| 23755:3 | 15 | R.QLAHQSN[+2204.772]STNIFFSPVSIATAFAMLSLGTK.A | HexNAc(4)Hex(5)NeuAc(2) | N[+2205] | 1796.1491 | 3 | 5386.4328 | 5386.4075 | 0 | 4.7 | 64 | Specific | 488.49 | 478.43 | 478.43 | 10.3 | 33 | >sp\|P01009\|A1AT_HUMAN Alpha-1-antitrypsin OS=Homo sapiens OX=9606 GN=SERPINA1 PE=1 SV=3 | 642 | 20201118_12833_1ng_glyco_hcd203040_compound15.27634.27634.3 | controllerType=0 controllerNumber=1 scan=27634 | 55.9138 |
| 23757:3 | 15 | R.QLAHQSN[+2204.772]STNIFFSPVSIATAFAMLSLGTK.A | HexNAc(4)Hex(5)NeuAc(2) | N[+2205] | 1796.1491 | 3 | 5386.4328 | 5386.4075 | 0 | 4.7 | 64 | Specific | 624.27 | 563 | 563 | 12.85 | 33 | >sp\|P01009\|A1AT_HUMAN Alpha-1-antitrypsin OS=Homo sapiens OX=9606 GN=SERPINA1 PE=1 SV=3 | 642 | 20201118_12833_1ng_glyco_hcd203040_compound15.27636.27636.3 | controllerType=0 controllerNumber=1 scan=27636 | 55.9204 |
| 23756:4 | 15 | R.QLAHQSN[+2204.772]STNIFFSPVSIATAFAMLSLGTK.A | HexNAc(4)Hex(5)NeuAc(2) | N[+2205] | 1347.3613 | 4 | 5386.4235 | 5386.4075 | 0 | 3 | 64 | Specific | 873.69 | 818.74 | 580.03 | 18.22 | 33 | >sp\|P01009\|A1AT_HUMAN Alpha-1-antitrypsin OS=Homo sapiens OX=9606 GN=SERPINA1 PE=1 SV=3 | 642 | 20201118_12833_1ng_glyco_hcd203040_compound15.27635.27635.4 | controllerType=0 controllerNumber=1 scan=27635 | 55.9185 |
| 23754:4 | 15 | R.QLAHQSN[+2204.772]STNIFFSPVSIATAFAMLSLGTK.A | HexNAc(4)Hex(5)NeuAc(2) | N[+2205] | 1347.3613 | 4 | 5386.4235 | 5386.4075 | 0 | 3 | 64 | Specific | 896.35 | 837.4 | 582.21 | 18.22 | 33 | >sp\|P01009\|A1AT_HUMAN Alpha-1-antitrypsin OS=Homo sapiens OX=9606 GN=SERPINA1 PE=1 SV=3 | 642 | 20201118_12833_1ng_glyco_hcd203040_compound15.27633.27633.4 | controllerType=0 controllerNumber=1 scan=27633 | 55.9123 |
| 22757:4 | 15 | R.QLAHQSN[+2204.772]STNIFFSPVSIATAFAM[+15.995]LSLGTK.A | HexNAc(4)Hex(5)NeuAc(2) | M[+16], N[+2205] | 1351.3561 | 4 | 5402.4025 | 5402.4024 | 0 | 0 | 64 | Specific | 342.21 | 342.21 | 342.21 | 7.94 | 33 | >sp\|P01009\|A1AT_HUMAN Alpha-1-antitrypsin OS=Homo sapiens OX=9606 GN=SERPINA1 PE=1 SV=3 | 642 | 20201118_12833_1ng_glyco_hcd203040_compound15.26598.26598.4 | controllerType=0 controllerNumber=1 scan=26598 | 53.9901 |
| 22759:4 | 15 | R.QLAHQSN[+2204.772]STNIFFSPVSIATAFAM[+15.995]LSLGTK.A | HexNAc(4)Hex(5)NeuAc(2) | M[+16], N[+2205] | 1351.3561 | 4 | 5402.4025 | 5402.4024 | 0 | 0 | 64 | Specific | 426.47 | 426.47 | 426.47 | 9.93 | 33 | >sp\|P01009\|A1AT_HUMAN Alpha-1-antitrypsin OS=Homo sapiens OX=9606 GN=SERPINA1 PE=1 SV=3 | 642 | 20201118_12833_1ng_glyco_hcd203040_compound15.26600.26600.4 | controllerType=0 controllerNumber=1 scan=26600 | 53.9936 |
| 19673:5 | 15 | K.YLGN[+2204.772]ATAIFFLPDEGKLQHLENELTHDIITK.F | HexNAc(4)Hex(5)NeuAc(2) | N[+2205] | 1149.9294 | 5 | 5745.6181 | 5745.6098 | 0 | 1.5 | 268 | Specific | 565.9 | 565.9 | 565.9 | 10.92 | 33 | >sp\|P01009\|A1AT_HUMAN Alpha-1-antitrypsin OS=Homo sapiens OX=9606 GN=SERPINA1 PE=1 SV=3 | 642 | 20201118_12833_1ng_glyco_hcd203040_compound15.23382.23382.5 | controllerType=0 controllerNumber=1 scan=23382 | 47.7436 |
| 19672:4 | 15 | K.YLGN[+2204.772]ATAIFFLPDEGKLQHLENELTHDIITK.F | HexNAc(4)Hex(5)NeuAc(2) | N[+2205] | 1437.1625 | 4 | 5745.6281 | 5745.6098 | 0 | 3.2 | 268 | Specific | 627.47 | 627.47 | 627.47 | 12.68 | 33 | >sp\|P01009\|A1AT_HUMAN Alpha-1-antitrypsin OS=Homo sapiens OX=9606 GN=SERPINA1 PE=1 SV=3 | 642 | 20201118_12833_1ng_glyco_hcd203040_compound15.23381.23381.4 | controllerType=0 controllerNumber=1 scan=23381 | 47.7423 |
| 19675:5 | 15 | K.YLGN[+2204.772]ATAIFFLPDEGKLQHLENELTHDIITK.F | HexNAc(4)Hex(5)NeuAc(2) | N[+2205] | 1149.9294 | 5 | 5745.6181 | 5745.6098 | 0 | 1.5 | 268 | Specific | 641.06 | 585.71 | 585.71 | 13.11 | 33 | >sp\|P01009\|A1AT_HUMAN Alpha-1-antitrypsin OS=Homo sapiens OX=9606 GN=SERPINA1 PE=1 SV=3 | 642 | 20201118_12833_1ng_glyco_hcd203040_compound15.23384.23384.5 | controllerType=0 controllerNumber=1 scan=23384 | 47.7483 |
| 19674:4 | 15 | K.YLGN[+2204.772]ATAIFFLPDEGKLQHLENELTHDIITK.F | HexNAc(4)Hex(5)NeuAc(2) | N[+2205] | 1437.1625 | 4 | 5745.6281 | 5745.6098 | 0 | 3.2 | 268 | Specific | 935.6 | 863.12 | 863.12 | 18.68 | 33 | >sp\|P01009\|A1AT_HUMAN Alpha-1-antitrypsin OS=Homo sapiens OX=9606 GN=SERPINA1 PE=1 SV=3 | 642 | 20201118_12833_1ng_glyco_hcd203040_compound15.23383.23383.4 | controllerType=0 controllerNumber=1 scan=23383 | 47.7464 |
| 09149:4 | 17 | K.[+42.011]QTDEIKDTRN[+2018.708]ESTQNC[+57.021]VVAEPEKMESSISSSSEEMSLS | HexNAc(3)Hex(6)Fuc(1)NeuAc(1) | .[42], C[+57], N[+2019] | 1613.6846 | 4 | 6451.7166 | 6451.6996 | 0 | 2.6 | 60 | Specific | 272.85 | 159.07 | 159.07 | 3.76 | 30 | >sp\|P47710\|CASA1_HUMAN Alpha-S1-casein OS=Homo sapiens OX=9606 GN=CSN1S1 PE=1 SV=1 | 989 | 20201118_12833_1ng_glyco_hcd203040_compound15.12384.12384.4 | controllerType=0 controllerNumber=1 scan=12384 | 27.2721 |
| 09147:4 | 17 | K.[+42.011]QTDEIKDTRN[+2018.708]ESTQNC[+57.021]VVAEPEKMESSISSSSEEMSLS | HexNAc(3)Hex(6)Fuc(1)NeuAc(1) | .[42], C[+57], N[+2019] | 1613.6846 | 4 | 6451.7166 | 6451.6996 | 0 | 2.6 | 60 | Specific | 321.46 | 275.63 | 275.63 | 6.62 | 30 | >sp\|P47710\|CASA1_HUMAN Alpha-S1-casein OS=Homo sapiens OX=9606 GN=CSN1S1 PE=1 SV=1 | 989 | 20201118_12833_1ng_glyco_hcd203040_compound15.12382.12382.4 | controllerType=0 controllerNumber=1 scan=12382 | 27.258 |
| 13645:4 | 19 | K.LKELPGVC[+57.021]N[+2205.793]ETMMALWEEC[+57.021]KPC[+57.021]LK.Q | HexNAc(4)Hex(5)Fuc(2)NeuAc(1) | C[+57]*3, N[+2206] | 1286.3043 | 4 | 5142.1955 | 5142.1776 | 0 | 3.5 | 95 | Specific | 557.37 | 557.37 | 557.37 | 10.3 | 24 | >sp\|P10909\|CLUS_HUMAN Clusterin OS=Homo sapiens OX=9606 GN=CLU PE=1 SV=1 | 1467 | 20201118_12833_1ng_glyco_hcd203040_compound15.17064.17064.4 | controllerType=0 controllerNumber=1 scan=17064 | 35.3356 |
| 13643:4 | 19 | K.LKELPGVC[+57.021]N[+2205.793]ETMMALWEEC[+57.021]KPC[+57.021]LK.Q | HexNAc(4)Hex(5)Fuc(2)NeuAc(1) | C[+57]*3, N[+2206] | 1286.3043 | 4 | 5142.1955 | 5142.1776 | 0 | 3.5 | 95 | Specific | 569.19 | 569.19 | 569.19 | 10.3 | 24 | >sp\|P10909\|CLUS_HUMAN Clusterin OS=Homo sapiens OX=9606 GN=CLU PE=1 SV=1 | 1467 | 20201118_12833_1ng_glyco_hcd203040_compound15.17062.17062.4 | controllerType=0 controllerNumber=1 scan=17062 | 35.3318 |
| 13616:4 | 19 | K.LKELPGVC[+57.021]N[+2205.793]ETMMALWEEC[+57.021]KPC[+57.021]LK.Q | HexNAc(4)Hex(5)Fuc(2)NeuAc(1) | C[+57]*3, N[+2206] | 1286.3037 | 4 | 5142.193 | 5142.1776 | 0 | 3 | 95 | Specific | 602.93 | 576.63 | 576.63 | 12.48 | 24 | >sp\|P10909\|CLUS_HUMAN Clusterin OS=Homo sapiens OX=9606 GN=CLU PE=1 SV=1 | 1467 | 20201118_12833_1ng_glyco_hcd203040_compound15.17034.17034.4 | controllerType=0 controllerNumber=1 scan=17034 | 35.284 |
| 13614:4 | 19 | K.LKELPGVC[+57.021]N[+2205.793]ETMMALWEEC[+57.021]KPC[+57.021]LK.Q | HexNAc(4)Hex(5)Fuc(2)NeuAc(1) | C[+57]*3, N[+2206] | 1286.3037 | 4 | 5142.193 | 5142.1776 | 0 | 3 | 95 | Specific | 666.57 | 659.46 | 659.46 | 13.48 | 24 | >sp\|P10909\|CLUS_HUMAN Clusterin OS=Homo sapiens OX=9606 GN=CLU PE=1 SV=1 | 1467 | 20201118_12833_1ng_glyco_hcd203040_compound15.17032.17032.4 | controllerType=0 controllerNumber=1 scan=17032 | 35.2799 |
| 13654:4 | 19 | K.LKELPGVC[+57.021]N[+2059.735]ETMMALWEEC[+57.021]KPC[+57.021]LK.Q | HexNAc(4)Hex(5)Fuc(1)NeuAc(1) | C[+57]*3, N[+2060] | 1249.7896 | 4 | 4996.1364 | 4996.1196 | 0 | 3.4 | 95 | Specific | 748.85 | 748.85 | 748.85 | 14.48 | 24 | >sp\|P10909\|CLUS_HUMAN Clusterin OS=Homo sapiens OX=9606 GN=CLU PE=1 SV=1 | 1467 | 20201118_12833_1ng_glyco_hcd203040_compound15.17073.17073.2 | controllerType=0 controllerNumber=1 scan=17073 | 35.3531 |
| 13656:4 | 19 | K.LKELPGVC[+57.021]N[+2059.735]ETMMALWEEC[+57.021]KPC[+57.021]LK.Q | HexNAc(4)Hex(5)Fuc(1)NeuAc(1) | C[+57]*3, N[+2060] | 1249.7896 | 4 | 4996.1364 | 4996.1196 | 0 | 3.4 | 95 | Specific | 841.37 | 744.43 | 744.43 | 16.48 | 24 | >sp\|P10909\|CLUS_HUMAN Clusterin OS=Homo sapiens OX=9606 GN=CLU PE=1 SV=1 | 1467 | 20201118_12833_1ng_glyco_hcd203040_compound15.17075.17075.2 | controllerType=0 controllerNumber=1 scan=17075 | 35.3574 |
| 23367:3 | 19 | K.MLN[+2205.793]TSSLLEQLNEQFNWVSR.L | HexNAc(4)Hex(5)Fuc(2)NeuAc(1) | N[+2206] | 1538.9991 | 3 | 4614.9827 | 4614.9796 | 0 | 0.7 | 352 | Specific | 233.31 | 233.31 | 233.31 | 5.7 | 24 | >sp\|P10909\|CLUS_HUMAN Clusterin OS=Homo sapiens OX=9606 GN=CLU PE=1 SV=1 | 1467 | 20201118_12833_1ng_glyco_hcd203040_compound15.27231.27231.3 | controllerType=0 controllerNumber=1 scan=27231 | 55.1769 |
| 23369:3 | 19 | K.MLN[+2205.793]TSSLLEQLNEQFNWVSR.L | HexNAc(4)Hex(5)Fuc(2)NeuAc(1) | N[+2206] | 1538.9991 | 3 | 4614.9827 | 4614.9796 | 0 | 0.7 | 352 | Specific | 334.25 | 318.27 | 318.27 | 6.75 | 24 | >sp\|P10909\|CLUS_HUMAN Clusterin OS=Homo sapiens OX=9606 GN=CLU PE=1 SV=1 | 1467 | 20201118_12833_1ng_glyco_hcd203040_compound15.27233.27233.3 | controllerType=0 controllerNumber=1 scan=27233 | 55.1808 |
| 22950:4 | 19 | K.MLN[+2350.830]TSSLLEQLNEQFNWVSR.L | HexNAc(4)Hex(5)Fuc(1)NeuAc(2) | N[+2351] | 1190.7664 | 4 | 4760.0438 | 4760.0171 | 0 | 5.6 | 352 | Specific | 480.54 | 380.25 | 380.25 | 9.31 | 24 | >sp\|P10909\|CLUS_HUMAN Clusterin OS=Homo sapiens OX=9606 GN=CLU PE=1 SV=1 | 1467 | 20201118_12833_1ng_glyco_hcd203040_compound15.26799.26799.4 | controllerType=0 controllerNumber=1 scan=26799 | 54.3802 |
| 22981:4 | 19 | K.MLN[+2350.830]TSSLLEQLNEQFNWVSR.L | HexNAc(4)Hex(5)Fuc(1)NeuAc(2) | N[+2351] | 1190.7662 | 4 | 4760.0428 | 4760.0171 | 0 | 5.4 | 352 | Specific | 633.6 | 411.62 | 411.62 | 11.86 | 24 | >sp\|P10909\|CLUS_HUMAN Clusterin OS=Homo sapiens OX=9606 GN=CLU PE=1 SV=1 | 1467 | 20201118_12833_1ng_glyco_hcd203040_compound15.26831.26831.4 | controllerType=0 controllerNumber=1 scan=26831 | 54.4425 |
| 22979:4 | 19 | K.MLN[+2350.830]TSSLLEQLNEQFNWVSR.L | HexNAc(4)Hex(5)Fuc(1)NeuAc(2) | N[+2351] | 1190.7662 | 4 | 4760.0428 | 4760.0171 | 0 | 5.4 | 352 | Specific | 636.76 | 636.76 | 636.76 | 11.86 | 24 | >sp\|P10909\|CLUS_HUMAN Clusterin OS=Homo sapiens OX=9606 GN=CLU PE=1 SV=1 | 1467 | 20201118_12833_1ng_glyco_hcd203040_compound15.26829.26829.4 | controllerType=0 controllerNumber=1 scan=26829 | 54.4382 |
| 22975:3 | 19 | K.MLN[+2350.830]TSSLLEQLNEQFNWVSR.L | HexNAc(4)Hex(5)Fuc(1)NeuAc(2) | N[+2351] | 1587.3542 | 3 | 4760.048 | 4760.0171 | 0 | 6.5 | 352 | Specific | 679.42 | 558.84 | 558.84 | 12.23 | 24 | >sp\|P10909\|CLUS_HUMAN Clusterin OS=Homo sapiens OX=9606 GN=CLU PE=1 SV=1 | 1467 | 20201118_12833_1ng_glyco_hcd203040_compound15.26825.26825.3 | controllerType=0 controllerNumber=1 scan=26825 | 54.4298 |
| 22977:3 | 19 | K.MLN[+2350.830]TSSLLEQLNEQFNWVSR.L | HexNAc(4)Hex(5)Fuc(1)NeuAc(2) | N[+2351] | 1587.3542 | 3 | 4760.048 | 4760.0171 | 0 | 6.5 | 352 | Specific | 723.35 | 467.22 | 467.22 | 13.23 | 24 | >sp\|P10909\|CLUS_HUMAN Clusterin OS=Homo sapiens OX=9606 GN=CLU PE=1 SV=1 | 1467 | 20201118_12833_1ng_glyco_hcd203040_compound15.26827.26827.3 | controllerType=0 controllerNumber=1 scan=26827 | 54.4336 |
| 22952:4 | 19 | K.MLN[+2350.830]TSSLLEQLNEQFNWVSR.L | HexNAc(4)Hex(5)Fuc(1)NeuAc(2) | N[+2351] | 1190.7664 | 4 | 4760.0438 | 4760.0171 | 0 | 5.6 | 352 | Specific | 779.11 | 585.83 | 585.83 | 14.86 | 24 | >sp\|P10909\|CLUS_HUMAN Clusterin OS=Homo sapiens OX=9606 GN=CLU PE=1 SV=1 | 1467 | 20201118_12833_1ng_glyco_hcd203040_compound15.26801.26801.4 | controllerType=0 controllerNumber=1 scan=26801 | 54.3846 |
| 18369:3 | 23 | R.C[+57.021]MWSSALNSLN[+1864.634]LSFAGLEQVPK.G | HexNAc(2)Hex(9) | C[+57], N[+1865] | 1439.6213 | 3 | 4316.8493 | 4316.8341 | 0 | 3.5 | 272 | Specific | 488.18 | 487.45 | 487.45 | 10.02 | 19 | >sp\|P08571\|CD14_HUMAN Monocyte differentiation antigen CD14 OS=Homo sapiens OX=9606 GN=CD14 PE=1 SV=2 | 1232 | 20201118_12833_1ng_glyco_hcd203040_compound15.22016.22016.3 | controllerType=0 controllerNumber=1 scan=22016 | 44.7983 |
| 18371:3 | 23 | R.C[+57.021]MWSSALNSLN[+1864.634]LSFAGLEQVPK.G | HexNAc(2)Hex(9) | C[+57], N[+1865] | 1439.6213 | 3 | 4316.8493 | 4316.8341 | 0 | 3.5 | 272 | Specific | 645.87 | 622.29 | 622.29 | 12.56 | 19 | >sp\|P08571\|CD14_HUMAN Monocyte differentiation antigen CD14 OS=Homo sapiens OX=9606 GN=CD14 PE=1 SV=2 | 1232 | 20201118_12833_1ng_glyco_hcd203040_compound15.22018.22018.3 | controllerType=0 controllerNumber=1 scan=22018 | 44.8019 |
| 09294:3 | 30 | R.IIVPLNNREN[+1460.529]ISDPTSPLR.T | HexNAc(4)Hex(4) | N[+1461] | 1203.5777 | 3 | 3608.7185 | 3608.7059 | 0 | 3.5 | 62 | Specific | 241.76 | 241.76 | 241.76 | 6.19 | 21 | >sp\|P01591\|IGJ_HUMAN Immunoglobulin J chain OS=Homo sapiens OX=9606 GN=JCHAIN PE=1 SV=4 | 1392 | 20201118_12833_1ng_glyco_hcd203040_compound15.12536.12536.3 | controllerType=0 controllerNumber=1 scan=12536 | 27.5355 |
| 09296:3 | 30 | R.IIVPLNNREN[+1460.529]ISDPTSPLR.T | HexNAc(4)Hex(4) | N[+1461] | 1203.5777 | 3 | 3608.7185 | 3608.7059 | 0 | 3.5 | 62 | Specific | 248.71 | 248.71 | 248.71 | 6.19 | 21 | >sp\|P01591\|IGJ_HUMAN Immunoglobulin J chain OS=Homo sapiens OX=9606 GN=JCHAIN PE=1 SV=4 | 1392 | 20201118_12833_1ng_glyco_hcd203040_compound15.12538.12538.3 | controllerType=0 controllerNumber=1 scan=12538 | 27.5395 |
| 11122:3 | 30 | R.IIVPLNNREN[+1694.603]ISDPTSPLR.T | HexNAc(3)Hex(4)Fuc(1)NeuAc(1) | N[+1695] | 1281.602 | 3 | 3842.7914 | 3842.7799 | 0 | 3 | 62 | Specific | 328.6 | 232.76 | 232.76 | 6.69 | 21 | >sp\|P01591\|IGJ_HUMAN Immunoglobulin J chain OS=Homo sapiens OX=9606 GN=JCHAIN PE=1 SV=4 | 1392 | 20201118_12833_1ng_glyco_hcd203040_compound15.14429.14429.3 | controllerType=0 controllerNumber=1 scan=14429 | 30.7755 |
| 12974:3 | 30 | R.IIVPLNNREN[+2204.772]ISDPTSPLR.T | HexNAc(4)Hex(5)NeuAc(2) | N[+2205] | 1451.6587 | 3 | 4352.9617 | 4352.9496 | 0 | 2.8 | 62 | Specific | 330.32 | 330.32 | 330.32 | 7.38 | 21 | >sp\|P01591\|IGJ_HUMAN Immunoglobulin J chain OS=Homo sapiens OX=9606 GN=JCHAIN PE=1 SV=4 | 1392 | 20201118_12833_1ng_glyco_hcd203040_compound15.16369.16369.3 | controllerType=0 controllerNumber=1 scan=16369 | 34.1058 |
| 10941:3 | 30 | R.IIVPLNNREN[+2059.735]ISDPTSPLR.T | HexNAc(4)Hex(5)Fuc(1)NeuAc(1) | N[+2060] | 1403.3115 | 3 | 4207.9198 | 4207.9121 | 0 | 1.8 | 62 | Specific | 348.21 | 348.21 | 348.21 | 7.32 | 21 | >sp\|P01591\|IGJ_HUMAN Immunoglobulin J chain OS=Homo sapiens OX=9606 GN=JCHAIN PE=1 SV=4 | 1392 | 20201118_12833_1ng_glyco_hcd203040_compound15.14241.14241.3 | controllerType=0 controllerNumber=1 scan=14241 | 30.384 |
| 09284:3 | 30 | R.IIVPLNNREN[+1663.608]ISDPTSPLR.T | HexNAc(5)Hex(4) | N[+1664] | 1271.2642 | 3 | 3811.7781 | 3811.7853 | 0 | -1.9 | 62 | Specific | 363.78 | 363.78 | 363.78 | 6.58 | 21 | >sp\|P01591\|IGJ_HUMAN Immunoglobulin J chain OS=Homo sapiens OX=9606 GN=JCHAIN PE=1 SV=4 | 1392 | 20201118_12833_1ng_glyco_hcd203040_compound15.12526.12526.3 | controllerType=0 controllerNumber=1 scan=12526 | 27.5188 |
| 11120:3 | 30 | R.IIVPLNNREN[+1694.603]ISDPTSPLR.T | HexNAc(3)Hex(4)Fuc(1)NeuAc(1) | N[+1695] | 1281.602 | 3 | 3842.7914 | 3842.7799 | 0 | 3 | 62 | Specific | 371.87 | 371.87 | 371.87 | 7.61 | 21 | >sp\|P01591\|IGJ_HUMAN Immunoglobulin J chain OS=Homo sapiens OX=9606 GN=JCHAIN PE=1 SV=4 | 1392 | 20201118_12833_1ng_glyco_hcd203040_compound15.14427.14427.3 | controllerType=0 controllerNumber=1 scan=14427 | 30.7695 |
| 09286:3 | 30 | R.IIVPLNNREN[+1663.608]ISDPTSPLR.T | HexNAc(5)Hex(4) | N[+1664] | 1271.2642 | 3 | 3811.7781 | 3811.7853 | 0 | -1.9 | 62 | Specific | 425.33 | 425.33 | 425.33 | 8.31 | 21 | >sp\|P01591\|IGJ_HUMAN Immunoglobulin J chain OS=Homo sapiens OX=9606 GN=JCHAIN PE=1 SV=4 | 1392 | 20201118_12833_1ng_glyco_hcd203040_compound15.12528.12528.3 | controllerType=0 controllerNumber=1 scan=12528 | 27.523 |
| 12976:3 | 30 | R.IIVPLNNREN[+2204.772]ISDPTSPLR.T | HexNAc(4)Hex(5)NeuAc(2) | N[+2205] | 1451.6587 | 3 | 4352.9617 | 4352.9496 | 0 | 2.8 | 62 | Specific | 441.53 | 409.03 | 409.03 | 9.36 | 21 | >sp\|P01591\|IGJ_HUMAN Immunoglobulin J chain OS=Homo sapiens OX=9606 GN=JCHAIN PE=1 SV=4 | 1392 | 20201118_12833_1ng_glyco_hcd203040_compound15.16371.16371.3 | controllerType=0 controllerNumber=1 scan=16371 | 34.111 |
| 10931:3 | 30 | R.IIVPLNNREN[+1913.677]ISDPTSPLR.T | HexNAc(4)Hex(5)NeuAc(1) | N[+1914] | 1354.6269 | 3 | 4061.8661 | 4061.8542 | 0 | 2.9 | 62 | Specific | 450.58 | 400.12 | 400.12 | 9.36 | 21 | >sp\|P01591\|IGJ_HUMAN Immunoglobulin J chain OS=Homo sapiens OX=9606 GN=JCHAIN PE=1 SV=4 | 1392 | 20201118_12833_1ng_glyco_hcd203040_compound15.14230.14230.3 | controllerType=0 controllerNumber=1 scan=14230 | 30.3433 |
| 10943:3 | 30 | R.IIVPLNNREN[+2059.735]ISDPTSPLR.T | HexNAc(4)Hex(5)Fuc(1)NeuAc(1) | N[+2060] | 1403.3115 | 3 | 4207.9198 | 4207.9121 | 0 | 1.8 | 62 | Specific | 458.58 | 414.51 | 414.51 | 9.33 | 21 | >sp\|P01591\|IGJ_HUMAN Immunoglobulin J chain OS=Homo sapiens OX=9606 GN=JCHAIN PE=1 SV=4 | 1392 | 20201118_12833_1ng_glyco_hcd203040_compound15.14243.14243.3 | controllerType=0 controllerNumber=1 scan=14243 | 30.3911 |
| 10929:3 | 30 | R.IIVPLNNREN[+1913.677]ISDPTSPLR.T | HexNAc(4)Hex(5)NeuAc(1) | N[+1914] | 1354.6269 | 3 | 4061.8661 | 4061.8542 | 0 | 2.9 | 62 | Specific | 600.31 | 567.24 | 567.24 | 12.13 | 21 | >sp\|P01591\|IGJ_HUMAN Immunoglobulin J chain OS=Homo sapiens OX=9606 GN=JCHAIN PE=1 SV=4 | 1392 | 20201118_12833_1ng_glyco_hcd203040_compound15.14228.14228.3 | controllerType=0 controllerNumber=1 scan=14228 | 30.3391 |
| 23310:4 | 57 | K.VFPLSLDSTPQDGNVVVAC[+57.021]LVQGFFPQEPLSVTWSESGQN[+2262.814]VTAI | HexNAc(5)Hex(5)Fuc(1)NeuAc(1) | C[+57], N[+2263] | 1760.5537 | 4 | 7039.1929 | 7039.1757 | 0 | 2.4 | 123 | Specific | 227.34 | 206.5 | 206.5 | 7.48 | 17 | >sp\|P0DOX2\|IGA2_HUMAN Immunoglobulin alpha-2 heavy chain OS=Homo sapiens OX=9606 PE=1 SV=2 | 631 | 20201118_12833_1ng_glyco_hcd203040_compound15.27172.27172.2 | controllerType=0 controllerNumber=1 scan=27172 | 55.0663 |
| 02688:2 | 57 | K.TPLTAN[+1501.555]ITK.S | HexNAc(5)Hex(3) | N[+1502] | 1230.5617 | 2 | 2460.1161 | 2460.1121 | 0 | 1.6 | 315 | Specific | 283.94 | 110.42 | 110.42 | 4.87 | 17 | >sp\|P0DOX2\|IGA2_HUMAN Immunoglobulin alpha-2 heavy chain OS=Homo sapiens OX=9606 PE=1 SV=2 | 631 | 20201118_12833_1ng_glyco_hcd203040_compound15.5674.5674.2 | controllerType=0 controllerNumber=1 scan=5674 | 16.0219 |
| 02634:2 | 57 | K.TPLTAN[+1444.534]ITK.S | HexNAc(4)Hex(3)Fuc(1) | N[+1445] | 1202.0507 | 2 | 2403.0941 | 2403.0906 | 0 | 1.5 | 315 | Specific | 363.05 | 156.71 | 156.71 | 5.74 | 17 | >sp\|P0DOX2\|IGA2_HUMAN Immunoglobulin alpha-2 heavy chain OS=Homo sapiens OX=9606 PE=1 SV=2 | 631 | 20201118_12833_1ng_glyco_hcd203040_compound15.5618.5618.2 | controllerType=0 controllerNumber=1 scan=5618 | 15.9233 |
| 02690:2 | 57 | K.TPLTAN[+1501.555]ITK.S | HexNAc(5)Hex(3) | N[+1502] | 1230.5617 | 2 | 2460.1161 | 2460.1121 | 0 | 1.6 | 315 | Specific | 369.97 | 336.43 | 336.43 | 7.31 | 17 | >sp\|P0DOX2\|IGA2_HUMAN Immunoglobulin alpha-2 heavy chain OS=Homo sapiens OX=9606 PE=1 SV=2 | 631 | 20201118_12833_1ng_glyco_hcd203040_compound15.5676.5676.2 | controllerType=0 controllerNumber=1 scan=5676 | 16.026 |
| 02621:3 | 57 | K.TPLTAN[+1809.666]ITK.S | HexNAc(5)Hex(4)Fuc(1) | N[+1810] | 923.4134 | 3 | 2768.2255 | 2768.2228 | 0 | 1 | 315 | Specific | 376.64 | 129.08 | 129.08 | 5.33 | 17 | >sp\|P0DOX2\|IGA2_HUMAN Immunoglobulin alpha-2 heavy chain OS=Homo sapiens OX=9606 PE=1 SV=2 | 631 | 20201118_12833_1ng_glyco_hcd203040_compound15.5605.5605.3 | controllerType=0 controllerNumber=1 scan=5605 | 15.8996 |
| 02623:3 | 57 | K.TPLTAN[+1809.666]ITK.S | HexNAc(5)Hex(4)Fuc(1) | N[+1810] | 923.4134 | 3 | 2768.2255 | 2768.2228 | 0 | 1 | 315 | Specific | 435.82 | 206.05 | 206.05 | 6.41 | 17 | >sp\|P0DOX2\|IGA2_HUMAN Immunoglobulin alpha-2 heavy chain OS=Homo sapiens OX=9606 PE=1 SV=2 | 631 | 20201118_12833_1ng_glyco_hcd203040_compound15.5607.5607.3 | controllerType=0 controllerNumber=1 scan=5607 | 15.9037 |
| 02699:3 | 57 | K.TPLTAN[+1647.613]ITK.S | HexNAc(5)Hex(3)Fuc(1) | N[+1648] | 869.397 | 3 | 2606.1765 | 2606.17 | 0 | 2.5 | 315 | Specific | 443.5 | 145.88 | 145.88 | 5.76 | 17 | >sp\|P0DOX2\|IGA2_HUMAN Immunoglobulin alpha-2 heavy chain OS=Homo sapiens OX=9606 PE=1 SV=2 | 631 | 20201118_12833_1ng_glyco_hcd203040_compound15.5686.5686.3 | controllerType=0 controllerNumber=1 scan=5686 | 16.0452 |
| 02636:2 | 57 | K.TPLTAN[+1444.534]ITK.S | HexNAc(4)Hex(3)Fuc(1) | N[+1445] | 1202.0507 | 2 | 2403.0941 | 2403.0906 | 0 | 1.5 | 315 | Specific | 449.18 | 246.97 | 246.97 | 6.85 | 17 | >sp\|P0DOX2\|IGA2_HUMAN Immunoglobulin alpha-2 heavy chain OS=Homo sapiens OX=9606 PE=1 SV=2 | 631 | 20201118_12833_1ng_glyco_hcd203040_compound15.5620.5620.2 | controllerType=0 controllerNumber=1 scan=5620 | 15.9267 |

Table S4. Cont.

| Query #:z | Protein Rank | Glycopeptide | Glycan | Modification Type(s) | Observed m/z | z | Observed (M+H) | Calc. mass (M+H) | Off-by-x error | Mass error (ppm) | StartPosition | Cleavage | Score | Delta | Delta Mod | \|Log Prob\| | # of unique peptides | Protein Name | Protein DB number | Comment | Scan # | Scan Time |
|---|---|---|---|---|---|---|---|---|---|---|---|---|---|---|---|---|---|---|---|---|---|---|
| 02624:2 | 57 | K.TPLTAN[+1809.666]ITK.S | HexNAc(5)Hex(4)Fuc(1) | N[+1810] | 1384.615 | 2 | 2768.2228 | 2768.2228 | 0 | 0 | 315 | Specific | 562.41 | 258.48 | 258.48 | 6.97 | 17 | >sp\|P0DOX2\|IGA2_HUMAN Immunoglobulin alpha-2 heavy chain OS=Homo sapiens OX=9606 PE=1 SV=2 | 631 | 20201118_12833_1ng_glyco_hcd203040_compound15.5608.5608.2 | controllerType=0 controllerNumber=1 scan=5608 | 15.9065 |
| 02701:3 | 57 | K.TPLTAN[+1647.613]ITK.S | HexNAc(5)Hex(3)Fuc(1) | N[+1648] | 869.397 | 3 | 2606.1765 | 2606.17 | 0 | 2.5 | 315 | Specific | 562.83 | 282.96 | 282.96 | 7.61 | 17 | >sp\|P0DOX2\|IGA2_HUMAN Immunoglobulin alpha-2 heavy chain OS=Homo sapiens OX=9606 PE=1 SV=2 | 631 | 20201118_12833_1ng_glyco_hcd203040_compound15.5688.5688.3 | controllerType=0 controllerNumber=1 scan=5688 | 16.0505 |
| 02677:2 | 57 | K.TPLTAN[+1647.613]ITK.S | HexNAc(5)Hex(3)Fuc(1) | N[+1648] | 1303.5915 | 2 | 2606.1757 | 2606.17 | 0 | 2.2 | 315 | Specific | 593.54 | 323.17 | 323.17 | 9.25 | 17 | >sp\|P0DOX2\|IGA2_HUMAN Immunoglobulin alpha-2 heavy chain OS=Homo sapiens OX=9606 PE=1 SV=2 | 631 | 20201118_12833_1ng_glyco_hcd203040_compound15.5663.5663.2 | controllerType=0 controllerNumber=1 scan=5663 | 16.0044 |
| 02622:2 | 57 | K.TPLTAN[+1809.666]ITK.S | HexNAc(5)Hex(4)Fuc(1) | N[+1810] | 1384.615 | 2 | 2768.2228 | 2768.2228 | 0 | 0 | 315 | Specific | 599.23 | 291.73 | 291.73 | 8.47 | 17 | >sp\|P0DOX2\|IGA2_HUMAN Immunoglobulin alpha-2 heavy chain OS=Homo sapiens OX=9606 PE=1 SV=2 | 631 | 20201118_12833_1ng_glyco_hcd203040_compound15.5606.5606.2 | controllerType=0 controllerNumber=1 scan=5606 | 15.9011 |
| 02675:2 | 57 | K.TPLTAN[+1647.613]ITK.S | HexNAc(5)Hex(3)Fuc(1) | N[+1648] | 1303.5915 | 2 | 2606.1757 | 2606.17 | 0 | 2.2 | 315 | Specific | 615.67 | 341.16 | 341.16 | 10.25 | 17 | >sp\|P0DOX2\|IGA2_HUMAN Immunoglobulin alpha-2 heavy chain OS=Homo sapiens OX=9606 PE=1 SV=2 | 631 | 20201118_12833_1ng_glyco_hcd203040_compound15.5661.5661.2 | controllerType=0 controllerNumber=1 scan=5661 | 15.9991 |
| 16183:3 | 64 | R.NYIVPILWLN[+1864.634]ETGTIGDEKANM[+15.995]FR.S | HexNAc(2)Hex(9) | M[+16], N[+1865] | 1559.0247 | 3 | 4675.0596 | 4675.0523 | 0 | 1.6 | 408 | Specific | 832.04 | 805.19 | 805.19 | 16.45 | 11 | >sp\|P16671\|CD36_HUMAN Platelet glycoprotein 4 OS=Homo sapiens OX=9606 GN=CD36 PE=1 SV=2 | 812 | 20201118_12833_1ng_glyco_hcd203040_compound15.19740.19740.3 | controllerType=0 controllerNumber=1 scan=19740 | 40.0651 |
| 16181:3 | 64 | R.NYIVPILWLN[+1864.634]ETGTIGDEKANM[+15.995]FR.S | HexNAc(2)Hex(9) | M[+16], N[+1865] | 1559.0247 | 3 | 4675.0596 | 4675.0523 | 0 | 1.6 | 408 | Specific | 836.83 | 836.83 | 836.83 | 16.45 | 11 | >sp\|P16671\|CD36_HUMAN Platelet glycoprotein 4 OS=Homo sapiens OX=9606 GN=CD36 PE=1 SV=2 | 812 | 20201118_12833_1ng_glyco_hcd203040_compound15.19738.19738.3 | controllerType=0 controllerNumber=1 scan=19738 | 40.06 |
| 24426:3 | 111 | R.AVN[+2204.772]ITSENLIDDVVSLIR.G | HexNAc(4)Hex(5)NeuAc(2) | N[+2205] | 1392.6266 | 3 | 4175.8654 | 4175.8481 | 0 | 4.1 | 413 | Specific | 475.83 | 428.08 | 428.08 | 9.11 | 6 | >sp\|P02748\|CO9_HUMAN Complement component C9 OS=Homo sapiens OX=9606 GN=C9 PE=1 SV=2 | 1547 | 20201118_12833_1ng_glyco_hcd203040_compound15.28331.28331.3 | controllerType=0 controllerNumber=1 scan=28331 | 57.2078 |
| 24428:3 | 111 | R.AVN[+2204.772]ITSENLIDDVVSLIR.G | HexNAc(4)Hex(5)NeuAc(2) | N[+2205] | 1392.6266 | 3 | 4175.8654 | 4175.8481 | 0 | 4.1 | 413 | Specific | 587.97 | 485.45 | 485.45 | 10.66 | 6 | >sp\|P02748\|CO9_HUMAN Complement component C9 OS=Homo sapiens OX=9606 GN=C9 PE=1 SV=2 | 1547 | 20201118_12833_1ng_glyco_hcd203040_compound15.28333.28333.3 | controllerType=0 controllerNumber=1 scan=28333 | 57.2117 |
| 17982:4 | 1 | R.RTAGWNVPIGTLRPFLN[+2716.983]WTGPPEPIEAAVAR.F | HexNAc(5)Hex(6)Fuc(3)NeuAc(1) | N[+2717] | 1526.7086 | 4 | 6103.8126 | 6103.795 | 0 | 2.9 | 140 | Specific | 209.12 | 209.12 | 209.12 | 5.46 | 135 | >sp\|P02788\|TRFL_HUMAN Lactotransferrin OS=Homo sapiens OX=9606 GN=LTF PE=1 SV=6 | 1270 | 20201114_13813_1ng_glyco_hcd203040_compound15.21494.21494.4 | controllerType=0 controllerNumber=1 scan=21494 | 43.3873 |
| 18381:4 | 1 | R.RTAGWNVPIGTLRPFLN[+2059.735]WTGPPEPIEAAVAR.F | HexNAc(4)Hex(5)Fuc(1)NeuAc(1) | N[+2060] | 1362.4026 | 4 | 5446.5886 | 5446.547 | 0 | 7.6 | 140 | Specific | 278.9 | 278.9 | 278.9 | 6.5 | 135 | >sp\|P02788\|TRFL_HUMAN Lactotransferrin OS=Homo sapiens OX=9606 GN=LTF PE=1 SV=6 | 1270 | 20201114_13813_1ng_glyco_hcd203040_compound15.21913.21913.4 | controllerType=0 controllerNumber=1 scan=21913 | 44.2062 |
| 17984:4 | 1 | R.RTAGWNVPIGTLRPFLN[+2716.983]WTGPPEPIEAAVAR.F | HexNAc(5)Hex(6)Fuc(3)NeuAc(1) | N[+2717] | 1526.7086 | 4 | 6103.8126 | 6103.795 | 0 | 2.9 | 140 | Specific | 294.26 | 294.26 | 294.26 | 7.53 | 135 | >sp\|P02788\|TRFL_HUMAN Lactotransferrin OS=Homo sapiens OX=9606 GN=LTF PE=1 SV=6 | 1270 | 20201114_13813_1ng_glyco_hcd203040_compound15.21496.21496.4 | controllerType=0 controllerNumber=1 scan=21496 | 43.3911 |
| 18383:4 | 1 | R.RTAGWNVPIGTLRPFLN[+2059.735]WTGPPEPIEAAVAR.F | HexNAc(4)Hex(5)Fuc(1)NeuAc(1) | N[+2060] | 1362.4026 | 4 | 5446.5886 | 5446.547 | 0 | 7.6 | 140 | Specific | 334.74 | 334.74 | 334.74 | 5.36 | 135 | >sp\|P02788\|TRFL_HUMAN Lactotransferrin OS=Homo sapiens OX=9606 GN=LTF PE=1 SV=6 | 1270 | 20201114_13813_1ng_glyco_hcd203040_compound15.21915.21915.4 | controllerType=0 controllerNumber=1 scan=21915 | 44.2113 |
| 17950:4 | 1 | R.RTAGWNVPIGTLRPFLN[+2570.925]WTGPPEPIEAAVAR.F | HexNAc(5)Hex(6)Fuc(2)NeuAc(1) | N[+2571] | 1490.193 | 4 | 5957.7502 | 5957.7371 | 0 | 2.2 | 140 | Specific | 461.53 | 410.38 | 410.38 | 9.73 | 135 | >sp\|P02788\|TRFL_HUMAN Lactotransferrin OS=Homo sapiens OX=9606 GN=LTF PE=1 SV=6 | 1270 | 20201114_13813_1ng_glyco_hcd203040_compound15.21461.21461.4 | controllerType=0 controllerNumber=1 scan=21461 | 43.3132 |
| 17074:4 | 1 | R.RTAGWNVPIGTLRPFLN[+1768.640]WTGPPEPIEAAVAR.F | HexNAc(4)Hex(5)Fuc(1) | N[+1769] | 1289.6205 | 4 | 5155.4601 | 5155.4516 | 0 | 1.7 | 140 | Specific | 504.14 | 504.14 | 504.14 | 10.02 | 135 | >sp\|P02788\|TRFL_HUMAN Lactotransferrin OS=Homo sapiens OX=9606 GN=LTF PE=1 SV=6 | 1270 | 20201114_13813_1ng_glyco_hcd203040_compound15.20547.20547.4 | controllerType=0 controllerNumber=1 scan=20547 | 41.4872 |
| 18040:4 | 1 | R.RTAGWNVPIGTLRPFLN[+2205.793]WTGPPEPIEAAVAR.F | HexNAc(4)Hex(5)Fuc(2)NeuAc(1) | N[+2206] | 1398.9089 | 4 | 5592.6139 | 5592.6049 | 0 | 1.6 | 140 | Specific | 564.16 | 520.26 | 520.26 | 10.27 | 135 | >sp\|P02788\|TRFL_HUMAN Lactotransferrin OS=Homo sapiens OX=9606 GN=LTF PE=1 SV=6 | 1270 | 20201114_13813_1ng_glyco_hcd203040_compound15.21557.21557.4 | controllerType=0 controllerNumber=1 scan=21557 | 43.5151 |
| 19339:3 | 1 | R.TAGWNVPIGTLRPFLN[+2205.793]WTGPPEPIEAAVAR.F | HexNAc(4)Hex(5)Fuc(2)NeuAc(1) | N[+2206] | 1812.857 | 3 | 5436.5565 | 5436.5038 | 0 | 9.7 | 141 | Specific | 209.69 | 209.69 | 209.69 | 4.76 | 135 | >sp\|P02788\|TRFL_HUMAN Lactotransferrin OS=Homo sapiens OX=9606 GN=LTF PE=1 SV=6 | 1270 | 20201114_13813_1ng_glyco_hcd203040_compound15.22917.22917.3 | controllerType=0 controllerNumber=1 scan=22917 | 46.3942 |
| 19277:4 | 1 | R.TAGWNVPIGTLRPFLN[+2424.867]WTGPPEPIEAAVAR.F | HexNAc(5)Hex(6)Fuc(1)NeuAc(1) | N[+2425] | 1414.6521 | 4 | 5655.5866 | 5655.5781 | 0 | 1.5 | 141 | Specific | 211.23 | 140.98 | 140.98 | 4.44 | 135 | >sp\|P02788\|TRFL_HUMAN Lactotransferrin OS=Homo sapiens OX=9606 GN=LTF PE=1 SV=6 | 1270 | 20201114_13813_1ng_glyco_hcd203040_compound15.22852.22852.4 | controllerType=0 controllerNumber=1 scan=22852 | 46.2744 |
| 19312:3 | 1 | R.TAGWNVPIGTLRPFLN[+2059.735]WTGPPEPIEAAVAR.F | HexNAc(4)Hex(5)Fuc(1)NeuAc(1) | N[+2060] | 1764.1497 | 3 | 5290.4347 | 5290.4459 | 0 | -2.1 | 141 | Specific | 240.84 | 240.84 | 240.84 | 5.96 | 135 | >sp\|P02788\|TRFL_HUMAN Lactotransferrin OS=Homo sapiens OX=9606 GN=LTF PE=1 SV=6 | 1270 | 20201114_13813_1ng_glyco_hcd203040_compound15.22889.22889.2 | controllerType=0 controllerNumber=1 scan=22889 | 46.3461 |
| 19269:3 | 1 | R.TAGWNVPIGTLRPFLN[+2205.793]WTGPPEPIEAAVAR.F | HexNAc(4)Hex(5)Fuc(2)NeuAc(1) | N[+2206] | 1812.8418 | 3 | 5436.5107 | 5436.5038 | 0 | 1.3 | 141 | Specific | 265.48 | 208.23 | 208.23 | 6.77 | 135 | >sp\|P02788\|TRFL_HUMAN Lactotransferrin OS=Homo sapiens OX=9606 GN=LTF PE=1 SV=6 | 1270 | 20201114_13813_1ng_glyco_hcd203040_compound15.22844.22844.3 | controllerType=0 controllerNumber=1 scan=22844 | 46.2513 |
| 19280:4 | 1 | R.TAGWNVPIGTLRPFLN[+2424.867]WTGPPEPIEAAVAR.F | HexNAc(5)Hex(6)Fuc(1)NeuAc(1) | N[+2425] | 1414.6521 | 4 | 5655.5866 | 5655.5781 | 0 | 1.5 | 141 | Specific | 270.37 | 182.77 | 182.77 | 5.19 | 135 | >sp\|P02788\|TRFL_HUMAN Lactotransferrin OS=Homo sapiens OX=9606 GN=LTF PE=1 SV=6 | 1270 | 20201114_13813_1ng_glyco_hcd203040_compound15.22855.22855.4 | controllerType=0 controllerNumber=1 scan=22855 | 46.2809 |
| 18225:3 | 1 | R.TAGWNVPIGTLRPFLN[+2278.809]WTGPPEPIEAAVAR.F | HexNAc(5)Hex(6)NeuAc(1) | N[+2279] | 1837.1797 | 3 | 5509.5244 | 5509.5201 | 0 | 0.8 | 141 | Specific | 271.67 | 90.07 | 90.07 | 3.84 | 135 | >sp\|P02788\|TRFL_HUMAN Lactotransferrin OS=Homo sapiens OX=9606 GN=LTF PE=1 SV=6 | 1270 | 20201114_13813_1ng_glyco_hcd203040_compound15.21749.21749.3 | controllerType=0 controllerNumber=1 scan=21749 | 43.866 |
| 19276:3 | 1 | R.TAGWNVPIGTLRPFLN[+2351.851]WTGPPEPIEAAVAR.F | HexNAc(4)Hex(5)Fuc(3)NeuAc(1) | N[+2352] | 1861.5266 | 3 | 5582.5652 | 5582.5617 | 0 | 0.6 | 141 | Specific | 298.65 | 294.69 | 294.69 | 7.55 | 135 | >sp\|P02788\|TRFL_HUMAN Lactotransferrin OS=Homo sapiens OX=9606 GN=LTF PE=1 SV=6 | 1270 | 20201114_13813_1ng_glyco_hcd203040_compound15.22851.22851.3 | controllerType=0 controllerNumber=1 scan=22851 | 46.2727 |
| 19273:3 | 1 | R.TAGWNVPIGTLRPFLN[+2351.851]WTGPPEPIEAAVAR.F | HexNAc(4)Hex(5)Fuc(3)NeuAc(1) | N[+2352] | 1861.5266 | 3 | 5582.5652 | 5582.5617 | 0 | 0.6 | 141 | Specific | 301.53 | 301.53 | 301.53 | 7.55 | 135 | >sp\|P02788\|TRFL_HUMAN Lactotransferrin OS=Homo sapiens OX=9606 GN=LTF PE=1 SV=6 | 1270 | 20201114_13813_1ng_glyco_hcd203040_compound15.22848.22848.3 | controllerType=0 controllerNumber=1 scan=22848 | 46.2561 |
| 19671:4 | 1 | R.TAGWNVPIGTLRPFLN[+2059.735]WTGPPEPIEAAVAR.F | HexNAc(4)Hex(5)Fuc(1)NeuAc(1) | N[+2060] | 1323.3684 | 4 | 5290.4518 | 5290.4459 | 0 | 1.1 | 141 | Specific | 307.43 | 307.43 | 307.43 | 7.51 | 135 | >sp\|P02788\|TRFL_HUMAN Lactotransferrin OS=Homo sapiens OX=9606 GN=LTF PE=1 SV=6 | 1270 | 20201114_13813_1ng_glyco_hcd203040_compound15.23267.23267.4 | controllerType=0 controllerNumber=1 scan=23267 | 47.0729 |
| 19272:4 | 1 | R.TAGWNVPIGTLRPFLN[+2351.851]WTGPPEPIEAAVAR.F | HexNAc(4)Hex(5)Fuc(3)NeuAc(1) | N[+2352] | 1396.3971 | 4 | 5582.5666 | 5582.5617 | 0 | 0.9 | 141 | Specific | 342.7 | 342.7 | 342.7 | 7.71 | 135 | >sp\|P02788\|TRFL_HUMAN Lactotransferrin OS=Homo sapiens OX=9606 GN=LTF PE=1 SV=6 | 1270 | 20201114_13813_1ng_glyco_hcd203040_compound15.22847.22847.4 | controllerType=0 controllerNumber=1 scan=22847 | 46.2547 |
| 19315:3 | 1 | R.TAGWNVPIGTLRPFLN[+2059.735]WTGPPEPIEAAVAR.F | HexNAc(4)Hex(5)Fuc(1)NeuAc(1) | N[+2060] | 1764.1497 | 3 | 5290.4347 | 5290.4459 | 0 | -2.1 | 141 | Specific | 353.96 | 332.59 | 332.59 | 7.09 | 135 | >sp\|P02788\|TRFL_HUMAN Lactotransferrin OS=Homo sapiens OX=9606 GN=LTF PE=1 SV=6 | 1270 | 20201114_13813_1ng_glyco_hcd203040_compound15.22892.22892.2 | controllerType=0 controllerNumber=1 scan=22892 | 46.3494 |
| 18222:3 | 1 | R.TAGWNVPIGTLRPFLN[+2278.809]WTGPPEPIEAAVAR.F | HexNAc(5)Hex(6)NeuAc(1) | N[+2279] | 1837.1797 | 3 | 5509.5244 | 5509.5201 | 0 | 0.8 | 141 | Specific | 363.64 | 198.48 | 198.48 | 7.24 | 135 | >sp\|P02788\|TRFL_HUMAN Lactotransferrin OS=Homo sapiens OX=9606 GN=LTF PE=1 SV=6 | 1270 | 20201114_13813_1ng_glyco_hcd203040_compound15.21746.21746.3 | controllerType=0 controllerNumber=1 scan=21746 | 43.8607 |
| 19116:3 | 1 | R.TAGWNVPIGTLRPFLN[+2716.983]WTGPPEPIEAAVAR.F | HexNAc(5)Hex(6)Fuc(3)NeuAc(1) | N[+2717] | 1983.2403 | 3 | 5947.7064 | 5947.6939 | 0 | 2.1 | 141 | Specific | 459.87 | 354.05 | 354.05 | 8.42 | 135 | >sp\|P02788\|TRFL_HUMAN Lactotransferrin OS=Homo sapiens OX=9606 GN=LTF PE=1 SV=6 | 1270 | 20201114_13813_1ng_glyco_hcd203040_compound15.22683.22683.3 | controllerType=0 controllerNumber=1 scan=22683 | 45.915 |
| 19086:4 | 1 | R.TAGWNVPIGTLRPFLN[+2716.983]WTGPPEPIEAAVAR.F | HexNAc(5)Hex(6)Fuc(3)NeuAc(1) | N[+2717] | 1487.6821 | 4 | 5947.7067 | 5947.6939 | 0 | 2.2 | 141 | Specific | 480.71 | 462.72 | 462.72 | 9.69 | 135 | >sp\|P02788\|TRFL_HUMAN Lactotransferrin OS=Homo sapiens OX=9606 GN=LTF PE=1 SV=6 | 1270 | 20201114_13813_1ng_glyco_hcd203040_compound15.22652.22652.2 | controllerType=0 controllerNumber=1 scan=22652 | 45.8581 |
| 19668:4 | 1 | R.TAGWNVPIGTLRPFLN[+2059.735]WTGPPEPIEAAVAR.F | HexNAc(4)Hex(5)Fuc(1)NeuAc(1) | N[+2060] | 1323.3684 | 4 | 5290.4518 | 5290.4459 | 0 | 1.1 | 141 | Specific | 481.22 | 416 | 416 | 9.71 | 135 | >sp\|P02788\|TRFL_HUMAN Lactotransferrin OS=Homo sapiens OX=9606 GN=LTF PE=1 SV=6 | 1270 | 20201114_13813_1ng_glyco_hcd203040_compound15.23264.23264.4 | controllerType=0 controllerNumber=1 scan=23264 | 47.0495 |
| 19099:4 | 1 | R.TAGWNVPIGTLRPFLN[+2716.983]WTGPPEPIEAAVAR.F | HexNAc(5)Hex(6)Fuc(3)NeuAc(1) | N[+2717] | 1487.6826 | 4 | 5947.7087 | 5947.6939 | 0 | 2.5 | 141 | Specific | 506.49 | 506.49 | 506.49 | 9.94 | 135 | >sp\|P02788\|TRFL_HUMAN Lactotransferrin OS=Homo sapiens OX=9606 GN=LTF PE=1 SV=6 | 1270 | 20201114_13813_1ng_glyco_hcd203040_compound15.22666.22666.4 | controllerType=0 controllerNumber=1 scan=22666 | 45.8847 |
| 18329:4 | 1 | R.TAGWNVPIGTLRPFLN[+1768.640]WTGPPEPIEAAVAR.F | HexNAc(4)Hex(5)Fuc(1) | N[+1769] | 1250.5964 | 4 | 4999.3639 | 4999.3504 | 0 | 2.7 | 141 | Specific | 540.31 | 509.56 | 509.56 | 10.06 | 135 | >sp\|P02788\|TRFL_HUMAN Lactotransferrin OS=Homo sapiens OX=9606 GN=LTF PE=1 SV=6 | 1270 | 20201114_13813_1ng_glyco_hcd203040_compound15.21859.21859.4 | controllerType=0 controllerNumber=1 scan=21859 | 44.0912 |
| 19104:4 | 1 | R.TAGWNVPIGTLRPFLN[+2863.041]WTGPPEPIEAAVAR.F | HexNAc(5)Hex(6)Fuc(4)NeuAc(1) | N[+2863] | 1524.1979 | 4 | 6093.7697 | 6093.7518 | 0 | 2.9 | 141 | Specific | 565.2 | 525.67 | 525.67 | 10.2 | 135 | >sp\|P02788\|TRFL_HUMAN Lactotransferrin OS=Homo sapiens OX=9606 GN=LTF PE=1 SV=6 | 1270 | 20201114_13813_1ng_glyco_hcd203040_compound15.22671.22671.4 | controllerType=0 controllerNumber=1 scan=22671 | 45.8949 |
| 19119:3 | 1 | R.TAGWNVPIGTLRPFLN[+2716.983]WTGPPEPIEAAVAR.F | HexNAc(5)Hex(6)Fuc(3)NeuAc(1) | N[+2717] | 1983.2403 | 3 | 5947.7064 | 5947.6939 | 0 | 2.1 | 141 | Specific | 583.67 | 528.67 | 528.67 | 11.44 | 135 | >sp\|P02788\|TRFL_HUMAN Lactotransferrin OS=Homo sapiens OX=9606 GN=LTF PE=1 SV=6 | 1270 | 20201114_13813_1ng_glyco_hcd203040_compound15.22686.22686.3 | controllerType=0 controllerNumber=1 scan=22686 | 45.9215 |
| 19083:4 | 1 | R.TAGWNVPIGTLRPFLN[+2716.983]WTGPPEPIEAAVAR.F | HexNAc(5)Hex(6)Fuc(3)NeuAc(1) | N[+2717] | 1487.6821 | 4 | 5947.7067 | 5947.6939 | 0 | 2.2 | 141 | Specific | 585.74 | 524.81 | 524.81 | 11.39 | 135 | >sp\|P02788\|TRFL_HUMAN Lactotransferrin OS=Homo sapiens OX=9606 GN=LTF PE=1 SV=6 | 1270 | 20201114_13813_1ng_glyco_hcd203040_compound15.22649.22649.2 | controllerType=0 controllerNumber=1 scan=22649 | 45.8526 |
| 19251:3 | 1 | R.TAGWNVPIGTLRPFLN[+2205.793]WTGPPEPIEAAVAR.F | HexNAc(4)Hex(5)Fuc(2)NeuAc(1) | N[+2206] | 1812.8433 | 3 | 5436.5155 | 5436.5038 | 0 | 2.2 | 141 | Specific | 594.41 | 492.32 | 492.32 | 11.44 | 135 | >sp\|P02788\|TRFL_HUMAN Lactotransferrin OS=Homo sapiens OX=9606 GN=LTF PE=1 SV=6 | 1270 | 20201114_13813_1ng_glyco_hcd203040_compound15.22825.22825.3 | controllerType=0 controllerNumber=1 scan=22825 | 46.2054 |
| 19249:3 | 1 | R.TAGWNVPIGTLRPFLN[+2205.793]WTGPPEPIEAAVAR.F | HexNAc(4)Hex(5)Fuc(2)NeuAc(1) | N[+2206] | 1812.8433 | 3 | 5436.5155 | 5436.5038 | 0 | 2.2 | 141 | Specific | 616.58 | 565.44 | 565.44 | 12.44 | 135 | >sp\|P02788\|TRFL_HUMAN Lactotransferrin OS=Homo sapiens OX=9606 GN=LTF PE=1 SV=6 | 1270 | 20201114_13813_1ng_glyco_hcd203040_compound15.22823.22823.3 | controllerType=0 controllerNumber=1 scan=22823 | 46.2002 |
| 19267:3 | 1 | R.TAGWNVPIGTLRPFLN[+2205.793]WTGPPEPIEAAVAR.F | HexNAc(4)Hex(5)Fuc(2)NeuAc(1) | N[+2206] | 1812.8418 | 3 | 5436.5107 | 5436.5038 | 0 | 1.3 | 141 | Specific | 629.57 | 579.71 | 579.71 | 12.46 | 135 | >sp\|P02788\|TRFL_HUMAN Lactotransferrin OS=Homo sapiens OX=9606 GN=LTF PE=1 SV=6 | 1270 | 20201114_13813_1ng_glyco_hcd203040_compound15.22842.22842.3 | controllerType=0 controllerNumber=1 scan=22842 | 46.2434 |
| 19341:3 | 1 | R.TAGWNVPIGTLRPFLN[+2205.793]WTGPPEPIEAAVAR.F | HexNAc(4)Hex(5)Fuc(2)NeuAc(1) | N[+2206] | 1812.857 | 3 | 5436.5565 | 5436.5038 | 0 | 9.7 | 141 | Specific | 653.93 | 588.39 | 588.39 | 12.73 | 135 | >sp\|P02788\|TRFL_HUMAN Lactotransferrin OS=Homo sapiens OX=9606 GN=LTF PE=1 SV=6 | 1270 | 20201114_13813_1ng_glyco_hcd203040_compound15.22919.22919.3 | controllerType=0 controllerNumber=1 scan=22919 | 46.3986 |
| 19275:4 | 1 | R.TAGWNVPIGTLRPFLN[+2351.851]WTGPPEPIEAAVAR.F | HexNAc(4)Hex(5)Fuc(3)NeuAc(1) | N[+2352] | 1396.3971 | 4 | 5582.5666 | 5582.5617 | 0 | 0.9 | 141 | Specific | 666.59 | 524.82 | 524.82 | 13.41 | 135 | >sp\|P02788\|TRFL_HUMAN Lactotransferrin OS=Homo sapiens OX=9606 GN=LTF PE=1 SV=6 | 1270 | 20201114_13813_1ng_glyco_hcd203040_compound15.22850.22850.4 | controllerType=0 controllerNumber=1 scan=22850 | 46.2711 |
| 19266:4 | 1 | R.TAGWNVPIGTLRPFLN[+2205.793]WTGPPEPIEAAVAR.F | HexNAc(4)Hex(5)Fuc(2)NeuAc(1) | N[+2206] | 1359.8831 | 4 | 5436.5104 | 5436.5038 | 0 | 1.2 | 141 | Specific | 670.1 | 629.9 | 629.9 | 13.41 | 135 | >sp\|P02788\|TRFL_HUMAN Lactotransferrin OS=Homo sapiens OX=9606 GN=LTF PE=1 SV=6 | 1270 | 20201114_13813_1ng_glyco_hcd203040_compound15.22841.22841.4 | controllerType=0 controllerNumber=1 scan=22841 | 46.242 |
| 19107:4 | 1 | R.TAGWNVPIGTLRPFLN[+2863.041]WTGPPEPIEAAVAR.F | HexNAc(5)Hex(6)Fuc(4)NeuAc(1) | N[+2863] | 1524.1979 | 4 | 6093.7697 | 6093.7518 | 0 | 2.9 | 141 | Specific | 702.13 | 580.38 | 580.38 | 14.39 | 135 | >sp\|P02788\|TRFL_HUMAN Lactotransferrin OS=Homo sapiens OX=9606 GN=LTF PE=1 SV=6 | 1270 | 20201114_13813_1ng_glyco_hcd203040_compound15.22674.22674.4 | controllerType=0 controllerNumber=1 scan=22674 | 45.9016 |
| 19271:3 | 1 | R.TAGWNVPIGTLRPFLN[+2059.735]WTGPPEPIEAAVAR.F | HexNAc(4)Hex(5)Fuc(1)NeuAc(1) | N[+2060] | 1764.1563 | 3 | 5290.4545 | 5290.4459 | 0 | 1.6 | 141 | Specific | 739.1 | 656.05 | 656.05 | 14.46 | 135 | >sp\|P02788\|TRFL_HUMAN Lactotransferrin OS=Homo sapiens OX=9606 GN=LTF PE=1 SV=6 | 1270 | 20201114_13813_1ng_glyco_hcd203040_compound15.22846.22846.3 | controllerType=0 controllerNumber=1 scan=22846 | 46.2536 |
| 18327:3 | 1 | R.TAGWNVPIGTLRPFLN[+1768.640]WTGPPEPIEAAVAR.F | HexNAc(4)Hex(5)Fuc(1) | N[+1769] | 1667.1256 | 3 | 4999.3622 | 4999.3504 | 0 | 2.3 | 141 | Specific | 749.12 | 631.35 | 631.35 | 14.44 | 135 | >sp\|P02788\|TRFL_HUMAN Lactotransferrin OS=Homo sapiens OX=9606 GN=LTF PE=1 SV=6 | 1270 | 20201114_13813_1ng_glyco_hcd203040_compound15.21857.21857.3 | controllerType=0 controllerNumber=1 scan=21857 | 44.089 |
| 19270:4 | 1 | R.TAGWNVPIGTLRPFLN[+2059.735]WTGPPEPIEAAVAR.F | HexNAc(4)Hex(5)Fuc(1)NeuAc(1) | N[+2060] | 1323.3688 | 4 | 5290.4533 | 5290.4459 | 0 | 1.4 | 141 | Specific | 751.37 | 673.26 | 673.26 | 15.41 | 135 | >sp\|P02788\|TRFL_HUMAN Lactotransferrin OS=Homo sapiens OX=9606 GN=LTF PE=1 SV=6 | 1270 | 20201114_13813_1ng_glyco_hcd203040_compound15.22845.22845.4 | controllerType=0 controllerNumber=1 scan=22845 | 46.2524 |
| 18325:3 | 1 | R.TAGWNVPIGTLRPFLN[+1768.640]WTGPPEPIEAAVAR.F | HexNAc(4)Hex(5)Fuc(1) | N[+1769] | 1667.1256 | 3 | 4999.3622 | 4999.3504 | 0 | 2.3 | 141 | Specific | 761.82 | 677.5 | 677.5 | 15.44 | 135 | >sp\|P02788\|TRFL_HUMAN Lactotransferrin OS=Homo sapiens OX=9606 GN=LTF PE=1 SV=6 | 1270 | 20201114_13813_1ng_glyco_hcd203040_compound15.21855.21855.3 | controllerType=0 controllerNumber=1 scan=21855 | 44.0833 |
| 19459:3 | 1 | R.TAGWNVPIGTLRPFLN[+2059.735]WTGPPEPIEAAVAR.F | HexNAc(4)Hex(5)Fuc(1)NeuAc(1) | N[+2060] | 1764.1544 | 3 | 5290.4486 | 5290.4459 | 0 | 0.5 | 141 | Specific | 771.88 | 692.31 | 692.31 | 15.14 | 135 | >sp\|P02788\|TRFL_HUMAN Lactotransferrin OS=Homo sapiens OX=9606 GN=LTF PE=1 SV=6 | 1270 | 20201114_13813_1ng_glyco_hcd203040_compound15.23047.23047.3 | controllerType=0 controllerNumber=1 scan=23047 | 46.6419 |
| 19268:4 | 1 | R.TAGWNVPIGTLRPFLN[+2205.793]WTGPPEPIEAAVAR.F | HexNAc(4)Hex(5)Fuc(2)NeuAc(1) | N[+2206] | 1359.8831 | 4 | 5436.5104 | 5436.5038 | 0 | 1.2 | 141 | Specific | 825.18 | 729.77 | 729.77 | 16.41 | 135 | >sp\|P02788\|TRFL_HUMAN Lactotransferrin OS=Homo sapiens OX=9606 GN=LTF PE=1 SV=6 | 1270 | 20201114_13813_1ng_glyco_hcd203040_compound15.22843.22843.4 | controllerType=0 controllerNumber=1 scan=22843 | 46.2487 |
| 19274:4 | 1 | R.TAGWNVPIGTLRPFLN[+2059.735]WTGPPEPIEAAVAR.F | HexNAc(4)Hex(5)Fuc(1)NeuAc(1) | N[+2060] | 1323.3688 | 4 | 5290.4533 | 5290.4459 | 0 | 1.4 | 141 | Specific | 871.29 | 738.85 | 738.85 | 17.41 | 135 | >sp\|P02788\|TRFL_HUMAN Lactotransferrin OS=Homo sapiens OX=9606 GN=LTF PE=1 SV=6 | 1270 | 20201114_13813_1ng_glyco_hcd203040_compound15.22849.22849.4 | controllerType=0 controllerNumber=1 scan=22849 | 46.2703 |
| 18332:4 | 1 | R.TAGWNVPIGTLRPFLN[+1768.640]WTGPPEPIEAAVAR.F | HexNAc(4)Hex(5)Fuc(1) | N[+1769] | 1250.5964 | 4 | 4999.3639 | 4999.3504 | 0 | 2.7 | 141 | Specific | 908.84 | 858.98 | 858.98 | 18.39 | 135 | >sp\|P02788\|TRFL_HUMAN Lactotransferrin OS=Homo sapiens OX=9606 GN=LTF PE=1 SV=6 | 1270 | 20201114_13813_1ng_glyco_hcd203040_compound15.21862.21862.4 | controllerType=0 controllerNumber=1 scan=21862 | 44.0951 |
| 16966:3 | 1 | R.TAGWNIPMGLLFN[+1768.640]QTGSC[+57.021]K.F | HexNAc(4)Hex(5)Fuc(1) | C[+57], N[+1769] | 1288.5602 | 3 | 3863.6662 | 3863.6495 | 0 | 4.3 | 485 | Specific | 225.72 | 174.75 | 174.75 | 3.45 | 135 | >sp\|P02788\|TRFL_HUMAN Lactotransferrin OS=Homo sapiens OX=9606 GN=LTF PE=1 SV=6 | 1270 | 20201114_13813_1ng_glyco_hcd203040_compound15.20435.20435.2 | controllerType=0 controllerNumber=1 scan=20435 | 41.2813 |
| 20783:3 | 1 | R.[+42.011]TAGWNIPMGLLFN[+2336.851]QTGSC[+57.021]K.F | HexNAc(6)Hex(6)Fuc(1) | .[42], C[+57], N[+2337] | 1491.9672 | 3 | 4473.8871 | 4473.8716 | 0 | 3.5 | 485 | Specific | 268.72 | 234.53 | 234.53 | 5.01 | 135 | >sp\|P02788\|TRFL_HUMAN Lactotransferrin OS=Homo sapiens OX=9606 GN=LTF PE=1 SV=6 | 1270 | 20201114_13813_1ng_glyco_hcd203040_compound15.24432.24432.3 | controllerType=0 controllerNumber=1 scan=24432 | 49.4833 |
| 16795:3 | 1 | R.TAGWNIPMGLLFN[+1606.587]QTGSC[+57.021]K.F | HexNAc(4)Hex(4)Fuc(1) | C[+57], N[+1607] | 1234.5405 | 3 | 3701.6068 | 3701.5966 | 0 | 2.8 | 485 | Specific | 294.39 | 294.39 | 294.39 | 7.16 | 135 | >sp\|P02788\|TRFL_HUMAN Lactotransferrin OS=Homo sapiens OX=9606 GN=LTF PE=1 SV=6 | 1270 | 20201114_13813_1ng_glyco_hcd203040_compound15.20257.20257.3 | controllerType=0 controllerNumber=1 scan=20257 | 40.923 |
| 18761:3 | 1 | R.TAGWNIPMGLLFN[+2205.793]QTGSC[+57.021]K.F | HexNAc(4)Hex(5)Fuc(2)NeuAc(1) | C[+57], N[+2206] | 1434.2731 | 3 | 4300.8048 | 4300.8028 | 0 | 0.5 | 485 | Specific | 312.9 | 273.48 | 273.48 | 5.21 | 135 | >sp\|P02788\|TRFL_HUMAN Lactotransferrin OS=Homo sapiens OX=9606 GN=LTF PE=1 SV=6 | 1270 | 20201114_13813_1ng_glyco_hcd203040_compound15.22313.22313.3 | controllerType=0 controllerNumber=1 scan=22313 | 45.09 |
| 16887:3 | 1 | R.TAGWNIPMGLLFN[+1768.640]QTGSC[+57.021]K.F | HexNAc(4)Hex(5)Fuc(1) | C[+57], N[+1769] | 1288.558 | 3 | 3863.6596 | 3863.6495 | 0 | 2.6 | 485 | Specific | 315.13 | 315.13 | 315.13 | 7.36 | 135 | >sp\|P02788\|TRFL_HUMAN Lactotransferrin OS=Homo sapiens OX=9606 GN=LTF PE=1 SV=6 | 1270 | 20201114_13813_1ng_glyco_hcd203040_compound15.20353.20353.3 | controllerType=0 controllerNumber=1 scan=20353 | 41.1195 |
| 18537:3 | 1 | R.TAGWNIPMGLLFN[+2424.867]QTGSC[+57.021]K.F | HexNAc(5)Hex(6)Fuc(1)NeuAc(1) | C[+57], N[+2425] | 1507.3003 | 3 | 4519.8865 | 4519.8771 | 0 | 2.1 | 485 | Specific | 362.01 | 261.66 | 261.66 | 7.39 | 135 | >sp\|P02788\|TRFL_HUMAN Lactotransferrin OS=Homo sapiens OX=9606 GN=LTF PE=1 SV=6 | 1270 | 20201114_13813_1ng_glyco_hcd203040_compound15.22077.22077.3 | controllerType=0 controllerNumber=1 scan=22077 | 44.5073 |
| 16871:3 | 1 | R.TAGWNIPMGLLFN[+1622.582]QTGSC[+57.021]K.F | HexNAc(4)Hex(5) | C[+57], N[+1623] | 1239.869 | 3 | 3717.5923 | 3717.5916 | 0 | 0.2 | 485 | Specific | 376.03 | 376.03 | 203.09 | 8.85 | 135 | >sp\|P02788\|TRFL_HUMAN Lactotransferrin OS=Homo sapiens OX=9606 GN=LTF PE=1 SV=6 | 1270 | 20201114_13813_1ng_glyco_hcd203040_compound15.20336.20336.3 | controllerType=0 controllerNumber=1 scan=20336 | 41.0798 |
| 16702:2 | 1 | R.TAGWNIPMGLLFN[+1768.640]QTGSC[+57.021]K.F | HexNAc(4)Hex(5)Fuc(1) | C[+57], N[+1769] | 1932.3374 | 2 | 3863.6675 | 3863.6495 | 0 | 4.7 | 485 | Specific | 379.51 | 317.78 | 317.78 | 7.39 | 135 | >sp\|P02788\|TRFL_HUMAN Lactotransferrin OS=Homo sapiens OX=9606 GN=LTF PE=1 SV=6 | 1270 | 20201114_13813_1ng_glyco_hcd203040_compound15.20160.20160.2 | controllerType=0 controllerNumber=1 scan=20160 | 40.7461 |
| 18535:3 | 1 | R.TAGWNIPMGLLFN[+2424.867]QTGSC[+57.021]K.F | HexNAc(5)Hex(6)Fuc(1)NeuAc(1) | C[+57], N[+2425] | 1507.3003 | 3 | 4519.8865 | 4519.8771 | 0 | 2.1 | 485 | Specific | 380.93 | 362.77 | 191.84 | 7.84 | 135 | >sp\|P02788\|TRFL_HUMAN Lactotransferrin OS=Homo sapiens OX=9606 GN=LTF PE=1 SV=6 | 1270 | 20201114_13813_1ng_glyco_hcd203040_compound15.22075.22075.3 | controllerType=0 controllerNumber=1 scan=22075 | 44.5038 |
| 16890:3 | 1 | R.TAGWNIPMGLLFN[+1768.640]QTGSC[+57.021]K.F | HexNAc(4)Hex(5)Fuc(1) | C[+57], N[+1769] | 1288.558 | 3 | 3863.6596 | 3863.6495 | 0 | 2.6 | 485 | Specific | 382.63 | 382.63 | 382.63 | 9.15 | 135 | >sp\|P02788\|TRFL_HUMAN Lactotransferrin OS=Homo sapiens OX=9606 GN=LTF PE=1 SV=6 | 1270 | 20201114_13813_1ng_glyco_hcd203040_compound15.20356.20356.3 | controllerType=0 controllerNumber=1 scan=20356 | 41.1237 |
| 18901:3 | 1 | R.TAGWNIPMGLLFN[+1710.598]QTGSC[+57.021]K.F | HexNAc(3)Hex(5)NeuAc(1) | C[+57], N[+1711] | 1269.2184 | 3 | 3805.6408 | 3805.6076 | 0 | 8.7 | 485 | Specific | 388.03 | 361.69 | 242.66 | 7.18 | 135 | >sp\|P02788\|TRFL_HUMAN Lactotransferrin OS=Homo sapiens OX=9606 GN=LTF PE=1 SV=6 | 1270 | 20201114_13813_1ng_glyco_hcd203040_compound15.22460.22460.3 | controllerType=0 controllerNumber=1 scan=22460 | 45.47 |
| 18903:3 | 1 | R.TAGWNIPMGLLFN[+1751.624]QTGSC[+57.021]K.F | HexNAc(4)Hex(4)NeuAc(1) | C[+57], N[+1752] | 1282.8945 | 3 | 3846.6688 | 3846.6341 | 0 | 9 | 485 | Specific | 439.53 | 439.53 | 439.53 | 8.73 | 135 | >sp\|P02788\|TRFL_HUMAN Lactotransferrin OS=Homo sapiens OX=9606 GN=LTF PE=1 SV=6 | 1270 | 20201114_13813_1ng_glyco_hcd203040_compound15.22462.22462.3 | controllerType=0 controllerNumber=1 scan=22462 | 45.4723 |
| 16797:3 | 1 | R.TAGWNIPMGLLFN[+1606.587]QTGSC[+57.021]K.F | HexNAc(4)Hex(4)Fuc(1) | C[+57], N[+1607] | 1234.5405 | 3 | 3701.6068 | 3701.5966 | 0 | 2.8 | 485 | Specific | 479.21 | 479.21 | 479.21 | 9.37 | 135 | >sp\|P02788\|TRFL_HUMAN Lactotransferrin OS=Homo sapiens OX=9606 GN=LTF PE=1 SV=6 | 1270 | 20201114_13813_1ng_glyco_hcd203040_compound15.20259.20259.3 | controllerType=0 controllerNumber=1 scan=20259 | 40.9271 |
| 18906:3 | 1 | R.TAGWNIPMGLLFN[+1751.624]QTGSC[+57.021]K.F | HexNAc(4)Hex(4)NeuAc(1) | C[+57], N[+1752] | 1282.8945 | 3 | 3846.6688 | 3846.6341 | 0 | 9 | 485 | Specific | 516.4 | 516.4 | 516.4 | 9.08 | 135 | >sp\|P02788\|TRFL_HUMAN Lactotransferrin OS=Homo sapiens OX=9606 GN=LTF PE=1 SV=6 | 1270 | 20201114_13813_1ng_glyco_hcd203040_compound15.22465.22465.3 | controllerType=0 controllerNumber=1 scan=22465 | [illegible] |
| 18844:3 | 1 | R.TAGWNIPMGLLFN[+1872.651]QTGSC[+57.021]K.F | HexNAc(3)Hex(6)NeuAc(1) | C[+57], N[+1873] | 1323.2319 | 3 | 3967.6811 | 3967.6604 | 0 | 5.2 | 485 | Specific | 549.32 | 525.53 | 224.91 | 9.42 | 135 | >sp\|P02788\|TRFL_HUMAN Lactotransferrin OS=Homo sapiens OX=9606 GN=LTF PE=1 SV=6 | 1270 | 20201114_13813_1ng_glyco_hcd203040_compound15.22401.22401.3 | controllerType=0 controllerNumber=1 scan=22401 | [illegible] |
| 16467:3 | 1 | R.TAGWNIPMGLLFN[+2279.830]QTGSC[+57.021]K.F | HexNAc(5)Hex(6)Fuc(2) | C[+57], N[+2280] | 1458.9557 | 3 | 4374.8527 | 4374.8396 | 0 | 3 | 485 | Specific | 587.43 | 577.82 | 350.23 | 11.07 | 135 | >sp\|P02788\|TRFL_HUMAN Lactotransferrin OS=Homo sapiens OX=9606 GN=LTF PE=1 SV=6 | 1270 | 20201114_13813_1ng_glyco_hcd203040_compound15.19914.19914.3 | controllerType=0 controllerNumber=1 scan=19914 | 40.2709 |
| 16704:2 | 1 | R.TAGWNIPMGLLFN[+1768.640]QTGSC[+57.021]K.F | HexNAc(4)Hex(5)Fuc(1) | C[+57], N[+1769] | 1932.3374 | 2 | 3863.6675 | 3863.6495 | 0 | 4.7 | 485 | Specific | 619.69 | 559.26 | 559.26 | 11.82 | 135 | >sp\|P02788\|TRFL_HUMAN Lactotransferrin OS=Homo sapiens OX=9606 GN=LTF PE=1 SV=6 | 1270 | 20201114_13813_1ng_glyco_hcd203040_compound15.20162.20162.2 | controllerType=0 controllerNumber=1 scan=20162 | 40.7508 |
| 18846:3 | 1 | R.TAGWNIPMGLLFN[+1872.651]QTGSC[+57.021]K.F | HexNAc(3)Hex(6)NeuAc(1) | C[+57], N[+1873] | 1323.2319 | 3 | 3967.6811 | 3967.6604 | 0 | 5.2 | 485 | Specific | 624.6 | 594.33 | 326.99 | 11.61 | 135 | >sp\|P02788\|TRFL_HUMAN Lactotransferrin OS=Homo sapiens OX=9606 GN=LTF PE=1 SV=6 | 1270 | 20201114_13813_1ng_glyco_hcd203040_compound15.22403.22403.3 | controllerType=0 controllerNumber=1 scan=22403 | 45.3586 |
| 20792:3 | 1 | R.TAGWNIPMGLLFN[+2350.830]QTGSC[+57.021]K.F | HexNAc(4)Hex(5)Fuc(1)NeuAc(2) | C[+57], N[+2351] | 1482.624 | 3 | 4445.8573 | 4445.8403 | 0 | 3.8 | 485 | Specific | 715.88 | 648.96 | 648.96 | 13.78 | 135 | >sp\|P02788\|TRFL_HUMAN Lactotransferrin OS=Homo sapiens OX=9606 GN=LTF PE=1 SV=6 | 1270 | 20201114_13813_1ng_glyco_hcd203040_compound15.24442.24442.3 | controllerType=0 controllerNumber=1 scan=24442 | 49.509 |
| 18904:3 | 1 | R.TAGWNIPMGLLFN[+1710.598]QTGSC[+57.021]K.F | HexNAc(3)Hex(5)NeuAc(1) | C[+57], N[+1711] | 1269.2184 | 3 | 3805.6408 | 3805.6076 | 0 | 8.7 | 485 | Specific | 721.07 | 545.82 | 390.51 | 13.41 | 135 | >sp\|P02788\|TRFL_HUMAN Lactotransferrin OS=Homo sapiens OX=9606 GN=LTF PE=1 SV=6 | 1270 | 20201114_13813_1ng_glyco_hcd203040_compound15.22463.22463.3 | controllerType=0 controllerNumber=1 scan=22463 | 45.4744 |
| 16499:3 | 1 | R.TAGWNIPMGLLFN[+2133.772]QTGSC[+57.021]K.F | HexNAc(5)Hex(6)Fuc(1) | C[+57], N[+2134] | 1410.2722 | 3 | 4228.8019 | 4228.7817 | 0 | 4.8 | 485 | Specific | 729.57 | 626.25 | 381.25 | 13.78 | 135 | >sp\|P02788\|TRFL_HUMAN Lactotransferrin OS=Homo sapiens OX=9606 GN=LTF PE=1 SV=6 | 1270 | 20201114_13813_1ng_glyco_hcd203040_compound15.19947.19947.3 | controllerType=0 controllerNumber=1 scan=19947 | 40.3395 |
| 17109:2 | 1 | R.TAGWNIPMGLLFN[+1216.423]QTGSC[+57.021]K.F | HexNAc(2)Hex(5) | C[+57], N[+1216] | 1656.2279 | 2 | 3311.4486 | 3311.4328 | 0 | 4.8 | 485 | Specific | 795.94 | 789.02 | 265.06 | 14.82 | 135 | >sp\|P02788\|TRFL_HUMAN Lactotransferrin OS=Homo sapiens OX=9606 GN=LTF PE=1 SV=6 | 1270 | 20201114_13813_1ng_glyco_hcd203040_compound15.20583.20583.2 | controllerType=0 controllerNumber=1 scan=20583 | 41.5588 |
| 18691:3 | 1 | R.TAGWNIPMGLLFN[+2059.735]QTGSC[+57.021]K.F | HexNAc(4)Hex(5)Fuc(1)NeuAc(1) | C[+57], N[+2060] | 1385.5882 | 3 | 4154.75 | 4154.7449 | 0 | 1.2 | 485 | Specific | 866.19 | 617.17 | 617.17 | 17.1 | 135 | >sp\|P02788\|TRFL_HUMAN Lactotransferrin OS=Homo sapiens OX=9606 GN=LTF PE=1 SV=6 | 1270 | 20201114_13813_1ng_glyco_hcd203040_compound15.22239.22239.3 | controllerType=0 controllerNumber=1 scan=22239 | 44.9167 |
| 20716:3 | 1 | R.TAGWNIPMGLLFN[+2350.830]QTGSC[+57.021]K.F | HexNAc(4)Hex(5)Fuc(1)NeuAc(2) | C[+57], N[+2351] | 1482.6188 | 3 | 4445.8419 | 4445.8403 | 0 | 0.4 | 485 | Specific | 876.71 | 731.65 | 731.65 | 16.77 | 135 | >sp\|P02788\|TRFL_HUMAN Lactotransferrin OS=Homo sapiens OX=9606 GN=LTF PE=1 SV=6 | 1270 | 20201114_13813_1ng_glyco_hcd203040_compound15.24361.24361.3 | controllerType=0 controllerNumber=1 scan=24361 | 49.3125 |
| 16496:3 | 1 | R.TAGWNIPMGLLFN[+2133.772]QTGSC[+57.021]K.F | HexNAc(5)Hex(6)Fuc(1) | C[+57], N[+2134] | 1410.2722 | 3 | 4228.8019 | 4228.7817 | 0 | 4.8 | 485 | Specific | 877.82 | 877.82 | 422.21 | 16.78 | 135 | >sp\|P02788\|TRFL_HUMAN Lactotransferrin OS=Homo sapiens OX=9606 GN=LTF PE=1 SV=6 | 1270 | 20201114_13813_1ng_glyco_hcd203040_compound15.19944.19944.3 | controllerType=0 controllerNumber=1 scan=19944 | 40.3343 |
| 17107:2 | 1 | R.TAGWNIPMGLLFN[+1216.423]QTGSC[+57.021]K.F | HexNAc(2)Hex(5) | C[+57], N[+1216] | 1656.2279 | 2 | 3311.4486 | 3311.4328 | 0 | 4.8 | 485 | Specific | 880.91 | 873.92 | 328.12 | 16.82 | 135 | >sp\|P02788\|TRFL_HUMAN Lactotransferrin OS=Homo sapiens OX=9606 GN=LTF PE=1 SV=6 | 1270 | 20201114_13813_1ng_glyco_hcd203040_compound15.20581.20581.2 | controllerType=0 controllerNumber=1 scan=20581 | 41.5539 |
| 20713:3 | 1 | R.TAGWNIPMGLLFN[+2350.830]QTGSC[+57.021]K.F | HexNAc(4)Hex(5)Fuc(1)NeuAc(2) | C[+57], N[+2351] | 1482.6188 | 3 | 4445.8419 | 4445.8403 | 0 | 0.4 | 485 | Specific | 883.99 | 766.02 | 766.02 | 16.77 | 135 | >sp\|P02788\|TRFL_HUMAN Lactotransferrin OS=Homo sapiens OX=9606 GN=LTF PE=1 SV=6 | 1270 | 20201114_13813_1ng_glyco_hcd203040_compound15.24358.24358.3 | controllerType=0 controllerNumber=1 scan=24358 | 49.3038 |
| 18638:3 | 1 | R.TAGWNIPMGLLFN[+2205.793]QTGSC[+57.021]K.F | HexNAc(4)Hex(5)Fuc(2)NeuAc(1) | C[+57], N[+2206] | 1434.2763 | 3 | 4300.8144 | 4300.8028 | 0 | 2.7 | 485 | Specific | 885.85 | 860.96 | 860.96 | 17.07 | 135 | >sp\|P02788\|TRFL_HUMAN Lactotransferrin OS=Homo sapiens OX=9606 GN=LTF PE=1 SV=6 | 1270 | 20201114_13813_1ng_glyco_hcd203040_compound15.22183.22183.3 | controllerType=0 controllerNumber=1 scan=22183 | 44.7769 |
| 16874:3 | 1 | R.TAGWNIPMGLLFN[+1622.582]QTGSC[+57.021]K.F | HexNAc(4)Hex(5) | C[+57], N[+1623] | 1239.869 | 3 | 3717.5923 | 3717.5916 | 0 | 0.2 | 485 | Specific | 891.75 | 859.27 | 426.69 | 16.77 | 135 | >sp\|P02788\|TRFL_HUMAN Lactotransferrin OS=Homo sapiens OX=9606 GN=LTF PE=1 SV=6 | 1270 | 20201114_13813_1ng_glyco_hcd203040_compound15.20339.20339.3 | controllerType=0 controllerNumber=1 scan=20339 | 41.0908 |
| 18636:3 | 1 | R.TAGWNIPMGLLFN[+2205.793]QTGSC[+57.021]K.F | HexNAc(4)Hex(5)Fuc(2)NeuAc(1) | C[+57], N[+2206] | 1434.2763 | 3 | 4300.8144 | 4300.8028 | 0 | 2.7 | 485 | Specific | 892.82 | 864.37 | 864.37 | 17.07 | 135 | >sp\|P02788\|TRFL_HUMAN Lactotransferrin OS=Homo sapiens OX=9606 GN=LTF PE=1 SV=6 | 1270 | 20201114_13813_1ng_glyco_hcd203040_compound15.22181.22181.3 | controllerType=0 controllerNumber=1 scan=22181 | 44.7645 |
| 18688:3 | 1 | R.TAGWNIPMGLLFN[+2059.735]QTGSC[+57.021]K.F | HexNAc(4)Hex(5)Fuc(1)NeuAc(1) | C[+57], N[+2060] | 1385.5882 | 3 | 4154.75 | 4154.7449 | 0 | 1.2 | 485 | Specific | 900.28 | 823.18 | 823.18 | 18.1 | 135 | >sp\|P02788\|TRFL_HUMAN Lactotransferrin OS=Homo sapiens OX=9606 GN=LTF PE=1 SV=6 | 1270 | 20201114_13813_1ng_glyco_hcd203040_compound15.22236.22236.3 | controllerType=0 controllerNumber=1 scan=22236 | 44.9025 |
| 20790:3 | 1 | R.TAGWNIPMGLLFN[+2350.830]QTGSC[+57.021]K.F | HexNAc(4)Hex(5)Fuc(1)NeuAc(2) | C[+57], N[+2351] | 1482.624 | 3 | 4445.8573 | 4445.8403 | 0 | 3.8 | 485 | Specific | 901.49 | 861.27 | 861.27 | 17.78 | 135 | >sp\|P02788\|TRFL_HUMAN Lactotransferrin OS=Homo sapiens OX=9606 GN=LTF PE=1 SV=6 | 1270 | 20201114_13813_1ng_glyco_hcd203040_compound15.24440.24440.3 | controllerType=0 controllerNumber=1 scan=24440 | 49.5017 |
| 16639:3 | 1 | R.TAGWNIPMGLLFN[+1768.640]QTGSC[+57.021]K.F | HexNAc(4)Hex(5)Fuc(1) | C[+57], N[+1769] | 1288.561 | 3 | 3863.6684 | 3863.6495 | 0 | 4.9 | 485 | Specific | 906.71 | 728.49 | 728.49 | 17.78 | 135 | >sp\|P02788\|TRFL_HUMAN Lactotransferrin OS=Homo sapiens OX=9606 GN=LTF PE=1 SV=6 | 1270 | 20201114_13813_1ng_glyco_hcd203040_compound15.20094.20094.3 | controllerType=0 controllerNumber=1 scan=20094 | 40.5995 |
| 18774:3 | 1 | R.TAGWNIPMGLLFN[+1913.677]QTGSC[+57.021]K.F | HexNAc(4)Hex(5)NeuAc(1) | C[+57], N[+1914] | 1336.9124 | 3 | 4008.7227 | 4008.687 | 0 | 8.9 | 485 | Specific | 908.51 | 908.51 | 908.51 | 17.41 | 135 | >sp\|P02788\|TRFL_HUMAN Lactotransferrin OS=Homo sapiens OX=9606 GN=LTF PE=1 SV=6 | 1270 | 20201114_13813_1ng_glyco_hcd203040_compound15.22327.22327.3 | controllerType=0 controllerNumber=1 scan=22327 | 45.1225 |
| 18646:3 | 1 | R.TAGWNIPMGLLFN[+2059.735]QTGSC[+57.021]K.F | HexNAc(4)Hex(5)Fuc(1)NeuAc(1) | C[+57], N[+2060] | 1385.5893 | 3 | 4154.7533 | 4154.7449 | 0 | 2 | 485 | Specific | 914.04 | 873.72 | 873.72 | 18.1 | 135 | >sp\|P02788\|TRFL_HUMAN Lactotransferrin OS=Homo sapiens OX=9606 GN=LTF PE=1 SV=6 | 1270 | 20201114_13813_1ng_glyco_hcd203040_compound15.22192.22192.3 | controllerType=0 controllerNumber=1 scan=22192 | 44.8075 |
| 16666:3 | 1 | R.TAGWNIPMGLLFN[+1768.640]QTGSC[+57.021]K.F | HexNAc(4)Hex(5)Fuc(1) | C[+57], N[+1769] | 1288.5619 | 3 | 3863.6713 | 3863.6495 | 0 | 5.6 | 485 | Specific | 931.12 | 931.12 | 931.12 | 17.61 | 135 | >sp\|P02788\|TRFL_HUMAN Lactotransferrin OS=Homo sapiens OX=9606 GN=LTF PE=1 SV=6 | 1270 | 20201114_13813_1ng_glyco_hcd203040_compound15.20123.20123.3 | controllerType=0 controllerNumber=1 scan=20123 | 40.6656 |
| 16664:3 | 1 | R.TAGWNIPMGLLFN[+1768.640]QTGSC[+57.021]K.F | HexNAc(4)Hex(5)Fuc(1) | C[+57], N[+1769] | 1288.5619 | 3 | 3863.6713 | 3863.6495 | 0 | 5.6 | 485 | Specific | 958.8 | 958.8 | 958.8 | 18.61 | 135 | >sp\|P02788\|TRFL_HUMAN Lactotransferrin OS=Homo sapiens OX=9606 GN=LTF PE=1 SV=6 | 1270 | 20201114_13813_1ng_glyco_hcd203040_compound15.20121.20121.3 | controllerType=0 controllerNumber=1 scan=20121 | 40.6524 |
| 16637:3 | 1 | R.TAGWNIPMGLLFN[+1768.640]QTGSC[+57.021]K.F | HexNAc(4)Hex(5)Fuc(1) | C[+57], N[+1769] | 1288.561 | 3 | 3863.6684 | 3863.6495 | 0 | 4.9 | 485 | Specific | 960.81 | 960.81 | 960.81 | 18.78 | 135 | >sp\|P02788\|TRFL_HUMAN Lactotransferrin OS=Homo sapiens OX=9606 GN=LTF PE=1 SV=6 | 1270 | 20201114_13813_1ng_glyco_hcd203040_compound15.20092.20092.3 | controllerType=0 controllerNumber=1 scan=20092 | 40.5939 |
| 18644:3 | 1 | R.TAGWNIPMGLLFN[+2059.735]QTGSC[+57.021]K.F | HexNAc(4)Hex(5)Fuc(1)NeuAc(1) | C[+57], N[+2060] | 1385.5893 | 3 | 4154.7533 | 4154.7449 | 0 | 2 | 485 | Specific | 961.27 | 812.25 | 812.25 | 19.1 | 135 | >sp\|P02788\|TRFL_HUMAN Lactotransferrin OS=Homo sapiens OX=9606 GN=LTF PE=1 SV=6 | 1270 | 20201114_13813_1ng_glyco_hcd203040_compound15.22190.22190.3 | controllerType=0 controllerNumber=1 scan=22190 | 44.7982 |
| 18731:3 | 1 | R.TAGWNIPMGLLFN[+2059.735]QTGSC[+57.021]K.F | HexNAc(4)Hex(5)Fuc(1)NeuAc(1) | C[+57], N[+2060] | 1385.5906 | 3 | 4154.7573 | 4154.7449 | 0 | 3 | 485 | Specific | 963.17 | 917.8 | 917.8 | 19.07 | 135 | >sp\|P02788\|TRFL_HUMAN Lactotransferrin OS=Homo sapiens OX=9606 GN=LTF PE=1 SV=6 | 1270 | 20201114_13813_1ng_glyco_hcd203040_compound15.22282.22282.3 | controllerType=0 controllerNumber=1 scan=22282 | 45.0192 |
| 18772:3 | 1 | R.TAGWNIPMGLLFN[+1913.677]QTGSC[+57.021]K.F | HexNAc(4)Hex(5)NeuAc(1) | C[+57], N[+1914] | 1336.9124 | 3 | 4008.7227 | 4008.687 | 0 | 8.9 | 485 | Specific | 973.96 | 966.5 | 966.5 | 18.41 | 135 | >sp\|P02788\|TRFL_HUMAN Lactotransferrin OS=Homo sapiens OX=9606 GN=LTF PE=1 SV=6 | 1270 | 20201114_13813_1ng_glyco_hcd203040_compound15.22325.22325.3 | controllerType=0 controllerNumber=1 scan=22325 | 45.1155 |
| 17059:3 | 1 | R.TAGWNIPMGLLFN[+1216.423]QTGSC[+57.021]K.F | HexNAc(2)Hex(5) | C[+57], N[+1216] | 1104.4846 | 3 | 3311.4394 | 3311.4328 | 0 | 2 | 485 | Specific | 992.36 | 964.29 | 459.71 | 19.1 | 135 | >sp\|P02788\|TRFL_HUMAN Lactotransferrin OS=Homo sapiens OX=9606 GN=LTF PE=1 SV=6 | 1270 | 20201114_13813_1ng_glyco_hcd203040_compound15.20531.20531.3 | controllerType=0 controllerNumber=1 scan=20531 | 41.4577 |
| 18729:3 | 1 | R.TAGWNIPMGLLFN[+2059.735]QTGSC[+57.021]K.F | HexNAc(4)Hex(5)Fuc(1)NeuAc(1) | C[+57], N[+2060] | 1385.5906 | 3 | 4154.7573 | 4154.7449 | 0 | 3 | 485 | Specific | 1013.95 | 879.83 | 879.83 | 20.07 | 135 | >sp\|P02788\|TRFL_HUMAN Lactotransferrin OS=Homo sapiens OX=9606 GN=LTF PE=1 SV=6 | 1270 | 20201114_13813_1ng_glyco_hcd203040_compound15.22280.22280.3 | controllerType=0 controllerNumber=1 scan=22280 | 45.012 |
| 17057:3 | 1 | R.TAGWNIPMGLLFN[+1216.423]QTGSC[+57.021]K.F | HexNAc(2)Hex(5) | C[+57], N[+1216] | 1104.4846 | 3 | 3311.4394 | 3311.4328 | 0 | 2 | 485 | Specific | 1017.67 | 1003.72 | 474.09 | 20.1 | 135 | >sp\|P02788\|TRFL_HUMAN Lactotransferrin OS=Homo sapiens OX=9606 GN=LTF PE=1 SV=6 | 1270 | 20201114_13813_1ng_glyco_hcd203040_compound15.20529.20529.3 | controllerType=0 controllerNumber=1 scan=20529 | 41.4525 |
| 17915:3 | 1 | R.TAGWNIPM[+15.995]GLLFN[+2059.735]QTGSC[+57.021]K.F | HexNAc(4)Hex(5)Fuc(1)NeuAc(1) | C[+57], M[+16], N[+2060] | 1390.9219 | 3 | 4170.7512 | 4170.7398 | 0 | 2.7 | 485 | Specific | 353.81 | 254.3 | 254.3 | 7.16 | 135 | >sp\|P02788\|TRFL_HUMAN Lactotransferrin OS=Homo sapiens OX=9606 GN=LTF PE=1 SV=6 | 1270 | 20201114_13813_1ng_glyco_hcd203040_compound15.21424.21424.3 | controllerType=0 controllerNumber=1 scan=21424 | 43.2353 |
| 17047:3 | 1 | R.TAGWNIPM[+15.995]GLLFN[+1913.677]QTGSC[+57.021]K.F | HexNAc(4)Hex(5)NeuAc(1) | C[+57], M[+16], N[+1914] | 1342.2316 | 3 | 4024.6803 | 4024.6819 | 0 | -0.4 | 485 | Specific | 417.98 | 399.97 | 399.97 | 8.88 | 135 | >sp\|P02788\|TRFL_HUMAN Lactotransferrin OS=Homo sapiens OX=9606 GN=LTF PE=1 SV=6 | 1270 | 20201114_13813_1ng_glyco_hcd203040_compound15.20519.20519.3 | controllerType=0 controllerNumber=1 scan=20519 | 41.4322 |
| 17962:3 | 1 | R.TAGWNIPM[+15.995]GLLFN[+2059.735]QTGSC[+57.021]K.F | HexNAc(4)Hex(5)Fuc(1)NeuAc(1) | C[+57], M[+16], N[+2060] | 1390.9233 | 3 | 4170.7553 | 4170.7398 | 0 | 3.7 | 485 | Specific | 448.92 | 355.85 | 355.85 | 6.73 | 135 | >sp\|P02788\|TRFL_HUMAN Lactotransferrin OS=Homo sapiens OX=9606 GN=LTF PE=1 SV=6 | 1270 | 20201114_13813_1ng_glyco_hcd203040_compound15.21473.21473.3 | controllerType=0 controllerNumber=1 scan=21473 | 43.3323 |
| 17049:3 | 1 | R.TAGWNIPM[+15.995]GLLFN[+1913.677]QTGSC[+57.021]K.F | HexNAc(4)Hex(5)NeuAc(1) | C[+57], M[+16], N[+1914] | 1342.2316 | 3 | 4024.6803 | 4024.6819 | 0 | -0.4 | 485 | Specific | 488.18 | 464.13 | 464.13 | 9.32 | 135 | >sp\|P02788\|TRFL_HUMAN Lactotransferrin OS=Homo sapiens OX=9606 GN=LTF PE=1 SV=6 | 1270 | 20201114_13813_1ng_glyco_hcd203040_compound15.20521.20521.3 | controllerType=0 controllerNumber=1 scan=20521 | 41.4358 |
| 17917:3 | 1 | R.TAGWNIPM[+15.995]GLLFN[+2059.735]QTGSC[+57.021]K.F | HexNAc(4)Hex(5)Fuc(1)NeuAc(1) | C[+57], M[+16], N[+2060] | 1390.9219 | 3 | 4170.7512 | 4170.7398 | 0 | 2.7 | 485 | Specific | 498.25 | 407.83 | 407.83 | 9.62 | 135 | >sp\|P02788\|TRFL_HUMAN Lactotransferrin OS=Homo sapiens OX=9606 GN=LTF PE=1 SV=6 | 1270 | 20201114_13813_1ng_glyco_hcd203040_compound15.21426.21426.3 | controllerType=0 controllerNumber=1 scan=21426 | 43.2392 |
| 17021:3 | 1 | R.TAGWNIPM[+15.995]GLLFN[+2059.735]QTGSC[+57.021]K.F | HexNAc(4)Hex(5)Fuc(1)NeuAc(1) | C[+57], M[+16], N[+2060] | 1390.9246 | 3 | 4170.7593 | 4170.7398 | 0 | 4.7 | 485 | Specific | 819.81 | 655.27 | 655.27 | 15.78 | 135 | >sp\|P02788\|TRFL_HUMAN Lactotransferrin OS=Homo sapiens OX=9606 GN=LTF PE=1 SV=6 | 1270 | 20201114_13813_1ng_glyco_hcd203040_compound15.20492.20492.3 | controllerType=0 controllerNumber=1 scan=20492 | 41.3844 |
| 17019:3 | 1 | R.TAGWNIPM[+15.995]GLLFN[+2059.735]QTGSC[+57.021]K.F | HexNAc(4)Hex(5)Fuc(1)NeuAc(1) | C[+57], M[+16], N[+2060] | 1390.9246 | 3 | 4170.7593 | 4170.7398 | 0 | 4.7 | 485 | Specific | 873.85 | 748.36 | 748.36 | 16.78 | 135 | >sp\|P02788\|TRFL_HUMAN Lactotransferrin OS=Homo sapiens OX=9606 GN=LTF PE=1 SV=6 | 1270 | 20201114_13813_1ng_glyco_hcd203040_compound15.20490.20490.3 | controllerType=0 controllerNumber=1 scan=20490 | 41.3781 |
| 18883:4 | 1 | R.TAGWNIPMGLLFN[+2205.793]QTGSC[+57.021]KFDEYFSQSC[+57.021]APGSDPR.S | HexNAc(4)Hex(5)Fuc(2)NeuAc(1) | C[+57]*2, N[+2206] | 1536.8976 | 4 | 6144.5685 | 6144.5499 | 0 | 3 | 485 | Specific | 667.8 | 658.46 | 658.46 | 12.69 | 135 | >sp\|P02788\|TRFL_HUMAN Lactotransferrin OS=Homo sapiens OX=9606 GN=LTF PE=1 SV=6 | 1270 | 20201114_13813_1ng_glyco_hcd203040_compound15.22441.22441.4 | controllerType=0 controllerNumber=1 scan=22441 | 45.4321 |
| 20258:4 | 1 | R.TAGWNIPMGLLFN[+2350.830]QTGSC[+57.021]KFDEYFSQSC[+57.021]APGSDPR.S | HexNAc(4)Hex(5)Fuc(1)NeuAc(2) | C[+57]*2, N[+2351] | 1573.1523 | 4 | 6289.5876 | 6289.5875 | 0 | 0 | 485 | Specific | 712.42 | 699.25 | 699.25 | 13.39 | 135 | >sp\|P02788\|TRFL_HUMAN Lactotransferrin OS=Homo sapiens OX=9606 GN=LTF PE=1 SV=6 | 1270 | 20201114_13813_1ng_glyco_hcd203040_compound15.23884.23884.4 | controllerType=0 controllerNumber=1 scan=23884 | 48.3732 |
| 20260:4 | 1 | R.TAGWNIPMGLLFN[+2350.830]QTGSC[+57.021]KFDEYFSQSC[+57.021]APGSDPR.S | HexNAc(4)Hex(5)Fuc(1)NeuAc(2) | C[+57]*2, N[+2351] | 1573.1523 | 4 | 6289.5876 | 6289.5875 | 0 | 0 | 485 | Specific | 819.71 | 785.6 | 785.6 | 15.39 | 135 | >sp\|P02788\|TRFL_HUMAN Lactotransferrin OS=Homo sapiens OX=9606 GN=LTF PE=1 SV=6 | 1270 | 20201114_13813_1ng_glyco_hcd203040_compound15.23886.23886.4 | controllerType=0 controllerNumber=1 scan=23886 | 48.3778 |
| 18979:4 | 1 | R.TAGWNIPMGLLFN[+1913.677]QTGSC[+57.021]KFDEYFSQSC[+57.021]APGSDPR.S | HexNAc(4)Hex(5)NeuAc(1) | C[+57]*2, N[+1914] | 1463.8665 | 4 | 5852.444 | 5852.4341 | 0 | 1.7 | 485 | Specific | 851.49 | 804.84 | 804.84 | 16.71 | 135 | >sp\|P02788\|TRFL_HUMAN Lactotransferrin OS=Homo sapiens OX=9606 GN=LTF PE=1 SV=6 | 1270 | 20201114_13813_1ng_glyco_hcd203040_compound15.22541.22541.4 | controllerType=0 controllerNumber=1 scan=22541 | 45.6436 |
| 18977:4 | 1 | R.TAGWNIPMGLLFN[+1913.677]QTGSC[+57.021]KFDEYFSQSC[+57.021]APGSDPR.S | HexNAc(4)Hex(5)NeuAc(1) | C[+57]*2, N[+1914] | 1463.8665 | 4 | 5852.444 | 5852.4341 | 0 | 1.7 | 485 | Specific | 856.57 | 778.5 | 778.5 | 16.71 | 135 | >sp\|P02788\|TRFL_HUMAN Lactotransferrin OS=Homo sapiens OX=9606 GN=LTF PE=1 SV=6 | 1270 | 20201114_13813_1ng_glyco_hcd203040_compound15.22539.22539.4 | controllerType=0 controllerNumber=1 scan=22539 | 45.6381 |
| 18885:4 | 1 | R.TAGWNIPMGLLFN[+2205.793]QTGSC[+57.021]KFDEYFSQSC[+57.021]APGSDPR.S | HexNAc(4)Hex(5)Fuc(2)NeuAc(1) | C[+57]*2, N[+2206] | 1536.8976 | 4 | 6144.5685 | 6144.5499 | 0 | 3 | 485 | Specific | 891.32 | 837.07 | 837.07 | 16.69 | 135 | >sp\|P02788\|TRFL_HUMAN Lactotransferrin OS=Homo sapiens OX=9606 GN=LTF PE=1 SV=6 | 1270 | 20201114_13813_1ng_glyco_hcd203040_compound15.22443.22443.4 | controllerType=0 controllerNumber=1 scan=22443 | 45.4363 |
| 08835:3 | 1 | R.N[+2205.793]GSDC[+57.021]PDKFC[+57.021]LFQSETK.N | HexNAc(4)Hex(5)Fuc(2)NeuAc(1) | C[+57]*2, N[+2206] | 1413.5627 | 3 | 4238.6735 | 4238.6667 | 0 | 1.6 | 642 | Specific | 210.26 | 79.55 | 79.55 | 3.25 | 135 | >sp\|P02788\|TRFL_HUMAN Lactotransferrin OS=Homo sapiens OX=9606 GN=LTF PE=1 SV=6 | 1270 | 20201114_13813_1ng_glyco_hcd203040_compound15.11941.11941.3 | controllerType=0 controllerNumber=1 scan=11941 | 26.4642 |
| 11284:3 | 1 | R.N[+2350.830]GSDC[+57.021]PDKFC[+57.021]LFQSETK.N | HexNAc(4)Hex(5)Fuc(1)NeuAc(2) | C[+57]*2, N[+2351] | 1461.9162 | 3 | 4383.734 | 4383.7043 | 0 | 6.8 | 642 | Specific | 383.42 | 331.22 | 331.22 | 6.32 | 135 | >sp\|P02788\|TRFL_HUMAN Lactotransferrin OS=Homo sapiens OX=9606 GN=LTF PE=1 SV=6 | 1270 | 20201114_13813_1ng_glyco_hcd203040_compound15.14484.14484.3 | controllerType=0 controllerNumber=1 scan=14484 | 30.7413 |
| 08833:3 | 1 | R.N[+2205.793]GSDC[+57.021]PDKFC[+57.021]LFQSETK.N | HexNAc(4)Hex(5)Fuc(2)NeuAc(1) | C[+57]*2, N[+2206] | 1413.5627 | 3 | 4238.6735 | 4238.6667 | 0 | 1.6 | 642 | Specific | 394.03 | 298.56 | 298.56 | 6.98 | 135 | >sp\|P02788\|TRFL_HUMAN Lactotransferrin OS=Homo sapiens OX=9606 GN=LTF PE=1 SV=6 | 1270 | 20201114_13813_1ng_glyco_hcd203040_compound15.11939.11939.3 | controllerType=0 controllerNumber=1 scan=11939 | 26.4597 |
| 11286:3 | 1 | R.N[+2350.830]GSDC[+57.021]PDKFC[+57.021]LFQSETK.N | HexNAc(4)Hex(5)Fuc(1)NeuAc(2) | C[+57]*2, N[+2351] | 1461.9162 | 3 | 4383.734 | 4383.7043 | 0 | 6.8 | 642 | Specific | 471.41 | 385.54 | 385.54 | 8.05 | 135 | >sp\|P02788\|TRFL_HUMAN Lactotransferrin OS=Homo sapiens OX=9606 GN=LTF PE=1 SV=6 | 1270 | 20201114_13813_1ng_glyco_hcd203040_compound15.14486.14486.3 | controllerType=0 controllerNumber=1 scan=14486 | 30.7455 |
| 15394:3 | 2 | R.LN[+1216.423]YSLPTGQWVGVQLPR.N | HexNAc(2)Hex(5) | N[+1216] | 1048.8278 | 3 | 3144.4688 | 3144.4617 | 0 | 2.3 | 1017 | Specific | 348.11 | 348.11 | 348.11 | 7.78 | 87 | >sp\|P24821\|TENA_HUMAN Tenascin OS=Homo sapiens OX=9606 GN=TNC PE=1 SV=3 | 486 | 20201114_13813_1ng_glyco_hcd203040_compound15.18769.18769.3 | controllerType=0 controllerNumber=1 scan=18769 | 38.1317 |
| 15393:3 | 2 | R.LN[+1216.423]YSLPTGQWVGVQLPR.N | HexNAc(2)Hex(5) | N[+1216] | 1048.8278 | 3 | 3144.4688 | 3144.4617 | 0 | 2.3 | 1017 | Specific | 685.45 | 685.45 | 685.45 | 12.99 | 87 | >sp\|P24821\|TENA_HUMAN Tenascin OS=Homo sapiens OX=9606 GN=TNC PE=1 SV=3 | 486 | 20201114_13813_1ng_glyco_hcd203040_compound15.18768.18768.3 | controllerType=0 controllerNumber=1 scan=18768 | 38.1273 |
| 09985:3 | 5 | K.TVLTPATNHMGN[+1378.476]VTFTIPANR.E | HexNAc(2)Hex(6) | N[+1378] | 1211.8837 | 3 | 3633.6366 | 3633.6358 | 0 | 0.2 | 74 | Specific | 340.54 | 340.54 | 161.01 | 7.11 | 86 | >sp\|P01024\|CO3_HUMAN Complement C3 OS=Homo sapiens OX=9606 GN=C3 PE=1 SV=2 | 492 | 20201114_13813_1ng_glyco_hcd203040_compound15.13134.13134.2 | controllerType=0 controllerNumber=1 scan=13134 | 28.4212 |
| 09925:3 | 5 | K.TVLTPATNHMGN[+1378.476]VTFTIPANR.E | HexNAc(2)Hex(6) | N[+1378] | 1211.8849 | 3 | 3633.6403 | 3633.6358 | 0 | 1.2 | 74 | Specific | 528.39 | 528.39 | 203.11 | 9.81 | 86 | >sp\|P01024\|CO3_HUMAN Complement C3 OS=Homo sapiens OX=9606 GN=C3 PE=1 SV=2 | 492 | 20201114_13813_1ng_glyco_hcd203040_compound15.13072.13072.3 | controllerType=0 controllerNumber=1 scan=13072 | 28.3181 |
| 09923:3 | 5 | K.TVLTPATNHMGN[+1378.476]VTFTIPANR.E | HexNAc(2)Hex(6) | N[+1378] | 1211.8849 | 3 | 3633.6403 | 3633.6358 | 0 | 1.2 | 74 | Specific | 531.79 | 531.79 | 266.74 | 9.81 | 86 | >sp\|P01024\|CO3_HUMAN Complement C3 OS=Homo sapiens OX=9606 GN=C3 PE=1 SV=2 | 492 | 20201114_13813_1ng_glyco_hcd203040_compound15.13070.13070.3 | controllerType=0 controllerNumber=1 scan=13070 | 28.3141 |
| 07397:3 | 6 | K.WEC[+57.021]KN[+2205.793]DTLLGIK.G | HexNAc(4)Hex(5)Fuc(2)NeuAc(1) | C[+57], N[+2206] | 1228.1885 | 3 | 3682.551 | 3682.5444 | 0 | 1.8 | 100 | Specific | 291.7 | 291.7 | 291.7 | 6.36 | 76 | >sp\|P22897\|MRC1_HUMAN Macrophage mannose receptor 1 OS=Homo sapiens OX=9606 GN=MRC1 PE=1 SV=1 | 1049 | 20201114_13813_1ng_glyco_hcd203040_compound15.10433.10433.3 | controllerType=0 controllerNumber=1 scan=10433 | 23.9085 |
| 07399:3 | 6 | K.WEC[+57.021]KN[+2205.793]DTLLGIK.G | HexNAc(4)Hex(5)Fuc(2)NeuAc(1) | C[+57], N[+2206] | 1228.1885 | 3 | 3682.551 | 3682.5444 | 0 | 1.8 | 100 | Specific | 349.65 | 316.9 | 316.9 | 6.85 | 76 | >sp\|P22897\|MRC1_HUMAN Macrophage mannose receptor 1 OS=Homo sapiens OX=9606 GN=MRC1 PE=1 SV=1 | 1049 | 20201114_13813_1ng_glyco_hcd203040_compound15.10435.10435.3 | controllerType=0 controllerNumber=1 scan=10435 | 23.912 |
| 07500:3 | 6 | K.WEC[+57.021]KN[+2059.735]DTLLGIK.G | HexNAc(4)Hex(5)Fuc(1)NeuAc(1) | C[+57], N[+2060] | 1179.5031 | 3 | 3536.4948 | 3536.4865 | 0 | 2.4 | 100 | Specific | 388.32 | 305.48 | 305.48 | 6.72 | 76 | >sp\|P22897\|MRC1_HUMAN Macrophage mannose receptor 1 OS=Homo sapiens OX=9606 GN=MRC1 PE=1 SV=1 | 1049 | 20201114_13813_1ng_glyco_hcd203040_compound15.10539.10539.3 | controllerType=0 controllerNumber=1 scan=10539 | 24.0858 |
| 07502:3 | 6 | K.WEC[+57.021]KN[+2059.735]DTLLGIK.G | HexNAc(4)Hex(5)Fuc(1)NeuAc(1) | C[+57], N[+2060] | 1179.5031 | 3 | 3536.4948 | 3536.4865 | 0 | 2.4 | 100 | Specific | 401.82 | 55.15 | 55.15 | 4.12 | 76 | >sp\|P22897\|MRC1_HUMAN Macrophage mannose receptor 1 OS=Homo sapiens OX=9606 GN=MRC1 PE=1 SV=1 | 1049 | 20201114_13813_1ng_glyco_hcd203040_compound15.10541.10541.3 | controllerType=0 controllerNumber=1 scan=10541 | 24.0902 |
| 08245:3 | 6 | K.TAHC[+57.021]N[+2059.735]ESFYFLC[+57.021]K.R | HexNAc(4)Hex(5)Fuc(1)NeuAc(1) | C[+57]*2, N[+2060] | 1246.1578 | 3 | 3736.4588 | 3736.4545 | 0 | 1.1 | 1201 | Specific | 388.7 | 226.39 | 226.39 | 6.63 | 76 | >sp\|P22897\|MRC1_HUMAN Macrophage mannose receptor 1 OS=Homo sapiens OX=9606 GN=MRC1 PE=1 SV=1 | 1049 | 20201114_13813_1ng_glyco_hcd203040_compound15.11316.11316.3 | controllerType=0 controllerNumber=1 scan=11316 | 25.3726 |
| 08247:3 | 6 | K.TAHC[+57.021]N[+2059.735]ESFYFLC[+57.021]K.R | HexNAc(4)Hex(5)Fuc(1)NeuAc(1) | C[+57]*2, N[+2060] | 1246.1578 | 3 | 3736.4588 | 3736.4545 | 0 | 1.1 | 1201 | Specific | 461.36 | 395.06 | 395.06 | 9.12 | 76 | >sp\|P22897\|MRC1_HUMAN Macrophage mannose receptor 1 OS=Homo sapiens OX=9606 GN=MRC1 PE=1 SV=1 | 1049 | 20201114_13813_1ng_glyco_hcd203040_compound15.11318.11318.3 | controllerType=0 controllerNumber=1 scan=11318 | 25.3765 |
| 08223:3 | 6 | K.TAHC[+57.021]N[+2059.735]ESFYFLC[+57.021]K.R | HexNAc(4)Hex(5)Fuc(1)NeuAc(1) | C[+57]*2, N[+2060] | 1246.1585 | 3 | 3736.461 | 3736.4545 | 0 | 1.7 | 1201 | Specific | 580.17 | 452.1 | 452.1 | 10.82 | 76 | >sp\|P22897\|MRC1_HUMAN Macrophage mannose receptor 1 OS=Homo sapiens OX=9606 GN=MRC1 PE=1 SV=1 | 1049 | 20201114_13813_1ng_glyco_hcd203040_compound15.11293.11293.3 | controllerType=0 controllerNumber=1 scan=11293 | 25.3346 |
| 08221:3 | 6 | K.TAHC[+57.021]N[+2059.735]ESFYFLC[+57.021]K.R | HexNAc(4)Hex(5)Fuc(1)NeuAc(1) | C[+57]*2, N[+2060] | 1246.1585 | 3 | 3736.461 | 3736.4545 | 0 | 1.7 | 1201 | Specific | 649.72 | 468.7 | 468.7 | 11.82 | 76 | >sp\|P22897\|MRC1_HUMAN Macrophage mannose receptor 1 OS=Homo sapiens OX=9606 GN=MRC1 PE=1 SV=1 | 1049 | 20201114_13813_1ng_glyco_hcd203040_compound15.11291.11291.3 | controllerType=0 controllerNumber=1 scan=11291 | 25.3307 |
| 19504:3 | 7 | R.GLSFDVSLEVSQGPGLLN[+2350.830]DTK.V | HexNAc(4)Hex(5)Fuc(1)NeuAc(2) | N[+2351] | 1509.6602 | 3 | 4526.966 | 4526.9436 | 0 | 5 | 118 | Specific | 667.75 | 609.28 | 609.28 | 12.78 | 64 | >sp\|P01833\|PIGR_HUMAN Polymeric immunoglobulin receptor OS=Homo sapiens OX=9606 GN=PIGR PE=1 SV=4 | 902 | 20201114_13813_1ng_glyco_hcd203040_compound15.23094.23094.3 | controllerType=0 controllerNumber=1 scan=23094 | 46.7433 |
| 17646:3 | 7 | R.GLSFDVSLEVSQGPGLLN[+2059.735]DTK.V | HexNAc(4)Hex(5)Fuc(1)NeuAc(1) | N[+2060] | 1412.6249 | 3 | 4235.8603 | 4235.8481 | 0 | 2.9 | 118 | Specific | 936.66 | 891.77 | 891.77 | 18.07 | 64 | >sp\|P01833\|PIGR_HUMAN Polymeric immunoglobulin receptor OS=Homo sapiens OX=9606 GN=PIGR PE=1 SV=4 | 902 | 20201114_13813_1ng_glyco_hcd203040_compound15.21144.21144.3 | controllerType=0 controllerNumber=1 scan=21144 | 42.6638 |
| 18858:3 | 7 | K.QIGLYPVLVIDSSGYVNPN[+1913.677]YTGR.I | HexNAc(4)Hex(5)NeuAc(1) | N[+1914] | 1480.3428 | 3 | 4439.0139 | 4438.9805 | 0 | 7.5 | 168 | Specific | 625 | 525.22 | 525.22 | 12.15 | 64 | >sp\|P01833\|PIGR_HUMAN Polymeric immunoglobulin receptor OS=Homo sapiens OX=9606 GN=PIGR PE=1 SV=4 | 902 | 20201114_13813_1ng_glyco_hcd203040_compound15.22415.22415.3 | controllerType=0 controllerNumber=1 scan=22415 | 45.3835 |
| 18856:3 | 7 | K.QIGLYPVLVIDSSGYVNPN[+1913.677]YTGR.I | HexNAc(4)Hex(5)NeuAc(1) | N[+1914] | 1480.3428 | 3 | 4439.0139 | 4438.9805 | 0 | 7.5 | 168 | Specific | 801.61 | 621.71 | 621.71 | 16.15 | 64 | >sp\|P01833\|PIGR_HUMAN Polymeric immunoglobulin receptor OS=Homo sapiens OX=9606 GN=PIGR PE=1 SV=4 | 902 | 20201114_13813_1ng_glyco_hcd203040_compound15.22413.22413.3 | controllerType=0 controllerNumber=1 scan=22413 | 45.3784 |
| 22675:3 | 7 | K.AQYEGRLSLLEEPGN[+1663.608]GTFTVILNQLTSR.D | HexNAc(5)Hex(4) | N[+1664] | 1590.7326 | 3 | 4770.1832 | 4770.2249 | 0 | -8.7 | 407 | Specific | 223.07 | 217.09 | 217.09 | 2.22 | 64 | >sp\|P01833\|PIGR_HUMAN Polymeric immunoglobulin receptor OS=Homo sapiens OX=9606 GN=PIGR PE=1 SV=4 | 902 | 20201114_13813_1ng_glyco_hcd203040_compound15.26425.26425.3 | controllerType=0 controllerNumber=1 scan=26425 | 53.274 |
| 22737:3 | 7 | R.[+42.011]LSLLEEPGN[+2336.851]GTFTVILNQLTSR.D | HexNAc(6)Hex(6)Fuc(1) | .[42], N[+2337] | 1594.3852 | 3 | 4781.141 | 4781.1542 | 0 | -2.8 | 413 | Specific | 259.64 | 239.32 | 239.32 | 5.37 | 64 | >sp\|P01833\|PIGR_HUMAN Polymeric immunoglobulin receptor OS=Homo sapiens OX=9606 GN=PIGR PE=1 SV=4 | 902 | 20201114_13813_1ng_glyco_hcd203040_compound15.26489.26489.2 | controllerType=0 controllerNumber=1 scan=26489 | 53.383 |
| 22738:3 | 7 | R.[+42.011]LSLLEEPGN[+2336.851]GTFTVILNQLTSR.D | HexNAc(6)Hex(6)Fuc(1) | .[42], N[+2337] | 1594.3852 | 3 | 4781.141 | 4781.1542 | 0 | -2.8 | 413 | Specific | 298.01 | 262.62 | 262.62 | 5.72 | 64 | >sp\|P01833\|PIGR_HUMAN Polymeric immunoglobulin receptor OS=Homo sapiens OX=9606 GN=PIGR PE=1 SV=4 | 902 | 20201114_13813_1ng_glyco_hcd203040_compound15.26490.26490.2 | controllerType=0 controllerNumber=1 scan=26490 | 53.3863 |
| 20448:3 | 7 | R.LSLLEEPGN[+2059.735]GTFTVILNQLTSR.D | HexNAc(4)Hex(5)Fuc(1)NeuAc(1) | N[+2060] | 1488.0153 | 3 | 4462.0314 | 4462.0275 | 0 | 0.9 | 413 | Specific | 334.57 | 334.57 | 334.57 | 7.39 | 64 | >sp\|P01833\|PIGR_HUMAN Polymeric immunoglobulin receptor OS=Homo sapiens OX=9606 GN=PIGR PE=1 SV=4 | 902 | 20201114_13813_1ng_glyco_hcd203040_compound15.24082.24082.3 | controllerType=0 controllerNumber=1 scan=24082 | 48.7668 |
| 28719:3 | 7 | R.[+42.011]LSLLEEPGN[+2351.851]GTFTVILNQLTSR.D | HexNAc(4)Hex(5)Fuc(3)NeuAc(1) | .[42], N[+2352] | 1599.3881 | 3 | 4796.1498 | 4796.1539 | 0 | -0.9 | 413 | Specific | 368.31 | 322.28 | 322.28 | 7.35 | 64 | >sp\|P01833\|PIGR_HUMAN Polymeric immunoglobulin receptor OS=Homo sapiens OX=9606 GN=PIGR PE=1 SV=4 | 902 | 20201114_13813_1ng_glyco_hcd203040_compound15.32826.32826.3 | controllerType=0 controllerNumber=1 scan=32826 | 64.9846 |
| 20467:3 | 7 | R.LSLLEEPGN[+2076.750]GTFTVILNQLTSR.D | HexNAc(4)Hex(6)Fuc(2) | N[+2077] | 1493.6905 | 3 | 4479.0569 | 4479.0428 | 0 | 3.1 | 413 | Specific | 418.69 | 368.11 | 368.11 | 7.86 | 64 | >sp\|P01833\|PIGR_HUMAN Polymeric immunoglobulin receptor OS=Homo sapiens OX=9606 GN=PIGR PE=1 SV=4 | 902 | 20201114_13813_1ng_glyco_hcd203040_compound15.24102.24102.3 | controllerType=0 controllerNumber=1 scan=24102 | 48.8018 |

Table S4. Cont.

| Query #:z | Protein Rank | Glycopeptide | Glycan | Modification Type(s) | Observed m/z | z | Observed (M+H) | Calc. mass (M+H) | Off-by-x error | Mass error (ppm) | StartPosition | Cleavage | Score | Delta | Delta Mod | \|Log Prob\| | # of unique peptides | Protein Name | Protein DB number | Comment | Scan # | Scan Time |
|---|---|---|---|---|---|---|---|---|---|---|---|---|---|---|---|---|---|---|---|---|---|---|
| 22893:3 | 7 | R.LSLLEEPGN[+2350.830]GTFTVILNQLTSR.D | HexNAc(4)Hex(5)Fuc(1)NeuAc(2) | N[+2351] | 1585.0508 | 3 | 4753.138 | 4753.1229 | 0 | 3.2 | 413 | Specific | 476.44 | 452.73 | 452.73 | 9.37 | 64 | >sp\|P01833\|PIGR_HUMAN Polymeric immunoglobulin receptor OS=Homo sapiens OX=9606 GN=PIGR PE=1 SV=4 | 902 | 20201114_13813_1ng_glyco_hcd203040_compound15.26652.26652.3 | controllerType=0 controllerNumber=1 scan=26652 | 53.6929 |
| 20282:3 | 7 | R.LSLLEEPGN[+2569.905]GTFTVILNQLTSR.D | HexNAc(5)Hex(6)NeuAc(2) | N[+2570] | 1658.0775 | 3 | 4972.2179 | 4972.1972 | 0 | 4.2 | 413 | Specific | 628.27 | 459.23 | 459.23 | 11.78 | 64 | >sp\|P01833\|PIGR_HUMAN Polymeric immunoglobulin receptor OS=Homo sapiens OX=9606 GN=PIGR PE=1 SV=4 | 902 | 20201114_13813_1ng_glyco_hcd203040_compound15.23908.23908.3 | controllerType=0 controllerNumber=1 scan=23908 | 48.4177 |
| 20450:3 | 7 | R.LSLLEEPGN[+2059.735]GTFTVILNQLTSR.D | HexNAc(4)Hex(5)Fuc(1)NeuAc(1) | N[+2060] | 1488.0153 | 3 | 4462.0314 | 4462.0275 | 0 | 0.9 | 413 | Specific | 725.79 | 553.84 | 553.84 | 14.1 | 64 | >sp\|P01833\|PIGR_HUMAN Polymeric immunoglobulin receptor OS=Homo sapiens OX=9606 GN=PIGR PE=1 SV=4 | 902 | 20201114_13813_1ng_glyco_hcd203040_compound15.24084.24084.3 | controllerType=0 controllerNumber=1 scan=24084 | 48.7708 |
| 20280:3 | 7 | R.LSLLEEPGN[+2569.905]GTFTVILNQLTSR.D | HexNAc(5)Hex(6)NeuAc(2) | N[+2570] | 1658.0775 | 3 | 4972.2179 | 4972.1972 | 0 | 4.2 | 413 | Specific | 883.12 | 651.28 | 651.28 | 16.78 | 64 | >sp\|P01833\|PIGR_HUMAN Polymeric immunoglobulin receptor OS=Homo sapiens OX=9606 GN=PIGR PE=1 SV=4 | 902 | 20201114_13813_1ng_glyco_hcd203040_compound15.23906.23906.3 | controllerType=0 controllerNumber=1 scan=23906 | 48.4133 |
| 17896:4 | 7 | K.IIEGEPNLKVPGN[+2350.830]VTAVLGETLK.V | HexNAc(4)Hex(5)Fuc(1)NeuAc(2) | N[+2351] | 1186.3088 | 4 | 4742.2132 | 4742.1797 | 0 | 7.1 | 457 | Specific | 299.53 | 298.78 | 298.78 | 6.69 | 64 | >sp\|P01833\|PIGR_HUMAN Polymeric immunoglobulin receptor OS=Homo sapiens OX=9606 GN=PIGR PE=1 SV=4 | 902 | 20201114_13813_1ng_glyco_hcd203040_compound15.21404.21404.4 | controllerType=0 controllerNumber=1 scan=21404 | 43.1916 |
| 15988:4 | 7 | K.IIEGEPNLKVPGN[+2059.735]VTAVLGETLK.V | HexNAc(4)Hex(5)Fuc(1)NeuAc(1) | N[+2060] | 1113.5276 | 4 | 4451.0886 | 4451.0843 | 0 | 1 | 457 | Specific | 367.35 | 309.27 | 309.27 | 7.85 | 64 | >sp\|P01833\|PIGR_HUMAN Polymeric immunoglobulin receptor OS=Homo sapiens OX=9606 GN=PIGR PE=1 SV=4 | 902 | 20201114_13813_1ng_glyco_hcd203040_compound15.19412.19412.4 | controllerType=0 controllerNumber=1 scan=19412 | 39.3301 |
| 15986:4 | 7 | K.IIEGEPNLKVPGN[+2059.735]VTAVLGETLK.V | HexNAc(4)Hex(5)Fuc(1)NeuAc(1) | N[+2060] | 1113.5276 | 4 | 4451.0886 | 4451.0843 | 0 | 1 | 457 | Specific | 418.17 | 326.68 | 326.68 | 6.91 | 64 | >sp\|P01833\|PIGR_HUMAN Polymeric immunoglobulin receptor OS=Homo sapiens OX=9606 GN=PIGR PE=1 SV=4 | 902 | 20201114_13813_1ng_glyco_hcd203040_compound15.19410.19410.4 | controllerType=0 controllerNumber=1 scan=19410 | 39.3251 |
| 17899:4 | 7 | K.IIEGEPNLKVPGN[+2350.830]VTAVLGETLK.V | HexNAc(4)Hex(5)Fuc(1)NeuAc(2) | N[+2351] | 1186.3088 | 4 | 4742.2132 | 4742.1797 | 0 | 7.1 | 457 | Specific | 521.54 | 430.98 | 430.98 | 9.27 | 64 | >sp\|P01833\|PIGR_HUMAN Polymeric immunoglobulin receptor OS=Homo sapiens OX=9606 GN=PIGR PE=1 SV=4 | 902 | 20201114_13813_1ng_glyco_hcd203040_compound15.21407.21407.4 | controllerType=0 controllerNumber=1 scan=21407 | 43.1958 |
| 17966:3 | 7 | K.IIEGEPNLKVPGN[+2350.830]VTAVLGETLK.V | HexNAc(4)Hex(5)Fuc(1)NeuAc(2) | N[+2351] | 1581.4013 | 3 | 4742.1894 | 4742.1797 | 0 | 2 | 457 | Specific | 525.92 | 435.16 | 435.16 | 10.08 | 64 | >sp\|P01833\|PIGR_HUMAN Polymeric immunoglobulin receptor OS=Homo sapiens OX=9606 GN=PIGR PE=1 SV=4 | 902 | 20201114_13813_1ng_glyco_hcd203040_compound15.21477.21477.3 | controllerType=0 controllerNumber=1 scan=21477 | 43.3382 |
| 15974:3 | 7 | K.IIEGEPNLKVPGN[+1913.677]VTAVLGETLK.V | HexNAc(4)Hex(5)NeuAc(1) | N[+1914] | 1435.6832 | 3 | 4305.035 | 4305.0264 | 0 | 2 | 457 | Specific | 545.4 | 545.4 | 545.4 | 10.22 | 64 | >sp\|P01833\|PIGR_HUMAN Polymeric immunoglobulin receptor OS=Homo sapiens OX=9606 GN=PIGR PE=1 SV=4 | 902 | 20201114_13813_1ng_glyco_hcd203040_compound15.19398.19398.3 | controllerType=0 controllerNumber=1 scan=19398 | 39.2979 |
| 15963:3 | 7 | K.IIEGEPNLKVPGN[+2205.793]VTAVLGETLK.V | HexNAc(4)Hex(5)Fuc(2)NeuAc(1) | N[+2206] | 1533.0593 | 3 | 4597.1632 | 4597.1422 | 0 | 4.6 | 457 | Specific | 669.64 | 669.64 | 669.64 | 13.09 | 64 | >sp\|P01833\|PIGR_HUMAN Polymeric immunoglobulin receptor OS=Homo sapiens OX=9606 GN=PIGR PE=1 SV=4 | 902 | 20201114_13813_1ng_glyco_hcd203040_compound15.19386.19386.3 | controllerType=0 controllerNumber=1 scan=19386 | 39.2763 |
| 15961:3 | 7 | K.IIEGEPNLKVPGN[+2205.793]VTAVLGETLK.V | HexNAc(4)Hex(5)Fuc(2)NeuAc(1) | N[+2206] | 1533.0593 | 3 | 4597.1632 | 4597.1422 | 0 | 4.6 | 457 | Specific | 1017.58 | 1005.56 | 1005.56 | 20.09 | 64 | >sp\|P01833\|PIGR_HUMAN Polymeric immunoglobulin receptor OS=Homo sapiens OX=9606 GN=PIGR PE=1 SV=4 | 902 | 20201114_13813_1ng_glyco_hcd203040_compound15.19384.19384.3 | controllerType=0 controllerNumber=1 scan=19384 | 39.2712 |
| 15549:5 | 7 | K.IIEGEPNLKVPGN[+2059.735]VTAVLGETLKVPC[+57.021]HFPC[+57.021]K.F | HexNAc(4)Hex(5)Fuc(1)NeuAc(1) | C[+57]*2, N[+2060] | 1096.1245 | 5 | 5476.5934 | 5476.5418 | 0 | 9.4 | 457 | Specific | 621.44 | 577.61 | 577.61 | 10.87 | 64 | >sp\|P01833\|PIGR_HUMAN Polymeric immunoglobulin receptor OS=Homo sapiens OX=9606 GN=PIGR PE=1 SV=4 | 902 | 20201114_13813_1ng_glyco_hcd203040_compound15.18931.18931.5 | controllerType=0 controllerNumber=1 scan=18931 | 38.4262 |
| 15547:5 | 7 | K.IIEGEPNLKVPGN[+2059.735]VTAVLGETLKVPC[+57.021]HFPC[+57.021]K.F | HexNAc(4)Hex(5)Fuc(1)NeuAc(1) | C[+57]*2, N[+2060] | 1096.1245 | 5 | 5476.5934 | 5476.5418 | 0 | 9.4 | 457 | Specific | 644.13 | 644.13 | 644.13 | 10.87 | 64 | >sp\|P01833\|PIGR_HUMAN Polymeric immunoglobulin receptor OS=Homo sapiens OX=9606 GN=PIGR PE=1 SV=4 | 902 | 20201114_13813_1ng_glyco_hcd203040_compound15.18929.18929.5 | controllerType=0 controllerNumber=1 scan=18929 | 38.421 |
| 15553:4 | 7 | K.IIEGEPNLKVPGN[+2059.735]VTAVLGETLKVPC[+57.021]HFPC[+57.021]K.F | HexNAc(4)Hex(5)Fuc(1)NeuAc(1) | C[+57]*2, N[+2060] | 1369.9038 | 4 | 5476.5934 | 5476.5418 | 0 | 9.4 | 457 | Specific | 677.39 | 677.39 | 677.39 | 11.19 | 64 | >sp\|P01833\|PIGR_HUMAN Polymeric immunoglobulin receptor OS=Homo sapiens OX=9606 GN=PIGR PE=1 SV=4 | 902 | 20201114_13813_1ng_glyco_hcd203040_compound15.18935.18935.4 | controllerType=0 controllerNumber=1 scan=18935 | 38.4358 |
| 15555:4 | 7 | K.IIEGEPNLKVPGN[+2059.735]VTAVLGETLKVPC[+57.021]HFPC[+57.021]K.F | HexNAc(4)Hex(5)Fuc(1)NeuAc(1) | C[+57]*2, N[+2060] | 1369.9038 | 4 | 5476.5934 | 5476.5418 | 0 | 9.4 | 457 | Specific | 689.03 | 670.88 | 670.88 | 11.19 | 64 | >sp\|P01833\|PIGR_HUMAN Polymeric immunoglobulin receptor OS=Homo sapiens OX=9606 GN=PIGR PE=1 SV=4 | 902 | 20201114_13813_1ng_glyco_hcd203040_compound15.18937.18937.4 | controllerType=0 controllerNumber=1 scan=18937 | 38.4408 |
| 13485:3 | 7 | K.VPGN[+2059.735]VTAVLGETLK.V | HexNAc(4)Hex(5)Fuc(1)NeuAc(1) | N[+2060] | 1153.1862 | 3 | 3457.5441 | 3457.5348 | 0 | 2.7 | 466 | Specific | 513.54 | 382.76 | 382.76 | 9.55 | 64 | >sp\|P01833\|PIGR_HUMAN Polymeric immunoglobulin receptor OS=Homo sapiens OX=9606 GN=PIGR PE=1 SV=4 | 902 | 20201114_13813_1ng_glyco_hcd203040_compound15.16784.16784.3 | controllerType=0 controllerNumber=1 scan=16784 | 34.6196 |
| 16516:3 | 7 | K.VPGN[+2350.830]VTAVLGETLK.V | HexNAc(4)Hex(5)Fuc(1)NeuAc(2) | N[+2351] | 1250.22 | 3 | 3748.6454 | 3748.6302 | 0 | 4.1 | 466 | Specific | 747.96 | 559.48 | 559.48 | 13.7 | 64 | >sp\|P01833\|PIGR_HUMAN Polymeric immunoglobulin receptor OS=Homo sapiens OX=9606 GN=PIGR PE=1 SV=4 | 902 | 20201114_13813_1ng_glyco_hcd203040_compound15.19965.19965.3 | controllerType=0 controllerNumber=1 scan=19965 | 40.3734 |
| 13487:3 | 7 | K.VPGN[+2059.735]VTAVLGETLK.V | HexNAc(4)Hex(5)Fuc(1)NeuAc(1) | N[+2060] | 1153.1862 | 3 | 3457.5441 | 3457.5348 | 0 | 2.7 | 466 | Specific | 748.92 | 567.2 | 567.2 | 13.99 | 64 | >sp\|P01833\|PIGR_HUMAN Polymeric immunoglobulin receptor OS=Homo sapiens OX=9606 GN=PIGR PE=1 SV=4 | 902 | 20201114_13813_1ng_glyco_hcd203040_compound15.16786.16786.3 | controllerType=0 controllerNumber=1 scan=16786 | 34.6239 |
| 13371:3 | 7 | K.VPGN[+2205.793]VTAVLGETLK.V | HexNAc(4)Hex(5)Fuc(2)NeuAc(1) | N[+2206] | 1201.8694 | 3 | 3603.5938 | 3603.5927 | 0 | 0.3 | 466 | Specific | 758.02 | 679.87 | 679.87 | 14.69 | 64 | >sp\|P01833\|PIGR_HUMAN Polymeric immunoglobulin receptor OS=Homo sapiens OX=9606 GN=PIGR PE=1 SV=4 | 902 | 20201114_13813_1ng_glyco_hcd203040_compound15.16665.16665.3 | controllerType=0 controllerNumber=1 scan=16665 | 34.4026 |
| 13369:3 | 7 | K.VPGN[+2205.793]VTAVLGETLK.V | HexNAc(4)Hex(5)Fuc(2)NeuAc(1) | N[+2206] | 1201.8694 | 3 | 3603.5938 | 3603.5927 | 0 | 0.3 | 466 | Specific | 825.15 | 769.72 | 769.72 | 15.69 | 64 | >sp\|P01833\|PIGR_HUMAN Polymeric immunoglobulin receptor OS=Homo sapiens OX=9606 GN=PIGR PE=1 SV=4 | 902 | 20201114_13813_1ng_glyco_hcd203040_compound15.16663.16663.3 | controllerType=0 controllerNumber=1 scan=16663 | 34.398 |
| 13532:4 | 7 | K.VPGN[+2059.735]VTAVLGETLKVPC[+57.021]HFPC[+57.021]K.F | HexNAc(4)Hex(5)Fuc(1)NeuAc(1) | C[+57]*2, N[+2060] | 1121.5092 | 4 | 4483.015 | 4482.9923 | 0 | 5.1 | 466 | Specific | 217.73 | 206.92 | 206.92 | 4.31 | 64 | >sp\|P01833\|PIGR_HUMAN Polymeric immunoglobulin receptor OS=Homo sapiens OX=9606 GN=PIGR PE=1 SV=4 | 902 | 20201114_13813_1ng_glyco_hcd203040_compound15.16832.16832.2 | controllerType=0 controllerNumber=1 scan=16832 | 34.7048 |
| 15568:3 | 7 | K.VPGN[+2351.851]VTAVLGETLKVPC[+57.021]HFPC[+57.021]K.F | HexNAc(4)Hex(5)Fuc(3)NeuAc(1) | C[+57]*2, N[+2352] | 1592.3765 | 3 | 4775.1149 | 4775.1081 | 0 | 1.4 | 466 | Specific | 278.9 | 278.9 | 278.9 | 5.72 | 64 | >sp\|P01833\|PIGR_HUMAN Polymeric immunoglobulin receptor OS=Homo sapiens OX=9606 GN=PIGR PE=1 SV=4 | 902 | 20201114_13813_1ng_glyco_hcd203040_compound15.18950.18950.3 | controllerType=0 controllerNumber=1 scan=18950 | 38.4607 |
| 13566:4 | 7 | K.VPGN[+2205.793]VTAVLGETLKVPC[+57.021]HFPC[+57.021]K.F | HexNAc(4)Hex(5)Fuc(2)NeuAc(1) | C[+57]*2, N[+2206] | 1158.0196 | 4 | 4629.0565 | 4629.0502 | 0 | 1.3 | 466 | Specific | 320.22 | 207.05 | 207.05 | 6.13 | 64 | >sp\|P01833\|PIGR_HUMAN Polymeric immunoglobulin receptor OS=Homo sapiens OX=9606 GN=PIGR PE=1 SV=4 | 902 | 20201114_13813_1ng_glyco_hcd203040_compound15.16868.16868.2 | controllerType=0 controllerNumber=1 scan=16868 | 34.7958 |
| 13438:4 | 7 | K.VPGN[+2205.793]VTAVLGETLKVPC[+57.021]HFPC[+57.021]K.F | HexNAc(4)Hex(5)Fuc(2)NeuAc(1) | C[+57]*2, N[+2206] | 1158.0231 | 4 | 4629.0706 | 4629.0502 | 0 | 4.4 | 466 | Specific | 357.88 | 357.88 | 357.88 | 7.06 | 64 | >sp\|P01833\|PIGR_HUMAN Polymeric immunoglobulin receptor OS=Homo sapiens OX=9606 GN=PIGR PE=1 SV=4 | 902 | 20201114_13813_1ng_glyco_hcd203040_compound15.16735.16735.3 | controllerType=0 controllerNumber=1 scan=16735 | 34.5328 |
| 13448:3 | 7 | K.VPGN[+2205.793]VTAVLGETLKVPC[+57.021]HFPC[+57.021]K.F | HexNAc(4)Hex(5)Fuc(2)NeuAc(1) | C[+57]*2, N[+2206] | 1543.7019 | 3 | 4629.0911 | 4629.0502 | 0 | 8.8 | 466 | Specific | 471.47 | 471.47 | 471.47 | 7.53 | 64 | >sp\|P01833\|PIGR_HUMAN Polymeric immunoglobulin receptor OS=Homo sapiens OX=9606 GN=PIGR PE=1 SV=4 | 902 | 20201114_13813_1ng_glyco_hcd203040_compound15.16745.16745.3 | controllerType=0 controllerNumber=1 scan=16745 | 34.5487 |
| 13535:4 | 7 | K.VPGN[+2205.793]VTAVLGETLKVPC[+57.021]HFPC[+57.021]K.F | HexNAc(4)Hex(5)Fuc(2)NeuAc(1) | C[+57]*2, N[+2206] | 1158.0241 | 4 | 4629.0745 | 4629.0502 | 0 | 5.3 | 466 | Specific | 548.89 | 519.29 | 519.29 | 8.92 | 64 | >sp\|P01833\|PIGR_HUMAN Polymeric immunoglobulin receptor OS=Homo sapiens OX=9606 GN=PIGR PE=1 SV=4 | 902 | 20201114_13813_1ng_glyco_hcd203040_compound15.16835.16835.2 | controllerType=0 controllerNumber=1 scan=16835 | 34.712 |
| 13450:3 | 7 | K.VPGN[+2205.793]VTAVLGETLKVPC[+57.021]HFPC[+57.021]K.F | HexNAc(4)Hex(5)Fuc(2)NeuAc(1) | C[+57]*2, N[+2206] | 1543.7019 | 3 | 4629.0911 | 4629.0502 | 0 | 8.8 | 466 | Specific | 774.46 | 726.39 | 726.39 | 13.23 | 64 | >sp\|P01833\|PIGR_HUMAN Polymeric immunoglobulin receptor OS=Homo sapiens OX=9606 GN=PIGR PE=1 SV=4 | 902 | 20201114_13813_1ng_glyco_hcd203040_compound15.16747.16747.3 | controllerType=0 controllerNumber=1 scan=16747 | 34.5527 |
| 13533:4 | 7 | K.VPGN[+2205.793]VTAVLGETLKVPC[+57.021]HFPC[+57.021]K.F | HexNAc(4)Hex(5)Fuc(2)NeuAc(1) | C[+57]*2, N[+2206] | 1158.0241 | 4 | 4629.0745 | 4629.0502 | 0 | 5.3 | 466 | Specific | 981.52 | 861.32 | 861.32 | 18.11 | 64 | >sp\|P01833\|PIGR_HUMAN Polymeric immunoglobulin receptor OS=Homo sapiens OX=9606 GN=PIGR PE=1 SV=4 | 902 | 20201114_13813_1ng_glyco_hcd203040_compound15.16833.16833.2 | controllerType=0 controllerNumber=1 scan=16833 | 34.7067 |
| 05728:3 | 7 | K.WN[+1548.545]NTGC[+57.021]QALPSQDEGPSK.A | HexNAc(3)Hex(4)NeuAc(1) | C[+57], N[+1549] | 1179.8217 | 3 | 3537.4506 | 3537.4215 | 0 | 8.2 | 498 | Specific | 416.86 | 375.84 | 375.84 | 8.52 | 64 | >sp\|P01833\|PIGR_HUMAN Polymeric immunoglobulin receptor OS=Homo sapiens OX=9606 GN=PIGR PE=1 SV=4 | 902 | 20201114_13813_1ng_glyco_hcd203040_compound15.8699.8699.3 | controllerType=0 controllerNumber=1 scan=8699 | 21.0034 |
| 05726:3 | 7 | K.WN[+1548.545]NTGC[+57.021]QALPSQDEGPSK.A | HexNAc(3)Hex(4)NeuAc(1) | C[+57], N[+1549] | 1179.8217 | 3 | 3537.4506 | 3537.4215 | 0 | 8.2 | 498 | Specific | 550.37 | 498.55 | 498.55 | 9.22 | 64 | >sp\|P01833\|PIGR_HUMAN Polymeric immunoglobulin receptor OS=Homo sapiens OX=9606 GN=PIGR PE=1 SV=4 | 902 | 20201114_13813_1ng_glyco_hcd203040_compound15.8697.8697.3 | controllerType=0 controllerNumber=1 scan=8697 | 21 |
| 07559:3 | 7 | K.WN[+2204.772]NTGC[+57.021]QALPSQDEGPSK.A | HexNAc(4)Hex(5)NeuAc(2) | C[+57], N[+2205] | 1398.558 | 3 | 4193.6596 | 4193.6491 | 0 | 2.5 | 498 | Specific | 622.46 | 520.02 | 520.02 | 12.07 | 64 | >sp\|P01833\|PIGR_HUMAN Polymeric immunoglobulin receptor OS=Homo sapiens OX=9606 GN=PIGR PE=1 SV=4 | 902 | 20201114_13813_1ng_glyco_hcd203040_compound15.10600.10600.3 | controllerType=0 controllerNumber=1 scan=10600 | 24.1864 |
| 05466:3 | 7 | K.WN[+2059.735]NTGC[+57.021]QALPSQDEGPSK.A | HexNAc(4)Hex(5)Fuc(1)NeuAc(1) | C[+57], N[+2060] | 1350.2123 | 3 | 4048.6225 | 4048.6116 | 0 | 2.7 | 498 | Specific | 623.46 | 544.93 | 544.93 | 12.07 | 64 | >sp\|P01833\|PIGR_HUMAN Polymeric immunoglobulin receptor OS=Homo sapiens OX=9606 GN=PIGR PE=1 SV=4 | 902 | 20201114_13813_1ng_glyco_hcd203040_compound15.8428.8428.3 | controllerType=0 controllerNumber=1 scan=8428 | 20.5489 |
| 05464:3 | 7 | K.WN[+2059.735]NTGC[+57.021]QALPSQDEGPSK.A | HexNAc(4)Hex(5)Fuc(1)NeuAc(1) | C[+57], N[+2060] | 1350.2123 | 3 | 4048.6225 | 4048.6116 | 0 | 2.7 | 498 | Specific | 697.2 | 620.73 | 620.73 | 13.07 | 64 | >sp\|P01833\|PIGR_HUMAN Polymeric immunoglobulin receptor OS=Homo sapiens OX=9606 GN=PIGR PE=1 SV=4 | 902 | 20201114_13813_1ng_glyco_hcd203040_compound15.8426.8426.3 | controllerType=0 controllerNumber=1 scan=8426 | 20.5451 |
| 07561:3 | 7 | K.WN[+2204.772]NTGC[+57.021]QALPSQDEGPSK.A | HexNAc(4)Hex(5)NeuAc(2) | C[+57], N[+2205] | 1398.558 | 3 | 4193.6596 | 4193.6491 | 0 | 2.5 | 498 | Specific | 740.01 | 639.7 | 639.7 | 14.07 | 64 | >sp\|P01833\|PIGR_HUMAN Polymeric immunoglobulin receptor OS=Homo sapiens OX=9606 GN=PIGR PE=1 SV=4 | 902 | 20201114_13813_1ng_glyco_hcd203040_compound15.10602.10602.3 | controllerType=0 controllerNumber=1 scan=10602 | 24.1903 |
| 07123:3 | 8 | R.FSDGLESN[+1864.634]SSTQFEVK.K | HexNAc(2)Hex(9) | N[+1865] | 1213.8242 | 3 | 3639.4579 | 3639.4472 | 0 | 3 | 219 | Specific | 219.81 | 219.81 | 219.81 | 5.02 | 62 | >sp\|P0C0L5\|CO4B_HUMAN Complement C4-B OS=Homo sapiens OX=9606 GN=C4B PE=1 SV=2 | 1424 | 20201114_13813_1ng_glyco_hcd203040_compound15.10150.10150.3 | controllerType=0 controllerNumber=1 scan=10150 | 23.4377 |
| 07125:3 | 8 | R.FSDGLESN[+1864.634]SSTQFEVK.K | HexNAc(2)Hex(9) | N[+1865] | 1213.8242 | 3 | 3639.4579 | 3639.4472 | 0 | 3 | 219 | Specific | 432.95 | 432.95 | 432.95 | 9.31 | 62 | >sp\|P0C0L5\|CO4B_HUMAN Complement C4-B OS=Homo sapiens OX=9606 GN=C4B PE=1 SV=2 | 1424 | 20201114_13813_1ng_glyco_hcd203040_compound15.10152.10152.3 | controllerType=0 controllerNumber=1 scan=10152 | 23.4417 |
| 17649:5 | 11 | R.LSLHRPALEDLLLGSEAN[+1524.534]LTC[+57.021]TLTGLR.D | HexNAc(2)Hex(6)Fuc(1) | C[+57], N[+1525] | 898.4307 | 5 | 4488.1246 | 4488.1319 | 0 | -1.6 | 127 | Specific | 266.16 | 262.63 | 262.63 | 4.57 | 44 | >sp\|P01876\|IGHA1_HUMAN Immunoglobulin heavy constant alpha 1 OS=Homo sapiens OX=9606 GN=IGHA1 PE=1 SV=2 | 255 | 20201114_13813_1ng_glyco_hcd203040_compound15.21147.21147.5 | controllerType=0 controllerNumber=1 scan=21147 | 42.6692 |
| 19730:4 | 11 | R.LSLHRPALEDLLLGSEAN[+2204.772]LTC[+57.021]TLTGLR.D | HexNAc(4)Hex(5)NeuAc(2) | C[+57], N[+2205] | 1292.8475 | 4 | 5168.3683 | 5168.3707 | 0 | -0.5 | 127 | Specific | 288.55 | 288.55 | 288.55 | 7.23 | 44 | >sp\|P01876\|IGHA1_HUMAN Immunoglobulin heavy constant alpha 1 OS=Homo sapiens OX=9606 GN=IGHA1 PE=1 SV=2 | 255 | 20201114_13813_1ng_glyco_hcd203040_compound15.23329.23329.2 | controllerType=0 controllerNumber=1 scan=23329 | 47.2005 |
| 17683:3 | 11 | R.LSLHRPALEDLLLGSEAN[+1095.397]LTC[+57.021]TLTGLR.D | HexNAc(3)Hex(3) | C[+57], N[+1095] | 1353.667 | 3 | 4058.9866 | 4058.9949 | 0 | -2 | 127 | Specific | 361.93 | 328.51 | 328.51 | 6.41 | 44 | >sp\|P01876\|IGHA1_HUMAN Immunoglobulin heavy constant alpha 1 OS=Homo sapiens OX=9606 GN=IGHA1 PE=1 SV=2 | 255 | 20201114_13813_1ng_glyco_hcd203040_compound15.21183.21183.3 | controllerType=0 controllerNumber=1 scan=21183 | 42.7631 |
| 17626:4 | 11 | R.LSLHRPALEDLLLGSEAN[+1501.555]LTC[+57.021]TLTGLR.D | HexNAc(5)Hex(3) | C[+57], N[+1502] | 1117.0477 | 4 | 4465.1688 | 4465.1536 | 0 | 3.4 | 127 | Specific | 470.86 | 470.86 | 470.86 | 9.73 | 44 | >sp\|P01876\|IGHA1_HUMAN Immunoglobulin heavy constant alpha 1 OS=Homo sapiens OX=9606 GN=IGHA1 PE=1 SV=2 | 255 | 20201114_13813_1ng_glyco_hcd203040_compound15.21124.21124.4 | controllerType=0 controllerNumber=1 scan=21124 | 42.6258 |
| 17685:3 | 11 | R.LSLHRPALEDLLLGSEAN[+1095.397]LTC[+57.021]TLTGLR.D | HexNAc(3)Hex(3) | C[+57], N[+1095] | 1353.667 | 3 | 4058.9866 | 4058.9949 | 0 | -2 | 127 | Specific | 488.38 | 431.78 | 431.78 | 8.38 | 44 | >sp\|P01876\|IGHA1_HUMAN Immunoglobulin heavy constant alpha 1 OS=Homo sapiens OX=9606 GN=IGHA1 PE=1 SV=2 | 255 | 20201114_13813_1ng_glyco_hcd203040_compound15.21185.21185.3 | controllerType=0 controllerNumber=1 scan=21185 | 42.7668 |
| 19732:4 | 11 | R.LSLHRPALEDLLLGSEAN[+2204.772]LTC[+57.021]TLTGLR.D | HexNAc(4)Hex(5)NeuAc(2) | C[+57], N[+2205] | 1292.8475 | 4 | 5168.3683 | 5168.3707 | 0 | -0.5 | 127 | Specific | 547.16 | 374.02 | 374.02 | 9.94 | 44 | >sp\|P01876\|IGHA1_HUMAN Immunoglobulin heavy constant alpha 1 OS=Homo sapiens OX=9606 GN=IGHA1 PE=1 SV=2 | 255 | 20201114_13813_1ng_glyco_hcd203040_compound15.23331.23331.2 | controllerType=0 controllerNumber=1 scan=23331 | 47.2047 |
| 17597:3 | 11 | R.LSLHRPALEDLLLGSEAN[+1622.582]LTC[+57.021]TLTGLR.D | HexNAc(4)Hex(5) | C[+57], N[+1623] | 1529.4026 | 3 | 4586.1932 | 4586.1799 | 0 | 2.9 | 127 | Specific | 575.7 | 516.17 | 516.17 | 10.76 | 44 | >sp\|P01876\|IGHA1_HUMAN Immunoglobulin heavy constant alpha 1 OS=Homo sapiens OX=9606 GN=IGHA1 PE=1 SV=2 | 255 | 20201114_13813_1ng_glyco_hcd203040_compound15.21093.21093.3 | controllerType=0 controllerNumber=1 scan=21093 | 42.5699 |
| 17554:4 | 11 | R.LSLHRPALEDLLLGSEAN[+1663.608]LTC[+57.021]TLTGLR.D | HexNAc(5)Hex(4) | C[+57], N[+1664] | 1157.5605 | 4 | 4627.22 | 4627.2064 | 0 | 2.9 | 127 | Specific | 613.75 | 613.75 | 613.75 | 12.44 | 44 | >sp\|P01876\|IGHA1_HUMAN Immunoglobulin heavy constant alpha 1 OS=Homo sapiens OX=9606 GN=IGHA1 PE=1 SV=2 | 255 | 20201114_13813_1ng_glyco_hcd203040_compound15.21049.21049.4 | controllerType=0 controllerNumber=1 scan=21049 | 42.4748 |
| 17490:4 | 11 | R.LSLHRPALEDLLLGSEAN[+1702.581]LTC[+57.021]TLTGLR.D | HexNAc(2)Hex(8) | C[+57], N[+1703] | 1167.3039 | 4 | 4666.1937 | 4666.1796 | 0 | 3 | 127 | Specific | 649.36 | 649.36 | 649.36 | 12.44 | 44 | >sp\|P01876\|IGHA1_HUMAN Immunoglobulin heavy constant alpha 1 OS=Homo sapiens OX=9606 GN=IGHA1 PE=1 SV=2 | 255 | 20201114_13813_1ng_glyco_hcd203040_compound15.20982.20982.4 | controllerType=0 controllerNumber=1 scan=20982 | 42.3281 |
| 17556:4 | 11 | R.LSLHRPALEDLLLGSEAN[+1663.608]LTC[+57.021]TLTGLR.D | HexNAc(5)Hex(4) | C[+57], N[+1664] | 1157.5605 | 4 | 4627.22 | 4627.2064 | 0 | 2.9 | 127 | Specific | 697.14 | 642.49 | 642.49 | 13.44 | 44 | >sp\|P01876\|IGHA1_HUMAN Immunoglobulin heavy constant alpha 1 OS=Homo sapiens OX=9606 GN=IGHA1 PE=1 SV=2 | 255 | 20201114_13813_1ng_glyco_hcd203040_compound15.21051.21051.4 | controllerType=0 controllerNumber=1 scan=21051 | 42.4826 |
| 17642:4 | 11 | R.LSLHRPALEDLLLGSEAN[+1298.476]LTC[+57.021]TLTGLR.D | HexNAc(4)Hex(3) | C[+57], N[+1298] | 1066.2775 | 4 | 4262.0882 | 4262.0742 | 0 | 3.3 | 127 | Specific | 821.22 | 804.34 | 804.34 | 16.44 | 44 | >sp\|P01876\|IGHA1_HUMAN Immunoglobulin heavy constant alpha 1 OS=Homo sapiens OX=9606 GN=IGHA1 PE=1 SV=2 | 255 | 20201114_13813_1ng_glyco_hcd203040_compound15.21140.21140.4 | controllerType=0 controllerNumber=1 scan=21140 | 42.6564 |
| 17639:4 | 11 | R.LSLHRPALEDLLLGSEAN[+1298.476]LTC[+57.021]TLTGLR.D | HexNAc(4)Hex(3) | C[+57], N[+1298] | 1066.2775 | 4 | 4262.0882 | 4262.0742 | 0 | 3.3 | 127 | Specific | 847.21 | 809.62 | 809.62 | 16.44 | 44 | >sp\|P01876\|IGHA1_HUMAN Immunoglobulin heavy constant alpha 1 OS=Homo sapiens OX=9606 GN=IGHA1 PE=1 SV=2 | 255 | 20201114_13813_1ng_glyco_hcd203040_compound15.21137.21137.4 | controllerType=0 controllerNumber=1 scan=21137 | 42.648 |
| 17488:4 | 11 | R.LSLHRPALEDLLLGSEAN[+1702.581]LTC[+57.021]TLTGLR.D | HexNAc(2)Hex(8) | C[+57], N[+1703] | 1167.3039 | 4 | 4666.1937 | 4666.1796 | 0 | 3 | 127 | Specific | 882.05 | 882.05 | 882.05 | 17.44 | 44 | >sp\|P01876\|IGHA1_HUMAN Immunoglobulin heavy constant alpha 1 OS=Homo sapiens OX=9606 GN=IGHA1 PE=1 SV=2 | 255 | 20201114_13813_1ng_glyco_hcd203040_compound15.20980.20980.4 | controllerType=0 controllerNumber=1 scan=20980 | 42.3207 |
| 17632:4 | 11 | R.LSLHRPALEDLLLGSEAN[+1216.423]LTC[+57.021]TLTGLR.D | HexNAc(2)Hex(5) | C[+57], N[+1216] | 1045.7632 | 4 | 4180.0311 | 4180.0211 | 0 | 2.4 | 127 | Specific | 939.77 | 939.77 | 939.77 | 18.44 | 44 | >sp\|P01876\|IGHA1_HUMAN Immunoglobulin heavy constant alpha 1 OS=Homo sapiens OX=9606 GN=IGHA1 PE=1 SV=2 | 255 | 20201114_13813_1ng_glyco_hcd203040_compound15.21130.21130.4 | controllerType=0 controllerNumber=1 scan=21130 | 42.6348 |
| 11083:3 | 11 | R.LAGKPTHVN[+1540.529]VSVVMAEVDGTC[+57.021]Y.- | HexNAc(2)Hex(7) | C[+57], N[+1541] | 1296.565 | 3 | 3887.6804 | 3887.6706 | 0 | 2.5 | 332 | CRagged | 243.66 | 243.66 | 210.34 | 6.58 | 44 | >sp\|P01876\|IGHA1_HUMAN Immunoglobulin heavy constant alpha 1 OS=Homo sapiens OX=9606 GN=IGHA1 PE=1 SV=2 | 255 | 20201114_13813_1ng_glyco_hcd203040_compound15.14274.14274.3 | controllerType=0 controllerNumber=1 scan=14274 | 30.3537 |
| 11217:3 | 11 | R.LAGKPTHVN[+1501.555]VSVVMAEVDGTC[+57.021]Y.- | HexNAc(5)Hex(3) | C[+57], N[+1502] | 1283.5711 | 3 | 3848.6988 | 3848.6974 | 0 | 0.3 | 332 | CRagged | 378.36 | 281.12 | 281.12 | 7.12 | 44 | >sp\|P01876\|IGHA1_HUMAN Immunoglobulin heavy constant alpha 1 OS=Homo sapiens OX=9606 GN=IGHA1 PE=1 SV=2 | 255 | 20201114_13813_1ng_glyco_hcd203040_compound15.14415.14415.3 | controllerType=0 controllerNumber=1 scan=14415 | 30.619 |
| 12867:3 | 11 | R.LAGKPTHVN[+1913.677]VSVVMAEVDGTC[+57.021]Y.- | HexNAc(4)Hex(5)NeuAc(1) | C[+57], N[+1914] | 1420.9476 | 3 | 4260.8281 | 4260.8191 | 0 | 2.1 | 332 | CRagged | 438.31 | 353.36 | 353.36 | 8.15 | 44 | >sp\|P01876\|IGHA1_HUMAN Immunoglobulin heavy constant alpha 1 OS=Homo sapiens OX=9606 GN=IGHA1 PE=1 SV=2 | 255 | 20201114_13813_1ng_glyco_hcd203040_compound15.16141.16141.3 | controllerType=0 controllerNumber=1 scan=16141 | 33.5252 |
| 12908:3 | 11 | R.LAGKPTHVN[+2059.735]VSVVMAEVDGTC[+57.021]Y.- | HexNAc(4)Hex(5)Fuc(1)NeuAc(1) | C[+57], N[+2060] | 1469.634 | 3 | 4406.8874 | 4406.877 | 0 | 2.3 | 332 | CRagged | 472.25 | 464.48 | 464.48 | 9.42 | 44 | >sp\|P01876\|IGHA1_HUMAN Immunoglobulin heavy constant alpha 1 OS=Homo sapiens OX=9606 GN=IGHA1 PE=1 SV=2 | 255 | 20201114_13813_1ng_glyco_hcd203040_compound15.16184.16184.3 | controllerType=0 controllerNumber=1 scan=16184 | 33.5998 |
| 12910:3 | 11 | R.LAGKPTHVN[+2059.735]VSVVMAEVDGTC[+57.021]Y.- | HexNAc(4)Hex(5)Fuc(1)NeuAc(1) | C[+57], N[+2060] | 1469.634 | 3 | 4406.8874 | 4406.877 | 0 | 2.3 | 332 | CRagged | 515.39 | 420.74 | 420.74 | 9.79 | 44 | >sp\|P01876\|IGHA1_HUMAN Immunoglobulin heavy constant alpha 1 OS=Homo sapiens OX=9606 GN=IGHA1 PE=1 SV=2 | 255 | 20201114_13813_1ng_glyco_hcd203040_compound15.16186.16186.3 | controllerType=0 controllerNumber=1 scan=16186 | 33.6035 |
| 12892:3 | 11 | R.LAGKPTHVN[+2059.735]VSVVMAEVDGTC[+57.021]Y.- | HexNAc(4)Hex(5)Fuc(1)NeuAc(1) | C[+57], N[+2060] | 1469.6357 | 3 | 4406.8925 | 4406.877 | 0 | 3.5 | 332 | CRagged | 519.84 | 452.21 | 452.21 | 9.79 | 44 | >sp\|P01876\|IGHA1_HUMAN Immunoglobulin heavy constant alpha 1 OS=Homo sapiens OX=9606 GN=IGHA1 PE=1 SV=2 | 255 | 20201114_13813_1ng_glyco_hcd203040_compound15.16167.16167.3 | controllerType=0 controllerNumber=1 scan=16167 | 33.5734 |
| 11219:3 | 11 | R.LAGKPTHVN[+1501.555]VSVVMAEVDGTC[+57.021]Y.- | HexNAc(5)Hex(3) | C[+57], N[+1502] | 1283.5711 | 3 | 3848.6988 | 3848.6974 | 0 | 0.3 | 332 | CRagged | 556.9 | 457.72 | 457.72 | 9.63 | 44 | >sp\|P01876\|IGHA1_HUMAN Immunoglobulin heavy constant alpha 1 OS=Homo sapiens OX=9606 GN=IGHA1 PE=1 SV=2 | 255 | 20201114_13813_1ng_glyco_hcd203040_compound15.14417.14417.3 | controllerType=0 controllerNumber=1 scan=14417 | 30.6227 |
| 12890:3 | 11 | R.LAGKPTHVN[+2059.735]VSVVMAEVDGTC[+57.021]Y.- | HexNAc(4)Hex(5)Fuc(1)NeuAc(1) | C[+57], N[+2060] | 1469.6357 | 3 | 4406.8925 | 4406.877 | 0 | 3.5 | 332 | CRagged | 587.61 | 587.61 | 587.61 | 11.12 | 44 | >sp\|P01876\|IGHA1_HUMAN Immunoglobulin heavy constant alpha 1 OS=Homo sapiens OX=9606 GN=IGHA1 PE=1 SV=2 | 255 | 20201114_13813_1ng_glyco_hcd203040_compound15.16165.16165.3 | controllerType=0 controllerNumber=1 scan=16165 | 33.5695 |
| 12858:3 | 11 | R.LAGKPTHVN[+2262.814]VSVVMAEVDGTC[+57.021]Y.- | HexNAc(5)Hex(5)Fuc(1)NeuAc(1) | C[+57], N[+2263] | 1537.327 | 3 | 4609.9663 | 4609.9564 | 0 | 2.2 | 332 | CRagged | 594.91 | 493.43 | 493.43 | 11.12 | 44 | >sp\|P01876\|IGHA1_HUMAN Immunoglobulin heavy constant alpha 1 OS=Homo sapiens OX=9606 GN=IGHA1 PE=1 SV=2 | 255 | 20201114_13813_1ng_glyco_hcd203040_compound15.16131.16131.3 | controllerType=0 controllerNumber=1 scan=16131 | 33.5063 |
| 11107:3 | 11 | R.LAGKPTHVN[+1809.666]VSVVMAEVDGTC[+57.021]Y.- | HexNAc(5)Hex(4)Fuc(1) | C[+57], N[+1810] | 1386.2872 | 3 | 4156.8469 | 4156.8081 | 0 | 9.3 | 332 | CRagged | 664.08 | 664.08 | 449.42 | 12.46 | 44 | >sp\|P01876\|IGHA1_HUMAN Immunoglobulin heavy constant alpha 1 OS=Homo sapiens OX=9606 GN=IGHA1 PE=1 SV=2 | 255 | 20201114_13813_1ng_glyco_hcd203040_compound15.14299.14299.3 | controllerType=0 controllerNumber=1 scan=14299 | 30.4083 |
| 12859:3 | 11 | R.LAGKPTHVN[+2262.814]VSVVMAEVDGTC[+57.021]Y.- | HexNAc(5)Hex(5)Fuc(1)NeuAc(1) | C[+57], N[+2263] | 1537.327 | 3 | 4609.9663 | 4609.9564 | 0 | 2.2 | 332 | CRagged | 686.73 | 576.84 | 576.84 | 13.12 | 44 | >sp\|P01876\|IGHA1_HUMAN Immunoglobulin heavy constant alpha 1 OS=Homo sapiens OX=9606 GN=IGHA1 PE=1 SV=2 | 255 | 20201114_13813_1ng_glyco_hcd203040_compound15.16132.16132.3 | controllerType=0 controllerNumber=1 scan=16132 | 33.5108 |
| 11298:3 | 11 | R.LAGKPTHVN[+1647.613]VSVVMAEVDGTC[+57.021]Y.- | HexNAc(5)Hex(3)Fuc(1) | C[+57], N[+1648] | 1332.2615 | 3 | 3994.77 | 3994.7553 | 0 | 3.7 | 332 | CRagged | 710.61 | 418.52 | 418.52 | 13.82 | 44 | >sp\|P01876\|IGHA1_HUMAN Immunoglobulin heavy constant alpha 1 OS=Homo sapiens OX=9606 GN=IGHA1 PE=1 SV=2 | 255 | 20201114_13813_1ng_glyco_hcd203040_compound15.14498.14498.3 | controllerType=0 controllerNumber=1 scan=14498 | 30.7648 |
| 12886:3 | 11 | R.LAGKPTHVN[+2262.814]VSVVMAEVDGTC[+57.021]Y.- | HexNAc(5)Hex(5)Fuc(1)NeuAc(1) | C[+57], N[+2263] | 1537.3255 | 3 | 4609.9619 | 4609.9564 | 0 | 1.2 | 332 | CRagged | 722.17 | 574.83 | 574.83 | 14.14 | 44 | >sp\|P01876\|IGHA1_HUMAN Immunoglobulin heavy constant alpha 1 OS=Homo sapiens OX=9606 GN=IGHA1 PE=1 SV=2 | 255 | 20201114_13813_1ng_glyco_hcd203040_compound15.16161.16161.3 | controllerType=0 controllerNumber=1 scan=16161 | 33.561 |
| 12883:3 | 11 | R.LAGKPTHVN[+2262.814]VSVVMAEVDGTC[+57.021]Y.- | HexNAc(5)Hex(5)Fuc(1)NeuAc(1) | C[+57], N[+2263] | 1537.3255 | 3 | 4609.9619 | 4609.9564 | 0 | 1.2 | 332 | CRagged | 737.49 | 534.09 | 534.09 | 14.14 | 44 | >sp\|P01876\|IGHA1_HUMAN Immunoglobulin heavy constant alpha 1 OS=Homo sapiens OX=9606 GN=IGHA1 PE=1 SV=2 | 255 | 20201114_13813_1ng_glyco_hcd203040_compound15.16158.16158.3 | controllerType=0 controllerNumber=1 scan=16158 | 33.5563 |
| 12885:3 | 11 | R.LAGKPTHVN[+1913.677]VSVVMAEVDGTC[+57.021]Y.- | HexNAc(4)Hex(5)NeuAc(1) | C[+57], N[+1914] | 1420.9463 | 3 | 4260.8245 | 4260.8191 | 0 | 1.3 | 332 | CRagged | 776.95 | 489.61 | 489.61 | 15.14 | 44 | >sp\|P01876\|IGHA1_HUMAN Immunoglobulin heavy constant alpha 1 OS=Homo sapiens OX=9606 GN=IGHA1 PE=1 SV=2 | 255 | 20201114_13813_1ng_glyco_hcd203040_compound15.16160.16160.3 | controllerType=0 controllerNumber=1 scan=16160 | 33.[illegible]59 |
| 11077:3 | 11 | R.LAGKPTHVN[+1702.581]VSVVMAEVDGTC[+57.021]Y.- | HexNAc(2)Hex(8) | C[+57], N[+1703] | 1350.5823 | 3 | 4049.7324 | 4049.7234 | 0 | 2.2 | 332 | CRagged | 820.5 | 820.5 | 532.12 | 16.12 | 44 | >sp\|P01876\|IGHA1_HUMAN Immunoglobulin heavy constant alpha 1 OS=Homo sapiens OX=9606 GN=IGHA1 PE=1 SV=2 | 255 | 20201114_13813_1ng_glyco_hcd203040_compound15.14268.14268.3 | controllerType=0 controllerNumber=1 scan=14268 | 30.3388 |
| 11079:3 | 11 | R.LAGKPTHVN[+1702.581]VSVVMAEVDGTC[+57.021]Y.- | HexNAc(2)Hex(8) | C[+57], N[+1703] | 1350.5823 | 3 | 4049.7324 | 4049.7234 | 0 | 2.2 | 332 | CRagged | 832.02 | 816.64 | 498.18 | 16.12 | 44 | >sp\|P01876\|IGHA1_HUMAN Immunoglobulin heavy constant alpha 1 OS=Homo sapiens OX=9606 GN=IGHA1 PE=1 SV=2 | 255 | 20201114_13813_1ng_glyco_hcd203040_compound15.14270.14270.3 | controllerType=0 controllerNumber=1 scan=14270 | 30.344 |
| 11215:3 | 11 | R.LAGKPTHVN[+1647.613]VSVVMAEVDGTC[+57.021]Y.- | HexNAc(5)Hex(3)Fuc(1) | C[+57], N[+1648] | 1332.2574 | 3 | 3994.7576 | 3994.7553 | 0 | 0.6 | 332 | CRagged | 847.34 | 782.38 | 782.38 | 15.82 | 44 | >sp\|P01876\|IGHA1_HUMAN Immunoglobulin heavy constant alpha 1 OS=Homo sapiens OX=9606 GN=IGHA1 PE=1 SV=2 | 255 | 20201114_13813_1ng_glyco_hcd203040_compound15.14412.14412.3 | controllerType=0 controllerNumber=1 scan=14412 | 30.6131 |
| 11213:3 | 11 | R.LAGKPTHVN[+1647.613]VSVVMAEVDGTC[+57.021]Y.- | HexNAc(5)Hex(3)Fuc(1) | C[+57], N[+1648] | 1332.2574 | 3 | 3994.7576 | 3994.7553 | 0 | 0.6 | 332 | CRagged | 855.21 | 855.21 | 855.21 | 16.82 | 44 | >sp\|P01876\|IGHA1_HUMAN Immunoglobulin heavy constant alpha 1 OS=Homo sapiens OX=9606 GN=IGHA1 PE=1 SV=2 | 255 | 20201114_13813_1ng_glyco_hcd203040_compound15.14410.14410.3 | controllerType=0 controllerNumber=1 scan=14410 | 30.6063 |
| 11081:3 | 11 | R.LAGKPTHVN[+1540.529]VSVVMAEVDGTC[+57.021]Y.- | HexNAc(2)Hex(7) | C[+57], N[+1541] | 1296.565 | 3 | 3887.6804 | 3887.6706 | 0 | 2.5 | 332 | CRagged | 855.37 | 855.37 | 538.1 | 17.12 | 44 | >sp\|P01876\|IGHA1_HUMAN Immunoglobulin heavy constant alpha 1 OS=Homo sapiens OX=9606 GN=IGHA1 PE=1 SV=2 | 255 | 20201114_13813_1ng_glyco_hcd203040_compound15.14272.14272.3 | controllerType=0 controllerNumber=1 scan=14272 | 30.3473 |
| 11103:3 | 11 | R.LAGKPTHVN[+1378.476]VSVVMAEVDGTC[+57.021]Y.- | HexNAc(2)Hex(6) | C[+57], N[+1378] | 1242.5563 | 3 | 3725.6544 | 3725.6178 | 0 | 9.8 | 332 | CRagged | 874.46 | 874.46 | 546.67 | 16.41 | 44 | >sp\|P01876\|IGHA1_HUMAN Immunoglobulin heavy constant alpha 1 OS=Homo sapiens OX=9606 GN=IGHA1 PE=1 SV=2 | 255 | 20201114_13813_1ng_glyco_hcd203040_compound15.14295.14295.3 | controllerType=0 controllerNumber=1 scan=14295 | 30.399 |
| 11105:3 | 11 | R.LAGKPTHVN[+1809.666]VSVVMAEVDGTC[+57.021]Y.- | HexNAc(5)Hex(4)Fuc(1) | C[+57], N[+1810] | 1386.2872 | 3 | 4156.8469 | 4156.8081 | 0 | 9.3 | 332 | CRagged | 880.78 | 753.51 | 596.02 | 16.46 | 44 | >sp\|P01876\|IGHA1_HUMAN Immunoglobulin heavy constant alpha 1 OS=Homo sapiens OX=9606 GN=IGHA1 PE=1 SV=2 | 255 | 20201114_13813_1ng_glyco_hcd203040_compound15.14297.14297.3 | controllerType=0 controllerNumber=1 scan=14297 | 30.4023 |
| 11140:3 | 11 | R.LAGKPTHVN[+1216.423]VSVVMAEVDGTC[+57.021]Y.- | HexNAc(2)Hex(5) | C[+57], N[+1216] | 1188.5371 | 3 | 3563.5967 | 3563.5649 | 0 | 8.9 | 332 | CRagged | 884.44 | 884.44 | 540.77 | 16.46 | 44 | >sp\|P01876\|IGHA1_HUMAN Immunoglobulin heavy constant alpha 1 OS=Homo sapiens OX=9606 GN=IGHA1 PE=1 SV=2 | 255 | 20201114_13813_1ng_glyco_hcd203040_compound15.14333.14333.3 | controllerType=0 controllerNumber=1 scan=14333 | 30.4737 |
| 11101:3 | 11 | R.LAGKPTHVN[+1378.476]VSVVMAEVDGTC[+57.021]Y.- | HexNAc(2)Hex(6) | C[+57], N[+1378] | 1242.5563 | 3 | 3725.6544 | 3725.6178 | 0 | 9.8 | 332 | CRagged | 923.9 | 923.9 | 601.32 | 17.41 | 44 | >sp\|P01876\|IGHA1_HUMAN Immunoglobulin heavy constant alpha 1 OS=Homo sapiens OX=9606 GN=IGHA1 PE=1 SV=2 | 255 | 20201114_13813_1ng_glyco_hcd203040_compound15.14293.14293.3 | controllerType=0 controllerNumber=1 scan=14293 | 30.3899 |
| 11142:3 | 11 | R.LAGKPTHVN[+1216.423]VSVVMAEVDGTC[+57.021]Y.- | HexNAc(2)Hex(5) | C[+57], N[+1216] | 1188.5371 | 3 | 3563.5967 | 3563.5649 | 0 | 8.9 | 332 | CRagged | 924.44 | 924.44 | 574.1 | 17.46 | 44 | >sp\|P01876\|IGHA1_HUMAN Immunoglobulin heavy constant alpha 1 OS=Homo sapiens OX=9606 GN=IGHA1 PE=1 SV=2 | 255 | 20201114_13813_1ng_glyco_hcd203040_compound15.14335.14335.3 | controllerType=0 controllerNumber=1 scan=14335 | 30.4811 |
| 05456:3 | 13 | R.LSPN[+2205.793]ASAEHLELR.W | HexNAc(4)Hex(5)Fuc(2)NeuAc(1) | N[+2206] | 1214.8548 | 3 | 3642.5498 | 3642.5421 | 0 | 2.1 | 52 | Specific | 522.01 | 457.7 | 457.7 | 9.47 | 32 | >sp\|Q13410\|BT1A1_HUMAN Butyrophilin subfamily 1 member A1 OS=Homo sapiens OX=9606 GN=BTN1A1 PE=1 SV=3 | 179 | 20201114_13813_1ng_glyco_hcd203040_compound15.8418.8418.3 | controllerType=0 controllerNumber=1 scan=8418 | 20.5318 |
| 07450:3 | 13 | R.LSPN[+2350.830]ASAEHLELR.W | HexNAc(4)Hex(5)Fuc(1)NeuAc(2) | N[+2351] | 1263.1999 | 3 | 3787.5851 | 3787.5796 | 0 | 1.5 | 52 | Specific | 541.37 | 432.27 | 432.27 | 9.49 | 32 | >sp\|Q13410\|BT1A1_HUMAN Butyrophilin subfamily 1 member A1 OS=Homo sapiens OX=9606 GN=BTN1A1 PE=1 SV=3 | 179 | 20201114_13813_1ng_glyco_hcd203040_compound15.10488.10488.3 | controllerType=0 controllerNumber=1 scan=10488 | 24.002 |
| 07448:3 | 13 | R.LSPN[+2350.830]ASAEHLELR.W | HexNAc(4)Hex(5)Fuc(1)NeuAc(2) | N[+2351] | 1263.1999 | 3 | 3787.5851 | 3787.5796 | 0 | 1.5 | 52 | Specific | 581.25 | 575.04 | 575.04 | 10.82 | 32 | >sp\|Q13410\|BT1A1_HUMAN Butyrophilin subfamily 1 member A1 OS=Homo sapiens OX=9606 GN=BTN1A1 PE=1 SV=3 | 179 | 20201114_13813_1ng_glyco_hcd203040_compound15.10486.10486.3 | controllerType=0 controllerNumber=1 scan=10486 | 23.9979 |
| 05458:3 | 13 | R.LSPN[+2205.793]ASAEHLELR.W | HexNAc(4)Hex(5)Fuc(2)NeuAc(1) | N[+2206] | 1214.8548 | 3 | 3642.5498 | 3642.5421 | 0 | 2.1 | 52 | Specific | 626.89 | 425.1 | 425.1 | 11.8 | 32 | >sp\|Q13410\|BT1A1_HUMAN Butyrophilin subfamily 1 member A1 OS=Homo sapiens OX=9606 GN=BTN1A1 PE=1 SV=3 | 179 | 20201114_13813_1ng_glyco_hcd203040_compound15.8420.8420.3 | controllerType=0 controllerNumber=1 scan=8420 | 20.536 |
| 17546:3 | 13 | K.N[+1913.677]VSC[+57.021]YIQNLLLGQEK.K | HexNAc(4)Hex(5)NeuAc(1) | C[+57], N[+1914] | 1231.5372 | 3 | 3692.5969 | 3692.5876 | 0 | 2.5 | 215 | Specific | 484.2 | 417.8 | 417.8 | 9.29 | 32 | >sp\|Q13410\|BT1A1_HUMAN Butyrophilin subfamily 1 member A1 OS=Homo sapiens OX=9606 GN=BTN1A1 PE=1 SV=3 | 179 | 20201114_13813_1ng_glyco_hcd203040_compound15.21040.21040.3 | controllerType=0 controllerNumber=1 scan=21040 | 42.4583 |
| 17544:3 | 13 | K.N[+1913.677]VSC[+57.021]YIQNLLLGQEK.K | HexNAc(4)Hex(5)NeuAc(1) | C[+57], N[+1914] | 1231.5372 | 3 | 3692.5969 | 3692.5876 | 0 | 2.5 | 215 | Specific | 555.74 | 475.4 | 475.4 | 9.8 | 32 | >sp\|Q13410\|BT1A1_HUMAN Butyrophilin subfamily 1 member A1 OS=Homo sapiens OX=9606 GN=BTN1A1 PE=1 SV=3 | 179 | 20201114_13813_1ng_glyco_hcd203040_compound15.21038.21038.3 | controllerType=0 controllerNumber=1 scan=21038 | 42.4544 |
| 24005:4 | 14 | R.QLAHQSN[+2351.851]STNIFFSPVSIATAFAMLSLGTK.A | HexNAc(4)Hex(5)Fuc(3)NeuAc(1) | N[+2352] | 1384.1211 | 4 | 5533.4626 | 5533.4858 | 0 | -4.2 | 64 | Specific | 270.26 | 270.26 | 270.26 | 6 | 34 | >sp\|P01009\|A1AT_HUMAN Alpha-1-antitrypsin OS=Homo sapiens OX=9606 GN=SERPINA1 PE=1 SV=3 | 642 | 20201114_13813_1ng_glyco_hcd203040_compound15.27814.27814.4 | controllerType=0 controllerNumber=1 scan=27814 | 55.8161 |
| 23976:3 | 14 | R.QLAHQSN[+2351.851]STNIFFSPVSIATAFAMLSLGTK.A | HexNAc(4)Hex(5)Fuc(3)NeuAc(1) | N[+2352] | 1845.1734 | 3 | 5533.5057 | 5533.4858 | 0 | 3.6 | 64 | Specific | 342.55 | 342.55 | 342.55 | 7.43 | 34 | >sp\|P01009\|A1AT_HUMAN Alpha-1-antitrypsin OS=Homo sapiens OX=9606 GN=SERPINA1 PE=1 SV=3 | 642 | 20201114_13813_1ng_glyco_hcd203040_compound15.27784.27784.3 | controllerType=0 controllerNumber=1 scan=27784 | 55.7572 |
| 24000:4 | 14 | R.QLAHQSN[+2204.772]STNIFFSPVSIATAFAMLSLGTK.A | HexNAc(4)Hex(5)NeuAc(2) | N[+2205] | 1347.3527 | 4 | 5386.3888 | 5386.4075 | 0 | -3.5 | 64 | Specific | 914.4 | 801.41 | 582.64 | 17.12 | 34 | >sp\|P01009\|A1AT_HUMAN Alpha-1-antitrypsin OS=Homo sapiens OX=9606 GN=SERPINA1 PE=1 SV=3 | 642 | 20201114_13813_1ng_glyco_hcd203040_compound15.27809.27809.4 | controllerType=0 controllerNumber=1 scan=27809 | 55.8046 |
| 24012:4 | 14 | R.QLAHQSN[+2204.772]STNIFFSPVSIATAFAMLSLGTK.A | HexNAc(4)Hex(5)NeuAc(2) | N[+2205] | 1347.3607 | 4 | 5386.4211 | 5386.4075 | 0 | 2.5 | 64 | Specific | 996.44 | 895.79 | 592.75 | 19.39 | 34 | >sp\|P01009\|A1AT_HUMAN Alpha-1-antitrypsin OS=Homo sapiens OX=9606 GN=SERPINA1 PE=1 SV=3 | 642 | 20201114_13813_1ng_glyco_hcd203040_compound15.27822.27822.4 | controllerType=0 controllerNumber=1 scan=27822 | 55.8312 |
| 22614:4 | 14 | K.ADTHDEILEGLNFN[+2861.000]LTEIPEAQIHEGFQELLR.T | HexNAc(5)Hex(6)NeuAc(3) | N[+2861] | 1638.964 | 4 | 6552.8341 | 6552.8239 | 0 | 1.6 | 94 | Specific | 941.52 | 780.14 | 780.14 | 18.31 | 34 | >sp\|P01009\|A1AT_HUMAN Alpha-1-antitrypsin OS=Homo sapiens OX=9606 GN=SERPINA1 PE=1 SV=3 | 642 | 20201114_13813_1ng_glyco_hcd203040_compound15.26361.26361.4 | controllerType=0 controllerNumber=1 scan=26361 | 53.1316 |
| 20144:3 | 14 | K.YLGN[+2204.772]ATAIFFLPDEGK.L | HexNAc(4)Hex(5)NeuAc(2) | N[+2205] | 1320.9067 | 3 | 3960.7055 | 3960.6677 | 0 | 9.5 | 268 | Specific | 585.52 | 572.01 | 572.01 | 10.33 | 34 | >sp\|P01009\|A1AT_HUMAN Alpha-1-antitrypsin OS=Homo sapiens OX=9606 GN=SERPINA1 PE=1 SV=3 | 642 | 20201114_13813_1ng_glyco_hcd203040_compound15.23765.23765.3 | controllerType=0 controllerNumber=1 scan=23765 | 48.1097 |
| 20146:3 | 14 | K.YLGN[+2204.772]ATAIFFLPDEGK.L | HexNAc(4)Hex(5)NeuAc(2) | N[+2205] | 1320.9067 | 3 | 3960.7055 | 3960.6677 | 0 | 9.5 | 268 | Specific | 820.09 | 784.68 | 784.68 | 15.33 | 34 | >sp\|P01009\|A1AT_HUMAN Alpha-1-antitrypsin OS=Homo sapiens OX=9606 GN=SERPINA1 PE=1 SV=3 | 642 | 20201114_13813_1ng_glyco_hcd203040_compound15.23767.23767.3 | controllerType=0 controllerNumber=1 scan=23767 | 48.114 |
| 13846:4 | 15 | K.LKELPGVC[+57.021]N[+2205.793]ETMMALWEEC[+57.021]KPC[+57.021]LK.Q | HexNAc(4)Hex(5)Fuc(2)NeuAc(1) | C[+57]*3, N[+2206] | 1286.3054 | 4 | 5142.1999 | 5142.1776 | 0 | 4.3 | 95 | Specific | 244.19 | 244.19 | 244.19 | 6.55 | 35 | >sp\|P10909\|CLUS_HUMAN Clusterin OS=Homo sapiens OX=9606 GN=CLU PE=1 SV=1 | 1467 | 20201114_13813_1ng_glyco_hcd203040_compound15.17158.17158.4 | controllerType=0 controllerNumber=1 scan=17158 | 35.2952 |
| 15609:4 | 15 | K.LKELPGVC[+57.021]N[+2350.830]ETMMALWEEC[+57.021]KPC[+57.021]LK.Q | HexNAc(4)Hex(5)Fuc(1)NeuAc(2) | C[+57]*3, N[+2351] | 1322.5588 | 4 | 5287.2134 | 5287.2151 | 0 | -0.3 | 95 | Specific | 291.01 | 268.02 | 268.02 | 7 | 35 | >sp\|P10909\|CLUS_HUMAN Clusterin OS=Homo sapiens OX=9606 GN=CLU PE=1 SV=1 | 1467 | 20201114_13813_1ng_glyco_hcd203040_compound15.18994.18994.4 | controllerType=0 controllerNumber=1 scan=18994 | 38.5446 |
| 15607:4 | 15 | K.LKELPGVC[+57.021]N[+2350.830]ETMMALWEEC[+57.021]KPC[+57.021]LK.Q | HexNAc(4)Hex(5)Fuc(1)NeuAc(2) | C[+57]*3, N[+2351] | 1322.5588 | 4 | 5287.2134 | 5287.2151 | 0 | -0.3 | 95 | Specific | 325.35 | 325.35 | 325.35 | 7.37 | 35 | >sp\|P10909\|CLUS_HUMAN Clusterin OS=Homo sapiens OX=9606 GN=CLU PE=1 SV=1 | 1467 | 20201114_13813_1ng_glyco_hcd203040_compound15.18992.18992.4 | controllerType=0 controllerNumber=1 scan=18992 | 38.5409 |
| 13889:3 | 15 | K.LKELPGVC[+57.021]N[+2059.735]ETMMALWEEC[+57.021]KPC[+57.021]LK.Q | HexNAc(4)Hex(5)Fuc(1)NeuAc(1) | C[+57]*3, N[+2060] | 1666.0572 | 3 | 4996.1571 | 4996.1196 | 0 | 7.5 | 95 | Specific | 523.34 | 523.34 | 523.34 | 8.64 | 35 | >sp\|P10909\|CLUS_HUMAN Clusterin OS=Homo sapiens OX=9606 GN=CLU PE=1 SV=1 | 1467 | 20201114_13813_1ng_glyco_hcd203040_compound15.17202.17202.3 | controllerType=0 controllerNumber=1 scan=17202 | 35.3745 |
| 13844:4 | 15 | K.LKELPGVC[+57.021]N[+2205.793]ETMMALWEEC[+57.021]KPC[+57.021]LK.Q | HexNAc(4)Hex(5)Fuc(2)NeuAc(1) | C[+57]*3, N[+2206] | 1286.3054 | 4 | 5142.1999 | 5142.1776 | 0 | 4.3 | 95 | Specific | 547.92 | 547.92 | 547.92 | 9.9 | 35 | >sp\|P10909\|CLUS_HUMAN Clusterin OS=Homo sapiens OX=9606 GN=CLU PE=1 SV=1 | 1467 | 20201114_13813_1ng_glyco_hcd203040_compound15.17156.17156.4 | controllerType=0 controllerNumber=1 scan=17156 | 35.2913 |
| 13891:3 | 15 | K.LKELPGVC[+57.021]N[+2059.735]ETMMALWEEC[+57.021]KPC[+57.021]LK.Q | HexNAc(4)Hex(5)Fuc(1)NeuAc(1) | C[+57]*3, N[+2060] | 1666.0572 | 3 | 4996.1571 | 4996.1196 | 0 | 7.5 | 95 | Specific | 570.27 | 570.27 | 570.27 | 8.78 | 35 | >sp\|P10909\|CLUS_HUMAN Clusterin OS=Homo sapiens OX=9606 GN=CLU PE=1 SV=1 | 1467 | 20201114_13813_1ng_glyco_hcd203040_compound15.17204.17204.3 | controllerType=0 controllerNumber=1 scan=17204 | 35.3787 |
| 15698:3 | 15 | K.ELPGVC[+57.021]N[+2059.735]ETMMALWEEC[+57.021]KPC[+57.021]LK.Q | HexNAc(4)Hex(5)Fuc(1)NeuAc(1) | C[+57]*3, N[+2060] | 1585.6492 | 3 | 4754.9331 | 4754.9406 | 0 | -1.6 | 97 | Specific | 570.77 | 495.26 | 495.26 | 8.99 | 35 | >sp\|P10909\|CLUS_HUMAN Clusterin OS=Homo sapiens OX=9606 GN=CLU PE=1 SV=1 | 1467 | 20201114_13813_1ng_glyco_hcd203040_compound15.19089.19089.3 | controllerType=0 controllerNumber=1 scan=19089 | 38.7336 |
| 15696:3 | 15 | K.ELPGVC[+57.021]N[+2059.735]ETMMALWEEC[+57.021]KPC[+57.021]LK.Q | HexNAc(4)Hex(5)Fuc(1)NeuAc(1) | C[+57]*3, N[+2060] | 1585.6492 | 3 | 4754.9331 | 4754.9406 | 0 | -1.6 | 97 | Specific | 695.45 | 541.12 | 541.12 | 12.18 | 35 | >sp\|P10909\|CLUS_HUMAN Clusterin OS=Homo sapiens OX=9606 GN=CLU PE=1 SV=1 | 1467 | 20201114_13813_1ng_glyco_hcd203040_compound15.19087.19087.3 | controllerType=0 controllerNumber=1 scan=19087 | 38.7284 |
| 00348:3 | 15 | R.EIRHN[+2059.735]STGC[+57.021]LR.M | HexNAc(4)Hex(5)Fuc(1)NeuAc(1) | C[+57], N[+2060] | 1134.8054 | 3 | 3402.4018 | 3402.3994 | 0 | 0.7 | 287 | Specific | 331.65 | 224.27 | 224.27 | 5.13 | 35 | >sp\|P10909\|CLUS_HUMAN Clusterin OS=Homo sapiens OX=9606 GN=CLU PE=1 SV=1 | 1467 | 20201114_13813_1ng_glyco_hcd203040_compound15.3109.3109.3 | controllerType=0 controllerNumber=1 scan=3109 | 11.2523 |
| 00350:3 | 15 | R.EIRHN[+2059.735]STGC[+57.021]LR.M | HexNAc(4)Hex(5)Fuc(1)NeuAc(1) | C[+57], N[+2060] | 1134.8054 | 3 | 3402.4018 | 3402.3994 | 0 | 0.7 | 287 | Specific | 402.14 | 250.98 | 250.98 | 5.69 | 35 | >sp\|P10909\|CLUS_HUMAN Clusterin OS=Homo sapiens OX=9606 GN=CLU PE=1 SV=1 | 1467 | 20201114_13813_1ng_glyco_hcd203040_compound15.3111.3111.3 | controllerType=0 controllerNumber=1 scan=3111 | 11.2573 |
| 23565:3 | 15 | K.MLN[+2205.793]TSSLLEQLNEQFNWVSR.L | HexNAc(4)Hex(5)Fuc(2)NeuAc(1) | N[+2206] | 1539.0009 | 3 | 4614.9881 | 4614.9796 | 0 | 1.9 | 352 | Specific | 470.46 | 425.73 | 425.73 | 9.39 | 35 | >sp\|P10909\|CLUS_HUMAN Clusterin OS=Homo sapiens OX=9606 GN=CLU PE=1 SV=1 | 1467 | 20201114_13813_1ng_glyco_hcd203040_compound15.27352.27352.3 | controllerType=0 controllerNumber=1 scan=27352 | 54.9759 |
| 23261:3 | 15 | K.MLN[+2350.830]TSSLLEQLNEQFNWVSR.L | HexNAc(4)Hex(5)Fuc(1)NeuAc(2) | N[+2351] | 1587.3423 | 3 | 4760.0125 | 4760.0171 | 0 | -1 | 352 | Specific | 492.53 | 354.58 | 354.58 | 8.01 | 35 | >sp\|P10909\|CLUS_HUMAN Clusterin OS=Homo sapiens OX=9606 GN=CLU PE=1 SV=1 | 1467 | 20201114_13813_1ng_glyco_hcd203040_compound15.27036.27036.3 | controllerType=0 controllerNumber=1 scan=27036 | 54.4076 |
| 20873:3 | 15 | K.MLN[+2059.735]TSSLLEQLNEQFNWVSR.L | HexNAc(4)Hex(5)Fuc(1)NeuAc(1) | N[+2060] | 1490.3239 | 3 | 4468.9572 | 4468.9217 | 0 | 7.9 | 352 | Specific | 520.28 | 511.09 | 511.09 | 9 | 35 | >sp\|P10909\|CLUS_HUMAN Clusterin OS=Homo sapiens OX=9606 GN=CLU PE=1 SV=1 | 1467 | 20201114_13813_1ng_glyco_hcd203040_compound15.24528.24528.3 | controllerType=0 controllerNumber=1 scan=24528 | 49.7632 |
| 20875:3 | 15 | K.MLN[+2059.735]TSSLLEQLNEQFNWVSR.L | HexNAc(4)Hex(5)Fuc(1)NeuAc(1) | N[+2060] | 1490.3239 | 3 | 4468.9572 | 4468.9217 | 0 | 7.9 | 352 | Specific | 914.68 | 646.25 | 646.25 | 17.33 | 35 | >sp\|P10909\|CLUS_HUMAN Clusterin OS=Homo sapiens OX=9606 GN=CLU PE=1 SV=1 | 1467 | 20201114_13813_1ng_glyco_hcd203040_compound15.24530.24530.3 | controllerType=0 controllerNumber=1 scan=24530 | 49.7666 |
| 02064:3 | 17 | K.N[+1216.423]NSIPDKQITASSSYK.T | HexNAc(2)Hex(5) | N[+1216] | 990.4404 | 3 | 2969.3066 | 2969.2991 | 0 | 2.5 | 238 | Specific | 280.46 | 280.46 | 280.46 | 6.29 | 31 | >sp\|Q08431\|MFGM_HUMAN Lactadherin OS=Homo sapiens OX=9606 GN=MFGE8 PE=1 SV=3 | 1064 | 20201114_13813_1ng_glyco_hcd203040_compound15.4895.4895.3 | controllerType=0 controllerNumber=1 scan=4895 | 14.4601 |
| 15613:3 | 18 | R.TLN[+2059.735]QSSDELQLSMGNAMFVK.E | HexNAc(4)Hex(5)Fuc(1)NeuAc(1) | N[+2060] | 1424.9399 | 3 | 4272.8051 | 4272.7926 | 0 | 2.9 | 125 | Specific | 297.2 | 297.2 | 297.2 | 7.16 | 28 | >sp\|P01011\|AACT_HUMAN Alpha-1-antichymotrypsin OS=Homo sapiens OX=9606 GN=SERPINA3 PE=1 SV=2 | 1314 | 20201114_13813_1ng_glyco_hcd203040_compound15.18998.18998.3 | controllerType=0 controllerNumber=1 scan=18998 | 38.551 |
| 15585:3 | 18 | R.TLN[+2059.735]QSSDELQLSMGNAMFVK.E | HexNAc(4)Hex(5)Fuc(1)NeuAc(1) | N[+2060] | 1424.9413 | 3 | 4272.8095 | 4272.7926 | 0 | 3.9 | 125 | Specific | 346.31 | 346.31 | 346.31 | 7.56 | 28 | >sp\|P01011\|AACT_HUMAN Alpha-1-antichymotrypsin OS=Homo sapiens OX=9606 GN=SERPINA3 PE=1 SV=2 | 1314 | 20201114_13813_1ng_glyco_hcd203040_compound15.18969.18969.3 | controllerType=0 controllerNumber=1 scan=18969 | 38.5018 |
| 02022:5 | 20 | R.EKQTDEIKDTRN[+2059.735]ESTQNC[+57.021]VVAEPEK.M | HexNAc(4)Hex(5)Fuc(1)NeuAc(1) | C[+57], N[+2060] | 1002.4328 | 5 | 5008.1349 | 5008.1251 | 0 | 2 | 58 | Specific | 304.97 | 257.85 | 257.85 | 6.64 | 25 | >sp\|P47710\|CASA1_HUMAN Alpha-S1-casein OS=Homo sapiens OX=9606 GN=CSN1S1 PE=1 SV=1 | 989 | 20201114_13813_1ng_glyco_hcd203040_compound15.4852.4852.5 | controllerType=0 controllerNumber=1 scan=4852 | 14.3834 |
| 02020:5 | 20 | R.EKQTDEIKDTRN[+2059.735]ESTQNC[+57.021]VVAEPEK.M | HexNAc(4)Hex(5)Fuc(1)NeuAc(1) | C[+57], N[+2060] | 1002.4328 | 5 | 5008.1349 | 5008.1251 | 0 | 2 | 58 | Specific | 352.25 | 352.25 | 352.25 | 7.5 | 25 | >sp\|P47710\|CASA1_HUMAN Alpha-S1-casein OS=Homo sapiens OX=9606 GN=CSN1S1 PE=1 SV=1 | 989 | 20201114_13813_1ng_glyco_hcd203040_compound15.4850.4850.5 | controllerType=0 controllerNumber=1 scan=4850 | 14.3797 |
| 02012:4 | 20 | R.EKQTDEIKDTRN[+2059.735]ESTQNC[+57.021]VVAEPEK.M | HexNAc(4)Hex(5)Fuc(1)NeuAc(1) | C[+57], N[+2060] | 1252.7885 | 4 | 5008.132 | 5008.1251 | 0 | 1.4 | 58 | Specific | 662.41 | 505.23 | 505.23 | 12.04 | 25 | >sp\|P47710\|CASA1_HUMAN Alpha-S1-casein OS=Homo sapiens OX=9606 GN=CSN1S1 PE=1 SV=1 | 989 | 20201114_13813_1ng_glyco_hcd203040_compound15.4842.4842.4 | controllerType=0 controllerNumber=1 scan=4842 | 14.3642 |
| 02014:4 | 20 | R.EKQTDEIKDTRN[+2059.735]ESTQNC[+57.021]VVAEPEK.M | HexNAc(4)Hex(5)Fuc(1)NeuAc(1) | C[+57], N[+2060] | 1252.7885 | 4 | 5008.132 | 5008.1251 | 0 | 1.4 | 58 | Specific | 670.52 | 536.12 | 536.12 | 12.04 | 25 | >sp\|P47710\|CASA1_HUMAN Alpha-S1-casein OS=Homo sapiens OX=9606 GN=CSN1S1 PE=1 SV=1 | 989 | 20201114_13813_1ng_glyco_hcd203040_compound15.4844.4844.4 | controllerType=0 controllerNumber=1 scan=4844 | 14.3691 |
| 09242:5 | 20 | K.QTDEIKDTRN[+2059.735]ESTQNC[+57.021]VVAEPEKMESSISSSSEEMSLSK.C | HexNAc(4)Hex(5)Fuc(1)NeuAc(1) | C[+57], N[+2060] | 1290.9498 | 5 | 6450.7201 | 6450.7156 | 0 | 0.7 | 60 | Specific | 743.76 | 550.69 | 550.69 | 13.52 | 25 | >sp\|P47710\|CASA1_HUMAN Alpha-S1-casein OS=Homo sapiens OX=9606 GN=CSN1S1 PE=1 SV=1 | 989 | 20201114_13813_1ng_glyco_hcd203040_compound15.12362.12362.5 | controllerType=0 controllerNumber=1 scan=12362 | 27.1587 |
| 09240:5 | 20 | K.QTDEIKDTRN[+2059.735]ESTQNC[+57.021]VVAEPEKMESSISSSSEEMSLSK.C | HexNAc(4)Hex(5)Fuc(1)NeuAc(1) | C[+57], N[+2060] | 1290.9498 | 5 | 6450.7201 | 6450.7156 | 0 | 0.7 | 60 | Specific | 799.04 | 621.1 | 621.1 | 14.52 | 25 | >sp\|P47710\|CASA1_HUMAN Alpha-S1-casein OS=Homo sapiens OX=9606 GN=CSN1S1 PE=1 SV=1 | 989 | 20201114_13813_1ng_glyco_hcd203040_compound15.12360.12360.5 | controllerType=0 controllerNumber=1 scan=12360 | 27.1539 |
| 08312:4 | 24 | K.[+42.011]SC[+57.021]QHN[+2018.708]GTMYQHGEIFSAHELFPSR.L | HexNAc(3)Hex(6)Fuc(1)NeuAc(1) | .[42], C[+57], N[+2019] | 1224.2561 | 4 | 4894.0028 | 4893.9759 | 0 | 5.5 | 110 | Specific | 232.96 | 232.96 | 232.96 | 6.26 | 21 | >sp\|Q6WN34\|CRDL2_HUMAN Chordin-like protein 2 OS=Homo sapiens OX=9606 GN=CHRDL2 PE=1 SV=1 | 1189 | 20201114_13813_1ng_glyco_hcd203040_compound15.11386.11386.4 | controllerType=0 controllerNumber=1 scan=11386 | 25.4925 |
| 08314:4 | 24 | K.[+42.011]SC[+57.021]QHN[+2018.708]GTMYQHGEIFSAHELFPSR.L | HexNAc(3)Hex(6)Fuc(1)NeuAc(1) | .[42], C[+57], N[+2019] | 1224.2561 | 4 | 4894.0028 | 4893.9759 | 0 | 5.5 | 110 | Specific | 234.66 | 199.12 | 199.12 | 4.48 | 21 | >sp\|Q6WN34\|CRDL2_HUMAN Chordin-like protein 2 OS=Homo sapiens OX=9606 GN=CHRDL2 PE=1 SV=1 | 1189 | 20201114_13813_1ng_glyco_hcd203040_compound15.11388.11388.4 | controllerType=0 controllerNumber=1 scan=11388 | 25.4964 |
| 14428:4 | 28 | K.MVSHHN[+2204.772]LTTGATLINEQWLLTTAK.N | HexNAc(4)Hex(5)NeuAc(2) | N[+2205] | 1221.8041 | 4 | 4884.1947 | 4884.1647 | 0 | 6.1 | 179 | Specific | 591.92 | 591.92 | 591.92 | 11.48 | 22 | >sp\|P00738\|HPT_HUMAN Haptoglobin OS=Homo sapiens OX=9606 GN=HP PE=1 SV=1 | 407 | 20201114_13813_1ng_glyco_hcd203040_compound15.17763.17763.2 | controllerType=0 controllerNumber=1 scan=17763 | 36.3429 |
| 14426:4 | 28 | K.MVSHHN[+2204.772]LTTGATLINEQWLLTTAK.N | HexNAc(4)Hex(5)NeuAc(2) | N[+2205] | 1221.8041 | 4 | 4884.1947 | 4884.1647 | 0 | 6.1 | 179 | Specific | 716.14 | 570.58 | 570.58 | 14.48 | 22 | >sp\|P00738\|HPT_HUMAN Haptoglobin OS=Homo sapiens OX=9606 GN=HP PE=1 SV=1 | 407 | 20201114_13813_1ng_glyco_hcd203040_compound15.17761.17761.2 | controllerType=0 controllerNumber=1 scan=17761 | 36.3384 |
| 01231:3 | 42 | K.HYTN[+1809.666]SSQDVTVPC[+57.021]R.V | HexNAc(5)Hex(4)Fuc(1) | C[+57], N[+1810] | 1158.4793 | 3 | 3473.4234 | 3473.4154 | 0 | 2.3 | 89 | Specific | 230.05 | 172.95 | 172.95 | 6.19 | 17 | >sp\|P01877\|IGHA2_HUMAN Immunoglobulin heavy constant alpha 2 OS=Homo sapiens OX=9606 GN=IGHA2 PE=1 SV=4 | 576 | 20201114_13813_1ng_glyco_hcd203040_compound15.4028.4028.3 | controllerType=0 controllerNumber=1 scan=4028 | 12.9278 |
| 01229:3 | 42 | K.HYTN[+1809.666]SSQDVTVPC[+57.021]R.V | HexNAc(5)Hex(4)Fuc(1) | C[+57], N[+1810] | 1158.4793 | 3 | 3473.4234 | 3473.4154 | 0 | 2.3 | 89 | Specific | 258.09 | 209.41 | 209.41 | 6.35 | 17 | >sp\|P01877\|IGHA2_HUMAN Immunoglobulin heavy constant alpha 2 OS=Homo sapiens OX=9606 GN=IGHA2 PE=1 SV=4 | 576 | 20201114_13813_1ng_glyco_hcd203040_compound15.4026.4026.3 | controllerType=0 controllerNumber=1 scan=4026 | 12.9238 |
| 01240:3 | 42 | K.HYTN[+1647.613]SSQDVTVPC[+57.021]R.V | HexNAc(5)Hex(3)Fuc(1) | C[+57], N[+1648] | 1104.4619 | 3 | 3311.3713 | 3311.3625 | 0 | 2.6 | 89 | Specific | 293.81 | 252.67 | 252.67 | 6.74 | 17 | >sp\|P01877\|IGHA2_HUMAN Immunoglobulin heavy constant alpha 2 OS=Homo sapiens OX=9606 GN=IGHA2 PE=1 SV=4 | 576 | 20201114_13813_1ng_glyco_hcd203040_compound15.4037.4037.2 | controllerType=0 controllerNumber=1 scan=4037 | 12.9428 |
| 01238:3 | 42 | K.HYTN[+1647.613]SSQDVTVPC[+57.021]R.V | HexNAc(5)Hex(3)Fuc(1) | C[+57], N[+1648] | 1104.4619 | 3 | 3311.3713 | 3311.3625 | 0 | 2.6 | 89 | Specific | 316.11 | 311.6 | 311.6 | 7.24 | 17 | >sp\|P01877\|IGHA2_HUMAN Immunoglobulin heavy constant alpha 2 OS=Homo sapiens OX=9606 GN=IGHA2 PE=1 SV=4 | 576 | 20201114_13813_1ng_glyco_hcd203040_compound15.4035.4035.2 | controllerType=0 controllerNumber=1 scan=4035 | 12.9388 |
| 01158:3 | 42 | K.HYTN[+1809.666]SSQDVTVPC[+57.021]R.V | HexNAc(5)Hex(4)Fuc(1) | C[+57], N[+1810] | 1158.4788 | 3 | 3473.4219 | 3473.4154 | 0 | 1.9 | 89 | Specific | 488.44 | 347.17 | 347.17 | 8.3 | 17 | >sp\|P01877\|IGHA2_HUMAN Immunoglobulin heavy constant alpha 2 OS=Homo sapiens OX=9606 GN=IGHA2 PE=1 SV=4 | 576 | 20201114_13813_1ng_glyco_hcd203040_compound15.3952.3952.3 | controllerType=0 controllerNumber=1 scan=3952 | 12.7915 |
| 01160:3 | 42 | K.HYTN[+1809.666]SSQDVTVPC[+57.021]R.V | HexNAc(5)Hex(4)Fuc(1) | C[+57], N[+1810] | 1158.4788 | 3 | 3473.4219 | 3473.4154 | 0 | 1.9 | 89 | Specific | 491.88 | 342.96 | 342.96 | 8.3 | 17 | >sp\|P01877\|IGHA2_HUMAN Immunoglobulin heavy constant alpha 2 OS=Homo sapiens OX=9606 GN=IGHA2 PE=1 SV=4 | 576 | 20201114_13813_1ng_glyco_hcd203040_compound15.3954.3954.3 | controllerType=0 controllerNumber=1 scan=3954 | 12.7973 |
| 01200:3 | 42 | K.HYTN[+1647.613]SSQDVTVPC[+57.021]R.V | HexNAc(5)Hex(3)Fuc(1) | C[+57], N[+1648] | 1104.4618 | 3 | 3311.3709 | 3311.3625 | 0 | 2.5 | 89 | Specific | 643.41 | 546.76 | 546.76 | 12.04 | 17 | >sp\|P01877\|IGHA2_HUMAN Immunoglobulin heavy constant alpha 2 OS=Homo sapiens OX=9606 GN=IGHA2 PE=1 SV=4 | 576 | 20201114_13813_1ng_glyco_hcd203040_compound15.3996.3996.3 | controllerType=0 controllerNumber=1 scan=3996 | 12.8699 |
| 01202:3 | 42 | K.HYTN[+1647.613]SSQDVTVPC[+57.021]R.V | HexNAc(5)Hex(3)Fuc(1) | C[+57], N[+1648] | 1104.4618 | 3 | 3311.3709 | 3311.3625 | 0 | 2.5 | 89 | Specific | 705.59 | 595.33 | 595.33 | 14.04 | 17 | >sp\|P01877\|IGHA2_HUMAN Immunoglobulin heavy constant alpha 2 OS=Homo sapiens OX=9606 GN=IGHA2 PE=1 SV=4 | 576 | 20201114_13813_1ng_glyco_hcd203040_compound15.3998.3998.3 | controllerType=0 controllerNumber=1 scan=3998 | 12.875 |
| 04613:3 | 42 | K.TPLTAN[+2262.814]ITK.S | HexNAc(5)Hex(5)Fuc(1)NeuAc(1) | N[+2263] | 1074.4619 | 3 | 3221.3713 | 3221.3711 | 0 | 0.1 | 200 | Specific | 213.62 | 148.19 | 148.19 | 3.86 | 17 | >sp\|P01877\|IGHA2_HUMAN Immunoglobulin heavy constant alpha 2 OS=Homo sapiens OX=9606 GN=IGHA2 PE=1 SV=4 | 576 | 20201114_13813_1ng_glyco_hcd203040_compound15.7545.7545.3 | controllerType=0 controllerNumber=1 scan=7545 | 19.069 |
| 02883:3 | 42 | K.TPLTAN[+1809.666]ITK.S | HexNAc(5)Hex(4)Fuc(1) | N[+1810] | 923.4151 | 3 | 2768.2308 | 2768.2228 | 0 | 2.9 | 200 | Specific | 284.19 | 119.59 | 119.59 | 5.05 | 17 | >sp\|P01877\|IGHA2_HUMAN Immunoglobulin heavy constant alpha 2 OS=Homo sapiens OX=9606 GN=IGHA2 PE=1 SV=4 | 576 | 20201114_13813_1ng_glyco_hcd203040_compound15.5745.5745.3 | controllerType=0 controllerNumber=1 scan=5745 | 15.9872 |
| 02901:2 | 42 | K.TPLTAN[+1444.534]ITK.S | HexNAc(4)Hex(3)Fuc(1) | N[+1445] | 1202.0538 | 2 | 2403.1002 | 2403.0906 | 0 | 4 | 200 | Specific | 378.53 | 199.38 | 199.38 | 6.26 | 17 | >sp\|P01877\|IGHA2_HUMAN Immunoglobulin heavy constant alpha 2 OS=Homo sapiens OX=9606 GN=IGHA2 PE=1 SV=4 | 576 | 20201114_13813_1ng_glyco_hcd203040_compound15.5763.5763.2 | controllerType=0 controllerNumber=1 scan=5763 | 16.0196 |
| 02903:2 | 42 | K.TPLTAN[+1444.534]ITK.S | HexNAc(4)Hex(3)Fuc(1) | N[+1445] | 1202.0538 | 2 | 2403.1002 | 2403.0906 | 0 | 4 | 200 | Specific | 424.96 | 242.71 | 242.71 | 6.6 | 17 | >sp\|P01877\|IGHA2_HUMAN Immunoglobulin heavy constant alpha 2 OS=Homo sapiens OX=9606 GN=IGHA2 PE=1 SV=4 | 576 | 20201114_13813_1ng_glyco_hcd203040_compound15.5765.5765.2 | controllerType=0 controllerNumber=1 scan=5765 | 16.0237 |
| 02948:3 | 42 | K.TPLTAN[+1647.613]ITK.S | HexNAc(5)Hex(3)Fuc(1) | N[+1648] | 869.3964 | 3 | 2606.1746 | 2606.17 | 0 | 1.8 | 200 | Specific | 473.32 | 218.34 | 218.34 | 6.8 | 17 | >sp\|P01877\|IGHA2_HUMAN Immunoglobulin heavy constant alpha 2 OS=Homo sapiens OX=9606 GN=IGHA2 PE=1 SV=4 | 576 | 20201114_13813_1ng_glyco_hcd203040_compound15.5812.5812.3 | controllerType=0 controllerNumber=1 scan=5812 | 16.11 |
| 02885:3 | 42 | K.TPLTAN[+1809.666]ITK.S | HexNAc(5)Hex(4)Fuc(1) | N[+1810] | 923.4151 | 3 | 2768.2308 | 2768.2228 | 0 | 2.9 | 200 | Specific | 489.21 | 249.19 | 249.19 | 7.29 | 17 | >sp\|P01877\|IGHA2_HUMAN Immunoglobulin heavy constant alpha 2 OS=Homo sapiens OX=9606 GN=IGHA2 PE=1 SV=4 | 576 | 20201114_13813_1ng_glyco_hcd203040_compound15.5747.5747.3 | controllerType=0 controllerNumber=1 scan=5747 | 15.991 |
| 02878:2 | 42 | K.TPLTAN[+1809.666]ITK.S | HexNAc(5)Hex(4)Fuc(1) | N[+1810] | 1384.6192 | 2 | 2768.2311 | 2768.2228 | 0 | 3 | 200 | Specific | 512.3 | 266.67 | 266.67 | 7.57 | 17 | >sp\|P01877\|IGHA2_HUMAN Immunoglobulin heavy constant alpha 2 OS=Homo sapiens OX=9606 GN=IGHA2 PE=1 SV=4 | 576 | 20201114_13813_1ng_glyco_hcd203040_compound15.5740.5740.2 | controllerType=0 controllerNumber=1 scan=5740 | 15.9778 |
| 02880:2 | 42 | K.TPLTAN[+1809.666]ITK.S | HexNAc(5)Hex(4)Fuc(1) | N[+1810] | 1384.6192 | 2 | 2768.2311 | 2768.2228 | 0 | 3 | 200 | Specific | 578.16 | 309.6 | 309.6 | 9.19 | 17 | >sp\|P01877\|IGHA2_HUMAN Immunoglobulin heavy constant alpha 2 OS=Homo sapiens OX=9606 GN=IGHA2 PE=1 SV=4 | 576 | 20201114_13813_1ng_glyco_hcd203040_compound15.5742.5742.2 | controllerType=0 controllerNumber=1 scan=5742 | 15.9823 |
| 02932:2 | 42 | K.TPLTAN[+1647.613]ITK.S | HexNAc(5)Hex(3)Fuc(1) | N[+1648] | 1303.5914 | 2 | 2606.1754 | 2606.17 | 0 | 2.1 | 200 | Specific | 594.96 | 345.7 | 345.7 | 9.48 | 17 | >sp\|P01877\|IGHA2_HUMAN Immunoglobulin heavy constant alpha 2 OS=Homo sapiens OX=9606 GN=IGHA2 PE=1 SV=4 | 576 | 20201114_13813_1ng_glyco_hcd203040_compound15.5796.5796.2 | controllerType=0 controllerNumber=1 scan=5796 | 16.0792 |
| 02934:2 | 42 | K.TPLTAN[+1647.613]ITK.S | HexNAc(5)Hex(3)Fuc(1) | N[+1648] | 1303.5914 | 2 | 2606.1754 | 2606.17 | 0 | 2.1 | 200 | Specific | 618 | 360.21 | 360.21 | 10.48 | 17 | >sp\|P01877\|IGHA2_HUMAN Immunoglobulin heavy constant alpha 2 OS=Homo sapiens OX=9606 GN=IGHA2 PE=1 SV=4 | 576 | 20201114_13813_1ng_glyco_hcd203040_compound15.5798.5798.2 | controllerType=0 controllerNumber=1 scan=5798 | 16.0847 |
| 10024:3 | 42 | R.MAGKPTHIN[+1540.529]VSVVMAEADGTC[+57.021]Y.- | HexNAc(2)Hex(7) | C[+57], N[+1541] | 1297.8793 | 3 | 3891.6234 | 3891.6114 | 0 | 3.1 | 319 | CRagged | 337.45 | 337.45 | 286.2 | 7.41 | 17 | >sp\|P01877\|IGHA2_HUMAN Immunoglobulin heavy constant alpha 2 OS=Homo sapiens OX=9606 GN=IGHA2 PE=1 SV=4 | 576 | 20201114_13813_1ng_glyco_hcd203040_compound15.13174.13174.3 | controllerType=0 controllerNumber=1 scan=13174 | 28.4857 |

Table S4. Cont.

| Query #:z | Protein Rank | Glycopeptide | Glycan | Modification Type(s) | Observed m/z | z | Observed (M+H) | Calc. mass (M+H) | Off-by-x error | Mass error (ppm) | StartPosition | Cleavage | Score | Delta | Delta Mod | \|Log Prob\| | # of unique peptides | Protein Name | Protein DB number | Comment | Scan # | Scan Time |
|---|---|---|---|---|---|---|---|---|---|---|---|---|---|---|---|---|---|---|---|---|---|---|
| 10057:3 | 42 | R.MAGKPTHIN[+1378.476]VSVVMAEADGTC[+57.021]Y.- | HexNAc(2)Hex(6) | C[+57], N[+1378] | 1243.8596 | 3 | 3729.5641 | 3729.5585 | 0 | 1.5 | 319 | CRagged | 364.67 | 364.67 | 241.02 | 7.93 | 17 | >sp\|P01877\|IGHA2_HUMAN Immunoglobulin heavy constant alpha 2 OS=Homo sapiens OX=9606 GN=IGHA2 PE=1 SV=4 | 576 | 20201114_13813_1ng_glyco_hcd203040_compound15.13209.13209.3 | controllerType=0 controllerNumber=1 scan=13209 | 28.5468 |
| 10022:3 | 42 | R.MAGKPTHIN[+1540.529]VSVVMAEADGTC[+57.021]Y.- | HexNAc(2)Hex(7) | C[+57], N[+1541] | 1297.8793 | 3 | 3891.6234 | 3891.6114 | 0 | 3.1 | 319 | CRagged | 369.09 | 369.09 | 208.59 | 7.9 | 17 | >sp\|P01877\|IGHA2_HUMAN Immunoglobulin heavy constant alpha 2 OS=Homo sapiens OX=9606 GN=IGHA2 PE=1 SV=4 | 576 | 20201114_13813_1ng_glyco_hcd203040_compound15.13172.13172.3 | controllerType=0 controllerNumber=1 scan=13172 | 28.482 |
| 10059:3 | 42 | R.MAGKPTHIN[+1378.476]VSVVMAEADGTC[+57.021]Y.- | HexNAc(2)Hex(6) | C[+57], N[+1378] | 1243.8596 | 3 | 3729.5641 | 3729.5585 | 0 | 1.5 | 319 | CRagged | 789.14 | 789.14 | 406.29 | 15.14 | 17 | >sp\|P01877\|IGHA2_HUMAN Immunoglobulin heavy constant alpha 2 OS=Homo sapiens OX=9606 GN=IGHA2 PE=1 SV=4 | 576 | 20201114_13813_1ng_glyco_hcd203040_compound15.13211.13211.3 | controllerType=0 controllerNumber=1 scan=13211 | 28.551 |
| 14998:3 | 44 | K.RNYIVPILWLN[+1864.634]ETGTIGDEK.A | HexNAc(2)Hex(9) | N[+1865] | 1399.3065 | 3 | 4195.9048 | 4195.8685 | 0 | 8.7 | 407 | Specific | 203.94 | 203.94 | 203.94 | 4.49 | 13 | >sp\|P16671\|CD36_HUMAN Platelet glycoprotein 4 OS=Homo sapiens OX=9606 GN=CD36 PE=1 SV=2 | 812 | 20201114_13813_1ng_glyco_hcd203040_compound15.18358.18358.3 | controllerType=0 controllerNumber=1 scan=18358 | 37.3783 |
| 15016:3 | 44 | K.RNYIVPILWLN[+1864.634]ETGTIGDEK.A | HexNAc(2)Hex(9) | N[+1865] | 1399.3014 | 3 | 4195.8898 | 4195.8685 | 0 | 5.1 | 407 | Specific | 536.87 | 536.87 | 536.87 | 9.49 | 13 | >sp\|P16671\|CD36_HUMAN Platelet glycoprotein 4 OS=Homo sapiens OX=9606 GN=CD36 PE=1 SV=2 | 812 | 20201114_13813_1ng_glyco_hcd203040_compound15.18377.18377.3 | controllerType=0 controllerNumber=1 scan=18377 | 37.4169 |
| 15018:3 | 44 | K.RNYIVPILWLN[+1864.634]ETGTIGDEK.A | HexNAc(2)Hex(9) | N[+1865] | 1399.3014 | 3 | 4195.8898 | 4195.8685 | 0 | 5.1 | 407 | Specific | 667.97 | 667.97 | 667.97 | 12.82 | 13 | >sp\|P16671\|CD36_HUMAN Platelet glycoprotein 4 OS=Homo sapiens OX=9606 GN=CD36 PE=1 SV=2 | 812 | 20201114_13813_1ng_glyco_hcd203040_compound15.18379.18379.3 | controllerType=0 controllerNumber=1 scan=18379 | 37.421 |
| 15000:3 | 44 | K.RNYIVPILWLN[+1864.634]ETGTIGDEK.A | HexNAc(2)Hex(9) | N[+1865] | 1399.3065 | 3 | 4195.9048 | 4195.8685 | 0 | 8.7 | 407 | Specific | 863.56 | 863.56 | 863.56 | 16.46 | 13 | >sp\|P16671\|CD36_HUMAN Platelet glycoprotein 4 OS=Homo sapiens OX=9606 GN=CD36 PE=1 SV=2 | 812 | 20201114_13813_1ng_glyco_hcd203040_compound15.18360.18360.3 | controllerType=0 controllerNumber=1 scan=18360 | 37.3829 |
| 13125:3 | 59 | R.IIVPLNNREN[+2351.851]ISDPTSPLR.T | HexNAc(4)Hex(5)Fuc(3)NeuAc(1) | N[+2352] | 1500.6745 | 3 | 4500.0089 | 4500.0279 | 0 | -4.2 | 62 | Specific | 210.27 | 210.27 | 210.27 | 3.54 | 12 | >sp\|P01591\|IGJ_HUMAN Immunoglobulin J chain OS=Homo sapiens OX=9606 GN=JCHAIN PE=1 SV=4 | 1392 | 20201114_13813_1ng_glyco_hcd203040_compound15.16411.16411.3 | controllerType=0 controllerNumber=1 scan=16411 | 33.9801 |
| 13123:3 | 59 | R.IIVPLNNREN[+2351.851]ISDPTSPLR.T | HexNAc(4)Hex(5)Fuc(3)NeuAc(1) | N[+2352] | 1500.6745 | 3 | 4500.0089 | 4500.0279 | 0 | -4.2 | 62 | Specific | 260.04 | 260.04 | 260.04 | 5.18 | 12 | >sp\|P01591\|IGJ_HUMAN Immunoglobulin J chain OS=Homo sapiens OX=9606 GN=JCHAIN PE=1 SV=4 | 1392 | 20201114_13813_1ng_glyco_hcd203040_compound15.16409.16409.3 | controllerType=0 controllerNumber=1 scan=16409 | 33.9764 |
| 09474:3 | 59 | R.IIVPLNNREN[+1809.666]ISDPTSPLR.T | HexNAc(5)Hex(4)Fuc(1) | N[+1810] | 1319.9546 | 3 | 3957.8494 | 3957.8432 | 0 | 1.6 | 62 | Specific | 281.77 | 238.26 | 238.26 | 6.16 | 12 | >sp\|P01591\|IGJ_HUMAN Immunoglobulin J chain OS=Homo sapiens OX=9606 GN=JCHAIN PE=1 SV=4 | 1392 | 20201114_13813_1ng_glyco_hcd203040_compound15.12604.12604.3 | controllerType=0 controllerNumber=1 scan=12604 | 27.5469 |
| 09445:3 | 59 | R.IIVPLNNREN[+1663.608]ISDPTSPLR.T | HexNAc(5)Hex(4) | N[+1664] | 1271.2705 | 3 | 3811.7969 | 3811.7853 | 0 | 3.1 | 62 | Specific | 332.31 | 319.04 | 319.04 | 6.86 | 12 | >sp\|P01591\|IGJ_HUMAN Immunoglobulin J chain OS=Homo sapiens OX=9606 GN=JCHAIN PE=1 SV=4 | 1392 | 20201114_13813_1ng_glyco_hcd203040_compound15.12574.12574.3 | controllerType=0 controllerNumber=1 scan=12574 | 27.4996 |
| 09447:3 | 59 | R.IIVPLNNREN[+1663.608]ISDPTSPLR.T | HexNAc(5)Hex(4) | N[+1664] | 1271.2705 | 3 | 3811.7969 | 3811.7853 | 0 | 3.1 | 62 | Specific | 382.3 | 341.15 | 341.15 | 7.13 | 12 | >sp\|P01591\|IGJ_HUMAN Immunoglobulin J chain OS=Homo sapiens OX=9606 GN=JCHAIN PE=1 SV=4 | 1392 | 20201114_13813_1ng_glyco_hcd203040_compound15.12576.12576.3 | controllerType=0 controllerNumber=1 scan=12576 | 27.5033 |
| 11170:3 | 59 | R.IIVPLNNREN[+1856.656]ISDPTSPLR.T | HexNAc(3)Hex(5)Fuc(1)NeuAc(1) | N[+1857] | 1335.6203 | 3 | 4004.8463 | 4004.8327 | 0 | 3.4 | 62 | Specific | 388.45 | 388.45 | 388.45 | 8.65 | 12 | >sp\|P01591\|IGJ_HUMAN Immunoglobulin J chain OS=Homo sapiens OX=9606 GN=JCHAIN PE=1 SV=4 | 1392 | 20201114_13813_1ng_glyco_hcd203040_compound15.14366.14366.3 | controllerType=0 controllerNumber=1 scan=14366 | 30.5348 |
| 11168:3 | 59 | R.IIVPLNNREN[+1856.656]ISDPTSPLR.T | HexNAc(3)Hex(5)Fuc(1)NeuAc(1) | N[+1857] | 1335.6203 | 3 | 4004.8463 | 4004.8327 | 0 | 3.4 | 62 | Specific | 468.37 | 424.08 | 424.08 | 8.86 | 12 | >sp\|P01591\|IGJ_HUMAN Immunoglobulin J chain OS=Homo sapiens OX=9606 GN=JCHAIN PE=1 SV=4 | 1392 | 20201114_13813_1ng_glyco_hcd203040_compound15.14364.14364.3 | controllerType=0 controllerNumber=1 scan=14364 | 30.5296 |
| 18368:3 | 2 | R.RTAGWNVPIGTLRPFLN[+2205.793]WTGPPEPIEAAVAR.F | HexNAc(4)Hex(5)Fuc(2)NeuAc(1) | N[+2206] | 1864.8805 | 3 | 5592.6268 | 5592.6049 | 0 | 3.9 | 140 | Specific | 261.99 | 250.24 | 250.24 | 5.03 | 125 | >sp\|P02788\|TRFL_HUMAN Lactotransferrin OS=Homo sapiens OX=9606 GN=LTF PE=1 SV=6 | 1270 | 20201116_13832_1ng_glyco_hcd203040_compound15.21896.21896.3 | controllerType=0 controllerNumber=1 scan=21896 | 43.5583 |
| 18364:5 | 2 | R.RTAGWNVPIGTLRPFLN[+2204.772]WTGPPEPIEAAVAR.F | HexNAc(4)Hex(5)NeuAc(2) | N[+2205] | 1119.1292 | 5 | 5591.6167 | 5591.5845 | 0 | 5.8 | 140 | Specific | 263.7 | 263.7 | 263.7 | 6.12 | 125 | >sp\|P02788\|TRFL_HUMAN Lactotransferrin OS=Homo sapiens OX=9606 GN=LTF PE=1 SV=6 | 1270 | 20201116_13832_1ng_glyco_hcd203040_compound15.21892.21892.5 | controllerType=0 controllerNumber=1 scan=21892 | 43.5513 |
| 18274:4 | 2 | R.RTAGWNVPIGTLRPFLN[+2570.925]WTGPPEPIEAAVAR.F | HexNAc(5)Hex(6)Fuc(2)NeuAc(1) | N[+2571] | 1490.1932 | 4 | 5957.7511 | 5957.7371 | 0 | 2.4 | 140 | Specific | 327.22 | 295.53 | 295.53 | 6.73 | 125 | >sp\|P02788\|TRFL_HUMAN Lactotransferrin OS=Homo sapiens OX=9606 GN=LTF PE=1 SV=6 | 1270 | 20201116_13832_1ng_glyco_hcd203040_compound15.21798.21798.4 | controllerType=0 controllerNumber=1 scan=21798 | 43.3541 |
| 18367:5 | 2 | R.RTAGWNVPIGTLRPFLN[+2204.772]WTGPPEPIEAAVAR.F | HexNAc(4)Hex(5)NeuAc(2) | N[+2205] | 1119.1292 | 5 | 5591.6167 | 5591.5845 | 0 | 5.8 | 140 | Specific | 338.16 | 338.16 | 338.16 | 6.46 | 125 | >sp\|P02788\|TRFL_HUMAN Lactotransferrin OS=Homo sapiens OX=9606 GN=LTF PE=1 SV=6 | 1270 | 20201116_13832_1ng_glyco_hcd203040_compound15.21895.21895.5 | controllerType=0 controllerNumber=1 scan=21895 | 43.5566 |
| 18276:4 | 2 | R.RTAGWNVPIGTLRPFLN[+2570.925]WTGPPEPIEAAVAR.F | HexNAc(5)Hex(6)Fuc(2)NeuAc(1) | N[+2571] | 1490.1932 | 4 | 5957.7511 | 5957.7371 | 0 | 2.4 | 140 | Specific | 431.39 | 411.13 | 411.13 | 9.59 | 125 | >sp\|P02788\|TRFL_HUMAN Lactotransferrin OS=Homo sapiens OX=9606 GN=LTF PE=1 SV=6 | 1270 | 20201116_13832_1ng_glyco_hcd203040_compound15.21800.21800.4 | controllerType=0 controllerNumber=1 scan=21800 | 43.3586 |
| 18353:4 | 2 | R.RTAGWNVPIGTLRPFLN[+2205.793]WTGPPEPIEAAVAR.F | HexNAc(4)Hex(5)Fuc(2)NeuAc(1) | N[+2206] | 1398.9119 | 4 | 5592.6256 | 5592.6049 | 0 | 3.7 | 140 | Specific | 487.99 | 377.8 | 377.8 | 9.64 | 125 | >sp\|P02788\|TRFL_HUMAN Lactotransferrin OS=Homo sapiens OX=9606 GN=LTF PE=1 SV=6 | 1270 | 20201116_13832_1ng_glyco_hcd203040_compound15.21881.21881.4 | controllerType=0 controllerNumber=1 scan=21881 | 43.527 |
| 19356:4 | 2 | R.RTAGWNVPIGTLRPFLN[+2350.830]WTGPPEPIEAAVAR.F | HexNAc(4)Hex(5)Fuc(1)NeuAc(2) | N[+2351] | 1435.1691 | 4 | 5737.6545 | 5737.6424 | 0 | 2.1 | 140 | Specific | 527.24 | 443.3 | 443.3 | 9.95 | 125 | >sp\|P02788\|TRFL_HUMAN Lactotransferrin OS=Homo sapiens OX=9606 GN=LTF PE=1 SV=6 | 1270 | 20201116_13832_1ng_glyco_hcd203040_compound15.22926.22926.4 | controllerType=0 controllerNumber=1 scan=22926 | 45.7733 |
| 18355:4 | 2 | R.RTAGWNVPIGTLRPFLN[+2205.793]WTGPPEPIEAAVAR.F | HexNAc(4)Hex(5)Fuc(2)NeuAc(1) | N[+2206] | 1398.9119 | 4 | 5592.6256 | 5592.6049 | 0 | 3.7 | 140 | Specific | 571.81 | 465.21 | 465.21 | 11.13 | 125 | >sp\|P02788\|TRFL_HUMAN Lactotransferrin OS=Homo sapiens OX=9606 GN=LTF PE=1 SV=6 | 1270 | 20201116_13832_1ng_glyco_hcd203040_compound15.21883.21883.4 | controllerType=0 controllerNumber=1 scan=21883 | 43.5322 |
| 19476:3 | 2 | R.TAGWNVPIGTLRPFLN[+2570.925]WTGPPEPIEAAVAR.F | HexNAc(5)Hex(6)Fuc(2)NeuAc(1) | N[+2571] | 1934.5554 | 3 | 5801.6516 | 5801.636 | 0 | 2.7 | 141 | Specific | 269.78 | 172.34 | 172.34 | 5.85 | 125 | >sp\|P02788\|TRFL_HUMAN Lactotransferrin OS=Homo sapiens OX=9606 GN=LTF PE=1 SV=6 | 1270 | 20201116_13832_1ng_glyco_hcd203040_compound15.23051.23051.3 | controllerType=0 controllerNumber=1 scan=23051 | 46.0348 |
| 18562:4 | 2 | R.[+42.011]TAGWNVPIGTLRPFLN[+2018.708]WTGPPEPIEAAVAR.F | HexNAc(3)Hex(6)Fuc(1)NeuAc(1) | .[42], N[+2019] | 1323.6245 | 4 | 5291.4762 | 5291.4299 | 0 | 8.8 | 141 | Specific | 275.61 | 176.22 | 176.22 | 3.59 | 125 | >sp\|P02788\|TRFL_HUMAN Lactotransferrin OS=Homo sapiens OX=9606 GN=LTF PE=1 SV=6 | 1270 | 20201116_13832_1ng_glyco_hcd203040_compound15.22098.22098.4 | controllerType=0 controllerNumber=1 scan=22098 | 43.9747 |
| 18559:4 | 2 | R.[+42.011]TAGWNVPIGTLRPFLN[+2018.708]WTGPPEPIEAAVAR.F | HexNAc(3)Hex(6)Fuc(1)NeuAc(1) | .[42], N[+2019] | 1323.6245 | 4 | 5291.4762 | 5291.4299 | 0 | 8.8 | 141 | Specific | 278.25 | 278.25 | 278.25 | 5 | 125 | >sp\|P02788\|TRFL_HUMAN Lactotransferrin OS=Homo sapiens OX=9606 GN=LTF PE=1 SV=6 | 1270 | 20201116_13832_1ng_glyco_hcd203040_compound15.22095.22095.4 | controllerType=0 controllerNumber=1 scan=22095 | 43.9654 |
| 19605:3 | 2 | R.TAGWNVPIGTLRPFLN[+2205.793]WTGPPEPIEAAVAR.F | HexNAc(4)Hex(5)Fuc(2)NeuAc(1) | N[+2206] | 1812.8468 | 3 | 5436.5257 | 5436.5038 | 0 | 4 | 141 | Specific | 300.72 | 300.72 | 300.72 | 6.79 | 125 | >sp\|P02788\|TRFL_HUMAN Lactotransferrin OS=Homo sapiens OX=9606 GN=LTF PE=1 SV=6 | 1270 | 20201116_13832_1ng_glyco_hcd203040_compound15.23186.23186.3 | controllerType=0 controllerNumber=1 scan=23186 | 46.3275 |
| 19604:4 | 2 | R.TAGWNVPIGTLRPFLN[+2205.793]WTGPPEPIEAAVAR.F | HexNAc(4)Hex(5)Fuc(2)NeuAc(1) | N[+2206] | 1359.8824 | 4 | 5436.508 | 5436.5038 | 0 | 0.8 | 141 | Specific | 304.98 | 304.98 | 304.98 | 6.5 | 125 | >sp\|P02788\|TRFL_HUMAN Lactotransferrin OS=Homo sapiens OX=9606 GN=LTF PE=1 SV=6 | 1270 | 20201116_13832_1ng_glyco_hcd203040_compound15.23185.23185.4 | controllerType=0 controllerNumber=1 scan=23185 | 46.3261 |
| 19479:3 | 2 | R.TAGWNVPIGTLRPFLN[+2570.925]WTGPPEPIEAAVAR.F | HexNAc(5)Hex(6)Fuc(2)NeuAc(1) | N[+2571] | 1934.5554 | 3 | 5801.6516 | 5801.636 | 0 | 2.7 | 141 | Specific | 312.56 | 205.35 | 205.35 | 5.89 | 125 | >sp\|P02788\|TRFL_HUMAN Lactotransferrin OS=Homo sapiens OX=9606 GN=LTF PE=1 SV=6 | 1270 | 20201116_13832_1ng_glyco_hcd203040_compound15.23054.23054.3 | controllerType=0 controllerNumber=1 scan=23054 | 46.0394 |
| 19611:5 | 2 | R.TAGWNVPIGTLRPFLN[+2205.793]WTGPPEPIEAAVAR.F | HexNAc(4)Hex(5)Fuc(2)NeuAc(1) | N[+2206] | 1088.1129 | 5 | 5436.5355 | 5436.5038 | 0 | 5.8 | 141 | Specific | 406.61 | 332.08 | 332.08 | 6.88 | 125 | >sp\|P02788\|TRFL_HUMAN Lactotransferrin OS=Homo sapiens OX=9606 GN=LTF PE=1 SV=6 | 1270 | 20201116_13832_1ng_glyco_hcd203040_compound15.23192.23192.5 | controllerType=0 controllerNumber=1 scan=23192 | 46.3358 |
| 18499:4 | 2 | R.TAGWNVPIGTLRPFLN[+2644.962]WTGPPEPIEAAVAR.F | HexNAc(6)Hex(7)Fuc(2) | N[+2645] | 1469.6779 | 4 | 5875.6896 | 5875.6727 | 0 | 2.9 | 141 | Specific | 518.72 | 487.18 | 487.18 | 9.62 | 125 | >sp\|P02788\|TRFL_HUMAN Lactotransferrin OS=Homo sapiens OX=9606 GN=LTF PE=1 SV=6 | 1270 | 20201116_13832_1ng_glyco_hcd203040_compound15.22032.22032.2 | controllerType=0 controllerNumber=1 scan=22032 | 43.8241 |
| 21023:5 | 2 | R.TAGWNVPIGTLRPFLN[+2351.851]WTGPPEPIEAAVAR.F | HexNAc(4)Hex(5)Fuc(3)NeuAc(1) | N[+2352] | 1117.3131 | 5 | 5582.5365 | 5582.5617 | 0 | -4.5 | 141 | Specific | 546.78 | 546.78 | 546.78 | 8.66 | 125 | >sp\|P02788\|TRFL_HUMAN Lactotransferrin OS=Homo sapiens OX=9606 GN=LTF PE=1 SV=6 | 1270 | 20201116_13832_1ng_glyco_hcd203040_compound15.24662.24662.5 | controllerType=0 controllerNumber=1 scan=24662 | 49.193 |
| 19613:5 | 2 | R.TAGWNVPIGTLRPFLN[+2205.793]WTGPPEPIEAAVAR.F | HexNAc(4)Hex(5)Fuc(2)NeuAc(1) | N[+2206] | 1088.1129 | 5 | 5436.5355 | 5436.5038 | 0 | 5.8 | 141 | Specific | 595.01 | 595.01 | 595.01 | 10.39 | 125 | >sp\|P02788\|TRFL_HUMAN Lactotransferrin OS=Homo sapiens OX=9606 GN=LTF PE=1 SV=6 | 1270 | 20201116_13832_1ng_glyco_hcd203040_compound15.23194.23194.5 | controllerType=0 controllerNumber=1 scan=23194 | 46.3401 |
| 19543:3 | 2 | R.TAGWNVPIGTLRPFLN[+2205.793]WTGPPEPIEAAVAR.F | HexNAc(4)Hex(5)Fuc(2)NeuAc(1) | N[+2206] | 1812.8465 | 3 | 5436.525 | 5436.5038 | 0 | 3.9 | 141 | Specific | 607.62 | 516.09 | 516.09 | 12 | 125 | >sp\|P02788\|TRFL_HUMAN Lactotransferrin OS=Homo sapiens OX=9606 GN=LTF PE=1 SV=6 | 1270 | 20201116_13832_1ng_glyco_hcd203040_compound15.23121.23121.3 | controllerType=0 controllerNumber=1 scan=23121 | 46.1778 |
| 21026:5 | 2 | R.TAGWNVPIGTLRPFLN[+2351.851]WTGPPEPIEAAVAR.F | HexNAc(4)Hex(5)Fuc(3)NeuAc(1) | N[+2352] | 1117.3131 | 5 | 5582.5365 | 5582.5617 | 0 | -4.5 | 141 | Specific | 613.65 | 583.36 | 583.36 | 10.79 | 125 | >sp\|P02788\|TRFL_HUMAN Lactotransferrin OS=Homo sapiens OX=9606 GN=LTF PE=1 SV=6 | 1270 | 20201116_13832_1ng_glyco_hcd203040_compound15.24665.24665.5 | controllerType=0 controllerNumber=1 scan=24665 | 49.1984 |
| 19545:3 | 2 | R.TAGWNVPIGTLRPFLN[+2205.793]WTGPPEPIEAAVAR.F | HexNAc(4)Hex(5)Fuc(2)NeuAc(1) | N[+2206] | 1812.8465 | 3 | 5436.525 | 5436.5038 | 0 | 3.9 | 141 | Specific | 618.9 | 524.04 | 524.04 | 12 | 125 | >sp\|P02788\|TRFL_HUMAN Lactotransferrin OS=Homo sapiens OX=9606 GN=LTF PE=1 SV=6 | 1270 | 20201116_13832_1ng_glyco_hcd203040_compound15.23123.23123.3 | controllerType=0 controllerNumber=1 scan=23123 | 46.1842 |
| 19607:3 | 2 | R.TAGWNVPIGTLRPFLN[+2205.793]WTGPPEPIEAAVAR.F | HexNAc(4)Hex(5)Fuc(2)NeuAc(1) | N[+2206] | 1812.8468 | 3 | 5436.5257 | 5436.5038 | 0 | 4 | 141 | Specific | 632.29 | 569.18 | 569.18 | 12 | 125 | >sp\|P02788\|TRFL_HUMAN Lactotransferrin OS=Homo sapiens OX=9606 GN=LTF PE=1 SV=6 | 1270 | 20201116_13832_1ng_glyco_hcd203040_compound15.23188.23188.3 | controllerType=0 controllerNumber=1 scan=23188 | 46.3313 |
| 18501:4 | 2 | R.TAGWNVPIGTLRPFLN[+2644.962]WTGPPEPIEAAVAR.F | HexNAc(6)Hex(7)Fuc(2) | N[+2645] | 1469.6779 | 4 | 5875.6896 | 5875.6727 | 0 | 2.9 | 141 | Specific | 643.22 | 586.16 | 586.16 | 11.98 | 125 | >sp\|P02788\|TRFL_HUMAN Lactotransferrin OS=Homo sapiens OX=9606 GN=LTF PE=1 SV=6 | 1270 | 20201116_13832_1ng_glyco_hcd203040_compound15.22034.22034.2 | controllerType=0 controllerNumber=1 scan=22034 | 43.8282 |
| 19606:4 | 2 | R.TAGWNVPIGTLRPFLN[+2205.793]WTGPPEPIEAAVAR.F | HexNAc(4)Hex(5)Fuc(2)NeuAc(1) | N[+2206] | 1359.8824 | 4 | 5436.508 | 5436.5038 | 0 | 0.8 | 141 | Specific | 710.74 | 586.28 | 586.28 | 13.7 | 125 | >sp\|P02788\|TRFL_HUMAN Lactotransferrin OS=Homo sapiens OX=9606 GN=LTF PE=1 SV=6 | 1270 | 20201116_13832_1ng_glyco_hcd203040_compound15.23187.23187.4 | controllerType=0 controllerNumber=1 scan=23187 | 46.3298 |
| 18604:4 | 2 | R.TAGWNVPIGTLRPFLN[+1768.640]WTGPPEPIEAAVAR.F | HexNAc(4)Hex(5)Fuc(1) | N[+1769] | 1250.5974 | 4 | 4999.3678 | 4999.3504 | 0 | 3.5 | 141 | Specific | 721.3 | 687.89 | 687.89 | 13.81 | 125 | >sp\|P02788\|TRFL_HUMAN Lactotransferrin OS=Homo sapiens OX=9606 GN=LTF PE=1 SV=6 | 1270 | 20201116_13832_1ng_glyco_hcd203040_compound15.22142.22142.4 | controllerType=0 controllerNumber=1 scan=22142 | 44.0657 |
| 18606:4 | 2 | R.TAGWNVPIGTLRPFLN[+1768.640]WTGPPEPIEAAVAR.F | HexNAc(4)Hex(5)Fuc(1) | N[+1769] | 1250.5974 | 4 | 4999.3678 | 4999.3504 | 0 | 3.5 | 141 | Specific | 793.31 | 627.97 | 627.97 | 14.81 | 125 | >sp\|P02788\|TRFL_HUMAN Lactotransferrin OS=Homo sapiens OX=9606 GN=LTF PE=1 SV=6 | 1270 | 20201116_13832_1ng_glyco_hcd203040_compound15.22144.22144.4 | controllerType=0 controllerNumber=1 scan=22144 | 44.0723 |
| 19565:5 | 2 | R.TAGWNVPIGTLRPFLN[+2205.793]WTGPPEPIEAAVAR.F | HexNAc(4)Hex(5)Fuc(2)NeuAc(1) | N[+2206] | 1088.11 | 5 | 5436.5208 | 5436.5038 | 0 | 3.1 | 141 | Specific | 873.25 | 707.06 | 707.06 | 17.29 | 125 | >sp\|P02788\|TRFL_HUMAN Lactotransferrin OS=Homo sapiens OX=9606 GN=LTF PE=1 SV=6 | 1270 | 20201116_13832_1ng_glyco_hcd203040_compound15.23144.23144.5 | controllerType=0 controllerNumber=1 scan=23144 | 46.2321 |
| 19567:5 | 2 | R.TAGWNVPIGTLRPFLN[+2205.793]WTGPPEPIEAAVAR.F | HexNAc(4)Hex(5)Fuc(2)NeuAc(1) | N[+2206] | 1088.11 | 5 | 5436.5208 | 5436.5038 | 0 | 3.1 | 141 | Specific | 921.16 | 739.97 | 739.97 | 18.29 | 125 | >sp\|P02788\|TRFL_HUMAN Lactotransferrin OS=Homo sapiens OX=9606 GN=LTF PE=1 SV=6 | 1270 | 20201116_13832_1ng_glyco_hcd203040_compound15.23146.23146.5 | controllerType=0 controllerNumber=1 scan=23146 | 46.2373 |
| 17052:4 | 2 | R.TAGWNIPMGLLFN[+1768.640]QTGSC[+57.021]K.F | HexNAc(4)Hex(5)Fuc(1) | C[+57], N[+1769] | 966.6722 | 4 | 3863.6668 | 3863.6495 | 0 | 4.5 | 485 | Specific | 375.36 | 336.52 | 336.52 | 6.56 | 125 | >sp\|P02788\|TRFL_HUMAN Lactotransferrin OS=Homo sapiens OX=9606 GN=LTF PE=1 SV=6 | 1270 | 20201116_13832_1ng_glyco_hcd203040_compound15.20526.20526.4 | controllerType=0 controllerNumber=1 scan=20526 | 40.7842 |
| 17266:3 | 2 | R.TAGWNIPMGLLFN[+1622.582]QTGSC[+57.021]K.F | HexNAc(4)Hex(5) | C[+57], N[+1623] | 1239.8774 | 3 | 3717.6176 | 3717.5916 | 0 | 7 | 485 | Specific | 403.69 | 403.69 | 178.39 | 8.12 | 125 | >sp\|P02788\|TRFL_HUMAN Lactotransferrin OS=Homo sapiens OX=9606 GN=LTF PE=1 SV=6 | 1270 | 20201116_13832_1ng_glyco_hcd203040_compound15.20748.20748.3 | controllerType=0 controllerNumber=1 scan=20748 | 41.215 |
| 17268:3 | 2 | R.TAGWNIPMGLLFN[+1622.582]QTGSC[+57.021]K.F | HexNAc(4)Hex(5) | C[+57], N[+1623] | 1239.8774 | 3 | 3717.6176 | 3717.5916 | 0 | 7 | 485 | Specific | 404.05 | 349.41 | 304.91 | 6.72 | 125 | >sp\|P02788\|TRFL_HUMAN Lactotransferrin OS=Homo sapiens OX=9606 GN=LTF PE=1 SV=6 | 1270 | 20201116_13832_1ng_glyco_hcd203040_compound15.20750.20750.3 | controllerType=0 controllerNumber=1 scan=20750 | 41.2199 |
| 17054:4 | 2 | R.TAGWNIPMGLLFN[+1768.640]QTGSC[+57.021]K.F | HexNAc(4)Hex(5)Fuc(1) | C[+57], N[+1769] | 966.6722 | 4 | 3863.6668 | 3863.6495 | 0 | 4.5 | 485 | Specific | 477.76 | 295 | 295 | 7.39 | 125 | >sp\|P02788\|TRFL_HUMAN Lactotransferrin OS=Homo sapiens OX=9606 GN=LTF PE=1 SV=6 | 1270 | 20201116_13832_1ng_glyco_hcd203040_compound15.20528.20528.4 | controllerType=0 controllerNumber=1 scan=20528 | 40.7885 |
| 19113:3 | 2 | R.TAGWNIPMGLLFN[+1913.677]QTGSC[+57.021]K.F | HexNAc(4)Hex(5)NeuAc(1) | C[+57], N[+1914] | 1336.9025 | 3 | 4008.693 | 4008.687 | 0 | 1.5 | 485 | Specific | 652.67 | 583.64 | 583.64 | 12.74 | 125 | >sp\|P02788\|TRFL_HUMAN Lactotransferrin OS=Homo sapiens OX=9606 GN=LTF PE=1 SV=6 | 1270 | 20201116_13832_1ng_glyco_hcd203040_compound15.22673.22673.3 | controllerType=0 controllerNumber=1 scan=22673 | 45.2295 |
| 19000:3 | 2 | R.TAGWNIPMGLLFN[+2059.735]QTGSC[+57.021]K.F | HexNAc(4)Hex(5)Fuc(1)NeuAc(1) | C[+57], N[+2060] | 1385.59 | 3 | 4154.7555 | 4154.7449 | 0 | 2.6 | 485 | Specific | 693.81 | 691.41 | 691.41 | 13.02 | 125 | >sp\|P02788\|TRFL_HUMAN Lactotransferrin OS=Homo sapiens OX=9606 GN=LTF PE=1 SV=6 | 1270 | 20201116_13832_1ng_glyco_hcd203040_compound15.22554.22554.3 | controllerType=0 controllerNumber=1 scan=22554 | 44.9291 |
| 18933:3 | 2 | R.TAGWNIPMGLLFN[+2205.793]QTGSC[+57.021]K.F | HexNAc(4)Hex(5)Fuc(2)NeuAc(1) | C[+57], N[+2206] | 1434.2754 | 3 | 4300.8118 | 4300.8028 | 0 | 2.1 | 485 | Specific | 736.41 | 698.18 | 698.18 | 14.02 | 125 | >sp\|P02788\|TRFL_HUMAN Lactotransferrin OS=Homo sapiens OX=9606 GN=LTF PE=1 SV=6 | 1270 | 20201116_13832_1ng_glyco_hcd203040_compound15.22484.22484.3 | controllerType=0 controllerNumber=1 scan=22484 | 44.797 |
| 18935:3 | 2 | R.TAGWNIPMGLLFN[+2205.793]QTGSC[+57.021]K.F | HexNAc(4)Hex(5)Fuc(2)NeuAc(1) | C[+57], N[+2206] | 1434.2754 | 3 | 4300.8118 | 4300.8028 | 0 | 2.1 | 485 | Specific | 845.19 | 834.55 | 834.55 | 16.02 | 125 | >sp\|P02788\|TRFL_HUMAN Lactotransferrin OS=Homo sapiens OX=9606 GN=LTF PE=1 SV=6 | 1270 | 20201116_13832_1ng_glyco_hcd203040_compound15.22486.22486.3 | controllerType=0 controllerNumber=1 scan=22486 | 44.8034 |
| 19116:3 | 2 | R.TAGWNIPMGLLFN[+1913.677]QTGSC[+57.021]K.F | HexNAc(4)Hex(5)NeuAc(1) | C[+57], N[+1914] | 1336.9025 | 3 | 4008.693 | 4008.687 | 0 | 1.5 | 485 | Specific | 847.24 | 682.48 | 682.48 | 15.74 | 125 | >sp\|P02788\|TRFL_HUMAN Lactotransferrin OS=Homo sapiens OX=9606 GN=LTF PE=1 SV=6 | 1270 | 20201116_13832_1ng_glyco_hcd203040_compound15.22676.22676.3 | controllerType=0 controllerNumber=1 scan=22676 | 45.2337 |
| 17007:3 | 2 | R.TAGWNIPMGLLFN[+1768.640]QTGSC[+57.021]K.F | HexNAc(4)Hex(5)Fuc(1) | C[+57], N[+1769] | 1288.5587 | 3 | 3863.6614 | 3863.6495 | 0 | 3.1 | 485 | Specific | 875.18 | 685.27 | 685.27 | 16.84 | 125 | >sp\|P02788\|TRFL_HUMAN Lactotransferrin OS=Homo sapiens OX=9606 GN=LTF PE=1 SV=6 | 1270 | 20201116_13832_1ng_glyco_hcd203040_compound15.20479.20479.3 | controllerType=0 controllerNumber=1 scan=20479 | 40.6802 |
| 16990:3 | 2 | R.TAGWNIPMGLLFN[+1768.640]QTGSC[+57.021]K.F | HexNAc(4)Hex(5)Fuc(1) | C[+57], N[+1769] | 1288.5596 | 3 | 3863.6643 | 3863.6495 | 0 | 3.8 | 485 | Specific | 918.21 | 762.57 | 762.57 | 17.84 | 125 | >sp\|P02788\|TRFL_HUMAN Lactotransferrin OS=Homo sapiens OX=9606 GN=LTF PE=1 SV=6 | 1270 | 20201116_13832_1ng_glyco_hcd203040_compound15.20461.20461.3 | controllerType=0 controllerNumber=1 scan=20461 | 40.642 |
| 16992:3 | 2 | R.TAGWNIPMGLLFN[+1768.640]QTGSC[+57.021]K.F | HexNAc(4)Hex(5)Fuc(1) | C[+57], N[+1769] | 1288.5596 | 3 | 3863.6643 | 3863.6495 | 0 | 3.8 | 485 | Specific | 924.4 | 703.57 | 703.57 | 17.84 | 125 | >sp\|P02788\|TRFL_HUMAN Lactotransferrin OS=Homo sapiens OX=9606 GN=LTF PE=1 SV=6 | 1270 | 20201116_13832_1ng_glyco_hcd203040_compound15.20463.20463.3 | controllerType=0 controllerNumber=1 scan=20463 | 40.6478 |
| 17004:3 | 2 | R.TAGWNIPMGLLFN[+1768.640]QTGSC[+57.021]K.F | HexNAc(4)Hex(5)Fuc(1) | C[+57], N[+1769] | 1288.5587 | 3 | 3863.6614 | 3863.6495 | 0 | 3.1 | 485 | Specific | 928.11 | 928.11 | 928.11 | 17.84 | 125 | >sp\|P02788\|TRFL_HUMAN Lactotransferrin OS=Homo sapiens OX=9606 GN=LTF PE=1 SV=6 | 1270 | 20201116_13832_1ng_glyco_hcd203040_compound15.20476.20476.3 | controllerType=0 controllerNumber=1 scan=20476 | 40.6757 |
| 19002:3 | 2 | R.TAGWNIPMGLLFN[+2059.735]QTGSC[+57.021]K.F | HexNAc(4)Hex(5)Fuc(1)NeuAc(1) | C[+57], N[+2060] | 1385.59 | 3 | 4154.7555 | 4154.7449 | 0 | 2.6 | 485 | Specific | 953.43 | 948.64 | 948.64 | 19.02 | 125 | >sp\|P02788\|TRFL_HUMAN Lactotransferrin OS=Homo sapiens OX=9606 GN=LTF PE=1 SV=6 | 1270 | 20201116_13832_1ng_glyco_hcd203040_compound15.22556.22556.3 | controllerType=0 controllerNumber=1 scan=22556 | 44.9333 |
| 18176:4 | 2 | R.[+42.011]TAGWNIPMGLLFN[+1872.651]QTGSC[+57.021]KFDEYFSQSC[+57.021]A | HexNAc(3)Hex(6)NeuAc(1) | .[42], C[+57]*2, N[+1873] | 1464.1229 | 4 | 5853.4699 | 5853.4181 | 0 | 8.8 | 485 | Specific | 232.87 | 232.87 | 232.87 | 5.07 | 125 | >sp\|P02788\|TRFL_HUMAN Lactotransferrin OS=Homo sapiens OX=9606 GN=LTF PE=1 SV=6 | 1270 | 20201116_13832_1ng_glyco_hcd203040_compound15.21697.21697.4 | controllerType=0 controllerNumber=1 scan=21697 | 43.1665 |
| 18140:4 | 2 | R.TAGWNIPMGLLFN[+2076.750]QTGSC[+57.021]KFDEYFSQSC[+57.021]APGSDPR.S | HexNAc(4)Hex(6)Fuc(2) | C[+57]*2, N[+2077] | 1504.6414 | 4 | 6015.5436 | 6015.5074 | 0 | 6 | 485 | Specific | 236.45 | 236.45 | 236.45 | 5.19 | 125 | >sp\|P02788\|TRFL_HUMAN Lactotransferrin OS=Homo sapiens OX=9606 GN=LTF PE=1 SV=6 | 1270 | 20201116_13832_1ng_glyco_hcd203040_compound15.21659.21659.4 | controllerType=0 controllerNumber=1 scan=21659 | 43.09 |
| 20743:4 | 2 | R.TAGWNIPMGLLFN[+2351.851]QTGSC[+57.021]KFDEYFSQSC[+57.021]APGSDPR.S | HexNAc(4)Hex(5)Fuc(3)NeuAc(1) | C[+57]*2, N[+2352] | 1573.4089 | 4 | 6290.6139 | 6290.6079 | 0 | 1 | 485 | Specific | 242.82 | 221.07 | 221.07 | 6.19 | 125 | >sp\|P02788\|TRFL_HUMAN Lactotransferrin OS=Homo sapiens OX=9606 GN=LTF PE=1 SV=6 | 1270 | 20201116_13832_1ng_glyco_hcd203040_compound15.24370.24370.4 | controllerType=0 controllerNumber=1 scan=24370 | 48.5643 |
| 19198:4 | 2 | R.TAGWNIPMGLLFN[+2076.750]QTGSC[+57.021]KFDEYFSQSC[+57.021]APGSDPR.S | HexNAc(4)Hex(6)Fuc(2) | C[+57]*2, N[+2077] | 1504.6399 | 4 | 6015.5378 | 6015.5074 | 0 | 5.1 | 485 | Specific | 252.59 | 252.59 | 252.59 | 6.06 | 125 | >sp\|P02788\|TRFL_HUMAN Lactotransferrin OS=Homo sapiens OX=9606 GN=LTF PE=1 SV=6 | 1270 | 20201116_13832_1ng_glyco_hcd203040_compound15.22762.22762.4 | controllerType=0 controllerNumber=1 scan=22762 | 45.4425 |
| 18138:4 | 2 | R.TAGWNIPMGLLFN[+2076.750]QTGSC[+57.021]KFDEYFSQSC[+57.021]APGSDPR.S | HexNAc(4)Hex(6)Fuc(2) | C[+57]*2, N[+2077] | 1504.6414 | 4 | 6015.5436 | 6015.5074 | 0 | 6 | 485 | Specific | 264.61 | 251.32 | 251.32 | 6.02 | 125 | >sp\|P02788\|TRFL_HUMAN Lactotransferrin OS=Homo sapiens OX=9606 GN=LTF PE=1 SV=6 | 1270 | 20201116_13832_1ng_glyco_hcd203040_compound15.21657.21657.4 | controllerType=0 controllerNumber=1 scan=21657 | 43.0867 |
| 18178:4 | 2 | R.[+42.011]TAGWNIPMGLLFN[+1872.651]QTGSC[+57.021]KFDEYFSQSC[+57.021]A | HexNAc(3)Hex(6)NeuAc(1) | .[42], C[+57]*2, N[+1873] | 1464.1229 | 4 | 5853.4699 | 5853.4181 | 0 | 8.8 | 485 | Specific | 265.63 | 250.09 | 250.09 | 4.19 | 125 | >sp\|P02788\|TRFL_HUMAN Lactotransferrin OS=Homo sapiens OX=9606 GN=LTF PE=1 SV=6 | 1270 | 20201116_13832_1ng_glyco_hcd203040_compound15.21699.21699.4 | controllerType=0 controllerNumber=1 scan=21699 | 43.1711 |
| 20745:4 | 2 | R.TAGWNIPMGLLFN[+2351.851]QTGSC[+57.021]KFDEYFSQSC[+57.021]APGSDPR.S | HexNAc(4)Hex(5)Fuc(3)NeuAc(1) | C[+57]*2, N[+2352] | 1573.4089 | 4 | 6290.6139 | 6290.6079 | 0 | 1 | 485 | Specific | 370.04 | 273.96 | 273.96 | 7.25 | 125 | >sp\|P02788\|TRFL_HUMAN Lactotransferrin OS=Homo sapiens OX=9606 GN=LTF PE=1 SV=6 | 1270 | 20201116_13832_1ng_glyco_hcd203040_compound15.24372.24372.4 | controllerType=0 controllerNumber=1 scan=24372 | 48.5684 |
| 18323:4 | 2 | R.TAGWNIPMGLLFN[+2076.750]QTGSC[+57.021]KFDEYFSQSC[+57.021]APGSDPR.S | HexNAc(4)Hex(6)Fuc(2) | C[+57]*2, N[+2077] | 1504.6295 | 4 | 6015.4962 | 6015.5074 | 0 | -1.8 | 485 | Specific | 390.15 | 390.15 | 390.15 | 8.17 | 125 | >sp\|P02788\|TRFL_HUMAN Lactotransferrin OS=Homo sapiens OX=9606 GN=LTF PE=1 SV=6 | 1270 | 20201116_13832_1ng_glyco_hcd203040_compound15.21849.21849.4 | controllerType=0 controllerNumber=1 scan=21849 | 43.4641 |
| 20708:4 | 2 | R.TAGWNIPMGLLFN[+2350.830]QTGSC[+57.021]KFDEYFSQSC[+57.021]APGSDPR.S | HexNAc(4)Hex(5)Fuc(1)NeuAc(2) | C[+57]*2, N[+2351] | 1573.1587 | 4 | 6289.6129 | 6289.5875 | 0 | 4.1 | 485 | Specific | 407.37 | 388.53 | 388.53 | 9.49 | 125 | >sp\|P02788\|TRFL_HUMAN Lactotransferrin OS=Homo sapiens OX=9606 GN=LTF PE=1 SV=6 | 1270 | 20201116_13832_1ng_glyco_hcd203040_compound15.24334.24334.4 | controllerType=0 controllerNumber=1 scan=24334 | 48.4883 |
| 18325:4 | 2 | R.TAGWNIPMGLLFN[+2076.750]QTGSC[+57.021]KFDEYFSQSC[+57.021]APGSDPR.S | HexNAc(4)Hex(6)Fuc(2) | C[+57]*2, N[+2077] | 1504.6295 | 4 | 6015.4962 | 6015.5074 | 0 | -1.8 | 485 | Specific | 413.65 | 393.99 | 393.99 | 8.34 | 125 | >sp\|P02788\|TRFL_HUMAN Lactotransferrin OS=Homo sapiens OX=9606 GN=LTF PE=1 SV=6 | 1270 | 20201116_13832_1ng_glyco_hcd203040_compound15.21851.21851.4 | controllerType=0 controllerNumber=1 scan=21851 | [illegible] |
| 18285:4 | 2 | R.TAGWNIPMGLLFN[+1606.587]QTGSC[+57.021]KFDEYFSQSC[+57.021]APGSDPR.S | HexNAc(4)Hex(4)Fuc(1) | C[+57]*2, N[+1607] | 1387.092 | 4 | 5545.3463 | 5545.3438 | 0 | 0.5 | 485 | Specific | 448.19 | 448.19 | 448.19 | 9.5 | 125 | >sp\|P02788\|TRFL_HUMAN Lactotransferrin OS=Homo sapiens OX=9606 GN=LTF PE=1 SV=6 | 1270 | 20201116_13832_1ng_glyco_hcd203040_compound15.21810.21810.4 | controllerType=0 controllerNumber=1 scan=21810 | 43.3866 |
| 19247:4 | 2 | R.TAGWNIPMGLLFN[+2076.750]QTGSC[+57.021]KFDEYFSQSC[+57.021]APGSDPR.S | HexNAc(4)Hex(6)Fuc(2) | C[+57]*2, N[+2077] | 1504.6415 | 4 | 6015.5441 | 6015.5074 | 0 | 6.1 | 485 | Specific | 488.45 | 488.45 | 488.45 | 8.93 | 125 | >sp\|P02788\|TRFL_HUMAN Lactotransferrin OS=Homo sapiens OX=9606 GN=LTF PE=1 SV=6 | 1270 | 20201116_13832_1ng_glyco_hcd203040_compound15.22813.22813.4 | controllerType=0 controllerNumber=1 scan=22813 | 45.5445 |
| 19271:5 | 2 | R.TAGWNIPMGLLFN[+2059.735]QTGSC[+57.021]KFDEYFSQSC[+57.021]APGSDPR.S | HexNAc(4)Hex(5)Fuc(1)NeuAc(1) | C[+57]*2, N[+2060] | 1200.5044 | 5 | 5998.4929 | 5998.492 | 0 | 0.1 | 485 | Specific | 512.32 | 512.32 | 512.32 | 9.53 | 125 | >sp\|P02788\|TRFL_HUMAN Lactotransferrin OS=Homo sapiens OX=9606 GN=LTF PE=1 SV=6 | 1270 | 20201116_13832_1ng_glyco_hcd203040_compound15.22838.22838.5 | controllerType=0 controllerNumber=1 scan=22838 | 45.5895 |
| 19291:3 | 2 | R.TAGWNIPMGLLFN[+1913.677]QTGSC[+57.021]KFDEYFSQSC[+57.021]APGSDPR.S | HexNAc(4)Hex(5)NeuAc(1) | C[+57]*2, N[+1914] | 1951.4871 | 3 | 5852.4469 | 5852.4341 | 0 | 2.2 | 485 | Specific | 518.32 | 437.59 | 437.59 | 10.33 | 125 | >sp\|P02788\|TRFL_HUMAN Lactotransferrin OS=Homo sapiens OX=9606 GN=LTF PE=1 SV=6 | 1270 | 20201116_13832_1ng_glyco_hcd203040_compound15.22859.22859.3 | controllerType=0 controllerNumber=1 scan=22859 | 45.6468 |
| 19218:4 | 2 | R.TAGWNIPMGLLFN[+2205.793]QTGSC[+57.021]KFDEYFSQSC[+57.021]APGSDPR.S | HexNAc(4)Hex(5)Fuc(2)NeuAc(1) | C[+57]*2, N[+2206] | 1536.8956 | 4 | 6144.5607 | 6144.5499 | 0 | 1.8 | 485 | Specific | 527.48 | 511 | 511 | 10.13 | 125 | >sp\|P02788\|TRFL_HUMAN Lactotransferrin OS=Homo sapiens OX=9606 GN=LTF PE=1 SV=6 | 1270 | 20201116_13832_1ng_glyco_hcd203040_compound15.22783.22783.4 | controllerType=0 controllerNumber=1 scan=22783 | 45.4855 |
| 18288:4 | 2 | R.TAGWNIPMGLLFN[+1606.587]QTGSC[+57.021]KFDEYFSQSC[+57.021]APGSDPR.S | HexNAc(4)Hex(4)Fuc(1) | C[+57]*2, N[+1607] | 1387.092 | 4 | 5545.3463 | 5545.3438 | 0 | 0.5 | 485 | Specific | 568.36 | 553.78 | 553.78 | 10.08 | 125 | >sp\|P02788\|TRFL_HUMAN Lactotransferrin OS=Homo sapiens OX=9606 GN=LTF PE=1 SV=6 | 1270 | 20201116_13832_1ng_glyco_hcd203040_compound15.21813.21813.4 | controllerType=0 controllerNumber=1 scan=21813 | 43.391 |
| 18270:3 | 2 | R.TAGWNIPMGLLFN[+1622.582]QTGSC[+57.021]KFDEYFSQSC[+57.021]APGSDPR.S | HexNAc(4)Hex(5) | C[+57]*2, N[+1623] | 1854.4563 | 3 | 5561.3542 | 5561.3387 | 0 | 2.8 | 485 | Specific | 571.21 | 507.35 | 162.72 | 10.55 | 125 | >sp\|P02788\|TRFL_HUMAN Lactotransferrin OS=Homo sapiens OX=9606 GN=LTF PE=1 SV=6 | 1270 | 20201116_13832_1ng_glyco_hcd203040_compound15.21794.21794.3 | controllerType=0 controllerNumber=1 scan=21794 | 43.34 |
| 18272:3 | 2 | R.TAGWNIPMGLLFN[+1622.582]QTGSC[+57.021]KFDEYFSQSC[+57.021]APGSDPR.S | HexNAc(4)Hex(5) | C[+57]*2, N[+1623] | 1854.4563 | 3 | 5561.3542 | 5561.3387 | 0 | 2.8 | 485 | Specific | 598.2 | 479.25 | 162.69 | 11.68 | 125 | >sp\|P02788\|TRFL_HUMAN Lactotransferrin OS=Homo sapiens OX=9606 GN=LTF PE=1 SV=6 | 1270 | 20201116_13832_1ng_glyco_hcd203040_compound15.21796.21796.3 | controllerType=0 controllerNumber=1 scan=21796 | 43.3436 |
| 18193:5 | 2 | R.TAGWNIPMGLLFN[+1768.640]QTGSC[+57.021]KFDEYFSQSC[+57.021]APGSDPR.S | HexNAc(4)Hex(5)Fuc(1) | C[+57]*2, N[+1769] | 1142.2861 | 5 | 5707.4016 | 5707.3966 | 0 | 0.9 | 485 | Specific | 634.49 | 634.49 | 634.49 | 12.7 | 125 | >sp\|P02788\|TRFL_HUMAN Lactotransferrin OS=Homo sapiens OX=9606 GN=LTF PE=1 SV=6 | 1270 | 20201116_13832_1ng_glyco_hcd203040_compound15.21714.21714.5 | controllerType=0 controllerNumber=1 scan=21714 | 43.1929 |
| 19273:5 | 2 | R.TAGWNIPMGLLFN[+2059.735]QTGSC[+57.021]KFDEYFSQSC[+57.021]APGSDPR.S | HexNAc(4)Hex(5)Fuc(1)NeuAc(1) | C[+57]*2, N[+2060] | 1200.5044 | 5 | 5998.4929 | 5998.492 | 0 | 0.1 | 485 | Specific | 674.54 | 674.54 | 674.54 | 13.02 | 125 | >sp\|P02788\|TRFL_HUMAN Lactotransferrin OS=Homo sapiens OX=9606 GN=LTF PE=1 SV=6 | 1270 | 20201116_13832_1ng_glyco_hcd203040_compound15.22840.22840.5 | controllerType=0 controllerNumber=1 scan=22840 | 45.5957 |
| 19299:5 | 2 | R.TAGWNIPMGLLFN[+1913.677]QTGSC[+57.021]KFDEYFSQSC[+57.021]APGSDPR.S | HexNAc(4)Hex(5)NeuAc(1) | C[+57]*2, N[+1914] | 1171.2988 | 5 | 5852.465 | 5852.4341 | 0 | 5.3 | 485 | Specific | 683.64 | 632.67 | 632.67 | 13.24 | 125 | >sp\|P02788\|TRFL_HUMAN Lactotransferrin OS=Homo sapiens OX=9606 GN=LTF PE=1 SV=6 | 1270 | 20201116_13832_1ng_glyco_hcd203040_compound15.22867.22867.5 | controllerType=0 controllerNumber=1 scan=22867 | 45.6625 |
| 18187:3 | 2 | R.TAGWNIPMGLLFN[+1768.640]QTGSC[+57.021]KFDEYFSQSC[+57.021]APGSDPR.S | HexNAc(4)Hex(5)Fuc(1) | C[+57]*2, N[+1769] | 1903.1403 | 3 | 5707.4065 | 5707.3966 | 0 | 1.7 | 485 | Specific | 693.41 | 601.78 | 601.78 | 13.68 | 125 | >sp\|P02788\|TRFL_HUMAN Lactotransferrin OS=Homo sapiens OX=9606 GN=LTF PE=1 SV=6 | 1270 | 20201116_13832_1ng_glyco_hcd203040_compound15.21708.21708.3 | controllerType=0 controllerNumber=1 scan=21708 | 43.1844 |
| 20711:4 | 2 | R.TAGWNIPMGLLFN[+2350.830]QTGSC[+57.021]KFDEYFSQSC[+57.021]APGSDPR.S | HexNAc(4)Hex(5)Fuc(1)NeuAc(2) | C[+57]*2, N[+2351] | 1573.1587 | 4 | 6289.6129 | 6289.5875 | 0 | 4.1 | 485 | Specific | 700.55 | 667.34 | 667.34 | 14.32 | 125 | >sp\|P02788\|TRFL_HUMAN Lactotransferrin OS=Homo sapiens OX=9606 GN=LTF PE=1 SV=6 | 1270 | 20201116_13832_1ng_glyco_hcd203040_compound15.24337.24337.4 | controllerType=0 controllerNumber=1 scan=24337 | 48.4933 |
| 19250:4 | 2 | R.TAGWNIPMGLLFN[+2076.750]QTGSC[+57.021]KFDEYFSQSC[+57.021]APGSDPR.S | HexNAc(4)Hex(6)Fuc(2) | C[+57]*2, N[+2077] | 1504.6415 | 4 | 6015.5441 | 6015.5074 | 0 | 6.1 | 485 | Specific | 718.32 | 718.32 | 718.32 | 13.42 | 125 | >sp\|P02788\|TRFL_HUMAN Lactotransferrin OS=Homo sapiens OX=9606 GN=LTF PE=1 SV=6 | 1270 | 20201116_13832_1ng_glyco_hcd203040_compound15.22816.22816.4 | controllerType=0 controllerNumber=1 scan=22816 | 45.5486 |
| 19294:3 | 2 | R.TAGWNIPMGLLFN[+1913.677]QTGSC[+57.021]KFDEYFSQSC[+57.021]APGSDPR.S | HexNAc(4)Hex(5)NeuAc(1) | C[+57]*2, N[+1914] | 1951.4871 | 3 | 5852.4469 | 5852.4341 | 0 | 2.2 | 485 | Specific | 725.23 | 568.94 | 568.94 | 14.68 | 125 | >sp\|P02788\|TRFL_HUMAN Lactotransferrin OS=Homo sapiens OX=9606 GN=LTF PE=1 SV=6 | 1270 | 20201116_13832_1ng_glyco_hcd203040_compound15.22862.22862.3 | controllerType=0 controllerNumber=1 scan=22862 | 45.6511 |
| 18233:4 | 2 | R.TAGWNIPMGLLFN[+1622.582]QTGSC[+57.021]KFDEYFSQSC[+57.021]APGSDPR.S | HexNAc(4)Hex(5) | C[+57]*2, N[+1623] | 1391.095 | 4 | 5561.3581 | 5561.3387 | 0 | 3.5 | 485 | Specific | 782.29 | 732.05 | 254.68 | 15.32 | 125 | >sp\|P02788\|TRFL_HUMAN Lactotransferrin OS=Homo sapiens OX=9606 GN=LTF PE=1 SV=6 | 1270 | 20201116_13832_1ng_glyco_hcd203040_compound15.21756.21756.4 | controllerType=0 controllerNumber=1 scan=21756 | 43.268 |
| 18175:4 | 2 | R.TAGWNIPMGLLFN[+1768.640]QTGSC[+57.021]KFDEYFSQSC[+57.021]APGSDPR.S | HexNAc(4)Hex(5)Fuc(1) | C[+57]*2, N[+1769] | 1427.6083 | 4 | 5707.4113 | 5707.3966 | 0 | 2.6 | 485 | Specific | 864.15 | 804.43 | 804.43 | 17.49 | 125 | >sp\|P02788\|TRFL_HUMAN Lactotransferrin OS=Homo sapiens OX=9606 GN=LTF PE=1 SV=6 | 1270 | 20201116_13832_1ng_glyco_hcd203040_compound15.21696.21696.4 | controllerType=0 controllerNumber=1 scan=21696 | 43.165 |
| 19194:4 | 2 | R.TAGWNIPMGLLFN[+2059.735]QTGSC[+57.021]KFDEYFSQSC[+57.021]APGSDPR.S | HexNAc(4)Hex(5)Fuc(1)NeuAc(1) | C[+57]*2, N[+2060] | 1500.3844 | 4 | 5998.5158 | 5998.492 | 0 | 4 | 485 | Specific | 912.73 | 886.51 | 886.51 | 18.32 | 125 | >sp\|P02788\|TRFL_HUMAN Lactotransferrin OS=Homo sapiens OX=9606 GN=LTF PE=1 SV=6 | 1270 | 20201116_13832_1ng_glyco_hcd203040_compound15.22758.22758.4 | controllerType=0 controllerNumber=1 scan=22758 | 45.4344 |
| 19301:5 | 2 | R.TAGWNIPMGLLFN[+1913.677]QTGSC[+57.021]KFDEYFSQSC[+57.021]APGSDPR.S | HexNAc(4)Hex(5)NeuAc(1) | C[+57]*2, N[+1914] | 1171.2988 | 5 | 5852.465 | 5852.4341 | 0 | 5.3 | 485 | Specific | 914.16 | 874.9 | 874.9 | 18.24 | 125 | >sp\|P02788\|TRFL_HUMAN Lactotransferrin OS=Homo sapiens OX=9606 GN=LTF PE=1 SV=6 | 1270 | 20201116_13832_1ng_glyco_hcd203040_compound15.22869.22869.5 | controllerType=0 controllerNumber=1 scan=22869 | 45.6666 |
| 19220:4 | 2 | R.TAGWNIPMGLLFN[+2205.793]QTGSC[+57.021]KFDEYFSQSC[+57.021]APGSDPR.S | HexNAc(4)Hex(5)Fuc(2)NeuAc(1) | C[+57]*2, N[+2206] | 1536.8956 | 4 | 6144.5607 | 6144.5499 | 0 | 1.8 | 485 | Specific | 914.94 | 873.88 | 873.88 | 18.49 | 125 | >sp\|P02788\|TRFL_HUMAN Lactotransferrin OS=Homo sapiens OX=9606 GN=LTF PE=1 SV=6 | 1270 | 20201116_13832_1ng_glyco_hcd203040_compound15.22785.22785.4 | controllerType=0 controllerNumber=1 scan=22785 | 45.4901 |
| 18177:4 | 2 | R.TAGWNIPMGLLFN[+1768.640]QTGSC[+57.021]KFDEYFSQSC[+57.021]APGSDPR.S | HexNAc(4)Hex(5)Fuc(1) | C[+57]*2, N[+1769] | 1427.6083 | 4 | 5707.4113 | 5707.3966 | 0 | 2.6 | 485 | Specific | 921.01 | 878.38 | 878.38 | 18.49 | 125 | >sp\|P02788\|TRFL_HUMAN Lactotransferrin OS=Homo sapiens OX=9606 GN=LTF PE=1 SV=6 | 1270 | 20201116_13832_1ng_glyco_hcd203040_compound15.21698.21698.4 | controllerType=0 controllerNumber=1 scan=21698 | 43.17 |
| 19196:4 | 2 | R.TAGWNIPMGLLFN[+2059.735]QTGSC[+57.021]KFDEYFSQSC[+57.021]APGSDPR.S | HexNAc(4)Hex(5)Fuc(1)NeuAc(1) | C[+57]*2, N[+2060] | 1500.3844 | 4 | 5998.5158 | 5998.492 | 0 | 4 | 485 | Specific | 935.78 | 901.4 | 901.4 | 18.32 | 125 | >sp\|P02788\|TRFL_HUMAN Lactotransferrin OS=Homo sapiens OX=9606 GN=LTF PE=1 SV=6 | 1270 | 20201116_13832_1ng_glyco_hcd203040_compound15.22760.22760.4 | controllerType=0 controllerNumber=1 scan=22760 | 45.4406 |
| 18184:3 | 2 | R.TAGWNIPMGLLFN[+1768.640]QTGSC[+57.021]KFDEYFSQSC[+57.021]APGSDPR.S | HexNAc(4)Hex(5)Fuc(1) | C[+57]*2, N[+1769] | 1903.1403 | 3 | 5707.4065 | 5707.3966 | 0 | 1.7 | 485 | Specific | 940.38 | 857.13 | 857.13 | 18.68 | 125 | >sp\|P02788\|TRFL_HUMAN Lactotransferrin OS=Homo sapiens OX=9606 GN=LTF PE=1 SV=6 | 1270 | 20201116_13832_1ng_glyco_hcd203040_compound15.21705.21705.3 | controllerType=0 controllerNumber=1 scan=21705 | 43.179 |
| 19272:4 | 2 | R.TAGWNIPMGLLFN[+1913.677]QTGSC[+57.021]KFDEYFSQSC[+57.021]APGSDPR.S | HexNAc(4)Hex(5)NeuAc(1) | C[+57]*2, N[+1914] | 1463.8683 | 4 | 5852.4513 | 5852.4341 | 0 | 2.9 | 485 | Specific | 941.98 | 871.56 | 871.56 | 18.49 | 125 | >sp\|P02788\|TRFL_HUMAN Lactotransferrin OS=Homo sapiens OX=9606 GN=LTF PE=1 SV=6 | 1270 | 20201116_13832_1ng_glyco_hcd203040_compound15.22839.22839.4 | controllerType=0 controllerNumber=1 scan=22839 | 45.5909 |
| 19274:4 | 2 | R.TAGWNIPMGLLFN[+1913.677]QTGSC[+57.021]KFDEYFSQSC[+57.021]APGSDPR.S | HexNAc(4)Hex(5)NeuAc(1) | C[+57]*2, N[+1914] | 1463.8683 | 4 | 5852.4513 | 5852.4341 | 0 | 2.9 | 485 | Specific | 951.2 | 897.38 | 897.38 | 19.49 | 125 | >sp\|P02788\|TRFL_HUMAN Lactotransferrin OS=Homo sapiens OX=9606 GN=LTF PE=1 SV=6 | 1270 | 20201116_13832_1ng_glyco_hcd203040_compound15.22841.22841.4 | controllerType=0 controllerNumber=1 scan=22841 | 45.5983 |
| 18235:4 | 2 | R.TAGWNIPMGLLFN[+1622.582]QTGSC[+57.021]KFDEYFSQSC[+57.021]APGSDPR.S | HexNAc(4)Hex(5) | C[+57]*2, N[+1623] | 1391.095 | 4 | 5561.3581 | 5561.3387 | 0 | 3.5 | 485 | Specific | 959.67 | 898.51 | 360.47 | 19.32 | 125 | >sp\|P02788\|TRFL_HUMAN Lactotransferrin OS=Homo sapiens OX=9606 GN=LTF PE=1 SV=6 | 1270 | 20201116_13832_1ng_glyco_hcd203040_compound15.21758.21758.4 | controllerType=0 controllerNumber=1 scan=21758 | 43.2729 |
| 18196:5 | 2 | R.TAGWNIPMGLLFN[+1768.640]QTGSC[+57.021]KFDEYFSQSC[+57.021]APGSDPR.S | HexNAc(4)Hex(5)Fuc(1) | C[+57]*2, N[+1769] | 1142.2861 | 5 | 5707.4016 | 5707.3966 | 0 | 0.9 | 485 | Specific | 1002.05 | 955.99 | 955.99 | 20.7 | 125 | >sp\|P02788\|TRFL_HUMAN Lactotransferrin OS=Homo sapiens OX=9606 GN=LTF PE=1 SV=6 | 1270 | 20201116_13832_1ng_glyco_hcd203040_compound15.21717.21717.5 | controllerType=0 controllerNumber=1 scan=21717 | 43.1972 |
| 18156:4 | 2 | R.TAGWNIPMGLLFN[+1768.640]QTGSC[+57.021]KFDEYFSQSC[+57.021]APGSDPR.S | HexNAc(4)Hex(5)Fuc(1) | C[+57]*2, N[+1769] | 1427.6095 | 4 | 5707.4162 | 5707.3966 | 0 | 3.4 | 485 | Specific | 1015.63 | 958.83 | 958.83 | 20.32 | 125 | >sp\|P02788\|TRFL_HUMAN Lactotransferrin OS=Homo sapiens OX=9606 GN=LTF PE=1 SV=6 | 1270 | 20201116_13832_1ng_glyco_hcd203040_compound15.21676.21676.4 | controllerType=0 controllerNumber=1 scan=21676 | 43.1271 |
| 18153:4 | 2 | R.TAGWNIPMGLLFN[+1768.640]QTGSC[+57.021]KFDEYFSQSC[+57.021]APGSDPR.S | HexNAc(4)Hex(5)Fuc(1) | C[+57]*2, N[+1769] | 1427.6095 | 4 | 5707.4162 | 5707.3966 | 0 | 3.4 | 485 | Specific | 1025.86 | 973.1 | 973.1 | 20.32 | 125 | >sp\|P02788\|TRFL_HUMAN Lactotransferrin OS=Homo sapiens OX=9606 GN=LTF PE=1 SV=6 | 1270 | 20201116_13832_1ng_glyco_hcd203040_compound15.21673.21673.4 | controllerType=0 controllerNumber=1 scan=21673 | 43.1193 |
| 13137:5 | 5 | K.KGN[+2059.735]TTLNSFVIPSESDVPTHC[+57.021]PSQWWPYAGHC[+57.021]YK.I | HexNAc(4)Hex(5)Fuc(1)NeuAc(1) | C[+57]*2, N[+2060] | 1205.5183 | 5 | 6023.5625 | 6023.5567 | 0 | 1 | 342 | Specific | 439.95 | 439.95 | 439.95 | 9.32 | 99 | >sp\|P22897\|MRC1_HUMAN Macrophage mannose receptor 1 OS=Homo sapiens OX=9606 GN=MRC1 PE=1 SV=1 | 1049 | 20201116_13832_1ng_glyco_hcd203040_compound15.16432.16432.5 | controllerType=0 controllerNumber=1 scan=16432 | 33.7298 |
| 13139:5 | 5 | K.KGN[+2059.735]TTLNSFVIPSESDVPTHC[+57.021]PSQWWPYAGHC[+57.021]YK.I | HexNAc(4)Hex(5)Fuc(1)NeuAc(1) | C[+57]*2, N[+2060] | 1205.5183 | 5 | 6023.5625 | 6023.5567 | 0 | 1 | 342 | Specific | 502.66 | 502.66 | 502.66 | 9.54 | 99 | >sp\|P22897\|MRC1_HUMAN Macrophage mannose receptor 1 OS=Homo sapiens OX=9606 GN=MRC1 PE=1 SV=1 | 1049 | 20201116_13832_1ng_glyco_hcd203040_compound15.16434.16434.5 | controllerType=0 controllerNumber=1 scan=16434 | 33.7356 |
| 19123:5 | 5 | K.[+42.011]HHFYC[+57.021]YMIGHTLSTFAEAN[+2018.708]QTC[+57.021]NNENAYL | HexNAc(3)Hex(6)Fuc(1)NeuAc(1) | .[42], C[+57]*2, N[+2019] | 1630.5193 | 5 | 8148.5675 | 8148.5353 | 0 | 4 | 511 | Specific | 263.58 | 263.58 | 263.58 | 4.68 | 99 | >sp\|P22897\|MRC1_HUMAN Macrophage mannose receptor 1 OS=Homo sapiens OX=9606 GN=MRC1 PE=1 SV=1 | 1049 | 20201116_13832_1ng_glyco_hcd203040_compound15.22683.22683.5 | controllerType=0 controllerNumber=1 scan=22683 | 45.2457 |
| 19126:5 | 5 | K.[+42.011]HHFYC[+57.021]YMIGHTLSTFAEAN[+2018.708]QTC[+57.021]NNENAYL | HexNAc(3)Hex(6)Fuc(1)NeuAc(1) | .[42], C[+57]*2, N[+2019] | 1630.5193 | 5 | 8148.5675 | 8148.5353 | 0 | 4 | 511 | Specific | 266.89 | 137.85 | 137.85 | 2.58 | 99 | >sp\|P22897\|MRC1_HUMAN Macrophage mannose receptor 1 OS=Homo sapiens OX=9606 GN=MRC1 PE=1 SV=1 | 1049 | 20201116_13832_1ng_glyco_hcd203040_compound15.22686.22686.5 | controllerType=0 controllerNumber=1 scan=22686 | 45.2558 |
| 16721:3 | 5 | R.VWIALNSN[+2059.735]LTDNQYTWTDKWR.V | HexNAc(4)Hex(5)Fuc(1)NeuAc(1) | N[+2060] | 1562.019 | 3 | 4684.0424 | 4684.0241 | 0 | 3.9 | 1153 | Specific | 229.69 | 118.76 | 118.76 | 3.84 | 99 | >sp\|P22897\|MRC1_HUMAN Macrophage mannose receptor 1 OS=Homo sapiens OX=9606 GN=MRC1 PE=1 SV=1 | 1049 | 20201116_13832_1ng_glyco_hcd203040_compound15.20181.20181.3 | controllerType=0 controllerNumber=1 scan=20181 | 40.1272 |
| 16719:3 | 5 | R.VWIALNSN[+2059.735]LTDNQYTWTDKWR.V | HexNAc(4)Hex(5)Fuc(1)NeuAc(1) | N[+2060] | 1562.019 | 3 | 4684.0424 | 4684.0241 | 0 | 3.9 | 1153 | Specific | 251.47 | 249.54 | 249.54 | 6 | 99 | >sp\|P22897\|MRC1_HUMAN Macrophage mannose receptor 1 OS=Homo sapiens OX=9606 GN=MRC1 PE=1 SV=1 | 1049 | 20201116_13832_1ng_glyco_hcd203040_compound15.20179.20179.3 | controllerType=0 controllerNumber=1 scan=20179 | 40.1218 |
| 16617:4 | 5 | R.VWIALNSN[+2205.793]LTDNQYTWTDKWR.V | HexNAc(4)Hex(5)Fuc(2)NeuAc(1) | N[+2206] | 1208.2759 | 4 | 4830.0819 | 4830.0821 | 0 | 0 | 1153 | Specific | 543.26 | 419.26 | 419.26 | 8.6 | 99 | >sp\|P22897\|MRC1_HUMAN Macrophage mannose receptor 1 OS=Homo sapiens OX=9606 GN=MRC1 PE=1 SV=1 | 1049 | 20201116_13832_1ng_glyco_hcd203040_compound15.20074.20074.4 | controllerType=0 controllerNumber=1 scan=20074 | 39.9488 |
| 16619:4 | 5 | R.VWIALNSN[+2205.793]LTDNQYTWTDKWR.V | HexNAc(4)Hex(5)Fuc(2)NeuAc(1) | N[+2206] | 1208.2759 | 4 | 4830.0819 | 4830.0821 | 0 | 0 | 1153 | Specific | 564.5 | 408.68 | 408.68 | 8.6 | 99 | >sp\|P22897\|MRC1_HUMAN Macrophage mannose receptor 1 OS=Homo sapiens OX=9606 GN=MRC1 PE=1 SV=1 | 1049 | 20201116_13832_1ng_glyco_hcd203040_compound15.20076.20076.4 | controllerType=0 controllerNumber=1 scan=20076 | 39.9527 |
| 08402:3 | 5 | K.TAHC[+57.021]N[+2059.735]ESFYFLC[+57.021]K.R | HexNAc(4)Hex(5)Fuc(1)NeuAc(1) | C[+57]*2, N[+2060] | 1246.168 | 3 | 3736.4895 | 3736.4545 | 0 | 9.4 | 1201 | Specific | 322.7 | 272.25 | 272.25 | 5.03 | 99 | >sp\|P22897\|MRC1_HUMAN Macrophage mannose receptor 1 OS=Homo sapiens OX=9606 GN=MRC1 PE=1 SV=1 | 1049 | 20201116_13832_1ng_glyco_hcd203040_compound15.11500.11500.3 | controllerType=0 controllerNumber=1 scan=11500 | 25.4977 |
| 08400:3 | 5 | K.TAHC[+57.021]N[+2059.735]ESFYFLC[+57.021]K.R | HexNAc(4)Hex(5)Fuc(1)NeuAc(1) | C[+57]*2, N[+2060] | 1246.168 | 3 | 3736.4895 | 3736.4545 | 0 | 9.4 | 1201 | Specific | 379.15 | 365.37 | 365.37 | 5.77 | 99 | >sp\|P22897\|MRC1_HUMAN Macrophage mannose receptor 1 OS=Homo sapiens OX=9606 GN=MRC1 PE=1 SV=1 | 1049 | 20201116_13832_1ng_glyco_hcd203040_compound15.11498.11498.3 | controllerType=0 controllerNumber=1 scan=11498 | 25.4936 |
| 08287:3 | 5 | K.TAHC[+57.021]N[+2205.793]ESFYFLC[+57.021]K.R | HexNAc(4)Hex(5)Fuc(2)NeuAc(1) | C[+57]*2, N[+2206] | 1294.8487 | 3 | 3882.5315 | 3882.5124 | 0 | 4.9 | 1201 | Specific | 379.2 | 273.08 | 273.08 | 6.26 | 99 | >sp\|P22897\|MRC1_HUMAN Macrophage mannose receptor 1 OS=Homo sapiens OX=9606 GN=MRC1 PE=1 SV=1 | 1049 | 20201116_13832_1ng_glyco_hcd203040_compound15.11381.11381.3 | controllerType=0 controllerNumber=1 scan=11381 | 25.3021 |
| 08257:3 | 5 | K.TAHC[+57.021]N[+2205.793]ESFYFLC[+57.021]K.R | HexNAc(4)Hex(5)Fuc(2)NeuAc(1) | C[+57]*2, N[+2206] | 1294.8461 | 3 | 3882.5238 | 3882.5124 | 0 | 2.9 | 1201 | Specific | 391.53 | 302.07 | 302.07 | 7.09 | 99 | >sp\|P22897\|MRC1_HUMAN Macrophage mannose receptor 1 OS=Homo sapiens OX=9606 GN=MRC1 PE=1 SV=1 | 1049 | 20201116_13832_1ng_glyco_hcd203040_compound15.11350.11350.3 | controllerType=0 controllerNumber=1 scan=11350 | 25.2519 |
| 08255:3 | 5 | K.TAHC[+57.021]N[+2205.793]ESFYFLC[+57.021]K.R | HexNAc(4)Hex(5)Fuc(2)NeuAc(1) | C[+57]*2, N[+2206] | 1294.8461 | 3 | 3882.5238 | 3882.5124 | 0 | 2.9 | 1201 | Specific | 412.75 | 358.33 | 358.33 | 7.67 | 99 | >sp\|P22897\|MRC1_HUMAN Macrophage mannose receptor 1 OS=Homo sapiens OX=9606 GN=MRC1 PE=1 SV=1 | 1049 | 20201116_13832_1ng_glyco_hcd203040_compound15.11348.11348.3 | controllerType=0 controllerNumber=1 scan=11348 | 25.2478 |
| 17989:4 | 6 | R.NIAAFGGDPNN[+1622.582]ITLFGESAGGASVSLQTLSPYNK.G | HexNAc(4)Hex(5) | N[+1623] | 1259.0776 | 4 | 5033.2887 | 5033.2679 | 0 | 4.1 | 197 | Specific | 305.08 | 305.08 | 305.08 | 7.21 | 56 | >sp\|P19835\|CEL_HUMAN Bile salt-activated lipase OS=Homo sapiens OX=9606 GN=CEL PE=1 SV=3 | 476 | 20201116_13832_1ng_glyco_hcd203040_compound15.21500.21500.4 | controllerType=0 controllerNumber=1 scan=21500 | 42.7312 |
| 19158:4 | 6 | R.NIAAFGGDPNN[+1913.677]ITLFGESAGGASVSLQTLSPYNK.G | HexNAc(4)Hex(5)NeuAc(1) | N[+1914] | 1331.8503 | 4 | 5324.3796 | 5324.3633 | 0 | 3 | 197 | Specific | 883.94 | 883.94 | 883.94 | 17.6 | 56 | >sp\|P19835\|CEL_HUMAN Bile salt-activated lipase OS=Homo sapiens OX=9606 GN=CEL PE=1 SV=3 | 476 | 20201116_13832_1ng_glyco_hcd203040_compound15.22720.22720.4 | controllerType=0 controllerNumber=1 scan=22720 | 45.3443 |
| 19164:3 | 6 | R.NIAAFGGDPNN[+1913.677]ITLFGESAGGASVSLQTLSPYNK.G | HexNAc(4)Hex(5)NeuAc(1) | N[+1914] | 1775.4658 | 3 | 5324.3828 | 5324.3633 | 0 | 3.7 | 197 | Specific | 977.04 | 925.33 | 925.33 | 18.94 | 56 | >sp\|P19835\|CEL_HUMAN Bile salt-activated lipase OS=Homo sapiens OX=9606 GN=CEL PE=1 SV=3 | 476 | 20201116_13832_1ng_glyco_hcd203040_compound15.22726.22726.3 | controllerType=0 controllerNumber=1 scan=22726 | 45.3542 |
| 19161:4 | 6 | R.NIAAFGGDPNN[+1913.677]ITLFGESAGGASVSLQTLSPYNK.G | HexNAc(4)Hex(5)NeuAc(1) | N[+1914] | 1331.8503 | 4 | 5324.3796 | 5324.3633 | 0 | 3 | 197 | Specific | 1000.97 | 1000.97 | 1000.97 | 20.6 | 56 | >sp\|P19835\|CEL_HUMAN Bile salt-activated lipase OS=Homo sapiens OX=9606 GN=CEL PE=1 SV=3 | 476 | 20201116_13832_1ng_glyco_hcd203040_compound15.22723.22723.4 | controllerType=0 controllerNumber=1 scan=22723 | 45.3491 |
| 19166:3 | 6 | R.NIAAFGGDPNN[+1913.677]ITLFGESAGGASVSLQTLSPYNK.G | HexNAc(4)Hex(5)NeuAc(1) | N[+1914] | 1775.4658 | 3 | 5324.3828 | 5324.3633 | 0 | 3.7 | 197 | Specific | 1004.06 | 977.01 | 977.01 | 19.94 | 56 | >sp\|P19835\|CEL_HUMAN Bile salt-activated lipase OS=Homo sapiens OX=9606 GN=CEL PE=1 SV=3 | 476 | 20201116_13832_1ng_glyco_hcd203040_compound15.22728.22728.3 | controllerType=0 controllerNumber=1 scan=22728 | 45.3601 |
| 20116:3 | 7 | R.GLSFDVSLEVSQGPGLLN[+2350.830]DTKVYTVDLGR.T | HexNAc(4)Hex(5)Fuc(1)NeuAc(2) | N[+2351] | 1810.82 | 3 | 5430.4455 | 5430.425 | 0 | 3.8 | 118 | Specific | 304.92 | 283.77 | 283.77 | 6.36 | 61 | >sp\|P01833\|PIGR_HUMAN Polymeric immunoglobulin receptor OS=Homo sapiens OX=9606 GN=PIGR PE=1 SV=4 | 902 | 20201116_13832_1ng_glyco_hcd203040_compound15.23718.23718.3 | controllerType=0 controllerNumber=1 scan=23718 | 47.2555 |
| 20114:3 | 7 | R.GLSFDVSLEVSQGPGLLN[+2350.830]DTKVYTVDLGR.T | HexNAc(4)Hex(5)Fuc(1)NeuAc(2) | N[+2351] | 1810.82 | 3 | 5430.4455 | 5430.425 | 0 | 3.8 | 118 | Specific | 316.78 | 281.22 | 281.22 | 6.52 | 61 | >sp\|P01833\|PIGR_HUMAN Polymeric immunoglobulin receptor OS=Homo sapiens OX=9606 GN=PIGR PE=1 SV=4 | 902 | 20201116_13832_1ng_glyco_hcd203040_compound15.23716.23716.3 | controllerType=0 controllerNumber=1 scan=23716 | 47.2521 |
| 18792:4 | 7 | R.GLSFDVSLEVSQGPGLLN[+2204.772]DTKVYTVDLGR.T | HexNAc(4)Hex(5)NeuAc(2) | N[+2205] | 1321.8562 | 4 | 5284.403 | 5284.3671 | 0 | 6.8 | 118 | Specific | 821.4 | 554.11 | 554.11 | 14.57 | 61 | >sp\|P01833\|PIGR_HUMAN Polymeric immunoglobulin receptor OS=Homo sapiens OX=9606 GN=PIGR PE=1 SV=4 | 902 | 20201116_13832_1ng_glyco_hcd203040_compound15.22337.22337.2 | controllerType=0 controllerNumber=1 scan=22337 | 44.4639 |
| 18790:4 | 7 | R.GLSFDVSLEVSQGPGLLN[+2204.772]DTKVYTVDLGR.T | HexNAc(4)Hex(5)NeuAc(2) | N[+2205] | 1321.8562 | 4 | 5284.403 | 5284.3671 | 0 | 6.8 | 118 | Specific | 931.05 | 797.17 | 797.17 | 16.57 | 61 | >sp\|P01833\|PIGR_HUMAN Polymeric immunoglobulin receptor OS=Homo sapiens OX=9606 GN=PIGR PE=1 SV=4 | 902 | 20201116_13832_1ng_glyco_hcd203040_compound15.22335.22335.2 | controllerType=0 controllerNumber=1 scan=22335 | 44.459 |
| 19136:4 | 7 | K.QIGLYPVLVIDSSGYVNPN[+2059.735]YTGR.I | HexNAc(4)Hex(5)Fuc(1)NeuAc(1) | N[+2060] | 1147.0191 | 4 | 4585.0545 | 4585.0384 | 0 | 3.5 | 168 | Specific | 591.44 | 525.77 | 525.77 | 11.58 | 61 | >sp\|P01833\|PIGR_HUMAN Polymeric immunoglobulin receptor OS=Homo sapiens OX=9606 GN=PIGR PE=1 SV=4 | 902 | 20201116_13832_1ng_glyco_hcd203040_compound15.22697.22697.4 | controllerType=0 controllerNumber=1 scan=22697 | 45.2823 |
| 19130:3 | 7 | K.QIGLYPVLVIDSSGYVNPN[+2059.735]YTGR.I | HexNAc(4)Hex(5)Fuc(1)NeuAc(1) | N[+2060] | 1529.0245 | 3 | 4585.0589 | 4585.0384 | 0 | 4.5 | 168 | Specific | 833.13 | 821.93 | 821.93 | 15.54 | 61 | >sp\|P01833\|PIGR_HUMAN Polymeric immunoglobulin receptor OS=Homo sapiens OX=9606 GN=PIGR PE=1 SV=4 | 902 | 20201116_13832_1ng_glyco_hcd203040_compound15.22691.22691.3 | controllerType=0 controllerNumber=1 scan=22691 | 45.2699 |
| 19139:4 | 7 | K.QIGLYPVLVIDSSGYVNPN[+2059.735]YTGR.I | HexNAc(4)Hex(5)Fuc(1)NeuAc(1) | N[+2060] | 1147.0191 | 4 | 4585.0545 | 4585.0384 | 0 | 3.5 | 168 | Specific | 894.92 | 785.07 | 785.07 | 17.58 | 61 | >sp\|P01833\|PIGR_HUMAN Polymeric immunoglobulin receptor OS=Homo sapiens OX=9606 GN=PIGR PE=1 SV=4 | 902 | 20201116_13832_1ng_glyco_hcd203040_compound15.22700.22700.4 | controllerType=0 controllerNumber=1 scan=22700 | 45.2955 |
| 19132:3 | 7 | K.QIGLYPVLVIDSSGYVNPN[+2059.735]YTGR.I | HexNAc(4)Hex(5)Fuc(1)NeuAc(1) | N[+2060] | 1529.0245 | 3 | 4585.0589 | 4585.0384 | 0 | 4.5 | 168 | Specific | 899.81 | 640.78 | 640.78 | 16.54 | 61 | >sp\|P01833\|PIGR_HUMAN Polymeric immunoglobulin receptor OS=Homo sapiens OX=9606 GN=PIGR PE=1 SV=4 | 902 | 20201116_13832_1ng_glyco_hcd203040_compound15.22693.22693.3 | controllerType=0 controllerNumber=1 scan=22693 | 45.2754 |
| 23296:4 | 7 | R.LSLLEEPGN[+2350.830]GTFTVILNQLTSR.D | HexNAc(4)Hex(5)Fuc(1)NeuAc(2) | N[+2351] | 1189.0385 | 4 | 4753.1322 | 4753.1229 | 0 | 1.9 | 413 | Specific | 228.77 | 228.77 | 228.77 | 6.73 | 61 | >sp\|P01833\|PIGR_HUMAN Polymeric immunoglobulin receptor OS=Homo sapiens OX=9606 GN=PIGR PE=1 SV=4 | 902 | 20201116_13832_1ng_glyco_hcd203040_compound15.27025.27025.4 | controllerType=0 controllerNumber=1 scan=27025 | 53.3099 |
| 20848:4 | 7 | R.LSLLEEPGN[+2204.772]GTFTVILNQLTSR.D | HexNAc(4)Hex(5)NeuAc(2) | N[+2205] | 1152.5317 | 4 | 4607.1048 | 4607.065 | 0 | 8.6 | 413 | Specific | 735.73 | 417.07 | 417.07 | 13.03 | 61 | >sp\|P01833\|PIGR_HUMAN Polymeric immunoglobulin receptor OS=Homo sapiens OX=9606 GN=PIGR PE=1 SV=4 | 902 | 20201116_13832_1ng_glyco_hcd203040_compound15.24480.24480.4 | controllerType=0 controllerNumber=1 scan=24480 | 48.8027 |
| 23294:4 | 7 | R.LSLLEEPGN[+2350.830]GTFTVILNQLTSR.D | HexNAc(4)Hex(5)Fuc(1)NeuAc(2) | N[+2351] | 1189.0385 | 4 | 4753.1322 | 4753.1229 | 0 | 1.9 | 413 | Specific | 807.84 | 610.1 | 610.1 | 16.75 | 61 | >sp\|P01833\|PIGR_HUMAN Polymeric immunoglobulin receptor OS=Homo sapiens OX=9606 GN=PIGR PE=1 SV=4 | 902 | 20201116_13832_1ng_glyco_hcd203040_compound15.27023.27023.4 | controllerType=0 controllerNumber=1 scan=27023 | 53.2987 |
| 20851:4 | 7 | R.LSLLEEPGN[+2204.772]GTFTVILNQLTSR.D | HexNAc(4)Hex(5)NeuAc(2) | N[+2205] | 1152.5317 | 4 | 4607.1048 | 4607.065 | 0 | 8.6 | 413 | Specific | 836.01 | 469.11 | 469.11 | 15.03 | 61 | >sp\|P01833\|PIGR_HUMAN Polymeric immunoglobulin receptor OS=Homo sapiens OX=9606 GN=PIGR PE=1 SV=4 | 902 | 20201116_13832_1ng_glyco_hcd203040_compound15.24483.24483.4 | controllerType=0 controllerNumber=1 scan=24483 | 48.8073 |
| 20845:3 | 7 | R.LSLLEEPGN[+2205.793]GTFTVILNQLTSR.D | HexNAc(4)Hex(5)Fuc(2)NeuAc(1) | N[+2206] | 1536.707 | 3 | 4608.1063 | 4608.0854 | 0 | 4.5 | 413 | Specific | 875.7 | 655.42 | 655.42 | 16.54 | 61 | >sp\|P01833\|PIGR_HUMAN Polymeric immunoglobulin receptor OS=Homo sapiens OX=9606 GN=PIGR PE=1 SV=4 | 902 | 20201116_13832_1ng_glyco_hcd203040_compound15.24477.24477.3 | controllerType=0 controllerNumber=1 scan=24477 | 48.797 |
| 23258:3 | 7 | R.LSLLEEPGN[+2350.830]GTFTVILNQLTSR.D | HexNAc(4)Hex(5)Fuc(1)NeuAc(2) | N[+2351] | 1585.0484 | 3 | 4753.1306 | 4753.1229 | 0 | 1.6 | 413 | Specific | 956.76 | 663.8 | 663.8 | 18.99 | 61 | >sp\|P01833\|PIGR_HUMAN Polymeric immunoglobulin receptor OS=Homo sapiens OX=9606 GN=PIGR PE=1 SV=4 | 902 | 20201116_13832_1ng_glyco_hcd203040_compound15.26985.26985.3 | controllerType=0 controllerNumber=1 scan=26985 | 53.2208 |
| 23268:3 | 7 | R.LSLLEEPGN[+2350.830]GTFTVILNQLTSR.D | HexNAc(4)Hex(5)Fuc(1)NeuAc(2) | N[+2351] | 1585.0511 | 3 | 4753.1387 | 4753.1229 | 0 | 3.3 | 413 | Specific | 964.21 | 721.08 | 721.08 | 19.09 | 61 | >sp\|P01833\|PIGR_HUMAN Polymeric immunoglobulin receptor OS=Homo sapiens OX=9606 GN=PIGR PE=1 SV=4 | 902 | 20201116_13832_1ng_glyco_hcd203040_compound15.26996.26996.3 | controllerType=0 controllerNumber=1 scan=26996 | 53.2446 |
| 20847:3 | 7 | R.LSLLEEPGN[+2205.793]GTFTVILNQLTSR.D | HexNAc(4)Hex(5)Fuc(2)NeuAc(1) | N[+2206] | 1536.707 | 3 | 4608.1063 | 4608.0854 | 0 | 4.5 | 413 | Specific | 965.21 | 703.01 | 703.01 | 18.54 | 61 | >sp\|P01833\|PIGR_HUMAN Polymeric immunoglobulin receptor OS=Homo sapiens OX=9606 GN=PIGR PE=1 SV=4 | 902 | 20201116_13832_1ng_glyco_hcd203040_compound15.24479.24479.3 | controllerType=0 controllerNumber=1 scan=24479 | 48.8012 |
| 23256:3 | 7 | R.LSLLEEPGN[+2350.830]GTFTVILNQLTSR.D | HexNAc(4)Hex(5)Fuc(1)NeuAc(2) | N[+2351] | 1585.0484 | 3 | 4753.1306 | 4753.1229 | 0 | 1.6 | 413 | Specific | 975.86 | 709.25 | 709.25 | 18.99 | 61 | >sp\|P01833\|PIGR_HUMAN Polymeric immunoglobulin receptor OS=Homo sapiens OX=9606 GN=PIGR PE=1 SV=4 | 902 | 20201116_13832_1ng_glyco_hcd203040_compound15.26983.26983.3 | controllerType=0 controllerNumber=1 scan=26983 | 53.2139 |
| 23266:3 | 7 | R.LSLLEEPGN[+2350.830]GTFTVILNQLTSR.D | HexNAc(4)Hex(5)Fuc(1)NeuAc(2) | N[+2351] | 1585.0511 | 3 | 4753.1387 | 4753.1229 | 0 | 3.3 | 413 | Specific | 1130.09 | 883.09 | 883.09 | 22.09 | 61 | >sp\|P01833\|PIGR_HUMAN Polymeric immunoglobulin receptor OS=Homo sapiens OX=9606 GN=PIGR PE=1 SV=4 | 902 | 20201116_13832_1ng_glyco_hcd203040_compound15.26994.26994.3 | controllerType=0 controllerNumber=1 scan=26994 | 53.2382 |
| 16228:3 | 7 | K.[+42.011]IIEGEPNLKVPGN[+2018.708]VTAVLGETLK.V | HexNAc(3)Hex(6)Fuc(1)NeuAc(1) | .[42], N[+2019] | 1484.6962 | 3 | 4452.074 | 4452.0683 | 0 | 1.3 | 457 | Specific | 206.92 | 206.92 | 206.92 | 4.65 | 61 | >sp\|P01833\|PIGR_HUMAN Polymeric immunoglobulin receptor OS=Homo sapiens OX=9606 GN=PIGR PE=1 SV=4 | 902 | 20201116_13832_1ng_glyco_hcd203040_compound15.19666.19666.3 | controllerType=0 controllerNumber=1 scan=19666 | 39.2879 |
| 14353:3 | 7 | K.IIEGEPNLKVPGN[+2059.735]VTAVLGETLK.V | HexNAc(4)Hex(5)Fuc(1)NeuAc(1) | N[+2060] | 1484.3747 | 3 | 4451.1095 | 4451.0843 | 0 | 5.7 | 457 | Specific | 218.48 | 218.48 | 218.48 | 4.22 | 61 | >sp\|P01833\|PIGR_HUMAN Polymeric immunoglobulin receptor OS=Homo sapiens OX=9606 GN=PIGR PE=1 SV=4 | 902 | 20201116_13832_1ng_glyco_hcd203040_compound15.17690.17690.3 | controllerType=0 controllerNumber=1 scan=17690 | 35.8296 |
| 14351:3 | 7 | K.IIEGEPNLKVPGN[+2059.735]VTAVLGETLK.V | HexNAc(4)Hex(5)Fuc(1)NeuAc(1) | N[+2060] | 1484.3747 | 3 | 4451.1095 | 4451.0843 | 0 | 5.7 | 457 | Specific | 258.1 | 258.1 | 258.1 | 6 | 61 | >sp\|P01833\|PIGR_HUMAN Polymeric immunoglobulin receptor OS=Homo sapiens OX=9606 GN=PIGR PE=1 SV=4 | 902 | 20201116_13832_1ng_glyco_hcd203040_compound15.17688.17688.3 | controllerType=0 controllerNumber=1 scan=17688 | 35.8245 |
| 16219:3 | 7 | K.IIEGEPNLKVPGN[+2204.772]VTAVLGETLK.V | HexNAc(4)Hex(5)NeuAc(2) | N[+2205] | 1532.7184 | 3 | 4596.1408 | 4596.1218 | 0 | 4.1 | 457 | Specific | 668.27 | 643.21 | 643.21 | 12.95 | 61 | >sp\|P01833\|PIGR_HUMAN Polymeric immunoglobulin receptor OS=Homo sapiens OX=9606 GN=PIGR PE=1 SV=4 | 902 | 20201116_13832_1ng_glyco_hcd203040_compound15.19657.19657.3 | controllerType=0 controllerNumber=1 scan=19657 | 39.2736 |

Table S4. Cont.

| Query #:z | Protein Rank | Glycopeptide | Glycan | Modification Type(s) | Observed m/z | z | Observed (M+H) | Calc. mass (M+H) | Off-by-x error | Mass error (ppm) | StartPosition | Cleavage | Score | Delta | Delta Mod | \|Log Prob\| | # of unique peptides | Protein Name | Protein DB number | Comment | Scan # | Scan Time |
|---|---|---|---|---|---|---|---|---|---|---|---|---|---|---|---|---|---|---|---|---|---|---|
| 16217:3 | 7 | K.IIEGEPNLKVPGN[+2204.772]VTAVLGETLK.V | HexNAc(4)Hex(5)NeuAc(2) | N[+2205] | 1532.7184 | 3 | 4596.1408 | 4596.1218 | 0 | 4.1 | 457 | Specific | 757.88 | 686.95 | 686.95 | 14.95 | 61 | >sp\|P01833\|PIGR_HUMAN Polymeric immunoglobulin receptor OS=Homo sapiens OX=9606 GN=PIGR PE=1 SV=4 | 902 | 20201116_13832_1ng_glyco_hcd203040_compound15.19655.19655.3 | controllerType=0 controllerNumber=1 scan=19655 | 39.2666 |
| 15702:5 | 7 | K.[+42.011]IIEGEPNLKVPGN[+2018.708]VTAVLGETLKVPC[+57.021]HFPC[+57.021] | HexNAc(3)Hex(6)Fuc(1)NeuAc(1) | .[42], C[+57]*2, N[+2019] | 1096.3176 | 5 | 5477.5589 | 5477.5258 | 0 | 6 | 457 | Specific | 220.28 | 187.92 | 187.92 | 3.83 | 61 | >sp\|P01833\|PIGR_HUMAN Polymeric immunoglobulin receptor OS=Homo sapiens OX=9606 GN=PIGR PE=1 SV=4 | 902 | 20201116_13832_1ng_glyco_hcd203040_compound15.19088.19088.5 | controllerType=0 controllerNumber=1 scan=19088 | 38.257 |
| 15633:4 | 7 | K.IIEGEPNLKVPGN[+2059.735]VTAVLGETLKVPC[+57.021]HFPC[+57.021]K.F | HexNAc(4)Hex(5)Fuc(1)NeuAc(1) | C[+57]*2, N[+2060] | 1369.8937 | 4 | 5476.5529 | 5476.5418 | 0 | 2 | 457 | Specific | 545.53 | 545.53 | 545.53 | 9.29 | 61 | >sp\|P01833\|PIGR_HUMAN Polymeric immunoglobulin receptor OS=Homo sapiens OX=9606 GN=PIGR PE=1 SV=4 | 902 | 20201116_13832_1ng_glyco_hcd203040_compound15.19016.19016.4 | controllerType=0 controllerNumber=1 scan=19016 | 38.1296 |
| 17263:4 | 7 | K.IIEGEPNLKVPGN[+2350.830]VTAVLGETLKVPC[+57.021]HFPC[+57.021]K.F | HexNAc(4)Hex(5)Fuc(1)NeuAc(2) | C[+57]*2, N[+2351] | 1442.671 | 4 | 5767.6623 | 5767.6372 | 0 | 4.3 | 457 | Specific | 669.19 | 624.12 | 624.12 | 12.25 | 61 | >sp\|P01833\|PIGR_HUMAN Polymeric immunoglobulin receptor OS=Homo sapiens OX=9606 GN=PIGR PE=1 SV=4 | 902 | 20201116_13832_1ng_glyco_hcd203040_compound15.20745.20745.4 | controllerType=0 controllerNumber=1 scan=20745 | 41.2102 |
| 15631:4 | 7 | K.IIEGEPNLKVPGN[+2059.735]VTAVLGETLKVPC[+57.021]HFPC[+57.021]K.F | HexNAc(4)Hex(5)Fuc(1)NeuAc(1) | C[+57]*2, N[+2060] | 1369.8937 | 4 | 5476.5529 | 5476.5418 | 0 | 2 | 457 | Specific | 673 | 575.84 | 575.84 | 12.42 | 61 | >sp\|P01833\|PIGR_HUMAN Polymeric immunoglobulin receptor OS=Homo sapiens OX=9606 GN=PIGR PE=1 SV=4 | 902 | 20201116_13832_1ng_glyco_hcd203040_compound15.19014.19014.4 | controllerType=0 controllerNumber=1 scan=19014 | 38.1258 |
| 17209:5 | 7 | K.IIEGEPNLKVPGN[+2350.830]VTAVLGETLKVPC[+57.021]HFPC[+57.021]K.F | HexNAc(4)Hex(5)Fuc(1)NeuAc(2) | C[+57]*2, N[+2351] | 1154.3369 | 5 | 5767.6555 | 5767.6372 | 0 | 3.2 | 457 | Specific | 764.16 | 675.37 | 675.37 | 14.75 | 61 | >sp\|P01833\|PIGR_HUMAN Polymeric immunoglobulin receptor OS=Homo sapiens OX=9606 GN=PIGR PE=1 SV=4 | 902 | 20201116_13832_1ng_glyco_hcd203040_compound15.20689.20689.5 | controllerType=0 controllerNumber=1 scan=20689 | 41.0922 |
| 17212:5 | 7 | K.IIEGEPNLKVPGN[+2350.830]VTAVLGETLKVPC[+57.021]HFPC[+57.021]K.F | HexNAc(4)Hex(5)Fuc(1)NeuAc(2) | C[+57]*2, N[+2351] | 1154.3369 | 5 | 5767.6555 | 5767.6372 | 0 | 3.2 | 457 | Specific | 827.43 | 789.22 | 789.22 | 15.75 | 61 | >sp\|P01833\|PIGR_HUMAN Polymeric immunoglobulin receptor OS=Homo sapiens OX=9606 GN=PIGR PE=1 SV=4 | 902 | 20201116_13832_1ng_glyco_hcd203040_compound15.20692.20692.5 | controllerType=0 controllerNumber=1 scan=20692 | 41.0973 |
| 17260:5 | 7 | K.IIEGEPNLKVPGN[+2350.830]VTAVLGETLKVPC[+57.021]HFPC[+57.021]K.F | HexNAc(4)Hex(5)Fuc(1)NeuAc(2) | C[+57]*2, N[+2351] | 1154.3373 | 5 | 5767.6573 | 5767.6372 | 0 | 3.5 | 457 | Specific | 863.04 | 649.64 | 649.64 | 16.75 | 61 | >sp\|P01833\|PIGR_HUMAN Polymeric immunoglobulin receptor OS=Homo sapiens OX=9606 GN=PIGR PE=1 SV=4 | 902 | 20201116_13832_1ng_glyco_hcd203040_compound15.20742.20742.5 | controllerType=0 controllerNumber=1 scan=20742 | 41.2038 |
| 17261:4 | 7 | K.IIEGEPNLKVPGN[+2350.830]VTAVLGETLKVPC[+57.021]HFPC[+57.021]K.F | HexNAc(4)Hex(5)Fuc(1)NeuAc(2) | C[+57]*2, N[+2351] | 1442.671 | 4 | 5767.6623 | 5767.6372 | 0 | 4.3 | 457 | Specific | 973.66 | 746.91 | 746.91 | 18.25 | 61 | >sp\|P01833\|PIGR_HUMAN Polymeric immunoglobulin receptor OS=Homo sapiens OX=9606 GN=PIGR PE=1 SV=4 | 902 | 20201116_13832_1ng_glyco_hcd203040_compound15.20743.20743.4 | controllerType=0 controllerNumber=1 scan=20743 | 41.2057 |
| 17258:5 | 7 | K.IIEGEPNLKVPGN[+2350.830]VTAVLGETLKVPC[+57.021]HFPC[+57.021]K.F | HexNAc(4)Hex(5)Fuc(1)NeuAc(2) | C[+57]*2, N[+2351] | 1154.3373 | 5 | 5767.6573 | 5767.6372 | 0 | 3.5 | 457 | Specific | 978.49 | 761.69 | 761.69 | 18.75 | 61 | >sp\|P01833\|PIGR_HUMAN Polymeric immunoglobulin receptor OS=Homo sapiens OX=9606 GN=PIGR PE=1 SV=4 | 902 | 20201116_13832_1ng_glyco_hcd203040_compound15.20740.20740.5 | controllerType=0 controllerNumber=1 scan=20740 | 41.198 |
| 13639:4 | 7 | K.VPGN[+2205.793]VTAVLGETLKVPC[+57.021]HFPC[+57.021]K.F | HexNAc(4)Hex(5)Fuc(2)NeuAc(1) | C[+57]*2, N[+2206] | 1158.0193 | 4 | 4629.0555 | 4629.0502 | 0 | 1.1 | 466 | Specific | 278.72 | 169.5 | 169.5 | 4.41 | 61 | >sp\|P01833\|PIGR_HUMAN Polymeric immunoglobulin receptor OS=Homo sapiens OX=9606 GN=PIGR PE=1 SV=4 | 902 | 20201116_13832_1ng_glyco_hcd203040_compound15.16951.16951.3 | controllerType=0 controllerNumber=1 scan=16951 | 34.5818 |
| 13708:4 | 7 | K.VPGN[+2205.793]VTAVLGETLKVPC[+57.021]HFPC[+57.021]K.F | HexNAc(4)Hex(5)Fuc(2)NeuAc(1) | C[+57]*2, N[+2206] | 1158.0241 | 4 | 4629.0745 | 4629.0502 | 0 | 5.3 | 466 | Specific | 325.91 | 325.91 | 325.91 | 6.47 | 61 | >sp\|P01833\|PIGR_HUMAN Polymeric immunoglobulin receptor OS=Homo sapiens OX=9606 GN=PIGR PE=1 SV=4 | 902 | 20201116_13832_1ng_glyco_hcd203040_compound15.17022.17022.4 | controllerType=0 controllerNumber=1 scan=17022 | 34.7016 |
| 13710:4 | 7 | K.VPGN[+2205.793]VTAVLGETLKVPC[+57.021]HFPC[+57.021]K.F | HexNAc(4)Hex(5)Fuc(2)NeuAc(1) | C[+57]*2, N[+2206] | 1158.0241 | 4 | 4629.0745 | 4629.0502 | 0 | 5.3 | 466 | Specific | 374.14 | 349.46 | 349.46 | 6.99 | 61 | >sp\|P01833\|PIGR_HUMAN Polymeric immunoglobulin receptor OS=Homo sapiens OX=9606 GN=PIGR PE=1 SV=4 | 902 | 20201116_13832_1ng_glyco_hcd203040_compound15.17024.17024.4 | controllerType=0 controllerNumber=1 scan=17024 | 34.7053 |
| 13641:4 | 7 | K.VPGN[+2205.793]VTAVLGETLKVPC[+57.021]HFPC[+57.021]K.F | HexNAc(4)Hex(5)Fuc(2)NeuAc(1) | C[+57]*2, N[+2206] | 1158.0193 | 4 | 4629.0555 | 4629.0502 | 0 | 1.1 | 466 | Specific | 381.65 | 346.67 | 346.67 | 6.31 | 61 | >sp\|P01833\|PIGR_HUMAN Polymeric immunoglobulin receptor OS=Homo sapiens OX=9606 GN=PIGR PE=1 SV=4 | 902 | 20201116_13832_1ng_glyco_hcd203040_compound15.16953.16953.3 | controllerType=0 controllerNumber=1 scan=16953 | 34.5867 |
| 05669:3 | 7 | K.WN[+1913.677]NTGC[+57.021]QALPSQDEGPSK.A | HexNAc(4)Hex(5)NeuAc(1) | C[+57], N[+1914] | 1301.5259 | 3 | 3902.5633 | 3902.5537 | 0 | 2.4 | 498 | Specific | 290.26 | 247.14 | 247.14 | 6.59 | 61 | >sp\|P01833\|PIGR_HUMAN Polymeric immunoglobulin receptor OS=Homo sapiens OX=9606 GN=PIGR PE=1 SV=4 | 902 | 20201116_13832_1ng_glyco_hcd203040_compound15.8672.8672.3 | controllerType=0 controllerNumber=1 scan=8672 | 20.8128 |
| 05671:3 | 7 | K.WN[+1913.677]NTGC[+57.021]QALPSQDEGPSK.A | HexNAc(4)Hex(5)NeuAc(1) | C[+57], N[+1914] | 1301.5259 | 3 | 3902.5633 | 3902.5537 | 0 | 2.4 | 498 | Specific | 302.02 | 259.36 | 259.36 | 6.72 | 61 | >sp\|P01833\|PIGR_HUMAN Polymeric immunoglobulin receptor OS=Homo sapiens OX=9606 GN=PIGR PE=1 SV=4 | 902 | 20201116_13832_1ng_glyco_hcd203040_compound15.8674.8674.3 | controllerType=0 controllerNumber=1 scan=8674 | 20.8168 |
| 05551:3 | 7 | K.WN[+2059.735]NTGC[+57.021]QALPSQDEGPSK.A | HexNAc(4)Hex(5)Fuc(1)NeuAc(1) | C[+57], N[+2060] | 1350.2137 | 3 | 4048.6265 | 4048.6116 | 0 | 3.7 | 498 | Specific | 416.33 | 365.9 | 365.9 | 7.61 | 61 | >sp\|P01833\|PIGR_HUMAN Polymeric immunoglobulin receptor OS=Homo sapiens OX=9606 GN=PIGR PE=1 SV=4 | 902 | 20201116_13832_1ng_glyco_hcd203040_compound15.8550.8550.3 | controllerType=0 controllerNumber=1 scan=8550 | 20.6086 |
| 05553:3 | 7 | K.WN[+2059.735]NTGC[+57.021]QALPSQDEGPSK.A | HexNAc(4)Hex(5)Fuc(1)NeuAc(1) | C[+57], N[+2060] | 1350.2137 | 3 | 4048.6265 | 4048.6116 | 0 | 3.7 | 498 | Specific | 422.02 | 361.14 | 361.14 | 7.61 | 61 | >sp\|P01833\|PIGR_HUMAN Polymeric immunoglobulin receptor OS=Homo sapiens OX=9606 GN=PIGR PE=1 SV=4 | 902 | 20201116_13832_1ng_glyco_hcd203040_compound15.8552.8552.3 | controllerType=0 controllerNumber=1 scan=8552 | 20.6122 |
| 18014:3 | 9 | R.LSLHRPALEDLLLGSEAN[+1298.476]LTC[+57.021]TLTGLR.D | HexNAc(4)Hex(3) | C[+57], N[+1298] | 1421.3679 | 3 | 4262.089 | 4262.0742 | 0 | 3.5 | 127 | Specific | 326.53 | 326.53 | 326.53 | 6.58 | 42 | >sp\|P01876\|IGHA1_HUMAN Immunoglobulin heavy constant alpha 1 OS=Homo sapiens OX=9606 GN=IGHA1 PE=1 SV=2 | 255 | 20201116_13832_1ng_glyco_hcd203040_compound15.21527.21527.3 | controllerType=0 controllerNumber=1 scan=21527 | 42.7978 |
| 17891:3 | 9 | R.LSLHRPALEDLLLGSEAN[+1864.634]LTC[+57.021]TLTGLR.D | HexNAc(2)Hex(9) | C[+57], N[+1865] | 1610.0816 | 3 | 4828.2302 | 4828.2324 | 0 | -0.5 | 127 | Specific | 336.15 | 336.15 | 336.15 | 5.8 | 42 | >sp\|P01876\|IGHA1_HUMAN Immunoglobulin heavy constant alpha 1 OS=Homo sapiens OX=9606 GN=IGHA1 PE=1 SV=2 | 255 | 20201116_13832_1ng_glyco_hcd203040_compound15.21398.21398.3 | controllerType=0 controllerNumber=1 scan=21398 | 42.4902 |
| 17842:4 | 9 | R.LSLHRPALEDLLLGSEAN[+568.212]LTC[+57.021]TLTGLR.D | HexNAc(2)Hex(1) | C[+57], N[+568] | 883.703 | 4 | 3531.7904 | 3531.8098 | 0 | -5.5 | 127 | Specific | 353.73 | 353.73 | 353.73 | 6.04 | 42 | >sp\|P01876\|IGHA1_HUMAN Immunoglobulin heavy constant alpha 1 OS=Homo sapiens OX=9606 GN=IGHA1 PE=1 SV=2 | 255 | 20201116_13832_1ng_glyco_hcd203040_compound15.21346.21346.2 | controllerType=0 controllerNumber=1 scan=21346 | 42.3604 |
| 18964:3 | 9 | R.LSLHRPALEDLLLGSEAN[+1751.624]LTC[+57.021]TLTGLR.D | HexNAc(4)Hex(4)NeuAc(1) | C[+57], N[+1752] | 1572.4175 | 3 | 4715.2381 | 4715.2225 | 0 | 3.3 | 127 | Specific | 420.37 | 420.37 | 420.37 | 8.67 | 42 | >sp\|P01876\|IGHA1_HUMAN Immunoglobulin heavy constant alpha 1 OS=Homo sapiens OX=9606 GN=IGHA1 PE=1 SV=2 | 255 | 20201116_13832_1ng_glyco_hcd203040_compound15.22516.22516.3 | controllerType=0 controllerNumber=1 scan=22516 | 44.8562 |
| 17894:3 | 9 | R.LSLHRPALEDLLLGSEAN[+1864.634]LTC[+57.021]TLTGLR.D | HexNAc(2)Hex(9) | C[+57], N[+1865] | 1610.0816 | 3 | 4828.2302 | 4828.2324 | 0 | -0.5 | 127 | Specific | 430.3 | 424.29 | 424.29 | 8.01 | 42 | >sp\|P01876\|IGHA1_HUMAN Immunoglobulin heavy constant alpha 1 OS=Homo sapiens OX=9606 GN=IGHA1 PE=1 SV=2 | 255 | 20201116_13832_1ng_glyco_hcd203040_compound15.21401.21401.3 | controllerType=0 controllerNumber=1 scan=21401 | 42.4945 |
| 17956:3 | 9 | R.LSLHRPALEDLLLGSEAN[+1501.555]LTC[+57.021]TLTGLR.D | HexNAc(5)Hex(3) | C[+57], N[+1502] | 1489.0607 | 3 | 4465.1676 | 4465.1536 | 0 | 3.1 | 127 | Specific | 464.22 | 464.22 | 464.22 | 9.1 | 42 | >sp\|P01876\|IGHA1_HUMAN Immunoglobulin heavy constant alpha 1 OS=Homo sapiens OX=9606 GN=IGHA1 PE=1 SV=2 | 255 | 20201116_13832_1ng_glyco_hcd203040_compound15.21466.21466.3 | controllerType=0 controllerNumber=1 scan=21466 | 42.6421 |
| 17843:4 | 9 | R.LSLHRPALEDLLLGSEAN[+568.212]LTC[+57.021]TLTGLR.D | HexNAc(2)Hex(1) | C[+57], N[+568] | 883.703 | 4 | 3531.7904 | 3531.8098 | 0 | -5.5 | 127 | Specific | 469.35 | 439.77 | 439.77 | 7.93 | 42 | >sp\|P01876\|IGHA1_HUMAN Immunoglobulin heavy constant alpha 1 OS=Homo sapiens OX=9606 GN=IGHA1 PE=1 SV=2 | 255 | 20201116_13832_1ng_glyco_hcd203040_compound15.21347.21347.2 | controllerType=0 controllerNumber=1 scan=21347 | 42.3641 |
| 17935:3 | 9 | R.LSLHRPALEDLLLGSEAN[+1622.582]LTC[+57.021]TLTGLR.D | HexNAc(4)Hex(5) | C[+57], N[+1623] | 1529.4013 | 3 | 4586.1894 | 4586.1799 | 0 | 2.1 | 127 | Specific | 503.28 | 493.88 | 493.88 | 9.18 | 42 | >sp\|P01876\|IGHA1_HUMAN Immunoglobulin heavy constant alpha 1 OS=Homo sapiens OX=9606 GN=IGHA1 PE=1 SV=2 | 255 | 20201116_13832_1ng_glyco_hcd203040_compound15.21444.21444.3 | controllerType=0 controllerNumber=1 scan=21444 | 42.5978 |
| 18017:3 | 9 | R.LSLHRPALEDLLLGSEAN[+1298.476]LTC[+57.021]TLTGLR.D | HexNAc(4)Hex(3) | C[+57], N[+1298] | 1421.3679 | 3 | 4262.089 | 4262.0742 | 0 | 3.5 | 127 | Specific | 519.41 | 501.26 | 501.26 | 9.14 | 42 | >sp\|P01876\|IGHA1_HUMAN Immunoglobulin heavy constant alpha 1 OS=Homo sapiens OX=9606 GN=IGHA1 PE=1 SV=2 | 255 | 20201116_13832_1ng_glyco_hcd203040_compound15.21530.21530.3 | controllerType=0 controllerNumber=1 scan=21530 | 42.8019 |
| 17985:5 | 9 | R.LSLHRPALEDLLLGSEAN[+1501.555]LTC[+57.021]TLTGLR.D | HexNAc(5)Hex(3) | C[+57], N[+1502] | 893.8403 | 5 | 4465.1724 | 4465.1536 | 0 | 4.2 | 127 | Specific | 538.55 | 538.55 | 538.55 | 9.45 | 42 | >sp\|P01876\|IGHA1_HUMAN Immunoglobulin heavy constant alpha 1 OS=Homo sapiens OX=9606 GN=IGHA1 PE=1 SV=2 | 255 | 20201116_13832_1ng_glyco_hcd203040_compound15.21496.21496.5 | controllerType=0 controllerNumber=1 scan=21496 | 42.7176 |
| 17880:3 | 9 | R.LSLHRPALEDLLLGSEAN[+1540.529]LTC[+57.021]TLTGLR.D | HexNAc(2)Hex(7) | C[+57], N[+1541] | 1502.051 | 3 | 4504.1383 | 4504.1268 | 0 | 2.6 | 127 | Specific | 552.79 | 520.4 | 520.4 | 9.54 | 42 | >sp\|P01876\|IGHA1_HUMAN Immunoglobulin heavy constant alpha 1 OS=Homo sapiens OX=9606 GN=IGHA1 PE=1 SV=2 | 255 | 20201116_13832_1ng_glyco_hcd203040_compound15.21386.21386.3 | controllerType=0 controllerNumber=1 scan=21386 | 42.464 |
| 17987:5 | 9 | R.LSLHRPALEDLLLGSEAN[+1501.555]LTC[+57.021]TLTGLR.D | HexNAc(5)Hex(3) | C[+57], N[+1502] | 893.8403 | 5 | 4465.1724 | 4465.1536 | 0 | 4.2 | 127 | Specific | 555.09 | 555.09 | 555.09 | 9.67 | 42 | >sp\|P01876\|IGHA1_HUMAN Immunoglobulin heavy constant alpha 1 OS=Homo sapiens OX=9606 GN=IGHA1 PE=1 SV=2 | 255 | 20201116_13832_1ng_glyco_hcd203040_compound15.21498.21498.5 | controllerType=0 controllerNumber=1 scan=21498 | 42.7254 |
| 18966:3 | 9 | R.LSLHRPALEDLLLGSEAN[+1751.624]LTC[+57.021]TLTGLR.D | HexNAc(4)Hex(4)NeuAc(1) | C[+57], N[+1752] | 1572.4175 | 3 | 4715.2381 | 4715.2225 | 0 | 3.3 | 127 | Specific | 567.18 | 567.18 | 567.18 | 9.36 | 42 | >sp\|P01876\|IGHA1_HUMAN Immunoglobulin heavy constant alpha 1 OS=Homo sapiens OX=9606 GN=IGHA1 PE=1 SV=2 | 255 | 20201116_13832_1ng_glyco_hcd203040_compound15.22518.22518.3 | controllerType=0 controllerNumber=1 scan=22518 | 44.8602 |
| 17914:3 | 9 | R.LSLHRPALEDLLLGSEAN[+1663.608]LTC[+57.021]TLTGLR.D | HexNAc(5)Hex(4) | C[+57], N[+1664] | 1543.0761 | 3 | 4627.2138 | 4627.2064 | 0 | 1.6 | 127 | Specific | 568.05 | 524.2 | 524.2 | 9.26 | 42 | >sp\|P01876\|IGHA1_HUMAN Immunoglobulin heavy constant alpha 1 OS=Homo sapiens OX=9606 GN=IGHA1 PE=1 SV=2 | 255 | 20201116_13832_1ng_glyco_hcd203040_compound15.21422.21422.3 | controllerType=0 controllerNumber=1 scan=21422 | 42.5513 |
| 17916:3 | 9 | R.LSLHRPALEDLLLGSEAN[+1663.608]LTC[+57.021]TLTGLR.D | HexNAc(5)Hex(4) | C[+57], N[+1664] | 1543.0761 | 3 | 4627.2138 | 4627.2064 | 0 | 1.6 | 127 | Specific | 575.97 | 531.63 | 531.63 | 10.4 | 42 | >sp\|P01876\|IGHA1_HUMAN Immunoglobulin heavy constant alpha 1 OS=Homo sapiens OX=9606 GN=IGHA1 PE=1 SV=2 | 255 | 20201116_13832_1ng_glyco_hcd203040_compound15.21424.21424.3 | controllerType=0 controllerNumber=1 scan=21424 | 42.5555 |
| 18874:4 | 9 | R.LSLHRPALEDLLLGSEAN[+2116.756]LTC[+57.021]TLTGLR.D | HexNAc(5)Hex(5)NeuAc(1) | C[+57], N[+2117] | 1270.8463 | 4 | 5080.3634 | 5080.3547 | 0 | 1.7 | 127 | Specific | 577.04 | 577.04 | 577.04 | 11.17 | 42 | >sp\|P01876\|IGHA1_HUMAN Immunoglobulin heavy constant alpha 1 OS=Homo sapiens OX=9606 GN=IGHA1 PE=1 SV=2 | 255 | 20201116_13832_1ng_glyco_hcd203040_compound15.22423.22423.4 | controllerType=0 controllerNumber=1 scan=22423 | 44.67 |
| 17927:4 | 9 | R.LSLHRPALEDLLLGSEAN[+1622.582]LTC[+57.021]TLTGLR.D | HexNAc(4)Hex(5) | C[+57], N[+1623] | 1147.3024 | 4 | 4586.1878 | 4586.1799 | 0 | 1.7 | 127 | Specific | 595.56 | 595.56 | 595.56 | 11.17 | 42 | >sp\|P01876\|IGHA1_HUMAN Immunoglobulin heavy constant alpha 1 OS=Homo sapiens OX=9606 GN=IGHA1 PE=1 SV=2 | 255 | 20201116_13832_1ng_glyco_hcd203040_compound15.21436.21436.4 | controllerType=0 controllerNumber=1 scan=21436 | 42.5814 |
| 18918:4 | 9 | R.LSLHRPALEDLLLGSEAN[+1872.651]LTC[+57.021]TLTGLR.D | HexNAc(3)Hex(6)NeuAc(1) | C[+57], N[+1873] | 1209.8206 | 4 | 4836.2606 | 4836.2487 | 0 | 2.4 | 127 | Specific | 600.56 | 600.56 | 600.56 | 12.17 | 42 | >sp\|P01876\|IGHA1_HUMAN Immunoglobulin heavy constant alpha 1 OS=Homo sapiens OX=9606 GN=IGHA1 PE=1 SV=2 | 255 | 20201116_13832_1ng_glyco_hcd203040_compound15.22468.22468.4 | controllerType=0 controllerNumber=1 scan=22468 | 44.7684 |
| 17887:3 | 9 | R.LSLHRPALEDLLLGSEAN[+1702.581]LTC[+57.021]TLTGLR.D | HexNAc(2)Hex(8) | C[+57], N[+1703] | 1556.0668 | 3 | 4666.1859 | 4666.1796 | 0 | 1.4 | 127 | Specific | 618.93 | 618.93 | 618.93 | 11.4 | 42 | >sp\|P01876\|IGHA1_HUMAN Immunoglobulin heavy constant alpha 1 OS=Homo sapiens OX=9606 GN=IGHA1 PE=1 SV=2 | 255 | 20201116_13832_1ng_glyco_hcd203040_compound15.21394.21394.3 | controllerType=0 controllerNumber=1 scan=21394 | 42.4821 |
| 17958:3 | 9 | R.LSLHRPALEDLLLGSEAN[+1501.555]LTC[+57.021]TLTGLR.D | HexNAc(5)Hex(3) | C[+57], N[+1502] | 1489.0607 | 3 | 4465.1676 | 4465.1536 | 0 | 3.1 | 127 | Specific | 619.06 | 565.77 | 565.77 | 11.67 | 42 | >sp\|P01876\|IGHA1_HUMAN Immunoglobulin heavy constant alpha 1 OS=Homo sapiens OX=9606 GN=IGHA1 PE=1 SV=2 | 255 | 20201116_13832_1ng_glyco_hcd203040_compound15.21468.21468.3 | controllerType=0 controllerNumber=1 scan=21468 | 42.6457 |
| 17938:3 | 9 | R.LSLHRPALEDLLLGSEAN[+1622.582]LTC[+57.021]TLTGLR.D | HexNAc(4)Hex(5) | C[+57], N[+1623] | 1529.4013 | 3 | 4586.1894 | 4586.1799 | 0 | 2.1 | 127 | Specific | 626.7 | 577.61 | 577.61 | 11.67 | 42 | >sp\|P01876\|IGHA1_HUMAN Immunoglobulin heavy constant alpha 1 OS=Homo sapiens OX=9606 GN=IGHA1 PE=1 SV=2 | 255 | 20201116_13832_1ng_glyco_hcd203040_compound15.21447.21447.3 | controllerType=0 controllerNumber=1 scan=21447 | 42.6037 |
| 17996:3 | 9 | R.LSLHRPALEDLLLGSEAN[+1216.423]LTC[+57.021]TLTGLR.D | HexNAc(2)Hex(5) | C[+57], N[+1216] | 1394.0175 | 3 | 4180.038 | 4180.0211 | 0 | 4 | 127 | Specific | 627.44 | 590.08 | 590.08 | 11.5 | 42 | >sp\|P01876\|IGHA1_HUMAN Immunoglobulin heavy constant alpha 1 OS=Homo sapiens OX=9606 GN=IGHA1 PE=1 SV=2 | 255 | 20201116_13832_1ng_glyco_hcd203040_compound15.21508.21508.3 | controllerType=0 controllerNumber=1 scan=21508 | 42.7486 |
| 17878:3 | 9 | R.LSLHRPALEDLLLGSEAN[+1540.529]LTC[+57.021]TLTGLR.D | HexNAc(2)Hex(7) | C[+57], N[+1541] | 1502.051 | 3 | 4504.1383 | 4504.1268 | 0 | 2.6 | 127 | Specific | 632.96 | 632.96 | 632.96 | 11.67 | 42 | >sp\|P01876\|IGHA1_HUMAN Immunoglobulin heavy constant alpha 1 OS=Homo sapiens OX=9606 GN=IGHA1 PE=1 SV=2 | 255 | 20201116_13832_1ng_glyco_hcd203040_compound15.21384.21384.3 | controllerType=0 controllerNumber=1 scan=21384 | 42.4572 |
| 17951:4 | 9 | R.LSLHRPALEDLLLGSEAN[+1378.476]LTC[+57.021]TLTGLR.D | HexNAc(2)Hex(6) | C[+57], N[+1378] | 1086.2771 | 4 | 4342.0867 | 4342.074 | 0 | 2.9 | 127 | Specific | 643.71 | 643.71 | 643.71 | 12.17 | 42 | >sp\|P01876\|IGHA1_HUMAN Immunoglobulin heavy constant alpha 1 OS=Homo sapiens OX=9606 GN=IGHA1 PE=1 SV=2 | 255 | 20201116_13832_1ng_glyco_hcd203040_compound15.21461.21461.4 | controllerType=0 controllerNumber=1 scan=21461 | 42.6332 |
| 17893:4 | 9 | R.LSLHRPALEDLLLGSEAN[+1663.608]LTC[+57.021]TLTGLR.D | HexNAc(5)Hex(4) | C[+57], N[+1664] | 1157.5594 | 4 | 4627.2156 | 4627.2064 | 0 | 2 | 127 | Specific | 656.88 | 656.88 | 656.88 | 13.17 | 42 | >sp\|P01876\|IGHA1_HUMAN Immunoglobulin heavy constant alpha 1 OS=Homo sapiens OX=9606 GN=IGHA1 PE=1 SV=2 | 255 | 20201116_13832_1ng_glyco_hcd203040_compound15.21400.21400.4 | controllerType=0 controllerNumber=1 scan=21400 | 42.4929 |
| 17931:4 | 9 | R.LSLHRPALEDLLLGSEAN[+1419.502]LTC[+57.021]TLTGLR.D | HexNAc(3)Hex(5) | C[+57], N[+1420] | 1096.5328 | 4 | 4383.1092 | 4383.1005 | 0 | 2 | 127 | Specific | 659.97 | 659.97 | 659.97 | 13.17 | 42 | >sp\|P01876\|IGHA1_HUMAN Immunoglobulin heavy constant alpha 1 OS=Homo sapiens OX=9606 GN=IGHA1 PE=1 SV=2 | 255 | 20201116_13832_1ng_glyco_hcd203040_compound15.21440.21440.4 | controllerType=0 controllerNumber=1 scan=21440 | 42.5895 |
| 17885:3 | 9 | R.LSLHRPALEDLLLGSEAN[+1702.581]LTC[+57.021]TLTGLR.D | HexNAc(2)Hex(8) | C[+57], N[+1703] | 1556.0668 | 3 | 4666.1859 | 4666.1796 | 0 | 1.4 | 127 | Specific | 674.3 | 674.3 | 674.3 | 12.4 | 42 | >sp\|P01876\|IGHA1_HUMAN Immunoglobulin heavy constant alpha 1 OS=Homo sapiens OX=9606 GN=IGHA1 PE=1 SV=2 | 255 | 20201116_13832_1ng_glyco_hcd203040_compound15.21392.21392.3 | controllerType=0 controllerNumber=1 scan=21392 | 42.4775 |
| 17994:3 | 9 | R.LSLHRPALEDLLLGSEAN[+1216.423]LTC[+57.021]TLTGLR.D | HexNAc(2)Hex(5) | C[+57], N[+1216] | 1394.0175 | 3 | 4180.038 | 4180.0211 | 0 | 4 | 127 | Specific | 681.43 | 659.8 | 659.8 | 12.5 | 42 | >sp\|P01876\|IGHA1_HUMAN Immunoglobulin heavy constant alpha 1 OS=Homo sapiens OX=9606 GN=IGHA1 PE=1 SV=2 | 255 | 20201116_13832_1ng_glyco_hcd203040_compound15.21506.21506.3 | controllerType=0 controllerNumber=1 scan=21506 | 42.7447 |
| 18038:4 | 9 | R.LSLHRPALEDLLLGSEAN[+1095.397]LTC[+57.021]TLTGLR.D | HexNAc(3)Hex(3) | C[+57], N[+1095] | 1015.5074 | 4 | 4059.0076 | 4058.9949 | 0 | 3.1 | 127 | Specific | 682.39 | 654.91 | 654.91 | 13 | 42 | >sp\|P01876\|IGHA1_HUMAN Immunoglobulin heavy constant alpha 1 OS=Homo sapiens OX=9606 GN=IGHA1 PE=1 SV=2 | 255 | 20201116_13832_1ng_glyco_hcd203040_compound15.21552.21552.4 | controllerType=0 controllerNumber=1 scan=21552 | 42.8632 |
| 17884:4 | 9 | R.LSLHRPALEDLLLGSEAN[+1864.634]LTC[+57.021]TLTGLR.D | HexNAc(2)Hex(9) | C[+57], N[+1865] | 1207.8154 | 4 | 4828.2396 | 4828.2324 | 0 | 1.5 | 127 | Specific | 684.22 | 684.22 | 684.22 | 12.9 | 42 | >sp\|P01876\|IGHA1_HUMAN Immunoglobulin heavy constant alpha 1 OS=Homo sapiens OX=9606 GN=IGHA1 PE=1 SV=2 | 255 | 20201116_13832_1ng_glyco_hcd203040_compound15.21391.21391.4 | controllerType=0 controllerNumber=1 scan=21391 | 42.4757 |
| 18041:4 | 9 | R.LSLHRPALEDLLLGSEAN[+1095.397]LTC[+57.021]TLTGLR.D | HexNAc(3)Hex(3) | C[+57], N[+1095] | 1015.5074 | 4 | 4059.0076 | 4058.9949 | 0 | 3.1 | 127 | Specific | 709.16 | 468.87 | 468.87 | 14 | 42 | >sp\|P01876\|IGHA1_HUMAN Immunoglobulin heavy constant alpha 1 OS=Homo sapiens OX=9606 GN=IGHA1 PE=1 SV=2 | 255 | 20201116_13832_1ng_glyco_hcd203040_compound15.21555.21555.4 | controllerType=0 controllerNumber=1 scan=21555 | 42.8777 |
| 17990:4 | 9 | R.LSLHRPALEDLLLGSEAN[+1216.423]LTC[+57.021]TLTGLR.D | HexNAc(2)Hex(5) | C[+57], N[+1216] | 1045.7648 | 4 | 4180.0374 | 4180.0211 | 0 | 3.9 | 127 | Specific | 735.37 | 735.37 | 735.37 | 14 | 42 | >sp\|P01876\|IGHA1_HUMAN Immunoglobulin heavy constant alpha 1 OS=Homo sapiens OX=9606 GN=IGHA1 PE=1 SV=2 | 255 | 20201116_13832_1ng_glyco_hcd203040_compound15.21502.21502.4 | controllerType=0 controllerNumber=1 scan=21502 | 42.7357 |
| 17896:4 | 9 | R.LSLHRPALEDLLLGSEAN[+1663.608]LTC[+57.021]TLTGLR.D | HexNAc(5)Hex(4) | C[+57], N[+1664] | 1157.5594 | 4 | 4627.2156 | 4627.2064 | 0 | 2 | 127 | Specific | 755.32 | 689.35 | 689.35 | 15.17 | 42 | >sp\|P01876\|IGHA1_HUMAN Immunoglobulin heavy constant alpha 1 OS=Homo sapiens OX=9606 GN=IGHA1 PE=1 SV=2 | 255 | 20201116_13832_1ng_glyco_hcd203040_compound15.21403.21403.4 | controllerType=0 controllerNumber=1 scan=21403 | 42.5062 |
| 17998:4 | 9 | R.LSLHRPALEDLLLGSEAN[+1298.476]LTC[+57.021]TLTGLR.D | HexNAc(4)Hex(3) | C[+57], N[+1298] | 1066.278 | 4 | 4262.0902 | 4262.0742 | 0 | 3.7 | 127 | Specific | 765.92 | 765.92 | 765.92 | 15 | 42 | >sp\|P01876\|IGHA1_HUMAN Immunoglobulin heavy constant alpha 1 OS=Homo sapiens OX=9606 GN=IGHA1 PE=1 SV=2 | 255 | 20201116_13832_1ng_glyco_hcd203040_compound15.21510.21510.4 | controllerType=0 controllerNumber=1 scan=21510 | 42.7533 |
| 18876:4 | 9 | R.LSLHRPALEDLLLGSEAN[+2116.756]LTC[+57.021]TLTGLR.D | HexNAc(5)Hex(5)NeuAc(1) | C[+57], N[+2117] | 1270.8463 | 4 | 5080.3634 | 5080.3547 | 0 | 1.7 | 127 | Specific | 781.65 | 704.81 | 704.81 | 15.17 | 42 | >sp\|P01876\|IGHA1_HUMAN Immunoglobulin heavy constant alpha 1 OS=Homo sapiens OX=9606 GN=IGHA1 PE=1 SV=2 | 255 | 20201116_13832_1ng_glyco_hcd203040_compound15.22425.22425.4 | controllerType=0 controllerNumber=1 scan=22425 | 44.6743 |
| 17947:4 | 9 | R.LSLHRPALEDLLLGSEAN[+1460.529]LTC[+57.021]TLTGLR.D | HexNAc(4)Hex(4) | C[+57], N[+1461] | 1106.7896 | 4 | 4424.1365 | 4424.1271 | 0 | 2.1 | 127 | Specific | 797.65 | 797.65 | 797.65 | 15.17 | 42 | >sp\|P01876\|IGHA1_HUMAN Immunoglobulin heavy constant alpha 1 OS=Homo sapiens OX=9606 GN=IGHA1 PE=1 SV=2 | 255 | 20201116_13832_1ng_glyco_hcd203040_compound15.21456.21456.4 | controllerType=0 controllerNumber=1 scan=21456 | 42.6213 |
| 17929:4 | 9 | R.LSLHRPALEDLLLGSEAN[+1622.582]LTC[+57.021]TLTGLR.D | HexNAc(4)Hex(5) | C[+57], N[+1623] | 1147.3024 | 4 | 4586.1878 | 4586.1799 | 0 | 1.7 | 127 | Specific | 802.15 | 665.92 | 665.92 | 16.17 | 42 | >sp\|P01876\|IGHA1_HUMAN Immunoglobulin heavy constant alpha 1 OS=Homo sapiens OX=9606 GN=IGHA1 PE=1 SV=2 | 255 | 20201116_13832_1ng_glyco_hcd203040_compound15.21438.21438.4 | controllerType=0 controllerNumber=1 scan=21438 | 42.5858 |
| 18919:4 | 9 | R.LSLHRPALEDLLLGSEAN[+1872.651]LTC[+57.021]TLTGLR.D | HexNAc(3)Hex(6)NeuAc(1) | C[+57], N[+1873] | 1209.8206 | 4 | 4836.2606 | 4836.2487 | 0 | 2.4 | 127 | Specific | 815.69 | 763.99 | 763.99 | 16.17 | 42 | >sp\|P01876\|IGHA1_HUMAN Immunoglobulin heavy constant alpha 1 OS=Homo sapiens OX=9606 GN=IGHA1 PE=1 SV=2 | 255 | 20201116_13832_1ng_glyco_hcd203040_compound15.22469.22469.4 | controllerType=0 controllerNumber=1 scan=22469 | 44.7733 |
| 17949:4 | 9 | R.LSLHRPALEDLLLGSEAN[+1460.529]LTC[+57.021]TLTGLR.D | HexNAc(4)Hex(4) | C[+57], N[+1461] | 1106.7896 | 4 | 4424.1365 | 4424.1271 | 0 | 2.1 | 127 | Specific | 827.09 | 736.41 | 736.41 | 16.17 | 42 | >sp\|P01876\|IGHA1_HUMAN Immunoglobulin heavy constant alpha 1 OS=Homo sapiens OX=9606 GN=IGHA1 PE=1 SV=2 | 255 | 20201116_13832_1ng_glyco_hcd203040_compound15.21458.21458.4 | controllerType=0 controllerNumber=1 scan=21458 | 42.6272 |
| 17933:4 | 9 | R.LSLHRPALEDLLLGSEAN[+1419.502]LTC[+57.021]TLTGLR.D | HexNAc(3)Hex(5) | C[+57], N[+1420] | 1096.5328 | 4 | 4383.1092 | 4383.1005 | 0 | 2 | 127 | Specific | 835.78 | 835.78 | 835.78 | 16.17 | 42 | >sp\|P01876\|IGHA1_HUMAN Immunoglobulin heavy constant alpha 1 OS=Homo sapiens OX=9606 GN=IGHA1 PE=1 SV=2 | 255 | 20201116_13832_1ng_glyco_hcd203040_compound15.21442.21442.4 | controllerType=0 controllerNumber=1 scan=21442 | 42.5933 |
| 17847:4 | 9 | R.LSLHRPALEDLLLGSEAN[+1702.581]LTC[+57.021]TLTGLR.D | HexNAc(2)Hex(8) | C[+57], N[+1703] | 1167.3035 | 4 | 4666.1922 | 4666.1796 | 0 | 2.7 | 127 | Specific | 857.43 | 857.43 | 857.43 | 17.17 | 42 | >sp\|P01876\|IGHA1_HUMAN Immunoglobulin heavy constant alpha 1 OS=Homo sapiens OX=9606 GN=IGHA1 PE=1 SV=2 | 255 | 20201116_13832_1ng_glyco_hcd203040_compound15.21352.21352.4 | controllerType=0 controllerNumber=1 scan=21352 | 42.3737 |
| 18001:4 | 9 | R.LSLHRPALEDLLLGSEAN[+1298.476]LTC[+57.021]TLTGLR.D | HexNAc(4)Hex(3) | C[+57], N[+1298] | 1066.278 | 4 | 4262.0902 | 4262.0742 | 0 | 3.7 | 127 | Specific | 919.16 | 866.92 | 866.92 | 18 | 42 | >sp\|P01876\|IGHA1_HUMAN Immunoglobulin heavy constant alpha 1 OS=Homo sapiens OX=9606 GN=IGHA1 PE=1 SV=2 | 255 | 20201116_13832_1ng_glyco_hcd203040_compound15.21513.21513.4 | controllerType=0 controllerNumber=1 scan=21513 | 42.7618 |
| 17869:4 | 9 | R.LSLHRPALEDLLLGSEAN[+1540.529]LTC[+57.021]TLTGLR.D | HexNAc(2)Hex(7) | C[+57], N[+1541] | 1126.7901 | 4 | 4504.1385 | 4504.1268 | 0 | 2.6 | 127 | Specific | 919.84 | 880.81 | 880.81 | 18.17 | 42 | >sp\|P01876\|IGHA1_HUMAN Immunoglobulin heavy constant alpha 1 OS=Homo sapiens OX=9606 GN=IGHA1 PE=1 SV=2 | 255 | 20201116_13832_1ng_glyco_hcd203040_compound15.21375.21375.4 | controllerType=0 controllerNumber=1 scan=21375 | 42.4201 |
| 17886:4 | 9 | R.LSLHRPALEDLLLGSEAN[+1864.634]LTC[+57.021]TLTGLR.D | HexNAc(2)Hex(9) | C[+57], N[+1865] | 1207.8154 | 4 | 4828.2396 | 4828.2324 | 0 | 1.5 | 127 | Specific | 951.89 | 951.89 | 951.89 | 18.9 | 42 | >sp\|P01876\|IGHA1_HUMAN Immunoglobulin heavy constant alpha 1 OS=Homo sapiens OX=9606 GN=IGHA1 PE=1 SV=2 | 255 | 20201116_13832_1ng_glyco_hcd203040_compound15.21393.21393.4 | controllerType=0 controllerNumber=1 scan=21393 | 42.4801 |
| 17992:4 | 9 | R.LSLHRPALEDLLLGSEAN[+1216.423]LTC[+57.021]TLTGLR.D | HexNAc(2)Hex(5) | C[+57], N[+1216] | 1045.7648 | 4 | 4180.0374 | 4180.0211 | 0 | 3.9 | 127 | Specific | 955.37 | 891.21 | 891.21 | 19 | 42 | >sp\|P01876\|IGHA1_HUMAN Immunoglobulin heavy constant alpha 1 OS=Homo sapiens OX=9606 GN=IGHA1 PE=1 SV=2 | 255 | 20201116_13832_1ng_glyco_hcd203040_compound15.21504.21504.4 | controllerType=0 controllerNumber=1 scan=21504 | 42.7404 |
| 17850:4 | 9 | R.LSLHRPALEDLLLGSEAN[+1702.581]LTC[+57.021]TLTGLR.D | HexNAc(2)Hex(8) | C[+57], N[+1703] | 1167.3035 | 4 | 4666.1922 | 4666.1796 | 0 | 2.7 | 127 | Specific | 967.05 | 967.05 | 967.05 | 19.17 | 42 | >sp\|P01876\|IGHA1_HUMAN Immunoglobulin heavy constant alpha 1 OS=Homo sapiens OX=9606 GN=IGHA1 PE=1 SV=2 | 255 | 20201116_13832_1ng_glyco_hcd203040_compound15.21355.21355.4 | controllerType=0 controllerNumber=1 scan=21355 | 42.3795 |
| 17871:4 | 9 | R.LSLHRPALEDLLLGSEAN[+1540.529]LTC[+57.021]TLTGLR.D | HexNAc(2)Hex(7) | C[+57], N[+1541] | 1126.7901 | 4 | 4504.1385 | 4504.1268 | 0 | 2.6 | 127 | Specific | 981.89 | 981.89 | 981.89 | 19.17 | 42 | >sp\|P01876\|IGHA1_HUMAN Immunoglobulin heavy constant alpha 1 OS=Homo sapiens OX=9606 GN=IGHA1 PE=1 SV=2 | 255 | 20201116_13832_1ng_glyco_hcd203040_compound15.21377.21377.4 | controllerType=0 controllerNumber=1 scan=21377 | [illegible] |
| 11445:3 | 9 | R.LAGKPTHVN[+1501.555]VSVVMAEVDGTC[+57.021]Y.- | HexNAc(5)Hex(3) | C[+57], N[+1502] | 1283.5738 | 3 | 3848.7068 | 3848.6974 | 0 | 2.4 | 332 | CRagged | 584.38 | 584.38 | 584.38 | 11.46 | 42 | >sp\|P01876\|IGHA1_HUMAN Immunoglobulin heavy constant alpha 1 OS=Homo sapiens OX=9606 GN=IGHA1 PE=1 SV=2 | 255 | 20201116_13832_1ng_glyco_hcd203040_compound15.14665.14665.3 | controllerType=0 controllerNumber=1 scan=14665 | [illegible] |
| 11443:3 | 9 | R.LAGKPTHVN[+1501.555]VSVVMAEVDGTC[+57.021]Y.- | HexNAc(5)Hex(3) | C[+57], N[+1502] | 1283.5738 | 3 | 3848.7068 | 3848.6974 | 0 | 2.4 | 332 | CRagged | 664.25 | 659.87 | 659.87 | 13.46 | 42 | >sp\|P01876\|IGHA1_HUMAN Immunoglobulin heavy constant alpha 1 OS=Homo sapiens OX=9606 GN=IGHA1 PE=1 SV=2 | 255 | 20201116_13832_1ng_glyco_hcd203040_compound15.14663.14663.3 | controllerType=0 controllerNumber=1 scan=14663 | 30.8037 |
| 11284:3 | 9 | R.LAGKPTHVN[+1540.529]VSVVMAEVDGTC[+57.021]Y.- | HexNAc(2)Hex(7) | C[+57], N[+1541] | 1296.5629 | 3 | 3887.6742 | 3887.6706 | 0 | 0.9 | 332 | CRagged | 728.81 | 559.58 | 473.08 | 14.19 | 42 | >sp\|P01876\|IGHA1_HUMAN Immunoglobulin heavy constant alpha 1 OS=Homo sapiens OX=9606 GN=IGHA1 PE=1 SV=2 | 255 | 20201116_13832_1ng_glyco_hcd203040_compound15.14497.14497.3 | controllerType=0 controllerNumber=1 scan=14497 | 30.4742 |
| 11289:3 | 9 | R.LAGKPTHVN[+1540.529]VSVVMAEVDGTC[+57.021]Y.- | HexNAc(2)Hex(7) | C[+57], N[+1541] | 1296.5644 | 3 | 3887.6786 | 3887.6706 | 0 | 2.1 | 332 | CRagged | 754.83 | 754.83 | 487.14 | 15.46 | 42 | >sp\|P01876\|IGHA1_HUMAN Immunoglobulin heavy constant alpha 1 OS=Homo sapiens OX=9606 GN=IGHA1 PE=1 SV=2 | 255 | 20201116_13832_1ng_glyco_hcd203040_compound15.14503.14503.3 | controllerType=0 controllerNumber=1 scan=14503 | 30.4865 |
| 11423:3 | 9 | R.LAGKPTHVN[+1647.613]VSVVMAEVDGTC[+57.021]Y.- | HexNAc(5)Hex(3)Fuc(1) | C[+57], N[+1648] | 1332.2596 | 3 | 3994.7642 | 3994.7553 | 0 | 2.2 | 332 | CRagged | 765.11 | 765.11 | 765.11 | 15.46 | 42 | >sp\|P01876\|IGHA1_HUMAN Immunoglobulin heavy constant alpha 1 OS=Homo sapiens OX=9606 GN=IGHA1 PE=1 SV=2 | 255 | 20201116_13832_1ng_glyco_hcd203040_compound15.14642.14642.3 | controllerType=0 controllerNumber=1 scan=14642 | 30.7668 |
| 11425:3 | 9 | R.LAGKPTHVN[+1647.613]VSVVMAEVDGTC[+57.021]Y.- | HexNAc(5)Hex(3)Fuc(1) | C[+57], N[+1648] | 1332.2596 | 3 | 3994.7642 | 3994.7553 | 0 | 2.2 | 332 | CRagged | 783.13 | 648.2 | 648.2 | 15.46 | 42 | >sp\|P01876\|IGHA1_HUMAN Immunoglobulin heavy constant alpha 1 OS=Homo sapiens OX=9606 GN=IGHA1 PE=1 SV=2 | 255 | 20201116_13832_1ng_glyco_hcd203040_compound15.14644.14644.3 | controllerType=0 controllerNumber=1 scan=14644 | 30.7719 |
| 11291:3 | 9 | R.LAGKPTHVN[+1540.529]VSVVMAEVDGTC[+57.021]Y.- | HexNAc(2)Hex(7) | C[+57], N[+1541] | 1296.5644 | 3 | 3887.6786 | 3887.6706 | 0 | 2.1 | 332 | CRagged | 791.59 | 754.22 | 462.31 | 15.46 | 42 | >sp\|P01876\|IGHA1_HUMAN Immunoglobulin heavy constant alpha 1 OS=Homo sapiens OX=9606 GN=IGHA1 PE=1 SV=2 | 255 | 20201116_13832_1ng_glyco_hcd203040_compound15.14505.14505.3 | controllerType=0 controllerNumber=1 scan=14505 | 30.5091 |
| 11282:3 | 9 | R.LAGKPTHVN[+1540.529]VSVVMAEVDGTC[+57.021]Y.- | HexNAc(2)Hex(7) | C[+57], N[+1541] | 1296.5629 | 3 | 3887.6742 | 3887.6706 | 0 | 0.9 | 332 | CRagged | 837.9 | 837.9 | 511.08 | 16.19 | 42 | >sp\|P01876\|IGHA1_HUMAN Immunoglobulin heavy constant alpha 1 OS=Homo sapiens OX=9606 GN=IGHA1 PE=1 SV=2 | 255 | 20201116_13832_1ng_glyco_hcd203040_compound15.14495.14495.3 | controllerType=0 controllerNumber=1 scan=14495 | 30.4695 |
| 11313:3 | 9 | R.LAGKPTHVN[+1378.476]VSVVMAEVDGTC[+57.021]Y.- | HexNAc(2)Hex(6) | C[+57], N[+1378] | 1242.5458 | 3 | 3725.6229 | 3725.6178 | 0 | 1.4 | 332 | CRagged | 878.9 | 834.85 | 549.76 | 17.19 | 42 | >sp\|P01876\|IGHA1_HUMAN Immunoglobulin heavy constant alpha 1 OS=Homo sapiens OX=9606 GN=IGHA1 PE=1 SV=2 | 255 | 20201116_13832_1ng_glyco_hcd203040_compound15.14528.14528.3 | controllerType=0 controllerNumber=1 scan=14528 | 30.5596 |
| 11369:3 | 9 | R.LAGKPTHVN[+1216.423]VSVVMAEVDGTC[+57.021]Y.- | HexNAc(2)Hex(5) | C[+57], N[+1216] | 1188.5286 | 3 | 3563.5713 | 3563.5649 | 0 | 1.8 | 332 | CRagged | 889.55 | 827.72 | 532.87 | 17.46 | 42 | >sp\|P01876\|IGHA1_HUMAN Immunoglobulin heavy constant alpha 1 OS=Homo sapiens OX=9606 GN=IGHA1 PE=1 SV=2 | 255 | 20201116_13832_1ng_glyco_hcd203040_compound15.14586.14586.3 | controllerType=0 controllerNumber=1 scan=14586 | 30.6691 |
| 11310:3 | 9 | R.LAGKPTHVN[+1378.476]VSVVMAEVDGTC[+57.021]Y.- | HexNAc(2)Hex(6) | C[+57], N[+1378] | 1242.5458 | 3 | 3725.6229 | 3725.6178 | 0 | 1.4 | 332 | CRagged | 894.85 | 894.85 | 557.94 | 17.19 | 42 | >sp\|P01876\|IGHA1_HUMAN Immunoglobulin heavy constant alpha 1 OS=Homo sapiens OX=9606 GN=IGHA1 PE=1 SV=2 | 255 | 20201116_13832_1ng_glyco_hcd203040_compound15.14525.14525.3 | controllerType=0 controllerNumber=1 scan=14525 | 30.5496 |
| 11367:3 | 9 | R.LAGKPTHVN[+1216.423]VSVVMAEVDGTC[+57.021]Y.- | HexNAc(2)Hex(5) | C[+57], N[+1216] | 1188.5286 | 3 | 3563.5713 | 3563.5649 | 0 | 1.8 | 332 | CRagged | 910.16 | 910.16 | 576.96 | 18.46 | 42 | >sp\|P01876\|IGHA1_HUMAN Immunoglobulin heavy constant alpha 1 OS=Homo sapiens OX=9606 GN=IGHA1 PE=1 SV=2 | 255 | 20201116_13832_1ng_glyco_hcd203040_compound15.14584.14584.3 | controllerType=0 controllerNumber=1 scan=14584 | 30.6632 |
| 07603:3 | 12 | R.LSPN[+2350.830]ASAEHLELR.W | HexNAc(4)Hex(5)Fuc(1)NeuAc(2) | N[+2351] | 1263.201 | 3 | 3787.5884 | 3787.5796 | 0 | 2.3 | 52 | Specific | 322.34 | 322.34 | 322.34 | 6.97 | 37 | >sp\|Q13410\|BT1A1_HUMAN Butyrophilin subfamily 1 member A1 OS=Homo sapiens OX=9606 GN=BTN1A1 PE=1 SV=3 | 179 | 20201116_13832_1ng_glyco_hcd203040_compound15.10672.10672.3 | controllerType=0 controllerNumber=1 scan=10672 | 24.125 |
| 07605:3 | 12 | R.LSPN[+2350.830]ASAEHLELR.W | HexNAc(4)Hex(5)Fuc(1)NeuAc(2) | N[+2351] | 1263.201 | 3 | 3787.5884 | 3787.5796 | 0 | 2.3 | 52 | Specific | 346.38 | 333.14 | 333.14 | 7.15 | 37 | >sp\|Q13410\|BT1A1_HUMAN Butyrophilin subfamily 1 member A1 OS=Homo sapiens OX=9606 GN=BTN1A1 PE=1 SV=3 | 179 | 20201116_13832_1ng_glyco_hcd203040_compound15.10674.10674.3 | controllerType=0 controllerNumber=1 scan=10674 | 24.1287 |
| 05545:3 | 12 | R.LSPN[+2205.793]ASAEHLELR.W | HexNAc(4)Hex(5)Fuc(2)NeuAc(1) | N[+2206] | 1214.8567 | 3 | 3642.5557 | 3642.5421 | 0 | 3.7 | 52 | Specific | 547.97 | 508.13 | 508.13 | 9.59 | 37 | >sp\|Q13410\|BT1A1_HUMAN Butyrophilin subfamily 1 member A1 OS=Homo sapiens OX=9606 GN=BTN1A1 PE=1 SV=3 | 179 | 20201116_13832_1ng_glyco_hcd203040_compound15.8544.8544.3 | controllerType=0 controllerNumber=1 scan=8544 | 20.5977 |
| 05547:3 | 12 | R.LSPN[+2205.793]ASAEHLELR.W | HexNAc(4)Hex(5)Fuc(2)NeuAc(1) | N[+2206] | 1214.8567 | 3 | 3642.5557 | 3642.5421 | 0 | 3.7 | 52 | Specific | 561.27 | 423.74 | 423.74 | 9.59 | 37 | >sp\|Q13410\|BT1A1_HUMAN Butyrophilin subfamily 1 member A1 OS=Homo sapiens OX=9606 GN=BTN1A1 PE=1 SV=3 | 179 | 20201116_13832_1ng_glyco_hcd203040_compound15.8546.8546.3 | controllerType=0 controllerNumber=1 scan=8546 | 20.6023 |
| 28268:3 | 19 | R.QLAHQSN[+2188.740]STNIFFSPVSIATAFAMLSLGTK.A | HexNAc(2)Hex(11) | N[+2189] | 1790.7975 | 3 | 5370.3779 | 5370.3749 | 0 | 0.6 | 64 | Specific | 359.17 | 339.46 | 339.46 | 7.15 | 30 | >sp\|P01009\|A1AT_HUMAN Alpha-1-antitrypsin OS=Homo sapiens OX=9606 GN=SERPINA1 PE=1 SV=3 | 642 | 20201116_13832_1ng_glyco_hcd203040_compound15.32184.32184.3 | controllerType=0 controllerNumber=1 scan=32184 | 62.5565 |
| 28270:3 | 19 | R.QLAHQSN[+2188.740]STNIFFSPVSIATAFAMLSLGTK.A | HexNAc(2)Hex(11) | N[+2189] | 1790.7975 | 3 | 5370.3779 | 5370.3749 | 0 | 0.6 | 64 | Specific | 375.16 | 304.55 | 304.55 | 7.09 | 30 | >sp\|P01009\|A1AT_HUMAN Alpha-1-antitrypsin OS=Homo sapiens OX=9606 GN=SERPINA1 PE=1 SV=3 | 642 | 20201116_13832_1ng_glyco_hcd203040_compound15.32186.32186.3 | controllerType=0 controllerNumber=1 scan=32186 | 62.5612 |
| 24700:3 | 19 | R.QLAHQSN[+2204.772]STNIFFSPVSIATAFAMLSLGTK.A | HexNAc(4)Hex(5)NeuAc(2) | N[+2205] | 1796.1462 | 3 | 5386.4241 | 5386.4075 | 0 | 3.1 | 64 | Specific | 534.69 | 533.73 | 533.73 | 9.81 | 30 | >sp\|P01009\|A1AT_HUMAN Alpha-1-antitrypsin OS=Homo sapiens OX=9606 GN=SERPINA1 PE=1 SV=3 | 642 | 20201116_13832_1ng_glyco_hcd203040_compound15.28481.28481.3 | controllerType=0 controllerNumber=1 scan=28481 | 55.9133 |
| 24702:3 | 19 | R.QLAHQSN[+2204.772]STNIFFSPVSIATAFAMLSLGTK.A | HexNAc(4)Hex(5)NeuAc(2) | N[+2205] | 1796.1462 | 3 | 5386.4241 | 5386.4075 | 0 | 3.1 | 64 | Specific | 649.47 | 595.59 | 595.59 | 12.17 | 30 | >sp\|P01009\|A1AT_HUMAN Alpha-1-antitrypsin OS=Homo sapiens OX=9606 GN=SERPINA1 PE=1 SV=3 | 642 | 20201116_13832_1ng_glyco_hcd203040_compound15.28483.28483.3 | controllerType=0 controllerNumber=1 scan=28483 | 55.919 |
| 24699:4 | 19 | R.QLAHQSN[+2204.772]STNIFFSPVSIATAFAMLSLGTK.A | HexNAc(4)Hex(5)NeuAc(2) | N[+2205] | 1347.3588 | 4 | 5386.4132 | 5386.4075 | 0 | 1.1 | 64 | Specific | 717.33 | 631.81 | 515.93 | 13.7 | 30 | >sp\|P01009\|A1AT_HUMAN Alpha-1-antitrypsin OS=Homo sapiens OX=9606 GN=SERPINA1 PE=1 SV=3 | 642 | 20201116_13832_1ng_glyco_hcd203040_compound15.28480.28480.4 | controllerType=0 controllerNumber=1 scan=28480 | 55.9119 |
| 24701:4 | 19 | R.QLAHQSN[+2204.772]STNIFFSPVSIATAFAMLSLGTK.A | HexNAc(4)Hex(5)NeuAc(2) | N[+2205] | 1347.3588 | 4 | 5386.4132 | 5386.4075 | 0 | 1.1 | 64 | Specific | 827.87 | 699.12 | 581 | 15.7 | 30 | >sp\|P01009\|A1AT_HUMAN Alpha-1-antitrypsin OS=Homo sapiens OX=9606 GN=SERPINA1 PE=1 SV=3 | 642 | 20201116_13832_1ng_glyco_hcd203040_compound15.28482.28482.4 | controllerType=0 controllerNumber=1 scan=28482 | 55.9171 |
| 20268:4 | 19 | K.YLGN[+2204.772]ATAIFFLPDEGKLQHLENELTHDIITK.F | HexNAc(4)Hex(5)NeuAc(2) | N[+2205] | 1437.1615 | 4 | 5745.6242 | 5745.6098 | 0 | 2.5 | 268 | Specific | 753.33 | 595.67 | 595.67 | 14.47 | 30 | >sp\|P01009\|A1AT_HUMAN Alpha-1-antitrypsin OS=Homo sapiens OX=9606 GN=SERPINA1 PE=1 SV=3 | 642 | 20201116_13832_1ng_glyco_hcd203040_compound15.23876.23876.6 | controllerType=0 controllerNumber=1 scan=23876 | 47.5656 |
| 20290:4 | 19 | K.YLGN[+2204.772]ATAIFFLPDEGKLQHLENELTHDIITK.F | HexNAc(4)Hex(5)NeuAc(2) | N[+2205] | 1437.1641 | 4 | 5745.6344 | 5745.6098 | 0 | 4.3 | 268 | Specific | 867.49 | 867.49 | 867.49 | 16.3 | 30 | >sp\|P01009\|A1AT_HUMAN Alpha-1-antitrypsin OS=Homo sapiens OX=9606 GN=SERPINA1 PE=1 SV=3 | 642 | 20201116_13832_1ng_glyco_hcd203040_compound15.23899.23899.4 | controllerType=0 controllerNumber=1 scan=23899 | 47.6251 |
| 20266:4 | 19 | K.YLGN[+2204.772]ATAIFFLPDEGKLQHLENELTHDIITK.F | HexNAc(4)Hex(5)NeuAc(2) | N[+2205] | 1437.1615 | 4 | 5745.6242 | 5745.6098 | 0 | 2.5 | 268 | Specific | 892.85 | 575.03 | 575.03 | 16.47 | 30 | >sp\|P01009\|A1AT_HUMAN Alpha-1-antitrypsin OS=Homo sapiens OX=9606 GN=SERPINA1 PE=1 SV=3 | 642 | 20201116_13832_1ng_glyco_hcd203040_compound15.23874.23874.6 | controllerType=0 controllerNumber=1 scan=23874 | 47.5595 |
| 20303:5 | 19 | K.YLGN[+2204.772]ATAIFFLPDEGKLQHLENELTHDIITK.F | HexNAc(4)Hex(5)NeuAc(2) | N[+2205] | 1149.932 | 5 | 5745.6309 | 5745.6098 | 0 | 3.7 | 268 | Specific | 925.74 | 713.92 | 713.92 | 17.8 | 30 | >sp\|P01009\|A1AT_HUMAN Alpha-1-antitrypsin OS=Homo sapiens OX=9606 GN=SERPINA1 PE=1 SV=3 | 642 | 20201116_13832_1ng_glyco_hcd203040_compound15.23913.23913.5 | controllerType=0 controllerNumber=1 scan=23913 | 47.6537 |
| 20300:5 | 19 | K.YLGN[+2204.772]ATAIFFLPDEGKLQHLENELTHDIITK.F | HexNAc(4)Hex(5)NeuAc(2) | N[+2205] | 1149.932 | 5 | 5745.6309 | 5745.6098 | 0 | 3.7 | 268 | Specific | 944.94 | 767.6 | 767.6 | 17.8 | 30 | >sp\|P01009\|A1AT_HUMAN Alpha-1-antitrypsin OS=Homo sapiens OX=9606 GN=SERPINA1 PE=1 SV=3 | 642 | 20201116_13832_1ng_glyco_hcd203040_compound15.23910.23910.5 | controllerType=0 controllerNumber=1 scan=23910 | 47.6485 |
| 20287:4 | 19 | K.YLGN[+2204.772]ATAIFFLPDEGKLQHLENELTHDIITK.F | HexNAc(4)Hex(5)NeuAc(2) | N[+2205] | 1437.1641 | 4 | 5745.6344 | 5745.6098 | 0 | 4.3 | 268 | Specific | 1248.48 | 1103.57 | 1103.57 | 23.3 | 30 | >sp\|P01009\|A1AT_HUMAN Alpha-1-antitrypsin OS=Homo sapiens OX=9606 GN=SERPINA1 PE=1 SV=3 | 642 | 20201116_13832_1ng_glyco_hcd203040_compound15.23896.23896.4 | controllerType=0 controllerNumber=1 scan=23896 | 47.6092 |
| 14121:4 | 25 | K.LKELPGVC[+57.021]N[+2059.735]ETMMALWEEC[+57.021]KPC[+57.021]LK.Q | HexNAc(4)Hex(5)Fuc(1)NeuAc(1) | C[+57]*3, N[+2060] | 1249.7894 | 4 | 4996.1359 | 4996.1196 | 0 | 3.3 | 95 | Specific | 684.1 | 684.1 | 684.1 | 12.95 | 21 | >sp\|P10909\|CLUS_HUMAN Clusterin OS=Homo sapiens OX=9606 GN=CLU PE=1 SV=1 | 1467 | 20201116_13832_1ng_glyco_hcd203040_compound15.17449.17449.4 | controllerType=0 controllerNumber=1 scan=17449 | 35.4111 |
| 14119:4 | 25 | K.LKELPGVC[+57.021]N[+2059.735]ETMMALWEEC[+57.021]KPC[+57.021]LK.Q | HexNAc(4)Hex(5)Fuc(1)NeuAc(1) | C[+57]*3, N[+2060] | 1249.7894 | 4 | 4996.1359 | 4996.1196 | 0 | 3.3 | 95 | Specific | 689.49 | 689.49 | 689.49 | 12.95 | 21 | >sp\|P10909\|CLUS_HUMAN Clusterin OS=Homo sapiens OX=9606 GN=CLU PE=1 SV=1 | 1467 | 20201116_13832_1ng_glyco_hcd203040_compound15.17447.17447.4 | controllerType=0 controllerNumber=1 scan=17447 | 35.4067 |
| 14037:4 | 25 | K.LKELPGVC[+57.021]N[+2205.793]ETMMALWEEC[+57.021]KPC[+57.021]LK.Q | HexNAc(4)Hex(5)Fuc(2)NeuAc(1) | C[+57]*3, N[+2206] | 1286.3037 | 4 | 5142.193 | 5142.1776 | 0 | 3 | 95 | Specific | 710.43 | 698.97 | 698.97 | 14.12 | 21 | >sp\|P10909\|CLUS_HUMAN Clusterin OS=Homo sapiens OX=9606 GN=CLU PE=1 SV=1 | 1467 | 20201116_13832_1ng_glyco_hcd203040_compound15.17362.17362.4 | controllerType=0 controllerNumber=1 scan=17362 | 35.2622 |
| 14039:4 | 25 | K.LKELPGVC[+57.021]N[+2205.793]ETMMALWEEC[+57.021]KPC[+57.021]LK.Q | HexNAc(4)Hex(5)Fuc(2)NeuAc(1) | C[+57]*3, N[+2206] | 1286.3037 | 4 | 5142.193 | 5142.1776 | 0 | 3 | 95 | Specific | 757.14 | 757.14 | 757.14 | 15.12 | 21 | >sp\|P10909\|CLUS_HUMAN Clusterin OS=Homo sapiens OX=9606 GN=CLU PE=1 SV=1 | 1467 | 20201116_13832_1ng_glyco_hcd203040_compound15.17364.17364.4 | controllerType=0 controllerNumber=1 scan=17364 | 35.2666 |
| 21335:4 | 25 | K.MLN[+2059.735]TSSLLEQLNEQFNWVSR.L | HexNAc(4)Hex(5)Fuc(1)NeuAc(1) | N[+2060] | 1117.9911 | 4 | 4468.9426 | 4468.9217 | 0 | 4.7 | 352 | Specific | 249.08 | 230.94 | 230.94 | 3.91 | 21 | >sp\|P10909\|CLUS_HUMAN Clusterin OS=Homo sapiens OX=9606 GN=CLU PE=1 SV=1 | 1467 | 20201116_13832_1ng_glyco_hcd203040_compound15.24986.24986.4 | controllerType=0 controllerNumber=1 scan=24986 | 49.825 |
| 21338:4 | 25 | K.MLN[+2059.735]TSSLLEQLNEQFNWVSR.L | HexNAc(4)Hex(5)Fuc(1)NeuAc(1) | N[+2060] | 1117.9911 | 4 | 4468.9426 | 4468.9217 | 0 | 4.7 | 352 | Specific | 264.59 | 252.92 | 252.92 | 4.49 | 21 | >sp\|P10909\|CLUS_HUMAN Clusterin OS=Homo sapiens OX=9606 GN=CLU PE=1 SV=1 | 1467 | 20201116_13832_1ng_glyco_hcd203040_compound15.24989.24989.4 | controllerType=0 controllerNumber=1 scan=24989 | 49.8318 |
| 23850:3 | 25 | K.MLN[+2350.830]TSSLLEQLNEQFNWVSR.L | HexNAc(4)Hex(5)Fuc(1)NeuAc(2) | N[+2351] | 1587.3517 | 3 | 4760.0407 | 4760.0171 | 0 | 5 | 352 | Specific | 757.77 | 566.43 | 566.43 | 14.29 | 21 | >sp\|P10909\|CLUS_HUMAN Clusterin OS=Homo sapiens OX=9606 GN=CLU PE=1 SV=1 | 1467 | 20201116_13832_1ng_glyco_hcd203040_compound15.27600.27600.3 | controllerType=0 controllerNumber=1 scan=27600 | 54.357 |
| 23848:3 | 25 | K.MLN[+2350.830]TSSLLEQLNEQFNWVSR.L | HexNAc(4)Hex(5)Fuc(1)NeuAc(2) | N[+2351] | 1587.3517 | 3 | 4760.0407 | 4760.0171 | 0 | 5 | 352 | Specific | 804.39 | 537.25 | 537.25 | 15.29 | 21 | >sp\|P10909\|CLUS_HUMAN Clusterin OS=Homo sapiens OX=9606 GN=CLU PE=1 SV=1 | 1467 | 20201116_13832_1ng_glyco_hcd203040_compound15.27598.27598.3 | controllerType=0 controllerNumber=1 scan=27598 | 54.3524 |
| 10233:3 | 25 | R.LAN[+2205.793]LTQGEDQYYLR.V | HexNAc(4)Hex(5)Fuc(2)NeuAc(1) | N[+2206] | 1297.2167 | 3 | 3889.6356 | 3889.6265 | 0 | 2.3 | 372 | Specific | 235.08 | 235.08 | 235.08 | 6 | 21 | >sp\|P10909\|CLUS_HUMAN Clusterin OS=Homo sapiens OX=9606 GN=CLU PE=1 SV=1 | 1467 | 20201116_13832_1ng_glyco_hcd203040_compound15.13411.13411.3 | controllerType=0 controllerNumber=1 scan=13411 | 28.6903 |
| 10235:3 | 25 | R.LAN[+2205.793]LTQGEDQYYLR.V | HexNAc(4)Hex(5)Fuc(2)NeuAc(1) | N[+2206] | 1297.2167 | 3 | 3889.6356 | 3889.6265 | 0 | 2.3 | 372 | Specific | 258.58 | 258.58 | 258.58 | 6.31 | 21 | >sp\|P10909\|CLUS_HUMAN Clusterin OS=Homo sapiens OX=9606 GN=CLU PE=1 SV=1 | 1467 | 20201116_13832_1ng_glyco_hcd203040_compound15.13413.13413.3 | controllerType=0 controllerNumber=1 scan=13413 | 28.6943 |
| 22511:4 | 36 | K.[+42.011]VFPLSLDSTPQDGNVVVAC[+57.021]LVQGFFPQEPLSVTWSESGQN[+1606 | HexNAc(4)Hex(4)Fuc(1) | .[42], C[+57], N[+1607] | 1606.9894 | 4 | 6424.9357 | 6424.9587 | 0 | -3.6 | 8 | Specific | 313.65 | 297.31 | 297.31 | 5.55 | 17 | >sp\|P01877\|IGHA2_HUMAN Immunoglobulin heavy constant alpha 2 OS=Homo sapiens OX=9606 GN=IGHA2 PE=1 SV=4 | 576 | 20201116_13832_1ng_glyco_hcd203040_compound15.26205.26205.4 | controllerType=0 controllerNumber=1 scan=26205 | 51.9281 |
| 22513:4 | 36 | K.[+42.011]VFPLSLDSTPQDGNVVVAC[+57.021]LVQGFFPQEPLSVTWSESGQN[+1606 | HexNAc(4)Hex(4)Fuc(1) | .[42], C[+57], N[+1607] | 1606.9894 | 4 | 6424.9357 | 6424.9587 | 0 | -3.6 | 8 | Specific | 331.19 | 297.75 | 297.75 | 5.03 | 17 | >sp\|P01877\|IGHA2_HUMAN Immunoglobulin heavy constant alpha 2 OS=Homo sapiens OX=9606 GN=IGHA2 PE=1 SV=4 | 576 | 20201116_13832_1ng_glyco_hcd203040_compound15.26207.26207.4 | controllerType=0 controllerNumber=1 scan=26207 | 51.9337 |
| 01268:3 | 36 | K.HYTN[+1647.613]SSQDVTVPC[+57.021]R.V | HexNAc(5)Hex(3)Fuc(1) | C[+57], N[+1648] | 1104.4629 | 3 | 3311.3742 | 3311.3625 | 0 | 3.5 | 89 | Specific | 327.61 | 301.87 | 301.87 | 6.67 | 17 | >sp\|P01877\|IGHA2_HUMAN Immunoglobulin heavy constant alpha 2 OS=Homo sapiens OX=9606 GN=IGHA2 PE=1 SV=4 | 576 | 20201116_13832_1ng_glyco_hcd203040_compound15.4111.4111.3 | controllerType=0 controllerNumber=1 scan=4111 | 13.0078 |
| 01266:3 | 36 | K.HYTN[+1647.613]SSQDVTVPC[+57.021]R.V | HexNAc(5)Hex(3)Fuc(1) | C[+57], N[+1648] | 1104.4629 | 3 | 3311.3742 | 3311.3625 | 0 | 3.5 | 89 | Specific | 397.71 | 311.69 | 311.69 | 6.91 | 17 | >sp\|P01877\|IGHA2_HUMAN Immunoglobulin heavy constant alpha 2 OS=Homo sapiens OX=9606 GN=IGHA2 PE=1 SV=4 | 576 | 20201116_13832_1ng_glyco_hcd203040_compound15.4109.4109.3 | controllerType=0 controllerNumber=1 scan=4109 | 13.0032 |
| 01243:3 | 36 | K.HYTN[+1647.613]SSQDVTVPC[+57.021]R.V | HexNAc(5)Hex(3)Fuc(1) | C[+57], N[+1648] | 1104.4621 | 3 | 3311.3716 | 3311.3625 | 0 | 2.7 | 89 | Specific | 404.32 | 345.91 | 345.91 | 7.67 | 17 | >sp\|P01877\|IGHA2_HUMAN Immunoglobulin heavy constant alpha 2 OS=Homo sapiens OX=9606 GN=IGHA2 PE=1 SV=4 | 576 | 20201116_13832_1ng_glyco_hcd203040_compound15.4085.4085.3 | controllerType=0 controllerNumber=1 scan=4085 | 12.9494 |
| 01245:3 | 36 | K.HYTN[+1647.613]SSQDVTVPC[+57.021]R.V | HexNAc(5)Hex(3)Fuc(1) | C[+57], N[+1648] | 1104.4621 | 3 | 3311.3716 | 3311.3625 | 0 | 2.7 | 89 | Specific | 521.27 | 408.33 | 408.33 | 9.54 | 17 | >sp\|P01877\|IGHA2_HUMAN Immunoglobulin heavy constant alpha 2 OS=Homo sapiens OX=9606 GN=IGHA2 PE=1 SV=4 | 576 | 20201116_13832_1ng_glyco_hcd203040_compound15.4087.4087.3 | controllerType=0 controllerNumber=1 scan=4087 | 12.9531 |
| 03265:4 | 36 | K.HYTN[+1647.613]SSQDVTVPC[+57.021]RVPPPPPC[+57.021]C[+57.021]HPR.L | HexNAc(5)Hex(3)Fuc(1) | C[+57]*3, N[+1648] | 1152.2482 | 4 | 4605.971 | 4605.9689 | 0 | 0.5 | 89 | Specific | 233.54 | 148.22 | 148.22 | 4.73 | 17 | >sp\|P01877\|IGHA2_HUMAN Immunoglobulin heavy constant alpha 2 OS=Homo sapiens OX=9606 GN=IGHA2 PE=1 SV=4 | 576 | 20201116_13832_1ng_glyco_hcd203040_compound15.6181.6181.4 | controllerType=0 controllerNumber=1 scan=6181 | 16.6497 |
| 03267:4 | 36 | K.HYTN[+1647.613]SSQDVTVPC[+57.021]RVPPPPPC[+57.021]C[+57.021]HPR.L | HexNAc(5)Hex(3)Fuc(1) | C[+57]*3, N[+1648] | 1152.2482 | 4 | 4605.971 | 4605.9689 | 0 | 0.5 | 89 | Specific | 236.59 | 236.59 | 236.59 | 5.65 | 17 | >sp\|P01877\|IGHA2_HUMAN Immunoglobulin heavy constant alpha 2 OS=Homo sapiens OX=9606 GN=IGHA2 PE=1 SV=4 | 576 | 20201116_13832_1ng_glyco_hcd203040_compound15.6183.6183.4 | controllerType=0 controllerNumber=1 scan=6183 | 16.6537 |
| 02905:3 | 36 | K.TPLTAN[+1809.666]ITK.S | HexNAc(5)Hex(4)Fuc(1) | N[+1810] | 923.4151 | 3 | 2768.2306 | 2768.2228 | 0 | 2.8 | 200 | Specific | 266.01 | 114 | 114 | 4.3 | 17 | >sp\|P01877\|IGHA2_HUMAN Immunoglobulin heavy constant alpha 2 OS=Homo sapiens OX=9606 GN=IGHA2 PE=1 SV=4 | 576 | 20201116_13832_1ng_glyco_hcd203040_compound15.5808.5808.3 | controllerType=0 controllerNumber=1 scan=5808 | 15.997 |
| 02943:2 | 36 | K.TPLTAN[+1444.534]ITK.S | HexNAc(4)Hex(3)Fuc(1) | N[+1445] | 1202.0533 | 2 | 2403.0993 | 2403.0906 | 0 | 3.6 | 200 | Specific | 311.46 | 87.93 | 87.93 | 4.33 | 17 | >sp\|P01877\|IGHA2_HUMAN Immunoglobulin heavy constant alpha 2 OS=Homo sapiens OX=9606 GN=IGHA2 PE=1 SV=4 | 576 | 20201116_13832_1ng_glyco_hcd203040_compound15.5847.5847.2 | controllerType=0 controllerNumber=1 scan=5847 | 16.0651 |
| 02945:2 | 36 | K.TPLTAN[+1444.534]ITK.S | HexNAc(4)Hex(3)Fuc(1) | N[+1445] | 1202.0533 | 2 | 2403.0993 | 2403.0906 | 0 | 3.6 | 200 | Specific | 312.59 | 312.59 | 312.59 | 6.27 | 17 | >sp\|P01877\|IGHA2_HUMAN Immunoglobulin heavy constant alpha 2 OS=Homo sapiens OX=9606 GN=IGHA2 PE=1 SV=4 | 576 | 20201116_13832_1ng_glyco_hcd203040_compound15.5849.5849.2 | controllerType=0 controllerNumber=1 scan=5849 | 16.069 |
| 02904:2 | 36 | K.TPLTAN[+1809.666]ITK.S | HexNAc(5)Hex(4)Fuc(1) | N[+1810] | 1384.6177 | 2 | 2768.2282 | 2768.2228 | 0 | 1.9 | 200 | Specific | 324.51 | 202.17 | 202.17 | 6.17 | 17 | >sp\|P01877\|IGHA2_HUMAN Immunoglobulin heavy constant alpha 2 OS=Homo sapiens OX=9606 GN=IGHA2 PE=1 SV=4 | 576 | 20201116_13832_1ng_glyco_hcd203040_compound15.5807.5807.2 | controllerType=0 controllerNumber=1 scan=5807 | 15.9945 |
| 02902:2 | 36 | K.TPLTAN[+1809.666]ITK.S | HexNAc(5)Hex(4)Fuc(1) | N[+1810] | 1384.6177 | 2 | 2768.2282 | 2768.2228 | 0 | 1.9 | 200 | Specific | 324.98 | 321.85 | 321.85 | 6.73 | 17 | >sp\|P01877\|IGHA2_HUMAN Immunoglobulin heavy constant alpha 2 OS=Homo sapiens OX=9606 GN=IGHA2 PE=1 SV=4 | 576 | 20201116_13832_1ng_glyco_hcd203040_compound15.5805.5805.2 | controllerType=0 controllerNumber=1 scan=5805 | 15.9904 |
| 02959:3 | 36 | K.TPLTAN[+1647.613]ITK.S | HexNAc(5)Hex(3)Fuc(1) | N[+1648] | 869.3984 | 3 | 2606.1807 | 2606.17 | 0 | 4.1 | 200 | Specific | 344.6 | 121.62 | 121.62 | 4.83 | 17 | >sp\|P01877\|IGHA2_HUMAN Immunoglobulin heavy constant alpha 2 OS=Homo sapiens OX=9606 GN=IGHA2 PE=1 SV=4 | 576 | 20201116_13832_1ng_glyco_hcd203040_compound15.5864.5864.3 | controllerType=0 controllerNumber=1 scan=5864 | 16.0971 |
| 02961:3 | 36 | K.TPLTAN[+1647.613]ITK.S | HexNAc(5)Hex(3)Fuc(1) | N[+1648] | 869.3984 | 3 | 2606.1807 | 2606.17 | 0 | 4.1 | 200 | Specific | 491.64 | 242.44 | 242.44 | 6.76 | 17 | >sp\|P01877\|IGHA2_HUMAN Immunoglobulin heavy constant alpha 2 OS=Homo sapiens OX=9606 GN=IGHA2 PE=1 SV=4 | 576 | 20201116_13832_1ng_glyco_hcd203040_compound15.5866.5866.3 | controllerType=0 controllerNumber=1 scan=5866 | 16.1011 |
| 02951:2 | 36 | K.TPLTAN[+1647.613]ITK.S | HexNAc(5)Hex(3)Fuc(1) | N[+1648] | 1303.5943 | 2 | 2606.1813 | 2606.17 | 0 | 4.3 | 200 | Specific | 569.07 | 254.36 | 254.36 | 7.31 | 17 | >sp\|P01877\|IGHA2_HUMAN Immunoglobulin heavy constant alpha 2 OS=Homo sapiens OX=9606 GN=IGHA2 PE=1 SV=4 | 576 | 20201116_13832_1ng_glyco_hcd203040_compound15.5856.5856.2 | controllerType=0 controllerNumber=1 scan=5856 | 16.0848 |
| 02949:2 | 36 | K.TPLTAN[+1647.613]ITK.S | HexNAc(5)Hex(3)Fuc(1) | N[+1648] | 1303.5943 | 2 | 2606.1813 | 2606.17 | 0 | 4.3 | 200 | Specific | 581.04 | 337.07 | 337.07 | 8.79 | 17 | >sp\|P01877\|IGHA2_HUMAN Immunoglobulin heavy constant alpha 2 OS=Homo sapiens OX=9606 GN=IGHA2 PE=1 SV=4 | 576 | 20201116_13832_1ng_glyco_hcd203040_compound15.5854.5854.2 | controllerType=0 controllerNumber=1 scan=5854 | 16.0799 |
| 10222:3 | 36 | R.MAGKPTHIN[+1378.476]VSVVMAEADGTC[+57.021]Y.- | HexNAc(2)Hex(6) | C[+57], N[+1378] | 1243.8619 | 3 | 3729.5711 | 3729.5585 | 0 | 3.4 | 319 | CRagged | 548.34 | 548.34 | 352.91 | 10.15 | 17 | >sp\|P01877\|IGHA2_HUMAN Immunoglobulin heavy constant alpha 2 OS=Homo sapiens OX=9606 GN=IGHA2 PE=1 SV=4 | 576 | 20201116_13832_1ng_glyco_hcd203040_compound15.13399.13399.3 | controllerType=0 controllerNumber=1 scan=13399 | 28.6666 |
| 10224:3 | 36 | R.MAGKPTHIN[+1378.476]VSVVMAEADGTC[+57.021]Y.- | HexNAc(2)Hex(6) | C[+57], N[+1378] | 1243.8619 | 3 | 3729.5711 | 3729.5585 | 0 | 3.4 | 319 | CRagged | 652.36 | 652.36 | 397.19 | 13.29 | 17 | >sp\|P01877\|IGHA2_HUMAN Immunoglobulin heavy constant alpha 2 OS=Homo sapiens OX=9606 GN=IGHA2 PE=1 SV=4 | 576 | 20201116_13832_1ng_glyco_hcd203040_compound15.13401.13401.3 | controllerType=0 controllerNumber=1 scan=13401 | 28.6719 |
| 17853:3 | 68 | R.NYIVPILWLN[+1864.634]ETGTIGDEKANMFR.S | HexNAc(2)Hex(9) | N[+1865] | 1553.6948 | 3 | 4659.0697 | 4659.0574 | 0 | 2.6 | 408 | Specific | 566.43 | 566.43 | 566.43 | 9.99 | 12 | >sp\|P16671\|CD36_HUMAN Platelet glycoprotein 4 OS=Homo sapiens OX=9606 GN=CD36 PE=1 SV=2 | 812 | 20201116_13832_1ng_glyco_hcd203040_compound15.21358.21358.3 | controllerType=0 controllerNumber=1 scan=21358 | 42.3849 |
| 17851:3 | 68 | R.NYIVPILWLN[+1864.634]ETGTIGDEKANMFR.S | HexNAc(2)Hex(9) | N[+1865] | 1553.6948 | 3 | 4659.0697 | 4659.0574 | 0 | 2.6 | 408 | Specific | 639.9 | 639.9 | 639.9 | 12.12 | 12 | >sp\|P16671\|CD36_HUMAN Platelet glycoprotein 4 OS=Homo sapiens OX=9606 GN=CD36 PE=1 SV=2 | 812 | 20201116_13832_1ng_glyco_hcd203040_compound15.21356.21356.3 | controllerType=0 controllerNumber=1 scan=21356 | 42.3812 |
| 06564:3 | 316 | R.[+42.011]VEDAYILTC[+57.021]N[+730.264]VSLEYEK.T | HexNAc(2)Hex(2) | .[42], C[+57], N[+730] | 940.0955 | 3 | 2818.2721 | 2818.2486 | 0 | 8.4 | 225 | Specific | 257.29 | 208.79 | 208.79 | 3.52 | 4 | >sp\|P40227\|TCPZ_HUMAN T-complex protein 1 subunit zeta OS=Homo sapiens OX=9606 GN=CCT6A PE=1 SV=3 | 1115 | 20201116_13832_1ng_glyco_hcd203040_compound15.9597.9597.3 | controllerType=0 controllerNumber=1 scan=9597 | 22.3352 |
| 06566:3 | 316 | R.[+42.011]VEDAYILTC[+57.021]N[+730.264]VSLEYEK.T | HexNAc(2)Hex(2) | .[42], C[+57], N[+730] | 940.0955 | 3 | 2818.2721 | 2818.2486 | 0 | 8.4 | 225 | Specific | 272.9 | 201.19 | 201.19 | 3.19 | 4 | >sp\|P40227\|TCPZ_HUMAN T-complex protein 1 subunit zeta OS=Homo sapiens OX=9606 GN=CCT6A PE=1 SV=3 | 1115 | 20201116_13832_1ng_glyco_hcd203040_compound15.9599.9599.3 | controllerType=0 controllerNumber=1 scan=9599 | 22.3395 |
| 20756:4 | 1 | R.RTAGWNVPIGTLRPFLN[+2350.830]WTGPPEPIEAAVAR.F | HexNAc(4)Hex(5)Fuc(1)NeuAc(2) | N[+2351] | 1435.1715 | 4 | 5737.6642 | 5737.6424 | 0 | 3.8 | 140 | Specific | 239.33 | 236.4 | 236.4 | 7.4 | 128 | >sp\|P02788\|TRFL_HUMAN Lactotransferrin OS=Homo sapiens OX=9606 GN=LTF PE=1 SV=6 | 1270 | 20201115_15212_1ng_glyco_hcd203040_compound15.23694.23694.4 | controllerType=0 controllerNumber=1 scan=23694 | 45.8294 |
| 21065:3 | 1 | R.[+42.011]RTAGWNVPIGTLRPFLN[+1864.634]WTGPPEPIEAAVAR.F | HexNAc(2)Hex(9) | .[42], N[+1865] | 1765.1642 | 3 | 5293.478 | 5293.4568 | 0 | 4 | 140 | Specific | 299.38 | 299.38 | 299.38 | 7.69 | 128 | >sp\|P02788\|TRFL_HUMAN Lactotransferrin OS=Homo sapiens OX=9606 GN=LTF PE=1 SV=6 | 1270 | 20201115_15212_1ng_glyco_hcd203040_compound15.24017.24017.3 | controllerType=0 controllerNumber=1 scan=24017 | 46.4904 |
| 19719:3 | 1 | R.RTAGWNVPIGTLRPFLN[+2059.735]WTGPPEPIEAAVAR.F | HexNAc(4)Hex(5)Fuc(1)NeuAc(1) | N[+2060] | 1816.1928 | 3 | 5446.564 | 5446.547 | 0 | 3.1 | 140 | Specific | 315.05 | 315.05 | 315.05 | 7.87 | 128 | >sp\|P02788\|TRFL_HUMAN Lactotransferrin OS=Homo sapiens OX=9606 GN=LTF PE=1 SV=6 | 1270 | 20201115_15212_1ng_glyco_hcd203040_compound15.22613.22613.3 | controllerType=0 controllerNumber=1 scan=22613 | 43.5676 |
| 19600:4 | 1 | R.RTAGWNVPIGTLRPFLN[+2570.925]WTGPPEPIEAAVAR.F | HexNAc(5)Hex(6)Fuc(2)NeuAc(1) | N[+2571] | 1490.1951 | 4 | 5957.7585 | 5957.7371 | 0 | 3.6 | 140 | Specific | 452 | 400.08 | 400.08 | 10.85 | 128 | >sp\|P02788\|TRFL_HUMAN Lactotransferrin OS=Homo sapiens OX=9606 GN=LTF PE=1 SV=6 | 1270 | 20201115_15212_1ng_glyco_hcd203040_compound15.22489.22489.4 | controllerType=0 controllerNumber=1 scan=22489 | 43.3119 |
| 19704:4 | 1 | R.RTAGWNVPIGTLRPFLN[+2350.830]WTGPPEPIEAAVAR.F | HexNAc(4)Hex(5)Fuc(1)NeuAc(2) | N[+2351] | 1435.1757 | 4 | 5737.6808 | 5737.6424 | 0 | 6.7 | 140 | Specific | 514.76 | 463.49 | 463.49 | 10.13 | 128 | >sp\|P02788\|TRFL_HUMAN Lactotransferrin OS=Homo sapiens OX=9606 GN=LTF PE=1 SV=6 | 1270 | 20201115_15212_1ng_glyco_hcd203040_compound15.22597.22597.2 | controllerType=0 controllerNumber=1 scan=22597 | 43.5364 |
| 19597:4 | 1 | R.RTAGWNVPIGTLRPFLN[+2570.925]WTGPPEPIEAAVAR.F | HexNAc(5)Hex(6)Fuc(2)NeuAc(1) | N[+2571] | 1490.1951 | 4 | 5957.7585 | 5957.7371 | 0 | 3.6 | 140 | Specific | 519.52 | 519.52 | 519.52 | 11.14 | 128 | >sp\|P02788\|TRFL_HUMAN Lactotransferrin OS=Homo sapiens OX=9606 GN=LTF PE=1 SV=6 | 1270 | 20201115_15212_1ng_glyco_hcd203040_compound15.22486.22486.4 | controllerType=0 controllerNumber=1 scan=22486 | 43.2926 |
| 18775:4 | 1 | R.RTAGWNVPIGTLRPFLN[+1768.640]WTGPPEPIEAAVAR.F | HexNAc(4)Hex(5)Fuc(1) | N[+1769] | 1289.6229 | 4 | 5155.4699 | 5155.4516 | 0 | 3.6 | 140 | Specific | 540.81 | 416.49 | 416.49 | 11.32 | 128 | >sp\|P02788\|TRFL_HUMAN Lactotransferrin OS=Homo sapiens OX=9606 GN=LTF PE=1 SV=6 | 1270 | 20201115_15212_1ng_glyco_hcd203040_compound15.21625.21625.4 | controllerType=0 controllerNumber=1 scan=21625 | 41.3504 |
| 20721:4 | 1 | R.RTAGWNVPIGTLRPFLN[+2350.830]WTGPPEPIEAAVAR.F | HexNAc(4)Hex(5)Fuc(1)NeuAc(2) | N[+2351] | 1435.1672 | 4 | 5737.6471 | 5737.6424 | 0 | 0.8 | 140 | Specific | 591.41 | 497.63 | 497.63 | 12.29 | 128 | >sp\|P02788\|TRFL_HUMAN Lactotransferrin OS=Homo sapiens OX=9606 GN=LTF PE=1 SV=6 | 1270 | 20201115_15212_1ng_glyco_hcd203040_compound15.23657.23657.2 | controllerType=0 controllerNumber=1 scan=23657 | 45.767 |
| 18778:4 | 1 | R.RTAGWNVPIGTLRPFLN[+1768.640]WTGPPEPIEAAVAR.F | HexNAc(4)Hex(5)Fuc(1) | N[+1769] | 1289.6229 | 4 | 5155.4699 | 5155.4516 | 0 | 3.6 | 140 | Specific | 592.07 | 497.58 | 497.58 | 12.84 | 128 | >sp\|P02788\|TRFL_HUMAN Lactotransferrin OS=Homo sapiens OX=9606 GN=LTF PE=1 SV=6 | 1270 | 20201115_15212_1ng_glyco_hcd203040_compound15.21628.21628.4 | controllerType=0 controllerNumber=1 scan=21628 | 41.356 |

Table S4. Cont.

| Query #:z | Protein Rank | Glycopeptide | Glycan | Modification Type(s) | Observed m/z | z | Observed (M+H) | Calc. mass (M+H) | Off-by-x error | Mass error (ppm) | StartPosition | Cleavage | Score | Delta | Delta Mod | \|Log Prob\| | # of unique peptides | Protein Name | Protein DB number | Comment | Scan # | Scan Time |
|---|---|---|---|---|---|---|---|---|---|---|---|---|---|---|---|---|---|---|---|---|---|---|
| 19707:4 | 1 | R.RTAGWNVPIGTLRPFLN[+2350.830]WTGPPEPIEAAVAR.F | HexNAc(4)Hex(5)Fuc(1)NeuAc(2) | N[+2351] | 1435.1757 | 4 | 5737.6808 | 5737.6424 | 0 | 6.7 | 140 | Specific | 616.88 | 539 | 539 | 12.83 | 128 | >sp\|P02788\|TRFL_HUMAN Lactotransferrin OS=Homo sapiens OX=9606 GN=LTF PE=1 SV=6 | 1270 | 20201115_15212_1ng_glyco_hcd203040_compound15.22600.22600.2 | controllerType=0 controllerNumber=1 scan=22600 | 43.542 |
| 20759:4 | 1 | R.RTAGWNVPIGTLRPFLN[+2350.830]WTGPPEPIEAAVAR.F | HexNAc(4)Hex(5)Fuc(1)NeuAc(2) | N[+2351] | 1435.1715 | 4 | 5737.6642 | 5737.6424 | 0 | 3.8 | 140 | Specific | 623.83 | 509.77 | 509.77 | 13.84 | 128 | >sp\|P02788\|TRFL_HUMAN Lactotransferrin OS=Homo sapiens OX=9606 GN=LTF PE=1 SV=6 | 1270 | 20201115_15212_1ng_glyco_hcd203040_compound15.23697.23697.4 | controllerType=0 controllerNumber=1 scan=23697 | 45.8333 |
| 20718:4 | 1 | R.RTAGWNVPIGTLRPFLN[+2350.830]WTGPPEPIEAAVAR.F | HexNAc(4)Hex(5)Fuc(1)NeuAc(2) | N[+2351] | 1435.1672 | 4 | 5737.6471 | 5737.6424 | 0 | 0.8 | 140 | Specific | 633.22 | 522.99 | 522.99 | 13.29 | 128 | >sp\|P02788\|TRFL_HUMAN Lactotransferrin OS=Homo sapiens OX=9606 GN=LTF PE=1 SV=6 | 1270 | 20201115_15212_1ng_glyco_hcd203040_compound15.23654.23654.2 | controllerType=0 controllerNumber=1 scan=23654 | 45.7573 |
| 22277:4 | 1 | R.[+42.011]TAGWNVPIGTLRPFLN[+2336.851]WTGPPEPIEAAVAR.F | HexNAc(6)Hex(6)Fuc(1) | .[42], N[+2337] | 1403.1436 | 4 | 5609.5524 | 5609.5726 | 0 | -3.6 | 141 | Specific | 305.73 | 305.73 | 305.73 | 6.6 | 128 | >sp\|P02788\|TRFL_HUMAN Lactotransferrin OS=Homo sapiens OX=9606 GN=LTF PE=1 SV=6 | 1270 | 20201115_15212_1ng_glyco_hcd203040_compound15.25287.25287.2 | controllerType=0 controllerNumber=1 scan=25287 | 49.2708 |
| 20963:3 | 1 | R.TAGWNVPIGTLRPFLN[+2205.793]WTGPPEPIEAAVAR.F | HexNAc(4)Hex(5)Fuc(2)NeuAc(1) | N[+2206] | 1812.843 | 3 | 5436.5144 | 5436.5038 | 0 | 2 | 141 | Specific | 349.12 | 349.12 | 349.12 | 8.89 | 128 | >sp\|P02788\|TRFL_HUMAN Lactotransferrin OS=Homo sapiens OX=9606 GN=LTF PE=1 SV=6 | 1270 | 20201115_15212_1ng_glyco_hcd203040_compound15.23911.23911.3 | controllerType=0 controllerNumber=1 scan=23911 | 46.3084 |
| 22279:4 | 1 | R.[+42.011]TAGWNVPIGTLRPFLN[+2336.851]WTGPPEPIEAAVAR.F | HexNAc(6)Hex(6)Fuc(1) | .[42], N[+2337] | 1403.1436 | 4 | 5609.5524 | 5609.5726 | 0 | -3.6 | 141 | Specific | 376.29 | 365.55 | 365.55 | 7.42 | 128 | >sp\|P02788\|TRFL_HUMAN Lactotransferrin OS=Homo sapiens OX=9606 GN=LTF PE=1 SV=6 | 1270 | 20201115_15212_1ng_glyco_hcd203040_compound15.25289.25289.2 | controllerType=0 controllerNumber=1 scan=25289 | 49.2819 |
| 21040:3 | 1 | R.[+42.011]TAGWNVPIGTLRPFLN[+2018.708]WTGPPEPIEAAVAR.F | HexNAc(3)Hex(6)Fuc(1)NeuAc(1) | .[42], N[+2019] | 1764.4952 | 3 | 5291.471 | 5291.4299 | 0 | 7.8 | 141 | Specific | 391.78 | 391.78 | 391.78 | 9.35 | 128 | >sp\|P02788\|TRFL_HUMAN Lactotransferrin OS=Homo sapiens OX=9606 GN=LTF PE=1 SV=6 | 1270 | 20201115_15212_1ng_glyco_hcd203040_compound15.23991.23991.3 | controllerType=0 controllerNumber=1 scan=23991 | 46.4554 |
| 22241:3 | 1 | R.TAGWNVPIGTLRPFLN[+2350.830]WTGPPEPIEAAVAR.F | HexNAc(4)Hex(5)Fuc(1)NeuAc(2) | N[+2351] | 1861.1915 | 3 | 5581.5599 | 5581.5413 | 0 | 3.3 | 141 | Specific | 482.42 | 305.02 | 305.02 | 9.14 | 128 | >sp\|P02788\|TRFL_HUMAN Lactotransferrin OS=Homo sapiens OX=9606 GN=LTF PE=1 SV=6 | 1270 | 20201115_15212_1ng_glyco_hcd203040_compound15.25249.25249.2 | controllerType=0 controllerNumber=1 scan=25249 | 49.1801 |
| 19908:3 | 1 | R.TAGWNVPIGTLRPFLN[+2279.830]WTGPPEPIEAAVAR.F | HexNAc(5)Hex(6)Fuc(2) | N[+2280] | 1837.5233 | 3 | 5510.5553 | 5510.5406 | 0 | 2.7 | 141 | Specific | 512.4 | 397.54 | 397.54 | 11.2 | 128 | >sp\|P02788\|TRFL_HUMAN Lactotransferrin OS=Homo sapiens OX=9606 GN=LTF PE=1 SV=6 | 1270 | 20201115_15212_1ng_glyco_hcd203040_compound15.22810.22810.3 | controllerType=0 controllerNumber=1 scan=22810 | 43.9735 |
| 20965:3 | 1 | R.TAGWNVPIGTLRPFLN[+2205.793]WTGPPEPIEAAVAR.F | HexNAc(4)Hex(5)Fuc(2)NeuAc(1) | N[+2206] | 1812.843 | 3 | 5436.5144 | 5436.5038 | 0 | 2 | 141 | Specific | 530.02 | 481.3 | 481.3 | 11.12 | 128 | >sp\|P02788\|TRFL_HUMAN Lactotransferrin OS=Homo sapiens OX=9606 GN=LTF PE=1 SV=6 | 1270 | 20201115_15212_1ng_glyco_hcd203040_compound15.23913.23913.3 | controllerType=0 controllerNumber=1 scan=23913 | 46.3122 |
| 22208:3 | 1 | R.TAGWNVPIGTLRPFLN[+2350.830]WTGPPEPIEAAVAR.F | HexNAc(4)Hex(5)Fuc(1)NeuAc(2) | N[+2351] | 1861.1914 | 3 | 5581.5596 | 5581.5413 | 0 | 3.3 | 141 | Specific | 534.55 | 338 | 338 | 9.74 | 128 | >sp\|P02788\|TRFL_HUMAN Lactotransferrin OS=Homo sapiens OX=9606 GN=LTF PE=1 SV=6 | 1270 | 20201115_15212_1ng_glyco_hcd203040_compound15.25215.25215.3 | controllerType=0 controllerNumber=1 scan=25215 | 49.1066 |
| 20922:3 | 1 | R.TAGWNVPIGTLRPFLN[+2351.851]WTGPPEPIEAAVAR.F | HexNAc(4)Hex(5)Fuc(3)NeuAc(1) | N[+2352] | 1861.5291 | 3 | 5582.5729 | 5582.5617 | 0 | 2 | 141 | Specific | 545.22 | 480.75 | 480.75 | 11.3 | 128 | >sp\|P02788\|TRFL_HUMAN Lactotransferrin OS=Homo sapiens OX=9606 GN=LTF PE=1 SV=6 | 1270 | 20201115_15212_1ng_glyco_hcd203040_compound15.23868.23868.3 | controllerType=0 controllerNumber=1 scan=23868 | 46.2114 |
| 22244:3 | 1 | R.TAGWNVPIGTLRPFLN[+2350.830]WTGPPEPIEAAVAR.F | HexNAc(4)Hex(5)Fuc(1)NeuAc(2) | N[+2351] | 1861.1915 | 3 | 5581.5599 | 5581.5413 | 0 | 3.3 | 141 | Specific | 566.21 | 400.91 | 400.91 | 11.63 | 128 | >sp\|P02788\|TRFL_HUMAN Lactotransferrin OS=Homo sapiens OX=9606 GN=LTF PE=1 SV=6 | 1270 | 20201115_15212_1ng_glyco_hcd203040_compound15.25252.25252.2 | controllerType=0 controllerNumber=1 scan=25252 | 49.1862 |
| 20925:3 | 1 | R.TAGWNVPIGTLRPFLN[+2351.851]WTGPPEPIEAAVAR.F | HexNAc(4)Hex(5)Fuc(3)NeuAc(1) | N[+2352] | 1861.5291 | 3 | 5582.5729 | 5582.5617 | 0 | 2 | 141 | Specific | 567.44 | 516.5 | 516.5 | 11.55 | 128 | >sp\|P02788\|TRFL_HUMAN Lactotransferrin OS=Homo sapiens OX=9606 GN=LTF PE=1 SV=6 | 1270 | 20201115_15212_1ng_glyco_hcd203040_compound15.23871.23871.3 | controllerType=0 controllerNumber=1 scan=23871 | 46.2174 |
| 20923:4 | 1 | R.TAGWNVPIGTLRPFLN[+2351.851]WTGPPEPIEAAVAR.F | HexNAc(4)Hex(5)Fuc(3)NeuAc(1) | N[+2352] | 1396.3997 | 4 | 5582.5768 | 5582.5617 | 0 | 2.7 | 141 | Specific | 617.83 | 566.51 | 566.51 | 13.92 | 128 | >sp\|P02788\|TRFL_HUMAN Lactotransferrin OS=Homo sapiens OX=9606 GN=LTF PE=1 SV=6 | 1270 | 20201115_15212_1ng_glyco_hcd203040_compound15.23869.23869.4 | controllerType=0 controllerNumber=1 scan=23869 | 46.2126 |
| 22206:3 | 1 | R.TAGWNVPIGTLRPFLN[+2350.830]WTGPPEPIEAAVAR.F | HexNAc(4)Hex(5)Fuc(1)NeuAc(2) | N[+2351] | 1861.1914 | 3 | 5581.5596 | 5581.5413 | 0 | 3.3 | 141 | Specific | 624.12 | 448.31 | 448.31 | 13.9 | 128 | >sp\|P02788\|TRFL_HUMAN Lactotransferrin OS=Homo sapiens OX=9606 GN=LTF PE=1 SV=6 | 1270 | 20201115_15212_1ng_glyco_hcd203040_compound15.25213.25213.3 | controllerType=0 controllerNumber=1 scan=25213 | 49.1 |
| 20964:5 | 1 | R.TAGWNVPIGTLRPFLN[+2205.793]WTGPPEPIEAAVAR.F | HexNAc(4)Hex(5)Fuc(2)NeuAc(1) | N[+2206] | 1088.1079 | 5 | 5436.5105 | 5436.5038 | 0 | 1.2 | 141 | Specific | 628.03 | 548.11 | 548.11 | 13.67 | 128 | >sp\|P02788\|TRFL_HUMAN Lactotransferrin OS=Homo sapiens OX=9606 GN=LTF PE=1 SV=6 | 1270 | 20201115_15212_1ng_glyco_hcd203040_compound15.23912.23912.5 | controllerType=0 controllerNumber=1 scan=23912 | 46.3099 |
| 19900:4 | 1 | R.TAGWNVPIGTLRPFLN[+2279.830]WTGPPEPIEAAVAR.F | HexNAc(5)Hex(6)Fuc(2) | N[+2280] | 1378.3953 | 4 | 5510.5592 | 5510.5406 | 0 | 3.4 | 141 | Specific | 656.09 | 618.02 | 618.02 | 14.92 | 128 | >sp\|P02788\|TRFL_HUMAN Lactotransferrin OS=Homo sapiens OX=9606 GN=LTF PE=1 SV=6 | 1270 | 20201115_15212_1ng_glyco_hcd203040_compound15.22802.22802.4 | controllerType=0 controllerNumber=1 scan=22802 | 43.9566 |
| 20915:4 | 1 | R.TAGWNVPIGTLRPFLN[+2205.793]WTGPPEPIEAAVAR.F | HexNAc(4)Hex(5)Fuc(2)NeuAc(1) | N[+2206] | 1359.8851 | 4 | 5436.5187 | 5436.5038 | 0 | 2.7 | 141 | Specific | 679.27 | 597.98 | 597.98 | 14.92 | 128 | >sp\|P02788\|TRFL_HUMAN Lactotransferrin OS=Homo sapiens OX=9606 GN=LTF PE=1 SV=6 | 1270 | 20201115_15212_1ng_glyco_hcd203040_compound15.23861.23861.4 | controllerType=0 controllerNumber=1 scan=23861 | 46.1975 |
| 20942:3 | 1 | R.TAGWNVPIGTLRPFLN[+2059.735]WTGPPEPIEAAVAR.F | HexNAc(4)Hex(5)Fuc(1)NeuAc(1) | N[+2060] | 1764.1607 | 3 | 5290.4676 | 5290.4459 | 0 | 4.1 | 141 | Specific | 681.87 | 550.37 | 550.37 | 14.84 | 128 | >sp\|P02788\|TRFL_HUMAN Lactotransferrin OS=Homo sapiens OX=9606 GN=LTF PE=1 SV=6 | 1270 | 20201115_15212_1ng_glyco_hcd203040_compound15.23889.23889.3 | controllerType=0 controllerNumber=1 scan=23889 | 46.2568 |
| 19903:4 | 1 | R.TAGWNVPIGTLRPFLN[+2279.830]WTGPPEPIEAAVAR.F | HexNAc(5)Hex(6)Fuc(2) | N[+2280] | 1378.3953 | 4 | 5510.5592 | 5510.5406 | 0 | 3.4 | 141 | Specific | 704.11 | 622.87 | 622.87 | 15.92 | 128 | >sp\|P02788\|TRFL_HUMAN Lactotransferrin OS=Homo sapiens OX=9606 GN=LTF PE=1 SV=6 | 1270 | 20201115_15212_1ng_glyco_hcd203040_compound15.22805.22805.4 | controllerType=0 controllerNumber=1 scan=22805 | 43.966 |
| 21156:4 | 1 | R.TAGWNVPIGTLRPFLN[+2059.735]WTGPPEPIEAAVAR.F | HexNAc(4)Hex(5)Fuc(1)NeuAc(1) | N[+2060] | 1323.3727 | 4 | 5290.4689 | 5290.4459 | 0 | 4.4 | 141 | Specific | 718.78 | 671.24 | 671.24 | 15.86 | 128 | >sp\|P02788\|TRFL_HUMAN Lactotransferrin OS=Homo sapiens OX=9606 GN=LTF PE=1 SV=6 | 1270 | 20201115_15212_1ng_glyco_hcd203040_compound15.24116.24116.2 | controllerType=0 controllerNumber=1 scan=24116 | 46.6589 |
| 20940:3 | 1 | R.TAGWNVPIGTLRPFLN[+2059.735]WTGPPEPIEAAVAR.F | HexNAc(4)Hex(5)Fuc(1)NeuAc(1) | N[+2060] | 1764.1607 | 3 | 5290.4676 | 5290.4459 | 0 | 4.1 | 141 | Specific | 755.26 | 648.52 | 648.52 | 16.84 | 128 | >sp\|P02788\|TRFL_HUMAN Lactotransferrin OS=Homo sapiens OX=9606 GN=LTF PE=1 SV=6 | 1270 | 20201115_15212_1ng_glyco_hcd203040_compound15.23887.23887.3 | controllerType=0 controllerNumber=1 scan=23887 | 46.2506 |
| 20986:4 | 1 | R.TAGWNVPIGTLRPFLN[+2059.735]WTGPPEPIEAAVAR.F | HexNAc(4)Hex(5)Fuc(1)NeuAc(1) | N[+2060] | 1323.371 | 4 | 5290.4621 | 5290.4459 | 0 | 3.1 | 141 | Specific | 786.25 | 712.37 | 712.37 | 16.92 | 128 | >sp\|P02788\|TRFL_HUMAN Lactotransferrin OS=Homo sapiens OX=9606 GN=LTF PE=1 SV=6 | 1270 | 20201115_15212_1ng_glyco_hcd203040_compound15.23935.23935.4 | controllerType=0 controllerNumber=1 scan=23935 | 46.3711 |
| 20926:4 | 1 | R.TAGWNVPIGTLRPFLN[+2351.851]WTGPPEPIEAAVAR.F | HexNAc(4)Hex(5)Fuc(3)NeuAc(1) | N[+2352] | 1396.3997 | 4 | 5582.5768 | 5582.5617 | 0 | 2.7 | 141 | Specific | 853.9 | 808.3 | 808.3 | 18.92 | 128 | >sp\|P02788\|TRFL_HUMAN Lactotransferrin OS=Homo sapiens OX=9606 GN=LTF PE=1 SV=6 | 1270 | 20201115_15212_1ng_glyco_hcd203040_compound15.23872.23872.4 | controllerType=0 controllerNumber=1 scan=23872 | 46.2193 |
| 21159:4 | 1 | R.TAGWNVPIGTLRPFLN[+2059.735]WTGPPEPIEAAVAR.F | HexNAc(4)Hex(5)Fuc(1)NeuAc(1) | N[+2060] | 1323.3727 | 4 | 5290.4689 | 5290.4459 | 0 | 4.4 | 141 | Specific | 876.64 | 694.91 | 694.91 | 18.86 | 128 | >sp\|P02788\|TRFL_HUMAN Lactotransferrin OS=Homo sapiens OX=9606 GN=LTF PE=1 SV=6 | 1270 | 20201115_15212_1ng_glyco_hcd203040_compound15.24119.24119.2 | controllerType=0 controllerNumber=1 scan=24119 | 46.666 |
| 19955:4 | 1 | R.TAGWNVPIGTLRPFLN[+1768.640]WTGPPEPIEAAVAR.F | HexNAc(4)Hex(5)Fuc(1) | N[+1769] | 1250.5977 | 4 | 4999.3688 | 4999.3504 | 0 | 3.7 | 141 | Specific | 884.55 | 813.13 | 813.13 | 18.86 | 128 | >sp\|P02788\|TRFL_HUMAN Lactotransferrin OS=Homo sapiens OX=9606 GN=LTF PE=1 SV=6 | 1270 | 20201115_15212_1ng_glyco_hcd203040_compound15.22859.22859.2 | controllerType=0 controllerNumber=1 scan=22859 | 44.0703 |
| 20917:4 | 1 | R.TAGWNVPIGTLRPFLN[+2205.793]WTGPPEPIEAAVAR.F | HexNAc(4)Hex(5)Fuc(2)NeuAc(1) | N[+2206] | 1359.8851 | 4 | 5436.5187 | 5436.5038 | 0 | 2.7 | 141 | Specific | 887.23 | 724.87 | 724.87 | 18.92 | 128 | >sp\|P02788\|TRFL_HUMAN Lactotransferrin OS=Homo sapiens OX=9606 GN=LTF PE=1 SV=6 | 1270 | 20201115_15212_1ng_glyco_hcd203040_compound15.23863.23863.4 | controllerType=0 controllerNumber=1 scan=23863 | 46.2045 |
| 20966:5 | 1 | R.TAGWNVPIGTLRPFLN[+2205.793]WTGPPEPIEAAVAR.F | HexNAc(4)Hex(5)Fuc(2)NeuAc(1) | N[+2206] | 1088.1079 | 5 | 5436.5105 | 5436.5038 | 0 | 1.2 | 141 | Specific | 918.97 | 711.41 | 711.41 | 19.67 | 128 | >sp\|P02788\|TRFL_HUMAN Lactotransferrin OS=Homo sapiens OX=9606 GN=LTF PE=1 SV=6 | 1270 | 20201115_15212_1ng_glyco_hcd203040_compound15.23914.23914.5 | controllerType=0 controllerNumber=1 scan=23914 | 46.3143 |
| 19958:4 | 1 | R.TAGWNVPIGTLRPFLN[+1768.640]WTGPPEPIEAAVAR.F | HexNAc(4)Hex(5)Fuc(1) | N[+1769] | 1250.5977 | 4 | 4999.3688 | 4999.3504 | 0 | 3.7 | 141 | Specific | 989.09 | 818.45 | 818.45 | 20.86 | 128 | >sp\|P02788\|TRFL_HUMAN Lactotransferrin OS=Homo sapiens OX=9606 GN=LTF PE=1 SV=6 | 1270 | 20201115_15212_1ng_glyco_hcd203040_compound15.22862.22862.2 | controllerType=0 controllerNumber=1 scan=22862 | 44.0759 |
| 18302:3 | 1 | R.TAGWNIPMGLLFN[+2278.809]QTGSC[+57.021]K.F | HexNAc(5)Hex(6)NeuAc(1) | C[+57], N[+2279] | 1458.6224 | 3 | 4373.8526 | 4373.8192 | 0 | 7.6 | 485 | Specific | 289.7 | 289.7 | 207.91 | 6.97 | 128 | >sp\|P02788\|TRFL_HUMAN Lactotransferrin OS=Homo sapiens OX=9606 GN=LTF PE=1 SV=6 | 1270 | 20201115_15212_1ng_glyco_hcd203040_compound15.21132.21132.3 | controllerType=0 controllerNumber=1 scan=21132 | 40.3712 |
| 18885:3 | 1 | R.TAGWNIPMGLLFN[+1216.423]QTGSC[+57.021]K.F | HexNAc(2)Hex(5) | C[+57], N[+1216] | 1104.4934 | 3 | 3311.4658 | 3311.4328 | 0 | 10 | 485 | Specific | 332.56 | 332.56 | 100.63 | 5.39 | 128 | >sp\|P02788\|TRFL_HUMAN Lactotransferrin OS=Homo sapiens OX=9606 GN=LTF PE=1 SV=6 | 1270 | 20201115_15212_1ng_glyco_hcd203040_compound15.21740.21740.3 | controllerType=0 controllerNumber=1 scan=21740 | 41.6055 |
| 20325:4 | 1 | R.TAGWNIPMGLLFN[+2205.793]QTGSC[+57.021]K.F | HexNAc(4)Hex(5)Fuc(2)NeuAc(1) | C[+57], N[+2206] | 1075.9591 | 4 | 4300.8148 | 4300.8028 | 0 | 2.8 | 485 | Specific | 360.74 | 300.76 | 300.76 | 8.88 | 128 | >sp\|P02788\|TRFL_HUMAN Lactotransferrin OS=Homo sapiens OX=9606 GN=LTF PE=1 SV=6 | 1270 | 20201115_15212_1ng_glyco_hcd203040_compound15.23244.23244.4 | controllerType=0 controllerNumber=1 scan=23244 | 44.8651 |
| 20411:3 | 1 | R.TAGWNIPMGLLFN[+1038.375]QTGSC[+57.021]K.F | HexNAc(2)Hex(3)Fuc(1) | C[+57], N[+1038] | 1045.1231 | 3 | 3133.3547 | 3133.3851 | 0 | -9.7 | 485 | Specific | 390.74 | 350.73 | 350.73 | 4.68 | 128 | >sp\|P02788\|TRFL_HUMAN Lactotransferrin OS=Homo sapiens OX=9606 GN=LTF PE=1 SV=6 | 1270 | 20201115_15212_1ng_glyco_hcd203040_compound15.23334.23334.4 | controllerType=0 controllerNumber=1 scan=23334 | 45.0821 |
| 22473:3 | 1 | R.TAGWNIPMGLLFN[+2204.772]QTGSC[+57.021]K.F | HexNAc(4)Hex(5)NeuAc(2) | C[+57], N[+2205] | 1433.9382 | 3 | 4299.7999 | 4299.7824 | 0 | 4.1 | 485 | Specific | 455.98 | 425.45 | 425.45 | 10.6 | 128 | >sp\|P02788\|TRFL_HUMAN Lactotransferrin OS=Homo sapiens OX=9606 GN=LTF PE=1 SV=6 | 1270 | 20201115_15212_1ng_glyco_hcd203040_compound15.25492.25492.3 | controllerType=0 controllerNumber=1 scan=25492 | 49.782 |
| 18798:3 | 1 | R.TAGWNIPMGLLFN[+1460.529]QTGSC[+57.021]K.F | HexNAc(4)Hex(4) | C[+57], N[+1461] | 1185.8561 | 3 | 3555.5539 | 3555.5387 | 0 | 4.3 | 485 | Specific | 582.59 | 557.13 | 236.11 | 12.59 | 128 | >sp\|P02788\|TRFL_HUMAN Lactotransferrin OS=Homo sapiens OX=9606 GN=LTF PE=1 SV=6 | 1270 | 20201115_15212_1ng_glyco_hcd203040_compound15.21649.21649.3 | controllerType=0 controllerNumber=1 scan=21649 | 41.4001 |
| 20327:4 | 1 | R.TAGWNIPMGLLFN[+2059.735]QTGSC[+57.021]K.F | HexNAc(4)Hex(5)Fuc(1)NeuAc(1) | C[+57], N[+2060] | 1039.4457 | 4 | 4154.7611 | 4154.7449 | 0 | 3.9 | 485 | Specific | 633.11 | 353.31 | 353.31 | 12.3 | 128 | >sp\|P02788\|TRFL_HUMAN Lactotransferrin OS=Homo sapiens OX=9606 GN=LTF PE=1 SV=6 | 1270 | 20201115_15212_1ng_glyco_hcd203040_compound15.23246.23246.4 | controllerType=0 controllerNumber=1 scan=23246 | 44.8739 |
| 18459:3 | 1 | R.TAGWNIPMGLLFN[+1768.640]QTGSC[+57.021]K.F | HexNAc(4)Hex(5)Fuc(1) | C[+57], N[+1769] | 1288.5591 | 3 | 3863.6629 | 3863.6495 | 0 | 3.5 | 485 | Specific | 644.64 | 644.64 | 644.64 | 13.59 | 128 | >sp\|P02788\|TRFL_HUMAN Lactotransferrin OS=Homo sapiens OX=9606 GN=LTF PE=1 SV=6 | 1270 | 20201115_15212_1ng_glyco_hcd203040_compound15.21296.21296.3 | controllerType=0 controllerNumber=1 scan=21296 | 40.6893 |
| 18754:3 | 1 | R.TAGWNIPMGLLFN[+1622.582]QTGSC[+57.021]K.F | HexNAc(4)Hex(5) | C[+57], N[+1623] | 1239.873 | 3 | 3717.6044 | 3717.5916 | 0 | 3.5 | 485 | Specific | 689.28 | 665.16 | 336.27 | 14.59 | 128 | >sp\|P02788\|TRFL_HUMAN Lactotransferrin OS=Homo sapiens OX=9606 GN=LTF PE=1 SV=6 | 1270 | 20201115_15212_1ng_glyco_hcd203040_compound15.21603.21603.3 | controllerType=0 controllerNumber=1 scan=21603 | 41.3144 |
| 20328:4 | 1 | R.TAGWNIPMGLLFN[+2205.793]QTGSC[+57.021]K.F | HexNAc(4)Hex(5)Fuc(2)NeuAc(1) | C[+57], N[+2206] | 1075.9591 | 4 | 4300.8148 | 4300.8028 | 0 | 2.8 | 485 | Specific | 699.46 | 404.23 | 404.23 | 15 | 128 | >sp\|P02788\|TRFL_HUMAN Lactotransferrin OS=Homo sapiens OX=9606 GN=LTF PE=1 SV=6 | 1270 | 20201115_15212_1ng_glyco_hcd203040_compound15.23247.23247.4 | controllerType=0 controllerNumber=1 scan=23247 | 44.8755 |
| 22305:4 | 1 | R.TAGWNIPMGLLFN[+2350.830]QTGSC[+57.021]K.F | HexNAc(4)Hex(5)Fuc(1)NeuAc(2) | C[+57], N[+2351] | 1112.2166 | 4 | 4445.8446 | 4445.8403 | 0 | 1 | 485 | Specific | 718.2 | 581.95 | 581.95 | 15.4 | 128 | >sp\|P02788\|TRFL_HUMAN Lactotransferrin OS=Homo sapiens OX=9606 GN=LTF PE=1 SV=6 | 1270 | 20201115_15212_1ng_glyco_hcd203040_compound15.25316.25316.4 | controllerType=0 controllerNumber=1 scan=25316 | 49.3308 |
| 18555:3 | 1 | R.TAGWNIPMGLLFN[+1606.587]QTGSC[+57.021]K.F | HexNAc(4)Hex(4)Fuc(1) | C[+57], N[+1607] | 1234.541 | 3 | 3701.6083 | 3701.5966 | 0 | 3.2 | 485 | Specific | 719.64 | 719.64 | 719.64 | 15.65 | 128 | >sp\|P02788\|TRFL_HUMAN Lactotransferrin OS=Homo sapiens OX=9606 GN=LTF PE=1 SV=6 | 1270 | 20201115_15212_1ng_glyco_hcd203040_compound15.21397.21397.3 | controllerType=0 controllerNumber=1 scan=21397 | 40.9074 |
| 20324:4 | 1 | R.TAGWNIPMGLLFN[+2059.735]QTGSC[+57.021]K.F | HexNAc(4)Hex(5)Fuc(1)NeuAc(1) | C[+57], N[+2060] | 1039.4457 | 4 | 4154.7611 | 4154.7449 | 0 | 3.9 | 485 | Specific | 755.74 | 509.41 | 509.41 | 16.94 | 128 | >sp\|P02788\|TRFL_HUMAN Lactotransferrin OS=Homo sapiens OX=9606 GN=LTF PE=1 SV=6 | 1270 | 20201115_15212_1ng_glyco_hcd203040_compound15.23243.23243.4 | controllerType=0 controllerNumber=1 scan=23243 | 44.8639 |
| 20559:3 | 1 | R.TAGWNIPMGLLFN[+1751.624]QTGSC[+57.021]K.F | HexNAc(4)Hex(4)NeuAc(1) | C[+57], N[+1752] | 1282.8863 | 3 | 3846.6443 | 3846.6341 | 0 | 2.6 | 485 | Specific | 793.23 | 760.79 | 760.79 | 16.65 | 128 | >sp\|P02788\|TRFL_HUMAN Lactotransferrin OS=Homo sapiens OX=9606 GN=LTF PE=1 SV=6 | 1270 | 20201115_15212_1ng_glyco_hcd203040_compound15.23489.23489.3 | controllerType=0 controllerNumber=1 scan=23489 | 45.4388 |
| 18888:3 | 1 | R.TAGWNIPMGLLFN[+1216.423]QTGSC[+57.021]K.F | HexNAc(2)Hex(5) | C[+57], N[+1216] | 1104.4934 | 3 | 3311.4658 | 3311.4328 | 0 | 10 | 485 | Specific | 795.12 | 770.66 | 321.64 | 15.46 | 128 | >sp\|P02788\|TRFL_HUMAN Lactotransferrin OS=Homo sapiens OX=9606 GN=LTF PE=1 SV=6 | 1270 | 20201115_15212_1ng_glyco_hcd203040_compound15.21743.21743.3 | controllerType=0 controllerNumber=1 scan=21743 | 41.6098 |
| 18801:3 | 1 | R.TAGWNIPMGLLFN[+1460.529]QTGSC[+57.021]K.F | HexNAc(4)Hex(4) | C[+57], N[+1461] | 1185.8561 | 3 | 3555.5539 | 3555.5387 | 0 | 4.3 | 485 | Specific | 807.41 | 775.5 | 396.16 | 17.59 | 128 | >sp\|P02788\|TRFL_HUMAN Lactotransferrin OS=Homo sapiens OX=9606 GN=LTF PE=1 SV=6 | 1270 | 20201115_15212_1ng_glyco_hcd203040_compound15.21652.21652.3 | controllerType=0 controllerNumber=1 scan=21652 | 41.4088 |
| 22476:3 | 1 | R.TAGWNIPMGLLFN[+2204.772]QTGSC[+57.021]K.F | HexNAc(4)Hex(5)NeuAc(2) | C[+57], N[+2205] | 1433.9382 | 3 | 4299.7999 | 4299.7824 | 0 | 4.1 | 485 | Specific | 820.65 | 750.43 | 750.43 | 17.59 | 128 | >sp\|P02788\|TRFL_HUMAN Lactotransferrin OS=Homo sapiens OX=9606 GN=LTF PE=1 SV=6 | 1270 | 20201115_15212_1ng_glyco_hcd203040_compound15.25495.25495.3 | controllerType=0 controllerNumber=1 scan=25495 | 49.7862 |
| 22308:4 | 1 | R.TAGWNIPMGLLFN[+2350.830]QTGSC[+57.021]K.F | HexNAc(4)Hex(5)Fuc(1)NeuAc(2) | C[+57], N[+2351] | 1112.2166 | 4 | 4445.8446 | 4445.8403 | 0 | 1 | 485 | Specific | 822.08 | 618.98 | 618.98 | 17.4 | 128 | >sp\|P02788\|TRFL_HUMAN Lactotransferrin OS=Homo sapiens OX=9606 GN=LTF PE=1 SV=6 | 1270 | 20201115_15212_1ng_glyco_hcd203040_compound15.25319.25319.4 | controllerType=0 controllerNumber=1 scan=25319 | 49.3363 |
| 20330:3 | 1 | R.TAGWNIPMGLLFN[+2059.735]QTGSC[+57.021]K.F | HexNAc(4)Hex(5)Fuc(1)NeuAc(1) | C[+57], N[+2060] | 1385.5917 | 3 | 4154.7606 | 4154.7449 | 0 | 3.8 | 485 | Specific | 827.46 | 805.2 | 805.2 | 17.59 | 128 | >sp\|P02788\|TRFL_HUMAN Lactotransferrin OS=Homo sapiens OX=9606 GN=LTF PE=1 SV=6 | 1270 | 20201115_15212_1ng_glyco_hcd203040_compound15.23250.23250.3 | controllerType=0 controllerNumber=1 scan=23250 | 44.8819 |
| 20465:4 | 1 | R.TAGWNIPMGLLFN[+1913.677]QTGSC[+57.021]K.F | HexNAc(4)Hex(5)NeuAc(1) | C[+57], N[+1914] | 1002.9294 | 4 | 4008.6956 | 4008.687 | 0 | 2.2 | 485 | Specific | 843.72 | 729.52 | 729.52 | 17.92 | 128 | >sp\|P02788\|TRFL_HUMAN Lactotransferrin OS=Homo sapiens OX=9606 GN=LTF PE=1 SV=6 | 1270 | 20201115_15212_1ng_glyco_hcd203040_compound15.23391.23391.4 | controllerType=0 controllerNumber=1 scan=23391 | 45.2149 |
| 20468:4 | 1 | R.TAGWNIPMGLLFN[+1913.677]QTGSC[+57.021]K.F | HexNAc(4)Hex(5)NeuAc(1) | C[+57], N[+1914] | 1002.9294 | 4 | 4008.6956 | 4008.687 | 0 | 2.2 | 485 | Specific | 847.2 | 847.2 | 847.2 | 17.92 | 128 | >sp\|P02788\|TRFL_HUMAN Lactotransferrin OS=Homo sapiens OX=9606 GN=LTF PE=1 SV=6 | 1270 | 20201115_15212_1ng_glyco_hcd203040_compound15.23394.23394.4 | controllerType=0 controllerNumber=1 scan=23394 | 45.2195 |
| 18852:2 | 1 | R.TAGWNIPMGLLFN[+1216.423]QTGSC[+57.021]K.F | HexNAc(2)Hex(5) | C[+57], N[+1216] | 1656.2262 | 2 | 3311.4452 | 3311.4328 | 0 | 3.7 | 485 | Specific | 873.32 | 838.93 | 291.67 | 18.57 | 128 | >sp\|P02788\|TRFL_HUMAN Lactotransferrin OS=Homo sapiens OX=9606 GN=LTF PE=1 SV=6 | 1270 | 20201115_15212_1ng_glyco_hcd203040_compound15.21705.21705.2 | controllerType=0 controllerNumber=1 scan=21705 | 41.5427 |
| 18539:3 | 1 | R.TAGWNIPMGLLFN[+1606.587]QTGSC[+57.021]K.F | HexNAc(4)Hex(4)Fuc(1) | C[+57], N[+1607] | 1234.5373 | 3 | 3701.5973 | 3701.5966 | 0 | 0.2 | 485 | Specific | 894.49 | 794.23 | 794.23 | 18.05 | 128 | >sp\|P02788\|TRFL_HUMAN Lactotransferrin OS=Homo sapiens OX=9606 GN=LTF PE=1 SV=6 | 1270 | 20201115_15212_1ng_glyco_hcd203040_compound15.21380.21380.3 | controllerType=0 controllerNumber=1 scan=21380 | 40.877 |
| 22297:3 | 1 | R.TAGWNIPMGLLFN[+2350.830]QTGSC[+57.021]K.F | HexNAc(4)Hex(5)Fuc(1)NeuAc(2) | C[+57], N[+2351] | 1482.6241 | 3 | 4445.8577 | 4445.8403 | 0 | 3.9 | 485 | Specific | 901.82 | 874.1 | 874.1 | 19.59 | 128 | >sp\|P02788\|TRFL_HUMAN Lactotransferrin OS=Homo sapiens OX=9606 GN=LTF PE=1 SV=6 | 1270 | 20201115_15212_1ng_glyco_hcd203040_compound15.25308.25308.3 | controllerType=0 controllerNumber=1 scan=25308 | 49.3205 |
| 18558:3 | 1 | R.TAGWNIPMGLLFN[+1606.587]QTGSC[+57.021]K.F | HexNAc(4)Hex(4)Fuc(1) | C[+57], N[+1607] | 1234.541 | 3 | 3701.6083 | 3701.5966 | 0 | 3.2 | 485 | Specific | 916.63 | 714.25 | 714.25 | 19.65 | 128 | >sp\|P02788\|TRFL_HUMAN Lactotransferrin OS=Homo sapiens OX=9606 GN=LTF PE=1 SV=6 | 1270 | 20201115_15212_1ng_glyco_hcd203040_compound15.21400.21400.3 | controllerType=0 controllerNumber=1 scan=21400 | 40.9117 |
| 20437:3 | 1 | R.TAGWNIPMGLLFN[+1913.677]QTGSC[+57.021]K.F | HexNAc(4)Hex(5)NeuAc(1) | C[+57], N[+1914] | 1336.905 | 3 | 4008.7003 | 4008.687 | 0 | 3.3 | 485 | Specific | 935.08 | 935.08 | 935.08 | 19.65 | 128 | >sp\|P02788\|TRFL_HUMAN Lactotransferrin OS=Homo sapiens OX=9606 GN=LTF PE=1 SV=6 | 1270 | 20201115_15212_1ng_glyco_hcd203040_compound15.23362.23362.3 | controllerType=0 controllerNumber=1 scan=23362 | 45.153 |
| 20266:3 | 1 | R.TAGWNIPMGLLFN[+2205.793]QTGSC[+57.021]K.F | HexNAc(4)Hex(5)Fuc(2)NeuAc(1) | C[+57], N[+2206] | 1434.277 | 3 | 4300.8166 | 4300.8028 | 0 | 3.2 | 485 | Specific | 935.64 | 931.86 | 931.86 | 19.65 | 128 | >sp\|P02788\|TRFL_HUMAN Lactotransferrin OS=Homo sapiens OX=9606 GN=LTF PE=1 SV=6 | 1270 | 20201115_15212_1ng_glyco_hcd203040_compound15.23183.23183.3 | controllerType=0 controllerNumber=1 scan=23183 | 44.7441 |
| 18462:3 | 1 | R.TAGWNIPMGLLFN[+1768.640]QTGSC[+57.021]K.F | HexNAc(4)Hex(5)Fuc(1) | C[+57], N[+1769] | 1288.5591 | 3 | 3863.6629 | 3863.6495 | 0 | 3.5 | 485 | Specific | 939.72 | 939.72 | 939.72 | 19.59 | 128 | >sp\|P02788\|TRFL_HUMAN Lactotransferrin OS=Homo sapiens OX=9606 GN=LTF PE=1 SV=6 | 1270 | 20201115_15212_1ng_glyco_hcd203040_compound15.21299.21299.3 | controllerType=0 controllerNumber=1 scan=21299 | 40.693 |
| 22300:3 | 1 | R.TAGWNIPMGLLFN[+2350.830]QTGSC[+57.021]K.F | HexNAc(4)Hex(5)Fuc(1)NeuAc(2) | C[+57], N[+2351] | 1482.6241 | 3 | 4445.8577 | 4445.8403 | 0 | 3.9 | 485 | Specific | 941.36 | 923.59 | 923.59 | 19.59 | 128 | >sp\|P02788\|TRFL_HUMAN Lactotransferrin OS=Homo sapiens OX=9606 GN=LTF PE=1 SV=6 | 1270 | 20201115_15212_1ng_glyco_hcd203040_compound15.25311.25311.3 | controllerType=0 controllerNumber=1 scan=25311 | 49.3252 |
| 20562:3 | 1 | R.TAGWNIPMGLLFN[+1751.624]QTGSC[+57.021]K.F | HexNAc(4)Hex(4)NeuAc(1) | C[+57], N[+1752] | 1282.8863 | 3 | 3846.6443 | 3846.6341 | 0 | 2.6 | 485 | Specific | 943.25 | 730.66 | 730.66 | 19.65 | 128 | >sp\|P02788\|TRFL_HUMAN Lactotransferrin OS=Homo sapiens OX=9606 GN=LTF PE=1 SV=6 | 1270 | 20201115_15212_1ng_glyco_hcd203040_compound15.23492.23492.3 | controllerType=0 controllerNumber=1 scan=23492 | 45.4468 |
| 20332:3 | 1 | R.TAGWNIPMGLLFN[+2059.735]QTGSC[+57.021]K.F | HexNAc(4)Hex(5)Fuc(1)NeuAc(1) | C[+57], N[+2060] | 1385.5917 | 3 | 4154.7606 | 4154.7449 | 0 | 3.8 | 485 | Specific | 946.9 | 942.56 | 942.56 | 19.59 | 128 | >sp\|P02788\|TRFL_HUMAN Lactotransferrin OS=Homo sapiens OX=9606 GN=LTF PE=1 SV=6 | 1270 | 20201115_15212_1ng_glyco_hcd203040_compound15.23252.23252.3 | controllerType=0 controllerNumber=1 scan=23252 | 44.8867 |
| 18426:3 | 1 | R.TAGWNIPMGLLFN[+1768.640]QTGSC[+57.021]K.F | HexNAc(4)Hex(5)Fuc(1) | C[+57], N[+1769] | 1288.5582 | 3 | 3863.6599 | 3863.6495 | 0 | 2.7 | 485 | Specific | 955.9 | 955.9 | 955.9 | 20.65 | 128 | >sp\|P02788\|TRFL_HUMAN Lactotransferrin OS=Homo sapiens OX=9606 GN=LTF PE=1 SV=6 | 1270 | 20201115_15212_1ng_glyco_hcd203040_compound15.21261.21261.3 | controllerType=0 controllerNumber=1 scan=21261 | 40.6154 |
| 18819:3 | 1 | R.TAGWNIPMGLLFN[+1216.423]QTGSC[+57.021]K.F | HexNAc(2)Hex(5) | C[+57], N[+1216] | 1104.4853 | 3 | 3311.4412 | 3311.4328 | 0 | 2.5 | 485 | Specific | 956.14 | 956.14 | 385.99 | 20.65 | 128 | >sp\|P02788\|TRFL_HUMAN Lactotransferrin OS=Homo sapiens OX=9606 GN=LTF PE=1 SV=6 | 1270 | 20201115_15212_1ng_glyco_hcd203040_compound15.21671.21671.3 | controllerType=0 controllerNumber=1 scan=21671 | 41.4579 |
| 18424:3 | 1 | R.TAGWNIPMGLLFN[+1768.640]QTGSC[+57.021]K.F | HexNAc(4)Hex(5)Fuc(1) | C[+57], N[+1769] | 1288.5582 | 3 | 3863.6599 | 3863.6495 | 0 | 2.7 | 485 | Specific | 957.65 | 957.65 | 957.65 | 20.65 | 128 | >sp\|P02788\|TRFL_HUMAN Lactotransferrin OS=Homo sapiens OX=9606 GN=LTF PE=1 SV=6 | 1270 | 20201115_15212_1ng_glyco_hcd203040_compound15.21259.21259.3 | controllerType=0 controllerNumber=1 scan=21259 | 40.6103 |
| 18850:2 | 1 | R.TAGWNIPMGLLFN[+1216.423]QTGSC[+57.021]K.F | HexNAc(2)Hex(5) | C[+57], N[+1216] | 1656.2262 | 2 | 3311.4452 | 3311.4328 | 0 | 3.7 | 485 | Specific | 966.03 | 944.91 | 348.54 | 20.57 | 128 | >sp\|P02788\|TRFL_HUMAN Lactotransferrin OS=Homo sapiens OX=9606 GN=LTF PE=1 SV=6 | 1270 | 20201115_15212_1ng_glyco_hcd203040_compound15.21703.21703.2 | controllerType=0 controllerNumber=1 scan=21703 | [illegible] |
| 20263:3 | 1 | R.TAGWNIPMGLLFN[+2205.793]QTGSC[+57.021]K.F | HexNAc(4)Hex(5)Fuc(2)NeuAc(1) | C[+57], N[+2206] | 1434.277 | 3 | 4300.8166 | 4300.8028 | 0 | 3.2 | 485 | Specific | 1006.03 | 893.71 | 893.71 | 21.65 | 128 | >sp\|P02788\|TRFL_HUMAN Lactotransferrin OS=Homo sapiens OX=9606 GN=LTF PE=1 SV=6 | 1270 | 20201115_15212_1ng_glyco_hcd203040_compound15.23180.23180.3 | controllerType=0 controllerNumber=1 scan=23180 | [illegible] |
| 18822:3 | 1 | R.TAGWNIPMGLLFN[+1216.423]QTGSC[+57.021]K.F | HexNAc(2)Hex(5) | C[+57], N[+1216] | 1104.4853 | 3 | 3311.4412 | 3311.4328 | 0 | 2.5 | 485 | Specific | 1043.89 | 869.05 | 482.13 | 21.65 | 128 | >sp\|P02788\|TRFL_HUMAN Lactotransferrin OS=Homo sapiens OX=9606 GN=LTF PE=1 SV=6 | 1270 | 20201115_15212_1ng_glyco_hcd203040_compound15.21674.21674.3 | controllerType=0 controllerNumber=1 scan=21674 | 41.4633 |
| 18840:3 | 1 | R.TAGWNIPM[+15.995]GLLFN[+2059.735]QTGSC[+57.021]K.F | HexNAc(4)Hex(5)Fuc(1)NeuAc(1) | C[+57], M[+16], N[+2060] | 1390.925 | 3 | 4170.7604 | 4170.7398 | 0 | 4.9 | 485 | Specific | 252.14 | 222.1 | 222.1 | 6.61 | 128 | >sp\|P02788\|TRFL_HUMAN Lactotransferrin OS=Homo sapiens OX=9606 GN=LTF PE=1 SV=6 | 1270 | 20201115_15212_1ng_glyco_hcd203040_compound15.21693.21693.3 | controllerType=0 controllerNumber=1 scan=21693 | 41.5106 |
| 18473:4 | 1 | R.TAGWNIPM[+15.995]GLLFN[+1751.624]QTGSC[+57.021]K.F | HexNAc(4)Hex(4)NeuAc(1) | C[+57], M[+16], N[+1752] | 966.4197 | 4 | 3862.6568 | 3862.6291 | 0 | 7.2 | 485 | Specific | 305.2 | 235.43 | 235.43 | 4.36 | 128 | >sp\|P02788\|TRFL_HUMAN Lactotransferrin OS=Homo sapiens OX=9606 GN=LTF PE=1 SV=6 | 1270 | 20201115_15212_1ng_glyco_hcd203040_compound15.21310.21310.4 | controllerType=0 controllerNumber=1 scan=21310 | 40.7098 |
| 18476:4 | 1 | R.TAGWNIPM[+15.995]GLLFN[+1751.624]QTGSC[+57.021]K.F | HexNAc(4)Hex(4)NeuAc(1) | C[+57], M[+16], N[+1752] | 966.4197 | 4 | 3862.6568 | 3862.6291 | 0 | 7.2 | 485 | Specific | 308.53 | 235.98 | 235.98 | 4.36 | 128 | >sp\|P02788\|TRFL_HUMAN Lactotransferrin OS=Homo sapiens OX=9606 GN=LTF PE=1 SV=6 | 1270 | 20201115_15212_1ng_glyco_hcd203040_compound15.21313.21313.4 | controllerType=0 controllerNumber=1 scan=21313 | 40.7154 |
| 18843:3 | 1 | R.TAGWNIPM[+15.995]GLLFN[+2059.735]QTGSC[+57.021]K.F | HexNAc(4)Hex(5)Fuc(1)NeuAc(1) | C[+57], M[+16], N[+2060] | 1390.925 | 3 | 4170.7604 | 4170.7398 | 0 | 4.9 | 485 | Specific | 719.46 | 579.42 | 579.42 | 15.04 | 128 | >sp\|P02788\|TRFL_HUMAN Lactotransferrin OS=Homo sapiens OX=9606 GN=LTF PE=1 SV=6 | 1270 | 20201115_15212_1ng_glyco_hcd203040_compound15.21696.21696.3 | controllerType=0 controllerNumber=1 scan=21696 | 41.5165 |
| 20674:4 | 1 | R.[+42.011]TAGWNIPMGLLFN[+2018.708]QTGSC[+57.021]KFDEYFSQSC[+57.021]A | HexNAc(3)Hex(6)Fuc(1)NeuAc(1) | .[42], C[+57]*2, N[+2019] | 1500.6324 | 4 | 5999.508 | 5999.4761 | 0 | 5.3 | 485 | Specific | 205.92 | 205.92 | 205.92 | 4.74 | 128 | >sp\|P02788\|TRFL_HUMAN Lactotransferrin OS=Homo sapiens OX=9606 GN=LTF PE=1 SV=6 | 1270 | 20201115_15212_1ng_glyco_hcd203040_compound15.23609.23609.4 | controllerType=0 controllerNumber=1 scan=23609 | 45.6773 |
| 20671:4 | 1 | R.[+42.011]TAGWNIPMGLLFN[+2018.708]QTGSC[+57.021]KFDEYFSQSC[+57.021]A | HexNAc(3)Hex(6)Fuc(1)NeuAc(1) | .[42], C[+57]*2, N[+2019] | 1500.6324 | 4 | 5999.508 | 5999.4761 | 0 | 5.3 | 485 | Specific | 321.56 | 321.56 | 321.56 | 6.94 | 128 | >sp\|P02788\|TRFL_HUMAN Lactotransferrin OS=Homo sapiens OX=9606 GN=LTF PE=1 SV=6 | 1270 | 20201115_15212_1ng_glyco_hcd203040_compound15.23606.23606.4 | controllerType=0 controllerNumber=1 scan=23606 | 45.6639 |
| 20588:4 | 1 | R.TAGWNIPMGLLFN[+2059.735]QTGSC[+57.021]KFDEYFSQSC[+57.021]APGSDPR.S | HexNAc(4)Hex(5)Fuc(1)NeuAc(1) | C[+57]*2, N[+2060] | 1500.3838 | 4 | 5998.5133 | 5998.492 | 0 | 3.6 | 485 | Specific | 713.97 | 707.98 | 707.98 | 14.91 | 128 | >sp\|P02788\|TRFL_HUMAN Lactotransferrin OS=Homo sapiens OX=9606 GN=LTF PE=1 SV=6 | 1270 | 20201115_15212_1ng_glyco_hcd203040_compound15.23520.23520.4 | controllerType=0 controllerNumber=1 scan=23520 | 45.5032 |
| 21865:4 | 1 | R.TAGWNIPMGLLFN[+2350.830]QTGSC[+57.021]KFDEYFSQSC[+57.021]APGSDPR.S | HexNAc(4)Hex(5)Fuc(1)NeuAc(2) | C[+57]*2, N[+2351] | 1573.1581 | 4 | 6289.6105 | 6289.5875 | 0 | 3.7 | 485 | Specific | 891.79 | 821.73 | 821.73 | 17.91 | 128 | >sp\|P02788\|TRFL_HUMAN Lactotransferrin OS=Homo sapiens OX=9606 GN=LTF PE=1 SV=6 | 1270 | 20201115_15212_1ng_glyco_hcd203040_compound15.24857.24857.4 | controllerType=0 controllerNumber=1 scan=24857 | 48.3357 |
| 20590:4 | 1 | R.TAGWNIPMGLLFN[+2059.735]QTGSC[+57.021]KFDEYFSQSC[+57.021]APGSDPR.S | HexNAc(4)Hex(5)Fuc(1)NeuAc(1) | C[+57]*2, N[+2060] | 1500.3838 | 4 | 5998.5133 | 5998.492 | 0 | 3.6 | 485 | Specific | 903.08 | 789.99 | 789.99 | 18.91 | 128 | >sp\|P02788\|TRFL_HUMAN Lactotransferrin OS=Homo sapiens OX=9606 GN=LTF PE=1 SV=6 | 1270 | 20201115_15212_1ng_glyco_hcd203040_compound15.23522.23522.4 | controllerType=0 controllerNumber=1 scan=23522 | 45.5078 |
| 20552:4 | 1 | R.TAGWNIPMGLLFN[+2205.793]QTGSC[+57.021]KFDEYFSQSC[+57.021]APGSDPR.S | HexNAc(4)Hex(5)Fuc(2)NeuAc(1) | C[+57]*2, N[+2206] | 1536.898 | 4 | 6144.57 | 6144.5499 | 0 | 3.3 | 485 | Specific | 934.14 | 850.66 | 850.66 | 18.97 | 128 | >sp\|P02788\|TRFL_HUMAN Lactotransferrin OS=Homo sapiens OX=9606 GN=LTF PE=1 SV=6 | 1270 | 20201115_15212_1ng_glyco_hcd203040_compound15.23482.23482.4 | controllerType=0 controllerNumber=1 scan=23482 | 45.4183 |
| 21868:4 | 1 | R.TAGWNIPMGLLFN[+2350.830]QTGSC[+57.021]KFDEYFSQSC[+57.021]APGSDPR.S | HexNAc(4)Hex(5)Fuc(1)NeuAc(2) | C[+57]*2, N[+2351] | 1573.1581 | 4 | 6289.6105 | 6289.5875 | 0 | 3.7 | 485 | Specific | 942.97 | 907.82 | 907.82 | 18.91 | 128 | >sp\|P02788\|TRFL_HUMAN Lactotransferrin OS=Homo sapiens OX=9606 GN=LTF PE=1 SV=6 | 1270 | 20201115_15212_1ng_glyco_hcd203040_compound15.24860.24860.4 | controllerType=0 controllerNumber=1 scan=24860 | 48.3408 |
| 20555:4 | 1 | R.TAGWNIPMGLLFN[+2205.793]QTGSC[+57.021]KFDEYFSQSC[+57.021]APGSDPR.S | HexNAc(4)Hex(5)Fuc(2)NeuAc(1) | C[+57]*2, N[+2206] | 1536.898 | 4 | 6144.57 | 6144.5499 | 0 | 3.3 | 485 | Specific | 1003.04 | 974.47 | 974.47 | 20.97 | 128 | >sp\|P02788\|TRFL_HUMAN Lactotransferrin OS=Homo sapiens OX=9606 GN=LTF PE=1 SV=6 | 1270 | 20201115_15212_1ng_glyco_hcd203040_compound15.23485.23485.4 | controllerType=0 controllerNumber=1 scan=23485 | 45.434 |
| 09990:3 | 1 | K.FGRN[+1501.555]GSDC[+57.021]PDKFC[+57.021]LFQSETK.N | HexNAc(5)Hex(3) | C[+57]*2, N[+1502] | 1298.8846 | 3 | 3894.6392 | 3894.6202 | 0 | 4.9 | 639 | Specific | 278.01 | 278.01 | 278.01 | 5.75 | 128 | >sp\|P02788\|TRFL_HUMAN Lactotransferrin OS=Homo sapiens OX=9606 GN=LTF PE=1 SV=6 | 1270 | 20201115_15212_1ng_glyco_hcd203040_compound15.12428.12428.3 | controllerType=0 controllerNumber=1 scan=12428 | 25.1555 |
| 09988:3 | 1 | K.FGRN[+1501.555]GSDC[+57.021]PDKFC[+57.021]LFQSETK.N | HexNAc(5)Hex(3) | C[+57]*2, N[+1502] | 1298.8846 | 3 | 3894.6392 | 3894.6202 | 0 | 4.9 | 639 | Specific | 387.15 | 387.15 | 387.15 | 8.32 | 128 | >sp\|P02788\|TRFL_HUMAN Lactotransferrin OS=Homo sapiens OX=9606 GN=LTF PE=1 SV=6 | 1270 | 20201115_15212_1ng_glyco_hcd203040_compound15.12426.12426.3 | controllerType=0 controllerNumber=1 scan=12426 | 25.1506 |
| 10818:3 | 1 | R.N[+2205.793]GSDC[+57.021]PDKFC[+57.021]LFQSETK.N | HexNAc(4)Hex(5)Fuc(2)NeuAc(1) | C[+57]*2, N[+2206] | 1413.5648 | 3 | 4238.6797 | 4238.6667 | 0 | 3.1 | 642 | Specific | 534.28 | 431.88 | 431.88 | 10.4 | 128 | >sp\|P02788\|TRFL_HUMAN Lactotransferrin OS=Homo sapiens OX=9606 GN=LTF PE=1 SV=6 | 1270 | 20201115_15212_1ng_glyco_hcd203040_compound15.13306.13306.3 | controllerType=0 controllerNumber=1 scan=13306 | 26.6199 |
| 10882:3 | 1 | R.N[+2059.735]GSDC[+57.021]PDKFC[+57.021]LFQSETK.N | HexNAc(4)Hex(5)Fuc(1)NeuAc(1) | C[+57]*2, N[+2060] | 1364.8781 | 3 | 4092.6198 | 4092.6088 | 0 | 2.7 | 642 | Specific | 566.2 | 372.34 | 372.34 | 9.12 | 128 | >sp\|P02788\|TRFL_HUMAN Lactotransferrin OS=Homo sapiens OX=9606 GN=LTF PE=1 SV=6 | 1270 | 20201115_15212_1ng_glyco_hcd203040_compound15.13372.13372.3 | controllerType=0 controllerNumber=1 scan=13372 | 26.7361 |
| 10816:3 | 1 | R.N[+2205.793]GSDC[+57.021]PDKFC[+57.021]LFQSETK.N | HexNAc(4)Hex(5)Fuc(2)NeuAc(1) | C[+57]*2, N[+2206] | 1413.5648 | 3 | 4238.6797 | 4238.6667 | 0 | 3.1 | 642 | Specific | 620.32 | 521.4 | 521.4 | 12.91 | 128 | >sp\|P02788\|TRFL_HUMAN Lactotransferrin OS=Homo sapiens OX=9606 GN=LTF PE=1 SV=6 | 1270 | 20201115_15212_1ng_glyco_hcd203040_compound15.13304.13304.3 | controllerType=0 controllerNumber=1 scan=13304 | 26.6154 |
| 10880:3 | 1 | R.N[+2059.735]GSDC[+57.021]PDKFC[+57.021]LFQSETK.N | HexNAc(4)Hex(5)Fuc(1)NeuAc(1) | C[+57]*2, N[+2060] | 1364.8781 | 3 | 4092.6198 | 4092.6088 | 0 | 2.7 | 642 | Specific | 763.79 | 660.55 | 660.55 | 15.91 | 128 | >sp\|P02788\|TRFL_HUMAN Lactotransferrin OS=Homo sapiens OX=9606 GN=LTF PE=1 SV=6 | 1270 | 20201115_15212_1ng_glyco_hcd203040_compound15.13370.13370.3 | controllerType=0 controllerNumber=1 scan=13370 | 26.7313 |
| 03347:3 | 2 | K.GPN[+2059.735]C[+57.021]SEPEC[+57.021]PGNC[+57.021]HLR.G | HexNAc(4)Hex(5)Fuc(1)NeuAc(1) | C[+57]*3, N[+2060] | 1315.1718 | 3 | 3943.5009 | 3943.4931 | 0 | 2 | 182 | Specific | 347.47 | 318.31 | 318.31 | 8.2 | 101 | >sp\|P24821\|TENA_HUMAN Tenascin OS=Homo sapiens OX=9606 GN=TNC PE=1 SV=3 | 486 | 20201115_15212_1ng_glyco_hcd203040_compound15.5534.5534.3 | controllerType=0 controllerNumber=1 scan=5534 | 13.7591 |
| 03349:3 | 2 | K.GPN[+2059.735]C[+57.021]SEPEC[+57.021]PGNC[+57.021]HLR.G | HexNAc(4)Hex(5)Fuc(1)NeuAc(1) | C[+57]*3, N[+2060] | 1315.1718 | 3 | 3943.5009 | 3943.4931 | 0 | 2 | 182 | Specific | 353 | 277.49 | 277.49 | 7.96 | 101 | >sp\|P24821\|TENA_HUMAN Tenascin OS=Homo sapiens OX=9606 GN=TNC PE=1 SV=3 | 486 | 20201115_15212_1ng_glyco_hcd203040_compound15.5536.5536.3 | controllerType=0 controllerNumber=1 scan=5536 | 13.7638 |
| 10778:3 | 2 | R.QTGLAPGQEYEISLHIVKN[+2059.735]NTR.G | HexNAc(4)Hex(5)Fuc(1)NeuAc(1) | N[+2060] | 1510.0179 | 3 | 4528.0391 | 4528.0241 | 0 | 3.3 | 770 | Specific | 414.5 | 201.47 | 201.47 | 6.83 | 101 | >sp\|P24821\|TENA_HUMAN Tenascin OS=Homo sapiens OX=9606 GN=TNC PE=1 SV=3 | 486 | 20201115_15212_1ng_glyco_hcd203040_compound15.13264.13264.3 | controllerType=0 controllerNumber=1 scan=13264 | 26.5469 |
| 10780:3 | 2 | R.QTGLAPGQEYEISLHIVKN[+2059.735]NTR.G | HexNAc(4)Hex(5)Fuc(1)NeuAc(1) | N[+2060] | 1510.0179 | 3 | 4528.0391 | 4528.0241 | 0 | 3.3 | 770 | Specific | 520.22 | 372.66 | 372.66 | 8.27 | 101 | >sp\|P24821\|TENA_HUMAN Tenascin OS=Homo sapiens OX=9606 GN=TNC PE=1 SV=3 | 486 | 20201115_15212_1ng_glyco_hcd203040_compound15.13266.13266.3 | controllerType=0 controllerNumber=1 scan=13266 | 26.5512 |
| 16987:3 | 2 | R.LN[+1702.581]YSLPTGQWVGVQLPR.N | HexNAc(2)Hex(8) | N[+1703] | 1210.881 | 3 | 3630.6286 | 3630.6202 | 0 | 2.3 | 1017 | Specific | 724.25 | 724.25 | 724.25 | 15.47 | 101 | >sp\|P24821\|TENA_HUMAN Tenascin OS=Homo sapiens OX=9606 GN=TNC PE=1 SV=3 | 486 | 20201115_15212_1ng_glyco_hcd203040_compound15.19726.19726.3 | controllerType=0 controllerNumber=1 scan=19726 | 37.7688 |
| 17088:3 | 2 | R.LN[+1378.476]YSLPTGQWVGVQLPR.N | HexNAc(2)Hex(6) | N[+1378] | 1102.846 | 3 | 3306.5234 | 3306.5145 | 0 | 2.7 | 1017 | Specific | 869.73 | 869.73 | 869.73 | 18.47 | 101 | >sp\|P24821\|TENA_HUMAN Tenascin OS=Homo sapiens OX=9606 GN=TNC PE=1 SV=3 | 486 | 20201115_15212_1ng_glyco_hcd203040_compound15.19831.19831.3 | controllerType=0 controllerNumber=1 scan=19831 | 37.9778 |
| 17026:3 | 2 | R.LN[+1540.529]YSLPTGQWVGVQLPR.N | HexNAc(2)Hex(7) | N[+1541] | 1156.8654 | 3 | 3468.5817 | 3468.5674 | 0 | 4.1 | 1017 | Specific | 896.19 | 882.69 | 882.69 | 18.41 | 101 | >sp\|P24821\|TENA_HUMAN Tenascin OS=Homo sapiens OX=9606 GN=TNC PE=1 SV=3 | 486 | 20201115_15212_1ng_glyco_hcd203040_compound15.19766.19766.3 | controllerType=0 controllerNumber=1 scan=19766 | 37.851 |
| 17091:3 | 2 | R.LN[+1378.476]YSLPTGQWVGVQLPR.N | HexNAc(2)Hex(6) | N[+1378] | 1102.846 | 3 | 3306.5234 | 3306.5145 | 0 | 2.7 | 1017 | Specific | 1021.18 | 829.12 | 829.12 | 21.47 | 101 | >sp\|P24821\|TENA_HUMAN Tenascin OS=Homo sapiens OX=9606 GN=TNC PE=1 SV=3 | 486 | 20201115_15212_1ng_glyco_hcd203040_compound15.19834.19834.3 | controllerType=0 controllerNumber=1 scan=19834 | 37.9828 |
| 17029:3 | 2 | R.LN[+1540.529]YSLPTGQWVGVQLPR.N | HexNAc(2)Hex(7) | N[+1541] | 1156.8654 | 3 | 3468.5817 | 3468.5674 | 0 | 4.1 | 1017 | Specific | 1118.82 | 916.75 | 916.75 | 23.41 | 101 | >sp\|P24821\|TENA_HUMAN Tenascin OS=Homo sapiens OX=9606 GN=TNC PE=1 SV=3 | 486 | 20201115_15212_1ng_glyco_hcd203040_compound15.19769.19769.3 | controllerType=0 controllerNumber=1 scan=19769 | 37.8566 |
| 11784:3 | 3 | K.TVLTPATNHMGN[+1378.476]VTFTIPANR.E | HexNAc(2)Hex(6) | N[+1378] | 1211.8875 | 3 | 3633.648 | 3633.6358 | 0 | 3.4 | 74 | Specific | 863.03 | 627.62 | 485.3 | 18.63 | 95 | >sp\|P01024\|CO3_HUMAN Complement C3 OS=Homo sapiens OX=9606 GN=C3 PE=1 SV=2 | 492 | 20201115_15212_1ng_glyco_hcd203040_compound15.14307.14307.3 | controllerType=0 controllerNumber=1 scan=14307 | 28.3002 |
| 11786:3 | 3 | K.TVLTPATNHMGN[+1378.476]VTFTIPANR.E | HexNAc(2)Hex(6) | N[+1378] | 1211.8875 | 3 | 3633.648 | 3633.6358 | 0 | 3.4 | 74 | Specific | 880.27 | 863.57 | 456.28 | 18.63 | 95 | >sp\|P01024\|CO3_HUMAN Complement C3 OS=Homo sapiens OX=9606 GN=C3 PE=1 SV=2 | 492 | 20201115_15212_1ng_glyco_hcd203040_compound15.14309.14309.3 | controllerType=0 controllerNumber=1 scan=14309 | 28.3059 |
| 09421:3 | 6 | K.WEC[+57.021]KN[+2059.735]DTLLGIK.G | HexNAc(4)Hex(5)Fuc(1)NeuAc(1) | C[+57], N[+2060] | 1179.504 | 3 | 3536.4973 | 3536.4865 | 0 | 3.1 | 100 | Specific | 484.92 | 189.31 | 189.31 | 7.31 | 79 | >sp\|P22897\|MRC1_HUMAN Macrophage mannose receptor 1 OS=Homo sapiens OX=9606 GN=MRC1 PE=1 SV=1 | 1049 | 20201115_15212_1ng_glyco_hcd203040_compound15.11830.11830.3 | controllerType=0 controllerNumber=1 scan=11830 | 24.1686 |
| 09423:3 | 6 | K.WEC[+57.021]KN[+2059.735]DTLLGIK.G | HexNAc(4)Hex(5)Fuc(1)NeuAc(1) | C[+57], N[+2060] | 1179.504 | 3 | 3536.4973 | 3536.4865 | 0 | 3.1 | 100 | Specific | 510.19 | 243.31 | 243.31 | 7.6 | 79 | >sp\|P22897\|MRC1_HUMAN Macrophage mannose receptor 1 OS=Homo sapiens OX=9606 GN=MRC1 PE=1 SV=1 | 1049 | 20201115_15212_1ng_glyco_hcd203040_compound15.11832.11832.3 | controllerType=0 controllerNumber=1 scan=11832 | 24.1724 |
| 14826:5 | 6 | K.KGN[+2059.735]TTLNSFVIPSESDVPTHC[+57.021]PSQWWPYAGHC[+57.021]YK.I | HexNAc(4)Hex(5)Fuc(1)NeuAc(1) | C[+57]*2, N[+2060] | 1205.5198 | 5 | 6023.5698 | 6023.5567 | 0 | 2.2 | 342 | Specific | 294.94 | 294.94 | 294.94 | 5.66 | 79 | >sp\|P22897\|MRC1_HUMAN Macrophage mannose receptor 1 OS=Homo sapiens OX=9606 GN=MRC1 PE=1 SV=1 | 1049 | 20201115_15212_1ng_glyco_hcd203040_compound15.17487.17487.3 | controllerType=0 controllerNumber=1 scan=17487 | 33.7103 |
| 14823:5 | 6 | K.KGN[+2059.735]TTLNSFVIPSESDVPTHC[+57.021]PSQWWPYAGHC[+57.021]YK.I | HexNAc(4)Hex(5)Fuc(1)NeuAc(1) | C[+57]*2, N[+2060] | 1205.5198 | 5 | 6023.5698 | 6023.5567 | 0 | 2.2 | 342 | Specific | 326.35 | 326.35 | 326.35 | 6.05 | 79 | >sp\|P22897\|MRC1_HUMAN Macrophage mannose receptor 1 OS=Homo sapiens OX=9606 GN=MRC1 PE=1 SV=1 | 1049 | 20201115_15212_1ng_glyco_hcd203040_compound15.17484.17484.3 | controllerType=0 controllerNumber=1 scan=17484 | 33.7015 |
| 16029:5 | 6 | K.KGN[+2351.851]TTLNSFVIPSESDVPTHC[+57.021]PSQWWPYAGHC[+57.021]YK.I | HexNAc(4)Hex(5)Fuc(3)NeuAc(1) | C[+57]*2, N[+2352] | 1263.9415 | 5 | 6315.6785 | 6315.6725 | 0 | 1 | 342 | Specific | 512 | 437.51 | 437.51 | 10.18 | 79 | >sp\|P22897\|MRC1_HUMAN Macrophage mannose receptor 1 OS=Homo sapiens OX=9606 GN=MRC1 PE=1 SV=1 | 1049 | 20201115_15212_1ng_glyco_hcd203040_compound15.18734.18734.5 | controllerType=0 controllerNumber=1 scan=18734 | 35.9993 |
| 16031:5 | 6 | K.KGN[+2351.851]TTLNSFVIPSESDVPTHC[+57.021]PSQWWPYAGHC[+57.021]YK.I | HexNAc(4)Hex(5)Fuc(3)NeuAc(1) | C[+57]*2, N[+2352] | 1263.9415 | 5 | 6315.6785 | 6315.6725 | 0 | 1 | 342 | Specific | 552.76 | 310.32 | 310.32 | 8.43 | 79 | >sp\|P22897\|MRC1_HUMAN Macrophage mannose receptor 1 OS=Homo sapiens OX=9606 GN=MRC1 PE=1 SV=1 | 1049 | 20201115_15212_1ng_glyco_hcd203040_compound15.18736.18736.5 | controllerType=0 controllerNumber=1 scan=18736 | 36.0037 |
| 21285:3 | 7 | R.GLSFDVSLEVSQGPGLLN[+2351.851]DTK.V | HexNAc(4)Hex(5)Fuc(3)NeuAc(1) | N[+2352] | 1509.9978 | 3 | 4527.9788 | 4527.964 | 0 | 3.3 | 118 | Specific | 396.84 | 308.81 | 308.81 | 8.44 | 68 | >sp\|P01833\|PIGR_HUMAN Polymeric immunoglobulin receptor OS=Homo sapiens OX=9606 GN=PIGR PE=1 SV=4 | 902 | 20201115_15212_1ng_glyco_hcd203040_compound15.24250.24250.3 | controllerType=0 controllerNumber=1 scan=24250 | 46.9288 |
| 19367:3 | 7 | R.GLSFDVSLEVSQGPGLLN[+2059.735]DTK.V | HexNAc(4)Hex(5)Fuc(1)NeuAc(1) | N[+2060] | 1412.6257 | 3 | 4235.8624 | 4235.8481 | 0 | 3.4 | 118 | Specific | 494.02 | 478.38 | 478.38 | 10.9 | 68 | >sp\|P01833\|PIGR_HUMAN Polymeric immunoglobulin receptor OS=Homo sapiens OX=9606 GN=PIGR PE=1 SV=4 | 902 | 20201115_15212_1ng_glyco_hcd203040_compound15.22244.22244.3 | controllerType=0 controllerNumber=1 scan=22244 | 42.711 |
| 21287:3 | 7 | R.GLSFDVSLEVSQGPGLLN[+2351.851]DTK.V | HexNAc(4)Hex(5)Fuc(3)NeuAc(1) | N[+2352] | 1509.9978 | 3 | 4527.9788 | 4527.964 | 0 | 3.3 | 118 | Specific | 495.28 | 276.05 | 276.05 | 8.78 | 68 | >sp\|P01833\|PIGR_HUMAN Polymeric immunoglobulin receptor OS=Homo sapiens OX=9606 GN=PIGR PE=1 SV=4 | 902 | 20201115_15212_1ng_glyco_hcd203040_compound15.24252.24252.3 | controllerType=0 controllerNumber=1 scan=24252 | 46.9363 |
| 19130:3 | 7 | R.GLSFDVSLEVSQGPGLLN[+2716.983]DTK.V | HexNAc(5)Hex(6)Fuc(3)NeuAc(1) | N[+2717] | 1631.7108 | 3 | 4893.1177 | 4893.0962 | 0 | 4.4 | 118 | Specific | 621.34 | 621.34 | 621.34 | 13.59 | 68 | >sp\|P01833\|PIGR_HUMAN Polymeric immunoglobulin receptor OS=Homo sapiens OX=9606 GN=PIGR PE=1 SV=4 | 902 | 20201115_15212_1ng_glyco_hcd203040_compound15.21995.21995.3 | controllerType=0 controllerNumber=1 scan=21995 | 42.1268 |
| 19133:3 | 7 | R.GLSFDVSLEVSQGPGLLN[+2716.983]DTK.V | HexNAc(5)Hex(6)Fuc(3)NeuAc(1) | N[+2717] | 1631.7108 | 3 | 4893.1177 | 4893.0962 | 0 | 4.4 | 118 | Specific | 740.55 | 649.76 | 649.76 | 15.59 | 68 | >sp\|P01833\|PIGR_HUMAN Polymeric immunoglobulin receptor OS=Homo sapiens OX=9606 GN=PIGR PE=1 SV=4 | 902 | 20201115_15212_1ng_glyco_hcd203040_compound15.21998.21998.3 | controllerType=0 controllerNumber=1 scan=21998 | 42.1341 |
| 21186:4 | 7 | R.GLSFDVSLEVSQGPGLLN[+2350.830]DTK.V | HexNAc(4)Hex(5)Fuc(1)NeuAc(2) | N[+2351] | 1132.4941 | 4 | 4526.9544 | 4526.9436 | 0 | 2.4 | 118 | Specific | 757.22 | 511.35 | 511.35 | 17 | 68 | >sp\|P01833\|PIGR_HUMAN Polymeric immunoglobulin receptor OS=Homo sapiens OX=9606 GN=PIGR PE=1 SV=4 | 902 | 20201115_15212_1ng_glyco_hcd203040_compound15.24147.24147.4 | controllerType=0 controllerNumber=1 scan=24147 | 46.7201 |
| 19370:3 | 7 | R.GLSFDVSLEVSQGPGLLN[+2059.735]DTK.V | HexNAc(4)Hex(5)Fuc(1)NeuAc(1) | N[+2060] | 1412.6257 | 3 | 4235.8624 | 4235.8481 | 0 | 3.4 | 118 | Specific | 1042.63 | 973.39 | 973.39 | 21.65 | 68 | >sp\|P01833\|PIGR_HUMAN Polymeric immunoglobulin receptor OS=Homo sapiens OX=9606 GN=PIGR PE=1 SV=4 | 902 | 20201115_15212_1ng_glyco_hcd203040_compound15.22247.22247.3 | controllerType=0 controllerNumber=1 scan=22247 | 42.7153 |
| 20497:4 | 7 | K.QIGLYPVLVIDSSGYVNPN[+2059.735]YTGR.I | HexNAc(4)Hex(5)Fuc(1)NeuAc(1) | N[+2060] | 1147.0192 | 4 | 4585.055 | 4585.0384 | 0 | 3.6 | 168 | Specific | 768.83 | 705.44 | 705.44 | 17.04 | 68 | >sp\|P01833\|PIGR_HUMAN Polymeric immunoglobulin receptor OS=Homo sapiens OX=9606 GN=PIGR PE=1 SV=4 | 902 | 20201115_15212_1ng_glyco_hcd203040_compound15.23424.23424.4 | controllerType=0 controllerNumber=1 scan=23424 | 45.2879 |
| 20500:4 | 7 | K.QIGLYPVLVIDSSGYVNPN[+2059.735]YTGR.I | HexNAc(4)Hex(5)Fuc(1)NeuAc(1) | N[+2060] | 1147.0192 | 4 | 4585.055 | 4585.0384 | 0 | 3.6 | 168 | Specific | 829.6 | 728.57 | 728.57 | 18.04 | 68 | >sp\|P01833\|PIGR_HUMAN Polymeric immunoglobulin receptor OS=Homo sapiens OX=9606 GN=PIGR PE=1 SV=4 | 902 | 20201115_15212_1ng_glyco_hcd203040_compound15.23427.23427.4 | controllerType=0 controllerNumber=1 scan=23427 | 45.2934 |
| 20482:3 | 7 | K.QIGLYPVLVIDSSGYVNPN[+2059.735]YTGR.I | HexNAc(4)Hex(5)Fuc(1)NeuAc(1) | N[+2060] | 1529.0242 | 3 | 4585.0581 | 4585.0384 | 0 | 4.3 | 168 | Specific | 841.07 | 715.01 | 715.01 | 17.69 | 68 | >sp\|P01833\|PIGR_HUMAN Polymeric immunoglobulin receptor OS=Homo sapiens OX=9606 GN=PIGR PE=1 SV=4 | 902 | 20201115_15212_1ng_glyco_hcd203040_compound15.23409.23409.3 | controllerType=0 controllerNumber=1 scan=23409 | 45.2558 |
| 20485:3 | 7 | K.QIGLYPVLVIDSSGYVNPN[+2059.735]YTGR.I | HexNAc(4)Hex(5)Fuc(1)NeuAc(1) | N[+2060] | 1529.0242 | 3 | 4585.0581 | 4585.0384 | 0 | 4.3 | 168 | Specific | 870.73 | 772.43 | 772.43 | 18.69 | 68 | >sp\|P01833\|PIGR_HUMAN Polymeric immunoglobulin receptor OS=Homo sapiens OX=9606 GN=PIGR PE=1 SV=4 | 902 | 20201115_15212_1ng_glyco_hcd203040_compound15.23412.23412.3 | controllerType=0 controllerNumber=1 scan=23412 | 45.2636 |
| 24238:4 | 7 | R.[+42.011]LSLLEEPGN[+2336.851]GTFTVILNQLTSR.D | HexNAc(6)Hex(6)Fuc(1) | .[42], N[+2337] | 1196.0406 | 4 | 4781.1405 | 4781.1542 | 0 | -2.9 | 413 | Specific | 249.41 | 154.68 | 154.68 | 3.26 | 68 | >sp\|P01833\|PIGR_HUMAN Polymeric immunoglobulin receptor OS=Homo sapiens OX=9606 GN=PIGR PE=1 SV=4 | 902 | 20201115_15212_1ng_glyco_hcd203040_compound15.27339.27339.4 | controllerType=0 controllerNumber=1 scan=27339 | 53.4718 |
| 24234:4 | 7 | R.[+42.011]LSLLEEPGN[+2336.851]GTFTVILNQLTSR.D | HexNAc(6)Hex(6)Fuc(1) | .[42], N[+2337] | 1196.043 | 4 | 4781.1502 | 4781.1542 | 0 | -0.8 | 413 | Specific | 265.71 | 188.05 | 188.05 | 4.18 | 68 | >sp\|P01833\|PIGR_HUMAN Polymeric immunoglobulin receptor OS=Homo sapiens OX=9606 GN=PIGR PE=1 SV=4 | 902 | 20201115_15212_1ng_glyco_hcd203040_compound15.27334.27334.4 | controllerType=0 controllerNumber=1 scan=27334 | 53.4569 |
| 24236:4 | 7 | R.[+42.011]LSLLEEPGN[+2336.851]GTFTVILNQLTSR.D | HexNAc(6)Hex(6)Fuc(1) | .[42], N[+2337] | 1196.0406 | 4 | 4781.1405 | 4781.1542 | 0 | -2.9 | 413 | Specific | 293.49 | 239.77 | 239.77 | 4.41 | 68 | >sp\|P01833\|PIGR_HUMAN Polymeric immunoglobulin receptor OS=Homo sapiens OX=9606 GN=PIGR PE=1 SV=4 | 902 | 20201115_15212_1ng_glyco_hcd203040_compound15.27337.27337.4 | controllerType=0 controllerNumber=1 scan=27337 | 53.4638 |
| 24232:4 | 7 | R.[+42.011]LSLLEEPGN[+2336.851]GTFTVILNQLTSR.D | HexNAc(6)Hex(6)Fuc(1) | .[42], N[+2337] | 1196.043 | 4 | 4781.1502 | 4781.1542 | 0 | -0.8 | 413 | Specific | 328.9 | 277.12 | 277.12 | 5.99 | 68 | >sp\|P01833\|PIGR_HUMAN Polymeric immunoglobulin receptor OS=Homo sapiens OX=9606 GN=PIGR PE=1 SV=4 | 902 | 20201115_15212_1ng_glyco_hcd203040_compound15.27332.27332.4 | controllerType=0 controllerNumber=1 scan=27332 | 53.4495 |
| 21964:3 | 7 | R.LSLLEEPGN[+2571.946]GTFTVILNQLTSR.D | HexNAc(5)Hex(6)Fuc(4) | N[+2572] | 1658.7386 | 3 | 4974.2013 | 4974.238 | 0 | -7.4 | 413 | Specific | 386.61 | 253.43 | 253.43 | 4.63 | 68 | >sp\|P01833\|PIGR_HUMAN Polymeric immunoglobulin receptor OS=Homo sapiens OX=9606 GN=PIGR PE=1 SV=4 | 902 | 20201115_15212_1ng_glyco_hcd203040_compound15.24959.24959.3 | controllerType=0 controllerNumber=1 scan=24959 | 48.5254 |
| 22066:3 | 7 | R.LSLLEEPGN[+2351.851]GTFTVILNQLTSR.D | HexNAc(4)Hex(5)Fuc(3)NeuAc(1) | N[+2352] | 1585.3875 | 3 | 4754.148 | 4754.1433 | 0 | 1 | 413 | Specific | 740.38 | 643.23 | 643.23 | 15.05 | 68 | >sp\|P01833\|PIGR_HUMAN Polymeric immunoglobulin receptor OS=Homo sapiens OX=9606 GN=PIGR PE=1 SV=4 | 902 | 20201115_15212_1ng_glyco_hcd203040_compound15.25065.25065.3 | controllerType=0 controllerNumber=1 scan=25065 | 48.7294 |
| 22069:3 | 7 | R.LSLLEEPGN[+2351.851]GTFTVILNQLTSR.D | HexNAc(4)Hex(5)Fuc(3)NeuAc(1) | N[+2352] | 1585.3875 | 3 | 4754.148 | 4754.1433 | 0 | 1 | 413 | Specific | 787.02 | 557.24 | 557.24 | 16.05 | 68 | >sp\|P01833\|PIGR_HUMAN Polymeric immunoglobulin receptor OS=Homo sapiens OX=9606 GN=PIGR PE=1 SV=4 | 902 | 20201115_15212_1ng_glyco_hcd203040_compound15.25068.25068.3 | controllerType=0 controllerNumber=1 scan=25068 | 48.7348 |
| 22067:4 | 7 | R.LSLLEEPGN[+2205.793]GTFTVILNQLTSR.D | HexNAc(4)Hex(5)Fuc(2)NeuAc(1) | N[+2206] | 1152.7824 | 4 | 4608.1077 | 4608.0854 | 0 | 4.8 | 413 | Specific | 890.55 | 696.43 | 696.43 | 18.39 | 68 | >sp\|P01833\|PIGR_HUMAN Polymeric immunoglobulin receptor OS=Homo sapiens OX=9606 GN=PIGR PE=1 SV=4 | 902 | 20201115_15212_1ng_glyco_hcd203040_compound15.25066.25066.3 | controllerType=0 controllerNumber=1 scan=25066 | 48.7308 |
| 22070:4 | 7 | R.LSLLEEPGN[+2205.793]GTFTVILNQLTSR.D | HexNAc(4)Hex(5)Fuc(2)NeuAc(1) | N[+2206] | 1152.7824 | 4 | 4608.1077 | 4608.0854 | 0 | 4.8 | 413 | Specific | 896.06 | 709.96 | 709.96 | 18.39 | 68 | >sp\|P01833\|PIGR_HUMAN Polymeric immunoglobulin receptor OS=Homo sapiens OX=9606 GN=PIGR PE=1 SV=4 | 902 | 20201115_15212_1ng_glyco_hcd203040_compound15.25069.25069.3 | controllerType=0 controllerNumber=1 scan=25069 | 48.7384 |
| 22136:4 | 7 | R.LSLLEEPGN[+2059.735]GTFTVILNQLTSR.D | HexNAc(4)Hex(5)Fuc(1)NeuAc(1) | N[+2060] | 1116.2666 | 4 | 4462.0447 | 4462.0275 | 0 | 3.9 | 413 | Specific | 938.42 | 676.16 | 676.16 | 19.94 | 68 | >sp\|P01833\|PIGR_HUMAN Polymeric immunoglobulin receptor OS=Homo sapiens OX=9606 GN=PIGR PE=1 SV=4 | 902 | 20201115_15212_1ng_glyco_hcd203040_compound15.25140.25140.4 | controllerType=0 controllerNumber=1 scan=25140 | 48.9386 |
| 22134:4 | 7 | R.LSLLEEPGN[+2059.735]GTFTVILNQLTSR.D | HexNAc(4)Hex(5)Fuc(1)NeuAc(1) | N[+2060] | 1116.2666 | 4 | 4462.0447 | 4462.0275 | 0 | 3.9 | 413 | Specific | 1075.15 | 857.29 | 857.29 | 22.94 | 68 | >sp\|P01833\|PIGR_HUMAN Polymeric immunoglobulin receptor OS=Homo sapiens OX=9606 GN=PIGR PE=1 SV=4 | 902 | 20201115_15212_1ng_glyco_hcd203040_compound15.25138.25138.4 | controllerType=0 controllerNumber=1 scan=25138 | 48.9326 |
| 19562:3 | 7 | K.IIEGEPNLKVPGN[+2350.830]VTAVLGETLK.V | HexNAc(4)Hex(5)Fuc(1)NeuAc(2) | N[+2351] | 1581.4131 | 3 | 4742.2249 | 4742.1797 | 0 | 9.5 | 457 | Specific | 501.89 | 501.89 | 501.89 | 9.24 | 68 | >sp\|P01833\|PIGR_HUMAN Polymeric immunoglobulin receptor OS=Homo sapiens OX=9606 GN=PIGR PE=1 SV=4 | 902 | 20201115_15212_1ng_glyco_hcd203040_compound15.22449.22449.2 | controllerType=0 controllerNumber=1 scan=22449 | 43.1968 |
| 19565:3 | 7 | K.IIEGEPNLKVPGN[+2350.830]VTAVLGETLK.V | HexNAc(4)Hex(5)Fuc(1)NeuAc(2) | N[+2351] | 1581.4131 | 3 | 4742.2249 | 4742.1797 | 0 | 9.5 | 457 | Specific | 525.56 | 373.18 | 373.18 | 7.77 | 68 | >sp\|P01833\|PIGR_HUMAN Polymeric immunoglobulin receptor OS=Homo sapiens OX=9606 GN=PIGR PE=1 SV=4 | 902 | 20201115_15212_1ng_glyco_hcd203040_compound15.22452.22452.2 | controllerType=0 controllerNumber=1 scan=22452 | 43.2031 |
| 17797:3 | 7 | K.IIEGEPNLKVPGN[+2205.793]VTAVLGETLK.V | HexNAc(4)Hex(5)Fuc(2)NeuAc(1) | N[+2206] | 1533.059 | 3 | 4597.1624 | 4597.1422 | 0 | 4.4 | 457 | Specific | 535.7 | 535.7 | 535.7 | 10.61 | 68 | >sp\|P01833\|PIGR_HUMAN Polymeric immunoglobulin receptor OS=Homo sapiens OX=9606 GN=PIGR PE=1 SV=4 | 902 | 20201115_15212_1ng_glyco_hcd203040_compound15.20604.20604.3 | controllerType=0 controllerNumber=1 scan=20604 | 39.4462 |
| 17800:3 | 7 | K.IIEGEPNLKVPGN[+2205.793]VTAVLGETLK.V | HexNAc(4)Hex(5)Fuc(2)NeuAc(1) | N[+2206] | 1533.059 | 3 | 4597.1624 | 4597.1422 | 0 | 4.4 | 457 | Specific | 565.46 | 565.46 | 565.46 | 10.86 | 68 | >sp\|P01833\|PIGR_HUMAN Polymeric immunoglobulin receptor OS=Homo sapiens OX=9606 GN=PIGR PE=1 SV=4 | 902 | 20201115_15212_1ng_glyco_hcd203040_compound15.20607.20607.3 | controllerType=0 controllerNumber=1 scan=20607 | 39.4504 |
| 17770:4 | 7 | K.IIEGEPNLKVPGN[+2059.735]VTAVLGETLK.V | HexNAc(4)Hex(5)Fuc(1)NeuAc(1) | N[+2060] | 1113.5297 | 4 | 4451.097 | 4451.0843 | 0 | 2.9 | 457 | Specific | 579.97 | 487.81 | 487.81 | 12.22 | 68 | >sp\|P01833\|PIGR_HUMAN Polymeric immunoglobulin receptor OS=Homo sapiens OX=9606 GN=PIGR PE=1 SV=4 | 902 | 20201115_15212_1ng_glyco_hcd203040_compound15.20575.20575.4 | controllerType=0 controllerNumber=1 scan=20575 | 39.3642 |
| 17767:4 | 7 | K.IIEGEPNLKVPGN[+2059.735]VTAVLGETLK.V | HexNAc(4)Hex(5)Fuc(1)NeuAc(1) | N[+2060] | 1113.5297 | 4 | 4451.097 | 4451.0843 | 0 | 2.9 | 457 | Specific | 604.97 | 604.97 | 604.97 | 13.22 | 68 | >sp\|P01833\|PIGR_HUMAN Polymeric immunoglobulin receptor OS=Homo sapiens OX=9606 GN=PIGR PE=1 SV=4 | 902 | 20201115_15212_1ng_glyco_hcd203040_compound15.20572.20572.4 | controllerType=0 controllerNumber=1 scan=20572 | 39.3585 |
| 19534:3 | 7 | K.IIEGEPNLKVPGN[+2350.830]VTAVLGETLK.V | HexNAc(4)Hex(5)Fuc(1)NeuAc(2) | N[+2351] | 1581.4031 | 3 | 4742.1948 | 4742.1797 | 0 | 3.2 | 457 | Specific | 747.24 | 632.12 | 632.12 | 15.19 | 68 | >sp\|P01833\|PIGR_HUMAN Polymeric immunoglobulin receptor OS=Homo sapiens OX=9606 GN=PIGR PE=1 SV=4 | 902 | 20201115_15212_1ng_glyco_hcd203040_compound15.22420.22420.2 | controllerType=0 controllerNumber=1 scan=22420 | 43.1463 |
| 19579:3 | 7 | K.IIEGEPNLKVPGN[+2350.830]VTAVLGETLK.V | HexNAc(4)Hex(5)Fuc(1)NeuAc(2) | N[+2351] | 1581.4135 | 3 | 4742.226 | 4742.1797 | 0 | 9.8 | 457 | Specific | 782.23 | 782.23 | 782.23 | 15 | 68 | >sp\|P01833\|PIGR_HUMAN Polymeric immunoglobulin receptor OS=Homo sapiens OX=9606 GN=PIGR PE=1 SV=4 | 902 | 20201115_15212_1ng_glyco_hcd203040_compound15.22467.22467.3 | controllerType=0 controllerNumber=1 scan=22467 | 43.2384 |
| 17666:3 | 7 | K.IIEGEPNLKVPGN[+2205.793]VTAVLGETLK.V | HexNAc(4)Hex(5)Fuc(2)NeuAc(1) | N[+2206] | 1533.0544 | 3 | 4597.1486 | 4597.1422 | 0 | 1.4 | 457 | Specific | 824.67 | 716.74 | 716.74 | 17.11 | 68 | >sp\|P01833\|PIGR_HUMAN Polymeric immunoglobulin receptor OS=Homo sapiens OX=9606 GN=PIGR PE=1 SV=4 | 902 | 20201115_15212_1ng_glyco_hcd203040_compound15.20466.20466.3 | controllerType=0 controllerNumber=1 scan=20466 | 39.1632 |
| 17729:3 | 7 | K.IIEGEPNLKVPGN[+2059.735]VTAVLGETLK.V | HexNAc(4)Hex(5)Fuc(1)NeuAc(1) | N[+2060] | 1484.3713 | 3 | 4451.0993 | 4451.0843 | 0 | 3.4 | 457 | Specific | 829.08 | 631.23 | 631.23 | 17.19 | 68 | >sp\|P01833\|PIGR_HUMAN Polymeric immunoglobulin receptor OS=Homo sapiens OX=9606 GN=PIGR PE=1 SV=4 | 902 | 20201115_15212_1ng_glyco_hcd203040_compound15.20533.20533.3 | controllerType=0 controllerNumber=1 scan=20533 | 39.2808 |
| 17684:3 | 7 | K.IIEGEPNLKVPGN[+2205.793]VTAVLGETLK.V | HexNAc(4)Hex(5)Fuc(2)NeuAc(1) | N[+2206] | 1533.0558 | 3 | 4597.153 | 4597.1422 | 0 | 2.3 | 457 | Specific | 868.39 | 859.1 | 859.1 | 18.19 | 68 | >sp\|P01833\|PIGR_HUMAN Polymeric immunoglobulin receptor OS=Homo sapiens OX=9606 GN=PIGR PE=1 SV=4 | 902 | 20201115_15212_1ng_glyco_hcd203040_compound15.20486.20486.3 | controllerType=0 controllerNumber=1 scan=20486 | 39.1997 |

Table S4. Cont.

| Query #:z | Protein Rank | Glycopeptide | Glycan | Modification Type(s) | Observed m/z | z | Observed (M+H) | Calc. mass (M+H) | Off-by-x error | Mass error (ppm) | StartPosition | Cleavage | Score | Delta | Delta Mod | \|Log Prob\| | # of unique peptides | Protein Name | Protein DB number | Comment | Scan # | Scan Time |
|---|---|---|---|---|---|---|---|---|---|---|---|---|---|---|---|---|---|---|---|---|---|---|
| 19577:3 | 7 | K.IIEGEPNLKVPGN[+2350.830]VTAVLGETLK.V | HexNAc(4)Hex(5)Fuc(1)NeuAc(2) | N[+2351] | 1581.4135 | 3 | 4742.226 | 4742.1797 | 0 | 9.8 | 457 | Specific | 982.93 | 897.75 | 897.75 | 19 | 68 | >sp\|P01833\|PIGR_HUMAN Polymeric immunoglobulin receptor OS=Homo sapiens OX=9606 GN=PIGR PE=1 SV=4 | 902 | 20201115_15212_1ng_glyco_hcd203040_compound15.22465.22465.3 | controllerType=0 controllerNumber=1 scan=22465 | 43.2283 |
| 19531:3 | 7 | K.IIEGEPNLKVPGN[+2350.830]VTAVLGETLK.V | HexNAc(4)Hex(5)Fuc(1)NeuAc(2) | N[+2351] | 1581.4031 | 3 | 4742.1948 | 4742.1797 | 0 | 3.2 | 457 | Specific | 1007.77 | 824.7 | 824.7 | 21.19 | 68 | >sp\|P01833\|PIGR_HUMAN Polymeric immunoglobulin receptor OS=Homo sapiens OX=9606 GN=PIGR PE=1 SV=4 | 902 | 20201115_15212_1ng_glyco_hcd203040_compound15.22417.22417.2 | controllerType=0 controllerNumber=1 scan=22417 | 43.1366 |
| 17726:3 | 7 | K.IIEGEPNLKVPGN[+2059.735]VTAVLGETLK.V | HexNAc(4)Hex(5)Fuc(1)NeuAc(1) | N[+2060] | 1484.3713 | 3 | 4451.0993 | 4451.0843 | 0 | 3.4 | 457 | Specific | 1020.87 | 926.54 | 926.54 | 21.19 | 68 | >sp\|P01833\|PIGR_HUMAN Polymeric immunoglobulin receptor OS=Homo sapiens OX=9606 GN=PIGR PE=1 SV=4 | 902 | 20201115_15212_1ng_glyco_hcd203040_compound15.20530.20530.3 | controllerType=0 controllerNumber=1 scan=20530 | 39.2742 |
| 17664:3 | 7 | K.IIEGEPNLKVPGN[+2205.793]VTAVLGETLK.V | HexNAc(4)Hex(5)Fuc(2)NeuAc(1) | N[+2206] | 1533.0544 | 3 | 4597.1486 | 4597.1422 | 0 | 1.4 | 457 | Specific | 1028.78 | 999.29 | 999.29 | 21.11 | 68 | >sp\|P01833\|PIGR_HUMAN Polymeric immunoglobulin receptor OS=Homo sapiens OX=9606 GN=PIGR PE=1 SV=4 | 902 | 20201115_15212_1ng_glyco_hcd203040_compound15.20464.20464.3 | controllerType=0 controllerNumber=1 scan=20464 | 39.1585 |
| 17271:4 | 7 | K.IIEGEPNLKVPGN[+2059.735]VTAVLGETLKVPC[+57.021]HFPC[+57.021]K.F | HexNAc(4)Hex(5)Fuc(1)NeuAc(1) | C[+57]*2, N[+2060] | 1369.8927 | 4 | 5476.549 | 5476.5418 | 0 | 1.3 | 457 | Specific | 590.41 | 529.51 | 529.51 | 11.4 | 68 | >sp\|P01833\|PIGR_HUMAN Polymeric immunoglobulin receptor OS=Homo sapiens OX=9606 GN=PIGR PE=1 SV=4 | 902 | 20201115_15212_1ng_glyco_hcd203040_compound15.20021.20021.4 | controllerType=0 controllerNumber=1 scan=20021 | 38.3534 |
| 17312:5 | 7 | K.IIEGEPNLKVPGN[+2205.793]VTAVLGETLKVPC[+57.021]HFPC[+57.021]K.F | HexNAc(4)Hex(5)Fuc(2)NeuAc(1) | C[+57]*2, N[+2206] | 1125.328 | 5 | 5622.6109 | 5622.5997 | 0 | 2 | 457 | Specific | 595.73 | 595.73 | 595.73 | 12.01 | 68 | >sp\|P01833\|PIGR_HUMAN Polymeric immunoglobulin receptor OS=Homo sapiens OX=9606 GN=PIGR PE=1 SV=4 | 902 | 20201115_15212_1ng_glyco_hcd203040_compound15.20064.20064.5 | controllerType=0 controllerNumber=1 scan=20064 | 38.4412 |
| 17323:4 | 7 | K.IIEGEPNLKVPGN[+2205.793]VTAVLGETLKVPC[+57.021]HFPC[+57.021]K.F | HexNAc(4)Hex(5)Fuc(2)NeuAc(1) | C[+57]*2, N[+2206] | 1406.4102 | 4 | 5622.619 | 5622.5997 | 0 | 3.4 | 457 | Specific | 610.4 | 610.4 | 610.4 | 12.42 | 68 | >sp\|P01833\|PIGR_HUMAN Polymeric immunoglobulin receptor OS=Homo sapiens OX=9606 GN=PIGR PE=1 SV=4 | 902 | 20201115_15212_1ng_glyco_hcd203040_compound15.20075.20075.4 | controllerType=0 controllerNumber=1 scan=20075 | 38.4609 |
| 17220:5 | 7 | K.IIEGEPNLKVPGN[+2059.735]VTAVLGETLKVPC[+57.021]HFPC[+57.021]K.F | HexNAc(4)Hex(5)Fuc(1)NeuAc(1) | C[+57]*2, N[+2060] | 1096.1167 | 5 | 5476.5544 | 5476.5418 | 0 | 2.3 | 457 | Specific | 682.09 | 660.42 | 660.42 | 14.01 | 68 | >sp\|P01833\|PIGR_HUMAN Polymeric immunoglobulin receptor OS=Homo sapiens OX=9606 GN=PIGR PE=1 SV=4 | 902 | 20201115_15212_1ng_glyco_hcd203040_compound15.19968.19968.5 | controllerType=0 controllerNumber=1 scan=19968 | 38.2468 |
| 17244:4 | 7 | K.IIEGEPNLKVPGN[+2204.772]VTAVLGETLKVPC[+57.021]HFPC[+57.021]K.F | HexNAc(4)Hex(5)NeuAc(2) | C[+57]*2, N[+2205] | 1406.1595 | 4 | 5621.6164 | 5621.5793 | 0 | 6.6 | 457 | Specific | 684.25 | 684.25 | 684.25 | 12.41 | 68 | >sp\|P01833\|PIGR_HUMAN Polymeric immunoglobulin receptor OS=Homo sapiens OX=9606 GN=PIGR PE=1 SV=4 | 902 | 20201115_15212_1ng_glyco_hcd203040_compound15.19993.19993.4 | controllerType=0 controllerNumber=1 scan=19993 | 38.2985 |
| 17321:4 | 7 | K.IIEGEPNLKVPGN[+2205.793]VTAVLGETLKVPC[+57.021]HFPC[+57.021]K.F | HexNAc(4)Hex(5)Fuc(2)NeuAc(1) | C[+57]*2, N[+2206] | 1406.4102 | 4 | 5622.619 | 5622.5997 | 0 | 3.4 | 457 | Specific | 700.76 | 606.1 | 606.1 | 14.42 | 68 | >sp\|P01833\|PIGR_HUMAN Polymeric immunoglobulin receptor OS=Homo sapiens OX=9606 GN=PIGR PE=1 SV=4 | 902 | 20201115_15212_1ng_glyco_hcd203040_compound15.20073.20073.4 | controllerType=0 controllerNumber=1 scan=20073 | 38.4563 |
| 18692:4 | 7 | K.IIEGEPNLKVPGN[+2350.830]VTAVLGETLKVPC[+57.021]HFPC[+57.021]K.F | HexNAc(4)Hex(5)Fuc(1)NeuAc(2) | C[+57]*2, N[+2351] | 1442.6711 | 4 | 5767.6628 | 5767.6372 | 0 | 4.4 | 457 | Specific | 738.87 | 738.87 | 738.87 | 14.42 | 68 | >sp\|P01833\|PIGR_HUMAN Polymeric immunoglobulin receptor OS=Homo sapiens OX=9606 GN=PIGR PE=1 SV=4 | 902 | 20201115_15212_1ng_glyco_hcd203040_compound15.21539.21539.4 | controllerType=0 controllerNumber=1 scan=21539 | 41.1977 |
| 17261:5 | 7 | K.IIEGEPNLKVPGN[+2059.735]VTAVLGETLKVPC[+57.021]HFPC[+57.021]K.F | HexNAc(4)Hex(5)Fuc(1)NeuAc(1) | C[+57]*2, N[+2060] | 1096.118 | 5 | 5476.5611 | 5476.5418 | 0 | 3.5 | 457 | Specific | 774.1 | 646.4 | 646.4 | 16.03 | 68 | >sp\|P01833\|PIGR_HUMAN Polymeric immunoglobulin receptor OS=Homo sapiens OX=9606 GN=PIGR PE=1 SV=4 | 902 | 20201115_15212_1ng_glyco_hcd203040_compound15.20011.20011.5 | controllerType=0 controllerNumber=1 scan=20011 | 38.334 |
| 17241:4 | 7 | K.IIEGEPNLKVPGN[+2204.772]VTAVLGETLKVPC[+57.021]HFPC[+57.021]K.F | HexNAc(4)Hex(5)NeuAc(2) | C[+57]*2, N[+2205] | 1406.1595 | 4 | 5621.6164 | 5621.5793 | 0 | 6.6 | 457 | Specific | 790.13 | 616.44 | 616.44 | 14.41 | 68 | >sp\|P01833\|PIGR_HUMAN Polymeric immunoglobulin receptor OS=Homo sapiens OX=9606 GN=PIGR PE=1 SV=4 | 902 | 20201115_15212_1ng_glyco_hcd203040_compound15.19990.19990.4 | controllerType=0 controllerNumber=1 scan=19990 | 38.2936 |
| 18689:4 | 7 | K.IIEGEPNLKVPGN[+2350.830]VTAVLGETLKVPC[+57.021]HFPC[+57.021]K.F | HexNAc(4)Hex(5)Fuc(1)NeuAc(2) | C[+57]*2, N[+2351] | 1442.6711 | 4 | 5767.6628 | 5767.6372 | 0 | 4.4 | 457 | Specific | 792.18 | 568.47 | 568.47 | 15.42 | 68 | >sp\|P01833\|PIGR_HUMAN Polymeric immunoglobulin receptor OS=Homo sapiens OX=9606 GN=PIGR PE=1 SV=4 | 902 | 20201115_15212_1ng_glyco_hcd203040_compound15.21536.21536.4 | controllerType=0 controllerNumber=1 scan=21536 | 41.1929 |
| 17223:5 | 7 | K.IIEGEPNLKVPGN[+2059.735]VTAVLGETLKVPC[+57.021]HFPC[+57.021]K.F | HexNAc(4)Hex(5)Fuc(1)NeuAc(1) | C[+57]*2, N[+2060] | 1096.1167 | 5 | 5476.5544 | 5476.5418 | 0 | 2.3 | 457 | Specific | 807.37 | 697.07 | 697.07 | 17.01 | 68 | >sp\|P01833\|PIGR_HUMAN Polymeric immunoglobulin receptor OS=Homo sapiens OX=9606 GN=PIGR PE=1 SV=4 | 902 | 20201115_15212_1ng_glyco_hcd203040_compound15.19971.19971.5 | controllerType=0 controllerNumber=1 scan=19971 | 38.255 |
| 17314:5 | 7 | K.IIEGEPNLKVPGN[+2205.793]VTAVLGETLKVPC[+57.021]HFPC[+57.021]K.F | HexNAc(4)Hex(5)Fuc(2)NeuAc(1) | C[+57]*2, N[+2206] | 1125.328 | 5 | 5622.6109 | 5622.5997 | 0 | 2 | 457 | Specific | 847.62 | 801.3 | 801.3 | 17.01 | 68 | >sp\|P01833\|PIGR_HUMAN Polymeric immunoglobulin receptor OS=Homo sapiens OX=9606 GN=PIGR PE=1 SV=4 | 902 | 20201115_15212_1ng_glyco_hcd203040_compound15.20066.20066.5 | controllerType=0 controllerNumber=1 scan=20066 | 38.4471 |
| 17264:5 | 7 | K.IIEGEPNLKVPGN[+2059.735]VTAVLGETLKVPC[+57.021]HFPC[+57.021]K.F | HexNAc(4)Hex(5)Fuc(1)NeuAc(1) | C[+57]*2, N[+2060] | 1096.118 | 5 | 5476.5611 | 5476.5418 | 0 | 3.5 | 457 | Specific | 868.68 | 721.5 | 721.5 | 18.03 | 68 | >sp\|P01833\|PIGR_HUMAN Polymeric immunoglobulin receptor OS=Homo sapiens OX=9606 GN=PIGR PE=1 SV=4 | 902 | 20201115_15212_1ng_glyco_hcd203040_compound15.20014.20014.5 | controllerType=0 controllerNumber=1 scan=20014 | 38.3391 |
| 18659:5 | 7 | K.IIEGEPNLKVPGN[+2350.830]VTAVLGETLKVPC[+57.021]HFPC[+57.021]K.F | HexNAc(4)Hex(5)Fuc(1)NeuAc(2) | C[+57]*2, N[+2351] | 1154.3373 | 5 | 5767.6573 | 5767.6372 | 0 | 3.5 | 457 | Specific | 906.57 | 725.9 | 725.9 | 19.03 | 68 | >sp\|P01833\|PIGR_HUMAN Polymeric immunoglobulin receptor OS=Homo sapiens OX=9606 GN=PIGR PE=1 SV=4 | 902 | 20201115_15212_1ng_glyco_hcd203040_compound15.21505.21505.5 | controllerType=0 controllerNumber=1 scan=21505 | 41.1309 |
| 18687:5 | 7 | K.IIEGEPNLKVPGN[+2350.830]VTAVLGETLKVPC[+57.021]HFPC[+57.021]K.F | HexNAc(4)Hex(5)Fuc(1)NeuAc(2) | C[+57]*2, N[+2351] | 1154.3369 | 5 | 5767.6555 | 5767.6372 | 0 | 3.2 | 457 | Specific | 922.89 | 738.3 | 738.3 | 19.09 | 68 | >sp\|P01833\|PIGR_HUMAN Polymeric immunoglobulin receptor OS=Homo sapiens OX=9606 GN=PIGR PE=1 SV=4 | 902 | 20201115_15212_1ng_glyco_hcd203040_compound15.21534.21534.5 | controllerType=0 controllerNumber=1 scan=21534 | 41.1903 |
| 18656:5 | 7 | K.IIEGEPNLKVPGN[+2350.830]VTAVLGETLKVPC[+57.021]HFPC[+57.021]K.F | HexNAc(4)Hex(5)Fuc(1)NeuAc(2) | C[+57]*2, N[+2351] | 1154.3373 | 5 | 5767.6573 | 5767.6372 | 0 | 3.5 | 457 | Specific | 1024.73 | 832.87 | 832.87 | 21.03 | 68 | >sp\|P01833\|PIGR_HUMAN Polymeric immunoglobulin receptor OS=Homo sapiens OX=9606 GN=PIGR PE=1 SV=4 | 902 | 20201115_15212_1ng_glyco_hcd203040_compound15.21502.21502.5 | controllerType=0 controllerNumber=1 scan=21502 | 41.125 |
| 18685:5 | 7 | K.IIEGEPNLKVPGN[+2350.830]VTAVLGETLKVPC[+57.021]HFPC[+57.021]K.F | HexNAc(4)Hex(5)Fuc(1)NeuAc(2) | C[+57]*2, N[+2351] | 1154.3369 | 5 | 5767.6555 | 5767.6372 | 0 | 3.2 | 457 | Specific | 1070.93 | 854.16 | 854.16 | 22.09 | 68 | >sp\|P01833\|PIGR_HUMAN Polymeric immunoglobulin receptor OS=Homo sapiens OX=9606 GN=PIGR PE=1 SV=4 | 902 | 20201115_15212_1ng_glyco_hcd203040_compound15.21532.21532.5 | controllerType=0 controllerNumber=1 scan=21532 | 41.184 |
| 17269:4 | 7 | K.IIEGEPNLKVPGN[+2059.735]VTAVLGETLKVPC[+57.021]HFPC[+57.021]K.F | HexNAc(4)Hex(5)Fuc(1)NeuAc(1) | C[+57]*2, N[+2060] | 1369.8927 | 4 | 5476.549 | 5476.5418 | 0 | 1.3 | 457 | Specific | 1094.79 | 953.11 | 953.11 | 21.4 | 68 | >sp\|P01833\|PIGR_HUMAN Polymeric immunoglobulin receptor OS=Homo sapiens OX=9606 GN=PIGR PE=1 SV=4 | 902 | 20201115_15212_1ng_glyco_hcd203040_compound15.20019.20019.4 | controllerType=0 controllerNumber=1 scan=20019 | 38.3458 |
| 15329:3 | 7 | K.VPGN[+2059.735]VTAVLGETLK.V | HexNAc(4)Hex(5)Fuc(1)NeuAc(1) | N[+2060] | 1153.1878 | 3 | 3457.5489 | 3457.5348 | 0 | 4.1 | 466 | Specific | 467.86 | 455.96 | 455.96 | 10.47 | 68 | >sp\|P01833\|PIGR_HUMAN Polymeric immunoglobulin receptor OS=Homo sapiens OX=9606 GN=PIGR PE=1 SV=4 | 902 | 20201115_15212_1ng_glyco_hcd203040_compound15.18009.18009.3 | controllerType=0 controllerNumber=1 scan=18009 | 34.7099 |
| 15225:3 | 7 | K.VPGN[+2205.793]VTAVLGETLK.V | HexNAc(4)Hex(5)Fuc(2)NeuAc(1) | N[+2206] | 1201.873 | 3 | 3603.6044 | 3603.5927 | 0 | 3.2 | 466 | Specific | 675.46 | 624.82 | 624.82 | 14.36 | 68 | >sp\|P01833\|PIGR_HUMAN Polymeric immunoglobulin receptor OS=Homo sapiens OX=9606 GN=PIGR PE=1 SV=4 | 902 | 20201115_15212_1ng_glyco_hcd203040_compound15.17901.17901.3 | controllerType=0 controllerNumber=1 scan=17901 | 34.4757 |
| 15213:3 | 7 | K.VPGN[+2205.793]VTAVLGETLK.V | HexNAc(4)Hex(5)Fuc(2)NeuAc(1) | N[+2206] | 1201.8746 | 3 | 3603.6092 | 3603.5927 | 0 | 4.6 | 466 | Specific | 694.56 | 624.14 | 624.14 | 13.75 | 68 | >sp\|P01833\|PIGR_HUMAN Polymeric immunoglobulin receptor OS=Homo sapiens OX=9606 GN=PIGR PE=1 SV=4 | 902 | 20201115_15212_1ng_glyco_hcd203040_compound15.17888.17888.3 | controllerType=0 controllerNumber=1 scan=17888 | 34.4421 |
| 15314:3 | 7 | K.VPGN[+2059.735]VTAVLGETLK.V | HexNAc(4)Hex(5)Fuc(1)NeuAc(1) | N[+2060] | 1153.1872 | 3 | 3457.547 | 3457.5348 | 0 | 3.5 | 466 | Specific | 731.41 | 446.61 | 446.61 | 15.3 | 68 | >sp\|P01833\|PIGR_HUMAN Polymeric immunoglobulin receptor OS=Homo sapiens OX=9606 GN=PIGR PE=1 SV=4 | 902 | 20201115_15212_1ng_glyco_hcd203040_compound15.17993.17993.3 | controllerType=0 controllerNumber=1 scan=17993 | 34.6844 |
| 18317:3 | 7 | K.VPGN[+2350.830]VTAVLGETLK.V | HexNAc(4)Hex(5)Fuc(1)NeuAc(2) | N[+2351] | 1250.2186 | 3 | 3748.6411 | 3748.6302 | 0 | 2.9 | 466 | Specific | 737.78 | 506.36 | 506.36 | 15.36 | 68 | >sp\|P01833\|PIGR_HUMAN Polymeric immunoglobulin receptor OS=Homo sapiens OX=9606 GN=PIGR PE=1 SV=4 | 902 | 20201115_15212_1ng_glyco_hcd203040_compound15.21148.21148.3 | controllerType=0 controllerNumber=1 scan=21148 | 40.406 |
| 15312:3 | 7 | K.VPGN[+2059.735]VTAVLGETLK.V | HexNAc(4)Hex(5)Fuc(1)NeuAc(1) | N[+2060] | 1153.1872 | 3 | 3457.547 | 3457.5348 | 0 | 3.5 | 466 | Specific | 738.31 | 695.17 | 695.17 | 15.3 | 68 | >sp\|P01833\|PIGR_HUMAN Polymeric immunoglobulin receptor OS=Homo sapiens OX=9606 GN=PIGR PE=1 SV=4 | 902 | 20201115_15212_1ng_glyco_hcd203040_compound15.17991.17991.3 | controllerType=0 controllerNumber=1 scan=17991 | 34.6799 |
| 15332:3 | 7 | K.VPGN[+2059.735]VTAVLGETLK.V | HexNAc(4)Hex(5)Fuc(1)NeuAc(1) | N[+2060] | 1153.1878 | 3 | 3457.5489 | 3457.5348 | 0 | 4.1 | 466 | Specific | 746.63 | 520.79 | 520.79 | 15.3 | 68 | >sp\|P01833\|PIGR_HUMAN Polymeric immunoglobulin receptor OS=Homo sapiens OX=9606 GN=PIGR PE=1 SV=4 | 902 | 20201115_15212_1ng_glyco_hcd203040_compound15.18012.18012.3 | controllerType=0 controllerNumber=1 scan=18012 | 34.7136 |
| 18314:3 | 7 | K.VPGN[+2350.830]VTAVLGETLK.V | HexNAc(4)Hex(5)Fuc(1)NeuAc(2) | N[+2351] | 1250.2186 | 3 | 3748.6411 | 3748.6302 | 0 | 2.9 | 466 | Specific | 763.59 | 499.48 | 499.48 | 16.36 | 68 | >sp\|P01833\|PIGR_HUMAN Polymeric immunoglobulin receptor OS=Homo sapiens OX=9606 GN=PIGR PE=1 SV=4 | 902 | 20201115_15212_1ng_glyco_hcd203040_compound15.21145.21145.3 | controllerType=0 controllerNumber=1 scan=21145 | 40.391 |
| 15215:3 | 7 | K.VPGN[+2205.793]VTAVLGETLK.V | HexNAc(4)Hex(5)Fuc(2)NeuAc(1) | N[+2206] | 1201.8746 | 3 | 3603.6092 | 3603.5927 | 0 | 4.6 | 466 | Specific | 787.78 | 633.27 | 633.27 | 15.75 | 68 | >sp\|P01833\|PIGR_HUMAN Polymeric immunoglobulin receptor OS=Homo sapiens OX=9606 GN=PIGR PE=1 SV=4 | 902 | 20201115_15212_1ng_glyco_hcd203040_compound15.17890.17890.3 | controllerType=0 controllerNumber=1 scan=17890 | 34.4478 |
| 15358:4 | 7 | K.VPGN[+2204.772]VTAVLGETLKVPC[+57.021]HFPC[+57.021]K.F | HexNAc(4)Hex(5)NeuAc(2) | C[+57]*2, N[+2205] | 1157.7708 | 4 | 4628.0613 | 4628.0298 | 0 | 6.8 | 466 | Specific | 324.28 | 247.44 | 247.44 | 6.18 | 68 | >sp\|P01833\|PIGR_HUMAN Polymeric immunoglobulin receptor OS=Homo sapiens OX=9606 GN=PIGR PE=1 SV=4 | 902 | 20201115_15212_1ng_glyco_hcd203040_compound15.18039.18039.4 | controllerType=0 controllerNumber=1 scan=18039 | 34.7597 |
| 15361:4 | 7 | K.VPGN[+2204.772]VTAVLGETLKVPC[+57.021]HFPC[+57.021]K.F | HexNAc(4)Hex(5)NeuAc(2) | C[+57]*2, N[+2205] | 1157.7708 | 4 | 4628.0613 | 4628.0298 | 0 | 6.8 | 466 | Specific | 449.11 | 330.23 | 330.23 | 5.45 | 68 | >sp\|P01833\|PIGR_HUMAN Polymeric immunoglobulin receptor OS=Homo sapiens OX=9606 GN=PIGR PE=1 SV=4 | 902 | 20201115_15212_1ng_glyco_hcd203040_compound15.18042.18042.4 | controllerType=0 controllerNumber=1 scan=18042 | 34.7654 |
| 17191:4 | 7 | K.VPGN[+2350.830]VTAVLGETLKVPC[+57.021]HFPC[+57.021]K.F | HexNAc(4)Hex(5)Fuc(1)NeuAc(2) | C[+57]*2, N[+2351] | 1194.2818 | 4 | 4774.1053 | 4774.0877 | 0 | 3.7 | 466 | Specific | 704.67 | 459.29 | 459.29 | 15.04 | 68 | >sp\|P01833\|PIGR_HUMAN Polymeric immunoglobulin receptor OS=Homo sapiens OX=9606 GN=PIGR PE=1 SV=4 | 902 | 20201115_15212_1ng_glyco_hcd203040_compound15.19938.19938.4 | controllerType=0 controllerNumber=1 scan=19938 | 38.1888 |
| 17270:4 | 7 | K.VPGN[+2350.830]VTAVLGETLKVPC[+57.021]HFPC[+57.021]K.F | HexNAc(4)Hex(5)Fuc(1)NeuAc(2) | C[+57]*2, N[+2351] | 1194.2812 | 4 | 4774.1029 | 4774.0877 | 0 | 3.2 | 466 | Specific | 792.81 | 698.74 | 698.74 | 16.09 | 68 | >sp\|P01833\|PIGR_HUMAN Polymeric immunoglobulin receptor OS=Homo sapiens OX=9606 GN=PIGR PE=1 SV=4 | 902 | 20201115_15212_1ng_glyco_hcd203040_compound15.20020.20020.4 | controllerType=0 controllerNumber=1 scan=20020 | 38.3491 |
| 17189:4 | 7 | K.VPGN[+2350.830]VTAVLGETLKVPC[+57.021]HFPC[+57.021]K.F | HexNAc(4)Hex(5)Fuc(1)NeuAc(2) | C[+57]*2, N[+2351] | 1194.2818 | 4 | 4774.1053 | 4774.0877 | 0 | 3.7 | 466 | Specific | 802.41 | 642.46 | 642.46 | 17.04 | 68 | >sp\|P01833\|PIGR_HUMAN Polymeric immunoglobulin receptor OS=Homo sapiens OX=9606 GN=PIGR PE=1 SV=4 | 902 | 20201115_15212_1ng_glyco_hcd203040_compound15.19936.19936.4 | controllerType=0 controllerNumber=1 scan=19936 | 38.1843 |
| 17267:4 | 7 | K.VPGN[+2350.830]VTAVLGETLKVPC[+57.021]HFPC[+57.021]K.F | HexNAc(4)Hex(5)Fuc(1)NeuAc(2) | C[+57]*2, N[+2351] | 1194.2812 | 4 | 4774.1029 | 4774.0877 | 0 | 3.2 | 466 | Specific | 828.96 | 585.62 | 585.62 | 17.09 | 68 | >sp\|P01833\|PIGR_HUMAN Polymeric immunoglobulin receptor OS=Homo sapiens OX=9606 GN=PIGR PE=1 SV=4 | 902 | 20201115_15212_1ng_glyco_hcd203040_compound15.20017.20017.4 | controllerType=0 controllerNumber=1 scan=20017 | 38.3432 |
| 07399:3 | 7 | K.WN[+1872.651]NTGC[+57.021]QALPSQDEGPSK.A | HexNAc(3)Hex(6)NeuAc(1) | C[+57], N[+1873] | 1287.8497 | 3 | 3861.5344 | 3861.5272 | 0 | 1.9 | 498 | Specific | 595.64 | 549.35 | 549.35 | 12.39 | 68 | >sp\|P01833\|PIGR_HUMAN Polymeric immunoglobulin receptor OS=Homo sapiens OX=9606 GN=PIGR PE=1 SV=4 | 902 | 20201115_15212_1ng_glyco_hcd203040_compound15.9733.9733.3 | controllerType=0 controllerNumber=1 scan=9733 | 20.7729 |
| 07510:3 | 7 | K.WN[+1710.598]NTGC[+57.021]QALPSQDEGPSK.A | HexNAc(3)Hex(5)NeuAc(1) | C[+57], N[+1711] | 1233.8328 | 3 | 3699.4839 | 3699.4743 | 0 | 2.6 | 498 | Specific | 636.25 | 578.1 | 578.1 | 13.47 | 68 | >sp\|P01833\|PIGR_HUMAN Polymeric immunoglobulin receptor OS=Homo sapiens OX=9606 GN=PIGR PE=1 SV=4 | 902 | 20201115_15212_1ng_glyco_hcd203040_compound15.9848.9848.3 | controllerType=0 controllerNumber=1 scan=9848 | 20.9584 |
| 07401:3 | 7 | K.WN[+1872.651]NTGC[+57.021]QALPSQDEGPSK.A | HexNAc(3)Hex(6)NeuAc(1) | C[+57], N[+1873] | 1287.8497 | 3 | 3861.5344 | 3861.5272 | 0 | 1.9 | 498 | Specific | 690.68 | 641.92 | 641.92 | 14.39 | 68 | >sp\|P01833\|PIGR_HUMAN Polymeric immunoglobulin receptor OS=Homo sapiens OX=9606 GN=PIGR PE=1 SV=4 | 902 | 20201115_15212_1ng_glyco_hcd203040_compound15.9735.9735.3 | controllerType=0 controllerNumber=1 scan=9735 | 20.7767 |
| 07513:3 | 7 | K.WN[+1710.598]NTGC[+57.021]QALPSQDEGPSK.A | HexNAc(3)Hex(5)NeuAc(1) | C[+57], N[+1711] | 1233.8328 | 3 | 3699.4839 | 3699.4743 | 0 | 2.6 | 498 | Specific | 702.11 | 627.73 | 627.73 | 15.47 | 68 | >sp\|P01833\|PIGR_HUMAN Polymeric immunoglobulin receptor OS=Homo sapiens OX=9606 GN=PIGR PE=1 SV=4 | 902 | 20201115_15212_1ng_glyco_hcd203040_compound15.9851.9851.3 | controllerType=0 controllerNumber=1 scan=9851 | 20.9626 |
| 07414:3 | 7 | K.WN[+1913.677]NTGC[+57.021]QALPSQDEGPSK.A | HexNAc(4)Hex(5)NeuAc(1) | C[+57], N[+1914] | 1301.5257 | 3 | 3902.5625 | 3902.5537 | 0 | 2.3 | 498 | Specific | 767.4 | 706.1 | 706.1 | 16.39 | 68 | >sp\|P01833\|PIGR_HUMAN Polymeric immunoglobulin receptor OS=Homo sapiens OX=9606 GN=PIGR PE=1 SV=4 | 902 | 20201115_15212_1ng_glyco_hcd203040_compound15.9749.9749.3 | controllerType=0 controllerNumber=1 scan=9749 | 20.801 |
| 07417:3 | 7 | K.WN[+1913.677]NTGC[+57.021]QALPSQDEGPSK.A | HexNAc(4)Hex(5)NeuAc(1) | C[+57], N[+1914] | 1301.5257 | 3 | 3902.5625 | 3902.5537 | 0 | 2.3 | 498 | Specific | 822.72 | 723.29 | 723.29 | 17.39 | 68 | >sp\|P01833\|PIGR_HUMAN Polymeric immunoglobulin receptor OS=Homo sapiens OX=9606 GN=PIGR PE=1 SV=4 | 902 | 20201115_15212_1ng_glyco_hcd203040_compound15.9752.9752.3 | controllerType=0 controllerNumber=1 scan=9752 | 20.8054 |
| 20338:3 | 9 | R.LSLHRPALEDLLLGSEAN[+1751.624]LTC[+57.021]TLTGLR.D | HexNAc(4)Hex(4)NeuAc(1) | C[+57], N[+1752] | 1572.4164 | 3 | 4715.2348 | 4715.2225 | 0 | 2.6 | 127 | Specific | 231.17 | 231.17 | 231.17 | 6.35 | 59 | >sp\|P01876\|IGHA1_HUMAN Immunoglobulin heavy constant alpha 1 OS=Homo sapiens OX=9606 GN=IGHA1 PE=1 SV=2 | 255 | 20201115_15212_1ng_glyco_hcd203040_compound15.23258.23258.3 | controllerType=0 controllerNumber=1 scan=23258 | 44.8937 |
| 19475:4 | 9 | R.LSLHRPALEDLLLGSEAN[+1622.582]LTC[+57.021]TLTGLR.D | HexNAc(4)Hex(5) | C[+57], N[+1623] | 1147.298 | 4 | 4586.1702 | 4586.1799 | 0 | -2.1 | 127 | Specific | 373.94 | 200.31 | 200.31 | 4.19 | 59 | >sp\|P01876\|IGHA1_HUMAN Immunoglobulin heavy constant alpha 1 OS=Homo sapiens OX=9606 GN=IGHA1 PE=1 SV=2 | 255 | 20201115_15212_1ng_glyco_hcd203040_compound15.22358.22358.2 | controllerType=0 controllerNumber=1 scan=22358 | 42.9875 |
| 20286:3 | 9 | R.LSLHRPALEDLLLGSEAN[+1872.651]LTC[+57.021]TLTGLR.D | HexNAc(3)Hex(6)NeuAc(1) | C[+57], N[+1873] | 1612.7608 | 3 | 4836.2678 | 4836.2487 | 0 | 3.9 | 127 | Specific | 376.56 | 375.8 | 375.8 | 8 | 59 | >sp\|P01876\|IGHA1_HUMAN Immunoglobulin heavy constant alpha 1 OS=Homo sapiens OX=9606 GN=IGHA1 PE=1 SV=2 | 255 | 20201115_15212_1ng_glyco_hcd203040_compound15.23204.23204.3 | controllerType=0 controllerNumber=1 scan=23204 | 44.7869 |
| 19387:3 | 9 | R.LSLHRPALEDLLLGSEAN[+1216.423]LTC[+57.021]TLTGLR.D | HexNAc(2)Hex(5) | C[+57], N[+1216] | 1394.0154 | 3 | 4180.0318 | 4180.0211 | 0 | 2.5 | 127 | Specific | 384.95 | 384.95 | 384.95 | 9.5 | 59 | >sp\|P01876\|IGHA1_HUMAN Immunoglobulin heavy constant alpha 1 OS=Homo sapiens OX=9606 GN=IGHA1 PE=1 SV=2 | 255 | 20201115_15212_1ng_glyco_hcd203040_compound15.22265.22265.3 | controllerType=0 controllerNumber=1 scan=22265 | 42.7576 |
| 20331:4 | 9 | R.LSLHRPALEDLLLGSEAN[+1751.624]LTC[+57.021]TLTGLR.D | HexNAc(4)Hex(4)NeuAc(1) | C[+57], N[+1752] | 1179.5651 | 4 | 4715.2386 | 4715.2225 | 0 | 3.4 | 127 | Specific | 439.91 | 439.91 | 439.91 | 10.34 | 59 | >sp\|P01876\|IGHA1_HUMAN Immunoglobulin heavy constant alpha 1 OS=Homo sapiens OX=9606 GN=IGHA1 PE=1 SV=2 | 255 | 20201115_15212_1ng_glyco_hcd203040_compound15.23251.23251.4 | controllerType=0 controllerNumber=1 scan=23251 | 44.8833 |
| 19309:4 | 9 | R.LSLHRPALEDLLLGSEAN[+1663.608]LTC[+57.021]TLTGLR.D | HexNAc(5)Hex(4) | C[+57], N[+1664] | 1157.559 | 4 | 4627.2142 | 4627.2064 | 0 | 1.7 | 127 | Specific | 484.04 | 484.04 | 484.04 | 10.48 | 59 | >sp\|P01876\|IGHA1_HUMAN Immunoglobulin heavy constant alpha 1 OS=Homo sapiens OX=9606 GN=IGHA1 PE=1 SV=2 | 255 | 20201115_15212_1ng_glyco_hcd203040_compound15.22184.22184.4 | controllerType=0 controllerNumber=1 scan=22184 | 42.5837 |
| 20329:4 | 9 | R.LSLHRPALEDLLLGSEAN[+1913.677]LTC[+57.021]TLTGLR.D | HexNAc(4)Hex(5)NeuAc(1) | C[+57], N[+1914] | 1220.0815 | 4 | 4877.304 | 4877.2753 | 0 | 5.9 | 127 | Specific | 493.76 | 388.08 | 388.08 | 9.56 | 59 | >sp\|P01876\|IGHA1_HUMAN Immunoglobulin heavy constant alpha 1 OS=Homo sapiens OX=9606 GN=IGHA1 PE=1 SV=2 | 255 | 20201115_15212_1ng_glyco_hcd203040_compound15.23248.23248.4 | controllerType=0 controllerNumber=1 scan=23248 | 44.8773 |
| 19365:5 | 9 | R.LSLHRPALEDLLLGSEAN[+1501.555]LTC[+57.021]TLTGLR.D | HexNAc(5)Hex(3) | C[+57], N[+1502] | 893.8391 | 5 | 4465.1663 | 4465.1536 | 0 | 2.8 | 127 | Specific | 496.13 | 496.13 | 496.13 | 10.66 | 59 | >sp\|P01876\|IGHA1_HUMAN Immunoglobulin heavy constant alpha 1 OS=Homo sapiens OX=9606 GN=IGHA1 PE=1 SV=2 | 255 | 20201115_15212_1ng_glyco_hcd203040_compound15.22242.22242.5 | controllerType=0 controllerNumber=1 scan=22242 | 42.7084 |
| 19310:3 | 9 | R.LSLHRPALEDLLLGSEAN[+1663.608]LTC[+57.021]TLTGLR.D | HexNAc(5)Hex(4) | C[+57], N[+1664] | 1543.0767 | 3 | 4627.2156 | 4627.2064 | 0 | 2 | 127 | Specific | 513.96 | 494.45 | 494.45 | 10 | 59 | >sp\|P01876\|IGHA1_HUMAN Immunoglobulin heavy constant alpha 1 OS=Homo sapiens OX=9606 GN=IGHA1 PE=1 SV=2 | 255 | 20201115_15212_1ng_glyco_hcd203040_compound15.22185.22185.3 | controllerType=0 controllerNumber=1 scan=22185 | 42.5851 |
| 19296:4 | 9 | R.LSLHRPALEDLLLGSEAN[+1581.555]LTC[+57.021]TLTGLR.D | HexNAc(3)Hex(6) | C[+57], N[+1582] | 1137.045 | 4 | 4545.158 | 4545.1533 | 0 | 1 | 127 | Specific | 515.9 | 515.9 | 515.9 | 10.08 | 59 | >sp\|P01876\|IGHA1_HUMAN Immunoglobulin heavy constant alpha 1 OS=Homo sapiens OX=9606 GN=IGHA1 PE=1 SV=2 | 255 | 20201115_15212_1ng_glyco_hcd203040_compound15.22170.22170.4 | controllerType=0 controllerNumber=1 scan=22170 | 42.5531 |
| 19345:3 | 9 | R.LSLHRPALEDLLLGSEAN[+1298.476]LTC[+57.021]TLTGLR.D | HexNAc(4)Hex(3) | C[+57], N[+1298] | 1421.369 | 3 | 4262.0923 | 4262.0742 | 0 | 4.2 | 127 | Specific | 538.39 | 531.55 | 531.55 | 10.2 | 59 | >sp\|P01876\|IGHA1_HUMAN Immunoglobulin heavy constant alpha 1 OS=Homo sapiens OX=9606 GN=IGHA1 PE=1 SV=2 | 255 | 20201115_15212_1ng_glyco_hcd203040_compound15.22221.22221.3 | controllerType=0 controllerNumber=1 scan=22221 | 42.6619 |
| 20340:3 | 9 | R.LSLHRPALEDLLLGSEAN[+1751.624]LTC[+57.021]TLTGLR.D | HexNAc(4)Hex(4)NeuAc(1) | C[+57], N[+1752] | 1572.4164 | 3 | 4715.2348 | 4715.2225 | 0 | 2.6 | 127 | Specific | 543.47 | 491.55 | 491.55 | 10.26 | 59 | >sp\|P01876\|IGHA1_HUMAN Immunoglobulin heavy constant alpha 1 OS=Homo sapiens OX=9606 GN=IGHA1 PE=1 SV=2 | 255 | 20201115_15212_1ng_glyco_hcd203040_compound15.23260.23260.3 | controllerType=0 controllerNumber=1 scan=23260 | 44.899 |
| 20246:5 | 9 | R.LSLHRPALEDLLLGSEAN[+2116.756]LTC[+57.021]TLTGLR.D | HexNAc(5)Hex(5)NeuAc(1) | C[+57], N[+2117] | 1016.8801 | 5 | 5080.3712 | 5080.3547 | 0 | 3.3 | 127 | Specific | 557.15 | 517.57 | 517.57 | 11.15 | 59 | >sp\|P01876\|IGHA1_HUMAN Immunoglobulin heavy constant alpha 1 OS=Homo sapiens OX=9606 GN=IGHA1 PE=1 SV=2 | 255 | 20201115_15212_1ng_glyco_hcd203040_compound15.23162.23162.5 | controllerType=0 controllerNumber=1 scan=23162 | 44.6987 |
| 19362:5 | 9 | R.LSLHRPALEDLLLGSEAN[+1501.555]LTC[+57.021]TLTGLR.D | HexNAc(5)Hex(3) | C[+57], N[+1502] | 893.8391 | 5 | 4465.1663 | 4465.1536 | 0 | 2.8 | 127 | Specific | 560.13 | 560.13 | 560.13 | 11.15 | 59 | >sp\|P01876\|IGHA1_HUMAN Immunoglobulin heavy constant alpha 1 OS=Homo sapiens OX=9606 GN=IGHA1 PE=1 SV=2 | 255 | 20201115_15212_1ng_glyco_hcd203040_compound15.22239.22239.5 | controllerType=0 controllerNumber=1 scan=22239 | 42.7009 |
| 20289:3 | 9 | R.LSLHRPALEDLLLGSEAN[+1872.651]LTC[+57.021]TLTGLR.D | HexNAc(3)Hex(6)NeuAc(1) | C[+57], N[+1873] | 1612.7608 | 3 | 4836.2678 | 4836.2487 | 0 | 3.9 | 127 | Specific | 561.31 | 505.64 | 505.64 | 10.45 | 59 | >sp\|P01876\|IGHA1_HUMAN Immunoglobulin heavy constant alpha 1 OS=Homo sapiens OX=9606 GN=IGHA1 PE=1 SV=2 | 255 | 20201115_15212_1ng_glyco_hcd203040_compound15.23207.23207.3 | controllerType=0 controllerNumber=1 scan=23207 | 44.7942 |
| 19348:3 | 9 | R.LSLHRPALEDLLLGSEAN[+1298.476]LTC[+57.021]TLTGLR.D | HexNAc(4)Hex(3) | C[+57], N[+1298] | 1421.369 | 3 | 4262.0923 | 4262.0742 | 0 | 4.2 | 127 | Specific | 562.89 | 515.11 | 515.11 | 10.45 | 59 | >sp\|P01876\|IGHA1_HUMAN Immunoglobulin heavy constant alpha 1 OS=Homo sapiens OX=9606 GN=IGHA1 PE=1 SV=2 | 255 | 20201115_15212_1ng_glyco_hcd203040_compound15.22224.22224.3 | controllerType=0 controllerNumber=1 scan=22224 | 42.6698 |
| 20308:3 | 9 | R.LSLHRPALEDLLLGSEAN[+1710.598]LTC[+57.021]TLTGLR.D | HexNAc(3)Hex(5)NeuAc(1) | C[+57], N[+1711] | 1558.7437 | 3 | 4674.2166 | 4674.1959 | 0 | 4.4 | 127 | Specific | 566.98 | 512.21 | 512.21 | 10.45 | 59 | >sp\|P01876\|IGHA1_HUMAN Immunoglobulin heavy constant alpha 1 OS=Homo sapiens OX=9606 GN=IGHA1 PE=1 SV=2 | 255 | 20201115_15212_1ng_glyco_hcd203040_compound15.23227.23227.3 | controllerType=0 controllerNumber=1 scan=23227 | 44.834 |
| 19358:3 | 9 | R.LSLHRPALEDLLLGSEAN[+1501.555]LTC[+57.021]TLTGLR.D | HexNAc(5)Hex(3) | C[+57], N[+1502] | 1489.0596 | 3 | 4465.1643 | 4465.1536 | 0 | 2.4 | 127 | Specific | 569.61 | 569.61 | 569.61 | 10.51 | 59 | >sp\|P01876\|IGHA1_HUMAN Immunoglobulin heavy constant alpha 1 OS=Homo sapiens OX=9606 GN=IGHA1 PE=1 SV=2 | 255 | 20201115_15212_1ng_glyco_hcd203040_compound15.22235.22235.3 | controllerType=0 controllerNumber=1 scan=22235 | 42.6945 |
| 19357:4 | 9 | R.LSLHRPALEDLLLGSEAN[+1501.555]LTC[+57.021]TLTGLR.D | HexNAc(5)Hex(3) | C[+57], N[+1502] | 1117.0489 | 4 | 4465.1737 | 4465.1536 | 0 | 4.5 | 127 | Specific | 583.38 | 583.38 | 583.38 | 12.33 | 59 | >sp\|P01876\|IGHA1_HUMAN Immunoglobulin heavy constant alpha 1 OS=Homo sapiens OX=9606 GN=IGHA1 PE=1 SV=2 | 255 | 20201115_15212_1ng_glyco_hcd203040_compound15.22234.22234.4 | controllerType=0 controllerNumber=1 scan=22234 | 42.6931 |
| 21413:4 | 9 | R.LSLHRPALEDLLLGSEAN[+2204.772]LTC[+57.021]TLTGLR.D | HexNAc(4)Hex(5)NeuAc(2) | C[+57], N[+2205] | 1292.8601 | 4 | 5168.4186 | 5168.3707 | 0 | 9.3 | 127 | Specific | 610.13 | 529.98 | 529.98 | 12.19 | 59 | >sp\|P01876\|IGHA1_HUMAN Immunoglobulin heavy constant alpha 1 OS=Homo sapiens OX=9606 GN=IGHA1 PE=1 SV=2 | 255 | 20201115_15212_1ng_glyco_hcd203040_compound15.24383.24383.4 | controllerType=0 controllerNumber=1 scan=24383 | 47.1741 |
| 19351:4 | 9 | R.LSLHRPALEDLLLGSEAN[+1647.613]LTC[+57.021]TLTGLR.D | HexNAc(5)Hex(3)Fuc(1) | C[+57], N[+1648] | 1153.5597 | 4 | 4611.2171 | 4611.2115 | 0 | 1.2 | 127 | Specific | 628.45 | 628.45 | 628.45 | 13.31 | 59 | >sp\|P01876\|IGHA1_HUMAN Immunoglobulin heavy constant alpha 1 OS=Homo sapiens OX=9606 GN=IGHA1 PE=1 SV=2 | 255 | 20201115_15212_1ng_glyco_hcd203040_compound15.22227.22227.4 | controllerType=0 controllerNumber=1 scan=22227 | 42.6732 |
| 19321:3 | 9 | R.LSLHRPALEDLLLGSEAN[+1460.529]LTC[+57.021]TLTGLR.D | HexNAc(4)Hex(4) | C[+57], N[+1461] | 1475.3837 | 3 | 4424.1366 | 4424.1271 | 0 | 2.2 | 127 | Specific | 638.28 | 638.28 | 638.28 | 12.7 | 59 | >sp\|P01876\|IGHA1_HUMAN Immunoglobulin heavy constant alpha 1 OS=Homo sapiens OX=9606 GN=IGHA1 PE=1 SV=2 | 255 | 20201115_15212_1ng_glyco_hcd203040_compound15.22196.22196.3 | controllerType=0 controllerNumber=1 scan=22196 | [illegible] |
| 19252:3 | 9 | R.LSLHRPALEDLLLGSEAN[+1540.529]LTC[+57.021]TLTGLR.D | HexNAc(2)Hex(7) | C[+57], N[+1541] | 1502.0524 | 3 | 4504.1427 | 4504.1268 | 0 | 3.5 | 127 | Specific | 639.73 | 635.53 | 635.53 | 12.72 | 59 | >sp\|P01876\|IGHA1_HUMAN Immunoglobulin heavy constant alpha 1 OS=Homo sapiens OX=9606 GN=IGHA1 PE=1 SV=2 | 255 | 20201115_15212_1ng_glyco_hcd203040_compound15.22124.22124.3 | controllerType=0 controllerNumber=1 scan=22124 | [illegible] |
| 19324:3 | 9 | R.LSLHRPALEDLLLGSEAN[+1460.529]LTC[+57.021]TLTGLR.D | HexNAc(4)Hex(4) | C[+57], N[+1461] | 1475.3837 | 3 | 4424.1366 | 4424.1271 | 0 | 2.2 | 127 | Specific | 647.57 | 604.59 | 604.59 | 12.7 | 59 | >sp\|P01876\|IGHA1_HUMAN Immunoglobulin heavy constant alpha 1 OS=Homo sapiens OX=9606 GN=IGHA1 PE=1 SV=2 | 255 | 20201115_15212_1ng_glyco_hcd203040_compound15.22199.22199.3 | controllerType=0 controllerNumber=1 scan=22199 | 42.6148 |
| 19219:3 | 9 | R.LSLHRPALEDLLLGSEAN[+1864.634]LTC[+57.021]TLTGLR.D | HexNAc(2)Hex(9) | C[+57], N[+1865] | 1610.0868 | 3 | 4828.246 | 4828.2324 | 0 | 2.8 | 127 | Specific | 649.36 | 602.37 | 602.37 | 12.78 | 59 | >sp\|P01876\|IGHA1_HUMAN Immunoglobulin heavy constant alpha 1 OS=Homo sapiens OX=9606 GN=IGHA1 PE=1 SV=2 | 255 | 20201115_15212_1ng_glyco_hcd203040_compound15.22090.22090.3 | controllerType=0 controllerNumber=1 scan=22090 | 42.357 |
| 20305:3 | 9 | R.LSLHRPALEDLLLGSEAN[+1710.598]LTC[+57.021]TLTGLR.D | HexNAc(3)Hex(5)NeuAc(1) | C[+57], N[+1711] | 1558.7437 | 3 | 4674.2166 | 4674.1959 | 0 | 4.4 | 127 | Specific | 652.65 | 577.03 | 577.03 | 13.72 | 59 | >sp\|P01876\|IGHA1_HUMAN Immunoglobulin heavy constant alpha 1 OS=Homo sapiens OX=9606 GN=IGHA1 PE=1 SV=2 | 255 | 20201115_15212_1ng_glyco_hcd203040_compound15.23224.23224.3 | controllerType=0 controllerNumber=1 scan=23224 | 44.8287 |
| 19360:3 | 9 | R.LSLHRPALEDLLLGSEAN[+1501.555]LTC[+57.021]TLTGLR.D | HexNAc(5)Hex(3) | C[+57], N[+1502] | 1489.0596 | 3 | 4465.1643 | 4465.1536 | 0 | 2.4 | 127 | Specific | 655.04 | 621.33 | 621.33 | 13.78 | 59 | >sp\|P01876\|IGHA1_HUMAN Immunoglobulin heavy constant alpha 1 OS=Homo sapiens OX=9606 GN=IGHA1 PE=1 SV=2 | 255 | 20201115_15212_1ng_glyco_hcd203040_compound15.22237.22237.3 | controllerType=0 controllerNumber=1 scan=22237 | 42.6987 |
| 19312:3 | 9 | R.LSLHRPALEDLLLGSEAN[+1663.608]LTC[+57.021]TLTGLR.D | HexNAc(5)Hex(4) | C[+57], N[+1664] | 1543.0767 | 3 | 4627.2156 | 4627.2064 | 0 | 2 | 127 | Specific | 660.72 | 605.37 | 605.37 | 13.7 | 59 | >sp\|P01876\|IGHA1_HUMAN Immunoglobulin heavy constant alpha 1 OS=Homo sapiens OX=9606 GN=IGHA1 PE=1 SV=2 | 255 | 20201115_15212_1ng_glyco_hcd203040_compound15.22187.22187.3 | controllerType=0 controllerNumber=1 scan=22187 | 42.589 |
| 19334:3 | 9 | R.LSLHRPALEDLLLGSEAN[+1378.476]LTC[+57.021]TLTGLR.D | HexNAc(2)Hex(6) | C[+57], N[+1378] | 1448.0319 | 3 | 4342.0812 | 4342.074 | 0 | 1.7 | 127 | Specific | 676.34 | 653.82 | 653.82 | 13.7 | 59 | >sp\|P01876\|IGHA1_HUMAN Immunoglobulin heavy constant alpha 1 OS=Homo sapiens OX=9606 GN=IGHA1 PE=1 SV=2 | 255 | 20201115_15212_1ng_glyco_hcd203040_compound15.22210.22210.3 | controllerType=0 controllerNumber=1 scan=22210 | 42.6389 |
| 19217:3 | 9 | R.LSLHRPALEDLLLGSEAN[+1864.634]LTC[+57.021]TLTGLR.D | HexNAc(2)Hex(9) | C[+57], N[+1865] | 1610.0868 | 3 | 4828.246 | 4828.2324 | 0 | 2.8 | 127 | Specific | 686.74 | 686.74 | 686.74 | 13.78 | 59 | >sp\|P01876\|IGHA1_HUMAN Immunoglobulin heavy constant alpha 1 OS=Homo sapiens OX=9606 GN=IGHA1 PE=1 SV=2 | 255 | 20201115_15212_1ng_glyco_hcd203040_compound15.22088.22088.3 | controllerType=0 controllerNumber=1 scan=22088 | 42.3518 |
| 19349:3 | 9 | R.LSLHRPALEDLLLGSEAN[+1257.449]LTC[+57.021]TLTGLR.D | HexNAc(3)Hex(4) | C[+57], N[+1257] | 1407.6907 | 3 | 4221.0576 | 4221.0477 | 0 | 2.4 | 127 | Specific | 689.55 | 689.55 | 689.55 | 13.7 | 59 | >sp\|P01876\|IGHA1_HUMAN Immunoglobulin heavy constant alpha 1 OS=Homo sapiens OX=9606 GN=IGHA1 PE=1 SV=2 | 255 | 20201115_15212_1ng_glyco_hcd203040_compound15.22225.22225.3 | controllerType=0 controllerNumber=1 scan=22225 | 42.6709 |
| 19336:3 | 9 | R.LSLHRPALEDLLLGSEAN[+1378.476]LTC[+57.021]TLTGLR.D | HexNAc(2)Hex(6) | C[+57], N[+1378] | 1448.0319 | 3 | 4342.0812 | 4342.074 | 0 | 1.7 | 127 | Specific | 696.65 | 696.65 | 696.65 | 13.7 | 59 | >sp\|P01876\|IGHA1_HUMAN Immunoglobulin heavy constant alpha 1 OS=Homo sapiens OX=9606 GN=IGHA1 PE=1 SV=2 | 255 | 20201115_15212_1ng_glyco_hcd203040_compound15.22212.22212.3 | controllerType=0 controllerNumber=1 scan=22212 | 42.6435 |
| 20301:4 | 9 | R.LSLHRPALEDLLLGSEAN[+1710.598]LTC[+57.021]TLTGLR.D | HexNAc(3)Hex(5)NeuAc(1) | C[+57], N[+1711] | 1169.3086 | 4 | 4674.2127 | 4674.1959 | 0 | 3.6 | 127 | Specific | 700.05 | 689.62 | 689.62 | 15.33 | 59 | >sp\|P01876\|IGHA1_HUMAN Immunoglobulin heavy constant alpha 1 OS=Homo sapiens OX=9606 GN=IGHA1 PE=1 SV=2 | 255 | 20201115_15212_1ng_glyco_hcd203040_compound15.23220.23220.4 | controllerType=0 controllerNumber=1 scan=23220 | 44.8198 |
| 19250:3 | 9 | R.LSLHRPALEDLLLGSEAN[+1540.529]LTC[+57.021]TLTGLR.D | HexNAc(2)Hex(7) | C[+57], N[+1541] | 1502.0524 | 3 | 4504.1427 | 4504.1268 | 0 | 3.5 | 127 | Specific | 708.23 | 643.49 | 643.49 | 14.72 | 59 | >sp\|P01876\|IGHA1_HUMAN Immunoglobulin heavy constant alpha 1 OS=Homo sapiens OX=9606 GN=IGHA1 PE=1 SV=2 | 255 | 20201115_15212_1ng_glyco_hcd203040_compound15.22122.22122.3 | controllerType=0 controllerNumber=1 scan=22122 | 42.4198 |
| 19389:3 | 9 | R.LSLHRPALEDLLLGSEAN[+1216.423]LTC[+57.021]TLTGLR.D | HexNAc(2)Hex(5) | C[+57], N[+1216] | 1394.0154 | 3 | 4180.0318 | 4180.0211 | 0 | 2.5 | 127 | Specific | 709 | 689.42 | 689.42 | 14.78 | 59 | >sp\|P01876\|IGHA1_HUMAN Immunoglobulin heavy constant alpha 1 OS=Homo sapiens OX=9606 GN=IGHA1 PE=1 SV=2 | 255 | 20201115_15212_1ng_glyco_hcd203040_compound15.22267.22267.3 | controllerType=0 controllerNumber=1 scan=22267 | 42.7609 |
| 19241:3 | 9 | R.LSLHRPALEDLLLGSEAN[+1702.581]LTC[+57.021]TLTGLR.D | HexNAc(2)Hex(8) | C[+57], N[+1703] | 1556.0704 | 3 | 4666.1966 | 4666.1796 | 0 | 3.6 | 127 | Specific | 724.73 | 724.73 | 724.73 | 14.72 | 59 | >sp\|P01876\|IGHA1_HUMAN Immunoglobulin heavy constant alpha 1 OS=Homo sapiens OX=9606 GN=IGHA1 PE=1 SV=2 | 255 | 20201115_15212_1ng_glyco_hcd203040_compound15.22113.22113.3 | controllerType=0 controllerNumber=1 scan=22113 | 42.4057 |
| 19314:4 | 9 | R.LSLHRPALEDLLLGSEAN[+1460.529]LTC[+57.021]TLTGLR.D | HexNAc(4)Hex(4) | C[+57], N[+1461] | 1106.7917 | 4 | 4424.1448 | 4424.1271 | 0 | 4 | 127 | Specific | 727.89 | 727.89 | 727.89 | 15.33 | 59 | >sp\|P01876\|IGHA1_HUMAN Immunoglobulin heavy constant alpha 1 OS=Homo sapiens OX=9606 GN=IGHA1 PE=1 SV=2 | 255 | 20201115_15212_1ng_glyco_hcd203040_compound15.22189.22189.4 | controllerType=0 controllerNumber=1 scan=22189 | 42.5921 |
| 21416:4 | 9 | R.LSLHRPALEDLLLGSEAN[+2204.772]LTC[+57.021]TLTGLR.D | HexNAc(4)Hex(5)NeuAc(2) | C[+57], N[+2205] | 1292.8601 | 4 | 5168.4186 | 5168.3707 | 0 | 9.3 | 127 | Specific | 738.04 | 721.45 | 721.45 | 14.19 | 59 | >sp\|P01876\|IGHA1_HUMAN Immunoglobulin heavy constant alpha 1 OS=Homo sapiens OX=9606 GN=IGHA1 PE=1 SV=2 | 255 | 20201115_15212_1ng_glyco_hcd203040_compound15.24386.24386.4 | controllerType=0 controllerNumber=1 scan=24386 | 47.1829 |
| 19415:4 | 9 | R.LSLHRPALEDLLLGSEAN[+1054.370]LTC[+57.021]TLTGLR.D | HexNAc(2)Hex(4) | C[+57], N[+1054] | 1005.2504 | 4 | 4017.9798 | 4017.9683 | 0 | 2.9 | 127 | Specific | 740.94 | 731.68 | 731.68 | 15.39 | 59 | >sp\|P01876\|IGHA1_HUMAN Immunoglobulin heavy constant alpha 1 OS=Homo sapiens OX=9606 GN=IGHA1 PE=1 SV=2 | 255 | 20201115_15212_1ng_glyco_hcd203040_compound15.22294.22294.4 | controllerType=0 controllerNumber=1 scan=22294 | 42.817 |
| 20249:5 | 9 | R.LSLHRPALEDLLLGSEAN[+2116.756]LTC[+57.021]TLTGLR.D | HexNAc(5)Hex(5)NeuAc(1) | C[+57], N[+2117] | 1016.8801 | 5 | 5080.3712 | 5080.3547 | 0 | 3.3 | 127 | Specific | 747.07 | 653.67 | 653.67 | 15.41 | 59 | >sp\|P01876\|IGHA1_HUMAN Immunoglobulin heavy constant alpha 1 OS=Homo sapiens OX=9606 GN=IGHA1 PE=1 SV=2 | 255 | 20201115_15212_1ng_glyco_hcd203040_compound15.23165.23165.5 | controllerType=0 controllerNumber=1 scan=23165 | 44.7054 |
| 19311:4 | 9 | R.LSLHRPALEDLLLGSEAN[+1663.608]LTC[+57.021]TLTGLR.D | HexNAc(5)Hex(4) | C[+57], N[+1664] | 1157.559 | 4 | 4627.2142 | 4627.2064 | 0 | 1.7 | 127 | Specific | 748.33 | 683.99 | 683.99 | 15.31 | 59 | >sp\|P01876\|IGHA1_HUMAN Immunoglobulin heavy constant alpha 1 OS=Homo sapiens OX=9606 GN=IGHA1 PE=1 SV=2 | 255 | 20201115_15212_1ng_glyco_hcd203040_compound15.22186.22186.4 | controllerType=0 controllerNumber=1 scan=22186 | 42.5879 |
| 19352:3 | 9 | R.LSLHRPALEDLLLGSEAN[+1257.449]LTC[+57.021]TLTGLR.D | HexNAc(3)Hex(4) | C[+57], N[+1257] | 1407.6907 | 3 | 4221.0576 | 4221.0477 | 0 | 2.4 | 127 | Specific | 770.08 | 727.58 | 727.58 | 15.7 | 59 | >sp\|P01876\|IGHA1_HUMAN Immunoglobulin heavy constant alpha 1 OS=Homo sapiens OX=9606 GN=IGHA1 PE=1 SV=2 | 255 | 20201115_15212_1ng_glyco_hcd203040_compound15.22228.22228.3 | controllerType=0 controllerNumber=1 scan=22228 | 42.676 |
| 20253:4 | 9 | R.LSLHRPALEDLLLGSEAN[+1872.651]LTC[+57.021]TLTGLR.D | HexNAc(3)Hex(6)NeuAc(1) | C[+57], N[+1873] | 1209.8221 | 4 | 4836.2664 | 4836.2487 | 0 | 3.7 | 127 | Specific | 774.53 | 684 | 684 | 16.33 | 59 | >sp\|P01876\|IGHA1_HUMAN Immunoglobulin heavy constant alpha 1 OS=Homo sapiens OX=9606 GN=IGHA1 PE=1 SV=2 | 255 | 20201115_15212_1ng_glyco_hcd203040_compound15.23170.23170.4 | controllerType=0 controllerNumber=1 scan=23170 | 44.7149 |
| 19297:4 | 9 | R.LSLHRPALEDLLLGSEAN[+1419.502]LTC[+57.021]TLTGLR.D | HexNAc(3)Hex(5) | C[+57], N[+1420] | 1096.5334 | 4 | 4383.1116 | 4383.1005 | 0 | 2.5 | 127 | Specific | 780.5 | 780.5 | 780.5 | 16.39 | 59 | >sp\|P01876\|IGHA1_HUMAN Immunoglobulin heavy constant alpha 1 OS=Homo sapiens OX=9606 GN=IGHA1 PE=1 SV=2 | 255 | 20201115_15212_1ng_glyco_hcd203040_compound15.22171.22171.4 | controllerType=0 controllerNumber=1 scan=22171 | 42.5542 |
| 19338:4 | 9 | R.LSLHRPALEDLLLGSEAN[+1298.476]LTC[+57.021]TLTGLR.D | HexNAc(4)Hex(3) | C[+57], N[+1298] | 1066.276 | 4 | 4262.0823 | 4262.0742 | 0 | 1.9 | 127 | Specific | 787.55 | 787.55 | 787.55 | 16.31 | 59 | >sp\|P01876\|IGHA1_HUMAN Immunoglobulin heavy constant alpha 1 OS=Homo sapiens OX=9606 GN=IGHA1 PE=1 SV=2 | 255 | 20201115_15212_1ng_glyco_hcd203040_compound15.22214.22214.4 | controllerType=0 controllerNumber=1 scan=22214 | 42.6471 |
| 19361:4 | 9 | R.LSLHRPALEDLLLGSEAN[+1216.423]LTC[+57.021]TLTGLR.D | HexNAc(2)Hex(5) | C[+57], N[+1216] | 1045.7631 | 4 | 4180.0306 | 4180.0211 | 0 | 2.3 | 127 | Specific | 794.92 | 794.92 | 794.92 | 16.39 | 59 | >sp\|P01876\|IGHA1_HUMAN Immunoglobulin heavy constant alpha 1 OS=Homo sapiens OX=9606 GN=IGHA1 PE=1 SV=2 | 255 | 20201115_15212_1ng_glyco_hcd203040_compound15.22238.22238.4 | controllerType=0 controllerNumber=1 scan=22238 | 42.6998 |
| 20333:4 | 9 | R.LSLHRPALEDLLLGSEAN[+1751.624]LTC[+57.021]TLTGLR.D | HexNAc(4)Hex(4)NeuAc(1) | C[+57], N[+1752] | 1179.5651 | 4 | 4715.2386 | 4715.2225 | 0 | 3.4 | 127 | Specific | 796.09 | 780.45 | 780.45 | 16.39 | 59 | >sp\|P01876\|IGHA1_HUMAN Immunoglobulin heavy constant alpha 1 OS=Homo sapiens OX=9606 GN=IGHA1 PE=1 SV=2 | 255 | 20201115_15212_1ng_glyco_hcd203040_compound15.23253.23253.4 | controllerType=0 controllerNumber=1 scan=23253 | 44.8877 |
| 19359:4 | 9 | R.LSLHRPALEDLLLGSEAN[+1501.555]LTC[+57.021]TLTGLR.D | HexNAc(5)Hex(3) | C[+57], N[+1502] | 1117.0489 | 4 | 4465.1737 | 4465.1536 | 0 | 4.5 | 127 | Specific | 817.22 | 813.46 | 813.46 | 17.33 | 59 | >sp\|P01876\|IGHA1_HUMAN Immunoglobulin heavy constant alpha 1 OS=Homo sapiens OX=9606 GN=IGHA1 PE=1 SV=2 | 255 | 20201115_15212_1ng_glyco_hcd203040_compound15.22236.22236.4 | controllerType=0 controllerNumber=1 scan=22236 | 42.6974 |
| 19354:4 | 9 | R.LSLHRPALEDLLLGSEAN[+1647.613]LTC[+57.021]TLTGLR.D | HexNAc(5)Hex(3)Fuc(1) | C[+57], N[+1648] | 1153.5597 | 4 | 4611.2171 | 4611.2115 | 0 | 1.2 | 127 | Specific | 840.68 | 756.53 | 756.53 | 17.31 | 59 | >sp\|P01876\|IGHA1_HUMAN Immunoglobulin heavy constant alpha 1 OS=Homo sapiens OX=9606 GN=IGHA1 PE=1 SV=2 | 255 | 20201115_15212_1ng_glyco_hcd203040_compound15.22230.22230.4 | controllerType=0 controllerNumber=1 scan=22230 | 42.6834 |
| 20303:4 | 9 | R.LSLHRPALEDLLLGSEAN[+1710.598]LTC[+57.021]TLTGLR.D | HexNAc(3)Hex(5)NeuAc(1) | C[+57], N[+1711] | 1169.3086 | 4 | 4674.2127 | 4674.1959 | 0 | 3.6 | 127 | Specific | 845.74 | 806.01 | 806.01 | 17.33 | 59 | >sp\|P01876\|IGHA1_HUMAN Immunoglobulin heavy constant alpha 1 OS=Homo sapiens OX=9606 GN=IGHA1 PE=1 SV=2 | 255 | 20201115_15212_1ng_glyco_hcd203040_compound15.23222.23222.4 | controllerType=0 controllerNumber=1 scan=23222 | 44.8244 |
| 19317:4 | 9 | R.LSLHRPALEDLLLGSEAN[+1460.529]LTC[+57.021]TLTGLR.D | HexNAc(4)Hex(4) | C[+57], N[+1461] | 1106.7917 | 4 | 4424.1448 | 4424.1271 | 0 | 4 | 127 | Specific | 852.87 | 731.01 | 731.01 | 18.33 | 59 | >sp\|P01876\|IGHA1_HUMAN Immunoglobulin heavy constant alpha 1 OS=Homo sapiens OX=9606 GN=IGHA1 PE=1 SV=2 | 255 | 20201115_15212_1ng_glyco_hcd203040_compound15.22192.22192.4 | controllerType=0 controllerNumber=1 scan=22192 | 42.6 |
| 20256:4 | 9 | R.LSLHRPALEDLLLGSEAN[+1872.651]LTC[+57.021]TLTGLR.D | HexNAc(3)Hex(6)NeuAc(1) | C[+57], N[+1873] | 1209.8221 | 4 | 4836.2664 | 4836.2487 | 0 | 3.7 | 127 | Specific | 874.29 | 874.29 | 874.29 | 18.33 | 59 | >sp\|P01876\|IGHA1_HUMAN Immunoglobulin heavy constant alpha 1 OS=Homo sapiens OX=9606 GN=IGHA1 PE=1 SV=2 | 255 | 20201115_15212_1ng_glyco_hcd203040_compound15.23173.23173.4 | controllerType=0 controllerNumber=1 scan=23173 | 44.7195 |
| 19240:4 | 9 | R.LSLHRPALEDLLLGSEAN[+1702.581]LTC[+57.021]TLTGLR.D | HexNAc(2)Hex(8) | C[+57], N[+1703] | 1167.3045 | 4 | 4666.1961 | 4666.1796 | 0 | 3.5 | 127 | Specific | 893.13 | 893.13 | 893.13 | 18.33 | 59 | >sp\|P01876\|IGHA1_HUMAN Immunoglobulin heavy constant alpha 1 OS=Homo sapiens OX=9606 GN=IGHA1 PE=1 SV=2 | 255 | 20201115_15212_1ng_glyco_hcd203040_compound15.22112.22112.4 | controllerType=0 controllerNumber=1 scan=22112 | 42.4042 |
| 19341:4 | 9 | R.LSLHRPALEDLLLGSEAN[+1298.476]LTC[+57.021]TLTGLR.D | HexNAc(4)Hex(3) | C[+57], N[+1298] | 1066.276 | 4 | 4262.0823 | 4262.0742 | 0 | 1.9 | 127 | Specific | 903.91 | 900.79 | 900.79 | 19.31 | 59 | >sp\|P01876\|IGHA1_HUMAN Immunoglobulin heavy constant alpha 1 OS=Homo sapiens OX=9606 GN=IGHA1 PE=1 SV=2 | 255 | 20201115_15212_1ng_glyco_hcd203040_compound15.22217.22217.4 | controllerType=0 controllerNumber=1 scan=22217 | 42.6539 |
| 19339:4 | 9 | R.LSLHRPALEDLLLGSEAN[+1257.449]LTC[+57.021]TLTGLR.D | HexNAc(3)Hex(4) | C[+57], N[+1257] | 1056.0192 | 4 | 4221.055 | 4221.0477 | 0 | 1.7 | 127 | Specific | 964.84 | 915.56 | 915.56 | 20.31 | 59 | >sp\|P01876\|IGHA1_HUMAN Immunoglobulin heavy constant alpha 1 OS=Homo sapiens OX=9606 GN=IGHA1 PE=1 SV=2 | 255 | 20201115_15212_1ng_glyco_hcd203040_compound15.22215.22215.4 | controllerType=0 controllerNumber=1 scan=22215 | 42.6483 |
| 19242:4 | 9 | R.LSLHRPALEDLLLGSEAN[+1702.581]LTC[+57.021]TLTGLR.D | HexNAc(2)Hex(8) | C[+57], N[+1703] | 1167.3045 | 4 | 4666.1961 | 4666.1796 | 0 | 3.5 | 127 | Specific | 972.78 | 972.78 | 972.78 | 20.33 | 59 | >sp\|P01876\|IGHA1_HUMAN Immunoglobulin heavy constant alpha 1 OS=Homo sapiens OX=9606 GN=IGHA1 PE=1 SV=2 | 255 | 20201115_15212_1ng_glyco_hcd203040_compound15.22114.22114.4 | controllerType=0 controllerNumber=1 scan=22114 | 42.4095 |
| 19210:4 | 9 | R.LSLHRPALEDLLLGSEAN[+1540.529]LTC[+57.021]TLTGLR.D | HexNAc(2)Hex(7) | C[+57], N[+1541] | 1126.7892 | 4 | 4504.1351 | 4504.1268 | 0 | 1.8 | 127 | Specific | 974.09 | 922.12 | 922.12 | 20.31 | 59 | >sp\|P01876\|IGHA1_HUMAN Immunoglobulin heavy constant alpha 1 OS=Homo sapiens OX=9606 GN=IGHA1 PE=1 SV=2 | 255 | 20201115_15212_1ng_glyco_hcd203040_compound15.22080.22080.2 | controllerType=0 controllerNumber=1 scan=22080 | 42.3288 |
| 19364:4 | 9 | R.LSLHRPALEDLLLGSEAN[+1216.423]LTC[+57.021]TLTGLR.D | HexNAc(2)Hex(5) | C[+57], N[+1216] | 1045.7631 | 4 | 4180.0306 | 4180.0211 | 0 | 2.3 | 127 | Specific | 990.3 | 898.68 | 898.68 | 20.39 | 59 | >sp\|P01876\|IGHA1_HUMAN Immunoglobulin heavy constant alpha 1 OS=Homo sapiens OX=9606 GN=IGHA1 PE=1 SV=2 | 255 | 20201115_15212_1ng_glyco_hcd203040_compound15.22241.22241.4 | controllerType=0 controllerNumber=1 scan=22241 | 42.7042 |
| 19418:4 | 9 | R.LSLHRPALEDLLLGSEAN[+1054.370]LTC[+57.021]TLTGLR.D | HexNAc(2)Hex(4) | C[+57], N[+1054] | 1005.2504 | 4 | 4017.9798 | 4017.9683 | 0 | 2.9 | 127 | Specific | 997.93 | 975.11 | 975.11 | 20.39 | 59 | >sp\|P01876\|IGHA1_HUMAN Immunoglobulin heavy constant alpha 1 OS=Homo sapiens OX=9606 GN=IGHA1 PE=1 SV=2 | 255 | 20201115_15212_1ng_glyco_hcd203040_compound15.22297.22297.4 | controllerType=0 controllerNumber=1 scan=22297 | 42.8213 |
| 19342:4 | 9 | R.LSLHRPALEDLLLGSEAN[+1257.449]LTC[+57.021]TLTGLR.D | HexNAc(3)Hex(4) | C[+57], N[+1257] | 1056.0192 | 4 | 4221.055 | 4221.0477 | 0 | 1.7 | 127 | Specific | 1075.77 | 1010.4 | 1010.4 | 22.31 | 59 | >sp\|P01876\|IGHA1_HUMAN Immunoglobulin heavy constant alpha 1 OS=Homo sapiens OX=9606 GN=IGHA1 PE=1 SV=2 | 255 | 20201115_15212_1ng_glyco_hcd203040_compound15.22218.22218.4 | controllerType=0 controllerNumber=1 scan=22218 | 42.6579 |
| 19212:4 | 9 | R.LSLHRPALEDLLLGSEAN[+1540.529]LTC[+57.021]TLTGLR.D | HexNAc(2)Hex(7) | C[+57], N[+1541] | 1126.7892 | 4 | 4504.1351 | 4504.1268 | 0 | 1.8 | 127 | Specific | 1094.75 | 1094.75 | 1094.75 | 22.31 | 59 | >sp\|P01876\|IGHA1_HUMAN Immunoglobulin heavy constant alpha 1 OS=Homo sapiens OX=9606 GN=IGHA1 PE=1 SV=2 | 255 | 20201115_15212_1ng_glyco_hcd203040_compound15.22082.22082.2 | controllerType=0 controllerNumber=1 scan=22082 | 42.3376 |
| 14783:3 | 9 | R.LAGKPTHVN[+2263.835]VSVVMAEVDGTC[+57.021].Y | HexNAc(5)Hex(5)Fuc(3) | C[+57], N[+2264] | 1483.3099 | 3 | 4447.9151 | 4447.9135 | 0 | 0.4 | 332 | CRagged | 237.42 | 191.34 | 191.34 | 6 | 59 | >sp\|P01876\|IGHA1_HUMAN Immunoglobulin heavy constant alpha 1 OS=Homo sapiens OX=9606 GN=IGHA1 PE=1 SV=2 | 255 | 20201115_15212_1ng_glyco_hcd203040_compound15.17443.17443.3 | controllerType=0 controllerNumber=1 scan=17443 | 33.6303 |
| 13046:4 | 9 | R.LAGKPTHVN[+1809.666]VSVVMAEVDGTC[+57.021]Y.- | HexNAc(5)Hex(4)Fuc(1) | C[+57], N[+1810] | 1039.957 | 4 | 4156.806 | 4156.8081 | 0 | -0.5 | 332 | CRagged | 306.81 | 224.07 | 224.07 | 4.79 | 59 | >sp\|P01876\|IGHA1_HUMAN Immunoglobulin heavy constant alpha 1 OS=Homo sapiens OX=9606 GN=IGHA1 PE=1 SV=2 | 255 | 20201115_15212_1ng_glyco_hcd203040_compound15.15614.15614.2 | controllerType=0 controllerNumber=1 scan=15614 | 30.6067 |
| 13111:4 | 9 | R.LAGKPTHVN[+1647.613]VSVVMAEVDGTC[+57.021]Y.- | HexNAc(5)Hex(3)Fuc(1) | C[+57], N[+1648] | 999.4469 | 4 | 3994.7657 | 3994.7553 | 0 | 2.6 | 332 | CRagged | 426.44 | 426.44 | 426.44 | 10.61 | 59 | >sp\|P01876\|IGHA1_HUMAN Immunoglobulin heavy constant alpha 1 OS=Homo sapiens OX=9606 GN=IGHA1 PE=1 SV=2 | 255 | 20201115_15212_1ng_glyco_hcd203040_compound15.15681.15681.4 | controllerType=0 controllerNumber=1 scan=15681 | 30.7174 |
| 12996:3 | 9 | R.LAGKPTHVN[+1971.719]VSVVMAEVDGTC[+57.021]Y.- | HexNAc(5)Hex(5)Fuc(1) | C[+57], N[+1972] | 1440.2959 | 3 | 4318.8731 | 4318.861 | 0 | 2.8 | 332 | CRagged | 517.06 | 513.76 | 329.06 | 10.93 | 59 | >sp\|P01876\|IGHA1_HUMAN Immunoglobulin heavy constant alpha 1 OS=Homo sapiens OX=9606 GN=IGHA1 PE=1 SV=2 | 255 | 20201115_15212_1ng_glyco_hcd203040_compound15.15562.15562.3 | controllerType=0 controllerNumber=1 scan=15562 | 30.5051 |
| 14789:4 | 9 | R.LAGKPTHVN[+2262.814]VSVVMAEVDGTC[+57.021]Y.- | HexNAc(5)Hex(5)Fuc(1)NeuAc(1) | C[+57], N[+2263] | 1153.2483 | 4 | 4609.9715 | 4609.9564 | 0 | 3.3 | 332 | CRagged | 536.56 | 426.22 | 426.22 | 11.13 | 59 | >sp\|P01876\|IGHA1_HUMAN Immunoglobulin heavy constant alpha 1 OS=Homo sapiens OX=9606 GN=IGHA1 PE=1 SV=2 | 255 | 20201115_15212_1ng_glyco_hcd203040_compound15.17449.17449.4 | controllerType=0 controllerNumber=1 scan=17449 | 33.6421 |
| 11897:3 | 9 | R.LAGKPTHVN[+1216.423]VSVVMAEVDGTC[+57.021]Y.- | HexNAc(2)Hex(5) | C[+57], N[+1216] | 1188.5292 | 3 | 3563.5731 | 3563.5649 | 0 | 2.3 | 332 | CRagged | 561.64 | 561.64 | 312.63 | 11.28 | 59 | >sp\|P01876\|IGHA1_HUMAN Immunoglobulin heavy constant alpha 1 OS=Homo sapiens OX=9606 GN=IGHA1 PE=1 SV=2 | 255 | 20201115_15212_1ng_glyco_hcd203040_compound15.14424.14424.3 | controllerType=0 controllerNumber=1 scan=14424 | 28.5246 |
| 13114:4 | 9 | R.LAGKPTHVN[+1647.613]VSVVMAEVDGTC[+57.021]Y.- | HexNAc(5)Hex(3)Fuc(1) | C[+57], N[+1648] | 999.4469 | 4 | 3994.7657 | 3994.7553 | 0 | 2.6 | 332 | CRagged | 566.16 | 353.88 | 353.88 | 9.74 | 59 | >sp\|P01876\|IGHA1_HUMAN Immunoglobulin heavy constant alpha 1 OS=Homo sapiens OX=9606 GN=IGHA1 PE=1 SV=2 | 255 | 20201115_15212_1ng_glyco_hcd203040_compound15.15684.15684.4 | controllerType=0 controllerNumber=1 scan=15684 | 30.7215 |
| 14927:3 | 9 | R.LAGKPTHVN[+1710.598]VSVVMAEVDGTC[+57.021]Y.- | HexNAc(3)Hex(5)NeuAc(1) | C[+57], N[+1711] | 1353.2561 | 3 | 4057.7537 | 4057.7397 | 0 | 3.5 | 332 | CRagged | 572.79 | 572.79 | 391.27 | 11.3 | 59 | >sp\|P01876\|IGHA1_HUMAN Immunoglobulin heavy constant alpha 1 OS=Homo sapiens OX=9606 GN=IGHA1 PE=1 SV=2 | 255 | 20201115_15212_1ng_glyco_hcd203040_compound15.17592.17592.3 | controllerType=0 controllerNumber=1 scan=17592 | 33.9212 |
| 12998:3 | 9 | R.LAGKPTHVN[+1971.719]VSVVMAEVDGTC[+57.021]Y.- | HexNAc(5)Hex(5)Fuc(1) | C[+57], N[+1972] | 1440.2959 | 3 | 4318.8731 | 4318.861 | 0 | 2.8 | 332 | CRagged | 579.05 | 513.83 | 363.09 | 12.63 | 59 | >sp\|P01876\|IGHA1_HUMAN Immunoglobulin heavy constant alpha 1 OS=Homo sapiens OX=9606 GN=IGHA1 PE=1 SV=2 | 255 | 20201115_15212_1ng_glyco_hcd203040_compound15.15564.15564.3 | controllerType=0 controllerNumber=1 scan=15564 | 30.5093 |
| 16754:3 | 9 | R.LAGKPTHVN[+2553.910]VSVVMAEVDGTC[+57.021]Y.- | HexNAc(5)Hex(5)Fuc(1)NeuAc(2) | C[+57], N[+2554] | 1634.3598 | 3 | 4901.0648 | 4901.0518 | 0 | 2.7 | 332 | CRagged | 602.3 | 465.9 | 465.9 | 13.63 | 59 | >sp\|P01876\|IGHA1_HUMAN Immunoglobulin heavy constant alpha 1 OS=Homo sapiens OX=9606 GN=IGHA1 PE=1 SV=2 | 255 | 20201115_15212_1ng_glyco_hcd203040_compound15.19484.19484.3 | controllerType=0 controllerNumber=1 scan=19484 | 37.3196 |
| 11900:3 | 9 | R.LAGKPTHVN[+1216.423]VSVVMAEVDGTC[+57.021]Y.- | HexNAc(2)Hex(5) | C[+57], N[+1216] | 1188.5292 | 3 | 3563.5731 | 3563.5649 | 0 | 2.3 | 332 | CRagged | 615.38 | 557.66 | 373.08 | 13.55 | 59 | >sp\|P01876\|IGHA1_HUMAN Immunoglobulin heavy constant alpha 1 OS=Homo sapiens OX=9606 GN=IGHA1 PE=1 SV=2 | 255 | 20201115_15212_1ng_glyco_hcd203040_compound15.14427.14427.3 | controllerType=0 controllerNumber=1 scan=14427 | 28.5299 |
| 14880:3 | 9 | R.LAGKPTHVN[+1872.651]VSVVMAEVDGTC[+57.021]Y.- | HexNAc(3)Hex(6)NeuAc(1) | C[+57], N[+1873] | 1407.2722 | 3 | 4219.8019 | 4219.7926 | 0 | 2.2 | 332 | CRagged | 645.38 | 623.14 | 399.86 | 13.55 | 59 | >sp\|P01876\|IGHA1_HUMAN Immunoglobulin heavy constant alpha 1 OS=Homo sapiens OX=9606 GN=IGHA1 PE=1 SV=2 | 255 | 20201115_15212_1ng_glyco_hcd203040_compound15.17543.17543.3 | controllerType=0 controllerNumber=1 scan=17543 | 33.8069 |
| 14930:3 | 9 | R.LAGKPTHVN[+1710.598]VSVVMAEVDGTC[+57.021]Y.- | HexNAc(3)Hex(5)NeuAc(1) | C[+57], N[+1711] | 1353.2561 | 3 | 4057.7537 | 4057.7397 | 0 | 3.5 | 332 | CRagged | 645.5 | 645.5 | 397.04 | 13.57 | 59 | >sp\|P01876\|IGHA1_HUMAN Immunoglobulin heavy constant alpha 1 OS=Homo sapiens OX=9606 GN=IGHA1 PE=1 SV=2 | 255 | 20201115_15212_1ng_glyco_hcd203040_compound15.17595.17595.3 | controllerType=0 controllerNumber=1 scan=17595 | 33.9273 |
| 16745:4 | 9 | R.LAGKPTHVN[+2553.910]VSVVMAEVDGTC[+57.021]Y.- | HexNAc(5)Hex(5)Fuc(1)NeuAc(2) | C[+57], N[+2554] | 1226.0212 | 4 | 4901.0628 | 4901.0518 | 0 | 2.2 | 332 | CRagged | 652.15 | 501.11 | 501.11 | 14.57 | 59 | >sp\|P01876\|IGHA1_HUMAN Immunoglobulin heavy constant alpha 1 OS=Homo sapiens OX=9606 GN=IGHA1 PE=1 SV=2 | 255 | 20201115_15212_1ng_glyco_hcd203040_compound15.19475.19475.4 | controllerType=0 controllerNumber=1 scan=19475 | 37.3036 |
| 14878:3 | 9 | R.LAGKPTHVN[+1872.651]VSVVMAEVDGTC[+57.021]Y.- | HexNAc(3)Hex(6)NeuAc(1) | C[+57], N[+1873] | 1407.2722 | 3 | 4219.8019 | 4219.7926 | 0 | 2.2 | 332 | CRagged | 665.23 | 665.23 | 457.18 | 14.55 | 59 | >sp\|P01876\|IGHA1_HUMAN Immunoglobulin heavy constant alpha 1 OS=Homo sapiens OX=9606 GN=IGHA1 PE=1 SV=2 | 255 | 20201115_15212_1ng_glyco_hcd203040_compound15.17541.17541.3 | controllerType=0 controllerNumber=1 scan=17541 | 33.8028 |
| 16743:4 | 9 | R.LAGKPTHVN[+2553.910]VSVVMAEVDGTC[+57.021]Y.- | HexNAc(5)Hex(5)Fuc(1)NeuAc(2) | C[+57], N[+2554] | 1226.0212 | 4 | 4901.0628 | 4901.0518 | 0 | 2.2 | 332 | CRagged | 710.96 | 534.21 | 534.21 | 15.57 | 59 | >sp\|P01876\|IGHA1_HUMAN Immunoglobulin heavy constant alpha 1 OS=Homo sapiens OX=9606 GN=IGHA1 PE=1 SV=2 | 255 | 20201115_15212_1ng_glyco_hcd203040_compound15.19473.19473.4 | controllerType=0 controllerNumber=1 scan=19473 | 37.2986 |
| 14791:4 | 9 | R.LAGKPTHVN[+2262.814]VSVVMAEVDGTC[+57.021]Y.- | HexNAc(5)Hex(5)Fuc(1)NeuAc(1) | C[+57], N[+2263] | 1153.2483 | 4 | 4609.9715 | 4609.9564 | 0 | 3.3 | 332 | CRagged | 720.29 | 592.31 | 592.31 | 15.65 | 59 | >sp\|P01876\|IGHA1_HUMAN Immunoglobulin heavy constant alpha 1 OS=Homo sapiens OX=9606 GN=IGHA1 PE=1 SV=2 | 255 | 20201115_15212_1ng_glyco_hcd203040_compound15.17451.17451.4 | controllerType=0 controllerNumber=1 scan=17451 | 33.6496 |
| 12982:3 | 9 | R.LAGKPTHVN[+1663.608]VSVVMAEVDGTC[+57.021]Y.- | HexNAc(5)Hex(4) | C[+57], N[+1664] | 1337.5903 | 3 | 4010.7562 | 4010.7502 | 0 | 1.5 | 332 | CRagged | 737.21 | 737.21 | 460.38 | 15.55 | 59 | >sp\|P01876\|IGHA1_HUMAN Immunoglobulin heavy constant alpha 1 OS=Homo sapiens OX=9606 GN=IGHA1 PE=1 SV=2 | 255 | 20201115_15212_1ng_glyco_hcd203040_compound15.15548.15548.3 | controllerType=0 controllerNumber=1 scan=15548 | 30.4824 |
| 14786:3 | 9 | R.LAGKPTHVN[+2262.814]VSVVMAEVDGTC[+57.021]Y.- | HexNAc(5)Hex(5)Fuc(1)NeuAc(1) | C[+57], N[+2263] | 1537.3284 | 3 | 4609.9707 | 4609.9564 | 0 | 3.1 | 332 | CRagged | 747.92 | 596.65 | 596.65 | 15.63 | 59 | >sp\|P01876\|IGHA1_HUMAN Immunoglobulin heavy constant alpha 1 OS=Homo sapiens OX=9606 GN=IGHA1 PE=1 SV=2 | 255 | 20201115_15212_1ng_glyco_hcd203040_compound15.17446.17446.3 | controllerType=0 controllerNumber=1 scan=17446 | 33.6377 |
| 14784:3 | 9 | R.LAGKPTHVN[+2262.814]VSVVMAEVDGTC[+57.021]Y.- | HexNAc(5)Hex(5)Fuc(1)NeuAc(1) | C[+57], N[+2263] | 1537.3284 | 3 | 4609.9707 | 4609.9564 | 0 | 3.1 | 332 | CRagged | 753.08 | 502.48 | 502.48 | 16.63 | 59 | >sp\|P01876\|IGHA1_HUMAN Immunoglobulin heavy constant alpha 1 OS=Homo sapiens OX=9606 GN=IGHA1 PE=1 SV=2 | 255 | 20201115_15212_1ng_glyco_hcd203040_compound15.17444.17444.3 | controllerType=0 controllerNumber=1 scan=17444 | 33.6317 |
| 13095:3 | 9 | R.LAGKPTHVN[+1501.555]VSVVMAEVDGTC[+57.021]Y.- | HexNAc(5)Hex(3) | C[+57], N[+1502] | 1283.574 | 3 | 3848.7075 | 3848.6974 | 0 | 2.6 | 332 | CRagged | 763.77 | 614.82 | 614.82 | 16.63 | 59 | >sp\|P01876\|IGHA1_HUMAN Immunoglobulin heavy constant alpha 1 OS=Homo sapiens OX=9606 GN=IGHA1 PE=1 SV=2 | 255 | 20201115_15212_1ng_glyco_hcd203040_compound15.15665.15665.3 | controllerType=0 controllerNumber=1 scan=15665 | 30.6946 |
| 16752:3 | 9 | R.LAGKPTHVN[+2553.910]VSVVMAEVDGTC[+57.021]Y.- | HexNAc(5)Hex(5)Fuc(1)NeuAc(2) | C[+57], N[+2554] | 1634.3598 | 3 | 4901.0648 | 4901.0518 | 0 | 2.7 | 332 | CRagged | 765.67 | 520.73 | 520.73 | 16.63 | 59 | >sp\|P01876\|IGHA1_HUMAN Immunoglobulin heavy constant alpha 1 OS=Homo sapiens OX=9606 GN=IGHA1 PE=1 SV=2 | 255 | 20201115_15212_1ng_glyco_hcd203040_compound15.19482.19482.3 | controllerType=0 controllerNumber=1 scan=19482 | 37.3152 |
| 12984:3 | 9 | R.LAGKPTHVN[+1663.608]VSVVMAEVDGTC[+57.021]Y.- | HexNAc(5)Hex(4) | C[+57], N[+1664] | 1337.5903 | 3 | 4010.7562 | 4010.7502 | 0 | 1.5 | 332 | CRagged | 775.36 | 714.71 | 480.76 | 16.55 | 59 | >sp\|P01876\|IGHA1_HUMAN Immunoglobulin heavy constant alpha 1 OS=Homo sapiens OX=9606 GN=IGHA1 PE=1 SV=2 | 255 | 20201115_15212_1ng_glyco_hcd203040_compound15.15550.15550.3 | controllerType=0 controllerNumber=1 scan=15550 | 30.4875 |
| 13099:3 | 9 | R.LAGKPTHVN[+1647.613]VSVVMAEVDGTC[+57.021]Y.- | HexNAc(5)Hex(3)Fuc(1) | C[+57], N[+1648] | 1332.2596 | 3 | 3994.7642 | 3994.7553 | 0 | 2.2 | 332 | CRagged | 828.58 | 665.99 | 665.99 | 17.55 | 59 | >sp\|P01876\|IGHA1_HUMAN Immunoglobulin heavy constant alpha 1 OS=Homo sapiens OX=9606 GN=IGHA1 PE=1 SV=2 | 255 | 20201115_15212_1ng_glyco_hcd203040_compound15.15669.15669.3 | controllerType=0 controllerNumber=1 scan=15669 | 30.7025 |
| 12988:3 | 9 | R.LAGKPTHVN[+1809.666]VSVVMAEVDGTC[+57.021]Y.- | HexNAc(5)Hex(4)Fuc(1) | C[+57], N[+1810] | 1386.2763 | 3 | 4156.8144 | 4156.8081 | 0 | 1.5 | 332 | CRagged | 829.07 | 687.12 | 541.29 | 17.55 | 59 | >sp\|P01876\|IGHA1_HUMAN Immunoglobulin heavy constant alpha 1 OS=Homo sapiens OX=9606 GN=IGHA1 PE=1 SV=2 | 255 | 20201115_15212_1ng_glyco_hcd203040_compound15.15554.15554.3 | controllerType=0 controllerNumber=1 scan=15554 | 30.4955 |
| 13097:3 | 9 | R.LAGKPTHVN[+1647.613]VSVVMAEVDGTC[+57.021]Y.- | HexNAc(5)Hex(3)Fuc(1) | C[+57], N[+1648] | 1332.2596 | 3 | 3994.7642 | 3994.7553 | 0 | 2.2 | 332 | CRagged | 847.86 | 654.73 | 654.73 | 17.55 | 59 | >sp\|P01876\|IGHA1_HUMAN Immunoglobulin heavy constant alpha 1 OS=Homo sapiens OX=9606 GN=IGHA1 PE=1 SV=2 | 255 | 20201115_15212_1ng_glyco_hcd203040_compound15.15667.15667.3 | controllerType=0 controllerNumber=1 scan=15667 | 30.6965 |
| 12935:3 | 9 | R.LAGKPTHVN[+1702.581]VSVVMAEVDGTC[+57.021]Y.- | HexNAc(2)Hex(8) | C[+57], N[+1703] | 1350.5834 | 3 | 4049.7357 | 4049.7234 | 0 | 3 | 332 | CRagged | 855.3 | 837.22 | 504.94 | 18.63 | 59 | >sp\|P01876\|IGHA1_HUMAN Immunoglobulin heavy constant alpha 1 OS=Homo sapiens OX=9606 GN=IGHA1 PE=1 SV=2 | 255 | 20201115_15212_1ng_glyco_hcd203040_compound15.15498.15498.3 | controllerType=0 controllerNumber=1 scan=15498 | 30.3453 |
| 12985:3 | 9 | R.LAGKPTHVN[+1809.666]VSVVMAEVDGTC[+57.021]Y.- | HexNAc(5)Hex(4)Fuc(1) | C[+57], N[+1810] | 1386.2763 | 3 | 4156.8144 | 4156.8081 | 0 | 1.5 | 332 | CRagged | 858.61 | 711.72 | 553.55 | 18.55 | 59 | >sp\|P01876\|IGHA1_HUMAN Immunoglobulin heavy constant alpha 1 OS=Homo sapiens OX=9606 GN=IGHA1 PE=1 SV=2 | 255 | 20201115_15212_1ng_glyco_hcd203040_compound15.15551.15551.3 | controllerType=0 controllerNumber=1 scan=15551 | 30.4893 |
| 13098:3 | 9 | R.LAGKPTHVN[+1501.555]VSVVMAEVDGTC[+57.021]Y.- | HexNAc(5)Hex(3) | C[+57], N[+1502] | 1283.574 | 3 | 3848.7075 | 3848.6974 | 0 | 2.6 | 332 | CRagged | 864.66 | 710.09 | 710.09 | 18.63 | 59 | >sp\|P01876\|IGHA1_HUMAN Immunoglobulin heavy constant alpha 1 OS=Homo sapiens OX=9606 GN=IGHA1 PE=1 SV=2 | 255 | 20201115_15212_1ng_glyco_hcd203040_compound15.15668.15668.3 | controllerType=0 controllerNumber=1 scan=15668 | 30.6988 |
| 12967:3 | 9 | R.LAGKPTHVN[+1378.476]VSVVMAEVDGTC[+57.021]Y.- | HexNAc(2)Hex(6) | C[+57], N[+1378] | 1242.5468 | 3 | 3725.6259 | 3725.6178 | 0 | 2.2 | 332 | CRagged | 866.1 | 814.13 | 545.27 | 18.55 | 59 | >sp\|P01876\|IGHA1_HUMAN Immunoglobulin heavy constant alpha 1 OS=Homo sapiens OX=9606 GN=IGHA1 PE=1 SV=2 | 255 | 20201115_15212_1ng_glyco_hcd203040_compound15.15532.15532.3 | controllerType=0 controllerNumber=1 scan=15532 | 30.4477 |
| 12956:3 | 9 | R.LAGKPTHVN[+1540.529]VSVVMAEVDGTC[+57.021]Y.- | HexNAc(2)Hex(7) | C[+57], N[+1541] | 1296.5659 | 3 | 3887.683 | 3887.6706 | 0 | 3.2 | 332 | CRagged | 891.05 | 828.96 | 524.51 | 18.63 | 59 | >sp\|P01876\|IGHA1_HUMAN Immunoglobulin heavy constant alpha 1 OS=Homo sapiens OX=9606 GN=IGHA1 PE=1 SV=2 | 255 | 20201115_15212_1ng_glyco_hcd203040_compound15.15520.15520.3 | controllerType=0 controllerNumber=1 scan=15520 | 30.4153 |

Table S4. Cont.

| Query #:z | Protein Rank | Glycopeptide | Glycan | Modification Type(s) | Observed m/z | z | Observed (M+H) | Calc. mass (M+H) | Off-by-x error | Mass error (ppm) | StartPosition | Cleavage | Score | Delta | Delta Mod | \|Log Prob\| | # of unique peptides | Protein Name | Protein DB number | Comment | Scan # | Scan Time |
|---|---|---|---|---|---|---|---|---|---|---|---|---|---|---|---|---|---|---|---|---|---|---|
| 13038:3 | 9 | R.LAGKPTHVN[+1216.423]VSVVMAEVDGTC[+57.021]Y.- | HexNAc(2)Hex(5) | C[+57], N[+1216] | 1188.5291 | 3 | 3563.5728 | 3563.5649 | 0 | 2.2 | 332 | CRagged | 903.6 | 903.6 | 536.03 | 19.55 | 59 | >sp\|P01876\|IGHA1_HUMAN Immunoglobulin heavy constant alpha 1 OS=Homo sapiens OX=9606 GN=IGHA1 PE=1 SV=2 | 255 | 20201115_15212_1ng_glyco_hcd203040_compound15.15606.15606.3 | controllerType=0 controllerNumber=1 scan=15606 | 30.595 |
| 12965:3 | 9 | R.LAGKPTHVN[+1378.476]VSVVMAEVDGTC[+57.021]Y.- | HexNAc(2)Hex(6) | C[+57], N[+1378] | 1242.5468 | 3 | 3725.6259 | 3725.6178 | 0 | 2.2 | 332 | CRagged | 904.67 | 800.41 | 539.74 | 19.55 | 59 | >sp\|P01876\|IGHA1_HUMAN Immunoglobulin heavy constant alpha 1 OS=Homo sapiens OX=9606 GN=IGHA1 PE=1 SV=2 | 255 | 20201115_15212_1ng_glyco_hcd203040_compound15.15530.15530.3 | controllerType=0 controllerNumber=1 scan=15530 | 30.4362 |
| 12953:3 | 9 | R.LAGKPTHVN[+1540.529]VSVVMAEVDGTC[+57.021]Y.- | HexNAc(2)Hex(7) | C[+57], N[+1541] | 1296.5659 | 3 | 3887.683 | 3887.6706 | 0 | 3.2 | 332 | CRagged | 904.88 | 904.88 | 593.75 | 19.63 | 59 | >sp\|P01876\|IGHA1_HUMAN Immunoglobulin heavy constant alpha 1 OS=Homo sapiens OX=9606 GN=IGHA1 PE=1 SV=2 | 255 | 20201115_15212_1ng_glyco_hcd203040_compound15.15517.15517.3 | controllerType=0 controllerNumber=1 scan=15517 | 30.3823 |
| 12933:3 | 9 | R.LAGKPTHVN[+1702.581]VSVVMAEVDGTC[+57.021]Y.- | HexNAc(2)Hex(8) | C[+57], N[+1703] | 1350.5834 | 3 | 4049.7357 | 4049.7234 | 0 | 3 | 332 | CRagged | 911.91 | 911.91 | 595.36 | 19.63 | 59 | >sp\|P01876\|IGHA1_HUMAN Immunoglobulin heavy constant alpha 1 OS=Homo sapiens OX=9606 GN=IGHA1 PE=1 SV=2 | 255 | 20201115_15212_1ng_glyco_hcd203040_compound15.15496.15496.3 | controllerType=0 controllerNumber=1 scan=15496 | 30.336 |
| 13041:3 | 9 | R.LAGKPTHVN[+1216.423]VSVVMAEVDGTC[+57.021]Y.- | HexNAc(2)Hex(5) | C[+57], N[+1216] | 1188.5291 | 3 | 3563.5728 | 3563.5649 | 0 | 2.2 | 332 | CRagged | 918.38 | 713.88 | 549.33 | 19.55 | 59 | >sp\|P01876\|IGHA1_HUMAN Immunoglobulin heavy constant alpha 1 OS=Homo sapiens OX=9606 GN=IGHA1 PE=1 SV=2 | 255 | 20201115_15212_1ng_glyco_hcd203040_compound15.15609.15609.3 | controllerType=0 controllerNumber=1 scan=15609 | 30.601 |
| 10948:3 | 9 | R.LAGKPTHVN[+1378.476]VSVVM[+15.995]AEVDGTC[+57.021]Y.- | HexNAc(2)Hex(6) | C[+57], M[+16], N[+1378] | 1247.8765 | 3 | 3741.615 | 3741.6127 | 0 | 0.6 | 332 | CRagged | 255.14 | 255.14 | 255.14 | 5.21 | 59 | >sp\|P01876\|IGHA1_HUMAN Immunoglobulin heavy constant alpha 1 OS=Homo sapiens OX=9606 GN=IGHA1 PE=1 SV=2 | 255 | 20201115_15212_1ng_glyco_hcd203040_compound15.13440.13440.3 | controllerType=0 controllerNumber=1 scan=13440 | 26.8536 |
| 18829:3 | 11 | K.RNIAAFGGDPNN[+1913.677]ITLFGESAGGASVSLQTLSPYNK.G | HexNAc(4)Hex(5)NeuAc(1) | N[+1914] | 1827.4972 | 3 | 5480.4769 | 5480.4644 | 0 | 2.3 | 196 | Specific | 439.21 | 280.6 | 280.6 | 8.24 | 49 | >sp\|P19835\|CEL_HUMAN Bile salt-activated lipase OS=Homo sapiens OX=9606 GN=CEL PE=1 SV=3 | 476 | 20201115_15212_1ng_glyco_hcd203040_compound15.21681.21681.3 | controllerType=0 controllerNumber=1 scan=21681 | 41.4794 |
| 18826:3 | 11 | K.RNIAAFGGDPNN[+1913.677]ITLFGESAGGASVSLQTLSPYNK.G | HexNAc(4)Hex(5)NeuAc(1) | N[+1914] | 1827.4972 | 3 | 5480.4769 | 5480.4644 | 0 | 2.3 | 196 | Specific | 451.59 | 350.97 | 350.97 | 8.77 | 49 | >sp\|P19835\|CEL_HUMAN Bile salt-activated lipase OS=Homo sapiens OX=9606 GN=CEL PE=1 SV=3 | 476 | 20201115_15212_1ng_glyco_hcd203040_compound15.21678.21678.3 | controllerType=0 controllerNumber=1 scan=21678 | 41.4686 |
| 20535:3 | 11 | R.NIAAFGGDPNN[+1913.677]ITLFGESAGGASVSLQTLSPYNK.G | HexNAc(4)Hex(5)NeuAc(1) | N[+1914] | 1775.4679 | 3 | 5324.389 | 5324.3633 | 0 | 4.8 | 197 | Specific | 893.93 | 859.76 | 859.76 | 17.89 | 49 | >sp\|P19835\|CEL_HUMAN Bile salt-activated lipase OS=Homo sapiens OX=9606 GN=CEL PE=1 SV=3 | 476 | 20201115_15212_1ng_glyco_hcd203040_compound15.23464.23464.3 | controllerType=0 controllerNumber=1 scan=23464 | 45.375 |
| 20538:3 | 11 | R.NIAAFGGDPNN[+1913.677]ITLFGESAGGASVSLQTLSPYNK.G | HexNAc(4)Hex(5)NeuAc(1) | N[+1914] | 1775.4679 | 3 | 5324.389 | 5324.3633 | 0 | 4.8 | 197 | Specific | 991.09 | 987.2 | 987.2 | 19.89 | 49 | >sp\|P19835\|CEL_HUMAN Bile salt-activated lipase OS=Homo sapiens OX=9606 GN=CEL PE=1 SV=3 | 476 | 20201115_15212_1ng_glyco_hcd203040_compound15.23467.23467.3 | controllerType=0 controllerNumber=1 scan=23467 | 45.3802 |
| 25181:5 | 13 | K.[+42.011]VFPLSLDSTPQDGNVVVAC[+57.021]LVQGFFPQEPLSVTWSESGQN[+2059 | HexNAc(4)Hex(5)Fuc(1)NeuAc(1) | .[42], C[+57], N[+2060] | 1376.4324 | 5 | 6878.1329 | 6878.1069 | 0 | 3.8 | 8 | Specific | 225.81 | 225.81 | 225.81 | 6.22 | 42 | >sp\|P01877\|IGHA2_HUMAN Immunoglobulin heavy constant alpha 2 OS=Homo sapiens OX=9606 GN=IGHA2 PE=1 SV=4 | 576 | 20201115_15212_1ng_glyco_hcd203040_compound15.28324.28324.5 | controllerType=0 controllerNumber=1 scan=28324 | 55.5307 |
| 25003:4 | 13 | K.VFPLSLDSTPQDGNVVVAC[+57.021]LVQGFFPQEPLSVTWSESGQN[+2117.777]VTAI | HexNAc(5)Hex(5)Fuc(2) | C[+57], N[+2118] | 1724.3059 | 4 | 6894.2017 | 6894.1382 | 0 | 9.2 | 8 | Specific | 227.07 | 227.07 | 227.07 | 7.12 | 42 | >sp\|P01877\|IGHA2_HUMAN Immunoglobulin heavy constant alpha 2 OS=Homo sapiens OX=9606 GN=IGHA2 PE=1 SV=4 | 576 | 20201115_15212_1ng_glyco_hcd203040_compound15.28139.28139.4 | controllerType=0 controllerNumber=1 scan=28139 | 55.1657 |
| 25005:4 | 13 | K.VFPLSLDSTPQDGNVVVAC[+57.021]LVQGFFPQEPLSVTWSESGQN[+2117.777]VTAI | HexNAc(5)Hex(5)Fuc(2) | C[+57], N[+2118] | 1724.3059 | 4 | 6894.2017 | 6894.1382 | 0 | 9.2 | 8 | Specific | 268.63 | 268.63 | 268.63 | 7.7 | 42 | >sp\|P01877\|IGHA2_HUMAN Immunoglobulin heavy constant alpha 2 OS=Homo sapiens OX=9606 GN=IGHA2 PE=1 SV=4 | 576 | 20201115_15212_1ng_glyco_hcd203040_compound15.28141.28141.4 | controllerType=0 controllerNumber=1 scan=28141 | 55.1694 |
| 27437:4 | 13 | K.VFPLSLDSTPQDGNVVVAC[+57.021]LVQGFFPQEPLSVTWSESGQN[+2350.830]VTAI | HexNAc(4)Hex(5)Fuc(1)NeuAc(2) | C[+57], N[+2351] | 1782.5565 | 4 | 7127.2041 | 7127.1917 | 0 | 1.7 | 8 | Specific | 334.7 | 334.7 | 334.7 | 9.74 | 42 | >sp\|P01877\|IGHA2_HUMAN Immunoglobulin heavy constant alpha 2 OS=Homo sapiens OX=9606 GN=IGHA2 PE=1 SV=4 | 576 | 20201115_15212_1ng_glyco_hcd203040_compound15.30675.30675.4 | controllerType=0 controllerNumber=1 scan=30675 | 59.9835 |
| 25183:5 | 13 | K.[+42.011]VFPLSLDSTPQDGNVVVAC[+57.021]LVQGFFPQEPLSVTWSESGQN[+2059 | HexNAc(4)Hex(5)Fuc(1)NeuAc(1) | .[42], C[+57], N[+2060] | 1376.4324 | 5 | 6878.1329 | 6878.1069 | 0 | 3.8 | 8 | Specific | 338.88 | 319.23 | 319.23 | 7.34 | 42 | >sp\|P01877\|IGHA2_HUMAN Immunoglobulin heavy constant alpha 2 OS=Homo sapiens OX=9606 GN=IGHA2 PE=1 SV=4 | 576 | 20201115_15212_1ng_glyco_hcd203040_compound15.28326.28326.5 | controllerType=0 controllerNumber=1 scan=28326 | 55.5377 |
| 25015:4 | 13 | K.VFPLSLDSTPQDGNVVVAC[+57.021]LVQGFFPQEPLSVTWSESGQN[+2059.735]VTAI | HexNAc(4)Hex(5)Fuc(1)NeuAc(1) | C[+57], N[+2060] | 1709.7874 | 4 | 6836.1279 | 6836.0963 | 0 | 4.6 | 8 | Specific | 344.64 | 344.64 | 344.64 | 9.21 | 42 | >sp\|P01877\|IGHA2_HUMAN Immunoglobulin heavy constant alpha 2 OS=Homo sapiens OX=9606 GN=IGHA2 PE=1 SV=4 | 576 | 20201115_15212_1ng_glyco_hcd203040_compound15.28151.28151.4 | controllerType=0 controllerNumber=1 scan=28151 | 55.1957 |
| 25017:4 | 13 | K.VFPLSLDSTPQDGNVVVAC[+57.021]LVQGFFPQEPLSVTWSESGQN[+2059.735]VTAI | HexNAc(4)Hex(5)Fuc(1)NeuAc(1) | C[+57], N[+2060] | 1709.7874 | 4 | 6836.1279 | 6836.0963 | 0 | 4.6 | 8 | Specific | 571.37 | 553.64 | 553.64 | 11.87 | 42 | >sp\|P01877\|IGHA2_HUMAN Immunoglobulin heavy constant alpha 2 OS=Homo sapiens OX=9606 GN=IGHA2 PE=1 SV=4 | 576 | 20201115_15212_1ng_glyco_hcd203040_compound15.28153.28153.4 | controllerType=0 controllerNumber=1 scan=28153 | 55.2013 |
| 03732:4 | 13 | K.HYTN[+2262.814]SSQDVTVPC[+57.021]R.V | HexNAc(5)Hex(5)Fuc(1)NeuAc(1) | C[+57], N[+2263] | 982.399 | 4 | 3926.574 | 3926.5636 | 0 | 2.7 | 89 | Specific | 325.71 | 211.76 | 211.76 | 7.31 | 42 | >sp\|P01877\|IGHA2_HUMAN Immunoglobulin heavy constant alpha 2 OS=Homo sapiens OX=9606 GN=IGHA2 PE=1 SV=4 | 576 | 20201115_15212_1ng_glyco_hcd203040_compound15.5939.5939.4 | controllerType=0 controllerNumber=1 scan=5939 | 14.4672 |
| 03734:4 | 13 | K.HYTN[+2262.814]SSQDVTVPC[+57.021]R.V | HexNAc(5)Hex(5)Fuc(1)NeuAc(1) | C[+57], N[+2263] | 982.399 | 4 | 3926.574 | 3926.5636 | 0 | 2.7 | 89 | Specific | 348.21 | 244.37 | 244.37 | 7.63 | 42 | >sp\|P01877\|IGHA2_HUMAN Immunoglobulin heavy constant alpha 2 OS=Homo sapiens OX=9606 GN=IGHA2 PE=1 SV=4 | 576 | 20201115_15212_1ng_glyco_hcd203040_compound15.5941.5941.4 | controllerType=0 controllerNumber=1 scan=5941 | 14.4719 |
| 02807:3 | 13 | K.HYTN[+1971.719]SSQDVTVPC[+57.021]R.V | HexNAc(5)Hex(5)Fuc(1) | C[+57], N[+1972] | 1212.4965 | 3 | 3635.475 | 3635.4682 | 0 | 1.9 | 89 | Specific | 395.44 | 208.4 | 208.4 | 7.47 | 42 | >sp\|P01877\|IGHA2_HUMAN Immunoglobulin heavy constant alpha 2 OS=Homo sapiens OX=9606 GN=IGHA2 PE=1 SV=4 | 576 | 20201115_15212_1ng_glyco_hcd203040_compound15.4975.4975.3 | controllerType=0 controllerNumber=1 scan=4975 | 12.8024 |
| 02891:2 | 13 | K.HYTN[+1647.613]SSQDVTVPC[+57.021]R.V | HexNAc(5)Hex(3)Fuc(1) | C[+57], N[+1648] | 1656.19 | 2 | 3311.3727 | 3311.3625 | 0 | 3.1 | 89 | Specific | 412.7 | 412.7 | 412.7 | 9.56 | 42 | >sp\|P01877\|IGHA2_HUMAN Immunoglobulin heavy constant alpha 2 OS=Homo sapiens OX=9606 GN=IGHA2 PE=1 SV=4 | 576 | 20201115_15212_1ng_glyco_hcd203040_compound15.5063.5063.2 | controllerType=0 controllerNumber=1 scan=5063 | 12.9634 |
| 02809:3 | 13 | K.HYTN[+1971.719]SSQDVTVPC[+57.021]R.V | HexNAc(5)Hex(5)Fuc(1) | C[+57], N[+1972] | 1212.4965 | 3 | 3635.475 | 3635.4682 | 0 | 1.9 | 89 | Specific | 422.87 | 277.67 | 277.67 | 8.09 | 42 | >sp\|P01877\|IGHA2_HUMAN Immunoglobulin heavy constant alpha 2 OS=Homo sapiens OX=9606 GN=IGHA2 PE=1 SV=4 | 576 | 20201115_15212_1ng_glyco_hcd203040_compound15.4977.4977.3 | controllerType=0 controllerNumber=1 scan=4977 | 12.8065 |
| 02893:2 | 13 | K.HYTN[+1647.613]SSQDVTVPC[+57.021]R.V | HexNAc(5)Hex(3)Fuc(1) | C[+57], N[+1648] | 1656.19 | 2 | 3311.3727 | 3311.3625 | 0 | 3.1 | 89 | Specific | 449.06 | 325.35 | 325.35 | 7.86 | 42 | >sp\|P01877\|IGHA2_HUMAN Immunoglobulin heavy constant alpha 2 OS=Homo sapiens OX=9606 GN=IGHA2 PE=1 SV=4 | 576 | 20201115_15212_1ng_glyco_hcd203040_compound15.5065.5065.2 | controllerType=0 controllerNumber=1 scan=5065 | 12.9668 |
| 02849:3 | 13 | K.HYTN[+1647.613]SSQDVTVPC[+57.021]R.V | HexNAc(5)Hex(3)Fuc(1) | C[+57], N[+1648] | 1104.4606 | 3 | 3311.3672 | 3311.3625 | 0 | 1.4 | 89 | Specific | 492.06 | 476 | 476 | 10.5 | 42 | >sp\|P01877\|IGHA2_HUMAN Immunoglobulin heavy constant alpha 2 OS=Homo sapiens OX=9606 GN=IGHA2 PE=1 SV=4 | 576 | 20201115_15212_1ng_glyco_hcd203040_compound15.5019.5019.2 | controllerType=0 controllerNumber=1 scan=5019 | 12.8826 |
| 02847:3 | 13 | K.HYTN[+1647.613]SSQDVTVPC[+57.021]R.V | HexNAc(5)Hex(3)Fuc(1) | C[+57], N[+1648] | 1104.4606 | 3 | 3311.3672 | 3311.3625 | 0 | 1.4 | 89 | Specific | 500.7 | 397.02 | 397.02 | 10.5 | 42 | >sp\|P01877\|IGHA2_HUMAN Immunoglobulin heavy constant alpha 2 OS=Homo sapiens OX=9606 GN=IGHA2 PE=1 SV=4 | 576 | 20201115_15212_1ng_glyco_hcd203040_compound15.5017.5017.2 | controllerType=0 controllerNumber=1 scan=5017 | 12.8752 |
| 03704:3 | 13 | K.HYTN[+2262.814]SSQDVTVPC[+57.021]R.V | HexNAc(5)Hex(5)Fuc(1)NeuAc(1) | C[+57], N[+2263] | 1309.5302 | 3 | 3926.5761 | 3926.5636 | 0 | 3.2 | 89 | Specific | 585.39 | 330.95 | 330.95 | 10.46 | 42 | >sp\|P01877\|IGHA2_HUMAN Immunoglobulin heavy constant alpha 2 OS=Homo sapiens OX=9606 GN=IGHA2 PE=1 SV=4 | 576 | 20201115_15212_1ng_glyco_hcd203040_compound15.5910.5910.3 | controllerType=0 controllerNumber=1 scan=5910 | 14.4143 |
| 03706:3 | 13 | K.HYTN[+2262.814]SSQDVTVPC[+57.021]R.V | HexNAc(5)Hex(5)Fuc(1)NeuAc(1) | C[+57], N[+2263] | 1309.5302 | 3 | 3926.5761 | 3926.5636 | 0 | 3.2 | 89 | Specific | 621.18 | 410.23 | 410.23 | 13.33 | 42 | >sp\|P01877\|IGHA2_HUMAN Immunoglobulin heavy constant alpha 2 OS=Homo sapiens OX=9606 GN=IGHA2 PE=1 SV=4 | 576 | 20201115_15212_1ng_glyco_hcd203040_compound15.5912.5912.3 | controllerType=0 controllerNumber=1 scan=5912 | 14.4181 |
| 02805:3 | 13 | K.HYTN[+1809.666]SSQDVTVPC[+57.021]R.V | HexNAc(5)Hex(4)Fuc(1) | C[+57], N[+1810] | 1158.4787 | 3 | 3473.4215 | 3473.4154 | 0 | 1.8 | 89 | Specific | 624.14 | 506.8 | 506.8 | 13.25 | 42 | >sp\|P01877\|IGHA2_HUMAN Immunoglobulin heavy constant alpha 2 OS=Homo sapiens OX=9606 GN=IGHA2 PE=1 SV=4 | 576 | 20201115_15212_1ng_glyco_hcd203040_compound15.4973.4973.3 | controllerType=0 controllerNumber=1 scan=4973 | 12.799 |
| 02834:3 | 13 | K.HYTN[+1647.613]SSQDVTVPC[+57.021]R.V | HexNAc(5)Hex(3)Fuc(1) | C[+57], N[+1648] | 1104.4601 | 3 | 3311.3658 | 3311.3625 | 0 | 1 | 89 | Specific | 677.9 | 464.84 | 464.84 | 13.73 | 42 | >sp\|P01877\|IGHA2_HUMAN Immunoglobulin heavy constant alpha 2 OS=Homo sapiens OX=9606 GN=IGHA2 PE=1 SV=4 | 576 | 20201115_15212_1ng_glyco_hcd203040_compound15.5003.5003.3 | controllerType=0 controllerNumber=1 scan=5003 | 12.8503 |
| 02803:3 | 13 | K.HYTN[+1809.666]SSQDVTVPC[+57.021]R.V | HexNAc(5)Hex(4)Fuc(1) | C[+57], N[+1810] | 1158.4787 | 3 | 3473.4215 | 3473.4154 | 0 | 1.8 | 89 | Specific | 684.5 | 637.29 | 637.29 | 14.25 | 42 | >sp\|P01877\|IGHA2_HUMAN Immunoglobulin heavy constant alpha 2 OS=Homo sapiens OX=9606 GN=IGHA2 PE=1 SV=4 | 576 | 20201115_15212_1ng_glyco_hcd203040_compound15.4971.4971.3 | controllerType=0 controllerNumber=1 scan=4971 | 12.7949 |
| 02832:3 | 13 | K.HYTN[+1647.613]SSQDVTVPC[+57.021]R.V | HexNAc(5)Hex(3)Fuc(1) | C[+57], N[+1648] | 1104.4601 | 3 | 3311.3658 | 3311.3625 | 0 | 1 | 89 | Specific | 735.24 | 670.42 | 670.42 | 14.73 | 42 | >sp\|P01877\|IGHA2_HUMAN Immunoglobulin heavy constant alpha 2 OS=Homo sapiens OX=9606 GN=IGHA2 PE=1 SV=4 | 576 | 20201115_15212_1ng_glyco_hcd203040_compound15.5001.5001.3 | controllerType=0 controllerNumber=1 scan=5001 | 12.8446 |
| 04822:4 | 13 | K.HYTN[+1647.613]SSQDVTVPC[+57.021]RVPPPPPC[+57.021]C[+57.021]HPR.L | HexNAc(5)Hex(3)Fuc(1) | C[+57]*3, N[+1648] | 1152.251 | 4 | 4605.9822 | 4605.9689 | 0 | 2.9 | 89 | Specific | 548.46 | 490.92 | 490.92 | 10.07 | 42 | >sp\|P01877\|IGHA2_HUMAN Immunoglobulin heavy constant alpha 2 OS=Homo sapiens OX=9606 GN=IGHA2 PE=1 SV=4 | 576 | 20201115_15212_1ng_glyco_hcd203040_compound15.7068.7068.4 | controllerType=0 controllerNumber=1 scan=7068 | 16.4751 |
| 04841:4 | 13 | K.HYTN[+1647.613]SSQDVTVPC[+57.021]RVPPPPPC[+57.021]C[+57.021]HPR.L | HexNAc(5)Hex(3)Fuc(1) | C[+57]*3, N[+1648] | 1152.2497 | 4 | 4605.9769 | 4605.9689 | 0 | 1.7 | 89 | Specific | 677.35 | 427.28 | 427.28 | 13.5 | 42 | >sp\|P01877\|IGHA2_HUMAN Immunoglobulin heavy constant alpha 2 OS=Homo sapiens OX=9606 GN=IGHA2 PE=1 SV=4 | 576 | 20201115_15212_1ng_glyco_hcd203040_compound15.7088.7088.4 | controllerType=0 controllerNumber=1 scan=7088 | 16.5131 |
| 04819:5 | 13 | K.HYTN[+1647.613]SSQDVTVPC[+57.021]RVPPPPPC[+57.021]C[+57.021]HPR.L | HexNAc(5)Hex(3)Fuc(1) | C[+57]*3, N[+1648] | 922.0017 | 5 | 4605.9795 | 4605.9689 | 0 | 2.3 | 89 | Specific | 744.32 | 491.79 | 491.79 | 15.11 | 42 | >sp\|P01877\|IGHA2_HUMAN Immunoglobulin heavy constant alpha 2 OS=Homo sapiens OX=9606 GN=IGHA2 PE=1 SV=4 | 576 | 20201115_15212_1ng_glyco_hcd203040_compound15.7065.7065.5 | controllerType=0 controllerNumber=1 scan=7065 | 16.4686 |
| 04824:4 | 13 | K.HYTN[+1647.613]SSQDVTVPC[+57.021]RVPPPPPC[+57.021]C[+57.021]HPR.L | HexNAc(5)Hex(3)Fuc(1) | C[+57]*3, N[+1648] | 1152.251 | 4 | 4605.9822 | 4605.9689 | 0 | 2.9 | 89 | Specific | 760.5 | 555.16 | 555.16 | 15.58 | 42 | >sp\|P01877\|IGHA2_HUMAN Immunoglobulin heavy constant alpha 2 OS=Homo sapiens OX=9606 GN=IGHA2 PE=1 SV=4 | 576 | 20201115_15212_1ng_glyco_hcd203040_compound15.7070.7070.4 | controllerType=0 controllerNumber=1 scan=7070 | 16.4792 |
| 04821:5 | 13 | K.HYTN[+1647.613]SSQDVTVPC[+57.021]RVPPPPPC[+57.021]C[+57.021]HPR.L | HexNAc(5)Hex(3)Fuc(1) | C[+57]*3, N[+1648] | 922.0017 | 5 | 4605.9795 | 4605.9689 | 0 | 2.3 | 89 | Specific | 799.09 | 557.07 | 557.07 | 16.11 | 42 | >sp\|P01877\|IGHA2_HUMAN Immunoglobulin heavy constant alpha 2 OS=Homo sapiens OX=9606 GN=IGHA2 PE=1 SV=4 | 576 | 20201115_15212_1ng_glyco_hcd203040_compound15.7067.7067.5 | controllerType=0 controllerNumber=1 scan=7067 | 16.4732 |
| 04843:4 | 13 | K.HYTN[+1647.613]SSQDVTVPC[+57.021]RVPPPPPC[+57.021]C[+57.021]HPR.L | HexNAc(5)Hex(3)Fuc(1) | C[+57]*3, N[+1648] | 1152.2497 | 4 | 4605.9769 | 4605.9689 | 0 | 1.7 | 89 | Specific | 807.1 | 566.41 | 566.41 | 16.5 | 42 | >sp\|P01877\|IGHA2_HUMAN Immunoglobulin heavy constant alpha 2 OS=Homo sapiens OX=9606 GN=IGHA2 PE=1 SV=4 | 576 | 20201115_15212_1ng_glyco_hcd203040_compound15.7090.7090.4 | controllerType=0 controllerNumber=1 scan=7090 | 16.5174 |
| 04544:2 | 13 | K.TPLTAN[+1971.719]ITK.S | HexNAc(5)Hex(5)Fuc(1) | N[+1972] | 1465.6482 | 2 | 2930.2892 | 2930.2757 | 0 | 4.6 | 200 | Specific | 251.82 | 212.08 | 212.08 | 6.07 | 42 | >sp\|P01877\|IGHA2_HUMAN Immunoglobulin heavy constant alpha 2 OS=Homo sapiens OX=9606 GN=IGHA2 PE=1 SV=4 | 576 | 20201115_15212_1ng_glyco_hcd203040_compound15.6779.6779.2 | controllerType=0 controllerNumber=1 scan=6779 | 15.9099 |
| 04546:2 | 13 | K.TPLTAN[+1971.719]ITK.S | HexNAc(5)Hex(5)Fuc(1) | N[+1972] | 1465.6482 | 2 | 2930.2892 | 2930.2757 | 0 | 4.6 | 200 | Specific | 254.33 | 184.21 | 184.21 | 5.89 | 42 | >sp\|P01877\|IGHA2_HUMAN Immunoglobulin heavy constant alpha 2 OS=Homo sapiens OX=9606 GN=IGHA2 PE=1 SV=4 | 576 | 20201115_15212_1ng_glyco_hcd203040_compound15.6781.6781.2 | controllerType=0 controllerNumber=1 scan=6781 | 15.9138 |
| 06395:2 | 13 | K.TPLTAN[+2059.735]ITK.S | HexNAc(4)Hex(5)Fuc(1)NeuAc(1) | N[+2060] | 1509.6504 | 2 | 3018.2936 | 3018.2917 | 0 | 0.6 | 200 | Specific | 271.91 | 127.23 | 127.23 | 5.2 | 42 | >sp\|P01877\|IGHA2_HUMAN Immunoglobulin heavy constant alpha 2 OS=Homo sapiens OX=9606 GN=IGHA2 PE=1 SV=4 | 576 | 20201115_15212_1ng_glyco_hcd203040_compound15.8696.8696.2 | controllerType=0 controllerNumber=1 scan=8696 | 19.1172 |
| 06409:3 | 13 | K.TPLTAN[+2263.835]ITK.S | HexNAc(5)Hex(5)Fuc(3) | N[+2264] | 1074.7993 | 3 | 3222.3835 | 3222.3915 | 0 | -2.5 | 200 | Specific | 294.8 | 176.82 | 176.82 | 5.14 | 42 | >sp\|P01877\|IGHA2_HUMAN Immunoglobulin heavy constant alpha 2 OS=Homo sapiens OX=9606 GN=IGHA2 PE=1 SV=4 | 576 | 20201115_15212_1ng_glyco_hcd203040_compound15.8711.8711.3 | controllerType=0 controllerNumber=1 scan=8711 | 19.142 |
| 06393:2 | 13 | K.TPLTAN[+2059.735]ITK.S | HexNAc(4)Hex(5)Fuc(1)NeuAc(1) | N[+2060] | 1509.6504 | 2 | 3018.2936 | 3018.2917 | 0 | 0.6 | 200 | Specific | 329.74 | 92.4 | 92.4 | 5.11 | 42 | >sp\|P01877\|IGHA2_HUMAN Immunoglobulin heavy constant alpha 2 OS=Homo sapiens OX=9606 GN=IGHA2 PE=1 SV=4 | 576 | 20201115_15212_1ng_glyco_hcd203040_compound15.8694.8694.2 | controllerType=0 controllerNumber=1 scan=8694 | 19.1141 |
| 04488:2 | 13 | K.TPLTAN[+1606.587]ITK.S | HexNAc(4)Hex(4)Fuc(1) | N[+1607] | 1283.0799 | 2 | 2565.1525 | 2565.1435 | 0 | 3.5 | 200 | Specific | 336.67 | 251.74 | 251.74 | 7.53 | 42 | >sp\|P01877\|IGHA2_HUMAN Immunoglobulin heavy constant alpha 2 OS=Homo sapiens OX=9606 GN=IGHA2 PE=1 SV=4 | 576 | 20201115_15212_1ng_glyco_hcd203040_compound15.6721.6721.2 | controllerType=0 controllerNumber=1 scan=6721 | 15.7999 |
| 04512:3 | 13 | K.TPLTAN[+1971.719]ITK.S | HexNAc(5)Hex(5)Fuc(1) | N[+1972] | 977.4325 | 3 | 2930.283 | 2930.2757 | 0 | 2.5 | 200 | Specific | 438.87 | 157.47 | 157.47 | 7.22 | 42 | >sp\|P01877\|IGHA2_HUMAN Immunoglobulin heavy constant alpha 2 OS=Homo sapiens OX=9606 GN=IGHA2 PE=1 SV=4 | 576 | 20201115_15212_1ng_glyco_hcd203040_compound15.6746.6746.3 | controllerType=0 controllerNumber=1 scan=6746 | 15.8505 |
| 04514:3 | 13 | K.TPLTAN[+1971.719]ITK.S | HexNAc(5)Hex(5)Fuc(1) | N[+1972] | 977.4325 | 3 | 2930.283 | 2930.2757 | 0 | 2.5 | 200 | Specific | 461.62 | 203.82 | 203.82 | 7.65 | 42 | >sp\|P01877\|IGHA2_HUMAN Immunoglobulin heavy constant alpha 2 OS=Homo sapiens OX=9606 GN=IGHA2 PE=1 SV=4 | 576 | 20201115_15212_1ng_glyco_hcd203040_compound15.6748.6748.3 | controllerType=0 controllerNumber=1 scan=6748 | 15.8547 |
| 04558:3 | 13 | K.TPLTAN[+1809.666]ITK.S | HexNAc(5)Hex(4)Fuc(1) | N[+1810] | 923.415 | 3 | 2768.2305 | 2768.2228 | 0 | 2.8 | 200 | Specific | 486.41 | 220.21 | 220.21 | 7.82 | 42 | >sp\|P01877\|IGHA2_HUMAN Immunoglobulin heavy constant alpha 2 OS=Homo sapiens OX=9606 GN=IGHA2 PE=1 SV=4 | 576 | 20201115_15212_1ng_glyco_hcd203040_compound15.6794.6794.3 | controllerType=0 controllerNumber=1 scan=6794 | 15.9382 |
| 04637:2 | 13 | K.TPLTAN[+1501.555]ITK.S | HexNAc(5)Hex(3) | N[+1502] | 1230.5633 | 2 | 2460.1193 | 2460.1121 | 0 | 2.9 | 200 | Specific | 510.46 | 259.14 | 259.14 | 8.12 | 42 | >sp\|P01877\|IGHA2_HUMAN Immunoglobulin heavy constant alpha 2 OS=Homo sapiens OX=9606 GN=IGHA2 PE=1 SV=4 | 576 | 20201115_15212_1ng_glyco_hcd203040_compound15.6876.6876.2 | controllerType=0 controllerNumber=1 scan=6876 | 16.128 |
| 04620:3 | 13 | K.TPLTAN[+1647.613]ITK.S | HexNAc(5)Hex(3)Fuc(1) | N[+1648] | 869.3974 | 3 | 2606.1777 | 2606.17 | 0 | 3 | 200 | Specific | 528.42 | 235.01 | 235.01 | 7.99 | 42 | >sp\|P01877\|IGHA2_HUMAN Immunoglobulin heavy constant alpha 2 OS=Homo sapiens OX=9606 GN=IGHA2 PE=1 SV=4 | 576 | 20201115_15212_1ng_glyco_hcd203040_compound15.6859.6859.3 | controllerType=0 controllerNumber=1 scan=6859 | 16.0906 |
| 04560:3 | 13 | K.TPLTAN[+1809.666]ITK.S | HexNAc(5)Hex(4)Fuc(1) | N[+1810] | 923.415 | 3 | 2768.2305 | 2768.2228 | 0 | 2.8 | 200 | Specific | 543.39 | 293.58 | 293.58 | 8.68 | 42 | >sp\|P01877\|IGHA2_HUMAN Immunoglobulin heavy constant alpha 2 OS=Homo sapiens OX=9606 GN=IGHA2 PE=1 SV=4 | 576 | 20201115_15212_1ng_glyco_hcd203040_compound15.6796.6796.3 | controllerType=0 controllerNumber=1 scan=6796 | 15.9436 |
| 04622:3 | 13 | K.TPLTAN[+1647.613]ITK.S | HexNAc(5)Hex(3)Fuc(1) | N[+1648] | 869.3974 | 3 | 2606.1777 | 2606.17 | 0 | 3 | 200 | Specific | 552 | 252.07 | 252.07 | 8.39 | 42 | >sp\|P01877\|IGHA2_HUMAN Immunoglobulin heavy constant alpha 2 OS=Homo sapiens OX=9606 GN=IGHA2 PE=1 SV=4 | 576 | 20201115_15212_1ng_glyco_hcd203040_compound15.6861.6861.3 | controllerType=0 controllerNumber=1 scan=6861 | 16.0967 |
| 04635:2 | 13 | K.TPLTAN[+1501.555]ITK.S | HexNAc(5)Hex(3) | N[+1502] | 1230.5633 | 2 | 2460.1193 | 2460.1121 | 0 | 2.9 | 200 | Specific | 567.18 | 386.94 | 386.94 | 10.84 | 42 | >sp\|P01877\|IGHA2_HUMAN Immunoglobulin heavy constant alpha 2 OS=Homo sapiens OX=9606 GN=IGHA2 PE=1 SV=4 | 576 | 20201115_15212_1ng_glyco_hcd203040_compound15.6874.6874.2 | controllerType=0 controllerNumber=1 scan=6874 | 16.1242 |
| 04667:2 | 13 | K.TPLTAN[+1241.454]ITK.S | HexNAc(3)Hex(3)Fuc(1) | N[+1241] | 1100.5114 | 2 | 2200.0155 | 2200.0113 | 0 | 1.9 | 200 | Specific | 590.78 | 348.09 | 348.09 | 10.39 | 42 | >sp\|P01877\|IGHA2_HUMAN Immunoglobulin heavy constant alpha 2 OS=Homo sapiens OX=9606 GN=IGHA2 PE=1 SV=4 | 576 | 20201115_15212_1ng_glyco_hcd203040_compound15.6907.6907.2 | controllerType=0 controllerNumber=1 scan=6907 | 16.1849 |
| 04669:2 | 13 | K.TPLTAN[+1241.454]ITK.S | HexNAc(3)Hex(3)Fuc(1) | N[+1241] | 1100.5114 | 2 | 2200.0155 | 2200.0113 | 0 | 1.9 | 200 | Specific | 596.75 | 340.86 | 340.86 | 10.39 | 42 | >sp\|P01877\|IGHA2_HUMAN Immunoglobulin heavy constant alpha 2 OS=Homo sapiens OX=9606 GN=IGHA2 PE=1 SV=4 | 576 | 20201115_15212_1ng_glyco_hcd203040_compound15.6909.6909.2 | controllerType=0 controllerNumber=1 scan=6909 | 16.1885 |
| 04575:2 | 13 | K.TPLTAN[+1444.534]ITK.S | HexNAc(4)Hex(3)Fuc(1) | N[+1445] | 1202.0528 | 2 | 2403.0983 | 2403.0906 | 0 | 3.2 | 200 | Specific | 614.21 | 350.28 | 350.28 | 11.47 | 42 | >sp\|P01877\|IGHA2_HUMAN Immunoglobulin heavy constant alpha 2 OS=Homo sapiens OX=9606 GN=IGHA2 PE=1 SV=4 | 576 | 20201115_15212_1ng_glyco_hcd203040_compound15.6811.6811.2 | controllerType=0 controllerNumber=1 scan=6811 | 15.9785 |
| 04611:2 | 13 | K.TPLTAN[+1647.613]ITK.S | HexNAc(5)Hex(3)Fuc(1) | N[+1648] | 1303.5938 | 2 | 2606.1803 | 2606.17 | 0 | 3.9 | 200 | Specific | 616.87 | 300.53 | 300.53 | 11.12 | 42 | >sp\|P01877\|IGHA2_HUMAN Immunoglobulin heavy constant alpha 2 OS=Homo sapiens OX=9606 GN=IGHA2 PE=1 SV=4 | 576 | 20201115_15212_1ng_glyco_hcd203040_compound15.6849.6849.2 | controllerType=0 controllerNumber=1 scan=6849 | 16.071 |
| 04609:2 | 13 | K.TPLTAN[+1647.613]ITK.S | HexNAc(5)Hex(3)Fuc(1) | N[+1648] | 1303.5938 | 2 | 2606.1803 | 2606.17 | 0 | 3.9 | 200 | Specific | 617.42 | 356.72 | 356.72 | 11.52 | 42 | >sp\|P01877\|IGHA2_HUMAN Immunoglobulin heavy constant alpha 2 OS=Homo sapiens OX=9606 GN=IGHA2 PE=1 SV=4 | 576 | 20201115_15212_1ng_glyco_hcd203040_compound15.6847.6847.2 | controllerType=0 controllerNumber=1 scan=6847 | 16.0632 |
| 04568:2 | 13 | K.TPLTAN[+1809.666]ITK.S | HexNAc(5)Hex(4)Fuc(1) | N[+1810] | 1384.6185 | 2 | 2768.2296 | 2768.2228 | 0 | 2.5 | 200 | Specific | 625.06 | 325.44 | 325.44 | 11.24 | 42 | >sp\|P01877\|IGHA2_HUMAN Immunoglobulin heavy constant alpha 2 OS=Homo sapiens OX=9606 GN=IGHA2 PE=1 SV=4 | 576 | 20201115_15212_1ng_glyco_hcd203040_compound15.6804.6804.2 | controllerType=0 controllerNumber=1 scan=6804 | 15.9619 |
| 04576:2 | 13 | K.TPLTAN[+1444.534]ITK.S | HexNAc(4)Hex(3)Fuc(1) | N[+1445] | 1202.0528 | 2 | 2403.0983 | 2403.0906 | 0 | 3.2 | 200 | Specific | 626.87 | 296.9 | 296.9 | 11.18 | 42 | >sp\|P01877\|IGHA2_HUMAN Immunoglobulin heavy constant alpha 2 OS=Homo sapiens OX=9606 GN=IGHA2 PE=1 SV=4 | 576 | 20201115_15212_1ng_glyco_hcd203040_compound15.6812.6812.2 | controllerType=0 controllerNumber=1 scan=6812 | 15.9825 |
| 04566:2 | 13 | K.TPLTAN[+1809.666]ITK.S | HexNAc(5)Hex(4)Fuc(1) | N[+1810] | 1384.6185 | 2 | 2768.2296 | 2768.2228 | 0 | 2.5 | 200 | Specific | 629.48 | 348.09 | 348.09 | 11.47 | 42 | >sp\|P01877\|IGHA2_HUMAN Immunoglobulin heavy constant alpha 2 OS=Homo sapiens OX=9606 GN=IGHA2 PE=1 SV=4 | 576 | 20201115_15212_1ng_glyco_hcd203040_compound15.6802.6802.2 | controllerType=0 controllerNumber=1 scan=6802 | 15.9541 |
| 12081:3 | 13 | R.MAGKPTHIN[+1647.613]VSVVMAEADGTC[+57.021]Y.- | HexNAc(5)Hex(3)Fuc(1) | C[+57], N[+1648] | 1333.5723 | 3 | 3998.7024 | 3998.6961 | 0 | 1.6 | 319 | CRagged | 401.89 | 401.89 | 401.89 | 10.38 | 42 | >sp\|P01877\|IGHA2_HUMAN Immunoglobulin heavy constant alpha 2 OS=Homo sapiens OX=9606 GN=IGHA2 PE=1 SV=4 | 576 | 20201115_15212_1ng_glyco_hcd203040_compound15.14615.14615.3 | controllerType=0 controllerNumber=1 scan=14615 | 28.8743 |
| 12083:3 | 13 | R.MAGKPTHIN[+1647.613]VSVVMAEADGTC[+57.021]Y.- | HexNAc(5)Hex(3)Fuc(1) | C[+57], N[+1648] | 1333.5723 | 3 | 3998.7024 | 3998.6961 | 0 | 1.6 | 319 | CRagged | 539.93 | 539.93 | 539.93 | 11.03 | 42 | >sp\|P01877\|IGHA2_HUMAN Immunoglobulin heavy constant alpha 2 OS=Homo sapiens OX=9606 GN=IGHA2 PE=1 SV=4 | 576 | 20201115_15212_1ng_glyco_hcd203040_compound15.14617.14617.3 | controllerType=0 controllerNumber=1 scan=14617 | 28.8779 |
| 12007:3 | 13 | R.MAGKPTHIN[+1216.423]VSVVMAEADGTC[+57.021]Y.- | HexNAc(2)Hex(5) | C[+57], N[+1216] | 1189.8448 | 3 | 3567.5198 | 3567.5057 | 0 | 3.9 | 319 | CRagged | 874.98 | 874.98 | 486.22 | 18.57 | 42 | >sp\|P01877\|IGHA2_HUMAN Immunoglobulin heavy constant alpha 2 OS=Homo sapiens OX=9606 GN=IGHA2 PE=1 SV=4 | 576 | 20201115_15212_1ng_glyco_hcd203040_compound15.14539.14539.3 | controllerType=0 controllerNumber=1 scan=14539 | 28.7466 |
| 11935:3 | 13 | R.MAGKPTHIN[+1540.529]VSVVMAEADGTC[+57.021]Y.- | HexNAc(2)Hex(7) | C[+57], N[+1541] | 1297.8797 | 3 | 3891.6245 | 3891.6114 | 0 | 3.4 | 319 | CRagged | 879.01 | 730.06 | 519.7 | 18.63 | 42 | >sp\|P01877\|IGHA2_HUMAN Immunoglobulin heavy constant alpha 2 OS=Homo sapiens OX=9606 GN=IGHA2 PE=1 SV=4 | 576 | 20201115_15212_1ng_glyco_hcd203040_compound15.14464.14464.3 | controllerType=0 controllerNumber=1 scan=14464 | 28.5959 |
| 11883:3 | 13 | R.MAGKPTHIN[+1702.581]VSVVMAEADGTC[+57.021]Y.- | HexNAc(2)Hex(8) | C[+57], N[+1703] | 1351.8954 | 3 | 4053.6718 | 4053.6642 | 0 | 1.9 | 319 | CRagged | 904.28 | 754.4 | 514.18 | 19.55 | 42 | >sp\|P01877\|IGHA2_HUMAN Immunoglobulin heavy constant alpha 2 OS=Homo sapiens OX=9606 GN=IGHA2 PE=1 SV=4 | 576 | 20201115_15212_1ng_glyco_hcd203040_compound15.14410.14410.3 | controllerType=0 controllerNumber=1 scan=14410 | 28.4953 |
| 11958:3 | 13 | R.MAGKPTHIN[+1378.476]VSVVMAEADGTC[+57.021]Y.- | HexNAc(2)Hex(6) | C[+57], N[+1378] | 1243.8614 | 3 | 3729.5696 | 3729.5585 | 0 | 3 | 319 | CRagged | 905.07 | 758.95 | 514.69 | 19.63 | 42 | >sp\|P01877\|IGHA2_HUMAN Immunoglobulin heavy constant alpha 2 OS=Homo sapiens OX=9606 GN=IGHA2 PE=1 SV=4 | 576 | 20201115_15212_1ng_glyco_hcd203040_compound15.14488.14488.3 | controllerType=0 controllerNumber=1 scan=14488 | 28.6475 |
| 11933:3 | 13 | R.MAGKPTHIN[+1540.529]VSVVMAEADGTC[+57.021]Y.- | HexNAc(2)Hex(7) | C[+57], N[+1541] | 1297.8797 | 3 | 3891.6245 | 3891.6114 | 0 | 3.4 | 319 | CRagged | 924.44 | 924.44 | 542.77 | 19.63 | 42 | >sp\|P01877\|IGHA2_HUMAN Immunoglobulin heavy constant alpha 2 OS=Homo sapiens OX=9606 GN=IGHA2 PE=1 SV=4 | 576 | 20201115_15212_1ng_glyco_hcd203040_compound15.14462.14462.3 | controllerType=0 controllerNumber=1 scan=14462 | 28.[illegible]908 |
| 12009:3 | 13 | R.MAGKPTHIN[+1216.423]VSVVMAEADGTC[+57.021]Y.- | HexNAc(2)Hex(5) | C[+57], N[+1216] | 1189.8448 | 3 | 3567.5198 | 3567.5057 | 0 | 3.9 | 319 | CRagged | 926.64 | 807.6 | 513.7 | 19.57 | 42 | >sp\|P01877\|IGHA2_HUMAN Immunoglobulin heavy constant alpha 2 OS=Homo sapiens OX=9606 GN=IGHA2 PE=1 SV=4 | 576 | 20201115_15212_1ng_glyco_hcd203040_compound15.14541.14541.3 | controllerType=0 controllerNumber=1 scan=14541 | 28.7527 |
| 11880:3 | 13 | R.MAGKPTHIN[+1702.581]VSVVMAEADGTC[+57.021]Y.- | HexNAc(2)Hex(8) | C[+57], N[+1703] | 1351.8954 | 3 | 4053.6718 | 4053.6642 | 0 | 1.9 | 319 | CRagged | 968.47 | 968.47 | 553.37 | 20.55 | 42 | >sp\|P01877\|IGHA2_HUMAN Immunoglobulin heavy constant alpha 2 OS=Homo sapiens OX=9606 GN=IGHA2 PE=1 SV=4 | 576 | 20201115_15212_1ng_glyco_hcd203040_compound15.14407.14407.3 | controllerType=0 controllerNumber=1 scan=14407 | 28.4881 |
| 11956:3 | 13 | R.MAGKPTHIN[+1378.476]VSVVMAEADGTC[+57.021]Y.- | HexNAc(2)Hex(6) | C[+57], N[+1378] | 1243.8614 | 3 | 3729.5696 | 3729.5585 | 0 | 3 | 319 | CRagged | 982.74 | 982.74 | 592.85 | 20.63 | 42 | >sp\|P01877\|IGHA2_HUMAN Immunoglobulin heavy constant alpha 2 OS=Homo sapiens OX=9606 GN=IGHA2 PE=1 SV=4 | 576 | 20201115_15212_1ng_glyco_hcd203040_compound15.14486.14486.3 | controllerType=0 controllerNumber=1 scan=14486 | 28.6398 |
| 07295:3 | 15 | R.LSPN[+2205.793]ASAEHLELR.W | HexNAc(4)Hex(5)Fuc(2)NeuAc(1) | N[+2206] | 1214.8559 | 3 | 3642.5531 | 3642.5421 | 0 | 3 | 52 | Specific | 491.69 | 345.57 | 345.57 | 8.93 | 33 | >sp\|Q13410\|BT1A1_HUMAN Butyrophilin subfamily 1 member A1 OS=Homo sapiens OX=9606 GN=BTN1A1 PE=1 SV=3 | 179 | 20201115_15212_1ng_glyco_hcd203040_compound15.9626.9626.3 | controllerType=0 controllerNumber=1 scan=9626 | 20.5992 |
| 07297:3 | 15 | R.LSPN[+2205.793]ASAEHLELR.W | HexNAc(4)Hex(5)Fuc(2)NeuAc(1) | N[+2206] | 1214.8559 | 3 | 3642.5531 | 3642.5421 | 0 | 3 | 52 | Specific | 532.71 | 380.74 | 380.74 | 10.63 | 33 | >sp\|Q13410\|BT1A1_HUMAN Butyrophilin subfamily 1 member A1 OS=Homo sapiens OX=9606 GN=BTN1A1 PE=1 SV=3 | 179 | 20201115_15212_1ng_glyco_hcd203040_compound15.9628.9628.3 | controllerType=0 controllerNumber=1 scan=9628 | 20.6032 |
| 25350:4 | 16 | R.[+42.011]QLAHQSN[+2190.793]STNIFFSPVSIATAFAMLSLGTK.A | HexNAc(6)Hex(6) | .[42], N[+2191] | 1354.3594 | 4 | 5414.4157 | 5414.4388 | 0 | -4.3 | 64 | Specific | 259.14 | 259.14 | 259.14 | 5.85 | 34 | >sp\|P01009\|A1AT_HUMAN Alpha-1-antitrypsin OS=Homo sapiens OX=9606 GN=SERPINA1 PE=1 SV=3 | 642 | 20201115_15212_1ng_glyco_hcd203040_compound15.28501.28501.4 | controllerType=0 controllerNumber=1 scan=28501 | 55.9128 |
| 25352:4 | 16 | R.[+42.011]QLAHQSN[+2190.793]STNIFFSPVSIATAFAMLSLGTK.A | HexNAc(6)Hex(6) | .[42], N[+2191] | 1354.3594 | 4 | 5414.4157 | 5414.4388 | 0 | -4.3 | 64 | Specific | 277.01 | 277.01 | 277.01 | 6.16 | 34 | >sp\|P01009\|A1AT_HUMAN Alpha-1-antitrypsin OS=Homo sapiens OX=9606 GN=SERPINA1 PE=1 SV=3 | 642 | 20201115_15212_1ng_glyco_hcd203040_compound15.28503.28503.4 | controllerType=0 controllerNumber=1 scan=28503 | 55.9187 |
| 25304:3 | 16 | R.QLAHQSN[+2204.772]STNIFFSPVSIATAFAMLSLGTK.A | HexNAc(4)Hex(5)NeuAc(2) | N[+2205] | 1796.1499 | 3 | 5386.435 | 5386.4075 | 0 | 5.1 | 64 | Specific | 790.69 | 750.9 | 750.9 | 16.29 | 34 | >sp\|P01009\|A1AT_HUMAN Alpha-1-antitrypsin OS=Homo sapiens OX=9606 GN=SERPINA1 PE=1 SV=3 | 642 | 20201115_15212_1ng_glyco_hcd203040_compound15.28453.28453.3 | controllerType=0 controllerNumber=1 scan=28453 | 55.8102 |
| 25306:3 | 16 | R.QLAHQSN[+2204.772]STNIFFSPVSIATAFAMLSLGTK.A | HexNAc(4)Hex(5)NeuAc(2) | N[+2205] | 1796.1499 | 3 | 5386.435 | 5386.4075 | 0 | 5.1 | 64 | Specific | 864.26 | 789.44 | 789.44 | 18.29 | 34 | >sp\|P01009\|A1AT_HUMAN Alpha-1-antitrypsin OS=Homo sapiens OX=9606 GN=SERPINA1 PE=1 SV=3 | 642 | 20201115_15212_1ng_glyco_hcd203040_compound15.28455.28455.3 | controllerType=0 controllerNumber=1 scan=28455 | 55.8153 |
| 25303:4 | 16 | R.QLAHQSN[+2204.772]STNIFFSPVSIATAFAMLSLGTK.A | HexNAc(4)Hex(5)NeuAc(2) | N[+2205] | 1347.3645 | 4 | 5386.4362 | 5386.4075 | 0 | 5.3 | 64 | Specific | 868.22 | 809.73 | 809.73 | 18.31 | 34 | >sp\|P01009\|A1AT_HUMAN Alpha-1-antitrypsin OS=Homo sapiens OX=9606 GN=SERPINA1 PE=1 SV=3 | 642 | 20201115_15212_1ng_glyco_hcd203040_compound15.28452.28452.4 | controllerType=0 controllerNumber=1 scan=28452 | 55.8083 |
| 25301:4 | 16 | R.QLAHQSN[+2204.772]STNIFFSPVSIATAFAMLSLGTK.A | HexNAc(4)Hex(5)NeuAc(2) | N[+2205] | 1347.3645 | 4 | 5386.4362 | 5386.4075 | 0 | 5.3 | 64 | Specific | 871.87 | 871.87 | 871.87 | 18.31 | 34 | >sp\|P01009\|A1AT_HUMAN Alpha-1-antitrypsin OS=Homo sapiens OX=9606 GN=SERPINA1 PE=1 SV=3 | 642 | 20201115_15212_1ng_glyco_hcd203040_compound15.28450.28450.4 | controllerType=0 controllerNumber=1 scan=28450 | 55.7889 |
| 24083:4 | 16 | K.ADTHDEILEGLNFN[+2862.021]LTEIPEAQIHEGFQELLR.T | HexNAc(5)Hex(6)Fuc(2)NeuAc(2) | N[+2862] | 1639.2218 | 4 | 6553.8654 | 6553.8443 | 0 | 3.2 | 94 | Specific | 689.81 | 594.19 | 594.19 | 14.9 | 34 | >sp\|P01009\|A1AT_HUMAN Alpha-1-antitrypsin OS=Homo sapiens OX=9606 GN=SERPINA1 PE=1 SV=3 | 642 | 20201115_15212_1ng_glyco_hcd203040_compound15.27176.27176.4 | controllerType=0 controllerNumber=1 scan=27176 | 53.1102 |
| 24086:4 | 16 | K.ADTHDEILEGLNFN[+2862.021]LTEIPEAQIHEGFQELLR.T | HexNAc(5)Hex(6)Fuc(2)NeuAc(2) | N[+2862] | 1639.2218 | 4 | 6553.8654 | 6553.8443 | 0 | 3.2 | 94 | Specific | 939.1 | 879.77 | 879.77 | 19.9 | 34 | >sp\|P01009\|A1AT_HUMAN Alpha-1-antitrypsin OS=Homo sapiens OX=9606 GN=SERPINA1 PE=1 SV=3 | 642 | 20201115_15212_1ng_glyco_hcd203040_compound15.27179.27179.4 | controllerType=0 controllerNumber=1 scan=27179 | 53.119 |
| 21796:3 | 16 | K.YLGN[+2204.772]ATAIFFLPDEGK.L | HexNAc(4)Hex(5)NeuAc(2) | N[+2205] | 1320.8974 | 3 | 3960.6776 | 3960.6677 | 0 | 2.5 | 268 | Specific | 465.77 | 465.77 | 465.77 | 10.48 | 34 | >sp\|P01009\|A1AT_HUMAN Alpha-1-antitrypsin OS=Homo sapiens OX=9606 GN=SERPINA1 PE=1 SV=3 | 642 | 20201115_15212_1ng_glyco_hcd203040_compound15.24785.24785.3 | controllerType=0 controllerNumber=1 scan=24785 | 48.1778 |
| 21799:3 | 16 | K.YLGN[+2204.772]ATAIFFLPDEGK.L | HexNAc(4)Hex(5)NeuAc(2) | N[+2205] | 1320.8974 | 3 | 3960.6776 | 3960.6677 | 0 | 2.5 | 268 | Specific | 650.97 | 524.84 | 524.84 | 14.47 | 34 | >sp\|P01009\|A1AT_HUMAN Alpha-1-antitrypsin OS=Homo sapiens OX=9606 GN=SERPINA1 PE=1 SV=3 | 642 | 20201115_15212_1ng_glyco_hcd203040_compound15.24788.24788.3 | controllerType=0 controllerNumber=1 scan=24788 | 48.1848 |
| 03695:3 | 17 | K.N[+1378.476]NSIPDKQITASSSYK.T | HexNAc(2)Hex(6) | N[+1378] | 1044.459 | 3 | 3131.3625 | 3131.352 | 0 | 3.4 | 238 | Specific | 220.59 | 117.99 | 117.99 | 4.24 | 30 | >sp\|Q08431\|MFGM_HUMAN Lactadherin OS=Homo sapiens OX=9606 GN=MFGE8 PE=1 SV=3 | 1064 | 20201115_15212_1ng_glyco_hcd203040_compound15.5900.5900.3 | controllerType=0 controllerNumber=1 scan=5900 | 14.3957 |
| 03625:3 | 17 | K.N[+1540.529]NSIPDKQITASSSYK.T | HexNAc(2)Hex(7) | N[+1541] | 1098.4755 | 3 | 3293.4119 | 3293.4048 | 0 | 2.2 | 238 | Specific | 309.2 | 215.65 | 215.65 | 6.72 | 30 | >sp\|Q08431\|MFGM_HUMAN Lactadherin OS=Homo sapiens OX=9606 GN=MFGE8 PE=1 SV=3 | 1064 | 20201115_15212_1ng_glyco_hcd203040_compound15.5828.5828.3 | controllerType=0 controllerNumber=1 scan=5828 | 14.2707 |
| 03627:3 | 17 | K.N[+1540.529]NSIPDKQITASSSYK.T | HexNAc(2)Hex(7) | N[+1541] | 1098.4755 | 3 | 3293.4119 | 3293.4048 | 0 | 2.2 | 238 | Specific | 370.24 | 189.97 | 189.97 | 6.97 | 30 | >sp\|Q08431\|MFGM_HUMAN Lactadherin OS=Homo sapiens OX=9606 GN=MFGE8 PE=1 SV=3 | 1064 | 20201115_15212_1ng_glyco_hcd203040_compound15.5830.5830.3 | controllerType=0 controllerNumber=1 scan=5830 | 14.2742 |
| 03763:3 | 17 | K.N[+1216.423]NSIPDKQITASSSYK.T | HexNAc(2)Hex(5) | N[+1216] | 990.4418 | 3 | 2969.3108 | 2969.2991 | 0 | 3.9 | 238 | Specific | 495.68 | 278.09 | 278.09 | 7.98 | 30 | >sp\|Q08431\|MFGM_HUMAN Lactadherin OS=Homo sapiens OX=9606 GN=MFGE8 PE=1 SV=3 | 1064 | 20201115_15212_1ng_glyco_hcd203040_compound15.5971.5971.3 | controllerType=0 controllerNumber=1 scan=5971 | 14.5225 |
| 03761:3 | 17 | K.N[+1216.423]NSIPDKQITASSSYK.T | HexNAc(2)Hex(5) | N[+1216] | 990.4418 | 3 | 2969.3108 | 2969.2991 | 0 | 3.9 | 238 | Specific | 526.38 | 345.11 | 345.11 | 8.51 | 30 | >sp\|Q08431\|MFGM_HUMAN Lactadherin OS=Homo sapiens OX=9606 GN=MFGE8 PE=1 SV=3 | 1064 | 20201115_15212_1ng_glyco_hcd203040_compound15.5969.5969.3 | controllerType=0 controllerNumber=1 scan=5969 | 14.5187 |
| 15789:4 | 18 | K.LKELPGVC[+57.021]N[+2059.735]ETMMALWEEC[+57.021]KPC[+57.021]LK.Q | HexNAc(4)Hex(5)Fuc(1)NeuAc(1) | C[+57]*3, N[+2060] | 1249.7944 | 4 | 4996.1559 | 4996.1196 | 0 | 7.3 | 95 | Specific | 348.85 | 343.39 | 343.39 | 7.13 | 35 | >sp\|P10909\|CLUS_HUMAN Clusterin OS=Homo sapiens OX=9606 GN=CLU PE=1 SV=1 | 1467 | 20201115_15212_1ng_glyco_hcd203040_compound15.18485.18485.4 | controllerType=0 controllerNumber=1 scan=18485 | 35.5682 |
| 15697:4 | 18 | K.LKELPGVC[+57.021]N[+2205.793]ETMMALWEEC[+57.021]KPC[+57.021]LK.Q | HexNAc(4)Hex(5)Fuc(2)NeuAc(1) | C[+57]*3, N[+2206] | 1286.3076 | 4 | 5142.2087 | 5142.1776 | 0 | 6 | 95 | Specific | 463.08 | 433.4 | 433.4 | 9.13 | 35 | >sp\|P10909\|CLUS_HUMAN Clusterin OS=Homo sapiens OX=9606 GN=CLU PE=1 SV=1 | 1467 | 20201115_15212_1ng_glyco_hcd203040_compound15.18390.18390.4 | controllerType=0 controllerNumber=1 scan=18390 | 35.4008 |
| 15700:4 | 18 | K.LKELPGVC[+57.021]N[+2205.793]ETMMALWEEC[+57.021]KPC[+57.021]LK.Q | HexNAc(4)Hex(5)Fuc(2)NeuAc(1) | C[+57]*3, N[+2206] | 1286.3076 | 4 | 5142.2087 | 5142.1776 | 0 | 6 | 95 | Specific | 614.14 | 530.07 | 530.07 | 12.12 | 35 | >sp\|P10909\|CLUS_HUMAN Clusterin OS=Homo sapiens OX=9606 GN=CLU PE=1 SV=1 | 1467 | 20201115_15212_1ng_glyco_hcd203040_compound15.18393.18393.4 | controllerType=0 controllerNumber=1 scan=18393 | 35.4053 |
| 15539:4 | 18 | K.LKELPGVC[+57.021]N[+2570.925]ETMMALWEEC[+57.021]KPC[+57.021]LK.Q | HexNAc(5)Hex(6)Fuc(2)NeuAc(1) | C[+57]*3, N[+2571] | 1377.5875 | 4 | 5507.3283 | 5507.3097 | 0 | 3.4 | 95 | Specific | 639.02 | 551.72 | 551.72 | 13.19 | 35 | >sp\|P10909\|CLUS_HUMAN Clusterin OS=Homo sapiens OX=9606 GN=CLU PE=1 SV=1 | 1467 | 20201115_15212_1ng_glyco_hcd203040_compound15.18226.18226.4 | controllerType=0 controllerNumber=1 scan=18226 | 35.0938 |
| 01961:3 | 18 | R.EIRHN[+2059.735]STGC[+57.021]LR.M | HexNAc(4)Hex(5)Fuc(1)NeuAc(1) | C[+57], N[+2060] | 1134.8075 | 3 | 3402.408 | 3402.3994 | 0 | 2.5 | 287 | Specific | 316.97 | 151.45 | 151.45 | 5.27 | 35 | >sp\|P10909\|CLUS_HUMAN Clusterin OS=Homo sapiens OX=9606 GN=CLU PE=1 SV=1 | 1467 | 20201115_15212_1ng_glyco_hcd203040_compound15.4095.4095.3 | controllerType=0 controllerNumber=1 scan=4095 | 11.2543 |
| 01963:3 | 18 | R.EIRHN[+2059.735]STGC[+57.021]LR.M | HexNAc(4)Hex(5)Fuc(1)NeuAc(1) | C[+57], N[+2060] | 1134.8075 | 3 | 3402.408 | 3402.3994 | 0 | 2.5 | 287 | Specific | 401.75 | 281.47 | 281.47 | 6.69 | 35 | >sp\|P10909\|CLUS_HUMAN Clusterin OS=Homo sapiens OX=9606 GN=CLU PE=1 SV=1 | 1467 | 20201115_15212_1ng_glyco_hcd203040_compound15.4097.4097.3 | controllerType=0 controllerNumber=1 scan=4097 | 11.2593 |
| 22462:3 | 18 | K.MLN[+2059.735]TSSLLEQLNEQFNWVSR.L | HexNAc(4)Hex(5)Fuc(1)NeuAc(1) | N[+2060] | 1490.3168 | 3 | 4468.9359 | 4468.9217 | 0 | 3.2 | 352 | Specific | 648.01 | 495.41 | 495.41 | 13.65 | 35 | >sp\|P10909\|CLUS_HUMAN Clusterin OS=Homo sapiens OX=9606 GN=CLU PE=1 SV=1 | 1467 | 20201115_15212_1ng_glyco_hcd203040_compound15.25481.25481.3 | controllerType=0 controllerNumber=1 scan=25481 | 49.7598 |
| 22744:3 | 18 | K.MLN[+1913.677]TSSLLEQLNEQFNWVSR.L | HexNAc(4)Hex(5)NeuAc(1) | N[+1914] | 1441.6334 | 3 | 4322.8855 | 4322.8637 | 0 | 5 | 352 | Specific | 661.95 | 617.62 | 617.62 | 14.04 | 35 | >sp\|P10909\|CLUS_HUMAN Clusterin OS=Homo sapiens OX=9606 GN=CLU PE=1 SV=1 | 1467 | 20201115_15212_1ng_glyco_hcd203040_compound15.25775.25775.3 | controllerType=0 controllerNumber=1 scan=25775 | 50.3572 |
| 22742:3 | 18 | K.MLN[+1913.677]TSSLLEQLNEQFNWVSR.L | HexNAc(4)Hex(5)NeuAc(1) | N[+1914] | 1441.6334 | 3 | 4322.8855 | 4322.8637 | 0 | 5 | 352 | Specific | 719.85 | 656.97 | 656.97 | 15.04 | 35 | >sp\|P10909\|CLUS_HUMAN Clusterin OS=Homo sapiens OX=9606 GN=CLU PE=1 SV=1 | 1467 | 20201115_15212_1ng_glyco_hcd203040_compound15.25773.25773.3 | controllerType=0 controllerNumber=1 scan=25773 | 50.3517 |
| 22465:3 | 18 | K.MLN[+2059.735]TSSLLEQLNEQFNWVSR.L | HexNAc(4)Hex(5)Fuc(1)NeuAc(1) | N[+2060] | 1490.3168 | 3 | 4468.9359 | 4468.9217 | 0 | 3.2 | 352 | Specific | 955.24 | 660.55 | 660.55 | 20.65 | 35 | >sp\|P10909\|CLUS_HUMAN Clusterin OS=Homo sapiens OX=9606 GN=CLU PE=1 SV=1 | 1467 | 20201115_15212_1ng_glyco_hcd203040_compound15.25484.25484.3 | controllerType=0 controllerNumber=1 scan=25484 | 49.7647 |
| 12041:3 | 18 | R.LAN[+2205.793]LTQGEDQYYLR.V | HexNAc(4)Hex(5)Fuc(2)NeuAc(1) | N[+2206] | 1297.2186 | 3 | 3889.6411 | 3889.6265 | 0 | 3.8 | 372 | Specific | 210.52 | 210.52 | 210.52 | 5.67 | 35 | >sp\|P10909\|CLUS_HUMAN Clusterin OS=Homo sapiens OX=9606 GN=CLU PE=1 SV=1 | 1467 | 20201115_15212_1ng_glyco_hcd203040_compound15.14574.14574.3 | controllerType=0 controllerNumber=1 scan=14574 | 28.8104 |
| 11785:3 | 18 | R.LAN[+2715.963]LTQGEDQYYLR.V | HexNAc(5)Hex(6)Fuc(1)NeuAc(2) | N[+2716] | 1467.2833 | 3 | 4399.8352 | 4399.7962 | 0 | 8.9 | 372 | Specific | 269.51 | 269.51 | 269.51 | 6.31 | 35 | >sp\|P10909\|CLUS_HUMAN Clusterin OS=Homo sapiens OX=9606 GN=CLU PE=1 SV=1 | 1467 | 20201115_15212_1ng_glyco_hcd203040_compound15.14308.14308.3 | controllerType=0 controllerNumber=1 scan=14308 | 28.3027 |
| 12043:3 | 18 | R.LAN[+2205.793]LTQGEDQYYLR.V | HexNAc(4)Hex(5)Fuc(2)NeuAc(1) | N[+2206] | 1297.2186 | 3 | 3889.6411 | 3889.6265 | 0 | 3.8 | 372 | Specific | 283.86 | 206.81 | 206.81 | 7.06 | 35 | >sp\|P10909\|CLUS_HUMAN Clusterin OS=Homo sapiens OX=9606 GN=CLU PE=1 SV=1 | 1467 | 20201115_15212_1ng_glyco_hcd203040_compound15.14576.14576.3 | controllerType=0 controllerNumber=1 scan=14576 | 28.8144 |
| 11787:3 | 18 | R.LAN[+2570.925]LTQGEDQYYLR.V | HexNAc(5)Hex(6)Fuc(2)NeuAc(1) | N[+2571] | 1418.93 | 3 | 4254.7754 | 4254.7587 | 0 | 3.9 | 372 | Specific | 293.83 | 293.83 | 293.83 | 7.75 | 35 | >sp\|P10909\|CLUS_HUMAN Clusterin OS=Homo sapiens OX=9606 GN=CLU PE=1 SV=1 | 1467 | 20201115_15212_1ng_glyco_hcd203040_compound15.14310.14310.3 | controllerType=0 controllerNumber=1 scan=14310 | 28.3078 |
| 14738:3 | 18 | R.LAN[+2350.830]LTQGEDQYYLR.V | HexNAc(4)Hex(5)Fuc(1)NeuAc(2) | N[+2351] | 1345.5621 | 3 | 4034.6717 | 4034.664 | 0 | 1.9 | 372 | Specific | 320.28 | 285.45 | 285.45 | 7.62 | 35 | >sp\|P10909\|CLUS_HUMAN Clusterin OS=Homo sapiens OX=9606 GN=CLU PE=1 SV=1 | 1467 | 20201115_15212_1ng_glyco_hcd203040_compound15.17396.17396.3 | controllerType=0 controllerNumber=1 scan=17396 | 33.5486 |
| 11789:3 | 18 | R.LAN[+2570.925]LTQGEDQYYLR.V | HexNAc(5)Hex(6)Fuc(2)NeuAc(1) | N[+2571] | 1418.93 | 3 | 4254.7754 | 4254.7587 | 0 | 3.9 | 372 | Specific | 443.29 | 390.36 | 390.36 | 10.25 | 35 | >sp\|P10909\|CLUS_HUMAN Clusterin OS=Homo sapiens OX=9606 GN=CLU PE=1 SV=1 | 1467 | 20201115_15212_1ng_glyco_hcd203040_compound15.14312.14312.3 | controllerType=0 controllerNumber=1 scan=14312 | 28.3112 |
| 14741:3 | 18 | R.LAN[+2350.830]LTQGEDQYYLR.V | HexNAc(4)Hex(5)Fuc(1)NeuAc(2) | N[+2351] | 1345.5621 | 3 | 4034.6717 | 4034.664 | 0 | 1.9 | 372 | Specific | 520.63 | 467.77 | 467.77 | 10.58 | 35 | >sp\|P10909\|CLUS_HUMAN Clusterin OS=Homo sapiens OX=9606 GN=CLU PE=1 SV=1 | 1467 | 20201115_15212_1ng_glyco_hcd203040_compound15.17399.17399.3 | controllerType=0 controllerNumber=1 scan=17399 | 33.5545 |
| 10257:5 | 20 | K.[+42.011]SC[+57.021]QHN[+2018.708]GTMYQHGEIFSAHELFPSR.L | HexNAc(3)Hex(6)Fuc(1)NeuAc(1) | .[42], C[+57], N[+2019] | 979.6074 | 5 | 4894.0081 | 4893.9759 | 0 | 6.6 | 110 | Specific | 274.22 | 165.4 | 165.4 | 4.42 | 23 | >sp\|Q6WN34\|CRDL2_HUMAN Chordin-like protein 2 OS=Homo sapiens OX=9606 GN=CHRDL2 PE=1 SV=1 | 1189 | 20201115_15212_1ng_glyco_hcd203040_compound15.12704.12704.2 | controllerType=0 controllerNumber=1 scan=12704 | 25.6 |
| 10255:5 | 20 | K.[+42.011]SC[+57.021]QHN[+2018.708]GTMYQHGEIFSAHELFPSR.L | HexNAc(3)Hex(6)Fuc(1)NeuAc(1) | .[42], C[+57], N[+2019] | 979.6074 | 5 | 4894.0081 | 4893.9759 | 0 | 6.6 | 110 | Specific | 295.77 | 292.63 | 292.63 | 7.11 | 23 | >sp\|Q6WN34\|CRDL2_HUMAN Chordin-like protein 2 OS=Homo sapiens OX=9606 GN=CHRDL2 PE=1 SV=1 | 1189 | 20201115_15212_1ng_glyco_hcd203040_compound15.12702.12702.2 | controllerType=0 controllerNumber=1 scan=12702 | 25.5965 |
| 11685:4 | 20 | K.SC[+57.021]QHN[+2351.851]GTMYQHGEIFSAHELFPSR.L | HexNAc(4)Hex(5)Fuc(3)NeuAc(1) | C[+57], N[+2352] | 1297.0303 | 4 | 5185.0994 | 5185.1077 | 0 | -1.6 | 110 | Specific | 719.78 | 719.78 | 719.78 | 13.61 | 23 | >sp\|Q6WN34\|CRDL2_HUMAN Chordin-like protein 2 OS=Homo sapiens OX=9606 GN=CHRDL2 PE=1 SV=1 | 1189 | 20201115_15212_1ng_glyco_hcd203040_compound15.14204.14204.4 | controllerType=0 controllerNumber=1 scan=14204 | 28.1236 |
| 11688:4 | 20 | K.SC[+57.021]QHN[+2351.851]GTMYQHGEIFSAHELFPSR.L | HexNAc(4)Hex(5)Fuc(3)NeuAc(1) | C[+57], N[+2352] | 1297.0303 | 4 | 5185.0994 | 5185.1077 | 0 | -1.6 | 110 | Specific | 766.82 | 744.46 | 744.46 | 14.61 | 23 | >sp\|Q6WN34\|CRDL2_HUMAN Chordin-like protein 2 OS=Homo sapiens OX=9606 GN=CHRDL2 PE=1 SV=1 | 1189 | 20201115_15212_1ng_glyco_hcd203040_compound15.14207.14207.4 | controllerType=0 controllerNumber=1 scan=14207 | 28.1369 |
| 10296:4 | 20 | K.SC[+57.021]QHN[+2059.735]GTMYQHGEIFSAHELFPSR.L | HexNAc(4)Hex(5)Fuc(1)NeuAc(1) | C[+57], N[+2060] | 1224.009 | 4 | 4893.014 | 4892.9919 | 0 | 4.5 | 110 | Specific | 773.26 | 773.26 | 773.26 | 15.51 | 23 | >sp\|Q6WN34\|CRDL2_HUMAN Chordin-like protein 2 OS=Homo sapiens OX=9606 GN=CHRDL2 PE=1 SV=1 | 1189 | 20201115_15212_1ng_glyco_hcd203040_compound15.12744.12744.4 | controllerType=0 controllerNumber=1 scan=12744 | 25.6626 |
| 10299:4 | 20 | K.SC[+57.021]QHN[+2059.735]GTMYQHGEIFSAHELFPSR.L | HexNAc(4)Hex(5)Fuc(1)NeuAc(1) | C[+57], N[+2060] | 1224.009 | 4 | 4893.014 | 4892.9919 | 0 | 4.5 | 110 | Specific | 858.83 | 855.66 | 855.66 | 17.51 | 23 | >sp\|Q6WN34\|CRDL2_HUMAN Chordin-like protein 2 OS=Homo sapiens OX=9606 GN=CHRDL2 PE=1 SV=1 | 1189 | 20201115_15212_1ng_glyco_hcd203040_compound15.12747.12747.4 | controllerType=0 controllerNumber=1 scan=12747 | 25.6678 |
| 16279:4 | 27 | K.MVSHHN[+2204.772]LTTGATLINEQWLLTTAK.N | HexNAc(4)Hex(5)NeuAc(2) | N[+2205] | 1221.8008 | 4 | 4884.1814 | 4884.1647 | 0 | 3.4 | 179 | Specific | 243.18 | 111.91 | 111.91 | 6.16 | 24 | >sp\|P00738\|HPT_HUMAN Haptoglobin OS=Homo sapiens OX=9606 GN=HP PE=1 SV=1 | 407 | 20201115_15212_1ng_glyco_hcd203040_compound15.18994.18994.4 | controllerType=0 controllerNumber=1 scan=18994 | 36.5013 |
| 16247:4 | 27 | K.MVSHHN[+2204.772]LTTGATLINEQWLLTTAK.N | HexNAc(4)Hex(5)NeuAc(2) | N[+2205] | 1221.8006 | 4 | 4884.1805 | 4884.1647 | 0 | 3.2 | 179 | Specific | 450.28 | 444.72 | 444.72 | 10.73 | 24 | >sp\|P00738\|HPT_HUMAN Haptoglobin OS=Homo sapiens OX=9606 GN=HP PE=1 SV=1 | 407 | 20201115_15212_1ng_glyco_hcd203040_compound15.18961.18961.3 | controllerType=0 controllerNumber=1 scan=18961 | 36.4386 |
| 16249:4 | 27 | K.MVSHHN[+2204.772]LTTGATLINEQWLLTTAK.N | HexNAc(4)Hex(5)NeuAc(2) | N[+2205] | 1221.8006 | 4 | 4884.1805 | 4884.1647 | 0 | 3.2 | 179 | Specific | 506.77 | 357.86 | 357.86 | 9.44 | 24 | >sp\|P00738\|HPT_HUMAN Haptoglobin OS=Homo sapiens OX=9606 GN=HP PE=1 SV=1 | 407 | 20201115_15212_1ng_glyco_hcd203040_compound15.18963.18963.3 | controllerType=0 controllerNumber=1 scan=18963 | 36.4426 |
| 15979:3 | 27 | K.VVLHPN[+2204.772]YSQVDIGLIK.L | HexNAc(4)Hex(5)NeuAc(2) | N[+2205] | 1333.9374 | 3 | 3999.7978 | 3999.7837 | 0 | 3.5 | 236 | Specific | 710.31 | 663.98 | 663.98 | 15.38 | 24 | >sp\|P00738\|HPT_HUMAN Haptoglobin OS=Homo sapiens OX=9606 GN=HP PE=1 SV=1 | 407 | 20201115_15212_1ng_glyco_hcd203040_compound15.18682.18682.3 | controllerType=0 controllerNumber=1 scan=18682 | 35.9091 |
| 15978:3 | 27 | K.VVLHPN[+2204.772]YSQVDIGLIK.L | HexNAc(4)Hex(5)NeuAc(2) | N[+2205] | 1333.9374 | 3 | 3999.7978 | 3999.7837 | 0 | 3.5 | 236 | Specific | 737.01 | 676.71 | 676.71 | 15.38 | 24 | >sp\|P00738\|HPT_HUMAN Haptoglobin OS=Homo sapiens OX=9606 GN=HP PE=1 SV=1 | 407 | 20201115_15212_1ng_glyco_hcd203040_compound15.18681.18681.3 | controllerType=0 controllerNumber=1 scan=18681 | 35.9049 |
| 17935:4 | 29 | R.FSLLGHASISC[+57.021]TVEN[+2350.830]ETIGVWRPSPPTC[+57.021]EK.I | HexNAc(4)Hex(5)Fuc(1)NeuAc(2) | C[+57]*2, N[+2351] | 1431.6188 | 4 | 5723.4533 | 5723.4655 | 0 | -2.1 | 207 | Specific | 574.96 | 415.87 | 415.87 | 10.12 | 21 | >sp\|P04003\|C4BPA_HUMAN C4b-binding protein alpha chain OS=Homo sapiens OX=9606 GN=C4BPA PE=1 SV=2 | 218 | 20201115_15212_1ng_glyco_hcd203040_compound15.20752.20752.4 | controllerType=0 controllerNumber=1 scan=20752 | 39.6955 |
| 17937:4 | 29 | R.FSLLGHASISC[+57.021]TVEN[+2350.830]ETIGVWRPSPPTC[+57.021]EK.I | HexNAc(4)Hex(5)Fuc(1)NeuAc(2) | C[+57]*2, N[+2351] | 1431.6188 | 4 | 5723.4533 | 5723.4655 | 0 | -2.1 | 207 | Specific | 614.56 | 534.13 | 534.13 | 12.38 | 21 | >sp\|P04003\|C4BPA_HUMAN C4b-binding protein alpha chain OS=Homo sapiens OX=9606 GN=C4BPA PE=1 SV=2 | 218 | 20201115_15212_1ng_glyco_hcd203040_compound15.20754.20754.4 | controllerType=0 controllerNumber=1 scan=20754 | 39.7023 |
| 03631:3 | 34 | R.LRN[+1216.423]VSWATGR.S | HexNAc(2)Hex(5) | N[+1216] | 792.6924 | 3 | 2376.0625 | 2376.0559 | 0 | 2.8 | 149 | Specific | 338.93 | 128.39 | 128.39 | 5.72 | 16 | >sp\|P08571\|CD14_HUMAN Monocyte differentiation antigen CD14 OS=Homo sapiens OX=9606 GN=CD14 PE=1 SV=2 | 1232 | 20201115_15212_1ng_glyco_hcd203040_compound15.5834.5834.3 | controllerType=0 controllerNumber=1 scan=5834 | 14.2816 |
| 26295:4 | 42 | R.QLDMLDLSN[+2205.793]NSLASVPEGLWASLGQPNWDMR.D | HexNAc(4)Hex(5)Fuc(2)NeuAc(1) | N[+2206] | 1416.6287 | 4 | 5663.493 | 5663.4443 | 0 | 8.6 | 261 | Specific | 243.52 | 243.52 | 243.52 | 6.57 | 13 | >sp\|P02750\|A2GL_HUMAN Leucine-rich alpha-2-glycoprotein OS=Homo sapiens OX=9606 GN=LRG1 PE=1 SV=2 | 518 | 20201115_15212_1ng_glyco_hcd203040_compound15.29488.29488.4 | controllerType=0 controllerNumber=1 scan=29488 | 57.7936 |
| 26297:4 | 42 | R.QLDMLDLSN[+2205.793]NSLASVPEGLWASLGQPNWDMR.D | HexNAc(4)Hex(5)Fuc(2)NeuAc(1) | N[+2206] | 1416.6287 | 4 | 5663.493 | 5663.4443 | 0 | 8.6 | 261 | Specific | 453.01 | 453.01 | 453.01 | 10.02 | 13 | >sp\|P02750\|A2GL_HUMAN Leucine-rich alpha-2-glycoprotein OS=Homo sapiens OX=9606 GN=LRG1 PE=1 SV=2 | 518 | 20201115_15212_1ng_glyco_hcd203040_compound15.29490.29490.4 | controllerType=0 controllerNumber=1 scan=29490 | 57.7972 |
| 14928:4 | 43 | R.IIVPLNNREN[+2350.830]ISDPTSPLR.T | HexNAc(4)Hex(5)Fuc(1)NeuAc(2) | N[+2351] | 1125.509 | 4 | 4499.0142 | 4499.0075 | 0 | 1.5 | 62 | Specific | 234.19 | 222.81 | 222.81 | 5.04 | 15 | >sp\|P01591\|IGJ_HUMAN Immunoglobulin J chain OS=Homo sapiens OX=9606 GN=JCHAIN PE=1 SV=4 | 1392 | 20201115_15212_1ng_glyco_hcd203040_compound15.17593.17593.4 | controllerType=0 controllerNumber=1 scan=17593 | 33.9227 |
| 11435:3 | 43 | R.IIVPLNNREN[+1647.613]ISDPTSPLR.T | HexNAc(5)Hex(3)Fuc(1) | N[+1648] | 1265.9395 | 3 | 3795.804 | 3795.7904 | 0 | 3.6 | 62 | Specific | 258.67 | 188.59 | 188.59 | 6.43 | 15 | >sp\|P01591\|IGJ_HUMAN Immunoglobulin J chain OS=Homo sapiens OX=9606 GN=JCHAIN PE=1 SV=4 | 1392 | 20201115_15212_1ng_glyco_hcd203040_compound15.13946.13946.3 | controllerType=0 controllerNumber=1 scan=13946 | 27.7231 |
| 13130:3 | 43 | R.IIVPLNNREN[+1694.603]ISDPTSPLR.T | HexNAc(3)Hex(4)Fuc(1)NeuAc(1) | N[+1695] | 1281.6022 | 3 | 3842.7921 | 3842.7799 | 0 | 3.2 | 62 | Specific | 291.19 | 226.53 | 226.53 | 7.04 | 15 | >sp\|P01591\|IGJ_HUMAN Immunoglobulin J chain OS=Homo sapiens OX=9606 GN=JCHAIN PE=1 SV=4 | 1392 | 20201115_15212_1ng_glyco_hcd203040_compound15.15701.15701.3 | controllerType=0 controllerNumber=1 scan=15701 | 30.7521 |
| 14931:4 | 43 | R.IIVPLNNREN[+2350.830]ISDPTSPLR.T | HexNAc(4)Hex(5)Fuc(1)NeuAc(2) | N[+2351] | 1125.509 | 4 | 4499.0142 | 4499.0075 | 0 | 1.5 | 62 | Specific | 304.56 | 245.45 | 245.45 | 7.2 | 15 | >sp\|P01591\|IGJ_HUMAN Immunoglobulin J chain OS=Homo sapiens OX=9606 GN=JCHAIN PE=1 SV=4 | 1392 | 20201115_15212_1ng_glyco_hcd203040_compound15.17596.17596.4 | controllerType=0 controllerNumber=1 scan=17596 | 33.9292 |
| 13132:3 | 43 | R.IIVPLNNREN[+1694.603]ISDPTSPLR.T | HexNAc(3)Hex(4)Fuc(1)NeuAc(1) | N[+1695] | 1281.6022 | 3 | 3842.7921 | 3842.7799 | 0 | 3.2 | 62 | Specific | 371.07 | 295.06 | 295.06 | 6.4 | 15 | >sp\|P01591\|IGJ_HUMAN Immunoglobulin J chain OS=Homo sapiens OX=9606 GN=JCHAIN PE=1 SV=4 | 1392 | 20201115_15212_1ng_glyco_hcd203040_compound15.15703.15703.3 | controllerType=0 controllerNumber=1 scan=15703 | 30.757 |
| 14926:3 | 43 | R.IIVPLNNREN[+2350.830]ISDPTSPLR.T | HexNAc(4)Hex(5)Fuc(1)NeuAc(2) | N[+2351] | 1500.3398 | 3 | 4499.0048 | 4499.0075 | 0 | -0.6 | 62 | Specific | 386.31 | 308.61 | 308.61 | 7.1 | 15 | >sp\|P01591\|IGJ_HUMAN Immunoglobulin J chain OS=Homo sapiens OX=9606 GN=JCHAIN PE=1 SV=4 | 1392 | 20201115_15212_1ng_glyco_hcd203040_compound15.17591.17591.3 | controllerType=0 controllerNumber=1 scan=17591 | 33.9201 |
| 11356:3 | 43 | R.IIVPLNNREN[+1663.608]ISDPTSPLR.T | HexNAc(5)Hex(4) | N[+1664] | 1271.2695 | 3 | 3811.7938 | 3811.7853 | 0 | 2.2 | 62 | Specific | 439.72 | 420.39 | 420.39 | 9.97 | 15 | >sp\|P01591\|IGJ_HUMAN Immunoglobulin J chain OS=Homo sapiens OX=9606 GN=JCHAIN PE=1 SV=4 | 1392 | 20201115_15212_1ng_glyco_hcd203040_compound15.13864.13864.3 | controllerType=0 controllerNumber=1 scan=13864 | 27.5835 |
| 11358:3 | 43 | R.IIVPLNNREN[+1663.608]ISDPTSPLR.T | HexNAc(5)Hex(4) | N[+1664] | 1271.2695 | 3 | 3811.7938 | 3811.7853 | 0 | 2.2 | 62 | Specific | 463.23 | 445.79 | 445.79 | 10.02 | 15 | >sp\|P01591\|IGJ_HUMAN Immunoglobulin J chain OS=Homo sapiens OX=9606 GN=JCHAIN PE=1 SV=4 | 1392 | 20201115_15212_1ng_glyco_hcd203040_compound15.13866.13866.3 | controllerType=0 controllerNumber=1 scan=13866 | 27.5869 |
| 14929:3 | 43 | R.IIVPLNNREN[+2350.830]ISDPTSPLR.T | HexNAc(4)Hex(5)Fuc(1)NeuAc(2) | N[+2351] | 1500.3398 | 3 | 4499.0048 | 4499.0075 | 0 | -0.6 | 62 | Specific | 520.17 | 504.03 | 504.03 | 9.62 | 15 | >sp\|P01591\|IGJ_HUMAN Immunoglobulin J chain OS=Homo sapiens OX=9606 GN=JCHAIN PE=1 SV=4 | 1392 | 20201115_15212_1ng_glyco_hcd203040_compound15.17594.17594.3 | controllerType=0 controllerNumber=1 scan=17594 | 33.9254 |
| 16135:7 | 106 | K.GNPEQTPVLKPEEEAPAPEVGASKPEGIDSRPETLHPGRPQPPAEEELC[+57.021]SGKPFI | HexNAc(4)Hex(5)Fuc(2)NeuAc(1) | C[+57], N[+2206] | 1382.5044 | 7 | 9671.4874 | 9671.4638 | 0 | 2.4 | 108 | Specific | 268.57 | 268.57 | 268.57 | 3.83 | 8 | >sp\|P04004\|VTNC_HUMAN Vitronectin OS=Homo sapiens OX=9606 GN=VTN PE=1 SV=1 | 219 | 20201115_15212_1ng_glyco_hcd203040_compound15.18844.18844.5 | controllerType=0 controllerNumber=1 scan=18844 | 36.2231 |
| 16133:7 | 106 | K.GNPEQTPVLKPEEEAPAPEVGASKPEGIDSRPETLHPGRPQPPAEEELC[+57.021]SGKPFI | HexNAc(4)Hex(5)Fuc(2)NeuAc(1) | C[+57], N[+2206] | 1382.5044 | 7 | 9671.4874 | 9671.4638 | 0 | 2.4 | 108 | Specific | 301.84 | 301.84 | 301.84 | 4.17 | 8 | >sp\|P04004\|VTNC_HUMAN Vitronectin OS=Homo sapiens OX=9606 GN=VTN PE=1 SV=1 | 219 | 20201115_15212_1ng_glyco_hcd203040_compound15.18842.18842.5 | controllerType=0 controllerNumber=1 scan=18842 | 36.2188 |
| 17768:4 | 1 | R.RTAGWNVPIGTLRPFLN[+2570.925]WTGPPEPIEAAVAR.F | HexNAc(5)Hex(6)Fuc(2)NeuAc(1) | N[+2571] | 1490.1969 | 4 | 5957.7658 | 5957.7371 | 0 | 4.8 | 140 | Specific | 220.13 | 220.13 | 220.13 | 4.04 | 160 | >sp\|P02788\|TRFL_HUMAN Lactotransferrin OS=Homo sapiens OX=9606 GN=LTF PE=1 SV=6 | 1270 | 20201119_15232_1ng_glyco_hcd203040_compound15.21422.21422.4 | controllerType=0 controllerNumber=1 scan=21422 | 43.3881 |
| 18897:4 | 1 | R.RTAGWNVPIGTLRPFLN[+2350.830]WTGPPEPIEAAVAR.F | HexNAc(4)Hex(5)Fuc(1)NeuAc(2) | N[+2351] | 1435.1732 | 4 | 5737.6711 | 5737.6424 | 0 | 5 | 140 | Specific | 222.9 | 222.9 | 222.9 | 5.18 | 160 | >sp\|P02788\|TRFL_HUMAN Lactotransferrin OS=Homo sapiens OX=9606 GN=LTF PE=1 SV=6 | 1270 | 20201119_15232_1ng_glyco_hcd203040_compound15.22598.22598.4 | controllerType=0 controllerNumber=1 scan=22598 | 45.8863 |
| 17872:3 | 1 | R.RTAGWNVPIGTLRPFLN[+2059.735]WTGPPEPIEAAVAR.F | HexNAc(4)Hex(5)Fuc(1)NeuAc(1) | N[+2060] | 1816.1944 | 3 | 5446.5687 | 5446.547 | 0 | 4 | 140 | Specific | 289.14 | 289.14 | 289.14 | 5.44 | 160 | >sp\|P02788\|TRFL_HUMAN Lactotransferrin OS=Homo sapiens OX=9606 GN=LTF PE=1 SV=6 | 1270 | 20201119_15232_1ng_glyco_hcd203040_compound15.21530.21530.3 | controllerType=0 controllerNumber=1 scan=21530 | 43.6177 |
| 17850:5 | 1 | R.RTAGWNVPIGTLRPFLN[+2205.793]WTGPPEPIEAAVAR.F | HexNAc(4)Hex(5)Fuc(2)NeuAc(1) | N[+2206] | 1119.3323 | 5 | 5592.6323 | 5592.6049 | 0 | 4.9 | 140 | Specific | 425.85 | 425.85 | 425.85 | 9.63 | 160 | >sp\|P02788\|TRFL_HUMAN Lactotransferrin OS=Homo sapiens OX=9606 GN=LTF PE=1 SV=6 | 1270 | 20201119_15232_1ng_glyco_hcd203040_compound15.21507.21507.5 | controllerType=0 controllerNumber=1 scan=21507 | 43.5624 |
| 17771:4 | 1 | R.RTAGWNVPIGTLRPFLN[+2570.925]WTGPPEPIEAAVAR.F | HexNAc(5)Hex(6)Fuc(2)NeuAc(1) | N[+2571] | 1490.1969 | 4 | 5957.7658 | 5957.7371 | 0 | 4.8 | 140 | Specific | 427.58 | 392.73 | 392.73 | 9.68 | 160 | >sp\|P02788\|TRFL_HUMAN Lactotransferrin OS=Homo sapiens OX=9606 GN=LTF PE=1 SV=6 | 1270 | 20201119_15232_1ng_glyco_hcd203040_compound15.21425.21425.4 | controllerType=0 controllerNumber=1 scan=21425 | 43.3957 |

Table S4. Cont.

| Query #:z | Protein Rank | Glycopeptide | Glycan | Modification Type(s) | Observed m/z | z | Observed (M+H) | Calc. mass (M+H) | Off-by-x error | Mass error (ppm) | StartPosition | Cleavage | Score | Delta | Delta Mod | \|Log Prob\| | # of unique peptides | Protein Name | Protein DB number | Comment | Scan # | Scan Time |
|---|---|---|---|---|---|---|---|---|---|---|---|---|---|---|---|---|---|---|---|---|---|---|
| 17870:5 | 1 | R.RTAGWNVPIGTLRPFLN[+2059.735]WTGPPEPIEAAVAR.F | HexNAc(4)Hex(5)Fuc(1)NeuAc(1) | N[+2060] | 1090.1189 | 5 | 5446.5654 | 5446.547 | 0 | 3.4 | 140 | Specific | 495.88 | 495.88 | 495.88 | 9.94 | 160 | >sp\|P02788\|TRFL_HUMAN Lactotransferrin OS=Homo sapiens OX=9606 GN=LTF PE=1 SV=6 | 1270 | 20201119_15232_1ng_glyco_hcd203040_compound15.21528.21528.5 | controllerType=0 controllerNumber=1 scan=21528 | 43.6132 |
| 17819:5 | 1 | R.RTAGWNVPIGTLRPFLN[+2205.793]WTGPPEPIEAAVAR.F | HexNAc(4)Hex(5)Fuc(2)NeuAc(1) | N[+2206] | 1119.3325 | 5 | 5592.6335 | 5592.6049 | 0 | 5.1 | 140 | Specific | 511.75 | 473.73 | 473.73 | 9.94 | 160 | >sp\|P02788\|TRFL_HUMAN Lactotransferrin OS=Homo sapiens OX=9606 GN=LTF PE=1 SV=6 | 1270 | 20201119_15232_1ng_glyco_hcd203040_compound15.21475.21475.3 | controllerType=0 controllerNumber=1 scan=21475 | 43.4863 |
| 16931:4 | 1 | R.RTAGWNVPIGTLRPFLN[+1768.640]WTGPPEPIEAAVAR.F | HexNAc(4)Hex(5)Fuc(1) | N[+1769] | 1289.6301 | 4 | 5155.4987 | 5155.4516 | 0 | 9.1 | 140 | Specific | 526.45 | 526.45 | 526.45 | 8.71 | 160 | >sp\|P02788\|TRFL_HUMAN Lactotransferrin OS=Homo sapiens OX=9606 GN=LTF PE=1 SV=6 | 1270 | 20201119_15232_1ng_glyco_hcd203040_compound15.20546.20546.4 | controllerType=0 controllerNumber=1 scan=20546 | 41.4457 |
| 18899:4 | 1 | R.RTAGWNVPIGTLRPFLN[+2350.830]WTGPPEPIEAAVAR.F | HexNAc(4)Hex(5)Fuc(1)NeuAc(2) | N[+2351] | 1435.1732 | 4 | 5737.6711 | 5737.6424 | 0 | 5 | 140 | Specific | 537.39 | 498.7 | 498.7 | 10.07 | 160 | >sp\|P02788\|TRFL_HUMAN Lactotransferrin OS=Homo sapiens OX=9606 GN=LTF PE=1 SV=6 | 1270 | 20201119_15232_1ng_glyco_hcd203040_compound15.22600.22600.4 | controllerType=0 controllerNumber=1 scan=22600 | 45.8903 |
| 16909:4 | 1 | R.RTAGWNVPIGTLRPFLN[+1768.640]WTGPPEPIEAAVAR.F | HexNAc(4)Hex(5)Fuc(1) | N[+1769] | 1289.6244 | 4 | 5155.4757 | 5155.4516 | 0 | 4.7 | 140 | Specific | 537.98 | 459.7 | 459.7 | 10.07 | 160 | >sp\|P02788\|TRFL_HUMAN Lactotransferrin OS=Homo sapiens OX=9606 GN=LTF PE=1 SV=6 | 1270 | 20201119_15232_1ng_glyco_hcd203040_compound15.20523.20523.4 | controllerType=0 controllerNumber=1 scan=20523 | 41.3918 |
| 16929:4 | 1 | R.RTAGWNVPIGTLRPFLN[+1768.640]WTGPPEPIEAAVAR.F | HexNAc(4)Hex(5)Fuc(1) | N[+1769] | 1289.6301 | 4 | 5155.4987 | 5155.4516 | 0 | 9.1 | 140 | Specific | 541.36 | 518.54 | 518.54 | 8.71 | 160 | >sp\|P02788\|TRFL_HUMAN Lactotransferrin OS=Homo sapiens OX=9606 GN=LTF PE=1 SV=6 | 1270 | 20201119_15232_1ng_glyco_hcd203040_compound15.20544.20544.4 | controllerType=0 controllerNumber=1 scan=20544 | 41.441 |
| 17822:5 | 1 | R.RTAGWNVPIGTLRPFLN[+2205.793]WTGPPEPIEAAVAR.F | HexNAc(4)Hex(5)Fuc(2)NeuAc(1) | N[+2206] | 1119.3325 | 5 | 5592.6335 | 5592.6049 | 0 | 5.1 | 140 | Specific | 548.38 | 519.15 | 519.15 | 10.01 | 160 | >sp\|P02788\|TRFL_HUMAN Lactotransferrin OS=Homo sapiens OX=9606 GN=LTF PE=1 SV=6 | 1270 | 20201119_15232_1ng_glyco_hcd203040_compound15.21478.21478.3 | controllerType=0 controllerNumber=1 scan=21478 | 43.493 |
| 17853:5 | 1 | R.RTAGWNVPIGTLRPFLN[+2205.793]WTGPPEPIEAAVAR.F | HexNAc(4)Hex(5)Fuc(2)NeuAc(1) | N[+2206] | 1119.3323 | 5 | 5592.6323 | 5592.6049 | 0 | 4.9 | 140 | Specific | 554.54 | 509.48 | 509.48 | 10.22 | 160 | >sp\|P02788\|TRFL_HUMAN Lactotransferrin OS=Homo sapiens OX=9606 GN=LTF PE=1 SV=6 | 1270 | 20201119_15232_1ng_glyco_hcd203040_compound15.21510.21510.5 | controllerType=0 controllerNumber=1 scan=21510 | 43.5702 |
| 17873:5 | 1 | R.RTAGWNVPIGTLRPFLN[+2059.735]WTGPPEPIEAAVAR.F | HexNAc(4)Hex(5)Fuc(1)NeuAc(1) | N[+2060] | 1090.1189 | 5 | 5446.5654 | 5446.547 | 0 | 3.4 | 140 | Specific | 563.84 | 556.64 | 556.64 | 10.45 | 160 | >sp\|P02788\|TRFL_HUMAN Lactotransferrin OS=Homo sapiens OX=9606 GN=LTF PE=1 SV=6 | 1270 | 20201119_15232_1ng_glyco_hcd203040_compound15.21531.21531.5 | controllerType=0 controllerNumber=1 scan=21531 | 43.6196 |
| 17842:4 | 1 | R.RTAGWNVPIGTLRPFLN[+2059.735]WTGPPEPIEAAVAR.F | HexNAc(4)Hex(5)Fuc(1)NeuAc(1) | N[+2060] | 1362.3978 | 4 | 5446.5695 | 5446.547 | 0 | 4.1 | 140 | Specific | 575.48 | 575.48 | 575.48 | 11.51 | 160 | >sp\|P02788\|TRFL_HUMAN Lactotransferrin OS=Homo sapiens OX=9606 GN=LTF PE=1 SV=6 | 1270 | 20201119_15232_1ng_glyco_hcd203040_compound15.21499.21499.4 | controllerType=0 controllerNumber=1 scan=21499 | 43.5437 |
| 16907:4 | 1 | R.RTAGWNVPIGTLRPFLN[+1768.640]WTGPPEPIEAAVAR.F | HexNAc(4)Hex(5)Fuc(1) | N[+1769] | 1289.6244 | 4 | 5155.4757 | 5155.4516 | 0 | 4.7 | 140 | Specific | 584.74 | 408.44 | 408.44 | 11.51 | 160 | >sp\|P02788\|TRFL_HUMAN Lactotransferrin OS=Homo sapiens OX=9606 GN=LTF PE=1 SV=6 | 1270 | 20201119_15232_1ng_glyco_hcd203040_compound15.20521.20521.4 | controllerType=0 controllerNumber=1 scan=20521 | 41.3823 |
| 18004:4 | 1 | R.RTAGWNVPIGTLRPFLN[+2059.735]WTGPPEPIEAAVAR.F | HexNAc(4)Hex(5)Fuc(1)NeuAc(1) | N[+2060] | 1362.3964 | 4 | 5446.5636 | 5446.547 | 0 | 3.1 | 140 | Specific | 588.98 | 452.48 | 452.48 | 11.73 | 160 | >sp\|P02788\|TRFL_HUMAN Lactotransferrin OS=Homo sapiens OX=9606 GN=LTF PE=1 SV=6 | 1270 | 20201119_15232_1ng_glyco_hcd203040_compound15.21667.21667.2 | controllerType=0 controllerNumber=1 scan=21667 | 43.8811 |
| 17845:4 | 1 | R.RTAGWNVPIGTLRPFLN[+2059.735]WTGPPEPIEAAVAR.F | HexNAc(4)Hex(5)Fuc(1)NeuAc(1) | N[+2060] | 1362.3978 | 4 | 5446.5695 | 5446.547 | 0 | 4.1 | 140 | Specific | 596.52 | 480.36 | 480.36 | 11.51 | 160 | >sp\|P02788\|TRFL_HUMAN Lactotransferrin OS=Homo sapiens OX=9606 GN=LTF PE=1 SV=6 | 1270 | 20201119_15232_1ng_glyco_hcd203040_compound15.21502.21502.4 | controllerType=0 controllerNumber=1 scan=21502 | 43.5516 |
| 17838:4 | 1 | R.RTAGWNVPIGTLRPFLN[+2205.793]WTGPPEPIEAAVAR.F | HexNAc(4)Hex(5)Fuc(2)NeuAc(1) | N[+2206] | 1398.9125 | 4 | 5592.6281 | 5592.6049 | 0 | 4.1 | 140 | Specific | 598.8 | 598.8 | 598.8 | 11.51 | 160 | >sp\|P02788\|TRFL_HUMAN Lactotransferrin OS=Homo sapiens OX=9606 GN=LTF PE=1 SV=6 | 1270 | 20201119_15232_1ng_glyco_hcd203040_compound15.21495.21495.4 | controllerType=0 controllerNumber=1 scan=21495 | 43.5352 |
| 17840:4 | 1 | R.RTAGWNVPIGTLRPFLN[+2205.793]WTGPPEPIEAAVAR.F | HexNAc(4)Hex(5)Fuc(2)NeuAc(1) | N[+2206] | 1398.9125 | 4 | 5592.6281 | 5592.6049 | 0 | 4.1 | 140 | Specific | 628 | 514 | 514 | 12.51 | 160 | >sp\|P02788\|TRFL_HUMAN Lactotransferrin OS=Homo sapiens OX=9606 GN=LTF PE=1 SV=6 | 1270 | 20201119_15232_1ng_glyco_hcd203040_compound15.21497.21497.4 | controllerType=0 controllerNumber=1 scan=21497 | 43.5409 |
| 18006:4 | 1 | R.RTAGWNVPIGTLRPFLN[+2059.735]WTGPPEPIEAAVAR.F | HexNAc(4)Hex(5)Fuc(1)NeuAc(1) | N[+2060] | 1362.3964 | 4 | 5446.5636 | 5446.547 | 0 | 3.1 | 140 | Specific | 645.07 | 534 | 534 | 12.73 | 160 | >sp\|P02788\|TRFL_HUMAN Lactotransferrin OS=Homo sapiens OX=9606 GN=LTF PE=1 SV=6 | 1270 | 20201119_15232_1ng_glyco_hcd203040_compound15.21669.21669.2 | controllerType=0 controllerNumber=1 scan=21669 | 43.8881 |
| 19192:3 | 1 | R.[+42.011]TAGWNVPIGTLRPFLN[+2018.708]WTGPPEPIEAAVAR.F | HexNAc(3)Hex(6)Fuc(1)NeuAc(1) | .[42], N[+2019] | 1764.4872 | 3 | 5291.447 | 5291.4299 | 0 | 3.2 | 141 | Specific | 214.69 | 190.61 | 190.61 | 5.02 | 160 | >sp\|P02788\|TRFL_HUMAN Lactotransferrin OS=Homo sapiens OX=9606 GN=LTF PE=1 SV=6 | 1270 | 20201119_15232_1ng_glyco_hcd203040_compound15.22905.22905.3 | controllerType=0 controllerNumber=1 scan=22905 | 46.4833 |
| 18082:4 | 1 | R.TAGWNVPIGTLRPFLN[+2279.830]WTGPPEPIEAAVAR.F | HexNAc(5)Hex(6)Fuc(2) | N[+2280] | 1378.3947 | 4 | 5510.5568 | 5510.5406 | 0 | 2.9 | 141 | Specific | 233.99 | 188.99 | 188.99 | 5.02 | 160 | >sp\|P02788\|TRFL_HUMAN Lactotransferrin OS=Homo sapiens OX=9606 GN=LTF PE=1 SV=6 | 1270 | 20201119_15232_1ng_glyco_hcd203040_compound15.21748.21748.4 | controllerType=0 controllerNumber=1 scan=21748 | 44.0564 |
| 19102:4 | 1 | R.[+42.011]TAGWNVPIGTLRPFLN[+2190.793]WTGPPEPIEAAVAR.F | HexNAc(6)Hex(6) | .[42], N[+2191] | 1366.6299 | 4 | 5463.4977 | 5463.5147 | 0 | -3.1 | 141 | Specific | 283.8 | 283.8 | 283.8 | 5.75 | 160 | >sp\|P02788\|TRFL_HUMAN Lactotransferrin OS=Homo sapiens OX=9606 GN=LTF PE=1 SV=6 | 1270 | 20201119_15232_1ng_glyco_hcd203040_compound15.22812.22812.4 | controllerType=0 controllerNumber=1 scan=22812 | 46.3238 |
| 19106:3 | 1 | R.[+42.011]TAGWNVPIGTLRPFLN[+2190.793]WTGPPEPIEAAVAR.F | HexNAc(6)Hex(6) | .[42], N[+2191] | 1821.8431 | 3 | 5463.5148 | 5463.5147 | 0 | 0 | 141 | Specific | 307.87 | 85.64 | 85.64 | 4.45 | 160 | >sp\|P02788\|TRFL_HUMAN Lactotransferrin OS=Homo sapiens OX=9606 GN=LTF PE=1 SV=6 | 1270 | 20201119_15232_1ng_glyco_hcd203040_compound15.22816.22816.3 | controllerType=0 controllerNumber=1 scan=22816 | 46.334 |
| 18050:4 | 1 | R.TAGWNVPIGTLRPFLN[+2425.887]WTGPPEPIEAAVAR.F | HexNAc(5)Hex(6)Fuc(3) | N[+2426] | 1414.9105 | 4 | 5656.6203 | 5656.5985 | 0 | 3.9 | 141 | Specific | 308.18 | 226.82 | 226.82 | 6.87 | 160 | >sp\|P02788\|TRFL_HUMAN Lactotransferrin OS=Homo sapiens OX=9606 GN=LTF PE=1 SV=6 | 1270 | 20201119_15232_1ng_glyco_hcd203040_compound15.21715.21715.4 | controllerType=0 controllerNumber=1 scan=21715 | 43.9903 |
| 19195:3 | 1 | R.[+42.011]TAGWNVPIGTLRPFLN[+2018.708]WTGPPEPIEAAVAR.F | HexNAc(3)Hex(6)Fuc(1)NeuAc(1) | .[42], N[+2019] | 1764.4872 | 3 | 5291.447 | 5291.4299 | 0 | 3.2 | 141 | Specific | 342.79 | 306.5 | 306.5 | 7.4 | 160 | >sp\|P02788\|TRFL_HUMAN Lactotransferrin OS=Homo sapiens OX=9606 GN=LTF PE=1 SV=6 | 1270 | 20201119_15232_1ng_glyco_hcd203040_compound15.22908.22908.3 | controllerType=0 controllerNumber=1 scan=22908 | 46.487 |
| 20519:4 | 1 | R.[+42.011]TAGWNVPIGTLRPFLN[+2336.851]WTGPPEPIEAAVAR.F | HexNAc(6)Hex(6)Fuc(1) | .[42], N[+2337] | 1403.1408 | 4 | 5609.5412 | 5609.5726 | 0 | -5.6 | 141 | Specific | 359.91 | 359.91 | 359.91 | 6.31 | 160 | >sp\|P02788\|TRFL_HUMAN Lactotransferrin OS=Homo sapiens OX=9606 GN=LTF PE=1 SV=6 | 1270 | 20201119_15232_1ng_glyco_hcd203040_compound15.24292.24292.4 | controllerType=0 controllerNumber=1 scan=24292 | 49.3139 |
| 18080:4 | 1 | R.TAGWNVPIGTLRPFLN[+2279.830]WTGPPEPIEAAVAR.F | HexNAc(5)Hex(6)Fuc(2) | N[+2280] | 1378.3947 | 4 | 5510.5568 | 5510.5406 | 0 | 2.9 | 141 | Specific | 362.38 | 361.99 | 361.99 | 7.93 | 160 | >sp\|P02788\|TRFL_HUMAN Lactotransferrin OS=Homo sapiens OX=9606 GN=LTF PE=1 SV=6 | 1270 | 20201119_15232_1ng_glyco_hcd203040_compound15.21746.21746.4 | controllerType=0 controllerNumber=1 scan=21746 | 44.0502 |
| 18960:3 | 1 | R.TAGWNVPIGTLRPFLN[+2569.905]WTGPPEPIEAAVAR.F | HexNAc(5)Hex(6)NeuAc(2) | N[+2570] | 1934.2203 | 3 | 5800.6463 | 5800.6156 | 0 | 5.3 | 141 | Specific | 363.1 | 166.17 | 166.17 | 5.85 | 160 | >sp\|P02788\|TRFL_HUMAN Lactotransferrin OS=Homo sapiens OX=9606 GN=LTF PE=1 SV=6 | 1270 | 20201119_15232_1ng_glyco_hcd203040_compound15.22663.22663.3 | controllerType=0 controllerNumber=1 scan=22663 | 46.007 |
| 19109:3 | 1 | R.[+42.011]TAGWNVPIGTLRPFLN[+2190.793]WTGPPEPIEAAVAR.F | HexNAc(6)Hex(6) | .[42], N[+2191] | 1821.8431 | 3 | 5463.5148 | 5463.5147 | 0 | 0 | 141 | Specific | 375.35 | 81.44 | 81.44 | 4.77 | 160 | >sp\|P02788\|TRFL_HUMAN Lactotransferrin OS=Homo sapiens OX=9606 GN=LTF PE=1 SV=6 | 1270 | 20201119_15232_1ng_glyco_hcd203040_compound15.22819.22819.3 | controllerType=0 controllerNumber=1 scan=22819 | 46.3383 |
| 19104:4 | 1 | R.[+42.011]TAGWNVPIGTLRPFLN[+2190.793]WTGPPEPIEAAVAR.F | HexNAc(6)Hex(6) | .[42], N[+2191] | 1366.6299 | 4 | 5463.4977 | 5463.5147 | 0 | -3.1 | 141 | Specific | 419.57 | 408.87 | 408.87 | 8.16 | 160 | >sp\|P02788\|TRFL_HUMAN Lactotransferrin OS=Homo sapiens OX=9606 GN=LTF PE=1 SV=6 | 1270 | 20201119_15232_1ng_glyco_hcd203040_compound15.22814.22814.4 | controllerType=0 controllerNumber=1 scan=22814 | 46.3284 |
| 18962:3 | 1 | R.TAGWNVPIGTLRPFLN[+2569.905]WTGPPEPIEAAVAR.F | HexNAc(5)Hex(6)NeuAc(2) | N[+2570] | 1934.2203 | 3 | 5800.6463 | 5800.6156 | 0 | 5.3 | 141 | Specific | 472.91 | 288.18 | 288.18 | 7.11 | 160 | >sp\|P02788\|TRFL_HUMAN Lactotransferrin OS=Homo sapiens OX=9606 GN=LTF PE=1 SV=6 | 1270 | 20201119_15232_1ng_glyco_hcd203040_compound15.22665.22665.3 | controllerType=0 controllerNumber=1 scan=22665 | 46.0157 |
| 20481:4 | 1 | R.TAGWNVPIGTLRPFLN[+2350.830]WTGPPEPIEAAVAR.F | HexNAc(4)Hex(5)Fuc(1)NeuAc(2) | N[+2351] | 1396.1434 | 4 | 5581.5519 | 5581.5413 | 0 | 1.9 | 141 | Specific | 506.74 | 356.6 | 356.6 | 8.47 | 160 | >sp\|P02788\|TRFL_HUMAN Lactotransferrin OS=Homo sapiens OX=9606 GN=LTF PE=1 SV=6 | 1270 | 20201119_15232_1ng_glyco_hcd203040_compound15.24253.24253.4 | controllerType=0 controllerNumber=1 scan=24253 | 49.2427 |
| 19067:4 | 1 | R.TAGWNVPIGTLRPFLN[+2424.867]WTGPPEPIEAAVAR.F | HexNAc(5)Hex(6)Fuc(1)NeuAc(1) | N[+2425] | 1414.6523 | 4 | 5655.5876 | 5655.5781 | 0 | 1.7 | 141 | Specific | 526.7 | 401.08 | 401.08 | 10.01 | 160 | >sp\|P02788\|TRFL_HUMAN Lactotransferrin OS=Homo sapiens OX=9606 GN=LTF PE=1 SV=6 | 1270 | 20201119_15232_1ng_glyco_hcd203040_compound15.22775.22775.4 | controllerType=0 controllerNumber=1 scan=22775 | 46.2325 |
| 19020:4 | 1 | R.TAGWNVPIGTLRPFLN[+2425.887]WTGPPEPIEAAVAR.F | HexNAc(5)Hex(6)Fuc(3) | N[+2426] | 1414.906 | 4 | 5656.6022 | 5656.5985 | 0 | 0.7 | 141 | Specific | 536.08 | 283.86 | 283.86 | 7.9 | 160 | >sp\|P02788\|TRFL_HUMAN Lactotransferrin OS=Homo sapiens OX=9606 GN=LTF PE=1 SV=6 | 1270 | 20201119_15232_1ng_glyco_hcd203040_compound15.22725.22725.4 | controllerType=0 controllerNumber=1 scan=22725 | 46.123 |
| 20454:3 | 1 | R.TAGWNVPIGTLRPFLN[+2350.830]WTGPPEPIEAAVAR.F | HexNAc(4)Hex(5)Fuc(1)NeuAc(2) | N[+2351] | 1861.1906 | 3 | 5581.5574 | 5581.5413 | 0 | 2.9 | 141 | Specific | 541.13 | 455.99 | 455.99 | 10.13 | 160 | >sp\|P02788\|TRFL_HUMAN Lactotransferrin OS=Homo sapiens OX=9606 GN=LTF PE=1 SV=6 | 1270 | 20201119_15232_1ng_glyco_hcd203040_compound15.24224.24224.3 | controllerType=0 controllerNumber=1 scan=24224 | 49.1814 |
| 19033:3 | 1 | R.TAGWNVPIGTLRPFLN[+2205.793]WTGPPEPIEAAVAR.F | HexNAc(4)Hex(5)Fuc(2)NeuAc(1) | N[+2206] | 1812.8471 | 3 | 5436.5268 | 5436.5038 | 0 | 4.2 | 141 | Specific | 550.12 | 534.09 | 534.09 | 10.11 | 160 | >sp\|P02788\|TRFL_HUMAN Lactotransferrin OS=Homo sapiens OX=9606 GN=LTF PE=1 SV=6 | 1270 | 20201119_15232_1ng_glyco_hcd203040_compound15.22740.22740.3 | controllerType=0 controllerNumber=1 scan=22740 | 46.1567 |
| 19107:3 | 1 | R.TAGWNVPIGTLRPFLN[+2059.735]WTGPPEPIEAAVAR.F | HexNAc(4)Hex(5)Fuc(1)NeuAc(1) | N[+2060] | 1764.1613 | 3 | 5290.4695 | 5290.4459 | 0 | 4.5 | 141 | Specific | 552.52 | 464.41 | 464.41 | 10.11 | 160 | >sp\|P02788\|TRFL_HUMAN Lactotransferrin OS=Homo sapiens OX=9606 GN=LTF PE=1 SV=6 | 1270 | 20201119_15232_1ng_glyco_hcd203040_compound15.22817.22817.2 | controllerType=0 controllerNumber=1 scan=22817 | 46.3352 |
| 19051:3 | 1 | R.TAGWNVPIGTLRPFLN[+2351.851]WTGPPEPIEAAVAR.F | HexNAc(4)Hex(5)Fuc(3)NeuAc(1) | N[+2352] | 1861.5354 | 3 | 5582.5915 | 5582.5617 | 0 | 5.3 | 141 | Specific | 577.17 | 497.33 | 497.33 | 10.8 | 160 | >sp\|P02788\|TRFL_HUMAN Lactotransferrin OS=Homo sapiens OX=9606 GN=LTF PE=1 SV=6 | 1270 | 20201119_15232_1ng_glyco_hcd203040_compound15.22758.22758.3 | controllerType=0 controllerNumber=1 scan=22758 | 46.1926 |
| 19048:3 | 1 | R.TAGWNVPIGTLRPFLN[+2351.851]WTGPPEPIEAAVAR.F | HexNAc(4)Hex(5)Fuc(3)NeuAc(1) | N[+2352] | 1861.5354 | 3 | 5582.5915 | 5582.5617 | 0 | 5.3 | 141 | Specific | 586.76 | 497.58 | 497.58 | 10.8 | 160 | >sp\|P02788\|TRFL_HUMAN Lactotransferrin OS=Homo sapiens OX=9606 GN=LTF PE=1 SV=6 | 1270 | 20201119_15232_1ng_glyco_hcd203040_compound15.22755.22755.3 | controllerType=0 controllerNumber=1 scan=22755 | 46.1844 |
| 19032:4 | 1 | R.TAGWNVPIGTLRPFLN[+2205.793]WTGPPEPIEAAVAR.F | HexNAc(4)Hex(5)Fuc(2)NeuAc(1) | N[+2206] | 1359.8867 | 4 | 5436.5251 | 5436.5038 | 0 | 3.9 | 141 | Specific | 589.12 | 475.82 | 475.82 | 11.52 | 160 | >sp\|P02788\|TRFL_HUMAN Lactotransferrin OS=Homo sapiens OX=9606 GN=LTF PE=1 SV=6 | 1270 | 20201119_15232_1ng_glyco_hcd203040_compound15.22739.22739.4 | controllerType=0 controllerNumber=1 scan=22739 | 46.1559 |
| 19023:4 | 1 | R.TAGWNVPIGTLRPFLN[+2425.887]WTGPPEPIEAAVAR.F | HexNAc(5)Hex(6)Fuc(3) | N[+2426] | 1414.906 | 4 | 5656.6022 | 5656.5985 | 0 | 0.7 | 141 | Specific | 603.62 | 368.08 | 368.08 | 10.84 | 160 | >sp\|P02788\|TRFL_HUMAN Lactotransferrin OS=Homo sapiens OX=9606 GN=LTF PE=1 SV=6 | 1270 | 20201119_15232_1ng_glyco_hcd203040_compound15.22728.22728.4 | controllerType=0 controllerNumber=1 scan=22728 | 46.1277 |
| 18953:4 | 1 | R.TAGWNVPIGTLRPFLN[+2570.925]WTGPPEPIEAAVAR.F | HexNAc(5)Hex(6)Fuc(2)NeuAc(1) | N[+2571] | 1451.1682 | 4 | 5801.651 | 5801.636 | 0 | 2.6 | 141 | Specific | 613.42 | 523.45 | 523.45 | 12.46 | 160 | >sp\|P02788\|TRFL_HUMAN Lactotransferrin OS=Homo sapiens OX=9606 GN=LTF PE=1 SV=6 | 1270 | 20201119_15232_1ng_glyco_hcd203040_compound15.22656.22656.4 | controllerType=0 controllerNumber=1 scan=22656 | 45.9929 |
| 19069:4 | 1 | R.TAGWNVPIGTLRPFLN[+2424.867]WTGPPEPIEAAVAR.F | HexNAc(5)Hex(6)Fuc(1)NeuAc(1) | N[+2425] | 1414.6523 | 4 | 5655.5876 | 5655.5781 | 0 | 1.7 | 141 | Specific | 644 | 428.83 | 428.83 | 12.46 | 160 | >sp\|P02788\|TRFL_HUMAN Lactotransferrin OS=Homo sapiens OX=9606 GN=LTF PE=1 SV=6 | 1270 | 20201119_15232_1ng_glyco_hcd203040_compound15.22777.22777.4 | controllerType=0 controllerNumber=1 scan=22777 | 46.2379 |
| 18088:4 | 1 | R.TAGWNVPIGTLRPFLN[+1768.640]WTGPPEPIEAAVAR.F | HexNAc(4)Hex(5)Fuc(1) | N[+1769] | 1250.6049 | 4 | 4999.3976 | 4999.3504 | 0 | 9.4 | 141 | Specific | 652.77 | 652.77 | 652.77 | 11.93 | 160 | >sp\|P02788\|TRFL_HUMAN Lactotransferrin OS=Homo sapiens OX=9606 GN=LTF PE=1 SV=6 | 1270 | 20201119_15232_1ng_glyco_hcd203040_compound15.21755.21755.4 | controllerType=0 controllerNumber=1 scan=21755 | 44.0707 |
| 20451:3 | 1 | R.TAGWNVPIGTLRPFLN[+2350.830]WTGPPEPIEAAVAR.F | HexNAc(4)Hex(5)Fuc(1)NeuAc(2) | N[+2351] | 1861.1906 | 3 | 5581.5574 | 5581.5413 | 0 | 2.9 | 141 | Specific | 658.71 | 495.13 | 495.13 | 13.57 | 160 | >sp\|P02788\|TRFL_HUMAN Lactotransferrin OS=Homo sapiens OX=9606 GN=LTF PE=1 SV=6 | 1270 | 20201119_15232_1ng_glyco_hcd203040_compound15.24221.24221.3 | controllerType=0 controllerNumber=1 scan=24221 | 49.1758 |
| 19036:3 | 1 | R.TAGWNVPIGTLRPFLN[+2205.793]WTGPPEPIEAAVAR.F | HexNAc(4)Hex(5)Fuc(2)NeuAc(1) | N[+2206] | 1812.8471 | 3 | 5436.5268 | 5436.5038 | 0 | 4.2 | 141 | Specific | 674.37 | 580.21 | 580.21 | 13.35 | 160 | >sp\|P02788\|TRFL_HUMAN Lactotransferrin OS=Homo sapiens OX=9606 GN=LTF PE=1 SV=6 | 1270 | 20201119_15232_1ng_glyco_hcd203040_compound15.22743.22743.3 | controllerType=0 controllerNumber=1 scan=22743 | 46.1641 |
| 19034:4 | 1 | R.TAGWNVPIGTLRPFLN[+2351.851]WTGPPEPIEAAVAR.F | HexNAc(4)Hex(5)Fuc(3)NeuAc(1) | N[+2352] | 1396.4019 | 4 | 5582.5856 | 5582.5617 | 0 | 4.3 | 141 | Specific | 683.29 | 634.14 | 634.14 | 13.29 | 160 | >sp\|P02788\|TRFL_HUMAN Lactotransferrin OS=Homo sapiens OX=9606 GN=LTF PE=1 SV=6 | 1270 | 20201119_15232_1ng_glyco_hcd203040_compound15.22741.22741.4 | controllerType=0 controllerNumber=1 scan=22741 | 46.1577 |
| 18955:4 | 1 | R.TAGWNVPIGTLRPFLN[+2570.925]WTGPPEPIEAAVAR.F | HexNAc(5)Hex(6)Fuc(2)NeuAc(1) | N[+2571] | 1451.1682 | 4 | 5801.651 | 5801.636 | 0 | 2.6 | 141 | Specific | 691.96 | 568.02 | 568.02 | 13.46 | 160 | >sp\|P02788\|TRFL_HUMAN Lactotransferrin OS=Homo sapiens OX=9606 GN=LTF PE=1 SV=6 | 1270 | 20201119_15232_1ng_glyco_hcd203040_compound15.22658.22658.4 | controllerType=0 controllerNumber=1 scan=22658 | 46.0013 |
| 19110:3 | 1 | R.TAGWNVPIGTLRPFLN[+2059.735]WTGPPEPIEAAVAR.F | HexNAc(4)Hex(5)Fuc(1)NeuAc(1) | N[+2060] | 1764.1613 | 3 | 5290.4695 | 5290.4459 | 0 | 4.5 | 141 | Specific | 728.96 | 646.85 | 646.85 | 14.35 | 160 | >sp\|P02788\|TRFL_HUMAN Lactotransferrin OS=Homo sapiens OX=9606 GN=LTF PE=1 SV=6 | 1270 | 20201119_15232_1ng_glyco_hcd203040_compound15.22820.22820.2 | controllerType=0 controllerNumber=1 scan=22820 | 46.3401 |
| 20482:5 | 1 | R.TAGWNVPIGTLRPFLN[+2350.830]WTGPPEPIEAAVAR.F | HexNAc(4)Hex(5)Fuc(1)NeuAc(2) | N[+2351] | 1117.1179 | 5 | 5581.5605 | 5581.5413 | 0 | 3.4 | 141 | Specific | 758.5 | 758.5 | 758.5 | 15.69 | 160 | >sp\|P02788\|TRFL_HUMAN Lactotransferrin OS=Homo sapiens OX=9606 GN=LTF PE=1 SV=6 | 1270 | 20201119_15232_1ng_glyco_hcd203040_compound15.24254.24254.5 | controllerType=0 controllerNumber=1 scan=24254 | 49.2441 |
| 18090:4 | 1 | R.TAGWNVPIGTLRPFLN[+1768.640]WTGPPEPIEAAVAR.F | HexNAc(4)Hex(5)Fuc(1) | N[+1769] | 1250.6049 | 4 | 4999.3976 | 4999.3504 | 0 | 9.4 | 141 | Specific | 789.59 | 739.96 | 739.96 | 13.93 | 160 | >sp\|P02788\|TRFL_HUMAN Lactotransferrin OS=Homo sapiens OX=9606 GN=LTF PE=1 SV=6 | 1270 | 20201119_15232_1ng_glyco_hcd203040_compound15.21757.21757.4 | controllerType=0 controllerNumber=1 scan=21757 | 44.0766 |
| 19037:4 | 1 | R.TAGWNVPIGTLRPFLN[+2351.851]WTGPPEPIEAAVAR.F | HexNAc(4)Hex(5)Fuc(3)NeuAc(1) | N[+2352] | 1396.4019 | 4 | 5582.5856 | 5582.5617 | 0 | 4.3 | 141 | Specific | 810.26 | 689.61 | 689.61 | 16.29 | 160 | >sp\|P02788\|TRFL_HUMAN Lactotransferrin OS=Homo sapiens OX=9606 GN=LTF PE=1 SV=6 | 1270 | 20201119_15232_1ng_glyco_hcd203040_compound15.22744.22744.4 | controllerType=0 controllerNumber=1 scan=22744 | 46.1653 |
| 19035:4 | 1 | R.TAGWNVPIGTLRPFLN[+2205.793]WTGPPEPIEAAVAR.F | HexNAc(4)Hex(5)Fuc(2)NeuAc(1) | N[+2206] | 1359.8867 | 4 | 5436.5251 | 5436.5038 | 0 | 3.9 | 141 | Specific | 823.66 | 691.11 | 691.11 | 16.52 | 160 | >sp\|P02788\|TRFL_HUMAN Lactotransferrin OS=Homo sapiens OX=9606 GN=LTF PE=1 SV=6 | 1270 | 20201119_15232_1ng_glyco_hcd203040_compound15.22742.22742.4 | controllerType=0 controllerNumber=1 scan=22742 | 46.1607 |
| 20484:5 | 1 | R.TAGWNVPIGTLRPFLN[+2350.830]WTGPPEPIEAAVAR.F | HexNAc(4)Hex(5)Fuc(1)NeuAc(2) | N[+2351] | 1117.1179 | 5 | 5581.5605 | 5581.5413 | 0 | 3.4 | 141 | Specific | 848.25 | 581.28 | 581.28 | 16.69 | 160 | >sp\|P02788\|TRFL_HUMAN Lactotransferrin OS=Homo sapiens OX=9606 GN=LTF PE=1 SV=6 | 1270 | 20201119_15232_1ng_glyco_hcd203040_compound15.24256.24256.5 | controllerType=0 controllerNumber=1 scan=24256 | 49.2492 |
| 20483:4 | 1 | R.TAGWNVPIGTLRPFLN[+2350.830]WTGPPEPIEAAVAR.F | HexNAc(4)Hex(5)Fuc(1)NeuAc(2) | N[+2351] | 1396.1434 | 4 | 5581.5519 | 5581.5413 | 0 | 1.9 | 141 | Specific | 859.59 | 681.44 | 681.44 | 17.46 | 160 | >sp\|P02788\|TRFL_HUMAN Lactotransferrin OS=Homo sapiens OX=9606 GN=LTF PE=1 SV=6 | 1270 | 20201119_15232_1ng_glyco_hcd203040_compound15.24255.24255.4 | controllerType=0 controllerNumber=1 scan=24255 | 49.2469 |
| 16991:3 | 1 | R.TAGWNIPMGLLFN[+1257.449]QTGSC[+57.021]K.F | HexNAc(3)Hex(4) | C[+57], N[+1257] | 1118.1587 | 3 | 3352.4615 | 3352.4594 | 0 | 0.7 | 485 | Specific | 358.7 | 316.77 | 201.08 | 5.8 | 160 | >sp\|P02788\|TRFL_HUMAN Lactotransferrin OS=Homo sapiens OX=9606 GN=LTF PE=1 SV=6 | 1270 | 20201119_15232_1ng_glyco_hcd203040_compound15.20608.20608.3 | controllerType=0 controllerNumber=1 scan=20608 | 41.571 |
| 16589:2 | 1 | R.TAGWNIPMGLLFN[+1768.640]QTGSC[+57.021]K.F | HexNAc(4)Hex(5)Fuc(1) | C[+57], N[+1769] | 1932.344 | 2 | 3863.6806 | 3863.6495 | 0 | 8.1 | 485 | Specific | 406.88 | 406.88 | 406.88 | 7.93 | 160 | >sp\|P02788\|TRFL_HUMAN Lactotransferrin OS=Homo sapiens OX=9606 GN=LTF PE=1 SV=6 | 1270 | 20201119_15232_1ng_glyco_hcd203040_compound15.20189.20189.2 | controllerType=0 controllerNumber=1 scan=20189 | 40.6846 |
| 16844:3 | 1 | R.TAGWNIPMGLLFN[+1606.587]QTGSC[+57.021]K.F | HexNAc(4)Hex(4)Fuc(1) | C[+57], N[+1607] | 1234.5391 | 3 | 3701.6028 | 3701.5966 | 0 | 1.7 | 485 | Specific | 424.29 | 424.29 | 424.29 | 9.07 | 160 | >sp\|P02788\|TRFL_HUMAN Lactotransferrin OS=Homo sapiens OX=9606 GN=LTF PE=1 SV=6 | 1270 | 20201119_15232_1ng_glyco_hcd203040_compound15.20455.20455.3 | controllerType=0 controllerNumber=1 scan=20455 | 41.2502 |
| 18474:3 | 1 | R.TAGWNIPMGLLFN[+2205.793]QTGSC[+57.021]K.F | HexNAc(4)Hex(5)Fuc(2)NeuAc(1) | C[+57], N[+2206] | 1434.2779 | 3 | 4300.8191 | 4300.8028 | 0 | 3.8 | 485 | Specific | 541.96 | 441.88 | 441.88 | 9.72 | 160 | >sp\|P02788\|TRFL_HUMAN Lactotransferrin OS=Homo sapiens OX=9606 GN=LTF PE=1 SV=6 | 1270 | 20201119_15232_1ng_glyco_hcd203040_compound15.22156.22156.3 | controllerType=0 controllerNumber=1 scan=22156 | 44.9098 |
| 16993:3 | 1 | R.TAGWNIPMGLLFN[+1257.449]QTGSC[+57.021]K.F | HexNAc(3)Hex(4) | C[+57], N[+1257] | 1118.1587 | 3 | 3352.4615 | 3352.4594 | 0 | 0.7 | 485 | Specific | 627.72 | 627.72 | 383.81 | 11.96 | 160 | >sp\|P02788\|TRFL_HUMAN Lactotransferrin OS=Homo sapiens OX=9606 GN=LTF PE=1 SV=6 | 1270 | 20201119_15232_1ng_glyco_hcd203040_compound15.20610.20610.3 | controllerType=0 controllerNumber=1 scan=20610 | 41.5745 |
| 16712:3 | 1 | R.TAGWNIPMGLLFN[+1622.582]QTGSC[+57.021]K.F | HexNAc(4)Hex(5) | C[+57], N[+1623] | 1239.8736 | 3 | 3717.6062 | 3717.5916 | 0 | 3.9 | 485 | Specific | 631.86 | 631.86 | 240.45 | 12.16 | 160 | >sp\|P02788\|TRFL_HUMAN Lactotransferrin OS=Homo sapiens OX=9606 GN=LTF PE=1 SV=6 | 1270 | 20201119_15232_1ng_glyco_hcd203040_compound15.20318.20318.3 | controllerType=0 controllerNumber=1 scan=20318 | 40.9694 |
| 20574:3 | 1 | R.TAGWNIPMGLLFN[+2350.830]QTGSC[+57.021]K.F | HexNAc(4)Hex(5)Fuc(1)NeuAc(2) | C[+57], N[+2351] | 1482.6232 | 3 | 4445.8551 | 4445.8403 | 0 | 3.3 | 485 | Specific | 698.88 | 571.47 | 571.47 | 13.16 | 160 | >sp\|P02788\|TRFL_HUMAN Lactotransferrin OS=Homo sapiens OX=9606 GN=LTF PE=1 SV=6 | 1270 | 20201119_15232_1ng_glyco_hcd203040_compound15.24349.24349.3 | controllerType=0 controllerNumber=1 scan=24349 | 49.4255 |
| 18621:4 | 1 | R.TAGWNIPMGLLFN[+1913.677]QTGSC[+57.021]K.F | HexNAc(4)Hex(5)NeuAc(1) | C[+57], N[+1914] | 1002.9309 | 4 | 4008.7017 | 4008.687 | 0 | 3.7 | 485 | Specific | 728.61 | 662.87 | 662.87 | 14.34 | 160 | >sp\|P02788\|TRFL_HUMAN Lactotransferrin OS=Homo sapiens OX=9606 GN=LTF PE=1 SV=6 | 1270 | 20201119_15232_1ng_glyco_hcd203040_compound15.22310.22310.4 | controllerType=0 controllerNumber=1 scan=22310 | 45.2812 |
| 16945:2 | 1 | R.TAGWNIPMGLLFN[+1216.423]QTGSC[+57.021]K.F | HexNAc(2)Hex(5) | C[+57], N[+1216] | 1656.2359 | 2 | 3311.4645 | 3311.4328 | 0 | 9.6 | 485 | Specific | 790.92 | 767.48 | 270.1 | 13.63 | 160 | >sp\|P02788\|TRFL_HUMAN Lactotransferrin OS=Homo sapiens OX=9606 GN=LTF PE=1 SV=6 | 1270 | 20201119_15232_1ng_glyco_hcd203040_compound15.20560.20560.2 | controllerType=0 controllerNumber=1 scan=20560 | 41.4756 |
| 16857:3 | 1 | R.TAGWNIPMGLLFN[+1460.529]QTGSC[+57.021]K.F | HexNAc(4)Hex(4) | C[+57], N[+1461] | 1185.8533 | 3 | 3555.5454 | 3555.5387 | 0 | 1.9 | 485 | Specific | 791.66 | 670.53 | 396.5 | 15.1 | 160 | >sp\|P02788\|TRFL_HUMAN Lactotransferrin OS=Homo sapiens OX=9606 GN=LTF PE=1 SV=6 | 1270 | 20201119_15232_1ng_glyco_hcd203040_compound15.20469.20469.3 | controllerType=0 controllerNumber=1 scan=20469 | 41.2754 |
| 16714:3 | 1 | R.TAGWNIPMGLLFN[+1622.582]QTGSC[+57.021]K.F | HexNAc(4)Hex(5) | C[+57], N[+1623] | 1239.8736 | 3 | 3717.6062 | 3717.5916 | 0 | 3.9 | 485 | Specific | 841.95 | 832.46 | 386.08 | 16.16 | 160 | >sp\|P02788\|TRFL_HUMAN Lactotransferrin OS=Homo sapiens OX=9606 GN=LTF PE=1 SV=6 | 1270 | 20201119_15232_1ng_glyco_hcd203040_compound15.20320.20320.3 | controllerType=0 controllerNumber=1 scan=20320 | 40.9735 |
| 18477:3 | 1 | R.TAGWNIPMGLLFN[+2205.793]QTGSC[+57.021]K.F | HexNAc(4)Hex(5)Fuc(2)NeuAc(1) | C[+57], N[+2206] | 1434.2779 | 3 | 4300.8191 | 4300.8028 | 0 | 3.8 | 485 | Specific | 854.29 | 786.17 | 786.17 | 17.16 | 160 | >sp\|P02788\|TRFL_HUMAN Lactotransferrin OS=Homo sapiens OX=9606 GN=LTF PE=1 SV=6 | 1270 | 20201119_15232_1ng_glyco_hcd203040_compound15.22159.22159.3 | controllerType=0 controllerNumber=1 scan=22159 | 44.9141 |
| 16943:2 | 1 | R.TAGWNIPMGLLFN[+1216.423]QTGSC[+57.021]K.F | HexNAc(2)Hex(5) | C[+57], N[+1216] | 1656.2359 | 2 | 3311.4645 | 3311.4328 | 0 | 9.6 | 485 | Specific | 855.67 | 855.67 | 321.87 | 15.63 | 160 | >sp\|P02788\|TRFL_HUMAN Lactotransferrin OS=Homo sapiens OX=9606 GN=LTF PE=1 SV=6 | 1270 | 20201119_15232_1ng_glyco_hcd203040_compound15.20558.20558.2 | controllerType=0 controllerNumber=1 scan=20558 | 41.4712 |
| 16859:3 | 1 | R.TAGWNIPMGLLFN[+1460.529]QTGSC[+57.021]K.F | HexNAc(4)Hex(4) | C[+57], N[+1461] | 1185.8533 | 3 | 3555.5454 | 3555.5387 | 0 | 1.9 | 485 | Specific | 857.57 | 855.42 | 399.64 | 17.1 | 160 | >sp\|P02788\|TRFL_HUMAN Lactotransferrin OS=Homo sapiens OX=9606 GN=LTF PE=1 SV=6 | 1270 | 20201119_15232_1ng_glyco_hcd203040_compound15.20471.20471.3 | controllerType=0 controllerNumber=1 scan=20471 | 41.2808 |
| 18618:4 | 1 | R.TAGWNIPMGLLFN[+1913.677]QTGSC[+57.021]K.F | HexNAc(4)Hex(5)NeuAc(1) | C[+57], N[+1914] | 1002.9309 | 4 | 4008.7017 | 4008.687 | 0 | 3.7 | 485 | Specific | 860.5 | 819.31 | 819.31 | 17.34 | 160 | >sp\|P02788\|TRFL_HUMAN Lactotransferrin OS=Homo sapiens OX=9606 GN=LTF PE=1 SV=6 | 1270 | 20201119_15232_1ng_glyco_hcd203040_compound15.22307.22307.4 | controllerType=0 controllerNumber=1 scan=22307 | 45.2667 |
| 16550:3 | 1 | R.TAGWNIPMGLLFN[+1768.640]QTGSC[+57.021]K.F | HexNAc(4)Hex(5)Fuc(1) | C[+57], N[+1769] | 1288.5599 | 3 | 3863.6651 | 3863.6495 | 0 | 4 | 485 | Specific | 876.43 | 876.43 | 876.43 | 16.94 | 160 | >sp\|P02788\|TRFL_HUMAN Lactotransferrin OS=Homo sapiens OX=9606 GN=LTF PE=1 SV=6 | 1270 | 20201119_15232_1ng_glyco_hcd203040_compound15.20149.20149.3 | controllerType=0 controllerNumber=1 scan=20149 | 40.6048 |
| 18461:3 | 1 | R.TAGWNIPMGLLFN[+2059.735]QTGSC[+57.021]K.F | HexNAc(4)Hex(5)Fuc(1)NeuAc(1) | C[+57], N[+2060] | 1385.5932 | 3 | 4154.765 | 4154.7449 | 0 | 4.9 | 485 | Specific | 878.33 | 862.49 | 862.49 | 16.94 | 160 | >sp\|P02788\|TRFL_HUMAN Lactotransferrin OS=Homo sapiens OX=9606 GN=LTF PE=1 SV=6 | 1270 | 20201119_15232_1ng_glyco_hcd203040_compound15.22143.22143.3 | controllerType=0 controllerNumber=1 scan=22143 | 44.8723 |
| 20577:3 | 1 | R.TAGWNIPMGLLFN[+2350.830]QTGSC[+57.021]K.F | HexNAc(4)Hex(5)Fuc(1)NeuAc(2) | C[+57], N[+2351] | 1482.6232 | 3 | 4445.8551 | 4445.8403 | 0 | 3.3 | 485 | Specific | 881.43 | 870.48 | 870.48 | 17.16 | 160 | >sp\|P02788\|TRFL_HUMAN Lactotransferrin OS=Homo sapiens OX=9606 GN=LTF PE=1 SV=6 | 1270 | 20201119_15232_1ng_glyco_hcd203040_compound15.24352.24352.3 | controllerType=0 controllerNumber=1 scan=24352 | 49.4309 |
| 16641:3 | 1 | R.TAGWNIPMGLLFN[+1606.587]QTGSC[+57.021]K.F | HexNAc(4)Hex(4)Fuc(1) | C[+57], N[+1607] | 1234.5395 | 3 | 3701.6039 | 3701.5966 | 0 | 2 | 485 | Specific | 885.34 | 831.68 | 831.68 | 17.1 | 160 | >sp\|P02788\|TRFL_HUMAN Lactotransferrin OS=Homo sapiens OX=9606 GN=LTF PE=1 SV=6 | 1270 | 20201119_15232_1ng_glyco_hcd203040_compound15.20244.20244.3 | controllerType=0 controllerNumber=1 scan=20244 | 40.8177 |
| 18463:3 | 1 | R.TAGWNIPMGLLFN[+2059.735]QTGSC[+57.021]K.F | HexNAc(4)Hex(5)Fuc(1)NeuAc(1) | C[+57], N[+2060] | 1385.5932 | 3 | 4154.765 | 4154.7449 | 0 | 4.9 | 485 | Specific | 899 | 866.73 | 866.73 | 16.94 | 160 | >sp\|P02788\|TRFL_HUMAN Lactotransferrin OS=Homo sapiens OX=9606 GN=LTF PE=1 SV=6 | 1270 | 20201119_15232_1ng_glyco_hcd203040_compound15.22145.22145.3 | controllerType=0 controllerNumber=1 scan=22145 | 44.8804 |
| 18594:3 | 1 | R.TAGWNIPMGLLFN[+1913.677]QTGSC[+57.021]K.F | HexNAc(4)Hex(5)NeuAc(1) | C[+57], N[+1914] | 1336.9039 | 3 | 4008.697 | 4008.687 | 0 | 2.5 | 485 | Specific | 903.43 | 734.98 | 734.98 | 18.1 | 160 | >sp\|P02788\|TRFL_HUMAN Lactotransferrin OS=Homo sapiens OX=9606 GN=LTF PE=1 SV=6 | 1270 | 20201119_15232_1ng_glyco_hcd203040_compound15.22282.22282.3 | controllerType=0 controllerNumber=1 scan=22282 | 45.2002 |
| 16523:3 | 1 | R.TAGWNIPMGLLFN[+1768.640]QTGSC[+57.021]K.F | HexNAc(4)Hex(5)Fuc(1) | C[+57], N[+1769] | 1288.5527 | 3 | 3863.6435 | 3863.6495 | 0 | -1.6 | 485 | Specific | 931.85 | 931.85 | 931.85 | 16.88 | 160 | >sp\|P02788\|TRFL_HUMAN Lactotransferrin OS=Homo sapiens OX=9606 GN=LTF PE=1 SV=6 | 1270 | 20201119_15232_1ng_glyco_hcd203040_compound15.20120.20120.3 | controllerType=0 controllerNumber=1 scan=20120 | 40.5498 |
| 16553:3 | 1 | R.TAGWNIPMGLLFN[+1768.640]QTGSC[+57.021]K.F | HexNAc(4)Hex(5)Fuc(1) | C[+57], N[+1769] | 1288.5599 | 3 | 3863.6651 | 3863.6495 | 0 | 4 | 485 | Specific | 931.88 | 931.88 | 931.88 | 17.94 | 160 | >sp\|P02788\|TRFL_HUMAN Lactotransferrin OS=Homo sapiens OX=9606 GN=LTF PE=1 SV=6 | 1270 | 20201119_15232_1ng_glyco_hcd203040_compound15.20152.20152.3 | controllerType=0 controllerNumber=1 scan=20152 | 40.6097 |
| 18592:3 | 1 | R.TAGWNIPMGLLFN[+1913.677]QTGSC[+57.021]K.F | HexNAc(4)Hex(5)NeuAc(1) | C[+57], N[+1914] | 1336.9039 | 3 | 4008.697 | 4008.687 | 0 | 2.5 | 485 | Specific | 951.66 | 945.82 | 945.82 | 19.1 | 160 | >sp\|P02788\|TRFL_HUMAN Lactotransferrin OS=Homo sapiens OX=9606 GN=LTF PE=1 SV=6 | 1270 | 20201119_15232_1ng_glyco_hcd203040_compound15.22280.22280.3 | controllerType=0 controllerNumber=1 scan=22280 | 45.1919 |
| 18704:3 | 1 | R.TAGWNIPMGLLFN[+1751.624]QTGSC[+57.021]K.F | HexNAc(4)Hex(4)NeuAc(1) | C[+57], N[+1752] | 1282.8867 | 3 | 3846.6454 | 3846.6341 | 0 | 2.9 | 485 | Specific | 971.53 | 971.53 | 971.53 | 19.16 | 160 | >sp\|P02788\|TRFL_HUMAN Lactotransferrin OS=Homo sapiens OX=9606 GN=LTF PE=1 SV=6 | 1270 | 20201119_15232_1ng_glyco_hcd203040_compound15.22397.22397.3 | controllerType=0 controllerNumber=1 scan=22397 | 45.4799 |
| 16643:3 | 1 | R.TAGWNIPMGLLFN[+1606.587]QTGSC[+57.021]K.F | HexNAc(4)Hex(4)Fuc(1) | C[+57], N[+1607] | 1234.5395 | 3 | 3701.6039 | 3701.5966 | 0 | 2 | 485 | Specific | 975.59 | 973.45 | 973.45 | 19.1 | 160 | >sp\|P02788\|TRFL_HUMAN Lactotransferrin OS=Homo sapiens OX=9606 GN=LTF PE=1 SV=6 | 1270 | 20201119_15232_1ng_glyco_hcd203040_compound15.20246.20246.3 | controllerType=0 controllerNumber=1 scan=20246 | 40.8224 |
| 16525:3 | 1 | R.TAGWNIPMGLLFN[+1768.640]QTGSC[+57.021]K.F | HexNAc(4)Hex(5)Fuc(1) | C[+57], N[+1769] | 1288.5527 | 3 | 3863.6435 | 3863.6495 | 0 | -1.6 | 485 | Specific | 985.2 | 985.2 | 985.2 | 17.88 | 160 | >sp\|P02788\|TRFL_HUMAN Lactotransferrin OS=Homo sapiens OX=9606 GN=LTF PE=1 SV=6 | 1270 | 20201119_15232_1ng_glyco_hcd203040_compound15.20122.20122.3 | controllerType=0 controllerNumber=1 scan=20122 | 40.5547 |
| 18701:3 | 1 | R.TAGWNIPMGLLFN[+1751.624]QTGSC[+57.021]K.F | HexNAc(4)Hex(4)NeuAc(1) | C[+57], N[+1752] | 1282.8867 | 3 | 3846.6454 | 3846.6341 | 0 | 2.9 | 485 | Specific | 1005.08 | 925.38 | 925.38 | 20.16 | 160 | >sp\|P02788\|TRFL_HUMAN Lactotransferrin OS=Homo sapiens OX=9606 GN=LTF PE=1 SV=6 | 1270 | 20201119_15232_1ng_glyco_hcd203040_compound15.22394.22394.3 | controllerType=0 controllerNumber=1 scan=22394 | 45.4711 |
| 16923:3 | 1 | R.TAGWNIPMGLLFN[+1216.423]QTGSC[+57.021]K.F | HexNAc(2)Hex(5) | C[+57], N[+1216] | 1104.4868 | 3 | 3311.446 | 3311.4328 | 0 | 4 | 485 | Specific | 1023.3 | 908.6 | 468.4 | 19.94 | 160 | >sp\|P02788\|TRFL_HUMAN Lactotransferrin OS=Homo sapiens OX=9606 GN=LTF PE=1 SV=6 | 1270 | 20201119_15232_1ng_glyco_hcd203040_compound15.20537.20537.3 | controllerType=0 controllerNumber=1 scan=20537 | 41.4191 |
| 16925:3 | 1 | R.TAGWNIPMGLLFN[+1216.423]QTGSC[+57.021]K.F | HexNAc(2)Hex(5) | C[+57], N[+1216] | 1104.4868 | 3 | 3311.446 | 3311.4328 | 0 | 4 | 485 | Specific | 1060.85 | 1060.85 | 477.76 | 20.94 | 160 | >sp\|P02788\|TRFL_HUMAN Lactotransferrin OS=Homo sapiens OX=9606 GN=LTF PE=1 SV=6 | 1270 | 20201119_15232_1ng_glyco_hcd203040_compound15.20539.20539.3 | controllerType=0 controllerNumber=1 scan=20539 | 41.4266 |
| 17147:3 | 1 | R.TAGWNIPM[+15.995]GLLFN[+1913.677]QTGSC[+57.021]K.F | HexNAc(4)Hex(5)NeuAc(1) | C[+57], M[+16], N[+1914] | 1342.237 | 3 | 4024.6964 | 4024.6819 | 0 | 3.6 | 485 | Specific | 295.92 | 295.92 | 295.92 | 6.73 | 160 | >sp\|P02788\|TRFL_HUMAN Lactotransferrin OS=Homo sapiens OX=9606 GN=LTF PE=1 SV=6 | 1270 | 20201119_15232_1ng_glyco_hcd203040_compound15.20772.20772.3 | controllerType=0 controllerNumber=1 scan=20772 | 41.8906 |
| 16864:3 | 1 | R.TAGWNIPM[+15.995]GLLFN[+2059.735]QTGSC[+57.021]K.F | HexNAc(4)Hex(5)Fuc(1)NeuAc(1) | C[+57], M[+16], N[+2060] | 1390.9228 | 3 | 4170.7538 | 4170.7398 | 0 | 3.4 | 485 | Specific | 339.68 | 297.46 | 297.46 | 6.88 | 160 | >sp\|P02788\|TRFL_HUMAN Lactotransferrin OS=Homo sapiens OX=9606 GN=LTF PE=1 SV=6 | 1270 | 20201119_15232_1ng_glyco_hcd203040_compound15.20476.20476.3 | controllerType=0 controllerNumber=1 scan=20476 | 41.2912 |
| 16694:3 | 1 | R.TAGWNIPM[+15.995]GLLFN[+2059.735]QTGSC[+57.021]K.F | HexNAc(4)Hex(5)Fuc(1)NeuAc(1) | C[+57], M[+16], N[+2060] | 1390.9222 | 3 | 4170.752 | 4170.7398 | 0 | 2.9 | 485 | Specific | 387.4 | 296.79 | 296.79 | 7.16 | 160 | >sp\|P02788\|TRFL_HUMAN Lactotransferrin OS=Homo sapiens OX=9606 GN=LTF PE=1 SV=6 | 1270 | 20201119_15232_1ng_glyco_hcd203040_compound15.20299.20299.3 | controllerType=0 controllerNumber=1 scan=20299 | 40.9307 |
| 17149:3 | 1 | R.TAGWNIPM[+15.995]GLLFN[+1913.677]QTGSC[+57.021]K.F | HexNAc(4)Hex(5)NeuAc(1) | C[+57], M[+16], N[+1914] | 1342.237 | 3 | 4024.6964 | 4024.6819 | 0 | 3.6 | 485 | Specific | 514.44 | 458.91 | 458.91 | 9.65 | 160 | >sp\|P02788\|TRFL_HUMAN Lactotransferrin OS=Homo sapiens OX=9606 GN=LTF PE=1 SV=6 | 1270 | 20201119_15232_1ng_glyco_hcd203040_compound15.20774.20774.3 | controllerType=0 controllerNumber=1 scan=20774 | 41.8943 |
| 16696:3 | 1 | R.TAGWNIPM[+15.995]GLLFN[+2059.735]QTGSC[+57.021]K.F | HexNAc(4)Hex(5)Fuc(1)NeuAc(1) | C[+57], M[+16], N[+2060] | 1390.9222 | 3 | 4170.752 | 4170.7398 | 0 | 2.9 | 485 | Specific | 589.99 | 435.17 | 435.17 | 11.16 | 160 | >sp\|P02788\|TRFL_HUMAN Lactotransferrin OS=Homo sapiens OX=9606 GN=LTF PE=1 SV=6 | 1270 | 20201119_15232_1ng_glyco_hcd203040_compound15.20301.20301.3 | controllerType=0 controllerNumber=1 scan=20301 | 40.9352 |
| 16866:3 | 1 | R.TAGWNIPM[+15.995]GLLFN[+2059.735]QTGSC[+57.021]K.F | HexNAc(4)Hex(5)Fuc(1)NeuAc(1) | C[+57], M[+16], N[+2060] | 1390.9228 | 3 | 4170.7538 | 4170.7398 | 0 | 3.4 | 485 | Specific | 720.56 | 631.96 | 631.96 | 14.16 | 160 | >sp\|P02788\|TRFL_HUMAN Lactotransferrin OS=Homo sapiens OX=9606 GN=LTF PE=1 SV=6 | 1270 | 20201119_15232_1ng_glyco_hcd203040_compound15.20478.20478.3 | controllerType=0 controllerNumber=1 scan=20478 | 41.2953 |
| 17783:3 | 1 | R.TAGWNIPMGLLFN[+1606.587]QTGSC[+57.021]KFDEYFSQSC[+57.021]APGSDPR.S | HexNAc(4)Hex(4)Fuc(1) | C[+57]*2, N[+1607] | 1849.1299 | 3 | 5545.3751 | 5545.3438 | 0 | 5.7 | 485 | Specific | 232.97 | 232.97 | 232.97 | 6.03 | 160 | >sp\|P02788\|TRFL_HUMAN Lactotransferrin OS=Homo sapiens OX=9606 GN=LTF PE=1 SV=6 | 1270 | 20201119_15232_1ng_glyco_hcd203040_compound15.21437.21437.3 | controllerType=0 controllerNumber=1 scan=21437 | 43.4192 |
| 18901:4 | 1 | R.TAGWNIPMGLLFN[+1751.624]QTGSC[+57.021]KFDEYFSQSC[+57.021]APGSDPR.S | HexNAc(4)Hex(4)NeuAc(1) | C[+57]*2, N[+1752] | 1423.3571 | 4 | 5690.4064 | 5690.3813 | 0 | 4.4 | 485 | Specific | 240.69 | 240.69 | 240.69 | 6.53 | 160 | >sp\|P02788\|TRFL_HUMAN Lactotransferrin OS=Homo sapiens OX=9606 GN=LTF PE=1 SV=6 | 1270 | 20201119_15232_1ng_glyco_hcd203040_compound15.22602.22602.4 | controllerType=0 controllerNumber=1 scan=22602 | 45.8933 |
| 17785:3 | 1 | R.TAGWNIPMGLLFN[+1606.587]QTGSC[+57.021]KFDEYFSQSC[+57.021]APGSDPR.S | HexNAc(4)Hex(4)Fuc(1) | C[+57]*2, N[+1607] | 1849.1299 | 3 | 5545.3751 | 5545.3438 | 0 | 5.7 | 485 | Specific | 313.31 | 290.18 | 290.18 | 6.68 | 160 | >sp\|P02788\|TRFL_HUMAN Lactotransferrin OS=Homo sapiens OX=9606 GN=LTF PE=1 SV=6 | 1270 | 20201119_15232_1ng_glyco_hcd203040_compound15.21439.21439.3 | controllerType=0 controllerNumber=1 scan=21439 | 43.4233 |
| 17734:4 | 1 | R.[+42.011]TAGWNIPMGLLFN[+1565.560]QTGSC[+57.021]KFDEYFSQSC[+57.021]A | HexNAc(3)Hex(5)Fuc(1) | .[42], C[+57]*2, N[+1566] | 1387.3462 | 4 | 5546.3629 | 5546.3278 | 0 | 6.3 | 485 | Specific | 339.67 | 339.67 | 339.67 | 5.6 | 160 | >sp\|P02788\|TRFL_HUMAN Lactotransferrin OS=Homo sapiens OX=9606 GN=LTF PE=1 SV=6 | 1270 | 20201119_15232_1ng_glyco_hcd203040_compound15.21386.21386.4 | controllerType=0 controllerNumber=1 scan=21386 | 43.273 |
| 18951:4 | 1 | R.TAGWNIPMGLLFN[+1548.545]QTGSC[+57.021]KFDEYFSQSC[+57.021]APGSDPR.S | HexNAc(3)Hex(4)NeuAc(1) | C[+57]*2, N[+1549] | 1372.5831 | 4 | 5487.3106 | 5487.3019 | 0 | 1.6 | 485 | Specific | 441.9 | 362.46 | 362.46 | 8.37 | 160 | >sp\|P02788\|TRFL_HUMAN Lactotransferrin OS=Homo sapiens OX=9606 GN=LTF PE=1 SV=6 | 1270 | 20201119_15232_1ng_glyco_hcd203040_compound15.22653.22653.4 | controllerType=0 controllerNumber=1 scan=22653 | 45.9869 |
| 20191:4 | 1 | R.TAGWNIPMGLLFN[+2204.772]QTGSC[+57.021]KFDEYFSQSC[+57.021]APGSDPR.S | HexNAc(4)Hex(5)NeuAc(2) | C[+57]*2, N[+2205] | 1536.6432 | 4 | 6143.5509 | 6143.5295 | 0 | 3.5 | 485 | Specific | 462.72 | 419.45 | 198.42 | 10.01 | 160 | >sp\|P02788\|TRFL_HUMAN Lactotransferrin OS=Homo sapiens OX=9606 GN=LTF PE=1 SV=6 | 1270 | 20201119_15232_1ng_glyco_hcd203040_compound15.23951.23951.4 | controllerType=0 controllerNumber=1 scan=23951 | 48.631 |
| 17732:4 | 1 | R.[+42.011]TAGWNIPMGLLFN[+1565.560]QTGSC[+57.021]KFDEYFSQSC[+57.021]A | HexNAc(3)Hex(5)Fuc(1) | .[42], C[+57]*2, N[+1566] | 1387.3462 | 4 | 5546.3629 | 5546.3278 | 0 | 6.3 | 485 | Specific | 554.3 | 554.3 | 554.3 | 9.73 | 160 | >sp\|P02788\|TRFL_HUMAN Lactotransferrin OS=Homo sapiens OX=9606 GN=LTF PE=1 SV=6 | 1270 | 20201119_15232_1ng_glyco_hcd203040_compound15.21384.21384.4 | controllerType=0 controllerNumber=1 scan=21384 | 43.2675 |
| 18904:4 | 1 | R.TAGWNIPMGLLFN[+1751.624]QTGSC[+57.021]KFDEYFSQSC[+57.021]APGSDPR.S | HexNAc(4)Hex(4)NeuAc(1) | C[+57]*2, N[+1752] | 1423.3571 | 4 | 5690.4064 | 5690.3813 | 0 | 4.4 | 485 | Specific | 585.38 | 548.58 | 548.58 | 11.51 | 160 | >sp\|P02788\|TRFL_HUMAN Lactotransferrin OS=Homo sapiens OX=9606 GN=LTF PE=1 SV=6 | 1270 | 20201119_15232_1ng_glyco_hcd203040_compound15.22605.22605.4 | controllerType=0 controllerNumber=1 scan=22605 | 45.8974 |
| 17843:4 | 1 | R.TAGWNIPMGLLFN[+1216.423]QTGSC[+57.021]KFDEYFSQSC[+57.021]APGSDPR.S | HexNAc(2)Hex(5) | C[+57]*2, N[+1216] | 1289.5564 | 4 | 5155.2038 | 5155.18 | 0 | 4.6 | 485 | Specific | 592.78 | 592.78 | 140.15 | 11.51 | 160 | >sp\|P02788\|TRFL_HUMAN Lactotransferrin OS=Homo sapiens OX=9606 GN=LTF PE=1 SV=6 | 1270 | 20201119_15232_1ng_glyco_hcd203040_compound15.21500.21500.4 | controllerType=0 controllerNumber=1 scan=21500 | 43.5448 |
| 17846:4 | 1 | R.TAGWNIPMGLLFN[+1216.423]QTGSC[+57.021]KFDEYFSQSC[+57.021]APGSDPR.S | HexNAc(2)Hex(5) | C[+57]*2, N[+1216] | 1289.5564 | 4 | 5155.2038 | 5155.18 | 0 | 4.6 | 485 | Specific | 617.53 | 601.81 | 231.07 | 12.51 | 160 | >sp\|P02788\|TRFL_HUMAN Lactotransferrin OS=Homo sapiens OX=9606 GN=LTF PE=1 SV=6 | 1270 | 20201119_15232_1ng_glyco_hcd203040_compound15.21503.21503.4 | controllerType=0 controllerNumber=1 scan=21503 | 43.5564 |
| 17839:3 | 1 | R.TAGWNIPMGLLFN[+1216.423]QTGSC[+57.021]KFDEYFSQSC[+57.021]APGSDPR.S | HexNAc(2)Hex(5) | C[+57]*2, N[+1216] | 1719.0717 | 3 | 5155.2007 | 5155.18 | 0 | 4 | 485 | Specific | 619.89 | 587.56 | 157.65 | 12.56 | 160 | >sp\|P02788\|TRFL_HUMAN Lactotransferrin OS=Homo sapiens OX=9606 GN=LTF PE=1 SV=6 | 1270 | 20201119_15232_1ng_glyco_hcd203040_compound15.21496.21496.3 | controllerType=0 controllerNumber=1 scan=21496 | 43.5366 |
| 20088:4 | 1 | R.TAGWNIPMGLLFN[+2350.830]QTGSC[+57.021]KFDEYFSQSC[+57.021]APGSDPR.S | HexNAc(4)Hex(5)Fuc(1)NeuAc(2) | C[+57]*2, N[+2351] | 1573.1582 | 4 | 6289.611 | 6289.5875 | 0 | 3.7 | 485 | Specific | 700.47 | 659.11 | 659.11 | 14.73 | 160 | >sp\|P02788\|TRFL_HUMAN Lactotransferrin OS=Homo sapiens OX=9606 GN=LTF PE=1 SV=6 | 1270 | 20201119_15232_1ng_glyco_hcd203040_compound15.23844.23844.4 | controllerType=0 controllerNumber=1 scan=23844 | 48.4181 |
| 18696:4 | 1 | R.TAGWNIPMGLLFN[+2205.793]QTGSC[+57.021]KFDEYFSQSC[+57.021]APGSDPR.S | HexNAc(4)Hex(5)Fuc(2)NeuAc(1) | C[+57]*2, N[+2206] | 1536.8976 | 4 | 6144.5685 | 6144.5499 | 0 | 3 | 485 | Specific | 701.38 | 631.79 | 631.79 | 14.73 | 160 | >sp\|P02788\|TRFL_HUMAN Lactotransferrin OS=Homo sapiens OX=9606 GN=LTF PE=1 SV=6 | 1270 | 20201119_15232_1ng_glyco_hcd203040_compound15.22389.22389.4 | controllerType=0 controllerNumber=1 scan=22389 | 45.4611 |
| 20193:4 | 1 | R.TAGWNIPMGLLFN[+2204.772]QTGSC[+57.021]KFDEYFSQSC[+57.021]APGSDPR.S | HexNAc(4)Hex(5)NeuAc(2) | C[+57]*2, N[+2205] | 1536.6432 | 4 | 6143.5509 | 6143.5295 | 0 | 3.5 | 485 | Specific | 772.96 | 768.31 | 419.21 | 15.73 | 160 | >sp\|P02788\|TRFL_HUMAN Lactotransferrin OS=Homo sapiens OX=9606 GN=LTF PE=1 SV=6 | 1270 | 20201119_15232_1ng_glyco_hcd203040_compound15.23953.23953.4 | controllerType=0 controllerNumber=1 scan=23953 | 48.6352 |
| 17728:4 | 1 | R.TAGWNIPMGLLFN[+1622.582]QTGSC[+57.021]KFDEYFSQSC[+57.021]APGSDPR.S | HexNAc(4)Hex(5) | C[+57]*2, N[+1623] | 1391.0966 | 4 | 5561.3644 | 5561.3387 | 0 | 4.6 | 485 | Specific | 778.54 | 694.93 | 212.75 | 15.51 | 160 | >sp\|P02788\|TRFL_HUMAN Lactotransferrin OS=Homo sapiens OX=9606 GN=LTF PE=1 SV=6 | 1270 | 20201119_15232_1ng_glyco_hcd203040_compound15.21380.21380.4 | controllerType=0 controllerNumber=1 scan=21380 | 43.2575 |
| 17841:3 | 1 | R.TAGWNIPMGLLFN[+1216.423]QTGSC[+57.021]KFDEYFSQSC[+57.021]APGSDPR.S | HexNAc(2)Hex(5) | C[+57]*2, N[+1216] | 1719.0717 | 3 | 5155.2007 | 5155.18 | 0 | 4 | 485 | Specific | 780.93 | 706.01 | 220.02 | 15.56 | 160 | >sp\|P02788\|TRFL_HUMAN Lactotransferrin OS=Homo sapiens OX=9606 GN=LTF PE=1 SV=6 | 1270 | 20201119_15232_1ng_glyco_hcd203040_compound15.21498.21498.3 | controllerType=0 controllerNumber=1 scan=21498 | 43.5418 |
| 18763:4 | 1 | R.TAGWNIPMGLLFN[+1913.677]QTGSC[+57.021]KFDEYFSQSC[+57.021]APGSDPR.S | HexNAc(4)Hex(5)NeuAc(1) | C[+57]*2, N[+1914] | 1463.8672 | 4 | 5852.4469 | 5852.4341 | 0 | 2.2 | 485 | Specific | 853.3 | 788.92 | 788.92 | 17.67 | 160 | >sp\|P02788\|TRFL_HUMAN Lactotransferrin OS=Homo sapiens OX=9606 GN=LTF PE=1 SV=6 | 1270 | 20201119_15232_1ng_glyco_hcd203040_compound15.22459.22459.4 | controllerType=0 controllerNumber=1 scan=22459 | 45.6158 |
| 18765:4 | 1 | R.TAGWNIPMGLLFN[+1913.677]QTGSC[+57.021]KFDEYFSQSC[+57.021]APGSDPR.S | HexNAc(4)Hex(5)NeuAc(1) | C[+57]*2, N[+1914] | 1463.8672 | 4 | 5852.4469 | 5852.4341 | 0 | 2.2 | 485 | Specific | 899.15 | 849.89 | 849.89 | 17.67 | 160 | >sp\|P02788\|TRFL_HUMAN Lactotransferrin OS=Homo sapiens OX=9606 GN=LTF PE=1 SV=6 | 1270 | 20201119_15232_1ng_glyco_hcd203040_compound15.22461.22461.4 | controllerType=0 controllerNumber=1 scan=22461 | 45.621 |
| 18695:4 | 1 | R.TAGWNIPMGLLFN[+2059.735]QTGSC[+57.021]KFDEYFSQSC[+57.021]APGSDPR.S | HexNAc(4)Hex(5)Fuc(1)NeuAc(1) | C[+57]*2, N[+2060] | 1500.3853 | 4 | 5998.5192 | 5998.492 | 0 | 4.5 | 485 | Specific | 902.12 | 848.12 | 848.12 | 18.51 | 160 | >sp\|P02788\|TRFL_HUMAN Lactotransferrin OS=Homo sapiens OX=9606 GN=LTF PE=1 SV=6 | 1270 | 20201119_15232_1ng_glyco_hcd203040_compound15.22388.22388.4 | controllerType=0 controllerNumber=1 scan=22388 | 45.4596 |
| 18692:4 | 1 | R.TAGWNIPMGLLFN[+2059.735]QTGSC[+57.021]KFDEYFSQSC[+57.021]APGSDPR.S | HexNAc(4)Hex(5)Fuc(1)NeuAc(1) | C[+57]*2, N[+2060] | 1500.3854 | 4 | 5998.5197 | 5998.492 | 0 | 4.6 | 485 | Specific | 934.46 | 903.23 | 903.23 | 18.51 | 160 | >sp\|P02788\|TRFL_HUMAN Lactotransferrin OS=Homo sapiens OX=9606 GN=LTF PE=1 SV=6 | 1270 | 20201119_15232_1ng_glyco_hcd203040_compound15.22384.22384.4 | controllerType=0 controllerNumber=1 scan=22384 | 45.4499 |
| 20091:4 | 1 | R.TAGWNIPMGLLFN[+2350.830]QTGSC[+57.021]KFDEYFSQSC[+57.021]APGSDPR.S | HexNAc(4)Hex(5)Fuc(1)NeuAc(2) | C[+57]*2, N[+2351] | 1573.1582 | 4 | 6289.611 | 6289.5875 | 0 | 3.7 | 485 | Specific | 936.02 | 893.96 | 893.96 | 18.73 | 160 | >sp\|P02788\|TRFL_HUMAN Lactotransferrin OS=Homo sapiens OX=9606 GN=LTF PE=1 SV=6 | 1270 | 20201119_15232_1ng_glyco_hcd203040_compound15.23847.23847.4 | controllerType=0 controllerNumber=1 scan=23847 | 48.4228 |
| 18697:4 | 1 | R.TAGWNIPMGLLFN[+2059.735]QTGSC[+57.021]KFDEYFSQSC[+57.021]APGSDPR.S | HexNAc(4)Hex(5)Fuc(1)NeuAc(1) | C[+57]*2, N[+2060] | 1500.3853 | 4 | 5998.5192 | 5998.492 | 0 | 4.5 | 485 | Specific | 937.74 | 907.45 | 907.45 | 18.51 | 160 | >sp\|P02788\|TRFL_HUMAN Lactotransferrin OS=Homo sapiens OX=9606 GN=LTF PE=1 SV=6 | 1270 | 20201119_15232_1ng_glyco_hcd203040_compound15.22390.22390.4 | controllerType=0 controllerNumber=1 scan=22390 | 45.466 |
| 18689:4 | 1 | R.TAGWNIPMGLLFN[+2059.735]QTGSC[+57.021]KFDEYFSQSC[+57.021]APGSDPR.S | HexNAc(4)Hex(5)Fuc(1)NeuAc(1) | C[+57]*2, N[+2060] | 1500.3854 | 4 | 5998.5197 | 5998.492 | 0 | 4.6 | 485 | Specific | 945.3 | 857.26 | 857.26 | 18.51 | 160 | >sp\|P02788\|TRFL_HUMAN Lactotransferrin OS=Homo sapiens OX=9606 GN=LTF PE=1 SV=6 | 1270 | 20201119_15232_1ng_glyco_hcd203040_compound15.22381.22381.4 | controllerType=0 controllerNumber=1 scan=22381 | 45.4402 |
| 18698:4 | 1 | R.TAGWNIPMGLLFN[+2205.793]QTGSC[+57.021]KFDEYFSQSC[+57.021]APGSDPR.S | HexNAc(4)Hex(5)Fuc(2)NeuAc(1) | C[+57]*2, N[+2206] | 1536.8976 | 4 | 6144.5685 | 6144.5499 | 0 | 3 | 485 | Specific | 952.27 | 921.82 | 921.82 | 19.73 | 160 | >sp\|P02788\|TRFL_HUMAN Lactotransferrin OS=Homo sapiens OX=9606 GN=LTF PE=1 SV=6 | 1270 | 20201119_15232_1ng_glyco_hcd203040_compound15.22391.22391.4 | controllerType=0 controllerNumber=1 scan=22391 | 45.4674 |
| 17730:4 | 1 | R.TAGWNIPMGLLFN[+1622.582]QTGSC[+57.021]KFDEYFSQSC[+57.021]APGSDPR.S | HexNAc(4)Hex(5) | C[+57]*2, N[+1623] | 1391.0966 | 4 | 5561.3644 | 5561.3387 | 0 | 4.6 | 485 | Specific | 980.61 | 915.94 | 346.17 | 19.51 | 160 | >sp\|P02788\|TRFL_HUMAN Lactotransferrin OS=Homo sapiens OX=9606 GN=LTF PE=1 SV=6 | 1270 | 20201119_15232_1ng_glyco_hcd203040_compound15.21382.21382.4 | controllerType=0 controllerNumber=1 scan=21382 | 43.2627 |
| 17729:4 | 1 | R.TAGWNIPMGLLFN[+1768.640]QTGSC[+57.021]KFDEYFSQSC[+57.021]APGSDPR.S | HexNAc(4)Hex(5)Fuc(1) | C[+57]*2, N[+1769] | 1427.6091 | 4 | 5707.4147 | 5707.3966 | 0 | 3.2 | 485 | Specific | 985.62 | 951.17 | 951.17 | 19.73 | 160 | >sp\|P02788\|TRFL_HUMAN Lactotransferrin OS=Homo sapiens OX=9606 GN=LTF PE=1 SV=6 | 1270 | 20201119_15232_1ng_glyco_hcd203040_compound15.21381.21381.4 | controllerType=0 controllerNumber=1 scan=21381 | 43.2588 |
| 17731:4 | 1 | R.TAGWNIPMGLLFN[+1768.640]QTGSC[+57.021]KFDEYFSQSC[+57.021]APGSDPR.S | HexNAc(4)Hex(5)Fuc(1) | C[+57]*2, N[+1769] | 1427.6091 | 4 | 5707.4147 | 5707.3966 | 0 | 3.2 | 485 | Specific | 991.78 | 857.49 | 857.49 | 19.73 | 160 | >sp\|P02788\|TRFL_HUMAN Lactotransferrin OS=Homo sapiens OX=9606 GN=LTF PE=1 SV=6 | 1270 | 20201119_15232_1ng_glyco_hcd203040_compound15.21383.21383.4 | controllerType=0 controllerNumber=1 scan=21383 | 43.2655 |
| 20107:4 | 1 | R.TAGWNIPM[+15.995]GLLFN[+2351.851]QTGSC[+57.021]KFDEYFSQSC[+57.021]A | HexNAc(4)Hex(5)Fuc(3)NeuAc(1) | C[+57]*2, M[+16], N[+2352] | 1577.4149 | 4 | 6306.6379 | 6306.6028 | 0 | 5.6 | 485 | Specific | 362.25 | 359.74 | 359.74 | 7.37 | 160 | >sp\|P02788\|TRFL_HUMAN Lactotransferrin OS=Homo sapiens OX=9606 GN=LTF PE=1 SV=6 | 1270 | 20201119_15232_1ng_glyco_hcd203040_compound15.23863.23863.4 | controllerType=0 controllerNumber=1 scan=23863 | 48.4548 |
| 20109:4 | 1 | R.TAGWNIPM[+15.995]GLLFN[+2351.851]QTGSC[+57.021]KFDEYFSQSC[+57.021]A | HexNAc(4)Hex(5)Fuc(3)NeuAc(1) | C[+57]*2, M[+16], N[+2352] | 1577.4149 | 4 | 6306.6379 | 6306.6028 | 0 | 5.6 | 485 | Specific | 444.42 | 364.54 | 364.54 | 7.66 | 160 | >sp\|P02788\|TRFL_HUMAN Lactotransferrin OS=Homo sapiens OX=9606 GN=LTF PE=1 SV=6 | 1270 | 20201119_15232_1ng_glyco_hcd203040_compound15.23865.23865.4 | controllerType=0 controllerNumber=1 scan=23865 | 48.4595 |
| 17830:4 | 1 | R.TAGWNIPM[+15.995]GLLFN[+2059.735]QTGSC[+57.021]KFDEYFSQSC[+57.021]A | HexNAc(4)Hex(5)Fuc(1)NeuAc(1) | C[+57]*2, M[+16], N[+2060] | 1504.3844 | 4 | 6014.5158 | 6014.487 | 0 | 4.8 | 485 | Specific | 479.02 | 413.5 | 413.5 | 9.77 | 160 | >sp\|P02788\|TRFL_HUMAN Lactotransferrin OS=Homo sapiens OX=9606 GN=LTF PE=1 SV=6 | 1270 | 20201119_15232_1ng_glyco_hcd203040_compound15.21486.21486.4 | controllerType=0 controllerNumber=1 scan=21486 | 43.5094 |
| 17832:4 | 1 | R.TAGWNIPM[+15.995]GLLFN[+2059.735]QTGSC[+57.021]KFDEYFSQSC[+57.021]A | HexNAc(4)Hex(5)Fuc(1)NeuAc(1) | C[+57]*2, M[+16], N[+2060] | 1504.3844 | 4 | 6014.5158 | 6014.487 | 0 | 4.8 | 485 | Specific | 690.75 | 610.35 | 610.35 | 13.51 | 160 | >sp\|P02788\|TRFL_HUMAN Lactotransferrin OS=Homo sapiens OX=9606 GN=LTF PE=1 SV=6 | 1270 | 20201119_15232_1ng_glyco_hcd203040_compound15.21488.21488.4 | controllerType=0 controllerNumber=1 scan=21488 | 43.5131 |
| 08729:3 | 1 | R.N[+2351.851]GSDC[+57.021]PDKFC[+57.021]LFQSETK.N | HexNAc(4)Hex(5)Fuc(3)NeuAc(1) | C[+57]*2, N[+2352] | 1462.2485 | 3 | 4384.7309 | 4384.7247 | 0 | 1.4 | 642 | Specific | 244.88 | 194.78 | 194.78 | 5.23 | 160 | >sp\|P02788\|TRFL_HUMAN Lactotransferrin OS=Homo sapiens OX=9606 GN=LTF PE=1 SV=6 | 1270 | 20201119_15232_1ng_glyco_hcd203040_compound15.11959.11959.3 | controllerType=0 controllerNumber=1 scan=11959 | 26.4812 |
| 11249:3 | 1 | R.N[+2351.851]GSDC[+57.021]PDKFC[+57.021]LFQSETK.N | HexNAc(4)Hex(5)Fuc(3)NeuAc(1) | C[+57]*2, N[+2352] | 1462.2441 | 3 | 4384.7177 | 4384.7247 | 0 | -1.6 | 642 | Specific | 260.24 | 214.69 | 214.69 | 4.42 | 160 | >sp\|P02788\|TRFL_HUMAN Lactotransferrin OS=Homo sapiens OX=9606 GN=LTF PE=1 SV=6 | 1270 | 20201119_15232_1ng_glyco_hcd203040_compound15.14571.14571.3 | controllerType=0 controllerNumber=1 scan=14571 | 30.9576 |
| 08801:3 | 1 | R.N[+2059.735]GSDC[+57.021]PDKFC[+57.021]LFQSETK.N | HexNAc(4)Hex(5)Fuc(1)NeuAc(1) | C[+57]*2, N[+2060] | 1364.8804 | 3 | 4092.6267 | 4092.6088 | 0 | 4.4 | 642 | Specific | 373.01 | 328.03 | 328.03 | 6.59 | 160 | >sp\|P02788\|TRFL_HUMAN Lactotransferrin OS=Homo sapiens OX=9606 GN=LTF PE=1 SV=6 | 1270 | 20201119_15232_1ng_glyco_hcd203040_compound15.12034.12034.3 | controllerType=0 controllerNumber=1 scan=12034 | 26.6117 |
| 08803:3 | 1 | R.N[+2059.735]GSDC[+57.021]PDKFC[+57.021]LFQSETK.N | HexNAc(4)Hex(5)Fuc(1)NeuAc(1) | C[+57]*2, N[+2060] | 1364.8804 | 3 | 4092.6267 | 4092.6088 | 0 | 4.4 | 642 | Specific | 417.89 | 308.89 | 308.89 | 6.63 | 160 | >sp\|P02788\|TRFL_HUMAN Lactotransferrin OS=Homo sapiens OX=9606 GN=LTF PE=1 SV=6 | 1270 | 20201119_15232_1ng_glyco_hcd203040_compound15.12036.12036.3 | controllerType=0 controllerNumber=1 scan=12036 | 26.6155 |
| 11197:3 | 1 | R.N[+2350.830]GSDC[+57.021]PDKFC[+57.021]LFQSETK.N | HexNAc(4)Hex(5)Fuc(1)NeuAc(2) | C[+57]*2, N[+2351] | 1461.91 | 3 | 4383.7154 | 4383.7043 | 0 | 2.5 | 642 | Specific | 423.71 | 334.67 | 334.67 | 6.84 | 160 | >sp\|P02788\|TRFL_HUMAN Lactotransferrin OS=Homo sapiens OX=9606 GN=LTF PE=1 SV=6 | 1270 | 20201119_15232_1ng_glyco_hcd203040_compound15.14517.14517.3 | controllerType=0 controllerNumber=1 scan=14517 | 30.8543 |
| 08831:3 | 1 | R.N[+2059.735]GSDC[+57.021]PDKFC[+57.021]LFQSETK.N | HexNAc(4)Hex(5)Fuc(1)NeuAc(1) | C[+57]*2, N[+2060] | 1364.877 | 3 | 4092.6165 | 4092.6088 | 0 | 1.9 | 642 | Specific | 501.86 | 400.5 | 400.5 | 9.16 | 160 | >sp\|P02788\|TRFL_HUMAN Lactotransferrin OS=Homo sapiens OX=9606 GN=LTF PE=1 SV=6 | 1270 | 20201119_15232_1ng_glyco_hcd203040_compound15.12065.12065.3 | controllerType=0 controllerNumber=1 scan=12065 | 26.6649 |
| 11199:3 | 1 | R.N[+2350.830]GSDC[+57.021]PDKFC[+57.021]LFQSETK.N | HexNAc(4)Hex(5)Fuc(1)NeuAc(2) | C[+57]*2, N[+2351] | 1461.91 | 3 | 4383.7154 | 4383.7043 | 0 | 2.5 | 642 | Specific | 521.38 | 400.07 | 400.07 | 9.16 | 160 | >sp\|P02788\|TRFL_HUMAN Lactotransferrin OS=Homo sapiens OX=9606 GN=LTF PE=1 SV=6 | 1270 | 20201119_15232_1ng_glyco_hcd203040_compound15.14519.14519.3 | controllerType=0 controllerNumber=1 scan=14519 | 30.861 |
| 08829:3 | 1 | R.N[+2059.735]GSDC[+57.021]PDKFC[+57.021]LFQSETK.N | HexNAc(4)Hex(5)Fuc(1)NeuAc(1) | C[+57]*2, N[+2060] | 1364.877 | 3 | 4092.6165 | 4092.6088 | 0 | 1.9 | 642 | Specific | 539.46 | 398.73 | 398.73 | 9.23 | 160 | >sp\|P02788\|TRFL_HUMAN Lactotransferrin OS=Homo sapiens OX=9606 GN=LTF PE=1 SV=6 | 1270 | 20201119_15232_1ng_glyco_hcd203040_compound15.12063.12063.3 | controllerType=0 controllerNumber=1 scan=12063 | 26.6607 |
| 08743:3 | 1 | R.N[+2205.793]GSDC[+57.021]PDKFC[+57.021]LFQSETK.N | HexNAc(4)Hex(5)Fuc(2)NeuAc(1) | C[+57]*2, N[+2206] | 1413.5645 | 3 | 4238.679 | 4238.6667 | 0 | 2.9 | 642 | Specific | 561.61 | 419.72 | 419.72 | 9.5 | 160 | >sp\|P02788\|TRFL_HUMAN Lactotransferrin OS=Homo sapiens OX=9606 GN=LTF PE=1 SV=6 | 1270 | 20201119_15232_1ng_glyco_hcd203040_compound15.11974.11974.3 | controllerType=0 controllerNumber=1 scan=11974 | 26.5092 |
| 08741:3 | 1 | R.N[+2205.793]GSDC[+57.021]PDKFC[+57.021]LFQSETK.N | HexNAc(4)Hex(5)Fuc(2)NeuAc(1) | C[+57]*2, N[+2206] | 1413.5645 | 3 | 4238.679 | 4238.6667 | 0 | 2.9 | 642 | Specific | 613.08 | 482.86 | 482.86 | 11.73 | 160 | >sp\|P02788\|TRFL_HUMAN Lactotransferrin OS=Homo sapiens OX=9606 GN=LTF PE=1 SV=6 | 1270 | 20201119_15232_1ng_glyco_hcd203040_compound15.11972.11972.3 | controllerType=0 controllerNumber=1 scan=11972 | 26.5041 |
| 17561:5 | 5 | R.LSLHRPALEDLLLGSEAN[+1524.534]LTC[+57.021]TLTGLR.D | HexNAc(2)Hex(6)Fuc(1) | C[+57], N[+1525] | 898.4373 | 5 | 4488.1573 | 4488.1319 | 0 | 5.7 | 127 | Specific | 249.94 | 166 | 166 | 3.54 | 60 | >sp\|P01876\|IGHA1_HUMAN Immunoglobulin heavy constant alpha 1 OS=Homo sapiens OX=9606 GN=IGHA1 PE=1 SV=2 | 255 | 20201119_15232_1ng_glyco_hcd203040_compound15.21204.21204.2 | controllerType=0 controllerNumber=1 scan=21204 | 42.8651 |
| 17625:4 | 5 | R.LSLHRPALEDLLLGSEAN[+1298.476]LTC[+57.021]TLTGLR.D | HexNAc(4)Hex(3) | C[+57], N[+1298] | 1066.2784 | 4 | 4262.0916 | 4262.0742 | 0 | 4.1 | 127 | Specific | 280.43 | 280.43 | 280.43 | 5.72 | 60 | >sp\|P01876\|IGHA1_HUMAN Immunoglobulin heavy constant alpha 1 OS=Homo sapiens OX=9606 GN=IGHA1 PE=1 SV=2 | 255 | 20201119_15232_1ng_glyco_hcd203040_compound15.21271.21271.3 | controllerType=0 controllerNumber=1 scan=21271 | 43.0187 |
| 18421:3 | 5 | R.LSLHRPALEDLLLGSEAN[+1913.677]LTC[+57.021]TLTGLR.D | HexNAc(4)Hex(5)NeuAc(1) | C[+57], N[+1914] | 1626.4369 | 3 | 4877.2963 | 4877.2753 | 0 | 4.3 | 127 | Specific | 342.64 | 342.64 | 342.64 | 6.66 | 60 | >sp\|P01876\|IGHA1_HUMAN Immunoglobulin heavy constant alpha 1 OS=Homo sapiens OX=9606 GN=IGHA1 PE=1 SV=2 | 255 | 20201119_15232_1ng_glyco_hcd203040_compound15.22101.22101.3 | controllerType=0 controllerNumber=1 scan=22101 | 44.788 |
| 18533:3 | 5 | R.LSLHRPALEDLLLGSEAN[+1548.545]LTC[+57.021]TLTGLR.D | HexNAc(3)Hex(4)NeuAc(1) | C[+57], N[+1549] | 1504.7227 | 3 | 4512.1536 | 4512.1431 | 0 | 2.3 | 127 | Specific | 357.13 | 357.13 | 357.13 | 7.35 | 60 | >sp\|P01876\|IGHA1_HUMAN Immunoglobulin heavy constant alpha 1 OS=Homo sapiens OX=9606 GN=IGHA1 PE=1 SV=2 | 255 | 20201119_15232_1ng_glyco_hcd203040_compound15.22218.22218.3 | controllerType=0 controllerNumber=1 scan=22218 | 45.0467 |
| 18530:3 | 5 | R.LSLHRPALEDLLLGSEAN[+1548.545]LTC[+57.021]TLTGLR.D | HexNAc(3)Hex(4)NeuAc(1) | C[+57], N[+1549] | 1504.7227 | 3 | 4512.1536 | 4512.1431 | 0 | 2.3 | 127 | Specific | 360.03 | 360.03 | 360.03 | 7.35 | 60 | >sp\|P01876\|IGHA1_HUMAN Immunoglobulin heavy constant alpha 1 OS=Homo sapiens OX=9606 GN=IGHA1 PE=1 SV=2 | 255 | 20201119_15232_1ng_glyco_hcd203040_compound15.22215.22215.3 | controllerType=0 controllerNumber=1 scan=22215 | 45.0388 |
| 17682:3 | 5 | R.LSLHRPALEDLLLGSEAN[+1501.555]LTC[+57.021]TLTGLR.D | HexNAc(5)Hex(3) | C[+57], N[+1502] | 1489.0638 | 3 | 4465.1768 | 4465.1536 | 0 | 5.2 | 127 | Specific | 481.27 | 481.27 | 481.27 | 9.04 | 60 | >sp\|P01876\|IGHA1_HUMAN Immunoglobulin heavy constant alpha 1 OS=Homo sapiens OX=9606 GN=IGHA1 PE=1 SV=2 | 255 | 20201119_15232_1ng_glyco_hcd203040_compound15.21331.21331.3 | controllerType=0 controllerNumber=1 scan=21331 | 43.1633 |

Table S4. Cont.

| Query #:z | Protein Rank | Glycopeptide | Glycan | Modification Type(s) | Observed m/z | z | Observed (M+H) | Calc. mass (M+H) | Off-by-x error | Mass error (ppm) | StartPosition | Cleavage | Score | Delta | Delta Mod | \|Log Prob\| | # of unique peptides | Protein Name | Protein DB number | Comment | Scan # | Scan Time |
|---|---|---|---|---|---|---|---|---|---|---|---|---|---|---|---|---|---|---|---|---|---|---|
| 17485:3 | 5 | R.LSLHRPALEDLLLGSEAN[+1663.608]LTC[+57.021]TLTGLR.D | HexNAc(5)Hex(4) | C[+57], N[+1664] | 1543.0799 | 3 | 4627.2251 | 4627.2064 | 0 | 4 | 127 | Specific | 488.28 | 488.28 | 488.28 | 9.04 | 60 | >sp\|P01876\|IGHA1_HUMAN Immunoglobulin heavy constant alpha 1 OS=Homo sapiens OX=9606 GN=IGHA1 PE=1 SV=2 | 255 | 20201119_15232_1ng_glyco_hcd203040_compound15.21125.21125.3 | controllerType=0 controllerNumber=1 scan=21125 | 42.6754 |
| 17628:4 | 5 | R.LSLHRPALEDLLLGSEAN[+1298.476]LTC[+57.021]TLTGLR.D | HexNAc(4)Hex(3) | C[+57], N[+1298] | 1066.2784 | 4 | 4262.0916 | 4262.0742 | 0 | 4.1 | 127 | Specific | 494.21 | 494.21 | 494.21 | 9.61 | 60 | >sp\|P01876\|IGHA1_HUMAN Immunoglobulin heavy constant alpha 1 OS=Homo sapiens OX=9606 GN=IGHA1 PE=1 SV=2 | 255 | 20201119_15232_1ng_glyco_hcd203040_compound15.21274.21274.3 | controllerType=0 controllerNumber=1 scan=21274 | 43.0246 |
| 17594:4 | 5 | R.LSLHRPALEDLLLGSEAN[+1298.476]LTC[+57.021]TLTGLR.D | HexNAc(4)Hex(3) | C[+57], N[+1298] | 1066.2762 | 4 | 4262.0828 | 4262.0742 | 0 | 2 | 127 | Specific | 542.6 | 400.82 | 400.82 | 10.07 | 60 | >sp\|P01876\|IGHA1_HUMAN Immunoglobulin heavy constant alpha 1 OS=Homo sapiens OX=9606 GN=IGHA1 PE=1 SV=2 | 255 | 20201119_15232_1ng_glyco_hcd203040_compound15.21239.21239.3 | controllerType=0 controllerNumber=1 scan=21239 | 42.9537 |
| 17684:3 | 5 | R.LSLHRPALEDLLLGSEAN[+1501.555]LTC[+57.021]TLTGLR.D | HexNAc(5)Hex(3) | C[+57], N[+1502] | 1489.0638 | 3 | 4465.1768 | 4465.1536 | 0 | 5.2 | 127 | Specific | 553.38 | 553.38 | 553.38 | 9.54 | 60 | >sp\|P01876\|IGHA1_HUMAN Immunoglobulin heavy constant alpha 1 OS=Homo sapiens OX=9606 GN=IGHA1 PE=1 SV=2 | 255 | 20201119_15232_1ng_glyco_hcd203040_compound15.21333.21333.3 | controllerType=0 controllerNumber=1 scan=21333 | 43.168 |
| 18410:4 | 5 | R.LSLHRPALEDLLLGSEAN[+1913.677]LTC[+57.021]TLTGLR.D | HexNAc(4)Hex(5)NeuAc(1) | C[+57], N[+1914] | 1220.0788 | 4 | 4877.2933 | 4877.2753 | 0 | 3.7 | 127 | Specific | 556.33 | 556.33 | 556.33 | 10.33 | 60 | >sp\|P01876\|IGHA1_HUMAN Immunoglobulin heavy constant alpha 1 OS=Homo sapiens OX=9606 GN=IGHA1 PE=1 SV=2 | 255 | 20201119_15232_1ng_glyco_hcd203040_compound15.22090.22090.4 | controllerType=0 controllerNumber=1 scan=22090 | 44.7649 |
| 18423:3 | 5 | R.LSLHRPALEDLLLGSEAN[+1913.677]LTC[+57.021]TLTGLR.D | HexNAc(4)Hex(5)NeuAc(1) | C[+57], N[+1914] | 1626.4369 | 3 | 4877.2963 | 4877.2753 | 0 | 4.3 | 127 | Specific | 591.93 | 484.32 | 484.32 | 10.78 | 60 | >sp\|P01876\|IGHA1_HUMAN Immunoglobulin heavy constant alpha 1 OS=Homo sapiens OX=9606 GN=IGHA1 PE=1 SV=2 | 255 | 20201119_15232_1ng_glyco_hcd203040_compound15.22103.22103.3 | controllerType=0 controllerNumber=1 scan=22103 | 44.7914 |
| 17427:3 | 5 | R.LSLHRPALEDLLLGSEAN[+1702.581]LTC[+57.021]TLTGLR.D | HexNAc(2)Hex(8) | C[+57], N[+1703] | 1556.0706 | 3 | 4666.1973 | 4666.1796 | 0 | 3.8 | 127 | Specific | 595.81 | 550.4 | 550.4 | 11 | 60 | >sp\|P01876\|IGHA1_HUMAN Immunoglobulin heavy constant alpha 1 OS=Homo sapiens OX=9606 GN=IGHA1 PE=1 SV=2 | 255 | 20201119_15232_1ng_glyco_hcd203040_compound15.21064.21064.3 | controllerType=0 controllerNumber=1 scan=21064 | 42.5094 |
| 17518:3 | 5 | R.LSLHRPALEDLLLGSEAN[+1501.555]LTC[+57.021]TLTGLR.D | HexNAc(5)Hex(3) | C[+57], N[+1502] | 1489.0632 | 3 | 4465.1749 | 4465.1536 | 0 | 4.8 | 127 | Specific | 600.89 | 600.89 | 600.89 | 11.78 | 60 | >sp\|P01876\|IGHA1_HUMAN Immunoglobulin heavy constant alpha 1 OS=Homo sapiens OX=9606 GN=IGHA1 PE=1 SV=2 | 255 | 20201119_15232_1ng_glyco_hcd203040_compound15.21160.21160.3 | controllerType=0 controllerNumber=1 scan=21160 | 42.7685 |
| 17549:4 | 5 | R.LSLHRPALEDLLLGSEAN[+1647.613]LTC[+57.021]TLTGLR.D | HexNAc(5)Hex(3)Fuc(1) | C[+57], N[+1648] | 1153.5683 | 4 | 4611.2513 | 4611.2115 | 0 | 8.6 | 127 | Specific | 607 | 607 | 607 | 11.31 | 60 | >sp\|P01876\|IGHA1_HUMAN Immunoglobulin heavy constant alpha 1 OS=Homo sapiens OX=9606 GN=IGHA1 PE=1 SV=2 | 255 | 20201119_15232_1ng_glyco_hcd203040_compound15.21192.21192.4 | controllerType=0 controllerNumber=1 scan=21192 | 42.8382 |
| 17403:3 | 5 | R.LSLHRPALEDLLLGSEAN[+1864.634]LTC[+57.021]TLTGLR.D | HexNAc(2)Hex(9) | C[+57], N[+1865] | 1610.091 | 3 | 4828.2584 | 4828.2324 | 0 | 5.4 | 127 | Specific | 609.19 | 564.35 | 564.35 | 11.23 | 60 | >sp\|P01876\|IGHA1_HUMAN Immunoglobulin heavy constant alpha 1 OS=Homo sapiens OX=9606 GN=IGHA1 PE=1 SV=2 | 255 | 20201119_15232_1ng_glyco_hcd203040_compound15.21039.21039.3 | controllerType=0 controllerNumber=1 scan=21039 | 42.4498 |
| 17566:3 | 5 | R.LSLHRPALEDLLLGSEAN[+1298.476]LTC[+57.021]TLTGLR.D | HexNAc(4)Hex(3) | C[+57], N[+1298] | 1421.3697 | 3 | 4262.0945 | 4262.0742 | 0 | 4.8 | 127 | Specific | 610.52 | 567.55 | 567.55 | 11.78 | 60 | >sp\|P01876\|IGHA1_HUMAN Immunoglobulin heavy constant alpha 1 OS=Homo sapiens OX=9606 GN=IGHA1 PE=1 SV=2 | 255 | 20201119_15232_1ng_glyco_hcd203040_compound15.21210.21210.3 | controllerType=0 controllerNumber=1 scan=21210 | 42.8757 |
| 17489:3 | 5 | R.LSLHRPALEDLLLGSEAN[+1622.582]LTC[+57.021]TLTGLR.D | HexNAc(4)Hex(5) | C[+57], N[+1623] | 1529.4047 | 3 | 4586.1996 | 4586.1799 | 0 | 4.3 | 127 | Specific | 624.88 | 517.83 | 517.83 | 11.78 | 60 | >sp\|P01876\|IGHA1_HUMAN Immunoglobulin heavy constant alpha 1 OS=Homo sapiens OX=9606 GN=IGHA1 PE=1 SV=2 | 255 | 20201119_15232_1ng_glyco_hcd203040_compound15.21129.21129.3 | controllerType=0 controllerNumber=1 scan=21129 | 42.6902 |
| 17429:3 | 5 | R.LSLHRPALEDLLLGSEAN[+1540.529]LTC[+57.021]TLTGLR.D | HexNAc(2)Hex(7) | C[+57], N[+1541] | 1502.0524 | 3 | 4504.1427 | 4504.1268 | 0 | 3.5 | 127 | Specific | 627 | 584.53 | 584.53 | 12 | 60 | >sp\|P01876\|IGHA1_HUMAN Immunoglobulin heavy constant alpha 1 OS=Homo sapiens OX=9606 GN=IGHA1 PE=1 SV=2 | 255 | 20201119_15232_1ng_glyco_hcd203040_compound15.21066.21066.3 | controllerType=0 controllerNumber=1 scan=21066 | 42.523 |
| 17488:3 | 5 | R.LSLHRPALEDLLLGSEAN[+1663.608]LTC[+57.021]TLTGLR.D | HexNAc(5)Hex(4) | C[+57], N[+1664] | 1543.0799 | 3 | 4627.2251 | 4627.2064 | 0 | 4 | 127 | Specific | 627.25 | 553.64 | 553.64 | 11.78 | 60 | >sp\|P01876\|IGHA1_HUMAN Immunoglobulin heavy constant alpha 1 OS=Homo sapiens OX=9606 GN=IGHA1 PE=1 SV=2 | 255 | 20201119_15232_1ng_glyco_hcd203040_compound15.21128.21128.3 | controllerType=0 controllerNumber=1 scan=21128 | 42.6884 |
| 17520:3 | 5 | R.LSLHRPALEDLLLGSEAN[+1501.555]LTC[+57.021]TLTGLR.D | HexNAc(5)Hex(3) | C[+57], N[+1502] | 1489.0632 | 3 | 4465.1749 | 4465.1536 | 0 | 4.8 | 127 | Specific | 632.83 | 587.94 | 587.94 | 11.78 | 60 | >sp\|P01876\|IGHA1_HUMAN Immunoglobulin heavy constant alpha 1 OS=Homo sapiens OX=9606 GN=IGHA1 PE=1 SV=2 | 255 | 20201119_15232_1ng_glyco_hcd203040_compound15.21162.21162.3 | controllerType=0 controllerNumber=1 scan=21162 | 42.7738 |
| 17486:3 | 5 | R.LSLHRPALEDLLLGSEAN[+1622.582]LTC[+57.021]TLTGLR.D | HexNAc(4)Hex(5) | C[+57], N[+1623] | 1529.4047 | 3 | 4586.1996 | 4586.1799 | 0 | 4.3 | 127 | Specific | 638.81 | 556.18 | 556.18 | 11.78 | 60 | >sp\|P01876\|IGHA1_HUMAN Immunoglobulin heavy constant alpha 1 OS=Homo sapiens OX=9606 GN=IGHA1 PE=1 SV=2 | 255 | 20201119_15232_1ng_glyco_hcd203040_compound15.21126.21126.3 | controllerType=0 controllerNumber=1 scan=21126 | 42.6766 |
| 18388:4 | 5 | R.LSLHRPALEDLLLGSEAN[+2116.756]LTC[+57.021]TLTGLR.D | HexNAc(5)Hex(5)NeuAc(1) | C[+57], N[+2117] | 1270.8491 | 4 | 5080.3747 | 5080.3547 | 0 | 3.9 | 127 | Specific | 638.89 | 589.91 | 589.91 | 12.57 | 60 | >sp\|P01876\|IGHA1_HUMAN Immunoglobulin heavy constant alpha 1 OS=Homo sapiens OX=9606 GN=IGHA1 PE=1 SV=2 | 255 | 20201119_15232_1ng_glyco_hcd203040_compound15.22067.22067.4 | controllerType=0 controllerNumber=1 scan=22067 | 44.7084 |
| 17569:3 | 5 | R.LSLHRPALEDLLLGSEAN[+1298.476]LTC[+57.021]TLTGLR.D | HexNAc(4)Hex(3) | C[+57], N[+1298] | 1421.3697 | 3 | 4262.0945 | 4262.0742 | 0 | 4.8 | 127 | Specific | 645.11 | 560.13 | 560.13 | 11.78 | 60 | >sp\|P01876\|IGHA1_HUMAN Immunoglobulin heavy constant alpha 1 OS=Homo sapiens OX=9606 GN=IGHA1 PE=1 SV=2 | 255 | 20201119_15232_1ng_glyco_hcd203040_compound15.21213.21213.3 | controllerType=0 controllerNumber=1 scan=21213 | 42.8875 |
| 17401:3 | 5 | R.LSLHRPALEDLLLGSEAN[+1864.634]LTC[+57.021]TLTGLR.D | HexNAc(2)Hex(9) | C[+57], N[+1865] | 1610.091 | 3 | 4828.2584 | 4828.2324 | 0 | 5.4 | 127 | Specific | 646.95 | 644.09 | 644.09 | 11.23 | 60 | >sp\|P01876\|IGHA1_HUMAN Immunoglobulin heavy constant alpha 1 OS=Homo sapiens OX=9606 GN=IGHA1 PE=1 SV=2 | 255 | 20201119_15232_1ng_glyco_hcd203040_compound15.21037.21037.3 | controllerType=0 controllerNumber=1 scan=21037 | 42.4448 |
| 17424:3 | 5 | R.LSLHRPALEDLLLGSEAN[+1702.581]LTC[+57.021]TLTGLR.D | HexNAc(2)Hex(8) | C[+57], N[+1703] | 1556.0706 | 3 | 4666.1973 | 4666.1796 | 0 | 3.8 | 127 | Specific | 648.87 | 641.52 | 641.52 | 12 | 60 | >sp\|P01876\|IGHA1_HUMAN Immunoglobulin heavy constant alpha 1 OS=Homo sapiens OX=9606 GN=IGHA1 PE=1 SV=2 | 255 | 20201119_15232_1ng_glyco_hcd203040_compound15.21061.21061.3 | controllerType=0 controllerNumber=1 scan=21061 | 42.5038 |
| 17522:5 | 5 | R.LSLHRPALEDLLLGSEAN[+1501.555]LTC[+57.021]TLTGLR.D | HexNAc(5)Hex(3) | C[+57], N[+1502] | 893.8406 | 5 | 4465.1739 | 4465.1536 | 0 | 4.5 | 127 | Specific | 658.69 | 658.69 | 658.69 | 13.29 | 60 | >sp\|P01876\|IGHA1_HUMAN Immunoglobulin heavy constant alpha 1 OS=Homo sapiens OX=9606 GN=IGHA1 PE=1 SV=2 | 255 | 20201119_15232_1ng_glyco_hcd203040_compound15.21164.21164.5 | controllerType=0 controllerNumber=1 scan=21164 | 42.7791 |
| 17438:4 | 5 | R.LSLHRPALEDLLLGSEAN[+1663.608]LTC[+57.021]TLTGLR.D | HexNAc(5)Hex(4) | C[+57], N[+1664] | 1157.5613 | 4 | 4627.2235 | 4627.2064 | 0 | 3.7 | 127 | Specific | 663.53 | 663.53 | 663.53 | 13.57 | 60 | >sp\|P01876\|IGHA1_HUMAN Immunoglobulin heavy constant alpha 1 OS=Homo sapiens OX=9606 GN=IGHA1 PE=1 SV=2 | 255 | 20201119_15232_1ng_glyco_hcd203040_compound15.21076.21076.4 | controllerType=0 controllerNumber=1 scan=21076 | 42.5574 |
| 17484:4 | 5 | R.LSLHRPALEDLLLGSEAN[+1501.555]LTC[+57.021]TLTGLR.D | HexNAc(5)Hex(3) | C[+57], N[+1502] | 1117.0484 | 4 | 4465.1717 | 4465.1536 | 0 | 4.1 | 127 | Specific | 663.56 | 663.56 | 663.56 | 13.35 | 60 | >sp\|P01876\|IGHA1_HUMAN Immunoglobulin heavy constant alpha 1 OS=Homo sapiens OX=9606 GN=IGHA1 PE=1 SV=2 | 255 | 20201119_15232_1ng_glyco_hcd203040_compound15.21124.21124.4 | controllerType=0 controllerNumber=1 scan=21124 | 42.674 |
| 17570:3 | 5 | R.LSLHRPALEDLLLGSEAN[+1095.397]LTC[+57.021]TLTGLR.D | HexNAc(3)Hex(3) | C[+57], N[+1095] | 1353.675 | 3 | 4059.0104 | 4058.9949 | 0 | 3.8 | 127 | Specific | 681.24 | 657.27 | 657.27 | 13 | 60 | >sp\|P01876\|IGHA1_HUMAN Immunoglobulin heavy constant alpha 1 OS=Homo sapiens OX=9606 GN=IGHA1 PE=1 SV=2 | 255 | 20201119_15232_1ng_glyco_hcd203040_compound15.21214.21214.3 | controllerType=0 controllerNumber=1 scan=21214 | 42.8948 |
| 17449:4 | 5 | R.LSLHRPALEDLLLGSEAN[+1622.582]LTC[+57.021]TLTGLR.D | HexNAc(4)Hex(5) | C[+57], N[+1623] | 1147.3042 | 4 | 4586.1951 | 4586.1799 | 0 | 3.3 | 127 | Specific | 684.56 | 676.26 | 676.26 | 13.57 | 60 | >sp\|P01876\|IGHA1_HUMAN Immunoglobulin heavy constant alpha 1 OS=Homo sapiens OX=9606 GN=IGHA1 PE=1 SV=2 | 255 | 20201119_15232_1ng_glyco_hcd203040_compound15.21087.21087.4 | controllerType=0 controllerNumber=1 scan=21087 | 42.5809 |
| 17568:3 | 5 | R.LSLHRPALEDLLLGSEAN[+1095.397]LTC[+57.021]TLTGLR.D | HexNAc(3)Hex(3) | C[+57], N[+1095] | 1353.675 | 3 | 4059.0104 | 4058.9949 | 0 | 3.8 | 127 | Specific | 687.22 | 687.22 | 687.22 | 13 | 60 | >sp\|P01876\|IGHA1_HUMAN Immunoglobulin heavy constant alpha 1 OS=Homo sapiens OX=9606 GN=IGHA1 PE=1 SV=2 | 255 | 20201119_15232_1ng_glyco_hcd203040_compound15.21212.21212.3 | controllerType=0 controllerNumber=1 scan=21212 | 42.8842 |
| 17525:5 | 5 | R.LSLHRPALEDLLLGSEAN[+1501.555]LTC[+57.021]TLTGLR.D | HexNAc(5)Hex(3) | C[+57], N[+1502] | 893.8406 | 5 | 4465.1739 | 4465.1536 | 0 | 4.5 | 127 | Specific | 702.95 | 702.95 | 702.95 | 14.29 | 60 | >sp\|P01876\|IGHA1_HUMAN Immunoglobulin heavy constant alpha 1 OS=Homo sapiens OX=9606 GN=IGHA1 PE=1 SV=2 | 255 | 20201119_15232_1ng_glyco_hcd203040_compound15.21167.21167.5 | controllerType=0 controllerNumber=1 scan=21167 | 42.7843 |
| 17505:3 | 5 | R.LSLHRPALEDLLLGSEAN[+1216.423]LTC[+57.021]TLTGLR.D | HexNAc(2)Hex(5) | C[+57], N[+1216] | 1394.0147 | 3 | 4180.0296 | 4180.0211 | 0 | 2 | 127 | Specific | 704.87 | 608.15 | 608.15 | 13.94 | 60 | >sp\|P01876\|IGHA1_HUMAN Immunoglobulin heavy constant alpha 1 OS=Homo sapiens OX=9606 GN=IGHA1 PE=1 SV=2 | 255 | 20201119_15232_1ng_glyco_hcd203040_compound15.21146.21146.3 | controllerType=0 controllerNumber=1 scan=21146 | 42.7339 |
| 17538:4 | 5 | R.LSLHRPALEDLLLGSEAN[+1647.613]LTC[+57.021]TLTGLR.D | HexNAc(5)Hex(3)Fuc(1) | C[+57], N[+1648] | 1153.5613 | 4 | 4611.2235 | 4611.2115 | 0 | 2.6 | 127 | Specific | 712.44 | 696.09 | 696.09 | 14.51 | 60 | >sp\|P01876\|IGHA1_HUMAN Immunoglobulin heavy constant alpha 1 OS=Homo sapiens OX=9606 GN=IGHA1 PE=1 SV=2 | 255 | 20201119_15232_1ng_glyco_hcd203040_compound15.21180.21180.4 | controllerType=0 controllerNumber=1 scan=21180 | 42.8107 |
| 17547:4 | 5 | R.LSLHRPALEDLLLGSEAN[+1647.613]LTC[+57.021]TLTGLR.D | HexNAc(5)Hex(3)Fuc(1) | C[+57], N[+1648] | 1153.5683 | 4 | 4611.2513 | 4611.2115 | 0 | 8.6 | 127 | Specific | 713.35 | 713.35 | 713.35 | 13.31 | 60 | >sp\|P01876\|IGHA1_HUMAN Immunoglobulin heavy constant alpha 1 OS=Homo sapiens OX=9606 GN=IGHA1 PE=1 SV=2 | 255 | 20201119_15232_1ng_glyco_hcd203040_compound15.21190.21190.4 | controllerType=0 controllerNumber=1 scan=21190 | 42.8328 |
| 18469:4 | 5 | R.LSLHRPALEDLLLGSEAN[+1710.598]LTC[+57.021]TLTGLR.D | HexNAc(3)Hex(5)NeuAc(1) | C[+57], N[+1711] | 1169.31 | 4 | 4674.2181 | 4674.1959 | 0 | 4.7 | 127 | Specific | 715.53 | 671.74 | 671.74 | 14.35 | 60 | >sp\|P01876\|IGHA1_HUMAN Immunoglobulin heavy constant alpha 1 OS=Homo sapiens OX=9606 GN=IGHA1 PE=1 SV=2 | 255 | 20201119_15232_1ng_glyco_hcd203040_compound15.22151.22151.4 | controllerType=0 controllerNumber=1 scan=22151 | 44.8997 |
| 17490:4 | 5 | R.LSLHRPALEDLLLGSEAN[+1419.502]LTC[+57.021]TLTGLR.D | HexNAc(3)Hex(5) | C[+57], N[+1420] | 1096.5347 | 4 | 4383.117 | 4383.1005 | 0 | 3.8 | 127 | Specific | 716.35 | 716.35 | 716.35 | 14.57 | 60 | >sp\|P01876\|IGHA1_HUMAN Immunoglobulin heavy constant alpha 1 OS=Homo sapiens OX=9606 GN=IGHA1 PE=1 SV=2 | 255 | 20201119_15232_1ng_glyco_hcd203040_compound15.21130.21130.4 | controllerType=0 controllerNumber=1 scan=21130 | 42.6921 |
| 17426:3 | 5 | R.LSLHRPALEDLLLGSEAN[+1540.529]LTC[+57.021]TLTGLR.D | HexNAc(2)Hex(7) | C[+57], N[+1541] | 1502.0524 | 3 | 4504.1427 | 4504.1268 | 0 | 3.5 | 127 | Specific | 722.13 | 662.22 | 662.22 | 14 | 60 | >sp\|P01876\|IGHA1_HUMAN Immunoglobulin heavy constant alpha 1 OS=Homo sapiens OX=9606 GN=IGHA1 PE=1 SV=2 | 255 | 20201119_15232_1ng_glyco_hcd203040_compound15.21063.21063.3 | controllerType=0 controllerNumber=1 scan=21063 | 42.506 |
| 17502:3 | 5 | R.LSLHRPALEDLLLGSEAN[+1216.423]LTC[+57.021]TLTGLR.D | HexNAc(2)Hex(5) | C[+57], N[+1216] | 1394.0147 | 3 | 4180.0296 | 4180.0211 | 0 | 2 | 127 | Specific | 743.77 | 650.11 | 650.11 | 13.94 | 60 | >sp\|P01876\|IGHA1_HUMAN Immunoglobulin heavy constant alpha 1 OS=Homo sapiens OX=9606 GN=IGHA1 PE=1 SV=2 | 255 | 20201119_15232_1ng_glyco_hcd203040_compound15.21143.21143.3 | controllerType=0 controllerNumber=1 scan=21143 | 42.7271 |
| 18424:4 | 5 | R.LSLHRPALEDLLLGSEAN[+1872.651]LTC[+57.021]TLTGLR.D | HexNAc(3)Hex(6)NeuAc(1) | C[+57], N[+1873] | 1209.8227 | 4 | 4836.2689 | 4836.2487 | 0 | 4.2 | 127 | Specific | 743.78 | 676.33 | 676.33 | 14.35 | 60 | >sp\|P01876\|IGHA1_HUMAN Immunoglobulin heavy constant alpha 1 OS=Homo sapiens OX=9606 GN=IGHA1 PE=1 SV=2 | 255 | 20201119_15232_1ng_glyco_hcd203040_compound15.22104.22104.4 | controllerType=0 controllerNumber=1 scan=22104 | 44.7934 |
| 17540:4 | 5 | R.LSLHRPALEDLLLGSEAN[+1647.613]LTC[+57.021]TLTGLR.D | HexNAc(5)Hex(3)Fuc(1) | C[+57], N[+1648] | 1153.5613 | 4 | 4611.2235 | 4611.2115 | 0 | 2.6 | 127 | Specific | 744.68 | 707.76 | 707.76 | 14.51 | 60 | >sp\|P01876\|IGHA1_HUMAN Immunoglobulin heavy constant alpha 1 OS=Homo sapiens OX=9606 GN=IGHA1 PE=1 SV=2 | 255 | 20201119_15232_1ng_glyco_hcd203040_compound15.21182.21182.4 | controllerType=0 controllerNumber=1 scan=21182 | 42.8158 |
| 17575:4 | 5 | R.LSLHRPALEDLLLGSEAN[+1054.370]LTC[+57.021]TLTGLR.D | HexNAc(2)Hex(4) | C[+57], N[+1054] | 1005.2507 | 4 | 4017.9808 | 4017.9683 | 0 | 3.1 | 127 | Specific | 746.31 | 746.31 | 746.31 | 14.57 | 60 | >sp\|P01876\|IGHA1_HUMAN Immunoglobulin heavy constant alpha 1 OS=Homo sapiens OX=9606 GN=IGHA1 PE=1 SV=2 | 255 | 20201119_15232_1ng_glyco_hcd203040_compound15.21219.21219.4 | controllerType=0 controllerNumber=1 scan=21219 | 42.9122 |
| 18391:4 | 5 | R.LSLHRPALEDLLLGSEAN[+2116.756]LTC[+57.021]TLTGLR.D | HexNAc(5)Hex(5)NeuAc(1) | C[+57], N[+2117] | 1270.8491 | 4 | 5080.3747 | 5080.3547 | 0 | 3.9 | 127 | Specific | 750.41 | 750.41 | 750.41 | 15.57 | 60 | >sp\|P01876\|IGHA1_HUMAN Immunoglobulin heavy constant alpha 1 OS=Homo sapiens OX=9606 GN=IGHA1 PE=1 SV=2 | 255 | 20201119_15232_1ng_glyco_hcd203040_compound15.22070.22070.4 | controllerType=0 controllerNumber=1 scan=22070 | 44.7137 |
| 18412:4 | 5 | R.LSLHRPALEDLLLGSEAN[+1913.677]LTC[+57.021]TLTGLR.D | HexNAc(4)Hex(5)NeuAc(1) | C[+57], N[+1914] | 1220.0788 | 4 | 4877.2933 | 4877.2753 | 0 | 3.7 | 127 | Specific | 756.63 | 721.12 | 721.12 | 15.57 | 60 | >sp\|P01876\|IGHA1_HUMAN Immunoglobulin heavy constant alpha 1 OS=Homo sapiens OX=9606 GN=IGHA1 PE=1 SV=2 | 255 | 20201119_15232_1ng_glyco_hcd203040_compound15.22092.22092.4 | controllerType=0 controllerNumber=1 scan=22092 | 44.7693 |
| 17494:4 | 5 | R.LSLHRPALEDLLLGSEAN[+1460.529]LTC[+57.021]TLTGLR.D | HexNAc(4)Hex(4) | C[+57], N[+1461] | 1106.7924 | 4 | 4424.1478 | 4424.1271 | 0 | 4.7 | 127 | Specific | 757.22 | 757.22 | 757.22 | 15.35 | 60 | >sp\|P01876\|IGHA1_HUMAN Immunoglobulin heavy constant alpha 1 OS=Homo sapiens OX=9606 GN=IGHA1 PE=1 SV=2 | 255 | 20201119_15232_1ng_glyco_hcd203040_compound15.21134.21134.4 | controllerType=0 controllerNumber=1 scan=21134 | 42.7015 |
| 17487:4 | 5 | R.LSLHRPALEDLLLGSEAN[+1501.555]LTC[+57.021]TLTGLR.D | HexNAc(5)Hex(3) | C[+57], N[+1502] | 1117.0484 | 4 | 4465.1717 | 4465.1536 | 0 | 4.1 | 127 | Specific | 770.74 | 767.75 | 767.75 | 15.35 | 60 | >sp\|P01876\|IGHA1_HUMAN Immunoglobulin heavy constant alpha 1 OS=Homo sapiens OX=9606 GN=IGHA1 PE=1 SV=2 | 255 | 20201119_15232_1ng_glyco_hcd203040_compound15.21127.21127.4 | controllerType=0 controllerNumber=1 scan=21127 | 42.6845 |
| 17509:4 | 5 | R.LSLHRPALEDLLLGSEAN[+1298.476]LTC[+57.021]TLTGLR.D | HexNAc(4)Hex(3) | C[+57], N[+1298] | 1066.2759 | 4 | 4262.0819 | 4262.0742 | 0 | 1.8 | 127 | Specific | 773.43 | 769.05 | 769.05 | 15.51 | 60 | >sp\|P01876\|IGHA1_HUMAN Immunoglobulin heavy constant alpha 1 OS=Homo sapiens OX=9606 GN=IGHA1 PE=1 SV=2 | 255 | 20201119_15232_1ng_glyco_hcd203040_compound15.21150.21150.4 | controllerType=0 controllerNumber=1 scan=21150 | 42.7432 |
| 17440:4 | 5 | R.LSLHRPALEDLLLGSEAN[+1663.608]LTC[+57.021]TLTGLR.D | HexNAc(5)Hex(4) | C[+57], N[+1664] | 1157.5613 | 4 | 4627.2235 | 4627.2064 | 0 | 3.7 | 127 | Specific | 778.77 | 764.57 | 764.57 | 15.57 | 60 | >sp\|P01876\|IGHA1_HUMAN Immunoglobulin heavy constant alpha 1 OS=Homo sapiens OX=9606 GN=IGHA1 PE=1 SV=2 | 255 | 20201119_15232_1ng_glyco_hcd203040_compound15.21078.21078.4 | controllerType=0 controllerNumber=1 scan=21078 | 42.5636 |
| 17451:4 | 5 | R.LSLHRPALEDLLLGSEAN[+1622.582]LTC[+57.021]TLTGLR.D | HexNAc(4)Hex(5) | C[+57], N[+1623] | 1147.3042 | 4 | 4586.1951 | 4586.1799 | 0 | 3.3 | 127 | Specific | 838.26 | 737.99 | 737.99 | 16.57 | 60 | >sp\|P01876\|IGHA1_HUMAN Immunoglobulin heavy constant alpha 1 OS=Homo sapiens OX=9606 GN=IGHA1 PE=1 SV=2 | 255 | 20201119_15232_1ng_glyco_hcd203040_compound15.21089.21089.4 | controllerType=0 controllerNumber=1 scan=21089 | 42.587 |
| 17496:4 | 5 | R.LSLHRPALEDLLLGSEAN[+1460.529]LTC[+57.021]TLTGLR.D | HexNAc(4)Hex(4) | C[+57], N[+1461] | 1106.7924 | 4 | 4424.1478 | 4424.1271 | 0 | 4.7 | 127 | Specific | 842.29 | 787.83 | 787.83 | 16.35 | 60 | >sp\|P01876\|IGHA1_HUMAN Immunoglobulin heavy constant alpha 1 OS=Homo sapiens OX=9606 GN=IGHA1 PE=1 SV=2 | 255 | 20201119_15232_1ng_glyco_hcd203040_compound15.21136.21136.4 | controllerType=0 controllerNumber=1 scan=21136 | 42.707 |
| 18427:4 | 5 | R.LSLHRPALEDLLLGSEAN[+1872.651]LTC[+57.021]TLTGLR.D | HexNAc(3)Hex(6)NeuAc(1) | C[+57], N[+1873] | 1209.8227 | 4 | 4836.2689 | 4836.2487 | 0 | 4.2 | 127 | Specific | 842.93 | 799.37 | 799.37 | 16.35 | 60 | >sp\|P01876\|IGHA1_HUMAN Immunoglobulin heavy constant alpha 1 OS=Homo sapiens OX=9606 GN=IGHA1 PE=1 SV=2 | 255 | 20201119_15232_1ng_glyco_hcd203040_compound15.22107.22107.4 | controllerType=0 controllerNumber=1 scan=22107 | 44.8004 |
| 18472:4 | 5 | R.LSLHRPALEDLLLGSEAN[+1710.598]LTC[+57.021]TLTGLR.D | HexNAc(3)Hex(5)NeuAc(1) | C[+57], N[+1711] | 1169.31 | 4 | 4674.2181 | 4674.1959 | 0 | 4.7 | 127 | Specific | 855.87 | 823.32 | 823.32 | 17.35 | 60 | >sp\|P01876\|IGHA1_HUMAN Immunoglobulin heavy constant alpha 1 OS=Homo sapiens OX=9606 GN=IGHA1 PE=1 SV=2 | 255 | 20201119_15232_1ng_glyco_hcd203040_compound15.22154.22154.4 | controllerType=0 controllerNumber=1 scan=22154 | 44.9048 |
| 17512:4 | 5 | R.LSLHRPALEDLLLGSEAN[+1298.476]LTC[+57.021]TLTGLR.D | HexNAc(4)Hex(3) | C[+57], N[+1298] | 1066.2759 | 4 | 4262.0819 | 4262.0742 | 0 | 1.8 | 127 | Specific | 878.68 | 868.69 | 868.69 | 17.51 | 60 | >sp\|P01876\|IGHA1_HUMAN Immunoglobulin heavy constant alpha 1 OS=Homo sapiens OX=9606 GN=IGHA1 PE=1 SV=2 | 255 | 20201119_15232_1ng_glyco_hcd203040_compound15.21153.21153.4 | controllerType=0 controllerNumber=1 scan=21153 | 42.7542 |
| 17425:4 | 5 | R.LSLHRPALEDLLLGSEAN[+1540.529]LTC[+57.021]TLTGLR.D | HexNAc(2)Hex(7) | C[+57], N[+1541] | 1126.7908 | 4 | 4504.1414 | 4504.1268 | 0 | 3.3 | 127 | Specific | 881.27 | 825.44 | 825.44 | 17.57 | 60 | >sp\|P01876\|IGHA1_HUMAN Immunoglobulin heavy constant alpha 1 OS=Homo sapiens OX=9606 GN=IGHA1 PE=1 SV=2 | 255 | 20201119_15232_1ng_glyco_hcd203040_compound15.21062.21062.4 | controllerType=0 controllerNumber=1 scan=21062 | 42.5049 |
| 17492:4 | 5 | R.LSLHRPALEDLLLGSEAN[+1419.502]LTC[+57.021]TLTGLR.D | HexNAc(3)Hex(5) | C[+57], N[+1420] | 1096.5347 | 4 | 4383.117 | 4383.1005 | 0 | 3.8 | 127 | Specific | 892.3 | 849.04 | 849.04 | 17.57 | 60 | >sp\|P01876\|IGHA1_HUMAN Immunoglobulin heavy constant alpha 1 OS=Homo sapiens OX=9606 GN=IGHA1 PE=1 SV=2 | 255 | 20201119_15232_1ng_glyco_hcd203040_compound15.21132.21132.4 | controllerType=0 controllerNumber=1 scan=21132 | 42.6966 |
| 17577:4 | 5 | R.LSLHRPALEDLLLGSEAN[+1054.370]LTC[+57.021]TLTGLR.D | HexNAc(2)Hex(4) | C[+57], N[+1054] | 1005.2507 | 4 | 4017.9808 | 4017.9683 | 0 | 3.1 | 127 | Specific | 908.82 | 868.44 | 868.44 | 18.57 | 60 | >sp\|P01876\|IGHA1_HUMAN Immunoglobulin heavy constant alpha 1 OS=Homo sapiens OX=9606 GN=IGHA1 PE=1 SV=2 | 255 | 20201119_15232_1ng_glyco_hcd203040_compound15.21221.21221.4 | controllerType=0 controllerNumber=1 scan=21221 | 42.9188 |
| 17498:4 | 5 | R.LSLHRPALEDLLLGSEAN[+1216.423]LTC[+57.021]TLTGLR.D | HexNAc(2)Hex(5) | C[+57], N[+1216] | 1045.7635 | 4 | 4180.0321 | 4180.0211 | 0 | 2.6 | 127 | Specific | 914.02 | 914.02 | 914.02 | 18.51 | 60 | >sp\|P01876\|IGHA1_HUMAN Immunoglobulin heavy constant alpha 1 OS=Homo sapiens OX=9606 GN=IGHA1 PE=1 SV=2 | 255 | 20201119_15232_1ng_glyco_hcd203040_compound15.21139.21139.4 | controllerType=0 controllerNumber=1 scan=21139 | 42.7154 |
| 17524:4 | 5 | R.LSLHRPALEDLLLGSEAN[+1257.449]LTC[+57.021]TLTGLR.D | HexNAc(3)Hex(4) | C[+57], N[+1257] | 1056.021 | 4 | 4221.0623 | 4221.0477 | 0 | 3.5 | 127 | Specific | 915.29 | 871.5 | 871.5 | 18.57 | 60 | >sp\|P01876\|IGHA1_HUMAN Immunoglobulin heavy constant alpha 1 OS=Homo sapiens OX=9606 GN=IGHA1 PE=1 SV=2 | 255 | 20201119_15232_1ng_glyco_hcd203040_compound15.21166.21166.4 | controllerType=0 controllerNumber=1 scan=21166 | 42.7813 |
| 17445:4 | 5 | R.LSLHRPALEDLLLGSEAN[+1378.476]LTC[+57.021]TLTGLR.D | HexNAc(2)Hex(6) | C[+57], N[+1378] | 1086.278 | 4 | 4342.0902 | 4342.074 | 0 | 3.7 | 127 | Specific | 921 | 843.17 | 843.17 | 18.57 | 60 | >sp\|P01876\|IGHA1_HUMAN Immunoglobulin heavy constant alpha 1 OS=Homo sapiens OX=9606 GN=IGHA1 PE=1 SV=2 | 255 | 20201119_15232_1ng_glyco_hcd203040_compound15.21083.21083.4 | controllerType=0 controllerNumber=1 scan=21083 | 42.5701 |
| 17420:4 | 5 | R.LSLHRPALEDLLLGSEAN[+1702.581]LTC[+57.021]TLTGLR.D | HexNAc(2)Hex(8) | C[+57], N[+1703] | 1167.3045 | 4 | 4666.1961 | 4666.1796 | 0 | 3.5 | 127 | Specific | 923.9 | 923.9 | 923.9 | 18.57 | 60 | >sp\|P01876\|IGHA1_HUMAN Immunoglobulin heavy constant alpha 1 OS=Homo sapiens OX=9606 GN=IGHA1 PE=1 SV=2 | 255 | 20201119_15232_1ng_glyco_hcd203040_compound15.21057.21057.4 | controllerType=0 controllerNumber=1 scan=21057 | 42.4915 |
| 17397:4 | 5 | R.LSLHRPALEDLLLGSEAN[+1864.634]LTC[+57.021]TLTGLR.D | HexNAc(2)Hex(9) | C[+57], N[+1865] | 1207.82 | 4 | 4828.2581 | 4828.2324 | 0 | 5.3 | 127 | Specific | 943.94 | 943.94 | 943.94 | 17.8 | 60 | >sp\|P01876\|IGHA1_HUMAN Immunoglobulin heavy constant alpha 1 OS=Homo sapiens OX=9606 GN=IGHA1 PE=1 SV=2 | 255 | 20201119_15232_1ng_glyco_hcd203040_compound15.21033.21033.4 | controllerType=0 controllerNumber=1 scan=21033 | 42.4343 |
| 17399:4 | 5 | R.LSLHRPALEDLLLGSEAN[+1864.634]LTC[+57.021]TLTGLR.D | HexNAc(2)Hex(9) | C[+57], N[+1865] | 1207.82 | 4 | 4828.2581 | 4828.2324 | 0 | 5.3 | 127 | Specific | 968.7 | 968.7 | 968.7 | 18.8 | 60 | >sp\|P01876\|IGHA1_HUMAN Immunoglobulin heavy constant alpha 1 OS=Homo sapiens OX=9606 GN=IGHA1 PE=1 SV=2 | 255 | 20201119_15232_1ng_glyco_hcd203040_compound15.21035.21035.4 | controllerType=0 controllerNumber=1 scan=21035 | 42.4403 |
| 17500:4 | 5 | R.LSLHRPALEDLLLGSEAN[+1216.423]LTC[+57.021]TLTGLR.D | HexNAc(2)Hex(5) | C[+57], N[+1216] | 1045.7635 | 4 | 4180.0321 | 4180.0211 | 0 | 2.6 | 127 | Specific | 974.03 | 974.03 | 974.03 | 19.51 | 60 | >sp\|P01876\|IGHA1_HUMAN Immunoglobulin heavy constant alpha 1 OS=Homo sapiens OX=9606 GN=IGHA1 PE=1 SV=2 | 255 | 20201119_15232_1ng_glyco_hcd203040_compound15.21141.21141.4 | controllerType=0 controllerNumber=1 scan=21141 | 42.7219 |
| 17422:4 | 5 | R.LSLHRPALEDLLLGSEAN[+1702.581]LTC[+57.021]TLTGLR.D | HexNAc(2)Hex(8) | C[+57], N[+1703] | 1167.3045 | 4 | 4666.1961 | 4666.1796 | 0 | 3.5 | 127 | Specific | 983.53 | 927.65 | 927.65 | 19.57 | 60 | >sp\|P01876\|IGHA1_HUMAN Immunoglobulin heavy constant alpha 1 OS=Homo sapiens OX=9606 GN=IGHA1 PE=1 SV=2 | 255 | 20201119_15232_1ng_glyco_hcd203040_compound15.21059.21059.4 | controllerType=0 controllerNumber=1 scan=21059 | 42.4963 |
| 17428:4 | 5 | R.LSLHRPALEDLLLGSEAN[+1540.529]LTC[+57.021]TLTGLR.D | HexNAc(2)Hex(7) | C[+57], N[+1541] | 1126.7908 | 4 | 4504.1414 | 4504.1268 | 0 | 3.3 | 127 | Specific | 987.8 | 932.85 | 932.85 | 19.57 | 60 | >sp\|P01876\|IGHA1_HUMAN Immunoglobulin heavy constant alpha 1 OS=Homo sapiens OX=9606 GN=IGHA1 PE=1 SV=2 | 255 | 20201119_15232_1ng_glyco_hcd203040_compound15.21065.21065.4 | controllerType=0 controllerNumber=1 scan=21065 | 42.5219 |
| 17448:4 | 5 | R.LSLHRPALEDLLLGSEAN[+1378.476]LTC[+57.021]TLTGLR.D | HexNAc(2)Hex(6) | C[+57], N[+1378] | 1086.278 | 4 | 4342.0902 | 4342.074 | 0 | 3.7 | 127 | Specific | 1023.42 | 1023.42 | 1023.42 | 20.57 | 60 | >sp\|P01876\|IGHA1_HUMAN Immunoglobulin heavy constant alpha 1 OS=Homo sapiens OX=9606 GN=IGHA1 PE=1 SV=2 | 255 | 20201119_15232_1ng_glyco_hcd203040_compound15.21086.21086.4 | controllerType=0 controllerNumber=1 scan=21086 | 42.5795 |
| 12833:3 | 5 | R.LAGKPTHVN[+2263.835]VSVVMAEVDGTC[+57.021].Y | HexNAc(5)Hex(5)Fuc(3) | C[+57], N[+2264] | 1483.3096 | 3 | 4447.9143 | 4447.9135 | 0 | 0.2 | 332 | CRagged | 214.45 | 192.39 | 192.39 | 2.46 | 60 | >sp\|P01876\|IGHA1_HUMAN Immunoglobulin heavy constant alpha 1 OS=Homo sapiens OX=9606 GN=IGHA1 PE=1 SV=2 | 255 | 20201119_15232_1ng_glyco_hcd203040_compound15.16240.16240.3 | controllerType=0 controllerNumber=1 scan=16240 | 33.7116 |
| 12831:3 | 5 | R.LAGKPTHVN[+2263.835]VSVVMAEVDGTC[+57.021].Y | HexNAc(5)Hex(5)Fuc(3) | C[+57], N[+2264] | 1483.3096 | 3 | 4447.9143 | 4447.9135 | 0 | 0.2 | 332 | CRagged | 253.2 | 252.31 | 252.31 | 5.41 | 60 | >sp\|P01876\|IGHA1_HUMAN Immunoglobulin heavy constant alpha 1 OS=Homo sapiens OX=9606 GN=IGHA1 PE=1 SV=2 | 255 | 20201119_15232_1ng_glyco_hcd203040_compound15.16238.16238.3 | controllerType=0 controllerNumber=1 scan=16238 | 33.6914 |
| 11198:4 | 5 | R.LAGKPTHVN[+1647.613]VSVVMAEVDGTC[+57.021]Y.- | HexNAc(5)Hex(3)Fuc(1) | C[+57], N[+1648] | 999.4455 | 4 | 3994.7601 | 3994.7553 | 0 | 1.2 | 332 | CRagged | 214.84 | 214.84 | 214.84 | 4.53 | 60 | >sp\|P01876\|IGHA1_HUMAN Immunoglobulin heavy constant alpha 1 OS=Homo sapiens OX=9606 GN=IGHA1 PE=1 SV=2 | 255 | 20201119_15232_1ng_glyco_hcd203040_compound15.14518.14518.4 | controllerType=0 controllerNumber=1 scan=14518 | 30.8558 |
| 11129:4 | 5 | R.LAGKPTHVN[+1809.666]VSVVMAEVDGTC[+57.021]Y.- | HexNAc(5)Hex(4)Fuc(1) | C[+57], N[+1810] | 1039.9595 | 4 | 4156.8162 | 4156.8081 | 0 | 1.9 | 332 | CRagged | 284.26 | 194.79 | 194.79 | 4.16 | 60 | >sp\|P01876\|IGHA1_HUMAN Immunoglobulin heavy constant alpha 1 OS=Homo sapiens OX=9606 GN=IGHA1 PE=1 SV=2 | 255 | 20201119_15232_1ng_glyco_hcd203040_compound15.14446.14446.4 | controllerType=0 controllerNumber=1 scan=14446 | 30.7105 |
| 11201:3 | 5 | R.LAGKPTHVN[+1216.423]VSVVMAEVDGTC[+57.021]Y.- | HexNAc(2)Hex(5) | C[+57], N[+1216] | 1188.5214 | 3 | 3563.5497 | 3563.5649 | 0 | -4.3 | 332 | CRagged | 287.15 | 287.15 | 245.06 | 5.59 | 60 | >sp\|P01876\|IGHA1_HUMAN Immunoglobulin heavy constant alpha 1 OS=Homo sapiens OX=9606 GN=IGHA1 PE=1 SV=2 | 255 | 20201119_15232_1ng_glyco_hcd203040_compound15.14521.14521.3 | controllerType=0 controllerNumber=1 scan=14521 | 30.8678 |
| 14825:3 | 5 | R.LAGKPTHVN[+2350.830]VSVVMAEVDGTC[+57.021]Y.- | HexNAc(4)Hex(5)Fuc(1)NeuAc(2) | C[+57], N[+2351] | 1566.6639 | 3 | 4697.9771 | 4697.9724 | 0 | 1 | 332 | CRagged | 348.14 | 221.85 | 221.85 | 6.8 | 60 | >sp\|P01876\|IGHA1_HUMAN Immunoglobulin heavy constant alpha 1 OS=Homo sapiens OX=9606 GN=IGHA1 PE=1 SV=2 | 255 | 20201119_15232_1ng_glyco_hcd203040_compound15.18303.18303.3 | controllerType=0 controllerNumber=1 scan=18303 | 37.3179 |
| 11250:3 | 5 | R.LAGKPTHVN[+1095.397]VSVVMAEVDGTC[+57.021]Y.- | HexNAc(3)Hex(3) | C[+57], N[+1095] | 1148.1895 | 3 | 3442.5539 | 3442.5387 | 0 | 4.4 | 332 | CRagged | 355.74 | 350.17 | 350.17 | 6.05 | 60 | >sp\|P01876\|IGHA1_HUMAN Immunoglobulin heavy constant alpha 1 OS=Homo sapiens OX=9606 GN=IGHA1 PE=1 SV=2 | 255 | 20201119_15232_1ng_glyco_hcd203040_compound15.14572.14572.3 | controllerType=0 controllerNumber=1 scan=14572 | 30.9596 |
| 11143:3 | 5 | R.LAGKPTHVN[+1809.666]VSVVMAEVDGTC[+57.021]Y.- | HexNAc(5)Hex(4)Fuc(1) | C[+57], N[+1810] | 1386.2812 | 3 | 4156.829 | 4156.8081 | 0 | 5 | 332 | CRagged | 373.11 | 373.11 | 311.08 | 7.88 | 60 | >sp\|P01876\|IGHA1_HUMAN Immunoglobulin heavy constant alpha 1 OS=Homo sapiens OX=9606 GN=IGHA1 PE=1 SV=2 | 255 | 20201119_15232_1ng_glyco_hcd203040_compound15.14461.14461.3 | controllerType=0 controllerNumber=1 scan=14461 | 30.7391 |
| 14832:4 | 5 | R.LAGKPTHVN[+2553.910]VSVVMAEVDGTC[+57.021]Y.- | HexNAc(5)Hex(5)Fuc(1)NeuAc(2) | C[+57], N[+2554] | 1226.023 | 4 | 4901.0701 | 4901.0518 | 0 | 3.7 | 332 | CRagged | 405.03 | 306.47 | 306.47 | 7.76 | 60 | >sp\|P01876\|IGHA1_HUMAN Immunoglobulin heavy constant alpha 1 OS=Homo sapiens OX=9606 GN=IGHA1 PE=1 SV=2 | 255 | 20201119_15232_1ng_glyco_hcd203040_compound15.18311.18311.4 | controllerType=0 controllerNumber=1 scan=18311 | 37.334 |
| 14823:3 | 5 | R.LAGKPTHVN[+2350.830]VSVVMAEVDGTC[+57.021]Y.- | HexNAc(4)Hex(5)Fuc(1)NeuAc(2) | C[+57], N[+2351] | 1566.6639 | 3 | 4697.9771 | 4697.9724 | 0 | 1 | 332 | CRagged | 414.98 | 351.21 | 351.21 | 7.99 | 60 | >sp\|P01876\|IGHA1_HUMAN Immunoglobulin heavy constant alpha 1 OS=Homo sapiens OX=9606 GN=IGHA1 PE=1 SV=2 | 255 | 20201119_15232_1ng_glyco_hcd203040_compound15.18301.18301.3 | controllerType=0 controllerNumber=1 scan=18301 | 37.3144 |
| 13009:3 | 5 | R.LAGKPTHVN[+1548.545]VSVVMAEVDGTC[+57.021]Y.- | HexNAc(3)Hex(4)NeuAc(1) | C[+57], N[+1549] | 1299.2358 | 3 | 3895.6928 | 3895.6869 | 0 | 1.5 | 332 | CRagged | 439.03 | 439.03 | 439.03 | 9.66 | 60 | >sp\|P01876\|IGHA1_HUMAN Immunoglobulin heavy constant alpha 1 OS=Homo sapiens OX=9606 GN=IGHA1 PE=1 SV=2 | 255 | 20201119_15232_1ng_glyco_hcd203040_compound15.16422.16422.3 | controllerType=0 controllerNumber=1 scan=16422 | 34.0372 |
| 11203:3 | 5 | R.LAGKPTHVN[+1216.423]VSVVMAEVDGTC[+57.021]Y.- | HexNAc(2)Hex(5) | C[+57], N[+1216] | 1188.5214 | 3 | 3563.5497 | 3563.5649 | 0 | -4.3 | 332 | CRagged | 439.95 | 414.2 | 284.59 | 8.19 | 60 | >sp\|P01876\|IGHA1_HUMAN Immunoglobulin heavy constant alpha 1 OS=Homo sapiens OX=9606 GN=IGHA1 PE=1 SV=2 | 255 | 20201119_15232_1ng_glyco_hcd203040_compound15.14523.14523.3 | controllerType=0 controllerNumber=1 scan=14523 | 30.8734 |
| 14834:4 | 5 | R.LAGKPTHVN[+2553.910]VSVVMAEVDGTC[+57.021]Y.- | HexNAc(5)Hex(5)Fuc(1)NeuAc(2) | C[+57], N[+2554] | 1226.023 | 4 | 4901.0701 | 4901.0518 | 0 | 3.7 | 332 | CRagged | 498.73 | 378.99 | 378.99 | 9.91 | 60 | >sp\|P01876\|IGHA1_HUMAN Immunoglobulin heavy constant alpha 1 OS=Homo sapiens OX=9606 GN=IGHA1 PE=1 SV=2 | 255 | 20201119_15232_1ng_glyco_hcd203040_compound15.18313.18313.4 | controllerType=0 controllerNumber=1 scan=18313 | 37.3378 |
| 13010:3 | 5 | R.LAGKPTHVN[+1548.545]VSVVMAEVDGTC[+57.021]Y.- | HexNAc(3)Hex(4)NeuAc(1) | C[+57], N[+1549] | 1299.2358 | 3 | 3895.6928 | 3895.6869 | 0 | 1.5 | 332 | CRagged | 583.11 | 583.11 | 583.11 | 11.49 | 60 | >sp\|P01876\|IGHA1_HUMAN Immunoglobulin heavy constant alpha 1 OS=Homo sapiens OX=9606 GN=IGHA1 PE=1 SV=2 | 255 | 20201119_15232_1ng_glyco_hcd203040_compound15.16423.16423.3 | controllerType=0 controllerNumber=1 scan=16423 | 34.0412 |
| 11145:3 | 5 | R.LAGKPTHVN[+1809.666]VSVVMAEVDGTC[+57.021]Y.- | HexNAc(5)Hex(4)Fuc(1) | C[+57], N[+1810] | 1386.2812 | 3 | 4156.829 | 4156.8081 | 0 | 5 | 332 | CRagged | 654.31 | 515.84 | 449.62 | 13.47 | 60 | >sp\|P01876\|IGHA1_HUMAN Immunoglobulin heavy constant alpha 1 OS=Homo sapiens OX=9606 GN=IGHA1 PE=1 SV=2 | 255 | 20201119_15232_1ng_glyco_hcd203040_compound15.14463.14463.3 | controllerType=0 controllerNumber=1 scan=14463 | 30.7428 |
| 11147:3 | 5 | R.LAGKPTHVN[+1501.555]VSVVMAEVDGTC[+57.021]Y.- | HexNAc(5)Hex(3) | C[+57], N[+1502] | 1283.5782 | 3 | 3848.72 | 3848.6974 | 0 | 5.9 | 332 | CRagged | 681.39 | 549.48 | 549.48 | 12.93 | 60 | >sp\|P01876\|IGHA1_HUMAN Immunoglobulin heavy constant alpha 1 OS=Homo sapiens OX=9606 GN=IGHA1 PE=1 SV=2 | 255 | 20201119_15232_1ng_glyco_hcd203040_compound15.14465.14465.3 | controllerType=0 controllerNumber=1 scan=14465 | 30.7459 |
| 11094:3 | 5 | R.LAGKPTHVN[+1378.476]VSVVMAEVDGTC[+57.021]Y.- | HexNAc(2)Hex(6) | C[+57], N[+1378] | 1242.5475 | 3 | 3725.6281 | 3725.6178 | 0 | 2.8 | 332 | CRagged | 693.29 | 693.29 | 449.73 | 13.7 | 60 | >sp\|P01876\|IGHA1_HUMAN Immunoglobulin heavy constant alpha 1 OS=Homo sapiens OX=9606 GN=IGHA1 PE=1 SV=2 | 255 | 20201119_15232_1ng_glyco_hcd203040_compound15.14410.14410.3 | controllerType=0 controllerNumber=1 scan=14410 | 30.6385 |
| 11066:3 | 5 | R.LAGKPTHVN[+1663.608]VSVVMAEVDGTC[+57.021]Y.- | HexNAc(5)Hex(4) | C[+57], N[+1664] | 1337.59 | 3 | 4010.7555 | 4010.7502 | 0 | 1.3 | 332 | CRagged | 749.92 | 675.33 | 499.05 | 14.49 | 60 | >sp\|P01876\|IGHA1_HUMAN Immunoglobulin heavy constant alpha 1 OS=Homo sapiens OX=9606 GN=IGHA1 PE=1 SV=2 | 255 | 20201119_15232_1ng_glyco_hcd203040_compound15.14380.14380.3 | controllerType=0 controllerNumber=1 scan=14380 | 30.5721 |
| 11064:3 | 5 | R.LAGKPTHVN[+1663.608]VSVVMAEVDGTC[+57.021]Y.- | HexNAc(5)Hex(4) | C[+57], N[+1664] | 1337.59 | 3 | 4010.7555 | 4010.7502 | 0 | 1.3 | 332 | CRagged | 776.39 | 776.39 | 490.9 | 15.49 | 60 | >sp\|P01876\|IGHA1_HUMAN Immunoglobulin heavy constant alpha 1 OS=Homo sapiens OX=9606 GN=IGHA1 PE=1 SV=2 | 255 | 20201119_15232_1ng_glyco_hcd203040_compound15.14378.14378.3 | controllerType=0 controllerNumber=1 scan=14378 | 30.5664 |
| 10967:3 | 5 | R.LAGKPTHVN[+1702.581]VSVVMAEVDGTC[+57.021]Y.- | HexNAc(2)Hex(8) | C[+57], N[+1703] | 1350.5817 | 3 | 4049.7306 | 4049.7234 | 0 | 1.8 | 332 | CRagged | 785.9 | 785.9 | 463.49 | 15.64 | 60 | >sp\|P01876\|IGHA1_HUMAN Immunoglobulin heavy constant alpha 1 OS=Homo sapiens OX=9606 GN=IGHA1 PE=1 SV=2 | 255 | 20201119_15232_1ng_glyco_hcd203040_compound15.14278.14278.3 | controllerType=0 controllerNumber=1 scan=14278 | 30.3841 |
| 11152:3 | 5 | R.LAGKPTHVN[+1647.613]VSVVMAEVDGTC[+57.021]Y.- | HexNAc(5)Hex(3)Fuc(1) | C[+57], N[+1648] | 1332.2642 | 3 | 3994.7781 | 3994.7553 | 0 | 5.7 | 332 | CRagged | 793.06 | 616.52 | 616.52 | 14.93 | 60 | >sp\|P01876\|IGHA1_HUMAN Immunoglobulin heavy constant alpha 1 OS=Homo sapiens OX=9606 GN=IGHA1 PE=1 SV=2 | 255 | 20201119_15232_1ng_glyco_hcd203040_compound15.14470.14470.3 | controllerType=0 controllerNumber=1 scan=14470 | 30.7581 |
| 11150:3 | 5 | R.LAGKPTHVN[+1501.555]VSVVMAEVDGTC[+57.021]Y.- | HexNAc(5)Hex(3) | C[+57], N[+1502] | 1283.5782 | 3 | 3848.72 | 3848.6974 | 0 | 5.9 | 332 | CRagged | 818.21 | 680.99 | 680.99 | 15.93 | 60 | >sp\|P01876\|IGHA1_HUMAN Immunoglobulin heavy constant alpha 1 OS=Homo sapiens OX=9606 GN=IGHA1 PE=1 SV=2 | 255 | 20201119_15232_1ng_glyco_hcd203040_compound15.14468.14468.3 | controllerType=0 controllerNumber=1 scan=14468 | 30.7507 |
| 11149:3 | 5 | R.LAGKPTHVN[+1647.613]VSVVMAEVDGTC[+57.021]Y.- | HexNAc(5)Hex(3)Fuc(1) | C[+57], N[+1648] | 1332.2642 | 3 | 3994.7781 | 3994.7553 | 0 | 5.7 | 332 | CRagged | 839.64 | 839.64 | 839.64 | 15.93 | 60 | >sp\|P01876\|IGHA1_HUMAN Immunoglobulin heavy constant alpha 1 OS=Homo sapiens OX=9606 GN=IGHA1 PE=1 SV=2 | 255 | 20201119_15232_1ng_glyco_hcd203040_compound15.14467.14467.3 | controllerType=0 controllerNumber=1 scan=14467 | 30.7482 |
| 10998:3 | 5 | R.LAGKPTHVN[+1540.529]VSVVMAEVDGTC[+57.021]Y.- | HexNAc(2)Hex(7) | C[+57], N[+1541] | 1296.5657 | 3 | 3887.6826 | 3887.6706 | 0 | 3.1 | 332 | CRagged | 842.44 | 842.44 | 552.37 | 16.7 | 60 | >sp\|P01876\|IGHA1_HUMAN Immunoglobulin heavy constant alpha 1 OS=Homo sapiens OX=9606 GN=IGHA1 PE=1 SV=2 | 255 | 20201119_15232_1ng_glyco_hcd203040_compound15.14310.14310.3 | controllerType=0 controllerNumber=1 scan=14310 | 30.444 |
| 11102:3 | 5 | R.LAGKPTHVN[+1216.423]VSVVMAEVDGTC[+57.021]Y.- | HexNAc(2)Hex(5) | C[+57], N[+1216] | 1188.5294 | 3 | 3563.5735 | 3563.5649 | 0 | 2.4 | 332 | CRagged | 867.03 | 845.16 | 536.07 | 17.64 | 60 | >sp\|P01876\|IGHA1_HUMAN Immunoglobulin heavy constant alpha 1 OS=Homo sapiens OX=9606 GN=IGHA1 PE=1 SV=2 | 255 | 20201119_15232_1ng_glyco_hcd203040_compound15.14418.14418.3 | controllerType=0 controllerNumber=1 scan=14418 | 30.6575 |
| 10965:3 | 5 | R.LAGKPTHVN[+1702.581]VSVVMAEVDGTC[+57.021]Y.- | HexNAc(2)Hex(8) | C[+57], N[+1703] | 1350.5817 | 3 | 4049.7306 | 4049.7234 | 0 | 1.8 | 332 | CRagged | 869.93 | 869.93 | 554.24 | 17.64 | 60 | >sp\|P01876\|IGHA1_HUMAN Immunoglobulin heavy constant alpha 1 OS=Homo sapiens OX=9606 GN=IGHA1 PE=1 SV=2 | 255 | 20201119_15232_1ng_glyco_hcd203040_compound15.14276.14276.3 | controllerType=0 controllerNumber=1 scan=14276 | 30.3776 |
| 11096:3 | 5 | R.LAGKPTHVN[+1378.476]VSVVMAEVDGTC[+57.021]Y.- | HexNAc(2)Hex(6) | C[+57], N[+1378] | 1242.5475 | 3 | 3725.6281 | 3725.6178 | 0 | 2.8 | 332 | CRagged | 877.67 | 815.45 | 526.09 | 17.7 | 60 | >sp\|P01876\|IGHA1_HUMAN Immunoglobulin heavy constant alpha 1 OS=Homo sapiens OX=9606 GN=IGHA1 PE=1 SV=2 | 255 | 20201119_15232_1ng_glyco_hcd203040_compound15.14412.14412.3 | controllerType=0 controllerNumber=1 scan=14412 | 30.6422 |
| 10996:3 | 5 | R.LAGKPTHVN[+1540.529]VSVVMAEVDGTC[+57.021]Y.- | HexNAc(2)Hex(7) | C[+57], N[+1541] | 1296.5657 | 3 | 3887.6826 | 3887.6706 | 0 | 3.1 | 332 | CRagged | 891.85 | 891.85 | 578.8 | 17.7 | 60 | >sp\|P01876\|IGHA1_HUMAN Immunoglobulin heavy constant alpha 1 OS=Homo sapiens OX=9606 GN=IGHA1 PE=1 SV=2 | 255 | 20201119_15232_1ng_glyco_hcd203040_compound15.14308.14308.3 | controllerType=0 controllerNumber=1 scan=14308 | 30.4376 |
| 11099:3 | 5 | R.LAGKPTHVN[+1216.423]VSVVMAEVDGTC[+57.021]Y.- | HexNAc(2)Hex(5) | C[+57], N[+1216] | 1188.5294 | 3 | 3563.5735 | 3563.5649 | 0 | 2.4 | 332 | CRagged | 922.96 | 922.96 | 567.44 | 18.64 | 60 | >sp\|P01876\|IGHA1_HUMAN Immunoglobulin heavy constant alpha 1 OS=Homo sapiens OX=9606 GN=IGHA1 PE=1 SV=2 | 255 | 20201119_15232_1ng_glyco_hcd203040_compound15.14415.14415.3 | controllerType=0 controllerNumber=1 scan=14415 | 30.6511 |
| 18711:3 | 6 | K.QIGLYPVLVIDSSGYVNPN[+1913.677]YTGR.I | HexNAc(4)Hex(5)NeuAc(1) | N[+1914] | 1480.3383 | 3 | 4439.0004 | 4438.9805 | 0 | 4.5 | 168 | Specific | 428.2 | 379.32 | 379.32 | 9.59 | 60 | >sp\|P01833\|PIGR_HUMAN Polymeric immunoglobulin receptor OS=Homo sapiens OX=9606 GN=PIGR PE=1 SV=4 | 902 | 20201119_15232_1ng_glyco_hcd203040_compound15.22404.22404.3 | controllerType=0 controllerNumber=1 scan=22404 | 45.498 |
| 18713:3 | 6 | K.QIGLYPVLVIDSSGYVNPN[+1913.677]YTGR.I | HexNAc(4)Hex(5)NeuAc(1) | N[+1914] | 1480.3383 | 3 | 4439.0004 | 4438.9805 | 0 | 4.5 | 168 | Specific | 477.77 | 396.65 | 396.65 | 9.68 | 60 | >sp\|P01833\|PIGR_HUMAN Polymeric immunoglobulin receptor OS=Homo sapiens OX=9606 GN=PIGR PE=1 SV=4 | 902 | 20201119_15232_1ng_glyco_hcd203040_compound15.22406.22406.3 | controllerType=0 controllerNumber=1 scan=22406 | 45.5014 |
| 18638:4 | 6 | K.QIGLYPVLVIDSSGYVNPN[+2059.735]YTGR.I | HexNAc(4)Hex(5)Fuc(1)NeuAc(1) | N[+2060] | 1147.0182 | 4 | 4585.0511 | 4585.0384 | 0 | 2.8 | 168 | Specific | 485.83 | 485.83 | 485.83 | 10.08 | 60 | >sp\|P01833\|PIGR_HUMAN Polymeric immunoglobulin receptor OS=Homo sapiens OX=9606 GN=PIGR PE=1 SV=4 | 902 | 20201119_15232_1ng_glyco_hcd203040_compound15.22328.22328.4 | controllerType=0 controllerNumber=1 scan=22328 | 45.331 |
| 18641:4 | 6 | K.QIGLYPVLVIDSSGYVNPN[+2059.735]YTGR.I | HexNAc(4)Hex(5)Fuc(1)NeuAc(1) | N[+2060] | 1147.0182 | 4 | 4585.0511 | 4585.0384 | 0 | 2.8 | 168 | Specific | 923.31 | 795.01 | 795.01 | 18.82 | 60 | >sp\|P01833\|PIGR_HUMAN Polymeric immunoglobulin receptor OS=Homo sapiens OX=9606 GN=PIGR PE=1 SV=4 | 902 | 20201119_15232_1ng_glyco_hcd203040_compound15.22331.22331.4 | controllerType=0 controllerNumber=1 scan=22331 | 45.3352 |
| 20321:4 | 6 | R.LSLLEEPGN[+2076.750]GTFTVILNQLTSR.D | HexNAc(4)Hex(6)Fuc(2) | N[+2077] | 1120.5216 | 4 | 4479.0648 | 4479.0428 | 0 | 4.9 | 413 | Specific | 255.7 | 239.13 | 239.13 | 5.18 | 60 | >sp\|P01833\|PIGR_HUMAN Polymeric immunoglobulin receptor OS=Homo sapiens OX=9606 GN=PIGR PE=1 SV=4 | 902 | 20201119_15232_1ng_glyco_hcd203040_compound15.24086.24086.4 | controllerType=0 controllerNumber=1 scan=24086 | 48.9088 |
| 20323:4 | 6 | R.LSLLEEPGN[+2076.750]GTFTVILNQLTSR.D | HexNAc(4)Hex(6)Fuc(2) | N[+2077] | 1120.5216 | 4 | 4479.0648 | 4479.0428 | 0 | 4.9 | 413 | Specific | 308.1 | 239.2 | 239.2 | 5.49 | 60 | >sp\|P01833\|PIGR_HUMAN Polymeric immunoglobulin receptor OS=Homo sapiens OX=9606 GN=PIGR PE=1 SV=4 | 902 | 20201119_15232_1ng_glyco_hcd203040_compound15.24088.24088.4 | controllerType=0 controllerNumber=1 scan=24088 | 48.9178 |
| 20286:4 | 6 | R.LSLLEEPGN[+2205.793]GTFTVILNQLTSR.D | HexNAc(4)Hex(5)Fuc(2)NeuAc(1) | N[+2206] | 1152.783 | 4 | 4608.1102 | 4608.0854 | 0 | 5.4 | 413 | Specific | 736.78 | 556.32 | 556.32 | 14.05 | 60 | >sp\|P01833\|PIGR_HUMAN Polymeric immunoglobulin receptor OS=Homo sapiens OX=9606 GN=PIGR PE=1 SV=4 | 902 | 20201119_15232_1ng_glyco_hcd203040_compound15.24050.24050.4 | controllerType=0 controllerNumber=1 scan=24050 | 48.8297 |
| 20267:4 | 6 | R.LSLLEEPGN[+2205.793]GTFTVILNQLTSR.D | HexNAc(4)Hex(5)Fuc(2)NeuAc(1) | N[+2206] | 1152.7842 | 4 | 4608.1151 | 4608.0854 | 0 | 6.4 | 413 | Specific | 875.33 | 688.48 | 688.48 | 16.77 | 60 | >sp\|P01833\|PIGR_HUMAN Polymeric immunoglobulin receptor OS=Homo sapiens OX=9606 GN=PIGR PE=1 SV=4 | 902 | 20201119_15232_1ng_glyco_hcd203040_compound15.24030.24030.3 | controllerType=0 controllerNumber=1 scan=24030 | 48.7928 |
| 20270:4 | 6 | R.LSLLEEPGN[+2205.793]GTFTVILNQLTSR.D | HexNAc(4)Hex(5)Fuc(2)NeuAc(1) | N[+2206] | 1152.7842 | 4 | 4608.1151 | 4608.0854 | 0 | 6.4 | 413 | Specific | 913.64 | 724.93 | 724.93 | 17.77 | 60 | >sp\|P01833\|PIGR_HUMAN Polymeric immunoglobulin receptor OS=Homo sapiens OX=9606 GN=PIGR PE=1 SV=4 | 902 | 20201119_15232_1ng_glyco_hcd203040_compound15.24033.24033.3 | controllerType=0 controllerNumber=1 scan=24033 | 48.7978 |
| 20264:3 | 6 | R.LSLLEEPGN[+2205.793]GTFTVILNQLTSR.D | HexNAc(4)Hex(5)Fuc(2)NeuAc(1) | N[+2206] | 1536.7073 | 3 | 4608.1074 | 4608.0854 | 0 | 4.8 | 413 | Specific | 936.12 | 677.68 | 677.68 | 18.42 | 60 | >sp\|P01833\|PIGR_HUMAN Polymeric immunoglobulin receptor OS=Homo sapiens OX=9606 GN=PIGR PE=1 SV=4 | 902 | 20201119_15232_1ng_glyco_hcd203040_compound15.24027.24027.3 | controllerType=0 controllerNumber=1 scan=24027 | 48.7867 |
| 22611:4 | 6 | R.LSLLEEPGN[+2350.830]GTFTVILNQLTSR.D | HexNAc(4)Hex(5)Fuc(1)NeuAc(2) | N[+2351] | 1189.0398 | 4 | 4753.1375 | 4753.1229 | 0 | 3.1 | 413 | Specific | 946.43 | 733.77 | 733.77 | 18.82 | 60 | >sp\|P01833\|PIGR_HUMAN Polymeric immunoglobulin receptor OS=Homo sapiens OX=9606 GN=PIGR PE=1 SV=4 | 902 | 20201119_15232_1ng_glyco_hcd203040_compound15.26474.26474.4 | controllerType=0 controllerNumber=1 scan=26474 | 53.3074 |
| 20288:4 | 6 | R.LSLLEEPGN[+2205.793]GTFTVILNQLTSR.D | HexNAc(4)Hex(5)Fuc(2)NeuAc(1) | N[+2206] | 1152.783 | 4 | 4608.1102 | 4608.0854 | 0 | 5.4 | 413 | Specific | 958.2 | 718.24 | 718.24 | 19.05 | 60 | >sp\|P01833\|PIGR_HUMAN Polymeric immunoglobulin receptor OS=Homo sapiens OX=9606 GN=PIGR PE=1 SV=4 | 902 | 20201119_15232_1ng_glyco_hcd203040_compound15.24052.24052.4 | controllerType=0 controllerNumber=1 scan=24052 | 48.8343 |
| 22606:3 | 6 | R.LSLLEEPGN[+2350.830]GTFTVILNQLTSR.D | HexNAc(4)Hex(5)Fuc(1)NeuAc(2) | N[+2351] | 1585.0496 | 3 | 4753.1343 | 4753.1229 | 0 | 2.4 | 413 | Specific | 958.82 | 672.71 | 672.71 | 19.59 | 60 | >sp\|P01833\|PIGR_HUMAN Polymeric immunoglobulin receptor OS=Homo sapiens OX=9606 GN=PIGR PE=1 SV=4 | 902 | 20201119_15232_1ng_glyco_hcd203040_compound15.26469.26469.3 | controllerType=0 controllerNumber=1 scan=26469 | 53.2928 |
| 22619:4 | 6 | R.LSLLEEPGN[+2350.830]GTFTVILNQLTSR.D | HexNAc(4)Hex(5)Fuc(1)NeuAc(2) | N[+2351] | 1189.0409 | 4 | 4753.1419 | 4753.1229 | 0 | 4 | 413 | Specific | 958.88 | 710.53 | 710.53 | 19.59 | 60 | >sp\|P01833\|PIGR_HUMAN Polymeric immunoglobulin receptor OS=Homo sapiens OX=9606 GN=PIGR PE=1 SV=4 | 902 | 20201119_15232_1ng_glyco_hcd203040_compound15.26483.26483.4 | controllerType=0 controllerNumber=1 scan=26483 | 53.333 |
| 20289:3 | 6 | R.LSLLEEPGN[+2059.735]GTFTVILNQLTSR.D | HexNAc(4)Hex(5)Fuc(1)NeuAc(1) | N[+2060] | 1488.0209 | 3 | 4462.0482 | 4462.0275 | 0 | 4.6 | 413 | Specific | 993.53 | 712.52 | 712.52 | 19.42 | 60 | >sp\|P01833\|PIGR_HUMAN Polymeric immunoglobulin receptor OS=Homo sapiens OX=9606 GN=PIGR PE=1 SV=4 | 902 | 20201119_15232_1ng_glyco_hcd203040_compound15.24053.24053.3 | controllerType=0 controllerNumber=1 scan=24053 | 48.8382 |
| 20261:3 | 6 | R.LSLLEEPGN[+2205.793]GTFTVILNQLTSR.D | HexNAc(4)Hex(5)Fuc(2)NeuAc(1) | N[+2206] | 1536.7073 | 3 | 4608.1074 | 4608.0854 | 0 | 4.8 | 413 | Specific | 1020.18 | 779.77 | 779.77 | 20.42 | 60 | >sp\|P01833\|PIGR_HUMAN Polymeric immunoglobulin receptor OS=Homo sapiens OX=9606 GN=PIGR PE=1 SV=4 | 902 | 20201119_15232_1ng_glyco_hcd203040_compound15.24024.24024.3 | controllerType=0 controllerNumber=1 scan=24024 | 48.7813 |
| 22617:4 | 6 | R.LSLLEEPGN[+2350.830]GTFTVILNQLTSR.D | HexNAc(4)Hex(5)Fuc(1)NeuAc(2) | N[+2351] | 1189.0409 | 4 | 4753.1419 | 4753.1229 | 0 | 4 | 413 | Specific | 1031.89 | 819.94 | 819.94 | 20.59 | 60 | >sp\|P01833\|PIGR_HUMAN Polymeric immunoglobulin receptor OS=Homo sapiens OX=9606 GN=PIGR PE=1 SV=4 | 902 | 20201119_15232_1ng_glyco_hcd203040_compound15.26481.26481.4 | controllerType=0 controllerNumber=1 scan=26481 | 53.3245 |
| 22609:4 | 6 | R.LSLLEEPGN[+2350.830]GTFTVILNQLTSR.D | HexNAc(4)Hex(5)Fuc(1)NeuAc(2) | N[+2351] | 1189.0398 | 4 | 4753.1375 | 4753.1229 | 0 | 3.1 | 413 | Specific | 1053.13 | 832.4 | 832.4 | 21.82 | 60 | >sp\|P01833\|PIGR_HUMAN Polymeric immunoglobulin receptor OS=Homo sapiens OX=9606 GN=PIGR PE=1 SV=4 | 902 | 20201119_15232_1ng_glyco_hcd203040_compound15.26472.26472.4 | controllerType=0 controllerNumber=1 scan=26472 | 53.2976 |
| 20287:3 | 6 | R.LSLLEEPGN[+2059.735]GTFTVILNQLTSR.D | HexNAc(4)Hex(5)Fuc(1)NeuAc(1) | N[+2060] | 1488.0209 | 3 | 4462.0482 | 4462.0275 | 0 | 4.6 | 413 | Specific | 1055.92 | 824.02 | 824.02 | 21.42 | 60 | >sp\|P01833\|PIGR_HUMAN Polymeric immunoglobulin receptor OS=Homo sapiens OX=9606 GN=PIGR PE=1 SV=4 | 902 | 20201119_15232_1ng_glyco_hcd203040_compound15.24051.24051.3 | controllerType=0 controllerNumber=1 scan=24051 | 48.8312 |
| 22604:3 | 6 | R.LSLLEEPGN[+2350.830]GTFTVILNQLTSR.D | HexNAc(4)Hex(5)Fuc(1)NeuAc(2) | N[+2351] | 1585.0496 | 3 | 4753.1343 | 4753.1229 | 0 | 2.4 | 413 | Specific | 1079.73 | 835.1 | 835.1 | 21.59 | 60 | >sp\|P01833\|PIGR_HUMAN Polymeric immunoglobulin receptor OS=Homo sapiens OX=9606 GN=PIGR PE=1 SV=4 | 902 | 20201119_15232_1ng_glyco_hcd203040_compound15.26467.26467.3 | controllerType=0 controllerNumber=1 scan=26467 | 53.2816 |
| 15778:3 | 6 | K.[+42.011]IIEGEPNLKVPGN[+2018.708]VTAVLGETLK.V | HexNAc(3)Hex(6)Fuc(1)NeuAc(1) | .[42], N[+2019] | 1484.7015 | 3 | 4452.0899 | 4452.0683 | 0 | 4.8 | 457 | Specific | 261.33 | 261.33 | 261.33 | 6.28 | 60 | >sp\|P01833\|PIGR_HUMAN Polymeric immunoglobulin receptor OS=Homo sapiens OX=9606 GN=PIGR PE=1 SV=4 | 902 | 20201119_15232_1ng_glyco_hcd203040_compound15.19337.19337.3 | controllerType=0 controllerNumber=1 scan=19337 | 39.194 |
| 15873:3 | 6 | K.IIEGEPNLKVPGN[+2205.793]VTAVLGETLK.V | HexNAc(4)Hex(5)Fuc(2)NeuAc(1) | N[+2206] | 1533.0569 | 3 | 4597.1561 | 4597.1422 | 0 | 3 | 457 | Specific | 269.48 | 269.48 | 269.48 | 6.5 | 60 | >sp\|P01833\|PIGR_HUMAN Polymeric immunoglobulin receptor OS=Homo sapiens OX=9606 GN=PIGR PE=1 SV=4 | 902 | 20201119_15232_1ng_glyco_hcd203040_compound15.19436.19436.3 | controllerType=0 controllerNumber=1 scan=19436 | 39.4054 |
| 15776:3 | 6 | K.[+42.011]IIEGEPNLKVPGN[+2018.708]VTAVLGETLK.V | HexNAc(3)Hex(6)Fuc(1)NeuAc(1) | .[42], N[+2019] | 1484.7015 | 3 | 4452.0899 | 4452.0683 | 0 | 4.8 | 457 | Specific | 374.83 | 374.83 | 374.83 | 7.4 | 60 | >sp\|P01833\|PIGR_HUMAN Polymeric immunoglobulin receptor OS=Homo sapiens OX=9606 GN=PIGR PE=1 SV=4 | 902 | 20201119_15232_1ng_glyco_hcd203040_compound15.19335.19335.3 | controllerType=0 controllerNumber=1 scan=19335 | 39.1886 |
| 15831:3 | 6 | K.IIEGEPNLKVPGN[+1913.677]VTAVLGETLK.V | HexNAc(4)Hex(5)NeuAc(1) | N[+1914] | 1435.691 | 3 | 4305.0584 | 4305.0264 | 0 | 7.4 | 457 | Specific | 382.12 | 310.79 | 310.79 | 6.27 | 60 | >sp\|P01833\|PIGR_HUMAN Polymeric immunoglobulin receptor OS=Homo sapiens OX=9606 GN=PIGR PE=1 SV=4 | 902 | 20201119_15232_1ng_glyco_hcd203040_compound15.19392.19392.3 | controllerType=0 controllerNumber=1 scan=19392 | 39.2934 |
| 15871:3 | 6 | K.IIEGEPNLKVPGN[+2205.793]VTAVLGETLK.V | HexNAc(4)Hex(5)Fuc(2)NeuAc(1) | N[+2206] | 1533.0569 | 3 | 4597.1561 | 4597.1422 | 0 | 3 | 457 | Specific | 426.46 | 426.46 | 426.46 | 9.38 | 60 | >sp\|P01833\|PIGR_HUMAN Polymeric immunoglobulin receptor OS=Homo sapiens OX=9606 GN=PIGR PE=1 SV=4 | 902 | 20201119_15232_1ng_glyco_hcd203040_compound15.19434.19434.3 | controllerType=0 controllerNumber=1 scan=19434 | 39.4014 |
| 15808:4 | 6 | K.IIEGEPNLKVPGN[+2205.793]VTAVLGETLK.V | HexNAc(4)Hex(5)Fuc(2)NeuAc(1) | N[+2206] | 1150.0456 | 4 | 4597.1605 | 4597.1422 | 0 | 4 | 457 | Specific | 440.22 | 440.22 | 440.22 | 9.11 | 60 | >sp\|P01833\|PIGR_HUMAN Polymeric immunoglobulin receptor OS=Homo sapiens OX=9606 GN=PIGR PE=1 SV=4 | 902 | 20201119_15232_1ng_glyco_hcd203040_compound15.19368.19368.4 | controllerType=0 controllerNumber=1 scan=19368 | 39.2508 |
| 15810:4 | 6 | K.IIEGEPNLKVPGN[+2205.793]VTAVLGETLK.V | HexNAc(4)Hex(5)Fuc(2)NeuAc(1) | N[+2206] | 1150.0456 | 4 | 4597.1605 | 4597.1422 | 0 | 4 | 457 | Specific | 442.13 | 417.22 | 417.22 | 9.11 | 60 | >sp\|P01833\|PIGR_HUMAN Polymeric immunoglobulin receptor OS=Homo sapiens OX=9606 GN=PIGR PE=1 SV=4 | 902 | 20201119_15232_1ng_glyco_hcd203040_compound15.19370.19370.4 | controllerType=0 controllerNumber=1 scan=19370 | 39.2546 |
| 15829:3 | 6 | K.IIEGEPNLKVPGN[+1913.677]VTAVLGETLK.V | HexNAc(4)Hex(5)NeuAc(1) | N[+1914] | 1435.691 | 3 | 4305.0584 | 4305.0264 | 0 | 7.4 | 457 | Specific | 683.27 | 683.27 | 683.27 | 12.17 | 60 | >sp\|P01833\|PIGR_HUMAN Polymeric immunoglobulin receptor OS=Homo sapiens OX=9606 GN=PIGR PE=1 SV=4 | 902 | 20201119_15232_1ng_glyco_hcd203040_compound15.19390.19390.3 | controllerType=0 controllerNumber=1 scan=19390 | 39.2898 |
| 15803:3 | 6 | K.IIEGEPNLKVPGN[+2059.735]VTAVLGETLK.V | HexNAc(4)Hex(5)Fuc(1)NeuAc(1) | N[+2060] | 1484.373 | 3 | 4451.1044 | 4451.0843 | 0 | 4.5 | 457 | Specific | 753.67 | 672.03 | 672.03 | 14.99 | 60 | >sp\|P01833\|PIGR_HUMAN Polymeric immunoglobulin receptor OS=Homo sapiens OX=9606 GN=PIGR PE=1 SV=4 | 902 | 20201119_15232_1ng_glyco_hcd203040_compound15.19363.19363.3 | controllerType=0 controllerNumber=1 scan=19363 | 39.2432 |
| 15801:3 | 6 | K.IIEGEPNLKVPGN[+2059.735]VTAVLGETLK.V | HexNAc(4)Hex(5)Fuc(1)NeuAc(1) | N[+2060] | 1484.373 | 3 | 4451.1044 | 4451.0843 | 0 | 4.5 | 457 | Specific | 936.26 | 853.43 | 853.43 | 17.99 | 60 | >sp\|P01833\|PIGR_HUMAN Polymeric immunoglobulin receptor OS=Homo sapiens OX=9606 GN=PIGR PE=1 SV=4 | 902 | 20201119_15232_1ng_glyco_hcd203040_compound15.19361.19361.3 | controllerType=0 controllerNumber=1 scan=19361 | 39.2352 |
| 15243:5 | 6 | K.IIEGEPNLKVPGN[+2059.735]VTAVLGETLKVPC[+57.021]HFPC[+57.021]K.F | HexNAc(4)Hex(5)Fuc(1)NeuAc(1) | C[+57]*2, N[+2060] | 1096.1182 | 5 | 5476.5617 | 5476.5418 | 0 | 3.6 | 457 | Specific | 250.28 | 250.28 | 250.28 | 5.62 | 60 | >sp\|P01833\|PIGR_HUMAN Polymeric immunoglobulin receptor OS=Homo sapiens OX=9606 GN=PIGR PE=1 SV=4 | 902 | 20201119_15232_1ng_glyco_hcd203040_compound15.18738.18738.3 | controllerType=0 controllerNumber=1 scan=18738 | 38.0987 |
| 15420:5 | 6 | K.IIEGEPNLKVPGN[+2059.735]VTAVLGETLKVPC[+57.021]HFPC[+57.021]K.F | HexNAc(4)Hex(5)Fuc(1)NeuAc(1) | C[+57]*2, N[+2060] | 1096.1172 | 5 | 5476.5568 | 5476.5418 | 0 | 2.7 | 457 | Specific | 350.2 | 350.2 | 350.2 | 7.53 | 60 | >sp\|P01833\|PIGR_HUMAN Polymeric immunoglobulin receptor OS=Homo sapiens OX=9606 GN=PIGR PE=1 SV=4 | 902 | 20201119_15232_1ng_glyco_hcd203040_compound15.18922.18922.2 | controllerType=0 controllerNumber=1 scan=18922 | 38.4095 |
| 15341:4 | 6 | K.IIEGEPNLKVPGN[+2059.735]VTAVLGETLKVPC[+57.021]HFPC[+57.021]K.F | HexNAc(4)Hex(5)Fuc(1)NeuAc(1) | C[+57]*2, N[+2060] | 1369.8918 | 4 | 5476.5456 | 5476.5418 | 0 | 0.7 | 457 | Specific | 369.02 | 369.02 | 369.02 | 6.81 | 60 | >sp\|P01833\|PIGR_HUMAN Polymeric immunoglobulin receptor OS=Homo sapiens OX=9606 GN=PIGR PE=1 SV=4 | 902 | 20201119_15232_1ng_glyco_hcd203040_compound15.18839.18839.2 | controllerType=0 controllerNumber=1 scan=18839 | 38.28 |
| 15245:5 | 6 | K.IIEGEPNLKVPGN[+2059.735]VTAVLGETLKVPC[+57.021]HFPC[+57.021]K.F | HexNAc(4)Hex(5)Fuc(1)NeuAc(1) | C[+57]*2, N[+2060] | 1096.1182 | 5 | 5476.5617 | 5476.5418 | 0 | 3.6 | 457 | Specific | 403.05 | 403.05 | 403.05 | 8.55 | 60 | >sp\|P01833\|PIGR_HUMAN Polymeric immunoglobulin receptor OS=Homo sapiens OX=9606 GN=PIGR PE=1 SV=4 | 902 | 20201119_15232_1ng_glyco_hcd203040_compound15.18740.18740.3 | controllerType=0 controllerNumber=1 scan=18740 | 38.1026 |
| 14056:5 | 6 | K.IIEGEPNLKVPGN[+2059.735]VTAVLGETLKVPC[+57.021]HFPC[+57.021]K.F | HexNAc(4)Hex(5)Fuc(1)NeuAc(1) | C[+57]*2, N[+2060] | 1096.1218 | 5 | 5476.58 | 5476.5418 | 0 | 7 | 457 | Specific | 454.57 | 454.57 | 454.57 | 8.4 | 60 | >sp\|P01833\|PIGR_HUMAN Polymeric immunoglobulin receptor OS=Homo sapiens OX=9606 GN=PIGR PE=1 SV=4 | 902 | 20201119_15232_1ng_glyco_hcd203040_compound15.17507.17507.5 | controllerType=0 controllerNumber=1 scan=17507 | 35.9455 |
| 14054:5 | 6 | K.IIEGEPNLKVPGN[+2059.735]VTAVLGETLKVPC[+57.021]HFPC[+57.021]K.F | HexNAc(4)Hex(5)Fuc(1)NeuAc(1) | C[+57]*2, N[+2060] | 1096.1218 | 5 | 5476.58 | 5476.5418 | 0 | 7 | 457 | Specific | 516.84 | 516.84 | 516.84 | 8.61 | 60 | >sp\|P01833\|PIGR_HUMAN Polymeric immunoglobulin receptor OS=Homo sapiens OX=9606 GN=PIGR PE=1 SV=4 | 902 | 20201119_15232_1ng_glyco_hcd203040_compound15.17505.17505.5 | controllerType=0 controllerNumber=1 scan=17505 | 35.9402 |
| 15253:4 | 6 | K.IIEGEPNLKVPGN[+2059.735]VTAVLGETLKVPC[+57.021]HFPC[+57.021]K.F | HexNAc(4)Hex(5)Fuc(1)NeuAc(1) | C[+57]*2, N[+2060] | 1369.8949 | 4 | 5476.5578 | 5476.5418 | 0 | 2.9 | 457 | Specific | 650.97 | 543.11 | 543.11 | 12.61 | 60 | >sp\|P01833\|PIGR_HUMAN Polymeric immunoglobulin receptor OS=Homo sapiens OX=9606 GN=PIGR PE=1 SV=4 | 902 | 20201119_15232_1ng_glyco_hcd203040_compound15.18748.18748.4 | controllerType=0 controllerNumber=1 scan=18748 | 38.1181 |
| 15344:4 | 6 | K.IIEGEPNLKVPGN[+2059.735]VTAVLGETLKVPC[+57.021]HFPC[+57.021]K.F | HexNAc(4)Hex(5)Fuc(1)NeuAc(1) | C[+57]*2, N[+2060] | 1369.8918 | 4 | 5476.5456 | 5476.5418 | 0 | 0.7 | 457 | Specific | 651.07 | 651.07 | 651.07 | 12.4 | 60 | >sp\|P01833\|PIGR_HUMAN Polymeric immunoglobulin receptor OS=Homo sapiens OX=9606 GN=PIGR PE=1 SV=4 | 902 | 20201119_15232_1ng_glyco_hcd203040_compound15.18842.18842.2 | controllerType=0 controllerNumber=1 scan=18842 | 38.2844 |
| 15390:4 | 6 | K.IIEGEPNLKVPGN[+2204.772]VTAVLGETLKVPC[+57.021]HFPC[+57.021]K.F | HexNAc(4)Hex(5)NeuAc(2) | C[+57]*2, N[+2205] | 1406.1593 | 4 | 5621.6154 | 5621.5793 | 0 | 6.4 | 457 | Specific | 676.12 | 676.12 | 676.12 | 11.56 | 60 | >sp\|P01833\|PIGR_HUMAN Polymeric immunoglobulin receptor OS=Homo sapiens OX=9606 GN=PIGR PE=1 SV=4 | 902 | 20201119_15232_1ng_glyco_hcd203040_compound15.18891.18891.4 | controllerType=0 controllerNumber=1 scan=18891 | 38.3605 |
| 15294:5 | 6 | K.IIEGEPNLKVPGN[+2205.793]VTAVLGETLKVPC[+57.021]HFPC[+57.021]K.F | HexNAc(4)Hex(5)Fuc(2)NeuAc(1) | C[+57]*2, N[+2206] | 1125.3278 | 5 | 5622.6097 | 5622.5997 | 0 | 1.8 | 457 | Specific | 690.92 | 690.92 | 690.92 | 13.12 | 60 | >sp\|P01833\|PIGR_HUMAN Polymeric immunoglobulin receptor OS=Homo sapiens OX=9606 GN=PIGR PE=1 SV=4 | 902 | 20201119_15232_1ng_glyco_hcd203040_compound15.18791.18791.5 | controllerType=0 controllerNumber=1 scan=18791 | 38.201 |
| 15423:5 | 6 | K.IIEGEPNLKVPGN[+2059.735]VTAVLGETLKVPC[+57.021]HFPC[+57.021]K.F | HexNAc(4)Hex(5)Fuc(1)NeuAc(1) | C[+57]*2, N[+2060] | 1096.1172 | 5 | 5476.5568 | 5476.5418 | 0 | 2.7 | 457 | Specific | 709.37 | 591.43 | 591.43 | 14.12 | 60 | >sp\|P01833\|PIGR_HUMAN Polymeric immunoglobulin receptor OS=Homo sapiens OX=9606 GN=PIGR PE=1 SV=4 | 902 | 20201119_15232_1ng_glyco_hcd203040_compound15.18925.18925.2 | controllerType=0 controllerNumber=1 scan=18925 | 38.4132 |
| 15297:4 | 6 | K.IIEGEPNLKVPGN[+2205.793]VTAVLGETLKVPC[+57.021]HFPC[+57.021]K.F | HexNAc(4)Hex(5)Fuc(2)NeuAc(1) | C[+57]*2, N[+2206] | 1406.4119 | 4 | 5622.6256 | 5622.5997 | 0 | 4.6 | 457 | Specific | 794.63 | 652.72 | 652.72 | 14.38 | 60 | >sp\|P01833\|PIGR_HUMAN Polymeric immunoglobulin receptor OS=Homo sapiens OX=9606 GN=PIGR PE=1 SV=4 | 902 | 20201119_15232_1ng_glyco_hcd203040_compound15.18794.18794.4 | controllerType=0 controllerNumber=1 scan=18794 | 38.2074 |
| 15302:5 | 6 | K.IIEGEPNLKVPGN[+2059.735]VTAVLGETLKVPC[+57.021]HFPC[+57.021]K.F | HexNAc(4)Hex(5)Fuc(1)NeuAc(1) | C[+57]*2, N[+2060] | 1096.1188 | 5 | 5476.5648 | 5476.5418 | 0 | 4.2 | 457 | Specific | 797.04 | 694.43 | 694.43 | 14.95 | 60 | >sp\|P01833\|PIGR_HUMAN Polymeric immunoglobulin receptor OS=Homo sapiens OX=9606 GN=PIGR PE=1 SV=4 | 902 | 20201119_15232_1ng_glyco_hcd203040_compound15.18799.18799.5 | controllerType=0 controllerNumber=1 scan=18799 | 38.2167 |
| 15273:5 | 6 | K.IIEGEPNLKVPGN[+2059.735]VTAVLGETLKVPC[+57.021]HFPC[+57.021]K.F | HexNAc(4)Hex(5)Fuc(1)NeuAc(1) | C[+57]*2, N[+2060] | 1096.118 | 5 | 5476.5611 | 5476.5418 | 0 | 3.5 | 457 | Specific | 820.48 | 750.2 | 750.2 | 16.18 | 60 | >sp\|P01833\|PIGR_HUMAN Polymeric immunoglobulin receptor OS=Homo sapiens OX=9606 GN=PIGR PE=1 SV=4 | 902 | 20201119_15232_1ng_glyco_hcd203040_compound15.18769.18769.5 | controllerType=0 controllerNumber=1 scan=18769 | 38.1583 |
| 15296:5 | 6 | K.IIEGEPNLKVPGN[+2205.793]VTAVLGETLKVPC[+57.021]HFPC[+57.021]K.F | HexNAc(4)Hex(5)Fuc(2)NeuAc(1) | C[+57]*2, N[+2206] | 1125.3278 | 5 | 5622.6097 | 5622.5997 | 0 | 1.8 | 457 | Specific | 824.5 | 711.28 | 711.28 | 16.12 | 60 | >sp\|P01833\|PIGR_HUMAN Polymeric immunoglobulin receptor OS=Homo sapiens OX=9606 GN=PIGR PE=1 SV=4 | 902 | 20201119_15232_1ng_glyco_hcd203040_compound15.18793.18793.5 | controllerType=0 controllerNumber=1 scan=18793 | 38.2046 |

Table S4. Cont.

| Query #:z | Protein Rank | Glycopeptide | Glycan | Modification Type(s) | Observed m/z | z | Observed (M+H) | Calc. mass (M+H) | Off-by-x error | Mass error (ppm) | StartPosition | Cleavage | Score | Delta | Delta Mod | \|Log Prob\| | # of unique peptides | Protein Name | Protein DB number | Comment | Scan # | Scan Time |
|---|---|---|---|---|---|---|---|---|---|---|---|---|---|---|---|---|---|---|---|---|---|---|
| 15387:4 | 6 | K.IIEGEPNLKVPGN[+2204.772]VTAVLGETLKVPC[+57.021]HFPC[+57.021]K.F | HexNAc(4)Hex(5)NeuAc(2) | C[+57]*2, N[+2205] | 1406.1593 | 4 | 5621.6154 | 5621.5793 | 0 | 6.4 | 457 | Specific | 842.42 | 685.84 | 685.84 | 14.56 | 60 | >sp\|P01833\|PIGR_HUMAN Polymeric immunoglobulin receptor OS=Homo sapiens OX=9606 GN=PIGR PE=1 SV=4 | 902 | 20201119_15232_1ng_glyco_hcd203040_compound15.18888.18888.4 | controllerType=0 controllerNumber=1 scan=18888 | 38.3551 |
| 15271:5 | 6 | K.IIEGEPNLKVPGN[+2059.735]VTAVLGETLKVPC[+57.021]HFPC[+57.021]K.F | HexNAc(4)Hex(5)Fuc(1)NeuAc(1) | C[+57]*2, N[+2060] | 1096.118 | 5 | 5476.5611 | 5476.5418 | 0 | 3.5 | 457 | Specific | 845.19 | 738.15 | 738.15 | 16.18 | 60 | >sp\|P01833\|PIGR_HUMAN Polymeric immunoglobulin receptor OS=Homo sapiens OX=9606 GN=PIGR PE=1 SV=4 | 902 | 20201119_15232_1ng_glyco_hcd203040_compound15.18767.18767.5 | controllerType=0 controllerNumber=1 scan=18767 | 38.1523 |
| 15306:4 | 6 | K.IIEGEPNLKVPGN[+2059.735]VTAVLGETLKVPC[+57.021]HFPC[+57.021]K.F | HexNAc(4)Hex(5)Fuc(1)NeuAc(1) | C[+57]*2, N[+2060] | 1369.8953 | 4 | 5476.5594 | 5476.5418 | 0 | 3.2 | 457 | Specific | 852.37 | 847.04 | 847.04 | 16.61 | 60 | >sp\|P01833\|PIGR_HUMAN Polymeric immunoglobulin receptor OS=Homo sapiens OX=9606 GN=PIGR PE=1 SV=4 | 902 | 20201119_15232_1ng_glyco_hcd203040_compound15.18803.18803.4 | controllerType=0 controllerNumber=1 scan=18803 | 38.2243 |
| 15251:4 | 6 | K.IIEGEPNLKVPGN[+2059.735]VTAVLGETLKVPC[+57.021]HFPC[+57.021]K.F | HexNAc(4)Hex(5)Fuc(1)NeuAc(1) | C[+57]*2, N[+2060] | 1369.8949 | 4 | 5476.5578 | 5476.5418 | 0 | 2.9 | 457 | Specific | 883.22 | 883.22 | 883.22 | 16.61 | 60 | >sp\|P01833\|PIGR_HUMAN Polymeric immunoglobulin receptor OS=Homo sapiens OX=9606 GN=PIGR PE=1 SV=4 | 902 | 20201119_15232_1ng_glyco_hcd203040_compound15.18746.18746.4 | controllerType=0 controllerNumber=1 scan=18746 | 38.1114 |
| 15304:4 | 6 | K.IIEGEPNLKVPGN[+2059.735]VTAVLGETLKVPC[+57.021]HFPC[+57.021]K.F | HexNAc(4)Hex(5)Fuc(1)NeuAc(1) | C[+57]*2, N[+2060] | 1369.8953 | 4 | 5476.5594 | 5476.5418 | 0 | 3.2 | 457 | Specific | 952.75 | 855.28 | 855.28 | 18.61 | 60 | >sp\|P01833\|PIGR_HUMAN Polymeric immunoglobulin receptor OS=Homo sapiens OX=9606 GN=PIGR PE=1 SV=4 | 902 | 20201119_15232_1ng_glyco_hcd203040_compound15.18801.18801.4 | controllerType=0 controllerNumber=1 scan=18801 | 38.2197 |
| 15300:5 | 6 | K.IIEGEPNLKVPGN[+2059.735]VTAVLGETLKVPC[+57.021]HFPC[+57.021]K.F | HexNAc(4)Hex(5)Fuc(1)NeuAc(1) | C[+57]*2, N[+2060] | 1096.1188 | 5 | 5476.5648 | 5476.5418 | 0 | 4.2 | 457 | Specific | 955.96 | 827.65 | 827.65 | 18.95 | 60 | >sp\|P01833\|PIGR_HUMAN Polymeric immunoglobulin receptor OS=Homo sapiens OX=9606 GN=PIGR PE=1 SV=4 | 902 | 20201119_15232_1ng_glyco_hcd203040_compound15.18797.18797.5 | controllerType=0 controllerNumber=1 scan=18797 | 38.2116 |
| 15295:4 | 6 | K.IIEGEPNLKVPGN[+2205.793]VTAVLGETLKVPC[+57.021]HFPC[+57.021]K.F | HexNAc(4)Hex(5)Fuc(2)NeuAc(1) | C[+57]*2, N[+2206] | 1406.4119 | 4 | 5622.6256 | 5622.5997 | 0 | 4.6 | 457 | Specific | 1092.19 | 977.43 | 977.43 | 20.38 | 60 | >sp\|P01833\|PIGR_HUMAN Polymeric immunoglobulin receptor OS=Homo sapiens OX=9606 GN=PIGR PE=1 SV=4 | 902 | 20201119_15232_1ng_glyco_hcd203040_compound15.18792.18792.4 | controllerType=0 controllerNumber=1 scan=18792 | 38.2019 |
| 13367:3 | 6 | K.VPGN[+2059.735]VTAVLGETLK.V | HexNAc(4)Hex(5)Fuc(1)NeuAc(1) | N[+2060] | 1153.1845 | 3 | 3457.539 | 3457.5348 | 0 | 1.2 | 466 | Specific | 325.49 | 308.45 | 308.45 | 6.72 | 60 | >sp\|P01833\|PIGR_HUMAN Polymeric immunoglobulin receptor OS=Homo sapiens OX=9606 GN=PIGR PE=1 SV=4 | 902 | 20201119_15232_1ng_glyco_hcd203040_compound15.16793.16793.3 | controllerType=0 controllerNumber=1 scan=16793 | 34.6954 |
| 13342:4 | 6 | K.VPGN[+2205.793]VTAVLGETLKVPC[+57.021]HFPC[+57.021]K.F | HexNAc(4)Hex(5)Fuc(2)NeuAc(1) | C[+57]*2, N[+2206] | 1158.0241 | 4 | 4629.0745 | 4629.0502 | 0 | 5.3 | 466 | Specific | 454.73 | 384.7 | 384.7 | 8.72 | 60 | >sp\|P01833\|PIGR_HUMAN Polymeric immunoglobulin receptor OS=Homo sapiens OX=9606 GN=PIGR PE=1 SV=4 | 902 | 20201119_15232_1ng_glyco_hcd203040_compound15.16767.16767.3 | controllerType=0 controllerNumber=1 scan=16767 | 34.6474 |
| 13237:4 | 6 | K.VPGN[+2205.793]VTAVLGETLKVPC[+57.021]HFPC[+57.021]K.F | HexNAc(4)Hex(5)Fuc(2)NeuAc(1) | C[+57]*2, N[+2206] | 1158.0207 | 4 | 4629.0609 | 4629.0502 | 0 | 2.3 | 466 | Specific | 521.62 | 378.53 | 378.53 | 9.64 | 60 | >sp\|P01833\|PIGR_HUMAN Polymeric immunoglobulin receptor OS=Homo sapiens OX=9606 GN=PIGR PE=1 SV=4 | 902 | 20201119_15232_1ng_glyco_hcd203040_compound15.16658.16658.4 | controllerType=0 controllerNumber=1 scan=16658 | 34.4414 |
| 13239:4 | 6 | K.VPGN[+2205.793]VTAVLGETLKVPC[+57.021]HFPC[+57.021]K.F | HexNAc(4)Hex(5)Fuc(2)NeuAc(1) | C[+57]*2, N[+2206] | 1158.0207 | 4 | 4629.0609 | 4629.0502 | 0 | 2.3 | 466 | Specific | 551.8 | 392.75 | 392.75 | 9.92 | 60 | >sp\|P01833\|PIGR_HUMAN Polymeric immunoglobulin receptor OS=Homo sapiens OX=9606 GN=PIGR PE=1 SV=4 | 902 | 20201119_15232_1ng_glyco_hcd203040_compound15.16660.16660.4 | controllerType=0 controllerNumber=1 scan=16660 | 34.4456 |
| 13278:4 | 6 | K.VPGN[+2205.793]VTAVLGETLKVPC[+57.021]HFPC[+57.021]K.F | HexNAc(4)Hex(5)Fuc(2)NeuAc(1) | C[+57]*2, N[+2206] | 1158.0201 | 4 | 4629.0584 | 4629.0502 | 0 | 1.8 | 466 | Specific | 561.67 | 305.59 | 305.59 | 8.07 | 60 | >sp\|P01833\|PIGR_HUMAN Polymeric immunoglobulin receptor OS=Homo sapiens OX=9606 GN=PIGR PE=1 SV=4 | 902 | 20201119_15232_1ng_glyco_hcd203040_compound15.16701.16701.4 | controllerType=0 controllerNumber=1 scan=16701 | 34.5185 |
| 13392:4 | 6 | K.VPGN[+2059.735]VTAVLGETLKVPC[+57.021]HFPC[+57.021]K.F | HexNAc(4)Hex(5)Fuc(1)NeuAc(1) | C[+57]*2, N[+2060] | 1121.5077 | 4 | 4483.0091 | 4482.9923 | 0 | 3.7 | 466 | Specific | 617.29 | 575.26 | 575.26 | 12.21 | 60 | >sp\|P01833\|PIGR_HUMAN Polymeric immunoglobulin receptor OS=Homo sapiens OX=9606 GN=PIGR PE=1 SV=4 | 902 | 20201119_15232_1ng_glyco_hcd203040_compound15.16819.16819.4 | controllerType=0 controllerNumber=1 scan=16819 | 34.7441 |
| 13390:4 | 6 | K.VPGN[+2059.735]VTAVLGETLKVPC[+57.021]HFPC[+57.021]K.F | HexNAc(4)Hex(5)Fuc(1)NeuAc(1) | C[+57]*2, N[+2060] | 1121.5077 | 4 | 4483.0091 | 4482.9923 | 0 | 3.7 | 466 | Specific | 630.53 | 630.53 | 630.53 | 12.21 | 60 | >sp\|P01833\|PIGR_HUMAN Polymeric immunoglobulin receptor OS=Homo sapiens OX=9606 GN=PIGR PE=1 SV=4 | 902 | 20201119_15232_1ng_glyco_hcd203040_compound15.16817.16817.4 | controllerType=0 controllerNumber=1 scan=16817 | 34.7401 |
| 13345:4 | 6 | K.VPGN[+2205.793]VTAVLGETLKVPC[+57.021]HFPC[+57.021]K.F | HexNAc(4)Hex(5)Fuc(2)NeuAc(1) | C[+57]*2, N[+2206] | 1158.0241 | 4 | 4629.0745 | 4629.0502 | 0 | 5.3 | 466 | Specific | 644.96 | 604.86 | 604.86 | 11.44 | 60 | >sp\|P01833\|PIGR_HUMAN Polymeric immunoglobulin receptor OS=Homo sapiens OX=9606 GN=PIGR PE=1 SV=4 | 902 | 20201119_15232_1ng_glyco_hcd203040_compound15.16770.16770.3 | controllerType=0 controllerNumber=1 scan=16770 | 34.6515 |
| 13276:4 | 6 | K.VPGN[+2205.793]VTAVLGETLKVPC[+57.021]HFPC[+57.021]K.F | HexNAc(4)Hex(5)Fuc(2)NeuAc(1) | C[+57]*2, N[+2206] | 1158.0201 | 4 | 4629.0584 | 4629.0502 | 0 | 1.8 | 466 | Specific | 659.38 | 560.85 | 560.85 | 13.16 | 60 | >sp\|P01833\|PIGR_HUMAN Polymeric immunoglobulin receptor OS=Homo sapiens OX=9606 GN=PIGR PE=1 SV=4 | 902 | 20201119_15232_1ng_glyco_hcd203040_compound15.16699.16699.4 | controllerType=0 controllerNumber=1 scan=16699 | 34.5126 |
| 05416:3 | 6 | K.WN[+1872.651]NTGC[+57.021]QALPSQDEGPSK.A | HexNAc(3)Hex(6)NeuAc(1) | C[+57], N[+1873] | 1287.853 | 3 | 3861.5443 | 3861.5272 | 0 | 4.4 | 498 | Specific | 371.89 | 348 | 348 | 7.02 | 60 | >sp\|P01833\|PIGR_HUMAN Polymeric immunoglobulin receptor OS=Homo sapiens OX=9606 GN=PIGR PE=1 SV=4 | 902 | 20201119_15232_1ng_glyco_hcd203040_compound15.8504.8504.3 | controllerType=0 controllerNumber=1 scan=8504 | 20.7728 |
| 05414:3 | 6 | K.WN[+1872.651]NTGC[+57.021]QALPSQDEGPSK.A | HexNAc(3)Hex(6)NeuAc(1) | C[+57], N[+1873] | 1287.853 | 3 | 3861.5443 | 3861.5272 | 0 | 4.4 | 498 | Specific | 381.8 | 367.1 | 367.1 | 7.43 | 60 | >sp\|P01833\|PIGR_HUMAN Polymeric immunoglobulin receptor OS=Homo sapiens OX=9606 GN=PIGR PE=1 SV=4 | 902 | 20201119_15232_1ng_glyco_hcd203040_compound15.8502.8502.3 | controllerType=0 controllerNumber=1 scan=8502 | 20.7687 |
| 05340:3 | 6 | K.WN[+2059.735]NTGC[+57.021]QALPSQDEGPSK.A | HexNAc(4)Hex(5)Fuc(1)NeuAc(1) | C[+57], N[+2060] | 1350.212 | 3 | 4048.6214 | 4048.6116 | 0 | 2.4 | 498 | Specific | 557.69 | 485.27 | 485.27 | 9.87 | 60 | >sp\|P01833\|PIGR_HUMAN Polymeric immunoglobulin receptor OS=Homo sapiens OX=9606 GN=PIGR PE=1 SV=4 | 902 | 20201119_15232_1ng_glyco_hcd203040_compound15.8426.8426.3 | controllerType=0 controllerNumber=1 scan=8426 | 20.6416 |
| 05338:3 | 6 | K.WN[+2059.735]NTGC[+57.021]QALPSQDEGPSK.A | HexNAc(4)Hex(5)Fuc(1)NeuAc(1) | C[+57], N[+2060] | 1350.212 | 3 | 4048.6214 | 4048.6116 | 0 | 2.4 | 498 | Specific | 573.19 | 507.08 | 507.08 | 9.87 | 60 | >sp\|P01833\|PIGR_HUMAN Polymeric immunoglobulin receptor OS=Homo sapiens OX=9606 GN=PIGR PE=1 SV=4 | 902 | 20201119_15232_1ng_glyco_hcd203040_compound15.8424.8424.3 | controllerType=0 controllerNumber=1 scan=8424 | 20.637 |
| 05394:3 | 6 | K.WN[+1913.677]NTGC[+57.021]QALPSQDEGPSK.A | HexNAc(4)Hex(5)NeuAc(1) | C[+57], N[+1914] | 1301.5256 | 3 | 3902.5622 | 3902.5537 | 0 | 2.2 | 498 | Specific | 668.68 | 597.61 | 597.61 | 13.1 | 60 | >sp\|P01833\|PIGR_HUMAN Polymeric immunoglobulin receptor OS=Homo sapiens OX=9606 GN=PIGR PE=1 SV=4 | 902 | 20201119_15232_1ng_glyco_hcd203040_compound15.8481.8481.3 | controllerType=0 controllerNumber=1 scan=8481 | 20.7328 |
| 05428:3 | 6 | K.WN[+1913.677]NTGC[+57.021]QALPSQDEGPSK.A | HexNAc(4)Hex(5)NeuAc(1) | C[+57], N[+1914] | 1301.5265 | 3 | 3902.5651 | 3902.5537 | 0 | 2.9 | 498 | Specific | 696.22 | 631.77 | 631.77 | 13.16 | 60 | >sp\|P01833\|PIGR_HUMAN Polymeric immunoglobulin receptor OS=Homo sapiens OX=9606 GN=PIGR PE=1 SV=4 | 902 | 20201119_15232_1ng_glyco_hcd203040_compound15.8517.8517.3 | controllerType=0 controllerNumber=1 scan=8517 | 20.7953 |
| 05395:3 | 6 | K.WN[+1913.677]NTGC[+57.021]QALPSQDEGPSK.A | HexNAc(4)Hex(5)NeuAc(1) | C[+57], N[+1914] | 1301.5256 | 3 | 3902.5622 | 3902.5537 | 0 | 2.2 | 498 | Specific | 758.48 | 675.03 | 675.03 | 15.1 | 60 | >sp\|P01833\|PIGR_HUMAN Polymeric immunoglobulin receptor OS=Homo sapiens OX=9606 GN=PIGR PE=1 SV=4 | 902 | 20201119_15232_1ng_glyco_hcd203040_compound15.8482.8482.3 | controllerType=0 controllerNumber=1 scan=8482 | 20.7365 |
| 05431:3 | 6 | K.WN[+1913.677]NTGC[+57.021]QALPSQDEGPSK.A | HexNAc(4)Hex(5)NeuAc(1) | C[+57], N[+1914] | 1301.5265 | 3 | 3902.5651 | 3902.5537 | 0 | 2.9 | 498 | Specific | 762.15 | 702.23 | 702.23 | 15.16 | 60 | >sp\|P01833\|PIGR_HUMAN Polymeric immunoglobulin receptor OS=Homo sapiens OX=9606 GN=PIGR PE=1 SV=4 | 902 | 20201119_15232_1ng_glyco_hcd203040_compound15.8520.8520.3 | controllerType=0 controllerNumber=1 scan=8520 | 20.8004 |
| 09666:3 | 7 | K.TVLTPATNHMGN[+1540.529]VTFTIPANR.E | HexNAc(2)Hex(7) | N[+1541] | 1265.9024 | 3 | 3795.6926 | 3795.6886 | 0 | 1.1 | 74 | Specific | 310.15 | 310.15 | 120.4 | 6.67 | 73 | >sp\|P01024\|CO3_HUMAN Complement C3 OS=Homo sapiens OX=9606 GN=C3 PE=1 SV=2 | 492 | 20201119_15232_1ng_glyco_hcd203040_compound15.12929.12929.3 | controllerType=0 controllerNumber=1 scan=12929 | 28.1157 |
| 09668:3 | 7 | K.TVLTPATNHMGN[+1540.529]VTFTIPANR.E | HexNAc(2)Hex(7) | N[+1541] | 1265.9024 | 3 | 3795.6926 | 3795.6886 | 0 | 1.1 | 74 | Specific | 413.59 | 362.76 | 135.18 | 6.16 | 73 | >sp\|P01024\|CO3_HUMAN Complement C3 OS=Homo sapiens OX=9606 GN=C3 PE=1 SV=2 | 492 | 20201119_15232_1ng_glyco_hcd203040_compound15.12931.12931.3 | controllerType=0 controllerNumber=1 scan=12931 | 28.1202 |
| 18656:4 | 8 | R.NIAAFGGDPNN[+1913.677]ITLFGESAGGASVSLQTLSPYNK.G | HexNAc(4)Hex(5)NeuAc(1) | N[+1914] | 1331.8495 | 4 | 5324.3761 | 5324.3633 | 0 | 2.4 | 197 | Specific | 1026.57 | 1004.51 | 1004.51 | 20.79 | 52 | >sp\|P19835\|CEL_HUMAN Bile salt-activated lipase OS=Homo sapiens OX=9606 GN=CEL PE=1 SV=3 | 476 | 20201119_15232_1ng_glyco_hcd203040_compound15.22347.22347.4 | controllerType=0 controllerNumber=1 scan=22347 | 45.3692 |
| 18659:4 | 8 | R.NIAAFGGDPNN[+1913.677]ITLFGESAGGASVSLQTLSPYNK.G | HexNAc(4)Hex(5)NeuAc(1) | N[+1914] | 1331.8495 | 4 | 5324.3761 | 5324.3633 | 0 | 2.4 | 197 | Specific | 1030.64 | 977.22 | 977.22 | 20.79 | 52 | >sp\|P19835\|CEL_HUMAN Bile salt-activated lipase OS=Homo sapiens OX=9606 GN=CEL PE=1 SV=3 | 476 | 20201119_15232_1ng_glyco_hcd203040_compound15.22350.22350.4 | controllerType=0 controllerNumber=1 scan=22350 | 45.3802 |
| 05285:3 | 12 | R.LSPN[+2205.793]ASAEHLELR.W | HexNAc(4)Hex(5)Fuc(2)NeuAc(1) | N[+2206] | 1214.8559 | 3 | 3642.5531 | 3642.5421 | 0 | 3 | 52 | Specific | 578.46 | 420.58 | 420.58 | 11.15 | 38 | >sp\|Q13410\|BT1A1_HUMAN Butyrophilin subfamily 1 member A1 OS=Homo sapiens OX=9606 GN=BTN1A1 PE=1 SV=3 | 179 | 20201119_15232_1ng_glyco_hcd203040_compound15.8369.8369.3 | controllerType=0 controllerNumber=1 scan=8369 | 20.5425 |
| 07367:3 | 12 | R.LSPN[+2350.830]ASAEHLELR.W | HexNAc(4)Hex(5)Fuc(1)NeuAc(2) | N[+2351] | 1263.2031 | 3 | 3787.5946 | 3787.5796 | 0 | 4 | 52 | Specific | 595.35 | 512.6 | 512.6 | 10.92 | 38 | >sp\|Q13410\|BT1A1_HUMAN Butyrophilin subfamily 1 member A1 OS=Homo sapiens OX=9606 GN=BTN1A1 PE=1 SV=3 | 179 | 20201119_15232_1ng_glyco_hcd203040_compound15.10525.10525.3 | controllerType=0 controllerNumber=1 scan=10525 | 24.0657 |
| 05283:3 | 12 | R.LSPN[+2205.793]ASAEHLELR.W | HexNAc(4)Hex(5)Fuc(2)NeuAc(1) | N[+2206] | 1214.8559 | 3 | 3642.5531 | 3642.5421 | 0 | 3 | 52 | Specific | 597.32 | 394.33 | 394.33 | 11.15 | 38 | >sp\|Q13410\|BT1A1_HUMAN Butyrophilin subfamily 1 member A1 OS=Homo sapiens OX=9606 GN=BTN1A1 PE=1 SV=3 | 179 | 20201119_15232_1ng_glyco_hcd203040_compound15.8367.8367.3 | controllerType=0 controllerNumber=1 scan=8367 | 20.5349 |
| 07365:3 | 12 | R.LSPN[+2350.830]ASAEHLELR.W | HexNAc(4)Hex(5)Fuc(1)NeuAc(2) | N[+2351] | 1263.2031 | 3 | 3787.5946 | 3787.5796 | 0 | 4 | 52 | Specific | 616.99 | 505.7 | 505.7 | 11.92 | 38 | >sp\|Q13410\|BT1A1_HUMAN Butyrophilin subfamily 1 member A1 OS=Homo sapiens OX=9606 GN=BTN1A1 PE=1 SV=3 | 179 | 20201119_15232_1ng_glyco_hcd203040_compound15.10523.10523.3 | controllerType=0 controllerNumber=1 scan=10523 | 24.0613 |
| 22757:4 | 15 | K.DIDGYGGIALPELIC[+57.021]TM[+15.995]FHTSGYDTQAIVENN[+2190.793]ESTEYG | HexNAc(6)Hex(6) | C[+57], M[+16], N[+2191] | 1805.2903 | 4 | 7218.1394 | 7218.1169 | 0 | 3.1 | 33 | Specific | 404.59 | 397.93 | 397.93 | 10.46 | 27 | >sp\|P00709\|LALBA_HUMAN Alpha-lactalbumin OS=Homo sapiens OX=9606 GN=LALBA PE=1 SV=1 | 468 | 20201119_15232_1ng_glyco_hcd203040_compound15.26627.26627.4 | controllerType=0 controllerNumber=1 scan=26627 | 53.6428 |
| 22759:4 | 15 | K.DIDGYGGIALPELIC[+57.021]TM[+15.995]FHTSGYDTQAIVENN[+2190.793]ESTEYG | HexNAc(6)Hex(6) | C[+57], M[+16], N[+2191] | 1805.2903 | 4 | 7218.1394 | 7218.1169 | 0 | 3.1 | 33 | Specific | 451.4 | 425.68 | 425.68 | 10.76 | 27 | >sp\|P00709\|LALBA_HUMAN Alpha-lactalbumin OS=Homo sapiens OX=9606 GN=LALBA PE=1 SV=1 | 468 | 20201119_15232_1ng_glyco_hcd203040_compound15.26629.26629.4 | controllerType=0 controllerNumber=1 scan=26629 | 53.6464 |
| 23521:4 | 16 | K.VFPLSLDSTPQDGNVVVAC[+57.021]LVQGFFPQEPLSVTWSESGQN[+2059.735]VTAI | HexNAc(4)Hex(5)Fuc(1)NeuAc(1) | C[+57], N[+2060] | 1709.7905 | 4 | 6836.1401 | 6836.0963 | 0 | 6.4 | 8 | Specific | 226.65 | 226.65 | 226.65 | 6.62 | 40 | >sp\|P01877\|IGHA2_HUMAN Immunoglobulin heavy constant alpha 2 OS=Homo sapiens OX=9606 GN=IGHA2 PE=1 SV=4 | 576 | 20201119_15232_1ng_glyco_hcd203040_compound15.27422.27422.4 | controllerType=0 controllerNumber=1 scan=27422 | 55.1968 |
| 23575:5 | 16 | K.VFPLSLDSTPQDGNVVVAC[+57.021]LVQGFFPQEPLSVTWSESGQN[+2262.814]VTAI | HexNAc(5)Hex(5)Fuc(1)NeuAc(1) | C[+57], N[+2263] | 1408.649 | 5 | 7039.2157 | 7039.1757 | 0 | 5.7 | 8 | Specific | 236.78 | 236.78 | 236.78 | 5.71 | 40 | >sp\|P01877\|IGHA2_HUMAN Immunoglobulin heavy constant alpha 2 OS=Homo sapiens OX=9606 GN=IGHA2 PE=1 SV=4 | 576 | 20201119_15232_1ng_glyco_hcd203040_compound15.27478.27478.5 | controllerType=0 controllerNumber=1 scan=27478 | 55.3065 |
| 23577:5 | 16 | K.VFPLSLDSTPQDGNVVVAC[+57.021]LVQGFFPQEPLSVTWSESGQN[+2262.814]VTAI | HexNAc(5)Hex(5)Fuc(1)NeuAc(1) | C[+57], N[+2263] | 1408.649 | 5 | 7039.2157 | 7039.1757 | 0 | 5.7 | 8 | Specific | 285.49 | 285.49 | 285.49 | 7.45 | 40 | >sp\|P01877\|IGHA2_HUMAN Immunoglobulin heavy constant alpha 2 OS=Homo sapiens OX=9606 GN=IGHA2 PE=1 SV=4 | 576 | 20201119_15232_1ng_glyco_hcd203040_compound15.27480.27480.5 | controllerType=0 controllerNumber=1 scan=27480 | 55.3117 |
| 23523:4 | 16 | K.VFPLSLDSTPQDGNVVVAC[+57.021]LVQGFFPQEPLSVTWSESGQN[+2059.735]VTAI | HexNAc(4)Hex(5)Fuc(1)NeuAc(1) | C[+57], N[+2060] | 1709.7905 | 4 | 6836.1401 | 6836.0963 | 0 | 6.4 | 8 | Specific | 367.85 | 367.85 | 367.85 | 8.02 | 40 | >sp\|P01877\|IGHA2_HUMAN Immunoglobulin heavy constant alpha 2 OS=Homo sapiens OX=9606 GN=IGHA2 PE=1 SV=4 | 576 | 20201119_15232_1ng_glyco_hcd203040_compound15.27424.27424.4 | controllerType=0 controllerNumber=1 scan=27424 | 55.202 |
| 21875:4 | 16 | K.VFPLSLDSTPQDGNVVVAC[+57.021]LVQGFFPQEPLSVTWSESGQN[+1647.613]VTAI | HexNAc(5)Hex(3)Fuc(1) | C[+57], N[+1648] | 1606.7571 | 4 | 6424.0065 | 6423.9746 | 0 | 5 | 8 | Specific | 500.79 | 500.79 | 500.79 | 10.92 | 40 | >sp\|P01877\|IGHA2_HUMAN Immunoglobulin heavy constant alpha 2 OS=Homo sapiens OX=9606 GN=IGHA2 PE=1 SV=4 | 576 | 20201119_15232_1ng_glyco_hcd203040_compound15.25699.25699.4 | controllerType=0 controllerNumber=1 scan=25699 | 51.9409 |
| 21910:4 | 16 | K.VFPLSLDSTPQDGNVVVAC[+57.021]LVQGFFPQEPLSVTWSESGQN[+1647.613]VTAI | HexNAc(5)Hex(3)Fuc(1) | C[+57], N[+1648] | 1606.7533 | 4 | 6423.9914 | 6423.9746 | 0 | 2.6 | 8 | Specific | 652.06 | 562.44 | 562.44 | 14.6 | 40 | >sp\|P01877\|IGHA2_HUMAN Immunoglobulin heavy constant alpha 2 OS=Homo sapiens OX=9606 GN=IGHA2 PE=1 SV=4 | 576 | 20201119_15232_1ng_glyco_hcd203040_compound15.25735.25735.4 | controllerType=0 controllerNumber=1 scan=25735 | 51.9906 |
| 21878:4 | 16 | K.VFPLSLDSTPQDGNVVVAC[+57.021]LVQGFFPQEPLSVTWSESGQN[+1647.613]VTAI | HexNAc(5)Hex(3)Fuc(1) | C[+57], N[+1648] | 1606.7571 | 4 | 6424.0065 | 6423.9746 | 0 | 5 | 8 | Specific | 895.1 | 895.1 | 895.1 | 18.43 | 40 | >sp\|P01877\|IGHA2_HUMAN Immunoglobulin heavy constant alpha 2 OS=Homo sapiens OX=9606 GN=IGHA2 PE=1 SV=4 | 576 | 20201119_15232_1ng_glyco_hcd203040_compound15.25702.25702.4 | controllerType=0 controllerNumber=1 scan=25702 | 51.9454 |
| 21913:4 | 16 | K.VFPLSLDSTPQDGNVVVAC[+57.021]LVQGFFPQEPLSVTWSESGQN[+1647.613]VTAI | HexNAc(5)Hex(3)Fuc(1) | C[+57], N[+1648] | 1606.7533 | 4 | 6423.9914 | 6423.9746 | 0 | 2.6 | 8 | Specific | 941.95 | 866.03 | 866.03 | 19.6 | 40 | >sp\|P01877\|IGHA2_HUMAN Immunoglobulin heavy constant alpha 2 OS=Homo sapiens OX=9606 GN=IGHA2 PE=1 SV=4 | 576 | 20201119_15232_1ng_glyco_hcd203040_compound15.25738.25738.4 | controllerType=0 controllerNumber=1 scan=25738 | 51.9987 |
| 01011:2 | 16 | K.HYTN[+1647.613]SSQDVTVPC[+57.021]R.V | HexNAc(5)Hex(3)Fuc(1) | C[+57], N[+1648] | 1656.1868 | 2 | 3311.3663 | 3311.3625 | 0 | 1.1 | 89 | Specific | 208.25 | 138.95 | 138.95 | 3.44 | 40 | >sp\|P01877\|IGHA2_HUMAN Immunoglobulin heavy constant alpha 2 OS=Homo sapiens OX=9606 GN=IGHA2 PE=1 SV=4 | 576 | 20201119_15232_1ng_glyco_hcd203040_compound15.3937.3937.2 | controllerType=0 controllerNumber=1 scan=3937 | 12.9819 |
| 01817:3 | 16 | K.HYTN[+2262.814]SSQDVTVPC[+57.021]R.V | HexNAc(5)Hex(5)Fuc(1)NeuAc(1) | C[+57], N[+2263] | 1309.5325 | 3 | 3926.583 | 3926.5636 | 0 | 4.9 | 89 | Specific | 244.93 | 177.39 | 177.39 | 5.62 | 40 | >sp\|P01877\|IGHA2_HUMAN Immunoglobulin heavy constant alpha 2 OS=Homo sapiens OX=9606 GN=IGHA2 PE=1 SV=4 | 576 | 20201119_15232_1ng_glyco_hcd203040_compound15.4775.4775.3 | controllerType=0 controllerNumber=1 scan=4775 | 14.45 |
| 01815:3 | 16 | K.HYTN[+2262.814]SSQDVTVPC[+57.021]R.V | HexNAc(5)Hex(5)Fuc(1)NeuAc(1) | C[+57], N[+2263] | 1309.5325 | 3 | 3926.583 | 3926.5636 | 0 | 4.9 | 89 | Specific | 300.93 | 166.21 | 166.21 | 5.73 | 40 | >sp\|P01877\|IGHA2_HUMAN Immunoglobulin heavy constant alpha 2 OS=Homo sapiens OX=9606 GN=IGHA2 PE=1 SV=4 | 576 | 20201119_15232_1ng_glyco_hcd203040_compound15.4773.4773.3 | controllerType=0 controllerNumber=1 scan=4773 | 14.4461 |
| 00937:3 | 16 | K.HYTN[+1809.666]SSQDVTVPC[+57.021]R.V | HexNAc(5)Hex(4)Fuc(1) | C[+57], N[+1810] | 1158.4808 | 3 | 3473.4278 | 3473.4154 | 0 | 3.6 | 89 | Specific | 510.85 | 326 | 326 | 7.83 | 40 | >sp\|P01877\|IGHA2_HUMAN Immunoglobulin heavy constant alpha 2 OS=Homo sapiens OX=9606 GN=IGHA2 PE=1 SV=4 | 576 | 20201119_15232_1ng_glyco_hcd203040_compound15.3860.3860.3 | controllerType=0 controllerNumber=1 scan=3860 | 12.8393 |
| 01039:3 | 16 | K.HYTN[+1647.613]SSQDVTVPC[+57.021]R.V | HexNAc(5)Hex(3)Fuc(1) | C[+57], N[+1648] | 1104.4621 | 3 | 3311.3716 | 3311.3625 | 0 | 2.7 | 89 | Specific | 537.22 | 404.98 | 404.98 | 9.64 | 40 | >sp\|P01877\|IGHA2_HUMAN Immunoglobulin heavy constant alpha 2 OS=Homo sapiens OX=9606 GN=IGHA2 PE=1 SV=4 | 576 | 20201119_15232_1ng_glyco_hcd203040_compound15.3966.3966.2 | controllerType=0 controllerNumber=1 scan=3966 | 13.0315 |
| 01041:3 | 16 | K.HYTN[+1647.613]SSQDVTVPC[+57.021]R.V | HexNAc(5)Hex(3)Fuc(1) | C[+57], N[+1648] | 1104.4621 | 3 | 3311.3716 | 3311.3625 | 0 | 2.7 | 89 | Specific | 549.22 | 549.22 | 549.22 | 9.64 | 40 | >sp\|P01877\|IGHA2_HUMAN Immunoglobulin heavy constant alpha 2 OS=Homo sapiens OX=9606 GN=IGHA2 PE=1 SV=4 | 576 | 20201119_15232_1ng_glyco_hcd203040_compound15.3968.3968.2 | controllerType=0 controllerNumber=1 scan=3968 | 13.035 |
| 00939:3 | 16 | K.HYTN[+1809.666]SSQDVTVPC[+57.021]R.V | HexNAc(5)Hex(4)Fuc(1) | C[+57], N[+1810] | 1158.4808 | 3 | 3473.4278 | 3473.4154 | 0 | 3.6 | 89 | Specific | 598.91 | 413.3 | 413.3 | 11.15 | 40 | >sp\|P01877\|IGHA2_HUMAN Immunoglobulin heavy constant alpha 2 OS=Homo sapiens OX=9606 GN=IGHA2 PE=1 SV=4 | 576 | 20201119_15232_1ng_glyco_hcd203040_compound15.3862.3862.3 | controllerType=0 controllerNumber=1 scan=3862 | 12.8442 |
| 00958:3 | 16 | K.HYTN[+1647.613]SSQDVTVPC[+57.021]R.V | HexNAc(5)Hex(3)Fuc(1) | C[+57], N[+1648] | 1104.461 | 3 | 3311.3683 | 3311.3625 | 0 | 1.8 | 89 | Specific | 760.16 | 744.04 | 744.04 | 15.09 | 40 | >sp\|P01877\|IGHA2_HUMAN Immunoglobulin heavy constant alpha 2 OS=Homo sapiens OX=9606 GN=IGHA2 PE=1 SV=4 | 576 | 20201119_15232_1ng_glyco_hcd203040_compound15.3882.3882.3 | controllerType=0 controllerNumber=1 scan=3882 | 12.8823 |
| 00960:3 | 16 | K.HYTN[+1647.613]SSQDVTVPC[+57.021]R.V | HexNAc(5)Hex(3)Fuc(1) | C[+57], N[+1648] | 1104.461 | 3 | 3311.3683 | 3311.3625 | 0 | 1.8 | 89 | Specific | 766.83 | 622.76 | 622.76 | 15.09 | 40 | >sp\|P01877\|IGHA2_HUMAN Immunoglobulin heavy constant alpha 2 OS=Homo sapiens OX=9606 GN=IGHA2 PE=1 SV=4 | 576 | 20201119_15232_1ng_glyco_hcd203040_compound15.3884.3884.3 | controllerType=0 controllerNumber=1 scan=3884 | 12.8896 |
| 03107:4 | 16 | K.HYTN[+1647.613]SSQDVTVPC[+57.021]RVPPPPPC[+57.021]C[+57.021]HPR.L | HexNAc(5)Hex(3)Fuc(1) | C[+57]*3, N[+1648] | 1152.2481 | 4 | 4605.9705 | 4605.9689 | 0 | 0.4 | 89 | Specific | 244.86 | 244.86 | 244.86 | 5.27 | 40 | >sp\|P01877\|IGHA2_HUMAN Immunoglobulin heavy constant alpha 2 OS=Homo sapiens OX=9606 GN=IGHA2 PE=1 SV=4 | 576 | 20201119_15232_1ng_glyco_hcd203040_compound15.6112.6112.2 | controllerType=0 controllerNumber=1 scan=6112 | 16.822 |
| 02931:4 | 16 | K.HYTN[+1647.613]SSQDVTVPC[+57.021]RVPPPPPC[+57.021]C[+57.021]HPR.L | HexNAc(5)Hex(3)Fuc(1) | C[+57]*3, N[+1648] | 1152.2515 | 4 | 4605.9842 | 4605.9689 | 0 | 3.3 | 89 | Specific | 588.58 | 451.08 | 451.08 | 10.45 | 40 | >sp\|P01877\|IGHA2_HUMAN Immunoglobulin heavy constant alpha 2 OS=Homo sapiens OX=9606 GN=IGHA2 PE=1 SV=4 | 576 | 20201119_15232_1ng_glyco_hcd203040_compound15.5930.5930.4 | controllerType=0 controllerNumber=1 scan=5930 | 16.5087 |
| 02898:5 | 16 | K.HYTN[+1647.613]SSQDVTVPC[+57.021]RVPPPPPC[+57.021]C[+57.021]HPR.L | HexNAc(5)Hex(3)Fuc(1) | C[+57]*3, N[+1648] | 922.0023 | 5 | 4605.9826 | 4605.9689 | 0 | 3 | 89 | Specific | 653.64 | 391.71 | 391.71 | 13.02 | 40 | >sp\|P01877\|IGHA2_HUMAN Immunoglobulin heavy constant alpha 2 OS=Homo sapiens OX=9606 GN=IGHA2 PE=1 SV=4 | 576 | 20201119_15232_1ng_glyco_hcd203040_compound15.5895.5895.5 | controllerType=0 controllerNumber=1 scan=5895 | 16.44 |
| 02928:5 | 16 | K.HYTN[+1647.613]SSQDVTVPC[+57.021]RVPPPPPC[+57.021]C[+57.021]HPR.L | HexNAc(5)Hex(3)Fuc(1) | C[+57]*3, N[+1648] | 922.0025 | 5 | 4605.9835 | 4605.9689 | 0 | 3.2 | 89 | Specific | 684.09 | 446.08 | 446.08 | 13.02 | 40 | >sp\|P01877\|IGHA2_HUMAN Immunoglobulin heavy constant alpha 2 OS=Homo sapiens OX=9606 GN=IGHA2 PE=1 SV=4 | 576 | 20201119_15232_1ng_glyco_hcd203040_compound15.5927.5927.5 | controllerType=0 controllerNumber=1 scan=5927 | 16.5014 |
| 02933:4 | 16 | K.HYTN[+1647.613]SSQDVTVPC[+57.021]RVPPPPPC[+57.021]C[+57.021]HPR.L | HexNAc(5)Hex(3)Fuc(1) | C[+57]*3, N[+1648] | 1152.2515 | 4 | 4605.9842 | 4605.9689 | 0 | 3.3 | 89 | Specific | 685.07 | 448.68 | 448.68 | 12.45 | 40 | >sp\|P01877\|IGHA2_HUMAN Immunoglobulin heavy constant alpha 2 OS=Homo sapiens OX=9606 GN=IGHA2 PE=1 SV=4 | 576 | 20201119_15232_1ng_glyco_hcd203040_compound15.5932.5932.4 | controllerType=0 controllerNumber=1 scan=5932 | 16.5127 |
| 02930:5 | 16 | K.HYTN[+1647.613]SSQDVTVPC[+57.021]RVPPPPPC[+57.021]C[+57.021]HPR.L | HexNAc(5)Hex(3)Fuc(1) | C[+57]*3, N[+1648] | 922.0025 | 5 | 4605.9835 | 4605.9689 | 0 | 3.2 | 89 | Specific | 759.79 | 615.77 | 615.77 | 15.02 | 40 | >sp\|P01877\|IGHA2_HUMAN Immunoglobulin heavy constant alpha 2 OS=Homo sapiens OX=9606 GN=IGHA2 PE=1 SV=4 | 576 | 20201119_15232_1ng_glyco_hcd203040_compound15.5929.5929.5 | controllerType=0 controllerNumber=1 scan=5929 | 16.5068 |
| 02900:5 | 16 | K.HYTN[+1647.613]SSQDVTVPC[+57.021]RVPPPPPC[+57.021]C[+57.021]HPR.L | HexNAc(5)Hex(3)Fuc(1) | C[+57]*3, N[+1648] | 922.0023 | 5 | 4605.9826 | 4605.9689 | 0 | 3 | 89 | Specific | 820.83 | 564.8 | 564.8 | 16.02 | 40 | >sp\|P01877\|IGHA2_HUMAN Immunoglobulin heavy constant alpha 2 OS=Homo sapiens OX=9606 GN=IGHA2 PE=1 SV=4 | 576 | 20201119_15232_1ng_glyco_hcd203040_compound15.5897.5897.5 | controllerType=0 controllerNumber=1 scan=5897 | 16.4478 |
| 04426:3 | 16 | K.TPLTAN[+2262.814]ITK.S | HexNAc(5)Hex(5)Fuc(1)NeuAc(1) | N[+2263] | 1074.4656 | 3 | 3221.3823 | 3221.3711 | 0 | 3.5 | 200 | Specific | 229.07 | 162.84 | 162.84 | 5.3 | 40 | >sp\|P01877\|IGHA2_HUMAN Immunoglobulin heavy constant alpha 2 OS=Homo sapiens OX=9606 GN=IGHA2 PE=1 SV=4 | 576 | 20201119_15232_1ng_glyco_hcd203040_compound15.7481.7481.3 | controllerType=0 controllerNumber=1 scan=7481 | 19.0754 |
| 02554:3 | 16 | K.TPLTAN[+1971.719]ITK.S | HexNAc(5)Hex(5)Fuc(1) | N[+1972] | 977.4327 | 3 | 2930.2836 | 2930.2757 | 0 | 2.7 | 200 | Specific | 301.08 | 198.18 | 198.18 | 5.81 | 40 | >sp\|P01877\|IGHA2_HUMAN Immunoglobulin heavy constant alpha 2 OS=Homo sapiens OX=9606 GN=IGHA2 PE=1 SV=4 | 576 | 20201119_15232_1ng_glyco_hcd203040_compound15.5539.5539.3 | controllerType=0 controllerNumber=1 scan=5539 | 15.7973 |
| 02679:2 | 16 | K.TPLTAN[+1501.555]ITK.S | HexNAc(5)Hex(3) | N[+1502] | 1230.5625 | 2 | 2460.1178 | 2460.1121 | 0 | 2.3 | 200 | Specific | 351.11 | 216.59 | 216.59 | 6.34 | 40 | >sp\|P01877\|IGHA2_HUMAN Immunoglobulin heavy constant alpha 2 OS=Homo sapiens OX=9606 GN=IGHA2 PE=1 SV=4 | 576 | 20201119_15232_1ng_glyco_hcd203040_compound15.5668.5668.2 | controllerType=0 controllerNumber=1 scan=5668 | 16.0325 |
| 02604:3 | 16 | K.TPLTAN[+1809.666]ITK.S | HexNAc(5)Hex(4)Fuc(1) | N[+1810] | 923.4147 | 3 | 2768.2296 | 2768.2228 | 0 | 2.4 | 200 | Specific | 361.46 | 94.56 | 94.56 | 4.92 | 40 | >sp\|P01877\|IGHA2_HUMAN Immunoglobulin heavy constant alpha 2 OS=Homo sapiens OX=9606 GN=IGHA2 PE=1 SV=4 | 576 | 20201119_15232_1ng_glyco_hcd203040_compound15.5591.5591.3 | controllerType=0 controllerNumber=1 scan=5591 | 15.8925 |
| 02680:2 | 16 | K.TPLTAN[+1501.555]ITK.S | HexNAc(5)Hex(3) | N[+1502] | 1230.5625 | 2 | 2460.1178 | 2460.1121 | 0 | 2.3 | 200 | Specific | 407.06 | 216.06 | 216.06 | 6.42 | 40 | >sp\|P01877\|IGHA2_HUMAN Immunoglobulin heavy constant alpha 2 OS=Homo sapiens OX=9606 GN=IGHA2 PE=1 SV=4 | 576 | 20201119_15232_1ng_glyco_hcd203040_compound15.5669.5669.2 | controllerType=0 controllerNumber=1 scan=5669 | 16.0358 |
| 02681:3 | 16 | K.TPLTAN[+1647.613]ITK.S | HexNAc(5)Hex(3)Fuc(1) | N[+1648] | 869.3976 | 3 | 2606.1781 | 2606.17 | 0 | 3.1 | 200 | Specific | 458.86 | 178.74 | 178.74 | 6.52 | 40 | >sp\|P01877\|IGHA2_HUMAN Immunoglobulin heavy constant alpha 2 OS=Homo sapiens OX=9606 GN=IGHA2 PE=1 SV=4 | 576 | 20201119_15232_1ng_glyco_hcd203040_compound15.5671.5671.3 | controllerType=0 controllerNumber=1 scan=5671 | 16.0404 |
| 02606:3 | 16 | K.TPLTAN[+1809.666]ITK.S | HexNAc(5)Hex(4)Fuc(1) | N[+1810] | 923.4147 | 3 | 2768.2296 | 2768.2228 | 0 | 2.4 | 200 | Specific | 491.74 | 255.83 | 255.83 | 7 | 40 | >sp\|P01877\|IGHA2_HUMAN Immunoglobulin heavy constant alpha 2 OS=Homo sapiens OX=9606 GN=IGHA2 PE=1 SV=4 | 576 | 20201119_15232_1ng_glyco_hcd203040_compound15.5593.5593.3 | controllerType=0 controllerNumber=1 scan=5593 | [illegible] |
| 02602:2 | 16 | K.TPLTAN[+1444.534]ITK.S | HexNAc(4)Hex(3)Fuc(1) | N[+1445] | 1202.0519 | 2 | 2403.0966 | 2403.0906 | 0 | 2.5 | 200 | Specific | 560.57 | 317.53 | 317.53 | 7.74 | 40 | >sp\|P01877\|IGHA2_HUMAN Immunoglobulin heavy constant alpha 2 OS=Homo sapiens OX=9606 GN=IGHA2 PE=1 SV=4 | 576 | 20201119_15232_1ng_glyco_hcd203040_compound15.5589.5589.2 | controllerType=0 controllerNumber=1 scan=5589 | [illegible] |
| 02683:3 | 16 | K.TPLTAN[+1647.613]ITK.S | HexNAc(5)Hex(3)Fuc(1) | N[+1648] | 869.3976 | 3 | 2606.1781 | 2606.17 | 0 | 3.1 | 200 | Specific | 563.06 | 266.81 | 266.81 | 7.57 | 40 | >sp\|P01877\|IGHA2_HUMAN Immunoglobulin heavy constant alpha 2 OS=Homo sapiens OX=9606 GN=IGHA2 PE=1 SV=4 | 576 | 20201119_15232_1ng_glyco_hcd203040_compound15.5673.5673.3 | controllerType=0 controllerNumber=1 scan=5673 | 16.0461 |
| 02600:2 | 16 | K.TPLTAN[+1444.534]ITK.S | HexNAc(4)Hex(3)Fuc(1) | N[+1445] | 1202.0519 | 2 | 2403.0966 | 2403.0906 | 0 | 2.5 | 200 | Specific | 563.84 | 310.1 | 310.1 | 7.74 | 40 | >sp\|P01877\|IGHA2_HUMAN Immunoglobulin heavy constant alpha 2 OS=Homo sapiens OX=9606 GN=IGHA2 PE=1 SV=4 | 576 | 20201119_15232_1ng_glyco_hcd203040_compound15.5587.5587.2 | controllerType=0 controllerNumber=1 scan=5587 | 15.8852 |
| 02605:2 | 16 | K.TPLTAN[+1809.666]ITK.S | HexNAc(5)Hex(4)Fuc(1) | N[+1810] | 1384.6176 | 2 | 2768.2279 | 2768.2228 | 0 | 1.8 | 200 | Specific | 579.49 | 330.92 | 330.92 | 9.02 | 40 | >sp\|P01877\|IGHA2_HUMAN Immunoglobulin heavy constant alpha 2 OS=Homo sapiens OX=9606 GN=IGHA2 PE=1 SV=4 | 576 | 20201119_15232_1ng_glyco_hcd203040_compound15.5592.5592.2 | controllerType=0 controllerNumber=1 scan=5592 | 15.894 |
| 02607:2 | 16 | K.TPLTAN[+1809.666]ITK.S | HexNAc(5)Hex(4)Fuc(1) | N[+1810] | 1384.6176 | 2 | 2768.2279 | 2768.2228 | 0 | 1.8 | 200 | Specific | 580.66 | 271.07 | 271.07 | 8.8 | 40 | >sp\|P01877\|IGHA2_HUMAN Immunoglobulin heavy constant alpha 2 OS=Homo sapiens OX=9606 GN=IGHA2 PE=1 SV=4 | 576 | 20201119_15232_1ng_glyco_hcd203040_compound15.5594.5594.2 | controllerType=0 controllerNumber=1 scan=5594 | 15.8998 |
| 02663:2 | 16 | K.TPLTAN[+1647.613]ITK.S | HexNAc(5)Hex(3)Fuc(1) | N[+1648] | 1303.5914 | 2 | 2606.1754 | 2606.17 | 0 | 2.1 | 200 | Specific | 602.12 | 299.69 | 299.69 | 9.86 | 40 | >sp\|P01877\|IGHA2_HUMAN Immunoglobulin heavy constant alpha 2 OS=Homo sapiens OX=9606 GN=IGHA2 PE=1 SV=4 | 576 | 20201119_15232_1ng_glyco_hcd203040_compound15.5652.5652.2 | controllerType=0 controllerNumber=1 scan=5652 | 16.0041 |
| 02661:2 | 16 | K.TPLTAN[+1647.613]ITK.S | HexNAc(5)Hex(3)Fuc(1) | N[+1648] | 1303.5914 | 2 | 2606.1754 | 2606.17 | 0 | 2.1 | 200 | Specific | 622.71 | 360.68 | 360.68 | 10.36 | 40 | >sp\|P01877\|IGHA2_HUMAN Immunoglobulin heavy constant alpha 2 OS=Homo sapiens OX=9606 GN=IGHA2 PE=1 SV=4 | 576 | 20201119_15232_1ng_glyco_hcd203040_compound15.5650.5650.2 | controllerType=0 controllerNumber=1 scan=5650 | 15.9983 |
| 10132:3 | 16 | R.MAGKPTHIN[+1501.555]VSVVMAEADGTC[+57.021]Y.- | HexNAc(5)Hex(3) | C[+57], N[+1502] | 1284.8868 | 3 | 3852.6458 | 3852.6382 | 0 | 2 | 319 | CRagged | 374.84 | 371.07 | 371.07 | 8.05 | 40 | >sp\|P01877\|IGHA2_HUMAN Immunoglobulin heavy constant alpha 2 OS=Homo sapiens OX=9606 GN=IGHA2 PE=1 SV=4 | 576 | 20201119_15232_1ng_glyco_hcd203040_compound15.13413.13413.3 | controllerType=0 controllerNumber=1 scan=13413 | 28.9373 |
| 10130:3 | 16 | R.MAGKPTHIN[+1501.555]VSVVMAEADGTC[+57.021]Y.- | HexNAc(5)Hex(3) | C[+57], N[+1502] | 1284.8868 | 3 | 3852.6458 | 3852.6382 | 0 | 2 | 319 | CRagged | 414.71 | 414.71 | 414.71 | 9.61 | 40 | >sp\|P01877\|IGHA2_HUMAN Immunoglobulin heavy constant alpha 2 OS=Homo sapiens OX=9606 GN=IGHA2 PE=1 SV=4 | 576 | 20201119_15232_1ng_glyco_hcd203040_compound15.13411.13411.3 | controllerType=0 controllerNumber=1 scan=13411 | 28.9332 |
| 10051:3 | 16 | R.MAGKPTHIN[+1216.423]VSVVMAEADGTC[+57.021]Y.- | HexNAc(2)Hex(5) | C[+57], N[+1216] | 1189.8411 | 3 | 3567.5088 | 3567.5057 | 0 | 0.9 | 319 | CRagged | 683.04 | 683.04 | 372.4 | 13.49 | 40 | >sp\|P01877\|IGHA2_HUMAN Immunoglobulin heavy constant alpha 2 OS=Homo sapiens OX=9606 GN=IGHA2 PE=1 SV=4 | 576 | 20201119_15232_1ng_glyco_hcd203040_compound15.13329.13329.3 | controllerType=0 controllerNumber=1 scan=13329 | 28.7907 |
| 09904:3 | 16 | R.MAGKPTHIN[+1702.581]VSVVMAEADGTC[+57.021]Y.- | HexNAc(2)Hex(8) | C[+57], N[+1703] | 1351.8967 | 3 | 4053.6754 | 4053.6642 | 0 | 2.8 | 319 | CRagged | 703.58 | 497.39 | 394.05 | 14.7 | 40 | >sp\|P01877\|IGHA2_HUMAN Immunoglobulin heavy constant alpha 2 OS=Homo sapiens OX=9606 GN=IGHA2 PE=1 SV=4 | 576 | 20201119_15232_1ng_glyco_hcd203040_compound15.13177.13177.3 | controllerType=0 controllerNumber=1 scan=13177 | 28.5222 |
| 09962:3 | 16 | R.MAGKPTHIN[+1378.476]VSVVMAEADGTC[+57.021]Y.- | HexNAc(2)Hex(6) | C[+57], N[+1378] | 1243.8633 | 3 | 3729.5755 | 3729.5585 | 0 | 4.5 | 319 | CRagged | 792.73 | 674.44 | 410.95 | 15.47 | 40 | >sp\|P01877\|IGHA2_HUMAN Immunoglobulin heavy constant alpha 2 OS=Homo sapiens OX=9606 GN=IGHA2 PE=1 SV=4 | 576 | 20201119_15232_1ng_glyco_hcd203040_compound15.13237.13237.3 | controllerType=0 controllerNumber=1 scan=13237 | 28.6313 |
| 10053:3 | 16 | R.MAGKPTHIN[+1216.423]VSVVMAEADGTC[+57.021]Y.- | HexNAc(2)Hex(5) | C[+57], N[+1216] | 1189.8411 | 3 | 3567.5088 | 3567.5057 | 0 | 0.9 | 319 | CRagged | 802.26 | 629.96 | 442.88 | 16.49 | 40 | >sp\|P01877\|IGHA2_HUMAN Immunoglobulin heavy constant alpha 2 OS=Homo sapiens OX=9606 GN=IGHA2 PE=1 SV=4 | 576 | 20201119_15232_1ng_glyco_hcd203040_compound15.13331.13331.3 | controllerType=0 controllerNumber=1 scan=13331 | 28.7952 |
| 09960:3 | 16 | R.MAGKPTHIN[+1378.476]VSVVMAEADGTC[+57.021]Y.- | HexNAc(2)Hex(6) | C[+57], N[+1378] | 1243.8633 | 3 | 3729.5755 | 3729.5585 | 0 | 4.5 | 319 | CRagged | 806.86 | 738.38 | 470.04 | 16.47 | 40 | >sp\|P01877\|IGHA2_HUMAN Immunoglobulin heavy constant alpha 2 OS=Homo sapiens OX=9606 GN=IGHA2 PE=1 SV=4 | 576 | 20201119_15232_1ng_glyco_hcd203040_compound15.13235.13235.3 | controllerType=0 controllerNumber=1 scan=13235 | 28.6263 |
| 09937:3 | 16 | R.MAGKPTHIN[+1540.529]VSVVMAEADGTC[+57.021]Y.- | HexNAc(2)Hex(7) | C[+57], N[+1541] | 1297.8793 | 3 | 3891.6234 | 3891.6114 | 0 | 3.1 | 319 | CRagged | 813.8 | 813.8 | 428.59 | 16.7 | 40 | >sp\|P01877\|IGHA2_HUMAN Immunoglobulin heavy constant alpha 2 OS=Homo sapiens OX=9606 GN=IGHA2 PE=1 SV=4 | 576 | 20201119_15232_1ng_glyco_hcd203040_compound15.13211.13211.3 | controllerType=0 controllerNumber=1 scan=13211 | 28.5826 |
| 09906:3 | 16 | R.MAGKPTHIN[+1702.581]VSVVMAEADGTC[+57.021]Y.- | HexNAc(2)Hex(8) | C[+57], N[+1703] | 1351.8967 | 3 | 4053.6754 | 4053.6642 | 0 | 2.8 | 319 | CRagged | 824.1 | 652.36 | 458.45 | 16.7 | 40 | >sp\|P01877\|IGHA2_HUMAN Immunoglobulin heavy constant alpha 2 OS=Homo sapiens OX=9606 GN=IGHA2 PE=1 SV=4 | 576 | 20201119_15232_1ng_glyco_hcd203040_compound15.13179.13179.3 | controllerType=0 controllerNumber=1 scan=13179 | 28.527 |
| 09934:3 | 16 | R.MAGKPTHIN[+1540.529]VSVVMAEADGTC[+57.021]Y.- | HexNAc(2)Hex(7) | C[+57], N[+1541] | 1297.8793 | 3 | 3891.6234 | 3891.6114 | 0 | 3.1 | 319 | CRagged | 875.14 | 875.14 | 524.96 | 17.7 | 40 | >sp\|P01877\|IGHA2_HUMAN Immunoglobulin heavy constant alpha 2 OS=Homo sapiens OX=9606 GN=IGHA2 PE=1 SV=4 | 576 | 20201119_15232_1ng_glyco_hcd203040_compound15.13208.13208.3 | controllerType=0 controllerNumber=1 scan=13208 | 28.5768 |
| 22732:4 | 18 | K.APDKNVIFSPLSISTALAFLSLGAHN[+2350.830]TTLTEILK.G | HexNAc(4)Hex(5)Fuc(1)NeuAc(2) | N[+2351] | 1484.2108 | 4 | 5933.8214 | 5933.8085 | 0 | 2.2 | 68 | Specific | 423.19 | 422.43 | 422.43 | 9.16 | 30 | >sp\|P01011\|AACT_HUMAN Alpha-1-antichymotrypsin OS=Homo sapiens OX=9606 GN=SERPINA3 PE=1 SV=2 | 1314 | 20201119_15232_1ng_glyco_hcd203040_compound15.26601.26601.4 | controllerType=0 controllerNumber=1 scan=26601 | 53.5912 |
| 22730:4 | 18 | K.APDKNVIFSPLSISTALAFLSLGAHN[+2350.830]TTLTEILK.G | HexNAc(4)Hex(5)Fuc(1)NeuAc(2) | N[+2351] | 1484.2108 | 4 | 5933.8214 | 5933.8085 | 0 | 2.2 | 68 | Specific | 614.92 | 614.92 | 614.92 | 12.19 | 30 | >sp\|P01011\|AACT_HUMAN Alpha-1-antichymotrypsin OS=Homo sapiens OX=9606 GN=SERPINA3 PE=1 SV=2 | 1314 | 20201119_15232_1ng_glyco_hcd203040_compound15.26599.26599.4 | controllerType=0 controllerNumber=1 scan=26599 | 53.5873 |
| 13739:4 | 19 | K.LKELPGVC[+57.021]N[+2204.772]ETMMALWEEC[+57.021]KPC[+57.021]LK.Q | HexNAc(4)Hex(5)NeuAc(2) | C[+57]*3, N[+2205] | 1286.052 | 4 | 5141.1862 | 5141.1572 | 0 | 5.6 | 95 | Specific | 279.38 | 236.36 | 236.36 | 4.91 | 31 | >sp\|P10909\|CLUS_HUMAN Clusterin OS=Homo sapiens OX=9606 GN=CLU PE=1 SV=1 | 1467 | 20201119_15232_1ng_glyco_hcd203040_compound15.17179.17179.4 | controllerType=0 controllerNumber=1 scan=17179 | 35.3857 |
| 13714:4 | 19 | K.LKELPGVC[+57.021]N[+2059.735]ETMMALWEEC[+57.021]KPC[+57.021]LK.Q | HexNAc(4)Hex(5)Fuc(1)NeuAc(1) | C[+57]*3, N[+2060] | 1249.7892 | 4 | 4996.1349 | 4996.1196 | 0 | 3.1 | 95 | Specific | 863.36 | 803.56 | 803.56 | 17.02 | 31 | >sp\|P10909\|CLUS_HUMAN Clusterin OS=Homo sapiens OX=9606 GN=CLU PE=1 SV=1 | 1467 | 20201119_15232_1ng_glyco_hcd203040_compound15.17153.17153.4 | controllerType=0 controllerNumber=1 scan=17153 | 35.3341 |
| 13711:4 | 19 | K.LKELPGVC[+57.021]N[+2059.735]ETMMALWEEC[+57.021]KPC[+57.021]LK.Q | HexNAc(4)Hex(5)Fuc(1)NeuAc(1) | C[+57]*3, N[+2060] | 1249.7892 | 4 | 4996.1349 | 4996.1196 | 0 | 3.1 | 95 | Specific | 941.75 | 941.75 | 941.75 | 18.02 | 31 | >sp\|P10909\|CLUS_HUMAN Clusterin OS=Homo sapiens OX=9606 GN=CLU PE=1 SV=1 | 1467 | 20201119_15232_1ng_glyco_hcd203040_compound15.17150.17150.4 | controllerType=0 controllerNumber=1 scan=17150 | 35.3276 |
| 20773:4 | 19 | K.MLN[+2059.735]TSSLLEQLNEQFNWVSR.L | HexNAc(4)Hex(5)Fuc(1)NeuAc(1) | N[+2060] | 1117.9899 | 4 | 4468.9378 | 4468.9217 | 0 | 3.6 | 352 | Specific | 283.47 | 186.4 | 186.4 | 4.92 | 31 | >sp\|P10909\|CLUS_HUMAN Clusterin OS=Homo sapiens OX=9606 GN=CLU PE=1 SV=1 | 1467 | 20201119_15232_1ng_glyco_hcd203040_compound15.24556.24556.4 | controllerType=0 controllerNumber=1 scan=24556 | 49.8764 |
| 23138:4 | 19 | K.MLN[+2350.830]TSSLLEQLNEQFNWVSR.L | HexNAc(4)Hex(5)Fuc(1)NeuAc(2) | N[+2351] | 1190.7633 | 4 | 4760.0316 | 4760.0171 | 0 | 3 | 352 | Specific | 392.59 | 325.83 | 325.83 | 7.97 | 31 | >sp\|P10909\|CLUS_HUMAN Clusterin OS=Homo sapiens OX=9606 GN=CLU PE=1 SV=1 | 1467 | 20201119_15232_1ng_glyco_hcd203040_compound15.27024.27024.4 | controllerType=0 controllerNumber=1 scan=27024 | 54.438 |
| 23183:4 | 19 | K.MLN[+2351.851]TSSLLEQLNEQFNWVSR.L | HexNAc(4)Hex(5)Fuc(3)NeuAc(1) | N[+2352] | 1191.0166 | 4 | 4761.0447 | 4761.0375 | 0 | 1.5 | 352 | Specific | 480.77 | 419.07 | 419.07 | 9.87 | 31 | >sp\|P10909\|CLUS_HUMAN Clusterin OS=Homo sapiens OX=9606 GN=CLU PE=1 SV=1 | 1467 | 20201119_15232_1ng_glyco_hcd203040_compound15.27071.27071.4 | controllerType=0 controllerNumber=1 scan=27071 | 54.5281 |
| 23185:4 | 19 | K.MLN[+2351.851]TSSLLEQLNEQFNWVSR.L | HexNAc(4)Hex(5)Fuc(3)NeuAc(1) | N[+2352] | 1191.0166 | 4 | 4761.0447 | 4761.0375 | 0 | 1.5 | 352 | Specific | 567.69 | 412.64 | 412.64 | 10.38 | 31 | >sp\|P10909\|CLUS_HUMAN Clusterin OS=Homo sapiens OX=9606 GN=CLU PE=1 SV=1 | 1467 | 20201119_15232_1ng_glyco_hcd203040_compound15.27073.27073.4 | controllerType=0 controllerNumber=1 scan=27073 | 54.5327 |
| 23136:4 | 19 | K.MLN[+2350.830]TSSLLEQLNEQFNWVSR.L | HexNAc(4)Hex(5)Fuc(1)NeuAc(2) | N[+2351] | 1190.7633 | 4 | 4760.0316 | 4760.0171 | 0 | 3 | 352 | Specific | 726.09 | 506.46 | 506.46 | 14.82 | 31 | >sp\|P10909\|CLUS_HUMAN Clusterin OS=Homo sapiens OX=9606 GN=CLU PE=1 SV=1 | 1467 | 20201119_15232_1ng_glyco_hcd203040_compound15.27022.27022.4 | controllerType=0 controllerNumber=1 scan=27022 | 54.4329 |
| 20729:4 | 19 | K.MLN[+2059.735]TSSLLEQLNEQFNWVSR.L | HexNAc(4)Hex(5)Fuc(1)NeuAc(1) | N[+2060] | 1117.9905 | 4 | 4468.9403 | 4468.9217 | 0 | 4.2 | 352 | Specific | 741.83 | 565.22 | 565.22 | 14.59 | 31 | >sp\|P10909\|CLUS_HUMAN Clusterin OS=Homo sapiens OX=9606 GN=CLU PE=1 SV=1 | 1467 | 20201119_15232_1ng_glyco_hcd203040_compound15.24510.24510.7 | controllerType=0 controllerNumber=1 scan=24510 | 49.7845 |
| 20731:4 | 19 | K.MLN[+2059.735]TSSLLEQLNEQFNWVSR.L | HexNAc(4)Hex(5)Fuc(1)NeuAc(1) | N[+2060] | 1117.9905 | 4 | 4468.9403 | 4468.9217 | 0 | 4.2 | 352 | Specific | 766.5 | 659.07 | 659.07 | 15.59 | 31 | >sp\|P10909\|CLUS_HUMAN Clusterin OS=Homo sapiens OX=9606 GN=CLU PE=1 SV=1 | 1467 | 20201119_15232_1ng_glyco_hcd203040_compound15.24512.24512.7 | controllerType=0 controllerNumber=1 scan=24512 | 49.7893 |
| 20716:3 | 19 | K.MLN[+2059.735]TSSLLEQLNEQFNWVSR.L | HexNAc(4)Hex(5)Fuc(1)NeuAc(1) | N[+2060] | 1490.3179 | 3 | 4468.9392 | 4468.9217 | 0 | 3.9 | 352 | Specific | 885.42 | 622.84 | 622.84 | 17.64 | 31 | >sp\|P10909\|CLUS_HUMAN Clusterin OS=Homo sapiens OX=9606 GN=CLU PE=1 SV=1 | 1467 | 20201119_15232_1ng_glyco_hcd203040_compound15.24497.24497.5 | controllerType=0 controllerNumber=1 scan=24497 | 49.7611 |
| 20713:3 | 19 | K.MLN[+2059.735]TSSLLEQLNEQFNWVSR.L | HexNAc(4)Hex(5)Fuc(1)NeuAc(1) | N[+2060] | 1490.3179 | 3 | 4468.9392 | 4468.9217 | 0 | 3.9 | 352 | Specific | 1012.01 | 699.76 | 699.76 | 20.64 | 31 | >sp\|P10909\|CLUS_HUMAN Clusterin OS=Homo sapiens OX=9606 GN=CLU PE=1 SV=1 | 1467 | 20201119_15232_1ng_glyco_hcd203040_compound15.24494.24494.5 | controllerType=0 controllerNumber=1 scan=24494 | 49.7551 |
| 12744:3 | 19 | R.LAN[+2350.830]LTQGEDQYYLR.V | HexNAc(4)Hex(5)Fuc(1)NeuAc(2) | N[+2351] | 1345.565 | 3 | 4034.6804 | 4034.664 | 0 | 4.1 | 372 | Specific | 471.76 | 429.34 | 429.34 | 9.15 | 31 | >sp\|P10909\|CLUS_HUMAN Clusterin OS=Homo sapiens OX=9606 GN=CLU PE=1 SV=1 | 1467 | 20201119_15232_1ng_glyco_hcd203040_compound15.16148.16148.3 | controllerType=0 controllerNumber=1 scan=16148 | 33.5422 |
| 12746:3 | 19 | R.LAN[+2350.830]LTQGEDQYYLR.V | HexNAc(4)Hex(5)Fuc(1)NeuAc(2) | N[+2351] | 1345.565 | 3 | 4034.6804 | 4034.664 | 0 | 4.1 | 372 | Specific | 566.73 | 479.6 | 479.6 | 9.63 | 31 | >sp\|P10909\|CLUS_HUMAN Clusterin OS=Homo sapiens OX=9606 GN=CLU PE=1 SV=1 | 1467 | 20201119_15232_1ng_glyco_hcd203040_compound15.16150.16150.3 | controllerType=0 controllerNumber=1 scan=16150 | 33.5459 |
| 23895:3 | 21 | R.QLAHQSN[+2204.772]STNIFFSPVSIATAFAMLSLGTK.A | HexNAc(4)Hex(5)NeuAc(2) | N[+2205] | 1796.1494 | 3 | 5386.4336 | 5386.4075 | 0 | 4.8 | 64 | Specific | 458.74 | 450.25 | 450.25 | 9.62 | 31 | >sp\|P01009\|A1AT_HUMAN Alpha-1-antitrypsin OS=Homo sapiens OX=9606 GN=SERPINA1 PE=1 SV=3 | 642 | 20201119_15232_1ng_glyco_hcd203040_compound15.27811.27811.3 | controllerType=0 controllerNumber=1 scan=27811 | 55.9474 |
| 23894:4 | 21 | R.QLAHQSN[+2204.772]STNIFFSPVSIATAFAMLSLGTK.A | HexNAc(4)Hex(5)NeuAc(2) | N[+2205] | 1347.3606 | 4 | 5386.4206 | 5386.4075 | 0 | 2.4 | 64 | Specific | 706.18 | 650.11 | 485.59 | 14.46 | 31 | >sp\|P01009\|A1AT_HUMAN Alpha-1-antitrypsin OS=Homo sapiens OX=9606 GN=SERPINA1 PE=1 SV=3 | 642 | 20201119_15232_1ng_glyco_hcd203040_compound15.27810.27810.4 | controllerType=0 controllerNumber=1 scan=27810 | 55.946 |
| 23897:3 | 21 | R.QLAHQSN[+2204.772]STNIFFSPVSIATAFAMLSLGTK.A | HexNAc(4)Hex(5)NeuAc(2) | N[+2205] | 1796.1494 | 3 | 5386.4336 | 5386.4075 | 0 | 4.8 | 64 | Specific | 712.51 | 707.58 | 707.58 | 14.35 | 31 | >sp\|P01009\|A1AT_HUMAN Alpha-1-antitrypsin OS=Homo sapiens OX=9606 GN=SERPINA1 PE=1 SV=3 | 642 | 20201119_15232_1ng_glyco_hcd203040_compound15.27813.27813.3 | controllerType=0 controllerNumber=1 scan=27813 | 55.9528 |
| 23896:4 | 21 | R.QLAHQSN[+2204.772]STNIFFSPVSIATAFAMLSLGTK.A | HexNAc(4)Hex(5)NeuAc(2) | N[+2205] | 1347.3606 | 4 | 5386.4206 | 5386.4075 | 0 | 2.4 | 64 | Specific | 941.33 | 786.67 | 632.24 | 18.46 | 31 | >sp\|P01009\|A1AT_HUMAN Alpha-1-antitrypsin OS=Homo sapiens OX=9606 GN=SERPINA1 PE=1 SV=3 | 642 | 20201119_15232_1ng_glyco_hcd203040_compound15.27812.27812.4 | controllerType=0 controllerNumber=1 scan=27812 | 55.9509 |
| 08270:4 | 24 | K.SC[+57.021]QHN[+2204.772]GTMYQHGEIFSAHELFPSR.L | HexNAc(4)Hex(5)NeuAc(2) | C[+57], N[+2205] | 1260.2728 | 4 | 5038.0695 | 5038.0294 | 0 | 8 | 110 | Specific | 418.54 | 415.9 | 415.9 | 7.65 | 25 | >sp\|Q6WN34\|CRDL2_HUMAN Chordin-like protein 2 OS=Homo sapiens OX=9606 GN=CHRDL2 PE=1 SV=1 | 1189 | 20201119_15232_1ng_glyco_hcd203040_compound15.11467.11467.2 | controllerType=0 controllerNumber=1 scan=11467 | 25.6389 |
| 08272:4 | 24 | K.SC[+57.021]QHN[+2204.772]GTMYQHGEIFSAHELFPSR.L | HexNAc(4)Hex(5)NeuAc(2) | C[+57], N[+2205] | 1260.2728 | 4 | 5038.0695 | 5038.0294 | 0 | 8 | 110 | Specific | 462.09 | 462.09 | 462.09 | 7.96 | 25 | >sp\|Q6WN34\|CRDL2_HUMAN Chordin-like protein 2 OS=Homo sapiens OX=9606 GN=CHRDL2 PE=1 SV=1 | 1189 | 20201119_15232_1ng_glyco_hcd203040_compound15.11469.11469.2 | controllerType=0 controllerNumber=1 scan=11469 | 25.6424 |
| 13965:2 | 33 | K.LC[+57.021]DN[+1866.688]LSTK.N | HexNAc(6)Hex(4) | C[+57], N[+1867] | 1409.0697 | 2 | 2817.1322 | 2817.1487 | 0 | -5.8 | 285 | Specific | 315.54 | 188.27 | 188.27 | 2.85 | 22 | >sp\|P02774\|VTDB_HUMAN Vitamin D-binding protein OS=Homo sapiens OX=9606 GN=GC PE=1 SV=2 | 1103 | 20201119_15232_1ng_glyco_hcd203040_compound15.17413.17413.2 | controllerType=0 controllerNumber=1 scan=17413 | 35.7824 |
| 13655:3 | 38 | R.SPYYN[+1913.677]VSDEISFHC[+57.021]YDGYTLR.G | HexNAc(4)Hex(5)NeuAc(1) | C[+57], N[+1914] | 1500.6083 | 3 | 4499.8104 | 4499.8012 | 0 | 2.1 | 118 | Specific | 249.22 | 249.22 | 249.22 | 6.65 | 19 | >sp\|P00751\|CFAB_HUMAN Complement factor B OS=Homo sapiens OX=9606 GN=CFB PE=1 SV=2 | 1353 | 20201119_15232_1ng_glyco_hcd203040_compound15.17092.17092.3 | controllerType=0 controllerNumber=1 scan=17092 | 35.2259 |
| 13657:3 | 38 | R.SPYYN[+1913.677]VSDEISFHC[+57.021]YDGYTLR.G | HexNAc(4)Hex(5)NeuAc(1) | C[+57], N[+1914] | 1500.6083 | 3 | 4499.8104 | 4499.8012 | 0 | 2.1 | 118 | Specific | 256.05 | 255.71 | 255.71 | 6.93 | 19 | >sp\|P00751\|CFAB_HUMAN Complement factor B OS=Homo sapiens OX=9606 GN=CFB PE=1 SV=2 | 1353 | 20201119_15232_1ng_glyco_hcd203040_compound15.17094.17094.3 | controllerType=0 controllerNumber=1 scan=17094 | 35.2299 |
| 09326:3 | 40 | R.IIVPLNNREN[+1663.608]ISDPTSPLR.T | HexNAc(5)Hex(4) | N[+1664] | 1271.2637 | 3 | 3811.7766 | 3811.7853 | 0 | -2.3 | 62 | Specific | 318.86 | 318.86 | 318.86 | 5.19 | 18 | >sp\|P01591\|IGJ_HUMAN Immunoglobulin J chain OS=Homo sapiens OX=9606 GN=JCHAIN PE=1 SV=4 | 1392 | 20201119_15232_1ng_glyco_hcd203040_compound15.12578.12578.3 | controllerType=0 controllerNumber=1 scan=12578 | 27.5391 |
| 09328:3 | 40 | R.IIVPLNNREN[+1663.608]ISDPTSPLR.T | HexNAc(5)Hex(4) | N[+1664] | 1271.2637 | 3 | 3811.7766 | 3811.7853 | 0 | -2.3 | 62 | Specific | 351.85 | 349.29 | 349.29 | 5.49 | 18 | >sp\|P01591\|IGJ_HUMAN Immunoglobulin J chain OS=Homo sapiens OX=9606 GN=JCHAIN PE=1 SV=4 | 1392 | 20201119_15232_1ng_glyco_hcd203040_compound15.12580.12580.3 | controllerType=0 controllerNumber=1 scan=12580 | 27.5438 |
| 09258:3 | 40 | R.IIVPLNNREN[+1809.666]ISDPTSPLR.T | HexNAc(5)Hex(4)Fuc(1) | N[+1810] | 1319.9561 | 3 | 3957.8538 | 3957.8432 | 0 | 2.7 | 62 | Specific | 421.09 | 357.67 | 357.67 | 7.17 | 18 | >sp\|P01591\|IGJ_HUMAN Immunoglobulin J chain OS=Homo sapiens OX=9606 GN=JCHAIN PE=1 SV=4 | 1392 | 20201119_15232_1ng_glyco_hcd203040_compound15.12507.12507.3 | controllerType=0 controllerNumber=1 scan=12507 | 27.4207 |
| 10941:3 | 40 | R.IIVPLNNREN[+2059.735]ISDPTSPLR.T | HexNAc(4)Hex(5)Fuc(1)NeuAc(1) | N[+2060] | 1403.3126 | 3 | 4207.9231 | 4207.9121 | 0 | 2.6 | 62 | Specific | 441.37 | 384.01 | 384.01 | 8.84 | 18 | >sp\|P01591\|IGJ_HUMAN Immunoglobulin J chain OS=Homo sapiens OX=9606 GN=JCHAIN PE=1 SV=4 | 1392 | 20201119_15232_1ng_glyco_hcd203040_compound15.14251.14251.3 | controllerType=0 controllerNumber=1 scan=14251 | 30.3302 |
| 10939:3 | 40 | R.IIVPLNNREN[+2059.735]ISDPTSPLR.T | HexNAc(4)Hex(5)Fuc(1)NeuAc(1) | N[+2060] | 1403.3126 | 3 | 4207.9231 | 4207.9121 | 0 | 2.6 | 62 | Specific | 457.95 | 442.84 | 442.84 | 8.95 | 18 | >sp\|P01591\|IGJ_HUMAN Immunoglobulin J chain OS=Homo sapiens OX=9606 GN=JCHAIN PE=1 SV=4 | 1392 | 20201119_15232_1ng_glyco_hcd203040_compound15.14249.14249.3 | controllerType=0 controllerNumber=1 scan=14249 | 30.3253 |
| 11034:3 | 40 | R.IIVPLNNREN[+1856.656]ISDPTSPLR.T | HexNAc(3)Hex(5)Fuc(1)NeuAc(1) | N[+1857] | 1335.617 | 3 | 4004.8364 | 4004.8327 | 0 | 0.9 | 62 | Specific | 483.95 | 483.95 | 483.95 | 8.79 | 18 | >sp\|P01591\|IGJ_HUMAN Immunoglobulin J chain OS=Homo sapiens OX=9606 GN=JCHAIN PE=1 SV=4 | 1392 | 20201119_15232_1ng_glyco_hcd203040_compound15.14347.14347.3 | controllerType=0 controllerNumber=1 scan=14347 | 30.5135 |
| 09260:3 | 40 | R.IIVPLNNREN[+1809.666]ISDPTSPLR.T | HexNAc(5)Hex(4)Fuc(1) | N[+1810] | 1319.9561 | 3 | 3957.8538 | 3957.8432 | 0 | 2.7 | 62 | Specific | 484.08 | 368.73 | 368.73 | 7.46 | 18 | >sp\|P01591\|IGJ_HUMAN Immunoglobulin J chain OS=Homo sapiens OX=9606 GN=JCHAIN PE=1 SV=4 | 1392 | 20201119_15232_1ng_glyco_hcd203040_compound15.12509.12509.3 | controllerType=0 controllerNumber=1 scan=12509 | 27.4247 |
| 11036:3 | 40 | R.IIVPLNNREN[+1856.656]ISDPTSPLR.T | HexNAc(3)Hex(5)Fuc(1)NeuAc(1) | N[+1857] | 1335.617 | 3 | 4004.8364 | 4004.8327 | 0 | 0.9 | 62 | Specific | 506.28 | 418.34 | 418.34 | 9.02 | 18 | >sp\|P01591\|IGJ_HUMAN Immunoglobulin J chain OS=Homo sapiens OX=9606 GN=JCHAIN PE=1 SV=4 | 1392 | 20201119_15232_1ng_glyco_hcd203040_compound15.14349.14349.3 | controllerType=0 controllerNumber=1 scan=14349 | 30.5186 |
| 10953:3 | 40 | R.IIVPLNNREN[+1913.677]ISDPTSPLR.T | HexNAc(4)Hex(5)NeuAc(1) | N[+1914] | 1354.6258 | 3 | 4061.8628 | 4061.8542 | 0 | 2.1 | 62 | Specific | 513.49 | 471.26 | 471.26 | 9.16 | 18 | >sp\|P01591\|IGJ_HUMAN Immunoglobulin J chain OS=Homo sapiens OX=9606 GN=JCHAIN PE=1 SV=4 | 1392 | 20201119_15232_1ng_glyco_hcd203040_compound15.14263.14263.3 | controllerType=0 controllerNumber=1 scan=14263 | 30.3492 |
| 10955:3 | 40 | R.IIVPLNNREN[+1913.677]ISDPTSPLR.T | HexNAc(4)Hex(5)NeuAc(1) | N[+1914] | 1354.6258 | 3 | 4061.8628 | 4061.8542 | 0 | 2.1 | 62 | Specific | 522.34 | 405.28 | 405.28 | 9.16 | 18 | >sp\|P01591\|IGJ_HUMAN Immunoglobulin J chain OS=Homo sapiens OX=9606 GN=JCHAIN PE=1 SV=4 | 1392 | 20201119_15232_1ng_glyco_hcd203040_compound15.14265.14265.3 | controllerType=0 controllerNumber=1 scan=14265 | 30.3535 |
| 16473:3 | 46 | K.AALAAFNAQNN[+2205.793]GSNFQLEEISR.A | HexNAc(4)Hex(5)Fuc(2)NeuAc(1) | N[+2206] | 1524.3276 | 3 | 4570.9682 | 4570.946 | 0 | 4.9 | 166 | Specific | 220.01 | 220.01 | 220.01 | 5.09 | 10 | >sp\|P02765\|FETUA_HUMAN Alpha-2-HS-glycoprotein OS=Homo sapiens OX=9606 GN=AHSG PE=1 SV=2 | 135 | 20201119_15232_1ng_glyco_hcd203040_compound15.20067.20067.3 | controllerType=0 controllerNumber=1 scan=20067 | 40.4578 |
| 16427:3 | 46 | K.AALAAFNAQNN[+2205.793]GSNFQLEEISR.A | HexNAc(4)Hex(5)Fuc(2)NeuAc(1) | N[+2206] | 1524.3202 | 3 | 4570.9462 | 4570.946 | 0 | 0.1 | 166 | Specific | 467.16 | 467.16 | 467.16 | 8.9 | 10 | >sp\|P02765\|FETUA_HUMAN Alpha-2-HS-glycoprotein OS=Homo sapiens OX=9606 GN=AHSG PE=1 SV=2 | 135 | 20201119_15232_1ng_glyco_hcd203040_compound15.20020.20020.3 | controllerType=0 controllerNumber=1 scan=20020 | 40.3808 |
| 16430:3 | 46 | K.AALAAFNAQNN[+2205.793]GSNFQLEEISR.A | HexNAc(4)Hex(5)Fuc(2)NeuAc(1) | N[+2206] | 1524.3202 | 3 | 4570.9462 | 4570.946 | 0 | 0.1 | 166 | Specific | 590.02 | 568.68 | 568.68 | 10.62 | 10 | >sp\|P02765\|FETUA_HUMAN Alpha-2-HS-glycoprotein OS=Homo sapiens OX=9606 GN=AHSG PE=1 SV=2 | 135 | 20201119_15232_1ng_glyco_hcd203040_compound15.20023.20023.3 | controllerType=0 controllerNumber=1 scan=20023 | 40.3855 |
| 18049:3 | 1 | R.RTAGWNVPIGTLRPFLN[+2205.793]WTGPPEPIEAAVAR.F | HexNAc(4)Hex(5)Fuc(2)NeuAc(1) | N[+2206] | 1864.8819 | 3 | 5592.6312 | 5592.6049 | 0 | 4.7 | 140 | Specific | 310.3 | 285.97 | 285.97 | 5.65 | 140 | >sp\|P02788\|TRFL_HUMAN Lactotransferrin OS=Homo sapiens OX=9606 GN=LTF PE=1 SV=6 | 1270 | 20201113_15413_1ng_glyco_hcd203040_compound15.21607.21607.3 | controllerType=0 controllerNumber=1 scan=21607 | 43.5914 |
| 19128:4 | 1 | R.RTAGWNVPIGTLRPFLN[+2350.830]WTGPPEPIEAAVAR.F | HexNAc(4)Hex(5)Fuc(1)NeuAc(2) | N[+2351] | 1435.172 | 4 | 5737.6662 | 5737.6424 | 0 | 4.1 | 140 | Specific | 407.19 | 407.19 | 407.19 | 9.77 | 140 | >sp\|P02788\|TRFL_HUMAN Lactotransferrin OS=Homo sapiens OX=9606 GN=LTF PE=1 SV=6 | 1270 | 20201113_15413_1ng_glyco_hcd203040_compound15.22735.22735.4 | controllerType=0 controllerNumber=1 scan=22735 | 45.9434 |
| 18050:4 | 1 | R.RTAGWNVPIGTLRPFLN[+2351.851]WTGPPEPIEAAVAR.F | HexNAc(4)Hex(5)Fuc(3)NeuAc(1) | N[+2352] | 1435.4274 | 4 | 5738.6877 | 5738.6628 | 0 | 4.3 | 140 | Specific | 410.74 | 290.38 | 290.38 | 7.93 | 140 | >sp\|P02788\|TRFL_HUMAN Lactotransferrin OS=Homo sapiens OX=9606 GN=LTF PE=1 SV=6 | 1270 | 20201113_15413_1ng_glyco_hcd203040_compound15.21608.21608.4 | controllerType=0 controllerNumber=1 scan=21608 | 43.593 |
| 19131:4 | 1 | R.RTAGWNVPIGTLRPFLN[+2350.830]WTGPPEPIEAAVAR.F | HexNAc(4)Hex(5)Fuc(1)NeuAc(2) | N[+2351] | 1435.172 | 4 | 5737.6662 | 5737.6424 | 0 | 4.1 | 140 | Specific | 467.99 | 467.99 | 467.99 | 9.79 | 140 | >sp\|P02788\|TRFL_HUMAN Lactotransferrin OS=Homo sapiens OX=9606 GN=LTF PE=1 SV=6 | 1270 | 20201113_15413_1ng_glyco_hcd203040_compound15.22738.22738.4 | controllerType=0 controllerNumber=1 scan=22738 | 45.9527 |
| 18052:4 | 1 | R.RTAGWNVPIGTLRPFLN[+2351.851]WTGPPEPIEAAVAR.F | HexNAc(4)Hex(5)Fuc(3)NeuAc(1) | N[+2352] | 1435.4274 | 4 | 5738.6877 | 5738.6628 | 0 | 4.3 | 140 | Specific | 480.93 | 341.39 | 341.39 | 8.44 | 140 | >sp\|P02788\|TRFL_HUMAN Lactotransferrin OS=Homo sapiens OX=9606 GN=LTF PE=1 SV=6 | 1270 | 20201113_15413_1ng_glyco_hcd203040_compound15.21610.21610.4 | controllerType=0 controllerNumber=1 scan=21610 | 43.5975 |
| 18055:4 | 1 | R.RTAGWNVPIGTLRPFLN[+2059.735]WTGPPEPIEAAVAR.F | HexNAc(4)Hex(5)Fuc(1)NeuAc(1) | N[+2060] | 1362.3984 | 4 | 5446.5719 | 5446.547 | 0 | 4.6 | 140 | Specific | 501.45 | 501.45 | 501.45 | 10.2 | 140 | >sp\|P02788\|TRFL_HUMAN Lactotransferrin OS=Homo sapiens OX=9606 GN=LTF PE=1 SV=6 | 1270 | 20201113_15413_1ng_glyco_hcd203040_compound15.21613.21613.4 | controllerType=0 controllerNumber=1 scan=21613 | 43.602 |
| 18038:4 | 1 | R.RTAGWNVPIGTLRPFLN[+2205.793]WTGPPEPIEAAVAR.F | HexNAc(4)Hex(5)Fuc(2)NeuAc(1) | N[+2206] | 1398.9136 | 4 | 5592.6325 | 5592.6049 | 0 | 4.9 | 140 | Specific | 539.18 | 529.28 | 529.28 | 10.33 | 140 | >sp\|P02788\|TRFL_HUMAN Lactotransferrin OS=Homo sapiens OX=9606 GN=LTF PE=1 SV=6 | 1270 | 20201113_15413_1ng_glyco_hcd203040_compound15.21596.21596.4 | controllerType=0 controllerNumber=1 scan=21596 | 43.5726 |
| 18023:4 | 1 | R.RTAGWNVPIGTLRPFLN[+2205.793]WTGPPEPIEAAVAR.F | HexNAc(4)Hex(5)Fuc(2)NeuAc(1) | N[+2206] | 1398.9083 | 4 | 5592.6115 | 5592.6049 | 0 | 1.2 | 140 | Specific | 553.1 | 551.13 | 551.13 | 10.34 | 140 | >sp\|P02788\|TRFL_HUMAN Lactotransferrin OS=Homo sapiens OX=9606 GN=LTF PE=1 SV=6 | 1270 | 20201113_15413_1ng_glyco_hcd203040_compound15.21580.21580.4 | controllerType=0 controllerNumber=1 scan=21580 | 43.5373 |
| 18040:4 | 1 | R.RTAGWNVPIGTLRPFLN[+2205.793]WTGPPEPIEAAVAR.F | HexNAc(4)Hex(5)Fuc(2)NeuAc(1) | N[+2206] | 1398.9136 | 4 | 5592.6325 | 5592.6049 | 0 | 4.9 | 140 | Specific | 620.54 | 526.21 | 526.21 | 12.72 | 140 | >sp\|P02788\|TRFL_HUMAN Lactotransferrin OS=Homo sapiens OX=9606 GN=LTF PE=1 SV=6 | 1270 | 20201113_15413_1ng_glyco_hcd203040_compound15.21598.21598.4 | controllerType=0 controllerNumber=1 scan=21598 | 43.5781 |
| 18026:4 | 1 | R.RTAGWNVPIGTLRPFLN[+2205.793]WTGPPEPIEAAVAR.F | HexNAc(4)Hex(5)Fuc(2)NeuAc(1) | N[+2206] | 1398.9083 | 4 | 5592.6115 | 5592.6049 | 0 | 1.2 | 140 | Specific | 632.85 | 532.82 | 532.82 | 12.54 | 140 | >sp\|P02788\|TRFL_HUMAN Lactotransferrin OS=Homo sapiens OX=9606 GN=LTF PE=1 SV=6 | 1270 | 20201113_15413_1ng_glyco_hcd203040_compound15.21583.21583.4 | controllerType=0 controllerNumber=1 scan=21583 | 43.5462 |
| 18058:4 | 1 | R.RTAGWNVPIGTLRPFLN[+2059.735]WTGPPEPIEAAVAR.F | HexNAc(4)Hex(5)Fuc(1)NeuAc(1) | N[+2060] | 1362.3984 | 4 | 5446.5719 | 5446.547 | 0 | 4.6 | 140 | Specific | 638.78 | 537.81 | 537.81 | 12.72 | 140 | >sp\|P02788\|TRFL_HUMAN Lactotransferrin OS=Homo sapiens OX=9606 GN=LTF PE=1 SV=6 | 1270 | 20201113_15413_1ng_glyco_hcd203040_compound15.21616.21616.4 | controllerType=0 controllerNumber=1 scan=21616 | 43.6088 |
| 19616:3 | 1 | R.[+42.011]TAGWNVPIGTLRPFLN[+2018.708]WTGPPEPIEAAVAR.F | HexNAc(3)Hex(6)Fuc(1)NeuAc(1) | .[42], N[+2019] | 1764.4929 | 3 | 5291.4641 | 5291.4299 | 0 | 6.5 | 141 | Specific | 203.67 | 203.67 | 203.67 | 4.33 | 140 | >sp\|P02788\|TRFL_HUMAN Lactotransferrin OS=Homo sapiens OX=9606 GN=LTF PE=1 SV=6 | 1270 | 20201113_15413_1ng_glyco_hcd203040_compound15.23246.23246.3 | controllerType=0 controllerNumber=1 scan=23246 | 46.9885 |
| 20772:5 | 1 | R.TAGWNVPIGTLRPFLN[+2351.851]WTGPPEPIEAAVAR.F | HexNAc(4)Hex(5)Fuc(3)NeuAc(1) | N[+2352] | 1117.3159 | 5 | 5582.5505 | 5582.5617 | 0 | -2 | 141 | Specific | 242.17 | 242.17 | 242.17 | 3.62 | 140 | >sp\|P02788\|TRFL_HUMAN Lactotransferrin OS=Homo sapiens OX=9606 GN=LTF PE=1 SV=6 | 1270 | 20201113_15413_1ng_glyco_hcd203040_compound15.24449.24449.5 | controllerType=0 controllerNumber=1 scan=24449 | 49.4149 |
| 19392:3 | 1 | R.TAGWNVPIGTLRPFLN[+2059.735]WTGPPEPIEAAVAR.F | HexNAc(4)Hex(5)Fuc(1)NeuAc(1) | N[+2060] | 1764.1646 | 3 | 5290.4792 | 5290.4459 | 0 | 6.3 | 141 | Specific | 264.5 | 264.5 | 264.5 | 6.32 | 140 | >sp\|P02788\|TRFL_HUMAN Lactotransferrin OS=Homo sapiens OX=9606 GN=LTF PE=1 SV=6 | 1270 | 20201113_15413_1ng_glyco_hcd203040_compound15.23010.23010.3 | controllerType=0 controllerNumber=1 scan=23010 | 46.4942 |
| 19275:3 | 1 | R.TAGWNVPIGTLRPFLN[+2424.867]WTGPPEPIEAAVAR.F | HexNAc(5)Hex(6)Fuc(1)NeuAc(1) | N[+2425] | 1885.8735 | 3 | 5655.6059 | 5655.5781 | 0 | 4.9 | 141 | Specific | 272.23 | 184.11 | 184.11 | 6.01 | 140 | >sp\|P02788\|TRFL_HUMAN Lactotransferrin OS=Homo sapiens OX=9606 GN=LTF PE=1 SV=6 | 1270 | 20201113_15413_1ng_glyco_hcd203040_compound15.22887.22887.3 | controllerType=0 controllerNumber=1 scan=22887 | 46.2275 |

Table S4. Cont.

| Query #:z | Protein Rank | Glycopeptide | Glycan | Modification Type(s) | Observed m/z | z | Observed (M+H) | Calc. mass (M+H) | Off-by-x error | Mass error (ppm) | StartPosition | Cleavage | Score | Delta | Delta Mod | \|Log Prob\| | # of unique peptides | Protein Name | Protein DB number | Comment | Scan # | Scan Time |
|---|---|---|---|---|---|---|---|---|---|---|---|---|---|---|---|---|---|---|---|---|---|---|
| 19223:5 | 1 | R.TAGWNVPIGTLRPFLN[+2569.905]WTGPPEPIEAAVAR.F | HexNAc(5)Hex(6)NeuAc(2) | N[+2570] | 1160.9355 | 5 | 5800.6486 | 5800.6156 | 0 | 5.7 | 141 | Specific | 273.74 | 273.74 | 273.74 | 6.25 | 140 | >sp\|P02788\|TRFL_HUMAN Lactotransferrin OS=Homo sapiens OX=9606 GN=LTF PE=1 SV=6 | 1270 | 20201113_15413_1ng_glyco_hcd203040_compound15.22833.22833.5 | controllerType=0 controllerNumber=1 scan=22833 | 46.1172 |
| 19342:3 | 1 | R.TAGWNVPIGTLRPFLN[+2205.793]WTGPPEPIEAAVAR.F | HexNAc(4)Hex(5)Fuc(2)NeuAc(1) | N[+2206] | 1812.8469 | 3 | 5436.5261 | 5436.5038 | 0 | 4.1 | 141 | Specific | 300.77 | 300.77 | 300.77 | 7.22 | 140 | >sp\|P02788\|TRFL_HUMAN Lactotransferrin OS=Homo sapiens OX=9606 GN=LTF PE=1 SV=6 | 1270 | 20201113_15413_1ng_glyco_hcd203040_compound15.22958.22958.2 | controllerType=0 controllerNumber=1 scan=22958 | 46.3872 |
| 20719:3 | 1 | R.[+42.011]TAGWNVPIGTLRPFLN[+2336.851]WTGPPEPIEAAVAR.F | HexNAc(6)Hex(6)Fuc(1) | .[42], N[+2337] | 1870.5254 | 3 | 5609.5615 | 5609.5726 | 0 | -2 | 141 | Specific | 301.33 | 295.56 | 295.56 | 6.02 | 140 | >sp\|P02788\|TRFL_HUMAN Lactotransferrin OS=Homo sapiens OX=9606 GN=LTF PE=1 SV=6 | 1270 | 20201113_15413_1ng_glyco_hcd203040_compound15.24394.24394.3 | controllerType=0 controllerNumber=1 scan=24394 | 49.3075 |
| 20718:4 | 1 | R.[+42.011]TAGWNVPIGTLRPFLN[+2336.851]WTGPPEPIEAAVAR.F | HexNAc(6)Hex(6)Fuc(1) | .[42], N[+2337] | 1403.1439 | 4 | 5609.5539 | 5609.5726 | 0 | -3.3 | 141 | Specific | 324.06 | 297.77 | 297.77 | 5.46 | 140 | >sp\|P02788\|TRFL_HUMAN Lactotransferrin OS=Homo sapiens OX=9606 GN=LTF PE=1 SV=6 | 1270 | 20201113_15413_1ng_glyco_hcd203040_compound15.24393.24393.4 | controllerType=0 controllerNumber=1 scan=24393 | 49.3064 |
| 19332:3 | 1 | R.TAGWNVPIGTLRPFLN[+2205.793]WTGPPEPIEAAVAR.F | HexNAc(4)Hex(5)Fuc(2)NeuAc(1) | N[+2206] | 1812.8464 | 3 | 5436.5246 | 5436.5038 | 0 | 3.8 | 141 | Specific | 330.76 | 295.03 | 295.03 | 7.37 | 140 | >sp\|P02788\|TRFL_HUMAN Lactotransferrin OS=Homo sapiens OX=9606 GN=LTF PE=1 SV=6 | 1270 | 20201113_15413_1ng_glyco_hcd203040_compound15.22947.22947.2 | controllerType=0 controllerNumber=1 scan=22947 | 46.3607 |
| 18279:3 | 1 | R.TAGWNVPIGTLRPFLN[+2279.830]WTGPPEPIEAAVAR.F | HexNAc(5)Hex(6)Fuc(2) | N[+2280] | 1837.525 | 3 | 5510.5604 | 5510.5406 | 0 | 3.6 | 141 | Specific | 372.05 | 288.28 | 288.28 | 7.53 | 140 | >sp\|P02788\|TRFL_HUMAN Lactotransferrin OS=Homo sapiens OX=9606 GN=LTF PE=1 SV=6 | 1270 | 20201113_15413_1ng_glyco_hcd203040_compound15.21848.21848.3 | controllerType=0 controllerNumber=1 scan=21848 | 44.0597 |
| 19208:3 | 1 | R.TAGWNVPIGTLRPFLN[+2570.925]WTGPPEPIEAAVAR.F | HexNAc(5)Hex(6)Fuc(2)NeuAc(1) | N[+2571] | 1934.557 | 3 | 5801.6564 | 5801.636 | 0 | 3.5 | 141 | Specific | 379.06 | 257.56 | 257.56 | 7.37 | 140 | >sp\|P02788\|TRFL_HUMAN Lactotransferrin OS=Homo sapiens OX=9606 GN=LTF PE=1 SV=6 | 1270 | 20201113_15413_1ng_glyco_hcd203040_compound15.22818.22818.3 | controllerType=0 controllerNumber=1 scan=22818 | 46.0882 |
| 18281:3 | 1 | R.TAGWNVPIGTLRPFLN[+2279.830]WTGPPEPIEAAVAR.F | HexNAc(5)Hex(6)Fuc(2) | N[+2280] | 1837.525 | 3 | 5510.5604 | 5510.5406 | 0 | 3.6 | 141 | Specific | 384.27 | 311.4 | 311.4 | 7.64 | 140 | >sp\|P02788\|TRFL_HUMAN Lactotransferrin OS=Homo sapiens OX=9606 GN=LTF PE=1 SV=6 | 1270 | 20201113_15413_1ng_glyco_hcd203040_compound15.21850.21850.3 | controllerType=0 controllerNumber=1 scan=21850 | 44.0637 |
| 20716:4 | 1 | R.[+42.011]TAGWNVPIGTLRPFLN[+2336.851]WTGPPEPIEAAVAR.F | HexNAc(6)Hex(6)Fuc(1) | .[42], N[+2337] | 1403.1439 | 4 | 5609.5539 | 5609.5726 | 0 | -3.3 | 141 | Specific | 386.32 | 386.32 | 386.32 | 7.6 | 140 | >sp\|P02788\|TRFL_HUMAN Lactotransferrin OS=Homo sapiens OX=9606 GN=LTF PE=1 SV=6 | 1270 | 20201113_15413_1ng_glyco_hcd203040_compound15.24391.24391.4 | controllerType=0 controllerNumber=1 scan=24391 | 49.3015 |
| 20555:4 | 1 | R.TAGWNVPIGTLRPFLN[+2862.021]WTGPPEPIEAAVAR.F | HexNAc(5)Hex(6)Fuc(2)NeuAc(2) | N[+2862] | 1523.9413 | 4 | 6092.7434 | 6092.7314 | 0 | 2 | 141 | Specific | 438.98 | 278.78 | 278.78 | 7.71 | 140 | >sp\|P02788\|TRFL_HUMAN Lactotransferrin OS=Homo sapiens OX=9606 GN=LTF PE=1 SV=6 | 1270 | 20201113_15413_1ng_glyco_hcd203040_compound15.24223.24223.4 | controllerType=0 controllerNumber=1 scan=24223 | 48.9571 |
| 19154:3 | 1 | R.TAGWNVPIGTLRPFLN[+2715.963]WTGPPEPIEAAVAR.F | HexNAc(5)Hex(6)Fuc(1)NeuAc(2) | N[+2716] | 1982.9071 | 3 | 5946.7067 | 5946.6735 | 0 | 5.6 | 141 | Specific | 465.65 | 298.64 | 298.64 | 7.16 | 140 | >sp\|P02788\|TRFL_HUMAN Lactotransferrin OS=Homo sapiens OX=9606 GN=LTF PE=1 SV=6 | 1270 | 20201113_15413_1ng_glyco_hcd203040_compound15.22762.22762.3 | controllerType=0 controllerNumber=1 scan=22762 | 45.9957 |
| 20725:3 | 1 | R.TAGWNVPIGTLRPFLN[+2350.830]WTGPPEPIEAAVAR.F | HexNAc(4)Hex(5)Fuc(1)NeuAc(2) | N[+2351] | 1861.1932 | 3 | 5581.565 | 5581.5413 | 0 | 4.3 | 141 | Specific | 473.12 | 315.87 | 315.87 | 7.67 | 140 | >sp\|P02788\|TRFL_HUMAN Lactotransferrin OS=Homo sapiens OX=9606 GN=LTF PE=1 SV=6 | 1270 | 20201113_15413_1ng_glyco_hcd203040_compound15.24400.24400.2 | controllerType=0 controllerNumber=1 scan=24400 | 49.3179 |
| 19225:5 | 1 | R.TAGWNVPIGTLRPFLN[+2569.905]WTGPPEPIEAAVAR.F | HexNAc(5)Hex(6)NeuAc(2) | N[+2570] | 1160.9355 | 5 | 5800.6486 | 5800.6156 | 0 | 5.7 | 141 | Specific | 520.5 | 412.62 | 412.62 | 9.33 | 140 | >sp\|P02788\|TRFL_HUMAN Lactotransferrin OS=Homo sapiens OX=9606 GN=LTF PE=1 SV=6 | 1270 | 20201113_15413_1ng_glyco_hcd203040_compound15.22835.22835.5 | controllerType=0 controllerNumber=1 scan=22835 | 46.1232 |
| 19211:3 | 1 | R.TAGWNVPIGTLRPFLN[+2570.925]WTGPPEPIEAAVAR.F | HexNAc(5)Hex(6)Fuc(2)NeuAc(1) | N[+2571] | 1934.557 | 3 | 5801.6564 | 5801.636 | 0 | 3.5 | 141 | Specific | 539.21 | 416.2 | 416.2 | 10.05 | 140 | >sp\|P02788\|TRFL_HUMAN Lactotransferrin OS=Homo sapiens OX=9606 GN=LTF PE=1 SV=6 | 1270 | 20201113_15413_1ng_glyco_hcd203040_compound15.22821.22821.3 | controllerType=0 controllerNumber=1 scan=22821 | 46.0955 |
| 20799:4 | 1 | R.TAGWNVPIGTLRPFLN[+2350.830]WTGPPEPIEAAVAR.F | HexNAc(4)Hex(5)Fuc(1)NeuAc(2) | N[+2351] | 1396.145 | 4 | 5581.5583 | 5581.5413 | 0 | 3 | 141 | Specific | 568.09 | 443.42 | 443.42 | 10.19 | 140 | >sp\|P02788\|TRFL_HUMAN Lactotransferrin OS=Homo sapiens OX=9606 GN=LTF PE=1 SV=6 | 1270 | 20201113_15413_1ng_glyco_hcd203040_compound15.24477.24477.4 | controllerType=0 controllerNumber=1 scan=24477 | 49.4691 |
| 18269:3 | 1 | R.TAGWNVPIGTLRPFLN[+2279.830]WTGPPEPIEAAVAR.F | HexNAc(5)Hex(6)Fuc(2) | N[+2280] | 1837.5244 | 3 | 5510.5586 | 5510.5406 | 0 | 3.3 | 141 | Specific | 576.25 | 487.07 | 487.07 | 11.44 | 140 | >sp\|P02788\|TRFL_HUMAN Lactotransferrin OS=Homo sapiens OX=9606 GN=LTF PE=1 SV=6 | 1270 | 20201113_15413_1ng_glyco_hcd203040_compound15.21837.21837.3 | controllerType=0 controllerNumber=1 scan=21837 | 44.0299 |
| 19127:4 | 1 | R.TAGWNVPIGTLRPFLN[+2863.041]WTGPPEPIEAAVAR.F | HexNAc(5)Hex(6)Fuc(4)NeuAc(1) | N[+2863] | 1524.1993 | 4 | 6093.7755 | 6093.7518 | 0 | 3.9 | 141 | Specific | 592.11 | 547.87 | 547.87 | 11.39 | 140 | >sp\|P02788\|TRFL_HUMAN Lactotransferrin OS=Homo sapiens OX=9606 GN=LTF PE=1 SV=6 | 1270 | 20201113_15413_1ng_glyco_hcd203040_compound15.22734.22734.4 | controllerType=0 controllerNumber=1 scan=22734 | 45.9422 |
| 19335:3 | 1 | R.TAGWNVPIGTLRPFLN[+2205.793]WTGPPEPIEAAVAR.F | HexNAc(4)Hex(5)Fuc(2)NeuAc(1) | N[+2206] | 1812.8464 | 3 | 5436.5246 | 5436.5038 | 0 | 3.8 | 141 | Specific | 609.32 | 605.06 | 605.06 | 12.44 | 140 | >sp\|P02788\|TRFL_HUMAN Lactotransferrin OS=Homo sapiens OX=9606 GN=LTF PE=1 SV=6 | 1270 | 20201113_15413_1ng_glyco_hcd203040_compound15.22950.22950.2 | controllerType=0 controllerNumber=1 scan=22950 | 46.365 |
| 19206:4 | 1 | R.TAGWNVPIGTLRPFLN[+2570.925]WTGPPEPIEAAVAR.F | HexNAc(5)Hex(6)Fuc(2)NeuAc(1) | N[+2571] | 1451.1708 | 4 | 5801.6613 | 5801.636 | 0 | 4.4 | 141 | Specific | 611.36 | 521.47 | 521.47 | 12.28 | 140 | >sp\|P02788\|TRFL_HUMAN Lactotransferrin OS=Homo sapiens OX=9606 GN=LTF PE=1 SV=6 | 1270 | 20201113_15413_1ng_glyco_hcd203040_compound15.22816.22816.4 | controllerType=0 controllerNumber=1 scan=22816 | 46.0859 |
| 18313:4 | 1 | R.TAGWNVPIGTLRPFLN[+1768.640]WTGPPEPIEAAVAR.F | HexNAc(4)Hex(5)Fuc(1) | N[+1769] | 1250.5951 | 4 | 4999.3586 | 4999.3504 | 0 | 1.6 | 141 | Specific | 614.24 | 559.48 | 559.48 | 12.34 | 140 | >sp\|P02788\|TRFL_HUMAN Lactotransferrin OS=Homo sapiens OX=9606 GN=LTF PE=1 SV=6 | 1270 | 20201113_15413_1ng_glyco_hcd203040_compound15.21883.21883.4 | controllerType=0 controllerNumber=1 scan=21883 | 44.1388 |
| 19252:4 | 1 | R.TAGWNVPIGTLRPFLN[+2424.867]WTGPPEPIEAAVAR.F | HexNAc(5)Hex(6)Fuc(1)NeuAc(1) | N[+2425] | 1414.6555 | 4 | 5655.6003 | 5655.5781 | 0 | 3.9 | 141 | Specific | 634.1 | 518.51 | 518.51 | 12.39 | 140 | >sp\|P02788\|TRFL_HUMAN Lactotransferrin OS=Homo sapiens OX=9606 GN=LTF PE=1 SV=6 | 1270 | 20201113_15413_1ng_glyco_hcd203040_compound15.22863.22863.4 | controllerType=0 controllerNumber=1 scan=22863 | 46.1689 |
| 18289:4 | 1 | R.TAGWNVPIGTLRPFLN[+2279.830]WTGPPEPIEAAVAR.F | HexNAc(5)Hex(6)Fuc(2) | N[+2280] | 1378.3955 | 4 | 5510.5602 | 5510.5406 | 0 | 3.6 | 141 | Specific | 701.56 | 594.17 | 594.17 | 14.39 | 140 | >sp\|P02788\|TRFL_HUMAN Lactotransferrin OS=Homo sapiens OX=9606 GN=LTF PE=1 SV=6 | 1270 | 20201113_15413_1ng_glyco_hcd203040_compound15.21858.21858.4 | controllerType=0 controllerNumber=1 scan=21858 | 44.0793 |
| 18308:3 | 1 | R.TAGWNVPIGTLRPFLN[+1768.640]WTGPPEPIEAAVAR.F | HexNAc(4)Hex(5)Fuc(1) | N[+1769] | 1667.1263 | 3 | 4999.3644 | 4999.3504 | 0 | 2.8 | 141 | Specific | 712.49 | 631.02 | 631.02 | 14.44 | 140 | >sp\|P02788\|TRFL_HUMAN Lactotransferrin OS=Homo sapiens OX=9606 GN=LTF PE=1 SV=6 | 1270 | 20201113_15413_1ng_glyco_hcd203040_compound15.21878.21878.3 | controllerType=0 controllerNumber=1 scan=21878 | 44.1232 |
| 19344:3 | 1 | R.TAGWNVPIGTLRPFLN[+2205.793]WTGPPEPIEAAVAR.F | HexNAc(4)Hex(5)Fuc(2)NeuAc(1) | N[+2206] | 1812.8469 | 3 | 5436.5261 | 5436.5038 | 0 | 4.1 | 141 | Specific | 716.21 | 659.64 | 659.64 | 14.33 | 140 | >sp\|P02788\|TRFL_HUMAN Lactotransferrin OS=Homo sapiens OX=9606 GN=LTF PE=1 SV=6 | 1270 | 20201113_15413_1ng_glyco_hcd203040_compound15.22960.22960.2 | controllerType=0 controllerNumber=1 scan=22960 | 46.391 |
| 18310:3 | 1 | R.TAGWNVPIGTLRPFLN[+1768.640]WTGPPEPIEAAVAR.F | HexNAc(4)Hex(5)Fuc(1) | N[+1769] | 1667.1263 | 3 | 4999.3644 | 4999.3504 | 0 | 2.8 | 141 | Specific | 725.02 | 613.93 | 613.93 | 14.44 | 140 | >sp\|P02788\|TRFL_HUMAN Lactotransferrin OS=Homo sapiens OX=9606 GN=LTF PE=1 SV=6 | 1270 | 20201113_15413_1ng_glyco_hcd203040_compound15.21880.21880.3 | controllerType=0 controllerNumber=1 scan=21880 | 44.13 |
| 18311:4 | 1 | R.TAGWNVPIGTLRPFLN[+1768.640]WTGPPEPIEAAVAR.F | HexNAc(4)Hex(5)Fuc(1) | N[+1769] | 1250.5951 | 4 | 4999.3586 | 4999.3504 | 0 | 1.6 | 141 | Specific | 732.68 | 694.77 | 694.77 | 14.34 | 140 | >sp\|P02788\|TRFL_HUMAN Lactotransferrin OS=Homo sapiens OX=9606 GN=LTF PE=1 SV=6 | 1270 | 20201113_15413_1ng_glyco_hcd203040_compound15.21881.21881.4 | controllerType=0 controllerNumber=1 scan=21881 | 44.1311 |
| 19130:4 | 1 | R.TAGWNVPIGTLRPFLN[+2863.041]WTGPPEPIEAAVAR.F | HexNAc(5)Hex(6)Fuc(4)NeuAc(1) | N[+2863] | 1524.1993 | 4 | 6093.7755 | 6093.7518 | 0 | 3.9 | 141 | Specific | 739 | 560.52 | 560.52 | 14.39 | 140 | >sp\|P02788\|TRFL_HUMAN Lactotransferrin OS=Homo sapiens OX=9606 GN=LTF PE=1 SV=6 | 1270 | 20201113_15413_1ng_glyco_hcd203040_compound15.22737.22737.4 | controllerType=0 controllerNumber=1 scan=22737 | 45.9475 |
| 19395:3 | 1 | R.TAGWNVPIGTLRPFLN[+2059.735]WTGPPEPIEAAVAR.F | HexNAc(4)Hex(5)Fuc(1)NeuAc(1) | N[+2060] | 1764.1646 | 3 | 5290.4792 | 5290.4459 | 0 | 6.3 | 141 | Specific | 739.47 | 695.19 | 695.19 | 13.93 | 140 | >sp\|P02788\|TRFL_HUMAN Lactotransferrin OS=Homo sapiens OX=9606 GN=LTF PE=1 SV=6 | 1270 | 20201113_15413_1ng_glyco_hcd203040_compound15.23013.23013.3 | controllerType=0 controllerNumber=1 scan=23013 | 46.4977 |
| 19209:4 | 1 | R.TAGWNVPIGTLRPFLN[+2570.925]WTGPPEPIEAAVAR.F | HexNAc(5)Hex(6)Fuc(2)NeuAc(1) | N[+2571] | 1451.1708 | 4 | 5801.6613 | 5801.636 | 0 | 4.4 | 141 | Specific | 789.27 | 685.77 | 685.77 | 15.28 | 140 | >sp\|P02788\|TRFL_HUMAN Lactotransferrin OS=Homo sapiens OX=9606 GN=LTF PE=1 SV=6 | 1270 | 20201113_15413_1ng_glyco_hcd203040_compound15.22819.22819.4 | controllerType=0 controllerNumber=1 scan=22819 | 46.0915 |
| 20801:4 | 1 | R.TAGWNVPIGTLRPFLN[+2350.830]WTGPPEPIEAAVAR.F | HexNAc(4)Hex(5)Fuc(1)NeuAc(2) | N[+2351] | 1396.145 | 4 | 5581.5583 | 5581.5413 | 0 | 3 | 141 | Specific | 823.91 | 547.42 | 547.42 | 16.39 | 140 | >sp\|P02788\|TRFL_HUMAN Lactotransferrin OS=Homo sapiens OX=9606 GN=LTF PE=1 SV=6 | 1270 | 20201113_15413_1ng_glyco_hcd203040_compound15.24479.24479.4 | controllerType=0 controllerNumber=1 scan=24479 | 49.4728 |
| 19254:4 | 1 | R.TAGWNVPIGTLRPFLN[+2424.867]WTGPPEPIEAAVAR.F | HexNAc(5)Hex(6)Fuc(1)NeuAc(1) | N[+2425] | 1414.6555 | 4 | 5655.6003 | 5655.5781 | 0 | 3.9 | 141 | Specific | 871.79 | 730.57 | 730.57 | 17.39 | 140 | >sp\|P02788\|TRFL_HUMAN Lactotransferrin OS=Homo sapiens OX=9606 GN=LTF PE=1 SV=6 | 1270 | 20201113_15413_1ng_glyco_hcd203040_compound15.22865.22865.4 | controllerType=0 controllerNumber=1 scan=22865 | 46.1735 |
| 20843:3 | 1 | R.[+42.011]TAGWNIPMGLLFN[+2336.851]QTGSC[+57.021]K.F | HexNAc(6)Hex(6)Fuc(1) | .[42], C[+57], N[+2337] | 1491.9505 | 3 | 4473.8369 | 4473.8716 | 0 | -7.7 | 485 | Specific | 259.61 | 227.68 | 227.68 | 4.88 | 140 | >sp\|P02788\|TRFL_HUMAN Lactotransferrin OS=Homo sapiens OX=9606 GN=LTF PE=1 SV=6 | 1270 | 20201113_15413_1ng_glyco_hcd203040_compound15.24522.24522.3 | controllerType=0 controllerNumber=1 scan=24522 | 49.5545 |
| 20846:3 | 1 | R.[+42.011]TAGWNIPMGLLFN[+2336.851]QTGSC[+57.021]K.F | HexNAc(6)Hex(6)Fuc(1) | .[42], C[+57], N[+2337] | 1491.9505 | 3 | 4473.8369 | 4473.8716 | 0 | -7.7 | 485 | Specific | 278.18 | 208.88 | 208.88 | 3.68 | 140 | >sp\|P02788\|TRFL_HUMAN Lactotransferrin OS=Homo sapiens OX=9606 GN=LTF PE=1 SV=6 | 1270 | 20201113_15413_1ng_glyco_hcd203040_compound15.24525.24525.3 | controllerType=0 controllerNumber=1 scan=24525 | 49.5598 |
| 18647:3 | 1 | R.TAGWNIPMGLLFN[+2351.851]QTGSC[+57.021]K.F | HexNAc(4)Hex(5)Fuc(3)NeuAc(1) | C[+57], N[+2352] | 1482.964 | 3 | 4446.8774 | 4446.8607 | 0 | 3.8 | 485 | Specific | 280.19 | 157.97 | 157.97 | 5.85 | 140 | >sp\|P02788\|TRFL_HUMAN Lactotransferrin OS=Homo sapiens OX=9606 GN=LTF PE=1 SV=6 | 1270 | 20201113_15413_1ng_glyco_hcd203040_compound15.22232.22232.3 | controllerType=0 controllerNumber=1 scan=22232 | 44.8822 |
| 18878:3 | 1 | R.TAGWNIPMGLLFN[+1872.651]QTGSC[+57.021]K.F | HexNAc(3)Hex(6)NeuAc(1) | C[+57], N[+1873] | 1323.2241 | 3 | 3967.6576 | 3967.6604 | 0 | -0.7 | 485 | Specific | 325.58 | 289.01 | 170.49 | 6.4 | 140 | >sp\|P02788\|TRFL_HUMAN Lactotransferrin OS=Homo sapiens OX=9606 GN=LTF PE=1 SV=6 | 1270 | 20201113_15413_1ng_glyco_hcd203040_compound15.22474.22474.3 | controllerType=0 controllerNumber=1 scan=22474 | 45.4573 |
| 18458:3 | 1 | R.TAGWNIPMGLLFN[+2570.925]QTGSC[+57.021]K.F | HexNAc(5)Hex(6)Fuc(2)NeuAc(1) | C[+57], N[+2571] | 1555.9979 | 3 | 4665.9791 | 4665.935 | 0 | 9.5 | 485 | Specific | 337.63 | 326.62 | 326.62 | 6.36 | 140 | >sp\|P02788\|TRFL_HUMAN Lactotransferrin OS=Homo sapiens OX=9606 GN=LTF PE=1 SV=6 | 1270 | 20201113_15413_1ng_glyco_hcd203040_compound15.22035.22035.3 | controllerType=0 controllerNumber=1 scan=22035 | 44.4341 |
| 18885:3 | 1 | R.TAGWNIPMGLLFN[+1710.598]QTGSC[+57.021]K.F | HexNAc(3)Hex(5)NeuAc(1) | C[+57], N[+1711] | 1269.2112 | 3 | 3805.6192 | 3805.6076 | 0 | 3 | 485 | Specific | 365.59 | 360.77 | 360.77 | 7.66 | 140 | >sp\|P02788\|TRFL_HUMAN Lactotransferrin OS=Homo sapiens OX=9606 GN=LTF PE=1 SV=6 | 1270 | 20201113_15413_1ng_glyco_hcd203040_compound15.22482.22482.3 | controllerType=0 controllerNumber=1 scan=22482 | 45.4782 |
| 18838:3 | 1 | R.TAGWNIPMGLLFN[+1694.603]QTGSC[+57.021]K.F | HexNAc(3)Hex(4)Fuc(1)NeuAc(1) | C[+57], N[+1695] | 1263.8768 | 3 | 3789.6157 | 3789.6127 | 0 | 0.8 | 485 | Specific | 409.76 | 409.76 | 409.76 | 8.94 | 140 | >sp\|P02788\|TRFL_HUMAN Lactotransferrin OS=Homo sapiens OX=9606 GN=LTF PE=1 SV=6 | 1270 | 20201113_15413_1ng_glyco_hcd203040_compound15.22432.22432.3 | controllerType=0 controllerNumber=1 scan=22432 | 45.3454 |
| 16553:3 | 1 | R.TAGWNIPMGLLFN[+2279.830]QTGSC[+57.021]K.F | HexNAc(5)Hex(6)Fuc(2) | C[+57], N[+2280] | 1458.9579 | 3 | 4374.8593 | 4374.8396 | 0 | 4.5 | 485 | Specific | 450.77 | 389.88 | 283.85 | 9.16 | 140 | >sp\|P02788\|TRFL_HUMAN Lactotransferrin OS=Homo sapiens OX=9606 GN=LTF PE=1 SV=6 | 1270 | 20201113_15413_1ng_glyco_hcd203040_compound15.20048.20048.3 | controllerType=0 controllerNumber=1 scan=20048 | 40.3982 |
| 18881:3 | 1 | R.TAGWNIPMGLLFN[+1872.651]QTGSC[+57.021]K.F | HexNAc(3)Hex(6)NeuAc(1) | C[+57], N[+1873] | 1323.2241 | 3 | 3967.6576 | 3967.6604 | 0 | -0.7 | 485 | Specific | 469.63 | 422.99 | 329.72 | 8.54 | 140 | >sp\|P02788\|TRFL_HUMAN Lactotransferrin OS=Homo sapiens OX=9606 GN=LTF PE=1 SV=6 | 1270 | 20201113_15413_1ng_glyco_hcd203040_compound15.22477.22477.3 | controllerType=0 controllerNumber=1 scan=22477 | 45.4653 |
| 18841:3 | 1 | R.TAGWNIPMGLLFN[+1694.603]QTGSC[+57.021]K.F | HexNAc(3)Hex(4)Fuc(1)NeuAc(1) | C[+57], N[+1695] | 1263.8768 | 3 | 3789.6157 | 3789.6127 | 0 | 0.8 | 485 | Specific | 480.33 | 331.45 | 331.45 | 7.62 | 140 | >sp\|P02788\|TRFL_HUMAN Lactotransferrin OS=Homo sapiens OX=9606 GN=LTF PE=1 SV=6 | 1270 | 20201113_15413_1ng_glyco_hcd203040_compound15.22435.22435.3 | controllerType=0 controllerNumber=1 scan=22435 | 45.3519 |
| 18726:3 | 1 | R.TAGWNIPMGLLFN[+1038.375]QTGSC[+57.021]K.F | HexNAc(2)Hex(3)Fuc(1) | C[+57], N[+1038] | 1045.1233 | 3 | 3133.3554 | 3133.3851 | 0 | -9.5 | 485 | Specific | 483.19 | 451.13 | 451.13 | 7.32 | 140 | >sp\|P02788\|TRFL_HUMAN Lactotransferrin OS=Homo sapiens OX=9606 GN=LTF PE=1 SV=6 | 1270 | 20201113_15413_1ng_glyco_hcd203040_compound15.22315.22315.2 | controllerType=0 controllerNumber=1 scan=22315 | 45.0792 |
| 16977:3 | 1 | R.TAGWNIPMGLLFN[+1622.582]QTGSC[+57.021]K.F | HexNAc(4)Hex(5) | C[+57], N[+1623] | 1239.8726 | 3 | 3717.6033 | 3717.5916 | 0 | 3.2 | 485 | Specific | 502.8 | 502.8 | 225.98 | 9.67 | 140 | >sp\|P02788\|TRFL_HUMAN Lactotransferrin OS=Homo sapiens OX=9606 GN=LTF PE=1 SV=6 | 1270 | 20201113_15413_1ng_glyco_hcd203040_compound15.20490.20490.3 | controllerType=0 controllerNumber=1 scan=20490 | 41.2698 |
| 18882:3 | 1 | R.TAGWNIPMGLLFN[+1751.624]QTGSC[+57.021]K.F | HexNAc(4)Hex(4)NeuAc(1) | C[+57], N[+1752] | 1282.8868 | 3 | 3846.6458 | 3846.6341 | 0 | 3 | 485 | Specific | 543.82 | 523.54 | 523.54 | 9.8 | 140 | >sp\|P02788\|TRFL_HUMAN Lactotransferrin OS=Homo sapiens OX=9606 GN=LTF PE=1 SV=6 | 1270 | 20201113_15413_1ng_glyco_hcd203040_compound15.22479.22479.3 | controllerType=0 controllerNumber=1 scan=22479 | 45.47 |
| 18728:3 | 1 | R.TAGWNIPMGLLFN[+1038.375]QTGSC[+57.021]K.F | HexNAc(2)Hex(3)Fuc(1) | C[+57], N[+1038] | 1045.1233 | 3 | 3133.3554 | 3133.3851 | 0 | -9.5 | 485 | Specific | 552.26 | 540.38 | 540.38 | 7.79 | 140 | >sp\|P02788\|TRFL_HUMAN Lactotransferrin OS=Homo sapiens OX=9606 GN=LTF PE=1 SV=6 | 1270 | 20201113_15413_1ng_glyco_hcd203040_compound15.22317.22317.2 | controllerType=0 controllerNumber=1 scan=22317 | 45.0849 |
| 20750:4 | 1 | R.TAGWNIPMGLLFN[+2350.830]QTGSC[+57.021]K.F | HexNAc(4)Hex(5)Fuc(1)NeuAc(2) | C[+57], N[+2351] | 1112.2179 | 4 | 4445.8499 | 4445.8403 | 0 | 2.2 | 485 | Specific | 588.92 | 550.99 | 550.99 | 11.12 | 140 | >sp\|P02788\|TRFL_HUMAN Lactotransferrin OS=Homo sapiens OX=9606 GN=LTF PE=1 SV=6 | 1270 | 20201113_15413_1ng_glyco_hcd203040_compound15.24426.24426.4 | controllerType=0 controllerNumber=1 scan=24426 | 49.3673 |
| 16555:3 | 1 | R.TAGWNIPMGLLFN[+2279.830]QTGSC[+57.021]K.F | HexNAc(5)Hex(6)Fuc(2) | C[+57], N[+2280] | 1458.9579 | 3 | 4374.8593 | 4374.8396 | 0 | 4.5 | 485 | Specific | 636.87 | 510.37 | 361.54 | 12.09 | 140 | >sp\|P02788\|TRFL_HUMAN Lactotransferrin OS=Homo sapiens OX=9606 GN=LTF PE=1 SV=6 | 1270 | 20201113_15413_1ng_glyco_hcd203040_compound15.20050.20050.3 | controllerType=0 controllerNumber=1 scan=20050 | 40.4019 |
| 20934:3 | 1 | R.TAGWNIPMGLLFN[+2204.772]QTGSC[+57.021]K.F | HexNAc(4)Hex(5)NeuAc(2) | C[+57], N[+2205] | 1433.9383 | 3 | 4299.8003 | 4299.7824 | 0 | 4.2 | 485 | Specific | 676.87 | 588.17 | 588.17 | 13.09 | 140 | >sp\|P02788\|TRFL_HUMAN Lactotransferrin OS=Homo sapiens OX=9606 GN=LTF PE=1 SV=6 | 1270 | 20201113_15413_1ng_glyco_hcd203040_compound15.24618.24618.3 | controllerType=0 controllerNumber=1 scan=24618 | 49.7881 |
| 18460:3 | 1 | R.TAGWNIPMGLLFN[+2570.925]QTGSC[+57.021]K.F | HexNAc(5)Hex(6)Fuc(2)NeuAc(1) | C[+57], N[+2571] | 1555.9979 | 3 | 4665.9791 | 4665.935 | 0 | 9.5 | 485 | Specific | 680.71 | 479 | 479 | 12.21 | 140 | >sp\|P02788\|TRFL_HUMAN Lactotransferrin OS=Homo sapiens OX=9606 GN=LTF PE=1 SV=6 | 1270 | 20201113_15413_1ng_glyco_hcd203040_compound15.22037.22037.3 | controllerType=0 controllerNumber=1 scan=22037 | 44.4378 |
| 16979:3 | 1 | R.TAGWNIPMGLLFN[+1622.582]QTGSC[+57.021]K.F | HexNAc(4)Hex(5) | C[+57], N[+1623] | 1239.8726 | 3 | 3717.6033 | 3717.5916 | 0 | 3.2 | 485 | Specific | 740.31 | 735.68 | 341.33 | 14.19 | 140 | >sp\|P02788\|TRFL_HUMAN Lactotransferrin OS=Homo sapiens OX=9606 GN=LTF PE=1 SV=6 | 1270 | 20201113_15413_1ng_glyco_hcd203040_compound15.20492.20492.3 | controllerType=0 controllerNumber=1 scan=20492 | 41.2732 |
| 20742:3 | 1 | R.TAGWNIPMGLLFN[+2350.830]QTGSC[+57.021]K.F | HexNAc(4)Hex(5)Fuc(1)NeuAc(2) | C[+57], N[+2351] | 1482.6231 | 3 | 4445.8548 | 4445.8403 | 0 | 3.3 | 485 | Specific | 751.1 | 728.67 | 728.67 | 15.19 | 140 | >sp\|P02788\|TRFL_HUMAN Lactotransferrin OS=Homo sapiens OX=9606 GN=LTF PE=1 SV=6 | 1270 | 20201113_15413_1ng_glyco_hcd203040_compound15.24418.24418.3 | controllerType=0 controllerNumber=1 scan=24418 | 49.3527 |
| 17153:3 | 1 | R.TAGWNIPMGLLFN[+1216.423]QTGSC[+57.021]K.F | HexNAc(2)Hex(5) | C[+57], N[+1216] | 1104.4849 | 3 | 3311.4401 | 3311.4328 | 0 | 2.2 | 485 | Specific | 798.59 | 798.59 | 267.93 | 15.15 | 140 | >sp\|P02788\|TRFL_HUMAN Lactotransferrin OS=Homo sapiens OX=9606 GN=LTF PE=1 SV=6 | 1270 | 20201113_15413_1ng_glyco_hcd203040_compound15.20673.20673.3 | controllerType=0 controllerNumber=1 scan=20673 | 41.6401 |
| 17208:3 | 1 | R.TAGWNIPMGLLFN[+1216.423]QTGSC[+57.021]K.F | HexNAc(2)Hex(5) | C[+57], N[+1216] | 1104.4873 | 3 | 3311.4474 | 3311.4328 | 0 | 4.4 | 485 | Specific | 806.89 | 801.54 | 293.95 | 16.09 | 140 | >sp\|P02788\|TRFL_HUMAN Lactotransferrin OS=Homo sapiens OX=9606 GN=LTF PE=1 SV=6 | 1270 | 20201113_15413_1ng_glyco_hcd203040_compound15.20730.20730.3 | controllerType=0 controllerNumber=1 scan=20730 | 41.7498 |
| 20747:4 | 1 | R.TAGWNIPMGLLFN[+2350.830]QTGSC[+57.021]K.F | HexNAc(4)Hex(5)Fuc(1)NeuAc(2) | C[+57], N[+2351] | 1112.2179 | 4 | 4445.8499 | 4445.8403 | 0 | 2.2 | 485 | Specific | 810.01 | 678.72 | 678.72 | 16.12 | 140 | >sp\|P02788\|TRFL_HUMAN Lactotransferrin OS=Homo sapiens OX=9606 GN=LTF PE=1 SV=6 | 1270 | 20201113_15413_1ng_glyco_hcd203040_compound15.24423.24423.4 | controllerType=0 controllerNumber=1 scan=24423 | 49.3613 |
| 16833:3 | 1 | R.TAGWNIPMGLLFN[+1606.587]QTGSC[+57.021]K.F | HexNAc(4)Hex(4)Fuc(1) | C[+57], N[+1607] | 1234.5405 | 3 | 3701.6068 | 3701.5966 | 0 | 2.8 | 485 | Specific | 819.75 | 819.75 | 819.75 | 16.19 | 140 | >sp\|P02788\|TRFL_HUMAN Lactotransferrin OS=Homo sapiens OX=9606 GN=LTF PE=1 SV=6 | 1270 | 20201113_15413_1ng_glyco_hcd203040_compound15.20340.20340.3 | controllerType=0 controllerNumber=1 scan=20340 | 40.9838 |
| 17162:2 | 1 | R.TAGWNIPMGLLFN[+1216.423]QTGSC[+57.021]K.F | HexNAc(2)Hex(5) | C[+57], N[+1216] | 1656.2263 | 2 | 3311.4453 | 3311.4328 | 0 | 3.8 | 485 | Specific | 863.6 | 783.71 | 302.15 | 17.24 | 140 | >sp\|P02788\|TRFL_HUMAN Lactotransferrin OS=Homo sapiens OX=9606 GN=LTF PE=1 SV=6 | 1270 | 20201113_15413_1ng_glyco_hcd203040_compound15.20682.20682.2 | controllerType=0 controllerNumber=1 scan=20682 | 41.6608 |
| 20683:3 | 1 | R.TAGWNIPMGLLFN[+2350.830]QTGSC[+57.021]K.F | HexNAc(4)Hex(5)Fuc(1)NeuAc(2) | C[+57], N[+2351] | 1482.6225 | 3 | 4445.8529 | 4445.8403 | 0 | 2.8 | 485 | Specific | 871.89 | 774.83 | 774.83 | 17.19 | 140 | >sp\|P02788\|TRFL_HUMAN Lactotransferrin OS=Homo sapiens OX=9606 GN=LTF PE=1 SV=6 | 1270 | 20201113_15413_1ng_glyco_hcd203040_compound15.24356.24356.3 | controllerType=0 controllerNumber=1 scan=24356 | 49.2263 |
| 17160:2 | 1 | R.TAGWNIPMGLLFN[+1216.423]QTGSC[+57.021]K.F | HexNAc(2)Hex(5) | C[+57], N[+1216] | 1656.2263 | 2 | 3311.4453 | 3311.4328 | 0 | 3.8 | 485 | Specific | 881.18 | 881.18 | 318.95 | 17.24 | 140 | >sp\|P02788\|TRFL_HUMAN Lactotransferrin OS=Homo sapiens OX=9606 GN=LTF PE=1 SV=6 | 1270 | 20201113_15413_1ng_glyco_hcd203040_compound15.20680.20680.2 | controllerType=0 controllerNumber=1 scan=20680 | 41.6564 |
| 16706:3 | 1 | R.TAGWNIPMGLLFN[+1768.640]QTGSC[+57.021]K.F | HexNAc(4)Hex(5)Fuc(1) | C[+57], N[+1769] | 1288.564 | 3 | 3863.6775 | 3863.6495 | 0 | 7.3 | 485 | Specific | 888.82 | 783.97 | 783.97 | 16.4 | 140 | >sp\|P02788\|TRFL_HUMAN Lactotransferrin OS=Homo sapiens OX=9606 GN=LTF PE=1 SV=6 | 1270 | 20201113_15413_1ng_glyco_hcd203040_compound15.20208.20208.3 | controllerType=0 controllerNumber=1 scan=20208 | 40.7178 |
| 20936:3 | 1 | R.TAGWNIPMGLLFN[+2204.772]QTGSC[+57.021]K.F | HexNAc(4)Hex(5)NeuAc(2) | C[+57], N[+2205] | 1433.9383 | 3 | 4299.8003 | 4299.7824 | 0 | 4.2 | 485 | Specific | 897.18 | 897.18 | 897.18 | 17.09 | 140 | >sp\|P02788\|TRFL_HUMAN Lactotransferrin OS=Homo sapiens OX=9606 GN=LTF PE=1 SV=6 | 1270 | 20201113_15413_1ng_glyco_hcd203040_compound15.24620.24620.3 | controllerType=0 controllerNumber=1 scan=24620 | 49.7928 |
| 20686:3 | 1 | R.TAGWNIPMGLLFN[+2350.830]QTGSC[+57.021]K.F | HexNAc(4)Hex(5)Fuc(1)NeuAc(2) | C[+57], N[+2351] | 1482.6225 | 3 | 4445.8529 | 4445.8403 | 0 | 2.8 | 485 | Specific | 897.81 | 856.03 | 856.03 | 17.19 | 140 | >sp\|P02788\|TRFL_HUMAN Lactotransferrin OS=Homo sapiens OX=9606 GN=LTF PE=1 SV=6 | 1270 | 20201113_15413_1ng_glyco_hcd203040_compound15.24359.24359.3 | controllerType=0 controllerNumber=1 scan=24359 | 49.2311 |
| 16727:3 | 1 | R.TAGWNIPMGLLFN[+1768.640]QTGSC[+57.021]K.F | HexNAc(4)Hex(5)Fuc(1) | C[+57], N[+1769] | 1288.5593 | 3 | 3863.6632 | 3863.6495 | 0 | 3.6 | 485 | Specific | 904.08 | 670.67 | 670.67 | 18.19 | 140 | >sp\|P02788\|TRFL_HUMAN Lactotransferrin OS=Homo sapiens OX=9606 GN=LTF PE=1 SV=6 | 1270 | 20201113_15413_1ng_glyco_hcd203040_compound15.20230.20230.3 | controllerType=0 controllerNumber=1 scan=20230 | 40.7653 |
| 18888:3 | 1 | R.TAGWNIPMGLLFN[+1710.598]QTGSC[+57.021]K.F | HexNAc(3)Hex(5)NeuAc(1) | C[+57], N[+1711] | 1269.2112 | 3 | 3805.6192 | 3805.6076 | 0 | 3 | 485 | Specific | 915.75 | 684.15 | 391.81 | 18.19 | 140 | >sp\|P02788\|TRFL_HUMAN Lactotransferrin OS=Homo sapiens OX=9606 GN=LTF PE=1 SV=6 | 1270 | 20201113_15413_1ng_glyco_hcd203040_compound15.22485.22485.3 | controllerType=0 controllerNumber=1 scan=22485 | 45.4837 |
| 20744:3 | 1 | R.TAGWNIPMGLLFN[+2350.830]QTGSC[+57.021]K.F | HexNAc(4)Hex(5)Fuc(1)NeuAc(2) | C[+57], N[+2351] | 1482.6231 | 3 | 4445.8548 | 4445.8403 | 0 | 3.3 | 485 | Specific | 928.52 | 905.77 | 905.77 | 18.19 | 140 | >sp\|P02788\|TRFL_HUMAN Lactotransferrin OS=Homo sapiens OX=9606 GN=LTF PE=1 SV=6 | 1270 | 20201113_15413_1ng_glyco_hcd203040_compound15.24420.24420.3 | controllerType=0 controllerNumber=1 scan=24420 | 49.3573 |
| 17210:3 | 1 | R.TAGWNIPMGLLFN[+1216.423]QTGSC[+57.021]K.F | HexNAc(2)Hex(5) | C[+57], N[+1216] | 1104.4873 | 3 | 3311.4474 | 3311.4328 | 0 | 4.4 | 485 | Specific | 934.08 | 746.64 | 406.65 | 18.09 | 140 | >sp\|P02788\|TRFL_HUMAN Lactotransferrin OS=Homo sapiens OX=9606 GN=LTF PE=1 SV=6 | 1270 | 20201113_15413_1ng_glyco_hcd203040_compound15.20732.20732.3 | controllerType=0 controllerNumber=1 scan=20732 | 41.7546 |
| 16709:3 | 1 | R.TAGWNIPMGLLFN[+1768.640]QTGSC[+57.021]K.F | HexNAc(4)Hex(5)Fuc(1) | C[+57], N[+1769] | 1288.564 | 3 | 3863.6775 | 3863.6495 | 0 | 7.3 | 485 | Specific | 934.76 | 934.76 | 934.76 | 17.4 | 140 | >sp\|P02788\|TRFL_HUMAN Lactotransferrin OS=Homo sapiens OX=9606 GN=LTF PE=1 SV=6 | 1270 | 20201113_15413_1ng_glyco_hcd203040_compound15.20211.20211.3 | controllerType=0 controllerNumber=1 scan=20211 | 40.7293 |
| 18634:3 | 1 | R.TAGWNIPMGLLFN[+2059.735]QTGSC[+57.021]K.F | HexNAc(4)Hex(5)Fuc(1)NeuAc(1) | C[+57], N[+2060] | 1385.5921 | 3 | 4154.7617 | 4154.7449 | 0 | 4.1 | 485 | Specific | 952.64 | 944.42 | 944.42 | 19.09 | 140 | >sp\|P02788\|TRFL_HUMAN Lactotransferrin OS=Homo sapiens OX=9606 GN=LTF PE=1 SV=6 | 1270 | 20201113_15413_1ng_glyco_hcd203040_compound15.22219.22219.3 | controllerType=0 controllerNumber=1 scan=22219 | 44.8468 |
| 16725:3 | 1 | R.TAGWNIPMGLLFN[+1768.640]QTGSC[+57.021]K.F | HexNAc(4)Hex(5)Fuc(1) | C[+57], N[+1769] | 1288.5593 | 3 | 3863.6632 | 3863.6495 | 0 | 3.6 | 485 | Specific | 954.43 | 954.43 | 954.43 | 19.19 | 140 | >sp\|P02788\|TRFL_HUMAN Lactotransferrin OS=Homo sapiens OX=9606 GN=LTF PE=1 SV=6 | 1270 | 20201113_15413_1ng_glyco_hcd203040_compound15.20228.20228.3 | controllerType=0 controllerNumber=1 scan=20228 | 40.7591 |
| 18584:3 | 1 | R.TAGWNIPMGLLFN[+2205.793]QTGSC[+57.021]K.F | HexNAc(4)Hex(5)Fuc(2)NeuAc(1) | C[+57], N[+2206] | 1434.2741 | 3 | 4300.8078 | 4300.8028 | 0 | 1.2 | 485 | Specific | 965.48 | 871.12 | 871.12 | 18.9 | 140 | >sp\|P02788\|TRFL_HUMAN Lactotransferrin OS=Homo sapiens OX=9606 GN=LTF PE=1 SV=6 | 1270 | 20201113_15413_1ng_glyco_hcd203040_compound15.22167.22167.3 | controllerType=0 controllerNumber=1 scan=22167 | 44.7316 |
| 16835:3 | 1 | R.TAGWNIPMGLLFN[+1606.587]QTGSC[+57.021]K.F | HexNAc(4)Hex(4)Fuc(1) | C[+57], N[+1607] | 1234.5405 | 3 | 3701.6068 | 3701.5966 | 0 | 2.8 | 485 | Specific | 971.59 | 953.75 | 953.75 | 19.19 | 140 | >sp\|P02788\|TRFL_HUMAN Lactotransferrin OS=Homo sapiens OX=9606 GN=LTF PE=1 SV=6 | 1270 | 20201113_15413_1ng_glyco_hcd203040_compound15.20342.20342.3 | controllerType=0 controllerNumber=1 scan=20342 | 40.9881 |
| 18884:3 | 1 | R.TAGWNIPMGLLFN[+1751.624]QTGSC[+57.021]K.F | HexNAc(4)Hex(4)NeuAc(1) | C[+57], N[+1752] | 1282.8868 | 3 | 3846.6458 | 3846.6341 | 0 | 3 | 485 | Specific | 977.53 | 803.57 | 803.57 | 19.19 | 140 | >sp\|P02788\|TRFL_HUMAN Lactotransferrin OS=Homo sapiens OX=9606 GN=LTF PE=1 SV=6 | 1270 | 20201113_15413_1ng_glyco_hcd203040_compound15.22481.22481.3 | controllerType=0 controllerNumber=1 scan=22481 | 45.4766 |
| 18631:3 | 1 | R.TAGWNIPMGLLFN[+2059.735]QTGSC[+57.021]K.F | HexNAc(4)Hex(5)Fuc(1)NeuAc(1) | C[+57], N[+2060] | 1385.5921 | 3 | 4154.7617 | 4154.7449 | 0 | 4.1 | 485 | Specific | 992.59 | 951.66 | 951.66 | 19.09 | 140 | >sp\|P02788\|TRFL_HUMAN Lactotransferrin OS=Homo sapiens OX=9606 GN=LTF PE=1 SV=6 | 1270 | 20201113_15413_1ng_glyco_hcd203040_compound15.22216.22216.3 | controllerType=0 controllerNumber=1 scan=22216 | 44.8327 |
| 18582:3 | 1 | R.TAGWNIPMGLLFN[+2205.793]QTGSC[+57.021]K.F | HexNAc(4)Hex(5)Fuc(2)NeuAc(1) | C[+57], N[+2206] | 1434.2741 | 3 | 4300.8078 | 4300.8028 | 0 | 1.2 | 485 | Specific | 1015.24 | 989.83 | 989.83 | 19.9 | 140 | >sp\|P02788\|TRFL_HUMAN Lactotransferrin OS=Homo sapiens OX=9606 GN=LTF PE=1 SV=6 | 1270 | 20201113_15413_1ng_glyco_hcd203040_compound15.22165.22165.3 | controllerType=0 controllerNumber=1 scan=22165 | 44.726 |
| 17155:3 | 1 | R.TAGWNIPMGLLFN[+1216.423]QTGSC[+57.021]K.F | HexNAc(2)Hex(5) | C[+57], N[+1216] | 1104.4849 | 3 | 3311.4401 | 3311.4328 | 0 | 2.2 | 485 | Specific | 1023.67 | 975.76 | 488.52 | 20.15 | 140 | >sp\|P02788\|TRFL_HUMAN Lactotransferrin OS=Homo sapiens OX=9606 GN=LTF PE=1 SV=6 | 1270 | 20201113_15413_1ng_glyco_hcd203040_compound15.20675.20675.3 | controllerType=0 controllerNumber=1 scan=20675 | 41.645 |
| 16968:3 | 1 | R.TAGWNIPM[+15.995]GLLFN[+2204.772]QTGSC[+57.021]K.F | HexNAc(4)Hex(5)NeuAc(2) | C[+57], M[+16], N[+2205] | 1439.2697 | 3 | 4315.7946 | 4315.7773 | 0 | 4 | 485 | Specific | 256.01 | 256.01 | 256.01 | 6.48 | 140 | >sp\|P02788\|TRFL_HUMAN Lactotransferrin OS=Homo sapiens OX=9606 GN=LTF PE=1 SV=6 | 1270 | 20201113_15413_1ng_glyco_hcd203040_compound15.20480.20480.4 | controllerType=0 controllerNumber=1 scan=20480 | 41.2511 |
| 17071:3 | 1 | R.TAGWNIPM[+15.995]GLLFN[+2204.772]QTGSC[+57.021]K.F | HexNAc(4)Hex(5)NeuAc(2) | C[+57], M[+16], N[+2205] | 1439.2735 | 3 | 4315.8059 | 4315.7773 | 0 | 6.6 | 485 | Specific | 293.3 | 273.54 | 273.54 | 5.91 | 140 | >sp\|P02788\|TRFL_HUMAN Lactotransferrin OS=Homo sapiens OX=9606 GN=LTF PE=1 SV=6 | 1270 | 20201113_15413_1ng_glyco_hcd203040_compound15.20587.20587.3 | controllerType=0 controllerNumber=1 scan=20587 | 41.4711 |
| 16970:3 | 1 | R.TAGWNIPM[+15.995]GLLFN[+2204.772]QTGSC[+57.021]K.F | HexNAc(4)Hex(5)NeuAc(2) | C[+57], M[+16], N[+2205] | 1439.2697 | 3 | 4315.7946 | 4315.7773 | 0 | 4 | 485 | Specific | 332.16 | 275.82 | 275.82 | 7.02 | 140 | >sp\|P02788\|TRFL_HUMAN Lactotransferrin OS=Homo sapiens OX=9606 GN=LTF PE=1 SV=6 | 1270 | 20201113_15413_1ng_glyco_hcd203040_compound15.20482.20482.4 | controllerType=0 controllerNumber=1 scan=20482 | 41.2552 |
| 17835:3 | 1 | R.TAGWNIPM[+15.995]GLLFN[+2059.735]QTGSC[+57.021]K.F | HexNAc(4)Hex(5)Fuc(1)NeuAc(1) | C[+57], M[+16], N[+2060] | 1390.9285 | 3 | 4170.771 | 4170.7398 | 0 | 7.5 | 485 | Specific | 342.99 | 232.15 | 232.15 | 5.98 | 140 | >sp\|P02788\|TRFL_HUMAN Lactotransferrin OS=Homo sapiens OX=9606 GN=LTF PE=1 SV=6 | 1270 | 20201113_15413_1ng_glyco_hcd203040_compound15.21384.21384.3 | controllerType=0 controllerNumber=1 scan=21384 | 43.1429 |
| 17298:3 | 1 | R.TAGWNIPM[+15.995]GLLFN[+1913.677]QTGSC[+57.021]K.F | HexNAc(4)Hex(5)NeuAc(1) | C[+57], M[+16], N[+1914] | 1342.2366 | 3 | 4024.6953 | 4024.6819 | 0 | 3.3 | 485 | Specific | 344.17 | 286.39 | 286.39 | 7.12 | 140 | >sp\|P02788\|TRFL_HUMAN Lactotransferrin OS=Homo sapiens OX=9606 GN=LTF PE=1 SV=6 | 1270 | 20201113_15413_1ng_glyco_hcd203040_compound15.20823.20823.3 | controllerType=0 controllerNumber=1 scan=20823 | 41.9282 |
| 17837:3 | 1 | R.TAGWNIPM[+15.995]GLLFN[+2059.735]QTGSC[+57.021]K.F | HexNAc(4)Hex(5)Fuc(1)NeuAc(1) | C[+57], M[+16], N[+2060] | 1390.9285 | 3 | 4170.771 | 4170.7398 | 0 | 7.5 | 485 | Specific | 485.92 | 224.11 | 224.11 | 6.34 | 140 | >sp\|P02788\|TRFL_HUMAN Lactotransferrin OS=Homo sapiens OX=9606 GN=LTF PE=1 SV=6 | 1270 | 20201113_15413_1ng_glyco_hcd203040_compound15.21386.21386.3 | controllerType=0 controllerNumber=1 scan=21386 | 43.1492 |
| 17360:3 | 1 | R.TAGWNIPM[+15.995]GLLFN[+2059.735]QTGSC[+57.021]K.F | HexNAc(4)Hex(5)Fuc(1)NeuAc(1) | C[+57], M[+16], N[+2060] | 1390.9236 | 3 | 4170.7564 | 4170.7398 | 0 | 4 | 485 | Specific | 500.48 | 380.86 | 380.86 | 9.56 | 140 | >sp\|P02788\|TRFL_HUMAN Lactotransferrin OS=Homo sapiens OX=9606 GN=LTF PE=1 SV=6 | 1270 | 20201113_15413_1ng_glyco_hcd203040_compound15.20888.20888.3 | controllerType=0 controllerNumber=1 scan=20888 | 42.0506 |
| 17362:3 | 1 | R.TAGWNIPM[+15.995]GLLFN[+2059.735]QTGSC[+57.021]K.F | HexNAc(4)Hex(5)Fuc(1)NeuAc(1) | C[+57], M[+16], N[+2060] | 1390.9236 | 3 | 4170.7564 | 4170.7398 | 0 | 4 | 485 | Specific | 603.43 | 535.61 | 535.61 | 12.09 | 140 | >sp\|P02788\|TRFL_HUMAN Lactotransferrin OS=Homo sapiens OX=9606 GN=LTF PE=1 SV=6 | 1270 | 20201113_15413_1ng_glyco_hcd203040_compound15.20890.20890.3 | controllerType=0 controllerNumber=1 scan=20890 | 42.0555 |
| 17221:3 | 1 | R.TAGWNIPM[+15.995]GLLFN[+2059.735]QTGSC[+57.021]K.F | HexNAc(4)Hex(5)Fuc(1)NeuAc(1) | C[+57], M[+16], N[+2060] | 1390.9222 | 3 | 4170.752 | 4170.7398 | 0 | 2.9 | 485 | Specific | 619.17 | 590.25 | 590.25 | 12.19 | 140 | >sp\|P02788\|TRFL_HUMAN Lactotransferrin OS=Homo sapiens OX=9606 GN=LTF PE=1 SV=6 | 1270 | 20201113_15413_1ng_glyco_hcd203040_compound15.20744.20744.3 | controllerType=0 controllerNumber=1 scan=20744 | 41.7783 |
| 16829:3 | 1 | R.TAGWNIPM[+15.995]GLLFN[+2059.735]QTGSC[+57.021]K.F | HexNAc(4)Hex(5)Fuc(1)NeuAc(1) | C[+57], M[+16], N[+2060] | 1390.9236 | 3 | 4170.7564 | 4170.7398 | 0 | 4 | 485 | Specific | 667.48 | 505.76 | 505.76 | 13.09 | 140 | >sp\|P02788\|TRFL_HUMAN Lactotransferrin OS=Homo sapiens OX=9606 GN=LTF PE=1 SV=6 | 1270 | 20201113_15413_1ng_glyco_hcd203040_compound15.20336.20336.3 | controllerType=0 controllerNumber=1 scan=20336 | 40.9751 |
| 16831:3 | 1 | R.TAGWNIPM[+15.995]GLLFN[+2059.735]QTGSC[+57.021]K.F | HexNAc(4)Hex(5)Fuc(1)NeuAc(1) | C[+57], M[+16], N[+2060] | 1390.9236 | 3 | 4170.7564 | 4170.7398 | 0 | 4 | 485 | Specific | 779.79 | 667.13 | 667.13 | 15.09 | 140 | >sp\|P02788\|TRFL_HUMAN Lactotransferrin OS=Homo sapiens OX=9606 GN=LTF PE=1 SV=6 | 1270 | 20201113_15413_1ng_glyco_hcd203040_compound15.20338.20338.3 | controllerType=0 controllerNumber=1 scan=20338 | 40.9805 |
| 17223:3 | 1 | R.TAGWNIPM[+15.995]GLLFN[+2059.735]QTGSC[+57.021]K.F | HexNAc(4)Hex(5)Fuc(1)NeuAc(1) | C[+57], M[+16], N[+2060] | 1390.9222 | 3 | 4170.752 | 4170.7398 | 0 | 2.9 | 485 | Specific | 810.42 | 714.11 | 714.11 | 16.19 | 140 | >sp\|P02788\|TRFL_HUMAN Lactotransferrin OS=Homo sapiens OX=9606 GN=LTF PE=1 SV=6 | 1270 | 20201113_15413_1ng_glyco_hcd203040_compound15.20746.20746.3 | controllerType=0 controllerNumber=1 scan=20746 | 41.7824 |
| 18914:4 | 1 | R.TAGWNIPMGLLFN[+2205.793]QTGSC[+57.021]KFDEYFSQSC[+57.021]APGSDPR.S | HexNAc(4)Hex(5)Fuc(2)NeuAc(1) | C[+57]*2, N[+2206] | 1536.8939 | 4 | 6144.5539 | 6144.5499 | 0 | 0.6 | 485 | Specific | 597.61 | 576.09 | 576.09 | 10.55 | 140 | >sp\|P02788\|TRFL_HUMAN Lactotransferrin OS=Homo sapiens OX=9606 GN=LTF PE=1 SV=6 | 1270 | 20201113_15413_1ng_glyco_hcd203040_compound15.22512.22512.4 | controllerType=0 controllerNumber=1 scan=22512 | 45.5285 |
| 18917:4 | 1 | R.TAGWNIPMGLLFN[+2205.793]QTGSC[+57.021]KFDEYFSQSC[+57.021]APGSDPR.S | HexNAc(4)Hex(5)Fuc(2)NeuAc(1) | C[+57]*2, N[+2206] | 1536.8939 | 4 | 6144.5539 | 6144.5499 | 0 | 0.6 | 485 | Specific | 687.3 | 642.28 | 642.28 | 12.55 | 140 | >sp\|P02788\|TRFL_HUMAN Lactotransferrin OS=Homo sapiens OX=9606 GN=LTF PE=1 SV=6 | 1270 | 20201113_15413_1ng_glyco_hcd203040_compound15.22515.22515.4 | controllerType=0 controllerNumber=1 scan=22515 | 45.5365 |
| 18913:4 | 1 | R.TAGWNIPMGLLFN[+2059.735]QTGSC[+57.021]KFDEYFSQSC[+57.021]APGSDPR.S | HexNAc(4)Hex(5)Fuc(1)NeuAc(1) | C[+57]*2, N[+2060] | 1500.3811 | 4 | 5998.5026 | 5998.492 | 0 | 1.8 | 485 | Specific | 744.28 | 690.83 | 690.83 | 13.79 | 140 | >sp\|P02788\|TRFL_HUMAN Lactotransferrin OS=Homo sapiens OX=9606 GN=LTF PE=1 SV=6 | 1270 | 20201113_15413_1ng_glyco_hcd203040_compound15.22511.22511.4 | controllerType=0 controllerNumber=1 scan=22511 | 45.5274 |
| 18916:4 | 1 | R.TAGWNIPMGLLFN[+2059.735]QTGSC[+57.021]KFDEYFSQSC[+57.021]APGSDPR.S | HexNAc(4)Hex(5)Fuc(1)NeuAc(1) | C[+57]*2, N[+2060] | 1500.3811 | 4 | 5998.5026 | 5998.492 | 0 | 1.8 | 485 | Specific | 866.1 | 843.69 | 843.69 | 16.79 | 140 | >sp\|P02788\|TRFL_HUMAN Lactotransferrin OS=Homo sapiens OX=9606 GN=LTF PE=1 SV=6 | 1270 | 20201113_15413_1ng_glyco_hcd203040_compound15.22514.22514.4 | controllerType=0 controllerNumber=1 scan=22514 | 45.5353 |
| 08801:3 | 1 | R.N[+2205.793]GSDC[+57.021]PDKFC[+57.021]LFQSETK.N | HexNAc(4)Hex(5)Fuc(2)NeuAc(1) | C[+57]*2, N[+2206] | 1413.5629 | 3 | 4238.6742 | 4238.6667 | 0 | 1.8 | 642 | Specific | 363.22 | 246.13 | 246.13 | 6.29 | 140 | >sp\|P02788\|TRFL_HUMAN Lactotransferrin OS=Homo sapiens OX=9606 GN=LTF PE=1 SV=6 | 1270 | 20201113_15413_1ng_glyco_hcd203040_compound15.11972.11972.3 | controllerType=0 controllerNumber=1 scan=11972 | 26.5566 |
| 08944:3 | 1 | R.N[+2059.735]GSDC[+57.021]PDKFC[+57.021]LFQSETK.N | HexNAc(4)Hex(5)Fuc(1)NeuAc(1) | C[+57]*2, N[+2060] | 1364.8791 | 3 | 4092.6227 | 4092.6088 | 0 | 3.4 | 642 | Specific | 387.89 | 312.73 | 312.73 | 6.81 | 140 | >sp\|P02788\|TRFL_HUMAN Lactotransferrin OS=Homo sapiens OX=9606 GN=LTF PE=1 SV=6 | 1270 | 20201113_15413_1ng_glyco_hcd203040_compound15.12121.12121.3 | controllerType=0 controllerNumber=1 scan=12121 | 26.8105 |
| 08803:3 | 1 | R.N[+2205.793]GSDC[+57.021]PDKFC[+57.021]LFQSETK.N | HexNAc(4)Hex(5)Fuc(2)NeuAc(1) | C[+57]*2, N[+2206] | 1413.5629 | 3 | 4238.6742 | 4238.6667 | 0 | 1.8 | 642 | Specific | 429.16 | 280.28 | 280.28 | 6.93 | 140 | >sp\|P02788\|TRFL_HUMAN Lactotransferrin OS=Homo sapiens OX=9606 GN=LTF PE=1 SV=6 | 1270 | 20201113_15413_1ng_glyco_hcd203040_compound15.11974.11974.3 | controllerType=0 controllerNumber=1 scan=11974 | 26.5607 |
| 08946:3 | 1 | R.N[+2059.735]GSDC[+57.021]PDKFC[+57.021]LFQSETK.N | HexNAc(4)Hex(5)Fuc(1)NeuAc(1) | C[+57]*2, N[+2060] | 1364.8791 | 3 | 4092.6227 | 4092.6088 | 0 | 3.4 | 642 | Specific | 435.41 | 264.12 | 264.12 | 6.82 | 140 | >sp\|P02788\|TRFL_HUMAN Lactotransferrin OS=Homo sapiens OX=9606 GN=LTF PE=1 SV=6 | 1270 | 20201113_15413_1ng_glyco_hcd203040_compound15.12123.12123.3 | controllerType=0 controllerNumber=1 scan=12123 | 26.8143 |
| 11283:3 | 1 | R.N[+2350.830]GSDC[+57.021]PDKFC[+57.021]LFQSETK.N | HexNAc(4)Hex(5)Fuc(1)NeuAc(2) | C[+57]*2, N[+2351] | 1461.9124 | 3 | 4383.7227 | 4383.7043 | 0 | 4.2 | 642 | Specific | 474.84 | 378.45 | 378.45 | 8.57 | 140 | >sp\|P02788\|TRFL_HUMAN Lactotransferrin OS=Homo sapiens OX=9606 GN=LTF PE=1 SV=6 | 1270 | 20201113_15413_1ng_glyco_hcd203040_compound15.14544.14544.3 | controllerType=0 controllerNumber=1 scan=14544 | 30.9188 |
| 08650:3 | 1 | R.N[+2570.925]GSDC[+57.021]PDKFC[+57.021]LFQSETK.N | HexNAc(5)Hex(6)Fuc(2)NeuAc(1) | C[+57]*2, N[+2571] | 1535.2769 | 3 | 4603.8162 | 4603.7989 | 0 | 3.7 | 642 | Specific | 499.59 | 443.29 | 443.29 | 8.94 | 140 | >sp\|P02788\|TRFL_HUMAN Lactotransferrin OS=Homo sapiens OX=9606 GN=LTF PE=1 SV=6 | 1270 | 20201113_15413_1ng_glyco_hcd203040_compound15.11810.11810.3 | controllerType=0 controllerNumber=1 scan=11810 | 26.275 |
| 08652:3 | 1 | R.N[+2570.925]GSDC[+57.021]PDKFC[+57.021]LFQSETK.N | HexNAc(5)Hex(6)Fuc(2)NeuAc(1) | C[+57]*2, N[+2571] | 1535.2769 | 3 | 4603.8162 | 4603.7989 | 0 | 3.7 | 642 | Specific | 529.01 | 458.49 | 458.49 | 9.22 | 140 | >sp\|P02788\|TRFL_HUMAN Lactotransferrin OS=Homo sapiens OX=9606 GN=LTF PE=1 SV=6 | 1270 | 20201113_15413_1ng_glyco_hcd203040_compound15.11812.11812.3 | controllerType=0 controllerNumber=1 scan=11812 | 26.2792 |
| 08827:3 | 1 | R.N[+2205.793]GSDC[+57.021]PDKFC[+57.021]LFQSETK.N | HexNAc(4)Hex(5)Fuc(2)NeuAc(1) | C[+57]*2, N[+2206] | 1413.5635 | 3 | 4238.676 | 4238.6667 | 0 | 2.2 | 642 | Specific | 556.37 | 399.98 | 399.98 | 9.36 | 140 | >sp\|P02788\|TRFL_HUMAN Lactotransferrin OS=Homo sapiens OX=9606 GN=LTF PE=1 SV=6 | 1270 | 20201113_15413_1ng_glyco_hcd203040_compound15.12000.12000.3 | controllerType=0 controllerNumber=1 scan=12000 | 26.608 |
| 11281:3 | 1 | R.N[+2350.830]GSDC[+57.021]PDKFC[+57.021]LFQSETK.N | HexNAc(4)Hex(5)Fuc(1)NeuAc(2) | C[+57]*2, N[+2351] | 1461.9124 | 3 | 4383.7227 | 4383.7043 | 0 | 4.2 | 642 | Specific | 598.51 | 542.75 | 542.75 | 10.5 | 140 | >sp\|P02788\|TRFL_HUMAN Lactotransferrin OS=Homo sapiens OX=9606 GN=LTF PE=1 SV=6 | 1270 | 20201113_15413_1ng_glyco_hcd203040_compound15.14542.14542.3 | controllerType=0 controllerNumber=1 scan=14542 | 30.914 |
| 08825:3 | 1 | R.N[+2205.793]GSDC[+57.021]PDKFC[+57.021]LFQSETK.N | HexNAc(4)Hex(5)Fuc(2)NeuAc(1) | C[+57]*2, N[+2206] | 1413.5635 | 3 | 4238.676 | 4238.6667 | 0 | 2.2 | 642 | Specific | 658.63 | 539.53 | 539.53 | 12.56 | 140 | >sp\|P02788\|TRFL_HUMAN Lactotransferrin OS=Homo sapiens OX=9606 GN=LTF PE=1 SV=6 | 1270 | 20201113_15413_1ng_glyco_hcd203040_compound15.11998.11998.3 | controllerType=0 controllerNumber=1 scan=11998 | 26.6034 |
| 07127:3 | 3 | R.FSDGLESN[+1864.634]SSTQFEVK.K | HexNAc(2)Hex(9) | N[+1865] | 1213.8234 | 3 | 3639.4557 | 3639.4472 | 0 | 2.4 | 219 | Specific | 449.86 | 449.86 | 449.86 | 9.35 | 88 | >sp\|P0C0L5\|CO4B_HUMAN Complement C4-B OS=Homo sapiens OX=9606 GN=C4B PE=1 SV=2 | 1424 | 20201113_15413_1ng_glyco_hcd203040_compound15.10224.10224.3 | controllerType=0 controllerNumber=1 scan=10224 | 23.6062 |
| 07125:3 | 3 | R.FSDGLESN[+1864.634]SSTQFEVK.K | HexNAc(2)Hex(9) | N[+1865] | 1213.8234 | 3 | 3639.4557 | 3639.4472 | 0 | 2.4 | 219 | Specific | 550.99 | 550.99 | 550.99 | 9.95 | 88 | >sp\|P0C0L5\|CO4B_HUMAN Complement C4-B OS=Homo sapiens OX=9606 GN=C4B PE=1 SV=2 | 1424 | 20201113_15413_1ng_glyco_hcd203040_compound15.10222.10222.3 | controllerType=0 controllerNumber=1 scan=10222 | 23.602 |
| 07079:3 | 3 | R.FSDGLESN[+1864.634]SSTQFEVK.K | HexNAc(2)Hex(9) | N[+1865] | 1213.8271 | 3 | 3639.4667 | 3639.4472 | 0 | 5.4 | 219 | Specific | 850.65 | 850.65 | 850.65 | 16.68 | 88 | >sp\|P0C0L5\|CO4B_HUMAN Complement C4-B OS=Homo sapiens OX=9606 GN=C4B PE=1 SV=2 | 1424 | 20201113_15413_1ng_glyco_hcd203040_compound15.10174.10174.3 | controllerType=0 controllerNumber=1 scan=10174 | 23.5179 |
| 07077:3 | 3 | R.FSDGLESN[+1864.634]SSTQFEVK.K | HexNAc(2)Hex(9) | N[+1865] | 1213.8271 | 3 | 3639.4667 | 3639.4472 | 0 | 5.4 | 219 | Specific | 856 | 856 | 856 | 16.68 | 88 | >sp\|P0C0L5\|CO4B_HUMAN Complement C4-B OS=Homo sapiens OX=9606 GN=C4B PE=1 SV=2 | 1424 | 20201113_15413_1ng_glyco_hcd203040_compound15.10172.10172.3 | controllerType=0 controllerNumber=1 scan=10172 | 23.5139 |
| 04620:3 | 3 | R.FSDGLESN[+1864.634]SSTQFEVKK.Y | HexNAc(2)Hex(9) | N[+1865] | 1256.525 | 3 | 3767.5603 | 3767.5421 | 0 | 4.8 | 219 | Specific | 229.93 | 229.93 | 229.93 | 5.48 | 88 | >sp\|P0C0L5\|CO4B_HUMAN Complement C4-B OS=Homo sapiens OX=9606 GN=C4B PE=1 SV=2 | 1424 | 20201113_15413_1ng_glyco_hcd203040_compound15.7623.7623.3 | controllerType=0 controllerNumber=1 scan=7623 | 19.3015 |
| 04618:3 | 3 | R.FSDGLESN[+1864.634]SSTQFEVKK.Y | HexNAc(2)Hex(9) | N[+1865] | 1256.525 | 3 | 3767.5603 | 3767.5421 | 0 | 4.8 | 219 | Specific | 241.4 | 241.4 | 241.4 | 5.48 | 88 | >sp\|P0C0L5\|CO4B_HUMAN Complement C4-B OS=Homo sapiens OX=9606 GN=C4B PE=1 SV=2 | 1424 | 20201113_15413_1ng_glyco_hcd203040_compound15.7621.7621.3 | controllerType=0 controllerNumber=1 scan=7621 | 19.2976 |
| 15443:2 | 5 | R.LN[+1216.423]YSLPTGQWVGVQLPR.N | HexNAc(2)Hex(5) | N[+1216] | 1572.7478 | 2 | 3144.4883 | 3144.4617 | 0 | 8.5 | 1017 | Specific | 206.39 | 206.39 | 206.39 | 4.09 | 83 | >sp\|P24821\|TENA_HUMAN Tenascin OS=Homo sapiens OX=9606 GN=TNC PE=1 SV=3 | 486 | 20201113_15413_1ng_glyco_hcd203040_compound15.18870.18870.2 | controllerType=0 controllerNumber=1 scan=18870 | 38.2955 |
| 15388:2 | 5 | R.LN[+1378.476]YSLPTGQWVGVQLPR.N | HexNAc(2)Hex(6) | N[+1378] | 1653.7729 | 2 | 3306.5385 | 3306.5145 | 0 | 7.2 | 1017 | Specific | 375.02 | 371.59 | 371.59 | 6.93 | 83 | >sp\|P24821\|TENA_HUMAN Tenascin OS=Homo sapiens OX=9606 GN=TNC PE=1 SV=3 | 486 | 20201113_15413_1ng_glyco_hcd203040_compound15.18813.18813.2 | controllerType=0 controllerNumber=1 scan=18813 | 38.1938 |
| 15445:2 | 5 | R.LN[+1216.423]YSLPTGQWVGVQLPR.N | HexNAc(2)Hex(5) | N[+1216] | 1572.7478 | 2 | 3144.4883 | 3144.4617 | 0 | 8.5 | 1017 | Specific | 484.05 | 484.05 | 484.05 | 8.73 | 83 | >sp\|P24821\|TENA_HUMAN Tenascin OS=Homo sapiens OX=9606 GN=TNC PE=1 SV=3 | 486 | 20201113_15413_1ng_glyco_hcd203040_compound15.18872.18872.2 | controllerType=0 controllerNumber=1 scan=18872 | 38.2994 |
| 09883:3 | 6 | K.TVLTPATNHMGN[+1378.476]VTFTIPANR.E | HexNAc(2)Hex(6) | N[+1378] | 1211.8881 | 3 | 3633.6498 | 3633.6358 | 0 | 3.9 | 74 | Specific | 568.24 | 421.87 | 297.92 | 10.07 | 87 | >sp\|P01024\|CO3_HUMAN Complement C3 OS=Homo sapiens OX=9606 GN=C3 PE=1 SV=2 | 492 | 20201113_15413_1ng_glyco_hcd203040_compound15.13094.13094.3 | controllerType=0 controllerNumber=1 scan=13094 | 28.407 |
| 09885:3 | 6 | K.TVLTPATNHMGN[+1378.476]VTFTIPANR.E | HexNAc(2)Hex(6) | N[+1378] | 1211.8881 | 3 | 3633.6498 | 3633.6358 | 0 | 3.9 | 74 | Specific | 587.05 | 534.61 | 235.96 | 11.27 | 87 | >sp\|P01024\|CO3_HUMAN Complement C3 OS=Homo sapiens OX=9606 GN=C3 PE=1 SV=2 | 492 | 20201113_15413_1ng_glyco_hcd203040_compound15.13096.13096.3 | controllerType=0 controllerNumber=1 scan=13096 | 28.4109 |
| 09830:3 | 6 | K.TVLTPATNHMGN[+1540.529]VTFTIPANR.E | HexNAc(2)Hex(7) | N[+1541] | 1265.9085 | 3 | 3795.711 | 3795.6886 | 0 | 5.9 | 74 | Specific | 702.68 | 663.68 | 350.62 | 13.75 | 87 | >sp\|P01024\|CO3_HUMAN Complement C3 OS=Homo sapiens OX=9606 GN=C3 PE=1 SV=2 | 492 | 20201113_15413_1ng_glyco_hcd203040_compound15.13039.13039.3 | controllerType=0 controllerNumber=1 scan=13039 | 28.3157 |
| 09828:3 | 6 | K.TVLTPATNHMGN[+1540.529]VTFTIPANR.E | HexNAc(2)Hex(7) | N[+1541] | 1265.9085 | 3 | 3795.711 | 3795.6886 | 0 | 5.9 | 74 | Specific | 747.14 | 747.14 | 406.92 | 13.75 | 87 | >sp\|P01024\|CO3_HUMAN Complement C3 OS=Homo sapiens OX=9606 GN=C3 PE=1 SV=2 | 492 | 20201113_15413_1ng_glyco_hcd203040_compound15.13037.13037.3 | controllerType=0 controllerNumber=1 scan=13037 | 28.3119 |
| 07386:3 | 7 | K.WEC[+57.021]KN[+2205.793]DTLLGIK.G | HexNAc(4)Hex(5)Fuc(2)NeuAc(1) | C[+57], N[+2206] | 1228.1921 | 3 | 3682.5617 | 3682.5444 | 0 | 4.7 | 100 | Specific | 219.6 | 112.61 | 112.61 | 3.08 | 83 | >sp\|P22897\|MRC1_HUMAN Macrophage mannose receptor 1 OS=Homo sapiens OX=9606 GN=MRC1 PE=1 SV=1 | 1049 | 20201113_15413_1ng_glyco_hcd203040_compound15.10492.10492.3 | controllerType=0 controllerNumber=1 scan=10492 | 24.0578 |
| 12863:4 | 7 | K.KGN[+2205.793]TTLNSFVIPSESDVPTHC[+57.021]PSQWWPYAGHC[+57.021]YK.I | HexNAc(4)Hex(5)Fuc(2)NeuAc(1) | C[+57]*2, N[+2206] | 1543.1597 | 4 | 6169.6169 | 6169.6146 | 0 | 0.4 | 342 | Specific | 712.43 | 452.27 | 452.27 | 13.86 | 83 | >sp\|P22897\|MRC1_HUMAN Macrophage mannose receptor 1 OS=Homo sapiens OX=9606 GN=MRC1 PE=1 SV=1 | 1049 | 20201113_15413_1ng_glyco_hcd203040_compound15.16198.16198.2 | controllerType=0 controllerNumber=1 scan=16198 | 33.7149 |
| 12860:4 | 7 | K.KGN[+2205.793]TTLNSFVIPSESDVPTHC[+57.021]PSQWWPYAGHC[+57.021]YK.I | HexNAc(4)Hex(5)Fuc(2)NeuAc(1) | C[+57]*2, N[+2206] | 1543.1597 | 4 | 6169.6169 | 6169.6146 | 0 | 0.4 | 342 | Specific | 826.16 | 826.16 | 826.16 | 15.86 | 83 | >sp\|P22897\|MRC1_HUMAN Macrophage mannose receptor 1 OS=Homo sapiens OX=9606 GN=MRC1 PE=1 SV=1 | 1049 | 20201113_15413_1ng_glyco_hcd203040_compound15.16195.16195.2 | controllerType=0 controllerNumber=1 scan=16195 | 33.7003 |
| 08216:3 | 7 | K.TAHC[+57.021]N[+2059.735]ESFYFLC[+57.021]K.R | HexNAc(4)Hex(5)Fuc(1)NeuAc(1) | C[+57]*2, N[+2060] | 1246.1603 | 3 | 3736.4665 | 3736.4545 | 0 | 3.2 | 1201 | Specific | 463.09 | 372.56 | 372.56 | 7.64 | 83 | >sp\|P22897\|MRC1_HUMAN Macrophage mannose receptor 1 OS=Homo sapiens OX=9606 GN=MRC1 PE=1 SV=1 | 1049 | 20201113_15413_1ng_glyco_hcd203040_compound15.11360.11360.3 | controllerType=0 controllerNumber=1 scan=11360 | 25.5223 |
| 08214:3 | 7 | K.TAHC[+57.021]N[+2059.735]ESFYFLC[+57.021]K.R | HexNAc(4)Hex(5)Fuc(1)NeuAc(1) | C[+57]*2, N[+2060] | 1246.1603 | 3 | 3736.4665 | 3736.4545 | 0 | 3.2 | 1201 | Specific | 484.28 | 402.57 | 402.57 | 9.35 | 83 | >sp\|P22897\|MRC1_HUMAN Macrophage mannose receptor 1 OS=Homo sapiens OX=9606 GN=MRC1 PE=1 SV=1 | 1049 | 20201113_15413_1ng_glyco_hcd203040_compound15.11358.11358.3 | controllerType=0 controllerNumber=1 scan=11358 | 25.5184 |
| 08194:3 | 7 | K.TAHC[+57.021]N[+2059.735]ESFYFLC[+57.021]K.R | HexNAc(4)Hex(5)Fuc(1)NeuAc(1) | C[+57]*2, N[+2060] | 1246.1591 | 3 | 3736.4628 | 3736.4545 | 0 | 2.2 | 1201 | Specific | 603.8 | 415.28 | 415.28 | 11.97 | 83 | >sp\|P22897\|MRC1_HUMAN Macrophage mannose receptor 1 OS=Homo sapiens OX=9606 GN=MRC1 PE=1 SV=1 | 1049 | 20201113_15413_1ng_glyco_hcd203040_compound15.11337.11337.3 | controllerType=0 controllerNumber=1 scan=11337 | 25.4816 |
| 08196:3 | 7 | K.TAHC[+57.021]N[+2059.735]ESFYFLC[+57.021]K.R | HexNAc(4)Hex(5)Fuc(1)NeuAc(1) | C[+57]*2, N[+2060] | 1246.1591 | 3 | 3736.4628 | 3736.4545 | 0 | 2.2 | 1201 | Specific | 657.6 | 562.82 | 562.82 | 12.97 | 83 | >sp\|P22897\|MRC1_HUMAN Macrophage mannose receptor 1 OS=Homo sapiens OX=9606 GN=MRC1 PE=1 SV=1 | 1049 | 20201113_15413_1ng_glyco_hcd203040_compound15.11339.11339.3 | controllerType=0 controllerNumber=1 scan=11339 | 25.4856 |
| 17099:3 | 9 | K.RNIAAFGGDPNN[+1913.677]ITLFGESAGGASVSLQTLSPYNK.G | HexNAc(4)Hex(5)NeuAc(1) | N[+1914] | 1827.5009 | 3 | 5480.4883 | 5480.4644 | 0 | 4.4 | 196 | Specific | 274.36 | 211.29 | 211.29 | 6.41 | 51 | >sp\|P19835\|CEL_HUMAN Bile salt-activated lipase OS=Homo sapiens OX=9606 GN=CEL PE=1 SV=3 | 476 | 20201113_15413_1ng_glyco_hcd203040_compound15.20616.20616.3 | controllerType=0 controllerNumber=1 scan=20616 | 41.5206 |
| 18899:3 | 9 | R.NIAAFGGDPNN[+1913.677]ITLFGESAGGASVSLQTLSPYNK.G | HexNAc(4)Hex(5)NeuAc(1) | N[+1914] | 1775.4647 | 3 | 5324.3795 | 5324.3633 | 0 | 3 | 197 | Specific | 233.47 | 233.47 | 233.47 | 6.76 | 51 | >sp\|P19835\|CEL_HUMAN Bile salt-activated lipase OS=Homo sapiens OX=9606 GN=CEL PE=1 SV=3 | 476 | 20201113_15413_1ng_glyco_hcd203040_compound15.22496.22496.3 | controllerType=0 controllerNumber=1 scan=22496 | 45.4995 |
| 18704:3 | 9 | R.NIAAFGGDPNN[+1702.581]ITLFGESAGGASVSLQTLSPYNK.G | HexNAc(2)Hex(8) | N[+1703] | 1705.0938 | 3 | 5113.267 | 5113.2676 | 0 | -0.1 | 197 | Specific | 497.74 | 497.74 | 497.74 | 9.39 | 51 | >sp\|P19835\|CEL_HUMAN Bile salt-activated lipase OS=Homo sapiens OX=9606 GN=CEL PE=1 SV=3 | 476 | 20201113_15413_1ng_glyco_hcd203040_compound15.22292.22292.3 | controllerType=0 controllerNumber=1 scan=22292 | 45.0138 |
| 18706:3 | 9 | R.NIAAFGGDPNN[+1702.581]ITLFGESAGGASVSLQTLSPYNK.G | HexNAc(2)Hex(8) | N[+1703] | 1705.0938 | 3 | 5113.267 | 5113.2676 | 0 | -0.1 | 197 | Specific | 551.68 | 551.68 | 551.68 | 9.86 | 51 | >sp\|P19835\|CEL_HUMAN Bile salt-activated lipase OS=Homo sapiens OX=9606 GN=CEL PE=1 SV=3 | 476 | 20201113_15413_1ng_glyco_hcd203040_compound15.22294.22294.3 | controllerType=0 controllerNumber=1 scan=22294 | 45.0178 |
| 18862:4 | 9 | R.NIAAFGGDPNN[+1913.677]ITLFGESAGGASVSLQTLSPYNK.G | HexNAc(4)Hex(5)NeuAc(1) | N[+1914] | 1331.8362 | 4 | 5324.3229 | 5324.3633 | 0 | -7.6 | 197 | Specific | 644.54 | 620.09 | 620.09 | 11.23 | 51 | >sp\|P19835\|CEL_HUMAN Bile salt-activated lipase OS=Homo sapiens OX=9606 GN=CEL PE=1 SV=3 | 476 | 20201113_15413_1ng_glyco_hcd203040_compound15.22458.22458.4 | controllerType=0 controllerNumber=1 scan=22458 | 45.4217 |
| 18865:4 | 9 | R.NIAAFGGDPNN[+1913.677]ITLFGESAGGASVSLQTLSPYNK.G | HexNAc(4)Hex(5)NeuAc(1) | N[+1914] | 1331.8362 | 4 | 5324.3229 | 5324.3633 | 0 | -7.6 | 197 | Specific | 767.26 | 653.09 | 653.09 | 14.23 | 51 | >sp\|P19835\|CEL_HUMAN Bile salt-activated lipase OS=Homo sapiens OX=9606 GN=CEL PE=1 SV=3 | 476 | 20201113_15413_1ng_glyco_hcd203040_compound15.22461.22461.4 | controllerType=0 controllerNumber=1 scan=22461 | 45.4265 |
| 18849:3 | 9 | R.NIAAFGGDPNN[+1913.677]ITLFGESAGGASVSLQTLSPYNK.G | HexNAc(4)Hex(5)NeuAc(1) | N[+1914] | 1775.4627 | 3 | 5324.3736 | 5324.3633 | 0 | 1.9 | 197 | Specific | 945.26 | 749.52 | 749.52 | 18.73 | 51 | >sp\|P19835\|CEL_HUMAN Bile salt-activated lipase OS=Homo sapiens OX=9606 GN=CEL PE=1 SV=3 | 476 | 20201113_15413_1ng_glyco_hcd203040_compound15.22444.22444.3 | controllerType=0 controllerNumber=1 scan=22444 | 45.3826 |
| 18847:3 | 9 | R.NIAAFGGDPNN[+1913.677]ITLFGESAGGASVSLQTLSPYNK.G | HexNAc(4)Hex(5)NeuAc(1) | N[+1914] | 1775.4627 | 3 | 5324.3736 | 5324.3633 | 0 | 1.9 | 197 | Specific | 1001.8 | 942.57 | 942.57 | 20.73 | 51 | >sp\|P19835\|CEL_HUMAN Bile salt-activated lipase OS=Homo sapiens OX=9606 GN=CEL PE=1 SV=3 | 476 | 20201113_15413_1ng_glyco_hcd203040_compound15.22442.22442.3 | controllerType=0 controllerNumber=1 scan=22442 | 45.365 |
| 19459:3 | 10 | R.GLSFDVSLEVSQGPGLLN[+2351.851]DTK.V | HexNAc(4)Hex(5)Fuc(3)NeuAc(1) | N[+2352] | 1509.9921 | 3 | 4527.9617 | 4527.964 | 0 | -0.5 | 118 | Specific | 561.8 | 548.11 | 548.11 | 9.29 | 51 | >sp\|P01833\|PIGR_HUMAN Polymeric immunoglobulin receptor OS=Homo sapiens OX=9606 GN=PIGR PE=1 SV=4 | 902 | 20201113_15413_1ng_glyco_hcd203040_compound15.23081.23081.3 | controllerType=0 controllerNumber=1 scan=23081 | 46.6185 |
| 20871:3 | 10 | K.QIGLYPVLVIDSSGYVNPN[+2350.830]YTGR.I | HexNAc(4)Hex(5)Fuc(1)NeuAc(2) | N[+2351] | 1626.0587 | 3 | 4876.1614 | 4876.1338 | 0 | 5.7 | 168 | Specific | 593.23 | 489.78 | 489.78 | 10.7 | 51 | >sp\|P01833\|PIGR_HUMAN Polymeric immunoglobulin receptor OS=Homo sapiens OX=9606 GN=PIGR PE=1 SV=4 | 902 | 20201113_15413_1ng_glyco_hcd203040_compound15.24551.24551.3 | controllerType=0 controllerNumber=1 scan=24551 | 49.6118 |
| 20872:3 | 10 | K.QIGLYPVLVIDSSGYVNPN[+2350.830]YTGR.I | HexNAc(4)Hex(5)Fuc(1)NeuAc(2) | N[+2351] | 1626.0587 | 3 | 4876.1614 | 4876.1338 | 0 | 5.7 | 168 | Specific | 637.97 | 589.11 | 589.11 | 11.7 | 51 | >sp\|P01833\|PIGR_HUMAN Polymeric immunoglobulin receptor OS=Homo sapiens OX=9606 GN=PIGR PE=1 SV=4 | 902 | 20201113_15413_1ng_glyco_hcd203040_compound15.24552.24552.3 | controllerType=0 controllerNumber=1 scan=24552 | 49.6154 |
| 28995:3 | 10 | R.[+42.011]LSLLEEPGN[+2351.851]GTFTVILNQLTSR.D | HexNAc(4)Hex(5)Fuc(3)NeuAc(1) | .[42], N[+2352] | 1599.3957 | 3 | 4796.1725 | 4796.1539 | 0 | 3.9 | 413 | Specific | 247.06 | 247.06 | 247.06 | 6.19 | 51 | >sp\|P01833\|PIGR_HUMAN Polymeric immunoglobulin receptor OS=Homo sapiens OX=9606 GN=PIGR PE=1 SV=4 | 902 | 20201113_15413_1ng_glyco_hcd203040_compound15.33017.33017.3 | controllerType=0 controllerNumber=1 scan=33017 | 64.9601 |
| 22926:3 | 10 | R.LSLLEEPGN[+2350.830]GTFTVILNQLTSR.D | HexNAc(4)Hex(5)Fuc(1)NeuAc(2) | N[+2351] | 1585.0495 | 3 | 4753.1339 | 4753.1229 | 0 | 2.3 | 413 | Specific | 277.48 | 277.48 | 277.48 | 6.84 | 51 | >sp\|P01833\|PIGR_HUMAN Polymeric immunoglobulin receptor OS=Homo sapiens OX=9606 GN=PIGR PE=1 SV=4 | 902 | 20201113_15413_1ng_glyco_hcd203040_compound15.26687.26687.3 | controllerType=0 controllerNumber=1 scan=26687 | 53.564 |
| 22928:3 | 10 | R.LSLLEEPGN[+2350.830]GTFTVILNQLTSR.D | HexNAc(4)Hex(5)Fuc(1)NeuAc(2) | N[+2351] | 1585.0495 | 3 | 4753.1339 | 4753.1229 | 0 | 2.3 | 413 | Specific | 707.12 | 567.81 | 567.81 | 14.17 | 51 | >sp\|P01833\|PIGR_HUMAN Polymeric immunoglobulin receptor OS=Homo sapiens OX=9606 GN=PIGR PE=1 SV=4 | 902 | 20201113_15413_1ng_glyco_hcd203040_compound15.26689.26689.3 | controllerType=0 controllerNumber=1 scan=26689 | 53.568 |
| 22748:4 | 10 | R.LSLLEEPGN[+2350.830]GTFTVILNQLTSR.D | HexNAc(4)Hex(5)Fuc(1)NeuAc(2) | N[+2351] | 1189.0425 | 4 | 4753.1483 | 4753.1229 | 0 | 5.3 | 413 | Specific | 855.28 | 657.78 | 657.78 | 16.68 | 51 | >sp\|P01833\|PIGR_HUMAN Polymeric immunoglobulin receptor OS=Homo sapiens OX=9606 GN=PIGR PE=1 SV=4 | 902 | 20201113_15413_1ng_glyco_hcd203040_compound15.26502.26502.4 | controllerType=0 controllerNumber=1 scan=26502 | 53.2095 |

Table S4. Cont.

| Query #:z | Protein Rank | Glycopeptide | Glycan | Modification Type(s) | Observed m/z | z | Observed (M+H) | Calc. mass (M+H) | Off-by-x error | Mass error (ppm) | StartPosition | Cleavage | Score | Delta | Delta Mod | \|Log Prob\| | # of unique peptides | Protein Name | Protein DB number | Comment | Scan # | Scan Time |
|---|---|---|---|---|---|---|---|---|---|---|---|---|---|---|---|---|---|---|---|---|---|---|
| 22718:3 | 10 | R.LSLLEEPGN[+2350.830]GTFTVILNQLTSR.D | HexNAc(4)Hex(5)Fuc(1)NeuAc(2) | N[+2351] | 1585.0483 | 3 | 4753.1303 | 4753.1229 | 0 | 1.5 | 413 | Specific | 952.43 | 690.41 | 690.41 | 19.17 | 51 | >sp\|P01833\|PIGR_HUMAN Polymeric immunoglobulin receptor OS=Homo sapiens OX=9606 GN=PIGR PE=1 SV=4 | 902 | 20201113_15413_1ng_glyco_hcd203040_compound15.26471.26471.3 | controllerType=0 controllerNumber=1 scan=26471 | 53.1494 |
| 20522:3 | 10 | R.LSLLEEPGN[+2059.735]GTFTVILNQLTSR.D | HexNAc(4)Hex(5)Fuc(1)NeuAc(1) | N[+2060] | 1488.0169 | 3 | 4462.0362 | 4462.0275 | 0 | 1.9 | 413 | Specific | 966.37 | 695.13 | 695.13 | 19.17 | 51 | >sp\|P01833\|PIGR_HUMAN Polymeric immunoglobulin receptor OS=Homo sapiens OX=9606 GN=PIGR PE=1 SV=4 | 902 | 20201113_15413_1ng_glyco_hcd203040_compound15.24189.24189.3 | controllerType=0 controllerNumber=1 scan=24189 | 48.8956 |
| 22746:4 | 10 | R.LSLLEEPGN[+2350.830]GTFTVILNQLTSR.D | HexNAc(4)Hex(5)Fuc(1)NeuAc(2) | N[+2351] | 1189.0425 | 4 | 4753.1483 | 4753.1229 | 0 | 5.3 | 413 | Specific | 1022.2 | 790.28 | 790.28 | 19.68 | 51 | >sp\|P01833\|PIGR_HUMAN Polymeric immunoglobulin receptor OS=Homo sapiens OX=9606 GN=PIGR PE=1 SV=4 | 902 | 20201113_15413_1ng_glyco_hcd203040_compound15.26500.26500.4 | controllerType=0 controllerNumber=1 scan=26500 | 53.2039 |
| 20520:3 | 10 | R.LSLLEEPGN[+2059.735]GTFTVILNQLTSR.D | HexNAc(4)Hex(5)Fuc(1)NeuAc(1) | N[+2060] | 1488.0169 | 3 | 4462.0362 | 4462.0275 | 0 | 1.9 | 413 | Specific | 1090.16 | 848.37 | 848.37 | 21.17 | 51 | >sp\|P01833\|PIGR_HUMAN Polymeric immunoglobulin receptor OS=Homo sapiens OX=9606 GN=PIGR PE=1 SV=4 | 902 | 20201113_15413_1ng_glyco_hcd203040_compound15.24187.24187.3 | controllerType=0 controllerNumber=1 scan=24187 | 48.8895 |
| 22716:3 | 10 | R.LSLLEEPGN[+2350.830]GTFTVILNQLTSR.D | HexNAc(4)Hex(5)Fuc(1)NeuAc(2) | N[+2351] | 1585.0483 | 3 | 4753.1303 | 4753.1229 | 0 | 1.5 | 413 | Specific | 1102.9 | 861.11 | 861.11 | 22.17 | 51 | >sp\|P01833\|PIGR_HUMAN Polymeric immunoglobulin receptor OS=Homo sapiens OX=9606 GN=PIGR PE=1 SV=4 | 902 | 20201113_15413_1ng_glyco_hcd203040_compound15.26469.26469.3 | controllerType=0 controllerNumber=1 scan=26469 | 53.1433 |
| 17938:3 | 10 | K.IIEGEPNLKVPGN[+2350.830]VTAVLGETLK.V | HexNAc(4)Hex(5)Fuc(1)NeuAc(2) | N[+2351] | 1581.4039 | 3 | 4742.197 | 4742.1797 | 0 | 3.7 | 457 | Specific | 220.03 | 220.03 | 220.03 | 4.31 | 51 | >sp\|P01833\|PIGR_HUMAN Polymeric immunoglobulin receptor OS=Homo sapiens OX=9606 GN=PIGR PE=1 SV=4 | 902 | 20201113_15413_1ng_glyco_hcd203040_compound15.21492.21492.3 | controllerType=0 controllerNumber=1 scan=21492 | 43.3647 |
| 15989:3 | 10 | K.[+42.011]IIEGEPNLKVPGN[+2018.708]VTAVLGETLK.V | HexNAc(3)Hex(6)Fuc(1)NeuAc(1) | .[42], N[+2019] | 1484.7068 | 3 | 4452.1058 | 4452.0683 | 0 | 8.4 | 457 | Specific | 320.87 | 320.87 | 320.87 | 5.67 | 51 | >sp\|P01833\|PIGR_HUMAN Polymeric immunoglobulin receptor OS=Homo sapiens OX=9606 GN=PIGR PE=1 SV=4 | 902 | 20201113_15413_1ng_glyco_hcd203040_compound15.19461.19461.3 | controllerType=0 controllerNumber=1 scan=19461 | 39.3696 |
| 16007:3 | 10 | K.IIEGEPNLKVPGN[+2205.793]VTAVLGETLK.V | HexNAc(4)Hex(5)Fuc(2)NeuAc(1) | N[+2206] | 1533.0555 | 3 | 4597.1519 | 4597.1422 | 0 | 2.1 | 457 | Specific | 386.45 | 386.45 | 386.45 | 8.51 | 51 | >sp\|P01833\|PIGR_HUMAN Polymeric immunoglobulin receptor OS=Homo sapiens OX=9606 GN=PIGR PE=1 SV=4 | 902 | 20201113_15413_1ng_glyco_hcd203040_compound15.19480.19480.3 | controllerType=0 controllerNumber=1 scan=19480 | 39.4066 |
| 17896:4 | 10 | K.IIEGEPNLKVPGN[+2350.830]VTAVLGETLK.V | HexNAc(4)Hex(5)Fuc(1)NeuAc(2) | N[+2351] | 1186.3083 | 4 | 4742.2113 | 4742.1797 | 0 | 6.7 | 457 | Specific | 473.01 | 388.41 | 388.41 | 7.86 | 51 | >sp\|P01833\|PIGR_HUMAN Polymeric immunoglobulin receptor OS=Homo sapiens OX=9606 GN=PIGR PE=1 SV=4 | 902 | 20201113_15413_1ng_glyco_hcd203040_compound15.21448.21448.4 | controllerType=0 controllerNumber=1 scan=21448 | 43.2723 |
| 17941:3 | 10 | K.IIEGEPNLKVPGN[+2350.830]VTAVLGETLK.V | HexNAc(4)Hex(5)Fuc(1)NeuAc(2) | N[+2351] | 1581.4039 | 3 | 4742.197 | 4742.1797 | 0 | 3.7 | 457 | Specific | 520.32 | 407.79 | 407.79 | 9.11 | 51 | >sp\|P01833\|PIGR_HUMAN Polymeric immunoglobulin receptor OS=Homo sapiens OX=9606 GN=PIGR PE=1 SV=4 | 902 | 20201113_15413_1ng_glyco_hcd203040_compound15.21495.21495.3 | controllerType=0 controllerNumber=1 scan=21495 | 43.3684 |
| 17894:4 | 10 | K.IIEGEPNLKVPGN[+2350.830]VTAVLGETLK.V | HexNAc(4)Hex(5)Fuc(1)NeuAc(2) | N[+2351] | 1186.3083 | 4 | 4742.2113 | 4742.1797 | 0 | 6.7 | 457 | Specific | 594.8 | 517.65 | 517.65 | 9.79 | 51 | >sp\|P01833\|PIGR_HUMAN Polymeric immunoglobulin receptor OS=Homo sapiens OX=9606 GN=PIGR PE=1 SV=4 | 902 | 20201113_15413_1ng_glyco_hcd203040_compound15.21446.21446.4 | controllerType=0 controllerNumber=1 scan=21446 | 43.2668 |
| 15956:3 | 10 | K.IIEGEPNLKVPGN[+2205.793]VTAVLGETLK.V | HexNAc(4)Hex(5)Fuc(2)NeuAc(1) | N[+2206] | 1533.056 | 3 | 4597.1534 | 4597.1422 | 0 | 2.4 | 457 | Specific | 840.55 | 760.26 | 760.26 | 15.58 | 51 | >sp\|P01833\|PIGR_HUMAN Polymeric immunoglobulin receptor OS=Homo sapiens OX=9606 GN=PIGR PE=1 SV=4 | 902 | 20201113_15413_1ng_glyco_hcd203040_compound15.19426.19426.3 | controllerType=0 controllerNumber=1 scan=19426 | 39.3019 |
| 16010:3 | 10 | K.IIEGEPNLKVPGN[+2205.793]VTAVLGETLK.V | HexNAc(4)Hex(5)Fuc(2)NeuAc(1) | N[+2206] | 1533.0555 | 3 | 4597.1519 | 4597.1422 | 0 | 2.1 | 457 | Specific | 909.52 | 817.17 | 817.17 | 17.58 | 51 | >sp\|P01833\|PIGR_HUMAN Polymeric immunoglobulin receptor OS=Homo sapiens OX=9606 GN=PIGR PE=1 SV=4 | 902 | 20201113_15413_1ng_glyco_hcd203040_compound15.19483.19483.3 | controllerType=0 controllerNumber=1 scan=19483 | 39.4107 |
| 15954:3 | 10 | K.IIEGEPNLKVPGN[+2205.793]VTAVLGETLK.V | HexNAc(4)Hex(5)Fuc(2)NeuAc(1) | N[+2206] | 1533.056 | 3 | 4597.1534 | 4597.1422 | 0 | 2.4 | 457 | Specific | 983.38 | 896.36 | 896.36 | 18.58 | 51 | >sp\|P01833\|PIGR_HUMAN Polymeric immunoglobulin receptor OS=Homo sapiens OX=9606 GN=PIGR PE=1 SV=4 | 902 | 20201113_15413_1ng_glyco_hcd203040_compound15.19424.19424.3 | controllerType=0 controllerNumber=1 scan=19424 | 39.2896 |
| 16585:2 | 10 | K.VPGN[+2350.830]VTAVLGETLK.V | HexNAc(4)Hex(5)Fuc(1)NeuAc(2) | N[+2351] | 1874.8278 | 2 | 3748.6484 | 3748.6302 | 0 | 4.9 | 466 | Specific | 288.07 | 63.59 | 63.59 | 4.34 | 51 | >sp\|P01833\|PIGR_HUMAN Polymeric immunoglobulin receptor OS=Homo sapiens OX=9606 GN=PIGR PE=1 SV=4 | 902 | 20201113_15413_1ng_glyco_hcd203040_compound15.20081.20081.2 | controllerType=0 controllerNumber=1 scan=20081 | 40.4642 |
| 16543:3 | 10 | K.VPGN[+2350.830]VTAVLGETLK.V | HexNAc(4)Hex(5)Fuc(1)NeuAc(2) | N[+2351] | 1250.218 | 3 | 3748.6393 | 3748.6302 | 0 | 2.4 | 466 | Specific | 770.6 | 597.36 | 597.36 | 14.92 | 51 | >sp\|P01833\|PIGR_HUMAN Polymeric immunoglobulin receptor OS=Homo sapiens OX=9606 GN=PIGR PE=1 SV=4 | 902 | 20201113_15413_1ng_glyco_hcd203040_compound15.20038.20038.3 | controllerType=0 controllerNumber=1 scan=20038 | 40.3812 |
| 13292:3 | 10 | K.VPGN[+2205.793]VTAVLGETLK.V | HexNAc(4)Hex(5)Fuc(2)NeuAc(1) | N[+2206] | 1201.8746 | 3 | 3603.6092 | 3603.5927 | 0 | 4.6 | 466 | Specific | 785.31 | 720.05 | 720.05 | 14.86 | 51 | >sp\|P01833\|PIGR_HUMAN Polymeric immunoglobulin receptor OS=Homo sapiens OX=9606 GN=PIGR PE=1 SV=4 | 902 | 20201113_15413_1ng_glyco_hcd203040_compound15.16643.16643.3 | controllerType=0 controllerNumber=1 scan=16643 | 34.4626 |
| 13294:3 | 10 | K.VPGN[+2205.793]VTAVLGETLK.V | HexNAc(4)Hex(5)Fuc(2)NeuAc(1) | N[+2206] | 1201.8746 | 3 | 3603.6092 | 3603.5927 | 0 | 4.6 | 466 | Specific | 807.13 | 733.45 | 733.45 | 15.86 | 51 | >sp\|P01833\|PIGR_HUMAN Polymeric immunoglobulin receptor OS=Homo sapiens OX=9606 GN=PIGR PE=1 SV=4 | 902 | 20201113_15413_1ng_glyco_hcd203040_compound15.16645.16645.3 | controllerType=0 controllerNumber=1 scan=16645 | 34.4675 |
| 16541:3 | 10 | K.VPGN[+2350.830]VTAVLGETLK.V | HexNAc(4)Hex(5)Fuc(1)NeuAc(2) | N[+2351] | 1250.218 | 3 | 3748.6393 | 3748.6302 | 0 | 2.4 | 466 | Specific | 838.55 | 605.72 | 605.72 | 15.92 | 51 | >sp\|P01833\|PIGR_HUMAN Polymeric immunoglobulin receptor OS=Homo sapiens OX=9606 GN=PIGR PE=1 SV=4 | 902 | 20201113_15413_1ng_glyco_hcd203040_compound15.20036.20036.3 | controllerType=0 controllerNumber=1 scan=20036 | 40.376 |
| 15437:4 | 10 | K.VPGN[+2350.830]VTAVLGETLKVPC[+57.021]HFPC[+57.021]K.F | HexNAc(4)Hex(5)Fuc(1)NeuAc(2) | C[+57]*2, N[+2351] | 1194.2825 | 4 | 4774.1082 | 4774.0877 | 0 | 4.3 | 466 | Specific | 330.68 | 289.74 | 289.74 | 6.45 | 51 | >sp\|P01833\|PIGR_HUMAN Polymeric immunoglobulin receptor OS=Homo sapiens OX=9606 GN=PIGR PE=1 SV=4 | 902 | 20201113_15413_1ng_glyco_hcd203040_compound15.18864.18864.4 | controllerType=0 controllerNumber=1 scan=18864 | 38.2846 |
| 15439:4 | 10 | K.VPGN[+2350.830]VTAVLGETLKVPC[+57.021]HFPC[+57.021]K.F | HexNAc(4)Hex(5)Fuc(1)NeuAc(2) | C[+57]*2, N[+2351] | 1194.2825 | 4 | 4774.1082 | 4774.0877 | 0 | 4.3 | 466 | Specific | 453.51 | 370.69 | 370.69 | 7.14 | 51 | >sp\|P01833\|PIGR_HUMAN Polymeric immunoglobulin receptor OS=Homo sapiens OX=9606 GN=PIGR PE=1 SV=4 | 902 | 20201113_15413_1ng_glyco_hcd203040_compound15.18866.18866.4 | controllerType=0 controllerNumber=1 scan=18866 | 38.2884 |
| 05611:3 | 10 | K.WN[+1710.598]NTGC[+57.021]QALPSQDEGPSK.A | HexNAc(3)Hex(5)NeuAc(1) | C[+57], N[+1711] | 1233.8326 | 3 | 3699.4832 | 3699.4743 | 0 | 2.4 | 498 | Specific | 310.13 | 254.34 | 254.34 | 6.76 | 51 | >sp\|P01833\|PIGR_HUMAN Polymeric immunoglobulin receptor OS=Homo sapiens OX=9606 GN=PIGR PE=1 SV=4 | 902 | 20201113_15413_1ng_glyco_hcd203040_compound15.8648.8648.3 | controllerType=0 controllerNumber=1 scan=8648 | 21.0045 |
| 05613:3 | 10 | K.WN[+1710.598]NTGC[+57.021]QALPSQDEGPSK.A | HexNAc(3)Hex(5)NeuAc(1) | C[+57], N[+1711] | 1233.8326 | 3 | 3699.4832 | 3699.4743 | 0 | 2.4 | 498 | Specific | 447.22 | 368.25 | 368.25 | 7.9 | 51 | >sp\|P01833\|PIGR_HUMAN Polymeric immunoglobulin receptor OS=Homo sapiens OX=9606 GN=PIGR PE=1 SV=4 | 902 | 20201113_15413_1ng_glyco_hcd203040_compound15.8650.8650.3 | controllerType=0 controllerNumber=1 scan=8650 | 21.0198 |
| 05470:3 | 10 | K.WN[+1872.651]NTGC[+57.021]QALPSQDEGPSK.A | HexNAc(3)Hex(6)NeuAc(1) | C[+57], N[+1873] | 1287.8539 | 3 | 3861.5473 | 3861.5272 | 0 | 5.2 | 498 | Specific | 530.69 | 506.78 | 506.78 | 9.29 | 51 | >sp\|P01833\|PIGR_HUMAN Polymeric immunoglobulin receptor OS=Homo sapiens OX=9606 GN=PIGR PE=1 SV=4 | 902 | 20201113_15413_1ng_glyco_hcd203040_compound15.8502.8502.3 | controllerType=0 controllerNumber=1 scan=8502 | 20.7634 |
| 05472:3 | 10 | K.WN[+1872.651]NTGC[+57.021]QALPSQDEGPSK.A | HexNAc(3)Hex(6)NeuAc(1) | C[+57], N[+1873] | 1287.8539 | 3 | 3861.5473 | 3861.5272 | 0 | 5.2 | 498 | Specific | 548.9 | 488.14 | 488.14 | 9.29 | 51 | >sp\|P01833\|PIGR_HUMAN Polymeric immunoglobulin receptor OS=Homo sapiens OX=9606 GN=PIGR PE=1 SV=4 | 902 | 20201113_15413_1ng_glyco_hcd203040_compound15.8504.8504.3 | controllerType=0 controllerNumber=1 scan=8504 | 20.7669 |
| 05522:3 | 10 | K.WN[+1913.677]NTGC[+57.021]QALPSQDEGPSK.A | HexNAc(4)Hex(5)NeuAc(1) | C[+57], N[+1914] | 1301.5275 | 3 | 3902.568 | 3902.5537 | 0 | 3.7 | 498 | Specific | 554.78 | 509.44 | 509.44 | 10 | 51 | >sp\|P01833\|PIGR_HUMAN Polymeric immunoglobulin receptor OS=Homo sapiens OX=9606 GN=PIGR PE=1 SV=4 | 902 | 20201113_15413_1ng_glyco_hcd203040_compound15.8556.8556.3 | controllerType=0 controllerNumber=1 scan=8556 | 20.8547 |
| 05524:3 | 10 | K.WN[+1913.677]NTGC[+57.021]QALPSQDEGPSK.A | HexNAc(4)Hex(5)NeuAc(1) | C[+57], N[+1914] | 1301.5275 | 3 | 3902.568 | 3902.5537 | 0 | 3.7 | 498 | Specific | 571.99 | 504.51 | 504.51 | 10 | 51 | >sp\|P01833\|PIGR_HUMAN Polymeric immunoglobulin receptor OS=Homo sapiens OX=9606 GN=PIGR PE=1 SV=4 | 902 | 20201113_15413_1ng_glyco_hcd203040_compound15.8558.8558.3 | controllerType=0 controllerNumber=1 scan=8558 | 20.8583 |
| 05434:3 | 10 | K.WN[+2059.735]NTGC[+57.021]QALPSQDEGPSK.A | HexNAc(4)Hex(5)Fuc(1)NeuAc(1) | C[+57], N[+2060] | 1350.2109 | 3 | 4048.6181 | 4048.6116 | 0 | 1.6 | 498 | Specific | 574.95 | 509.47 | 509.47 | 9.95 | 51 | >sp\|P01833\|PIGR_HUMAN Polymeric immunoglobulin receptor OS=Homo sapiens OX=9606 GN=PIGR PE=1 SV=4 | 902 | 20201113_15413_1ng_glyco_hcd203040_compound15.8465.8465.3 | controllerType=0 controllerNumber=1 scan=8465 | 20.7025 |
| 05477:3 | 10 | K.WN[+1913.677]NTGC[+57.021]QALPSQDEGPSK.A | HexNAc(4)Hex(5)NeuAc(1) | C[+57], N[+1914] | 1301.5297 | 3 | 3902.5746 | 3902.5537 | 0 | 5.4 | 498 | Specific | 589.92 | 523.08 | 523.08 | 10.68 | 51 | >sp\|P01833\|PIGR_HUMAN Polymeric immunoglobulin receptor OS=Homo sapiens OX=9606 GN=PIGR PE=1 SV=4 | 902 | 20201113_15413_1ng_glyco_hcd203040_compound15.8509.8509.3 | controllerType=0 controllerNumber=1 scan=8509 | 20.7756 |
| 05564:3 | 10 | K.WN[+1710.598]NTGC[+57.021]QALPSQDEGPSK.A | HexNAc(3)Hex(5)NeuAc(1) | C[+57], N[+1711] | 1233.832 | 3 | 3699.4813 | 3699.4743 | 0 | 1.9 | 498 | Specific | 600.26 | 536.66 | 536.66 | 12.15 | 51 | >sp\|P01833\|PIGR_HUMAN Polymeric immunoglobulin receptor OS=Homo sapiens OX=9606 GN=PIGR PE=1 SV=4 | 902 | 20201113_15413_1ng_glyco_hcd203040_compound15.8599.8599.3 | controllerType=0 controllerNumber=1 scan=8599 | 20.9244 |
| 05562:3 | 10 | K.WN[+1710.598]NTGC[+57.021]QALPSQDEGPSK.A | HexNAc(3)Hex(5)NeuAc(1) | C[+57], N[+1711] | 1233.832 | 3 | 3699.4813 | 3699.4743 | 0 | 1.9 | 498 | Specific | 601.82 | 548.43 | 548.43 | 12.15 | 51 | >sp\|P01833\|PIGR_HUMAN Polymeric immunoglobulin receptor OS=Homo sapiens OX=9606 GN=PIGR PE=1 SV=4 | 902 | 20201113_15413_1ng_glyco_hcd203040_compound15.8597.8597.3 | controllerType=0 controllerNumber=1 scan=8597 | 20.9207 |
| 05475:3 | 10 | K.WN[+1913.677]NTGC[+57.021]QALPSQDEGPSK.A | HexNAc(4)Hex(5)NeuAc(1) | C[+57], N[+1914] | 1301.5297 | 3 | 3902.5746 | 3902.5537 | 0 | 5.4 | 498 | Specific | 613.52 | 551.02 | 551.02 | 11.68 | 51 | >sp\|P01833\|PIGR_HUMAN Polymeric immunoglobulin receptor OS=Homo sapiens OX=9606 GN=PIGR PE=1 SV=4 | 902 | 20201113_15413_1ng_glyco_hcd203040_compound15.8507.8507.3 | controllerType=0 controllerNumber=1 scan=8507 | 20.7719 |
| 05436:3 | 10 | K.WN[+2059.735]NTGC[+57.021]QALPSQDEGPSK.A | HexNAc(4)Hex(5)Fuc(1)NeuAc(1) | C[+57], N[+2060] | 1350.2109 | 3 | 4048.6181 | 4048.6116 | 0 | 1.6 | 498 | Specific | 632.02 | 563.9 | 563.9 | 12.15 | 51 | >sp\|P01833\|PIGR_HUMAN Polymeric immunoglobulin receptor OS=Homo sapiens OX=9606 GN=PIGR PE=1 SV=4 | 902 | 20201113_15413_1ng_glyco_hcd203040_compound15.8467.8467.3 | controllerType=0 controllerNumber=1 scan=8467 | 20.7063 |
| 18656:3 | 11 | R.LSLHRPALEDLLLGSEAN[+1751.624]LTC[+57.021]TLTGLR.D | HexNAc(4)Hex(4)NeuAc(1) | C[+57], N[+1752] | 1572.4258 | 3 | 4715.263 | 4715.2225 | 0 | 8.6 | 127 | Specific | 381.59 | 381.59 | 381.59 | 7.83 | 56 | >sp\|P01876\|IGHA1_HUMAN Immunoglobulin heavy constant alpha 1 OS=Homo sapiens OX=9606 GN=IGHA1 PE=1 SV=2 | 255 | 20201113_15413_1ng_glyco_hcd203040_compound15.22242.22242.3 | controllerType=0 controllerNumber=1 scan=22242 | 44.8999 |
| 17558:3 | 11 | R.LSLHRPALEDLLLGSEAN[+1702.581]LTC[+57.021]TLTGLR.D | HexNAc(2)Hex(8) | C[+57], N[+1703] | 1556.0723 | 3 | 4666.2024 | 4666.1796 | 0 | 4.9 | 127 | Specific | 396.86 | 396.86 | 396.86 | 8.58 | 56 | >sp\|P01876\|IGHA1_HUMAN Immunoglobulin heavy constant alpha 1 OS=Homo sapiens OX=9606 GN=IGHA1 PE=1 SV=2 | 255 | 20201113_15413_1ng_glyco_hcd203040_compound15.21094.21094.3 | controllerType=0 controllerNumber=1 scan=21094 | 42.4637 |
| 17576:4 | 11 | R.LSLHRPALEDLLLGSEAN[+1622.582]LTC[+57.021]TLTGLR.D | HexNAc(4)Hex(5) | C[+57], N[+1623] | 1147.3033 | 4 | 4586.1912 | 4586.1799 | 0 | 2.5 | 127 | Specific | 444.9 | 369.73 | 369.73 | 8.14 | 56 | >sp\|P01876\|IGHA1_HUMAN Immunoglobulin heavy constant alpha 1 OS=Homo sapiens OX=9606 GN=IGHA1 PE=1 SV=2 | 255 | 20201113_15413_1ng_glyco_hcd203040_compound15.21113.21113.4 | controllerType=0 controllerNumber=1 scan=21113 | 42.5051 |
| 17569:4 | 11 | R.LSLHRPALEDLLLGSEAN[+1663.608]LTC[+57.021]TLTGLR.D | HexNAc(5)Hex(4) | C[+57], N[+1664] | 1157.5602 | 4 | 4627.2191 | 4627.2064 | 0 | 2.7 | 127 | Specific | 448.08 | 448.08 | 448.08 | 9.65 | 56 | >sp\|P01876\|IGHA1_HUMAN Immunoglobulin heavy constant alpha 1 OS=Homo sapiens OX=9606 GN=IGHA1 PE=1 SV=2 | 255 | 20201113_15413_1ng_glyco_hcd203040_compound15.21106.21106.4 | controllerType=0 controllerNumber=1 scan=21106 | 42.4875 |
| 17703:4 | 11 | R.LSLHRPALEDLLLGSEAN[+1216.423]LTC[+57.021]TLTGLR.D | HexNAc(2)Hex(5) | C[+57], N[+1216] | 1045.7671 | 4 | 4180.0467 | 4180.0211 | 0 | 6.1 | 127 | Specific | 489.26 | 479.94 | 479.94 | 9.26 | 56 | >sp\|P01876\|IGHA1_HUMAN Immunoglobulin heavy constant alpha 1 OS=Homo sapiens OX=9606 GN=IGHA1 PE=1 SV=2 | 255 | 20201113_15413_1ng_glyco_hcd203040_compound15.21247.21247.3 | controllerType=0 controllerNumber=1 scan=21247 | 42.8521 |
| 17731:4 | 11 | R.LSLHRPALEDLLLGSEAN[+1095.397]LTC[+57.021]TLTGLR.D | HexNAc(3)Hex(3) | C[+57], N[+1095] | 1015.5109 | 4 | 4059.0218 | 4058.9949 | 0 | 6.6 | 127 | Specific | 511.12 | 511.12 | 511.12 | 9.13 | 56 | >sp\|P01876\|IGHA1_HUMAN Immunoglobulin heavy constant alpha 1 OS=Homo sapiens OX=9606 GN=IGHA1 PE=1 SV=2 | 255 | 20201113_15413_1ng_glyco_hcd203040_compound15.21276.21276.4 | controllerType=0 controllerNumber=1 scan=21276 | 42.9112 |
| 18659:3 | 11 | R.LSLHRPALEDLLLGSEAN[+1751.624]LTC[+57.021]TLTGLR.D | HexNAc(4)Hex(4)NeuAc(1) | C[+57], N[+1752] | 1572.4258 | 3 | 4715.263 | 4715.2225 | 0 | 8.6 | 127 | Specific | 580.86 | 580.86 | 580.86 | 9.91 | 56 | >sp\|P01876\|IGHA1_HUMAN Immunoglobulin heavy constant alpha 1 OS=Homo sapiens OX=9606 GN=IGHA1 PE=1 SV=2 | 255 | 20201113_15413_1ng_glyco_hcd203040_compound15.22245.22245.3 | controllerType=0 controllerNumber=1 scan=22245 | 44.9067 |
| 18571:4 | 11 | R.LSLHRPALEDLLLGSEAN[+2116.756]LTC[+57.021]TLTGLR.D | HexNAc(5)Hex(5)NeuAc(1) | C[+57], N[+2117] | 1270.8434 | 4 | 5080.3517 | 5080.3547 | 0 | -0.6 | 127 | Specific | 599.72 | 496.28 | 496.28 | 10.72 | 56 | >sp\|P01876\|IGHA1_HUMAN Immunoglobulin heavy constant alpha 1 OS=Homo sapiens OX=9606 GN=IGHA1 PE=1 SV=2 | 255 | 20201113_15413_1ng_glyco_hcd203040_compound15.22153.22153.4 | controllerType=0 controllerNumber=1 scan=22153 | 44.6951 |
| 17573:3 | 11 | R.LSLHRPALEDLLLGSEAN[+1540.529]LTC[+57.021]TLTGLR.D | HexNAc(2)Hex(7) | C[+57], N[+1541] | 1502.0524 | 3 | 4504.1427 | 4504.1268 | 0 | 3.5 | 127 | Specific | 603.95 | 603.95 | 603.95 | 11.76 | 56 | >sp\|P01876\|IGHA1_HUMAN Immunoglobulin heavy constant alpha 1 OS=Homo sapiens OX=9606 GN=IGHA1 PE=1 SV=2 | 255 | 20201113_15413_1ng_glyco_hcd203040_compound15.21110.21110.3 | controllerType=0 controllerNumber=1 scan=21110 | 42.4999 |
| 17648:3 | 11 | R.LSLHRPALEDLLLGSEAN[+1460.529]LTC[+57.021]TLTGLR.D | HexNAc(4)Hex(4) | C[+57], N[+1461] | 1475.3897 | 3 | 4424.1546 | 4424.1271 | 0 | 6.2 | 127 | Specific | 606.99 | 590.64 | 590.64 | 11.25 | 56 | >sp\|P01876\|IGHA1_HUMAN Immunoglobulin heavy constant alpha 1 OS=Homo sapiens OX=9606 GN=IGHA1 PE=1 SV=2 | 255 | 20201113_15413_1ng_glyco_hcd203040_compound15.21189.21189.3 | controllerType=0 controllerNumber=1 scan=21189 | 42.695 |
| 17578:4 | 11 | R.LSLHRPALEDLLLGSEAN[+1622.582]LTC[+57.021]TLTGLR.D | HexNAc(4)Hex(5) | C[+57], N[+1623] | 1147.3033 | 4 | 4586.1912 | 4586.1799 | 0 | 2.5 | 127 | Specific | 625.54 | 368.6 | 368.6 | 10.94 | 56 | >sp\|P01876\|IGHA1_HUMAN Immunoglobulin heavy constant alpha 1 OS=Homo sapiens OX=9606 GN=IGHA1 PE=1 SV=2 | 255 | 20201113_15413_1ng_glyco_hcd203040_compound15.21115.21115.4 | controllerType=0 controllerNumber=1 scan=21115 | 42.5127 |
| 17614:3 | 11 | R.LSLHRPALEDLLLGSEAN[+1501.555]LTC[+57.021]TLTGLR.D | HexNAc(5)Hex(3) | C[+57], N[+1502] | 1489.0602 | 3 | 4465.1662 | 4465.1536 | 0 | 2.8 | 127 | Specific | 641.84 | 618.63 | 618.63 | 11.76 | 56 | >sp\|P01876\|IGHA1_HUMAN Immunoglobulin heavy constant alpha 1 OS=Homo sapiens OX=9606 GN=IGHA1 PE=1 SV=2 | 255 | 20201113_15413_1ng_glyco_hcd203040_compound15.21153.21153.3 | controllerType=0 controllerNumber=1 scan=21153 | 42.6072 |
| 17574:3 | 11 | R.LSLHRPALEDLLLGSEAN[+1702.581]LTC[+57.021]TLTGLR.D | HexNAc(2)Hex(8) | C[+57], N[+1703] | 1556.0685 | 3 | 4666.1911 | 4666.1796 | 0 | 2.5 | 127 | Specific | 643.37 | 618.3 | 618.3 | 11.71 | 56 | >sp\|P01876\|IGHA1_HUMAN Immunoglobulin heavy constant alpha 1 OS=Homo sapiens OX=9606 GN=IGHA1 PE=1 SV=2 | 255 | 20201113_15413_1ng_glyco_hcd203040_compound15.21111.21111.3 | controllerType=0 controllerNumber=1 scan=21111 | 42.5018 |
| 17582:3 | 11 | R.LSLHRPALEDLLLGSEAN[+1663.608]LTC[+57.021]TLTGLR.D | HexNAc(5)Hex(4) | C[+57], N[+1664] | 1543.0788 | 3 | 4627.2218 | 4627.2064 | 0 | 3.3 | 127 | Specific | 647.37 | 584.21 | 584.21 | 11.76 | 56 | >sp\|P01876\|IGHA1_HUMAN Immunoglobulin heavy constant alpha 1 OS=Homo sapiens OX=9606 GN=IGHA1 PE=1 SV=2 | 255 | 20201113_15413_1ng_glyco_hcd203040_compound15.21119.21119.3 | controllerType=0 controllerNumber=1 scan=21119 | 42.5196 |
| 17650:3 | 11 | R.LSLHRPALEDLLLGSEAN[+1460.529]LTC[+57.021]TLTGLR.D | HexNAc(4)Hex(4) | C[+57], N[+1461] | 1475.3897 | 3 | 4424.1546 | 4424.1271 | 0 | 6.2 | 127 | Specific | 647.97 | 593.56 | 593.56 | 11.25 | 56 | >sp\|P01876\|IGHA1_HUMAN Immunoglobulin heavy constant alpha 1 OS=Homo sapiens OX=9606 GN=IGHA1 PE=1 SV=2 | 255 | 20201113_15413_1ng_glyco_hcd203040_compound15.21191.21191.3 | controllerType=0 controllerNumber=1 scan=21191 | 42.7073 |
| 17595:3 | 11 | R.LSLHRPALEDLLLGSEAN[+1378.476]LTC[+57.021]TLTGLR.D | HexNAc(2)Hex(6) | C[+57], N[+1378] | 1448.0367 | 3 | 4342.0955 | 4342.074 | 0 | 5 | 127 | Specific | 650.6 | 595.78 | 595.78 | 12.66 | 56 | >sp\|P01876\|IGHA1_HUMAN Immunoglobulin heavy constant alpha 1 OS=Homo sapiens OX=9606 GN=IGHA1 PE=1 SV=2 | 255 | 20201113_15413_1ng_glyco_hcd203040_compound15.21133.21133.3 | controllerType=0 controllerNumber=1 scan=21133 | 42.5564 |
| 17611:3 | 11 | R.LSLHRPALEDLLLGSEAN[+1501.555]LTC[+57.021]TLTGLR.D | HexNAc(5)Hex(3) | C[+57], N[+1502] | 1489.0602 | 3 | 4465.1662 | 4465.1536 | 0 | 2.8 | 127 | Specific | 662.07 | 662.07 | 662.07 | 12.76 | 56 | >sp\|P01876\|IGHA1_HUMAN Immunoglobulin heavy constant alpha 1 OS=Homo sapiens OX=9606 GN=IGHA1 PE=1 SV=2 | 255 | 20201113_15413_1ng_glyco_hcd203040_compound15.21150.21150.3 | controllerType=0 controllerNumber=1 scan=21150 | 42.5923 |
| 17564:4 | 11 | R.LSLHRPALEDLLLGSEAN[+1702.581]LTC[+57.021]TLTGLR.D | HexNAc(2)Hex(8) | C[+57], N[+1703] | 1167.3035 | 4 | 4666.1922 | 4666.1796 | 0 | 2.7 | 127 | Specific | 665.84 | 665.84 | 665.84 | 13.44 | 56 | >sp\|P01876\|IGHA1_HUMAN Immunoglobulin heavy constant alpha 1 OS=Homo sapiens OX=9606 GN=IGHA1 PE=1 SV=2 | 255 | 20201113_15413_1ng_glyco_hcd203040_compound15.21101.21101.4 | controllerType=0 controllerNumber=1 scan=21101 | 42.4785 |
| 17561:3 | 11 | R.LSLHRPALEDLLLGSEAN[+1702.581]LTC[+57.021]TLTGLR.D | HexNAc(2)Hex(8) | C[+57], N[+1703] | 1556.0723 | 3 | 4666.2024 | 4666.1796 | 0 | 4.9 | 127 | Specific | 666.41 | 666.41 | 666.41 | 12.66 | 56 | >sp\|P01876\|IGHA1_HUMAN Immunoglobulin heavy constant alpha 1 OS=Homo sapiens OX=9606 GN=IGHA1 PE=1 SV=2 | 255 | 20201113_15413_1ng_glyco_hcd203040_compound15.21097.21097.3 | controllerType=0 controllerNumber=1 scan=21097 | 42.4709 |
| 17570:3 | 11 | R.LSLHRPALEDLLLGSEAN[+1540.529]LTC[+57.021]TLTGLR.D | HexNAc(2)Hex(7) | C[+57], N[+1541] | 1502.0524 | 3 | 4504.1427 | 4504.1268 | 0 | 3.5 | 127 | Specific | 717.66 | 650.15 | 650.15 | 13.76 | 56 | >sp\|P01876\|IGHA1_HUMAN Immunoglobulin heavy constant alpha 1 OS=Homo sapiens OX=9606 GN=IGHA1 PE=1 SV=2 | 255 | 20201113_15413_1ng_glyco_hcd203040_compound15.21107.21107.3 | controllerType=0 controllerNumber=1 scan=21107 | 42.4887 |
| 17571:3 | 11 | R.LSLHRPALEDLLLGSEAN[+1702.581]LTC[+57.021]TLTGLR.D | HexNAc(2)Hex(8) | C[+57], N[+1703] | 1556.0685 | 3 | 4666.1911 | 4666.1796 | 0 | 2.5 | 127 | Specific | 720.86 | 720.86 | 720.86 | 13.71 | 56 | >sp\|P01876\|IGHA1_HUMAN Immunoglobulin heavy constant alpha 1 OS=Homo sapiens OX=9606 GN=IGHA1 PE=1 SV=2 | 255 | 20201113_15413_1ng_glyco_hcd203040_compound15.21108.21108.3 | controllerType=0 controllerNumber=1 scan=21108 | 42.4899 |
| 17610:4 | 11 | R.LSLHRPALEDLLLGSEAN[+1501.555]LTC[+57.021]TLTGLR.D | HexNAc(5)Hex(3) | C[+57], N[+1502] | 1117.0462 | 4 | 4465.1629 | 4465.1536 | 0 | 2.1 | 127 | Specific | 731.62 | 731.62 | 731.62 | 14.39 | 56 | >sp\|P01876\|IGHA1_HUMAN Immunoglobulin heavy constant alpha 1 OS=Homo sapiens OX=9606 GN=IGHA1 PE=1 SV=2 | 255 | 20201113_15413_1ng_glyco_hcd203040_compound15.21149.21149.4 | controllerType=0 controllerNumber=1 scan=21149 | 42.5912 |
| 17638:3 | 11 | R.LSLHRPALEDLLLGSEAN[+1216.423]LTC[+57.021]TLTGLR.D | HexNAc(2)Hex(5) | C[+57], N[+1216] | 1394.0156 | 3 | 4180.0321 | 4180.0211 | 0 | 2.6 | 127 | Specific | 733.55 | 725.28 | 725.28 | 13.71 | 56 | >sp\|P01876\|IGHA1_HUMAN Immunoglobulin heavy constant alpha 1 OS=Homo sapiens OX=9606 GN=IGHA1 PE=1 SV=2 | 255 | 20201113_15413_1ng_glyco_hcd203040_compound15.21178.21178.3 | controllerType=0 controllerNumber=1 scan=21178 | 42.6693 |
| 17593:3 | 11 | R.LSLHRPALEDLLLGSEAN[+1378.476]LTC[+57.021]TLTGLR.D | HexNAc(2)Hex(6) | C[+57], N[+1378] | 1448.0367 | 3 | 4342.0955 | 4342.074 | 0 | 5 | 127 | Specific | 752.14 | 681.11 | 681.11 | 14.66 | 56 | >sp\|P01876\|IGHA1_HUMAN Immunoglobulin heavy constant alpha 1 OS=Homo sapiens OX=9606 GN=IGHA1 PE=1 SV=2 | 255 | 20201113_15413_1ng_glyco_hcd203040_compound15.21131.21131.3 | controllerType=0 controllerNumber=1 scan=21131 | 42.548 |
| 17635:3 | 11 | R.LSLHRPALEDLLLGSEAN[+1216.423]LTC[+57.021]TLTGLR.D | HexNAc(2)Hex(5) | C[+57], N[+1216] | 1394.0156 | 3 | 4180.0321 | 4180.0211 | 0 | 2.6 | 127 | Specific | 771.98 | 700.97 | 700.97 | 14.71 | 56 | >sp\|P01876\|IGHA1_HUMAN Immunoglobulin heavy constant alpha 1 OS=Homo sapiens OX=9606 GN=IGHA1 PE=1 SV=2 | 255 | 20201113_15413_1ng_glyco_hcd203040_compound15.21175.21175.3 | controllerType=0 controllerNumber=1 scan=21175 | 42.6598 |
| 17572:4 | 11 | R.LSLHRPALEDLLLGSEAN[+1663.608]LTC[+57.021]TLTGLR.D | HexNAc(5)Hex(4) | C[+57], N[+1664] | 1157.5602 | 4 | 4627.2191 | 4627.2064 | 0 | 2.7 | 127 | Specific | 784.56 | 757.3 | 757.3 | 15.44 | 56 | >sp\|P01876\|IGHA1_HUMAN Immunoglobulin heavy constant alpha 1 OS=Homo sapiens OX=9606 GN=IGHA1 PE=1 SV=2 | 255 | 20201113_15413_1ng_glyco_hcd203040_compound15.21109.21109.4 | controllerType=0 controllerNumber=1 scan=21109 | 42.4923 |
| 17613:4 | 11 | R.LSLHRPALEDLLLGSEAN[+1501.555]LTC[+57.021]TLTGLR.D | HexNAc(5)Hex(3) | C[+57], N[+1502] | 1117.0462 | 4 | 4465.1629 | 4465.1536 | 0 | 2.1 | 127 | Specific | 817.65 | 817.65 | 817.65 | 16.39 | 56 | >sp\|P01876\|IGHA1_HUMAN Immunoglobulin heavy constant alpha 1 OS=Homo sapiens OX=9606 GN=IGHA1 PE=1 SV=2 | 255 | 20201113_15413_1ng_glyco_hcd203040_compound15.21152.21152.4 | controllerType=0 controllerNumber=1 scan=21152 | 42.6053 |
| 17563:4 | 11 | R.LSLHRPALEDLLLGSEAN[+1540.529]LTC[+57.021]TLTGLR.D | HexNAc(2)Hex(7) | C[+57], N[+1541] | 1126.7901 | 4 | 4504.1385 | 4504.1268 | 0 | 2.6 | 127 | Specific | 856.57 | 807.43 | 807.43 | 17.39 | 56 | >sp\|P01876\|IGHA1_HUMAN Immunoglobulin heavy constant alpha 1 OS=Homo sapiens OX=9606 GN=IGHA1 PE=1 SV=2 | 255 | 20201113_15413_1ng_glyco_hcd203040_compound15.21100.21100.4 | controllerType=0 controllerNumber=1 scan=21100 | 42.4768 |
| 17605:4 | 11 | R.LSLHRPALEDLLLGSEAN[+1378.476]LTC[+57.021]TLTGLR.D | HexNAc(2)Hex(6) | C[+57], N[+1378] | 1086.2774 | 4 | 4342.0877 | 4342.074 | 0 | 3.2 | 127 | Specific | 875.32 | 875.32 | 875.32 | 17.44 | 56 | >sp\|P01876\|IGHA1_HUMAN Immunoglobulin heavy constant alpha 1 OS=Homo sapiens OX=9606 GN=IGHA1 PE=1 SV=2 | 255 | 20201113_15413_1ng_glyco_hcd203040_compound15.21144.21144.4 | controllerType=0 controllerNumber=1 scan=21144 | 42.5787 |
| 17626:4 | 11 | R.LSLHRPALEDLLLGSEAN[+1216.423]LTC[+57.021]TLTGLR.D | HexNAc(2)Hex(5) | C[+57], N[+1216] | 1045.7633 | 4 | 4180.0316 | 4180.0211 | 0 | 2.5 | 127 | Specific | 889.33 | 889.33 | 889.33 | 17.39 | 56 | >sp\|P01876\|IGHA1_HUMAN Immunoglobulin heavy constant alpha 1 OS=Homo sapiens OX=9606 GN=IGHA1 PE=1 SV=2 | 255 | 20201113_15413_1ng_glyco_hcd203040_compound15.21166.21166.4 | controllerType=0 controllerNumber=1 scan=21166 | 42.6324 |
| 17629:4 | 11 | R.LSLHRPALEDLLLGSEAN[+1216.423]LTC[+57.021]TLTGLR.D | HexNAc(2)Hex(5) | C[+57], N[+1216] | 1045.7633 | 4 | 4180.0316 | 4180.0211 | 0 | 2.5 | 127 | Specific | 892.09 | 892.09 | 892.09 | 17.39 | 56 | >sp\|P01876\|IGHA1_HUMAN Immunoglobulin heavy constant alpha 1 OS=Homo sapiens OX=9606 GN=IGHA1 PE=1 SV=2 | 255 | 20201113_15413_1ng_glyco_hcd203040_compound15.21169.21169.4 | controllerType=0 controllerNumber=1 scan=21169 | 42.6462 |
| 17506:4 | 11 | R.LSLHRPALEDLLLGSEAN[+1540.529]LTC[+57.021]TLTGLR.D | HexNAc(2)Hex(7) | C[+57], N[+1541] | 1126.7909 | 4 | 4504.1419 | 4504.1268 | 0 | 3.4 | 127 | Specific | 949.37 | 865.68 | 865.68 | 18.44 | 56 | >sp\|P01876\|IGHA1_HUMAN Immunoglobulin heavy constant alpha 1 OS=Homo sapiens OX=9606 GN=IGHA1 PE=1 SV=2 | 255 | 20201113_15413_1ng_glyco_hcd203040_compound15.21040.21040.2 | controllerType=0 controllerNumber=1 scan=21040 | 42.[illegible]04 |
| 17509:4 | 11 | R.LSLHRPALEDLLLGSEAN[+1540.529]LTC[+57.021]TLTGLR.D | HexNAc(2)Hex(7) | C[+57], N[+1541] | 1126.7909 | 4 | 4504.1419 | 4504.1268 | 0 | 3.4 | 127 | Specific | 961.54 | 961.54 | 961.54 | 19.44 | 56 | >sp\|P01876\|IGHA1_HUMAN Immunoglobulin heavy constant alpha 1 OS=Homo sapiens OX=9606 GN=IGHA1 PE=1 SV=2 | 255 | 20201113_15413_1ng_glyco_hcd203040_compound15.21043.21043.2 | controllerType=0 controllerNumber=1 scan=21043 | 42.[illegible]86 |
| 17607:4 | 11 | R.LSLHRPALEDLLLGSEAN[+1378.476]LTC[+57.021]TLTGLR.D | HexNAc(2)Hex(6) | C[+57], N[+1378] | 1086.2774 | 4 | 4342.0877 | 4342.074 | 0 | 3.2 | 127 | Specific | 972.96 | 972.96 | 972.96 | 19.44 | 56 | >sp\|P01876\|IGHA1_HUMAN Immunoglobulin heavy constant alpha 1 OS=Homo sapiens OX=9606 GN=IGHA1 PE=1 SV=2 | 255 | 20201113_15413_1ng_glyco_hcd203040_compound15.21146.21146.4 | controllerType=0 controllerNumber=1 scan=21146 | 42.5837 |
| 17566:4 | 11 | R.LSLHRPALEDLLLGSEAN[+1702.581]LTC[+57.021]TLTGLR.D | HexNAc(2)Hex(8) | C[+57], N[+1703] | 1167.3035 | 4 | 4666.1922 | 4666.1796 | 0 | 2.7 | 127 | Specific | 999.27 | 968.45 | 968.45 | 19.44 | 56 | >sp\|P01876\|IGHA1_HUMAN Immunoglobulin heavy constant alpha 1 OS=Homo sapiens OX=9606 GN=IGHA1 PE=1 SV=2 | 255 | 20201113_15413_1ng_glyco_hcd203040_compound15.21103.21103.4 | controllerType=0 controllerNumber=1 scan=21103 | 42.4834 |
| 17565:4 | 11 | R.LSLHRPALEDLLLGSEAN[+1540.529]LTC[+57.021]TLTGLR.D | HexNAc(2)Hex(7) | C[+57], N[+1541] | 1126.7901 | 4 | 4504.1385 | 4504.1268 | 0 | 2.6 | 127 | Specific | 1070.6 | 1070.6 | 1070.6 | 21.39 | 56 | >sp\|P01876\|IGHA1_HUMAN Immunoglobulin heavy constant alpha 1 OS=Homo sapiens OX=9606 GN=IGHA1 PE=1 SV=2 | 255 | 20201113_15413_1ng_glyco_hcd203040_compound15.21102.21102.4 | controllerType=0 controllerNumber=1 scan=21102 | 42.4817 |
| 17555:4 | 11 | R.LSLHRPALEDLLLGSEAN[+1540.529]LTC[+57.021]TLTGLR.D | HexNAc(2)Hex(7) | C[+57], N[+1541] | 1126.7923 | 4 | 4504.1473 | 4504.1268 | 0 | 4.6 | 127 | Specific | 1073.94 | 1073.94 | 1073.94 | 21.33 | 56 | >sp\|P01876\|IGHA1_HUMAN Immunoglobulin heavy constant alpha 1 OS=Homo sapiens OX=9606 GN=IGHA1 PE=1 SV=2 | 255 | 20201113_15413_1ng_glyco_hcd203040_compound15.21091.21091.2 | controllerType=0 controllerNumber=1 scan=21091 | 42.4596 |
| 12960:3 | 11 | R.LAGKPTHVN[+2262.814]VSVVMAEVDGTC[+57.021].Y | HexNAc(5)Hex(5)Fuc(1)NeuAc(1) | C[+57], N[+2263] | 1482.9741 | 3 | 4446.9076 | 4446.8931 | 0 | 3.3 | 332 | CRagged | 210.21 | 138.65 | 138.65 | 3.13 | 56 | >sp\|P01876\|IGHA1_HUMAN Immunoglobulin heavy constant alpha 1 OS=Homo sapiens OX=9606 GN=IGHA1 PE=1 SV=2 | 255 | 20201113_15413_1ng_glyco_hcd203040_compound15.16299.16299.3 | controllerType=0 controllerNumber=1 scan=16299 | 33.8832 |
| 12835:3 | 11 | R.LAGKPTHVN[+2263.835]VSVVMAEVDGTC[+57.021].Y | HexNAc(5)Hex(5)Fuc(3) | C[+57], N[+2264] | 1483.3091 | 3 | 4447.9129 | 4447.9135 | 0 | -0.1 | 332 | CRagged | 233.28 | 233.28 | 233.28 | 3.72 | 56 | >sp\|P01876\|IGHA1_HUMAN Immunoglobulin heavy constant alpha 1 OS=Homo sapiens OX=9606 GN=IGHA1 PE=1 SV=2 | 255 | 20201113_15413_1ng_glyco_hcd203040_compound15.16169.16169.3 | controllerType=0 controllerNumber=1 scan=16169 | 33.6391 |
| 12833:3 | 11 | R.LAGKPTHVN[+2263.835]VSVVMAEVDGTC[+57.021].Y | HexNAc(5)Hex(5)Fuc(3) | C[+57], N[+2264] | 1483.3091 | 3 | 4447.9129 | 4447.9135 | 0 | -0.1 | 332 | CRagged | 263.59 | 237.1 | 237.1 | 5.5 | 56 | >sp\|P01876\|IGHA1_HUMAN Immunoglobulin heavy constant alpha 1 OS=Homo sapiens OX=9606 GN=IGHA1 PE=1 SV=2 | 255 | 20201113_15413_1ng_glyco_hcd203040_compound15.16167.16167.3 | controllerType=0 controllerNumber=1 scan=16167 | 33.6335 |
| 13027:3 | 11 | R.LAGKPTHVN[+1710.598]VSVVMAEVDGTC[+57.021]Y.- | HexNAc(3)Hex(5)NeuAc(1) | C[+57], N[+1711] | 1353.2557 | 3 | 4057.7525 | 4057.7397 | 0 | 3.1 | 332 | CRagged | 233.06 | 233.06 | 233.06 | 6.24 | 56 | >sp\|P01876\|IGHA1_HUMAN Immunoglobulin heavy constant alpha 1 OS=Homo sapiens OX=9606 GN=IGHA1 PE=1 SV=2 | 255 | 20201113_15413_1ng_glyco_hcd203040_compound15.16368.16368.3 | controllerType=0 controllerNumber=1 scan=16368 | 34.002 |
| 12857:3 | 11 | R.LAGKPTHVN[+2117.777]VSVVMAEVDGTC[+57.021]Y.- | HexNAc(5)Hex(5)Fuc(2) | C[+57], N[+2118] | 1488.9756 | 3 | 4464.9124 | 4464.9189 | 0 | -1.5 | 332 | CRagged | 281.61 | 281.61 | 281.61 | 5.74 | 56 | >sp\|P01876\|IGHA1_HUMAN Immunoglobulin heavy constant alpha 1 OS=Homo sapiens OX=9606 GN=IGHA1 PE=1 SV=2 | 255 | 20201113_15413_1ng_glyco_hcd203040_compound15.16192.16192.3 | controllerType=0 controllerNumber=1 scan=16192 | 33.6958 |
| 12897:3 | 11 | R.LAGKPTHVN[+1913.677]VSVVMAEVDGTC[+57.021]Y.- | HexNAc(4)Hex(5)NeuAc(1) | C[+57], N[+1914] | 1420.951 | 3 | 4260.8384 | 4260.8191 | 0 | 4.5 | 332 | CRagged | 292.3 | 265.28 | 265.28 | 6.66 | 56 | >sp\|P01876\|IGHA1_HUMAN Immunoglobulin heavy constant alpha 1 OS=Homo sapiens OX=9606 GN=IGHA1 PE=1 SV=2 | 255 | 20201113_15413_1ng_glyco_hcd203040_compound15.16234.16234.3 | controllerType=0 controllerNumber=1 scan=16234 | 33.7793 |
| 11270:3 | 11 | R.LAGKPTHVN[+1241.454]VSVVMAEVDGTC[+57.021]Y.- | HexNAc(3)Hex(3)Fuc(1) | C[+57], N[+1241] | 1196.8773 | 3 | 3588.6172 | 3588.5966 | 0 | 5.7 | 332 | CRagged | 317.09 | 317.09 | 317.09 | 6.64 | 56 | >sp\|P01876\|IGHA1_HUMAN Immunoglobulin heavy constant alpha 1 OS=Homo sapiens OX=9606 GN=IGHA1 PE=1 SV=2 | 255 | 20201113_15413_1ng_glyco_hcd203040_compound15.14530.14530.3 | controllerType=0 controllerNumber=1 scan=14530 | 30.8912 |
| 13047:3 | 11 | R.LAGKPTHVN[+1548.545]VSVVMAEVDGTC[+57.021]Y.- | HexNAc(3)Hex(4)NeuAc(1) | C[+57], N[+1549] | 1299.2372 | 3 | 3895.6972 | 3895.6869 | 0 | 2.6 | 332 | CRagged | 339.28 | 339.28 | 339.28 | 7.36 | 56 | >sp\|P01876\|IGHA1_HUMAN Immunoglobulin heavy constant alpha 1 OS=Homo sapiens OX=9606 GN=IGHA1 PE=1 SV=2 | 255 | 20201113_15413_1ng_glyco_hcd203040_compound15.16389.16389.3 | controllerType=0 controllerNumber=1 scan=16389 | 34.0359 |
| 14932:3 | 11 | R.LAGKPTHVN[+2553.910]VSVVMAEVDGTC[+57.021]Y.- | HexNAc(5)Hex(5)Fuc(1)NeuAc(2) | C[+57], N[+2554] | 1634.3578 | 3 | 4901.059 | 4901.0518 | 0 | 1.5 | 332 | CRagged | 339.62 | 262.73 | 262.73 | 6.98 | 56 | >sp\|P01876\|IGHA1_HUMAN Immunoglobulin heavy constant alpha 1 OS=Homo sapiens OX=9606 GN=IGHA1 PE=1 SV=2 | 255 | 20201113_15413_1ng_glyco_hcd203040_compound15.18341.18341.3 | controllerType=0 controllerNumber=1 scan=18341 | 37.3556 |
| 11071:3 | 11 | R.LAGKPTHVN[+1825.661]VSVVMAEVDGTC[+57.021]Y.- | HexNAc(5)Hex(5) | C[+57], N[+1826] | 1391.6067 | 3 | 4172.8057 | 4172.8031 | 0 | 0.6 | 332 | CRagged | 354.38 | 288.14 | 288.14 | 7.05 | 56 | >sp\|P01876\|IGHA1_HUMAN Immunoglobulin heavy constant alpha 1 OS=Homo sapiens OX=9606 GN=IGHA1 PE=1 SV=2 | 255 | 20201113_15413_1ng_glyco_hcd203040_compound15.14323.14323.3 | controllerType=0 controllerNumber=1 scan=14323 | 30.5038 |
| 12970:3 | 11 | R.LAGKPTHVN[+1710.598]VSVVMAEVDGTC[+57.021]Y.- | HexNAc(3)Hex(5)NeuAc(1) | C[+57], N[+1711] | 1353.2523 | 3 | 4057.7422 | 4057.7397 | 0 | 0.6 | 332 | CRagged | 362.71 | 341.97 | 277.22 | 7.27 | 56 | >sp\|P01876\|IGHA1_HUMAN Immunoglobulin heavy constant alpha 1 OS=Homo sapiens OX=9606 GN=IGHA1 PE=1 SV=2 | 255 | 20201113_15413_1ng_glyco_hcd203040_compound15.16309.16309.3 | controllerType=0 controllerNumber=1 scan=16309 | 33.9012 |
| 09965:3 | 11 | R.LAGKPTHVN[+1216.423]VSVVMAEVDGTC[+57.021]Y.- | HexNAc(2)Hex(5) | C[+57], N[+1216] | 1188.5295 | 3 | 3563.5739 | 3563.5649 | 0 | 2.5 | 332 | CRagged | 366.61 | 366.61 | 176.58 | 7.68 | 56 | >sp\|P01876\|IGHA1_HUMAN Immunoglobulin heavy constant alpha 1 OS=Homo sapiens OX=9606 GN=IGHA1 PE=1 SV=2 | 255 | 20201113_15413_1ng_glyco_hcd203040_compound15.13179.13179.3 | controllerType=0 controllerNumber=1 scan=13179 | 28.5528 |
| 09925:3 | 11 | R.LAGKPTHVN[+1378.476]VSVVMAEVDGTC[+57.021]Y.- | HexNAc(2)Hex(6) | C[+57], N[+1378] | 1242.5485 | 3 | 3725.631 | 3725.6178 | 0 | 3.6 | 332 | CRagged | 368.77 | 368.77 | 292.45 | 7.73 | 56 | >sp\|P01876\|IGHA1_HUMAN Immunoglobulin heavy constant alpha 1 OS=Homo sapiens OX=9606 GN=IGHA1 PE=1 SV=2 | 255 | 20201113_15413_1ng_glyco_hcd203040_compound15.13138.13138.3 | controllerType=0 controllerNumber=1 scan=13138 | 28.4817 |
| 14930:3 | 11 | R.LAGKPTHVN[+2553.910]VSVVMAEVDGTC[+57.021]Y.- | HexNAc(5)Hex(5)Fuc(1)NeuAc(2) | C[+57], N[+2554] | 1634.3578 | 3 | 4901.059 | 4901.0518 | 0 | 1.5 | 332 | CRagged | 369.19 | 294.25 | 294.25 | 7.3 | 56 | >sp\|P01876\|IGHA1_HUMAN Immunoglobulin heavy constant alpha 1 OS=Homo sapiens OX=9606 GN=IGHA1 PE=1 SV=2 | 255 | 20201113_15413_1ng_glyco_hcd203040_compound15.18339.18339.3 | controllerType=0 controllerNumber=1 scan=18339 | 37.3516 |
| 14976:3 | 11 | R.LAGKPTHVN[+2204.772]VSVVMAEVDGTC[+57.021]Y.- | HexNAc(4)Hex(5)NeuAc(2) | C[+57], N[+2205] | 1517.9859 | 3 | 4551.9431 | 4551.9145 | 0 | 6.3 | 332 | CRagged | 383.64 | 360.89 | 360.89 | 7.22 | 56 | >sp\|P01876\|IGHA1_HUMAN Immunoglobulin heavy constant alpha 1 OS=Homo sapiens OX=9606 GN=IGHA1 PE=1 SV=2 | 255 | 20201113_15413_1ng_glyco_hcd203040_compound15.18386.18386.3 | controllerType=0 controllerNumber=1 scan=18386 | 37.4292 |
| 12894:3 | 11 | R.LAGKPTHVN[+1913.677]VSVVMAEVDGTC[+57.021]Y.- | HexNAc(4)Hex(5)NeuAc(1) | C[+57], N[+1914] | 1420.951 | 3 | 4260.8384 | 4260.8191 | 0 | 4.5 | 332 | CRagged | 414.1 | 242.91 | 242.91 | 7.01 | 56 | >sp\|P01876\|IGHA1_HUMAN Immunoglobulin heavy constant alpha 1 OS=Homo sapiens OX=9606 GN=IGHA1 PE=1 SV=2 | 255 | 20201113_15413_1ng_glyco_hcd203040_compound15.16231.16231.3 | controllerType=0 controllerNumber=1 scan=16231 | 33.775 |
| 12855:3 | 11 | R.LAGKPTHVN[+2117.777]VSVVMAEVDGTC[+57.021]Y.- | HexNAc(5)Hex(5)Fuc(2) | C[+57], N[+2118] | 1488.9756 | 3 | 4464.9124 | 4464.9189 | 0 | -1.5 | 332 | CRagged | 417.94 | 417.94 | 417.94 | 8.11 | 56 | >sp\|P01876\|IGHA1_HUMAN Immunoglobulin heavy constant alpha 1 OS=Homo sapiens OX=9606 GN=IGHA1 PE=1 SV=2 | 255 | 20201113_15413_1ng_glyco_hcd203040_compound15.16190.16190.3 | controllerType=0 controllerNumber=1 scan=16190 | 33.69 |
| 12972:3 | 11 | R.LAGKPTHVN[+1710.598]VSVVMAEVDGTC[+57.021]Y.- | HexNAc(3)Hex(5)NeuAc(1) | C[+57], N[+1711] | 1353.2523 | 3 | 4057.7422 | 4057.7397 | 0 | 0.6 | 332 | CRagged | 420.28 | 380.17 | 258.79 | 9.01 | 56 | >sp\|P01876\|IGHA1_HUMAN Immunoglobulin heavy constant alpha 1 OS=Homo sapiens OX=9606 GN=IGHA1 PE=1 SV=2 | 255 | 20201113_15413_1ng_glyco_hcd203040_compound15.16311.16311.3 | controllerType=0 controllerNumber=1 scan=16311 | 33.9048 |
| 14920:4 | 11 | R.LAGKPTHVN[+2553.910]VSVVMAEVDGTC[+57.021]Y.- | HexNAc(5)Hex(5)Fuc(1)NeuAc(2) | C[+57], N[+2554] | 1226.0208 | 4 | 4901.0613 | 4901.0518 | 0 | 1.9 | 332 | CRagged | 433.61 | 333.39 | 333.39 | 7.76 | 56 | >sp\|P01876\|IGHA1_HUMAN Immunoglobulin heavy constant alpha 1 OS=Homo sapiens OX=9606 GN=IGHA1 PE=1 SV=2 | 255 | 20201113_15413_1ng_glyco_hcd203040_compound15.18328.18328.2 | controllerType=0 controllerNumber=1 scan=18328 | 37.3327 |
| 14865:3 | 11 | R.LAGKPTHVN[+2350.830]VSVVMAEVDGTC[+57.021]Y.- | HexNAc(4)Hex(5)Fuc(1)NeuAc(2) | C[+57], N[+2351] | 1566.6523 | 3 | 4697.9423 | 4697.9724 | 0 | -6.4 | 332 | CRagged | 436.66 | 407.11 | 407.11 | 8.09 | 56 | >sp\|P01876\|IGHA1_HUMAN Immunoglobulin heavy constant alpha 1 OS=Homo sapiens OX=9606 GN=IGHA1 PE=1 SV=2 | 255 | 20201113_15413_1ng_glyco_hcd203040_compound15.18270.18270.3 | controllerType=0 controllerNumber=1 scan=18270 | 37.2237 |
| 12854:4 | 11 | R.LAGKPTHVN[+2262.814]VSVVMAEVDGTC[+57.021]Y.- | HexNAc(5)Hex(5)Fuc(1)NeuAc(1) | C[+57], N[+2263] | 1153.2471 | 4 | 4609.9666 | 4609.9564 | 0 | 2.2 | 332 | CRagged | 438.81 | 438.81 | 438.81 | 9.37 | 56 | >sp\|P01876\|IGHA1_HUMAN Immunoglobulin heavy constant alpha 1 OS=Homo sapiens OX=9606 GN=IGHA1 PE=1 SV=2 | 255 | 20201113_15413_1ng_glyco_hcd203040_compound15.16189.16189.4 | controllerType=0 controllerNumber=1 scan=16189 | 33.6885 |
| 09967:3 | 11 | R.LAGKPTHVN[+1216.423]VSVVMAEVDGTC[+57.021]Y.- | HexNAc(2)Hex(5) | C[+57], N[+1216] | 1188.5295 | 3 | 3563.5739 | 3563.5649 | 0 | 2.5 | 332 | CRagged | 447.65 | 447.65 | 241.84 | 9.42 | 56 | >sp\|P01876\|IGHA1_HUMAN Immunoglobulin heavy constant alpha 1 OS=Homo sapiens OX=9606 GN=IGHA1 PE=1 SV=2 | 255 | 20201113_15413_1ng_glyco_hcd203040_compound15.13181.13181.3 | controllerType=0 controllerNumber=1 scan=13181 | 28.5566 |
| 13049:3 | 11 | R.LAGKPTHVN[+1548.545]VSVVMAEVDGTC[+57.021]Y.- | HexNAc(3)Hex(4)NeuAc(1) | C[+57], N[+1549] | 1299.2372 | 3 | 3895.6972 | 3895.6869 | 0 | 2.6 | 332 | CRagged | 466.31 | 333.33 | 333.33 | 7.67 | 56 | >sp\|P01876\|IGHA1_HUMAN Immunoglobulin heavy constant alpha 1 OS=Homo sapiens OX=9606 GN=IGHA1 PE=1 SV=2 | 255 | 20201113_15413_1ng_glyco_hcd203040_compound15.16391.16391.3 | controllerType=0 controllerNumber=1 scan=16391 | 34.0395 |
| 14867:3 | 11 | R.LAGKPTHVN[+2350.830]VSVVMAEVDGTC[+57.021]Y.- | HexNAc(4)Hex(5)Fuc(1)NeuAc(2) | C[+57], N[+2351] | 1566.6523 | 3 | 4697.9423 | 4697.9724 | 0 | -6.4 | 332 | CRagged | 481.9 | 387.84 | 387.84 | 8.22 | 56 | >sp\|P01876\|IGHA1_HUMAN Immunoglobulin heavy constant alpha 1 OS=Homo sapiens OX=9606 GN=IGHA1 PE=1 SV=2 | 255 | 20201113_15413_1ng_glyco_hcd203040_compound15.18272.18272.3 | controllerType=0 controllerNumber=1 scan=18272 | 37.2275 |
| 12856:4 | 11 | R.LAGKPTHVN[+2262.814]VSVVMAEVDGTC[+57.021]Y.- | HexNAc(5)Hex(5)Fuc(1)NeuAc(1) | C[+57], N[+2263] | 1153.2471 | 4 | 4609.9666 | 4609.9564 | 0 | 2.2 | 332 | CRagged | 512.24 | 512.24 | 512.24 | 9.64 | 56 | >sp\|P01876\|IGHA1_HUMAN Immunoglobulin heavy constant alpha 1 OS=Homo sapiens OX=9606 GN=IGHA1 PE=1 SV=2 | 255 | 20201113_15413_1ng_glyco_hcd203040_compound15.16191.16191.4 | controllerType=0 controllerNumber=1 scan=16191 | 33.6935 |
| 14998:3 | 11 | R.LAGKPTHVN[+2204.772]VSVVMAEVDGTC[+57.021]Y.- | HexNAc(4)Hex(5)NeuAc(2) | C[+57], N[+2205] | 1517.9786 | 3 | 4551.9212 | 4551.9145 | 0 | 1.5 | 332 | CRagged | 516.65 | 499.92 | 448.27 | 9.69 | 56 | >sp\|P01876\|IGHA1_HUMAN Immunoglobulin heavy constant alpha 1 OS=Homo sapiens OX=9606 GN=IGHA1 PE=1 SV=2 | 255 | 20201113_15413_1ng_glyco_hcd203040_compound15.18409.18409.3 | controllerType=0 controllerNumber=1 scan=18409 | 37.4703 |
| 09927:3 | 11 | R.LAGKPTHVN[+1378.476]VSVVMAEVDGTC[+57.021]Y.- | HexNAc(2)Hex(6) | C[+57], N[+1378] | 1242.5485 | 3 | 3725.631 | 3725.6178 | 0 | 3.6 | 332 | CRagged | 517.92 | 449.38 | 321.55 | 9.74 | 56 | >sp\|P01876\|IGHA1_HUMAN Immunoglobulin heavy constant alpha 1 OS=Homo sapiens OX=9606 GN=IGHA1 PE=1 SV=2 | 255 | 20201113_15413_1ng_glyco_hcd203040_compound15.13140.13140.3 | controllerType=0 controllerNumber=1 scan=13140 | 28.4853 |
| 11070:3 | 11 | R.LAGKPTHVN[+1825.661]VSVVMAEVDGTC[+57.021]Y.- | HexNAc(5)Hex(5) | C[+57], N[+1826] | 1391.6067 | 3 | 4172.8057 | 4172.8031 | 0 | 0.6 | 332 | CRagged | 527.53 | 341.71 | 341.71 | 7.96 | 56 | >sp\|P01876\|IGHA1_HUMAN Immunoglobulin heavy constant alpha 1 OS=Homo sapiens OX=9606 GN=IGHA1 PE=1 SV=2 | 255 | 20201113_15413_1ng_glyco_hcd203040_compound15.14322.14322.3 | controllerType=0 controllerNumber=1 scan=14322 | 30.5004 |
| 15000:3 | 11 | R.LAGKPTHVN[+2204.772]VSVVMAEVDGTC[+57.021]Y.- | HexNAc(4)Hex(5)NeuAc(2) | C[+57], N[+2205] | 1517.9786 | 3 | 4551.9212 | 4551.9145 | 0 | 1.5 | 332 | CRagged | 540.4 | 377.55 | 377.55 | 9.82 | 56 | >sp\|P01876\|IGHA1_HUMAN Immunoglobulin heavy constant alpha 1 OS=Homo sapiens OX=9606 GN=IGHA1 PE=1 SV=2 | 255 | 20201113_15413_1ng_glyco_hcd203040_compound15.18411.18411.3 | controllerType=0 controllerNumber=1 scan=18411 | 37.474 |
| 12834:3 | 11 | R.LAGKPTHVN[+2262.814]VSVVMAEVDGTC[+57.021]Y.- | HexNAc(5)Hex(5)Fuc(1)NeuAc(1) | C[+57], N[+2263] | 1537.3259 | 3 | 4609.963 | 4609.9564 | 0 | 1.4 | 332 | CRagged | 585.49 | 411.64 | 411.64 | 10.97 | 56 | >sp\|P01876\|IGHA1_HUMAN Immunoglobulin heavy constant alpha 1 OS=Homo sapiens OX=9606 GN=IGHA1 PE=1 SV=2 | 255 | 20201113_15413_1ng_glyco_hcd203040_compound15.16168.16168.3 | controllerType=0 controllerNumber=1 scan=16168 | 33.6368 |
| 11226:3 | 11 | R.LAGKPTHVN[+1501.555]VSVVMAEVDGTC[+57.021]Y.- | HexNAc(5)Hex(3) | C[+57], N[+1502] | 1283.5749 | 3 | 3848.7101 | 3848.6974 | 0 | 3.3 | 332 | CRagged | 614.12 | 614.12 | 614.12 | 12.27 | 56 | >sp\|P01876\|IGHA1_HUMAN Immunoglobulin heavy constant alpha 1 OS=Homo sapiens OX=9606 GN=IGHA1 PE=1 SV=2 | 255 | 20201113_15413_1ng_glyco_hcd203040_compound15.14485.14485.3 | controllerType=0 controllerNumber=1 scan=14485 | 30.8084 |
| 11228:3 | 11 | R.LAGKPTHVN[+1501.555]VSVVMAEVDGTC[+57.021]Y.- | HexNAc(5)Hex(3) | C[+57], N[+1502] | 1283.5749 | 3 | 3848.7101 | 3848.6974 | 0 | 3.3 | 332 | CRagged | 753.82 | 612.23 | 612.23 | 15.27 | 56 | >sp\|P01876\|IGHA1_HUMAN Immunoglobulin heavy constant alpha 1 OS=Homo sapiens OX=9606 GN=IGHA1 PE=1 SV=2 | 255 | 20201113_15413_1ng_glyco_hcd203040_compound15.14487.14487.3 | controllerType=0 controllerNumber=1 scan=14487 | 30.8128 |
| 11014:3 | 11 | R.LAGKPTHVN[+1702.581]VSVVMAEVDGTC[+57.021]Y.- | HexNAc(2)Hex(8) | C[+57], N[+1703] | 1350.5811 | 3 | 4049.7288 | 4049.7234 | 0 | 1.3 | 332 | CRagged | 756.88 | 732.44 | 447.62 | 14.97 | 56 | >sp\|P01876\|IGHA1_HUMAN Immunoglobulin heavy constant alpha 1 OS=Homo sapiens OX=9606 GN=IGHA1 PE=1 SV=2 | 255 | 20201113_15413_1ng_glyco_hcd203040_compound15.14264.14264.3 | controllerType=0 controllerNumber=1 scan=14264 | 30.3994 |
| 11085:3 | 11 | R.LAGKPTHVN[+1809.666]VSVVMAEVDGTC[+57.021]Y.- | HexNAc(5)Hex(4)Fuc(1) | C[+57], N[+1810] | 1386.2772 | 3 | 4156.8169 | 4156.8081 | 0 | 2.1 | 332 | CRagged | 766.46 | 766.46 | 494.81 | 15.22 | 56 | >sp\|P01876\|IGHA1_HUMAN Immunoglobulin heavy constant alpha 1 OS=Homo sapiens OX=9606 GN=IGHA1 PE=1 SV=2 | 255 | 20201113_15413_1ng_glyco_hcd203040_compound15.14338.14338.3 | controllerType=0 controllerNumber=1 scan=14338 | 30.5371 |
| 11077:3 | 11 | R.LAGKPTHVN[+1663.608]VSVVMAEVDGTC[+57.021]Y.- | HexNAc(5)Hex(4) | C[+57], N[+1664] | 1337.5914 | 3 | 4010.7595 | 4010.7502 | 0 | 2.3 | 332 | CRagged | 767.03 | 725.8 | 461.02 | 15.22 | 56 | >sp\|P01876\|IGHA1_HUMAN Immunoglobulin heavy constant alpha 1 OS=Homo sapiens OX=9606 GN=IGHA1 PE=1 SV=2 | 255 | 20201113_15413_1ng_glyco_hcd203040_compound15.14330.14330.3 | controllerType=0 controllerNumber=1 scan=14330 | 30.5216 |
| 11208:3 | 11 | R.LAGKPTHVN[+1647.613]VSVVMAEVDGTC[+57.021]Y.- | HexNAc(5)Hex(3)Fuc(1) | C[+57], N[+1648] | 1332.2573 | 3 | 3994.7572 | 3994.7553 | 0 | 0.5 | 332 | CRagged | 790.02 | 628.32 | 628.32 | 14.97 | 56 | >sp\|P01876\|IGHA1_HUMAN Immunoglobulin heavy constant alpha 1 OS=Homo sapiens OX=9606 GN=IGHA1 PE=1 SV=2 | 255 | 20201113_15413_1ng_glyco_hcd203040_compound15.14466.14466.3 | controllerType=0 controllerNumber=1 scan=14466 | 30.7638 |
| 11012:3 | 11 | R.LAGKPTHVN[+1702.581]VSVVMAEVDGTC[+57.021]Y.- | HexNAc(2)Hex(8) | C[+57], N[+1703] | 1350.5811 | 3 | 4049.7288 | 4049.7234 | 0 | 1.3 | 332 | CRagged | 809.91 | 809.91 | 491.65 | 15.97 | 56 | >sp\|P01876\|IGHA1_HUMAN Immunoglobulin heavy constant alpha 1 OS=Homo sapiens OX=9606 GN=IGHA1 PE=1 SV=2 | 255 | 20201113_15413_1ng_glyco_hcd203040_compound15.14262.14262.3 | controllerType=0 controllerNumber=1 scan=14262 | 30.3937 |
| 12832:3 | 11 | R.LAGKPTHVN[+2262.814]VSVVMAEVDGTC[+57.021]Y.- | HexNAc(5)Hex(5)Fuc(1)NeuAc(1) | C[+57], N[+2263] | 1537.3259 | 3 | 4609.963 | 4609.9564 | 0 | 1.4 | 332 | CRagged | 818.99 | 559.57 | 559.57 | 15.97 | 56 | >sp\|P01876\|IGHA1_HUMAN Immunoglobulin heavy constant alpha 1 OS=Homo sapiens OX=9606 GN=IGHA1 PE=1 SV=2 | 255 | 20201113_15413_1ng_glyco_hcd203040_compound15.16166.16166.3 | controllerType=0 controllerNumber=1 scan=16166 | 33.6321 |
| 11206:3 | 11 | R.LAGKPTHVN[+1647.613]VSVVMAEVDGTC[+57.021]Y.- | HexNAc(5)Hex(3)Fuc(1) | C[+57], N[+1648] | 1332.2573 | 3 | 3994.7572 | 3994.7553 | 0 | 0.5 | 332 | CRagged | 828.66 | 814.68 | 814.68 | 15.97 | 56 | >sp\|P01876\|IGHA1_HUMAN Immunoglobulin heavy constant alpha 1 OS=Homo sapiens OX=9606 GN=IGHA1 PE=1 SV=2 | 255 | 20201113_15413_1ng_glyco_hcd203040_compound15.14464.14464.3 | controllerType=0 controllerNumber=1 scan=14464 | 30.7594 |
| 11083:3 | 11 | R.LAGKPTHVN[+1809.666]VSVVMAEVDGTC[+57.021]Y.- | HexNAc(5)Hex(4)Fuc(1) | C[+57], N[+1810] | 1386.2772 | 3 | 4156.8169 | 4156.8081 | 0 | 2.1 | 332 | CRagged | 832.27 | 832.27 | 542.53 | 16.22 | 56 | >sp\|P01876\|IGHA1_HUMAN Immunoglobulin heavy constant alpha 1 OS=Homo sapiens OX=9606 GN=IGHA1 PE=1 SV=2 | 255 | 20201113_15413_1ng_glyco_hcd203040_compound15.14336.14336.3 | controllerType=0 controllerNumber=1 scan=14336 | 30.5307 |
| 11075:3 | 11 | R.LAGKPTHVN[+1663.608]VSVVMAEVDGTC[+57.021]Y.- | HexNAc(5)Hex(4) | C[+57], N[+1664] | 1337.5914 | 3 | 4010.7595 | 4010.7502 | 0 | 2.3 | 332 | CRagged | 832.41 | 705.99 | 534.11 | 16.22 | 56 | >sp\|P01876\|IGHA1_HUMAN Immunoglobulin heavy constant alpha 1 OS=Homo sapiens OX=9606 GN=IGHA1 PE=1 SV=2 | 255 | 20201113_15413_1ng_glyco_hcd203040_compound15.14328.14328.3 | controllerType=0 controllerNumber=1 scan=14328 | 30.5161 |
| 11131:3 | 11 | R.LAGKPTHVN[+1216.423]VSVVMAEVDGTC[+57.021]Y.- | HexNAc(2)Hex(5) | C[+57], N[+1216] | 1188.5292 | 3 | 3563.5731 | 3563.5649 | 0 | 2.3 | 332 | CRagged | 853.79 | 817.79 | 548.61 | 17.22 | 56 | >sp\|P01876\|IGHA1_HUMAN Immunoglobulin heavy constant alpha 1 OS=Homo sapiens OX=9606 GN=IGHA1 PE=1 SV=2 | 255 | 20201113_15413_1ng_glyco_hcd203040_compound15.14386.14386.3 | controllerType=0 controllerNumber=1 scan=14386 | 30.6205 |
| 11048:3 | 11 | R.LAGKPTHVN[+1540.529]VSVVMAEVDGTC[+57.021]Y.- | HexNAc(2)Hex(7) | C[+57], N[+1541] | 1296.565 | 3 | 3887.6804 | 3887.6706 | 0 | 2.5 | 332 | CRagged | 859.24 | 780.25 | 510.97 | 17.22 | 56 | >sp\|P01876\|IGHA1_HUMAN Immunoglobulin heavy constant alpha 1 OS=Homo sapiens OX=9606 GN=IGHA1 PE=1 SV=2 | 255 | 20201113_15413_1ng_glyco_hcd203040_compound15.14300.14300.3 | controllerType=0 controllerNumber=1 scan=14300 | 30.4633 |
| 11072:3 | 11 | R.LAGKPTHVN[+1378.476]VSVVMAEVDGTC[+57.021]Y.- | HexNAc(2)Hex(6) | C[+57], N[+1378] | 1242.5469 | 3 | 3725.6262 | 3725.6178 | 0 | 2.3 | 332 | CRagged | 889.21 | 889.21 | 570.02 | 17.22 | 56 | >sp\|P01876\|IGHA1_HUMAN Immunoglobulin heavy constant alpha 1 OS=Homo sapiens OX=9606 GN=IGHA1 PE=1 SV=2 | 255 | 20201113_15413_1ng_glyco_hcd203040_compound15.14325.14325.3 | controllerType=0 controllerNumber=1 scan=14325 | 30.5083 |
| 11074:3 | 11 | R.LAGKPTHVN[+1378.476]VSVVMAEVDGTC[+57.021]Y.- | HexNAc(2)Hex(6) | C[+57], N[+1378] | 1242.5469 | 3 | 3725.6262 | 3725.6178 | 0 | 2.3 | 332 | CRagged | 897.18 | 829.87 | 553.63 | 17.22 | 56 | >sp\|P01876\|IGHA1_HUMAN Immunoglobulin heavy constant alpha 1 OS=Homo sapiens OX=9606 GN=IGHA1 PE=1 SV=2 | 255 | 20201113_15413_1ng_glyco_hcd203040_compound15.14327.14327.3 | controllerType=0 controllerNumber=1 scan=14327 | 30.5152 |
| 11046:3 | 11 | R.LAGKPTHVN[+1540.529]VSVVMAEVDGTC[+57.021]Y.- | HexNAc(2)Hex(7) | C[+57], N[+1541] | 1296.565 | 3 | 3887.6804 | 3887.6706 | 0 | 2.5 | 332 | CRagged | 902.87 | 902.87 | 583.53 | 18.22 | 56 | >sp\|P01876\|IGHA1_HUMAN Immunoglobulin heavy constant alpha 1 OS=Homo sapiens OX=9606 GN=IGHA1 PE=1 SV=2 | 255 | 20201113_15413_1ng_glyco_hcd203040_compound15.14298.14298.3 | controllerType=0 controllerNumber=1 scan=14298 | 30.4568 |
| 11128:3 | 11 | R.LAGKPTHVN[+1216.423]VSVVMAEVDGTC[+57.021]Y.- | HexNAc(2)Hex(5) | C[+57], N[+1216] | 1188.5292 | 3 | 3563.5731 | 3563.5649 | 0 | 2.3 | 332 | CRagged | 907.6 | 907.6 | 567.78 | 18.22 | 56 | >sp\|P01876\|IGHA1_HUMAN Immunoglobulin heavy constant alpha 1 OS=Homo sapiens OX=9606 GN=IGHA1 PE=1 SV=2 | 255 | 20201113_15413_1ng_glyco_hcd203040_compound15.14383.14383.3 | controllerType=0 controllerNumber=1 scan=14383 | 30.6139 |
| 07442:3 | 12 | R.LSPN[+2350.830]ASAEHLELR.W | HexNAc(4)Hex(5)Fuc(1)NeuAc(2) | N[+2351] | 1263.2014 | 3 | 3787.5895 | 3787.5796 | 0 | 2.6 | 52 | Specific | 466.72 | 466.72 | 466.72 | 9.04 | 34 | >sp\|Q13410\|BT1A1_HUMAN Butyrophilin subfamily 1 member A1 OS=Homo sapiens OX=9606 GN=BTN1A1 PE=1 SV=3 | 179 | 20201113_15413_1ng_glyco_hcd203040_compound15.10550.10550.3 | controllerType=0 controllerNumber=1 scan=10550 | 24.1565 |
| 05393:3 | 12 | R.LSPN[+2205.793]ASAEHLELR.W | HexNAc(4)Hex(5)Fuc(2)NeuAc(1) | N[+2206] | 1214.8563 | 3 | 3642.5542 | 3642.5421 | 0 | 3.3 | 52 | Specific | 533.23 | 493.48 | 493.48 | 9.63 | 34 | >sp\|Q13410\|BT1A1_HUMAN Butyrophilin subfamily 1 member A1 OS=Homo sapiens OX=9606 GN=BTN1A1 PE=1 SV=3 | 179 | 20201113_15413_1ng_glyco_hcd203040_compound15.8422.8422.3 | controllerType=0 controllerNumber=1 scan=8422 | 20.6272 |
| 07444:3 | 12 | R.LSPN[+2350.830]ASAEHLELR.W | HexNAc(4)Hex(5)Fuc(1)NeuAc(2) | N[+2351] | 1263.2014 | 3 | 3787.5895 | 3787.5796 | 0 | 2.6 | 52 | Specific | 539.65 | 449.29 | 449.29 | 9.58 | 34 | >sp\|Q13410\|BT1A1_HUMAN Butyrophilin subfamily 1 member A1 OS=Homo sapiens OX=9606 GN=BTN1A1 PE=1 SV=3 | 179 | 20201113_15413_1ng_glyco_hcd203040_compound15.10552.10552.3 | controllerType=0 controllerNumber=1 scan=10552 | 24.1609 |
| 05396:3 | 12 | R.LSPN[+2205.793]ASAEHLELR.W | HexNAc(4)Hex(5)Fuc(2)NeuAc(1) | N[+2206] | 1214.8563 | 3 | 3642.5542 | 3642.5421 | 0 | 3.3 | 52 | Specific | 619.44 | 403.49 | 403.49 | 12.02 | 34 | >sp\|Q13410\|BT1A1_HUMAN Butyrophilin subfamily 1 member A1 OS=Homo sapiens OX=9606 GN=BTN1A1 PE=1 SV=3 | 179 | 20201113_15413_1ng_glyco_hcd203040_compound15.8425.8425.3 | controllerType=0 controllerNumber=1 scan=8425 | 20.633 |
| 17527:3 | 12 | K.N[+2059.735]VSC[+57.021]YIQNLLLGQEK.K | HexNAc(4)Hex(5)Fuc(1)NeuAc(1) | C[+57], N[+2060] | 1280.226 | 3 | 3838.6634 | 3838.6455 | 0 | 4.7 | 215 | Specific | 418.05 | 266.61 | 266.61 | 6.91 | 34 | >sp\|Q13410\|BT1A1_HUMAN Butyrophilin subfamily 1 member A1 OS=Homo sapiens OX=9606 GN=BTN1A1 PE=1 SV=3 | 179 | 20201113_15413_1ng_glyco_hcd203040_compound15.21062.21062.3 | controllerType=0 controllerNumber=1 scan=21062 | 42.4038 |
| 22009:4 | 14 | K.[+42.011]VFPLSLDSTPQDGNVVVAC[+57.021]LVQGFFPQEPLSVTWSESGQN[+1768 | HexNAc(4)Hex(5)Fuc(1) | .[42], C[+57], N[+1769] | 1647.5195 | 4 | 6587.0563 | 6587.0115 | 0 | 6.8 | 8 | Specific | 291.02 | 214.24 | 214.24 | 4.94 | 38 | >sp\|P01877\|IGHA2_HUMAN Immunoglobulin heavy constant alpha 2 OS=Homo sapiens OX=9606 GN=IGHA2 PE=1 SV=4 | 576 | 20201113_15413_1ng_glyco_hcd203040_compound15.25732.25732.2 | controllerType=0 controllerNumber=1 scan=25732 | 51.8104 |
| 23820:4 | 14 | K.VFPLSLDSTPQDGNVVVAC[+57.021]LVQGFFPQEPLSVTWSESGQN[+2117.777]VTAI | HexNAc(5)Hex(5)Fuc(2) | C[+57], N[+2118] | 1724.2954 | 4 | 6894.1597 | 6894.1382 | 0 | 3.1 | 8 | Specific | 340.96 | 340.96 | 340.96 | 8.36 | 38 | >sp\|P01877\|IGHA2_HUMAN Immunoglobulin heavy constant alpha 2 OS=Homo sapiens OX=9606 GN=IGHA2 PE=1 SV=4 | 576 | 20201113_15413_1ng_glyco_hcd203040_compound15.27616.27616.4 | controllerType=0 controllerNumber=1 scan=27616 | 55.3264 |
| 22007:4 | 14 | K.[+42.011]VFPLSLDSTPQDGNVVVAC[+57.021]LVQGFFPQEPLSVTWSESGQN[+1768 | HexNAc(4)Hex(5)Fuc(1) | .[42], C[+57], N[+1769] | 1647.5195 | 4 | 6587.0563 | 6587.0115 | 0 | 6.8 | 8 | Specific | 353.51 | 353.51 | 353.51 | 6.83 | 38 | >sp\|P01877\|IGHA2_HUMAN Immunoglobulin heavy constant alpha 2 OS=Homo sapiens OX=9606 GN=IGHA2 PE=1 SV=4 | 576 | 20201113_15413_1ng_glyco_hcd203040_compound15.25730.25730.2 | controllerType=0 controllerNumber=1 scan=25730 | 51.8043 |
| 23733:5 | 14 | K.VFPLSLDSTPQDGNVVVAC[+57.021]LVQGFFPQEPLSVTWSESGQN[+2262.814]VTAI | HexNAc(5)Hex(5)Fuc(1)NeuAc(1) | C[+57], N[+2263] | 1408.6546 | 5 | 7039.2438 | 7039.1757 | 0 | 9.7 | 8 | Specific | 545.41 | 545.41 | 545.41 | 9.83 | 38 | >sp\|P01877\|IGHA2_HUMAN Immunoglobulin heavy constant alpha 2 OS=Homo sapiens OX=9606 GN=IGHA2 PE=1 SV=4 | 576 | 20201113_15413_1ng_glyco_hcd203040_compound15.27525.27525.5 | controllerType=0 controllerNumber=1 scan=27525 | 55.1336 |
| 22032:4 | 14 | K.VFPLSLDSTPQDGNVVVAC[+57.021]LVQGFFPQEPLSVTWSESGQN[+1809.666]VTAI | HexNAc(5)Hex(4)Fuc(1) | C[+57], N[+1810] | 1647.272 | 4 | 6586.0661 | 6586.0275 | 0 | 5.9 | 8 | Specific | 692.32 | 616.89 | 616.89 | 13.69 | 38 | >sp\|P01877\|IGHA2_HUMAN Immunoglobulin heavy constant alpha 2 OS=Homo sapiens OX=9606 GN=IGHA2 PE=1 SV=4 | 576 | 20201113_15413_1ng_glyco_hcd203040_compound15.25756.25756.4 | controllerType=0 controllerNumber=1 scan=25756 | 51.8584 |
| 22034:4 | 14 | K.VFPLSLDSTPQDGNVVVAC[+57.021]LVQGFFPQEPLSVTWSESGQN[+1809.666]VTAI | HexNAc(5)Hex(4)Fuc(1) | C[+57], N[+1810] | 1647.272 | 4 | 6586.0661 | 6586.0275 | 0 | 5.9 | 8 | Specific | 710.71 | 680.95 | 680.95 | 14.69 | 38 | >sp\|P01877\|IGHA2_HUMAN Immunoglobulin heavy constant alpha 2 OS=Homo sapiens OX=9606 GN=IGHA2 PE=1 SV=4 | 576 | 20201113_15413_1ng_glyco_hcd203040_compound15.25758.25758.4 | controllerType=0 controllerNumber=1 scan=25758 | 51.8651 |
| 01103:3 | 14 | K.HYTN[+1647.613]SSQDVTVPC[+57.021]R.V | HexNAc(5)Hex(3)Fuc(1) | C[+57], N[+1648] | 1104.4623 | 3 | 3311.3724 | 3311.3625 | 0 | 3 | 89 | Specific | 292.79 | 292.79 | 292.79 | 6.69 | 38 | >sp\|P01877\|IGHA2_HUMAN Immunoglobulin heavy constant alpha 2 OS=Homo sapiens OX=9606 GN=IGHA2 PE=1 SV=4 | 576 | 20201113_15413_1ng_glyco_hcd203040_compound15.3974.3974.2 | controllerType=0 controllerNumber=1 scan=3974 | 12.983 |
| 01031:3 | 14 | K.HYTN[+1971.719]SSQDVTVPC[+57.021]R.V | HexNAc(5)Hex(5)Fuc(1) | C[+57], N[+1972] | 1212.4982 | 3 | 3635.4801 | 3635.4682 | 0 | 3.3 | 89 | Specific | 355 | 273.36 | 273.36 | 6.94 | 38 | >sp\|P01877\|IGHA2_HUMAN Immunoglobulin heavy constant alpha 2 OS=Homo sapiens OX=9606 GN=IGHA2 PE=1 SV=4 | 576 | 20201113_15413_1ng_glyco_hcd203040_compound15.3899.3899.3 | controllerType=0 controllerNumber=1 scan=3899 | 12.8423 |
| 01106:3 | 14 | K.HYTN[+1647.613]SSQDVTVPC[+57.021]R.V | HexNAc(5)Hex(3)Fuc(1) | C[+57], N[+1648] | 1104.4623 | 3 | 3311.3724 | 3311.3625 | 0 | 3 | 89 | Specific | 367.23 | 367.23 | 367.23 | 7.49 | 38 | >sp\|P01877\|IGHA2_HUMAN Immunoglobulin heavy constant alpha 2 OS=Homo sapiens OX=9606 GN=IGHA2 PE=1 SV=4 | 576 | 20201113_15413_1ng_glyco_hcd203040_compound15.3977.3977.2 | controllerType=0 controllerNumber=1 scan=3977 | 12.9888 |
| 01029:3 | 14 | K.HYTN[+1971.719]SSQDVTVPC[+57.021]R.V | HexNAc(5)Hex(5)Fuc(1) | C[+57], N[+1972] | 1212.4982 | 3 | 3635.4801 | 3635.4682 | 0 | 3.3 | 89 | Specific | 420.79 | 284.81 | 284.81 | 7.23 | 38 | >sp\|P01877\|IGHA2_HUMAN Immunoglobulin heavy constant alpha 2 OS=Homo sapiens OX=9606 GN=IGHA2 PE=1 SV=4 | 576 | 20201113_15413_1ng_glyco_hcd203040_compound15.3897.3897.3 | controllerType=0 controllerNumber=1 scan=3897 | 12.8383 |
| 01032:3 | 14 | K.HYTN[+1809.666]SSQDVTVPC[+57.021]R.V | HexNAc(5)Hex(4)Fuc(1) | C[+57], N[+1810] | 1158.4815 | 3 | 3473.43 | 3473.4154 | 0 | 4.2 | 89 | Specific | 509.25 | 448.69 | 448.69 | 9.39 | 38 | >sp\|P01877\|IGHA2_HUMAN Immunoglobulin heavy constant alpha 2 OS=Homo sapiens OX=9606 GN=IGHA2 PE=1 SV=4 | 576 | 20201113_15413_1ng_glyco_hcd203040_compound15.3900.3900.3 | controllerType=0 controllerNumber=1 scan=3900 | 12.8443 |
| 01034:3 | 14 | K.HYTN[+1809.666]SSQDVTVPC[+57.021]R.V | HexNAc(5)Hex(4)Fuc(1) | C[+57], N[+1810] | 1158.4815 | 3 | 3473.43 | 3473.4154 | 0 | 4.2 | 89 | Specific | 633.51 | 532.58 | 532.58 | 11.91 | 38 | >sp\|P01877\|IGHA2_HUMAN Immunoglobulin heavy constant alpha 2 OS=Homo sapiens OX=9606 GN=IGHA2 PE=1 SV=4 | 576 | 20201113_15413_1ng_glyco_hcd203040_compound15.3902.3902.3 | controllerType=0 controllerNumber=1 scan=3902 | 12.8481 |
| 01064:3 | 14 | K.HYTN[+1647.613]SSQDVTVPC[+57.021]R.V | HexNAc(5)Hex(3)Fuc(1) | C[+57], N[+1648] | 1104.4612 | 3 | 3311.3691 | 3311.3625 | 0 | 2 | 89 | Specific | 707.88 | 592.6 | 592.6 | 13.97 | 38 | >sp\|P01877\|IGHA2_HUMAN Immunoglobulin heavy constant alpha 2 OS=Homo sapiens OX=9606 GN=IGHA2 PE=1 SV=4 | 576 | 20201113_15413_1ng_glyco_hcd203040_compound15.3934.3934.3 | controllerType=0 controllerNumber=1 scan=3934 | 12.903 |

Table S4. Cont.

| Query #:z | Protein Rank | Glycopeptide | Glycan | Modification Type(s) | Observed m/z | z | Observed (M+H) | Calc. mass (M+H) | Off-by-x error | Mass error (ppm) | StartPosition | Cleavage | Score | Delta | Delta Mod | \|Log Prob\| | # of unique peptides | Protein Name | Protein DB number | Comment | Scan # | Scan Time |
|---|---|---|---|---|---|---|---|---|---|---|---|---|---|---|---|---|---|---|---|---|---|---|
| 01066:3 | 14 | K.HYTN[+1647.613]SSQDVTVPC[+57.021]R.V | HexNAc(5)Hex(3)Fuc(1) | C[+57], N[+1648] | 1104.4612 | 3 | 3311.3691 | 3311.3625 | 0 | 2 | 89 | Specific | 756.24 | 610.06 | 610.06 | 14.97 | 38 | >sp\|P01877\|IGHA2_HUMAN Immunoglobulin heavy constant alpha 2 OS=Homo sapiens OX=9606 GN=IGHA2 PE=1 SV=4 | 576 | 20201113_15413_1ng_glyco_hcd203040_compound15.3936.3936.3 | controllerType=0 controllerNumber=1 scan=3936 | 12.9083 |
| 02697:2 | 14 | K.TPLTAN[+1606.587]ITK.S | HexNAc(4)Hex(4)Fuc(1) | N[+1607] | 1283.0778 | 2 | 2565.1483 | 2565.1435 | 0 | 1.9 | 200 | Specific | 210.86 | 61.95 | 61.95 | 3.14 | 38 | >sp\|P01877\|IGHA2_HUMAN Immunoglobulin heavy constant alpha 2 OS=Homo sapiens OX=9606 GN=IGHA2 PE=1 SV=4 | 576 | 20201113_15413_1ng_glyco_hcd203040_compound15.5629.5629.2 | controllerType=0 controllerNumber=1 scan=5629 | 15.9419 |
| 02690:3 | 14 | K.TPLTAN[+1971.719]ITK.S | HexNAc(5)Hex(5)Fuc(1) | N[+1972] | 977.4326 | 3 | 2930.2832 | 2930.2757 | 0 | 2.6 | 200 | Specific | 226.17 | 202.98 | 202.98 | 5.47 | 38 | >sp\|P01877\|IGHA2_HUMAN Immunoglobulin heavy constant alpha 2 OS=Homo sapiens OX=9606 GN=IGHA2 PE=1 SV=4 | 576 | 20201113_15413_1ng_glyco_hcd203040_compound15.5622.5622.3 | controllerType=0 controllerNumber=1 scan=5622 | 15.9293 |
| 02692:3 | 14 | K.TPLTAN[+1971.719]ITK.S | HexNAc(5)Hex(5)Fuc(1) | N[+1972] | 977.4326 | 3 | 2930.2832 | 2930.2757 | 0 | 2.6 | 200 | Specific | 319.62 | 187.83 | 187.83 | 5.6 | 38 | >sp\|P01877\|IGHA2_HUMAN Immunoglobulin heavy constant alpha 2 OS=Homo sapiens OX=9606 GN=IGHA2 PE=1 SV=4 | 576 | 20201113_15413_1ng_glyco_hcd203040_compound15.5624.5624.3 | controllerType=0 controllerNumber=1 scan=5624 | 15.9339 |
| 02741:3 | 14 | K.TPLTAN[+1809.666]ITK.S | HexNAc(5)Hex(4)Fuc(1) | N[+1810] | 923.4158 | 3 | 2768.2328 | 2768.2228 | 0 | 3.6 | 200 | Specific | 353.43 | 89.29 | 89.29 | 4.86 | 38 | >sp\|P01877\|IGHA2_HUMAN Immunoglobulin heavy constant alpha 2 OS=Homo sapiens OX=9606 GN=IGHA2 PE=1 SV=4 | 576 | 20201113_15413_1ng_glyco_hcd203040_compound15.5675.5675.3 | controllerType=0 controllerNumber=1 scan=5675 | 16.026 |
| 02817:2 | 14 | K.TPLTAN[+1501.555]ITK.S | HexNAc(5)Hex(3) | N[+1502] | 1230.5635 | 2 | 2460.1198 | 2460.1121 | 0 | 3.1 | 200 | Specific | 359.81 | 359.81 | 359.81 | 7.26 | 38 | >sp\|P01877\|IGHA2_HUMAN Immunoglobulin heavy constant alpha 2 OS=Homo sapiens OX=9606 GN=IGHA2 PE=1 SV=4 | 576 | 20201113_15413_1ng_glyco_hcd203040_compound15.5754.5754.2 | controllerType=0 controllerNumber=1 scan=5754 | 16.1684 |
| 02813:3 | 14 | K.TPLTAN[+1647.613]ITK.S | HexNAc(5)Hex(3)Fuc(1) | N[+1648] | 869.3974 | 3 | 2606.1777 | 2606.17 | 0 | 3 | 200 | Specific | 371.56 | 165.14 | 165.14 | 5.82 | 38 | >sp\|P01877\|IGHA2_HUMAN Immunoglobulin heavy constant alpha 2 OS=Homo sapiens OX=9606 GN=IGHA2 PE=1 SV=4 | 576 | 20201113_15413_1ng_glyco_hcd203040_compound15.5750.5750.3 | controllerType=0 controllerNumber=1 scan=5750 | 16.1609 |
| 04513:3 | 14 | K.TPLTAN[+2262.814]ITK.S | HexNAc(5)Hex(5)Fuc(1)NeuAc(1) | N[+2263] | 1074.465 | 3 | 3221.3804 | 3221.3711 | 0 | 2.9 | 200 | Specific | 382.61 | 304.18 | 304.18 | 6.94 | 38 | >sp\|P01877\|IGHA2_HUMAN Immunoglobulin heavy constant alpha 2 OS=Homo sapiens OX=9606 GN=IGHA2 PE=1 SV=4 | 576 | 20201113_15413_1ng_glyco_hcd203040_compound15.7513.7513.3 | controllerType=0 controllerNumber=1 scan=7513 | 19.1224 |
| 02819:2 | 14 | K.TPLTAN[+1501.555]ITK.S | HexNAc(5)Hex(3) | N[+1502] | 1230.5635 | 2 | 2460.1198 | 2460.1121 | 0 | 3.1 | 200 | Specific | 445.63 | 255.39 | 255.39 | 7 | 38 | >sp\|P01877\|IGHA2_HUMAN Immunoglobulin heavy constant alpha 2 OS=Homo sapiens OX=9606 GN=IGHA2 PE=1 SV=4 | 576 | 20201113_15413_1ng_glyco_hcd203040_compound15.5756.5756.2 | controllerType=0 controllerNumber=1 scan=5756 | 16.1725 |
| 02743:3 | 14 | K.TPLTAN[+1809.666]ITK.S | HexNAc(5)Hex(4)Fuc(1) | N[+1810] | 923.4158 | 3 | 2768.2328 | 2768.2228 | 0 | 3.6 | 200 | Specific | 519.44 | 258.69 | 258.69 | 7.23 | 38 | >sp\|P01877\|IGHA2_HUMAN Immunoglobulin heavy constant alpha 2 OS=Homo sapiens OX=9606 GN=IGHA2 PE=1 SV=4 | 576 | 20201113_15413_1ng_glyco_hcd203040_compound15.5677.5677.3 | controllerType=0 controllerNumber=1 scan=5677 | 16.0309 |
| 02749:2 | 14 | K.TPLTAN[+1444.534]ITK.S | HexNAc(4)Hex(3)Fuc(1) | N[+1445] | 1202.0541 | 2 | 2403.101 | 2403.0906 | 0 | 4.3 | 200 | Specific | 519.77 | 332.86 | 332.86 | 7.55 | 38 | >sp\|P01877\|IGHA2_HUMAN Immunoglobulin heavy constant alpha 2 OS=Homo sapiens OX=9606 GN=IGHA2 PE=1 SV=4 | 576 | 20201113_15413_1ng_glyco_hcd203040_compound15.5683.5683.2 | controllerType=0 controllerNumber=1 scan=5683 | 16.0402 |
| 02815:3 | 14 | K.TPLTAN[+1647.613]ITK.S | HexNAc(5)Hex(3)Fuc(1) | N[+1648] | 869.3974 | 3 | 2606.1777 | 2606.17 | 0 | 3 | 200 | Specific | 521.94 | 248.23 | 248.23 | 7.04 | 38 | >sp\|P01877\|IGHA2_HUMAN Immunoglobulin heavy constant alpha 2 OS=Homo sapiens OX=9606 GN=IGHA2 PE=1 SV=4 | 576 | 20201113_15413_1ng_glyco_hcd203040_compound15.5752.5752.3 | controllerType=0 controllerNumber=1 scan=5752 | 16.1653 |
| 02736:2 | 14 | K.TPLTAN[+1809.666]ITK.S | HexNAc(5)Hex(4)Fuc(1) | N[+1810] | 1384.6192 | 2 | 2768.2311 | 2768.2228 | 0 | 3 | 200 | Specific | 571.3 | 275.7 | 275.7 | 7.77 | 38 | >sp\|P01877\|IGHA2_HUMAN Immunoglobulin heavy constant alpha 2 OS=Homo sapiens OX=9606 GN=IGHA2 PE=1 SV=4 | 576 | 20201113_15413_1ng_glyco_hcd203040_compound15.5670.5670.2 | controllerType=0 controllerNumber=1 scan=5670 | 16.0175 |
| 02734:2 | 14 | K.TPLTAN[+1809.666]ITK.S | HexNAc(5)Hex(4)Fuc(1) | N[+1810] | 1384.6192 | 2 | 2768.2311 | 2768.2228 | 0 | 3 | 200 | Specific | 572.61 | 319.06 | 319.06 | 7.87 | 38 | >sp\|P01877\|IGHA2_HUMAN Immunoglobulin heavy constant alpha 2 OS=Homo sapiens OX=9606 GN=IGHA2 PE=1 SV=4 | 576 | 20201113_15413_1ng_glyco_hcd203040_compound15.5668.5668.2 | controllerType=0 controllerNumber=1 scan=5668 | 16.0132 |
| 02751:2 | 14 | K.TPLTAN[+1444.534]ITK.S | HexNAc(4)Hex(3)Fuc(1) | N[+1445] | 1202.0541 | 2 | 2403.101 | 2403.0906 | 0 | 4.3 | 200 | Specific | 575.89 | 314.01 | 314.01 | 8.96 | 38 | >sp\|P01877\|IGHA2_HUMAN Immunoglobulin heavy constant alpha 2 OS=Homo sapiens OX=9606 GN=IGHA2 PE=1 SV=4 | 576 | 20201113_15413_1ng_glyco_hcd203040_compound15.5685.5685.2 | controllerType=0 controllerNumber=1 scan=5685 | 16.044 |
| 02800:2 | 14 | K.TPLTAN[+1647.613]ITK.S | HexNAc(5)Hex(3)Fuc(1) | N[+1648] | 1303.5909 | 2 | 2606.1744 | 2606.17 | 0 | 1.7 | 200 | Specific | 591.45 | 311 | 311 | 9.02 | 38 | >sp\|P01877\|IGHA2_HUMAN Immunoglobulin heavy constant alpha 2 OS=Homo sapiens OX=9606 GN=IGHA2 PE=1 SV=4 | 576 | 20201113_15413_1ng_glyco_hcd203040_compound15.5736.5736.2 | controllerType=0 controllerNumber=1 scan=5736 | 16.1354 |
| 02798:2 | 14 | K.TPLTAN[+1647.613]ITK.S | HexNAc(5)Hex(3)Fuc(1) | N[+1648] | 1303.5909 | 2 | 2606.1744 | 2606.17 | 0 | 1.7 | 200 | Specific | 603.71 | 334 | 334 | 10.13 | 38 | >sp\|P01877\|IGHA2_HUMAN Immunoglobulin heavy constant alpha 2 OS=Homo sapiens OX=9606 GN=IGHA2 PE=1 SV=4 | 576 | 20201113_15413_1ng_glyco_hcd203040_compound15.5734.5734.2 | controllerType=0 controllerNumber=1 scan=5734 | 16.1285 |
| 10137:3 | 14 | R.MAGKPTHIN[+1501.555]VSVVMAEADGTC[+57.021]Y.- | HexNAc(5)Hex(3) | C[+57], N[+1502] | 1284.8875 | 3 | 3852.648 | 3852.6382 | 0 | 2.5 | 319 | CRagged | 222.81 | 191.95 | 191.95 | 4.57 | 38 | >sp\|P01877\|IGHA2_HUMAN Immunoglobulin heavy constant alpha 2 OS=Homo sapiens OX=9606 GN=IGHA2 PE=1 SV=4 | 576 | 20201113_15413_1ng_glyco_hcd203040_compound15.13358.13358.3 | controllerType=0 controllerNumber=1 scan=13358 | 28.8989 |
| 10053:3 | 14 | R.MAGKPTHIN[+1809.666]VSVVMAEADGTC[+57.021]Y.- | HexNAc(5)Hex(4)Fuc(1) | C[+57], N[+1810] | 1387.5928 | 3 | 4160.7639 | 4160.7489 | 0 | 3.6 | 319 | CRagged | 259.07 | 259.07 | 247.88 | 6.66 | 38 | >sp\|P01877\|IGHA2_HUMAN Immunoglobulin heavy constant alpha 2 OS=Homo sapiens OX=9606 GN=IGHA2 PE=1 SV=4 | 576 | 20201113_15413_1ng_glyco_hcd203040_compound15.13271.13271.3 | controllerType=0 controllerNumber=1 scan=13271 | 28.7478 |
| 11738:3 | 14 | R.MAGKPTHIN[+1913.677]VSVVMAEADGTC[+57.021]Y.- | HexNAc(4)Hex(5)NeuAc(1) | C[+57], N[+1914] | 1422.2647 | 3 | 4264.7796 | 4264.7599 | 0 | 4.6 | 319 | CRagged | 279.55 | 279.55 | 279.55 | 6.83 | 38 | >sp\|P01877\|IGHA2_HUMAN Immunoglobulin heavy constant alpha 2 OS=Homo sapiens OX=9606 GN=IGHA2 PE=1 SV=4 | 576 | 20201113_15413_1ng_glyco_hcd203040_compound15.15015.15015.3 | controllerType=0 controllerNumber=1 scan=15015 | 31.71 |
| 10139:3 | 14 | R.MAGKPTHIN[+1501.555]VSVVMAEADGTC[+57.021]Y.- | HexNAc(5)Hex(3) | C[+57], N[+1502] | 1284.8875 | 3 | 3852.648 | 3852.6382 | 0 | 2.5 | 319 | CRagged | 312.1 | 312.1 | 312.1 | 7.1 | 38 | >sp\|P01877\|IGHA2_HUMAN Immunoglobulin heavy constant alpha 2 OS=Homo sapiens OX=9606 GN=IGHA2 PE=1 SV=4 | 576 | 20201113_15413_1ng_glyco_hcd203040_compound15.13360.13360.3 | controllerType=0 controllerNumber=1 scan=13360 | 28.9029 |
| 10051:3 | 14 | R.MAGKPTHIN[+1809.666]VSVVMAEADGTC[+57.021]Y.- | HexNAc(5)Hex(4)Fuc(1) | C[+57], N[+1810] | 1387.5928 | 3 | 4160.7639 | 4160.7489 | 0 | 3.6 | 319 | CRagged | 351.11 | 351.11 | 307.56 | 7.73 | 38 | >sp\|P01877\|IGHA2_HUMAN Immunoglobulin heavy constant alpha 2 OS=Homo sapiens OX=9606 GN=IGHA2 PE=1 SV=4 | 576 | 20201113_15413_1ng_glyco_hcd203040_compound15.13269.13269.3 | controllerType=0 controllerNumber=1 scan=13269 | 28.7418 |
| 10000:3 | 14 | R.MAGKPTHIN[+1540.529]VSVVMAEADGTC[+57.021]Y.- | HexNAc(2)Hex(7) | C[+57], N[+1541] | 1297.8802 | 3 | 3891.626 | 3891.6114 | 0 | 3.8 | 319 | CRagged | 673.46 | 673.46 | 411.56 | 13.27 | 38 | >sp\|P01877\|IGHA2_HUMAN Immunoglobulin heavy constant alpha 2 OS=Homo sapiens OX=9606 GN=IGHA2 PE=1 SV=4 | 576 | 20201113_15413_1ng_glyco_hcd203040_compound15.13216.13216.3 | controllerType=0 controllerNumber=1 scan=13216 | 28.6367 |
| 09952:3 | 14 | R.MAGKPTHIN[+1702.581]VSVVMAEADGTC[+57.021]Y.- | HexNAc(2)Hex(8) | C[+57], N[+1703] | 1351.897 | 3 | 4053.6765 | 4053.6642 | 0 | 3 | 319 | CRagged | 723.54 | 723.54 | 423.43 | 14.27 | 38 | >sp\|P01877\|IGHA2_HUMAN Immunoglobulin heavy constant alpha 2 OS=Homo sapiens OX=9606 GN=IGHA2 PE=1 SV=4 | 576 | 20201113_15413_1ng_glyco_hcd203040_compound15.13166.13166.3 | controllerType=0 controllerNumber=1 scan=13166 | 28.5307 |
| 09974:3 | 14 | R.MAGKPTHIN[+1540.529]VSVVMAEADGTC[+57.021]Y.- | HexNAc(2)Hex(7) | C[+57], N[+1541] | 1297.8797 | 3 | 3891.6245 | 3891.6114 | 0 | 3.4 | 319 | CRagged | 833.68 | 669.47 | 466.85 | 16.27 | 38 | >sp\|P01877\|IGHA2_HUMAN Immunoglobulin heavy constant alpha 2 OS=Homo sapiens OX=9606 GN=IGHA2 PE=1 SV=4 | 576 | 20201113_15413_1ng_glyco_hcd203040_compound15.13188.13188.3 | controllerType=0 controllerNumber=1 scan=13188 | 28.5695 |
| 09954:3 | 14 | R.MAGKPTHIN[+1702.581]VSVVMAEADGTC[+57.021]Y.- | HexNAc(2)Hex(8) | C[+57], N[+1703] | 1351.897 | 3 | 4053.6765 | 4053.6642 | 0 | 3 | 319 | CRagged | 838.66 | 706.82 | 450.1 | 16.27 | 38 | >sp\|P01877\|IGHA2_HUMAN Immunoglobulin heavy constant alpha 2 OS=Homo sapiens OX=9606 GN=IGHA2 PE=1 SV=4 | 576 | 20201113_15413_1ng_glyco_hcd203040_compound15.13168.13168.3 | controllerType=0 controllerNumber=1 scan=13168 | 28.5358 |
| 09972:3 | 14 | R.MAGKPTHIN[+1540.529]VSVVMAEADGTC[+57.021]Y.- | HexNAc(2)Hex(7) | C[+57], N[+1541] | 1297.8797 | 3 | 3891.6245 | 3891.6114 | 0 | 3.4 | 319 | CRagged | 865.8 | 689.41 | 497.9 | 17.27 | 38 | >sp\|P01877\|IGHA2_HUMAN Immunoglobulin heavy constant alpha 2 OS=Homo sapiens OX=9606 GN=IGHA2 PE=1 SV=4 | 576 | 20201113_15413_1ng_glyco_hcd203040_compound15.13186.13186.3 | controllerType=0 controllerNumber=1 scan=13186 | 28.5646 |
| 10003:3 | 14 | R.MAGKPTHIN[+1540.529]VSVVMAEADGTC[+57.021]Y.- | HexNAc(2)Hex(7) | C[+57], N[+1541] | 1297.8802 | 3 | 3891.626 | 3891.6114 | 0 | 3.8 | 319 | CRagged | 884.88 | 884.88 | 482.02 | 17.27 | 38 | >sp\|P01877\|IGHA2_HUMAN Immunoglobulin heavy constant alpha 2 OS=Homo sapiens OX=9606 GN=IGHA2 PE=1 SV=4 | 576 | 20201113_15413_1ng_glyco_hcd203040_compound15.13219.13219.3 | controllerType=0 controllerNumber=1 scan=13219 | 28.6437 |
| 10024:3 | 14 | R.MAGKPTHIN[+1378.476]VSVVMAEADGTC[+57.021]Y.- | HexNAc(2)Hex(6) | C[+57], N[+1378] | 1243.8626 | 3 | 3729.5733 | 3729.5585 | 0 | 3.9 | 319 | CRagged | 904.43 | 795.8 | 528.06 | 18.16 | 38 | >sp\|P01877\|IGHA2_HUMAN Immunoglobulin heavy constant alpha 2 OS=Homo sapiens OX=9606 GN=IGHA2 PE=1 SV=4 | 576 | 20201113_15413_1ng_glyco_hcd203040_compound15.13241.13241.3 | controllerType=0 controllerNumber=1 scan=13241 | 28.6915 |
| 10074:3 | 14 | R.MAGKPTHIN[+1216.423]VSVVMAEADGTC[+57.021]Y.- | HexNAc(2)Hex(5) | C[+57], N[+1216] | 1189.8441 | 3 | 3567.5176 | 3567.5057 | 0 | 3.3 | 319 | CRagged | 924.48 | 903.93 | 538.9 | 18.27 | 38 | >sp\|P01877\|IGHA2_HUMAN Immunoglobulin heavy constant alpha 2 OS=Homo sapiens OX=9606 GN=IGHA2 PE=1 SV=4 | 576 | 20201113_15413_1ng_glyco_hcd203040_compound15.13293.13293.3 | controllerType=0 controllerNumber=1 scan=13293 | 28.7862 |
| 10076:3 | 14 | R.MAGKPTHIN[+1216.423]VSVVMAEADGTC[+57.021]Y.- | HexNAc(2)Hex(5) | C[+57], N[+1216] | 1189.8441 | 3 | 3567.5176 | 3567.5057 | 0 | 3.3 | 319 | CRagged | 952.42 | 952.42 | 582.31 | 19.27 | 38 | >sp\|P01877\|IGHA2_HUMAN Immunoglobulin heavy constant alpha 2 OS=Homo sapiens OX=9606 GN=IGHA2 PE=1 SV=4 | 576 | 20201113_15413_1ng_glyco_hcd203040_compound15.13295.13295.3 | controllerType=0 controllerNumber=1 scan=13295 | 28.7922 |
| 10022:3 | 14 | R.MAGKPTHIN[+1378.476]VSVVMAEADGTC[+57.021]Y.- | HexNAc(2)Hex(6) | C[+57], N[+1378] | 1243.8626 | 3 | 3729.5733 | 3729.5585 | 0 | 3.9 | 319 | CRagged | 1016.86 | 1013.97 | 616.1 | 20.16 | 38 | >sp\|P01877\|IGHA2_HUMAN Immunoglobulin heavy constant alpha 2 OS=Homo sapiens OX=9606 GN=IGHA2 PE=1 SV=4 | 576 | 20201113_15413_1ng_glyco_hcd203040_compound15.13239.13239.3 | controllerType=0 controllerNumber=1 scan=13239 | 28.6833 |
| 24170:4 | 15 | R.QLAHQSN[+2204.772]STNIFFSPVSIATAFAMLSLGTK.A | HexNAc(4)Hex(5)NeuAc(2) | N[+2205] | 1347.36 | 4 | 5386.4181 | 5386.4075 | 0 | 2 | 64 | Specific | 215.63 | 215.63 | 215.63 | 5.01 | 33 | >sp\|P01009\|A1AT_HUMAN Alpha-1-antitrypsin OS=Homo sapiens OX=9606 GN=SERPINA1 PE=1 SV=3 | 642 | 20201113_15413_1ng_glyco_hcd203040_compound15.27980.27980.4 | controllerType=0 controllerNumber=1 scan=27980 | 55.9758 |
| 27004:4 | 15 | R.QLAHQSN[+2861.000]STNIFFSPVSIATAFAMLSLGTK.A | HexNAc(5)Hex(6)NeuAc(3) | N[+2861] | 1511.4205 | 4 | 6042.6603 | 6042.6351 | 0 | 4.2 | 64 | Specific | 345.71 | 345.71 | 345.71 | 7.43 | 33 | >sp\|P01009\|A1AT_HUMAN Alpha-1-antitrypsin OS=Homo sapiens OX=9606 GN=SERPINA1 PE=1 SV=3 | 642 | 20201113_15413_1ng_glyco_hcd203040_compound15.30936.30936.4 | controllerType=0 controllerNumber=1 scan=30936 | 61.2512 |
| 27006:4 | 15 | R.QLAHQSN[+2861.000]STNIFFSPVSIATAFAMLSLGTK.A | HexNAc(5)Hex(6)NeuAc(3) | N[+2861] | 1511.4205 | 4 | 6042.6603 | 6042.6351 | 0 | 4.2 | 64 | Specific | 349.87 | 349.87 | 349.87 | 7.43 | 33 | >sp\|P01009\|A1AT_HUMAN Alpha-1-antitrypsin OS=Homo sapiens OX=9606 GN=SERPINA1 PE=1 SV=3 | 642 | 20201113_15413_1ng_glyco_hcd203040_compound15.30938.30938.4 | controllerType=0 controllerNumber=1 scan=30938 | 61.2547 |
| 24105:4 | 15 | R.QLAHQSN[+2204.772]STNIFFSPVSIATAFAMLSLGTK.A | HexNAc(4)Hex(5)NeuAc(2) | N[+2205] | 1347.3588 | 4 | 5386.4132 | 5386.4075 | 0 | 1.1 | 64 | Specific | 863.41 | 797.98 | 553.2 | 17.1 | 33 | >sp\|P01009\|A1AT_HUMAN Alpha-1-antitrypsin OS=Homo sapiens OX=9606 GN=SERPINA1 PE=1 SV=3 | 642 | 20201113_15413_1ng_glyco_hcd203040_compound15.27912.27912.4 | controllerType=0 controllerNumber=1 scan=27912 | 55.8546 |
| 24172:4 | 15 | R.QLAHQSN[+2204.772]STNIFFSPVSIATAFAMLSLGTK.A | HexNAc(4)Hex(5)NeuAc(2) | N[+2205] | 1347.36 | 4 | 5386.4181 | 5386.4075 | 0 | 2 | 64 | Specific | 867.44 | 813.19 | 565.25 | 17.34 | 33 | >sp\|P01009\|A1AT_HUMAN Alpha-1-antitrypsin OS=Homo sapiens OX=9606 GN=SERPINA1 PE=1 SV=3 | 642 | 20201113_15413_1ng_glyco_hcd203040_compound15.27982.27982.4 | controllerType=0 controllerNumber=1 scan=27982 | 55.9794 |
| 24107:4 | 15 | R.QLAHQSN[+2204.772]STNIFFSPVSIATAFAMLSLGTK.A | HexNAc(4)Hex(5)NeuAc(2) | N[+2205] | 1347.3588 | 4 | 5386.4132 | 5386.4075 | 0 | 1.1 | 64 | Specific | 882.21 | 829.84 | 583.78 | 17.1 | 33 | >sp\|P01009\|A1AT_HUMAN Alpha-1-antitrypsin OS=Homo sapiens OX=9606 GN=SERPINA1 PE=1 SV=3 | 642 | 20201113_15413_1ng_glyco_hcd203040_compound15.27914.27914.4 | controllerType=0 controllerNumber=1 scan=27914 | 55.8605 |
| 13890:4 | 19 | K.LKELPGVC[+57.021]N[+2059.735]ETMMALWEEC[+57.021]KPC[+57.021]LK.Q | HexNAc(4)Hex(5)Fuc(1)NeuAc(1) | C[+57]*3, N[+2060] | 1249.7903 | 4 | 4996.1393 | 4996.1196 | 0 | 3.9 | 95 | Specific | 225.68 | 225.68 | 225.68 | 5.6 | 27 | >sp\|P10909\|CLUS_HUMAN Clusterin OS=Homo sapiens OX=9606 GN=CLU PE=1 SV=1 | 1467 | 20201113_15413_1ng_glyco_hcd203040_compound15.17262.17262.4 | controllerType=0 controllerNumber=1 scan=17262 | 35.5446 |
| 13826:4 | 19 | K.LKELPGVC[+57.021]N[+2204.772]ETMMALWEEC[+57.021]KPC[+57.021]LK.Q | HexNAc(4)Hex(5)NeuAc(2) | C[+57]*3, N[+2205] | 1286.0512 | 4 | 5141.1828 | 5141.1572 | 0 | 5 | 95 | Specific | 429.81 | 429.81 | 429.81 | 8.82 | 27 | >sp\|P10909\|CLUS_HUMAN Clusterin OS=Homo sapiens OX=9606 GN=CLU PE=1 SV=1 | 1467 | 20201113_15413_1ng_glyco_hcd203040_compound15.17196.17196.4 | controllerType=0 controllerNumber=1 scan=17196 | 35.4323 |
| 13893:4 | 19 | K.LKELPGVC[+57.021]N[+2059.735]ETMMALWEEC[+57.021]KPC[+57.021]LK.Q | HexNAc(4)Hex(5)Fuc(1)NeuAc(1) | C[+57]*3, N[+2060] | 1249.7903 | 4 | 4996.1393 | 4996.1196 | 0 | 3.9 | 95 | Specific | 634.18 | 515.76 | 515.76 | 11.62 | 27 | >sp\|P10909\|CLUS_HUMAN Clusterin OS=Homo sapiens OX=9606 GN=CLU PE=1 SV=1 | 1467 | 20201113_15413_1ng_glyco_hcd203040_compound15.17265.17265.4 | controllerType=0 controllerNumber=1 scan=17265 | 35.5486 |
| 13828:4 | 19 | K.LKELPGVC[+57.021]N[+2204.772]ETMMALWEEC[+57.021]KPC[+57.021]LK.Q | HexNAc(4)Hex(5)NeuAc(2) | C[+57]*3, N[+2205] | 1286.0512 | 4 | 5141.1828 | 5141.1572 | 0 | 5 | 95 | Specific | 650.79 | 529.24 | 529.24 | 12.62 | 27 | >sp\|P10909\|CLUS_HUMAN Clusterin OS=Homo sapiens OX=9606 GN=CLU PE=1 SV=1 | 1467 | 20201113_15413_1ng_glyco_hcd203040_compound15.17198.17198.4 | controllerType=0 controllerNumber=1 scan=17198 | 35.4364 |
| 17653:3 | 19 | K.ELPGVC[+57.021]N[+2350.830]ETMMALWEEC[+57.021]KPC[+57.021]LK.Q | HexNAc(4)Hex(5)Fuc(1)NeuAc(2) | C[+57]*3, N[+2351] | 1682.6947 | 3 | 5046.0694 | 5046.036 | 0 | 6.6 | 97 | Specific | 425.5 | 368 | 368 | 7.23 | 27 | >sp\|P10909\|CLUS_HUMAN Clusterin OS=Homo sapiens OX=9606 GN=CLU PE=1 SV=1 | 1467 | 20201113_15413_1ng_glyco_hcd203040_compound15.21194.21194.3 | controllerType=0 controllerNumber=1 scan=21194 | 42.7139 |
| 17651:3 | 19 | K.ELPGVC[+57.021]N[+2350.830]ETMMALWEEC[+57.021]KPC[+57.021]LK.Q | HexNAc(4)Hex(5)Fuc(1)NeuAc(2) | C[+57]*3, N[+2351] | 1682.6947 | 3 | 5046.0694 | 5046.036 | 0 | 6.6 | 97 | Specific | 454.14 | 429.34 | 429.34 | 8.54 | 27 | >sp\|P10909\|CLUS_HUMAN Clusterin OS=Homo sapiens OX=9606 GN=CLU PE=1 SV=1 | 1467 | 20201113_15413_1ng_glyco_hcd203040_compound15.21192.21192.3 | controllerType=0 controllerNumber=1 scan=21192 | 42.7092 |
| 23354:4 | 19 | K.MLN[+2350.830]TSSLLEQLNEQFNWVSR.L | HexNAc(4)Hex(5)Fuc(1)NeuAc(2) | N[+2351] | 1190.7627 | 4 | 4760.0291 | 4760.0171 | 0 | 2.5 | 352 | Specific | 326.39 | 254.21 | 254.21 | 5.73 | 27 | >sp\|P10909\|CLUS_HUMAN Clusterin OS=Homo sapiens OX=9606 GN=CLU PE=1 SV=1 | 1467 | 20201113_15413_1ng_glyco_hcd203040_compound15.27132.27132.4 | controllerType=0 controllerNumber=1 scan=27132 | 54.4264 |
| 23351:4 | 19 | K.MLN[+2350.830]TSSLLEQLNEQFNWVSR.L | HexNAc(4)Hex(5)Fuc(1)NeuAc(2) | N[+2351] | 1190.7627 | 4 | 4760.0291 | 4760.0171 | 0 | 2.5 | 352 | Specific | 370.6 | 327.71 | 327.71 | 7.45 | 27 | >sp\|P10909\|CLUS_HUMAN Clusterin OS=Homo sapiens OX=9606 GN=CLU PE=1 SV=1 | 1467 | 20201113_15413_1ng_glyco_hcd203040_compound15.27129.27129.4 | controllerType=0 controllerNumber=1 scan=27129 | 54.4193 |
| 20968:4 | 19 | K.MLN[+2059.735]TSSLLEQLNEQFNWVSR.L | HexNAc(4)Hex(5)Fuc(1)NeuAc(1) | N[+2060] | 1117.9924 | 4 | 4468.9476 | 4468.9217 | 0 | 5.8 | 352 | Specific | 524.51 | 395.25 | 395.25 | 9.16 | 27 | >sp\|P10909\|CLUS_HUMAN Clusterin OS=Homo sapiens OX=9606 GN=CLU PE=1 SV=1 | 1467 | 20201113_15413_1ng_glyco_hcd203040_compound15.24653.24653.4 | controllerType=0 controllerNumber=1 scan=24653 | 49.8617 |
| 20971:4 | 19 | K.MLN[+2059.735]TSSLLEQLNEQFNWVSR.L | HexNAc(4)Hex(5)Fuc(1)NeuAc(1) | N[+2060] | 1117.9924 | 4 | 4468.9476 | 4468.9217 | 0 | 5.8 | 352 | Specific | 605.67 | 550.92 | 550.92 | 11.68 | 27 | >sp\|P10909\|CLUS_HUMAN Clusterin OS=Homo sapiens OX=9606 GN=CLU PE=1 SV=1 | 1467 | 20201113_15413_1ng_glyco_hcd203040_compound15.24656.24656.4 | controllerType=0 controllerNumber=1 scan=24656 | 49.8672 |
| 20960:3 | 19 | K.MLN[+2059.735]TSSLLEQLNEQFNWVSR.L | HexNAc(4)Hex(5)Fuc(1)NeuAc(1) | N[+2060] | 1490.3198 | 3 | 4468.9447 | 4468.9217 | 0 | 5.2 | 352 | Specific | 626.19 | 511.59 | 511.59 | 11.7 | 27 | >sp\|P10909\|CLUS_HUMAN Clusterin OS=Homo sapiens OX=9606 GN=CLU PE=1 SV=1 | 1467 | 20201113_15413_1ng_glyco_hcd203040_compound15.24645.24645.3 | controllerType=0 controllerNumber=1 scan=24645 | 49.8464 |
| 23346:3 | 19 | K.MLN[+2350.830]TSSLLEQLNEQFNWVSR.L | HexNAc(4)Hex(5)Fuc(1)NeuAc(2) | N[+2351] | 1587.3486 | 3 | 4760.0312 | 4760.0171 | 0 | 3 | 352 | Specific | 680.82 | 547.02 | 547.02 | 13.22 | 27 | >sp\|P10909\|CLUS_HUMAN Clusterin OS=Homo sapiens OX=9606 GN=CLU PE=1 SV=1 | 1467 | 20201113_15413_1ng_glyco_hcd203040_compound15.27124.27124.3 | controllerType=0 controllerNumber=1 scan=27124 | 54.4113 |
| 23348:3 | 19 | K.MLN[+2350.830]TSSLLEQLNEQFNWVSR.L | HexNAc(4)Hex(5)Fuc(1)NeuAc(2) | N[+2351] | 1587.3486 | 3 | 4760.0312 | 4760.0171 | 0 | 3 | 352 | Specific | 863.75 | 621.13 | 621.13 | 17.22 | 27 | >sp\|P10909\|CLUS_HUMAN Clusterin OS=Homo sapiens OX=9606 GN=CLU PE=1 SV=1 | 1467 | 20201113_15413_1ng_glyco_hcd203040_compound15.27126.27126.3 | controllerType=0 controllerNumber=1 scan=27126 | 54.4152 |
| 20912:3 | 19 | K.MLN[+2205.793]TSSLLEQLNEQFNWVSR.L | HexNAc(4)Hex(5)Fuc(2)NeuAc(1) | N[+2206] | 1539.0068 | 3 | 4615.0058 | 4614.9796 | 0 | 5.7 | 352 | Specific | 900.58 | 636.46 | 636.46 | 17.7 | 27 | >sp\|P10909\|CLUS_HUMAN Clusterin OS=Homo sapiens OX=9606 GN=CLU PE=1 SV=1 | 1467 | 20201113_15413_1ng_glyco_hcd203040_compound15.24594.24594.3 | controllerType=0 controllerNumber=1 scan=24594 | 49.722 |
| 20910:3 | 19 | K.MLN[+2205.793]TSSLLEQLNEQFNWVSR.L | HexNAc(4)Hex(5)Fuc(2)NeuAc(1) | N[+2206] | 1539.0068 | 3 | 4615.0058 | 4614.9796 | 0 | 5.7 | 352 | Specific | 976.48 | 718.98 | 718.98 | 18.7 | 27 | >sp\|P10909\|CLUS_HUMAN Clusterin OS=Homo sapiens OX=9606 GN=CLU PE=1 SV=1 | 1467 | 20201113_15413_1ng_glyco_hcd203040_compound15.24592.24592.3 | controllerType=0 controllerNumber=1 scan=24592 | 49.715 |
| 20962:3 | 19 | K.MLN[+2059.735]TSSLLEQLNEQFNWVSR.L | HexNAc(4)Hex(5)Fuc(1)NeuAc(1) | N[+2060] | 1490.3198 | 3 | 4468.9447 | 4468.9217 | 0 | 5.2 | 352 | Specific | 983.76 | 725.03 | 725.03 | 18.7 | 27 | >sp\|P10909\|CLUS_HUMAN Clusterin OS=Homo sapiens OX=9606 GN=CLU PE=1 SV=1 | 1467 | 20201113_15413_1ng_glyco_hcd203040_compound15.24647.24647.3 | controllerType=0 controllerNumber=1 scan=24647 | 49.85 |
| 22709:3 | 19 | K.M[+15.995]LN[+2350.830]TSSLLEQLNEQFNWVSR.L | HexNAc(4)Hex(5)Fuc(1)NeuAc(2) | M[+16], N[+2351] | 1592.6878 | 3 | 4776.0489 | 4776.012 | 0 | 7.7 | 352 | Specific | 359.7 | 214.07 | 214.07 | 5.88 | 27 | >sp\|P10909\|CLUS_HUMAN Clusterin OS=Homo sapiens OX=9606 GN=CLU PE=1 SV=1 | 1467 | 20201113_15413_1ng_glyco_hcd203040_compound15.26461.26461.3 | controllerType=0 controllerNumber=1 scan=26461 | 53.1279 |
| 20492:3 | 19 | K.M[+15.995]LN[+2059.735]TSSLLEQLNEQFNWVSR.L | HexNAc(4)Hex(5)Fuc(1)NeuAc(1) | M[+16], N[+2060] | 1495.649 | 3 | 4484.9324 | 4484.9166 | 0 | 3.5 | 352 | Specific | 447.41 | 374.88 | 374.88 | 7.97 | 27 | >sp\|P10909\|CLUS_HUMAN Clusterin OS=Homo sapiens OX=9606 GN=CLU PE=1 SV=1 | 1467 | 20201113_15413_1ng_glyco_hcd203040_compound15.24157.24157.3 | controllerType=0 controllerNumber=1 scan=24157 | 48.8269 |
| 20455:3 | 19 | K.M[+15.995]LN[+2059.735]TSSLLEQLNEQFNWVSR.L | HexNAc(4)Hex(5)Fuc(1)NeuAc(1) | M[+16], N[+2060] | 1495.6529 | 3 | 4484.9441 | 4484.9166 | 0 | 6.1 | 352 | Specific | 549.2 | 416.31 | 416.31 | 9.31 | 27 | >sp\|P10909\|CLUS_HUMAN Clusterin OS=Homo sapiens OX=9606 GN=CLU PE=1 SV=1 | 1467 | 20201113_15413_1ng_glyco_hcd203040_compound15.24119.24119.3 | controllerType=0 controllerNumber=1 scan=24119 | 48.7508 |
| 20493:3 | 19 | K.M[+15.995]LN[+2059.735]TSSLLEQLNEQFNWVSR.L | HexNAc(4)Hex(5)Fuc(1)NeuAc(1) | M[+16], N[+2060] | 1495.649 | 3 | 4484.9324 | 4484.9166 | 0 | 3.5 | 352 | Specific | 553.84 | 406.3 | 406.3 | 10.02 | 27 | >sp\|P10909\|CLUS_HUMAN Clusterin OS=Homo sapiens OX=9606 GN=CLU PE=1 SV=1 | 1467 | 20201113_15413_1ng_glyco_hcd203040_compound15.24158.24158.3 | controllerType=0 controllerNumber=1 scan=24158 | 48.8306 |
| 20457:3 | 19 | K.M[+15.995]LN[+2059.735]TSSLLEQLNEQFNWVSR.L | HexNAc(4)Hex(5)Fuc(1)NeuAc(1) | M[+16], N[+2060] | 1495.6529 | 3 | 4484.9441 | 4484.9166 | 0 | 6.1 | 352 | Specific | 657.23 | 375.34 | 375.34 | 12.7 | 27 | >sp\|P10909\|CLUS_HUMAN Clusterin OS=Homo sapiens OX=9606 GN=CLU PE=1 SV=1 | 1467 | 20201113_15413_1ng_glyco_hcd203040_compound15.24121.24121.3 | controllerType=0 controllerNumber=1 scan=24121 | 48.7546 |
| 22697:4 | 21 | K.DIDGYGGIALPELIC[+57.021]TMFHTSGYDTQAIVENN[+1686.586]ESTEYGLFQISNK. | HexNAc(2)Hex(7)Fuc(1) | C[+57], N[+1687] | 1675.2335 | 4 | 6697.9121 | 6697.9153 | 0 | -0.5 | 33 | Specific | 357.26 | 357.26 | 357.26 | 7.97 | 25 | >sp\|P00709\|LALBA_HUMAN Alpha-lactalbumin OS=Homo sapiens OX=9606 GN=LALBA PE=1 SV=1 | 468 | 20201113_15413_1ng_glyco_hcd203040_compound15.26449.26449.3 | controllerType=0 controllerNumber=1 scan=26449 | 53.1044 |
| 24151:4 | 21 | K.DIDGYGGIALPELIC[+57.021]TMFHTSGYDTQAIVENN[+2336.851]ESTEYGLFQISNK. | HexNAc(6)Hex(6)Fuc(1) | C[+57], N[+2337] | 1837.7965 | 4 | 7348.1641 | 7348.1799 | 0 | -2.2 | 33 | Specific | 487.71 | 470.1 | 16.8 | 9.37 | 25 | >sp\|P00709\|LALBA_HUMAN Alpha-lactalbumin OS=Homo sapiens OX=9606 GN=LALBA PE=1 SV=1 | 468 | 20201113_15413_1ng_glyco_hcd203040_compound15.27960.27960.2 | controllerType=0 controllerNumber=1 scan=27960 | 55.9393 |
| 22699:4 | 21 | K.DIDGYGGIALPELIC[+57.021]TMFHTSGYDTQAIVENN[+1686.586]ESTEYGLFQISNK. | HexNAc(2)Hex(7)Fuc(1) | C[+57], N[+1687] | 1675.2335 | 4 | 6697.9121 | 6697.9153 | 0 | -0.5 | 33 | Specific | 513.1 | 513.1 | 513.1 | 9.98 | 25 | >sp\|P00709\|LALBA_HUMAN Alpha-lactalbumin OS=Homo sapiens OX=9606 GN=LALBA PE=1 SV=1 | 468 | 20201113_15413_1ng_glyco_hcd203040_compound15.26451.26451.3 | controllerType=0 controllerNumber=1 scan=26451 | 53.108 |
| 24153:4 | 21 | K.DIDGYGGIALPELIC[+57.021]TMFHTSGYDTQAIVENN[+2336.851]ESTEYGLFQISNK. | HexNAc(6)Hex(6)Fuc(1) | C[+57], N[+2337] | 1837.7965 | 4 | 7348.1641 | 7348.1799 | 0 | -2.2 | 33 | Specific | 544.13 | 489.95 | 16.8 | 9.64 | 25 | >sp\|P00709\|LALBA_HUMAN Alpha-lactalbumin OS=Homo sapiens OX=9606 GN=LALBA PE=1 SV=1 | 468 | 20201113_15413_1ng_glyco_hcd203040_compound15.27962.27962.2 | controllerType=0 controllerNumber=1 scan=27962 | 55.9431 |
| 09814:4 | 26 | K.SC[+57.021]QHN[+2351.851]GTMYQHGEIFSAHELFPSR.L | HexNAc(4)Hex(5)Fuc(3)NeuAc(1) | C[+57], N[+2352] | 1297.0335 | 4 | 5185.112 | 5185.1077 | 0 | 0.8 | 110 | Specific | 269.99 | 269.99 | 269.99 | 5.78 | 23 | >sp\|Q6WN34\|CRDL2_HUMAN Chordin-like protein 2 OS=Homo sapiens OX=9606 GN=CHRDL2 PE=1 SV=1 | 1189 | 20201113_15413_1ng_glyco_hcd203040_compound15.13022.13022.2 | controllerType=0 controllerNumber=1 scan=13022 | 28.2862 |
| 09824:4 | 26 | K.SC[+57.021]QHN[+2350.830]GTMYQHGEIFSAHELFPSR.L | HexNAc(4)Hex(5)Fuc(1)NeuAc(2) | C[+57], N[+2351] | 1296.7858 | 4 | 5184.1212 | 5184.0873 | 0 | 6.6 | 110 | Specific | 274.2 | 274.2 | 274.2 | 5.29 | 23 | >sp\|Q6WN34\|CRDL2_HUMAN Chordin-like protein 2 OS=Homo sapiens OX=9606 GN=CHRDL2 PE=1 SV=1 | 1189 | 20201113_15413_1ng_glyco_hcd203040_compound15.13033.13033.2 | controllerType=0 controllerNumber=1 scan=13033 | [illegible] |
| 09826:4 | 26 | K.SC[+57.021]QHN[+2350.830]GTMYQHGEIFSAHELFPSR.L | HexNAc(4)Hex(5)Fuc(1)NeuAc(2) | C[+57], N[+2351] | 1296.7858 | 4 | 5184.1212 | 5184.0873 | 0 | 6.6 | 110 | Specific | 309.34 | 258.27 | 258.27 | 5.51 | 23 | >sp\|Q6WN34\|CRDL2_HUMAN Chordin-like protein 2 OS=Homo sapiens OX=9606 GN=CHRDL2 PE=1 SV=1 | 1189 | 20201113_15413_1ng_glyco_hcd203040_compound15.13035.13035.2 | controllerType=0 controllerNumber=1 scan=13035 | [illegible] |
| 09816:4 | 26 | K.SC[+57.021]QHN[+2351.851]GTMYQHGEIFSAHELFPSR.L | HexNAc(4)Hex(5)Fuc(3)NeuAc(1) | C[+57], N[+2352] | 1297.0335 | 4 | 5185.112 | 5185.1077 | 0 | 0.8 | 110 | Specific | 353.39 | 351.56 | 351.56 | 6.85 | 23 | >sp\|Q6WN34\|CRDL2_HUMAN Chordin-like protein 2 OS=Homo sapiens OX=9606 GN=CHRDL2 PE=1 SV=1 | 1189 | 20201113_15413_1ng_glyco_hcd203040_compound15.13024.13024.2 | controllerType=0 controllerNumber=1 scan=13024 | 28.2896 |
| 08289:4 | 26 | K.SC[+57.021]QHN[+2204.772]GTMYQHGEIFSAHELFPSR.L | HexNAc(4)Hex(5)NeuAc(2) | C[+57], N[+2205] | 1260.2693 | 4 | 5038.0553 | 5038.0294 | 0 | 5.2 | 110 | Specific | 404.19 | 404.19 | 404.19 | 8.21 | 23 | >sp\|Q6WN34\|CRDL2_HUMAN Chordin-like protein 2 OS=Homo sapiens OX=9606 GN=CHRDL2 PE=1 SV=1 | 1189 | 20201113_15413_1ng_glyco_hcd203040_compound15.11436.11436.4 | controllerType=0 controllerNumber=1 scan=11436 | 25.651 |
| 09765:5 | 26 | K.SC[+57.021]QHN[+2351.851]GTMYQHGEIFSAHELFPSR.L | HexNAc(4)Hex(5)Fuc(3)NeuAc(1) | C[+57], N[+2352] | 1037.8267 | 5 | 5185.1044 | 5185.1077 | 0 | -0.6 | 110 | Specific | 405.37 | 379.98 | 379.98 | 8.68 | 23 | >sp\|Q6WN34\|CRDL2_HUMAN Chordin-like protein 2 OS=Homo sapiens OX=9606 GN=CHRDL2 PE=1 SV=1 | 1189 | 20201113_15413_1ng_glyco_hcd203040_compound15.12971.12971.5 | controllerType=0 controllerNumber=1 scan=12971 | 28.2051 |
| 08287:4 | 26 | K.SC[+57.021]QHN[+2204.772]GTMYQHGEIFSAHELFPSR.L | HexNAc(4)Hex(5)NeuAc(2) | C[+57], N[+2205] | 1260.2693 | 4 | 5038.0553 | 5038.0294 | 0 | 5.2 | 110 | Specific | 456.81 | 456.81 | 456.81 | 8.24 | 23 | >sp\|Q6WN34\|CRDL2_HUMAN Chordin-like protein 2 OS=Homo sapiens OX=9606 GN=CHRDL2 PE=1 SV=1 | 1189 | 20201113_15413_1ng_glyco_hcd203040_compound15.11434.11434.4 | controllerType=0 controllerNumber=1 scan=11434 | 25.6475 |
| 09703:4 | 26 | K.SC[+57.021]QHN[+2350.830]GTMYQHGEIFSAHELFPSR.L | HexNAc(4)Hex(5)Fuc(1)NeuAc(2) | C[+57], N[+2351] | 1296.7811 | 4 | 5184.1027 | 5184.0873 | 0 | 3 | 110 | Specific | 502.29 | 429.43 | 429.43 | 9.16 | 23 | >sp\|Q6WN34\|CRDL2_HUMAN Chordin-like protein 2 OS=Homo sapiens OX=9606 GN=CHRDL2 PE=1 SV=1 | 1189 | 20201113_15413_1ng_glyco_hcd203040_compound15.12907.12907.4 | controllerType=0 controllerNumber=1 scan=12907 | 28.0987 |
| 09705:4 | 26 | K.SC[+57.021]QHN[+2350.830]GTMYQHGEIFSAHELFPSR.L | HexNAc(4)Hex(5)Fuc(1)NeuAc(2) | C[+57], N[+2351] | 1296.7811 | 4 | 5184.1027 | 5184.0873 | 0 | 3 | 110 | Specific | 553.64 | 458.22 | 458.22 | 9.48 | 23 | >sp\|Q6WN34\|CRDL2_HUMAN Chordin-like protein 2 OS=Homo sapiens OX=9606 GN=CHRDL2 PE=1 SV=1 | 1189 | 20201113_15413_1ng_glyco_hcd203040_compound15.12909.12909.4 | controllerType=0 controllerNumber=1 scan=12909 | 28.1023 |
| 01913:5 | 31 | R.EKQTDEIKDTRN[+2059.735]ESTQNC[+57.021]VVAEPEK.M | HexNAc(4)Hex(5)Fuc(1)NeuAc(1) | C[+57], N[+2060] | 1002.4354 | 5 | 5008.148 | 5008.1251 | 0 | 4.6 | 58 | Specific | 299.53 | 299.53 | 299.53 | 5.75 | 26 | >sp\|P47710\|CASA1_HUMAN Alpha-S1-casein OS=Homo sapiens OX=9606 GN=CSN1S1 PE=1 SV=1 | 989 | 20201113_15413_1ng_glyco_hcd203040_compound15.4816.4816.5 | controllerType=0 controllerNumber=1 scan=4816 | 14.4974 |
| 01911:5 | 31 | R.EKQTDEIKDTRN[+2059.735]ESTQNC[+57.021]VVAEPEK.M | HexNAc(4)Hex(5)Fuc(1)NeuAc(1) | C[+57], N[+2060] | 1002.4354 | 5 | 5008.148 | 5008.1251 | 0 | 4.6 | 58 | Specific | 309.84 | 309.84 | 309.84 | 5.96 | 26 | >sp\|P47710\|CASA1_HUMAN Alpha-S1-casein OS=Homo sapiens OX=9606 GN=CSN1S1 PE=1 SV=1 | 989 | 20201113_15413_1ng_glyco_hcd203040_compound15.4814.4814.5 | controllerType=0 controllerNumber=1 scan=4814 | 14.4935 |
| 01887:4 | 31 | R.EKQTDEIKDTRN[+2059.735]ESTQNC[+57.021]VVAEPEK.M | HexNAc(4)Hex(5)Fuc(1)NeuAc(1) | C[+57], N[+2060] | 1252.792 | 4 | 5008.1462 | 5008.1251 | 0 | 4.2 | 58 | Specific | 367.98 | 367.98 | 367.98 | 5.86 | 26 | >sp\|P47710\|CASA1_HUMAN Alpha-S1-casein OS=Homo sapiens OX=9606 GN=CSN1S1 PE=1 SV=1 | 989 | 20201113_15413_1ng_glyco_hcd203040_compound15.4789.4789.4 | controllerType=0 controllerNumber=1 scan=4789 | 14.4438 |
| 01889:4 | 31 | R.EKQTDEIKDTRN[+2059.735]ESTQNC[+57.021]VVAEPEK.M | HexNAc(4)Hex(5)Fuc(1)NeuAc(1) | C[+57], N[+2060] | 1252.792 | 4 | 5008.1462 | 5008.1251 | 0 | 4.2 | 58 | Specific | 523.03 | 296.33 | 296.33 | 6.04 | 26 | >sp\|P47710\|CASA1_HUMAN Alpha-S1-casein OS=Homo sapiens OX=9606 GN=CSN1S1 PE=1 SV=1 | 989 | 20201113_15413_1ng_glyco_hcd203040_compound15.4791.4791.4 | controllerType=0 controllerNumber=1 scan=4791 | 14.4483 |
| 09199:5 | 31 | K.[+42.011]QTDEIKDTRN[+2018.708]ESTQNC[+57.021]VVAEPEKMESSISSSSEEMSLS | HexNAc(3)Hex(6)Fuc(1)NeuAc(1) | .[42], C[+57], N[+2019] | 1291.1503 | 5 | 6451.7222 | 6451.6996 | 0 | 3.5 | 60 | Specific | 221.63 | 221.63 | 221.63 | 3.76 | 26 | >sp\|P47710\|CASA1_HUMAN Alpha-S1-casein OS=Homo sapiens OX=9606 GN=CSN1S1 PE=1 SV=1 | 989 | 20201113_15413_1ng_glyco_hcd203040_compound15.12384.12384.3 | controllerType=0 controllerNumber=1 scan=12384 | 27.2476 |
| 09214:4 | 31 | K.[+42.011]QTDEIKDTRN[+2018.708]ESTQNC[+57.021]VVAEPEKMESSISSSSEEMSLS | HexNAc(3)Hex(6)Fuc(1)NeuAc(1) | .[42], C[+57], N[+2019] | 1613.6903 | 4 | 6451.7394 | 6451.6996 | 0 | 6.2 | 60 | Specific | 231.93 | 197.02 | 197.02 | 3.44 | 26 | >sp\|P47710\|CASA1_HUMAN Alpha-S1-casein OS=Homo sapiens OX=9606 GN=CSN1S1 PE=1 SV=1 | 989 | 20201113_15413_1ng_glyco_hcd203040_compound15.12400.12400.2 | controllerType=0 controllerNumber=1 scan=12400 | 27.2749 |
| 09247:4 | 31 | K.QTDEIKDTRN[+2059.735]ESTQNC[+57.021]VVAEPEKMESSISSSSEEMSLSK.C | HexNAc(4)Hex(5)Fuc(1)NeuAc(1) | C[+57], N[+2060] | 1613.4387 | 4 | 6450.7331 | 6450.7156 | 0 | 2.7 | 60 | Specific | 312.57 | 228.69 | 228.69 | 4.73 | 26 | >sp\|P47710\|CASA1_HUMAN Alpha-S1-casein OS=Homo sapiens OX=9606 GN=CSN1S1 PE=1 SV=1 | 989 | 20201113_15413_1ng_glyco_hcd203040_compound15.12435.12435.2 | controllerType=0 controllerNumber=1 scan=12435 | 27.3338 |
| 09244:4 | 31 | K.QTDEIKDTRN[+2059.735]ESTQNC[+57.021]VVAEPEKMESSISSSSEEMSLSK.C | HexNAc(4)Hex(5)Fuc(1)NeuAc(1) | C[+57], N[+2060] | 1613.4387 | 4 | 6450.7331 | 6450.7156 | 0 | 2.7 | 60 | Specific | 324.1 | 231.2 | 231.2 | 4.73 | 26 | >sp\|P47710\|CASA1_HUMAN Alpha-S1-casein OS=Homo sapiens OX=9606 GN=CSN1S1 PE=1 SV=1 | 989 | 20201113_15413_1ng_glyco_hcd203040_compound15.12432.12432.2 | controllerType=0 controllerNumber=1 scan=12432 | 27.3292 |
| 09223:5 | 31 | K.QTDEIKDTRN[+2059.735]ESTQNC[+57.021]VVAEPEKMESSISSSSEEMSLSK.C | HexNAc(4)Hex(5)Fuc(1)NeuAc(1) | C[+57], N[+2060] | 1290.9464 | 5 | 6450.703 | 6450.7156 | 0 | -2 | 60 | Specific | 423.51 | 386.88 | 386.88 | 6.87 | 26 | >sp\|P47710\|CASA1_HUMAN Alpha-S1-casein OS=Homo sapiens OX=9606 GN=CSN1S1 PE=1 SV=1 | 989 | 20201113_15413_1ng_glyco_hcd203040_compound15.12409.12409.3 | controllerType=0 controllerNumber=1 scan=12409 | 27.2892 |
| 09225:5 | 31 | K.QTDEIKDTRN[+2059.735]ESTQNC[+57.021]VVAEPEKMESSISSSSEEMSLSK.C | HexNAc(4)Hex(5)Fuc(1)NeuAc(1) | C[+57], N[+2060] | 1290.9464 | 5 | 6450.703 | 6450.7156 | 0 | -2 | 60 | Specific | 449.38 | 387.05 | 387.05 | 7.03 | 26 | >sp\|P47710\|CASA1_HUMAN Alpha-S1-casein OS=Homo sapiens OX=9606 GN=CSN1S1 PE=1 SV=1 | 989 | 20201113_15413_1ng_glyco_hcd203040_compound15.12411.12411.3 | controllerType=0 controllerNumber=1 scan=12411 | 27.293 |
| 14179:3 | 34 | K.VVLHPN[+2204.772]YSQVDIGLIK.L | HexNAc(4)Hex(5)NeuAc(2) | N[+2205] | 1333.939 | 3 | 3999.8025 | 3999.7837 | 0 | 4.7 | 236 | Specific | 373.42 | 373.42 | 373.42 | 7.6 | 20 | >sp\|P00738\|HPT_HUMAN Haptoglobin OS=Homo sapiens OX=9606 GN=HP PE=1 SV=1 | 407 | 20201113_15413_1ng_glyco_hcd203040_compound15.17561.17561.3 | controllerType=0 controllerNumber=1 scan=17561 | 36.0427 |
| 14181:3 | 34 | K.VVLHPN[+2204.772]YSQVDIGLIK.L | HexNAc(4)Hex(5)NeuAc(2) | N[+2205] | 1333.939 | 3 | 3999.8025 | 3999.7837 | 0 | 4.7 | 236 | Specific | 388.55 | 388.55 | 388.55 | 9.06 | 20 | >sp\|P00738\|HPT_HUMAN Haptoglobin OS=Homo sapiens OX=9606 GN=HP PE=1 SV=1 | 407 | 20201113_15413_1ng_glyco_hcd203040_compound15.17563.17563.3 | controllerType=0 controllerNumber=1 scan=17563 | 36.0474 |
| 14154:3 | 34 | K.VVLHPN[+2204.772]YSQVDIGLIK.L | HexNAc(4)Hex(5)NeuAc(2) | N[+2205] | 1333.9341 | 3 | 3999.7879 | 3999.7837 | 0 | 1 | 236 | Specific | 579.27 | 579.27 | 579.27 | 10.95 | 20 | >sp\|P00738\|HPT_HUMAN Haptoglobin OS=Homo sapiens OX=9606 GN=HP PE=1 SV=1 | 407 | 20201113_15413_1ng_glyco_hcd203040_compound15.17535.17535.3 | controllerType=0 controllerNumber=1 scan=17535 | 35.9956 |
| 14156:3 | 34 | K.VVLHPN[+2204.772]YSQVDIGLIK.L | HexNAc(4)Hex(5)NeuAc(2) | N[+2205] | 1333.9341 | 3 | 3999.7879 | 3999.7837 | 0 | 1 | 236 | Specific | 626.18 | 609.09 | 609.09 | 11.95 | 20 | >sp\|P00738\|HPT_HUMAN Haptoglobin OS=Homo sapiens OX=9606 GN=HP PE=1 SV=1 | 407 | 20201113_15413_1ng_glyco_hcd203040_compound15.17537.17537.3 | controllerType=0 controllerNumber=1 scan=17537 | 35.9991 |
| 15011:3 | 46 | K.RNYIVPILWLN[+1864.634]ETGTIGDEK.A | HexNAc(2)Hex(9) | N[+1865] | 1399.2961 | 3 | 4195.8737 | 4195.8685 | 0 | 1.2 | 407 | Specific | 591.76 | 591.76 | 591.76 | 10.97 | 16 | >sp\|P16671\|CD36_HUMAN Platelet glycoprotein 4 OS=Homo sapiens OX=9606 GN=CD36 PE=1 SV=2 | 812 | 20201113_15413_1ng_glyco_hcd203040_compound15.18422.18422.3 | controllerType=0 controllerNumber=1 scan=18422 | 37.4944 |
| 15009:3 | 46 | K.RNYIVPILWLN[+1864.634]ETGTIGDEK.A | HexNAc(2)Hex(9) | N[+1865] | 1399.2961 | 3 | 4195.8737 | 4195.8685 | 0 | 1.2 | 407 | Specific | 696.77 | 664.03 | 664.03 | 12.97 | 16 | >sp\|P16671\|CD36_HUMAN Platelet glycoprotein 4 OS=Homo sapiens OX=9606 GN=CD36 PE=1 SV=2 | 812 | 20201113_15413_1ng_glyco_hcd203040_compound15.18420.18420.3 | controllerType=0 controllerNumber=1 scan=18420 | 37.4906 |
| 16504:3 | 46 | R.NYIVPILWLN[+1864.634]ETGTIGDEKANM[+15.995]FR.S | HexNAc(2)Hex(9) | M[+16], N[+1865] | 1559.0325 | 3 | 4675.083 | 4675.0523 | 0 | 6.6 | 408 | Specific | 221.66 | 221.66 | 221.66 | 3.68 | 16 | >sp\|P16671\|CD36_HUMAN Platelet glycoprotein 4 OS=Homo sapiens OX=9606 GN=CD36 PE=1 SV=2 | 812 | 20201113_15413_1ng_glyco_hcd203040_compound15.19997.19997.3 | controllerType=0 controllerNumber=1 scan=19997 | 40.3029 |
| 09387:3 | 51 | R.IIVPLNNREN[+1809.666]ISDPTSPLR.T | HexNAc(5)Hex(4)Fuc(1) | N[+1810] | 1319.9572 | 3 | 3957.8571 | 3957.8432 | 0 | 3.5 | 62 | Specific | 318.4 | 280.87 | 280.87 | 6.39 | 18 | >sp\|P01591\|IGJ_HUMAN Immunoglobulin J chain OS=Homo sapiens OX=9606 GN=JCHAIN PE=1 SV=4 | 1392 | 20201113_15413_1ng_glyco_hcd203040_compound15.12581.12581.3 | controllerType=0 controllerNumber=1 scan=12581 | 27.564 |
| 09415:3 | 51 | R.IIVPLNNREN[+1809.666]ISDPTSPLR.T | HexNAc(5)Hex(4)Fuc(1) | N[+1810] | 1319.9562 | 3 | 3957.8541 | 3957.8432 | 0 | 2.8 | 62 | Specific | 352.32 | 276.98 | 276.98 | 6.7 | 18 | >sp\|P01591\|IGJ_HUMAN Immunoglobulin J chain OS=Homo sapiens OX=9606 GN=JCHAIN PE=1 SV=4 | 1392 | 20201113_15413_1ng_glyco_hcd203040_compound15.12610.12610.3 | controllerType=0 controllerNumber=1 scan=12610 | 27.6123 |
| 11167:3 | 51 | R.IIVPLNNREN[+1856.656]ISDPTSPLR.T | HexNAc(3)Hex(5)Fuc(1)NeuAc(1) | N[+1857] | 1335.6187 | 3 | 4004.8416 | 4004.8327 | 0 | 2.2 | 62 | Specific | 361.4 | 361.4 | 361.4 | 7.03 | 18 | >sp\|P01591\|IGJ_HUMAN Immunoglobulin J chain OS=Homo sapiens OX=9606 GN=JCHAIN PE=1 SV=4 | 1392 | 20201113_15413_1ng_glyco_hcd203040_compound15.14423.14423.3 | controllerType=0 controllerNumber=1 scan=14423 | 30.6832 |
| 13023:3 | 51 | R.IIVPLNNREN[+2350.830]ISDPTSPLR.T | HexNAc(4)Hex(5)Fuc(1)NeuAc(2) | N[+2351] | 1500.3464 | 3 | 4499.0246 | 4499.0075 | 0 | 3.8 | 62 | Specific | 366.31 | 366.31 | 366.31 | 7.08 | 18 | >sp\|P01591\|IGJ_HUMAN Immunoglobulin J chain OS=Homo sapiens OX=9606 GN=JCHAIN PE=1 SV=4 | 1392 | 20201113_15413_1ng_glyco_hcd203040_compound15.16364.16364.3 | controllerType=0 controllerNumber=1 scan=16364 | 33.9946 |
| 13021:3 | 51 | R.IIVPLNNREN[+2350.830]ISDPTSPLR.T | HexNAc(4)Hex(5)Fuc(1)NeuAc(2) | N[+2351] | 1500.3464 | 3 | 4499.0246 | 4499.0075 | 0 | 3.8 | 62 | Specific | 369.87 | 369.87 | 369.87 | 7.08 | 18 | >sp\|P01591\|IGJ_HUMAN Immunoglobulin J chain OS=Homo sapiens OX=9606 GN=JCHAIN PE=1 SV=4 | 1392 | 20201113_15413_1ng_glyco_hcd203040_compound15.16362.16362.3 | controllerType=0 controllerNumber=1 scan=16362 | 33.9904 |
| 09413:3 | 51 | R.IIVPLNNREN[+1809.666]ISDPTSPLR.T | HexNAc(5)Hex(4)Fuc(1) | N[+1810] | 1319.9562 | 3 | 3957.8541 | 3957.8432 | 0 | 2.8 | 62 | Specific | 378.9 | 282.99 | 282.99 | 6.7 | 18 | >sp\|P01591\|IGJ_HUMAN Immunoglobulin J chain OS=Homo sapiens OX=9606 GN=JCHAIN PE=1 SV=4 | 1392 | 20201113_15413_1ng_glyco_hcd203040_compound15.12608.12608.3 | controllerType=0 controllerNumber=1 scan=12608 | 27.6084 |
| 09389:3 | 51 | R.IIVPLNNREN[+1809.666]ISDPTSPLR.T | HexNAc(5)Hex(4)Fuc(1) | N[+1810] | 1319.9572 | 3 | 3957.8571 | 3957.8432 | 0 | 3.5 | 62 | Specific | 417.33 | 339.81 | 339.81 | 7.04 | 18 | >sp\|P01591\|IGJ_HUMAN Immunoglobulin J chain OS=Homo sapiens OX=9606 GN=JCHAIN PE=1 SV=4 | 1392 | 20201113_15413_1ng_glyco_hcd203040_compound15.12583.12583.3 | controllerType=0 controllerNumber=1 scan=12583 | 27.5678 |
| 11170:3 | 51 | R.IIVPLNNREN[+1856.656]ISDPTSPLR.T | HexNAc(3)Hex(5)Fuc(1)NeuAc(1) | N[+1857] | 1335.6187 | 3 | 4004.8416 | 4004.8327 | 0 | 2.2 | 62 | Specific | 521.26 | 521.26 | 521.26 | 9.04 | 18 | >sp\|P01591\|IGJ_HUMAN Immunoglobulin J chain OS=Homo sapiens OX=9606 GN=JCHAIN PE=1 SV=4 | 1392 | 20201113_15413_1ng_glyco_hcd203040_compound15.14426.14426.3 | controllerType=0 controllerNumber=1 scan=14426 | 30.6876 |
| 06366:3 | 480 | R.[+42.011]VEDAYILTC[+57.021]N[+730.264]VSLEYEK.T | HexNAc(2)Hex(2) | .[42], C[+57], N[+730] | 940.0954 | 3 | 2818.2715 | 2818.2486 | 0 | 8.2 | 225 | Specific | 230.97 | 204.21 | 204.21 | 3.07 | 3 | >sp\|P40227\|TCPZ_HUMAN T-complex protein 1 subunit zeta OS=Homo sapiens OX=9606 GN=CCT6A PE=1 SV=3 | 1115 | 20201113_15413_1ng_glyco_hcd203040_compound15.9428.9428.3 | controllerType=0 controllerNumber=1 scan=9428 | 22.3014 |
| 18773:5 | 1 | R.RTAGWNVPIGTLRPFLN[+2350.830]WTGPPEPIEAAVAR.F | HexNAc(4)Hex(5)Fuc(1)NeuAc(2) | N[+2351] | 1148.3396 | 5 | 5737.6689 | 5737.6424 | 0 | 4.6 | 140 | Specific | 313.13 | 313.13 | 313.13 | 6.63 | 122 | >sp\|P02788\|TRFL_HUMAN Lactotransferrin OS=Homo sapiens OX=9606 GN=LTF PE=1 SV=6 | 1270 | 20201119_15432_1ng_glyco_hcd203040_compound15.22477.22477.5 | controllerType=0 controllerNumber=1 scan=22477 | 45.8745 |
| 17740:5 | 1 | R.RTAGWNVPIGTLRPFLN[+2204.772]WTGPPEPIEAAVAR.F | HexNAc(4)Hex(5)NeuAc(2) | N[+2205] | 1119.1265 | 5 | 5591.6032 | 5591.5845 | 0 | 3.4 | 140 | Specific | 364.76 | 364.76 | 364.76 | 7.41 | 122 | >sp\|P02788\|TRFL_HUMAN Lactotransferrin OS=Homo sapiens OX=9606 GN=LTF PE=1 SV=6 | 1270 | 20201119_15432_1ng_glyco_hcd203040_compound15.21399.21399.5 | controllerType=0 controllerNumber=1 scan=21399 | 43.5784 |
| 17743:5 | 1 | R.RTAGWNVPIGTLRPFLN[+2204.772]WTGPPEPIEAAVAR.F | HexNAc(4)Hex(5)NeuAc(2) | N[+2205] | 1119.1265 | 5 | 5591.6032 | 5591.5845 | 0 | 3.4 | 140 | Specific | 368.15 | 368.15 | 368.15 | 7.41 | 122 | >sp\|P02788\|TRFL_HUMAN Lactotransferrin OS=Homo sapiens OX=9606 GN=LTF PE=1 SV=6 | 1270 | 20201119_15432_1ng_glyco_hcd203040_compound15.21402.21402.5 | controllerType=0 controllerNumber=1 scan=21402 | 43.5887 |
| 18767:4 | 1 | R.RTAGWNVPIGTLRPFLN[+2350.830]WTGPPEPIEAAVAR.F | HexNAc(4)Hex(5)Fuc(1)NeuAc(2) | N[+2351] | 1435.1735 | 4 | 5737.672 | 5737.6424 | 0 | 5.2 | 140 | Specific | 430.1 | 352.16 | 352.16 | 7.77 | 122 | >sp\|P02788\|TRFL_HUMAN Lactotransferrin OS=Homo sapiens OX=9606 GN=LTF PE=1 SV=6 | 1270 | 20201119_15432_1ng_glyco_hcd203040_compound15.22471.22471.4 | controllerType=0 controllerNumber=1 scan=22471 | 45.8645 |
| 17611:4 | 1 | R.RTAGWNVPIGTLRPFLN[+2570.925]WTGPPEPIEAAVAR.F | HexNAc(5)Hex(6)Fuc(2)NeuAc(1) | N[+2571] | 1490.1948 | 4 | 5957.7575 | 5957.7371 | 0 | 3.4 | 140 | Specific | 483.52 | 457.69 | 457.69 | 9.46 | 122 | >sp\|P02788\|TRFL_HUMAN Lactotransferrin OS=Homo sapiens OX=9606 GN=LTF PE=1 SV=6 | 1270 | 20201119_15432_1ng_glyco_hcd203040_compound15.21265.21265.4 | controllerType=0 controllerNumber=1 scan=21265 | 43.3151 |
| 17736:4 | 1 | R.RTAGWNVPIGTLRPFLN[+2205.793]WTGPPEPIEAAVAR.F | HexNAc(4)Hex(5)Fuc(2)NeuAc(1) | N[+2206] | 1398.9067 | 4 | 5592.6051 | 5592.6049 | 0 | 0 | 140 | Specific | 515.97 | 500.67 | 500.67 | 8.61 | 122 | >sp\|P02788\|TRFL_HUMAN Lactotransferrin OS=Homo sapiens OX=9606 GN=LTF PE=1 SV=6 | 1270 | 20201119_15432_1ng_glyco_hcd203040_compound15.21395.21395.4 | controllerType=0 controllerNumber=1 scan=21395 | 43.5742 |
| 17734:4 | 1 | R.RTAGWNVPIGTLRPFLN[+2205.793]WTGPPEPIEAAVAR.F | HexNAc(4)Hex(5)Fuc(2)NeuAc(1) | N[+2206] | 1398.9067 | 4 | 5592.6051 | 5592.6049 | 0 | 0 | 140 | Specific | 516.06 | 509.79 | 509.79 | 8.61 | 122 | >sp\|P02788\|TRFL_HUMAN Lactotransferrin OS=Homo sapiens OX=9606 GN=LTF PE=1 SV=6 | 1270 | 20201119_15432_1ng_glyco_hcd203040_compound15.21393.21393.4 | controllerType=0 controllerNumber=1 scan=21393 | 43.5696 |
| 17613:4 | 1 | R.RTAGWNVPIGTLRPFLN[+2570.925]WTGPPEPIEAAVAR.F | HexNAc(5)Hex(6)Fuc(2)NeuAc(1) | N[+2571] | 1490.1948 | 4 | 5957.7575 | 5957.7371 | 0 | 3.4 | 140 | Specific | 529.18 | 436.6 | 436.6 | 9.71 | 122 | >sp\|P02788\|TRFL_HUMAN Lactotransferrin OS=Homo sapiens OX=9606 GN=LTF PE=1 SV=6 | 1270 | 20201119_15432_1ng_glyco_hcd203040_compound15.21267.21267.4 | controllerType=0 controllerNumber=1 scan=21267 | 43.3203 |
| 18770:4 | 1 | R.RTAGWNVPIGTLRPFLN[+2350.830]WTGPPEPIEAAVAR.F | HexNAc(4)Hex(5)Fuc(1)NeuAc(2) | N[+2351] | 1435.1735 | 4 | 5737.672 | 5737.6424 | 0 | 5.2 | 140 | Specific | 638.05 | 514.91 | 514.91 | 12 | 122 | >sp\|P02788\|TRFL_HUMAN Lactotransferrin OS=Homo sapiens OX=9606 GN=LTF PE=1 SV=6 | 1270 | 20201119_15432_1ng_glyco_hcd203040_compound15.22474.22474.4 | controllerType=0 controllerNumber=1 scan=22474 | 45.869 |
| 18899:5 | 1 | R.TAGWNVPIGTLRPFLN[+2570.925]WTGPPEPIEAAVAR.F | HexNAc(5)Hex(6)Fuc(2)NeuAc(1) | N[+2571] | 1161.1383 | 5 | 5801.6624 | 5801.636 | 0 | 4.6 | 141 | Specific | 214.1 | 188.44 | 188.44 | 3.8 | 122 | >sp\|P02788\|TRFL_HUMAN Lactotransferrin OS=Homo sapiens OX=9606 GN=LTF PE=1 SV=6 | 1270 | 20201119_15432_1ng_glyco_hcd203040_compound15.22608.22608.5 | controllerType=0 controllerNumber=1 scan=22608 | 46.1609 |
| 18857:3 | 1 | R.TAGWNVPIGTLRPFLN[+2570.925]WTGPPEPIEAAVAR.F | HexNAc(5)Hex(6)Fuc(2)NeuAc(1) | N[+2571] | 1934.5575 | 3 | 5801.6578 | 5801.636 | 0 | 3.8 | 141 | Specific | 216.57 | 139.68 | 139.68 | 4.49 | 122 | >sp\|P02788\|TRFL_HUMAN Lactotransferrin OS=Homo sapiens OX=9606 GN=LTF PE=1 SV=6 | 1270 | 20201119_15432_1ng_glyco_hcd203040_compound15.22565.22565.3 | controllerType=0 controllerNumber=1 scan=22565 | 46.0713 |
| 17910:3 | 1 | R.TAGWNVPIGTLRPFLN[+2425.887]WTGPPEPIEAAVAR.F | HexNAc(5)Hex(6)Fuc(3) | N[+2426] | 1886.2119 | 3 | 5656.6211 | 5656.5985 | 0 | 4 | 141 | Specific | 223.43 | 178.25 | 178.25 | 5.18 | 122 | >sp\|P02788\|TRFL_HUMAN Lactotransferrin OS=Homo sapiens OX=9606 GN=LTF PE=1 SV=6 | 1270 | 20201119_15432_1ng_glyco_hcd203040_compound15.21576.21576.3 | controllerType=0 controllerNumber=1 scan=21576 | 43.9282 |
| 17945:4 | 1 | R.TAGWNVPIGTLRPFLN[+2279.830]WTGPPEPIEAAVAR.F | HexNAc(5)Hex(6)Fuc(2) | N[+2280] | 1378.3981 | 4 | 5510.5705 | 5510.5406 | 0 | 5.4 | 141 | Specific | 235.72 | 235.72 | 235.72 | 5.61 | 122 | >sp\|P02788\|TRFL_HUMAN Lactotransferrin OS=Homo sapiens OX=9606 GN=LTF PE=1 SV=6 | 1270 | 20201119_15432_1ng_glyco_hcd203040_compound15.21613.21613.4 | controllerType=0 controllerNumber=1 scan=21613 | 44.0129 |
| 17912:3 | 1 | R.TAGWNVPIGTLRPFLN[+2425.887]WTGPPEPIEAAVAR.F | HexNAc(5)Hex(6)Fuc(3) | N[+2426] | 1886.2119 | 3 | 5656.6211 | 5656.5985 | 0 | 4 | 141 | Specific | 255.56 | 128.17 | 128.17 | 5.78 | 122 | >sp\|P02788\|TRFL_HUMAN Lactotransferrin OS=Homo sapiens OX=9606 GN=LTF PE=1 SV=6 | 1270 | 20201119_15432_1ng_glyco_hcd203040_compound15.21578.21578.3 | controllerType=0 controllerNumber=1 scan=21578 | 43.9334 |
| 17987:3 | 1 | R.TAGWNVPIGTLRPFLN[+1768.640]WTGPPEPIEAAVAR.F | HexNAc(4)Hex(5)Fuc(1) | N[+1769] | 1667.1237 | 3 | 4999.3567 | 4999.3504 | 0 | 1.2 | 141 | Specific | 278.87 | 250.41 | 250.41 | 6.56 | 122 | >sp\|P02788\|TRFL_HUMAN Lactotransferrin OS=Homo sapiens OX=9606 GN=LTF PE=1 SV=6 | 1270 | 20201119_15432_1ng_glyco_hcd203040_compound15.21657.21657.3 | controllerType=0 controllerNumber=1 scan=21657 | 44.1053 |
| 18929:4 | 1 | R.TAGWNVPIGTLRPFLN[+2424.867]WTGPPEPIEAAVAR.F | HexNAc(5)Hex(6)Fuc(1)NeuAc(1) | N[+2425] | 1414.6553 | 4 | 5655.5993 | 5655.5781 | 0 | 3.8 | 141 | Specific | 332.68 | 220.38 | 220.38 | 6.94 | 122 | >sp\|P02788\|TRFL_HUMAN Lactotransferrin OS=Homo sapiens OX=9606 GN=LTF PE=1 SV=6 | 1270 | 20201119_15432_1ng_glyco_hcd203040_compound15.22640.22640.4 | controllerType=0 controllerNumber=1 scan=22640 | 46.2408 |
| 18941:3 | 1 | R.TAGWNVPIGTLRPFLN[+2205.793]WTGPPEPIEAAVAR.F | HexNAc(4)Hex(5)Fuc(2)NeuAc(1) | N[+2206] | 1812.847 | 3 | 5436.5265 | 5436.5038 | 0 | 4.2 | 141 | Specific | 342.2 | 342.2 | 342.2 | 7.74 | 122 | >sp\|P02788\|TRFL_HUMAN Lactotransferrin OS=Homo sapiens OX=9606 GN=LTF PE=1 SV=6 | 1270 | 20201119_15432_1ng_glyco_hcd203040_compound15.22653.22653.3 | controllerType=0 controllerNumber=1 scan=22653 | 46.2721 |
| 18946:3 | 1 | R.TAGWNVPIGTLRPFLN[+2351.851]WTGPPEPIEAAVAR.F | HexNAc(4)Hex(5)Fuc(3)NeuAc(1) | N[+2352] | 1861.531 | 3 | 5582.5784 | 5582.5617 | 0 | 3 | 141 | Specific | 357.26 | 350.69 | 350.69 | 8.04 | 122 | >sp\|P02788\|TRFL_HUMAN Lactotransferrin OS=Homo sapiens OX=9606 GN=LTF PE=1 SV=6 | 1270 | 20201119_15432_1ng_glyco_hcd203040_compound15.22658.22658.3 | controllerType=0 controllerNumber=1 scan=22658 | 46.2801 |
| 19341:4 | 1 | R.TAGWNVPIGTLRPFLN[+2205.793]WTGPPEPIEAAVAR.F | HexNAc(4)Hex(5)Fuc(2)NeuAc(1) | N[+2206] | 1359.8857 | 4 | 5436.5212 | 5436.5038 | 0 | 3.2 | 141 | Specific | 388.47 | 365.66 | 365.66 | 8.12 | 122 | >sp\|P02788\|TRFL_HUMAN Lactotransferrin OS=Homo sapiens OX=9606 GN=LTF PE=1 SV=6 | 1270 | 20201119_15432_1ng_glyco_hcd203040_compound15.23073.23073.4 | controllerType=0 controllerNumber=1 scan=23073 | 47.1512 |
| 18945:4 | 1 | R.TAGWNVPIGTLRPFLN[+2351.851]WTGPPEPIEAAVAR.F | HexNAc(4)Hex(5)Fuc(3)NeuAc(1) | N[+2352] | 1396.3992 | 4 | 5582.5749 | 5582.5617 | 0 | 2.4 | 141 | Specific | 405.76 | 405.76 | 405.76 | 9.41 | 122 | >sp\|P02788\|TRFL_HUMAN Lactotransferrin OS=Homo sapiens OX=9606 GN=LTF PE=1 SV=6 | 1270 | 20201119_15432_1ng_glyco_hcd203040_compound15.22657.22657.4 | controllerType=0 controllerNumber=1 scan=22657 | 46.2789 |
| 18860:3 | 1 | R.TAGWNVPIGTLRPFLN[+2570.925]WTGPPEPIEAAVAR.F | HexNAc(5)Hex(6)Fuc(2)NeuAc(1) | N[+2571] | 1934.5575 | 3 | 5801.6578 | 5801.636 | 0 | 3.8 | 141 | Specific | 452.46 | 330.35 | 330.35 | 8.26 | 122 | >sp\|P02788\|TRFL_HUMAN Lactotransferrin OS=Homo sapiens OX=9606 GN=LTF PE=1 SV=6 | 1270 | 20201119_15432_1ng_glyco_hcd203040_compound15.22568.22568.3 | controllerType=0 controllerNumber=1 scan=22568 | 46.075 |
| 18949:3 | 1 | R.TAGWNVPIGTLRPFLN[+2351.851]WTGPPEPIEAAVAR.F | HexNAc(4)Hex(5)Fuc(3)NeuAc(1) | N[+2352] | 1861.531 | 3 | 5582.5784 | 5582.5617 | 0 | 3 | 141 | Specific | 458.94 | 403.14 | 403.14 | 9.83 | 122 | >sp\|P02788\|TRFL_HUMAN Lactotransferrin OS=Homo sapiens OX=9606 GN=LTF PE=1 SV=6 | 1270 | 20201119_15432_1ng_glyco_hcd203040_compound15.22661.22661.3 | controllerType=0 controllerNumber=1 scan=22661 | 46.2879 |
| 19202:4 | 1 | R.TAGWNVPIGTLRPFLN[+2059.735]WTGPPEPIEAAVAR.F | HexNAc(4)Hex(5)Fuc(1)NeuAc(1) | N[+2060] | 1323.376 | 4 | 5290.4821 | 5290.4459 | 0 | 6.8 | 141 | Specific | 462.61 | 441.73 | 441.73 | 8.65 | 122 | >sp\|P02788\|TRFL_HUMAN Lactotransferrin OS=Homo sapiens OX=9606 GN=LTF PE=1 SV=6 | 1270 | 20201119_15432_1ng_glyco_hcd203040_compound15.22928.22928.4 | controllerType=0 controllerNumber=1 scan=22928 | 46.8455 |
| 17948:4 | 1 | R.TAGWNVPIGTLRPFLN[+2279.830]WTGPPEPIEAAVAR.F | HexNAc(5)Hex(6)Fuc(2) | N[+2280] | 1378.3981 | 4 | 5510.5705 | 5510.5406 | 0 | 5.4 | 141 | Specific | 483.92 | 385.08 | 385.08 | 9.07 | 122 | >sp\|P02788\|TRFL_HUMAN Lactotransferrin OS=Homo sapiens OX=9606 GN=LTF PE=1 SV=6 | 1270 | 20201119_15432_1ng_glyco_hcd203040_compound15.21616.21616.4 | controllerType=0 controllerNumber=1 scan=21616 | 44.0256 |
| 18852:4 | 1 | R.TAGWNVPIGTLRPFLN[+2570.925]WTGPPEPIEAAVAR.F | HexNAc(5)Hex(6)Fuc(2)NeuAc(1) | N[+2571] | 1451.1702 | 4 | 5801.6588 | 5801.636 | 0 | 3.9 | 141 | Specific | 493.71 | 393.98 | 393.98 | 9.83 | 122 | >sp\|P02788\|TRFL_HUMAN Lactotransferrin OS=Homo sapiens OX=9606 GN=LTF PE=1 SV=6 | 1270 | 20201119_15432_1ng_glyco_hcd203040_compound15.22560.22560.4 | controllerType=0 controllerNumber=1 scan=22560 | 46.0632 |
| 18931:4 | 1 | R.TAGWNVPIGTLRPFLN[+2424.867]WTGPPEPIEAAVAR.F | HexNAc(5)Hex(6)Fuc(1)NeuAc(1) | N[+2425] | 1414.6553 | 4 | 5655.5993 | 5655.5781 | 0 | 3.8 | 141 | Specific | 549.96 | 417.07 | 417.07 | 10.08 | 122 | >sp\|P02788\|TRFL_HUMAN Lactotransferrin OS=Homo sapiens OX=9606 GN=LTF PE=1 SV=6 | 1270 | 20201119_15432_1ng_glyco_hcd203040_compound15.22642.22642.4 | controllerType=0 controllerNumber=1 scan=22642 | 46.246 |
| 18794:4 | 1 | R.TAGWNVPIGTLRPFLN[+2863.041]WTGPPEPIEAAVAR.F | HexNAc(5)Hex(6)Fuc(4)NeuAc(1) | N[+2863] | 1524.2006 | 4 | 6093.7804 | 6093.7518 | 0 | 4.7 | 141 | Specific | 561.6 | 529.81 | 529.81 | 10.19 | 122 | >sp\|P02788\|TRFL_HUMAN Lactotransferrin OS=Homo sapiens OX=9606 GN=LTF PE=1 SV=6 | 1270 | 20201119_15432_1ng_glyco_hcd203040_compound15.22499.22499.4 | controllerType=0 controllerNumber=1 scan=22499 | 45.9283 |
| 18943:3 | 1 | R.TAGWNVPIGTLRPFLN[+2205.793]WTGPPEPIEAAVAR.F | HexNAc(4)Hex(5)Fuc(2)NeuAc(1) | N[+2206] | 1812.847 | 3 | 5436.5265 | 5436.5038 | 0 | 4.2 | 141 | Specific | 577.46 | 478.33 | 478.33 | 11.5 | 122 | >sp\|P02788\|TRFL_HUMAN Lactotransferrin OS=Homo sapiens OX=9606 GN=LTF PE=1 SV=6 | 1270 | 20201119_15432_1ng_glyco_hcd203040_compound15.22655.22655.3 | controllerType=0 controllerNumber=1 scan=22655 | 46.2767 |
| 19343:4 | 1 | R.TAGWNVPIGTLRPFLN[+2205.793]WTGPPEPIEAAVAR.F | HexNAc(4)Hex(5)Fuc(2)NeuAc(1) | N[+2206] | 1359.8857 | 4 | 5436.5212 | 5436.5038 | 0 | 3.2 | 141 | Specific | 577.51 | 442.12 | 442.12 | 11.49 | 122 | >sp\|P02788\|TRFL_HUMAN Lactotransferrin OS=Homo sapiens OX=9606 GN=LTF PE=1 SV=6 | 1270 | 20201119_15432_1ng_glyco_hcd203040_compound15.23075.23075.4 | controllerType=0 controllerNumber=1 scan=23075 | 47.1578 |
| 18787:4 | 1 | R.TAGWNVPIGTLRPFLN[+2716.983]WTGPPEPIEAAVAR.F | HexNAc(5)Hex(6)Fuc(3)NeuAc(1) | N[+2717] | 1487.6876 | 4 | 5947.7287 | 5947.6939 | 0 | 5.9 | 141 | Specific | 578.12 | 518.41 | 518.41 | 10.74 | 122 | >sp\|P02788\|TRFL_HUMAN Lactotransferrin OS=Homo sapiens OX=9606 GN=LTF PE=1 SV=6 | 1270 | 20201119_15432_1ng_glyco_hcd203040_compound15.22492.22492.4 | controllerType=0 controllerNumber=1 scan=22492 | 45.9127 |
| 17988:4 | 1 | R.TAGWNVPIGTLRPFLN[+1768.640]WTGPPEPIEAAVAR.F | HexNAc(4)Hex(5)Fuc(1) | N[+1769] | 1250.5935 | 4 | 4999.3522 | 4999.3504 | 0 | 0.4 | 141 | Specific | 579.19 | 522.46 | 522.46 | 10.49 | 122 | >sp\|P02788\|TRFL_HUMAN Lactotransferrin OS=Homo sapiens OX=9606 GN=LTF PE=1 SV=6 | 1270 | 20201119_15432_1ng_glyco_hcd203040_compound15.21658.21658.4 | controllerType=0 controllerNumber=1 scan=21658 | 44.1067 |
| 17989:3 | 1 | R.TAGWNVPIGTLRPFLN[+1768.640]WTGPPEPIEAAVAR.F | HexNAc(4)Hex(5)Fuc(1) | N[+1769] | 1667.1237 | 3 | 4999.3567 | 4999.3504 | 0 | 1.2 | 141 | Specific | 661.05 | 575.4 | 575.4 | 13.31 | 122 | >sp\|P02788\|TRFL_HUMAN Lactotransferrin OS=Homo sapiens OX=9606 GN=LTF PE=1 SV=6 | 1270 | 20201119_15432_1ng_glyco_hcd203040_compound15.21659.21659.3 | controllerType=0 controllerNumber=1 scan=21659 | 44.109 |
| 18797:4 | 1 | R.TAGWNVPIGTLRPFLN[+2863.041]WTGPPEPIEAAVAR.F | HexNAc(5)Hex(6)Fuc(4)NeuAc(1) | N[+2863] | 1524.2006 | 4 | 6093.7804 | 6093.7518 | 0 | 4.7 | 141 | Specific | 689.56 | 560.39 | 560.39 | 13.37 | 122 | >sp\|P02788\|TRFL_HUMAN Lactotransferrin OS=Homo sapiens OX=9606 GN=LTF PE=1 SV=6 | 1270 | 20201119_15432_1ng_glyco_hcd203040_compound15.22502.22502.4 | controllerType=0 controllerNumber=1 scan=22502 | 45.9367 |
| 18937:3 | 1 | R.TAGWNVPIGTLRPFLN[+2059.735]WTGPPEPIEAAVAR.F | HexNAc(4)Hex(5)Fuc(1)NeuAc(1) | N[+2060] | 1764.1629 | 3 | 5290.4742 | 5290.4459 | 0 | 5.4 | 141 | Specific | 695.24 | 628.95 | 628.95 | 12.87 | 122 | >sp\|P02788\|TRFL_HUMAN Lactotransferrin OS=Homo sapiens OX=9606 GN=LTF PE=1 SV=6 | 1270 | 20201119_15432_1ng_glyco_hcd203040_compound15.22648.22648.3 | controllerType=0 controllerNumber=1 scan=22648 | 46.2594 |
| 20226:4 | 1 | R.TAGWNVPIGTLRPFLN[+2350.830]WTGPPEPIEAAVAR.F | HexNAc(4)Hex(5)Fuc(1)NeuAc(2) | N[+2351] | 1396.1453 | 4 | 5581.5592 | 5581.5413 | 0 | 3.2 | 141 | Specific | 700.93 | 575.99 | 575.99 | 14.49 | 122 | >sp\|P02788\|TRFL_HUMAN Lactotransferrin OS=Homo sapiens OX=9606 GN=LTF PE=1 SV=6 | 1270 | 20201119_15432_1ng_glyco_hcd203040_compound15.23998.23998.4 | controllerType=0 controllerNumber=1 scan=23998 | 49.1145 |
| 18939:3 | 1 | R.TAGWNVPIGTLRPFLN[+2059.735]WTGPPEPIEAAVAR.F | HexNAc(4)Hex(5)Fuc(1)NeuAc(1) | N[+2060] | 1764.1629 | 3 | 5290.4742 | 5290.4459 | 0 | 5.4 | 141 | Specific | 707.09 | 618.08 | 618.08 | 13.87 | 122 | >sp\|P02788\|TRFL_HUMAN Lactotransferrin OS=Homo sapiens OX=9606 GN=LTF PE=1 SV=6 | 1270 | 20201119_15432_1ng_glyco_hcd203040_compound15.22650.22650.3 | controllerType=0 controllerNumber=1 scan=22650 | 46.2668 |
| 19205:4 | 1 | R.TAGWNVPIGTLRPFLN[+2059.735]WTGPPEPIEAAVAR.F | HexNAc(4)Hex(5)Fuc(1)NeuAc(1) | N[+2060] | 1323.376 | 4 | 5290.4821 | 5290.4459 | 0 | 6.8 | 141 | Specific | 709.69 | 662.65 | 662.65 | 13.45 | 122 | >sp\|P02788\|TRFL_HUMAN Lactotransferrin OS=Homo sapiens OX=9606 GN=LTF PE=1 SV=6 | 1270 | 20201119_15432_1ng_glyco_hcd203040_compound15.22931.22931.4 | controllerType=0 controllerNumber=1 scan=22931 | 46.851 |

Table S4. Cont.

| Query #:z | Protein Rank | Glycopeptide | Glycan | Modification Type(s) | Observed m/z | z | Observed (M+H) | Calc. mass (M+H) | Off-by-x error | Mass error (ppm) | StartPosition | Cleavage | Score | Delta | Delta Mod | \|Log Prob\| | # of unique peptides | Protein Name | Protein DB number | Comment | Scan # | Scan Time |
|---|---|---|---|---|---|---|---|---|---|---|---|---|---|---|---|---|---|---|---|---|---|---|
| 18995:3 | 1 | R.TAGWNVPIGTLRPFLN[+2059.735]WTGPPEPIEAAVAR.F | HexNAc(4)Hex(5)Fuc(1)NeuAc(1) | N[+2060] | 1764.1574 | 3 | 5290.4577 | 5290.4459 | 0 | 2.2 | 141 | Specific | 725.6 | 633.97 | 633.97 | 14.63 | 122 | >sp\|P02788\|TRFL_HUMAN Lactotransferrin OS=Homo sapiens OX=9606 GN=LTF PE=1 SV=6 | 1270 | 20201119_15432_1ng_glyco_hcd203040_compound15.22709.22709.3 | controllerType=0 controllerNumber=1 scan=22709 | 46.3931 |
| 18940:4 | 1 | R.TAGWNVPIGTLRPFLN[+2205.793]WTGPPEPIEAAVAR.F | HexNAc(4)Hex(5)Fuc(2)NeuAc(1) | N[+2206] | 1359.8878 | 4 | 5436.5295 | 5436.5038 | 0 | 4.7 | 141 | Specific | 727.12 | 657.7 | 657.7 | 14.37 | 122 | >sp\|P02788\|TRFL_HUMAN Lactotransferrin OS=Homo sapiens OX=9606 GN=LTF PE=1 SV=6 | 1270 | 20201119_15432_1ng_glyco_hcd203040_compound15.22652.22652.4 | controllerType=0 controllerNumber=1 scan=22652 | 46.2707 |
| 18790:4 | 1 | R.TAGWNVPIGTLRPFLN[+2716.983]WTGPPEPIEAAVAR.F | HexNAc(5)Hex(6)Fuc(3)NeuAc(1) | N[+2717] | 1487.6876 | 4 | 5947.7287 | 5947.6939 | 0 | 5.9 | 141 | Specific | 743.07 | 621.67 | 621.67 | 13.74 | 122 | >sp\|P02788\|TRFL_HUMAN Lactotransferrin OS=Homo sapiens OX=9606 GN=LTF PE=1 SV=6 | 1270 | 20201119_15432_1ng_glyco_hcd203040_compound15.22495.22495.4 | controllerType=0 controllerNumber=1 scan=22495 | 45.9193 |
| 18854:4 | 1 | R.TAGWNVPIGTLRPFLN[+2570.925]WTGPPEPIEAAVAR.F | HexNAc(5)Hex(6)Fuc(2)NeuAc(1) | N[+2571] | 1451.1702 | 4 | 5801.6588 | 5801.636 | 0 | 3.9 | 141 | Specific | 764.74 | 628.03 | 628.03 | 15.49 | 122 | >sp\|P02788\|TRFL_HUMAN Lactotransferrin OS=Homo sapiens OX=9606 GN=LTF PE=1 SV=6 | 1270 | 20201119_15432_1ng_glyco_hcd203040_compound15.22562.22562.4 | controllerType=0 controllerNumber=1 scan=22562 | 46.0678 |
| 18992:3 | 1 | R.TAGWNVPIGTLRPFLN[+2059.735]WTGPPEPIEAAVAR.F | HexNAc(4)Hex(5)Fuc(1)NeuAc(1) | N[+2060] | 1764.1574 | 3 | 5290.4577 | 5290.4459 | 0 | 2.2 | 141 | Specific | 766.77 | 642.82 | 642.82 | 15.63 | 122 | >sp\|P02788\|TRFL_HUMAN Lactotransferrin OS=Homo sapiens OX=9606 GN=LTF PE=1 SV=6 | 1270 | 20201119_15432_1ng_glyco_hcd203040_compound15.22706.22706.3 | controllerType=0 controllerNumber=1 scan=22706 | 46.3817 |
| 20228:4 | 1 | R.TAGWNVPIGTLRPFLN[+2350.830]WTGPPEPIEAAVAR.F | HexNAc(4)Hex(5)Fuc(1)NeuAc(2) | N[+2351] | 1396.1453 | 4 | 5581.5592 | 5581.5413 | 0 | 3.2 | 141 | Specific | 817.97 | 661.81 | 661.81 | 16.49 | 122 | >sp\|P02788\|TRFL_HUMAN Lactotransferrin OS=Homo sapiens OX=9606 GN=LTF PE=1 SV=6 | 1270 | 20201119_15432_1ng_glyco_hcd203040_compound15.24000.24000.4 | controllerType=0 controllerNumber=1 scan=24000 | 49.1221 |
| 17990:4 | 1 | R.TAGWNVPIGTLRPFLN[+1768.640]WTGPPEPIEAAVAR.F | HexNAc(4)Hex(5)Fuc(1) | N[+1769] | 1250.5935 | 4 | 4999.3522 | 4999.3504 | 0 | 0.4 | 141 | Specific | 832.98 | 724.59 | 724.59 | 15.49 | 122 | >sp\|P02788\|TRFL_HUMAN Lactotransferrin OS=Homo sapiens OX=9606 GN=LTF PE=1 SV=6 | 1270 | 20201119_15432_1ng_glyco_hcd203040_compound15.21660.21660.4 | controllerType=0 controllerNumber=1 scan=21660 | 44.1111 |
| 18942:4 | 1 | R.TAGWNVPIGTLRPFLN[+2205.793]WTGPPEPIEAAVAR.F | HexNAc(4)Hex(5)Fuc(2)NeuAc(1) | N[+2206] | 1359.8878 | 4 | 5436.5295 | 5436.5038 | 0 | 4.7 | 141 | Specific | 841.96 | 695.45 | 695.45 | 16.37 | 122 | >sp\|P02788\|TRFL_HUMAN Lactotransferrin OS=Homo sapiens OX=9606 GN=LTF PE=1 SV=6 | 1270 | 20201119_15432_1ng_glyco_hcd203040_compound15.22654.22654.4 | controllerType=0 controllerNumber=1 scan=22654 | 46.2755 |
| 18948:4 | 1 | R.TAGWNVPIGTLRPFLN[+2351.851]WTGPPEPIEAAVAR.F | HexNAc(4)Hex(5)Fuc(3)NeuAc(1) | N[+2352] | 1396.3992 | 4 | 5582.5749 | 5582.5617 | 0 | 2.4 | 141 | Specific | 878.84 | 744.46 | 744.46 | 17.49 | 122 | >sp\|P02788\|TRFL_HUMAN Lactotransferrin OS=Homo sapiens OX=9606 GN=LTF PE=1 SV=6 | 1270 | 20201119_15432_1ng_glyco_hcd203040_compound15.22660.22660.4 | controllerType=0 controllerNumber=1 scan=22660 | 46.2862 |
| 18944:5 | 1 | R.TAGWNVPIGTLRPFLN[+2205.793]WTGPPEPIEAAVAR.F | HexNAc(4)Hex(5)Fuc(2)NeuAc(1) | N[+2206] | 1088.1102 | 5 | 5436.5221 | 5436.5038 | 0 | 3.4 | 141 | Specific | 904.95 | 785.6 | 785.6 | 18.76 | 122 | >sp\|P02788\|TRFL_HUMAN Lactotransferrin OS=Homo sapiens OX=9606 GN=LTF PE=1 SV=6 | 1270 | 20201119_15432_1ng_glyco_hcd203040_compound15.22656.22656.5 | controllerType=0 controllerNumber=1 scan=22656 | 46.2778 |
| 18947:5 | 1 | R.TAGWNVPIGTLRPFLN[+2205.793]WTGPPEPIEAAVAR.F | HexNAc(4)Hex(5)Fuc(2)NeuAc(1) | N[+2206] | 1088.1102 | 5 | 5436.5221 | 5436.5038 | 0 | 3.4 | 141 | Specific | 930.01 | 784.48 | 784.48 | 18.76 | 122 | >sp\|P02788\|TRFL_HUMAN Lactotransferrin OS=Homo sapiens OX=9606 GN=LTF PE=1 SV=6 | 1270 | 20201119_15432_1ng_glyco_hcd203040_compound15.22659.22659.5 | controllerType=0 controllerNumber=1 scan=22659 | 46.2841 |
| 16194:3 | 1 | R.TAGWNIPMGLLFN[+2279.830]QTGSC[+57.021]K.F | HexNAc(5)Hex(6)Fuc(2) | C[+57], N[+2280] | 1458.9656 | 3 | 4374.8823 | 4374.8396 | 0 | 9.8 | 485 | Specific | 212.65 | 212.65 | 212.65 | 3.49 | 122 | >sp\|P02788\|TRFL_HUMAN Lactotransferrin OS=Homo sapiens OX=9606 GN=LTF PE=1 SV=6 | 1270 | 20201119_15432_1ng_glyco_hcd203040_compound15.19782.19782.3 | controllerType=0 controllerNumber=1 scan=19782 | 40.2076 |
| 18502:4 | 1 | R.[+42.011]TAGWNIPMGLLFN[+1872.651]QTGSC[+57.021]K.F | HexNAc(3)Hex(6)NeuAc(1) | .[42], C[+57], N[+1873] | 1003.1781 | 4 | 4009.6905 | 4009.671 | 0 | 4.9 | 485 | Specific | 216.01 | 162.46 | 162.46 | 3.44 | 122 | >sp\|P02788\|TRFL_HUMAN Lactotransferrin OS=Homo sapiens OX=9606 GN=LTF PE=1 SV=6 | 1270 | 20201119_15432_1ng_glyco_hcd203040_compound15.22195.22195.4 | controllerType=0 controllerNumber=1 scan=22195 | 45.2971 |
| 18504:4 | 1 | R.[+42.011]TAGWNIPMGLLFN[+1872.651]QTGSC[+57.021]K.F | HexNAc(3)Hex(6)NeuAc(1) | .[42], C[+57], N[+1873] | 1003.1781 | 4 | 4009.6905 | 4009.671 | 0 | 4.9 | 485 | Specific | 249.82 | 178.35 | 178.35 | 4.39 | 122 | >sp\|P02788\|TRFL_HUMAN Lactotransferrin OS=Homo sapiens OX=9606 GN=LTF PE=1 SV=6 | 1270 | 20201119_15432_1ng_glyco_hcd203040_compound15.22197.22197.4 | controllerType=0 controllerNumber=1 scan=22197 | 45.3019 |
| 16196:3 | 1 | R.TAGWNIPMGLLFN[+2279.830]QTGSC[+57.021]K.F | HexNAc(5)Hex(6)Fuc(2) | C[+57], N[+2280] | 1458.9656 | 3 | 4374.8823 | 4374.8396 | 0 | 9.8 | 485 | Specific | 367.82 | 367.82 | 299.06 | 5.96 | 122 | >sp\|P02788\|TRFL_HUMAN Lactotransferrin OS=Homo sapiens OX=9606 GN=LTF PE=1 SV=6 | 1270 | 20201119_15432_1ng_glyco_hcd203040_compound15.19784.19784.3 | controllerType=0 controllerNumber=1 scan=19784 | 40.2115 |
| 18332:4 | 1 | R.TAGWNIPMGLLFN[+2205.793]QTGSC[+57.021]K.F | HexNAc(4)Hex(5)Fuc(2)NeuAc(1) | C[+57], N[+2206] | 1075.9637 | 4 | 4300.833 | 4300.8028 | 0 | 7 | 485 | Specific | 475.75 | 472.22 | 472.22 | 8.58 | 122 | >sp\|P02788\|TRFL_HUMAN Lactotransferrin OS=Homo sapiens OX=9606 GN=LTF PE=1 SV=6 | 1270 | 20201119_15432_1ng_glyco_hcd203040_compound15.22017.22017.4 | controllerType=0 controllerNumber=1 scan=22017 | 44.8504 |
| 16449:2 | 1 | R.TAGWNIPMGLLFN[+1768.640]QTGSC[+57.021]K.F | HexNAc(4)Hex(5)Fuc(1) | C[+57], N[+1769] | 1932.3376 | 2 | 3863.6679 | 3863.6495 | 0 | 4.8 | 485 | Specific | 518.93 | 419.67 | 419.67 | 9.52 | 122 | >sp\|P02788\|TRFL_HUMAN Lactotransferrin OS=Homo sapiens OX=9606 GN=LTF PE=1 SV=6 | 1270 | 20201119_15432_1ng_glyco_hcd203040_compound15.20048.20048.2 | controllerType=0 controllerNumber=1 scan=20048 | 40.7121 |
| 16724:3 | 1 | R.TAGWNIPMGLLFN[+1460.529]QTGSC[+57.021]K.F | HexNAc(4)Hex(4) | C[+57], N[+1461] | 1185.8565 | 3 | 3555.555 | 3555.5387 | 0 | 4.6 | 485 | Specific | 543.27 | 543.27 | 212.97 | 9.49 | 122 | >sp\|P02788\|TRFL_HUMAN Lactotransferrin OS=Homo sapiens OX=9606 GN=LTF PE=1 SV=6 | 1270 | 20201119_15432_1ng_glyco_hcd203040_compound15.20336.20336.3 | controllerType=0 controllerNumber=1 scan=20336 | 41.3389 |
| 20372:4 | 1 | R.TAGWNIPMGLLFN[+2350.830]QTGSC[+57.021]K.F | HexNAc(4)Hex(5)Fuc(1)NeuAc(2) | C[+57], N[+2351] | 1112.2194 | 4 | 4445.8558 | 4445.8403 | 0 | 3.5 | 485 | Specific | 570.37 | 471 | 471 | 10.11 | 122 | >sp\|P02788\|TRFL_HUMAN Lactotransferrin OS=Homo sapiens OX=9606 GN=LTF PE=1 SV=6 | 1270 | 20201119_15432_1ng_glyco_hcd203040_compound15.24150.24150.4 | controllerType=0 controllerNumber=1 scan=24150 | 49.44 |
| 20375:4 | 1 | R.TAGWNIPMGLLFN[+2350.830]QTGSC[+57.021]K.F | HexNAc(4)Hex(5)Fuc(1)NeuAc(2) | C[+57], N[+2351] | 1112.2194 | 4 | 4445.8558 | 4445.8403 | 0 | 3.5 | 485 | Specific | 607.56 | 486.52 | 486.52 | 12.29 | 122 | >sp\|P02788\|TRFL_HUMAN Lactotransferrin OS=Homo sapiens OX=9606 GN=LTF PE=1 SV=6 | 1270 | 20201119_15432_1ng_glyco_hcd203040_compound15.24153.24153.4 | controllerType=0 controllerNumber=1 scan=24153 | 49.4504 |
| 16448:2 | 1 | R.TAGWNIPMGLLFN[+1768.640]QTGSC[+57.021]K.F | HexNAc(4)Hex(5)Fuc(1) | C[+57], N[+1769] | 1932.3376 | 2 | 3863.6679 | 3863.6495 | 0 | 4.8 | 485 | Specific | 617.36 | 586.63 | 586.63 | 12.04 | 122 | >sp\|P02788\|TRFL_HUMAN Lactotransferrin OS=Homo sapiens OX=9606 GN=LTF PE=1 SV=6 | 1270 | 20201119_15432_1ng_glyco_hcd203040_compound15.20047.20047.2 | controllerType=0 controllerNumber=1 scan=20047 | 40.7088 |
| 16421:2 | 1 | R.TAGWNIPMGLLFN[+1768.640]QTGSC[+57.021]K.F | HexNAc(4)Hex(5)Fuc(1) | C[+57], N[+1769] | 1932.3341 | 2 | 3863.6609 | 3863.6495 | 0 | 3 | 485 | Specific | 635.11 | 634.69 | 634.69 | 12.16 | 122 | >sp\|P02788\|TRFL_HUMAN Lactotransferrin OS=Homo sapiens OX=9606 GN=LTF PE=1 SV=6 | 1270 | 20201119_15432_1ng_glyco_hcd203040_compound15.20019.20019.2 | controllerType=0 controllerNumber=1 scan=20019 | 40.6505 |
| 16423:2 | 1 | R.TAGWNIPMGLLFN[+1768.640]QTGSC[+57.021]K.F | HexNAc(4)Hex(5)Fuc(1) | C[+57], N[+1769] | 1932.3341 | 2 | 3863.6609 | 3863.6495 | 0 | 3 | 485 | Specific | 665.06 | 582.86 | 582.86 | 13.16 | 122 | >sp\|P02788\|TRFL_HUMAN Lactotransferrin OS=Homo sapiens OX=9606 GN=LTF PE=1 SV=6 | 1270 | 20201119_15432_1ng_glyco_hcd203040_compound15.20021.20021.2 | controllerType=0 controllerNumber=1 scan=20021 | 40.6543 |
| 16727:3 | 1 | R.TAGWNIPMGLLFN[+1460.529]QTGSC[+57.021]K.F | HexNAc(4)Hex(4) | C[+57], N[+1461] | 1185.8565 | 3 | 3555.555 | 3555.5387 | 0 | 4.6 | 485 | Specific | 733.5 | 658.45 | 399.38 | 13.9 | 122 | >sp\|P02788\|TRFL_HUMAN Lactotransferrin OS=Homo sapiens OX=9606 GN=LTF PE=1 SV=6 | 1270 | 20201119_15432_1ng_glyco_hcd203040_compound15.20339.20339.3 | controllerType=0 controllerNumber=1 scan=20339 | 41.3442 |
| 18334:4 | 1 | R.TAGWNIPMGLLFN[+2205.793]QTGSC[+57.021]K.F | HexNAc(4)Hex(5)Fuc(2)NeuAc(1) | C[+57], N[+2206] | 1075.9637 | 4 | 4300.833 | 4300.8028 | 0 | 7 | 485 | Specific | 754.38 | 565.99 | 565.99 | 14.25 | 122 | >sp\|P02788\|TRFL_HUMAN Lactotransferrin OS=Homo sapiens OX=9606 GN=LTF PE=1 SV=6 | 1270 | 20201119_15432_1ng_glyco_hcd203040_compound15.22019.22019.4 | controllerType=0 controllerNumber=1 scan=22019 | 44.8541 |
| 16595:3 | 1 | R.TAGWNIPMGLLFN[+1622.582]QTGSC[+57.021]K.F | HexNAc(4)Hex(5) | C[+57], N[+1623] | 1239.8724 | 3 | 3717.6026 | 3717.5916 | 0 | 3 | 485 | Specific | 755.36 | 755.36 | 310.55 | 15.03 | 122 | >sp\|P02788\|TRFL_HUMAN Lactotransferrin OS=Homo sapiens OX=9606 GN=LTF PE=1 SV=6 | 1270 | 20201119_15432_1ng_glyco_hcd203040_compound15.20201.20201.3 | controllerType=0 controllerNumber=1 scan=20201 | 41.0478 |
| 18587:3 | 1 | R.TAGWNIPMGLLFN[+1751.624]QTGSC[+57.021]K.F | HexNAc(4)Hex(4)NeuAc(1) | C[+57], N[+1752] | 1282.8893 | 3 | 3846.6535 | 3846.6341 | 0 | 5 | 485 | Specific | 847.49 | 767.41 | 767.41 | 15.9 | 122 | >sp\|P02788\|TRFL_HUMAN Lactotransferrin OS=Homo sapiens OX=9606 GN=LTF PE=1 SV=6 | 1270 | 20201119_15432_1ng_glyco_hcd203040_compound15.22284.22284.3 | controllerType=0 controllerNumber=1 scan=22284 | 45.4801 |
| 18278:3 | 1 | R.TAGWNIPMGLLFN[+2205.793]QTGSC[+57.021]K.F | HexNAc(4)Hex(5)Fuc(2)NeuAc(1) | C[+57], N[+2206] | 1434.278 | 3 | 4300.8195 | 4300.8028 | 0 | 3.9 | 485 | Specific | 862.22 | 860.91 | 860.91 | 17.03 | 122 | >sp\|P02788\|TRFL_HUMAN Lactotransferrin OS=Homo sapiens OX=9606 GN=LTF PE=1 SV=6 | 1270 | 20201119_15432_1ng_glyco_hcd203040_compound15.21961.21961.3 | controllerType=0 controllerNumber=1 scan=21961 | 44.7297 |
| 16598:3 | 1 | R.TAGWNIPMGLLFN[+1622.582]QTGSC[+57.021]K.F | HexNAc(4)Hex(5) | C[+57], N[+1623] | 1239.8724 | 3 | 3717.6026 | 3717.5916 | 0 | 3 | 485 | Specific | 881.52 | 856.72 | 434.72 | 17.03 | 122 | >sp\|P02788\|TRFL_HUMAN Lactotransferrin OS=Homo sapiens OX=9606 GN=LTF PE=1 SV=6 | 1270 | 20201119_15432_1ng_glyco_hcd203040_compound15.20204.20204.3 | controllerType=0 controllerNumber=1 scan=20204 | 41.0524 |
| 16785:3 | 1 | R.TAGWNIPMGLLFN[+1216.423]QTGSC[+57.021]K.F | HexNAc(2)Hex(5) | C[+57], N[+1216] | 1104.4878 | 3 | 3311.4489 | 3311.4328 | 0 | 4.9 | 485 | Specific | 901.88 | 901.88 | 362.19 | 17.9 | 122 | >sp\|P02788\|TRFL_HUMAN Lactotransferrin OS=Homo sapiens OX=9606 GN=LTF PE=1 SV=6 | 1270 | 20201119_15432_1ng_glyco_hcd203040_compound15.20400.20400.3 | controllerType=0 controllerNumber=1 scan=20400 | 41.4817 |
| 18589:3 | 1 | R.TAGWNIPMGLLFN[+1751.624]QTGSC[+57.021]K.F | HexNAc(4)Hex(4)NeuAc(1) | C[+57], N[+1752] | 1282.8893 | 3 | 3846.6535 | 3846.6341 | 0 | 5 | 485 | Specific | 923.01 | 837.54 | 837.54 | 17.9 | 122 | >sp\|P02788\|TRFL_HUMAN Lactotransferrin OS=Homo sapiens OX=9606 GN=LTF PE=1 SV=6 | 1270 | 20201119_15432_1ng_glyco_hcd203040_compound15.22286.22286.3 | controllerType=0 controllerNumber=1 scan=22286 | 45.4849 |
| 18467:3 | 1 | R.TAGWNIPMGLLFN[+1913.677]QTGSC[+57.021]K.F | HexNAc(4)Hex(5)NeuAc(1) | C[+57], N[+1914] | 1336.903 | 3 | 4008.6945 | 4008.687 | 0 | 1.9 | 485 | Specific | 923.53 | 914.78 | 914.78 | 18.03 | 122 | >sp\|P02788\|TRFL_HUMAN Lactotransferrin OS=Homo sapiens OX=9606 GN=LTF PE=1 SV=6 | 1270 | 20201119_15432_1ng_glyco_hcd203040_compound15.22158.22158.3 | controllerType=0 controllerNumber=1 scan=22158 | 45.2042 |
| 16402:3 | 1 | R.TAGWNIPMGLLFN[+1768.640]QTGSC[+57.021]K.F | HexNAc(4)Hex(5)Fuc(1) | C[+57], N[+1769] | 1288.5582 | 3 | 3863.6599 | 3863.6495 | 0 | 2.7 | 485 | Specific | 934.37 | 934.37 | 934.37 | 18.03 | 122 | >sp\|P02788\|TRFL_HUMAN Lactotransferrin OS=Homo sapiens OX=9606 GN=LTF PE=1 SV=6 | 1270 | 20201119_15432_1ng_glyco_hcd203040_compound15.20000.20000.3 | controllerType=0 controllerNumber=1 scan=20000 | 40.6174 |
| 18464:3 | 1 | R.TAGWNIPMGLLFN[+1913.677]QTGSC[+57.021]K.F | HexNAc(4)Hex(5)NeuAc(1) | C[+57], N[+1914] | 1336.903 | 3 | 4008.6945 | 4008.687 | 0 | 1.9 | 485 | Specific | 948.54 | 865.38 | 865.38 | 18.03 | 122 | >sp\|P02788\|TRFL_HUMAN Lactotransferrin OS=Homo sapiens OX=9606 GN=LTF PE=1 SV=6 | 1270 | 20201119_15432_1ng_glyco_hcd203040_compound15.22155.22155.3 | controllerType=0 controllerNumber=1 scan=22155 | 45.1849 |
| 16787:3 | 1 | R.TAGWNIPMGLLFN[+1216.423]QTGSC[+57.021]K.F | HexNAc(2)Hex(5) | C[+57], N[+1216] | 1104.4878 | 3 | 3311.4489 | 3311.4328 | 0 | 4.9 | 485 | Specific | 970.37 | 970.37 | 459.5 | 18.9 | 122 | >sp\|P02788\|TRFL_HUMAN Lactotransferrin OS=Homo sapiens OX=9606 GN=LTF PE=1 SV=6 | 1270 | 20201119_15432_1ng_glyco_hcd203040_compound15.20402.20402.3 | controllerType=0 controllerNumber=1 scan=20402 | 41.4863 |
| 16498:3 | 1 | R.TAGWNIPMGLLFN[+1606.587]QTGSC[+57.021]K.F | HexNAc(4)Hex(4)Fuc(1) | C[+57], N[+1607] | 1234.5389 | 3 | 3701.6021 | 3701.5966 | 0 | 1.5 | 485 | Specific | 972.32 | 972.32 | 972.32 | 18.7 | 122 | >sp\|P02788\|TRFL_HUMAN Lactotransferrin OS=Homo sapiens OX=9606 GN=LTF PE=1 SV=6 | 1270 | 20201119_15432_1ng_glyco_hcd203040_compound15.20100.20100.3 | controllerType=0 controllerNumber=1 scan=20100 | 40.8346 |
| 16501:3 | 1 | R.TAGWNIPMGLLFN[+1606.587]QTGSC[+57.021]K.F | HexNAc(4)Hex(4)Fuc(1) | C[+57], N[+1607] | 1234.5389 | 3 | 3701.6021 | 3701.5966 | 0 | 1.5 | 485 | Specific | 973.55 | 732.45 | 732.45 | 18.7 | 122 | >sp\|P02788\|TRFL_HUMAN Lactotransferrin OS=Homo sapiens OX=9606 GN=LTF PE=1 SV=6 | 1270 | 20201119_15432_1ng_glyco_hcd203040_compound15.20103.20103.3 | controllerType=0 controllerNumber=1 scan=20103 | 40.8414 |
| 16400:3 | 1 | R.TAGWNIPMGLLFN[+1768.640]QTGSC[+57.021]K.F | HexNAc(4)Hex(5)Fuc(1) | C[+57], N[+1769] | 1288.5582 | 3 | 3863.6599 | 3863.6495 | 0 | 2.7 | 485 | Specific | 985.72 | 985.72 | 985.72 | 19.03 | 122 | >sp\|P02788\|TRFL_HUMAN Lactotransferrin OS=Homo sapiens OX=9606 GN=LTF PE=1 SV=6 | 1270 | 20201119_15432_1ng_glyco_hcd203040_compound15.19998.19998.3 | controllerType=0 controllerNumber=1 scan=19998 | 40.6121 |
| 18275:3 | 1 | R.TAGWNIPMGLLFN[+2205.793]QTGSC[+57.021]K.F | HexNAc(4)Hex(5)Fuc(2)NeuAc(1) | C[+57], N[+2206] | 1434.278 | 3 | 4300.8195 | 4300.8028 | 0 | 3.9 | 485 | Specific | 1029.58 | 924.16 | 924.16 | 20.03 | 122 | >sp\|P02788\|TRFL_HUMAN Lactotransferrin OS=Homo sapiens OX=9606 GN=LTF PE=1 SV=6 | 1270 | 20201119_15432_1ng_glyco_hcd203040_compound15.21958.21958.3 | controllerType=0 controllerNumber=1 scan=21958 | 44.7236 |
| 16772:3 | 1 | R.TAGWNIPM[+15.995]GLLFN[+2059.735]QTGSC[+57.021]K.F | HexNAc(4)Hex(5)Fuc(1)NeuAc(1) | C[+57], M[+16], N[+2060] | 1390.9261 | 3 | 4170.7637 | 4170.7398 | 0 | 5.7 | 485 | Specific | 271.7 | 210.92 | 210.92 | 4.25 | 122 | >sp\|P02788\|TRFL_HUMAN Lactotransferrin OS=Homo sapiens OX=9606 GN=LTF PE=1 SV=6 | 1270 | 20201119_15432_1ng_glyco_hcd203040_compound15.20386.20386.3 | controllerType=0 controllerNumber=1 scan=20386 | 41.4381 |
| 16774:3 | 1 | R.TAGWNIPM[+15.995]GLLFN[+2059.735]QTGSC[+57.021]K.F | HexNAc(4)Hex(5)Fuc(1)NeuAc(1) | C[+57], M[+16], N[+2060] | 1390.9261 | 3 | 4170.7637 | 4170.7398 | 0 | 5.7 | 485 | Specific | 511.87 | 403.9 | 403.9 | 8.76 | 122 | >sp\|P02788\|TRFL_HUMAN Lactotransferrin OS=Homo sapiens OX=9606 GN=LTF PE=1 SV=6 | 1270 | 20201119_15432_1ng_glyco_hcd203040_compound15.20388.20388.3 | controllerType=0 controllerNumber=1 scan=20388 | 41.4474 |
| 16649:3 | 1 | R.TAGWNIPM[+15.995]GLLFN[+2059.735]QTGSC[+57.021]K.F | HexNAc(4)Hex(5)Fuc(1)NeuAc(1) | C[+57], M[+16], N[+2060] | 1390.923 | 3 | 4170.7545 | 4170.7398 | 0 | 3.5 | 485 | Specific | 581.52 | 493.97 | 493.97 | 11.03 | 122 | >sp\|P02788\|TRFL_HUMAN Lactotransferrin OS=Homo sapiens OX=9606 GN=LTF PE=1 SV=6 | 1270 | 20201119_15432_1ng_glyco_hcd203040_compound15.20258.20258.3 | controllerType=0 controllerNumber=1 scan=20258 | 41.162 |
| 16651:3 | 1 | R.TAGWNIPM[+15.995]GLLFN[+2059.735]QTGSC[+57.021]K.F | HexNAc(4)Hex(5)Fuc(1)NeuAc(1) | C[+57], M[+16], N[+2060] | 1390.923 | 3 | 4170.7545 | 4170.7398 | 0 | 3.5 | 485 | Specific | 776.9 | 561.58 | 561.58 | 15.03 | 122 | >sp\|P02788\|TRFL_HUMAN Lactotransferrin OS=Homo sapiens OX=9606 GN=LTF PE=1 SV=6 | 1270 | 20201119_15432_1ng_glyco_hcd203040_compound15.20260.20260.3 | controllerType=0 controllerNumber=1 scan=20260 | 41.1666 |
| 17526:4 | 1 | R.[+42.011]TAGWNIPMGLLFN[+1872.651]QTGSC[+57.021]KFDEYFSQSC[+57.021]A | HexNAc(3)Hex(6)NeuAc(1) | .[42], C[+57]*2, N[+1873] | 1464.1237 | 4 | 5853.4728 | 5853.4181 | 0 | 9.3 | 485 | Specific | 210.25 | 210.25 | 210.25 | 3.42 | 122 | >sp\|P02788\|TRFL_HUMAN Lactotransferrin OS=Homo sapiens OX=9606 GN=LTF PE=1 SV=6 | 1270 | 20201119_15432_1ng_glyco_hcd203040_compound15.21177.21177.4 | controllerType=0 controllerNumber=1 scan=21177 | 43.1525 |
| 18603:4 | 1 | R.TAGWNIPMGLLFN[+2205.793]QTGSC[+57.021]KFDEYFSQSC[+57.021]APGSDPR.S | HexNAc(4)Hex(5)Fuc(2)NeuAc(1) | C[+57]*2, N[+2206] | 1536.9001 | 4 | 6144.5788 | 6144.5499 | 0 | 4.7 | 485 | Specific | 246.82 | 237.75 | 237.75 | 5.67 | 122 | >sp\|P02788\|TRFL_HUMAN Lactotransferrin OS=Homo sapiens OX=9606 GN=LTF PE=1 SV=6 | 1270 | 20201119_15432_1ng_glyco_hcd203040_compound15.22300.22300.3 | controllerType=0 controllerNumber=1 scan=22300 | 45.5214 |
| 17532:3 | 1 | R.TAGWNIPMGLLFN[+1768.640]QTGSC[+57.021]KFDEYFSQSC[+57.021]APGSDPR.S | HexNAc(4)Hex(5)Fuc(1) | C[+57]*2, N[+1769] | 1903.1427 | 3 | 5707.4134 | 5707.3966 | 0 | 2.9 | 485 | Specific | 404.32 | 338.77 | 338.77 | 7.41 | 122 | >sp\|P02788\|TRFL_HUMAN Lactotransferrin OS=Homo sapiens OX=9606 GN=LTF PE=1 SV=6 | 1270 | 20201119_15432_1ng_glyco_hcd203040_compound15.21183.21183.3 | controllerType=0 controllerNumber=1 scan=21183 | 43.16 |
| 17641:3 | 1 | R.TAGWNIPMGLLFN[+1606.587]QTGSC[+57.021]KFDEYFSQSC[+57.021]APGSDPR.S | HexNAc(4)Hex(4)Fuc(1) | C[+57]*2, N[+1607] | 1849.1286 | 3 | 5545.3713 | 5545.3438 | 0 | 5 | 485 | Specific | 411.15 | 385.14 | 385.14 | 8.85 | 122 | >sp\|P02788\|TRFL_HUMAN Lactotransferrin OS=Homo sapiens OX=9606 GN=LTF PE=1 SV=6 | 1270 | 20201119_15432_1ng_glyco_hcd203040_compound15.21296.21296.3 | controllerType=0 controllerNumber=1 scan=21296 | 43.3735 |
| 18586:4 | 1 | R.TAGWNIPMGLLFN[+2076.750]QTGSC[+57.021]KFDEYFSQSC[+57.021]APGSDPR.S | HexNAc(4)Hex(6)Fuc(2) | C[+57]*2, N[+2077] | 1504.6423 | 4 | 6015.5475 | 6015.5074 | 0 | 6.7 | 485 | Specific | 429.81 | 429.81 | 429.81 | 8.03 | 122 | >sp\|P02788\|TRFL_HUMAN Lactotransferrin OS=Homo sapiens OX=9606 GN=LTF PE=1 SV=6 | 1270 | 20201119_15432_1ng_glyco_hcd203040_compound15.22283.22283.4 | controllerType=0 controllerNumber=1 scan=22283 | 45.4786 |
| 18606:4 | 1 | R.TAGWNIPMGLLFN[+2205.793]QTGSC[+57.021]KFDEYFSQSC[+57.021]APGSDPR.S | HexNAc(4)Hex(5)Fuc(2)NeuAc(1) | C[+57]*2, N[+2206] | 1536.9001 | 4 | 6144.5788 | 6144.5499 | 0 | 4.7 | 485 | Specific | 450.8 | 420.11 | 420.11 | 9 | 122 | >sp\|P02788\|TRFL_HUMAN Lactotransferrin OS=Homo sapiens OX=9606 GN=LTF PE=1 SV=6 | 1270 | 20201119_15432_1ng_glyco_hcd203040_compound15.22303.22303.4 | controllerType=0 controllerNumber=1 scan=22303 | 45.5281 |
| 17634:4 | 1 | R.TAGWNIPMGLLFN[+1606.587]QTGSC[+57.021]KFDEYFSQSC[+57.021]APGSDPR.S | HexNAc(4)Hex(4)Fuc(1) | C[+57]*2, N[+1607] | 1387.0957 | 4 | 5545.361 | 5545.3438 | 0 | 3.1 | 485 | Specific | 508.26 | 507.48 | 507.48 | 9.4 | 122 | >sp\|P02788\|TRFL_HUMAN Lactotransferrin OS=Homo sapiens OX=9606 GN=LTF PE=1 SV=6 | 1270 | 20201119_15432_1ng_glyco_hcd203040_compound15.21289.21289.4 | controllerType=0 controllerNumber=1 scan=21289 | 43.3611 |
| 17600:3 | 1 | R.TAGWNIPMGLLFN[+1606.587]QTGSC[+57.021]KFDEYFSQSC[+57.021]APGSDPR.S | HexNAc(4)Hex(4)Fuc(1) | C[+57]*2, N[+1607] | 1849.1237 | 3 | 5545.3567 | 5545.3438 | 0 | 2.3 | 485 | Specific | 607.01 | 570.7 | 570.7 | 12.05 | 122 | >sp\|P02788\|TRFL_HUMAN Lactotransferrin OS=Homo sapiens OX=9606 GN=LTF PE=1 SV=6 | 1270 | 20201119_15432_1ng_glyco_hcd203040_compound15.21253.21253.3 | controllerType=0 controllerNumber=1 scan=21253 | 43.2883 |
| 17523:4 | 1 | R.TAGWNIPMGLLFN[+1768.640]QTGSC[+57.021]KFDEYFSQSC[+57.021]APGSDPR.S | HexNAc(4)Hex(5)Fuc(1) | C[+57]*2, N[+1769] | 1427.6094 | 4 | 5707.4157 | 5707.3966 | 0 | 3.3 | 485 | Specific | 633.09 | 617 | 617 | 11.92 | 122 | >sp\|P02788\|TRFL_HUMAN Lactotransferrin OS=Homo sapiens OX=9606 GN=LTF PE=1 SV=6 | 1270 | 20201119_15432_1ng_glyco_hcd203040_compound15.21174.21174.4 | controllerType=0 controllerNumber=1 scan=21174 | 43.1476 |
| 17535:3 | 1 | R.TAGWNIPMGLLFN[+1768.640]QTGSC[+57.021]KFDEYFSQSC[+57.021]APGSDPR.S | HexNAc(4)Hex(5)Fuc(1) | C[+57]*2, N[+1769] | 1903.1427 | 3 | 5707.4134 | 5707.3966 | 0 | 2.9 | 485 | Specific | 649.88 | 521.27 | 521.27 | 12.05 | 122 | >sp\|P02788\|TRFL_HUMAN Lactotransferrin OS=Homo sapiens OX=9606 GN=LTF PE=1 SV=6 | 1270 | 20201119_15432_1ng_glyco_hcd203040_compound15.21186.21186.3 | controllerType=0 controllerNumber=1 scan=21186 | 43.1637 |
| 17602:3 | 1 | R.TAGWNIPMGLLFN[+1606.587]QTGSC[+57.021]KFDEYFSQSC[+57.021]APGSDPR.S | HexNAc(4)Hex(4)Fuc(1) | C[+57]*2, N[+1607] | 1849.1237 | 3 | 5545.3567 | 5545.3438 | 0 | 2.3 | 485 | Specific | 669.67 | 601.42 | 601.42 | 13.05 | 122 | >sp\|P02788\|TRFL_HUMAN Lactotransferrin OS=Homo sapiens OX=9606 GN=LTF PE=1 SV=6 | 1270 | 20201119_15432_1ng_glyco_hcd203040_compound15.21255.21255.3 | controllerType=0 controllerNumber=1 scan=21255 | 43.2921 |
| 18633:4 | 1 | R.TAGWNIPMGLLFN[+2059.735]QTGSC[+57.021]KFDEYFSQSC[+57.021]APGSDPR.S | HexNAc(4)Hex(5)Fuc(1)NeuAc(1) | C[+57]*2, N[+2060] | 1500.3822 | 4 | 5998.507 | 5998.492 | 0 | 2.5 | 485 | Specific | 700.8 | 678.36 | 678.36 | 13.92 | 122 | >sp\|P02788\|TRFL_HUMAN Lactotransferrin OS=Homo sapiens OX=9606 GN=LTF PE=1 SV=6 | 1270 | 20201119_15432_1ng_glyco_hcd203040_compound15.22332.22332.4 | controllerType=0 controllerNumber=1 scan=22332 | 45.5842 |
| 18562:4 | 1 | R.TAGWNIPMGLLFN[+2205.793]QTGSC[+57.021]KFDEYFSQSC[+57.021]APGSDPR.S | HexNAc(4)Hex(5)Fuc(2)NeuAc(1) | C[+57]*2, N[+2206] | 1536.9028 | 4 | 6144.5895 | 6144.5499 | 0 | 6.4 | 485 | Specific | 798.56 | 766.57 | 766.57 | 14.16 | 122 | >sp\|P02788\|TRFL_HUMAN Lactotransferrin OS=Homo sapiens OX=9606 GN=LTF PE=1 SV=6 | 1270 | 20201119_15432_1ng_glyco_hcd203040_compound15.22258.22258.4 | controllerType=0 controllerNumber=1 scan=22258 | 45.4243 |
| 17637:4 | 1 | R.TAGWNIPMGLLFN[+1606.587]QTGSC[+57.021]KFDEYFSQSC[+57.021]APGSDPR.S | HexNAc(4)Hex(4)Fuc(1) | C[+57]*2, N[+1607] | 1387.0957 | 4 | 5545.361 | 5545.3438 | 0 | 3.1 | 485 | Specific | 854.54 | 842.85 | 842.85 | 16.92 | 122 | >sp\|P02788\|TRFL_HUMAN Lactotransferrin OS=Homo sapiens OX=9606 GN=LTF PE=1 SV=6 | 1270 | 20201119_15432_1ng_glyco_hcd203040_compound15.21292.21292.4 | controllerType=0 controllerNumber=1 scan=21292 | 43.3662 |
| 19876:4 | 1 | R.TAGWNIPMGLLFN[+2350.830]QTGSC[+57.021]KFDEYFSQSC[+57.021]APGSDPR.S | HexNAc(4)Hex(5)Fuc(1)NeuAc(2) | C[+57]*2, N[+2351] | 1573.1588 | 4 | 6289.6134 | 6289.5875 | 0 | 4.1 | 485 | Specific | 899.33 | 840.64 | 840.64 | 16.8 | 122 | >sp\|P02788\|TRFL_HUMAN Lactotransferrin OS=Homo sapiens OX=9606 GN=LTF PE=1 SV=6 | 1270 | 20201119_15432_1ng_glyco_hcd203040_compound15.23633.23633.4 | controllerType=0 controllerNumber=1 scan=23633 | 48.3588 |
| 17525:4 | 1 | R.TAGWNIPMGLLFN[+1768.640]QTGSC[+57.021]KFDEYFSQSC[+57.021]APGSDPR.S | HexNAc(4)Hex(5)Fuc(1) | C[+57]*2, N[+1769] | 1427.6094 | 4 | 5707.4157 | 5707.3966 | 0 | 3.3 | 485 | Specific | 920.27 | 864.63 | 864.63 | 17.92 | 122 | >sp\|P02788\|TRFL_HUMAN Lactotransferrin OS=Homo sapiens OX=9606 GN=LTF PE=1 SV=6 | 1270 | 20201119_15432_1ng_glyco_hcd203040_compound15.21176.21176.4 | controllerType=0 controllerNumber=1 scan=21176 | 43.1516 |
| 18564:4 | 1 | R.TAGWNIPMGLLFN[+2205.793]QTGSC[+57.021]KFDEYFSQSC[+57.021]APGSDPR.S | HexNAc(4)Hex(5)Fuc(2)NeuAc(1) | C[+57]*2, N[+2206] | 1536.9028 | 4 | 6144.5895 | 6144.5499 | 0 | 6.4 | 485 | Specific | 928.26 | 914.68 | 914.68 | 17.16 | 122 | >sp\|P02788\|TRFL_HUMAN Lactotransferrin OS=Homo sapiens OX=9606 GN=LTF PE=1 SV=6 | 1270 | 20201119_15432_1ng_glyco_hcd203040_compound15.22260.22260.4 | controllerType=0 controllerNumber=1 scan=22260 | 45.429 |
| 18635:4 | 1 | R.TAGWNIPMGLLFN[+2059.735]QTGSC[+57.021]KFDEYFSQSC[+57.021]APGSDPR.S | HexNAc(4)Hex(5)Fuc(1)NeuAc(1) | C[+57]*2, N[+2060] | 1500.3822 | 4 | 5998.507 | 5998.492 | 0 | 2.5 | 485 | Specific | 938.45 | 899.54 | 899.54 | 17.92 | 122 | >sp\|P02788\|TRFL_HUMAN Lactotransferrin OS=Homo sapiens OX=9606 GN=LTF PE=1 SV=6 | 1270 | 20201119_15432_1ng_glyco_hcd203040_compound15.22334.22334.4 | controllerType=0 controllerNumber=1 scan=22334 | 45.5883 |
| 18573:4 | 1 | R.TAGWNIPMGLLFN[+2059.735]QTGSC[+57.021]KFDEYFSQSC[+57.021]APGSDPR.S | HexNAc(4)Hex(5)Fuc(1)NeuAc(1) | C[+57]*2, N[+2060] | 1500.3892 | 4 | 5998.5348 | 5998.492 | 0 | 7.1 | 485 | Specific | 947.18 | 900.75 | 900.75 | 16.87 | 122 | >sp\|P02788\|TRFL_HUMAN Lactotransferrin OS=Homo sapiens OX=9606 GN=LTF PE=1 SV=6 | 1270 | 20201119_15432_1ng_glyco_hcd203040_compound15.22269.22269.4 | controllerType=0 controllerNumber=1 scan=22269 | 45.4476 |
| 18576:4 | 1 | R.TAGWNIPMGLLFN[+2059.735]QTGSC[+57.021]KFDEYFSQSC[+57.021]APGSDPR.S | HexNAc(4)Hex(5)Fuc(1)NeuAc(1) | C[+57]*2, N[+2060] | 1500.3892 | 4 | 5998.5348 | 5998.492 | 0 | 7.1 | 485 | Specific | 953.24 | 887.75 | 887.75 | 17.87 | 122 | >sp\|P02788\|TRFL_HUMAN Lactotransferrin OS=Homo sapiens OX=9606 GN=LTF PE=1 SV=6 | 1270 | 20201119_15432_1ng_glyco_hcd203040_compound15.22272.22272.4 | controllerType=0 controllerNumber=1 scan=22272 | 45.4558 |
| 19878:4 | 1 | R.TAGWNIPMGLLFN[+2350.830]QTGSC[+57.021]KFDEYFSQSC[+57.021]APGSDPR.S | HexNAc(4)Hex(5)Fuc(1)NeuAc(2) | C[+57]*2, N[+2351] | 1573.1588 | 4 | 6289.6134 | 6289.5875 | 0 | 4.1 | 485 | Specific | 963.74 | 930.81 | 930.81 | 18.8 | 122 | >sp\|P02788\|TRFL_HUMAN Lactotransferrin OS=Homo sapiens OX=9606 GN=LTF PE=1 SV=6 | 1270 | 20201119_15432_1ng_glyco_hcd203040_compound15.23635.23635.4 | controllerType=0 controllerNumber=1 scan=23635 | 48.3645 |
| 19890:4 | 1 | R.TAGWNIPM[+15.995]GLLFN[+2351.851]QTGSC[+57.021]KFDEYFSQSC[+57.021]A | HexNAc(4)Hex(5)Fuc(3)NeuAc(1) | C[+57]*2, M[+16], N[+2352] | 1577.4157 | 4 | 6306.6408 | 6306.6028 | 0 | 6 | 485 | Specific | 216.48 | 202.72 | 202.72 | 3.95 | 122 | >sp\|P02788\|TRFL_HUMAN Lactotransferrin OS=Homo sapiens OX=9606 GN=LTF PE=1 SV=6 | 1270 | 20201119_15432_1ng_glyco_hcd203040_compound15.23647.23647.4 | controllerType=0 controllerNumber=1 scan=23647 | [illegible] |
| 08806:3 | 1 | R.N[+2059.735]GSDC[+57.021]PDKFC[+57.021]LFQSETK.N | HexNAc(4)Hex(5)Fuc(1)NeuAc(1) | C[+57]*2, N[+2060] | 1364.879 | 3 | 4092.6223 | 4092.6088 | 0 | 3.3 | 642 | Specific | 371.3 | 311.18 | 311.18 | 6.73 | 122 | >sp\|P02788\|TRFL_HUMAN Lactotransferrin OS=Homo sapiens OX=9606 GN=LTF PE=1 SV=6 | 1270 | 20201119_15432_1ng_glyco_hcd203040_compound15.12053.12053.3 | controllerType=0 controllerNumber=1 scan=12053 | [illegible] |
| 08808:3 | 1 | R.N[+2059.735]GSDC[+57.021]PDKFC[+57.021]LFQSETK.N | HexNAc(4)Hex(5)Fuc(1)NeuAc(1) | C[+57]*2, N[+2060] | 1364.879 | 3 | 4092.6223 | 4092.6088 | 0 | 3.3 | 642 | Specific | 398.17 | 267.74 | 267.74 | 6.65 | 122 | >sp\|P02788\|TRFL_HUMAN Lactotransferrin OS=Homo sapiens OX=9606 GN=LTF PE=1 SV=6 | 1270 | 20201119_15432_1ng_glyco_hcd203040_compound15.12055.12055.3 | controllerType=0 controllerNumber=1 scan=12055 | 26.7257 |
| 11199:3 | 1 | R.N[+2350.830]GSDC[+57.021]PDKFC[+57.021]LFQSETK.N | HexNAc(4)Hex(5)Fuc(1)NeuAc(2) | C[+57]*2, N[+2351] | 1461.904 | 3 | 4383.6974 | 4383.7043 | 0 | -1.6 | 642 | Specific | 404.75 | 332.33 | 332.33 | 5.77 | 122 | >sp\|P02788\|TRFL_HUMAN Lactotransferrin OS=Homo sapiens OX=9606 GN=LTF PE=1 SV=6 | 1270 | 20201119_15432_1ng_glyco_hcd203040_compound15.14530.14530.3 | controllerType=0 controllerNumber=1 scan=14530 | 30.9442 |
| 11201:3 | 1 | R.N[+2350.830]GSDC[+57.021]PDKFC[+57.021]LFQSETK.N | HexNAc(4)Hex(5)Fuc(1)NeuAc(2) | C[+57]*2, N[+2351] | 1461.904 | 3 | 4383.6974 | 4383.7043 | 0 | -1.6 | 642 | Specific | 423.86 | 283.31 | 283.31 | 5.27 | 122 | >sp\|P02788\|TRFL_HUMAN Lactotransferrin OS=Homo sapiens OX=9606 GN=LTF PE=1 SV=6 | 1270 | 20201119_15432_1ng_glyco_hcd203040_compound15.14532.14532.3 | controllerType=0 controllerNumber=1 scan=14532 | 30.948 |
| 08681:3 | 1 | R.N[+2205.793]GSDC[+57.021]PDKFC[+57.021]LFQSETK.N | HexNAc(4)Hex(5)Fuc(2)NeuAc(1) | C[+57]*2, N[+2206] | 1413.564 | 3 | 4238.6775 | 4238.6667 | 0 | 2.5 | 642 | Specific | 545.54 | 400.3 | 400.3 | 9.35 | 122 | >sp\|P02788\|TRFL_HUMAN Lactotransferrin OS=Homo sapiens OX=9606 GN=LTF PE=1 SV=6 | 1270 | 20201119_15432_1ng_glyco_hcd203040_compound15.11923.11923.3 | controllerType=0 controllerNumber=1 scan=11923 | 26.5032 |
| 08678:3 | 1 | R.N[+2205.793]GSDC[+57.021]PDKFC[+57.021]LFQSETK.N | HexNAc(4)Hex(5)Fuc(2)NeuAc(1) | C[+57]*2, N[+2206] | 1413.564 | 3 | 4238.6775 | 4238.6667 | 0 | 2.5 | 642 | Specific | 573.23 | 442.37 | 442.37 | 9.58 | 122 | >sp\|P02788\|TRFL_HUMAN Lactotransferrin OS=Homo sapiens OX=9606 GN=LTF PE=1 SV=6 | 1270 | 20201119_15432_1ng_glyco_hcd203040_compound15.11920.11920.3 | controllerType=0 controllerNumber=1 scan=11920 | 26.4978 |
| 04426:3 | 4 | R.FSDGLESN[+1864.634]SSTQFEVKK.Y | HexNAc(2)Hex(9) | N[+1865] | 1256.5231 | 3 | 3767.5548 | 3767.5421 | 0 | 3.4 | 219 | Specific | 428.91 | 286.01 | 286.01 | 6.86 | 90 | >sp\|P0C0L4\|CO4A_HUMAN Complement C4-A OS=Homo sapiens OX=9606 GN=C4A PE=1 SV=2 | 1235 | 20201119_15432_1ng_glyco_hcd203040_compound15.7500.7500.3 | controllerType=0 controllerNumber=1 scan=7500 | 19.1574 |
| 04424:3 | 4 | R.FSDGLESN[+1864.634]SSTQFEVKK.Y | HexNAc(2)Hex(9) | N[+1865] | 1256.5231 | 3 | 3767.5548 | 3767.5421 | 0 | 3.4 | 219 | Specific | 591.42 | 450.16 | 450.16 | 10.76 | 90 | >sp\|P0C0L4\|CO4A_HUMAN Complement C4-A OS=Homo sapiens OX=9606 GN=C4A PE=1 SV=2 | 1235 | 20201119_15432_1ng_glyco_hcd203040_compound15.7498.7498.3 | controllerType=0 controllerNumber=1 scan=7498 | 19.1535 |
| 07402:3 | 5 | K.WEC[+57.021]KN[+2059.735]DTLLGIK.G | HexNAc(4)Hex(5)Fuc(1)NeuAc(1) | C[+57], N[+2060] | 1179.503 | 3 | 3536.4944 | 3536.4865 | 0 | 2.2 | 100 | Specific | 222.17 | 43.27 | 43.27 | 2.06 | 84 | >sp\|P22897\|MRC1_HUMAN Macrophage mannose receptor 1 OS=Homo sapiens OX=9606 GN=MRC1 PE=1 SV=1 | 1049 | 20201119_15432_1ng_glyco_hcd203040_compound15.10579.10579.3 | controllerType=0 controllerNumber=1 scan=10579 | 24.2136 |
| 07257:3 | 5 | K.WEC[+57.021]KN[+2205.793]DTLLGIK.G | HexNAc(4)Hex(5)Fuc(2)NeuAc(1) | C[+57], N[+2206] | 1228.1898 | 3 | 3682.5547 | 3682.5444 | 0 | 2.8 | 100 | Specific | 309.24 | 219.32 | 219.32 | 5.93 | 84 | >sp\|P22897\|MRC1_HUMAN Macrophage mannose receptor 1 OS=Homo sapiens OX=9606 GN=MRC1 PE=1 SV=1 | 1049 | 20201119_15432_1ng_glyco_hcd203040_compound15.10429.10429.3 | controllerType=0 controllerNumber=1 scan=10429 | 23.9639 |
| 12817:4 | 5 | K.KGN[+2059.735]TTLNSFVIPSESDVPTHC[+57.021]PSQWWPYAGHC[+57.021]YK.I | HexNAc(4)Hex(5)Fuc(1)NeuAc(1) | C[+57]*2, N[+2060] | 1506.6475 | 4 | 6023.568 | 6023.5567 | 0 | 1.9 | 342 | Specific | 464.4 | 464.4 | 464.4 | 8.78 | 84 | >sp\|P22897\|MRC1_HUMAN Macrophage mannose receptor 1 OS=Homo sapiens OX=9606 GN=MRC1 PE=1 SV=1 | 1049 | 20201119_15432_1ng_glyco_hcd203040_compound15.16230.16230.4 | controllerType=0 controllerNumber=1 scan=16230 | 33.7955 |
| 12819:4 | 5 | K.KGN[+2059.735]TTLNSFVIPSESDVPTHC[+57.021]PSQWWPYAGHC[+57.021]YK.I | HexNAc(4)Hex(5)Fuc(1)NeuAc(1) | C[+57]*2, N[+2060] | 1506.6475 | 4 | 6023.568 | 6023.5567 | 0 | 1.9 | 342 | Specific | 511.31 | 374.57 | 374.57 | 7.67 | 84 | >sp\|P22897\|MRC1_HUMAN Macrophage mannose receptor 1 OS=Homo sapiens OX=9606 GN=MRC1 PE=1 SV=1 | 1049 | 20201119_15432_1ng_glyco_hcd203040_compound15.16232.16232.4 | controllerType=0 controllerNumber=1 scan=16232 | 33.7995 |
| 12703:4 | 5 | K.KGN[+2205.793]TTLNSFVIPSESDVPTHC[+57.021]PSQWWPYAGHC[+57.021]YK.I | HexNAc(4)Hex(5)Fuc(2)NeuAc(1) | C[+57]*2, N[+2206] | 1543.165 | 4 | 6169.6383 | 6169.6146 | 0 | 3.8 | 342 | Specific | 576.11 | 519.55 | 519.55 | 10.58 | 84 | >sp\|P22897\|MRC1_HUMAN Macrophage mannose receptor 1 OS=Homo sapiens OX=9606 GN=MRC1 PE=1 SV=1 | 1049 | 20201119_15432_1ng_glyco_hcd203040_compound15.16112.16112.2 | controllerType=0 controllerNumber=1 scan=16112 | 33.5896 |
| 12705:4 | 5 | K.KGN[+2205.793]TTLNSFVIPSESDVPTHC[+57.021]PSQWWPYAGHC[+57.021]YK.I | HexNAc(4)Hex(5)Fuc(2)NeuAc(1) | C[+57]*2, N[+2206] | 1543.165 | 4 | 6169.6383 | 6169.6146 | 0 | 3.8 | 342 | Specific | 737.22 | 484.44 | 484.44 | 13.58 | 84 | >sp\|P22897\|MRC1_HUMAN Macrophage mannose receptor 1 OS=Homo sapiens OX=9606 GN=MRC1 PE=1 SV=1 | 1049 | 20201119_15432_1ng_glyco_hcd203040_compound15.16114.16114.2 | controllerType=0 controllerNumber=1 scan=16114 | 33.5936 |
| 18527:4 | 8 | K.QIGLYPVLVIDSSGYVNPN[+2076.750]YTGR.I | HexNAc(4)Hex(6)Fuc(2) | N[+2077] | 1151.2759 | 4 | 4602.0819 | 4602.0537 | 0 | 6.1 | 168 | Specific | 213.77 | 211.8 | 211.8 | 3.79 | 58 | >sp\|P01833\|PIGR_HUMAN Polymeric immunoglobulin receptor OS=Homo sapiens OX=9606 GN=PIGR PE=1 SV=4 | 902 | 20201119_15432_1ng_glyco_hcd203040_compound15.22221.22221.4 | controllerType=0 controllerNumber=1 scan=22221 | 45.3537 |
| 18529:4 | 8 | K.QIGLYPVLVIDSSGYVNPN[+2076.750]YTGR.I | HexNAc(4)Hex(6)Fuc(2) | N[+2077] | 1151.2759 | 4 | 4602.0819 | 4602.0537 | 0 | 6.1 | 168 | Specific | 373.79 | 313.21 | 313.21 | 5.93 | 58 | >sp\|P01833\|PIGR_HUMAN Polymeric immunoglobulin receptor OS=Homo sapiens OX=9606 GN=PIGR PE=1 SV=4 | 902 | 20201119_15432_1ng_glyco_hcd203040_compound15.22223.22223.4 | controllerType=0 controllerNumber=1 scan=22223 | 45.3577 |
| 18478:3 | 8 | K.QIGLYPVLVIDSSGYVNPN[+2205.793]YTGR.I | HexNAc(4)Hex(5)Fuc(2)NeuAc(1) | N[+2206] | 1577.707 | 3 | 4731.1063 | 4731.0963 | 0 | 2.1 | 168 | Specific | 498.77 | 498.77 | 498.77 | 9.89 | 58 | >sp\|P01833\|PIGR_HUMAN Polymeric immunoglobulin receptor OS=Homo sapiens OX=9606 GN=PIGR PE=1 SV=4 | 902 | 20201119_15432_1ng_glyco_hcd203040_compound15.22170.22170.3 | controllerType=0 controllerNumber=1 scan=22170 | 45.2419 |
| 18480:3 | 8 | K.QIGLYPVLVIDSSGYVNPN[+2205.793]YTGR.I | HexNAc(4)Hex(5)Fuc(2)NeuAc(1) | N[+2206] | 1577.707 | 3 | 4731.1063 | 4731.0963 | 0 | 2.1 | 168 | Specific | 551.19 | 546.68 | 546.68 | 10.37 | 58 | >sp\|P01833\|PIGR_HUMAN Polymeric immunoglobulin receptor OS=Homo sapiens OX=9606 GN=PIGR PE=1 SV=4 | 902 | 20201119_15432_1ng_glyco_hcd203040_compound15.22172.22172.3 | controllerType=0 controllerNumber=1 scan=22172 | 45.2456 |
| 18489:3 | 8 | K.QIGLYPVLVIDSSGYVNPN[+2059.735]YTGR.I | HexNAc(4)Hex(5)Fuc(1)NeuAc(1) | N[+2060] | 1529.0195 | 3 | 4585.0438 | 4585.0384 | 0 | 1.2 | 168 | Specific | 935.38 | 849.5 | 849.5 | 18.23 | 58 | >sp\|P01833\|PIGR_HUMAN Polymeric immunoglobulin receptor OS=Homo sapiens OX=9606 GN=PIGR PE=1 SV=4 | 902 | 20201119_15432_1ng_glyco_hcd203040_compound15.22182.22182.3 | controllerType=0 controllerNumber=1 scan=22182 | 45.2697 |
| 18487:3 | 8 | K.QIGLYPVLVIDSSGYVNPN[+2059.735]YTGR.I | HexNAc(4)Hex(5)Fuc(1)NeuAc(1) | N[+2060] | 1529.0195 | 3 | 4585.0438 | 4585.0384 | 0 | 1.2 | 168 | Specific | 950.85 | 839.8 | 839.8 | 19.23 | 58 | >sp\|P01833\|PIGR_HUMAN Polymeric immunoglobulin receptor OS=Homo sapiens OX=9606 GN=PIGR PE=1 SV=4 | 902 | 20201119_15432_1ng_glyco_hcd203040_compound15.22180.22180.3 | controllerType=0 controllerNumber=1 scan=22180 | 45.2617 |
| 22471:4 | 8 | K.AQYEGRLSLLEEPGN[+1663.608]GTFTVILNQLTSR.D | HexNAc(5)Hex(4) | N[+1664] | 1193.3017 | 4 | 4770.1849 | 4770.2249 | 0 | -8.4 | 407 | Specific | 241.49 | 208.86 | 208.86 | 4.2 | 58 | >sp\|P01833\|PIGR_HUMAN Polymeric immunoglobulin receptor OS=Homo sapiens OX=9606 GN=PIGR PE=1 SV=4 | 902 | 20201119_15432_1ng_glyco_hcd203040_compound15.26335.26335.4 | controllerType=0 controllerNumber=1 scan=26335 | 53.3724 |
| 22469:4 | 8 | K.AQYEGRLSLLEEPGN[+1663.608]GTFTVILNQLTSR.D | HexNAc(5)Hex(4) | N[+1664] | 1193.3017 | 4 | 4770.1849 | 4770.2249 | 0 | -8.4 | 407 | Specific | 292.29 | 279.27 | 279.27 | 4.65 | 58 | >sp\|P01833\|PIGR_HUMAN Polymeric immunoglobulin receptor OS=Homo sapiens OX=9606 GN=PIGR PE=1 SV=4 | 902 | 20201119_15432_1ng_glyco_hcd203040_compound15.26333.26333.4 | controllerType=0 controllerNumber=1 scan=26333 | 53.3683 |
| 22677:3 | 8 | R.LSLLEEPGN[+2351.851]GTFTVILNQLTSR.D | HexNAc(4)Hex(5)Fuc(3)NeuAc(1) | N[+2352] | 1585.3827 | 3 | 4754.1337 | 4754.1433 | 0 | -2 | 413 | Specific | 353.76 | 330.3 | 330.3 | 6.18 | 58 | >sp\|P01833\|PIGR_HUMAN Polymeric immunoglobulin receptor OS=Homo sapiens OX=9606 GN=PIGR PE=1 SV=4 | 902 | 20201119_15432_1ng_glyco_hcd203040_compound15.26548.26548.3 | controllerType=0 controllerNumber=1 scan=26548 | 53.7714 |
| 22678:3 | 8 | R.LSLLEEPGN[+2351.851]GTFTVILNQLTSR.D | HexNAc(4)Hex(5)Fuc(3)NeuAc(1) | N[+2352] | 1585.3827 | 3 | 4754.1337 | 4754.1433 | 0 | -2 | 413 | Specific | 467.71 | 467.71 | 467.71 | 8.13 | 58 | >sp\|P01833\|PIGR_HUMAN Polymeric immunoglobulin receptor OS=Homo sapiens OX=9606 GN=PIGR PE=1 SV=4 | 902 | 20201119_15432_1ng_glyco_hcd203040_compound15.26549.26549.3 | controllerType=0 controllerNumber=1 scan=26549 | 53.7746 |
| 22546:3 | 8 | R.LSLLEEPGN[+2350.830]GTFTVILNQLTSR.D | HexNAc(4)Hex(5)Fuc(1)NeuAc(2) | N[+2351] | 1585.0508 | 3 | 4753.138 | 4753.1229 | 0 | 3.2 | 413 | Specific | 648.03 | 560.91 | 560.91 | 12.55 | 58 | >sp\|P01833\|PIGR_HUMAN Polymeric immunoglobulin receptor OS=Homo sapiens OX=9606 GN=PIGR PE=1 SV=4 | 902 | 20201119_15432_1ng_glyco_hcd203040_compound15.26413.26413.3 | controllerType=0 controllerNumber=1 scan=26413 | 53.5149 |
| 22548:3 | 8 | R.LSLLEEPGN[+2350.830]GTFTVILNQLTSR.D | HexNAc(4)Hex(5)Fuc(1)NeuAc(2) | N[+2351] | 1585.0508 | 3 | 4753.138 | 4753.1229 | 0 | 3.2 | 413 | Specific | 715.18 | 454.34 | 454.34 | 14.55 | 58 | >sp\|P01833\|PIGR_HUMAN Polymeric immunoglobulin receptor OS=Homo sapiens OX=9606 GN=PIGR PE=1 SV=4 | 902 | 20201119_15432_1ng_glyco_hcd203040_compound15.26415.26415.3 | controllerType=0 controllerNumber=1 scan=26415 | 53.5186 |
| 22447:4 | 8 | R.LSLLEEPGN[+2350.830]GTFTVILNQLTSR.D | HexNAc(4)Hex(5)Fuc(1)NeuAc(2) | N[+2351] | 1189.0412 | 4 | 4753.1429 | 4753.1229 | 0 | 4.2 | 413 | Specific | 928.16 | 705.26 | 705.26 | 18.69 | 58 | >sp\|P01833\|PIGR_HUMAN Polymeric immunoglobulin receptor OS=Homo sapiens OX=9606 GN=PIGR PE=1 SV=4 | 902 | 20201119_15432_1ng_glyco_hcd203040_compound15.26310.26310.4 | controllerType=0 controllerNumber=1 scan=26310 | 53.3174 |
| 20077:4 | 8 | R.LSLLEEPGN[+2205.793]GTFTVILNQLTSR.D | HexNAc(4)Hex(5)Fuc(2)NeuAc(1) | N[+2206] | 1152.7792 | 4 | 4608.095 | 4608.0854 | 0 | 2.1 | 413 | Specific | 972.26 | 736.71 | 736.71 | 19.81 | 58 | >sp\|P01833\|PIGR_HUMAN Polymeric immunoglobulin receptor OS=Homo sapiens OX=9606 GN=PIGR PE=1 SV=4 | 902 | 20201119_15432_1ng_glyco_hcd203040_compound15.23842.23842.4 | controllerType=0 controllerNumber=1 scan=23842 | 48.7886 |
| 20059:3 | 8 | R.LSLLEEPGN[+2205.793]GTFTVILNQLTSR.D | HexNAc(4)Hex(5)Fuc(2)NeuAc(1) | N[+2206] | 1536.7053 | 3 | 4608.1012 | 4608.0854 | 0 | 3.4 | 413 | Specific | 985.96 | 733.39 | 733.39 | 19.55 | 58 | >sp\|P01833\|PIGR_HUMAN Polymeric immunoglobulin receptor OS=Homo sapiens OX=9606 GN=PIGR PE=1 SV=4 | 902 | 20201119_15432_1ng_glyco_hcd203040_compound15.23824.23824.3 | controllerType=0 controllerNumber=1 scan=23824 | 48.7496 |
| 22445:4 | 8 | R.LSLLEEPGN[+2350.830]GTFTVILNQLTSR.D | HexNAc(4)Hex(5)Fuc(1)NeuAc(2) | N[+2351] | 1189.0412 | 4 | 4753.1429 | 4753.1229 | 0 | 4.2 | 413 | Specific | 1057.42 | 839.9 | 839.9 | 21.69 | 58 | >sp\|P01833\|PIGR_HUMAN Polymeric immunoglobulin receptor OS=Homo sapiens OX=9606 GN=PIGR PE=1 SV=4 | 902 | 20201119_15432_1ng_glyco_hcd203040_compound15.26308.26308.4 | controllerType=0 controllerNumber=1 scan=26308 | 53.3111 |
| 20075:4 | 8 | R.LSLLEEPGN[+2205.793]GTFTVILNQLTSR.D | HexNAc(4)Hex(5)Fuc(2)NeuAc(1) | N[+2206] | 1152.7792 | 4 | 4608.095 | 4608.0854 | 0 | 2.1 | 413 | Specific | 1061.92 | 834.36 | 834.36 | 21.81 | 58 | >sp\|P01833\|PIGR_HUMAN Polymeric immunoglobulin receptor OS=Homo sapiens OX=9606 GN=PIGR PE=1 SV=4 | 902 | 20201119_15432_1ng_glyco_hcd203040_compound15.23840.23840.4 | controllerType=0 controllerNumber=1 scan=23840 | 48.7818 |
| 20057:3 | 8 | R.LSLLEEPGN[+2205.793]GTFTVILNQLTSR.D | HexNAc(4)Hex(5)Fuc(2)NeuAc(1) | N[+2206] | 1536.7053 | 3 | 4608.1012 | 4608.0854 | 0 | 3.4 | 413 | Specific | 1064.54 | 823.55 | 823.55 | 21.55 | 58 | >sp\|P01833\|PIGR_HUMAN Polymeric immunoglobulin receptor OS=Homo sapiens OX=9606 GN=PIGR PE=1 SV=4 | 902 | 20201119_15432_1ng_glyco_hcd203040_compound15.23822.23822.3 | controllerType=0 controllerNumber=1 scan=23822 | 48.7429 |
| 15664:3 | 8 | K.[+42.011]IIEGEPNLKVPGN[+2018.708]VTAVLGETLK.V | HexNAc(3)Hex(6)Fuc(1)NeuAc(1) | .[42], N[+2019] | 1484.7043 | 3 | 4452.0983 | 4452.0683 | 0 | 6.7 | 457 | Specific | 266.16 | 266.16 | 266.16 | 5.49 | 58 | >sp\|P01833\|PIGR_HUMAN Polymeric immunoglobulin receptor OS=Homo sapiens OX=9606 GN=PIGR PE=1 SV=4 | 902 | 20201119_15432_1ng_glyco_hcd203040_compound15.19226.19226.3 | controllerType=0 controllerNumber=1 scan=19226 | 39.2158 |
| 15666:3 | 8 | K.[+42.011]IIEGEPNLKVPGN[+2018.708]VTAVLGETLK.V | HexNAc(3)Hex(6)Fuc(1)NeuAc(1) | .[42], N[+2019] | 1484.7043 | 3 | 4452.0983 | 4452.0683 | 0 | 6.7 | 457 | Specific | 275.56 | 268.19 | 268.19 | 5.49 | 58 | >sp\|P01833\|PIGR_HUMAN Polymeric immunoglobulin receptor OS=Homo sapiens OX=9606 GN=PIGR PE=1 SV=4 | 902 | 20201119_15432_1ng_glyco_hcd203040_compound15.19228.19228.3 | controllerType=0 controllerNumber=1 scan=19228 | 39.2203 |
| 15726:3 | 8 | K.IIEGEPNLKVPGN[+2059.735]VTAVLGETLK.V | HexNAc(4)Hex(5)Fuc(1)NeuAc(1) | N[+2060] | 1484.3716 | 3 | 4451.1004 | 4451.0843 | 0 | 3.6 | 457 | Specific | 616.62 | 616.62 | 616.62 | 12.28 | 58 | >sp\|P01833\|PIGR_HUMAN Polymeric immunoglobulin receptor OS=Homo sapiens OX=9606 GN=PIGR PE=1 SV=4 | 902 | 20201119_15432_1ng_glyco_hcd203040_compound15.19291.19291.3 | controllerType=0 controllerNumber=1 scan=19291 | 39.3339 |
| 15728:3 | 8 | K.IIEGEPNLKVPGN[+2059.735]VTAVLGETLK.V | HexNAc(4)Hex(5)Fuc(1)NeuAc(1) | N[+2060] | 1484.3716 | 3 | 4451.1004 | 4451.0843 | 0 | 3.6 | 457 | Specific | 752.7 | 651.36 | 651.36 | 15.28 | 58 | >sp\|P01833\|PIGR_HUMAN Polymeric immunoglobulin receptor OS=Homo sapiens OX=9606 GN=PIGR PE=1 SV=4 | 902 | 20201119_15432_1ng_glyco_hcd203040_compound15.19293.19293.3 | controllerType=0 controllerNumber=1 scan=19293 | 39.3373 |
| 15637:3 | 8 | K.IIEGEPNLKVPGN[+2205.793]VTAVLGETLK.V | HexNAc(4)Hex(5)Fuc(2)NeuAc(1) | N[+2206] | 1533.0551 | 3 | 4597.1508 | 4597.1422 | 0 | 1.9 | 457 | Specific | 835.53 | 743.81 | 743.81 | 16.28 | 58 | >sp\|P01833\|PIGR_HUMAN Polymeric immunoglobulin receptor OS=Homo sapiens OX=9606 GN=PIGR PE=1 SV=4 | 902 | 20201119_15432_1ng_glyco_hcd203040_compound15.19198.19198.3 | controllerType=0 controllerNumber=1 scan=19198 | 39.1679 |
| 15635:3 | 8 | K.IIEGEPNLKVPGN[+2205.793]VTAVLGETLK.V | HexNAc(4)Hex(5)Fuc(2)NeuAc(1) | N[+2206] | 1533.0551 | 3 | 4597.1508 | 4597.1422 | 0 | 1.9 | 457 | Specific | 1056.56 | 973.34 | 973.34 | 21.28 | 58 | >sp\|P01833\|PIGR_HUMAN Polymeric immunoglobulin receptor OS=Homo sapiens OX=9606 GN=PIGR PE=1 SV=4 | 902 | 20201119_15432_1ng_glyco_hcd203040_compound15.19196.19196.3 | controllerType=0 controllerNumber=1 scan=19196 | 39.1623 |
| 15208:5 | 8 | K.IIEGEPNLKVPGN[+2059.735]VTAVLGETLKVPC[+57.021]HFPC[+57.021]K.F | HexNAc(4)Hex(5)Fuc(1)NeuAc(1) | C[+57]*2, N[+2060] | 1096.1239 | 5 | 5476.5904 | 5476.5418 | 0 | 8.9 | 457 | Specific | 551.01 | 551.01 | 551.01 | 8.3 | 58 | >sp\|P01833\|PIGR_HUMAN Polymeric immunoglobulin receptor OS=Homo sapiens OX=9606 GN=PIGR PE=1 SV=4 | 902 | 20201119_15432_1ng_glyco_hcd203040_compound15.18712.18712.5 | controllerType=0 controllerNumber=1 scan=18712 | 38.2939 |
| 16661:4 | 8 | K.IIEGEPNLKVPGN[+2350.830]VTAVLGETLKVPC[+57.021]HFPC[+57.021]K.F | HexNAc(4)Hex(5)Fuc(1)NeuAc(2) | C[+57]*2, N[+2351] | 1442.6721 | 4 | 5767.6667 | 5767.6372 | 0 | 5.1 | 457 | Specific | 605.49 | 526.02 | 526.02 | 11.01 | 58 | >sp\|P01833\|PIGR_HUMAN Polymeric immunoglobulin receptor OS=Homo sapiens OX=9606 GN=PIGR PE=1 SV=4 | 902 | 20201119_15432_1ng_glyco_hcd203040_compound15.20270.20270.4 | controllerType=0 controllerNumber=1 scan=20270 | 41.192 |
| 15247:5 | 8 | K.IIEGEPNLKVPGN[+2205.793]VTAVLGETLKVPC[+57.021]HFPC[+57.021]K.F | HexNAc(4)Hex(5)Fuc(2)NeuAc(1) | C[+57]*2, N[+2206] | 1125.327 | 5 | 5622.606 | 5622.5997 | 0 | 1.1 | 457 | Specific | 607.67 | 571.8 | 571.8 | 11.35 | 58 | >sp\|P01833\|PIGR_HUMAN Polymeric immunoglobulin receptor OS=Homo sapiens OX=9606 GN=PIGR PE=1 SV=4 | 902 | 20201119_15432_1ng_glyco_hcd203040_compound15.18753.18753.5 | controllerType=0 controllerNumber=1 scan=18753 | 38.374 |
| 16671:4 | 8 | K.IIEGEPNLKVPGN[+2350.830]VTAVLGETLKVPC[+57.021]HFPC[+57.021]K.F | HexNAc(4)Hex(5)Fuc(1)NeuAc(2) | C[+57]*2, N[+2351] | 1442.6704 | 4 | 5767.6598 | 5767.6372 | 0 | 3.9 | 457 | Specific | 619.71 | 574.15 | 574.15 | 11.13 | 58 | >sp\|P01833\|PIGR_HUMAN Polymeric immunoglobulin receptor OS=Homo sapiens OX=9606 GN=PIGR PE=1 SV=4 | 902 | 20201119_15432_1ng_glyco_hcd203040_compound15.20281.20281.4 | controllerType=0 controllerNumber=1 scan=20281 | 41.2196 |
| 15249:5 | 8 | K.IIEGEPNLKVPGN[+2205.793]VTAVLGETLKVPC[+57.021]HFPC[+57.021]K.F | HexNAc(4)Hex(5)Fuc(2)NeuAc(1) | C[+57]*2, N[+2206] | 1125.327 | 5 | 5622.606 | 5622.5997 | 0 | 1.1 | 457 | Specific | 640.54 | 640.54 | 640.54 | 11.35 | 58 | >sp\|P01833\|PIGR_HUMAN Polymeric immunoglobulin receptor OS=Homo sapiens OX=9606 GN=PIGR PE=1 SV=4 | 902 | 20201119_15432_1ng_glyco_hcd203040_compound15.18755.18755.5 | controllerType=0 controllerNumber=1 scan=18755 | 38.3776 |
| 15257:4 | 8 | K.IIEGEPNLKVPGN[+2205.793]VTAVLGETLKVPC[+57.021]HFPC[+57.021]K.F | HexNAc(4)Hex(5)Fuc(2)NeuAc(1) | C[+57]*2, N[+2206] | 1406.4102 | 4 | 5622.619 | 5622.5997 | 0 | 3.4 | 457 | Specific | 641.82 | 613.47 | 613.47 | 11.13 | 58 | >sp\|P01833\|PIGR_HUMAN Polymeric immunoglobulin receptor OS=Homo sapiens OX=9606 GN=PIGR PE=1 SV=4 | 902 | 20201119_15432_1ng_glyco_hcd203040_compound15.18763.18763.4 | controllerType=0 controllerNumber=1 scan=18763 | 38.3923 |
| 15254:4 | 8 | K.IIEGEPNLKVPGN[+2205.793]VTAVLGETLKVPC[+57.021]HFPC[+57.021]K.F | HexNAc(4)Hex(5)Fuc(2)NeuAc(1) | C[+57]*2, N[+2206] | 1406.4102 | 4 | 5622.619 | 5622.5997 | 0 | 3.4 | 457 | Specific | 711.93 | 680.29 | 680.29 | 13.13 | 58 | >sp\|P01833\|PIGR_HUMAN Polymeric immunoglobulin receptor OS=Homo sapiens OX=9606 GN=PIGR PE=1 SV=4 | 902 | 20201119_15432_1ng_glyco_hcd203040_compound15.18760.18760.4 | controllerType=0 controllerNumber=1 scan=18760 | 38.3872 |
| 15211:5 | 8 | K.IIEGEPNLKVPGN[+2059.735]VTAVLGETLKVPC[+57.021]HFPC[+57.021]K.F | HexNAc(4)Hex(5)Fuc(1)NeuAc(1) | C[+57]*2, N[+2060] | 1096.1239 | 5 | 5476.5904 | 5476.5418 | 0 | 8.9 | 457 | Specific | 773.49 | 773.49 | 773.49 | 13.48 | 58 | >sp\|P01833\|PIGR_HUMAN Polymeric immunoglobulin receptor OS=Homo sapiens OX=9606 GN=PIGR PE=1 SV=4 | 902 | 20201119_15432_1ng_glyco_hcd203040_compound15.18715.18715.5 | controllerType=0 controllerNumber=1 scan=18715 | 38.2986 |
| 16669:4 | 8 | K.IIEGEPNLKVPGN[+2350.830]VTAVLGETLKVPC[+57.021]HFPC[+57.021]K.F | HexNAc(4)Hex(5)Fuc(1)NeuAc(2) | C[+57]*2, N[+2351] | 1442.6704 | 4 | 5767.6598 | 5767.6372 | 0 | 3.9 | 457 | Specific | 833.46 | 618.15 | 618.15 | 15.13 | 58 | >sp\|P01833\|PIGR_HUMAN Polymeric immunoglobulin receptor OS=Homo sapiens OX=9606 GN=PIGR PE=1 SV=4 | 902 | 20201119_15432_1ng_glyco_hcd203040_compound15.20279.20279.4 | controllerType=0 controllerNumber=1 scan=20279 | 41.2152 |
| 16653:5 | 8 | K.IIEGEPNLKVPGN[+2350.830]VTAVLGETLKVPC[+57.021]HFPC[+57.021]K.F | HexNAc(4)Hex(5)Fuc(1)NeuAc(2) | C[+57]*2, N[+2351] | 1154.3374 | 5 | 5767.6579 | 5767.6372 | 0 | 3.6 | 457 | Specific | 903.85 | 665.01 | 665.01 | 17.67 | 58 | >sp\|P01833\|PIGR_HUMAN Polymeric immunoglobulin receptor OS=Homo sapiens OX=9606 GN=PIGR PE=1 SV=4 | 902 | 20201119_15432_1ng_glyco_hcd203040_compound15.20262.20262.5 | controllerType=0 controllerNumber=1 scan=20262 | 41.1699 |
| 16659:4 | 8 | K.IIEGEPNLKVPGN[+2350.830]VTAVLGETLKVPC[+57.021]HFPC[+57.021]K.F | HexNAc(4)Hex(5)Fuc(1)NeuAc(2) | C[+57]*2, N[+2351] | 1442.6721 | 4 | 5767.6667 | 5767.6372 | 0 | 5.1 | 457 | Specific | 937.63 | 806.7 | 806.7 | 17.01 | 58 | >sp\|P01833\|PIGR_HUMAN Polymeric immunoglobulin receptor OS=Homo sapiens OX=9606 GN=PIGR PE=1 SV=4 | 902 | 20201119_15432_1ng_glyco_hcd203040_compound15.20268.20268.4 | controllerType=0 controllerNumber=1 scan=20268 | 41.187 |
| 16655:5 | 8 | K.IIEGEPNLKVPGN[+2350.830]VTAVLGETLKVPC[+57.021]HFPC[+57.021]K.F | HexNAc(4)Hex(5)Fuc(1)NeuAc(2) | C[+57]*2, N[+2351] | 1154.3374 | 5 | 5767.6579 | 5767.6372 | 0 | 3.6 | 457 | Specific | 972.33 | 812.34 | 812.34 | 18.67 | 58 | >sp\|P01833\|PIGR_HUMAN Polymeric immunoglobulin receptor OS=Homo sapiens OX=9606 GN=PIGR PE=1 SV=4 | 902 | 20201119_15432_1ng_glyco_hcd203040_compound15.20264.20264.5 | controllerType=0 controllerNumber=1 scan=20264 | 41.1769 |
| 13165:3 | 8 | K.VPGN[+2205.793]VTAVLGETLK.V | HexNAc(4)Hex(5)Fuc(2)NeuAc(1) | N[+2206] | 1201.8726 | 3 | 3603.6033 | 3603.5927 | 0 | 2.9 | 466 | Specific | 643.75 | 566.19 | 566.19 | 11.93 | 58 | >sp\|P01833\|PIGR_HUMAN Polymeric immunoglobulin receptor OS=Homo sapiens OX=9606 GN=PIGR PE=1 SV=4 | 902 | 20201119_15432_1ng_glyco_hcd203040_compound15.16590.16590.3 | controllerType=0 controllerNumber=1 scan=16590 | 34.439 |
| 13167:3 | 8 | K.VPGN[+2205.793]VTAVLGETLK.V | HexNAc(4)Hex(5)Fuc(2)NeuAc(1) | N[+2206] | 1201.8726 | 3 | 3603.6033 | 3603.5927 | 0 | 2.9 | 466 | Specific | 677.2 | 561.53 | 561.53 | 12.93 | 58 | >sp\|P01833\|PIGR_HUMAN Polymeric immunoglobulin receptor OS=Homo sapiens OX=9606 GN=PIGR PE=1 SV=4 | 902 | 20201119_15432_1ng_glyco_hcd203040_compound15.16592.16592.3 | controllerType=0 controllerNumber=1 scan=16592 | 34.4432 |
| 13292:4 | 8 | K.VPGN[+2205.793]VTAVLGETLKVPC[+57.021]HFPC[+57.021]K.F | HexNAc(4)Hex(5)Fuc(2)NeuAc(1) | C[+57]*2, N[+2206] | 1158.0226 | 4 | 4629.0687 | 4629.0502 | 0 | 4 | 466 | Specific | 528.08 | 381.19 | 381.19 | 9.87 | 58 | >sp\|P01833\|PIGR_HUMAN Polymeric immunoglobulin receptor OS=Homo sapiens OX=9606 GN=PIGR PE=1 SV=4 | 902 | 20201119_15432_1ng_glyco_hcd203040_compound15.16722.16722.3 | controllerType=0 controllerNumber=1 scan=16722 | 34.6917 |
| 13294:4 | 8 | K.VPGN[+2205.793]VTAVLGETLKVPC[+57.021]HFPC[+57.021]K.F | HexNAc(4)Hex(5)Fuc(2)NeuAc(1) | C[+57]*2, N[+2206] | 1158.0226 | 4 | 4629.0687 | 4629.0502 | 0 | 4 | 466 | Specific | 638.53 | 364.78 | 364.78 | 10.89 | 58 | >sp\|P01833\|PIGR_HUMAN Polymeric immunoglobulin receptor OS=Homo sapiens OX=9606 GN=PIGR PE=1 SV=4 | 902 | 20201119_15432_1ng_glyco_hcd203040_compound15.16724.16724.3 | controllerType=0 controllerNumber=1 scan=16724 | 34.6957 |
| 13202:4 | 8 | K.VPGN[+2205.793]VTAVLGETLKVPC[+57.021]HFPC[+57.021]K.F | HexNAc(4)Hex(5)Fuc(2)NeuAc(1) | C[+57]*2, N[+2206] | 1158.0226 | 4 | 4629.0687 | 4629.0502 | 0 | 4 | 466 | Specific | 701.22 | 357.66 | 357.66 | 12.89 | 58 | >sp\|P01833\|PIGR_HUMAN Polymeric immunoglobulin receptor OS=Homo sapiens OX=9606 GN=PIGR PE=1 SV=4 | 902 | 20201119_15432_1ng_glyco_hcd203040_compound15.16629.16629.4 | controllerType=0 controllerNumber=1 scan=16629 | 34.505 |
| 13200:4 | 8 | K.VPGN[+2205.793]VTAVLGETLKVPC[+57.021]HFPC[+57.021]K.F | HexNAc(4)Hex(5)Fuc(2)NeuAc(1) | C[+57]*2, N[+2206] | 1158.0226 | 4 | 4629.0687 | 4629.0502 | 0 | 4 | 466 | Specific | 853.53 | 478.62 | 478.62 | 17.28 | 58 | >sp\|P01833\|PIGR_HUMAN Polymeric immunoglobulin receptor OS=Homo sapiens OX=9606 GN=PIGR PE=1 SV=4 | 902 | 20201119_15432_1ng_glyco_hcd203040_compound15.16627.16627.4 | controllerType=0 controllerNumber=1 scan=16627 | 34.4995 |
| 05391:3 | 8 | K.WN[+1872.651]NTGC[+57.021]QALPSQDEGPSK.A | HexNAc(3)Hex(6)NeuAc(1) | C[+57], N[+1873] | 1287.8508 | 3 | 3861.5377 | 3861.5272 | 0 | 2.7 | 498 | Specific | 323.94 | 305.45 | 305.45 | 6.79 | 58 | >sp\|P01833\|PIGR_HUMAN Polymeric immunoglobulin receptor OS=Homo sapiens OX=9606 GN=PIGR PE=1 SV=4 | 902 | 20201119_15432_1ng_glyco_hcd203040_compound15.8498.8498.3 | controllerType=0 controllerNumber=1 scan=8498 | 20.8177 |
| 05369:3 | 8 | K.WN[+1872.651]NTGC[+57.021]QALPSQDEGPSK.A | HexNAc(3)Hex(6)NeuAc(1) | C[+57], N[+1873] | 1287.8515 | 3 | 3861.5399 | 3861.5272 | 0 | 3.3 | 498 | Specific | 351.14 | 329.4 | 329.4 | 7.27 | 58 | >sp\|P01833\|PIGR_HUMAN Polymeric immunoglobulin receptor OS=Homo sapiens OX=9606 GN=PIGR PE=1 SV=4 | 902 | 20201119_15432_1ng_glyco_hcd203040_compound15.8475.8475.3 | controllerType=0 controllerNumber=1 scan=8475 | 20.7777 |
| 05393:3 | 8 | K.WN[+1872.651]NTGC[+57.021]QALPSQDEGPSK.A | HexNAc(3)Hex(6)NeuAc(1) | C[+57], N[+1873] | 1287.8508 | 3 | 3861.5377 | 3861.5272 | 0 | 2.7 | 498 | Specific | 431.45 | 399.5 | 399.5 | 9.18 | 58 | >sp\|P01833\|PIGR_HUMAN Polymeric immunoglobulin receptor OS=Homo sapiens OX=9606 GN=PIGR PE=1 SV=4 | 902 | 20201119_15432_1ng_glyco_hcd203040_compound15.8500.8500.3 | controllerType=0 controllerNumber=1 scan=8500 | 20.8216 |
| 05371:3 | 8 | K.WN[+1872.651]NTGC[+57.021]QALPSQDEGPSK.A | HexNAc(3)Hex(6)NeuAc(1) | C[+57], N[+1873] | 1287.8515 | 3 | 3861.5399 | 3861.5272 | 0 | 3.3 | 498 | Specific | 434.74 | 408.13 | 408.13 | 9.19 | 58 | >sp\|P01833\|PIGR_HUMAN Polymeric immunoglobulin receptor OS=Homo sapiens OX=9606 GN=PIGR PE=1 SV=4 | 902 | 20201119_15432_1ng_glyco_hcd203040_compound15.8477.8477.3 | controllerType=0 controllerNumber=1 scan=8477 | 20.7817 |
| 05390:3 | 8 | K.WN[+1913.677]NTGC[+57.021]QALPSQDEGPSK.A | HexNAc(4)Hex(5)NeuAc(1) | C[+57], N[+1914] | 1301.5269 | 3 | 3902.5662 | 3902.5537 | 0 | 3.2 | 498 | Specific | 615.26 | 553.05 | 553.05 | 12.03 | 58 | >sp\|P01833\|PIGR_HUMAN Polymeric immunoglobulin receptor OS=Homo sapiens OX=9606 GN=PIGR PE=1 SV=4 | 902 | 20201119_15432_1ng_glyco_hcd203040_compound15.8497.8497.3 | controllerType=0 controllerNumber=1 scan=8497 | 20.8157 |
| 05388:3 | 8 | K.WN[+1913.677]NTGC[+57.021]QALPSQDEGPSK.A | HexNAc(4)Hex(5)NeuAc(1) | C[+57], N[+1914] | 1301.5269 | 3 | 3902.5662 | 3902.5537 | 0 | 3.2 | 498 | Specific | 710.45 | 642.88 | 642.88 | 14.03 | 58 | >sp\|P01833\|PIGR_HUMAN Polymeric immunoglobulin receptor OS=Homo sapiens OX=9606 GN=PIGR PE=1 SV=4 | 902 | 20201119_15432_1ng_glyco_hcd203040_compound15.8495.8495.3 | controllerType=0 controllerNumber=1 scan=8495 | 20.812 |
| 23480:4 | 9 | K.VFPLSLDSTPQDGNVVVAC[+57.021]LVQGFFPQEPLSVTWSESGQN[+2263.835]VTAI | HexNAc(5)Hex(5)Fuc(3) | C[+57], N[+2264] | 1760.8083 | 4 | 7040.2114 | 7040.1961 | 0 | 2.2 | 8 | Specific | 232.85 | 223.64 | 223.64 | 7.08 | 60 | >sp\|P01877\|IGHA2_HUMAN Immunoglobulin heavy constant alpha 2 OS=Homo sapiens OX=9606 GN=IGHA2 PE=1 SV=4 | 576 | 20201119_15432_1ng_glyco_hcd203040_compound15.27384.27384.4 | controllerType=0 controllerNumber=1 scan=27384 | 55.3379 |
| 23482:4 | 9 | K.VFPLSLDSTPQDGNVVVAC[+57.021]LVQGFFPQEPLSVTWSESGQN[+2263.835]VTAI | HexNAc(5)Hex(5)Fuc(3) | C[+57], N[+2264] | 1760.8083 | 4 | 7040.2114 | 7040.1961 | 0 | 2.2 | 8 | Specific | 233.17 | 230.39 | 230.39 | 7.11 | 60 | >sp\|P01877\|IGHA2_HUMAN Immunoglobulin heavy constant alpha 2 OS=Homo sapiens OX=9606 GN=IGHA2 PE=1 SV=4 | 576 | 20201119_15432_1ng_glyco_hcd203040_compound15.27386.27386.4 | controllerType=0 controllerNumber=1 scan=27386 | 55.3417 |
| 21968:4 | 9 | K.VFPLSLDSTPQDGNVVVAC[+57.021]LVQGFFPQEPLSVTWSESGQN[+1647.613]VTAI | HexNAc(5)Hex(3)Fuc(1) | C[+57], N[+1648] | 1606.7501 | 4 | 6423.9787 | 6423.9746 | 0 | 0.6 | 8 | Specific | 518 | 518 | 518 | 10.4 | 60 | >sp\|P01877\|IGHA2_HUMAN Immunoglobulin heavy constant alpha 2 OS=Homo sapiens OX=9606 GN=IGHA2 PE=1 SV=4 | 576 | 20201119_15432_1ng_glyco_hcd203040_compound15.25813.25813.5 | controllerType=0 controllerNumber=1 scan=25813 | 52.3994 |
| 21970:4 | 9 | K.VFPLSLDSTPQDGNVVVAC[+57.021]LVQGFFPQEPLSVTWSESGQN[+1647.613]VTAI | HexNAc(5)Hex(3)Fuc(1) | C[+57], N[+1648] | 1606.7501 | 4 | 6423.9787 | 6423.9746 | 0 | 0.6 | 8 | Specific | 603.97 | 603.97 | 603.97 | 12.92 | 60 | >sp\|P01877\|IGHA2_HUMAN Immunoglobulin heavy constant alpha 2 OS=Homo sapiens OX=9606 GN=IGHA2 PE=1 SV=4 | 576 | 20201119_15432_1ng_glyco_hcd203040_compound15.25815.25815.5 | controllerType=0 controllerNumber=1 scan=25815 | 52.4038 |
| 00919:3 | 9 | K.HYTN[+1971.719]SSQDVTVPC[+57.021]R.V | HexNAc(5)Hex(5)Fuc(1) | C[+57], N[+1972] | 1212.5002 | 3 | 3635.486 | 3635.4682 | 0 | 4.9 | 89 | Specific | 278.48 | 193.32 | 193.32 | 5.8 | 60 | >sp\|P01877\|IGHA2_HUMAN Immunoglobulin heavy constant alpha 2 OS=Homo sapiens OX=9606 GN=IGHA2 PE=1 SV=4 | 576 | 20201119_15432_1ng_glyco_hcd203040_compound15.3860.3860.3 | controllerType=0 controllerNumber=1 scan=3860 | 12.8883 |
| 01026:3 | 9 | K.HYTN[+1647.613]SSQDVTVPC[+57.021]R.V | HexNAc(5)Hex(3)Fuc(1) | C[+57], N[+1648] | 1104.4624 | 3 | 3311.3727 | 3311.3625 | 0 | 3.1 | 89 | Specific | 332.5 | 274.84 | 274.84 | 6.74 | 60 | >sp\|P01877\|IGHA2_HUMAN Immunoglobulin heavy constant alpha 2 OS=Homo sapiens OX=9606 GN=IGHA2 PE=1 SV=4 | 576 | 20201119_15432_1ng_glyco_hcd203040_compound15.3971.3971.2 | controllerType=0 controllerNumber=1 scan=3971 | 13.0932 |
| 01788:3 | 9 | K.HYTN[+2262.814]SSQDVTVPC[+57.021]R.V | HexNAc(5)Hex(5)Fuc(1)NeuAc(1) | C[+57], N[+2263] | 1309.5313 | 3 | 3926.5794 | 3926.5636 | 0 | 4 | 89 | Specific | 344.82 | 233.17 | 233.17 | 6.56 | 60 | >sp\|P01877\|IGHA2_HUMAN Immunoglobulin heavy constant alpha 2 OS=Homo sapiens OX=9606 GN=IGHA2 PE=1 SV=4 | 576 | 20201119_15432_1ng_glyco_hcd203040_compound15.4764.4764.3 | controllerType=0 controllerNumber=1 scan=4764 | 14.4981 |
| 01790:3 | 9 | K.HYTN[+2262.814]SSQDVTVPC[+57.021]R.V | HexNAc(5)Hex(5)Fuc(1)NeuAc(1) | C[+57], N[+2263] | 1309.5313 | 3 | 3926.5794 | 3926.5636 | 0 | 4 | 89 | Specific | 360.65 | 244.2 | 244.2 | 6.57 | 60 | >sp\|P01877\|IGHA2_HUMAN Immunoglobulin heavy constant alpha 2 OS=Homo sapiens OX=9606 GN=IGHA2 PE=1 SV=4 | 576 | 20201119_15432_1ng_glyco_hcd203040_compound15.4766.4766.3 | controllerType=0 controllerNumber=1 scan=4766 | 14.5019 |

Table S4. Cont.

| Query #:z | Protein Rank | Glycopeptide | Glycan | Modification Type(s) | Observed m/z | z | Observed (M+H) | Calc. mass (M+H) | Off-by-x error | Mass error (ppm) | StartPosition | Cleavage | Score | Delta | Delta Mod | \|Log Prob\| | # of unique peptides | Protein Name | Protein DB number | Comment | Scan # | Scan Time |
|---|---|---|---|---|---|---|---|---|---|---|---|---|---|---|---|---|---|---|---|---|---|---|
| 01024:3 | 9 | K.HYTN[+1647.613]SSQDVTVPC[+57.021]R.V | HexNAc(5)Hex(3)Fuc(1) | C[+57], N[+1648] | 1104.4624 | 3 | 3311.3727 | 3311.3625 | 0 | 3.1 | 89 | Specific | 374.17 | 297.25 | 297.25 | 6.82 | 60 | >sp\|P01877\|IGHA2_HUMAN Immunoglobulin heavy constant alpha 2 OS=Homo sapiens OX=9606 GN=IGHA2 PE=1 SV=4 | 576 | 20201119_15432_1ng_glyco_hcd203040_compound15.3969.3969.2 | controllerType=0 controllerNumber=1 scan=3969 | 13.0887 |
| 00901:3 | 9 | K.HYTN[+1809.666]SSQDVTVPC[+57.021]R.V | HexNAc(5)Hex(4)Fuc(1) | C[+57], N[+1810] | 1158.4813 | 3 | 3473.4292 | 3473.4154 | 0 | 4 | 89 | Specific | 404.79 | 272.56 | 272.56 | 6.87 | 60 | >sp\|P01877\|IGHA2_HUMAN Immunoglobulin heavy constant alpha 2 OS=Homo sapiens OX=9606 GN=IGHA2 PE=1 SV=4 | 576 | 20201119_15432_1ng_glyco_hcd203040_compound15.3841.3841.3 | controllerType=0 controllerNumber=1 scan=3841 | 12.8531 |
| 00903:3 | 9 | K.HYTN[+1809.666]SSQDVTVPC[+57.021]R.V | HexNAc(5)Hex(4)Fuc(1) | C[+57], N[+1810] | 1158.4813 | 3 | 3473.4292 | 3473.4154 | 0 | 4 | 89 | Specific | 511.8 | 334.16 | 334.16 | 7.99 | 60 | >sp\|P01877\|IGHA2_HUMAN Immunoglobulin heavy constant alpha 2 OS=Homo sapiens OX=9606 GN=IGHA2 PE=1 SV=4 | 576 | 20201119_15432_1ng_glyco_hcd203040_compound15.3843.3843.3 | controllerType=0 controllerNumber=1 scan=3843 | 12.8567 |
| 00918:3 | 9 | K.HYTN[+1647.613]SSQDVTVPC[+57.021]R.V | HexNAc(5)Hex(3)Fuc(1) | C[+57], N[+1648] | 1104.4634 | 3 | 3311.3757 | 3311.3625 | 0 | 4 | 89 | Specific | 559.26 | 387.41 | 387.41 | 9.89 | 60 | >sp\|P01877\|IGHA2_HUMAN Immunoglobulin heavy constant alpha 2 OS=Homo sapiens OX=9606 GN=IGHA2 PE=1 SV=4 | 576 | 20201119_15432_1ng_glyco_hcd203040_compound15.3859.3859.3 | controllerType=0 controllerNumber=1 scan=3859 | 12.8868 |
| 00920:3 | 9 | K.HYTN[+1647.613]SSQDVTVPC[+57.021]R.V | HexNAc(5)Hex(3)Fuc(1) | C[+57], N[+1648] | 1104.4634 | 3 | 3311.3757 | 3311.3625 | 0 | 4 | 89 | Specific | 629.51 | 540.18 | 540.18 | 12.07 | 60 | >sp\|P01877\|IGHA2_HUMAN Immunoglobulin heavy constant alpha 2 OS=Homo sapiens OX=9606 GN=IGHA2 PE=1 SV=4 | 576 | 20201119_15432_1ng_glyco_hcd203040_compound15.3861.3861.3 | controllerType=0 controllerNumber=1 scan=3861 | 12.8906 |
| 02905:4 | 9 | K.HYTN[+1647.613]SSQDVTVPC[+57.021]RVPPPPPC[+57.021]C[+57.021]HPR.L | HexNAc(5)Hex(3)Fuc(1) | C[+57]*3, N[+1648] | 1152.2526 | 4 | 4605.9886 | 4605.9689 | 0 | 4.3 | 89 | Specific | 516.68 | 391.85 | 391.85 | 8.99 | 60 | >sp\|P01877\|IGHA2_HUMAN Immunoglobulin heavy constant alpha 2 OS=Homo sapiens OX=9606 GN=IGHA2 PE=1 SV=4 | 576 | 20201119_15432_1ng_glyco_hcd203040_compound15.5922.5922.4 | controllerType=0 controllerNumber=1 scan=5922 | 16.5379 |
| 02903:4 | 9 | K.HYTN[+1647.613]SSQDVTVPC[+57.021]RVPPPPPC[+57.021]C[+57.021]HPR.L | HexNAc(5)Hex(3)Fuc(1) | C[+57]*3, N[+1648] | 1152.2526 | 4 | 4605.9886 | 4605.9689 | 0 | 4.3 | 89 | Specific | 516.71 | 516.71 | 516.71 | 8.99 | 60 | >sp\|P01877\|IGHA2_HUMAN Immunoglobulin heavy constant alpha 2 OS=Homo sapiens OX=9606 GN=IGHA2 PE=1 SV=4 | 576 | 20201119_15432_1ng_glyco_hcd203040_compound15.5920.5920.4 | controllerType=0 controllerNumber=1 scan=5920 | 16.5334 |
| 02941:4 | 9 | K.HYTN[+1647.613]SSQDVTVPC[+57.021]RVPPPPPC[+57.021]C[+57.021]HPR.L | HexNAc(5)Hex(3)Fuc(1) | C[+57]*3, N[+1648] | 1152.2515 | 4 | 4605.9842 | 4605.9689 | 0 | 3.3 | 89 | Specific | 533.99 | 284.84 | 284.84 | 7.15 | 60 | >sp\|P01877\|IGHA2_HUMAN Immunoglobulin heavy constant alpha 2 OS=Homo sapiens OX=9606 GN=IGHA2 PE=1 SV=4 | 576 | 20201119_15432_1ng_glyco_hcd203040_compound15.5960.5960.4 | controllerType=0 controllerNumber=1 scan=5960 | 16.6095 |
| 02943:4 | 9 | K.HYTN[+1647.613]SSQDVTVPC[+57.021]RVPPPPPC[+57.021]C[+57.021]HPR.L | HexNAc(5)Hex(3)Fuc(1) | C[+57]*3, N[+1648] | 1152.2515 | 4 | 4605.9842 | 4605.9689 | 0 | 3.3 | 89 | Specific | 567.32 | 407.11 | 407.11 | 9.45 | 60 | >sp\|P01877\|IGHA2_HUMAN Immunoglobulin heavy constant alpha 2 OS=Homo sapiens OX=9606 GN=IGHA2 PE=1 SV=4 | 576 | 20201119_15432_1ng_glyco_hcd203040_compound15.5962.5962.4 | controllerType=0 controllerNumber=1 scan=5962 | 16.6133 |
| 02937:5 | 9 | K.HYTN[+1647.613]SSQDVTVPC[+57.021]RVPPPPPC[+57.021]C[+57.021]HPR.L | HexNAc(5)Hex(3)Fuc(1) | C[+57]*3, N[+1648] | 922.0028 | 5 | 4605.9847 | 4605.9689 | 0 | 3.4 | 89 | Specific | 634.31 | 373.5 | 373.5 | 10.77 | 60 | >sp\|P01877\|IGHA2_HUMAN Immunoglobulin heavy constant alpha 2 OS=Homo sapiens OX=9606 GN=IGHA2 PE=1 SV=4 | 576 | 20201119_15432_1ng_glyco_hcd203040_compound15.5956.5956.5 | controllerType=0 controllerNumber=1 scan=5956 | 16.6019 |
| 02939:5 | 9 | K.HYTN[+1647.613]SSQDVTVPC[+57.021]RVPPPPPC[+57.021]C[+57.021]HPR.L | HexNAc(5)Hex(3)Fuc(1) | C[+57]*3, N[+1648] | 922.0028 | 5 | 4605.9847 | 4605.9689 | 0 | 3.4 | 89 | Specific | 740.19 | 619.63 | 619.63 | 14.17 | 60 | >sp\|P01877\|IGHA2_HUMAN Immunoglobulin heavy constant alpha 2 OS=Homo sapiens OX=9606 GN=IGHA2 PE=1 SV=4 | 576 | 20201119_15432_1ng_glyco_hcd203040_compound15.5958.5958.5 | controllerType=0 controllerNumber=1 scan=5958 | 16.6062 |
| 18295:4 | 9 | R.LSLHRPALEDLLLGSEAN[+1913.677]LTC[+57.021]TLTGLR.D | HexNAc(4)Hex(5)NeuAc(1) | C[+57], N[+1914] | 1220.0813 | 4 | 4877.3035 | 4877.2753 | 0 | 5.8 | 114 | Specific | 223.16 | 223.16 | 223.16 | 4.61 | 60 | >sp\|P01877\|IGHA2_HUMAN Immunoglobulin heavy constant alpha 2 OS=Homo sapiens OX=9606 GN=IGHA2 PE=1 SV=4 | 576 | 20201119_15432_1ng_glyco_hcd203040_compound15.21979.21979.4 | controllerType=0 controllerNumber=1 scan=21979 | 44.7729 |
| 17378:4 | 9 | R.LSLHRPALEDLLLGSEAN[+1663.608]LTC[+57.021]TLTGLR.D | HexNAc(5)Hex(4) | C[+57], N[+1664] | 1157.5606 | 4 | 4627.2205 | 4627.2064 | 0 | 3 | 114 | Specific | 283.51 | 283.51 | 283.51 | 6.95 | 60 | >sp\|P01877\|IGHA2_HUMAN Immunoglobulin heavy constant alpha 2 OS=Homo sapiens OX=9606 GN=IGHA2 PE=1 SV=4 | 576 | 20201119_15432_1ng_glyco_hcd203040_compound15.21021.21021.4 | controllerType=0 controllerNumber=1 scan=21021 | 42.7823 |
| 17335:4 | 9 | R.LSLHRPALEDLLLGSEAN[+1825.661]LTC[+57.021]TLTGLR.D | HexNAc(5)Hex(5) | C[+57], N[+1826] | 1198.0757 | 4 | 4789.2811 | 4789.2592 | 0 | 4.6 | 114 | Specific | 322.75 | 322.75 | 322.75 | 7.27 | 60 | >sp\|P01877\|IGHA2_HUMAN Immunoglobulin heavy constant alpha 2 OS=Homo sapiens OX=9606 GN=IGHA2 PE=1 SV=4 | 576 | 20201119_15432_1ng_glyco_hcd203040_compound15.20975.20975.4 | controllerType=0 controllerNumber=1 scan=20975 | 42.6535 |
| 17286:5 | 9 | R.LSLHRPALEDLLLGSEAN[+1864.634]LTC[+57.021]TLTGLR.D | HexNAc(2)Hex(9) | C[+57], N[+1865] | 966.455 | 5 | 4828.2461 | 4828.2324 | 0 | 2.8 | 114 | Specific | 394 | 394 | 394 | 8.52 | 60 | >sp\|P01877\|IGHA2_HUMAN Immunoglobulin heavy constant alpha 2 OS=Homo sapiens OX=9606 GN=IGHA2 PE=1 SV=4 | 576 | 20201119_15432_1ng_glyco_hcd203040_compound15.20924.20924.5 | controllerType=0 controllerNumber=1 scan=20924 | 42.5392 |
| 17337:4 | 9 | R.LSLHRPALEDLLLGSEAN[+1825.661]LTC[+57.021]TLTGLR.D | HexNAc(5)Hex(5) | C[+57], N[+1826] | 1198.0757 | 4 | 4789.2811 | 4789.2592 | 0 | 4.6 | 114 | Specific | 488.49 | 362.27 | 362.27 | 8.45 | 60 | >sp\|P01877\|IGHA2_HUMAN Immunoglobulin heavy constant alpha 2 OS=Homo sapiens OX=9606 GN=IGHA2 PE=1 SV=4 | 576 | 20201119_15432_1ng_glyco_hcd203040_compound15.20977.20977.4 | controllerType=0 controllerNumber=1 scan=20977 | 42.6572 |
| 17361:3 | 9 | R.LSLHRPALEDLLLGSEAN[+1622.582]LTC[+57.021]TLTGLR.D | HexNAc(4)Hex(5) | C[+57], N[+1623] | 1529.4033 | 3 | 4586.1952 | 4586.1799 | 0 | 3.3 | 114 | Specific | 504.28 | 479.09 | 479.09 | 9.57 | 60 | >sp\|P01877\|IGHA2_HUMAN Immunoglobulin heavy constant alpha 2 OS=Homo sapiens OX=9606 GN=IGHA2 PE=1 SV=4 | 576 | 20201119_15432_1ng_glyco_hcd203040_compound15.21003.21003.3 | controllerType=0 controllerNumber=1 scan=21003 | 42.7376 |
| 18328:4 | 9 | R.LSLHRPALEDLLLGSEAN[+1872.651]LTC[+57.021]TLTGLR.D | HexNAc(3)Hex(6)NeuAc(1) | C[+57], N[+1873] | 1209.8195 | 4 | 4836.2562 | 4836.2487 | 0 | 1.5 | 114 | Specific | 530.33 | 440.69 | 440.69 | 9.89 | 60 | >sp\|P01877\|IGHA2_HUMAN Immunoglobulin heavy constant alpha 2 OS=Homo sapiens OX=9606 GN=IGHA2 PE=1 SV=4 | 576 | 20201119_15432_1ng_glyco_hcd203040_compound15.22013.22013.4 | controllerType=0 controllerNumber=1 scan=22013 | 44.8404 |
| 18356:3 | 9 | R.LSLHRPALEDLLLGSEAN[+1751.624]LTC[+57.021]TLTGLR.D | HexNAc(4)Hex(4)NeuAc(1) | C[+57], N[+1752] | 1572.4192 | 3 | 4715.2432 | 4715.2225 | 0 | 4.4 | 114 | Specific | 546.08 | 546.08 | 546.08 | 9.55 | 60 | >sp\|P01877\|IGHA2_HUMAN Immunoglobulin heavy constant alpha 2 OS=Homo sapiens OX=9606 GN=IGHA2 PE=1 SV=4 | 576 | 20201119_15432_1ng_glyco_hcd203040_compound15.22042.22042.3 | controllerType=0 controllerNumber=1 scan=22042 | 44.9036 |
| 17383:3 | 9 | R.LSLHRPALEDLLLGSEAN[+1460.529]LTC[+57.021]TLTGLR.D | HexNAc(4)Hex(4) | C[+57], N[+1461] | 1475.3874 | 3 | 4424.1476 | 4424.1271 | 0 | 4.6 | 114 | Specific | 553.3 | 553.3 | 553.3 | 9.78 | 60 | >sp\|P01877\|IGHA2_HUMAN Immunoglobulin heavy constant alpha 2 OS=Homo sapiens OX=9606 GN=IGHA2 PE=1 SV=4 | 576 | 20201119_15432_1ng_glyco_hcd203040_compound15.21026.21026.3 | controllerType=0 controllerNumber=1 scan=21026 | 42.7927 |
| 17295:3 | 9 | R.LSLHRPALEDLLLGSEAN[+1702.581]LTC[+57.021]TLTGLR.D | HexNAc(2)Hex(8) | C[+57], N[+1703] | 1556.0694 | 3 | 4666.1936 | 4666.1796 | 0 | 3 | 114 | Specific | 593.57 | 593.57 | 593.57 | 11.09 | 60 | >sp\|P01877\|IGHA2_HUMAN Immunoglobulin heavy constant alpha 2 OS=Homo sapiens OX=9606 GN=IGHA2 PE=1 SV=4 | 576 | 20201119_15432_1ng_glyco_hcd203040_compound15.20934.20934.3 | controllerType=0 controllerNumber=1 scan=20934 | 42.5596 |
| 18359:3 | 9 | R.LSLHRPALEDLLLGSEAN[+1751.624]LTC[+57.021]TLTGLR.D | HexNAc(4)Hex(4)NeuAc(1) | C[+57], N[+1752] | 1572.4192 | 3 | 4715.2432 | 4715.2225 | 0 | 4.4 | 114 | Specific | 601.04 | 601.04 | 601.04 | 11.96 | 60 | >sp\|P01877\|IGHA2_HUMAN Immunoglobulin heavy constant alpha 2 OS=Homo sapiens OX=9606 GN=IGHA2 PE=1 SV=4 | 576 | 20201119_15432_1ng_glyco_hcd203040_compound15.22045.22045.3 | controllerType=0 controllerNumber=1 scan=22045 | 44.9092 |
| 17297:3 | 9 | R.LSLHRPALEDLLLGSEAN[+1702.581]LTC[+57.021]TLTGLR.D | HexNAc(2)Hex(8) | C[+57], N[+1703] | 1556.0694 | 3 | 4666.1936 | 4666.1796 | 0 | 3 | 114 | Specific | 611.02 | 565.15 | 565.15 | 12.09 | 60 | >sp\|P01877\|IGHA2_HUMAN Immunoglobulin heavy constant alpha 2 OS=Homo sapiens OX=9606 GN=IGHA2 PE=1 SV=4 | 576 | 20201119_15432_1ng_glyco_hcd203040_compound15.20936.20936.3 | controllerType=0 controllerNumber=1 scan=20936 | 42.5649 |
| 18307:3 | 9 | R.LSLHRPALEDLLLGSEAN[+1913.677]LTC[+57.021]TLTGLR.D | HexNAc(4)Hex(5)NeuAc(1) | C[+57], N[+1914] | 1626.4412 | 3 | 4877.3091 | 4877.2753 | 0 | 6.9 | 114 | Specific | 611.18 | 531.97 | 531.97 | 11.04 | 60 | >sp\|P01877\|IGHA2_HUMAN Immunoglobulin heavy constant alpha 2 OS=Homo sapiens OX=9606 GN=IGHA2 PE=1 SV=4 | 576 | 20201119_15432_1ng_glyco_hcd203040_compound15.21991.21991.3 | controllerType=0 controllerNumber=1 scan=21991 | 44.7898 |
| 17370:3 | 9 | R.LSLHRPALEDLLLGSEAN[+1501.555]LTC[+57.021]TLTGLR.D | HexNAc(5)Hex(3) | C[+57], N[+1502] | 1489.0604 | 3 | 4465.1665 | 4465.1536 | 0 | 2.9 | 114 | Specific | 613.26 | 572.45 | 572.45 | 12.08 | 60 | >sp\|P01877\|IGHA2_HUMAN Immunoglobulin heavy constant alpha 2 OS=Homo sapiens OX=9606 GN=IGHA2 PE=1 SV=4 | 576 | 20201119_15432_1ng_glyco_hcd203040_compound15.21012.21012.3 | controllerType=0 controllerNumber=1 scan=21012 | 42.7625 |
| 17386:3 | 9 | R.LSLHRPALEDLLLGSEAN[+1460.529]LTC[+57.021]TLTGLR.D | HexNAc(4)Hex(4) | C[+57], N[+1461] | 1475.3874 | 3 | 4424.1476 | 4424.1271 | 0 | 4.6 | 114 | Specific | 623.71 | 504.32 | 504.32 | 11.96 | 60 | >sp\|P01877\|IGHA2_HUMAN Immunoglobulin heavy constant alpha 2 OS=Homo sapiens OX=9606 GN=IGHA2 PE=1 SV=4 | 576 | 20201119_15432_1ng_glyco_hcd203040_compound15.21029.21029.3 | controllerType=0 controllerNumber=1 scan=21029 | 42.8022 |
| 17325:3 | 9 | R.LSLHRPALEDLLLGSEAN[+1540.529]LTC[+57.021]TLTGLR.D | HexNAc(2)Hex(7) | C[+57], N[+1541] | 1502.0551 | 3 | 4504.1508 | 4504.1268 | 0 | 5.3 | 114 | Specific | 625.23 | 611.2 | 611.2 | 11.33 | 60 | >sp\|P01877\|IGHA2_HUMAN Immunoglobulin heavy constant alpha 2 OS=Homo sapiens OX=9606 GN=IGHA2 PE=1 SV=4 | 576 | 20201119_15432_1ng_glyco_hcd203040_compound15.20965.20965.3 | controllerType=0 controllerNumber=1 scan=20965 | 42.6296 |
| 17367:3 | 9 | R.LSLHRPALEDLLLGSEAN[+1501.555]LTC[+57.021]TLTGLR.D | HexNAc(5)Hex(3) | C[+57], N[+1502] | 1489.0604 | 3 | 4465.1665 | 4465.1536 | 0 | 2.9 | 114 | Specific | 632.71 | 632.71 | 632.71 | 12.08 | 60 | >sp\|P01877\|IGHA2_HUMAN Immunoglobulin heavy constant alpha 2 OS=Homo sapiens OX=9606 GN=IGHA2 PE=1 SV=4 | 576 | 20201119_15432_1ng_glyco_hcd203040_compound15.21009.21009.3 | controllerType=0 controllerNumber=1 scan=21009 | 42.7514 |
| 17273:3 | 9 | R.LSLHRPALEDLLLGSEAN[+1864.634]LTC[+57.021]TLTGLR.D | HexNAc(2)Hex(9) | C[+57], N[+1865] | 1610.0878 | 3 | 4828.2489 | 4828.2324 | 0 | 3.4 | 114 | Specific | 639.63 | 639.63 | 639.63 | 12.09 | 60 | >sp\|P01877\|IGHA2_HUMAN Immunoglobulin heavy constant alpha 2 OS=Homo sapiens OX=9606 GN=IGHA2 PE=1 SV=4 | 576 | 20201119_15432_1ng_glyco_hcd203040_compound15.20911.20911.3 | controllerType=0 controllerNumber=1 scan=20911 | 42.5078 |
| 17364:3 | 9 | R.LSLHRPALEDLLLGSEAN[+1622.582]LTC[+57.021]TLTGLR.D | HexNAc(4)Hex(5) | C[+57], N[+1623] | 1529.4033 | 3 | 4586.1952 | 4586.1799 | 0 | 3.3 | 114 | Specific | 643.56 | 562.13 | 562.13 | 12.09 | 60 | >sp\|P01877\|IGHA2_HUMAN Immunoglobulin heavy constant alpha 2 OS=Homo sapiens OX=9606 GN=IGHA2 PE=1 SV=4 | 576 | 20201119_15432_1ng_glyco_hcd203040_compound15.21006.21006.3 | controllerType=0 controllerNumber=1 scan=21006 | 42.7471 |
| 17322:3 | 9 | R.LSLHRPALEDLLLGSEAN[+1540.529]LTC[+57.021]TLTGLR.D | HexNAc(2)Hex(7) | C[+57], N[+1541] | 1502.0551 | 3 | 4504.1508 | 4504.1268 | 0 | 5.3 | 114 | Specific | 672.42 | 605.68 | 605.68 | 12.33 | 60 | >sp\|P01877\|IGHA2_HUMAN Immunoglobulin heavy constant alpha 2 OS=Homo sapiens OX=9606 GN=IGHA2 PE=1 SV=4 | 576 | 20201119_15432_1ng_glyco_hcd203040_compound15.20962.20962.3 | controllerType=0 controllerNumber=1 scan=20962 | 42.6237 |
| 17328:4 | 9 | R.LSLHRPALEDLLLGSEAN[+1622.582]LTC[+57.021]TLTGLR.D | HexNAc(4)Hex(5) | C[+57], N[+1623] | 1147.3067 | 4 | 4586.2049 | 4586.1799 | 0 | 5.5 | 114 | Specific | 674.34 | 674.34 | 674.34 | 12.87 | 60 | >sp\|P01877\|IGHA2_HUMAN Immunoglobulin heavy constant alpha 2 OS=Homo sapiens OX=9606 GN=IGHA2 PE=1 SV=4 | 576 | 20201119_15432_1ng_glyco_hcd203040_compound15.20968.20968.4 | controllerType=0 controllerNumber=1 scan=20968 | 42.6368 |
| 17389:4 | 9 | R.LSLHRPALEDLLLGSEAN[+1298.476]LTC[+57.021]TLTGLR.D | HexNAc(4)Hex(3) | C[+57], N[+1298] | 1066.2771 | 4 | 4262.0867 | 4262.0742 | 0 | 2.9 | 114 | Specific | 675.44 | 675.44 | 675.44 | 13.63 | 60 | >sp\|P01877\|IGHA2_HUMAN Immunoglobulin heavy constant alpha 2 OS=Homo sapiens OX=9606 GN=IGHA2 PE=1 SV=4 | 576 | 20201119_15432_1ng_glyco_hcd203040_compound15.21032.21032.4 | controllerType=0 controllerNumber=1 scan=21032 | 42.8071 |
| 17376:3 | 9 | R.LSLHRPALEDLLLGSEAN[+1216.423]LTC[+57.021]TLTGLR.D | HexNAc(2)Hex(5) | C[+57], N[+1216] | 1394.0158 | 3 | 4180.0329 | 4180.0211 | 0 | 2.8 | 114 | Specific | 681.22 | 586.02 | 586.02 | 13.08 | 60 | >sp\|P01877\|IGHA2_HUMAN Immunoglobulin heavy constant alpha 2 OS=Homo sapiens OX=9606 GN=IGHA2 PE=1 SV=4 | 576 | 20201119_15432_1ng_glyco_hcd203040_compound15.21018.21018.3 | controllerType=0 controllerNumber=1 scan=21018 | 42.7762 |
| 17271:3 | 9 | R.LSLHRPALEDLLLGSEAN[+1864.634]LTC[+57.021]TLTGLR.D | HexNAc(2)Hex(9) | C[+57], N[+1865] | 1610.0878 | 3 | 4828.2489 | 4828.2324 | 0 | 3.4 | 114 | Specific | 698.64 | 698.64 | 698.64 | 13.09 | 60 | >sp\|P01877\|IGHA2_HUMAN Immunoglobulin heavy constant alpha 2 OS=Homo sapiens OX=9606 GN=IGHA2 PE=1 SV=4 | 576 | 20201119_15432_1ng_glyco_hcd203040_compound15.20909.20909.3 | controllerType=0 controllerNumber=1 scan=20909 | 42.5005 |
| 17340:3 | 9 | R.LSLHRPALEDLLLGSEAN[+1378.476]LTC[+57.021]TLTGLR.D | HexNAc(2)Hex(6) | C[+57], N[+1378] | 1448.0334 | 3 | 4342.0856 | 4342.074 | 0 | 2.7 | 114 | Specific | 699.52 | 645.01 | 645.01 | 13.08 | 60 | >sp\|P01877\|IGHA2_HUMAN Immunoglobulin heavy constant alpha 2 OS=Homo sapiens OX=9606 GN=IGHA2 PE=1 SV=4 | 576 | 20201119_15432_1ng_glyco_hcd203040_compound15.20981.20981.3 | controllerType=0 controllerNumber=1 scan=20981 | 42.6688 |
| 17356:4 | 9 | R.LSLHRPALEDLLLGSEAN[+1501.555]LTC[+57.021]TLTGLR.D | HexNAc(5)Hex(3) | C[+57], N[+1502] | 1117.0489 | 4 | 4465.1737 | 4465.1536 | 0 | 4.5 | 114 | Specific | 713.06 | 713.06 | 713.06 | 14.5 | 60 | >sp\|P01877\|IGHA2_HUMAN Immunoglobulin heavy constant alpha 2 OS=Homo sapiens OX=9606 GN=IGHA2 PE=1 SV=4 | 576 | 20201119_15432_1ng_glyco_hcd203040_compound15.20998.20998.4 | controllerType=0 controllerNumber=1 scan=20998 | 42.7255 |
| 17342:3 | 9 | R.LSLHRPALEDLLLGSEAN[+1378.476]LTC[+57.021]TLTGLR.D | HexNAc(2)Hex(6) | C[+57], N[+1378] | 1448.0334 | 3 | 4342.0856 | 4342.074 | 0 | 2.7 | 114 | Specific | 721.79 | 631.88 | 631.88 | 14.08 | 60 | >sp\|P01877\|IGHA2_HUMAN Immunoglobulin heavy constant alpha 2 OS=Homo sapiens OX=9606 GN=IGHA2 PE=1 SV=4 | 576 | 20201119_15432_1ng_glyco_hcd203040_compound15.20983.20983.3 | controllerType=0 controllerNumber=1 scan=20983 | 42.6769 |
| 18298:4 | 9 | R.LSLHRPALEDLLLGSEAN[+1913.677]LTC[+57.021]TLTGLR.D | HexNAc(4)Hex(5)NeuAc(1) | C[+57], N[+1914] | 1220.0813 | 4 | 4877.3035 | 4877.2753 | 0 | 5.8 | 114 | Specific | 752.33 | 725.02 | 725.02 | 14.87 | 60 | >sp\|P01877\|IGHA2_HUMAN Immunoglobulin heavy constant alpha 2 OS=Homo sapiens OX=9606 GN=IGHA2 PE=1 SV=4 | 576 | 20201119_15432_1ng_glyco_hcd203040_compound15.21982.21982.4 | controllerType=0 controllerNumber=1 scan=21982 | 44.776 |
| 17344:4 | 9 | R.LSLHRPALEDLLLGSEAN[+1419.502]LTC[+57.021]TLTGLR.D | HexNAc(3)Hex(5) | C[+57], N[+1420] | 1096.5331 | 4 | 4383.1107 | 4383.1005 | 0 | 2.3 | 114 | Specific | 758.01 | 758.01 | 758.01 | 15.63 | 60 | >sp\|P01877\|IGHA2_HUMAN Immunoglobulin heavy constant alpha 2 OS=Homo sapiens OX=9606 GN=IGHA2 PE=1 SV=4 | 576 | 20201119_15432_1ng_glyco_hcd203040_compound15.20985.20985.4 | controllerType=0 controllerNumber=1 scan=20985 | 42.6793 |
| 17373:3 | 9 | R.LSLHRPALEDLLLGSEAN[+1216.423]LTC[+57.021]TLTGLR.D | HexNAc(2)Hex(5) | C[+57], N[+1216] | 1394.0158 | 3 | 4180.0329 | 4180.0211 | 0 | 2.8 | 114 | Specific | 760.76 | 695.22 | 695.22 | 15.08 | 60 | >sp\|P01877\|IGHA2_HUMAN Immunoglobulin heavy constant alpha 2 OS=Homo sapiens OX=9606 GN=IGHA2 PE=1 SV=4 | 576 | 20201119_15432_1ng_glyco_hcd203040_compound15.21015.21015.3 | controllerType=0 controllerNumber=1 scan=21015 | 42.7664 |
| 17366:4 | 9 | R.LSLHRPALEDLLLGSEAN[+1460.529]LTC[+57.021]TLTGLR.D | HexNAc(4)Hex(4) | C[+57], N[+1461] | 1106.7936 | 4 | 4424.1527 | 4424.1271 | 0 | 5.8 | 114 | Specific | 761.23 | 761.23 | 761.23 | 14.87 | 60 | >sp\|P01877\|IGHA2_HUMAN Immunoglobulin heavy constant alpha 2 OS=Homo sapiens OX=9606 GN=IGHA2 PE=1 SV=4 | 576 | 20201119_15432_1ng_glyco_hcd203040_compound15.21008.21008.4 | controllerType=0 controllerNumber=1 scan=21008 | 42.7503 |
| 17358:4 | 9 | R.LSLHRPALEDLLLGSEAN[+1501.555]LTC[+57.021]TLTGLR.D | HexNAc(5)Hex(3) | C[+57], N[+1502] | 1117.0489 | 4 | 4465.1737 | 4465.1536 | 0 | 4.5 | 114 | Specific | 787.47 | 787.47 | 787.47 | 15.5 | 60 | >sp\|P01877\|IGHA2_HUMAN Immunoglobulin heavy constant alpha 2 OS=Homo sapiens OX=9606 GN=IGHA2 PE=1 SV=4 | 576 | 20201119_15432_1ng_glyco_hcd203040_compound15.21000.21000.4 | controllerType=0 controllerNumber=1 scan=21000 | 42.7342 |
| 17380:4 | 9 | R.LSLHRPALEDLLLGSEAN[+1663.608]LTC[+57.021]TLTGLR.D | HexNAc(5)Hex(4) | C[+57], N[+1664] | 1157.5606 | 4 | 4627.2205 | 4627.2064 | 0 | 3 | 114 | Specific | 793.08 | 755.51 | 755.51 | 15.63 | 60 | >sp\|P01877\|IGHA2_HUMAN Immunoglobulin heavy constant alpha 2 OS=Homo sapiens OX=9606 GN=IGHA2 PE=1 SV=4 | 576 | 20201119_15432_1ng_glyco_hcd203040_compound15.21023.21023.4 | controllerType=0 controllerNumber=1 scan=21023 | 42.7862 |
| 17331:4 | 9 | R.LSLHRPALEDLLLGSEAN[+1622.582]LTC[+57.021]TLTGLR.D | HexNAc(4)Hex(5) | C[+57], N[+1623] | 1147.3067 | 4 | 4586.2049 | 4586.1799 | 0 | 5.5 | 114 | Specific | 840.94 | 733.67 | 733.67 | 15.87 | 60 | >sp\|P01877\|IGHA2_HUMAN Immunoglobulin heavy constant alpha 2 OS=Homo sapiens OX=9606 GN=IGHA2 PE=1 SV=4 | 576 | 20201119_15432_1ng_glyco_hcd203040_compound15.20971.20971.4 | controllerType=0 controllerNumber=1 scan=20971 | 42.6471 |
| 18330:4 | 9 | R.LSLHRPALEDLLLGSEAN[+1872.651]LTC[+57.021]TLTGLR.D | HexNAc(3)Hex(6)NeuAc(1) | C[+57], N[+1873] | 1209.8195 | 4 | 4836.2562 | 4836.2487 | 0 | 1.5 | 114 | Specific | 852.82 | 840.79 | 840.79 | 17.31 | 60 | >sp\|P01877\|IGHA2_HUMAN Immunoglobulin heavy constant alpha 2 OS=Homo sapiens OX=9606 GN=IGHA2 PE=1 SV=4 | 576 | 20201119_15432_1ng_glyco_hcd203040_compound15.22015.22015.4 | controllerType=0 controllerNumber=1 scan=22015 | 44.8472 |
| 17360:4 | 9 | R.LSLHRPALEDLLLGSEAN[+1216.423]LTC[+57.021]TLTGLR.D | HexNAc(2)Hex(5) | C[+57], N[+1216] | 1045.7638 | 4 | 4180.0335 | 4180.0211 | 0 | 3 | 114 | Specific | 856.03 | 788.1 | 788.1 | 17.63 | 60 | >sp\|P01877\|IGHA2_HUMAN Immunoglobulin heavy constant alpha 2 OS=Homo sapiens OX=9606 GN=IGHA2 PE=1 SV=4 | 576 | 20201119_15432_1ng_glyco_hcd203040_compound15.21002.21002.4 | controllerType=0 controllerNumber=1 scan=21002 | 42.7365 |
| 17318:4 | 9 | R.LSLHRPALEDLLLGSEAN[+1540.529]LTC[+57.021]TLTGLR.D | HexNAc(2)Hex(7) | C[+57], N[+1541] | 1126.7931 | 4 | 4504.1507 | 4504.1268 | 0 | 5.3 | 114 | Specific | 857.38 | 820.21 | 820.21 | 16.87 | 60 | >sp\|P01877\|IGHA2_HUMAN Immunoglobulin heavy constant alpha 2 OS=Homo sapiens OX=9606 GN=IGHA2 PE=1 SV=4 | 576 | 20201119_15432_1ng_glyco_hcd203040_compound15.20958.20958.4 | controllerType=0 controllerNumber=1 scan=20958 | 42.6113 |
| 17369:4 | 9 | R.LSLHRPALEDLLLGSEAN[+1460.529]LTC[+57.021]TLTGLR.D | HexNAc(4)Hex(4) | C[+57], N[+1461] | 1106.7936 | 4 | 4424.1527 | 4424.1271 | 0 | 5.8 | 114 | Specific | 859.11 | 734.99 | 734.99 | 16.87 | 60 | >sp\|P01877\|IGHA2_HUMAN Immunoglobulin heavy constant alpha 2 OS=Homo sapiens OX=9606 GN=IGHA2 PE=1 SV=4 | 576 | 20201119_15432_1ng_glyco_hcd203040_compound15.21011.21011.4 | controllerType=0 controllerNumber=1 scan=21011 | 42.7561 |
| 17438:4 | 9 | R.LSLHRPALEDLLLGSEAN[+1095.397]LTC[+57.021]TLTGLR.D | HexNAc(3)Hex(3) | C[+57], N[+1095] | 1015.5074 | 4 | 4059.0079 | 4058.9949 | 0 | 3.2 | 114 | Specific | 865.84 | 865.84 | 865.84 | 17.63 | 60 | >sp\|P01877\|IGHA2_HUMAN Immunoglobulin heavy constant alpha 2 OS=Homo sapiens OX=9606 GN=IGHA2 PE=1 SV=4 | 576 | 20201119_15432_1ng_glyco_hcd203040_compound15.21083.21083.4 | controllerType=0 controllerNumber=1 scan=21083 | 42.9299 |
| 17339:4 | 9 | R.LSLHRPALEDLLLGSEAN[+1378.476]LTC[+57.021]TLTGLR.D | HexNAc(2)Hex(6) | C[+57], N[+1378] | 1086.2768 | 4 | 4342.0853 | 4342.074 | 0 | 2.6 | 114 | Specific | 904.97 | 820.12 | 820.12 | 18.63 | 60 | >sp\|P01877\|IGHA2_HUMAN Immunoglobulin heavy constant alpha 2 OS=Homo sapiens OX=9606 GN=IGHA2 PE=1 SV=4 | 576 | 20201119_15432_1ng_glyco_hcd203040_compound15.20980.20980.4 | controllerType=0 controllerNumber=1 scan=20980 | 42.6673 |
| 17294:4 | 9 | R.LSLHRPALEDLLLGSEAN[+1702.581]LTC[+57.021]TLTGLR.D | HexNAc(2)Hex(8) | C[+57], N[+1703] | 1167.3042 | 4 | 4666.1951 | 4666.1796 | 0 | 3.3 | 114 | Specific | 911.12 | 911.12 | 911.12 | 18.63 | 60 | >sp\|P01877\|IGHA2_HUMAN Immunoglobulin heavy constant alpha 2 OS=Homo sapiens OX=9606 GN=IGHA2 PE=1 SV=4 | 576 | 20201119_15432_1ng_glyco_hcd203040_compound15.20933.20933.4 | controllerType=0 controllerNumber=1 scan=20933 | 42.5581 |
| 17270:4 | 9 | R.LSLHRPALEDLLLGSEAN[+1864.634]LTC[+57.021]TLTGLR.D | HexNAc(2)Hex(9) | C[+57], N[+1865] | 1207.8182 | 4 | 4828.2508 | 4828.2324 | 0 | 3.8 | 114 | Specific | 924.39 | 924.39 | 924.39 | 18.63 | 60 | >sp\|P01877\|IGHA2_HUMAN Immunoglobulin heavy constant alpha 2 OS=Homo sapiens OX=9606 GN=IGHA2 PE=1 SV=4 | 576 | 20201119_15432_1ng_glyco_hcd203040_compound15.20908.20908.4 | controllerType=0 controllerNumber=1 scan=20908 | 42.4991 |
| 17411:4 | 9 | R.LSLHRPALEDLLLGSEAN[+1257.449]LTC[+57.021]TLTGLR.D | HexNAc(3)Hex(4) | C[+57], N[+1257] | 1056.0202 | 4 | 4221.0589 | 4221.0477 | 0 | 2.7 | 114 | Specific | 926.28 | 871.1 | 871.1 | 18.63 | 60 | >sp\|P01877\|IGHA2_HUMAN Immunoglobulin heavy constant alpha 2 OS=Homo sapiens OX=9606 GN=IGHA2 PE=1 SV=4 | 576 | 20201119_15432_1ng_glyco_hcd203040_compound15.21055.21055.4 | controllerType=0 controllerNumber=1 scan=21055 | 42.8574 |
| 17363:4 | 9 | R.LSLHRPALEDLLLGSEAN[+1216.423]LTC[+57.021]TLTGLR.D | HexNAc(2)Hex(5) | C[+57], N[+1216] | 1045.7638 | 4 | 4180.0335 | 4180.0211 | 0 | 3 | 114 | Specific | 927.36 | 927.36 | 927.36 | 18.63 | 60 | >sp\|P01877\|IGHA2_HUMAN Immunoglobulin heavy constant alpha 2 OS=Homo sapiens OX=9606 GN=IGHA2 PE=1 SV=4 | 576 | 20201119_15432_1ng_glyco_hcd203040_compound15.21005.21005.4 | controllerType=0 controllerNumber=1 scan=21005 | 42.7463 |
| 17440:4 | 9 | R.LSLHRPALEDLLLGSEAN[+1095.397]LTC[+57.021]TLTGLR.D | HexNAc(3)Hex(3) | C[+57], N[+1095] | 1015.5074 | 4 | 4059.0079 | 4058.9949 | 0 | 3.2 | 114 | Specific | 943.03 | 928.99 | 928.99 | 18.63 | 60 | >sp\|P01877\|IGHA2_HUMAN Immunoglobulin heavy constant alpha 2 OS=Homo sapiens OX=9606 GN=IGHA2 PE=1 SV=4 | 576 | 20201119_15432_1ng_glyco_hcd203040_compound15.21085.21085.4 | controllerType=0 controllerNumber=1 scan=21085 | 42.9369 |
| 17346:4 | 9 | R.LSLHRPALEDLLLGSEAN[+1419.502]LTC[+57.021]TLTGLR.D | HexNAc(3)Hex(5) | C[+57], N[+1420] | 1096.5331 | 4 | 4383.1107 | 4383.1005 | 0 | 2.3 | 114 | Specific | 953.23 | 937 | 937 | 19.63 | 60 | >sp\|P01877\|IGHA2_HUMAN Immunoglobulin heavy constant alpha 2 OS=Homo sapiens OX=9606 GN=IGHA2 PE=1 SV=4 | 576 | 20201119_15432_1ng_glyco_hcd203040_compound15.20987.20987.4 | controllerType=0 controllerNumber=1 scan=20987 | 42.6863 |
| 17296:4 | 9 | R.LSLHRPALEDLLLGSEAN[+1702.581]LTC[+57.021]TLTGLR.D | HexNAc(2)Hex(8) | C[+57], N[+1703] | 1167.3042 | 4 | 4666.1951 | 4666.1796 | 0 | 3.3 | 114 | Specific | 992.92 | 964.83 | 964.83 | 19.63 | 60 | >sp\|P01877\|IGHA2_HUMAN Immunoglobulin heavy constant alpha 2 OS=Homo sapiens OX=9606 GN=IGHA2 PE=1 SV=4 | 576 | 20201119_15432_1ng_glyco_hcd203040_compound15.20935.20935.4 | controllerType=0 controllerNumber=1 scan=20935 | 42.5632 |
| 17341:4 | 9 | R.LSLHRPALEDLLLGSEAN[+1378.476]LTC[+57.021]TLTGLR.D | HexNAc(2)Hex(6) | C[+57], N[+1378] | 1086.2768 | 4 | 4342.0853 | 4342.074 | 0 | 2.6 | 114 | Specific | 1003.37 | 973.12 | 973.12 | 20.63 | 60 | >sp\|P01877\|IGHA2_HUMAN Immunoglobulin heavy constant alpha 2 OS=Homo sapiens OX=9606 GN=IGHA2 PE=1 SV=4 | 576 | 20201119_15432_1ng_glyco_hcd203040_compound15.20982.20982.4 | controllerType=0 controllerNumber=1 scan=20982 | 42.6741 |
| 17320:4 | 9 | R.LSLHRPALEDLLLGSEAN[+1540.529]LTC[+57.021]TLTGLR.D | HexNAc(2)Hex(7) | C[+57], N[+1541] | 1126.7931 | 4 | 4504.1507 | 4504.1268 | 0 | 5.3 | 114 | Specific | 1003.73 | 964.03 | 964.03 | 19.87 | 60 | >sp\|P01877\|IGHA2_HUMAN Immunoglobulin heavy constant alpha 2 OS=Homo sapiens OX=9606 GN=IGHA2 PE=1 SV=4 | 576 | 20201119_15432_1ng_glyco_hcd203040_compound15.20960.20960.4 | controllerType=0 controllerNumber=1 scan=20960 | 42.6169 |
| 17272:4 | 9 | R.LSLHRPALEDLLLGSEAN[+1864.634]LTC[+57.021]TLTGLR.D | HexNAc(2)Hex(9) | C[+57], N[+1865] | 1207.8182 | 4 | 4828.2508 | 4828.2324 | 0 | 3.8 | 114 | Specific | 1086.89 | 1058.76 | 1058.76 | 21.63 | 60 | >sp\|P01877\|IGHA2_HUMAN Immunoglobulin heavy constant alpha 2 OS=Homo sapiens OX=9606 GN=IGHA2 PE=1 SV=4 | 576 | 20201119_15432_1ng_glyco_hcd203040_compound15.20910.20910.4 | controllerType=0 controllerNumber=1 scan=20910 | 42.5053 |
| 17413:4 | 9 | R.LSLHRPALEDLLLGSEAN[+1257.449]LTC[+57.021]TLTGLR.D | HexNAc(3)Hex(4) | C[+57], N[+1257] | 1056.0202 | 4 | 4221.0589 | 4221.0477 | 0 | 2.7 | 114 | Specific | 1117.89 | 1057.37 | 1057.37 | 22.63 | 60 | >sp\|P01877\|IGHA2_HUMAN Immunoglobulin heavy constant alpha 2 OS=Homo sapiens OX=9606 GN=IGHA2 PE=1 SV=4 | 576 | 20201119_15432_1ng_glyco_hcd203040_compound15.21057.21057.4 | controllerType=0 controllerNumber=1 scan=21057 | 42.8626 |
| 04396:3 | 9 | K.TPLTAN[+2262.814]ITK.S | HexNAc(5)Hex(5)Fuc(1)NeuAc(1) | N[+2263] | 1074.4655 | 3 | 3221.3819 | 3221.3711 | 0 | 3.4 | 200 | Specific | 243.17 | 114.3 | 114.3 | 4.35 | 60 | >sp\|P01877\|IGHA2_HUMAN Immunoglobulin heavy constant alpha 2 OS=Homo sapiens OX=9606 GN=IGHA2 PE=1 SV=4 | 576 | 20201119_15432_1ng_glyco_hcd203040_compound15.7469.7469.3 | controllerType=0 controllerNumber=1 scan=7469 | 19.1055 |
| 02508:3 | 9 | K.TPLTAN[+1971.719]ITK.S | HexNAc(5)Hex(5)Fuc(1) | N[+1972] | 977.4315 | 3 | 2930.2799 | 2930.2757 | 0 | 1.4 | 200 | Specific | 244.34 | 146.1 | 146.1 | 4.35 | 60 | >sp\|P01877\|IGHA2_HUMAN Immunoglobulin heavy constant alpha 2 OS=Homo sapiens OX=9606 GN=IGHA2 PE=1 SV=4 | 576 | 20201119_15432_1ng_glyco_hcd203040_compound15.5510.5510.3 | controllerType=0 controllerNumber=1 scan=5510 | 15.7998 |
| 02548:3 | 9 | K.TPLTAN[+1809.666]ITK.S | HexNAc(5)Hex(4)Fuc(1) | N[+1810] | 923.4162 | 3 | 2768.2341 | 2768.2228 | 0 | 4.1 | 200 | Specific | 274.87 | 123.91 | 123.91 | 4.53 | 60 | >sp\|P01877\|IGHA2_HUMAN Immunoglobulin heavy constant alpha 2 OS=Homo sapiens OX=9606 GN=IGHA2 PE=1 SV=4 | 576 | 20201119_15432_1ng_glyco_hcd203040_compound15.5553.5553.3 | controllerType=0 controllerNumber=1 scan=5553 | 15.8752 |
| 02572:2 | 9 | K.TPLTAN[+1606.587]ITK.S | HexNAc(4)Hex(4)Fuc(1) | N[+1607] | 1283.0829 | 2 | 2565.1586 | 2565.1435 | 0 | 5.9 | 200 | Specific | 320.55 | 225.66 | 225.66 | 5.36 | 60 | >sp\|P01877\|IGHA2_HUMAN Immunoglobulin heavy constant alpha 2 OS=Homo sapiens OX=9606 GN=IGHA2 PE=1 SV=4 | 576 | 20201119_15432_1ng_glyco_hcd203040_compound15.5577.5577.2 | controllerType=0 controllerNumber=1 scan=5577 | 15.9169 |
| 02646:2 | 9 | K.TPLTAN[+1501.555]ITK.S | HexNAc(5)Hex(3) | N[+1502] | 1230.5632 | 2 | 2460.119 | 2460.1121 | 0 | 2.8 | 200 | Specific | 351.48 | 136.21 | 136.21 | 5.41 | 60 | >sp\|P01877\|IGHA2_HUMAN Immunoglobulin heavy constant alpha 2 OS=Homo sapiens OX=9606 GN=IGHA2 PE=1 SV=4 | 576 | 20201119_15432_1ng_glyco_hcd203040_compound15.5654.5654.2 | controllerType=0 controllerNumber=1 scan=5654 | 16.0649 |
| 02648:2 | 9 | K.TPLTAN[+1501.555]ITK.S | HexNAc(5)Hex(3) | N[+1502] | 1230.5632 | 2 | 2460.119 | 2460.1121 | 0 | 2.8 | 200 | Specific | 404.68 | 186.62 | 186.62 | 6.22 | 60 | >sp\|P01877\|IGHA2_HUMAN Immunoglobulin heavy constant alpha 2 OS=Homo sapiens OX=9606 GN=IGHA2 PE=1 SV=4 | 576 | 20201119_15432_1ng_glyco_hcd203040_compound15.5656.5656.2 | controllerType=0 controllerNumber=1 scan=5656 | 16.069 |
| 02581:2 | 9 | K.TPLTAN[+1444.534]ITK.S | HexNAc(4)Hex(3)Fuc(1) | N[+1445] | 1202.0534 | 2 | 2403.0995 | 2403.0906 | 0 | 3.7 | 200 | Specific | 419.37 | 238.23 | 238.23 | 6.43 | 60 | >sp\|P01877\|IGHA2_HUMAN Immunoglobulin heavy constant alpha 2 OS=Homo sapiens OX=9606 GN=IGHA2 PE=1 SV=4 | 576 | 20201119_15432_1ng_glyco_hcd203040_compound15.5587.5587.2 | controllerType=0 controllerNumber=1 scan=5587 | 15.937 |
| 02642:3 | 9 | K.TPLTAN[+1647.613]ITK.S | HexNAc(5)Hex(3)Fuc(1) | N[+1648] | 869.3976 | 3 | 2606.1781 | 2606.17 | 0 | 3.1 | 200 | Specific | 464.62 | 182.61 | 182.61 | 6.36 | 60 | >sp\|P01877\|IGHA2_HUMAN Immunoglobulin heavy constant alpha 2 OS=Homo sapiens OX=9606 GN=IGHA2 PE=1 SV=4 | 576 | 20201119_15432_1ng_glyco_hcd203040_compound15.5650.5650.3 | controllerType=0 controllerNumber=1 scan=5650 | 16.0561 |
| 02550:3 | 9 | K.TPLTAN[+1809.666]ITK.S | HexNAc(5)Hex(4)Fuc(1) | N[+1810] | 923.4162 | 3 | 2768.2341 | 2768.2228 | 0 | 4.1 | 200 | Specific | 514.99 | 238.77 | 238.77 | 6.86 | 60 | >sp\|P01877\|IGHA2_HUMAN Immunoglobulin heavy constant alpha 2 OS=Homo sapiens OX=9606 GN=IGHA2 PE=1 SV=4 | 576 | 20201119_15432_1ng_glyco_hcd203040_compound15.5555.5555.3 | controllerType=0 controllerNumber=1 scan=5555 | 15.8791 |
| 02583:2 | 9 | K.TPLTAN[+1444.534]ITK.S | HexNAc(4)Hex(3)Fuc(1) | N[+1445] | 1202.0534 | 2 | 2403.0995 | 2403.0906 | 0 | 3.7 | 200 | Specific | 534.68 | 293.82 | 293.82 | 7.35 | 60 | >sp\|P01877\|IGHA2_HUMAN Immunoglobulin heavy constant alpha 2 OS=Homo sapiens OX=9606 GN=IGHA2 PE=1 SV=4 | 576 | 20201119_15432_1ng_glyco_hcd203040_compound15.5589.5589.2 | controllerType=0 controllerNumber=1 scan=5589 | 15.9405 |
| 02559:2 | 9 | K.TPLTAN[+1809.666]ITK.S | HexNAc(5)Hex(4)Fuc(1) | N[+1810] | 1384.621 | 2 | 2768.2348 | 2768.2228 | 0 | 4.3 | 200 | Specific | 540.44 | 283.57 | 283.57 | 7.22 | 60 | >sp\|P01877\|IGHA2_HUMAN Immunoglobulin heavy constant alpha 2 OS=Homo sapiens OX=9606 GN=IGHA2 PE=1 SV=4 | 576 | 20201119_15432_1ng_glyco_hcd203040_compound15.5564.5564.2 | controllerType=0 controllerNumber=1 scan=5564 | 15.8946 |
| 02557:2 | 9 | K.TPLTAN[+1809.666]ITK.S | HexNAc(5)Hex(4)Fuc(1) | N[+1810] | 1384.621 | 2 | 2768.2348 | 2768.2228 | 0 | 4.3 | 200 | Specific | 568.68 | 359.5 | 359.5 | 8.12 | 60 | >sp\|P01877\|IGHA2_HUMAN Immunoglobulin heavy constant alpha 2 OS=Homo sapiens OX=9606 GN=IGHA2 PE=1 SV=4 | 576 | 20201119_15432_1ng_glyco_hcd203040_compound15.5562.5562.2 | controllerType=0 controllerNumber=1 scan=5562 | 15.8906 |
| 02644:3 | 9 | K.TPLTAN[+1647.613]ITK.S | HexNAc(5)Hex(3)Fuc(1) | N[+1648] | 869.3976 | 3 | 2606.1781 | 2606.17 | 0 | 3.1 | 200 | Specific | 575.36 | 276.57 | 276.57 | 8.62 | 60 | >sp\|P01877\|IGHA2_HUMAN Immunoglobulin heavy constant alpha 2 OS=Homo sapiens OX=9606 GN=IGHA2 PE=1 SV=4 | 576 | 20201119_15432_1ng_glyco_hcd203040_compound15.5652.5652.3 | controllerType=0 controllerNumber=1 scan=5652 | 16.0615 |
| 02634:2 | 9 | K.TPLTAN[+1647.613]ITK.S | HexNAc(5)Hex(3)Fuc(1) | N[+1648] | 1303.5925 | 2 | 2606.1776 | 2606.17 | 0 | 2.9 | 200 | Specific | 597.08 | 296.85 | 296.85 | 8.76 | 60 | >sp\|P01877\|IGHA2_HUMAN Immunoglobulin heavy constant alpha 2 OS=Homo sapiens OX=9606 GN=IGHA2 PE=1 SV=4 | 576 | 20201119_15432_1ng_glyco_hcd203040_compound15.5642.5642.2 | controllerType=0 controllerNumber=1 scan=5642 | 16.0408 |
| 02632:2 | 9 | K.TPLTAN[+1647.613]ITK.S | HexNAc(5)Hex(3)Fuc(1) | N[+1648] | 1303.5925 | 2 | 2606.1776 | 2606.17 | 0 | 2.9 | 200 | Specific | 615.27 | 358.9 | 358.9 | 10.43 | 60 | >sp\|P01877\|IGHA2_HUMAN Immunoglobulin heavy constant alpha 2 OS=Homo sapiens OX=9606 GN=IGHA2 PE=1 SV=4 | 576 | 20201119_15432_1ng_glyco_hcd203040_compound15.5640.5640.2 | controllerType=0 controllerNumber=1 scan=5640 | 16.0341 |
| 10027:3 | 9 | R.MAGKPTHIN[+1809.666]VSVVMAEADGTC[+57.021]Y.- | HexNAc(5)Hex(4)Fuc(1) | C[+57], N[+1810] | 1387.5926 | 3 | 4160.7632 | 4160.7489 | 0 | 3.4 | 319 | CRagged | 232.86 | 232.86 | 181.61 | 6.56 | 60 | >sp\|P01877\|IGHA2_HUMAN Immunoglobulin heavy constant alpha 2 OS=Homo sapiens OX=9606 GN=IGHA2 PE=1 SV=4 | 576 | 20201119_15432_1ng_glyco_hcd203040_compound15.13316.13316.3 | controllerType=0 controllerNumber=1 scan=13316 | 28.8456 |
| 10002:3 | 9 | R.MAGKPTHIN[+1663.608]VSVVMAEADGTC[+57.021]Y.- | HexNAc(5)Hex(4) | C[+57], N[+1664] | 1338.9068 | 3 | 4014.7058 | 4014.691 | 0 | 3.7 | 319 | CRagged | 270.47 | 263.75 | 263.75 | 6.93 | 60 | >sp\|P01877\|IGHA2_HUMAN Immunoglobulin heavy constant alpha 2 OS=Homo sapiens OX=9606 GN=IGHA2 PE=1 SV=4 | 576 | 20201119_15432_1ng_glyco_hcd203040_compound15.13290.13290.3 | controllerType=0 controllerNumber=1 scan=13290 | 28.7961 |
| 10029:3 | 9 | R.MAGKPTHIN[+1809.666]VSVVMAEADGTC[+57.021]Y.- | HexNAc(5)Hex(4)Fuc(1) | C[+57], N[+1810] | 1387.5926 | 3 | 4160.7632 | 4160.7489 | 0 | 3.4 | 319 | CRagged | 278.35 | 278.23 | 278.23 | 7 | 60 | >sp\|P01877\|IGHA2_HUMAN Immunoglobulin heavy constant alpha 2 OS=Homo sapiens OX=9606 GN=IGHA2 PE=1 SV=4 | 576 | 20201119_15432_1ng_glyco_hcd203040_compound15.13318.13318.3 | controllerType=0 controllerNumber=1 scan=13318 | 28.8495 |
| 10000:3 | 9 | R.MAGKPTHIN[+1663.608]VSVVMAEADGTC[+57.021]Y.- | HexNAc(5)Hex(4) | C[+57], N[+1664] | 1338.9068 | 3 | 4014.7058 | 4014.691 | 0 | 3.7 | 319 | CRagged | 283.01 | 283.01 | 218.61 | 7 | 60 | >sp\|P01877\|IGHA2_HUMAN Immunoglobulin heavy constant alpha 2 OS=Homo sapiens OX=9606 GN=IGHA2 PE=1 SV=4 | 576 | 20201119_15432_1ng_glyco_hcd203040_compound15.13288.13288.3 | controllerType=0 controllerNumber=1 scan=13288 | 28.7921 |
| 10106:3 | 9 | R.MAGKPTHIN[+1501.555]VSVVMAEADGTC[+57.021]Y.- | HexNAc(5)Hex(3) | C[+57], N[+1502] | 1284.888 | 3 | 3852.6494 | 3852.6382 | 0 | 2.9 | 319 | CRagged | 430.11 | 430.11 | 430.11 | 9.84 | 60 | >sp\|P01877\|IGHA2_HUMAN Immunoglobulin heavy constant alpha 2 OS=Homo sapiens OX=9606 GN=IGHA2 PE=1 SV=4 | 576 | 20201119_15432_1ng_glyco_hcd203040_compound15.13398.13398.3 | controllerType=0 controllerNumber=1 scan=13398 | 28.9886 |
| 10104:3 | 9 | R.MAGKPTHIN[+1501.555]VSVVMAEADGTC[+57.021]Y.- | HexNAc(5)Hex(3) | C[+57], N[+1502] | 1284.888 | 3 | 3852.6494 | 3852.6382 | 0 | 2.9 | 319 | CRagged | 439.4 | 439.4 | 439.4 | 9.84 | 60 | >sp\|P01877\|IGHA2_HUMAN Immunoglobulin heavy constant alpha 2 OS=Homo sapiens OX=9606 GN=IGHA2 PE=1 SV=4 | 576 | 20201119_15432_1ng_glyco_hcd203040_compound15.13396.13396.3 | controllerType=0 controllerNumber=1 scan=13396 | 28.9851 |
| 10079:3 | 9 | R.MAGKPTHIN[+1501.555]VSVVMAEADGTC[+57.021]Y.- | HexNAc(5)Hex(3) | C[+57], N[+1502] | 1284.8887 | 3 | 3852.6516 | 3852.6382 | 0 | 3.5 | 319 | CRagged | 519.12 | 512.92 | 512.92 | 10.17 | 60 | >sp\|P01877\|IGHA2_HUMAN Immunoglobulin heavy constant alpha 2 OS=Homo sapiens OX=9606 GN=IGHA2 PE=1 SV=4 | 576 | 20201119_15432_1ng_glyco_hcd203040_compound15.13370.13370.3 | controllerType=0 controllerNumber=1 scan=13370 | 28.941 |
| 10081:3 | 9 | R.MAGKPTHIN[+1501.555]VSVVMAEADGTC[+57.021]Y.- | HexNAc(5)Hex(3) | C[+57], N[+1502] | 1284.8887 | 3 | 3852.6516 | 3852.6382 | 0 | 3.5 | 319 | CRagged | 520.66 | 504.7 | 504.7 | 10.17 | 60 | >sp\|P01877\|IGHA2_HUMAN Immunoglobulin heavy constant alpha 2 OS=Homo sapiens OX=9606 GN=IGHA2 PE=1 SV=4 | 576 | 20201119_15432_1ng_glyco_hcd203040_compound15.13372.13372.3 | controllerType=0 controllerNumber=1 scan=13372 | 28.9446 |
| 09902:3 | 9 | R.MAGKPTHIN[+1702.581]VSVVMAEADGTC[+57.021]Y.- | HexNAc(2)Hex(8) | C[+57], N[+1703] | 1351.8967 | 3 | 4053.6754 | 4053.6642 | 0 | 2.8 | 319 | CRagged | 827.48 | 827.48 | 455.23 | 16.69 | 60 | >sp\|P01877\|IGHA2_HUMAN Immunoglobulin heavy constant alpha 2 OS=Homo sapiens OX=9606 GN=IGHA2 PE=1 SV=4 | 576 | 20201119_15432_1ng_glyco_hcd203040_compound15.13187.13187.3 | controllerType=0 controllerNumber=1 scan=13187 | 28.6133 |
| 09904:3 | 9 | R.MAGKPTHIN[+1702.581]VSVVMAEADGTC[+57.021]Y.- | HexNAc(2)Hex(8) | C[+57], N[+1703] | 1351.8967 | 3 | 4053.6754 | 4053.6642 | 0 | 2.8 | 319 | CRagged | 828.27 | 652.25 | 458.18 | 16.69 | 60 | >sp\|P01877\|IGHA2_HUMAN Immunoglobulin heavy constant alpha 2 OS=Homo sapiens OX=9606 GN=IGHA2 PE=1 SV=4 | 576 | 20201119_15432_1ng_glyco_hcd203040_compound15.13189.13189.3 | controllerType=0 controllerNumber=1 scan=13189 | 28.6179 |
| 10026:3 | 9 | R.MAGKPTHIN[+1216.423]VSVVMAEADGTC[+57.021]Y.- | HexNAc(2)Hex(5) | C[+57], N[+1216] | 1189.8451 | 3 | 3567.5209 | 3567.5057 | 0 | 4.3 | 319 | CRagged | 833.4 | 833.4 | 462.58 | 16.56 | 60 | >sp\|P01877\|IGHA2_HUMAN Immunoglobulin heavy constant alpha 2 OS=Homo sapiens OX=9606 GN=IGHA2 PE=1 SV=4 | 576 | 20201119_15432_1ng_glyco_hcd203040_compound15.13315.13315.3 | controllerType=0 controllerNumber=1 scan=13315 | 28.8437 |
| 10024:3 | 9 | R.MAGKPTHIN[+1216.423]VSVVMAEADGTC[+57.021]Y.- | HexNAc(2)Hex(5) | C[+57], N[+1216] | 1189.8451 | 3 | 3567.5209 | 3567.5057 | 0 | 4.3 | 319 | CRagged | 838.3 | 838.3 | 441.4 | 16.56 | 60 | >sp\|P01877\|IGHA2_HUMAN Immunoglobulin heavy constant alpha 2 OS=Homo sapiens OX=9606 GN=IGHA2 PE=1 SV=4 | 576 | 20201119_15432_1ng_glyco_hcd203040_compound15.13313.13313.3 | controllerType=0 controllerNumber=1 scan=13313 | 28.8381 |
| 09960:3 | 9 | R.MAGKPTHIN[+1378.476]VSVVMAEADGTC[+57.021]Y.- | HexNAc(2)Hex(6) | C[+57], N[+1378] | 1243.8619 | 3 | 3729.5711 | 3729.5585 | 0 | 3.4 | 319 | CRagged | 917.4 | 800.49 | 521.73 | 18.69 | 60 | >sp\|P01877\|IGHA2_HUMAN Immunoglobulin heavy constant alpha 2 OS=Homo sapiens OX=9606 GN=IGHA2 PE=1 SV=4 | 576 | 20201119_15432_1ng_glyco_hcd203040_compound15.13247.13247.3 | controllerType=0 controllerNumber=1 scan=13247 | 28.7231 |
| 09958:3 | 9 | R.MAGKPTHIN[+1378.476]VSVVMAEADGTC[+57.021]Y.- | HexNAc(2)Hex(6) | C[+57], N[+1378] | 1243.8619 | 3 | 3729.5711 | 3729.5585 | 0 | 3.4 | 319 | CRagged | 965.81 | 965.81 | 565.49 | 19.69 | 60 | >sp\|P01877\|IGHA2_HUMAN Immunoglobulin heavy constant alpha 2 OS=Homo sapiens OX=9606 GN=IGHA2 PE=1 SV=4 | 576 | 20201119_15432_1ng_glyco_hcd203040_compound15.13245.13245.3 | controllerType=0 controllerNumber=1 scan=13245 | 28.7174 |
| 16793:3 | 11 | K.RNIAAFGGDPNN[+1913.677]ITLFGESAGGASVSLQTLSPYNK.G | HexNAc(4)Hex(5)NeuAc(1) | N[+1914] | 1827.503 | 3 | 5480.4945 | 5480.4644 | 0 | 5.5 | 196 | Specific | 223.58 | 155.73 | 155.73 | 3.98 | 45 | >sp\|P19835\|CEL_HUMAN Bile salt-activated lipase OS=Homo sapiens OX=9606 GN=CEL PE=1 SV=3 | 476 | 20201119_15432_1ng_glyco_hcd203040_compound15.20408.20408.3 | controllerType=0 controllerNumber=1 scan=20408 | 41.4956 |
| 18563:4 | 11 | R.NIAAFGGDPNN[+1913.677]ITLFGESAGGASVSLQTLSPYNK.G | HexNAc(4)Hex(5)NeuAc(1) | N[+1914] | 1331.8534 | 4 | 5324.3918 | 5324.3633 | 0 | 5.3 | 197 | Specific | 574.2 | 574.2 | 574.2 | 9.32 | 45 | >sp\|P19835\|CEL_HUMAN Bile salt-activated lipase OS=Homo sapiens OX=9606 GN=CEL PE=1 SV=3 | 476 | 20201119_15432_1ng_glyco_hcd203040_compound15.22259.22259.4 | controllerType=0 controllerNumber=1 scan=22259 | 45.4257 |
| 18566:3 | 11 | R.NIAAFGGDPNN[+1913.677]ITLFGESAGGASVSLQTLSPYNK.G | HexNAc(4)Hex(5)NeuAc(1) | N[+1914] | 1775.4715 | 3 | 5324.4 | 5324.3633 | 0 | 6.9 | 197 | Specific | 619.66 | 619.66 | 619.66 | 10.94 | 45 | >sp\|P19835\|CEL_HUMAN Bile salt-activated lipase OS=Homo sapiens OX=9606 GN=CEL PE=1 SV=3 | 476 | 20201119_15432_1ng_glyco_hcd203040_compound15.22262.22262.3 | controllerType=0 controllerNumber=1 scan=22262 | 45.4322 |
| 18565:4 | 11 | R.NIAAFGGDPNN[+1913.677]ITLFGESAGGASVSLQTLSPYNK.G | HexNAc(4)Hex(5)NeuAc(1) | N[+1914] | 1331.8534 | 4 | 5324.3918 | 5324.3633 | 0 | 5.3 | 197 | Specific | 929.21 | 929.21 | 929.21 | 17.5 | 45 | >sp\|P19835\|CEL_HUMAN Bile salt-activated lipase OS=Homo sapiens OX=9606 GN=CEL PE=1 SV=3 | 476 | 20201119_15432_1ng_glyco_hcd203040_compound15.22261.22261.4 | controllerType=0 controllerNumber=1 scan=22261 | 45.4303 |
| 18569:3 | 11 | R.NIAAFGGDPNN[+1913.677]ITLFGESAGGASVSLQTLSPYNK.G | HexNAc(4)Hex(5)NeuAc(1) | N[+1914] | 1775.4715 | 3 | 5324.4 | 5324.3633 | 0 | 6.9 | 197 | Specific | 1048.8 | 1030.44 | 1030.44 | 18.94 | 45 | >sp\|P19835\|CEL_HUMAN Bile salt-activated lipase OS=Homo sapiens OX=9606 GN=CEL PE=1 SV=3 | 476 | 20201119_15432_1ng_glyco_hcd203040_compound15.22265.22265.3 | controllerType=0 controllerNumber=1 scan=22265 | 45.4357 |
| 07362:3 | 13 | R.LSPN[+2350.830]ASAEHLELR.W | HexNAc(4)Hex(5)Fuc(1)NeuAc(2) | N[+2351] | 1263.2021 | 3 | 3787.5917 | 3787.5796 | 0 | 3.2 | 52 | Specific | 555.24 | 523.5 | 523.5 | 9.89 | 43 | >sp\|Q13410\|BT1A1_HUMAN Butyrophilin subfamily 1 member A1 OS=Homo sapiens OX=9606 GN=BTN1A1 PE=1 SV=3 | 179 | 20201119_15432_1ng_glyco_hcd203040_compound15.10538.10538.3 | controllerType=0 controllerNumber=1 scan=10538 | 24.1473 |
| 05269:3 | 13 | R.LSPN[+2205.793]ASAEHLELR.W | HexNAc(4)Hex(5)Fuc(2)NeuAc(1) | N[+2206] | 1214.8561 | 3 | 3642.5539 | 3642.5421 | 0 | 3.2 | 52 | Specific | 570.42 | 529.1 | 529.1 | 9.89 | 43 | >sp\|Q13410\|BT1A1_HUMAN Butyrophilin subfamily 1 member A1 OS=Homo sapiens OX=9606 GN=BTN1A1 PE=1 SV=3 | 179 | 20201119_15432_1ng_glyco_hcd203040_compound15.8372.8372.3 | controllerType=0 controllerNumber=1 scan=8372 | 20.5945 |
| 07364:3 | 13 | R.LSPN[+2350.830]ASAEHLELR.W | HexNAc(4)Hex(5)Fuc(1)NeuAc(2) | N[+2351] | 1263.2021 | 3 | 3787.5917 | 3787.5796 | 0 | 3.2 | 52 | Specific | 575.3 | 543.57 | 543.57 | 11.07 | 43 | >sp\|Q13410\|BT1A1_HUMAN Butyrophilin subfamily 1 member A1 OS=Homo sapiens OX=9606 GN=BTN1A1 PE=1 SV=3 | 179 | 20201119_15432_1ng_glyco_hcd203040_compound15.10540.10540.3 | controllerType=0 controllerNumber=1 scan=10540 | 24.1515 |
| 05271:3 | 13 | R.LSPN[+2205.793]ASAEHLELR.W | HexNAc(4)Hex(5)Fuc(2)NeuAc(1) | N[+2206] | 1214.8561 | 3 | 3642.5539 | 3642.5421 | 0 | 3.2 | 52 | Specific | 596.39 | 456.31 | 456.31 | 11.07 | 43 | >sp\|Q13410\|BT1A1_HUMAN Butyrophilin subfamily 1 member A1 OS=Homo sapiens OX=9606 GN=BTN1A1 PE=1 SV=3 | 179 | 20201119_15432_1ng_glyco_hcd203040_compound15.8374.8374.3 | controllerType=0 controllerNumber=1 scan=8374 | 20.6016 |
| 19319:3 | 13 | K.N[+2204.772]VSC[+57.021]YIQNLLLGQEK.K | HexNAc(4)Hex(5)NeuAc(2) | C[+57], N[+2205] | 1328.5677 | 3 | 3983.6885 | 3983.683 | 0 | 1.4 | 215 | Specific | 209.6 | 179.67 | 179.67 | 3.43 | 43 | >sp\|Q13410\|BT1A1_HUMAN Butyrophilin subfamily 1 member A1 OS=Homo sapiens OX=9606 GN=BTN1A1 PE=1 SV=3 | 179 | 20201119_15432_1ng_glyco_hcd203040_compound15.23050.23050.3 | controllerType=0 controllerNumber=1 scan=23050 | 47.106 |
| 17202:3 | 13 | K.N[+2059.735]VSC[+57.021]YIQNLLLGQEK.K | HexNAc(4)Hex(5)Fuc(1)NeuAc(1) | C[+57], N[+2060] | 1280.223 | 3 | 3838.6543 | 3838.6455 | 0 | 2.3 | 215 | Specific | 450.54 | 384.04 | 384.04 | 9.13 | 43 | >sp\|Q13410\|BT1A1_HUMAN Butyrophilin subfamily 1 member A1 OS=Homo sapiens OX=9606 GN=BTN1A1 PE=1 SV=3 | 179 | 20201119_15432_1ng_glyco_hcd203040_compound15.20836.20836.2 | controllerType=0 controllerNumber=1 scan=20836 | 42.3328 |
| 17230:3 | 13 | K.N[+2059.735]VSC[+57.021]YIQNLLLGQEK.K | HexNAc(4)Hex(5)Fuc(1)NeuAc(1) | C[+57], N[+2060] | 1280.2244 | 3 | 3838.6587 | 3838.6455 | 0 | 3.4 | 215 | Specific | 452.19 | 364.8 | 364.8 | 7.74 | 43 | >sp\|Q13410\|BT1A1_HUMAN Butyrophilin subfamily 1 member A1 OS=Homo sapiens OX=9606 GN=BTN1A1 PE=1 SV=3 | 179 | 20201119_15432_1ng_glyco_hcd203040_compound15.20866.20866.3 | controllerType=0 controllerNumber=1 scan=20866 | 42.4054 |
| 17228:3 | 13 | K.N[+2059.735]VSC[+57.021]YIQNLLLGQEK.K | HexNAc(4)Hex(5)Fuc(1)NeuAc(1) | C[+57], N[+2060] | 1280.2244 | 3 | 3838.6587 | 3838.6455 | 0 | 3.4 | 215 | Specific | 463.82 | 315.55 | 315.55 | 7.29 | 43 | >sp\|Q13410\|BT1A1_HUMAN Butyrophilin subfamily 1 member A1 OS=Homo sapiens OX=9606 GN=BTN1A1 PE=1 SV=3 | 179 | 20201119_15432_1ng_glyco_hcd203040_compound15.20864.20864.3 | controllerType=0 controllerNumber=1 scan=20864 | 42.4015 |
| 17204:3 | 13 | K.N[+2059.735]VSC[+57.021]YIQNLLLGQEK.K | HexNAc(4)Hex(5)Fuc(1)NeuAc(1) | C[+57], N[+2060] | 1280.223 | 3 | 3838.6543 | 3838.6455 | 0 | 2.3 | 215 | Specific | 484.63 | 367.98 | 367.98 | 7.88 | 43 | >sp\|Q13410\|BT1A1_HUMAN Butyrophilin subfamily 1 member A1 OS=Homo sapiens OX=9606 GN=BTN1A1 PE=1 SV=3 | 179 | 20201119_15432_1ng_glyco_hcd203040_compound15.20838.20838.2 | controllerType=0 controllerNumber=1 scan=20838 | 42.3379 |
| 13684:4 | 15 | K.LKELPGVC[+57.021]N[+2059.735]ETMMALWEEC[+57.021]KPC[+57.021]LK.Q | HexNAc(4)Hex(5)Fuc(1)NeuAc(1) | C[+57]*3, N[+2060] | 1249.79 | 4 | 4996.1383 | 4996.1196 | 0 | 3.7 | 95 | Specific | 233.59 | 233.59 | 233.59 | 6.05 | 32 | >sp\|P10909\|CLUS_HUMAN Clusterin OS=Homo sapiens OX=9606 GN=CLU PE=1 SV=1 | 1467 | 20201119_15432_1ng_glyco_hcd203040_compound15.17129.17129.4 | controllerType=0 controllerNumber=1 scan=17129 | 35.4307 |
| 15333:4 | 15 | K.LKELPGVC[+57.021]N[+2350.830]ETMMALWEEC[+57.021]KPC[+57.021]LK.Q | HexNAc(4)Hex(5)Fuc(1)NeuAc(2) | C[+57]*3, N[+2351] | 1322.5621 | 4 | 5287.2267 | 5287.2151 | 0 | 2.2 | 95 | Specific | 626.02 | 543.51 | 543.51 | 12.17 | 32 | >sp\|P10909\|CLUS_HUMAN Clusterin OS=Homo sapiens OX=9606 GN=CLU PE=1 SV=1 | 1467 | 20201119_15432_1ng_glyco_hcd203040_compound15.18843.18843.2 | controllerType=0 controllerNumber=1 scan=18843 | 38.5181 |
| 15335:4 | 15 | K.LKELPGVC[+57.021]N[+2350.830]ETMMALWEEC[+57.021]KPC[+57.021]LK.Q | HexNAc(4)Hex(5)Fuc(1)NeuAc(2) | C[+57]*3, N[+2351] | 1322.5621 | 4 | 5287.2267 | 5287.2151 | 0 | 2.2 | 95 | Specific | 729.18 | 609.07 | 609.07 | 14.17 | 32 | >sp\|P10909\|CLUS_HUMAN Clusterin OS=Homo sapiens OX=9606 GN=CLU PE=1 SV=1 | 1467 | 20201119_15432_1ng_glyco_hcd203040_compound15.18845.18845.2 | controllerType=0 controllerNumber=1 scan=18845 | 38.522 |
| 13600:4 | 15 | K.LKELPGVC[+57.021]N[+2205.793]ETMMALWEEC[+57.021]KPC[+57.021]LK.Q | HexNAc(4)Hex(5)Fuc(2)NeuAc(1) | C[+57]*3, N[+2206] | 1286.3029 | 4 | 5142.1896 | 5142.1776 | 0 | 2.3 | 95 | Specific | 815.16 | 815.16 | 815.16 | 16.17 | 32 | >sp\|P10909\|CLUS_HUMAN Clusterin OS=Homo sapiens OX=9606 GN=CLU PE=1 SV=1 | 1467 | 20201119_15432_1ng_glyco_hcd203040_compound15.17042.17042.4 | controllerType=0 controllerNumber=1 scan=17042 | 35.276 |
| 13682:4 | 15 | K.LKELPGVC[+57.021]N[+2059.735]ETMMALWEEC[+57.021]KPC[+57.021]LK.Q | HexNAc(4)Hex(5)Fuc(1)NeuAc(1) | C[+57]*3, N[+2060] | 1249.79 | 4 | 4996.1383 | 4996.1196 | 0 | 3.7 | 95 | Specific | 869.03 | 869.03 | 869.03 | 17.17 | 32 | >sp\|P10909\|CLUS_HUMAN Clusterin OS=Homo sapiens OX=9606 GN=CLU PE=1 SV=1 | 1467 | 20201119_15432_1ng_glyco_hcd203040_compound15.17127.17127.4 | controllerType=0 controllerNumber=1 scan=17127 | 35.426 |
| 13598:4 | 15 | K.LKELPGVC[+57.021]N[+2205.793]ETMMALWEEC[+57.021]KPC[+57.021]LK.Q | HexNAc(4)Hex(5)Fuc(2)NeuAc(1) | C[+57]*3, N[+2206] | 1286.3029 | 4 | 5142.1896 | 5142.1776 | 0 | 2.3 | 95 | Specific | 1022.5 | 879.28 | 879.28 | 20.17 | 32 | >sp\|P10909\|CLUS_HUMAN Clusterin OS=Homo sapiens OX=9606 GN=CLU PE=1 SV=1 | 1467 | 20201119_15432_1ng_glyco_hcd203040_compound15.17040.17040.4 | controllerType=0 controllerNumber=1 scan=17040 | 35.2707 |
| 23083:3 | 15 | K.[+42.011]MLN[+2336.851]TSSLLEQLNEQFNWVSR.L | HexNAc(6)Hex(6)Fuc(1) | .[42], N[+2337] | 1596.6795 | 3 | 4788.0239 | 4788.0484 | 0 | -5.1 | 352 | Specific | 213.6 | 197.16 | 197.16 | 3.55 | 32 | >sp\|P10909\|CLUS_HUMAN Clusterin OS=Homo sapiens OX=9606 GN=CLU PE=1 SV=1 | 1467 | 20201119_15432_1ng_glyco_hcd203040_compound15.26971.26971.3 | controllerType=0 controllerNumber=1 scan=26971 | 54.5815 |
| 20505:3 | 15 | K.MLN[+2205.793]TSSLLEQLNEQFNWVSR.L | HexNAc(4)Hex(5)Fuc(2)NeuAc(1) | N[+2206] | 1539.0046 | 3 | 4614.9992 | 4614.9796 | 0 | 4.2 | 352 | Specific | 584.56 | 534.86 | 534.86 | 11.43 | 32 | >sp\|P10909\|CLUS_HUMAN Clusterin OS=Homo sapiens OX=9606 GN=CLU PE=1 SV=1 | 1467 | 20201119_15432_1ng_glyco_hcd203040_compound15.24290.24290.3 | controllerType=0 controllerNumber=1 scan=24290 | 49.7838 |
| 20507:3 | 15 | K.MLN[+2205.793]TSSLLEQLNEQFNWVSR.L | HexNAc(4)Hex(5)Fuc(2)NeuAc(1) | N[+2206] | 1539.0046 | 3 | 4614.9992 | 4614.9796 | 0 | 4.2 | 352 | Specific | 686.5 | 615.41 | 615.41 | 13.43 | 32 | >sp\|P10909\|CLUS_HUMAN Clusterin OS=Homo sapiens OX=9606 GN=CLU PE=1 SV=1 | 1467 | 20201119_15432_1ng_glyco_hcd203040_compound15.24292.24292.3 | controllerType=0 controllerNumber=1 scan=24292 | 49.7884 |
| 22992:3 | 15 | K.MLN[+2350.830]TSSLLEQLNEQFNWVSR.L | HexNAc(4)Hex(5)Fuc(1)NeuAc(2) | N[+2351] | 1587.352 | 3 | 4760.0414 | 4760.0171 | 0 | 5.1 | 352 | Specific | 826.19 | 561.66 | 561.66 | 16.43 | 32 | >sp\|P10909\|CLUS_HUMAN Clusterin OS=Homo sapiens OX=9606 GN=CLU PE=1 SV=1 | 1467 | 20201119_15432_1ng_glyco_hcd203040_compound15.26876.26876.3 | controllerType=0 controllerNumber=1 scan=26876 | 54.4003 |
| 22994:3 | 15 | K.MLN[+2350.830]TSSLLEQLNEQFNWVSR.L | HexNAc(4)Hex(5)Fuc(1)NeuAc(2) | N[+2351] | 1587.352 | 3 | 4760.0414 | 4760.0171 | 0 | 5.1 | 352 | Specific | 934.76 | 634.63 | 634.63 | 18.43 | 32 | >sp\|P10909\|CLUS_HUMAN Clusterin OS=Homo sapiens OX=9606 GN=CLU PE=1 SV=1 | 1467 | 20201119_15432_1ng_glyco_hcd203040_compound15.26878.26878.3 | controllerType=0 controllerNumber=1 scan=26878 | 54.4045 |
| 12698:3 | 15 | R.LAN[+2350.830]LTQGEDQYYLR.V | HexNAc(4)Hex(5)Fuc(1)NeuAc(2) | N[+2351] | 1345.5622 | 3 | 4034.672 | 4034.664 | 0 | 2 | 372 | Specific | 310.37 | 285.39 | 285.39 | 6.48 | 32 | >sp\|P10909\|CLUS_HUMAN Clusterin OS=Homo sapiens OX=9606 GN=CLU PE=1 SV=1 | 1467 | 20201119_15432_1ng_glyco_hcd203040_compound15.16107.16107.3 | controllerType=0 controllerNumber=1 scan=16107 | 33.5811 |
| 12684:3 | 15 | R.LAN[+2350.830]LTQGEDQYYLR.V | HexNAc(4)Hex(5)Fuc(1)NeuAc(2) | N[+2351] | 1345.5615 | 3 | 4034.6698 | 4034.664 | 0 | 1.4 | 372 | Specific | 315.81 | 281.89 | 281.89 | 6.16 | 32 | >sp\|P10909\|CLUS_HUMAN Clusterin OS=Homo sapiens OX=9606 GN=CLU PE=1 SV=1 | 1467 | 20201119_15432_1ng_glyco_hcd203040_compound15.16092.16092.3 | controllerType=0 controllerNumber=1 scan=16092 | 33.5549 |
| 12725:3 | 15 | R.LAN[+2350.830]LTQGEDQYYLR.V | HexNAc(4)Hex(5)Fuc(1)NeuAc(2) | N[+2351] | 1345.5668 | 3 | 4034.6859 | 4034.664 | 0 | 5.4 | 372 | Specific | 329.32 | 253.45 | 253.45 | 5.85 | 32 | >sp\|P10909\|CLUS_HUMAN Clusterin OS=Homo sapiens OX=9606 GN=CLU PE=1 SV=1 | 1467 | 20201119_15432_1ng_glyco_hcd203040_compound15.16135.16135.3 | controllerType=0 controllerNumber=1 scan=16135 | 33.6293 |
| 12686:3 | 15 | R.LAN[+2350.830]LTQGEDQYYLR.V | HexNAc(4)Hex(5)Fuc(1)NeuAc(2) | N[+2351] | 1345.5615 | 3 | 4034.6698 | 4034.664 | 0 | 1.4 | 372 | Specific | 387.3 | 379.38 | 379.38 | 8.63 | 32 | >sp\|P10909\|CLUS_HUMAN Clusterin OS=Homo sapiens OX=9606 GN=CLU PE=1 SV=1 | 1467 | 20201119_15432_1ng_glyco_hcd203040_compound15.16094.16094.3 | controllerType=0 controllerNumber=1 scan=16094 | 33.5585 |
| 12700:3 | 15 | R.LAN[+2350.830]LTQGEDQYYLR.V | HexNAc(4)Hex(5)Fuc(1)NeuAc(2) | N[+2351] | 1345.5622 | 3 | 4034.672 | 4034.664 | 0 | 2 | 372 | Specific | 406.19 | 265.71 | 265.71 | 6.73 | 32 | >sp\|P10909\|CLUS_HUMAN Clusterin OS=Homo sapiens OX=9606 GN=CLU PE=1 SV=1 | 1467 | 20201119_15432_1ng_glyco_hcd203040_compound15.16109.16109.3 | controllerType=0 controllerNumber=1 scan=16109 | 33.5847 |
| 11082:3 | 17 | R.LAGKPTHVN[+1971.719]VSVVMAEVDGTC[+57.021]Y.- | HexNAc(5)Hex(5)Fuc(1) | C[+57], N[+1972] | 1440.2973 | 3 | 4318.8773 | 4318.861 | 0 | 3.8 | 332 | CRagged | 348.58 | 335.47 | 287.42 | 7.92 | 24 | >sp\|P01876\|IGHA1_HUMAN Immunoglobulin heavy constant alpha 1 OS=Homo sapiens OX=9606 GN=IGHA1 PE=1 SV=2 | 255 | 20201119_15432_1ng_glyco_hcd203040_compound15.14408.14408.3 | controllerType=0 controllerNumber=1 scan=14408 | 30.6834 |
| 12938:3 | 17 | R.LAGKPTHVN[+1710.598]VSVVMAEVDGTC[+57.021]Y.- | HexNAc(3)Hex(5)NeuAc(1) | C[+57], N[+1711] | 1353.2551 | 3 | 4057.7506 | 4057.7397 | 0 | 2.7 | 332 | CRagged | 379.34 | 355.54 | 283.86 | 8.31 | 24 | >sp\|P01876\|IGHA1_HUMAN Immunoglobulin heavy constant alpha 1 OS=Homo sapiens OX=9606 GN=IGHA1 PE=1 SV=2 | 255 | 20201119_15432_1ng_glyco_hcd203040_compound15.16355.16355.3 | controllerType=0 controllerNumber=1 scan=16355 | 34.0084 |
| 11084:3 | 17 | R.LAGKPTHVN[+1971.719]VSVVMAEVDGTC[+57.021]Y.- | HexNAc(5)Hex(5)Fuc(1) | C[+57], N[+1972] | 1440.2973 | 3 | 4318.8773 | 4318.861 | 0 | 3.8 | 332 | CRagged | 388.05 | 363.81 | 312.25 | 8.31 | 24 | >sp\|P01876\|IGHA1_HUMAN Immunoglobulin heavy constant alpha 1 OS=Homo sapiens OX=9606 GN=IGHA1 PE=1 SV=2 | 255 | 20201119_15432_1ng_glyco_hcd203040_compound15.14410.14410.3 | controllerType=0 controllerNumber=1 scan=14410 | 30.6906 |
| 12936:3 | 17 | R.LAGKPTHVN[+1710.598]VSVVMAEVDGTC[+57.021]Y.- | HexNAc(3)Hex(5)NeuAc(1) | C[+57], N[+1711] | 1353.2551 | 3 | 4057.7506 | 4057.7397 | 0 | 2.7 | 332 | CRagged | 392.77 | 392.77 | 315.96 | 9.71 | 24 | >sp\|P01876\|IGHA1_HUMAN Immunoglobulin heavy constant alpha 1 OS=Homo sapiens OX=9606 GN=IGHA1 PE=1 SV=2 | 255 | 20201119_15432_1ng_glyco_hcd203040_compound15.16353.16353.3 | controllerType=0 controllerNumber=1 scan=16353 | 34.0049 |
| 11075:3 | 17 | R.LAGKPTHVN[+1663.608]VSVVMAEVDGTC[+57.021]Y.- | HexNAc(5)Hex(4) | C[+57], N[+1664] | 1337.5956 | 3 | 4010.7724 | 4010.7502 | 0 | 5.5 | 332 | CRagged | 658.74 | 658.74 | 429.45 | 12.93 | 24 | >sp\|P01876\|IGHA1_HUMAN Immunoglobulin heavy constant alpha 1 OS=Homo sapiens OX=9606 GN=IGHA1 PE=1 SV=2 | 255 | 20201119_15432_1ng_glyco_hcd203040_compound15.14401.14401.3 | controllerType=0 controllerNumber=1 scan=14401 | 30.6695 |
| 11121:3 | 17 | R.LAGKPTHVN[+1501.555]VSVVMAEVDGTC[+57.021]Y.- | HexNAc(5)Hex(3) | C[+57], N[+1502] | 1283.5756 | 3 | 3848.7123 | 3848.6974 | 0 | 3.9 | 332 | CRagged | 719.82 | 719.82 | 719.82 | 14.69 | 24 | >sp\|P01876\|IGHA1_HUMAN Immunoglobulin heavy constant alpha 1 OS=Homo sapiens OX=9606 GN=IGHA1 PE=1 SV=2 | 255 | 20201119_15432_1ng_glyco_hcd203040_compound15.14449.14449.3 | controllerType=0 controllerNumber=1 scan=14449 | 30.7747 |
| 11101:3 | 17 | R.LAGKPTHVN[+1809.666]VSVVMAEVDGTC[+57.021]Y.- | HexNAc(5)Hex(4)Fuc(1) | C[+57], N[+1810] | 1386.2773 | 3 | 4156.8173 | 4156.8081 | 0 | 2.2 | 332 | CRagged | 756.58 | 632.16 | 482.23 | 15.69 | 24 | >sp\|P01876\|IGHA1_HUMAN Immunoglobulin heavy constant alpha 1 OS=Homo sapiens OX=9606 GN=IGHA1 PE=1 SV=2 | 255 | 20201119_15432_1ng_glyco_hcd203040_compound15.14428.14428.3 | controllerType=0 controllerNumber=1 scan=14428 | 30.7305 |
| 11078:3 | 17 | R.LAGKPTHVN[+1663.608]VSVVMAEVDGTC[+57.021]Y.- | HexNAc(5)Hex(4) | C[+57], N[+1664] | 1337.5956 | 3 | 4010.7724 | 4010.7502 | 0 | 5.5 | 332 | CRagged | 800.78 | 800.78 | 488.32 | 15.93 | 24 | >sp\|P01876\|IGHA1_HUMAN Immunoglobulin heavy constant alpha 1 OS=Homo sapiens OX=9606 GN=IGHA1 PE=1 SV=2 | 255 | 20201119_15432_1ng_glyco_hcd203040_compound15.14404.14404.3 | controllerType=0 controllerNumber=1 scan=14404 | 30.6768 |
| 11123:3 | 17 | R.LAGKPTHVN[+1501.555]VSVVMAEVDGTC[+57.021]Y.- | HexNAc(5)Hex(3) | C[+57], N[+1502] | 1283.5756 | 3 | 3848.7123 | 3848.6974 | 0 | 3.9 | 332 | CRagged | 820.05 | 820.05 | 820.05 | 16.69 | 24 | >sp\|P01876\|IGHA1_HUMAN Immunoglobulin heavy constant alpha 1 OS=Homo sapiens OX=9606 GN=IGHA1 PE=1 SV=2 | 255 | 20201119_15432_1ng_glyco_hcd203040_compound15.14451.14451.3 | controllerType=0 controllerNumber=1 scan=14451 | 30.7799 |
| 10962:3 | 17 | R.LAGKPTHVN[+1702.581]VSVVMAEVDGTC[+57.021]Y.- | HexNAc(2)Hex(8) | C[+57], N[+1703] | 1350.5837 | 3 | 4049.7365 | 4049.7234 | 0 | 3.2 | 332 | CRagged | 827 | 775.65 | 500.04 | 16.69 | 24 | >sp\|P01876\|IGHA1_HUMAN Immunoglobulin heavy constant alpha 1 OS=Homo sapiens OX=9606 GN=IGHA1 PE=1 SV=2 | 255 | 20201119_15432_1ng_glyco_hcd203040_compound15.14284.14284.3 | controllerType=0 controllerNumber=1 scan=14284 | 30.4637 |
| 11145:3 | 17 | R.LAGKPTHVN[+1647.613]VSVVMAEVDGTC[+57.021]Y.- | HexNAc(5)Hex(3)Fuc(1) | C[+57], N[+1648] | 1332.2615 | 3 | 3994.77 | 3994.7553 | 0 | 3.7 | 332 | CRagged | 831.78 | 663.54 | 663.54 | 16.69 | 24 | >sp\|P01876\|IGHA1_HUMAN Immunoglobulin heavy constant alpha 1 OS=Homo sapiens OX=9606 GN=IGHA1 PE=1 SV=2 | 255 | 20201119_15432_1ng_glyco_hcd203040_compound15.14473.14473.3 | controllerType=0 controllerNumber=1 scan=14473 | 30.8321 |
| 11048:3 | 17 | R.LAGKPTHVN[+1378.476]VSVVMAEVDGTC[+57.021]Y.- | HexNAc(2)Hex(6) | C[+57], N[+1378] | 1242.5474 | 3 | 3725.6277 | 3725.6178 | 0 | 2.7 | 332 | CRagged | 851.5 | 830.97 | 533.58 | 17.69 | 24 | >sp\|P01876\|IGHA1_HUMAN Immunoglobulin heavy constant alpha 1 OS=Homo sapiens OX=9606 GN=IGHA1 PE=1 SV=2 | 255 | 20201119_15432_1ng_glyco_hcd203040_compound15.14373.14373.3 | controllerType=0 controllerNumber=1 scan=14373 | 30.6211 |
| 10959:3 | 17 | R.LAGKPTHVN[+1702.581]VSVVMAEVDGTC[+57.021]Y.- | HexNAc(2)Hex(8) | C[+57], N[+1703] | 1350.5837 | 3 | 4049.7365 | 4049.7234 | 0 | 3.2 | 332 | CRagged | 867.66 | 867.66 | 554.39 | 17.69 | 24 | >sp\|P01876\|IGHA1_HUMAN Immunoglobulin heavy constant alpha 1 OS=Homo sapiens OX=9606 GN=IGHA1 PE=1 SV=2 | 255 | 20201119_15432_1ng_glyco_hcd203040_compound15.14281.14281.3 | controllerType=0 controllerNumber=1 scan=14281 | 30.4577 |
| 11102:3 | 17 | R.LAGKPTHVN[+1216.423]VSVVMAEVDGTC[+57.021]Y.- | HexNAc(2)Hex(5) | C[+57], N[+1216] | 1188.5292 | 3 | 3563.5731 | 3563.5649 | 0 | 2.3 | 332 | CRagged | 889.92 | 889.92 | 548.73 | 17.69 | 24 | >sp\|P01876\|IGHA1_HUMAN Immunoglobulin heavy constant alpha 1 OS=Homo sapiens OX=9606 GN=IGHA1 PE=1 SV=2 | 255 | 20201119_15432_1ng_glyco_hcd203040_compound15.14429.14429.3 | controllerType=0 controllerNumber=1 scan=14429 | 30.7319 |

Table S4. Cont.

| Query #:z | Protein Rank | Glycopeptide | Glycan | Modification Type(s) | Observed m/z | z | Observed (M+H) | Calc. mass (M+H) | Off-by-x error | Mass error (ppm) | StartPosition | Cleavage | Score | Delta | Delta Mod | \|Log Prob\| | # of unique peptides | Protein Name | Protein DB number | Comment | Scan # | Scan Time |
|---|---|---|---|---|---|---|---|---|---|---|---|---|---|---|---|---|---|---|---|---|---|---|
| 11045:3 | 17 | R.LAGKPTHVN[+1378.476]VSVVMAEVDGTC[+57.021]Y.- | HexNAc(2)Hex(6) | C[+57], N[+1378] | 1242.5474 | 3 | 3725.6277 | 3725.6178 | 0 | 2.7 | 332 | CRagged | 896.69 | 896.69 | 558.24 | 17.69 | 24 | >sp\|P01876\|IGHA1_HUMAN Immunoglobulin heavy constant alpha 1 OS=Homo sapiens OX=9606 GN=IGHA1 PE=1 SV=2 | 255 | 20201119_15432_1ng_glyco_hcd203040_compound15.14370.14370.3 | controllerType=0 controllerNumber=1 scan=14370 | 30.6144 |
| 11105:3 | 17 | R.LAGKPTHVN[+1216.423]VSVVMAEVDGTC[+57.021]Y.- | HexNAc(2)Hex(5) | C[+57], N[+1216] | 1188.5292 | 3 | 3563.5731 | 3563.5649 | 0 | 2.3 | 332 | CRagged | 899.33 | 852.86 | 567.41 | 17.69 | 24 | >sp\|P01876\|IGHA1_HUMAN Immunoglobulin heavy constant alpha 1 OS=Homo sapiens OX=9606 GN=IGHA1 PE=1 SV=2 | 255 | 20201119_15432_1ng_glyco_hcd203040_compound15.14432.14432.3 | controllerType=0 controllerNumber=1 scan=14432 | 30.7419 |
| 11099:3 | 17 | R.LAGKPTHVN[+1809.666]VSVVMAEVDGTC[+57.021]Y.- | HexNAc(5)Hex(4)Fuc(1) | C[+57], N[+1810] | 1386.2773 | 3 | 4156.8173 | 4156.8081 | 0 | 2.2 | 332 | CRagged | 922.62 | 728 | 623.4 | 18.69 | 24 | >sp\|P01876\|IGHA1_HUMAN Immunoglobulin heavy constant alpha 1 OS=Homo sapiens OX=9606 GN=IGHA1 PE=1 SV=2 | 255 | 20201119_15432_1ng_glyco_hcd203040_compound15.14426.14426.3 | controllerType=0 controllerNumber=1 scan=14426 | 30.7232 |
| 11143:3 | 17 | R.LAGKPTHVN[+1647.613]VSVVMAEVDGTC[+57.021]Y.- | HexNAc(5)Hex(3)Fuc(1) | C[+57], N[+1648] | 1332.2615 | 3 | 3994.77 | 3994.7553 | 0 | 3.7 | 332 | CRagged | 955.57 | 785.65 | 785.65 | 19.69 | 24 | >sp\|P01876\|IGHA1_HUMAN Immunoglobulin heavy constant alpha 1 OS=Homo sapiens OX=9606 GN=IGHA1 PE=1 SV=2 | 255 | 20201119_15432_1ng_glyco_hcd203040_compound15.14471.14471.3 | controllerType=0 controllerNumber=1 scan=14471 | 30.8139 |
| 23735:4 | 18 | R.QLAHQSN[+2351.851]STNIFFSPVSIATAFAMLSLGTK.A | HexNAc(4)Hex(5)Fuc(3)NeuAc(1) | N[+2352] | 1384.1287 | 4 | 5533.4928 | 5533.4858 | 0 | 1.3 | 64 | Specific | 384.81 | 384.81 | 384.81 | 9.19 | 33 | >sp\|P01009\|A1AT_HUMAN Alpha-1-antitrypsin OS=Homo sapiens OX=9606 GN=SERPINA1 PE=1 SV=3 | 642 | 20201119_15432_1ng_glyco_hcd203040_compound15.27649.27649.4 | controllerType=0 controllerNumber=1 scan=27649 | 55.8039 |
| 23737:4 | 18 | R.QLAHQSN[+2351.851]STNIFFSPVSIATAFAMLSLGTK.A | HexNAc(4)Hex(5)Fuc(3)NeuAc(1) | N[+2352] | 1384.1287 | 4 | 5533.4928 | 5533.4858 | 0 | 1.3 | 64 | Specific | 471.51 | 438.73 | 438.73 | 9.37 | 33 | >sp\|P01009\|A1AT_HUMAN Alpha-1-antitrypsin OS=Homo sapiens OX=9606 GN=SERPINA1 PE=1 SV=3 | 642 | 20201119_15432_1ng_glyco_hcd203040_compound15.27651.27651.4 | controllerType=0 controllerNumber=1 scan=27651 | 55.8075 |
| 22338:4 | 18 | K.ADTHDEILEGLNFN[+2861.000]LTEIPEAQIHEGFQELLR.T | HexNAc(5)Hex(6)NeuAc(3) | N[+2861] | 1638.9756 | 4 | 6552.8805 | 6552.8239 | 0 | 8.6 | 94 | Specific | 343.64 | 343.64 | 343.64 | 6.16 | 33 | >sp\|P01009\|A1AT_HUMAN Alpha-1-antitrypsin OS=Homo sapiens OX=9606 GN=SERPINA1 PE=1 SV=3 | 642 | 20201119_15432_1ng_glyco_hcd203040_compound15.26196.26196.4 | controllerType=0 controllerNumber=1 scan=26196 | 53.0906 |
| 22340:4 | 18 | K.ADTHDEILEGLNFN[+2861.000]LTEIPEAQIHEGFQELLR.T | HexNAc(5)Hex(6)NeuAc(3) | N[+2861] | 1638.9756 | 4 | 6552.8805 | 6552.8239 | 0 | 8.6 | 94 | Specific | 406.22 | 406.22 | 406.22 | 7.85 | 33 | >sp\|P01009\|A1AT_HUMAN Alpha-1-antitrypsin OS=Homo sapiens OX=9606 GN=SERPINA1 PE=1 SV=3 | 642 | 20201119_15432_1ng_glyco_hcd203040_compound15.26198.26198.4 | controllerType=0 controllerNumber=1 scan=26198 | 53.0977 |
| 19606:5 | 18 | K.YLGN[+2204.772]ATAIFFLPDEGKLQHLENELTHDIITK.F | HexNAc(4)Hex(5)NeuAc(2) | N[+2205] | 1149.9314 | 5 | 5745.6279 | 5745.6098 | 0 | 3.2 | 268 | Specific | 633.34 | 607.84 | 607.84 | 11.72 | 33 | >sp\|P01009\|A1AT_HUMAN Alpha-1-antitrypsin OS=Homo sapiens OX=9606 GN=SERPINA1 PE=1 SV=3 | 642 | 20201119_15432_1ng_glyco_hcd203040_compound15.23351.23351.5 | controllerType=0 controllerNumber=1 scan=23351 | 47.7638 |
| 19609:5 | 18 | K.YLGN[+2204.772]ATAIFFLPDEGKLQHLENELTHDIITK.F | HexNAc(4)Hex(5)NeuAc(2) | N[+2205] | 1149.9314 | 5 | 5745.6279 | 5745.6098 | 0 | 3.2 | 268 | Specific | 738.97 | 695.53 | 695.53 | 13.72 | 33 | >sp\|P01009\|A1AT_HUMAN Alpha-1-antitrypsin OS=Homo sapiens OX=9606 GN=SERPINA1 PE=1 SV=3 | 642 | 20201119_15432_1ng_glyco_hcd203040_compound15.23354.23354.5 | controllerType=0 controllerNumber=1 scan=23354 | 47.7689 |
| 19582:4 | 18 | K.YLGN[+2204.772]ATAIFFLPDEGKLQHLENELTHDIITK.F | HexNAc(4)Hex(5)NeuAc(2) | N[+2205] | 1437.1624 | 4 | 5745.6276 | 5745.6098 | 0 | 3.1 | 268 | Specific | 974.2 | 947.8 | 947.8 | 18.18 | 33 | >sp\|P01009\|A1AT_HUMAN Alpha-1-antitrypsin OS=Homo sapiens OX=9606 GN=SERPINA1 PE=1 SV=3 | 642 | 20201119_15432_1ng_glyco_hcd203040_compound15.23326.23326.4 | controllerType=0 controllerNumber=1 scan=23326 | 47.7142 |
| 19580:4 | 18 | K.YLGN[+2204.772]ATAIFFLPDEGKLQHLENELTHDIITK.F | HexNAc(4)Hex(5)NeuAc(2) | N[+2205] | 1437.1624 | 4 | 5745.6276 | 5745.6098 | 0 | 3.1 | 268 | Specific | 1179.14 | 1055.6 | 1055.6 | 22.18 | 33 | >sp\|P01009\|A1AT_HUMAN Alpha-1-antitrypsin OS=Homo sapiens OX=9606 GN=SERPINA1 PE=1 SV=3 | 642 | 20201119_15432_1ng_glyco_hcd203040_compound15.23324.23324.4 | controllerType=0 controllerNumber=1 scan=23324 | 47.7095 |
| 09090:5 | 24 | K.[+42.011]QTDEIKDTRN[+2018.708]ESTQNC[+57.021]VVAEPEKMESSISSSSEEMSLS | HexNAc(3)Hex(6)Fuc(1)NeuAc(1) | .[42], C[+57], N[+2019] | 1291.1528 | 5 | 6451.7351 | 6451.6996 | 0 | 5.5 | 60 | Specific | 231.89 | 195.14 | 195.14 | 3.23 | 31 | >sp\|P47710\|CASA1_HUMAN Alpha-S1-casein OS=Homo sapiens OX=9606 GN=CSN1S1 PE=1 SV=1 | 989 | 20201119_15432_1ng_glyco_hcd203040_compound15.12347.12347.3 | controllerType=0 controllerNumber=1 scan=12347 | 27.23 |
| 09088:5 | 24 | K.[+42.011]QTDEIKDTRN[+2018.708]ESTQNC[+57.021]VVAEPEKMESSISSSSEEMSLS | HexNAc(3)Hex(6)Fuc(1)NeuAc(1) | .[42], C[+57], N[+2019] | 1291.1528 | 5 | 6451.7351 | 6451.6996 | 0 | 5.5 | 60 | Specific | 287.57 | 275.56 | 275.56 | 4.56 | 31 | >sp\|P47710\|CASA1_HUMAN Alpha-S1-casein OS=Homo sapiens OX=9606 GN=CSN1S1 PE=1 SV=1 | 989 | 20201119_15432_1ng_glyco_hcd203040_compound15.12345.12345.3 | controllerType=0 controllerNumber=1 scan=12345 | 27.2263 |
| 09262:3 | 40 | R.IIVPLNNREN[+1809.666]ISDPTSPLR.T | HexNAc(5)Hex(4)Fuc(1) | N[+1810] | 1319.9583 | 3 | 3957.8604 | 3957.8432 | 0 | 4.3 | 62 | Specific | 316.15 | 240.52 | 240.52 | 5.93 | 16 | >sp\|P01591\|IGJ_HUMAN Immunoglobulin J chain OS=Homo sapiens OX=9606 GN=JCHAIN PE=1 SV=4 | 1392 | 20201119_15432_1ng_glyco_hcd203040_compound15.12525.12525.3 | controllerType=0 controllerNumber=1 scan=12525 | 27.5077 |
| 09259:3 | 40 | R.IIVPLNNREN[+1809.666]ISDPTSPLR.T | HexNAc(5)Hex(4)Fuc(1) | N[+1810] | 1319.9583 | 3 | 3957.8604 | 3957.8432 | 0 | 4.3 | 62 | Specific | 404.55 | 338.34 | 338.34 | 6.99 | 16 | >sp\|P01591\|IGJ_HUMAN Immunoglobulin J chain OS=Homo sapiens OX=9606 GN=JCHAIN PE=1 SV=4 | 1392 | 20201119_15432_1ng_glyco_hcd203040_compound15.12522.12522.3 | controllerType=0 controllerNumber=1 scan=12522 | 27.503 |
| 12971:3 | 40 | R.IIVPLNNREN[+2204.772]ISDPTSPLR.T | HexNAc(4)Hex(5)NeuAc(2) | N[+2205] | 1451.6614 | 3 | 4352.9697 | 4352.9496 | 0 | 4.6 | 62 | Specific | 409.81 | 407.55 | 407.55 | 8.56 | 16 | >sp\|P01591\|IGJ_HUMAN Immunoglobulin J chain OS=Homo sapiens OX=9606 GN=JCHAIN PE=1 SV=4 | 1392 | 20201119_15432_1ng_glyco_hcd203040_compound15.16389.16389.3 | controllerType=0 controllerNumber=1 scan=16389 | 34.0702 |
| 12969:3 | 40 | R.IIVPLNNREN[+2204.772]ISDPTSPLR.T | HexNAc(4)Hex(5)NeuAc(2) | N[+2205] | 1451.6614 | 3 | 4352.9697 | 4352.9496 | 0 | 4.6 | 62 | Specific | 431.09 | 431.09 | 431.09 | 8.79 | 16 | >sp\|P01591\|IGJ_HUMAN Immunoglobulin J chain OS=Homo sapiens OX=9606 GN=JCHAIN PE=1 SV=4 | 1392 | 20201119_15432_1ng_glyco_hcd203040_compound15.16387.16387.3 | controllerType=0 controllerNumber=1 scan=16387 | 34.0651 |
| 11008:3 | 40 | R.IIVPLNNREN[+1856.656]ISDPTSPLR.T | HexNAc(3)Hex(5)Fuc(1)NeuAc(1) | N[+1857] | 1335.6182 | 3 | 4004.8401 | 4004.8327 | 0 | 1.8 | 62 | Specific | 482.36 | 482.36 | 482.36 | 9.1 | 16 | >sp\|P01591\|IGJ_HUMAN Immunoglobulin J chain OS=Homo sapiens OX=9606 GN=JCHAIN PE=1 SV=4 | 1392 | 20201119_15432_1ng_glyco_hcd203040_compound15.14331.14331.3 | controllerType=0 controllerNumber=1 scan=14331 | 30.5372 |
| 09271:3 | 40 | R.IIVPLNNREN[+1663.608]ISDPTSPLR.T | HexNAc(5)Hex(4) | N[+1664] | 1271.2711 | 3 | 3811.7986 | 3811.7853 | 0 | 3.5 | 62 | Specific | 488.2 | 483.17 | 483.17 | 9.1 | 16 | >sp\|P01591\|IGJ_HUMAN Immunoglobulin J chain OS=Homo sapiens OX=9606 GN=JCHAIN PE=1 SV=4 | 1392 | 20201119_15432_1ng_glyco_hcd203040_compound15.12534.12534.3 | controllerType=0 controllerNumber=1 scan=12534 | 27.5234 |
| 09269:3 | 40 | R.IIVPLNNREN[+1663.608]ISDPTSPLR.T | HexNAc(5)Hex(4) | N[+1664] | 1271.2711 | 3 | 3811.7986 | 3811.7853 | 0 | 3.5 | 62 | Specific | 490.43 | 455.24 | 455.24 | 9.1 | 16 | >sp\|P01591\|IGJ_HUMAN Immunoglobulin J chain OS=Homo sapiens OX=9606 GN=JCHAIN PE=1 SV=4 | 1392 | 20201119_15432_1ng_glyco_hcd203040_compound15.12532.12532.3 | controllerType=0 controllerNumber=1 scan=12532 | 27.5185 |
| 11010:3 | 40 | R.IIVPLNNREN[+1856.656]ISDPTSPLR.T | HexNAc(3)Hex(5)Fuc(1)NeuAc(1) | N[+1857] | 1335.6182 | 3 | 4004.8401 | 4004.8327 | 0 | 1.8 | 62 | Specific | 530.87 | 428.94 | 428.94 | 9.35 | 16 | >sp\|P01591\|IGJ_HUMAN Immunoglobulin J chain OS=Homo sapiens OX=9606 GN=JCHAIN PE=1 SV=4 | 1392 | 20201119_15432_1ng_glyco_hcd203040_compound15.14333.14333.3 | controllerType=0 controllerNumber=1 scan=14333 | 30.5424 |
| 06301:3 | 406 | R.[+42.011]VEDAYILTC[+57.021]N[+730.264]VSLEYEK.T | HexNAc(2)Hex(2) | .[42], C[+57], N[+730] | 940.0955 | 3 | 2818.2719 | 2818.2486 | 0 | 8.3 | 225 | Specific | 257.66 | 124.14 | 124.14 | 2.01 | 3 | >sp\|P40227\|TCPZ_HUMAN T-complex protein 1 subunit zeta OS=Homo sapiens OX=9606 GN=CCT6A PE=1 SV=3 | 1115 | 20201119_15432_1ng_glyco_hcd203040_compound15.9438.9438.3 | controllerType=0 controllerNumber=1 scan=9438 | 22.3441 |
| 06303:3 | 406 | R.[+42.011]VEDAYILTC[+57.021]N[+730.264]VSLEYEK.T | HexNAc(2)Hex(2) | .[42], C[+57], N[+730] | 940.0955 | 3 | 2818.2719 | 2818.2486 | 0 | 8.3 | 225 | Specific | 285.05 | 132.56 | 132.56 | 2.63 | 3 | >sp\|P40227\|TCPZ_HUMAN T-complex protein 1 subunit zeta OS=Homo sapiens OX=9606 GN=CCT6A PE=1 SV=3 | 1115 | 20201119_15432_1ng_glyco_hcd203040_compound15.9440.9440.3 | controllerType=0 controllerNumber=1 scan=9440 | 22.3484 |
| 18500:3 | 1 | R.RTAGWNVPIGTLRPFLN[+1930.692]WTGPPEPIEAAVAR.F | HexNAc(4)Hex(6)Fuc(1) | N[+1931] | 1773.1607 | 3 | 5317.4676 | 5317.5044 | 0 | -6.9 | 140 | Specific | 228.88 | 228.88 | 228.88 | 3.55 | 138 | >sp\|P02788\|TRFL_HUMAN Lactotransferrin OS=Homo sapiens OX=9606 GN=LTF PE=1 SV=6 | 1270 | 20201119_16913_1ng_glyco_hcd203040_compound15.22193.22193.3 | controllerType=0 controllerNumber=1 scan=22193 | 46.4169 |
| 17572:4 | 1 | R.RTAGWNVPIGTLRPFLN[+2059.735]WTGPPEPIEAAVAR.F | HexNAc(4)Hex(5)Fuc(1)NeuAc(1) | N[+2060] | 1362.3976 | 4 | 5446.5685 | 5446.547 | 0 | 4 | 140 | Specific | 274.64 | 169.07 | 169.07 | 5.93 | 138 | >sp\|P02788\|TRFL_HUMAN Lactotransferrin OS=Homo sapiens OX=9606 GN=LTF PE=1 SV=6 | 1270 | 20201119_16913_1ng_glyco_hcd203040_compound15.21222.21222.4 | controllerType=0 controllerNumber=1 scan=21222 | 44.2146 |
| 18558:3 | 1 | R.[+42.011]RTAGWNVPIGTLRPFLN[+1864.634]WTGPPEPIEAAVAR.F | HexNAc(2)Hex(9) | .[42], N[+1865] | 1765.1626 | 3 | 5293.4732 | 5293.4568 | 0 | 3.1 | 140 | Specific | 332.07 | 325.26 | 325.26 | 6.63 | 138 | >sp\|P02788\|TRFL_HUMAN Lactotransferrin OS=Homo sapiens OX=9606 GN=LTF PE=1 SV=6 | 1270 | 20201119_16913_1ng_glyco_hcd203040_compound15.22253.22253.3 | controllerType=0 controllerNumber=1 scan=22253 | 46.4972 |
| 17279:5 | 1 | R.RTAGWNVPIGTLRPFLN[+2059.735]WTGPPEPIEAAVAR.F | HexNAc(4)Hex(5)Fuc(1)NeuAc(1) | N[+2060] | 1090.1177 | 5 | 5446.5593 | 5446.547 | 0 | 2.3 | 140 | Specific | 372.83 | 372.83 | 372.83 | 7.81 | 138 | >sp\|P02788\|TRFL_HUMAN Lactotransferrin OS=Homo sapiens OX=9606 GN=LTF PE=1 SV=6 | 1270 | 20201119_16913_1ng_glyco_hcd203040_compound15.20918.20918.5 | controllerType=0 controllerNumber=1 scan=20918 | 43.6316 |
| 17574:4 | 1 | R.RTAGWNVPIGTLRPFLN[+2059.735]WTGPPEPIEAAVAR.F | HexNAc(4)Hex(5)Fuc(1)NeuAc(1) | N[+2060] | 1362.3976 | 4 | 5446.5685 | 5446.547 | 0 | 4 | 140 | Specific | 430.61 | 430.61 | 430.61 | 9.6 | 138 | >sp\|P02788\|TRFL_HUMAN Lactotransferrin OS=Homo sapiens OX=9606 GN=LTF PE=1 SV=6 | 1270 | 20201119_16913_1ng_glyco_hcd203040_compound15.21224.21224.4 | controllerType=0 controllerNumber=1 scan=21224 | 44.2215 |
| 17253:4 | 1 | R.RTAGWNVPIGTLRPFLN[+2204.772]WTGPPEPIEAAVAR.F | HexNAc(4)Hex(5)NeuAc(2) | N[+2205] | 1398.6597 | 4 | 5591.6169 | 5591.5845 | 0 | 5.8 | 140 | Specific | 452.26 | 452.26 | 452.26 | 8.98 | 138 | >sp\|P02788\|TRFL_HUMAN Lactotransferrin OS=Homo sapiens OX=9606 GN=LTF PE=1 SV=6 | 1270 | 20201119_16913_1ng_glyco_hcd203040_compound15.20891.20891.4 | controllerType=0 controllerNumber=1 scan=20891 | 43.5621 |
| 17144:4 | 1 | R.RTAGWNVPIGTLRPFLN[+2570.925]WTGPPEPIEAAVAR.F | HexNAc(5)Hex(6)Fuc(2)NeuAc(1) | N[+2571] | 1490.1963 | 4 | 5957.7633 | 5957.7371 | 0 | 4.4 | 140 | Specific | 494.07 | 494.07 | 494.07 | 9.32 | 138 | >sp\|P02788\|TRFL_HUMAN Lactotransferrin OS=Homo sapiens OX=9606 GN=LTF PE=1 SV=6 | 1270 | 20201119_16913_1ng_glyco_hcd203040_compound15.20777.20777.4 | controllerType=0 controllerNumber=1 scan=20777 | 43.2991 |
| 17266:4 | 1 | R.RTAGWNVPIGTLRPFLN[+2059.735]WTGPPEPIEAAVAR.F | HexNAc(4)Hex(5)Fuc(1)NeuAc(1) | N[+2060] | 1362.3958 | 4 | 5446.5612 | 5446.547 | 0 | 2.6 | 140 | Specific | 496.48 | 396.87 | 396.87 | 9.62 | 138 | >sp\|P02788\|TRFL_HUMAN Lactotransferrin OS=Homo sapiens OX=9606 GN=LTF PE=1 SV=6 | 1270 | 20201119_16913_1ng_glyco_hcd203040_compound15.20905.20905.4 | controllerType=0 controllerNumber=1 scan=20905 | 43.6059 |
| 17240:4 | 1 | R.RTAGWNVPIGTLRPFLN[+2351.851]WTGPPEPIEAAVAR.F | HexNAc(4)Hex(5)Fuc(3)NeuAc(1) | N[+2352] | 1435.4303 | 4 | 5738.6994 | 5738.6628 | 0 | 6.4 | 140 | Specific | 499.46 | 396.55 | 396.55 | 8.98 | 138 | >sp\|P02788\|TRFL_HUMAN Lactotransferrin OS=Homo sapiens OX=9606 GN=LTF PE=1 SV=6 | 1270 | 20201119_16913_1ng_glyco_hcd203040_compound15.20877.20877.4 | controllerType=0 controllerNumber=1 scan=20877 | 43.5263 |
| 17142:4 | 1 | R.RTAGWNVPIGTLRPFLN[+2570.925]WTGPPEPIEAAVAR.F | HexNAc(5)Hex(6)Fuc(2)NeuAc(1) | N[+2571] | 1490.1963 | 4 | 5957.7633 | 5957.7371 | 0 | 4.4 | 140 | Specific | 501.25 | 458.24 | 458.24 | 9.55 | 138 | >sp\|P02788\|TRFL_HUMAN Lactotransferrin OS=Homo sapiens OX=9606 GN=LTF PE=1 SV=6 | 1270 | 20201119_16913_1ng_glyco_hcd203040_compound15.20775.20775.4 | controllerType=0 controllerNumber=1 scan=20775 | 43.2925 |
| 17175:4 | 1 | R.RTAGWNVPIGTLRPFLN[+2424.867]WTGPPEPIEAAVAR.F | HexNAc(5)Hex(6)Fuc(1)NeuAc(1) | N[+2425] | 1453.6793 | 4 | 5811.6955 | 5811.6792 | 0 | 2.8 | 140 | Specific | 502.12 | 502.12 | 502.12 | 9.92 | 138 | >sp\|P02788\|TRFL_HUMAN Lactotransferrin OS=Homo sapiens OX=9606 GN=LTF PE=1 SV=6 | 1270 | 20201119_16913_1ng_glyco_hcd203040_compound15.20809.20809.4 | controllerType=0 controllerNumber=1 scan=20809 | 43.3764 |
| 17282:5 | 1 | R.RTAGWNVPIGTLRPFLN[+2059.735]WTGPPEPIEAAVAR.F | HexNAc(4)Hex(5)Fuc(1)NeuAc(1) | N[+2060] | 1090.1177 | 5 | 5446.5593 | 5446.547 | 0 | 2.3 | 140 | Specific | 511.55 | 462.62 | 462.62 | 9.76 | 138 | >sp\|P02788\|TRFL_HUMAN Lactotransferrin OS=Homo sapiens OX=9606 GN=LTF PE=1 SV=6 | 1270 | 20201119_16913_1ng_glyco_hcd203040_compound15.20921.20921.5 | controllerType=0 controllerNumber=1 scan=20921 | 43.6438 |
| 17251:4 | 1 | R.RTAGWNVPIGTLRPFLN[+2204.772]WTGPPEPIEAAVAR.F | HexNAc(4)Hex(5)NeuAc(2) | N[+2205] | 1398.6597 | 4 | 5591.6169 | 5591.5845 | 0 | 5.8 | 140 | Specific | 512.74 | 398 | 398 | 9.21 | 138 | >sp\|P02788\|TRFL_HUMAN Lactotransferrin OS=Homo sapiens OX=9606 GN=LTF PE=1 SV=6 | 1270 | 20201119_16913_1ng_glyco_hcd203040_compound15.20889.20889.4 | controllerType=0 controllerNumber=1 scan=20889 | 43.5561 |
| 17224:4 | 1 | R.RTAGWNVPIGTLRPFLN[+2205.793]WTGPPEPIEAAVAR.F | HexNAc(4)Hex(5)Fuc(2)NeuAc(1) | N[+2206] | 1398.9109 | 4 | 5592.6217 | 5592.6049 | 0 | 3 | 140 | Specific | 562.27 | 426.63 | 426.63 | 10.21 | 138 | >sp\|P02788\|TRFL_HUMAN Lactotransferrin OS=Homo sapiens OX=9606 GN=LTF PE=1 SV=6 | 1270 | 20201119_16913_1ng_glyco_hcd203040_compound15.20860.20860.4 | controllerType=0 controllerNumber=1 scan=20860 | 43.4893 |
| 17242:4 | 1 | R.RTAGWNVPIGTLRPFLN[+2351.851]WTGPPEPIEAAVAR.F | HexNAc(4)Hex(5)Fuc(3)NeuAc(1) | N[+2352] | 1435.4303 | 4 | 5738.6994 | 5738.6628 | 0 | 6.4 | 140 | Specific | 568.87 | 384.65 | 384.65 | 9.5 | 138 | >sp\|P02788\|TRFL_HUMAN Lactotransferrin OS=Homo sapiens OX=9606 GN=LTF PE=1 SV=6 | 1270 | 20201119_16913_1ng_glyco_hcd203040_compound15.20879.20879.4 | controllerType=0 controllerNumber=1 scan=20879 | 43.534 |
| 17226:4 | 1 | R.RTAGWNVPIGTLRPFLN[+2205.793]WTGPPEPIEAAVAR.F | HexNAc(4)Hex(5)Fuc(2)NeuAc(1) | N[+2206] | 1398.9109 | 4 | 5592.6217 | 5592.6049 | 0 | 3 | 140 | Specific | 589.63 | 527.85 | 527.85 | 11.46 | 138 | >sp\|P02788\|TRFL_HUMAN Lactotransferrin OS=Homo sapiens OX=9606 GN=LTF PE=1 SV=6 | 1270 | 20201119_16913_1ng_glyco_hcd203040_compound15.20862.20862.4 | controllerType=0 controllerNumber=1 scan=20862 | 43.498 |
| 16343:4 | 1 | R.RTAGWNVPIGTLRPFLN[+1768.640]WTGPPEPIEAAVAR.F | HexNAc(4)Hex(5)Fuc(1) | N[+1769] | 1289.6246 | 4 | 5155.4767 | 5155.4516 | 0 | 4.9 | 140 | Specific | 596.99 | 563.07 | 563.07 | 11.1 | 138 | >sp\|P02788\|TRFL_HUMAN Lactotransferrin OS=Homo sapiens OX=9606 GN=LTF PE=1 SV=6 | 1270 | 20201119_16913_1ng_glyco_hcd203040_compound15.19939.19939.4 | controllerType=0 controllerNumber=1 scan=19939 | 41.3449 |
| 16345:4 | 1 | R.RTAGWNVPIGTLRPFLN[+1768.640]WTGPPEPIEAAVAR.F | HexNAc(4)Hex(5)Fuc(1) | N[+1769] | 1289.6246 | 4 | 5155.4767 | 5155.4516 | 0 | 4.9 | 140 | Specific | 644.23 | 520.84 | 520.84 | 12.1 | 138 | >sp\|P02788\|TRFL_HUMAN Lactotransferrin OS=Homo sapiens OX=9606 GN=LTF PE=1 SV=6 | 1270 | 20201119_16913_1ng_glyco_hcd203040_compound15.19941.19941.4 | controllerType=0 controllerNumber=1 scan=19941 | 41.357 |
| 17268:4 | 1 | R.RTAGWNVPIGTLRPFLN[+2059.735]WTGPPEPIEAAVAR.F | HexNAc(4)Hex(5)Fuc(1)NeuAc(1) | N[+2060] | 1362.3958 | 4 | 5446.5612 | 5446.547 | 0 | 2.6 | 140 | Specific | 697.48 | 558.04 | 558.04 | 13.4 | 138 | >sp\|P02788\|TRFL_HUMAN Lactotransferrin OS=Homo sapiens OX=9606 GN=LTF PE=1 SV=6 | 1270 | 20201119_16913_1ng_glyco_hcd203040_compound15.20907.20907.4 | controllerType=0 controllerNumber=1 scan=20907 | 43.612 |
| 17178:4 | 1 | R.RTAGWNVPIGTLRPFLN[+2424.867]WTGPPEPIEAAVAR.F | HexNAc(5)Hex(6)Fuc(1)NeuAc(1) | N[+2425] | 1453.6793 | 4 | 5811.6955 | 5811.6792 | 0 | 2.8 | 140 | Specific | 719.43 | 557.55 | 557.55 | 14.46 | 138 | >sp\|P02788\|TRFL_HUMAN Lactotransferrin OS=Homo sapiens OX=9606 GN=LTF PE=1 SV=6 | 1270 | 20201119_16913_1ng_glyco_hcd203040_compound15.20812.20812.4 | controllerType=0 controllerNumber=1 scan=20812 | 43.3872 |
| 19804:4 | 1 | R.[+42.011]TAGWNVPIGTLRPFLN[+2336.851]WTGPPEPIEAAVAR.F | HexNAc(6)Hex(6)Fuc(1) | .[42], N[+2337] | 1403.1445 | 4 | 5609.5563 | 5609.5726 | 0 | -2.9 | 141 | Specific | 202.72 | 202.72 | 202.72 | 3.78 | 138 | >sp\|P02788\|TRFL_HUMAN Lactotransferrin OS=Homo sapiens OX=9606 GN=LTF PE=1 SV=6 | 1270 | 20201119_16913_1ng_glyco_hcd203040_compound15.23558.23558.4 | controllerType=0 controllerNumber=1 scan=23558 | 49.4435 |
| 18335:3 | 1 | R.TAGWNVPIGTLRPFLN[+2424.867]WTGPPEPIEAAVAR.F | HexNAc(5)Hex(6)Fuc(1)NeuAc(1) | N[+2425] | 1885.8735 | 3 | 5655.6059 | 5655.5781 | 0 | 4.9 | 141 | Specific | 236.22 | 149.64 | 149.64 | 5.33 | 138 | >sp\|P02788\|TRFL_HUMAN Lactotransferrin OS=Homo sapiens OX=9606 GN=LTF PE=1 SV=6 | 1270 | 20201119_16913_1ng_glyco_hcd203040_compound15.22021.22021.3 | controllerType=0 controllerNumber=1 scan=22021 | 46.0713 |
| 19705:4 | 1 | R.[+42.011]TAGWNVPIGTLRPFLN[+2336.851]WTGPPEPIEAAVAR.F | HexNAc(6)Hex(6)Fuc(1) | .[42], N[+2337] | 1403.1382 | 4 | 5609.5309 | 5609.5726 | 0 | -7.4 | 141 | Specific | 237.75 | 237.75 | 237.75 | 3.26 | 138 | >sp\|P02788\|TRFL_HUMAN Lactotransferrin OS=Homo sapiens OX=9606 GN=LTF PE=1 SV=6 | 1270 | 20201119_16913_1ng_glyco_hcd203040_compound15.23453.23453.3 | controllerType=0 controllerNumber=1 scan=23453 | 49.2008 |
| 17491:4 | 1 | R.TAGWNVPIGTLRPFLN[+2279.830]WTGPPEPIEAAVAR.F | HexNAc(5)Hex(6)Fuc(2) | N[+2280] | 1378.3887 | 4 | 5510.5329 | 5510.5406 | 0 | -1.4 | 141 | Specific | 319.04 | 319.04 | 319.04 | 5.85 | 138 | >sp\|P02788\|TRFL_HUMAN Lactotransferrin OS=Homo sapiens OX=9606 GN=LTF PE=1 SV=6 | 1270 | 20201119_16913_1ng_glyco_hcd203040_compound15.21138.21138.4 | controllerType=0 controllerNumber=1 scan=21138 | 44.0381 |
| 18443:3 | 1 | R.TAGWNVPIGTLRPFLN[+2205.793]WTGPPEPIEAAVAR.F | HexNAc(4)Hex(5)Fuc(2)NeuAc(1) | N[+2206] | 1812.8433 | 3 | 5436.5155 | 5436.5038 | 0 | 2.2 | 141 | Specific | 349.79 | 293.87 | 293.87 | 7.32 | 138 | >sp\|P02788\|TRFL_HUMAN Lactotransferrin OS=Homo sapiens OX=9606 GN=LTF PE=1 SV=6 | 1270 | 20201119_16913_1ng_glyco_hcd203040_compound15.22134.22134.3 | controllerType=0 controllerNumber=1 scan=22134 | 46.3041 |
| 18338:3 | 1 | R.TAGWNVPIGTLRPFLN[+2424.867]WTGPPEPIEAAVAR.F | HexNAc(5)Hex(6)Fuc(1)NeuAc(1) | N[+2425] | 1885.8735 | 3 | 5655.6059 | 5655.5781 | 0 | 4.9 | 141 | Specific | 363.58 | 235.29 | 235.29 | 7.07 | 138 | >sp\|P02788\|TRFL_HUMAN Lactotransferrin OS=Homo sapiens OX=9606 GN=LTF PE=1 SV=6 | 1270 | 20201119_16913_1ng_glyco_hcd203040_compound15.22024.22024.3 | controllerType=0 controllerNumber=1 scan=22024 | 46.08 |
| 19806:4 | 1 | R.[+42.011]TAGWNVPIGTLRPFLN[+2336.851]WTGPPEPIEAAVAR.F | HexNAc(6)Hex(6)Fuc(1) | .[42], N[+2337] | 1403.1445 | 4 | 5609.5563 | 5609.5726 | 0 | -2.9 | 141 | Specific | 381.21 | 381.21 | 381.21 | 8.1 | 138 | >sp\|P02788\|TRFL_HUMAN Lactotransferrin OS=Homo sapiens OX=9606 GN=LTF PE=1 SV=6 | 1270 | 20201119_16913_1ng_glyco_hcd203040_compound15.23560.23560.4 | controllerType=0 controllerNumber=1 scan=23560 | 49.448 |
| 18329:5 | 1 | R.TAGWNVPIGTLRPFLN[+2570.925]WTGPPEPIEAAVAR.F | HexNAc(5)Hex(6)Fuc(2)NeuAc(1) | N[+2571] | 1161.1356 | 5 | 5801.649 | 5801.636 | 0 | 2.2 | 141 | Specific | 419.41 | 394.34 | 394.34 | 9.72 | 138 | >sp\|P02788\|TRFL_HUMAN Lactotransferrin OS=Homo sapiens OX=9606 GN=LTF PE=1 SV=6 | 1270 | 20201119_16913_1ng_glyco_hcd203040_compound15.22015.22015.5 | controllerType=0 controllerNumber=1 scan=22015 | 46.0545 |
| 18320:3 | 1 | R.TAGWNVPIGTLRPFLN[+2570.925]WTGPPEPIEAAVAR.F | HexNAc(5)Hex(6)Fuc(2)NeuAc(1) | N[+2571] | 1934.5578 | 3 | 5801.6589 | 5801.636 | 0 | 4 | 141 | Specific | 456.66 | 318.01 | 318.01 | 8.03 | 138 | >sp\|P02788\|TRFL_HUMAN Lactotransferrin OS=Homo sapiens OX=9606 GN=LTF PE=1 SV=6 | 1270 | 20201119_16913_1ng_glyco_hcd203040_compound15.22006.22006.3 | controllerType=0 controllerNumber=1 scan=22006 | 46.0351 |
| 19700:3 | 1 | R.TAGWNVPIGTLRPFLN[+2350.830]WTGPPEPIEAAVAR.F | HexNAc(4)Hex(5)Fuc(1)NeuAc(2) | N[+2351] | 1861.1883 | 3 | 5581.5504 | 5581.5413 | 0 | 1.6 | 141 | Specific | 457.66 | 250.55 | 250.55 | 7.68 | 138 | >sp\|P02788\|TRFL_HUMAN Lactotransferrin OS=Homo sapiens OX=9606 GN=LTF PE=1 SV=6 | 1270 | 20201119_16913_1ng_glyco_hcd203040_compound15.23448.23448.6 | controllerType=0 controllerNumber=1 scan=23448 | 49.1886 |
| 19697:3 | 1 | R.TAGWNVPIGTLRPFLN[+2350.830]WTGPPEPIEAAVAR.F | HexNAc(4)Hex(5)Fuc(1)NeuAc(2) | N[+2351] | 1861.1883 | 3 | 5581.5504 | 5581.5413 | 0 | 1.6 | 141 | Specific | 465.78 | 324.49 | 324.49 | 7.97 | 138 | >sp\|P02788\|TRFL_HUMAN Lactotransferrin OS=Homo sapiens OX=9606 GN=LTF PE=1 SV=6 | 1270 | 20201119_16913_1ng_glyco_hcd203040_compound15.23445.23445.6 | controllerType=0 controllerNumber=1 scan=23445 | 49.1833 |
| 18317:4 | 1 | R.TAGWNVPIGTLRPFLN[+2570.925]WTGPPEPIEAAVAR.F | HexNAc(5)Hex(6)Fuc(2)NeuAc(1) | N[+2571] | 1451.1711 | 4 | 5801.6628 | 5801.636 | 0 | 4.6 | 141 | Specific | 483.65 | 385.11 | 385.11 | 9.42 | 138 | >sp\|P02788\|TRFL_HUMAN Lactotransferrin OS=Homo sapiens OX=9606 GN=LTF PE=1 SV=6 | 1270 | 20201119_16913_1ng_glyco_hcd203040_compound15.22003.22003.4 | controllerType=0 controllerNumber=1 scan=22003 | 46.0293 |
| 18309:5 | 1 | R.TAGWNVPIGTLRPFLN[+2570.925]WTGPPEPIEAAVAR.F | HexNAc(5)Hex(6)Fuc(2)NeuAc(1) | N[+2571] | 1161.1365 | 5 | 5801.6533 | 5801.636 | 0 | 3 | 141 | Specific | 517.93 | 480.27 | 480.27 | 10.16 | 138 | >sp\|P02788\|TRFL_HUMAN Lactotransferrin OS=Homo sapiens OX=9606 GN=LTF PE=1 SV=6 | 1270 | 20201119_16913_1ng_glyco_hcd203040_compound15.21994.21994.5 | controllerType=0 controllerNumber=1 scan=21994 | 45.9945 |
| 18261:3 | 1 | R.TAGWNVPIGTLRPFLN[+2716.983]WTGPPEPIEAAVAR.F | HexNAc(5)Hex(6)Fuc(3)NeuAc(1) | N[+2717] | 1983.2389 | 3 | 5947.702 | 5947.6939 | 0 | 1.4 | 141 | Specific | 537.98 | 467.48 | 467.48 | 9.81 | 138 | >sp\|P02788\|TRFL_HUMAN Lactotransferrin OS=Homo sapiens OX=9606 GN=LTF PE=1 SV=6 | 1270 | 20201119_16913_1ng_glyco_hcd203040_compound15.21944.21944.3 | controllerType=0 controllerNumber=1 scan=21944 | 45.8782 |
| 18439:4 | 1 | R.TAGWNVPIGTLRPFLN[+2205.793]WTGPPEPIEAAVAR.F | HexNAc(4)Hex(5)Fuc(2)NeuAc(1) | N[+2206] | 1359.884 | 4 | 5436.5143 | 5436.5038 | 0 | 1.9 | 141 | Specific | 538.41 | 535.33 | 535.33 | 9.98 | 138 | >sp\|P02788\|TRFL_HUMAN Lactotransferrin OS=Homo sapiens OX=9606 GN=LTF PE=1 SV=6 | 1270 | 20201119_16913_1ng_glyco_hcd203040_compound15.22130.22130.4 | controllerType=0 controllerNumber=1 scan=22130 | 46.2963 |
| 17493:4 | 1 | R.TAGWNVPIGTLRPFLN[+2279.830]WTGPPEPIEAAVAR.F | HexNAc(5)Hex(6)Fuc(2) | N[+2280] | 1378.3887 | 4 | 5510.5329 | 5510.5406 | 0 | -1.4 | 141 | Specific | 539.49 | 400.52 | 400.52 | 8.61 | 138 | >sp\|P02788\|TRFL_HUMAN Lactotransferrin OS=Homo sapiens OX=9606 GN=LTF PE=1 SV=6 | 1270 | 20201119_16913_1ng_glyco_hcd203040_compound15.21140.21140.4 | controllerType=0 controllerNumber=1 scan=21140 | 44.0445 |
| 18395:5 | 1 | R.TAGWNVPIGTLRPFLN[+2205.793]WTGPPEPIEAAVAR.F | HexNAc(4)Hex(5)Fuc(2)NeuAc(1) | N[+2206] | 1088.1096 | 5 | 5436.519 | 5436.5038 | 0 | 2.8 | 141 | Specific | 543.06 | 440.6 | 440.6 | 10.18 | 138 | >sp\|P02788\|TRFL_HUMAN Lactotransferrin OS=Homo sapiens OX=9606 GN=LTF PE=1 SV=6 | 1270 | 20201119_16913_1ng_glyco_hcd203040_compound15.22084.22084.4 | controllerType=0 controllerNumber=1 scan=22084 | 46.1961 |
| 18239:4 | 1 | R.TAGWNVPIGTLRPFLN[+2936.057]WTGPPEPIEAAVAR.F | HexNAc(6)Hex(7)Fuc(2)NeuAc(1) | N[+2936] | 1542.4487 | 4 | 6166.7731 | 6166.7682 | 0 | 0.8 | 141 | Specific | 553.91 | 504.85 | 504.85 | 9.99 | 138 | >sp\|P02788\|TRFL_HUMAN Lactotransferrin OS=Homo sapiens OX=9606 GN=LTF PE=1 SV=6 | 1270 | 20201119_16913_1ng_glyco_hcd203040_compound15.21921.21921.4 | controllerType=0 controllerNumber=1 scan=21921 | 45.8205 |
| 18454:5 | 1 | R.TAGWNVPIGTLRPFLN[+2205.793]WTGPPEPIEAAVAR.F | HexNAc(4)Hex(5)Fuc(2)NeuAc(1) | N[+2206] | 1088.1094 | 5 | 5436.5178 | 5436.5038 | 0 | 2.6 | 141 | Specific | 560.41 | 541.76 | 541.76 | 10.39 | 138 | >sp\|P02788\|TRFL_HUMAN Lactotransferrin OS=Homo sapiens OX=9606 GN=LTF PE=1 SV=6 | 1270 | 20201119_16913_1ng_glyco_hcd203040_compound15.22145.22145.5 | controllerType=0 controllerNumber=1 scan=22145 | 46.3261 |
| 17499:3 | 1 | R.TAGWNVPIGTLRPFLN[+1768.640]WTGPPEPIEAAVAR.F | HexNAc(4)Hex(5)Fuc(1) | N[+1769] | 1667.1286 | 3 | 4999.3713 | 4999.3504 | 0 | 4.2 | 141 | Specific | 605.05 | 568.87 | 568.87 | 12.3 | 138 | >sp\|P02788\|TRFL_HUMAN Lactotransferrin OS=Homo sapiens OX=9606 GN=LTF PE=1 SV=6 | 1270 | 20201119_16913_1ng_glyco_hcd203040_compound15.21147.21147.3 | controllerType=0 controllerNumber=1 scan=21147 | [illegible] |
| 18263:3 | 1 | R.TAGWNVPIGTLRPFLN[+2716.983]WTGPPEPIEAAVAR.F | HexNAc(5)Hex(6)Fuc(3)NeuAc(1) | N[+2717] | 1983.2389 | 3 | 5947.702 | 5947.6939 | 0 | 1.4 | 141 | Specific | 607.47 | 532.46 | 532.46 | 12.33 | 138 | >sp\|P02788\|TRFL_HUMAN Lactotransferrin OS=Homo sapiens OX=9606 GN=LTF PE=1 SV=6 | 1270 | 20201119_16913_1ng_glyco_hcd203040_compound15.21946.21946.3 | controllerType=0 controllerNumber=1 scan=21946 | [illegible] |
| 19679:3 | 1 | R.TAGWNVPIGTLRPFLN[+2350.830]WTGPPEPIEAAVAR.F | HexNAc(4)Hex(5)Fuc(1)NeuAc(2) | N[+2351] | 1861.1916 | 3 | 5581.5603 | 5581.5413 | 0 | 3.4 | 141 | Specific | 622 | 442.07 | 442.07 | 12.66 | 138 | >sp\|P02788\|TRFL_HUMAN Lactotransferrin OS=Homo sapiens OX=9606 GN=LTF PE=1 SV=6 | 1270 | 20201119_16913_1ng_glyco_hcd203040_compound15.23426.23426.3 | controllerType=0 controllerNumber=1 scan=23426 | [illegible] |
| 18322:4 | 1 | R.TAGWNVPIGTLRPFLN[+2424.867]WTGPPEPIEAAVAR.F | HexNAc(5)Hex(6)Fuc(1)NeuAc(1) | N[+2425] | 1414.6548 | 4 | 5655.5973 | 5655.5781 | 0 | 3.4 | 141 | Specific | 632.65 | 530.37 | 530.37 | 12.57 | 138 | >sp\|P02788\|TRFL_HUMAN Lactotransferrin OS=Homo sapiens OX=9606 GN=LTF PE=1 SV=6 | 1270 | 20201119_16913_1ng_glyco_hcd203040_compound15.22008.22008.4 | controllerType=0 controllerNumber=1 scan=22008 | [illegible] |
| 18445:3 | 1 | R.TAGWNVPIGTLRPFLN[+2059.735]WTGPPEPIEAAVAR.F | HexNAc(4)Hex(5)Fuc(1)NeuAc(1) | N[+2060] | 1764.1584 | 3 | 5290.4607 | 5290.4459 | 0 | 2.8 | 141 | Specific | 634.1 | 511.29 | 511.29 | 12.66 | 138 | >sp\|P02788\|TRFL_HUMAN Lactotransferrin OS=Homo sapiens OX=9606 GN=LTF PE=1 SV=6 | 1270 | 20201119_16913_1ng_glyco_hcd203040_compound15.22136.22136.3 | controllerType=0 controllerNumber=1 scan=22136 | 46.3079 |
| 18425:3 | 1 | R.TAGWNVPIGTLRPFLN[+2205.793]WTGPPEPIEAAVAR.F | HexNAc(4)Hex(5)Fuc(2)NeuAc(1) | N[+2206] | 1812.8466 | 3 | 5436.5254 | 5436.5038 | 0 | 4 | 141 | Specific | 641.21 | 557.72 | 557.72 | 12.66 | 138 | >sp\|P02788\|TRFL_HUMAN Lactotransferrin OS=Homo sapiens OX=9606 GN=LTF PE=1 SV=6 | 1270 | 20201119_16913_1ng_glyco_hcd203040_compound15.22115.22115.2 | controllerType=0 controllerNumber=1 scan=22115 | 46.2661 |
| 19682:3 | 1 | R.TAGWNVPIGTLRPFLN[+2350.830]WTGPPEPIEAAVAR.F | HexNAc(4)Hex(5)Fuc(1)NeuAc(2) | N[+2351] | 1861.1916 | 3 | 5581.5603 | 5581.5413 | 0 | 3.4 | 141 | Specific | 645.55 | 552.01 | 552.01 | 12.66 | 138 | >sp\|P02788\|TRFL_HUMAN Lactotransferrin OS=Homo sapiens OX=9606 GN=LTF PE=1 SV=6 | 1270 | 20201119_16913_1ng_glyco_hcd203040_compound15.23429.23429.3 | controllerType=0 controllerNumber=1 scan=23429 | 49.1507 |
| 18311:5 | 1 | R.TAGWNVPIGTLRPFLN[+2570.925]WTGPPEPIEAAVAR.F | HexNAc(5)Hex(6)Fuc(2)NeuAc(1) | N[+2571] | 1161.1365 | 5 | 5801.6533 | 5801.636 | 0 | 3 | 141 | Specific | 648.87 | 543.57 | 543.57 | 12.71 | 138 | >sp\|P02788\|TRFL_HUMAN Lactotransferrin OS=Homo sapiens OX=9606 GN=LTF PE=1 SV=6 | 1270 | 20201119_16913_1ng_glyco_hcd203040_compound15.21996.21996.5 | controllerType=0 controllerNumber=1 scan=21996 | 46.0023 |
| 18442:3 | 1 | R.TAGWNVPIGTLRPFLN[+2059.735]WTGPPEPIEAAVAR.F | HexNAc(4)Hex(5)Fuc(1)NeuAc(1) | N[+2060] | 1764.1584 | 3 | 5290.4607 | 5290.4459 | 0 | 2.8 | 141 | Specific | 656.09 | 570.14 | 570.14 | 13.66 | 138 | >sp\|P02788\|TRFL_HUMAN Lactotransferrin OS=Homo sapiens OX=9606 GN=LTF PE=1 SV=6 | 1270 | 20201119_16913_1ng_glyco_hcd203040_compound15.22133.22133.3 | controllerType=0 controllerNumber=1 scan=22133 | 46.303 |
| 19671:4 | 1 | R.TAGWNVPIGTLRPFLN[+2350.830]WTGPPEPIEAAVAR.F | HexNAc(4)Hex(5)Fuc(1)NeuAc(2) | N[+2351] | 1396.1455 | 4 | 5581.5602 | 5581.5413 | 0 | 3.4 | 141 | Specific | 676.96 | 545.53 | 545.53 | 13.57 | 138 | >sp\|P02788\|TRFL_HUMAN Lactotransferrin OS=Homo sapiens OX=9606 GN=LTF PE=1 SV=6 | 1270 | 20201119_16913_1ng_glyco_hcd203040_compound15.23418.23418.4 | controllerType=0 controllerNumber=1 scan=23418 | 49.12 |
| 18446:3 | 1 | R.TAGWNVPIGTLRPFLN[+2205.793]WTGPPEPIEAAVAR.F | HexNAc(4)Hex(5)Fuc(2)NeuAc(1) | N[+2206] | 1812.8433 | 3 | 5436.5155 | 5436.5038 | 0 | 2.2 | 141 | Specific | 684.89 | 623.11 | 623.11 | 13.6 | 138 | >sp\|P02788\|TRFL_HUMAN Lactotransferrin OS=Homo sapiens OX=9606 GN=LTF PE=1 SV=6 | 1270 | 20201119_16913_1ng_glyco_hcd203040_compound15.22137.22137.3 | controllerType=0 controllerNumber=1 scan=22137 | 46.3092 |
| 18241:4 | 1 | R.TAGWNVPIGTLRPFLN[+2936.057]WTGPPEPIEAAVAR.F | HexNAc(6)Hex(7)Fuc(2)NeuAc(1) | N[+2936] | 1542.4487 | 4 | 6166.7731 | 6166.7682 | 0 | 0.8 | 141 | Specific | 691.65 | 594.21 | 594.21 | 13.24 | 138 | >sp\|P02788\|TRFL_HUMAN Lactotransferrin OS=Homo sapiens OX=9606 GN=LTF PE=1 SV=6 | 1270 | 20201119_16913_1ng_glyco_hcd203040_compound15.21923.21923.4 | controllerType=0 controllerNumber=1 scan=21923 | 45.8275 |
| 18428:3 | 1 | R.TAGWNVPIGTLRPFLN[+2205.793]WTGPPEPIEAAVAR.F | HexNAc(4)Hex(5)Fuc(2)NeuAc(1) | N[+2206] | 1812.8466 | 3 | 5436.5254 | 5436.5038 | 0 | 4 | 141 | Specific | 693.36 | 595.79 | 595.79 | 13.66 | 138 | >sp\|P02788\|TRFL_HUMAN Lactotransferrin OS=Homo sapiens OX=9606 GN=LTF PE=1 SV=6 | 1270 | 20201119_16913_1ng_glyco_hcd203040_compound15.22118.22118.2 | controllerType=0 controllerNumber=1 scan=22118 | 46.2704 |
| 18319:4 | 1 | R.TAGWNVPIGTLRPFLN[+2570.925]WTGPPEPIEAAVAR.F | HexNAc(5)Hex(6)Fuc(2)NeuAc(1) | N[+2571] | 1451.1711 | 4 | 5801.6628 | 5801.636 | 0 | 4.6 | 141 | Specific | 703.42 | 591.91 | 591.91 | 14.21 | 138 | >sp\|P02788\|TRFL_HUMAN Lactotransferrin OS=Homo sapiens OX=9606 GN=LTF PE=1 SV=6 | 1270 | 20201119_16913_1ng_glyco_hcd203040_compound15.22005.22005.4 | controllerType=0 controllerNumber=1 scan=22005 | 46.0343 |
| 18438:4 | 1 | R.TAGWNVPIGTLRPFLN[+2059.735]WTGPPEPIEAAVAR.F | HexNAc(4)Hex(5)Fuc(1)NeuAc(1) | N[+2060] | 1323.3718 | 4 | 5290.4655 | 5290.4459 | 0 | 3.7 | 141 | Specific | 709.05 | 674.4 | 674.4 | 14.57 | 138 | >sp\|P02788\|TRFL_HUMAN Lactotransferrin OS=Homo sapiens OX=9606 GN=LTF PE=1 SV=6 | 1270 | 20201119_16913_1ng_glyco_hcd203040_compound15.22129.22129.4 | controllerType=0 controllerNumber=1 scan=22129 | 46.2949 |
| 17501:3 | 1 | R.TAGWNVPIGTLRPFLN[+1768.640]WTGPPEPIEAAVAR.F | HexNAc(4)Hex(5)Fuc(1) | N[+1769] | 1667.1286 | 3 | 4999.3713 | 4999.3504 | 0 | 4.2 | 141 | Specific | 719.29 | 617.82 | 617.82 | 14.3 | 138 | >sp\|P02788\|TRFL_HUMAN Lactotransferrin OS=Homo sapiens OX=9606 GN=LTF PE=1 SV=6 | 1270 | 20201119_16913_1ng_glyco_hcd203040_compound15.21149.21149.3 | controllerType=0 controllerNumber=1 scan=21149 | 44.0643 |
| 18325:4 | 1 | R.TAGWNVPIGTLRPFLN[+2424.867]WTGPPEPIEAAVAR.F | HexNAc(5)Hex(6)Fuc(1)NeuAc(1) | N[+2425] | 1414.6548 | 4 | 5655.5973 | 5655.5781 | 0 | 3.4 | 141 | Specific | 745.24 | 633.78 | 633.78 | 14.57 | 138 | >sp\|P02788\|TRFL_HUMAN Lactotransferrin OS=Homo sapiens OX=9606 GN=LTF PE=1 SV=6 | 1270 | 20201119_16913_1ng_glyco_hcd203040_compound15.22011.22011.4 | controllerType=0 controllerNumber=1 scan=22011 | 46.0501 |
| 17470:4 | 1 | R.TAGWNVPIGTLRPFLN[+1768.640]WTGPPEPIEAAVAR.F | HexNAc(4)Hex(5)Fuc(1) | N[+1769] | 1250.5995 | 4 | 4999.3761 | 4999.3504 | 0 | 5.1 | 141 | Specific | 781.8 | 717.33 | 717.33 | 15.21 | 138 | >sp\|P02788\|TRFL_HUMAN Lactotransferrin OS=Homo sapiens OX=9606 GN=LTF PE=1 SV=6 | 1270 | 20201119_16913_1ng_glyco_hcd203040_compound15.21116.21116.3 | controllerType=0 controllerNumber=1 scan=21116 | 43.9892 |
| 17500:4 | 1 | R.TAGWNVPIGTLRPFLN[+1768.640]WTGPPEPIEAAVAR.F | HexNAc(4)Hex(5)Fuc(1) | N[+1769] | 1250.5983 | 4 | 4999.3712 | 4999.3504 | 0 | 4.2 | 141 | Specific | 787.59 | 749.03 | 749.03 | 15.21 | 138 | >sp\|P02788\|TRFL_HUMAN Lactotransferrin OS=Homo sapiens OX=9606 GN=LTF PE=1 SV=6 | 1270 | 20201119_16913_1ng_glyco_hcd203040_compound15.21148.21148.4 | controllerType=0 controllerNumber=1 scan=21148 | 44.0623 |
| 18457:5 | 1 | R.TAGWNVPIGTLRPFLN[+2205.793]WTGPPEPIEAAVAR.F | HexNAc(4)Hex(5)Fuc(2)NeuAc(1) | N[+2206] | 1088.1094 | 5 | 5436.5178 | 5436.5038 | 0 | 2.6 | 141 | Specific | 825.78 | 685.31 | 685.31 | 16.65 | 138 | >sp\|P02788\|TRFL_HUMAN Lactotransferrin OS=Homo sapiens OX=9606 GN=LTF PE=1 SV=6 | 1270 | 20201119_16913_1ng_glyco_hcd203040_compound15.22148.22148.5 | controllerType=0 controllerNumber=1 scan=22148 | 46.3314 |
| 19673:4 | 1 | R.TAGWNVPIGTLRPFLN[+2350.830]WTGPPEPIEAAVAR.F | HexNAc(4)Hex(5)Fuc(1)NeuAc(2) | N[+2351] | 1396.1455 | 4 | 5581.5602 | 5581.5413 | 0 | 3.4 | 141 | Specific | 866.44 | 622.21 | 622.21 | 17.57 | 138 | >sp\|P02788\|TRFL_HUMAN Lactotransferrin OS=Homo sapiens OX=9606 GN=LTF PE=1 SV=6 | 1270 | 20201119_16913_1ng_glyco_hcd203040_compound15.23420.23420.4 | controllerType=0 controllerNumber=1 scan=23420 | 49.1298 |
| 18440:4 | 1 | R.TAGWNVPIGTLRPFLN[+2059.735]WTGPPEPIEAAVAR.F | HexNAc(4)Hex(5)Fuc(1)NeuAc(1) | N[+2060] | 1323.3718 | 4 | 5290.4655 | 5290.4459 | 0 | 3.7 | 141 | Specific | 867.02 | 691.2 | 691.2 | 17.57 | 138 | >sp\|P02788\|TRFL_HUMAN Lactotransferrin OS=Homo sapiens OX=9606 GN=LTF PE=1 SV=6 | 1270 | 20201119_16913_1ng_glyco_hcd203040_compound15.22131.22131.4 | controllerType=0 controllerNumber=1 scan=22131 | 46.3006 |
| 18455:5 | 1 | R.TAGWNVPIGTLRPFLN[+2059.735]WTGPPEPIEAAVAR.F | HexNAc(4)Hex(5)Fuc(1)NeuAc(1) | N[+2060] | 1058.8975 | 5 | 5290.4582 | 5290.4459 | 0 | 2.3 | 141 | Specific | 875.05 | 823.39 | 823.39 | 17.65 | 138 | >sp\|P02788\|TRFL_HUMAN Lactotransferrin OS=Homo sapiens OX=9606 GN=LTF PE=1 SV=6 | 1270 | 20201119_16913_1ng_glyco_hcd203040_compound15.22146.22146.5 | controllerType=0 controllerNumber=1 scan=22146 | 46.3272 |
| 18441:4 | 1 | R.TAGWNVPIGTLRPFLN[+2205.793]WTGPPEPIEAAVAR.F | HexNAc(4)Hex(5)Fuc(2)NeuAc(1) | N[+2206] | 1359.884 | 4 | 5436.5143 | 5436.5038 | 0 | 1.9 | 141 | Specific | 888.05 | 714.05 | 714.05 | 17.51 | 138 | >sp\|P02788\|TRFL_HUMAN Lactotransferrin OS=Homo sapiens OX=9606 GN=LTF PE=1 SV=6 | 1270 | 20201119_16913_1ng_glyco_hcd203040_compound15.22132.22132.4 | controllerType=0 controllerNumber=1 scan=22132 | 46.3019 |
| 17473:4 | 1 | R.TAGWNVPIGTLRPFLN[+1768.640]WTGPPEPIEAAVAR.F | HexNAc(4)Hex(5)Fuc(1) | N[+1769] | 1250.5995 | 4 | 4999.3761 | 4999.3504 | 0 | 5.1 | 141 | Specific | 917.77 | 713.76 | 713.76 | 18.21 | 138 | >sp\|P02788\|TRFL_HUMAN Lactotransferrin OS=Homo sapiens OX=9606 GN=LTF PE=1 SV=6 | 1270 | 20201119_16913_1ng_glyco_hcd203040_compound15.21119.21119.3 | controllerType=0 controllerNumber=1 scan=21119 | 43.996 |
| 17502:4 | 1 | R.TAGWNVPIGTLRPFLN[+1768.640]WTGPPEPIEAAVAR.F | HexNAc(4)Hex(5)Fuc(1) | N[+1769] | 1250.5983 | 4 | 4999.3712 | 4999.3504 | 0 | 4.2 | 141 | Specific | 922.45 | 820.58 | 820.58 | 18.21 | 138 | >sp\|P02788\|TRFL_HUMAN Lactotransferrin OS=Homo sapiens OX=9606 GN=LTF PE=1 SV=6 | 1270 | 20201119_16913_1ng_glyco_hcd203040_compound15.21150.21150.4 | controllerType=0 controllerNumber=1 scan=21150 | 44.0664 |
| 18458:5 | 1 | R.TAGWNVPIGTLRPFLN[+2059.735]WTGPPEPIEAAVAR.F | HexNAc(4)Hex(5)Fuc(1)NeuAc(1) | N[+2060] | 1058.8975 | 5 | 5290.4582 | 5290.4459 | 0 | 2.3 | 141 | Specific | 925.95 | 796.9 | 796.9 | 18.65 | 138 | >sp\|P02788\|TRFL_HUMAN Lactotransferrin OS=Homo sapiens OX=9606 GN=LTF PE=1 SV=6 | 1270 | 20201119_16913_1ng_glyco_hcd203040_compound15.22149.22149.5 | controllerType=0 controllerNumber=1 scan=22149 | 46.3346 |
| 18398:5 | 1 | R.TAGWNVPIGTLRPFLN[+2205.793]WTGPPEPIEAAVAR.F | HexNAc(4)Hex(5)Fuc(2)NeuAc(1) | N[+2206] | 1088.1096 | 5 | 5436.519 | 5436.5038 | 0 | 2.8 | 141 | Specific | 970.11 | 743.17 | 743.17 | 19.71 | 138 | >sp\|P02788\|TRFL_HUMAN Lactotransferrin OS=Homo sapiens OX=9606 GN=LTF PE=1 SV=6 | 1270 | 20201119_16913_1ng_glyco_hcd203040_compound15.22087.22087.4 | controllerType=0 controllerNumber=1 scan=22087 | 46.2062 |
| 18199:4 | 1 | R.TAGWNVPIGTLRPFLN[+3227.153]WTGPPEPIEAAVAR.F | HexNAc(6)Hex(7)Fuc(2)NeuAc(2) | N[+3227] | 1615.2255 | 4 | 6457.88 | 6457.8636 | 0 | 2.5 | 141 | Specific | 992.56 | 921.62 | 921.62 | 19.51 | 138 | >sp\|P02788\|TRFL_HUMAN Lactotransferrin OS=Homo sapiens OX=9606 GN=LTF PE=1 SV=6 | 1270 | 20201119_16913_1ng_glyco_hcd203040_compound15.21878.21878.2 | controllerType=0 controllerNumber=1 scan=21878 | 45.728 |
| 18196:4 | 1 | R.TAGWNVPIGTLRPFLN[+3227.153]WTGPPEPIEAAVAR.F | HexNAc(6)Hex(7)Fuc(2)NeuAc(2) | N[+3227] | 1615.2255 | 4 | 6457.88 | 6457.8636 | 0 | 2.5 | 141 | Specific | 1030.15 | 990.69 | 990.69 | 20.51 | 138 | >sp\|P02788\|TRFL_HUMAN Lactotransferrin OS=Homo sapiens OX=9606 GN=LTF PE=1 SV=6 | 1270 | 20201119_16913_1ng_glyco_hcd203040_compound15.21875.21875.2 | controllerType=0 controllerNumber=1 scan=21875 | 45.7183 |
| 24570:3 | 1 | R.[+42.011]TAGWNIPMGLLFN[+2351.851]QTGSC[+57.021]K.F | HexNAc(4)Hex(5)Fuc(3)NeuAc(1) | .[42], C[+57], N[+2352] | 1496.9614 | 3 | 4488.8695 | 4488.8713 | 0 | -0.4 | 485 | Specific | 231.73 | 211.1 | 211.1 | 5.17 | 138 | >sp\|P02788\|TRFL_HUMAN Lactotransferrin OS=Homo sapiens OX=9606 GN=LTF PE=1 SV=6 | 1270 | 20201119_16913_1ng_glyco_hcd203040_compound15.28535.28535.3 | controllerType=0 controllerNumber=1 scan=28535 | 59.8773 |
| 24572:3 | 1 | R.[+42.011]TAGWNIPMGLLFN[+2351.851]QTGSC[+57.021]K.F | HexNAc(4)Hex(5)Fuc(3)NeuAc(1) | .[42], C[+57], N[+2352] | 1496.9614 | 3 | 4488.8695 | 4488.8713 | 0 | -0.4 | 485 | Specific | 272.31 | 164.95 | 164.95 | 5.01 | 138 | >sp\|P02788\|TRFL_HUMAN Lactotransferrin OS=Homo sapiens OX=9606 GN=LTF PE=1 SV=6 | 1270 | 20201119_16913_1ng_glyco_hcd203040_compound15.28537.28537.3 | controllerType=0 controllerNumber=1 scan=28537 | 59.8833 |
| 15888:3 | 1 | R.TAGWNIPMGLLFN[+2279.830]QTGSC[+57.021]K.F | HexNAc(5)Hex(6)Fuc(2) | C[+57], N[+2280] | 1458.9638 | 3 | 4374.8769 | 4374.8396 | 0 | 8.5 | 485 | Specific | 366.47 | 366.47 | 277.44 | 6.57 | 138 | >sp\|P02788\|TRFL_HUMAN Lactotransferrin OS=Homo sapiens OX=9606 GN=LTF PE=1 SV=6 | 1270 | 20201119_16913_1ng_glyco_hcd203040_compound15.19464.19464.3 | controllerType=0 controllerNumber=1 scan=19464 | 40.336 |
| 18142:3 | 1 | R.TAGWNIPMGLLFN[+1548.545]QTGSC[+57.021]K.F | HexNAc(3)Hex(4)NeuAc(1) | C[+57], N[+1549] | 1215.1945 | 3 | 3643.569 | 3643.5548 | 0 | 3.9 | 485 | Specific | 428.73 | 400.18 | 400.18 | 9.55 | 138 | >sp\|P02788\|TRFL_HUMAN Lactotransferrin OS=Homo sapiens OX=9606 GN=LTF PE=1 SV=6 | 1270 | 20201119_16913_1ng_glyco_hcd203040_compound15.21819.21819.3 | controllerType=0 controllerNumber=1 scan=21819 | 45.5878 |
| 17878:4 | 1 | R.TAGWNIPMGLLFN[+2059.735]QTGSC[+57.021]K.F | HexNAc(4)Hex(5)Fuc(1)NeuAc(1) | C[+57], N[+2060] | 1039.4488 | 4 | 4154.7733 | 4154.7449 | 0 | 6.8 | 485 | Specific | 477.34 | 363.7 | 363.7 | 7.36 | 138 | >sp\|P02788\|TRFL_HUMAN Lactotransferrin OS=Homo sapiens OX=9606 GN=LTF PE=1 SV=6 | 1270 | 20201119_16913_1ng_glyco_hcd203040_compound15.21542.21542.4 | controllerType=0 controllerNumber=1 scan=21542 | 44.8743 |
| 17827:3 | 1 | R.TAGWNIPMGLLFN[+2351.851]QTGSC[+57.021]K.F | HexNAc(4)Hex(5)Fuc(3)NeuAc(1) | C[+57], N[+2352] | 1482.9676 | 3 | 4446.8882 | 4446.8607 | 0 | 6.2 | 485 | Specific | 491.13 | 396.51 | 396.51 | 8.93 | 138 | >sp\|P02788\|TRFL_HUMAN Lactotransferrin OS=Homo sapiens OX=9606 GN=LTF PE=1 SV=6 | 1270 | 20201119_16913_1ng_glyco_hcd203040_compound15.21489.21489.3 | controllerType=0 controllerNumber=1 scan=21489 | 44.7413 |
| 17881:4 | 1 | R.TAGWNIPMGLLFN[+2059.735]QTGSC[+57.021]K.F | HexNAc(4)Hex(5)Fuc(1)NeuAc(1) | C[+57], N[+2060] | 1039.4488 | 4 | 4154.7733 | 4154.7449 | 0 | 6.8 | 485 | Specific | 506.68 | 232.96 | 232.96 | 6.87 | 138 | >sp\|P02788\|TRFL_HUMAN Lactotransferrin OS=Homo sapiens OX=9606 GN=LTF PE=1 SV=6 | 1270 | 20201119_16913_1ng_glyco_hcd203040_compound15.21545.21545.4 | controllerType=0 controllerNumber=1 scan=21545 | 44.8993 |
| 15890:3 | 1 | R.TAGWNIPMGLLFN[+2279.830]QTGSC[+57.021]K.F | HexNAc(5)Hex(6)Fuc(2) | C[+57], N[+2280] | 1458.9638 | 3 | 4374.8769 | 4374.8396 | 0 | 8.5 | 485 | Specific | 508.32 | 508.32 | 357.93 | 8.53 | 138 | >sp\|P02788\|TRFL_HUMAN Lactotransferrin OS=Homo sapiens OX=9606 GN=LTF PE=1 SV=6 | 1270 | 20201119_16913_1ng_glyco_hcd203040_compound15.19466.19466.3 | controllerType=0 controllerNumber=1 scan=19466 | 40.3395 |
| 18145:3 | 1 | R.TAGWNIPMGLLFN[+1548.545]QTGSC[+57.021]K.F | HexNAc(3)Hex(4)NeuAc(1) | C[+57], N[+1549] | 1215.1945 | 3 | 3643.569 | 3643.5548 | 0 | 3.9 | 485 | Specific | 645.17 | 630.24 | 630.24 | 12.41 | 138 | >sp\|P02788\|TRFL_HUMAN Lactotransferrin OS=Homo sapiens OX=9606 GN=LTF PE=1 SV=6 | 1270 | 20201119_16913_1ng_glyco_hcd203040_compound15.21822.21822.3 | controllerType=0 controllerNumber=1 scan=21822 | 45.5934 |
| 18031:3 | 1 | R.TAGWNIPMGLLFN[+1694.603]QTGSC[+57.021]K.F | HexNAc(3)Hex(4)Fuc(1)NeuAc(1) | C[+57], N[+1695] | 1263.8823 | 3 | 3789.6322 | 3789.6127 | 0 | 5.2 | 485 | Specific | 664.7 | 629.99 | 629.99 | 13.05 | 138 | >sp\|P02788\|TRFL_HUMAN Lactotransferrin OS=Homo sapiens OX=9606 GN=LTF PE=1 SV=6 | 1270 | 20201119_16913_1ng_glyco_hcd203040_compound15.21703.21703.3 | controllerType=0 controllerNumber=1 scan=21703 | 45.3333 |
| 18027:3 | 1 | R.TAGWNIPMGLLFN[+1872.651]QTGSC[+57.021]K.F | HexNAc(3)Hex(6)NeuAc(1) | C[+57], N[+1873] | 1323.2287 | 3 | 3967.6715 | 3967.6604 | 0 | 2.8 | 485 | Specific | 760.19 | 730.78 | 301.41 | 15.41 | 138 | >sp\|P02788\|TRFL_HUMAN Lactotransferrin OS=Homo sapiens OX=9606 GN=LTF PE=1 SV=6 | 1270 | 20201119_16913_1ng_glyco_hcd203040_compound15.21699.21699.3 | controllerType=0 controllerNumber=1 scan=21699 | 45.3167 |
| 19781:4 | 1 | R.TAGWNIPMGLLFN[+2350.830]QTGSC[+57.021]K.F | HexNAc(4)Hex(5)Fuc(1)NeuAc(2) | C[+57], N[+2351] | 1112.2193 | 4 | 4445.8553 | 4445.8403 | 0 | 3.4 | 485 | Specific | 775.69 | 639.6 | 639.6 | 15.55 | 138 | >sp\|P02788\|TRFL_HUMAN Lactotransferrin OS=Homo sapiens OX=9606 GN=LTF PE=1 SV=6 | 1270 | 20201119_16913_1ng_glyco_hcd203040_compound15.23534.23534.4 | controllerType=0 controllerNumber=1 scan=23534 | 49.3917 |
| 18021:4 | 1 | R.TAGWNIPMGLLFN[+1913.677]QTGSC[+57.021]K.F | HexNAc(4)Hex(5)NeuAc(1) | C[+57], N[+1914] | 1002.9307 | 4 | 4008.701 | 4008.687 | 0 | 3.5 | 485 | Specific | 856.34 | 787.51 | 787.51 | 17.55 | 138 | >sp\|P02788\|TRFL_HUMAN Lactotransferrin OS=Homo sapiens OX=9606 GN=LTF PE=1 SV=6 | 1270 | 20201119_16913_1ng_glyco_hcd203040_compound15.21692.21692.4 | controllerType=0 controllerNumber=1 scan=21692 | 45.3019 |
| 19783:4 | 1 | R.TAGWNIPMGLLFN[+2350.830]QTGSC[+57.021]K.F | HexNAc(4)Hex(5)Fuc(1)NeuAc(2) | C[+57], N[+2351] | 1112.2193 | 4 | 4445.8553 | 4445.8403 | 0 | 3.4 | 485 | Specific | 857.28 | 628.76 | 628.76 | 17.55 | 138 | >sp\|P02788\|TRFL_HUMAN Lactotransferrin OS=Homo sapiens OX=9606 GN=LTF PE=1 SV=6 | 1270 | 20201119_16913_1ng_glyco_hcd203040_compound15.23536.23536.4 | controllerType=0 controllerNumber=1 scan=23536 | 49.3979 |
| 19951:3 | 1 | R.TAGWNIPMGLLFN[+2204.772]QTGSC[+57.021]K.F | HexNAc(4)Hex(5)NeuAc(2) | C[+57], N[+2205] | 1433.9378 | 3 | 4299.7989 | 4299.7824 | 0 | 3.8 | 485 | Specific | 863.81 | 830.89 | 830.89 | 17.41 | 138 | >sp\|P02788\|TRFL_HUMAN Lactotransferrin OS=Homo sapiens OX=9606 GN=LTF PE=1 SV=6 | 1270 | 20201119_16913_1ng_glyco_hcd203040_compound15.23711.23711.3 | controllerType=0 controllerNumber=1 scan=23711 | 49.8044 |
| 18033:3 | 1 | R.TAGWNIPMGLLFN[+1694.603]QTGSC[+57.021]K.F | HexNAc(3)Hex(4)Fuc(1)NeuAc(1) | C[+57], N[+1695] | 1263.8823 | 3 | 3789.6322 | 3789.6127 | 0 | 5.2 | 485 | Specific | 885.35 | 848.41 | 848.41 | 17.05 | 138 | >sp\|P02788\|TRFL_HUMAN Lactotransferrin OS=Homo sapiens OX=9606 GN=LTF PE=1 SV=6 | 1270 | 20201119_16913_1ng_glyco_hcd203040_compound15.21705.21705.3 | controllerType=0 controllerNumber=1 scan=21705 | 45.3408 |
| 16025:3 | 1 | R.TAGWNIPMGLLFN[+1768.640]QTGSC[+57.021]K.F | HexNAc(4)Hex(5)Fuc(1) | C[+57], N[+1769] | 1288.5611 | 3 | 3863.6687 | 3863.6495 | 0 | 5 | 485 | Specific | 885.35 | 885.35 | 885.35 | 17.05 | 138 | >sp\|P02788\|TRFL_HUMAN Lactotransferrin OS=Homo sapiens OX=9606 GN=LTF PE=1 SV=6 | 1270 | 20201119_16913_1ng_glyco_hcd203040_compound15.19608.19608.3 | controllerType=0 controllerNumber=1 scan=19608 | 40.6313 |
| 18018:4 | 1 | R.TAGWNIPMGLLFN[+1913.677]QTGSC[+57.021]K.F | HexNAc(4)Hex(5)NeuAc(1) | C[+57], N[+1914] | 1002.9307 | 4 | 4008.701 | 4008.687 | 0 | 3.5 | 485 | Specific | 886.53 | 849.49 | 849.49 | 17.55 | 138 | >sp\|P02788\|TRFL_HUMAN Lactotransferrin OS=Homo sapiens OX=9606 GN=LTF PE=1 SV=6 | 1270 | 20201119_16913_1ng_glyco_hcd203040_compound15.21689.21689.4 | controllerType=0 controllerNumber=1 scan=21689 | 45.2938 |
| 19782:3 | 1 | R.TAGWNIPMGLLFN[+2350.830]QTGSC[+57.021]K.F | HexNAc(4)Hex(5)Fuc(1)NeuAc(2) | C[+57], N[+2351] | 1482.6252 | 3 | 4445.861 | 4445.8403 | 0 | 4.7 | 485 | Specific | 893.35 | 862.24 | 862.24 | 17.05 | 138 | >sp\|P02788\|TRFL_HUMAN Lactotransferrin OS=Homo sapiens OX=9606 GN=LTF PE=1 SV=6 | 1270 | 20201119_16913_1ng_glyco_hcd203040_compound15.23535.23535.3 | controllerType=0 controllerNumber=1 scan=23535 | 49.3954 |
| 19954:3 | 1 | R.TAGWNIPMGLLFN[+2204.772]QTGSC[+57.021]K.F | HexNAc(4)Hex(5)NeuAc(2) | C[+57], N[+2205] | 1433.9378 | 3 | 4299.7989 | 4299.7824 | 0 | 3.8 | 485 | Specific | 899.54 | 827.42 | 827.42 | 17.41 | 138 | >sp\|P02788\|TRFL_HUMAN Lactotransferrin OS=Homo sapiens OX=9606 GN=LTF PE=1 SV=6 | 1270 | 20201119_16913_1ng_glyco_hcd203040_compound15.23714.23714.3 | controllerType=0 controllerNumber=1 scan=23714 | 49.8094 |
| 17975:3 | 1 | R.TAGWNIPMGLLFN[+1913.677]QTGSC[+57.021]K.F | HexNAc(4)Hex(5)NeuAc(1) | C[+57], N[+1914] | 1336.9128 | 3 | 4008.7238 | 4008.687 | 0 | 9.2 | 485 | Specific | 903.83 | 862.61 | 862.61 | 17 | 138 | >sp\|P02788\|TRFL_HUMAN Lactotransferrin OS=Homo sapiens OX=9606 GN=LTF PE=1 SV=6 | 1270 | 20201119_16913_1ng_glyco_hcd203040_compound15.21644.21644.3 | controllerType=0 controllerNumber=1 scan=21644 | 45.1905 |
| 17830:3 | 1 | R.TAGWNIPMGLLFN[+2351.851]QTGSC[+57.021]K.F | HexNAc(4)Hex(5)Fuc(3)NeuAc(1) | C[+57], N[+2352] | 1482.9676 | 3 | 4446.8882 | 4446.8607 | 0 | 6.2 | 485 | Specific | 913.05 | 733.6 | 733.6 | 17.71 | 138 | >sp\|P02788\|TRFL_HUMAN Lactotransferrin OS=Homo sapiens OX=9606 GN=LTF PE=1 SV=6 | 1270 | 20201119_16913_1ng_glyco_hcd203040_compound15.21492.21492.3 | controllerType=0 controllerNumber=1 scan=21492 | 44.7452 |
| 17831:3 | 1 | R.TAGWNIPMGLLFN[+2205.793]QTGSC[+57.021]K.F | HexNAc(4)Hex(5)Fuc(2)NeuAc(1) | C[+57], N[+2206] | 1434.2818 | 3 | 4300.8308 | 4300.8028 | 0 | 6.5 | 485 | Specific | 914.39 | 781.92 | 781.92 | 17.42 | 138 | >sp\|P02788\|TRFL_HUMAN Lactotransferrin OS=Homo sapiens OX=9606 GN=LTF PE=1 SV=6 | 1270 | 20201119_16913_1ng_glyco_hcd203040_compound15.21493.21493.3 | controllerType=0 controllerNumber=1 scan=21493 | 44.7499 |
| 16401:2 | 1 | R.TAGWNIPMGLLFN[+1216.423]QTGSC[+57.021]K.F | HexNAc(2)Hex(5) | C[+57], N[+1216] | 1656.2232 | 2 | 3311.4391 | 3311.4328 | 0 | 1.9 | 485 | Specific | 926.53 | 823.5 | 293.41 | 18.44 | 138 | >sp\|P02788\|TRFL_HUMAN Lactotransferrin OS=Homo sapiens OX=9606 GN=LTF PE=1 SV=6 | 1270 | 20201119_16913_1ng_glyco_hcd203040_compound15.20000.20000.2 | controllerType=0 controllerNumber=1 scan=20000 | 41.5018 |
| 16028:3 | 1 | R.TAGWNIPMGLLFN[+1768.640]QTGSC[+57.021]K.F | HexNAc(4)Hex(5)Fuc(1) | C[+57], N[+1769] | 1288.5611 | 3 | 3863.6687 | 3863.6495 | 0 | 5 | 485 | Specific | 937.31 | 709.21 | 709.21 | 18.05 | 138 | >sp\|P02788\|TRFL_HUMAN Lactotransferrin OS=Homo sapiens OX=9606 GN=LTF PE=1 SV=6 | 1270 | 20201119_16913_1ng_glyco_hcd203040_compound15.19611.19611.3 | controllerType=0 controllerNumber=1 scan=19611 | 40.6352 |
| 17850:3 | 1 | R.TAGWNIPMGLLFN[+2059.735]QTGSC[+57.021]K.F | HexNAc(4)Hex(5)Fuc(1)NeuAc(1) | C[+57], N[+2060] | 1385.5948 | 3 | 4154.7698 | 4154.7449 | 0 | 6 | 485 | Specific | 942.38 | 918.85 | 918.85 | 17.71 | 138 | >sp\|P02788\|TRFL_HUMAN Lactotransferrin OS=Homo sapiens OX=9606 GN=LTF PE=1 SV=6 | 1270 | 20201119_16913_1ng_glyco_hcd203040_compound15.21513.21513.3 | controllerType=0 controllerNumber=1 scan=21513 | 44.7985 |
| 19780:3 | 1 | R.TAGWNIPMGLLFN[+2350.830]QTGSC[+57.021]K.F | HexNAc(4)Hex(5)Fuc(1)NeuAc(2) | C[+57], N[+2351] | 1482.6252 | 3 | 4445.861 | 4445.8403 | 0 | 4.7 | 485 | Specific | 950.76 | 918.33 | 918.33 | 19.05 | 138 | >sp\|P02788\|TRFL_HUMAN Lactotransferrin OS=Homo sapiens OX=9606 GN=LTF PE=1 SV=6 | 1270 | 20201119_16913_1ng_glyco_hcd203040_compound15.23533.23533.3 | controllerType=0 controllerNumber=1 scan=23533 | 49.3903 |
| 16399:2 | 1 | R.TAGWNIPMGLLFN[+1216.423]QTGSC[+57.021]K.F | HexNAc(2)Hex(5) | C[+57], N[+1216] | 1656.2232 | 2 | 3311.4391 | 3311.4328 | 0 | 1.9 | 485 | Specific | 956.06 | 949.3 | 362.51 | 19.44 | 138 | >sp\|P02788\|TRFL_HUMAN Lactotransferrin OS=Homo sapiens OX=9606 GN=LTF PE=1 SV=6 | 1270 | 20201119_16913_1ng_glyco_hcd203040_compound15.19998.19998.2 | controllerType=0 controllerNumber=1 scan=19998 | 41.4963 |
| 17973:3 | 1 | R.TAGWNIPMGLLFN[+1913.677]QTGSC[+57.021]K.F | HexNAc(4)Hex(5)NeuAc(1) | C[+57], N[+1914] | 1336.9128 | 3 | 4008.7238 | 4008.687 | 0 | 9.2 | 485 | Specific | 957.4 | 955.69 | 955.69 | 18 | 138 | >sp\|P02788\|TRFL_HUMAN Lactotransferrin OS=Homo sapiens OX=9606 GN=LTF PE=1 SV=6 | 1270 | 20201119_16913_1ng_glyco_hcd203040_compound15.21642.21642.3 | controllerType=0 controllerNumber=1 scan=21642 | 45.1824 |
| 17829:3 | 1 | R.TAGWNIPMGLLFN[+2205.793]QTGSC[+57.021]K.F | HexNAc(4)Hex(5)Fuc(2)NeuAc(1) | C[+57], N[+2206] | 1434.2818 | 3 | 4300.8308 | 4300.8028 | 0 | 6.5 | 485 | Specific | 965.91 | 881.88 | 881.88 | 18.42 | 138 | >sp\|P02788\|TRFL_HUMAN Lactotransferrin OS=Homo sapiens OX=9606 GN=LTF PE=1 SV=6 | 1270 | 20201119_16913_1ng_glyco_hcd203040_compound15.21491.21491.3 | controllerType=0 controllerNumber=1 scan=21491 | 44.7435 |
| 18030:3 | 1 | R.TAGWNIPMGLLFN[+1872.651]QTGSC[+57.021]K.F | HexNAc(3)Hex(6)NeuAc(1) | C[+57], N[+1873] | 1323.2287 | 3 | 3967.6715 | 3967.6604 | 0 | 2.8 | 485 | Specific | 967.43 | 899.03 | 441.07 | 19.41 | 138 | >sp\|P02788\|TRFL_HUMAN Lactotransferrin OS=Homo sapiens OX=9606 GN=LTF PE=1 SV=6 | 1270 | 20201119_16913_1ng_glyco_hcd203040_compound15.21702.21702.3 | controllerType=0 controllerNumber=1 scan=21702 | 45.3314 |
| 17848:3 | 1 | R.TAGWNIPMGLLFN[+2059.735]QTGSC[+57.021]K.F | HexNAc(4)Hex(5)Fuc(1)NeuAc(1) | C[+57], N[+2060] | 1385.5948 | 3 | 4154.7698 | 4154.7449 | 0 | 6 | 485 | Specific | 988.63 | 973 | 973 | 18.71 | 138 | >sp\|P02788\|TRFL_HUMAN Lactotransferrin OS=Homo sapiens OX=9606 GN=LTF PE=1 SV=6 | 1270 | 20201119_16913_1ng_glyco_hcd203040_compound15.21511.21511.3 | controllerType=0 controllerNumber=1 scan=21511 | 44.7861 |
| 16363:3 | 1 | R.TAGWNIPMGLLFN[+1216.423]QTGSC[+57.021]K.F | HexNAc(2)Hex(5) | C[+57], N[+1216] | 1104.4868 | 3 | 3311.446 | 3311.4328 | 0 | 4 | 485 | Specific | 1024.47 | 1024.47 | 477.88 | 20.05 | 138 | >sp\|P02788\|TRFL_HUMAN Lactotransferrin OS=Homo sapiens OX=9606 GN=LTF PE=1 SV=6 | 1270 | 20201119_16913_1ng_glyco_hcd203040_compound15.19960.19960.3 | controllerType=0 controllerNumber=1 scan=19960 | 41.4051 |
| 16366:3 | 1 | R.TAGWNIPMGLLFN[+1216.423]QTGSC[+57.021]K.F | HexNAc(2)Hex(5) | C[+57], N[+1216] | 1104.4868 | 3 | 3311.446 | 3311.4328 | 0 | 4 | 485 | Specific | 1042.31 | 1042.31 | 478.34 | 20.05 | 138 | >sp\|P02788\|TRFL_HUMAN Lactotransferrin OS=Homo sapiens OX=9606 GN=LTF PE=1 SV=6 | 1270 | 20201119_16913_1ng_glyco_hcd203040_compound15.19963.19963.3 | controllerType=0 controllerNumber=1 scan=19963 | 41.4131 |
| 15710:3 | 1 | R.TAGWNIPM[+15.995]GLLFN[+2059.735]QTGSC[+57.021]K.F | HexNAc(4)Hex(5)Fuc(1)NeuAc(1) | C[+57], M[+16], N[+2060] | 1390.9222 | 3 | 4170.752 | 4170.7398 | 0 | 2.9 | 485 | Specific | 388.82 | 369.14 | 369.14 | 7.88 | 138 | >sp\|P02788\|TRFL_HUMAN Lactotransferrin OS=Homo sapiens OX=9606 GN=LTF PE=1 SV=6 | 1270 | 20201119_16913_1ng_glyco_hcd203040_compound15.19279.19279.3 | controllerType=0 controllerNumber=1 scan=19279 | 39.9508 |
| 15713:3 | 1 | R.TAGWNIPM[+15.995]GLLFN[+2059.735]QTGSC[+57.021]K.F | HexNAc(4)Hex(5)Fuc(1)NeuAc(1) | C[+57], M[+16], N[+2060] | 1390.9222 | 3 | 4170.752 | 4170.7398 | 0 | 2.9 | 485 | Specific | 602.2 | 489.65 | 489.65 | 12.41 | 138 | >sp\|P02788\|TRFL_HUMAN Lactotransferrin OS=Homo sapiens OX=9606 GN=LTF PE=1 SV=6 | 1270 | 20201119_16913_1ng_glyco_hcd203040_compound15.19282.19282.3 | controllerType=0 controllerNumber=1 scan=19282 | 39.9567 |
| 18170:4 | 1 | R.[+42.011]TAGWNIPMGLLFN[+1872.651]QTGSC[+57.021]KFDEYFSQSC[+57.021]A | HexNAc(3)Hex(6)NeuAc(1) | .[42], C[+57]*2, N[+1873] | 1464.1224 | 4 | 5853.4679 | 5853.4181 | 0 | 8.5 | 485 | Specific | 337.56 | 290.49 | 290.49 | 5.2 | 138 | >sp\|P02788\|TRFL_HUMAN Lactotransferrin OS=Homo sapiens OX=9606 GN=LTF PE=1 SV=6 | 1270 | 20201119_16913_1ng_glyco_hcd203040_compound15.21848.21848.4 | controllerType=0 controllerNumber=1 scan=21848 | 45.6642 |
| 18168:4 | 1 | R.[+42.011]TAGWNIPMGLLFN[+1872.651]QTGSC[+57.021]KFDEYFSQSC[+57.021]A | HexNAc(3)Hex(6)NeuAc(1) | .[42], C[+57]*2, N[+1873] | 1464.1224 | 4 | 5853.4679 | 5853.4181 | 0 | 8.5 | 485 | Specific | 514.98 | 514.98 | 514.98 | 7.93 | 138 | >sp\|P02788\|TRFL_HUMAN Lactotransferrin OS=Homo sapiens OX=9606 GN=LTF PE=1 SV=6 | 1270 | 20201119_16913_1ng_glyco_hcd203040_compound15.21846.21846.4 | controllerType=0 controllerNumber=1 scan=21846 | 45.6588 |
| 18180:4 | 1 | R.TAGWNIPMGLLFN[+2059.735]QTGSC[+57.021]KFDEYFSQSC[+57.021]APGSDPR.S | HexNAc(4)Hex(5)Fuc(1)NeuAc(1) | C[+57]*2, N[+2060] | 1500.3821 | 4 | 5998.5065 | 5998.492 | 0 | 2.4 | 485 | Specific | 638.59 | 604.44 | 604.44 | 11.75 | 138 | >sp\|P02788\|TRFL_HUMAN Lactotransferrin OS=Homo sapiens OX=9606 GN=LTF PE=1 SV=6 | 1270 | 20201119_16913_1ng_glyco_hcd203040_compound15.21859.21859.4 | controllerType=0 controllerNumber=1 scan=21859 | 45.6869 |
| 18118:4 | 1 | R.TAGWNIPMGLLFN[+2076.750]QTGSC[+57.021]KFDEYFSQSC[+57.021]APGSDPR.S | HexNAc(4)Hex(6)Fuc(2) | C[+57]*2, N[+2077] | 1504.6412 | 4 | 6015.5431 | 6015.5074 | 0 | 5.9 | 485 | Specific | 643.4 | 643.4 | 643.4 | 11.11 | 138 | >sp\|P02788\|TRFL_HUMAN Lactotransferrin OS=Homo sapiens OX=9606 GN=LTF PE=1 SV=6 | 1270 | 20201119_16913_1ng_glyco_hcd203040_compound15.21794.21794.4 | controllerType=0 controllerNumber=1 scan=21794 | 45.5323 |
| 18120:4 | 1 | R.TAGWNIPMGLLFN[+2076.750]QTGSC[+57.021]KFDEYFSQSC[+57.021]APGSDPR.S | HexNAc(4)Hex(6)Fuc(2) | C[+57]*2, N[+2077] | 1504.6412 | 4 | 6015.5431 | 6015.5074 | 0 | 5.9 | 485 | Specific | 794.28 | 754.44 | 754.44 | 14.11 | 138 | >sp\|P02788\|TRFL_HUMAN Lactotransferrin OS=Homo sapiens OX=9606 GN=LTF PE=1 SV=6 | 1270 | 20201119_16913_1ng_glyco_hcd203040_compound15.21796.21796.4 | controllerType=0 controllerNumber=1 scan=21796 | 45.5375 |
| 19365:4 | 1 | R.TAGWNIPMGLLFN[+2350.830]QTGSC[+57.021]KFDEYFSQSC[+57.021]APGSDPR.S | HexNAc(4)Hex(5)Fuc(1)NeuAc(2) | C[+57]*2, N[+2351] | 1573.1615 | 4 | 6289.6242 | 6289.5875 | 0 | 5.8 | 485 | Specific | 837.66 | 675.89 | 675.89 | 15.11 | 138 | >sp\|P02788\|TRFL_HUMAN Lactotransferrin OS=Homo sapiens OX=9606 GN=LTF PE=1 SV=6 | 1270 | 20201119_16913_1ng_glyco_hcd203040_compound15.23098.23098.4 | controllerType=0 controllerNumber=1 scan=23098 | 48.3984 |

Table S4. Cont.

| Query #:z | Protein Rank | Glycopeptide | Glycan | Modification Type(s) | Observed m/z | z | Observed (M+H) | Calc. mass (M+H) | Off-by-x error | Mass error (ppm) | StartPosition | Cleavage | Score | Delta | Delta Mod | \|Log Prob\| | # of unique peptides | Protein Name | Protein DB number | Comment | Scan # | Scan Time |
|---|---|---|---|---|---|---|---|---|---|---|---|---|---|---|---|---|---|---|---|---|---|---|
| 18183:4 | 1 | R.TAGWNIPMGLLFN[+2059.735]QTGSC[+57.021]KFDEYFSQSC[+57.021]APGSDPR. | HexNAc(4)Hex(5)Fuc(1)NeuAc(1) | C[+57]*2, N[+2060] | 1500.3821 | 4 | 5998.5065 | 5998.492 | 0 | 2.4 | 485 | Specific | 865.8 | 825.43 | 825.43 | 16.75 | 138 | >sp\|P02788\|TRFL_HUMAN Lactotransferrin OS=Homo sapiens OX=9606 GN=LTF PE=1 SV=6 | 1270 | 20201119_16913_1ng_glyco_hcd203040_compound15.21862.21862.4 | controllerType=0 controllerNumber=1 scan=21862 | 45.6909 |
| 19368:4 | 1 | R.TAGWNIPMGLLFN[+2350.830]QTGSC[+57.021]KFDEYFSQSC[+57.021]APGSDPR. | HexNAc(4)Hex(5)Fuc(1)NeuAc(2) | C[+57]*2, N[+2351] | 1573.1615 | 4 | 6289.6242 | 6289.5875 | 0 | 5.8 | 485 | Specific | 876.5 | 834.59 | 834.59 | 16.11 | 138 | >sp\|P02788\|TRFL_HUMAN Lactotransferrin OS=Homo sapiens OX=9606 GN=LTF PE=1 SV=6 | 1270 | 20201119_16913_1ng_glyco_hcd203040_compound15.23101.23101.4 | controllerType=0 controllerNumber=1 scan=23101 | 48.4039 |
| 08599:3 | 1 | R.N[+2571.946]GSDC[+57.021]PDKFC[+57.021]LFQSETK.N | HexNAc(5)Hex(6)Fuc(4) | C[+57]*2, N[+2572] | 1535.6031 | 3 | 4604.7947 | 4604.8193 | 0 | -5.3 | 642 | Specific | 318.67 | 255.63 | 255.63 | 4.63 | 138 | >sp\|P02788\|TRFL_HUMAN Lactotransferrin OS=Homo sapiens OX=9606 GN=LTF PE=1 SV=6 | 1270 | 20201119_16913_1ng_glyco_hcd203040_compound15.11848.11848.3 | controllerType=0 controllerNumber=1 scan=11848 | 26.2393 |
| 08717:3 | 1 | R.N[+2205.793]GSDC[+57.021]PDKFC[+57.021]LFQSETK.N | HexNAc(4)Hex(5)Fuc(2)NeuAc(1) | C[+57]*2, N[+2206] | 1413.5656 | 3 | 4238.6823 | 4238.6667 | 0 | 3.7 | 642 | Specific | 405.12 | 300.27 | 300.27 | 7.04 | 138 | >sp\|P02788\|TRFL_HUMAN Lactotransferrin OS=Homo sapiens OX=9606 GN=LTF PE=1 SV=6 | 1270 | 20201119_16913_1ng_glyco_hcd203040_compound15.11978.11978.3 | controllerType=0 controllerNumber=1 scan=11978 | 26.4804 |
| 08596:3 | 1 | R.N[+2571.946]GSDC[+57.021]PDKFC[+57.021]LFQSETK.N | HexNAc(5)Hex(6)Fuc(4) | C[+57]*2, N[+2572] | 1535.6031 | 3 | 4604.7947 | 4604.8193 | 0 | -5.3 | 642 | Specific | 412.87 | 387.74 | 387.74 | 7.24 | 138 | >sp\|P02788\|TRFL_HUMAN Lactotransferrin OS=Homo sapiens OX=9606 GN=LTF PE=1 SV=6 | 1270 | 20201119_16913_1ng_glyco_hcd203040_compound15.11845.11845.3 | controllerType=0 controllerNumber=1 scan=11845 | 26.2345 |
| 11119:3 | 1 | R.N[+2350.830]GSDC[+57.021]PDKFC[+57.021]LFQSETK.N | HexNAc(4)Hex(5)Fuc(1)NeuAc(2) | C[+57]*2, N[+2351] | 1461.9139 | 3 | 4383.7271 | 4383.7043 | 0 | 5.2 | 642 | Specific | 514.32 | 426.99 | 426.99 | 8.69 | 138 | >sp\|P02788\|TRFL_HUMAN Lactotransferrin OS=Homo sapiens OX=9606 GN=LTF PE=1 SV=6 | 1270 | 20201119_16913_1ng_glyco_hcd203040_compound15.14474.14474.3 | controllerType=0 controllerNumber=1 scan=14474 | 30.8768 |
| 08822:3 | 1 | R.N[+2059.735]GSDC[+57.021]PDKFC[+57.021]LFQSETK.N | HexNAc(4)Hex(5)Fuc(1)NeuAc(1) | C[+57]*2, N[+2060] | 1364.8793 | 3 | 4092.6233 | 4092.6088 | 0 | 3.5 | 642 | Specific | 519 | 438.75 | 438.75 | 9.4 | 138 | >sp\|P02788\|TRFL_HUMAN Lactotransferrin OS=Homo sapiens OX=9606 GN=LTF PE=1 SV=6 | 1270 | 20201119_16913_1ng_glyco_hcd203040_compound15.12087.12087.3 | controllerType=0 controllerNumber=1 scan=12087 | 26.6664 |
| 08719:3 | 1 | R.N[+2205.793]GSDC[+57.021]PDKFC[+57.021]LFQSETK.N | HexNAc(4)Hex(5)Fuc(2)NeuAc(1) | C[+57]*2, N[+2206] | 1413.5656 | 3 | 4238.6823 | 4238.6667 | 0 | 3.7 | 642 | Specific | 538.82 | 338.36 | 338.36 | 7.84 | 138 | >sp\|P02788\|TRFL_HUMAN Lactotransferrin OS=Homo sapiens OX=9606 GN=LTF PE=1 SV=6 | 1270 | 20201119_16913_1ng_glyco_hcd203040_compound15.11980.11980.3 | controllerType=0 controllerNumber=1 scan=11980 | 26.4844 |
| 08825:3 | 1 | R.N[+2059.735]GSDC[+57.021]PDKFC[+57.021]LFQSETK.N | HexNAc(4)Hex(5)Fuc(1)NeuAc(1) | C[+57]*2, N[+2060] | 1364.8793 | 3 | 4092.6233 | 4092.6088 | 0 | 3.5 | 642 | Specific | 545.85 | 422.9 | 422.9 | 9.42 | 138 | >sp\|P02788\|TRFL_HUMAN Lactotransferrin OS=Homo sapiens OX=9606 GN=LTF PE=1 SV=6 | 1270 | 20201119_16913_1ng_glyco_hcd203040_compound15.12090.12090.3 | controllerType=0 controllerNumber=1 scan=12090 | 26.6706 |
| 11117:3 | 1 | R.N[+2350.830]GSDC[+57.021]PDKFC[+57.021]LFQSETK.N | HexNAc(4)Hex(5)Fuc(1)NeuAc(2) | C[+57]*2, N[+2351] | 1461.9139 | 3 | 4383.7271 | 4383.7043 | 0 | 5.2 | 642 | Specific | 677.84 | 587.22 | 587.22 | 12.24 | 138 | >sp\|P02788\|TRFL_HUMAN Lactotransferrin OS=Homo sapiens OX=9606 GN=LTF PE=1 SV=6 | 1270 | 20201119_16913_1ng_glyco_hcd203040_compound15.14472.14472.3 | controllerType=0 controllerNumber=1 scan=14472 | 30.8717 |
| 07839:3 | 2 | R.GN[+2059.735]FSTEGC[+57.021]GC[+57.021]VC[+57.021]EPGWK.G | HexNAc(4)Hex(5)Fuc(1)NeuAc(1) | C[+57]*3, N[+2060] | 1335.5098 | 3 | 4004.5149 | 4004.5023 | 0 | 3.2 | 165 | Specific | 538.22 | 476.56 | 476.56 | 9.89 | 120 | >sp\|P24821\|TENA_HUMAN Tenascin OS=Homo sapiens OX=9606 GN=TNC PE=1 SV=3 | 486 | 20201119_16913_1ng_glyco_hcd203040_compound15.11048.11048.3 | controllerType=0 controllerNumber=1 scan=11048 | 24.8754 |
| 07841:3 | 2 | R.GN[+2059.735]FSTEGC[+57.021]GC[+57.021]VC[+57.021]EPGWK.G | HexNAc(4)Hex(5)Fuc(1)NeuAc(1) | C[+57]*3, N[+2060] | 1335.5098 | 3 | 4004.5149 | 4004.5023 | 0 | 3.2 | 165 | Specific | 569.65 | 466.01 | 466.01 | 10.16 | 120 | >sp\|P24821\|TENA_HUMAN Tenascin OS=Homo sapiens OX=9606 GN=TNC PE=1 SV=3 | 486 | 20201119_16913_1ng_glyco_hcd203040_compound15.11050.11050.3 | controllerType=0 controllerNumber=1 scan=11050 | 24.879 |
| 01429:3 | 2 | K.GPN[+1913.677]C[+57.021]SEPEC[+57.021]PGNC[+57.021]HLR.G | HexNAc(4)Hex(5)NeuAc(1) | C[+57]*3, N[+1914] | 1266.4863 | 3 | 3797.4442 | 3797.4352 | 0 | 2.4 | 182 | Specific | 267.99 | 242.26 | 242.26 | 6.49 | 120 | >sp\|P24821\|TENA_HUMAN Tenascin OS=Homo sapiens OX=9606 GN=TNC PE=1 SV=3 | 486 | 20201119_16913_1ng_glyco_hcd203040_compound15.4386.4386.3 | controllerType=0 controllerNumber=1 scan=4386 | 13.7556 |
| 01431:3 | 2 | K.GPN[+1913.677]C[+57.021]SEPEC[+57.021]PGNC[+57.021]HLR.G | HexNAc(4)Hex(5)NeuAc(1) | C[+57]*3, N[+1914] | 1266.4863 | 3 | 3797.4442 | 3797.4352 | 0 | 2.4 | 182 | Specific | 289.81 | 233.6 | 233.6 | 6.56 | 120 | >sp\|P24821\|TENA_HUMAN Tenascin OS=Homo sapiens OX=9606 GN=TNC PE=1 SV=3 | 486 | 20201119_16913_1ng_glyco_hcd203040_compound15.4388.4388.3 | controllerType=0 controllerNumber=1 scan=4388 | 13.7595 |
| 01393:3 | 2 | K.GPN[+2059.735]C[+57.021]SEPEC[+57.021]PGNC[+57.021]HLR.G | HexNAc(4)Hex(5)Fuc(1)NeuAc(1) | C[+57]*3, N[+2060] | 1315.1725 | 3 | 3943.5031 | 3943.4931 | 0 | 2.5 | 182 | Specific | 421.99 | 370.69 | 370.69 | 7.96 | 120 | >sp\|P24821\|TENA_HUMAN Tenascin OS=Homo sapiens OX=9606 GN=TNC PE=1 SV=3 | 486 | 20201119_16913_1ng_glyco_hcd203040_compound15.4349.4349.3 | controllerType=0 controllerNumber=1 scan=4349 | 13.6911 |
| 01395:3 | 2 | K.GPN[+2059.735]C[+57.021]SEPEC[+57.021]PGNC[+57.021]HLR.G | HexNAc(4)Hex(5)Fuc(1)NeuAc(1) | C[+57]*3, N[+2060] | 1315.1725 | 3 | 3943.5031 | 3943.4931 | 0 | 2.5 | 182 | Specific | 483.01 | 388.09 | 388.09 | 9.52 | 120 | >sp\|P24821\|TENA_HUMAN Tenascin OS=Homo sapiens OX=9606 GN=TNC PE=1 SV=3 | 486 | 20201119_16913_1ng_glyco_hcd203040_compound15.4351.4351.3 | controllerType=0 controllerNumber=1 scan=4351 | 13.6959 |
| 03683:5 | 2 | R.C[+57.021]IN[+2059.735]GTC[+57.021]YC[+57.021]EEGFTGEDC[+57.021]GKPTC[ | HexNAc(4)Hex(5)Fuc(1)NeuAc(1) | C[+57]*6, N[+2060] | 1144.6392 | 5 | 5719.1667 | 5719.1633 | 0 | 0.6 | 325 | Specific | 305.04 | 305.04 | 305.04 | 6.85 | 120 | >sp\|P24821\|TENA_HUMAN Tenascin OS=Homo sapiens OX=9606 GN=TNC PE=1 SV=3 | 486 | 20201119_16913_1ng_glyco_hcd203040_compound15.6732.6732.5 | controllerType=0 controllerNumber=1 scan=6732 | 17.7914 |
| 03666:4 | 2 | R.C[+57.021]IN[+2059.735]GTC[+57.021]YC[+57.021]EEGFTGEDC[+57.021]GKPTC[ | HexNAc(4)Hex(5)Fuc(1)NeuAc(1) | C[+57]*6, N[+2060] | 1430.5501 | 4 | 5719.1784 | 5719.1633 | 0 | 2.6 | 325 | Specific | 311.87 | 311.87 | 311.87 | 6.43 | 120 | >sp\|P24821\|TENA_HUMAN Tenascin OS=Homo sapiens OX=9606 GN=TNC PE=1 SV=3 | 486 | 20201119_16913_1ng_glyco_hcd203040_compound15.6714.6714.4 | controllerType=0 controllerNumber=1 scan=6714 | 17.7619 |
| 03659:5 | 2 | R.C[+57.021]IN[+2059.735]GTC[+57.021]YC[+57.021]EEGFTGEDC[+57.021]GKPTC[ | HexNAc(4)Hex(5)Fuc(1)NeuAc(1) | C[+57]*6, N[+2060] | 1144.6406 | 5 | 5719.174 | 5719.1633 | 0 | 1.9 | 325 | Specific | 331.76 | 331.76 | 331.76 | 7.53 | 120 | >sp\|P24821\|TENA_HUMAN Tenascin OS=Homo sapiens OX=9606 GN=TNC PE=1 SV=3 | 486 | 20201119_16913_1ng_glyco_hcd203040_compound15.6707.6707.5 | controllerType=0 controllerNumber=1 scan=6707 | 17.7505 |
| 03664:4 | 2 | R.C[+57.021]IN[+2059.735]GTC[+57.021]YC[+57.021]EEGFTGEDC[+57.021]GKPTC[ | HexNAc(4)Hex(5)Fuc(1)NeuAc(1) | C[+57]*6, N[+2060] | 1430.5501 | 4 | 5719.1784 | 5719.1633 | 0 | 2.6 | 325 | Specific | 337.23 | 337.23 | 337.23 | 6.85 | 120 | >sp\|P24821\|TENA_HUMAN Tenascin OS=Homo sapiens OX=9606 GN=TNC PE=1 SV=3 | 486 | 20201119_16913_1ng_glyco_hcd203040_compound15.6712.6712.4 | controllerType=0 controllerNumber=1 scan=6712 | 17.7585 |
| 03657:5 | 2 | R.C[+57.021]IN[+2059.735]GTC[+57.021]YC[+57.021]EEGFTGEDC[+57.021]GKPTC[ | HexNAc(4)Hex(5)Fuc(1)NeuAc(1) | C[+57]*6, N[+2060] | 1144.6406 | 5 | 5719.174 | 5719.1633 | 0 | 1.9 | 325 | Specific | 370.66 | 370.66 | 370.66 | 7.9 | 120 | >sp\|P24821\|TENA_HUMAN Tenascin OS=Homo sapiens OX=9606 GN=TNC PE=1 SV=3 | 486 | 20201119_16913_1ng_glyco_hcd203040_compound15.6705.6705.5 | controllerType=0 controllerNumber=1 scan=6705 | 17.7468 |
| 03685:5 | 2 | R.C[+57.021]IN[+2059.735]GTC[+57.021]YC[+57.021]EEGFTGEDC[+57.021]GKPTC[ | HexNAc(4)Hex(5)Fuc(1)NeuAc(1) | C[+57]*6, N[+2060] | 1144.6392 | 5 | 5719.1667 | 5719.1633 | 0 | 0.6 | 325 | Specific | 395.26 | 395.26 | 395.26 | 9.02 | 120 | >sp\|P24821\|TENA_HUMAN Tenascin OS=Homo sapiens OX=9606 GN=TNC PE=1 SV=3 | 486 | 20201119_16913_1ng_glyco_hcd203040_compound15.6734.6734.5 | controllerType=0 controllerNumber=1 scan=6734 | 17.7963 |
| 08699:4 | 2 | R.QTGLAPGQEYEISLHIVKN[+2059.735]NTR.G | HexNAc(4)Hex(5)Fuc(1)NeuAc(1) | N[+2060] | 1132.7646 | 4 | 4528.0364 | 4528.0241 | 0 | 2.7 | 770 | Specific | 540.42 | 351.83 | 351.83 | 8.07 | 120 | >sp\|P24821\|TENA_HUMAN Tenascin OS=Homo sapiens OX=9606 GN=TNC PE=1 SV=3 | 486 | 20201119_16913_1ng_glyco_hcd203040_compound15.11959.11959.2 | controllerType=0 controllerNumber=1 scan=11959 | 26.4407 |
| 08701:4 | 2 | R.QTGLAPGQEYEISLHIVKN[+2059.735]NTR.G | HexNAc(4)Hex(5)Fuc(1)NeuAc(1) | N[+2060] | 1132.7646 | 4 | 4528.0364 | 4528.0241 | 0 | 2.7 | 770 | Specific | 560 | 402.76 | 402.76 | 9.76 | 120 | >sp\|P24821\|TENA_HUMAN Tenascin OS=Homo sapiens OX=9606 GN=TNC PE=1 SV=3 | 486 | 20201119_16913_1ng_glyco_hcd203040_compound15.11961.11961.2 | controllerType=0 controllerNumber=1 scan=11961 | 26.4463 |
| 14888:2 | 2 | R.LN[+1216.423]YSLPTGQWVGVQLPR.N | HexNAc(2)Hex(5) | N[+1216] | 1572.7383 | 2 | 3144.4694 | 3144.4617 | 0 | 2.4 | 1017 | Specific | 458.89 | 358.61 | 358.61 | 8.25 | 120 | >sp\|P24821\|TENA_HUMAN Tenascin OS=Homo sapiens OX=9606 GN=TNC PE=1 SV=3 | 486 | 20201119_16913_1ng_glyco_hcd203040_compound15.18401.18401.2 | controllerType=0 controllerNumber=1 scan=18401 | 38.2347 |
| 14758:3 | 2 | R.LN[+1540.529]YSLPTGQWVGVQLPR.N | HexNAc(2)Hex(7) | N[+1541] | 1156.8635 | 3 | 3468.5758 | 3468.5674 | 0 | 2.4 | 1017 | Specific | 487.59 | 487.59 | 487.59 | 9.57 | 120 | >sp\|P24821\|TENA_HUMAN Tenascin OS=Homo sapiens OX=9606 GN=TNC PE=1 SV=3 | 486 | 20201119_16913_1ng_glyco_hcd203040_compound15.18266.18266.3 | controllerType=0 controllerNumber=1 scan=18266 | 37.9477 |
| 14890:2 | 2 | R.LN[+1216.423]YSLPTGQWVGVQLPR.N | HexNAc(2)Hex(5) | N[+1216] | 1572.7383 | 2 | 3144.4694 | 3144.4617 | 0 | 2.4 | 1017 | Specific | 570.37 | 420.97 | 420.97 | 10.19 | 120 | >sp\|P24821\|TENA_HUMAN Tenascin OS=Homo sapiens OX=9606 GN=TNC PE=1 SV=3 | 486 | 20201119_16913_1ng_glyco_hcd203040_compound15.18403.18403.2 | controllerType=0 controllerNumber=1 scan=18403 | 38.2401 |
| 14653:2 | 2 | R.LN[+1702.581]YSLPTGQWVGVQLPR.N | HexNAc(2)Hex(8) | N[+1703] | 1815.8208 | 2 | 3630.6343 | 3630.6202 | 0 | 3.9 | 1017 | Specific | 598.09 | 508.83 | 508.83 | 11.51 | 120 | >sp\|P24821\|TENA_HUMAN Tenascin OS=Homo sapiens OX=9606 GN=TNC PE=1 SV=3 | 486 | 20201119_16913_1ng_glyco_hcd203040_compound15.18157.18157.2 | controllerType=0 controllerNumber=1 scan=18157 | 37.7473 |
| 14651:2 | 2 | R.LN[+1702.581]YSLPTGQWVGVQLPR.N | HexNAc(2)Hex(8) | N[+1703] | 1815.8208 | 2 | 3630.6343 | 3630.6202 | 0 | 3.9 | 1017 | Specific | 658.98 | 658.98 | 658.98 | 13.51 | 120 | >sp\|P24821\|TENA_HUMAN Tenascin OS=Homo sapiens OX=9606 GN=TNC PE=1 SV=3 | 486 | 20201119_16913_1ng_glyco_hcd203040_compound15.18155.18155.2 | controllerType=0 controllerNumber=1 scan=18155 | 37.7421 |
| 14761:3 | 2 | R.LN[+1540.529]YSLPTGQWVGVQLPR.N | HexNAc(2)Hex(7) | N[+1541] | 1156.8635 | 3 | 3468.5758 | 3468.5674 | 0 | 2.4 | 1017 | Specific | 739.86 | 609 | 609 | 14.35 | 120 | >sp\|P24821\|TENA_HUMAN Tenascin OS=Homo sapiens OX=9606 GN=TNC PE=1 SV=3 | 486 | 20201119_16913_1ng_glyco_hcd203040_compound15.18269.18269.3 | controllerType=0 controllerNumber=1 scan=18269 | 37.9518 |
| 14765:2 | 2 | R.LN[+1378.476]YSLPTGQWVGVQLPR.N | HexNAc(2)Hex(6) | N[+1378] | 1653.7651 | 2 | 3306.5228 | 3306.5145 | 0 | 2.5 | 1017 | Specific | 823.44 | 626.55 | 626.55 | 16.44 | 120 | >sp\|P24821\|TENA_HUMAN Tenascin OS=Homo sapiens OX=9606 GN=TNC PE=1 SV=3 | 486 | 20201119_16913_1ng_glyco_hcd203040_compound15.18273.18273.2 | controllerType=0 controllerNumber=1 scan=18273 | 37.9601 |
| 14811:2 | 2 | R.LN[+1216.423]YSLPTGQWVGVQLPR.N | HexNAc(2)Hex(5) | N[+1216] | 1572.7403 | 2 | 3144.4733 | 3144.4617 | 0 | 3.7 | 1017 | Specific | 842.44 | 781.57 | 781.57 | 16.51 | 120 | >sp\|P24821\|TENA_HUMAN Tenascin OS=Homo sapiens OX=9606 GN=TNC PE=1 SV=3 | 486 | 20201119_16913_1ng_glyco_hcd203040_compound15.18321.18321.2 | controllerType=0 controllerNumber=1 scan=18321 | 38.0603 |
| 14763:2 | 2 | R.LN[+1378.476]YSLPTGQWVGVQLPR.N | HexNAc(2)Hex(6) | N[+1378] | 1653.7651 | 2 | 3306.5228 | 3306.5145 | 0 | 2.5 | 1017 | Specific | 859.67 | 745.28 | 745.28 | 17.44 | 120 | >sp\|P24821\|TENA_HUMAN Tenascin OS=Homo sapiens OX=9606 GN=TNC PE=1 SV=3 | 486 | 20201119_16913_1ng_glyco_hcd203040_compound15.18271.18271.2 | controllerType=0 controllerNumber=1 scan=18271 | 37.9551 |
| 14616:3 | 2 | R.LN[+1702.581]YSLPTGQWVGVQLPR.N | HexNAc(2)Hex(8) | N[+1703] | 1210.8796 | 3 | 3630.6242 | 3630.6202 | 0 | 1.1 | 1017 | Specific | 866.94 | 841.44 | 841.44 | 17.09 | 120 | >sp\|P24821\|TENA_HUMAN Tenascin OS=Homo sapiens OX=9606 GN=TNC PE=1 SV=3 | 486 | 20201119_16913_1ng_glyco_hcd203040_compound15.18119.18119.3 | controllerType=0 controllerNumber=1 scan=18119 | 37.6748 |
| 14809:2 | 2 | R.LN[+1216.423]YSLPTGQWVGVQLPR.N | HexNAc(2)Hex(5) | N[+1216] | 1572.7403 | 2 | 3144.4733 | 3144.4617 | 0 | 3.7 | 1017 | Specific | 901.62 | 781.58 | 781.58 | 18.51 | 120 | >sp\|P24821\|TENA_HUMAN Tenascin OS=Homo sapiens OX=9606 GN=TNC PE=1 SV=3 | 486 | 20201119_16913_1ng_glyco_hcd203040_compound15.18319.18319.2 | controllerType=0 controllerNumber=1 scan=18319 | 38.0531 |
| 14679:3 | 2 | R.LN[+1540.529]YSLPTGQWVGVQLPR.N | HexNAc(2)Hex(7) | N[+1541] | 1156.8647 | 3 | 3468.5795 | 3468.5674 | 0 | 3.5 | 1017 | Specific | 955.83 | 919.4 | 919.4 | 19.41 | 120 | >sp\|P24821\|TENA_HUMAN Tenascin OS=Homo sapiens OX=9606 GN=TNC PE=1 SV=3 | 486 | 20201119_16913_1ng_glyco_hcd203040_compound15.18184.18184.3 | controllerType=0 controllerNumber=1 scan=18184 | 37.7902 |
| 14827:3 | 2 | R.LN[+1216.423]YSLPTGQWVGVQLPR.N | HexNAc(2)Hex(5) | N[+1216] | 1048.829 | 3 | 3144.4725 | 3144.4617 | 0 | 3.4 | 1017 | Specific | 1012.83 | 1012.83 | 1012.83 | 20.41 | 120 | >sp\|P24821\|TENA_HUMAN Tenascin OS=Homo sapiens OX=9606 GN=TNC PE=1 SV=3 | 486 | 20201119_16913_1ng_glyco_hcd203040_compound15.18338.18338.3 | controllerType=0 controllerNumber=1 scan=18338 | 38.0936 |
| 14743:3 | 2 | R.LN[+1378.476]YSLPTGQWVGVQLPR.N | HexNAc(2)Hex(6) | N[+1378] | 1102.847 | 3 | 3306.5263 | 3306.5145 | 0 | 3.6 | 1017 | Specific | 1030.55 | 944.03 | 944.03 | 20.41 | 120 | >sp\|P24821\|TENA_HUMAN Tenascin OS=Homo sapiens OX=9606 GN=TNC PE=1 SV=3 | 486 | 20201119_16913_1ng_glyco_hcd203040_compound15.18250.18250.3 | controllerType=0 controllerNumber=1 scan=18250 | 37.9106 |
| 14682:3 | 2 | R.LN[+1540.529]YSLPTGQWVGVQLPR.N | HexNAc(2)Hex(7) | N[+1541] | 1156.8647 | 3 | 3468.5795 | 3468.5674 | 0 | 3.5 | 1017 | Specific | 1048.61 | 990.61 | 990.61 | 20.41 | 120 | >sp\|P24821\|TENA_HUMAN Tenascin OS=Homo sapiens OX=9606 GN=TNC PE=1 SV=3 | 486 | 20201119_16913_1ng_glyco_hcd203040_compound15.18187.18187.3 | controllerType=0 controllerNumber=1 scan=18187 | 37.7951 |
| 14745:3 | 2 | R.LN[+1378.476]YSLPTGQWVGVQLPR.N | HexNAc(2)Hex(6) | N[+1378] | 1102.847 | 3 | 3306.5263 | 3306.5145 | 0 | 3.6 | 1017 | Specific | 1052.72 | 866.6 | 866.6 | 21.41 | 120 | >sp\|P24821\|TENA_HUMAN Tenascin OS=Homo sapiens OX=9606 GN=TNC PE=1 SV=3 | 486 | 20201119_16913_1ng_glyco_hcd203040_compound15.18252.18252.3 | controllerType=0 controllerNumber=1 scan=18252 | 37.9162 |
| 14619:3 | 2 | R.LN[+1702.581]YSLPTGQWVGVQLPR.N | HexNAc(2)Hex(8) | N[+1703] | 1210.8796 | 3 | 3630.6242 | 3630.6202 | 0 | 1.1 | 1017 | Specific | 1053.38 | 841.44 | 841.44 | 21.09 | 120 | >sp\|P24821\|TENA_HUMAN Tenascin OS=Homo sapiens OX=9606 GN=TNC PE=1 SV=3 | 486 | 20201119_16913_1ng_glyco_hcd203040_compound15.18122.18122.3 | controllerType=0 controllerNumber=1 scan=18122 | 37.6829 |
| 14829:3 | 2 | R.LN[+1216.423]YSLPTGQWVGVQLPR.N | HexNAc(2)Hex(5) | N[+1216] | 1048.829 | 3 | 3144.4725 | 3144.4617 | 0 | 3.4 | 1017 | Specific | 1113.65 | 1113.65 | 1113.65 | 22.41 | 120 | >sp\|P24821\|TENA_HUMAN Tenascin OS=Homo sapiens OX=9606 GN=TNC PE=1 SV=3 | 486 | 20201119_16913_1ng_glyco_hcd203040_compound15.18340.18340.3 | controllerType=0 controllerNumber=1 scan=18340 | 38.0984 |
| 03598:3 | 2 | R.N[+2059.735]TTSYVLR.G | HexNAc(4)Hex(5)Fuc(1)NeuAc(1) | N[+2060] | 1005.0878 | 3 | 3013.2488 | 3013.24 | 0 | 2.9 | 1034 | Specific | 221.66 | 136.25 | 136.25 | 3.94 | 120 | >sp\|P24821\|TENA_HUMAN Tenascin OS=Homo sapiens OX=9606 GN=TNC PE=1 SV=3 | 486 | 20201119_16913_1ng_glyco_hcd203040_compound15.6644.6644.3 | controllerType=0 controllerNumber=1 scan=6644 | 17.6462 |
| 03612:2 | 2 | R.N[+2059.735]TTSYVLR.G | HexNAc(4)Hex(5)Fuc(1)NeuAc(1) | N[+2060] | 1507.1276 | 2 | 3013.2479 | 3013.24 | 0 | 2.6 | 1034 | Specific | 292.04 | 158.84 | 158.84 | 5.57 | 120 | >sp\|P24821\|TENA_HUMAN Tenascin OS=Homo sapiens OX=9606 GN=TNC PE=1 SV=3 | 486 | 20201119_16913_1ng_glyco_hcd203040_compound15.6658.6658.2 | controllerType=0 controllerNumber=1 scan=6658 | 17.6691 |
| 03600:3 | 2 | R.N[+2059.735]TTSYVLR.G | HexNAc(4)Hex(5)Fuc(1)NeuAc(1) | N[+2060] | 1005.0878 | 3 | 3013.2488 | 3013.24 | 0 | 2.9 | 1034 | Specific | 320.11 | 233.71 | 233.71 | 6.42 | 120 | >sp\|P24821\|TENA_HUMAN Tenascin OS=Homo sapiens OX=9606 GN=TNC PE=1 SV=3 | 486 | 20201119_16913_1ng_glyco_hcd203040_compound15.6646.6646.3 | controllerType=0 controllerNumber=1 scan=6646 | 17.6501 |
| 07479:3 | 2 | R.N[+2059.735]LTVPGSLR.A | HexNAc(4)Hex(5)Fuc(1)NeuAc(1) | N[+2060] | 1006.1016 | 3 | 3016.2902 | 3016.2873 | 0 | 1 | 1119 | Specific | 219.84 | 108.1 | 108.1 | 2.64 | 120 | >sp\|P24821\|TENA_HUMAN Tenascin OS=Homo sapiens OX=9606 GN=TNC PE=1 SV=3 | 486 | 20201119_16913_1ng_glyco_hcd203040_compound15.10672.10672.3 | controllerType=0 controllerNumber=1 scan=10672 | 24.2424 |
| 07514:2 | 2 | R.N[+2059.735]LTVPGSLR.A | HexNAc(4)Hex(5)Fuc(1)NeuAc(1) | N[+2060] | 1508.6535 | 2 | 3016.2997 | 3016.2873 | 0 | 4.1 | 1119 | Specific | 271.66 | 58.69 | 58.69 | 3.89 | 120 | >sp\|P24821\|TENA_HUMAN Tenascin OS=Homo sapiens OX=9606 GN=TNC PE=1 SV=3 | 486 | 20201119_16913_1ng_glyco_hcd203040_compound15.10708.10708.2 | controllerType=0 controllerNumber=1 scan=10708 | 24.3006 |
| 22950:4 | 2 | R.GVTQDFSTTPLSVEVLTEEVPDMGN[+2350.830]LTVTEVSWDALR.L | HexNAc(4)Hex(5)Fuc(1)NeuAc(2) | N[+2351] | 1597.4617 | 4 | 6386.825 | 6386.8047 | 0 | 3.2 | 1237 | Specific | 590.38 | 590.38 | 590.38 | 12.06 | 120 | >sp\|P24821\|TENA_HUMAN Tenascin OS=Homo sapiens OX=9606 GN=TNC PE=1 SV=3 | 486 | 20201119_16913_1ng_glyco_hcd203040_compound15.26848.26848.4 | controllerType=0 controllerNumber=1 scan=26848 | 56.5706 |
| 22952:4 | 2 | R.GVTQDFSTTPLSVEVLTEEVPDMGN[+2350.830]LTVTEVSWDALR.L | HexNAc(4)Hex(5)Fuc(1)NeuAc(2) | N[+2351] | 1597.4617 | 4 | 6386.825 | 6386.8047 | 0 | 3.2 | 1237 | Specific | 798.6 | 747.54 | 747.54 | 16.06 | 120 | >sp\|P24821\|TENA_HUMAN Tenascin OS=Homo sapiens OX=9606 GN=TNC PE=1 SV=3 | 486 | 20201119_16913_1ng_glyco_hcd203040_compound15.26850.26850.4 | controllerType=0 controllerNumber=1 scan=26850 | 56.5775 |
| 10788:3 | 2 | R.LLETVEYN[+2059.735]ISGAER.T | HexNAc(4)Hex(5)Fuc(1)NeuAc(1) | N[+2060] | 1218.5209 | 3 | 3653.5482 | 3653.5468 | 0 | 0.4 | 1478 | Specific | 229.69 | 163.45 | 163.45 | 5.27 | 120 | >sp\|P24821\|TENA_HUMAN Tenascin OS=Homo sapiens OX=9606 GN=TNC PE=1 SV=3 | 486 | 20201119_16913_1ng_glyco_hcd203040_compound15.14126.14126.3 | controllerType=0 controllerNumber=1 scan=14126 | 30.1277 |
| 10790:3 | 2 | R.LLETVEYN[+2059.735]ISGAER.T | HexNAc(4)Hex(5)Fuc(1)NeuAc(1) | N[+2060] | 1218.5209 | 3 | 3653.5482 | 3653.5468 | 0 | 0.4 | 1478 | Specific | 476.59 | 387.03 | 387.03 | 9.17 | 120 | >sp\|P24821\|TENA_HUMAN Tenascin OS=Homo sapiens OX=9606 GN=TNC PE=1 SV=3 | 486 | 20201119_16913_1ng_glyco_hcd203040_compound15.14128.14128.3 | controllerType=0 controllerNumber=1 scan=14128 | 30.1328 |
| 09617:3 | 3 | K.TVLTPATNHMGN[+1702.581]VTFTIPANR.E | HexNAc(2)Hex(8) | N[+1703] | 1319.9232 | 3 | 3957.755 | 3957.7414 | 0 | 3.4 | 74 | Specific | 653.63 | 653.63 | 304.32 | 13.51 | 111 | >sp\|P01024\|CO3_HUMAN Complement C3 OS=Homo sapiens OX=9606 GN=C3 PE=1 SV=2 | 492 | 20201119_16913_1ng_glyco_hcd203040_compound15.12912.12912.3 | controllerType=0 controllerNumber=1 scan=12912 | 28.0495 |
| 09615:3 | 3 | K.TVLTPATNHMGN[+1702.581]VTFTIPANR.E | HexNAc(2)Hex(8) | N[+1703] | 1319.9232 | 3 | 3957.755 | 3957.7414 | 0 | 3.4 | 74 | Specific | 719.43 | 719.43 | 413.03 | 14.51 | 111 | >sp\|P01024\|CO3_HUMAN Complement C3 OS=Homo sapiens OX=9606 GN=C3 PE=1 SV=2 | 492 | 20201119_16913_1ng_glyco_hcd203040_compound15.12910.12910.3 | controllerType=0 controllerNumber=1 scan=12910 | 28.0457 |
| 09665:3 | 3 | K.TVLTPATNHMGN[+1540.529]VTFTIPANR.E | HexNAc(2)Hex(7) | N[+1541] | 1265.9028 | 3 | 3795.6937 | 3795.6886 | 0 | 1.4 | 74 | Specific | 963.54 | 780.15 | 469.73 | 19.18 | 111 | >sp\|P01024\|CO3_HUMAN Complement C3 OS=Homo sapiens OX=9606 GN=C3 PE=1 SV=2 | 492 | 20201119_16913_1ng_glyco_hcd203040_compound15.12963.12963.3 | controllerType=0 controllerNumber=1 scan=12963 | 28.1358 |
| 09682:3 | 3 | K.TVLTPATNHMGN[+1378.476]VTFTIPANR.E | HexNAc(2)Hex(6) | N[+1378] | 1211.8867 | 3 | 3633.6454 | 3633.6358 | 0 | 2.6 | 74 | Specific | 970.2 | 849.38 | 513.92 | 19.44 | 111 | >sp\|P01024\|CO3_HUMAN Complement C3 OS=Homo sapiens OX=9606 GN=C3 PE=1 SV=2 | 492 | 20201119_16913_1ng_glyco_hcd203040_compound15.12981.12981.3 | controllerType=0 controllerNumber=1 scan=12981 | 28.1696 |
| 09663:3 | 3 | K.TVLTPATNHMGN[+1540.529]VTFTIPANR.E | HexNAc(2)Hex(7) | N[+1541] | 1265.9028 | 3 | 3795.6937 | 3795.6886 | 0 | 1.4 | 74 | Specific | 1046.31 | 999.91 | 569.14 | 20.18 | 111 | >sp\|P01024\|CO3_HUMAN Complement C3 OS=Homo sapiens OX=9606 GN=C3 PE=1 SV=2 | 492 | 20201119_16913_1ng_glyco_hcd203040_compound15.12961.12961.3 | controllerType=0 controllerNumber=1 scan=12961 | 28.1296 |
| 09680:3 | 3 | K.TVLTPATNHMGN[+1378.476]VTFTIPANR.E | HexNAc(2)Hex(6) | N[+1378] | 1211.8867 | 3 | 3633.6454 | 3633.6358 | 0 | 2.6 | 74 | Specific | 1127.59 | 1022.01 | 617.96 | 22.44 | 111 | >sp\|P01024\|CO3_HUMAN Complement C3 OS=Homo sapiens OX=9606 GN=C3 PE=1 SV=2 | 492 | 20201119_16913_1ng_glyco_hcd203040_compound15.12979.12979.3 | controllerType=0 controllerNumber=1 scan=12979 | 28.1642 |
| 06992:2 | 4 | R.FSDGLESN[+1864.634]SSTQFEVK.K | HexNAc(2)Hex(9) | N[+1865] | 1820.2352 | 2 | 3639.4631 | 3639.4472 | 0 | 4.4 | 219 | Specific | 320.4 | 320.4 | 320.4 | 6.71 | 79 | >sp\|P0C0L5\|CO4B_HUMAN Complement C4-B OS=Homo sapiens OX=9606 GN=C4B PE=1 SV=2 | 1424 | 20201119_16913_1ng_glyco_hcd203040_compound15.10168.10168.2 | controllerType=0 controllerNumber=1 scan=10168 | 23.4099 |
| 06993:2 | 4 | R.FSDGLESN[+1864.634]SSTQFEVK.K | HexNAc(2)Hex(9) | N[+1865] | 1820.2352 | 2 | 3639.4631 | 3639.4472 | 0 | 4.4 | 219 | Specific | 381 | 354.16 | 354.16 | 7.47 | 79 | >sp\|P0C0L5\|CO4B_HUMAN Complement C4-B OS=Homo sapiens OX=9606 GN=C4B PE=1 SV=2 | 1424 | 20201119_16913_1ng_glyco_hcd203040_compound15.10169.10169.2 | controllerType=0 controllerNumber=1 scan=10169 | 23.4136 |
| 06999:3 | 4 | R.FSDGLESN[+1864.634]SSTQFEVK.K | HexNAc(2)Hex(9) | N[+1865] | 1213.8253 | 3 | 3639.4612 | 3639.4472 | 0 | 3.9 | 219 | Specific | 719.51 | 676.95 | 676.95 | 14.27 | 79 | >sp\|P0C0L5\|CO4B_HUMAN Complement C4-B OS=Homo sapiens OX=9606 GN=C4B PE=1 SV=2 | 1424 | 20201119_16913_1ng_glyco_hcd203040_compound15.10176.10176.3 | controllerType=0 controllerNumber=1 scan=10176 | 23.4266 |
| 07001:3 | 4 | R.FSDGLESN[+1864.634]SSTQFEVK.K | HexNAc(2)Hex(9) | N[+1865] | 1213.8253 | 3 | 3639.4612 | 3639.4472 | 0 | 3.9 | 219 | Specific | 732.66 | 732.66 | 732.66 | 14.27 | 79 | >sp\|P0C0L5\|CO4B_HUMAN Complement C4-B OS=Homo sapiens OX=9606 GN=C4B PE=1 SV=2 | 1424 | 20201119_16913_1ng_glyco_hcd203040_compound15.10178.10178.3 | controllerType=0 controllerNumber=1 scan=10178 | 23.4305 |
| 04474:3 | 4 | R.FSDGLESN[+1864.634]SSTQFEVKK.Y | HexNAc(2)Hex(9) | N[+1865] | 1256.522 | 3 | 3767.5515 | 3767.5421 | 0 | 2.5 | 219 | Specific | 437.3 | 437.3 | 437.3 | 9.02 | 79 | >sp\|P0C0L5\|CO4B_HUMAN Complement C4-B OS=Homo sapiens OX=9606 GN=C4B PE=1 SV=2 | 1424 | 20201119_16913_1ng_glyco_hcd203040_compound15.7557.7557.3 | controllerType=0 controllerNumber=1 scan=7557 | 19.1135 |
| 04472:3 | 4 | R.FSDGLESN[+1864.634]SSTQFEVKK.Y | HexNAc(2)Hex(9) | N[+1865] | 1256.522 | 3 | 3767.5515 | 3767.5421 | 0 | 2.5 | 219 | Specific | 562.76 | 334.65 | 334.65 | 8.05 | 79 | >sp\|P0C0L5\|CO4B_HUMAN Complement C4-B OS=Homo sapiens OX=9606 GN=C4B PE=1 SV=2 | 1424 | 20201119_16913_1ng_glyco_hcd203040_compound15.7555.7555.3 | controllerType=0 controllerNumber=1 scan=7555 | 19.1098 |
| 16893:3 | 6 | R.GLSFDVSLEVSQGPGLLN[+2205.793]DTK.V | HexNAc(4)Hex(5)Fuc(2)NeuAc(1) | N[+2206] | 1461.3107 | 3 | 4381.9176 | 4381.906 | 0 | 2.6 | 118 | Specific | 261.97 | 223.28 | 223.28 | 6.12 | 82 | >sp\|P01833\|PIGR_HUMAN Polymeric immunoglobulin receptor OS=Homo sapiens OX=9606 GN=PIGR PE=1 SV=4 | 902 | 20201119_16913_1ng_glyco_hcd203040_compound15.20513.20513.2 | controllerType=0 controllerNumber=1 scan=20513 | 42.5[illegible]53 |
| 16771:3 | 6 | R.GLSFDVSLEVSQGPGLLN[+2570.925]DTK.V | HexNAc(5)Hex(6)Fuc(2)NeuAc(1) | N[+2571] | 1583.0256 | 3 | 4747.0622 | 4747.0382 | 0 | 5 | 118 | Specific | 299.5 | 255.04 | 255.04 | 6.33 | 82 | >sp\|P01833\|PIGR_HUMAN Polymeric immunoglobulin receptor OS=Homo sapiens OX=9606 GN=PIGR PE=1 SV=4 | 902 | 20201119_16913_1ng_glyco_hcd203040_compound15.20385.20385.3 | controllerType=0 controllerNumber=1 scan=20385 | 42.2[illegible]08 |
| 16652:3 | 6 | R.GLSFDVSLEVSQGPGLLN[+2716.983]DTK.V | HexNAc(5)Hex(6)Fuc(3)NeuAc(1) | N[+2717] | 1631.7125 | 3 | 4893.1228 | 4893.0962 | 0 | 5.5 | 118 | Specific | 312.14 | 292.67 | 292.67 | 6.29 | 82 | >sp\|P01833\|PIGR_HUMAN Polymeric immunoglobulin receptor OS=Homo sapiens OX=9606 GN=PIGR PE=1 SV=4 | 902 | 20201119_16913_1ng_glyco_hcd203040_compound15.20261.20261.3 | controllerType=0 controllerNumber=1 scan=20261 | 42.0[illegible]32 |
| 16697:3 | 6 | R.GLSFDVSLEVSQGPGLLN[+2571.946]DTK.V | HexNAc(5)Hex(6)Fuc(4) | N[+2572] | 1583.3556 | 3 | 4748.0524 | 4748.0586 | 0 | -1.3 | 118 | Specific | 346.03 | 236.33 | 236.33 | 5.55 | 82 | >sp\|P01833\|PIGR_HUMAN Polymeric immunoglobulin receptor OS=Homo sapiens OX=9606 GN=PIGR PE=1 SV=4 | 902 | 20201119_16913_1ng_glyco_hcd203040_compound15.20308.20308.4 | controllerType=0 controllerNumber=1 scan=20308 | [illegible] |
| 16699:3 | 6 | R.GLSFDVSLEVSQGPGLLN[+2571.946]DTK.V | HexNAc(5)Hex(6)Fuc(4) | N[+2572] | 1583.3556 | 3 | 4748.0524 | 4748.0586 | 0 | -1.3 | 118 | Specific | 568.19 | 381.07 | 381.07 | 8.73 | 82 | >sp\|P01833\|PIGR_HUMAN Polymeric immunoglobulin receptor OS=Homo sapiens OX=9606 GN=PIGR PE=1 SV=4 | 902 | 20201119_16913_1ng_glyco_hcd203040_compound15.20310.20310.4 | controllerType=0 controllerNumber=1 scan=20310 | 42.1441 |
| 16654:3 | 6 | R.GLSFDVSLEVSQGPGLLN[+2716.983]DTK.V | HexNAc(5)Hex(6)Fuc(3)NeuAc(1) | N[+2717] | 1631.7125 | 3 | 4893.1228 | 4893.0962 | 0 | 5.5 | 118 | Specific | 601.97 | 552.99 | 552.99 | 11.71 | 82 | >sp\|P01833\|PIGR_HUMAN Polymeric immunoglobulin receptor OS=Homo sapiens OX=9606 GN=PIGR PE=1 SV=4 | 902 | 20201119_16913_1ng_glyco_hcd203040_compound15.20263.20263.3 | controllerType=0 controllerNumber=1 scan=20263 | 42.0368 |
| 16774:3 | 6 | R.GLSFDVSLEVSQGPGLLN[+2570.925]DTK.V | HexNAc(5)Hex(6)Fuc(2)NeuAc(1) | N[+2571] | 1583.0256 | 3 | 4747.0622 | 4747.0382 | 0 | 5 | 118 | Specific | 753.18 | 661.18 | 661.18 | 15.05 | 82 | >sp\|P01833\|PIGR_HUMAN Polymeric immunoglobulin receptor OS=Homo sapiens OX=9606 GN=PIGR PE=1 SV=4 | 902 | 20201119_16913_1ng_glyco_hcd203040_compound15.20388.20388.3 | controllerType=0 controllerNumber=1 scan=20388 | 42.2872 |
| 18723:3 | 6 | R.GLSFDVSLEVSQGPGLLN[+2350.830]DTK.V | HexNAc(4)Hex(5)Fuc(1)NeuAc(2) | N[+2351] | 1509.6578 | 3 | 4526.9588 | 4526.9436 | 0 | 3.4 | 118 | Specific | 788.59 | 709.78 | 709.78 | 15.41 | 82 | >sp\|P01833\|PIGR_HUMAN Polymeric immunoglobulin receptor OS=Homo sapiens OX=9606 GN=PIGR PE=1 SV=4 | 902 | 20201119_16913_1ng_glyco_hcd203040_compound15.22427.22427.3 | controllerType=0 controllerNumber=1 scan=22427 | 46.8983 |
| 18644:3 | 6 | R.GLSFDVSLEVSQGPGLLN[+2350.830]DTK.V | HexNAc(4)Hex(5)Fuc(1)NeuAc(2) | N[+2351] | 1509.658 | 3 | 4526.9595 | 4526.9436 | 0 | 3.5 | 118 | Specific | 957.31 | 924.02 | 924.02 | 19.41 | 82 | >sp\|P01833\|PIGR_HUMAN Polymeric immunoglobulin receptor OS=Homo sapiens OX=9606 GN=PIGR PE=1 SV=4 | 902 | 20201119_16913_1ng_glyco_hcd203040_compound15.22344.22344.3 | controllerType=0 controllerNumber=1 scan=22344 | 46.7316 |
| 18641:3 | 6 | R.GLSFDVSLEVSQGPGLLN[+2350.830]DTK.V | HexNAc(4)Hex(5)Fuc(1)NeuAc(2) | N[+2351] | 1509.658 | 3 | 4526.9595 | 4526.9436 | 0 | 3.5 | 118 | Specific | 1073.16 | 1009.46 | 1009.46 | 21.41 | 82 | >sp\|P01833\|PIGR_HUMAN Polymeric immunoglobulin receptor OS=Homo sapiens OX=9606 GN=PIGR PE=1 SV=4 | 902 | 20201119_16913_1ng_glyco_hcd203040_compound15.22341.22341.3 | controllerType=0 controllerNumber=1 scan=22341 | 46.72 |
| 19901:3 | 6 | K.QIGLYPVLVIDSSGYVNPN[+2350.830]YTGR.I | HexNAc(4)Hex(5)Fuc(1)NeuAc(2) | N[+2351] | 1626.053 | 3 | 4876.1446 | 4876.1338 | 0 | 2.2 | 168 | Specific | 339.43 | 339.43 | 339.43 | 7.55 | 82 | >sp\|P01833\|PIGR_HUMAN Polymeric immunoglobulin receptor OS=Homo sapiens OX=9606 GN=PIGR PE=1 SV=4 | 902 | 20201119_16913_1ng_glyco_hcd203040_compound15.23659.23659.3 | controllerType=0 controllerNumber=1 scan=23659 | 49.6869 |
| 19982:4 | 6 | K.QIGLYPVLVIDSSGYVNPN[+2204.772]YTGR.I | HexNAc(4)Hex(5)NeuAc(2) | N[+2205] | 1183.2791 | 4 | 4730.0946 | 4730.0759 | 0 | 3.9 | 168 | Specific | 732.34 | 681.21 | 681.21 | 14.26 | 82 | >sp\|P01833\|PIGR_HUMAN Polymeric immunoglobulin receptor OS=Homo sapiens OX=9606 GN=PIGR PE=1 SV=4 | 902 | 20201119_16913_1ng_glyco_hcd203040_compound15.23744.23744.4 | controllerType=0 controllerNumber=1 scan=23744 | 49.8908 |
| 19904:3 | 6 | K.QIGLYPVLVIDSSGYVNPN[+2350.830]YTGR.I | HexNAc(4)Hex(5)Fuc(1)NeuAc(2) | N[+2351] | 1626.053 | 3 | 4876.1446 | 4876.1338 | 0 | 2.2 | 168 | Specific | 766.58 | 557.05 | 557.05 | 15.43 | 82 | >sp\|P01833\|PIGR_HUMAN Polymeric immunoglobulin receptor OS=Homo sapiens OX=9606 GN=PIGR PE=1 SV=4 | 902 | 20201119_16913_1ng_glyco_hcd203040_compound15.23662.23662.3 | controllerType=0 controllerNumber=1 scan=23662 | 49.691 |
| 19980:4 | 6 | K.QIGLYPVLVIDSSGYVNPN[+2204.772]YTGR.I | HexNAc(4)Hex(5)NeuAc(2) | N[+2205] | 1183.2791 | 4 | 4730.0946 | 4730.0759 | 0 | 3.9 | 168 | Specific | 781.36 | 669.69 | 669.69 | 15.26 | 82 | >sp\|P01833\|PIGR_HUMAN Polymeric immunoglobulin receptor OS=Homo sapiens OX=9606 GN=PIGR PE=1 SV=4 | 902 | 20201119_16913_1ng_glyco_hcd203040_compound15.23742.23742.4 | controllerType=0 controllerNumber=1 scan=23742 | 49.8836 |
| 18061:3 | 6 | K.QIGLYPVLVIDSSGYVNPN[+1913.677]YTGR.I | HexNAc(4)Hex(5)NeuAc(1) | N[+1914] | 1480.3372 | 3 | 4438.9971 | 4438.9805 | 0 | 3.7 | 168 | Specific | 842.64 | 808.87 | 808.87 | 16.49 | 82 | >sp\|P01833\|PIGR_HUMAN Polymeric immunoglobulin receptor OS=Homo sapiens OX=9606 GN=PIGR PE=1 SV=4 | 902 | 20201119_16913_1ng_glyco_hcd203040_compound15.21735.21735.3 | controllerType=0 controllerNumber=1 scan=21735 | 45.4135 |
| 18063:3 | 6 | K.QIGLYPVLVIDSSGYVNPN[+1913.677]YTGR.I | HexNAc(4)Hex(5)NeuAc(1) | N[+1914] | 1480.3372 | 3 | 4438.9971 | 4438.9805 | 0 | 3.7 | 168 | Specific | 887.67 | 803.62 | 803.62 | 17.49 | 82 | >sp\|P01833\|PIGR_HUMAN Polymeric immunoglobulin receptor OS=Homo sapiens OX=9606 GN=PIGR PE=1 SV=4 | 902 | 20201119_16913_1ng_glyco_hcd203040_compound15.21737.21737.3 | controllerType=0 controllerNumber=1 scan=21737 | 45.4215 |
| 18001:3 | 6 | K.QIGLYPVLVIDSSGYVNPN[+2059.735]YTGR.I | HexNAc(4)Hex(5)Fuc(1)NeuAc(1) | N[+2060] | 1529.025 | 3 | 4585.0603 | 4585.0384 | 0 | 4.8 | 168 | Specific | 899.36 | 792 | 792 | 17.12 | 82 | >sp\|P01833\|PIGR_HUMAN Polymeric immunoglobulin receptor OS=Homo sapiens OX=9606 GN=PIGR PE=1 SV=4 | 902 | 20201119_16913_1ng_glyco_hcd203040_compound15.21672.21672.3 | controllerType=0 controllerNumber=1 scan=21672 | 45.2598 |
| 18004:3 | 6 | K.QIGLYPVLVIDSSGYVNPN[+2059.735]YTGR.I | HexNAc(4)Hex(5)Fuc(1)NeuAc(1) | N[+2060] | 1529.025 | 3 | 4585.0603 | 4585.0384 | 0 | 4.8 | 168 | Specific | 931.8 | 853.46 | 853.46 | 18.12 | 82 | >sp\|P01833\|PIGR_HUMAN Polymeric immunoglobulin receptor OS=Homo sapiens OX=9606 GN=PIGR PE=1 SV=4 | 902 | 20201119_16913_1ng_glyco_hcd203040_compound15.21675.21675.3 | controllerType=0 controllerNumber=1 scan=21675 | 45.2661 |
| 22256:3 | 6 | R.LSLLEEPGN[+2351.851]GTFTVILNQLTSR.D | HexNAc(4)Hex(5)Fuc(3)NeuAc(1) | N[+2352] | 1585.3829 | 3 | 4754.1341 | 4754.1433 | 0 | -1.9 | 413 | Specific | 214.3 | 185.37 | 185.37 | 3.72 | 82 | >sp\|P01833\|PIGR_HUMAN Polymeric immunoglobulin receptor OS=Homo sapiens OX=9606 GN=PIGR PE=1 SV=4 | 902 | 20201119_16913_1ng_glyco_hcd203040_compound15.26123.26123.3 | controllerType=0 controllerNumber=1 scan=26123 | 55.0253 |
| 19607:4 | 6 | R.LSLLEEPGN[+2076.750]GTFTVILNQLTSR.D | HexNAc(4)Hex(6)Fuc(2) | N[+2077] | 1120.5256 | 4 | 4479.0804 | 4479.0428 | 0 | 8.4 | 413 | Specific | 244.85 | 200.31 | 200.31 | 3.44 | 82 | >sp\|P01833\|PIGR_HUMAN Polymeric immunoglobulin receptor OS=Homo sapiens OX=9606 GN=PIGR PE=1 SV=4 | 902 | 20201119_16913_1ng_glyco_hcd203040_compound15.23351.23351.4 | controllerType=0 controllerNumber=1 scan=23351 | 48.9723 |
| 19606:3 | 6 | R.LSLLEEPGN[+2205.793]GTFTVILNQLTSR.D | HexNAc(4)Hex(5)Fuc(2)NeuAc(1) | N[+2206] | 1536.7101 | 3 | 4608.1159 | 4608.0854 | 0 | 6.6 | 413 | Specific | 258.01 | 258.01 | 258.01 | 5.7 | 82 | >sp\|P01833\|PIGR_HUMAN Polymeric immunoglobulin receptor OS=Homo sapiens OX=9606 GN=PIGR PE=1 SV=4 | 902 | 20201119_16913_1ng_glyco_hcd203040_compound15.23350.23350.3 | controllerType=0 controllerNumber=1 scan=23350 | 48.9708 |
| 19590:4 | 6 | R.LSLLEEPGN[+2076.750]GTFTVILNQLTSR.D | HexNAc(4)Hex(6)Fuc(2) | N[+2077] | 1120.5218 | 4 | 4479.0653 | 4479.0428 | 0 | 5 | 413 | Specific | 274.53 | 216.27 | 216.27 | 6.03 | 82 | >sp\|P01833\|PIGR_HUMAN Polymeric immunoglobulin receptor OS=Homo sapiens OX=9606 GN=PIGR PE=1 SV=4 | 902 | 20201119_16913_1ng_glyco_hcd203040_compound15.23333.23333.4 | controllerType=0 controllerNumber=1 scan=23333 | 48.9137 |
| 19435:3 | 6 | R.LSLLEEPGN[+2570.925]GTFTVILNQLTSR.D | HexNAc(5)Hex(6)Fuc(2)NeuAc(1) | N[+2571] | 1658.4021 | 3 | 4973.1916 | 4973.2176 | 0 | -5.2 | 413 | Specific | 285.19 | 204.19 | 204.19 | 4.58 | 82 | >sp\|P01833\|PIGR_HUMAN Polymeric immunoglobulin receptor OS=Homo sapiens OX=9606 GN=PIGR PE=1 SV=4 | 902 | 20201119_16913_1ng_glyco_hcd203040_compound15.23172.23172.4 | controllerType=0 controllerNumber=1 scan=23172 | 48.571 |
| 24491:3 | 6 | R.LSLLEEPGN[+3007.058]GTFTVILNQLTSR.D | HexNAc(5)Hex(6)Fuc(1)NeuAc(3) | N[+3007] | 1803.7974 | 3 | 5409.3776 | 5409.3505 | 0 | 5 | 413 | Specific | 344.54 | 185.87 | 185.87 | 6.35 | 82 | >sp\|P01833\|PIGR_HUMAN Polymeric immunoglobulin receptor OS=Homo sapiens OX=9606 GN=PIGR PE=1 SV=4 | 902 | 20201119_16913_1ng_glyco_hcd203040_compound15.28452.28452.3 | controllerType=0 controllerNumber=1 scan=28452 | 59.7181 |
| 24398:4 | 6 | R.LSLLEEPGN[+3007.058]GTFTVILNQLTSR.D | HexNAc(5)Hex(6)Fuc(1)NeuAc(3) | N[+3007] | 1353.0963 | 4 | 5409.3634 | 5409.3505 | 0 | 2.4 | 413 | Specific | 355.05 | 334.29 | 334.29 | 7.9 | 82 | >sp\|P01833\|PIGR_HUMAN Polymeric immunoglobulin receptor OS=Homo sapiens OX=9606 GN=PIGR PE=1 SV=4 | 902 | 20201119_16913_1ng_glyco_hcd203040_compound15.28355.28355.2 | controllerType=0 controllerNumber=1 scan=28355 | 59.5366 |
| 19438:3 | 6 | R.LSLLEEPGN[+2570.925]GTFTVILNQLTSR.D | HexNAc(5)Hex(6)Fuc(2)NeuAc(1) | N[+2571] | 1658.4021 | 3 | 4973.1916 | 4973.2176 | 0 | -5.2 | 413 | Specific | 375.01 | 260.4 | 260.4 | 5.48 | 82 | >sp\|P01833\|PIGR_HUMAN Polymeric immunoglobulin receptor OS=Homo sapiens OX=9606 GN=PIGR PE=1 SV=4 | 902 | 20201119_16913_1ng_glyco_hcd203040_compound15.23175.23175.4 | controllerType=0 controllerNumber=1 scan=23175 | 48.5761 |
| 21597:3 | 6 | R.[+42.011]LSLLEEPGN[+2336.851]GTFTVILNQLTSR.D | HexNAc(6)Hex(6)Fuc(1) | .[42], N[+2337] | 1594.3797 | 3 | 4781.1245 | 4781.1542 | 0 | -6.2 | 413 | Specific | 383 | 346.2 | 346.2 | 6.08 | 82 | >sp\|P01833\|PIGR_HUMAN Polymeric immunoglobulin receptor OS=Homo sapiens OX=9606 GN=PIGR PE=1 SV=4 | 902 | 20201119_16913_1ng_glyco_hcd203040_compound15.25435.25435.3 | controllerType=0 controllerNumber=1 scan=25435 | 53.5076 |
| 19593:4 | 6 | R.LSLLEEPGN[+2076.750]GTFTVILNQLTSR.D | HexNAc(4)Hex(6)Fuc(2) | N[+2077] | 1120.5218 | 4 | 4479.0653 | 4479.0428 | 0 | 5 | 413 | Specific | 412.12 | 315.95 | 315.95 | 6.34 | 82 | >sp\|P01833\|PIGR_HUMAN Polymeric immunoglobulin receptor OS=Homo sapiens OX=9606 GN=PIGR PE=1 SV=4 | 902 | 20201119_16913_1ng_glyco_hcd203040_compound15.23336.23336.4 | controllerType=0 controllerNumber=1 scan=23336 | 48.9196 |
| 19574:3 | 6 | R.LSLLEEPGN[+2076.750]GTFTVILNQLTSR.D | HexNAc(4)Hex(6)Fuc(2) | N[+2077] | 1493.6946 | 3 | 4479.0694 | 4479.0428 | 0 | 5.9 | 413 | Specific | 424.04 | 357.06 | 357.06 | 7.44 | 82 | >sp\|P01833\|PIGR_HUMAN Polymeric immunoglobulin receptor OS=Homo sapiens OX=9606 GN=PIGR PE=1 SV=4 | 902 | 20201119_16913_1ng_glyco_hcd203040_compound15.23316.23316.3 | controllerType=0 controllerNumber=1 scan=23316 | 48.8822 |
| 19573:3 | 6 | R.LSLLEEPGN[+2076.750]GTFTVILNQLTSR.D | HexNAc(4)Hex(6)Fuc(2) | N[+2077] | 1493.6946 | 3 | 4479.0694 | 4479.0428 | 0 | 5.9 | 413 | Specific | 434.5 | 368.8 | 368.8 | 7.5 | 82 | >sp\|P01833\|PIGR_HUMAN Polymeric immunoglobulin receptor OS=Homo sapiens OX=9606 GN=PIGR PE=1 SV=4 | 902 | 20201119_16913_1ng_glyco_hcd203040_compound15.23315.23315.3 | controllerType=0 controllerNumber=1 scan=23315 | 48.877 |
| 27340:3 | 6 | R.[+42.011]LSLLEEPGN[+2351.851]GTFTVILNQLTSR.D | HexNAc(4)Hex(5)Fuc(3)NeuAc(1) | .[42], N[+2352] | 1599.3831 | 3 | 4796.1348 | 4796.1539 | 0 | -4 | 413 | Specific | 441.46 | 386.74 | 386.74 | 7.85 | 82 | >sp\|P01833\|PIGR_HUMAN Polymeric immunoglobulin receptor OS=Homo sapiens OX=9606 GN=PIGR PE=1 SV=4 | 902 | 20201119_16913_1ng_glyco_hcd203040_compound15.31425.31425.3 | controllerType=0 controllerNumber=1 scan=31425 | 65.3204 |
| 27338:3 | 6 | R.[+42.011]LSLLEEPGN[+2351.851]GTFTVILNQLTSR.D | HexNAc(4)Hex(5)Fuc(3)NeuAc(1) | .[42], N[+2352] | 1599.3831 | 3 | 4796.1348 | 4796.1539 | 0 | -4 | 413 | Specific | 441.66 | 411.65 | 411.65 | 7.85 | 82 | >sp\|P01833\|PIGR_HUMAN Polymeric immunoglobulin receptor OS=Homo sapiens OX=9606 GN=PIGR PE=1 SV=4 | 902 | 20201119_16913_1ng_glyco_hcd203040_compound15.31423.31423.3 | controllerType=0 controllerNumber=1 scan=31423 | 65.316 |
| 27352:3 | 6 | R.[+42.011]LSLLEEPGN[+2351.851]GTFTVILNQLTSR.D | HexNAc(4)Hex(5)Fuc(3)NeuAc(1) | .[42], N[+2352] | 1599.386 | 3 | 4796.1436 | 4796.1539 | 0 | -2.1 | 413 | Specific | 446.12 | 400.98 | 400.98 | 8.19 | 82 | >sp\|P01833\|PIGR_HUMAN Polymeric immunoglobulin receptor OS=Homo sapiens OX=9606 GN=PIGR PE=1 SV=4 | 902 | 20201119_16913_1ng_glyco_hcd203040_compound15.31438.31438.3 | controllerType=0 controllerNumber=1 scan=31438 | 65.3458 |
| 27354:3 | 6 | R.[+42.011]LSLLEEPGN[+2351.851]GTFTVILNQLTSR.D | HexNAc(4)Hex(5)Fuc(3)NeuAc(1) | .[42], N[+2352] | 1599.386 | 3 | 4796.1436 | 4796.1539 | 0 | -2.1 | 413 | Specific | 467.77 | 278.38 | 278.38 | 6.3 | 82 | >sp\|P01833\|PIGR_HUMAN Polymeric immunoglobulin receptor OS=Homo sapiens OX=9606 GN=PIGR PE=1 SV=4 | 902 | 20201119_16913_1ng_glyco_hcd203040_compound15.31440.31440.3 | controllerType=0 controllerNumber=1 scan=31440 | 65.3509 |
| 21595:3 | 6 | R.[+42.011]LSLLEEPGN[+2336.851]GTFTVILNQLTSR.D | HexNAc(6)Hex(6)Fuc(1) | .[42], N[+2337] | 1594.3797 | 3 | 4781.1245 | 4781.1542 | 0 | -6.2 | 413 | Specific | 472.06 | 472.06 | 472.06 | 7.99 | 82 | >sp\|P01833\|PIGR_HUMAN Polymeric immunoglobulin receptor OS=Homo sapiens OX=9606 GN=PIGR PE=1 SV=4 | 902 | 20201119_16913_1ng_glyco_hcd203040_compound15.25433.25433.3 | controllerType=0 controllerNumber=1 scan=25433 | 53.5028 |
| 24400:4 | 6 | R.LSLLEEPGN[+3007.058]GTFTVILNQLTSR.D | HexNAc(5)Hex(6)Fuc(1)NeuAc(3) | N[+3007] | 1353.0963 | 4 | 5409.3634 | 5409.3505 | 0 | 2.4 | 413 | Specific | 520.06 | 379.28 | 379.28 | 10.02 | 82 | >sp\|P01833\|PIGR_HUMAN Polymeric immunoglobulin receptor OS=Homo sapiens OX=9606 GN=PIGR PE=1 SV=4 | 902 | 20201119_16913_1ng_glyco_hcd203040_compound15.28357.28357.2 | controllerType=0 controllerNumber=1 scan=28357 | 59.5412 |
| 19609:3 | 6 | R.LSLLEEPGN[+2205.793]GTFTVILNQLTSR.D | HexNAc(4)Hex(5)Fuc(2)NeuAc(1) | N[+2206] | 1536.7101 | 3 | 4608.1159 | 4608.0854 | 0 | 6.6 | 413 | Specific | 710.32 | 472.75 | 472.75 | 13.49 | 82 | >sp\|P01833\|PIGR_HUMAN Polymeric immunoglobulin receptor OS=Homo sapiens OX=9606 GN=PIGR PE=1 SV=4 | 902 | 20201119_16913_1ng_glyco_hcd203040_compound15.23353.23353.3 | controllerType=0 controllerNumber=1 scan=23353 | 48.9769 |
| 19652:3 | 6 | R.LSLLEEPGN[+2059.735]GTFTVILNQLTSR.D | HexNAc(4)Hex(5)Fuc(1)NeuAc(1) | N[+2060] | 1488.0179 | 3 | 4462.0391 | 4462.0275 | 0 | 2.6 | 413 | Specific | 747.48 | 613.33 | 613.33 | 14.43 | 82 | >sp\|P01833\|PIGR_HUMAN Polymeric immunoglobulin receptor OS=Homo sapiens OX=9606 GN=PIGR PE=1 SV=4 | 902 | 20201119_16913_1ng_glyco_hcd203040_compound15.23398.23398.3 | controllerType=0 controllerNumber=1 scan=23398 | 49.0815 |
| 19505:4 | 6 | R.LSLLEEPGN[+2205.793]GTFTVILNQLTSR.D | HexNAc(4)Hex(5)Fuc(2)NeuAc(1) | N[+2206] | 1152.782 | 4 | 4608.1063 | 4608.0854 | 0 | 4.5 | 413 | Specific | 822.66 | 655.61 | 655.61 | 16.26 | 82 | >sp\|P01833\|PIGR_HUMAN Polymeric immunoglobulin receptor OS=Homo sapiens OX=9606 GN=PIGR PE=1 SV=4 | 902 | 20201119_16913_1ng_glyco_hcd203040_compound15.23245.23245.3 | controllerType=0 controllerNumber=1 scan=23245 | 48.7256 |
| 21733:3 | 6 | R.LSLLEEPGN[+2350.830]GTFTVILNQLTSR.D | HexNAc(4)Hex(5)Fuc(1)NeuAc(2) | N[+2351] | 1585.0606 | 3 | 4753.1673 | 4753.1229 | 0 | 9.3 | 413 | Specific | 838.89 | 618.4 | 618.4 | 15.07 | 82 | >sp\|P01833\|PIGR_HUMAN Polymeric immunoglobulin receptor OS=Homo sapiens OX=9606 GN=PIGR PE=1 SV=4 | 902 | 20201119_16913_1ng_glyco_hcd203040_compound15.25577.25577.3 | controllerType=0 controllerNumber=1 scan=25577 | 53.8198 |
| 19576:4 | 6 | R.LSLLEEPGN[+2059.735]GTFTVILNQLTSR.D | HexNAc(4)Hex(5)Fuc(1)NeuAc(1) | N[+2060] | 1116.2662 | 4 | 4462.0428 | 4462.0275 | 0 | 3.4 | 413 | Specific | 863.84 | 693.15 | 693.15 | 17.63 | 82 | >sp\|P01833\|PIGR_HUMAN Polymeric immunoglobulin receptor OS=Homo sapiens OX=9606 GN=PIGR PE=1 SV=4 | 902 | 20201119_16913_1ng_glyco_hcd203040_compound15.23319.23319.4 | controllerType=0 controllerNumber=1 scan=23319 | 48.8881 |
| 19508:4 | 6 | R.LSLLEEPGN[+2205.793]GTFTVILNQLTSR.D | HexNAc(4)Hex(5)Fuc(2)NeuAc(1) | N[+2206] | 1152.782 | 4 | 4608.1063 | 4608.0854 | 0 | 4.5 | 413 | Specific | 871.41 | 678.59 | 678.59 | 17.26 | 82 | >sp\|P01833\|PIGR_HUMAN Polymeric immunoglobulin receptor OS=Homo sapiens OX=9606 GN=PIGR PE=1 SV=4 | 902 | 20201119_16913_1ng_glyco_hcd203040_compound15.23248.23248.3 | controllerType=0 controllerNumber=1 scan=23248 | 48.7362 |
| 21735:3 | 6 | R.LSLLEEPGN[+2350.830]GTFTVILNQLTSR.D | HexNAc(4)Hex(5)Fuc(1)NeuAc(2) | N[+2351] | 1585.0606 | 3 | 4753.1673 | 4753.1229 | 0 | 9.3 | 413 | Specific | 873.56 | 657.66 | 657.66 | 16.07 | 82 | >sp\|P01833\|PIGR_HUMAN Polymeric immunoglobulin receptor OS=Homo sapiens OX=9606 GN=PIGR PE=1 SV=4 | 902 | 20201119_16913_1ng_glyco_hcd203040_compound15.25579.25579.3 | controllerType=0 controllerNumber=1 scan=25579 | 53.8251 |
| 19484:3 | 6 | R.LSLLEEPGN[+2205.793]GTFTVILNQLTSR.D | HexNAc(4)Hex(5)Fuc(2)NeuAc(1) | N[+2206] | 1536.7112 | 3 | 4608.1192 | 4608.0854 | 0 | 7.3 | 413 | Specific | 893.82 | 630.18 | 630.18 | 16.49 | 82 | >sp\|P01833\|PIGR_HUMAN Polymeric immunoglobulin receptor OS=Homo sapiens OX=9606 GN=PIGR PE=1 SV=4 | 902 | 20201119_16913_1ng_glyco_hcd203040_compound15.23223.23223.4 | controllerType=0 controllerNumber=1 scan=23223 | 48.682 |
| 19655:3 | 6 | R.LSLLEEPGN[+2059.735]GTFTVILNQLTSR.D | HexNAc(4)Hex(5)Fuc(1)NeuAc(1) | N[+2060] | 1488.0179 | 3 | 4462.0391 | 4462.0275 | 0 | 2.6 | 413 | Specific | 930.92 | 657.1 | 657.1 | 18.43 | 82 | >sp\|P01833\|PIGR_HUMAN Polymeric immunoglobulin receptor OS=Homo sapiens OX=9606 GN=PIGR PE=1 SV=4 | 902 | 20201119_16913_1ng_glyco_hcd203040_compound15.23401.23401.3 | controllerType=0 controllerNumber=1 scan=23401 | 49.0857 |
| 21499:3 | 6 | R.LSLLEEPGN[+2350.830]GTFTVILNQLTSR.D | HexNAc(4)Hex(5)Fuc(1)NeuAc(2) | N[+2351] | 1585.0517 | 3 | 4753.1405 | 4753.1229 | 0 | 3.7 | 413 | Specific | 943.52 | 692.21 | 692.21 | 18.49 | 82 | >sp\|P01833\|PIGR_HUMAN Polymeric immunoglobulin receptor OS=Homo sapiens OX=9606 GN=PIGR PE=1 SV=4 | 902 | 20201119_16913_1ng_glyco_hcd203040_compound15.25332.25332.3 | controllerType=0 controllerNumber=1 scan=25332 | 53.2533 |
| 19481:3 | 6 | R.LSLLEEPGN[+2205.793]GTFTVILNQLTSR.D | HexNAc(4)Hex(5)Fuc(2)NeuAc(1) | N[+2206] | 1536.7112 | 3 | 4608.1192 | 4608.0854 | 0 | 7.3 | 413 | Specific | 980.17 | 877.44 | 877.44 | 18.49 | 82 | >sp\|P01833\|PIGR_HUMAN Polymeric immunoglobulin receptor OS=Homo sapiens OX=9606 GN=PIGR PE=1 SV=4 | 902 | 20201119_16913_1ng_glyco_hcd203040_compound15.23220.23220.4 | controllerType=0 controllerNumber=1 scan=23220 | 48.676 |
| 19575:3 | 6 | R.LSLLEEPGN[+2059.735]GTFTVILNQLTSR.D | HexNAc(4)Hex(5)Fuc(1)NeuAc(1) | N[+2060] | 1488.0184 | 3 | 4462.0405 | 4462.0275 | 0 | 2.9 | 413 | Specific | 1063.54 | 995.94 | 995.94 | 21.49 | 82 | >sp\|P01833\|PIGR_HUMAN Polymeric immunoglobulin receptor OS=Homo sapiens OX=9606 GN=PIGR PE=1 SV=4 | 902 | 20201119_16913_1ng_glyco_hcd203040_compound15.23318.23318.3 | controllerType=0 controllerNumber=1 scan=23318 | 48.8867 |
| 21497:3 | 6 | R.LSLLEEPGN[+2350.830]GTFTVILNQLTSR.D | HexNAc(4)Hex(5)Fuc(1)NeuAc(2) | N[+2351] | 1585.0517 | 3 | 4753.1405 | 4753.1229 | 0 | 3.7 | 413 | Specific | 1114.49 | 866.19 | 866.19 | 22.49 | 82 | >sp\|P01833\|PIGR_HUMAN Polymeric immunoglobulin receptor OS=Homo sapiens OX=9606 GN=PIGR PE=1 SV=4 | 902 | 20201119_16913_1ng_glyco_hcd203040_compound15.25330.25330.3 | controllerType=0 controllerNumber=1 scan=25330 | 53.2459 |
| 15337:4 | 6 | K.IIEGEPNLKVPGN[+2205.793]VTAVLGETLK.V | HexNAc(4)Hex(5)Fuc(2)NeuAc(1) | N[+2206] | 1150.0475 | 4 | 4597.1683 | 4597.1422 | 0 | 5.7 | 457 | Specific | 222.73 | 222.73 | 222.73 | 3.79 | 82 | >sp\|P01833\|PIGR_HUMAN Polymeric immunoglobulin receptor OS=Homo sapiens OX=9606 GN=PIGR PE=1 SV=4 | 902 | 20201119_16913_1ng_glyco_hcd203040_compound15.18889.18889.4 | controllerType=0 controllerNumber=1 scan=18889 | 39.2168 |
| 17174:3 | 6 | K.[+42.011]IIEGEPNLKVPGN[+2336.851]VTAVLGETLK.V | HexNAc(6)Hex(6)Fuc(1) | .[42], N[+2337] | 1590.7347 | 3 | 4770.1895 | 4770.211 | 0 | -4.5 | 457 | Specific | 304.86 | 277.76 | 277.76 | 4.84 | 82 | >sp\|P01833\|PIGR_HUMAN Polymeric immunoglobulin receptor OS=Homo sapiens OX=9606 GN=PIGR PE=1 SV=4 | 902 | 20201119_16913_1ng_glyco_hcd203040_compound15.20808.20808.3 | controllerType=0 controllerNumber=1 scan=20808 | 43.3742 |
| 17171:3 | 6 | K.[+42.011]IIEGEPNLKVPGN[+2336.851]VTAVLGETLK.V | HexNAc(6)Hex(6)Fuc(1) | .[42], N[+2337] | 1590.7347 | 3 | 4770.1895 | 4770.211 | 0 | -4.5 | 457 | Specific | 373.07 | 349.1 | 349.1 | 5.58 | 82 | >sp\|P01833\|PIGR_HUMAN Polymeric immunoglobulin receptor OS=Homo sapiens OX=9606 GN=PIGR PE=1 SV=4 | 902 | 20201119_16913_1ng_glyco_hcd203040_compound15.20805.20805.3 | controllerType=0 controllerNumber=1 scan=20805 | 43.3637 |
| 15288:3 | 6 | K.IIEGEPNLKVPGN[+2205.793]VTAVLGETLK.V | HexNAc(4)Hex(5)Fuc(2)NeuAc(1) | N[+2206] | 1533.056 | 3 | 4597.1533 | 4597.1422 | 0 | 2.4 | 457 | Specific | 404.41 | 404.41 | 404.41 | 9.03 | 82 | >sp\|P01833\|PIGR_HUMAN Polymeric immunoglobulin receptor OS=Homo sapiens OX=9606 GN=PIGR PE=1 SV=4 | 902 | 20201119_16913_1ng_glyco_hcd203040_compound15.18837.18837.3 | controllerType=0 controllerNumber=1 scan=18837 | 39.1327 |
| 15350:4 | 6 | K.IIEGEPNLKVPGN[+2059.735]VTAVLGETLK.V | HexNAc(4)Hex(5)Fuc(1)NeuAc(1) | N[+2060] | 1113.5319 | 4 | 4451.1058 | 4451.0843 | 0 | 4.8 | 457 | Specific | 462.93 | 462.93 | 462.93 | 8.78 | 82 | >sp\|P01833\|PIGR_HUMAN Polymeric immunoglobulin receptor OS=Homo sapiens OX=9606 GN=PIGR PE=1 SV=4 | 902 | 20201119_16913_1ng_glyco_hcd203040_compound15.18902.18902.3 | controllerType=0 controllerNumber=1 scan=18902 | 39.2401 |
| 17119:3 | 6 | K.IIEGEPNLKVPGN[+2350.830]VTAVLGETLK.V | HexNAc(4)Hex(5)Fuc(1)NeuAc(2) | N[+2351] | 1581.4034 | 3 | 4742.1956 | 4742.1797 | 0 | 3.3 | 457 | Specific | 680.58 | 680.58 | 680.58 | 13.02 | 82 | >sp\|P01833\|PIGR_HUMAN Polymeric immunoglobulin receptor OS=Homo sapiens OX=9606 GN=PIGR PE=1 SV=4 | 902 | 20201119_16913_1ng_glyco_hcd203040_compound15.20751.20751.4 | controllerType=0 controllerNumber=1 scan=20751 | 43.2414 |
| 15379:4 | 6 | K.IIEGEPNLKVPGN[+2059.735]VTAVLGETLK.V | HexNAc(4)Hex(5)Fuc(1)NeuAc(1) | N[+2060] | 1113.529 | 4 | 4451.0941 | 4451.0843 | 0 | 2.2 | 457 | Specific | 690.84 | 690.84 | 690.84 | 12.86 | 82 | >sp\|P01833\|PIGR_HUMAN Polymeric immunoglobulin receptor OS=Homo sapiens OX=9606 GN=PIGR PE=1 SV=4 | 902 | 20201119_16913_1ng_glyco_hcd203040_compound15.18932.18932.4 | controllerType=0 controllerNumber=1 scan=18932 | 39.298 |
| 15353:4 | 6 | K.IIEGEPNLKVPGN[+2059.735]VTAVLGETLK.V | HexNAc(4)Hex(5)Fuc(1)NeuAc(1) | N[+2060] | 1113.5319 | 4 | 4451.1058 | 4451.0843 | 0 | 4.8 | 457 | Specific | 696.46 | 675.3 | 675.3 | 12.56 | 82 | >sp\|P01833\|PIGR_HUMAN Polymeric immunoglobulin receptor OS=Homo sapiens OX=9606 GN=PIGR PE=1 SV=4 | 902 | 20201119_16913_1ng_glyco_hcd203040_compound15.18905.18905.3 | controllerType=0 controllerNumber=1 scan=18905 | 39.2446 |
| 15339:4 | 6 | K.IIEGEPNLKVPGN[+2205.793]VTAVLGETLK.V | HexNAc(4)Hex(5)Fuc(2)NeuAc(1) | N[+2206] | 1150.0475 | 4 | 4597.1683 | 4597.1422 | 0 | 5.7 | 457 | Specific | 723.97 | 723.97 | 723.97 | 13.22 | 82 | >sp\|P01833\|PIGR_HUMAN Polymeric immunoglobulin receptor OS=Homo sapiens OX=9606 GN=PIGR PE=1 SV=4 | 902 | 20201119_16913_1ng_glyco_hcd203040_compound15.18891.18891.4 | controllerType=0 controllerNumber=1 scan=18891 | 39.222 |
| 17094:3 | 6 | K.IIEGEPNLKVPGN[+2350.830]VTAVLGETLK.V | HexNAc(4)Hex(5)Fuc(1)NeuAc(2) | N[+2351] | 1581.4011 | 3 | 4742.1886 | 4742.1797 | 0 | 1.9 | 457 | Specific | 773.01 | 743.76 | 743.76 | 14.96 | 82 | >sp\|P01833\|PIGR_HUMAN Polymeric immunoglobulin receptor OS=Homo sapiens OX=9606 GN=PIGR PE=1 SV=4 | 902 | 20201119_16913_1ng_glyco_hcd203040_compound15.20724.20724.3 | controllerType=0 controllerNumber=1 scan=20724 | 43.1766 |
| 15291:3 | 6 | K.IIEGEPNLKVPGN[+2205.793]VTAVLGETLK.V | HexNAc(4)Hex(5)Fuc(2)NeuAc(1) | N[+2206] | 1533.056 | 3 | 4597.1533 | 4597.1422 | 0 | 2.4 | 457 | Specific | 781.42 | 701.87 | 701.87 | 14.96 | 82 | >sp\|P01833\|PIGR_HUMAN Polymeric immunoglobulin receptor OS=Homo sapiens OX=9606 GN=PIGR PE=1 SV=4 | 902 | 20201119_16913_1ng_glyco_hcd203040_compound15.18840.18840.3 | controllerType=0 controllerNumber=1 scan=18840 | 39.1363 |
| 15338:3 | 6 | K.IIEGEPNLKVPGN[+2059.735]VTAVLGETLK.V | HexNAc(4)Hex(5)Fuc(1)NeuAc(1) | N[+2060] | 1484.3746 | 3 | 4451.1092 | 4451.0843 | 0 | 5.6 | 457 | Specific | 840.22 | 767.78 | 767.78 | 15.31 | 82 | >sp\|P01833\|PIGR_HUMAN Polymeric immunoglobulin receptor OS=Homo sapiens OX=9606 GN=PIGR PE=1 SV=4 | 902 | 20201119_16913_1ng_glyco_hcd203040_compound15.18890.18890.3 | controllerType=0 controllerNumber=1 scan=18890 | 39.2198 |
| 17104:4 | 6 | K.IIEGEPNLKVPGN[+2350.830]VTAVLGETLK.V | HexNAc(4)Hex(5)Fuc(1)NeuAc(2) | N[+2351] | 1186.3031 | 4 | 4742.1907 | 4742.1797 | 0 | 2.3 | 457 | Specific | 849.95 | 797.27 | 797.27 | 15.86 | 82 | >sp\|P01833\|PIGR_HUMAN Polymeric immunoglobulin receptor OS=Homo sapiens OX=9606 GN=PIGR PE=1 SV=4 | 902 | 20201119_16913_1ng_glyco_hcd203040_compound15.20735.20735.4 | controllerType=0 controllerNumber=1 scan=20735 | 43.2047 |
| 15329:3 | 6 | K.IIEGEPNLKVPGN[+2205.793]VTAVLGETLK.V | HexNAc(4)Hex(5)Fuc(2)NeuAc(1) | N[+2206] | 1533.0604 | 3 | 4597.1665 | 4597.1422 | 0 | 5.3 | 457 | Specific | 860.06 | 841.26 | 841.26 | 16.31 | 82 | >sp\|P01833\|PIGR_HUMAN Polymeric immunoglobulin receptor OS=Homo sapiens OX=9606 GN=PIGR PE=1 SV=4 | 902 | 20201119_16913_1ng_glyco_hcd203040_compound15.18881.18881.3 | controllerType=0 controllerNumber=1 scan=18881 | 39.2046 |
| 15331:3 | 6 | K.IIEGEPNLKVPGN[+2205.793]VTAVLGETLK.V | HexNAc(4)Hex(5)Fuc(2)NeuAc(1) | N[+2206] | 1533.0604 | 3 | 4597.1665 | 4597.1422 | 0 | 5.3 | 457 | Specific | 862.08 | 842.1 | 842.1 | 16.31 | 82 | >sp\|P01833\|PIGR_HUMAN Polymeric immunoglobulin receptor OS=Homo sapiens OX=9606 GN=PIGR PE=1 SV=4 | 902 | 20201119_16913_1ng_glyco_hcd203040_compound15.18883.18883.3 | controllerType=0 controllerNumber=1 scan=18883 | 39.2092 |
| 17122:3 | 6 | K.IIEGEPNLKVPGN[+2350.830]VTAVLGETLK.V | HexNAc(4)Hex(5)Fuc(1)NeuAc(2) | N[+2351] | 1581.4034 | 3 | 4742.1956 | 4742.1797 | 0 | 3.3 | 457 | Specific | 874.16 | 704.34 | 704.34 | 17.02 | 82 | >sp\|P01833\|PIGR_HUMAN Polymeric immunoglobulin receptor OS=Homo sapiens OX=9606 GN=PIGR PE=1 SV=4 | 902 | 20201119_16913_1ng_glyco_hcd203040_compound15.20754.20754.4 | controllerType=0 controllerNumber=1 scan=20754 | 43.246 |
| 17092:3 | 6 | K.IIEGEPNLKVPGN[+2350.830]VTAVLGETLK.V | HexNAc(4)Hex(5)Fuc(1)NeuAc(2) | N[+2351] | 1581.4011 | 3 | 4742.1886 | 4742.1797 | 0 | 1.9 | 457 | Specific | 957.7 | 885.49 | 885.49 | 18.96 | 82 | >sp\|P01833\|PIGR_HUMAN Polymeric immunoglobulin receptor OS=Homo sapiens OX=9606 GN=PIGR PE=1 SV=4 | 902 | 20201119_16913_1ng_glyco_hcd203040_compound15.20722.20722.3 | controllerType=0 controllerNumber=1 scan=20722 | 43.1668 |
| 15335:3 | 6 | K.IIEGEPNLKVPGN[+2059.735]VTAVLGETLK.V | HexNAc(4)Hex(5)Fuc(1)NeuAc(1) | N[+2060] | 1484.3746 | 3 | 4451.1092 | 4451.0843 | 0 | 5.6 | 457 | Specific | 1070.22 | 831.56 | 831.56 | 20.31 | 82 | >sp\|P01833\|PIGR_HUMAN Polymeric immunoglobulin receptor OS=Homo sapiens OX=9606 GN=PIGR PE=1 SV=4 | 902 | 20201119_16913_1ng_glyco_hcd203040_compound15.18887.18887.3 | controllerType=0 controllerNumber=1 scan=18887 | 39.2145 |
| 17101:4 | 6 | K.IIEGEPNLKVPGN[+2350.830]VTAVLGETLK.V | HexNAc(4)Hex(5)Fuc(1)NeuAc(2) | N[+2351] | 1186.3031 | 4 | 4742.1907 | 4742.1797 | 0 | 2.3 | 457 | Specific | 1079.81 | 1027.34 | 1027.34 | 20.86 | 82 | >sp\|P01833\|PIGR_HUMAN Polymeric immunoglobulin receptor OS=Homo sapiens OX=9606 GN=PIGR PE=1 SV=4 | 902 | 20201119_16913_1ng_glyco_hcd203040_compound15.20732.20732.4 | controllerType=0 controllerNumber=1 scan=20732 | 43.1895 |
| 16291:5 | 6 | K.IIEGEPNLKVPGN[+2350.830]VTAVLGETLKVPC[+57.021]HFPC[+57.021]K.F | HexNAc(4)Hex(5)Fuc(1)NeuAc(2) | C[+57]*2, N[+2351] | 1154.3402 | 5 | 5767.672 | 5767.6372 | 0 | 6 | 457 | Specific | 662.02 | 588.99 | 588.99 | 11.89 | 82 | >sp\|P01833\|PIGR_HUMAN Polymeric immunoglobulin receptor OS=Homo sapiens OX=9606 GN=PIGR PE=1 SV=4 | 902 | 20201119_16913_1ng_glyco_hcd203040_compound15.19885.19885.5 | controllerType=0 controllerNumber=1 scan=19885 | 41.2276 |
| 14964:4 | 6 | K.IIEGEPNLKVPGN[+2205.793]VTAVLGETLKVPC[+57.021]HFPC[+57.021]K.F | HexNAc(4)Hex(5)Fuc(2)NeuAc(1) | C[+57]*2, N[+2206] | 1406.4154 | 4 | 5622.6398 | 5622.5997 | 0 | 7.1 | 457 | Specific | 662.72 | 662.72 | 662.72 | 10.92 | 82 | >sp\|P01833\|PIGR_HUMAN Polymeric immunoglobulin receptor OS=Homo sapiens OX=9606 GN=PIGR PE=1 SV=4 | 902 | 20201119_16913_1ng_glyco_hcd203040_compound15.18480.18480.2 | controllerType=0 controllerNumber=1 scan=18480 | 38.3933 |
| 16293:4 | 6 | K.IIEGEPNLKVPGN[+2350.830]VTAVLGETLKVPC[+57.021]HFPC[+57.021]K.F | HexNAc(4)Hex(5)Fuc(1)NeuAc(2) | C[+57]*2, N[+2351] | 1442.6747 | 4 | 5767.6769 | 5767.6372 | 0 | 6.9 | 457 | Specific | 676.3 | 676.3 | 676.3 | 10.92 | 82 | >sp\|P01833\|PIGR_HUMAN Polymeric immunoglobulin receptor OS=Homo sapiens OX=9606 GN=PIGR PE=1 SV=4 | 902 | 20201119_16913_1ng_glyco_hcd203040_compound15.19887.19887.4 | controllerType=0 controllerNumber=1 scan=19887 | 41.2323 |
| 14966:4 | 6 | K.IIEGEPNLKVPGN[+2205.793]VTAVLGETLKVPC[+57.021]HFPC[+57.021]K.F | HexNAc(4)Hex(5)Fuc(2)NeuAc(1) | C[+57]*2, N[+2206] | 1406.4154 | 4 | 5622.6398 | 5622.5997 | 0 | 7.1 | 457 | Specific | 708.97 | 708.97 | 708.97 | 11.92 | 82 | >sp\|P01833\|PIGR_HUMAN Polymeric immunoglobulin receptor OS=Homo sapiens OX=9606 GN=PIGR PE=1 SV=4 | 902 | 20201119_16913_1ng_glyco_hcd203040_compound15.18482.18482.2 | controllerType=0 controllerNumber=1 scan=18482 | 38.3996 |

Table S4. Cont.

| Query #:z | Protein Rank | Glycopeptide | Glycan | Modification Type(s) | Observed m/z | z | Observed (M+H) | Calc. mass (M+H) | Off-by-x error | Mass error (ppm) | StartPosition | Cleavage | Score | Delta | Delta Mod | \|Log Prob\| | # of unique peptides | Protein Name | Protein DB number | Comment | Scan # | Scan Time |
|---|---|---|---|---|---|---|---|---|---|---|---|---|---|---|---|---|---|---|---|---|---|---|
| 16279:5 | 6 | K.IIEGEPNLKVPGN[+2350.830]VTAVLGETLKVPC[+57.021]HFPC[+57.021]K.F | HexNAc(4)Hex(5)Fuc(1)NeuAc(2) | C[+57]*2, N[+2351] | 1154.337 | 5 | 5767.6561 | 5767.6372 | 0 | 3.3 | 457 | Specific | 741.3 | 583.69 | 583.69 | 13.6 | 82 | >sp\|P01833\|PIGR_HUMAN Polymeric immunoglobulin receptor OS=Homo sapiens OX=9606 GN=PIGR PE=1 SV=4 | 902 | 20201119_16913_1ng_glyco_hcd203040_compound15.19872.19872.5 | controllerType=0 controllerNumber=1 scan=19872 | 41.2026 |
| 16294:5 | 6 | K.IIEGEPNLKVPGN[+2350.830]VTAVLGETLKVPC[+57.021]HFPC[+57.021]K.F | HexNAc(4)Hex(5)Fuc(1)NeuAc(2) | C[+57]*2, N[+2351] | 1154.3402 | 5 | 5767.672 | 5767.6372 | 0 | 6 | 457 | Specific | 768.81 | 621.86 | 621.86 | 13.89 | 82 | >sp\|P01833\|PIGR_HUMAN Polymeric immunoglobulin receptor OS=Homo sapiens OX=9606 GN=PIGR PE=1 SV=4 | 902 | 20201119_16913_1ng_glyco_hcd203040_compound15.19888.19888.5 | controllerType=0 controllerNumber=1 scan=19888 | 41.2342 |
| 14907:4 | 6 | K.IIEGEPNLKVPGN[+2205.793]VTAVLGETLKVPC[+57.021]HFPC[+57.021]K.F | HexNAc(4)Hex(5)Fuc(2)NeuAc(1) | C[+57]*2, N[+2206] | 1406.4128 | 4 | 5622.6296 | 5622.5997 | 0 | 5.3 | 457 | Specific | 796.35 | 796.35 | 796.35 | 13.21 | 82 | >sp\|P01833\|PIGR_HUMAN Polymeric immunoglobulin receptor OS=Homo sapiens OX=9606 GN=PIGR PE=1 SV=4 | 902 | 20201119_16913_1ng_glyco_hcd203040_compound15.18421.18421.4 | controllerType=0 controllerNumber=1 scan=18421 | 38.2769 |
| 14902:5 | 6 | K.IIEGEPNLKVPGN[+2205.793]VTAVLGETLKVPC[+57.021]HFPC[+57.021]K.F | HexNAc(4)Hex(5)Fuc(2)NeuAc(1) | C[+57]*2, N[+2206] | 1125.3315 | 5 | 5622.6286 | 5622.5997 | 0 | 5.1 | 457 | Specific | 884.05 | 809.93 | 809.93 | 16.23 | 82 | >sp\|P01833\|PIGR_HUMAN Polymeric immunoglobulin receptor OS=Homo sapiens OX=9606 GN=PIGR PE=1 SV=4 | 902 | 20201119_16913_1ng_glyco_hcd203040_compound15.18416.18416.5 | controllerType=0 controllerNumber=1 scan=18416 | 38.2663 |
| 16277:5 | 6 | K.IIEGEPNLKVPGN[+2350.830]VTAVLGETLKVPC[+57.021]HFPC[+57.021]K.F | HexNAc(4)Hex(5)Fuc(1)NeuAc(2) | C[+57]*2, N[+2351] | 1154.337 | 5 | 5767.6561 | 5767.6372 | 0 | 3.3 | 457 | Specific | 919.53 | 678.97 | 678.97 | 17.6 | 82 | >sp\|P01833\|PIGR_HUMAN Polymeric immunoglobulin receptor OS=Homo sapiens OX=9606 GN=PIGR PE=1 SV=4 | 902 | 20201119_16913_1ng_glyco_hcd203040_compound15.19870.19870.5 | controllerType=0 controllerNumber=1 scan=19870 | 41.1953 |
| 14899:5 | 6 | K.IIEGEPNLKVPGN[+2205.793]VTAVLGETLKVPC[+57.021]HFPC[+57.021]K.F | HexNAc(4)Hex(5)Fuc(2)NeuAc(1) | C[+57]*2, N[+2206] | 1125.3315 | 5 | 5622.6286 | 5622.5997 | 0 | 5.1 | 457 | Specific | 964.44 | 916.48 | 916.48 | 18.23 | 82 | >sp\|P01833\|PIGR_HUMAN Polymeric immunoglobulin receptor OS=Homo sapiens OX=9606 GN=PIGR PE=1 SV=4 | 902 | 20201119_16913_1ng_glyco_hcd203040_compound15.18413.18413.5 | controllerType=0 controllerNumber=1 scan=18413 | 38.2597 |
| 16290:4 | 6 | K.IIEGEPNLKVPGN[+2350.830]VTAVLGETLKVPC[+57.021]HFPC[+57.021]K.F | HexNAc(4)Hex(5)Fuc(1)NeuAc(2) | C[+57]*2, N[+2351] | 1442.6747 | 4 | 5767.6769 | 5767.6372 | 0 | 6.9 | 457 | Specific | 1016.55 | 731.15 | 731.15 | 17.92 | 82 | >sp\|P01833\|PIGR_HUMAN Polymeric immunoglobulin receptor OS=Homo sapiens OX=9606 GN=PIGR PE=1 SV=4 | 902 | 20201119_16913_1ng_glyco_hcd203040_compound15.19884.19884.4 | controllerType=0 controllerNumber=1 scan=19884 | 41.2265 |
| 16278:4 | 6 | K.IIEGEPNLKVPGN[+2350.830]VTAVLGETLKVPC[+57.021]HFPC[+57.021]K.F | HexNAc(4)Hex(5)Fuc(1)NeuAc(2) | C[+57]*2, N[+2351] | 1442.6707 | 4 | 5767.6608 | 5767.6372 | 0 | 4.1 | 457 | Specific | 1022.21 | 777.19 | 777.19 | 18.55 | 82 | >sp\|P01833\|PIGR_HUMAN Polymeric immunoglobulin receptor OS=Homo sapiens OX=9606 GN=PIGR PE=1 SV=4 | 902 | 20201119_16913_1ng_glyco_hcd203040_compound15.19871.19871.4 | controllerType=0 controllerNumber=1 scan=19871 | 41.1968 |
| 14904:4 | 6 | K.IIEGEPNLKVPGN[+2205.793]VTAVLGETLKVPC[+57.021]HFPC[+57.021]K.F | HexNAc(4)Hex(5)Fuc(2)NeuAc(1) | C[+57]*2, N[+2206] | 1406.4128 | 4 | 5622.6296 | 5622.5997 | 0 | 5.3 | 457 | Specific | 1064.5 | 929.73 | 929.73 | 19.21 | 82 | >sp\|P01833\|PIGR_HUMAN Polymeric immunoglobulin receptor OS=Homo sapiens OX=9606 GN=PIGR PE=1 SV=4 | 902 | 20201119_16913_1ng_glyco_hcd203040_compound15.18418.18418.4 | controllerType=0 controllerNumber=1 scan=18418 | 38.271 |
| 16035:3 | 6 | K.VPGN[+2350.830]VTAVLGETLK.V | HexNAc(4)Hex(5)Fuc(1)NeuAc(2) | N[+2351] | 1250.2215 | 3 | 3748.6499 | 3748.6302 | 0 | 5.3 | 466 | Specific | 401.13 | 277.91 | 277.91 | 6.66 | 82 | >sp\|P01833\|PIGR_HUMAN Polymeric immunoglobulin receptor OS=Homo sapiens OX=9606 GN=PIGR PE=1 SV=4 | 902 | 20201119_16913_1ng_glyco_hcd203040_compound15.19618.19618.3 | controllerType=0 controllerNumber=1 scan=19618 | 40.6473 |
| 12944:3 | 6 | K.VPGN[+2351.851]VTAVLGETLK.V | HexNAc(4)Hex(5)Fuc(3)NeuAc(1) | N[+2352] | 1250.5611 | 3 | 3749.6687 | 3749.6506 | 0 | 4.8 | 466 | Specific | 539.11 | 501.43 | 501.43 | 9.38 | 82 | >sp\|P01833\|PIGR_HUMAN Polymeric immunoglobulin receptor OS=Homo sapiens OX=9606 GN=PIGR PE=1 SV=4 | 902 | 20201119_16913_1ng_glyco_hcd203040_compound15.16379.16379.3 | controllerType=0 controllerNumber=1 scan=16379 | 34.2931 |
| 12942:3 | 6 | K.VPGN[+2351.851]VTAVLGETLK.V | HexNAc(4)Hex(5)Fuc(3)NeuAc(1) | N[+2352] | 1250.5611 | 3 | 3749.6687 | 3749.6506 | 0 | 4.8 | 466 | Specific | 571.34 | 571.34 | 571.34 | 9.66 | 82 | >sp\|P01833\|PIGR_HUMAN Polymeric immunoglobulin receptor OS=Homo sapiens OX=9606 GN=PIGR PE=1 SV=4 | 902 | 20201119_16913_1ng_glyco_hcd203040_compound15.16377.16377.3 | controllerType=0 controllerNumber=1 scan=16377 | 34.289 |
| 13004:3 | 6 | K.VPGN[+2205.793]VTAVLGETLK.V | HexNAc(4)Hex(5)Fuc(2)NeuAc(1) | N[+2206] | 1201.8729 | 3 | 3603.604 | 3603.5927 | 0 | 3.1 | 466 | Specific | 647.28 | 519.15 | 519.15 | 12.27 | 82 | >sp\|P01833\|PIGR_HUMAN Polymeric immunoglobulin receptor OS=Homo sapiens OX=9606 GN=PIGR PE=1 SV=4 | 902 | 20201119_16913_1ng_glyco_hcd203040_compound15.16442.16442.3 | controllerType=0 controllerNumber=1 scan=16442 | 34.423 |
| 15971:3 | 6 | K.VPGN[+2350.830]VTAVLGETLK.V | HexNAc(4)Hex(5)Fuc(1)NeuAc(2) | N[+2351] | 1250.2193 | 3 | 3748.6433 | 3748.6302 | 0 | 3.5 | 466 | Specific | 697.04 | 517.26 | 517.26 | 13.27 | 82 | >sp\|P01833\|PIGR_HUMAN Polymeric immunoglobulin receptor OS=Homo sapiens OX=9606 GN=PIGR PE=1 SV=4 | 902 | 20201119_16913_1ng_glyco_hcd203040_compound15.19551.19551.3 | controllerType=0 controllerNumber=1 scan=19551 | 40.5116 |
| 15974:3 | 6 | K.VPGN[+2350.830]VTAVLGETLK.V | HexNAc(4)Hex(5)Fuc(1)NeuAc(2) | N[+2351] | 1250.2193 | 3 | 3748.6433 | 3748.6302 | 0 | 3.5 | 466 | Specific | 733.06 | 547.72 | 547.72 | 14.27 | 82 | >sp\|P01833\|PIGR_HUMAN Polymeric immunoglobulin receptor OS=Homo sapiens OX=9606 GN=PIGR PE=1 SV=4 | 902 | 20201119_16913_1ng_glyco_hcd203040_compound15.19554.19554.3 | controllerType=0 controllerNumber=1 scan=19554 | 40.5173 |
| 13001:3 | 6 | K.VPGN[+2205.793]VTAVLGETLK.V | HexNAc(4)Hex(5)Fuc(2)NeuAc(1) | N[+2206] | 1201.8729 | 3 | 3603.604 | 3603.5927 | 0 | 3.1 | 466 | Specific | 758.35 | 738.75 | 738.75 | 15.27 | 82 | >sp\|P01833\|PIGR_HUMAN Polymeric immunoglobulin receptor OS=Homo sapiens OX=9606 GN=PIGR PE=1 SV=4 | 902 | 20201119_16913_1ng_glyco_hcd203040_compound15.16439.16439.3 | controllerType=0 controllerNumber=1 scan=16439 | 34.417 |
| 13087:3 | 6 | K.VPGN[+2059.735]VTAVLGETLK.V | HexNAc(4)Hex(5)Fuc(1)NeuAc(1) | N[+2060] | 1153.1865 | 3 | 3457.5448 | 3457.5348 | 0 | 2.9 | 466 | Specific | 790.45 | 523.72 | 523.72 | 15.27 | 82 | >sp\|P01833\|PIGR_HUMAN Polymeric immunoglobulin receptor OS=Homo sapiens OX=9606 GN=PIGR PE=1 SV=4 | 902 | 20201119_16913_1ng_glyco_hcd203040_compound15.16529.16529.3 | controllerType=0 controllerNumber=1 scan=16529 | 34.6272 |
| 13084:3 | 6 | K.VPGN[+2059.735]VTAVLGETLK.V | HexNAc(4)Hex(5)Fuc(1)NeuAc(1) | N[+2060] | 1153.1865 | 3 | 3457.5448 | 3457.5348 | 0 | 2.9 | 466 | Specific | 807.18 | 613.12 | 613.12 | 16.27 | 82 | >sp\|P01833\|PIGR_HUMAN Polymeric immunoglobulin receptor OS=Homo sapiens OX=9606 GN=PIGR PE=1 SV=4 | 902 | 20201119_16913_1ng_glyco_hcd203040_compound15.16526.16526.3 | controllerType=0 controllerNumber=1 scan=16526 | 34.6192 |
| 14986:3 | 6 | K.VPGN[+2350.830]VTAVLGETLKVPC[+57.021]HFPC[+57.021]K.F | HexNAc(4)Hex(5)Fuc(1)NeuAc(2) | C[+57]*2, N[+2351] | 1592.0436 | 3 | 4774.1164 | 4774.0877 | 0 | 6 | 466 | Specific | 339.59 | 222.71 | 222.71 | 3.38 | 82 | >sp\|P01833\|PIGR_HUMAN Polymeric immunoglobulin receptor OS=Homo sapiens OX=9606 GN=PIGR PE=1 SV=4 | 902 | 20201119_16913_1ng_glyco_hcd203040_compound15.18503.18503.3 | controllerType=0 controllerNumber=1 scan=18503 | 38.4421 |
| 14984:3 | 6 | K.VPGN[+2350.830]VTAVLGETLKVPC[+57.021]HFPC[+57.021]K.F | HexNAc(4)Hex(5)Fuc(1)NeuAc(2) | C[+57]*2, N[+2351] | 1592.0436 | 3 | 4774.1164 | 4774.0877 | 0 | 6 | 466 | Specific | 404.49 | 303.87 | 303.87 | 5.85 | 82 | >sp\|P01833\|PIGR_HUMAN Polymeric immunoglobulin receptor OS=Homo sapiens OX=9606 GN=PIGR PE=1 SV=4 | 902 | 20201119_16913_1ng_glyco_hcd203040_compound15.18501.18501.3 | controllerType=0 controllerNumber=1 scan=18501 | 38.4385 |
| 14976:4 | 6 | K.VPGN[+2350.830]VTAVLGETLKVPC[+57.021]HFPC[+57.021]K.F | HexNAc(4)Hex(5)Fuc(1)NeuAc(2) | C[+57]*2, N[+2351] | 1194.2791 | 4 | 4774.0946 | 4774.0877 | 0 | 1.4 | 466 | Specific | 535.3 | 279.01 | 279.01 | 7.18 | 82 | >sp\|P01833\|PIGR_HUMAN Polymeric immunoglobulin receptor OS=Homo sapiens OX=9606 GN=PIGR PE=1 SV=4 | 902 | 20201119_16913_1ng_glyco_hcd203040_compound15.18493.18493.4 | controllerType=0 controllerNumber=1 scan=18493 | 38.421 |
| 13167:4 | 6 | K.VPGN[+2059.735]VTAVLGETLKVPC[+57.021]HFPC[+57.021]K.F | HexNAc(4)Hex(5)Fuc(1)NeuAc(1) | C[+57]*2, N[+2060] | 1121.5057 | 4 | 4483.0008 | 4482.9923 | 0 | 1.9 | 466 | Specific | 633.42 | 542.96 | 542.96 | 11.96 | 82 | >sp\|P01833\|PIGR_HUMAN Polymeric immunoglobulin receptor OS=Homo sapiens OX=9606 GN=PIGR PE=1 SV=4 | 902 | 20201119_16913_1ng_glyco_hcd203040_compound15.16612.16612.4 | controllerType=0 controllerNumber=1 scan=16612 | 34.8129 |
| 13165:4 | 6 | K.VPGN[+2059.735]VTAVLGETLKVPC[+57.021]HFPC[+57.021]K.F | HexNAc(4)Hex(5)Fuc(1)NeuAc(1) | C[+57]*2, N[+2060] | 1121.5057 | 4 | 4483.0008 | 4482.9923 | 0 | 1.9 | 466 | Specific | 642.38 | 527.6 | 527.6 | 11.96 | 82 | >sp\|P01833\|PIGR_HUMAN Polymeric immunoglobulin receptor OS=Homo sapiens OX=9606 GN=PIGR PE=1 SV=4 | 902 | 20201119_16913_1ng_glyco_hcd203040_compound15.16610.16610.4 | controllerType=0 controllerNumber=1 scan=16610 | 34.8084 |
| 13032:4 | 6 | K.VPGN[+2205.793]VTAVLGETLKVPC[+57.021]HFPC[+57.021]K.F | HexNAc(4)Hex(5)Fuc(2)NeuAc(1) | C[+57]*2, N[+2206] | 1158.024 | 4 | 4629.074 | 4629.0502 | 0 | 5.1 | 466 | Specific | 655.61 | 630.11 | 630.11 | 12.65 | 82 | >sp\|P01833\|PIGR_HUMAN Polymeric immunoglobulin receptor OS=Homo sapiens OX=9606 GN=PIGR PE=1 SV=4 | 902 | 20201119_16913_1ng_glyco_hcd203040_compound15.16471.16471.4 | controllerType=0 controllerNumber=1 scan=16471 | 34.4995 |
| 14974:4 | 6 | K.VPGN[+2350.830]VTAVLGETLKVPC[+57.021]HFPC[+57.021]K.F | HexNAc(4)Hex(5)Fuc(1)NeuAc(2) | C[+57]*2, N[+2351] | 1194.2791 | 4 | 4774.0946 | 4774.0877 | 0 | 1.4 | 466 | Specific | 727.08 | 606.64 | 606.64 | 13.69 | 82 | >sp\|P01833\|PIGR_HUMAN Polymeric immunoglobulin receptor OS=Homo sapiens OX=9606 GN=PIGR PE=1 SV=4 | 902 | 20201119_16913_1ng_glyco_hcd203040_compound15.18491.18491.4 | controllerType=0 controllerNumber=1 scan=18491 | 38.4161 |
| 14885:3 | 6 | K.VPGN[+2350.830]VTAVLGETLKVPC[+57.021]HFPC[+57.021]K.F | HexNAc(4)Hex(5)Fuc(1)NeuAc(2) | C[+57]*2, N[+2351] | 1592.0474 | 3 | 4774.1277 | 4774.0877 | 0 | 8.4 | 466 | Specific | 727.31 | 628.77 | 628.77 | 12 | 82 | >sp\|P01833\|PIGR_HUMAN Polymeric immunoglobulin receptor OS=Homo sapiens OX=9606 GN=PIGR PE=1 SV=4 | 902 | 20201119_16913_1ng_glyco_hcd203040_compound15.18398.18398.3 | controllerType=0 controllerNumber=1 scan=18398 | 38.2301 |
| 13029:4 | 6 | K.VPGN[+2205.793]VTAVLGETLKVPC[+57.021]HFPC[+57.021]K.F | HexNAc(4)Hex(5)Fuc(2)NeuAc(1) | C[+57]*2, N[+2206] | 1158.024 | 4 | 4629.074 | 4629.0502 | 0 | 5.1 | 466 | Specific | 769.24 | 624.17 | 624.17 | 14.65 | 82 | >sp\|P01833\|PIGR_HUMAN Polymeric immunoglobulin receptor OS=Homo sapiens OX=9606 GN=PIGR PE=1 SV=4 | 902 | 20201119_16913_1ng_glyco_hcd203040_compound15.16468.16468.4 | controllerType=0 controllerNumber=1 scan=16468 | 34.4954 |
| 14839:4 | 6 | K.VPGN[+2351.851]VTAVLGETLKVPC[+57.021]HFPC[+57.021]K.F | HexNAc(4)Hex(5)Fuc(3)NeuAc(1) | C[+57]*2, N[+2352] | 1194.5326 | 4 | 4775.1087 | 4775.1081 | 0 | 0.1 | 466 | Specific | 788.07 | 788.07 | 788.07 | 14.14 | 82 | >sp\|P01833\|PIGR_HUMAN Polymeric immunoglobulin receptor OS=Homo sapiens OX=9606 GN=PIGR PE=1 SV=4 | 902 | 20201119_16913_1ng_glyco_hcd203040_compound15.18350.18350.4 | controllerType=0 controllerNumber=1 scan=18350 | 38.1277 |
| 14883:3 | 6 | K.VPGN[+2350.830]VTAVLGETLKVPC[+57.021]HFPC[+57.021]K.F | HexNAc(4)Hex(5)Fuc(1)NeuAc(2) | C[+57]*2, N[+2351] | 1592.0474 | 3 | 4774.1277 | 4774.0877 | 0 | 8.4 | 466 | Specific | 895.23 | 844.8 | 844.8 | 15 | 82 | >sp\|P01833\|PIGR_HUMAN Polymeric immunoglobulin receptor OS=Homo sapiens OX=9606 GN=PIGR PE=1 SV=4 | 902 | 20201119_16913_1ng_glyco_hcd203040_compound15.18396.18396.3 | controllerType=0 controllerNumber=1 scan=18396 | 38.2221 |
| 14837:4 | 6 | K.VPGN[+2351.851]VTAVLGETLKVPC[+57.021]HFPC[+57.021]K.F | HexNAc(4)Hex(5)Fuc(3)NeuAc(1) | C[+57]*2, N[+2352] | 1194.5326 | 4 | 4775.1087 | 4775.1081 | 0 | 0.1 | 466 | Specific | 955.2 | 704.83 | 704.83 | 18.14 | 82 | >sp\|P01833\|PIGR_HUMAN Polymeric immunoglobulin receptor OS=Homo sapiens OX=9606 GN=PIGR PE=1 SV=4 | 902 | 20201119_16913_1ng_glyco_hcd203040_compound15.18348.18348.4 | controllerType=0 controllerNumber=1 scan=18348 | 38.1174 |
| 05469:3 | 6 | K.WN[+2059.735]NTGC[+57.021]QALPSQDEGPSK.A | HexNAc(4)Hex(5)Fuc(1)NeuAc(1) | C[+57], N[+2060] | 1350.2106 | 3 | 4048.6173 | 4048.6116 | 0 | 1.4 | 498 | Specific | 213.1 | 155.72 | 155.72 | 4.3 | 82 | >sp\|P01833\|PIGR_HUMAN Polymeric immunoglobulin receptor OS=Homo sapiens OX=9606 GN=PIGR PE=1 SV=4 | 902 | 20201119_16913_1ng_glyco_hcd203040_compound15.8585.8585.3 | controllerType=0 controllerNumber=1 scan=8585 | 20.8154 |
| 03847:3 | 6 | K.WN[+1216.423]NTGC[+57.021]QALPSQDEGPSK.A | HexNAc(2)Hex(5) | C[+57], N[+1216] | 1069.1099 | 3 | 3205.3151 | 3205.2995 | 0 | 4.9 | 498 | Specific | 233.74 | 233.74 | 233.74 | 6.09 | 82 | >sp\|P01833\|PIGR_HUMAN Polymeric immunoglobulin receptor OS=Homo sapiens OX=9606 GN=PIGR PE=1 SV=4 | 902 | 20201119_16913_1ng_glyco_hcd203040_compound15.6901.6901.3 | controllerType=0 controllerNumber=1 scan=6901 | 18.0721 |
| 05345:3 | 6 | K.WN[+2205.793]NTGC[+57.021]QALPSQDEGPSK.A | HexNAc(4)Hex(5)Fuc(2)NeuAc(1) | C[+57], N[+2206] | 1398.8954 | 3 | 4194.6718 | 4194.6695 | 0 | 0.5 | 498 | Specific | 239.48 | 200.68 | 200.68 | 5.79 | 82 | >sp\|P01833\|PIGR_HUMAN Polymeric immunoglobulin receptor OS=Homo sapiens OX=9606 GN=PIGR PE=1 SV=4 | 902 | 20201119_16913_1ng_glyco_hcd203040_compound15.8457.8457.3 | controllerType=0 controllerNumber=1 scan=8457 | 20.6009 |
| 05347:3 | 6 | K.WN[+2205.793]NTGC[+57.021]QALPSQDEGPSK.A | HexNAc(4)Hex(5)Fuc(2)NeuAc(1) | C[+57], N[+2206] | 1398.8954 | 3 | 4194.6718 | 4194.6695 | 0 | 0.5 | 498 | Specific | 264.77 | 214.12 | 214.12 | 5.86 | 82 | >sp\|P01833\|PIGR_HUMAN Polymeric immunoglobulin receptor OS=Homo sapiens OX=9606 GN=PIGR PE=1 SV=4 | 902 | 20201119_16913_1ng_glyco_hcd203040_compound15.8459.8459.3 | controllerType=0 controllerNumber=1 scan=8459 | 20.6049 |
| 03850:3 | 6 | K.WN[+1216.423]NTGC[+57.021]QALPSQDEGPSK.A | HexNAc(2)Hex(5) | C[+57], N[+1216] | 1069.1099 | 3 | 3205.3151 | 3205.2995 | 0 | 4.9 | 498 | Specific | 363.44 | 361.6 | 361.6 | 7.55 | 82 | >sp\|P01833\|PIGR_HUMAN Polymeric immunoglobulin receptor OS=Homo sapiens OX=9606 GN=PIGR PE=1 SV=4 | 902 | 20201119_16913_1ng_glyco_hcd203040_compound15.6904.6904.3 | controllerType=0 controllerNumber=1 scan=6904 | 18.0766 |
| 07552:3 | 6 | K.WN[+2204.772]NTGC[+57.021]QALPSQDEGPSK.A | HexNAc(4)Hex(5)NeuAc(2) | C[+57], N[+2205] | 1398.5573 | 3 | 4193.6574 | 4193.6491 | 0 | 2 | 498 | Specific | 422.62 | 365.5 | 365.5 | 8.01 | 82 | >sp\|P01833\|PIGR_HUMAN Polymeric immunoglobulin receptor OS=Homo sapiens OX=9606 GN=PIGR PE=1 SV=4 | 902 | 20201119_16913_1ng_glyco_hcd203040_compound15.10748.10748.3 | controllerType=0 controllerNumber=1 scan=10748 | 24.367 |
| 05632:3 | 6 | K.WN[+1548.545]NTGC[+57.021]QALPSQDEGPSK.A | HexNAc(3)Hex(4)NeuAc(1) | C[+57], N[+1549] | 1179.814 | 3 | 3537.4275 | 3537.4215 | 0 | 1.7 | 498 | Specific | 423.53 | 372.6 | 372.6 | 8.01 | 82 | >sp\|P01833\|PIGR_HUMAN Polymeric immunoglobulin receptor OS=Homo sapiens OX=9606 GN=PIGR PE=1 SV=4 | 902 | 20201119_16913_1ng_glyco_hcd203040_compound15.8754.8754.3 | controllerType=0 controllerNumber=1 scan=8754 | 21.1008 |
| 05634:3 | 6 | K.WN[+1548.545]NTGC[+57.021]QALPSQDEGPSK.A | HexNAc(3)Hex(4)NeuAc(1) | C[+57], N[+1549] | 1179.814 | 3 | 3537.4275 | 3537.4215 | 0 | 1.7 | 498 | Specific | 564.03 | 473.76 | 473.76 | 10.1 | 82 | >sp\|P01833\|PIGR_HUMAN Polymeric immunoglobulin receptor OS=Homo sapiens OX=9606 GN=PIGR PE=1 SV=4 | 902 | 20201119_16913_1ng_glyco_hcd203040_compound15.8756.8756.3 | controllerType=0 controllerNumber=1 scan=8756 | 21.1043 |
| 07554:3 | 6 | K.WN[+2204.772]NTGC[+57.021]QALPSQDEGPSK.A | HexNAc(4)Hex(5)NeuAc(2) | C[+57], N[+2205] | 1398.5573 | 3 | 4193.6574 | 4193.6491 | 0 | 2 | 498 | Specific | 647.68 | 581.78 | 581.78 | 12.35 | 82 | >sp\|P01833\|PIGR_HUMAN Polymeric immunoglobulin receptor OS=Homo sapiens OX=9606 GN=PIGR PE=1 SV=4 | 902 | 20201119_16913_1ng_glyco_hcd203040_compound15.10750.10750.3 | controllerType=0 controllerNumber=1 scan=10750 | 24.3702 |
| 07544:3 | 6 | K.WN[+2204.772]NTGC[+57.021]QALPSQDEGPSK.A | HexNAc(4)Hex(5)NeuAc(2) | C[+57], N[+2205] | 1398.5608 | 3 | 4193.668 | 4193.6491 | 0 | 4.5 | 498 | Specific | 648.89 | 586.88 | 586.88 | 12.05 | 82 | >sp\|P01833\|PIGR_HUMAN Polymeric immunoglobulin receptor OS=Homo sapiens OX=9606 GN=PIGR PE=1 SV=4 | 902 | 20201119_16913_1ng_glyco_hcd203040_compound15.10739.10739.3 | controllerType=0 controllerNumber=1 scan=10739 | 24.35 |
| 07546:3 | 6 | K.WN[+2204.772]NTGC[+57.021]QALPSQDEGPSK.A | HexNAc(4)Hex(5)NeuAc(2) | C[+57], N[+2205] | 1398.5608 | 3 | 4193.668 | 4193.6491 | 0 | 4.5 | 498 | Specific | 707.37 | 637.55 | 637.55 | 14.05 | 82 | >sp\|P01833\|PIGR_HUMAN Polymeric immunoglobulin receptor OS=Homo sapiens OX=9606 GN=PIGR PE=1 SV=4 | 902 | 20201119_16913_1ng_glyco_hcd203040_compound15.10741.10741.3 | controllerType=0 controllerNumber=1 scan=10741 | 24.354 |
| 05337:3 | 6 | K.WN[+2059.735]NTGC[+57.021]QALPSQDEGPSK.A | HexNAc(4)Hex(5)Fuc(1)NeuAc(1) | C[+57], N[+2060] | 1350.21 | 3 | 4048.6155 | 4048.6116 | 0 | 1 | 498 | Specific | 749.09 | 664.93 | 664.93 | 14.09 | 82 | >sp\|P01833\|PIGR_HUMAN Polymeric immunoglobulin receptor OS=Homo sapiens OX=9606 GN=PIGR PE=1 SV=4 | 902 | 20201119_16913_1ng_glyco_hcd203040_compound15.8449.8449.3 | controllerType=0 controllerNumber=1 scan=8449 | 20.5874 |
| 05339:3 | 6 | K.WN[+2059.735]NTGC[+57.021]QALPSQDEGPSK.A | HexNAc(4)Hex(5)Fuc(1)NeuAc(1) | C[+57], N[+2060] | 1350.21 | 3 | 4048.6155 | 4048.6116 | 0 | 1 | 498 | Specific | 751.73 | 676.23 | 676.23 | 15.09 | 82 | >sp\|P01833\|PIGR_HUMAN Polymeric immunoglobulin receptor OS=Homo sapiens OX=9606 GN=PIGR PE=1 SV=4 | 902 | 20201119_16913_1ng_glyco_hcd203040_compound15.8451.8451.3 | controllerType=0 controllerNumber=1 scan=8451 | 20.5915 |
| 05419:3 | 6 | K.WN[+1872.651]NTGC[+57.021]QALPSQDEGPSK.A | HexNAc(3)Hex(6)NeuAc(1) | C[+57], N[+1873] | 1287.8495 | 3 | 3861.5341 | 3861.5272 | 0 | 1.8 | 498 | Specific | 768.66 | 706.14 | 706.14 | 15.35 | 82 | >sp\|P01833\|PIGR_HUMAN Polymeric immunoglobulin receptor OS=Homo sapiens OX=9606 GN=PIGR PE=1 SV=4 | 902 | 20201119_16913_1ng_glyco_hcd203040_compound15.8533.8533.3 | controllerType=0 controllerNumber=1 scan=8533 | 20.7283 |
| 05417:3 | 6 | K.WN[+1872.651]NTGC[+57.021]QALPSQDEGPSK.A | HexNAc(3)Hex(6)NeuAc(1) | C[+57], N[+1873] | 1287.8495 | 3 | 3861.5341 | 3861.5272 | 0 | 1.8 | 498 | Specific | 793.31 | 728.93 | 728.93 | 15.35 | 82 | >sp\|P01833\|PIGR_HUMAN Polymeric immunoglobulin receptor OS=Homo sapiens OX=9606 GN=PIGR PE=1 SV=4 | 902 | 20201119_16913_1ng_glyco_hcd203040_compound15.8531.8531.3 | controllerType=0 controllerNumber=1 scan=8531 | 20.7241 |
| 05516:3 | 6 | K.WN[+1710.598]NTGC[+57.021]QALPSQDEGPSK.A | HexNAc(3)Hex(5)NeuAc(1) | C[+57], N[+1711] | 1233.8348 | 3 | 3699.4898 | 3699.4743 | 0 | 4.2 | 498 | Specific | 812.42 | 699.99 | 699.99 | 16.05 | 82 | >sp\|P01833\|PIGR_HUMAN Polymeric immunoglobulin receptor OS=Homo sapiens OX=9606 GN=PIGR PE=1 SV=4 | 902 | 20201119_16913_1ng_glyco_hcd203040_compound15.8634.8634.3 | controllerType=0 controllerNumber=1 scan=8634 | 20.8981 |
| 05459:3 | 6 | K.WN[+1913.677]NTGC[+57.021]QALPSQDEGPSK.A | HexNAc(4)Hex(5)NeuAc(1) | C[+57], N[+1914] | 1301.5267 | 3 | 3902.5655 | 3902.5537 | 0 | 3 | 498 | Specific | 828.6 | 736.93 | 736.93 | 16.41 | 82 | >sp\|P01833\|PIGR_HUMAN Polymeric immunoglobulin receptor OS=Homo sapiens OX=9606 GN=PIGR PE=1 SV=4 | 902 | 20201119_16913_1ng_glyco_hcd203040_compound15.8575.8575.3 | controllerType=0 controllerNumber=1 scan=8575 | 20.7999 |
| 05457:3 | 6 | K.WN[+1913.677]NTGC[+57.021]QALPSQDEGPSK.A | HexNAc(4)Hex(5)NeuAc(1) | C[+57], N[+1914] | 1301.5267 | 3 | 3902.5655 | 3902.5537 | 0 | 3 | 498 | Specific | 878.07 | 823.93 | 823.93 | 17.41 | 82 | >sp\|P01833\|PIGR_HUMAN Polymeric immunoglobulin receptor OS=Homo sapiens OX=9606 GN=PIGR PE=1 SV=4 | 902 | 20201119_16913_1ng_glyco_hcd203040_compound15.8573.8573.3 | controllerType=0 controllerNumber=1 scan=8573 | 20.7956 |
| 05514:3 | 6 | K.WN[+1710.598]NTGC[+57.021]QALPSQDEGPSK.A | HexNAc(3)Hex(5)NeuAc(1) | C[+57], N[+1711] | 1233.8348 | 3 | 3699.4898 | 3699.4743 | 0 | 4.2 | 498 | Specific | 943.01 | 849.72 | 849.72 | 18.05 | 82 | >sp\|P01833\|PIGR_HUMAN Polymeric immunoglobulin receptor OS=Homo sapiens OX=9606 GN=PIGR PE=1 SV=4 | 902 | 20201119_16913_1ng_glyco_hcd203040_compound15.8632.8632.3 | controllerType=0 controllerNumber=1 scan=8632 | 20.8931 |
| 07425:3 | 8 | K.WEC[+57.021]KN[+2059.735]DTLLGIK.G | HexNAc(4)Hex(5)Fuc(1)NeuAc(1) | C[+57], N[+2060] | 1179.5031 | 3 | 3536.4948 | 3536.4865 | 0 | 2.4 | 100 | Specific | 377 | 172.43 | 172.43 | 5.49 | 70 | >sp\|P22897\|MRC1_HUMAN Macrophage mannose receptor 1 OS=Homo sapiens OX=9606 GN=MRC1 PE=1 SV=1 | 1049 | 20201119_16913_1ng_glyco_hcd203040_compound15.10616.10616.3 | controllerType=0 controllerNumber=1 scan=10616 | 24.1491 |
| 07427:3 | 8 | K.WEC[+57.021]KN[+2059.735]DTLLGIK.G | HexNAc(4)Hex(5)Fuc(1)NeuAc(1) | C[+57], N[+2060] | 1179.5031 | 3 | 3536.4948 | 3536.4865 | 0 | 2.4 | 100 | Specific | 479.79 | 385.95 | 385.95 | 8.68 | 70 | >sp\|P22897\|MRC1_HUMAN Macrophage mannose receptor 1 OS=Homo sapiens OX=9606 GN=MRC1 PE=1 SV=1 | 1049 | 20201119_16913_1ng_glyco_hcd203040_compound15.10618.10618.3 | controllerType=0 controllerNumber=1 scan=10618 | 24.1528 |
| 08094:3 | 8 | K.[+42.011]TAHC[+57.021]N[+2018.708]ESFYFLC[+57.021]K.R | HexNAc(3)Hex(6)Fuc(1)NeuAc(1) | .[42], C[+57]*2, N[+2019] | 1246.49 | 3 | 3737.4554 | 3737.4385 | 0 | 4.5 | 1201 | Specific | 200.4 | 102.93 | 102.93 | 3.49 | 70 | >sp\|P22897\|MRC1_HUMAN Macrophage mannose receptor 1 OS=Homo sapiens OX=9606 GN=MRC1 PE=1 SV=1 | 1049 | 20201119_16913_1ng_glyco_hcd203040_compound15.11314.11314.3 | controllerType=0 controllerNumber=1 scan=11314 | 25.3439 |
| 08096:3 | 8 | K.[+42.011]TAHC[+57.021]N[+2018.708]ESFYFLC[+57.021]K.R | HexNAc(3)Hex(6)Fuc(1)NeuAc(1) | .[42], C[+57]*2, N[+2019] | 1246.49 | 3 | 3737.4554 | 3737.4385 | 0 | 4.5 | 1201 | Specific | 206.22 | 79.12 | 79.12 | 3.46 | 70 | >sp\|P22897\|MRC1_HUMAN Macrophage mannose receptor 1 OS=Homo sapiens OX=9606 GN=MRC1 PE=1 SV=1 | 1049 | 20201119_16913_1ng_glyco_hcd203040_compound15.11316.11316.3 | controllerType=0 controllerNumber=1 scan=11316 | 25.3477 |
| 16970:3 | 10 | R.LSLHRPALEDLLLGSEAN[+1460.529]LTC[+57.021]TLTGLR.D | HexNAc(4)Hex(4) | C[+57], N[+1461] | 1475.3945 | 3 | 4424.1688 | 4424.1271 | 0 | 9.4 | 127 | Specific | 365.6 | 365.5 | 365.5 | 6.06 | 55 | >sp\|P01876\|IGHA1_HUMAN Immunoglobulin heavy constant alpha 1 OS=Homo sapiens OX=9606 GN=IGHA1 PE=1 SV=2 | 255 | 20201119_16913_1ng_glyco_hcd203040_compound15.20594.20594.3 | controllerType=0 controllerNumber=1 scan=20594 | 42.8242 |
| 16930:4 | 10 | R.LSLHRPALEDLLLGSEAN[+1216.423]LTC[+57.021]TLTGLR.D | HexNAc(2)Hex(5) | C[+57], N[+1216] | 1045.7673 | 4 | 4180.0472 | 4180.0211 | 0 | 6.2 | 127 | Specific | 409.1 | 409.1 | 409.1 | 9.03 | 55 | >sp\|P01876\|IGHA1_HUMAN Immunoglobulin heavy constant alpha 1 OS=Homo sapiens OX=9606 GN=IGHA1 PE=1 SV=2 | 255 | 20201119_16913_1ng_glyco_hcd203040_compound15.20552.20552.4 | controllerType=0 controllerNumber=1 scan=20552 | 42.7014 |
| 16988:3 | 10 | R.LSLHRPALEDLLLGSEAN[+1298.476]LTC[+57.021]TLTGLR.D | HexNAc(4)Hex(3) | C[+57], N[+1298] | 1421.3766 | 3 | 4262.1154 | 4262.0742 | 0 | 9.7 | 127 | Specific | 434.79 | 431.54 | 431.54 | 7.7 | 55 | >sp\|P01876\|IGHA1_HUMAN Immunoglobulin heavy constant alpha 1 OS=Homo sapiens OX=9606 GN=IGHA1 PE=1 SV=2 | 255 | 20201119_16913_1ng_glyco_hcd203040_compound15.20613.20613.3 | controllerType=0 controllerNumber=1 scan=20613 | 42.894 |
| 16962:3 | 10 | R.LSLHRPALEDLLLGSEAN[+1298.476]LTC[+57.021]TLTGLR.D | HexNAc(4)Hex(3) | C[+57], N[+1298] | 1421.3659 | 3 | 4262.0832 | 4262.0742 | 0 | 2.1 | 127 | Specific | 490.65 | 481.64 | 481.64 | 9.14 | 55 | >sp\|P01876\|IGHA1_HUMAN Immunoglobulin heavy constant alpha 1 OS=Homo sapiens OX=9606 GN=IGHA1 PE=1 SV=2 | 255 | 20201119_16913_1ng_glyco_hcd203040_compound15.20585.20585.2 | controllerType=0 controllerNumber=1 scan=20585 | 42.8022 |
| 16985:3 | 10 | R.LSLHRPALEDLLLGSEAN[+1298.476]LTC[+57.021]TLTGLR.D | HexNAc(4)Hex(3) | C[+57], N[+1298] | 1421.3766 | 3 | 4262.1154 | 4262.0742 | 0 | 9.7 | 127 | Specific | 531.63 | 531.63 | 531.63 | 8.04 | 55 | >sp\|P01876\|IGHA1_HUMAN Immunoglobulin heavy constant alpha 1 OS=Homo sapiens OX=9606 GN=IGHA1 PE=1 SV=2 | 255 | 20201119_16913_1ng_glyco_hcd203040_compound15.20610.20610.3 | controllerType=0 controllerNumber=1 scan=20610 | 42.8734 |
| 17038:3 | 10 | R.LSLHRPALEDLLLGSEAN[+1501.555]LTC[+57.021]TLTGLR.D | HexNAc(5)Hex(3) | C[+57], N[+1502] | 1489.0668 | 3 | 4465.1859 | 4465.1536 | 0 | 7.2 | 127 | Specific | 556.75 | 556.75 | 556.75 | 8.73 | 55 | >sp\|P01876\|IGHA1_HUMAN Immunoglobulin heavy constant alpha 1 OS=Homo sapiens OX=9606 GN=IGHA1 PE=1 SV=2 | 255 | 20201119_16913_1ng_glyco_hcd203040_compound15.20665.20665.3 | controllerType=0 controllerNumber=1 scan=20665 | 43.0226 |
| 16959:3 | 10 | R.LSLHRPALEDLLLGSEAN[+1298.476]LTC[+57.021]TLTGLR.D | HexNAc(4)Hex(3) | C[+57], N[+1298] | 1421.3659 | 3 | 4262.0832 | 4262.0742 | 0 | 2.1 | 127 | Specific | 580.02 | 534.98 | 534.98 | 10.92 | 55 | >sp\|P01876\|IGHA1_HUMAN Immunoglobulin heavy constant alpha 1 OS=Homo sapiens OX=9606 GN=IGHA1 PE=1 SV=2 | 255 | 20201119_16913_1ng_glyco_hcd203040_compound15.20582.20582.2 | controllerType=0 controllerNumber=1 scan=20582 | 42.791 |
| 16973:3 | 10 | R.LSLHRPALEDLLLGSEAN[+1460.529]LTC[+57.021]TLTGLR.D | HexNAc(4)Hex(4) | C[+57], N[+1461] | 1475.3945 | 3 | 4424.1688 | 4424.1271 | 0 | 9.4 | 127 | Specific | 600.53 | 600.53 | 600.53 | 10.56 | 55 | >sp\|P01876\|IGHA1_HUMAN Immunoglobulin heavy constant alpha 1 OS=Homo sapiens OX=9606 GN=IGHA1 PE=1 SV=2 | 255 | 20201119_16913_1ng_glyco_hcd203040_compound15.20597.20597.3 | controllerType=0 controllerNumber=1 scan=20597 | 42.8356 |
| 17899:3 | 10 | R.LSLHRPALEDLLLGSEAN[+1751.624]LTC[+57.021]TLTGLR.D | HexNAc(4)Hex(4)NeuAc(1) | C[+57], N[+1752] | 1572.4207 | 3 | 4715.2476 | 4715.2225 | 0 | 5.3 | 127 | Specific | 608.68 | 608.68 | 608.68 | 11.27 | 55 | >sp\|P01876\|IGHA1_HUMAN Immunoglobulin heavy constant alpha 1 OS=Homo sapiens OX=9606 GN=IGHA1 PE=1 SV=2 | 255 | 20201119_16913_1ng_glyco_hcd203040_compound15.21564.21564.3 | controllerType=0 controllerNumber=1 scan=21564 | 44.9428 |
| 17894:4 | 10 | R.LSLHRPALEDLLLGSEAN[+1751.624]LTC[+57.021]TLTGLR.D | HexNAc(4)Hex(4)NeuAc(1) | C[+57], N[+1752] | 1179.5658 | 4 | 4715.2415 | 4715.2225 | 0 | 4 | 127 | Specific | 623.47 | 622.01 | 622.01 | 12.3 | 55 | >sp\|P01876\|IGHA1_HUMAN Immunoglobulin heavy constant alpha 1 OS=Homo sapiens OX=9606 GN=IGHA1 PE=1 SV=2 | 255 | 20201119_16913_1ng_glyco_hcd203040_compound15.21559.21559.4 | controllerType=0 controllerNumber=1 scan=21559 | 44.9334 |
| 16945:3 | 10 | R.LSLHRPALEDLLLGSEAN[+1460.529]LTC[+57.021]TLTGLR.D | HexNAc(4)Hex(4) | C[+57], N[+1461] | 1475.3926 | 3 | 4424.1634 | 4424.1271 | 0 | 8.2 | 127 | Specific | 645.44 | 566.24 | 566.24 | 10.64 | 55 | >sp\|P01876\|IGHA1_HUMAN Immunoglobulin heavy constant alpha 1 OS=Homo sapiens OX=9606 GN=IGHA1 PE=1 SV=2 | 255 | 20201119_16913_1ng_glyco_hcd203040_compound15.20567.20567.3 | controllerType=0 controllerNumber=1 scan=20567 | [illegible] |
| 16939:3 | 10 | R.LSLHRPALEDLLLGSEAN[+1378.476]LTC[+57.021]TLTGLR.D | HexNAc(2)Hex(6) | C[+57], N[+1378] | 1448.0388 | 3 | 4342.1017 | 4342.074 | 0 | 6.4 | 127 | Specific | 646.39 | 613.25 | 613.25 | 11.27 | 55 | >sp\|P01876\|IGHA1_HUMAN Immunoglobulin heavy constant alpha 1 OS=Homo sapiens OX=9606 GN=IGHA1 PE=1 SV=2 | 255 | 20201119_16913_1ng_glyco_hcd203040_compound15.20561.20561.3 | controllerType=0 controllerNumber=1 scan=20561 | [illegible] |
| 16937:4 | 10 | R.LSLHRPALEDLLLGSEAN[+1419.502]LTC[+57.021]TLTGLR.D | HexNAc(3)Hex(5) | C[+57], N[+1420] | 1096.5407 | 4 | 4383.1409 | 4383.1005 | 0 | 9.2 | 127 | Specific | 647.65 | 647.65 | 647.65 | 11.24 | 55 | >sp\|P01876\|IGHA1_HUMAN Immunoglobulin heavy constant alpha 1 OS=Homo sapiens OX=9606 GN=IGHA1 PE=1 SV=2 | 255 | 20201119_16913_1ng_glyco_hcd203040_compound15.20559.20559.4 | controllerType=0 controllerNumber=1 scan=20559 | [illegible] |
| 17036:3 | 10 | R.LSLHRPALEDLLLGSEAN[+1501.555]LTC[+57.021]TLTGLR.D | HexNAc(5)Hex(3) | C[+57], N[+1502] | 1489.0668 | 3 | 4465.1859 | 4465.1536 | 0 | 7.2 | 127 | Specific | 647.73 | 647.73 | 647.73 | 10.99 | 55 | >sp\|P01876\|IGHA1_HUMAN Immunoglobulin heavy constant alpha 1 OS=Homo sapiens OX=9606 GN=IGHA1 PE=1 SV=2 | 255 | 20201119_16913_1ng_glyco_hcd203040_compound15.20663.20663.3 | controllerType=0 controllerNumber=1 scan=20663 | [illegible] |
| 16950:4 | 10 | R.LSLHRPALEDLLLGSEAN[+1501.555]LTC[+57.021]TLTGLR.D | HexNAc(5)Hex(3) | C[+57], N[+1502] | 1117.0455 | 4 | 4465.16 | 4465.1536 | 0 | 1.4 | 127 | Specific | 651.46 | 651.46 | 651.46 | 13.33 | 55 | >sp\|P01876\|IGHA1_HUMAN Immunoglobulin heavy constant alpha 1 OS=Homo sapiens OX=9606 GN=IGHA1 PE=1 SV=2 | 255 | 20201119_16913_1ng_glyco_hcd203040_compound15.20573.20573.4 | controllerType=0 controllerNumber=1 scan=20573 | 42.7584 |
| 16943:3 | 10 | R.LSLHRPALEDLLLGSEAN[+1460.529]LTC[+57.021]TLTGLR.D | HexNAc(4)Hex(4) | C[+57], N[+1461] | 1475.3926 | 3 | 4424.1634 | 4424.1271 | 0 | 8.2 | 127 | Specific | 663.24 | 663.24 | 663.24 | 11.64 | 55 | >sp\|P01876\|IGHA1_HUMAN Immunoglobulin heavy constant alpha 1 OS=Homo sapiens OX=9606 GN=IGHA1 PE=1 SV=2 | 255 | 20201119_16913_1ng_glyco_hcd203040_compound15.20565.20565.3 | controllerType=0 controllerNumber=1 scan=20565 | 42.7305 |
| 16957:3 | 10 | R.LSLHRPALEDLLLGSEAN[+1216.423]LTC[+57.021]TLTGLR.D | HexNAc(2)Hex(5) | C[+57], N[+1216] | 1394.0139 | 3 | 4180.027 | 4180.0211 | 0 | 1.4 | 127 | Specific | 673.81 | 654.5 | 654.5 | 12.65 | 55 | >sp\|P01876\|IGHA1_HUMAN Immunoglobulin heavy constant alpha 1 OS=Homo sapiens OX=9606 GN=IGHA1 PE=1 SV=2 | 255 | 20201119_16913_1ng_glyco_hcd203040_compound15.20580.20580.3 | controllerType=0 controllerNumber=1 scan=20580 | 42.7833 |
| 17891:4 | 10 | R.LSLHRPALEDLLLGSEAN[+1751.624]LTC[+57.021]TLTGLR.D | HexNAc(4)Hex(4)NeuAc(1) | C[+57], N[+1752] | 1179.5658 | 4 | 4715.2415 | 4715.2225 | 0 | 4 | 127 | Specific | 685.16 | 685.16 | 685.16 | 13.3 | 55 | >sp\|P01876\|IGHA1_HUMAN Immunoglobulin heavy constant alpha 1 OS=Homo sapiens OX=9606 GN=IGHA1 PE=1 SV=2 | 255 | 20201119_16913_1ng_glyco_hcd203040_compound15.21556.21556.4 | controllerType=0 controllerNumber=1 scan=21556 | 44.9254 |
| 16894:4 | 10 | R.LSLHRPALEDLLLGSEAN[+1663.608]LTC[+57.021]TLTGLR.D | HexNAc(5)Hex(4) | C[+57], N[+1664] | 1157.559 | 4 | 4627.2142 | 4627.2064 | 0 | 1.7 | 127 | Specific | 686.8 | 685.06 | 685.06 | 13.6 | 55 | >sp\|P01876\|IGHA1_HUMAN Immunoglobulin heavy constant alpha 1 OS=Homo sapiens OX=9606 GN=IGHA1 PE=1 SV=2 | 255 | 20201119_16913_1ng_glyco_hcd203040_compound15.20514.20514.4 | controllerType=0 controllerNumber=1 scan=20514 | 42.5963 |
| 16952:4 | 10 | R.LSLHRPALEDLLLGSEAN[+1501.555]LTC[+57.021]TLTGLR.D | HexNAc(5)Hex(3) | C[+57], N[+1502] | 1117.0455 | 4 | 4465.16 | 4465.1536 | 0 | 1.4 | 127 | Specific | 693.21 | 689.66 | 689.66 | 13.33 | 55 | >sp\|P01876\|IGHA1_HUMAN Immunoglobulin heavy constant alpha 1 OS=Homo sapiens OX=9606 GN=IGHA1 PE=1 SV=2 | 255 | 20201119_16913_1ng_glyco_hcd203040_compound15.20575.20575.4 | controllerType=0 controllerNumber=1 scan=20575 | 42.7655 |
| 16897:4 | 10 | R.LSLHRPALEDLLLGSEAN[+1663.608]LTC[+57.021]TLTGLR.D | HexNAc(5)Hex(4) | C[+57], N[+1664] | 1157.559 | 4 | 4627.2142 | 4627.2064 | 0 | 1.7 | 127 | Specific | 703.47 | 675.12 | 675.12 | 14.6 | 55 | >sp\|P01876\|IGHA1_HUMAN Immunoglobulin heavy constant alpha 1 OS=Homo sapiens OX=9606 GN=IGHA1 PE=1 SV=2 | 255 | 20201119_16913_1ng_glyco_hcd203040_compound15.20517.20517.4 | controllerType=0 controllerNumber=1 scan=20517 | 42.6114 |
| 17902:3 | 10 | R.LSLHRPALEDLLLGSEAN[+1751.624]LTC[+57.021]TLTGLR.D | HexNAc(4)Hex(4)NeuAc(1) | C[+57], N[+1752] | 1572.4207 | 3 | 4715.2476 | 4715.2225 | 0 | 5.3 | 127 | Specific | 719.07 | 695.38 | 695.38 | 13.27 | 55 | >sp\|P01876\|IGHA1_HUMAN Immunoglobulin heavy constant alpha 1 OS=Homo sapiens OX=9606 GN=IGHA1 PE=1 SV=2 | 255 | 20201119_16913_1ng_glyco_hcd203040_compound15.21567.21567.3 | controllerType=0 controllerNumber=1 scan=21567 | 44.9552 |
| 16974:4 | 10 | R.LSLHRPALEDLLLGSEAN[+1298.476]LTC[+57.021]TLTGLR.D | HexNAc(4)Hex(3) | C[+57], N[+1298] | 1066.2818 | 4 | 4262.1053 | 4262.0742 | 0 | 7.3 | 127 | Specific | 738.75 | 738.75 | 738.75 | 13.67 | 55 | >sp\|P01876\|IGHA1_HUMAN Immunoglobulin heavy constant alpha 1 OS=Homo sapiens OX=9606 GN=IGHA1 PE=1 SV=2 | 255 | 20201119_16913_1ng_glyco_hcd203040_compound15.20598.20598.4 | controllerType=0 controllerNumber=1 scan=20598 | 42.8375 |
| 17886:4 | 10 | R.LSLHRPALEDLLLGSEAN[+1710.598]LTC[+57.021]TLTGLR.D | HexNAc(3)Hex(5)NeuAc(1) | C[+57], N[+1711] | 1169.3097 | 4 | 4674.2171 | 4674.1959 | 0 | 4.5 | 127 | Specific | 761.31 | 761.31 | 761.31 | 15.3 | 55 | >sp\|P01876\|IGHA1_HUMAN Immunoglobulin heavy constant alpha 1 OS=Homo sapiens OX=9606 GN=IGHA1 PE=1 SV=2 | 255 | 20201119_16913_1ng_glyco_hcd203040_compound15.21551.21551.4 | controllerType=0 controllerNumber=1 scan=21551 | 44.9087 |
| 16940:4 | 10 | R.LSLHRPALEDLLLGSEAN[+1419.502]LTC[+57.021]TLTGLR.D | HexNAc(3)Hex(5) | C[+57], N[+1420] | 1096.5407 | 4 | 4383.1409 | 4383.1005 | 0 | 9.2 | 127 | Specific | 764.87 | 691.02 | 691.02 | 14.24 | 55 | >sp\|P01876\|IGHA1_HUMAN Immunoglobulin heavy constant alpha 1 OS=Homo sapiens OX=9606 GN=IGHA1 PE=1 SV=2 | 255 | 20201119_16913_1ng_glyco_hcd203040_compound15.20562.20562.4 | controllerType=0 controllerNumber=1 scan=20562 | 42.7255 |
| 16954:3 | 10 | R.LSLHRPALEDLLLGSEAN[+1216.423]LTC[+57.021]TLTGLR.D | HexNAc(2)Hex(5) | C[+57], N[+1216] | 1394.0139 | 3 | 4180.027 | 4180.0211 | 0 | 1.4 | 127 | Specific | 765.58 | 704.32 | 704.32 | 14.65 | 55 | >sp\|P01876\|IGHA1_HUMAN Immunoglobulin heavy constant alpha 1 OS=Homo sapiens OX=9606 GN=IGHA1 PE=1 SV=2 | 255 | 20201119_16913_1ng_glyco_hcd203040_compound15.20577.20577.3 | controllerType=0 controllerNumber=1 scan=20577 | 42.7728 |
| 17889:4 | 10 | R.LSLHRPALEDLLLGSEAN[+1710.598]LTC[+57.021]TLTGLR.D | HexNAc(3)Hex(5)NeuAc(1) | C[+57], N[+1711] | 1169.3097 | 4 | 4674.2171 | 4674.1959 | 0 | 4.5 | 127 | Specific | 826.94 | 791.1 | 791.1 | 16.3 | 55 | >sp\|P01876\|IGHA1_HUMAN Immunoglobulin heavy constant alpha 1 OS=Homo sapiens OX=9606 GN=IGHA1 PE=1 SV=2 | 255 | 20201119_16913_1ng_glyco_hcd203040_compound15.21554.21554.4 | controllerType=0 controllerNumber=1 scan=21554 | 44.9224 |
| 16867:4 | 10 | R.LSLHRPALEDLLLGSEAN[+1702.581]LTC[+57.021]TLTGLR.D | HexNAc(2)Hex(8) | C[+57], N[+1703] | 1167.305 | 4 | 4666.1981 | 4666.1796 | 0 | 4 | 127 | Specific | 873.93 | 873.93 | 873.93 | 17.66 | 55 | >sp\|P01876\|IGHA1_HUMAN Immunoglobulin heavy constant alpha 1 OS=Homo sapiens OX=9606 GN=IGHA1 PE=1 SV=2 | 255 | 20201119_16913_1ng_glyco_hcd203040_compound15.20485.20485.4 | controllerType=0 controllerNumber=1 scan=20485 | 42.4872 |
| 16977:4 | 10 | R.LSLHRPALEDLLLGSEAN[+1298.476]LTC[+57.021]TLTGLR.D | HexNAc(4)Hex(3) | C[+57], N[+1298] | 1066.2818 | 4 | 4262.1053 | 4262.0742 | 0 | 7.3 | 127 | Specific | 899.5 | 873.09 | 873.09 | 16.67 | 55 | >sp\|P01876\|IGHA1_HUMAN Immunoglobulin heavy constant alpha 1 OS=Homo sapiens OX=9606 GN=IGHA1 PE=1 SV=2 | 255 | 20201119_16913_1ng_glyco_hcd203040_compound15.20601.20601.4 | controllerType=0 controllerNumber=1 scan=20601 | 42.8446 |
| 16969:4 | 10 | R.LSLHRPALEDLLLGSEAN[+1257.449]LTC[+57.021]TLTGLR.D | HexNAc(3)Hex(4) | C[+57], N[+1257] | 1056.0237 | 4 | 4221.0731 | 4221.0477 | 0 | 6 | 127 | Specific | 905.2 | 859.6 | 859.6 | 17.96 | 55 | >sp\|P01876\|IGHA1_HUMAN Immunoglobulin heavy constant alpha 1 OS=Homo sapiens OX=9606 GN=IGHA1 PE=1 SV=2 | 255 | 20201119_16913_1ng_glyco_hcd203040_compound15.20593.20593.4 | controllerType=0 controllerNumber=1 scan=20593 | 42.8231 |
| 16889:4 | 10 | R.LSLHRPALEDLLLGSEAN[+1540.529]LTC[+57.021]TLTGLR.D | HexNAc(2)Hex(7) | C[+57], N[+1541] | 1126.7902 | 4 | 4504.139 | 4504.1268 | 0 | 2.7 | 127 | Specific | 922.24 | 922.24 | 922.24 | 18.6 | 55 | >sp\|P01876\|IGHA1_HUMAN Immunoglobulin heavy constant alpha 1 OS=Homo sapiens OX=9606 GN=IGHA1 PE=1 SV=2 | 255 | 20201119_16913_1ng_glyco_hcd203040_compound15.20508.20508.4 | controllerType=0 controllerNumber=1 scan=20508 | 42.5864 |
| 16888:4 | 10 | R.LSLHRPALEDLLLGSEAN[+1540.529]LTC[+57.021]TLTGLR.D | HexNAc(2)Hex(7) | C[+57], N[+1541] | 1126.7902 | 4 | 4504.139 | 4504.1268 | 0 | 2.7 | 127 | Specific | 935.77 | 827.3 | 827.3 | 18.6 | 55 | >sp\|P01876\|IGHA1_HUMAN Immunoglobulin heavy constant alpha 1 OS=Homo sapiens OX=9606 GN=IGHA1 PE=1 SV=2 | 255 | 20201119_16913_1ng_glyco_hcd203040_compound15.20507.20507.4 | controllerType=0 controllerNumber=1 scan=20507 | 42.577 |
| 16869:4 | 10 | R.LSLHRPALEDLLLGSEAN[+1702.581]LTC[+57.021]TLTGLR.D | HexNAc(2)Hex(8) | C[+57], N[+1703] | 1167.305 | 4 | 4666.1981 | 4666.1796 | 0 | 4 | 127 | Specific | 1016.51 | 1002.57 | 1002.57 | 20.66 | 55 | >sp\|P01876\|IGHA1_HUMAN Immunoglobulin heavy constant alpha 1 OS=Homo sapiens OX=9606 GN=IGHA1 PE=1 SV=2 | 255 | 20201119_16913_1ng_glyco_hcd203040_compound15.20487.20487.4 | controllerType=0 controllerNumber=1 scan=20487 | 42.5041 |
| 16972:4 | 10 | R.LSLHRPALEDLLLGSEAN[+1257.449]LTC[+57.021]TLTGLR.D | HexNAc(3)Hex(4) | C[+57], N[+1257] | 1056.0237 | 4 | 4221.0731 | 4221.0477 | 0 | 6 | 127 | Specific | 1062.93 | 1003.91 | 1003.91 | 20.96 | 55 | >sp\|P01876\|IGHA1_HUMAN Immunoglobulin heavy constant alpha 1 OS=Homo sapiens OX=9606 GN=IGHA1 PE=1 SV=2 | 255 | 20201119_16913_1ng_glyco_hcd203040_compound15.20596.20596.4 | controllerType=0 controllerNumber=1 scan=20596 | 42.8337 |
| 12764:3 | 10 | R.LAGKPTHVN[+2263.835]VSVVMAEVDGTC[+57.021].Y | HexNAc(5)Hex(5)Fuc(3) | C[+57], N[+2264] | 1483.3105 | 3 | 4447.9169 | 4447.9135 | 0 | 0.8 | 332 | CRagged | 217.14 | 182.89 | 182.89 | 3.12 | 55 | >sp\|P01876\|IGHA1_HUMAN Immunoglobulin heavy constant alpha 1 OS=Homo sapiens OX=9606 GN=IGHA1 PE=1 SV=2 | 255 | 20201119_16913_1ng_glyco_hcd203040_compound15.16191.16191.3 | controllerType=0 controllerNumber=1 scan=16191 | 33.939 |
| 12761:3 | 10 | R.LAGKPTHVN[+2263.835]VSVVMAEVDGTC[+57.021].Y | HexNAc(5)Hex(5)Fuc(3) | C[+57], N[+2264] | 1483.3105 | 3 | 4447.9169 | 4447.9135 | 0 | 0.8 | 332 | CRagged | 243.04 | 237.18 | 237.18 | 5.55 | 55 | >sp\|P01876\|IGHA1_HUMAN Immunoglobulin heavy constant alpha 1 OS=Homo sapiens OX=9606 GN=IGHA1 PE=1 SV=2 | 255 | 20201119_16913_1ng_glyco_hcd203040_compound15.16188.16188.3 | controllerType=0 controllerNumber=1 scan=16188 | 33.9337 |
| 12731:4 | 10 | R.LAGKPTHVN[+2263.835]VSVVMAEVDGTC[+57.021]Y.- | HexNAc(5)Hex(5)Fuc(3) | C[+57], N[+2264] | 1153.4955 | 4 | 4610.9603 | 4610.9768 | 0 | -3.6 | 332 | CRagged | 202.96 | 202.96 | 202.96 | 3.42 | 55 | >sp\|P01876\|IGHA1_HUMAN Immunoglobulin heavy constant alpha 1 OS=Homo sapiens OX=9606 GN=IGHA1 PE=1 SV=2 | 255 | 20201119_16913_1ng_glyco_hcd203040_compound15.16157.16157.4 | controllerType=0 controllerNumber=1 scan=16157 | 33.8857 |
| 11105:2 | 10 | R.LAGKPTHVN[+1647.613]VSVVMAEVDGTC[+57.021]Y.- | HexNAc(5)Hex(3)Fuc(1) | C[+57], N[+1648] | 1997.8911 | 2 | 3994.7749 | 3994.7553 | 0 | 4.9 | 332 | CRagged | 239.06 | 239.06 | 239.06 | 5.58 | 55 | >sp\|P01876\|IGHA1_HUMAN Immunoglobulin heavy constant alpha 1 OS=Homo sapiens OX=9606 GN=IGHA1 PE=1 SV=2 | 255 | 20201119_16913_1ng_glyco_hcd203040_compound15.14458.14458.2 | controllerType=0 controllerNumber=1 scan=14458 | 30.8417 |
| 11058:4 | 10 | R.LAGKPTHVN[+1809.666]VSVVMAEVDGTC[+57.021]Y.- | HexNAc(5)Hex(4)Fuc(1) | C[+57], N[+1810] | 1039.9593 | 4 | 4156.8153 | 4156.8081 | 0 | 1.7 | 332 | CRagged | 247.42 | 200.78 | 200.78 | 4.5 | 55 | >sp\|P01876\|IGHA1_HUMAN Immunoglobulin heavy constant alpha 1 OS=Homo sapiens OX=9606 GN=IGHA1 PE=1 SV=2 | 255 | 20201119_16913_1ng_glyco_hcd203040_compound15.14409.14409.4 | controllerType=0 controllerNumber=1 scan=14409 | 30.7357 |
| 09941:3 | 10 | R.LAGKPTHVN[+1647.613]VSVVMAEVDGTC[+57.021]Y.- | HexNAc(5)Hex(3)Fuc(1) | C[+57], N[+1648] | 1332.2595 | 3 | 3994.7638 | 3994.7553 | 0 | 2.1 | 332 | CRagged | 376.68 | 376.68 | 376.68 | 9.4 | 55 | >sp\|P01876\|IGHA1_HUMAN Immunoglobulin heavy constant alpha 1 OS=Homo sapiens OX=9606 GN=IGHA1 PE=1 SV=2 | 255 | 20201119_16913_1ng_glyco_hcd203040_compound15.13249.13249.3 | controllerType=0 controllerNumber=1 scan=13249 | 28.6305 |
| 09855:3 | 10 | R.LAGKPTHVN[+1809.666]VSVVMAEVDGTC[+57.021]Y.- | HexNAc(5)Hex(4)Fuc(1) | C[+57], N[+1810] | 1386.2756 | 3 | 4156.8122 | 4156.8081 | 0 | 1 | 332 | CRagged | 418.34 | 302.71 | 294.77 | 7.47 | 55 | >sp\|P01876\|IGHA1_HUMAN Immunoglobulin heavy constant alpha 1 OS=Homo sapiens OX=9606 GN=IGHA1 PE=1 SV=2 | 255 | 20201119_16913_1ng_glyco_hcd203040_compound15.13160.13160.3 | controllerType=0 controllerNumber=1 scan=13160 | 28.4718 |
| 09857:3 | 10 | R.LAGKPTHVN[+1809.666]VSVVMAEVDGTC[+57.021]Y.- | HexNAc(5)Hex(4)Fuc(1) | C[+57], N[+1810] | 1386.2756 | 3 | 4156.8122 | 4156.8081 | 0 | 1 | 332 | CRagged | 419 | 419 | 322.09 | 9.32 | 55 | >sp\|P01876\|IGHA1_HUMAN Immunoglobulin heavy constant alpha 1 OS=Homo sapiens OX=9606 GN=IGHA1 PE=1 SV=2 | 255 | 20201119_16913_1ng_glyco_hcd203040_compound15.13162.13162.3 | controllerType=0 controllerNumber=1 scan=13162 | 28.4759 |
| 09943:3 | 10 | R.LAGKPTHVN[+1647.613]VSVVMAEVDGTC[+57.021]Y.- | HexNAc(5)Hex(3)Fuc(1) | C[+57], N[+1648] | 1332.2595 | 3 | 3994.7638 | 3994.7553 | 0 | 2.1 | 332 | CRagged | 444.01 | 444.01 | 444.01 | 9.65 | 55 | >sp\|P01876\|IGHA1_HUMAN Immunoglobulin heavy constant alpha 1 OS=Homo sapiens OX=9606 GN=IGHA1 PE=1 SV=2 | 255 | 20201119_16913_1ng_glyco_hcd203040_compound15.13251.13251.3 | controllerType=0 controllerNumber=1 scan=13251 | 28.6422 |
| 14552:3 | 10 | R.LAGKPTHVN[+2204.772]VSVVMAEVDGTC[+57.021]Y.- | HexNAc(4)Hex(5)NeuAc(2) | C[+57], N[+2205] | 1517.9778 | 3 | 4551.919 | 4551.9145 | 0 | 1 | 332 | CRagged | 453.83 | 408.98 | 408.98 | 9.47 | 55 | >sp\|P01876\|IGHA1_HUMAN Immunoglobulin heavy constant alpha 1 OS=Homo sapiens OX=9606 GN=IGHA1 PE=1 SV=2 | 255 | 20201119_16913_1ng_glyco_hcd203040_compound15.18052.18052.3 | controllerType=0 controllerNumber=1 scan=18052 | 37.5543 |
| 11023:3 | 10 | R.LAGKPTHVN[+1809.666]VSVVMAEVDGTC[+57.021]Y.- | HexNAc(5)Hex(4)Fuc(1) | C[+57], N[+1810] | 1386.2765 | 3 | 4156.8151 | 4156.8081 | 0 | 1.7 | 332 | CRagged | 455.04 | 441.78 | 217.25 | 9.74 | 55 | >sp\|P01876\|IGHA1_HUMAN Immunoglobulin heavy constant alpha 1 OS=Homo sapiens OX=9606 GN=IGHA1 PE=1 SV=2 | 255 | 20201119_16913_1ng_glyco_hcd203040_compound15.14372.14372.2 | controllerType=0 controllerNumber=1 scan=14372 | 30.6315 |
| 10947:3 | 10 | R.LAGKPTHVN[+1663.608]VSVVMAEVDGTC[+57.021]Y.- | HexNAc(5)Hex(4) | C[+57], N[+1664] | 1337.5933 | 3 | 4010.7654 | 4010.7502 | 0 | 3.8 | 332 | CRagged | 464.12 | 464.12 | 371.34 | 9.8 | 55 | >sp\|P01876\|IGHA1_HUMAN Immunoglobulin heavy constant alpha 1 OS=Homo sapiens OX=9606 GN=IGHA1 PE=1 SV=2 | 255 | 20201119_16913_1ng_glyco_hcd203040_compound15.14291.14291.3 | controllerType=0 controllerNumber=1 scan=14291 | 30.4505 |
| 14554:3 | 10 | R.LAGKPTHVN[+2204.772]VSVVMAEVDGTC[+57.021]Y.- | HexNAc(4)Hex(5)NeuAc(2) | C[+57], N[+2205] | 1517.9778 | 3 | 4551.919 | 4551.9145 | 0 | 1 | 332 | CRagged | 477.68 | 379.38 | 379.38 | 9.47 | 55 | >sp\|P01876\|IGHA1_HUMAN Immunoglobulin heavy constant alpha 1 OS=Homo sapiens OX=9606 GN=IGHA1 PE=1 SV=2 | 255 | 20201119_16913_1ng_glyco_hcd203040_compound15.18054.18054.3 | controllerType=0 controllerNumber=1 scan=18054 | 37.558 |
| 11021:2 | 10 | R.LAGKPTHVN[+1378.476]VSVVMAEVDGTC[+57.021]Y.- | HexNAc(2)Hex(6) | C[+57], N[+1378] | 1863.3197 | 2 | 3725.6321 | 3725.6178 | 0 | 3.9 | 332 | CRagged | 484.02 | 431.67 | 237.32 | 9.12 | 55 | >sp\|P01876\|IGHA1_HUMAN Immunoglobulin heavy constant alpha 1 OS=Homo sapiens OX=9606 GN=IGHA1 PE=1 SV=2 | 255 | 20201119_16913_1ng_glyco_hcd203040_compound15.14370.14370.2 | controllerType=0 controllerNumber=1 scan=14370 | 30.6284 |
| 12715:3 | 10 | R.LAGKPTHVN[+2059.735]VSVVMAEVDGTC[+57.021]Y.- | HexNAc(4)Hex(5)Fuc(1)NeuAc(1) | C[+57], N[+2060] | 1469.6358 | 3 | 4406.8928 | 4406.877 | 0 | 3.6 | 332 | CRagged | 495.83 | 495.83 | 495.83 | 9.8 | 55 | >sp\|P01876\|IGHA1_HUMAN Immunoglobulin heavy constant alpha 1 OS=Homo sapiens OX=9606 GN=IGHA1 PE=1 SV=2 | 255 | 20201119_16913_1ng_glyco_hcd203040_compound15.16140.16140.3 | controllerType=0 controllerNumber=1 scan=16140 | 33.8494 |
| 12717:3 | 10 | R.LAGKPTHVN[+2059.735]VSVVMAEVDGTC[+57.021]Y.- | HexNAc(4)Hex(5)Fuc(1)NeuAc(1) | C[+57], N[+2060] | 1469.6358 | 3 | 4406.8928 | 4406.877 | 0 | 3.6 | 332 | CRagged | 568.39 | 568.39 | 568.39 | 10.32 | 55 | >sp\|P01876\|IGHA1_HUMAN Immunoglobulin heavy constant alpha 1 OS=Homo sapiens OX=9606 GN=IGHA1 PE=1 SV=2 | 255 | 20201119_16913_1ng_glyco_hcd203040_compound15.16142.16142.3 | controllerType=0 controllerNumber=1 scan=16142 | 33.8567 |
| 12777:3 | 10 | R.LAGKPTHVN[+1710.598]VSVVMAEVDGTC[+57.021]Y.- | HexNAc(3)Hex(5)NeuAc(1) | C[+57], N[+1711] | 1353.2571 | 3 | 4057.7569 | 4057.7397 | 0 | 4.2 | 332 | CRagged | 576.08 | 523.08 | 339.8 | 11.22 | 55 | >sp\|P01876\|IGHA1_HUMAN Immunoglobulin heavy constant alpha 1 OS=Homo sapiens OX=9606 GN=IGHA1 PE=1 SV=2 | 255 | 20201119_16913_1ng_glyco_hcd203040_compound15.16204.16204.3 | controllerType=0 controllerNumber=1 scan=16204 | 33.9563 |
| 09842:3 | 10 | R.LAGKPTHVN[+1378.476]VSVVMAEVDGTC[+57.021]Y.- | HexNAc(2)Hex(6) | C[+57], N[+1378] | 1242.5456 | 3 | 3725.6222 | 3725.6178 | 0 | 1.2 | 332 | CRagged | 585.52 | 497.42 | 353.61 | 11.25 | 55 | >sp\|P01876\|IGHA1_HUMAN Immunoglobulin heavy constant alpha 1 OS=Homo sapiens OX=9606 GN=IGHA1 PE=1 SV=2 | 255 | 20201119_16913_1ng_glyco_hcd203040_compound15.13146.13146.3 | controllerType=0 controllerNumber=1 scan=13146 | 28.4465 |
| 09840:3 | 10 | R.LAGKPTHVN[+1378.476]VSVVMAEVDGTC[+57.021]Y.- | HexNAc(2)Hex(6) | C[+57], N[+1378] | 1242.5456 | 3 | 3725.6222 | 3725.6178 | 0 | 1.2 | 332 | CRagged | 587.7 | 587.7 | 359.92 | 11.25 | 55 | >sp\|P01876\|IGHA1_HUMAN Immunoglobulin heavy constant alpha 1 OS=Homo sapiens OX=9606 GN=IGHA1 PE=1 SV=2 | 255 | 20201119_16913_1ng_glyco_hcd203040_compound15.13144.13144.3 | controllerType=0 controllerNumber=1 scan=13144 | 28.4425 |
| 11019:2 | 10 | R.LAGKPTHVN[+1378.476]VSVVMAEVDGTC[+57.021]Y.- | HexNAc(2)Hex(6) | C[+57], N[+1378] | 1863.3197 | 2 | 3725.6321 | 3725.6178 | 0 | 3.9 | 332 | CRagged | 595.08 | 595.08 | 314.27 | 10.9 | 55 | >sp\|P01876\|IGHA1_HUMAN Immunoglobulin heavy constant alpha 1 OS=Homo sapiens OX=9606 GN=IGHA1 PE=1 SV=2 | 255 | 20201119_16913_1ng_glyco_hcd203040_compound15.14368.14368.2 | controllerType=0 controllerNumber=1 scan=14368 | 30.6217 |
| 11026:3 | 10 | R.LAGKPTHVN[+1809.666]VSVVMAEVDGTC[+57.021]Y.- | HexNAc(5)Hex(4)Fuc(1) | C[+57], N[+1810] | 1386.2765 | 3 | 4156.8151 | 4156.8081 | 0 | 1.7 | 332 | CRagged | 615.25 | 555.54 | 327.6 | 12.52 | 55 | >sp\|P01876\|IGHA1_HUMAN Immunoglobulin heavy constant alpha 1 OS=Homo sapiens OX=9606 GN=IGHA1 PE=1 SV=2 | 255 | 20201119_16913_1ng_glyco_hcd203040_compound15.14375.14375.2 | controllerType=0 controllerNumber=1 scan=14375 | 30.6433 |
| 12774:3 | 10 | R.LAGKPTHVN[+1710.598]VSVVMAEVDGTC[+57.021]Y.- | HexNAc(3)Hex(5)NeuAc(1) | C[+57], N[+1711] | 1353.2571 | 3 | 4057.7569 | 4057.7397 | 0 | 4.2 | 332 | CRagged | 632.16 | 632.16 | 444.79 | 12.22 | 55 | >sp\|P01876\|IGHA1_HUMAN Immunoglobulin heavy constant alpha 1 OS=Homo sapiens OX=9606 GN=IGHA1 PE=1 SV=2 | 255 | 20201119_16913_1ng_glyco_hcd203040_compound15.16201.16201.3 | controllerType=0 controllerNumber=1 scan=16201 | 33.9526 |
| 09879:3 | 10 | R.LAGKPTHVN[+1216.423]VSVVMAEVDGTC[+57.021]Y.- | HexNAc(2)Hex(5) | C[+57], N[+1216] | 1188.5308 | 3 | 3563.5779 | 3563.5649 | 0 | 3.6 | 332 | CRagged | 632.22 | 632.22 | 370.06 | 12.58 | 55 | >sp\|P01876\|IGHA1_HUMAN Immunoglobulin heavy constant alpha 1 OS=Homo sapiens OX=9606 GN=IGHA1 PE=1 SV=2 | 255 | 20201119_16913_1ng_glyco_hcd203040_compound15.13185.13185.3 | controllerType=0 controllerNumber=1 scan=13185 | 28.5168 |
| 10922:3 | 10 | R.LAGKPTHVN[+1702.581]VSVVMAEVDGTC[+57.021]Y.- | HexNAc(2)Hex(8) | C[+57], N[+1703] | 1350.5829 | 3 | 4049.7343 | 4049.7234 | 0 | 2.7 | 332 | CRagged | 657.06 | 657.06 | 471.57 | 13.52 | 55 | >sp\|P01876\|IGHA1_HUMAN Immunoglobulin heavy constant alpha 1 OS=Homo sapiens OX=9606 GN=IGHA1 PE=1 SV=2 | 255 | 20201119_16913_1ng_glyco_hcd203040_compound15.14266.14266.3 | controllerType=0 controllerNumber=1 scan=14266 | 30.4046 |
| 09881:3 | 10 | R.LAGKPTHVN[+1216.423]VSVVMAEVDGTC[+57.021]Y.- | HexNAc(2)Hex(5) | C[+57], N[+1216] | 1188.5308 | 3 | 3563.5779 | 3563.5649 | 0 | 3.6 | 332 | CRagged | 708.59 | 708.59 | 403.64 | 14.58 | 55 | >sp\|P01876\|IGHA1_HUMAN Immunoglobulin heavy constant alpha 1 OS=Homo sapiens OX=9606 GN=IGHA1 PE=1 SV=2 | 255 | 20201119_16913_1ng_glyco_hcd203040_compound15.13187.13187.3 | controllerType=0 controllerNumber=1 scan=13187 | 28.5217 |
| 12647:3 | 10 | R.LAGKPTHVN[+2262.814]VSVVMAEVDGTC[+57.021]Y.- | HexNAc(5)Hex(5)Fuc(1)NeuAc(1) | C[+57], N[+2263] | 1537.3262 | 3 | 4609.9641 | 4609.9564 | 0 | 1.7 | 332 | CRagged | 742.34 | 602.66 | 602.66 | 14.52 | 55 | >sp\|P01876\|IGHA1_HUMAN Immunoglobulin heavy constant alpha 1 OS=Homo sapiens OX=9606 GN=IGHA1 PE=1 SV=2 | 255 | 20201119_16913_1ng_glyco_hcd203040_compound15.16069.16069.3 | controllerType=0 controllerNumber=1 scan=16069 | 33.687 |
| 12653:3 | 10 | R.LAGKPTHVN[+1913.677]VSVVMAEVDGTC[+57.021]Y.- | HexNAc(4)Hex(5)NeuAc(1) | C[+57], N[+1914] | 1420.9463 | 3 | 4260.8245 | 4260.8191 | 0 | 1.3 | 332 | CRagged | 758.91 | 514.84 | 514.84 | 15.25 | 55 | >sp\|P01876\|IGHA1_HUMAN Immunoglobulin heavy constant alpha 1 OS=Homo sapiens OX=9606 GN=IGHA1 PE=1 SV=2 | 255 | 20201119_16913_1ng_glyco_hcd203040_compound15.16075.16075.3 | controllerType=0 controllerNumber=1 scan=16075 | 33.6978 |
| 10945:3 | 10 | R.LAGKPTHVN[+1663.608]VSVVMAEVDGTC[+57.021]Y.- | HexNAc(5)Hex(4) | C[+57], N[+1664] | 1337.5933 | 3 | 4010.7654 | 4010.7502 | 0 | 3.8 | 332 | CRagged | 777.06 | 777.06 | 479.49 | 15.58 | 55 | >sp\|P01876\|IGHA1_HUMAN Immunoglobulin heavy constant alpha 1 OS=Homo sapiens OX=9606 GN=IGHA1 PE=1 SV=2 | 255 | 20201119_16913_1ng_glyco_hcd203040_compound15.14289.14289.3 | controllerType=0 controllerNumber=1 scan=14289 | 30.4453 |
| 11063:3 | 10 | R.LAGKPTHVN[+1647.613]VSVVMAEVDGTC[+57.021]Y.- | HexNAc(5)Hex(3)Fuc(1) | C[+57], N[+1648] | 1332.263 | 3 | 3994.7744 | 3994.7553 | 0 | 4.8 | 332 | CRagged | 783.69 | 705.93 | 705.93 | 15.22 | 55 | >sp\|P01876\|IGHA1_HUMAN Immunoglobulin heavy constant alpha 1 OS=Homo sapiens OX=9606 GN=IGHA1 PE=1 SV=2 | 255 | 20201119_16913_1ng_glyco_hcd203040_compound15.14415.14415.3 | controllerType=0 controllerNumber=1 scan=14415 | 30.752 |
| 11066:3 | 10 | R.LAGKPTHVN[+1647.613]VSVVMAEVDGTC[+57.021]Y.- | HexNAc(5)Hex(3)Fuc(1) | C[+57], N[+1648] | 1332.263 | 3 | 3994.7744 | 3994.7553 | 0 | 4.8 | 332 | CRagged | 812.68 | 768.21 | 768.21 | 16.22 | 55 | >sp\|P01876\|IGHA1_HUMAN Immunoglobulin heavy constant alpha 1 OS=Homo sapiens OX=9606 GN=IGHA1 PE=1 SV=2 | 255 | 20201119_16913_1ng_glyco_hcd203040_compound15.14418.14418.3 | controllerType=0 controllerNumber=1 scan=14418 | 30.758 |
| 11090:3 | 10 | R.LAGKPTHVN[+1501.555]VSVVMAEVDGTC[+57.021]Y.- | HexNAc(5)Hex(3) | C[+57], N[+1502] | 1283.577 | 3 | 3848.7164 | 3848.6974 | 0 | 4.9 | 332 | CRagged | 821.4 | 821.4 | 821.4 | 16.22 | 55 | >sp\|P01876\|IGHA1_HUMAN Immunoglobulin heavy constant alpha 1 OS=Homo sapiens OX=9606 GN=IGHA1 PE=1 SV=2 | 255 | 20201119_16913_1ng_glyco_hcd203040_compound15.14443.14443.3 | controllerType=0 controllerNumber=1 scan=14443 | 30.809 |
| 10956:3 | 10 | R.LAGKPTHVN[+1663.608]VSVVMAEVDGTC[+57.021]Y.- | HexNAc(5)Hex(4) | C[+57], N[+1664] | 1337.5935 | 3 | 4010.7659 | 4010.7502 | 0 | 3.9 | 332 | CRagged | 822.59 | 822.59 | 545.75 | 16.58 | 55 | >sp\|P01876\|IGHA1_HUMAN Immunoglobulin heavy constant alpha 1 OS=Homo sapiens OX=9606 GN=IGHA1 PE=1 SV=2 | 255 | 20201119_16913_1ng_glyco_hcd203040_compound15.14301.14301.3 | controllerType=0 controllerNumber=1 scan=14301 | 30.4729 |
| 10990:3 | 10 | R.LAGKPTHVN[+1809.666]VSVVMAEVDGTC[+57.021]Y.- | HexNAc(5)Hex(4)Fuc(1) | C[+57], N[+1810] | 1386.2795 | 3 | 4156.8239 | 4156.8081 | 0 | 3.8 | 332 | CRagged | 827.39 | 779.42 | 539.56 | 16.58 | 55 | >sp\|P01876\|IGHA1_HUMAN Immunoglobulin heavy constant alpha 1 OS=Homo sapiens OX=9606 GN=IGHA1 PE=1 SV=2 | 255 | 20201119_16913_1ng_glyco_hcd203040_compound15.14337.14337.3 | controllerType=0 controllerNumber=1 scan=14337 | 30.5523 |
| 11002:3 | 10 | R.LAGKPTHVN[+1378.476]VSVVMAEVDGTC[+57.021]Y.- | HexNAc(2)Hex(6) | C[+57], N[+1378] | 1242.5483 | 3 | 3725.6303 | 3725.6178 | 0 | 3.4 | 332 | CRagged | 831.94 | 815.53 | 530.53 | 16.58 | 55 | >sp\|P01876\|IGHA1_HUMAN Immunoglobulin heavy constant alpha 1 OS=Homo sapiens OX=9606 GN=IGHA1 PE=1 SV=2 | 255 | 20201119_16913_1ng_glyco_hcd203040_compound15.14350.14350.3 | controllerType=0 controllerNumber=1 scan=14350 | 30.5818 |
| 12645:3 | 10 | R.LAGKPTHVN[+2262.814]VSVVMAEVDGTC[+57.021]Y.- | HexNAc(5)Hex(5)Fuc(1)NeuAc(1) | C[+57], N[+2263] | 1537.3262 | 3 | 4609.9641 | 4609.9564 | 0 | 1.7 | 332 | CRagged | 850.35 | 598.13 | 598.13 | 17.52 | 55 | >sp\|P01876\|IGHA1_HUMAN Immunoglobulin heavy constant alpha 1 OS=Homo sapiens OX=9606 GN=IGHA1 PE=1 SV=2 | 255 | 20201119_16913_1ng_glyco_hcd203040_compound15.16067.16067.3 | controllerType=0 controllerNumber=1 scan=16067 | 33.6811 |
| 12650:3 | 10 | R.LAGKPTHVN[+1913.677]VSVVMAEVDGTC[+57.021]Y.- | HexNAc(4)Hex(5)NeuAc(1) | C[+57], N[+1914] | 1420.9463 | 3 | 4260.8245 | 4260.8191 | 0 | 1.3 | 332 | CRagged | 852.63 | 539.51 | 539.51 | 17.25 | 55 | >sp\|P01876\|IGHA1_HUMAN Immunoglobulin heavy constant alpha 1 OS=Homo sapiens OX=9606 GN=IGHA1 PE=1 SV=2 | 255 | 20201119_16913_1ng_glyco_hcd203040_compound15.16072.16072.3 | controllerType=0 controllerNumber=1 scan=16072 | 33.6918 |
| 11012:3 | 10 | R.LAGKPTHVN[+1216.423]VSVVMAEVDGTC[+57.021]Y.- | HexNAc(2)Hex(5) | C[+57], N[+1216] | 1188.5289 | 3 | 3563.572 | 3563.5649 | 0 | 2 | 332 | CRagged | 875.24 | 875.24 | 502.36 | 17.52 | 55 | >sp\|P01876\|IGHA1_HUMAN Immunoglobulin heavy constant alpha 1 OS=Homo sapiens OX=9606 GN=IGHA1 PE=1 SV=2 | 255 | 20201119_16913_1ng_glyco_hcd203040_compound15.14361.14361.3 | controllerType=0 controllerNumber=1 scan=14361 | 30.6023 |
| 10928:3 | 10 | R.LAGKPTHVN[+1540.529]VSVVMAEVDGTC[+57.021]Y.- | HexNAc(2)Hex(7) | C[+57], N[+1541] | 1296.5649 | 3 | 3887.6801 | 3887.6706 | 0 | 2.4 | 332 | CRagged | 878.06 | 734.65 | 553.03 | 17.52 | 55 | >sp\|P01876\|IGHA1_HUMAN Immunoglobulin heavy constant alpha 1 OS=Homo sapiens OX=9606 GN=IGHA1 PE=1 SV=2 | 255 | 20201119_16913_1ng_glyco_hcd203040_compound15.14272.14272.3 | controllerType=0 controllerNumber=1 scan=14272 | 30.4136 |
| 10931:3 | 10 | R.LAGKPTHVN[+1540.529]VSVVMAEVDGTC[+57.021]Y.- | HexNAc(2)Hex(7) | C[+57], N[+1541] | 1296.5649 | 3 | 3887.6801 | 3887.6706 | 0 | 2.4 | 332 | CRagged | 878.45 | 814.47 | 522.47 | 17.52 | 55 | >sp\|P01876\|IGHA1_HUMAN Immunoglobulin heavy constant alpha 1 OS=Homo sapiens OX=9606 GN=IGHA1 PE=1 SV=2 | 255 | 20201119_16913_1ng_glyco_hcd203040_compound15.14275.14275.3 | controllerType=0 controllerNumber=1 scan=14275 | 30.421 |
| 10954:3 | 10 | R.LAGKPTHVN[+1663.608]VSVVMAEVDGTC[+57.021]Y.- | HexNAc(5)Hex(4) | C[+57], N[+1664] | 1337.5935 | 3 | 4010.7659 | 4010.7502 | 0 | 3.9 | 332 | CRagged | 878.94 | 714.64 | 576.77 | 17.58 | 55 | >sp\|P01876\|IGHA1_HUMAN Immunoglobulin heavy constant alpha 1 OS=Homo sapiens OX=9606 GN=IGHA1 PE=1 SV=2 | 255 | 20201119_16913_1ng_glyco_hcd203040_compound15.14299.14299.3 | controllerType=0 controllerNumber=1 scan=14299 | 30.4649 |
| 10924:3 | 10 | R.LAGKPTHVN[+1702.581]VSVVMAEVDGTC[+57.021]Y.- | HexNAc(2)Hex(8) | C[+57], N[+1703] | 1350.5829 | 3 | 4049.7343 | 4049.7234 | 0 | 2.7 | 332 | CRagged | 884.73 | 865.15 | 532.67 | 17.52 | 55 | >sp\|P01876\|IGHA1_HUMAN Immunoglobulin heavy constant alpha 1 OS=Homo sapiens OX=9606 GN=IGHA1 PE=1 SV=2 | 255 | 20201119_16913_1ng_glyco_hcd203040_compound15.14268.14268.3 | controllerType=0 controllerNumber=1 scan=14268 | 30.4087 |
| 10999:3 | 10 | R.LAGKPTHVN[+1378.476]VSVVMAEVDGTC[+57.021]Y.- | HexNAc(2)Hex(6) | C[+57], N[+1378] | 1242.5483 | 3 | 3725.6303 | 3725.6178 | 0 | 3.4 | 332 | CRagged | 899.05 | 899.05 | 550.27 | 17.58 | 55 | >sp\|P01876\|IGHA1_HUMAN Immunoglobulin heavy constant alpha 1 OS=Homo sapiens OX=9606 GN=IGHA1 PE=1 SV=2 | 255 | 20201119_16913_1ng_glyco_hcd203040_compound15.14347.14347.3 | controllerType=0 controllerNumber=1 scan=14347 | 30.5748 |
| 10988:3 | 10 | R.LAGKPTHVN[+1809.666]VSVVMAEVDGTC[+57.021]Y.- | HexNAc(5)Hex(4)Fuc(1) | C[+57], N[+1810] | 1386.2795 | 3 | 4156.8239 | 4156.8081 | 0 | 3.8 | 332 | CRagged | 919.15 | 919.15 | 610.15 | 18.58 | 55 | >sp\|P01876\|IGHA1_HUMAN Immunoglobulin heavy constant alpha 1 OS=Homo sapiens OX=9606 GN=IGHA1 PE=1 SV=2 | 255 | 20201119_16913_1ng_glyco_hcd203040_compound15.14335.14335.3 | controllerType=0 controllerNumber=1 scan=14335 | 30.5391 |
| 11015:3 | 10 | R.LAGKPTHVN[+1216.423]VSVVMAEVDGTC[+57.021]Y.- | HexNAc(2)Hex(5) | C[+57], N[+1216] | 1188.5289 | 3 | 3563.572 | 3563.5649 | 0 | 2 | 332 | CRagged | 940.97 | 928.39 | 551.17 | 18.52 | 55 | >sp\|P01876\|IGHA1_HUMAN Immunoglobulin heavy constant alpha 1 OS=Homo sapiens OX=9606 GN=IGHA1 PE=1 SV=2 | 255 | 20201119_16913_1ng_glyco_hcd203040_compound15.14364.14364.3 | controllerType=0 controllerNumber=1 scan=14364 | 30.6096 |
| 08895:3 | 10 | R.LAGKPTHVN[+1216.423]VSVVM[+15.995]AEVDGTC[+57.021]Y.- | HexNAc(2)Hex(5) | C[+57], M[+16], N[+1216] | 1193.8604 | 3 | 3579.5667 | 3579.5599 | 0 | 1.9 | 332 | CRagged | 311.12 | 311.12 | 311.12 | 7.23 | 55 | >sp\|P01876\|IGHA1_HUMAN Immunoglobulin heavy constant alpha 1 OS=Homo sapiens OX=9606 GN=IGHA1 PE=1 SV=2 | 255 | 20201119_16913_1ng_glyco_hcd203040_compound15.12163.12163.3 | controllerType=0 controllerNumber=1 scan=12163 | 26.7948 |
| 18065:3 | 11 | R.NIAAFGGDPNN[+1913.677]ITLFGESAGGASVSLQTLSPYNK.G | HexNAc(4)Hex(5)NeuAc(1) | N[+1914] | 1775.4676 | 3 | 5324.3883 | 5324.3633 | 0 | 4.7 | 197 | Specific | 847.85 | 847.85 | 847.85 | 16.01 | 45 | >sp\|P19835\|CEL_HUMAN Bile salt-activated lipase OS=Homo sapiens OX=9606 GN=CEL PE=1 SV=3 | 476 | 20201119_16913_1ng_glyco_hcd203040_compound15.21739.21739.3 | controllerType=0 controllerNumber=1 scan=21739 | 45.4244 |
| 18062:4 | 11 | R.NIAAFGGDPNN[+1913.677]ITLFGESAGGASVSLQTLSPYNK.G | HexNAc(4)Hex(5)NeuAc(1) | N[+1914] | 1331.851 | 4 | 5324.382 | 5324.3633 | 0 | 3.5 | 197 | Specific | 945.56 | 930.59 | 930.59 | 18.51 | 45 | >sp\|P19835\|CEL_HUMAN Bile salt-activated lipase OS=Homo sapiens OX=9606 GN=CEL PE=1 SV=3 | 476 | 20201119_16913_1ng_glyco_hcd203040_compound15.21736.21736.4 | controllerType=0 controllerNumber=1 scan=21736 | 45.4185 |
| 18060:4 | 11 | R.NIAAFGGDPNN[+1913.677]ITLFGESAGGASVSLQTLSPYNK.G | HexNAc(4)Hex(5)NeuAc(1) | N[+1914] | 1331.851 | 4 | 5324.382 | 5324.3633 | 0 | 3.5 | 197 | Specific | 969 | 945.13 | 945.13 | 19.51 | 45 | >sp\|P19835\|CEL_HUMAN Bile salt-activated lipase OS=Homo sapiens OX=9606 GN=CEL PE=1 SV=3 | 476 | 20201119_16913_1ng_glyco_hcd203040_compound15.21734.21734.4 | controllerType=0 controllerNumber=1 scan=21734 | 45.4121 |

Table S4. Cont.

| Query #:z | Protein Rank | Glycopeptide | Glycan | Modification Type(s) | Observed m/z | z | Observed (M+H) | Calc. mass (M+H) | Off-by-x error | Mass error (ppm) | StartPosition | Cleavage | Score | Delta | Delta Mod | \|Log Prob\| | # of unique peptides | Protein Name | Protein DB number | Comment | Scan # | Scan Time |
|---|---|---|---|---|---|---|---|---|---|---|---|---|---|---|---|---|---|---|---|---|---|---|
| 18068:3 | 11 | R.NIAAFGGDPNN[+1913.677]ITLFGESAGGASVSLQTLSPYNK.G | HexNAc(4)Hex(5)NeuAc(1) | N[+1914] | 1775.4676 | 3 | 5324.3883 | 5324.3633 | 0 | 4.7 | 197 | Specific | 996.12 | 975.04 | 975.04 | 19.01 | 45 | >sp\|P19835\|CEL_HUMAN Bile salt-activated lipase OS=Homo sapiens OX=9606 GN=CEL PE=1 SV=3 | 476 | 20201119_16913_1ng_glyco_hcd203040_compound15.21742.21742.3 | controllerType=0 controllerNumber=1 scan=21742 | 45.4292 |
| 19596:9 | 13 | K.EPLTTSLAPTLTSELSTSQAETSTPRTETTMSPLTN[+1362.481]TTTSQGTTRC[+57.021]( | HexNAc(2)Hex(5)Fuc(1) | C[+57]*2, N[+1362] | 1098.6108 | 9 | 9879.4388 | 9879.4033 | 0 | 3.6 | 1739 | Specific | 261.7 | 261.7 | 261.7 | 3.33 | 48 | >sp\|Q9HC84\|MUC5B_HUMAN Mucin-5B OS=Homo sapiens OX=9606 GN=MUC5B PE=1 SV=3 | 677 | 20201119_16913_1ng_glyco_hcd203040_compound15.23339.23339.3 | controllerType=0 controllerNumber=1 scan=23339 | 48.9293 |
| 19599:9 | 13 | K.EPLTTSLAPTLTSELSTSQAETSTPRTETTMSPLTN[+1362.481]TTTSQGTTRC[+57.021]( | HexNAc(2)Hex(5)Fuc(1) | C[+57]*2, N[+1362] | 1098.6108 | 9 | 9879.4388 | 9879.4033 | 0 | 3.6 | 1739 | Specific | 304.46 | 304.46 | 304.46 | 3.57 | 48 | >sp\|Q9HC84\|MUC5B_HUMAN Mucin-5B OS=Homo sapiens OX=9606 GN=MUC5B PE=1 SV=3 | 677 | 20201119_16913_1ng_glyco_hcd203040_compound15.23342.23342.3 | controllerType=0 controllerNumber=1 scan=23342 | 48.9529 |
| 13372:4 | 14 | K.LKELPGVC[+57.021]N[+2424.867]ETMMALWEEC[+57.021]KPC[+57.021]LK.Q | HexNAc(5)Hex(6)Fuc(1)NeuAc(1) | C[+57]*3, N[+2425] | 1341.0704 | 4 | 5361.2599 | 5361.2518 | 0 | 1.5 | 95 | Specific | 380.88 | 322.63 | 257.28 | 6.72 | 48 | >sp\|P10909\|CLUS_HUMAN Clusterin OS=Homo sapiens OX=9606 GN=CLU PE=1 SV=1 | 1467 | 20201119_16913_1ng_glyco_hcd203040_compound15.16825.16825.4 | controllerType=0 controllerNumber=1 scan=16825 | 35.2447 |
| 13374:4 | 14 | K.LKELPGVC[+57.021]N[+2424.867]ETMMALWEEC[+57.021]KPC[+57.021]LK.Q | HexNAc(5)Hex(6)Fuc(1)NeuAc(1) | C[+57]*3, N[+2425] | 1341.0704 | 4 | 5361.2599 | 5361.2518 | 0 | 1.5 | 95 | Specific | 396.26 | 389.37 | 300.09 | 8.58 | 48 | >sp\|P10909\|CLUS_HUMAN Clusterin OS=Homo sapiens OX=9606 GN=CLU PE=1 SV=1 | 1467 | 20201119_16913_1ng_glyco_hcd203040_compound15.16827.16827.4 | controllerType=0 controllerNumber=1 scan=16827 | 35.2486 |
| 13468:3 | 14 | K.LKELPGVC[+57.021]N[+2059.735]ETMMALWEEC[+57.021]KPC[+57.021]LK.Q | HexNAc(4)Hex(5)Fuc(1)NeuAc(1) | C[+57]*3, N[+2060] | 1666.0522 | 3 | 4996.1421 | 4996.1196 | 0 | 4.5 | 95 | Specific | 406.36 | 406.36 | 406.36 | 8.04 | 48 | >sp\|P10909\|CLUS_HUMAN Clusterin OS=Homo sapiens OX=9606 GN=CLU PE=1 SV=1 | 1467 | 20201119_16913_1ng_glyco_hcd203040_compound15.16925.16925.3 | controllerType=0 controllerNumber=1 scan=16925 | 35.431 |
| 15062:3 | 14 | K.LKELPGVC[+57.021]N[+2350.830]ETMMALWEEC[+57.021]KPC[+57.021]LK.Q | HexNAc(4)Hex(5)Fuc(1)NeuAc(2) | C[+57]*3, N[+2351] | 1763.0889 | 3 | 5287.2521 | 5287.2151 | 0 | 7 | 95 | Specific | 497.01 | 497.01 | 497.01 | 7.56 | 48 | >sp\|P10909\|CLUS_HUMAN Clusterin OS=Homo sapiens OX=9606 GN=CLU PE=1 SV=1 | 1467 | 20201119_16913_1ng_glyco_hcd203040_compound15.18582.18582.3 | controllerType=0 controllerNumber=1 scan=18582 | 38.6032 |
| 15060:3 | 14 | K.LKELPGVC[+57.021]N[+2350.830]ETMMALWEEC[+57.021]KPC[+57.021]LK.Q | HexNAc(4)Hex(5)Fuc(1)NeuAc(2) | C[+57]*3, N[+2351] | 1763.0889 | 3 | 5287.2521 | 5287.2151 | 0 | 7 | 95 | Specific | 527.79 | 487.58 | 487.58 | 7.81 | 48 | >sp\|P10909\|CLUS_HUMAN Clusterin OS=Homo sapiens OX=9606 GN=CLU PE=1 SV=1 | 1467 | 20201119_16913_1ng_glyco_hcd203040_compound15.18580.18580.3 | controllerType=0 controllerNumber=1 scan=18580 | 38.5967 |
| 13471:3 | 14 | K.LKELPGVC[+57.021]N[+2059.735]ETMMALWEEC[+57.021]KPC[+57.021]LK.Q | HexNAc(4)Hex(5)Fuc(1)NeuAc(1) | C[+57]*3, N[+2060] | 1666.0522 | 3 | 4996.1421 | 4996.1196 | 0 | 4.5 | 95 | Specific | 564.36 | 543.84 | 543.84 | 8.72 | 48 | >sp\|P10909\|CLUS_HUMAN Clusterin OS=Homo sapiens OX=9606 GN=CLU PE=1 SV=1 | 1467 | 20201119_16913_1ng_glyco_hcd203040_compound15.16928.16928.3 | controllerType=0 controllerNumber=1 scan=16928 | 35.4354 |
| 15051:4 | 14 | K.LKELPGVC[+57.021]N[+2351.851]ETMMALWEEC[+57.021]KPC[+57.021]LK.Q | HexNAc(4)Hex(5)Fuc(3)NeuAc(1) | C[+57]*3, N[+2352] | 1322.8109 | 4 | 5288.2218 | 5288.2355 | 0 | -2.6 | 95 | Specific | 564.77 | 553.15 | 553.15 | 8.4 | 48 | >sp\|P10909\|CLUS_HUMAN Clusterin OS=Homo sapiens OX=9606 GN=CLU PE=1 SV=1 | 1467 | 20201119_16913_1ng_glyco_hcd203040_compound15.18571.18571.4 | controllerType=0 controllerNumber=1 scan=18571 | 38.5816 |
| 13283:4 | 14 | K.LKELPGVC[+57.021]N[+2571.946]ETMMALWEEC[+57.021]KPC[+57.021]LK.Q | HexNAc(5)Hex(6)Fuc(4) | C[+57]*3, N[+2572] | 1377.8336 | 4 | 5508.3127 | 5508.3302 | 0 | -3.2 | 95 | Specific | 602.62 | 602.62 | 602.62 | 10.66 | 48 | >sp\|P10909\|CLUS_HUMAN Clusterin OS=Homo sapiens OX=9606 GN=CLU PE=1 SV=1 | 1467 | 20201119_16913_1ng_glyco_hcd203040_compound15.16733.16733.4 | controllerType=0 controllerNumber=1 scan=16733 | 35.0807 |
| 13281:4 | 14 | K.LKELPGVC[+57.021]N[+2571.946]ETMMALWEEC[+57.021]KPC[+57.021]LK.Q | HexNAc(5)Hex(6)Fuc(4) | C[+57]*3, N[+2572] | 1377.8336 | 4 | 5508.3127 | 5508.3302 | 0 | -3.2 | 95 | Specific | 678.66 | 678.66 | 678.66 | 11.66 | 48 | >sp\|P10909\|CLUS_HUMAN Clusterin OS=Homo sapiens OX=9606 GN=CLU PE=1 SV=1 | 1467 | 20201119_16913_1ng_glyco_hcd203040_compound15.16731.16731.4 | controllerType=0 controllerNumber=1 scan=16731 | 35.0767 |
| 15053:4 | 14 | K.LKELPGVC[+57.021]N[+2351.851]ETMMALWEEC[+57.021]KPC[+57.021]LK.Q | HexNAc(4)Hex(5)Fuc(3)NeuAc(1) | C[+57]*3, N[+2352] | 1322.8109 | 4 | 5288.2218 | 5288.2355 | 0 | -2.6 | 95 | Specific | 834.28 | 824.39 | 824.39 | 14.66 | 48 | >sp\|P10909\|CLUS_HUMAN Clusterin OS=Homo sapiens OX=9606 GN=CLU PE=1 SV=1 | 1467 | 20201119_16913_1ng_glyco_hcd203040_compound15.18573.18573.4 | controllerType=0 controllerNumber=1 scan=18573 | 38.5851 |
| 13384:4 | 14 | K.LKELPGVC[+57.021]N[+2205.793]ETMMALWEEC[+57.021]KPC[+57.021]LK.Q | HexNAc(4)Hex(5)Fuc(2)NeuAc(1) | C[+57]*3, N[+2206] | 1286.3037 | 4 | 5142.193 | 5142.1776 | 0 | 3 | 95 | Specific | 905.55 | 905.55 | 905.55 | 18.02 | 48 | >sp\|P10909\|CLUS_HUMAN Clusterin OS=Homo sapiens OX=9606 GN=CLU PE=1 SV=1 | 1467 | 20201119_16913_1ng_glyco_hcd203040_compound15.16838.16838.4 | controllerType=0 controllerNumber=1 scan=16838 | 35.27 |
| 13356:4 | 14 | K.LKELPGVC[+57.021]N[+2351.851]ETMMALWEEC[+57.021]KPC[+57.021]LK.Q | HexNAc(4)Hex(5)Fuc(3)NeuAc(1) | C[+57]*3, N[+2352] | 1322.8173 | 4 | 5288.2472 | 5288.2355 | 0 | 2.2 | 95 | Specific | 906.82 | 906.82 | 906.82 | 17.96 | 48 | >sp\|P10909\|CLUS_HUMAN Clusterin OS=Homo sapiens OX=9606 GN=CLU PE=1 SV=1 | 1467 | 20201119_16913_1ng_glyco_hcd203040_compound15.16809.16809.4 | controllerType=0 controllerNumber=1 scan=16809 | 35.2144 |
| 13419:4 | 14 | K.LKELPGVC[+57.021]N[+2059.735]ETMMALWEEC[+57.021]KPC[+57.021]LK.Q | HexNAc(4)Hex(5)Fuc(1)NeuAc(1) | C[+57]*3, N[+2060] | 1249.7893 | 4 | 4996.1354 | 4996.1196 | 0 | 3.2 | 95 | Specific | 944.23 | 873.95 | 873.95 | 18.02 | 48 | >sp\|P10909\|CLUS_HUMAN Clusterin OS=Homo sapiens OX=9606 GN=CLU PE=1 SV=1 | 1467 | 20201119_16913_1ng_glyco_hcd203040_compound15.16874.16874.4 | controllerType=0 controllerNumber=1 scan=16874 | 35.335 |
| 13416:4 | 14 | K.LKELPGVC[+57.021]N[+2059.735]ETMMALWEEC[+57.021]KPC[+57.021]LK.Q | HexNAc(4)Hex(5)Fuc(1)NeuAc(1) | C[+57]*3, N[+2060] | 1249.7893 | 4 | 4996.1354 | 4996.1196 | 0 | 3.2 | 95 | Specific | 1069.4 | 952.16 | 952.16 | 21.02 | 48 | >sp\|P10909\|CLUS_HUMAN Clusterin OS=Homo sapiens OX=9606 GN=CLU PE=1 SV=1 | 1467 | 20201119_16913_1ng_glyco_hcd203040_compound15.16871.16871.4 | controllerType=0 controllerNumber=1 scan=16871 | 35.3268 |
| 13381:4 | 14 | K.LKELPGVC[+57.021]N[+2205.793]ETMMALWEEC[+57.021]KPC[+57.021]LK.Q | HexNAc(4)Hex(5)Fuc(2)NeuAc(1) | C[+57]*3, N[+2206] | 1286.3037 | 4 | 5142.193 | 5142.1776 | 0 | 3 | 95 | Specific | 1069.7 | 942.47 | 942.47 | 21.02 | 48 | >sp\|P10909\|CLUS_HUMAN Clusterin OS=Homo sapiens OX=9606 GN=CLU PE=1 SV=1 | 1467 | 20201119_16913_1ng_glyco_hcd203040_compound15.16835.16835.4 | controllerType=0 controllerNumber=1 scan=16835 | 35.2642 |
| 16961:3 | 14 | K.ELPGVC[+57.021]N[+2350.830]ETMMALWEEC[+57.021]KPC[+57.021]LK.Q | HexNAc(4)Hex(5)Fuc(1)NeuAc(2) | C[+57]*3, N[+2351] | 1682.6956 | 3 | 5046.0722 | 5046.036 | 0 | 7.2 | 97 | Specific | 227.6 | 222.53 | 222.53 | 5.22 | 48 | >sp\|P10909\|CLUS_HUMAN Clusterin OS=Homo sapiens OX=9606 GN=CLU PE=1 SV=1 | 1467 | 20201119_16913_1ng_glyco_hcd203040_compound15.20584.20584.3 | controllerType=0 controllerNumber=1 scan=20584 | 42.7942 |
| 15146:3 | 14 | K.ELPGVC[+57.021]N[+2059.735]ETMMALWEEC[+57.021]KPC[+57.021]LK.Q | HexNAc(4)Hex(5)Fuc(1)NeuAc(1) | C[+57]*3, N[+2060] | 1585.6503 | 3 | 4754.9364 | 4754.9406 | 0 | -0.9 | 97 | Specific | 504.78 | 495.91 | 495.91 | 9.16 | 48 | >sp\|P10909\|CLUS_HUMAN Clusterin OS=Homo sapiens OX=9606 GN=CLU PE=1 SV=1 | 1467 | 20201119_16913_1ng_glyco_hcd203040_compound15.18670.18670.3 | controllerType=0 controllerNumber=1 scan=18670 | 38.7976 |
| 15143:3 | 14 | K.ELPGVC[+57.021]N[+2059.735]ETMMALWEEC[+57.021]KPC[+57.021]LK.Q | HexNAc(4)Hex(5)Fuc(1)NeuAc(1) | C[+57]*3, N[+2060] | 1585.6503 | 3 | 4754.9364 | 4754.9406 | 0 | -0.9 | 97 | Specific | 661.15 | 595.77 | 595.77 | 12.7 | 48 | >sp\|P10909\|CLUS_HUMAN Clusterin OS=Homo sapiens OX=9606 GN=CLU PE=1 SV=1 | 1467 | 20201119_16913_1ng_glyco_hcd203040_compound15.18667.18667.3 | controllerType=0 controllerNumber=1 scan=18667 | 38.7911 |
| 19264:3 | 14 | R.QLEEFLN[+2059.735]QSSPFYFWMNGDR.I | HexNAc(4)Hex(5)Fuc(1)NeuAc(1) | N[+2060] | 1523.2932 | 3 | 4567.865 | 4567.8638 | 0 | 0.3 | 139 | Specific | 680.92 | 615.92 | 615.92 | 13.09 | 48 | >sp\|P10909\|CLUS_HUMAN Clusterin OS=Homo sapiens OX=9606 GN=CLU PE=1 SV=1 | 1467 | 20201119_16913_1ng_glyco_hcd203040_compound15.22992.22992.3 | controllerType=0 controllerNumber=1 scan=22992 | 48.1436 |
| 19267:3 | 14 | R.QLEEFLN[+2059.735]QSSPFYFWMNGDR.I | HexNAc(4)Hex(5)Fuc(1)NeuAc(1) | N[+2060] | 1523.2932 | 3 | 4567.865 | 4567.8638 | 0 | 0.3 | 139 | Specific | 843.84 | 719.12 | 719.12 | 16.09 | 48 | >sp\|P10909\|CLUS_HUMAN Clusterin OS=Homo sapiens OX=9606 GN=CLU PE=1 SV=1 | 1467 | 20201119_16913_1ng_glyco_hcd203040_compound15.22995.22995.3 | controllerType=0 controllerNumber=1 scan=22995 | 48.1478 |
| 22000:4 | 14 | R.QLEEFLN[+2351.851]QSSPFYFWMNGDRIDSLLENDR.Q | HexNAc(4)Hex(5)Fuc(3)NeuAc(1) | N[+2352] | 1479.6246 | 4 | 5915.4767 | 5915.5044 | 0 | -4.7 | 139 | Specific | 233.08 | 232.26 | 232.26 | 4.3 | 48 | >sp\|P10909\|CLUS_HUMAN Clusterin OS=Homo sapiens OX=9606 GN=CLU PE=1 SV=1 | 1467 | 20201119_16913_1ng_glyco_hcd203040_compound15.25856.25856.3 | controllerType=0 controllerNumber=1 scan=25856 | 54.4745 |
| 22003:4 | 14 | R.QLEEFLN[+2351.851]QSSPFYFWMNGDRIDSLLENDR.Q | HexNAc(4)Hex(5)Fuc(3)NeuAc(1) | N[+2352] | 1479.6246 | 4 | 5915.4767 | 5915.5044 | 0 | -4.7 | 139 | Specific | 498.77 | 415.91 | 415.91 | 7.45 | 48 | >sp\|P10909\|CLUS_HUMAN Clusterin OS=Homo sapiens OX=9606 GN=CLU PE=1 SV=1 | 1467 | 20201119_16913_1ng_glyco_hcd203040_compound15.25859.25859.3 | controllerType=0 controllerNumber=1 scan=25859 | 54.4828 |
| 00130:3 | 14 | R.EIRHN[+2350.830]STGC[+57.021]LR.M | HexNAc(4)Hex(5)Fuc(1)NeuAc(2) | C[+57], N[+2351] | 1231.8416 | 3 | 3693.5103 | 3693.4948 | 0 | 4.2 | 287 | Specific | 206.68 | 132.32 | 132.32 | 3.11 | 48 | >sp\|P10909\|CLUS_HUMAN Clusterin OS=Homo sapiens OX=9606 GN=CLU PE=1 SV=1 | 1467 | 20201119_16913_1ng_glyco_hcd203040_compound15.3033.3033.3 | controllerType=0 controllerNumber=1 scan=3033 | 11.3548 |
| 00088:4 | 14 | R.EIRHN[+2424.867]STGC[+57.021]LR.M | HexNAc(5)Hex(6)Fuc(1)NeuAc(1) | C[+57], N[+2425] | 942.6406 | 4 | 3767.5406 | 3767.5316 | 0 | 2.4 | 287 | Specific | 230.75 | 156.78 | 156.78 | 4.8 | 48 | >sp\|P10909\|CLUS_HUMAN Clusterin OS=Homo sapiens OX=9606 GN=CLU PE=1 SV=1 | 1467 | 20201119_16913_1ng_glyco_hcd203040_compound15.2989.2989.4 | controllerType=0 controllerNumber=1 scan=2989 | 11.2651 |
| 22128:3 | 14 | K.[+42.011]MLN[+2336.851]TSSLLEQLNEQFNWVSR.L | HexNAc(6)Hex(6)Fuc(1) | .[42], N[+2337] | 1596.6839 | 3 | 4788.0371 | 4788.0484 | 0 | -2.3 | 352 | Specific | 241.22 | 125.77 | 125.77 | 3.62 | 48 | >sp\|P10909\|CLUS_HUMAN Clusterin OS=Homo sapiens OX=9606 GN=CLU PE=1 SV=1 | 1467 | 20201119_16913_1ng_glyco_hcd203040_compound15.25990.25990.3 | controllerType=0 controllerNumber=1 scan=25990 | 54.7603 |
| 22126:3 | 14 | K.[+42.011]MLN[+2336.851]TSSLLEQLNEQFNWVSR.L | HexNAc(6)Hex(6)Fuc(1) | .[42], N[+2337] | 1596.6839 | 3 | 4788.0371 | 4788.0484 | 0 | -2.3 | 352 | Specific | 314.31 | 269.99 | 269.99 | 5.5 | 48 | >sp\|P10909\|CLUS_HUMAN Clusterin OS=Homo sapiens OX=9606 GN=CLU PE=1 SV=1 | 1467 | 20201119_16913_1ng_glyco_hcd203040_compound15.25988.25988.3 | controllerType=0 controllerNumber=1 scan=25988 | 54.7552 |
| 21985:3 | 14 | K.MLN[+2350.830]TSSLLEQLNEQFNWVSR.L | HexNAc(4)Hex(5)Fuc(1)NeuAc(2) | N[+2351] | 1587.3524 | 3 | 4760.0425 | 4760.0171 | 0 | 5.3 | 352 | Specific | 899.22 | 640.75 | 640.75 | 16.71 | 48 | >sp\|P10909\|CLUS_HUMAN Clusterin OS=Homo sapiens OX=9606 GN=CLU PE=1 SV=1 | 1467 | 20201119_16913_1ng_glyco_hcd203040_compound15.25840.25840.3 | controllerType=0 controllerNumber=1 scan=25840 | 54.42 |
| 21986:4 | 14 | K.MLN[+2350.830]TSSLLEQLNEQFNWVSR.L | HexNAc(4)Hex(5)Fuc(1)NeuAc(2) | N[+2351] | 1190.7664 | 4 | 4760.0438 | 4760.0171 | 0 | 5.6 | 352 | Specific | 941.75 | 684.25 | 684.25 | 17.85 | 48 | >sp\|P10909\|CLUS_HUMAN Clusterin OS=Homo sapiens OX=9606 GN=CLU PE=1 SV=1 | 1467 | 20201119_16913_1ng_glyco_hcd203040_compound15.25841.25841.4 | controllerType=0 controllerNumber=1 scan=25841 | 54.4243 |
| 19942:3 | 14 | K.MLN[+2059.735]TSSLLEQLNEQFNWVSR.L | HexNAc(4)Hex(5)Fuc(1)NeuAc(1) | N[+2060] | 1490.3191 | 3 | 4468.9429 | 4468.9217 | 0 | 4.8 | 352 | Specific | 955.7 | 690.03 | 690.03 | 19.05 | 48 | >sp\|P10909\|CLUS_HUMAN Clusterin OS=Homo sapiens OX=9606 GN=CLU PE=1 SV=1 | 1467 | 20201119_16913_1ng_glyco_hcd203040_compound15.23702.23702.3 | controllerType=0 controllerNumber=1 scan=23702 | 49.7881 |
| 19947:4 | 14 | K.MLN[+2059.735]TSSLLEQLNEQFNWVSR.L | HexNAc(4)Hex(5)Fuc(1)NeuAc(1) | N[+2060] | 1117.9906 | 4 | 4468.9407 | 4468.9217 | 0 | 4.3 | 352 | Specific | 1019.91 | 745.21 | 745.21 | 20.19 | 48 | >sp\|P10909\|CLUS_HUMAN Clusterin OS=Homo sapiens OX=9606 GN=CLU PE=1 SV=1 | 1467 | 20201119_16913_1ng_glyco_hcd203040_compound15.23707.23707.4 | controllerType=0 controllerNumber=1 scan=23707 | 49.7985 |
| 21983:3 | 14 | K.MLN[+2350.830]TSSLLEQLNEQFNWVSR.L | HexNAc(4)Hex(5)Fuc(1)NeuAc(2) | N[+2351] | 1587.3524 | 3 | 4760.0425 | 4760.0171 | 0 | 5.3 | 352 | Specific | 1038.2 | 772.08 | 772.08 | 19.71 | 48 | >sp\|P10909\|CLUS_HUMAN Clusterin OS=Homo sapiens OX=9606 GN=CLU PE=1 SV=1 | 1467 | 20201119_16913_1ng_glyco_hcd203040_compound15.25838.25838.3 | controllerType=0 controllerNumber=1 scan=25838 | 54.4146 |
| 21984:4 | 14 | K.MLN[+2350.830]TSSLLEQLNEQFNWVSR.L | HexNAc(4)Hex(5)Fuc(1)NeuAc(2) | N[+2351] | 1190.7664 | 4 | 4760.0438 | 4760.0171 | 0 | 5.6 | 352 | Specific | 1050.43 | 793.81 | 793.81 | 20.85 | 48 | >sp\|P10909\|CLUS_HUMAN Clusterin OS=Homo sapiens OX=9606 GN=CLU PE=1 SV=1 | 1467 | 20201119_16913_1ng_glyco_hcd203040_compound15.25839.25839.4 | controllerType=0 controllerNumber=1 scan=25839 | 54.4163 |
| 19944:4 | 14 | K.MLN[+2059.735]TSSLLEQLNEQFNWVSR.L | HexNAc(4)Hex(5)Fuc(1)NeuAc(1) | N[+2060] | 1117.9906 | 4 | 4468.9407 | 4468.9217 | 0 | 4.3 | 352 | Specific | 1056.9 | 827.46 | 827.46 | 21.19 | 48 | >sp\|P10909\|CLUS_HUMAN Clusterin OS=Homo sapiens OX=9606 GN=CLU PE=1 SV=1 | 1467 | 20201119_16913_1ng_glyco_hcd203040_compound15.23704.23704.4 | controllerType=0 controllerNumber=1 scan=23704 | 49.7929 |
| 19939:3 | 14 | K.MLN[+2059.735]TSSLLEQLNEQFNWVSR.L | HexNAc(4)Hex(5)Fuc(1)NeuAc(1) | N[+2060] | 1490.3191 | 3 | 4468.9429 | 4468.9217 | 0 | 4.8 | 352 | Specific | 1078.5 | 858.14 | 858.14 | 21.05 | 48 | >sp\|P10909\|CLUS_HUMAN Clusterin OS=Homo sapiens OX=9606 GN=CLU PE=1 SV=1 | 1467 | 20201119_16913_1ng_glyco_hcd203040_compound15.23699.23699.3 | controllerType=0 controllerNumber=1 scan=23699 | 49.7837 |
| 19533:3 | 14 | K.M[+15.995]LN[+2059.735]TSSLLEQLNEQFNWVSR.L | HexNAc(4)Hex(5)Fuc(1)NeuAc(1) | M[+16], N[+2060] | 1495.6531 | 3 | 4484.9448 | 4484.9166 | 0 | 6.3 | 352 | Specific | 700.23 | 575.76 | 575.76 | 13.71 | 48 | >sp\|P10909\|CLUS_HUMAN Clusterin OS=Homo sapiens OX=9606 GN=CLU PE=1 SV=1 | 1467 | 20201119_16913_1ng_glyco_hcd203040_compound15.23274.23274.3 | controllerType=0 controllerNumber=1 scan=23274 | 48.7905 |
| 19535:3 | 14 | K.M[+15.995]LN[+2059.735]TSSLLEQLNEQFNWVSR.L | HexNAc(4)Hex(5)Fuc(1)NeuAc(1) | M[+16], N[+2060] | 1495.6531 | 3 | 4484.9448 | 4484.9166 | 0 | 6.3 | 352 | Specific | 839.89 | 779.5 | 779.5 | 15.71 | 48 | >sp\|P10909\|CLUS_HUMAN Clusterin OS=Homo sapiens OX=9606 GN=CLU PE=1 SV=1 | 1467 | 20201119_16913_1ng_glyco_hcd203040_compound15.23276.23276.3 | controllerType=0 controllerNumber=1 scan=23276 | 48.7947 |
| 10007:3 | 14 | R.LAN[+2205.793]LTQGEDQYYLR.V | HexNAc(4)Hex(5)Fuc(2)NeuAc(1) | N[+2206] | 1297.2166 | 3 | 3889.6353 | 3889.6265 | 0 | 2.3 | 372 | Specific | 320.39 | 295.96 | 295.96 | 6.8 | 48 | >sp\|P10909\|CLUS_HUMAN Clusterin OS=Homo sapiens OX=9606 GN=CLU PE=1 SV=1 | 1467 | 20201119_16913_1ng_glyco_hcd203040_compound15.13318.13318.3 | controllerType=0 controllerNumber=1 scan=13318 | 28.774 |
| 10009:3 | 14 | R.LAN[+2205.793]LTQGEDQYYLR.V | HexNAc(4)Hex(5)Fuc(2)NeuAc(1) | N[+2206] | 1297.2166 | 3 | 3889.6353 | 3889.6265 | 0 | 2.3 | 372 | Specific | 421.26 | 354.7 | 354.7 | 7.87 | 48 | >sp\|P10909\|CLUS_HUMAN Clusterin OS=Homo sapiens OX=9606 GN=CLU PE=1 SV=1 | 1467 | 20201119_16913_1ng_glyco_hcd203040_compound15.13320.13320.3 | controllerType=0 controllerNumber=1 scan=13320 | 28.7778 |
| 22690:3 | 16 | R.[+42.011]QLAHQSN[+2190.793]STNIFFSPVSIATAFAMLSLGTK.A | HexNAc(6)Hex(6) | .[42], N[+2191] | 1805.4809 | 3 | 5414.4282 | 5414.4388 | 0 | -2 | 64 | Specific | 217.27 | 217.27 | 217.27 | 3.8 | 41 | >sp\|P01009\|A1AT_HUMAN Alpha-1-antitrypsin OS=Homo sapiens OX=9606 GN=SERPINA1 PE=1 SV=3 | 642 | 20201119_16913_1ng_glyco_hcd203040_compound15.26577.26577.3 | controllerType=0 controllerNumber=1 scan=26577 | 56.0207 |
| 25178:4 | 16 | R.QLAHQSN[+2861.000]STNIFFSPVSIATAFAMLSLGTK.A | HexNAc(5)Hex(6)NeuAc(3) | N[+2861] | 1511.4258 | 4 | 6042.6813 | 6042.6351 | 0 | 7.6 | 64 | Specific | 296.2 | 296.2 | 296.2 | 6 | 41 | >sp\|P01009\|A1AT_HUMAN Alpha-1-antitrypsin OS=Homo sapiens OX=9606 GN=SERPINA1 PE=1 SV=3 | 642 | 20201119_16913_1ng_glyco_hcd203040_compound15.29165.29165.4 | controllerType=0 controllerNumber=1 scan=29165 | 61.0622 |
| 25176:4 | 16 | R.QLAHQSN[+2861.000]STNIFFSPVSIATAFAMLSLGTK.A | HexNAc(5)Hex(6)NeuAc(3) | N[+2861] | 1511.4258 | 4 | 6042.6813 | 6042.6351 | 0 | 7.6 | 64 | Specific | 300.71 | 270.16 | 270.16 | 6.02 | 41 | >sp\|P01009\|A1AT_HUMAN Alpha-1-antitrypsin OS=Homo sapiens OX=9606 GN=SERPINA1 PE=1 SV=3 | 642 | 20201119_16913_1ng_glyco_hcd203040_compound15.29163.29163.4 | controllerType=0 controllerNumber=1 scan=29163 | 61.058 |
| 25267:4 | 16 | R.QLAHQSN[+2862.021]STNIFFSPVSIATAFAMLSLGTK.A | HexNAc(5)Hex(6)Fuc(2)NeuAc(2) | N[+2862] | 1511.6716 | 4 | 6043.6647 | 6043.6555 | 0 | 1.5 | 64 | Specific | 330.4 | 274.82 | 274.82 | 7.09 | 41 | >sp\|P01009\|A1AT_HUMAN Alpha-1-antitrypsin OS=Homo sapiens OX=9606 GN=SERPINA1 PE=1 SV=3 | 642 | 20201119_16913_1ng_glyco_hcd203040_compound15.29258.29258.4 | controllerType=0 controllerNumber=1 scan=29258 | 61.2415 |
| 22726:4 | 16 | R.QLAHQSN[+2350.830]STNIFFSPVSIATAFAMLSLGTK.A | HexNAc(4)Hex(5)Fuc(1)NeuAc(2) | N[+2351] | 1383.8711 | 4 | 5532.4626 | 5532.4654 | 0 | -0.5 | 64 | Specific | 392.58 | 392.58 | 392.58 | 8.58 | 41 | >sp\|P01009\|A1AT_HUMAN Alpha-1-antitrypsin OS=Homo sapiens OX=9606 GN=SERPINA1 PE=1 SV=3 | 642 | 20201119_16913_1ng_glyco_hcd203040_compound15.26614.26614.4 | controllerType=0 controllerNumber=1 scan=26614 | 56.1063 |
| 25236:4 | 16 | R.QLAHQSN[+3007.058]STNIFFSPVSIATAFAMLSLGTK.A | HexNAc(5)Hex(6)Fuc(1)NeuAc(3) | N[+3007] | 1547.9327 | 4 | 6188.7091 | 6188.693 | 0 | 2.6 | 64 | Specific | 397.26 | 397.26 | 397.26 | 9.39 | 41 | >sp\|P01009\|A1AT_HUMAN Alpha-1-antitrypsin OS=Homo sapiens OX=9606 GN=SERPINA1 PE=1 SV=3 | 642 | 20201119_16913_1ng_glyco_hcd203040_compound15.29226.29226.4 | controllerType=0 controllerNumber=1 scan=29226 | 61.1817 |
| 25265:4 | 16 | R.QLAHQSN[+2862.021]STNIFFSPVSIATAFAMLSLGTK.A | HexNAc(5)Hex(6)Fuc(2)NeuAc(2) | N[+2862] | 1511.6716 | 4 | 6043.6647 | 6043.6555 | 0 | 1.5 | 64 | Specific | 397.7 | 395.06 | 395.06 | 9.39 | 41 | >sp\|P01009\|A1AT_HUMAN Alpha-1-antitrypsin OS=Homo sapiens OX=9606 GN=SERPINA1 PE=1 SV=3 | 642 | 20201119_16913_1ng_glyco_hcd203040_compound15.29256.29256.4 | controllerType=0 controllerNumber=1 scan=29256 | 61.2372 |
| 22728:4 | 16 | R.QLAHQSN[+2350.830]STNIFFSPVSIATAFAMLSLGTK.A | HexNAc(4)Hex(5)Fuc(1)NeuAc(2) | N[+2351] | 1383.8711 | 4 | 5532.4626 | 5532.4654 | 0 | -0.5 | 64 | Specific | 430.53 | 388.16 | 388.16 | 8.83 | 41 | >sp\|P01009\|A1AT_HUMAN Alpha-1-antitrypsin OS=Homo sapiens OX=9606 GN=SERPINA1 PE=1 SV=3 | 642 | 20201119_16913_1ng_glyco_hcd203040_compound15.26616.26616.4 | controllerType=0 controllerNumber=1 scan=26616 | 56.1106 |
| 22593:3 | 16 | R.QLAHQSN[+2350.830]STNIFFSPVSIATAFAMLSLGTK.A | HexNAc(4)Hex(5)Fuc(1)NeuAc(2) | N[+2351] | 1844.8304 | 3 | 5532.4767 | 5532.4654 | 0 | 2 | 64 | Specific | 496.58 | 478.06 | 478.06 | 9.82 | 41 | >sp\|P01009\|A1AT_HUMAN Alpha-1-antitrypsin OS=Homo sapiens OX=9606 GN=SERPINA1 PE=1 SV=3 | 642 | 20201119_16913_1ng_glyco_hcd203040_compound15.26475.26475.3 | controllerType=0 controllerNumber=1 scan=26475 | 55.7804 |
| 25314:4 | 16 | R.QLAHQSN[+2861.000]STNIFFSPVSIATAFAMLSLGTK.A | HexNAc(5)Hex(6)NeuAc(3) | N[+2861] | 1511.4232 | 4 | 6042.6711 | 6042.6351 | 0 | 5.9 | 64 | Specific | 498.62 | 458.53 | 458.53 | 9.08 | 41 | >sp\|P01009\|A1AT_HUMAN Alpha-1-antitrypsin OS=Homo sapiens OX=9606 GN=SERPINA1 PE=1 SV=3 | 642 | 20201119_16913_1ng_glyco_hcd203040_compound15.29307.29307.4 | controllerType=0 controllerNumber=1 scan=29307 | 61.3354 |
| 25316:4 | 16 | R.QLAHQSN[+2861.000]STNIFFSPVSIATAFAMLSLGTK.A | HexNAc(5)Hex(6)NeuAc(3) | N[+2861] | 1511.4232 | 4 | 6042.6711 | 6042.6351 | 0 | 5.9 | 64 | Specific | 559.99 | 552.15 | 552.15 | 9.61 | 41 | >sp\|P01009\|A1AT_HUMAN Alpha-1-antitrypsin OS=Homo sapiens OX=9606 GN=SERPINA1 PE=1 SV=3 | 642 | 20201119_16913_1ng_glyco_hcd203040_compound15.29309.29309.4 | controllerType=0 controllerNumber=1 scan=29309 | 61.3392 |
| 25238:4 | 16 | R.QLAHQSN[+3007.058]STNIFFSPVSIATAFAMLSLGTK.A | HexNAc(5)Hex(6)Fuc(1)NeuAc(3) | N[+3007] | 1547.9327 | 4 | 6188.7091 | 6188.693 | 0 | 2.6 | 64 | Specific | 563.73 | 515.09 | 515.09 | 10.25 | 41 | >sp\|P01009\|A1AT_HUMAN Alpha-1-antitrypsin OS=Homo sapiens OX=9606 GN=SERPINA1 PE=1 SV=3 | 642 | 20201119_16913_1ng_glyco_hcd203040_compound15.29228.29228.4 | controllerType=0 controllerNumber=1 scan=29228 | 61.186 |
| 22595:3 | 16 | R.QLAHQSN[+2350.830]STNIFFSPVSIATAFAMLSLGTK.A | HexNAc(4)Hex(5)Fuc(1)NeuAc(2) | N[+2351] | 1844.8304 | 3 | 5532.4767 | 5532.4654 | 0 | 2 | 64 | Specific | 565.14 | 474.98 | 474.98 | 10.34 | 41 | >sp\|P01009\|A1AT_HUMAN Alpha-1-antitrypsin OS=Homo sapiens OX=9606 GN=SERPINA1 PE=1 SV=3 | 642 | 20201119_16913_1ng_glyco_hcd203040_compound15.26477.26477.3 | controllerType=0 controllerNumber=1 scan=26477 | 55.7842 |
| 22643:3 | 16 | R.QLAHQSN[+2204.772]STNIFFSPVSIATAFAMLSLGTK.A | HexNAc(4)Hex(5)NeuAc(2) | N[+2205] | 1796.1473 | 3 | 5386.4274 | 5386.4075 | 0 | 3.7 | 64 | Specific | 814.67 | 747.68 | 747.68 | 16.66 | 41 | >sp\|P01009\|A1AT_HUMAN Alpha-1-antitrypsin OS=Homo sapiens OX=9606 GN=SERPINA1 PE=1 SV=3 | 642 | 20201119_16913_1ng_glyco_hcd203040_compound15.26528.26528.3 | controllerType=0 controllerNumber=1 scan=26528 | 55.9156 |
| 22598:4 | 16 | R.QLAHQSN[+2350.830]STNIFFSPVSIATAFAMLSLGTK.A | HexNAc(4)Hex(5)Fuc(1)NeuAc(2) | N[+2351] | 1383.8778 | 4 | 5532.4894 | 5532.4654 | 0 | 4.3 | 64 | Specific | 854.95 | 854.22 | 854.22 | 17.21 | 41 | >sp\|P01009\|A1AT_HUMAN Alpha-1-antitrypsin OS=Homo sapiens OX=9606 GN=SERPINA1 PE=1 SV=3 | 642 | 20201119_16913_1ng_glyco_hcd203040_compound15.26480.26480.5 | controllerType=0 controllerNumber=1 scan=26480 | 55.7901 |
| 22600:4 | 16 | R.QLAHQSN[+2350.830]STNIFFSPVSIATAFAMLSLGTK.A | HexNAc(4)Hex(5)Fuc(1)NeuAc(2) | N[+2351] | 1383.8778 | 4 | 5532.4894 | 5532.4654 | 0 | 4.3 | 64 | Specific | 861.71 | 800.47 | 800.47 | 17.21 | 41 | >sp\|P01009\|A1AT_HUMAN Alpha-1-antitrypsin OS=Homo sapiens OX=9606 GN=SERPINA1 PE=1 SV=3 | 642 | 20201119_16913_1ng_glyco_hcd203040_compound15.26482.26482.5 | controllerType=0 controllerNumber=1 scan=26482 | 55.7961 |
| 22641:3 | 16 | R.QLAHQSN[+2204.772]STNIFFSPVSIATAFAMLSLGTK.A | HexNAc(4)Hex(5)NeuAc(2) | N[+2205] | 1796.1473 | 3 | 5386.4274 | 5386.4075 | 0 | 3.7 | 64 | Specific | 862.03 | 824.15 | 824.15 | 17.66 | 41 | >sp\|P01009\|A1AT_HUMAN Alpha-1-antitrypsin OS=Homo sapiens OX=9606 GN=SERPINA1 PE=1 SV=3 | 642 | 20201119_16913_1ng_glyco_hcd203040_compound15.26526.26526.3 | controllerType=0 controllerNumber=1 scan=26526 | 55.9086 |
| 22642:4 | 16 | R.QLAHQSN[+2204.772]STNIFFSPVSIATAFAMLSLGTK.A | HexNAc(4)Hex(5)NeuAc(2) | N[+2205] | 1347.3608 | 4 | 5386.4215 | 5386.4075 | 0 | 2.6 | 64 | Specific | 953.51 | 917.37 | 608.71 | 19.51 | 41 | >sp\|P01009\|A1AT_HUMAN Alpha-1-antitrypsin OS=Homo sapiens OX=9606 GN=SERPINA1 PE=1 SV=3 | 642 | 20201119_16913_1ng_glyco_hcd203040_compound15.26527.26527.4 | controllerType=0 controllerNumber=1 scan=26527 | 55.9132 |
| 22640:4 | 16 | R.QLAHQSN[+2204.772]STNIFFSPVSIATAFAMLSLGTK.A | HexNAc(4)Hex(5)NeuAc(2) | N[+2205] | 1347.3608 | 4 | 5386.4215 | 5386.4075 | 0 | 2.6 | 64 | Specific | 958.16 | 947.08 | 611.89 | 19.51 | 41 | >sp\|P01009\|A1AT_HUMAN Alpha-1-antitrypsin OS=Homo sapiens OX=9606 GN=SERPINA1 PE=1 SV=3 | 642 | 20201119_16913_1ng_glyco_hcd203040_compound15.26525.26525.4 | controllerType=0 controllerNumber=1 scan=26525 | 55.9072 |
| 21480:4 | 16 | K.ADTHDEILEGLNFN[+2861.000]LTEIPEAQIHEGFQELLR.T | HexNAc(5)Hex(6)NeuAc(3) | N[+2861] | 1638.9679 | 4 | 6552.8498 | 6552.8239 | 0 | 3.9 | 94 | Specific | 876.31 | 590.93 | 590.93 | 17.46 | 41 | >sp\|P01009\|A1AT_HUMAN Alpha-1-antitrypsin OS=Homo sapiens OX=9606 GN=SERPINA1 PE=1 SV=3 | 642 | 20201119_16913_1ng_glyco_hcd203040_compound15.25312.25312.4 | controllerType=0 controllerNumber=1 scan=25312 | 53.2046 |
| 21482:4 | 16 | K.ADTHDEILEGLNFN[+2861.000]LTEIPEAQIHEGFQELLR.T | HexNAc(5)Hex(6)NeuAc(3) | N[+2861] | 1638.9679 | 4 | 6552.8498 | 6552.8239 | 0 | 3.9 | 94 | Specific | 958.21 | 774.44 | 774.44 | 19.46 | 41 | >sp\|P01009\|A1AT_HUMAN Alpha-1-antitrypsin OS=Homo sapiens OX=9606 GN=SERPINA1 PE=1 SV=3 | 642 | 20201119_16913_1ng_glyco_hcd203040_compound15.25314.25314.4 | controllerType=0 controllerNumber=1 scan=25314 | 53.2089 |
| 19318:3 | 16 | K.YLGN[+2204.772]ATAIFFLPDEGK.L | HexNAc(4)Hex(5)NeuAc(2) | N[+2205] | 1320.897 | 3 | 3960.6765 | 3960.6677 | 0 | 2.2 | 268 | Specific | 474.2 | 399.94 | 399.94 | 9.43 | 41 | >sp\|P01009\|A1AT_HUMAN Alpha-1-antitrypsin OS=Homo sapiens OX=9606 GN=SERPINA1 PE=1 SV=3 | 642 | 20201119_16913_1ng_glyco_hcd203040_compound15.23049.23049.3 | controllerType=0 controllerNumber=1 scan=23049 | 48.2911 |
| 19296:3 | 16 | K.YLGN[+2204.772]ATAIFFLPDEGK.L | HexNAc(4)Hex(5)NeuAc(2) | N[+2205] | 1320.9001 | 3 | 3960.6857 | 3960.6677 | 0 | 4.6 | 268 | Specific | 696.48 | 639.59 | 639.59 | 12.91 | 41 | >sp\|P01009\|A1AT_HUMAN Alpha-1-antitrypsin OS=Homo sapiens OX=9606 GN=SERPINA1 PE=1 SV=3 | 642 | 20201119_16913_1ng_glyco_hcd203040_compound15.23026.23026.3 | controllerType=0 controllerNumber=1 scan=23026 | 48.[illegible]389 |
| 19321:3 | 16 | K.YLGN[+2204.772]ATAIFFLPDEGK.L | HexNAc(4)Hex(5)NeuAc(2) | N[+2205] | 1320.897 | 3 | 3960.6765 | 3960.6677 | 0 | 2.2 | 268 | Specific | 767.15 | 663.75 | 663.75 | 15.21 | 41 | >sp\|P01009\|A1AT_HUMAN Alpha-1-antitrypsin OS=Homo sapiens OX=9606 GN=SERPINA1 PE=1 SV=3 | 642 | 20201119_16913_1ng_glyco_hcd203040_compound15.23052.23052.3 | controllerType=0 controllerNumber=1 scan=23052 | 48.[illegible]964 |
| 19299:3 | 16 | K.YLGN[+2204.772]ATAIFFLPDEGK.L | HexNAc(4)Hex(5)NeuAc(2) | N[+2205] | 1320.9001 | 3 | 3960.6857 | 3960.6677 | 0 | 4.6 | 268 | Specific | 882.23 | 769.16 | 769.16 | 16.91 | 41 | >sp\|P01009\|A1AT_HUMAN Alpha-1-antitrypsin OS=Homo sapiens OX=9606 GN=SERPINA1 PE=1 SV=3 | 642 | 20201119_16913_1ng_glyco_hcd203040_compound15.23029.23029.3 | controllerType=0 controllerNumber=1 scan=23029 | 48.[illegible]433 |
| 19105:4 | 16 | K.[+42.011]YLGN[+2190.793]ATAIFFLPDEGKLQHLENELTHDIITK.F | HexNAc(6)Hex(6) | .[42], N[+2191] | 1444.1569 | 4 | 5773.6056 | 5773.6411 | 0 | -6.1 | 268 | Specific | 229.61 | 229.61 | 229.61 | 3.66 | 41 | >sp\|P01009\|A1AT_HUMAN Alpha-1-antitrypsin OS=Homo sapiens OX=9606 GN=SERPINA1 PE=1 SV=3 | 642 | 20201119_16913_1ng_glyco_hcd203040_compound15.22826.22826.4 | controllerType=0 controllerNumber=1 scan=22826 | [illegible] |
| 19108:4 | 16 | K.[+42.011]YLGN[+2190.793]ATAIFFLPDEGKLQHLENELTHDIITK.F | HexNAc(6)Hex(6) | .[42], N[+2191] | 1444.1569 | 4 | 5773.6056 | 5773.6411 | 0 | -6.1 | 268 | Specific | 240.73 | 208.86 | 208.86 | 2.07 | 41 | >sp\|P01009\|A1AT_HUMAN Alpha-1-antitrypsin OS=Homo sapiens OX=9606 GN=SERPINA1 PE=1 SV=3 | 642 | 20201119_16913_1ng_glyco_hcd203040_compound15.22829.22829.4 | controllerType=0 controllerNumber=1 scan=22829 | 47.7728 |
| 20317:4 | 17 | K.APDKNVIFSPLSISTALAFLSLGAHN[+2205.793]TTLTEILK.G | HexNAc(4)Hex(5)Fuc(2)NeuAc(1) | N[+2206] | 1447.9584 | 4 | 5788.8117 | 5788.771 | 0 | 7 | 68 | Specific | 383.69 | 383.69 | 383.69 | 7.79 | 39 | >sp\|P01011\|AACT_HUMAN Alpha-1-antichymotrypsin OS=Homo sapiens OX=9606 GN=SERPINA3 PE=1 SV=2 | 1314 | 20201119_16913_1ng_glyco_hcd203040_compound15.24093.24093.4 | controllerType=0 controllerNumber=1 scan=24093 | 50.6737 |
| 21578:5 | 17 | K.APDKNVIFSPLSISTALAFLSLGAHN[+2350.830]TTLTEILK.G | HexNAc(4)Hex(5)Fuc(1)NeuAc(2) | N[+2351] | 1187.5712 | 5 | 5933.8267 | 5933.8085 | 0 | 3.1 | 68 | Specific | 453.45 | 406.73 | 406.73 | 9.03 | 39 | >sp\|P01011\|AACT_HUMAN Alpha-1-antichymotrypsin OS=Homo sapiens OX=9606 GN=SERPINA3 PE=1 SV=2 | 1314 | 20201119_16913_1ng_glyco_hcd203040_compound15.25415.25415.4 | controllerType=0 controllerNumber=1 scan=25415 | 53.4679 |
| 20314:4 | 17 | K.APDKNVIFSPLSISTALAFLSLGAHN[+2205.793]TTLTEILK.G | HexNAc(4)Hex(5)Fuc(2)NeuAc(1) | N[+2206] | 1447.9584 | 4 | 5788.8117 | 5788.771 | 0 | 7 | 68 | Specific | 472.1 | 472.1 | 472.1 | 8.13 | 39 | >sp\|P01011\|AACT_HUMAN Alpha-1-antichymotrypsin OS=Homo sapiens OX=9606 GN=SERPINA3 PE=1 SV=2 | 1314 | 20201119_16913_1ng_glyco_hcd203040_compound15.24090.24090.4 | controllerType=0 controllerNumber=1 scan=24090 | 50.6688 |
| 20385:4 | 17 | K.APDKNVIFSPLSISTALAFLSLGAHN[+2059.735]TTLTEILK.G | HexNAc(4)Hex(5)Fuc(1)NeuAc(1) | N[+2060] | 1411.4423 | 4 | 5642.7474 | 5642.7131 | 0 | 6.1 | 68 | Specific | 1181.05 | 1177.49 | 1177.49 | 22.2 | 39 | >sp\|P01011\|AACT_HUMAN Alpha-1-antichymotrypsin OS=Homo sapiens OX=9606 GN=SERPINA3 PE=1 SV=2 | 1314 | 20201119_16913_1ng_glyco_hcd203040_compound15.24164.24164.4 | controllerType=0 controllerNumber=1 scan=24164 | 50.8389 |
| 23162:3 | 17 | K.NVIFSPLSISTALAFLSLGAHN[+2350.830]TTLTEILK.G | HexNAc(4)Hex(5)Fuc(1)NeuAc(2) | N[+2351] | 1841.5454 | 3 | 5522.6216 | 5522.5967 | 0 | 4.5 | 72 | Specific | 276.96 | 276.96 | 276.96 | 6.72 | 39 | >sp\|P01011\|AACT_HUMAN Alpha-1-antichymotrypsin OS=Homo sapiens OX=9606 GN=SERPINA3 PE=1 SV=2 | 1314 | 20201119_16913_1ng_glyco_hcd203040_compound15.27069.27069.3 | controllerType=0 controllerNumber=1 scan=27069 | 57.0164 |
| 23165:5 | 17 | K.NVIFSPLSISTALAFLSLGAHN[+2350.830]TTLTEILK.G | HexNAc(4)Hex(5)Fuc(1)NeuAc(2) | N[+2351] | 1105.3289 | 5 | 5522.6152 | 5522.5967 | 0 | 3.3 | 72 | Specific | 297.78 | 280.11 | 280.11 | 7.13 | 39 | >sp\|P01011\|AACT_HUMAN Alpha-1-antichymotrypsin OS=Homo sapiens OX=9606 GN=SERPINA3 PE=1 SV=2 | 1314 | 20201119_16913_1ng_glyco_hcd203040_compound15.27072.27072.5 | controllerType=0 controllerNumber=1 scan=27072 | 57.0211 |
| 23159:4 | 17 | K.NVIFSPLSISTALAFLSLGAHN[+2350.830]TTLTEILK.G | HexNAc(4)Hex(5)Fuc(1)NeuAc(2) | N[+2351] | 1381.41 | 4 | 5522.6183 | 5522.5967 | 0 | 3.9 | 72 | Specific | 466.35 | 466.35 | 466.35 | 9.79 | 39 | >sp\|P01011\|AACT_HUMAN Alpha-1-antichymotrypsin OS=Homo sapiens OX=9606 GN=SERPINA3 PE=1 SV=2 | 1314 | 20201119_16913_1ng_glyco_hcd203040_compound15.27066.27066.4 | controllerType=0 controllerNumber=1 scan=27066 | 57.01 |
| 23160:3 | 17 | K.NVIFSPLSISTALAFLSLGAHN[+2350.830]TTLTEILK.G | HexNAc(4)Hex(5)Fuc(1)NeuAc(2) | N[+2351] | 1841.5454 | 3 | 5522.6216 | 5522.5967 | 0 | 4.5 | 72 | Specific | 467.23 | 467.23 | 467.23 | 9.52 | 39 | >sp\|P01011\|AACT_HUMAN Alpha-1-antichymotrypsin OS=Homo sapiens OX=9606 GN=SERPINA3 PE=1 SV=2 | 1314 | 20201119_16913_1ng_glyco_hcd203040_compound15.27067.27067.3 | controllerType=0 controllerNumber=1 scan=27067 | 57.0113 |
| 23161:4 | 17 | K.NVIFSPLSISTALAFLSLGAHN[+2350.830]TTLTEILK.G | HexNAc(4)Hex(5)Fuc(1)NeuAc(2) | N[+2351] | 1381.41 | 4 | 5522.6183 | 5522.5967 | 0 | 3.9 | 72 | Specific | 764.5 | 764.5 | 764.5 | 15.57 | 39 | >sp\|P01011\|AACT_HUMAN Alpha-1-antichymotrypsin OS=Homo sapiens OX=9606 GN=SERPINA3 PE=1 SV=2 | 1314 | 20201119_16913_1ng_glyco_hcd203040_compound15.27068.27068.4 | controllerType=0 controllerNumber=1 scan=27068 | 57.0145 |
| 14815:4 | 17 | K.GLKFN[+2350.830]LTETSEAEIHQSFQHLLR.T | HexNAc(4)Hex(5)Fuc(1)NeuAc(2) | N[+2351] | 1263.0681 | 4 | 5049.2506 | 5049.2251 | 0 | 5.1 | 102 | Specific | 534.09 | 257.76 | 257.76 | 6.32 | 39 | >sp\|P01011\|AACT_HUMAN Alpha-1-antichymotrypsin OS=Homo sapiens OX=9606 GN=SERPINA3 PE=1 SV=2 | 1314 | 20201119_16913_1ng_glyco_hcd203040_compound15.18325.18325.4 | controllerType=0 controllerNumber=1 scan=18325 | 38.0672 |
| 14813:4 | 17 | K.GLKFN[+2350.830]LTETSEAEIHQSFQHLLR.T | HexNAc(4)Hex(5)Fuc(1)NeuAc(2) | N[+2351] | 1263.0681 | 4 | 5049.2506 | 5049.2251 | 0 | 5.1 | 102 | Specific | 560.09 | 321.67 | 321.67 | 6.86 | 39 | >sp\|P01011\|AACT_HUMAN Alpha-1-antichymotrypsin OS=Homo sapiens OX=9606 GN=SERPINA3 PE=1 SV=2 | 1314 | 20201119_16913_1ng_glyco_hcd203040_compound15.18323.18323.4 | controllerType=0 controllerNumber=1 scan=18323 | 38.0634 |
| 15011:3 | 17 | R.TLN[+2205.793]QSSDELQLSMGNAMFVK.E | HexNAc(4)Hex(5)Fuc(2)NeuAc(1) | N[+2206] | 1473.6231 | 3 | 4418.8548 | 4418.8505 | 0 | 1 | 125 | Specific | 249.78 | 141.25 | 141.25 | 5.12 | 39 | >sp\|P01011\|AACT_HUMAN Alpha-1-antichymotrypsin OS=Homo sapiens OX=9606 GN=SERPINA3 PE=1 SV=2 | 1314 | 20201119_16913_1ng_glyco_hcd203040_compound15.18529.18529.3 | controllerType=0 controllerNumber=1 scan=18529 | 38.4954 |
| 15013:3 | 17 | R.TLN[+2205.793]QSSDELQLSMGNAMFVK.E | HexNAc(4)Hex(5)Fuc(2)NeuAc(1) | N[+2206] | 1473.6231 | 3 | 4418.8548 | 4418.8505 | 0 | 1 | 125 | Specific | 378.39 | 258.86 | 258.86 | 6.85 | 39 | >sp\|P01011\|AACT_HUMAN Alpha-1-antichymotrypsin OS=Homo sapiens OX=9606 GN=SERPINA3 PE=1 SV=2 | 1314 | 20201119_16913_1ng_glyco_hcd203040_compound15.18531.18531.3 | controllerType=0 controllerNumber=1 scan=18531 | 38.5016 |
| 21170:4 | 17 | K.YTGN[+2861.000]ASALFILPDQDKMEEVEAMLLPETLKR.W | HexNAc(5)Hex(6)NeuAc(3) | N[+2861] | 1596.7151 | 4 | 6383.8385 | 6383.7859 | 0 | 8.2 | 268 | Specific | 234.43 | 204.88 | 204.88 | 3.21 | 39 | >sp\|P01011\|AACT_HUMAN Alpha-1-antichymotrypsin OS=Homo sapiens OX=9606 GN=SERPINA3 PE=1 SV=2 | 1314 | 20201119_16913_1ng_glyco_hcd203040_compound15.24988.24988.4 | controllerType=0 controllerNumber=1 scan=24988 | 52.5245 |
| 20768:4 | 18 | K.VFPLSLDSTPQDGNVVVAC[+57.021]LVQGFFPQEPLSVTWSESGQN[+1971.719]VTAI | HexNAc(5)Hex(5)Fuc(1) | C[+57], N[+1972] | 1687.7824 | 4 | 6748.1079 | 6748.0803 | 0 | 4.1 | 8 | Specific | 254.75 | 228.78 | 228.78 | 7.02 | 41 | >sp\|P01877\|IGHA2_HUMAN Immunoglobulin heavy constant alpha 2 OS=Homo sapiens OX=9606 GN=IGHA2 PE=1 SV=4 | 576 | 20201119_16913_1ng_glyco_hcd203040_compound15.24566.24566.2 | controllerType=0 controllerNumber=1 scan=24566 | 51.7398 |
| 20770:4 | 18 | K.VFPLSLDSTPQDGNVVVAC[+57.021]LVQGFFPQEPLSVTWSESGQN[+1971.719]VTAI | HexNAc(5)Hex(5)Fuc(1) | C[+57], N[+1972] | 1687.7824 | 4 | 6748.1079 | 6748.0803 | 0 | 4.1 | 8 | Specific | 376.15 | 342.71 | 342.71 | 8.14 | 41 | >sp\|P01877\|IGHA2_HUMAN Immunoglobulin heavy constant alpha 2 OS=Homo sapiens OX=9606 GN=IGHA2 PE=1 SV=4 | 576 | 20201119_16913_1ng_glyco_hcd203040_compound15.24568.24568.2 | controllerType=0 controllerNumber=1 scan=24568 | 51.7437 |
| 20800:4 | 18 | K.VFPLSLDSTPQDGNVVVAC[+57.021]LVQGFFPQEPLSVTWSESGQN[+1647.613]VTAI | HexNAc(5)Hex(3)Fuc(1) | C[+57], N[+1648] | 1606.7529 | 4 | 6423.9899 | 6423.9746 | 0 | 2.4 | 8 | Specific | 579.49 | 579.49 | 579.49 | 12.14 | 41 | >sp\|P01877\|IGHA2_HUMAN Immunoglobulin heavy constant alpha 2 OS=Homo sapiens OX=9606 GN=IGHA2 PE=1 SV=4 | 576 | 20201119_16913_1ng_glyco_hcd203040_compound15.24599.24599.4 | controllerType=0 controllerNumber=1 scan=24599 | 51.7961 |
| 22358:4 | 18 | K.VFPLSLDSTPQDGNVVVAC[+57.021]LVQGFFPQEPLSVTWSESGQN[+2262.814]VTAI | HexNAc(5)Hex(5)Fuc(1)NeuAc(1) | C[+57], N[+2263] | 1760.5575 | 4 | 7039.208 | 7039.1757 | 0 | 4.6 | 8 | Specific | 629.6 | 580.76 | 580.76 | 12.83 | 41 | >sp\|P01877\|IGHA2_HUMAN Immunoglobulin heavy constant alpha 2 OS=Homo sapiens OX=9606 GN=IGHA2 PE=1 SV=4 | 576 | 20201119_16913_1ng_glyco_hcd203040_compound15.26231.26231.4 | controllerType=0 controllerNumber=1 scan=26231 | 55.2924 |
| 22360:4 | 18 | K.VFPLSLDSTPQDGNVVVAC[+57.021]LVQGFFPQEPLSVTWSESGQN[+2262.814]VTAI | HexNAc(5)Hex(5)Fuc(1)NeuAc(1) | C[+57], N[+2263] | 1760.5575 | 4 | 7039.208 | 7039.1757 | 0 | 4.6 | 8 | Specific | 833.28 | 779.63 | 779.63 | 16.83 | 41 | >sp\|P01877\|IGHA2_HUMAN Immunoglobulin heavy constant alpha 2 OS=Homo sapiens OX=9606 GN=IGHA2 PE=1 SV=4 | 576 | 20201119_16913_1ng_glyco_hcd203040_compound15.26233.26233.4 | controllerType=0 controllerNumber=1 scan=26233 | 55.2973 |
| 20802:4 | 18 | K.VFPLSLDSTPQDGNVVVAC[+57.021]LVQGFFPQEPLSVTWSESGQN[+1647.613]VTAI | HexNAc(5)Hex(3)Fuc(1) | C[+57], N[+1648] | 1606.7529 | 4 | 6423.9899 | 6423.9746 | 0 | 2.4 | 8 | Specific | 936.13 | 752.88 | 752.88 | 19.14 | 41 | >sp\|P01877\|IGHA2_HUMAN Immunoglobulin heavy constant alpha 2 OS=Homo sapiens OX=9606 GN=IGHA2 PE=1 SV=4 | 576 | 20201119_16913_1ng_glyco_hcd203040_compound15.24601.24601.4 | controllerType=0 controllerNumber=1 scan=24601 | 51.8013 |
| 00862:3 | 18 | K.HYTN[+1606.587]SSQDVTVPC[+57.021]R.V | HexNAc(4)Hex(4)Fuc(1) | C[+57], N[+1607] | 1090.7859 | 3 | 3270.3432 | 3270.336 | 0 | 2.2 | 89 | Specific | 207.53 | 207.53 | 207.53 | 4.8 | 41 | >sp\|P01877\|IGHA2_HUMAN Immunoglobulin heavy constant alpha 2 OS=Homo sapiens OX=9606 GN=IGHA2 PE=1 SV=4 | 576 | 20201119_16913_1ng_glyco_hcd203040_compound15.3798.3798.3 | controllerType=0 controllerNumber=1 scan=3798 | 12.7448 |
| 01860:3 | 18 | K.HYTN[+2262.814]SSQDVTVPC[+57.021]R.V | HexNAc(5)Hex(5)Fuc(1)NeuAc(1) | C[+57], N[+2263] | 1309.5316 | 3 | 3926.5801 | 3926.5636 | 0 | 4.2 | 89 | Specific | 221.78 | 94.98 | 94.98 | 3.46 | 41 | >sp\|P01877\|IGHA2_HUMAN Immunoglobulin heavy constant alpha 2 OS=Homo sapiens OX=9606 GN=IGHA2 PE=1 SV=4 | 576 | 20201119_16913_1ng_glyco_hcd203040_compound15.4841.4841.3 | controllerType=0 controllerNumber=1 scan=4841 | 14.5585 |
| 01751:3 | 18 | K.HYTN[+2059.735]SSQDVTVPC[+57.021]R.V | HexNAc(4)Hex(5)Fuc(1)NeuAc(1) | C[+57], N[+2060] | 1241.836 | 3 | 3723.4934 | 3723.4842 | 0 | 2.5 | 89 | Specific | 227.47 | 172.35 | 172.35 | 5.63 | 41 | >sp\|P01877\|IGHA2_HUMAN Immunoglobulin heavy constant alpha 2 OS=Homo sapiens OX=9606 GN=IGHA2 PE=1 SV=4 | 576 | 20201119_16913_1ng_glyco_hcd203040_compound15.4728.4728.3 | controllerType=0 controllerNumber=1 scan=4728 | 14.3556 |
| 01732:3 | 18 | K.HYTN[+2059.735]SSQDVTVPC[+57.021]R.V | HexNAc(4)Hex(5)Fuc(1)NeuAc(1) | C[+57], N[+2060] | 1241.8365 | 3 | 3723.4949 | 3723.4842 | 0 | 2.9 | 89 | Specific | 255.79 | 193.88 | 193.88 | 6.22 | 41 | >sp\|P01877\|IGHA2_HUMAN Immunoglobulin heavy constant alpha 2 OS=Homo sapiens OX=9606 GN=IGHA2 PE=1 SV=4 | 576 | 20201119_16913_1ng_glyco_hcd203040_compound15.4708.4708.3 | controllerType=0 controllerNumber=1 scan=4708 | 14.3174 |
| 00860:3 | 18 | K.HYTN[+1606.587]SSQDVTVPC[+57.021]R.V | HexNAc(4)Hex(4)Fuc(1) | C[+57], N[+1607] | 1090.7859 | 3 | 3270.3432 | 3270.336 | 0 | 2.2 | 89 | Specific | 287.81 | 189.48 | 189.48 | 6.23 | 41 | >sp\|P01877\|IGHA2_HUMAN Immunoglobulin heavy constant alpha 2 OS=Homo sapiens OX=9606 GN=IGHA2 PE=1 SV=4 | 576 | 20201119_16913_1ng_glyco_hcd203040_compound15.3796.3796.3 | controllerType=0 controllerNumber=1 scan=3796 | 12.7416 |
| 01749:3 | 18 | K.HYTN[+2059.735]SSQDVTVPC[+57.021]R.V | HexNAc(4)Hex(5)Fuc(1)NeuAc(1) | C[+57], N[+2060] | 1241.836 | 3 | 3723.4934 | 3723.4842 | 0 | 2.5 | 89 | Specific | 306.29 | 223.66 | 223.66 | 6.31 | 41 | >sp\|P01877\|IGHA2_HUMAN Immunoglobulin heavy constant alpha 2 OS=Homo sapiens OX=9606 GN=IGHA2 PE=1 SV=4 | 576 | 20201119_16913_1ng_glyco_hcd203040_compound15.4726.4726.3 | controllerType=0 controllerNumber=1 scan=4726 | 14.3509 |
| 00989:2 | 18 | K.HYTN[+1647.613]SSQDVTVPC[+57.021]R.V | HexNAc(5)Hex(3)Fuc(1) | C[+57], N[+1648] | 1656.19 | 2 | 3311.3727 | 3311.3625 | 0 | 3.1 | 89 | Specific | 311 | 220.63 | 220.63 | 5.69 | 41 | >sp\|P01877\|IGHA2_HUMAN Immunoglobulin heavy constant alpha 2 OS=Homo sapiens OX=9606 GN=IGHA2 PE=1 SV=4 | 576 | 20201119_16913_1ng_glyco_hcd203040_compound15.3930.3930.2 | controllerType=0 controllerNumber=1 scan=3930 | 12.9714 |
| 00892:3 | 18 | K.HYTN[+1971.719]SSQDVTVPC[+57.021]R.V | HexNAc(5)Hex(5)Fuc(1) | C[+57], N[+1972] | 1212.4974 | 3 | 3635.4776 | 3635.4682 | 0 | 2.6 | 89 | Specific | 339.18 | 298.46 | 298.46 | 7.03 | 41 | >sp\|P01877\|IGHA2_HUMAN Immunoglobulin heavy constant alpha 2 OS=Homo sapiens OX=9606 GN=IGHA2 PE=1 SV=4 | 576 | 20201119_16913_1ng_glyco_hcd203040_compound15.3829.3829.3 | controllerType=0 controllerNumber=1 scan=3829 | 12.7985 |
| 00894:3 | 18 | K.HYTN[+1971.719]SSQDVTVPC[+57.021]R.V | HexNAc(5)Hex(5)Fuc(1) | C[+57], N[+1972] | 1212.4974 | 3 | 3635.4776 | 3635.4682 | 0 | 2.6 | 89 | Specific | 350.58 | 230.75 | 230.75 | 7.08 | 41 | >sp\|P01877\|IGHA2_HUMAN Immunoglobulin heavy constant alpha 2 OS=Homo sapiens OX=9606 GN=IGHA2 PE=1 SV=4 | 576 | 20201119_16913_1ng_glyco_hcd203040_compound15.3831.3831.3 | controllerType=0 controllerNumber=1 scan=3831 | 12.8025 |
| 01730:3 | 18 | K.HYTN[+2059.735]SSQDVTVPC[+57.021]R.V | HexNAc(4)Hex(5)Fuc(1)NeuAc(1) | C[+57], N[+2060] | 1241.8365 | 3 | 3723.4949 | 3723.4842 | 0 | 2.9 | 89 | Specific | 366.92 | 219.14 | 219.14 | 6.72 | 41 | >sp\|P01877\|IGHA2_HUMAN Immunoglobulin heavy constant alpha 2 OS=Homo sapiens OX=9606 GN=IGHA2 PE=1 SV=4 | 576 | 20201119_16913_1ng_glyco_hcd203040_compound15.4706.4706.3 | controllerType=0 controllerNumber=1 scan=4706 | 14.3129 |
| 00908:3 | 18 | K.HYTN[+1444.534]SSQDVTVPC[+57.021]R.V | HexNAc(4)Hex(3)Fuc(1) | C[+57], N[+1445] | 1036.7679 | 3 | 3108.289 | 3108.2832 | 0 | 1.9 | 89 | Specific | 460.96 | 360.78 | 360.78 | 8.11 | 41 | >sp\|P01877\|IGHA2_HUMAN Immunoglobulin heavy constant alpha 2 OS=Homo sapiens OX=9606 GN=IGHA2 PE=1 SV=4 | 576 | 20201119_16913_1ng_glyco_hcd203040_compound15.3845.3845.3 | controllerType=0 controllerNumber=1 scan=3845 | 12.8249 |
| 00910:3 | 18 | K.HYTN[+1444.534]SSQDVTVPC[+57.021]R.V | HexNAc(4)Hex(3)Fuc(1) | C[+57], N[+1445] | 1036.7679 | 3 | 3108.289 | 3108.2832 | 0 | 1.9 | 89 | Specific | 514.75 | 270.61 | 270.61 | 7.64 | 41 | >sp\|P01877\|IGHA2_HUMAN Immunoglobulin heavy constant alpha 2 OS=Homo sapiens OX=9606 GN=IGHA2 PE=1 SV=4 | 576 | 20201119_16913_1ng_glyco_hcd203040_compound15.3847.3847.3 | controllerType=0 controllerNumber=1 scan=3847 | 12.8289 |
| 00914:3 | 18 | K.HYTN[+1809.666]SSQDVTVPC[+57.021]R.V | HexNAc(5)Hex(4)Fuc(1) | C[+57], N[+1810] | 1158.4819 | 3 | 3473.4311 | 3473.4154 | 0 | 4.5 | 89 | Specific | 604.48 | 532.87 | 532.87 | 12 | 41 | >sp\|P01877\|IGHA2_HUMAN Immunoglobulin heavy constant alpha 2 OS=Homo sapiens OX=9606 GN=IGHA2 PE=1 SV=4 | 576 | 20201119_16913_1ng_glyco_hcd203040_compound15.3852.3852.3 | controllerType=0 controllerNumber=1 scan=3852 | 12.8384 |
| 00916:3 | 18 | K.HYTN[+1809.666]SSQDVTVPC[+57.021]R.V | HexNAc(5)Hex(4)Fuc(1) | C[+57], N[+1810] | 1158.4819 | 3 | 3473.4311 | 3473.4154 | 0 | 4.5 | 89 | Specific | 659.6 | 565.45 | 565.45 | 13 | 41 | >sp\|P01877\|IGHA2_HUMAN Immunoglobulin heavy constant alpha 2 OS=Homo sapiens OX=9606 GN=IGHA2 PE=1 SV=4 | 576 | 20201119_16913_1ng_glyco_hcd203040_compound15.3854.3854.3 | controllerType=0 controllerNumber=1 scan=3854 | 12.8428 |
| 00943:3 | 18 | K.HYTN[+1647.613]SSQDVTVPC[+57.021]R.V | HexNAc(5)Hex(3)Fuc(1) | C[+57], N[+1648] | 1104.4616 | 3 | 3311.3702 | 3311.3625 | 0 | 2.3 | 89 | Specific | 664.27 | 564.05 | 564.05 | 13.3 | 41 | >sp\|P01877\|IGHA2_HUMAN Immunoglobulin heavy constant alpha 2 OS=Homo sapiens OX=9606 GN=IGHA2 PE=1 SV=4 | 576 | 20201119_16913_1ng_glyco_hcd203040_compound15.3882.3882.3 | controllerType=0 controllerNumber=1 scan=3882 | 12.8901 |
| 00945:3 | 18 | K.HYTN[+1647.613]SSQDVTVPC[+57.021]R.V | HexNAc(5)Hex(3)Fuc(1) | C[+57], N[+1648] | 1104.4616 | 3 | 3311.3702 | 3311.3625 | 0 | 2.3 | 89 | Specific | 674.66 | 608.36 | 608.36 | 13.3 | 41 | >sp\|P01877\|IGHA2_HUMAN Immunoglobulin heavy constant alpha 2 OS=Homo sapiens OX=9606 GN=IGHA2 PE=1 SV=4 | 576 | 20201119_16913_1ng_glyco_hcd203040_compound15.3884.3884.3 | controllerType=0 controllerNumber=1 scan=3884 | 12.8955 |
| 02962:4 | 18 | K.HYTN[+1647.613]SSQDVTVPC[+57.021]RVPPPPPC[+57.021]C[+57.021]HPR.L | HexNAc(5)Hex(3)Fuc(1) | C[+57]*3, N[+1648] | 1152.2514 | 4 | 4605.9837 | 4605.9689 | 0 | 3.2 | 89 | Specific | 221.97 | 221.97 | 221.97 | 3.89 | 41 | >sp\|P01877\|IGHA2_HUMAN Immunoglobulin heavy constant alpha 2 OS=Homo sapiens OX=9606 GN=IGHA2 PE=1 SV=4 | 576 | 20201119_16913_1ng_glyco_hcd203040_compound15.5981.5981.4 | controllerType=0 controllerNumber=1 scan=5981 | 16.5356 |
| 02963:4 | 18 | K.HYTN[+1647.613]SSQDVTVPC[+57.021]RVPPPPPC[+57.021]C[+57.021]HPR.L | HexNAc(5)Hex(3)Fuc(1) | C[+57]*3, N[+1648] | 1152.2514 | 4 | 4605.9837 | 4605.9689 | 0 | 3.2 | 89 | Specific | 273.62 | 236.97 | 236.97 | 5.52 | 41 | >sp\|P01877\|IGHA2_HUMAN Immunoglobulin heavy constant alpha 2 OS=Homo sapiens OX=9606 GN=IGHA2 PE=1 SV=4 | 576 | 20201119_16913_1ng_glyco_hcd203040_compound15.5982.5982.4 | controllerType=0 controllerNumber=1 scan=5982 | 16.5394 |
| 04426:3 | 18 | K.TPLTAN[+2262.814]ITK.S | HexNAc(5)Hex(5)Fuc(1)NeuAc(1) | N[+2263] | 1074.4645 | 3 | 3221.379 | 3221.3711 | 0 | 2.4 | 200 | Specific | 211.76 | 200.05 | 200.05 | 4.52 | 41 | >sp\|P01877\|IGHA2_HUMAN Immunoglobulin heavy constant alpha 2 OS=Homo sapiens OX=9606 GN=IGHA2 PE=1 SV=4 | 576 | 20201119_16913_1ng_glyco_hcd203040_compound15.7507.7507.3 | controllerType=0 controllerNumber=1 scan=7507 | 19.0323 |
| 02543:2 | 18 | K.TPLTAN[+1606.587]ITK.S | HexNAc(4)Hex(4)Fuc(1) | N[+1607] | 1283.079 | 2 | 2565.1508 | 2565.1435 | 0 | 2.8 | 200 | Specific | 213.12 | 56.53 | 56.53 | 3.27 | 41 | >sp\|P01877\|IGHA2_HUMAN Immunoglobulin heavy constant alpha 2 OS=Homo sapiens OX=9606 GN=IGHA2 PE=1 SV=4 | 576 | 20201119_16913_1ng_glyco_hcd203040_compound15.5547.5547.2 | controllerType=0 controllerNumber=1 scan=5547 | 15.7575 |
| 02564:2 | 18 | K.TPLTAN[+1971.719]ITK.S | HexNAc(5)Hex(5)Fuc(1) | N[+1972] | 1465.6468 | 2 | 2930.2863 | 2930.2757 | 0 | 3.6 | 200 | Specific | 264.9 | 239.17 | 239.17 | 6.27 | 41 | >sp\|P01877\|IGHA2_HUMAN Immunoglobulin heavy constant alpha 2 OS=Homo sapiens OX=9606 GN=IGHA2 PE=1 SV=4 | 576 | 20201119_16913_1ng_glyco_hcd203040_compound15.5569.5569.2 | controllerType=0 controllerNumber=1 scan=5569 | 15.7982 |
| 04429:3 | 18 | K.TPLTAN[+2262.814]ITK.S | HexNAc(5)Hex(5)Fuc(1)NeuAc(1) | N[+2263] | 1074.4645 | 3 | 3221.379 | 3221.3711 | 0 | 2.4 | 200 | Specific | 276.72 | 120.67 | 120.67 | 4.6 | 41 | >sp\|P01877\|IGHA2_HUMAN Immunoglobulin heavy constant alpha 2 OS=Homo sapiens OX=9606 GN=IGHA2 PE=1 SV=4 | 576 | 20201119_16913_1ng_glyco_hcd203040_compound15.7510.7510.3 | controllerType=0 controllerNumber=1 scan=7510 | 19.0371 |
| 02552:3 | 18 | K.TPLTAN[+1971.719]ITK.S | HexNAc(5)Hex(5)Fuc(1) | N[+1972] | 977.4334 | 3 | 2930.2856 | 2930.2757 | 0 | 3.4 | 200 | Specific | 301.54 | 208.63 | 208.63 | 6 | 41 | >sp\|P01877\|IGHA2_HUMAN Immunoglobulin heavy constant alpha 2 OS=Homo sapiens OX=9606 GN=IGHA2 PE=1 SV=4 | 576 | 20201119_16913_1ng_glyco_hcd203040_compound15.5557.5557.3 | controllerType=0 controllerNumber=1 scan=5557 | 15.7772 |
| 02566:2 | 18 | K.TPLTAN[+1971.719]ITK.S | HexNAc(5)Hex(5)Fuc(1) | N[+1972] | 1465.6468 | 2 | 2930.2863 | 2930.2757 | 0 | 3.6 | 200 | Specific | 321.71 | 269.65 | 269.65 | 6.53 | 41 | >sp\|P01877\|IGHA2_HUMAN Immunoglobulin heavy constant alpha 2 OS=Homo sapiens OX=9606 GN=IGHA2 PE=1 SV=4 | 576 | 20201119_16913_1ng_glyco_hcd203040_compound15.5571.5571.2 | controllerType=0 controllerNumber=1 scan=5571 | 15.8022 |
| 02554:3 | 18 | K.TPLTAN[+1971.719]ITK.S | HexNAc(5)Hex(5)Fuc(1) | N[+1972] | 977.4334 | 3 | 2930.2856 | 2930.2757 | 0 | 3.4 | 200 | Specific | 364.41 | 182.19 | 182.19 | 6.43 | 41 | >sp\|P01877\|IGHA2_HUMAN Immunoglobulin heavy constant alpha 2 OS=Homo sapiens OX=9606 GN=IGHA2 PE=1 SV=4 | 576 | 20201119_16913_1ng_glyco_hcd203040_compound15.5559.5559.3 | controllerType=0 controllerNumber=1 scan=5559 | 15.781 |
| 02672:2 | 18 | K.TPLTAN[+1501.555]ITK.S | HexNAc(5)Hex(3) | N[+1502] | 1230.5612 | 2 | 2460.1151 | 2460.1121 | 0 | 1.2 | 200 | Specific | 364.49 | 205.09 | 205.09 | 6.12 | 41 | >sp\|P01877\|IGHA2_HUMAN Immunoglobulin heavy constant alpha 2 OS=Homo sapiens OX=9606 GN=IGHA2 PE=1 SV=4 | 576 | 20201119_16913_1ng_glyco_hcd203040_compound15.5681.5681.2 | controllerType=0 controllerNumber=1 scan=5681 | 16.009 |
| 02674:2 | 18 | K.TPLTAN[+1501.555]ITK.S | HexNAc(5)Hex(3) | N[+1502] | 1230.5612 | 2 | 2460.1151 | 2460.1121 | 0 | 1.2 | 200 | Specific | 457.06 | 256.29 | 256.29 | 6.86 | 41 | >sp\|P01877\|IGHA2_HUMAN Immunoglobulin heavy constant alpha 2 OS=Homo sapiens OX=9606 GN=IGHA2 PE=1 SV=4 | 576 | 20201119_16913_1ng_glyco_hcd203040_compound15.5683.5683.2 | controllerType=0 controllerNumber=1 scan=5683 | 16.02 |
| 02678:3 | 18 | K.TPLTAN[+1647.613]ITK.S | HexNAc(5)Hex(3)Fuc(1) | N[+1648] | 869.3972 | 3 | 2606.177 | 2606.17 | 0 | 2.7 | 200 | Specific | 457.85 | 164.89 | 164.89 | 6.29 | 41 | >sp\|P01877\|IGHA2_HUMAN Immunoglobulin heavy constant alpha 2 OS=Homo sapiens OX=9606 GN=IGHA2 PE=1 SV=4 | 576 | 20201119_16913_1ng_glyco_hcd203040_compound15.5688.5688.3 | controllerType=0 controllerNumber=1 scan=5688 | 16.0302 |
| 02605:3 | 18 | K.TPLTAN[+1809.666]ITK.S | HexNAc(5)Hex(4)Fuc(1) | N[+1810] | 923.4142 | 3 | 2768.2281 | 2768.2228 | 0 | 1.9 | 200 | Specific | 502.47 | 262.36 | 262.36 | 7.27 | 41 | >sp\|P01877\|IGHA2_HUMAN Immunoglobulin heavy constant alpha 2 OS=Homo sapiens OX=9606 GN=IGHA2 PE=1 SV=4 | 576 | 20201119_16913_1ng_glyco_hcd203040_compound15.5612.5612.3 | controllerType=0 controllerNumber=1 scan=5612 | 15.8848 |
| 02603:3 | 18 | K.TPLTAN[+1809.666]ITK.S | HexNAc(5)Hex(4)Fuc(1) | N[+1810] | 923.4142 | 3 | 2768.2281 | 2768.2228 | 0 | 1.9 | 200 | Specific | 507.83 | 181 | 181 | 6.91 | 41 | >sp\|P01877\|IGHA2_HUMAN Immunoglobulin heavy constant alpha 2 OS=Homo sapiens OX=9606 GN=IGHA2 PE=1 SV=4 | 576 | 20201119_16913_1ng_glyco_hcd203040_compound15.5610.5610.3 | controllerType=0 controllerNumber=1 scan=5610 | 15.8797 |
| 02613:2 | 18 | K.TPLTAN[+1444.534]ITK.S | HexNAc(4)Hex(3)Fuc(1) | N[+1445] | 1202.052 | 2 | 2403.0968 | 2403.0906 | 0 | 2.6 | 200 | Specific | 539.33 | 322.18 | 322.18 | 7.65 | 41 | >sp\|P01877\|IGHA2_HUMAN Immunoglobulin heavy constant alpha 2 OS=Homo sapiens OX=9606 GN=IGHA2 PE=1 SV=4 | 576 | 20201119_16913_1ng_glyco_hcd203040_compound15.5620.5620.2 | controllerType=0 controllerNumber=1 scan=5620 | 15.8969 |
| 02615:2 | 18 | K.TPLTAN[+1444.534]ITK.S | HexNAc(4)Hex(3)Fuc(1) | N[+1445] | 1202.052 | 2 | 2403.0968 | 2403.0906 | 0 | 2.6 | 200 | Specific | 563.74 | 311.76 | 311.76 | 7.92 | 41 | >sp\|P01877\|IGHA2_HUMAN Immunoglobulin heavy constant alpha 2 OS=Homo sapiens OX=9606 GN=IGHA2 PE=1 SV=4 | 576 | 20201119_16913_1ng_glyco_hcd203040_compound15.5622.5622.2 | controllerType=0 controllerNumber=1 scan=5622 | 15.9006 |
| 02680:3 | 18 | K.TPLTAN[+1647.613]ITK.S | HexNAc(5)Hex(3)Fuc(1) | N[+1648] | 869.3972 | 3 | 2606.177 | 2606.17 | 0 | 2.7 | 200 | Specific | 586.31 | 291.74 | 291.74 | 8.96 | 41 | >sp\|P01877\|IGHA2_HUMAN Immunoglobulin heavy constant alpha 2 OS=Homo sapiens OX=9606 GN=IGHA2 PE=1 SV=4 | 576 | 20201119_16913_1ng_glyco_hcd203040_compound15.5690.5690.3 | controllerType=0 controllerNumber=1 scan=5690 | 16.0389 |
| 02659:2 | 18 | K.TPLTAN[+1647.613]ITK.S | HexNAc(5)Hex(3)Fuc(1) | N[+1648] | 1303.5917 | 2 | 2606.1762 | 2606.17 | 0 | 2.4 | 200 | Specific | 602.82 | 307.32 | 307.32 | 10.17 | 41 | >sp\|P01877\|IGHA2_HUMAN Immunoglobulin heavy constant alpha 2 OS=Homo sapiens OX=9606 GN=IGHA2 PE=1 SV=4 | 576 | 20201119_16913_1ng_glyco_hcd203040_compound15.5668.5668.2 | controllerType=0 controllerNumber=1 scan=5668 | 15.9869 |
| 02602:2 | 18 | K.TPLTAN[+1809.666]ITK.S | HexNAc(5)Hex(4)Fuc(1) | N[+1810] | 1384.6181 | 2 | 2768.2289 | 2768.2228 | 0 | 2.2 | 200 | Specific | 618.9 | 349.87 | 349.87 | 10.45 | 41 | >sp\|P01877\|IGHA2_HUMAN Immunoglobulin heavy constant alpha 2 OS=Homo sapiens OX=9606 GN=IGHA2 PE=1 SV=4 | 576 | 20201119_16913_1ng_glyco_hcd203040_compound15.5609.5609.2 | controllerType=0 controllerNumber=1 scan=5609 | 15.8782 |
| 02657:2 | 18 | K.TPLTAN[+1647.613]ITK.S | HexNAc(5)Hex(3)Fuc(1) | N[+1648] | 1303.5917 | 2 | 2606.1762 | 2606.17 | 0 | 2.4 | 200 | Specific | 620.49 | 365.08 | 365.08 | 10.61 | 41 | >sp\|P01877\|IGHA2_HUMAN Immunoglobulin heavy constant alpha 2 OS=Homo sapiens OX=9606 GN=IGHA2 PE=1 SV=4 | 576 | 20201119_16913_1ng_glyco_hcd203040_compound15.5666.5666.2 | controllerType=0 controllerNumber=1 scan=5666 | 15.9806 |
| 02604:2 | 18 | K.TPLTAN[+1809.666]ITK.S | HexNAc(5)Hex(4)Fuc(1) | N[+1810] | 1384.6181 | 2 | 2768.2289 | 2768.2228 | 0 | 2.2 | 200 | Specific | 623.53 | 343.05 | 343.05 | 10.45 | 41 | >sp\|P01877\|IGHA2_HUMAN Immunoglobulin heavy constant alpha 2 OS=Homo sapiens OX=9606 GN=IGHA2 PE=1 SV=4 | 576 | 20201119_16913_1ng_glyco_hcd203040_compound15.5611.5611.2 | controllerType=0 controllerNumber=1 scan=5611 | 15.8829 |
| 10085:3 | 18 | R.MAGKPTHIN[+1501.555]VSVVMAEADGTC[+57.021]Y.- | HexNAc(5)Hex(3) | C[+57], N[+1502] | 1284.8871 | 3 | 3852.6469 | 3852.6382 | 0 | 2.3 | 319 | CRagged | 401.8 | 364.47 | 364.47 | 8.17 | 41 | >sp\|P01877\|IGHA2_HUMAN Immunoglobulin heavy constant alpha 2 OS=Homo sapiens OX=9606 GN=IGHA2 PE=1 SV=4 | 576 | 20201119_16913_1ng_glyco_hcd203040_compound15.13398.13398.3 | controllerType=0 controllerNumber=1 scan=13398 | 28.9054 |
| 10091:3 | 18 | R.MAGKPTHIN[+1647.613]VSVVMAEADGTC[+57.021]Y.- | HexNAc(5)Hex(3)Fuc(1) | C[+57], N[+1648] | 1333.5722 | 3 | 3998.702 | 3998.6961 | 0 | 1.5 | 319 | CRagged | 537.27 | 537.27 | 537.27 | 9.99 | 41 | >sp\|P01877\|IGHA2_HUMAN Immunoglobulin heavy constant alpha 2 OS=Homo sapiens OX=9606 GN=IGHA2 PE=1 SV=4 | 576 | 20201119_16913_1ng_glyco_hcd203040_compound15.13404.13404.3 | controllerType=0 controllerNumber=1 scan=13404 | 28.9148 |
| 10083:3 | 18 | R.MAGKPTHIN[+1501.555]VSVVMAEADGTC[+57.021]Y.- | HexNAc(5)Hex(3) | C[+57], N[+1502] | 1284.8871 | 3 | 3852.6469 | 3852.6382 | 0 | 2.3 | 319 | CRagged | 549.86 | 544.34 | 544.34 | 9.99 | 41 | >sp\|P01877\|IGHA2_HUMAN Immunoglobulin heavy constant alpha 2 OS=Homo sapiens OX=9606 GN=IGHA2 PE=1 SV=4 | 576 | 20201119_16913_1ng_glyco_hcd203040_compound15.13396.13396.3 | controllerType=0 controllerNumber=1 scan=13396 | 28.9016 |
| 10093:3 | 18 | R.MAGKPTHIN[+1647.613]VSVVMAEADGTC[+57.021]Y.- | HexNAc(5)Hex(3)Fuc(1) | C[+57], N[+1648] | 1333.5722 | 3 | 3998.702 | 3998.6961 | 0 | 1.5 | 319 | CRagged | 563.95 | 563.95 | 563.95 | 10.26 | 41 | >sp\|P01877\|IGHA2_HUMAN Immunoglobulin heavy constant alpha 2 OS=Homo sapiens OX=9606 GN=IGHA2 PE=1 SV=4 | 576 | 20201119_16913_1ng_glyco_hcd203040_compound15.13406.13406.3 | controllerType=0 controllerNumber=1 scan=13406 | 28.9197 |
| 10042:3 | 18 | R.MAGKPTHIN[+1216.423]VSVVMAEADGTC[+57.021]Y.- | HexNAc(2)Hex(5) | C[+57], N[+1216] | 1189.8428 | 3 | 3567.5139 | 3567.5057 | 0 | 2.3 | 319 | CRagged | 758.36 | 758.36 | 375.13 | 15.52 | 41 | >sp\|P01877\|IGHA2_HUMAN Immunoglobulin heavy constant alpha 2 OS=Homo sapiens OX=9606 GN=IGHA2 PE=1 SV=4 | 576 | 20201119_16913_1ng_glyco_hcd203040_compound15.13354.13354.3 | controllerType=0 controllerNumber=1 scan=13354 | 28.8336 |
| 09918:3 | 18 | R.MAGKPTHIN[+1540.529]VSVVMAEADGTC[+57.021]Y.- | HexNAc(2)Hex(7) | C[+57], N[+1541] | 1297.8791 | 3 | 3891.6227 | 3891.6114 | 0 | 2.9 | 319 | CRagged | 801.44 | 801.44 | 431.11 | 16.58 | 41 | >sp\|P01877\|IGHA2_HUMAN Immunoglobulin heavy constant alpha 2 OS=Homo sapiens OX=9606 GN=IGHA2 PE=1 SV=4 | 576 | 20201119_16913_1ng_glyco_hcd203040_compound15.13225.13225.3 | controllerType=0 controllerNumber=1 scan=13225 | 28.5884 |
| 09920:3 | 18 | R.MAGKPTHIN[+1540.529]VSVVMAEADGTC[+57.021]Y.- | HexNAc(2)Hex(7) | C[+57], N[+1541] | 1297.8791 | 3 | 3891.6227 | 3891.6114 | 0 | 2.9 | 319 | CRagged | 830.04 | 830.04 | 461.9 | 16.58 | 41 | >sp\|P01877\|IGHA2_HUMAN Immunoglobulin heavy constant alpha 2 OS=Homo sapiens OX=9606 GN=IGHA2 PE=1 SV=4 | 576 | 20201119_16913_1ng_glyco_hcd203040_compound15.13227.13227.3 | controllerType=0 controllerNumber=1 scan=13227 | 28.5936 |
| 09886:3 | 18 | R.MAGKPTHIN[+1702.581]VSVVMAEADGTC[+57.021]Y.- | HexNAc(2)Hex(8) | C[+57], N[+1703] | 1351.8967 | 3 | 4053.6754 | 4053.6642 | 0 | 2.8 | 319 | CRagged | 846.55 | 846.55 | 468.32 | 16.58 | 41 | >sp\|P01877\|IGHA2_HUMAN Immunoglobulin heavy constant alpha 2 OS=Homo sapiens OX=9606 GN=IGHA2 PE=1 SV=4 | 576 | 20201119_16913_1ng_glyco_hcd203040_compound15.13192.13192.3 | controllerType=0 controllerNumber=1 scan=13192 | 28.5296 |
| 09958:3 | 18 | R.MAGKPTHIN[+1378.476]VSVVMAEADGTC[+57.021]Y.- | HexNAc(2)Hex(6) | C[+57], N[+1378] | 1243.8607 | 3 | 3729.5674 | 3729.5585 | 0 | 2.4 | 319 | CRagged | 846.69 | 692.06 | 489.16 | 16.52 | 41 | >sp\|P01877\|IGHA2_HUMAN Immunoglobulin heavy constant alpha 2 OS=Homo sapiens OX=9606 GN=IGHA2 PE=1 SV=4 | 576 | 20201119_16913_1ng_glyco_hcd203040_compound15.13267.13267.3 | controllerType=0 controllerNumber=1 scan=13267 | 28.6834 |
| 10045:3 | 18 | R.MAGKPTHIN[+1216.423]VSVVMAEADGTC[+57.021]Y.- | HexNAc(2)Hex(5) | C[+57], N[+1216] | 1189.8428 | 3 | 3567.5139 | 3567.5057 | 0 | 2.3 | 319 | CRagged | 878.88 | 715.96 | 480.71 | 17.52 | 41 | >sp\|P01877\|IGHA2_HUMAN Immunoglobulin heavy constant alpha 2 OS=Homo sapiens OX=9606 GN=IGHA2 PE=1 SV=4 | 576 | 20201119_16913_1ng_glyco_hcd203040_compound15.13357.13357.3 | controllerType=0 controllerNumber=1 scan=13357 | 28.8385 |

Table S4. Cont.

| Query #:z | Protein Rank | Glycopeptide | Glycan | Modification Type(s) | Observed m/z | z | Observed (M+H) | Calc. mass (M+H) | Off-by-x error | Mass error (ppm) | StartPosition | Cleavage | Score | Delta | Delta Mod | \|Log Prob\| | # of unique peptides | Protein Name | Protein DB number | Comment | Scan # | Scan Time |
|---|---|---|---|---|---|---|---|---|---|---|---|---|---|---|---|---|---|---|---|---|---|---|
| 09888:3 | 18 | R.MAGKPTHIN[+1702.581]VSVVMAEADGTC[+57.021]Y.- | HexNAc(2)Hex(8) | C[+57], N[+1703] | 1351.8967 | 3 | 4053.6754 | 4053.6642 | 0 | 2.8 | 319 | CRagged | 892.45 | 704.91 | 485.27 | 17.58 | 41 | >sp\|P01877\|IGHA2_HUMAN Immunoglobulin heavy constant alpha 2 OS=Homo sapiens OX=9606 GN=IGHA2 PE=1 SV=4 | 576 | 20201119_16913_1ng_glyco_hcd203040_compound15.13194.13194.3 | controllerType=0 controllerNumber=1 scan=13194 | 28.5347 |
| 09955:3 | 18 | R.MAGKPTHIN[+1378.476]VSVVMAEADGTC[+57.021]Y.- | HexNAc(2)Hex(6) | C[+57], N[+1378] | 1243.8607 | 3 | 3729.5674 | 3729.5585 | 0 | 2.4 | 319 | CRagged | 969.77 | 819.99 | 550.59 | 19.52 | 41 | >sp\|P01877\|IGHA2_HUMAN Immunoglobulin heavy constant alpha 2 OS=Homo sapiens OX=9606 GN=IGHA2 PE=1 SV=4 | 576 | 20201119_16913_1ng_glyco_hcd203040_compound15.13264.13264.3 | controllerType=0 controllerNumber=1 scan=13264 | 28.673 |
| 05824:3 | 19 | K.KLLN[+2059.735]LTVR.I | HexNAc(4)Hex(5)Fuc(1)NeuAc(1) | N[+2060] | 1006.1262 | 3 | 3016.3642 | 3016.3601 | 0 | 1.4 | 133 | Specific | 210.2 | 210.2 | 210.2 | 3.66 | 36 | >sp\|Q6UX06\|OLFM4_HUMAN Olfactomedin-4 OS=Homo sapiens OX=9606 GN=OLFM4 PE=1 SV=1 | 624 | 20201119_16913_1ng_glyco_hcd203040_compound15.8952.8952.3 | controllerType=0 controllerNumber=1 scan=8952 | 21.4207 |
| 05822:3 | 19 | K.KLLN[+2059.735]LTVR.I | HexNAc(4)Hex(5)Fuc(1)NeuAc(1) | N[+2060] | 1006.1262 | 3 | 3016.3642 | 3016.3601 | 0 | 1.4 | 133 | Specific | 237.91 | 237.91 | 237.91 | 5.8 | 36 | >sp\|Q6UX06\|OLFM4_HUMAN Olfactomedin-4 OS=Homo sapiens OX=9606 GN=OLFM4 PE=1 SV=1 | 624 | 20201119_16913_1ng_glyco_hcd203040_compound15.8950.8950.3 | controllerType=0 controllerNumber=1 scan=8950 | 21.4171 |
| 08767:5 | 19 | K.LKEC[+57.021]EASKDQNTPVVHPPPTPGSC[+57.021]GHGGVVN[+2059.735]ISKPS | HexNAc(4)Hex(5)Fuc(1)NeuAc(1) | C[+57]*2, N[+2060] | 1348.0273 | 5 | 6736.1072 | 6736.081 | 0 | 3.9 | 223 | Specific | 338 | 338 | 338 | 4.91 | 36 | >sp\|Q6UX06\|OLFM4_HUMAN Olfactomedin-4 OS=Homo sapiens OX=9606 GN=OLFM4 PE=1 SV=1 | 624 | 20201119_16913_1ng_glyco_hcd203040_compound15.12030.12030.5 | controllerType=0 controllerNumber=1 scan=12030 | 26.5694 |
| 08765:5 | 19 | K.LKEC[+57.021]EASKDQNTPVVHPPPTPGSC[+57.021]GHGGVVN[+2059.735]ISKPS | HexNAc(4)Hex(5)Fuc(1)NeuAc(1) | C[+57]*2, N[+2060] | 1348.0273 | 5 | 6736.1072 | 6736.081 | 0 | 3.9 | 223 | Specific | 342.59 | 342.59 | 342.59 | 6.05 | 36 | >sp\|Q6UX06\|OLFM4_HUMAN Olfactomedin-4 OS=Homo sapiens OX=9606 GN=OLFM4 PE=1 SV=1 | 624 | 20201119_16913_1ng_glyco_hcd203040_compound15.12028.12028.5 | controllerType=0 controllerNumber=1 scan=12028 | 26.5657 |
| 08792:6 | 19 | K.LKEC[+57.021]EASKDQNTPVVHPPPTPGSC[+57.021]GHGGVVN[+2059.735]ISKPS | HexNAc(4)Hex(5)Fuc(1)NeuAc(1) | C[+57]*2, N[+2060] | 1123.5223 | 6 | 6736.0973 | 6736.081 | 0 | 2.4 | 223 | Specific | 521.7 | 521.7 | 521.7 | 8.99 | 36 | >sp\|Q6UX06\|OLFM4_HUMAN Olfactomedin-4 OS=Homo sapiens OX=9606 GN=OLFM4 PE=1 SV=1 | 624 | 20201119_16913_1ng_glyco_hcd203040_compound15.12056.12056.6 | controllerType=0 controllerNumber=1 scan=12056 | 26.6162 |
| 08794:6 | 19 | K.LKEC[+57.021]EASKDQNTPVVHPPPTPGSC[+57.021]GHGGVVN[+2059.735]ISKPS | HexNAc(4)Hex(5)Fuc(1)NeuAc(1) | C[+57]*2, N[+2060] | 1123.5223 | 6 | 6736.0973 | 6736.081 | 0 | 2.4 | 223 | Specific | 669.76 | 660.16 | 660.16 | 12.54 | 36 | >sp\|Q6UX06\|OLFM4_HUMAN Olfactomedin-4 OS=Homo sapiens OX=9606 GN=OLFM4 PE=1 SV=1 | 624 | 20201119_16913_1ng_glyco_hcd203040_compound15.12058.12058.6 | controllerType=0 controllerNumber=1 scan=12058 | 26.6202 |
| 08712:6 | 19 | K.LKEC[+57.021]EASKDQNTPVVHPPPTPGSC[+57.021]GHGGVVN[+2059.735]ISKPS | HexNAc(4)Hex(5)Fuc(1)NeuAc(1) | C[+57]*2, N[+2060] | 1123.5229 | 6 | 6736.1009 | 6736.081 | 0 | 3 | 223 | Specific | 960.21 | 816.6 | 816.6 | 18.6 | 36 | >sp\|Q6UX06\|OLFM4_HUMAN Olfactomedin-4 OS=Homo sapiens OX=9606 GN=OLFM4 PE=1 SV=1 | 624 | 20201119_16913_1ng_glyco_hcd203040_compound15.11973.11973.6 | controllerType=0 controllerNumber=1 scan=11973 | 26.473 |
| 08710:6 | 19 | K.LKEC[+57.021]EASKDQNTPVVHPPPTPGSC[+57.021]GHGGVVN[+2059.735]ISKPS | HexNAc(4)Hex(5)Fuc(1)NeuAc(1) | C[+57]*2, N[+2060] | 1123.5229 | 6 | 6736.1009 | 6736.081 | 0 | 3 | 223 | Specific | 1032.54 | 935.97 | 935.97 | 19.6 | 36 | >sp\|Q6UX06\|OLFM4_HUMAN Olfactomedin-4 OS=Homo sapiens OX=9606 GN=OLFM4 PE=1 SV=1 | 624 | 20201119_16913_1ng_glyco_hcd203040_compound15.11971.11971.6 | controllerType=0 controllerNumber=1 scan=11971 | 26.4669 |
| 11196:3 | 19 | R.VN[+1216.423]LTTNTIAVTQTLPNAAYNNR.F | HexNAc(2)Hex(5) | N[+1216] | 1202.565 | 3 | 3605.6804 | 3605.6699 | 0 | 2.9 | 351 | Specific | 464.92 | 464.92 | 464.92 | 9.71 | 36 | >sp\|Q6UX06\|OLFM4_HUMAN Olfactomedin-4 OS=Homo sapiens OX=9606 GN=OLFM4 PE=1 SV=1 | 624 | 20201119_16913_1ng_glyco_hcd203040_compound15.14555.14555.3 | controllerType=0 controllerNumber=1 scan=14555 | 31.0335 |
| 11274:3 | 19 | R.VN[+1054.370]LTTNTIAVTQTLPNAAYNNR.F | HexNAc(2)Hex(4) | N[+1054] | 1148.5467 | 3 | 3443.6254 | 3443.6171 | 0 | 2.4 | 351 | Specific | 512.26 | 512.26 | 512.26 | 9.88 | 36 | >sp\|Q6UX06\|OLFM4_HUMAN Olfactomedin-4 OS=Homo sapiens OX=9606 GN=OLFM4 PE=1 SV=1 | 624 | 20201119_16913_1ng_glyco_hcd203040_compound15.14636.14636.3 | controllerType=0 controllerNumber=1 scan=14636 | 31.1738 |
| 11277:3 | 19 | R.VN[+1054.370]LTTNTIAVTQTLPNAAYNNR.F | HexNAc(2)Hex(4) | N[+1054] | 1148.5467 | 3 | 3443.6254 | 3443.6171 | 0 | 2.4 | 351 | Specific | 726.02 | 726.02 | 726.02 | 14.43 | 36 | >sp\|Q6UX06\|OLFM4_HUMAN Olfactomedin-4 OS=Homo sapiens OX=9606 GN=OLFM4 PE=1 SV=1 | 624 | 20201119_16913_1ng_glyco_hcd203040_compound15.14639.14639.3 | controllerType=0 controllerNumber=1 scan=14639 | 31.1783 |
| 11133:3 | 19 | R.VN[+1216.423]LTTNTIAVTQTLPNAAYNNR.F | HexNAc(2)Hex(5) | N[+1216] | 1202.5654 | 3 | 3605.6816 | 3605.6699 | 0 | 3.3 | 351 | Specific | 771.15 | 771.15 | 771.15 | 15.49 | 36 | >sp\|Q6UX06\|OLFM4_HUMAN Olfactomedin-4 OS=Homo sapiens OX=9606 GN=OLFM4 PE=1 SV=1 | 624 | 20201119_16913_1ng_glyco_hcd203040_compound15.14488.14488.3 | controllerType=0 controllerNumber=1 scan=14488 | 30.9033 |
| 11159:2 | 19 | R.VN[+1216.423]LTTNTIAVTQTLPNAAYNNR.F | HexNAc(2)Hex(5) | N[+1216] | 1803.3416 | 2 | 3605.676 | 3605.6699 | 0 | 1.7 | 351 | Specific | 946.58 | 847.37 | 847.37 | 18.52 | 36 | >sp\|Q6UX06\|OLFM4_HUMAN Olfactomedin-4 OS=Homo sapiens OX=9606 GN=OLFM4 PE=1 SV=1 | 624 | 20201119_16913_1ng_glyco_hcd203040_compound15.14516.14516.2 | controllerType=0 controllerNumber=1 scan=14516 | 30.9568 |
| 11135:3 | 19 | R.VN[+1216.423]LTTNTIAVTQTLPNAAYNNR.F | HexNAc(2)Hex(5) | N[+1216] | 1202.5654 | 3 | 3605.6816 | 3605.6699 | 0 | 3.3 | 351 | Specific | 1011.43 | 1011.43 | 1011.43 | 20.49 | 36 | >sp\|Q6UX06\|OLFM4_HUMAN Olfactomedin-4 OS=Homo sapiens OX=9606 GN=OLFM4 PE=1 SV=1 | 624 | 20201119_16913_1ng_glyco_hcd203040_compound15.14490.14490.3 | controllerType=0 controllerNumber=1 scan=14490 | 30.9097 |
| 11198:3 | 19 | R.VN[+1216.423]LTTNTIAVTQTLPNAAYNNR.F | HexNAc(2)Hex(5) | N[+1216] | 1202.565 | 3 | 3605.6804 | 3605.6699 | 0 | 2.9 | 351 | Specific | 1023.82 | 956.53 | 956.53 | 20.49 | 36 | >sp\|Q6UX06\|OLFM4_HUMAN Olfactomedin-4 OS=Homo sapiens OX=9606 GN=OLFM4 PE=1 SV=1 | 624 | 20201119_16913_1ng_glyco_hcd203040_compound15.14557.14557.3 | controllerType=0 controllerNumber=1 scan=14557 | 31.0366 |
| 11156:2 | 19 | R.VN[+1216.423]LTTNTIAVTQTLPNAAYNNR.F | HexNAc(2)Hex(5) | N[+1216] | 1803.3416 | 2 | 3605.676 | 3605.6699 | 0 | 1.7 | 351 | Specific | 1042.11 | 1004.25 | 1004.25 | 20.52 | 36 | >sp\|Q6UX06\|OLFM4_HUMAN Olfactomedin-4 OS=Homo sapiens OX=9606 GN=OLFM4 PE=1 SV=1 | 624 | 20201119_16913_1ng_glyco_hcd203040_compound15.14513.14513.2 | controllerType=0 controllerNumber=1 scan=14513 | 30.9503 |
| 05312:3 | 20 | R.LSPN[+2205.793]ASAEHLELR.W | HexNAc(4)Hex(5)Fuc(2)NeuAc(1) | N[+2206] | 1214.8559 | 3 | 3642.5531 | 3642.5421 | 0 | 3 | 52 | Specific | 379.07 | 94.98 | 94.98 | 5.46 | 32 | >sp\|Q13410\|BT1A1_HUMAN Butyrophilin subfamily 1 member A1 OS=Homo sapiens OX=9606 GN=BTN1A1 PE=1 SV=3 | 179 | 20201119_16913_1ng_glyco_hcd203040_compound15.8423.8423.3 | controllerType=0 controllerNumber=1 scan=8423 | 20.543 |
| 05314:3 | 20 | R.LSPN[+2205.793]ASAEHLELR.W | HexNAc(4)Hex(5)Fuc(2)NeuAc(1) | N[+2206] | 1214.8559 | 3 | 3642.5531 | 3642.5421 | 0 | 3 | 52 | Specific | 428.52 | 294.62 | 294.62 | 7.52 | 32 | >sp\|Q13410\|BT1A1_HUMAN Butyrophilin subfamily 1 member A1 OS=Homo sapiens OX=9606 GN=BTN1A1 PE=1 SV=3 | 179 | 20201119_16913_1ng_glyco_hcd203040_compound15.8425.8425.3 | controllerType=0 controllerNumber=1 scan=8425 | 20.5468 |
| 07379:3 | 20 | R.LSPN[+2350.830]ASAEHLELR.W | HexNAc(4)Hex(5)Fuc(1)NeuAc(2) | N[+2351] | 1263.2005 | 3 | 3787.5869 | 3787.5796 | 0 | 1.9 | 52 | Specific | 634.35 | 593.82 | 593.82 | 12.3 | 32 | >sp\|Q13410\|BT1A1_HUMAN Butyrophilin subfamily 1 member A1 OS=Homo sapiens OX=9606 GN=BTN1A1 PE=1 SV=3 | 179 | 20201119_16913_1ng_glyco_hcd203040_compound15.10568.10568.3 | controllerType=0 controllerNumber=1 scan=10568 | 24.0661 |
| 07378:3 | 20 | R.LSPN[+2350.830]ASAEHLELR.W | HexNAc(4)Hex(5)Fuc(1)NeuAc(2) | N[+2351] | 1263.2005 | 3 | 3787.5869 | 3787.5796 | 0 | 1.9 | 52 | Specific | 752.85 | 629.13 | 629.13 | 15.3 | 32 | >sp\|Q13410\|BT1A1_HUMAN Butyrophilin subfamily 1 member A1 OS=Homo sapiens OX=9606 GN=BTN1A1 PE=1 SV=3 | 179 | 20201119_16913_1ng_glyco_hcd203040_compound15.10567.10567.3 | controllerType=0 controllerNumber=1 scan=10567 | 24.0619 |
| 16829:3 | 20 | K.N[+2059.735]VSC[+57.021]YIQNLLLGQEK.K | HexNAc(4)Hex(5)Fuc(1)NeuAc(1) | C[+57], N[+2060] | 1280.2221 | 3 | 3838.6518 | 3838.6455 | 0 | 1.6 | 215 | Specific | 225.12 | 86.13 | 86.13 | 3.6 | 32 | >sp\|Q13410\|BT1A1_HUMAN Butyrophilin subfamily 1 member A1 OS=Homo sapiens OX=9606 GN=BTN1A1 PE=1 SV=3 | 179 | 20201119_16913_1ng_glyco_hcd203040_compound15.20445.20445.3 | controllerType=0 controllerNumber=1 scan=20445 | 42.3956 |
| 18798:3 | 20 | K.N[+2204.772]VSC[+57.021]YIQNLLLGQEK.K | HexNAc(4)Hex(5)NeuAc(2) | C[+57], N[+2205] | 1328.5712 | 3 | 3983.6991 | 3983.683 | 0 | 4 | 215 | Specific | 280.81 | 103.18 | 103.18 | 4.58 | 32 | >sp\|Q13410\|BT1A1_HUMAN Butyrophilin subfamily 1 member A1 OS=Homo sapiens OX=9606 GN=BTN1A1 PE=1 SV=3 | 179 | 20201119_16913_1ng_glyco_hcd203040_compound15.22506.22506.3 | controllerType=0 controllerNumber=1 scan=22506 | 47.0933 |
| 18801:3 | 20 | K.N[+2204.772]VSC[+57.021]YIQNLLLGQEK.K | HexNAc(4)Hex(5)NeuAc(2) | C[+57], N[+2205] | 1328.5712 | 3 | 3983.6991 | 3983.683 | 0 | 4 | 215 | Specific | 282.34 | 152.59 | 152.59 | 4.76 | 32 | >sp\|Q13410\|BT1A1_HUMAN Butyrophilin subfamily 1 member A1 OS=Homo sapiens OX=9606 GN=BTN1A1 PE=1 SV=3 | 179 | 20201119_16913_1ng_glyco_hcd203040_compound15.22509.22509.3 | controllerType=0 controllerNumber=1 scan=22509 | 47.1051 |
| 01698:3 | 21 | K.N[+1540.529]NSIPDKQITASSSYK.T | HexNAc(2)Hex(7) | N[+1541] | 1098.4738 | 3 | 3293.4068 | 3293.4048 | 0 | 0.6 | 238 | Specific | 201.64 | 201.64 | 201.64 | 4.01 | 31 | >sp\|Q08431\|MFGM_HUMAN Lactadherin OS=Homo sapiens OX=9606 GN=MFGE8 PE=1 SV=3 | 1064 | 20201119_16913_1ng_glyco_hcd203040_compound15.4672.4672.3 | controllerType=0 controllerNumber=1 scan=4672 | 14.2519 |
| 01760:3 | 21 | K.N[+1378.476]NSIPDKQITASSSYK.T | HexNAc(2)Hex(6) | N[+1378] | 1044.4593 | 3 | 3131.3632 | 3131.352 | 0 | 3.6 | 238 | Specific | 226.31 | 226.31 | 226.31 | 5.85 | 31 | >sp\|Q08431\|MFGM_HUMAN Lactadherin OS=Homo sapiens OX=9606 GN=MFGE8 PE=1 SV=3 | 1064 | 20201119_16913_1ng_glyco_hcd203040_compound15.4737.4737.3 | controllerType=0 controllerNumber=1 scan=4737 | 14.3723 |
| 01700:3 | 21 | K.N[+1540.529]NSIPDKQITASSSYK.T | HexNAc(2)Hex(7) | N[+1541] | 1098.4738 | 3 | 3293.4068 | 3293.4048 | 0 | 0.6 | 238 | Specific | 263.73 | 263.73 | 263.73 | 5.68 | 31 | >sp\|Q08431\|MFGM_HUMAN Lactadherin OS=Homo sapiens OX=9606 GN=MFGE8 PE=1 SV=3 | 1064 | 20201119_16913_1ng_glyco_hcd203040_compound15.4674.4674.3 | controllerType=0 controllerNumber=1 scan=4674 | 14.2565 |
| 01831:3 | 21 | K.N[+1216.423]NSIPDKQITASSSYK.T | HexNAc(2)Hex(5) | N[+1216] | 990.4415 | 3 | 2969.3099 | 2969.2991 | 0 | 3.6 | 238 | Specific | 512.15 | 295.72 | 295.72 | 7.28 | 31 | >sp\|Q08431\|MFGM_HUMAN Lactadherin OS=Homo sapiens OX=9606 GN=MFGE8 PE=1 SV=3 | 1064 | 20201119_16913_1ng_glyco_hcd203040_compound15.4811.4811.3 | controllerType=0 controllerNumber=1 scan=4811 | 14.5077 |
| 01829:3 | 21 | K.N[+1216.423]NSIPDKQITASSSYK.T | HexNAc(2)Hex(5) | N[+1216] | 990.4415 | 3 | 2969.3099 | 2969.2991 | 0 | 3.6 | 238 | Specific | 570.82 | 483.48 | 483.48 | 9.55 | 31 | >sp\|Q08431\|MFGM_HUMAN Lactadherin OS=Homo sapiens OX=9606 GN=MFGE8 PE=1 SV=3 | 1064 | 20201119_16913_1ng_glyco_hcd203040_compound15.4809.4809.3 | controllerType=0 controllerNumber=1 scan=4809 | 14.5034 |
| 09139:4 | 26 | K.QTDEIKDTRN[+2059.735]ESTQNC[+57.021]VVAEPEKMESSISSSSEEMSLSK.C | HexNAc(4)Hex(5)Fuc(1)NeuAc(1) | C[+57], N[+2060] | 1613.436 | 4 | 6450.7223 | 6450.7156 | 0 | 1 | 60 | Specific | 376.72 | 247.61 | 247.61 | 5.48 | 24 | >sp\|P47710\|CASA1_HUMAN Alpha-S1-casein OS=Homo sapiens OX=9606 GN=CSN1S1 PE=1 SV=1 | 989 | 20201119_16913_1ng_glyco_hcd203040_compound15.12417.12417.2 | controllerType=0 controllerNumber=1 scan=12417 | 27.2343 |
| 09138:4 | 26 | K.QTDEIKDTRN[+2059.735]ESTQNC[+57.021]VVAEPEKMESSISSSSEEMSLSK.C | HexNAc(4)Hex(5)Fuc(1)NeuAc(1) | C[+57], N[+2060] | 1613.436 | 4 | 6450.7223 | 6450.7156 | 0 | 1 | 60 | Specific | 396.64 | 297.06 | 297.06 | 5.64 | 24 | >sp\|P47710\|CASA1_HUMAN Alpha-S1-casein OS=Homo sapiens OX=9606 GN=CSN1S1 PE=1 SV=1 | 989 | 20201119_16913_1ng_glyco_hcd203040_compound15.12416.12416.2 | controllerType=0 controllerNumber=1 scan=12416 | 27.2308 |
| 08245:5 | 35 | K.[+42.011]SC[+57.021]QHN[+2018.708]GTMYQHGEIFSAHELFPSR.L | HexNAc(3)Hex(6)Fuc(1)NeuAc(1) | .[42], C[+57], N[+2019] | 979.6086 | 5 | 4894.0139 | 4893.9759 | 0 | 7.8 | 110 | Specific | 218.47 | 149.12 | 149.12 | 3.61 | 21 | >sp\|Q6WN34\|CRDL2_HUMAN Chordin-like protein 2 OS=Homo sapiens OX=9606 GN=CHRDL2 PE=1 SV=1 | 1189 | 20201119_16913_1ng_glyco_hcd203040_compound15.11472.11472.5 | controllerType=0 controllerNumber=1 scan=11472 | 25.609 |
| 08246:5 | 35 | K.[+42.011]SC[+57.021]QHN[+2018.708]GTMYQHGEIFSAHELFPSR.L | HexNAc(3)Hex(6)Fuc(1)NeuAc(1) | .[42], C[+57], N[+2019] | 979.6086 | 5 | 4894.0139 | 4893.9759 | 0 | 7.8 | 110 | Specific | 229.46 | 198.97 | 198.97 | 4.18 | 21 | >sp\|Q6WN34\|CRDL2_HUMAN Chordin-like protein 2 OS=Homo sapiens OX=9606 GN=CHRDL2 PE=1 SV=1 | 1189 | 20201119_16913_1ng_glyco_hcd203040_compound15.11473.11473.5 | controllerType=0 controllerNumber=1 scan=11473 | 25.6128 |
| 09739:4 | 35 | K.SC[+57.021]QHN[+2350.830]GTMYQHGEIFSAHELFPSR.L | HexNAc(4)Hex(5)Fuc(1)NeuAc(2) | C[+57], N[+2351] | 1296.7828 | 4 | 5184.1095 | 5184.0873 | 0 | 4.3 | 110 | Specific | 551.62 | 551.62 | 551.62 | 9.28 | 21 | >sp\|Q6WN34\|CRDL2_HUMAN Chordin-like protein 2 OS=Homo sapiens OX=9606 GN=CHRDL2 PE=1 SV=1 | 1189 | 20201119_16913_1ng_glyco_hcd203040_compound15.13040.13040.4 | controllerType=0 controllerNumber=1 scan=13040 | 28.2665 |
| 09736:4 | 35 | K.SC[+57.021]QHN[+2350.830]GTMYQHGEIFSAHELFPSR.L | HexNAc(4)Hex(5)Fuc(1)NeuAc(2) | C[+57], N[+2351] | 1296.7828 | 4 | 5184.1095 | 5184.0873 | 0 | 4.3 | 110 | Specific | 567.59 | 567.59 | 567.59 | 9.28 | 21 | >sp\|Q6WN34\|CRDL2_HUMAN Chordin-like protein 2 OS=Homo sapiens OX=9606 GN=CHRDL2 PE=1 SV=1 | 1189 | 20201119_16913_1ng_glyco_hcd203040_compound15.13037.13037.4 | controllerType=0 controllerNumber=1 scan=13037 | 28.2627 |
| 09373:3 | 58 | R.IIVPLNNREN[+1647.613]ISDPTSPLR.T | HexNAc(5)Hex(3)Fuc(1) | N[+1648] | 1265.9367 | 3 | 3795.7956 | 3795.7904 | 0 | 1.4 | 62 | Specific | 332.38 | 283.38 | 283.38 | 6.34 | 16 | >sp\|P01591\|IGJ_HUMAN Immunoglobulin J chain OS=Homo sapiens OX=9606 GN=JCHAIN PE=1 SV=4 | 1392 | 20201119_16913_1ng_glyco_hcd203040_compound15.12660.12660.3 | controllerType=0 controllerNumber=1 scan=12660 | 27.6371 |
| 09291:3 | 58 | R.IIVPLNNREN[+1809.666]ISDPTSPLR.T | HexNAc(5)Hex(4)Fuc(1) | N[+1810] | 1319.9564 | 3 | 3957.8545 | 3957.8432 | 0 | 2.9 | 62 | Specific | 361.57 | 276.69 | 276.69 | 6.88 | 16 | >sp\|P01591\|IGJ_HUMAN Immunoglobulin J chain OS=Homo sapiens OX=9606 GN=JCHAIN PE=1 SV=4 | 1392 | 20201119_16913_1ng_glyco_hcd203040_compound15.12576.12576.3 | controllerType=0 controllerNumber=1 scan=12576 | 27.4955 |
| 09374:3 | 58 | R.IIVPLNNREN[+1647.613]ISDPTSPLR.T | HexNAc(5)Hex(3)Fuc(1) | N[+1648] | 1265.9367 | 3 | 3795.7956 | 3795.7904 | 0 | 1.4 | 62 | Specific | 365.7 | 271.48 | 271.48 | 6.41 | 16 | >sp\|P01591\|IGJ_HUMAN Immunoglobulin J chain OS=Homo sapiens OX=9606 GN=JCHAIN PE=1 SV=4 | 1392 | 20201119_16913_1ng_glyco_hcd203040_compound15.12661.12661.3 | controllerType=0 controllerNumber=1 scan=12661 | 27.6406 |
| 09293:3 | 58 | R.IIVPLNNREN[+1809.666]ISDPTSPLR.T | HexNAc(5)Hex(4)Fuc(1) | N[+1810] | 1319.9564 | 3 | 3957.8545 | 3957.8432 | 0 | 2.9 | 62 | Specific | 414.19 | 315.87 | 315.87 | 7.16 | 16 | >sp\|P01591\|IGJ_HUMAN Immunoglobulin J chain OS=Homo sapiens OX=9606 GN=JCHAIN PE=1 SV=4 | 1392 | 20201119_16913_1ng_glyco_hcd203040_compound15.12578.12578.3 | controllerType=0 controllerNumber=1 scan=12578 | 27.4995 |
| 09299:3 | 58 | R.IIVPLNNREN[+1663.608]ISDPTSPLR.T | HexNAc(5)Hex(4) | N[+1664] | 1271.2703 | 3 | 3811.7964 | 3811.7853 | 0 | 2.9 | 62 | Specific | 460.73 | 455.77 | 455.77 | 9.16 | 16 | >sp\|P01591\|IGJ_HUMAN Immunoglobulin J chain OS=Homo sapiens OX=9606 GN=JCHAIN PE=1 SV=4 | 1392 | 20201119_16913_1ng_glyco_hcd203040_compound15.12584.12584.3 | controllerType=0 controllerNumber=1 scan=12584 | 27.5096 |
| 09302:3 | 58 | R.IIVPLNNREN[+1663.608]ISDPTSPLR.T | HexNAc(5)Hex(4) | N[+1664] | 1271.2703 | 3 | 3811.7964 | 3811.7853 | 0 | 2.9 | 62 | Specific | 474.17 | 474.06 | 474.06 | 9.16 | 16 | >sp\|P01591\|IGJ_HUMAN Immunoglobulin J chain OS=Homo sapiens OX=9606 GN=JCHAIN PE=1 SV=4 | 1392 | 20201119_16913_1ng_glyco_hcd203040_compound15.12587.12587.3 | controllerType=0 controllerNumber=1 scan=12587 | 27.5145 |
| 10995:3 | 58 | R.IIVPLNNREN[+1856.656]ISDPTSPLR.T | HexNAc(3)Hex(5)Fuc(1)NeuAc(1) | N[+1857] | 1335.6148 | 3 | 4004.8299 | 4004.8327 | 0 | -0.7 | 62 | Specific | 496.99 | 393.41 | 393.41 | 8.29 | 16 | >sp\|P01591\|IGJ_HUMAN Immunoglobulin J chain OS=Homo sapiens OX=9606 GN=JCHAIN PE=1 SV=4 | 1392 | 20201119_16913_1ng_glyco_hcd203040_compound15.14342.14342.3 | controllerType=0 controllerNumber=1 scan=14342 | 30.5663 |
| 10993:3 | 58 | R.IIVPLNNREN[+1856.656]ISDPTSPLR.T | HexNAc(3)Hex(5)Fuc(1)NeuAc(1) | N[+1857] | 1335.6148 | 3 | 4004.8299 | 4004.8327 | 0 | -0.7 | 62 | Specific | 522.24 | 522.24 | 522.24 | 8.52 | 16 | >sp\|P01591\|IGJ_HUMAN Immunoglobulin J chain OS=Homo sapiens OX=9606 GN=JCHAIN PE=1 SV=4 | 1392 | 20201119_16913_1ng_glyco_hcd203040_compound15.14340.14340.3 | controllerType=0 controllerNumber=1 scan=14340 | 30.5571 |
| 11006:3 | 58 | R.IIVPLNNREN[+1856.656]ISDPTSPLR.T | HexNAc(3)Hex(5)Fuc(1)NeuAc(1) | N[+1857] | 1335.6204 | 3 | 4004.8467 | 4004.8327 | 0 | 3.5 | 62 | Specific | 522.92 | 479.85 | 479.85 | 9.4 | 16 | >sp\|P01591\|IGJ_HUMAN Immunoglobulin J chain OS=Homo sapiens OX=9606 GN=JCHAIN PE=1 SV=4 | 1392 | 20201119_16913_1ng_glyco_hcd203040_compound15.14354.14354.3 | controllerType=0 controllerNumber=1 scan=14354 | 30.5877 |
| 11008:3 | 58 | R.IIVPLNNREN[+1856.656]ISDPTSPLR.T | HexNAc(3)Hex(5)Fuc(1)NeuAc(1) | N[+1857] | 1335.6204 | 3 | 4004.8467 | 4004.8327 | 0 | 3.5 | 62 | Specific | 541.35 | 505.27 | 505.27 | 9.42 | 16 | >sp\|P01591\|IGJ_HUMAN Immunoglobulin J chain OS=Homo sapiens OX=9606 GN=JCHAIN PE=1 SV=4 | 1392 | 20201119_16913_1ng_glyco_hcd203040_compound15.14356.14356.3 | controllerType=0 controllerNumber=1 scan=14356 | 30.593 |
| 21436:3 | 68 | R.[+42.011]QLDMLDLSN[+1872.651]NSLASVPEGLWASLGQPNWDMR.D | HexNAc(3)Hex(6)NeuAc(1) | .[42], N[+1873] | 1791.46 | 3 | 5372.3656 | 5372.3125 | 0 | 9.9 | 261 | Specific | 216.74 | 216.74 | 216.74 | 3.59 | 11 | >sp\|P02750\|A2GL_HUMAN Leucine-rich alpha-2-glycoprotein OS=Homo sapiens OX=9606 GN=LRG1 PE=1 SV=2 | 518 | 20201119_16913_1ng_glyco_hcd203040_compound15.25266.25266.3 | controllerType=0 controllerNumber=1 scan=25266 | 53.1076 |
| 21438:3 | 68 | R.[+42.011]QLDMLDLSN[+1872.651]NSLASVPEGLWASLGQPNWDMR.D | HexNAc(3)Hex(6)NeuAc(1) | .[42], N[+1873] | 1791.46 | 3 | 5372.3656 | 5372.3125 | 0 | 9.9 | 261 | Specific | 262.07 | 254.06 | 254.06 | 5.16 | 11 | >sp\|P02750\|A2GL_HUMAN Leucine-rich alpha-2-glycoprotein OS=Homo sapiens OX=9606 GN=LRG1 PE=1 SV=2 | 518 | 20201119_16913_1ng_glyco_hcd203040_compound15.25268.25268.3 | controllerType=0 controllerNumber=1 scan=25268 | 53.1116 |
| 10444:3 | 102 | R.ADGTVNQIEGEATPVN[+1216.423]LTEPAKLEVK.F | HexNAc(2)Hex(5) | N[+1216] | 1313.9548 | 3 | 3939.8498 | 3939.8326 | 0 | 4.3 | 83 | Specific | 264.04 | 264.04 | 264.04 | 5.86 | 9 | >sp\|P05090\|APOD_HUMAN Apolipoprotein D OS=Homo sapiens OX=9606 GN=APOD PE=1 SV=1 | 594 | 20201119_16913_1ng_glyco_hcd203040_compound15.13770.13770.3 | controllerType=0 controllerNumber=1 scan=13770 | 29.5163 |
| 10446:3 | 102 | R.ADGTVNQIEGEATPVN[+1216.423]LTEPAKLEVK.F | HexNAc(2)Hex(5) | N[+1216] | 1313.9548 | 3 | 3939.8498 | 3939.8326 | 0 | 4.3 | 83 | Specific | 345.87 | 345.87 | 345.87 | 6.78 | 9 | >sp\|P05090\|APOD_HUMAN Apolipoprotein D OS=Homo sapiens OX=9606 GN=APOD PE=1 SV=1 | 594 | 20201119_16913_1ng_glyco_hcd203040_compound15.13772.13772.3 | controllerType=0 controllerNumber=1 scan=13772 | 29.5203 |
| 18349:5 | 1 | R.RTAGWNVPIGTLRPFLN[+2205.793]WTGPPEPIEAAVAR.F | HexNAc(4)Hex(5)Fuc(2)NeuAc(1) | N[+2206] | 1119.3342 | 5 | 5592.642 | 5592.6049 | 0 | 6.6 | 140 | Specific | 200.18 | 200.18 | 200.18 | 3.13 | 111 | >sp\|P02788\|TRFL_HUMAN Lactotransferrin OS=Homo sapiens OX=9606 GN=LTF PE=1 SV=6 | 1270 | 20201114_16933_1ng_glyco_hcd203040_compound15.21684.21684.4 | controllerType=0 controllerNumber=1 scan=21684 | 43.5282 |
| 17362:4 | 1 | R.RTAGWNVPIGTLRPFLN[+1913.677]WTGPPEPIEAAVAR.F | HexNAc(4)Hex(5)NeuAc(1) | N[+1914] | 1325.882 | 4 | 5300.506 | 5300.4891 | 0 | 3.2 | 140 | Specific | 215.2 | 215.2 | 215.2 | 5.36 | 111 | >sp\|P02788\|TRFL_HUMAN Lactotransferrin OS=Homo sapiens OX=9606 GN=LTF PE=1 SV=6 | 1270 | 20201114_16933_1ng_glyco_hcd203040_compound15.20653.20653.4 | controllerType=0 controllerNumber=1 scan=20653 | 41.4046 |
| 18351:5 | 1 | R.RTAGWNVPIGTLRPFLN[+2205.793]WTGPPEPIEAAVAR.F | HexNAc(4)Hex(5)Fuc(2)NeuAc(1) | N[+2206] | 1119.3342 | 5 | 5592.642 | 5592.6049 | 0 | 6.6 | 140 | Specific | 260.49 | 260.49 | 260.49 | 4.18 | 111 | >sp\|P02788\|TRFL_HUMAN Lactotransferrin OS=Homo sapiens OX=9606 GN=LTF PE=1 SV=6 | 1270 | 20201114_16933_1ng_glyco_hcd203040_compound15.21686.21686.4 | controllerType=0 controllerNumber=1 scan=21686 | 43.5329 |
| 19548:4 | 1 | R.RTAGWNVPIGTLRPFLN[+1930.692]WTGPPEPIEAAVAR.F | HexNAc(4)Hex(6)Fuc(1) | N[+1931] | 1330.1365 | 4 | 5317.5241 | 5317.5044 | 0 | 3.7 | 140 | Specific | 302.92 | 302.92 | 302.92 | 7.27 | 111 | >sp\|P02788\|TRFL_HUMAN Lactotransferrin OS=Homo sapiens OX=9606 GN=LTF PE=1 SV=6 | 1270 | 20201114_16933_1ng_glyco_hcd203040_compound15.22938.22938.2 | controllerType=0 controllerNumber=1 scan=22938 | 46.2921 |
| 17348:4 | 1 | R.RTAGWNVPIGTLRPFLN[+1768.640]WTGPPEPIEAAVAR.F | HexNAc(4)Hex(5)Fuc(1) | N[+1769] | 1289.6217 | 4 | 5155.465 | 5155.4516 | 0 | 2.6 | 140 | Specific | 309.72 | 279.61 | 279.61 | 7.06 | 111 | >sp\|P02788\|TRFL_HUMAN Lactotransferrin OS=Homo sapiens OX=9606 GN=LTF PE=1 SV=6 | 1270 | 20201114_16933_1ng_glyco_hcd203040_compound15.20638.20638.4 | controllerType=0 controllerNumber=1 scan=20638 | 41.3684 |
| 19551:4 | 1 | R.RTAGWNVPIGTLRPFLN[+1930.692]WTGPPEPIEAAVAR.F | HexNAc(4)Hex(6)Fuc(1) | N[+1931] | 1330.1365 | 4 | 5317.5241 | 5317.5044 | 0 | 3.7 | 140 | Specific | 337.87 | 337.87 | 337.87 | 6.51 | 111 | >sp\|P02788\|TRFL_HUMAN Lactotransferrin OS=Homo sapiens OX=9606 GN=LTF PE=1 SV=6 | 1270 | 20201114_16933_1ng_glyco_hcd203040_compound15.22941.22941.2 | controllerType=0 controllerNumber=1 scan=22941 | 46.3004 |
| 20902:4 | 1 | R.[+42.011]TAGWNVPIGTLRPFLN[+2336.851]WTGPPEPIEAAVAR.F | HexNAc(6)Hex(6)Fuc(1) | .[42], N[+2337] | 1403.1405 | 4 | 5609.5402 | 5609.5726 | 0 | -5.8 | 141 | Specific | 216.94 | 190.14 | 190.14 | 3.58 | 111 | >sp\|P02788\|TRFL_HUMAN Lactotransferrin OS=Homo sapiens OX=9606 GN=LTF PE=1 SV=6 | 1270 | 20201114_16933_1ng_glyco_hcd203040_compound15.24361.24361.4 | controllerType=0 controllerNumber=1 scan=24361 | 49.2827 |
| 21007:4 | 1 | R.TAGWNVPIGTLRPFLN[+2350.830]WTGPPEPIEAAVAR.F | HexNAc(4)Hex(5)Fuc(1)NeuAc(2) | N[+2351] | 1396.1428 | 4 | 5581.5495 | 5581.5413 | 0 | 1.5 | 141 | Specific | 265.36 | 265.36 | 265.36 | 6.7 | 111 | >sp\|P02788\|TRFL_HUMAN Lactotransferrin OS=Homo sapiens OX=9606 GN=LTF PE=1 SV=6 | 1270 | 20201114_16933_1ng_glyco_hcd203040_compound15.24469.24469.4 | controllerType=0 controllerNumber=1 scan=24469 | 49.4777 |
| 20949:4 | 1 | R.TAGWNVPIGTLRPFLN[+2350.830]WTGPPEPIEAAVAR.F | HexNAc(4)Hex(5)Fuc(1)NeuAc(2) | N[+2351] | 1396.1426 | 4 | 5581.5485 | 5581.5413 | 0 | 1.3 | 141 | Specific | 353.76 | 166.7 | 166.7 | 6.34 | 111 | >sp\|P02788\|TRFL_HUMAN Lactotransferrin OS=Homo sapiens OX=9606 GN=LTF PE=1 SV=6 | 1270 | 20201114_16933_1ng_glyco_hcd203040_compound15.24409.24409.3 | controllerType=0 controllerNumber=1 scan=24409 | 49.3627 |
| 20904:4 | 1 | R.[+42.011]TAGWNVPIGTLRPFLN[+2336.851]WTGPPEPIEAAVAR.F | HexNAc(6)Hex(6)Fuc(1) | .[42], N[+2337] | 1403.1405 | 4 | 5609.5402 | 5609.5726 | 0 | -5.8 | 141 | Specific | 365.85 | 356.68 | 356.68 | 6.4 | 111 | >sp\|P02788\|TRFL_HUMAN Lactotransferrin OS=Homo sapiens OX=9606 GN=LTF PE=1 SV=6 | 1270 | 20201114_16933_1ng_glyco_hcd203040_compound15.24363.24363.4 | controllerType=0 controllerNumber=1 scan=24363 | 49.287 |
| 20864:3 | 1 | R.TAGWNVPIGTLRPFLN[+2350.830]WTGPPEPIEAAVAR.F | HexNAc(4)Hex(5)Fuc(1)NeuAc(2) | N[+2351] | 1861.196 | 3 | 5581.5735 | 5581.5413 | 0 | 5.8 | 141 | Specific | 379.78 | 221.55 | 221.55 | 7.06 | 111 | >sp\|P02788\|TRFL_HUMAN Lactotransferrin OS=Homo sapiens OX=9606 GN=LTF PE=1 SV=6 | 1270 | 20201114_16933_1ng_glyco_hcd203040_compound15.24321.24321.3 | controllerType=0 controllerNumber=1 scan=24321 | 49.1857 |
| 21008:4 | 1 | R.TAGWNVPIGTLRPFLN[+2350.830]WTGPPEPIEAAVAR.F | HexNAc(4)Hex(5)Fuc(1)NeuAc(2) | N[+2351] | 1396.1428 | 4 | 5581.5495 | 5581.5413 | 0 | 1.5 | 141 | Specific | 381.38 | 381.38 | 381.38 | 9.35 | 111 | >sp\|P02788\|TRFL_HUMAN Lactotransferrin OS=Homo sapiens OX=9606 GN=LTF PE=1 SV=6 | 1270 | 20201114_16933_1ng_glyco_hcd203040_compound15.24470.24470.4 | controllerType=0 controllerNumber=1 scan=24470 | 49.4812 |
| 20946:4 | 1 | R.TAGWNVPIGTLRPFLN[+2350.830]WTGPPEPIEAAVAR.F | HexNAc(4)Hex(5)Fuc(1)NeuAc(2) | N[+2351] | 1396.1426 | 4 | 5581.5485 | 5581.5413 | 0 | 1.3 | 141 | Specific | 514.69 | 326.85 | 326.85 | 8.35 | 111 | >sp\|P02788\|TRFL_HUMAN Lactotransferrin OS=Homo sapiens OX=9606 GN=LTF PE=1 SV=6 | 1270 | 20201114_16933_1ng_glyco_hcd203040_compound15.24406.24406.3 | controllerType=0 controllerNumber=1 scan=24406 | 49.3582 |
| 19710:3 | 1 | R.TAGWNVPIGTLRPFLN[+2059.735]WTGPPEPIEAAVAR.F | HexNAc(4)Hex(5)Fuc(1)NeuAc(1) | N[+2060] | 1764.1541 | 3 | 5290.4479 | 5290.4459 | 0 | 0.4 | 141 | Specific | 543.17 | 499.59 | 499.59 | 10.03 | 111 | >sp\|P02788\|TRFL_HUMAN Lactotransferrin OS=Homo sapiens OX=9606 GN=LTF PE=1 SV=6 | 1270 | 20201114_16933_1ng_glyco_hcd203040_compound15.23111.23111.3 | controllerType=0 controllerNumber=1 scan=23111 | [illegible] |
| 19712:3 | 1 | R.TAGWNVPIGTLRPFLN[+2059.735]WTGPPEPIEAAVAR.F | HexNAc(4)Hex(5)Fuc(1)NeuAc(1) | N[+2060] | 1764.1541 | 3 | 5290.4479 | 5290.4459 | 0 | 0.4 | 141 | Specific | 570.4 | 520.93 | 520.93 | 10.03 | 111 | >sp\|P02788\|TRFL_HUMAN Lactotransferrin OS=Homo sapiens OX=9606 GN=LTF PE=1 SV=6 | 1270 | 20201114_16933_1ng_glyco_hcd203040_compound15.23113.23113.3 | controllerType=0 controllerNumber=1 scan=23113 | [illegible] |
| 20867:3 | 1 | R.TAGWNVPIGTLRPFLN[+2350.830]WTGPPEPIEAAVAR.F | HexNAc(4)Hex(5)Fuc(1)NeuAc(2) | N[+2351] | 1861.196 | 3 | 5581.5735 | 5581.5413 | 0 | 5.8 | 141 | Specific | 599.88 | 574.7 | 574.7 | 11.08 | 111 | >sp\|P02788\|TRFL_HUMAN Lactotransferrin OS=Homo sapiens OX=9606 GN=LTF PE=1 SV=6 | 1270 | 20201114_16933_1ng_glyco_hcd203040_compound15.24324.24324.3 | controllerType=0 controllerNumber=1 scan=24324 | [illegible] |
| 19575:3 | 1 | R.TAGWNVPIGTLRPFLN[+2059.735]WTGPPEPIEAAVAR.F | HexNAc(4)Hex(5)Fuc(1)NeuAc(1) | N[+2060] | 1764.1565 | 3 | 5290.4548 | 5290.4459 | 0 | 1.7 | 141 | Specific | 648.15 | 600.73 | 600.73 | 12.38 | 111 | >sp\|P02788\|TRFL_HUMAN Lactotransferrin OS=Homo sapiens OX=9606 GN=LTF PE=1 SV=6 | 1270 | 20201114_16933_1ng_glyco_hcd203040_compound15.22966.22966.2 | controllerType=0 controllerNumber=1 scan=22966 | [illegible] |
| 19513:4 | 1 | R.TAGWNVPIGTLRPFLN[+2205.793]WTGPPEPIEAAVAR.F | HexNAc(4)Hex(5)Fuc(2)NeuAc(1) | N[+2206] | 1359.8839 | 4 | 5436.5138 | 5436.5038 | 0 | 1.8 | 141 | Specific | 690.23 | 620.97 | 620.97 | 13.39 | 111 | >sp\|P02788\|TRFL_HUMAN Lactotransferrin OS=Homo sapiens OX=9606 GN=LTF PE=1 SV=6 | 1270 | 20201114_16933_1ng_glyco_hcd203040_compound15.22901.22901.4 | controllerType=0 controllerNumber=1 scan=22901 | 46.1826 |
| 19514:3 | 1 | R.TAGWNVPIGTLRPFLN[+2205.793]WTGPPEPIEAAVAR.F | HexNAc(4)Hex(5)Fuc(2)NeuAc(1) | N[+2206] | 1812.8421 | 3 | 5436.5118 | 5436.5038 | 0 | 1.5 | 141 | Specific | 697.48 | 583.64 | 583.64 | 13.18 | 111 | >sp\|P02788\|TRFL_HUMAN Lactotransferrin OS=Homo sapiens OX=9606 GN=LTF PE=1 SV=6 | 1270 | 20201114_16933_1ng_glyco_hcd203040_compound15.22902.22902.3 | controllerType=0 controllerNumber=1 scan=22902 | 46.1891 |
| 19572:3 | 1 | R.TAGWNVPIGTLRPFLN[+2059.735]WTGPPEPIEAAVAR.F | HexNAc(4)Hex(5)Fuc(1)NeuAc(1) | N[+2060] | 1764.1565 | 3 | 5290.4548 | 5290.4459 | 0 | 1.7 | 141 | Specific | 730.7 | 681.87 | 681.87 | 14.38 | 111 | >sp\|P02788\|TRFL_HUMAN Lactotransferrin OS=Homo sapiens OX=9606 GN=LTF PE=1 SV=6 | 1270 | 20201114_16933_1ng_glyco_hcd203040_compound15.22963.22963.2 | controllerType=0 controllerNumber=1 scan=22963 | 46.3518 |
| 19512:3 | 1 | R.TAGWNVPIGTLRPFLN[+2205.793]WTGPPEPIEAAVAR.F | HexNAc(4)Hex(5)Fuc(2)NeuAc(1) | N[+2206] | 1812.8421 | 3 | 5436.5118 | 5436.5038 | 0 | 1.5 | 141 | Specific | 748.1 | 627.55 | 627.55 | 14.18 | 111 | >sp\|P02788\|TRFL_HUMAN Lactotransferrin OS=Homo sapiens OX=9606 GN=LTF PE=1 SV=6 | 1270 | 20201114_16933_1ng_glyco_hcd203040_compound15.22900.22900.3 | controllerType=0 controllerNumber=1 scan=22900 | 46.1811 |
| 20941:4 | 1 | R.TAGWNVPIGTLRPFLN[+2350.830]WTGPPEPIEAAVAR.F | HexNAc(4)Hex(5)Fuc(1)NeuAc(2) | N[+2351] | 1396.1426 | 4 | 5581.5485 | 5581.5413 | 0 | 1.3 | 141 | Specific | 800.8 | 657.62 | 657.62 | 16.19 | 111 | >sp\|P02788\|TRFL_HUMAN Lactotransferrin OS=Homo sapiens OX=9606 GN=LTF PE=1 SV=6 | 1270 | 20201114_16933_1ng_glyco_hcd203040_compound15.24401.24401.2 | controllerType=0 controllerNumber=1 scan=24401 | 49.3508 |
| 19515:4 | 1 | R.TAGWNVPIGTLRPFLN[+2205.793]WTGPPEPIEAAVAR.F | HexNAc(4)Hex(5)Fuc(2)NeuAc(1) | N[+2206] | 1359.8839 | 4 | 5436.5138 | 5436.5038 | 0 | 1.8 | 141 | Specific | 822.81 | 720.77 | 720.77 | 16.39 | 111 | >sp\|P02788\|TRFL_HUMAN Lactotransferrin OS=Homo sapiens OX=9606 GN=LTF PE=1 SV=6 | 1270 | 20201114_16933_1ng_glyco_hcd203040_compound15.22903.22903.4 | controllerType=0 controllerNumber=1 scan=22903 | 46.2036 |
| 18945:3 | 1 | R.TAGWNVPIGTLRPFLN[+349.137]WTGPPEPIEAAVAR.F | HexNAc(1)Fuc(1) | N[+349] | 1193.9573 | 3 | 3579.8574 | 3579.8482 | 0 | 2.6 | 141 | Specific | 1016.2 | 1005.6 | 1005.6 | 20.38 | 111 | >sp\|P02788\|TRFL_HUMAN Lactotransferrin OS=Homo sapiens OX=9606 GN=LTF PE=1 SV=6 | 1270 | 20201114_16933_1ng_glyco_hcd203040_compound15.22306.22306.3 | controllerType=0 controllerNumber=1 scan=22306 | 44.8108 |
| 18947:3 | 1 | R.TAGWNVPIGTLRPFLN[+349.137]WTGPPEPIEAAVAR.F | HexNAc(1)Fuc(1) | N[+349] | 1193.9573 | 3 | 3579.8574 | 3579.8482 | 0 | 2.6 | 141 | Specific | 1033.43 | 1005.16 | 1005.16 | 20.38 | 111 | >sp\|P02788\|TRFL_HUMAN Lactotransferrin OS=Homo sapiens OX=9606 GN=LTF PE=1 SV=6 | 1270 | 20201114_16933_1ng_glyco_hcd203040_compound15.22308.22308.3 | controllerType=0 controllerNumber=1 scan=22308 | 44.8185 |
| 20982:3 | 1 | R.TAGWNIPMGLLFN[+2350.830]QTGSC[+57.021]K.F | HexNAc(4)Hex(5)Fuc(1)NeuAc(2) | C[+57], N[+2351] | 1482.6242 | 3 | 4445.8581 | 4445.8403 | 0 | 4 | 485 | Specific | 389.93 | 377.83 | 377.83 | 9.46 | 111 | >sp\|P02788\|TRFL_HUMAN Lactotransferrin OS=Homo sapiens OX=9606 GN=LTF PE=1 SV=6 | 1270 | 20201114_16933_1ng_glyco_hcd203040_compound15.24444.24444.3 | controllerType=0 controllerNumber=1 scan=24444 | 49.4318 |
| 19179:3 | 1 | R.TAGWNIPMGLLFN[+1751.624]QTGSC[+57.021]K.F | HexNAc(4)Hex(4)NeuAc(1) | C[+57], N[+1752] | 1282.8865 | 3 | 3846.645 | 3846.6341 | 0 | 2.8 | 485 | Specific | 444.74 | 401.48 | 401.48 | 9.73 | 111 | >sp\|P02788\|TRFL_HUMAN Lactotransferrin OS=Homo sapiens OX=9606 GN=LTF PE=1 SV=6 | 1270 | 20201114_16933_1ng_glyco_hcd203040_compound15.22552.22552.3 | controllerType=0 controllerNumber=1 scan=22552 | 45.4397 |
| 19181:3 | 1 | R.TAGWNIPMGLLFN[+1751.624]QTGSC[+57.021]K.F | HexNAc(4)Hex(4)NeuAc(1) | C[+57], N[+1752] | 1282.8865 | 3 | 3846.645 | 3846.6341 | 0 | 2.8 | 485 | Specific | 585.14 | 435.4 | 435.4 | 11.39 | 111 | >sp\|P02788\|TRFL_HUMAN Lactotransferrin OS=Homo sapiens OX=9606 GN=LTF PE=1 SV=6 | 1270 | 20201114_16933_1ng_glyco_hcd203040_compound15.22554.22554.3 | controllerType=0 controllerNumber=1 scan=22554 | 45.4459 |
| 17173:3 | 1 | R.TAGWNIPMGLLFN[+1622.582]QTGSC[+57.021]K.F | HexNAc(4)Hex(5) | C[+57], N[+1623] | 1239.8735 | 3 | 3717.6059 | 3717.5916 | 0 | 3.8 | 485 | Specific | 638.37 | 638.37 | 289.4 | 12.3 | 111 | >sp\|P02788\|TRFL_HUMAN Lactotransferrin OS=Homo sapiens OX=9606 GN=LTF PE=1 SV=6 | 1270 | 20201114_16933_1ng_glyco_hcd203040_compound15.20456.20456.3 | controllerType=0 controllerNumber=1 scan=20456 | 41.0067 |
| 18001:2 | 1 | R.TAGWNIPMGLLFN[+349.137]QTGSC[+57.021]K.F | HexNAc(1)Fuc(1) | C[+57], N[+349] | 1222.5778 | 2 | 2444.1483 | 2444.1472 | 0 | 0.5 | 485 | Specific | 695.28 | 625.8 | 625.8 | 13.19 | 111 | >sp\|P02788\|TRFL_HUMAN Lactotransferrin OS=Homo sapiens OX=9606 GN=LTF PE=1 SV=6 | 1270 | 20201114_16933_1ng_glyco_hcd203040_compound15.21322.21322.2 | controllerType=0 controllerNumber=1 scan=21322 | 42.8111 |
| 17175:3 | 1 | R.TAGWNIPMGLLFN[+1622.582]QTGSC[+57.021]K.F | HexNAc(4)Hex(5) | C[+57], N[+1623] | 1239.8735 | 3 | 3717.6059 | 3717.5916 | 0 | 3.8 | 485 | Specific | 779.14 | 751.45 | 358.46 | 15.3 | 111 | >sp\|P02788\|TRFL_HUMAN Lactotransferrin OS=Homo sapiens OX=9606 GN=LTF PE=1 SV=6 | 1270 | 20201114_16933_1ng_glyco_hcd203040_compound15.20458.20458.3 | controllerType=0 controllerNumber=1 scan=20458 | 41.0116 |
| 18003:2 | 1 | R.TAGWNIPMGLLFN[+349.137]QTGSC[+57.021]K.F | HexNAc(1)Fuc(1) | C[+57], N[+349] | 1222.5778 | 2 | 2444.1483 | 2444.1472 | 0 | 0.5 | 485 | Specific | 821.27 | 737.69 | 737.69 | 16.19 | 111 | >sp\|P02788\|TRFL_HUMAN Lactotransferrin OS=Homo sapiens OX=9606 GN=LTF PE=1 SV=6 | 1270 | 20201114_16933_1ng_glyco_hcd203040_compound15.21324.21324.2 | controllerType=0 controllerNumber=1 scan=21324 | 42.815 |
| 17189:3 | 1 | R.TAGWNIPMGLLFN[+1622.582]QTGSC[+57.021]K.F | HexNAc(4)Hex(5) | C[+57], N[+1623] | 1239.871 | 3 | 3717.5985 | 3717.5916 | 0 | 1.9 | 485 | Specific | 832.11 | 832.11 | 372.21 | 16.39 | 111 | >sp\|P02788\|TRFL_HUMAN Lactotransferrin OS=Homo sapiens OX=9606 GN=LTF PE=1 SV=6 | 1270 | 20201114_16933_1ng_glyco_hcd203040_compound15.20473.20473.3 | controllerType=0 controllerNumber=1 scan=20473 | 41.0387 |
| 17191:3 | 1 | R.TAGWNIPMGLLFN[+1622.582]QTGSC[+57.021]K.F | HexNAc(4)Hex(5) | C[+57], N[+1623] | 1239.871 | 3 | 3717.5985 | 3717.5916 | 0 | 1.9 | 485 | Specific | 871.92 | 696.05 | 405.07 | 17.39 | 111 | >sp\|P02788\|TRFL_HUMAN Lactotransferrin OS=Homo sapiens OX=9606 GN=LTF PE=1 SV=6 | 1270 | 20201114_16933_1ng_glyco_hcd203040_compound15.20475.20475.3 | controllerType=0 controllerNumber=1 scan=20475 | 41.0431 |
| 20984:3 | 1 | R.TAGWNIPMGLLFN[+2350.830]QTGSC[+57.021]K.F | HexNAc(4)Hex(5)Fuc(1)NeuAc(2) | C[+57], N[+2351] | 1482.6242 | 3 | 4445.8581 | 4445.8403 | 0 | 4 | 485 | Specific | 885.3 | 873.69 | 873.69 | 17.3 | 111 | >sp\|P02788\|TRFL_HUMAN Lactotransferrin OS=Homo sapiens OX=9606 GN=LTF PE=1 SV=6 | 1270 | 20201114_16933_1ng_glyco_hcd203040_compound15.24446.24446.3 | controllerType=0 controllerNumber=1 scan=24446 | 49.4355 |
| 17383:3 | 1 | R.TAGWNIPMGLLFN[+1216.423]QTGSC[+57.021]K.F | HexNAc(2)Hex(5) | C[+57], N[+1216] | 1104.4857 | 3 | 3311.4427 | 3311.4328 | 0 | 3 | 485 | Specific | 889.66 | 889.66 | 367.49 | 17.3 | 111 | >sp\|P02788\|TRFL_HUMAN Lactotransferrin OS=Homo sapiens OX=9606 GN=LTF PE=1 SV=6 | 1270 | 20201114_16933_1ng_glyco_hcd203040_compound15.20675.20675.3 | controllerType=0 controllerNumber=1 scan=20675 | 41.4534 |
| 19100:3 | 1 | R.TAGWNIPMGLLFN[+1913.677]QTGSC[+57.021]K.F | HexNAc(4)Hex(5)NeuAc(1) | C[+57], N[+1914] | 1336.9128 | 3 | 4008.7238 | 4008.687 | 0 | 9.2 | 485 | Specific | 891.11 | 891.11 | 891.11 | 16.54 | 111 | >sp\|P02788\|TRFL_HUMAN Lactotransferrin OS=Homo sapiens OX=9606 GN=LTF PE=1 SV=6 | 1270 | 20201114_16933_1ng_glyco_hcd203040_compound15.22469.22469.3 | controllerType=0 controllerNumber=1 scan=22469 | 45.2307 |
| 16969:3 | 1 | R.TAGWNIPMGLLFN[+1768.640]QTGSC[+57.021]K.F | HexNAc(4)Hex(5)Fuc(1) | C[+57], N[+1769] | 1288.5585 | 3 | 3863.661 | 3863.6495 | 0 | 3 | 485 | Specific | 927.81 | 692.44 | 692.44 | 18.3 | 111 | >sp\|P02788\|TRFL_HUMAN Lactotransferrin OS=Homo sapiens OX=9606 GN=LTF PE=1 SV=6 | 1270 | 20201114_16933_1ng_glyco_hcd203040_compound15.20243.20243.3 | controllerType=0 controllerNumber=1 scan=20243 | 40.613 |
| 16967:3 | 1 | R.TAGWNIPMGLLFN[+1768.640]QTGSC[+57.021]K.F | HexNAc(4)Hex(5)Fuc(1) | C[+57], N[+1769] | 1288.5585 | 3 | 3863.661 | 3863.6495 | 0 | 3 | 485 | Specific | 933.16 | 933.16 | 933.16 | 18.3 | 111 | >sp\|P02788\|TRFL_HUMAN Lactotransferrin OS=Homo sapiens OX=9606 GN=LTF PE=1 SV=6 | 1270 | 20201114_16933_1ng_glyco_hcd203040_compound15.20241.20241.3 | controllerType=0 controllerNumber=1 scan=20241 | 40.6069 |
| 18921:3 | 1 | R.TAGWNIPMGLLFN[+2059.735]QTGSC[+57.021]K.F | HexNAc(4)Hex(5)Fuc(1)NeuAc(1) | C[+57], N[+2060] | 1385.5982 | 3 | 4154.7801 | 4154.7449 | 0 | 8.5 | 485 | Specific | 933.39 | 933.39 | 933.39 | 17.7 | 111 | >sp\|P02788\|TRFL_HUMAN Lactotransferrin OS=Homo sapiens OX=9606 GN=LTF PE=1 SV=6 | 1270 | 20201114_16933_1ng_glyco_hcd203040_compound15.22281.22281.3 | controllerType=0 controllerNumber=1 scan=22281 | 44.7629 |
| 18900:3 | 1 | R.TAGWNIPMGLLFN[+2205.793]QTGSC[+57.021]K.F | HexNAc(4)Hex(5)Fuc(2)NeuAc(1) | C[+57], N[+2206] | 1434.2746 | 3 | 4300.8092 | 4300.8028 | 0 | 1.5 | 485 | Specific | 936.18 | 788.33 | 788.33 | 18.39 | 111 | >sp\|P02788\|TRFL_HUMAN Lactotransferrin OS=Homo sapiens OX=9606 GN=LTF PE=1 SV=6 | 1270 | 20201114_16933_1ng_glyco_hcd203040_compound15.22258.22258.3 | controllerType=0 controllerNumber=1 scan=22258 | 44.6999 |
| 19098:3 | 1 | R.TAGWNIPMGLLFN[+1913.677]QTGSC[+57.021]K.F | HexNAc(4)Hex(5)NeuAc(1) | C[+57], N[+1914] | 1336.9128 | 3 | 4008.7238 | 4008.687 | 0 | 9.2 | 485 | Specific | 959.22 | 959.22 | 959.22 | 18.54 | 111 | >sp\|P02788\|TRFL_HUMAN Lactotransferrin OS=Homo sapiens OX=9606 GN=LTF PE=1 SV=6 | 1270 | 20201114_16933_1ng_glyco_hcd203040_compound15.22467.22467.3 | controllerType=0 controllerNumber=1 scan=22467 | 45.2235 |
| 18919:3 | 1 | R.TAGWNIPMGLLFN[+2059.735]QTGSC[+57.021]K.F | HexNAc(4)Hex(5)Fuc(1)NeuAc(1) | C[+57], N[+2060] | 1385.5982 | 3 | 4154.7801 | 4154.7449 | 0 | 8.5 | 485 | Specific | 961.71 | 961.71 | 961.71 | 18.7 | 111 | >sp\|P02788\|TRFL_HUMAN Lactotransferrin OS=Homo sapiens OX=9606 GN=LTF PE=1 SV=6 | 1270 | 20201114_16933_1ng_glyco_hcd203040_compound15.22279.22279.3 | controllerType=0 controllerNumber=1 scan=22279 | 44.7556 |
| 17385:3 | 1 | R.TAGWNIPMGLLFN[+1216.423]QTGSC[+57.021]K.F | HexNAc(2)Hex(5) | C[+57], N[+1216] | 1104.4857 | 3 | 3311.4427 | 3311.4328 | 0 | 3 | 485 | Specific | 967.88 | 825.13 | 454.61 | 19.3 | 111 | >sp\|P02788\|TRFL_HUMAN Lactotransferrin OS=Homo sapiens OX=9606 GN=LTF PE=1 SV=6 | 1270 | 20201114_16933_1ng_glyco_hcd203040_compound15.20677.20677.3 | controllerType=0 controllerNumber=1 scan=20677 | 41.4586 |
| 18898:3 | 1 | R.TAGWNIPMGLLFN[+2205.793]QTGSC[+57.021]K.F | HexNAc(4)Hex(5)Fuc(2)NeuAc(1) | C[+57], N[+2206] | 1434.2746 | 3 | 4300.8092 | 4300.8028 | 0 | 1.5 | 485 | Specific | 997.7 | 903.55 | 903.55 | 19.39 | 111 | >sp\|P02788\|TRFL_HUMAN Lactotransferrin OS=Homo sapiens OX=9606 GN=LTF PE=1 SV=6 | 1270 | 20201114_16933_1ng_glyco_hcd203040_compound15.22256.22256.3 | controllerType=0 controllerNumber=1 scan=22256 | 44.6893 |
| 17038:3 | 1 | R.TAGWNIPM[+15.995]GLLFN[+2059.735]QTGSC[+57.021]K.F | HexNAc(4)Hex(5)Fuc(1)NeuAc(1) | C[+57], M[+16], N[+2060] | 1390.916 | 3 | 4170.7334 | 4170.7398 | 0 | -1.5 | 485 | Specific | 564.38 | 398.9 | 398.9 | 9.02 | 111 | >sp\|P02788\|TRFL_HUMAN Lactotransferrin OS=Homo sapiens OX=9606 GN=LTF PE=1 SV=6 | 1270 | 20201114_16933_1ng_glyco_hcd203040_compound15.20315.20315.3 | controllerType=0 controllerNumber=1 scan=20315 | 40.741 |
| 17245:3 | 1 | R.TAGWNIPM[+15.995]GLLFN[+2059.735]QTGSC[+57.021]K.F | HexNAc(4)Hex(5)Fuc(1)NeuAc(1) | C[+57], M[+16], N[+2060] | 1390.9279 | 3 | 4170.7692 | 4170.7398 | 0 | 7 | 485 | Specific | 583.61 | 430.62 | 430.62 | 10.68 | 111 | >sp\|P02788\|TRFL_HUMAN Lactotransferrin OS=Homo sapiens OX=9606 GN=LTF PE=1 SV=6 | 1270 | 20201114_16933_1ng_glyco_hcd203040_compound15.20531.20531.3 | controllerType=0 controllerNumber=1 scan=20531 | 41.1503 |
| 17588:3 | 1 | R.TAGWNIPM[+15.995]GLLFN[+2059.735]QTGSC[+57.021]K.F | HexNAc(4)Hex(5)Fuc(1)NeuAc(1) | C[+57], M[+16], N[+2060] | 1390.9199 | 3 | 4170.745 | 4170.7398 | 0 | 1.3 | 485 | Specific | 659.28 | 500.42 | 500.42 | 13.2 | 111 | >sp\|P02788\|TRFL_HUMAN Lactotransferrin OS=Homo sapiens OX=9606 GN=LTF PE=1 SV=6 | 1270 | 20201114_16933_1ng_glyco_hcd203040_compound15.20888.20888.3 | controllerType=0 controllerNumber=1 scan=20888 | 41.8628 |
| 18178:4 | 1 | R.TAGWNIPMGLLFN[+1768.640]QTGSC[+57.021]KFDEYFSQSC[+57.021]APGSDPR. | HexNAc(4)Hex(5)Fuc(1) | C[+57]*2, N[+1769] | 1427.6124 | 4 | 5707.4279 | 5707.3966 | 0 | 5.5 | 485 | Specific | 465.46 | 444.65 | 444.65 | 9.02 | 111 | >sp\|P02788\|TRFL_HUMAN Lactotransferrin OS=Homo sapiens OX=9606 GN=LTF PE=1 SV=6 | 1270 | 20201114_16933_1ng_glyco_hcd203040_compound15.21506.21506.2 | controllerType=0 controllerNumber=1 scan=21506 | 43.1523 |
| 20581:4 | 1 | R.TAGWNIPMGLLFN[+2204.772]QTGSC[+57.021]KFDEYFSQSC[+57.021]APGSDPR. | HexNAc(4)Hex(5)NeuAc(2) | C[+57]*2, N[+2205] | 1536.6478 | 4 | 6143.5694 | 6143.5295 | 0 | 6.5 | 485 | Specific | 478.94 | 470.17 | 470.17 | 8.61 | 111 | >sp\|P02788\|TRFL_HUMAN Lactotransferrin OS=Homo sapiens OX=9606 GN=LTF PE=1 SV=6 | 1270 | 20201114_16933_1ng_glyco_hcd203040_compound15.24025.24025.4 | controllerType=0 controllerNumber=1 scan=24025 | 48.5425 |
| 19253:3 | 1 | R.TAGWNIPMGLLFN[+1913.677]QTGSC[+57.021]KFDEYFSQSC[+57.021]APGSDPR. | HexNAc(4)Hex(5)NeuAc(1) | C[+57]*2, N[+1914] | 1951.4898 | 3 | 5852.4549 | 5852.4341 | 0 | 3.6 | 485 | Specific | 552.91 | 444.69 | 444.69 | 9.67 | 111 | >sp\|P02788\|TRFL_HUMAN Lactotransferrin OS=Homo sapiens OX=9606 GN=LTF PE=1 SV=6 | 1270 | 20201114_16933_1ng_glyco_hcd203040_compound15.22629.22629.3 | controllerType=0 controllerNumber=1 scan=22629 | 45.6048 |
| 20583:4 | 1 | R.TAGWNIPMGLLFN[+2204.772]QTGSC[+57.021]KFDEYFSQSC[+57.021]APGSDPR. | HexNAc(4)Hex(5)NeuAc(2) | C[+57]*2, N[+2205] | 1536.6478 | 4 | 6143.5694 | 6143.5295 | 0 | 6.5 | 485 | Specific | 585.28 | 575.76 | 575.76 | 10.22 | 111 | >sp\|P02788\|TRFL_HUMAN Lactotransferrin OS=Homo sapiens OX=9606 GN=LTF PE=1 SV=6 | 1270 | 20201114_16933_1ng_glyco_hcd203040_compound15.24027.24027.4 | controllerType=0 controllerNumber=1 scan=24027 | 48.5484 |
| 19256:3 | 1 | R.TAGWNIPMGLLFN[+1913.677]QTGSC[+57.021]KFDEYFSQSC[+57.021]APGSDPR. | HexNAc(4)Hex(5)NeuAc(1) | C[+57]*2, N[+1914] | 1951.4898 | 3 | 5852.4549 | 5852.4341 | 0 | 3.6 | 485 | Specific | 705.75 | 557.14 | 557.14 | 13.83 | 111 | >sp\|P02788\|TRFL_HUMAN Lactotransferrin OS=Homo sapiens OX=9606 GN=LTF PE=1 SV=6 | 1270 | 20201114_16933_1ng_glyco_hcd203040_compound15.22632.22632.3 | controllerType=0 controllerNumber=1 scan=22632 | 45.6258 |
| 19158:4 | 1 | R.TAGWNIPMGLLFN[+2205.793]QTGSC[+57.021]KFDEYFSQSC[+57.021]APGSDPR. | HexNAc(4)Hex(5)Fuc(2)NeuAc(1) | C[+57]*2, N[+2206] | 1536.8987 | 4 | 6144.5729 | 6144.5499 | 0 | 3.7 | 485 | Specific | 710.1 | 692.66 | 692.66 | 13.84 | 111 | >sp\|P02788\|TRFL_HUMAN Lactotransferrin OS=Homo sapiens OX=9606 GN=LTF PE=1 SV=6 | 1270 | 20201114_16933_1ng_glyco_hcd203040_compound15.22530.22530.4 | controllerType=0 controllerNumber=1 scan=22530 | 45.3902 |
| 20456:4 | 1 | R.TAGWNIPMGLLFN[+2350.830]QTGSC[+57.021]KFDEYFSQSC[+57.021]APGSDPR. | HexNAc(4)Hex(5)Fuc(1)NeuAc(2) | C[+57]*2, N[+2351] | 1573.1537 | 4 | 6289.5929 | 6289.5875 | 0 | 0.9 | 485 | Specific | 730.15 | 670.98 | 670.98 | 13.73 | 111 | >sp\|P02788\|TRFL_HUMAN Lactotransferrin OS=Homo sapiens OX=9606 GN=LTF PE=1 SV=6 | 1270 | 20201114_16933_1ng_glyco_hcd203040_compound15.23893.23893.2 | controllerType=0 controllerNumber=1 scan=23893 | 48.2747 |
| 19174:4 | 1 | R.TAGWNIPMGLLFN[+2059.735]QTGSC[+57.021]KFDEYFSQSC[+57.021]APGSDPR. | HexNAc(4)Hex(5)Fuc(1)NeuAc(1) | C[+57]*2, N[+2060] | 1500.3829 | 4 | 5998.5099 | 5998.492 | 0 | 3 | 485 | Specific | 801.07 | 748.08 | 748.08 | 15.84 | 111 | >sp\|P02788\|TRFL_HUMAN Lactotransferrin OS=Homo sapiens OX=9606 GN=LTF PE=1 SV=6 | 1270 | 20201114_16933_1ng_glyco_hcd203040_compound15.22547.22547.4 | controllerType=0 controllerNumber=1 scan=22547 | 45.4312 |
| 20468:4 | 1 | R.TAGWNIPMGLLFN[+2350.830]QTGSC[+57.021]KFDEYFSQSC[+57.021]APGSDPR. | HexNAc(4)Hex(5)Fuc(1)NeuAc(2) | C[+57]*2, N[+2351] | 1573.158 | 4 | 6289.61 | 6289.5875 | 0 | 3.6 | 485 | Specific | 835.79 | 796.81 | 796.81 | 15.84 | 111 | >sp\|P02788\|TRFL_HUMAN Lactotransferrin OS=Homo sapiens OX=9606 GN=LTF PE=1 SV=6 | 1270 | 20201114_16933_1ng_glyco_hcd203040_compound15.23906.23906.4 | controllerType=0 controllerNumber=1 scan=23906 | 48.3034 |
| 19176:4 | 1 | R.TAGWNIPMGLLFN[+2059.735]QTGSC[+57.021]KFDEYFSQSC[+57.021]APGSDPR. | HexNAc(4)Hex(5)Fuc(1)NeuAc(1) | C[+57]*2, N[+2060] | 1500.3829 | 4 | 5998.5099 | 5998.492 | 0 | 3 | 485 | Specific | 860.04 | 820.61 | 820.61 | 16.84 | 111 | >sp\|P02788\|TRFL_HUMAN Lactotransferrin OS=Homo sapiens OX=9606 GN=LTF PE=1 SV=6 | 1270 | 20201114_16933_1ng_glyco_hcd203040_compound15.22549.22549.4 | controllerType=0 controllerNumber=1 scan=22549 | 45.4362 |
| 19244:4 | 1 | R.TAGWNIPMGLLFN[+1913.677]QTGSC[+57.021]KFDEYFSQSC[+57.021]APGSDPR. | HexNAc(4)Hex(5)NeuAc(1) | C[+57]*2, N[+1914] | 1463.8712 | 4 | 5852.463 | 5852.4341 | 0 | 4.9 | 485 | Specific | 890.02 | 803.32 | 803.32 | 16.63 | 111 | >sp\|P02788\|TRFL_HUMAN Lactotransferrin OS=Homo sapiens OX=9606 GN=LTF PE=1 SV=6 | 1270 | 20201114_16933_1ng_glyco_hcd203040_compound15.22620.22620.4 | controllerType=0 controllerNumber=1 scan=22620 | 45.5876 |
| 19156:4 | 1 | R.TAGWNIPMGLLFN[+2205.793]QTGSC[+57.021]KFDEYFSQSC[+57.021]APGSDPR. | HexNAc(4)Hex(5)Fuc(2)NeuAc(1) | C[+57]*2, N[+2206] | 1536.8987 | 4 | 6144.5729 | 6144.5499 | 0 | 3.7 | 485 | Specific | 899.16 | 839.28 | 839.28 | 16.84 | 111 | >sp\|P02788\|TRFL_HUMAN Lactotransferrin OS=Homo sapiens OX=9606 GN=LTF PE=1 SV=6 | 1270 | 20201114_16933_1ng_glyco_hcd203040_compound15.22528.22528.4 | controllerType=0 controllerNumber=1 scan=22528 | 45.383 |
| 20458:4 | 1 | R.TAGWNIPMGLLFN[+2350.830]QTGSC[+57.021]KFDEYFSQSC[+57.021]APGSDPR. | HexNAc(4)Hex(5)Fuc(1)NeuAc(2) | C[+57]*2, N[+2351] | 1573.1537 | 4 | 6289.5929 | 6289.5875 | 0 | 0.9 | 485 | Specific | 917.95 | 866.13 | 866.13 | 17.73 | 111 | >sp\|P02788\|TRFL_HUMAN Lactotransferrin OS=Homo sapiens OX=9606 GN=LTF PE=1 SV=6 | 1270 | 20201114_16933_1ng_glyco_hcd203040_compound15.23895.23895.2 | controllerType=0 controllerNumber=1 scan=23895 | 48.2818 |
| 19240:4 | 1 | R.TAGWNIPMGLLFN[+1913.677]QTGSC[+57.021]KFDEYFSQSC[+57.021]APGSDPR. | HexNAc(4)Hex(5)NeuAc(1) | C[+57]*2, N[+1914] | 1463.8771 | 4 | 5852.4865 | 5852.4341 | 0 | 8.9 | 485 | Specific | 927.29 | 927.29 | 927.29 | 17.08 | 111 | >sp\|P02788\|TRFL_HUMAN Lactotransferrin OS=Homo sapiens OX=9606 GN=LTF PE=1 SV=6 | 1270 | 20201114_16933_1ng_glyco_hcd203040_compound15.22615.22615.4 | controllerType=0 controllerNumber=1 scan=22615 | 45.5761 |
| 19246:4 | 1 | R.TAGWNIPMGLLFN[+1913.677]QTGSC[+57.021]KFDEYFSQSC[+57.021]APGSDPR. | HexNAc(4)Hex(5)NeuAc(1) | C[+57]*2, N[+1914] | 1463.8712 | 4 | 5852.463 | 5852.4341 | 0 | 4.9 | 485 | Specific | 933.65 | 898.77 | 898.77 | 17.63 | 111 | >sp\|P02788\|TRFL_HUMAN Lactotransferrin OS=Homo sapiens OX=9606 GN=LTF PE=1 SV=6 | 1270 | 20201114_16933_1ng_glyco_hcd203040_compound15.22622.22622.4 | controllerType=0 controllerNumber=1 scan=22622 | 45.5934 |
| 20466:4 | 1 | R.TAGWNIPMGLLFN[+2350.830]QTGSC[+57.021]KFDEYFSQSC[+57.021]APGSDPR. | HexNAc(4)Hex(5)Fuc(1)NeuAc(2) | C[+57]*2, N[+2351] | 1573.158 | 4 | 6289.61 | 6289.5875 | 0 | 3.6 | 485 | Specific | 936.83 | 807.22 | 807.22 | 17.84 | 111 | >sp\|P02788\|TRFL_HUMAN Lactotransferrin OS=Homo sapiens OX=9606 GN=LTF PE=1 SV=6 | 1270 | 20201114_16933_1ng_glyco_hcd203040_compound15.23904.23904.4 | controllerType=0 controllerNumber=1 scan=23904 | 48.2972 |
| 19242:4 | 1 | R.TAGWNIPMGLLFN[+1913.677]QTGSC[+57.021]KFDEYFSQSC[+57.021]APGSDPR. | HexNAc(4)Hex(5)NeuAc(1) | C[+57]*2, N[+1914] | 1463.8771 | 4 | 5852.4865 | 5852.4341 | 0 | 8.9 | 485 | Specific | 950.16 | 950.16 | 950.16 | 18.08 | 111 | >sp\|P02788\|TRFL_HUMAN Lactotransferrin OS=Homo sapiens OX=9606 GN=LTF PE=1 SV=6 | 1270 | 20201114_16933_1ng_glyco_hcd203040_compound15.22617.22617.4 | controllerType=0 controllerNumber=1 scan=22617 | 45.5815 |
| 18181:4 | 1 | R.TAGWNIPMGLLFN[+1768.640]QTGSC[+57.021]KFDEYFSQSC[+57.021]APGSDPR. | HexNAc(4)Hex(5)Fuc(1) | C[+57]*2, N[+1769] | 1427.6124 | 4 | 5707.4279 | 5707.3966 | 0 | 5.5 | 485 | Specific | 996.34 | 947.33 | 947.33 | 18.63 | 111 | >sp\|P02788\|TRFL_HUMAN Lactotransferrin OS=Homo sapiens OX=9606 GN=LTF PE=1 SV=6 | 1270 | 20201114_16933_1ng_glyco_hcd203040_compound15.21509.21509.2 | controllerType=0 controllerNumber=1 scan=21509 | 43.156 |
| 09229:3 | 1 | R.N[+2059.735]GSDC[+57.021]PDKFC[+57.021]LFQSETK.N | HexNAc(4)Hex(5)Fuc(1)NeuAc(1) | C[+57]*2, N[+2060] | 1364.8788 | 3 | 4092.622 | 4092.6088 | 0 | 3.2 | 642 | Specific | 253.16 | 186.71 | 186.71 | 5.52 | 111 | >sp\|P02788\|TRFL_HUMAN Lactotransferrin OS=Homo sapiens OX=9606 GN=LTF PE=1 SV=6 | 1270 | 20201114_16933_1ng_glyco_hcd203040_compound15.12167.12167.3 | controllerType=0 controllerNumber=1 scan=12167 | 26.6869 |
| 09231:3 | 1 | R.N[+2059.735]GSDC[+57.021]PDKFC[+57.021]LFQSETK.N | HexNAc(4)Hex(5)Fuc(1)NeuAc(1) | C[+57]*2, N[+2060] | 1364.8788 | 3 | 4092.622 | 4092.6088 | 0 | 3.2 | 642 | Specific | 277.21 | 163.66 | 163.66 | 5.08 | 111 | >sp\|P02788\|TRFL_HUMAN Lactotransferrin OS=Homo sapiens OX=9606 GN=LTF PE=1 SV=6 | 1270 | 20201114_16933_1ng_glyco_hcd203040_compound15.12169.12169.3 | controllerType=0 controllerNumber=1 scan=12169 | 26.6904 |
| 09218:3 | 1 | R.N[+2059.735]GSDC[+57.021]PDKFC[+57.021]LFQSETK.N | HexNAc(4)Hex(5)Fuc(1)NeuAc(1) | C[+57]*2, N[+2060] | 1364.8784 | 3 | 4092.6205 | 4092.6088 | 0 | 2.9 | 642 | Specific | 297.53 | 134.28 | 134.28 | 4.92 | 111 | >sp\|P02788\|TRFL_HUMAN Lactotransferrin OS=Homo sapiens OX=9606 GN=LTF PE=1 SV=6 | 1270 | 20201114_16933_1ng_glyco_hcd203040_compound15.12155.12155.3 | controllerType=0 controllerNumber=1 scan=12155 | 26.6653 |
| 09097:3 | 1 | R.N[+2205.793]GSDC[+57.021]PDKFC[+57.021]LFQSETK.N | HexNAc(4)Hex(5)Fuc(2)NeuAc(1) | C[+57]*2, N[+2206] | 1413.5651 | 3 | 4238.6808 | 4238.6667 | 0 | 3.3 | 642 | Specific | 330.73 | 179.02 | 179.02 | 5.99 | 111 | >sp\|P02788\|TRFL_HUMAN Lactotransferrin OS=Homo sapiens OX=9606 GN=LTF PE=1 SV=6 | 1270 | 20201114_16933_1ng_glyco_hcd203040_compound15.12030.12030.3 | controllerType=0 controllerNumber=1 scan=12030 | 26.4576 |
| 09107:3 | 1 | R.N[+2205.793]GSDC[+57.021]PDKFC[+57.021]LFQSETK.N | HexNAc(4)Hex(5)Fuc(2)NeuAc(1) | C[+57]*2, N[+2206] | 1413.5632 | 3 | 4238.6749 | 4238.6667 | 0 | 1.9 | 642 | Specific | 349.33 | 209.01 | 209.01 | 6.53 | 111 | >sp\|P02788\|TRFL_HUMAN Lactotransferrin OS=Homo sapiens OX=9606 GN=LTF PE=1 SV=6 | 1270 | 20201114_16933_1ng_glyco_hcd203040_compound15.12041.12041.3 | controllerType=0 controllerNumber=1 scan=12041 | 26.4793 |
| 09099:3 | 1 | R.N[+2205.793]GSDC[+57.021]PDKFC[+57.021]LFQSETK.N | HexNAc(4)Hex(5)Fuc(2)NeuAc(1) | C[+57]*2, N[+2206] | 1413.5651 | 3 | 4238.6808 | 4238.6667 | 0 | 3.3 | 642 | Specific | 394.68 | 258 | 258 | 6.76 | 111 | >sp\|P02788\|TRFL_HUMAN Lactotransferrin OS=Homo sapiens OX=9606 GN=LTF PE=1 SV=6 | 1270 | 20201114_16933_1ng_glyco_hcd203040_compound15.12032.12032.3 | controllerType=0 controllerNumber=1 scan=12032 | 26.4619 |
| 09109:3 | 1 | R.N[+2205.793]GSDC[+57.021]PDKFC[+57.021]LFQSETK.N | HexNAc(4)Hex(5)Fuc(2)NeuAc(1) | C[+57]*2, N[+2206] | 1413.5632 | 3 | 4238.6749 | 4238.6667 | 0 | 1.9 | 642 | Specific | 462.37 | 370.59 | 370.59 | 7.67 | 111 | >sp\|P02788\|TRFL_HUMAN Lactotransferrin OS=Homo sapiens OX=9606 GN=LTF PE=1 SV=6 | 1270 | 20201114_16933_1ng_glyco_hcd203040_compound15.12043.12043.3 | controllerType=0 controllerNumber=1 scan=12043 | 26.4831 |
| 07360:3 | 4 | R.FSDGLESN[+1864.634]SSTQFEVK.K | HexNAc(2)Hex(9) | N[+1865] | 1213.8218 | 3 | 3639.451 | 3639.4472 | 0 | 1 | 219 | Specific | 497.63 | 497.63 | 497.63 | 9.59 | 76 | >sp\|P0C0L4\|CO4A_HUMAN Complement C4-A OS=Homo sapiens OX=9606 GN=C4A PE=1 SV=2 | 1235 | 20201114_16933_1ng_glyco_hcd203040_compound15.10218.10218.3 | controllerType=0 controllerNumber=1 scan=10218 | 23.4334 |
| 07362:3 | 4 | R.FSDGLESN[+1864.634]SSTQFEVK.K | HexNAc(2)Hex(9) | N[+1865] | 1213.8218 | 3 | 3639.451 | 3639.4472 | 0 | 1 | 219 | Specific | 498.75 | 498.75 | 498.75 | 9.59 | 76 | >sp\|P0C0L4\|CO4A_HUMAN Complement C4-A OS=Homo sapiens OX=9606 GN=C4A PE=1 SV=2 | 1235 | 20201114_16933_1ng_glyco_hcd203040_compound15.10220.10220.3 | controllerType=0 controllerNumber=1 scan=10220 | 23.4375 |
| 07635:3 | 5 | K.WEC[+57.021]KN[+2205.793]DTLLGIK.G | HexNAc(4)Hex(5)Fuc(2)NeuAc(1) | C[+57], N[+2206] | 1228.1905 | 3 | 3682.5569 | 3682.5444 | 0 | 3.4 | 100 | Specific | 268.95 | 268.95 | 268.95 | 5.79 | 78 | >sp\|P22897\|MRC1_HUMAN Macrophage mannose receptor 1 OS=Homo sapiens OX=9606 GN=MRC1 PE=1 SV=1 | 1049 | 20201114_16933_1ng_glyco_hcd203040_compound15.10503.10503.3 | controllerType=0 controllerNumber=1 scan=10503 | 23.9121 |
| 07746:3 | 5 | K.WEC[+57.021]KN[+2059.735]DTLLGIK.G | HexNAc(4)Hex(5)Fuc(1)NeuAc(1) | C[+57], N[+2060] | 1179.5053 | 3 | 3536.5014 | 3536.4865 | 0 | 4.2 | 100 | Specific | 289.76 | 289.76 | 289.76 | 6.27 | 78 | >sp\|P22897\|MRC1_HUMAN Macrophage mannose receptor 1 OS=Homo sapiens OX=9606 GN=MRC1 PE=1 SV=1 | 1049 | 20201114_16933_1ng_glyco_hcd203040_compound15.10618.10618.3 | controllerType=0 controllerNumber=1 scan=10618 | 24.1062 |
| 14414:5 | 5 | K.KGN[+2351.851]TTLNSFVIPSESDVPTHC[+57.021]PSQWWPYAGHC[+57.021]YK.I | HexNAc(4)Hex(5)Fuc(3)NeuAc(1) | C[+57]*2, N[+2352] | 1263.9406 | 5 | 6315.6737 | 6315.6725 | 0 | 0.2 | 342 | Specific | 381.18 | 203.47 | 203.47 | 7.3 | 78 | >sp\|P22897\|MRC1_HUMAN Macrophage mannose receptor 1 OS=Homo sapiens OX=9606 GN=MRC1 PE=1 SV=1 | 1049 | 20201114_16933_1ng_glyco_hcd203040_compound15.17566.17566.5 | controllerType=0 controllerNumber=1 scan=17566 | 35.922 |
| 14412:5 | 5 | K.KGN[+2351.851]TTLNSFVIPSESDVPTHC[+57.021]PSQWWPYAGHC[+57.021]YK.I | HexNAc(4)Hex(5)Fuc(3)NeuAc(1) | C[+57]*2, N[+2352] | 1263.9406 | 5 | 6315.6737 | 6315.6725 | 0 | 0.2 | 342 | Specific | 483.13 | 390.74 | 390.74 | 9.71 | 78 | >sp\|P22897\|MRC1_HUMAN Macrophage mannose receptor 1 OS=Homo sapiens OX=9606 GN=MRC1 PE=1 SV=1 | 1049 | 20201114_16933_1ng_glyco_hcd203040_compound15.17564.17564.5 | controllerType=0 controllerNumber=1 scan=17564 | 35.9183 |
| 13080:5 | 5 | K.KGN[+2204.772]TTLNSFVIPSESDVPTHC[+57.021]PSQWWPYAGHC[+57.021]YK.I | HexNAc(4)Hex(5)NeuAc(2) | C[+57]*2, N[+2205] | 1234.5305 | 5 | 6168.6235 | 6168.5942 | 0 | 4.7 | 342 | Specific | 614.3 | 614.3 | 614.3 | 12.21 | 78 | >sp\|P22897\|MRC1_HUMAN Macrophage mannose receptor 1 OS=Homo sapiens OX=9606 GN=MRC1 PE=1 SV=1 | 1049 | 20201114_16933_1ng_glyco_hcd203040_compound15.16182.16182.5 | controllerType=0 controllerNumber=1 scan=16182 | 33.5113 |
| 13078:5 | 5 | K.KGN[+2204.772]TTLNSFVIPSESDVPTHC[+57.021]PSQWWPYAGHC[+57.021]YK.I | HexNAc(4)Hex(5)NeuAc(2) | C[+57]*2, N[+2205] | 1234.5305 | 5 | 6168.6235 | 6168.5942 | 0 | 4.7 | 342 | Specific | 801.19 | 680.19 | 680.19 | 16.21 | 78 | >sp\|P22897\|MRC1_HUMAN Macrophage mannose receptor 1 OS=Homo sapiens OX=9606 GN=MRC1 PE=1 SV=1 | 1049 | 20201114_16933_1ng_glyco_hcd203040_compound15.16180.16180.5 | controllerType=0 controllerNumber=1 scan=16180 | 33.5068 |
| 08384:3 | 5 | K.TAHC[+57.021]N[+2205.793]ESFYFLC[+57.021]K.R | HexNAc(4)Hex(5)Fuc(2)NeuAc(1) | C[+57]*2, N[+2206] | 1294.8487 | 3 | 3882.5315 | 3882.5124 | 0 | 4.9 | 1201 | Specific | 242.21 | 169.93 | 169.93 | 5.21 | 78 | >sp\|P22897\|MRC1_HUMAN Macrophage mannose receptor 1 OS=Homo sapiens OX=9606 GN=MRC1 PE=1 SV=1 | 1049 | 20201114_16933_1ng_glyco_hcd203040_compound15.11281.11281.3 | controllerType=0 controllerNumber=1 scan=11281 | 25.1914 |
| 10167:3 | 7 | K.TVLTPATNHMGN[+1378.476]VTFTIPANR.E | HexNAc(2)Hex(6) | N[+1378] | 1211.8929 | 3 | 3633.6641 | 3633.6358 | 0 | 7.8 | 74 | Specific | 329.97 | 329.97 | 175.56 | 7.12 | 84 | >sp\|P01024\|CO3_HUMAN Complement C3 OS=Homo sapiens OX=9606 GN=C3 PE=1 SV=2 | 492 | 20201114_16933_1ng_glyco_hcd203040_compound15.13140.13140.3 | controllerType=0 controllerNumber=1 scan=13140 | 28.3379 |
| 20683:4 | 8 | R.NIAAFGGDPNN[+2204.772]ITLFGESAGGASVSLQTLSPYNK.G | HexNAc(4)Hex(5)NeuAc(2) | N[+2205] | 1404.634 | 4 | 5615.5143 | 5615.4587 | 0 | 9.9 | 197 | Specific | 333.84 | 333.84 | 333.84 | 7.39 | 49 | >sp\|P19835\|CEL_HUMAN Bile salt-activated lipase OS=Homo sapiens OX=9606 GN=CEL PE=1 SV=3 | 476 | 20201114_16933_1ng_glyco_hcd203040_compound15.24132.24132.4 | controllerType=0 controllerNumber=1 scan=24132 | 48.7588 |
| 20681:4 | 8 | R.NIAAFGGDPNN[+2204.772]ITLFGESAGGASVSLQTLSPYNK.G | HexNAc(4)Hex(5)NeuAc(2) | N[+2205] | 1404.634 | 4 | 5615.5143 | 5615.4587 | 0 | 9.9 | 197 | Specific | 405.93 | 405.93 | 405.93 | 9.09 | 49 | >sp\|P19835\|CEL_HUMAN Bile salt-activated lipase OS=Homo sapiens OX=9606 GN=CEL PE=1 SV=3 | 476 | 20201114_16933_1ng_glyco_hcd203040_compound15.24130.24130.4 | controllerType=0 controllerNumber=1 scan=24130 | 48.7551 |

Table S4. Cont.

| Query #:z | Protein Rank | Glycopeptide | Glycan | Modification Type(s) | Observed m/z | z | Observed (M+H) | Calc. mass (M+H) | Off-by-x error | Mass error (ppm) | StartPosition | Cleavage | Score | Delta | Delta Mod | \|Log Prob\| | # of unique peptides | Protein Name | Protein DB number | Comment | Scan # | Scan Time |
|---|---|---|---|---|---|---|---|---|---|---|---|---|---|---|---|---|---|---|---|---|---|---|
| 19146:4 | 8 | R.NIAAFGGDPNN[+1913.677]ITLFGESAGGASVSLQTLSPYNK.G | HexNAc(4)Hex(5)NeuAc(1) | N[+1914] | 1331.8539 | 4 | 5324.3937 | 5324.3633 | 0 | 5.7 | 197 | Specific | 561.73 | 561.73 | 561.73 | 10.36 | 49 | >sp\|P19835\|CEL_HUMAN Bile salt-activated lipase OS=Homo sapiens OX=9606 GN=CEL PE=1 SV=3 | 476 | 20201114_16933_1ng_glyco_hcd203040_compound15.22517.22517.4 | controllerType=0 controllerNumber=1 scan=22517 | 45.3582 |
| 19143:4 | 8 | R.NIAAFGGDPNN[+1913.677]ITLFGESAGGASVSLQTLSPYNK.G | HexNAc(4)Hex(5)NeuAc(1) | N[+1914] | 1331.8539 | 4 | 5324.3937 | 5324.3633 | 0 | 5.7 | 197 | Specific | 752.24 | 723.79 | 723.79 | 15.52 | 49 | >sp\|P19835\|CEL_HUMAN Bile salt-activated lipase OS=Homo sapiens OX=9606 GN=CEL PE=1 SV=3 | 476 | 20201114_16933_1ng_glyco_hcd203040_compound15.22514.22514.4 | controllerType=0 controllerNumber=1 scan=22514 | 45.3534 |
| 23050:3 | 9 | R.LSLLEEPGN[+2350.830]GTFTVILNQLTSR.D | HexNAc(4)Hex(5)Fuc(1)NeuAc(2) | N[+2351] | 1585.0524 | 3 | 4753.1426 | 4753.1229 | 0 | 4.2 | 413 | Specific | 347.1 | 347.1 | 347.1 | 7.99 | 54 | >sp\|P01833\|PIGR_HUMAN Polymeric immunoglobulin receptor OS=Homo sapiens OX=9606 GN=PIGR PE=1 SV=4 | 902 | 20201114_16933_1ng_glyco_hcd203040_compound15.26618.26618.3 | controllerType=0 controllerNumber=1 scan=26618 | 53.4667 |
| 20744:3 | 9 | R.LSLLEEPGN[+2059.735]GTFTVILNQLTSR.D | HexNAc(4)Hex(5)Fuc(1)NeuAc(1) | N[+2060] | 1488.0179 | 3 | 4462.0391 | 4462.0275 | 0 | 2.6 | 413 | Specific | 437.79 | 382.25 | 382.25 | 9.73 | 54 | >sp\|P01833\|PIGR_HUMAN Polymeric immunoglobulin receptor OS=Homo sapiens OX=9606 GN=PIGR PE=1 SV=4 | 902 | 20201114_16933_1ng_glyco_hcd203040_compound15.24196.24196.3 | controllerType=0 controllerNumber=1 scan=24196 | 48.9139 |
| 23052:3 | 9 | R.LSLLEEPGN[+2350.830]GTFTVILNQLTSR.D | HexNAc(4)Hex(5)Fuc(1)NeuAc(2) | N[+2351] | 1585.0524 | 3 | 4753.1426 | 4753.1229 | 0 | 4.2 | 413 | Specific | 668.85 | 433.86 | 433.86 | 13.3 | 54 | >sp\|P01833\|PIGR_HUMAN Polymeric immunoglobulin receptor OS=Homo sapiens OX=9606 GN=PIGR PE=1 SV=4 | 902 | 20201114_16933_1ng_glyco_hcd203040_compound15.26620.26620.3 | controllerType=0 controllerNumber=1 scan=26620 | 53.4714 |
| 16227:4 | 9 | K.IIEGEPNLKVPGN[+2204.772]VTAVLGETLK.V | HexNAc(4)Hex(5)NeuAc(2) | N[+2205] | 1149.7967 | 4 | 4596.1649 | 4596.1218 | 0 | 9.4 | 457 | Specific | 333.39 | 333.39 | 333.39 | 6.54 | 54 | >sp\|P01833\|PIGR_HUMAN Polymeric immunoglobulin receptor OS=Homo sapiens OX=9606 GN=PIGR PE=1 SV=4 | 902 | 20201114_16933_1ng_glyco_hcd203040_compound15.19469.19469.4 | controllerType=0 controllerNumber=1 scan=19469 | 39.2217 |
| 16228:4 | 9 | K.IIEGEPNLKVPGN[+2204.772]VTAVLGETLK.V | HexNAc(4)Hex(5)NeuAc(2) | N[+2205] | 1149.7967 | 4 | 4596.1649 | 4596.1218 | 0 | 9.4 | 457 | Specific | 419.97 | 419.97 | 419.97 | 8.24 | 54 | >sp\|P01833\|PIGR_HUMAN Polymeric immunoglobulin receptor OS=Homo sapiens OX=9606 GN=PIGR PE=1 SV=4 | 902 | 20201114_16933_1ng_glyco_hcd203040_compound15.19470.19470.4 | controllerType=0 controllerNumber=1 scan=19470 | 39.2255 |
| 16304:3 | 9 | K.IIEGEPNLKVPGN[+2059.735]VTAVLGETLK.V | HexNAc(4)Hex(5)Fuc(1)NeuAc(1) | N[+2060] | 1484.3715 | 3 | 4451.1 | 4451.0843 | 0 | 3.5 | 457 | Specific | 492.37 | 492.37 | 492.37 | 9.4 | 54 | >sp\|P01833\|PIGR_HUMAN Polymeric immunoglobulin receptor OS=Homo sapiens OX=9606 GN=PIGR PE=1 SV=4 | 902 | 20201114_16933_1ng_glyco_hcd203040_compound15.19549.19549.3 | controllerType=0 controllerNumber=1 scan=19549 | 39.3713 |
| 16204:3 | 9 | K.IIEGEPNLKVPGN[+2059.735]VTAVLGETLK.V | HexNAc(4)Hex(5)Fuc(1)NeuAc(1) | N[+2060] | 1484.3707 | 3 | 4451.0974 | 4451.0843 | 0 | 3 | 457 | Specific | 604.24 | 604.24 | 604.24 | 11.86 | 54 | >sp\|P01833\|PIGR_HUMAN Polymeric immunoglobulin receptor OS=Homo sapiens OX=9606 GN=PIGR PE=1 SV=4 | 902 | 20201114_16933_1ng_glyco_hcd203040_compound15.19446.19446.3 | controllerType=0 controllerNumber=1 scan=19446 | 39.1849 |
| 18165:3 | 9 | K.IIEGEPNLKVPGN[+2350.830]VTAVLGETLK.V | HexNAc(4)Hex(5)Fuc(1)NeuAc(2) | N[+2351] | 1581.3979 | 3 | 4742.1791 | 4742.1797 | 0 | -0.1 | 457 | Specific | 609.92 | 452.45 | 452.45 | 11.37 | 54 | >sp\|P01833\|PIGR_HUMAN Polymeric immunoglobulin receptor OS=Homo sapiens OX=9606 GN=PIGR PE=1 SV=4 | 902 | 20201114_16933_1ng_glyco_hcd203040_compound15.21492.21492.3 | controllerType=0 controllerNumber=1 scan=21492 | 43.1281 |
| 16206:3 | 9 | K.IIEGEPNLKVPGN[+2059.735]VTAVLGETLK.V | HexNAc(4)Hex(5)Fuc(1)NeuAc(1) | N[+2060] | 1484.3707 | 3 | 4451.0974 | 4451.0843 | 0 | 3 | 457 | Specific | 684.14 | 661.04 | 661.04 | 12.86 | 54 | >sp\|P01833\|PIGR_HUMAN Polymeric immunoglobulin receptor OS=Homo sapiens OX=9606 GN=PIGR PE=1 SV=4 | 902 | 20201114_16933_1ng_glyco_hcd203040_compound15.19448.19448.3 | controllerType=0 controllerNumber=1 scan=19448 | 39.1886 |
| 16187:3 | 9 | K.IIEGEPNLKVPGN[+2205.793]VTAVLGETLK.V | HexNAc(4)Hex(5)Fuc(2)NeuAc(1) | N[+2206] | 1533.0557 | 3 | 4597.1526 | 4597.1422 | 0 | 2.3 | 457 | Specific | 731.35 | 731.35 | 731.35 | 13.96 | 54 | >sp\|P01833\|PIGR_HUMAN Polymeric immunoglobulin receptor OS=Homo sapiens OX=9606 GN=PIGR PE=1 SV=4 | 902 | 20201114_16933_1ng_glyco_hcd203040_compound15.19428.19428.3 | controllerType=0 controllerNumber=1 scan=19428 | 39.151 |
| 16178:3 | 9 | K.IIEGEPNLKVPGN[+2205.793]VTAVLGETLK.V | HexNAc(4)Hex(5)Fuc(2)NeuAc(1) | N[+2206] | 1533.0516 | 3 | 4597.1402 | 4597.1422 | 0 | -0.4 | 457 | Specific | 741.07 | 567.94 | 567.94 | 13.37 | 54 | >sp\|P01833\|PIGR_HUMAN Polymeric immunoglobulin receptor OS=Homo sapiens OX=9606 GN=PIGR PE=1 SV=4 | 902 | 20201114_16933_1ng_glyco_hcd203040_compound15.19417.19417.3 | controllerType=0 controllerNumber=1 scan=19417 | 39.1281 |
| 16189:3 | 9 | K.IIEGEPNLKVPGN[+2205.793]VTAVLGETLK.V | HexNAc(4)Hex(5)Fuc(2)NeuAc(1) | N[+2206] | 1533.0557 | 3 | 4597.1526 | 4597.1422 | 0 | 2.3 | 457 | Specific | 776.91 | 691.37 | 691.37 | 14.96 | 54 | >sp\|P01833\|PIGR_HUMAN Polymeric immunoglobulin receptor OS=Homo sapiens OX=9606 GN=PIGR PE=1 SV=4 | 902 | 20201114_16933_1ng_glyco_hcd203040_compound15.19430.19430.3 | controllerType=0 controllerNumber=1 scan=19430 | 39.155 |
| 16259:3 | 9 | K.IIEGEPNLKVPGN[+2059.735]VTAVLGETLK.V | HexNAc(4)Hex(5)Fuc(1)NeuAc(1) | N[+2060] | 1484.3671 | 3 | 4451.0868 | 4451.0843 | 0 | 0.6 | 457 | Specific | 779.59 | 481.99 | 481.99 | 14.76 | 54 | >sp\|P01833\|PIGR_HUMAN Polymeric immunoglobulin receptor OS=Homo sapiens OX=9606 GN=PIGR PE=1 SV=4 | 902 | 20201114_16933_1ng_glyco_hcd203040_compound15.19503.19503.3 | controllerType=0 controllerNumber=1 scan=19503 | 39.2921 |
| 16186:3 | 9 | K.IIEGEPNLKVPGN[+2205.793]VTAVLGETLK.V | HexNAc(4)Hex(5)Fuc(2)NeuAc(1) | N[+2206] | 1533.0529 | 3 | 4597.1442 | 4597.1422 | 0 | 0.4 | 457 | Specific | 827.57 | 763.1 | 763.1 | 15.76 | 54 | >sp\|P01833\|PIGR_HUMAN Polymeric immunoglobulin receptor OS=Homo sapiens OX=9606 GN=PIGR PE=1 SV=4 | 902 | 20201114_16933_1ng_glyco_hcd203040_compound15.19426.19426.3 | controllerType=0 controllerNumber=1 scan=19426 | 39.1467 |
| 18163:3 | 9 | K.IIEGEPNLKVPGN[+2350.830]VTAVLGETLK.V | HexNAc(4)Hex(5)Fuc(1)NeuAc(2) | N[+2351] | 1581.3979 | 3 | 4742.1791 | 4742.1797 | 0 | -0.1 | 457 | Specific | 937.74 | 708.43 | 708.43 | 17.37 | 54 | >sp\|P01833\|PIGR_HUMAN Polymeric immunoglobulin receptor OS=Homo sapiens OX=9606 GN=PIGR PE=1 SV=4 | 902 | 20201114_16933_1ng_glyco_hcd203040_compound15.21490.21490.3 | controllerType=0 controllerNumber=1 scan=21490 | 43.1212 |
| 16175:3 | 9 | K.IIEGEPNLKVPGN[+2205.793]VTAVLGETLK.V | HexNAc(4)Hex(5)Fuc(2)NeuAc(1) | N[+2206] | 1533.0516 | 3 | 4597.1402 | 4597.1422 | 0 | -0.4 | 457 | Specific | 993.6 | 925.48 | 925.48 | 18.37 | 54 | >sp\|P01833\|PIGR_HUMAN Polymeric immunoglobulin receptor OS=Homo sapiens OX=9606 GN=PIGR PE=1 SV=4 | 902 | 20201114_16933_1ng_glyco_hcd203040_compound15.19414.19414.3 | controllerType=0 controllerNumber=1 scan=19414 | 39.1221 |
| 16257:3 | 9 | K.IIEGEPNLKVPGN[+2059.735]VTAVLGETLK.V | HexNAc(4)Hex(5)Fuc(1)NeuAc(1) | N[+2060] | 1484.3671 | 3 | 4451.0868 | 4451.0843 | 0 | 0.6 | 457 | Specific | 994.84 | 891.02 | 891.02 | 18.76 | 54 | >sp\|P01833\|PIGR_HUMAN Polymeric immunoglobulin receptor OS=Homo sapiens OX=9606 GN=PIGR PE=1 SV=4 | 902 | 20201114_16933_1ng_glyco_hcd203040_compound15.19501.19501.3 | controllerType=0 controllerNumber=1 scan=19501 | 39.287 |
| 16184:3 | 9 | K.IIEGEPNLKVPGN[+2205.793]VTAVLGETLK.V | HexNAc(4)Hex(5)Fuc(2)NeuAc(1) | N[+2206] | 1533.0529 | 3 | 4597.1442 | 4597.1422 | 0 | 0.4 | 457 | Specific | 997.19 | 902.44 | 902.44 | 18.76 | 54 | >sp\|P01833\|PIGR_HUMAN Polymeric immunoglobulin receptor OS=Homo sapiens OX=9606 GN=PIGR PE=1 SV=4 | 902 | 20201114_16933_1ng_glyco_hcd203040_compound15.19424.19424.3 | controllerType=0 controllerNumber=1 scan=19424 | 39.1412 |
| 15805:4 | 9 | K.IIEGEPNLKVPGN[+2059.735]VTAVLGETLKVPC[+57.021]HFPC[+57.021]K.F | HexNAc(4)Hex(5)Fuc(1)NeuAc(1) | C[+57]*2, N[+2060] | 1369.8983 | 4 | 5476.5714 | 5476.5418 | 0 | 5.4 | 457 | Specific | 216.76 | 216.76 | 216.76 | 3.8 | 54 | >sp\|P01833\|PIGR_HUMAN Polymeric immunoglobulin receptor OS=Homo sapiens OX=9606 GN=PIGR PE=1 SV=4 | 902 | 20201114_16933_1ng_glyco_hcd203040_compound15.19009.19009.4 | controllerType=0 controllerNumber=1 scan=19009 | 38.4069 |
| 15813:4 | 9 | K.IIEGEPNLKVPGN[+2059.735]VTAVLGETLKVPC[+57.021]HFPC[+57.021]K.F | HexNAc(4)Hex(5)Fuc(1)NeuAc(1) | C[+57]*2, N[+2060] | 1369.8999 | 4 | 5476.5778 | 5476.5418 | 0 | 6.6 | 457 | Specific | 218.23 | 218.23 | 218.23 | 3.47 | 54 | >sp\|P01833\|PIGR_HUMAN Polymeric immunoglobulin receptor OS=Homo sapiens OX=9606 GN=PIGR PE=1 SV=4 | 902 | 20201114_16933_1ng_glyco_hcd203040_compound15.19018.19018.2 | controllerType=0 controllerNumber=1 scan=19018 | 38.4219 |
| 15807:4 | 9 | K.IIEGEPNLKVPGN[+2059.735]VTAVLGETLKVPC[+57.021]HFPC[+57.021]K.F | HexNAc(4)Hex(5)Fuc(1)NeuAc(1) | C[+57]*2, N[+2060] | 1369.8983 | 4 | 5476.5714 | 5476.5418 | 0 | 5.4 | 457 | Specific | 519.87 | 519.87 | 519.87 | 8.32 | 54 | >sp\|P01833\|PIGR_HUMAN Polymeric immunoglobulin receptor OS=Homo sapiens OX=9606 GN=PIGR PE=1 SV=4 | 902 | 20201114_16933_1ng_glyco_hcd203040_compound15.19011.19011.4 | controllerType=0 controllerNumber=1 scan=19011 | 38.411 |
| 15793:5 | 9 | K.IIEGEPNLKVPGN[+2205.793]VTAVLGETLKVPC[+57.021]HFPC[+57.021]K.F | HexNAc(4)Hex(5)Fuc(2)NeuAc(1) | C[+57]*2, N[+2206] | 1125.33 | 5 | 5622.6209 | 5622.5997 | 0 | 3.8 | 457 | Specific | 573.92 | 573.92 | 573.92 | 10.49 | 54 | >sp\|P01833\|PIGR_HUMAN Polymeric immunoglobulin receptor OS=Homo sapiens OX=9606 GN=PIGR PE=1 SV=4 | 902 | 20201114_16933_1ng_glyco_hcd203040_compound15.18997.18997.5 | controllerType=0 controllerNumber=1 scan=18997 | 38.3871 |
| 15816:4 | 9 | K.IIEGEPNLKVPGN[+2059.735]VTAVLGETLKVPC[+57.021]HFPC[+57.021]K.F | HexNAc(4)Hex(5)Fuc(1)NeuAc(1) | C[+57]*2, N[+2060] | 1369.8999 | 4 | 5476.5778 | 5476.5418 | 0 | 6.6 | 457 | Specific | 585.09 | 585.09 | 585.09 | 9.24 | 54 | >sp\|P01833\|PIGR_HUMAN Polymeric immunoglobulin receptor OS=Homo sapiens OX=9606 GN=PIGR PE=1 SV=4 | 902 | 20201114_16933_1ng_glyco_hcd203040_compound15.19021.19021.2 | controllerType=0 controllerNumber=1 scan=19021 | 38.4261 |
| 15798:4 | 9 | K.IIEGEPNLKVPGN[+2205.793]VTAVLGETLKVPC[+57.021]HFPC[+57.021]K.F | HexNAc(4)Hex(5)Fuc(2)NeuAc(1) | C[+57]*2, N[+2206] | 1406.4106 | 4 | 5622.6206 | 5622.5997 | 0 | 3.7 | 457 | Specific | 612.55 | 560.4 | 560.4 | 10.86 | 54 | >sp\|P01833\|PIGR_HUMAN Polymeric immunoglobulin receptor OS=Homo sapiens OX=9606 GN=PIGR PE=1 SV=4 | 902 | 20201114_16933_1ng_glyco_hcd203040_compound15.19002.19002.4 | controllerType=0 controllerNumber=1 scan=19002 | 38.3968 |
| 15791:5 | 9 | K.IIEGEPNLKVPGN[+2205.793]VTAVLGETLKVPC[+57.021]HFPC[+57.021]K.F | HexNAc(4)Hex(5)Fuc(2)NeuAc(1) | C[+57]*2, N[+2206] | 1125.33 | 5 | 5622.6209 | 5622.5997 | 0 | 3.8 | 457 | Specific | 648.91 | 607.89 | 607.89 | 11.49 | 54 | >sp\|P01833\|PIGR_HUMAN Polymeric immunoglobulin receptor OS=Homo sapiens OX=9606 GN=PIGR PE=1 SV=4 | 902 | 20201114_16933_1ng_glyco_hcd203040_compound15.18995.18995.5 | controllerType=0 controllerNumber=1 scan=18995 | 38.3829 |
| 17297:4 | 9 | K.IIEGEPNLKVPGN[+2350.830]VTAVLGETLKVPC[+57.021]HFPC[+57.021]K.F | HexNAc(4)Hex(5)Fuc(1)NeuAc(2) | C[+57]*2, N[+2351] | 1442.6693 | 4 | 5767.6554 | 5767.6372 | 0 | 3.2 | 457 | Specific | 722.53 | 606.91 | 606.91 | 12.86 | 54 | >sp\|P01833\|PIGR_HUMAN Polymeric immunoglobulin receptor OS=Homo sapiens OX=9606 GN=PIGR PE=1 SV=4 | 902 | 20201114_16933_1ng_glyco_hcd203040_compound15.20585.20585.4 | controllerType=0 controllerNumber=1 scan=20585 | 41.2469 |
| 15796:4 | 9 | K.IIEGEPNLKVPGN[+2205.793]VTAVLGETLKVPC[+57.021]HFPC[+57.021]K.F | HexNAc(4)Hex(5)Fuc(2)NeuAc(1) | C[+57]*2, N[+2206] | 1406.4106 | 4 | 5622.6206 | 5622.5997 | 0 | 3.7 | 457 | Specific | 1072.08 | 975.17 | 975.17 | 19.86 | 54 | >sp\|P01833\|PIGR_HUMAN Polymeric immunoglobulin receptor OS=Homo sapiens OX=9606 GN=PIGR PE=1 SV=4 | 902 | 20201114_16933_1ng_glyco_hcd203040_compound15.19000.19000.4 | controllerType=0 controllerNumber=1 scan=19000 | 38.3915 |
| 13572:3 | 9 | K.VPGN[+2205.793]VTAVLGETLK.V | HexNAc(4)Hex(5)Fuc(2)NeuAc(1) | N[+2206] | 1201.8709 | 3 | 3603.5982 | 3603.5927 | 0 | 1.5 | 466 | Specific | 530.86 | 485.55 | 485.55 | 9.91 | 54 | >sp\|P01833\|PIGR_HUMAN Polymeric immunoglobulin receptor OS=Homo sapiens OX=9606 GN=PIGR PE=1 SV=4 | 902 | 20201114_16933_1ng_glyco_hcd203040_compound15.16694.16694.3 | controllerType=0 controllerNumber=1 scan=16694 | 34.411 |
| 13727:3 | 9 | K.VPGN[+2059.735]VTAVLGETLK.V | HexNAc(4)Hex(5)Fuc(1)NeuAc(1) | N[+2060] | 1153.1855 | 3 | 3457.5419 | 3457.5348 | 0 | 2.1 | 466 | Specific | 536.07 | 382.35 | 382.35 | 9.91 | 54 | >sp\|P01833\|PIGR_HUMAN Polymeric immunoglobulin receptor OS=Homo sapiens OX=9606 GN=PIGR PE=1 SV=4 | 902 | 20201114_16933_1ng_glyco_hcd203040_compound15.16854.16854.3 | controllerType=0 controllerNumber=1 scan=16854 | 34.6807 |
| 13574:3 | 9 | K.VPGN[+2205.793]VTAVLGETLK.V | HexNAc(4)Hex(5)Fuc(2)NeuAc(1) | N[+2206] | 1201.8709 | 3 | 3603.5982 | 3603.5927 | 0 | 1.5 | 466 | Specific | 610.65 | 507.58 | 507.58 | 12.24 | 54 | >sp\|P01833\|PIGR_HUMAN Polymeric immunoglobulin receptor OS=Homo sapiens OX=9606 GN=PIGR PE=1 SV=4 | 902 | 20201114_16933_1ng_glyco_hcd203040_compound15.16696.16696.3 | controllerType=0 controllerNumber=1 scan=16696 | 34.415 |
| 15694:3 | 9 | K.VPGN[+2351.851]VTAVLGETLKVPC[+57.021]HFPC[+57.021]K.F | HexNAc(4)Hex(5)Fuc(3)NeuAc(1) | C[+57]*2, N[+2352] | 1592.3679 | 3 | 4775.089 | 4775.1081 | 0 | -4 | 466 | Specific | 363.05 | 363.05 | 363.05 | 5.37 | 54 | >sp\|P01833\|PIGR_HUMAN Polymeric immunoglobulin receptor OS=Homo sapiens OX=9606 GN=PIGR PE=1 SV=4 | 902 | 20201114_16933_1ng_glyco_hcd203040_compound15.18893.18893.3 | controllerType=0 controllerNumber=1 scan=18893 | 38.197 |
| 13624:4 | 9 | K.VPGN[+2204.772]VTAVLGETLKVPC[+57.021]HFPC[+57.021]K.F | HexNAc(4)Hex(5)NeuAc(2) | C[+57]*2, N[+2205] | 1157.7649 | 4 | 4628.0379 | 4628.0298 | 0 | 1.7 | 466 | Specific | 441.87 | 280.48 | 280.48 | 7.18 | 54 | >sp\|P01833\|PIGR_HUMAN Polymeric immunoglobulin receptor OS=Homo sapiens OX=9606 GN=PIGR PE=1 SV=4 | 902 | 20201114_16933_1ng_glyco_hcd203040_compound15.16748.16748.3 | controllerType=0 controllerNumber=1 scan=16748 | 34.5053 |
| 13626:4 | 9 | K.VPGN[+2204.772]VTAVLGETLKVPC[+57.021]HFPC[+57.021]K.F | HexNAc(4)Hex(5)NeuAc(2) | C[+57]*2, N[+2205] | 1157.7649 | 4 | 4628.0379 | 4628.0298 | 0 | 1.7 | 466 | Specific | 751.04 | 589.98 | 589.98 | 14.79 | 54 | >sp\|P01833\|PIGR_HUMAN Polymeric immunoglobulin receptor OS=Homo sapiens OX=9606 GN=PIGR PE=1 SV=4 | 902 | 20201114_16933_1ng_glyco_hcd203040_compound15.16750.16750.3 | controllerType=0 controllerNumber=1 scan=16750 | 34.5091 |
| 05930:3 | 9 | K.WN[+1710.598]NTGC[+57.021]QALPSQDEGPSK.A | HexNAc(3)Hex(5)NeuAc(1) | C[+57], N[+1711] | 1233.8309 | 3 | 3699.478 | 3699.4743 | 0 | 1 | 498 | Specific | 338.92 | 284.84 | 284.84 | 7.19 | 54 | >sp\|P01833\|PIGR_HUMAN Polymeric immunoglobulin receptor OS=Homo sapiens OX=9606 GN=PIGR PE=1 SV=4 | 902 | 20201114_16933_1ng_glyco_hcd203040_compound15.8730.8730.3 | controllerType=0 controllerNumber=1 scan=8730 | 20.9016 |
| 05932:3 | 9 | K.WN[+1710.598]NTGC[+57.021]QALPSQDEGPSK.A | HexNAc(3)Hex(5)NeuAc(1) | C[+57], N[+1711] | 1233.8309 | 3 | 3699.478 | 3699.4743 | 0 | 1 | 498 | Specific | 435 | 376.56 | 376.56 | 9.53 | 54 | >sp\|P01833\|PIGR_HUMAN Polymeric immunoglobulin receptor OS=Homo sapiens OX=9606 GN=PIGR PE=1 SV=4 | 902 | 20201114_16933_1ng_glyco_hcd203040_compound15.8732.8732.3 | controllerType=0 controllerNumber=1 scan=8732 | 20.9057 |
| 05864:3 | 9 | K.WN[+1913.677]NTGC[+57.021]QALPSQDEGPSK.A | HexNAc(4)Hex(5)NeuAc(1) | C[+57], N[+1914] | 1301.5341 | 3 | 3902.5878 | 3902.5537 | 0 | 8.7 | 498 | Specific | 448.24 | 430.49 | 430.49 | 8.87 | 54 | >sp\|P01833\|PIGR_HUMAN Polymeric immunoglobulin receptor OS=Homo sapiens OX=9606 GN=PIGR PE=1 SV=4 | 902 | 20201114_16933_1ng_glyco_hcd203040_compound15.8661.8661.3 | controllerType=0 controllerNumber=1 scan=8661 | 20.7818 |
| 05866:3 | 9 | K.WN[+1913.677]NTGC[+57.021]QALPSQDEGPSK.A | HexNAc(4)Hex(5)NeuAc(1) | C[+57], N[+1914] | 1301.5341 | 3 | 3902.5878 | 3902.5537 | 0 | 8.7 | 498 | Specific | 474.64 | 466.73 | 466.73 | 8.93 | 54 | >sp\|P01833\|PIGR_HUMAN Polymeric immunoglobulin receptor OS=Homo sapiens OX=9606 GN=PIGR PE=1 SV=4 | 902 | 20201114_16933_1ng_glyco_hcd203040_compound15.8663.8663.3 | controllerType=0 controllerNumber=1 scan=8663 | 20.7858 |
| 07804:3 | 9 | K.WN[+2204.772]NTGC[+57.021]QALPSQDEGPSK.A | HexNAc(4)Hex(5)NeuAc(2) | C[+57], N[+2205] | 1398.5627 | 3 | 4193.6735 | 4193.6491 | 0 | 5.8 | 498 | Specific | 518.42 | 414.4 | 414.4 | 9.34 | 54 | >sp\|P01833\|PIGR_HUMAN Polymeric immunoglobulin receptor OS=Homo sapiens OX=9606 GN=PIGR PE=1 SV=4 | 902 | 20201114_16933_1ng_glyco_hcd203040_compound15.10678.10678.3 | controllerType=0 controllerNumber=1 scan=10678 | 24.2048 |
| 07806:3 | 9 | K.WN[+2204.772]NTGC[+57.021]QALPSQDEGPSK.A | HexNAc(4)Hex(5)NeuAc(2) | C[+57], N[+2205] | 1398.5627 | 3 | 4193.6735 | 4193.6491 | 0 | 5.8 | 498 | Specific | 641.14 | 555.52 | 555.52 | 11.68 | 54 | >sp\|P01833\|PIGR_HUMAN Polymeric immunoglobulin receptor OS=Homo sapiens OX=9606 GN=PIGR PE=1 SV=4 | 902 | 20201114_16933_1ng_glyco_hcd203040_compound15.10680.10680.3 | controllerType=0 controllerNumber=1 scan=10680 | 24.2091 |
| 17819:3 | 11 | R.LSLHRPALEDLLLGSEAN[+1864.634]LTC[+57.021]TLTGLR.D | HexNAc(2)Hex(9) | C[+57], N[+1865] | 1610.0859 | 3 | 4828.2431 | 4828.2324 | 0 | 2.2 | 127 | Specific | 207.94 | 207.94 | 207.94 | 5.01 | 40 | >sp\|P01876\|IGHA1_HUMAN Immunoglobulin heavy constant alpha 1 OS=Homo sapiens OX=9606 GN=IGHA1 PE=1 SV=2 | 255 | 20201114_16933_1ng_glyco_hcd203040_compound15.21131.21131.3 | controllerType=0 controllerNumber=1 scan=21131 | 42.3833 |
| 17817:3 | 11 | R.LSLHRPALEDLLLGSEAN[+1864.634]LTC[+57.021]TLTGLR.D | HexNAc(2)Hex(9) | C[+57], N[+1865] | 1610.0859 | 3 | 4828.2431 | 4828.2324 | 0 | 2.2 | 127 | Specific | 306.7 | 306.7 | 306.7 | 6.92 | 40 | >sp\|P01876\|IGHA1_HUMAN Immunoglobulin heavy constant alpha 1 OS=Homo sapiens OX=9606 GN=IGHA1 PE=1 SV=2 | 255 | 20201114_16933_1ng_glyco_hcd203040_compound15.21129.21129.3 | controllerType=0 controllerNumber=1 scan=21129 | 42.3799 |
| 17802:3 | 11 | R.LSLHRPALEDLLLGSEAN[+1702.581]LTC[+57.021]TLTGLR.D | HexNAc(2)Hex(8) | C[+57], N[+1703] | 1556.0684 | 3 | 4666.1907 | 4666.1796 | 0 | 2.4 | 127 | Specific | 414.17 | 414.17 | 414.17 | 9.06 | 40 | >sp\|P01876\|IGHA1_HUMAN Immunoglobulin heavy constant alpha 1 OS=Homo sapiens OX=9606 GN=IGHA1 PE=1 SV=2 | 255 | 20201114_16933_1ng_glyco_hcd203040_compound15.21113.21113.3 | controllerType=0 controllerNumber=1 scan=21113 | 42.338 |
| 17939:3 | 11 | R.LSLHRPALEDLLLGSEAN[+1257.449]LTC[+57.021]TLTGLR.D | HexNAc(3)Hex(4) | C[+57], N[+1257] | 1407.701 | 3 | 4221.0884 | 4221.0477 | 0 | 9.7 | 127 | Specific | 448.08 | 448.08 | 448.08 | 8.41 | 40 | >sp\|P01876\|IGHA1_HUMAN Immunoglobulin heavy constant alpha 1 OS=Homo sapiens OX=9606 GN=IGHA1 PE=1 SV=2 | 255 | 20201114_16933_1ng_glyco_hcd203040_compound15.21256.21256.3 | controllerType=0 controllerNumber=1 scan=21256 | 42.6597 |
| 17880:3 | 11 | R.LSLHRPALEDLLLGSEAN[+1378.476]LTC[+57.021]TLTGLR.D | HexNAc(2)Hex(6) | C[+57], N[+1378] | 1448.0352 | 3 | 4342.0911 | 4342.074 | 0 | 3.9 | 127 | Specific | 457.03 | 457.03 | 457.03 | 9.17 | 40 | >sp\|P01876\|IGHA1_HUMAN Immunoglobulin heavy constant alpha 1 OS=Homo sapiens OX=9606 GN=IGHA1 PE=1 SV=2 | 255 | 20201114_16933_1ng_glyco_hcd203040_compound15.21195.21195.3 | controllerType=0 controllerNumber=1 scan=21195 | 42.5379 |
| 17860:3 | 11 | R.LSLHRPALEDLLLGSEAN[+1540.529]LTC[+57.021]TLTGLR.D | HexNAc(2)Hex(7) | C[+57], N[+1541] | 1502.0488 | 3 | 4504.1317 | 4504.1268 | 0 | 1.1 | 127 | Specific | 504.58 | 504.58 | 504.58 | 9.27 | 40 | >sp\|P01876\|IGHA1_HUMAN Immunoglobulin heavy constant alpha 1 OS=Homo sapiens OX=9606 GN=IGHA1 PE=1 SV=2 | 255 | 20201114_16933_1ng_glyco_hcd203040_compound15.21174.21174.3 | controllerType=0 controllerNumber=1 scan=21174 | 42.4854 |
| 17847:3 | 11 | R.LSLHRPALEDLLLGSEAN[+1702.581]LTC[+57.021]TLTGLR.D | HexNAc(2)Hex(8) | C[+57], N[+1703] | 1556.075 | 3 | 4666.2105 | 4666.1796 | 0 | 6.6 | 127 | Specific | 514.02 | 514.02 | 514.02 | 8.75 | 40 | >sp\|P01876\|IGHA1_HUMAN Immunoglobulin heavy constant alpha 1 OS=Homo sapiens OX=9606 GN=IGHA1 PE=1 SV=2 | 255 | 20201114_16933_1ng_glyco_hcd203040_compound15.21160.21160.3 | controllerType=0 controllerNumber=1 scan=21160 | 42.446 |
| 17875:3 | 11 | R.LSLHRPALEDLLLGSEAN[+1501.555]LTC[+57.021]TLTGLR.D | HexNAc(5)Hex(3) | C[+57], N[+1502] | 1489.06 | 3 | 4465.1654 | 4465.1536 | 0 | 2.6 | 127 | Specific | 553.81 | 544.56 | 544.56 | 9.78 | 40 | >sp\|P01876\|IGHA1_HUMAN Immunoglobulin heavy constant alpha 1 OS=Homo sapiens OX=9606 GN=IGHA1 PE=1 SV=2 | 255 | 20201114_16933_1ng_glyco_hcd203040_compound15.21190.21190.3 | controllerType=0 controllerNumber=1 scan=21190 | 42.5266 |
| 17804:3 | 11 | R.LSLHRPALEDLLLGSEAN[+1702.581]LTC[+57.021]TLTGLR.D | HexNAc(2)Hex(8) | C[+57], N[+1703] | 1556.0684 | 3 | 4666.1907 | 4666.1796 | 0 | 2.4 | 127 | Specific | 566.02 | 537.78 | 537.78 | 9.78 | 40 | >sp\|P01876\|IGHA1_HUMAN Immunoglobulin heavy constant alpha 1 OS=Homo sapiens OX=9606 GN=IGHA1 PE=1 SV=2 | 255 | 20201114_16933_1ng_glyco_hcd203040_compound15.21115.21115.3 | controllerType=0 controllerNumber=1 scan=21115 | 42.3417 |
| 17852:4 | 11 | R.LSLHRPALEDLLLGSEAN[+1663.608]LTC[+57.021]TLTGLR.D | HexNAc(5)Hex(4) | C[+57], N[+1664] | 1157.5605 | 4 | 4627.22 | 4627.2064 | 0 | 2.9 | 127 | Specific | 578.25 | 578.25 | 578.25 | 11.46 | 40 | >sp\|P01876\|IGHA1_HUMAN Immunoglobulin heavy constant alpha 1 OS=Homo sapiens OX=9606 GN=IGHA1 PE=1 SV=2 | 255 | 20201114_16933_1ng_glyco_hcd203040_compound15.21166.21166.4 | controllerType=0 controllerNumber=1 scan=21166 | 42.4594 |
| 17856:4 | 11 | R.LSLHRPALEDLLLGSEAN[+1540.529]LTC[+57.021]TLTGLR.D | HexNAc(2)Hex(7) | C[+57], N[+1541] | 1126.7893 | 4 | 4504.1356 | 4504.1268 | 0 | 2 | 127 | Specific | 632.11 | 632.11 | 632.11 | 12.55 | 40 | >sp\|P01876\|IGHA1_HUMAN Immunoglobulin heavy constant alpha 1 OS=Homo sapiens OX=9606 GN=IGHA1 PE=1 SV=2 | 255 | 20201114_16933_1ng_glyco_hcd203040_compound15.21170.21170.4 | controllerType=0 controllerNumber=1 scan=21170 | 42.4762 |
| 17941:3 | 11 | R.LSLHRPALEDLLLGSEAN[+1257.449]LTC[+57.021]TLTGLR.D | HexNAc(3)Hex(4) | C[+57], N[+1257] | 1407.701 | 3 | 4221.0884 | 4221.0477 | 0 | 9.7 | 127 | Specific | 640.81 | 608.01 | 608.01 | 11.07 | 40 | >sp\|P01876\|IGHA1_HUMAN Immunoglobulin heavy constant alpha 1 OS=Homo sapiens OX=9606 GN=IGHA1 PE=1 SV=2 | 255 | 20201114_16933_1ng_glyco_hcd203040_compound15.21258.21258.3 | controllerType=0 controllerNumber=1 scan=21258 | 42.6641 |
| 17877:3 | 11 | R.LSLHRPALEDLLLGSEAN[+1501.555]LTC[+57.021]TLTGLR.D | HexNAc(5)Hex(3) | C[+57], N[+1502] | 1489.06 | 3 | 4465.1654 | 4465.1536 | 0 | 2.6 | 127 | Specific | 676.73 | 602.47 | 602.47 | 12.93 | 40 | >sp\|P01876\|IGHA1_HUMAN Immunoglobulin heavy constant alpha 1 OS=Homo sapiens OX=9606 GN=IGHA1 PE=1 SV=2 | 255 | 20201114_16933_1ng_glyco_hcd203040_compound15.21192.21192.3 | controllerType=0 controllerNumber=1 scan=21192 | 42.531 |
| 17864:4 | 11 | R.LSLHRPALEDLLLGSEAN[+1501.555]LTC[+57.021]TLTGLR.D | HexNAc(5)Hex(3) | C[+57], N[+1502] | 1117.0458 | 4 | 4465.1614 | 4465.1536 | 0 | 1.8 | 127 | Specific | 695.58 | 695.58 | 695.58 | 13.55 | 40 | >sp\|P01876\|IGHA1_HUMAN Immunoglobulin heavy constant alpha 1 OS=Homo sapiens OX=9606 GN=IGHA1 PE=1 SV=2 | 255 | 20201114_16933_1ng_glyco_hcd203040_compound15.21178.21178.4 | controllerType=0 controllerNumber=1 scan=21178 | 42.4967 |
| 17862:4 | 11 | R.LSLHRPALEDLLLGSEAN[+1501.555]LTC[+57.021]TLTGLR.D | HexNAc(5)Hex(3) | C[+57], N[+1502] | 1117.0458 | 4 | 4465.1614 | 4465.1536 | 0 | 1.8 | 127 | Specific | 708.58 | 708.58 | 708.58 | 14.55 | 40 | >sp\|P01876\|IGHA1_HUMAN Immunoglobulin heavy constant alpha 1 OS=Homo sapiens OX=9606 GN=IGHA1 PE=1 SV=2 | 255 | 20201114_16933_1ng_glyco_hcd203040_compound15.21176.21176.4 | controllerType=0 controllerNumber=1 scan=21176 | 42.4892 |
| 17883:4 | 11 | R.LSLHRPALEDLLLGSEAN[+1216.423]LTC[+57.021]TLTGLR.D | HexNAc(2)Hex(5) | C[+57], N[+1216] | 1045.7631 | 4 | 4180.0306 | 4180.0211 | 0 | 2.3 | 127 | Specific | 735.48 | 735.48 | 735.48 | 14.55 | 40 | >sp\|P01876\|IGHA1_HUMAN Immunoglobulin heavy constant alpha 1 OS=Homo sapiens OX=9606 GN=IGHA1 PE=1 SV=2 | 255 | 20201114_16933_1ng_glyco_hcd203040_compound15.21198.21198.4 | controllerType=0 controllerNumber=1 scan=21198 | 42.5427 |
| 17858:3 | 11 | R.LSLHRPALEDLLLGSEAN[+1540.529]LTC[+57.021]TLTGLR.D | HexNAc(2)Hex(7) | C[+57], N[+1541] | 1502.0488 | 3 | 4504.1317 | 4504.1268 | 0 | 1.1 | 127 | Specific | 770.29 | 696.13 | 696.13 | 14.73 | 40 | >sp\|P01876\|IGHA1_HUMAN Immunoglobulin heavy constant alpha 1 OS=Homo sapiens OX=9606 GN=IGHA1 PE=1 SV=2 | 255 | 20201114_16933_1ng_glyco_hcd203040_compound15.21172.21172.3 | controllerType=0 controllerNumber=1 scan=21172 | 42.4795 |
| 17853:4 | 11 | R.LSLHRPALEDLLLGSEAN[+1540.529]LTC[+57.021]TLTGLR.D | HexNAc(2)Hex(7) | C[+57], N[+1541] | 1126.7893 | 4 | 4504.1356 | 4504.1268 | 0 | 2 | 127 | Specific | 808.14 | 808.14 | 808.14 | 16.55 | 40 | >sp\|P01876\|IGHA1_HUMAN Immunoglobulin heavy constant alpha 1 OS=Homo sapiens OX=9606 GN=IGHA1 PE=1 SV=2 | 255 | 20201114_16933_1ng_glyco_hcd203040_compound15.21167.21167.4 | controllerType=0 controllerNumber=1 scan=21167 | 42.4608 |
| 17904:4 | 11 | R.LSLHRPALEDLLLGSEAN[+1257.449]LTC[+57.021]TLTGLR.D | HexNAc(3)Hex(4) | C[+57], N[+1257] | 1056.0275 | 4 | 4221.0882 | 4221.0477 | 0 | 9.6 | 127 | Specific | 830.86 | 830.86 | 830.86 | 15.7 | 40 | >sp\|P01876\|IGHA1_HUMAN Immunoglobulin heavy constant alpha 1 OS=Homo sapiens OX=9606 GN=IGHA1 PE=1 SV=2 | 255 | 20201114_16933_1ng_glyco_hcd203040_compound15.21220.21220.4 | controllerType=0 controllerNumber=1 scan=21220 | 42.5856 |
| 17855:4 | 11 | R.LSLHRPALEDLLLGSEAN[+1663.608]LTC[+57.021]TLTGLR.D | HexNAc(5)Hex(4) | C[+57], N[+1664] | 1157.5605 | 4 | 4627.22 | 4627.2064 | 0 | 2.9 | 127 | Specific | 832.44 | 798.93 | 798.93 | 16.46 | 40 | >sp\|P01876\|IGHA1_HUMAN Immunoglobulin heavy constant alpha 1 OS=Homo sapiens OX=9606 GN=IGHA1 PE=1 SV=2 | 255 | 20201114_16933_1ng_glyco_hcd203040_compound15.21169.21169.4 | controllerType=0 controllerNumber=1 scan=21169 | 42.4743 |
| 17785:4 | 11 | R.LSLHRPALEDLLLGSEAN[+1864.634]LTC[+57.021]TLTGLR.D | HexNAc(2)Hex(9) | C[+57], N[+1865] | 1207.815 | 4 | 4828.2381 | 4828.2324 | 0 | 1.2 | 127 | Specific | 906.79 | 906.79 | 906.79 | 18.36 | 40 | >sp\|P01876\|IGHA1_HUMAN Immunoglobulin heavy constant alpha 1 OS=Homo sapiens OX=9606 GN=IGHA1 PE=1 SV=2 | 255 | 20201114_16933_1ng_glyco_hcd203040_compound15.21096.21096.4 | controllerType=0 controllerNumber=1 scan=21096 | 42.3045 |
| 17885:4 | 11 | R.LSLHRPALEDLLLGSEAN[+1216.423]LTC[+57.021]TLTGLR.D | HexNAc(2)Hex(5) | C[+57], N[+1216] | 1045.7631 | 4 | 4180.0306 | 4180.0211 | 0 | 2.3 | 127 | Specific | 1014.15 | 986.72 | 986.72 | 20.55 | 40 | >sp\|P01876\|IGHA1_HUMAN Immunoglobulin heavy constant alpha 1 OS=Homo sapiens OX=9606 GN=IGHA1 PE=1 SV=2 | 255 | 20201114_16933_1ng_glyco_hcd203040_compound15.21200.21200.4 | controllerType=0 controllerNumber=1 scan=21200 | 42.5481 |
| 17907:4 | 11 | R.LSLHRPALEDLLLGSEAN[+1257.449]LTC[+57.021]TLTGLR.D | HexNAc(3)Hex(4) | C[+57], N[+1257] | 1056.0275 | 4 | 4221.0882 | 4221.0477 | 0 | 9.6 | 127 | Specific | 1050.41 | 1046.62 | 1046.62 | 20.7 | 40 | >sp\|P01876\|IGHA1_HUMAN Immunoglobulin heavy constant alpha 1 OS=Homo sapiens OX=9606 GN=IGHA1 PE=1 SV=2 | 255 | 20201114_16933_1ng_glyco_hcd203040_compound15.21223.21223.4 | controllerType=0 controllerNumber=1 scan=21223 | [illegible] |
| 13277:3 | 11 | R.LAGKPTHVN[+1548.545]VSVVMAEVDGTC[+57.021]Y.- | HexNAc(3)Hex(4)NeuAc(1) | C[+57], N[+1549] | 1299.2405 | 3 | 3895.7071 | 3895.6869 | 0 | 5.2 | 332 | CRagged | 230 | 63.12 | 63.12 | 4.76 | 40 | >sp\|P01876\|IGHA1_HUMAN Immunoglobulin heavy constant alpha 1 OS=Homo sapiens OX=9606 GN=IGHA1 PE=1 SV=2 | 255 | 20201114_16933_1ng_glyco_hcd203040_compound15.16388.16388.3 | controllerType=0 controllerNumber=1 scan=16388 | [illegible] |
| 11403:3 | 11 | R.LAGKPTHVN[+1809.666]VSVVMAEVDGTC[+57.021]Y.- | HexNAc(5)Hex(4)Fuc(1) | C[+57], N[+1810] | 1386.2773 | 3 | 4156.8173 | 4156.8081 | 0 | 2.2 | 332 | CRagged | 313.29 | 100.11 | 100.11 | 5.23 | 40 | >sp\|P01876\|IGHA1_HUMAN Immunoglobulin heavy constant alpha 1 OS=Homo sapiens OX=9606 GN=IGHA1 PE=1 SV=2 | 255 | 20201114_16933_1ng_glyco_hcd203040_compound15.14420.14420.3 | controllerType=0 controllerNumber=1 scan=14420 | [illegible] |
| 11326:3 | 11 | R.LAGKPTHVN[+1702.581]VSVVMAEVDGTC[+57.021]Y.- | HexNAc(2)Hex(8) | C[+57], N[+1703] | 1350.5829 | 3 | 4049.7343 | 4049.7234 | 0 | 2.7 | 332 | CRagged | 315.04 | 268.39 | 268.39 | 7.38 | 40 | >sp\|P01876\|IGHA1_HUMAN Immunoglobulin heavy constant alpha 1 OS=Homo sapiens OX=9606 GN=IGHA1 PE=1 SV=2 | 255 | 20201114_16933_1ng_glyco_hcd203040_compound15.14340.14340.3 | controllerType=0 controllerNumber=1 scan=14340 | [illegible] |
| 11435:3 | 11 | R.LAGKPTHVN[+1501.555]VSVVMAEVDGTC[+57.021]Y.- | HexNAc(5)Hex(3) | C[+57], N[+1502] | 1283.5732 | 3 | 3848.705 | 3848.6974 | 0 | 2 | 332 | CRagged | 396.67 | 390.18 | 390.18 | 9.54 | 40 | >sp\|P01876\|IGHA1_HUMAN Immunoglobulin heavy constant alpha 1 OS=Homo sapiens OX=9606 GN=IGHA1 PE=1 SV=2 | 255 | 20201114_16933_1ng_glyco_hcd203040_compound15.14454.14454.3 | controllerType=0 controllerNumber=1 scan=14454 | 30.5665 |
| 11428:3 | 11 | R.LAGKPTHVN[+1647.613]VSVVMAEVDGTC[+57.021]Y.- | HexNAc(5)Hex(3)Fuc(1) | C[+57], N[+1648] | 1332.2588 | 3 | 3994.762 | 3994.7553 | 0 | 1.7 | 332 | CRagged | 526.11 | 526.11 | 526.11 | 10.05 | 40 | >sp\|P01876\|IGHA1_HUMAN Immunoglobulin heavy constant alpha 1 OS=Homo sapiens OX=9606 GN=IGHA1 PE=1 SV=2 | 255 | 20201114_16933_1ng_glyco_hcd203040_compound15.14447.14447.3 | controllerType=0 controllerNumber=1 scan=14447 | 30.5533 |
| 13118:3 | 11 | R.LAGKPTHVN[+2262.814]VSVVMAEVDGTC[+57.021]Y.- | HexNAc(5)Hex(5)Fuc(1)NeuAc(1) | C[+57], N[+2263] | 1537.3252 | 3 | 4609.961 | 4609.9564 | 0 | 1 | 332 | CRagged | 538.02 | 488.23 | 488.23 | 9.85 | 40 | >sp\|P01876\|IGHA1_HUMAN Immunoglobulin heavy constant alpha 1 OS=Homo sapiens OX=9606 GN=IGHA1 PE=1 SV=2 | 255 | 20201114_16933_1ng_glyco_hcd203040_compound15.16221.16221.3 | controllerType=0 controllerNumber=1 scan=16221 | 33.5841 |
| 11370:3 | 11 | R.LAGKPTHVN[+1378.476]VSVVMAEVDGTC[+57.021]Y.- | HexNAc(2)Hex(6) | C[+57], N[+1378] | 1242.5468 | 3 | 3725.6259 | 3725.6178 | 0 | 2.2 | 332 | CRagged | 596.68 | 596.68 | 435.09 | 11.38 | 40 | >sp\|P01876\|IGHA1_HUMAN Immunoglobulin heavy constant alpha 1 OS=Homo sapiens OX=9606 GN=IGHA1 PE=1 SV=2 | 255 | 20201114_16933_1ng_glyco_hcd203040_compound15.14386.14386.3 | controllerType=0 controllerNumber=1 scan=14386 | 30.4459 |
| 11401:3 | 11 | R.LAGKPTHVN[+1809.666]VSVVMAEVDGTC[+57.021]Y.- | HexNAc(5)Hex(4)Fuc(1) | C[+57], N[+1810] | 1386.2773 | 3 | 4156.8173 | 4156.8081 | 0 | 2.2 | 332 | CRagged | 691.74 | 691.74 | 441.13 | 13.38 | 40 | >sp\|P01876\|IGHA1_HUMAN Immunoglobulin heavy constant alpha 1 OS=Homo sapiens OX=9606 GN=IGHA1 PE=1 SV=2 | 255 | 20201114_16933_1ng_glyco_hcd203040_compound15.14418.14418.3 | controllerType=0 controllerNumber=1 scan=14418 | 30.5022 |
| 11430:3 | 11 | R.LAGKPTHVN[+1216.423]VSVVMAEVDGTC[+57.021]Y.- | HexNAc(2)Hex(5) | C[+57], N[+1216] | 1188.5289 | 3 | 3563.572 | 3563.5649 | 0 | 2 | 332 | CRagged | 693.72 | 693.72 | 449.78 | 13.38 | 40 | >sp\|P01876\|IGHA1_HUMAN Immunoglobulin heavy constant alpha 1 OS=Homo sapiens OX=9606 GN=IGHA1 PE=1 SV=2 | 255 | 20201114_16933_1ng_glyco_hcd203040_compound15.14449.14449.3 | controllerType=0 controllerNumber=1 scan=14449 | 30.5554 |
| 11350:3 | 11 | R.LAGKPTHVN[+1663.608]VSVVMAEVDGTC[+57.021]Y.- | HexNAc(5)Hex(4) | C[+57], N[+1664] | 1337.5916 | 3 | 4010.7603 | 4010.7502 | 0 | 2.5 | 332 | CRagged | 707.16 | 658.38 | 413.41 | 14.38 | 40 | >sp\|P01876\|IGHA1_HUMAN Immunoglobulin heavy constant alpha 1 OS=Homo sapiens OX=9606 GN=IGHA1 PE=1 SV=2 | 255 | 20201114_16933_1ng_glyco_hcd203040_compound15.14365.14365.3 | controllerType=0 controllerNumber=1 scan=14365 | 30.403 |
| 13116:3 | 11 | R.LAGKPTHVN[+2262.814]VSVVMAEVDGTC[+57.021]Y.- | HexNAc(5)Hex(5)Fuc(1)NeuAc(1) | C[+57], N[+2263] | 1537.3252 | 3 | 4609.961 | 4609.9564 | 0 | 1 | 332 | CRagged | 743.68 | 536.3 | 536.3 | 14.19 | 40 | >sp\|P01876\|IGHA1_HUMAN Immunoglobulin heavy constant alpha 1 OS=Homo sapiens OX=9606 GN=IGHA1 PE=1 SV=2 | 255 | 20201114_16933_1ng_glyco_hcd203040_compound15.16219.16219.3 | controllerType=0 controllerNumber=1 scan=16219 | 33.5803 |
| 11433:3 | 11 | R.LAGKPTHVN[+1501.555]VSVVMAEVDGTC[+57.021]Y.- | HexNAc(5)Hex(3) | C[+57], N[+1502] | 1283.5732 | 3 | 3848.705 | 3848.6974 | 0 | 2 | 332 | CRagged | 757.15 | 637.14 | 637.14 | 15.38 | 40 | >sp\|P01876\|IGHA1_HUMAN Immunoglobulin heavy constant alpha 1 OS=Homo sapiens OX=9606 GN=IGHA1 PE=1 SV=2 | 255 | 20201114_16933_1ng_glyco_hcd203040_compound15.14452.14452.3 | controllerType=0 controllerNumber=1 scan=14452 | 30.562 |
| 11347:3 | 11 | R.LAGKPTHVN[+1663.608]VSVVMAEVDGTC[+57.021]Y.- | HexNAc(5)Hex(4) | C[+57], N[+1664] | 1337.5916 | 3 | 4010.7603 | 4010.7502 | 0 | 2.5 | 332 | CRagged | 782.32 | 782.32 | 497.82 | 15.38 | 40 | >sp\|P01876\|IGHA1_HUMAN Immunoglobulin heavy constant alpha 1 OS=Homo sapiens OX=9606 GN=IGHA1 PE=1 SV=2 | 255 | 20201114_16933_1ng_glyco_hcd203040_compound15.14362.14362.3 | controllerType=0 controllerNumber=1 scan=14362 | 30.3959 |
| 11324:3 | 11 | R.LAGKPTHVN[+1702.581]VSVVMAEVDGTC[+57.021]Y.- | HexNAc(2)Hex(8) | C[+57], N[+1703] | 1350.5829 | 3 | 4049.7343 | 4049.7234 | 0 | 2.7 | 332 | CRagged | 812.02 | 812.02 | 522.96 | 16.38 | 40 | >sp\|P01876\|IGHA1_HUMAN Immunoglobulin heavy constant alpha 1 OS=Homo sapiens OX=9606 GN=IGHA1 PE=1 SV=2 | 255 | 20201114_16933_1ng_glyco_hcd203040_compound15.14338.14338.3 | controllerType=0 controllerNumber=1 scan=14338 | 30.3518 |
| 11431:3 | 11 | R.LAGKPTHVN[+1647.613]VSVVMAEVDGTC[+57.021]Y.- | HexNAc(5)Hex(3)Fuc(1) | C[+57], N[+1648] | 1332.2588 | 3 | 3994.762 | 3994.7553 | 0 | 1.7 | 332 | CRagged | 842.68 | 755.31 | 755.31 | 16.38 | 40 | >sp\|P01876\|IGHA1_HUMAN Immunoglobulin heavy constant alpha 1 OS=Homo sapiens OX=9606 GN=IGHA1 PE=1 SV=2 | 255 | 20201114_16933_1ng_glyco_hcd203040_compound15.14450.14450.3 | controllerType=0 controllerNumber=1 scan=14450 | 30.5571 |
| 11340:3 | 11 | R.LAGKPTHVN[+1663.608]VSVVMAEVDGTC[+57.021]Y.- | HexNAc(5)Hex(4) | C[+57], N[+1664] | 1337.5909 | 3 | 4010.7581 | 4010.7502 | 0 | 2 | 332 | CRagged | 844.21 | 844.21 | 536.96 | 16.38 | 40 | >sp\|P01876\|IGHA1_HUMAN Immunoglobulin heavy constant alpha 1 OS=Homo sapiens OX=9606 GN=IGHA1 PE=1 SV=2 | 255 | 20201114_16933_1ng_glyco_hcd203040_compound15.14354.14354.3 | controllerType=0 controllerNumber=1 scan=14354 | 30.3809 |
| 11338:3 | 11 | R.LAGKPTHVN[+1663.608]VSVVMAEVDGTC[+57.021]Y.- | HexNAc(5)Hex(4) | C[+57], N[+1664] | 1337.5909 | 3 | 4010.7581 | 4010.7502 | 0 | 2 | 332 | CRagged | 844.4 | 844.4 | 530.59 | 16.38 | 40 | >sp\|P01876\|IGHA1_HUMAN Immunoglobulin heavy constant alpha 1 OS=Homo sapiens OX=9606 GN=IGHA1 PE=1 SV=2 | 255 | 20201114_16933_1ng_glyco_hcd203040_compound15.14352.14352.3 | controllerType=0 controllerNumber=1 scan=14352 | 30.3761 |
| 11373:3 | 11 | R.LAGKPTHVN[+1378.476]VSVVMAEVDGTC[+57.021]Y.- | HexNAc(2)Hex(6) | C[+57], N[+1378] | 1242.5468 | 3 | 3725.6259 | 3725.6178 | 0 | 2.2 | 332 | CRagged | 849.07 | 801.17 | 517.36 | 16.38 | 40 | >sp\|P01876\|IGHA1_HUMAN Immunoglobulin heavy constant alpha 1 OS=Homo sapiens OX=9606 GN=IGHA1 PE=1 SV=2 | 255 | 20201114_16933_1ng_glyco_hcd203040_compound15.14389.14389.3 | controllerType=0 controllerNumber=1 scan=14389 | 30.4501 |
| 11432:3 | 11 | R.LAGKPTHVN[+1216.423]VSVVMAEVDGTC[+57.021]Y.- | HexNAc(2)Hex(5) | C[+57], N[+1216] | 1188.5289 | 3 | 3563.572 | 3563.5649 | 0 | 2 | 332 | CRagged | 899.25 | 856.74 | 570.49 | 17.38 | 40 | >sp\|P01876\|IGHA1_HUMAN Immunoglobulin heavy constant alpha 1 OS=Homo sapiens OX=9606 GN=IGHA1 PE=1 SV=2 | 255 | 20201114_16933_1ng_glyco_hcd203040_compound15.14451.14451.3 | controllerType=0 controllerNumber=1 scan=14451 | 30.5602 |
| 11382:3 | 11 | R.LAGKPTHVN[+1216.423]VSVVMAEVDGTC[+57.021]Y.- | HexNAc(2)Hex(5) | C[+57], N[+1216] | 1188.53 | 3 | 3563.5753 | 3563.5649 | 0 | 2.9 | 332 | CRagged | 926.27 | 874.44 | 536.6 | 18.29 | 40 | >sp\|P01876\|IGHA1_HUMAN Immunoglobulin heavy constant alpha 1 OS=Homo sapiens OX=9606 GN=IGHA1 PE=1 SV=2 | 255 | 20201114_16933_1ng_glyco_hcd203040_compound15.14398.14398.3 | controllerType=0 controllerNumber=1 scan=14398 | 30.4679 |
| 11380:3 | 11 | R.LAGKPTHVN[+1216.423]VSVVMAEVDGTC[+57.021]Y.- | HexNAc(2)Hex(5) | C[+57], N[+1216] | 1188.53 | 3 | 3563.5753 | 3563.5649 | 0 | 2.9 | 332 | CRagged | 938.07 | 938.07 | 556.72 | 18.29 | 40 | >sp\|P01876\|IGHA1_HUMAN Immunoglobulin heavy constant alpha 1 OS=Homo sapiens OX=9606 GN=IGHA1 PE=1 SV=2 | 255 | 20201114_16933_1ng_glyco_hcd203040_compound15.14396.14396.3 | controllerType=0 controllerNumber=1 scan=14396 | 30.461 |
| 07686:3 | 13 | R.LSPN[+2350.830]ASAEHLELR.W | HexNAc(4)Hex(5)Fuc(1)NeuAc(2) | N[+2351] | 1263.2005 | 3 | 3787.5869 | 3787.5796 | 0 | 1.9 | 52 | Specific | 619.71 | 520.23 | 520.23 | 11.97 | 33 | >sp\|Q13410\|BT1A1_HUMAN Butyrophilin subfamily 1 member A1 OS=Homo sapiens OX=9606 GN=BTN1A1 PE=1 SV=3 | 179 | 20201114_16933_1ng_glyco_hcd203040_compound15.10556.10556.3 | controllerType=0 controllerNumber=1 scan=10556 | 24.0058 |
| 07684:3 | 13 | R.LSPN[+2350.830]ASAEHLELR.W | HexNAc(4)Hex(5)Fuc(1)NeuAc(2) | N[+2351] | 1263.2005 | 3 | 3787.5869 | 3787.5796 | 0 | 1.9 | 52 | Specific | 653.48 | 628.21 | 628.21 | 12.97 | 33 | >sp\|Q13410\|BT1A1_HUMAN Butyrophilin subfamily 1 member A1 OS=Homo sapiens OX=9606 GN=BTN1A1 PE=1 SV=3 | 179 | 20201114_16933_1ng_glyco_hcd203040_compound15.10554.10554.3 | controllerType=0 controllerNumber=1 scan=10554 | 24.0017 |
| 15872:4 | 16 | K.LKELPGVC[+57.021]N[+2350.830]ETMMALWEEC[+57.021]KPC[+57.021]LK.Q | HexNAc(4)Hex(5)Fuc(1)NeuAc(2) | C[+57]*3, N[+2351] | 1322.5632 | 4 | 5287.2311 | 5287.2151 | 0 | 3 | 95 | Specific | 248.86 | 248.86 | 248.86 | 6.35 | 30 | >sp\|P10909\|CLUS_HUMAN Clusterin OS=Homo sapiens OX=9606 GN=CLU PE=1 SV=1 | 1467 | 20201114_16933_1ng_glyco_hcd203040_compound15.19080.19080.4 | controllerType=0 controllerNumber=1 scan=19080 | 38.5196 |
| 14113:3 | 16 | K.LKELPGVC[+57.021]N[+2059.735]ETMMALWEEC[+57.021]KPC[+57.021]LK.Q | HexNAc(4)Hex(5)Fuc(1)NeuAc(1) | C[+57]*3, N[+2060] | 1666.05 | 3 | 4996.1355 | 4996.1196 | 0 | 3.2 | 95 | Specific | 262.85 | 255.02 | 255.02 | 5.72 | 30 | >sp\|P10909\|CLUS_HUMAN Clusterin OS=Homo sapiens OX=9606 GN=CLU PE=1 SV=1 | 1467 | 20201114_16933_1ng_glyco_hcd203040_compound15.17254.17254.3 | controllerType=0 controllerNumber=1 scan=17254 | 35.3857 |
| 14092:3 | 16 | K.LKELPGVC[+57.021]N[+2059.735]ETMMALWEEC[+57.021]KPC[+57.021]LK.Q | HexNAc(4)Hex(5)Fuc(1)NeuAc(1) | C[+57]*3, N[+2060] | 1666.046 | 3 | 4996.1234 | 4996.1196 | 0 | 0.8 | 95 | Specific | 289.64 | 289.64 | 289.64 | 6.13 | 30 | >sp\|P10909\|CLUS_HUMAN Clusterin OS=Homo sapiens OX=9606 GN=CLU PE=1 SV=1 | 1467 | 20201114_16933_1ng_glyco_hcd203040_compound15.17232.17232.3 | controllerType=0 controllerNumber=1 scan=17232 | 35.344 |
| 14076:4 | 16 | K.LKELPGVC[+57.021]N[+2205.793]ETMMALWEEC[+57.021]KPC[+57.021]LK.Q | HexNAc(4)Hex(5)Fuc(2)NeuAc(1) | C[+57]*3, N[+2206] | 1286.3036 | 4 | 5142.1925 | 5142.1776 | 0 | 2.9 | 95 | Specific | 310.95 | 310.95 | 310.95 | 6.95 | 30 | >sp\|P10909\|CLUS_HUMAN Clusterin OS=Homo sapiens OX=9606 GN=CLU PE=1 SV=1 | 1467 | 20201114_16933_1ng_glyco_hcd203040_compound15.17216.17216.4 | controllerType=0 controllerNumber=1 scan=17216 | 35.3071 |
| 15870:4 | 16 | K.LKELPGVC[+57.021]N[+2350.830]ETMMALWEEC[+57.021]KPC[+57.021]LK.Q | HexNAc(4)Hex(5)Fuc(1)NeuAc(2) | C[+57]*3, N[+2351] | 1322.5632 | 4 | 5287.2311 | 5287.2151 | 0 | 3 | 95 | Specific | 320.08 | 320.08 | 320.08 | 7.29 | 30 | >sp\|P10909\|CLUS_HUMAN Clusterin OS=Homo sapiens OX=9606 GN=CLU PE=1 SV=1 | 1467 | 20201114_16933_1ng_glyco_hcd203040_compound15.19078.19078.4 | controllerType=0 controllerNumber=1 scan=19078 | 38.5161 |
| 14111:3 | 16 | K.LKELPGVC[+57.021]N[+2059.735]ETMMALWEEC[+57.021]KPC[+57.021]LK.Q | HexNAc(4)Hex(5)Fuc(1)NeuAc(1) | C[+57]*3, N[+2060] | 1666.05 | 3 | 4996.1355 | 4996.1196 | 0 | 3.2 | 95 | Specific | 493.89 | 493.89 | 493.89 | 8.78 | 30 | >sp\|P10909\|CLUS_HUMAN Clusterin OS=Homo sapiens OX=9606 GN=CLU PE=1 SV=1 | 1467 | 20201114_16933_1ng_glyco_hcd203040_compound15.17252.17252.3 | controllerType=0 controllerNumber=1 scan=17252 | 35.3821 |
| 14078:4 | 16 | K.LKELPGVC[+57.021]N[+2205.793]ETMMALWEEC[+57.021]KPC[+57.021]LK.Q | HexNAc(4)Hex(5)Fuc(2)NeuAc(1) | C[+57]*3, N[+2206] | 1286.3036 | 4 | 5142.1925 | 5142.1776 | 0 | 2.9 | 95 | Specific | 575.77 | 575.77 | 575.77 | 10.96 | 30 | >sp\|P10909\|CLUS_HUMAN Clusterin OS=Homo sapiens OX=9606 GN=CLU PE=1 SV=1 | 1467 | 20201114_16933_1ng_glyco_hcd203040_compound15.17218.17218.4 | controllerType=0 controllerNumber=1 scan=17218 | 35.3112 |
| 14064:4 | 16 | K.LKELPGVC[+57.021]N[+2059.735]ETMMALWEEC[+57.021]KPC[+57.021]LK.Q | HexNAc(4)Hex(5)Fuc(1)NeuAc(1) | C[+57]*3, N[+2060] | 1249.7892 | 4 | 4996.1349 | 4996.1196 | 0 | 3.1 | 95 | Specific | 787.84 | 756.07 | 756.07 | 14.86 | 30 | >sp\|P10909\|CLUS_HUMAN Clusterin OS=Homo sapiens OX=9606 GN=CLU PE=1 SV=1 | 1467 | 20201114_16933_1ng_glyco_hcd203040_compound15.17203.17203.4 | controllerType=0 controllerNumber=1 scan=17203 | 35.2835 |
| 14062:4 | 16 | K.LKELPGVC[+57.021]N[+2059.735]ETMMALWEEC[+57.021]KPC[+57.021]LK.Q | HexNAc(4)Hex(5)Fuc(1)NeuAc(1) | C[+57]*3, N[+2060] | 1249.7892 | 4 | 4996.1349 | 4996.1196 | 0 | 3.1 | 95 | Specific | 924.67 | 817.65 | 817.65 | 17.86 | 30 | >sp\|P10909\|CLUS_HUMAN Clusterin OS=Homo sapiens OX=9606 GN=CLU PE=1 SV=1 | 1467 | 20201114_16933_1ng_glyco_hcd203040_compound15.17201.17201.4 | controllerType=0 controllerNumber=1 scan=17201 | 35.2789 |
| 21107:3 | 16 | K.MLN[+2059.735]TSSLLEQLNEQFNWVSR.L | HexNAc(4)Hex(5)Fuc(1)NeuAc(1) | N[+2060] | 1490.3155 | 3 | 4468.9319 | 4468.9217 | 0 | 2.3 | 352 | Specific | 736.53 | 524.34 | 524.34 | 14.39 | 30 | >sp\|P10909\|CLUS_HUMAN Clusterin OS=Homo sapiens OX=9606 GN=CLU PE=1 SV=1 | 1467 | 20201114_16933_1ng_glyco_hcd203040_compound15.24575.24575.3 | controllerType=0 controllerNumber=1 scan=24575 | 49.7339 |
| 21105:3 | 16 | K.MLN[+2059.735]TSSLLEQLNEQFNWVSR.L | HexNAc(4)Hex(5)Fuc(1)NeuAc(1) | N[+2060] | 1490.3155 | 3 | 4468.9319 | 4468.9217 | 0 | 2.3 | 352 | Specific | 1023.51 | 789.29 | 789.29 | 20.39 | 30 | >sp\|P10909\|CLUS_HUMAN Clusterin OS=Homo sapiens OX=9606 GN=CLU PE=1 SV=1 | 1467 | 20201114_16933_1ng_glyco_hcd203040_compound15.24573.24573.3 | controllerType=0 controllerNumber=1 scan=24573 | 49.7287 |
| 20640:3 | 16 | K.M[+15.995]LN[+2059.735]TSSLLEQLNEQFNWVSR.L | HexNAc(4)Hex(5)Fuc(1)NeuAc(1) | M[+16], N[+2060] | 1495.6478 | 3 | 4484.9287 | 4484.9166 | 0 | 2.7 | 352 | Specific | 264.83 | 264.83 | 264.83 | 6.91 | 30 | >sp\|P10909\|CLUS_HUMAN Clusterin OS=Homo sapiens OX=9606 GN=CLU PE=1 SV=1 | 1467 | 20201114_16933_1ng_glyco_hcd203040_compound15.24087.24087.3 | controllerType=0 controllerNumber=1 scan=24087 | 48.6639 |
| 10351:3 | 16 | R.LAN[+2205.793]LTQGEDQYYLR.V | HexNAc(4)Hex(5)Fuc(2)NeuAc(1) | N[+2206] | 1297.2155 | 3 | 3889.632 | 3889.6265 | 0 | 1.4 | 372 | Specific | 281.64 | 281.64 | 281.64 | 6.56 | 30 | >sp\|P10909\|CLUS_HUMAN Clusterin OS=Homo sapiens OX=9606 GN=CLU PE=1 SV=1 | 1467 | 20201114_16933_1ng_glyco_hcd203040_compound15.13331.13331.3 | controllerType=0 controllerNumber=1 scan=13331 | 28.6715 |
| 10354:3 | 16 | R.LAN[+2205.793]LTQGEDQYYLR.V | HexNAc(4)Hex(5)Fuc(2)NeuAc(1) | N[+2206] | 1297.2155 | 3 | 3889.632 | 3889.6265 | 0 | 1.4 | 372 | Specific | 478.34 | 434.36 | 434.36 | 9.44 | 30 | >sp\|P10909\|CLUS_HUMAN Clusterin OS=Homo sapiens OX=9606 GN=CLU PE=1 SV=1 | 1467 | 20201114_16933_1ng_glyco_hcd203040_compound15.13334.13334.3 | controllerType=0 controllerNumber=1 scan=13334 | 28.6757 |
| 24265:3 | 19 | R.QLAHQSN[+2204.772]STNIFFSPVSIATAFAMLSLGTK.A | HexNAc(4)Hex(5)NeuAc(2) | N[+2205] | 1796.1445 | 3 | 5386.4189 | 5386.4075 | 0 | 2.1 | 64 | Specific | 569.09 | 568.46 | 497.83 | 10.23 | 29 | >sp\|P01009\|A1AT_HUMAN Alpha-1-antitrypsin OS=Homo sapiens OX=9606 GN=SERPINA1 PE=1 SV=3 | 642 | 20201114_16933_1ng_glyco_hcd203040_compound15.27887.27887.3 | controllerType=0 controllerNumber=1 scan=27887 | 55.8022 |
| 24268:3 | 19 | R.QLAHQSN[+2204.772]STNIFFSPVSIATAFAMLSLGTK.A | HexNAc(4)Hex(5)NeuAc(2) | N[+2205] | 1796.1445 | 3 | 5386.4189 | 5386.4075 | 0 | 2.1 | 64 | Specific | 778.96 | 702.54 | 468.5 | 15.38 | 29 | >sp\|P01009\|A1AT_HUMAN Alpha-1-antitrypsin OS=Homo sapiens OX=9606 GN=SERPINA1 PE=1 SV=3 | 642 | 20201114_16933_1ng_glyco_hcd203040_compound15.27890.27890.3 | controllerType=0 controllerNumber=1 scan=27890 | 55.8071 |
| 24257:4 | 19 | R.QLAHQSN[+2204.772]STNIFFSPVSIATAFAMLSLGTK.A | HexNAc(4)Hex(5)NeuAc(2) | N[+2205] | 1347.3602 | 4 | 5386.4191 | 5386.4075 | 0 | 2.2 | 64 | Specific | 875.14 | 806.83 | 598.74 | 17.39 | 29 | >sp\|P01009\|A1AT_HUMAN Alpha-1-antitrypsin OS=Homo sapiens OX=9606 GN=SERPINA1 PE=1 SV=3 | 642 | 20201114_16933_1ng_glyco_hcd203040_compound15.27879.27879.4 | controllerType=0 controllerNumber=1 scan=27879 | 55.7865 |
| 24259:4 | 19 | R.QLAHQSN[+2204.772]STNIFFSPVSIATAFAMLSLGTK.A | HexNAc(4)Hex(5)NeuAc(2) | N[+2205] | 1347.3602 | 4 | 5386.4191 | 5386.4075 | 0 | 2.2 | 64 | Specific | 953.48 | 841.34 | 609.02 | 19.39 | 29 | >sp\|P01009\|A1AT_HUMAN Alpha-1-antitrypsin OS=Homo sapiens OX=9606 GN=SERPINA1 PE=1 SV=3 | 642 | 20201114_16933_1ng_glyco_hcd203040_compound15.27881.27881.4 | controllerType=0 controllerNumber=1 scan=27881 | 55.7926 |
| 20186:4 | 19 | K.YLGN[+2204.772]ATAIFFLPDEGKLQHLENELTHDIITK.F | HexNAc(4)Hex(5)NeuAc(2) | N[+2205] | 1437.1594 | 4 | 5745.6159 | 5745.6098 | 0 | 1.1 | 268 | Specific | 717.54 | 625.55 | 625.55 | 12.96 | 29 | >sp\|P01009\|A1AT_HUMAN Alpha-1-antitrypsin OS=Homo sapiens OX=9606 GN=SERPINA1 PE=1 SV=3 | 642 | 20201114_16933_1ng_glyco_hcd203040_compound15.23608.23608.4 | controllerType=0 controllerNumber=1 scan=23608 | 47.6612 |
| 20184:4 | 19 | K.YLGN[+2204.772]ATAIFFLPDEGKLQHLENELTHDIITK.F | HexNAc(4)Hex(5)NeuAc(2) | N[+2205] | 1437.1594 | 4 | 5745.6159 | 5745.6098 | 0 | 1.1 | 268 | Specific | 1250.27 | 1146.72 | 1146.72 | 23.96 | 29 | >sp\|P01009\|A1AT_HUMAN Alpha-1-antitrypsin OS=Homo sapiens OX=9606 GN=SERPINA1 PE=1 SV=3 | 642 | 20201114_16933_1ng_glyco_hcd203040_compound15.23606.23606.4 | controllerType=0 controllerNumber=1 scan=23606 | 47.6545 |
| 09502:4 | 23 | K.QTDEIKDTRN[+2059.735]ESTQNC[+57.021]VVAEPEKMESSISSSSEEMSLSK.C | HexNAc(4)Hex(5)Fuc(1)NeuAc(1) | C[+57], N[+2060] | 1613.4362 | 4 | 6450.7228 | 6450.7156 | 0 | 1.1 | 60 | Specific | 305.29 | 248.95 | 248.95 | 5.71 | 26 | >sp\|P47710\|CASA1_HUMAN Alpha-S1-casein OS=Homo sapiens OX=9606 GN=CSN1S1 PE=1 SV=1 | 989 | 20201114_16933_1ng_glyco_hcd203040_compound15.12449.12449.2 | controllerType=0 controllerNumber=1 scan=12449 | 27.153 |
| 09515:4 | 23 | K.QTDEIKDTRN[+2059.735]ESTQNC[+57.021]VVAEPEKMESSISSSSEEMSLSK.C | HexNAc(4)Hex(5)Fuc(1)NeuAc(1) | C[+57], N[+2060] | 1613.436 | 4 | 6450.7223 | 6450.7156 | 0 | 1 | 60 | Specific | 400.56 | 303.8 | 303.8 | 6.56 | 26 | >sp\|P47710\|CASA1_HUMAN Alpha-S1-casein OS=Homo sapiens OX=9606 GN=CSN1S1 PE=1 SV=1 | 989 | 20201114_16933_1ng_glyco_hcd203040_compound15.12463.12463.4 | controllerType=0 controllerNumber=1 scan=12463 | 27.1776 |
| 09504:4 | 23 | K.QTDEIKDTRN[+2059.735]ESTQNC[+57.021]VVAEPEKMESSISSSSEEMSLSK.C | HexNAc(4)Hex(5)Fuc(1)NeuAc(1) | C[+57], N[+2060] | 1613.4362 | 4 | 6450.7228 | 6450.7156 | 0 | 1.1 | 60 | Specific | 476.26 | 319.94 | 319.94 | 6.95 | 26 | >sp\|P47710\|CASA1_HUMAN Alpha-S1-casein OS=Homo sapiens OX=9606 GN=CSN1S1 PE=1 SV=1 | 989 | 20201114_16933_1ng_glyco_hcd203040_compound15.12451.12451.2 | controllerType=0 controllerNumber=1 scan=12451 | 27.1566 |
| 09517:4 | 23 | K.QTDEIKDTRN[+2059.735]ESTQNC[+57.021]VVAEPEKMESSISSSSEEMSLSK.C | HexNAc(4)Hex(5)Fuc(1)NeuAc(1) | C[+57], N[+2060] | 1613.436 | 4 | 6450.7223 | 6450.7156 | 0 | 1 | 60 | Specific | 602.74 | 371.27 | 371.27 | 9.7 | 26 | >sp\|P47710\|CASA1_HUMAN Alpha-S1-casein OS=Homo sapiens OX=9606 GN=CSN1S1 PE=1 SV=1 | 989 | 20201114_16933_1ng_glyco_hcd203040_compound15.12465.12465.4 | controllerType=0 controllerNumber=1 scan=12465 | 27.1816 |
| 23072:4 | 25 | K.APDKNVIFSPLSISTALAFLSLGAHN[+2350.830]TTLTEILK.G | HexNAc(4)Hex(5)Fuc(1)NeuAc(2) | N[+2351] | 1484.2157 | 4 | 5933.841 | 5933.8085 | 0 | 5.5 | 68 | Specific | 410.32 | 410.32 | 410.32 | 8.75 | 24 | >sp\|P01011\|AACT_HUMAN Alpha-1-antichymotrypsin OS=Homo sapiens OX=9606 GN=SERPINA3 PE=1 SV=2 | 1314 | 20201114_16933_1ng_glyco_hcd203040_compound15.26644.26644.4 | controllerType=0 controllerNumber=1 scan=26644 | 53.5269 |
| 23070:4 | 25 | K.APDKNVIFSPLSISTALAFLSLGAHN[+2350.830]TTLTEILK.G | HexNAc(4)Hex(5)Fuc(1)NeuAc(2) | N[+2351] | 1484.2157 | 4 | 5933.841 | 5933.8085 | 0 | 5.5 | 68 | Specific | 468.28 | 468.28 | 468.28 | 9.02 | 24 | >sp\|P01011\|AACT_HUMAN Alpha-1-antichymotrypsin OS=Homo sapiens OX=9606 GN=SERPINA3 PE=1 SV=2 | 1314 | 20201114_16933_1ng_glyco_hcd203040_compound15.26642.26642.4 | controllerType=0 controllerNumber=1 scan=26642 | 53.523 |
| 08546:5 | 26 | K.SC[+57.021]QHN[+2205.793]GTMYQHGEIFSAHELFPSR.L | HexNAc(4)Hex(5)Fuc(2)NeuAc(1) | C[+57], N[+2206] | 1008.6243 | 5 | 5039.0924 | 5039.0498 | 0 | 8.5 | 110 | Specific | 311.33 | 311.33 | 311.33 | 6.84 | 19 | >sp\|Q6WN34\|CRDL2_HUMAN Chordin-like protein 2 OS=Homo sapiens OX=9606 GN=CHRDL2 PE=1 SV=1 | 1189 | 20201114_16933_1ng_glyco_hcd203040_compound15.11449.11449.5 | controllerType=0 controllerNumber=1 scan=11449 | 25.469 |
| 08549:4 | 26 | K.SC[+57.021]QHN[+2205.793]GTMYQHGEIFSAHELFPSR.L | HexNAc(4)Hex(5)Fuc(2)NeuAc(1) | C[+57], N[+2206] | 1260.5286 | 4 | 5039.0926 | 5039.0498 | 0 | 8.5 | 110 | Specific | 468.66 | 468.66 | 468.66 | 8.63 | 19 | >sp\|Q6WN34\|CRDL2_HUMAN Chordin-like protein 2 OS=Homo sapiens OX=9606 GN=CHRDL2 PE=1 SV=1 | 1189 | 20201114_16933_1ng_glyco_hcd203040_compound15.11452.11452.4 | controllerType=0 controllerNumber=1 scan=11452 | 25.4749 |
| 01563:3 | 29 | K.HYTN[+1647.613]SSQDVTVPC[+57.021]R.V | HexNAc(5)Hex(3)Fuc(1) | C[+57], N[+1648] | 1104.4622 | 3 | 3311.372 | 3311.3625 | 0 | 2.9 | 89 | Specific | 297.17 | 280.52 | 280.52 | 6.82 | 20 | >sp\|P01877\|IGHA2_HUMAN Immunoglobulin heavy constant alpha 2 OS=Homo sapiens OX=9606 GN=IGHA2 PE=1 SV=4 | 576 | 20201114_16933_1ng_glyco_hcd203040_compound15.4190.4190.2 | controllerType=0 controllerNumber=1 scan=4190 | 12.9641 |
| 01561:3 | 29 | K.HYTN[+1647.613]SSQDVTVPC[+57.021]R.V | HexNAc(5)Hex(3)Fuc(1) | C[+57], N[+1648] | 1104.4622 | 3 | 3311.372 | 3311.3625 | 0 | 2.9 | 89 | Specific | 342.94 | 263.27 | 263.27 | 7.26 | 20 | >sp\|P01877\|IGHA2_HUMAN Immunoglobulin heavy constant alpha 2 OS=Homo sapiens OX=9606 GN=IGHA2 PE=1 SV=4 | 576 | 20201114_16933_1ng_glyco_hcd203040_compound15.4188.4188.2 | controllerType=0 controllerNumber=1 scan=4188 | 12.9592 |
| 01524:3 | 29 | K.HYTN[+1809.666]SSQDVTVPC[+57.021]R.V | HexNAc(5)Hex(4)Fuc(1) | C[+57], N[+1810] | 1158.4793 | 3 | 3473.4234 | 3473.4154 | 0 | 2.3 | 89 | Specific | 369.49 | 255.75 | 255.75 | 7.32 | 20 | >sp\|P01877\|IGHA2_HUMAN Immunoglobulin heavy constant alpha 2 OS=Homo sapiens OX=9606 GN=IGHA2 PE=1 SV=4 | 576 | 20201114_16933_1ng_glyco_hcd203040_compound15.4150.4150.3 | controllerType=0 controllerNumber=1 scan=4150 | 12.8917 |
| 01522:3 | 29 | K.HYTN[+1809.666]SSQDVTVPC[+57.021]R.V | HexNAc(5)Hex(4)Fuc(1) | C[+57], N[+1810] | 1158.4793 | 3 | 3473.4234 | 3473.4154 | 0 | 2.3 | 89 | Specific | 434.82 | 320.84 | 320.84 | 8.06 | 20 | >sp\|P01877\|IGHA2_HUMAN Immunoglobulin heavy constant alpha 2 OS=Homo sapiens OX=9606 GN=IGHA2 PE=1 SV=4 | 576 | 20201114_16933_1ng_glyco_hcd203040_compound15.4148.4148.3 | controllerType=0 controllerNumber=1 scan=4148 | 12.8877 |
| 01508:3 | 29 | K.HYTN[+1647.613]SSQDVTVPC[+57.021]R.V | HexNAc(5)Hex(3)Fuc(1) | C[+57], N[+1648] | 1104.4617 | 3 | 3311.3705 | 3311.3625 | 0 | 2.4 | 89 | Specific | 475.78 | 424.32 | 424.32 | 9.63 | 20 | >sp\|P01877\|IGHA2_HUMAN Immunoglobulin heavy constant alpha 2 OS=Homo sapiens OX=9606 GN=IGHA2 PE=1 SV=4 | 576 | 20201114_16933_1ng_glyco_hcd203040_compound15.4133.4133.2 | controllerType=0 controllerNumber=1 scan=4133 | 12.8535 |
| 01496:3 | 29 | K.HYTN[+1809.666]SSQDVTVPC[+57.021]R.V | HexNAc(5)Hex(4)Fuc(1) | C[+57], N[+1810] | 1158.4791 | 3 | 3473.4226 | 3473.4154 | 0 | 2.1 | 89 | Specific | 491.14 | 356.31 | 356.31 | 8.4 | 20 | >sp\|P01877\|IGHA2_HUMAN Immunoglobulin heavy constant alpha 2 OS=Homo sapiens OX=9606 GN=IGHA2 PE=1 SV=4 | 576 | 20201114_16933_1ng_glyco_hcd203040_compound15.4121.4121.3 | controllerType=0 controllerNumber=1 scan=4121 | 12.8305 |
| 01510:3 | 29 | K.HYTN[+1647.613]SSQDVTVPC[+57.021]R.V | HexNAc(5)Hex(3)Fuc(1) | C[+57], N[+1648] | 1104.4617 | 3 | 3311.3705 | 3311.3625 | 0 | 2.4 | 89 | Specific | 518.48 | 308.57 | 308.57 | 8.26 | 20 | >sp\|P01877\|IGHA2_HUMAN Immunoglobulin heavy constant alpha 2 OS=Homo sapiens OX=9606 GN=IGHA2 PE=1 SV=4 | 576 | 20201114_16933_1ng_glyco_hcd203040_compound15.4135.4135.2 | controllerType=0 controllerNumber=1 scan=4135 | 12.8612 |
| 01498:3 | 29 | K.HYTN[+1809.666]SSQDVTVPC[+57.021]R.V | HexNAc(5)Hex(4)Fuc(1) | C[+57], N[+1810] | 1158.4791 | 3 | 3473.4226 | 3473.4154 | 0 | 2.1 | 89 | Specific | 543.91 | 346.82 | 346.82 | 8.71 | 20 | >sp\|P01877\|IGHA2_HUMAN Immunoglobulin heavy constant alpha 2 OS=Homo sapiens OX=9606 GN=IGHA2 PE=1 SV=4 | 576 | 20201114_16933_1ng_glyco_hcd203040_compound15.4123.4123.3 | controllerType=0 controllerNumber=1 scan=4123 | 12.8361 |
| 01489:3 | 29 | K.HYTN[+1647.613]SSQDVTVPC[+57.021]R.V | HexNAc(5)Hex(3)Fuc(1) | C[+57], N[+1648] | 1104.4619 | 3 | 3311.3713 | 3311.3625 | 0 | 2.6 | 89 | Specific | 627.27 | 492.25 | 492.25 | 12.24 | 20 | >sp\|P01877\|IGHA2_HUMAN Immunoglobulin heavy constant alpha 2 OS=Homo sapiens OX=9606 GN=IGHA2 PE=1 SV=4 | 576 | 20201114_16933_1ng_glyco_hcd203040_compound15.4113.4113.3 | controllerType=0 controllerNumber=1 scan=4113 | 12.8047 |
| 01491:3 | 29 | K.HYTN[+1647.613]SSQDVTVPC[+57.021]R.V | HexNAc(5)Hex(3)Fuc(1) | C[+57], N[+1648] | 1104.4619 | 3 | 3311.3713 | 3311.3625 | 0 | 2.6 | 89 | Specific | 683.62 | 575.6 | 575.6 | 13.24 | 20 | >sp\|P01877\|IGHA2_HUMAN Immunoglobulin heavy constant alpha 2 OS=Homo sapiens OX=9606 GN=IGHA2 PE=1 SV=4 | 576 | 20201114_16933_1ng_glyco_hcd203040_compound15.4115.4115.3 | controllerType=0 controllerNumber=1 scan=4115 | 12.8184 |
| 01519:3 | 29 | K.HYTN[+1647.613]SSQDVTVPC[+57.021]R.V | HexNAc(5)Hex(3)Fuc(1) | C[+57], N[+1648] | 1104.4615 | 3 | 3311.3698 | 3311.3625 | 0 | 2.2 | 89 | Specific | 762.43 | 762.43 | 762.43 | 15.24 | 20 | >sp\|P01877\|IGHA2_HUMAN Immunoglobulin heavy constant alpha 2 OS=Homo sapiens OX=9606 GN=IGHA2 PE=1 SV=4 | 576 | 20201114_16933_1ng_glyco_hcd203040_compound15.4145.4145.3 | controllerType=0 controllerNumber=1 scan=4145 | 12.8801 |
| 01521:3 | 29 | K.HYTN[+1647.613]SSQDVTVPC[+57.021]R.V | HexNAc(5)Hex(3)Fuc(1) | C[+57], N[+1648] | 1104.4615 | 3 | 3311.3698 | 3311.3625 | 0 | 2.2 | 89 | Specific | 780.17 | 592.94 | 592.94 | 15.24 | 20 | >sp\|P01877\|IGHA2_HUMAN Immunoglobulin heavy constant alpha 2 OS=Homo sapiens OX=9606 GN=IGHA2 PE=1 SV=4 | 576 | 20201114_16933_1ng_glyco_hcd203040_compound15.4147.4147.3 | controllerType=0 controllerNumber=1 scan=4147 | 12.8858 |
| 03482:5 | 29 | K.HYTN[+1647.613]SSQDVTVPC[+57.021]RVPPPPPC[+57.021]C[+57.021]HPR.L | HexNAc(5)Hex(3)Fuc(1) | C[+57]*3, N[+1648] | 921.9997 | 5 | 4605.9695 | 4605.9689 | 0 | 0.1 | 89 | Specific | 339.21 | 306.16 | 306.16 | 7.06 | 20 | >sp\|P01877\|IGHA2_HUMAN Immunoglobulin heavy constant alpha 2 OS=Homo sapiens OX=9606 GN=IGHA2 PE=1 SV=4 | 576 | 20201114_16933_1ng_glyco_hcd203040_compound15.6185.6185.5 | controllerType=0 controllerNumber=1 scan=6185 | 16.5352 |
| 03480:5 | 29 | K.HYTN[+1647.613]SSQDVTVPC[+57.021]RVPPPPPC[+57.021]C[+57.021]HPR.L | HexNAc(5)Hex(3)Fuc(1) | C[+57]*3, N[+1648] | 921.9997 | 5 | 4605.9695 | 4605.9689 | 0 | 0.1 | 89 | Specific | 349.69 | 162.63 | 162.63 | 5.91 | 20 | >sp\|P01877\|IGHA2_HUMAN Immunoglobulin heavy constant alpha 2 OS=Homo sapiens OX=9606 GN=IGHA2 PE=1 SV=4 | 576 | 20201114_16933_1ng_glyco_hcd203040_compound15.6183.6183.5 | controllerType=0 controllerNumber=1 scan=6183 | 16.5318 |
| 03182:2 | 29 | K.TPLTAN[+1444.534]ITK.S | HexNAc(4)Hex(3)Fuc(1) | N[+1445] | 1202.0524 | 2 | 2403.0975 | 2403.0906 | 0 | 2.9 | 200 | Specific | 219.97 | 33.31 | 33.31 | 3.24 | 20 | >sp\|P01877\|IGHA2_HUMAN Immunoglobulin heavy constant alpha 2 OS=Homo sapiens OX=9606 GN=IGHA2 PE=1 SV=4 | 576 | 20201114_16933_1ng_glyco_hcd203040_compound15.5872.5872.2 | controllerType=0 controllerNumber=1 scan=5872 | 15.9668 |
| 03231:2 | 29 | K.TPLTAN[+1501.555]ITK.S | HexNAc(5)Hex(3) | N[+1502] | 1230.5647 | 2 | 2460.1222 | 2460.1121 | 0 | 4.1 | 200 | Specific | 243.37 | 101.85 | 101.85 | 4.56 | 20 | >sp\|P01877\|IGHA2_HUMAN Immunoglobulin heavy constant alpha 2 OS=Homo sapiens OX=9606 GN=IGHA2 PE=1 SV=4 | 576 | 20201114_16933_1ng_glyco_hcd203040_compound15.5923.5923.2 | controllerType=0 controllerNumber=1 scan=5923 | 16.0588 |
| 03251:3 | 29 | K.TPLTAN[+1647.613]ITK.S | HexNAc(5)Hex(3)Fuc(1) | N[+1648] | 869.3981 | 3 | 2606.1797 | 2606.17 | 0 | 3.7 | 200 | Specific | 312.62 | 277.8 | 277.8 | 6.56 | 20 | >sp\|P01877\|IGHA2_HUMAN Immunoglobulin heavy constant alpha 2 OS=Homo sapiens OX=9606 GN=IGHA2 PE=1 SV=4 | 576 | 20201114_16933_1ng_glyco_hcd203040_compound15.5944.5944.3 | controllerType=0 controllerNumber=1 scan=5944 | 16.0979 |
| 03184:2 | 29 | K.TPLTAN[+1444.534]ITK.S | HexNAc(4)Hex(3)Fuc(1) | N[+1445] | 1202.0524 | 2 | 2403.0975 | 2403.0906 | 0 | 2.9 | 200 | Specific | 354.81 | 169.85 | 169.85 | 6.02 | 20 | >sp\|P01877\|IGHA2_HUMAN Immunoglobulin heavy constant alpha 2 OS=Homo sapiens OX=9606 GN=IGHA2 PE=1 SV=4 | 576 | 20201114_16933_1ng_glyco_hcd203040_compound15.5874.5874.2 | controllerType=0 controllerNumber=1 scan=5874 | 15.9714 |
| 03176:2 | 29 | K.TPLTAN[+1809.666]ITK.S | HexNAc(5)Hex(4)Fuc(1) | N[+1810] | 1384.6176 | 2 | 2768.2279 | 2768.2228 | 0 | 1.8 | 200 | Specific | 425.81 | 217.98 | 217.98 | 6.92 | 20 | >sp\|P01877\|IGHA2_HUMAN Immunoglobulin heavy constant alpha 2 OS=Homo sapiens OX=9606 GN=IGHA2 PE=1 SV=4 | 576 | 20201114_16933_1ng_glyco_hcd203040_compound15.5866.5866.2 | controllerType=0 controllerNumber=1 scan=5866 | 15.9561 |
| 03210:2 | 29 | K.TPLTAN[+1647.613]ITK.S | HexNAc(5)Hex(3)Fuc(1) | N[+1648] | 1303.5931 | 2 | 2606.1788 | 2606.17 | 0 | 3.4 | 200 | Specific | 450.14 | 258.95 | 258.95 | 7.26 | 20 | >sp\|P01877\|IGHA2_HUMAN Immunoglobulin heavy constant alpha 2 OS=Homo sapiens OX=9606 GN=IGHA2 PE=1 SV=4 | 576 | 20201114_16933_1ng_glyco_hcd203040_compound15.5902.5902.2 | controllerType=0 controllerNumber=1 scan=5902 | 16.0239 |

Table S4. Cont.

| Query #:z | Protein Rank | Glycopeptide | Glycan | Modification Type(s) | Observed m/z | z | Observed (M+H) | Calc. mass (M+H) | Off-by-x error | Mass error (ppm) | StartPosition | Cleavage | Score | Delta | Delta Mod | \|Log Prob\| | # of unique peptides | Protein Name | Protein DB number | Comment | Scan # | Scan Time |
|---|---|---|---|---|---|---|---|---|---|---|---|---|---|---|---|---|---|---|---|---|---|---|
| 03212:2 | 29 | K.TPLTAN[+1647.613]ITK.S | HexNAc(5)Hex(3)Fuc(1) | N[+1648] | 1303.5931 | 2 | 2606.1788 | 2606.17 | 0 | 3.4 | 200 | Specific | 485.65 | 264.46 | 264.46 | 7.32 | 20 | >sp\|P01877\|IGHA2_HUMAN Immunoglobulin heavy constant alpha 2 OS=Homo sapiens OX=9606 GN=IGHA2 PE=1 SV=4 | 576 | 20201114_16933_1ng_glyco_hcd203040_compound15.5904.5904.2 | controllerType=0 controllerNumber=1 scan=5904 | 16.0276 |
| 03178:2 | 29 | K.TPLTAN[+1809.666]ITK.S | HexNAc(5)Hex(4)Fuc(1) | N[+1810] | 1384.6176 | 2 | 2768.2279 | 2768.2228 | 0 | 1.8 | 200 | Specific | 517.36 | 276.98 | 276.98 | 7.69 | 20 | >sp\|P01877\|IGHA2_HUMAN Immunoglobulin heavy constant alpha 2 OS=Homo sapiens OX=9606 GN=IGHA2 PE=1 SV=4 | 576 | 20201114_16933_1ng_glyco_hcd203040_compound15.5868.5868.2 | controllerType=0 controllerNumber=1 scan=5868 | 15.9597 |
| 03253:3 | 29 | K.TPLTAN[+1647.613]ITK.S | HexNAc(5)Hex(3)Fuc(1) | N[+1648] | 869.3981 | 3 | 2606.1797 | 2606.17 | 0 | 3.7 | 200 | Specific | 540.68 | 263.73 | 263.73 | 7.6 | 20 | >sp\|P01877\|IGHA2_HUMAN Immunoglobulin heavy constant alpha 2 OS=Homo sapiens OX=9606 GN=IGHA2 PE=1 SV=4 | 576 | 20201114_16933_1ng_glyco_hcd203040_compound15.5946.5946.3 | controllerType=0 controllerNumber=1 scan=5946 | 16.1017 |
| 10276:3 | 29 | R.MAGKPTHIN[+1540.529]VSVVMAEADGTC[+57.021]Y.- | HexNAc(2)Hex(7) | C[+57], N[+1541] | 1297.8806 | 3 | 3891.6271 | 3891.6114 | 0 | 4 | 319 | CRagged | 220.56 | 220.56 | 220.56 | 5.37 | 20 | >sp\|P01877\|IGHA2_HUMAN Immunoglobulin heavy constant alpha 2 OS=Homo sapiens OX=9606 GN=IGHA2 PE=1 SV=4 | 576 | 20201114_16933_1ng_glyco_hcd203040_compound15.13254.13254.3 | controllerType=0 controllerNumber=1 scan=13254 | 28.538 |
| 10198:3 | 29 | R.MAGKPTHIN[+1702.581]VSVVMAEADGTC[+57.021]Y.- | HexNAc(2)Hex(8) | C[+57], N[+1703] | 1351.8904 | 3 | 4053.6568 | 4053.6642 | 0 | -1.8 | 319 | CRagged | 538.98 | 538.98 | 335.64 | 8.82 | 20 | >sp\|P01877\|IGHA2_HUMAN Immunoglobulin heavy constant alpha 2 OS=Homo sapiens OX=9606 GN=IGHA2 PE=1 SV=4 | 576 | 20201114_16933_1ng_glyco_hcd203040_compound15.13173.13173.3 | controllerType=0 controllerNumber=1 scan=13173 | 28.3966 |
| 10200:3 | 29 | R.MAGKPTHIN[+1702.581]VSVVMAEADGTC[+57.021]Y.- | HexNAc(2)Hex(8) | C[+57], N[+1703] | 1351.8904 | 3 | 4053.6568 | 4053.6642 | 0 | -1.8 | 319 | CRagged | 573.28 | 573.28 | 318.54 | 10.16 | 20 | >sp\|P01877\|IGHA2_HUMAN Immunoglobulin heavy constant alpha 2 OS=Homo sapiens OX=9606 GN=IGHA2 PE=1 SV=4 | 576 | 20201114_16933_1ng_glyco_hcd203040_compound15.13175.13175.3 | controllerType=0 controllerNumber=1 scan=13175 | 28.4005 |
| 10339:3 | 29 | R.MAGKPTHIN[+1216.423]VSVVMAEADGTC[+57.021]Y.- | HexNAc(2)Hex(5) | C[+57], N[+1216] | 1189.8428 | 3 | 3567.5139 | 3567.5057 | 0 | 2.3 | 319 | CRagged | 608.11 | 608.11 | 283.75 | 12.38 | 20 | >sp\|P01877\|IGHA2_HUMAN Immunoglobulin heavy constant alpha 2 OS=Homo sapiens OX=9606 GN=IGHA2 PE=1 SV=4 | 576 | 20201114_16933_1ng_glyco_hcd203040_compound15.13319.13319.3 | controllerType=0 controllerNumber=1 scan=13319 | 28.6501 |
| 10337:3 | 29 | R.MAGKPTHIN[+1216.423]VSVVMAEADGTC[+57.021]Y.- | HexNAc(2)Hex(5) | C[+57], N[+1216] | 1189.8428 | 3 | 3567.5139 | 3567.5057 | 0 | 2.3 | 319 | CRagged | 697.19 | 697.14 | 392.04 | 13.38 | 20 | >sp\|P01877\|IGHA2_HUMAN Immunoglobulin heavy constant alpha 2 OS=Homo sapiens OX=9606 GN=IGHA2 PE=1 SV=4 | 576 | 20201114_16933_1ng_glyco_hcd203040_compound15.13317.13317.3 | controllerType=0 controllerNumber=1 scan=13317 | 28.6459 |
| 10303:3 | 29 | R.MAGKPTHIN[+1378.476]VSVVMAEADGTC[+57.021]Y.- | HexNAc(2)Hex(6) | C[+57], N[+1378] | 1243.8608 | 3 | 3729.5678 | 3729.5585 | 0 | 2.5 | 319 | CRagged | 796.07 | 780.27 | 444.63 | 15.38 | 20 | >sp\|P01877\|IGHA2_HUMAN Immunoglobulin heavy constant alpha 2 OS=Homo sapiens OX=9606 GN=IGHA2 PE=1 SV=4 | 576 | 20201114_16933_1ng_glyco_hcd203040_compound15.13282.13282.3 | controllerType=0 controllerNumber=1 scan=13282 | 28.5874 |
| 10305:3 | 29 | R.MAGKPTHIN[+1378.476]VSVVMAEADGTC[+57.021]Y.- | HexNAc(2)Hex(6) | C[+57], N[+1378] | 1243.8608 | 3 | 3729.5678 | 3729.5585 | 0 | 2.5 | 319 | CRagged | 820.53 | 820.53 | 420.2 | 16.38 | 20 | >sp\|P01877\|IGHA2_HUMAN Immunoglobulin heavy constant alpha 2 OS=Homo sapiens OX=9606 GN=IGHA2 PE=1 SV=4 | 576 | 20201114_16933_1ng_glyco_hcd203040_compound15.13284.13284.3 | controllerType=0 controllerNumber=1 scan=13284 | 28.5917 |
| 16803:3 | 63 | K.AALAAFNAQNN[+2204.772]GSNFQLEEISR.A | HexNAc(4)Hex(5)NeuAc(2) | N[+2205] | 1523.988 | 3 | 4569.9494 | 4569.9256 | 0 | 5.2 | 166 | Specific | 374.11 | 315.62 | 315.62 | 7.74 | 9 | >sp\|P02765\|FETUA_HUMAN Alpha-2-HS-glycoprotein OS=Homo sapiens OX=9606 GN=AHSG PE=1 SV=2 | 135 | 20201114_16933_1ng_glyco_hcd203040_compound15.20070.20070.3 | controllerType=0 controllerNumber=1 scan=20070 | 40.2896 |
| 09717:3 | 65 | R.IIVPLNNREN[+1809.666]ISDPTSPLR.T | HexNAc(5)Hex(4)Fuc(1) | N[+1810] | 1319.9562 | 3 | 3957.8541 | 3957.8432 | 0 | 2.8 | 62 | Specific | 254.38 | 178.8 | 178.8 | 5.76 | 12 | >sp\|P01591\|IGJ_HUMAN Immunoglobulin J chain OS=Homo sapiens OX=9606 GN=JCHAIN PE=1 SV=4 | 1392 | 20201114_16933_1ng_glyco_hcd203040_compound15.12674.12674.3 | controllerType=0 controllerNumber=1 scan=12674 | 27.5527 |
| 14546:7 | 115 | K.GNPEQTPVLKPEEEAPAPEVGASKPEGIDSRPETLHPGRPQPPAEEELC[+57.021]SGKPFI | HexNAc(4)Hex(5)Fuc(2)NeuAc(1) | C[+57], N[+2206] | 1382.5039 | 7 | 9671.484 | 9671.4638 | 0 | 2.1 | 108 | Specific | 296.04 | 296.04 | 296.04 | 5.77 | 8 | >sp\|P04004\|VTNC_HUMAN Vitronectin OS=Homo sapiens OX=9606 GN=VTN PE=1 SV=1 | 219 | 20201114_16933_1ng_glyco_hcd203040_compound15.17704.17704.3 | controllerType=0 controllerNumber=1 scan=17704 | 36.1553 |
| 01513:3 | 1 | K.GPN[+1913.677]C[+57.021]SEPEC[+57.021]PGNC[+57.021]HLR.G | HexNAc(4)Hex(5)NeuAc(1) | C[+57]*3, N[+1914] | 1266.4854 | 3 | 3797.4417 | 3797.4352 | 0 | 1.7 | 182 | Specific | 436.66 | 349.45 | 349.45 | 8.4 | 152 | >sp\|P24821\|TENA_HUMAN Tenascin OS=Homo sapiens OX=9606 GN=TNC PE=1 SV=3 | 486 | 20201116_fqg_colostrum_1ng_glyco_hcd203040_compound15.4409.4409.3 | controllerType=0 controllerNumber=1 scan=4409 | 13.7383 |
| 01511:3 | 1 | K.GPN[+1913.677]C[+57.021]SEPEC[+57.021]PGNC[+57.021]HLR.G | HexNAc(4)Hex(5)NeuAc(1) | C[+57]*3, N[+1914] | 1266.4854 | 3 | 3797.4417 | 3797.4352 | 0 | 1.7 | 182 | Specific | 505.4 | 449.37 | 449.37 | 10.23 | 152 | >sp\|P24821\|TENA_HUMAN Tenascin OS=Homo sapiens OX=9606 GN=TNC PE=1 SV=3 | 486 | 20201116_fqg_colostrum_1ng_glyco_hcd203040_compound15.4407.4407.3 | controllerType=0 controllerNumber=1 scan=4407 | 13.7343 |
| 00733:3 | 1 | K.GPN[+1216.423]C[+57.021]SEPEC[+57.021]PGNC[+57.021]HLR.G | HexNAc(2)Hex(5) | C[+57]*3, N[+1216] | 1034.0683 | 3 | 3100.1902 | 3100.181 | 0 | 3 | 182 | Specific | 595.3 | 582.05 | 582.05 | 11.91 | 152 | >sp\|P24821\|TENA_HUMAN Tenascin OS=Homo sapiens OX=9606 GN=TNC PE=1 SV=3 | 486 | 20201116_fqg_colostrum_1ng_glyco_hcd203040_compound15.3599.3599.3 | controllerType=0 controllerNumber=1 scan=3599 | 12.2188 |
| 00731:3 | 1 | K.GPN[+1216.423]C[+57.021]SEPEC[+57.021]PGNC[+57.021]HLR.G | HexNAc(2)Hex(5) | C[+57]*3, N[+1216] | 1034.0683 | 3 | 3100.1902 | 3100.181 | 0 | 3 | 182 | Specific | 701.91 | 701.91 | 701.91 | 14.91 | 152 | >sp\|P24821\|TENA_HUMAN Tenascin OS=Homo sapiens OX=9606 GN=TNC PE=1 SV=3 | 486 | 20201116_fqg_colostrum_1ng_glyco_hcd203040_compound15.3597.3597.3 | controllerType=0 controllerNumber=1 scan=3597 | 12.2153 |
| 01532:3 | 1 | K.GPN[+2059.735]C[+57.021]SEPEC[+57.021]PGNC[+57.021]HLR.G | HexNAc(4)Hex(5)Fuc(1)NeuAc(1) | C[+57]*3, N[+2060] | 1315.1736 | 3 | 3943.5064 | 3943.4931 | 0 | 3.4 | 182 | Specific | 710.68 | 598.77 | 598.77 | 14.91 | 152 | >sp\|P24821\|TENA_HUMAN Tenascin OS=Homo sapiens OX=9606 GN=TNC PE=1 SV=3 | 486 | 20201116_fqg_colostrum_1ng_glyco_hcd203040_compound15.4429.4429.3 | controllerType=0 controllerNumber=1 scan=4429 | 13.7876 |
| 01530:3 | 1 | K.GPN[+2059.735]C[+57.021]SEPEC[+57.021]PGNC[+57.021]HLR.G | HexNAc(4)Hex(5)Fuc(1)NeuAc(1) | C[+57]*3, N[+2060] | 1315.1736 | 3 | 3943.5064 | 3943.4931 | 0 | 3.4 | 182 | Specific | 756.19 | 685.09 | 685.09 | 15.91 | 152 | >sp\|P24821\|TENA_HUMAN Tenascin OS=Homo sapiens OX=9606 GN=TNC PE=1 SV=3 | 486 | 20201116_fqg_colostrum_1ng_glyco_hcd203040_compound15.4427.4427.3 | controllerType=0 controllerNumber=1 scan=4427 | 13.7719 |
| 00710:3 | 1 | K.GPN[+1216.423]C[+57.021]SEPEC[+57.021]PGNC[+57.021]HLR.G | HexNAc(2)Hex(5) | C[+57]*3, N[+1216] | 1034.0688 | 3 | 3100.1917 | 3100.181 | 0 | 3.4 | 182 | Specific | 819.57 | 819.57 | 819.57 | 16.91 | 152 | >sp\|P24821\|TENA_HUMAN Tenascin OS=Homo sapiens OX=9606 GN=TNC PE=1 SV=3 | 486 | 20201116_fqg_colostrum_1ng_glyco_hcd203040_compound15.3575.3575.3 | controllerType=0 controllerNumber=1 scan=3575 | 12.1758 |
| 00712:3 | 1 | K.GPN[+1216.423]C[+57.021]SEPEC[+57.021]PGNC[+57.021]HLR.G | HexNAc(2)Hex(5) | C[+57]*3, N[+1216] | 1034.0688 | 3 | 3100.1917 | 3100.181 | 0 | 3.4 | 182 | Specific | 841.02 | 790.78 | 790.78 | 16.91 | 152 | >sp\|P24821\|TENA_HUMAN Tenascin OS=Homo sapiens OX=9606 GN=TNC PE=1 SV=3 | 486 | 20201116_fqg_colostrum_1ng_glyco_hcd203040_compound15.3577.3577.3 | controllerType=0 controllerNumber=1 scan=3577 | 12.1798 |
| 03598:4 | 1 | R.C[+57.021]IN[+2059.735]GTC[+57.021]YC[+57.021]EEGFTGEDC[+57.021]GKPTC[ | HexNAc(4)Hex(5)Fuc(1)NeuAc(1) | C[+57]*6, N[+2060] | 1430.549 | 4 | 5719.174 | 5719.1633 | 0 | 1.9 | 325 | Specific | 320 | 320 | 320 | 8.01 | 152 | >sp\|P24821\|TENA_HUMAN Tenascin OS=Homo sapiens OX=9606 GN=TNC PE=1 SV=3 | 486 | 20201116_fqg_colostrum_1ng_glyco_hcd203040_compound15.6570.6570.4 | controllerType=0 controllerNumber=1 scan=6570 | 17.6996 |
| 03601:4 | 1 | R.C[+57.021]IN[+2059.735]GTC[+57.021]YC[+57.021]EEGFTGEDC[+57.021]GKPTC[ | HexNAc(4)Hex(5)Fuc(1)NeuAc(1) | C[+57]*6, N[+2060] | 1430.549 | 4 | 5719.174 | 5719.1633 | 0 | 1.9 | 325 | Specific | 420.92 | 420.92 | 420.92 | 10.08 | 152 | >sp\|P24821\|TENA_HUMAN Tenascin OS=Homo sapiens OX=9606 GN=TNC PE=1 SV=3 | 486 | 20201116_fqg_colostrum_1ng_glyco_hcd203040_compound15.6573.6573.4 | controllerType=0 controllerNumber=1 scan=6573 | 17.7035 |
| 03529:5 | 1 | R.C[+57.021]IN[+2351.851]GTC[+57.021]YC[+57.021]EEGFTGEDC[+57.021]GKPTC[ | HexNAc(4)Hex(5)Fuc(3)NeuAc(1) | C[+57]*6, N[+2352] | 1203.0642 | 5 | 6011.292 | 6011.2791 | 0 | 2.1 | 325 | Specific | 477 | 477 | 477 | 10.91 | 152 | >sp\|P24821\|TENA_HUMAN Tenascin OS=Homo sapiens OX=9606 GN=TNC PE=1 SV=3 | 486 | 20201116_fqg_colostrum_1ng_glyco_hcd203040_compound15.6499.6499.5 | controllerType=0 controllerNumber=1 scan=6499 | 17.5895 |
| 03532:5 | 1 | R.C[+57.021]IN[+2351.851]GTC[+57.021]YC[+57.021]EEGFTGEDC[+57.021]GKPTC[ | HexNAc(4)Hex(5)Fuc(3)NeuAc(1) | C[+57]*6, N[+2352] | 1203.0642 | 5 | 6011.292 | 6011.2791 | 0 | 2.1 | 325 | Specific | 479.25 | 479.25 | 479.25 | 10.91 | 152 | >sp\|P24821\|TENA_HUMAN Tenascin OS=Homo sapiens OX=9606 GN=TNC PE=1 SV=3 | 486 | 20201116_fqg_colostrum_1ng_glyco_hcd203040_compound15.6502.6502.5 | controllerType=0 controllerNumber=1 scan=6502 | 17.594 |
| 08889:3 | 1 | R.QTGLAPGQEYEISLHIVKN[+2059.735]NTR.G | HexNAc(4)Hex(5)Fuc(1)NeuAc(1) | N[+2060] | 1510.0173 | 3 | 4528.0373 | 4528.0241 | 0 | 2.9 | 770 | Specific | 543.22 | 343.26 | 343.26 | 8.05 | 152 | >sp\|P24821\|TENA_HUMAN Tenascin OS=Homo sapiens OX=9606 GN=TNC PE=1 SV=3 | 486 | 20201116_fqg_colostrum_1ng_glyco_hcd203040_compound15.12075.12075.3 | controllerType=0 controllerNumber=1 scan=12075 | 26.4715 |
| 08887:3 | 1 | R.QTGLAPGQEYEISLHIVKN[+2059.735]NTR.G | HexNAc(4)Hex(5)Fuc(1)NeuAc(1) | N[+2060] | 1510.0173 | 3 | 4528.0373 | 4528.0241 | 0 | 2.9 | 770 | Specific | 743.82 | 424.08 | 424.08 | 14.05 | 152 | >sp\|P24821\|TENA_HUMAN Tenascin OS=Homo sapiens OX=9606 GN=TNC PE=1 SV=3 | 486 | 20201116_fqg_colostrum_1ng_glyco_hcd203040_compound15.12073.12073.3 | controllerType=0 controllerNumber=1 scan=12073 | 26.4659 |
| 15761:3 | 1 | R.LN[+1540.529]YSLPTGQWVGVQLPR.N | HexNAc(2)Hex(7) | N[+1541] | 1156.8599 | 3 | 3468.5651 | 3468.5674 | 0 | -0.6 | 1017 | Specific | 222.47 | 222.47 | 222.47 | 3.85 | 152 | >sp\|P24821\|TENA_HUMAN Tenascin OS=Homo sapiens OX=9606 GN=TNC PE=1 SV=3 | 486 | 20201116_fqg_colostrum_1ng_glyco_hcd203040_compound15.19183.19183.3 | controllerType=0 controllerNumber=1 scan=19183 | 37.8916 |
| 15815:3 | 1 | R.LN[+1378.476]YSLPTGQWVGVQLPR.N | HexNAc(2)Hex(6) | N[+1378] | 1102.8457 | 3 | 3306.5226 | 3306.5145 | 0 | 2.4 | 1017 | Specific | 233.96 | 233.96 | 233.96 | 3.61 | 152 | >sp\|P24821\|TENA_HUMAN Tenascin OS=Homo sapiens OX=9606 GN=TNC PE=1 SV=3 | 486 | 20201116_fqg_colostrum_1ng_glyco_hcd203040_compound15.19239.19239.3 | controllerType=0 controllerNumber=1 scan=19239 | 37.9846 |
| 15764:3 | 1 | R.LN[+1540.529]YSLPTGQWVGVQLPR.N | HexNAc(2)Hex(7) | N[+1541] | 1156.8599 | 3 | 3468.5651 | 3468.5674 | 0 | -0.6 | 1017 | Specific | 324.31 | 201.07 | 201.07 | 4.15 | 152 | >sp\|P24821\|TENA_HUMAN Tenascin OS=Homo sapiens OX=9606 GN=TNC PE=1 SV=3 | 486 | 20201116_fqg_colostrum_1ng_glyco_hcd203040_compound15.19186.19186.3 | controllerType=0 controllerNumber=1 scan=19186 | 37.8983 |
| 15649:3 | 1 | R.LN[+1702.581]YSLPTGQWVGVQLPR.N | HexNAc(2)Hex(8) | N[+1703] | 1210.8799 | 3 | 3630.6253 | 3630.6202 | 0 | 1.4 | 1017 | Specific | 603.39 | 595.69 | 595.69 | 13 | 152 | >sp\|P24821\|TENA_HUMAN Tenascin OS=Homo sapiens OX=9606 GN=TNC PE=1 SV=3 | 486 | 20201116_fqg_colostrum_1ng_glyco_hcd203040_compound15.19068.19068.3 | controllerType=0 controllerNumber=1 scan=19068 | 37.7 |
| 15827:2 | 1 | R.LN[+1378.476]YSLPTGQWVGVQLPR.N | HexNAc(2)Hex(6) | N[+1378] | 1653.7685 | 2 | 3306.5297 | 3306.5145 | 0 | 4.6 | 1017 | Specific | 668.9 | 584.34 | 584.34 | 13.47 | 152 | >sp\|P24821\|TENA_HUMAN Tenascin OS=Homo sapiens OX=9606 GN=TNC PE=1 SV=3 | 486 | 20201116_fqg_colostrum_1ng_glyco_hcd203040_compound15.19251.19251.2 | controllerType=0 controllerNumber=1 scan=19251 | 38.0086 |
| 15652:3 | 1 | R.LN[+1702.581]YSLPTGQWVGVQLPR.N | HexNAc(2)Hex(8) | N[+1703] | 1210.8799 | 3 | 3630.6253 | 3630.6202 | 0 | 1.4 | 1017 | Specific | 806.49 | 668.33 | 668.33 | 17 | 152 | >sp\|P24821\|TENA_HUMAN Tenascin OS=Homo sapiens OX=9606 GN=TNC PE=1 SV=3 | 486 | 20201116_fqg_colostrum_1ng_glyco_hcd203040_compound15.19071.19071.3 | controllerType=0 controllerNumber=1 scan=19071 | 37.7043 |
| 15830:2 | 1 | R.LN[+1378.476]YSLPTGQWVGVQLPR.N | HexNAc(2)Hex(6) | N[+1378] | 1653.7685 | 2 | 3306.5297 | 3306.5145 | 0 | 4.6 | 1017 | Specific | 814.04 | 736.91 | 736.91 | 16.47 | 152 | >sp\|P24821\|TENA_HUMAN Tenascin OS=Homo sapiens OX=9606 GN=TNC PE=1 SV=3 | 486 | 20201116_fqg_colostrum_1ng_glyco_hcd203040_compound15.19254.19254.2 | controllerType=0 controllerNumber=1 scan=19254 | 38.0146 |
| 15832:2 | 1 | R.LN[+1216.423]YSLPTGQWVGVQLPR.N | HexNAc(2)Hex(5) | N[+1216] | 1572.7384 | 2 | 3144.4696 | 3144.4617 | 0 | 2.5 | 1017 | Specific | 876.02 | 740.44 | 740.44 | 18.02 | 152 | >sp\|P24821\|TENA_HUMAN Tenascin OS=Homo sapiens OX=9606 GN=TNC PE=1 SV=3 | 486 | 20201116_fqg_colostrum_1ng_glyco_hcd203040_compound15.19256.19256.2 | controllerType=0 controllerNumber=1 scan=19256 | 38.0225 |
| 15831:2 | 1 | R.LN[+1216.423]YSLPTGQWVGVQLPR.N | HexNAc(2)Hex(5) | N[+1216] | 1572.7384 | 2 | 3144.4696 | 3144.4617 | 0 | 2.5 | 1017 | Specific | 913.31 | 913.31 | 913.31 | 19.02 | 152 | >sp\|P24821\|TENA_HUMAN Tenascin OS=Homo sapiens OX=9606 GN=TNC PE=1 SV=3 | 486 | 20201116_fqg_colostrum_1ng_glyco_hcd203040_compound15.19255.19255.2 | controllerType=0 controllerNumber=1 scan=19255 | 38.0164 |
| 15835:3 | 1 | R.LN[+1216.423]YSLPTGQWVGVQLPR.N | HexNAc(2)Hex(5) | N[+1216] | 1048.8283 | 3 | 3144.4703 | 3144.4617 | 0 | 2.7 | 1017 | Specific | 938.89 | 935.49 | 935.49 | 19.24 | 152 | >sp\|P24821\|TENA_HUMAN Tenascin OS=Homo sapiens OX=9606 GN=TNC PE=1 SV=3 | 486 | 20201116_fqg_colostrum_1ng_glyco_hcd203040_compound15.19260.19260.3 | controllerType=0 controllerNumber=1 scan=19260 | 38.0294 |
| 15838:3 | 1 | R.LN[+1216.423]YSLPTGQWVGVQLPR.N | HexNAc(2)Hex(5) | N[+1216] | 1048.8283 | 3 | 3144.4703 | 3144.4617 | 0 | 2.7 | 1017 | Specific | 1074.75 | 922.12 | 922.12 | 22.24 | 152 | >sp\|P24821\|TENA_HUMAN Tenascin OS=Homo sapiens OX=9606 GN=TNC PE=1 SV=3 | 486 | 20201116_fqg_colostrum_1ng_glyco_hcd203040_compound15.19263.19263.3 | controllerType=0 controllerNumber=1 scan=19263 | 38.0339 |
| 01842:3 | 1 | R.N[+1768.640]TTSYVLR.G | HexNAc(4)Hex(5)Fuc(1) | N[+1769] | 908.0559 | 3 | 2722.1531 | 2722.1446 | 0 | 3.1 | 1034 | Specific | 208.64 | 158.63 | 158.63 | 4.47 | 152 | >sp\|P24821\|TENA_HUMAN Tenascin OS=Homo sapiens OX=9606 GN=TNC PE=1 SV=3 | 486 | 20201116_fqg_colostrum_1ng_glyco_hcd203040_compound15.4751.4751.3 | controllerType=0 controllerNumber=1 scan=4751 | 14.3675 |
| 03552:3 | 1 | R.N[+2059.735]TTSYVLR.G | HexNAc(4)Hex(5)Fuc(1)NeuAc(1) | N[+2060] | 1005.0887 | 3 | 3013.2516 | 3013.24 | 0 | 3.8 | 1034 | Specific | 215.3 | 136 | 136 | 4.35 | 152 | >sp\|P24821\|TENA_HUMAN Tenascin OS=Homo sapiens OX=9606 GN=TNC PE=1 SV=3 | 486 | 20201116_fqg_colostrum_1ng_glyco_hcd203040_compound15.6523.6523.3 | controllerType=0 controllerNumber=1 scan=6523 | 17.6266 |
| 01826:2 | 1 | R.N[+1768.640]TTSYVLR.G | HexNAc(4)Hex(5)Fuc(1) | N[+1769] | 1361.5774 | 2 | 2722.1476 | 2722.1446 | 0 | 1.1 | 1034 | Specific | 247.18 | 143.11 | 143.11 | 5.43 | 152 | >sp\|P24821\|TENA_HUMAN Tenascin OS=Homo sapiens OX=9606 GN=TNC PE=1 SV=3 | 486 | 20201116_fqg_colostrum_1ng_glyco_hcd203040_compound15.4734.4734.2 | controllerType=0 controllerNumber=1 scan=4734 | 14.3349 |
| 01844:3 | 1 | R.N[+1768.640]TTSYVLR.G | HexNAc(4)Hex(5)Fuc(1) | N[+1769] | 908.0559 | 3 | 2722.1531 | 2722.1446 | 0 | 3.1 | 1034 | Specific | 294.86 | 234.54 | 234.54 | 6.96 | 152 | >sp\|P24821\|TENA_HUMAN Tenascin OS=Homo sapiens OX=9606 GN=TNC PE=1 SV=3 | 486 | 20201116_fqg_colostrum_1ng_glyco_hcd203040_compound15.4753.4753.3 | controllerType=0 controllerNumber=1 scan=4753 | 14.3713 |
| 03555:3 | 1 | R.N[+2059.735]TTSYVLR.G | HexNAc(4)Hex(5)Fuc(1)NeuAc(1) | N[+2060] | 1005.0887 | 3 | 3013.2516 | 3013.24 | 0 | 3.8 | 1034 | Specific | 324.42 | 216.55 | 216.55 | 7.03 | 152 | >sp\|P24821\|TENA_HUMAN Tenascin OS=Homo sapiens OX=9606 GN=TNC PE=1 SV=3 | 486 | 20201116_fqg_colostrum_1ng_glyco_hcd203040_compound15.6526.6526.3 | controllerType=0 controllerNumber=1 scan=6526 | 17.631 |
| 01860:2 | 1 | R.N[+1216.423]TTSYVLR.G | HexNAc(2)Hex(5) | N[+1216] | 1085.4707 | 2 | 2169.9342 | 2169.9279 | 0 | 2.9 | 1034 | Specific | 332.38 | 246.89 | 246.89 | 6.89 | 152 | >sp\|P24821\|TENA_HUMAN Tenascin OS=Homo sapiens OX=9606 GN=TNC PE=1 SV=3 | 486 | 20201116_fqg_colostrum_1ng_glyco_hcd203040_compound15.4770.4770.2 | controllerType=0 controllerNumber=1 scan=4770 | 14.4072 |
| 03558:2 | 1 | R.N[+2059.735]TTSYVLR.G | HexNAc(4)Hex(5)Fuc(1)NeuAc(1) | N[+2060] | 1507.1291 | 2 | 3013.2509 | 3013.24 | 0 | 3.6 | 1034 | Specific | 339.53 | 212.57 | 212.57 | 6.81 | 152 | >sp\|P24821\|TENA_HUMAN Tenascin OS=Homo sapiens OX=9606 GN=TNC PE=1 SV=3 | 486 | 20201116_fqg_colostrum_1ng_glyco_hcd203040_compound15.6529.6529.2 | controllerType=0 controllerNumber=1 scan=6529 | 17.6346 |
| 01829:2 | 1 | R.N[+1768.640]TTSYVLR.G | HexNAc(4)Hex(5)Fuc(1) | N[+1769] | 1361.5774 | 2 | 2722.1476 | 2722.1446 | 0 | 1.1 | 1034 | Specific | 384.99 | 257.42 | 257.42 | 7.09 | 152 | >sp\|P24821\|TENA_HUMAN Tenascin OS=Homo sapiens OX=9606 GN=TNC PE=1 SV=3 | 486 | 20201116_fqg_colostrum_1ng_glyco_hcd203040_compound15.4737.4737.2 | controllerType=0 controllerNumber=1 scan=4737 | 14.34 |
| 03560:2 | 1 | R.N[+2059.735]TTSYVLR.G | HexNAc(4)Hex(5)Fuc(1)NeuAc(1) | N[+2060] | 1507.1291 | 2 | 3013.2509 | 3013.24 | 0 | 3.6 | 1034 | Specific | 404.11 | 264.79 | 264.79 | 7.58 | 152 | >sp\|P24821\|TENA_HUMAN Tenascin OS=Homo sapiens OX=9606 GN=TNC PE=1 SV=3 | 486 | 20201116_fqg_colostrum_1ng_glyco_hcd203040_compound15.6531.6531.2 | controllerType=0 controllerNumber=1 scan=6531 | 17.6407 |
| 01862:2 | 1 | R.N[+1216.423]TTSYVLR.G | HexNAc(2)Hex(5) | N[+1216] | 1085.4707 | 2 | 2169.9342 | 2169.9279 | 0 | 2.9 | 1034 | Specific | 460.17 | 366.26 | 366.26 | 8.44 | 152 | >sp\|P24821\|TENA_HUMAN Tenascin OS=Homo sapiens OX=9606 GN=TNC PE=1 SV=3 | 486 | 20201116_fqg_colostrum_1ng_glyco_hcd203040_compound15.4772.4772.2 | controllerType=0 controllerNumber=1 scan=4772 | 14.4116 |
| 20637:5 | 1 | K.ASTEQAPELENLTVTEVGWDGLRLN[+2059.735]WTAADQAYEHFIIQVQEANKVEAAR | HexNAc(4)Hex(5)Fuc(1)NeuAc(1) | N[+2060] | 1535.1113 | 5 | 7671.5271 | 7671.5078 | 0 | 2.5 | 1069 | Specific | 470.95 | 470.95 | 9.46 | 10.28 | 152 | >sp\|P24821\|TENA_HUMAN Tenascin OS=Homo sapiens OX=9606 GN=TNC PE=1 SV=3 | 486 | 20201116_fqg_colostrum_1ng_glyco_hcd203040_compound15.24249.24249.5 | controllerType=0 controllerNumber=1 scan=24249 | 47.4072 |
| 20640:5 | 1 | K.ASTEQAPELENLTVTEVGWDGLRLN[+2059.735]WTAADQAYEHFIIQVQEANKVEAAR | HexNAc(4)Hex(5)Fuc(1)NeuAc(1) | N[+2060] | 1535.1113 | 5 | 7671.5271 | 7671.5078 | 0 | 2.5 | 1069 | Specific | 880.14 | 866.1 | 2.61 | 18.26 | 152 | >sp\|P24821\|TENA_HUMAN Tenascin OS=Homo sapiens OX=9606 GN=TNC PE=1 SV=3 | 486 | 20201116_fqg_colostrum_1ng_glyco_hcd203040_compound15.24252.24252.5 | controllerType=0 controllerNumber=1 scan=24252 | 47.4115 |
| 07567:2 | 1 | R.N[+2059.735]LTVPGSLR.A | HexNAc(4)Hex(5)Fuc(1)NeuAc(1) | N[+2060] | 1508.6533 | 2 | 3016.2993 | 3016.2873 | 0 | 4 | 1119 | Specific | 317.05 | 255.42 | 255.42 | 6.46 | 152 | >sp\|P24821\|TENA_HUMAN Tenascin OS=Homo sapiens OX=9606 GN=TNC PE=1 SV=3 | 486 | 20201116_fqg_colostrum_1ng_glyco_hcd203040_compound15.10707.10707.2 | controllerType=0 controllerNumber=1 scan=10707 | 24.2297 |
| 07569:2 | 1 | R.N[+2059.735]LTVPGSLR.A | HexNAc(4)Hex(5)Fuc(1)NeuAc(1) | N[+2060] | 1508.6533 | 2 | 3016.2993 | 3016.2873 | 0 | 4 | 1119 | Specific | 366.39 | 211.68 | 211.68 | 6.65 | 152 | >sp\|P24821\|TENA_HUMAN Tenascin OS=Homo sapiens OX=9606 GN=TNC PE=1 SV=3 | 486 | 20201116_fqg_colostrum_1ng_glyco_hcd203040_compound15.10709.10709.2 | controllerType=0 controllerNumber=1 scan=10709 | 24.2352 |
| 09893:3 | 2 | K.TVLTPATNHMGN[+1540.529]VTFTIPANR.E | HexNAc(2)Hex(7) | N[+1541] | 1265.904 | 3 | 3795.6974 | 3795.6886 | 0 | 2.3 | 74 | Specific | 613.56 | 613.56 | 326.56 | 13.42 | 110 | >sp\|P01024\|CO3_HUMAN Complement C3 OS=Homo sapiens OX=9606 GN=C3 PE=1 SV=2 | 492 | 20201116_fqg_colostrum_1ng_glyco_hcd203040_compound15.13115.13115.3 | controllerType=0 controllerNumber=1 scan=13115 | 28.1436 |
| 09896:3 | 2 | K.TVLTPATNHMGN[+1540.529]VTFTIPANR.E | HexNAc(2)Hex(7) | N[+1541] | 1265.904 | 3 | 3795.6974 | 3795.6886 | 0 | 2.3 | 74 | Specific | 618.56 | 584 | 321.13 | 13.42 | 110 | >sp\|P01024\|CO3_HUMAN Complement C3 OS=Homo sapiens OX=9606 GN=C3 PE=1 SV=2 | 492 | 20201116_fqg_colostrum_1ng_glyco_hcd203040_compound15.13118.13118.3 | controllerType=0 controllerNumber=1 scan=13118 | 28.1481 |
| 09921:3 | 2 | K.TVLTPATNHMGN[+1378.476]VTFTIPANR.E | HexNAc(2)Hex(6) | N[+1378] | 1211.8863 | 3 | 3633.6443 | 3633.6358 | 0 | 2.3 | 74 | Specific | 764.02 | 764.02 | 402.17 | 16.42 | 110 | >sp\|P01024\|CO3_HUMAN Complement C3 OS=Homo sapiens OX=9606 GN=C3 PE=1 SV=2 | 492 | 20201116_fqg_colostrum_1ng_glyco_hcd203040_compound15.13144.13144.3 | controllerType=0 controllerNumber=1 scan=13144 | 28.194 |
| 09856:3 | 2 | K.TVLTPATNHMGN[+1540.529]VTFTIPANR.E | HexNAc(2)Hex(7) | N[+1541] | 1265.9072 | 3 | 3795.7069 | 3795.6886 | 0 | 4.8 | 74 | Specific | 851.87 | 800.74 | 455.85 | 17.87 | 110 | >sp\|P01024\|CO3_HUMAN Complement C3 OS=Homo sapiens OX=9606 GN=C3 PE=1 SV=2 | 492 | 20201116_fqg_colostrum_1ng_glyco_hcd203040_compound15.13077.13077.3 | controllerType=0 controllerNumber=1 scan=13077 | 28.0811 |
| 09853:3 | 2 | K.TVLTPATNHMGN[+1540.529]VTFTIPANR.E | HexNAc(2)Hex(7) | N[+1541] | 1265.9072 | 3 | 3795.7069 | 3795.6886 | 0 | 4.8 | 74 | Specific | 851.92 | 851.92 | 473.89 | 17.87 | 110 | >sp\|P01024\|CO3_HUMAN Complement C3 OS=Homo sapiens OX=9606 GN=C3 PE=1 SV=2 | 492 | 20201116_fqg_colostrum_1ng_glyco_hcd203040_compound15.13074.13074.3 | controllerType=0 controllerNumber=1 scan=13074 | 28.0766 |
| 09924:3 | 2 | K.TVLTPATNHMGN[+1378.476]VTFTIPANR.E | HexNAc(2)Hex(6) | N[+1378] | 1211.8863 | 3 | 3633.6443 | 3633.6358 | 0 | 2.3 | 74 | Specific | 862.13 | 847.13 | 447.4 | 18.42 | 110 | >sp\|P01024\|CO3_HUMAN Complement C3 OS=Homo sapiens OX=9606 GN=C3 PE=1 SV=2 | 492 | 20201116_fqg_colostrum_1ng_glyco_hcd203040_compound15.13147.13147.3 | controllerType=0 controllerNumber=1 scan=13147 | 28.1988 |
| 24215:4 | 3 | A.[+42.011]SPTSPKVFPLSLDSTPQDGNVVVAC[+57.021]LVQGFFPQEPLSVTWSESGQN | HexNAc(3)Hex(5) | .[42], C[+57], N[+1420] | 1709.5371 | 4 | 6835.1265 | 6835.1864 | 0 | -8.8 | 2 | NRagged | 225.08 | 207.3 | 207.3 | 3.32 | 120 | >sp\|P01877\|IGHA2_HUMAN Immunoglobulin heavy constant alpha 2 OS=Homo sapiens OX=9606 GN=IGHA2 PE=1 SV=4 | 576 | 20201116_fqg_colostrum_1ng_glyco_hcd203040_compound15.27981.27981.4 | controllerType=0 controllerNumber=1 scan=27981 | 55.3963 |
| 26259:4 | 3 | A.[+42.011]SPTSPKVFPLSLDSTPQDGNVVVAC[+57.021]LVQGFFPQEPLSVTWSESGQN | HexNAc(4)Hex(5)NeuAc(1) | .[42], C[+57], N[+1914] | 1833.0826 | 4 | 7329.3086 | 7329.3612 | 0 | -7.2 | 2 | NRagged | 262.92 | 231.08 | 231.08 | 4.51 | 120 | >sp\|P01877\|IGHA2_HUMAN Immunoglobulin heavy constant alpha 2 OS=Homo sapiens OX=9606 GN=IGHA2 PE=1 SV=4 | 576 | 20201116_fqg_colostrum_1ng_glyco_hcd203040_compound15.30118.30118.4 | controllerType=0 controllerNumber=1 scan=30118 | 60.0663 |
| 26262:4 | 3 | A.[+42.011]SPTSPKVFPLSLDSTPQDGNVVVAC[+57.021]LVQGFFPQEPLSVTWSESGQN | HexNAc(4)Hex(5)NeuAc(1) | .[42], C[+57], N[+1914] | 1833.0826 | 4 | 7329.3086 | 7329.3612 | 0 | -7.2 | 2 | NRagged | 275.32 | 229.76 | 229.76 | 4.51 | 120 | >sp\|P01877\|IGHA2_HUMAN Immunoglobulin heavy constant alpha 2 OS=Homo sapiens OX=9606 GN=IGHA2 PE=1 SV=4 | 576 | 20201116_fqg_colostrum_1ng_glyco_hcd203040_compound15.30121.30121.4 | controllerType=0 controllerNumber=1 scan=30121 | 60.0777 |
| 24217:4 | 3 | A.[+42.011]SPTSPKVFPLSLDSTPQDGNVVVAC[+57.021]LVQGFFPQEPLSVTWSESGQN | HexNAc(3)Hex(5) | .[42], C[+57], N[+1420] | 1709.5371 | 4 | 6835.1265 | 6835.1864 | 0 | -8.8 | 2 | NRagged | 293.87 | 268.33 | 268.33 | 3.93 | 120 | >sp\|P01877\|IGHA2_HUMAN Immunoglobulin heavy constant alpha 2 OS=Homo sapiens OX=9606 GN=IGHA2 PE=1 SV=4 | 576 | 20201116_fqg_colostrum_1ng_glyco_hcd203040_compound15.27983.27983.4 | controllerType=0 controllerNumber=1 scan=27983 | 55.4007 |
| 24734:4 | 3 | A.[+42.011]SPTSPKVFPLSLDSTPQDGNVVVAC[+57.021]LVQGFFPQEPLSVTWSESGQN | HexNAc(3)Hex(5) | .[42], C[+57], N[+1420] | 1709.5376 | 4 | 6835.1284 | 6835.1864 | 0 | -8.5 | 2 | NRagged | 315.88 | 315.88 | 315.88 | 4.88 | 120 | >sp\|P01877\|IGHA2_HUMAN Immunoglobulin heavy constant alpha 2 OS=Homo sapiens OX=9606 GN=IGHA2 PE=1 SV=4 | 576 | 20201116_fqg_colostrum_1ng_glyco_hcd203040_compound15.28523.28523.4 | controllerType=0 controllerNumber=1 scan=28523 | 56.6228 |
| 24732:4 | 3 | A.[+42.011]SPTSPKVFPLSLDSTPQDGNVVVAC[+57.021]LVQGFFPQEPLSVTWSESGQN | HexNAc(3)Hex(5) | .[42], C[+57], N[+1420] | 1709.5376 | 4 | 6835.1284 | 6835.1864 | 0 | -8.5 | 2 | NRagged | 366.22 | 306.69 | 306.69 | 5.69 | 120 | >sp\|P01877\|IGHA2_HUMAN Immunoglobulin heavy constant alpha 2 OS=Homo sapiens OX=9606 GN=IGHA2 PE=1 SV=4 | 576 | 20201116_fqg_colostrum_1ng_glyco_hcd203040_compound15.28521.28521.4 | controllerType=0 controllerNumber=1 scan=28521 | [illegible] |
| 24386:4 | 3 | K.VFPLSLDSTPQDGNVVVAC[+57.021]LVQGFFPQEPLSVTWSESGQN[+2059.735]VTAI | HexNAc(4)Hex(5)Fuc(1)NeuAc(1) | C[+57], N[+2060] | 1709.7824 | 4 | 6836.1079 | 6836.0963 | 0 | 1.7 | 8 | Specific | 202.08 | 202.08 | 202.08 | 6.15 | 120 | >sp\|P01877\|IGHA2_HUMAN Immunoglobulin heavy constant alpha 2 OS=Homo sapiens OX=9606 GN=IGHA2 PE=1 SV=4 | 576 | 20201116_fqg_colostrum_1ng_glyco_hcd203040_compound15.28159.28159.4 | controllerType=0 controllerNumber=1 scan=28159 | [illegible] |
| 24077:4 | 3 | K.VFPLSLDSTPQDGNVVVAC[+57.021]LVQGFFPQEPLSVTWSESGQN[+2076.750]VTAI | HexNAc(4)Hex(6)Fuc(2) | C[+57], N[+2077] | 1714.0454 | 4 | 6853.1597 | 6853.1116 | 0 | 7 | 8 | Specific | 207.73 | 207.73 | 207.73 | 5.16 | 120 | >sp\|P01877\|IGHA2_HUMAN Immunoglobulin heavy constant alpha 2 OS=Homo sapiens OX=9606 GN=IGHA2 PE=1 SV=4 | 576 | 20201116_fqg_colostrum_1ng_glyco_hcd203040_compound15.27836.27836.4 | controllerType=0 controllerNumber=1 scan=27836 | [illegible] |
| 26103:4 | 3 | K.VFPLSLDSTPQDGNVVVAC[+57.021]LVQGFFPQEPLSVTWSESGQN[+2570.925]VTAI | HexNAc(5)Hex(6)Fuc(2)NeuAc(1) | C[+57], N[+2571] | 1837.5832 | 4 | 7347.311 | 7347.2864 | 0 | 3.3 | 8 | Specific | 211.17 | 211.17 | 211.17 | 7.4 | 120 | >sp\|P01877\|IGHA2_HUMAN Immunoglobulin heavy constant alpha 2 OS=Homo sapiens OX=9606 GN=IGHA2 PE=1 SV=4 | 576 | 20201116_fqg_colostrum_1ng_glyco_hcd203040_compound15.29955.29955.4 | controllerType=0 controllerNumber=1 scan=29955 | [illegible] |
| 27366:4 | 3 | K.VFPLSLDSTPQDGNVVVAC[+57.021]LVQGFFPQEPLSVTWSESGQN[+2350.830]VTAI | HexNAc(4)Hex(5)Fuc(1)NeuAc(2) | C[+57], N[+2351] | 1782.5615 | 4 | 7127.2241 | 7127.1917 | 0 | 4.5 | 8 | Specific | 214 | 214 | 214 | 6.95 | 120 | >sp\|P01877\|IGHA2_HUMAN Immunoglobulin heavy constant alpha 2 OS=Homo sapiens OX=9606 GN=IGHA2 PE=1 SV=4 | 576 | 20201116_fqg_colostrum_1ng_glyco_hcd203040_compound15.31271.31271.4 | controllerType=0 controllerNumber=1 scan=31271 | 62.4287 |
| 24893:5 | 3 | K.VFPLSLDSTPQDGNVVVAC[+57.021]LVQGFFPQEPLSVTWSESGQN[+2117.777]VTAI | HexNAc(5)Hex(5)Fuc(2) | C[+57], N[+2118] | 1379.6379 | 5 | 6894.1602 | 6894.1382 | 0 | 3.2 | 8 | Specific | 229.01 | 229.01 | 229.01 | 6.79 | 120 | >sp\|P01877\|IGHA2_HUMAN Immunoglobulin heavy constant alpha 2 OS=Homo sapiens OX=9606 GN=IGHA2 PE=1 SV=4 | 576 | 20201116_fqg_colostrum_1ng_glyco_hcd203040_compound15.28689.28689.5 | controllerType=0 controllerNumber=1 scan=28689 | 56.986 |
| 26288:7 | 3 | K.VFPLSLDSTPQDGNVVVAC[+57.021]LVQGFFPQEPLSVTWSESGQN[+2553.910]VTAI | HexNAc(5)Hex(5)Fuc(1)NeuAc(2) | C[+57], N[+2554] | 1048.0516 | 7 | 7330.3175 | 7330.2711 | 0 | 6.3 | 8 | Specific | 251.42 | 251.42 | 251.42 | 6.02 | 120 | >sp\|P01877\|IGHA2_HUMAN Immunoglobulin heavy constant alpha 2 OS=Homo sapiens OX=9606 GN=IGHA2 PE=1 SV=4 | 576 | 20201116_fqg_colostrum_1ng_glyco_hcd203040_compound15.30148.30148.7 | controllerType=0 controllerNumber=1 scan=30148 | 60.1332 |
| 22887:7 | 3 | K.VFPLSLDSTPQDGNVVVAC[+57.021]LVQGFFPQEPLSVTWSESGQN[+1809.666]VTAI | HexNAc(5)Hex(4)Fuc(1) | C[+57], N[+1810] | 941.7282 | 7 | 6586.0537 | 6586.0275 | 0 | 4 | 8 | Specific | 252.09 | 252.09 | 252.09 | 5.71 | 120 | >sp\|P01877\|IGHA2_HUMAN Immunoglobulin heavy constant alpha 2 OS=Homo sapiens OX=9606 GN=IGHA2 PE=1 SV=4 | 576 | 20201116_fqg_colostrum_1ng_glyco_hcd203040_compound15.26591.26591.7 | controllerType=0 controllerNumber=1 scan=26591 | 52.2465 |
| 27289:4 | 3 | K.VFPLSLDSTPQDGNVVVAC[+57.021]LVQGFFPQEPLSVTWSESGQN[+2553.910]VTAI | HexNAc(5)Hex(5)Fuc(1)NeuAc(2) | C[+57], N[+2554] | 1833.3368 | 4 | 7330.3252 | 7330.2711 | 0 | 7.4 | 8 | Specific | 254.26 | 254.26 | 254.26 | 7.77 | 120 | >sp\|P01877\|IGHA2_HUMAN Immunoglobulin heavy constant alpha 2 OS=Homo sapiens OX=9606 GN=IGHA2 PE=1 SV=4 | 576 | 20201116_fqg_colostrum_1ng_glyco_hcd203040_compound15.31191.31191.4 | controllerType=0 controllerNumber=1 scan=31191 | 62.2714 |
| 22889:7 | 3 | K.VFPLSLDSTPQDGNVVVAC[+57.021]LVQGFFPQEPLSVTWSESGQN[+1809.666]VTAI | HexNAc(5)Hex(4)Fuc(1) | C[+57], N[+1810] | 941.7282 | 7 | 6586.0537 | 6586.0275 | 0 | 4 | 8 | Specific | 279.2 | 279.2 | 279.2 | 6.35 | 120 | >sp\|P01877\|IGHA2_HUMAN Immunoglobulin heavy constant alpha 2 OS=Homo sapiens OX=9606 GN=IGHA2 PE=1 SV=4 | 576 | 20201116_fqg_colostrum_1ng_glyco_hcd203040_compound15.26593.26593.7 | controllerType=0 controllerNumber=1 scan=26593 | 52.253 |
| 26290:7 | 3 | K.VFPLSLDSTPQDGNVVVAC[+57.021]LVQGFFPQEPLSVTWSESGQN[+2553.910]VTAI | HexNAc(5)Hex(5)Fuc(1)NeuAc(2) | C[+57], N[+2554] | 1048.0516 | 7 | 7330.3175 | 7330.2711 | 0 | 6.3 | 8 | Specific | 283.94 | 283.94 | 283.94 | 6.33 | 120 | >sp\|P01877\|IGHA2_HUMAN Immunoglobulin heavy constant alpha 2 OS=Homo sapiens OX=9606 GN=IGHA2 PE=1 SV=4 | 576 | 20201116_fqg_colostrum_1ng_glyco_hcd203040_compound15.30150.30150.7 | controllerType=0 controllerNumber=1 scan=30150 | 60.1383 |
| 24729:4 | 3 | K.VFPLSLDSTPQDGNVVVAC[+57.021]LVQGFFPQEPLSVTWSESGQN[+2076.750]VTAI | HexNAc(4)Hex(6)Fuc(2) | C[+57], N[+2077] | 1714.0418 | 4 | 6853.1455 | 6853.1116 | 0 | 4.9 | 8 | Specific | 284.65 | 284.65 | 284.65 | 8.89 | 120 | >sp\|P01877\|IGHA2_HUMAN Immunoglobulin heavy constant alpha 2 OS=Homo sapiens OX=9606 GN=IGHA2 PE=1 SV=4 | 576 | 20201116_fqg_colostrum_1ng_glyco_hcd203040_compound15.28518.28518.4 | controllerType=0 controllerNumber=1 scan=28518 | 56.6113 |
| 26132:6 | 3 | K.VFPLSLDSTPQDGNVVVAC[+57.021]LVQGFFPQEPLSVTWSESGQN[+2350.830]VTAI | HexNAc(4)Hex(5)Fuc(1)NeuAc(2) | C[+57], N[+2351] | 1188.7055 | 6 | 7127.1966 | 7127.1917 | 0 | 0.7 | 8 | Specific | 310.42 | 310.42 | 310.42 | 9.6 | 120 | >sp\|P01877\|IGHA2_HUMAN Immunoglobulin heavy constant alpha 2 OS=Homo sapiens OX=9606 GN=IGHA2 PE=1 SV=4 | 576 | 20201116_fqg_colostrum_1ng_glyco_hcd203040_compound15.29985.29985.6 | controllerType=0 controllerNumber=1 scan=29985 | 59.7623 |
| 26278:5 | 3 | K.VFPLSLDSTPQDGNVVVAC[+57.021]LVQGFFPQEPLSVTWSESGQN[+2570.925]VTAI | HexNAc(5)Hex(6)Fuc(2)NeuAc(1) | C[+57], N[+2571] | 1470.2701 | 5 | 7347.3212 | 7347.2864 | 0 | 4.7 | 8 | Specific | 315.74 | 315.74 | 315.74 | 9.28 | 120 | >sp\|P01877\|IGHA2_HUMAN Immunoglobulin heavy constant alpha 2 OS=Homo sapiens OX=9606 GN=IGHA2 PE=1 SV=4 | 576 | 20201116_fqg_colostrum_1ng_glyco_hcd203040_compound15.30138.30138.5 | controllerType=0 controllerNumber=1 scan=30138 | 60.1131 |
| 22688:5 | 3 | K.VFPLSLDSTPQDGNVVVAC[+57.021]LVQGFFPQEPLSVTWSESGQN[+1647.613]VTAI | HexNAc(5)Hex(3)Fuc(1) | C[+57], N[+1648] | 1285.6126 | 5 | 6424.0337 | 6423.9746 | 0 | 9.2 | 8 | Specific | 331.81 | 331.81 | 331.81 | 7.34 | 120 | >sp\|P01877\|IGHA2_HUMAN Immunoglobulin heavy constant alpha 2 OS=Homo sapiens OX=9606 GN=IGHA2 PE=1 SV=4 | 576 | 20201116_fqg_colostrum_1ng_glyco_hcd203040_compound15.26385.26385.5 | controllerType=0 controllerNumber=1 scan=26385 | 51.8143 |
| 22533:4 | 3 | K.VFPLSLDSTPQDGNVVVAC[+57.021]LVQGFFPQEPLSVTWSESGQN[+1663.608]VTAI | HexNAc(5)Hex(4) | C[+57], N[+1664] | 1610.7594 | 4 | 6440.0158 | 6439.9696 | 0 | 7.2 | 8 | Specific | 333.75 | 295.17 | 295.17 | 8.63 | 120 | >sp\|P01877\|IGHA2_HUMAN Immunoglobulin heavy constant alpha 2 OS=Homo sapiens OX=9606 GN=IGHA2 PE=1 SV=4 | 576 | 20201116_fqg_colostrum_1ng_glyco_hcd203040_compound15.26223.26223.4 | controllerType=0 controllerNumber=1 scan=26223 | 51.5121 |
| 22830:4 | 3 | K.VFPLSLDSTPQDGNVVVAC[+57.021]LVQGFFPQEPLSVTWSESGQN[+1809.666]VTAI | HexNAc(5)Hex(4)Fuc(1) | C[+57], N[+1810] | 1647.2671 | 4 | 6586.0465 | 6586.0275 | 0 | 2.9 | 8 | Specific | 347.69 | 347.69 | 347.69 | 10.2 | 120 | >sp\|P01877\|IGHA2_HUMAN Immunoglobulin heavy constant alpha 2 OS=Homo sapiens OX=9606 GN=IGHA2 PE=1 SV=4 | 576 | 20201116_fqg_colostrum_1ng_glyco_hcd203040_compound15.26532.26532.4 | controllerType=0 controllerNumber=1 scan=26532 | 52.1282 |
| 23279:5 | 3 | K.VFPLSLDSTPQDGNVVVAC[+57.021]LVQGFFPQEPLSVTWSESGQN[+1809.666]VTAI | HexNAc(5)Hex(4)Fuc(1) | C[+57], N[+1810] | 1318.0157 | 5 | 6586.0496 | 6586.0275 | 0 | 3.4 | 8 | Specific | 370.29 | 370.29 | 370.29 | 10.39 | 120 | >sp\|P01877\|IGHA2_HUMAN Immunoglobulin heavy constant alpha 2 OS=Homo sapiens OX=9606 GN=IGHA2 PE=1 SV=4 | 576 | 20201116_fqg_colostrum_1ng_glyco_hcd203040_compound15.27002.27002.5 | controllerType=0 controllerNumber=1 scan=27002 | 53.1292 |
| 23221:4 | 3 | K.VFPLSLDSTPQDGNVVVAC[+57.021]LVQGFFPQEPLSVTWSESGQN[+1864.634]VTAI | HexNAc(2)Hex(9) | C[+57], N[+1865] | 1661.0136 | 4 | 6641.0324 | 6640.9956 | 0 | 5.5 | 8 | Specific | 379.51 | 379.51 | 379.51 | 9.73 | 120 | >sp\|P01877\|IGHA2_HUMAN Immunoglobulin heavy constant alpha 2 OS=Homo sapiens OX=9606 GN=IGHA2 PE=1 SV=4 | 576 | 20201116_fqg_colostrum_1ng_glyco_hcd203040_compound15.26940.26940.4 | controllerType=0 controllerNumber=1 scan=26940 | 52.9923 |
| 23356:4 | 3 | K.VFPLSLDSTPQDGNVVVAC[+57.021]LVQGFFPQEPLSVTWSESGQN[+1622.582]VTAI | HexNAc(4)Hex(5) | C[+57], N[+1623] | 1600.5042 | 4 | 6398.9948 | 6398.943 | 0 | 8.1 | 8 | Specific | 382.25 | 339.21 | 339.21 | 6.72 | 120 | >sp\|P01877\|IGHA2_HUMAN Immunoglobulin heavy constant alpha 2 OS=Homo sapiens OX=9606 GN=IGHA2 PE=1 SV=4 | 576 | 20201116_fqg_colostrum_1ng_glyco_hcd203040_compound15.27082.27082.4 | controllerType=0 controllerNumber=1 scan=27082 | 53.2995 |
| 27291:4 | 3 | K.VFPLSLDSTPQDGNVVVAC[+57.021]LVQGFFPQEPLSVTWSESGQN[+2553.910]VTAI | HexNAc(5)Hex(5)Fuc(1)NeuAc(2) | C[+57], N[+2554] | 1833.3368 | 4 | 7330.3252 | 7330.2711 | 0 | 7.4 | 8 | Specific | 386.59 | 313.49 | 313.49 | 9.03 | 120 | >sp\|P01877\|IGHA2_HUMAN Immunoglobulin heavy constant alpha 2 OS=Homo sapiens OX=9606 GN=IGHA2 PE=1 SV=4 | 576 | 20201116_fqg_colostrum_1ng_glyco_hcd203040_compound15.31193.31193.4 | controllerType=0 controllerNumber=1 scan=31193 | 62.2758 |
| 24756:4 | 3 | K.VFPLSLDSTPQDGNVVVAC[+57.021]LVQGFFPQEPLSVTWSESGQN[+2117.777]VTAI | HexNAc(5)Hex(5)Fuc(2) | C[+57], N[+2118] | 1724.3004 | 4 | 6894.1797 | 6894.1382 | 0 | 6 | 8 | Specific | 389.23 | 389.23 | 389.23 | 11.04 | 120 | >sp\|P01877\|IGHA2_HUMAN Immunoglobulin heavy constant alpha 2 OS=Homo sapiens OX=9606 GN=IGHA2 PE=1 SV=4 | 576 | 20201116_fqg_colostrum_1ng_glyco_hcd203040_compound15.28546.28546.4 | controllerType=0 controllerNumber=1 scan=28546 | 56.6647 |
| 24862:5 | 3 | K.VFPLSLDSTPQDGNVVVAC[+57.021]LVQGFFPQEPLSVTWSESGQN[+2059.735]VTAI | HexNAc(4)Hex(5)Fuc(1)NeuAc(1) | C[+57], N[+2060] | 1368.0326 | 5 | 6836.1341 | 6836.0963 | 0 | 5.5 | 8 | Specific | 391.58 | 391.58 | 391.58 | 11.04 | 120 | >sp\|P01877\|IGHA2_HUMAN Immunoglobulin heavy constant alpha 2 OS=Homo sapiens OX=9606 GN=IGHA2 PE=1 SV=4 | 576 | 20201116_fqg_colostrum_1ng_glyco_hcd203040_compound15.28657.28657.5 | controllerType=0 controllerNumber=1 scan=28657 | 56.9225 |
| 23224:4 | 3 | K.VFPLSLDSTPQDGNVVVAC[+57.021]LVQGFFPQEPLSVTWSESGQN[+1864.634]VTAI | HexNAc(2)Hex(9) | C[+57], N[+1865] | 1661.0136 | 4 | 6641.0324 | 6640.9956 | 0 | 5.5 | 8 | Specific | 396.53 | 306.36 | 306.36 | 7 | 120 | >sp\|P01877\|IGHA2_HUMAN Immunoglobulin heavy constant alpha 2 OS=Homo sapiens OX=9606 GN=IGHA2 PE=1 SV=4 | 576 | 20201116_fqg_colostrum_1ng_glyco_hcd203040_compound15.26943.26943.4 | controllerType=0 controllerNumber=1 scan=26943 | 52.9978 |
| 26112:6 | 3 | K.VFPLSLDSTPQDGNVVVAC[+57.021]LVQGFFPQEPLSVTWSESGQN[+2553.910]VTAI | HexNAc(5)Hex(5)Fuc(1)NeuAc(2) | C[+57], N[+2554] | 1222.5549 | 6 | 7330.2928 | 7330.2711 | 0 | 3 | 8 | Specific | 399.15 | 399.15 | 399.15 | 11.7 | 120 | >sp\|P01877\|IGHA2_HUMAN Immunoglobulin heavy constant alpha 2 OS=Homo sapiens OX=9606 GN=IGHA2 PE=1 SV=4 | 576 | 20201116_fqg_colostrum_1ng_glyco_hcd203040_compound15.29964.29964.6 | controllerType=0 controllerNumber=1 scan=29964 | 59.7192 |
| 27118:4 | 3 | K.VFPLSLDSTPQDGNVVVAC[+57.021]LVQGFFPQEPLSVTWSESGQN[+2350.830]VTAI | HexNAc(4)Hex(5)Fuc(1)NeuAc(2) | C[+57], N[+2351] | 1782.5548 | 4 | 7127.1973 | 7127.1917 | 0 | 0.8 | 8 | Specific | 412.41 | 412.41 | 412.41 | 11.67 | 120 | >sp\|P01877\|IGHA2_HUMAN Immunoglobulin heavy constant alpha 2 OS=Homo sapiens OX=9606 GN=IGHA2 PE=1 SV=4 | 576 | 20201116_fqg_colostrum_1ng_glyco_hcd203040_compound15.31013.31013.4 | controllerType=0 controllerNumber=1 scan=31013 | 61.9123 |
| 24388:4 | 3 | K.VFPLSLDSTPQDGNVVVAC[+57.021]LVQGFFPQEPLSVTWSESGQN[+2059.735]VTAI | HexNAc(4)Hex(5)Fuc(1)NeuAc(1) | C[+57], N[+2060] | 1709.7824 | 4 | 6836.1079 | 6836.0963 | 0 | 1.7 | 8 | Specific | 414.2 | 381.59 | 381.59 | 11.91 | 120 | >sp\|P01877\|IGHA2_HUMAN Immunoglobulin heavy constant alpha 2 OS=Homo sapiens OX=9606 GN=IGHA2 PE=1 SV=4 | 576 | 20201116_fqg_colostrum_1ng_glyco_hcd203040_compound15.28161.28161.4 | controllerType=0 controllerNumber=1 scan=28161 | 55.8 |
| 24200:5 | 3 | K.VFPLSLDSTPQDGNVVVAC[+57.021]LVQGFFPQEPLSVTWSESGQN[+2117.777]VTAI | HexNAc(5)Hex(5)Fuc(2) | C[+57], N[+2118] | 1379.6376 | 5 | 6894.159 | 6894.1382 | 0 | 3 | 8 | Specific | 414.89 | 414.89 | 414.89 | 11.81 | 120 | >sp\|P01877\|IGHA2_HUMAN Immunoglobulin heavy constant alpha 2 OS=Homo sapiens OX=9606 GN=IGHA2 PE=1 SV=4 | 576 | 20201116_fqg_colostrum_1ng_glyco_hcd203040_compound15.27965.27965.5 | controllerType=0 controllerNumber=1 scan=27965 | 55.3584 |
| 23296:5 | 3 | K.VFPLSLDSTPQDGNVVVAC[+57.021]LVQGFFPQEPLSVTWSESGQN[+1809.666]VTAI | HexNAc(5)Hex(4)Fuc(1) | C[+57], N[+1810] | 1318.0166 | 5 | 6586.0539 | 6586.0275 | 0 | 4 | 8 | Specific | 416.92 | 416.92 | 416.92 | 11.36 | 120 | >sp\|P01877\|IGHA2_HUMAN Immunoglobulin heavy constant alpha 2 OS=Homo sapiens OX=9606 GN=IGHA2 PE=1 SV=4 | 576 | 20201116_fqg_colostrum_1ng_glyco_hcd203040_compound15.27020.27020.5 | controllerType=0 controllerNumber=1 scan=27020 | 53.1768 |
| 24218:5 | 3 | K.VFPLSLDSTPQDGNVVVAC[+57.021]LVQGFFPQEPLSVTWSESGQN[+2059.735]VTAI | HexNAc(4)Hex(5)Fuc(1)NeuAc(1) | C[+57], N[+2060] | 1368.0303 | 5 | 6836.1225 | 6836.0963 | 0 | 3.8 | 8 | Specific | 418.34 | 418.34 | 418.34 | 11.81 | 120 | >sp\|P01877\|IGHA2_HUMAN Immunoglobulin heavy constant alpha 2 OS=Homo sapiens OX=9606 GN=IGHA2 PE=1 SV=4 | 576 | 20201116_fqg_colostrum_1ng_glyco_hcd203040_compound15.27984.27984.5 | controllerType=0 controllerNumber=1 scan=27984 | 55.402 |
| 24954:5 | 3 | K.VFPLSLDSTPQDGNVVVAC[+57.021]LVQGFFPQEPLSVTWSESGQN[+1694.603]VTAI | HexNAc(3)Hex(4)Fuc(1)NeuAc(1) | C[+57], N[+1695] | 1295.0071 | 5 | 6471.0063 | 6470.9641 | 0 | 6.5 | 8 | Specific | 434.33 | 434.33 | 434.33 | 10.89 | 120 | >sp\|P01877\|IGHA2_HUMAN Immunoglobulin heavy constant alpha 2 OS=Homo sapiens OX=9606 GN=IGHA2 PE=1 SV=4 | 576 | 20201116_fqg_colostrum_1ng_glyco_hcd203040_compound15.28752.28752.5 | controllerType=0 controllerNumber=1 scan=28752 | 57.1162 |
| 22536:4 | 3 | K.VFPLSLDSTPQDGNVVVAC[+57.021]LVQGFFPQEPLSVTWSESGQN[+1663.608]VTAI | HexNAc(5)Hex(4) | C[+57], N[+1664] | 1610.7594 | 4 | 6440.0158 | 6439.9696 | 0 | 7.2 | 8 | Specific | 455.53 | 414.44 | 414.44 | 9.59 | 120 | >sp\|P01877\|IGHA2_HUMAN Immunoglobulin heavy constant alpha 2 OS=Homo sapiens OX=9606 GN=IGHA2 PE=1 SV=4 | 576 | 20201116_fqg_colostrum_1ng_glyco_hcd203040_compound15.26226.26226.4 | controllerType=0 controllerNumber=1 scan=26226 | 51.5183 |
| 26350:4 | 3 | K.VFPLSLDSTPQDGNVVVAC[+57.021]LVQGFFPQEPLSVTWSESGQN[+2570.925]VTAI | HexNAc(5)Hex(6)Fuc(2)NeuAc(1) | C[+57], N[+2571] | 1837.5885 | 4 | 7347.332 | 7347.2864 | 0 | 6.2 | 8 | Specific | 456.68 | 362.81 | 362.81 | 9.91 | 120 | >sp\|P01877\|IGHA2_HUMAN Immunoglobulin heavy constant alpha 2 OS=Homo sapiens OX=9606 GN=IGHA2 PE=1 SV=4 | 576 | 20201116_fqg_colostrum_1ng_glyco_hcd203040_compound15.30213.30213.4 | controllerType=0 controllerNumber=1 scan=30213 | 60.268 |
| 26134:6 | 3 | K.VFPLSLDSTPQDGNVVVAC[+57.021]LVQGFFPQEPLSVTWSESGQN[+2350.830]VTAI | HexNAc(4)Hex(5)Fuc(1)NeuAc(2) | C[+57], N[+2351] | 1188.7055 | 6 | 7127.1966 | 7127.1917 | 0 | 0.7 | 8 | Specific | 463.71 | 420.26 | 420.26 | 11.86 | 120 | >sp\|P01877\|IGHA2_HUMAN Immunoglobulin heavy constant alpha 2 OS=Homo sapiens OX=9606 GN=IGHA2 PE=1 SV=4 | 576 | 20201116_fqg_colostrum_1ng_glyco_hcd203040_compound15.29987.29987.6 | controllerType=0 controllerNumber=1 scan=29987 | 59.7677 |
| 27259:4 | 3 | K.VFPLSLDSTPQDGNVVVAC[+57.021]LVQGFFPQEPLSVTWSESGQN[+2350.830]VTAI | HexNAc(4)Hex(5)Fuc(1)NeuAc(2) | C[+57], N[+2351] | 1782.557 | 4 | 7127.2061 | 7127.1917 | 0 | 2 | 8 | Specific | 469.42 | 446.69 | 446.69 | 12.1 | 120 | >sp\|P01877\|IGHA2_HUMAN Immunoglobulin heavy constant alpha 2 OS=Homo sapiens OX=9606 GN=IGHA2 PE=1 SV=4 | 576 | 20201116_fqg_colostrum_1ng_glyco_hcd203040_compound15.31160.31160.4 | controllerType=0 controllerNumber=1 scan=31160 | 62.2012 |
| 22867:5 | 3 | K.VFPLSLDSTPQDGNVVVAC[+57.021]LVQGFFPQEPLSVTWSESGQN[+1622.582]VTAI | HexNAc(4)Hex(5) | C[+57], N[+1623] | 1280.6033 | 5 | 6398.9873 | 6398.943 | 0 | 6.9 | 8 | Specific | 469.69 | 446.55 | 446.55 | 11.1 | 120 | >sp\|P01877\|IGHA2_HUMAN Immunoglobulin heavy constant alpha 2 OS=Homo sapiens OX=9606 GN=IGHA2 PE=1 SV=4 | 576 | 20201116_fqg_colostrum_1ng_glyco_hcd203040_compound15.26571.26571.5 | controllerType=0 controllerNumber=1 scan=26571 | 52.2089 |
| 27120:4 | 3 | K.VFPLSLDSTPQDGNVVVAC[+57.021]LVQGFFPQEPLSVTWSESGQN[+2350.830]VTAI | HexNAc(4)Hex(5)Fuc(1)NeuAc(2) | C[+57], N[+2351] | 1782.5548 | 4 | 7127.1973 | 7127.1917 | 0 | 0.8 | 8 | Specific | 480.72 | 425.07 | 425.07 | 11.94 | 120 | >sp\|P01877\|IGHA2_HUMAN Immunoglobulin heavy constant alpha 2 OS=Homo sapiens OX=9606 GN=IGHA2 PE=1 SV=4 | 576 | 20201116_fqg_colostrum_1ng_glyco_hcd203040_compound15.31015.31015.4 | controllerType=0 controllerNumber=1 scan=31015 | 61.9164 |
| 24957:5 | 3 | K.VFPLSLDSTPQDGNVVVAC[+57.021]LVQGFFPQEPLSVTWSESGQN[+1694.603]VTAI | HexNAc(3)Hex(4)Fuc(1)NeuAc(1) | C[+57], N[+1695] | 1295.0071 | 5 | 6471.0063 | 6470.9641 | 0 | 6.5 | 8 | Specific | 510.85 | 510.85 | 510.85 | 11.3 | 120 | >sp\|P01877\|IGHA2_HUMAN Immunoglobulin heavy constant alpha 2 OS=Homo sapiens OX=9606 GN=IGHA2 PE=1 SV=4 | 576 | 20201116_fqg_colostrum_1ng_glyco_hcd203040_compound15.28755.28755.5 | controllerType=0 controllerNumber=1 scan=28755 | 57.1238 |
| 24108:4 | 3 | K.VFPLSLDSTPQDGNVVVAC[+57.021]LVQGFFPQEPLSVTWSESGQN[+2279.830]VTAI | HexNAc(5)Hex(6)Fuc(2) | C[+57], N[+2280] | 1764.8144 | 4 | 7056.2358 | 7056.191 | 0 | 6.4 | 8 | Specific | 524.78 | 515.33 | 515.33 | 11.41 | 120 | >sp\|P01877\|IGHA2_HUMAN Immunoglobulin heavy constant alpha 2 OS=Homo sapiens OX=9606 GN=IGHA2 PE=1 SV=4 | 576 | 20201116_fqg_colostrum_1ng_glyco_hcd203040_compound15.27869.27869.4 | controllerType=0 controllerNumber=1 scan=27869 | 55.128 |
| 24889:4 | 3 | K.VFPLSLDSTPQDGNVVVAC[+57.021]LVQGFFPQEPLSVTWSESGQN[+2117.777]VTAI | HexNAc(5)Hex(5)Fuc(2) | C[+57], N[+2118] | 1724.2954 | 4 | 6894.1597 | 6894.1382 | 0 | 3.1 | 8 | Specific | 541.2 | 460.84 | 460.84 | 12.4 | 120 | >sp\|P01877\|IGHA2_HUMAN Immunoglobulin heavy constant alpha 2 OS=Homo sapiens OX=9606 GN=IGHA2 PE=1 SV=4 | 576 | 20201116_fqg_colostrum_1ng_glyco_hcd203040_compound15.28685.28685.4 | controllerType=0 controllerNumber=1 scan=28685 | 56.9785 |
| 24057:6 | 3 | K.VFPLSLDSTPQDGNVVVAC[+57.021]LVQGFFPQEPLSVTWSESGQN[+2263.835]VTAI | HexNAc(5)Hex(5)Fuc(3) | C[+57], N[+2264] | 1174.2087 | 6 | 7040.2157 | 7040.1961 | 0 | 2.8 | 8 | Specific | 543.11 | 493.21 | 493.21 | 12.4 | 120 | >sp\|P01877\|IGHA2_HUMAN Immunoglobulin heavy constant alpha 2 OS=Homo sapiens OX=9606 GN=IGHA2 PE=1 SV=4 | 576 | 20201116_fqg_colostrum_1ng_glyco_hcd203040_compound15.27815.27815.2 | controllerType=0 controllerNumber=1 scan=27815 | 54.9899 |
| 26283:6 | 3 | K.VFPLSLDSTPQDGNVVVAC[+57.021]LVQGFFPQEPLSVTWSESGQN[+2350.830]VTAI | HexNAc(4)Hex(5)Fuc(1)NeuAc(2) | C[+57], N[+2351] | 1188.7114 | 6 | 7127.2318 | 7127.1917 | 0 | 5.6 | 8 | Specific | 554.81 | 554.81 | 554.81 | 11.74 | 120 | >sp\|P01877\|IGHA2_HUMAN Immunoglobulin heavy constant alpha 2 OS=Homo sapiens OX=9606 GN=IGHA2 PE=1 SV=4 | 576 | 20201116_fqg_colostrum_1ng_glyco_hcd203040_compound15.30143.30143.6 | controllerType=0 controllerNumber=1 scan=30143 | 60.1236 |
| 24059:6 | 3 | K.VFPLSLDSTPQDGNVVVAC[+57.021]LVQGFFPQEPLSVTWSESGQN[+2263.835]VTAI | HexNAc(5)Hex(5)Fuc(3) | C[+57], N[+2264] | 1174.2087 | 6 | 7040.2157 | 7040.1961 | 0 | 2.8 | 8 | Specific | 555.59 | 555.59 | 555.59 | 12.4 | 120 | >sp\|P01877\|IGHA2_HUMAN Immunoglobulin heavy constant alpha 2 OS=Homo sapiens OX=9606 GN=IGHA2 PE=1 SV=4 | 576 | 20201116_fqg_colostrum_1ng_glyco_hcd203040_compound15.27817.27817.2 | controllerType=0 controllerNumber=1 scan=27817 | 54.9967 |
| 22691:5 | 3 | K.VFPLSLDSTPQDGNVVVAC[+57.021]LVQGFFPQEPLSVTWSESGQN[+1647.613]VTAI | HexNAc(5)Hex(3)Fuc(1) | C[+57], N[+1648] | 1285.6126 | 5 | 6424.0337 | 6423.9746 | 0 | 9.2 | 8 | Specific | 584.16 | 584.16 | 584.16 | 12.88 | 120 | >sp\|P01877\|IGHA2_HUMAN Immunoglobulin heavy constant alpha 2 OS=Homo sapiens OX=9606 GN=IGHA2 PE=1 SV=4 | 576 | 20201116_fqg_colostrum_1ng_glyco_hcd203040_compound15.26388.26388.5 | controllerType=0 controllerNumber=1 scan=26388 | 51.8327 |
| 26114:6 | 3 | K.VFPLSLDSTPQDGNVVVAC[+57.021]LVQGFFPQEPLSVTWSESGQN[+2553.910]VTAI | HexNAc(5)Hex(5)Fuc(1)NeuAc(2) | C[+57], N[+2554] | 1222.5549 | 6 | 7330.2928 | 7330.2711 | 0 | 3 | 8 | Specific | 603.15 | 561.25 | 561.25 | 14.97 | 120 | >sp\|P01877\|IGHA2_HUMAN Immunoglobulin heavy constant alpha 2 OS=Homo sapiens OX=9606 GN=IGHA2 PE=1 SV=4 | 576 | 20201116_fqg_colostrum_1ng_glyco_hcd203040_compound15.29966.29966.6 | controllerType=0 controllerNumber=1 scan=29966 | 59.725 |
| 22705:5 | 3 | K.VFPLSLDSTPQDGNVVVAC[+57.021]LVQGFFPQEPLSVTWSESGQN[+1971.719]VTAI | HexNAc(5)Hex(5)Fuc(1) | C[+57], N[+1972] | 1350.4252 | 5 | 6748.097 | 6748.0803 | 0 | 2.5 | 8 | Specific | 608.74 | 608.74 | 608.74 | 15.08 | 120 | >sp\|P01877\|IGHA2_HUMAN Immunoglobulin heavy constant alpha 2 OS=Homo sapiens OX=9606 GN=IGHA2 PE=1 SV=4 | 576 | 20201116_fqg_colostrum_1ng_glyco_hcd203040_compound15.26403.26403.5 | controllerType=0 controllerNumber=1 scan=26403 | 51.8704 |
| 22707:5 | 3 | K.VFPLSLDSTPQDGNVVVAC[+57.021]LVQGFFPQEPLSVTWSESGQN[+1971.719]VTAI | HexNAc(5)Hex(5)Fuc(1) | C[+57], N[+1972] | 1350.4252 | 5 | 6748.097 | 6748.0803 | 0 | 2.5 | 8 | Specific | 610.51 | 610.51 | 610.51 | 15.08 | 120 | >sp\|P01877\|IGHA2_HUMAN Immunoglobulin heavy constant alpha 2 OS=Homo sapiens OX=9606 GN=IGHA2 PE=1 SV=4 | 576 | 20201116_fqg_colostrum_1ng_glyco_hcd203040_compound15.26405.26405.5 | controllerType=0 controllerNumber=1 scan=26405 | 51.8827 |
| 22978:4 | 3 | K.VFPLSLDSTPQDGNVVVAC[+57.021]LVQGFFPQEPLSVTWSESGQN[+1038.375]VTAI | HexNAc(2)Hex(3)Fuc(1) | C[+57], N[+1038] | 1454.4465 | 4 | 5814.7643 | 5814.7365 | 0 | 4.8 | 8 | Specific | 616.02 | 616.02 | 616.02 | 14.53 | 120 | >sp\|P01877\|IGHA2_HUMAN Immunoglobulin heavy constant alpha 2 OS=Homo sapiens OX=9606 GN=IGHA2 PE=1 SV=4 | 576 | 20201116_fqg_colostrum_1ng_glyco_hcd203040_compound15.26686.26686.4 | controllerType=0 controllerNumber=1 scan=26686 | 52.417 |
| 26258:5 | 3 | K.VFPLSLDSTPQDGNVVVAC[+57.021]LVQGFFPQEPLSVTWSESGQN[+2553.910]VTAI | HexNAc(5)Hex(5)Fuc(1)NeuAc(2) | C[+57], N[+2554] | 1466.8682 | 5 | 7330.3119 | 7330.2711 | 0 | 5.6 | 8 | Specific | 632.88 | 627.93 | 627.93 | 14.31 | 120 | >sp\|P01877\|IGHA2_HUMAN Immunoglobulin heavy constant alpha 2 OS=Homo sapiens OX=9606 GN=IGHA2 PE=1 SV=4 | 576 | 20201116_fqg_colostrum_1ng_glyco_hcd203040_compound15.30117.30117.5 | controllerType=0 controllerNumber=1 scan=30117 | 60.0652 |
| 24036:5 | 3 | K.VFPLSLDSTPQDGNVVVAC[+57.021]LVQGFFPQEPLSVTWSESGQN[+2262.814]VTAI | HexNAc(5)Hex(5)Fuc(1)NeuAc(1) | C[+57], N[+2263] | 1408.6458 | 5 | 7039.1999 | 7039.1757 | 0 | 3.4 | 8 | Specific | 634.38 | 634.38 | 634.38 | 14.97 | 120 | >sp\|P01877\|IGHA2_HUMAN Immunoglobulin heavy constant alpha 2 OS=Homo sapiens OX=9606 GN=IGHA2 PE=1 SV=4 | 576 | 20201116_fqg_colostrum_1ng_glyco_hcd203040_compound15.27793.27793.5 | controllerType=0 controllerNumber=1 scan=27793 | 54.9426 |
| 26180:4 | 3 | K.VFPLSLDSTPQDGNVVVAC[+57.021]LVQGFFPQEPLSVTWSESGQN[+2553.910]VTAI | HexNAc(5)Hex(5)Fuc(1)NeuAc(2) | C[+57], N[+2554] | 1833.3297 | 4 | 7330.2969 | 7330.2711 | 0 | 3.5 | 8 | Specific | 637.63 | 566.84 | 566.84 | 14.97 | 120 | >sp\|P01877\|IGHA2_HUMAN Immunoglobulin heavy constant alpha 2 OS=Homo sapiens OX=9606 GN=IGHA2 PE=1 SV=4 | 576 | 20201116_fqg_colostrum_1ng_glyco_hcd203040_compound15.30035.30035.4 | controllerType=0 controllerNumber=1 scan=30035 | 59.8968 |
| 24651:4 | 3 | K.VFPLSLDSTPQDGNVVVAC[+57.021]LVQGFFPQEPLSVTWSESGQN[+2262.814]VTAI | HexNAc(5)Hex(5)Fuc(1)NeuAc(1) | C[+57], N[+2263] | 1760.554 | 4 | 7039.1943 | 7039.1757 | 0 | 2.6 | 8 | Specific | 640.25 | 599.65 | 599.65 | 15.08 | 120 | >sp\|P01877\|IGHA2_HUMAN Immunoglobulin heavy constant alpha 2 OS=Homo sapiens OX=9606 GN=IGHA2 PE=1 SV=4 | 576 | 20201116_fqg_colostrum_1ng_glyco_hcd203040_compound15.28436.28436.4 | controllerType=0 controllerNumber=1 scan=28436 | 56.4177 |
| 23254:4 | 3 | K.VFPLSLDSTPQDGNVVVAC[+57.021]LVQGFFPQEPLSVTWSESGQN[+1809.666]VTAI | HexNAc(5)Hex(4)Fuc(1) | C[+57], N[+1810] | 1647.269 | 4 | 6586.0544 | 6586.0275 | 0 | 4.1 | 8 | Specific | 647.35 | 630.69 | 630.69 | 14.53 | 120 | >sp\|P01877\|IGHA2_HUMAN Immunoglobulin heavy constant alpha 2 OS=Homo sapiens OX=9606 GN=IGHA2 PE=1 SV=4 | 576 | 20201116_fqg_colostrum_1ng_glyco_hcd203040_compound15.26976.26976.4 | controllerType=0 controllerNumber=1 scan=26976 | 53.0702 |
| 26285:6 | 3 | K.VFPLSLDSTPQDGNVVVAC[+57.021]LVQGFFPQEPLSVTWSESGQN[+2350.830]VTAI | HexNAc(4)Hex(5)Fuc(1)NeuAc(2) | C[+57], N[+2351] | 1188.7114 | 6 | 7127.2318 | 7127.1917 | 0 | 5.6 | 8 | Specific | 650.89 | 647.12 | 647.12 | 15.31 | 120 | >sp\|P01877\|IGHA2_HUMAN Immunoglobulin heavy constant alpha 2 OS=Homo sapiens OX=9606 GN=IGHA2 PE=1 SV=4 | 576 | 20201116_fqg_colostrum_1ng_glyco_hcd203040_compound15.30145.30145.6 | controllerType=0 controllerNumber=1 scan=30145 | 60.1284 |
| 23469:4 | 3 | K.VFPLSLDSTPQDGNVVVAC[+57.021]LVQGFFPQEPLSVTWSESGQN[+1444.534]VTAI | HexNAc(4)Hex(3)Fuc(1) | C[+57], N[+1445] | 1555.9858 | 4 | 6220.9215 | 6220.8953 | 0 | 4.2 | 8 | Specific | 653.76 | 613.47 | 613.47 | 15.53 | 120 | >sp\|P01877\|IGHA2_HUMAN Immunoglobulin heavy constant alpha 2 OS=Homo sapiens OX=9606 GN=IGHA2 PE=1 SV=4 | 576 | 20201116_fqg_colostrum_1ng_glyco_hcd203040_compound15.27200.27200.4 | controllerType=0 controllerNumber=1 scan=27200 | 53.552 |
| 27261:4 | 3 | K.VFPLSLDSTPQDGNVVVAC[+57.021]LVQGFFPQEPLSVTWSESGQN[+2350.830]VTAI | HexNAc(4)Hex(5)Fuc(1)NeuAc(2) | C[+57], N[+2351] | 1782.557 | 4 | 7127.2061 | 7127.1917 | 0 | 2 | 8 | Specific | 656.22 | 573.84 | 573.84 | 16.08 | 120 | >sp\|P01877\|IGHA2_HUMAN Immunoglobulin heavy constant alpha 2 OS=Homo sapiens OX=9606 GN=IGHA2 PE=1 SV=4 | 576 | 20201116_fqg_colostrum_1ng_glyco_hcd203040_compound15.31162.31162.4 | controllerType=0 controllerNumber=1 scan=31162 | 62.209 |
| 22710:5 | 3 | K.VFPLSLDSTPQDGNVVVAC[+57.021]LVQGFFPQEPLSVTWSESGQN[+1606.587]VTAI | HexNAc(4)Hex(4)Fuc(1) | C[+57], N[+1607] | 1277.3978 | 5 | 6382.96 | 6382.9481 | 0 | 1.9 | 8 | Specific | 658.06 | 634.51 | 634.51 | 16.08 | 120 | >sp\|P01877\|IGHA2_HUMAN Immunoglobulin heavy constant alpha 2 OS=Homo sapiens OX=9606 GN=IGHA2 PE=1 SV=4 | 576 | 20201116_fqg_colostrum_1ng_glyco_hcd203040_compound15.26408.26408.5 | controllerType=0 controllerNumber=1 scan=26408 | 51.8872 |
| 26178:4 | 3 | K.VFPLSLDSTPQDGNVVVAC[+57.021]LVQGFFPQEPLSVTWSESGQN[+2553.910]VTAI | HexNAc(5)Hex(5)Fuc(1)NeuAc(2) | C[+57], N[+2554] | 1833.3297 | 4 | 7330.2969 | 7330.2711 | 0 | 3.5 | 8 | Specific | 659.55 | 629.76 | 629.76 | 15.97 | 120 | >sp\|P01877\|IGHA2_HUMAN Immunoglobulin heavy constant alpha 2 OS=Homo sapiens OX=9606 GN=IGHA2 PE=1 SV=4 | 576 | 20201116_fqg_colostrum_1ng_glyco_hcd203040_compound15.30033.30033.4 | controllerType=0 controllerNumber=1 scan=30033 | 59.8904 |
| 22713:5 | 3 | K.VFPLSLDSTPQDGNVVVAC[+57.021]LVQGFFPQEPLSVTWSESGQN[+1606.587]VTAI | HexNAc(4)Hex(4)Fuc(1) | C[+57], N[+1607] | 1277.3978 | 5 | 6382.96 | 6382.9481 | 0 | 1.9 | 8 | Specific | 666.82 | 623.53 | 623.53 | 16.08 | 120 | >sp\|P01877\|IGHA2_HUMAN Immunoglobulin heavy constant alpha 2 OS=Homo sapiens OX=9606 GN=IGHA2 PE=1 SV=4 | 576 | 20201116_fqg_colostrum_1ng_glyco_hcd203040_compound15.26411.26411.5 | controllerType=0 controllerNumber=1 scan=26411 | 51.9068 |
| 24991:4 | 3 | K.VFPLSLDSTPQDGNVVVAC[+57.021]LVQGFFPQEPLSVTWSESGQN[+1694.603]VTAI | HexNAc(3)Hex(4)Fuc(1)NeuAc(1) | C[+57], N[+1695] | 1618.4961 | 4 | 6470.9626 | 6470.9641 | 0 | -0.2 | 8 | Specific | 677.6 | 661.57 | 661.57 | 15.34 | 120 | >sp\|P01877\|IGHA2_HUMAN Immunoglobulin heavy constant alpha 2 OS=Homo sapiens OX=9606 GN=IGHA2 PE=1 SV=4 | 576 | 20201116_fqg_colostrum_1ng_glyco_hcd203040_compound15.28791.28791.4 | controllerType=0 controllerNumber=1 scan=28791 | 57.2039 |
| 26171:5 | 3 | K.VFPLSLDSTPQDGNVVVAC[+57.021]LVQGFFPQEPLSVTWSESGQN[+2553.910]VTAI | HexNAc(5)Hex(5)Fuc(1)NeuAc(2) | C[+57], N[+2554] | 1466.8685 | 5 | 7330.3132 | 7330.2711 | 0 | 5.7 | 8 | Specific | 680.85 | 674.24 | 674.24 | 15.31 | 120 | >sp\|P01877\|IGHA2_HUMAN Immunoglobulin heavy constant alpha 2 OS=Homo sapiens OX=9606 GN=IGHA2 PE=1 SV=4 | 576 | 20201116_fqg_colostrum_1ng_glyco_hcd203040_compound15.30026.30026.5 | controllerType=0 controllerNumber=1 scan=30026 | 59.8659 |
| 24066:4 | 3 | K.VFPLSLDSTPQDGNVVVAC[+57.021]LVQGFFPQEPLSVTWSESGQN[+2262.814]VTAI | HexNAc(5)Hex(5)Fuc(1)NeuAc(1) | C[+57], N[+2263] | 1760.5605 | 4 | 7039.2202 | 7039.1757 | 0 | 6.3 | 8 | Specific | 683.7 | 664.12 | 664.12 | 15.08 | 120 | >sp\|P01877\|IGHA2_HUMAN Immunoglobulin heavy constant alpha 2 OS=Homo sapiens OX=9606 GN=IGHA2 PE=1 SV=4 | 576 | 20201116_fqg_colostrum_1ng_glyco_hcd203040_compound15.27825.27825.4 | controllerType=0 controllerNumber=1 scan=27825 | 55.0132 |
| 24710:4 | 3 | K.VFPLSLDSTPQDGNVVVAC[+57.021]LVQGFFPQEPLSVTWSESGQN[+2262.814]VTAI | HexNAc(5)Hex(5)Fuc(1)NeuAc(1) | C[+57], N[+2263] | 1760.5503 | 4 | 7039.1792 | 7039.1757 | 0 | 0.5 | 8 | Specific | 684.48 | 676.09 | 676.09 | 15.84 | 120 | >sp\|P01877\|IGHA2_HUMAN Immunoglobulin heavy constant alpha 2 OS=Homo sapiens OX=9606 GN=IGHA2 PE=1 SV=4 | 576 | 20201116_fqg_colostrum_1ng_glyco_hcd203040_compound15.28498.28498.4 | controllerType=0 controllerNumber=1 scan=28498 | 56.5546 |
| 23433:5 | 3 | K.VFPLSLDSTPQDGNVVVAC[+57.021]LVQGFFPQEPLSVTWSESGQN[+1444.534]VTAI | HexNAc(4)Hex(3)Fuc(1) | C[+57], N[+1445] | 1244.9895 | 5 | 6220.9184 | 6220.8953 | 0 | 3.7 | 8 | Specific | 685.22 | 669.55 | 669.55 | 15.97 | 120 | >sp\|P01877\|IGHA2_HUMAN Immunoglobulin heavy constant alpha 2 OS=Homo sapiens OX=9606 GN=IGHA2 PE=1 SV=4 | 576 | 20201116_fqg_colostrum_1ng_glyco_hcd203040_compound15.27162.27162.5 | controllerType=0 controllerNumber=1 scan=27162 | 53.4603 |
| 26148:4 | 3 | K.VFPLSLDSTPQDGNVVVAC[+57.021]LVQGFFPQEPLSVTWSESGQN[+2350.830]VTAI | HexNAc(4)Hex(5)Fuc(1)NeuAc(2) | C[+57], N[+2351] | 1782.5621 | 4 | 7127.2266 | 7127.1917 | 0 | 4.9 | 8 | Specific | 688.8 | 650.52 | 650.52 | 15.53 | 120 | >sp\|P01877\|IGHA2_HUMAN Immunoglobulin heavy constant alpha 2 OS=Homo sapiens OX=9606 GN=IGHA2 PE=1 SV=4 | 576 | 20201116_fqg_colostrum_1ng_glyco_hcd203040_compound15.30002.30002.4 | controllerType=0 controllerNumber=1 scan=30002 | 59.7985 |
| 24370:4 | 3 | K.VFPLSLDSTPQDGNVVVAC[+57.021]LVQGFFPQEPLSVTWSESGQN[+1694.603]VTAI | HexNAc(3)Hex(4)Fuc(1)NeuAc(1) | C[+57], N[+1695] | 1618.5013 | 4 | 6470.9836 | 6470.9641 | 0 | 3 | 8 | Specific | 693.87 | 645.88 | 645.88 | 15.97 | 120 | >sp\|P01877\|IGHA2_HUMAN Immunoglobulin heavy constant alpha 2 OS=Homo sapiens OX=9606 GN=IGHA2 PE=1 SV=4 | 576 | 20201116_fqg_colostrum_1ng_glyco_hcd203040_compound15.28143.28143.4 | controllerType=0 controllerNumber=1 scan=28143 | 55.7618 |
| 24069:5 | 3 | K.VFPLSLDSTPQDGNVVVAC[+57.021]LVQGFFPQEPLSVTWSESGQN[+2262.814]VTAI | HexNAc(5)Hex(5)Fuc(1)NeuAc(1) | C[+57], N[+2263] | 1408.6487 | 5 | 7039.2145 | 7039.1757 | 0 | 5.5 | 8 | Specific | 703.5 | 697.31 | 697.31 | 16.31 | 120 | >sp\|P01877\|IGHA2_HUMAN Immunoglobulin heavy constant alpha 2 OS=Homo sapiens OX=9606 GN=IGHA2 PE=1 SV=4 | 576 | 20201116_fqg_colostrum_1ng_glyco_hcd203040_compound15.27828.27828.5 | controllerType=0 controllerNumber=1 scan=27828 | 55.021 |
| 26169:5 | 3 | K.VFPLSLDSTPQDGNVVVAC[+57.021]LVQGFFPQEPLSVTWSESGQN[+2553.910]VTAI | HexNAc(5)Hex(5)Fuc(1)NeuAc(2) | C[+57], N[+2554] | 1466.8685 | 5 | 7330.3132 | 7330.2711 | 0 | 5.7 | 8 | Specific | 705.34 | 705.34 | 705.34 | 16.31 | 120 | >sp\|P01877\|IGHA2_HUMAN Immunoglobulin heavy constant alpha 2 OS=Homo sapiens OX=9606 GN=IGHA2 PE=1 SV=4 | 576 | 20201116_fqg_colostrum_1ng_glyco_hcd203040_compound15.30024.30024.5 | controllerType=0 controllerNumber=1 scan=30024 | 59.8481 |
| 22812:5 | 3 | K.VFPLSLDSTPQDGNVVVAC[+57.021]LVQGFFPQEPLSVTWSESGQN[+1444.534]VTAI | HexNAc(4)Hex(3)Fuc(1) | C[+57], N[+1445] | 1244.9877 | 5 | 6220.9093 | 6220.8953 | 0 | 2.2 | 8 | Specific | 707.31 | 707.31 | 707.31 | 17.08 | 120 | >sp\|P01877\|IGHA2_HUMAN Immunoglobulin heavy constant alpha 2 OS=Homo sapiens OX=9606 GN=IGHA2 PE=1 SV=4 | 576 | 20201116_fqg_colostrum_1ng_glyco_hcd203040_compound15.26514.26514.5 | controllerType=0 controllerNumber=1 scan=26514 | 52.0927 |
| 24068:4 | 3 | K.VFPLSLDSTPQDGNVVVAC[+57.021]LVQGFFPQEPLSVTWSESGQN[+2262.814]VTAI | HexNAc(5)Hex(5)Fuc(1)NeuAc(1) | C[+57], N[+2263] | 1760.5605 | 4 | 7039.2202 | 7039.1757 | 0 | 6.3 | 8 | Specific | 716.65 | 685.48 | 685.48 | 16.08 | 120 | >sp\|P01877\|IGHA2_HUMAN Immunoglobulin heavy constant alpha 2 OS=Homo sapiens OX=9606 GN=IGHA2 PE=1 SV=4 | 576 | 20201116_fqg_colostrum_1ng_glyco_hcd203040_compound15.27827.27827.4 | controllerType=0 controllerNumber=1 scan=27827 | 55.0199 |
| 26174:6 | 3 | K.VFPLSLDSTPQDGNVVVAC[+57.021]LVQGFFPQEPLSVTWSESGQN[+2553.910]VTAI | HexNAc(5)Hex(5)Fuc(1)NeuAc(2) | C[+57], N[+2554] | 1222.5582 | 6 | 7330.3126 | 7330.2711 | 0 | 5.7 | 8 | Specific | 717.73 | 692.99 | 692.99 | 16.31 | 120 | >sp\|P01877\|IGHA2_HUMAN Immunoglobulin heavy constant alpha 2 OS=Homo sapiens OX=9606 GN=IGHA2 PE=1 SV=4 | 576 | 20201116_fqg_colostrum_1ng_glyco_hcd203040_compound15.30029.30029.6 | controllerType=0 controllerNumber=1 scan=30029 | 59.876 |
| 22829:6 | 3 | K.VFPLSLDSTPQDGNVVVAC[+57.021]LVQGFFPQEPLSVTWSESGQN[+1444.534]VTAI | HexNAc(4)Hex(3)Fuc(1) | C[+57], N[+1445] | 1037.6571 | 6 | 6220.9065 | 6220.8953 | 0 | 1.8 | 8 | Specific | 725.83 | 725.83 | 725.83 | 17.08 | 120 | >sp\|P01877\|IGHA2_HUMAN Immunoglobulin heavy constant alpha 2 OS=Homo sapiens OX=9606 GN=IGHA2 PE=1 SV=4 | 576 | 20201116_fqg_colostrum_1ng_glyco_hcd203040_compound15.26531.26531.6 | controllerType=0 controllerNumber=1 scan=26531 | 52.1267 |

Table S4. Cont.

<table>
<tr><th>Query #:z</th><th>Protein Rank</th><th>Glycopeptide</th><th>Glycan</th><th>Modification Type(s)</th><th>Observed m/z</th><th>z</th><th>Observed (M+H)</th><th>Calc. mass (M+H)</th><th>Off-by-x error</th><th>Mass error (ppm)</th><th>StartPosition</th><th>Cleavage</th><th>Score</th><th>Delta</th><th>Delta Mod</th><th>|Log Prob|</th><th># of unique peptides</th><th>Protein Name</th><th>Protein DB number</th><th>Comment</th><th>Scan #</th><th>Scan Time</th></tr>
<tr><td>24653:4</td><td>3</td><td>K.VFPLSLDSTPQDGNVVVAC[+57.021]LVQGFFPQEPLSVTWSESGQN[+2262.814]VTAI</td><td>HexNAc(5)Hex(5)Fuc(1)NeuAc(1)</td><td>C[+57], N[+2263]</td><td>1760.554</td><td>4</td><td>7039.1943</td><td>7039.1757</td><td>0</td><td>2.6</td><td>8</td><td>Specific</td><td>727.83</td><td>685.46</td><td>685.46</td><td>17.08</td><td>120</td><td>>sp|P01877|IGHA2_HUMAN Immunoglobulin heavy constant alpha 2 OS=Homo sapiens OX=9606 GN=IGHA2 PE=1 SV=4</td><td>576</td><td>20201116_fqg_colostrum_1ng_glyco_hcd203040_compound15.28438.28438.4</td><td>controllerType=0 controllerNumber=1 scan=28438</td><td>56.4243</td></tr>
<tr><td>23349:4</td><td>3</td><td>K.VFPLSLDSTPQDGNVVVAC[+57.021]LVQGFFPQEPLSVTWSESGQN[+1647.613]VTAI</td><td>HexNAc(5)Hex(3)Fuc(1)</td><td>C[+57], N[+1648]</td><td>1606.7545</td><td>4</td><td>6423.9962</td><td>6423.9746</td><td>0</td><td>3.4</td><td>8</td><td>Specific</td><td>734.29</td><td>726.68</td><td>726.68</td><td>16.97</td><td>120</td><td>>sp|P01877|IGHA2_HUMAN Immunoglobulin heavy constant alpha 2 OS=Homo sapiens OX=9606 GN=IGHA2 PE=1 SV=4</td><td>576</td><td>20201116_fqg_colostrum_1ng_glyco_hcd203040_compound15.27075.27075.4</td><td>controllerType=0 controllerNumber=1 scan=27075</td><td>53.287</td></tr>
<tr><td>24080:6</td><td>3</td><td>K.VFPLSLDSTPQDGNVVVAC[+57.021]LVQGFFPQEPLSVTWSESGQN[+2262.814]VTAI</td><td>HexNAc(5)Hex(5)Fuc(1)NeuAc(1)</td><td>C[+57], N[+2263]</td><td>1174.0412</td><td>6</td><td>7039.2108</td><td>7039.1757</td><td>0</td><td>5</td><td>8</td><td>Specific</td><td>737.68</td><td>703.41</td><td>703.41</td><td>16.53</td><td>120</td><td>>sp|P01877|IGHA2_HUMAN Immunoglobulin heavy constant alpha 2 OS=Homo sapiens OX=9606 GN=IGHA2 PE=1 SV=4</td><td>576</td><td>20201116_fqg_colostrum_1ng_glyco_hcd203040_compound15.27839.27839.6</td><td>controllerType=0 controllerNumber=1 scan=27839</td><td>55.0514</td></tr>
<tr><td>24082:6</td><td>3</td><td>K.VFPLSLDSTPQDGNVVVAC[+57.021]LVQGFFPQEPLSVTWSESGQN[+2262.814]VTAI</td><td>HexNAc(5)Hex(5)Fuc(1)NeuAc(1)</td><td>C[+57], N[+2263]</td><td>1174.0412</td><td>6</td><td>7039.2108</td><td>7039.1757</td><td>0</td><td>5</td><td>8</td><td>Specific</td><td>743.12</td><td>741.07</td><td>741.07</td><td>16.53</td><td>120</td><td>>sp|P01877|IGHA2_HUMAN Immunoglobulin heavy constant alpha 2 OS=Homo sapiens OX=9606 GN=IGHA2 PE=1 SV=4</td><td>576</td><td>20201116_fqg_colostrum_1ng_glyco_hcd203040_compound15.27841.27841.6</td><td>controllerType=0 controllerNumber=1 scan=27841</td><td>55.06</td></tr>
<tr><td>22704:4</td><td>3</td><td>K.VFPLSLDSTPQDGNVVVAC[+57.021]LVQGFFPQEPLSVTWSESGQN[+1971.719]VTAI</td><td>HexNAc(5)Hex(5)Fuc(1)</td><td>C[+57], N[+1972]</td><td>1687.7811</td><td>4</td><td>6748.1025</td><td>6748.0803</td><td>0</td><td>3.3</td><td>8</td><td>Specific</td><td>752.42</td><td>752.42</td><td>752.42</td><td>17.97</td><td>120</td><td>>sp|P01877|IGHA2_HUMAN Immunoglobulin heavy constant alpha 2 OS=Homo sapiens OX=9606 GN=IGHA2 PE=1 SV=4</td><td>576</td><td>20201116_fqg_colostrum_1ng_glyco_hcd203040_compound15.26402.26402.4</td><td>controllerType=0 controllerNumber=1 scan=26402</td><td>51.8689</td></tr>
<tr><td>22706:4</td><td>3</td><td>K.VFPLSLDSTPQDGNVVVAC[+57.021]LVQGFFPQEPLSVTWSESGQN[+1971.719]VTAI</td><td>HexNAc(5)Hex(5)Fuc(1)</td><td>C[+57], N[+1972]</td><td>1687.7811</td><td>4</td><td>6748.1025</td><td>6748.0803</td><td>0</td><td>3.3</td><td>8</td><td>Specific</td><td>754.8</td><td>630.67</td><td>630.67</td><td>17.97</td><td>120</td><td>>sp|P01877|IGHA2_HUMAN Immunoglobulin heavy constant alpha 2 OS=Homo sapiens OX=9606 GN=IGHA2 PE=1 SV=4</td><td>576</td><td>20201116_fqg_colostrum_1ng_glyco_hcd203040_compound15.26404.26404.4</td><td>controllerType=0 controllerNumber=1 scan=26404</td><td>51.8799</td></tr>
<tr><td>22708:4</td><td>3</td><td>K.VFPLSLDSTPQDGNVVVAC[+57.021]LVQGFFPQEPLSVTWSESGQN[+1606.587]VTAI</td><td>HexNAc(4)Hex(4)Fuc(1)</td><td>C[+57], N[+1607]</td><td>1596.497</td><td>4</td><td>6382.966</td><td>6382.9481</td><td>0</td><td>2.8</td><td>8</td><td>Specific</td><td>754.91</td><td>727.97</td><td>727.97</td><td>18.08</td><td>120</td><td>>sp|P01877|IGHA2_HUMAN Immunoglobulin heavy constant alpha 2 OS=Homo sapiens OX=9606 GN=IGHA2 PE=1 SV=4</td><td>576</td><td>20201116_fqg_colostrum_1ng_glyco_hcd203040_compound15.26406.26406.4</td><td>controllerType=0 controllerNumber=1 scan=26406</td><td>51.8841</td></tr>
<tr><td>22832:6</td><td>3</td><td>K.VFPLSLDSTPQDGNVVVAC[+57.021]LVQGFFPQEPLSVTWSESGQN[+1444.534]VTAI</td><td>HexNAc(4)Hex(3)Fuc(1)</td><td>C[+57], N[+1445]</td><td>1037.6571</td><td>6</td><td>6220.9065</td><td>6220.8953</td><td>0</td><td>1.8</td><td>8</td><td>Specific</td><td>756.31</td><td>756.31</td><td>756.31</td><td>18.08</td><td>120</td><td>>sp|P01877|IGHA2_HUMAN Immunoglobulin heavy constant alpha 2 OS=Homo sapiens OX=9606 GN=IGHA2 PE=1 SV=4</td><td>576</td><td>20201116_fqg_colostrum_1ng_glyco_hcd203040_compound15.26534.26534.6</td><td>controllerType=0 controllerNumber=1 scan=26534</td><td>52.1355</td></tr>
<tr><td>23270:5</td><td>3</td><td>K.VFPLSLDSTPQDGNVVVAC[+57.021]LVQGFFPQEPLSVTWSESGQN[+1606.587]VTAI</td><td>HexNAc(4)Hex(4)Fuc(1)</td><td>C[+57], N[+1607]</td><td>1277.4001</td><td>5</td><td>6382.9716</td><td>6382.9481</td><td>0</td><td>3.7</td><td>8</td><td>Specific</td><td>769.16</td><td>719.26</td><td>719.26</td><td>17.97</td><td>120</td><td>>sp|P01877|IGHA2_HUMAN Immunoglobulin heavy constant alpha 2 OS=Homo sapiens OX=9606 GN=IGHA2 PE=1 SV=4</td><td>576</td><td>20201116_fqg_colostrum_1ng_glyco_hcd203040_compound15.26992.26992.5</td><td>controllerType=0 controllerNumber=1 scan=26992</td><td>53.108</td></tr>
<tr><td>22833:4</td><td>3</td><td>K.VFPLSLDSTPQDGNVVVAC[+57.021]LVQGFFPQEPLSVTWSESGQN[+1809.666]VTAI</td><td>HexNAc(5)Hex(4)Fuc(1)</td><td>C[+57], N[+1810]</td><td>1647.2671</td><td>4</td><td>6586.0465</td><td>6586.0275</td><td>0</td><td>2.9</td><td>8</td><td>Specific</td><td>770.87</td><td>716.92</td><td>716.92</td><td>17.97</td><td>120</td><td>>sp|P01877|IGHA2_HUMAN Immunoglobulin heavy constant alpha 2 OS=Homo sapiens OX=9606 GN=IGHA2 PE=1 SV=4</td><td>576</td><td>20201116_fqg_colostrum_1ng_glyco_hcd203040_compound15.26535.26535.4</td><td>controllerType=0 controllerNumber=1 scan=26535</td><td>52.139</td></tr>
<tr><td>22779:4</td><td>3</td><td>K.VFPLSLDSTPQDGNVVVAC[+57.021]LVQGFFPQEPLSVTWSESGQN[+1444.534]VTAI</td><td>HexNAc(4)Hex(3)Fuc(1)</td><td>C[+57], N[+1445]</td><td>1555.986</td><td>4</td><td>6220.922</td><td>6220.8953</td><td>0</td><td>4.3</td><td>8</td><td>Specific</td><td>773.22</td><td>723.5</td><td>723.5</td><td>17.53</td><td>120</td><td>>sp|P01877|IGHA2_HUMAN Immunoglobulin heavy constant alpha 2 OS=Homo sapiens OX=9606 GN=IGHA2 PE=1 SV=4</td><td>576</td><td>20201116_fqg_colostrum_1ng_glyco_hcd203040_compound15.26480.26480.4</td><td>controllerType=0 controllerNumber=1 scan=26480</td><td>52.0385</td></tr>
<tr><td>23282:5</td><td>3</td><td>K.VFPLSLDSTPQDGNVVVAC[+57.021]LVQGFFPQEPLSVTWSESGQN[+1809.666]VTAI</td><td>HexNAc(5)Hex(4)Fuc(1)</td><td>C[+57], N[+1810]</td><td>1318.0157</td><td>5</td><td>6586.0496</td><td>6586.0275</td><td>0</td><td>3.4</td><td>8</td><td>Specific</td><td>778.82</td><td>778.82</td><td>778.82</td><td>17.97</td><td>120</td><td>>sp|P01877|IGHA2_HUMAN Immunoglobulin heavy constant alpha 2 OS=Homo sapiens OX=9606 GN=IGHA2 PE=1 SV=4</td><td>576</td><td>20201116_fqg_colostrum_1ng_glyco_hcd203040_compound15.27005.27005.5</td><td>controllerType=0 controllerNumber=1 scan=27005</td><td>53.1336</td></tr>
<tr><td>23298:5</td><td>3</td><td>K.VFPLSLDSTPQDGNVVVAC[+57.021]LVQGFFPQEPLSVTWSESGQN[+1809.666]VTAI</td><td>HexNAc(5)Hex(4)Fuc(1)</td><td>C[+57], N[+1810]</td><td>1318.0166</td><td>5</td><td>6586.0539</td><td>6586.0275</td><td>0</td><td>4</td><td>8</td><td>Specific</td><td>787.52</td><td>787.52</td><td>787.52</td><td>17.53</td><td>120</td><td>>sp|P01877|IGHA2_HUMAN Immunoglobulin heavy constant alpha 2 OS=Homo sapiens OX=9606 GN=IGHA2 PE=1 SV=4</td><td>576</td><td>20201116_fqg_colostrum_1ng_glyco_hcd203040_compound15.27022.27022.5</td><td>controllerType=0 controllerNumber=1 scan=27022</td><td>53.1816</td></tr>
<tr><td>26150:4</td><td>3</td><td>K.VFPLSLDSTPQDGNVVVAC[+57.021]LVQGFFPQEPLSVTWSESGQN[+2350.830]VTAI</td><td>HexNAc(4)Hex(5)Fuc(1)NeuAc(2)</td><td>C[+57], N[+2351]</td><td>1782.5621</td><td>4</td><td>7127.2266</td><td>7127.1917</td><td>0</td><td>4.9</td><td>8</td><td>Specific</td><td>788.02</td><td>731.01</td><td>731.01</td><td>17.53</td><td>120</td><td>>sp|P01877|IGHA2_HUMAN Immunoglobulin heavy constant alpha 2 OS=Homo sapiens OX=9606 GN=IGHA2 PE=1 SV=4</td><td>576</td><td>20201116_fqg_colostrum_1ng_glyco_hcd203040_compound15.30004.30004.4</td><td>controllerType=0 controllerNumber=1 scan=30004</td><td>59.8059</td></tr>
<tr><td>24038:5</td><td>3</td><td>K.VFPLSLDSTPQDGNVVVAC[+57.021]LVQGFFPQEPLSVTWSESGQN[+2262.814]VTAI</td><td>HexNAc(5)Hex(5)Fuc(1)NeuAc(1)</td><td>C[+57], N[+2263]</td><td>1408.6458</td><td>5</td><td>7039.1999</td><td>7039.1757</td><td>0</td><td>3.4</td><td>8</td><td>Specific</td><td>791.5</td><td>752.17</td><td>752.17</td><td>17.97</td><td>120</td><td>>sp|P01877|IGHA2_HUMAN Immunoglobulin heavy constant alpha 2 OS=Homo sapiens OX=9606 GN=IGHA2 PE=1 SV=4</td><td>576</td><td>20201116_fqg_colostrum_1ng_glyco_hcd203040_compound15.27795.27795.5</td><td>controllerType=0 controllerNumber=1 scan=27795</td><td>54.9499</td></tr>
<tr><td>24712:4</td><td>3</td><td>K.VFPLSLDSTPQDGNVVVAC[+57.021]LVQGFFPQEPLSVTWSESGQN[+2262.814]VTAI</td><td>HexNAc(5)Hex(5)Fuc(1)NeuAc(1)</td><td>C[+57], N[+2263]</td><td>1760.5503</td><td>4</td><td>7039.1792</td><td>7039.1757</td><td>0</td><td>0.5</td><td>8</td><td>Specific</td><td>791.74</td><td>771.38</td><td>771.38</td><td>17.84</td><td>120</td><td>>sp|P01877|IGHA2_HUMAN Immunoglobulin heavy constant alpha 2 OS=Homo sapiens OX=9606 GN=IGHA2 PE=1 SV=4</td><td>576</td><td>20201116_fqg_colostrum_1ng_glyco_hcd203040_compound15.28500.28500.4</td><td>controllerType=0 controllerNumber=1 scan=28500</td><td>56.5631</td></tr>
<tr><td>23259:4</td><td>3</td><td>K.VFPLSLDSTPQDGNVVVAC[+57.021]LVQGFFPQEPLSVTWSESGQN[+1606.587]VTAI</td><td>HexNAc(4)Hex(4)Fuc(1)</td><td>C[+57], N[+1607]</td><td>1596.4956</td><td>4</td><td>6382.9606</td><td>6382.9481</td><td>0</td><td>2</td><td>8</td><td>Specific</td><td>793.91</td><td>750.77</td><td>750.77</td><td>18.08</td><td>120</td><td>>sp|P01877|IGHA2_HUMAN Immunoglobulin heavy constant alpha 2 OS=Homo sapiens OX=9606 GN=IGHA2 PE=1 SV=4</td><td>576</td><td>20201116_fqg_colostrum_1ng_glyco_hcd203040_compound15.26981.26981.4</td><td>controllerType=0 controllerNumber=1 scan=26981</td><td>53.0817</td></tr>
<tr><td>24071:5</td><td>3</td><td>K.VFPLSLDSTPQDGNVVVAC[+57.021]LVQGFFPQEPLSVTWSESGQN[+2262.814]VTAI</td><td>HexNAc(5)Hex(5)Fuc(1)NeuAc(1)</td><td>C[+57], N[+2263]</td><td>1408.6487</td><td>5</td><td>7039.2145</td><td>7039.1757</td><td>0</td><td>5.5</td><td>8</td><td>Specific</td><td>794.33</td><td>794.33</td><td>794.33</td><td>17.31</td><td>120</td><td>>sp|P01877|IGHA2_HUMAN Immunoglobulin heavy constant alpha 2 OS=Homo sapiens OX=9606 GN=IGHA2 PE=1 SV=4</td><td>576</td><td>20201116_fqg_colostrum_1ng_glyco_hcd203040_compound15.27830.27830.5</td><td>controllerType=0 controllerNumber=1 scan=27830</td><td>55.0297</td></tr>
<tr><td>26176:6</td><td>3</td><td>K.VFPLSLDSTPQDGNVVVAC[+57.021]LVQGFFPQEPLSVTWSESGQN[+2553.910]VTAI</td><td>HexNAc(5)Hex(5)Fuc(1)NeuAc(2)</td><td>C[+57], N[+2554]</td><td>1222.5582</td><td>6</td><td>7330.3126</td><td>7330.2711</td><td>0</td><td>5.7</td><td>8</td><td>Specific</td><td>805.96</td><td>728.6</td><td>728.6</td><td>18.31</td><td>120</td><td>>sp|P01877|IGHA2_HUMAN Immunoglobulin heavy constant alpha 2 OS=Homo sapiens OX=9606 GN=IGHA2 PE=1 SV=4</td><td>576</td><td>20201116_fqg_colostrum_1ng_glyco_hcd203040_compound15.30031.30031.6</td><td>controllerType=0 controllerNumber=1 scan=30031</td><td>59.8871</td></tr>
<tr><td>22790:5</td><td>3</td><td>K.VFPLSLDSTPQDGNVVVAC[+57.021]LVQGFFPQEPLSVTWSESGQN[+1606.587]VTAI</td><td>HexNAc(4)Hex(4)Fuc(1)</td><td>C[+57], N[+1607]</td><td>1277.4009</td><td>5</td><td>6382.9753</td><td>6382.9481</td><td>0</td><td>4.3</td><td>8</td><td>Specific</td><td>810.56</td><td>769.51</td><td>769.51</td><td>18.53</td><td>120</td><td>>sp|P01877|IGHA2_HUMAN Immunoglobulin heavy constant alpha 2 OS=Homo sapiens OX=9606 GN=IGHA2 PE=1 SV=4</td><td>576</td><td>20201116_fqg_colostrum_1ng_glyco_hcd203040_compound15.26491.26491.5</td><td>controllerType=0 controllerNumber=1 scan=26491</td><td>52.058</td></tr>
<tr><td>26261:5</td><td>3</td><td>K.VFPLSLDSTPQDGNVVVAC[+57.021]LVQGFFPQEPLSVTWSESGQN[+2553.910]VTAI</td><td>HexNAc(5)Hex(5)Fuc(1)NeuAc(2)</td><td>C[+57], N[+2554]</td><td>1466.8682</td><td>5</td><td>7330.3119</td><td>7330.2711</td><td>0</td><td>5.6</td><td>8</td><td>Specific</td><td>844.78</td><td>844.78</td><td>844.78</td><td>18.31</td><td>120</td><td>>sp|P01877|IGHA2_HUMAN Immunoglobulin heavy constant alpha 2 OS=Homo sapiens OX=9606 GN=IGHA2 PE=1 SV=4</td><td>576</td><td>20201116_fqg_colostrum_1ng_glyco_hcd203040_compound15.30120.30120.5</td><td>controllerType=0 controllerNumber=1 scan=30120</td><td>60.0736</td></tr>
<tr><td>22815:5</td><td>3</td><td>K.VFPLSLDSTPQDGNVVVAC[+57.021]LVQGFFPQEPLSVTWSESGQN[+1444.534]VTAI</td><td>HexNAc(4)Hex(3)Fuc(1)</td><td>C[+57], N[+1445]</td><td>1244.9877</td><td>5</td><td>6220.9093</td><td>6220.8953</td><td>0</td><td>2.2</td><td>8</td><td>Specific</td><td>848.32</td><td>772.4</td><td>772.4</td><td>19.08</td><td>120</td><td>>sp|P01877|IGHA2_HUMAN Immunoglobulin heavy constant alpha 2 OS=Homo sapiens OX=9606 GN=IGHA2 PE=1 SV=4</td><td>576</td><td>20201116_fqg_colostrum_1ng_glyco_hcd203040_compound15.26517.26517.5</td><td>controllerType=0 controllerNumber=1 scan=26517</td><td>52.0983</td></tr>
<tr><td>23436:5</td><td>3</td><td>K.VFPLSLDSTPQDGNVVVAC[+57.021]LVQGFFPQEPLSVTWSESGQN[+1444.534]VTAI</td><td>HexNAc(4)Hex(3)Fuc(1)</td><td>C[+57], N[+1445]</td><td>1244.9895</td><td>5</td><td>6220.9184</td><td>6220.8953</td><td>0</td><td>3.7</td><td>8</td><td>Specific</td><td>872.1</td><td>823.54</td><td>823.54</td><td>19.97</td><td>120</td><td>>sp|P01877|IGHA2_HUMAN Immunoglobulin heavy constant alpha 2 OS=Homo sapiens OX=9606 GN=IGHA2 PE=1 SV=4</td><td>576</td><td>20201116_fqg_colostrum_1ng_glyco_hcd203040_compound15.27165.27165.5</td><td>controllerType=0 controllerNumber=1 scan=27165</td><td>53.4683</td></tr>
<tr><td>24993:4</td><td>3</td><td>K.VFPLSLDSTPQDGNVVVAC[+57.021]LVQGFFPQEPLSVTWSESGQN[+1694.603]VTAI</td><td>HexNAc(3)Hex(4)Fuc(1)NeuAc(1)</td><td>C[+57], N[+1695]</td><td>1618.4961</td><td>4</td><td>6470.9626</td><td>6470.9641</td><td>0</td><td>-0.2</td><td>8</td><td>Specific</td><td>875.29</td><td>793.58</td><td>793.58</td><td>19.34</td><td>120</td><td>>sp|P01877|IGHA2_HUMAN Immunoglobulin heavy constant alpha 2 OS=Homo sapiens OX=9606 GN=IGHA2 PE=1 SV=4</td><td>576</td><td>20201116_fqg_colostrum_1ng_glyco_hcd203040_compound15.28793.28793.4</td><td>controllerType=0 controllerNumber=1 scan=28793</td><td>57.2124</td></tr>
<tr><td>22863:4</td><td>3</td><td>K.VFPLSLDSTPQDGNVVVAC[+57.021]LVQGFFPQEPLSVTWSESGQN[+1038.375]VTAI</td><td>HexNAc(2)Hex(3)Fuc(1)</td><td>C[+57], N[+1038]</td><td>1454.4446</td><td>4</td><td>5814.7565</td><td>5814.7365</td><td>0</td><td>3.4</td><td>8</td><td>Specific</td><td>875.44</td><td>875.44</td><td>875.44</td><td>19.97</td><td>120</td><td>>sp|P01877|IGHA2_HUMAN Immunoglobulin heavy constant alpha 2 OS=Homo sapiens OX=9606 GN=IGHA2 PE=1 SV=4</td><td>576</td><td>20201116_fqg_colostrum_1ng_glyco_hcd203040_compound15.26566.26566.4</td><td>controllerType=0 controllerNumber=1 scan=26566</td><td>52.1995</td></tr>
<tr><td>23257:4</td><td>3</td><td>K.VFPLSLDSTPQDGNVVVAC[+57.021]LVQGFFPQEPLSVTWSESGQN[+1809.666]VTAI</td><td>HexNAc(5)Hex(4)Fuc(1)</td><td>C[+57], N[+1810]</td><td>1647.269</td><td>4</td><td>6586.0544</td><td>6586.0275</td><td>0</td><td>4.1</td><td>8</td><td>Specific</td><td>877.17</td><td>877.17</td><td>877.17</td><td>19.53</td><td>120</td><td>>sp|P01877|IGHA2_HUMAN Immunoglobulin heavy constant alpha 2 OS=Homo sapiens OX=9606 GN=IGHA2 PE=1 SV=4</td><td>576</td><td>20201116_fqg_colostrum_1ng_glyco_hcd203040_compound15.26979.26979.4</td><td>controllerType=0 controllerNumber=1 scan=26979</td><td>53.0792</td></tr>
<tr><td>22711:4</td><td>3</td><td>K.VFPLSLDSTPQDGNVVVAC[+57.021]LVQGFFPQEPLSVTWSESGQN[+1606.587]VTAI</td><td>HexNAc(4)Hex(4)Fuc(1)</td><td>C[+57], N[+1607]</td><td>1596.497</td><td>4</td><td>6382.966</td><td>6382.9481</td><td>0</td><td>2.8</td><td>8</td><td>Specific</td><td>880.95</td><td>859.2</td><td>859.2</td><td>20.08</td><td>120</td><td>>sp|P01877|IGHA2_HUMAN Immunoglobulin heavy constant alpha 2 OS=Homo sapiens OX=9606 GN=IGHA2 PE=1 SV=4</td><td>576</td><td>20201116_fqg_colostrum_1ng_glyco_hcd203040_compound15.26409.26409.4</td><td>controllerType=0 controllerNumber=1 scan=26409</td><td>51.9022</td></tr>
<tr><td>24221:5</td><td>3</td><td>K.VFPLSLDSTPQDGNVVVAC[+57.021]LVQGFFPQEPLSVTWSESGQN[+2059.735]VTAI</td><td>HexNAc(4)Hex(5)Fuc(1)NeuAc(1)</td><td>C[+57], N[+2060]</td><td>1368.0303</td><td>5</td><td>6836.1225</td><td>6836.0963</td><td>0</td><td>3.8</td><td>8</td><td>Specific</td><td>891.6</td><td>891.6</td><td>891.6</td><td>19.97</td><td>120</td><td>>sp|P01877|IGHA2_HUMAN Immunoglobulin heavy constant alpha 2 OS=Homo sapiens OX=9606 GN=IGHA2 PE=1 SV=4</td><td>576</td><td>20201116_fqg_colostrum_1ng_glyco_hcd203040_compound15.27987.27987.5</td><td>controllerType=0 controllerNumber=1 scan=27987</td><td>55.4068</td></tr>
<tr><td>23351:4</td><td>3</td><td>K.VFPLSLDSTPQDGNVVVAC[+57.021]LVQGFFPQEPLSVTWSESGQN[+1647.613]VTAI</td><td>HexNAc(5)Hex(3)Fuc(1)</td><td>C[+57], N[+1648]</td><td>1606.7545</td><td>4</td><td>6423.9962</td><td>6423.9746</td><td>0</td><td>3.4</td><td>8</td><td>Specific</td><td>892.13</td><td>892.13</td><td>892.13</td><td>19.97</td><td>120</td><td>>sp|P01877|IGHA2_HUMAN Immunoglobulin heavy constant alpha 2 OS=Homo sapiens OX=9606 GN=IGHA2 PE=1 SV=4</td><td>576</td><td>20201116_fqg_colostrum_1ng_glyco_hcd203040_compound15.27077.27077.4</td><td>controllerType=0 controllerNumber=1 scan=27077</td><td>53.2935</td></tr>
<tr><td>22860:4</td><td>3</td><td>K.VFPLSLDSTPQDGNVVVAC[+57.021]LVQGFFPQEPLSVTWSESGQN[+1038.375]VTAI</td><td>HexNAc(2)Hex(3)Fuc(1)</td><td>C[+57], N[+1038]</td><td>1454.4446</td><td>4</td><td>5814.7565</td><td>5814.7365</td><td>0</td><td>3.4</td><td>8</td><td>Specific</td><td>898.01</td><td>880.25</td><td>880.25</td><td>19.97</td><td>120</td><td>>sp|P01877|IGHA2_HUMAN Immunoglobulin heavy constant alpha 2 OS=Homo sapiens OX=9606 GN=IGHA2 PE=1 SV=4</td><td>576</td><td>20201116_fqg_colostrum_1ng_glyco_hcd203040_compound15.26563.26563.4</td><td>controllerType=0 controllerNumber=1 scan=26563</td><td>52.1904</td></tr>
<tr><td>23262:4</td><td>3</td><td>K.VFPLSLDSTPQDGNVVVAC[+57.021]LVQGFFPQEPLSVTWSESGQN[+1606.587]VTAI</td><td>HexNAc(4)Hex(4)Fuc(1)</td><td>C[+57], N[+1607]</td><td>1596.4956</td><td>4</td><td>6382.9606</td><td>6382.9481</td><td>0</td><td>2</td><td>8</td><td>Specific</td><td>903.19</td><td>903.19</td><td>903.19</td><td>21.08</td><td>120</td><td>>sp|P01877|IGHA2_HUMAN Immunoglobulin heavy constant alpha 2 OS=Homo sapiens OX=9606 GN=IGHA2 PE=1 SV=4</td><td>576</td><td>20201116_fqg_colostrum_1ng_glyco_hcd203040_compound15.26984.26984.4</td><td>controllerType=0 controllerNumber=1 scan=26984</td><td>53.0938</td></tr>
<tr><td>24372:4</td><td>3</td><td>K.VFPLSLDSTPQDGNVVVAC[+57.021]LVQGFFPQEPLSVTWSESGQN[+1694.603]VTAI</td><td>HexNAc(3)Hex(4)Fuc(1)NeuAc(1)</td><td>C[+57], N[+1695]</td><td>1618.5013</td><td>4</td><td>6470.9836</td><td>6470.9641</td><td>0</td><td>3</td><td>8</td><td>Specific</td><td>911.35</td><td>848.07</td><td>848.07</td><td>20.97</td><td>120</td><td>>sp|P01877|IGHA2_HUMAN Immunoglobulin heavy constant alpha 2 OS=Homo sapiens OX=9606 GN=IGHA2 PE=1 SV=4</td><td>576</td><td>20201116_fqg_colostrum_1ng_glyco_hcd203040_compound15.28145.28145.4</td><td>controllerType=0 controllerNumber=1 scan=28145</td><td>55.7678</td></tr>
<tr><td>23471:4</td><td>3</td><td>K.VFPLSLDSTPQDGNVVVAC[+57.021]LVQGFFPQEPLSVTWSESGQN[+1444.534]VTAI</td><td>HexNAc(4)Hex(3)Fuc(1)</td><td>C[+57], N[+1445]</td><td>1555.9858</td><td>4</td><td>6220.9215</td><td>6220.8953</td><td>0</td><td>4.2</td><td>8</td><td>Specific</td><td>937.1</td><td>925.57</td><td>925.57</td><td>20.53</td><td>120</td><td>>sp|P01877|IGHA2_HUMAN Immunoglobulin heavy constant alpha 2 OS=Homo sapiens OX=9606 GN=IGHA2 PE=1 SV=4</td><td>576</td><td>20201116_fqg_colostrum_1ng_glyco_hcd203040_compound15.27202.27202.4</td><td>controllerType=0 controllerNumber=1 scan=27202</td><td>53.5583</td></tr>
<tr><td>22980:4</td><td>3</td><td>K.VFPLSLDSTPQDGNVVVAC[+57.021]LVQGFFPQEPLSVTWSESGQN[+1038.375]VTAI</td><td>HexNAc(2)Hex(3)Fuc(1)</td><td>C[+57], N[+1038]</td><td>1454.4465</td><td>4</td><td>5814.7643</td><td>5814.7365</td><td>0</td><td>4.8</td><td>8</td><td>Specific</td><td>952.53</td><td>944.87</td><td>944.87</td><td>21.53</td><td>120</td><td>>sp|P01877|IGHA2_HUMAN Immunoglobulin heavy constant alpha 2 OS=Homo sapiens OX=9606 GN=IGHA2 PE=1 SV=4</td><td>576</td><td>20201116_fqg_colostrum_1ng_glyco_hcd203040_compound15.26688.26688.4</td><td>controllerType=0 controllerNumber=1 scan=26688</td><td>52.4239</td></tr>
<tr><td>22781:4</td><td>3</td><td>K.VFPLSLDSTPQDGNVVVAC[+57.021]LVQGFFPQEPLSVTWSESGQN[+1444.534]VTAI</td><td>HexNAc(4)Hex(3)Fuc(1)</td><td>C[+57], N[+1445]</td><td>1555.986</td><td>4</td><td>6220.922</td><td>6220.8953</td><td>0</td><td>4.3</td><td>8</td><td>Specific</td><td>996.46</td><td>950.59</td><td>950.59</td><td>21.53</td><td>120</td><td>>sp|P01877|IGHA2_HUMAN Immunoglobulin heavy constant alpha 2 OS=Homo sapiens OX=9606 GN=IGHA2 PE=1 SV=4</td><td>576</td><td>20201116_fqg_colostrum_1ng_glyco_hcd203040_compound15.26482.26482.4</td><td>controllerType=0 controllerNumber=1 scan=26482</td><td>52.0457</td></tr>
<tr><td>01101:4</td><td>3</td><td>K.HYTN[+1647.613]SSQDVTVPC[+57.021]R.V</td><td>HexNAc(5)Hex(3)Fuc(1)</td><td>C[+57], N[+1648]</td><td>828.5987</td><td>4</td><td>3311.3728</td><td>3311.3625</td><td>0</td><td>3.1</td><td>89</td><td>Specific</td><td>314.71</td><td>285.51</td><td>285.51</td><td>5.22</td><td>120</td><td>>sp|P01877|IGHA2_HUMAN Immunoglobulin heavy constant alpha 2 OS=Homo sapiens OX=9606 GN=IGHA2 PE=1 SV=4</td><td>576</td><td>20201116_fqg_colostrum_1ng_glyco_hcd203040_compound15.3982.3982.4</td><td>controllerType=0 controllerNumber=1 scan=3982</td><td>12.979</td></tr>
<tr><td>01103:4</td><td>3</td><td>K.HYTN[+1647.613]SSQDVTVPC[+57.021]R.V</td><td>HexNAc(5)Hex(3)Fuc(1)</td><td>C[+57], N[+1648]</td><td>828.5987</td><td>4</td><td>3311.3728</td><td>3311.3625</td><td>0</td><td>3.1</td><td>89</td><td>Specific</td><td>389.6</td><td>334.74</td><td>334.74</td><td>5.81</td><td>120</td><td>>sp|P01877|IGHA2_HUMAN Immunoglobulin heavy constant alpha 2 OS=Homo sapiens OX=9606 GN=IGHA2 PE=1 SV=4</td><td>576</td><td>20201116_fqg_colostrum_1ng_glyco_hcd203040_compound15.3984.3984.4</td><td>controllerType=0 controllerNumber=1 scan=3984</td><td>12.9846</td></tr>
<tr><td>01920:4</td><td>3</td><td>K.HYTN[+2262.814]SSQDVTVPC[+57.021]R.V</td><td>HexNAc(5)Hex(5)Fuc(1)NeuAc(1)</td><td>C[+57], N[+2263]</td><td>982.3981</td><td>4</td><td>3926.5704</td><td>3926.5636</td><td>0</td><td>1.7</td><td>89</td><td>Specific</td><td>403.7</td><td>266.7</td><td>266.7</td><td>8.05</td><td>120</td><td>>sp|P01877|IGHA2_HUMAN Immunoglobulin heavy constant alpha 2 OS=Homo sapiens OX=9606 GN=IGHA2 PE=1 SV=4</td><td>576</td><td>20201116_fqg_colostrum_1ng_glyco_hcd203040_compound15.4832.4832.4</td><td>controllerType=0 controllerNumber=1 scan=4832</td><td>14.528</td></tr>
<tr><td>01835:3</td><td>3</td><td>K.HYTN[+2059.735]SSQDVTVPC[+57.021]R.V</td><td>HexNAc(4)Hex(5)Fuc(1)NeuAc(1)</td><td>C[+57], N[+2060]</td><td>1241.8362</td><td>3</td><td>3723.4942</td><td>3723.4842</td><td>0</td><td>2.7</td><td>89</td><td>Specific</td><td>428.93</td><td>286.63</td><td>286.63</td><td>7.89</td><td>120</td><td>>sp|P01877|IGHA2_HUMAN Immunoglobulin heavy constant alpha 2 OS=Homo sapiens OX=9606 GN=IGHA2 PE=1 SV=4</td><td>576</td><td>20201116_fqg_colostrum_1ng_glyco_hcd203040_compound15.4744.4744.3</td><td>controllerType=0 controllerNumber=1 scan=4744</td><td>14.3557</td></tr>
<tr><td>01922:4</td><td>3</td><td>K.HYTN[+2262.814]SSQDVTVPC[+57.021]R.V</td><td>HexNAc(5)Hex(5)Fuc(1)NeuAc(1)</td><td>C[+57], N[+2263]</td><td>982.3981</td><td>4</td><td>3926.5704</td><td>3926.5636</td><td>0</td><td>1.7</td><td>89</td><td>Specific</td><td>430.42</td><td>286.31</td><td>286.31</td><td>8.11</td><td>120</td><td>>sp|P01877|IGHA2_HUMAN Immunoglobulin heavy constant alpha 2 OS=Homo sapiens OX=9606 GN=IGHA2 PE=1 SV=4</td><td>576</td><td>20201116_fqg_colostrum_1ng_glyco_hcd203040_compound15.4834.4834.4</td><td>controllerType=0 controllerNumber=1 scan=4834</td><td>14.5316</td></tr>
<tr><td>00923:3</td><td>3</td><td>K.HYTN[+1216.423]SSQDVTVPC[+57.021]R.V</td><td>HexNAc(2)Hex(5)</td><td>C[+57], N[+1216]</td><td>960.7316</td><td>3</td><td>2880.1802</td><td>2880.1722</td><td>0</td><td>2.8</td><td>89</td><td>Specific</td><td>440.58</td><td>360.49</td><td>360.49</td><td>8.42</td><td>120</td><td>>sp|P01877|IGHA2_HUMAN Immunoglobulin heavy constant alpha 2 OS=Homo sapiens OX=9606 GN=IGHA2 PE=1 SV=4</td><td>576</td><td>20201116_fqg_colostrum_1ng_glyco_hcd203040_compound15.3796.3796.3</td><td>controllerType=0 controllerNumber=1 scan=3796</td><td>12.5703</td></tr>
<tr><td>01520:3</td><td>3</td><td>K.HYTN[+1606.587]SSQDVTVPC[+57.021]R.V</td><td>HexNAc(4)Hex(4)Fuc(1)</td><td>C[+57], N[+1607]</td><td>1090.7856</td><td>3</td><td>3270.3421</td><td>3270.336</td><td>0</td><td>1.9</td><td>89</td><td>Specific</td><td>463.26</td><td>290.23</td><td>290.23</td><td>8.1</td><td>120</td><td>>sp|P01877|IGHA2_HUMAN Immunoglobulin heavy constant alpha 2 OS=Homo sapiens OX=9606 GN=IGHA2 PE=1 SV=4</td><td>576</td><td>20201116_fqg_colostrum_1ng_glyco_hcd203040_compound15.4416.4416.3</td><td>controllerType=0 controllerNumber=1 scan=4416</td><td>13.7505</td></tr>
<tr><td>00994:3</td><td>3</td><td>K.HYTN[+1930.692]SSQDVTVPC[+57.021]R.V</td><td>HexNAc(4)Hex(6)Fuc(1)</td><td>C[+57], N[+1931]</td><td>1198.8212</td><td>3</td><td>3594.4491</td><td>3594.4416</td><td>0</td><td>2.1</td><td>89</td><td>Specific</td><td>467.13</td><td>360.25</td><td>360.25</td><td>8.69</td><td>120</td><td>>sp|P01877|IGHA2_HUMAN Immunoglobulin heavy constant alpha 2 OS=Homo sapiens OX=9606 GN=IGHA2 PE=1 SV=4</td><td>576</td><td>20201116_fqg_colostrum_1ng_glyco_hcd203040_compound15.3870.3870.3</td><td>controllerType=0 controllerNumber=1 scan=3870</td><td>12.7153</td></tr>
<tr><td>03004:3</td><td>3</td><td>K.HYTN[+2350.830]SSQDVTVPC[+57.021]R.V</td><td>HexNAc(4)Hex(5)Fuc(1)NeuAc(2)</td><td>C[+57], N[+2351]</td><td>1338.8676</td><td>3</td><td>4014.5883</td><td>4014.5796</td><td>0</td><td>2.2</td><td>89</td><td>Specific</td><td>474.29</td><td>243.3</td><td>243.3</td><td>7.76</td><td>120</td><td>>sp|P01877|IGHA2_HUMAN Immunoglobulin heavy constant alpha 2 OS=Homo sapiens OX=9606 GN=IGHA2 PE=1 SV=4</td><td>576</td><td>20201116_fqg_colostrum_1ng_glyco_hcd203040_compound15.5955.5955.3</td><td>controllerType=0 controllerNumber=1 scan=5955</td><td>16.5907</td></tr>
<tr><td>00913:3</td><td>3</td><td>K.HYTN[+1768.640]SSQDVTVPC[+57.021]R.V</td><td>HexNAc(4)Hex(5)Fuc(1)</td><td>C[+57], N[+1769]</td><td>1144.8036</td><td>3</td><td>3432.3963</td><td>3432.3888</td><td>0</td><td>2.2</td><td>89</td><td>Specific</td><td>474.33</td><td>288.8</td><td>288.8</td><td>8.1</td><td>120</td><td>>sp|P01877|IGHA2_HUMAN Immunoglobulin heavy constant alpha 2 OS=Homo sapiens OX=9606 GN=IGHA2 PE=1 SV=4</td><td>576</td><td>20201116_fqg_colostrum_1ng_glyco_hcd203040_compound15.3786.3786.3</td><td>controllerType=0 controllerNumber=1 scan=3786</td><td>12.5532</td></tr>
<tr><td>01518:3</td><td>3</td><td>K.HYTN[+1606.587]SSQDVTVPC[+57.021]R.V</td><td>HexNAc(4)Hex(4)Fuc(1)</td><td>C[+57], N[+1607]</td><td>1090.7856</td><td>3</td><td>3270.3421</td><td>3270.336</td><td>0</td><td>1.9</td><td>89</td><td>Specific</td><td>474.34</td><td>288.11</td><td>288.11</td><td>8.1</td><td>120</td><td>>sp|P01877|IGHA2_HUMAN Immunoglobulin heavy constant alpha 2 OS=Homo sapiens OX=9606 GN=IGHA2 PE=1 SV=4</td><td>576</td><td>20201116_fqg_colostrum_1ng_glyco_hcd203040_compound15.4414.4414.3</td><td>controllerType=0 controllerNumber=1 scan=4414</td><td>13.7463</td></tr>
<tr><td>00992:3</td><td>3</td><td>K.HYTN[+1930.692]SSQDVTVPC[+57.021]R.V</td><td>HexNAc(4)Hex(6)Fuc(1)</td><td>C[+57], N[+1931]</td><td>1198.8212</td><td>3</td><td>3594.4491</td><td>3594.4416</td><td>0</td><td>2.1</td><td>89</td><td>Specific</td><td>479.12</td><td>376.24</td><td>376.24</td><td>8.69</td><td>120</td><td>>sp|P01877|IGHA2_HUMAN Immunoglobulin heavy constant alpha 2 OS=Homo sapiens OX=9606 GN=IGHA2 PE=1 SV=4</td><td>576</td><td>20201116_fqg_colostrum_1ng_glyco_hcd203040_compound15.3868.3868.3</td><td>controllerType=0 controllerNumber=1 scan=3868</td><td>12.7117</td></tr>
<tr><td>01679:3</td><td>3</td><td>K.HYTN[+1444.534]SSQDVTVPC[+57.021]R.V</td><td>HexNAc(4)Hex(3)Fuc(1)</td><td>C[+57], N[+1445]</td><td>1036.7688</td><td>3</td><td>3108.2919</td><td>3108.2832</td><td>0</td><td>2.8</td><td>89</td><td>Specific</td><td>481.55</td><td>452.83</td><td>452.83</td><td>9.92</td><td>120</td><td>>sp|P01877|IGHA2_HUMAN Immunoglobulin heavy constant alpha 2 OS=Homo sapiens OX=9606 GN=IGHA2 PE=1 SV=4</td><td>576</td><td>20201116_fqg_colostrum_1ng_glyco_hcd203040_compound15.4582.4582.3</td><td>controllerType=0 controllerNumber=1 scan=4582</td><td>14.0535</td></tr>
<tr><td>03001:3</td><td>3</td><td>K.HYTN[+2350.830]SSQDVTVPC[+57.021]R.V</td><td>HexNAc(4)Hex(5)Fuc(1)NeuAc(2)</td><td>C[+57], N[+2351]</td><td>1338.8676</td><td>3</td><td>4014.5883</td><td>4014.5796</td><td>0</td><td>2.2</td><td>89</td><td>Specific</td><td>488.21</td><td>397.66</td><td>397.66</td><td>10.03</td><td>120</td><td>>sp|P01877|IGHA2_HUMAN Immunoglobulin heavy constant alpha 2 OS=Homo sapiens OX=9606 GN=IGHA2 PE=1 SV=4</td><td>576</td><td>20201116_fqg_colostrum_1ng_glyco_hcd203040_compound15.5952.5952.3</td><td>controllerType=0 controllerNumber=1 scan=5952</td><td>16.5861</td></tr>
<tr><td>00925:3</td><td>3</td><td>K.HYTN[+1216.423]SSQDVTVPC[+57.021]R.V</td><td>HexNAc(2)Hex(5)</td><td>C[+57], N[+1216]</td><td>960.7316</td><td>3</td><td>2880.1802</td><td>2880.1722</td><td>0</td><td>2.8</td><td>89</td><td>Specific</td><td>496.29</td><td>406.97</td><td>406.97</td><td>9.92</td><td>120</td><td>>sp|P01877|IGHA2_HUMAN Immunoglobulin heavy constant alpha 2 OS=Homo sapiens OX=9606 GN=IGHA2 PE=1 SV=4</td><td>576</td><td>20201116_fqg_colostrum_1ng_glyco_hcd203040_compound15.3798.3798.3</td><td>controllerType=0 controllerNumber=1 scan=3798</td><td>12.574</td></tr>
<tr><td>01055:2</td><td>3</td><td>K.HYTN[+1809.666]SSQDVTVPC[+57.021]R.V</td><td>HexNAc(5)Hex(4)Fuc(1)</td><td>C[+57], N[+1810]</td><td>1737.2175</td><td>2</td><td>3473.4277</td><td>3473.4154</td><td>0</td><td>3.6</td><td>89</td><td>Specific</td><td>498.25</td><td>457.03</td><td>457.03</td><td>9.3</td><td>120</td><td>>sp|P01877|IGHA2_HUMAN Immunoglobulin heavy constant alpha 2 OS=Homo sapiens OX=9606 GN=IGHA2 PE=1 SV=4</td><td>576</td><td>20201116_fqg_colostrum_1ng_glyco_hcd203040_compound15.3934.3934.2</td><td>controllerType=0 controllerNumber=1 scan=3934</td><td>12.8799</td></tr>
<tr><td>01909:3</td><td>3</td><td>K.HYTN[+2262.814]SSQDVTVPC[+57.021]R.V</td><td>HexNAc(5)Hex(5)Fuc(1)NeuAc(1)</td><td>C[+57], N[+2263]</td><td>1309.529</td><td>3</td><td>3926.5724</td><td>3926.5636</td><td>0</td><td>2.2</td><td>89</td><td>Specific</td><td>514.79</td><td>285.14</td><td>285.14</td><td>8.3</td><td>120</td><td>>sp|P01877|IGHA2_HUMAN Immunoglobulin heavy constant alpha 2 OS=Homo sapiens OX=9606 GN=IGHA2 PE=1 SV=4</td><td>576</td><td>20201116_fqg_colostrum_1ng_glyco_hcd203040_compound15.4821.4821.3</td><td>controllerType=0 controllerNumber=1 scan=4821</td><td>14.509</td></tr>
<tr><td>01054:2</td><td>3</td><td>K.HYTN[+1606.587]SSQDVTVPC[+57.021]R.V</td><td>HexNAc(4)Hex(4)Fuc(1)</td><td>C[+57], N[+1607]</td><td>1635.6784</td><td>2</td><td>3270.3495</td><td>3270.336</td><td>0</td><td>4.1</td><td>89</td><td>Specific</td><td>517.3</td><td>455.45</td><td>455.45</td><td>8.97</td><td>120</td><td>>sp|P01877|IGHA2_HUMAN Immunoglobulin heavy constant alpha 2 OS=Homo sapiens OX=9606 GN=IGHA2 PE=1 SV=4</td><td>576</td><td>20201116_fqg_colostrum_1ng_glyco_hcd203040_compound15.3933.3933.2</td><td>controllerType=0 controllerNumber=1 scan=3933</td><td>12.8784</td></tr>
<tr><td>01537:3</td><td>3</td><td>K.HYTN[+1444.534]SSQDVTVPC[+57.021]R.V</td><td>HexNAc(4)Hex(3)Fuc(1)</td><td>C[+57], N[+1445]</td><td>1036.7688</td><td>3</td><td>3108.2919</td><td>3108.2832</td><td>0</td><td>2.8</td><td>89</td><td>Specific</td><td>550.7</td><td>391.58</td><td>391.58</td><td>10.25</td><td>120</td><td>>sp|P01877|IGHA2_HUMAN Immunoglobulin heavy constant alpha 2 OS=Homo sapiens OX=9606 GN=IGHA2 PE=1 SV=4</td><td>576</td><td>20201116_fqg_colostrum_1ng_glyco_hcd203040_compound15.4434.4434.3</td><td>controllerType=0 controllerNumber=1 scan=4434</td><td>13.7962</td></tr>
<tr><td>01681:3</td><td>3</td><td>K.HYTN[+1444.534]SSQDVTVPC[+57.021]R.V</td><td>HexNAc(4)Hex(3)Fuc(1)</td><td>C[+57], N[+1445]</td><td>1036.7688</td><td>3</td><td>3108.2919</td><td>3108.2832</td><td>0</td><td>2.8</td><td>89</td><td>Specific</td><td>579.05</td><td>419.09</td><td>419.09</td><td>10.44</td><td>120</td><td>>sp|P01877|IGHA2_HUMAN Immunoglobulin heavy constant alpha 2 OS=Homo sapiens OX=9606 GN=IGHA2 PE=1 SV=4</td><td>576</td><td>20201116_fqg_colostrum_1ng_glyco_hcd203040_compound15.4584.4584.3</td><td>controllerType=0 controllerNumber=1 scan=4584</td><td>14.0587</td></tr>
<tr><td>01056:2</td><td>3</td><td>K.HYTN[+1606.587]SSQDVTVPC[+57.021]R.V</td><td>HexNAc(4)Hex(4)Fuc(1)</td><td>C[+57], N[+1607]</td><td>1635.6784</td><td>2</td><td>3270.3495</td><td>3270.336</td><td>0</td><td>4.1</td><td>89</td><td>Specific</td><td>583.88</td><td>433.68</td><td>433.68</td><td>10.76</td><td>120</td><td>>sp|P01877|IGHA2_HUMAN Immunoglobulin heavy constant alpha 2 OS=Homo sapiens OX=9606 GN=IGHA2 PE=1 SV=4</td><td>576</td><td>20201116_fqg_colostrum_1ng_glyco_hcd203040_compound15.3935.3935.2</td><td>controllerType=0 controllerNumber=1 scan=3935</td><td>12.8821</td></tr>
<tr><td>00915:3</td><td>3</td><td>K.HYTN[+1768.640]SSQDVTVPC[+57.021]R.V</td><td>HexNAc(4)Hex(5)Fuc(1)</td><td>C[+57], N[+1769]</td><td>1144.8036</td><td>3</td><td>3432.3963</td><td>3432.3888</td><td>0</td><td>2.2</td><td>89</td><td>Specific</td><td>594.52</td><td>412.8</td><td>412.8</td><td>11.93</td><td>120</td><td>>sp|P01877|IGHA2_HUMAN Immunoglobulin heavy constant alpha 2 OS=Homo sapiens OX=9606 GN=IGHA2 PE=1 SV=4</td><td>576</td><td>20201116_fqg_colostrum_1ng_glyco_hcd203040_compound15.3788.3788.3</td><td>controllerType=0 controllerNumber=1 scan=3788</td><td>12.5569</td></tr>
<tr><td>01057:2</td><td>3</td><td>K.HYTN[+1809.666]SSQDVTVPC[+57.021]R.V</td><td>HexNAc(5)Hex(4)Fuc(1)</td><td>C[+57], N[+1810]</td><td>1737.2175</td><td>2</td><td>3473.4277</td><td>3473.4154</td><td>0</td><td>3.6</td><td>89</td><td>Specific</td><td>615.31</td><td>469.12</td><td>469.12</td><td>12.2</td><td>120</td><td>>sp|P01877|IGHA2_HUMAN Immunoglobulin heavy constant alpha 2 OS=Homo sapiens OX=9606 GN=IGHA2 PE=1 SV=4</td><td>576</td><td>20201116_fqg_colostrum_1ng_glyco_hcd203040_compound15.3936.3936.2</td><td>controllerType=0 controllerNumber=1 scan=3936</td><td>12.8849</td></tr>
<tr><td>01277:3</td><td>3</td><td>K.HYTN[+876.322]SSQDVTVPC[+57.021]R.V</td><td>HexNAc(2)Hex(2)Fuc(1)</td><td>C[+57], N[+876]</td><td>847.3639</td><td>3</td><td>2540.0772</td><td>2540.0716</td><td>0</td><td>2.2</td><td>89</td><td>Specific</td><td>664.16</td><td>517.95</td><td>517.95</td><td>13.93</td><td>120</td><td>>sp|P01877|IGHA2_HUMAN Immunoglobulin heavy constant alpha 2 OS=Homo sapiens OX=9606 GN=IGHA2 PE=1 SV=4</td><td>576</td><td>20201116_fqg_colostrum_1ng_glyco_hcd203040_compound15.4164.4164.3</td><td>controllerType=0 controllerNumber=1 scan=4164</td><td>13.3048</td></tr>
<tr><td>01010:3</td><td>3</td><td>K.HYTN[+1971.719]SSQDVTVPC[+57.021]R.V</td><td>HexNAc(5)Hex(5)Fuc(1)</td><td>C[+57], N[+1972]</td><td>1212.4968</td><td>3</td><td>3635.4757</td><td>3635.4682</td><td>0</td><td>2.1</td><td>89</td><td>Specific</td><td>665.94</td><td>568.4</td><td>568.4</td><td>13.93</td><td>120</td><td>>sp|P01877|IGHA2_HUMAN Immunoglobulin heavy constant alpha 2 OS=Homo sapiens OX=9606 GN=IGHA2 PE=1 SV=4</td><td>576</td><td>20201116_fqg_colostrum_1ng_glyco_hcd203040_compound15.3887.3887.3</td><td>controllerType=0 controllerNumber=1 scan=3887</td><td>12.7539</td></tr>
<tr><td>01837:3</td><td>3</td><td>K.HYTN[+2059.735]SSQDVTVPC[+57.021]R.V</td><td>HexNAc(4)Hex(5)Fuc(1)NeuAc(1)</td><td>C[+57], N[+2060]</td><td>1241.8362</td><td>3</td><td>3723.4942</td><td>3723.4842</td><td>0</td><td>2.7</td><td>89</td><td>Specific</td><td>668.2</td><td>532.01</td><td>532.01</td><td>13.93</td><td>120</td><td>>sp|P01877|IGHA2_HUMAN Immunoglobulin heavy constant alpha 2 OS=Homo sapiens OX=9606 GN=IGHA2 PE=1 SV=4</td><td>576</td><td>20201116_fqg_colostrum_1ng_glyco_hcd203040_compound15.4746.4746.3</td><td>controllerType=0 controllerNumber=1 scan=4746</td><td>14.3599</td></tr>
<tr><td>01071:2</td><td>3</td><td>K.HYTN[+1647.613]SSQDVTVPC[+57.021]R.V</td><td>HexNAc(5)Hex(3)Fuc(1)</td><td>C[+57], N[+1648]</td><td>1656.188</td><td>2</td><td>3311.3688</td><td>3311.3625</td><td>0</td><td>1.9</td><td>89</td><td>Specific</td><td>673.29</td><td>621.71</td><td>621.71</td><td>13.31</td><td>120</td><td>>sp|P01877|IGHA2_HUMAN Immunoglobulin heavy constant alpha 2 OS=Homo sapiens OX=9606 GN=IGHA2 PE=1 SV=4</td><td>576</td><td>20201116_fqg_colostrum_1ng_glyco_hcd203040_compound15.3951.3951.2</td><td>controllerType=0 controllerNumber=1 scan=3951</td><td>12.9157</td></tr>
<tr><td>01008:3</td><td>3</td><td>K.HYTN[+1971.719]SSQDVTVPC[+57.021]R.V</td><td>HexNAc(5)Hex(5)Fuc(1)</td><td>C[+57], N[+1972]</td><td>1212.4968</td><td>3</td><td>3635.4757</td><td>3635.4682</td><td>0</td><td>2.1</td><td>89</td><td>Specific</td><td>676.03</td><td>585.37</td><td>585.37</td><td>13.93</td><td>120</td><td>>sp|P01877|IGHA2_HUMAN Immunoglobulin heavy constant alpha 2 OS=Homo sapiens OX=9606 GN=IGHA2 PE=1 SV=4</td><td>576</td><td>20201116_fqg_colostrum_1ng_glyco_hcd203040_compound15.3885.3885.3</td><td>controllerType=0 controllerNumber=1 scan=3885</td><td>12.7478</td></tr>
<tr><td>01066:3</td><td>3</td><td>K.HYTN[+1647.613]SSQDVTVPC[+57.021]R.V</td><td>HexNAc(5)Hex(3)Fuc(1)</td><td>C[+57], N[+1648]</td><td>1104.4621</td><td>3</td><td>3311.3716</td><td>3311.3625</td><td>0</td><td>2.7</td><td>89</td><td>Specific</td><td>681.69</td><td>557.63</td><td>557.63</td><td>13.93</td><td>120</td><td>>sp|P01877|IGHA2_HUMAN Immunoglobulin heavy constant alpha 2 OS=Homo sapiens OX=9606 GN=IGHA2 PE=1 SV=4</td><td>576</td><td>20201116_fqg_colostrum_1ng_glyco_hcd203040_compound15.3946.3946.3</td><td>controllerType=0 controllerNumber=1 scan=3946</td><td>[illegible]</td></tr>
<tr><td>00962:3</td><td>3</td><td>K.HYTN[+1403.507]SSQDVTVPC[+57.021]R.V</td><td>HexNAc(3)Hex(4)Fuc(1)</td><td>C[+57], N[+1404]</td><td>1023.0927</td><td>3</td><td>3067.2637</td><td>3067.2566</td><td>0</td><td>2.3</td><td>89</td><td>Specific</td><td>687.41</td><td>633.16</td><td>633.16</td><td>13.93</td><td>120</td><td>>sp|P01877|IGHA2_HUMAN Immunoglobulin heavy constant alpha 2 OS=Homo sapiens OX=9606 GN=IGHA2 PE=1 SV=4</td><td>576</td><td>20201116_fqg_colostrum_1ng_glyco_hcd203040_compound15.3837.3837.3</td><td>controllerType=0 controllerNumber=1 scan=3837</td><td>[illegible]</td></tr>
<tr><td>01015:3</td><td>3</td><td>K.HYTN[+1809.666]SSQDVTVPC[+57.021]R.V</td><td>HexNAc(5)Hex(4)Fuc(1)</td><td>C[+57], N[+1810]</td><td>1158.4788</td><td>3</td><td>3473.4219</td><td>3473.4154</td><td>0</td><td>1.9</td><td>89</td><td>Specific</td><td>691.45</td><td>575.42</td><td>575.42</td><td>13.93</td><td>120</td><td>>sp|P01877|IGHA2_HUMAN Immunoglobulin heavy constant alpha 2 OS=Homo sapiens OX=9606 GN=IGHA2 PE=1 SV=4</td><td>576</td><td>20201116_fqg_colostrum_1ng_glyco_hcd203040_compound15.3892.3892.3</td><td>controllerType=0 controllerNumber=1 scan=3892</td><td>[illegible]</td></tr>
<tr><td>01275:3</td><td>3</td><td>K.HYTN[+876.322]SSQDVTVPC[+57.021]R.V</td><td>HexNAc(2)Hex(2)Fuc(1)</td><td>C[+57], N[+876]</td><td>847.3639</td><td>3</td><td>2540.0772</td><td>2540.0716</td><td>0</td><td>2.2</td><td>89</td><td>Specific</td><td>692.85</td><td>573.93</td><td>573.93</td><td>13.93</td><td>120</td><td>>sp|P01877|IGHA2_HUMAN Immunoglobulin heavy constant alpha 2 OS=Homo sapiens OX=9606 GN=IGHA2 PE=1 SV=4</td><td>576</td><td>20201116_fqg_colostrum_1ng_glyco_hcd203040_compound15.4162.4162.3</td><td>controllerType=0 controllerNumber=1 scan=4162</td><td>[illegible]</td></tr>
<tr><td>00983:3</td><td>3</td><td>K.HYTN[+1444.534]SSQDVTVPC[+57.021]R.V</td><td>HexNAc(4)Hex(3)Fuc(1)</td><td>C[+57], N[+1445]</td><td>1036.7681</td><td>3</td><td>3108.2897</td><td>3108.2832</td><td>0</td><td>2.1</td><td>89</td><td>Specific</td><td>706.81</td><td>632.63</td><td>632.63</td><td>14.93</td><td>120</td><td>>sp|P01877|IGHA2_HUMAN Immunoglobulin heavy constant alpha 2 OS=Homo sapiens OX=9606 GN=IGHA2 PE=1 SV=4</td><td>576</td><td>20201116_fqg_colostrum_1ng_glyco_hcd203040_compound15.3859.3859.3</td><td>controllerType=0 controllerNumber=1 scan=3859</td><td>12.6936</td></tr>
<tr><td>01064:3</td><td>3</td><td>K.HYTN[+1647.613]SSQDVTVPC[+57.021]R.V</td><td>HexNAc(5)Hex(3)Fuc(1)</td><td>C[+57], N[+1648]</td><td>1104.4621</td><td>3</td><td>3311.3716</td><td>3311.3625</td><td>0</td><td>2.7</td><td>89</td><td>Specific</td><td>710.2</td><td>553.81</td><td>553.81</td><td>14.93</td><td>120</td><td>>sp|P01877|IGHA2_HUMAN Immunoglobulin heavy constant alpha 2 OS=Homo sapiens OX=9606 GN=IGHA2 PE=1 SV=4</td><td>576</td><td>20201116_fqg_colostrum_1ng_glyco_hcd203040_compound15.3944.3944.3</td><td>controllerType=0 controllerNumber=1 scan=3944</td><td>12.9012</td></tr>
<tr><td>01534:3</td><td>3</td><td>K.HYTN[+1444.534]SSQDVTVPC[+57.021]R.V</td><td>HexNAc(4)Hex(3)Fuc(1)</td><td>C[+57], N[+1445]</td><td>1036.7688</td><td>3</td><td>3108.2919</td><td>3108.2832</td><td>0</td><td>2.8</td><td>89</td><td>Specific</td><td>713.37</td><td>697.06</td><td>697.06</td><td>14.82</td><td>120</td><td>>sp|P01877|IGHA2_HUMAN Immunoglobulin heavy constant alpha 2 OS=Homo sapiens OX=9606 GN=IGHA2 PE=1 SV=4</td><td>576</td><td>20201116_fqg_colostrum_1ng_glyco_hcd203040_compound15.4431.4431.3</td><td>controllerType=0 controllerNumber=1 scan=4431</td><td>13.791</td></tr>
<tr><td>01074:2</td><td>3</td><td>K.HYTN[+1647.613]SSQDVTVPC[+57.021]R.V</td><td>HexNAc(5)Hex(3)Fuc(1)</td><td>C[+57], N[+1648]</td><td>1656.188</td><td>2</td><td>3311.3688</td><td>3311.3625</td><td>0</td><td>1.9</td><td>89</td><td>Specific</td><td>715.02</td><td>574.54</td><td>574.54</td><td>14.31</td><td>120</td><td>>sp|P01877|IGHA2_HUMAN Immunoglobulin heavy constant alpha 2 OS=Homo sapiens OX=9606 GN=IGHA2 PE=1 SV=4</td><td>576</td><td>20201116_fqg_colostrum_1ng_glyco_hcd203040_compound15.3954.3954.2</td><td>controllerType=0 controllerNumber=1 scan=3954</td><td>12.9235</td></tr>
<tr><td>01025:2</td><td>3</td><td>K.HYTN[+1444.534]SSQDVTVPC[+57.021]R.V</td><td>HexNAc(4)Hex(3)Fuc(1)</td><td>C[+57], N[+1445]</td><td>1554.6492</td><td>2</td><td>3108.2911</td><td>3108.2832</td><td>0</td><td>2.6</td><td>89</td><td>Specific</td><td>720.92</td><td>595.79</td><td>595.79</td><td>14.31</td><td>120</td><td>>sp|P01877|IGHA2_HUMAN Immunoglobulin heavy constant alpha 2 OS=Homo sapiens OX=9606 GN=IGHA2 PE=1 SV=4</td><td>576</td><td>20201116_fqg_colostrum_1ng_glyco_hcd203040_compound15.3903.3903.2</td><td>controllerType=0 controllerNumber=1 scan=3903</td><td>12.8086</td></tr>
<tr><td>01012:3</td><td>3</td><td>K.HYTN[+1809.666]SSQDVTVPC[+57.021]R.V</td><td>HexNAc(5)Hex(4)Fuc(1)</td><td>C[+57], N[+1810]</td><td>1158.4788</td><td>3</td><td>3473.4219</td><td>3473.4154</td><td>0</td><td>1.9</td><td>89</td><td>Specific</td><td>724.17</td><td>681.83</td><td>681.83</td><td>14.93</td><td>120</td><td>>sp|P01877|IGHA2_HUMAN Immunoglobulin heavy constant alpha 2 OS=Homo sapiens OX=9606 GN=IGHA2 PE=1 SV=4</td><td>576</td><td>20201116_fqg_colostrum_1ng_glyco_hcd203040_compound15.3889.3889.3</td><td>controllerType=0 controllerNumber=1 scan=3889</td><td>12.7563</td></tr>
<tr><td>01911:3</td><td>3</td><td>K.HYTN[+2262.814]SSQDVTVPC[+57.021]R.V</td><td>HexNAc(5)Hex(5)Fuc(1)NeuAc(1)</td><td>C[+57], N[+2263]</td><td>1309.529</td><td>3</td><td>3926.5724</td><td>3926.5636</td><td>0</td><td>2.2</td><td>89</td><td>Specific</td><td>726.75</td><td>496.95</td><td>496.95</td><td>14.93</td><td>120</td><td>>sp|P01877|IGHA2_HUMAN Immunoglobulin heavy constant alpha 2 OS=Homo sapiens OX=9606 GN=IGHA2 PE=1 SV=4</td><td>576</td><td>20201116_fqg_colostrum_1ng_glyco_hcd203040_compound15.4823.4823.3</td><td>controllerType=0 controllerNumber=1 scan=4823</td><td>14.5128</td></tr>
<tr><td>00947:3</td><td>3</td><td>K.HYTN[+1606.587]SSQDVTVPC[+57.021]R.V</td><td>HexNAc(4)Hex(4)Fuc(1)</td><td>C[+57], N[+1607]</td><td>1090.7858</td><td>3</td><td>3270.3428</td><td>3270.336</td><td>0</td><td>2.1</td><td>89</td><td>Specific</td><td>732.68</td><td>574.62</td><td>574.62</td><td>14.93</td><td>120</td><td>>sp|P01877|IGHA2_HUMAN Immunoglobulin heavy constant alpha 2 OS=Homo sapiens OX=9606 GN=IGHA2 PE=1 SV=4</td><td>576</td><td>20201116_fqg_colostrum_1ng_glyco_hcd203040_compound15.3821.3821.3</td><td>controllerType=0 controllerNumber=1 scan=3821</td><td>12.6204</td></tr>
<tr><td>01105:2</td><td>3</td><td>K.HYTN[+876.322]SSQDVTVPC[+57.021]R.V</td><td>HexNAc(2)Hex(2)Fuc(1)</td><td>C[+57], N[+876]</td><td>1270.543</td><td>2</td><td>2540.0787</td><td>2540.0716</td><td>0</td><td>2.8</td><td>89</td><td>Specific</td><td>752.15</td><td>704.21</td><td>704.21</td><td>15.2</td><td>120</td><td>>sp|P01877|IGHA2_HUMAN Immunoglobulin heavy constant alpha 2 OS=Homo sapiens OX=9606 GN=IGHA2 PE=1 SV=4</td><td>576</td><td>20201116_fqg_colostrum_1ng_glyco_hcd203040_compound15.3986.3986.2</td><td>controllerType=0 controllerNumber=1 scan=3986</td><td>12.9879</td></tr>
<tr><td>01023:2</td><td>3</td><td>K.HYTN[+1444.534]SSQDVTVPC[+57.021]R.V</td><td>HexNAc(4)Hex(3)Fuc(1)</td><td>C[+57], N[+1445]</td><td>1554.6492</td><td>2</td><td>3108.2911</td><td>3108.2832</td><td>0</td><td>2.6</td><td>89</td><td>Specific</td><td>763.69</td><td>696.98</td><td>696.98</td><td>15.31</td><td>120</td><td>>sp|P01877|IGHA2_HUMAN Immunoglobulin heavy constant alpha 2 OS=Homo sapiens OX=9606 GN=IGHA2 PE=1 SV=4</td><td>576</td><td>20201116_fqg_colostrum_1ng_glyco_hcd203040_compound15.3901.3901.2</td><td>controllerType=0 controllerNumber=1 scan=3901</td><td>12.8018</td></tr>
<tr><td>00985:3</td><td>3</td><td>K.HYTN[+1444.534]SSQDVTVPC[+57.021]R.V</td><td>HexNAc(4)Hex(3)Fuc(1)</td><td>C[+57], N[+1445]</td><td>1036.7681</td><td>3</td><td>3108.2897</td><td>3108.2832</td><td>0</td><td>2.1</td><td>89</td><td>Specific</td><td>771.52</td><td>622.45</td><td>622.45</td><td>15.93</td><td>120</td><td>>sp|P01877|IGHA2_HUMAN Immunoglobulin heavy constant alpha 2 OS=Homo sapiens OX=9606 GN=IGHA2 PE=1 SV=4</td><td>576</td><td>20201116_fqg_colostrum_1ng_glyco_hcd203040_compound15.3861.3861.3</td><td>controllerType=0 controllerNumber=1 scan=3861</td><td>12.7019</td></tr>
<tr><td>01892:3</td><td>3</td><td>K.HYTN[+1694.603]SSQDVTVPC[+57.021]R.V</td><td>HexNAc(3)Hex(4)Fuc(1)NeuAc(1)</td><td>C[+57], N[+1695]</td><td>1120.125</td><td>3</td><td>3358.3605</td><td>3358.352</td><td>0</td><td>2.5</td><td>89</td><td>Specific</td><td>780.52</td><td>667.05</td><td>667.05</td><td>15.93</td><td>120</td><td>>sp|P01877|IGHA2_HUMAN Immunoglobulin heavy constant alpha 2 OS=Homo sapiens OX=9606 GN=IGHA2 PE=1 SV=4</td><td>576</td><td>20201116_fqg_colostrum_1ng_glyco_hcd203040_compound15.4803.4803.3</td><td>controllerType=0 controllerNumber=1 scan=4803</td><td>14.4734</td></tr>
<tr><td>01107:2</td><td>3</td><td>K.HYTN[+876.322]SSQDVTVPC[+57.021]R.V</td><td>HexNAc(2)Hex(2)Fuc(1)</td><td>C[+57], N[+876]</td><td>1270.543</td><td>2</td><td>2540.0787</td><td>2540.0716</td><td>0</td><td>2.8</td><td>89</td><td>Specific</td><td>787.91</td><td>663.27</td><td>663.27</td><td>15.2</td><td>120</td><td>>sp|P01877|IGHA2_HUMAN Immunoglobulin heavy constant alpha 2 OS=Homo sapiens OX=9606 GN=IGHA2 PE=1 SV=4</td><td>576</td><td>20201116_fqg_colostrum_1ng_glyco_hcd203040_compound15.3988.3988.2</td><td>controllerType=0 controllerNumber=1 scan=3988</td><td>12.9939</td></tr>
<tr><td>01047:2</td><td>3</td><td>K.HYTN[+1241.454]SSQDVTVPC[+57.021]R.V</td><td>HexNAc(3)Hex(3)Fuc(1)</td><td>C[+57], N[+1241]</td><td>1453.1087</td><td>2</td><td>2905.2101</td><td>2905.2038</td><td>0</td><td>2.2</td><td>89</td><td>Specific</td><td>788.97</td><td>666.2</td><td>666.2</td><td>15.31</td><td>120</td><td>>sp|P01877|IGHA2_HUMAN Immunoglobulin heavy constant alpha 2 OS=Homo sapiens OX=9606 GN=IGHA2 PE=1 SV=4</td><td>576</td><td>20201116_fqg_colostrum_1ng_glyco_hcd203040_compound15.3926.3926.2</td><td>controllerType=0 controllerNumber=1 scan=3926</td><td>12.8606</td></tr>
<tr><td>00945:3</td><td>3</td><td>K.HYTN[+1606.587]SSQDVTVPC[+57.021]R.V</td><td>HexNAc(4)Hex(4)Fuc(1)</td><td>C[+57], N[+1607]</td><td>1090.7858</td><td>3</td><td>3270.3428</td><td>3270.336</td><td>0</td><td>2.1</td><td>89</td><td>Specific</td><td>792.87</td><td>639.7</td><td>639.7</td><td>15.93</td><td>120</td><td>>sp|P01877|IGHA2_HUMAN Immunoglobulin heavy constant alpha 2 OS=Homo sapiens OX=9606 GN=IGHA2 PE=1 SV=4</td><td>576</td><td>20201116_fqg_colostrum_1ng_glyco_hcd203040_compound15.3819.3819.3</td><td>controllerType=0 controllerNumber=1 scan=3819</td><td>12.6152</td></tr>
<tr><td>01011:3</td><td>3</td><td>K.HYTN[+1241.454]SSQDVTVPC[+57.021]R.V</td><td>HexNAc(3)Hex(3)Fuc(1)</td><td>C[+57], N[+1241]</td><td>969.0747</td><td>3</td><td>2905.2095</td><td>2905.2038</td><td>0</td><td>2</td><td>89</td><td>Specific</td><td>800.37</td><td>725.4</td><td>725.4</td><td>16.93</td><td>120</td><td>>sp|P01877|IGHA2_HUMAN Immunoglobulin heavy constant alpha 2 OS=Homo sapiens OX=9606 GN=IGHA2 PE=1 SV=4</td><td>576</td><td>20201116_fqg_colostrum_1ng_glyco_hcd203040_compound15.3888.3888.3</td><td>controllerType=0 controllerNumber=1 scan=3888</td><td>12.7549</td></tr>
<tr><td>01045:2</td><td>3</td><td>K.HYTN[+1241.454]SSQDVTVPC[+57.021]R.V</td><td>HexNAc(3)Hex(3)Fuc(1)</td><td>C[+57], N[+1241]</td><td>1453.1087</td><td>2</td><td>2905.2101</td><td>2905.2038</td><td>0</td><td>2.2</td><td>89</td><td>Specific</td><td>801.14</td><td>696.81</td><td>696.81</td><td>16.31</td><td>120</td><td>>sp|P01877|IGHA2_HUMAN Immunoglobulin heavy constant alpha 2 OS=Homo sapiens OX=9606 GN=IGHA2 PE=1 SV=4</td><td>576</td><td>20201116_fqg_colostrum_1ng_glyco_hcd203040_compound15.3924.3924.2</td><td>controllerType=0 controllerNumber=1 scan=3924</td><td>12.8556</td></tr>
<tr><td>01037:2</td><td>3</td><td>K.HYTN[+1038.375]SSQDVTVPC[+57.021]R.V</td><td>HexNAc(2)Hex(3)Fuc(1)</td><td>C[+57], N[+1038]</td><td>1351.5691</td><td>2</td><td>2702.131</td><td>2702.1244</td><td>0</td><td>2.4</td><td>89</td><td>Specific</td><td>803.76</td><td>728.57</td><td>728.57</td><td>16.31</td><td>120</td><td>>sp|P01877|IGHA2_HUMAN Immunoglobulin heavy constant alpha 2 OS=Homo sapiens OX=9606 GN=IGHA2 PE=1 SV=4</td><td>576</td><td>20201116_fqg_colostrum_1ng_glyco_hcd203040_compound15.3915.3915.2</td><td>controllerType=0 controllerNumber=1 scan=3915</td><td>12.8388</td></tr>
<tr><td>00960:3</td><td>3</td><td>K.HYTN[+1403.507]SSQDVTVPC[+57.021]R.V</td><td>HexNAc(3)Hex(4)Fuc(1)</td><td>C[+57], N[+1404]</td><td>1023.0927</td><td>3</td><td>3067.2637</td><td>3067.2566</td><td>0</td><td>2.3</td><td>89</td><td>Specific</td><td>805.43</td><td>792.87</td><td>792.87</td><td>16.93</td><td>120</td><td>>sp|P01877|IGHA2_HUMAN Immunoglobulin heavy constant alpha 2 OS=Homo sapiens OX=9606 GN=IGHA2 PE=1 SV=4</td><td>576</td><td>20201116_fqg_colostrum_1ng_glyco_hcd203040_compound15.3835.3835.3</td><td>controllerType=0 controllerNumber=1 scan=3835</td><td>12.6478</td></tr>
<tr><td>01890:3</td><td>3</td><td>K.HYTN[+1694.603]SSQDVTVPC[+57.021]R.V</td><td>HexNAc(3)Hex(4)Fuc(1)NeuAc(1)</td><td>C[+57], N[+1695]</td><td>1120.125</td><td>3</td><td>3358.3605</td><td>3358.352</td><td>0</td><td>2.5</td><td>89</td><td>Specific</td><td>810.89</td><td>601.97</td><td>601.97</td><td>16.93</td><td>120</td><td>>sp|P01877|IGHA2_HUMAN Immunoglobulin heavy constant alpha 2 OS=Homo sapiens OX=9606 GN=IGHA2 PE=1 SV=4</td><td>576</td><td>20201116_fqg_colostrum_1ng_glyco_hcd203040_compound15.4801.4801.3</td><td>controllerType=0 controllerNumber=1 scan=4801</td><td>14.4678</td></tr>
<tr><td>01035:2</td><td>3</td><td>K.HYTN[+1038.375]SSQDVTVPC[+57.021]R.V</td><td>HexNAc(2)Hex(3)Fuc(1)</td><td>C[+57], N[+1038]</td><td>1351.5691</td><td>2</td><td>2702.131</td><td>2702.1244</td><td>0</td><td>2.4</td><td>89</td><td>Specific</td><td>815.74</td><td>811.51</td><td>811.51</td><td>16.31</td><td>120</td><td>>sp|P01877|IGHA2_HUMAN Immunoglobulin heavy constant alpha 2 OS=Homo sapiens OX=9606 GN=IGHA2 PE=1 SV=4</td><td>576</td><td>20201116_fqg_colostrum_1ng_glyco_hcd203040_compound15.3913.3913.2</td><td>controllerType=0 controllerNumber=1 scan=3913</td><td>12.833</td></tr>
<tr><td>01014:3</td><td>3</td><td>K.HYTN[+1241.454]SSQDVTVPC[+57.021]R.V</td><td>HexNAc(3)Hex(3)Fuc(1)</td><td>C[+57], N[+1241]</td><td>969.0747</td><td>3</td><td>2905.2095</td><td>2905.2038</td><td>0</td><td>2</td><td>89</td><td>Specific</td><td>820.64</td><td>644.3</td><td>644.3</td><td>16.93</td><td>120</td><td>>sp|P01877|IGHA2_HUMAN Immunoglobulin heavy constant alpha 2 OS=Homo sapiens OX=9606 GN=IGHA2 PE=1 SV=4</td><td>576</td><td>20201116_fqg_colostrum_1ng_glyco_hcd203040_compound15.3891.3891.3</td><td>controllerType=0 controllerNumber=1 scan=3891</td><td>12.7726</td></tr>
<tr><td>01094:3</td><td>3</td><td>K.HYTN[+876.322]SSQDVTVPC[+57.021]R.V</td><td>HexNAc(2)Hex(2)Fuc(1)</td><td>C[+57], N[+876]</td><td>847.3643</td><td>3</td><td>2540.0785</td><td>2540.0716</td><td>0</td><td>2.7</td><td>89</td><td>Specific</td><td>856.36</td><td>856.36</td><td>856.36</td><td>17.93</td><td>120</td><td>>sp|P01877|IGHA2_HUMAN Immunoglobulin heavy constant alpha 2 OS=Homo sapiens OX=9606 GN=IGHA2 PE=1 SV=4</td><td>576</td><td>20201116_fqg_colostrum_1ng_glyco_hcd203040_compound15.3975.3975.3</td><td>controllerType=0 controllerNumber=1 scan=3975</td><td>12.9657</td></tr>
<tr><td>01097:3</td><td>3</td><td>K.HYTN[+876.322]SSQDVTVPC[+57.021]R.V</td><td>HexNAc(2)Hex(2)Fuc(1)</td><td>C[+57], N[+876]</td><td>847.3643</td><td>3</td><td>2540.0785</td><td>2540.0716</td><td>0</td><td>2.7</td><td>89</td><td>Specific</td><td>871.33</td><td>723.45</td><td>723.45</td><td>17.93</td><td>120</td><td>>sp|P01877|IGHA2_HUMAN Immunoglobulin heavy constant alpha 2 OS=Homo sapiens OX=9606 GN=IGHA2 PE=1 SV=4</td><td>576</td><td>20201116_fqg_colostrum_1ng_glyco_hcd203040_compound15.3978.3978.3</td><td>controllerType=0 controllerNumber=1 scan=3978</td><td>12.9716</td></tr>
<tr><td>01016:3</td><td>3</td><td>K.HYTN[+1038.375]SSQDVTVPC[+57.021]R.V</td><td>HexNAc(2)Hex(3)Fuc(1)</td><td>C[+57], N[+1038]</td><td>901.3816</td><td>3</td><td>2702.1301</td><td>2702.1244</td><td>0</td><td>2.1</td><td>89</td><td>Specific</td><td>895.51</td><td>748.95</td><td>748.95</td><td>17.93</td><td>120</td><td>>sp|P01877|IGHA2_HUMAN Immunoglobulin heavy constant alpha 2 OS=Homo sapiens OX=9606 GN=IGHA2 PE=1 SV=4</td><td>576</td><td>20201116_fqg_colostrum_1ng_glyco_hcd203040_compound15.3893.3893.3</td><td>controllerType=0 controllerNumber=1 scan=3893</td><td>12.7852</td></tr>
<tr><td>01013:3</td><td>3</td><td>K.HYTN[+1038.375]SSQDVTVPC[+57.021]R.V</td><td>HexNAc(2)Hex(3)Fuc(1)</td><td>C[+57], N[+1038]</td><td>901.3816</td><td>3</td><td>2702.1301</td><td>2702.1244</td><td>0</td><td>2.1</td><td>89</td><td>Specific</td><td>897.84</td><td>897.84</td><td>897.84</td><td>17.93</td><td>120</td><td>>sp|P01877|IGHA2_HUMAN Immunoglobulin heavy constant alpha 2 OS=Homo sapiens OX=9606 GN=IGHA2 PE=1 SV=4</td><td>576</td><td>20201116_fqg_colostrum_1ng_glyco_hcd203040_compound15.3890.3890.3</td><td>controllerType=0 controllerNumber=1 scan=3890</td><td>12.7576</td></tr>
<tr><td>02931:5</td><td>3</td><td>K.HYTN[+1809.666]SSQDVTVPC[+57.021]RVPPPPPC[+57.021]C[+57.021]HPR.L</td><td>HexNAc(5)Hex(4)Fuc(1)</td><td>C[+57]*3, N[+1810]</td><td>954.4123</td><td>5</td><td>4768.0325</td><td>4768.0217</td><td>0</td><td>2.3</td><td>89</td><td>Specific</td><td>295.69</td><td>244.49</td><td>244.49</td><td>6.79</td><td>120</td><td>>sp|P01877|IGHA2_HUMAN Immunoglobulin heavy constant alpha 2 OS=Homo sapiens OX=9606 GN=IGHA2 PE=1 SV=4</td><td>576</td><td>20201116_fqg_colostrum_1ng_glyco_hcd203040_compound15.5880.5880.5</td><td>controllerType=0 controllerNumber=1 scan=5880</td><td>16.4547</td></tr>
<tr><td>02933:5</td><td>3</td><td>K.HYTN[+1809.666]SSQDVTVPC[+57.021]RVPPPPPC[+57.021]C[+57.021]HPR.L</td><td>HexNAc(5)Hex(4)Fuc(1)</td><td>C[+57]*3, N[+1810]</td><td>954.4123</td><td>5</td><td>4768.0325</td><td>4768.0217</td><td>0</td><td>2.3</td><td>89</td><td>Specific</td><td>412</td><td>318.57</td><td>318.57</td><td>7.96</td><td>120</td><td>>sp|P01877|IGHA2_HUMAN Immunoglobulin heavy constant alpha 2 OS=Homo sapiens OX=9606 GN=IGHA2 PE=1 SV=4</td><td>576</td><td>20201116_fqg_colostrum_1ng_glyco_hcd203040_compound15.5882.5882.5</td><td>controllerType=0 controllerNumber=1 scan=5882</td><td>16.4594</td></tr>
<tr><td>03647:4</td><td>3</td><td>K.HYTN[+2263.835]SSQDVTVPC[+57.021]RVPPPPPC[+57.021]C[+57.021]HPR.L</td><td>HexNAc(5)Hex(5)Fuc(3)</td><td>C[+57]*3, N[+2264]</td><td>1306.2983</td><td>4</td><td>5222.1714</td><td>5222.1903</td><td>0</td><td>-3.6</td><td>89</td><td>Specific</td><td>455.31</td><td>373.91</td><td>373.91</td><td>6.18</td><td>120</td><td>>sp|P01877|IGHA2_HUMAN Immunoglobulin heavy constant alpha 2 OS=Homo sapiens OX=9606 GN=IGHA2 PE=1 SV=4</td><td>576</td><td>20201116_fqg_colostrum_1ng_glyco_hcd203040_compound15.6620.6620.4</td><td>controllerType=0 controllerNumber=1 scan=6620</td><td>17.7711</td></tr>
<tr><td>03645:4</td><td>3</td><td>K.HYTN[+2263.835]SSQDVTVPC[+57.021]RVPPPPPC[+57.021]C[+57.021]HPR.L</td><td>HexNAc(5)Hex(5)Fuc(3)</td><td>C[+57]*3, N[+2264]</td><td>1306.2983</td><td>4</td><td>5222.1714</td><td>5222.1903</td><td>0</td><td>-3.6</td><td>89</td><td>Specific</td><td>472.24</td><td>472.24</td><td>472.24</td><td>7.6</td><td>120</td><td>>sp|P01877|IGHA2_HUMAN Immunoglobulin heavy constant alpha 2 OS=Homo sapiens OX=9606 GN=IGHA2 PE=1 SV=4</td><td>576</td><td>20201116_fqg_colostrum_1ng_glyco_hcd203040_compound15.6618.6618.4</td><td>controllerType=0 controllerNumber=1 scan=6618</td><td>17.7674</td></tr>
<tr><td>03610:4</td><td>3</td><td>K.HYTN[+2262.814]SSQDVTVPC[+57.021]RVPPPPPC[+57.021]C[+57.021]HPR.L</td><td>HexNAc(5)Hex(5)Fuc(1)NeuAc(1)</td><td>C[+57]*3, N[+2263]</td><td>1306.0492</td><td>4</td><td>5221.175</td><td>5221.1699</td><td>0</td><td>1</td><td>89</td><td>Specific</td><td>499.81</td><td>412.32</td><td>412.32</td><td>9.01</td><td>120</td><td>>sp|P01877|IGHA2_HUMAN Immunoglobulin heavy constant alpha 2 OS=Homo sapiens OX=9606 GN=IGHA2 PE=1 SV=4</td><td>576</td><td>20201116_fqg_colostrum_1ng_glyco_hcd203040_compound15.6582.6582.4</td><td>controllerType=0 controllerNumber=1 scan=6582</td><td>17.7177</td></tr>
<tr><td>03613:4</td><td>3</td><td>K.HYTN[+2262.814]SSQDVTVPC[+57.021]RVPPPPPC[+57.021]C[+57.021]HPR.L</td><td>HexNAc(5)Hex(5)Fuc(1)NeuAc(1)</td><td>C[+57]*3, N[+2263]</td><td>1306.0492</td><td>4</td><td>5221.175</td><td>5221.1699</td><td>0</td><td>1</td><td>89</td><td>Specific</td><td>673.08</td><td>432.11</td><td>432.11</td><td>12.91</td><td>120</td><td>>sp|P01877|IGHA2_HUMAN Immunoglobulin heavy constant alpha 2 OS=Homo sapiens OX=9606 GN=IGHA2 PE=1 SV=4</td><td>576</td><td>20201116_fqg_colostrum_1ng_glyco_hcd203040_compound15.6585.6585.4</td><td>controllerType=0 controllerNumber=1 scan=6585</td><td>17.722</td></tr>
<tr><td>02929:4</td><td>3</td><td>K.HYTN[+1809.666]SSQDVTVPC[+57.021]RVPPPPPC[+57.021]C[+57.021]HPR.L</td><td>HexNAc(5)Hex(4)Fuc(1)</td><td>C[+57]*3, N[+1810]</td><td>1192.7632</td><td>4</td><td>4768.0311</td><td>4768.0217</td><td>0</td><td>2</td><td>89</td><td>Specific</td><td>728.74</td><td>516.16</td><td>516.16</td><td>14.15</td><td>120</td><td>>sp|P01877|IGHA2_HUMAN Immunoglobulin heavy constant alpha 2 OS=Homo sapiens OX=9606 GN=IGHA2 PE=1 SV=4</td><td>576</td><td>20201116_fqg_colostrum_1ng_glyco_hcd203040_compound15.5878.5878.4</td><td>controllerType=0 controllerNumber=1 scan=5878</td><td>16.4524</td></tr>
<tr><td>02958:5</td><td>3</td><td>K.HYTN[+1647.613]SSQDVTVPC[+57.021]RVPPPPPC[+57.021]C[+57.021]HPR.L</td><td>HexNAc(5)Hex(3)Fuc(1)</td><td>C[+57]*3, N[+1648]</td><td>922.0022</td><td>5</td><td>4605.982</td><td>4605.9689</td><td>0</td><td>2.9</td><td>89</td><td>Specific</td><td>767.77</td><td>487.59</td><td>487.59</td><td>15.67</td><td>120</td><td>>sp|P01877|IGHA2_HUMAN Immunoglobulin heavy constant alpha 2 OS=Homo sapiens OX=9606 GN=IGHA2 PE=1 SV=4</td><td>576</td><td>20201116_fqg_colostrum_1ng_glyco_hcd203040_compound15.5908.5908.5</td><td>controllerType=0 controllerNumber=1 scan=5908</td><td>16.5055</td></tr>
<tr><td>02953:3</td><td>3</td><td>K.HYTN[+1444.534]SSQDVTVPC[+57.021]RVPPPPPC[+57.021]C[+57.021]HPR.L</td><td>HexNAc(4)Hex(3)Fuc(1)</td><td>C[+57]*3, N[+1445]</td><td>1468.3045</td><td>3</td><td>4402.8989</td><td>4402.8895</td><td>0</td><td>2.2</td><td>89</td><td>Specific</td><td>771.3</td><td>675.63</td><td>675.63</td><td>14.63</td><td>120</td><td>>sp|P01877|IGHA2_HUMAN Immunoglobulin heavy constant alpha 2 OS=Homo sapiens OX=9606 GN=IGHA2 PE=1 SV=4</td><td>576</td><td>20201116_fqg_colostrum_1ng_glyco_hcd203040_compound15.5902.5902.3</td><td>controllerType=0 controllerNumber=1 scan=5902</td><td>16.4912</td></tr>
<tr><td>03510:5</td><td>3</td><td>K.HYTN[+2262.814]SSQDVTVPC[+57.021]RVPPPPPC[+57.021]C[+57.021]HPR.L</td><td>HexNAc(5)Hex(5)Fuc(1)NeuAc(1)</td><td>C[+57]*3, N[+2263]</td><td>1045.0418</td><td>5</td><td>5221.1796</td><td>5221.1699</td><td>0</td><td>1.9</td><td>89</td><td>Specific</td><td>778.33</td><td>584.89</td><td>584.89</td><td>15.77</td><td>120</td><td>>sp|P01877|IGHA2_HUMAN Immunoglobulin heavy constant alpha 2 OS=Homo sapiens OX=9606 GN=IGHA2 PE=1 SV=4</td><td>576</td><td>20201116_fqg_colostrum_1ng_glyco_hcd203040_compound15.6479.6479.5</td><td>controllerType=0 controllerNumber=1 scan=6479</td><td>17.5558</td></tr>
<tr><td>03513:5</td><td>3</td><td>K.HYTN[+2262.814]SSQDVTVPC[+57.021]RVPPPPPC[+57.021]C[+57.021]HPR.L</td><td>HexNAc(5)Hex(5)Fuc(1)NeuAc(1)</td><td>C[+57]*3, N[+2263]</td><td>1045.0418</td><td>5</td><td>5221.1796</td><td>5221.1699</td><td>0</td><td>1.9</td><td>89</td><td>Specific</td><td>791.09</td><td>544.99</td><td>544.99</td><td>15.77</td><td>120</td><td>>sp|P01877|IGHA2_HUMAN Immunoglobulin heavy constant alpha 2 OS=Homo sapiens OX=9606 GN=IGHA2 PE=1 SV=4</td><td>576</td><td>20201116_fqg_colostrum_1ng_glyco_hcd203040_compound15.6482.6482.5</td><td>controllerType=0 controllerNumber=1 scan=6482</td><td>17.562</td></tr>
<tr><td>02960:5</td><td>3</td><td>K.HYTN[+1647.613]SSQDVTVPC[+57.021]RVPPPPPC[+57.021]C[+57.021]HPR.L</td><td>HexNAc(5)Hex(3)Fuc(1)</td><td>C[+57]*3, N[+1648]</td><td>922.0022</td><td>5</td><td>4605.982</td><td>4605.9689</td><td>0</td><td>2.9</td><td>89</td><td>Specific</td><td>794.41</td><td>530.38</td><td>530.38</td><td>15.67</td><td>120</td><td>>sp|P01877|IGHA2_HUMAN Immunoglobulin heavy constant alpha 2 OS=Homo sapiens OX=9606 GN=IGHA2 PE=1 SV=4</td><td>576</td><td>20201116_fqg_colostrum_1ng_glyco_hcd203040_compound15.5910.5910.5</td><td>controllerType=0 controllerNumber=1 scan=5910</td><td>16.5097</td></tr>
<tr><td>03281:5</td><td>3</td><td>K.HYTN[+1444.534]SSQDVTVPC[+57.021]RVPPPPPC[+57.021]C[+57.021]HPR.L</td><td>HexNAc(4)Hex(3)Fuc(1)</td><td>C[+57]*3, N[+1445]</td><td>881.3861</td><td>5</td><td>4402.9012</td><td>4402.8895</td><td>0</td><td>2.7</td><td>89</td><td>Specific</td><td>803.69</td><td>733.6</td><td>733.6</td><td>16.77</td><td>120</td><td>>sp|P01877|IGHA2_HUMAN Immunoglobulin heavy constant alpha 2 OS=Homo sapiens OX=9606 GN=IGHA2 PE=1 SV=4</td><td>576</td><td>20201116_fqg_colostrum_1ng_glyco_hcd203040_compound15.6242.6242.5</td><td>controllerType=0 controllerNumber=1 scan=6242</td><td>17.1325</td></tr>
<tr><td>02902:4</td><td>3</td><td>K.HYTN[+1809.666]SSQDVTVPC[+57.021]RVPPPPPC[+57.021]C[+57.021]HPR.L</td><td>HexNAc(5)Hex(4)Fuc(1)</td><td>C[+57]*3, N[+1810]</td><td>1192.7652</td><td>4</td><td>4768.0389</td><td>4768.0217</td><td>0</td><td>3.6</td><td>89</td><td>Specific</td><td>827.7</td><td>751.17</td><td>751.17</td><td>16.05</td><td>120</td><td>>sp|P01877|IGHA2_HUMAN Immunoglobulin heavy constant alpha 2 OS=Homo sapiens OX=9606 GN=IGHA2 PE=1 SV=4</td><td>576</td><td>20201116_fqg_colostrum_1ng_glyco_hcd203040_compound15.5850.5850.4</td><td>controllerType=0 controllerNumber=1 scan=5850</td><td>16.4012</td></tr>
<tr><td>02932:4</td><td>3</td><td>K.HYTN[+1809.666]SSQDVTVPC[+57.021]RVPPPPPC[+57.021]C[+57.021]HPR.L</td><td>HexNAc(5)Hex(4)Fuc(1)</td><td>C[+57]*3, N[+1810]</td><td>1192.7632</td><td>4</td><td>4768.0311</td><td>4768.0217</td><td>0</td><td>2</td><td>89</td><td>Specific</td><td>852.28</td><td>578.31</td><td>578.31</td><td>17.15</td><td>120</td><td>>sp|P01877|IGHA2_HUMAN Immunoglobulin heavy constant alpha 2 OS=Homo sapiens OX=9606 GN=IGHA2 PE=1 SV=4</td><td>576</td><td>20201116_fqg_colostrum_1ng_glyco_hcd203040_compound15.5881.5881.4</td><td>controllerType=0 controllerNumber=1 scan=5881</td><td>16.4566</td></tr>
<tr><td>02955:3</td><td>3</td><td>K.HYTN[+1444.534]SSQDVTVPC[+57.021]RVPPPPPC[+57.021]C[+57.021]HPR.L</td><td>HexNAc(4)Hex(3)Fuc(1)</td><td>C[+57]*3, N[+1445]</td><td>1468.3045</td><td>3</td><td>4402.8989</td><td>4402.8895</td><td>0</td><td>2.2</td><td>89</td><td>Specific</td><td>859.72</td><td>672.29</td><td>672.29</td><td>16.63</td><td>120</td><td>>sp|P01877|IGHA2_HUMAN Immunoglobulin heavy constant alpha 2 OS=Homo sapiens OX=9606 GN=IGHA2 PE=1 SV=4</td><td>576</td><td>20201116_fqg_colostrum_1ng_glyco_hcd203040_compound15.5904.5904.3</td><td>controllerType=0 controllerNumber=1 scan=5904</td><td>16.4979</td></tr>
<tr><td>02905:4</td><td>3</td><td>K.HYTN[+1809.666]SSQDVTVPC[+57.021]RVPPPPPC[+57.021]C[+57.021]HPR.L</td><td>HexNAc(5)Hex(4)Fuc(1)</td><td>C[+57]*3, N[+1810]</td><td>1192.7652</td><td>4</td><td>4768.0389</td><td>4768.0217</td><td>0</td><td>3.6</td><td>89</td><td>Specific</td><td>886.84</td><td>616.69</td><td>616.69</td><td>17.05</td><td>120</td><td>>sp|P01877|IGHA2_HUMAN Immunoglobulin heavy constant alpha 2 OS=Homo sapiens OX=9606 GN=IGHA2 PE=1 SV=4</td><td>576</td><td>20201116_fqg_colostrum_1ng_glyco_hcd203040_compound15.5853.5853.4</td><td>controllerType=0 controllerNumber=1 scan=5853</td><td>16.4063</td></tr>
<tr><td>02959:4</td><td>3</td><td>K.HYTN[+1647.613]SSQDVTVPC[+57.021]RVPPPPPC[+57.021]C[+57.021]HPR.L</td><td>HexNAc(5)Hex(3)Fuc(1)</td><td>C[+57]*3, N[+1648]</td><td>1152.251</td><td>4</td><td>4605.9822</td><td>4605.9689</td><td>0</td><td>2.9</td><td>89</td><td>Specific</td><td>900.95</td><td>641.58</td><td>641.58</td><td>18.05</td><td>120</td><td>>sp|P01877|IGHA2_HUMAN Immunoglobulin heavy constant alpha 2 OS=Homo sapiens OX=9606 GN=IGHA2 PE=1 SV=4</td><td>576</td><td>20201116_fqg_colostrum_1ng_glyco_hcd203040_compound15.5909.5909.4</td><td>controllerType=0 controllerNumber=1 scan=5909</td><td>16.5079</td></tr>
<tr><td>02898:5</td><td>3</td><td>K.HYTN[+1444.534]SSQDVTVPC[+57.021]RVPPPPPC[+57.021]C[+57.021]HPR.L</td><td>HexNAc(4)Hex(3)Fuc(1)</td><td>C[+57]*3, N[+1445]</td><td>881.3861</td><td>5</td><td>4402.9012</td><td>4402.8895</td><td>0</td><td>2.7</td><td>89</td><td>Specific</td><td>914.17</td><td>816.59</td><td>816.59</td><td>18.77</td><td>120</td><td>>sp|P01877|IGHA2_HUMAN Immunoglobulin heavy constant alpha 2 OS=Homo sapiens OX=9606 GN=IGHA2 PE=1 SV=4</td><td>576</td><td>20201116_fqg_colostrum_1ng_glyco_hcd203040_compound15.5846.5846.5</td><td>controllerType=0 controllerNumber=1 scan=5846</td><td>16.3928</td></tr>
<tr><td>02970:4</td><td>3</td><td>K.HYTN[+1038.375]SSQDVTVPC[+57.021]RVPPPPPC[+57.021]C[+57.021]HPR.L</td><td>HexNAc(2)Hex(3)Fuc(1)</td><td>C[+57]*3, N[+1038]</td><td>999.9413</td><td>4</td><td>3996.7435</td><td>3996.7307</td><td>0</td><td>3.2</td><td>89</td><td>Specific</td><td>948.5</td><td>948.5</td><td>948.5</td><td>18.05</td><td>120</td><td>>sp|P01877|IGHA2_HUMAN Immunoglobulin heavy constant alpha 2 OS=Homo sapiens OX=9606 GN=IGHA2 PE=1 SV=4</td><td>576</td><td>20201116_fqg_colostrum_1ng_glyco_hcd203040_compound15.5920.5920.4</td><td>controllerType=0 controllerNumber=1 scan=5920</td><td>16.5219</td></tr>
<tr><td>02977:5</td><td>3</td><td>K.HYTN[+1038.375]SSQDVTVPC[+57.021]RVPPPPPC[+57.021]C[+57.021]HPR.L</td><td>HexNAc(2)Hex(3)Fuc(1)</td><td>C[+57]*3, N[+1038]</td><td>800.1534</td><td>5</td><td>3996.7379</td><td>3996.7307</td><td>0</td><td>1.8</td><td>89</td><td>Specific</td><td>972.95</td><td>972.95</td><td>972.95</td><td>19.77</td><td>120</td><td>>sp|P01877|IGHA2_HUMAN Immunoglobulin heavy constant alpha 2 OS=Homo sapiens OX=9606 GN=IGHA2 PE=1 SV=4</td><td>576</td><td>20201116_fqg_colostrum_1ng_glyco_hcd203040_compound15.5927.5927.5</td><td>controllerType=0 controllerNumber=1 scan=5927</td><td>16.535</td></tr>
<tr><td>02867:4</td><td>3</td><td>K.HYTN[+1606.587]SSQDVTVPC[+57.021]RVPPPPPC[+57.021]C[+57.021]HPR.L</td><td>HexNAc(4)Hex(4)Fuc(1)</td><td>C[+57]*3, N[+1607]</td><td>1141.9943</td><td>4</td><td>4564.9554</td><td>4564.9423</td><td>0</td><td>2.9</td><td>89</td><td>Specific</td><td>984.02</td><td>759.77</td><td>759.77</td><td>19.05</td><td>120</td><td>>sp|P01877|IGHA2_HUMAN Immunoglobulin heavy constant alpha 2 OS=Homo sapiens OX=9606 GN=IGHA2 PE=1 SV=4</td><td>576</td><td>20201116_fqg_colostrum_1ng_glyco_hcd203040_compound15.5813.5813.2</td><td>controllerType=0 controllerNumber=1 scan=5813</td><td>16.3298</td></tr>
<tr><td>03283:5</td><td>3</td><td>K.HYTN[+1444.534]SSQDVTVPC[+57.021]RVPPPPPC[+57.021]C[+57.021]HPR.L</td><td>HexNAc(4)Hex(3)Fuc(1)</td><td>C[+57]*3, N[+1445]</td><td>881.3861</td><td>5</td><td>4402.9012</td><td>4402.8895</td><td>0</td><td>2.7</td><td>89</td><td>Specific</td><td>1007.67</td><td>813.25</td><td>813.25</td><td>20.77</td><td>120</td><td>>sp|P01877|IGHA2_HUMAN Immunoglobulin heavy constant alpha 2 OS=Homo sapiens OX=9606 GN=IGHA2 PE=1 SV=4</td><td>576</td><td>20201116_fqg_colostrum_1ng_glyco_hcd203040_compound15.6244.6244.5</td><td>controllerType=0 controllerNumber=1 scan=6244</td><td>17.1377</td></tr>
<tr><td>02957:4</td><td>3</td><td>K.HYTN[+1647.613]SSQDVTVPC[+57.021]RVPPPPPC[+57.021]C[+57.021]HPR.L</td><td>HexNAc(5)Hex(3)Fuc(1)</td><td>C[+57]*3, N[+1648]</td><td>1152.251</td><td>4</td><td>4605.9822</td><td>4605.9689</td><td>0</td><td>2.9</td><td>89</td><td>Specific</td><td>1020.41</td><td>900.11</td><td>900.11</td><td>20.05</td><td>120</td><td>>sp|P01877|IGHA2_HUMAN Immunoglobulin heavy constant alpha 2 OS=Homo sapiens OX=9606 GN=IGHA2 PE=1 SV=4</td><td>576</td><td>20201116_fqg_colostrum_1ng_glyco_hcd203040_compound15.5907.5907.4</td><td>controllerType=0 controllerNumber=1 scan=5907</td><td>16.5038</td></tr>
<tr><td>02899:4</td><td>3</td><td>K.HYTN[+1444.534]SSQDVTVPC[+57.021]RVPPPPPC[+57.021]C[+57.021]HPR.L</td><td>HexNAc(4)Hex(3)Fuc(1)</td><td>C[+57]*3, N[+1445]</td><td>1101.4806</td><td>4</td><td>4402.9007</td><td>4402.8895</td><td>0</td><td>2.5</td><td>89</td><td>Specific</td><td>1021.53</td><td>841.85</td><td>841.85</td><td>20.15</td><td>120</td><td>>sp|P01877|IGHA2_HUMAN Immunoglobulin heavy constant alpha 2 OS=Homo sapiens OX=9606 GN=IGHA2 PE=1 SV=4</td><td>576</td><td>20201116_fqg_colostrum_1ng_glyco_hcd203040_compound15.5847.5847.4</td><td>controllerType=0 controllerNumber=1 scan=5847</td><td>16.3942</td></tr>
<tr><td>03280:4</td><td>3</td><td>K.HYTN[+1444.534]SSQDVTVPC[+57.021]RVPPPPPC[+57.021]C[+57.021]HPR.L</td><td>HexNAc(4)Hex(3)Fuc(1)</td><td>C[+57]*3, N[+1445]</td><td>1101.4795</td><td>4</td><td>4402.8963</td><td>4402.8895</td><td>0</td><td>1.6</td><td>89</td><td>Specific</td><td>1022.5</td><td>867.72</td><td>867.72</td><td>19.91</td><td>120</td><td>>sp|P01877|IGHA2_HUMAN Immunoglobulin heavy constant alpha 2 OS=Homo sapiens OX=9606 GN=IGHA2 PE=1 SV=4</td><td>576</td><td>20201116_fqg_colostrum_1ng_glyco_hcd203040_compound15.6241.6241.4</td><td>controllerType=0 controllerNumber=1 scan=6241</td><td>17.131</td></tr>
<tr><td>02901:4</td><td>3</td><td>K.HYTN[+1444.534]SSQDVTVPC[+57.021]RVPPPPPC[+57.021]C[+57.021]HPR.L</td><td>HexNAc(4)Hex(3)Fuc(1)</td><td>C[+57]*3, N[+1445]</td><td>1101.4806</td><td>4</td><td>4402.9007</td><td>4402.8895</td><td>0</td><td>2.5</td><td>89</td><td>Specific</td><td>1036.62</td><td>726.84</td><td>726.84</td><td>20.15</td><td>120</td><td>>sp|P01877|IGHA2_HUMAN Immunoglobulin heavy constant alpha 2 OS=Homo sapiens OX=9606 GN=IGHA2 PE=1 SV=4</td><td>576</td><td>20201116_fqg_colostrum_1ng_glyco_hcd203040_compound15.5849.5849.4</td><td>controllerType=0 controllerNumber=1 scan=5849</td><td>16.4</td></tr>
<tr><td>03282:4</td><td>3</td><td>K.HYTN[+1444.534]SSQDVTVPC[+57.021]RVPPPPPC[+57.021]C[+57.021]HPR.L</td><td>HexNAc(4)Hex(3)Fuc(1)</td><td>C[+57]*3, N[+1445]</td><td>1101.4795</td><td>4</td><td>4402.8963</td><td>4402.8895</td><td>0</td><td>1.6</td><td>89</td><td>Specific</td><td>1041</td><td>738.92</td><td>738.92</td><td>19.91</td><td>120</td><td>>sp|P01877|IGHA2_HUMAN Immunoglobulin heavy constant alpha 2 OS=Homo sapiens OX=9606 GN=IGHA2 PE=1 SV=4</td><td>576</td><td>20201116_fqg_colostrum_1ng_glyco_hcd203040_compound15.6243.6243.4</td><td>controllerType=0 controllerNumber=1 scan=6243</td><td>17.1358</td></tr>
<tr><td>02900:5</td><td>3</td><td>K.HYTN[+1444.534]SSQDVTVPC[+57.021]RVPPPPPC[+57.021]C[+57.021]HPR.L</td><td>HexNAc(4)Hex(3)Fuc(1)</td><td>C[+57]*3, N[+1445]</td><td>881.3861</td><td>5</td><td>4402.9012</td><td>4402.8895</td><td>0</td><td>2.7</td><td>89</td><td>Specific</td><td>1060.04</td><td>739.51</td><td>739.51</td><td>21.77</td><td>120</td><td>>sp|P01877|IGHA2_HUMAN Immunoglobulin heavy constant alpha 2 OS=Homo sapiens OX=9606 GN=IGHA2 PE=1 SV=4</td><td>576</td><td>20201116_fqg_colostrum_1ng_glyco_hcd203040_compound15.5848.5848.5</td><td>controllerType=0 controllerNumber=1 scan=5848</td><td>16.3979</td></tr>
<tr><td>02980:5</td><td>3</td><td>K.HYTN[+1038.375]SSQDVTVPC[+57.021]RVPPPPPC[+57.021]C[+57.021]HPR.L</td><td>HexNAc(2)Hex(3)Fuc(1)</td><td>C[+57]*3, N[+1038]</td><td>800.1534</td><td>5</td><td>3996.7379</td><td>3996.7307</td><td>0</td><td>1.8</td><td>89</td><td>Specific</td><td>1072.34</td><td>1066.94</td><td>1066.94</td><td>21.77</td><td>120</td><td>>sp|P01877|IGHA2_HUMAN Immunoglobulin heavy constant alpha 2 OS=Homo sapiens OX=9606 GN=IGHA2 PE=1 SV=4</td><td>576</td><td>20201116_fqg_colostrum_1ng_glyco_hcd203040_compound15.5930.5930.5</td><td>controllerType=0 controllerNumber=1 scan=5930</td><td>16.5415</td></tr>
<tr><td>02973:4</td><td>3</td><td>K.HYTN[+1038.375]SSQDVTVPC[+57.021]RVPPPPPC[+57.021]C[+57.021]HPR.L</td><td>HexNAc(2)Hex(3)Fuc(1)</td><td>C[+57]*3, N[+1038]</td><td>999.9413</td><td>4</td><td>3996.7435</td><td>3996.7307</td><td>0</td><td>3.2</td><td>89</td><td>Specific</td><td>1080.6</td><td>820.13</td><td>820.13</td><td>21.05</td><td>120</td><td>>sp|P01877|IGHA2_HUMAN Immunoglobulin heavy constant alpha 2 OS=Homo sapiens OX=9606 GN=IGHA2 PE=1 SV=4</td><td>576</td><td>20201116_fqg_colostrum_1ng_glyco_hcd203040_compound15.5923.5923.4</td><td>controllerType=0 controllerNumber=1 scan=5923</td><td>16.529</td></tr>
<tr><td>02865:4</td><td>3</td><td>K.HYTN[+1606.587]SSQDVTVPC[+57.021]RVPPPPPC[+57.021]C[+57.021]HPR.L</td><td>HexNAc(4)Hex(4)Fuc(1)</td><td>C[+57]*3, N[+1607]</td><td>1141.9943</td><td>4</td><td>4564.9554</td><td>4564.9423</td><td>0</td><td>2.9</td><td>89</td><td>Specific</td><td>1099.74</td><td>982.15</td><td>982.15</td><td>21.05</td><td>120</td><td>>sp|P01877|IGHA2_HUMAN Immunoglobulin heavy constant alpha 2 OS=Homo sapiens OX=9606 GN=IGHA2 PE=1 SV=4</td><td>576</td><td>20201116_fqg_colostrum_1ng_glyco_hcd203040_compound15.5811.5811.2</td><td>controllerType=0 controllerNumber=1 scan=5811</td><td>16.3217</td></tr>
<tr><td>19261:5</td><td>3</td><td>R.LSLHRPALEDLLLGSEAN[+1501.555]LTC[+57.021]TLTGLRDASGATFTWTPSSGK.S</td><td>HexNAc(5)Hex(3)</td><td>C[+57], N[+1502]</td><td>1192.5772</td><td>5</td><td>5958.8566</td><td>5958.8323</td><td>0</td><td>4.1</td><td>114</td><td>Specific</td><td>397.47</td><td>397.47</td><td>397.47</td><td>8.45</td><td>120</td><td>>sp|P01877|IGHA2_HUMAN Immunoglobulin heavy constant alpha 2 OS=Homo sapiens OX=9606 GN=IGHA2 PE=1 SV=4</td><td>576</td><td>20201116_fqg_colostrum_1ng_glyco_hcd203040_compound15.22816.22816.5</td><td>controllerType=0 controllerNumber=1 scan=22816</td><td>44.6161</td></tr>
<tr><td>19264:5</td><td>3</td><td>R.LSLHRPALEDLLLGSEAN[+1501.555]LTC[+57.021]TLTGLRDASGATFTWTPSSGK.S</td><td>HexNAc(5)Hex(3)</td><td>C[+57], N[+1502]</td><td>1192.5772</td><td>5</td><td>5958.8566</td><td>5958.8323</td><td>0</td><td>4.1</td><td>114</td><td>Specific</td><td>715.18</td><td>670.13</td><td>670.13</td><td>13.72</td><td>120</td><td>>sp|P01877|IGHA2_HUMAN Immunoglobulin heavy constant alpha 2 OS=Homo sapiens OX=9606 GN=IGHA2 PE=1 SV=4</td><td>576</td><td>20201116_fqg_colostrum_1ng_glyco_hcd203040_compound15.22819.22819.5</td><td>controllerType=0 controllerNumber=1 scan=22819</td><td>44.6236</td></tr>
<tr><td>02543:3</td><td>3</td><td>K.TPLTAN[+1768.640]ITK.S</td><td>HexNAc(4)Hex(5)Fuc(1)</td><td>N[+1769]</td><td>909.7395</td><td>3</td><td>2727.204</td><td>2727.1963</td><td>0</td><td>2.8</td><td>200</td><td>Specific</td><td>229.3</td><td>94.4</td><td>94.4</td><td>3.7</td><td>120</td><td>>sp|P01877|IGHA2_HUMAN Immunoglobulin heavy constant alpha 2 OS=Homo sapiens OX=9606 GN=IGHA2 PE=1 SV=4</td><td>576</td><td>20201116_fqg_colostrum_1ng_glyco_hcd203040_compound15.5476.5476.3</td><td>controllerType=0 controllerNumber=1 scan=5476</td><td>15.6365</td></tr>
<tr><td>07163:3</td><td>3</td><td>K.TPLTAN[+2350.830]ITK.S</td><td>HexNAc(4)Hex(5)Fuc(1)NeuAc(2)</td><td>N[+2351]</td><td>1103.8031</td><td>3</td><td>3309.3948</td><td>3309.3871</td><td>0</td><td>2.3</td><td>200</td><td>Specific</td><td>254.23</td><td>77.42</td><td>77.42</td><td>5.25</td><td>120</td><td>>sp|P01877|IGHA2_HUMAN Immunoglobulin heavy constant alpha 2 OS=Homo sapiens OX=9606 GN=IGHA2 PE=1 SV=4</td><td>576</td><td>20201116_fqg_colostrum_1ng_glyco_hcd203040_compound15.10289.10289.3</td><td>controllerType=0 controllerNumber=1 scan=10289</td><td>23.5879</td></tr>
<tr><td>02764:2</td><td>3</td><td>K.TPLTAN[+1444.534]ITK.S</td><td>HexNAc(4)Hex(3)Fuc(1)</td><td>N[+1445]</td><td>1202.0512</td><td>2</td><td>2403.0951</td><td>2403.0906</td><td>0</td><td>1.9</td><td>200</td><td>Specific</td><td>262.86</td><td>106.19</td><td>106.19</td><td>5.33</td><td>120</td><td>>sp|P01877|IGHA2_HUMAN Immunoglobulin heavy constant alpha 2 OS=Homo sapiens OX=9606 GN=IGHA2 PE=1 SV=4</td><td>576</td><td>20201116_fqg_colostrum_1ng_glyco_hcd203040_compound15.5706.5706.2</td><td>controllerType=0 controllerNumber=1 scan=5706</td><td>16.12</td></tr>
<tr><td>07165:3</td><td>3</td><td>K.TPLTAN[+2350.830]ITK.S</td><td>HexNAc(4)Hex(5)Fuc(1)NeuAc(2)</td><td>N[+2351]</td><td>1103.8031</td><td>3</td><td>3309.3948</td><td>3309.3871</td><td>0</td><td>2.3</td><td>200</td><td>Specific</td><td>283.28</td><td>242.5</td><td>242.5</td><td>5.26</td><td>120</td><td>>sp|P01877|IGHA2_HUMAN Immunoglobulin heavy constant alpha 2 OS=Homo sapiens OX=9606 GN=IGHA2 PE=1 SV=4</td><td>576</td><td>20201116_fqg_colostrum_1ng_glyco_hcd203040_compound15.10291.10291.3</td><td>controllerType=0 controllerNumber=1 scan=10291</td><td>23.5918</td></tr>
<tr><td>04413:3</td><td>3</td><td>K.TPLTAN[+2262.814]ITK.S</td><td>HexNAc(5)Hex(5)Fuc(1)NeuAc(1)</td><td>N[+2263]</td><td>1074.4643</td><td>3</td><td>3221.3782</td><td>3221.3711</td><td>0</td><td>2.2</td><td>200</td><td>Specific</td><td>323.82</td><td>132.48</td><td>132.48</td><td>6.14</td><td>120</td><td>>sp|P01877|IGHA2_HUMAN Immunoglobulin heavy constant alpha 2 OS=Homo sapiens OX=9606 GN=IGHA2 PE=1 SV=4</td><td>576</td><td>20201116_fqg_colostrum_1ng_glyco_hcd203040_compound15.7429.7429.3</td><td>controllerType=0 controllerNumber=1 scan=7429</td><td>19.0302</td></tr>
<tr><td>02573:3</td><td>3</td><td>K.TPLTAN[+1971.719]ITK.S</td><td>HexNAc(5)Hex(5)Fuc(1)</td><td>N[+1972]</td><td>977.4322</td><td>3</td><td>2930.2819</td><td>2930.2757</td><td>0</td><td>2.1</td><td>200</td><td>Specific</td><td>509.42</td><td>236.71</td><td>236.71</td><td>8.08</td><td>120</td><td>>sp|P01877|IGHA2_HUMAN Immunoglobulin heavy constant alpha 2 OS=Homo sapiens OX=9606 GN=IGHA2 PE=1 SV=4</td><td>576</td><td>20201116_fqg_colostrum_1ng_glyco_hcd203040_compound15.5508.5508.3</td><td>controllerType=0 controllerNumber=1 scan=5508</td><td>15.6978</td></tr>
<tr><td>02784:2</td><td>3</td><td>K.TPLTAN[+1809.666]ITK.S</td><td>HexNAc(5)Hex(4)Fuc(1)</td><td>N[+1810]</td><td>1384.6171</td><td>2</td><td>2768.2269</td><td>2768.2228</td><td>0</td><td>1.5</td><td>200</td><td>Specific</td><td>511.23</td><td>306.9</td><td>306.9</td><td>8.12</td><td>120</td><td>>sp|P01877|IGHA2_HUMAN Immunoglobulin heavy constant alpha 2 OS=Homo sapiens OX=9606 GN=IGHA2 PE=1 SV=4</td><td>576</td><td>20201116_fqg_colostrum_1ng_glyco_hcd203040_compound15.5727.5727.2</td><td>controllerType=0 controllerNumber=1 scan=5727</td><td>16.158</td></tr>
<tr><td>02611:3</td><td>3</td><td>K.TPLTAN[+1809.666]ITK.S</td><td>HexNAc(5)Hex(4)Fuc(1)</td><td>N[+1810]</td><td>923.4137</td><td>3</td><td>2768.2264</td><td>2768.2228</td><td>0</td><td>1.3</td><td>200</td><td>Specific</td><td>513.15</td><td>232.05</td><td>232.05</td><td>7.85</td><td>120</td><td>>sp|P01877|IGHA2_HUMAN Immunoglobulin heavy constant alpha 2 OS=Homo sapiens OX=9606 GN=IGHA2 PE=1 SV=4</td><td>576</td><td>20201116_fqg_colostrum_1ng_glyco_hcd203040_compound15.5548.5548.3</td><td>controllerType=0 controllerNumber=1 scan=5548</td><td>15.8089</td></tr>
</table>

Table S4. Cont.

| Query #:z | Protein Rank | Glycopeptide | Glycan | Modification Type(s) | Observed m/z | z | Observed (M+H) | Calc. mass (M+H) | Off-by-x error | Mass error (ppm) | StartPosition | Cleavage | Score | Delta | Delta Mod | \|Log Prob\| | # of unique peptides | Protein Name | Protein DB number | Comment | Scan # | Scan Time |
|---|---|---|---|---|---|---|---|---|---|---|---|---|---|---|---|---|---|---|---|---|---|---|
| 02678:3 | 3 | K.TPLTAN[+1647.613]ITK.S | HexNAc(5)Hex(3)Fuc(1) | N[+1648] | 869.3964 | 3 | 2606.1746 | 2606.17 | 0 | 1.8 | 200 | Specific | 532.19 | 228.75 | 228.75 | 8.19 | 120 | >sp\|P01877\|IGHA2_HUMAN Immunoglobulin heavy constant alpha 2 OS=Homo sapiens OX=9606 GN=IGHA2 PE=1 SV=4 | 576 | 20201116_fqg_colostrum_1ng_glyco_hcd203040_compound15.5618.5618.3 | controllerType=0 controllerNumber=1 scan=5618 | 15.965 |
| 02576:3 | 3 | K.TPLTAN[+1971.719]ITK.S | HexNAc(5)Hex(5)Fuc(1) | N[+1972] | 977.4322 | 3 | 2930.2819 | 2930.2757 | 0 | 2.1 | 200 | Specific | 535.88 | 298.73 | 298.73 | 8.49 | 120 | >sp\|P01877\|IGHA2_HUMAN Immunoglobulin heavy constant alpha 2 OS=Homo sapiens OX=9606 GN=IGHA2 PE=1 SV=4 | 576 | 20201116_fqg_colostrum_1ng_glyco_hcd203040_compound15.5511.5511.3 | controllerType=0 controllerNumber=1 scan=5511 | 15.7049 |
| 02681:3 | 3 | K.TPLTAN[+1647.613]ITK.S | HexNAc(5)Hex(3)Fuc(1) | N[+1648] | 869.3964 | 3 | 2606.1746 | 2606.17 | 0 | 1.8 | 200 | Specific | 544.8 | 274.97 | 274.97 | 8.5 | 120 | >sp\|P01877\|IGHA2_HUMAN Immunoglobulin heavy constant alpha 2 OS=Homo sapiens OX=9606 GN=IGHA2 PE=1 SV=4 | 576 | 20201116_fqg_colostrum_1ng_glyco_hcd203040_compound15.5621.5621.3 | controllerType=0 controllerNumber=1 scan=5621 | 15.9715 |
| 02782:2 | 3 | K.TPLTAN[+1809.666]ITK.S | HexNAc(5)Hex(4)Fuc(1) | N[+1810] | 1384.6171 | 2 | 2768.2269 | 2768.2228 | 0 | 1.5 | 200 | Specific | 546.18 | 295.76 | 295.76 | 8.13 | 120 | >sp\|P01877\|IGHA2_HUMAN Immunoglobulin heavy constant alpha 2 OS=Homo sapiens OX=9606 GN=IGHA2 PE=1 SV=4 | 576 | 20201116_fqg_colostrum_1ng_glyco_hcd203040_compound15.5725.5725.2 | controllerType=0 controllerNumber=1 scan=5725 | 16.1535 |
| 02884:2 | 3 | K.TPLTAN[+876.322]ITK.S | HexNAc(2)Hex(2)Fuc(1) | N[+876] | 917.9456 | 2 | 1834.8838 | 1834.8791 | 0 | 2.6 | 200 | Specific | 562.92 | 309.96 | 309.96 | 8.76 | 120 | >sp\|P01877\|IGHA2_HUMAN Immunoglobulin heavy constant alpha 2 OS=Homo sapiens OX=9606 GN=IGHA2 PE=1 SV=4 | 576 | 20201116_fqg_colostrum_1ng_glyco_hcd203040_compound15.5831.5831.2 | controllerType=0 controllerNumber=1 scan=5831 | 16.361 |
| 02582:2 | 3 | K.TPLTAN[+1971.719]ITK.S | HexNAc(5)Hex(5)Fuc(1) | N[+1972] | 1465.6453 | 2 | 2930.2833 | 2930.2757 | 0 | 2.6 | 200 | Specific | 565.94 | 289.98 | 289.98 | 8.56 | 120 | >sp\|P01877\|IGHA2_HUMAN Immunoglobulin heavy constant alpha 2 OS=Homo sapiens OX=9606 GN=IGHA2 PE=1 SV=4 | 576 | 20201116_fqg_colostrum_1ng_glyco_hcd203040_compound15.5517.5517.2 | controllerType=0 controllerNumber=1 scan=5517 | 15.7199 |
| 02493:2 | 3 | K.TPLTAN[+1768.640]ITK.S | HexNAc(4)Hex(5)Fuc(1) | N[+1769] | 1364.1058 | 2 | 2727.2042 | 2727.1963 | 0 | 2.9 | 200 | Specific | 569.57 | 318.19 | 318.19 | 8.65 | 120 | >sp\|P01877\|IGHA2_HUMAN Immunoglobulin heavy constant alpha 2 OS=Homo sapiens OX=9606 GN=IGHA2 PE=1 SV=4 | 576 | 20201116_fqg_colostrum_1ng_glyco_hcd203040_compound15.5425.5425.2 | controllerType=0 controllerNumber=1 scan=5425 | 15.5451 |
| 02746:2 | 3 | K.TPLTAN[+1647.613]ITK.S | HexNAc(5)Hex(3)Fuc(1) | N[+1648] | 1303.5909 | 2 | 2606.1744 | 2606.17 | 0 | 1.7 | 200 | Specific | 571 | 359.38 | 359.38 | 8.98 | 120 | >sp\|P01877\|IGHA2_HUMAN Immunoglobulin heavy constant alpha 2 OS=Homo sapiens OX=9606 GN=IGHA2 PE=1 SV=4 | 576 | 20201116_fqg_colostrum_1ng_glyco_hcd203040_compound15.5688.5688.2 | controllerType=0 controllerNumber=1 scan=5688 | 16.0911 |
| 04385:2 | 3 | K.TPLTAN[+2059.735]ITK.S | HexNAc(4)Hex(5)Fuc(1)NeuAc(1) | N[+2060] | 1509.6542 | 2 | 3018.3012 | 3018.2917 | 0 | 3.1 | 200 | Specific | 575.9 | 313.02 | 313.02 | 8.65 | 120 | >sp\|P01877\|IGHA2_HUMAN Immunoglobulin heavy constant alpha 2 OS=Homo sapiens OX=9606 GN=IGHA2 PE=1 SV=4 | 576 | 20201116_fqg_colostrum_1ng_glyco_hcd203040_compound15.7400.7400.2 | controllerType=0 controllerNumber=1 scan=7400 | 18.9816 |
| 04415:3 | 3 | K.TPLTAN[+2262.814]ITK.S | HexNAc(5)Hex(5)Fuc(1)NeuAc(1) | N[+2263] | 1074.4643 | 3 | 3221.3782 | 3221.3711 | 0 | 2.2 | 200 | Specific | 577.21 | 309.74 | 309.74 | 8.98 | 120 | >sp\|P01877\|IGHA2_HUMAN Immunoglobulin heavy constant alpha 2 OS=Homo sapiens OX=9606 GN=IGHA2 PE=1 SV=4 | 576 | 20201116_fqg_colostrum_1ng_glyco_hcd203040_compound15.7431.7431.3 | controllerType=0 controllerNumber=1 scan=7431 | 19.0343 |
| 04361:3 | 3 | K.TPLTAN[+2059.735]ITK.S | HexNAc(4)Hex(5)Fuc(1)NeuAc(1) | N[+2060] | 1006.7711 | 3 | 3018.2989 | 3018.2917 | 0 | 2.4 | 200 | Specific | 578.56 | 266.2 | 266.2 | 8.69 | 120 | >sp\|P01877\|IGHA2_HUMAN Immunoglobulin heavy constant alpha 2 OS=Homo sapiens OX=9606 GN=IGHA2 PE=1 SV=4 | 576 | 20201116_fqg_colostrum_1ng_glyco_hcd203040_compound15.7375.7375.3 | controllerType=0 controllerNumber=1 scan=7375 | 18.9292 |
| 02613:3 | 3 | K.TPLTAN[+1809.666]ITK.S | HexNAc(5)Hex(4)Fuc(1) | N[+1810] | 923.4137 | 3 | 2768.2264 | 2768.2228 | 0 | 1.3 | 200 | Specific | 584.95 | 288.16 | 288.16 | 9.93 | 120 | >sp\|P01877\|IGHA2_HUMAN Immunoglobulin heavy constant alpha 2 OS=Homo sapiens OX=9606 GN=IGHA2 PE=1 SV=4 | 576 | 20201116_fqg_colostrum_1ng_glyco_hcd203040_compound15.5550.5550.3 | controllerType=0 controllerNumber=1 scan=5550 | 15.8165 |
| 02618:2 | 3 | K.TPLTAN[+1809.666]ITK.S | HexNAc(5)Hex(4)Fuc(1) | N[+1810] | 1384.6171 | 2 | 2768.2269 | 2768.2228 | 0 | 1.5 | 200 | Specific | 591.19 | 320.56 | 320.56 | 9.97 | 120 | >sp\|P01877\|IGHA2_HUMAN Immunoglobulin heavy constant alpha 2 OS=Homo sapiens OX=9606 GN=IGHA2 PE=1 SV=4 | 576 | 20201116_fqg_colostrum_1ng_glyco_hcd203040_compound15.5555.5555.2 | controllerType=0 controllerNumber=1 scan=5555 | 15.8275 |
| 02723:2 | 3 | K.TPLTAN[+1444.534]ITK.S | HexNAc(4)Hex(3)Fuc(1) | N[+1445] | 1202.0509 | 2 | 2403.0946 | 2403.0906 | 0 | 1.7 | 200 | Specific | 595.91 | 337.7 | 337.7 | 10.21 | 120 | >sp\|P01877\|IGHA2_HUMAN Immunoglobulin heavy constant alpha 2 OS=Homo sapiens OX=9606 GN=IGHA2 PE=1 SV=4 | 576 | 20201116_fqg_colostrum_1ng_glyco_hcd203040_compound15.5664.5664.2 | controllerType=0 controllerNumber=1 scan=5664 | 16.0536 |
| 02668:2 | 3 | K.TPLTAN[+1809.666]ITK.S | HexNAc(5)Hex(4)Fuc(1) | N[+1810] | 1384.6183 | 2 | 2768.2294 | 2768.2228 | 0 | 2.4 | 200 | Specific | 603.74 | 335.44 | 335.44 | 11.21 | 120 | >sp\|P01877\|IGHA2_HUMAN Immunoglobulin heavy constant alpha 2 OS=Homo sapiens OX=9606 GN=IGHA2 PE=1 SV=4 | 576 | 20201116_fqg_colostrum_1ng_glyco_hcd203040_compound15.5607.5607.2 | controllerType=0 controllerNumber=1 scan=5607 | 15.9415 |
| 02628:2 | 3 | K.TPLTAN[+1663.608]ITK.S | HexNAc(5)Hex(4) | N[+1664] | 1311.5878 | 2 | 2622.1683 | 2622.1649 | 0 | 1.3 | 200 | Specific | 610.36 | 360.56 | 360.56 | 11.29 | 120 | >sp\|P01877\|IGHA2_HUMAN Immunoglobulin heavy constant alpha 2 OS=Homo sapiens OX=9606 GN=IGHA2 PE=1 SV=4 | 576 | 20201116_fqg_colostrum_1ng_glyco_hcd203040_compound15.5565.5565.2 | controllerType=0 controllerNumber=1 scan=5565 | 15.8495 |
| 04425:2 | 3 | K.TPLTAN[+2262.814]ITK.S | HexNAc(5)Hex(5)Fuc(1)NeuAc(1) | N[+2263] | 1611.1917 | 2 | 3221.3761 | 3221.3711 | 0 | 1.6 | 200 | Specific | 612.91 | 312.38 | 312.38 | 10.9 | 120 | >sp\|P01877\|IGHA2_HUMAN Immunoglobulin heavy constant alpha 2 OS=Homo sapiens OX=9606 GN=IGHA2 PE=1 SV=4 | 576 | 20201116_fqg_colostrum_1ng_glyco_hcd203040_compound15.7441.7441.2 | controllerType=0 controllerNumber=1 scan=7441 | 19.0458 |
| 02585:2 | 3 | K.TPLTAN[+1971.719]ITK.S | HexNAc(5)Hex(5)Fuc(1) | N[+1972] | 1465.6453 | 2 | 2930.2833 | 2930.2757 | 0 | 2.6 | 200 | Specific | 613.12 | 334.85 | 334.85 | 11.21 | 120 | >sp\|P01877\|IGHA2_HUMAN Immunoglobulin heavy constant alpha 2 OS=Homo sapiens OX=9606 GN=IGHA2 PE=1 SV=4 | 576 | 20201116_fqg_colostrum_1ng_glyco_hcd203040_compound15.5520.5520.2 | controllerType=0 controllerNumber=1 scan=5520 | 15.7281 |
| 02530:2 | 3 | K.TPLTAN[+1216.423]ITK.S | HexNAc(2)Hex(5) | N[+1216] | 1087.9954 | 2 | 2174.9835 | 2174.9796 | 0 | 1.8 | 200 | Specific | 613.54 | 311.01 | 311.01 | 11.14 | 120 | >sp\|P01877\|IGHA2_HUMAN Immunoglobulin heavy constant alpha 2 OS=Homo sapiens OX=9606 GN=IGHA2 PE=1 SV=4 | 576 | 20201116_fqg_colostrum_1ng_glyco_hcd203040_compound15.5463.5463.2 | controllerType=0 controllerNumber=1 scan=5463 | 15.6123 |
| 02528:2 | 3 | K.TPLTAN[+1216.423]ITK.S | HexNAc(2)Hex(5) | N[+1216] | 1087.9954 | 2 | 2174.9835 | 2174.9796 | 0 | 1.8 | 200 | Specific | 615.44 | 346.39 | 346.39 | 11.36 | 120 | >sp\|P01877\|IGHA2_HUMAN Immunoglobulin heavy constant alpha 2 OS=Homo sapiens OX=9606 GN=IGHA2 PE=1 SV=4 | 576 | 20201116_fqg_colostrum_1ng_glyco_hcd203040_compound15.5461.5461.2 | controllerType=0 controllerNumber=1 scan=5461 | 15.6066 |
| 02748:2 | 3 | K.TPLTAN[+1647.613]ITK.S | HexNAc(5)Hex(3)Fuc(1) | N[+1648] | 1303.5909 | 2 | 2606.1744 | 2606.17 | 0 | 1.7 | 200 | Specific | 616.69 | 323.34 | 323.34 | 11.21 | 120 | >sp\|P01877\|IGHA2_HUMAN Immunoglobulin heavy constant alpha 2 OS=Homo sapiens OX=9606 GN=IGHA2 PE=1 SV=4 | 576 | 20201116_fqg_colostrum_1ng_glyco_hcd203040_compound15.5690.5690.2 | controllerType=0 controllerNumber=1 scan=5690 | 16.0967 |
| 02612:3 | 3 | K.TPLTAN[+1606.587]ITK.S | HexNAc(4)Hex(4)Fuc(1) | N[+1607] | 855.7207 | 3 | 2565.1475 | 2565.1435 | 0 | 1.6 | 200 | Specific | 619.05 | 282.64 | 282.64 | 10.93 | 120 | >sp\|P01877\|IGHA2_HUMAN Immunoglobulin heavy constant alpha 2 OS=Homo sapiens OX=9606 GN=IGHA2 PE=1 SV=4 | 576 | 20201116_fqg_colostrum_1ng_glyco_hcd203040_compound15.5549.5549.3 | controllerType=0 controllerNumber=1 scan=5549 | 15.8118 |
| 02496:2 | 3 | K.TPLTAN[+1768.640]ITK.S | HexNAc(4)Hex(5)Fuc(1) | N[+1769] | 1364.1058 | 2 | 2727.2042 | 2727.1963 | 0 | 2.9 | 200 | Specific | 620.26 | 329.53 | 329.53 | 11.11 | 120 | >sp\|P01877\|IGHA2_HUMAN Immunoglobulin heavy constant alpha 2 OS=Homo sapiens OX=9606 GN=IGHA2 PE=1 SV=4 | 576 | 20201116_fqg_colostrum_1ng_glyco_hcd203040_compound15.5428.5428.2 | controllerType=0 controllerNumber=1 scan=5428 | 15.5507 |
| 02696:3 | 3 | K.TPLTAN[+1444.534]ITK.S | HexNAc(4)Hex(3)Fuc(1) | N[+1445] | 801.7036 | 3 | 2403.0962 | 2403.0906 | 0 | 2.3 | 200 | Specific | 620.89 | 290.4 | 290.4 | 11.16 | 120 | >sp\|P01877\|IGHA2_HUMAN Immunoglobulin heavy constant alpha 2 OS=Homo sapiens OX=9606 GN=IGHA2 PE=1 SV=4 | 576 | 20201116_fqg_colostrum_1ng_glyco_hcd203040_compound15.5636.5636.3 | controllerType=0 controllerNumber=1 scan=5636 | 15.9989 |
| 04428:2 | 3 | K.TPLTAN[+2262.814]ITK.S | HexNAc(5)Hex(5)Fuc(1)NeuAc(1) | N[+2263] | 1611.1917 | 2 | 3221.3761 | 3221.3711 | 0 | 1.6 | 200 | Specific | 621.78 | 294.86 | 294.86 | 10.71 | 120 | >sp\|P01877\|IGHA2_HUMAN Immunoglobulin heavy constant alpha 2 OS=Homo sapiens OX=9606 GN=IGHA2 PE=1 SV=4 | 576 | 20201116_fqg_colostrum_1ng_glyco_hcd203040_compound15.7444.7444.2 | controllerType=0 controllerNumber=1 scan=7444 | 19.0555 |
| 04387:2 | 3 | K.TPLTAN[+2059.735]ITK.S | HexNAc(4)Hex(5)Fuc(1)NeuAc(1) | N[+2060] | 1509.6542 | 2 | 3018.3012 | 3018.2917 | 0 | 3.1 | 200 | Specific | 624.28 | 343.92 | 343.92 | 11.26 | 120 | >sp\|P01877\|IGHA2_HUMAN Immunoglobulin heavy constant alpha 2 OS=Homo sapiens OX=9606 GN=IGHA2 PE=1 SV=4 | 576 | 20201116_fqg_colostrum_1ng_glyco_hcd203040_compound15.7402.7402.2 | controllerType=0 controllerNumber=1 scan=7402 | 18.987 |
| 02564:2 | 3 | K.TPLTAN[+1606.587]ITK.S | HexNAc(4)Hex(4)Fuc(1) | N[+1607] | 1283.078 | 2 | 2565.1488 | 2565.1435 | 0 | 2.1 | 200 | Specific | 624.74 | 343.69 | 343.69 | 11.36 | 120 | >sp\|P01877\|IGHA2_HUMAN Immunoglobulin heavy constant alpha 2 OS=Homo sapiens OX=9606 GN=IGHA2 PE=1 SV=4 | 576 | 20201116_fqg_colostrum_1ng_glyco_hcd203040_compound15.5498.5498.2 | controllerType=0 controllerNumber=1 scan=5498 | 15.6796 |
| 02609:3 | 3 | K.TPLTAN[+1606.587]ITK.S | HexNAc(4)Hex(4)Fuc(1) | N[+1607] | 855.7207 | 3 | 2565.1475 | 2565.1435 | 0 | 1.6 | 200 | Specific | 627.99 | 289.41 | 289.41 | 10.93 | 120 | >sp\|P01877\|IGHA2_HUMAN Immunoglobulin heavy constant alpha 2 OS=Homo sapiens OX=9606 GN=IGHA2 PE=1 SV=4 | 576 | 20201116_fqg_colostrum_1ng_glyco_hcd203040_compound15.5546.5546.3 | controllerType=0 controllerNumber=1 scan=5546 | 15.8069 |
| 04627:2 | 3 | K.TPLTAN[+1694.603]ITK.S | HexNAc(3)Hex(4)Fuc(1)NeuAc(1) | N[+1695] | 1327.0856 | 2 | 2653.164 | 2653.1595 | 0 | 1.7 | 200 | Specific | 628.95 | 328.62 | 328.62 | 11.21 | 120 | >sp\|P01877\|IGHA2_HUMAN Immunoglobulin heavy constant alpha 2 OS=Homo sapiens OX=9606 GN=IGHA2 PE=1 SV=4 | 576 | 20201116_fqg_colostrum_1ng_glyco_hcd203040_compound15.7650.7650.2 | controllerType=0 controllerNumber=1 scan=7650 | 19.3959 |
| 02672:2 | 3 | K.TPLTAN[+1647.613]ITK.S | HexNAc(5)Hex(3)Fuc(1) | N[+1648] | 1303.5916 | 2 | 2606.1759 | 2606.17 | 0 | 2.3 | 200 | Specific | 630.04 | 351.89 | 351.89 | 11.36 | 120 | >sp\|P01877\|IGHA2_HUMAN Immunoglobulin heavy constant alpha 2 OS=Homo sapiens OX=9606 GN=IGHA2 PE=1 SV=4 | 576 | 20201116_fqg_colostrum_1ng_glyco_hcd203040_compound15.5611.5611.2 | controllerType=0 controllerNumber=1 scan=5611 | 15.9519 |
| 04364:3 | 3 | K.TPLTAN[+2059.735]ITK.S | HexNAc(4)Hex(5)Fuc(1)NeuAc(1) | N[+2060] | 1006.7711 | 3 | 3018.2989 | 3018.2917 | 0 | 2.4 | 200 | Specific | 631.98 | 303.7 | 303.7 | 11.36 | 120 | >sp\|P01877\|IGHA2_HUMAN Immunoglobulin heavy constant alpha 2 OS=Homo sapiens OX=9606 GN=IGHA2 PE=1 SV=4 | 576 | 20201116_fqg_colostrum_1ng_glyco_hcd203040_compound15.7378.7378.3 | controllerType=0 controllerNumber=1 scan=7378 | 18.94 |
| 02767:2 | 3 | K.TPLTAN[+1444.534]ITK.S | HexNAc(4)Hex(3)Fuc(1) | N[+1445] | 1202.0512 | 2 | 2403.0951 | 2403.0906 | 0 | 1.9 | 200 | Specific | 637.62 | 290.22 | 290.22 | 10.94 | 120 | >sp\|P01877\|IGHA2_HUMAN Immunoglobulin heavy constant alpha 2 OS=Homo sapiens OX=9606 GN=IGHA2 PE=1 SV=4 | 576 | 20201116_fqg_colostrum_1ng_glyco_hcd203040_compound15.5709.5709.2 | controllerType=0 controllerNumber=1 scan=5709 | 16.1234 |
| 02615:2 | 3 | K.TPLTAN[+1809.666]ITK.S | HexNAc(5)Hex(4)Fuc(1) | N[+1810] | 1384.6171 | 2 | 2768.2269 | 2768.2228 | 0 | 1.5 | 200 | Specific | 642.54 | 353.55 | 353.55 | 11.13 | 120 | >sp\|P01877\|IGHA2_HUMAN Immunoglobulin heavy constant alpha 2 OS=Homo sapiens OX=9606 GN=IGHA2 PE=1 SV=4 | 576 | 20201116_fqg_colostrum_1ng_glyco_hcd203040_compound15.5552.5552.2 | controllerType=0 controllerNumber=1 scan=5552 | 15.8185 |
| 02675:2 | 3 | K.TPLTAN[+1647.613]ITK.S | HexNAc(5)Hex(3)Fuc(1) | N[+1648] | 1303.5916 | 2 | 2606.1759 | 2606.17 | 0 | 2.3 | 200 | Specific | 642.57 | 345.83 | 345.83 | 11.36 | 120 | >sp\|P01877\|IGHA2_HUMAN Immunoglobulin heavy constant alpha 2 OS=Homo sapiens OX=9606 GN=IGHA2 PE=1 SV=4 | 576 | 20201116_fqg_colostrum_1ng_glyco_hcd203040_compound15.5614.5614.2 | controllerType=0 controllerNumber=1 scan=5614 | 15.9569 |
| 02562:2 | 3 | K.TPLTAN[+1606.587]ITK.S | HexNAc(4)Hex(4)Fuc(1) | N[+1607] | 1283.078 | 2 | 2565.1488 | 2565.1435 | 0 | 2.1 | 200 | Specific | 646.41 | 382.95 | 382.95 | 12.79 | 120 | >sp\|P01877\|IGHA2_HUMAN Immunoglobulin heavy constant alpha 2 OS=Homo sapiens OX=9606 GN=IGHA2 PE=1 SV=4 | 576 | 20201116_fqg_colostrum_1ng_glyco_hcd203040_compound15.5496.5496.2 | controllerType=0 controllerNumber=1 scan=5496 | 15.6733 |
| 02662:2 | 3 | K.TPLTAN[+1501.555]ITK.S | HexNAc(5)Hex(3) | N[+1502] | 1230.5621 | 2 | 2460.1168 | 2460.1121 | 0 | 1.9 | 200 | Specific | 649.5 | 377.95 | 377.95 | 11.53 | 120 | >sp\|P01877\|IGHA2_HUMAN Immunoglobulin heavy constant alpha 2 OS=Homo sapiens OX=9606 GN=IGHA2 PE=1 SV=4 | 576 | 20201116_fqg_colostrum_1ng_glyco_hcd203040_compound15.5601.5601.2 | controllerType=0 controllerNumber=1 scan=5601 | 15.9286 |
| 02626:2 | 3 | K.TPLTAN[+1663.608]ITK.S | HexNAc(5)Hex(4) | N[+1664] | 1311.5878 | 2 | 2622.1683 | 2622.1649 | 0 | 1.3 | 200 | Specific | 649.93 | 375.2 | 375.2 | 11.29 | 120 | >sp\|P01877\|IGHA2_HUMAN Immunoglobulin heavy constant alpha 2 OS=Homo sapiens OX=9606 GN=IGHA2 PE=1 SV=4 | 576 | 20201116_fqg_colostrum_1ng_glyco_hcd203040_compound15.5563.5563.2 | controllerType=0 controllerNumber=1 scan=5563 | 15.8445 |
| 02882:2 | 3 | K.TPLTAN[+876.322]ITK.S | HexNAc(2)Hex(2)Fuc(1) | N[+876] | 917.9456 | 2 | 1834.8838 | 1834.8791 | 0 | 2.6 | 200 | Specific | 650.87 | 360.64 | 360.64 | 12.53 | 120 | >sp\|P01877\|IGHA2_HUMAN Immunoglobulin heavy constant alpha 2 OS=Homo sapiens OX=9606 GN=IGHA2 PE=1 SV=4 | 576 | 20201116_fqg_colostrum_1ng_glyco_hcd203040_compound15.5829.5829.2 | controllerType=0 controllerNumber=1 scan=5829 | 16.3559 |
| 02665:2 | 3 | K.TPLTAN[+1501.555]ITK.S | HexNAc(5)Hex(3) | N[+1502] | 1230.5621 | 2 | 2460.1168 | 2460.1121 | 0 | 1.9 | 200 | Specific | 655.66 | 316.39 | 316.39 | 12.14 | 120 | >sp\|P01877\|IGHA2_HUMAN Immunoglobulin heavy constant alpha 2 OS=Homo sapiens OX=9606 GN=IGHA2 PE=1 SV=4 | 576 | 20201116_fqg_colostrum_1ng_glyco_hcd203040_compound15.5604.5604.2 | controllerType=0 controllerNumber=1 scan=5604 | 15.9347 |
| 02726:2 | 3 | K.TPLTAN[+1444.534]ITK.S | HexNAc(4)Hex(3)Fuc(1) | N[+1445] | 1202.0509 | 2 | 2403.0946 | 2403.0906 | 0 | 1.7 | 200 | Specific | 655.82 | 306.23 | 306.23 | 12.14 | 120 | >sp\|P01877\|IGHA2_HUMAN Immunoglobulin heavy constant alpha 2 OS=Homo sapiens OX=9606 GN=IGHA2 PE=1 SV=4 | 576 | 20201116_fqg_colostrum_1ng_glyco_hcd203040_compound15.5667.5667.2 | controllerType=0 controllerNumber=1 scan=5667 | 16.059 |
| 02586:2 | 3 | K.TPLTAN[+1403.507]ITK.S | HexNAc(3)Hex(4)Fuc(1) | N[+1404] | 1181.5385 | 2 | 2362.0697 | 2362.0641 | 0 | 2.4 | 200 | Specific | 657.95 | 327.55 | 327.55 | 12.21 | 120 | >sp\|P01877\|IGHA2_HUMAN Immunoglobulin heavy constant alpha 2 OS=Homo sapiens OX=9606 GN=IGHA2 PE=1 SV=4 | 576 | 20201116_fqg_colostrum_1ng_glyco_hcd203040_compound15.5521.5521.2 | controllerType=0 controllerNumber=1 scan=5521 | 15.7343 |
| 02698:3 | 3 | K.TPLTAN[+1444.534]ITK.S | HexNAc(4)Hex(3)Fuc(1) | N[+1445] | 801.7036 | 3 | 2403.0962 | 2403.0906 | 0 | 2.3 | 200 | Specific | 659.44 | 314.44 | 314.44 | 12.36 | 120 | >sp\|P01877\|IGHA2_HUMAN Immunoglobulin heavy constant alpha 2 OS=Homo sapiens OX=9606 GN=IGHA2 PE=1 SV=4 | 576 | 20201116_fqg_colostrum_1ng_glyco_hcd203040_compound15.5638.5638.3 | controllerType=0 controllerNumber=1 scan=5638 | 16.0054 |
| 02584:2 | 3 | K.TPLTAN[+1403.507]ITK.S | HexNAc(3)Hex(4)Fuc(1) | N[+1404] | 1181.5385 | 2 | 2362.0697 | 2362.0641 | 0 | 2.4 | 200 | Specific | 661.03 | 364.15 | 364.15 | 12.53 | 120 | >sp\|P01877\|IGHA2_HUMAN Immunoglobulin heavy constant alpha 2 OS=Homo sapiens OX=9606 GN=IGHA2 PE=1 SV=4 | 576 | 20201116_fqg_colostrum_1ng_glyco_hcd203040_compound15.5519.5519.2 | controllerType=0 controllerNumber=1 scan=5519 | 15.7225 |
| 02658:2 | 3 | K.TPLTAN[+1298.476]ITK.S | HexNAc(4)Hex(3) | N[+1298] | 1129.0227 | 2 | 2257.0382 | 2257.0327 | 0 | 2.4 | 200 | Specific | 669.51 | 355 | 355 | 12.36 | 120 | >sp\|P01877\|IGHA2_HUMAN Immunoglobulin heavy constant alpha 2 OS=Homo sapiens OX=9606 GN=IGHA2 PE=1 SV=4 | 576 | 20201116_fqg_colostrum_1ng_glyco_hcd203040_compound15.5597.5597.2 | controllerType=0 controllerNumber=1 scan=5597 | 15.9158 |
| 02617:2 | 3 | K.TPLTAN[+1444.534]ITK.S | HexNAc(4)Hex(3)Fuc(1) | N[+1445] | 1202.0511 | 2 | 2403.0949 | 2403.0906 | 0 | 1.8 | 200 | Specific | 669.93 | 359.33 | 359.33 | 12.36 | 120 | >sp\|P01877\|IGHA2_HUMAN Immunoglobulin heavy constant alpha 2 OS=Homo sapiens OX=9606 GN=IGHA2 PE=1 SV=4 | 576 | 20201116_fqg_colostrum_1ng_glyco_hcd203040_compound15.5554.5554.2 | controllerType=0 controllerNumber=1 scan=5554 | 15.8208 |
| 04625:2 | 3 | K.TPLTAN[+1694.603]ITK.S | HexNAc(3)Hex(4)Fuc(1)NeuAc(1) | N[+1695] | 1327.0856 | 2 | 2653.164 | 2653.1595 | 0 | 1.7 | 200 | Specific | 671.57 | 380.74 | 380.74 | 13.79 | 120 | >sp\|P01877\|IGHA2_HUMAN Immunoglobulin heavy constant alpha 2 OS=Homo sapiens OX=9606 GN=IGHA2 PE=1 SV=4 | 576 | 20201116_fqg_colostrum_1ng_glyco_hcd203040_compound15.7648.7648.2 | controllerType=0 controllerNumber=1 scan=7648 | 19.3901 |
| 02660:2 | 3 | K.TPLTAN[+1298.476]ITK.S | HexNAc(4)Hex(3) | N[+1298] | 1129.0227 | 2 | 2257.0382 | 2257.0327 | 0 | 2.4 | 200 | Specific | 674.52 | 316.33 | 316.33 | 12.14 | 120 | >sp\|P01877\|IGHA2_HUMAN Immunoglobulin heavy constant alpha 2 OS=Homo sapiens OX=9606 GN=IGHA2 PE=1 SV=4 | 576 | 20201116_fqg_colostrum_1ng_glyco_hcd203040_compound15.5599.5599.2 | controllerType=0 controllerNumber=1 scan=5599 | 15.9212 |
| 02667:2 | 3 | K.TPLTAN[+1241.454]ITK.S | HexNAc(3)Hex(3)Fuc(1) | N[+1241] | 1100.5116 | 2 | 2200.016 | 2200.0113 | 0 | 2.2 | 200 | Specific | 677.23 | 377.71 | 377.71 | 12.53 | 120 | >sp\|P01877\|IGHA2_HUMAN Immunoglobulin heavy constant alpha 2 OS=Homo sapiens OX=9606 GN=IGHA2 PE=1 SV=4 | 576 | 20201116_fqg_colostrum_1ng_glyco_hcd203040_compound15.5606.5606.2 | controllerType=0 controllerNumber=1 scan=5606 | 15.9404 |
| 02619:2 | 3 | K.TPLTAN[+1444.534]ITK.S | HexNAc(4)Hex(3)Fuc(1) | N[+1445] | 1202.0511 | 2 | 2403.0949 | 2403.0906 | 0 | 1.8 | 200 | Specific | 678.37 | 331.3 | 331.3 | 12.21 | 120 | >sp\|P01877\|IGHA2_HUMAN Immunoglobulin heavy constant alpha 2 OS=Homo sapiens OX=9606 GN=IGHA2 PE=1 SV=4 | 576 | 20201116_fqg_colostrum_1ng_glyco_hcd203040_compound15.5556.5556.2 | controllerType=0 controllerNumber=1 scan=5556 | 15.8328 |
| 02710:2 | 3 | K.TPLTAN[+1038.375]ITK.S | HexNAc(2)Hex(3)Fuc(1) | N[+1038] | 998.9709 | 2 | 1996.9346 | 1996.9319 | 0 | 1.3 | 200 | Specific | 687.49 | 332.34 | 332.34 | 11.97 | 120 | >sp\|P01877\|IGHA2_HUMAN Immunoglobulin heavy constant alpha 2 OS=Homo sapiens OX=9606 GN=IGHA2 PE=1 SV=4 | 576 | 20201116_fqg_colostrum_1ng_glyco_hcd203040_compound15.5651.5651.2 | controllerType=0 controllerNumber=1 scan=5651 | 16.0333 |
| 02707:2 | 3 | K.TPLTAN[+1038.375]ITK.S | HexNAc(2)Hex(3)Fuc(1) | N[+1038] | 998.9709 | 2 | 1996.9346 | 1996.9319 | 0 | 1.3 | 200 | Specific | 701.21 | 405.15 | 405.15 | 14.55 | 120 | >sp\|P01877\|IGHA2_HUMAN Immunoglobulin heavy constant alpha 2 OS=Homo sapiens OX=9606 GN=IGHA2 PE=1 SV=4 | 576 | 20201116_fqg_colostrum_1ng_glyco_hcd203040_compound15.5648.5648.2 | controllerType=0 controllerNumber=1 scan=5648 | 16.0237 |
| 07754:3 | 3 | R.MAGKPTHIN[+1540.529]VSVVMAEADGTC[+57.021].Y | HexNAc(2)Hex(7) | C[+57], N[+1541] | 1243.5265 | 3 | 3728.5651 | 3728.548 | 0 | 4.6 | 319 | CRagged | 365.72 | 352.7 | 232.96 | 8.12 | 120 | >sp\|P01877\|IGHA2_HUMAN Immunoglobulin heavy constant alpha 2 OS=Homo sapiens OX=9606 GN=IGHA2 PE=1 SV=4 | 576 | 20201116_fqg_colostrum_1ng_glyco_hcd203040_compound15.10901.10901.3 | controllerType=0 controllerNumber=1 scan=10901 | 24.5564 |
| 07756:3 | 3 | R.MAGKPTHIN[+1540.529]VSVVMAEADGTC[+57.021].Y | HexNAc(2)Hex(7) | C[+57], N[+1541] | 1243.5265 | 3 | 3728.5651 | 3728.548 | 0 | 4.6 | 319 | CRagged | 430.09 | 378.66 | 325.23 | 8.42 | 120 | >sp\|P01877\|IGHA2_HUMAN Immunoglobulin heavy constant alpha 2 OS=Homo sapiens OX=9606 GN=IGHA2 PE=1 SV=4 | 576 | 20201116_fqg_colostrum_1ng_glyco_hcd203040_compound15.10903.10903.3 | controllerType=0 controllerNumber=1 scan=10903 | 24.5603 |
| 08927:3 | 3 | R.MAGKPTHIN[+1216.423]VSVVMAEADGTC[+57.021].Y | HexNAc(2)Hex(5) | C[+57], N[+1216] | 1135.4877 | 3 | 3404.4485 | 3404.4424 | 0 | 1.8 | 319 | CRagged | 516.86 | 516.86 | 314.27 | 10.64 | 120 | >sp\|P01877\|IGHA2_HUMAN Immunoglobulin heavy constant alpha 2 OS=Homo sapiens OX=9606 GN=IGHA2 PE=1 SV=4 | 576 | 20201116_fqg_colostrum_1ng_glyco_hcd203040_compound15.12114.12114.3 | controllerType=0 controllerNumber=1 scan=12114 | 26.5337 |
| 08930:3 | 3 | R.MAGKPTHIN[+1216.423]VSVVMAEADGTC[+57.021].Y | HexNAc(2)Hex(5) | C[+57], N[+1216] | 1135.4877 | 3 | 3404.4485 | 3404.4424 | 0 | 1.8 | 319 | CRagged | 825.66 | 625.7 | 511.45 | 17.42 | 120 | >sp\|P01877\|IGHA2_HUMAN Immunoglobulin heavy constant alpha 2 OS=Homo sapiens OX=9606 GN=IGHA2 PE=1 SV=4 | 576 | 20201116_fqg_colostrum_1ng_glyco_hcd203040_compound15.12117.12117.3 | controllerType=0 controllerNumber=1 scan=12117 | 26.5379 |
| 07846:3 | 3 | R.MAGKPTHIN[+1216.423]VSVVMAEADGTC[+57.021].Y | HexNAc(2)Hex(5) | C[+57], N[+1216] | 1135.4895 | 3 | 3404.454 | 3404.4424 | 0 | 3.4 | 319 | CRagged | 859.2 | 669.04 | 515.16 | 18.31 | 120 | >sp\|P01877\|IGHA2_HUMAN Immunoglobulin heavy constant alpha 2 OS=Homo sapiens OX=9606 GN=IGHA2 PE=1 SV=4 | 576 | 20201116_fqg_colostrum_1ng_glyco_hcd203040_compound15.10997.10997.3 | controllerType=0 controllerNumber=1 scan=10997 | 24.7386 |
| 07843:3 | 3 | R.MAGKPTHIN[+1216.423]VSVVMAEADGTC[+57.021].Y | HexNAc(2)Hex(5) | C[+57], N[+1216] | 1135.4895 | 3 | 3404.454 | 3404.4424 | 0 | 3.4 | 319 | CRagged | 898.92 | 777.55 | 516.08 | 18.31 | 120 | >sp\|P01877\|IGHA2_HUMAN Immunoglobulin heavy constant alpha 2 OS=Homo sapiens OX=9606 GN=IGHA2 PE=1 SV=4 | 576 | 20201116_fqg_colostrum_1ng_glyco_hcd203040_compound15.10994.10994.3 | controllerType=0 controllerNumber=1 scan=10994 | 24.7318 |
| 07072:3 | 3 | R.M[+15.995]AGKPTHIN[+1378.476]VSVVMAEADGTC[+57.021].Y | HexNAc(2)Hex(6) | C[+57], M[+16], N[+1378] | 1194.8381 | 3 | 3582.4997 | 3582.4901 | 0 | 2.7 | 319 | CRagged | 540.01 | 479.7 | 105.19 | 10.85 | 120 | >sp\|P01877\|IGHA2_HUMAN Immunoglobulin heavy constant alpha 2 OS=Homo sapiens OX=9606 GN=IGHA2 PE=1 SV=4 | 576 | 20201116_fqg_colostrum_1ng_glyco_hcd203040_compound15.10195.10195.3 | controllerType=0 controllerNumber=1 scan=10195 | 23.4457 |
| 07117:3 | 3 | R.M[+15.995]AGKPTHIN[+1216.423]VSVVMAEADGTC[+57.021].Y | HexNAc(2)Hex(5) | C[+57], M[+16], N[+1216] | 1140.821 | 3 | 3420.4483 | 3420.4373 | 0 | 3.2 | 319 | CRagged | 562 | 562 | 221.95 | 10.93 | 120 | >sp\|P01877\|IGHA2_HUMAN Immunoglobulin heavy constant alpha 2 OS=Homo sapiens OX=9606 GN=IGHA2 PE=1 SV=4 | 576 | 20201116_fqg_colostrum_1ng_glyco_hcd203040_compound15.10242.10242.3 | controllerType=0 controllerNumber=1 scan=10242 | 23.5201 |
| 07069:3 | 3 | R.M[+15.995]AGKPTHIN[+1378.476]VSVVMAEADGTC[+57.021].Y | HexNAc(2)Hex(6) | C[+57], M[+16], N[+1378] | 1194.8381 | 3 | 3582.4997 | 3582.4901 | 0 | 2.7 | 319 | CRagged | 604.92 | 604.92 | 106.59 | 13.42 | 120 | >sp\|P01877\|IGHA2_HUMAN Immunoglobulin heavy constant alpha 2 OS=Homo sapiens OX=9606 GN=IGHA2 PE=1 SV=4 | 576 | 20201116_fqg_colostrum_1ng_glyco_hcd203040_compound15.10192.10192.3 | controllerType=0 controllerNumber=1 scan=10192 | 23.4386 |
| 07120:3 | 3 | R.M[+15.995]AGKPTHIN[+1216.423]VSVVMAEADGTC[+57.021].Y | HexNAc(2)Hex(5) | C[+57], M[+16], N[+1216] | 1140.821 | 3 | 3420.4483 | 3420.4373 | 0 | 3.2 | 319 | CRagged | 699.13 | 682.72 | 253.58 | 14.31 | 120 | >sp\|P01877\|IGHA2_HUMAN Immunoglobulin heavy constant alpha 2 OS=Homo sapiens OX=9606 GN=IGHA2 PE=1 SV=4 | 576 | 20201116_fqg_colostrum_1ng_glyco_hcd203040_compound15.10245.10245.3 | controllerType=0 controllerNumber=1 scan=10245 | 23.5248 |
| 10212:3 | 3 | R.MAGKPTHIN[+1501.555]VSVVMAEADGTC[+57.021]Y.- | HexNAc(5)Hex(3) | C[+57], N[+1502] | 1284.8882 | 3 | 3852.6502 | 3852.6382 | 0 | 3.1 | 319 | CRagged | 218.65 | 218.65 | 218.65 | 5.56 | 120 | >sp\|P01877\|IGHA2_HUMAN Immunoglobulin heavy constant alpha 2 OS=Homo sapiens OX=9606 GN=IGHA2 PE=1 SV=4 | 576 | 20201116_fqg_colostrum_1ng_glyco_hcd203040_compound15.13446.13446.3 | controllerType=0 controllerNumber=1 scan=13446 | 28.7742 |
| 12194:3 | 3 | R.MAGKPTHIN[+1548.545]VSVVMAEADGTC[+57.021]Y.- | HexNAc(3)Hex(4)NeuAc(1) | C[+57], N[+1549] | 1300.5496 | 3 | 3899.6343 | 3899.6277 | 0 | 1.7 | 319 | CRagged | 274.95 | 238.73 | 238.73 | 7.24 | 120 | >sp\|P01877\|IGHA2_HUMAN Immunoglobulin heavy constant alpha 2 OS=Homo sapiens OX=9606 GN=IGHA2 PE=1 SV=4 | 576 | 20201116_fqg_colostrum_1ng_glyco_hcd203040_compound15.15494.15494.3 | controllerType=0 controllerNumber=1 scan=15494 | 31.9836 |
| 07793:3 | 3 | R.MAGKPTHIN[+1216.423]VSVVMAEADGTC[+57.021]Y.- | HexNAc(2)Hex(5) | C[+57], N[+1216] | 1189.8333 | 3 | 3567.4854 | 3567.5057 | 0 | -5.7 | 319 | CRagged | 309.26 | 309.26 | 166.83 | 6.45 | 120 | >sp\|P01877\|IGHA2_HUMAN Immunoglobulin heavy constant alpha 2 OS=Homo sapiens OX=9606 GN=IGHA2 PE=1 SV=4 | 576 | 20201116_fqg_colostrum_1ng_glyco_hcd203040_compound15.10942.10942.3 | controllerType=0 controllerNumber=1 scan=10942 | [illegible] |
| 10195:2 | 3 | R.MAGKPTHIN[+1216.423]VSVVMAEADGTC[+57.021]Y.- | HexNAc(2)Hex(5) | C[+57], N[+1216] | 1784.2604 | 2 | 3567.5135 | 3567.5057 | 0 | 2.2 | 319 | CRagged | 326.95 | 259.19 | 167.04 | 4.94 | 120 | >sp\|P01877\|IGHA2_HUMAN Immunoglobulin heavy constant alpha 2 OS=Homo sapiens OX=9606 GN=IGHA2 PE=1 SV=4 | 576 | 20201116_fqg_colostrum_1ng_glyco_hcd203040_compound15.13428.13428.2 | controllerType=0 controllerNumber=1 scan=13428 | [illegible] |
| 10210:3 | 3 | R.MAGKPTHIN[+1501.555]VSVVMAEADGTC[+57.021]Y.- | HexNAc(5)Hex(3) | C[+57], N[+1502] | 1284.8882 | 3 | 3852.6502 | 3852.6382 | 0 | 3.1 | 319 | CRagged | 543.21 | 528.49 | 528.49 | 10.56 | 120 | >sp\|P01877\|IGHA2_HUMAN Immunoglobulin heavy constant alpha 2 OS=Homo sapiens OX=9606 GN=IGHA2 PE=1 SV=4 | 576 | 20201116_fqg_colostrum_1ng_glyco_hcd203040_compound15.13444.13444.3 | controllerType=0 controllerNumber=1 scan=13444 | [illegible] |
| 10072:3 | 3 | R.MAGKPTHIN[+1540.529]VSVVMAEADGTC[+57.021]Y.- | HexNAc(2)Hex(7) | C[+57], N[+1541] | 1297.8771 | 3 | 3891.6168 | 3891.6114 | 0 | 1.4 | 319 | CRagged | 692.82 | 692.82 | 391.8 | 14 | 120 | >sp\|P01877\|IGHA2_HUMAN Immunoglobulin heavy constant alpha 2 OS=Homo sapiens OX=9606 GN=IGHA2 PE=1 SV=4 | 576 | 20201116_fqg_colostrum_1ng_glyco_hcd203040_compound15.13301.13301.3 | controllerType=0 controllerNumber=1 scan=13301 | [illegible] |
| 12197:3 | 3 | R.MAGKPTHIN[+1548.545]VSVVMAEADGTC[+57.021]Y.- | HexNAc(3)Hex(4)NeuAc(1) | C[+57], N[+1549] | 1300.5496 | 3 | 3899.6343 | 3899.6277 | 0 | 1.7 | 319 | CRagged | 694.51 | 625.02 | 625.02 | 14.24 | 120 | >sp\|P01877\|IGHA2_HUMAN Immunoglobulin heavy constant alpha 2 OS=Homo sapiens OX=9606 GN=IGHA2 PE=1 SV=4 | 576 | 20201116_fqg_colostrum_1ng_glyco_hcd203040_compound15.15497.15497.3 | controllerType=0 controllerNumber=1 scan=15497 | 31.9872 |
| 10208:3 | 3 | R.MAGKPTHIN[+1054.370]VSVVMAEADGTC[+57.021]Y.- | HexNAc(2)Hex(4) | C[+57], N[+1054] | 1135.8245 | 3 | 3405.4589 | 3405.4529 | 0 | 1.8 | 319 | CRagged | 728.43 | 728.43 | 335.36 | 15.24 | 120 | >sp\|P01877\|IGHA2_HUMAN Immunoglobulin heavy constant alpha 2 OS=Homo sapiens OX=9606 GN=IGHA2 PE=1 SV=4 | 576 | 20201116_fqg_colostrum_1ng_glyco_hcd203040_compound15.13442.13442.3 | controllerType=0 controllerNumber=1 scan=13442 | 28.7664 |
| 09274:3 | 3 | R.MAGKPTHIN[+1378.476]VSVVMAEADGTC[+57.021]Y.- | HexNAc(2)Hex(6) | C[+57], N[+1378] | 1243.8631 | 3 | 3729.5747 | 3729.5585 | 0 | 4.3 | 319 | CRagged | 762.88 | 702.15 | 384.19 | 15.69 | 120 | >sp\|P01877\|IGHA2_HUMAN Immunoglobulin heavy constant alpha 2 OS=Homo sapiens OX=9606 GN=IGHA2 PE=1 SV=4 | 576 | 20201116_fqg_colostrum_1ng_glyco_hcd203040_compound15.12472.12472.3 | controllerType=0 controllerNumber=1 scan=12472 | 27.0668 |
| 10118:3 | 3 | R.MAGKPTHIN[+1809.666]VSVVMAEADGTC[+57.021]Y.- | HexNAc(5)Hex(4)Fuc(1) | C[+57], N[+1810] | 1387.5884 | 3 | 4160.7508 | 4160.7489 | 0 | 0.4 | 319 | CRagged | 803.6 | 755.98 | 443.92 | 16.5 | 120 | >sp\|P01877\|IGHA2_HUMAN Immunoglobulin heavy constant alpha 2 OS=Homo sapiens OX=9606 GN=IGHA2 PE=1 SV=4 | 576 | 20201116_fqg_colostrum_1ng_glyco_hcd203040_compound15.13348.13348.3 | controllerType=0 controllerNumber=1 scan=13348 | 28.5891 |
| 10115:3 | 3 | R.MAGKPTHIN[+1809.666]VSVVMAEADGTC[+57.021]Y.- | HexNAc(5)Hex(4)Fuc(1) | C[+57], N[+1810] | 1387.5884 | 3 | 4160.7508 | 4160.7489 | 0 | 0.4 | 319 | CRagged | 827.81 | 736.73 | 424.66 | 16.5 | 120 | >sp\|P01877\|IGHA2_HUMAN Immunoglobulin heavy constant alpha 2 OS=Homo sapiens OX=9606 GN=IGHA2 PE=1 SV=4 | 576 | 20201116_fqg_colostrum_1ng_glyco_hcd203040_compound15.13345.13345.3 | controllerType=0 controllerNumber=1 scan=13345 | 28.583 |
| 09276:3 | 3 | R.MAGKPTHIN[+1378.476]VSVVMAEADGTC[+57.021]Y.- | HexNAc(2)Hex(6) | C[+57], N[+1378] | 1243.8631 | 3 | 3729.5747 | 3729.5585 | 0 | 4.3 | 319 | CRagged | 852.22 | 852.22 | 462.87 | 17.69 | 120 | >sp\|P01877\|IGHA2_HUMAN Immunoglobulin heavy constant alpha 2 OS=Homo sapiens OX=9606 GN=IGHA2 PE=1 SV=4 | 576 | 20201116_fqg_colostrum_1ng_glyco_hcd203040_compound15.12474.12474.3 | controllerType=0 controllerNumber=1 scan=12474 | 27.0741 |
| 10011:3 | 3 | R.MAGKPTHIN[+1702.581]VSVVMAEADGTC[+57.021]Y.- | HexNAc(2)Hex(8) | C[+57], N[+1703] | 1351.8965 | 3 | 4053.6751 | 4053.6642 | 0 | 2.7 | 319 | CRagged | 864.66 | 720.82 | 484.53 | 18.24 | 120 | >sp\|P01877\|IGHA2_HUMAN Immunoglobulin heavy constant alpha 2 OS=Homo sapiens OX=9606 GN=IGHA2 PE=1 SV=4 | 576 | 20201116_fqg_colostrum_1ng_glyco_hcd203040_compound15.13237.13237.3 | controllerType=0 controllerNumber=1 scan=13237 | 28.3823 |
| 10075:3 | 3 | R.MAGKPTHIN[+1540.529]VSVVMAEADGTC[+57.021]Y.- | HexNAc(2)Hex(7) | C[+57], N[+1541] | 1297.8771 | 3 | 3891.6168 | 3891.6114 | 0 | 1.4 | 319 | CRagged | 891.57 | 711.4 | 500.91 | 18 | 120 | >sp\|P01877\|IGHA2_HUMAN Immunoglobulin heavy constant alpha 2 OS=Homo sapiens OX=9606 GN=IGHA2 PE=1 SV=4 | 576 | 20201116_fqg_colostrum_1ng_glyco_hcd203040_compound15.13304.13304.3 | controllerType=0 controllerNumber=1 scan=13304 | 28.5072 |
| 10064:3 | 3 | R.MAGKPTHIN[+1378.476]VSVVMAEADGTC[+57.021]Y.- | HexNAc(2)Hex(6) | C[+57], N[+1378] | 1243.86 | 3 | 3729.5656 | 3729.5585 | 0 | 1.9 | 319 | CRagged | 898.31 | 898.31 | 518.02 | 18.24 | 120 | >sp\|P01877\|IGHA2_HUMAN Immunoglobulin heavy constant alpha 2 OS=Homo sapiens OX=9606 GN=IGHA2 PE=1 SV=4 | 576 | 20201116_fqg_colostrum_1ng_glyco_hcd203040_compound15.13292.13292.3 | controllerType=0 controllerNumber=1 scan=13292 | 28.4853 |
| 09356:3 | 3 | R.MAGKPTHIN[+1216.423]VSVVMAEADGTC[+57.021]Y.- | HexNAc(2)Hex(5) | C[+57], N[+1216] | 1189.8466 | 3 | 3567.5253 | 3567.5057 | 0 | 5.5 | 319 | CRagged | 907.11 | 907.11 | 485.61 | 18.47 | 120 | >sp\|P01877\|IGHA2_HUMAN Immunoglobulin heavy constant alpha 2 OS=Homo sapiens OX=9606 GN=IGHA2 PE=1 SV=4 | 576 | 20201116_fqg_colostrum_1ng_glyco_hcd203040_compound15.12557.12557.3 | controllerType=0 controllerNumber=1 scan=12557 | 27.2107 |
| 10008:3 | 3 | R.MAGKPTHIN[+1702.581]VSVVMAEADGTC[+57.021]Y.- | HexNAc(2)Hex(8) | C[+57], N[+1703] | 1351.8965 | 3 | 4053.6751 | 4053.6642 | 0 | 2.7 | 319 | CRagged | 929.11 | 794.11 | 490.03 | 19.24 | 120 | >sp\|P01877\|IGHA2_HUMAN Immunoglobulin heavy constant alpha 2 OS=Homo sapiens OX=9606 GN=IGHA2 PE=1 SV=4 | 576 | 20201116_fqg_colostrum_1ng_glyco_hcd203040_compound15.13234.13234.3 | controllerType=0 controllerNumber=1 scan=13234 | 28.3684 |
| 10060:3 | 3 | R.MAGKPTHIN[+1378.476]VSVVMAEADGTC[+57.021]Y.- | HexNAc(2)Hex(6) | C[+57], N[+1378] | 1243.86 | 3 | 3729.5656 | 3729.5585 | 0 | 1.9 | 319 | CRagged | 953.06 | 796.42 | 557.42 | 20.24 | 120 | >sp\|P01877\|IGHA2_HUMAN Immunoglobulin heavy constant alpha 2 OS=Homo sapiens OX=9606 GN=IGHA2 PE=1 SV=4 | 576 | 20201116_fqg_colostrum_1ng_glyco_hcd203040_compound15.13288.13288.3 | controllerType=0 controllerNumber=1 scan=13288 | 28.4731 |
| 09359:3 | 3 | R.MAGKPTHIN[+1216.423]VSVVMAEADGTC[+57.021]Y.- | HexNAc(2)Hex(5) | C[+57], N[+1216] | 1189.8466 | 3 | 3567.5253 | 3567.5057 | 0 | 5.5 | 319 | CRagged | 956.11 | 956.11 | 537.6 | 19.47 | 120 | >sp\|P01877\|IGHA2_HUMAN Immunoglobulin heavy constant alpha 2 OS=Homo sapiens OX=9606 GN=IGHA2 PE=1 SV=4 | 576 | 20201116_fqg_colostrum_1ng_glyco_hcd203040_compound15.12560.12560.3 | controllerType=0 controllerNumber=1 scan=12560 | 27.2189 |
| 10126:3 | 3 | R.MAGKPTHIN[+1216.423]VSVVMAEADGTC[+57.021]Y.- | HexNAc(2)Hex(5) | C[+57], N[+1216] | 1189.8428 | 3 | 3567.5139 | 3567.5057 | 0 | 2.3 | 319 | CRagged | 959.42 | 805.6 | 549.44 | 20.24 | 120 | >sp\|P01877\|IGHA2_HUMAN Immunoglobulin heavy constant alpha 2 OS=Homo sapiens OX=9606 GN=IGHA2 PE=1 SV=4 | 576 | 20201116_fqg_colostrum_1ng_glyco_hcd203040_compound15.13357.13357.3 | controllerType=0 controllerNumber=1 scan=13357 | 28.6047 |
| 10128:3 | 3 | R.MAGKPTHIN[+1216.423]VSVVMAEADGTC[+57.021]Y.- | HexNAc(2)Hex(5) | C[+57], N[+1216] | 1189.8428 | 3 | 3567.5139 | 3567.5057 | 0 | 2.3 | 319 | CRagged | 964.58 | 791.67 | 565.43 | 20.24 | 120 | >sp\|P01877\|IGHA2_HUMAN Immunoglobulin heavy constant alpha 2 OS=Homo sapiens OX=9606 GN=IGHA2 PE=1 SV=4 | 576 | 20201116_fqg_colostrum_1ng_glyco_hcd203040_compound15.13359.13359.3 | controllerType=0 controllerNumber=1 scan=13359 | 28.6109 |
| 09710:3 | 3 | R.M[+15.995]AGKPTHIN[+1054.370]VSVVMAEADGTC[+57.021]Y.- | HexNAc(2)Hex(4) | C[+57], M[+16], N[+1054] | 1141.156 | 3 | 3421.4536 | 3421.4478 | 0 | 1.7 | 319 | CRagged | 487.65 | 482.98 | 243.5 | 10.34 | 120 | >sp\|P01877\|IGHA2_HUMAN Immunoglobulin heavy constant alpha 2 OS=Homo sapiens OX=9606 GN=IGHA2 PE=1 SV=4 | 576 | 20201116_fqg_colostrum_1ng_glyco_hcd203040_compound15.12924.12924.3 | controllerType=0 controllerNumber=1 scan=12924 | 27.8458 |
| 09670:3 | 3 | R.M[+15.995]AGKPTHIN[+1216.423]VSVVMAEADGTC[+57.021]Y.- | HexNAc(2)Hex(5) | C[+57], M[+16], N[+1216] | 1195.175 | 3 | 3583.5104 | 3583.5006 | 0 | 2.7 | 319 | CRagged | 679.79 | 679.79 | 271.5 | 14.24 | 120 | >sp\|P01877\|IGHA2_HUMAN Immunoglobulin heavy constant alpha 2 OS=Homo sapiens OX=9606 GN=IGHA2 PE=1 SV=4 | 576 | 20201116_fqg_colostrum_1ng_glyco_hcd203040_compound15.12883.12883.3 | controllerType=0 controllerNumber=1 scan=12883 | 27.785 |
| 09713:3 | 3 | R.M[+15.995]AGKPTHIN[+1054.370]VSVVMAEADGTC[+57.021]Y.- | HexNAc(2)Hex(4) | C[+57], M[+16], N[+1054] | 1141.156 | 3 | 3421.4536 | 3421.4478 | 0 | 1.7 | 319 | CRagged | 696.04 | 682.53 | 256.65 | 14.24 | 120 | >sp\|P01877\|IGHA2_HUMAN Immunoglobulin heavy constant alpha 2 OS=Homo sapiens OX=9606 GN=IGHA2 PE=1 SV=4 | 576 | 20201116_fqg_colostrum_1ng_glyco_hcd203040_compound15.12927.12927.3 | controllerType=0 controllerNumber=1 scan=12927 | 27.8583 |
| 09533:3 | 3 | R.M[+15.995]AGKPTHIN[+1378.476]VSVVMAEADGTC[+57.021]Y.- | HexNAc(2)Hex(6) | C[+57], M[+16], N[+1378] | 1249.1917 | 3 | 3745.5606 | 3745.5535 | 0 | 1.9 | 319 | CRagged | 729.4 | 653.42 | 267.14 | 15.24 | 120 | >sp\|P01877\|IGHA2_HUMAN Immunoglobulin heavy constant alpha 2 OS=Homo sapiens OX=9606 GN=IGHA2 PE=1 SV=4 | 576 | 20201116_fqg_colostrum_1ng_glyco_hcd203040_compound15.12740.12740.3 | controllerType=0 controllerNumber=1 scan=12740 | 27.5556 |
| 09536:3 | 3 | R.M[+15.995]AGKPTHIN[+1378.476]VSVVMAEADGTC[+57.021]Y.- | HexNAc(2)Hex(6) | C[+57], M[+16], N[+1378] | 1249.1917 | 3 | 3745.5606 | 3745.5535 | 0 | 1.9 | 319 | CRagged | 762.49 | 762.49 | 267.51 | 16.24 | 120 | >sp\|P01877\|IGHA2_HUMAN Immunoglobulin heavy constant alpha 2 OS=Homo sapiens OX=9606 GN=IGHA2 PE=1 SV=4 | 576 | 20201116_fqg_colostrum_1ng_glyco_hcd203040_compound15.12743.12743.3 | controllerType=0 controllerNumber=1 scan=12743 | 27.5687 |
| 18366:4 | 4 | R.LSLHRPALEDLLLGSEAN[+1298.476]LTC[+57.021]TLTGLR.D | HexNAc(4)Hex(3) | C[+57], N[+1298] | 1066.277 | 4 | 4262.0863 | 4262.0742 | 0 | 2.8 | 127 | Specific | 209.36 | 209.36 | 209.36 | 3.42 | 104 | >sp\|P01876\|IGHA1_HUMAN Immunoglobulin heavy constant alpha 1 OS=Homo sapiens OX=9606 GN=IGHA1 PE=1 SV=2 | 255 | 20201116_fqg_colostrum_1ng_glyco_hcd203040_compound15.21885.21885.3 | controllerType=0 controllerNumber=1 scan=21885 | 42.7706 |
| 18319:3 | 4 | R.[+42.011]LSLHRPALEDLLLGSEAN[+892.317]LTC[+57.021]TLTGLR.D | HexNAc(2)Hex(3) | .[42], C[+57], N[+892] | 1299.9915 | 3 | 3897.9599 | 3897.926 | 0 | 8.7 | 127 | Specific | 211.05 | 211.05 | 211.05 | 2.26 | 104 | >sp\|P01876\|IGHA1_HUMAN Immunoglobulin heavy constant alpha 1 OS=Homo sapiens OX=9606 GN=IGHA1 PE=1 SV=2 | 255 | 20201116_fqg_colostrum_1ng_glyco_hcd203040_compound15.21836.21836.3 | controllerType=0 controllerNumber=1 scan=21836 | 42.6749 |
| 26036:4 | 4 | R.LSLHRPALEDLLLGSEAN[+1501.555]LTC[+57.021]TLTGLR.D | HexNAc(5)Hex(3) | C[+57], N[+1502] | 1117.0487 | 4 | 4465.1732 | 4465.1536 | 0 | 4.4 | 127 | Specific | 237.14 | 237.14 | 237.14 | 6.56 | 104 | >sp\|P01876\|IGHA1_HUMAN Immunoglobulin heavy constant alpha 1 OS=Homo sapiens OX=9606 GN=IGHA1 PE=1 SV=2 | 255 | 20201116_fqg_colostrum_1ng_glyco_hcd203040_compound15.29884.29884.4 | controllerType=0 controllerNumber=1 scan=29884 | 59.5585 |
| 19389:5 | 4 | R.LSLHRPALEDLLLGSEAN[+1710.598]LTC[+57.021]TLTGLR.D | HexNAc(3)Hex(5)NeuAc(1) | C[+57], N[+1711] | 935.6481 | 5 | 4674.2114 | 4674.1959 | 0 | 3.3 | 127 | Specific | 248.73 | 248.73 | 248.73 | 4.96 | 104 | >sp\|P01876\|IGHA1_HUMAN Immunoglobulin heavy constant alpha 1 OS=Homo sapiens OX=9606 GN=IGHA1 PE=1 SV=2 | 255 | 20201116_fqg_colostrum_1ng_glyco_hcd203040_compound15.22949.22949.5 | controllerType=0 controllerNumber=1 scan=22949 | 44.8848 |
| 28496:4 | 4 | R.LSLHRPALEDLLLGSEAN[+1501.555]LTC[+57.021]TLTGLR.D | HexNAc(5)Hex(3) | C[+57], N[+1502] | 1117.047 | 4 | 4465.1663 | 4465.1536 | 0 | 2.9 | 127 | Specific | 249.13 | 249.13 | 249.13 | 7.12 | 104 | >sp\|P01876\|IGHA1_HUMAN Immunoglobulin heavy constant alpha 1 OS=Homo sapiens OX=9606 GN=IGHA1 PE=1 SV=2 | 255 | 20201116_fqg_colostrum_1ng_glyco_hcd203040_compound15.32450.32450.4 | controllerType=0 controllerNumber=1 scan=32450 | 64.7199 |
| 37675:4 | 4 | R.LSLHRPALEDLLLGSEAN[+1216.423]LTC[+57.021]TLTGLR.D | HexNAc(2)Hex(5) | C[+57], N[+1216] | 1045.7653 | 4 | 4180.0394 | 4180.0211 | 0 | 4.4 | 127 | Specific | 255.44 | 255.44 | 255.44 | 6.67 | 104 | >sp\|P01876\|IGHA1_HUMAN Immunoglobulin heavy constant alpha 1 OS=Homo sapiens OX=9606 GN=IGHA1 PE=1 SV=2 | 255 | 20201116_fqg_colostrum_1ng_glyco_hcd203040_compound15.44104.44104.2 | controllerType=0 controllerNumber=1 scan=44104 | 90.2645 |
| 18322:3 | 4 | R.[+42.011]LSLHRPALEDLLLGSEAN[+892.317]LTC[+57.021]TLTGLR.D | HexNAc(2)Hex(3) | .[42], C[+57], N[+892] | 1299.9915 | 3 | 3897.9599 | 3897.926 | 0 | 8.7 | 127 | Specific | 260.37 | 235.69 | 235.69 | 3.74 | 104 | >sp\|P01876\|IGHA1_HUMAN Immunoglobulin heavy constant alpha 1 OS=Homo sapiens OX=9606 GN=IGHA1 PE=1 SV=2 | 255 | 20201116_fqg_colostrum_1ng_glyco_hcd203040_compound15.21839.21839.3 | controllerType=0 controllerNumber=1 scan=21839 | 42.6801 |
| 18345:3 | 4 | R.LSLHRPALEDLLLGSEAN[+1501.555]LTC[+57.021]TLTGLR.D | HexNAc(5)Hex(3) | C[+57], N[+1502] | 1489.0607 | 3 | 4465.1676 | 4465.1536 | 0 | 3.1 | 127 | Specific | 262 | 262 | 262 | 6.88 | 104 | >sp\|P01876\|IGHA1_HUMAN Immunoglobulin heavy constant alpha 1 OS=Homo sapiens OX=9606 GN=IGHA1 PE=1 SV=2 | 255 | 20201116_fqg_colostrum_1ng_glyco_hcd203040_compound15.21863.21863.2 | controllerType=0 controllerNumber=1 scan=21863 | 42.7289 |
| 39149:4 | 4 | R.LSLHRPALEDLLLGSEAN[+1501.555]LTC[+57.021]TLTGLR.D | HexNAc(5)Hex(3) | C[+57], N[+1502] | 1117.0456 | 4 | 4465.1605 | 4465.1536 | 0 | 1.5 | 127 | Specific | 264.12 | 264.12 | 264.12 | 7.37 | 104 | >sp\|P01876\|IGHA1_HUMAN Immunoglobulin heavy constant alpha 1 OS=Homo sapiens OX=9606 GN=IGHA1 PE=1 SV=2 | 255 | 20201116_fqg_colostrum_1ng_glyco_hcd203040_compound15.45640.45640.4 | controllerType=0 controllerNumber=1 scan=45640 | 93.1186 |
| 27441:4 | 4 | R.LSLHRPALEDLLLGSEAN[+1501.555]LTC[+57.021]TLTGLR.D | HexNAc(5)Hex(3) | C[+57], N[+1502] | 1117.0472 | 4 | 4465.1668 | 4465.1536 | 0 | 3 | 127 | Specific | 268.27 | 268.27 | 268.27 | 7.5 | 104 | >sp\|P01876\|IGHA1_HUMAN Immunoglobulin heavy constant alpha 1 OS=Homo sapiens OX=9606 GN=IGHA1 PE=1 SV=2 | 255 | 20201116_fqg_colostrum_1ng_glyco_hcd203040_compound15.31349.31349.2 | controllerType=0 controllerNumber=1 scan=31349 | 62.5865 |
| 37476:4 | 4 | R.LSLHRPALEDLLLGSEAN[+1216.423]LTC[+57.021]TLTGLR.D | HexNAc(2)Hex(5) | C[+57], N[+1216] | 1045.7622 | 4 | 4180.0272 | 4180.0211 | 0 | 1.4 | 127 | Specific | 268.52 | 268.52 | 268.52 | 7.37 | 104 | >sp\|P01876\|IGHA1_HUMAN Immunoglobulin heavy constant alpha 1 OS=Homo sapiens OX=9606 GN=IGHA1 PE=1 SV=2 | 255 | 20201116_fqg_colostrum_1ng_glyco_hcd203040_compound15.43869.43869.4 | controllerType=0 controllerNumber=1 scan=43869 | 89.7981 |
| 37176:4 | 4 | R.LSLHRPALEDLLLGSEAN[+1216.423]LTC[+57.021]TLTGLR.D | HexNAc(2)Hex(5) | C[+57], N[+1216] | 1045.763 | 4 | 4180.0301 | 4180.0211 | 0 | 2.1 | 127 | Specific | 268.81 | 268.81 | 268.81 | 7.61 | 104 | >sp\|P01876\|IGHA1_HUMAN Immunoglobulin heavy constant alpha 1 OS=Homo sapiens OX=9606 GN=IGHA1 PE=1 SV=2 | 255 | 20201116_fqg_colostrum_1ng_glyco_hcd203040_compound15.43465.43465.2 | controllerType=0 controllerNumber=1 scan=43465 | 88.8681 |
| 19324:5 | 4 | R.LSLHRPALEDLLLGSEAN[+1872.651]LTC[+57.021]TLTGLR.D | HexNAc(3)Hex(6)NeuAc(1) | C[+57], N[+1873] | 968.0589 | 5 | 4836.2655 | 4836.2487 | 0 | 3.5 | 127 | Specific | 270.8 | 270.8 | 270.8 | 7.73 | 104 | >sp\|P01876\|IGHA1_HUMAN Immunoglobulin heavy constant alpha 1 OS=Homo sapiens OX=9606 GN=IGHA1 PE=1 SV=2 | 255 | 20201116_fqg_colostrum_1ng_glyco_hcd203040_compound15.22882.22882.5 | controllerType=0 controllerNumber=1 scan=22882 | 44.7603 |
| 38424:4 | 4 | R.LSLHRPALEDLLLGSEAN[+1216.423]LTC[+57.021]TLTGLR.D | HexNAc(2)Hex(5) | C[+57], N[+1216] | 1045.7651 | 4 | 4180.0384 | 4180.0211 | 0 | 4.1 | 127 | Specific | 271.9 | 271.9 | 271.9 | 7.06 | 104 | >sp\|P01876\|IGHA1_HUMAN Immunoglobulin heavy constant alpha 1 OS=Homo sapiens OX=9606 GN=IGHA1 PE=1 SV=2 | 255 | 20201116_fqg_colostrum_1ng_glyco_hcd203040_compound15.44886.44886.4 | controllerType=0 controllerNumber=1 scan=44886 | 91.7133 |
| 18121:5 | 4 | R.LSLHRPALEDLLLGSEAN[+1864.634]LTC[+57.021]TLTGLR.D | HexNAc(2)Hex(9) | C[+57], N[+1865] | 966.4547 | 5 | 4828.2443 | 4828.2324 | 0 | 2.5 | 127 | Specific | 273.05 | 273.05 | 273.05 | 6.48 | 104 | >sp\|P01876\|IGHA1_HUMAN Immunoglobulin heavy constant alpha 1 OS=Homo sapiens OX=9606 GN=IGHA1 PE=1 SV=2 | 255 | 20201116_fqg_colostrum_1ng_glyco_hcd203040_compound15.21630.21630.5 | controllerType=0 controllerNumber=1 scan=21630 | 42.2423 |
| 26896:4 | 4 | R.LSLHRPALEDLLLGSEAN[+1501.555]LTC[+57.021]TLTGLR.D | HexNAc(5)Hex(3) | C[+57], N[+1502] | 1117.0444 | 4 | 4465.1556 | 4465.1536 | 0 | 0.4 | 127 | Specific | 273.31 | 273.31 | 273.31 | 6.87 | 104 | >sp\|P01876\|IGHA1_HUMAN Immunoglobulin heavy constant alpha 1 OS=Homo sapiens OX=9606 GN=IGHA1 PE=1 SV=2 | 255 | 20201116_fqg_colostrum_1ng_glyco_hcd203040_compound15.30781.30781.2 | controllerType=0 controllerNumber=1 scan=30781 | 61.4319 |
| 20963:4 | 4 | R.[+42.011]LSLHRPALEDLLLGSEAN[+1955.724]LTC[+57.021]TLTGLR.D | HexNAc(5)Hex(4)Fuc(2) | .[42], C[+57], N[+1956] | 1241.0834 | 4 | 4961.3117 | 4961.3328 | 0 | -4.3 | 127 | Specific | 280.66 | 182.81 | 182.81 | 3.32 | 104 | >sp\|P01876\|IGHA1_HUMAN Immunoglobulin heavy constant alpha 1 OS=Homo sapiens OX=9606 GN=IGHA1 PE=1 SV=2 | 255 | 20201116_fqg_colostrum_1ng_glyco_hcd203040_compound15.24587.24587.4 | controllerType=0 controllerNumber=1 scan=24587 | 48.072 |
| 28486:4 | 4 | R.LSLHRPALEDLLLGSEAN[+1501.555]LTC[+57.021]TLTGLR.D | HexNAc(5)Hex(3) | C[+57], N[+1502] | 1117.0486 | 4 | 4465.1727 | 4465.1536 | 0 | 4.3 | 127 | Specific | 281.39 | 281.39 | 281.39 | 7.21 | 104 | >sp\|P01876\|IGHA1_HUMAN Immunoglobulin heavy constant alpha 1 OS=Homo sapiens OX=9606 GN=IGHA1 PE=1 SV=2 | 255 | 20201116_fqg_colostrum_1ng_glyco_hcd203040_compound15.32439.32439.4 | controllerType=0 controllerNumber=1 scan=32439 | 64.6978 |
| 18157:4 | 4 | R.LSLHRPALEDLLLGSEAN[+1825.661]LTC[+57.021]TLTGLR.D | HexNAc(5)Hex(5) | C[+57], N[+1826] | 1198.0737 | 4 | 4789.2728 | 4789.2592 | 0 | 2.8 | 127 | Specific | 286.92 | 286.92 | 286.92 | 7.66 | 104 | >sp\|P01876\|IGHA1_HUMAN Immunoglobulin heavy constant alpha 1 OS=Homo sapiens OX=9606 GN=IGHA1 PE=1 SV=2 | 255 | 20201116_fqg_colostrum_1ng_glyco_hcd203040_compound15.21668.21668.4 | controllerType=0 controllerNumber=1 scan=21668 | 42.3279 |
| 18265:4 | 4 | R.LSLHRPALEDLLLGSEAN[+1622.582]LTC[+57.021]TLTGLR.D | HexNAc(4)Hex(5) | C[+57], N[+1623] | 1147.3039 | 4 | 4586.1937 | 4586.1799 | 0 | 3 | 127 | Specific | 289.18 | 289.18 | 289.18 | 7.66 | 104 | >sp\|P01876\|IGHA1_HUMAN Immunoglobulin heavy constant alpha 1 OS=Homo sapiens OX=9606 GN=IGHA1 PE=1 SV=2 | 255 | 20201116_fqg_colostrum_1ng_glyco_hcd203040_compound15.21780.21780.2 | controllerType=0 controllerNumber=1 scan=21780 | 42.5605 |
| 37644:4 | 4 | R.LSLHRPALEDLLLGSEAN[+1216.423]LTC[+57.021]TLTGLR.D | HexNAc(2)Hex(5) | C[+57], N[+1216] | 1045.7608 | 4 | 4180.0213 | 4180.0211 | 0 | 0 | 127 | Specific | 293.08 | 293.08 | 293.08 | 7.03 | 104 | >sp\|P01876\|IGHA1_HUMAN Immunoglobulin heavy constant alpha 1 OS=Homo sapiens OX=9606 GN=IGHA1 PE=1 SV=2 | 255 | 20201116_fqg_colostrum_1ng_glyco_hcd203040_compound15.44071.44071.2 | controllerType=0 controllerNumber=1 scan=44071 | 90.203 |
| 26806:4 | 4 | R.LSLHRPALEDLLLGSEAN[+1501.555]LTC[+57.021]TLTGLR.D | HexNAc(5)Hex(3) | C[+57], N[+1502] | 1117.0484 | 4 | 4465.1717 | 4465.1536 | 0 | 4.1 | 127 | Specific | 293.3 | 293.3 | 293.3 | 7.21 | 104 | >sp\|P01876\|IGHA1_HUMAN Immunoglobulin heavy constant alpha 1 OS=Homo sapiens OX=9606 GN=IGHA1 PE=1 SV=2 | 255 | 20201116_fqg_colostrum_1ng_glyco_hcd203040_compound15.30687.30687.4 | controllerType=0 controllerNumber=1 scan=30687 | 61.2423 |
| 37216:4 | 4 | R.LSLHRPALEDLLLGSEAN[+1501.555]LTC[+57.021]TLTGLR.D | HexNAc(5)Hex(3) | C[+57], N[+1502] | 1117.0451 | 4 | 4465.1585 | 4465.1536 | 0 | 1.1 | 127 | Specific | 299.54 | 268.73 | 268.73 | 7.43 | 104 | >sp\|P01876\|IGHA1_HUMAN Immunoglobulin heavy constant alpha 1 OS=Homo sapiens OX=9606 GN=IGHA1 PE=1 SV=2 | 255 | 20201116_fqg_colostrum_1ng_glyco_hcd203040_compound15.43512.43512.4 | controllerType=0 controllerNumber=1 scan=43512 | 88.9664 |
| 37049:4 | 4 | R.LSLHRPALEDLLLGSEAN[+1501.555]LTC[+57.021]TLTGLR.D | HexNAc(5)Hex(3) | C[+57], N[+1502] | 1117.0466 | 4 | 4465.1644 | 4465.1536 | 0 | 2.4 | 127 | Specific | 315.86 | 315.86 | 315.86 | 8.16 | 104 | >sp\|P01876\|IGHA1_HUMAN Immunoglobulin heavy constant alpha 1 OS=Homo sapiens OX=9606 GN=IGHA1 PE=1 SV=2 | 255 | 20201116_fqg_colostrum_1ng_glyco_hcd203040_compound15.43304.43304.4 | controllerType=0 controllerNumber=1 scan=43304 | 88.5162 |
| 19373:4 | 4 | R.LSLHRPALEDLLLGSEAN[+1710.598]LTC[+57.021]TLTGLR.D | HexNAc(3)Hex(5)NeuAc(1) | C[+57], N[+1711] | 1169.3078 | 4 | 4674.2093 | 4674.1959 | 0 | 2.9 | 127 | Specific | 317.2 | 317.2 | 317.2 | 8.05 | 104 | >sp\|P01876\|IGHA1_HUMAN Immunoglobulin heavy constant alpha 1 OS=Homo sapiens OX=9606 GN=IGHA1 PE=1 SV=2 | 255 | 20201116_fqg_colostrum_1ng_glyco_hcd203040_compound15.22933.22933.4 | controllerType=0 controllerNumber=1 scan=22933 | 44.8577 |
| 28498:4 | 4 | R.LSLHRPALEDLLLGSEAN[+1501.555]LTC[+57.021]TLTGLR.D | HexNAc(5)Hex(3) | C[+57], N[+1502] | 1117.047 | 4 | 4465.1663 | 4465.1536 | 0 | 2.9 | 127 | Specific | 320.17 | 189.73 | 189.73 | 7.2 | 104 | >sp\|P01876\|IGHA1_HUMAN Immunoglobulin heavy constant alpha 1 OS=Homo sapiens OX=9606 GN=IGHA1 PE=1 SV=2 | 255 | 20201116_fqg_colostrum_1ng_glyco_hcd203040_compound15.32452.32452.4 | controllerType=0 controllerNumber=1 scan=32452 | 64.7233 |
| 37838:4 | 4 | R.LSLHRPALEDLLLGSEAN[+1216.423]LTC[+57.021]TLTGLR.D | HexNAc(2)Hex(5) | C[+57], N[+1216] | 1045.7648 | 4 | 4180.0374 | 4180.0211 | 0 | 3.9 | 127 | Specific | 325.03 | 325.03 | 325.03 | 7.67 | 104 | >sp\|P01876\|IGHA1_HUMAN Immunoglobulin heavy constant alpha 1 OS=Homo sapiens OX=9606 GN=IGHA1 PE=1 SV=2 | 255 | 20201116_fqg_colostrum_1ng_glyco_hcd203040_compound15.44275.44275.4 | controllerType=0 controllerNumber=1 scan=44275 | 90.5658 |
| 18173:4 | 4 | R.LSLHRPALEDLLLGSEAN[+1663.608]LTC[+57.021]TLTGLR.D | HexNAc(5)Hex(4) | C[+57], N[+1664] | 1157.5599 | 4 | 4627.2176 | 4627.2064 | 0 | 2.4 | 127 | Specific | 330.29 | 330.29 | 330.29 | 8.22 | 104 | >sp\|P01876\|IGHA1_HUMAN Immunoglobulin heavy constant alpha 1 OS=Homo sapiens OX=9606 GN=IGHA1 PE=1 SV=2 | 255 | 20201116_fqg_colostrum_1ng_glyco_hcd203040_compound15.21685.21685.4 | controllerType=0 controllerNumber=1 scan=21685 | 42.3715 |
| 25928:4 | 4 | R.LSLHRPALEDLLLGSEAN[+1501.555]LTC[+57.021]TLTGLR.D | HexNAc(5)Hex(3) | C[+57], N[+1502] | 1117.0474 | 4 | 4465.1678 | 4465.1536 | 0 | 3.2 | 127 | Specific | 340.37 | 340.37 | 340.37 | 8.52 | 104 | >sp\|P01876\|IGHA1_HUMAN Immunoglobulin heavy constant alpha 1 OS=Homo sapiens OX=9606 GN=IGHA1 PE=1 SV=2 | 255 | 20201116_fqg_colostrum_1ng_glyco_hcd203040_compound15.29772.29772.2 | controllerType=0 controllerNumber=1 scan=29772 | 59.331 |
| 26756:4 | 4 | R.LSLHRPALEDLLLGSEAN[+1501.555]LTC[+57.021]TLTGLR.D | HexNAc(5)Hex(3) | C[+57], N[+1502] | 1117.0464 | 4 | 4465.1639 | 4465.1536 | 0 | 2.3 | 127 | Specific | 345.6 | 266.7 | 266.7 | 8.11 | 104 | >sp\|P01876\|IGHA1_HUMAN Immunoglobulin heavy constant alpha 1 OS=Homo sapiens OX=9606 GN=IGHA1 PE=1 SV=2 | 255 | 20201116_fqg_colostrum_1ng_glyco_hcd203040_compound15.30635.30635.4 | controllerType=0 controllerNumber=1 scan=30635 | 61.1308 |
| 18349:5 | 4 | R.LSLHRPALEDLLLGSEAN[+1054.370]LTC[+57.021]TLTGLR.D | HexNAc(2)Hex(4) | C[+57], N[+1054] | 804.4003 | 5 | 4017.9723 | 4017.9683 | 0 | 1 | 127 | Specific | 346.38 | 346.38 | 346.38 | 6.72 | 104 | >sp\|P01876\|IGHA1_HUMAN Immunoglobulin heavy constant alpha 1 OS=Homo sapiens OX=9606 GN=IGHA1 PE=1 SV=2 | 255 | 20201116_fqg_colostrum_1ng_glyco_hcd203040_compound15.21867.21867.5 | controllerType=0 controllerNumber=1 scan=21867 | 42.7361 |
| 29058:4 | 4 | R.LSLHRPALEDLLLGSEAN[+1216.423]LTC[+57.021]TLTGLR.D | HexNAc(2)Hex(5) | C[+57], N[+1216] | 1045.7627 | 4 | 4180.0291 | 4180.0211 | 0 | 1.9 | 127 | Specific | 349.89 | 349.89 | 349.89 | 8.62 | 104 | >sp\|P01876\|IGHA1_HUMAN Immunoglobulin heavy constant alpha 1 OS=Homo sapiens OX=9606 GN=IGHA1 PE=1 SV=2 | 255 | 20201116_fqg_colostrum_1ng_glyco_hcd203040_compound15.33034.33034.3 | controllerType=0 controllerNumber=1 scan=33034 | 65.8278 |
| 24681:4 | 4 | R.LSLHRPALEDLLLGSEAN[+1501.555]LTC[+57.021]TLTGLR.D | HexNAc(5)Hex(3) | C[+57], N[+1502] | 1117.0455 | 4 | 4465.16 | 4465.1536 | 0 | 1.4 | 127 | Specific | 350.57 | 350.57 | 350.57 | 8.39 | 104 | >sp\|P01876\|IGHA1_HUMAN Immunoglobulin heavy constant alpha 1 OS=Homo sapiens OX=9606 GN=IGHA1 PE=1 SV=2 | 255 | 20201116_fqg_colostrum_1ng_glyco_hcd203040_compound15.28467.28467.4 | controllerType=0 controllerNumber=1 scan=28467 | 56.4874 |
| 37377:4 | 4 | R.LSLHRPALEDLLLGSEAN[+1216.423]LTC[+57.021]TLTGLR.D | HexNAc(2)Hex(5) | C[+57], N[+1216] | 1045.7615 | 4 | 4180.0242 | 4180.0211 | 0 | 0.7 | 127 | Specific | 353.48 | 353.48 | 353.48 | 8.39 | 104 | >sp\|P01876\|IGHA1_HUMAN Immunoglobulin heavy constant alpha 1 OS=Homo sapiens OX=9606 GN=IGHA1 PE=1 SV=2 | 255 | 20201116_fqg_colostrum_1ng_glyco_hcd203040_compound15.43737.43737.4 | controllerType=0 controllerNumber=1 scan=43737 | 89.4999 |
| 29829:4 | 4 | R.LSLHRPALEDLLLGSEAN[+1501.555]LTC[+57.021]TLTGLR.D | HexNAc(5)Hex(3) | C[+57], N[+1502] | 1117.0466 | 4 | 4465.1644 | 4465.1536 | 0 | 2.4 | 127 | Specific | 375.81 | 375.81 | 375.81 | 8.82 | 104 | >sp\|P01876\|IGHA1_HUMAN Immunoglobulin heavy constant alpha 1 OS=Homo sapiens OX=9606 GN=IGHA1 PE=1 SV=2 | 255 | 20201116_fqg_colostrum_1ng_glyco_hcd203040_compound15.33837.33837.2 | controllerType=0 controllerNumber=1 scan=33837 | 67.3159 |
| 20966:4 | 4 | R.[+42.011]LSLHRPALEDLLLGSEAN[+1955.724]LTC[+57.021]TLTGLR.D | HexNAc(5)Hex(4)Fuc(2) | .[42], C[+57], N[+1956] | 1241.0834 | 4 | 4961.3117 | 4961.3328 | 0 | -4.3 | 127 | Specific | 377.46 | 306.81 | 306.81 | 5.16 | 104 | >sp\|P01876\|IGHA1_HUMAN Immunoglobulin heavy constant alpha 1 OS=Homo sapiens OX=9606 GN=IGHA1 PE=1 SV=2 | 255 | 20201116_fqg_colostrum_1ng_glyco_hcd203040_compound15.24590.24590.4 | controllerType=0 controllerNumber=1 scan=24590 | 48.081 |
| 18536:4 | 4 | R.LSLHRPALEDLLLGSEAN[+1444.534]LTC[+57.021]TLTGLR.D | HexNAc(4)Hex(3)Fuc(1) | C[+57], N[+1445] | 1102.7919 | 4 | 4408.1458 | 4408.1321 | 0 | 3.1 | 127 | Specific | 382.67 | 305.15 | 305.15 | 6.49 | 104 | >sp\|P01876\|IGHA1_HUMAN Immunoglobulin heavy constant alpha 1 OS=Homo sapiens OX=9606 GN=IGHA1 PE=1 SV=2 | 255 | 20201116_fqg_colostrum_1ng_glyco_hcd203040_compound15.22063.22063.4 | controllerType=0 controllerNumber=1 scan=22063 | 43.1859 |
| 37008:4 | 4 | R.LSLHRPALEDLLLGSEAN[+1216.423]LTC[+57.021]TLTGLR.D | HexNAc(2)Hex(5) | C[+57], N[+1216] | 1045.7612 | 4 | 4180.0228 | 4180.0211 | 0 | 0.4 | 127 | Specific | 383.17 | 383.17 | 383.17 | 9.39 | 104 | >sp\|P01876\|IGHA1_HUMAN Immunoglobulin heavy constant alpha 1 OS=Homo sapiens OX=9606 GN=IGHA1 PE=1 SV=2 | 255 | 20201116_fqg_colostrum_1ng_glyco_hcd203040_compound15.43261.43261.2 | controllerType=0 controllerNumber=1 scan=43261 | 88.4387 |
| 29802:4 | 4 | R.LSLHRPALEDLLLGSEAN[+1216.423]LTC[+57.021]TLTGLR.D | HexNAc(2)Hex(5) | C[+57], N[+1216] | 1045.7658 | 4 | 4180.0413 | 4180.0211 | 0 | 4.8 | 127 | Specific | 384.73 | 384.73 | 384.73 | 9.58 | 104 | >sp\|P01876\|IGHA1_HUMAN Immunoglobulin heavy constant alpha 1 OS=Homo sapiens OX=9606 GN=IGHA1 PE=1 SV=2 | 255 | 20201116_fqg_colostrum_1ng_glyco_hcd203040_compound15.33809.33809.4 | controllerType=0 controllerNumber=1 scan=33809 | 67.2616 |
| 37075:4 | 4 | R.LSLHRPALEDLLLGSEAN[+1216.423]LTC[+57.021]TLTGLR.D | HexNAc(2)Hex(5) | C[+57], N[+1216] | 1045.7631 | 4 | 4180.0306 | 4180.0211 | 0 | 2.3 | 127 | Specific | 392.2 | 392.2 | 392.2 | 10.13 | 104 | >sp\|P01876\|IGHA1_HUMAN Immunoglobulin heavy constant alpha 1 OS=Homo sapiens OX=9606 GN=IGHA1 PE=1 SV=2 | 255 | 20201116_fqg_colostrum_1ng_glyco_hcd203040_compound15.43332.43332.4 | controllerType=0 controllerNumber=1 scan=43332 | 88.5685 |
| 18357:3 | 4 | R.LSLHRPALEDLLLGSEAN[+1298.476]LTC[+57.021]TLTGLR.D | HexNAc(4)Hex(3) | C[+57], N[+1298] | 1421.3683 | 3 | 4262.0905 | 4262.0742 | 0 | 3.8 | 127 | Specific | 396.39 | 396.39 | 396.39 | 9.4 | 104 | >sp\|P01876\|IGHA1_HUMAN Immunoglobulin heavy constant alpha 1 OS=Homo sapiens OX=9606 GN=IGHA1 PE=1 SV=2 | 255 | 20201116_fqg_colostrum_1ng_glyco_hcd203040_compound15.21876.21876.2 | controllerType=0 controllerNumber=1 scan=21876 | 42.7528 |
| 28488:4 | 4 | R.LSLHRPALEDLLLGSEAN[+1501.555]LTC[+57.021]TLTGLR.D | HexNAc(5)Hex(3) | C[+57], N[+1502] | 1117.0486 | 4 | 4465.1727 | 4465.1536 | 0 | 4.3 | 127 | Specific | 401.65 | 278.88 | 278.88 | 7.75 | 104 | >sp\|P01876\|IGHA1_HUMAN Immunoglobulin heavy constant alpha 1 OS=Homo sapiens OX=9606 GN=IGHA1 PE=1 SV=2 | 255 | 20201116_fqg_colostrum_1ng_glyco_hcd203040_compound15.32441.32441.4 | controllerType=0 controllerNumber=1 scan=32441 | 64.7019 |
| 29804:4 | 4 | R.LSLHRPALEDLLLGSEAN[+1216.423]LTC[+57.021]TLTGLR.D | HexNAc(2)Hex(5) | C[+57], N[+1216] | 1045.7658 | 4 | 4180.0413 | 4180.0211 | 0 | 4.8 | 127 | Specific | 431.4 | 431.4 | 431.4 | 9.66 | 104 | >sp\|P01876\|IGHA1_HUMAN Immunoglobulin heavy constant alpha 1 OS=Homo sapiens OX=9606 GN=IGHA1 PE=1 SV=2 | 255 | 20201116_fqg_colostrum_1ng_glyco_hcd203040_compound15.33811.33811.4 | controllerType=0 controllerNumber=1 scan=33811 | 67.2652 |
| 26758:4 | 4 | R.LSLHRPALEDLLLGSEAN[+1501.555]LTC[+57.021]TLTGLR.D | HexNAc(5)Hex(3) | C[+57], N[+1502] | 1117.0464 | 4 | 4465.1639 | 4465.1536 | 0 | 2.3 | 127 | Specific | 432.02 | 326.56 | 326.56 | 8.63 | 104 | >sp\|P01876\|IGHA1_HUMAN Immunoglobulin heavy constant alpha 1 OS=Homo sapiens OX=9606 GN=IGHA1 PE=1 SV=2 | 255 | 20201116_fqg_colostrum_1ng_glyco_hcd203040_compound15.30637.30637.4 | controllerType=0 controllerNumber=1 scan=30637 | 61.1344 |
| 18502:4 | 4 | R.LSLHRPALEDLLLGSEAN[+892.317]LTC[+57.021]TLTGLR.D | HexNAc(2)Hex(3) | C[+57], N[+892] | 964.7371 | 4 | 3855.9266 | 3855.9155 | 0 | 2.9 | 127 | Specific | 437.73 | 437.73 | 437.73 | 10.11 | 104 | >sp\|P01876\|IGHA1_HUMAN Immunoglobulin heavy constant alpha 1 OS=Homo sapiens OX=9606 GN=IGHA1 PE=1 SV=2 | 255 | 20201116_fqg_colostrum_1ng_glyco_hcd203040_compound15.22028.22028.4 | controllerType=0 controllerNumber=1 scan=22028 | 43.1238 |
| 19317:4 | 4 | R.LSLHRPALEDLLLGSEAN[+1872.651]LTC[+57.021]TLTGLR.D | HexNAc(3)Hex(6)NeuAc(1) | C[+57], N[+1873] | 1209.8217 | 4 | 4836.265 | 4836.2487 | 0 | 3.4 | 127 | Specific | 438.36 | 438.36 | 438.36 | 10.11 | 104 | >sp\|P01876\|IGHA1_HUMAN Immunoglobulin heavy constant alpha 1 OS=Homo sapiens OX=9606 GN=IGHA1 PE=1 SV=2 | 255 | 20201116_fqg_colostrum_1ng_glyco_hcd203040_compound15.22875.22875.4 | controllerType=0 controllerNumber=1 scan=22875 | 44.7493 |
| 18360:3 | 4 | R.LSLHRPALEDLLLGSEAN[+1298.476]LTC[+57.021]TLTGLR.D | HexNAc(4)Hex(3) | C[+57], N[+1298] | 1421.3683 | 3 | 4262.0905 | 4262.0742 | 0 | 3.8 | 127 | Specific | 440.58 | 433.4 | 433.4 | 9.53 | 104 | >sp\|P01876\|IGHA1_HUMAN Immunoglobulin heavy constant alpha 1 OS=Homo sapiens OX=9606 GN=IGHA1 PE=1 SV=2 | 255 | 20201116_fqg_colostrum_1ng_glyco_hcd203040_compound15.21879.21879.2 | controllerType=0 controllerNumber=1 scan=21879 | 42.7575 |
| 37178:4 | 4 | R.LSLHRPALEDLLLGSEAN[+1216.423]LTC[+57.021]TLTGLR.D | HexNAc(2)Hex(5) | C[+57], N[+1216] | 1045.763 | 4 | 4180.0301 | 4180.0211 | 0 | 2.1 | 127 | Specific | 441.23 | 441.23 | 441.23 | 10.26 | 104 | >sp\|P01876\|IGHA1_HUMAN Immunoglobulin heavy constant alpha 1 OS=Homo sapiens OX=9606 GN=IGHA1 PE=1 SV=2 | 255 | 20201116_fqg_colostrum_1ng_glyco_hcd203040_compound15.43467.43467.2 | controllerType=0 controllerNumber=1 scan=43467 | 88.8722 |

Table S4. Cont.

| Query #:z | Protein Rank | Glycopeptide | Glycan | Modification Type(s) | Observed m/z | z | Observed (M+H) | Calc. mass (M+H) | Off-by-x error | Mass error (ppm) | StartPosition | Cleavage | Score | Delta | Delta Mod | \|Log Prob\| | # of unique peptides | Protein Name | Protein DB number | Comment | Scan # | Scan Time |
|---|---|---|---|---|---|---|---|---|---|---|---|---|---|---|---|---|---|---|---|---|---|---|
| 38426:4 | 4 | R.LSLHRPALEDLLLGSEAN[+1216.423]LTC[+57.021]TLTGLR.D | HexNAc(2)Hex(5) | C[+57], N[+1216] | 1045.7651 | 4 | 4180.0384 | 4180.0211 | 0 | 4.1 | 127 | Specific | 443.95 | 443.95 | 443.95 | 9.71 | 104 | >sp\|P01876\|IGHA1_HUMAN Immunoglobulin heavy constant alpha 1 OS=Homo sapiens OX=9606 GN=IGHA1 PE=1 SV=2 | 255 | 20201116_fqg_colostrum_1ng_glyco_hcd203040_compound15.44888.44888.4 | controllerType=0 controllerNumber=1 scan=44888 | 91.7168 |
| 26898:4 | 4 | R.LSLHRPALEDLLLGSEAN[+1501.555]LTC[+57.021]TLTGLR.D | HexNAc(5)Hex(3) | C[+57], N[+1502] | 1117.0444 | 4 | 4465.1556 | 4465.1536 | 0 | 0.4 | 127 | Specific | 445.91 | 445.91 | 445.91 | 9.52 | 104 | >sp\|P01876\|IGHA1_HUMAN Immunoglobulin heavy constant alpha 1 OS=Homo sapiens OX=9606 GN=IGHA1 PE=1 SV=2 | 255 | 20201116_fqg_colostrum_1ng_glyco_hcd203040_compound15.30783.30783.2 | controllerType=0 controllerNumber=1 scan=30783 | 61.4369 |
| 27443:4 | 4 | R.LSLHRPALEDLLLGSEAN[+1501.555]LTC[+57.021]TLTGLR.D | HexNAc(5)Hex(3) | C[+57], N[+1502] | 1117.0472 | 4 | 4465.1668 | 4465.1536 | 0 | 3 | 127 | Specific | 469.55 | 443.55 | 443.55 | 10.32 | 104 | >sp\|P01876\|IGHA1_HUMAN Immunoglobulin heavy constant alpha 1 OS=Homo sapiens OX=9606 GN=IGHA1 PE=1 SV=2 | 255 | 20201116_fqg_colostrum_1ng_glyco_hcd203040_compound15.31351.31351.2 | controllerType=0 controllerNumber=1 scan=31351 | 62.59 |
| 19392:5 | 4 | R.LSLHRPALEDLLLGSEAN[+1710.598]LTC[+57.021]TLTGLR.D | HexNAc(3)Hex(5)NeuAc(1) | C[+57], N[+1711] | 935.6481 | 5 | 4674.2114 | 4674.1959 | 0 | 3.3 | 127 | Specific | 489.44 | 405.23 | 405.23 | 8.73 | 104 | >sp\|P01876\|IGHA1_HUMAN Immunoglobulin heavy constant alpha 1 OS=Homo sapiens OX=9606 GN=IGHA1 PE=1 SV=2 | 255 | 20201116_fqg_colostrum_1ng_glyco_hcd203040_compound15.22952.22952.5 | controllerType=0 controllerNumber=1 scan=22952 | 44.8941 |
| 26038:4 | 4 | R.LSLHRPALEDLLLGSEAN[+1501.555]LTC[+57.021]TLTGLR.D | HexNAc(5)Hex(3) | C[+57], N[+1502] | 1117.0487 | 4 | 4465.1732 | 4465.1536 | 0 | 4.4 | 127 | Specific | 491.49 | 466.33 | 466.33 | 9.95 | 104 | >sp\|P01876\|IGHA1_HUMAN Immunoglobulin heavy constant alpha 1 OS=Homo sapiens OX=9606 GN=IGHA1 PE=1 SV=2 | 255 | 20201116_fqg_colostrum_1ng_glyco_hcd203040_compound15.29886.29886.4 | controllerType=0 controllerNumber=1 scan=29886 | 59.5622 |
| 25930:4 | 4 | R.LSLHRPALEDLLLGSEAN[+1501.555]LTC[+57.021]TLTGLR.D | HexNAc(5)Hex(3) | C[+57], N[+1502] | 1117.0474 | 4 | 4465.1678 | 4465.1536 | 0 | 3.2 | 127 | Specific | 495.45 | 495.45 | 495.45 | 10.4 | 104 | >sp\|P01876\|IGHA1_HUMAN Immunoglobulin heavy constant alpha 1 OS=Homo sapiens OX=9606 GN=IGHA1 PE=1 SV=2 | 255 | 20201116_fqg_colostrum_1ng_glyco_hcd203040_compound15.29774.29774.2 | controllerType=0 controllerNumber=1 scan=29774 | 59.3359 |
| 18235:3 | 4 | R.LSLHRPALEDLLLGSEAN[+1257.449]LTC[+57.021]TLTGLR.D | HexNAc(3)Hex(4) | C[+57], N[+1257] | 1407.6909 | 3 | 4221.058 | 4221.0477 | 0 | 2.4 | 127 | Specific | 519.56 | 518.26 | 518.26 | 10 | 104 | >sp\|P01876\|IGHA1_HUMAN Immunoglobulin heavy constant alpha 1 OS=Homo sapiens OX=9606 GN=IGHA1 PE=1 SV=2 | 255 | 20201116_fqg_colostrum_1ng_glyco_hcd203040_compound15.21749.21749.6 | controllerType=0 controllerNumber=1 scan=21749 | 42.4977 |
| 18351:3 | 4 | R.LSLHRPALEDLLLGSEAN[+1054.370]LTC[+57.021]TLTGLR.D | HexNAc(2)Hex(4) | C[+57], N[+1054] | 1339.9965 | 3 | 4017.975 | 4017.9683 | 0 | 1.7 | 127 | Specific | 525.18 | 525.18 | 525.18 | 10.11 | 104 | >sp\|P01876\|IGHA1_HUMAN Immunoglobulin heavy constant alpha 1 OS=Homo sapiens OX=9606 GN=IGHA1 PE=1 SV=2 | 255 | 20201116_fqg_colostrum_1ng_glyco_hcd203040_compound15.21869.21869.3 | controllerType=0 controllerNumber=1 scan=21869 | 42.7384 |
| 18174:4 | 4 | R.LSLHRPALEDLLLGSEAN[+1540.529]LTC[+57.021]TLTGLR.D | HexNAc(2)Hex(7) | C[+57], N[+1541] | 1126.7885 | 4 | 4504.1322 | 4504.1268 | 0 | 1.2 | 127 | Specific | 533.32 | 533.32 | 533.32 | 10.49 | 104 | >sp\|P01876\|IGHA1_HUMAN Immunoglobulin heavy constant alpha 1 OS=Homo sapiens OX=9606 GN=IGHA1 PE=1 SV=2 | 255 | 20201116_fqg_colostrum_1ng_glyco_hcd203040_compound15.21686.21686.4 | controllerType=0 controllerNumber=1 scan=21686 | 42.373 |
| 19251:3 | 4 | R.LSLHRPALEDLLLGSEAN[+2116.756]LTC[+57.021]TLTGLR.D | HexNAc(5)Hex(5)NeuAc(1) | C[+57], N[+2117] | 1694.1275 | 3 | 5080.368 | 5080.3547 | 0 | 2.6 | 127 | Specific | 546.67 | 532.47 | 532.47 | 10.21 | 104 | >sp\|P01876\|IGHA1_HUMAN Immunoglobulin heavy constant alpha 1 OS=Homo sapiens OX=9606 GN=IGHA1 PE=1 SV=2 | 255 | 20201116_fqg_colostrum_1ng_glyco_hcd203040_compound15.22806.22806.3 | controllerType=0 controllerNumber=1 scan=22806 | 44.5966 |
| 37077:4 | 4 | R.LSLHRPALEDLLLGSEAN[+1216.423]LTC[+57.021]TLTGLR.D | HexNAc(2)Hex(5) | C[+57], N[+1216] | 1045.7631 | 4 | 4180.0306 | 4180.0211 | 0 | 2.3 | 127 | Specific | 547.22 | 547.22 | 547.22 | 10.83 | 104 | >sp\|P01876\|IGHA1_HUMAN Immunoglobulin heavy constant alpha 1 OS=Homo sapiens OX=9606 GN=IGHA1 PE=1 SV=2 | 255 | 20201116_fqg_colostrum_1ng_glyco_hcd203040_compound15.43334.43334.4 | controllerType=0 controllerNumber=1 scan=43334 | 88.5722 |
| 18192:3 | 4 | R.LSLHRPALEDLLLGSEAN[+1501.555]LTC[+57.021]TLTGLR.D | HexNAc(5)Hex(3) | C[+57], N[+1502] | 1489.0607 | 3 | 4465.1676 | 4465.1536 | 0 | 3.1 | 127 | Specific | 553.94 | 553.94 | 553.94 | 10.1 | 104 | >sp\|P01876\|IGHA1_HUMAN Immunoglobulin heavy constant alpha 1 OS=Homo sapiens OX=9606 GN=IGHA1 PE=1 SV=2 | 255 | 20201116_fqg_colostrum_1ng_glyco_hcd203040_compound15.21704.21704.6 | controllerType=0 controllerNumber=1 scan=21704 | 42.4059 |
| 18124:5 | 4 | R.LSLHRPALEDLLLGSEAN[+1864.634]LTC[+57.021]TLTGLR.D | HexNAc(2)Hex(9) | C[+57], N[+1865] | 966.4547 | 5 | 4828.2443 | 4828.2324 | 0 | 2.5 | 127 | Specific | 554.4 | 554.4 | 554.4 | 11.05 | 104 | >sp\|P01876\|IGHA1_HUMAN Immunoglobulin heavy constant alpha 1 OS=Homo sapiens OX=9606 GN=IGHA1 PE=1 SV=2 | 255 | 20201116_fqg_colostrum_1ng_glyco_hcd203040_compound15.21633.21633.5 | controllerType=0 controllerNumber=1 scan=21633 | 42.2551 |
| 18276:4 | 4 | R.LSLHRPALEDLLLGSEAN[+1216.423]LTC[+57.021]TLTGLR.D | HexNAc(2)Hex(5) | C[+57], N[+1216] | 1045.763 | 4 | 4180.0301 | 4180.0211 | 0 | 2.1 | 127 | Specific | 554.5 | 554.5 | 554.5 | 10.83 | 104 | >sp\|P01876\|IGHA1_HUMAN Immunoglobulin heavy constant alpha 1 OS=Homo sapiens OX=9606 GN=IGHA1 PE=1 SV=2 | 255 | 20201116_fqg_colostrum_1ng_glyco_hcd203040_compound15.21791.21791.2 | controllerType=0 controllerNumber=1 scan=21791 | 42.5798 |
| 18189:3 | 4 | R.LSLHRPALEDLLLGSEAN[+1501.555]LTC[+57.021]TLTGLR.D | HexNAc(5)Hex(3) | C[+57], N[+1502] | 1489.0607 | 3 | 4465.1676 | 4465.1536 | 0 | 3.1 | 127 | Specific | 560.58 | 560.58 | 560.58 | 10.29 | 104 | >sp\|P01876\|IGHA1_HUMAN Immunoglobulin heavy constant alpha 1 OS=Homo sapiens OX=9606 GN=IGHA1 PE=1 SV=2 | 255 | 20201116_fqg_colostrum_1ng_glyco_hcd203040_compound15.21701.21701.6 | controllerType=0 controllerNumber=1 scan=21701 | 42.3992 |
| 24684:4 | 4 | R.LSLHRPALEDLLLGSEAN[+1501.555]LTC[+57.021]TLTGLR.D | HexNAc(5)Hex(3) | C[+57], N[+1502] | 1117.0455 | 4 | 4465.16 | 4465.1536 | 0 | 1.4 | 127 | Specific | 561.2 | 481.09 | 481.09 | 10.78 | 104 | >sp\|P01876\|IGHA1_HUMAN Immunoglobulin heavy constant alpha 1 OS=Homo sapiens OX=9606 GN=IGHA1 PE=1 SV=2 | 255 | 20201116_fqg_colostrum_1ng_glyco_hcd203040_compound15.28470.28470.4 | controllerType=0 controllerNumber=1 scan=28470 | 56.4954 |
| 18190:5 | 4 | R.LSLHRPALEDLLLGSEAN[+1378.476]LTC[+57.021]TLTGLR.D | HexNAc(2)Hex(6) | C[+57], N[+1378] | 869.2216 | 5 | 4342.0791 | 4342.074 | 0 | 1.2 | 127 | Specific | 580.03 | 580.03 | 580.03 | 12.39 | 104 | >sp\|P01876\|IGHA1_HUMAN Immunoglobulin heavy constant alpha 1 OS=Homo sapiens OX=9606 GN=IGHA1 PE=1 SV=2 | 255 | 20201116_fqg_colostrum_1ng_glyco_hcd203040_compound15.21702.21702.5 | controllerType=0 controllerNumber=1 scan=21702 | 42.4003 |
| 29060:4 | 4 | R.LSLHRPALEDLLLGSEAN[+1216.423]LTC[+57.021]TLTGLR.D | HexNAc(2)Hex(5) | C[+57], N[+1216] | 1045.7627 | 4 | 4180.0291 | 4180.0211 | 0 | 1.9 | 127 | Specific | 580.85 | 450.87 | 450.87 | 12.4 | 104 | >sp\|P01876\|IGHA1_HUMAN Immunoglobulin heavy constant alpha 1 OS=Homo sapiens OX=9606 GN=IGHA1 PE=1 SV=2 | 255 | 20201116_fqg_colostrum_1ng_glyco_hcd203040_compound15.33036.33036.3 | controllerType=0 controllerNumber=1 scan=33036 | 65.8314 |
| 18132:4 | 4 | R.LSLHRPALEDLLLGSEAN[+1864.634]LTC[+57.021]TLTGLR.D | HexNAc(2)Hex(9) | C[+57], N[+1865] | 1207.8161 | 4 | 4828.2425 | 4828.2324 | 0 | 2.1 | 127 | Specific | 586.18 | 586.18 | 586.18 | 12.4 | 104 | >sp\|P01876\|IGHA1_HUMAN Immunoglobulin heavy constant alpha 1 OS=Homo sapiens OX=9606 GN=IGHA1 PE=1 SV=2 | 255 | 20201116_fqg_colostrum_1ng_glyco_hcd203040_compound15.21642.21642.4 | controllerType=0 controllerNumber=1 scan=21642 | 42.2731 |
| 18167:4 | 4 | R.LSLHRPALEDLLLGSEAN[+1581.555]LTC[+57.021]TLTGLR.D | HexNAc(3)Hex(6) | C[+57], N[+1582] | 1137.0468 | 4 | 4545.1654 | 4545.1533 | 0 | 2.6 | 127 | Specific | 589.21 | 589.21 | 589.21 | 12.4 | 104 | >sp\|P01876\|IGHA1_HUMAN Immunoglobulin heavy constant alpha 1 OS=Homo sapiens OX=9606 GN=IGHA1 PE=1 SV=2 | 255 | 20201116_fqg_colostrum_1ng_glyco_hcd203040_compound15.21678.21678.4 | controllerType=0 controllerNumber=1 scan=21678 | 42.3577 |
| 19246:4 | 4 | R.LSLHRPALEDLLLGSEAN[+2116.756]LTC[+57.021]TLTGLR.D | HexNAc(5)Hex(5)NeuAc(1) | C[+57], N[+2117] | 1270.8464 | 4 | 5080.3639 | 5080.3547 | 0 | 1.8 | 127 | Specific | 594.74 | 594.74 | 594.74 | 12.4 | 104 | >sp\|P01876\|IGHA1_HUMAN Immunoglobulin heavy constant alpha 1 OS=Homo sapiens OX=9606 GN=IGHA1 PE=1 SV=2 | 255 | 20201116_fqg_colostrum_1ng_glyco_hcd203040_compound15.22801.22801.4 | controllerType=0 controllerNumber=1 scan=22801 | 44.5869 |
| 19380:3 | 4 | R.LSLHRPALEDLLLGSEAN[+1710.598]LTC[+57.021]TLTGLR.D | HexNAc(3)Hex(5)NeuAc(1) | C[+57], N[+1711] | 1558.7396 | 3 | 4674.2041 | 4674.1959 | 0 | 1.8 | 127 | Specific | 608.45 | 540.68 | 540.68 | 12.78 | 104 | >sp\|P01876\|IGHA1_HUMAN Immunoglobulin heavy constant alpha 1 OS=Homo sapiens OX=9606 GN=IGHA1 PE=1 SV=2 | 255 | 20201116_fqg_colostrum_1ng_glyco_hcd203040_compound15.22940.22940.3 | controllerType=0 controllerNumber=1 scan=22940 | 44.8698 |
| 18196:3 | 4 | R.LSLHRPALEDLLLGSEAN[+1460.529]LTC[+57.021]TLTGLR.D | HexNAc(4)Hex(4) | C[+57], N[+1461] | 1475.3849 | 3 | 4424.1403 | 4424.1271 | 0 | 3 | 127 | Specific | 612.96 | 528.2 | 528.2 | 12.68 | 104 | >sp\|P01876\|IGHA1_HUMAN Immunoglobulin heavy constant alpha 1 OS=Homo sapiens OX=9606 GN=IGHA1 PE=1 SV=2 | 255 | 20201116_fqg_colostrum_1ng_glyco_hcd203040_compound15.21708.21708.6 | controllerType=0 controllerNumber=1 scan=21708 | 42.4118 |
| 19233:3 | 4 | R.LSLHRPALEDLLLGSEAN[+1913.677]LTC[+57.021]TLTGLR.D | HexNAc(4)Hex(5)NeuAc(1) | C[+57], N[+1914] | 1626.4351 | 3 | 4877.2908 | 4877.2753 | 0 | 3.2 | 127 | Specific | 618.72 | 618.72 | 618.72 | 12.68 | 104 | >sp\|P01876\|IGHA1_HUMAN Immunoglobulin heavy constant alpha 1 OS=Homo sapiens OX=9606 GN=IGHA1 PE=1 SV=2 | 255 | 20201116_fqg_colostrum_1ng_glyco_hcd203040_compound15.22787.22787.3 | controllerType=0 controllerNumber=1 scan=22787 | 44.5591 |
| 19384:3 | 4 | R.LSLHRPALEDLLLGSEAN[+1548.545]LTC[+57.021]TLTGLR.D | HexNAc(3)Hex(4)NeuAc(1) | C[+57], N[+1549] | 1504.7236 | 3 | 4512.1562 | 4512.1431 | 0 | 2.9 | 127 | Specific | 618.78 | 618.78 | 618.78 | 12.68 | 104 | >sp\|P01876\|IGHA1_HUMAN Immunoglobulin heavy constant alpha 1 OS=Homo sapiens OX=9606 GN=IGHA1 PE=1 SV=2 | 255 | 20201116_fqg_colostrum_1ng_glyco_hcd203040_compound15.22944.22944.3 | controllerType=0 controllerNumber=1 scan=22944 | 44.8744 |
| 18076:3 | 4 | R.LSLHRPALEDLLLGSEAN[+1864.634]LTC[+57.021]TLTGLR.D | HexNAc(2)Hex(9) | C[+57], N[+1865] | 1610.0845 | 3 | 4828.239 | 4828.2324 | 0 | 1.4 | 127 | Specific | 621.24 | 577.63 | 577.63 | 12.54 | 104 | >sp\|P01876\|IGHA1_HUMAN Immunoglobulin heavy constant alpha 1 OS=Homo sapiens OX=9606 GN=IGHA1 PE=1 SV=2 | 255 | 20201116_fqg_colostrum_1ng_glyco_hcd203040_compound15.21583.21583.3 | controllerType=0 controllerNumber=1 scan=21583 | 42.1597 |
| 37010:4 | 4 | R.LSLHRPALEDLLLGSEAN[+1216.423]LTC[+57.021]TLTGLR.D | HexNAc(2)Hex(5) | C[+57], N[+1216] | 1045.7612 | 4 | 4180.0228 | 4180.0211 | 0 | 0.4 | 127 | Specific | 621.93 | 620.58 | 620.58 | 12.67 | 104 | >sp\|P01876\|IGHA1_HUMAN Immunoglobulin heavy constant alpha 1 OS=Homo sapiens OX=9606 GN=IGHA1 PE=1 SV=2 | 255 | 20201116_fqg_colostrum_1ng_glyco_hcd203040_compound15.43263.43263.2 | controllerType=0 controllerNumber=1 scan=43263 | 88.4424 |
| 18177:3 | 4 | R.LSLHRPALEDLLLGSEAN[+1663.608]LTC[+57.021]TLTGLR.D | HexNAc(5)Hex(4) | C[+57], N[+1664] | 1543.0788 | 3 | 4627.2218 | 4627.2064 | 0 | 3.3 | 127 | Specific | 625.47 | 590.13 | 590.13 | 12.68 | 104 | >sp\|P01876\|IGHA1_HUMAN Immunoglobulin heavy constant alpha 1 OS=Homo sapiens OX=9606 GN=IGHA1 PE=1 SV=2 | 255 | 20201116_fqg_colostrum_1ng_glyco_hcd203040_compound15.21689.21689.3 | controllerType=0 controllerNumber=1 scan=21689 | 42.3774 |
| 19322:3 | 4 | R.LSLHRPALEDLLLGSEAN[+1751.624]LTC[+57.021]TLTGLR.D | HexNAc(4)Hex(4)NeuAc(1) | C[+57], N[+1752] | 1572.4175 | 3 | 4715.2381 | 4715.2225 | 0 | 3.3 | 127 | Specific | 625.75 | 625.75 | 625.75 | 12.68 | 104 | >sp\|P01876\|IGHA1_HUMAN Immunoglobulin heavy constant alpha 1 OS=Homo sapiens OX=9606 GN=IGHA1 PE=1 SV=2 | 255 | 20201116_fqg_colostrum_1ng_glyco_hcd203040_compound15.22880.22880.3 | controllerType=0 controllerNumber=1 scan=22880 | 44.758 |
| 19254:3 | 4 | R.LSLHRPALEDLLLGSEAN[+2116.756]LTC[+57.021]TLTGLR.D | HexNAc(5)Hex(5)NeuAc(1) | C[+57], N[+2117] | 1694.1275 | 3 | 5080.368 | 5080.3547 | 0 | 2.6 | 127 | Specific | 627.3 | 610.53 | 610.53 | 12.78 | 104 | >sp\|P01876\|IGHA1_HUMAN Immunoglobulin heavy constant alpha 1 OS=Homo sapiens OX=9606 GN=IGHA1 PE=1 SV=2 | 255 | 20201116_fqg_colostrum_1ng_glyco_hcd203040_compound15.22809.22809.3 | controllerType=0 controllerNumber=1 scan=22809 | 44.6043 |
| 18201:4 | 4 | R.LSLHRPALEDLLLGSEAN[+1460.529]LTC[+57.021]TLTGLR.D | HexNAc(4)Hex(4) | C[+57], N[+1461] | 1106.7919 | 4 | 4424.1458 | 4424.1271 | 0 | 4.2 | 127 | Specific | 630.44 | 604.24 | 604.24 | 12.85 | 104 | >sp\|P01876\|IGHA1_HUMAN Immunoglobulin heavy constant alpha 1 OS=Homo sapiens OX=9606 GN=IGHA1 PE=1 SV=2 | 255 | 20201116_fqg_colostrum_1ng_glyco_hcd203040_compound15.21714.21714.4 | controllerType=0 controllerNumber=1 scan=21714 | 42.4252 |
| 19325:3 | 4 | R.LSLHRPALEDLLLGSEAN[+1751.624]LTC[+57.021]TLTGLR.D | HexNAc(4)Hex(4)NeuAc(1) | C[+57], N[+1752] | 1572.4175 | 3 | 4715.2381 | 4715.2225 | 0 | 3.3 | 127 | Specific | 638.27 | 638.27 | 638.27 | 12.68 | 104 | >sp\|P01876\|IGHA1_HUMAN Immunoglobulin heavy constant alpha 1 OS=Homo sapiens OX=9606 GN=IGHA1 PE=1 SV=2 | 255 | 20201116_fqg_colostrum_1ng_glyco_hcd203040_compound15.22883.22883.3 | controllerType=0 controllerNumber=1 scan=22883 | 44.7633 |
| 19236:3 | 4 | R.LSLHRPALEDLLLGSEAN[+1913.677]LTC[+57.021]TLTGLR.D | HexNAc(4)Hex(5)NeuAc(1) | C[+57], N[+1914] | 1626.4351 | 3 | 4877.2908 | 4877.2753 | 0 | 3.2 | 127 | Specific | 643.63 | 591.12 | 591.12 | 12.68 | 104 | >sp\|P01876\|IGHA1_HUMAN Immunoglobulin heavy constant alpha 1 OS=Homo sapiens OX=9606 GN=IGHA1 PE=1 SV=2 | 255 | 20201116_fqg_colostrum_1ng_glyco_hcd203040_compound15.22790.22790.3 | controllerType=0 controllerNumber=1 scan=22790 | 44.5673 |
| 18096:3 | 4 | R.LSLHRPALEDLLLGSEAN[+1864.634]LTC[+57.021]TLTGLR.D | HexNAc(2)Hex(9) | C[+57], N[+1865] | 1610.0848 | 3 | 4828.2398 | 4828.2324 | 0 | 1.5 | 127 | Specific | 644.1 | 644.1 | 644.1 | 12.54 | 104 | >sp\|P01876\|IGHA1_HUMAN Immunoglobulin heavy constant alpha 1 OS=Homo sapiens OX=9606 GN=IGHA1 PE=1 SV=2 | 255 | 20201116_fqg_colostrum_1ng_glyco_hcd203040_compound15.21604.21604.3 | controllerType=0 controllerNumber=1 scan=21604 | 42.1941 |
| 19282:3 | 4 | R.LSLHRPALEDLLLGSEAN[+1872.651]LTC[+57.021]TLTGLR.D | HexNAc(3)Hex(6)NeuAc(1) | C[+57], N[+1873] | 1612.7581 | 3 | 4836.2598 | 4836.2487 | 0 | 2.3 | 127 | Specific | 649.76 | 649.76 | 649.76 | 12.78 | 104 | >sp\|P01876\|IGHA1_HUMAN Immunoglobulin heavy constant alpha 1 OS=Homo sapiens OX=9606 GN=IGHA1 PE=1 SV=2 | 255 | 20201116_fqg_colostrum_1ng_glyco_hcd203040_compound15.22838.22838.3 | controllerType=0 controllerNumber=1 scan=22838 | 44.6602 |
| 18348:3 | 4 | R.LSLHRPALEDLLLGSEAN[+1501.555]LTC[+57.021]TLTGLR.D | HexNAc(5)Hex(3) | C[+57], N[+1502] | 1489.0607 | 3 | 4465.1676 | 4465.1536 | 0 | 3.1 | 127 | Specific | 652.85 | 652.85 | 652.85 | 13.68 | 104 | >sp\|P01876\|IGHA1_HUMAN Immunoglobulin heavy constant alpha 1 OS=Homo sapiens OX=9606 GN=IGHA1 PE=1 SV=2 | 255 | 20201116_fqg_colostrum_1ng_glyco_hcd203040_compound15.21866.21866.2 | controllerType=0 controllerNumber=1 scan=21866 | 42.735 |
| 18320:5 | 4 | R.LSLHRPALEDLLLGSEAN[+1298.476]LTC[+57.021]TLTGLR.D | HexNAc(4)Hex(3) | C[+57], N[+1298] | 853.2224 | 5 | 4262.0828 | 4262.0742 | 0 | 2 | 127 | Specific | 654.27 | 654.27 | 654.27 | 14.62 | 104 | >sp\|P01876\|IGHA1_HUMAN Immunoglobulin heavy constant alpha 1 OS=Homo sapiens OX=9606 GN=IGHA1 PE=1 SV=2 | 255 | 20201116_fqg_colostrum_1ng_glyco_hcd203040_compound15.21837.21837.5 | controllerType=0 controllerNumber=1 scan=21837 | 42.6761 |
| 18180:3 | 4 | R.LSLHRPALEDLLLGSEAN[+1663.608]LTC[+57.021]TLTGLR.D | HexNAc(5)Hex(4) | C[+57], N[+1664] | 1543.0788 | 3 | 4627.2218 | 4627.2064 | 0 | 3.3 | 127 | Specific | 659.22 | 626.29 | 626.29 | 13.68 | 104 | >sp\|P01876\|IGHA1_HUMAN Immunoglobulin heavy constant alpha 1 OS=Homo sapiens OX=9606 GN=IGHA1 PE=1 SV=2 | 255 | 20201116_fqg_colostrum_1ng_glyco_hcd203040_compound15.21692.21692.3 | controllerType=0 controllerNumber=1 scan=21692 | 42.3819 |
| 18199:3 | 4 | R.LSLHRPALEDLLLGSEAN[+1460.529]LTC[+57.021]TLTGLR.D | HexNAc(4)Hex(4) | C[+57], N[+1461] | 1475.3849 | 3 | 4424.1403 | 4424.1271 | 0 | 3 | 127 | Specific | 659.77 | 553.29 | 553.29 | 13.68 | 104 | >sp\|P01876\|IGHA1_HUMAN Immunoglobulin heavy constant alpha 1 OS=Homo sapiens OX=9606 GN=IGHA1 PE=1 SV=2 | 255 | 20201116_fqg_colostrum_1ng_glyco_hcd203040_compound15.21711.21711.6 | controllerType=0 controllerNumber=1 scan=21711 | 42.4208 |
| 19386:3 | 4 | R.LSLHRPALEDLLLGSEAN[+1548.545]LTC[+57.021]TLTGLR.D | HexNAc(3)Hex(4)NeuAc(1) | C[+57], N[+1549] | 1504.7236 | 3 | 4512.1562 | 4512.1431 | 0 | 2.9 | 127 | Specific | 660.09 | 647.77 | 647.77 | 13.68 | 104 | >sp\|P01876\|IGHA1_HUMAN Immunoglobulin heavy constant alpha 1 OS=Homo sapiens OX=9606 GN=IGHA1 PE=1 SV=2 | 255 | 20201116_fqg_colostrum_1ng_glyco_hcd203040_compound15.22946.22946.3 | controllerType=0 controllerNumber=1 scan=22946 | 44.8816 |
| 19221:4 | 4 | R.LSLHRPALEDLLLGSEAN[+1913.677]LTC[+57.021]TLTGLR.D | HexNAc(4)Hex(5)NeuAc(1) | C[+57], N[+1914] | 1220.0769 | 4 | 4877.286 | 4877.2753 | 0 | 2.2 | 127 | Specific | 662.62 | 662.62 | 662.62 | 14.4 | 104 | >sp\|P01876\|IGHA1_HUMAN Immunoglobulin heavy constant alpha 1 OS=Homo sapiens OX=9606 GN=IGHA1 PE=1 SV=2 | 255 | 20201116_fqg_colostrum_1ng_glyco_hcd203040_compound15.22775.22775.4 | controllerType=0 controllerNumber=1 scan=22775 | 44.5395 |
| 19279:3 | 4 | R.LSLHRPALEDLLLGSEAN[+1872.651]LTC[+57.021]TLTGLR.D | HexNAc(3)Hex(6)NeuAc(1) | C[+57], N[+1873] | 1612.7581 | 3 | 4836.2598 | 4836.2487 | 0 | 2.3 | 127 | Specific | 667.31 | 594.23 | 594.23 | 13.78 | 104 | >sp\|P01876\|IGHA1_HUMAN Immunoglobulin heavy constant alpha 1 OS=Homo sapiens OX=9606 GN=IGHA1 PE=1 SV=2 | 255 | 20201116_fqg_colostrum_1ng_glyco_hcd203040_compound15.22835.22835.3 | controllerType=0 controllerNumber=1 scan=22835 | 44.6536 |
| 18208:3 | 4 | R.LSLHRPALEDLLLGSEAN[+1460.529]LTC[+57.021]TLTGLR.D | HexNAc(4)Hex(4) | C[+57], N[+1461] | 1475.3867 | 3 | 4424.1454 | 4424.1271 | 0 | 4.1 | 127 | Specific | 676.1 | 590.64 | 590.64 | 13.23 | 104 | >sp\|P01876\|IGHA1_HUMAN Immunoglobulin heavy constant alpha 1 OS=Homo sapiens OX=9606 GN=IGHA1 PE=1 SV=2 | 255 | 20201116_fqg_colostrum_1ng_glyco_hcd203040_compound15.21721.21721.3 | controllerType=0 controllerNumber=1 scan=21721 | 42.4401 |
| 18505:4 | 4 | R.LSLHRPALEDLLLGSEAN[+892.317]LTC[+57.021]TLTGLR.D | HexNAc(2)Hex(3) | C[+57], N[+892] | 964.7371 | 4 | 3855.9266 | 3855.9155 | 0 | 2.9 | 127 | Specific | 678.74 | 678.74 | 678.74 | 14.3 | 104 | >sp\|P01876\|IGHA1_HUMAN Immunoglobulin heavy constant alpha 1 OS=Homo sapiens OX=9606 GN=IGHA1 PE=1 SV=2 | 255 | 20201116_fqg_colostrum_1ng_glyco_hcd203040_compound15.22031.22031.4 | controllerType=0 controllerNumber=1 scan=22031 | 43.1284 |
| 19347:5 | 4 | R.LSLHRPALEDLLLGSEAN[+1751.624]LTC[+57.021]TLTGLR.D | HexNAc(4)Hex(4)NeuAc(1) | C[+57], N[+1752] | 943.8521 | 5 | 4715.2313 | 4715.2225 | 0 | 1.9 | 127 | Specific | 678.78 | 678.78 | 678.78 | 14.62 | 104 | >sp\|P01876\|IGHA1_HUMAN Immunoglobulin heavy constant alpha 1 OS=Homo sapiens OX=9606 GN=IGHA1 PE=1 SV=2 | 255 | 20201116_fqg_colostrum_1ng_glyco_hcd203040_compound15.22906.22906.5 | controllerType=0 controllerNumber=1 scan=22906 | 44.8035 |
| 18205:3 | 4 | R.LSLHRPALEDLLLGSEAN[+1460.529]LTC[+57.021]TLTGLR.D | HexNAc(4)Hex(4) | C[+57], N[+1461] | 1475.3867 | 3 | 4424.1454 | 4424.1271 | 0 | 4.1 | 127 | Specific | 684.57 | 620.69 | 620.69 | 13.23 | 104 | >sp\|P01876\|IGHA1_HUMAN Immunoglobulin heavy constant alpha 1 OS=Homo sapiens OX=9606 GN=IGHA1 PE=1 SV=2 | 255 | 20201116_fqg_colostrum_1ng_glyco_hcd203040_compound15.21718.21718.3 | controllerType=0 controllerNumber=1 scan=21718 | 42.4353 |
| 18094:3 | 4 | R.LSLHRPALEDLLLGSEAN[+1864.634]LTC[+57.021]TLTGLR.D | HexNAc(2)Hex(9) | C[+57], N[+1865] | 1610.0848 | 3 | 4828.2398 | 4828.2324 | 0 | 1.5 | 127 | Specific | 688.56 | 680.46 | 680.46 | 13.54 | 104 | >sp\|P01876\|IGHA1_HUMAN Immunoglobulin heavy constant alpha 1 OS=Homo sapiens OX=9606 GN=IGHA1 PE=1 SV=2 | 255 | 20201116_fqg_colostrum_1ng_glyco_hcd203040_compound15.21602.21602.3 | controllerType=0 controllerNumber=1 scan=21602 | 42.1887 |
| 18140:3 | 4 | R.LSLHRPALEDLLLGSEAN[+1540.529]LTC[+57.021]TLTGLR.D | HexNAc(2)Hex(7) | C[+57], N[+1541] | 1502.0507 | 3 | 4504.1376 | 4504.1268 | 0 | 2.4 | 127 | Specific | 689.58 | 613.56 | 613.56 | 13.78 | 104 | >sp\|P01876\|IGHA1_HUMAN Immunoglobulin heavy constant alpha 1 OS=Homo sapiens OX=9606 GN=IGHA1 PE=1 SV=2 | 255 | 20201116_fqg_colostrum_1ng_glyco_hcd203040_compound15.21650.21650.3 | controllerType=0 controllerNumber=1 scan=21650 | 42.2922 |
| 19318:4 | 4 | R.LSLHRPALEDLLLGSEAN[+1751.624]LTC[+57.021]TLTGLR.D | HexNAc(4)Hex(4)NeuAc(1) | C[+57], N[+1752] | 1179.5644 | 4 | 4715.2357 | 4715.2225 | 0 | 2.8 | 127 | Specific | 690.26 | 648.11 | 648.11 | 14.3 | 104 | >sp\|P01876\|IGHA1_HUMAN Immunoglobulin heavy constant alpha 1 OS=Homo sapiens OX=9606 GN=IGHA1 PE=1 SV=2 | 255 | 20201116_fqg_colostrum_1ng_glyco_hcd203040_compound15.22876.22876.4 | controllerType=0 controllerNumber=1 scan=22876 | 44.7507 |
| 18267:3 | 4 | R.LSLHRPALEDLLLGSEAN[+1257.449]LTC[+57.021]TLTGLR.D | HexNAc(3)Hex(4) | C[+57], N[+1257] | 1407.6901 | 3 | 4221.0558 | 4221.0477 | 0 | 1.9 | 127 | Specific | 697.01 | 594.18 | 594.18 | 13.78 | 104 | >sp\|P01876\|IGHA1_HUMAN Immunoglobulin heavy constant alpha 1 OS=Homo sapiens OX=9606 GN=IGHA1 PE=1 SV=2 | 255 | 20201116_fqg_colostrum_1ng_glyco_hcd203040_compound15.21782.21782.3 | controllerType=0 controllerNumber=1 scan=21782 | 42.5656 |
| 18354:3 | 4 | R.LSLHRPALEDLLLGSEAN[+1054.370]LTC[+57.021]TLTGLR.D | HexNAc(2)Hex(4) | C[+57], N[+1054] | 1339.9965 | 3 | 4017.975 | 4017.9683 | 0 | 1.7 | 127 | Specific | 698.63 | 657.78 | 657.78 | 13.78 | 104 | >sp\|P01876\|IGHA1_HUMAN Immunoglobulin heavy constant alpha 1 OS=Homo sapiens OX=9606 GN=IGHA1 PE=1 SV=2 | 255 | 20201116_fqg_colostrum_1ng_glyco_hcd203040_compound15.21872.21872.3 | controllerType=0 controllerNumber=1 scan=21872 | 42.7459 |
| 18162:3 | 4 | R.LSLHRPALEDLLLGSEAN[+1622.582]LTC[+57.021]TLTGLR.D | HexNAc(4)Hex(5) | C[+57], N[+1623] | 1529.4007 | 3 | 4586.1875 | 4586.1799 | 0 | 1.7 | 127 | Specific | 702.48 | 658.41 | 658.41 | 14.78 | 104 | >sp\|P01876\|IGHA1_HUMAN Immunoglobulin heavy constant alpha 1 OS=Homo sapiens OX=9606 GN=IGHA1 PE=1 SV=2 | 255 | 20201116_fqg_colostrum_1ng_glyco_hcd203040_compound15.21673.21673.3 | controllerType=0 controllerNumber=1 scan=21673 | 42.3416 |
| 18268:4 | 4 | R.LSLHRPALEDLLLGSEAN[+1622.582]LTC[+57.021]TLTGLR.D | HexNAc(4)Hex(5) | C[+57], N[+1623] | 1147.3039 | 4 | 4586.1937 | 4586.1799 | 0 | 3 | 127 | Specific | 702.69 | 580.94 | 580.94 | 15.3 | 104 | >sp\|P01876\|IGHA1_HUMAN Immunoglobulin heavy constant alpha 1 OS=Homo sapiens OX=9606 GN=IGHA1 PE=1 SV=2 | 255 | 20201116_fqg_colostrum_1ng_glyco_hcd203040_compound15.21783.21783.2 | controllerType=0 controllerNumber=1 scan=21783 | 42.5669 |
| 18213:3 | 4 | R.LSLHRPALEDLLLGSEAN[+1298.476]LTC[+57.021]TLTGLR.D | HexNAc(4)Hex(3) | C[+57], N[+1298] | 1421.3658 | 3 | 4262.0828 | 4262.0742 | 0 | 2 | 127 | Specific | 706.49 | 623.83 | 623.83 | 14.78 | 104 | >sp\|P01876\|IGHA1_HUMAN Immunoglobulin heavy constant alpha 1 OS=Homo sapiens OX=9606 GN=IGHA1 PE=1 SV=2 | 255 | 20201116_fqg_colostrum_1ng_glyco_hcd203040_compound15.21726.21726.3 | controllerType=0 controllerNumber=1 scan=21726 | 42.4498 |
| 18264:3 | 4 | R.LSLHRPALEDLLLGSEAN[+1257.449]LTC[+57.021]TLTGLR.D | HexNAc(3)Hex(4) | C[+57], N[+1257] | 1407.6901 | 3 | 4221.0558 | 4221.0477 | 0 | 1.9 | 127 | Specific | 711.55 | 654.88 | 654.88 | 14.78 | 104 | >sp\|P01876\|IGHA1_HUMAN Immunoglobulin heavy constant alpha 1 OS=Homo sapiens OX=9606 GN=IGHA1 PE=1 SV=2 | 255 | 20201116_fqg_colostrum_1ng_glyco_hcd203040_compound15.21779.21779.3 | controllerType=0 controllerNumber=1 scan=21779 | 42.5593 |
| 18165:3 | 4 | R.LSLHRPALEDLLLGSEAN[+1622.582]LTC[+57.021]TLTGLR.D | HexNAc(4)Hex(5) | C[+57], N[+1623] | 1529.4007 | 3 | 4586.1875 | 4586.1799 | 0 | 1.7 | 127 | Specific | 712.83 | 622.29 | 622.29 | 14.78 | 104 | >sp\|P01876\|IGHA1_HUMAN Immunoglobulin heavy constant alpha 1 OS=Homo sapiens OX=9606 GN=IGHA1 PE=1 SV=2 | 255 | 20201116_fqg_colostrum_1ng_glyco_hcd203040_compound15.21676.21676.3 | controllerType=0 controllerNumber=1 scan=21676 | 42.3558 |
| 18119:3 | 4 | R.LSLHRPALEDLLLGSEAN[+1702.581]LTC[+57.021]TLTGLR.D | HexNAc(2)Hex(8) | C[+57], N[+1703] | 1556.0681 | 3 | 4666.1896 | 4666.1796 | 0 | 2.1 | 127 | Specific | 717.74 | 687.83 | 687.83 | 14.78 | 104 | >sp\|P01876\|IGHA1_HUMAN Immunoglobulin heavy constant alpha 1 OS=Homo sapiens OX=9606 GN=IGHA1 PE=1 SV=2 | 255 | 20201116_fqg_colostrum_1ng_glyco_hcd203040_compound15.21628.21628.3 | controllerType=0 controllerNumber=1 scan=21628 | 42.2401 |
| 19247:5 | 4 | R.LSLHRPALEDLLLGSEAN[+2116.756]LTC[+57.021]TLTGLR.D | HexNAc(5)Hex(5)NeuAc(1) | C[+57], N[+2117] | 1016.8779 | 5 | 5080.3606 | 5080.3547 | 0 | 1.2 | 127 | Specific | 718.59 | 718.59 | 718.59 | 15.39 | 104 | >sp\|P01876\|IGHA1_HUMAN Immunoglobulin heavy constant alpha 1 OS=Homo sapiens OX=9606 GN=IGHA1 PE=1 SV=2 | 255 | 20201116_fqg_colostrum_1ng_glyco_hcd203040_compound15.22802.22802.5 | controllerType=0 controllerNumber=1 scan=22802 | 44.5883 |
| 17724:4 | 4 | R.LSLHRPALEDLLLGSEAN[+1501.555]LTC[+57.021]TLTGLR.D | HexNAc(5)Hex(3) | C[+57], N[+1502] | 1117.0447 | 4 | 4465.1571 | 4465.1536 | 0 | 0.8 | 127 | Specific | 720.02 | 720.02 | 720.02 | 15.17 | 104 | >sp\|P01876\|IGHA1_HUMAN Immunoglobulin heavy constant alpha 1 OS=Homo sapiens OX=9606 GN=IGHA1 PE=1 SV=2 | 255 | 20201116_fqg_colostrum_1ng_glyco_hcd203040_compound15.21217.21217.2 | controllerType=0 controllerNumber=1 scan=21217 | 41.4301 |
| 18160:4 | 4 | R.LSLHRPALEDLLLGSEAN[+1825.661]LTC[+57.021]TLTGLR.D | HexNAc(5)Hex(5) | C[+57], N[+1826] | 1198.0737 | 4 | 4789.2728 | 4789.2592 | 0 | 2.8 | 127 | Specific | 720.91 | 610.46 | 610.46 | 15.3 | 104 | >sp\|P01876\|IGHA1_HUMAN Immunoglobulin heavy constant alpha 1 OS=Homo sapiens OX=9606 GN=IGHA1 PE=1 SV=2 | 255 | 20201116_fqg_colostrum_1ng_glyco_hcd203040_compound15.21671.21671.4 | controllerType=0 controllerNumber=1 scan=21671 | 42.3393 |
| 18224:4 | 4 | R.LSLHRPALEDLLLGSEAN[+1501.555]LTC[+57.021]TLTGLR.D | HexNAc(5)Hex(3) | C[+57], N[+1502] | 1117.0472 | 4 | 4465.1668 | 4465.1536 | 0 | 3 | 127 | Specific | 722.88 | 722.88 | 722.88 | 15.3 | 104 | >sp\|P01876\|IGHA1_HUMAN Immunoglobulin heavy constant alpha 1 OS=Homo sapiens OX=9606 GN=IGHA1 PE=1 SV=2 | 255 | 20201116_fqg_colostrum_1ng_glyco_hcd203040_compound15.21738.21738.4 | controllerType=0 controllerNumber=1 scan=21738 | 42.4789 |
| 18159:3 | 4 | R.LSLHRPALEDLLLGSEAN[+1378.476]LTC[+57.021]TLTGLR.D | HexNAc(2)Hex(6) | C[+57], N[+1378] | 1448.0335 | 3 | 4342.086 | 4342.074 | 0 | 2.8 | 127 | Specific | 723.31 | 688.17 | 688.17 | 14.68 | 104 | >sp\|P01876\|IGHA1_HUMAN Immunoglobulin heavy constant alpha 1 OS=Homo sapiens OX=9606 GN=IGHA1 PE=1 SV=2 | 255 | 20201116_fqg_colostrum_1ng_glyco_hcd203040_compound15.21670.21670.3 | controllerType=0 controllerNumber=1 scan=21670 | 42.3386 |
| 18073:3 | 4 | R.LSLHRPALEDLLLGSEAN[+1864.634]LTC[+57.021]TLTGLR.D | HexNAc(2)Hex(9) | C[+57], N[+1865] | 1610.0845 | 3 | 4828.239 | 4828.2324 | 0 | 1.4 | 127 | Specific | 724.1 | 716.93 | 716.93 | 14.54 | 104 | >sp\|P01876\|IGHA1_HUMAN Immunoglobulin heavy constant alpha 1 OS=Homo sapiens OX=9606 GN=IGHA1 PE=1 SV=2 | 255 | 20201116_fqg_colostrum_1ng_glyco_hcd203040_compound15.21580.21580.3 | controllerType=0 controllerNumber=1 scan=21580 | 42.1513 |
| 18151:4 | 4 | R.LSLHRPALEDLLLGSEAN[+1622.582]LTC[+57.021]TLTGLR.D | HexNAc(4)Hex(5) | C[+57], N[+1623] | 1147.3033 | 4 | 4586.1912 | 4586.1799 | 0 | 2.5 | 127 | Specific | 727.69 | 724.18 | 724.18 | 15.4 | 104 | >sp\|P01876\|IGHA1_HUMAN Immunoglobulin heavy constant alpha 1 OS=Homo sapiens OX=9606 GN=IGHA1 PE=1 SV=2 | 255 | 20201116_fqg_colostrum_1ng_glyco_hcd203040_compound15.21662.21662.4 | controllerType=0 controllerNumber=1 scan=21662 | [illegible] |
| 18229:5 | 4 | R.LSLHRPALEDLLLGSEAN[+1501.555]LTC[+57.021]TLTGLR.D | HexNAc(5)Hex(3) | C[+57], N[+1502] | 893.8388 | 5 | 4465.165 | 4465.1536 | 0 | 2.6 | 127 | Specific | 728.68 | 728.68 | 728.68 | 15.62 | 104 | >sp\|P01876\|IGHA1_HUMAN Immunoglobulin heavy constant alpha 1 OS=Homo sapiens OX=9606 GN=IGHA1 PE=1 SV=2 | 255 | 20201116_fqg_colostrum_1ng_glyco_hcd203040_compound15.21743.21743.5 | controllerType=0 controllerNumber=1 scan=21743 | [illegible] |
| 18217:5 | 4 | R.LSLHRPALEDLLLGSEAN[+1216.423]LTC[+57.021]TLTGLR.D | HexNAc(2)Hex(5) | C[+57], N[+1216] | 836.8141 | 5 | 4180.0414 | 4180.0211 | 0 | 4.9 | 127 | Specific | 737.33 | 737.33 | 737.33 | 15.07 | 104 | >sp\|P01876\|IGHA1_HUMAN Immunoglobulin heavy constant alpha 1 OS=Homo sapiens OX=9606 GN=IGHA1 PE=1 SV=2 | 255 | 20201116_fqg_colostrum_1ng_glyco_hcd203040_compound15.21730.21730.5 | controllerType=0 controllerNumber=1 scan=21730 | [illegible] |
| 19248:4 | 4 | R.LSLHRPALEDLLLGSEAN[+2116.756]LTC[+57.021]TLTGLR.D | HexNAc(5)Hex(5)NeuAc(1) | C[+57], N[+2117] | 1270.8464 | 4 | 5080.3639 | 5080.3547 | 0 | 1.8 | 127 | Specific | 745.58 | 664.44 | 664.44 | 15.4 | 104 | >sp\|P01876\|IGHA1_HUMAN Immunoglobulin heavy constant alpha 1 OS=Homo sapiens OX=9606 GN=IGHA1 PE=1 SV=2 | 255 | 20201116_fqg_colostrum_1ng_glyco_hcd203040_compound15.22803.22803.4 | controllerType=0 controllerNumber=1 scan=22803 | [illegible] |
| 18232:5 | 4 | R.LSLHRPALEDLLLGSEAN[+1501.555]LTC[+57.021]TLTGLR.D | HexNAc(5)Hex(3) | C[+57], N[+1502] | 893.8388 | 5 | 4465.165 | 4465.1536 | 0 | 2.6 | 127 | Specific | 748.93 | 748.93 | 748.93 | 15.62 | 104 | >sp\|P01876\|IGHA1_HUMAN Immunoglobulin heavy constant alpha 1 OS=Homo sapiens OX=9606 GN=IGHA1 PE=1 SV=2 | 255 | 20201116_fqg_colostrum_1ng_glyco_hcd203040_compound15.21746.21746.5 | controllerType=0 controllerNumber=1 scan=21746 | 42.4934 |
| 18210:3 | 4 | R.LSLHRPALEDLLLGSEAN[+1298.476]LTC[+57.021]TLTGLR.D | HexNAc(4)Hex(3) | C[+57], N[+1298] | 1421.3658 | 3 | 4262.0828 | 4262.0742 | 0 | 2 | 127 | Specific | 749.08 | 749.08 | 749.08 | 14.78 | 104 | >sp\|P01876\|IGHA1_HUMAN Immunoglobulin heavy constant alpha 1 OS=Homo sapiens OX=9606 GN=IGHA1 PE=1 SV=2 | 255 | 20201116_fqg_colostrum_1ng_glyco_hcd203040_compound15.21723.21723.3 | controllerType=0 controllerNumber=1 scan=21723 | 42.4432 |
| 19327:5 | 4 | R.LSLHRPALEDLLLGSEAN[+1872.651]LTC[+57.021]TLTGLR.D | HexNAc(3)Hex(6)NeuAc(1) | C[+57], N[+1873] | 968.0589 | 5 | 4836.2655 | 4836.2487 | 0 | 3.5 | 127 | Specific | 750.93 | 704.54 | 704.54 | 16.52 | 104 | >sp\|P01876\|IGHA1_HUMAN Immunoglobulin heavy constant alpha 1 OS=Homo sapiens OX=9606 GN=IGHA1 PE=1 SV=2 | 255 | 20201116_fqg_colostrum_1ng_glyco_hcd203040_compound15.22885.22885.5 | controllerType=0 controllerNumber=1 scan=22885 | 44.7683 |
| 17727:4 | 4 | R.LSLHRPALEDLLLGSEAN[+1501.555]LTC[+57.021]TLTGLR.D | HexNAc(5)Hex(3) | C[+57], N[+1502] | 1117.0447 | 4 | 4465.1571 | 4465.1536 | 0 | 0.8 | 127 | Specific | 753.86 | 511.4 | 511.4 | 16.17 | 104 | >sp\|P01876\|IGHA1_HUMAN Immunoglobulin heavy constant alpha 1 OS=Homo sapiens OX=9606 GN=IGHA1 PE=1 SV=2 | 255 | 20201116_fqg_colostrum_1ng_glyco_hcd203040_compound15.21220.21220.2 | controllerType=0 controllerNumber=1 scan=21220 | 41.4372 |
| 18352:5 | 4 | R.LSLHRPALEDLLLGSEAN[+1054.370]LTC[+57.021]TLTGLR.D | HexNAc(2)Hex(4) | C[+57], N[+1054] | 804.4003 | 5 | 4017.9723 | 4017.9683 | 0 | 1 | 127 | Specific | 760.29 | 675.43 | 675.43 | 16.39 | 104 | >sp\|P01876\|IGHA1_HUMAN Immunoglobulin heavy constant alpha 1 OS=Homo sapiens OX=9606 GN=IGHA1 PE=1 SV=2 | 255 | 20201116_fqg_colostrum_1ng_glyco_hcd203040_compound15.21870.21870.5 | controllerType=0 controllerNumber=1 scan=21870 | 42.7414 |
| 18137:3 | 4 | R.LSLHRPALEDLLLGSEAN[+1540.529]LTC[+57.021]TLTGLR.D | HexNAc(2)Hex(7) | C[+57], N[+1541] | 1502.0507 | 3 | 4504.1376 | 4504.1268 | 0 | 2.4 | 127 | Specific | 762.85 | 692.54 | 692.54 | 15.78 | 104 | >sp\|P01876\|IGHA1_HUMAN Immunoglobulin heavy constant alpha 1 OS=Homo sapiens OX=9606 GN=IGHA1 PE=1 SV=2 | 255 | 20201116_fqg_colostrum_1ng_glyco_hcd203040_compound15.21647.21647.3 | controllerType=0 controllerNumber=1 scan=21647 | 42.2833 |
| 18262:3 | 4 | R.LSLHRPALEDLLLGSEAN[+1054.370]LTC[+57.021]TLTGLR.D | HexNAc(2)Hex(4) | C[+57], N[+1054] | 1339.9958 | 3 | 4017.9728 | 4017.9683 | 0 | 1.1 | 127 | Specific | 765.04 | 765.04 | 765.04 | 15.54 | 104 | >sp\|P01876\|IGHA1_HUMAN Immunoglobulin heavy constant alpha 1 OS=Homo sapiens OX=9606 GN=IGHA1 PE=1 SV=2 | 255 | 20201116_fqg_colostrum_1ng_glyco_hcd203040_compound15.21777.21777.3 | controllerType=0 controllerNumber=1 scan=21777 | 42.5571 |
| 18100:4 | 4 | R.LSLHRPALEDLLLGSEAN[+1540.529]LTC[+57.021]TLTGLR.D | HexNAc(2)Hex(7) | C[+57], N[+1541] | 1126.7881 | 4 | 4504.1307 | 4504.1268 | 0 | 0.9 | 127 | Specific | 766.07 | 673.26 | 673.26 | 16.17 | 104 | >sp\|P01876\|IGHA1_HUMAN Immunoglobulin heavy constant alpha 1 OS=Homo sapiens OX=9606 GN=IGHA1 PE=1 SV=2 | 255 | 20201116_fqg_colostrum_1ng_glyco_hcd203040_compound15.21608.21608.2 | controllerType=0 controllerNumber=1 scan=21608 | 42.1991 |
| 19249:5 | 4 | R.LSLHRPALEDLLLGSEAN[+2116.756]LTC[+57.021]TLTGLR.D | HexNAc(5)Hex(5)NeuAc(1) | C[+57], N[+2117] | 1016.8779 | 5 | 5080.3606 | 5080.3547 | 0 | 1.2 | 127 | Specific | 766.55 | 739.65 | 739.65 | 16.39 | 104 | >sp\|P01876\|IGHA1_HUMAN Immunoglobulin heavy constant alpha 1 OS=Homo sapiens OX=9606 GN=IGHA1 PE=1 SV=2 | 255 | 20201116_fqg_colostrum_1ng_glyco_hcd203040_compound15.22804.22804.5 | controllerType=0 controllerNumber=1 scan=22804 | 44.5943 |
| 19272:5 | 4 | R.LSLHRPALEDLLLGSEAN[+1913.677]LTC[+57.021]TLTGLR.D | HexNAc(4)Hex(5)NeuAc(1) | C[+57], N[+1914] | 976.2617 | 5 | 4877.2796 | 4877.2753 | 0 | 0.9 | 127 | Specific | 772.26 | 772.26 | 772.26 | 16.39 | 104 | >sp\|P01876\|IGHA1_HUMAN Immunoglobulin heavy constant alpha 1 OS=Homo sapiens OX=9606 GN=IGHA1 PE=1 SV=2 | 255 | 20201116_fqg_colostrum_1ng_glyco_hcd203040_compound15.22828.22828.5 | controllerType=0 controllerNumber=1 scan=22828 | 44.6399 |
| 19320:4 | 4 | R.LSLHRPALEDLLLGSEAN[+1751.624]LTC[+57.021]TLTGLR.D | HexNAc(4)Hex(4)NeuAc(1) | C[+57], N[+1752] | 1179.5644 | 4 | 4715.2357 | 4715.2225 | 0 | 2.8 | 127 | Specific | 773.14 | 667.1 | 667.1 | 16.3 | 104 | >sp\|P01876\|IGHA1_HUMAN Immunoglobulin heavy constant alpha 1 OS=Homo sapiens OX=9606 GN=IGHA1 PE=1 SV=2 | 255 | 20201116_fqg_colostrum_1ng_glyco_hcd203040_compound15.22878.22878.4 | controllerType=0 controllerNumber=1 scan=22878 | 44.7557 |
| 18259:3 | 4 | R.LSLHRPALEDLLLGSEAN[+1054.370]LTC[+57.021]TLTGLR.D | HexNAc(2)Hex(4) | C[+57], N[+1054] | 1339.9958 | 3 | 4017.9728 | 4017.9683 | 0 | 1.1 | 127 | Specific | 778.56 | 737.64 | 737.64 | 15.54 | 104 | >sp\|P01876\|IGHA1_HUMAN Immunoglobulin heavy constant alpha 1 OS=Homo sapiens OX=9606 GN=IGHA1 PE=1 SV=2 | 255 | 20201116_fqg_colostrum_1ng_glyco_hcd203040_compound15.21774.21774.3 | controllerType=0 controllerNumber=1 scan=21774 | 42.5479 |
| 18175:4 | 4 | R.LSLHRPALEDLLLGSEAN[+1663.608]LTC[+57.021]TLTGLR.D | HexNAc(5)Hex(4) | C[+57], N[+1664] | 1157.5599 | 4 | 4627.2176 | 4627.2064 | 0 | 2.4 | 127 | Specific | 782.41 | 730.89 | 730.89 | 16.4 | 104 | >sp\|P01876\|IGHA1_HUMAN Immunoglobulin heavy constant alpha 1 OS=Homo sapiens OX=9606 GN=IGHA1 PE=1 SV=2 | 255 | 20201116_fqg_colostrum_1ng_glyco_hcd203040_compound15.21687.21687.4 | controllerType=0 controllerNumber=1 scan=21687 | 42.3754 |
| 19349:5 | 4 | R.LSLHRPALEDLLLGSEAN[+1751.624]LTC[+57.021]TLTGLR.D | HexNAc(4)Hex(4)NeuAc(1) | C[+57], N[+1752] | 943.8521 | 5 | 4715.2313 | 4715.2225 | 0 | 1.9 | 127 | Specific | 787.53 | 787.53 | 787.53 | 16.62 | 104 | >sp\|P01876\|IGHA1_HUMAN Immunoglobulin heavy constant alpha 1 OS=Homo sapiens OX=9606 GN=IGHA1 PE=1 SV=2 | 255 | 20201116_fqg_colostrum_1ng_glyco_hcd203040_compound15.22908.22908.5 | controllerType=0 controllerNumber=1 scan=22908 | 44.8075 |
| 18156:3 | 4 | R.LSLHRPALEDLLLGSEAN[+1378.476]LTC[+57.021]TLTGLR.D | HexNAc(2)Hex(6) | C[+57], N[+1378] | 1448.0335 | 3 | 4342.086 | 4342.074 | 0 | 2.8 | 127 | Specific | 788.74 | 733.73 | 733.73 | 15.68 | 104 | >sp\|P01876\|IGHA1_HUMAN Immunoglobulin heavy constant alpha 1 OS=Homo sapiens OX=9606 GN=IGHA1 PE=1 SV=2 | 255 | 20201116_fqg_colostrum_1ng_glyco_hcd203040_compound15.21667.21667.3 | controllerType=0 controllerNumber=1 scan=21667 | 42.3268 |
| 18161:3 | 4 | R.LSLHRPALEDLLLGSEAN[+1216.423]LTC[+57.021]TLTGLR.D | HexNAc(2)Hex(5) | C[+57], N[+1216] | 1394.0142 | 3 | 4180.0281 | 4180.0211 | 0 | 1.7 | 127 | Specific | 788.86 | 717.87 | 717.87 | 15.78 | 104 | >sp\|P01876\|IGHA1_HUMAN Immunoglobulin heavy constant alpha 1 OS=Homo sapiens OX=9606 GN=IGHA1 PE=1 SV=2 | 255 | 20201116_fqg_colostrum_1ng_glyco_hcd203040_compound15.21672.21672.3 | controllerType=0 controllerNumber=1 scan=21672 | 42.3404 |
| 18116:3 | 4 | R.LSLHRPALEDLLLGSEAN[+1702.581]LTC[+57.021]TLTGLR.D | HexNAc(2)Hex(8) | C[+57], N[+1703] | 1556.0681 | 3 | 4666.1896 | 4666.1796 | 0 | 2.1 | 127 | Specific | 790.22 | 790.22 | 790.22 | 15.78 | 104 | >sp\|P01876\|IGHA1_HUMAN Immunoglobulin heavy constant alpha 1 OS=Homo sapiens OX=9606 GN=IGHA1 PE=1 SV=2 | 255 | 20201116_fqg_colostrum_1ng_glyco_hcd203040_compound15.21625.21625.3 | controllerType=0 controllerNumber=1 scan=21625 | 42.2254 |
| 18133:4 | 4 | R.LSLHRPALEDLLLGSEAN[+1622.582]LTC[+57.021]TLTGLR.D | HexNAc(4)Hex(5) | C[+57], N[+1623] | 1147.3024 | 4 | 4586.1878 | 4586.1799 | 0 | 1.7 | 127 | Specific | 790.64 | 775.93 | 775.93 | 16.4 | 104 | >sp\|P01876\|IGHA1_HUMAN Immunoglobulin heavy constant alpha 1 OS=Homo sapiens OX=9606 GN=IGHA1 PE=1 SV=2 | 255 | 20201116_fqg_colostrum_1ng_glyco_hcd203040_compound15.21643.21643.2 | controllerType=0 controllerNumber=1 scan=21643 | 42.2745 |
| 19376:4 | 4 | R.LSLHRPALEDLLLGSEAN[+1548.545]LTC[+57.021]TLTGLR.D | HexNAc(3)Hex(4)NeuAc(1) | C[+57], N[+1549] | 1128.7945 | 4 | 4512.1561 | 4512.1431 | 0 | 2.9 | 127 | Specific | 790.73 | 790.73 | 790.73 | 16.3 | 104 | >sp\|P01876\|IGHA1_HUMAN Immunoglobulin heavy constant alpha 1 OS=Homo sapiens OX=9606 GN=IGHA1 PE=1 SV=2 | 255 | 20201116_fqg_colostrum_1ng_glyco_hcd203040_compound15.22936.22936.4 | controllerType=0 controllerNumber=1 scan=22936 | 44.8624 |
| 19224:4 | 4 | R.LSLHRPALEDLLLGSEAN[+1913.677]LTC[+57.021]TLTGLR.D | HexNAc(4)Hex(5)NeuAc(1) | C[+57], N[+1914] | 1220.0769 | 4 | 4877.286 | 4877.2753 | 0 | 2.2 | 127 | Specific | 793.92 | 761.67 | 761.67 | 16.4 | 104 | >sp\|P01876\|IGHA1_HUMAN Immunoglobulin heavy constant alpha 1 OS=Homo sapiens OX=9606 GN=IGHA1 PE=1 SV=2 | 255 | 20201116_fqg_colostrum_1ng_glyco_hcd203040_compound15.22778.22778.4 | controllerType=0 controllerNumber=1 scan=22778 | 44.5444 |
| 18253:4 | 4 | R.LSLHRPALEDLLLGSEAN[+1298.476]LTC[+57.021]TLTGLR.D | HexNAc(4)Hex(3) | C[+57], N[+1298] | 1066.2758 | 4 | 4262.0814 | 4262.0742 | 0 | 1.7 | 127 | Specific | 795.15 | 795.15 | 795.15 | 16.4 | 104 | >sp\|P01876\|IGHA1_HUMAN Immunoglobulin heavy constant alpha 1 OS=Homo sapiens OX=9606 GN=IGHA1 PE=1 SV=2 | 255 | 20201116_fqg_colostrum_1ng_glyco_hcd203040_compound15.21768.21768.4 | controllerType=0 controllerNumber=1 scan=21768 | 42.5347 |
| 18164:3 | 4 | R.LSLHRPALEDLLLGSEAN[+1216.423]LTC[+57.021]TLTGLR.D | HexNAc(2)Hex(5) | C[+57], N[+1216] | 1394.0142 | 3 | 4180.0281 | 4180.0211 | 0 | 1.7 | 127 | Specific | 801.55 | 766.82 | 766.82 | 16.78 | 104 | >sp\|P01876\|IGHA1_HUMAN Immunoglobulin heavy constant alpha 1 OS=Homo sapiens OX=9606 GN=IGHA1 PE=1 SV=2 | 255 | 20201116_fqg_colostrum_1ng_glyco_hcd203040_compound15.21675.21675.3 | controllerType=0 controllerNumber=1 scan=21675 | 42.3501 |
| 18336:3 | 4 | R.LSLHRPALEDLLLGSEAN[+892.317]LTC[+57.021]TLTGLR.D | HexNAc(2)Hex(3) | C[+57], N[+892] | 1285.9791 | 3 | 3855.9226 | 3855.9155 | 0 | 1.9 | 127 | Specific | 812.97 | 812.97 | 812.97 | 16.78 | 104 | >sp\|P01876\|IGHA1_HUMAN Immunoglobulin heavy constant alpha 1 OS=Homo sapiens OX=9606 GN=IGHA1 PE=1 SV=2 | 255 | 20201116_fqg_colostrum_1ng_glyco_hcd203040_compound15.21854.21854.3 | controllerType=0 controllerNumber=1 scan=21854 | 42.7118 |
| 18323:5 | 4 | R.LSLHRPALEDLLLGSEAN[+1298.476]LTC[+57.021]TLTGLR.D | HexNAc(4)Hex(3) | C[+57], N[+1298] | 853.2224 | 5 | 4262.0828 | 4262.0742 | 0 | 2 | 127 | Specific | 814.75 | 814.75 | 814.75 | 17.62 | 104 | >sp\|P01876\|IGHA1_HUMAN Immunoglobulin heavy constant alpha 1 OS=Homo sapiens OX=9606 GN=IGHA1 PE=1 SV=2 | 255 | 20201116_fqg_colostrum_1ng_glyco_hcd203040_compound15.21840.21840.5 | controllerType=0 controllerNumber=1 scan=21840 | 42.6818 |
| 18339:3 | 4 | R.LSLHRPALEDLLLGSEAN[+892.317]LTC[+57.021]TLTGLR.D | HexNAc(2)Hex(3) | C[+57], N[+892] | 1285.9791 | 3 | 3855.9226 | 3855.9155 | 0 | 1.9 | 127 | Specific | 826.64 | 826.64 | 826.64 | 16.78 | 104 | >sp\|P01876\|IGHA1_HUMAN Immunoglobulin heavy constant alpha 1 OS=Homo sapiens OX=9606 GN=IGHA1 PE=1 SV=2 | 255 | 20201116_fqg_colostrum_1ng_glyco_hcd203040_compound15.21857.21857.3 | controllerType=0 controllerNumber=1 scan=21857 | 42.7221 |
| 18169:4 | 4 | R.LSLHRPALEDLLLGSEAN[+1419.502]LTC[+57.021]TLTGLR.D | HexNAc(3)Hex(5) | C[+57], N[+1420] | 1096.5328 | 4 | 4383.1092 | 4383.1005 | 0 | 2 | 127 | Specific | 840.63 | 805.89 | 805.89 | 17.4 | 104 | >sp\|P01876\|IGHA1_HUMAN Immunoglobulin heavy constant alpha 1 OS=Homo sapiens OX=9606 GN=IGHA1 PE=1 SV=2 | 255 | 20201116_fqg_colostrum_1ng_glyco_hcd203040_compound15.21680.21680.4 | controllerType=0 controllerNumber=1 scan=21680 | 42.36 |
| 18226:4 | 4 | R.LSLHRPALEDLLLGSEAN[+1501.555]LTC[+57.021]TLTGLR.D | HexNAc(5)Hex(3) | C[+57], N[+1502] | 1117.0472 | 4 | 4465.1668 | 4465.1536 | 0 | 3 | 127 | Specific | 842.35 | 842.35 | 842.35 | 17.3 | 104 | >sp\|P01876\|IGHA1_HUMAN Immunoglobulin heavy constant alpha 1 OS=Homo sapiens OX=9606 GN=IGHA1 PE=1 SV=2 | 255 | 20201116_fqg_colostrum_1ng_glyco_hcd203040_compound15.21740.21740.4 | controllerType=0 controllerNumber=1 scan=21740 | 42.4834 |
| 19375:4 | 4 | R.LSLHRPALEDLLLGSEAN[+1710.598]LTC[+57.021]TLTGLR.D | HexNAc(3)Hex(5)NeuAc(1) | C[+57], N[+1711] | 1169.3078 | 4 | 4674.2093 | 4674.1959 | 0 | 2.9 | 127 | Specific | 845.45 | 807.05 | 807.05 | 17.3 | 104 | >sp\|P01876\|IGHA1_HUMAN Immunoglobulin heavy constant alpha 1 OS=Homo sapiens OX=9606 GN=IGHA1 PE=1 SV=2 | 255 | 20201116_fqg_colostrum_1ng_glyco_hcd203040_compound15.22935.22935.4 | controllerType=0 controllerNumber=1 scan=22935 | 44.8615 |
| 18331:4 | 4 | R.LSLHRPALEDLLLGSEAN[+892.317]LTC[+57.021]TLTGLR.D | HexNAc(2)Hex(3) | C[+57], N[+892] | 964.736 | 4 | 3855.9222 | 3855.9155 | 0 | 1.7 | 127 | Specific | 852.21 | 852.21 | 852.21 | 18.4 | 104 | >sp\|P01876\|IGHA1_HUMAN Immunoglobulin heavy constant alpha 1 OS=Homo sapiens OX=9606 GN=IGHA1 PE=1 SV=2 | 255 | 20201116_fqg_colostrum_1ng_glyco_hcd203040_compound15.21849.21849.4 | controllerType=0 controllerNumber=1 scan=21849 | 42.6983 |
| 19319:4 | 4 | R.LSLHRPALEDLLLGSEAN[+1872.651]LTC[+57.021]TLTGLR.D | HexNAc(3)Hex(6)NeuAc(1) | C[+57], N[+1873] | 1209.8217 | 4 | 4836.265 | 4836.2487 | 0 | 3.4 | 127 | Specific | 855.68 | 841.64 | 841.64 | 18.3 | 104 | >sp\|P01876\|IGHA1_HUMAN Immunoglobulin heavy constant alpha 1 OS=Homo sapiens OX=9606 GN=IGHA1 PE=1 SV=2 | 255 | 20201116_fqg_colostrum_1ng_glyco_hcd203040_compound15.22877.22877.4 | controllerType=0 controllerNumber=1 scan=22877 | 44.7534 |
| 18110:4 | 4 | R.LSLHRPALEDLLLGSEAN[+1702.581]LTC[+57.021]TLTGLR.D | HexNAc(2)Hex(8) | C[+57], N[+1703] | 1167.302 | 4 | 4666.1863 | 4666.1796 | 0 | 1.4 | 127 | Specific | 861.14 | 861.14 | 861.14 | 18.17 | 104 | >sp\|P01876\|IGHA1_HUMAN Immunoglobulin heavy constant alpha 1 OS=Homo sapiens OX=9606 GN=IGHA1 PE=1 SV=2 | 255 | 20201116_fqg_colostrum_1ng_glyco_hcd203040_compound15.21619.21619.4 | controllerType=0 controllerNumber=1 scan=21619 | 42.2165 |
| 18153:4 | 4 | R.LSLHRPALEDLLLGSEAN[+1622.582]LTC[+57.021]TLTGLR.D | HexNAc(4)Hex(5) | C[+57], N[+1623] | 1147.3033 | 4 | 4586.1912 | 4586.1799 | 0 | 2.5 | 127 | Specific | 864.57 | 814.14 | 814.14 | 18.4 | 104 | >sp\|P01876\|IGHA1_HUMAN Immunoglobulin heavy constant alpha 1 OS=Homo sapiens OX=9606 GN=IGHA1 PE=1 SV=2 | 255 | 20201116_fqg_colostrum_1ng_glyco_hcd203040_compound15.21664.21664.4 | controllerType=0 controllerNumber=1 scan=21664 | 42.3211 |
| 18152:4 | 4 | R.LSLHRPALEDLLLGSEAN[+1378.476]LTC[+57.021]TLTGLR.D | HexNAc(2)Hex(6) | C[+57], N[+1378] | 1086.2767 | 4 | 4342.0848 | 4342.074 | 0 | 2.5 | 127 | Specific | 867.44 | 789.49 | 789.49 | 18.4 | 104 | >sp\|P01876\|IGHA1_HUMAN Immunoglobulin heavy constant alpha 1 OS=Homo sapiens OX=9606 GN=IGHA1 PE=1 SV=2 | 255 | 20201116_fqg_colostrum_1ng_glyco_hcd203040_compound15.21663.21663.4 | controllerType=0 controllerNumber=1 scan=21663 | 42.3177 |
| 18203:4 | 4 | R.LSLHRPALEDLLLGSEAN[+1460.529]LTC[+57.021]TLTGLR.D | HexNAc(4)Hex(4) | C[+57], N[+1461] | 1106.7919 | 4 | 4424.1458 | 4424.1271 | 0 | 4.2 | 127 | Specific | 874.71 | 776.16 | 776.16 | 17.85 | 104 | >sp\|P01876\|IGHA1_HUMAN Immunoglobulin heavy constant alpha 1 OS=Homo sapiens OX=9606 GN=IGHA1 PE=1 SV=2 | 255 | 20201116_fqg_colostrum_1ng_glyco_hcd203040_compound15.21716.21716.4 | controllerType=0 controllerNumber=1 scan=21716 | 42.4293 |
| 18193:5 | 4 | R.LSLHRPALEDLLLGSEAN[+1378.476]LTC[+57.021]TLTGLR.D | HexNAc(2)Hex(6) | C[+57], N[+1378] | 869.2216 | 5 | 4342.0791 | 4342.074 | 0 | 1.2 | 127 | Specific | 878.36 | 594.74 | 594.74 | 18.39 | 104 | >sp\|P01876\|IGHA1_HUMAN Immunoglobulin heavy constant alpha 1 OS=Homo sapiens OX=9606 GN=IGHA1 PE=1 SV=2 | 255 | 20201116_fqg_colostrum_1ng_glyco_hcd203040_compound15.21705.21705.5 | controllerType=0 controllerNumber=1 scan=21705 | 42.4074 |
| 18142:4 | 4 | R.LSLHRPALEDLLLGSEAN[+1378.476]LTC[+57.021]TLTGLR.D | HexNAc(2)Hex(6) | C[+57], N[+1378] | 1086.2757 | 4 | 4342.0809 | 4342.074 | 0 | 1.6 | 127 | Specific | 915.16 | 822.12 | 822.12 | 19.4 | 104 | >sp\|P01876\|IGHA1_HUMAN Immunoglobulin heavy constant alpha 1 OS=Homo sapiens OX=9606 GN=IGHA1 PE=1 SV=2 | 255 | 20201116_fqg_colostrum_1ng_glyco_hcd203040_compound15.21652.21652.4 | controllerType=0 controllerNumber=1 scan=21652 | 42.2941 |
| 18135:4 | 4 | R.LSLHRPALEDLLLGSEAN[+1622.582]LTC[+57.021]TLTGLR.D | HexNAc(4)Hex(5) | C[+57], N[+1623] | 1147.3024 | 4 | 4586.1878 | 4586.1799 | 0 | 1.7 | 127 | Specific | 919.99 | 870.49 | 870.49 | 19.4 | 104 | >sp\|P01876\|IGHA1_HUMAN Immunoglobulin heavy constant alpha 1 OS=Homo sapiens OX=9606 GN=IGHA1 PE=1 SV=2 | 255 | 20201116_fqg_colostrum_1ng_glyco_hcd203040_compound15.21645.21645.2 | controllerType=0 controllerNumber=1 scan=21645 | 42.2815 |
| 18255:4 | 4 | R.LSLHRPALEDLLLGSEAN[+1298.476]LTC[+57.021]TLTGLR.D | HexNAc(4)Hex(3) | C[+57], N[+1298] | 1066.2758 | 4 | 4262.0814 | 4262.0742 | 0 | 1.7 | 127 | Specific | 923.72 | 862.19 | 862.19 | 19.4 | 104 | >sp\|P01876\|IGHA1_HUMAN Immunoglobulin heavy constant alpha 1 OS=Homo sapiens OX=9606 GN=IGHA1 PE=1 SV=2 | 255 | 20201116_fqg_colostrum_1ng_glyco_hcd203040_compound15.21770.21770.4 | controllerType=0 controllerNumber=1 scan=21770 | 42.5406 |
| 18115:4 | 4 | R.LSLHRPALEDLLLGSEAN[+1540.529]LTC[+57.021]TLTGLR.D | HexNAc(2)Hex(7) | C[+57], N[+1541] | 1126.7884 | 4 | 4504.1317 | 4504.1268 | 0 | 1.1 | 127 | Specific | 924.98 | 811.06 | 811.06 | 19.17 | 104 | >sp\|P01876\|IGHA1_HUMAN Immunoglobulin heavy constant alpha 1 OS=Homo sapiens OX=9606 GN=IGHA1 PE=1 SV=2 | 255 | 20201116_fqg_colostrum_1ng_glyco_hcd203040_compound15.21624.21624.4 | controllerType=0 controllerNumber=1 scan=21624 | 42.2242 |
| 18286:4 | 4 | R.LSLHRPALEDLLLGSEAN[+1257.449]LTC[+57.021]TLTGLR.D | HexNAc(3)Hex(4) | C[+57], N[+1257] | 1056.0191 | 4 | 4221.0545 | 4221.0477 | 0 | 1.6 | 127 | Specific | 926.62 | 909.47 | 909.47 | 19.17 | 104 | >sp\|P01876\|IGHA1_HUMAN Immunoglobulin heavy constant alpha 1 OS=Homo sapiens OX=9606 GN=IGHA1 PE=1 SV=2 | 255 | 20201116_fqg_colostrum_1ng_glyco_hcd203040_compound15.21802.21802.3 | controllerType=0 controllerNumber=1 scan=21802 | 42.6077 |
| 18212:4 | 4 | R.LSLHRPALEDLLLGSEAN[+1257.449]LTC[+57.021]TLTGLR.D | HexNAc(3)Hex(4) | C[+57], N[+1257] | 1056.0187 | 4 | 4221.053 | 4221.0477 | 0 | 1.3 | 127 | Specific | 927.85 | 907.15 | 907.15 | 19.17 | 104 | >sp\|P01876\|IGHA1_HUMAN Immunoglobulin heavy constant alpha 1 OS=Homo sapiens OX=9606 GN=IGHA1 PE=1 SV=2 | 255 | 20201116_fqg_colostrum_1ng_glyco_hcd203040_compound15.21725.21725.4 | controllerType=0 controllerNumber=1 scan=21725 | 42.4455 |
| 18155:4 | 4 | R.LSLHRPALEDLLLGSEAN[+1216.423]LTC[+57.021]TLTGLR.D | HexNAc(2)Hex(5) | C[+57], N[+1216] | 1045.7631 | 4 | 4180.0306 | 4180.0211 | 0 | 2.3 | 127 | Specific | 934.01 | 830.03 | 830.03 | 19.4 | 104 | >sp\|P01876\|IGHA1_HUMAN Immunoglobulin heavy constant alpha 1 OS=Homo sapiens OX=9606 GN=IGHA1 PE=1 SV=2 | 255 | 20201116_fqg_colostrum_1ng_glyco_hcd203040_compound15.21666.21666.4 | controllerType=0 controllerNumber=1 scan=21666 | 42.3256 |
| 19379:4 | 4 | R.LSLHRPALEDLLLGSEAN[+1548.545]LTC[+57.021]TLTGLR.D | HexNAc(3)Hex(4)NeuAc(1) | C[+57], N[+1549] | 1128.7945 | 4 | 4512.1561 | 4512.1431 | 0 | 2.9 | 127 | Specific | 935.29 | 918.22 | 918.22 | 19.3 | 104 | >sp\|P01876\|IGHA1_HUMAN Immunoglobulin heavy constant alpha 1 OS=Homo sapiens OX=9606 GN=IGHA1 PE=1 SV=2 | 255 | 20201116_fqg_colostrum_1ng_glyco_hcd203040_compound15.22939.22939.4 | controllerType=0 controllerNumber=1 scan=22939 | 44.8674 |
| 18170:4 | 4 | R.LSLHRPALEDLLLGSEAN[+1581.555]LTC[+57.021]TLTGLR.D | HexNAc(3)Hex(6) | C[+57], N[+1582] | 1137.0468 | 4 | 4545.1654 | 4545.1533 | 0 | 2.6 | 127 | Specific | 938 | 908.85 | 908.85 | 19.4 | 104 | >sp\|P01876\|IGHA1_HUMAN Immunoglobulin heavy constant alpha 1 OS=Homo sapiens OX=9606 GN=IGHA1 PE=1 SV=2 | 255 | 20201116_fqg_colostrum_1ng_glyco_hcd203040_compound15.21681.21681.4 | controllerType=0 controllerNumber=1 scan=21681 | 42.3616 |
| 18254:4 | 4 | R.LSLHRPALEDLLLGSEAN[+1054.370]LTC[+57.021]TLTGLR.D | HexNAc(2)Hex(4) | C[+57], N[+1054] | 1005.2494 | 4 | 4017.9759 | 4017.9683 | 0 | 1.9 | 127 | Specific | 939.66 | 877.03 | 877.03 | 19.4 | 104 | >sp\|P01876\|IGHA1_HUMAN Immunoglobulin heavy constant alpha 1 OS=Homo sapiens OX=9606 GN=IGHA1 PE=1 SV=2 | 255 | 20201116_fqg_colostrum_1ng_glyco_hcd203040_compound15.21769.21769.4 | controllerType=0 controllerNumber=1 scan=21769 | 42.5362 |
| 18220:5 | 4 | R.LSLHRPALEDLLLGSEAN[+1216.423]LTC[+57.021]TLTGLR.D | HexNAc(2)Hex(5) | C[+57], N[+1216] | 836.8141 | 5 | 4180.0414 | 4180.0211 | 0 | 4.9 | 127 | Specific | 950 | 692.32 | 692.32 | 20.07 | 104 | >sp\|P01876\|IGHA1_HUMAN Immunoglobulin heavy constant alpha 1 OS=Homo sapiens OX=9606 GN=IGHA1 PE=1 SV=2 | 255 | 20201116_fqg_colostrum_1ng_glyco_hcd203040_compound15.21733.21733.5 | controllerType=0 controllerNumber=1 scan=21733 | 42.4682 |
| 18172:4 | 4 | R.LSLHRPALEDLLLGSEAN[+1419.502]LTC[+57.021]TLTGLR.D | HexNAc(3)Hex(5) | C[+57], N[+1420] | 1096.5328 | 4 | 4383.1092 | 4383.1005 | 0 | 2 | 127 | Specific | 978.91 | 959.68 | 959.68 | 20.4 | 104 | >sp\|P01876\|IGHA1_HUMAN Immunoglobulin heavy constant alpha 1 OS=Homo sapiens OX=9606 GN=IGHA1 PE=1 SV=2 | 255 | 20201116_fqg_colostrum_1ng_glyco_hcd203040_compound15.21683.21683.4 | controllerType=0 controllerNumber=1 scan=21683 | 42.3678 |
| 18333:4 | 4 | R.LSLHRPALEDLLLGSEAN[+892.317]LTC[+57.021]TLTGLR.D | HexNAc(2)Hex(3) | C[+57], N[+892] | 964.736 | 4 | 3855.9222 | 3855.9155 | 0 | 1.7 | 127 | Specific | 989.45 | 989.45 | 989.45 | 20.4 | 104 | >sp\|P01876\|IGHA1_HUMAN Immunoglobulin heavy constant alpha 1 OS=Homo sapiens OX=9606 GN=IGHA1 PE=1 SV=2 | 255 | 20201116_fqg_colostrum_1ng_glyco_hcd203040_compound15.21851.21851.4 | controllerType=0 controllerNumber=1 scan=21851 | 42.7084 |
| 18158:4 | 4 | R.LSLHRPALEDLLLGSEAN[+1216.423]LTC[+57.021]TLTGLR.D | HexNAc(2)Hex(5) | C[+57], N[+1216] | 1045.7631 | 4 | 4180.0306 | 4180.0211 | 0 | 2.3 | 127 | Specific | 992.88 | 992.88 | 992.88 | 20.4 | 104 | >sp\|P01876\|IGHA1_HUMAN Immunoglobulin heavy constant alpha 1 OS=Homo sapiens OX=9606 GN=IGHA1 PE=1 SV=2 | 255 | 20201116_fqg_colostrum_1ng_glyco_hcd203040_compound15.21669.21669.4 | controllerType=0 controllerNumber=1 scan=21669 | 42.3334 |
| 18154:4 | 4 | R.LSLHRPALEDLLLGSEAN[+1378.476]LTC[+57.021]TLTGLR.D | HexNAc(2)Hex(6) | C[+57], N[+1378] | 1086.2767 | 4 | 4342.0848 | 4342.074 | 0 | 2.5 | 127 | Specific | 1009.35 | 1009.35 | 1009.35 | 21.4 | 104 | >sp\|P01876\|IGHA1_HUMAN Immunoglobulin heavy constant alpha 1 OS=Homo sapiens OX=9606 GN=IGHA1 PE=1 SV=2 | 255 | 20201116_fqg_colostrum_1ng_glyco_hcd203040_compound15.21665.21665.4 | controllerType=0 controllerNumber=1 scan=21665 | 42.3245 |
| 18118:4 | 4 | R.LSLHRPALEDLLLGSEAN[+1540.529]LTC[+57.021]TLTGLR.D | HexNAc(2)Hex(7) | C[+57], N[+1541] | 1126.7884 | 4 | 4504.1317 | 4504.1268 | 0 | 1.1 | 127 | Specific | 1019.77 | 1019.77 | 1019.77 | 21.17 | 104 | >sp\|P01876\|IGHA1_HUMAN Immunoglobulin heavy constant alpha 1 OS=Homo sapiens OX=9606 GN=IGHA1 PE=1 SV=2 | 255 | 20201116_fqg_colostrum_1ng_glyco_hcd203040_compound15.21627.21627.4 | controllerType=0 controllerNumber=1 scan=21627 | 42.2313 |
| 18256:4 | 4 | R.LSLHRPALEDLLLGSEAN[+1054.370]LTC[+57.021]TLTGLR.D | HexNAc(2)Hex(4) | C[+57], N[+1054] | 1005.2494 | 4 | 4017.9759 | 4017.9683 | 0 | 1.9 | 127 | Specific | 1022.52 | 1022.52 | 1022.52 | 21.4 | 104 | >sp\|P01876\|IGHA1_HUMAN Immunoglobulin heavy constant alpha 1 OS=Homo sapiens OX=9606 GN=IGHA1 PE=1 SV=2 | 255 | 20201116_fqg_colostrum_1ng_glyco_hcd203040_compound15.21771.21771.4 | controllerType=0 controllerNumber=1 scan=21771 | 42.5445 |
| 18279:4 | 4 | R.LSLHRPALEDLLLGSEAN[+1216.423]LTC[+57.021]TLTGLR.D | HexNAc(2)Hex(5) | C[+57], N[+1216] | 1045.763 | 4 | 4180.0301 | 4180.0211 | 0 | 2.1 | 127 | Specific | 1025.1 | 1017.22 | 1017.22 | 21.4 | 104 | >sp\|P01876\|IGHA1_HUMAN Immunoglobulin heavy constant alpha 1 OS=Homo sapiens OX=9606 GN=IGHA1 PE=1 SV=2 | 255 | 20201116_fqg_colostrum_1ng_glyco_hcd203040_compound15.21794.21794.2 | controllerType=0 controllerNumber=1 scan=21794 | 42.586 |
| 18145:4 | 4 | R.LSLHRPALEDLLLGSEAN[+1378.476]LTC[+57.021]TLTGLR.D | HexNAc(2)Hex(6) | C[+57], N[+1378] | 1086.2757 | 4 | 4342.0809 | 4342.074 | 0 | 1.6 | 127 | Specific | 1030.95 | 1030.95 | 1030.95 | 21.4 | 104 | >sp\|P01876\|IGHA1_HUMAN Immunoglobulin heavy constant alpha 1 OS=Homo sapiens OX=9606 GN=IGHA1 PE=1 SV=2 | 255 | 20201116_fqg_colostrum_1ng_glyco_hcd203040_compound15.21655.21655.4 | controllerType=0 controllerNumber=1 scan=21655 | 42.3029 |
| 18112:4 | 4 | R.LSLHRPALEDLLLGSEAN[+1702.581]LTC[+57.021]TLTGLR.D | HexNAc(2)Hex(8) | C[+57], N[+1703] | 1167.302 | 4 | 4666.1863 | 4666.1796 | 0 | 1.4 | 127 | Specific | 1048.48 | 1047.44 | 1047.44 | 21.17 | 104 | >sp\|P01876\|IGHA1_HUMAN Immunoglobulin heavy constant alpha 1 OS=Homo sapiens OX=9606 GN=IGHA1 PE=1 SV=2 | 255 | 20201116_fqg_colostrum_1ng_glyco_hcd203040_compound15.21621.21621.4 | controllerType=0 controllerNumber=1 scan=21621 | 42.2213 |
| 18176:4 | 4 | R.LSLHRPALEDLLLGSEAN[+1540.529]LTC[+57.021]TLTGLR.D | HexNAc(2)Hex(7) | C[+57], N[+1541] | 1126.7885 | 4 | 4504.1322 | 4504.1268 | 0 | 1.2 | 127 | Specific | 1056.3 | 955.49 | 955.49 | 22.17 | 104 | >sp\|P01876\|IGHA1_HUMAN Immunoglobulin heavy constant alpha 1 OS=Homo sapiens OX=9606 GN=IGHA1 PE=1 SV=2 | 255 | 20201116_fqg_colostrum_1ng_glyco_hcd203040_compound15.21688.21688.4 | controllerType=0 controllerNumber=1 scan=21688 | 42.3763 |
| 18289:4 | 4 | R.LSLHRPALEDLLLGSEAN[+1257.449]LTC[+57.021]TLTGLR.D | HexNAc(3)Hex(4) | C[+57], N[+1257] | 1056.0191 | 4 | 4221.0545 | 4221.0477 | 0 | 1.6 | 127 | Specific | 1067.31 | 1054.79 | 1054.79 | 22.17 | 104 | >sp\|P01876\|IGHA1_HUMAN Immunoglobulin heavy constant alpha 1 OS=Homo sapiens OX=9606 GN=IGHA1 PE=1 SV=2 | 255 | 20201116_fqg_colostrum_1ng_glyco_hcd203040_compound15.21805.21805.3 | controllerType=0 controllerNumber=1 scan=21805 | 42.6169 |
| 18134:4 | 4 | R.LSLHRPALEDLLLGSEAN[+1864.634]LTC[+57.021]TLTGLR.D | HexNAc(2)Hex(9) | C[+57], N[+1865] | 1207.8161 | 4 | 4828.2425 | 4828.2324 | 0 | 2.1 | 127 | Specific | 1067.37 | 1064.23 | 1064.23 | 22.4 | 104 | >sp\|P01876\|IGHA1_HUMAN Immunoglobulin heavy constant alpha 1 OS=Homo sapiens OX=9606 GN=IGHA1 PE=1 SV=2 | 255 | 20201116_fqg_colostrum_1ng_glyco_hcd203040_compound15.21644.21644.4 | controllerType=0 controllerNumber=1 scan=21644 | 42.2768 |
| 18215:4 | 4 | R.LSLHRPALEDLLLGSEAN[+1257.449]LTC[+57.021]TLTGLR.D | HexNAc(3)Hex(4) | C[+57], N[+1257] | 1056.0187 | 4 | 4221.053 | 4221.0477 | 0 | 1.3 | 127 | Specific | 1089.34 | 1064.28 | 1064.28 | 22.17 | 104 | >sp\|P01876\|IGHA1_HUMAN Immunoglobulin heavy constant alpha 1 OS=Homo sapiens OX=9606 GN=IGHA1 PE=1 SV=2 | 255 | 20201116_fqg_colostrum_1ng_glyco_hcd203040_compound15.21728.21728.4 | controllerType=0 controllerNumber=1 scan=21728 | 42.4561 |
| 10204:3 | 4 | R.LAGKPTHVN[+1444.534]VSVVMAEVDGTC[+57.021].Y | HexNAc(4)Hex(3)Fuc(1) | C[+57], N[+1445] | 1210.2128 | 3 | 3628.6239 | 3628.6126 | 0 | 3.1 | 332 | CRagged | 248.37 | 178.83 | 178.83 | 4.55 | 104 | >sp\|P01876\|IGHA1_HUMAN Immunoglobulin heavy constant alpha 1 OS=Homo sapiens OX=9606 GN=IGHA1 PE=1 SV=2 | 255 | 20201116_fqg_colostrum_1ng_glyco_hcd203040_compound15.13437.13437.3 | controllerType=0 controllerNumber=1 scan=13437 | 28.758 |
| 08686:3 | 4 | R.LAGKPTHVN[+1054.370]VSVVMAEVDGTC[+57.021].Y | HexNAc(2)Hex(4) | C[+57], N[+1054] | 1080.1576 | 3 | 3238.4584 | 3238.4488 | 0 | 3 | 332 | CRagged | 278.56 | 278.56 | 114.19 | 7.52 | 104 | >sp\|P01876\|IGHA1_HUMAN Immunoglobulin heavy constant alpha 1 OS=Homo sapiens OX=9606 GN=IGHA1 PE=1 SV=2 | 255 | 20201116_fqg_colostrum_1ng_glyco_hcd203040_compound15.11864.11864.3 | controllerType=0 controllerNumber=1 scan=11864 | 26.1104 |
| 08654:3 | 4 | R.LAGKPTHVN[+1444.534]VSVVMAEVDGTC[+57.021].Y | HexNAc(4)Hex(3)Fuc(1) | C[+57], N[+1445] | 1210.2093 | 3 | 3628.6133 | 3628.6126 | 0 | 0.2 | 332 | CRagged | 437.07 | 437.07 | 437.07 | 9.49 | 104 | >sp\|P01876\|IGHA1_HUMAN Immunoglobulin heavy constant alpha 1 OS=Homo sapiens OX=9606 GN=IGHA1 PE=1 SV=2 | 255 | 20201116_fqg_colostrum_1ng_glyco_hcd203040_compound15.11831.11831.3 | controllerType=0 controllerNumber=1 scan=11831 | 26.0568 |
| 10131:3 | 4 | R.LAGKPTHVN[+1216.423]VSVVMAEVDGTC[+57.021].Y | HexNAc(2)Hex(5) | C[+57], N[+1216] | 1134.1754 | 3 | 3400.5118 | 3400.5016 | 0 | 3 | 332 | CRagged | 494.46 | 454.03 | 271.78 | 10.41 | 104 | >sp\|P01876\|IGHA1_HUMAN Immunoglobulin heavy constant alpha 1 OS=Homo sapiens OX=9606 GN=IGHA1 PE=1 SV=2 | 255 | 20201116_fqg_colostrum_1ng_glyco_hcd203040_compound15.13362.13362.3 | controllerType=0 controllerNumber=1 scan=13362 | 28.614 |
| 08596:3 | 4 | R.LAGKPTHVN[+1216.423]VSVVMAEVDGTC[+57.021].Y | HexNAc(2)Hex(5) | C[+57], N[+1216] | 1134.1747 | 3 | 3400.5096 | 3400.5016 | 0 | 2.3 | 332 | CRagged | 515.8 | 308.19 | 308.19 | 8.99 | 104 | >sp\|P01876\|IGHA1_HUMAN Immunoglobulin heavy constant alpha 1 OS=Homo sapiens OX=9606 GN=IGHA1 PE=1 SV=2 | 255 | 20201116_fqg_colostrum_1ng_glyco_hcd203040_compound15.11771.11771.3 | controllerType=0 controllerNumber=1 scan=11771 | 25.9523 |
| 10056:3 | 4 | R.LAGKPTHVN[+1809.666]VSVVMAEVDGTC[+57.021].Y | HexNAc(5)Hex(4)Fuc(1) | C[+57], N[+1810] | 1331.9267 | 3 | 3993.7655 | 3993.7448 | 0 | 5.2 | 332 | CRagged | 520.54 | 376.92 | 222.26 | 8.72 | 104 | >sp\|P01876\|IGHA1_HUMAN Immunoglobulin heavy constant alpha 1 OS=Homo sapiens OX=9606 GN=IGHA1 PE=1 SV=2 | 255 | 20201116_fqg_colostrum_1ng_glyco_hcd203040_compound15.13284.13284.3 | controllerType=0 controllerNumber=1 scan=13284 | 28.4666 |
| 10059:3 | 4 | R.LAGKPTHVN[+1809.666]VSVVMAEVDGTC[+57.021].Y | HexNAc(5)Hex(4)Fuc(1) | C[+57], N[+1810] | 1331.9267 | 3 | 3993.7655 | 3993.7448 | 0 | 5.2 | 332 | CRagged | 535.97 | 381.86 | 287.16 | 9.98 | 104 | >sp\|P01876\|IGHA1_HUMAN Immunoglobulin heavy constant alpha 1 OS=Homo sapiens OX=9606 GN=IGHA1 PE=1 SV=2 | 255 | 20201116_fqg_colostrum_1ng_glyco_hcd203040_compound15.13287.13287.3 | controllerType=0 controllerNumber=1 scan=13287 | 28.4715 |
| 08594:3 | 4 | R.LAGKPTHVN[+1216.423]VSVVMAEVDGTC[+57.021].Y | HexNAc(2)Hex(5) | C[+57], N[+1216] | 1134.1747 | 3 | 3400.5096 | 3400.5016 | 0 | 2.3 | 332 | CRagged | 567.09 | 510.36 | 390.6 | 11.03 | 104 | >sp\|P01876\|IGHA1_HUMAN Immunoglobulin heavy constant alpha 1 OS=Homo sapiens OX=9606 GN=IGHA1 PE=1 SV=2 | 255 | 20201116_fqg_colostrum_1ng_glyco_hcd203040_compound15.11769.11769.3 | controllerType=0 controllerNumber=1 scan=11769 | 25.9446 |
| 10107:3 | 4 | R.LAGKPTHVN[+1606.587]VSVVMAEVDGTC[+57.021].Y | HexNAc(4)Hex(4)Fuc(1) | C[+57], N[+1607] | 1264.2291 | 3 | 3790.6726 | 3790.6654 | 0 | 1.9 | 332 | CRagged | 618.92 | 437.61 | 437.61 | 13.42 | 104 | >sp\|P01876\|IGHA1_HUMAN Immunoglobulin heavy constant alpha 1 OS=Homo sapiens OX=9606 GN=IGHA1 PE=1 SV=2 | 255 | 20201116_fqg_colostrum_1ng_glyco_hcd203040_compound15.13337.13337.3 | controllerType=0 controllerNumber=1 scan=13337 | 28.5655 |

Table S4. Cont.

| Query #:z | Protein Rank | Glycopeptide | Glycan | Modification Type(s) | Observed m/z | z | Observed (M+H) | Calc. mass (M+H) | Off-by-x error | Mass error (ppm) | StartPosition | Cleavage | Score | Delta | Delta Mod | \|Log Prob\| | # of unique peptides | Protein Name | Protein DB number | Comment | Scan # | Scan Time |
|---|---|---|---|---|---|---|---|---|---|---|---|---|---|---|---|---|---|---|---|---|---|---|
| 10134:3 | 4 | R.LAGKPTHVN[+1216.423]VSVVMAEVDGTC[+57.021].Y | HexNAc(2)Hex(5) | C[+57], N[+1216] | 1134.1754 | 3 | 3400.5118 | 3400.5016 | 0 | 3 | 332 | CRagged | 619.71 | 532.42 | 315.53 | 13.31 | 104 | >sp\|P01876\|IGHA1_HUMAN Immunoglobulin heavy constant alpha 1 OS=Homo sapiens OX=9606 GN=IGHA1 PE=1 SV=2 | 255 | 20201116_fqg_colostrum_1ng_glyco_hcd203040_compound15.13365.13365.3 | controllerType=0 controllerNumber=1 scan=13365 | 28.6209 |
| 08849:3 | 4 | R.LAGKPTHVN[+1216.423]VSVVMAEVDGTC[+57.021].Y | HexNAc(2)Hex(5) | C[+57], N[+1216] | 1134.1756 | 3 | 3400.5121 | 3400.5016 | 0 | 3.1 | 332 | CRagged | 636.51 | 561.52 | 381.38 | 13.31 | 104 | >sp\|P01876\|IGHA1_HUMAN Immunoglobulin heavy constant alpha 1 OS=Homo sapiens OX=9606 GN=IGHA1 PE=1 SV=2 | 255 | 20201116_fqg_colostrum_1ng_glyco_hcd203040_compound15.12033.12033.3 | controllerType=0 controllerNumber=1 scan=12033 | 26.4017 |
| 08852:3 | 4 | R.LAGKPTHVN[+1216.423]VSVVMAEVDGTC[+57.021].Y | HexNAc(2)Hex(5) | C[+57], N[+1216] | 1134.1756 | 3 | 3400.5121 | 3400.5016 | 0 | 3.1 | 332 | CRagged | 674.74 | 647.41 | 438.01 | 14.31 | 104 | >sp\|P01876\|IGHA1_HUMAN Immunoglobulin heavy constant alpha 1 OS=Homo sapiens OX=9606 GN=IGHA1 PE=1 SV=2 | 255 | 20201116_fqg_colostrum_1ng_glyco_hcd203040_compound15.12036.12036.3 | controllerType=0 controllerNumber=1 scan=12036 | 26.4077 |
| 08689:3 | 4 | R.LAGKPTHVN[+1054.370]VSVVMAEVDGTC[+57.021].Y | HexNAc(2)Hex(4) | C[+57], N[+1054] | 1080.1576 | 3 | 3238.4584 | 3238.4488 | 0 | 3 | 332 | CRagged | 695.66 | 576.77 | 425.62 | 14.31 | 104 | >sp\|P01876\|IGHA1_HUMAN Immunoglobulin heavy constant alpha 1 OS=Homo sapiens OX=9606 GN=IGHA1 PE=1 SV=2 | 255 | 20201116_fqg_colostrum_1ng_glyco_hcd203040_compound15.11867.11867.3 | controllerType=0 controllerNumber=1 scan=11867 | 26.1148 |
| 08657:3 | 4 | R.LAGKPTHVN[+1444.534]VSVVMAEVDGTC[+57.021].Y | HexNAc(4)Hex(3)Fuc(1) | C[+57], N[+1445] | 1210.2093 | 3 | 3628.6133 | 3628.6126 | 0 | 0.2 | 332 | CRagged | 713.6 | 568.92 | 568.92 | 14.68 | 104 | >sp\|P01876\|IGHA1_HUMAN Immunoglobulin heavy constant alpha 1 OS=Homo sapiens OX=9606 GN=IGHA1 PE=1 SV=2 | 255 | 20201116_fqg_colostrum_1ng_glyco_hcd203040_compound15.11834.11834.3 | controllerType=0 controllerNumber=1 scan=11834 | 26.0609 |
| 13189:4 | 4 | R.LAGKPTHVN[+2262.814]VSVVMAEVDGTC[+57.021]Y.- | HexNAc(5)Hex(5)Fuc(1)NeuAc(1) | C[+57], N[+2263] | 1153.2481 | 4 | 4609.9705 | 4609.9564 | 0 | 3.1 | 332 | CRagged | 213.16 | 213.16 | 213.16 | 5.78 | 104 | >sp\|P01876\|IGHA1_HUMAN Immunoglobulin heavy constant alpha 1 OS=Homo sapiens OX=9606 GN=IGHA1 PE=1 SV=2 | 255 | 20201116_fqg_colostrum_1ng_glyco_hcd203040_compound15.16520.16520.4 | controllerType=0 controllerNumber=1 scan=16520 | 33.5529 |
| 10203:3 | 4 | R.LAGKPTHVN[+892.317]VSVVMAEVDGTC[+57.021]Y.- | HexNAc(2)Hex(3) | C[+57], N[+892] | 1080.4901 | 3 | 3239.4559 | 3239.4593 | 0 | -1.1 | 332 | CRagged | 342.46 | 342.46 | 218.04 | 5.38 | 104 | >sp\|P01876\|IGHA1_HUMAN Immunoglobulin heavy constant alpha 1 OS=Homo sapiens OX=9606 GN=IGHA1 PE=1 SV=2 | 255 | 20201116_fqg_colostrum_1ng_glyco_hcd203040_compound15.13436.13436.3 | controllerType=0 controllerNumber=1 scan=13436 | 28.7552 |
| 11348:3 | 4 | R.LAGKPTHVN[+1054.370]VSVVMAEVDGTC[+57.021]Y.- | HexNAc(2)Hex(4) | C[+57], N[+1054] | 1134.5114 | 3 | 3401.5196 | 3401.5121 | 0 | 2.2 | 332 | CRagged | 344.05 | 344.05 | 186.72 | 8.46 | 104 | >sp\|P01876\|IGHA1_HUMAN Immunoglobulin heavy constant alpha 1 OS=Homo sapiens OX=9606 GN=IGHA1 PE=1 SV=2 | 255 | 20201116_fqg_colostrum_1ng_glyco_hcd203040_compound15.14618.14618.3 | controllerType=0 controllerNumber=1 scan=14618 | 30.6372 |
| 11160:3 | 4 | R.LAGKPTHVN[+1663.608]VSVVMAEVDGTC[+57.021]Y.- | HexNAc(5)Hex(4) | C[+57], N[+1664] | 1337.5898 | 3 | 4010.7548 | 4010.7502 | 0 | 1.1 | 332 | CRagged | 402.21 | 402.21 | 290.8 | 9.84 | 104 | >sp\|P01876\|IGHA1_HUMAN Immunoglobulin heavy constant alpha 1 OS=Homo sapiens OX=9606 GN=IGHA1 PE=1 SV=2 | 255 | 20201116_fqg_colostrum_1ng_glyco_hcd203040_compound15.14424.14424.3 | controllerType=0 controllerNumber=1 scan=14424 | 30.3043 |
| 11155:3 | 4 | R.LAGKPTHVN[+1825.661]VSVVMAEVDGTC[+57.021]Y.- | HexNAc(5)Hex(5) | C[+57], N[+1826] | 1391.6075 | 3 | 4172.8079 | 4172.8031 | 0 | 1.2 | 332 | CRagged | 405.85 | 398.3 | 244.64 | 9.84 | 104 | >sp\|P01876\|IGHA1_HUMAN Immunoglobulin heavy constant alpha 1 OS=Homo sapiens OX=9606 GN=IGHA1 PE=1 SV=2 | 255 | 20201116_fqg_colostrum_1ng_glyco_hcd203040_compound15.14418.14418.3 | controllerType=0 controllerNumber=1 scan=14418 | 30.2939 |
| 15579:3 | 4 | R.LAGKPTHVN[+2204.772]VSVVMAEVDGTC[+57.021]Y.- | HexNAc(4)Hex(5)NeuAc(2) | C[+57], N[+2205] | 1517.9843 | 3 | 4551.9383 | 4551.9145 | 0 | 5.2 | 332 | CRagged | 487.98 | 363.97 | 363.97 | 8.31 | 104 | >sp\|P01876\|IGHA1_HUMAN Immunoglobulin heavy constant alpha 1 OS=Homo sapiens OX=9606 GN=IGHA1 PE=1 SV=2 | 255 | 20201116_fqg_colostrum_1ng_glyco_hcd203040_compound15.18994.18994.3 | controllerType=0 controllerNumber=1 scan=18994 | 37.5494 |
| 15576:3 | 4 | R.LAGKPTHVN[+2204.772]VSVVMAEVDGTC[+57.021]Y.- | HexNAc(4)Hex(5)NeuAc(2) | C[+57], N[+2205] | 1517.9843 | 3 | 4551.9383 | 4551.9145 | 0 | 5.2 | 332 | CRagged | 502.42 | 342.27 | 342.27 | 8.26 | 104 | >sp\|P01876\|IGHA1_HUMAN Immunoglobulin heavy constant alpha 1 OS=Homo sapiens OX=9606 GN=IGHA1 PE=1 SV=2 | 255 | 20201116_fqg_colostrum_1ng_glyco_hcd203040_compound15.18991.18991.3 | controllerType=0 controllerNumber=1 scan=18991 | 37.5378 |
| 15418:3 | 4 | R.LAGKPTHVN[+2350.830]VSVVMAEVDGTC[+57.021]Y.- | HexNAc(4)Hex(5)Fuc(1)NeuAc(2) | C[+57], N[+2351] | 1566.6667 | 3 | 4697.9855 | 4697.9724 | 0 | 2.8 | 332 | CRagged | 520.84 | 276.9 | 276.9 | 8.63 | 104 | >sp\|P01876\|IGHA1_HUMAN Immunoglobulin heavy constant alpha 1 OS=Homo sapiens OX=9606 GN=IGHA1 PE=1 SV=2 | 255 | 20201116_fqg_colostrum_1ng_glyco_hcd203040_compound15.18826.18826.3 | controllerType=0 controllerNumber=1 scan=18826 | 37.2891 |
| 13258:3 | 4 | R.LAGKPTHVN[+2262.814]VSVVMAEVDGTC[+57.021]Y.- | HexNAc(5)Hex(5)Fuc(1)NeuAc(1) | C[+57], N[+2263] | 1537.3237 | 3 | 4609.9564 | 4609.9564 | 0 | 0 | 332 | CRagged | 522.56 | 522.56 | 522.56 | 9.83 | 104 | >sp\|P01876\|IGHA1_HUMAN Immunoglobulin heavy constant alpha 1 OS=Homo sapiens OX=9606 GN=IGHA1 PE=1 SV=2 | 255 | 20201116_fqg_colostrum_1ng_glyco_hcd203040_compound15.16591.16591.3 | controllerType=0 controllerNumber=1 scan=16591 | 33.6556 |
| 09984:3 | 4 | R.LAGKPTHVN[+1809.666]VSVVMAEVDGTC[+57.021]Y.- | HexNAc(5)Hex(4)Fuc(1) | C[+57], N[+1810] | 1386.2779 | 3 | 4156.8191 | 4156.8081 | 0 | 2.6 | 332 | CRagged | 560.78 | 560.78 | 362.18 | 10.86 | 104 | >sp\|P01876\|IGHA1_HUMAN Immunoglobulin heavy constant alpha 1 OS=Homo sapiens OX=9606 GN=IGHA1 PE=1 SV=2 | 255 | 20201116_fqg_colostrum_1ng_glyco_hcd203040_compound15.13209.13209.3 | controllerType=0 controllerNumber=1 scan=13209 | 28.3138 |
| 11200:3 | 4 | R.LAGKPTHVN[+1606.587]VSVVMAEVDGTC[+57.021]Y.- | HexNAc(4)Hex(4)Fuc(1) | C[+57], N[+1607] | 1318.5818 | 3 | 3953.731 | 3953.7288 | 0 | 0.6 | 332 | CRagged | 568.71 | 568.71 | 568.71 | 10.62 | 104 | >sp\|P01876\|IGHA1_HUMAN Immunoglobulin heavy constant alpha 1 OS=Homo sapiens OX=9606 GN=IGHA1 PE=1 SV=2 | 255 | 20201116_fqg_colostrum_1ng_glyco_hcd203040_compound15.14465.14465.3 | controllerType=0 controllerNumber=1 scan=14465 | 30.3679 |
| 09985:3 | 4 | R.LAGKPTHVN[+1606.587]VSVVMAEVDGTC[+57.021]Y.- | HexNAc(4)Hex(4)Fuc(1) | C[+57], N[+1607] | 1318.5826 | 3 | 3953.7332 | 3953.7288 | 0 | 1.1 | 332 | CRagged | 578.68 | 556.25 | 556.25 | 10.62 | 104 | >sp\|P01876\|IGHA1_HUMAN Immunoglobulin heavy constant alpha 1 OS=Homo sapiens OX=9606 GN=IGHA1 PE=1 SV=2 | 255 | 20201116_fqg_colostrum_1ng_glyco_hcd203040_compound15.13210.13210.3 | controllerType=0 controllerNumber=1 scan=13210 | 28.3149 |
| 13192:4 | 4 | R.LAGKPTHVN[+2262.814]VSVVMAEVDGTC[+57.021]Y.- | HexNAc(5)Hex(5)Fuc(1)NeuAc(1) | C[+57], N[+2263] | 1153.2481 | 4 | 4609.9705 | 4609.9564 | 0 | 3.1 | 332 | CRagged | 578.94 | 550.13 | 550.13 | 10.97 | 104 | >sp\|P01876\|IGHA1_HUMAN Immunoglobulin heavy constant alpha 1 OS=Homo sapiens OX=9606 GN=IGHA1 PE=1 SV=2 | 255 | 20201116_fqg_colostrum_1ng_glyco_hcd203040_compound15.16523.16523.4 | controllerType=0 controllerNumber=1 scan=16523 | 33.56 |
| 13243:3 | 4 | R.LAGKPTHVN[+2059.735]VSVVMAEVDGTC[+57.021]Y.- | HexNAc(4)Hex(5)Fuc(1)NeuAc(1) | C[+57], N[+2060] | 1469.638 | 3 | 4406.8994 | 4406.877 | 0 | 5.1 | 332 | CRagged | 602.71 | 601.4 | 601.4 | 12.47 | 104 | >sp\|P01876\|IGHA1_HUMAN Immunoglobulin heavy constant alpha 1 OS=Homo sapiens OX=9606 GN=IGHA1 PE=1 SV=2 | 255 | 20201116_fqg_colostrum_1ng_glyco_hcd203040_compound15.16575.16575.3 | controllerType=0 controllerNumber=1 scan=16575 | 33.6303 |
| 11165:3 | 4 | R.LAGKPTHVN[+1809.666]VSVVMAEVDGTC[+57.021]Y.- | HexNAc(5)Hex(4)Fuc(1) | C[+57], N[+1810] | 1386.2751 | 3 | 4156.8107 | 4156.8081 | 0 | 0.6 | 332 | CRagged | 604.3 | 604.3 | 379.99 | 13 | 104 | >sp\|P01876\|IGHA1_HUMAN Immunoglobulin heavy constant alpha 1 OS=Homo sapiens OX=9606 GN=IGHA1 PE=1 SV=2 | 255 | 20201116_fqg_colostrum_1ng_glyco_hcd203040_compound15.14429.14429.3 | controllerType=0 controllerNumber=1 scan=14429 | 30.3102 |
| 11175:2 | 4 | R.LAGKPTHVN[+1378.476]VSVVMAEVDGTC[+57.021]Y.- | HexNAc(2)Hex(6) | C[+57], N[+1378] | 1863.3127 | 2 | 3725.6181 | 3725.6178 | 0 | 0.1 | 332 | CRagged | 639.92 | 639.92 | 306.92 | 11.88 | 104 | >sp\|P01876\|IGHA1_HUMAN Immunoglobulin heavy constant alpha 1 OS=Homo sapiens OX=9606 GN=IGHA1 PE=1 SV=2 | 255 | 20201116_fqg_colostrum_1ng_glyco_hcd203040_compound15.14439.14439.2 | controllerType=0 controllerNumber=1 scan=14439 | 30.3256 |
| 11242:2 | 4 | R.LAGKPTHVN[+1216.423]VSVVMAEVDGTC[+57.021]Y.- | HexNAc(2)Hex(5) | C[+57], N[+1216] | 1782.2917 | 2 | 3563.5761 | 3563.5649 | 0 | 3.1 | 332 | CRagged | 648.89 | 648.89 | 306.3 | 12.52 | 104 | >sp\|P01876\|IGHA1_HUMAN Immunoglobulin heavy constant alpha 1 OS=Homo sapiens OX=9606 GN=IGHA1 PE=1 SV=2 | 255 | 20201116_fqg_colostrum_1ng_glyco_hcd203040_compound15.14508.14508.2 | controllerType=0 controllerNumber=1 scan=14508 | 30.4307 |
| 13185:3 | 4 | R.LAGKPTHVN[+2262.814]VSVVMAEVDGTC[+57.021]Y.- | HexNAc(5)Hex(5)Fuc(1)NeuAc(1) | C[+57], N[+2263] | 1537.3281 | 3 | 4609.9696 | 4609.9564 | 0 | 2.9 | 332 | CRagged | 665.5 | 465.29 | 465.29 | 14.14 | 104 | >sp\|P01876\|IGHA1_HUMAN Immunoglobulin heavy constant alpha 1 OS=Homo sapiens OX=9606 GN=IGHA1 PE=1 SV=2 | 255 | 20201116_fqg_colostrum_1ng_glyco_hcd203040_compound15.16516.16516.3 | controllerType=0 controllerNumber=1 scan=16516 | 33.5459 |
| 11386:3 | 4 | R.LAGKPTHVN[+1216.423]VSVVMAEVDGTC[+57.021]Y.- | HexNAc(2)Hex(5) | C[+57], N[+1216] | 1188.5267 | 3 | 3563.5655 | 3563.5649 | 0 | 0.1 | 332 | CRagged | 665.77 | 610.35 | 354.28 | 13.5 | 104 | >sp\|P01876\|IGHA1_HUMAN Immunoglobulin heavy constant alpha 1 OS=Homo sapiens OX=9606 GN=IGHA1 PE=1 SV=2 | 255 | 20201116_fqg_colostrum_1ng_glyco_hcd203040_compound15.14657.14657.3 | controllerType=0 controllerNumber=1 scan=14657 | 30.7039 |
| 11167:3 | 4 | R.LAGKPTHVN[+1216.423]VSVVMAEVDGTC[+57.021]Y.- | HexNAc(2)Hex(5) | C[+57], N[+1216] | 1188.5278 | 3 | 3563.5687 | 3563.5649 | 0 | 1.1 | 332 | CRagged | 667.89 | 667.89 | 322.69 | 14 | 104 | >sp\|P01876\|IGHA1_HUMAN Immunoglobulin heavy constant alpha 1 OS=Homo sapiens OX=9606 GN=IGHA1 PE=1 SV=2 | 255 | 20201116_fqg_colostrum_1ng_glyco_hcd203040_compound15.14431.14431.3 | controllerType=0 controllerNumber=1 scan=14431 | 30.3125 |
| 15421:3 | 4 | R.LAGKPTHVN[+2350.830]VSVVMAEVDGTC[+57.021]Y.- | HexNAc(4)Hex(5)Fuc(1)NeuAc(2) | C[+57], N[+2351] | 1566.6667 | 3 | 4697.9855 | 4697.9724 | 0 | 2.8 | 332 | CRagged | 672.47 | 584.04 | 584.04 | 14.24 | 104 | >sp\|P01876\|IGHA1_HUMAN Immunoglobulin heavy constant alpha 1 OS=Homo sapiens OX=9606 GN=IGHA1 PE=1 SV=2 | 255 | 20201116_fqg_colostrum_1ng_glyco_hcd203040_compound15.18829.18829.3 | controllerType=0 controllerNumber=1 scan=18829 | 37.2936 |
| 11311:3 | 4 | R.LAGKPTHVN[+1444.534]VSVVMAEVDGTC[+57.021]Y.- | HexNAc(4)Hex(3)Fuc(1) | C[+57], N[+1445] | 1264.565 | 3 | 3791.6804 | 3791.6759 | 0 | 1.2 | 332 | CRagged | 672.67 | 672.67 | 672.67 | 14 | 104 | >sp\|P01876\|IGHA1_HUMAN Immunoglobulin heavy constant alpha 1 OS=Homo sapiens OX=9606 GN=IGHA1 PE=1 SV=2 | 255 | 20201116_fqg_colostrum_1ng_glyco_hcd203040_compound15.14580.14580.3 | controllerType=0 controllerNumber=1 scan=14580 | 30.5704 |
| 13187:3 | 4 | R.LAGKPTHVN[+2262.814]VSVVMAEVDGTC[+57.021]Y.- | HexNAc(5)Hex(5)Fuc(1)NeuAc(1) | C[+57], N[+2263] | 1537.3281 | 3 | 4609.9696 | 4609.9564 | 0 | 2.9 | 332 | CRagged | 678.52 | 548.16 | 548.16 | 14.14 | 104 | >sp\|P01876\|IGHA1_HUMAN Immunoglobulin heavy constant alpha 1 OS=Homo sapiens OX=9606 GN=IGHA1 PE=1 SV=2 | 255 | 20201116_fqg_colostrum_1ng_glyco_hcd203040_compound15.16518.16518.3 | controllerType=0 controllerNumber=1 scan=16518 | 33.5507 |
| 10052:3 | 4 | R.LAGKPTHVN[+1501.555]VSVVMAEVDGTC[+57.021]Y.- | HexNAc(5)Hex(3) | C[+57], N[+1502] | 1283.5728 | 3 | 3848.7039 | 3848.6974 | 0 | 1.7 | 332 | CRagged | 692.97 | 625.34 | 625.34 | 14.24 | 104 | >sp\|P01876\|IGHA1_HUMAN Immunoglobulin heavy constant alpha 1 OS=Homo sapiens OX=9606 GN=IGHA1 PE=1 SV=2 | 255 | 20201116_fqg_colostrum_1ng_glyco_hcd203040_compound15.13280.13280.3 | controllerType=0 controllerNumber=1 scan=13280 | 28.4604 |
| 09946:3 | 4 | R.LAGKPTHVN[+1663.608]VSVVMAEVDGTC[+57.021]Y.- | HexNAc(5)Hex(4) | C[+57], N[+1664] | 1337.5904 | 3 | 4010.7566 | 4010.7502 | 0 | 1.6 | 332 | CRagged | 698.35 | 681.64 | 432.59 | 14 | 104 | >sp\|P01876\|IGHA1_HUMAN Immunoglobulin heavy constant alpha 1 OS=Homo sapiens OX=9606 GN=IGHA1 PE=1 SV=2 | 255 | 20201116_fqg_colostrum_1ng_glyco_hcd203040_compound15.13170.13170.3 | controllerType=0 controllerNumber=1 scan=13170 | 28.2376 |
| 11132:3 | 4 | R.LAGKPTHVN[+1663.608]VSVVMAEVDGTC[+57.021]Y.- | HexNAc(5)Hex(4) | C[+57], N[+1664] | 1337.5886 | 3 | 4010.7511 | 4010.7502 | 0 | 0.2 | 332 | CRagged | 712.31 | 712.31 | 421.34 | 14.5 | 104 | >sp\|P01876\|IGHA1_HUMAN Immunoglobulin heavy constant alpha 1 OS=Homo sapiens OX=9606 GN=IGHA1 PE=1 SV=2 | 255 | 20201116_fqg_colostrum_1ng_glyco_hcd203040_compound15.14395.14395.3 | controllerType=0 controllerNumber=1 scan=14395 | 30.2596 |
| 11365:3 | 4 | R.LAGKPTHVN[+1501.555]VSVVMAEVDGTC[+57.021]Y.- | HexNAc(5)Hex(3) | C[+57], N[+1502] | 1283.5762 | 3 | 3848.7141 | 3848.6974 | 0 | 4.3 | 332 | CRagged | 713.1 | 581.77 | 581.77 | 14.69 | 104 | >sp\|P01876\|IGHA1_HUMAN Immunoglobulin heavy constant alpha 1 OS=Homo sapiens OX=9606 GN=IGHA1 PE=1 SV=2 | 255 | 20201116_fqg_colostrum_1ng_glyco_hcd203040_compound15.14635.14635.3 | controllerType=0 controllerNumber=1 scan=14635 | 30.6634 |
| 13261:3 | 4 | R.LAGKPTHVN[+2262.814]VSVVMAEVDGTC[+57.021]Y.- | HexNAc(5)Hex(5)Fuc(1)NeuAc(1) | C[+57], N[+2263] | 1537.3237 | 3 | 4609.9564 | 4609.9564 | 0 | 0 | 332 | CRagged | 716.43 | 566.02 | 566.02 | 14.5 | 104 | >sp\|P01876\|IGHA1_HUMAN Immunoglobulin heavy constant alpha 1 OS=Homo sapiens OX=9606 GN=IGHA1 PE=1 SV=2 | 255 | 20201116_fqg_colostrum_1ng_glyco_hcd203040_compound15.16594.16594.3 | controllerType=0 controllerNumber=1 scan=16594 | 33.6596 |
| 11287:3 | 4 | R.LAGKPTHVN[+1647.613]VSVVMAEVDGTC[+57.021]Y.- | HexNAc(5)Hex(3)Fuc(1) | C[+57], N[+1648] | 1332.2592 | 3 | 3994.7631 | 3994.7553 | 0 | 1.9 | 332 | CRagged | 721.46 | 719.53 | 719.53 | 15.24 | 104 | >sp\|P01876\|IGHA1_HUMAN Immunoglobulin heavy constant alpha 1 OS=Homo sapiens OX=9606 GN=IGHA1 PE=1 SV=2 | 255 | 20201116_fqg_colostrum_1ng_glyco_hcd203040_compound15.14555.14555.3 | controllerType=0 controllerNumber=1 scan=14555 | 30.5153 |
| 13246:3 | 4 | R.LAGKPTHVN[+2059.735]VSVVMAEVDGTC[+57.021]Y.- | HexNAc(4)Hex(5)Fuc(1)NeuAc(1) | C[+57], N[+2060] | 1469.638 | 3 | 4406.8994 | 4406.877 | 0 | 5.1 | 332 | CRagged | 722.6 | 623.31 | 623.31 | 14.47 | 104 | >sp\|P01876\|IGHA1_HUMAN Immunoglobulin heavy constant alpha 1 OS=Homo sapiens OX=9606 GN=IGHA1 PE=1 SV=2 | 255 | 20201116_fqg_colostrum_1ng_glyco_hcd203040_compound15.16578.16578.3 | controllerType=0 controllerNumber=1 scan=16578 | 33.6347 |
| 11235:3 | 4 | R.LAGKPTHVN[+1501.555]VSVVMAEVDGTC[+57.021]Y.- | HexNAc(5)Hex(3) | C[+57], N[+1502] | 1283.5733 | 3 | 3848.7053 | 3848.6974 | 0 | 2.1 | 332 | CRagged | 725.48 | 725.48 | 725.48 | 15.24 | 104 | >sp\|P01876\|IGHA1_HUMAN Immunoglobulin heavy constant alpha 1 OS=Homo sapiens OX=9606 GN=IGHA1 PE=1 SV=2 | 255 | 20201116_fqg_colostrum_1ng_glyco_hcd203040_compound15.14501.14501.3 | controllerType=0 controllerNumber=1 scan=14501 | 30.4211 |
| 11134:3 | 4 | R.LAGKPTHVN[+1540.529]VSVVMAEVDGTC[+57.021]Y.- | HexNAc(2)Hex(7) | C[+57], N[+1541] | 1296.5626 | 3 | 3887.6731 | 3887.6706 | 0 | 0.7 | 332 | CRagged | 732.33 | 732.33 | 403.68 | 15 | 104 | >sp\|P01876\|IGHA1_HUMAN Immunoglobulin heavy constant alpha 1 OS=Homo sapiens OX=9606 GN=IGHA1 PE=1 SV=2 | 255 | 20201116_fqg_colostrum_1ng_glyco_hcd203040_compound15.14397.14397.3 | controllerType=0 controllerNumber=1 scan=14397 | 30.2619 |
| 10055:3 | 4 | R.LAGKPTHVN[+1501.555]VSVVMAEVDGTC[+57.021]Y.- | HexNAc(5)Hex(3) | C[+57], N[+1502] | 1283.5728 | 3 | 3848.7039 | 3848.6974 | 0 | 1.7 | 332 | CRagged | 732.56 | 626.9 | 626.9 | 15.24 | 104 | >sp\|P01876\|IGHA1_HUMAN Immunoglobulin heavy constant alpha 1 OS=Homo sapiens OX=9606 GN=IGHA1 PE=1 SV=2 | 255 | 20201116_fqg_colostrum_1ng_glyco_hcd203040_compound15.13283.13283.3 | controllerType=0 controllerNumber=1 scan=13283 | 28.4651 |
| 11178:2 | 4 | R.LAGKPTHVN[+1378.476]VSVVMAEVDGTC[+57.021]Y.- | HexNAc(2)Hex(6) | C[+57], N[+1378] | 1863.3127 | 2 | 3725.6181 | 3725.6178 | 0 | 0.1 | 332 | CRagged | 736.55 | 649.97 | 268.44 | 13.88 | 104 | >sp\|P01876\|IGHA1_HUMAN Immunoglobulin heavy constant alpha 1 OS=Homo sapiens OX=9606 GN=IGHA1 PE=1 SV=2 | 255 | 20201116_fqg_colostrum_1ng_glyco_hcd203040_compound15.14442.14442.2 | controllerType=0 controllerNumber=1 scan=14442 | 30.3294 |
| 09949:3 | 4 | R.LAGKPTHVN[+1663.608]VSVVMAEVDGTC[+57.021]Y.- | HexNAc(5)Hex(4) | C[+57], N[+1664] | 1337.5904 | 3 | 4010.7566 | 4010.7502 | 0 | 1.6 | 332 | CRagged | 752.89 | 587.76 | 466.34 | 16 | 104 | >sp\|P01876\|IGHA1_HUMAN Immunoglobulin heavy constant alpha 1 OS=Homo sapiens OX=9606 GN=IGHA1 PE=1 SV=2 | 255 | 20201116_fqg_colostrum_1ng_glyco_hcd203040_compound15.13173.13173.3 | controllerType=0 controllerNumber=1 scan=13173 | 28.2426 |
| 11163:3 | 4 | R.LAGKPTHVN[+1663.608]VSVVMAEVDGTC[+57.021]Y.- | HexNAc(5)Hex(4) | C[+57], N[+1664] | 1337.5898 | 3 | 4010.7548 | 4010.7502 | 0 | 1.1 | 332 | CRagged | 761.93 | 595.59 | 493.2 | 16 | 104 | >sp\|P01876\|IGHA1_HUMAN Immunoglobulin heavy constant alpha 1 OS=Homo sapiens OX=9606 GN=IGHA1 PE=1 SV=2 | 255 | 20201116_fqg_colostrum_1ng_glyco_hcd203040_compound15.14427.14427.3 | controllerType=0 controllerNumber=1 scan=14427 | 30.3081 |
| 09971:3 | 4 | R.LAGKPTHVN[+1378.476]VSVVMAEVDGTC[+57.021]Y.- | HexNAc(2)Hex(6) | C[+57], N[+1378] | 1242.5475 | 3 | 3725.6281 | 3725.6178 | 0 | 2.8 | 332 | CRagged | 769.92 | 706.07 | 443.46 | 16.14 | 104 | >sp\|P01876\|IGHA1_HUMAN Immunoglobulin heavy constant alpha 1 OS=Homo sapiens OX=9606 GN=IGHA1 PE=1 SV=2 | 255 | 20201116_fqg_colostrum_1ng_glyco_hcd203040_compound15.13196.13196.3 | controllerType=0 controllerNumber=1 scan=13196 | 28.2846 |
| 13332:3 | 4 | R.LAGKPTHVN[+1710.598]VSVVMAEVDGTC[+57.021]Y.- | HexNAc(3)Hex(5)NeuAc(1) | C[+57], N[+1711] | 1353.2558 | 3 | 4057.7528 | 4057.7397 | 0 | 3.2 | 332 | CRagged | 771.32 | 670.06 | 405.33 | 16.14 | 104 | >sp\|P01876\|IGHA1_HUMAN Immunoglobulin heavy constant alpha 1 OS=Homo sapiens OX=9606 GN=IGHA1 PE=1 SV=2 | 255 | 20201116_fqg_colostrum_1ng_glyco_hcd203040_compound15.16667.16667.3 | controllerType=0 controllerNumber=1 scan=16667 | 33.7783 |
| 11168:3 | 4 | R.LAGKPTHVN[+1809.666]VSVVMAEVDGTC[+57.021]Y.- | HexNAc(5)Hex(4)Fuc(1) | C[+57], N[+1810] | 1386.2751 | 3 | 4156.8107 | 4156.8081 | 0 | 0.6 | 332 | CRagged | 775.12 | 748.92 | 483.1 | 16 | 104 | >sp\|P01876\|IGHA1_HUMAN Immunoglobulin heavy constant alpha 1 OS=Homo sapiens OX=9606 GN=IGHA1 PE=1 SV=2 | 255 | 20201116_fqg_colostrum_1ng_glyco_hcd203040_compound15.14432.14432.3 | controllerType=0 controllerNumber=1 scan=14432 | 30.3146 |
| 11170:3 | 4 | R.LAGKPTHVN[+1216.423]VSVVMAEVDGTC[+57.021]Y.- | HexNAc(2)Hex(5) | C[+57], N[+1216] | 1188.5278 | 3 | 3563.5687 | 3563.5649 | 0 | 1.1 | 332 | CRagged | 777.95 | 668.43 | 444.2 | 16 | 104 | >sp\|P01876\|IGHA1_HUMAN Immunoglobulin heavy constant alpha 1 OS=Homo sapiens OX=9606 GN=IGHA1 PE=1 SV=2 | 255 | 20201116_fqg_colostrum_1ng_glyco_hcd203040_compound15.14434.14434.3 | controllerType=0 controllerNumber=1 scan=14434 | 30.3199 |
| 13334:3 | 4 | R.LAGKPTHVN[+1710.598]VSVVMAEVDGTC[+57.021]Y.- | HexNAc(3)Hex(5)NeuAc(1) | C[+57], N[+1711] | 1353.2558 | 3 | 4057.7528 | 4057.7397 | 0 | 3.2 | 332 | CRagged | 785.41 | 716.37 | 461.62 | 16.14 | 104 | >sp\|P01876\|IGHA1_HUMAN Immunoglobulin heavy constant alpha 1 OS=Homo sapiens OX=9606 GN=IGHA1 PE=1 SV=2 | 255 | 20201116_fqg_colostrum_1ng_glyco_hcd203040_compound15.16669.16669.3 | controllerType=0 controllerNumber=1 scan=16669 | 33.7848 |
| 11245:2 | 4 | R.LAGKPTHVN[+1216.423]VSVVMAEVDGTC[+57.021]Y.- | HexNAc(2)Hex(5) | C[+57], N[+1216] | 1782.2917 | 2 | 3563.5761 | 3563.5649 | 0 | 3.1 | 332 | CRagged | 794.27 | 709.07 | 286.92 | 15.52 | 104 | >sp\|P01876\|IGHA1_HUMAN Immunoglobulin heavy constant alpha 1 OS=Homo sapiens OX=9606 GN=IGHA1 PE=1 SV=2 | 255 | 20201116_fqg_colostrum_1ng_glyco_hcd203040_compound15.14511.14511.2 | controllerType=0 controllerNumber=1 scan=14511 | 30.4359 |
| 11389:3 | 4 | R.LAGKPTHVN[+1216.423]VSVVMAEVDGTC[+57.021]Y.- | HexNAc(2)Hex(5) | C[+57], N[+1216] | 1188.5267 | 3 | 3563.5655 | 3563.5649 | 0 | 0.1 | 332 | CRagged | 801.45 | 767.77 | 492.88 | 16.5 | 104 | >sp\|P01876\|IGHA1_HUMAN Immunoglobulin heavy constant alpha 1 OS=Homo sapiens OX=9606 GN=IGHA1 PE=1 SV=2 | 255 | 20201116_fqg_colostrum_1ng_glyco_hcd203040_compound15.14660.14660.3 | controllerType=0 controllerNumber=1 scan=14660 | 30.7091 |
| 11203:3 | 4 | R.LAGKPTHVN[+1606.587]VSVVMAEVDGTC[+57.021]Y.- | HexNAc(4)Hex(4)Fuc(1) | C[+57], N[+1607] | 1318.5818 | 3 | 3953.731 | 3953.7288 | 0 | 0.6 | 332 | CRagged | 802.99 | 795.8 | 795.8 | 17 | 104 | >sp\|P01876\|IGHA1_HUMAN Immunoglobulin heavy constant alpha 1 OS=Homo sapiens OX=9606 GN=IGHA1 PE=1 SV=2 | 255 | 20201116_fqg_colostrum_1ng_glyco_hcd203040_compound15.14468.14468.3 | controllerType=0 controllerNumber=1 scan=14468 | 30.3724 |
| 11238:3 | 4 | R.LAGKPTHVN[+1501.555]VSVVMAEVDGTC[+57.021]Y.- | HexNAc(5)Hex(3) | C[+57], N[+1502] | 1283.5733 | 3 | 3848.7053 | 3848.6974 | 0 | 2.1 | 332 | CRagged | 806.7 | 806.7 | 806.7 | 17.24 | 104 | >sp\|P01876\|IGHA1_HUMAN Immunoglobulin heavy constant alpha 1 OS=Homo sapiens OX=9606 GN=IGHA1 PE=1 SV=2 | 255 | 20201116_fqg_colostrum_1ng_glyco_hcd203040_compound15.14504.14504.3 | controllerType=0 controllerNumber=1 scan=14504 | 30.4264 |
| 09987:3 | 4 | R.LAGKPTHVN[+1809.666]VSVVMAEVDGTC[+57.021]Y.- | HexNAc(5)Hex(4)Fuc(1) | C[+57], N[+1810] | 1386.2779 | 3 | 4156.8191 | 4156.8081 | 0 | 2.6 | 332 | CRagged | 819.11 | 761.58 | 541.64 | 17.24 | 104 | >sp\|P01876\|IGHA1_HUMAN Immunoglobulin heavy constant alpha 1 OS=Homo sapiens OX=9606 GN=IGHA1 PE=1 SV=2 | 255 | 20201116_fqg_colostrum_1ng_glyco_hcd203040_compound15.13212.13212.3 | controllerType=0 controllerNumber=1 scan=13212 | 28.3179 |
| 09988:3 | 4 | R.LAGKPTHVN[+1606.587]VSVVMAEVDGTC[+57.021]Y.- | HexNAc(4)Hex(4)Fuc(1) | C[+57], N[+1607] | 1318.5826 | 3 | 3953.7332 | 3953.7288 | 0 | 1.1 | 332 | CRagged | 824.93 | 657.48 | 657.48 | 17 | 104 | >sp\|P01876\|IGHA1_HUMAN Immunoglobulin heavy constant alpha 1 OS=Homo sapiens OX=9606 GN=IGHA1 PE=1 SV=2 | 255 | 20201116_fqg_colostrum_1ng_glyco_hcd203040_compound15.13213.13213.3 | controllerType=0 controllerNumber=1 scan=13213 | 28.3199 |
| 09964:3 | 4 | R.LAGKPTHVN[+1216.423]VSVVMAEVDGTC[+57.021]Y.- | HexNAc(2)Hex(5) | C[+57], N[+1216] | 1188.5286 | 3 | 3563.5713 | 3563.5649 | 0 | 1.8 | 332 | CRagged | 826.47 | 789.8 | 405.53 | 17.24 | 104 | >sp\|P01876\|IGHA1_HUMAN Immunoglobulin heavy constant alpha 1 OS=Homo sapiens OX=9606 GN=IGHA1 PE=1 SV=2 | 255 | 20201116_fqg_colostrum_1ng_glyco_hcd203040_compound15.13188.13188.2 | controllerType=0 controllerNumber=1 scan=13188 | 28.2674 |
| 11257:3 | 4 | R.LAGKPTHVN[+1647.613]VSVVMAEVDGTC[+57.021]Y.- | HexNAc(5)Hex(3)Fuc(1) | C[+57], N[+1648] | 1332.2597 | 3 | 3994.7645 | 3994.7553 | 0 | 2.3 | 332 | CRagged | 828.63 | 828.63 | 828.63 | 17.24 | 104 | >sp\|P01876\|IGHA1_HUMAN Immunoglobulin heavy constant alpha 1 OS=Homo sapiens OX=9606 GN=IGHA1 PE=1 SV=2 | 255 | 20201116_fqg_colostrum_1ng_glyco_hcd203040_compound15.14524.14524.3 | controllerType=0 controllerNumber=1 scan=14524 | 30.4624 |
| 11351:3 | 4 | R.LAGKPTHVN[+1054.370]VSVVMAEVDGTC[+57.021]Y.- | HexNAc(2)Hex(4) | C[+57], N[+1054] | 1134.5114 | 3 | 3401.5196 | 3401.5121 | 0 | 2.2 | 332 | CRagged | 834.55 | 689.77 | 468.87 | 17.24 | 104 | >sp\|P01876\|IGHA1_HUMAN Immunoglobulin heavy constant alpha 1 OS=Homo sapiens OX=9606 GN=IGHA1 PE=1 SV=2 | 255 | 20201116_fqg_colostrum_1ng_glyco_hcd203040_compound15.14621.14621.3 | controllerType=0 controllerNumber=1 scan=14621 | 30.6413 |
| 11129:3 | 4 | R.LAGKPTHVN[+1378.476]VSVVMAEVDGTC[+57.021]Y.- | HexNAc(2)Hex(6) | C[+57], N[+1378] | 1242.545 | 3 | 3725.6204 | 3725.6178 | 0 | 0.7 | 332 | CRagged | 846.57 | 846.57 | 511.1 | 17 | 104 | >sp\|P01876\|IGHA1_HUMAN Immunoglobulin heavy constant alpha 1 OS=Homo sapiens OX=9606 GN=IGHA1 PE=1 SV=2 | 255 | 20201116_fqg_colostrum_1ng_glyco_hcd203040_compound15.14392.14392.3 | controllerType=0 controllerNumber=1 scan=14392 | 30.2522 |
| 11135:3 | 4 | R.LAGKPTHVN[+1663.608]VSVVMAEVDGTC[+57.021]Y.- | HexNAc(5)Hex(4) | C[+57], N[+1664] | 1337.5886 | 3 | 4010.7511 | 4010.7502 | 0 | 0.2 | 332 | CRagged | 847.5 | 845.76 | 528.14 | 16.5 | 104 | >sp\|P01876\|IGHA1_HUMAN Immunoglobulin heavy constant alpha 1 OS=Homo sapiens OX=9606 GN=IGHA1 PE=1 SV=2 | 255 | 20201116_fqg_colostrum_1ng_glyco_hcd203040_compound15.14398.14398.3 | controllerType=0 controllerNumber=1 scan=14398 | 30.2653 |
| 09993:3 | 4 | R.LAGKPTHVN[+1216.423]VSVVMAEVDGTC[+57.021]Y.- | HexNAc(2)Hex(5) | C[+57], N[+1216] | 1188.5294 | 3 | 3563.5735 | 3563.5649 | 0 | 2.4 | 332 | CRagged | 862.04 | 852.72 | 523.89 | 18.24 | 104 | >sp\|P01876\|IGHA1_HUMAN Immunoglobulin heavy constant alpha 1 OS=Homo sapiens OX=9606 GN=IGHA1 PE=1 SV=2 | 255 | 20201116_fqg_colostrum_1ng_glyco_hcd203040_compound15.13218.13218.3 | controllerType=0 controllerNumber=1 scan=13218 | 28.3316 |
| 11131:3 | 4 | R.LAGKPTHVN[+1378.476]VSVVMAEVDGTC[+57.021]Y.- | HexNAc(2)Hex(6) | C[+57], N[+1378] | 1242.545 | 3 | 3725.6204 | 3725.6178 | 0 | 0.7 | 332 | CRagged | 869.27 | 849.12 | 528.71 | 18 | 104 | >sp\|P01876\|IGHA1_HUMAN Immunoglobulin heavy constant alpha 1 OS=Homo sapiens OX=9606 GN=IGHA1 PE=1 SV=2 | 255 | 20201116_fqg_colostrum_1ng_glyco_hcd203040_compound15.14394.14394.3 | controllerType=0 controllerNumber=1 scan=14394 | 30.2589 |
| 11313:3 | 4 | R.LAGKPTHVN[+1444.534]VSVVMAEVDGTC[+57.021]Y.- | HexNAc(4)Hex(3)Fuc(1) | C[+57], N[+1445] | 1264.565 | 3 | 3791.6804 | 3791.6759 | 0 | 1.2 | 332 | CRagged | 870.71 | 749.06 | 749.06 | 18 | 104 | >sp\|P01876\|IGHA1_HUMAN Immunoglobulin heavy constant alpha 1 OS=Homo sapiens OX=9606 GN=IGHA1 PE=1 SV=2 | 255 | 20201116_fqg_colostrum_1ng_glyco_hcd203040_compound15.14582.14582.3 | controllerType=0 controllerNumber=1 scan=14582 | 30.5743 |
| 09990:3 | 4 | R.LAGKPTHVN[+1216.423]VSVVMAEVDGTC[+57.021]Y.- | HexNAc(2)Hex(5) | C[+57], N[+1216] | 1188.5294 | 3 | 3563.5735 | 3563.5649 | 0 | 2.4 | 332 | CRagged | 878.34 | 878.34 | 492.08 | 18.24 | 104 | >sp\|P01876\|IGHA1_HUMAN Immunoglobulin heavy constant alpha 1 OS=Homo sapiens OX=9606 GN=IGHA1 PE=1 SV=2 | 255 | 20201116_fqg_colostrum_1ng_glyco_hcd203040_compound15.13215.13215.3 | controllerType=0 controllerNumber=1 scan=13215 | 28.3229 |
| 11289:3 | 4 | R.LAGKPTHVN[+1647.613]VSVVMAEVDGTC[+57.021]Y.- | HexNAc(5)Hex(3)Fuc(1) | C[+57], N[+1648] | 1332.2592 | 3 | 3994.7631 | 3994.7553 | 0 | 1.9 | 332 | CRagged | 878.77 | 577.06 | 577.06 | 18.24 | 104 | >sp\|P01876\|IGHA1_HUMAN Immunoglobulin heavy constant alpha 1 OS=Homo sapiens OX=9606 GN=IGHA1 PE=1 SV=2 | 255 | 20201116_fqg_colostrum_1ng_glyco_hcd203040_compound15.14557.14557.3 | controllerType=0 controllerNumber=1 scan=14557 | 30.5197 |
| 11259:3 | 4 | R.LAGKPTHVN[+1647.613]VSVVMAEVDGTC[+57.021]Y.- | HexNAc(5)Hex(3)Fuc(1) | C[+57], N[+1648] | 1332.2597 | 3 | 3994.7645 | 3994.7553 | 0 | 2.3 | 332 | CRagged | 879.23 | 834.71 | 834.71 | 18.24 | 104 | >sp\|P01876\|IGHA1_HUMAN Immunoglobulin heavy constant alpha 1 OS=Homo sapiens OX=9606 GN=IGHA1 PE=1 SV=2 | 255 | 20201116_fqg_colostrum_1ng_glyco_hcd203040_compound15.14526.14526.3 | controllerType=0 controllerNumber=1 scan=14526 | 30.4679 |
| 09967:3 | 4 | R.LAGKPTHVN[+1216.423]VSVVMAEVDGTC[+57.021]Y.- | HexNAc(2)Hex(5) | C[+57], N[+1216] | 1188.5286 | 3 | 3563.5713 | 3563.5649 | 0 | 1.8 | 332 | CRagged | 880.1 | 855.82 | 532.35 | 18.24 | 104 | >sp\|P01876\|IGHA1_HUMAN Immunoglobulin heavy constant alpha 1 OS=Homo sapiens OX=9606 GN=IGHA1 PE=1 SV=2 | 255 | 20201116_fqg_colostrum_1ng_glyco_hcd203040_compound15.13191.13191.2 | controllerType=0 controllerNumber=1 scan=13191 | [illegible] |
| 11112:3 | 4 | R.LAGKPTHVN[+1702.581]VSVVMAEVDGTC[+57.021]Y.- | HexNAc(2)Hex(8) | C[+57], N[+1703] | 1350.5817 | 3 | 4049.7306 | 4049.7234 | 0 | 1.8 | 332 | CRagged | 893.92 | 859.2 | 537.91 | 18.24 | 104 | >sp\|P01876\|IGHA1_HUMAN Immunoglobulin heavy constant alpha 1 OS=Homo sapiens OX=9606 GN=IGHA1 PE=1 SV=2 | 255 | 20201116_fqg_colostrum_1ng_glyco_hcd203040_compound15.14374.14374.3 | controllerType=0 controllerNumber=1 scan=14374 | [illegible] |
| 09974:3 | 4 | R.LAGKPTHVN[+1378.476]VSVVMAEVDGTC[+57.021]Y.- | HexNAc(2)Hex(6) | C[+57], N[+1378] | 1242.5475 | 3 | 3725.6281 | 3725.6178 | 0 | 2.8 | 332 | CRagged | 898.05 | 846.51 | 543.74 | 18.14 | 104 | >sp\|P01876\|IGHA1_HUMAN Immunoglobulin heavy constant alpha 1 OS=Homo sapiens OX=9606 GN=IGHA1 PE=1 SV=2 | 255 | 20201116_fqg_colostrum_1ng_glyco_hcd203040_compound15.13199.13199.3 | controllerType=0 controllerNumber=1 scan=13199 | [illegible] |
| 11136:3 | 4 | R.LAGKPTHVN[+1540.529]VSVVMAEVDGTC[+57.021]Y.- | HexNAc(2)Hex(7) | C[+57], N[+1541] | 1296.5626 | 3 | 3887.6731 | 3887.6706 | 0 | 0.7 | 332 | CRagged | 904.92 | 792.84 | 514.3 | 19 | 104 | >sp\|P01876\|IGHA1_HUMAN Immunoglobulin heavy constant alpha 1 OS=Homo sapiens OX=9606 GN=IGHA1 PE=1 SV=2 | 255 | 20201116_fqg_colostrum_1ng_glyco_hcd203040_compound15.14399.14399.3 | controllerType=0 controllerNumber=1 scan=14399 | [illegible] |
| 11109:3 | 4 | R.LAGKPTHVN[+1702.581]VSVVMAEVDGTC[+57.021]Y.- | HexNAc(2)Hex(8) | C[+57], N[+1703] | 1350.5817 | 3 | 4049.7306 | 4049.7234 | 0 | 1.8 | 332 | CRagged | 948.55 | 948.55 | 551.03 | 19.24 | 104 | >sp\|P01876\|IGHA1_HUMAN Immunoglobulin heavy constant alpha 1 OS=Homo sapiens OX=9606 GN=IGHA1 PE=1 SV=2 | 255 | 20201116_fqg_colostrum_1ng_glyco_hcd203040_compound15.14371.14371.3 | controllerType=0 controllerNumber=1 scan=14371 | 30.2117 |
| 08545:3 | 4 | R.LAGKPTHVN[+1216.423]VSVVM[+15.995]AEVDGTC[+57.021]Y.- | HexNAc(2)Hex(5) | C[+57], M[+16], N[+1216] | 1193.8575 | 3 | 3579.5579 | 3579.5599 | 0 | -0.6 | 332 | CRagged | 677.01 | 661.77 | 661.77 | 13.5 | 104 | >sp\|P01876\|IGHA1_HUMAN Immunoglobulin heavy constant alpha 1 OS=Homo sapiens OX=9606 GN=IGHA1 PE=1 SV=2 | 255 | 20201116_fqg_colostrum_1ng_glyco_hcd203040_compound15.11718.11718.3 | controllerType=0 controllerNumber=1 scan=11718 | 25.8676 |
| 08507:3 | 4 | R.LAGKPTHVN[+1216.423]VSVVM[+15.995]AEVDGTC[+57.021]Y.- | HexNAc(2)Hex(5) | C[+57], M[+16], N[+1216] | 1193.8616 | 3 | 3579.5703 | 3579.5599 | 0 | 2.9 | 332 | CRagged | 680.57 | 680.57 | 680.57 | 14.14 | 104 | >sp\|P01876\|IGHA1_HUMAN Immunoglobulin heavy constant alpha 1 OS=Homo sapiens OX=9606 GN=IGHA1 PE=1 SV=2 | 255 | 20201116_fqg_colostrum_1ng_glyco_hcd203040_compound15.11679.11679.3 | controllerType=0 controllerNumber=1 scan=11679 | 25.7906 |
| 08510:3 | 4 | R.LAGKPTHVN[+1216.423]VSVVM[+15.995]AEVDGTC[+57.021]Y.- | HexNAc(2)Hex(5) | C[+57], M[+16], N[+1216] | 1193.8616 | 3 | 3579.5703 | 3579.5599 | 0 | 2.9 | 332 | CRagged | 808.29 | 649.38 | 649.38 | 17.14 | 104 | >sp\|P01876\|IGHA1_HUMAN Immunoglobulin heavy constant alpha 1 OS=Homo sapiens OX=9606 GN=IGHA1 PE=1 SV=2 | 255 | 20201116_fqg_colostrum_1ng_glyco_hcd203040_compound15.11682.11682.3 | controllerType=0 controllerNumber=1 scan=11682 | 25.7957 |
| 18552:4 | 5 | R.RTAGWNVPIGTLRPFLN[+2715.963]WTGPPEPIEAAVAR.F | HexNAc(5)Hex(6)Fuc(1)NeuAc(2) | N[+2716] | 1526.4578 | 4 | 6102.8092 | 6102.7746 | 0 | 5.7 | 140 | Specific | 457.07 | 360.6 | 360.6 | 8.72 | 91 | >sp\|P02788\|TRFL_HUMAN Lactotransferrin OS=Homo sapiens OX=9606 GN=LTF PE=1 SV=6 | 1270 | 20201116_fqg_colostrum_1ng_glyco_hcd203040_compound15.22080.22080.4 | controllerType=0 controllerNumber=1 scan=22080 | 43.2167 |
| 18555:4 | 5 | R.RTAGWNVPIGTLRPFLN[+2715.963]WTGPPEPIEAAVAR.F | HexNAc(5)Hex(6)Fuc(1)NeuAc(2) | N[+2716] | 1526.4578 | 4 | 6102.8092 | 6102.7746 | 0 | 5.7 | 140 | Specific | 520.28 | 456.75 | 456.75 | 10.45 | 91 | >sp\|P02788\|TRFL_HUMAN Lactotransferrin OS=Homo sapiens OX=9606 GN=LTF PE=1 SV=6 | 1270 | 20201116_fqg_colostrum_1ng_glyco_hcd203040_compound15.22083.22083.4 | controllerType=0 controllerNumber=1 scan=22083 | 43.225 |
| 21535:3 | 5 | R.[+42.011]TAGWNVPIGTLRPFLN[+2336.851]WTGPPEPIEAAVAR.F | HexNAc(6)Hex(6)Fuc(1) | .[42], N[+2337] | 1870.5256 | 3 | 5609.5622 | 5609.5726 | 0 | -1.8 | 141 | Specific | 245.1 | 236 | 236 | 6.37 | 91 | >sp\|P02788\|TRFL_HUMAN Lactotransferrin OS=Homo sapiens OX=9606 GN=LTF PE=1 SV=6 | 1270 | 20201116_fqg_colostrum_1ng_glyco_hcd203040_compound15.25183.25183.3 | controllerType=0 controllerNumber=1 scan=25183 | 49.2931 |
| 20116:3 | 5 | R.TAGWNVPIGTLRPFLN[+2205.793]WTGPPEPIEAAVAR.F | HexNAc(4)Hex(5)Fuc(2)NeuAc(1) | N[+2206] | 1812.8431 | 3 | 5436.5148 | 5436.5038 | 0 | 2 | 141 | Specific | 273.96 | 273.96 | 273.96 | 8.1 | 91 | >sp\|P02788\|TRFL_HUMAN Lactotransferrin OS=Homo sapiens OX=9606 GN=LTF PE=1 SV=6 | 1270 | 20201116_fqg_colostrum_1ng_glyco_hcd203040_compound15.23709.23709.2 | controllerType=0 controllerNumber=1 scan=23709 | 46.3741 |
| 19991:4 | 5 | R.TAGWNVPIGTLRPFLN[+2570.925]WTGPPEPIEAAVAR.F | HexNAc(5)Hex(6)Fuc(2)NeuAc(1) | N[+2571] | 1451.173 | 4 | 5801.6701 | 5801.636 | 0 | 5.9 | 141 | Specific | 310.35 | 271.39 | 271.39 | 7.81 | 91 | >sp\|P02788\|TRFL_HUMAN Lactotransferrin OS=Homo sapiens OX=9606 GN=LTF PE=1 SV=6 | 1270 | 20201116_fqg_colostrum_1ng_glyco_hcd203040_compound15.23580.23580.4 | controllerType=0 controllerNumber=1 scan=23580 | 46.1307 |
| 19934:3 | 5 | R.TAGWNVPIGTLRPFLN[+2570.925]WTGPPEPIEAAVAR.F | HexNAc(5)Hex(6)Fuc(2)NeuAc(1) | N[+2571] | 1934.5564 | 3 | 5801.6545 | 5801.636 | 0 | 3.2 | 141 | Specific | 328.52 | 195.53 | 195.53 | 7.69 | 91 | >sp\|P02788\|TRFL_HUMAN Lactotransferrin OS=Homo sapiens OX=9606 GN=LTF PE=1 SV=6 | 1270 | 20201116_fqg_colostrum_1ng_glyco_hcd203040_compound15.23520.23520.3 | controllerType=0 controllerNumber=1 scan=23520 | 46.008 |
| 19907:5 | 5 | R.TAGWNVPIGTLRPFLN[+2716.983]WTGPPEPIEAAVAR.F | HexNAc(5)Hex(6)Fuc(3)NeuAc(1) | N[+2717] | 1190.3477 | 5 | 5947.7092 | 5947.6939 | 0 | 2.6 | 141 | Specific | 351.76 | 351.76 | 351.76 | 9.54 | 91 | >sp\|P02788\|TRFL_HUMAN Lactotransferrin OS=Homo sapiens OX=9606 GN=LTF PE=1 SV=6 | 1270 | 20201116_fqg_colostrum_1ng_glyco_hcd203040_compound15.23492.23492.5 | controllerType=0 controllerNumber=1 scan=23492 | 45.9484 |
| 20099:4 | 5 | R.TAGWNVPIGTLRPFLN[+2076.750]WTGPPEPIEAAVAR.F | HexNAc(4)Hex(6)Fuc(2) | N[+2077] | 1327.6257 | 4 | 5307.4811 | 5307.4612 | 0 | 3.8 | 141 | Specific | 355.39 | 355.39 | 355.39 | 9.23 | 91 | >sp\|P02788\|TRFL_HUMAN Lactotransferrin OS=Homo sapiens OX=9606 GN=LTF PE=1 SV=6 | 1270 | 20201116_fqg_colostrum_1ng_glyco_hcd203040_compound15.23692.23692.4 | controllerType=0 controllerNumber=1 scan=23692 | 46.349 |
| 20050:3 | 5 | R.TAGWNVPIGTLRPFLN[+2205.793]WTGPPEPIEAAVAR.F | HexNAc(4)Hex(5)Fuc(2)NeuAc(1) | N[+2206] | 1812.8446 | 3 | 5436.5192 | 5436.5038 | 0 | 2.8 | 141 | Specific | 376.5 | 337.61 | 337.61 | 8.88 | 91 | >sp\|P02788\|TRFL_HUMAN Lactotransferrin OS=Homo sapiens OX=9606 GN=LTF PE=1 SV=6 | 1270 | 20201116_fqg_colostrum_1ng_glyco_hcd203040_compound15.23641.23641.3 | controllerType=0 controllerNumber=1 scan=23641 | 46.2513 |
| 20043:3 | 5 | R.TAGWNVPIGTLRPFLN[+2351.851]WTGPPEPIEAAVAR.F | HexNAc(4)Hex(5)Fuc(3)NeuAc(1) | N[+2352] | 1861.5289 | 3 | 5582.5721 | 5582.5617 | 0 | 1.9 | 141 | Specific | 434.09 | 381.59 | 381.59 | 10.7 | 91 | >sp\|P02788\|TRFL_HUMAN Lactotransferrin OS=Homo sapiens OX=9606 GN=LTF PE=1 SV=6 | 1270 | 20201116_fqg_colostrum_1ng_glyco_hcd203040_compound15.23634.23634.3 | controllerType=0 controllerNumber=1 scan=23634 | 46.2339 |
| 20042:4 | 5 | R.TAGWNVPIGTLRPFLN[+2351.851]WTGPPEPIEAAVAR.F | HexNAc(4)Hex(5)Fuc(3)NeuAc(1) | N[+2352] | 1396.3994 | 4 | 5582.5758 | 5582.5617 | 0 | 2.5 | 141 | Specific | 484.89 | 444.37 | 444.37 | 11.21 | 91 | >sp\|P02788\|TRFL_HUMAN Lactotransferrin OS=Homo sapiens OX=9606 GN=LTF PE=1 SV=6 | 1270 | 20201116_fqg_colostrum_1ng_glyco_hcd203040_compound15.23633.23633.4 | controllerType=0 controllerNumber=1 scan=23633 | 46.2324 |
| 19937:3 | 5 | R.TAGWNVPIGTLRPFLN[+2570.925]WTGPPEPIEAAVAR.F | HexNAc(5)Hex(6)Fuc(2)NeuAc(1) | N[+2571] | 1934.5564 | 3 | 5801.6545 | 5801.636 | 0 | 3.2 | 141 | Specific | 511.38 | 379.25 | 379.25 | 9.74 | 91 | >sp\|P02788\|TRFL_HUMAN Lactotransferrin OS=Homo sapiens OX=9606 GN=LTF PE=1 SV=6 | 1270 | 20201116_fqg_colostrum_1ng_glyco_hcd203040_compound15.23523.23523.3 | controllerType=0 controllerNumber=1 scan=23523 | 46.0188 |
| 19874:4 | 5 | R.TAGWNVPIGTLRPFLN[+2716.983]WTGPPEPIEAAVAR.F | HexNAc(5)Hex(6)Fuc(3)NeuAc(1) | N[+2717] | 1487.6812 | 4 | 5947.7028 | 5947.6939 | 0 | 1.5 | 141 | Specific | 545.64 | 487.51 | 487.51 | 11.3 | 91 | >sp\|P02788\|TRFL_HUMAN Lactotransferrin OS=Homo sapiens OX=9606 GN=LTF PE=1 SV=6 | 1270 | 20201116_fqg_colostrum_1ng_glyco_hcd203040_compound15.23458.23458.4 | controllerType=0 controllerNumber=1 scan=23458 | 45.8907 |
| 19052:3 | 5 | R.TAGWNVPIGTLRPFLN[+1768.640]WTGPPEPIEAAVAR.F | HexNAc(4)Hex(5)Fuc(1) | N[+1769] | 1667.1259 | 3 | 4999.3633 | 4999.3504 | 0 | 2.6 | 141 | Specific | 548.16 | 311.19 | 311.19 | 8.32 | 91 | >sp\|P02788\|TRFL_HUMAN Lactotransferrin OS=Homo sapiens OX=9606 GN=LTF PE=1 SV=6 | 1270 | 20201116_fqg_colostrum_1ng_glyco_hcd203040_compound15.22597.22597.3 | controllerType=0 controllerNumber=1 scan=22597 | 44.1534 |
| 20045:3 | 5 | R.TAGWNVPIGTLRPFLN[+2351.851]WTGPPEPIEAAVAR.F | HexNAc(4)Hex(5)Fuc(3)NeuAc(1) | N[+2352] | 1861.5289 | 3 | 5582.5721 | 5582.5617 | 0 | 1.9 | 141 | Specific | 562.25 | 520.9 | 520.9 | 11.5 | 91 | >sp\|P02788\|TRFL_HUMAN Lactotransferrin OS=Homo sapiens OX=9606 GN=LTF PE=1 SV=6 | 1270 | 20201116_fqg_colostrum_1ng_glyco_hcd203040_compound15.23636.23636.3 | controllerType=0 controllerNumber=1 scan=23636 | 46.2377 |
| 21468:3 | 5 | R.TAGWNVPIGTLRPFLN[+2350.830]WTGPPEPIEAAVAR.F | HexNAc(4)Hex(5)Fuc(1)NeuAc(2) | N[+2351] | 1861.1906 | 3 | 5581.5574 | 5581.5413 | 0 | 2.9 | 141 | Specific | 586.13 | 495.72 | 495.72 | 12.78 | 91 | >sp\|P02788\|TRFL_HUMAN Lactotransferrin OS=Homo sapiens OX=9606 GN=LTF PE=1 SV=6 | 1270 | 20201116_fqg_colostrum_1ng_glyco_hcd203040_compound15.25113.25113.3 | controllerType=0 controllerNumber=1 scan=25113 | 49.1387 |
| 19048:4 | 5 | R.TAGWNVPIGTLRPFLN[+1768.640]WTGPPEPIEAAVAR.F | HexNAc(4)Hex(5)Fuc(1) | N[+1769] | 1250.5979 | 4 | 4999.3698 | 4999.3504 | 0 | 3.9 | 141 | Specific | 593.91 | 521.85 | 521.85 | 13 | 91 | >sp\|P02788\|TRFL_HUMAN Lactotransferrin OS=Homo sapiens OX=9606 GN=LTF PE=1 SV=6 | 1270 | 20201116_fqg_colostrum_1ng_glyco_hcd203040_compound15.22593.22593.4 | controllerType=0 controllerNumber=1 scan=22593 | 44.1477 |
| 19886:4 | 5 | R.TAGWNVPIGTLRPFLN[+2716.983]WTGPPEPIEAAVAR.F | HexNAc(5)Hex(6)Fuc(3)NeuAc(1) | N[+2717] | 1487.6835 | 4 | 5947.7121 | 5947.6939 | 0 | 3.1 | 141 | Specific | 594.74 | 549.87 | 549.87 | 13 | 91 | >sp\|P02788\|TRFL_HUMAN Lactotransferrin OS=Homo sapiens OX=9606 GN=LTF PE=1 SV=6 | 1270 | 20201116_fqg_colostrum_1ng_glyco_hcd203040_compound15.23471.23471.4 | controllerType=0 controllerNumber=1 scan=23471 | 45.9141 |
| 21465:3 | 5 | R.TAGWNVPIGTLRPFLN[+2350.830]WTGPPEPIEAAVAR.F | HexNAc(4)Hex(5)Fuc(1)NeuAc(2) | N[+2351] | 1861.1906 | 3 | 5581.5574 | 5581.5413 | 0 | 2.9 | 141 | Specific | 598.93 | 415.46 | 415.46 | 12.78 | 91 | >sp\|P02788\|TRFL_HUMAN Lactotransferrin OS=Homo sapiens OX=9606 GN=LTF PE=1 SV=6 | 1270 | 20201116_fqg_colostrum_1ng_glyco_hcd203040_compound15.25110.25110.3 | controllerType=0 controllerNumber=1 scan=25110 | 49.1285 |
| 20118:3 | 5 | R.TAGWNVPIGTLRPFLN[+2205.793]WTGPPEPIEAAVAR.F | HexNAc(4)Hex(5)Fuc(2)NeuAc(1) | N[+2206] | 1812.8431 | 3 | 5436.5148 | 5436.5038 | 0 | 2 | 141 | Specific | 615.1 | 521.84 | 521.84 | 13.89 | 91 | >sp\|P02788\|TRFL_HUMAN Lactotransferrin OS=Homo sapiens OX=9606 GN=LTF PE=1 SV=6 | 1270 | 20201116_fqg_colostrum_1ng_glyco_hcd203040_compound15.23711.23711.2 | controllerType=0 controllerNumber=1 scan=23711 | 46.3773 |
| 20102:4 | 5 | R.TAGWNVPIGTLRPFLN[+2076.750]WTGPPEPIEAAVAR.F | HexNAc(4)Hex(6)Fuc(2) | N[+2077] | 1327.6257 | 4 | 5307.4811 | 5307.4612 | 0 | 3.8 | 141 | Specific | 619.36 | 528.59 | 528.59 | 14 | 91 | >sp\|P02788\|TRFL_HUMAN Lactotransferrin OS=Homo sapiens OX=9606 GN=LTF PE=1 SV=6 | 1270 | 20201116_fqg_colostrum_1ng_glyco_hcd203040_compound15.23695.23695.4 | controllerType=0 controllerNumber=1 scan=23695 | 46.3538 |
| 19994:4 | 5 | R.TAGWNVPIGTLRPFLN[+2570.925]WTGPPEPIEAAVAR.F | HexNAc(5)Hex(6)Fuc(2)NeuAc(1) | N[+2571] | 1451.173 | 4 | 5801.6701 | 5801.636 | 0 | 5.9 | 141 | Specific | 649.72 | 476.57 | 476.57 | 13.34 | 91 | >sp\|P02788\|TRFL_HUMAN Lactotransferrin OS=Homo sapiens OX=9606 GN=LTF PE=1 SV=6 | 1270 | 20201116_fqg_colostrum_1ng_glyco_hcd203040_compound15.23583.23583.4 | controllerType=0 controllerNumber=1 scan=23583 | 46.134 |
| 19910:5 | 5 | R.TAGWNVPIGTLRPFLN[+2716.983]WTGPPEPIEAAVAR.F | HexNAc(5)Hex(6)Fuc(3)NeuAc(1) | N[+2717] | 1190.3477 | 5 | 5947.7092 | 5947.6939 | 0 | 2.6 | 141 | Specific | 653.31 | 611.62 | 611.62 | 15.32 | 91 | >sp\|P02788\|TRFL_HUMAN Lactotransferrin OS=Homo sapiens OX=9606 GN=LTF PE=1 SV=6 | 1270 | 20201116_fqg_colostrum_1ng_glyco_hcd203040_compound15.23495.23495.5 | controllerType=0 controllerNumber=1 scan=23495 | 45.9531 |
| 20142:3 | 5 | R.TAGWNVPIGTLRPFLN[+2059.735]WTGPPEPIEAAVAR.F | HexNAc(4)Hex(5)Fuc(1)NeuAc(1) | N[+2060] | 1764.159 | 3 | 5290.4625 | 5290.4459 | 0 | 3.1 | 141 | Specific | 665.84 | 579.2 | 579.2 | 14.78 | 91 | >sp\|P02788\|TRFL_HUMAN Lactotransferrin OS=Homo sapiens OX=9606 GN=LTF PE=1 SV=6 | 1270 | 20201116_fqg_colostrum_1ng_glyco_hcd203040_compound15.23736.23736.2 | controllerType=0 controllerNumber=1 scan=23736 | 46.4301 |
| 20047:3 | 5 | R.TAGWNVPIGTLRPFLN[+2205.793]WTGPPEPIEAAVAR.F | HexNAc(4)Hex(5)Fuc(2)NeuAc(1) | N[+2206] | 1812.8446 | 3 | 5436.5192 | 5436.5038 | 0 | 2.8 | 141 | Specific | 665.96 | 590.42 | 590.42 | 14.78 | 91 | >sp\|P02788\|TRFL_HUMAN Lactotransferrin OS=Homo sapiens OX=9606 GN=LTF PE=1 SV=6 | 1270 | 20201116_fqg_colostrum_1ng_glyco_hcd203040_compound15.23638.23638.3 | controllerType=0 controllerNumber=1 scan=23638 | 46.24 |
| 20073:3 | 5 | R.TAGWNVPIGTLRPFLN[+2059.735]WTGPPEPIEAAVAR.F | HexNAc(4)Hex(5)Fuc(1)NeuAc(1) | N[+2060] | 1764.1591 | 3 | 5290.4629 | 5290.4459 | 0 | 3.2 | 141 | Specific | 677.18 | 629.32 | 629.32 | 14.78 | 91 | >sp\|P02788\|TRFL_HUMAN Lactotransferrin OS=Homo sapiens OX=9606 GN=LTF PE=1 SV=6 | 1270 | 20201116_fqg_colostrum_1ng_glyco_hcd203040_compound15.23665.23665.3 | controllerType=0 controllerNumber=1 scan=23665 | 46.2953 |
| 19877:4 | 5 | R.TAGWNVPIGTLRPFLN[+2716.983]WTGPPEPIEAAVAR.F | HexNAc(5)Hex(6)Fuc(3)NeuAc(1) | N[+2717] | 1487.6812 | 4 | 5947.7028 | 5947.6939 | 0 | 1.5 | 141 | Specific | 690.06 | 601.16 | 601.16 | 14.87 | 91 | >sp\|P02788\|TRFL_HUMAN Lactotransferrin OS=Homo sapiens OX=9606 GN=LTF PE=1 SV=6 | 1270 | 20201116_fqg_colostrum_1ng_glyco_hcd203040_compound15.23461.23461.4 | controllerType=0 controllerNumber=1 scan=23461 | 45.8971 |
| 20046:4 | 5 | R.TAGWNVPIGTLRPFLN[+2205.793]WTGPPEPIEAAVAR.F | HexNAc(4)Hex(5)Fuc(2)NeuAc(1) | N[+2206] | 1359.8842 | 4 | 5436.5148 | 5436.5038 | 0 | 2 | 141 | Specific | 691.49 | 655 | 655 | 15.11 | 91 | >sp\|P02788\|TRFL_HUMAN Lactotransferrin OS=Homo sapiens OX=9606 GN=LTF PE=1 SV=6 | 1270 | 20201116_fqg_colostrum_1ng_glyco_hcd203040_compound15.23637.23637.4 | controllerType=0 controllerNumber=1 scan=23637 | 46.2388 |
| 20069:4 | 5 | R.TAGWNVPIGTLRPFLN[+2059.735]WTGPPEPIEAAVAR.F | HexNAc(4)Hex(5)Fuc(1)NeuAc(1) | N[+2060] | 1323.3713 | 4 | 5290.4635 | 5290.4459 | 0 | 3.3 | 141 | Specific | 696.34 | 636.12 | 636.12 | 15 | 91 | >sp\|P02788\|TRFL_HUMAN Lactotransferrin OS=Homo sapiens OX=9606 GN=LTF PE=1 SV=6 | 1270 | 20201116_fqg_colostrum_1ng_glyco_hcd203040_compound15.23661.23661.4 | controllerType=0 controllerNumber=1 scan=23661 | 46.2861 |
| 20076:3 | 5 | R.TAGWNVPIGTLRPFLN[+2059.735]WTGPPEPIEAAVAR.F | HexNAc(4)Hex(5)Fuc(1)NeuAc(1) | N[+2060] | 1764.1591 | 3 | 5290.4629 | 5290.4459 | 0 | 3.2 | 141 | Specific | 701.76 | 577.19 | 577.19 | 15.78 | 91 | >sp\|P02788\|TRFL_HUMAN Lactotransferrin OS=Homo sapiens OX=9606 GN=LTF PE=1 SV=6 | 1270 | 20201116_fqg_colostrum_1ng_glyco_hcd203040_compound15.23668.23668.3 | controllerType=0 controllerNumber=1 scan=23668 | 46.3007 |
| 19889:4 | 5 | R.TAGWNVPIGTLRPFLN[+2716.983]WTGPPEPIEAAVAR.F | HexNAc(5)Hex(6)Fuc(3)NeuAc(1) | N[+2717] | 1487.6835 | 4 | 5947.7121 | 5947.6939 | 0 | 3.1 | 141 | Specific | 732.47 | 599.62 | 599.62 | 16 | 91 | >sp\|P02788\|TRFL_HUMAN Lactotransferrin OS=Homo sapiens OX=9606 GN=LTF PE=1 SV=6 | 1270 | 20201116_fqg_colostrum_1ng_glyco_hcd203040_compound15.23474.23474.4 | controllerType=0 controllerNumber=1 scan=23474 | 45.9196 |
| 20044:4 | 5 | R.TAGWNVPIGTLRPFLN[+2351.851]WTGPPEPIEAAVAR.F | HexNAc(4)Hex(5)Fuc(3)NeuAc(1) | N[+2352] | 1396.3994 | 4 | 5582.5758 | 5582.5617 | 0 | 2.5 | 141 | Specific | 790.04 | 658.57 | 658.57 | 17.11 | 91 | >sp\|P02788\|TRFL_HUMAN Lactotransferrin OS=Homo sapiens OX=9606 GN=LTF PE=1 SV=6 | 1270 | 20201116_fqg_colostrum_1ng_glyco_hcd203040_compound15.23635.23635.4 | controllerType=0 controllerNumber=1 scan=23635 | 46.2365 |
| 20071:4 | 5 | R.TAGWNVPIGTLRPFLN[+2059.735]WTGPPEPIEAAVAR.F | HexNAc(4)Hex(5)Fuc(1)NeuAc(1) | N[+2060] | 1323.3713 | 4 | 5290.4635 | 5290.4459 | 0 | 3.3 | 141 | Specific | 846.39 | 712.09 | 712.09 | 18 | 91 | >sp\|P02788\|TRFL_HUMAN Lactotransferrin OS=Homo sapiens OX=9606 GN=LTF PE=1 SV=6 | 1270 | 20201116_fqg_colostrum_1ng_glyco_hcd203040_compound15.23663.23663.4 | controllerType=0 controllerNumber=1 scan=23663 | 46.2922 |
| 20049:4 | 5 | R.TAGWNVPIGTLRPFLN[+2205.793]WTGPPEPIEAAVAR.F | HexNAc(4)Hex(5)Fuc(2)NeuAc(1) | N[+2206] | 1359.8842 | 4 | 5436.5148 | 5436.5038 | 0 | 2 | 141 | Specific | 853.11 | 797.64 | 797.64 | 19.11 | 91 | >sp\|P02788\|TRFL_HUMAN Lactotransferrin OS=Homo sapiens OX=9606 GN=LTF PE=1 SV=6 | 1270 | 20201116_fqg_colostrum_1ng_glyco_hcd203040_compound15.23640.23640.4 | controllerType=0 controllerNumber=1 scan=23640 | 46.244 |
| 20091:5 | 5 | R.TAGWNVPIGTLRPFLN[+2059.735]WTGPPEPIEAAVAR.F | HexNAc(4)Hex(5)Fuc(1)NeuAc(1) | N[+2060] | 1058.8969 | 5 | 5290.4552 | 5290.4459 | 0 | 1.8 | 141 | Specific | 897.06 | 822.69 | 822.69 | 19.32 | 91 | >sp\|P02788\|TRFL_HUMAN Lactotransferrin OS=Homo sapiens OX=9606 GN=LTF PE=1 SV=6 | 1270 | 20201116_fqg_colostrum_1ng_glyco_hcd203040_compound15.23683.23683.5 | controllerType=0 controllerNumber=1 scan=23683 | 46.3302 |
| 21493:5 | 5 | R.TAGWNVPIGTLRPFLN[+2350.830]WTGPPEPIEAAVAR.F | HexNAc(4)Hex(5)Fuc(1)NeuAc(2) | N[+2351] | 1117.1174 | 5 | 5581.5579 | 5581.5413 | 0 | 3 | 141 | Specific | 945.9 | 812.54 | 812.54 | 20.21 | 91 | >sp\|P02788\|TRFL_HUMAN Lactotransferrin OS=Homo sapiens OX=9606 GN=LTF PE=1 SV=6 | 1270 | 20201116_fqg_colostrum_1ng_glyco_hcd203040_compound15.25139.25139.5 | controllerType=0 controllerNumber=1 scan=25139 | 49.1811 |
| 20094:5 | 5 | R.TAGWNVPIGTLRPFLN[+2059.735]WTGPPEPIEAAVAR.F | HexNAc(4)Hex(5)Fuc(1)NeuAc(1) | N[+2060] | 1058.8969 | 5 | 5290.4552 | 5290.4459 | 0 | 1.8 | 141 | Specific | 955.48 | 796 | 796 | 21.32 | 91 | >sp\|P02788\|TRFL_HUMAN Lactotransferrin OS=Homo sapiens OX=9606 GN=LTF PE=1 SV=6 | 1270 | 20201116_fqg_colostrum_1ng_glyco_hcd203040_compound15.23686.23686.5 | controllerType=0 controllerNumber=1 scan=23686 | 46.3374 |
| 21586:4 | 5 | R.TAGWNIPMGLLFN[+2350.830]QTGSC[+57.021]K.F | HexNAc(4)Hex(5)Fuc(1)NeuAc(2) | C[+57], N[+2351] | 1112.2183 | 4 | 4445.8514 | 4445.8403 | 0 | 2.5 | 485 | Specific | 304.69 | 263.76 | 263.76 | 6.96 | 91 | >sp\|P02788\|TRFL_HUMAN Lactotransferrin OS=Homo sapiens OX=9606 GN=LTF PE=1 SV=6 | 1270 | 20201116_fqg_colostrum_1ng_glyco_hcd203040_compound15.25236.25236.4 | controllerType=0 controllerNumber=1 scan=25236 | 49.3955 |
| 19454:3 | 5 | R.TAGWNIPMGLLFN[+2059.735]QTGSC[+57.021]K.F | HexNAc(4)Hex(5)Fuc(1)NeuAc(1) | C[+57], N[+2060] | 1385.5917 | 3 | 4154.7606 | 4154.7449 | 0 | 3.8 | 485 | Specific | 341.77 | 305.19 | 305.19 | 8.53 | 91 | >sp\|P02788\|TRFL_HUMAN Lactotransferrin OS=Homo sapiens OX=9606 GN=LTF PE=1 SV=6 | 1270 | 20201116_fqg_colostrum_1ng_glyco_hcd203040_compound15.23017.23017.3 | controllerType=0 controllerNumber=1 scan=23017 | 45.0237 |
| 17798:2 | 5 | R.TAGWNIPMGLLFN[+1216.423]QTGSC[+57.021]K.F | HexNAc(2)Hex(5) | C[+57], N[+1216] | 1656.2305 | 2 | 3311.4537 | 3311.4328 | 0 | 6.3 | 485 | Specific | 672.11 | 672.11 | 224.02 | 13.42 | 91 | >sp\|P02788\|TRFL_HUMAN Lactotransferrin OS=Homo sapiens OX=9606 GN=LTF PE=1 SV=6 | 1270 | 20201116_fqg_colostrum_1ng_glyco_hcd203040_compound15.21294.21294.2 | controllerType=0 controllerNumber=1 scan=21294 | 41.5871 |
| 17735:3 | 5 | R.TAGWNIPMGLLFN[+1216.423]QTGSC[+57.021]K.F | HexNAc(2)Hex(5) | C[+57], N[+1216] | 1104.4857 | 3 | 3311.4427 | 3311.4328 | 0 | 3 | 485 | Specific | 731.06 | 731.06 | 260.41 | 15.53 | 91 | >sp\|P02788\|TRFL_HUMAN Lactotransferrin OS=Homo sapiens OX=9606 GN=LTF PE=1 SV=6 | 1270 | 20201116_fqg_colostrum_1ng_glyco_hcd203040_compound15.21229.21229.3 | controllerType=0 controllerNumber=1 scan=21229 | 41.4532 |
| 21562:3 | 5 | R.TAGWNIPMGLLFN[+2350.830]QTGSC[+57.021]K.F | HexNAc(4)Hex(5)Fuc(1)NeuAc(2) | C[+57], N[+2351] | 1482.6238 | 3 | 4445.857 | 4445.8403 | 0 | 3.7 | 485 | Specific | 897.75 | 864.38 | 864.38 | 18.53 | 91 | >sp\|P02788\|TRFL_HUMAN Lactotransferrin OS=Homo sapiens OX=9606 GN=LTF PE=1 SV=6 | 1270 | 20201116_fqg_colostrum_1ng_glyco_hcd203040_compound15.25211.25211.3 | controllerType=0 controllerNumber=1 scan=25211 | 49.3451 |
| 19352:3 | 5 | R.TAGWNIPMGLLFN[+2059.735]QTGSC[+57.021]K.F | HexNAc(4)Hex(5)Fuc(1)NeuAc(1) | C[+57], N[+2060] | 1385.5897 | 3 | 4154.7544 | 4154.7449 | 0 | 2.3 | 485 | Specific | 907.55 | 815.93 | 815.93 | 19.64 | 91 | >sp\|P02788\|TRFL_HUMAN Lactotransferrin OS=Homo sapiens OX=9606 GN=LTF PE=1 SV=6 | 1270 | 20201116_fqg_colostrum_1ng_glyco_hcd203040_compound15.22911.22911.3 | controllerType=0 controllerNumber=1 scan=22911 | 44.8114 |
| 19355:3 | 5 | R.TAGWNIPMGLLFN[+2059.735]QTGSC[+57.021]K.F | HexNAc(4)Hex(5)Fuc(1)NeuAc(1) | C[+57], N[+2060] | 1385.5897 | 3 | 4154.7544 | 4154.7449 | 0 | 2.3 | 485 | Specific | 928.79 | 837.87 | 837.87 | 19.64 | 91 | >sp\|P02788\|TRFL_HUMAN Lactotransferrin OS=Homo sapiens OX=9606 GN=LTF PE=1 SV=6 | 1270 | 20201116_fqg_colostrum_1ng_glyco_hcd203040_compound15.22914.22914.3 | controllerType=0 controllerNumber=1 scan=22914 | 44.8185 |
| 19456:3 | 5 | R.TAGWNIPMGLLFN[+2059.735]QTGSC[+57.021]K.F | HexNAc(4)Hex(5)Fuc(1)NeuAc(1) | C[+57], N[+2060] | 1385.5917 | 3 | 4154.7606 | 4154.7449 | 0 | 3.8 | 485 | Specific | 934.04 | 704.83 | 704.83 | 19.53 | 91 | >sp\|P02788\|TRFL_HUMAN Lactotransferrin OS=Homo sapiens OX=9606 GN=LTF PE=1 SV=6 | 1270 | 20201116_fqg_colostrum_1ng_glyco_hcd203040_compound15.23019.23019.3 | controllerType=0 controllerNumber=1 scan=23019 | 45.0276 |
| 21557:3 | 5 | R.TAGWNIPMGLLFN[+2350.830]QTGSC[+57.021]K.F | HexNAc(4)Hex(5)Fuc(1)NeuAc(2) | C[+57], N[+2351] | 1482.6243 | 3 | 4445.8584 | 4445.8403 | 0 | 4.1 | 485 | Specific | 936.18 | 935.99 | 935.99 | 19.09 | 91 | >sp\|P02788\|TRFL_HUMAN Lactotransferrin OS=Homo sapiens OX=9606 GN=LTF PE=1 SV=6 | 1270 | 20201116_fqg_colostrum_1ng_glyco_hcd203040_compound15.25205.25205.3 | controllerType=0 controllerNumber=1 scan=25205 | 49.3325 |
| 21554:3 | 5 | R.TAGWNIPMGLLFN[+2350.830]QTGSC[+57.021]K.F | HexNAc(4)Hex(5)Fuc(1)NeuAc(2) | C[+57], N[+2351] | 1482.6243 | 3 | 4445.8584 | 4445.8403 | 0 | 4.1 | 485 | Specific | 936.67 | 813.28 | 813.28 | 19.09 | 91 | >sp\|P02788\|TRFL_HUMAN Lactotransferrin OS=Homo sapiens OX=9606 GN=LTF PE=1 SV=6 | 1270 | 20201116_fqg_colostrum_1ng_glyco_hcd203040_compound15.25202.25202.3 | controllerType=0 controllerNumber=1 scan=25202 | 49.3251 |
| 21560:3 | 5 | R.TAGWNIPMGLLFN[+2350.830]QTGSC[+57.021]K.F | HexNAc(4)Hex(5)Fuc(1)NeuAc(2) | C[+57], N[+2351] | 1482.6238 | 3 | 4445.857 | 4445.8403 | 0 | 3.7 | 485 | Specific | 947.93 | 901.97 | 901.97 | 19.53 | 91 | >sp\|P02788\|TRFL_HUMAN Lactotransferrin OS=Homo sapiens OX=9606 GN=LTF PE=1 SV=6 | 1270 | 20201116_fqg_colostrum_1ng_glyco_hcd203040_compound15.25209.25209.3 | controllerType=0 controllerNumber=1 scan=25209 | 49.34 |
| 17738:3 | 5 | R.TAGWNIPMGLLFN[+1216.423]QTGSC[+57.021]K.F | HexNAc(2)Hex(5) | C[+57], N[+1216] | 1104.4857 | 3 | 3311.4427 | 3311.4328 | 0 | 3 | 485 | Specific | 1047.8 | 1006.52 | 489.84 | 21.53 | 91 | >sp\|P02788\|TRFL_HUMAN Lactotransferrin OS=Homo sapiens OX=9606 GN=LTF PE=1 SV=6 | 1270 | 20201116_fqg_colostrum_1ng_glyco_hcd203040_compound15.21232.21232.3 | controllerType=0 controllerNumber=1 scan=21232 | 41.4647 |
| 18527:4 | 5 | R.TAGWNIPMGLLFN[+1768.640]QTGSC[+57.021]KFDEYFSQSC[+57.021]APGSDPR.S | HexNAc(4)Hex(5)Fuc(1) | C[+57]*2, N[+1769] | 1427.6094 | 4 | 5707.4157 | 5707.3966 | 0 | 3.3 | 485 | Specific | 488.48 | 472 | 472 | 9.98 | 91 | >sp\|P02788\|TRFL_HUMAN Lactotransferrin OS=Homo sapiens OX=9606 GN=LTF PE=1 SV=6 | 1270 | 20201116_fqg_colostrum_1ng_glyco_hcd203040_compound15.22054.22054.4 | controllerType=0 controllerNumber=1 scan=22054 | 43.1673 |
| 21101:4 | 5 | R.TAGWNIPMGLLFN[+2350.830]QTGSC[+57.021]KFDEYFSQSC[+57.021]APGSDPR.S | HexNAc(4)Hex(5)Fuc(1)NeuAc(2) | C[+57]*2, N[+2351] | 1573.1565 | 4 | 6289.6042 | 6289.5875 | 0 | 2.7 | 485 | Specific | 723.58 | 701.7 | 701.7 | 14.98 | 91 | >sp\|P02788\|TRFL_HUMAN Lactotransferrin OS=Homo sapiens OX=9606 GN=LTF PE=1 SV=6 | 1270 | 20201116_fqg_colostrum_1ng_glyco_hcd203040_compound15.24732.24732.4 | controllerType=0 controllerNumber=1 scan=24732 | 48.3917 |
| 21078:4 | 5 | R.TAGWNIPMGLLFN[+2350.830]QTGSC[+57.021]KFDEYFSQSC[+57.021]APGSDPR.S | HexNAc(4)Hex(5)Fuc(1)NeuAc(2) | C[+57]*2, N[+2351] | 1573.1591 | 4 | 6289.6144 | 6289.5875 | 0 | 4.3 | 485 | Specific | 831.08 | 772.26 | 772.26 | 16.43 | 91 | >sp\|P02788\|TRFL_HUMAN Lactotransferrin OS=Homo sapiens OX=9606 GN=LTF PE=1 SV=6 | 1270 | 20201116_fqg_colostrum_1ng_glyco_hcd203040_compound15.24708.24708.4 | controllerType=0 controllerNumber=1 scan=24708 | 48.3446 |
| 18530:4 | 5 | R.TAGWNIPMGLLFN[+1768.640]QTGSC[+57.021]KFDEYFSQSC[+57.021]APGSDPR.S | HexNAc(4)Hex(5)Fuc(1) | C[+57]*2, N[+1769] | 1427.6094 | 4 | 5707.4157 | 5707.3966 | 0 | 3.3 | 485 | Specific | 862.62 | 778.86 | 778.86 | 17.88 | 91 | >sp\|P02788\|TRFL_HUMAN Lactotransferrin OS=Homo sapiens OX=9606 GN=LTF PE=1 SV=6 | 1270 | 20201116_fqg_colostrum_1ng_glyco_hcd203040_compound15.22057.22057.4 | controllerType=0 controllerNumber=1 scan=22057 | 43.1771 |
| 21081:4 | 5 | R.TAGWNIPMGLLFN[+2350.830]QTGSC[+57.021]KFDEYFSQSC[+57.021]APGSDPR.S | HexNAc(4)Hex(5)Fuc(1)NeuAc(2) | C[+57]*2, N[+2351] | 1573.1591 | 4 | 6289.6144 | 6289.5875 | 0 | 4.3 | 485 | Specific | 936.94 | 882.91 | 882.91 | 18.43 | 91 | >sp\|P02788\|TRFL_HUMAN Lactotransferrin OS=Homo sapiens OX=9606 GN=LTF PE=1 SV=6 | 1270 | 20201116_fqg_colostrum_1ng_glyco_hcd203040_compound15.24711.24711.4 | controllerType=0 controllerNumber=1 scan=24711 | 48.3502 |
| 21103:4 | 5 | R.TAGWNIPMGLLFN[+2350.830]QTGSC[+57.021]KFDEYFSQSC[+57.021]APGSDPR.S | HexNAc(4)Hex(5)Fuc(1)NeuAc(2) | C[+57]*2, N[+2351] | 1573.1565 | 4 | 6289.6042 | 6289.5875 | 0 | 2.7 | 485 | Specific | 940.76 | 899.73 | 899.73 | 18.98 | 91 | >sp\|P02788\|TRFL_HUMAN Lactotransferrin OS=Homo sapiens OX=9606 GN=LTF PE=1 SV=6 | 1270 | 20201116_fqg_colostrum_1ng_glyco_hcd203040_compound15.24734.24734.4 | controllerType=0 controllerNumber=1 scan=24734 | 48.3955 |
| 18298:3 | 7 | R.GLSFDVSLEVSQGPGLLN[+2351.851]DTK.V | HexNAc(4)Hex(5)Fuc(3)NeuAc(1) | N[+2352] | 1509.9987 | 3 | 4527.9816 | 4527.964 | 0 | 3.9 | 118 | Specific | 348.2 | 255.97 | 255.97 | 8 | 77 | >sp\|P01833\|PIGR_HUMAN Polymeric immunoglobulin receptor OS=Homo sapiens OX=9606 GN=PIGR PE=1 SV=4 | 902 | 20201116_fqg_colostrum_1ng_glyco_hcd203040_compound15.21814.21814.3 | controllerType=0 controllerNumber=1 scan=21814 | 42.6296 |
| 18301:3 | 7 | R.GLSFDVSLEVSQGPGLLN[+2351.851]DTK.V | HexNAc(4)Hex(5)Fuc(3)NeuAc(1) | N[+2352] | 1509.9987 | 3 | 4527.9816 | 4527.964 | 0 | 3.9 | 118 | Specific | 720.28 | 554.73 | 554.73 | 15.53 | 77 | >sp\|P01833\|PIGR_HUMAN Polymeric immunoglobulin receptor OS=Homo sapiens OX=9606 GN=PIGR PE=1 SV=4 | 902 | 20201116_fqg_colostrum_1ng_glyco_hcd203040_compound15.21817.21817.3 | controllerType=0 controllerNumber=1 scan=21817 | 42.6377 |
| 19555:4 | 7 | K.QIGLYPVLVIDSSGYVNPN[+2205.793]YTGR.I | HexNAc(4)Hex(5)Fuc(2)NeuAc(1) | N[+2206] | 1183.5319 | 4 | 4731.1058 | 4731.0963 | 0 | 2 | 168 | Specific | 229.19 | 197.8 | 197.8 | 4.92 | 77 | >sp\|P01833\|PIGR_HUMAN Polymeric immunoglobulin receptor OS=Homo sapiens OX=9606 GN=PIGR PE=1 SV=4 | 902 | 20201116_fqg_colostrum_1ng_glyco_hcd203040_compound15.23122.23122.4 | controllerType=0 controllerNumber=1 scan=23122 | 45.2374 |
| 18947:3 | 7 | K.QIGLYPVLVIDSSGYVNPN[+1257.449]YTGR.I | HexNAc(3)Hex(4) | N[+1257] | 1261.5861 | 3 | 3782.7438 | 3782.7529 | 0 | -2.4 | 168 | Specific | 471.81 | 471.81 | 471.81 | 9.09 | 77 | >sp\|P01833\|PIGR_HUMAN Polymeric immunoglobulin receptor OS=Homo sapiens OX=9606 GN=PIGR PE=1 SV=4 | 902 | 20201116_fqg_colostrum_1ng_glyco_hcd203040_compound15.22489.22489.3 | controllerType=0 controllerNumber=1 scan=22489 | 43.9548 |
| 19558:4 | 7 | K.QIGLYPVLVIDSSGYVNPN[+2205.793]YTGR.I | HexNAc(4)Hex(5)Fuc(2)NeuAc(1) | N[+2206] | 1183.5319 | 4 | 4731.1058 | 4731.0963 | 0 | 2 | 168 | Specific | 630.93 | 465.21 | 465.21 | 13.67 | 77 | >sp\|P01833\|PIGR_HUMAN Polymeric immunoglobulin receptor OS=Homo sapiens OX=9606 GN=PIGR PE=1 SV=4 | 902 | 20201116_fqg_colostrum_1ng_glyco_hcd203040_compound15.23125.23125.4 | controllerType=0 controllerNumber=1 scan=23125 | 45.2445 |
| 21777:4 | 7 | K.QIGLYPVLVIDSSGYVNPN[+2204.772]YTGR.I | HexNAc(4)Hex(5)NeuAc(2) | N[+2205] | 1183.2779 | 4 | 4730.0897 | 4730.0759 | 0 | 2.9 | 168 | Specific | 721.6 | 556.57 | 556.57 | 15.57 | 77 | >sp\|P01833\|PIGR_HUMAN Polymeric immunoglobulin receptor OS=Homo sapiens OX=9606 GN=PIGR PE=1 SV=4 | 902 | 20201116_fqg_colostrum_1ng_glyco_hcd203040_compound15.25435.25435.4 | controllerType=0 controllerNumber=1 scan=25435 | 49.8401 |
| 19554:3 | 7 | K.QIGLYPVLVIDSSGYVNPN[+2205.793]YTGR.I | HexNAc(4)Hex(5)Fuc(2)NeuAc(1) | N[+2206] | 1577.709 | 3 | 4731.1126 | 4731.0963 | 0 | 3.4 | 168 | Specific | 725.59 | 678.66 | 678.66 | 15.36 | 77 | >sp\|P01833\|PIGR_HUMAN Polymeric immunoglobulin receptor OS=Homo sapiens OX=9606 GN=PIGR PE=1 SV=4 | 902 | 20201116_fqg_colostrum_1ng_glyco_hcd203040_compound15.23121.23121.3 | controllerType=0 controllerNumber=1 scan=23121 | 45.2363 |
| 18950:3 | 7 | K.QIGLYPVLVIDSSGYVNPN[+1257.449]YTGR.I | HexNAc(3)Hex(4) | N[+1257] | 1261.5861 | 3 | 3782.7438 | 3782.7529 | 0 | -2.4 | 168 | Specific | 762.71 | 628 | 628 | 15.07 | 77 | >sp\|P01833\|PIGR_HUMAN Polymeric immunoglobulin receptor OS=Homo sapiens OX=9606 GN=PIGR PE=1 SV=4 | 902 | 20201116_fqg_colostrum_1ng_glyco_hcd203040_compound15.22492.22492.3 | controllerType=0 controllerNumber=1 scan=22492 | 43.9595 |
| 21779:4 | 7 | K.QIGLYPVLVIDSSGYVNPN[+2204.772]YTGR.I | HexNAc(4)Hex(5)NeuAc(2) | N[+2205] | 1183.2779 | 4 | 4730.0897 | 4730.0759 | 0 | 2.9 | 168 | Specific | 845.24 | 638.16 | 638.16 | 17.57 | 77 | >sp\|P01833\|PIGR_HUMAN Polymeric immunoglobulin receptor OS=Homo sapiens OX=9606 GN=PIGR PE=1 SV=4 | 902 | 20201116_fqg_colostrum_1ng_glyco_hcd203040_compound15.25437.25437.4 | controllerType=0 controllerNumber=1 scan=25437 | 49.8457 |

Table S4. Cont.

| Query #:z | Protein Rank | Glycopeptide | Glycan | Modification Type(s) | Observed m/z | z | Observed (M+H) | Calc. mass (M+H) | Off-by-x error | Mass error (ppm) | StartPosition | Cleavage | Score | Delta | Delta Mod | \|Log Prob\| | # of unique peptides | Protein Name | Protein DB number | Comment | Scan # | Scan Time |
|---|---|---|---|---|---|---|---|---|---|---|---|---|---|---|---|---|---|---|---|---|---|---|
| 19557:3 | 7 | K.QIGLYPVLVIDSSGYVNPN[+2205.793]YTGR.I | HexNAc(4)Hex(5)Fuc(2)NeuAc(1) | N[+2206] | 1577.709 | 3 | 4731.1126 | 4731.0963 | 0 | 3.4 | 168 | Specific | 869.92 | 832.69 | 832.69 | 18.36 | 77 | >sp\|P01833\|PIGR_HUMAN Polymeric immunoglobulin receptor OS=Homo sapiens OX=9606 GN=PIGR PE=1 SV=4 | 902 | 20201116_fqg_colostrum_1ng_glyco_hcd203040_compound15.23124.23124.3 | controllerType=0 controllerNumber=1 scan=23124 | 45.2434 |
| 21295:3 | 7 | K.[+42.011]AQYEGRLSLLEEPGN[+1622.582]GTFTVILNQLTSR.D | HexNAc(4)Hex(5) | .[42], N[+1623] | 1591.0655 | 3 | 4771.1819 | 4771.2089 | 0 | -5.7 | 407 | Specific | 300.87 | 270.69 | 270.69 | 3.77 | 77 | >sp\|P01833\|PIGR_HUMAN Polymeric immunoglobulin receptor OS=Homo sapiens OX=9606 GN=PIGR PE=1 SV=4 | 902 | 20201116_fqg_colostrum_1ng_glyco_hcd203040_compound15.24933.24933.3 | controllerType=0 controllerNumber=1 scan=24933 | 48.7517 |
| 21292:3 | 7 | K.[+42.011]AQYEGRLSLLEEPGN[+1622.582]GTFTVILNQLTSR.D | HexNAc(4)Hex(5) | .[42], N[+1623] | 1591.0655 | 3 | 4771.1819 | 4771.2089 | 0 | -5.7 | 407 | Specific | 327.58 | 327.58 | 327.58 | 6.21 | 77 | >sp\|P01833\|PIGR_HUMAN Polymeric immunoglobulin receptor OS=Homo sapiens OX=9606 GN=PIGR PE=1 SV=4 | 902 | 20201116_fqg_colostrum_1ng_glyco_hcd203040_compound15.24930.24930.3 | controllerType=0 controllerNumber=1 scan=24930 | 48.7452 |
| 28853:4 | 7 | R.[+42.011]LSLLEEPGN[+2351.851]GTFTVILNQLTSR.D | HexNAc(4)Hex(5)Fuc(3)NeuAc(1) | .[42], N[+2352] | 1199.7953 | 4 | 4796.1595 | 4796.1539 | 0 | 1.2 | 413 | Specific | 247.55 | 247.55 | 247.55 | 5.9 | 77 | >sp\|P01833\|PIGR_HUMAN Polymeric immunoglobulin receptor OS=Homo sapiens OX=9606 GN=PIGR PE=1 SV=4 | 902 | 20201116_fqg_colostrum_1ng_glyco_hcd203040_compound15.32822.32822.4 | controllerType=0 controllerNumber=1 scan=32822 | 65.434 |
| 28855:4 | 7 | R.[+42.011]LSLLEEPGN[+2351.851]GTFTVILNQLTSR.D | HexNAc(4)Hex(5)Fuc(3)NeuAc(1) | .[42], N[+2352] | 1199.7953 | 4 | 4796.1595 | 4796.1539 | 0 | 1.2 | 413 | Specific | 288.23 | 271.37 | 271.37 | 6.35 | 77 | >sp\|P01833\|PIGR_HUMAN Polymeric immunoglobulin receptor OS=Homo sapiens OX=9606 GN=PIGR PE=1 SV=4 | 902 | 20201116_fqg_colostrum_1ng_glyco_hcd203040_compound15.32824.32824.4 | controllerType=0 controllerNumber=1 scan=32824 | 65.438 |
| 26060:4 | 7 | R.LSLLEEPGN[+3008.078]GTFTVILNQLTSR.D | HexNAc(5)Hex(6)Fuc(3)NeuAc(2) | N[+3008] | 1353.3558 | 4 | 5410.4015 | 5410.3709 | 0 | 5.7 | 413 | Specific | 296.7 | 293.08 | 293.08 | 7.26 | 77 | >sp\|P01833\|PIGR_HUMAN Polymeric immunoglobulin receptor OS=Homo sapiens OX=9606 GN=PIGR PE=1 SV=4 | 902 | 20201116_fqg_colostrum_1ng_glyco_hcd203040_compound15.29909.29909.4 | controllerType=0 controllerNumber=1 scan=29909 | 59.6098 |
| 21121:4 | 7 | R.LSLLEEPGN[+2715.963]GTFTVILNQLTSR.D | HexNAc(5)Hex(6)Fuc(1)NeuAc(2) | N[+2716] | 1280.3313 | 4 | 5118.3034 | 5118.2551 | 0 | 9.4 | 413 | Specific | 333.69 | 292.68 | 292.68 | 7.02 | 77 | >sp\|P01833\|PIGR_HUMAN Polymeric immunoglobulin receptor OS=Homo sapiens OX=9606 GN=PIGR PE=1 SV=4 | 902 | 20201116_fqg_colostrum_1ng_glyco_hcd203040_compound15.24752.24752.2 | controllerType=0 controllerNumber=1 scan=24752 | 48.4276 |
| 25960:4 | 7 | R.LSLLEEPGN[+3153.116]GTFTVILNQLTSR.D | HexNAc(5)Hex(6)Fuc(2)NeuAc(3) | N[+3153] | 1389.6057 | 4 | 5555.401 | 5555.4084 | 0 | -1.3 | 413 | Specific | 386.35 | 319.81 | 319.81 | 7.56 | 77 | >sp\|P01833\|PIGR_HUMAN Polymeric immunoglobulin receptor OS=Homo sapiens OX=9606 GN=PIGR PE=1 SV=4 | 902 | 20201116_fqg_colostrum_1ng_glyco_hcd203040_compound15.29805.29805.4 | controllerType=0 controllerNumber=1 scan=29805 | 59.399 |
| 28791:4 | 7 | R.[+42.011]LSLLEEPGN[+2350.830]GTFTVILNQLTSR.D | HexNAc(4)Hex(5)Fuc(1)NeuAc(2) | .[42], N[+2351] | 1199.5426 | 4 | 4795.1488 | 4795.1335 | 0 | 3.2 | 413 | Specific | 397.53 | 380.29 | 380.29 | 10.29 | 77 | >sp\|P01833\|PIGR_HUMAN Polymeric immunoglobulin receptor OS=Homo sapiens OX=9606 GN=PIGR PE=1 SV=4 | 902 | 20201116_fqg_colostrum_1ng_glyco_hcd203040_compound15.32757.32757.3 | controllerType=0 controllerNumber=1 scan=32757 | 65.3095 |
| 28793:4 | 7 | R.[+42.011]LSLLEEPGN[+2350.830]GTFTVILNQLTSR.D | HexNAc(4)Hex(5)Fuc(1)NeuAc(2) | .[42], N[+2351] | 1199.5426 | 4 | 4795.1488 | 4795.1335 | 0 | 3.2 | 413 | Specific | 454.33 | 352.67 | 352.67 | 9 | 77 | >sp\|P01833\|PIGR_HUMAN Polymeric immunoglobulin receptor OS=Homo sapiens OX=9606 GN=PIGR PE=1 SV=4 | 902 | 20201116_fqg_colostrum_1ng_glyco_hcd203040_compound15.32759.32759.3 | controllerType=0 controllerNumber=1 scan=32759 | 65.3135 |
| 21393:3 | 7 | R.LSLLEEPGN[+2204.772]GTFTVILNQLTSR.D | HexNAc(4)Hex(5)NeuAc(2) | N[+2205] | 1536.373 | 3 | 4607.1044 | 4607.065 | 0 | 8.5 | 413 | Specific | 483.33 | 234.07 | 234.07 | 7.22 | 77 | >sp\|P01833\|PIGR_HUMAN Polymeric immunoglobulin receptor OS=Homo sapiens OX=9606 GN=PIGR PE=1 SV=4 | 902 | 20201116_fqg_colostrum_1ng_glyco_hcd203040_compound15.25035.25035.3 | controllerType=0 controllerNumber=1 scan=25035 | 48.9579 |
| 25962:4 | 7 | R.LSLLEEPGN[+3153.116]GTFTVILNQLTSR.D | HexNAc(5)Hex(6)Fuc(2)NeuAc(3) | N[+3153] | 1389.6057 | 4 | 5555.401 | 5555.4084 | 0 | -1.3 | 413 | Specific | 486.4 | 391.01 | 391.01 | 9.58 | 77 | >sp\|P01833\|PIGR_HUMAN Polymeric immunoglobulin receptor OS=Homo sapiens OX=9606 GN=PIGR PE=1 SV=4 | 902 | 20201116_fqg_colostrum_1ng_glyco_hcd203040_compound15.29807.29807.4 | controllerType=0 controllerNumber=1 scan=29807 | 59.4038 |
| 26062:4 | 7 | R.LSLLEEPGN[+3008.078]GTFTVILNQLTSR.D | HexNAc(5)Hex(6)Fuc(3)NeuAc(2) | N[+3008] | 1353.3558 | 4 | 5410.4015 | 5410.3709 | 0 | 5.7 | 413 | Specific | 492.71 | 492.71 | 492.71 | 10 | 77 | >sp\|P01833\|PIGR_HUMAN Polymeric immunoglobulin receptor OS=Homo sapiens OX=9606 GN=PIGR PE=1 SV=4 | 902 | 20201116_fqg_colostrum_1ng_glyco_hcd203040_compound15.29911.29911.4 | controllerType=0 controllerNumber=1 scan=29911 | 59.6133 |
| 28830:3 | 7 | R.[+42.011]LSLLEEPGN[+2351.851]GTFTVILNQLTSR.D | HexNAc(4)Hex(5)Fuc(3)NeuAc(1) | .[42], N[+2352] | 1599.3874 | 3 | 4796.1476 | 4796.1539 | 0 | -1.3 | 413 | Specific | 497.01 | 449.33 | 449.33 | 9.37 | 77 | >sp\|P01833\|PIGR_HUMAN Polymeric immunoglobulin receptor OS=Homo sapiens OX=9606 GN=PIGR PE=1 SV=4 | 902 | 20201116_fqg_colostrum_1ng_glyco_hcd203040_compound15.32798.32798.3 | controllerType=0 controllerNumber=1 scan=32798 | 65.3866 |
| 28832:3 | 7 | R.[+42.011]LSLLEEPGN[+2351.851]GTFTVILNQLTSR.D | HexNAc(4)Hex(5)Fuc(3)NeuAc(1) | .[42], N[+2352] | 1599.3874 | 3 | 4796.1476 | 4796.1539 | 0 | -1.3 | 413 | Specific | 503.62 | 445.37 | 445.37 | 9.48 | 77 | >sp\|P01833\|PIGR_HUMAN Polymeric immunoglobulin receptor OS=Homo sapiens OX=9606 GN=PIGR PE=1 SV=4 | 902 | 20201116_fqg_colostrum_1ng_glyco_hcd203040_compound15.32800.32800.3 | controllerType=0 controllerNumber=1 scan=32800 | 65.3907 |
| 26014:3 | 7 | R.LSLLEEPGN[+3007.058]GTFTVILNQLTSR.D | HexNAc(5)Hex(6)Fuc(1)NeuAc(3) | N[+3007] | 1803.7961 | 3 | 5409.3738 | 5409.3505 | 0 | 4.3 | 413 | Specific | 510.31 | 463.85 | 463.85 | 10.13 | 77 | >sp\|P01833\|PIGR_HUMAN Polymeric immunoglobulin receptor OS=Homo sapiens OX=9606 GN=PIGR PE=1 SV=4 | 902 | 20201116_fqg_colostrum_1ng_glyco_hcd203040_compound15.29861.29861.3 | controllerType=0 controllerNumber=1 scan=29861 | 59.5104 |
| 26012:3 | 7 | R.LSLLEEPGN[+3007.058]GTFTVILNQLTSR.D | HexNAc(5)Hex(6)Fuc(1)NeuAc(3) | N[+3007] | 1803.7961 | 3 | 5409.3738 | 5409.3505 | 0 | 4.3 | 413 | Specific | 514.92 | 514.92 | 514.92 | 10.13 | 77 | >sp\|P01833\|PIGR_HUMAN Polymeric immunoglobulin receptor OS=Homo sapiens OX=9606 GN=PIGR PE=1 SV=4 | 902 | 20201116_fqg_colostrum_1ng_glyco_hcd203040_compound15.29859.29859.3 | controllerType=0 controllerNumber=1 scan=29859 | 59.5069 |
| 23428:3 | 7 | R.LSLLEEPGN[+2350.830]GTFTVILNQLTSR.D | HexNAc(4)Hex(5)Fuc(1)NeuAc(2) | N[+2351] | 1585.0535 | 3 | 4753.146 | 4753.1229 | 0 | 4.9 | 413 | Specific | 603.97 | 526.3 | 526.3 | 12.91 | 77 | >sp\|P01833\|PIGR_HUMAN Polymeric immunoglobulin receptor OS=Homo sapiens OX=9606 GN=PIGR PE=1 SV=4 | 902 | 20201116_fqg_colostrum_1ng_glyco_hcd203040_compound15.27157.27157.3 | controllerType=0 controllerNumber=1 scan=27157 | 53.4512 |
| 21124:4 | 7 | R.LSLLEEPGN[+2715.963]GTFTVILNQLTSR.D | HexNAc(5)Hex(6)Fuc(1)NeuAc(2) | N[+2716] | 1280.3313 | 4 | 5118.3034 | 5118.2551 | 0 | 9.4 | 413 | Specific | 674.72 | 531.19 | 531.19 | 13.47 | 77 | >sp\|P01833\|PIGR_HUMAN Polymeric immunoglobulin receptor OS=Homo sapiens OX=9606 GN=PIGR PE=1 SV=4 | 902 | 20201116_fqg_colostrum_1ng_glyco_hcd203040_compound15.24755.24755.2 | controllerType=0 controllerNumber=1 scan=24755 | 48.432 |
| 23438:4 | 7 | R.LSLLEEPGN[+2350.830]GTFTVILNQLTSR.D | HexNAc(4)Hex(5)Fuc(1)NeuAc(2) | N[+2351] | 1189.0419 | 4 | 4753.1458 | 4753.1229 | 0 | 4.8 | 413 | Specific | 698.37 | 617.52 | 617.52 | 14.12 | 77 | >sp\|P01833\|PIGR_HUMAN Polymeric immunoglobulin receptor OS=Homo sapiens OX=9606 GN=PIGR PE=1 SV=4 | 902 | 20201116_fqg_colostrum_1ng_glyco_hcd203040_compound15.27167.27167.4 | controllerType=0 controllerNumber=1 scan=27167 | 53.4718 |
| 21367:4 | 7 | R.LSLLEEPGN[+2205.793]GTFTVILNQLTSR.D | HexNAc(4)Hex(5)Fuc(2)NeuAc(1) | N[+2206] | 1152.7807 | 4 | 4608.1009 | 4608.0854 | 0 | 3.4 | 413 | Specific | 713.61 | 611.35 | 611.35 | 15.57 | 77 | >sp\|P01833\|PIGR_HUMAN Polymeric immunoglobulin receptor OS=Homo sapiens OX=9606 GN=PIGR PE=1 SV=4 | 902 | 20201116_fqg_colostrum_1ng_glyco_hcd203040_compound15.25008.25008.4 | controllerType=0 controllerNumber=1 scan=25008 | 48.9019 |
| 21370:4 | 7 | R.LSLLEEPGN[+2205.793]GTFTVILNQLTSR.D | HexNAc(4)Hex(5)Fuc(2)NeuAc(1) | N[+2206] | 1152.7807 | 4 | 4608.1009 | 4608.0854 | 0 | 3.4 | 413 | Specific | 742.33 | 495.05 | 495.05 | 15.57 | 77 | >sp\|P01833\|PIGR_HUMAN Polymeric immunoglobulin receptor OS=Homo sapiens OX=9606 GN=PIGR PE=1 SV=4 | 902 | 20201116_fqg_colostrum_1ng_glyco_hcd203040_compound15.25011.25011.4 | controllerType=0 controllerNumber=1 scan=25011 | 48.9076 |
| 23453:3 | 7 | R.LSLLEEPGN[+2496.888]GTFTVILNQLTSR.D | HexNAc(4)Hex(5)Fuc(2)NeuAc(2) | N[+2497] | 1633.7389 | 3 | 4899.2023 | 4899.1808 | 0 | 4.4 | 413 | Specific | 761.34 | 699 | 699 | 15.91 | 77 | >sp\|P01833\|PIGR_HUMAN Polymeric immunoglobulin receptor OS=Homo sapiens OX=9606 GN=PIGR PE=1 SV=4 | 902 | 20201116_fqg_colostrum_1ng_glyco_hcd203040_compound15.27183.27183.3 | controllerType=0 controllerNumber=1 scan=27183 | 53.5127 |
| 21395:3 | 7 | R.LSLLEEPGN[+2204.772]GTFTVILNQLTSR.D | HexNAc(4)Hex(5)NeuAc(2) | N[+2205] | 1536.373 | 3 | 4607.1044 | 4607.065 | 0 | 8.5 | 413 | Specific | 815.96 | 411.1 | 411.1 | 16.26 | 77 | >sp\|P01833\|PIGR_HUMAN Polymeric immunoglobulin receptor OS=Homo sapiens OX=9606 GN=PIGR PE=1 SV=4 | 902 | 20201116_fqg_colostrum_1ng_glyco_hcd203040_compound15.25037.25037.3 | controllerType=0 controllerNumber=1 scan=25037 | 48.9616 |
| 23456:3 | 7 | R.LSLLEEPGN[+2496.888]GTFTVILNQLTSR.D | HexNAc(4)Hex(5)Fuc(2)NeuAc(2) | N[+2497] | 1633.7389 | 3 | 4899.2023 | 4899.1808 | 0 | 4.4 | 413 | Specific | 845.86 | 729.45 | 729.45 | 16.91 | 77 | >sp\|P01833\|PIGR_HUMAN Polymeric immunoglobulin receptor OS=Homo sapiens OX=9606 GN=PIGR PE=1 SV=4 | 902 | 20201116_fqg_colostrum_1ng_glyco_hcd203040_compound15.27186.27186.3 | controllerType=0 controllerNumber=1 scan=27186 | 53.5178 |
| 23441:4 | 7 | R.LSLLEEPGN[+2350.830]GTFTVILNQLTSR.D | HexNAc(4)Hex(5)Fuc(1)NeuAc(2) | N[+2351] | 1189.0419 | 4 | 4753.1458 | 4753.1229 | 0 | 4.8 | 413 | Specific | 872.7 | 664.65 | 664.65 | 18.12 | 77 | >sp\|P01833\|PIGR_HUMAN Polymeric immunoglobulin receptor OS=Homo sapiens OX=9606 GN=PIGR PE=1 SV=4 | 902 | 20201116_fqg_colostrum_1ng_glyco_hcd203040_compound15.27170.27170.4 | controllerType=0 controllerNumber=1 scan=27170 | 53.4767 |
| 23430:3 | 7 | R.LSLLEEPGN[+2350.830]GTFTVILNQLTSR.D | HexNAc(4)Hex(5)Fuc(1)NeuAc(2) | N[+2351] | 1585.0535 | 3 | 4753.146 | 4753.1229 | 0 | 4.9 | 413 | Specific | 937 | 654.03 | 654.03 | 18.91 | 77 | >sp\|P01833\|PIGR_HUMAN Polymeric immunoglobulin receptor OS=Homo sapiens OX=9606 GN=PIGR PE=1 SV=4 | 902 | 20201116_fqg_colostrum_1ng_glyco_hcd203040_compound15.27159.27159.3 | controllerType=0 controllerNumber=1 scan=27159 | 53.4552 |
| 21331:3 | 7 | R.LSLLEEPGN[+2059.735]GTFTVILNQLTSR.D | HexNAc(4)Hex(5)Fuc(1)NeuAc(1) | N[+2060] | 1488.0182 | 3 | 4462.0402 | 4462.0275 | 0 | 2.8 | 413 | Specific | 976.82 | 733.85 | 733.85 | 20.36 | 77 | >sp\|P01833\|PIGR_HUMAN Polymeric immunoglobulin receptor OS=Homo sapiens OX=9606 GN=PIGR PE=1 SV=4 | 902 | 20201116_fqg_colostrum_1ng_glyco_hcd203040_compound15.24970.24970.3 | controllerType=0 controllerNumber=1 scan=24970 | 48.8367 |
| 21283:4 | 7 | R.LSLLEEPGN[+2351.851]GTFTVILNQLTSR.D | HexNAc(4)Hex(5)Fuc(3)NeuAc(1) | N[+2352] | 1189.2931 | 4 | 4754.1507 | 4754.1433 | 0 | 1.6 | 413 | Specific | 985.62 | 830.54 | 830.54 | 20.43 | 77 | >sp\|P01833\|PIGR_HUMAN Polymeric immunoglobulin receptor OS=Homo sapiens OX=9606 GN=PIGR PE=1 SV=4 | 902 | 20201116_fqg_colostrum_1ng_glyco_hcd203040_compound15.24920.24920.4 | controllerType=0 controllerNumber=1 scan=24920 | 48.7263 |
| 21254:3 | 7 | R.LSLLEEPGN[+2351.851]GTFTVILNQLTSR.D | HexNAc(4)Hex(5)Fuc(3)NeuAc(1) | N[+2352] | 1585.3924 | 3 | 4754.1626 | 4754.1433 | 0 | 4.1 | 413 | Specific | 985.67 | 725.76 | 725.76 | 19.91 | 77 | >sp\|P01833\|PIGR_HUMAN Polymeric immunoglobulin receptor OS=Homo sapiens OX=9606 GN=PIGR PE=1 SV=4 | 902 | 20201116_fqg_colostrum_1ng_glyco_hcd203040_compound15.24890.24890.3 | controllerType=0 controllerNumber=1 scan=24890 | 48.6612 |
| 21329:3 | 7 | R.LSLLEEPGN[+2059.735]GTFTVILNQLTSR.D | HexNAc(4)Hex(5)Fuc(1)NeuAc(1) | N[+2060] | 1488.0182 | 3 | 4462.0402 | 4462.0275 | 0 | 2.8 | 413 | Specific | 1061.77 | 827.06 | 827.06 | 22.36 | 77 | >sp\|P01833\|PIGR_HUMAN Polymeric immunoglobulin receptor OS=Homo sapiens OX=9606 GN=PIGR PE=1 SV=4 | 902 | 20201116_fqg_colostrum_1ng_glyco_hcd203040_compound15.24968.24968.3 | controllerType=0 controllerNumber=1 scan=24968 | 48.8294 |
| 21251:3 | 7 | R.LSLLEEPGN[+2351.851]GTFTVILNQLTSR.D | HexNAc(4)Hex(5)Fuc(3)NeuAc(1) | N[+2352] | 1585.3924 | 3 | 4754.1626 | 4754.1433 | 0 | 4.1 | 413 | Specific | 1116.62 | 946.34 | 946.34 | 22.91 | 77 | >sp\|P01833\|PIGR_HUMAN Polymeric immunoglobulin receptor OS=Homo sapiens OX=9606 GN=PIGR PE=1 SV=4 | 902 | 20201116_fqg_colostrum_1ng_glyco_hcd203040_compound15.24887.24887.3 | controllerType=0 controllerNumber=1 scan=24887 | 48.6548 |
| 21280:4 | 7 | R.LSLLEEPGN[+2351.851]GTFTVILNQLTSR.D | HexNAc(4)Hex(5)Fuc(3)NeuAc(1) | N[+2352] | 1189.2931 | 4 | 4754.1507 | 4754.1433 | 0 | 1.6 | 413 | Specific | 1148.48 | 1061.14 | 1061.14 | 23.43 | 77 | >sp\|P01833\|PIGR_HUMAN Polymeric immunoglobulin receptor OS=Homo sapiens OX=9606 GN=PIGR PE=1 SV=4 | 902 | 20201116_fqg_colostrum_1ng_glyco_hcd203040_compound15.24917.24917.4 | controllerType=0 controllerNumber=1 scan=24917 | 48.7181 |
| 14457:3 | 7 | K.IIEGEPNLKVPGN[+2205.793]VTAVLGETLK.V | HexNAc(4)Hex(5)Fuc(2)NeuAc(1) | N[+2206] | 1533.0597 | 3 | 4597.1647 | 4597.1422 | 0 | 4.9 | 457 | Specific | 741.53 | 725.53 | 725.53 | 14.22 | 77 | >sp\|P01833\|PIGR_HUMAN Polymeric immunoglobulin receptor OS=Homo sapiens OX=9606 GN=PIGR PE=1 SV=4 | 902 | 20201116_fqg_colostrum_1ng_glyco_hcd203040_compound15.17834.17834.3 | controllerType=0 controllerNumber=1 scan=17834 | 35.6648 |
| 18547:3 | 7 | K.IIEGEPNLKVPGN[+2350.830]VTAVLGETLK.V | HexNAc(4)Hex(5)Fuc(1)NeuAc(2) | N[+2351] | 1581.4025 | 3 | 4742.193 | 4742.1797 | 0 | 2.8 | 457 | Specific | 774.87 | 597.92 | 597.92 | 15.67 | 77 | >sp\|P01833\|PIGR_HUMAN Polymeric immunoglobulin receptor OS=Homo sapiens OX=9606 GN=PIGR PE=1 SV=4 | 902 | 20201116_fqg_colostrum_1ng_glyco_hcd203040_compound15.22075.22075.3 | controllerType=0 controllerNumber=1 scan=22075 | 43.211 |
| 16524:3 | 7 | K.IIEGEPNLKVPGN[+2205.793]VTAVLGETLK.V | HexNAc(4)Hex(5)Fuc(2)NeuAc(1) | N[+2206] | 1533.0577 | 3 | 4597.1585 | 4597.1422 | 0 | 3.5 | 457 | Specific | 776.92 | 776.92 | 776.92 | 15.67 | 77 | >sp\|P01833\|PIGR_HUMAN Polymeric immunoglobulin receptor OS=Homo sapiens OX=9606 GN=PIGR PE=1 SV=4 | 902 | 20201116_fqg_colostrum_1ng_glyco_hcd203040_compound15.19974.19974.3 | controllerType=0 controllerNumber=1 scan=19974 | 39.2268 |
| 16543:3 | 7 | K.IIEGEPNLKVPGN[+2059.735]VTAVLGETLK.V | HexNAc(4)Hex(5)Fuc(1)NeuAc(1) | N[+2060] | 1484.3709 | 3 | 4451.0982 | 4451.0843 | 0 | 3.1 | 457 | Specific | 807.81 | 616.7 | 616.7 | 16.67 | 77 | >sp\|P01833\|PIGR_HUMAN Polymeric immunoglobulin receptor OS=Homo sapiens OX=9606 GN=PIGR PE=1 SV=4 | 902 | 20201116_fqg_colostrum_1ng_glyco_hcd203040_compound15.19993.19993.3 | controllerType=0 controllerNumber=1 scan=19993 | 39.2557 |
| 16527:3 | 7 | K.IIEGEPNLKVPGN[+2205.793]VTAVLGETLK.V | HexNAc(4)Hex(5)Fuc(2)NeuAc(1) | N[+2206] | 1533.0577 | 3 | 4597.1585 | 4597.1422 | 0 | 3.5 | 457 | Specific | 846.26 | 747.45 | 747.45 | 16.67 | 77 | >sp\|P01833\|PIGR_HUMAN Polymeric immunoglobulin receptor OS=Homo sapiens OX=9606 GN=PIGR PE=1 SV=4 | 902 | 20201116_fqg_colostrum_1ng_glyco_hcd203040_compound15.19977.19977.3 | controllerType=0 controllerNumber=1 scan=19977 | 39.2347 |
| 14455:3 | 7 | K.IIEGEPNLKVPGN[+2205.793]VTAVLGETLK.V | HexNAc(4)Hex(5)Fuc(2)NeuAc(1) | N[+2206] | 1533.0597 | 3 | 4597.1647 | 4597.1422 | 0 | 4.9 | 457 | Specific | 904.46 | 888.73 | 888.73 | 18.22 | 77 | >sp\|P01833\|PIGR_HUMAN Polymeric immunoglobulin receptor OS=Homo sapiens OX=9606 GN=PIGR PE=1 SV=4 | 902 | 20201116_fqg_colostrum_1ng_glyco_hcd203040_compound15.17832.17832.3 | controllerType=0 controllerNumber=1 scan=17832 | 35.6574 |
| 18545:3 | 7 | K.IIEGEPNLKVPGN[+2350.830]VTAVLGETLK.V | HexNAc(4)Hex(5)Fuc(1)NeuAc(2) | N[+2351] | 1581.4025 | 3 | 4742.193 | 4742.1797 | 0 | 2.8 | 457 | Specific | 929.88 | 875.62 | 875.62 | 18.67 | 77 | >sp\|P01833\|PIGR_HUMAN Polymeric immunoglobulin receptor OS=Homo sapiens OX=9606 GN=PIGR PE=1 SV=4 | 902 | 20201116_fqg_colostrum_1ng_glyco_hcd203040_compound15.22073.22073.3 | controllerType=0 controllerNumber=1 scan=22073 | 43.2068 |
| 16540:3 | 7 | K.IIEGEPNLKVPGN[+2059.735]VTAVLGETLK.V | HexNAc(4)Hex(5)Fuc(1)NeuAc(1) | N[+2060] | 1484.3709 | 3 | 4451.0982 | 4451.0843 | 0 | 3.1 | 457 | Specific | 1012.08 | 916.26 | 916.26 | 20.67 | 77 | >sp\|P01833\|PIGR_HUMAN Polymeric immunoglobulin receptor OS=Homo sapiens OX=9606 GN=PIGR PE=1 SV=4 | 902 | 20201116_fqg_colostrum_1ng_glyco_hcd203040_compound15.19990.19990.3 | controllerType=0 controllerNumber=1 scan=19990 | 39.2507 |
| 16021:4 | 7 | K.[+42.011]IIEGEPNLKVPGN[+2018.708]VTAVLGETLKVPC[+57.021]HFPC[+57.021] | HexNAc(3)Hex(6)Fuc(1)NeuAc(1) | .[42], C[+57]*2, N[+2019] | 1370.1499 | 4 | 5477.5778 | 5477.5258 | 0 | 9.5 | 457 | Specific | 201.68 | 161.34 | 161.34 | 3.29 | 77 | >sp\|P01833\|PIGR_HUMAN Polymeric immunoglobulin receptor OS=Homo sapiens OX=9606 GN=PIGR PE=1 SV=4 | 902 | 20201116_fqg_colostrum_1ng_glyco_hcd203040_compound15.19452.19452.4 | controllerType=0 controllerNumber=1 scan=19452 | 38.3592 |
| 15985:4 | 7 | K.IIEGEPNLKVPGN[+2205.793]VTAVLGETLKVPC[+57.021]HFPC[+57.021]K.F | HexNAc(4)Hex(5)Fuc(2)NeuAc(1) | C[+57]*2, N[+2206] | 1406.4105 | 4 | 5622.6203 | 5622.5997 | 0 | 3.7 | 457 | Specific | 304.49 | 304.49 | 304.49 | 7 | 77 | >sp\|P01833\|PIGR_HUMAN Polymeric immunoglobulin receptor OS=Homo sapiens OX=9606 GN=PIGR PE=1 SV=4 | 902 | 20201116_fqg_colostrum_1ng_glyco_hcd203040_compound15.19415.19415.3 | controllerType=0 controllerNumber=1 scan=19415 | 38.3013 |
| 16102:5 | 7 | K.IIEGEPNLKVPGN[+2204.772]VTAVLGETLKVPC[+57.021]HFPC[+57.021]K.F | HexNAc(4)Hex(5)NeuAc(2) | C[+57]*2, N[+2205] | 1125.1277 | 5 | 5621.6093 | 5621.5793 | 0 | 5.3 | 457 | Specific | 487.01 | 323.54 | 323.54 | 7.72 | 77 | >sp\|P01833\|PIGR_HUMAN Polymeric immunoglobulin receptor OS=Homo sapiens OX=9606 GN=PIGR PE=1 SV=4 | 902 | 20201116_fqg_colostrum_1ng_glyco_hcd203040_compound15.19536.19536.5 | controllerType=0 controllerNumber=1 scan=19536 | 38.5128 |
| 16105:5 | 7 | K.IIEGEPNLKVPGN[+2204.772]VTAVLGETLKVPC[+57.021]HFPC[+57.021]K.F | HexNAc(4)Hex(5)NeuAc(2) | C[+57]*2, N[+2205] | 1125.1277 | 5 | 5621.6093 | 5621.5793 | 0 | 5.3 | 457 | Specific | 623.73 | 383.18 | 383.18 | 12.2 | 77 | >sp\|P01833\|PIGR_HUMAN Polymeric immunoglobulin receptor OS=Homo sapiens OX=9606 GN=PIGR PE=1 SV=4 | 902 | 20201116_fqg_colostrum_1ng_glyco_hcd203040_compound15.19539.19539.5 | controllerType=0 controllerNumber=1 scan=19539 | 38.5171 |
| 16002:4 | 7 | K.IIEGEPNLKVPGN[+2059.735]VTAVLGETLKVPC[+57.021]HFPC[+57.021]K.F | HexNAc(4)Hex(5)Fuc(1)NeuAc(1) | C[+57]*2, N[+2060] | 1369.8939 | 4 | 5476.5539 | 5476.5418 | 0 | 2.2 | 457 | Specific | 634.24 | 577.68 | 577.68 | 12.35 | 77 | >sp\|P01833\|PIGR_HUMAN Polymeric immunoglobulin receptor OS=Homo sapiens OX=9606 GN=PIGR PE=1 SV=4 | 902 | 20201116_fqg_colostrum_1ng_glyco_hcd203040_compound15.19432.19432.2 | controllerType=0 controllerNumber=1 scan=19432 | 38.3257 |
| 15881:4 | 7 | K.IIEGEPNLKVPGN[+2204.772]VTAVLGETLKVPC[+57.021]HFPC[+57.021]K.F | HexNAc(4)Hex(5)NeuAc(2) | C[+57]*2, N[+2205] | 1406.1603 | 4 | 5621.6193 | 5621.5793 | 0 | 7.1 | 457 | Specific | 706.17 | 479.79 | 479.79 | 13.35 | 77 | >sp\|P01833\|PIGR_HUMAN Polymeric immunoglobulin receptor OS=Homo sapiens OX=9606 GN=PIGR PE=1 SV=4 | 902 | 20201116_fqg_colostrum_1ng_glyco_hcd203040_compound15.19307.19307.2 | controllerType=0 controllerNumber=1 scan=19307 | 38.112 |
| 15988:4 | 7 | K.IIEGEPNLKVPGN[+2205.793]VTAVLGETLKVPC[+57.021]HFPC[+57.021]K.F | HexNAc(4)Hex(5)Fuc(2)NeuAc(1) | C[+57]*2, N[+2206] | 1406.4105 | 4 | 5622.6203 | 5622.5997 | 0 | 3.7 | 457 | Specific | 722.99 | 657.33 | 657.33 | 14.24 | 77 | >sp\|P01833\|PIGR_HUMAN Polymeric immunoglobulin receptor OS=Homo sapiens OX=9606 GN=PIGR PE=1 SV=4 | 902 | 20201116_fqg_colostrum_1ng_glyco_hcd203040_compound15.19418.19418.3 | controllerType=0 controllerNumber=1 scan=19418 | 38.3051 |
| 15866:5 | 7 | K.IIEGEPNLKVPGN[+2351.851]VTAVLGETLKVPC[+57.021]HFPC[+57.021]K.F | HexNAc(4)Hex(5)Fuc(3)NeuAc(1) | C[+57]*2, N[+2352] | 1154.54 | 5 | 5768.6711 | 5768.6576 | 0 | 2.3 | 457 | Specific | 733.68 | 688.04 | 688.04 | 14.97 | 77 | >sp\|P01833\|PIGR_HUMAN Polymeric immunoglobulin receptor OS=Homo sapiens OX=9606 GN=PIGR PE=1 SV=4 | 902 | 20201116_fqg_colostrum_1ng_glyco_hcd203040_compound15.19292.19292.5 | controllerType=0 controllerNumber=1 scan=19292 | 38.0875 |
| 17566:5 | 7 | K.IIEGEPNLKVPGN[+2350.830]VTAVLGETLKVPC[+57.021]HFPC[+57.021]K.F | HexNAc(4)Hex(5)Fuc(1)NeuAc(2) | C[+57]*2, N[+2351] | 1154.3359 | 5 | 5767.6506 | 5767.6372 | 0 | 2.3 | 457 | Specific | 787.39 | 553.32 | 553.32 | 15.97 | 77 | >sp\|P01833\|PIGR_HUMAN Polymeric immunoglobulin receptor OS=Homo sapiens OX=9606 GN=PIGR PE=1 SV=4 | 902 | 20201116_fqg_colostrum_1ng_glyco_hcd203040_compound15.21054.21054.5 | controllerType=0 controllerNumber=1 scan=21054 | 41.1171 |
| 15898:4 | 7 | K.IIEGEPNLKVPGN[+2351.851]VTAVLGETLKVPC[+57.021]HFPC[+57.021]K.F | HexNAc(4)Hex(5)Fuc(3)NeuAc(1) | C[+57]*2, N[+2352] | 1442.9247 | 4 | 5768.6769 | 5768.6576 | 0 | 3.3 | 457 | Specific | 799.38 | 687.95 | 687.95 | 15.24 | 77 | >sp\|P01833\|PIGR_HUMAN Polymeric immunoglobulin receptor OS=Homo sapiens OX=9606 GN=PIGR PE=1 SV=4 | 902 | 20201116_fqg_colostrum_1ng_glyco_hcd203040_compound15.19325.19325.4 | controllerType=0 controllerNumber=1 scan=19325 | 38.1461 |
| 17568:5 | 7 | K.IIEGEPNLKVPGN[+2350.830]VTAVLGETLKVPC[+57.021]HFPC[+57.021]K.F | HexNAc(4)Hex(5)Fuc(1)NeuAc(2) | C[+57]*2, N[+2351] | 1154.3359 | 5 | 5767.6506 | 5767.6372 | 0 | 2.3 | 457 | Specific | 802.79 | 625.98 | 625.98 | 16.97 | 77 | >sp\|P01833\|PIGR_HUMAN Polymeric immunoglobulin receptor OS=Homo sapiens OX=9606 GN=PIGR PE=1 SV=4 | 902 | 20201116_fqg_colostrum_1ng_glyco_hcd203040_compound15.21056.21056.5 | controllerType=0 controllerNumber=1 scan=21056 | 41.1219 |
| 16004:4 | 7 | K.IIEGEPNLKVPGN[+2059.735]VTAVLGETLKVPC[+57.021]HFPC[+57.021]K.F | HexNAc(4)Hex(5)Fuc(1)NeuAc(1) | C[+57]*2, N[+2060] | 1369.8939 | 4 | 5476.5539 | 5476.5418 | 0 | 2.2 | 457 | Specific | 856.71 | 740.72 | 740.72 | 17.35 | 77 | >sp\|P01833\|PIGR_HUMAN Polymeric immunoglobulin receptor OS=Homo sapiens OX=9606 GN=PIGR PE=1 SV=4 | 902 | 20201116_fqg_colostrum_1ng_glyco_hcd203040_compound15.19434.19434.2 | controllerType=0 controllerNumber=1 scan=19434 | 38.3307 |
| 15895:4 | 7 | K.IIEGEPNLKVPGN[+2351.851]VTAVLGETLKVPC[+57.021]HFPC[+57.021]K.F | HexNAc(4)Hex(5)Fuc(3)NeuAc(1) | C[+57]*2, N[+2352] | 1442.9247 | 4 | 5768.6769 | 5768.6576 | 0 | 3.3 | 457 | Specific | 991.55 | 957.35 | 957.35 | 19.24 | 77 | >sp\|P01833\|PIGR_HUMAN Polymeric immunoglobulin receptor OS=Homo sapiens OX=9606 GN=PIGR PE=1 SV=4 | 902 | 20201116_fqg_colostrum_1ng_glyco_hcd203040_compound15.19322.19322.4 | controllerType=0 controllerNumber=1 scan=19322 | 38.1403 |
| 15878:4 | 7 | K.IIEGEPNLKVPGN[+2204.772]VTAVLGETLKVPC[+57.021]HFPC[+57.021]K.F | HexNAc(4)Hex(5)NeuAc(2) | C[+57]*2, N[+2205] | 1406.1603 | 4 | 5621.6193 | 5621.5793 | 0 | 7.1 | 457 | Specific | 1036.79 | 954.9 | 954.9 | 19.35 | 77 | >sp\|P01833\|PIGR_HUMAN Polymeric immunoglobulin receptor OS=Homo sapiens OX=9606 GN=PIGR PE=1 SV=4 | 902 | 20201116_fqg_colostrum_1ng_glyco_hcd203040_compound15.19304.19304.2 | controllerType=0 controllerNumber=1 scan=19304 | 38.1058 |
| 15863:5 | 7 | K.IIEGEPNLKVPGN[+2351.851]VTAVLGETLKVPC[+57.021]HFPC[+57.021]K.F | HexNAc(4)Hex(5)Fuc(3)NeuAc(1) | C[+57]*2, N[+2352] | 1154.54 | 5 | 5768.6711 | 5768.6576 | 0 | 2.3 | 457 | Specific | 1117.45 | 1104.25 | 1104.25 | 22.97 | 77 | >sp\|P01833\|PIGR_HUMAN Polymeric immunoglobulin receptor OS=Homo sapiens OX=9606 GN=PIGR PE=1 SV=4 | 902 | 20201116_fqg_colostrum_1ng_glyco_hcd203040_compound15.19289.19289.5 | controllerType=0 controllerNumber=1 scan=19289 | 38.0818 |
| 17517:5 | 7 | K.IIEGEPNLKVPGN[+2350.830]VTAVLGETLKVPC[+57.021]HFPC[+57.021]K.F | HexNAc(4)Hex(5)Fuc(1)NeuAc(2) | C[+57]*2, N[+2351] | 1154.3366 | 5 | 5767.6536 | 5767.6372 | 0 | 2.8 | 457 | Specific | 1138.88 | 836.84 | 836.84 | 22.86 | 77 | >sp\|P01833\|PIGR_HUMAN Polymeric immunoglobulin receptor OS=Homo sapiens OX=9606 GN=PIGR PE=1 SV=4 | 902 | 20201116_fqg_colostrum_1ng_glyco_hcd203040_compound15.21002.21002.5 | controllerType=0 controllerNumber=1 scan=21002 | 40.9933 |
| 05365:3 | 7 | K.WN[+2204.772]NTGC[+57.021]QALPSQDEGPSK.A | HexNAc(4)Hex(5)NeuAc(2) | C[+57], N[+2205] | 1398.5657 | 3 | 4193.6826 | 4193.6491 | 0 | 8 | 498 | Specific | 361.7 | 289.22 | 289.22 | 6.94 | 77 | >sp\|P01833\|PIGR_HUMAN Polymeric immunoglobulin receptor OS=Homo sapiens OX=9606 GN=PIGR PE=1 SV=4 | 902 | 20201116_fqg_colostrum_1ng_glyco_hcd203040_compound15.8412.8412.3 | controllerType=0 controllerNumber=1 scan=8412 | 20.6572 |
| 05367:3 | 7 | K.WN[+2204.772]NTGC[+57.021]QALPSQDEGPSK.A | HexNAc(4)Hex(5)NeuAc(2) | C[+57], N[+2205] | 1398.5657 | 3 | 4193.6826 | 4193.6491 | 0 | 8 | 498 | Specific | 454.82 | 403.13 | 403.13 | 8.96 | 77 | >sp\|P01833\|PIGR_HUMAN Polymeric immunoglobulin receptor OS=Homo sapiens OX=9606 GN=PIGR PE=1 SV=4 | 902 | 20201116_fqg_colostrum_1ng_glyco_hcd203040_compound15.8414.8414.3 | controllerType=0 controllerNumber=1 scan=8414 | 20.6634 |
| 05398:3 | 7 | K.WN[+2059.735]NTGC[+57.021]QALPSQDEGPSK.A | HexNAc(4)Hex(5)Fuc(1)NeuAc(1) | C[+57], N[+2060] | 1350.212 | 3 | 4048.6214 | 4048.6116 | 0 | 2.4 | 498 | Specific | 535.34 | 455.73 | 455.73 | 10.56 | 77 | >sp\|P01833\|PIGR_HUMAN Polymeric immunoglobulin receptor OS=Homo sapiens OX=9606 GN=PIGR PE=1 SV=4 | 902 | 20201116_fqg_colostrum_1ng_glyco_hcd203040_compound15.8446.8446.3 | controllerType=0 controllerNumber=1 scan=8446 | 20.7169 |
| 07622:3 | 7 | K.WN[+2204.772]NTGC[+57.021]QALPSQDEGPSK.A | HexNAc(4)Hex(5)NeuAc(2) | C[+57], N[+2205] | 1398.5567 | 3 | 4193.6555 | 4193.6491 | 0 | 1.5 | 498 | Specific | 639.01 | 568.15 | 568.15 | 13.24 | 77 | >sp\|P01833\|PIGR_HUMAN Polymeric immunoglobulin receptor OS=Homo sapiens OX=9606 GN=PIGR PE=1 SV=4 | 902 | 20201116_fqg_colostrum_1ng_glyco_hcd203040_compound15.10765.10765.3 | controllerType=0 controllerNumber=1 scan=10765 | 24.3227 |
| 05401:3 | 7 | K.WN[+2059.735]NTGC[+57.021]QALPSQDEGPSK.A | HexNAc(4)Hex(5)Fuc(1)NeuAc(1) | C[+57], N[+2060] | 1350.212 | 3 | 4048.6214 | 4048.6116 | 0 | 2.4 | 498 | Specific | 672.42 | 586.91 | 586.91 | 14.24 | 77 | >sp\|P01833\|PIGR_HUMAN Polymeric immunoglobulin receptor OS=Homo sapiens OX=9606 GN=PIGR PE=1 SV=4 | 902 | 20201116_fqg_colostrum_1ng_glyco_hcd203040_compound15.8449.8449.3 | controllerType=0 controllerNumber=1 scan=8449 | 20.7244 |
| 07625:3 | 7 | K.WN[+2204.772]NTGC[+57.021]QALPSQDEGPSK.A | HexNAc(4)Hex(5)NeuAc(2) | C[+57], N[+2205] | 1398.5567 | 3 | 4193.6555 | 4193.6491 | 0 | 1.5 | 498 | Specific | 781 | 651.91 | 651.91 | 16.24 | 77 | >sp\|P01833\|PIGR_HUMAN Polymeric immunoglobulin receptor OS=Homo sapiens OX=9606 GN=PIGR PE=1 SV=4 | 902 | 20201116_fqg_colostrum_1ng_glyco_hcd203040_compound15.10768.10768.3 | controllerType=0 controllerNumber=1 scan=10768 | 24.327 |
| 05392:3 | 7 | K.WN[+1913.677]NTGC[+57.021]QALPSQDEGPSK.A | HexNAc(4)Hex(5)NeuAc(1) | C[+57], N[+1914] | 1301.5252 | 3 | 3902.5611 | 3902.5537 | 0 | 1.9 | 498 | Specific | 818.55 | 736.05 | 736.05 | 17.24 | 77 | >sp\|P01833\|PIGR_HUMAN Polymeric immunoglobulin receptor OS=Homo sapiens OX=9606 GN=PIGR PE=1 SV=4 | 902 | 20201116_fqg_colostrum_1ng_glyco_hcd203040_compound15.8440.8440.3 | controllerType=0 controllerNumber=1 scan=8440 | 20.7069 |
| 05393:3 | 7 | K.WN[+1872.651]NTGC[+57.021]QALPSQDEGPSK.A | HexNAc(3)Hex(6)NeuAc(1) | C[+57], N[+1873] | 1287.8499 | 3 | 3861.5352 | 3861.5272 | 0 | 2.1 | 498 | Specific | 824.72 | 740 | 740 | 17.24 | 77 | >sp\|P01833\|PIGR_HUMAN Polymeric immunoglobulin receptor OS=Homo sapiens OX=9606 GN=PIGR PE=1 SV=4 | 902 | 20201116_fqg_colostrum_1ng_glyco_hcd203040_compound15.8441.8441.3 | controllerType=0 controllerNumber=1 scan=8441 | 20.7099 |
| 05389:3 | 7 | K.WN[+1913.677]NTGC[+57.021]QALPSQDEGPSK.A | HexNAc(4)Hex(5)NeuAc(1) | C[+57], N[+1914] | 1301.5252 | 3 | 3902.5611 | 3902.5537 | 0 | 1.9 | 498 | Specific | 826.78 | 772.31 | 772.31 | 17.24 | 77 | >sp\|P01833\|PIGR_HUMAN Polymeric immunoglobulin receptor OS=Homo sapiens OX=9606 GN=PIGR PE=1 SV=4 | 902 | 20201116_fqg_colostrum_1ng_glyco_hcd203040_compound15.8437.8437.3 | controllerType=0 controllerNumber=1 scan=8437 | 20.7024 |
| 05391:3 | 7 | K.WN[+1872.651]NTGC[+57.021]QALPSQDEGPSK.A | HexNAc(3)Hex(6)NeuAc(1) | C[+57], N[+1873] | 1287.8499 | 3 | 3861.5352 | 3861.5272 | 0 | 2.1 | 498 | Specific | 830.46 | 788.25 | 788.25 | 17.24 | 77 | >sp\|P01833\|PIGR_HUMAN Polymeric immunoglobulin receptor OS=Homo sapiens OX=9606 GN=PIGR PE=1 SV=4 | 902 | 20201116_fqg_colostrum_1ng_glyco_hcd203040_compound15.8439.8439.3 | controllerType=0 controllerNumber=1 scan=8439 | 20.7046 |
| 24418:5 | 11 | R.QLAHQSN[+2204.772]STNIFFSPVSIATAFAMLSLGTK.A | HexNAc(4)Hex(5)NeuAc(2) | N[+2205] | 1078.0925 | 5 | 5386.4336 | 5386.4075 | 0 | 4.8 | 64 | Specific | 221.19 | 198.56 | 198.56 | 5.01 | 39 | >sp\|P01009\|A1AT_HUMAN Alpha-1-antitrypsin OS=Homo sapiens OX=9606 GN=SERPINA1 PE=1 SV=3 | 642 | 20201116_fqg_colostrum_1ng_glyco_hcd203040_compound15.28193.28193.5 | controllerType=0 controllerNumber=1 scan=28193 | 55.8768 |
| 24465:4 | 11 | R.[+42.011]QLAHQSN[+2190.793]STNIFFSPVSIATAFAMLSLGTK.A | HexNAc(6)Hex(6) | .[42], N[+2191] | 1354.3602 | 4 | 5414.4191 | 5414.4388 | 0 | -3.6 | 64 | Specific | 395.66 | 395.66 | 395.66 | 9.26 | 39 | >sp\|P01009\|A1AT_HUMAN Alpha-1-antitrypsin OS=Homo sapiens OX=9606 GN=SERPINA1 PE=1 SV=3 | 642 | 20201116_fqg_colostrum_1ng_glyco_hcd203040_compound15.28242.28242.4 | controllerType=0 controllerNumber=1 scan=28242 | 55.9928 |
| 24468:4 | 11 | R.[+42.011]QLAHQSN[+2190.793]STNIFFSPVSIATAFAMLSLGTK.A | HexNAc(6)Hex(6) | .[42], N[+2191] | 1354.3602 | 4 | 5414.4191 | 5414.4388 | 0 | -3.6 | 64 | Specific | 444.49 | 416.95 | 416.95 | 9.39 | 39 | >sp\|P01009\|A1AT_HUMAN Alpha-1-antitrypsin OS=Homo sapiens OX=9606 GN=SERPINA1 PE=1 SV=3 | 642 | 20201116_fqg_colostrum_1ng_glyco_hcd203040_compound15.28245.28245.4 | controllerType=0 controllerNumber=1 scan=28245 | 55.9979 |
| 24413:3 | 11 | R.QLAHQSN[+2204.772]STNIFFSPVSIATAFAMLSLGTK.A | HexNAc(4)Hex(5)NeuAc(2) | N[+2205] | 1796.1485 | 3 | 5386.431 | 5386.4075 | 0 | 4.4 | 64 | Specific | 852.81 | 820.21 | 820.21 | 18.34 | 39 | >sp\|P01009\|A1AT_HUMAN Alpha-1-antitrypsin OS=Homo sapiens OX=9606 GN=SERPINA1 PE=1 SV=3 | 642 | 20201116_fqg_colostrum_1ng_glyco_hcd203040_compound15.28188.28188.3 | controllerType=0 controllerNumber=1 scan=28188 | 55.8682 |
| 24393:3 | 11 | R.QLAHQSN[+2204.772]STNIFFSPVSIATAFAMLSLGTK.A | HexNAc(4)Hex(5)NeuAc(2) | N[+2205] | 1796.145 | 3 | 5386.4204 | 5386.4075 | 0 | 2.4 | 64 | Specific | 860.67 | 803.09 | 590.18 | 18.89 | 39 | >sp\|P01009\|A1AT_HUMAN Alpha-1-antitrypsin OS=Homo sapiens OX=9606 GN=SERPINA1 PE=1 SV=3 | 642 | 20201116_fqg_colostrum_1ng_glyco_hcd203040_compound15.28166.28166.3 | controllerType=0 controllerNumber=1 scan=28166 | 55.8125 |
| 24396:4 | 11 | R.QLAHQSN[+2204.772]STNIFFSPVSIATAFAMLSLGTK.A | HexNAc(4)Hex(5)NeuAc(2) | N[+2205] | 1347.3627 | 4 | 5386.4289 | 5386.4075 | 0 | 4 | 64 | Specific | 887.18 | 839.56 | 839.56 | 19 | 39 | >sp\|P01009\|A1AT_HUMAN Alpha-1-antitrypsin OS=Homo sapiens OX=9606 GN=SERPINA1 PE=1 SV=3 | 642 | 20201116_fqg_colostrum_1ng_glyco_hcd203040_compound15.28170.28170.4 | controllerType=0 controllerNumber=1 scan=28170 | 55.8239 |
| 24415:3 | 11 | R.QLAHQSN[+2204.772]STNIFFSPVSIATAFAMLSLGTK.A | HexNAc(4)Hex(5)NeuAc(2) | N[+2205] | 1796.1485 | 3 | 5386.431 | 5386.4075 | 0 | 4.4 | 64 | Specific | 900 | 828.71 | 828.71 | 19.34 | 39 | >sp\|P01009\|A1AT_HUMAN Alpha-1-antitrypsin OS=Homo sapiens OX=9606 GN=SERPINA1 PE=1 SV=3 | 642 | 20201116_fqg_colostrum_1ng_glyco_hcd203040_compound15.28190.28190.3 | controllerType=0 controllerNumber=1 scan=28190 | 55.8731 |
| 24394:4 | 11 | R.QLAHQSN[+2204.772]STNIFFSPVSIATAFAMLSLGTK.A | HexNAc(4)Hex(5)NeuAc(2) | N[+2205] | 1347.3627 | 4 | 5386.4289 | 5386.4075 | 0 | 4 | 64 | Specific | 962.57 | 958.45 | 958.45 | 21 | 39 | >sp\|P01009\|A1AT_HUMAN Alpha-1-antitrypsin OS=Homo sapiens OX=9606 GN=SERPINA1 PE=1 SV=3 | 642 | 20201116_fqg_colostrum_1ng_glyco_hcd203040_compound15.28168.28168.4 | controllerType=0 controllerNumber=1 scan=28168 | 55.817 |
| 24391:3 | 11 | R.QLAHQSN[+2204.772]STNIFFSPVSIATAFAMLSLGTK.A | HexNAc(4)Hex(5)NeuAc(2) | N[+2205] | 1796.145 | 3 | 5386.4204 | 5386.4075 | 0 | 2.4 | 64 | Specific | 978.87 | 935.25 | 649.1 | 20.89 | 39 | >sp\|P01009\|A1AT_HUMAN Alpha-1-antitrypsin OS=Homo sapiens OX=9606 GN=SERPINA1 PE=1 SV=3 | 642 | 20201116_fqg_colostrum_1ng_glyco_hcd203040_compound15.28164.28164.3 | controllerType=0 controllerNumber=1 scan=28164 | 55.8048 |
| 23292:4 | 11 | K.ADTHDEILEGLNFN[+3007.058]LTEIPEAQIHEGFQELLR.T | HexNAc(5)Hex(6)Fuc(1)NeuAc(3) | N[+3007] | 1675.4743 | 4 | 6698.8755 | 6698.8818 | 0 | -0.9 | 94 | Specific | 465.03 | 372.56 | 372.56 | 8.46 | 39 | >sp\|P01009\|A1AT_HUMAN Alpha-1-antitrypsin OS=Homo sapiens OX=9606 GN=SERPINA1 PE=1 SV=3 | 642 | 20201116_fqg_colostrum_1ng_glyco_hcd203040_compound15.27015.27015.3 | controllerType=0 controllerNumber=1 scan=27015 | 53.1664 |
| 23294:4 | 11 | K.ADTHDEILEGLNFN[+3007.058]LTEIPEAQIHEGFQELLR.T | HexNAc(5)Hex(6)Fuc(1)NeuAc(3) | N[+3007] | 1675.4743 | 4 | 6698.8755 | 6698.8818 | 0 | -0.9 | 94 | Specific | 622.26 | 494.07 | 494.07 | 12.69 | 39 | >sp\|P01009\|A1AT_HUMAN Alpha-1-antitrypsin OS=Homo sapiens OX=9606 GN=SERPINA1 PE=1 SV=3 | 642 | 20201116_fqg_colostrum_1ng_glyco_hcd203040_compound15.27017.27017.3 | controllerType=0 controllerNumber=1 scan=27017 | 53.1709 |
| 21860:5 | 11 | K.ADTHDEILEGLNFN[+2204.772]LTEIPEAQIHEGFQELLR.T | HexNAc(4)Hex(5)NeuAc(2) | N[+2205] | 1180.1273 | 5 | 5896.6075 | 5896.5963 | 0 | 1.9 | 94 | Specific | 740.21 | 576.17 | 576.17 | 16.11 | 39 | >sp\|P01009\|A1AT_HUMAN Alpha-1-antitrypsin OS=Homo sapiens OX=9606 GN=SERPINA1 PE=1 SV=3 | 642 | 20201116_fqg_colostrum_1ng_glyco_hcd203040_compound15.25521.25521.5 | controllerType=0 controllerNumber=1 scan=25521 | 50.0366 |
| 21863:5 | 11 | K.ADTHDEILEGLNFN[+2204.772]LTEIPEAQIHEGFQELLR.T | HexNAc(4)Hex(5)NeuAc(2) | N[+2205] | 1180.1273 | 5 | 5896.6075 | 5896.5963 | 0 | 1.9 | 94 | Specific | 898.84 | 703 | 703 | 19.11 | 39 | >sp\|P01009\|A1AT_HUMAN Alpha-1-antitrypsin OS=Homo sapiens OX=9606 GN=SERPINA1 PE=1 SV=3 | 642 | 20201116_fqg_colostrum_1ng_glyco_hcd203040_compound15.25524.25524.5 | controllerType=0 controllerNumber=1 scan=25524 | 50.043 |
| 20740:4 | 11 | K.YLGN[+2204.772]ATAIFFLPDEGKLQHLENELTHDIITK.F | HexNAc(4)Hex(5)NeuAc(2) | N[+2205] | 1437.1616 | 4 | 5745.6247 | 5745.6098 | 0 | 2.6 | 268 | Specific | 935.17 | 850.57 | 850.57 | 18.8 | 39 | >sp\|P01009\|A1AT_HUMAN Alpha-1-antitrypsin OS=Homo sapiens OX=9606 GN=SERPINA1 PE=1 SV=3 | 642 | 20201116_fqg_colostrum_1ng_glyco_hcd203040_compound15.24356.24356.4 | controllerType=0 controllerNumber=1 scan=24356 | 47.6261 |
| 20737:4 | 11 | K.YLGN[+2204.772]ATAIFFLPDEGKLQHLENELTHDIITK.F | HexNAc(4)Hex(5)NeuAc(2) | N[+2205] | 1437.1616 | 4 | 5745.6247 | 5745.6098 | 0 | 2.6 | 268 | Specific | 1254.5 | 1149.54 | 1149.54 | 25.8 | 39 | >sp\|P01009\|A1AT_HUMAN Alpha-1-antitrypsin OS=Homo sapiens OX=9606 GN=SERPINA1 PE=1 SV=3 | 642 | 20201116_fqg_colostrum_1ng_glyco_hcd203040_compound15.24353.24353.4 | controllerType=0 controllerNumber=1 scan=24353 | 47.6129 |
| 22060:4 | 13 | K.APDKNVIFSPLSISTALAFLSLGAHN[+2570.925]TTLTEILK.G | HexNAc(5)Hex(6)Fuc(2)NeuAc(1) | N[+2571] | 1539.2362 | 4 | 6153.923 | 6153.9032 | 0 | 3.2 | 68 | Specific | 260.42 | 260.06 | 260.06 | 4.73 | 32 | >sp\|P01011\|AACT_HUMAN Alpha-1-antichymotrypsin OS=Homo sapiens OX=9606 GN=SERPINA3 PE=1 SV=2 | 1314 | 20201116_fqg_colostrum_1ng_glyco_hcd203040_compound15.25730.25730.2 | controllerType=0 controllerNumber=1 scan=25730 | 50.5103 |
| 00834:2 | 20 | R.SYN[+1241.454]DSVDPR.I | HexNAc(3)Hex(3)Fuc(1) | N[+1241] | 1147.4638 | 2 | 2293.9203 | 2293.9188 | 0 | 0.6 | 481 | Specific | 298.92 | 250.96 | 250.96 | 6.57 | 34 | >sp\|P05164\|PERM_HUMAN Myeloperoxidase OS=Homo sapiens OX=9606 GN=MPO PE=1 SV=1 | 12 | 20201116_fqg_colostrum_1ng_glyco_hcd203040_compound15.3704.3704.2 | controllerType=0 controllerNumber=1 scan=3704 | 12.4067 |
| 00837:2 | 20 | R.SYN[+1241.454]DSVDPR.I | HexNAc(3)Hex(3)Fuc(1) | N[+1241] | 1147.4638 | 2 | 2293.9203 | 2293.9188 | 0 | 0.6 | 481 | Specific | 314.5 | 195.48 | 195.48 | 6.47 | 34 | >sp\|P05164\|PERM_HUMAN Myeloperoxidase OS=Homo sapiens OX=9606 GN=MPO PE=1 SV=1 | 12 | 20201116_fqg_colostrum_1ng_glyco_hcd203040_compound15.3707.3707.2 | controllerType=0 controllerNumber=1 scan=3707 | 12.4109 |
| 00821:2 | 20 | R.SYN[+1241.454]DSVDPR.I | HexNAc(3)Hex(3)Fuc(1) | N[+1241] | 1147.4659 | 2 | 2293.9244 | 2293.9188 | 0 | 2.5 | 481 | Specific | 343.32 | 275.63 | 275.63 | 7.5 | 34 | >sp\|P05164\|PERM_HUMAN Myeloperoxidase OS=Homo sapiens OX=9606 GN=MPO PE=1 SV=1 | 12 | 20201116_fqg_colostrum_1ng_glyco_hcd203040_compound15.3690.3690.2 | controllerType=0 controllerNumber=1 scan=3690 | 12.3782 |
| 00852:2 | 20 | R.SYN[+1038.375]DSVDPR.I | HexNAc(2)Hex(3)Fuc(1) | N[+1038] | 1045.9257 | 2 | 2090.8441 | 2090.8395 | 0 | 2.2 | 481 | Specific | 365.12 | 221.3 | 221.3 | 7.32 | 34 | >sp\|P05164\|PERM_HUMAN Myeloperoxidase OS=Homo sapiens OX=9606 GN=MPO PE=1 SV=1 | 12 | 20201116_fqg_colostrum_1ng_glyco_hcd203040_compound15.3723.3723.2 | controllerType=0 controllerNumber=1 scan=3723 | 12.4423 |
| 00823:2 | 20 | R.SYN[+1241.454]DSVDPR.I | HexNAc(3)Hex(3)Fuc(1) | N[+1241] | 1147.4659 | 2 | 2293.9244 | 2293.9188 | 0 | 2.5 | 481 | Specific | 380.72 | 290.39 | 290.39 | 7.67 | 34 | >sp\|P05164\|PERM_HUMAN Myeloperoxidase OS=Homo sapiens OX=9606 GN=MPO PE=1 SV=1 | 12 | 20201116_fqg_colostrum_1ng_glyco_hcd203040_compound15.3692.3692.2 | controllerType=0 controllerNumber=1 scan=3692 | 12.3833 |
| 00854:2 | 20 | R.SYN[+1038.375]DSVDPR.I | HexNAc(2)Hex(3)Fuc(1) | N[+1038] | 1045.9257 | 2 | 2090.8441 | 2090.8395 | 0 | 2.2 | 481 | Specific | 405.48 | 358.94 | 358.94 | 8.2 | 34 | >sp\|P05164\|PERM_HUMAN Myeloperoxidase OS=Homo sapiens OX=9606 GN=MPO PE=1 SV=1 | 12 | 20201116_fqg_colostrum_1ng_glyco_hcd203040_compound15.3725.3725.2 | controllerType=0 controllerNumber=1 scan=3725 | 12.4465 |
| 01798:3 | 21 | K.N[+1540.529]NSIPDKQITASSSYK.T | HexNAc(2)Hex(7) | N[+1541] | 1098.4763 | 3 | 3293.4145 | 3293.4048 | 0 | 2.9 | 238 | Specific | 392.05 | 358.38 | 358.38 | 7.74 | 27 | >sp\|Q08431\|MFGM_HUMAN Lactadherin OS=Homo sapiens OX=9606 GN=MFGE8 PE=1 SV=3 | 1064 | 20201116_fqg_colostrum_1ng_glyco_hcd203040_compound15.4705.4705.3 | controllerType=0 controllerNumber=1 scan=4705 | 14.2815 |
| 01796:3 | 21 | K.N[+1540.529]NSIPDKQITASSSYK.T | HexNAc(2)Hex(7) | N[+1541] | 1098.4763 | 3 | 3293.4145 | 3293.4048 | 0 | 2.9 | 238 | Specific | 446.19 | 267.43 | 267.43 | 7.36 | 27 | >sp\|Q08431\|MFGM_HUMAN Lactadherin OS=Homo sapiens OX=9606 GN=MFGE8 PE=1 SV=3 | 1064 | 20201116_fqg_colostrum_1ng_glyco_hcd203040_compound15.4703.4703.3 | controllerType=0 controllerNumber=1 scan=4703 | 14.278 |
| 01888:3 | 21 | K.N[+1216.423]NSIPDKQITASSSYK.T | HexNAc(2)Hex(5) | N[+1216] | 990.4403 | 3 | 2969.3063 | 2969.2991 | 0 | 2.4 | 238 | Specific | 739.61 | 523.23 | 523.23 | 14.55 | 27 | >sp\|Q08431\|MFGM_HUMAN Lactadherin OS=Homo sapiens OX=9606 GN=MFGE8 PE=1 SV=3 | 1064 | 20201116_fqg_colostrum_1ng_glyco_hcd203040_compound15.4799.4799.3 | controllerType=0 controllerNumber=1 scan=4799 | 14.4646 |
| 01886:3 | 21 | K.N[+1216.423]NSIPDKQITASSSYK.T | HexNAc(2)Hex(5) | N[+1216] | 990.4403 | 3 | 2969.3063 | 2969.2991 | 0 | 2.4 | 238 | Specific | 888.41 | 654.34 | 654.34 | 17.55 | 27 | >sp\|Q08431\|MFGM_HUMAN Lactadherin OS=Homo sapiens OX=9606 GN=MFGE8 PE=1 SV=3 | 1064 | 20201116_fqg_colostrum_1ng_glyco_hcd203040_compound15.4797.4797.3 | controllerType=0 controllerNumber=1 scan=4797 | 14.4597 |
| 13490:4 | 22 | R.IIVPLNNREN[+2205.793]ISDPTSPLR.T | HexNAc(4)Hex(5)Fuc(2)NeuAc(1) | N[+2206] | 1089.2444 | 4 | 4353.9559 | 4353.97 | 0 | -3.2 | 62 | Specific | 260.23 | 202.98 | 202.98 | 2.55 | 29 | >sp\|P01591\|IGJ_HUMAN Immunoglobulin J chain OS=Homo sapiens OX=9606 GN=JCHAIN PE=1 SV=4 | 1392 | 20201116_fqg_colostrum_1ng_glyco_hcd203040_compound15.16832.16832.2 | controllerType=0 controllerNumber=1 scan=16832 | 34.0489 |
| 09515:3 | 22 | R.IIVPLNNREN[+1768.640]ISDPTSPLR.T | HexNAc(4)Hex(5)Fuc(1) | N[+1769] | 1306.2781 | 3 | 3916.8198 | 3916.8167 | 0 | 0.8 | 62 | Specific | 361.01 | 294.76 | 294.76 | 7.54 | 29 | >sp\|P01591\|IGJ_HUMAN Immunoglobulin J chain OS=Homo sapiens OX=9606 GN=JCHAIN PE=1 SV=4 | 1392 | 20201116_fqg_colostrum_1ng_glyco_hcd203040_compound15.12722.12722.3 | controllerType=0 controllerNumber=1 scan=12722 | 27.5287 |
| 09483:3 | 22 | R.IIVPLNNREN[+1663.608]ISDPTSPLR.T | HexNAc(5)Hex(4) | N[+1664] | 1271.2687 | 3 | 3811.7916 | 3811.7853 | 0 | 1.7 | 62 | Specific | 394.87 | 382.28 | 382.28 | 9.68 | 29 | >sp\|P01591\|IGJ_HUMAN Immunoglobulin J chain OS=Homo sapiens OX=9606 GN=JCHAIN PE=1 SV=4 | 1392 | 20201116_fqg_colostrum_1ng_glyco_hcd203040_compound15.12689.12689.3 | controllerType=0 controllerNumber=1 scan=12689 | 27.4683 |
| 09548:3 | 22 | R.IIVPLNNREN[+1647.613]ISDPTSPLR.T | HexNAc(5)Hex(3)Fuc(1) | N[+1648] | 1265.9341 | 3 | 3795.7879 | 3795.7904 | 0 | -0.7 | 62 | Specific | 433.54 | 349.52 | 349.52 | 7.6 | 29 | >sp\|P01591\|IGJ_HUMAN Immunoglobulin J chain OS=Homo sapiens OX=9606 GN=JCHAIN PE=1 SV=4 | 1392 | 20201116_fqg_colostrum_1ng_glyco_hcd203040_compound15.12756.12756.3 | controllerType=0 controllerNumber=1 scan=12756 | 27.5872 |
| 09697:3 | 22 | R.IIVPLNNREN[+1038.375]ISDPTSPLR.T | HexNAc(2)Hex(3)Fuc(1) | N[+1038] | 1062.8556 | 3 | 3186.5523 | 3186.5523 | 0 | 0 | 62 | Specific | 437.57 | 437.57 | 437.57 | 9.02 | 29 | >sp\|P01591\|IGJ_HUMAN Immunoglobulin J chain OS=Homo sapiens OX=9606 GN=JCHAIN PE=1 SV=4 | 1392 | 20201116_fqg_colostrum_1ng_glyco_hcd203040_compound15.12911.12911.3 | controllerType=0 controllerNumber=1 scan=12911 | 27.8286 |
| 09425:3 | 22 | R.IIVPLNNREN[+1809.666]ISDPTSPLR.T | HexNAc(5)Hex(4)Fuc(1) | N[+1810] | 1319.9556 | 3 | 3957.8523 | 3957.8432 | 0 | 2.3 | 62 | Specific | 472.03 | 373.02 | 373.02 | 8.71 | 29 | >sp\|P01591\|IGJ_HUMAN Immunoglobulin J chain OS=Homo sapiens OX=9606 GN=JCHAIN PE=1 SV=4 | 1392 | 20201116_fqg_colostrum_1ng_glyco_hcd203040_compound15.12629.12629.3 | controllerType=0 controllerNumber=1 scan=12629 | 27.359 |
| 09518:3 | 22 | R.IIVPLNNREN[+1768.640]ISDPTSPLR.T | HexNAc(4)Hex(5)Fuc(1) | N[+1769] | 1306.2781 | 3 | 3916.8198 | 3916.8167 | 0 | 0.8 | 62 | Specific | 474.35 | 394.8 | 394.8 | 9.73 | 29 | >sp\|P01591\|IGJ_HUMAN Immunoglobulin J chain OS=Homo sapiens OX=9606 GN=JCHAIN PE=1 SV=4 | 1392 | 20201116_fqg_colostrum_1ng_glyco_hcd203040_compound15.12725.12725.3 | controllerType=0 controllerNumber=1 scan=12725 | 27.5329 |
| 13394:3 | 22 | R.IIVPLNNREN[+2350.830]ISDPTSPLR.T | HexNAc(4)Hex(5)Fuc(1)NeuAc(2) | N[+2351] | 1500.3417 | 3 | 4499.0106 | 4499.0075 | 0 | 0.7 | 62 | Specific | 478.43 | 470.26 | 470.26 | 9.73 | 29 | >sp\|P01591\|IGJ_HUMAN Immunoglobulin J chain OS=Homo sapiens OX=9606 GN=JCHAIN PE=1 SV=4 | 1392 | 20201116_fqg_colostrum_1ng_glyco_hcd203040_compound15.16731.16731.3 | controllerType=0 controllerNumber=1 scan=16731 | 33.88 |
| 09551:3 | 22 | R.IIVPLNNREN[+1647.613]ISDPTSPLR.T | HexNAc(5)Hex(3)Fuc(1) | N[+1648] | 1265.9341 | 3 | 3795.7879 | 3795.7904 | 0 | -0.7 | 62 | Specific | 483.76 | 423.72 | 423.72 | 9.31 | 29 | >sp\|P01591\|IGJ_HUMAN Immunoglobulin J chain OS=Homo sapiens OX=9606 GN=JCHAIN PE=1 SV=4 | 1392 | 20201116_fqg_colostrum_1ng_glyco_hcd203040_compound15.12759.12759.3 | controllerType=0 controllerNumber=1 scan=12759 | 27.5962 |
| 11250:3 | 22 | R.IIVPLNNREN[+1856.656]ISDPTSPLR.T | HexNAc(3)Hex(5)Fuc(1)NeuAc(1) | N[+1857] | 1335.6177 | 3 | 4004.8386 | 4004.8327 | 0 | 1.5 | 62 | Specific | 507.12 | 418.34 | 418.34 | 9.93 | 29 | >sp\|P01591\|IGJ_HUMAN Immunoglobulin J chain OS=Homo sapiens OX=9606 GN=JCHAIN PE=1 SV=4 | 1392 | 20201116_fqg_colostrum_1ng_glyco_hcd203040_compound15.14516.14516.3 | controllerType=0 controllerNumber=1 scan=14516 | 30.4416 |
| 11253:3 | 22 | R.IIVPLNNREN[+1856.656]ISDPTSPLR.T | HexNAc(3)Hex(5)Fuc(1)NeuAc(1) | N[+1857] | 1335.6177 | 3 | 4004.8386 | 4004.8327 | 0 | 1.5 | 62 | Specific | 509.98 | 509.98 | 509.98 | 9.93 | 29 | >sp\|P01591\|IGJ_HUMAN Immunoglobulin J chain OS=Homo sapiens OX=9606 GN=JCHAIN PE=1 SV=4 | 1392 | 20201116_fqg_colostrum_1ng_glyco_hcd203040_compound15.14519.14519.3 | controllerType=0 controllerNumber=1 scan=14519 | 30.4469 |
| 13397:3 | 22 | R.IIVPLNNREN[+2350.830]ISDPTSPLR.T | HexNAc(4)Hex(5)Fuc(1)NeuAc(2) | N[+2351] | 1500.3417 | 3 | 4499.0106 | 4499.0075 | 0 | 0.7 | 62 | Specific | 516.88 | 449.67 | 449.67 | 9.93 | 29 | >sp\|P01591\|IGJ_HUMAN Immunoglobulin J chain OS=Homo sapiens OX=9606 GN=JCHAIN PE=1 SV=4 | 1392 | 20201116_fqg_colostrum_1ng_glyco_hcd203040_compound15.16734.16734.3 | controllerType=0 controllerNumber=1 scan=16734 | 33.8856 |
| 09479:3 | 22 | R.IIVPLNNREN[+1565.560]ISDPTSPLR.T | HexNAc(3)Hex(5)Fuc(1) | N[+1566] | 1238.5858 | 3 | 3713.7427 | 3713.7373 | 0 | 1.5 | 62 | Specific | 521.93 | 478.1 | 478.1 | 10.04 | 29 | >sp\|P01591\|IGJ_HUMAN Immunoglobulin J chain OS=Homo sapiens OX=9606 GN=JCHAIN PE=1 SV=4 | 1392 | 20201116_fqg_colostrum_1ng_glyco_hcd203040_compound15.12684.12684.3 | controllerType=0 controllerNumber=1 scan=12684 | 27.4538 |
| 09495:3 | 22 | R.IIVPLNNREN[+1257.449]ISDPTSPLR.T | HexNAc(3)Hex(4) | N[+1257] | 1135.882 | 3 | 3405.6314 | 3405.6266 | 0 | 1.4 | 62 | Specific | 532.44 | 528.49 | 528.49 | 10.04 | 29 | >sp\|P01591\|IGJ_HUMAN Immunoglobulin J chain OS=Homo sapiens OX=9606 GN=JCHAIN PE=1 SV=4 | 1392 | 20201116_fqg_colostrum_1ng_glyco_hcd203040_compound15.12701.12701.3 | controllerType=0 controllerNumber=1 scan=12701 | 27.4857 |
| 11320:3 | 22 | R.IIVPLNNREN[+1694.603]ISDPTSPLR.T | HexNAc(3)Hex(4)Fuc(1)NeuAc(1) | N[+1695] | 1281.6016 | 3 | 3842.7903 | 3842.7799 | 0 | 2.7 | 62 | Specific | 541.31 | 541.31 | 541.31 | 10.38 | 29 | >sp\|P01591\|IGJ_HUMAN Immunoglobulin J chain OS=Homo sapiens OX=9606 GN=JCHAIN PE=1 SV=4 | 1392 | 20201116_fqg_colostrum_1ng_glyco_hcd203040_compound15.14589.14589.3 | controllerType=0 controllerNumber=1 scan=14589 | 30.589 |
| 09481:3 | 22 | R.IIVPLNNREN[+1565.560]ISDPTSPLR.T | HexNAc(3)Hex(5)Fuc(1) | N[+1566] | 1238.5858 | 3 | 3713.7427 | 3713.7373 | 0 | 1.5 | 62 | Specific | 542.36 | 475.25 | 475.25 | 10.14 | 29 | >sp\|P01591\|IGJ_HUMAN Immunoglobulin J chain OS=Homo sapiens OX=9606 GN=JCHAIN PE=1 SV=4 | 1392 | 20201116_fqg_colostrum_1ng_glyco_hcd203040_compound15.12686.12686.3 | controllerType=0 controllerNumber=1 scan=12686 | 27.458 |
| 09540:3 | 22 | R.IIVPLNNREN[+1501.555]ISDPTSPLR.T | HexNAc(5)Hex(3) | N[+1502] | 1217.251 | 3 | 3649.7385 | 3649.7325 | 0 | 1.7 | 62 | Specific | 547.47 | 538.71 | 538.71 | 10.38 | 29 | >sp\|P01591\|IGJ_HUMAN Immunoglobulin J chain OS=Homo sapiens OX=9606 GN=JCHAIN PE=1 SV=4 | 1392 | 20201116_fqg_colostrum_1ng_glyco_hcd203040_compound15.12748.12748.3 | controllerType=0 controllerNumber=1 scan=12748 | 27.577 |
| 09492:3 | 22 | R.IIVPLNNREN[+1257.449]ISDPTSPLR.T | HexNAc(3)Hex(4) | N[+1257] | 1135.882 | 3 | 3405.6314 | 3405.6266 | 0 | 1.4 | 62 | Specific | 548.95 | 511.88 | 511.88 | 10.14 | 29 | >sp\|P01591\|IGJ_HUMAN Immunoglobulin J chain OS=Homo sapiens OX=9606 GN=JCHAIN PE=1 SV=4 | 1392 | 20201116_fqg_colostrum_1ng_glyco_hcd203040_compound15.12698.12698.3 | controllerType=0 controllerNumber=1 scan=12698 | 27.4815 |
| 09428:3 | 22 | R.IIVPLNNREN[+1809.666]ISDPTSPLR.T | HexNAc(5)Hex(4)Fuc(1) | N[+1810] | 1319.9556 | 3 | 3957.8523 | 3957.8432 | 0 | 2.3 | 62 | Specific | 559.35 | 465.15 | 465.15 | 10.38 | 29 | >sp\|P01591\|IGJ_HUMAN Immunoglobulin J chain OS=Homo sapiens OX=9606 GN=JCHAIN PE=1 SV=4 | 1392 | 20201116_fqg_colostrum_1ng_glyco_hcd203040_compound15.12632.12632.3 | controllerType=0 controllerNumber=1 scan=12632 | 27.3638 |
| 09457:3 | 22 | R.IIVPLNNREN[+1460.529]ISDPTSPLR.T | HexNAc(4)Hex(4) | N[+1461] | 1203.5753 | 3 | 3608.7112 | 3608.7059 | 0 | 1.5 | 62 | Specific | 560.95 | 544.18 | 544.18 | 10.33 | 29 | >sp\|P01591\|IGJ_HUMAN Immunoglobulin J chain OS=Homo sapiens OX=9606 GN=JCHAIN PE=1 SV=4 | 1392 | 20201116_fqg_colostrum_1ng_glyco_hcd203040_compound15.12662.12662.3 | controllerType=0 controllerNumber=1 scan=12662 | 27.4194 |
| 09543:3 | 22 | R.IIVPLNNREN[+1501.555]ISDPTSPLR.T | HexNAc(5)Hex(3) | N[+1502] | 1217.251 | 3 | 3649.7385 | 3649.7325 | 0 | 1.7 | 62 | Specific | 566.18 | 566.18 | 566.18 | 10.57 | 29 | >sp\|P01591\|IGJ_HUMAN Immunoglobulin J chain OS=Homo sapiens OX=9606 GN=JCHAIN PE=1 SV=4 | 1392 | 20201116_fqg_colostrum_1ng_glyco_hcd203040_compound15.12751.12751.3 | controllerType=0 controllerNumber=1 scan=12751 | 27.5815 |
| 11317:3 | 22 | R.IIVPLNNREN[+1694.603]ISDPTSPLR.T | HexNAc(3)Hex(4)Fuc(1)NeuAc(1) | N[+1695] | 1281.6016 | 3 | 3842.7903 | 3842.7799 | 0 | 2.7 | 62 | Specific | 567.41 | 567.41 | 567.41 | 10.57 | 29 | >sp\|P01591\|IGJ_HUMAN Immunoglobulin J chain OS=Homo sapiens OX=9606 GN=JCHAIN PE=1 SV=4 | 1392 | 20201116_fqg_colostrum_1ng_glyco_hcd203040_compound15.14586.14586.3 | controllerType=0 controllerNumber=1 scan=14586 | 30.5825 |
| 09700:3 | 22 | R.IIVPLNNREN[+1038.375]ISDPTSPLR.T | HexNAc(2)Hex(3)Fuc(1) | N[+1038] | 1062.8556 | 3 | 3186.5523 | 3186.5523 | 0 | 0 | 62 | Specific | 578.3 | 558.48 | 558.48 | 9.83 | 29 | >sp\|P01591\|IGJ_HUMAN Immunoglobulin J chain OS=Homo sapiens OX=9606 GN=JCHAIN PE=1 SV=4 | 1392 | 20201116_fqg_colostrum_1ng_glyco_hcd203040_compound15.12914.12914.3 | controllerType=0 controllerNumber=1 scan=12914 | 27.8327 |
| 09460:3 | 22 | R.IIVPLNNREN[+1460.529]ISDPTSPLR.T | HexNAc(4)Hex(4) | N[+1461] | 1203.5753 | 3 | 3608.7112 | 3608.7059 | 0 | 1.5 | 62 | Specific | 608.95 | 576.72 | 576.72 | 12.71 | 29 | >sp\|P01591\|IGJ_HUMAN Immunoglobulin J chain OS=Homo sapiens OX=9606 GN=JCHAIN PE=1 SV=4 | 1392 | 20201116_fqg_colostrum_1ng_glyco_hcd203040_compound15.12665.12665.3 | controllerType=0 controllerNumber=1 scan=12665 | 27.4242 |
| 09485:3 | 22 | R.IIVPLNNREN[+1663.608]ISDPTSPLR.T | HexNAc(5)Hex(4) | N[+1664] | 1271.2687 | 3 | 3811.7916 | 3811.7853 | 0 | 1.7 | 62 | Specific | 616.19 | 596.65 | 596.65 | 12.95 | 29 | >sp\|P01591\|IGJ_HUMAN Immunoglobulin J chain OS=Homo sapiens OX=9606 GN=JCHAIN PE=1 SV=4 | 1392 | 20201116_fqg_colostrum_1ng_glyco_hcd203040_compound15.12691.12691.3 | controllerType=0 controllerNumber=1 scan=12691 | 27.4725 |
| 04697:2 | 22 | R.EN[+1663.608]ISDPTSPLR.T | HexNAc(5)Hex(4) | N[+1664] | 1446.6192 | 2 | 2892.2311 | 2892.225 | 0 | 2.1 | 70 | Specific | 357.8 | 261.58 | 261.58 | 7.5 | 29 | >sp\|P01591\|IGJ_HUMAN Immunoglobulin J chain OS=Homo sapiens OX=9606 GN=JCHAIN PE=1 SV=4 | 1392 | 20201116_fqg_colostrum_1ng_glyco_hcd203040_compound15.7722.7722.2 | controllerType=0 controllerNumber=1 scan=7722 | 19.5151 |
| 04700:2 | 22 | R.EN[+1663.608]ISDPTSPLR.T | HexNAc(5)Hex(4) | N[+1664] | 1446.6192 | 2 | 2892.2311 | 2892.225 | 0 | 2.1 | 70 | Specific | 412.95 | 319.45 | 319.45 | 7.97 | 29 | >sp\|P01591\|IGJ_HUMAN Immunoglobulin J chain OS=Homo sapiens OX=9606 GN=JCHAIN PE=1 SV=4 | 1392 | 20201116_fqg_colostrum_1ng_glyco_hcd203040_compound15.7725.7725.2 | controllerType=0 controllerNumber=1 scan=7725 | 19.5196 |
| 06699:2 | 22 | R.EN[+1913.677]ISDPTSPLR.T | HexNAc(4)Hex(5)NeuAc(1) | N[+1914] | 1571.6547 | 2 | 3142.3021 | 3142.2938 | 0 | 2.6 | 70 | Specific | 422.06 | 285.82 | 285.82 | 7.75 | 29 | >sp\|P01591\|IGJ_HUMAN Immunoglobulin J chain OS=Homo sapiens OX=9606 GN=JCHAIN PE=1 SV=4 | 1392 | 20201116_fqg_colostrum_1ng_glyco_hcd203040_compound15.9808.9808.2 | controllerType=0 controllerNumber=1 scan=9808 | 22.787 |
| 06700:2 | 22 | R.EN[+1913.677]ISDPTSPLR.T | HexNAc(4)Hex(5)NeuAc(1) | N[+1914] | 1571.6547 | 2 | 3142.3021 | 3142.2938 | 0 | 2.6 | 70 | Specific | 475.03 | 305.83 | 305.83 | 8.16 | 29 | >sp\|P01591\|IGJ_HUMAN Immunoglobulin J chain OS=Homo sapiens OX=9606 GN=JCHAIN PE=1 SV=4 | 1392 | 20201116_fqg_colostrum_1ng_glyco_hcd203040_compound15.9809.9809.2 | controllerType=0 controllerNumber=1 scan=9809 | 22.7908 |
| 23898:3 | 23 | K.MLN[+2351.851]TSSLLEQLNEQFNWVSR.L | HexNAc(4)Hex(5)Fuc(3)NeuAc(1) | N[+2352] | 1587.6825 | 3 | 4761.0329 | 4761.0375 | 0 | -1 | 352 | Specific | 246.17 | 234.48 | 234.48 | 6.31 | 26 | >sp\|P10909\|CLUS_HUMAN Clusterin OS=Homo sapiens OX=9606 GN=CLU PE=1 SV=1 | 1467 | 20201116_fqg_colostrum_1ng_glyco_hcd203040_compound15.27649.27649.3 | controllerType=0 controllerNumber=1 scan=27649 | 54.6143 |
| 23786:4 | 23 | K.MLN[+2350.830]TSSLLEQLNEQFNWVSR.L | HexNAc(4)Hex(5)Fuc(1)NeuAc(2) | N[+2351] | 1190.7641 | 4 | 4760.0345 | 4760.0171 | 0 | 3.7 | 352 | Specific | 246.89 | 198.37 | 198.37 | 5.5 | 26 | >sp\|P10909\|CLUS_HUMAN Clusterin OS=Homo sapiens OX=9606 GN=CLU PE=1 SV=1 | 1467 | 20201116_fqg_colostrum_1ng_glyco_hcd203040_compound15.27532.27532.3 | controllerType=0 controllerNumber=1 scan=27532 | 54.3325 |
| 21975:3 | 23 | K.MLN[+1913.677]TSSLLEQLNEQFNWVSR.L | HexNAc(4)Hex(5)NeuAc(1) | N[+1914] | 1441.6342 | 3 | 4322.8881 | 4322.8637 | 0 | 5.6 | 352 | Specific | 664.71 | 621.83 | 621.83 | 13.87 | 26 | >sp\|P10909\|CLUS_HUMAN Clusterin OS=Homo sapiens OX=9606 GN=CLU PE=1 SV=1 | 1467 | 20201116_fqg_colostrum_1ng_glyco_hcd203040_compound15.25642.25642.3 | controllerType=0 controllerNumber=1 scan=25642 | 50.3178 |
| 21978:3 | 23 | K.MLN[+1913.677]TSSLLEQLNEQFNWVSR.L | HexNAc(4)Hex(5)NeuAc(1) | N[+1914] | 1441.6342 | 3 | 4322.8881 | 4322.8637 | 0 | 5.6 | 352 | Specific | 923.95 | 761.49 | 761.49 | 18.87 | 26 | >sp\|P10909\|CLUS_HUMAN Clusterin OS=Homo sapiens OX=9606 GN=CLU PE=1 SV=1 | 1467 | 20201116_fqg_colostrum_1ng_glyco_hcd203040_compound15.25645.25645.3 | controllerType=0 controllerNumber=1 scan=25645 | 50.3233 |
| 21738:3 | 23 | K.MLN[+2059.735]TSSLLEQLNEQFNWVSR.L | HexNAc(4)Hex(5)Fuc(1)NeuAc(1) | N[+2060] | 1490.3182 | 3 | 4468.94 | 4468.9217 | 0 | 4.1 | 352 | Specific | 969.21 | 695.74 | 695.74 | 20.09 | 26 | >sp\|P10909\|CLUS_HUMAN Clusterin OS=Homo sapiens OX=9606 GN=CLU PE=1 SV=1 | 1467 | 20201116_fqg_colostrum_1ng_glyco_hcd203040_compound15.25394.25394.3 | controllerType=0 controllerNumber=1 scan=25394 | 49.7585 |
| 21735:3 | 23 | K.MLN[+2059.735]TSSLLEQLNEQFNWVSR.L | HexNAc(4)Hex(5)Fuc(1)NeuAc(1) | N[+2060] | 1490.3182 | 3 | 4468.94 | 4468.9217 | 0 | 4.1 | 352 | Specific | 1054.64 | 769.89 | 769.89 | 22.09 | 26 | >sp\|P10909\|CLUS_HUMAN Clusterin OS=Homo sapiens OX=9606 GN=CLU PE=1 SV=1 | 1467 | 20201116_fqg_colostrum_1ng_glyco_hcd203040_compound15.25391.25391.3 | controllerType=0 controllerNumber=1 scan=25391 | 49.7519 |
| 03506:2 | 24 | K.AAIPSALDTN[+1216.423]SSK.S | HexNAc(2)Hex(5) | N[+1216] | 1246.0466 | 2 | 2491.0858 | 2491.0815 | 0 | 1.7 | 542 | Specific | 546.95 | 470.93 | 470.93 | 10.36 | 20 | >sp\|Q08380\|LG3BP_HUMAN Galectin-3-binding protein OS=Homo sapiens OX=9606 GN=LGALS3BP PE=1 SV=1 | 1562 | 20201116_fqg_colostrum_1ng_glyco_hcd203040_compound15.6475.6475.2 | controllerType=0 controllerNumber=1 scan=6475 | 17.5502 |
| 03503:2 | 24 | K.AAIPSALDTN[+1216.423]SSK.S | HexNAc(2)Hex(5) | N[+1216] | 1246.0466 | 2 | 2491.0858 | 2491.0815 | 0 | 1.7 | 542 | Specific | 615.07 | 471.31 | 471.31 | 12.93 | 20 | >sp\|Q08380\|LG3BP_HUMAN Galectin-3-binding protein OS=Homo sapiens OX=9606 GN=LGALS3BP PE=1 SV=1 | 1562 | 20201116_fqg_colostrum_1ng_glyco_hcd203040_compound15.6472.6472.2 | controllerType=0 controllerNumber=1 scan=6472 | 17.5453 |
| 22928:5 | 25 | R.SDHGSSISC[+57.021]QPPAEIPGYLPADTVHLAVEFFN[+2350.830]LTHLPANLLQGASI | HexNAc(4)Hex(5)Fuc(1)NeuAc(2) | C[+57], N[+2351] | 1450.8671 | 5 | 7250.3064 | 7250.2714 | 0 | 4.8 | 48 | Specific | 533.55 | 533.55 | 533.55 | 11.43 | 18 | >sp\|P02750\|A2GL_HUMAN Leucine-rich alpha-2-glycoprotein OS=Homo sapiens OX=9606 GN=LRG1 PE=1 SV=2 | 518 | 20201116_fqg_colostrum_1ng_glyco_hcd203040_compound15.26634.26634.5 | controllerType=0 controllerNumber=1 scan=26634 | 52.3267 |
| 21708:5 | 25 | R.SDHGSSISC[+57.021]QPPAEIPGYLPADTVHLAVEFFN[+2570.925]LTHLPANLLQGASI | HexNAc(5)Hex(6)Fuc(2)NeuAc(1) | C[+57], N[+2571] | 1494.8858 | 5 | 7470.3998 | 7470.3661 | 0 | 4.5 | 48 | Specific | 694.26 | 694.26 | 694.26 | 15.1 | 18 | >sp\|P02750\|A2GL_HUMAN Leucine-rich alpha-2-glycoprotein OS=Homo sapiens OX=9606 GN=LRG1 PE=1 SV=2 | 518 | 20201116_fqg_colostrum_1ng_glyco_hcd203040_compound15.25363.25363.5 | controllerType=0 controllerNumber=1 scan=25363 | 49.6975 |
| 22931:5 | 25 | R.SDHGSSISC[+57.021]QPPAEIPGYLPADTVHLAVEFFN[+2350.830]LTHLPANLLQGASI | HexNAc(4)Hex(5)Fuc(1)NeuAc(2) | C[+57], N[+2351] | 1450.8671 | 5 | 7250.3064 | 7250.2714 | 0 | 4.8 | 48 | Specific | 827.09 | 738.23 | 738.23 | 18.1 | 18 | >sp\|P02750\|A2GL_HUMAN Leucine-rich alpha-2-glycoprotein OS=Homo sapiens OX=9606 GN=LRG1 PE=1 SV=2 | 518 | 20201116_fqg_colostrum_1ng_glyco_hcd203040_compound15.26637.26637.5 | controllerType=0 controllerNumber=1 scan=26637 | 52.3322 |
| 21705:5 | 25 | R.SDHGSSISC[+57.021]QPPAEIPGYLPADTVHLAVEFFN[+2570.925]LTHLPANLLQGASI | HexNAc(5)Hex(6)Fuc(2)NeuAc(1) | C[+57], N[+2571] | 1494.8858 | 5 | 7470.3998 | 7470.3661 | 0 | 4.5 | 48 | Specific | 914.34 | 914.34 | 914.34 | 20.1 | 18 | >sp\|P02750\|A2GL_HUMAN Leucine-rich alpha-2-glycoprotein OS=Homo sapiens OX=9606 GN=LRG1 PE=1 SV=2 | 518 | 20201116_fqg_colostrum_1ng_glyco_hcd203040_compound15.25360.25360.5 | controllerType=0 controllerNumber=1 scan=25360 | 49.6912 |
| 23216:4 | 25 | R.QLDMLDLSN[+1913.677]NSLASVPEGLWASLGQPNWDMR.D | HexNAc(4)Hex(5)NeuAc(1) | N[+1914] | 1343.592 | 4 | 5371.3463 | 5371.3285 | 0 | 3.3 | 261 | Specific | 878.86 | 845.12 | 845.12 | 19.21 | 18 | >sp\|P02750\|A2GL_HUMAN Leucine-rich alpha-2-glycoprotein OS=Homo sapiens OX=9606 GN=LRG1 PE=1 SV=2 | 518 | 20201116_fqg_colostrum_1ng_glyco_hcd203040_compound15.26935.26935.2 | controllerType=0 controllerNumber=1 scan=26935 | 52.9759 |

Table S4. Cont.

| Query #:z | Protein Rank | Glycopeptide | Glycan | Modification Type(s) | Observed m/z | z | Observed (M+H) | Calc. mass (M+H) | Off-by-x error | Mass error (ppm) | StartPosition | Cleavage | Score | Delta | Delta Mod | \|Log Prob\| | # of unique peptides | Protein Name | Protein DB number | Comment | Scan # | Scan Time |
|---|---|---|---|---|---|---|---|---|---|---|---|---|---|---|---|---|---|---|---|---|---|---|
| 25248:4 | 25 | R.QLDMLDLSN[+2204.772]NSLASVPEGLWASLGQPNWDMR.D | HexNAc(4)Hex(5)NeuAc(2) | N[+2205] | 1416.3679 | 4 | 5662.4499 | 5662.4239 | 0 | 4.6 | 261 | Specific | 895.94 | 895.94 | 895.94 | 18.77 | 18 | >sp\|P02750\|A2GL_HUMAN Leucine-rich alpha-2-glycoprotein OS=Homo sapiens OX=9606 GN=LRG1 PE=1 SV=2 | 518 | 20201116_fqg_colostrum_1ng_glyco_hcd203040_compound15.29061.29061.4 | controllerType=0 controllerNumber=1 scan=29061 | 57.8221 |
| 25250:4 | 25 | R.QLDMLDLSN[+2204.772]NSLASVPEGLWASLGQPNWDMR.D | HexNAc(4)Hex(5)NeuAc(2) | N[+2205] | 1416.3679 | 4 | 5662.4499 | 5662.4239 | 0 | 4.6 | 261 | Specific | 1058.46 | 1020.25 | 1020.25 | 22.77 | 18 | >sp\|P02750\|A2GL_HUMAN Leucine-rich alpha-2-glycoprotein OS=Homo sapiens OX=9606 GN=LRG1 PE=1 SV=2 | 518 | 20201116_fqg_colostrum_1ng_glyco_hcd203040_compound15.29063.29063.4 | controllerType=0 controllerNumber=1 scan=29063 | 57.8284 |
| 23219:4 | 25 | R.QLDMLDLSN[+1913.677]NSLASVPEGLWASLGQPNWDMR.D | HexNAc(4)Hex(5)NeuAc(1) | N[+1914] | 1343.592 | 4 | 5371.3463 | 5371.3285 | 0 | 3.3 | 261 | Specific | 1134.89 | 1051.05 | 1051.05 | 24.21 | 18 | >sp\|P02750\|A2GL_HUMAN Leucine-rich alpha-2-glycoprotein OS=Homo sapiens OX=9606 GN=LRG1 PE=1 SV=2 | 518 | 20201116_fqg_colostrum_1ng_glyco_hcd203040_compound15.26938.26938.2 | controllerType=0 controllerNumber=1 scan=26938 | 52.9889 |
| 01770:3 | 35 | R.LRN[+1216.423]VSWATGR.S | HexNAc(2)Hex(5) | N[+1216] | 792.6922 | 3 | 2376.0621 | 2376.0559 | 0 | 2.6 | 149 | Specific | 313.34 | 128.51 | 128.51 | 5.47 | 14 | >sp\|P08571\|CD14_HUMAN Monocyte differentiation antigen CD14 OS=Homo sapiens OX=9606 GN=CD14 PE=1 SV=2 | 1232 | 20201116_fqg_colostrum_1ng_glyco_hcd203040_compound15.4676.4676.3 | controllerType=0 controllerNumber=1 scan=4676 | 14.229 |
| 01772:3 | 35 | R.LRN[+1216.423]VSWATGR.S | HexNAc(2)Hex(5) | N[+1216] | 792.6922 | 3 | 2376.0621 | 2376.0559 | 0 | 2.6 | 149 | Specific | 446.98 | 241.39 | 241.39 | 6.99 | 14 | >sp\|P08571\|CD14_HUMAN Monocyte differentiation antigen CD14 OS=Homo sapiens OX=9606 GN=CD14 PE=1 SV=2 | 1232 | 20201116_fqg_colostrum_1ng_glyco_hcd203040_compound15.4678.4678.3 | controllerType=0 controllerNumber=1 scan=4678 | 14.2327 |
| 14528:2 | 44 | K.LC[+57.021]DN[+1866.688]LSTK.N | HexNAc(6)Hex(4) | C[+57], N[+1867] | 1409.0776 | 2 | 2817.1478 | 2817.1487 | 0 | -0.3 | 285 | Specific | 262.76 | 200.82 | 200.82 | 3.37 | 17 | >sp\|P02774\|VTDB_HUMAN Vitamin D-binding protein OS=Homo sapiens OX=9606 GN=GC PE=1 SV=2 | 1103 | 20201116_fqg_colostrum_1ng_glyco_hcd203040_compound15.17907.17907.2 | controllerType=0 controllerNumber=1 scan=17907 | 35.7891 |
| 01896:4 | 57 | R.EKQTDEIKDTRN[+2059.735]ESTQNC[+57.021]VVAEPEK.M | HexNAc(4)Hex(5)Fuc(1)NeuAc(1) | C[+57], N[+2060] | 1252.7902 | 4 | 5008.1388 | 5008.1251 | 0 | 2.7 | 58 | Specific | 392.14 | 392.14 | 392.14 | 8.14 | 12 | >sp\|P47710\|CASA1_HUMAN Alpha-S1-casein OS=Homo sapiens OX=9606 GN=CSN1S1 PE=1 SV=1 | 989 | 20201116_fqg_colostrum_1ng_glyco_hcd203040_compound15.4807.4807.4 | controllerType=0 controllerNumber=1 scan=4807 | 14.4802 |
| 01898:4 | 57 | R.EKQTDEIKDTRN[+2059.735]ESTQNC[+57.021]VVAEPEK.M | HexNAc(4)Hex(5)Fuc(1)NeuAc(1) | C[+57], N[+2060] | 1252.7902 | 4 | 5008.1388 | 5008.1251 | 0 | 2.7 | 58 | Specific | 512.33 | 512.33 | 512.33 | 8.64 | 12 | >sp\|P47710\|CASA1_HUMAN Alpha-S1-casein OS=Homo sapiens OX=9606 GN=CSN1S1 PE=1 SV=1 | 989 | 20201116_fqg_colostrum_1ng_glyco_hcd203040_compound15.4809.4809.4 | controllerType=0 controllerNumber=1 scan=4809 | 14.4855 |
| 01762:2 | 71 | R.VVLGAHN[+730.264]LSR.R | HexNAc(2)Hex(2) | N[+730] | 898.4471 | 2 | 1795.8869 | 1795.8807 | 0 | 3.4 | 82 | Specific | 307.04 | 307.04 | 307.04 | 6.82 | 9 | >sp\|P08246\|ELNE_HUMAN Neutrophil elastase OS=Homo sapiens OX=9606 GN=ELANE PE=1 SV=1 | 1269 | 20201116_fqg_colostrum_1ng_glyco_hcd203040_compound15.4668.4668.2 | controllerType=0 controllerNumber=1 scan=4668 | 14.2112 |
| 01764:2 | 71 | R.VVLGAHN[+730.264]LSR.R | HexNAc(2)Hex(2) | N[+730] | 898.4471 | 2 | 1795.8869 | 1795.8807 | 0 | 3.4 | 82 | Specific | 390.49 | 359.54 | 359.54 | 4.98 | 9 | >sp\|P08246\|ELNE_HUMAN Neutrophil elastase OS=Homo sapiens OX=9606 GN=ELANE PE=1 SV=1 | 1269 | 20201116_fqg_colostrum_1ng_glyco_hcd203040_compound15.4670.4670.2 | controllerType=0 controllerNumber=1 scan=4670 | 14.216 |
| 01716:3 | 71 | R.VVLGAHN[+730.264]LSR.R | HexNAc(2)Hex(2) | N[+730] | 599.2993 | 3 | 1795.8834 | 1795.8807 | 0 | 1.5 | 82 | Specific | 504.55 | 504.55 | 504.55 | 9.77 | 9 | >sp\|P08246\|ELNE_HUMAN Neutrophil elastase OS=Homo sapiens OX=9606 GN=ELANE PE=1 SV=1 | 1269 | 20201116_fqg_colostrum_1ng_glyco_hcd203040_compound15.4620.4620.2 | controllerType=0 controllerNumber=1 scan=4620 | 14.1215 |
| 01744:3 | 71 | R.VVLGAHN[+730.264]LSR.R | HexNAc(2)Hex(2) | N[+730] | 599.2995 | 3 | 1795.8839 | 1795.8807 | 0 | 1.8 | 82 | Specific | 516.05 | 516.05 | 516.05 | 10 | 9 | >sp\|P08246\|ELNE_HUMAN Neutrophil elastase OS=Homo sapiens OX=9606 GN=ELANE PE=1 SV=1 | 1269 | 20201116_fqg_colostrum_1ng_glyco_hcd203040_compound15.4649.4649.3 | controllerType=0 controllerNumber=1 scan=4649 | 14.1792 |
| 01714:3 | 71 | R.VVLGAHN[+730.264]LSR.R | HexNAc(2)Hex(2) | N[+730] | 599.2993 | 3 | 1795.8834 | 1795.8807 | 0 | 1.5 | 82 | Specific | 544.48 | 544.48 | 544.48 | 9.98 | 9 | >sp\|P08246\|ELNE_HUMAN Neutrophil elastase OS=Homo sapiens OX=9606 GN=ELANE PE=1 SV=1 | 1269 | 20201116_fqg_colostrum_1ng_glyco_hcd203040_compound15.4618.4618.2 | controllerType=0 controllerNumber=1 scan=4618 | 14.1161 |
| 01742:3 | 71 | R.VVLGAHN[+730.264]LSR.R | HexNAc(2)Hex(2) | N[+730] | 599.2995 | 3 | 1795.8839 | 1795.8807 | 0 | 1.8 | 82 | Specific | 574.55 | 574.55 | 574.55 | 10.4 | 9 | >sp\|P08246\|ELNE_HUMAN Neutrophil elastase OS=Homo sapiens OX=9606 GN=ELANE PE=1 SV=1 | 1269 | 20201116_fqg_colostrum_1ng_glyco_hcd203040_compound15.4647.4647.3 | controllerType=0 controllerNumber=1 scan=4647 | 14.1732 |
| 20000:3 | 71 | R.GIASVLQELN[+876.322]VTVVTSLC[+57.021]R.R | HexNAc(2)Hex(2)Fuc(1) | C[+57], N[+876] | 979.1552 | 3 | 2935.451 | 2935.4439 | 0 | 2.4 | 164 | Specific | 267.92 | 267.92 | 267.92 | 4.86 | 9 | >sp\|P08246\|ELNE_HUMAN Neutrophil elastase OS=Homo sapiens OX=9606 GN=ELANE PE=1 SV=1 | 1269 | 20201116_fqg_colostrum_1ng_glyco_hcd203040_compound15.23589.23589.2 | controllerType=0 controllerNumber=1 scan=23589 | 46.1483 |
| 19997:3 | 71 | R.GIASVLQELN[+876.322]VTVVTSLC[+57.021]R.R | HexNAc(2)Hex(2)Fuc(1) | C[+57], N[+876] | 979.1552 | 3 | 2935.451 | 2935.4439 | 0 | 2.4 | 164 | Specific | 485.81 | 467.29 | 467.29 | 9.39 | 9 | >sp\|P08246\|ELNE_HUMAN Neutrophil elastase OS=Homo sapiens OX=9606 GN=ELANE PE=1 SV=1 | 1269 | 20201116_fqg_colostrum_1ng_glyco_hcd203040_compound15.23586.23586.2 | controllerType=0 controllerNumber=1 scan=23586 | 46.1393 |
| 19974:2 | 71 | R.GIASVLQELN[+876.322]VTVVTSLC[+57.021]R.R | HexNAc(2)Hex(2)Fuc(1) | C[+57], N[+876] | 1468.2275 | 2 | 2935.4476 | 2935.4439 | 0 | 1.3 | 164 | Specific | 1009.53 | 1009.53 | 1009.53 | 21.18 | 9 | >sp\|P08246\|ELNE_HUMAN Neutrophil elastase OS=Homo sapiens OX=9606 GN=ELANE PE=1 SV=1 | 1269 | 20201116_fqg_colostrum_1ng_glyco_hcd203040_compound15.23562.23562.2 | controllerType=0 controllerNumber=1 scan=23562 | 46.0882 |
| 19976:2 | 71 | R.GIASVLQELN[+876.322]VTVVTSLC[+57.021]R.R | HexNAc(2)Hex(2)Fuc(1) | C[+57], N[+876] | 1468.2275 | 2 | 2935.4476 | 2935.4439 | 0 | 1.3 | 164 | Specific | 1049.99 | 994.23 | 994.23 | 21.18 | 9 | >sp\|P08246\|ELNE_HUMAN Neutrophil elastase OS=Homo sapiens OX=9606 GN=ELANE PE=1 SV=1 | 1269 | 20201116_fqg_colostrum_1ng_glyco_hcd203040_compound15.23564.23564.2 | controllerType=0 controllerNumber=1 scan=23564 | 46.0963 |
| 19965:3 | 71 | R.GIASVLQELN[+876.322]VTVVTSLC[+57.021]R.R | HexNAc(2)Hex(2)Fuc(1) | C[+57], N[+876] | 979.155 | 3 | 2935.4504 | 2935.4439 | 0 | 2.2 | 164 | Specific | 1062.78 | 1062.78 | 1062.78 | 22.64 | 9 | >sp\|P08246\|ELNE_HUMAN Neutrophil elastase OS=Homo sapiens OX=9606 GN=ELANE PE=1 SV=1 | 1269 | 20201116_fqg_colostrum_1ng_glyco_hcd203040_compound15.23553.23553.3 | controllerType=0 controllerNumber=1 scan=23553 | 46.073 |
| 19967:3 | 71 | R.GIASVLQELN[+876.322]VTVVTSLC[+57.021]R.R | HexNAc(2)Hex(2)Fuc(1) | C[+57], N[+876] | 979.155 | 3 | 2935.4504 | 2935.4439 | 0 | 2.2 | 164 | Specific | 1178.48 | 1178.48 | 1178.48 | 24.64 | 9 | >sp\|P08246\|ELNE_HUMAN Neutrophil elastase OS=Homo sapiens OX=9606 GN=ELANE PE=1 SV=1 | 1269 | 20201116_fqg_colostrum_1ng_glyco_hcd203040_compound15.23555.23555.3 | controllerType=0 controllerNumber=1 scan=23555 | 46.0788 |
| 08896:3 | 75 | R.FVN[+714.269]VTVTPEDQC[+57.021]RPNNVC[+57.021]TGVLTR.R | HexNAc(2)Hex(1)Fuc(1) | C[+57]*2, N[+714] | 1164.2147 | 3 | 3490.6294 | 3490.62 | 0 | 2.7 | 169 | Specific | 688.11 | 662.03 | 662.03 | 14.24 | 9 | >sp\|P20160\|CAP7_HUMAN Azurocidin OS=Homo sapiens OX=9606 GN=AZU1 PE=1 SV=3 | 850 | 20201116_fqg_colostrum_1ng_glyco_hcd203040_compound15.12082.12082.3 | controllerType=0 controllerNumber=1 scan=12082 | 26.484 |
| 08899:3 | 75 | R.FVN[+714.269]VTVTPEDQC[+57.021]RPNNVC[+57.021]TGVLTR.R | HexNAc(2)Hex(1)Fuc(1) | C[+57]*2, N[+714] | 1164.2147 | 3 | 3490.6294 | 3490.62 | 0 | 2.7 | 169 | Specific | 710.06 | 662.04 | 662.04 | 15.24 | 9 | >sp\|P20160\|CAP7_HUMAN Azurocidin OS=Homo sapiens OX=9606 GN=AZU1 PE=1 SV=3 | 850 | 20201116_fqg_colostrum_1ng_glyco_hcd203040_compound15.12085.12085.3 | controllerType=0 controllerNumber=1 scan=12085 | 26.488 |
| 08759:3 | 75 | R.FVN[+876.322]VTVTPEDQC[+57.021]RPNNVC[+57.021]TGVLTR.R | HexNAc(2)Hex(2)Fuc(1) | C[+57]*2, N[+876] | 1218.2314 | 3 | 3652.6796 | 3652.6728 | 0 | 1.9 | 169 | Specific | 967.04 | 928.8 | 928.8 | 20.24 | 9 | >sp\|P20160\|CAP7_HUMAN Azurocidin OS=Homo sapiens OX=9606 GN=AZU1 PE=1 SV=3 | 850 | 20201116_fqg_colostrum_1ng_glyco_hcd203040_compound15.11940.11940.3 | controllerType=0 controllerNumber=1 scan=11940 | 26.2284 |
| 08745:3 | 75 | R.FVN[+876.322]VTVTPEDQC[+57.021]RPNNVC[+57.021]TGVLTR.R | HexNAc(2)Hex(2)Fuc(1) | C[+57]*2, N[+876] | 1218.2313 | 3 | 3652.6793 | 3652.6728 | 0 | 1.8 | 169 | Specific | 987.76 | 987.76 | 987.76 | 20.24 | 9 | >sp\|P20160\|CAP7_HUMAN Azurocidin OS=Homo sapiens OX=9606 GN=AZU1 PE=1 SV=3 | 850 | 20201116_fqg_colostrum_1ng_glyco_hcd203040_compound15.11925.11925.3 | controllerType=0 controllerNumber=1 scan=11925 | 26.201 |
| 08762:3 | 75 | R.FVN[+876.322]VTVTPEDQC[+57.021]RPNNVC[+57.021]TGVLTR.R | HexNAc(2)Hex(2)Fuc(1) | C[+57]*2, N[+876] | 1218.2314 | 3 | 3652.6796 | 3652.6728 | 0 | 1.9 | 169 | Specific | 1017.56 | 1015.96 | 1015.96 | 21.24 | 9 | >sp\|P20160\|CAP7_HUMAN Azurocidin OS=Homo sapiens OX=9606 GN=AZU1 PE=1 SV=3 | 850 | 20201116_fqg_colostrum_1ng_glyco_hcd203040_compound15.11943.11943.3 | controllerType=0 controllerNumber=1 scan=11943 | 26.2332 |
| 08748:3 | 75 | R.FVN[+876.322]VTVTPEDQC[+57.021]RPNNVC[+57.021]TGVLTR.R | HexNAc(2)Hex(2)Fuc(1) | C[+57]*2, N[+876] | 1218.2313 | 3 | 3652.6793 | 3652.6728 | 0 | 1.8 | 169 | Specific | 1040.15 | 1038.94 | 1038.94 | 21.24 | 9 | >sp\|P20160\|CAP7_HUMAN Azurocidin OS=Homo sapiens OX=9606 GN=AZU1 PE=1 SV=3 | 850 | 20201116_fqg_colostrum_1ng_glyco_hcd203040_compound15.11928.11928.3 | controllerType=0 controllerNumber=1 scan=11928 | 26.2064 |
| 00874:4 | 78 | K.TKPREEQFN[+1444.534]STFR.V | HexNAc(4)Hex(3)Fuc(1) | N[+1445] | 771.8451 | 4 | 3084.3584 | 3084.3526 | 0 | 1.9 | 168 | Specific | 252.86 | 182.56 | 182.56 | 4.63 | 12 | >sp\|P01859\|IGHG2_HUMAN Immunoglobulin heavy constant gamma 2 OS=Homo sapiens OX=9606 GN=IGHG2 PE=1 SV=2 | 387 | 20201116_fqg_colostrum_1ng_glyco_hcd203040_compound15.3744.3744.2 | controllerType=0 controllerNumber=1 scan=3744 | 12.4796 |
| 19226:5 | 88 | R.VGAHDPPAQVLQELN[+876.322]VTVVTFFC[+57.021]RPHNIC[+57.021]TFVPR.R | HexNAc(2)Hex(2)Fuc(1) | C[+57]*2, N[+876] | 959.8704 | 5 | 4795.3231 | 4795.3117 | 0 | 2.4 | 160 | Specific | 958.55 | 958.55 | 958.55 | 20.23 | 7 | >sp\|P24158\|PRTN3_HUMAN Myeloblastin OS=Homo sapiens OX=9606 GN=PRTN3 PE=1 SV=3 | 931 | 20201116_fqg_colostrum_1ng_glyco_hcd203040_compound15.22780.22780.5 | controllerType=0 controllerNumber=1 scan=22780 | 44.5494 |
| 19229:5 | 88 | R.VGAHDPPAQVLQELN[+876.322]VTVVTFFC[+57.021]RPHNIC[+57.021]TFVPR.R | HexNAc(2)Hex(2)Fuc(1) | C[+57]*2, N[+876] | 959.8704 | 5 | 4795.3231 | 4795.3117 | 0 | 2.4 | 160 | Specific | 1162.08 | 1162.08 | 1162.08 | 24.23 | 7 | >sp\|P24158\|PRTN3_HUMAN Myeloblastin OS=Homo sapiens OX=9606 GN=PRTN3 PE=1 SV=3 | 931 | 20201116_fqg_colostrum_1ng_glyco_hcd203040_compound15.22783.22783.5 | controllerType=0 controllerNumber=1 scan=22783 | 44.5547 |
| 16908:4 | 2 | R.RTAGWNVPIGTLRPFLN[+1768.640]WTGPPEPIEAAVAR.F | HexNAc(4)Hex(5)Fuc(1) | N[+1769] | 1289.6213 | 4 | 5155.4635 | 5155.4516 | 0 | 2.3 | 140 | Specific | 271.63 | 243.32 | 243.32 | 6.3 | 108 | >sp\|P02788\|TRFL_HUMAN Lactotransferrin OS=Homo sapiens OX=9606 GN=LTF PE=1 SV=6 | 1270 | 20201117_fqg_milk_1ng_glyco_hcd203040_compound15.20462.20462.4 | controllerType=0 controllerNumber=1 scan=20462 | 41.4066 |
| 17861:4 | 2 | R.RTAGWNVPIGTLRPFLN[+2059.735]WTGPPEPIEAAVAR.F | HexNAc(4)Hex(5)Fuc(1)NeuAc(1) | N[+2060] | 1362.3961 | 4 | 5446.5627 | 5446.547 | 0 | 2.9 | 140 | Specific | 533.45 | 533.45 | 533.45 | 10.16 | 108 | >sp\|P02788\|TRFL_HUMAN Lactotransferrin OS=Homo sapiens OX=9606 GN=LTF PE=1 SV=6 | 1270 | 20201117_fqg_milk_1ng_glyco_hcd203040_compound15.21457.21457.4 | controllerType=0 controllerNumber=1 scan=21457 | 43.5595 |
| 17863:4 | 2 | R.RTAGWNVPIGTLRPFLN[+2059.735]WTGPPEPIEAAVAR.F | HexNAc(4)Hex(5)Fuc(1)NeuAc(1) | N[+2060] | 1362.3961 | 4 | 5446.5627 | 5446.547 | 0 | 2.9 | 140 | Specific | 643.06 | 515.69 | 515.69 | 12.7 | 108 | >sp\|P02788\|TRFL_HUMAN Lactotransferrin OS=Homo sapiens OX=9606 GN=LTF PE=1 SV=6 | 1270 | 20201117_fqg_milk_1ng_glyco_hcd203040_compound15.21459.21459.4 | controllerType=0 controllerNumber=1 scan=21459 | 43.5702 |
| 17840:4 | 2 | R.RTAGWNVPIGTLRPFLN[+2205.793]WTGPPEPIEAAVAR.F | HexNAc(4)Hex(5)Fuc(2)NeuAc(1) | N[+2206] | 1398.9102 | 4 | 5592.6188 | 5592.6049 | 0 | 2.5 | 140 | Specific | 650.21 | 475.88 | 475.88 | 13.7 | 108 | >sp\|P02788\|TRFL_HUMAN Lactotransferrin OS=Homo sapiens OX=9606 GN=LTF PE=1 SV=6 | 1270 | 20201117_fqg_milk_1ng_glyco_hcd203040_compound15.21434.21434.2 | controllerType=0 controllerNumber=1 scan=21434 | 43.4924 |
| 17842:4 | 2 | R.RTAGWNVPIGTLRPFLN[+2205.793]WTGPPEPIEAAVAR.F | HexNAc(4)Hex(5)Fuc(2)NeuAc(1) | N[+2206] | 1398.9102 | 4 | 5592.6188 | 5592.6049 | 0 | 2.5 | 140 | Specific | 663.74 | 551.99 | 551.99 | 13.7 | 108 | >sp\|P02788\|TRFL_HUMAN Lactotransferrin OS=Homo sapiens OX=9606 GN=LTF PE=1 SV=6 | 1270 | 20201117_fqg_milk_1ng_glyco_hcd203040_compound15.21436.21436.2 | controllerType=0 controllerNumber=1 scan=21436 | 43.5026 |
| 18102:3 | 2 | R.TAGWNVPIGTLRPFLN[+2059.735]WTGPPEPIEAAVAR.F | HexNAc(4)Hex(5)Fuc(1)NeuAc(1) | N[+2060] | 1764.1639 | 3 | 5290.4772 | 5290.4459 | 0 | 5.9 | 141 | Specific | 216.77 | 216.77 | 216.77 | 3.57 | 108 | >sp\|P02788\|TRFL_HUMAN Lactotransferrin OS=Homo sapiens OX=9606 GN=LTF PE=1 SV=6 | 1270 | 20201117_fqg_milk_1ng_glyco_hcd203040_compound15.21707.21707.3 | controllerType=0 controllerNumber=1 scan=21707 | 44.0731 |
| 19053:4 | 2 | R.TAGWNVPIGTLRPFLN[+2569.905]WTGPPEPIEAAVAR.F | HexNAc(5)Hex(6)NeuAc(2) | N[+2570] | 1450.9216 | 4 | 5800.6647 | 5800.6156 | 0 | 8.5 | 141 | Specific | 254.65 | 213.98 | 213.98 | 4.92 | 108 | >sp\|P02788\|TRFL_HUMAN Lactotransferrin OS=Homo sapiens OX=9606 GN=LTF PE=1 SV=6 | 1270 | 20201117_fqg_milk_1ng_glyco_hcd203040_compound15.22699.22699.2 | controllerType=0 controllerNumber=1 scan=22699 | 46.2129 |
| 18073:3 | 2 | R.[+42.011]TAGWNVPIGTLRPFLN[+1872.651]WTGPPEPIEAAVAR.F | HexNAc(3)Hex(6)NeuAc(1) | .[42], N[+1873] | 1715.8116 | 3 | 5145.4203 | 5145.372 | 0 | 9.4 | 141 | Specific | 289.09 | 178.44 | 178.44 | 3.74 | 108 | >sp\|P02788\|TRFL_HUMAN Lactotransferrin OS=Homo sapiens OX=9606 GN=LTF PE=1 SV=6 | 1270 | 20201117_fqg_milk_1ng_glyco_hcd203040_compound15.21677.21677.3 | controllerType=0 controllerNumber=1 scan=21677 | 43.9998 |
| 18098:4 | 2 | R.[+42.011]TAGWNVPIGTLRPFLN[+2018.708]WTGPPEPIEAAVAR.F | HexNAc(3)Hex(6)Fuc(1)NeuAc(1) | .[42], N[+2019] | 1323.6241 | 4 | 5291.4748 | 5291.4299 | 0 | 8.5 | 141 | Specific | 333.14 | 267.34 | 267.34 | 5.74 | 108 | >sp\|P02788\|TRFL_HUMAN Lactotransferrin OS=Homo sapiens OX=9606 GN=LTF PE=1 SV=6 | 1270 | 20201117_fqg_milk_1ng_glyco_hcd203040_compound15.21703.21703.4 | controllerType=0 controllerNumber=1 scan=21703 | 44.0664 |
| 18071:3 | 2 | R.[+42.011]TAGWNVPIGTLRPFLN[+1872.651]WTGPPEPIEAAVAR.F | HexNAc(3)Hex(6)NeuAc(1) | .[42], N[+1873] | 1715.8116 | 3 | 5145.4203 | 5145.372 | 0 | 9.4 | 141 | Specific | 344.92 | 257.95 | 257.95 | 5.71 | 108 | >sp\|P02788\|TRFL_HUMAN Lactotransferrin OS=Homo sapiens OX=9606 GN=LTF PE=1 SV=6 | 1270 | 20201117_fqg_milk_1ng_glyco_hcd203040_compound15.21675.21675.3 | controllerType=0 controllerNumber=1 scan=21675 | 43.9936 |
| 18169:3 | 2 | R.TAGWNVPIGTLRPFLN[+1768.640]WTGPPEPIEAAVAR.F | HexNAc(4)Hex(5)Fuc(1) | N[+1769] | 1667.1256 | 3 | 4999.3622 | 4999.3504 | 0 | 2.4 | 141 | Specific | 396.82 | 258.89 | 258.89 | 7.6 | 108 | >sp\|P02788\|TRFL_HUMAN Lactotransferrin OS=Homo sapiens OX=9606 GN=LTF PE=1 SV=6 | 1270 | 20201117_fqg_milk_1ng_glyco_hcd203040_compound15.21777.21777.3 | controllerType=0 controllerNumber=1 scan=21777 | 44.2234 |
| 18172:3 | 2 | R.TAGWNVPIGTLRPFLN[+1768.640]WTGPPEPIEAAVAR.F | HexNAc(4)Hex(5)Fuc(1) | N[+1769] | 1667.1256 | 3 | 4999.3622 | 4999.3504 | 0 | 2.4 | 141 | Specific | 398.99 | 167.45 | 167.45 | 5.55 | 108 | >sp\|P02788\|TRFL_HUMAN Lactotransferrin OS=Homo sapiens OX=9606 GN=LTF PE=1 SV=6 | 1270 | 20201117_fqg_milk_1ng_glyco_hcd203040_compound15.21780.21780.3 | controllerType=0 controllerNumber=1 scan=21780 | 44.2306 |
| 19056:4 | 2 | R.TAGWNVPIGTLRPFLN[+2569.905]WTGPPEPIEAAVAR.F | HexNAc(5)Hex(6)NeuAc(2) | N[+2570] | 1450.9216 | 4 | 5800.6647 | 5800.6156 | 0 | 8.5 | 141 | Specific | 490.94 | 375.99 | 375.99 | 8.29 | 108 | >sp\|P02788\|TRFL_HUMAN Lactotransferrin OS=Homo sapiens OX=9606 GN=LTF PE=1 SV=6 | 1270 | 20201117_fqg_milk_1ng_glyco_hcd203040_compound15.22702.22702.2 | controllerType=0 controllerNumber=1 scan=22702 | 46.2211 |
| 19074:3 | 2 | R.TAGWNVPIGTLRPFLN[+2205.793]WTGPPEPIEAAVAR.F | HexNAc(4)Hex(5)Fuc(2)NeuAc(1) | N[+2206] | 1812.8448 | 3 | 5436.5199 | 5436.5038 | 0 | 3 | 141 | Specific | 532.3 | 481.24 | 481.24 | 10.16 | 108 | >sp\|P02788\|TRFL_HUMAN Lactotransferrin OS=Homo sapiens OX=9606 GN=LTF PE=1 SV=6 | 1270 | 20201117_fqg_milk_1ng_glyco_hcd203040_compound15.22721.22721.3 | controllerType=0 controllerNumber=1 scan=22721 | 46.2612 |
| 19063:3 | 2 | R.TAGWNVPIGTLRPFLN[+2350.830]WTGPPEPIEAAVAR.F | HexNAc(4)Hex(5)Fuc(1)NeuAc(2) | N[+2351] | 1861.1952 | 3 | 5581.5709 | 5581.5413 | 0 | 5.3 | 141 | Specific | 544.59 | 360.75 | 360.75 | 7.61 | 108 | >sp\|P02788\|TRFL_HUMAN Lactotransferrin OS=Homo sapiens OX=9606 GN=LTF PE=1 SV=6 | 1270 | 20201117_fqg_milk_1ng_glyco_hcd203040_compound15.22709.22709.3 | controllerType=0 controllerNumber=1 scan=22709 | 46.2346 |
| 19078:4 | 2 | R.TAGWNVPIGTLRPFLN[+2351.851]WTGPPEPIEAAVAR.F | HexNAc(4)Hex(5)Fuc(3)NeuAc(1) | N[+2352] | 1396.3994 | 4 | 5582.5758 | 5582.5617 | 0 | 2.5 | 141 | Specific | 550.18 | 510.78 | 510.78 | 10.26 | 108 | >sp\|P02788\|TRFL_HUMAN Lactotransferrin OS=Homo sapiens OX=9606 GN=LTF PE=1 SV=6 | 1270 | 20201117_fqg_milk_1ng_glyco_hcd203040_compound15.22725.22725.4 | controllerType=0 controllerNumber=1 scan=22725 | 46.2696 |
| 19048:4 | 2 | R.TAGWNVPIGTLRPFLN[+2424.867]WTGPPEPIEAAVAR.F | HexNAc(5)Hex(6)Fuc(1)NeuAc(1) | N[+2425] | 1414.6525 | 4 | 5655.588 | 5655.5781 | 0 | 1.8 | 141 | Specific | 589.78 | 474.74 | 474.74 | 11.43 | 108 | >sp\|P02788\|TRFL_HUMAN Lactotransferrin OS=Homo sapiens OX=9606 GN=LTF PE=1 SV=6 | 1270 | 20201117_fqg_milk_1ng_glyco_hcd203040_compound15.22694.22694.4 | controllerType=0 controllerNumber=1 scan=22694 | 46.2041 |
| 19065:3 | 2 | R.TAGWNVPIGTLRPFLN[+2350.830]WTGPPEPIEAAVAR.F | HexNAc(4)Hex(5)Fuc(1)NeuAc(2) | N[+2351] | 1861.1952 | 3 | 5581.5709 | 5581.5413 | 0 | 5.3 | 141 | Specific | 592.49 | 413.25 | 413.25 | 10.81 | 108 | >sp\|P02788\|TRFL_HUMAN Lactotransferrin OS=Homo sapiens OX=9606 GN=LTF PE=1 SV=6 | 1270 | 20201117_fqg_milk_1ng_glyco_hcd203040_compound15.22711.22711.3 | controllerType=0 controllerNumber=1 scan=22711 | 46.2387 |
| 18957:4 | 2 | R.TAGWNVPIGTLRPFLN[+2570.925]WTGPPEPIEAAVAR.F | HexNAc(5)Hex(6)Fuc(1)NeuAc(1) | N[+2571] | 1451.1694 | 4 | 5801.6559 | 5801.636 | 0 | 3.4 | 141 | Specific | 617.85 | 507.34 | 507.34 | 12.11 | 108 | >sp\|P02788\|TRFL_HUMAN Lactotransferrin OS=Homo sapiens OX=9606 GN=LTF PE=1 SV=6 | 1270 | 20201117_fqg_milk_1ng_glyco_hcd203040_compound15.22599.22599.4 | controllerType=0 controllerNumber=1 scan=22599 | 46.0151 |
| 19050:4 | 2 | R.TAGWNVPIGTLRPFLN[+2424.867]WTGPPEPIEAAVAR.F | HexNAc(5)Hex(6)Fuc(1)NeuAc(1) | N[+2425] | 1414.6525 | 4 | 5655.588 | 5655.5781 | 0 | 1.8 | 141 | Specific | 618.39 | 419.24 | 419.24 | 12.43 | 108 | >sp\|P02788\|TRFL_HUMAN Lactotransferrin OS=Homo sapiens OX=9606 GN=LTF PE=1 SV=6 | 1270 | 20201117_fqg_milk_1ng_glyco_hcd203040_compound15.22696.22696.4 | controllerType=0 controllerNumber=1 scan=22696 | 46.2087 |
| 19073:4 | 2 | R.TAGWNVPIGTLRPFLN[+2205.793]WTGPPEPIEAAVAR.F | HexNAc(4)Hex(5)Fuc(2)NeuAc(1) | N[+2206] | 1359.8864 | 4 | 5436.5236 | 5436.5038 | 0 | 3.6 | 141 | Specific | 683.79 | 585.48 | 585.48 | 13.11 | 108 | >sp\|P02788\|TRFL_HUMAN Lactotransferrin OS=Homo sapiens OX=9606 GN=LTF PE=1 SV=6 | 1270 | 20201117_fqg_milk_1ng_glyco_hcd203040_compound15.22720.22720.4 | controllerType=0 controllerNumber=1 scan=22720 | 46.2597 |
| 19076:3 | 2 | R.TAGWNVPIGTLRPFLN[+2205.793]WTGPPEPIEAAVAR.F | HexNAc(4)Hex(5)Fuc(2)NeuAc(1) | N[+2206] | 1812.8448 | 3 | 5436.5199 | 5436.5038 | 0 | 3 | 141 | Specific | 714.1 | 616.94 | 616.94 | 14.7 | 108 | >sp\|P02788\|TRFL_HUMAN Lactotransferrin OS=Homo sapiens OX=9606 GN=LTF PE=1 SV=6 | 1270 | 20201117_fqg_milk_1ng_glyco_hcd203040_compound15.22723.22723.3 | controllerType=0 controllerNumber=1 scan=22723 | 46.2673 |
| 19080:3 | 2 | R.TAGWNVPIGTLRPFLN[+2059.735]WTGPPEPIEAAVAR.F | HexNAc(4)Hex(5)Fuc(1)NeuAc(1) | N[+2060] | 1764.16 | 3 | 5290.4654 | 5290.4459 | 0 | 3.7 | 141 | Specific | 716.38 | 611.08 | 611.08 | 14.29 | 108 | >sp\|P02788\|TRFL_HUMAN Lactotransferrin OS=Homo sapiens OX=9606 GN=LTF PE=1 SV=6 | 1270 | 20201117_fqg_milk_1ng_glyco_hcd203040_compound15.22727.22727.3 | controllerType=0 controllerNumber=1 scan=22727 | 46.2754 |
| 19077:3 | 2 | R.TAGWNVPIGTLRPFLN[+2059.735]WTGPPEPIEAAVAR.F | HexNAc(4)Hex(5)Fuc(1)NeuAc(1) | N[+2060] | 1764.16 | 3 | 5290.4654 | 5290.4459 | 0 | 3.7 | 141 | Specific | 716.74 | 621.45 | 621.45 | 14.29 | 108 | >sp\|P02788\|TRFL_HUMAN Lactotransferrin OS=Homo sapiens OX=9606 GN=LTF PE=1 SV=6 | 1270 | 20201117_fqg_milk_1ng_glyco_hcd203040_compound15.22724.22724.3 | controllerType=0 controllerNumber=1 scan=22724 | 46.2684 |
| 19099:5 | 2 | R.TAGWNVPIGTLRPFLN[+2059.735]WTGPPEPIEAAVAR.F | HexNAc(4)Hex(5)Fuc(1)NeuAc(1) | N[+2060] | 1058.898 | 5 | 5290.4607 | 5290.4459 | 0 | 2.8 | 141 | Specific | 721.44 | 627.8 | 627.8 | 15.14 | 108 | >sp\|P02788\|TRFL_HUMAN Lactotransferrin OS=Homo sapiens OX=9606 GN=LTF PE=1 SV=6 | 1270 | 20201117_fqg_milk_1ng_glyco_hcd203040_compound15.22747.22747.5 | controllerType=0 controllerNumber=1 scan=22747 | 46.3176 |
| 20412:4 | 2 | R.TAGWNVPIGTLRPFLN[+2350.830]WTGPPEPIEAAVAR.F | HexNAc(4)Hex(5)Fuc(1)NeuAc(2) | N[+2351] | 1396.1432 | 4 | 5581.5509 | 5581.5413 | 0 | 1.7 | 141 | Specific | 733.75 | 606.99 | 606.99 | 14.43 | 108 | >sp\|P02788\|TRFL_HUMAN Lactotransferrin OS=Homo sapiens OX=9606 GN=LTF PE=1 SV=6 | 1270 | 20201117_fqg_milk_1ng_glyco_hcd203040_compound15.24123.24123.4 | controllerType=0 controllerNumber=1 scan=24123 | 49.1319 |
| 19102:5 | 2 | R.TAGWNVPIGTLRPFLN[+2059.735]WTGPPEPIEAAVAR.F | HexNAc(4)Hex(5)Fuc(1)NeuAc(1) | N[+2060] | 1058.898 | 5 | 5290.4607 | 5290.4459 | 0 | 2.8 | 141 | Specific | 756.96 | 642.36 | 642.36 | 16.14 | 108 | >sp\|P02788\|TRFL_HUMAN Lactotransferrin OS=Homo sapiens OX=9606 GN=LTF PE=1 SV=6 | 1270 | 20201117_fqg_milk_1ng_glyco_hcd203040_compound15.22750.22750.5 | controllerType=0 controllerNumber=1 scan=22750 | 46.3263 |
| 20415:4 | 2 | R.TAGWNVPIGTLRPFLN[+2350.830]WTGPPEPIEAAVAR.F | HexNAc(4)Hex(5)Fuc(1)NeuAc(2) | N[+2351] | 1396.1432 | 4 | 5581.5509 | 5581.5413 | 0 | 1.7 | 141 | Specific | 762.4 | 590.87 | 590.87 | 15.43 | 108 | >sp\|P02788\|TRFL_HUMAN Lactotransferrin OS=Homo sapiens OX=9606 GN=LTF PE=1 SV=6 | 1270 | 20201117_fqg_milk_1ng_glyco_hcd203040_compound15.24126.24126.4 | controllerType=0 controllerNumber=1 scan=24126 | 49.1382 |
| 18959:4 | 2 | R.TAGWNVPIGTLRPFLN[+2570.925]WTGPPEPIEAAVAR.F | HexNAc(5)Hex(6)Fuc(1)NeuAc(1) | N[+2571] | 1451.1694 | 4 | 5801.6559 | 5801.636 | 0 | 3.4 | 141 | Specific | 763.6 | 536.06 | 536.06 | 15.11 | 108 | >sp\|P02788\|TRFL_HUMAN Lactotransferrin OS=Homo sapiens OX=9606 GN=LTF PE=1 SV=6 | 1270 | 20201117_fqg_milk_1ng_glyco_hcd203040_compound15.22601.22601.4 | controllerType=0 controllerNumber=1 scan=22601 | 46.0246 |
| 19075:4 | 2 | R.TAGWNVPIGTLRPFLN[+2205.793]WTGPPEPIEAAVAR.F | HexNAc(4)Hex(5)Fuc(2)NeuAc(1) | N[+2206] | 1359.8864 | 4 | 5436.5236 | 5436.5038 | 0 | 3.6 | 141 | Specific | 796 | 687.4 | 687.4 | 15.11 | 108 | >sp\|P02788\|TRFL_HUMAN Lactotransferrin OS=Homo sapiens OX=9606 GN=LTF PE=1 SV=6 | 1270 | 20201117_fqg_milk_1ng_glyco_hcd203040_compound15.22722.22722.4 | controllerType=0 controllerNumber=1 scan=22722 | 46.2656 |
| 19079:5 | 2 | R.TAGWNVPIGTLRPFLN[+2205.793]WTGPPEPIEAAVAR.F | HexNAc(4)Hex(5)Fuc(2)NeuAc(1) | N[+2206] | 1088.1097 | 5 | 5436.5196 | 5436.5038 | 0 | 2.9 | 141 | Specific | 867.14 | 703.58 | 703.58 | 16.14 | 108 | >sp\|P02788\|TRFL_HUMAN Lactotransferrin OS=Homo sapiens OX=9606 GN=LTF PE=1 SV=6 | 1270 | 20201117_fqg_milk_1ng_glyco_hcd203040_compound15.22726.22726.5 | controllerType=0 controllerNumber=1 scan=22726 | 46.2707 |
| 19081:4 | 2 | R.TAGWNVPIGTLRPFLN[+2351.851]WTGPPEPIEAAVAR.F | HexNAc(4)Hex(5)Fuc(3)NeuAc(1) | N[+2352] | 1396.3994 | 4 | 5582.5758 | 5582.5617 | 0 | 2.5 | 141 | Specific | 874.67 | 740.77 | 740.77 | 17.51 | 108 | >sp\|P02788\|TRFL_HUMAN Lactotransferrin OS=Homo sapiens OX=9606 GN=LTF PE=1 SV=6 | 1270 | 20201117_fqg_milk_1ng_glyco_hcd203040_compound15.22728.22728.4 | controllerType=0 controllerNumber=1 scan=22728 | 46.2792 |
| 19082:5 | 2 | R.TAGWNVPIGTLRPFLN[+2205.793]WTGPPEPIEAAVAR.F | HexNAc(4)Hex(5)Fuc(2)NeuAc(1) | N[+2206] | 1088.1097 | 5 | 5436.5196 | 5436.5038 | 0 | 2.9 | 141 | Specific | 903.63 | 707.91 | 707.91 | 19.14 | 108 | >sp\|P02788\|TRFL_HUMAN Lactotransferrin OS=Homo sapiens OX=9606 GN=LTF PE=1 SV=6 | 1270 | 20201117_fqg_milk_1ng_glyco_hcd203040_compound15.22729.22729.5 | controllerType=0 controllerNumber=1 scan=22729 | 46.2811 |
| 18680:3 | 2 | R.TAGWNIPMGLLFN[+1913.677]QTGSC[+57.021]K.F | HexNAc(4)Hex(5)NeuAc(1) | C[+57], N[+1914] | 1336.9061 | 3 | 4008.7036 | 4008.687 | 0 | 4.2 | 485 | Specific | 240.25 | 192.75 | 192.75 | 5.53 | 108 | >sp\|P02788\|TRFL_HUMAN Lactotransferrin OS=Homo sapiens OX=9606 GN=LTF PE=1 SV=6 | 1270 | 20201117_fqg_milk_1ng_glyco_hcd203040_compound15.22311.22311.4 | controllerType=0 controllerNumber=1 scan=22311 | 45.4428 |
| 16767:3 | 2 | R.TAGWNIPMGLLFN[+1768.640]QTGSC[+57.021]K.F | HexNAc(4)Hex(5)Fuc(1) | C[+57], N[+1769] | 1288.5552 | 3 | 3863.6512 | 3863.6495 | 0 | 1.5 | 485 | Specific | 327.94 | 327.94 | 327.94 | 7.07 | 108 | >sp\|P02788\|TRFL_HUMAN Lactotransferrin OS=Homo sapiens OX=9606 GN=LTF PE=1 SV=6 | 1270 | 20201117_fqg_milk_1ng_glyco_hcd203040_compound15.20315.20315.3 | controllerType=0 controllerNumber=1 scan=20315 | [illegible] |
| 18698:3 | 2 | R.TAGWNIPMGLLFN[+1751.624]QTGSC[+57.021]K.F | HexNAc(4)Hex(4)NeuAc(1) | C[+57], N[+1752] | 1282.8852 | 3 | 3846.641 | 3846.6341 | 0 | 1.8 | 485 | Specific | 372.59 | 372.59 | 372.59 | 7.34 | 108 | >sp\|P02788\|TRFL_HUMAN Lactotransferrin OS=Homo sapiens OX=9606 GN=LTF PE=1 SV=6 | 1270 | 20201117_fqg_milk_1ng_glyco_hcd203040_compound15.22330.22330.3 | controllerType=0 controllerNumber=1 scan=22330 | [illegible] |
| 16756:3 | 2 | R.TAGWNIPMGLLFN[+1606.587]QTGSC[+57.021]K.F | HexNAc(4)Hex(4)Fuc(1) | C[+57], N[+1607] | 1234.544 | 3 | 3701.6175 | 3701.5966 | 0 | 5.6 | 485 | Specific | 375.85 | 375.85 | 375.85 | 8.2 | 108 | >sp\|P02788\|TRFL_HUMAN Lactotransferrin OS=Homo sapiens OX=9606 GN=LTF PE=1 SV=6 | 1270 | 20201117_fqg_milk_1ng_glyco_hcd203040_compound15.20304.20304.3 | controllerType=0 controllerNumber=1 scan=20304 | [illegible] |
| 16770:3 | 2 | R.TAGWNIPMGLLFN[+1768.640]QTGSC[+57.021]K.F | HexNAc(4)Hex(5)Fuc(1) | C[+57], N[+1769] | 1288.5566 | 3 | 3863.6552 | 3863.6495 | 0 | 1.5 | 485 | Specific | 451.32 | 442.66 | 442.66 | 9.27 | 108 | >sp\|P02788\|TRFL_HUMAN Lactotransferrin OS=Homo sapiens OX=9606 GN=LTF PE=1 SV=6 | 1270 | 20201117_fqg_milk_1ng_glyco_hcd203040_compound15.20318.20318.3 | controllerType=0 controllerNumber=1 scan=20318 | [illegible] |
| 16604:4 | 2 | R.TAGWNIPMGLLFN[+1768.640]QTGSC[+57.021]K.F | HexNAc(4)Hex(5)Fuc(1) | C[+57], N[+1769] | 966.6698 | 4 | 3863.6575 | 3863.6495 | 0 | 2.1 | 485 | Specific | 487.02 | 484.04 | 484.04 | 10.05 | 108 | >sp\|P02788\|TRFL_HUMAN Lactotransferrin OS=Homo sapiens OX=9606 GN=LTF PE=1 SV=6 | 1270 | 20201117_fqg_milk_1ng_glyco_hcd203040_compound15.20145.20145.4 | controllerType=0 controllerNumber=1 scan=20145 | 40.7327 |
| 16871:3 | 2 | R.TAGWNIPMGLLFN[+1460.529]QTGSC[+57.021]K.F | HexNAc(4)Hex(4) | C[+57], N[+1461] | 1185.8535 | 3 | 3555.5458 | 3555.5387 | 0 | 2 | 485 | Specific | 532.73 | 479.88 | 250.35 | 9.51 | 108 | >sp\|P02788\|TRFL_HUMAN Lactotransferrin OS=Homo sapiens OX=9606 GN=LTF PE=1 SV=6 | 1270 | 20201117_fqg_milk_1ng_glyco_hcd203040_compound15.20423.20423.3 | controllerType=0 controllerNumber=1 scan=20423 | 41.3157 |
| 16758:3 | 2 | R.TAGWNIPMGLLFN[+1606.587]QTGSC[+57.021]K.F | HexNAc(4)Hex(4)Fuc(1) | C[+57], N[+1607] | 1234.544 | 3 | 3701.6175 | 3701.5966 | 0 | 5.6 | 485 | Specific | 599.37 | 538.97 | 538.97 | 8.74 | 108 | >sp\|P02788\|TRFL_HUMAN Lactotransferrin OS=Homo sapiens OX=9606 GN=LTF PE=1 SV=6 | 1270 | 20201117_fqg_milk_1ng_glyco_hcd203040_compound15.20306.20306.3 | controllerType=0 controllerNumber=1 scan=20306 | 41.0881 |
| 18682:3 | 2 | R.TAGWNIPMGLLFN[+1913.677]QTGSC[+57.021]K.F | HexNAc(4)Hex(5)NeuAc(1) | C[+57], N[+1914] | 1336.9061 | 3 | 4008.7036 | 4008.687 | 0 | 4.2 | 485 | Specific | 638.49 | 521.39 | 521.39 | 11.73 | 108 | >sp\|P02788\|TRFL_HUMAN Lactotransferrin OS=Homo sapiens OX=9606 GN=LTF PE=1 SV=6 | 1270 | 20201117_fqg_milk_1ng_glyco_hcd203040_compound15.22313.22313.4 | controllerType=0 controllerNumber=1 scan=22313 | 45.4461 |
| 16777:3 | 2 | R.TAGWNIPMGLLFN[+1622.582]QTGSC[+57.021]K.F | HexNAc(4)Hex(5) | C[+57], N[+1623] | 1239.8716 | 3 | 3717.6004 | 3717.5916 | 0 | 2.4 | 485 | Specific | 643.16 | 282.32 | 282.32 | 12.14 | 108 | >sp\|P02788\|TRFL_HUMAN Lactotransferrin OS=Homo sapiens OX=9606 GN=LTF PE=1 SV=6 | 1270 | 20201117_fqg_milk_1ng_glyco_hcd203040_compound15.20326.20326.3 | controllerType=0 controllerNumber=1 scan=20326 | 41.124 |
| 16606:4 | 2 | R.TAGWNIPMGLLFN[+1768.640]QTGSC[+57.021]K.F | HexNAc(4)Hex(5)Fuc(1) | C[+57], N[+1769] | 966.6698 | 4 | 3863.6575 | 3863.6495 | 0 | 2.1 | 485 | Specific | 662.92 | 456.24 | 456.24 | 13.67 | 108 | >sp\|P02788\|TRFL_HUMAN Lactotransferrin OS=Homo sapiens OX=9606 GN=LTF PE=1 SV=6 | 1270 | 20201117_fqg_milk_1ng_glyco_hcd203040_compound15.20147.20147.4 | controllerType=0 controllerNumber=1 scan=20147 | 40.7387 |
| 16873:3 | 2 | R.TAGWNIPMGLLFN[+1460.529]QTGSC[+57.021]K.F | HexNAc(4)Hex(4) | C[+57], N[+1461] | 1185.8535 | 3 | 3555.5458 | 3555.5387 | 0 | 2 | 485 | Specific | 686.86 | 664.79 | 343.32 | 13.05 | 108 | >sp\|P02788\|TRFL_HUMAN Lactotransferrin OS=Homo sapiens OX=9606 GN=LTF PE=1 SV=6 | 1270 | 20201117_fqg_milk_1ng_glyco_hcd203040_compound15.20425.20425.3 | controllerType=0 controllerNumber=1 scan=20425 | 41.3211 |
| 18701:3 | 2 | R.TAGWNIPMGLLFN[+1751.624]QTGSC[+57.021]K.F | HexNAc(4)Hex(4)NeuAc(1) | C[+57], N[+1752] | 1282.8852 | 3 | 3846.641 | 3846.6341 | 0 | 1.8 | 485 | Specific | 815.35 | 772.91 | 772.91 | 16.05 | 108 | >sp\|P02788\|TRFL_HUMAN Lactotransferrin OS=Homo sapiens OX=9606 GN=LTF PE=1 SV=6 | 1270 | 20201117_fqg_milk_1ng_glyco_hcd203040_compound15.22333.22333.3 | controllerType=0 controllerNumber=1 scan=22333 | 45.4865 |
| 16664:3 | 2 | R.TAGWNIPMGLLFN[+1606.587]QTGSC[+57.021]K.F | HexNAc(4)Hex(4)Fuc(1) | C[+57], N[+1607] | 1234.5391 | 3 | 3701.6028 | 3701.5966 | 0 | 1.7 | 485 | Specific | 828.71 | 828.71 | 828.71 | 16.05 | 108 | >sp\|P02788\|TRFL_HUMAN Lactotransferrin OS=Homo sapiens OX=9606 GN=LTF PE=1 SV=6 | 1270 | 20201117_fqg_milk_1ng_glyco_hcd203040_compound15.20208.20208.3 | controllerType=0 controllerNumber=1 scan=20208 | 40.8655 |
| 18608:4 | 2 | R.TAGWNIPMGLLFN[+1913.677]QTGSC[+57.021]K.F | HexNAc(4)Hex(5)NeuAc(1) | C[+57], N[+1914] | 1002.9308 | 4 | 4008.7012 | 4008.687 | 0 | 3.6 | 485 | Specific | 879.63 | 843.68 | 843.68 | 17.35 | 108 | >sp\|P02788\|TRFL_HUMAN Lactotransferrin OS=Homo sapiens OX=9606 GN=LTF PE=1 SV=6 | 1270 | 20201117_fqg_milk_1ng_glyco_hcd203040_compound15.22235.22235.4 | controllerType=0 controllerNumber=1 scan=22235 | 45.2517 |
| 18441:3 | 2 | R.TAGWNIPMGLLFN[+2059.735]QTGSC[+57.021]K.F | HexNAc(4)Hex(5)Fuc(1)NeuAc(1) | C[+57], N[+2060] | 1385.59 | 3 | 4154.7555 | 4154.7449 | 0 | 2.6 | 485 | Specific | 901.04 | 836.61 | 836.61 | 18.14 | 108 | >sp\|P02788\|TRFL_HUMAN Lactotransferrin OS=Homo sapiens OX=9606 GN=LTF PE=1 SV=6 | 1270 | 20201117_fqg_milk_1ng_glyco_hcd203040_compound15.22060.22060.3 | controllerType=0 controllerNumber=1 scan=22060 | 44.8251 |
| 18404:3 | 2 | R.TAGWNIPMGLLFN[+2205.793]QTGSC[+57.021]K.F | HexNAc(4)Hex(5)Fuc(2)NeuAc(1) | C[+57], N[+2206] | 1434.2762 | 3 | 4300.814 | 4300.8028 | 0 | 2.6 | 485 | Specific | 901.56 | 889.17 | 889.17 | 18.14 | 108 | >sp\|P02788\|TRFL_HUMAN Lactotransferrin OS=Homo sapiens OX=9606 GN=LTF PE=1 SV=6 | 1270 | 20201117_fqg_milk_1ng_glyco_hcd203040_compound15.22021.22021.3 | controllerType=0 controllerNumber=1 scan=22021 | 44.7363 |
| 16544:3 | 2 | R.TAGWNIPMGLLFN[+1768.640]QTGSC[+57.021]K.F | HexNAc(4)Hex(5)Fuc(1) | C[+57], N[+1769] | 1288.5585 | 3 | 3863.661 | 3863.6495 | 0 | 3 | 485 | Specific | 902.35 | 690.32 | 690.32 | 18.14 | 108 | >sp\|P02788\|TRFL_HUMAN Lactotransferrin OS=Homo sapiens OX=9606 GN=LTF PE=1 SV=6 | 1270 | 20201117_fqg_milk_1ng_glyco_hcd203040_compound15.20083.20083.3 | controllerType=0 controllerNumber=1 scan=20083 | 40.6035 |
| 16667:3 | 2 | R.TAGWNIPMGLLFN[+1606.587]QTGSC[+57.021]K.F | HexNAc(4)Hex(4)Fuc(1) | C[+57], N[+1607] | 1234.5391 | 3 | 3701.6028 | 3701.5966 | 0 | 1.7 | 485 | Specific | 906.49 | 745.54 | 745.54 | 18.14 | 108 | >sp\|P02788\|TRFL_HUMAN Lactotransferrin OS=Homo sapiens OX=9606 GN=LTF PE=1 SV=6 | 1270 | 20201117_fqg_milk_1ng_glyco_hcd203040_compound15.20211.20211.3 | controllerType=0 controllerNumber=1 scan=20211 | 40.8866 |
| 18575:3 | 2 | R.TAGWNIPMGLLFN[+1913.677]QTGSC[+57.021]K.F | HexNAc(4)Hex(5)NeuAc(1) | C[+57], N[+1914] | 1336.9037 | 3 | 4008.6967 | 4008.687 | 0 | 2.4 | 485 | Specific | 920.68 | 913.15 | 913.15 | 18.14 | 108 | >sp\|P02788\|TRFL_HUMAN Lactotransferrin OS=Homo sapiens OX=9606 GN=LTF PE=1 SV=6 | 1270 | 20201117_fqg_milk_1ng_glyco_hcd203040_compound15.22200.22200.3 | controllerType=0 controllerNumber=1 scan=22200 | 45.1619 |
| 16780:3 | 2 | R.TAGWNIPMGLLFN[+1622.582]QTGSC[+57.021]K.F | HexNAc(4)Hex(5) | C[+57], N[+1623] | 1239.8716 | 3 | 3717.6004 | 3717.5916 | 0 | 2.4 | 485 | Specific | 922.39 | 805.1 | 454.67 | 18.14 | 108 | >sp\|P02788\|TRFL_HUMAN Lactotransferrin OS=Homo sapiens OX=9606 GN=LTF PE=1 SV=6 | 1270 | 20201117_fqg_milk_1ng_glyco_hcd203040_compound15.20329.20329.3 | controllerType=0 controllerNumber=1 scan=20329 | 41.1299 |
| 16939:3 | 2 | R.TAGWNIPMGLLFN[+1216.423]QTGSC[+57.021]K.F | HexNAc(2)Hex(5) | C[+57], N[+1216] | 1104.4855 | 3 | 3311.442 | 3311.4328 | 0 | 2.8 | 485 | Specific | 925.85 | 925.85 | 385.46 | 18.14 | 108 | >sp\|P02788\|TRFL_HUMAN Lactotransferrin OS=Homo sapiens OX=9606 GN=LTF PE=1 SV=6 | 1270 | 20201117_fqg_milk_1ng_glyco_hcd203040_compound15.20495.20495.3 | controllerType=0 controllerNumber=1 scan=20495 | 41.4804 |
| 18611:4 | 2 | R.TAGWNIPMGLLFN[+1913.677]QTGSC[+57.021]K.F | HexNAc(4)Hex(5)NeuAc(1) | C[+57], N[+1914] | 1002.9308 | 4 | 4008.7012 | 4008.687 | 0 | 3.6 | 485 | Specific | 927.11 | 878.91 | 878.91 | 18.35 | 108 | >sp\|P02788\|TRFL_HUMAN Lactotransferrin OS=Homo sapiens OX=9606 GN=LTF PE=1 SV=6 | 1270 | 20201117_fqg_milk_1ng_glyco_hcd203040_compound15.22238.22238.4 | controllerType=0 controllerNumber=1 scan=22238 | 45.2568 |
| 18401:3 | 2 | R.TAGWNIPMGLLFN[+2205.793]QTGSC[+57.021]K.F | HexNAc(4)Hex(5)Fuc(2)NeuAc(1) | C[+57], N[+2206] | 1434.2762 | 3 | 4300.814 | 4300.8028 | 0 | 2.6 | 485 | Specific | 927.7 | 788.8 | 788.8 | 18.14 | 108 | >sp\|P02788\|TRFL_HUMAN Lactotransferrin OS=Homo sapiens OX=9606 GN=LTF PE=1 SV=6 | 1270 | 20201117_fqg_milk_1ng_glyco_hcd203040_compound15.22018.22018.3 | controllerType=0 controllerNumber=1 scan=22018 | 44.7299 |
| 18439:3 | 2 | R.TAGWNIPMGLLFN[+2059.735]QTGSC[+57.021]K.F | HexNAc(4)Hex(5)Fuc(1)NeuAc(1) | C[+57], N[+2060] | 1385.59 | 3 | 4154.7555 | 4154.7449 | 0 | 2.6 | 485 | Specific | 929.96 | 839.43 | 839.43 | 18.14 | 108 | >sp\|P02788\|TRFL_HUMAN Lactotransferrin OS=Homo sapiens OX=9606 GN=LTF PE=1 SV=6 | 1270 | 20201117_fqg_milk_1ng_glyco_hcd203040_compound15.22058.22058.3 | controllerType=0 controllerNumber=1 scan=22058 | 44.8174 |
| 16542:3 | 2 | R.TAGWNIPMGLLFN[+1768.640]QTGSC[+57.021]K.F | HexNAc(4)Hex(5)Fuc(1) | C[+57], N[+1769] | 1288.5585 | 3 | 3863.661 | 3863.6495 | 0 | 3 | 485 | Specific | 938.56 | 938.56 | 938.56 | 18.14 | 108 | >sp\|P02788\|TRFL_HUMAN Lactotransferrin OS=Homo sapiens OX=9606 GN=LTF PE=1 SV=6 | 1270 | 20201117_fqg_milk_1ng_glyco_hcd203040_compound15.20081.20081.3 | controllerType=0 controllerNumber=1 scan=20081 | 40.5965 |
| 18572:3 | 2 | R.TAGWNIPMGLLFN[+1913.677]QTGSC[+57.021]K.F | HexNAc(4)Hex(5)NeuAc(1) | C[+57], N[+1914] | 1336.9037 | 3 | 4008.6967 | 4008.687 | 0 | 2.4 | 485 | Specific | 956.58 | 936.38 | 936.38 | 19.14 | 108 | >sp\|P02788\|TRFL_HUMAN Lactotransferrin OS=Homo sapiens OX=9606 GN=LTF PE=1 SV=6 | 1270 | 20201117_fqg_milk_1ng_glyco_hcd203040_compound15.22197.22197.3 | controllerType=0 controllerNumber=1 scan=22197 | 45.1514 |
| 16941:3 | 2 | R.TAGWNIPMGLLFN[+1216.423]QTGSC[+57.021]K.F | HexNAc(2)Hex(5) | C[+57], N[+1216] | 1104.4855 | 3 | 3311.442 | 3311.4328 | 0 | 2.8 | 485 | Specific | 1046.53 | 1039.04 | 490.51 | 20.14 | 108 | >sp\|P02788\|TRFL_HUMAN Lactotransferrin OS=Homo sapiens OX=9606 GN=LTF PE=1 SV=6 | 1270 | 20201117_fqg_milk_1ng_glyco_hcd203040_compound15.20497.20497.3 | controllerType=0 controllerNumber=1 scan=20497 | 41.4903 |
| 16786:3 | 2 | R.TAGWNIPM[+15.995]GLLFN[+2059.735]QTGSC[+57.021]K.F | HexNAc(4)Hex(5)Fuc(1)NeuAc(1) | C[+57], M[+16], N[+2060] | 1390.9229 | 3 | 4170.7542 | 4170.7398 | 0 | 3.4 | 485 | Specific | 663.68 | 530.58 | 530.58 | 12.73 | 108 | >sp\|P02788\|TRFL_HUMAN Lactotransferrin OS=Homo sapiens OX=9606 GN=LTF PE=1 SV=6 | 1270 | 20201117_fqg_milk_1ng_glyco_hcd203040_compound15.20335.20335.3 | controllerType=0 controllerNumber=1 scan=20335 | 41.142 |
| 16788:3 | 2 | R.TAGWNIPM[+15.995]GLLFN[+2059.735]QTGSC[+57.021]K.F | HexNAc(4)Hex(5)Fuc(1)NeuAc(1) | C[+57], M[+16], N[+2060] | 1390.9229 | 3 | 4170.7542 | 4170.7398 | 0 | 3.4 | 485 | Specific | 741 | 688.38 | 688.38 | 13.73 | 108 | >sp\|P02788\|TRFL_HUMAN Lactotransferrin OS=Homo sapiens OX=9606 GN=LTF PE=1 SV=6 | 1270 | 20201117_fqg_milk_1ng_glyco_hcd203040_compound15.20337.20337.3 | controllerType=0 controllerNumber=1 scan=20337 | 41.1463 |
| 18825:4 | 2 | R.TAGWNIPMGLLFN[+1913.677]QTGSC[+57.021]KFDEYFSQSC[+57.021]APGSDPR.S | HexNAc(4)Hex(5)NeuAc(1) | C[+57]*2, N[+1914] | 1463.8649 | 4 | 5852.4377 | 5852.4341 | 0 | 0.6 | 485 | Specific | 478.38 | 415.92 | 415.92 | 9.52 | 108 | >sp\|P02788\|TRFL_HUMAN Lactotransferrin OS=Homo sapiens OX=9606 GN=LTF PE=1 SV=6 | 1270 | 20201117_fqg_milk_1ng_glyco_hcd203040_compound15.22462.22462.4 | controllerType=0 controllerNumber=1 scan=22462 | 45.7471 |
| 18827:4 | 2 | R.TAGWNIPMGLLFN[+1913.677]QTGSC[+57.021]KFDEYFSQSC[+57.021]APGSDPR.S | HexNAc(4)Hex(5)NeuAc(1) | C[+57]*2, N[+1914] | 1463.8649 | 4 | 5852.4377 | 5852.4341 | 0 | 0.6 | 485 | Specific | 614.12 | 511.53 | 511.53 | 12.14 | 108 | >sp\|P02788\|TRFL_HUMAN Lactotransferrin OS=Homo sapiens OX=9606 GN=LTF PE=1 SV=6 | 1270 | 20201117_fqg_milk_1ng_glyco_hcd203040_compound15.22464.22464.4 | controllerType=0 controllerNumber=1 scan=22464 | 45.7515 |
| 17690:4 | 2 | R.TAGWNIPMGLLFN[+1768.640]QTGSC[+57.021]KFDEYFSQSC[+57.021]APGSDPR.S | HexNAc(4)Hex(5)Fuc(1) | C[+57]*2, N[+1769] | 1427.6079 | 4 | 5707.4098 | 5707.3966 | 0 | 2.3 | 485 | Specific | 707.17 | 648.09 | 648.09 | 14.62 | 108 | >sp\|P02788\|TRFL_HUMAN Lactotransferrin OS=Homo sapiens OX=9606 GN=LTF PE=1 SV=6 | 1270 | 20201117_fqg_milk_1ng_glyco_hcd203040_compound15.21279.21279.4 | controllerType=0 controllerNumber=1 scan=21279 | 43.1951 |
| 18692:4 | 2 | R.TAGWNIPMGLLFN[+2059.735]QTGSC[+57.021]KFDEYFSQSC[+57.021]APGSDPR.S | HexNAc(4)Hex(5)Fuc(1)NeuAc(1) | C[+57]*2, N[+2060] | 1500.3831 | 4 | 5998.5104 | 5998.492 | 0 | 3.1 | 485 | Specific | 795.23 | 752.11 | 752.11 | 15.62 | 108 | >sp\|P02788\|TRFL_HUMAN Lactotransferrin OS=Homo sapiens OX=9606 GN=LTF PE=1 SV=6 | 1270 | 20201117_fqg_milk_1ng_glyco_hcd203040_compound15.22324.22324.4 | controllerType=0 controllerNumber=1 scan=22324 | 45.4728 |
| 17692:4 | 2 | R.TAGWNIPMGLLFN[+1768.640]QTGSC[+57.021]KFDEYFSQSC[+57.021]APGSDPR.S | HexNAc(4)Hex(5)Fuc(1) | C[+57]*2, N[+1769] | 1427.6079 | 4 | 5707.4098 | 5707.3966 | 0 | 2.3 | 485 | Specific | 832.5 | 754.5 | 754.5 | 16.05 | 108 | >sp\|P02788\|TRFL_HUMAN Lactotransferrin OS=Homo sapiens OX=9606 GN=LTF PE=1 SV=6 | 1270 | 20201117_fqg_milk_1ng_glyco_hcd203040_compound15.21281.21281.4 | controllerType=0 controllerNumber=1 scan=21281 | 43.1994 |
| 18695:4 | 2 | R.TAGWNIPMGLLFN[+2059.735]QTGSC[+57.021]KFDEYFSQSC[+57.021]APGSDPR.S | HexNAc(4)Hex(5)Fuc(1)NeuAc(1) | C[+57]*2, N[+2060] | 1500.3831 | 4 | 5998.5104 | 5998.492 | 0 | 3.1 | 485 | Specific | 913.46 | 863.07 | 863.07 | 18.62 | 108 | >sp\|P02788\|TRFL_HUMAN Lactotransferrin OS=Homo sapiens OX=9606 GN=LTF PE=1 SV=6 | 1270 | 20201117_fqg_milk_1ng_glyco_hcd203040_compound15.22327.22327.4 | controllerType=0 controllerNumber=1 scan=22327 | 45.478 |
| 11264:3 | 2 | R.N[+2350.830]GSDC[+57.021]PDKFC[+57.021]LFQSETK.N | HexNAc(4)Hex(5)Fuc(1)NeuAc(2) | C[+57]*2, N[+2351] | 1461.9105 | 3 | 4383.7168 | 4383.7043 | 0 | 2.9 | 642 | Specific | 350.83 | 271.9 | 271.9 | 6.71 | 108 | >sp\|P02788\|TRFL_HUMAN Lactotransferrin OS=Homo sapiens OX=9606 GN=LTF PE=1 SV=6 | 1270 | 20201117_fqg_milk_1ng_glyco_hcd203040_compound15.14540.14540.3 | controllerType=0 controllerNumber=1 scan=14540 | 30.8273 |
| 11262:3 | 2 | R.N[+2350.830]GSDC[+57.021]PDKFC[+57.021]LFQSETK.N | HexNAc(4)Hex(5)Fuc(1)NeuAc(2) | C[+57]*2, N[+2351] | 1461.9105 | 3 | 4383.7168 | 4383.7043 | 0 | 2.9 | 642 | Specific | 391.69 | 340.32 | 340.32 | 7.04 | 108 | >sp\|P02788\|TRFL_HUMAN Lactotransferrin OS=Homo sapiens OX=9606 GN=LTF PE=1 SV=6 | 1270 | 20201117_fqg_milk_1ng_glyco_hcd203040_compound15.14538.14538.3 | controllerType=0 controllerNumber=1 scan=14538 | 30.8218 |
| 08866:3 | 2 | R.N[+2059.735]GSDC[+57.021]PDKFC[+57.021]LFQSETK.N | HexNAc(4)Hex(5)Fuc(1)NeuAc(1) | C[+57]*2, N[+2060] | 1364.8799 | 3 | 4092.6253 | 4092.6088 | 0 | 4 | 642 | Specific | 431.25 | 368.4 | 368.4 | 6.83 | 108 | >sp\|P02788\|TRFL_HUMAN Lactotransferrin OS=Homo sapiens OX=9606 GN=LTF PE=1 SV=6 | 1270 | 20201117_fqg_milk_1ng_glyco_hcd203040_compound15.12062.12062.3 | controllerType=0 controllerNumber=1 scan=12062 | 26.6341 |
| 08753:3 | 2 | R.N[+2205.793]GSDC[+57.021]PDKFC[+57.021]LFQSETK.N | HexNAc(4)Hex(5)Fuc(2)NeuAc(1) | C[+57]*2, N[+2206] | 1413.5615 | 3 | 4238.6698 | 4238.6667 | 0 | 0.7 | 642 | Specific | 460.34 | 330.72 | 330.72 | 7.2 | 108 | >sp\|P02788\|TRFL_HUMAN Lactotransferrin OS=Homo sapiens OX=9606 GN=LTF PE=1 SV=6 | 1270 | 20201117_fqg_milk_1ng_glyco_hcd203040_compound15.11943.11943.3 | controllerType=0 controllerNumber=1 scan=11943 | 26.4102 |
| 08869:3 | 2 | R.N[+2059.735]GSDC[+57.021]PDKFC[+57.021]LFQSETK.N | HexNAc(4)Hex(5)Fuc(1)NeuAc(1) | C[+57]*2, N[+2060] | 1364.8799 | 3 | 4092.6253 | 4092.6088 | 0 | 4 | 642 | Specific | 483.11 | 350.35 | 350.35 | 7.07 | 108 | >sp\|P02788\|TRFL_HUMAN Lactotransferrin OS=Homo sapiens OX=9606 GN=LTF PE=1 SV=6 | 1270 | 20201117_fqg_milk_1ng_glyco_hcd203040_compound15.12064.12064.3 | controllerType=0 controllerNumber=1 scan=12064 | 26.638 |
| 08755:3 | 2 | R.N[+2205.793]GSDC[+57.021]PDKFC[+57.021]LFQSETK.N | HexNAc(4)Hex(5)Fuc(2)NeuAc(1) | C[+57]*2, N[+2206] | 1413.5615 | 3 | 4238.6698 | 4238.6667 | 0 | 0.7 | 642 | Specific | 516.23 | 344.45 | 344.45 | 7.44 | 108 | >sp\|P02788\|TRFL_HUMAN Lactotransferrin OS=Homo sapiens OX=9606 GN=LTF PE=1 SV=6 | 1270 | 20201117_fqg_milk_1ng_glyco_hcd203040_compound15.11945.11945.2 | controllerType=0 controllerNumber=1 scan=11945 | 26.414 |
| 08792:3 | 2 | R.N[+2205.793]GSDC[+57.021]PDKFC[+57.021]LFQSETK.N | HexNAc(4)Hex(5)Fuc(2)NeuAc(1) | C[+57]*2, N[+2206] | 1413.5654 | 3 | 4238.6815 | 4238.6667 | 0 | 3.5 | 642 | Specific | 538 | 437.16 | 437.16 | 8.81 | 108 | >sp\|P02788\|TRFL_HUMAN Lactotransferrin OS=Homo sapiens OX=9606 GN=LTF PE=1 SV=6 | 1270 | 20201117_fqg_milk_1ng_glyco_hcd203040_compound15.11985.11985.3 | controllerType=0 controllerNumber=1 scan=11985 | 26.5051 |
| 08794:3 | 2 | R.N[+2205.793]GSDC[+57.021]PDKFC[+57.021]LFQSETK.N | HexNAc(4)Hex(5)Fuc(2)NeuAc(1) | C[+57]*2, N[+2206] | 1413.5654 | 3 | 4238.6815 | 4238.6667 | 0 | 3.5 | 642 | Specific | 562.78 | 451.89 | 451.89 | 9.1 | 108 | >sp\|P02788\|TRFL_HUMAN Lactotransferrin OS=Homo sapiens OX=9606 GN=LTF PE=1 SV=6 | 1270 | 20201117_fqg_milk_1ng_glyco_hcd203040_compound15.11987.11987.3 | controllerType=0 controllerNumber=1 scan=11987 | 26.5099 |
| 12805:4 | 5 | K.KGN[+2204.772]TTLNSFVIPSESDVPTHC[+57.021]PSQWWPYAGHC[+57.021]YK.I | HexNAc(4)Hex(5)NeuAc(2) | C[+57]*2, N[+2205] | 1542.9066 | 4 | 6168.6046 | 6168.5942 | 0 | 1.7 | 342 | Specific | 208.86 | 208.86 | 208.86 | 4.63 | 67 | >sp\|P22897\|MRC1_HUMAN Macrophage mannose receptor 1 OS=Homo sapiens OX=9606 GN=MRC1 PE=1 SV=1 | 1049 | 20201117_fqg_milk_1ng_glyco_hcd203040_compound15.16165.16165.2 | controllerType=0 controllerNumber=1 scan=16165 | 33.5456 |
| 12802:4 | 5 | K.KGN[+2204.772]TTLNSFVIPSESDVPTHC[+57.021]PSQWWPYAGHC[+57.021]YK.I | HexNAc(4)Hex(5)NeuAc(2) | C[+57]*2, N[+2205] | 1542.9066 | 4 | 6168.6046 | 6168.5942 | 0 | 1.7 | 342 | Specific | 259.66 | 259.66 | 259.66 | 7.01 | 67 | >sp\|P22897\|MRC1_HUMAN Macrophage mannose receptor 1 OS=Homo sapiens OX=9606 GN=MRC1 PE=1 SV=1 | 1049 | 20201117_fqg_milk_1ng_glyco_hcd203040_compound15.16162.16162.2 | controllerType=0 controllerNumber=1 scan=16162 | 33.5407 |
| 12830:4 | 5 | K.KGN[+2205.793]TTLNSFVIPSESDVPTHC[+57.021]PSQWWPYAGHC[+57.021]YK.I | HexNAc(4)Hex(5)Fuc(2)NeuAc(1) | C[+57]*2, N[+2206] | 1543.1609 | 4 | 6169.6217 | 6169.6146 | 0 | 1.2 | 342 | Specific | 580.98 | 580.98 | 580.98 | 11.7 | 67 | >sp\|P22897\|MRC1_HUMAN Macrophage mannose receptor 1 OS=Homo sapiens OX=9606 GN=MRC1 PE=1 SV=1 | 1049 | 20201117_fqg_milk_1ng_glyco_hcd203040_compound15.16191.16191.4 | controllerType=0 controllerNumber=1 scan=16191 | 33.5927 |
| 12828:4 | 5 | K.KGN[+2205.793]TTLNSFVIPSESDVPTHC[+57.021]PSQWWPYAGHC[+57.021]YK.I | HexNAc(4)Hex(5)Fuc(2)NeuAc(1) | C[+57]*2, N[+2206] | 1543.1609 | 4 | 6169.6217 | 6169.6146 | 0 | 1.2 | 342 | Specific | 629.42 | 491.63 | 491.63 | 12.54 | 67 | >sp\|P22897\|MRC1_HUMAN Macrophage mannose receptor 1 OS=Homo sapiens OX=9606 GN=MRC1 PE=1 SV=1 | 1049 | 20201117_fqg_milk_1ng_glyco_hcd203040_compound15.16189.16189.4 | controllerType=0 controllerNumber=1 scan=16189 | 33.5886 |
| 16176:3 | 5 | R.VWIALNSN[+2205.793]LTDNQYTWTDKWR.V | HexNAc(4)Hex(5)Fuc(2)NeuAc(1) | N[+2206] | 1610.7037 | 3 | 4830.0965 | 4830.0821 | 0 | 3 | 1153 | Specific | 653.62 | 497.52 | 497.52 | 12.76 | 67 | >sp\|P22897\|MRC1_HUMAN Macrophage mannose receptor 1 OS=Homo sapiens OX=9606 GN=MRC1 PE=1 SV=1 | 1049 | 20201117_fqg_milk_1ng_glyco_hcd203040_compound15.19699.19699.3 | controllerType=0 controllerNumber=1 scan=19699 | 39.8988 |
| 16174:3 | 5 | R.VWIALNSN[+2205.793]LTDNQYTWTDKWR.V | HexNAc(4)Hex(5)Fuc(2)NeuAc(1) | N[+2206] | 1610.7037 | 3 | 4830.0965 | 4830.0821 | 0 | 3 | 1153 | Specific | 772.87 | 566.52 | 566.52 | 14.76 | 67 | >sp\|P22897\|MRC1_HUMAN Macrophage mannose receptor 1 OS=Homo sapiens OX=9606 GN=MRC1 PE=1 SV=1 | 1049 | 20201117_fqg_milk_1ng_glyco_hcd203040_compound15.19697.19697.3 | controllerType=0 controllerNumber=1 scan=19697 | 39.8937 |
| 19292:3 | 6 | R.GLSFDVSLEVSQGPGLLN[+2350.830]DTK.V | HexNAc(4)Hex(5)Fuc(1)NeuAc(2) | N[+2351] | 1509.6557 | 3 | 4526.9525 | 4526.9436 | 0 | 2 | 118 | Specific | 822.16 | 539.57 | 539.57 | 16.05 | 57 | >sp\|P01833\|PIGR_HUMAN Polymeric immunoglobulin receptor OS=Homo sapiens OX=9606 GN=PIGR PE=1 SV=4 | 902 | 20201117_fqg_milk_1ng_glyco_hcd203040_compound15.22954.22954.3 | controllerType=0 controllerNumber=1 scan=22954 | 46.6733 |
| 19294:3 | 6 | R.GLSFDVSLEVSQGPGLLN[+2350.830]DTK.V | HexNAc(4)Hex(5)Fuc(1)NeuAc(2) | N[+2351] | 1509.6557 | 3 | 4526.9525 | 4526.9436 | 0 | 2 | 118 | Specific | 1046.5 | 1007.64 | 1007.64 | 20.05 | 57 | >sp\|P01833\|PIGR_HUMAN Polymeric immunoglobulin receptor OS=Homo sapiens OX=9606 GN=PIGR PE=1 SV=4 | 902 | 20201117_fqg_milk_1ng_glyco_hcd203040_compound15.22956.22956.3 | controllerType=0 controllerNumber=1 scan=22956 | 46.6796 |
| 18667:3 | 6 | K.QIGLYPVLVIDSSGYVNPN[+2059.735]YTGR.I | HexNAc(4)Hex(5)Fuc(1)NeuAc(1) | N[+2060] | 1529.0256 | 3 | 4585.0622 | 4585.0384 | 0 | 5.2 | 168 | Specific | 422.29 | 371.55 | 371.55 | 7.19 | 57 | >sp\|P01833\|PIGR_HUMAN Polymeric immunoglobulin receptor OS=Homo sapiens OX=9606 GN=PIGR PE=1 SV=4 | 902 | 20201117_fqg_milk_1ng_glyco_hcd203040_compound15.22298.22298.3 | controllerType=0 controllerNumber=1 scan=22298 | 45.4138 |
| 18654:3 | 6 | K.QIGLYPVLVIDSSGYVNPN[+2059.735]YTGR.I | HexNAc(4)Hex(5)Fuc(1)NeuAc(1) | N[+2060] | 1529.0295 | 3 | 4585.0739 | 4585.0384 | 0 | 7.7 | 168 | Specific | 545.7 | 545.45 | 545.45 | 8.46 | 57 | >sp\|P01833\|PIGR_HUMAN Polymeric immunoglobulin receptor OS=Homo sapiens OX=9606 GN=PIGR PE=1 SV=4 | 902 | 20201117_fqg_milk_1ng_glyco_hcd203040_compound15.22284.22284.3 | controllerType=0 controllerNumber=1 scan=22284 | 45.3606 |
| 18669:3 | 6 | K.QIGLYPVLVIDSSGYVNPN[+2059.735]YTGR.I | HexNAc(4)Hex(5)Fuc(1)NeuAc(1) | N[+2060] | 1529.0256 | 3 | 4585.0622 | 4585.0384 | 0 | 5.2 | 168 | Specific | 588.53 | 430.26 | 430.26 | 10.72 | 57 | >sp\|P01833\|PIGR_HUMAN Polymeric immunoglobulin receptor OS=Homo sapiens OX=9606 GN=PIGR PE=1 SV=4 | 902 | 20201117_fqg_milk_1ng_glyco_hcd203040_compound15.22300.22300.3 | controllerType=0 controllerNumber=1 scan=22300 | 45.4179 |
| 18657:3 | 6 | K.QIGLYPVLVIDSSGYVNPN[+2059.735]YTGR.I | HexNAc(4)Hex(5)Fuc(1)NeuAc(1) | N[+2060] | 1529.0295 | 3 | 4585.0739 | 4585.0384 | 0 | 7.7 | 168 | Specific | 777.72 | 727.83 | 727.83 | 14 | 57 | >sp\|P01833\|PIGR_HUMAN Polymeric immunoglobulin receptor OS=Homo sapiens OX=9606 GN=PIGR PE=1 SV=4 | 902 | 20201117_fqg_milk_1ng_glyco_hcd203040_compound15.22287.22287.3 | controllerType=0 controllerNumber=1 scan=22287 | 45.3767 |
| 24526:3 | 6 | R.[+42.011]LSLLEEPGN[+2205.793]GTFTVILNQLTSR.D | HexNAc(4)Hex(5)Fuc(2)NeuAc(1) | .[42], N[+2206] | 1550.7042 | 3 | 4650.0979 | 4650.096 | 0 | 0.4 | 413 | Specific | 226.62 | 226.62 | 226.62 | 4.8 | 57 | >sp\|P01833\|PIGR_HUMAN Polymeric immunoglobulin receptor OS=Homo sapiens OX=9606 GN=PIGR PE=1 SV=4 | 902 | 20201117_fqg_milk_1ng_glyco_hcd203040_compound15.28401.28401.3 | controllerType=0 controllerNumber=1 scan=28401 | 56.964 |
| 20308:4 | 6 | R.LSLLEEPGN[+2076.750]GTFTVILNQLTSR.D | HexNAc(4)Hex(6)Fuc(2) | N[+2077] | 1120.5223 | 4 | 4479.0672 | 4479.0428 | 0 | 5.4 | 413 | Specific | 244.12 | 227.55 | 227.55 | 6.19 | 57 | >sp\|P01833\|PIGR_HUMAN Polymeric immunoglobulin receptor OS=Homo sapiens OX=9606 GN=PIGR PE=1 SV=4 | 902 | 20201117_fqg_milk_1ng_glyco_hcd203040_compound15.24013.24013.4 | controllerType=0 controllerNumber=1 scan=24013 | 48.8578 |
| 22743:3 | 6 | R.[+42.011]LSLLEEPGN[+2336.851]GTFTVILNQLTSR.D | HexNAc(6)Hex(6)Fuc(1) | .[42], N[+2337] | 1594.3848 | 3 | 4781.1399 | 4781.1542 | 0 | -3 | 413 | Specific | 318.08 | 277.39 | 277.39 | 4.96 | 57 | >sp\|P01833\|PIGR_HUMAN Polymeric immunoglobulin receptor OS=Homo sapiens OX=9606 GN=PIGR PE=1 SV=4 | 902 | 20201117_fqg_milk_1ng_glyco_hcd203040_compound15.26547.26547.3 | controllerType=0 controllerNumber=1 scan=26547 | 53.5206 |
| 24528:3 | 6 | R.[+42.011]LSLLEEPGN[+2205.793]GTFTVILNQLTSR.D | HexNAc(4)Hex(5)Fuc(2)NeuAc(1) | .[42], N[+2206] | 1550.7042 | 3 | 4650.0979 | 4650.096 | 0 | 0.4 | 413 | Specific | 328.93 | 303.58 | 303.58 | 7.08 | 57 | >sp\|P01833\|PIGR_HUMAN Polymeric immunoglobulin receptor OS=Homo sapiens OX=9606 GN=PIGR PE=1 SV=4 | 902 | 20201117_fqg_milk_1ng_glyco_hcd203040_compound15.28403.28403.3 | controllerType=0 controllerNumber=1 scan=28403 | 56.9681 |
| 22741:3 | 6 | R.[+42.011]LSLLEEPGN[+2336.851]GTFTVILNQLTSR.D | HexNAc(6)Hex(6)Fuc(1) | .[42], N[+2337] | 1594.3848 | 3 | 4781.1399 | 4781.1542 | 0 | -3 | 413 | Specific | 357.38 | 328.79 | 328.79 | 6.35 | 57 | >sp\|P01833\|PIGR_HUMAN Polymeric immunoglobulin receptor OS=Homo sapiens OX=9606 GN=PIGR PE=1 SV=4 | 902 | 20201117_fqg_milk_1ng_glyco_hcd203040_compound15.26545.26545.3 | controllerType=0 controllerNumber=1 scan=26545 | 53.5138 |
| 20310:4 | 6 | R.LSLLEEPGN[+2076.750]GTFTVILNQLTSR.D | HexNAc(4)Hex(6)Fuc(2) | N[+2077] | 1120.5223 | 4 | 4479.0672 | 4479.0428 | 0 | 5.4 | 413 | Specific | 633.99 | 530.73 | 530.73 | 12.34 | 57 | >sp\|P01833\|PIGR_HUMAN Polymeric immunoglobulin receptor OS=Homo sapiens OX=9606 GN=PIGR PE=1 SV=4 | 902 | 20201117_fqg_milk_1ng_glyco_hcd203040_compound15.24015.24015.4 | controllerType=0 controllerNumber=1 scan=24015 | 48.866 |
| 20262:3 | 6 | R.LSLLEEPGN[+2205.793]GTFTVILNQLTSR.D | HexNAc(4)Hex(5)Fuc(2)NeuAc(1) | N[+2206] | 1536.7048 | 3 | 4608.0998 | 4608.0854 | 0 | 3.1 | 413 | Specific | 937.08 | 670.45 | 670.45 | 18.61 | 57 | >sp\|P01833\|PIGR_HUMAN Polymeric immunoglobulin receptor OS=Homo sapiens OX=9606 GN=PIGR PE=1 SV=4 | 902 | 20201117_fqg_milk_1ng_glyco_hcd203040_compound15.23965.23965.3 | controllerType=0 controllerNumber=1 scan=23965 | 48.7592 |
| 20269:4 | 6 | R.LSLLEEPGN[+2205.793]GTFTVILNQLTSR.D | HexNAc(4)Hex(5)Fuc(2)NeuAc(1) | N[+2206] | 1152.7795 | 4 | 4608.096 | 4608.0854 | 0 | 2.3 | 413 | Specific | 974.84 | 761.29 | 761.29 | 20.23 | 57 | >sp\|P01833\|PIGR_HUMAN Polymeric immunoglobulin receptor OS=Homo sapiens OX=9606 GN=PIGR PE=1 SV=4 | 902 | 20201117_fqg_milk_1ng_glyco_hcd203040_compound15.23972.23972.4 | controllerType=0 controllerNumber=1 scan=23972 | 48.7706 |
| 20260:3 | 6 | R.LSLLEEPGN[+2205.793]GTFTVILNQLTSR.D | HexNAc(4)Hex(5)Fuc(2)NeuAc(1) | N[+2206] | 1536.7048 | 3 | 4608.0998 | 4608.0854 | 0 | 3.1 | 413 | Specific | 1038.61 | 797.49 | 797.49 | 20.61 | 57 | >sp\|P01833\|PIGR_HUMAN Polymeric immunoglobulin receptor OS=Homo sapiens OX=9606 GN=PIGR PE=1 SV=4 | 902 | 20201117_fqg_milk_1ng_glyco_hcd203040_compound15.23963.23963.3 | controllerType=0 controllerNumber=1 scan=23963 | 48.754 |
| 20267:4 | 6 | R.LSLLEEPGN[+2205.793]GTFTVILNQLTSR.D | HexNAc(4)Hex(5)Fuc(2)NeuAc(1) | N[+2206] | 1152.7795 | 4 | 4608.096 | 4608.0854 | 0 | 2.3 | 413 | Specific | 1057.75 | 845.12 | 845.12 | 22.23 | 57 | >sp\|P01833\|PIGR_HUMAN Polymeric immunoglobulin receptor OS=Homo sapiens OX=9606 GN=PIGR PE=1 SV=4 | 902 | 20201117_fqg_milk_1ng_glyco_hcd203040_compound15.23970.23970.4 | controllerType=0 controllerNumber=1 scan=23970 | 48.7652 |
| 15795:4 | 6 | K.IIEGEPNLKVPGN[+2205.793]VTAVLGETLK.V | HexNAc(4)Hex(5)Fuc(2)NeuAc(1) | N[+2206] | 1150.0413 | 4 | 4597.1434 | 4597.1422 | 0 | 0.3 | 457 | Specific | 731.58 | 652.51 | 652.51 | 13.57 | 57 | >sp\|P01833\|PIGR_HUMAN Polymeric immunoglobulin receptor OS=Homo sapiens OX=9606 GN=PIGR PE=1 SV=4 | 902 | 20201117_fqg_milk_1ng_glyco_hcd203040_compound15.19297.19297.2 | controllerType=0 controllerNumber=1 scan=19297 | 39.2027 |
| 15781:3 | 6 | K.IIEGEPNLKVPGN[+2205.793]VTAVLGETLK.V | HexNAc(4)Hex(5)Fuc(2)NeuAc(1) | N[+2206] | 1533.0557 | 3 | 4597.1526 | 4597.1422 | 0 | 2.3 | 457 | Specific | 815.5 | 815.5 | 815.5 | 16.23 | 57 | >sp\|P01833\|PIGR_HUMAN Polymeric immunoglobulin receptor OS=Homo sapiens OX=9606 GN=PIGR PE=1 SV=4 | 902 | 20201117_fqg_milk_1ng_glyco_hcd203040_compound15.19282.19282.3 | controllerType=0 controllerNumber=1 scan=19282 | 39.172 |
| 15763:3 | 6 | K.IIEGEPNLKVPGN[+2205.793]VTAVLGETLK.V | HexNAc(4)Hex(5)Fuc(2)NeuAc(1) | N[+2206] | 1533.0546 | 3 | 4597.1493 | 4597.1422 | 0 | 1.5 | 457 | Specific | 836.11 | 742.71 | 742.71 | 16.15 | 57 | >sp\|P01833\|PIGR_HUMAN Polymeric immunoglobulin receptor OS=Homo sapiens OX=9606 GN=PIGR PE=1 SV=4 | 902 | 20201117_fqg_milk_1ng_glyco_hcd203040_compound15.19265.19265.3 | controllerType=0 controllerNumber=1 scan=19265 | 39.137 |
| 15792:4 | 6 | K.IIEGEPNLKVPGN[+2205.793]VTAVLGETLK.V | HexNAc(4)Hex(5)Fuc(2)NeuAc(1) | N[+2206] | 1150.0413 | 4 | 4597.1434 | 4597.1422 | 0 | 0.3 | 457 | Specific | 859.05 | 859.05 | 859.05 | 16.57 | 57 | >sp\|P01833\|PIGR_HUMAN Polymeric immunoglobulin receptor OS=Homo sapiens OX=9606 GN=PIGR PE=1 SV=4 | 902 | 20201117_fqg_milk_1ng_glyco_hcd203040_compound15.19294.19294.2 | controllerType=0 controllerNumber=1 scan=19294 | 39.197 |
| 15771:3 | 6 | K.IIEGEPNLKVPGN[+2205.793]VTAVLGETLK.V | HexNAc(4)Hex(5)Fuc(2)NeuAc(1) | N[+2206] | 1533.0514 | 3 | 4597.1398 | 4597.1422 | 0 | -0.5 | 457 | Specific | 860.49 | 761.72 | 761.72 | 16.24 | 57 | >sp\|P01833\|PIGR_HUMAN Polymeric immunoglobulin receptor OS=Homo sapiens OX=9606 GN=PIGR PE=1 SV=4 | 902 | 20201117_fqg_milk_1ng_glyco_hcd203040_compound15.19271.19271.3 | controllerType=0 controllerNumber=1 scan=19271 | 39.1509 |
| 15765:3 | 6 | K.IIEGEPNLKVPGN[+2205.793]VTAVLGETLK.V | HexNAc(4)Hex(5)Fuc(2)NeuAc(1) | N[+2206] | 1533.0546 | 3 | 4597.1493 | 4597.1422 | 0 | 1.5 | 457 | Specific | 927.8 | 863.88 | 863.88 | 18.15 | 57 | >sp\|P01833\|PIGR_HUMAN Polymeric immunoglobulin receptor OS=Homo sapiens OX=9606 GN=PIGR PE=1 SV=4 | 902 | 20201117_fqg_milk_1ng_glyco_hcd203040_compound15.19267.19267.3 | controllerType=0 controllerNumber=1 scan=19267 | 39.1412 |
| 15768:3 | 6 | K.IIEGEPNLKVPGN[+2205.793]VTAVLGETLK.V | HexNAc(4)Hex(5)Fuc(2)NeuAc(1) | N[+2206] | 1533.0514 | 3 | 4597.1398 | 4597.1422 | 0 | -0.5 | 457 | Specific | 945.85 | 872.83 | 872.83 | 17.24 | 57 | >sp\|P01833\|PIGR_HUMAN Polymeric immunoglobulin receptor OS=Homo sapiens OX=9606 GN=PIGR PE=1 SV=4 | 902 | 20201117_fqg_milk_1ng_glyco_hcd203040_compound15.19268.19268.3 | controllerType=0 controllerNumber=1 scan=19268 | 39.1456 |

Table S4. Cont.

| Query #:z | Protein Rank | Glycopeptide | Glycan | Modification Type(s) | Observed m/z | z | Observed (M+H) | Calc. mass (M+H) | Off-by-x error | Mass error (ppm) | StartPosition | Cleavage | Score | Delta | Delta Mod | \|Log Prob\| | # of unique peptides | Protein Name | Protein DB number | Comment | Scan # | Scan Time |
|---|---|---|---|---|---|---|---|---|---|---|---|---|---|---|---|---|---|---|---|---|---|---|
| 15778:3 | 6 | K.IIEGEPNLKVPGN[+2205.793]VTAVLGETLK.V | HexNAc(4)Hex(5)Fuc(2)NeuAc(1) | N[+2206] | 1533.0557 | 3 | 4597.1526 | 4597.1422 | 0 | 2.3 | 457 | Specific | 1003.27 | 928.13 | 928.13 | 20.23 | 57 | >sp\|P01833\|PIGR_HUMAN Polymeric immunoglobulin receptor OS=Homo sapiens OX=9606 GN=PIGR PE=1 SV=4 | 902 | 20201117_fqg_milk_1ng_glyco_hcd203040_compound15.19279.19279.3 | controllerType=0 controllerNumber=1 scan=19279 | 39.1668 |
| 15383:4 | 6 | K.IIEGEPNLKVPGN[+2059.735]VTAVLGETLKVPC[+57.021]HFPC[+57.021]K.F | HexNAc(4)Hex(5)Fuc(1)NeuAc(1) | C[+57]*2, N[+2060] | 1369.889 | 4 | 5476.5343 | 5476.5418 | 0 | -1.4 | 457 | Specific | 424.74 | 424.74 | 424.74 | 7.48 | 57 | >sp\|P01833\|PIGR_HUMAN Polymeric immunoglobulin receptor OS=Homo sapiens OX=9606 GN=PIGR PE=1 SV=4 | 902 | 20201117_fqg_milk_1ng_glyco_hcd203040_compound15.18838.18838.4 | controllerType=0 controllerNumber=1 scan=18838 | 38.3747 |
| 15379:5 | 6 | K.IIEGEPNLKVPGN[+2059.735]VTAVLGETLKVPC[+57.021]HFPC[+57.021]K.F | HexNAc(4)Hex(5)Fuc(1)NeuAc(1) | C[+57]*2, N[+2060] | 1096.1174 | 5 | 5476.5581 | 5476.5418 | 0 | 3 | 457 | Specific | 494.72 | 415.22 | 415.22 | 9.36 | 57 | >sp\|P01833\|PIGR_HUMAN Polymeric immunoglobulin receptor OS=Homo sapiens OX=9606 GN=PIGR PE=1 SV=4 | 902 | 20201117_fqg_milk_1ng_glyco_hcd203040_compound15.18834.18834.5 | controllerType=0 controllerNumber=1 scan=18834 | 38.3664 |
| 15324:4 | 6 | K.IIEGEPNLKVPGN[+2205.793]VTAVLGETLKVPC[+57.021]HFPC[+57.021]K.F | HexNAc(4)Hex(5)Fuc(2)NeuAc(1) | C[+57]*2, N[+2206] | 1406.4108 | 4 | 5622.6212 | 5622.5997 | 0 | 3.8 | 457 | Specific | 533.62 | 526.69 | 526.69 | 8.38 | 57 | >sp\|P01833\|PIGR_HUMAN Polymeric immunoglobulin receptor OS=Homo sapiens OX=9606 GN=PIGR PE=1 SV=4 | 902 | 20201117_fqg_milk_1ng_glyco_hcd203040_compound15.18777.18777.4 | controllerType=0 controllerNumber=1 scan=18777 | 38.2642 |
| 15345:5 | 6 | K.IIEGEPNLKVPGN[+2059.735]VTAVLGETLKVPC[+57.021]HFPC[+57.021]K.F | HexNAc(4)Hex(5)Fuc(1)NeuAc(1) | C[+57]*2, N[+2060] | 1096.1183 | 5 | 5476.5623 | 5476.5418 | 0 | 3.7 | 457 | Specific | 565.88 | 457.36 | 457.36 | 9.25 | 57 | >sp\|P01833\|PIGR_HUMAN Polymeric immunoglobulin receptor OS=Homo sapiens OX=9606 GN=PIGR PE=1 SV=4 | 902 | 20201117_fqg_milk_1ng_glyco_hcd203040_compound15.18799.18799.5 | controllerType=0 controllerNumber=1 scan=18799 | 38.3028 |
| 15380:4 | 6 | K.IIEGEPNLKVPGN[+2059.735]VTAVLGETLKVPC[+57.021]HFPC[+57.021]K.F | HexNAc(4)Hex(5)Fuc(1)NeuAc(1) | C[+57]*2, N[+2060] | 1369.889 | 4 | 5476.5343 | 5476.5418 | 0 | -1.4 | 457 | Specific | 603.62 | 603.62 | 603.62 | 10.34 | 57 | >sp\|P01833\|PIGR_HUMAN Polymeric immunoglobulin receptor OS=Homo sapiens OX=9606 GN=PIGR PE=1 SV=4 | 902 | 20201117_fqg_milk_1ng_glyco_hcd203040_compound15.18835.18835.4 | controllerType=0 controllerNumber=1 scan=18835 | 38.3676 |
| 15326:4 | 6 | K.IIEGEPNLKVPGN[+2205.793]VTAVLGETLKVPC[+57.021]HFPC[+57.021]K.F | HexNAc(4)Hex(5)Fuc(2)NeuAc(1) | C[+57]*2, N[+2206] | 1406.4108 | 4 | 5622.6212 | 5622.5997 | 0 | 3.8 | 457 | Specific | 616.91 | 616.91 | 616.91 | 10.93 | 57 | >sp\|P01833\|PIGR_HUMAN Polymeric immunoglobulin receptor OS=Homo sapiens OX=9606 GN=PIGR PE=1 SV=4 | 902 | 20201117_fqg_milk_1ng_glyco_hcd203040_compound15.18779.18779.4 | controllerType=0 controllerNumber=1 scan=18779 | 38.2686 |
| 15382:5 | 6 | K.IIEGEPNLKVPGN[+2059.735]VTAVLGETLKVPC[+57.021]HFPC[+57.021]K.F | HexNAc(4)Hex(5)Fuc(1)NeuAc(1) | C[+57]*2, N[+2060] | 1096.1174 | 5 | 5476.5581 | 5476.5418 | 0 | 3 | 457 | Specific | 628.97 | 573.23 | 573.23 | 11.91 | 57 | >sp\|P01833\|PIGR_HUMAN Polymeric immunoglobulin receptor OS=Homo sapiens OX=9606 GN=PIGR PE=1 SV=4 | 902 | 20201117_fqg_milk_1ng_glyco_hcd203040_compound15.18837.18837.5 | controllerType=0 controllerNumber=1 scan=18837 | 38.3728 |
| 15347:5 | 6 | K.IIEGEPNLKVPGN[+2059.735]VTAVLGETLKVPC[+57.021]HFPC[+57.021]K.F | HexNAc(4)Hex(5)Fuc(1)NeuAc(1) | C[+57]*2, N[+2060] | 1096.1183 | 5 | 5476.5623 | 5476.5418 | 0 | 3.7 | 457 | Specific | 715.62 | 613.03 | 613.03 | 13.51 | 57 | >sp\|P01833\|PIGR_HUMAN Polymeric immunoglobulin receptor OS=Homo sapiens OX=9606 GN=PIGR PE=1 SV=4 | 902 | 20201117_fqg_milk_1ng_glyco_hcd203040_compound15.18801.18801.5 | controllerType=0 controllerNumber=1 scan=18801 | 38.308 |
| 15315:5 | 6 | K.IIEGEPNLKVPGN[+2205.793]VTAVLGETLKVPC[+57.021]HFPC[+57.021]K.F | HexNAc(4)Hex(5)Fuc(2)NeuAc(1) | C[+57]*2, N[+2206] | 1125.3289 | 5 | 5622.6152 | 5622.5997 | 0 | 2.8 | 457 | Specific | 766.39 | 671.88 | 671.88 | 14.91 | 57 | >sp\|P01833\|PIGR_HUMAN Polymeric immunoglobulin receptor OS=Homo sapiens OX=9606 GN=PIGR PE=1 SV=4 | 902 | 20201117_fqg_milk_1ng_glyco_hcd203040_compound15.18768.18768.5 | controllerType=0 controllerNumber=1 scan=18768 | 38.2494 |
| 15313:5 | 6 | K.IIEGEPNLKVPGN[+2205.793]VTAVLGETLKVPC[+57.021]HFPC[+57.021]K.F | HexNAc(4)Hex(5)Fuc(2)NeuAc(1) | C[+57]*2, N[+2206] | 1125.3289 | 5 | 5622.6152 | 5622.5997 | 0 | 2.8 | 457 | Specific | 838.22 | 734.76 | 734.76 | 15.91 | 57 | >sp\|P01833\|PIGR_HUMAN Polymeric immunoglobulin receptor OS=Homo sapiens OX=9606 GN=PIGR PE=1 SV=4 | 902 | 20201117_fqg_milk_1ng_glyco_hcd203040_compound15.18766.18766.5 | controllerType=0 controllerNumber=1 scan=18766 | 38.2432 |
| 15290:5 | 6 | K.IIEGEPNLKVPGN[+2205.793]VTAVLGETLKVPC[+57.021]HFPC[+57.021]K.F | HexNAc(4)Hex(5)Fuc(2)NeuAc(1) | C[+57]*2, N[+2206] | 1125.3269 | 5 | 5622.6054 | 5622.5997 | 0 | 1 | 457 | Specific | 862.85 | 831.19 | 831.19 | 16.82 | 57 | >sp\|P01833\|PIGR_HUMAN Polymeric immunoglobulin receptor OS=Homo sapiens OX=9606 GN=PIGR PE=1 SV=4 | 902 | 20201117_fqg_milk_1ng_glyco_hcd203040_compound15.18742.18742.5 | controllerType=0 controllerNumber=1 scan=18742 | 38.1958 |
| 15288:5 | 6 | K.IIEGEPNLKVPGN[+2205.793]VTAVLGETLKVPC[+57.021]HFPC[+57.021]K.F | HexNAc(4)Hex(5)Fuc(2)NeuAc(1) | C[+57]*2, N[+2206] | 1125.3269 | 5 | 5622.6054 | 5622.5997 | 0 | 1 | 457 | Specific | 990.72 | 925.58 | 925.58 | 18.82 | 57 | >sp\|P01833\|PIGR_HUMAN Polymeric immunoglobulin receptor OS=Homo sapiens OX=9606 GN=PIGR PE=1 SV=4 | 902 | 20201117_fqg_milk_1ng_glyco_hcd203040_compound15.18740.18740.5 | controllerType=0 controllerNumber=1 scan=18740 | 38.19 |
| 16444:3 | 6 | K.VPGN[+2350.830]VTAVLGETLK.V | HexNAc(4)Hex(5)Fuc(1)NeuAc(2) | N[+2351] | 1250.2179 | 3 | 3748.6391 | 3748.6302 | 0 | 2.4 | 466 | Specific | 290.78 | 234.19 | 234.19 | 6.08 | 57 | >sp\|P01833\|PIGR_HUMAN Polymeric immunoglobulin receptor OS=Homo sapiens OX=9606 GN=PIGR PE=1 SV=4 | 902 | 20201117_fqg_milk_1ng_glyco_hcd203040_compound15.19978.19978.3 | controllerType=0 controllerNumber=1 scan=19978 | 40.4077 |
| 16446:3 | 6 | K.VPGN[+2350.830]VTAVLGETLK.V | HexNAc(4)Hex(5)Fuc(1)NeuAc(2) | N[+2351] | 1250.2179 | 3 | 3748.6391 | 3748.6302 | 0 | 2.4 | 466 | Specific | 316.56 | 237.07 | 237.07 | 6.39 | 57 | >sp\|P01833\|PIGR_HUMAN Polymeric immunoglobulin receptor OS=Homo sapiens OX=9606 GN=PIGR PE=1 SV=4 | 902 | 20201117_fqg_milk_1ng_glyco_hcd203040_compound15.19980.19980.3 | controllerType=0 controllerNumber=1 scan=19980 | 40.4137 |
| 13263:3 | 6 | K.VPGN[+2205.793]VTAVLGETLK.V | HexNAc(4)Hex(5)Fuc(2)NeuAc(1) | N[+2206] | 1201.8726 | 3 | 3603.6033 | 3603.5927 | 0 | 2.9 | 466 | Specific | 706.41 | 439.08 | 439.08 | 14.06 | 57 | >sp\|P01833\|PIGR_HUMAN Polymeric immunoglobulin receptor OS=Homo sapiens OX=9606 GN=PIGR PE=1 SV=4 | 902 | 20201117_fqg_milk_1ng_glyco_hcd203040_compound15.16640.16640.3 | controllerType=0 controllerNumber=1 scan=16640 | 34.4029 |
| 13266:3 | 6 | K.VPGN[+2205.793]VTAVLGETLK.V | HexNAc(4)Hex(5)Fuc(2)NeuAc(1) | N[+2206] | 1201.8726 | 3 | 3603.6033 | 3603.5927 | 0 | 2.9 | 466 | Specific | 720.1 | 443.8 | 443.8 | 14.06 | 57 | >sp\|P01833\|PIGR_HUMAN Polymeric immunoglobulin receptor OS=Homo sapiens OX=9606 GN=PIGR PE=1 SV=4 | 902 | 20201117_fqg_milk_1ng_glyco_hcd203040_compound15.16643.16643.3 | controllerType=0 controllerNumber=1 scan=16643 | 34.4102 |
| 13365:3 | 6 | K.VPGN[+2059.735]VTAVLGETLK.V | HexNAc(4)Hex(5)Fuc(1)NeuAc(1) | N[+2060] | 1153.1857 | 3 | 3457.5426 | 3457.5348 | 0 | 2.3 | 466 | Specific | 746.95 | 504.54 | 504.54 | 14.06 | 57 | >sp\|P01833\|PIGR_HUMAN Polymeric immunoglobulin receptor OS=Homo sapiens OX=9606 GN=PIGR PE=1 SV=4 | 902 | 20201117_fqg_milk_1ng_glyco_hcd203040_compound15.16746.16746.3 | controllerType=0 controllerNumber=1 scan=16746 | 34.614 |
| 13362:3 | 6 | K.VPGN[+2059.735]VTAVLGETLK.V | HexNAc(4)Hex(5)Fuc(1)NeuAc(1) | N[+2060] | 1153.1857 | 3 | 3457.5426 | 3457.5348 | 0 | 2.3 | 466 | Specific | 750.93 | 706.15 | 706.15 | 15.06 | 57 | >sp\|P01833\|PIGR_HUMAN Polymeric immunoglobulin receptor OS=Homo sapiens OX=9606 GN=PIGR PE=1 SV=4 | 902 | 20201117_fqg_milk_1ng_glyco_hcd203040_compound15.16743.16743.3 | controllerType=0 controllerNumber=1 scan=16743 | 34.6085 |
| 13397:4 | 6 | K.VPGN[+2205.793]VTAVLGETLKVPC[+57.021]HFPC[+57.021]K.F | HexNAc(4)Hex(5)Fuc(2)NeuAc(1) | C[+57]*2, N[+2206] | 1158.0212 | 4 | 4629.0628 | 4629.0502 | 0 | 2.7 | 466 | Specific | 450.47 | 450.47 | 450.47 | 9.45 | 57 | >sp\|P01833\|PIGR_HUMAN Polymeric immunoglobulin receptor OS=Homo sapiens OX=9606 GN=PIGR PE=1 SV=4 | 902 | 20201117_fqg_milk_1ng_glyco_hcd203040_compound15.16779.16779.4 | controllerType=0 controllerNumber=1 scan=16779 | 34.6705 |
| 13399:4 | 6 | K.VPGN[+2205.793]VTAVLGETLKVPC[+57.021]HFPC[+57.021]K.F | HexNAc(4)Hex(5)Fuc(2)NeuAc(1) | C[+57]*2, N[+2206] | 1158.0212 | 4 | 4629.0628 | 4629.0502 | 0 | 2.7 | 466 | Specific | 547.69 | 487.11 | 487.11 | 9.98 | 57 | >sp\|P01833\|PIGR_HUMAN Polymeric immunoglobulin receptor OS=Homo sapiens OX=9606 GN=PIGR PE=1 SV=4 | 902 | 20201117_fqg_milk_1ng_glyco_hcd203040_compound15.16781.16781.4 | controllerType=0 controllerNumber=1 scan=16781 | 34.6768 |
| 13411:4 | 6 | K.VPGN[+2059.735]VTAVLGETLKVPC[+57.021]HFPC[+57.021]K.F | HexNAc(4)Hex(5)Fuc(1)NeuAc(1) | C[+57]*2, N[+2060] | 1121.505 | 4 | 4482.9984 | 4482.9923 | 0 | 1.3 | 466 | Specific | 587.02 | 355.41 | 355.41 | 9.48 | 57 | >sp\|P01833\|PIGR_HUMAN Polymeric immunoglobulin receptor OS=Homo sapiens OX=9606 GN=PIGR PE=1 SV=4 | 902 | 20201117_fqg_milk_1ng_glyco_hcd203040_compound15.16793.16793.4 | controllerType=0 controllerNumber=1 scan=16793 | 34.6978 |
| 13413:4 | 6 | K.VPGN[+2059.735]VTAVLGETLKVPC[+57.021]HFPC[+57.021]K.F | HexNAc(4)Hex(5)Fuc(1)NeuAc(1) | C[+57]*2, N[+2060] | 1121.505 | 4 | 4482.9984 | 4482.9923 | 0 | 1.3 | 466 | Specific | 612.76 | 533.04 | 533.04 | 12.15 | 57 | >sp\|P01833\|PIGR_HUMAN Polymeric immunoglobulin receptor OS=Homo sapiens OX=9606 GN=PIGR PE=1 SV=4 | 902 | 20201117_fqg_milk_1ng_glyco_hcd203040_compound15.16795.16795.4 | controllerType=0 controllerNumber=1 scan=16795 | 34.7022 |
| 05346:3 | 6 | K.WN[+2059.735]NTGC[+57.021]QALPSQDEGPSK.A | HexNAc(4)Hex(5)Fuc(1)NeuAc(1) | C[+57], N[+2060] | 1350.2126 | 3 | 4048.6232 | 4048.6116 | 0 | 2.9 | 498 | Specific | 543.31 | 463.41 | 463.41 | 9.59 | 57 | >sp\|P01833\|PIGR_HUMAN Polymeric immunoglobulin receptor OS=Homo sapiens OX=9606 GN=PIGR PE=1 SV=4 | 902 | 20201117_fqg_milk_1ng_glyco_hcd203040_compound15.8395.8395.3 | controllerType=0 controllerNumber=1 scan=8395 | 20.5037 |
| 05348:3 | 6 | K.WN[+2059.735]NTGC[+57.021]QALPSQDEGPSK.A | HexNAc(4)Hex(5)Fuc(1)NeuAc(1) | C[+57], N[+2060] | 1350.2126 | 3 | 4048.6232 | 4048.6116 | 0 | 2.9 | 498 | Specific | 652.98 | 566.99 | 566.99 | 13.14 | 57 | >sp\|P01833\|PIGR_HUMAN Polymeric immunoglobulin receptor OS=Homo sapiens OX=9606 GN=PIGR PE=1 SV=4 | 902 | 20201117_fqg_milk_1ng_glyco_hcd203040_compound15.8397.8397.3 | controllerType=0 controllerNumber=1 scan=8397 | 20.5075 |
| 05443:3 | 6 | K.WN[+1913.677]NTGC[+57.021]QALPSQDEGPSK.A | HexNAc(4)Hex(5)NeuAc(1) | C[+57], N[+1914] | 1301.5254 | 3 | 3902.5618 | 3902.5537 | 0 | 2.1 | 498 | Specific | 716.99 | 627.85 | 627.85 | 14.05 | 57 | >sp\|P01833\|PIGR_HUMAN Polymeric immunoglobulin receptor OS=Homo sapiens OX=9606 GN=PIGR PE=1 SV=4 | 902 | 20201117_fqg_milk_1ng_glyco_hcd203040_compound15.8496.8496.3 | controllerType=0 controllerNumber=1 scan=8496 | 20.696 |
| 05382:3 | 6 | K.WN[+2059.735]NTGC[+57.021]QALPSQDEGPSK.A | HexNAc(4)Hex(5)Fuc(1)NeuAc(1) | C[+57], N[+2060] | 1350.2122 | 3 | 4048.6221 | 4048.6116 | 0 | 2.6 | 498 | Specific | 731.66 | 656.51 | 656.51 | 14.14 | 57 | >sp\|P01833\|PIGR_HUMAN Polymeric immunoglobulin receptor OS=Homo sapiens OX=9606 GN=PIGR PE=1 SV=4 | 902 | 20201117_fqg_milk_1ng_glyco_hcd203040_compound15.8433.8433.3 | controllerType=0 controllerNumber=1 scan=8433 | 20.5845 |
| 05441:3 | 6 | K.WN[+1913.677]NTGC[+57.021]QALPSQDEGPSK.A | HexNAc(4)Hex(5)NeuAc(1) | C[+57], N[+1914] | 1301.5254 | 3 | 3902.5618 | 3902.5537 | 0 | 2.1 | 498 | Specific | 739.79 | 650.36 | 650.36 | 14.05 | 57 | >sp\|P01833\|PIGR_HUMAN Polymeric immunoglobulin receptor OS=Homo sapiens OX=9606 GN=PIGR PE=1 SV=4 | 902 | 20201117_fqg_milk_1ng_glyco_hcd203040_compound15.8494.8494.3 | controllerType=0 controllerNumber=1 scan=8494 | 20.6898 |
| 05384:3 | 6 | K.WN[+2059.735]NTGC[+57.021]QALPSQDEGPSK.A | HexNAc(4)Hex(5)Fuc(1)NeuAc(1) | C[+57], N[+2060] | 1350.2122 | 3 | 4048.6221 | 4048.6116 | 0 | 2.6 | 498 | Specific | 784.95 | 693.1 | 693.1 | 15.14 | 57 | >sp\|P01833\|PIGR_HUMAN Polymeric immunoglobulin receptor OS=Homo sapiens OX=9606 GN=PIGR PE=1 SV=4 | 902 | 20201117_fqg_milk_1ng_glyco_hcd203040_compound15.8435.8435.3 | controllerType=0 controllerNumber=1 scan=8435 | 20.59 |
| 16950:3 | 7 | K.RNIAAFGGDPNN[+1913.677]ITLFGESAGGASVSLQTLSPYNK.G | HexNAc(4)Hex(5)NeuAc(1) | N[+1914] | 1827.4978 | 3 | 5480.4787 | 5480.4644 | 0 | 2.6 | 196 | Specific | 217.35 | 170.74 | 170.74 | 5.76 | 47 | >sp\|P19835\|CEL_HUMAN Bile salt-activated lipase OS=Homo sapiens OX=9606 GN=CEL PE=1 SV=3 | 476 | 20201117_fqg_milk_1ng_glyco_hcd203040_compound15.20506.20506.3 | controllerType=0 controllerNumber=1 scan=20506 | 41.507 |
| 16952:3 | 7 | K.RNIAAFGGDPNN[+1913.677]ITLFGESAGGASVSLQTLSPYNK.G | HexNAc(4)Hex(5)NeuAc(1) | N[+1914] | 1827.4978 | 3 | 5480.4787 | 5480.4644 | 0 | 2.6 | 196 | Specific | 221.9 | 118.05 | 118.05 | 4.94 | 47 | >sp\|P19835\|CEL_HUMAN Bile salt-activated lipase OS=Homo sapiens OX=9606 GN=CEL PE=1 SV=3 | 476 | 20201117_fqg_milk_1ng_glyco_hcd203040_compound15.20508.20508.3 | controllerType=0 controllerNumber=1 scan=20508 | 41.5108 |
| 16925:4 | 7 | K.RNIAAFGGDPNN[+1913.677]ITLFGESAGGASVSLQTLSPYNK.G | HexNAc(4)Hex(5)NeuAc(1) | N[+1914] | 1370.8754 | 4 | 5480.4796 | 5480.4644 | 0 | 2.8 | 196 | Specific | 905.95 | 783.78 | 783.78 | 19.18 | 47 | >sp\|P19835\|CEL_HUMAN Bile salt-activated lipase OS=Homo sapiens OX=9606 GN=CEL PE=1 SV=3 | 476 | 20201117_fqg_milk_1ng_glyco_hcd203040_compound15.20480.20480.4 | controllerType=0 controllerNumber=1 scan=20480 | 41.4489 |
| 16927:4 | 7 | K.RNIAAFGGDPNN[+1913.677]ITLFGESAGGASVSLQTLSPYNK.G | HexNAc(4)Hex(5)NeuAc(1) | N[+1914] | 1370.8754 | 4 | 5480.4796 | 5480.4644 | 0 | 2.8 | 196 | Specific | 1042.38 | 952.83 | 952.83 | 21.18 | 47 | >sp\|P19835\|CEL_HUMAN Bile salt-activated lipase OS=Homo sapiens OX=9606 GN=CEL PE=1 SV=3 | 476 | 20201117_fqg_milk_1ng_glyco_hcd203040_compound15.20482.20482.4 | controllerType=0 controllerNumber=1 scan=20482 | 41.4561 |
| 18693:3 | 7 | R.NIAAFGGDPNN[+1913.677]ITLFGESAGGASVSLQTLSPYNK.G | HexNAc(4)Hex(5)NeuAc(1) | N[+1914] | 1775.4714 | 3 | 5324.3996 | 5324.3633 | 0 | 6.8 | 197 | Specific | 277.63 | 277.63 | 277.63 | 6.41 | 47 | >sp\|P19835\|CEL_HUMAN Bile salt-activated lipase OS=Homo sapiens OX=9606 GN=CEL PE=1 SV=3 | 476 | 20201117_fqg_milk_1ng_glyco_hcd203040_compound15.22325.22325.3 | controllerType=0 controllerNumber=1 scan=22325 | 45.474 |
| 18696:3 | 7 | R.NIAAFGGDPNN[+1913.677]ITLFGESAGGASVSLQTLSPYNK.G | HexNAc(4)Hex(5)NeuAc(1) | N[+1914] | 1775.4714 | 3 | 5324.3996 | 5324.3633 | 0 | 6.8 | 197 | Specific | 704.32 | 686.12 | 686.12 | 13.92 | 47 | >sp\|P19835\|CEL_HUMAN Bile salt-activated lipase OS=Homo sapiens OX=9606 GN=CEL PE=1 SV=3 | 476 | 20201117_fqg_milk_1ng_glyco_hcd203040_compound15.22328.22328.3 | controllerType=0 controllerNumber=1 scan=22328 | 45.4791 |
| 17498:3 | 8 | R.LSLHRPALEDLLLGSEAN[+1460.529]LTC[+57.021]TLTGLR.D | HexNAc(4)Hex(4) | C[+57], N[+1461] | 1475.3859 | 3 | 4424.1432 | 4424.1271 | 0 | 3.7 | 127 | Specific | 381.11 | 381.11 | 381.11 | 8.25 | 47 | >sp\|P01876\|IGHA1_HUMAN Immunoglobulin heavy constant alpha 1 OS=Homo sapiens OX=9606 GN=IGHA1 PE=1 SV=2 | 255 | 20201117_fqg_milk_1ng_glyco_hcd203040_compound15.21079.21079.3 | controllerType=0 controllerNumber=1 scan=21079 | 42.7692 |
| 17500:3 | 8 | R.LSLHRPALEDLLLGSEAN[+1460.529]LTC[+57.021]TLTGLR.D | HexNAc(4)Hex(4) | C[+57], N[+1461] | 1475.3859 | 3 | 4424.1432 | 4424.1271 | 0 | 3.7 | 127 | Specific | 454.82 | 448.03 | 448.03 | 8.51 | 47 | >sp\|P01876\|IGHA1_HUMAN Immunoglobulin heavy constant alpha 1 OS=Homo sapiens OX=9606 GN=IGHA1 PE=1 SV=2 | 255 | 20201117_fqg_milk_1ng_glyco_hcd203040_compound15.21081.21081.3 | controllerType=0 controllerNumber=1 scan=21081 | 42.7752 |
| 19566:4 | 8 | R.LSLHRPALEDLLLGSEAN[+2204.772]LTC[+57.021]TLTGLR.D | HexNAc(4)Hex(5)NeuAc(2) | C[+57], N[+2205] | 1292.8508 | 4 | 5168.3815 | 5168.3707 | 0 | 2.1 | 127 | Specific | 456.52 | 356.54 | 356.54 | 7.75 | 47 | >sp\|P01876\|IGHA1_HUMAN Immunoglobulin heavy constant alpha 1 OS=Homo sapiens OX=9606 GN=IGHA1 PE=1 SV=2 | 255 | 20201117_fqg_milk_1ng_glyco_hcd203040_compound15.23238.23238.2 | controllerType=0 controllerNumber=1 scan=23238 | 47.2391 |
| 17399:4 | 8 | R.LSLHRPALEDLLLGSEAN[+1378.476]LTC[+57.021]TLTGLR.D | HexNAc(2)Hex(6) | C[+57], N[+1378] | 1086.2775 | 4 | 4342.0882 | 4342.074 | 0 | 3.3 | 127 | Specific | 459.4 | 459.4 | 459.4 | 9.5 | 47 | >sp\|P01876\|IGHA1_HUMAN Immunoglobulin heavy constant alpha 1 OS=Homo sapiens OX=9606 GN=IGHA1 PE=1 SV=2 | 255 | 20201117_fqg_milk_1ng_glyco_hcd203040_compound15.20975.20975.4 | controllerType=0 controllerNumber=1 scan=20975 | 42.5167 |
| 17458:4 | 8 | R.LSLHRPALEDLLLGSEAN[+1581.555]LTC[+57.021]TLTGLR.D | HexNAc(3)Hex(6) | C[+57], N[+1582] | 1137.0448 | 4 | 4545.1574 | 4545.1533 | 0 | 0.9 | 127 | Specific | 483.24 | 477.92 | 477.92 | 9.57 | 47 | >sp\|P01876\|IGHA1_HUMAN Immunoglobulin heavy constant alpha 1 OS=Homo sapiens OX=9606 GN=IGHA1 PE=1 SV=2 | 255 | 20201117_fqg_milk_1ng_glyco_hcd203040_compound15.21037.21037.4 | controllerType=0 controllerNumber=1 scan=21037 | 42.6724 |
| 17460:4 | 8 | R.LSLHRPALEDLLLGSEAN[+1581.555]LTC[+57.021]TLTGLR.D | HexNAc(3)Hex(6) | C[+57], N[+1582] | 1137.0448 | 4 | 4545.1574 | 4545.1533 | 0 | 0.9 | 127 | Specific | 493.82 | 364.44 | 364.44 | 6.73 | 47 | >sp\|P01876\|IGHA1_HUMAN Immunoglobulin heavy constant alpha 1 OS=Homo sapiens OX=9606 GN=IGHA1 PE=1 SV=2 | 255 | 20201117_fqg_milk_1ng_glyco_hcd203040_compound15.21039.21039.4 | controllerType=0 controllerNumber=1 scan=21039 | 42.6816 |
| 18385:3 | 8 | R.LSLHRPALEDLLLGSEAN[+1913.677]LTC[+57.021]TLTGLR.D | HexNAc(4)Hex(5)NeuAc(1) | C[+57], N[+1914] | 1626.4323 | 3 | 4877.2824 | 4877.2753 | 0 | 1.5 | 127 | Specific | 494.43 | 471.41 | 471.41 | 9.07 | 47 | >sp\|P01876\|IGHA1_HUMAN Immunoglobulin heavy constant alpha 1 OS=Homo sapiens OX=9606 GN=IGHA1 PE=1 SV=2 | 255 | 20201117_fqg_milk_1ng_glyco_hcd203040_compound15.22001.22001.3 | controllerType=0 controllerNumber=1 scan=22001 | 44.6898 |
| 17534:3 | 8 | R.LSLHRPALEDLLLGSEAN[+1054.370]LTC[+57.021]TLTGLR.D | HexNAc(2)Hex(4) | C[+57], N[+1054] | 1339.9971 | 3 | 4017.9768 | 4017.9683 | 0 | 2.1 | 127 | Specific | 569.69 | 527.9 | 527.9 | 9.45 | 47 | >sp\|P01876\|IGHA1_HUMAN Immunoglobulin heavy constant alpha 1 OS=Homo sapiens OX=9606 GN=IGHA1 PE=1 SV=2 | 255 | 20201117_fqg_milk_1ng_glyco_hcd203040_compound15.21116.21116.3 | controllerType=0 controllerNumber=1 scan=21116 | 42.846 |
| 17488:3 | 8 | R.LSLHRPALEDLLLGSEAN[+1257.449]LTC[+57.021]TLTGLR.D | HexNAc(3)Hex(4) | C[+57], N[+1257] | 1407.6921 | 3 | 4221.0617 | 4221.0477 | 0 | 3.3 | 127 | Specific | 570.7 | 546.22 | 546.22 | 9.45 | 47 | >sp\|P01876\|IGHA1_HUMAN Immunoglobulin heavy constant alpha 1 OS=Homo sapiens OX=9606 GN=IGHA1 PE=1 SV=2 | 255 | 20201117_fqg_milk_1ng_glyco_hcd203040_compound15.21068.21068.3 | controllerType=0 controllerNumber=1 scan=21068 | 42.7453 |
| 17436:3 | 8 | R.LSLHRPALEDLLLGSEAN[+1622.582]LTC[+57.021]TLTGLR.D | HexNAc(4)Hex(5) | C[+57], N[+1623] | 1529.4024 | 3 | 4586.1926 | 4586.1799 | 0 | 2.8 | 127 | Specific | 589.11 | 536.65 | 536.65 | 10.7 | 47 | >sp\|P01876\|IGHA1_HUMAN Immunoglobulin heavy constant alpha 1 OS=Homo sapiens OX=9606 GN=IGHA1 PE=1 SV=2 | 255 | 20201117_fqg_milk_1ng_glyco_hcd203040_compound15.21014.21014.3 | controllerType=0 controllerNumber=1 scan=21014 | 42.6121 |
| 17331:3 | 8 | R.LSLHRPALEDLLLGSEAN[+1864.634]LTC[+57.021]TLTGLR.D | HexNAc(2)Hex(9) | C[+57], N[+1865] | 1610.0862 | 3 | 4828.2442 | 4828.2324 | 0 | 2.4 | 127 | Specific | 594.08 | 545.24 | 545.24 | 10.7 | 47 | >sp\|P01876\|IGHA1_HUMAN Immunoglobulin heavy constant alpha 1 OS=Homo sapiens OX=9606 GN=IGHA1 PE=1 SV=2 | 255 | 20201117_fqg_milk_1ng_glyco_hcd203040_compound15.20904.20904.3 | controllerType=0 controllerNumber=1 scan=20904 | 42.3604 |
| 19568:4 | 8 | R.LSLHRPALEDLLLGSEAN[+2204.772]LTC[+57.021]TLTGLR.D | HexNAc(4)Hex(5)NeuAc(2) | C[+57], N[+2205] | 1292.8508 | 4 | 5168.3815 | 5168.3707 | 0 | 2.1 | 127 | Specific | 609.01 | 594.49 | 594.49 | 12.2 | 47 | >sp\|P01876\|IGHA1_HUMAN Immunoglobulin heavy constant alpha 1 OS=Homo sapiens OX=9606 GN=IGHA1 PE=1 SV=2 | 255 | 20201117_fqg_milk_1ng_glyco_hcd203040_compound15.23240.23240.2 | controllerType=0 controllerNumber=1 scan=23240 | 47.2447 |
| 17450:3 | 8 | R.LSLHRPALEDLLLGSEAN[+1501.555]LTC[+57.021]TLTGLR.D | HexNAc(5)Hex(3) | C[+57], N[+1502] | 1489.0596 | 3 | 4465.1643 | 4465.1536 | 0 | 2.4 | 127 | Specific | 617.01 | 555.82 | 555.82 | 11.7 | 47 | >sp\|P01876\|IGHA1_HUMAN Immunoglobulin heavy constant alpha 1 OS=Homo sapiens OX=9606 GN=IGHA1 PE=1 SV=2 | 255 | 20201117_fqg_milk_1ng_glyco_hcd203040_compound15.21029.21029.3 | controllerType=0 controllerNumber=1 scan=21029 | 42.6588 |
| 17490:3 | 8 | R.LSLHRPALEDLLLGSEAN[+1257.449]LTC[+57.021]TLTGLR.D | HexNAc(3)Hex(4) | C[+57], N[+1257] | 1407.6921 | 3 | 4221.0617 | 4221.0477 | 0 | 3.3 | 127 | Specific | 622.54 | 548.74 | 548.74 | 11.7 | 47 | >sp\|P01876\|IGHA1_HUMAN Immunoglobulin heavy constant alpha 1 OS=Homo sapiens OX=9606 GN=IGHA1 PE=1 SV=2 | 255 | 20201117_fqg_milk_1ng_glyco_hcd203040_compound15.21070.21070.3 | controllerType=0 controllerNumber=1 scan=21070 | 42.75 |
| 17439:3 | 8 | R.LSLHRPALEDLLLGSEAN[+1622.582]LTC[+57.021]TLTGLR.D | HexNAc(4)Hex(5) | C[+57], N[+1623] | 1529.4024 | 3 | 4586.1926 | 4586.1799 | 0 | 2.8 | 127 | Specific | 632.3 | 526.85 | 526.85 | 11.7 | 47 | >sp\|P01876\|IGHA1_HUMAN Immunoglobulin heavy constant alpha 1 OS=Homo sapiens OX=9606 GN=IGHA1 PE=1 SV=2 | 255 | 20201117_fqg_milk_1ng_glyco_hcd203040_compound15.21017.21017.3 | controllerType=0 controllerNumber=1 scan=21017 | 42.6219 |
| 17377:3 | 8 | R.LSLHRPALEDLLLGSEAN[+1540.529]LTC[+57.021]TLTGLR.D | HexNAc(2)Hex(7) | C[+57], N[+1541] | 1502.0499 | 3 | 4504.135 | 4504.1268 | 0 | 1.8 | 127 | Specific | 635.7 | 611.74 | 611.74 | 11.62 | 47 | >sp\|P01876\|IGHA1_HUMAN Immunoglobulin heavy constant alpha 1 OS=Homo sapiens OX=9606 GN=IGHA1 PE=1 SV=2 | 255 | 20201117_fqg_milk_1ng_glyco_hcd203040_compound15.20952.20952.3 | controllerType=0 controllerNumber=1 scan=20952 | 42.4742 |
| 17329:3 | 8 | R.LSLHRPALEDLLLGSEAN[+1864.634]LTC[+57.021]TLTGLR.D | HexNAc(2)Hex(9) | C[+57], N[+1865] | 1610.0862 | 3 | 4828.2442 | 4828.2324 | 0 | 2.4 | 127 | Specific | 637.19 | 628.52 | 628.52 | 11.7 | 47 | >sp\|P01876\|IGHA1_HUMAN Immunoglobulin heavy constant alpha 1 OS=Homo sapiens OX=9606 GN=IGHA1 PE=1 SV=2 | 255 | 20201117_fqg_milk_1ng_glyco_hcd203040_compound15.20902.20902.3 | controllerType=0 controllerNumber=1 scan=20902 | 42.3554 |
| 17448:3 | 8 | R.LSLHRPALEDLLLGSEAN[+1501.555]LTC[+57.021]TLTGLR.D | HexNAc(5)Hex(3) | C[+57], N[+1502] | 1489.0596 | 3 | 4465.1643 | 4465.1536 | 0 | 2.4 | 127 | Specific | 638.16 | 638.16 | 638.16 | 11.7 | 47 | >sp\|P01876\|IGHA1_HUMAN Immunoglobulin heavy constant alpha 1 OS=Homo sapiens OX=9606 GN=IGHA1 PE=1 SV=2 | 255 | 20201117_fqg_milk_1ng_glyco_hcd203040_compound15.21027.21027.3 | controllerType=0 controllerNumber=1 scan=21027 | 42.6498 |
| 18387:3 | 8 | R.LSLHRPALEDLLLGSEAN[+1913.677]LTC[+57.021]TLTGLR.D | HexNAc(4)Hex(5)NeuAc(1) | C[+57], N[+1914] | 1626.4323 | 3 | 4877.2824 | 4877.2753 | 0 | 1.5 | 127 | Specific | 640.61 | 568.52 | 568.52 | 11.62 | 47 | >sp\|P01876\|IGHA1_HUMAN Immunoglobulin heavy constant alpha 1 OS=Homo sapiens OX=9606 GN=IGHA1 PE=1 SV=2 | 255 | 20201117_fqg_milk_1ng_glyco_hcd203040_compound15.22003.22003.3 | controllerType=0 controllerNumber=1 scan=22003 | 44.6955 |
| 17536:3 | 8 | R.LSLHRPALEDLLLGSEAN[+1054.370]LTC[+57.021]TLTGLR.D | HexNAc(2)Hex(4) | C[+57], N[+1054] | 1339.9971 | 3 | 4017.9768 | 4017.9683 | 0 | 2.1 | 127 | Specific | 647.97 | 609.08 | 609.08 | 11.7 | 47 | >sp\|P01876\|IGHA1_HUMAN Immunoglobulin heavy constant alpha 1 OS=Homo sapiens OX=9606 GN=IGHA1 PE=1 SV=2 | 255 | 20201117_fqg_milk_1ng_glyco_hcd203040_compound15.21118.21118.3 | controllerType=0 controllerNumber=1 scan=21118 | 42.8531 |
| 17350:3 | 8 | R.LSLHRPALEDLLLGSEAN[+1702.581]LTC[+57.021]TLTGLR.D | HexNAc(2)Hex(8) | C[+57], N[+1703] | 1556.0678 | 3 | 4666.1889 | 4666.1796 | 0 | 2 | 127 | Specific | 654.15 | 654.15 | 654.15 | 12.62 | 47 | >sp\|P01876\|IGHA1_HUMAN Immunoglobulin heavy constant alpha 1 OS=Homo sapiens OX=9606 GN=IGHA1 PE=1 SV=2 | 255 | 20201117_fqg_milk_1ng_glyco_hcd203040_compound15.20924.20924.3 | controllerType=0 controllerNumber=1 scan=20924 | 42.4082 |
| 17413:3 | 8 | R.LSLHRPALEDLLLGSEAN[+1378.476]LTC[+57.021]TLTGLR.D | HexNAc(2)Hex(6) | C[+57], N[+1378] | 1448.0343 | 3 | 4342.0883 | 4342.074 | 0 | 3.3 | 127 | Specific | 654.55 | 623.18 | 623.18 | 12.7 | 47 | >sp\|P01876\|IGHA1_HUMAN Immunoglobulin heavy constant alpha 1 OS=Homo sapiens OX=9606 GN=IGHA1 PE=1 SV=2 | 255 | 20201117_fqg_milk_1ng_glyco_hcd203040_compound15.20989.20989.3 | controllerType=0 controllerNumber=1 scan=20989 | 42.5504 |
| 17425:4 | 8 | R.LSLHRPALEDLLLGSEAN[+1622.582]LTC[+57.021]TLTGLR.D | HexNAc(4)Hex(5) | C[+57], N[+1623] | 1147.3036 | 4 | 4586.1927 | 4586.1799 | 0 | 2.8 | 127 | Specific | 661.11 | 661.11 | 661.11 | 13.28 | 47 | >sp\|P01876\|IGHA1_HUMAN Immunoglobulin heavy constant alpha 1 OS=Homo sapiens OX=9606 GN=IGHA1 PE=1 SV=2 | 255 | 20201117_fqg_milk_1ng_glyco_hcd203040_compound15.21003.21003.4 | controllerType=0 controllerNumber=1 scan=21003 | 42.5866 |
| 17528:3 | 8 | R.LSLHRPALEDLLLGSEAN[+1095.397]LTC[+57.021]TLTGLR.D | HexNAc(3)Hex(3) | C[+57], N[+1095] | 1353.6728 | 3 | 4059.0038 | 4058.9949 | 0 | 2.2 | 127 | Specific | 661.46 | 661.46 | 661.46 | 12.7 | 47 | >sp\|P01876\|IGHA1_HUMAN Immunoglobulin heavy constant alpha 1 OS=Homo sapiens OX=9606 GN=IGHA1 PE=1 SV=2 | 255 | 20201117_fqg_milk_1ng_glyco_hcd203040_compound15.21110.21110.3 | controllerType=0 controllerNumber=1 scan=21110 | 42.8347 |
| 17447:4 | 8 | R.LSLHRPALEDLLLGSEAN[+1501.555]LTC[+57.021]TLTGLR.D | HexNAc(5)Hex(3) | C[+57], N[+1502] | 1117.0485 | 4 | 4465.1722 | 4465.1536 | 0 | 4.2 | 127 | Specific | 679.58 | 679.58 | 679.58 | 12.88 | 47 | >sp\|P01876\|IGHA1_HUMAN Immunoglobulin heavy constant alpha 1 OS=Homo sapiens OX=9606 GN=IGHA1 PE=1 SV=2 | 255 | 20201117_fqg_milk_1ng_glyco_hcd203040_compound15.21026.21026.4 | controllerType=0 controllerNumber=1 scan=21026 | 42.6483 |
| 17531:3 | 8 | R.LSLHRPALEDLLLGSEAN[+1095.397]LTC[+57.021]TLTGLR.D | HexNAc(3)Hex(3) | C[+57], N[+1095] | 1353.6728 | 3 | 4059.0038 | 4058.9949 | 0 | 2.2 | 127 | Specific | 687.28 | 653 | 653 | 12.7 | 47 | >sp\|P01876\|IGHA1_HUMAN Immunoglobulin heavy constant alpha 1 OS=Homo sapiens OX=9606 GN=IGHA1 PE=1 SV=2 | 255 | 20201117_fqg_milk_1ng_glyco_hcd203040_compound15.21113.21113.3 | controllerType=0 controllerNumber=1 scan=21113 | 42.841 |
| 17440:3 | 8 | R.LSLHRPALEDLLLGSEAN[+1216.423]LTC[+57.021]TLTGLR.D | HexNAc(2)Hex(5) | C[+57], N[+1216] | 1394.0152 | 3 | 4180.031 | 4180.0211 | 0 | 2.4 | 127 | Specific | 694.12 | 609.67 | 609.67 | 12.7 | 47 | >sp\|P01876\|IGHA1_HUMAN Immunoglobulin heavy constant alpha 1 OS=Homo sapiens OX=9606 GN=IGHA1 PE=1 SV=2 | 255 | 20201117_fqg_milk_1ng_glyco_hcd203040_compound15.21018.21018.3 | controllerType=0 controllerNumber=1 scan=21018 | 42.6238 |
| 17430:4 | 8 | R.LSLHRPALEDLLLGSEAN[+1460.529]LTC[+57.021]TLTGLR.D | HexNAc(4)Hex(4) | C[+57], N[+1461] | 1106.7897 | 4 | 4424.137 | 4424.1271 | 0 | 2.3 | 127 | Specific | 701.33 | 674.51 | 674.51 | 14.28 | 47 | >sp\|P01876\|IGHA1_HUMAN Immunoglobulin heavy constant alpha 1 OS=Homo sapiens OX=9606 GN=IGHA1 PE=1 SV=2 | 255 | 20201117_fqg_milk_1ng_glyco_hcd203040_compound15.21008.21008.4 | controllerType=0 controllerNumber=1 scan=21008 | 42.5955 |
| 17426:4 | 8 | R.LSLHRPALEDLLLGSEAN[+1419.502]LTC[+57.021]TLTGLR.D | HexNAc(3)Hex(5) | C[+57], N[+1420] | 1096.5335 | 4 | 4383.1121 | 4383.1005 | 0 | 2.7 | 127 | Specific | 701.8 | 701.8 | 701.8 | 14.28 | 47 | >sp\|P01876\|IGHA1_HUMAN Immunoglobulin heavy constant alpha 1 OS=Homo sapiens OX=9606 GN=IGHA1 PE=1 SV=2 | 255 | 20201117_fqg_milk_1ng_glyco_hcd203040_compound15.21004.21004.4 | controllerType=0 controllerNumber=1 scan=21004 | [illegible] |
| 17348:3 | 8 | R.LSLHRPALEDLLLGSEAN[+1702.581]LTC[+57.021]TLTGLR.D | HexNAc(2)Hex(8) | C[+57], N[+1703] | 1556.0678 | 3 | 4666.1889 | 4666.1796 | 0 | 2 | 127 | Specific | 702.94 | 702.16 | 702.16 | 13.62 | 47 | >sp\|P01876\|IGHA1_HUMAN Immunoglobulin heavy constant alpha 1 OS=Homo sapiens OX=9606 GN=IGHA1 PE=1 SV=2 | 255 | 20201117_fqg_milk_1ng_glyco_hcd203040_compound15.20922.20922.3 | controllerType=0 controllerNumber=1 scan=20922 | [illegible] |
| 17375:3 | 8 | R.LSLHRPALEDLLLGSEAN[+1540.529]LTC[+57.021]TLTGLR.D | HexNAc(2)Hex(7) | C[+57], N[+1541] | 1502.0499 | 3 | 4504.135 | 4504.1268 | 0 | 1.8 | 127 | Specific | 713.19 | 637.08 | 637.08 | 13.62 | 47 | >sp\|P01876\|IGHA1_HUMAN Immunoglobulin heavy constant alpha 1 OS=Homo sapiens OX=9606 GN=IGHA1 PE=1 SV=2 | 255 | 20201117_fqg_milk_1ng_glyco_hcd203040_compound15.20950.20950.3 | controllerType=0 controllerNumber=1 scan=20950 | [illegible] |
| 17473:5 | 8 | R.LSLHRPALEDLLLGSEAN[+1501.555]LTC[+57.021]TLTGLR.D | HexNAc(5)Hex(3) | C[+57], N[+1502] | 893.8384 | 5 | 4465.1629 | 4465.1536 | 0 | 2.1 | 127 | Specific | 723.74 | 723.74 | 723.74 | 14.01 | 47 | >sp\|P01876\|IGHA1_HUMAN Immunoglobulin heavy constant alpha 1 OS=Homo sapiens OX=9606 GN=IGHA1 PE=1 SV=2 | 255 | 20201117_fqg_milk_1ng_glyco_hcd203040_compound15.21053.21053.5 | controllerType=0 controllerNumber=1 scan=21053 | [illegible] |
| 17411:3 | 8 | R.LSLHRPALEDLLLGSEAN[+1378.476]LTC[+57.021]TLTGLR.D | HexNAc(2)Hex(6) | C[+57], N[+1378] | 1448.0343 | 3 | 4342.0883 | 4342.074 | 0 | 3.3 | 127 | Specific | 729.74 | 657.31 | 657.31 | 13.7 | 47 | >sp\|P01876\|IGHA1_HUMAN Immunoglobulin heavy constant alpha 1 OS=Homo sapiens OX=9606 GN=IGHA1 PE=1 SV=2 | 255 | 20201117_fqg_milk_1ng_glyco_hcd203040_compound15.20987.20987.3 | controllerType=0 controllerNumber=1 scan=20987 | 42.539 |
| 17356:5 | 8 | R.LSLHRPALEDLLLGSEAN[+1864.634]LTC[+57.021]TLTGLR.D | HexNAc(2)Hex(9) | C[+57], N[+1865] | 966.4543 | 5 | 4828.2424 | 4828.2324 | 0 | 2.1 | 127 | Specific | 734.1 | 734.1 | 734.1 | 14.01 | 47 | >sp\|P01876\|IGHA1_HUMAN Immunoglobulin heavy constant alpha 1 OS=Homo sapiens OX=9606 GN=IGHA1 PE=1 SV=2 | 255 | 20201117_fqg_milk_1ng_glyco_hcd203040_compound15.20930.20930.5 | controllerType=0 controllerNumber=1 scan=20930 | 42.419 |
| 17474:4 | 8 | R.LSLHRPALEDLLLGSEAN[+1663.608]LTC[+57.021]TLTGLR.D | HexNAc(5)Hex(4) | C[+57], N[+1664] | 1157.5601 | 4 | 4627.2186 | 4627.2064 | 0 | 2.6 | 127 | Specific | 746.43 | 659.34 | 659.34 | 14.28 | 47 | >sp\|P01876\|IGHA1_HUMAN Immunoglobulin heavy constant alpha 1 OS=Homo sapiens OX=9606 GN=IGHA1 PE=1 SV=2 | 255 | 20201117_fqg_milk_1ng_glyco_hcd203040_compound15.21054.21054.4 | controllerType=0 controllerNumber=1 scan=21054 | 42.7127 |
| 17476:5 | 8 | R.LSLHRPALEDLLLGSEAN[+1501.555]LTC[+57.021]TLTGLR.D | HexNAc(5)Hex(3) | C[+57], N[+1502] | 893.8384 | 5 | 4465.1629 | 4465.1536 | 0 | 2.1 | 127 | Specific | 750.42 | 738.42 | 738.42 | 15.01 | 47 | >sp\|P01876\|IGHA1_HUMAN Immunoglobulin heavy constant alpha 1 OS=Homo sapiens OX=9606 GN=IGHA1 PE=1 SV=2 | 255 | 20201117_fqg_milk_1ng_glyco_hcd203040_compound15.21056.21056.5 | controllerType=0 controllerNumber=1 scan=21056 | 42.7185 |
| 17437:3 | 8 | R.LSLHRPALEDLLLGSEAN[+1216.423]LTC[+57.021]TLTGLR.D | HexNAc(2)Hex(5) | C[+57], N[+1216] | 1394.0152 | 3 | 4180.031 | 4180.0211 | 0 | 2.4 | 127 | Specific | 756.65 | 695.81 | 695.81 | 14.7 | 47 | >sp\|P01876\|IGHA1_HUMAN Immunoglobulin heavy constant alpha 1 OS=Homo sapiens OX=9606 GN=IGHA1 PE=1 SV=2 | 255 | 20201117_fqg_milk_1ng_glyco_hcd203040_compound15.21015.21015.3 | controllerType=0 controllerNumber=1 scan=21015 | 42.6133 |
| 17468:4 | 8 | R.LSLHRPALEDLLLGSEAN[+1298.476]LTC[+57.021]TLTGLR.D | HexNAc(4)Hex(3) | C[+57], N[+1298] | 1066.2764 | 4 | 4262.0838 | 4262.0742 | 0 | 2.2 | 127 | Specific | 771.53 | 771.53 | 771.53 | 15.28 | 47 | >sp\|P01876\|IGHA1_HUMAN Immunoglobulin heavy constant alpha 1 OS=Homo sapiens OX=9606 GN=IGHA1 PE=1 SV=2 | 255 | 20201117_fqg_milk_1ng_glyco_hcd203040_compound15.21048.21048.4 | controllerType=0 controllerNumber=1 scan=21048 | 42.7018 |
| 17518:4 | 8 | R.LSLHRPALEDLLLGSEAN[+1095.397]LTC[+57.021]TLTGLR.D | HexNAc(3)Hex(3) | C[+57], N[+1095] | 1015.5065 | 4 | 4059.0042 | 4058.9949 | 0 | 2.3 | 127 | Specific | 777.52 | 777.52 | 777.52 | 15.28 | 47 | >sp\|P01876\|IGHA1_HUMAN Immunoglobulin heavy constant alpha 1 OS=Homo sapiens OX=9606 GN=IGHA1 PE=1 SV=2 | 255 | 20201117_fqg_milk_1ng_glyco_hcd203040_compound15.21100.21100.4 | controllerType=0 controllerNumber=1 scan=21100 | 42.8133 |
| 17449:4 | 8 | R.LSLHRPALEDLLLGSEAN[+1501.555]LTC[+57.021]TLTGLR.D | HexNAc(5)Hex(3) | C[+57], N[+1502] | 1117.0485 | 4 | 4465.1722 | 4465.1536 | 0 | 4.2 | 127 | Specific | 778.6 | 774.64 | 774.64 | 14.88 | 47 | >sp\|P01876\|IGHA1_HUMAN Immunoglobulin heavy constant alpha 1 OS=Homo sapiens OX=9606 GN=IGHA1 PE=1 SV=2 | 255 | 20201117_fqg_milk_1ng_glyco_hcd203040_compound15.21028.21028.4 | controllerType=0 controllerNumber=1 scan=21028 | 42.6548 |
| 17527:4 | 8 | R.LSLHRPALEDLLLGSEAN[+1054.370]LTC[+57.021]TLTGLR.D | HexNAc(2)Hex(4) | C[+57], N[+1054] | 1005.2497 | 4 | 4017.9769 | 4017.9683 | 0 | 2.1 | 127 | Specific | 780.23 | 780.23 | 780.23 | 15.28 | 47 | >sp\|P01876\|IGHA1_HUMAN Immunoglobulin heavy constant alpha 1 OS=Homo sapiens OX=9606 GN=IGHA1 PE=1 SV=2 | 255 | 20201117_fqg_milk_1ng_glyco_hcd203040_compound15.21109.21109.4 | controllerType=0 controllerNumber=1 scan=21109 | 42.8336 |
| 17427:4 | 8 | R.LSLHRPALEDLLLGSEAN[+1622.582]LTC[+57.021]TLTGLR.D | HexNAc(4)Hex(5) | C[+57], N[+1623] | 1147.3036 | 4 | 4586.1927 | 4586.1799 | 0 | 2.8 | 127 | Specific | 783.79 | 715.45 | 715.45 | 15.28 | 47 | >sp\|P01876\|IGHA1_HUMAN Immunoglobulin heavy constant alpha 1 OS=Homo sapiens OX=9606 GN=IGHA1 PE=1 SV=2 | 255 | 20201117_fqg_milk_1ng_glyco_hcd203040_compound15.21005.21005.4 | controllerType=0 controllerNumber=1 scan=21005 | 42.5915 |
| 17358:5 | 8 | R.LSLHRPALEDLLLGSEAN[+1864.634]LTC[+57.021]TLTGLR.D | HexNAc(2)Hex(9) | C[+57], N[+1865] | 966.4543 | 5 | 4828.2424 | 4828.2324 | 0 | 2.1 | 127 | Specific | 827.19 | 827.19 | 827.19 | 16.01 | 47 | >sp\|P01876\|IGHA1_HUMAN Immunoglobulin heavy constant alpha 1 OS=Homo sapiens OX=9606 GN=IGHA1 PE=1 SV=2 | 255 | 20201117_fqg_milk_1ng_glyco_hcd203040_compound15.20932.20932.5 | controllerType=0 controllerNumber=1 scan=20932 | 42.4247 |
| 17470:4 | 8 | R.LSLHRPALEDLLLGSEAN[+1298.476]LTC[+57.021]TLTGLR.D | HexNAc(4)Hex(3) | C[+57], N[+1298] | 1066.2764 | 4 | 4262.0838 | 4262.0742 | 0 | 2.2 | 127 | Specific | 846.96 | 832.53 | 832.53 | 16.28 | 47 | >sp\|P01876\|IGHA1_HUMAN Immunoglobulin heavy constant alpha 1 OS=Homo sapiens OX=9606 GN=IGHA1 PE=1 SV=2 | 255 | 20201117_fqg_milk_1ng_glyco_hcd203040_compound15.21050.21050.4 | controllerType=0 controllerNumber=1 scan=21050 | 42.7079 |
| 17433:4 | 8 | R.LSLHRPALEDLLLGSEAN[+1460.529]LTC[+57.021]TLTGLR.D | HexNAc(4)Hex(4) | C[+57], N[+1461] | 1106.7897 | 4 | 4424.137 | 4424.1271 | 0 | 2.3 | 127 | Specific | 855.45 | 752.71 | 752.71 | 17.28 | 47 | >sp\|P01876\|IGHA1_HUMAN Immunoglobulin heavy constant alpha 1 OS=Homo sapiens OX=9606 GN=IGHA1 PE=1 SV=2 | 255 | 20201117_fqg_milk_1ng_glyco_hcd203040_compound15.21011.21011.4 | controllerType=0 controllerNumber=1 scan=21011 | 42.6083 |
| 17371:4 | 8 | R.LSLHRPALEDLLLGSEAN[+1540.529]LTC[+57.021]TLTGLR.D | HexNAc(2)Hex(7) | C[+57], N[+1541] | 1126.7892 | 4 | 4504.1351 | 4504.1268 | 0 | 1.8 | 127 | Specific | 857.54 | 813.49 | 813.49 | 17.2 | 47 | >sp\|P01876\|IGHA1_HUMAN Immunoglobulin heavy constant alpha 1 OS=Homo sapiens OX=9606 GN=IGHA1 PE=1 SV=2 | 255 | 20201117_fqg_milk_1ng_glyco_hcd203040_compound15.20946.20946.4 | controllerType=0 controllerNumber=1 scan=20946 | 42.4553 |
| 17428:4 | 8 | R.LSLHRPALEDLLLGSEAN[+1419.502]LTC[+57.021]TLTGLR.D | HexNAc(3)Hex(5) | C[+57], N[+1420] | 1096.5335 | 4 | 4383.1121 | 4383.1005 | 0 | 2.7 | 127 | Specific | 875.9 | 753.71 | 753.71 | 17.28 | 47 | >sp\|P01876\|IGHA1_HUMAN Immunoglobulin heavy constant alpha 1 OS=Homo sapiens OX=9606 GN=IGHA1 PE=1 SV=2 | 255 | 20201117_fqg_milk_1ng_glyco_hcd203040_compound15.21006.21006.4 | controllerType=0 controllerNumber=1 scan=21006 | 42.5933 |
| 17347:4 | 8 | R.LSLHRPALEDLLLGSEAN[+1702.581]LTC[+57.021]TLTGLR.D | HexNAc(2)Hex(8) | C[+57], N[+1703] | 1167.3025 | 4 | 4666.1883 | 4666.1796 | 0 | 1.9 | 127 | Specific | 885.17 | 885.17 | 885.17 | 17.2 | 47 | >sp\|P01876\|IGHA1_HUMAN Immunoglobulin heavy constant alpha 1 OS=Homo sapiens OX=9606 GN=IGHA1 PE=1 SV=2 | 255 | 20201117_fqg_milk_1ng_glyco_hcd203040_compound15.20921.20921.4 | controllerType=0 controllerNumber=1 scan=20921 | 42.3993 |
| 17520:4 | 8 | R.LSLHRPALEDLLLGSEAN[+1095.397]LTC[+57.021]TLTGLR.D | HexNAc(3)Hex(3) | C[+57], N[+1095] | 1015.5065 | 4 | 4059.0042 | 4058.9949 | 0 | 2.3 | 127 | Specific | 896.72 | 896.72 | 896.72 | 17.28 | 47 | >sp\|P01876\|IGHA1_HUMAN Immunoglobulin heavy constant alpha 1 OS=Homo sapiens OX=9606 GN=IGHA1 PE=1 SV=2 | 255 | 20201117_fqg_milk_1ng_glyco_hcd203040_compound15.21102.21102.4 | controllerType=0 controllerNumber=1 scan=21102 | 42.8194 |
| 17457:4 | 8 | R.LSLHRPALEDLLLGSEAN[+1257.449]LTC[+57.021]TLTGLR.D | HexNAc(3)Hex(4) | C[+57], N[+1257] | 1056.0185 | 4 | 4221.0521 | 4221.0477 | 0 | 1 | 127 | Specific | 924.52 | 861.25 | 861.25 | 18.2 | 47 | >sp\|P01876\|IGHA1_HUMAN Immunoglobulin heavy constant alpha 1 OS=Homo sapiens OX=9606 GN=IGHA1 PE=1 SV=2 | 255 | 20201117_fqg_milk_1ng_glyco_hcd203040_compound15.21036.21036.4 | controllerType=0 controllerNumber=1 scan=21036 | 42.671 |
| 17431:4 | 8 | R.LSLHRPALEDLLLGSEAN[+1216.423]LTC[+57.021]TLTGLR.D | HexNAc(2)Hex(5) | C[+57], N[+1216] | 1045.7632 | 4 | 4180.0311 | 4180.0211 | 0 | 2.4 | 127 | Specific | 943.49 | 902.87 | 902.87 | 18.28 | 47 | >sp\|P01876\|IGHA1_HUMAN Immunoglobulin heavy constant alpha 1 OS=Homo sapiens OX=9606 GN=IGHA1 PE=1 SV=2 | 255 | 20201117_fqg_milk_1ng_glyco_hcd203040_compound15.21009.21009.4 | controllerType=0 controllerNumber=1 scan=21009 | 42.5967 |
| 17326:4 | 8 | R.LSLHRPALEDLLLGSEAN[+1864.634]LTC[+57.021]TLTGLR.D | HexNAc(2)Hex(9) | C[+57], N[+1865] | 1207.8162 | 4 | 4828.243 | 4828.2324 | 0 | 2.2 | 127 | Specific | 944.62 | 944.62 | 944.62 | 18.28 | 47 | >sp\|P01876\|IGHA1_HUMAN Immunoglobulin heavy constant alpha 1 OS=Homo sapiens OX=9606 GN=IGHA1 PE=1 SV=2 | 255 | 20201117_fqg_milk_1ng_glyco_hcd203040_compound15.20899.20899.4 | controllerType=0 controllerNumber=1 scan=20899 | 42.3489 |
| 17328:4 | 8 | R.LSLHRPALEDLLLGSEAN[+1864.634]LTC[+57.021]TLTGLR.D | HexNAc(2)Hex(9) | C[+57], N[+1865] | 1207.8162 | 4 | 4828.243 | 4828.2324 | 0 | 2.2 | 127 | Specific | 968.18 | 922.14 | 922.14 | 19.28 | 47 | >sp\|P01876\|IGHA1_HUMAN Immunoglobulin heavy constant alpha 1 OS=Homo sapiens OX=9606 GN=IGHA1 PE=1 SV=2 | 255 | 20201117_fqg_milk_1ng_glyco_hcd203040_compound15.20901.20901.4 | controllerType=0 controllerNumber=1 scan=20901 | 42.3546 |
| 17530:4 | 8 | R.LSLHRPALEDLLLGSEAN[+1054.370]LTC[+57.021]TLTGLR.D | HexNAc(2)Hex(4) | C[+57], N[+1054] | 1005.2497 | 4 | 4017.9769 | 4017.9683 | 0 | 2.1 | 127 | Specific | 981.74 | 981.74 | 981.74 | 19.28 | 47 | >sp\|P01876\|IGHA1_HUMAN Immunoglobulin heavy constant alpha 1 OS=Homo sapiens OX=9606 GN=IGHA1 PE=1 SV=2 | 255 | 20201117_fqg_milk_1ng_glyco_hcd203040_compound15.21112.21112.4 | controllerType=0 controllerNumber=1 scan=21112 | 42.8385 |
| 17434:4 | 8 | R.LSLHRPALEDLLLGSEAN[+1216.423]LTC[+57.021]TLTGLR.D | HexNAc(2)Hex(5) | C[+57], N[+1216] | 1045.7632 | 4 | 4180.0311 | 4180.0211 | 0 | 2.4 | 127 | Specific | 988.88 | 965.92 | 965.92 | 19.28 | 47 | >sp\|P01876\|IGHA1_HUMAN Immunoglobulin heavy constant alpha 1 OS=Homo sapiens OX=9606 GN=IGHA1 PE=1 SV=2 | 255 | 20201117_fqg_milk_1ng_glyco_hcd203040_compound15.21012.21012.4 | controllerType=0 controllerNumber=1 scan=21012 | 42.6096 |
| 17349:4 | 8 | R.LSLHRPALEDLLLGSEAN[+1702.581]LTC[+57.021]TLTGLR.D | HexNAc(2)Hex(8) | C[+57], N[+1703] | 1167.3025 | 4 | 4666.1883 | 4666.1796 | 0 | 1.9 | 127 | Specific | 1027.56 | 950.71 | 950.71 | 20.2 | 47 | >sp\|P01876\|IGHA1_HUMAN Immunoglobulin heavy constant alpha 1 OS=Homo sapiens OX=9606 GN=IGHA1 PE=1 SV=2 | 255 | 20201117_fqg_milk_1ng_glyco_hcd203040_compound15.20923.20923.4 | controllerType=0 controllerNumber=1 scan=20923 | 42.4052 |
| 17402:4 | 8 | R.LSLHRPALEDLLLGSEAN[+1378.476]LTC[+57.021]TLTGLR.D | HexNAc(2)Hex(6) | C[+57], N[+1378] | 1086.2775 | 4 | 4342.0882 | 4342.074 | 0 | 3.3 | 127 | Specific | 1036.45 | 1017.6 | 1017.6 | 20.28 | 47 | >sp\|P01876\|IGHA1_HUMAN Immunoglobulin heavy constant alpha 1 OS=Homo sapiens OX=9606 GN=IGHA1 PE=1 SV=2 | 255 | 20201117_fqg_milk_1ng_glyco_hcd203040_compound15.20978.20978.4 | controllerType=0 controllerNumber=1 scan=20978 | 42.5212 |
| 17373:4 | 8 | R.LSLHRPALEDLLLGSEAN[+1540.529]LTC[+57.021]TLTGLR.D | HexNAc(2)Hex(7) | C[+57], N[+1541] | 1126.7892 | 4 | 4504.1351 | 4504.1268 | 0 | 1.8 | 127 | Specific | 1041.55 | 977.74 | 977.74 | 20.2 | 47 | >sp\|P01876\|IGHA1_HUMAN Immunoglobulin heavy constant alpha 1 OS=Homo sapiens OX=9606 GN=IGHA1 PE=1 SV=2 | 255 | 20201117_fqg_milk_1ng_glyco_hcd203040_compound15.20948.20948.4 | controllerType=0 controllerNumber=1 scan=20948 | 42.4641 |
| 17459:4 | 8 | R.LSLHRPALEDLLLGSEAN[+1257.449]LTC[+57.021]TLTGLR.D | HexNAc(3)Hex(4) | C[+57], N[+1257] | 1056.0185 | 4 | 4221.0521 | 4221.0477 | 0 | 1 | 127 | Specific | 1081.59 | 1081.59 | 1081.59 | 21.2 | 47 | >sp\|P01876\|IGHA1_HUMAN Immunoglobulin heavy constant alpha 1 OS=Homo sapiens OX=9606 GN=IGHA1 PE=1 SV=2 | 255 | 20201117_fqg_milk_1ng_glyco_hcd203040_compound15.21038.21038.4 | controllerType=0 controllerNumber=1 scan=21038 | 42.6799 |
| 12960:3 | 8 | R.LAGKPTHVN[+2263.835]VSVVMAEVDGTC[+57.021].Y | HexNAc(5)Hex(5)Fuc(3) | C[+57], N[+2264] | 1483.3106 | 3 | 4447.9173 | 4447.9135 | 0 | 0.9 | 332 | CRagged | 225.05 | 225.05 | 225.05 | 4.3 | 47 | >sp\|P01876\|IGHA1_HUMAN Immunoglobulin heavy constant alpha 1 OS=Homo sapiens OX=9606 GN=IGHA1 PE=1 SV=2 | 255 | 20201117_fqg_milk_1ng_glyco_hcd203040_compound15.16326.16326.3 | controllerType=0 controllerNumber=1 scan=16326 | 33.8306 |
| 12983:3 | 8 | R.LAGKPTHVN[+1710.598]VSVVMAEVDGTC[+57.021]Y.- | HexNAc(3)Hex(5)NeuAc(1) | C[+57], N[+1711] | 1353.2549 | 3 | 4057.7503 | 4057.7397 | 0 | 2.6 | 332 | CRagged | 232.9 | 214.69 | 214.69 | 5.14 | 47 | >sp\|P01876\|IGHA1_HUMAN Immunoglobulin heavy constant alpha 1 OS=Homo sapiens OX=9606 GN=IGHA1 PE=1 SV=2 | 255 | 20201117_fqg_milk_1ng_glyco_hcd203040_compound15.16349.16349.3 | controllerType=0 controllerNumber=1 scan=16349 | 33.8686 |
| 11174:4 | 8 | R.LAGKPTHVN[+1809.666]VSVVMAEVDGTC[+57.021]Y.- | HexNAc(5)Hex(4)Fuc(1) | C[+57], N[+1810] | 1039.961 | 4 | 4156.8221 | 4156.8081 | 0 | 3.4 | 332 | CRagged | 260.78 | 183.48 | 183.48 | 4.45 | 47 | >sp\|P01876\|IGHA1_HUMAN Immunoglobulin heavy constant alpha 1 OS=Homo sapiens OX=9606 GN=IGHA1 PE=1 SV=2 | 255 | 20201117_fqg_milk_1ng_glyco_hcd203040_compound15.14447.14447.4 | controllerType=0 controllerNumber=1 scan=14447 | 30.6561 |
| 11131:4 | 8 | R.LAGKPTHVN[+1809.666]VSVVMAEVDGTC[+57.021]Y.- | HexNAc(5)Hex(4)Fuc(1) | C[+57], N[+1810] | 1039.9583 | 4 | 4156.8113 | 4156.8081 | 0 | 0.8 | 332 | CRagged | 270.87 | 194.57 | 194.57 | 5.39 | 47 | >sp\|P01876\|IGHA1_HUMAN Immunoglobulin heavy constant alpha 1 OS=Homo sapiens OX=9606 GN=IGHA1 PE=1 SV=2 | 255 | 20201117_fqg_milk_1ng_glyco_hcd203040_compound15.14403.14403.4 | controllerType=0 controllerNumber=1 scan=14403 | 30.5752 |
| 14843:3 | 8 | R.LAGKPTHVN[+2553.910]VSVVMAEVDGTC[+57.021]Y.- | HexNAc(5)Hex(5)Fuc(1)NeuAc(2) | C[+57], N[+2554] | 1634.3591 | 3 | 4901.0626 | 4901.0518 | 0 | 2.2 | 332 | CRagged | 324.37 | 295.64 | 295.64 | 7.74 | 47 | >sp\|P01876\|IGHA1_HUMAN Immunoglobulin heavy constant alpha 1 OS=Homo sapiens OX=9606 GN=IGHA1 PE=1 SV=2 | 255 | 20201117_fqg_milk_1ng_glyco_hcd203040_compound15.18278.18278.3 | controllerType=0 controllerNumber=1 scan=18278 | 37.3292 |
| 11156:3 | 8 | R.LAGKPTHVN[+1809.666]VSVVMAEVDGTC[+57.021]Y.- | HexNAc(5)Hex(4)Fuc(1) | C[+57], N[+1810] | 1386.2769 | 3 | 4156.8162 | 4156.8081 | 0 | 1.9 | 332 | CRagged | 386.62 | 386.62 | 300.07 | 9.66 | 47 | >sp\|P01876\|IGHA1_HUMAN Immunoglobulin heavy constant alpha 1 OS=Homo sapiens OX=9606 GN=IGHA1 PE=1 SV=2 | 255 | 20201117_fqg_milk_1ng_glyco_hcd203040_compound15.14429.14429.3 | controllerType=0 controllerNumber=1 scan=14429 | 30.6197 |
| 12864:4 | 8 | R.LAGKPTHVN[+2262.814]VSVVMAEVDGTC[+57.021]Y.- | HexNAc(5)Hex(5)Fuc(1)NeuAc(1) | C[+57], N[+2263] | 1153.2469 | 4 | 4609.9656 | 4609.9564 | 0 | 2 | 332 | CRagged | 393.38 | 393.38 | 393.38 | 8.79 | 47 | >sp\|P01876\|IGHA1_HUMAN Immunoglobulin heavy constant alpha 1 OS=Homo sapiens OX=9606 GN=IGHA1 PE=1 SV=2 | 255 | 20201117_fqg_milk_1ng_glyco_hcd203040_compound15.16226.16226.4 | controllerType=0 controllerNumber=1 scan=16226 | 33.654 |
| 11105:3 | 8 | R.LAGKPTHVN[+1663.608]VSVVMAEVDGTC[+57.021]Y.- | HexNAc(5)Hex(4) | C[+57], N[+1664] | 1337.5884 | 3 | 4010.7508 | 4010.7502 | 0 | 0.1 | 332 | CRagged | 430.15 | 430.15 | 261.14 | 9.45 | 47 | >sp\|P01876\|IGHA1_HUMAN Immunoglobulin heavy constant alpha 1 OS=Homo sapiens OX=9606 GN=IGHA1 PE=1 SV=2 | 255 | 20201117_fqg_milk_1ng_glyco_hcd203040_compound15.14376.14376.3 | controllerType=0 controllerNumber=1 scan=14376 | 30.5311 |
| 12862:4 | 8 | R.LAGKPTHVN[+2262.814]VSVVMAEVDGTC[+57.021]Y.- | HexNAc(5)Hex(5)Fuc(1)NeuAc(1) | C[+57], N[+2263] | 1153.2469 | 4 | 4609.9656 | 4609.9564 | 0 | 2 | 332 | CRagged | 439.64 | 439.64 | 439.64 | 9.65 | 47 | >sp\|P01876\|IGHA1_HUMAN Immunoglobulin heavy constant alpha 1 OS=Homo sapiens OX=9606 GN=IGHA1 PE=1 SV=2 | 255 | 20201117_fqg_milk_1ng_glyco_hcd203040_compound15.16224.16224.4 | controllerType=0 controllerNumber=1 scan=16224 | 33.6479 |
| 11128:4 | 8 | R.LAGKPTHVN[+1809.666]VSVVMAEVDGTC[+57.021]Y.- | HexNAc(5)Hex(4)Fuc(1) | C[+57], N[+1810] | 1039.9583 | 4 | 4156.8113 | 4156.8081 | 0 | 0.8 | 332 | CRagged | 496.39 | 326.79 | 326.79 | 8.29 | 47 | >sp\|P01876\|IGHA1_HUMAN Immunoglobulin heavy constant alpha 1 OS=Homo sapiens OX=9606 GN=IGHA1 PE=1 SV=2 | 255 | 20201117_fqg_milk_1ng_glyco_hcd203040_compound15.14400.14400.4 | controllerType=0 controllerNumber=1 scan=14400 | 30.5698 |
| 11211:4 | 8 | R.LAGKPTHVN[+1647.613]VSVVMAEVDGTC[+57.021]Y.- | HexNAc(5)Hex(3)Fuc(1) | C[+57], N[+1648] | 999.4464 | 4 | 3994.7637 | 3994.7553 | 0 | 2.1 | 332 | CRagged | 581 | 366.5 | 366.5 | 9.86 | 47 | >sp\|P01876\|IGHA1_HUMAN Immunoglobulin heavy constant alpha 1 OS=Homo sapiens OX=9606 GN=IGHA1 PE=1 SV=2 | 255 | 20201117_fqg_milk_1ng_glyco_hcd203040_compound15.14486.14486.4 | controllerType=0 controllerNumber=1 scan=14486 | 30.7328 |
| 11027:3 | 8 | R.LAGKPTHVN[+1702.581]VSVVMAEVDGTC[+57.021]Y.- | HexNAc(2)Hex(8) | C[+57], N[+1703] | 1350.5829 | 3 | 4049.7343 | 4049.7234 | 0 | 2.7 | 332 | CRagged | 645.03 | 645.03 | 365.55 | 12.79 | 47 | >sp\|P01876\|IGHA1_HUMAN Immunoglobulin heavy constant alpha 1 OS=Homo sapiens OX=9606 GN=IGHA1 PE=1 SV=2 | 255 | 20201117_fqg_milk_1ng_glyco_hcd203040_compound15.14295.14295.3 | controllerType=0 controllerNumber=1 scan=14295 | 30.3625 |
| 11214:4 | 8 | R.LAGKPTHVN[+1647.613]VSVVMAEVDGTC[+57.021]Y.- | HexNAc(5)Hex(3)Fuc(1) | C[+57], N[+1648] | 999.4464 | 4 | 3994.7637 | 3994.7553 | 0 | 2.1 | 332 | CRagged | 653.29 | 624.34 | 624.34 | 13.52 | 47 | >sp\|P01876\|IGHA1_HUMAN Immunoglobulin heavy constant alpha 1 OS=Homo sapiens OX=9606 GN=IGHA1 PE=1 SV=2 | 255 | 20201117_fqg_milk_1ng_glyco_hcd203040_compound15.14489.14489.4 | controllerType=0 controllerNumber=1 scan=14489 | 30.7378 |
| 11155:3 | 8 | R.LAGKPTHVN[+1216.423]VSVVMAEVDGTC[+57.021]Y.- | HexNAc(2)Hex(5) | C[+57], N[+1216] | 1188.5291 | 3 | 3563.5728 | 3563.5649 | 0 | 2.2 | 332 | CRagged | 714.92 | 714.92 | 440.92 | 14.79 | 47 | >sp\|P01876\|IGHA1_HUMAN Immunoglobulin heavy constant alpha 1 OS=Homo sapiens OX=9606 GN=IGHA1 PE=1 SV=2 | 255 | 20201117_fqg_milk_1ng_glyco_hcd203040_compound15.14428.14428.3 | controllerType=0 controllerNumber=1 scan=14428 | 30.6179 |
| 11108:3 | 8 | R.LAGKPTHVN[+1663.608]VSVVMAEVDGTC[+57.021]Y.- | HexNAc(5)Hex(4) | C[+57], N[+1664] | 1337.5884 | 3 | 4010.7508 | 4010.7502 | 0 | 0.1 | 332 | CRagged | 745.91 | 564.87 | 493.65 | 14.32 | 47 | >sp\|P01876\|IGHA1_HUMAN Immunoglobulin heavy constant alpha 1 OS=Homo sapiens OX=9606 GN=IGHA1 PE=1 SV=2 | 255 | 20201117_fqg_milk_1ng_glyco_hcd203040_compound15.14379.14379.3 | controllerType=0 controllerNumber=1 scan=14379 | 30.5353 |
| 11209:3 | 8 | R.LAGKPTHVN[+1647.613]VSVVMAEVDGTC[+57.021]Y.- | HexNAc(5)Hex(3)Fuc(1) | C[+57], N[+1648] | 1332.2601 | 3 | 3994.7656 | 3994.7553 | 0 | 2.6 | 332 | CRagged | 761.64 | 761.64 | 761.64 | 15.79 | 47 | >sp\|P01876\|IGHA1_HUMAN Immunoglobulin heavy constant alpha 1 OS=Homo sapiens OX=9606 GN=IGHA1 PE=1 SV=2 | 255 | 20201117_fqg_milk_1ng_glyco_hcd203040_compound15.14484.14484.3 | controllerType=0 controllerNumber=1 scan=14484 | 30.7306 |
| 11158:3 | 8 | R.LAGKPTHVN[+1809.666]VSVVMAEVDGTC[+57.021]Y.- | HexNAc(5)Hex(4)Fuc(1) | C[+57], N[+1810] | 1386.2769 | 3 | 4156.8162 | 4156.8081 | 0 | 1.9 | 332 | CRagged | 771.08 | 629.31 | 495.28 | 15.71 | 47 | >sp\|P01876\|IGHA1_HUMAN Immunoglobulin heavy constant alpha 1 OS=Homo sapiens OX=9606 GN=IGHA1 PE=1 SV=2 | 255 | 20201117_fqg_milk_1ng_glyco_hcd203040_compound15.14431.14431.3 | controllerType=0 controllerNumber=1 scan=14431 | 30.6239 |
| 11207:3 | 8 | R.LAGKPTHVN[+1647.613]VSVVMAEVDGTC[+57.021]Y.- | HexNAc(5)Hex(3)Fuc(1) | C[+57], N[+1648] | 1332.2601 | 3 | 3994.7656 | 3994.7553 | 0 | 2.6 | 332 | CRagged | 784.08 | 784.08 | 784.08 | 15.79 | 47 | >sp\|P01876\|IGHA1_HUMAN Immunoglobulin heavy constant alpha 1 OS=Homo sapiens OX=9606 GN=IGHA1 PE=1 SV=2 | 255 | 20201117_fqg_milk_1ng_glyco_hcd203040_compound15.14482.14482.3 | controllerType=0 controllerNumber=1 scan=14482 | 30.7256 |
| 11029:3 | 8 | R.LAGKPTHVN[+1702.581]VSVVMAEVDGTC[+57.021]Y.- | HexNAc(2)Hex(8) | C[+57], N[+1703] | 1350.5829 | 3 | 4049.7343 | 4049.7234 | 0 | 2.7 | 332 | CRagged | 784.4 | 731.21 | 451.91 | 15.79 | 47 | >sp\|P01876\|IGHA1_HUMAN Immunoglobulin heavy constant alpha 1 OS=Homo sapiens OX=9606 GN=IGHA1 PE=1 SV=2 | 255 | 20201117_fqg_milk_1ng_glyco_hcd203040_compound15.14297.14297.3 | controllerType=0 controllerNumber=1 scan=14297 | 30.3682 |
| 11169:3 | 8 | R.LAGKPTHVN[+1501.555]VSVVMAEVDGTC[+57.021]Y.- | HexNAc(5)Hex(3) | C[+57], N[+1502] | 1283.5732 | 3 | 3848.705 | 3848.6974 | 0 | 2 | 332 | CRagged | 787.61 | 665.44 | 665.44 | 15.71 | 47 | >sp\|P01876\|IGHA1_HUMAN Immunoglobulin heavy constant alpha 1 OS=Homo sapiens OX=9606 GN=IGHA1 PE=1 SV=2 | 255 | 20201117_fqg_milk_1ng_glyco_hcd203040_compound15.14442.14442.3 | controllerType=0 controllerNumber=1 scan=14442 | 30.6487 |
| 11083:3 | 8 | R.LAGKPTHVN[+1378.476]VSVVMAEVDGTC[+57.021]Y.- | HexNAc(2)Hex(6) | C[+57], N[+1378] | 1242.5468 | 3 | 3725.6259 | 3725.6178 | 0 | 2.2 | 332 | CRagged | 852.14 | 852.14 | 513.91 | 17.79 | 47 | >sp\|P01876\|IGHA1_HUMAN Immunoglobulin heavy constant alpha 1 OS=Homo sapiens OX=9606 GN=IGHA1 PE=1 SV=2 | 255 | 20201117_fqg_milk_1ng_glyco_hcd203040_compound15.14353.14353.3 | controllerType=0 controllerNumber=1 scan=14353 | 30.477 |
| 11167:3 | 8 | R.LAGKPTHVN[+1501.555]VSVVMAEVDGTC[+57.021]Y.- | HexNAc(5)Hex(3) | C[+57], N[+1502] | 1283.5732 | 3 | 3848.705 | 3848.6974 | 0 | 2 | 332 | CRagged | 857.36 | 857.36 | 857.36 | 17.71 | 47 | >sp\|P01876\|IGHA1_HUMAN Immunoglobulin heavy constant alpha 1 OS=Homo sapiens OX=9606 GN=IGHA1 PE=1 SV=2 | 255 | 20201117_fqg_milk_1ng_glyco_hcd203040_compound15.14440.14440.3 | controllerType=0 controllerNumber=1 scan=14440 | 30.6423 |
| 11086:3 | 8 | R.LAGKPTHVN[+1378.476]VSVVMAEVDGTC[+57.021]Y.- | HexNAc(2)Hex(6) | C[+57], N[+1378] | 1242.5468 | 3 | 3725.6259 | 3725.6178 | 0 | 2.2 | 332 | CRagged | 887.1 | 840.16 | 541.95 | 17.79 | 47 | >sp\|P01876\|IGHA1_HUMAN Immunoglobulin heavy constant alpha 1 OS=Homo sapiens OX=9606 GN=IGHA1 PE=1 SV=2 | 255 | 20201117_fqg_milk_1ng_glyco_hcd203040_compound15.14356.14356.3 | controllerType=0 controllerNumber=1 scan=14356 | 30.4872 |
| 11157:3 | 8 | R.LAGKPTHVN[+1216.423]VSVVMAEVDGTC[+57.021]Y.- | HexNAc(2)Hex(5) | C[+57], N[+1216] | 1188.5291 | 3 | 3563.5728 | 3563.5649 | 0 | 2.2 | 332 | CRagged | 895.9 | 879.35 | 537.03 | 17.79 | 47 | >sp\|P01876\|IGHA1_HUMAN Immunoglobulin heavy constant alpha 1 OS=Homo sapiens OX=9606 GN=IGHA1 PE=1 SV=2 | 255 | 20201117_fqg_milk_1ng_glyco_hcd203040_compound15.14430.14430.3 | controllerType=0 controllerNumber=1 scan=14430 | 30.6224 |
| 21872:4 | 11 | K.VFPLSLDSTPQDGNVVVAC[+57.021]LVQGFFPQEPLSVTWSESGQN[+1809.666]VTAI | HexNAc(5)Hex(4)Fuc(1) | C[+57], N[+1810] | 1647.2642 | 4 | 6586.0348 | 6586.0275 | 0 | 1.1 | 8 | Specific | 701.1 | 633.52 | 633.52 | 15.94 | 45 | >sp\|P01877\|IGHA2_HUMAN Immunoglobulin heavy constant alpha 2 OS=Homo sapiens OX=9606 GN=IGHA2 PE=1 SV=4 | 576 | 20201117_fqg_milk_1ng_glyco_hcd203040_compound15.25637.25637.4 | controllerType=0 controllerNumber=1 scan=25637 | 51.9239 |
| 21874:4 | 11 | K.VFPLSLDSTPQDGNVVVAC[+57.021]LVQGFFPQEPLSVTWSESGQN[+1809.666]VTAI | HexNAc(5)Hex(4)Fuc(1) | C[+57], N[+1810] | 1647.2642 | 4 | 6586.0348 | 6586.0275 | 0 | 1.1 | 8 | Specific | 759.35 | 683.51 | 683.51 | 16.94 | 45 | >sp\|P01877\|IGHA2_HUMAN Immunoglobulin heavy constant alpha 2 OS=Homo sapiens OX=9606 GN=IGHA2 PE=1 SV=4 | 576 | 20201117_fqg_milk_1ng_glyco_hcd203040_compound15.25639.25639.4 | controllerType=0 controllerNumber=1 scan=25639 | 51.9294 |
| 21846:4 | 11 | K.VFPLSLDSTPQDGNVVVAC[+57.021]LVQGFFPQEPLSVTWSESGQN[+1647.613]VTAI | HexNAc(5)Hex(3)Fuc(1) | C[+57], N[+1648] | 1606.7543 | 4 | 6423.9953 | 6423.9746 | 0 | 3.2 | 8 | Specific | 793.7 | 712.7 | 712.7 | 17.02 | 45 | >sp\|P01877\|IGHA2_HUMAN Immunoglobulin heavy constant alpha 2 OS=Homo sapiens OX=9606 GN=IGHA2 PE=1 SV=4 | 576 | 20201117_fqg_milk_1ng_glyco_hcd203040_compound15.25610.25610.4 | controllerType=0 controllerNumber=1 scan=25610 | 51.8722 |
| 21848:4 | 11 | K.VFPLSLDSTPQDGNVVVAC[+57.021]LVQGFFPQEPLSVTWSESGQN[+1647.613]VTAI | HexNAc(5)Hex(3)Fuc(1) | C[+57], N[+1648] | 1606.7543 | 4 | 6423.9953 | 6423.9746 | 0 | 3.2 | 8 | Specific | 963.53 | 830.02 | 830.02 | 21.02 | 45 | >sp\|P01877\|IGHA2_HUMAN Immunoglobulin heavy constant alpha 2 OS=Homo sapiens OX=9606 GN=IGHA2 PE=1 SV=4 | 576 | 20201117_fqg_milk_1ng_glyco_hcd203040_compound15.25612.25612.4 | controllerType=0 controllerNumber=1 scan=25612 | 51.8794 |
| 00991:3 | 11 | K.HYTN[+1971.719]SSQDVTVPC[+57.021]R.V | HexNAc(5)Hex(5)Fuc(1) | C[+57], N[+1972] | 1212.4982 | 3 | 3635.4801 | 3635.4682 | 0 | 3.3 | 89 | Specific | 240.09 | 135.69 | 135.69 | 5.24 | 45 | >sp\|P01877\|IGHA2_HUMAN Immunoglobulin heavy constant alpha 2 OS=Homo sapiens OX=9606 GN=IGHA2 PE=1 SV=4 | 576 | 20201117_fqg_milk_1ng_glyco_hcd203040_compound15.3882.3882.3 | controllerType=0 controllerNumber=1 scan=3882 | 12.8138 |
| 00977:3 | 11 | K.HYTN[+1444.534]SSQDVTVPC[+57.021]R.V | HexNAc(4)Hex(3)Fuc(1) | C[+57], N[+1445] | 1036.7693 | 3 | 3108.2934 | 3108.2832 | 0 | 3.3 | 89 | Specific | 301.73 | 199.47 | 199.47 | 6.22 | 45 | >sp\|P01877\|IGHA2_HUMAN Immunoglobulin heavy constant alpha 2 OS=Homo sapiens OX=9606 GN=IGHA2 PE=1 SV=4 | 576 | 20201117_fqg_milk_1ng_glyco_hcd203040_compound15.3867.3867.3 | controllerType=0 controllerNumber=1 scan=3867 | 12.7853 |
| 00993:3 | 11 | K.HYTN[+1971.719]SSQDVTVPC[+57.021]R.V | HexNAc(5)Hex(5)Fuc(1) | C[+57], N[+1972] | 1212.4982 | 3 | 3635.4801 | 3635.4682 | 0 | 3.3 | 89 | Specific | 302.27 | 241.04 | 241.04 | 6.57 | 45 | >sp\|P01877\|IGHA2_HUMAN Immunoglobulin heavy constant alpha 2 OS=Homo sapiens OX=9606 GN=IGHA2 PE=1 SV=4 | 576 | 20201117_fqg_milk_1ng_glyco_hcd203040_compound15.3884.3884.3 | controllerType=0 controllerNumber=1 scan=3884 | 12.8301 |
| 00975:3 | 11 | K.HYTN[+1444.534]SSQDVTVPC[+57.021]R.V | HexNAc(4)Hex(3)Fuc(1) | C[+57], N[+1445] | 1036.7693 | 3 | 3108.2934 | 3108.2832 | 0 | 3.3 | 89 | Specific | 302.47 | 302.47 | 302.47 | 6.88 | 45 | >sp\|P01877\|IGHA2_HUMAN Immunoglobulin heavy constant alpha 2 OS=Homo sapiens OX=9606 GN=IGHA2 PE=1 SV=4 | 576 | 20201117_fqg_milk_1ng_glyco_hcd203040_compound15.3865.3865.3 | controllerType=0 controllerNumber=1 scan=3865 | 12.7807 |
| 01776:3 | 11 | K.HYTN[+2059.735]SSQDVTVPC[+57.021]R.V | HexNAc(4)Hex(5)Fuc(1)NeuAc(1) | C[+57], N[+2060] | 1241.8356 | 3 | 3723.4923 | 3723.4842 | 0 | 2.2 | 89 | Specific | 311.36 | 224.55 | 224.55 | 6.58 | 45 | >sp\|P01877\|IGHA2_HUMAN Immunoglobulin heavy constant alpha 2 OS=Homo sapiens OX=9606 GN=IGHA2 PE=1 SV=4 | 576 | 20201117_fqg_milk_1ng_glyco_hcd203040_compound15.4697.4697.3 | controllerType=0 controllerNumber=1 scan=4697 | 14.2651 |
| 00979:3 | 11 | K.HYTN[+1971.719]SSQDVTVPC[+57.021]R.V | HexNAc(5)Hex(5)Fuc(1) | C[+57], N[+1972] | 1212.497 | 3 | 3635.4765 | 3635.4682 | 0 | 2.3 | 89 | Specific | 313.13 | 243.52 | 243.52 | 6.88 | 45 | >sp\|P01877\|IGHA2_HUMAN Immunoglobulin heavy constant alpha 2 OS=Homo sapiens OX=9606 GN=IGHA2 PE=1 SV=4 | 576 | 20201117_fqg_milk_1ng_glyco_hcd203040_compound15.3869.3869.3 | controllerType=0 controllerNumber=1 scan=3869 | 12.7886 |
| 01786:3 | 11 | K.HYTN[+2059.735]SSQDVTVPC[+57.021]R.V | HexNAc(4)Hex(5)Fuc(1)NeuAc(1) | C[+57], N[+2060] | 1241.8354 | 3 | 3723.4916 | 3723.4842 | 0 | 2 | 89 | Specific | 345.73 | 208.52 | 208.52 | 6.56 | 45 | >sp\|P01877\|IGHA2_HUMAN Immunoglobulin heavy constant alpha 2 OS=Homo sapiens OX=9606 GN=IGHA2 PE=1 SV=4 | 576 | 20201117_fqg_milk_1ng_glyco_hcd203040_compound15.4708.4708.3 | controllerType=0 controllerNumber=1 scan=4708 | 14.2883 |
| 01788:3 | 11 | K.HYTN[+2059.735]SSQDVTVPC[+57.021]R.V | HexNAc(4)Hex(5)Fuc(1)NeuAc(1) | C[+57], N[+2060] | 1241.8354 | 3 | 3723.4916 | 3723.4842 | 0 | 2 | 89 | Specific | 349.29 | 240.1 | 240.1 | 6.61 | 45 | >sp\|P01877\|IGHA2_HUMAN Immunoglobulin heavy constant alpha 2 OS=Homo sapiens OX=9606 GN=IGHA2 PE=1 SV=4 | 576 | 20201117_fqg_milk_1ng_glyco_hcd203040_compound15.4710.4710.3 | controllerType=0 controllerNumber=1 scan=4710 | 14.2923 |
| 01844:3 | 11 | K.HYTN[+2262.814]SSQDVTVPC[+57.021]R.V | HexNAc(5)Hex(5)Fuc(1)NeuAc(1) | C[+57], N[+2263] | 1309.53 | 3 | 3926.5753 | 3926.5636 | 0 | 3 | 89 | Specific | 393.6 | 186.32 | 186.32 | 6.57 | 45 | >sp\|P01877\|IGHA2_HUMAN Immunoglobulin heavy constant alpha 2 OS=Homo sapiens OX=9606 GN=IGHA2 PE=1 SV=4 | 576 | 20201117_fqg_milk_1ng_glyco_hcd203040_compound15.4768.4768.3 | controllerType=0 controllerNumber=1 scan=4768 | 14.393 |
| 00981:3 | 11 | K.HYTN[+1971.719]SSQDVTVPC[+57.021]R.V | HexNAc(5)Hex(5)Fuc(1) | C[+57], N[+1972] | 1212.497 | 3 | 3635.4765 | 3635.4682 | 0 | 2.3 | 89 | Specific | 400.18 | 227.12 | 227.12 | 6.84 | 45 | >sp\|P01877\|IGHA2_HUMAN Immunoglobulin heavy constant alpha 2 OS=Homo sapiens OX=9606 GN=IGHA2 PE=1 SV=4 | 576 | 20201117_fqg_milk_1ng_glyco_hcd203040_compound15.3871.3871.3 | controllerType=0 controllerNumber=1 scan=3871 | 12.7933 |
| 01842:3 | 11 | K.HYTN[+2262.814]SSQDVTVPC[+57.021]R.V | HexNAc(5)Hex(5)Fuc(1)NeuAc(1) | C[+57], N[+2263] | 1309.53 | 3 | 3926.5753 | 3926.5636 | 0 | 3 | 89 | Specific | 415.02 | 271.29 | 271.29 | 7.5 | 45 | >sp\|P01877\|IGHA2_HUMAN Immunoglobulin heavy constant alpha 2 OS=Homo sapiens OX=9606 GN=IGHA2 PE=1 SV=4 | 576 | 20201117_fqg_milk_1ng_glyco_hcd203040_compound15.4766.4766.3 | controllerType=0 controllerNumber=1 scan=4766 | 14.3889 |
| 01774:3 | 11 | K.HYTN[+2059.735]SSQDVTVPC[+57.021]R.V | HexNAc(4)Hex(5)Fuc(1)NeuAc(1) | C[+57], N[+2060] | 1241.8356 | 3 | 3723.4923 | 3723.4842 | 0 | 2.2 | 89 | Specific | 431.28 | 252.68 | 252.68 | 7.3 | 45 | >sp\|P01877\|IGHA2_HUMAN Immunoglobulin heavy constant alpha 2 OS=Homo sapiens OX=9606 GN=IGHA2 PE=1 SV=4 | 576 | 20201117_fqg_milk_1ng_glyco_hcd203040_compound15.4695.4695.3 | controllerType=0 controllerNumber=1 scan=4695 | 14.2611 |
| 01008:3 | 11 | K.HYTN[+1647.613]SSQDVTVPC[+57.021]R.V | HexNAc(5)Hex(3)Fuc(1) | C[+57], N[+1648] | 1104.4605 | 3 | 3311.3669 | 3311.3625 | 0 | 1.3 | 89 | Specific | 570.63 | 363.46 | 363.46 | 8.24 | 45 | >sp\|P01877\|IGHA2_HUMAN Immunoglobulin heavy constant alpha 2 OS=Homo sapiens OX=9606 GN=IGHA2 PE=1 SV=4 | 576 | 20201117_fqg_milk_1ng_glyco_hcd203040_compound15.3900.3900.2 | controllerType=0 controllerNumber=1 scan=3900 | 12.8643 |

Table S4. Cont.

| Query #:z | Protein Rank | Glycopeptide | Glycan | Modification Type(s) | Observed m/z | z | Observed (M+H) | Calc. mass (M+H) | Off-by-x error | Mass error (ppm) | StartPosition | Cleavage | Score | Delta | Delta Mod | \|Log Prob\| | # of unique peptides | Protein Name | Protein DB number | Comment | Scan # | Scan Time |
|---|---|---|---|---|---|---|---|---|---|---|---|---|---|---|---|---|---|---|---|---|---|---|
| 00974:3 | 11 | K.HYTN[+1809.666]SSQDVTVPC[+57.021]R.V | HexNAc(5)Hex(4)Fuc(1) | C[+57], N[+1810] | 1158.4791 | 3 | 3473.4226 | 3473.4154 | 0 | 2.1 | 89 | Specific | 574.76 | 357.62 | 357.62 | 9.58 | 45 | >sp\|P01877\|IGHA2_HUMAN Immunoglobulin heavy constant alpha 2 OS=Homo sapiens OX=9606 GN=IGHA2 PE=1 SV=4 | 576 | 20201117_fqg_milk_1ng_glyco_hcd203040_compound15.3864.3864.3 | controllerType=0 controllerNumber=1 scan=3864 | 12.7793 |
| 00976:3 | 11 | K.HYTN[+1809.666]SSQDVTVPC[+57.021]R.V | HexNAc(5)Hex(4)Fuc(1) | C[+57], N[+1810] | 1158.4791 | 3 | 3473.4226 | 3473.4154 | 0 | 2.1 | 89 | Specific | 581.25 | 501.54 | 501.54 | 11.25 | 45 | >sp\|P01877\|IGHA2_HUMAN Immunoglobulin heavy constant alpha 2 OS=Homo sapiens OX=9606 GN=IGHA2 PE=1 SV=4 | 576 | 20201117_fqg_milk_1ng_glyco_hcd203040_compound15.3866.3866.3 | controllerType=0 controllerNumber=1 scan=3866 | 12.7831 |
| 01010:3 | 11 | K.HYTN[+1647.613]SSQDVTVPC[+57.021]R.V | HexNAc(5)Hex(3)Fuc(1) | C[+57], N[+1648] | 1104.4605 | 3 | 3311.3669 | 3311.3625 | 0 | 1.3 | 89 | Specific | 678.91 | 459.78 | 459.78 | 13.16 | 45 | >sp\|P01877\|IGHA2_HUMAN Immunoglobulin heavy constant alpha 2 OS=Homo sapiens OX=9606 GN=IGHA2 PE=1 SV=4 | 576 | 20201117_fqg_milk_1ng_glyco_hcd203040_compound15.3902.3902.2 | controllerType=0 controllerNumber=1 scan=3902 | 12.8739 |
| 00995:3 | 11 | K.HYTN[+1647.613]SSQDVTVPC[+57.021]R.V | HexNAc(5)Hex(3)Fuc(1) | C[+57], N[+1648] | 1104.4615 | 3 | 3311.3698 | 3311.3625 | 0 | 2.2 | 89 | Specific | 736.76 | 561.94 | 561.94 | 14.25 | 45 | >sp\|P01877\|IGHA2_HUMAN Immunoglobulin heavy constant alpha 2 OS=Homo sapiens OX=9606 GN=IGHA2 PE=1 SV=4 | 576 | 20201117_fqg_milk_1ng_glyco_hcd203040_compound15.3886.3886.3 | controllerType=0 controllerNumber=1 scan=3886 | 12.8334 |
| 00997:3 | 11 | K.HYTN[+1647.613]SSQDVTVPC[+57.021]R.V | HexNAc(5)Hex(3)Fuc(1) | C[+57], N[+1648] | 1104.4615 | 3 | 3311.3698 | 3311.3625 | 0 | 2.2 | 89 | Specific | 750.53 | 558.6 | 558.6 | 15.25 | 45 | >sp\|P01877\|IGHA2_HUMAN Immunoglobulin heavy constant alpha 2 OS=Homo sapiens OX=9606 GN=IGHA2 PE=1 SV=4 | 576 | 20201117_fqg_milk_1ng_glyco_hcd203040_compound15.3888.3888.3 | controllerType=0 controllerNumber=1 scan=3888 | 12.842 |
| 03132:5 | 11 | K.HYTN[+1647.613]SSQDVTVPC[+57.021]RVPPPPPC[+57.021]C[+57.021]HPR.L | HexNAc(5)Hex(3)Fuc(1) | C[+57]*3, N[+1648] | 922.0018 | 5 | 4605.9799 | 4605.9689 | 0 | 2.4 | 89 | Specific | 228.67 | 214.06 | 214.06 | 6.08 | 45 | >sp\|P01877\|IGHA2_HUMAN Immunoglobulin heavy constant alpha 2 OS=Homo sapiens OX=9606 GN=IGHA2 PE=1 SV=4 | 576 | 20201117_fqg_milk_1ng_glyco_hcd203040_compound15.6104.6104.5 | controllerType=0 controllerNumber=1 scan=6104 | 16.7309 |
| 02691:3 | 11 | K.TPLTAN[+1809.666]ITK.S | HexNAc(5)Hex(4)Fuc(1) | N[+1810] | 923.4142 | 3 | 2768.2281 | 2768.2228 | 0 | 1.9 | 200 | Specific | 401.49 | 298.68 | 298.68 | 6.93 | 45 | >sp\|P01877\|IGHA2_HUMAN Immunoglobulin heavy constant alpha 2 OS=Homo sapiens OX=9606 GN=IGHA2 PE=1 SV=4 | 576 | 20201117_fqg_milk_1ng_glyco_hcd203040_compound15.5648.5648.3 | controllerType=0 controllerNumber=1 scan=5648 | 15.9312 |
| 02747:3 | 11 | K.TPLTAN[+1647.613]ITK.S | HexNAc(5)Hex(3)Fuc(1) | N[+1648] | 869.3976 | 3 | 2606.1783 | 2606.17 | 0 | 3.2 | 200 | Specific | 471.18 | 178.62 | 178.62 | 6.53 | 45 | >sp\|P01877\|IGHA2_HUMAN Immunoglobulin heavy constant alpha 2 OS=Homo sapiens OX=9606 GN=IGHA2 PE=1 SV=4 | 576 | 20201117_fqg_milk_1ng_glyco_hcd203040_compound15.5706.5706.3 | controllerType=0 controllerNumber=1 scan=5706 | 16.0367 |
| 02727:2 | 11 | K.TPLTAN[+1501.555]ITK.S | HexNAc(5)Hex(3) | N[+1502] | 1230.5621 | 2 | 2460.1168 | 2460.1121 | 0 | 1.9 | 200 | Specific | 474.61 | 287.95 | 287.95 | 7.4 | 45 | >sp\|P01877\|IGHA2_HUMAN Immunoglobulin heavy constant alpha 2 OS=Homo sapiens OX=9606 GN=IGHA2 PE=1 SV=4 | 576 | 20201117_fqg_milk_1ng_glyco_hcd203040_compound15.5685.5685.2 | controllerType=0 controllerNumber=1 scan=5685 | 15.9963 |
| 02665:2 | 11 | K.TPLTAN[+1444.534]ITK.S | HexNAc(4)Hex(3)Fuc(1) | N[+1445] | 1202.0512 | 2 | 2403.0951 | 2403.0906 | 0 | 1.9 | 200 | Specific | 481.25 | 290.47 | 290.47 | 7.4 | 45 | >sp\|P01877\|IGHA2_HUMAN Immunoglobulin heavy constant alpha 2 OS=Homo sapiens OX=9606 GN=IGHA2 PE=1 SV=4 | 576 | 20201117_fqg_milk_1ng_glyco_hcd203040_compound15.5621.5621.2 | controllerType=0 controllerNumber=1 scan=5621 | 15.8817 |
| 02794:2 | 11 | K.TPLTAN[+1241.454]ITK.S | HexNAc(3)Hex(3)Fuc(1) | N[+1241] | 1100.5115 | 2 | 2200.0158 | 2200.0113 | 0 | 2 | 200 | Specific | 481.95 | 246.53 | 246.53 | 7.29 | 45 | >sp\|P01877\|IGHA2_HUMAN Immunoglobulin heavy constant alpha 2 OS=Homo sapiens OX=9606 GN=IGHA2 PE=1 SV=4 | 576 | 20201117_fqg_milk_1ng_glyco_hcd203040_compound15.5754.5754.2 | controllerType=0 controllerNumber=1 scan=5754 | 16.1247 |
| 02693:3 | 11 | K.TPLTAN[+1809.666]ITK.S | HexNAc(5)Hex(4)Fuc(1) | N[+1810] | 923.4142 | 3 | 2768.2281 | 2768.2228 | 0 | 1.9 | 200 | Specific | 529.2 | 262.2 | 262.2 | 7.1 | 45 | >sp\|P01877\|IGHA2_HUMAN Immunoglobulin heavy constant alpha 2 OS=Homo sapiens OX=9606 GN=IGHA2 PE=1 SV=4 | 576 | 20201117_fqg_milk_1ng_glyco_hcd203040_compound15.5650.5650.3 | controllerType=0 controllerNumber=1 scan=5650 | 15.9352 |
| 02729:2 | 11 | K.TPLTAN[+1501.555]ITK.S | HexNAc(5)Hex(3) | N[+1502] | 1230.5621 | 2 | 2460.1168 | 2460.1121 | 0 | 1.9 | 200 | Specific | 541.17 | 284.18 | 284.18 | 7.48 | 45 | >sp\|P01877\|IGHA2_HUMAN Immunoglobulin heavy constant alpha 2 OS=Homo sapiens OX=9606 GN=IGHA2 PE=1 SV=4 | 576 | 20201117_fqg_milk_1ng_glyco_hcd203040_compound15.5687.5687.2 | controllerType=0 controllerNumber=1 scan=5687 | 16 |
| 02792:2 | 11 | K.TPLTAN[+1241.454]ITK.S | HexNAc(3)Hex(3)Fuc(1) | N[+1241] | 1100.5115 | 2 | 2200.0158 | 2200.0113 | 0 | 2 | 200 | Specific | 568.87 | 352.54 | 352.54 | 8.07 | 45 | >sp\|P01877\|IGHA2_HUMAN Immunoglobulin heavy constant alpha 2 OS=Homo sapiens OX=9606 GN=IGHA2 PE=1 SV=4 | 576 | 20201117_fqg_milk_1ng_glyco_hcd203040_compound15.5752.5752.2 | controllerType=0 controllerNumber=1 scan=5752 | 16.1211 |
| 02667:2 | 11 | K.TPLTAN[+1444.534]ITK.S | HexNAc(4)Hex(3)Fuc(1) | N[+1445] | 1202.0512 | 2 | 2403.0951 | 2403.0906 | 0 | 1.9 | 200 | Specific | 577.54 | 306.32 | 306.32 | 9.15 | 45 | >sp\|P01877\|IGHA2_HUMAN Immunoglobulin heavy constant alpha 2 OS=Homo sapiens OX=9606 GN=IGHA2 PE=1 SV=4 | 576 | 20201117_fqg_milk_1ng_glyco_hcd203040_compound15.5623.5623.2 | controllerType=0 controllerNumber=1 scan=5623 | 15.8854 |
| 02687:2 | 11 | K.TPLTAN[+1809.666]ITK.S | HexNAc(5)Hex(4)Fuc(1) | N[+1810] | 1384.6161 | 2 | 2768.225 | 2768.2228 | 0 | 0.8 | 200 | Specific | 581.71 | 343.02 | 343.02 | 9.22 | 45 | >sp\|P01877\|IGHA2_HUMAN Immunoglobulin heavy constant alpha 2 OS=Homo sapiens OX=9606 GN=IGHA2 PE=1 SV=4 | 576 | 20201117_fqg_milk_1ng_glyco_hcd203040_compound15.5643.5643.2 | controllerType=0 controllerNumber=1 scan=5643 | 15.9203 |
| 02689:2 | 11 | K.TPLTAN[+1809.666]ITK.S | HexNAc(5)Hex(4)Fuc(1) | N[+1810] | 1384.6161 | 2 | 2768.225 | 2768.2228 | 0 | 0.8 | 200 | Specific | 592.84 | 347.76 | 347.76 | 9.24 | 45 | >sp\|P01877\|IGHA2_HUMAN Immunoglobulin heavy constant alpha 2 OS=Homo sapiens OX=9606 GN=IGHA2 PE=1 SV=4 | 576 | 20201117_fqg_milk_1ng_glyco_hcd203040_compound15.5645.5645.2 | controllerType=0 controllerNumber=1 scan=5645 | 15.9252 |
| 02732:2 | 11 | K.TPLTAN[+1647.613]ITK.S | HexNAc(5)Hex(3)Fuc(1) | N[+1648] | 1303.5921 | 2 | 2606.1769 | 2606.17 | 0 | 2.6 | 200 | Specific | 598.43 | 329.93 | 329.93 | 9.31 | 45 | >sp\|P01877\|IGHA2_HUMAN Immunoglobulin heavy constant alpha 2 OS=Homo sapiens OX=9606 GN=IGHA2 PE=1 SV=4 | 576 | 20201117_fqg_milk_1ng_glyco_hcd203040_compound15.5690.5690.2 | controllerType=0 controllerNumber=1 scan=5690 | 16.0078 |
| 02749:3 | 11 | K.TPLTAN[+1647.613]ITK.S | HexNAc(5)Hex(3)Fuc(1) | N[+1648] | 869.3976 | 3 | 2606.1783 | 2606.17 | 0 | 3.2 | 200 | Specific | 604.78 | 290.59 | 290.59 | 9.93 | 45 | >sp\|P01877\|IGHA2_HUMAN Immunoglobulin heavy constant alpha 2 OS=Homo sapiens OX=9606 GN=IGHA2 PE=1 SV=4 | 576 | 20201117_fqg_milk_1ng_glyco_hcd203040_compound15.5708.5708.3 | controllerType=0 controllerNumber=1 scan=5708 | 16.0424 |
| 02730:2 | 11 | K.TPLTAN[+1647.613]ITK.S | HexNAc(5)Hex(3)Fuc(1) | N[+1648] | 1303.5921 | 2 | 2606.1769 | 2606.17 | 0 | 2.6 | 200 | Specific | 612.32 | 344.06 | 344.06 | 10.31 | 45 | >sp\|P01877\|IGHA2_HUMAN Immunoglobulin heavy constant alpha 2 OS=Homo sapiens OX=9606 GN=IGHA2 PE=1 SV=4 | 576 | 20201117_fqg_milk_1ng_glyco_hcd203040_compound15.5688.5688.2 | controllerType=0 controllerNumber=1 scan=5688 | 16.0019 |
| 10045:3 | 11 | R.MAGKPTHIN[+1663.608]VSVVMAEADGTC[+57.021]Y.- | HexNAc(5)Hex(4) | C[+57], N[+1664] | 1338.9053 | 3 | 4014.7014 | 4014.691 | 0 | 2.6 | 319 | CRagged | 294.68 | 289.86 | 263.64 | 7.31 | 45 | >sp\|P01877\|IGHA2_HUMAN Immunoglobulin heavy constant alpha 2 OS=Homo sapiens OX=9606 GN=IGHA2 PE=1 SV=4 | 576 | 20201117_fqg_milk_1ng_glyco_hcd203040_compound15.13280.13280.3 | controllerType=0 controllerNumber=1 scan=13280 | 28.6806 |
| 10133:3 | 11 | R.MAGKPTHIN[+1647.613]VSVVMAEADGTC[+57.021]Y.- | HexNAc(5)Hex(3)Fuc(1) | C[+57], N[+1648] | 1333.5745 | 3 | 3998.709 | 3998.6961 | 0 | 3.2 | 319 | CRagged | 456.34 | 456.34 | 456.34 | 10.01 | 45 | >sp\|P01877\|IGHA2_HUMAN Immunoglobulin heavy constant alpha 2 OS=Homo sapiens OX=9606 GN=IGHA2 PE=1 SV=4 | 576 | 20201117_fqg_milk_1ng_glyco_hcd203040_compound15.13371.13371.3 | controllerType=0 controllerNumber=1 scan=13371 | 28.8422 |
| 10169:3 | 11 | R.MAGKPTHIN[+1095.397]VSVVMAEADGTC[+57.021]Y.- | HexNAc(3)Hex(3) | C[+57], N[+1095] | 1149.5007 | 3 | 3446.4874 | 3446.4794 | 0 | 2.3 | 319 | CRagged | 476.76 | 295.95 | 295.95 | 8.41 | 45 | >sp\|P01877\|IGHA2_HUMAN Immunoglobulin heavy constant alpha 2 OS=Homo sapiens OX=9606 GN=IGHA2 PE=1 SV=4 | 576 | 20201117_fqg_milk_1ng_glyco_hcd203040_compound15.13408.13408.3 | controllerType=0 controllerNumber=1 scan=13408 | 28.9076 |
| 10148:3 | 11 | R.MAGKPTHIN[+1501.555]VSVVMAEADGTC[+57.021]Y.- | HexNAc(5)Hex(3) | C[+57], N[+1502] | 1284.8882 | 3 | 3852.6502 | 3852.6382 | 0 | 3.1 | 319 | CRagged | 496.05 | 496.05 | 496.05 | 10.24 | 45 | >sp\|P01877\|IGHA2_HUMAN Immunoglobulin heavy constant alpha 2 OS=Homo sapiens OX=9606 GN=IGHA2 PE=1 SV=4 | 576 | 20201117_fqg_milk_1ng_glyco_hcd203040_compound15.13387.13387.3 | controllerType=0 controllerNumber=1 scan=13387 | 28.874 |
| 10146:3 | 11 | R.MAGKPTHIN[+1501.555]VSVVMAEADGTC[+57.021]Y.- | HexNAc(5)Hex(3) | C[+57], N[+1502] | 1284.8882 | 3 | 3852.6502 | 3852.6382 | 0 | 3.1 | 319 | CRagged | 519.86 | 519.86 | 519.86 | 10.24 | 45 | >sp\|P01877\|IGHA2_HUMAN Immunoglobulin heavy constant alpha 2 OS=Homo sapiens OX=9606 GN=IGHA2 PE=1 SV=4 | 576 | 20201117_fqg_milk_1ng_glyco_hcd203040_compound15.13385.13385.3 | controllerType=0 controllerNumber=1 scan=13385 | 28.8686 |
| 10167:3 | 11 | R.MAGKPTHIN[+1095.397]VSVVMAEADGTC[+57.021]Y.- | HexNAc(3)Hex(3) | C[+57], N[+1095] | 1149.5007 | 3 | 3446.4874 | 3446.4794 | 0 | 2.3 | 319 | CRagged | 575.13 | 460.3 | 460.3 | 11.79 | 45 | >sp\|P01877\|IGHA2_HUMAN Immunoglobulin heavy constant alpha 2 OS=Homo sapiens OX=9606 GN=IGHA2 PE=1 SV=4 | 576 | 20201117_fqg_milk_1ng_glyco_hcd203040_compound15.13406.13406.3 | controllerType=0 controllerNumber=1 scan=13406 | 28.9024 |
| 10135:3 | 11 | R.MAGKPTHIN[+1647.613]VSVVMAEADGTC[+57.021]Y.- | HexNAc(5)Hex(3)Fuc(1) | C[+57], N[+1648] | 1333.5745 | 3 | 3998.709 | 3998.6961 | 0 | 3.2 | 319 | CRagged | 594.14 | 594.14 | 594.14 | 11.79 | 45 | >sp\|P01877\|IGHA2_HUMAN Immunoglobulin heavy constant alpha 2 OS=Homo sapiens OX=9606 GN=IGHA2 PE=1 SV=4 | 576 | 20201117_fqg_milk_1ng_glyco_hcd203040_compound15.13373.13373.3 | controllerType=0 controllerNumber=1 scan=13373 | 28.8458 |
| 09964:3 | 11 | R.MAGKPTHIN[+1702.581]VSVVMAEADGTC[+57.021]Y.- | HexNAc(2)Hex(8) | C[+57], N[+1703] | 1351.8962 | 3 | 4053.674 | 4053.6642 | 0 | 2.4 | 319 | CRagged | 732.1 | 548.32 | 431.82 | 14.79 | 45 | >sp\|P01877\|IGHA2_HUMAN Immunoglobulin heavy constant alpha 2 OS=Homo sapiens OX=9606 GN=IGHA2 PE=1 SV=4 | 576 | 20201117_fqg_milk_1ng_glyco_hcd203040_compound15.13196.13196.3 | controllerType=0 controllerNumber=1 scan=13196 | 28.5016 |
| 10047:3 | 11 | R.MAGKPTHIN[+1216.423]VSVVMAEADGTC[+57.021]Y.- | HexNAc(2)Hex(5) | C[+57], N[+1216] | 1189.8437 | 3 | 3567.5165 | 3567.5057 | 0 | 3 | 319 | CRagged | 756.2 | 678.58 | 421.31 | 15.79 | 45 | >sp\|P01877\|IGHA2_HUMAN Immunoglobulin heavy constant alpha 2 OS=Homo sapiens OX=9606 GN=IGHA2 PE=1 SV=4 | 576 | 20201117_fqg_milk_1ng_glyco_hcd203040_compound15.13282.13282.3 | controllerType=0 controllerNumber=1 scan=13282 | 28.684 |
| 09969:3 | 11 | R.MAGKPTHIN[+1540.529]VSVVMAEADGTC[+57.021]Y.- | HexNAc(2)Hex(7) | C[+57], N[+1541] | 1297.8787 | 3 | 3891.6216 | 3891.6114 | 0 | 2.6 | 319 | CRagged | 810.12 | 752.82 | 444.29 | 16.79 | 45 | >sp\|P01877\|IGHA2_HUMAN Immunoglobulin heavy constant alpha 2 OS=Homo sapiens OX=9606 GN=IGHA2 PE=1 SV=4 | 576 | 20201117_fqg_milk_1ng_glyco_hcd203040_compound15.13201.13201.3 | controllerType=0 controllerNumber=1 scan=13201 | 28.5113 |
| 10006:3 | 11 | R.MAGKPTHIN[+1378.476]VSVVMAEADGTC[+57.021]Y.- | HexNAc(2)Hex(6) | C[+57], N[+1378] | 1243.8599 | 3 | 3729.5652 | 3729.5585 | 0 | 1.8 | 319 | CRagged | 826.4 | 704.88 | 448.71 | 16.71 | 45 | >sp\|P01877\|IGHA2_HUMAN Immunoglobulin heavy constant alpha 2 OS=Homo sapiens OX=9606 GN=IGHA2 PE=1 SV=4 | 576 | 20201117_fqg_milk_1ng_glyco_hcd203040_compound15.13239.13239.3 | controllerType=0 controllerNumber=1 scan=13239 | 28.5795 |
| 09971:3 | 11 | R.MAGKPTHIN[+1540.529]VSVVMAEADGTC[+57.021]Y.- | HexNAc(2)Hex(7) | C[+57], N[+1541] | 1297.8787 | 3 | 3891.6216 | 3891.6114 | 0 | 2.6 | 319 | CRagged | 858.92 | 858.92 | 480.76 | 17.79 | 45 | >sp\|P01877\|IGHA2_HUMAN Immunoglobulin heavy constant alpha 2 OS=Homo sapiens OX=9606 GN=IGHA2 PE=1 SV=4 | 576 | 20201117_fqg_milk_1ng_glyco_hcd203040_compound15.13203.13203.3 | controllerType=0 controllerNumber=1 scan=13203 | 28.5175 |
| 10003:3 | 11 | R.MAGKPTHIN[+1378.476]VSVVMAEADGTC[+57.021]Y.- | HexNAc(2)Hex(6) | C[+57], N[+1378] | 1243.8599 | 3 | 3729.5652 | 3729.5585 | 0 | 1.8 | 319 | CRagged | 863.17 | 818.28 | 477.22 | 17.71 | 45 | >sp\|P01877\|IGHA2_HUMAN Immunoglobulin heavy constant alpha 2 OS=Homo sapiens OX=9606 GN=IGHA2 PE=1 SV=4 | 576 | 20201117_fqg_milk_1ng_glyco_hcd203040_compound15.13236.13236.3 | controllerType=0 controllerNumber=1 scan=13236 | 28.5724 |
| 09966:3 | 11 | R.MAGKPTHIN[+1702.581]VSVVMAEADGTC[+57.021]Y.- | HexNAc(2)Hex(8) | C[+57], N[+1703] | 1351.8962 | 3 | 4053.674 | 4053.6642 | 0 | 2.4 | 319 | CRagged | 869.21 | 700.27 | 484.73 | 17.79 | 45 | >sp\|P01877\|IGHA2_HUMAN Immunoglobulin heavy constant alpha 2 OS=Homo sapiens OX=9606 GN=IGHA2 PE=1 SV=4 | 576 | 20201117_fqg_milk_1ng_glyco_hcd203040_compound15.13198.13198.3 | controllerType=0 controllerNumber=1 scan=13198 | 28.5065 |
| 10049:3 | 11 | R.MAGKPTHIN[+1216.423]VSVVMAEADGTC[+57.021]Y.- | HexNAc(2)Hex(5) | C[+57], N[+1216] | 1189.8437 | 3 | 3567.5165 | 3567.5057 | 0 | 3 | 319 | CRagged | 883.9 | 883.9 | 487.91 | 17.79 | 45 | >sp\|P01877\|IGHA2_HUMAN Immunoglobulin heavy constant alpha 2 OS=Homo sapiens OX=9606 GN=IGHA2 PE=1 SV=4 | 576 | 20201117_fqg_milk_1ng_glyco_hcd203040_compound15.13284.13284.3 | controllerType=0 controllerNumber=1 scan=13284 | 28.6919 |
| 09586:3 | 11 | R.M[+15.995]AGKPTHIN[+1378.476]VSVVMAEADGTC[+57.021]Y.- | HexNAc(2)Hex(6) | C[+57], M[+16], N[+1378] | 1249.1944 | 3 | 3745.5686 | 3745.5535 | 0 | 4.1 | 319 | CRagged | 223.66 | 223.66 | 45.76 | 5.05 | 45 | >sp\|P01877\|IGHA2_HUMAN Immunoglobulin heavy constant alpha 2 OS=Homo sapiens OX=9606 GN=IGHA2 PE=1 SV=4 | 576 | 20201117_fqg_milk_1ng_glyco_hcd203040_compound15.12805.12805.3 | controllerType=0 controllerNumber=1 scan=12805 | 27.8602 |
| 05360:3 | 13 | R.LSPN[+2205.793]ASAEHLELR.W | HexNAc(4)Hex(5)Fuc(2)NeuAc(1) | N[+2206] | 1214.8556 | 3 | 3642.5524 | 3642.5421 | 0 | 2.8 | 52 | Specific | 573.63 | 374.4 | 374.4 | 11.25 | 31 | >sp\|Q13410\|BT1A1_HUMAN Butyrophilin subfamily 1 member A1 OS=Homo sapiens OX=9606 GN=BTN1A1 PE=1 SV=3 | 179 | 20201117_fqg_milk_1ng_glyco_hcd203040_compound15.8410.8410.3 | controllerType=0 controllerNumber=1 scan=8410 | 20.53 |
| 05362:3 | 13 | R.LSPN[+2205.793]ASAEHLELR.W | HexNAc(4)Hex(5)Fuc(2)NeuAc(1) | N[+2206] | 1214.8556 | 3 | 3642.5524 | 3642.5421 | 0 | 2.8 | 52 | Specific | 584.45 | 416.59 | 416.59 | 11.25 | 31 | >sp\|Q13410\|BT1A1_HUMAN Butyrophilin subfamily 1 member A1 OS=Homo sapiens OX=9606 GN=BTN1A1 PE=1 SV=3 | 179 | 20201117_fqg_milk_1ng_glyco_hcd203040_compound15.8412.8412.3 | controllerType=0 controllerNumber=1 scan=8412 | 20.5483 |
| 07450:3 | 13 | R.LSPN[+2350.830]ASAEHLELR.W | HexNAc(4)Hex(5)Fuc(1)NeuAc(2) | N[+2351] | 1263.2022 | 3 | 3787.5921 | 3787.5796 | 0 | 3.3 | 52 | Specific | 590.67 | 576.04 | 576.04 | 11.25 | 31 | >sp\|Q13410\|BT1A1_HUMAN Butyrophilin subfamily 1 member A1 OS=Homo sapiens OX=9606 GN=BTN1A1 PE=1 SV=3 | 179 | 20201117_fqg_milk_1ng_glyco_hcd203040_compound15.10575.10575.3 | controllerType=0 controllerNumber=1 scan=10575 | 24.0839 |
| 07448:3 | 13 | R.LSPN[+2350.830]ASAEHLELR.W | HexNAc(4)Hex(5)Fuc(1)NeuAc(2) | N[+2351] | 1263.2022 | 3 | 3787.5921 | 3787.5796 | 0 | 3.3 | 52 | Specific | 598.41 | 484.06 | 484.06 | 11.25 | 31 | >sp\|Q13410\|BT1A1_HUMAN Butyrophilin subfamily 1 member A1 OS=Homo sapiens OX=9606 GN=BTN1A1 PE=1 SV=3 | 179 | 20201117_fqg_milk_1ng_glyco_hcd203040_compound15.10573.10573.3 | controllerType=0 controllerNumber=1 scan=10573 | 24.08 |
| 17302:3 | 13 | K.N[+2059.735]VSC[+57.021]YIQNLLLGQEK.K | HexNAc(4)Hex(5)Fuc(1)NeuAc(1) | C[+57], N[+2060] | 1280.2231 | 3 | 3838.6547 | 3838.6455 | 0 | 2.4 | 215 | Specific | 514.01 | 473.24 | 473.24 | 9.51 | 31 | >sp\|Q13410\|BT1A1_HUMAN Butyrophilin subfamily 1 member A1 OS=Homo sapiens OX=9606 GN=BTN1A1 PE=1 SV=3 | 179 | 20201117_fqg_milk_1ng_glyco_hcd203040_compound15.20874.20874.3 | controllerType=0 controllerNumber=1 scan=20874 | 42.2951 |
| 17304:3 | 13 | K.N[+2059.735]VSC[+57.021]YIQNLLLGQEK.K | HexNAc(4)Hex(5)Fuc(1)NeuAc(1) | C[+57], N[+2060] | 1280.2231 | 3 | 3838.6547 | 3838.6455 | 0 | 2.4 | 215 | Specific | 623.52 | 455.59 | 455.59 | 12.06 | 31 | >sp\|Q13410\|BT1A1_HUMAN Butyrophilin subfamily 1 member A1 OS=Homo sapiens OX=9606 GN=BTN1A1 PE=1 SV=3 | 179 | 20201117_fqg_milk_1ng_glyco_hcd203040_compound15.20876.20876.3 | controllerType=0 controllerNumber=1 scan=20876 | 42.2989 |
| 24052:4 | 15 | R.[+42.011]QLAHQSN[+2190.793]STNIFFSPVSIATAFAMLSLGTK.A | HexNAc(6)Hex(6) | .[42], N[+2191] | 1354.3522 | 4 | 5414.3869 | 5414.4388 | 0 | -9.6 | 64 | Specific | 239.46 | 239.46 | 239.46 | 4.63 | 32 | >sp\|P01009\|A1AT_HUMAN Alpha-1-antitrypsin OS=Homo sapiens OX=9606 GN=SERPINA1 PE=1 SV=3 | 642 | 20201117_fqg_milk_1ng_glyco_hcd203040_compound15.27910.27910.4 | controllerType=0 controllerNumber=1 scan=27910 | 56.0662 |
| 24050:4 | 15 | R.[+42.011]QLAHQSN[+2190.793]STNIFFSPVSIATAFAMLSLGTK.A | HexNAc(6)Hex(6) | .[42], N[+2191] | 1354.3522 | 4 | 5414.3869 | 5414.4388 | 0 | -9.6 | 64 | Specific | 261.82 | 261.82 | 261.82 | 5.07 | 32 | >sp\|P01009\|A1AT_HUMAN Alpha-1-antitrypsin OS=Homo sapiens OX=9606 GN=SERPINA1 PE=1 SV=3 | 642 | 20201117_fqg_milk_1ng_glyco_hcd203040_compound15.27908.27908.4 | controllerType=0 controllerNumber=1 scan=27908 | 56.0627 |
| 23939:3 | 15 | R.QLAHQSN[+2204.772]STNIFFSPVSIATAFAMLSLGTK.A | HexNAc(4)Hex(5)NeuAc(2) | N[+2205] | 1796.1508 | 3 | 5386.438 | 5386.4075 | 0 | 5.7 | 64 | Specific | 607.39 | 607.39 | 607.39 | 11.81 | 32 | >sp\|P01009\|A1AT_HUMAN Alpha-1-antitrypsin OS=Homo sapiens OX=9606 GN=SERPINA1 PE=1 SV=3 | 642 | 20201117_fqg_milk_1ng_glyco_hcd203040_compound15.27792.27792.3 | controllerType=0 controllerNumber=1 scan=27792 | 55.8484 |
| 23941:3 | 15 | R.QLAHQSN[+2204.772]STNIFFSPVSIATAFAMLSLGTK.A | HexNAc(4)Hex(5)NeuAc(2) | N[+2205] | 1796.1508 | 3 | 5386.438 | 5386.4075 | 0 | 5.7 | 64 | Specific | 700.63 | 638.11 | 638.11 | 13.81 | 32 | >sp\|P01009\|A1AT_HUMAN Alpha-1-antitrypsin OS=Homo sapiens OX=9606 GN=SERPINA1 PE=1 SV=3 | 642 | 20201117_fqg_milk_1ng_glyco_hcd203040_compound15.27794.27794.3 | controllerType=0 controllerNumber=1 scan=27794 | 55.8538 |
| 23933:4 | 15 | R.QLAHQSN[+2204.772]STNIFFSPVSIATAFAMLSLGTK.A | HexNAc(4)Hex(5)NeuAc(2) | N[+2205] | 1347.3574 | 4 | 5386.4079 | 5386.4075 | 0 | 0.1 | 64 | Specific | 837.24 | 721.05 | 538.9 | 16.04 | 32 | >sp\|P01009\|A1AT_HUMAN Alpha-1-antitrypsin OS=Homo sapiens OX=9606 GN=SERPINA1 PE=1 SV=3 | 642 | 20201117_fqg_milk_1ng_glyco_hcd203040_compound15.27785.27785.4 | controllerType=0 controllerNumber=1 scan=27785 | 55.8332 |
| 23931:4 | 15 | R.QLAHQSN[+2204.772]STNIFFSPVSIATAFAMLSLGTK.A | HexNAc(4)Hex(5)NeuAc(2) | N[+2205] | 1347.3574 | 4 | 5386.4079 | 5386.4075 | 0 | 0.1 | 64 | Specific | 857.06 | 809.38 | 496.81 | 17.04 | 32 | >sp\|P01009\|A1AT_HUMAN Alpha-1-antitrypsin OS=Homo sapiens OX=9606 GN=SERPINA1 PE=1 SV=3 | 642 | 20201117_fqg_milk_1ng_glyco_hcd203040_compound15.27783.27783.4 | controllerType=0 controllerNumber=1 scan=27783 | 55.8266 |
| 20052:3 | 15 | K.YLGN[+2204.772]ATAIFFLPDEGK.L | HexNAc(4)Hex(5)NeuAc(2) | N[+2205] | 1320.8932 | 3 | 3960.6652 | 3960.6677 | 0 | -0.6 | 268 | Specific | 264.36 | 153.67 | 153.67 | 3.41 | 32 | >sp\|P01009\|A1AT_HUMAN Alpha-1-antitrypsin OS=Homo sapiens OX=9606 GN=SERPINA1 PE=1 SV=3 | 642 | 20201117_fqg_milk_1ng_glyco_hcd203040_compound15.23745.23745.3 | controllerType=0 controllerNumber=1 scan=23745 | 48.2768 |
| 20050:3 | 15 | K.YLGN[+2204.772]ATAIFFLPDEGK.L | HexNAc(4)Hex(5)NeuAc(2) | N[+2205] | 1320.8932 | 3 | 3960.6652 | 3960.6677 | 0 | -0.6 | 268 | Specific | 266.35 | 193.6 | 193.6 | 4.35 | 32 | >sp\|P01009\|A1AT_HUMAN Alpha-1-antitrypsin OS=Homo sapiens OX=9606 GN=SERPINA1 PE=1 SV=3 | 642 | 20201117_fqg_milk_1ng_glyco_hcd203040_compound15.23743.23743.3 | controllerType=0 controllerNumber=1 scan=23743 | 48.2732 |
| 19783:4 | 15 | K.YLGN[+2204.772]ATAIFFLPDEGKLQHLENELTHDIITK.F | HexNAc(4)Hex(5)NeuAc(2) | N[+2205] | 1437.1641 | 4 | 5745.6344 | 5745.6098 | 0 | 4.3 | 268 | Specific | 654.13 | 516.73 | 516.73 | 12.15 | 32 | >sp\|P01009\|A1AT_HUMAN Alpha-1-antitrypsin OS=Homo sapiens OX=9606 GN=SERPINA1 PE=1 SV=3 | 642 | 20201117_fqg_milk_1ng_glyco_hcd203040_compound15.23466.23466.4 | controllerType=0 controllerNumber=1 scan=23466 | 47.738 |
| 19781:4 | 15 | K.YLGN[+2204.772]ATAIFFLPDEGKLQHLENELTHDIITK.F | HexNAc(4)Hex(5)NeuAc(2) | N[+2205] | 1437.1641 | 4 | 5745.6344 | 5745.6098 | 0 | 4.3 | 268 | Specific | 771.2 | 703.29 | 703.29 | 14.15 | 32 | >sp\|P01009\|A1AT_HUMAN Alpha-1-antitrypsin OS=Homo sapiens OX=9606 GN=SERPINA1 PE=1 SV=3 | 642 | 20201117_fqg_milk_1ng_glyco_hcd203040_compound15.23464.23464.4 | controllerType=0 controllerNumber=1 scan=23464 | 47.7315 |
| 02299:3 | 22 | K.QTDEIKDTRN[+2059.735]ESTQNC[+57.021]VVAEPEK.M | HexNAc(4)Hex(5)Fuc(1)NeuAc(1) | C[+57], N[+2060] | 1584.3346 | 3 | 4750.9892 | 4750.9876 | 0 | 0.4 | 60 | Specific | 232.51 | 232.51 | 232.51 | 4.81 | 26 | >sp\|P47710\|CASA1_HUMAN Alpha-S1-casein OS=Homo sapiens OX=9606 GN=CSN1S1 PE=1 SV=1 | 989 | 20201117_fqg_milk_1ng_glyco_hcd203040_compound15.5239.5239.3 | controllerType=0 controllerNumber=1 scan=5239 | 15.2133 |
| 08282:5 | 25 | K.SC[+57.021]QHN[+2204.772]GTMYQHGEIFSAHELFPSR.L | HexNAc(4)Hex(5)NeuAc(2) | C[+57], N[+2205] | 1008.4183 | 5 | 5038.0623 | 5038.0294 | 0 | 6.5 | 110 | Specific | 432.59 | 353.03 | 353.03 | 7 | 24 | >sp\|Q6WN34\|CRDL2_HUMAN Chordin-like protein 2 OS=Homo sapiens OX=9606 GN=CHRDL2 PE=1 SV=1 | 1189 | 20201117_fqg_milk_1ng_glyco_hcd203040_compound15.11446.11446.5 | controllerType=0 controllerNumber=1 scan=11446 | 25.563 |
| 08272:4 | 25 | K.SC[+57.021]QHN[+2205.793]GTMYQHGEIFSAHELFPSR.L | HexNAc(4)Hex(5)Fuc(2)NeuAc(1) | C[+57], N[+2206] | 1260.5223 | 4 | 5039.0675 | 5039.0498 | 0 | 3.5 | 110 | Specific | 452.24 | 452.24 | 452.24 | 9.03 | 24 | >sp\|Q6WN34\|CRDL2_HUMAN Chordin-like protein 2 OS=Homo sapiens OX=9606 GN=CHRDL2 PE=1 SV=1 | 1189 | 20201117_fqg_milk_1ng_glyco_hcd203040_compound15.11435.11435.4 | controllerType=0 controllerNumber=1 scan=11435 | 25.5437 |
| 08284:5 | 25 | K.SC[+57.021]QHN[+2204.772]GTMYQHGEIFSAHELFPSR.L | HexNAc(4)Hex(5)NeuAc(2) | C[+57], N[+2205] | 1008.4183 | 5 | 5038.0623 | 5038.0294 | 0 | 6.5 | 110 | Specific | 462.69 | 386.52 | 386.52 | 8.74 | 24 | >sp\|Q6WN34\|CRDL2_HUMAN Chordin-like protein 2 OS=Homo sapiens OX=9606 GN=CHRDL2 PE=1 SV=1 | 1189 | 20201117_fqg_milk_1ng_glyco_hcd203040_compound15.11448.11448.5 | controllerType=0 controllerNumber=1 scan=11448 | 25.5671 |
| 08315:4 | 25 | K.SC[+57.021]QHN[+2059.735]GTMYQHGEIFSAHELFPSR.L | HexNAc(4)Hex(5)Fuc(1)NeuAc(1) | C[+57], N[+2060] | 1224.007 | 4 | 4893.0062 | 4892.9919 | 0 | 2.9 | 110 | Specific | 543.39 | 543.39 | 543.39 | 9.67 | 24 | >sp\|Q6WN34\|CRDL2_HUMAN Chordin-like protein 2 OS=Homo sapiens OX=9606 GN=CHRDL2 PE=1 SV=1 | 1189 | 20201117_fqg_milk_1ng_glyco_hcd203040_compound15.11480.11480.4 | controllerType=0 controllerNumber=1 scan=11480 | 25.6184 |
| 08274:4 | 25 | K.SC[+57.021]QHN[+2205.793]GTMYQHGEIFSAHELFPSR.L | HexNAc(4)Hex(5)Fuc(2)NeuAc(1) | C[+57], N[+2206] | 1260.5223 | 4 | 5039.0675 | 5039.0498 | 0 | 3.5 | 110 | Specific | 559.87 | 531.3 | 531.3 | 9.55 | 24 | >sp\|Q6WN34\|CRDL2_HUMAN Chordin-like protein 2 OS=Homo sapiens OX=9606 GN=CHRDL2 PE=1 SV=1 | 1189 | 20201117_fqg_milk_1ng_glyco_hcd203040_compound15.11437.11437.4 | controllerType=0 controllerNumber=1 scan=11437 | 25.5475 |
| 08313:4 | 25 | K.SC[+57.021]QHN[+2059.735]GTMYQHGEIFSAHELFPSR.L | HexNAc(4)Hex(5)Fuc(1)NeuAc(1) | C[+57], N[+2060] | 1224.007 | 4 | 4893.0062 | 4892.9919 | 0 | 2.9 | 110 | Specific | 581.1 | 581.1 | 581.1 | 11.21 | 24 | >sp\|Q6WN34\|CRDL2_HUMAN Chordin-like protein 2 OS=Homo sapiens OX=9606 GN=CHRDL2 PE=1 SV=1 | 1189 | 20201117_fqg_milk_1ng_glyco_hcd203040_compound15.11478.11478.4 | controllerType=0 controllerNumber=1 scan=11478 | 25.6146 |
| 18406:3 | 26 | R.C[+57.021]MWSSALNSLN[+1864.634]LSFAGLEQVPK.G | HexNAc(2)Hex(9) | C[+57], N[+1865] | 1439.6182 | 3 | 4316.8401 | 4316.8341 | 0 | 1.4 | 272 | Specific | 900.96 | 805.57 | 805.57 | 18.52 | 19 | >sp\|P08571\|CD14_HUMAN Monocyte differentiation antigen CD14 OS=Homo sapiens OX=9606 GN=CD14 PE=1 SV=2 | 1232 | 20201117_fqg_milk_1ng_glyco_hcd203040_compound15.22023.22023.3 | controllerType=0 controllerNumber=1 scan=22023 | 44.74 |
| 18408:3 | 26 | R.C[+57.021]MWSSALNSLN[+1864.634]LSFAGLEQVPK.G | HexNAc(2)Hex(9) | C[+57], N[+1865] | 1439.6182 | 3 | 4316.8401 | 4316.8341 | 0 | 1.4 | 272 | Specific | 1142.87 | 1032.17 | 1032.17 | 22.52 | 19 | >sp\|P08571\|CD14_HUMAN Monocyte differentiation antigen CD14 OS=Homo sapiens OX=9606 GN=CD14 PE=1 SV=2 | 1232 | 20201117_fqg_milk_1ng_glyco_hcd203040_compound15.22025.22025.3 | controllerType=0 controllerNumber=1 scan=22025 | 44.7443 |
| 13729:4 | 30 | K.LKELPGVC[+57.021]N[+2205.793]ETMMALWEEC[+57.021]KPC[+57.021]LK.Q | HexNAc(4)Hex(5)Fuc(2)NeuAc(1) | C[+57]*3, N[+2206] | 1286.3048 | 4 | 5142.1974 | 5142.1776 | 0 | 3.9 | 95 | Specific | 617.07 | 514 | 514 | 11.83 | 24 | >sp\|P10909\|CLUS_HUMAN Clusterin OS=Homo sapiens OX=9606 GN=CLU PE=1 SV=1 | 1467 | 20201117_fqg_milk_1ng_glyco_hcd203040_compound15.17123.17123.4 | controllerType=0 controllerNumber=1 scan=17123 | 35.292 |
| 13727:4 | 30 | K.LKELPGVC[+57.021]N[+2205.793]ETMMALWEEC[+57.021]KPC[+57.021]LK.Q | HexNAc(4)Hex(5)Fuc(2)NeuAc(1) | C[+57]*3, N[+2206] | 1286.3048 | 4 | 5142.1974 | 5142.1776 | 0 | 3.9 | 95 | Specific | 636.41 | 636.41 | 636.41 | 11.83 | 24 | >sp\|P10909\|CLUS_HUMAN Clusterin OS=Homo sapiens OX=9606 GN=CLU PE=1 SV=1 | 1467 | 20201117_fqg_milk_1ng_glyco_hcd203040_compound15.17121.17121.4 | controllerType=0 controllerNumber=1 scan=17121 | 35.2874 |
| 20722:3 | 30 | K.[+42.011]MLN[+2018.708]TSSLLEQLNEQFNWVSR.L | HexNAc(3)Hex(6)Fuc(1)NeuAc(1) | .[42], N[+2019] | 1490.6507 | 3 | 4469.9375 | 4469.9057 | 0 | 7.1 | 352 | Specific | 336.16 | 311.07 | 311.07 | 5.94 | 24 | >sp\|P10909\|CLUS_HUMAN Clusterin OS=Homo sapiens OX=9606 GN=CLU PE=1 SV=1 | 1467 | 20201117_fqg_milk_1ng_glyco_hcd203040_compound15.24444.24444.5 | controllerType=0 controllerNumber=1 scan=24444 | [illegible] |
| 20719:3 | 30 | K.[+42.011]MLN[+2018.708]TSSLLEQLNEQFNWVSR.L | HexNAc(3)Hex(6)Fuc(1)NeuAc(1) | .[42], N[+2019] | 1490.6507 | 3 | 4469.9375 | 4469.9057 | 0 | 7.1 | 352 | Specific | 341.77 | 315.87 | 315.87 | 5.94 | 24 | >sp\|P10909\|CLUS_HUMAN Clusterin OS=Homo sapiens OX=9606 GN=CLU PE=1 SV=1 | 1467 | 20201117_fqg_milk_1ng_glyco_hcd203040_compound15.24441.24441.5 | controllerType=0 controllerNumber=1 scan=24441 | [illegible] |
| 09395:3 | 45 | R.IIVPLNNREN[+1663.608]ISDPTSPLR.T | HexNAc(5)Hex(4) | N[+1664] | 1271.2698 | 3 | 3811.7949 | 3811.7853 | 0 | 2.5 | 62 | Specific | 358.49 | 349.49 | 349.49 | 6.92 | 17 | >sp\|P01591\|IGJ_HUMAN Immunoglobulin J chain OS=Homo sapiens OX=9606 GN=JCHAIN PE=1 SV=4 | 1392 | 20201117_fqg_milk_1ng_glyco_hcd203040_compound15.12608.12608.3 | controllerType=0 controllerNumber=1 scan=12608 | [illegible] |
| 09397:3 | 45 | R.IIVPLNNREN[+1663.608]ISDPTSPLR.T | HexNAc(5)Hex(4) | N[+1664] | 1271.2698 | 3 | 3811.7949 | 3811.7853 | 0 | 2.5 | 62 | Specific | 359.25 | 359.25 | 359.25 | 6.92 | 17 | >sp\|P01591\|IGJ_HUMAN Immunoglobulin J chain OS=Homo sapiens OX=9606 GN=JCHAIN PE=1 SV=4 | 1392 | 20201117_fqg_milk_1ng_glyco_hcd203040_compound15.12610.12610.3 | controllerType=0 controllerNumber=1 scan=12610 | [illegible] |
| 09354:3 | 45 | R.IIVPLNNREN[+1460.529]ISDPTSPLR.T | HexNAc(4)Hex(4) | N[+1461] | 1203.5761 | 3 | 3608.7138 | 3608.7059 | 0 | 2.2 | 62 | Specific | 360.37 | 355.94 | 355.94 | 7.05 | 17 | >sp\|P01591\|IGJ_HUMAN Immunoglobulin J chain OS=Homo sapiens OX=9606 GN=JCHAIN PE=1 SV=4 | 1392 | 20201117_fqg_milk_1ng_glyco_hcd203040_compound15.12565.12565.3 | controllerType=0 controllerNumber=1 scan=12565 | 27.4654 |
| 09356:3 | 45 | R.IIVPLNNREN[+1460.529]ISDPTSPLR.T | HexNAc(4)Hex(4) | N[+1461] | 1203.5761 | 3 | 3608.7138 | 3608.7059 | 0 | 2.2 | 62 | Specific | 386.72 | 360.23 | 360.23 | 7.07 | 17 | >sp\|P01591\|IGJ_HUMAN Immunoglobulin J chain OS=Homo sapiens OX=9606 GN=JCHAIN PE=1 SV=4 | 1392 | 20201117_fqg_milk_1ng_glyco_hcd203040_compound15.12567.12567.3 | controllerType=0 controllerNumber=1 scan=12567 | 27.4706 |
| 11012:3 | 45 | R.IIVPLNNREN[+1913.677]ISDPTSPLR.T | HexNAc(4)Hex(5)NeuAc(1) | N[+1914] | 1354.6266 | 3 | 4061.8654 | 4061.8542 | 0 | 2.8 | 62 | Specific | 536.33 | 481.7 | 481.7 | 9.22 | 17 | >sp\|P01591\|IGJ_HUMAN Immunoglobulin J chain OS=Homo sapiens OX=9606 GN=JCHAIN PE=1 SV=4 | 1392 | 20201117_fqg_milk_1ng_glyco_hcd203040_compound15.14279.14279.3 | controllerType=0 controllerNumber=1 scan=14279 | 30.3367 |
| 11015:3 | 45 | R.IIVPLNNREN[+1913.677]ISDPTSPLR.T | HexNAc(4)Hex(5)NeuAc(1) | N[+1914] | 1354.6266 | 3 | 4061.8654 | 4061.8542 | 0 | 2.8 | 62 | Specific | 542.59 | 511.44 | 511.44 | 9.22 | 17 | >sp\|P01591\|IGJ_HUMAN Immunoglobulin J chain OS=Homo sapiens OX=9606 GN=JCHAIN PE=1 SV=4 | 1392 | 20201117_fqg_milk_1ng_glyco_hcd203040_compound15.14282.14282.3 | controllerType=0 controllerNumber=1 scan=14282 | 30.3411 |
| 14343:4 | 55 | K.MVSHHN[+2204.772]LTTGATLINEQWLLTTAK.N | HexNAc(4)Hex(5)NeuAc(2) | N[+2205] | 1221.8022 | 4 | 4884.1868 | 4884.1647 | 0 | 4.5 | 179 | Specific | 222.92 | 222.92 | 222.92 | 5.05 | 16 | >sp\|P00738\|HPT_HUMAN Haptoglobin OS=Homo sapiens OX=9606 GN=HP PE=1 SV=1 | 407 | 20201117_fqg_milk_1ng_glyco_hcd203040_compound15.17761.17761.2 | controllerType=0 controllerNumber=1 scan=17761 | 36.4488 |
| 14341:4 | 55 | K.MVSHHN[+2204.772]LTTGATLINEQWLLTTAK.N | HexNAc(4)Hex(5)NeuAc(2) | N[+2205] | 1221.8022 | 4 | 4884.1868 | 4884.1647 | 0 | 4.5 | 179 | Specific | 252.83 | 252.83 | 252.83 | 6.54 | 16 | >sp\|P00738\|HPT_HUMAN Haptoglobin OS=Homo sapiens OX=9606 GN=HP PE=1 SV=1 | 407 | 20201117_fqg_milk_1ng_glyco_hcd203040_compound15.17759.17759.2 | controllerType=0 controllerNumber=1 scan=17759 | 36.4447 |
| 14085:3 | 55 | K.VVLHPN[+2204.772]YSQVDIGLIK.L | HexNAc(4)Hex(5)NeuAc(2) | N[+2205] | 1333.938 | 3 | 3999.7996 | 3999.7837 | 0 | 4 | 236 | Specific | 441.79 | 398.6 | 398.6 | 9.06 | 16 | >sp\|P00738\|HPT_HUMAN Haptoglobin OS=Homo sapiens OX=9606 GN=HP PE=1 SV=1 | 407 | 20201117_fqg_milk_1ng_glyco_hcd203040_compound15.17491.17491.3 | controllerType=0 controllerNumber=1 scan=17491 | 35.9376 |
| 14083:3 | 55 | K.VVLHPN[+2204.772]YSQVDIGLIK.L | HexNAc(4)Hex(5)NeuAc(2) | N[+2205] | 1333.938 | 3 | 3999.7996 | 3999.7837 | 0 | 4 | 236 | Specific | 491.16 | 481.99 | 481.99 | 9.21 | 16 | >sp\|P00738\|HPT_HUMAN Haptoglobin OS=Homo sapiens OX=9606 GN=HP PE=1 SV=1 | 407 | 20201117_fqg_milk_1ng_glyco_hcd203040_compound15.17489.17489.3 | controllerType=0 controllerNumber=1 scan=17489 | 35.9339 |
| 16435:3 | 60 | K.AALAAFNAQNN[+2204.772]GSNFQLEEISR.A | HexNAc(4)Hex(5)NeuAc(2) | N[+2205] | 1523.9809 | 3 | 4569.9281 | 4569.9256 | 0 | 0.6 | 166 | Specific | 227.76 | 176.54 | 176.54 | 5.71 | 10 | >sp\|P02765\|FETUA_HUMAN Alpha-2-HS-glycoprotein OS=Homo sapiens OX=9606 GN=AHSG PE=1 SV=2 | 135 | 20201117_fqg_milk_1ng_glyco_hcd203040_compound15.19968.19968.3 | controllerType=0 controllerNumber=1 scan=19968 | 40.3902 |
| 16437:3 | 60 | K.AALAAFNAQNN[+2204.772]GSNFQLEEISR.A | HexNAc(4)Hex(5)NeuAc(2) | N[+2205] | 1523.9809 | 3 | 4569.9281 | 4569.9256 | 0 | 0.6 | 166 | Specific | 486.68 | 425.32 | 425.32 | 9.5 | 10 | >sp\|P02765\|FETUA_HUMAN Alpha-2-HS-glycoprotein OS=Homo sapiens OX=9606 GN=AHSG PE=1 SV=2 | 135 | 20201117_fqg_milk_1ng_glyco_hcd203040_compound15.19970.19970.3 | controllerType=0 controllerNumber=1 scan=19970 | 40.3942 |
| 06362:3 | 462 | R.[+42.011]VEDAYILTC[+57.021]N[+730.264]VSLEYEK.T | HexNAc(2)Hex(2) | .[42], C[+57], N[+730] | 940.094 | 3 | 2818.2675 | 2818.2486 | 0 | 6.7 | 225 | Specific | 289.52 | 216.64 | 216.64 | 2.49 | 2 | >sp\|P40227\|TCPZ_HUMAN T-complex protein 1 subunit zeta OS=Homo sapiens OX=9606 GN=CCT6A PE=1 SV=3 | 1115 | 20201117_fqg_milk_1ng_glyco_hcd203040_compound15.9445.9445.3 | controllerType=0 controllerNumber=1 scan=9445 | 22.2342 |
| 01515:3 | 1 | K.GPN[+2059.735]C[+57.021]SEPEC[+57.021]PGNC[+57.021]HLR.G | HexNAc(4)Hex(5)Fuc(1)NeuAc(1) | C[+57]*3, N[+2060] | 1315.1736 | 3 | 3943.5064 | 3943.4931 | 0 | 3.4 | 182 | Specific | 454.48 | 412.4 | 412.4 | 9.46 | 156 | >sp\|P24821\|TENA_HUMAN Tenascin OS=Homo sapiens OX=9606 GN=TNC PE=1 SV=3 | 486 | 20201118_lss_colostrum_1ng_glyco_hcd203040_compound15.4461.4461.3 | controllerType=0 controllerNumber=1 scan=4461 | 13.8298 |
| 04759:5 | 1 | R.C[+57.021]IN[+2350.830]GTC[+57.021]YC[+57.021]EEGFTGEDC[+57.021]GKPTC[ | HexNAc(4)Hex(5)Fuc(1)NeuAc(2) | C[+57]*6, N[+2351] | 1202.8591 | 5 | 6010.2666 | 6010.2587 | 0 | 1.3 | 325 | Specific | 204.05 | 204.05 | 204.05 | 5.27 | 156 | >sp\|P24821\|TENA_HUMAN Tenascin OS=Homo sapiens OX=9606 GN=TNC PE=1 SV=3 | 486 | 20201118_lss_colostrum_1ng_glyco_hcd203040_compound15.7821.7821.5 | controllerType=0 controllerNumber=1 scan=7821 | 19.463 |
| 04750:4 | 1 | R.C[+57.021]IN[+2350.830]GTC[+57.021]YC[+57.021]EEGFTGEDC[+57.021]GKPTC[ | HexNAc(4)Hex(5)Fuc(1)NeuAc(2) | C[+57]*6, N[+2351] | 1503.3206 | 4 | 6010.2604 | 6010.2587 | 0 | 0.3 | 325 | Specific | 313.34 | 313.34 | 313.34 | 6.28 | 156 | >sp\|P24821\|TENA_HUMAN Tenascin OS=Homo sapiens OX=9606 GN=TNC PE=1 SV=3 | 486 | 20201118_lss_colostrum_1ng_glyco_hcd203040_compound15.7812.7812.4 | controllerType=0 controllerNumber=1 scan=7812 | 19.4485 |
| 03747:5 | 1 | R.C[+57.021]IN[+2059.735]GTC[+57.021]YC[+57.021]EEGFTGEDC[+57.021]GKPTC[ | HexNAc(4)Hex(5)Fuc(1)NeuAc(1) | C[+57]*6, N[+2060] | 1144.6417 | 5 | 5719.1795 | 5719.1633 | 0 | 2.8 | 325 | Specific | 424.05 | 424.05 | 424.05 | 9.89 | 156 | >sp\|P24821\|TENA_HUMAN Tenascin OS=Homo sapiens OX=9606 GN=TNC PE=1 SV=3 | 486 | 20201118_lss_colostrum_1ng_glyco_hcd203040_compound15.6773.6773.5 | controllerType=0 controllerNumber=1 scan=6773 | 17.7679 |
| 03743:4 | 1 | R.C[+57.021]IN[+2059.735]GTC[+57.021]YC[+57.021]EEGFTGEDC[+57.021]GKPTC[ | HexNAc(4)Hex(5)Fuc(1)NeuAc(1) | C[+57]*6, N[+2060] | 1430.5507 | 4 | 5719.1808 | 5719.1633 | 0 | 3.1 | 325 | Specific | 482.51 | 482.51 | 482.51 | 9.45 | 156 | >sp\|P24821\|TENA_HUMAN Tenascin OS=Homo sapiens OX=9606 GN=TNC PE=1 SV=3 | 486 | 20201118_lss_colostrum_1ng_glyco_hcd203040_compound15.6769.6769.4 | controllerType=0 controllerNumber=1 scan=6769 | 17.7608 |
| 03745:4 | 1 | R.C[+57.021]IN[+2059.735]GTC[+57.021]YC[+57.021]EEGFTGEDC[+57.021]GKPTC[ | HexNAc(4)Hex(5)Fuc(1)NeuAc(1) | C[+57]*6, N[+2060] | 1430.5507 | 4 | 5719.1808 | 5719.1633 | 0 | 3.1 | 325 | Specific | 546.09 | 546.09 | 546.09 | 9.68 | 156 | >sp\|P24821\|TENA_HUMAN Tenascin OS=Homo sapiens OX=9606 GN=TNC PE=1 SV=3 | 486 | 20201118_lss_colostrum_1ng_glyco_hcd203040_compound15.6771.6771.4 | controllerType=0 controllerNumber=1 scan=6771 | 17.7645 |
| 03749:5 | 1 | R.C[+57.021]IN[+2059.735]GTC[+57.021]YC[+57.021]EEGFTGEDC[+57.021]GKPTC[ | HexNAc(4)Hex(5)Fuc(1)NeuAc(1) | C[+57]*6, N[+2060] | 1144.6417 | 5 | 5719.1795 | 5719.1633 | 0 | 2.8 | 325 | Specific | 609.08 | 609.08 | 609.08 | 12.9 | 156 | >sp\|P24821\|TENA_HUMAN Tenascin OS=Homo sapiens OX=9606 GN=TNC PE=1 SV=3 | 486 | 20201118_lss_colostrum_1ng_glyco_hcd203040_compound15.6775.6775.5 | controllerType=0 controllerNumber=1 scan=6775 | 17.7716 |
| 08911:4 | 1 | R.QTGLAPGQEYEISLHIVKN[+2205.793]NTR.G | HexNAc(4)Hex(5)Fuc(2)NeuAc(1) | N[+2206] | 1169.2776 | 4 | 4674.0887 | 4674.0821 | 0 | 1.4 | 770 | Specific | 208.79 | 133.7 | 133.7 | 3.25 | 156 | >sp\|P24821\|TENA_HUMAN Tenascin OS=Homo sapiens OX=9606 GN=TNC PE=1 SV=3 | 486 | 20201118_lss_colostrum_1ng_glyco_hcd203040_compound15.12126.12126.4 | controllerType=0 controllerNumber=1 scan=12126 | 26.5669 |
| 08837:3 | 1 | R.QTGLAPGQEYEISLHIVKN[+2204.772]NTR.G | HexNAc(4)Hex(5)NeuAc(2) | N[+2205] | 1558.3644 | 3 | 4673.0788 | 4673.0617 | 0 | 3.7 | 770 | Specific | 316.99 | 99.25 | 99.25 | 4.54 | 156 | >sp\|P24821\|TENA_HUMAN Tenascin OS=Homo sapiens OX=9606 GN=TNC PE=1 SV=3 | 486 | 20201118_lss_colostrum_1ng_glyco_hcd203040_compound15.12049.12049.3 | controllerType=0 controllerNumber=1 scan=12049 | 26.4302 |
| 08812:3 | 1 | R.QTGLAPGQEYEISLHIVKN[+2351.851]NTR.G | HexNAc(4)Hex(5)Fuc(3)NeuAc(1) | N[+2352] | 1607.3879 | 3 | 4820.1491 | 4820.14 | 0 | 1.9 | 770 | Specific | 357.83 | 328.74 | 328.74 | 6.61 | 156 | >sp\|P24821\|TENA_HUMAN Tenascin OS=Homo sapiens OX=9606 GN=TNC PE=1 SV=3 | 486 | 20201118_lss_colostrum_1ng_glyco_hcd203040_compound15.12023.12023.3 | controllerType=0 controllerNumber=1 scan=12023 | 26.3842 |
| 08839:3 | 1 | R.QTGLAPGQEYEISLHIVKN[+2204.772]NTR.G | HexNAc(4)Hex(5)NeuAc(2) | N[+2205] | 1558.3644 | 3 | 4673.0788 | 4673.0617 | 0 | 3.7 | 770 | Specific | 377.03 | 221.51 | 221.51 | 6.11 | 156 | >sp\|P24821\|TENA_HUMAN Tenascin OS=Homo sapiens OX=9606 GN=TNC PE=1 SV=3 | 486 | 20201118_lss_colostrum_1ng_glyco_hcd203040_compound15.12051.12051.3 | controllerType=0 controllerNumber=1 scan=12051 | 26.4341 |
| 08987:4 | 1 | R.QTGLAPGQEYEISLHIVKN[+1913.677]NTR.G | HexNAc(4)Hex(5)NeuAc(1) | N[+1914] | 1096.2483 | 4 | 4381.9715 | 4381.9662 | 0 | 1.2 | 770 | Specific | 405.36 | 318.57 | 318.57 | 7.35 | 156 | >sp\|P24821\|TENA_HUMAN Tenascin OS=Homo sapiens OX=9606 GN=TNC PE=1 SV=3 | 486 | 20201118_lss_colostrum_1ng_glyco_hcd203040_compound15.12205.12205.4 | controllerType=0 controllerNumber=1 scan=12205 | 26.7101 |
| 08799:4 | 1 | R.QTGLAPGQEYEISLHIVKN[+2350.830]NTR.G | HexNAc(4)Hex(5)Fuc(1)NeuAc(2) | N[+2351] | 1205.5414 | 4 | 4819.1439 | 4819.1196 | 0 | 5 | 770 | Specific | 425.11 | 328.53 | 328.53 | 6.15 | 156 | >sp\|P24821\|TENA_HUMAN Tenascin OS=Homo sapiens OX=9606 GN=TNC PE=1 SV=3 | 486 | 20201118_lss_colostrum_1ng_glyco_hcd203040_compound15.12009.12009.4 | controllerType=0 controllerNumber=1 scan=12009 | 26.3604 |
| 08797:4 | 1 | R.QTGLAPGQEYEISLHIVKN[+2350.830]NTR.G | HexNAc(4)Hex(5)Fuc(1)NeuAc(2) | N[+2351] | 1205.5414 | 4 | 4819.1439 | 4819.1196 | 0 | 5 | 770 | Specific | 450.87 | 364.48 | 364.48 | 7.45 | 156 | >sp\|P24821\|TENA_HUMAN Tenascin OS=Homo sapiens OX=9606 GN=TNC PE=1 SV=3 | 486 | 20201118_lss_colostrum_1ng_glyco_hcd203040_compound15.12007.12007.4 | controllerType=0 controllerNumber=1 scan=12007 | 26.3555 |
| 08989:4 | 1 | R.QTGLAPGQEYEISLHIVKN[+1913.677]NTR.G | HexNAc(4)Hex(5)NeuAc(1) | N[+1914] | 1096.2483 | 4 | 4381.9715 | 4381.9662 | 0 | 1.2 | 770 | Specific | 455.38 | 401.3 | 401.3 | 9.46 | 156 | >sp\|P24821\|TENA_HUMAN Tenascin OS=Homo sapiens OX=9606 GN=TNC PE=1 SV=3 | 486 | 20201118_lss_colostrum_1ng_glyco_hcd203040_compound15.12207.12207.4 | controllerType=0 controllerNumber=1 scan=12207 | 26.7146 |
| 08835:4 | 1 | R.QTGLAPGQEYEISLHIVKN[+2059.735]NTR.G | HexNAc(4)Hex(5)Fuc(1)NeuAc(1) | N[+2060] | 1132.7641 | 4 | 4528.0345 | 4528.0241 | 0 | 2.3 | 770 | Specific | 569.99 | 418.8 | 418.8 | 10.19 | 156 | >sp\|P24821\|TENA_HUMAN Tenascin OS=Homo sapiens OX=9606 GN=TNC PE=1 SV=3 | 486 | 20201118_lss_colostrum_1ng_glyco_hcd203040_compound15.12047.12047.2 | controllerType=0 controllerNumber=1 scan=12047 | 26.4269 |
| 08833:4 | 1 | R.QTGLAPGQEYEISLHIVKN[+2059.735]NTR.G | HexNAc(4)Hex(5)Fuc(1)NeuAc(1) | N[+2060] | 1132.7641 | 4 | 4528.0345 | 4528.0241 | 0 | 2.3 | 770 | Specific | 609.45 | 440.09 | 440.09 | 12.52 | 156 | >sp\|P24821\|TENA_HUMAN Tenascin OS=Homo sapiens OX=9606 GN=TNC PE=1 SV=3 | 486 | 20201118_lss_colostrum_1ng_glyco_hcd203040_compound15.12045.12045.2 | controllerType=0 controllerNumber=1 scan=12045 | 26.422 |
| 08859:3 | 1 | R.QTGLAPGQEYEISLHIVKN[+2059.735]NTR.G | HexNAc(4)Hex(5)Fuc(1)NeuAc(1) | N[+2060] | 1510.0181 | 3 | 4528.0398 | 4528.0241 | 0 | 3.5 | 770 | Specific | 673.05 | 398.05 | 398.05 | 12.7 | 156 | >sp\|P24821\|TENA_HUMAN Tenascin OS=Homo sapiens OX=9606 GN=TNC PE=1 SV=3 | 486 | 20201118_lss_colostrum_1ng_glyco_hcd203040_compound15.12072.12072.3 | controllerType=0 controllerNumber=1 scan=12072 | 26.4737 |
| 08857:3 | 1 | R.QTGLAPGQEYEISLHIVKN[+2059.735]NTR.G | HexNAc(4)Hex(5)Fuc(1)NeuAc(1) | N[+2060] | 1510.0181 | 3 | 4528.0398 | 4528.0241 | 0 | 3.5 | 770 | Specific | 769.25 | 553.77 | 553.77 | 14.7 | 156 | >sp\|P24821\|TENA_HUMAN Tenascin OS=Homo sapiens OX=9606 GN=TNC PE=1 SV=3 | 486 | 20201118_lss_colostrum_1ng_glyco_hcd203040_compound15.12070.12070.3 | controllerType=0 controllerNumber=1 scan=12070 | 26.4681 |
| 14830:2 | 1 | R.LN[+1216.423]YSLPTGQWVGVQLPR.N | HexNAc(2)Hex(5) | N[+1216] | 1572.7392 | 2 | 3144.4711 | 3144.4617 | 0 | 3 | 1017 | Specific | 849.62 | 769.34 | 769.34 | 16.56 | 156 | >sp\|P24821\|TENA_HUMAN Tenascin OS=Homo sapiens OX=9606 GN=TNC PE=1 SV=3 | 486 | 20201118_lss_colostrum_1ng_glyco_hcd203040_compound15.18272.18272.2 | controllerType=0 controllerNumber=1 scan=18272 | 38.079 |
| 14827:2 | 1 | R.LN[+1216.423]YSLPTGQWVGVQLPR.N | HexNAc(2)Hex(5) | N[+1216] | 1572.7392 | 2 | 3144.4711 | 3144.4617 | 0 | 3 | 1017 | Specific | 874.9 | 840.28 | 840.28 | 17.56 | 156 | >sp\|P24821\|TENA_HUMAN Tenascin OS=Homo sapiens OX=9606 GN=TNC PE=1 SV=3 | 486 | 20201118_lss_colostrum_1ng_glyco_hcd203040_compound15.18269.18269.2 | controllerType=0 controllerNumber=1 scan=18269 | 38.0749 |
| 14629:3 | 1 | R.LN[+1702.581]YSLPTGQWVGVQLPR.N | HexNAc(2)Hex(8) | N[+1703] | 1210.8775 | 3 | 3630.6179 | 3630.6202 | 0 | -0.6 | 1017 | Specific | 875.95 | 806.09 | 806.09 | 16.87 | 156 | >sp\|P24821\|TENA_HUMAN Tenascin OS=Homo sapiens OX=9606 GN=TNC PE=1 SV=3 | 486 | 20201118_lss_colostrum_1ng_glyco_hcd203040_compound15.18063.18063.2 | controllerType=0 controllerNumber=1 scan=18063 | 37.6445 |
| 14669:3 | 1 | R.LN[+1702.581]YSLPTGQWVGVQLPR.N | HexNAc(2)Hex(8) | N[+1703] | 1210.8807 | 3 | 3630.6275 | 3630.6202 | 0 | 2 | 1017 | Specific | 897.77 | 839.84 | 839.84 | 17.49 | 156 | >sp\|P24821\|TENA_HUMAN Tenascin OS=Homo sapiens OX=9606 GN=TNC PE=1 SV=3 | 486 | 20201118_lss_colostrum_1ng_glyco_hcd203040_compound15.18105.18105.3 | controllerType=0 controllerNumber=1 scan=18105 | 37.7411 |
| 14758:3 | 1 | R.LN[+1378.476]YSLPTGQWVGVQLPR.N | HexNAc(2)Hex(6) | N[+1378] | 1102.8463 | 3 | 3306.5245 | 3306.5145 | 0 | 3 | 1017 | Specific | 1009.51 | 1009.51 | 1009.51 | 20.49 | 156 | >sp\|P24821\|TENA_HUMAN Tenascin OS=Homo sapiens OX=9606 GN=TNC PE=1 SV=3 | 486 | 20201118_lss_colostrum_1ng_glyco_hcd203040_compound15.18197.18197.3 | controllerType=0 controllerNumber=1 scan=18197 | 37.9184 |
| 14631:3 | 1 | R.LN[+1702.581]YSLPTGQWVGVQLPR.N | HexNAc(2)Hex(8) | N[+1703] | 1210.8775 | 3 | 3630.6179 | 3630.6202 | 0 | -0.6 | 1017 | Specific | 1031.44 | 832.33 | 832.33 | 19.87 | 156 | >sp\|P24821\|TENA_HUMAN Tenascin OS=Homo sapiens OX=9606 GN=TNC PE=1 SV=3 | 486 | 20201118_lss_colostrum_1ng_glyco_hcd203040_compound15.18065.18065.2 | controllerType=0 controllerNumber=1 scan=18065 | 37.6583 |
| 14671:3 | 1 | R.LN[+1702.581]YSLPTGQWVGVQLPR.N | HexNAc(2)Hex(8) | N[+1703] | 1210.8807 | 3 | 3630.6275 | 3630.6202 | 0 | 2 | 1017 | Specific | 1067.17 | 887.45 | 887.45 | 21.49 | 156 | >sp\|P24821\|TENA_HUMAN Tenascin OS=Homo sapiens OX=9606 GN=TNC PE=1 SV=3 | 486 | 20201118_lss_colostrum_1ng_glyco_hcd203040_compound15.18107.18107.3 | controllerType=0 controllerNumber=1 scan=18107 | 37.7449 |
| 14811:3 | 1 | R.LN[+1216.423]YSLPTGQWVGVQLPR.N | HexNAc(2)Hex(5) | N[+1216] | 1048.8297 | 3 | 3144.4747 | 3144.4617 | 0 | 4.1 | 1017 | Specific | 1078.58 | 1078.58 | 1078.58 | 21.29 | 156 | >sp\|P24821\|TENA_HUMAN Tenascin OS=Homo sapiens OX=9606 GN=TNC PE=1 SV=3 | 486 | 20201118_lss_colostrum_1ng_glyco_hcd203040_compound15.18252.18252.3 | controllerType=0 controllerNumber=1 scan=18252 | 38.0415 |
| 14814:3 | 1 | R.LN[+1216.423]YSLPTGQWVGVQLPR.N | HexNAc(2)Hex(5) | N[+1216] | 1048.8297 | 3 | 3144.4747 | 3144.4617 | 0 | 4.1 | 1017 | Specific | 1100.59 | 938.48 | 938.48 | 22.29 | 156 | >sp\|P24821\|TENA_HUMAN Tenascin OS=Homo sapiens OX=9606 GN=TNC PE=1 SV=3 | 486 | 20201118_lss_colostrum_1ng_glyco_hcd203040_compound15.18255.18255.3 | controllerType=0 controllerNumber=1 scan=18255 | 38.0483 |
| 14761:3 | 1 | R.LN[+1378.476]YSLPTGQWVGVQLPR.N | HexNAc(2)Hex(6) | N[+1378] | 1102.8463 | 3 | 3306.5245 | 3306.5145 | 0 | 3 | 1017 | Specific | 1112.99 | 923.65 | 923.65 | 22.49 | 156 | >sp\|P24821\|TENA_HUMAN Tenascin OS=Homo sapiens OX=9606 GN=TNC PE=1 SV=3 | 486 | 20201118_lss_colostrum_1ng_glyco_hcd203040_compound15.18200.18200.3 | controllerType=0 controllerNumber=1 scan=18200 | 37.9243 |
| 01858:2 | 1 | R.N[+1216.423]TTSYVLR.G | HexNAc(2)Hex(5) | N[+1216] | 1085.4709 | 2 | 2169.9345 | 2169.9279 | 0 | 3 | 1034 | Specific | 241.24 | 152.87 | 152.87 | 5.21 | 156 | >sp\|P24821\|TENA_HUMAN Tenascin OS=Homo sapiens OX=9606 GN=TNC PE=1 SV=3 | 486 | 20201118_lss_colostrum_1ng_glyco_hcd203040_compound15.4818.4818.2 | controllerType=0 controllerNumber=1 scan=4818 | 14.4329 |
| 03709:3 | 1 | R.N[+2059.735]TTSYVLR.G | HexNAc(4)Hex(5)Fuc(1)NeuAc(1) | N[+2060] | 1005.0885 | 3 | 3013.2509 | 3013.24 | 0 | 3.6 | 1034 | Specific | 334.46 | 216.98 | 216.98 | 6.27 | 156 | >sp\|P24821\|TENA_HUMAN Tenascin OS=Homo sapiens OX=9606 GN=TNC PE=1 SV=3 | 486 | 20201118_lss_colostrum_1ng_glyco_hcd203040_compound15.6734.6734.3 | controllerType=0 controllerNumber=1 scan=6734 | 17.7041 |
| 03687:2 | 1 | R.N[+2059.735]TTSYVLR.G | HexNAc(4)Hex(5)Fuc(1)NeuAc(1) | N[+2060] | 1507.1275 | 2 | 3013.2477 | 3013.24 | 0 | 2.6 | 1034 | Specific | 357.77 | 193.65 | 193.65 | 6.44 | 156 | >sp\|P24821\|TENA_HUMAN Tenascin OS=Homo sapiens OX=9606 GN=TNC PE=1 SV=3 | 486 | 20201118_lss_colostrum_1ng_glyco_hcd203040_compound15.6711.6711.2 | controllerType=0 controllerNumber=1 scan=6711 | 17.6643 |
| 01860:2 | 1 | R.N[+1216.423]TTSYVLR.G | HexNAc(2)Hex(5) | N[+1216] | 1085.4709 | 2 | 2169.9345 | 2169.9279 | 0 | 3 | 1034 | Specific | 365.36 | 259.22 | 259.22 | 6.93 | 156 | >sp\|P24821\|TENA_HUMAN Tenascin OS=Homo sapiens OX=9606 GN=TNC PE=1 SV=3 | 486 | 20201118_lss_colostrum_1ng_glyco_hcd203040_compound15.4820.4820.2 | controllerType=0 controllerNumber=1 scan=4820 | 14.437 |
| 07608:3 | 1 | R.N[+2059.735]LTVPGSLR.A | HexNAc(4)Hex(5)Fuc(1)NeuAc(1) | N[+2060] | 1006.1009 | 3 | 3016.288 | 3016.2873 | 0 | 0.2 | 1119 | Specific | 213.97 | 129.31 | 129.31 | 3.31 | 156 | >sp\|P24821\|TENA_HUMAN Tenascin OS=Homo sapiens OX=9606 GN=TNC PE=1 SV=3 | 486 | 20201118_lss_colostrum_1ng_glyco_hcd203040_compound15.10769.10769.3 | controllerType=0 controllerNumber=1 scan=10769 | 24.2821 |
| 07610:3 | 1 | R.N[+2059.735]LTVPGSLR.A | HexNAc(4)Hex(5)Fuc(1)NeuAc(1) | N[+2060] | 1006.1009 | 3 | 3016.288 | 3016.2873 | 0 | 0.2 | 1119 | Specific | 234.11 | 90.85 | 90.85 | 3.44 | 156 | >sp\|P24821\|TENA_HUMAN Tenascin OS=Homo sapiens OX=9606 GN=TNC PE=1 SV=3 | 486 | 20201118_lss_colostrum_1ng_glyco_hcd203040_compound15.10771.10771.3 | controllerType=0 controllerNumber=1 scan=10771 | 24.2861 |
| 09860:3 | 2 | K.TVLTPATNHMGN[+1378.476]VTFTIPANR.E | HexNAc(2)Hex(6) | N[+1378] | 1211.8865 | 3 | 3633.645 | 3633.6358 | 0 | 2.5 | 74 | Specific | 489.79 | 489.79 | 225.4 | 9.92 | 148 | >sp\|P01024\|CO3_HUMAN Complement C3 OS=Homo sapiens OX=9606 GN=C3 PE=1 SV=2 | 492 | 20201118_lss_colostrum_1ng_glyco_hcd203040_compound15.13112.13112.3 | controllerType=0 controllerNumber=1 scan=13112 | 28.326 |
| 09857:3 | 2 | K.TVLTPATNHMGN[+1378.476]VTFTIPANR.E | HexNAc(2)Hex(6) | N[+1378] | 1211.8865 | 3 | 3633.645 | 3633.6358 | 0 | 2.5 | 74 | Specific | 670.59 | 670.59 | 390.9 | 13.7 | 148 | >sp\|P01024\|CO3_HUMAN Complement C3 OS=Homo sapiens OX=9606 GN=C3 PE=1 SV=2 | 492 | 20201118_lss_colostrum_1ng_glyco_hcd203040_compound15.13109.13109.3 | controllerType=0 controllerNumber=1 scan=13109 | 28.3163 |
| 09806:3 | 2 | K.TVLTPATNHMGN[+1540.529]VTFTIPANR.E | HexNAc(2)Hex(7) | N[+1541] | 1265.9036 | 3 | 3795.6963 | 3795.6886 | 0 | 2 | 74 | Specific | 704.28 | 487.58 | 382.68 | 14.7 | 148 | >sp\|P01024\|CO3_HUMAN Complement C3 OS=Homo sapiens OX=9606 GN=C3 PE=1 SV=2 | 492 | 20201118_lss_colostrum_1ng_glyco_hcd203040_compound15.13056.13056.3 | controllerType=0 controllerNumber=1 scan=13056 | 28.206 |
| 09753:3 | 2 | K.TVLTPATNHMGN[+1540.529]VTFTIPANR.E | HexNAc(2)Hex(7) | N[+1541] | 1265.9048 | 3 | 3795.7 | 3795.6886 | 0 | 3 | 74 | Specific | 893.08 | 797.06 | 422.28 | 17.7 | 148 | >sp\|P01024\|CO3_HUMAN Complement C3 OS=Homo sapiens OX=9606 GN=C3 PE=1 SV=2 | 492 | 20201118_lss_colostrum_1ng_glyco_hcd203040_compound15.13000.13000.3 | controllerType=0 controllerNumber=1 scan=13000 | 28.103 |
| 09808:3 | 2 | K.TVLTPATNHMGN[+1540.529]VTFTIPANR.E | HexNAc(2)Hex(7) | N[+1541] | 1265.9036 | 3 | 3795.6963 | 3795.6886 | 0 | 2 | 74 | Specific | 898.81 | 798.61 | 442.45 | 17.7 | 148 | >sp\|P01024\|CO3_HUMAN Complement C3 OS=Homo sapiens OX=9606 GN=C3 PE=1 SV=2 | 492 | 20201118_lss_colostrum_1ng_glyco_hcd203040_compound15.13058.13058.3 | controllerType=0 controllerNumber=1 scan=13058 | 28.2098 |
| 09820:3 | 2 | K.TVLTPATNHMGN[+1378.476]VTFTIPANR.E | HexNAc(2)Hex(6) | N[+1378] | 1211.8863 | 3 | 3633.6443 | 3633.6358 | 0 | 2.3 | 74 | Specific | 979.24 | 952.79 | 539.19 | 19.7 | 148 | >sp\|P01024\|CO3_HUMAN Complement C3 OS=Homo sapiens OX=9606 GN=C3 PE=1 SV=2 | 492 | 20201118_lss_colostrum_1ng_glyco_hcd203040_compound15.13070.13070.3 | controllerType=0 controllerNumber=1 scan=13070 | 28.2332 |
| 09750:3 | 2 | K.TVLTPATNHMGN[+1540.529]VTFTIPANR.E | HexNAc(2)Hex(7) | N[+1541] | 1265.9048 | 3 | 3795.7 | 3795.6886 | 0 | 3 | 74 | Specific | 1104.42 | 1042.69 | 580.92 | 22.7 | 148 | >sp\|P01024\|CO3_HUMAN Complement C3 OS=Homo sapiens OX=9606 GN=C3 PE=1 SV=2 | 492 | 20201118_lss_colostrum_1ng_glyco_hcd203040_compound15.12997.12997.3 | controllerType=0 controllerNumber=1 scan=12997 | 28.0959 |
| 09818:3 | 2 | K.TVLTPATNHMGN[+1378.476]VTFTIPANR.E | HexNAc(2)Hex(6) | N[+1378] | 1211.8863 | 3 | 3633.6443 | 3633.6358 | 0 | 2.3 | 74 | Specific | 1178.35 | 1134.91 | 635.89 | 23.7 | 148 | >sp\|P01024\|CO3_HUMAN Complement C3 OS=Homo sapiens OX=9606 GN=C3 PE=1 SV=2 | 492 | 20201118_lss_colostrum_1ng_glyco_hcd203040_compound15.13068.13068.3 | controllerType=0 controllerNumber=1 scan=13068 | 28.2276 |
| 18783:3 | 3 | R.RTAGWNVPIGTLRPFLN[+1930.692]WTGPPEPIEAAVAR.F | HexNAc(4)Hex(6)Fuc(1) | N[+1931] | 1773.1572 | 3 | 5317.457 | 5317.5044 | 0 | -8.9 | 140 | Specific | 282.71 | 282.71 | 282.71 | 4.78 | 146 | >sp\|P02788\|TRFL_HUMAN Lactotransferrin OS=Homo sapiens OX=9606 GN=LTF PE=1 SV=6 | 1270 | 20201118_lss_colostrum_1ng_glyco_hcd203040_compound15.22390.22390.3 | controllerType=0 controllerNumber=1 scan=22390 | 46.5384 |
| 16477:4 | 3 | R.RTAGWNVPIGTLRPFLN[+1768.640]WTGPPEPIEAAVAR.F | HexNAc(4)Hex(5)Fuc(1) | N[+1769] | 1289.6222 | 4 | 5155.467 | 5155.4516 | 0 | 3 | 140 | Specific | 535.12 | 383.02 | 383.02 | 10.3 | 146 | >sp\|P02788\|TRFL_HUMAN Lactotransferrin OS=Homo sapiens OX=9606 GN=LTF PE=1 SV=6 | 1270 | 20201118_lss_colostrum_1ng_glyco_hcd203040_compound15.19987.19987.4 | controllerType=0 controllerNumber=1 scan=19987 | 41.3879 |
| 17303:4 | 3 | R.RTAGWNVPIGTLRPFLN[+2716.983]WTGPPEPIEAAVAR.F | HexNAc(5)Hex(6)Fuc(3)NeuAc(1) | N[+2717] | 1526.7092 | 4 | 6103.8151 | 6103.795 | 0 | 3.3 | 140 | Specific | 558.38 | 427.19 | 427.19 | 10.36 | 146 | >sp\|P02788\|TRFL_HUMAN Lactotransferrin OS=Homo sapiens OX=9606 GN=LTF PE=1 SV=6 | 1270 | 20201118_lss_colostrum_1ng_glyco_hcd203040_compound15.20847.20847.4 | controllerType=0 controllerNumber=1 scan=20847 | 43.2472 |
| 17321:4 | 3 | R.RTAGWNVPIGTLRPFLN[+2570.925]WTGPPEPIEAAVAR.F | HexNAc(5)Hex(6)Fuc(2)NeuAc(1) | N[+2571] | 1490.1936 | 4 | 5957.7526 | 5957.7371 | 0 | 2.6 | 140 | Specific | 563.37 | 461.51 | 461.51 | 10.56 | 146 | >sp\|P02788\|TRFL_HUMAN Lactotransferrin OS=Homo sapiens OX=9606 GN=LTF PE=1 SV=6 | 1270 | 20201118_lss_colostrum_1ng_glyco_hcd203040_compound15.20866.20866.4 | controllerType=0 controllerNumber=1 scan=20866 | 43.3009 |
| 17319:4 | 3 | R.RTAGWNVPIGTLRPFLN[+2570.925]WTGPPEPIEAAVAR.F | HexNAc(5)Hex(6)Fuc(2)NeuAc(1) | N[+2571] | 1490.1936 | 4 | 5957.7526 | 5957.7371 | 0 | 2.6 | 140 | Specific | 566.59 | 551.19 | 551.19 | 10.56 | 146 | >sp\|P02788\|TRFL_HUMAN Lactotransferrin OS=Homo sapiens OX=9606 GN=LTF PE=1 SV=6 | 1270 | 20201118_lss_colostrum_1ng_glyco_hcd203040_compound15.20864.20864.4 | controllerType=0 controllerNumber=1 scan=20864 | 43.295 |
| 17440:4 | 3 | R.RTAGWNVPIGTLRPFLN[+2059.735]WTGPPEPIEAAVAR.F | HexNAc(4)Hex(5)Fuc(1)NeuAc(1) | N[+2060] | 1362.3956 | 4 | 5446.5607 | 5446.547 | 0 | 2.5 | 140 | Specific | 569.17 | 444.08 | 444.08 | 10.56 | 146 | >sp\|P02788\|TRFL_HUMAN Lactotransferrin OS=Homo sapiens OX=9606 GN=LTF PE=1 SV=6 | 1270 | 20201118_lss_colostrum_1ng_glyco_hcd203040_compound15.20989.20989.4 | controllerType=0 controllerNumber=1 scan=20989 | 43.5291 |
| 17488:4 | 3 | R.RTAGWNVPIGTLRPFLN[+2351.851]WTGPPEPIEAAVAR.F | HexNAc(4)Hex(5)Fuc(3)NeuAc(1) | N[+2352] | 1435.4238 | 4 | 5738.6735 | 5738.6628 | 0 | 1.9 | 140 | Specific | 579.26 | 354.98 | 354.98 | 10.03 | 146 | >sp\|P02788\|TRFL_HUMAN Lactotransferrin OS=Homo sapiens OX=9606 GN=LTF PE=1 SV=6 | 1270 | 20201118_lss_colostrum_1ng_glyco_hcd203040_compound15.21039.21039.4 | controllerType=0 controllerNumber=1 scan=21039 | 43.6473 |
| 17306:4 | 3 | R.RTAGWNVPIGTLRPFLN[+2716.983]WTGPPEPIEAAVAR.F | HexNAc(5)Hex(6)Fuc(3)NeuAc(1) | N[+2717] | 1526.7092 | 4 | 6103.8151 | 6103.795 | 0 | 3.3 | 140 | Specific | 581.92 | 482.46 | 482.46 | 11.7 | 146 | >sp\|P02788\|TRFL_HUMAN Lactotransferrin OS=Homo sapiens OX=9606 GN=LTF PE=1 SV=6 | 1270 | 20201118_lss_colostrum_1ng_glyco_hcd203040_compound15.20850.20850.4 | controllerType=0 controllerNumber=1 scan=20850 | 43.2571 |
| 17442:4 | 3 | R.RTAGWNVPIGTLRPFLN[+2059.735]WTGPPEPIEAAVAR.F | HexNAc(4)Hex(5)Fuc(1)NeuAc(1) | N[+2060] | 1362.3956 | 4 | 5446.5607 | 5446.547 | 0 | 2.5 | 140 | Specific | 622.03 | 530.41 | 530.41 | 12.9 | 146 | >sp\|P02788\|TRFL_HUMAN Lactotransferrin OS=Homo sapiens OX=9606 GN=LTF PE=1 SV=6 | 1270 | 20201118_lss_colostrum_1ng_glyco_hcd203040_compound15.20991.20991.4 | controllerType=0 controllerNumber=1 scan=20991 | 43.5372 |
| 16479:4 | 3 | R.RTAGWNVPIGTLRPFLN[+1768.640]WTGPPEPIEAAVAR.F | HexNAc(4)Hex(5)Fuc(1) | N[+1769] | 1289.6222 | 4 | 5155.467 | 5155.4516 | 0 | 3 | 140 | Specific | 646.26 | 574.79 | 574.79 | 12.9 | 146 | >sp\|P02788\|TRFL_HUMAN Lactotransferrin OS=Homo sapiens OX=9606 GN=LTF PE=1 SV=6 | 1270 | 20201118_lss_colostrum_1ng_glyco_hcd203040_compound15.19989.19989.4 | controllerType=0 controllerNumber=1 scan=19989 | 41.3931 |
| 19992:4 | 3 | R.[+42.011]TAGWNVPIGTLRPFLN[+2336.851]WTGPPEPIEAAVAR.F | HexNAc(6)Hex(6)Fuc(1) | .[42], N[+2337] | 1403.1434 | 4 | 5609.5519 | 5609.5726 | 0 | -3.7 | 141 | Specific | 239.81 | 222.57 | 222.57 | 5.24 | 146 | >sp\|P02788\|TRFL_HUMAN Lactotransferrin OS=Homo sapiens OX=9606 GN=LTF PE=1 SV=6 | 1270 | 20201118_lss_colostrum_1ng_glyco_hcd203040_compound15.23651.23651.4 | controllerType=0 controllerNumber=1 scan=23651 | 49.3681 |
| 20006:5 | 3 | R.[+42.011]TAGWNVPIGTLRPFLN[+2336.851]WTGPPEPIEAAVAR.F | HexNAc(6)Hex(6)Fuc(1) | .[42], N[+2337] | 1122.7166 | 5 | 5609.5537 | 5609.5726 | 0 | -3.4 | 141 | Specific | 269.63 | 256.99 | 256.99 | 4.55 | 146 | >sp\|P02788\|TRFL_HUMAN Lactotransferrin OS=Homo sapiens OX=9606 GN=LTF PE=1 SV=6 | 1270 | 20201118_lss_colostrum_1ng_glyco_hcd203040_compound15.23665.23665.5 | controllerType=0 controllerNumber=1 scan=23665 | 49.3913 |
| 17676:4 | 3 | R.[+42.011]TAGWNVPIGTLRPFLN[+1872.651]WTGPPEPIEAAVAR.F | HexNAc(3)Hex(6)NeuAc(1) | .[42], N[+1873] | 1287.1064 | 4 | 5145.404 | 5145.372 | 0 | 6.2 | 141 | Specific | 285.41 | 244.95 | 244.95 | 5.94 | 146 | >sp\|P02788\|TRFL_HUMAN Lactotransferrin OS=Homo sapiens OX=9606 GN=LTF PE=1 SV=6 | 1270 | 20201118_lss_colostrum_1ng_glyco_hcd203040_compound15.21236.21236.4 | controllerType=0 controllerNumber=1 scan=21236 | 44.0614 |
| 19994:4 | 3 | R.[+42.011]TAGWNVPIGTLRPFLN[+2336.851]WTGPPEPIEAAVAR.F | HexNAc(6)Hex(6)Fuc(1) | .[42], N[+2337] | 1403.1434 | 4 | 5609.5519 | 5609.5726 | 0 | -3.7 | 141 | Specific | 320.05 | 320.05 | 320.05 | 6.09 | 146 | >sp\|P02788\|TRFL_HUMAN Lactotransferrin OS=Homo sapiens OX=9606 GN=LTF PE=1 SV=6 | 1270 | 20201118_lss_colostrum_1ng_glyco_hcd203040_compound15.23653.23653.4 | controllerType=0 controllerNumber=1 scan=23653 | 49.3722 |

Table S4. Cont.

| Query #:z | Protein Rank | Glycopeptide | Glycan | Modification Type(s) | Observed m/z | z | Observed (M+H) | Calc. mass (M+H) | Off-by-x error | Mass error (ppm) | StartPosition | Cleavage | Score | Delta | Delta Mod | \|Log Prob\| | # of unique peptides | Protein Name | Protein DB number | Comment | Scan # | Scan Time |
|---|---|---|---|---|---|---|---|---|---|---|---|---|---|---|---|---|---|---|---|---|---|---|
| 18537:4 | 3 | R.TAGWNVPIGTLRPFLN[+2863.041]WTGPPEPIEAAVAR.F | HexNAc(5)Hex(6)Fuc(4)NeuAc(1) | N[+2863] | 1524.1964 | 4 | 6093.7638 | 6093.7518 | 0 | 2 | 141 | Specific | 330.49 | 330.49 | 330.49 | 7.46 | 146 | >sp\|P02788\|TRFL_HUMAN Lactotransferrin OS=Homo sapiens OX=9606 GN=LTF PE=1 SV=6 | 1270 | 20201118_lss_colostrum_1ng_glyco_hcd203040_compound15.22134.22134.4 | controllerType=0 controllerNumber=1 scan=22134 | 45.9902 |
| 18612:4 | 3 | R.TAGWNVPIGTLRPFLN[+2716.983]WTGPPEPIEAAVAR.F | HexNAc(5)Hex(6)Fuc(3)NeuAc(1) | N[+2717] | 1487.683 | 4 | 5947.7101 | 5947.6939 | 0 | 2.7 | 141 | Specific | 405.97 | 401.6 | 401.6 | 9.74 | 146 | >sp\|P02788\|TRFL_HUMAN Lactotransferrin OS=Homo sapiens OX=9606 GN=LTF PE=1 SV=6 | 1270 | 20201118_lss_colostrum_1ng_glyco_hcd203040_compound15.22212.22212.4 | controllerType=0 controllerNumber=1 scan=22212 | 46.163 |
| 18523:3 | 3 | R.TAGWNVPIGTLRPFLN[+2716.983]WTGPPEPIEAAVAR.F | HexNAc(5)Hex(6)Fuc(3)NeuAc(1) | N[+2717] | 1983.2429 | 3 | 5947.7141 | 5947.6939 | 0 | 3.4 | 141 | Specific | 407.46 | 314.19 | 314.19 | 7.68 | 146 | >sp\|P02788\|TRFL_HUMAN Lactotransferrin OS=Homo sapiens OX=9606 GN=LTF PE=1 SV=6 | 1270 | 20201118_lss_colostrum_1ng_glyco_hcd203040_compound15.22119.22119.3 | controllerType=0 controllerNumber=1 scan=22119 | 45.9587 |
| 19991:4 | 3 | R.TAGWNVPIGTLRPFLN[+2350.830]WTGPPEPIEAAVAR.F | HexNAc(4)Hex(5)Fuc(1)NeuAc(2) | N[+2351] | 1396.1455 | 4 | 5581.5602 | 5581.5413 | 0 | 3.4 | 141 | Specific | 407.91 | 285.08 | 285.08 | 7.43 | 146 | >sp\|P02788\|TRFL_HUMAN Lactotransferrin OS=Homo sapiens OX=9606 GN=LTF PE=1 SV=6 | 1270 | 20201118_lss_colostrum_1ng_glyco_hcd203040_compound15.23650.23650.4 | controllerType=0 controllerNumber=1 scan=23650 | 49.3667 |
| 17635:3 | 3 | R.TAGWNVPIGTLRPFLN[+2425.887]WTGPPEPIEAAVAR.F | HexNAc(5)Hex(6)Fuc(3) | N[+2426] | 1886.2091 | 3 | 5656.6127 | 5656.5985 | 0 | 2.5 | 141 | Specific | 418.25 | 207.88 | 207.88 | 7.42 | 146 | >sp\|P02788\|TRFL_HUMAN Lactotransferrin OS=Homo sapiens OX=9606 GN=LTF PE=1 SV=6 | 1270 | 20201118_lss_colostrum_1ng_glyco_hcd203040_compound15.21191.21191.3 | controllerType=0 controllerNumber=1 scan=21191 | 43.9325 |
| 18789:4 | 3 | R.TAGWNVPIGTLRPFLN[+2204.772]WTGPPEPIEAAVAR.F | HexNAc(4)Hex(5)NeuAc(2) | N[+2205] | 1359.6355 | 4 | 5435.5202 | 5435.4834 | 0 | 6.8 | 141 | Specific | 440.27 | 322.4 | 322.4 | 6.84 | 146 | >sp\|P02788\|TRFL_HUMAN Lactotransferrin OS=Homo sapiens OX=9606 GN=LTF PE=1 SV=6 | 1270 | 20201118_lss_colostrum_1ng_glyco_hcd203040_compound15.22396.22396.4 | controllerType=0 controllerNumber=1 scan=22396 | 46.5519 |
| 18681:3 | 3 | R.TAGWNVPIGTLRPFLN[+2205.793]WTGPPEPIEAAVAR.F | HexNAc(4)Hex(5)Fuc(2)NeuAc(1) | N[+2206] | 1812.8452 | 3 | 5436.521 | 5436.5038 | 0 | 3.2 | 141 | Specific | 449.5 | 397.27 | 397.27 | 9.99 | 146 | >sp\|P02788\|TRFL_HUMAN Lactotransferrin OS=Homo sapiens OX=9606 GN=LTF PE=1 SV=6 | 1270 | 20201118_lss_colostrum_1ng_glyco_hcd203040_compound15.22283.22283.2 | controllerType=0 controllerNumber=1 scan=22283 | 46.3124 |
| 18526:3 | 3 | R.TAGWNVPIGTLRPFLN[+2716.983]WTGPPEPIEAAVAR.F | HexNAc(5)Hex(6)Fuc(3)NeuAc(1) | N[+2717] | 1983.2429 | 3 | 5947.7141 | 5947.6939 | 0 | 3.4 | 141 | Specific | 473.58 | 416.13 | 416.13 | 9.88 | 146 | >sp\|P02788\|TRFL_HUMAN Lactotransferrin OS=Homo sapiens OX=9606 GN=LTF PE=1 SV=6 | 1270 | 20201118_lss_colostrum_1ng_glyco_hcd203040_compound15.22122.22122.3 | controllerType=0 controllerNumber=1 scan=22122 | 45.963 |
| 20001:3 | 3 | R.TAGWNVPIGTLRPFLN[+2350.830]WTGPPEPIEAAVAR.F | HexNAc(4)Hex(5)Fuc(1)NeuAc(2) | N[+2351] | 1861.1912 | 3 | 5581.5592 | 5581.5413 | 0 | 3.2 | 141 | Specific | 510.34 | 345.03 | 345.03 | 8.41 | 146 | >sp\|P02788\|TRFL_HUMAN Lactotransferrin OS=Homo sapiens OX=9606 GN=LTF PE=1 SV=6 | 1270 | 20201118_lss_colostrum_1ng_glyco_hcd203040_compound15.23660.23660.3 | controllerType=0 controllerNumber=1 scan=23660 | 49.3829 |
| 17679:3 | 3 | R.TAGWNVPIGTLRPFLN[+1913.677]WTGPPEPIEAAVAR.F | HexNAc(4)Hex(5)NeuAc(1) | N[+1914] | 1715.4781 | 3 | 5144.4198 | 5144.388 | 0 | 6.2 | 141 | Specific | 515.83 | 441.56 | 441.56 | 9.24 | 146 | >sp\|P02788\|TRFL_HUMAN Lactotransferrin OS=Homo sapiens OX=9606 GN=LTF PE=1 SV=6 | 1270 | 20201118_lss_colostrum_1ng_glyco_hcd203040_compound15.21239.21239.3 | controllerType=0 controllerNumber=1 scan=21239 | 44.087 |
| 18646:3 | 3 | R.TAGWNVPIGTLRPFLN[+2351.851]WTGPPEPIEAAVAR.F | HexNAc(4)Hex(5)Fuc(3)NeuAc(1) | N[+2352] | 1861.5299 | 3 | 5582.5751 | 5582.5617 | 0 | 2.4 | 141 | Specific | 523.56 | 471.78 | 471.78 | 10.29 | 146 | >sp\|P02788\|TRFL_HUMAN Lactotransferrin OS=Homo sapiens OX=9606 GN=LTF PE=1 SV=6 | 1270 | 20201118_lss_colostrum_1ng_glyco_hcd203040_compound15.22247.22247.3 | controllerType=0 controllerNumber=1 scan=22247 | 46.2373 |
| 18649:3 | 3 | R.TAGWNVPIGTLRPFLN[+2351.851]WTGPPEPIEAAVAR.F | HexNAc(4)Hex(5)Fuc(3)NeuAc(1) | N[+2352] | 1861.5299 | 3 | 5582.5751 | 5582.5617 | 0 | 2.4 | 141 | Specific | 554.35 | 464.45 | 464.45 | 10.3 | 146 | >sp\|P02788\|TRFL_HUMAN Lactotransferrin OS=Homo sapiens OX=9606 GN=LTF PE=1 SV=6 | 1270 | 20201118_lss_colostrum_1ng_glyco_hcd203040_compound15.22250.22250.3 | controllerType=0 controllerNumber=1 scan=22250 | 46.2421 |
| 17677:3 | 3 | R.TAGWNVPIGTLRPFLN[+1913.677]WTGPPEPIEAAVAR.F | HexNAc(4)Hex(5)NeuAc(1) | N[+1914] | 1715.4781 | 3 | 5144.4198 | 5144.388 | 0 | 6.2 | 141 | Specific | 575.97 | 529.18 | 529.18 | 9.51 | 146 | >sp\|P02788\|TRFL_HUMAN Lactotransferrin OS=Homo sapiens OX=9606 GN=LTF PE=1 SV=6 | 1270 | 20201118_lss_colostrum_1ng_glyco_hcd203040_compound15.21237.21237.3 | controllerType=0 controllerNumber=1 scan=21237 | 44.0631 |
| 18655:3 | 3 | R.TAGWNVPIGTLRPFLN[+2205.793]WTGPPEPIEAAVAR.F | HexNAc(4)Hex(5)Fuc(2)NeuAc(1) | N[+2206] | 1812.8444 | 3 | 5436.5188 | 5436.5038 | 0 | 2.8 | 141 | Specific | 586.29 | 540.18 | 540.18 | 11.9 | 146 | >sp\|P02788\|TRFL_HUMAN Lactotransferrin OS=Homo sapiens OX=9606 GN=LTF PE=1 SV=6 | 1270 | 20201118_lss_colostrum_1ng_glyco_hcd203040_compound15.22256.22256.3 | controllerType=0 controllerNumber=1 scan=22256 | 46.2597 |
| 18706:4 | 3 | R.TAGWNVPIGTLRPFLN[+2076.750]WTGPPEPIEAAVAR.F | HexNAc(4)Hex(6)Fuc(2) | N[+2077] | 1327.6256 | 4 | 5307.4806 | 5307.4612 | 0 | 3.7 | 141 | Specific | 592.06 | 525.95 | 525.95 | 11.63 | 146 | >sp\|P02788\|TRFL_HUMAN Lactotransferrin OS=Homo sapiens OX=9606 GN=LTF PE=1 SV=6 | 1270 | 20201118_lss_colostrum_1ng_glyco_hcd203040_compound15.22310.22310.4 | controllerType=0 controllerNumber=1 scan=22310 | 46.3697 |
| 20011:5 | 3 | R.TAGWNVPIGTLRPFLN[+2350.830]WTGPPEPIEAAVAR.F | HexNAc(4)Hex(5)Fuc(1)NeuAc(2) | N[+2351] | 1117.1185 | 5 | 5581.5636 | 5581.5413 | 0 | 4 | 141 | Specific | 619.85 | 599.53 | 599.53 | 12.87 | 146 | >sp\|P02788\|TRFL_HUMAN Lactotransferrin OS=Homo sapiens OX=9606 GN=LTF PE=1 SV=6 | 1270 | 20201118_lss_colostrum_1ng_glyco_hcd203040_compound15.23670.23670.5 | controllerType=0 controllerNumber=1 scan=23670 | 49.3986 |
| 18635:4 | 3 | R.TAGWNVPIGTLRPFLN[+2351.851]WTGPPEPIEAAVAR.F | HexNAc(4)Hex(5)Fuc(3)NeuAc(1) | N[+2352] | 1396.3992 | 4 | 5582.5749 | 5582.5617 | 0 | 2.4 | 141 | Specific | 640.26 | 577.29 | 577.29 | 12.83 | 146 | >sp\|P02788\|TRFL_HUMAN Lactotransferrin OS=Homo sapiens OX=9606 GN=LTF PE=1 SV=6 | 1270 | 20201118_lss_colostrum_1ng_glyco_hcd203040_compound15.22236.22236.4 | controllerType=0 controllerNumber=1 scan=22236 | 46.2155 |
| 18524:4 | 3 | R.TAGWNVPIGTLRPFLN[+2570.925]WTGPPEPIEAAVAR.F | HexNAc(5)Hex(6)Fuc(2)NeuAc(1) | N[+2571] | 1451.1687 | 4 | 5801.653 | 5801.636 | 0 | 2.9 | 141 | Specific | 644.98 | 548.26 | 548.26 | 12.83 | 146 | >sp\|P02788\|TRFL_HUMAN Lactotransferrin OS=Homo sapiens OX=9606 GN=LTF PE=1 SV=6 | 1270 | 20201118_lss_colostrum_1ng_glyco_hcd203040_compound15.22120.22120.4 | controllerType=0 controllerNumber=1 scan=22120 | 45.9598 |
| 18708:4 | 3 | R.TAGWNVPIGTLRPFLN[+2076.750]WTGPPEPIEAAVAR.F | HexNAc(4)Hex(6)Fuc(2) | N[+2077] | 1327.6256 | 4 | 5307.4806 | 5307.4612 | 0 | 3.7 | 141 | Specific | 660.08 | 590.38 | 590.38 | 13.63 | 146 | >sp\|P02788\|TRFL_HUMAN Lactotransferrin OS=Homo sapiens OX=9606 GN=LTF PE=1 SV=6 | 1270 | 20201118_lss_colostrum_1ng_glyco_hcd203040_compound15.22312.22312.4 | controllerType=0 controllerNumber=1 scan=22312 | 46.3764 |
| 17688:3 | 3 | R.TAGWNVPIGTLRPFLN[+1768.640]WTGPPEPIEAAVAR.F | HexNAc(4)Hex(5)Fuc(1) | N[+1769] | 1667.1274 | 3 | 4999.3677 | 4999.3504 | 0 | 3.4 | 141 | Specific | 664.04 | 566.85 | 566.85 | 13.7 | 146 | >sp\|P02788\|TRFL_HUMAN Lactotransferrin OS=Homo sapiens OX=9606 GN=LTF PE=1 SV=6 | 1270 | 20201118_lss_colostrum_1ng_glyco_hcd203040_compound15.21249.21249.3 | controllerType=0 controllerNumber=1 scan=21249 | 44.1197 |
| 18540:4 | 3 | R.TAGWNVPIGTLRPFLN[+2863.041]WTGPPEPIEAAVAR.F | HexNAc(5)Hex(6)Fuc(4)NeuAc(1) | N[+2863] | 1524.1964 | 4 | 6093.7638 | 6093.7518 | 0 | 2 | 141 | Specific | 671.19 | 519.21 | 519.21 | 13.83 | 146 | >sp\|P02788\|TRFL_HUMAN Lactotransferrin OS=Homo sapiens OX=9606 GN=LTF PE=1 SV=6 | 1270 | 20201118_lss_colostrum_1ng_glyco_hcd203040_compound15.22137.22137.4 | controllerType=0 controllerNumber=1 scan=22137 | 45.994 |
| 17686:3 | 3 | R.TAGWNVPIGTLRPFLN[+1768.640]WTGPPEPIEAAVAR.F | HexNAc(4)Hex(5)Fuc(1) | N[+1769] | 1667.1274 | 3 | 4999.3677 | 4999.3504 | 0 | 3.4 | 141 | Specific | 677.78 | 605.83 | 605.83 | 13.7 | 146 | >sp\|P02788\|TRFL_HUMAN Lactotransferrin OS=Homo sapiens OX=9606 GN=LTF PE=1 SV=6 | 1270 | 20201118_lss_colostrum_1ng_glyco_hcd203040_compound15.21247.21247.3 | controllerType=0 controllerNumber=1 scan=21247 | 44.1153 |
| 17685:4 | 3 | R.TAGWNVPIGTLRPFLN[+1768.640]WTGPPEPIEAAVAR.F | HexNAc(4)Hex(5)Fuc(1) | N[+1769] | 1250.5977 | 4 | 4999.3688 | 4999.3504 | 0 | 3.7 | 141 | Specific | 680.35 | 641.79 | 641.79 | 13.63 | 146 | >sp\|P02788\|TRFL_HUMAN Lactotransferrin OS=Homo sapiens OX=9606 GN=LTF PE=1 SV=6 | 1270 | 20201118_lss_colostrum_1ng_glyco_hcd203040_compound15.21246.21246.4 | controllerType=0 controllerNumber=1 scan=21246 | 44.1135 |
| 18652:3 | 3 | R.TAGWNVPIGTLRPFLN[+2205.793]WTGPPEPIEAAVAR.F | HexNAc(4)Hex(5)Fuc(2)NeuAc(1) | N[+2206] | 1812.8444 | 3 | 5436.5188 | 5436.5038 | 0 | 2.8 | 141 | Specific | 680.59 | 635.9 | 635.9 | 13.9 | 146 | >sp\|P02788\|TRFL_HUMAN Lactotransferrin OS=Homo sapiens OX=9606 GN=LTF PE=1 SV=6 | 1270 | 20201118_lss_colostrum_1ng_glyco_hcd203040_compound15.22253.22253.3 | controllerType=0 controllerNumber=1 scan=22253 | 46.2529 |
| 18683:3 | 3 | R.TAGWNVPIGTLRPFLN[+2205.793]WTGPPEPIEAAVAR.F | HexNAc(4)Hex(5)Fuc(2)NeuAc(1) | N[+2206] | 1812.8452 | 3 | 5436.521 | 5436.5038 | 0 | 3.2 | 141 | Specific | 684.44 | 601.32 | 601.32 | 13.9 | 146 | >sp\|P02788\|TRFL_HUMAN Lactotransferrin OS=Homo sapiens OX=9606 GN=LTF PE=1 SV=6 | 1270 | 20201118_lss_colostrum_1ng_glyco_hcd203040_compound15.22285.22285.2 | controllerType=0 controllerNumber=1 scan=22285 | 46.3158 |
| 18743:5 | 3 | R.TAGWNVPIGTLRPFLN[+2205.793]WTGPPEPIEAAVAR.F | HexNAc(4)Hex(5)Fuc(2)NeuAc(1) | N[+2206] | 1088.1105 | 5 | 5436.5233 | 5436.5038 | 0 | 3.6 | 141 | Specific | 693.04 | 488.25 | 488.25 | 13.87 | 146 | >sp\|P02788\|TRFL_HUMAN Lactotransferrin OS=Homo sapiens OX=9606 GN=LTF PE=1 SV=6 | 1270 | 20201118_lss_colostrum_1ng_glyco_hcd203040_compound15.22348.22348.5 | controllerType=0 controllerNumber=1 scan=22348 | 46.4548 |
| 18657:4 | 3 | R.TAGWNVPIGTLRPFLN[+2059.735]WTGPPEPIEAAVAR.F | HexNAc(4)Hex(5)Fuc(1)NeuAc(1) | N[+2060] | 1323.3715 | 4 | 5290.464 | 5290.4459 | 0 | 3.4 | 141 | Specific | 699.03 | 643.86 | 643.86 | 13.63 | 146 | >sp\|P02788\|TRFL_HUMAN Lactotransferrin OS=Homo sapiens OX=9606 GN=LTF PE=1 SV=6 | 1270 | 20201118_lss_colostrum_1ng_glyco_hcd203040_compound15.22259.22259.4 | controllerType=0 controllerNumber=1 scan=22259 | 46.2681 |
| 18615:4 | 3 | R.TAGWNVPIGTLRPFLN[+2716.983]WTGPPEPIEAAVAR.F | HexNAc(5)Hex(6)Fuc(3)NeuAc(1) | N[+2717] | 1487.683 | 4 | 5947.7101 | 5947.6939 | 0 | 2.7 | 141 | Specific | 699.49 | 589.71 | 589.71 | 13.83 | 146 | >sp\|P02788\|TRFL_HUMAN Lactotransferrin OS=Homo sapiens OX=9606 GN=LTF PE=1 SV=6 | 1270 | 20201118_lss_colostrum_1ng_glyco_hcd203040_compound15.22215.22215.4 | controllerType=0 controllerNumber=1 scan=22215 | 46.1672 |
| 18647:4 | 3 | R.TAGWNVPIGTLRPFLN[+2205.793]WTGPPEPIEAAVAR.F | HexNAc(4)Hex(5)Fuc(2)NeuAc(1) | N[+2206] | 1359.8846 | 4 | 5436.5168 | 5436.5038 | 0 | 2.4 | 141 | Specific | 721.16 | 648.57 | 648.57 | 14.83 | 146 | >sp\|P02788\|TRFL_HUMAN Lactotransferrin OS=Homo sapiens OX=9606 GN=LTF PE=1 SV=6 | 1270 | 20201118_lss_colostrum_1ng_glyco_hcd203040_compound15.22248.22248.4 | controllerType=0 controllerNumber=1 scan=22248 | 46.2385 |
| 18769:3 | 3 | R.TAGWNVPIGTLRPFLN[+2059.735]WTGPPEPIEAAVAR.F | HexNAc(4)Hex(5)Fuc(1)NeuAc(1) | N[+2060] | 1764.1558 | 3 | 5290.453 | 5290.4459 | 0 | 1.3 | 141 | Specific | 727.93 | 642.6 | 642.6 | 14.74 | 146 | >sp\|P02788\|TRFL_HUMAN Lactotransferrin OS=Homo sapiens OX=9606 GN=LTF PE=1 SV=6 | 1270 | 20201118_lss_colostrum_1ng_glyco_hcd203040_compound15.22375.22375.2 | controllerType=0 controllerNumber=1 scan=22375 | 46.508 |
| 18527:4 | 3 | R.TAGWNVPIGTLRPFLN[+2570.925]WTGPPEPIEAAVAR.F | HexNAc(5)Hex(6)Fuc(2)NeuAc(1) | N[+2571] | 1451.1687 | 4 | 5801.653 | 5801.636 | 0 | 2.9 | 141 | Specific | 745.66 | 613.23 | 613.23 | 14.83 | 146 | >sp\|P02788\|TRFL_HUMAN Lactotransferrin OS=Homo sapiens OX=9606 GN=LTF PE=1 SV=6 | 1270 | 20201118_lss_colostrum_1ng_glyco_hcd203040_compound15.22123.22123.4 | controllerType=0 controllerNumber=1 scan=22123 | 45.9691 |
| 18659:4 | 3 | R.TAGWNVPIGTLRPFLN[+2059.735]WTGPPEPIEAAVAR.F | HexNAc(4)Hex(5)Fuc(1)NeuAc(1) | N[+2060] | 1323.3715 | 4 | 5290.464 | 5290.4459 | 0 | 3.4 | 141 | Specific | 763.15 | 634.57 | 634.57 | 15.63 | 146 | >sp\|P02788\|TRFL_HUMAN Lactotransferrin OS=Homo sapiens OX=9606 GN=LTF PE=1 SV=6 | 1270 | 20201118_lss_colostrum_1ng_glyco_hcd203040_compound15.22261.22261.4 | controllerType=0 controllerNumber=1 scan=22261 | 46.2755 |
| 18792:4 | 3 | R.TAGWNVPIGTLRPFLN[+2204.772]WTGPPEPIEAAVAR.F | HexNAc(4)Hex(5)NeuAc(2) | N[+2205] | 1359.6355 | 4 | 5435.5202 | 5435.4834 | 0 | 6.8 | 141 | Specific | 773.22 | 510.91 | 510.91 | 14.78 | 146 | >sp\|P02788\|TRFL_HUMAN Lactotransferrin OS=Homo sapiens OX=9606 GN=LTF PE=1 SV=6 | 1270 | 20201118_lss_colostrum_1ng_glyco_hcd203040_compound15.22399.22399.4 | controllerType=0 controllerNumber=1 scan=22399 | 46.5561 |
| 19993:4 | 3 | R.TAGWNVPIGTLRPFLN[+2350.830]WTGPPEPIEAAVAR.F | HexNAc(4)Hex(5)Fuc(1)NeuAc(2) | N[+2351] | 1396.1455 | 4 | 5581.5602 | 5581.5413 | 0 | 3.4 | 141 | Specific | 786.53 | 526.79 | 526.79 | 15.63 | 146 | >sp\|P02788\|TRFL_HUMAN Lactotransferrin OS=Homo sapiens OX=9606 GN=LTF PE=1 SV=6 | 1270 | 20201118_lss_colostrum_1ng_glyco_hcd203040_compound15.23652.23652.4 | controllerType=0 controllerNumber=1 scan=23652 | 49.3708 |
| 18637:4 | 3 | R.TAGWNVPIGTLRPFLN[+2351.851]WTGPPEPIEAAVAR.F | HexNAc(4)Hex(5)Fuc(3)NeuAc(1) | N[+2352] | 1396.3992 | 4 | 5582.5749 | 5582.5617 | 0 | 2.4 | 141 | Specific | 795.16 | 697.87 | 697.87 | 15.83 | 146 | >sp\|P02788\|TRFL_HUMAN Lactotransferrin OS=Homo sapiens OX=9606 GN=LTF PE=1 SV=6 | 1270 | 20201118_lss_colostrum_1ng_glyco_hcd203040_compound15.22238.22238.4 | controllerType=0 controllerNumber=1 scan=22238 | 46.2214 |
| 18698:5 | 3 | R.TAGWNVPIGTLRPFLN[+2076.750]WTGPPEPIEAAVAR.F | HexNAc(4)Hex(6)Fuc(2) | N[+2077] | 1062.3025 | 5 | 5307.4834 | 5307.4612 | 0 | 4.2 | 141 | Specific | 802.1 | 740.63 | 740.63 | 16.87 | 146 | >sp\|P02788\|TRFL_HUMAN Lactotransferrin OS=Homo sapiens OX=9606 GN=LTF PE=1 SV=6 | 1270 | 20201118_lss_colostrum_1ng_glyco_hcd203040_compound15.22301.22301.5 | controllerType=0 controllerNumber=1 scan=22301 | 46.354 |
| 18650:4 | 3 | R.TAGWNVPIGTLRPFLN[+2205.793]WTGPPEPIEAAVAR.F | HexNAc(4)Hex(5)Fuc(2)NeuAc(1) | N[+2206] | 1359.8846 | 4 | 5436.5168 | 5436.5038 | 0 | 2.4 | 141 | Specific | 806.03 | 728.62 | 728.62 | 16.83 | 146 | >sp\|P02788\|TRFL_HUMAN Lactotransferrin OS=Homo sapiens OX=9606 GN=LTF PE=1 SV=6 | 1270 | 20201118_lss_colostrum_1ng_glyco_hcd203040_compound15.22251.22251.4 | controllerType=0 controllerNumber=1 scan=22251 | 46.2511 |
| 17687:4 | 3 | R.TAGWNVPIGTLRPFLN[+1768.640]WTGPPEPIEAAVAR.F | HexNAc(4)Hex(5)Fuc(1) | N[+1769] | 1250.5977 | 4 | 4999.3688 | 4999.3504 | 0 | 3.7 | 141 | Specific | 878.02 | 713.18 | 713.18 | 17.63 | 146 | >sp\|P02788\|TRFL_HUMAN Lactotransferrin OS=Homo sapiens OX=9606 GN=LTF PE=1 SV=6 | 1270 | 20201118_lss_colostrum_1ng_glyco_hcd203040_compound15.21248.21248.4 | controllerType=0 controllerNumber=1 scan=21248 | 44.1179 |
| 18695:5 | 3 | R.TAGWNVPIGTLRPFLN[+2076.750]WTGPPEPIEAAVAR.F | HexNAc(4)Hex(6)Fuc(2) | N[+2077] | 1062.3025 | 5 | 5307.4834 | 5307.4612 | 0 | 4.2 | 141 | Specific | 880.05 | 845.5 | 845.5 | 17.87 | 146 | >sp\|P02788\|TRFL_HUMAN Lactotransferrin OS=Homo sapiens OX=9606 GN=LTF PE=1 SV=6 | 1270 | 20201118_lss_colostrum_1ng_glyco_hcd203040_compound15.22298.22298.5 | controllerType=0 controllerNumber=1 scan=22298 | 46.3428 |
| 16695:3 | 3 | R.TAGWNIPMGLLFN[+2076.750]QTGSC[+57.021]K.F | HexNAc(4)Hex(6)Fuc(2) | C[+57], N[+2077] | 1391.2563 | 3 | 4171.7543 | 4171.7602 | 0 | -1.4 | 485 | Specific | 268.37 | 200.33 | 200.33 | 3.88 | 146 | >sp\|P02788\|TRFL_HUMAN Lactotransferrin OS=Homo sapiens OX=9606 GN=LTF PE=1 SV=6 | 1270 | 20201118_lss_colostrum_1ng_glyco_hcd203040_compound15.20213.20213.3 | controllerType=0 controllerNumber=1 scan=20213 | 41.8547 |
| 16694:3 | 3 | R.TAGWNIPMGLLFN[+2076.750]QTGSC[+57.021]K.F | HexNAc(4)Hex(6)Fuc(2) | C[+57], N[+2077] | 1391.2563 | 3 | 4171.7543 | 4171.7602 | 0 | -1.4 | 485 | Specific | 318.73 | 248.3 | 248.3 | 6.01 | 146 | >sp\|P02788\|TRFL_HUMAN Lactotransferrin OS=Homo sapiens OX=9606 GN=LTF PE=1 SV=6 | 1270 | 20201118_lss_colostrum_1ng_glyco_hcd203040_compound15.20212.20212.3 | controllerType=0 controllerNumber=1 scan=20212 | 41.851 |
| 18064:4 | 3 | R.TAGWNIPMGLLFN[+2059.735]QTGSC[+57.021]K.F | HexNAc(4)Hex(5)Fuc(1)NeuAc(1) | C[+57], N[+2060] | 1039.444 | 4 | 4154.7542 | 4154.7449 | 0 | 2.2 | 485 | Specific | 358.19 | 327.76 | 327.76 | 7.74 | 146 | >sp\|P02788\|TRFL_HUMAN Lactotransferrin OS=Homo sapiens OX=9606 GN=LTF PE=1 SV=6 | 1270 | 20201118_lss_colostrum_1ng_glyco_hcd203040_compound15.21642.21642.4 | controllerType=0 controllerNumber=1 scan=21642 | 44.9424 |
| 18001:3 | 3 | R.TAGWNIPMGLLFN[+2205.793]QTGSC[+57.021]K.F | HexNAc(4)Hex(5)Fuc(2)NeuAc(1) | C[+57], N[+2206] | 1434.2773 | 3 | 4300.8173 | 4300.8028 | 0 | 3.4 | 485 | Specific | 591.32 | 572.56 | 572.56 | 11.43 | 146 | >sp\|P02788\|TRFL_HUMAN Lactotransferrin OS=Homo sapiens OX=9606 GN=LTF PE=1 SV=6 | 1270 | 20201118_lss_colostrum_1ng_glyco_hcd203040_compound15.21576.21576.3 | controllerType=0 controllerNumber=1 scan=21576 | 44.7953 |
| 20217:3 | 3 | R.TAGWNIPMGLLFN[+2204.772]QTGSC[+57.021]K.F | HexNAc(4)Hex(5)NeuAc(2) | C[+57], N[+2205] | 1433.9404 | 3 | 4299.8065 | 4299.7824 | 0 | 5.6 | 485 | Specific | 623.87 | 614.21 | 614.21 | 11.97 | 146 | >sp\|P02788\|TRFL_HUMAN Lactotransferrin OS=Homo sapiens OX=9606 GN=LTF PE=1 SV=6 | 1270 | 20201118_lss_colostrum_1ng_glyco_hcd203040_compound15.23886.23886.3 | controllerType=0 controllerNumber=1 scan=23886 | 49.9229 |
| 18066:4 | 3 | R.TAGWNIPMGLLFN[+2059.735]QTGSC[+57.021]K.F | HexNAc(4)Hex(5)Fuc(1)NeuAc(1) | C[+57], N[+2060] | 1039.444 | 4 | 4154.7542 | 4154.7449 | 0 | 2.2 | 485 | Specific | 631.55 | 348.12 | 348.12 | 11.16 | 146 | >sp\|P02788\|TRFL_HUMAN Lactotransferrin OS=Homo sapiens OX=9606 GN=LTF PE=1 SV=6 | 1270 | 20201118_lss_colostrum_1ng_glyco_hcd203040_compound15.21644.21644.4 | controllerType=0 controllerNumber=1 scan=21644 | 44.9487 |
| 18209:4 | 3 | R.TAGWNIPMGLLFN[+1913.677]QTGSC[+57.021]K.F | HexNAc(4)Hex(5)NeuAc(1) | C[+57], N[+1914] | 1002.9294 | 4 | 4008.6956 | 4008.687 | 0 | 2.2 | 485 | Specific | 650.99 | 606.9 | 606.9 | 13.87 | 146 | >sp\|P02788\|TRFL_HUMAN Lactotransferrin OS=Homo sapiens OX=9606 GN=LTF PE=1 SV=6 | 1270 | 20201118_lss_colostrum_1ng_glyco_hcd203040_compound15.21793.21793.4 | controllerType=0 controllerNumber=1 scan=21793 | 45.3055 |
| 16153:3 | 3 | R.TAGWNIPMGLLFN[+1768.640]QTGSC[+57.021]K.F | HexNAc(4)Hex(5)Fuc(1) | C[+57], N[+1769] | 1288.5585 | 3 | 3863.661 | 3863.6495 | 0 | 3 | 485 | Specific | 669.19 | 669.19 | 669.19 | 13.63 | 146 | >sp\|P02788\|TRFL_HUMAN Lactotransferrin OS=Homo sapiens OX=9606 GN=LTF PE=1 SV=6 | 1270 | 20201118_lss_colostrum_1ng_glyco_hcd203040_compound15.19650.19650.3 | controllerType=0 controllerNumber=1 scan=19650 | 40.691 |
| 20219:3 | 3 | R.TAGWNIPMGLLFN[+2204.772]QTGSC[+57.021]K.F | HexNAc(4)Hex(5)NeuAc(2) | C[+57], N[+2205] | 1433.9404 | 3 | 4299.8065 | 4299.7824 | 0 | 5.6 | 485 | Specific | 722.94 | 559.51 | 559.51 | 13.97 | 146 | >sp\|P02788\|TRFL_HUMAN Lactotransferrin OS=Homo sapiens OX=9606 GN=LTF PE=1 SV=6 | 1270 | 20201118_lss_colostrum_1ng_glyco_hcd203040_compound15.23888.23888.3 | controllerType=0 controllerNumber=1 scan=23888 | 49.9276 |
| 20017:4 | 3 | R.TAGWNIPMGLLFN[+2351.851]QTGSC[+57.021]K.F | HexNAc(4)Hex(5)Fuc(3)NeuAc(1) | C[+57], N[+2352] | 1112.4685 | 4 | 4446.8524 | 4446.8607 | 0 | -1.9 | 485 | Specific | 769.63 | 559.96 | 559.96 | 14.8 | 146 | >sp\|P02788\|TRFL_HUMAN Lactotransferrin OS=Homo sapiens OX=9606 GN=LTF PE=1 SV=6 | 1270 | 20201118_lss_colostrum_1ng_glyco_hcd203040_compound15.23676.23676.4 | controllerType=0 controllerNumber=1 scan=23676 | 49.4125 |
| 20014:4 | 3 | R.TAGWNIPMGLLFN[+2351.851]QTGSC[+57.021]K.F | HexNAc(4)Hex(5)Fuc(3)NeuAc(1) | C[+57], N[+2352] | 1112.4685 | 4 | 4446.8524 | 4446.8607 | 0 | -1.9 | 485 | Specific | 785.44 | 701.89 | 701.89 | 14.8 | 146 | >sp\|P02788\|TRFL_HUMAN Lactotransferrin OS=Homo sapiens OX=9606 GN=LTF PE=1 SV=6 | 1270 | 20201118_lss_colostrum_1ng_glyco_hcd203040_compound15.23673.23673.4 | controllerType=0 controllerNumber=1 scan=23673 | 49.405 |
| 18212:4 | 3 | R.TAGWNIPMGLLFN[+1913.677]QTGSC[+57.021]K.F | HexNAc(4)Hex(5)NeuAc(1) | C[+57], N[+1914] | 1002.9294 | 4 | 4008.6956 | 4008.687 | 0 | 2.2 | 485 | Specific | 822.02 | 654.75 | 654.75 | 16.87 | 146 | >sp\|P02788\|TRFL_HUMAN Lactotransferrin OS=Homo sapiens OX=9606 GN=LTF PE=1 SV=6 | 1270 | 20201118_lss_colostrum_1ng_glyco_hcd203040_compound15.21796.21796.4 | controllerType=0 controllerNumber=1 scan=21796 | 45.3105 |
| 18004:3 | 3 | R.TAGWNIPMGLLFN[+2205.793]QTGSC[+57.021]K.F | HexNAc(4)Hex(5)Fuc(2)NeuAc(1) | C[+57], N[+2206] | 1434.2773 | 3 | 4300.8173 | 4300.8028 | 0 | 3.4 | 485 | Specific | 863.62 | 823.59 | 823.59 | 17.43 | 146 | >sp\|P02788\|TRFL_HUMAN Lactotransferrin OS=Homo sapiens OX=9606 GN=LTF PE=1 SV=6 | 1270 | 20201118_lss_colostrum_1ng_glyco_hcd203040_compound15.21579.21579.3 | controllerType=0 controllerNumber=1 scan=21579 | 44.7996 |
| 16573:2 | 3 | R.TAGWNIPMGLLFN[+1216.423]QTGSC[+57.021]K.F | HexNAc(2)Hex(5) | C[+57], N[+1216] | 1656.2236 | 2 | 3311.4399 | 3311.4328 | 0 | 2.1 | 485 | Specific | 866.25 | 785.22 | 267.25 | 17.7 | 146 | >sp\|P02788\|TRFL_HUMAN Lactotransferrin OS=Homo sapiens OX=9606 GN=LTF PE=1 SV=6 | 1270 | 20201118_lss_colostrum_1ng_glyco_hcd203040_compound15.20087.20087.2 | controllerType=0 controllerNumber=1 scan=20087 | 41.6075 |
| 16155:3 | 3 | R.TAGWNIPMGLLFN[+1768.640]QTGSC[+57.021]K.F | HexNAc(4)Hex(5)Fuc(1) | C[+57], N[+1769] | 1288.5585 | 3 | 3863.661 | 3863.6495 | 0 | 3 | 485 | Specific | 894.08 | 661.49 | 661.49 | 17.63 | 146 | >sp\|P02788\|TRFL_HUMAN Lactotransferrin OS=Homo sapiens OX=9606 GN=LTF PE=1 SV=6 | 1270 | 20201118_lss_colostrum_1ng_glyco_hcd203040_compound15.19652.19652.3 | controllerType=0 controllerNumber=1 scan=19652 | 40.6954 |
| 16571:2 | 3 | R.TAGWNIPMGLLFN[+1216.423]QTGSC[+57.021]K.F | HexNAc(2)Hex(5) | C[+57], N[+1216] | 1656.2236 | 2 | 3311.4399 | 3311.4328 | 0 | 2.1 | 485 | Specific | 905.62 | 905.62 | 318.19 | 18.7 | 146 | >sp\|P02788\|TRFL_HUMAN Lactotransferrin OS=Homo sapiens OX=9606 GN=LTF PE=1 SV=6 | 1270 | 20201118_lss_colostrum_1ng_glyco_hcd203040_compound15.20085.20085.2 | controllerType=0 controllerNumber=1 scan=20085 | 41.6029 |
| 16524:3 | 3 | R.TAGWNIPMGLLFN[+1216.423]QTGSC[+57.021]K.F | HexNAc(2)Hex(5) | C[+57], N[+1216] | 1104.4864 | 3 | 3311.4445 | 3311.4328 | 0 | 3.5 | 485 | Specific | 907.25 | 907.25 | 389.17 | 18.43 | 146 | >sp\|P02788\|TRFL_HUMAN Lactotransferrin OS=Homo sapiens OX=9606 GN=LTF PE=1 SV=6 | 1270 | 20201118_lss_colostrum_1ng_glyco_hcd203040_compound15.20036.20036.3 | controllerType=0 controllerNumber=1 scan=20036 | 41.4945 |
| 18178:3 | 3 | R.TAGWNIPMGLLFN[+1913.677]QTGSC[+57.021]K.F | HexNAc(4)Hex(5)NeuAc(1) | C[+57], N[+1914] | 1336.9034 | 3 | 4008.6956 | 4008.687 | 0 | 2.1 | 485 | Specific | 908.18 | 908.18 | 908.18 | 18.63 | 146 | >sp\|P02788\|TRFL_HUMAN Lactotransferrin OS=Homo sapiens OX=9606 GN=LTF PE=1 SV=6 | 1270 | 20201118_lss_colostrum_1ng_glyco_hcd203040_compound15.21761.21761.3 | controllerType=0 controllerNumber=1 scan=21761 | 45.2186 |
| 19997:3 | 3 | R.TAGWNIPMGLLFN[+2350.830]QTGSC[+57.021]K.F | HexNAc(4)Hex(5)Fuc(1)NeuAc(2) | C[+57], N[+2351] | 1482.6243 | 3 | 4445.8584 | 4445.8403 | 0 | 4.1 | 485 | Specific | 916 | 870.85 | 870.85 | 18.43 | 146 | >sp\|P02788\|TRFL_HUMAN Lactotransferrin OS=Homo sapiens OX=9606 GN=LTF PE=1 SV=6 | 1270 | 20201118_lss_colostrum_1ng_glyco_hcd203040_compound15.23656.23656.3 | controllerType=0 controllerNumber=1 scan=23656 | 49.3756 |
| 20000:3 | 3 | R.TAGWNIPMGLLFN[+2350.830]QTGSC[+57.021]K.F | HexNAc(4)Hex(5)Fuc(1)NeuAc(2) | C[+57], N[+2351] | 1482.6243 | 3 | 4445.8584 | 4445.8403 | 0 | 4.1 | 485 | Specific | 918.15 | 886.42 | 886.42 | 18.43 | 146 | >sp\|P02788\|TRFL_HUMAN Lactotransferrin OS=Homo sapiens OX=9606 GN=LTF PE=1 SV=6 | 1270 | 20201118_lss_colostrum_1ng_glyco_hcd203040_compound15.23659.23659.3 | controllerType=0 controllerNumber=1 scan=23659 | 49.3818 |
| 18015:3 | 3 | R.TAGWNIPMGLLFN[+2059.735]QTGSC[+57.021]K.F | HexNAc(4)Hex(5)Fuc(1)NeuAc(1) | C[+57], N[+2060] | 1385.5928 | 3 | 4154.7639 | 4154.7449 | 0 | 4.6 | 485 | Specific | 932.41 | 817.12 | 817.12 | 17.97 | 146 | >sp\|P02788\|TRFL_HUMAN Lactotransferrin OS=Homo sapiens OX=9606 GN=LTF PE=1 SV=6 | 1270 | 20201118_lss_colostrum_1ng_glyco_hcd203040_compound15.21590.21590.3 | controllerType=0 controllerNumber=1 scan=21590 | 44.821 |
| 18012:3 | 3 | R.TAGWNIPMGLLFN[+2059.735]QTGSC[+57.021]K.F | HexNAc(4)Hex(5)Fuc(1)NeuAc(1) | C[+57], N[+2060] | 1385.5928 | 3 | 4154.7639 | 4154.7449 | 0 | 4.6 | 485 | Specific | 950.86 | 815.33 | 815.33 | 18.97 | 146 | >sp\|P02788\|TRFL_HUMAN Lactotransferrin OS=Homo sapiens OX=9606 GN=LTF PE=1 SV=6 | 1270 | 20201118_lss_colostrum_1ng_glyco_hcd203040_compound15.21587.21587.3 | controllerType=0 controllerNumber=1 scan=21587 | 44.8127 |
| 18176:3 | 3 | R.TAGWNIPMGLLFN[+1913.677]QTGSC[+57.021]K.F | HexNAc(4)Hex(5)NeuAc(1) | C[+57], N[+1914] | 1336.9034 | 3 | 4008.6956 | 4008.687 | 0 | 2.1 | 485 | Specific | 960.38 | 941.53 | 941.53 | 19.63 | 146 | >sp\|P02788\|TRFL_HUMAN Lactotransferrin OS=Homo sapiens OX=9606 GN=LTF PE=1 SV=6 | 1270 | 20201118_lss_colostrum_1ng_glyco_hcd203040_compound15.21759.21759.3 | controllerType=0 controllerNumber=1 scan=21759 | 45.2137 |
| 16527:3 | 3 | R.TAGWNIPMGLLFN[+1216.423]QTGSC[+57.021]K.F | HexNAc(2)Hex(5) | C[+57], N[+1216] | 1104.4864 | 3 | 3311.4445 | 3311.4328 | 0 | 3.5 | 485 | Specific | 1056.71 | 885.54 | 505.2 | 21.43 | 146 | >sp\|P02788\|TRFL_HUMAN Lactotransferrin OS=Homo sapiens OX=9606 GN=LTF PE=1 SV=6 | 1270 | 20201118_lss_colostrum_1ng_glyco_hcd203040_compound15.20039.20039.3 | controllerType=0 controllerNumber=1 scan=20039 | 41.4998 |
| 19699:4 | 3 | R.TAGWNIPMGLLFN[+2204.772]QTGSC[+57.021]KFDEYFSQSC[+57.021]APGSDPR.S | HexNAc(4)Hex(5)NeuAc(2) | C[+57]*2, N[+2205] | 1536.6334 | 4 | 6143.5119 | 6143.5295 | 0 | -2.9 | 485 | Specific | 243.54 | 211.33 | 211.33 | 4.59 | 146 | >sp\|P02788\|TRFL_HUMAN Lactotransferrin OS=Homo sapiens OX=9606 GN=LTF PE=1 SV=6 | 1270 | 20201118_lss_colostrum_1ng_glyco_hcd203040_compound15.23345.23345.4 | controllerType=0 controllerNumber=1 scan=23345 | [illegible] |
| 19696:4 | 3 | R.TAGWNIPMGLLFN[+2204.772]QTGSC[+57.021]KFDEYFSQSC[+57.021]APGSDPR.S | HexNAc(4)Hex(5)NeuAc(2) | C[+57]*2, N[+2205] | 1536.6334 | 4 | 6143.5119 | 6143.5295 | 0 | -2.9 | 485 | Specific | 327.47 | 253.59 | 164.82 | 5.24 | 146 | >sp\|P02788\|TRFL_HUMAN Lactotransferrin OS=Homo sapiens OX=9606 GN=LTF PE=1 SV=6 | 1270 | 20201118_lss_colostrum_1ng_glyco_hcd203040_compound15.23342.23342.4 | controllerType=0 controllerNumber=1 scan=23342 | [illegible] |
| 18244:4 | 3 | R.TAGWNIPMGLLFN[+2205.793]QTGSC[+57.021]KFDEYFSQSC[+57.021]APGSDPR.S | HexNAc(4)Hex(5)Fuc(2)NeuAc(1) | C[+57]*2, N[+2206] | 1536.9033 | 4 | 6144.5915 | 6144.5499 | 0 | 6.8 | 485 | Specific | 355.29 | 348.75 | 348.75 | 5.27 | 146 | >sp\|P02788\|TRFL_HUMAN Lactotransferrin OS=Homo sapiens OX=9606 GN=LTF PE=1 SV=6 | 1270 | 20201118_lss_colostrum_1ng_glyco_hcd203040_compound15.21829.21829.3 | controllerType=0 controllerNumber=1 scan=21829 | [illegible] |
| 18489:4 | 3 | R.TAGWNIPMGLLFN[+1913.677]QTGSC[+57.021]KFDEYFSQSC[+57.021]APGSDPR.S | HexNAc(4)Hex(5)NeuAc(1) | C[+57]*2, N[+1914] | 1463.8683 | 4 | 5852.4513 | 5852.4341 | 0 | 2.9 | 485 | Specific | 363.72 | 310.76 | 310.76 | 6.95 | 146 | >sp\|P02788\|TRFL_HUMAN Lactotransferrin OS=Homo sapiens OX=9606 GN=LTF PE=1 SV=6 | 1270 | 20201118_lss_colostrum_1ng_glyco_hcd203040_compound15.22084.22084.4 | controllerType=0 controllerNumber=1 scan=22084 | [illegible] |
| 18289:5 | 3 | R.TAGWNIPMGLLFN[+2204.772]QTGSC[+57.021]KFDEYFSQSC[+57.021]APGSDPR.S | HexNAc(4)Hex(5)NeuAc(2) | C[+57]*2, N[+2205] | 1229.5188 | 5 | 6143.5649 | 6143.5295 | 0 | 5.8 | 485 | Specific | 388.52 | 388.52 | 388.52 | 8.59 | 146 | >sp\|P02788\|TRFL_HUMAN Lactotransferrin OS=Homo sapiens OX=9606 GN=LTF PE=1 SV=6 | 1270 | 20201118_lss_colostrum_1ng_glyco_hcd203040_compound15.21876.21876.5 | controllerType=0 controllerNumber=1 scan=21876 | 45.4935 |
| 18241:4 | 3 | R.TAGWNIPMGLLFN[+2205.793]QTGSC[+57.021]KFDEYFSQSC[+57.021]APGSDPR.S | HexNAc(4)Hex(5)Fuc(2)NeuAc(1) | C[+57]*2, N[+2206] | 1536.9033 | 4 | 6144.5915 | 6144.5499 | 0 | 6.8 | 485 | Specific | 417.49 | 405.56 | 405.56 | 8.01 | 146 | >sp\|P02788\|TRFL_HUMAN Lactotransferrin OS=Homo sapiens OX=9606 GN=LTF PE=1 SV=6 | 1270 | 20201118_lss_colostrum_1ng_glyco_hcd203040_compound15.21826.21826.3 | controllerType=0 controllerNumber=1 scan=21826 | 45.3867 |
| 17454:4 | 3 | R.TAGWNIPMGLLFN[+1216.423]QTGSC[+57.021]KFDEYFSQSC[+57.021]APGSDPR.S | HexNAc(2)Hex(5) | C[+57]*2, N[+1216] | 1289.5559 | 4 | 5155.2018 | 5155.18 | 0 | 4.2 | 485 | Specific | 463.19 | 463.19 | 115.77 | 9.04 | 146 | >sp\|P02788\|TRFL_HUMAN Lactotransferrin OS=Homo sapiens OX=9606 GN=LTF PE=1 SV=6 | 1270 | 20201118_lss_colostrum_1ng_glyco_hcd203040_compound15.21004.21004.4 | controllerType=0 controllerNumber=1 scan=21004 | 43.5668 |
| 18292:5 | 3 | R.TAGWNIPMGLLFN[+2204.772]QTGSC[+57.021]KFDEYFSQSC[+57.021]APGSDPR.S | HexNAc(4)Hex(5)NeuAc(2) | C[+57]*2, N[+2205] | 1229.5188 | 5 | 6143.5649 | 6143.5295 | 0 | 5.8 | 485 | Specific | 518.85 | 489.52 | 489.52 | 9.13 | 146 | >sp\|P02788\|TRFL_HUMAN Lactotransferrin OS=Homo sapiens OX=9606 GN=LTF PE=1 SV=6 | 1270 | 20201118_lss_colostrum_1ng_glyco_hcd203040_compound15.21879.21879.5 | controllerType=0 controllerNumber=1 scan=21879 | 45.5015 |
| 17299:4 | 3 | R.TAGWNIPMGLLFN[+1768.640]QTGSC[+57.021]KFDEYFSQSC[+57.021]APGSDPR.S | HexNAc(4)Hex(5)Fuc(1) | C[+57]*2, N[+1769] | 1427.61 | 4 | 5707.4181 | 5707.3966 | 0 | 3.8 | 485 | Specific | 551.25 | 536.44 | 536.44 | 9.35 | 146 | >sp\|P02788\|TRFL_HUMAN Lactotransferrin OS=Homo sapiens OX=9606 GN=LTF PE=1 SV=6 | 1270 | 20201118_lss_colostrum_1ng_glyco_hcd203040_compound15.20843.20843.4 | controllerType=0 controllerNumber=1 scan=20843 | 43.2392 |
| 17457:4 | 3 | R.TAGWNIPMGLLFN[+1216.423]QTGSC[+57.021]KFDEYFSQSC[+57.021]APGSDPR.S | HexNAc(2)Hex(5) | C[+57]*2, N[+1216] | 1289.5559 | 4 | 5155.2018 | 5155.18 | 0 | 4.2 | 485 | Specific | 555.68 | 555.68 | 253.06 | 9.61 | 146 | >sp\|P02788\|TRFL_HUMAN Lactotransferrin OS=Homo sapiens OX=9606 GN=LTF PE=1 SV=6 | 1270 | 20201118_lss_colostrum_1ng_glyco_hcd203040_compound15.21007.21007.4 | controllerType=0 controllerNumber=1 scan=21007 | 43.5784 |
| 18296:4 | 3 | R.TAGWNIPMGLLFN[+2059.735]QTGSC[+57.021]KFDEYFSQSC[+57.021]APGSDPR.S | HexNAc(4)Hex(5)Fuc(1)NeuAc(1) | C[+57]*2, N[+2060] | 1500.3822 | 4 | 5998.507 | 5998.492 | 0 | 2.5 | 485 | Specific | 632.41 | 602.67 | 602.67 | 12.15 | 146 | >sp\|P02788\|TRFL_HUMAN Lactotransferrin OS=Homo sapiens OX=9606 GN=LTF PE=1 SV=6 | 1270 | 20201118_lss_colostrum_1ng_glyco_hcd203040_compound15.21884.21884.4 | controllerType=0 controllerNumber=1 scan=21884 | 45.5118 |
| 17453:3 | 3 | R.TAGWNIPMGLLFN[+1216.423]QTGSC[+57.021]KFDEYFSQSC[+57.021]APGSDPR.S | HexNAc(2)Hex(5) | C[+57]*2, N[+1216] | 1719.0721 | 3 | 5155.2018 | 5155.18 | 0 | 4.2 | 485 | Specific | 633.28 | 586.04 | 129.18 | 12.02 | 146 | >sp\|P02788\|TRFL_HUMAN Lactotransferrin OS=Homo sapiens OX=9606 GN=LTF PE=1 SV=6 | 1270 | 20201118_lss_colostrum_1ng_glyco_hcd203040_compound15.21003.21003.3 | controllerType=0 controllerNumber=1 scan=21003 | 43.5657 |
| 19578:5 | 3 | R.TAGWNIPMGLLFN[+2350.830]QTGSC[+57.021]KFDEYFSQSC[+57.021]APGSDPR.S | HexNAc(4)Hex(5)Fuc(1)NeuAc(2) | C[+57]*2, N[+2351] | 1258.7316 | 5 | 6289.6287 | 6289.5875 | 0 | 6.6 | 485 | Specific | 688.16 | 676.66 | 676.66 | 12.34 | 146 | >sp\|P02788\|TRFL_HUMAN Lactotransferrin OS=Homo sapiens OX=9606 GN=LTF PE=1 SV=6 | 1270 | 20201118_lss_colostrum_1ng_glyco_hcd203040_compound15.23219.23219.5 | controllerType=0 controllerNumber=1 scan=23219 | 48.4084 |
| 18258:4 | 3 | R.TAGWNIPMGLLFN[+2351.851]QTGSC[+57.021]KFDEYFSQSC[+57.021]APGSDPR.S | HexNAc(4)Hex(5)Fuc(3)NeuAc(1) | C[+57]*2, N[+2352] | 1573.4119 | 4 | 6290.6256 | 6290.6079 | 0 | 2.8 | 485 | Specific | 710 | 707.68 | 707.68 | 14.15 | 146 | >sp\|P02788\|TRFL_HUMAN Lactotransferrin OS=Homo sapiens OX=9606 GN=LTF PE=1 SV=6 | 1270 | 20201118_lss_colostrum_1ng_glyco_hcd203040_compound15.21844.21844.4 | controllerType=0 controllerNumber=1 scan=21844 | 45.4271 |
| 18268:4 | 3 | R.TAGWNIPMGLLFN[+2205.793]QTGSC[+57.021]KFDEYFSQSC[+57.021]APGSDPR.S | HexNAc(4)Hex(5)Fuc(2)NeuAc(1) | C[+57]*2, N[+2206] | 1536.897 | 4 | 6144.5661 | 6144.5499 | 0 | 2.6 | 485 | Specific | 758.34 | 709.37 | 709.37 | 15.15 | 146 | >sp\|P02788\|TRFL_HUMAN Lactotransferrin OS=Homo sapiens OX=9606 GN=LTF PE=1 SV=6 | 1270 | 20201118_lss_colostrum_1ng_glyco_hcd203040_compound15.21855.21855.4 | controllerType=0 controllerNumber=1 scan=21855 | 45.4573 |
| 19593:4 | 3 | R.TAGWNIPMGLLFN[+2350.830]QTGSC[+57.021]KFDEYFSQSC[+57.021]APGSDPR.S | HexNAc(4)Hex(5)Fuc(1)NeuAc(2) | C[+57]*2, N[+2351] | 1573.157 | 4 | 6289.6061 | 6289.5875 | 0 | 3 | 485 | Specific | 797.11 | 740.14 | 740.14 | 15.15 | 146 | >sp\|P02788\|TRFL_HUMAN Lactotransferrin OS=Homo sapiens OX=9606 GN=LTF PE=1 SV=6 | 1270 | 20201118_lss_colostrum_1ng_glyco_hcd203040_compound15.23235.23235.4 | controllerType=0 controllerNumber=1 scan=23235 | 48.4423 |
| 17456:3 | 3 | R.TAGWNIPMGLLFN[+1216.423]QTGSC[+57.021]KFDEYFSQSC[+57.021]APGSDPR.S | HexNAc(2)Hex(5) | C[+57]*2, N[+1216] | 1719.0721 | 3 | 5155.2018 | 5155.18 | 0 | 4.2 | 485 | Specific | 817.84 | 782.13 | 210.19 | 16.02 | 146 | >sp\|P02788\|TRFL_HUMAN Lactotransferrin OS=Homo sapiens OX=9606 GN=LTF PE=1 SV=6 | 1270 | 20201118_lss_colostrum_1ng_glyco_hcd203040_compound15.21006.21006.3 | controllerType=0 controllerNumber=1 scan=21006 | 43.5765 |
| 18492:4 | 3 | R.TAGWNIPMGLLFN[+1913.677]QTGSC[+57.021]KFDEYFSQSC[+57.021]APGSDPR.S | HexNAc(4)Hex(5)NeuAc(1) | C[+57]*2, N[+1914] | 1463.8683 | 4 | 5852.4513 | 5852.4341 | 0 | 2.9 | 485 | Specific | 823.26 | 729.3 | 729.3 | 16.15 | 146 | >sp\|P02788\|TRFL_HUMAN Lactotransferrin OS=Homo sapiens OX=9606 GN=LTF PE=1 SV=6 | 1270 | 20201118_lss_colostrum_1ng_glyco_hcd203040_compound15.22087.22087.4 | controllerType=0 controllerNumber=1 scan=22087 | 45.8903 |
| 18261:4 | 3 | R.TAGWNIPMGLLFN[+2351.851]QTGSC[+57.021]KFDEYFSQSC[+57.021]APGSDPR.S | HexNAc(4)Hex(5)Fuc(3)NeuAc(1) | C[+57]*2, N[+2352] | 1573.4119 | 4 | 6290.6256 | 6290.6079 | 0 | 2.8 | 485 | Specific | 831.23 | 819.38 | 819.38 | 16.15 | 146 | >sp\|P02788\|TRFL_HUMAN Lactotransferrin OS=Homo sapiens OX=9606 GN=LTF PE=1 SV=6 | 1270 | 20201118_lss_colostrum_1ng_glyco_hcd203040_compound15.21847.21847.4 | controllerType=0 controllerNumber=1 scan=21847 | 45.4365 |
| 18299:4 | 3 | R.TAGWNIPMGLLFN[+2059.735]QTGSC[+57.021]KFDEYFSQSC[+57.021]APGSDPR.S | HexNAc(4)Hex(5)Fuc(1)NeuAc(1) | C[+57]*2, N[+2060] | 1500.3822 | 4 | 5998.507 | 5998.492 | 0 | 2.5 | 485 | Specific | 851.22 | 802.82 | 802.82 | 17.15 | 146 | >sp\|P02788\|TRFL_HUMAN Lactotransferrin OS=Homo sapiens OX=9606 GN=LTF PE=1 SV=6 | 1270 | 20201118_lss_colostrum_1ng_glyco_hcd203040_compound15.21887.21887.4 | controllerType=0 controllerNumber=1 scan=21887 | 45.5157 |
| 18370:4 | 3 | R.TAGWNIPMGLLFN[+1913.677]QTGSC[+57.021]KFDEYFSQSC[+57.021]APGSDPR.S | HexNAc(4)Hex(5)NeuAc(1) | C[+57]*2, N[+1914] | 1463.8679 | 4 | 5852.4499 | 5852.4341 | 0 | 2.7 | 485 | Specific | 861.06 | 753.33 | 753.33 | 17.15 | 146 | >sp\|P02788\|TRFL_HUMAN Lactotransferrin OS=Homo sapiens OX=9606 GN=LTF PE=1 SV=6 | 1270 | 20201118_lss_colostrum_1ng_glyco_hcd203040_compound15.21960.21960.2 | controllerType=0 controllerNumber=1 scan=21960 | 45.6525 |
| 18270:4 | 3 | R.TAGWNIPMGLLFN[+2205.793]QTGSC[+57.021]KFDEYFSQSC[+57.021]APGSDPR.S | HexNAc(4)Hex(5)Fuc(2)NeuAc(1) | C[+57]*2, N[+2206] | 1536.897 | 4 | 6144.5661 | 6144.5499 | 0 | 2.6 | 485 | Specific | 869.74 | 846.44 | 846.44 | 17.15 | 146 | >sp\|P02788\|TRFL_HUMAN Lactotransferrin OS=Homo sapiens OX=9606 GN=LTF PE=1 SV=6 | 1270 | 20201118_lss_colostrum_1ng_glyco_hcd203040_compound15.21857.21857.4 | controllerType=0 controllerNumber=1 scan=21857 | 45.4621 |
| 18373:4 | 3 | R.TAGWNIPMGLLFN[+1913.677]QTGSC[+57.021]KFDEYFSQSC[+57.021]APGSDPR.S | HexNAc(4)Hex(5)NeuAc(1) | C[+57]*2, N[+1914] | 1463.8679 | 4 | 5852.4499 | 5852.4341 | 0 | 2.7 | 485 | Specific | 899.41 | 785.9 | 785.9 | 17.15 | 146 | >sp\|P02788\|TRFL_HUMAN Lactotransferrin OS=Homo sapiens OX=9606 GN=LTF PE=1 SV=6 | 1270 | 20201118_lss_colostrum_1ng_glyco_hcd203040_compound15.21963.21963.2 | controllerType=0 controllerNumber=1 scan=21963 | 45.6655 |
| 19581:5 | 3 | R.TAGWNIPMGLLFN[+2350.830]QTGSC[+57.021]KFDEYFSQSC[+57.021]APGSDPR.S | HexNAc(4)Hex(5)Fuc(1)NeuAc(2) | C[+57]*2, N[+2351] | 1258.7316 | 5 | 6289.6287 | 6289.5875 | 0 | 6.6 | 485 | Specific | 914.26 | 876.61 | 876.61 | 17.34 | 146 | >sp\|P02788\|TRFL_HUMAN Lactotransferrin OS=Homo sapiens OX=9606 GN=LTF PE=1 SV=6 | 1270 | 20201118_lss_colostrum_1ng_glyco_hcd203040_compound15.23222.23222.5 | controllerType=0 controllerNumber=1 scan=23222 | 48.4163 |
| 17301:4 | 3 | R.TAGWNIPMGLLFN[+1768.640]QTGSC[+57.021]KFDEYFSQSC[+57.021]APGSDPR.S | HexNAc(4)Hex(5)Fuc(1) | C[+57]*2, N[+1769] | 1427.61 | 4 | 5707.4181 | 5707.3966 | 0 | 3.8 | 485 | Specific | 919.17 | 855.17 | 855.17 | 17.95 | 146 | >sp\|P02788\|TRFL_HUMAN Lactotransferrin OS=Homo sapiens OX=9606 GN=LTF PE=1 SV=6 | 1270 | 20201118_lss_colostrum_1ng_glyco_hcd203040_compound15.20845.20845.4 | controllerType=0 controllerNumber=1 scan=20845 | 43.2441 |
| 19595:4 | 3 | R.TAGWNIPMGLLFN[+2350.830]QTGSC[+57.021]KFDEYFSQSC[+57.021]APGSDPR.S | HexNAc(4)Hex(5)Fuc(1)NeuAc(2) | C[+57]*2, N[+2351] | 1573.157 | 4 | 6289.6061 | 6289.5875 | 0 | 3 | 485 | Specific | 950.27 | 915.98 | 915.98 | 19.15 | 146 | >sp\|P02788\|TRFL_HUMAN Lactotransferrin OS=Homo sapiens OX=9606 GN=LTF PE=1 SV=6 | 1270 | 20201118_lss_colostrum_1ng_glyco_hcd203040_compound15.23237.23237.4 | controllerType=0 controllerNumber=1 scan=23237 | 48.4468 |
| 17240:4 | 3 | R.TAGWNIPM[+15.995]GLLFN[+2116.756]QTGSC[+57.021]KFDEYFSQSC[+57.021]A | HexNAc(5)Hex(5)NeuAc(1) | C[+57]*2, M[+16], N[+2117] | 1518.6382 | 4 | 6071.5309 | 6071.5084 | 0 | 3.7 | 485 | Specific | 315.06 | 289.84 | 289.84 | 6.52 | 146 | >sp\|P02788\|TRFL_HUMAN Lactotransferrin OS=Homo sapiens OX=9606 GN=LTF PE=1 SV=6 | 1270 | 20201118_lss_colostrum_1ng_glyco_hcd203040_compound15.20781.20781.4 | controllerType=0 controllerNumber=1 scan=20781 | 43.1109 |
| 17300:3 | 3 | R.TAGWNIPM[+15.995]GLLFN[+1751.624]QTGSC[+57.021]KFDEYFSQSC[+57.021]A | HexNAc(4)Hex(4)NeuAc(1) | C[+57]*2, M[+16], N[+1752] | 1902.8099 | 3 | 5706.4151 | 5706.3762 | 0 | 6.8 | 485 | Specific | 321.17 | 290.57 | 290.57 | 5.74 | 146 | >sp\|P02788\|TRFL_HUMAN Lactotransferrin OS=Homo sapiens OX=9606 GN=LTF PE=1 SV=6 | 1270 | 20201118_lss_colostrum_1ng_glyco_hcd203040_compound15.20844.20844.3 | controllerType=0 controllerNumber=1 scan=20844 | 43.2407 |
| 17243:4 | 3 | R.TAGWNIPM[+15.995]GLLFN[+2116.756]QTGSC[+57.021]KFDEYFSQSC[+57.021]A | HexNAc(5)Hex(5)NeuAc(1) | C[+57]*2, M[+16], N[+2117] | 1518.6382 | 4 | 6071.5309 | 6071.5084 | 0 | 3.7 | 485 | Specific | 354.01 | 271.28 | 271.28 | 6.51 | 146 | >sp\|P02788\|TRFL_HUMAN Lactotransferrin OS=Homo sapiens OX=9606 GN=LTF PE=1 SV=6 | 1270 | 20201118_lss_colostrum_1ng_glyco_hcd203040_compound15.20784.20784.4 | controllerType=0 controllerNumber=1 scan=20784 | 43.115 |
| 17302:3 | 3 | R.TAGWNIPM[+15.995]GLLFN[+1751.624]QTGSC[+57.021]KFDEYFSQSC[+57.021]A | HexNAc(4)Hex(4)NeuAc(1) | C[+57]*2, M[+16], N[+1752] | 1902.8099 | 3 | 5706.4151 | 5706.3762 | 0 | 6.8 | 485 | Specific | 371.63 | 202.18 | 202.18 | 4.8 | 146 | >sp\|P02788\|TRFL_HUMAN Lactotransferrin OS=Homo sapiens OX=9606 GN=LTF PE=1 SV=6 | 1270 | 20201118_lss_colostrum_1ng_glyco_hcd203040_compound15.20846.20846.3 | controllerType=0 controllerNumber=1 scan=20846 | 43.2453 |
| 08061:3 | 3 | K.FGRN[+1501.555]GSDC[+57.021]PDKFC[+57.021]LFQSETK.N | HexNAc(5)Hex(3) | C[+57]*2, N[+1502] | 1298.8847 | 3 | 3894.6395 | 3894.6202 | 0 | 5 | 639 | Specific | 298.8 | 298.8 | 298.8 | 5.25 | 146 | >sp\|P02788\|TRFL_HUMAN Lactotransferrin OS=Homo sapiens OX=9606 GN=LTF PE=1 SV=6 | 1270 | 20201118_lss_colostrum_1ng_glyco_hcd203040_compound15.11239.11239.3 | controllerType=0 controllerNumber=1 scan=11239 | 25.0597 |
| 08975:3 | 3 | R.N[+2059.735]GSDC[+57.021]PDKFC[+57.021]LFQSETK.N | HexNAc(4)Hex(5)Fuc(1)NeuAc(1) | C[+57]*2, N[+2060] | 1364.8798 | 3 | 4092.6249 | 4092.6088 | 0 | 3.9 | 642 | Specific | 451.89 | 409.14 | 409.14 | 8.99 | 146 | >sp\|P02788\|TRFL_HUMAN Lactotransferrin OS=Homo sapiens OX=9606 GN=LTF PE=1 SV=6 | 1270 | 20201118_lss_colostrum_1ng_glyco_hcd203040_compound15.12192.12192.3 | controllerType=0 controllerNumber=1 scan=12192 | 26.6845 |
| 08977:3 | 3 | R.N[+2059.735]GSDC[+57.021]PDKFC[+57.021]LFQSETK.N | HexNAc(4)Hex(5)Fuc(1)NeuAc(1) | C[+57]*2, N[+2060] | 1364.8798 | 3 | 4092.6249 | 4092.6088 | 0 | 3.9 | 642 | Specific | 569.89 | 451.06 | 451.06 | 9.56 | 146 | >sp\|P02788\|TRFL_HUMAN Lactotransferrin OS=Homo sapiens OX=9606 GN=LTF PE=1 SV=6 | 1270 | 20201118_lss_colostrum_1ng_glyco_hcd203040_compound15.12194.12194.3 | controllerType=0 controllerNumber=1 scan=12194 | 26.6901 |
| 04625:3 | 4 | R.FSDGLESN[+1864.634]SSTQFEVKK.Y | HexNAc(2)Hex(9) | N[+1865] | 1256.5189 | 3 | 3767.5421 | 3767.5421 | 0 | 0 | 219 | Specific | 235.38 | 235.38 | 235.38 | 5.24 | 91 | >sp\|P0C0L4\|CO4A_HUMAN Complement C4-A OS=Homo sapiens OX=9606 GN=C4A PE=1 SV=2 | 1235 | 20201118_lss_colostrum_1ng_glyco_hcd203040_compound15.7683.7683.3 | controllerType=0 controllerNumber=1 scan=7683 | 19.2372 |
| 17036:3 | 5 | R.GLSFDVSLEVSQGPGLLN[+2350.830]DTK.V | HexNAc(4)Hex(5)Fuc(1)NeuAc(2) | N[+2351] | 1509.666 | 3 | 4526.9833 | 4526.9436 | 0 | 8.8 | 118 | Specific | 442.67 | 404.25 | 404.25 | 6.71 | 97 | >sp\|P01833\|PIGR_HUMAN Polymeric immunoglobulin receptor OS=Homo sapiens OX=9606 GN=PIGR PE=1 SV=4 | 902 | 20201118_lss_colostrum_1ng_glyco_hcd203040_compound15.20568.20568.3 | controllerType=0 controllerNumber=1 scan=20568 | 42.6265 |
| 16873:3 | 5 | R.GLSFDVSLEVSQGPGLLN[+2571.946]DTK.V | HexNAc(5)Hex(6)Fuc(4) | N[+2572] | 1583.3571 | 3 | 4748.0568 | 4748.0586 | 0 | -0.4 | 118 | Specific | 772.35 | 710.52 | 710.52 | 15 | 97 | >sp\|P01833\|PIGR_HUMAN Polymeric immunoglobulin receptor OS=Homo sapiens OX=9606 GN=PIGR PE=1 SV=4 | 902 | 20201118_lss_colostrum_1ng_glyco_hcd203040_compound15.20398.20398.3 | controllerType=0 controllerNumber=1 scan=20398 | 42.2435 |
| 16875:3 | 5 | R.GLSFDVSLEVSQGPGLLN[+2571.946]DTK.V | HexNAc(5)Hex(6)Fuc(4) | N[+2572] | 1583.3571 | 3 | 4748.0568 | 4748.0586 | 0 | -0.4 | 118 | Specific | 779.72 | 684.31 | 684.31 | 15 | 97 | >sp\|P01833\|PIGR_HUMAN Polymeric immunoglobulin receptor OS=Homo sapiens OX=9606 GN=PIGR PE=1 SV=4 | 902 | 20201118_lss_colostrum_1ng_glyco_hcd203040_compound15.20400.20400.3 | controllerType=0 controllerNumber=1 scan=20400 | 42.2492 |
| 18866:4 | 5 | R.GLSFDVSLEVSQGPGLLN[+2350.830]DTK.V | HexNAc(4)Hex(5)Fuc(1)NeuAc(2) | N[+2351] | 1132.4954 | 4 | 4526.9598 | 4526.9436 | 0 | 3.6 | 118 | Specific | 889.5 | 652.51 | 652.51 | 17.67 | 97 | >sp\|P01833\|PIGR_HUMAN Polymeric immunoglobulin receptor OS=Homo sapiens OX=9606 GN=PIGR PE=1 SV=4 | 902 | 20201118_lss_colostrum_1ng_glyco_hcd203040_compound15.22476.22476.4 | controllerType=0 controllerNumber=1 scan=22476 | 46.7175 |
| 18869:4 | 5 | R.GLSFDVSLEVSQGPGLLN[+2350.830]DTK.V | HexNAc(4)Hex(5)Fuc(1)NeuAc(2) | N[+2351] | 1132.4954 | 4 | 4526.9598 | 4526.9436 | 0 | 3.6 | 118 | Specific | 892.87 | 869.32 | 869.32 | 17.67 | 97 | >sp\|P01833\|PIGR_HUMAN Polymeric immunoglobulin receptor OS=Homo sapiens OX=9606 GN=PIGR PE=1 SV=4 | 902 | 20201118_lss_colostrum_1ng_glyco_hcd203040_compound15.22479.22479.4 | controllerType=0 controllerNumber=1 scan=22479 | 46.7268 |
| 18860:3 | 5 | R.GLSFDVSLEVSQGPGLLN[+2350.830]DTK.V | HexNAc(4)Hex(5)Fuc(1)NeuAc(2) | N[+2351] | 1509.6596 | 3 | 4526.9643 | 4526.9436 | 0 | 4.6 | 118 | Specific | 974.77 | 796.9 | 796.9 | 18.97 | 97 | >sp\|P01833\|PIGR_HUMAN Polymeric immunoglobulin receptor OS=Homo sapiens OX=9606 GN=PIGR PE=1 SV=4 | 902 | 20201118_lss_colostrum_1ng_glyco_hcd203040_compound15.22470.22470.3 | controllerType=0 controllerNumber=1 scan=22470 | 46.7024 |
| 18862:3 | 5 | R.GLSFDVSLEVSQGPGLLN[+2350.830]DTK.V | HexNAc(4)Hex(5)Fuc(1)NeuAc(2) | N[+2351] | 1509.6596 | 3 | 4526.9643 | 4526.9436 | 0 | 4.6 | 118 | Specific | 1025.17 | 1002.12 | 1002.12 | 19.97 | 97 | >sp\|P01833\|PIGR_HUMAN Polymeric immunoglobulin receptor OS=Homo sapiens OX=9606 GN=PIGR PE=1 SV=4 | 902 | 20201118_lss_colostrum_1ng_glyco_hcd203040_compound15.22472.22472.3 | controllerType=0 controllerNumber=1 scan=22472 | 46.7079 |
| 22124:3 | 5 | K.QIGLYPVLVIDSSGYVNPN[+1913.677]YTGR.I | HexNAc(4)Hex(5)NeuAc(1) | N[+1914] | 1480.336 | 3 | 4438.9934 | 4438.9805 | 0 | 2.9 | 168 | Specific | 236.08 | 236.08 | 236.08 | 5.91 | 97 | >sp\|P01833\|PIGR_HUMAN Polymeric immunoglobulin receptor OS=Homo sapiens OX=9606 GN=PIGR PE=1 SV=4 | 902 | 20201118_lss_colostrum_1ng_glyco_hcd203040_compound15.25879.25879.3 | controllerType=0 controllerNumber=1 scan=25879 | 54.3819 |
| 21982:3 | 5 | K.QIGLYPVLVIDSSGYVNPN[+2059.735]YTGR.I | HexNAc(4)Hex(5)Fuc(1)NeuAc(1) | N[+2060] | 1529.025 | 3 | 4585.0603 | 4585.0384 | 0 | 4.8 | 168 | Specific | 236.37 | 219.13 | 219.13 | 5.27 | 97 | >sp\|P01833\|PIGR_HUMAN Polymeric immunoglobulin receptor OS=Homo sapiens OX=9606 GN=PIGR PE=1 SV=4 | 902 | 20201118_lss_colostrum_1ng_glyco_hcd203040_compound15.25731.25731.3 | controllerType=0 controllerNumber=1 scan=25731 | 54.0493 |
| 22126:3 | 5 | K.QIGLYPVLVIDSSGYVNPN[+1913.677]YTGR.I | HexNAc(4)Hex(5)NeuAc(1) | N[+1914] | 1480.336 | 3 | 4438.9934 | 4438.9805 | 0 | 2.9 | 168 | Specific | 284.74 | 284.74 | 284.74 | 6.13 | 97 | >sp\|P01833\|PIGR_HUMAN Polymeric immunoglobulin receptor OS=Homo sapiens OX=9606 GN=PIGR PE=1 SV=4 | 902 | 20201118_lss_colostrum_1ng_glyco_hcd203040_compound15.25881.25881.3 | controllerType=0 controllerNumber=1 scan=25881 | 54.3876 |
| 18208:3 | 5 | K.QIGLYPVLVIDSSGYVNPN[+2076.750]YTGR.I | HexNAc(4)Hex(6)Fuc(2) | N[+2077] | 1534.6979 | 3 | 4602.0792 | 4602.0537 | 0 | 5.5 | 168 | Specific | 287.21 | 231.57 | 231.57 | 5.19 | 97 | >sp\|P01833\|PIGR_HUMAN Polymeric immunoglobulin receptor OS=Homo sapiens OX=9606 GN=PIGR PE=1 SV=4 | 902 | 20201118_lss_colostrum_1ng_glyco_hcd203040_compound15.21792.21792.3 | controllerType=0 controllerNumber=1 scan=21792 | 45.3045 |
| 20208:5 | 5 | K.QIGLYPVLVIDSSGYVNPN[+2204.772]YTGR.I | HexNAc(4)Hex(5)NeuAc(2) | N[+2205] | 946.8241 | 5 | 4730.0912 | 4730.0759 | 0 | 3.2 | 168 | Specific | 353.17 | 296.4 | 296.4 | 7.66 | 97 | >sp\|P01833\|PIGR_HUMAN Polymeric immunoglobulin receptor OS=Homo sapiens OX=9606 GN=PIGR PE=1 SV=4 | 902 | 20201118_lss_colostrum_1ng_glyco_hcd203040_compound15.23876.23876.5 | controllerType=0 controllerNumber=1 scan=23876 | 49.8996 |
| 18200:4 | 5 | K.QIGLYPVLVIDSSGYVNPN[+2076.750]YTGR.I | HexNAc(4)Hex(6)Fuc(2) | N[+2077] | 1151.2747 | 4 | 4602.077 | 4602.0537 | 0 | 5.1 | 168 | Specific | 370.03 | 370.03 | 370.03 | 7.8 | 97 | >sp\|P01833\|PIGR_HUMAN Polymeric immunoglobulin receptor OS=Homo sapiens OX=9606 GN=PIGR PE=1 SV=4 | 902 | 20201118_lss_colostrum_1ng_glyco_hcd203040_compound15.21784.21784.4 | controllerType=0 controllerNumber=1 scan=21784 | 45.2888 |
| 18202:4 | 5 | K.QIGLYPVLVIDSSGYVNPN[+2076.750]YTGR.I | HexNAc(4)Hex(6)Fuc(2) | N[+2077] | 1151.2747 | 4 | 4602.077 | 4602.0537 | 0 | 5.1 | 168 | Specific | 375.51 | 340.74 | 340.74 | 7.6 | 97 | >sp\|P01833\|PIGR_HUMAN Polymeric immunoglobulin receptor OS=Homo sapiens OX=9606 GN=PIGR PE=1 SV=4 | 902 | 20201118_lss_colostrum_1ng_glyco_hcd203040_compound15.21786.21786.4 | controllerType=0 controllerNumber=1 scan=21786 | 45.2929 |
| 20143:4 | 5 | K.QIGLYPVLVIDSSGYVNPN[+2350.830]YTGR.I | HexNAc(4)Hex(5)Fuc(1)NeuAc(2) | N[+2351] | 1219.7962 | 4 | 4876.1629 | 4876.1338 | 0 | 6 | 168 | Specific | 456.33 | 385.9 | 385.9 | 9.21 | 97 | >sp\|P01833\|PIGR_HUMAN Polymeric immunoglobulin receptor OS=Homo sapiens OX=9606 GN=PIGR PE=1 SV=4 | 902 | 20201118_lss_colostrum_1ng_glyco_hcd203040_compound15.23808.23808.4 | controllerType=0 controllerNumber=1 scan=23808 | 49.7429 |
| 20210:5 | 5 | K.QIGLYPVLVIDSSGYVNPN[+2204.772]YTGR.I | HexNAc(4)Hex(5)NeuAc(2) | N[+2205] | 946.8241 | 5 | 4730.0912 | 4730.0759 | 0 | 3.2 | 168 | Specific | 521.59 | 498.77 | 498.77 | 10.37 | 97 | >sp\|P01833\|PIGR_HUMAN Polymeric immunoglobulin receptor OS=Homo sapiens OX=9606 GN=PIGR PE=1 SV=4 | 902 | 20201118_lss_colostrum_1ng_glyco_hcd203040_compound15.23878.23878.5 | controllerType=0 controllerNumber=1 scan=23878 | 49.9037 |
| 20146:4 | 5 | K.QIGLYPVLVIDSSGYVNPN[+2350.830]YTGR.I | HexNAc(4)Hex(5)Fuc(1)NeuAc(2) | N[+2351] | 1219.7962 | 4 | 4876.1629 | 4876.1338 | 0 | 6 | 168 | Specific | 687.7 | 568.3 | 568.3 | 13.12 | 97 | >sp\|P01833\|PIGR_HUMAN Polymeric immunoglobulin receptor OS=Homo sapiens OX=9606 GN=PIGR PE=1 SV=4 | 902 | 20201118_lss_colostrum_1ng_glyco_hcd203040_compound15.23811.23811.4 | controllerType=0 controllerNumber=1 scan=23811 | 49.7484 |
| 20190:4 | 5 | K.QIGLYPVLVIDSSGYVNPN[+2204.772]YTGR.I | HexNAc(4)Hex(5)NeuAc(2) | N[+2205] | 1183.2784 | 4 | 4730.0916 | 4730.0759 | 0 | 3.3 | 168 | Specific | 754.44 | 574.59 | 574.59 | 15.97 | 97 | >sp\|P01833\|PIGR_HUMAN Polymeric immunoglobulin receptor OS=Homo sapiens OX=9606 GN=PIGR PE=1 SV=4 | 902 | 20201118_lss_colostrum_1ng_glyco_hcd203040_compound15.23858.23858.4 | controllerType=0 controllerNumber=1 scan=23858 | 49.8571 |
| 18251:3 | 5 | K.QIGLYPVLVIDSSGYVNPN[+1913.677]YTGR.I | HexNAc(4)Hex(5)NeuAc(1) | N[+1914] | 1480.3354 | 3 | 4438.9916 | 4438.9805 | 0 | 2.5 | 168 | Specific | 804.4 | 763.65 | 763.65 | 16.93 | 97 | >sp\|P01833\|PIGR_HUMAN Polymeric immunoglobulin receptor OS=Homo sapiens OX=9606 GN=PIGR PE=1 SV=4 | 902 | 20201118_lss_colostrum_1ng_glyco_hcd203040_compound15.21837.21837.3 | controllerType=0 controllerNumber=1 scan=21837 | 45.4146 |
| 18267:4 | 5 | K.QIGLYPVLVIDSSGYVNPN[+1913.677]YTGR.I | HexNAc(4)Hex(5)NeuAc(1) | N[+1914] | 1110.5035 | 4 | 4438.992 | 4438.9805 | 0 | 2.6 | 168 | Specific | 814.55 | 690.55 | 690.55 | 17.17 | 97 | >sp\|P01833\|PIGR_HUMAN Polymeric immunoglobulin receptor OS=Homo sapiens OX=9606 GN=PIGR PE=1 SV=4 | 902 | 20201118_lss_colostrum_1ng_glyco_hcd203040_compound15.21853.21853.4 | controllerType=0 controllerNumber=1 scan=21853 | 45.4533 |
| 18203:4 | 5 | K.QIGLYPVLVIDSSGYVNPN[+2059.735]YTGR.I | HexNAc(4)Hex(5)Fuc(1)NeuAc(1) | N[+2060] | 1147.0185 | 4 | 4585.0521 | 4585.0384 | 0 | 3 | 168 | Specific | 869.26 | 840.38 | 840.38 | 18.17 | 97 | >sp\|P01833\|PIGR_HUMAN Polymeric immunoglobulin receptor OS=Homo sapiens OX=9606 GN=PIGR PE=1 SV=4 | 902 | 20201118_lss_colostrum_1ng_glyco_hcd203040_compound15.21787.21787.4 | controllerType=0 controllerNumber=1 scan=21787 | 45.2965 |
| 18201:4 | 5 | K.QIGLYPVLVIDSSGYVNPN[+2059.735]YTGR.I | HexNAc(4)Hex(5)Fuc(1)NeuAc(1) | N[+2060] | 1147.0185 | 4 | 4585.0521 | 4585.0384 | 0 | 3 | 168 | Specific | 869.64 | 795.31 | 795.31 | 18.17 | 97 | >sp\|P01833\|PIGR_HUMAN Polymeric immunoglobulin receptor OS=Homo sapiens OX=9606 GN=PIGR PE=1 SV=4 | 902 | 20201118_lss_colostrum_1ng_glyco_hcd203040_compound15.21785.21785.4 | controllerType=0 controllerNumber=1 scan=21785 | 45.2903 |
| 18264:4 | 5 | K.QIGLYPVLVIDSSGYVNPN[+1913.677]YTGR.I | HexNAc(4)Hex(5)NeuAc(1) | N[+1914] | 1110.5035 | 4 | 4438.992 | 4438.9805 | 0 | 2.6 | 168 | Specific | 872.8 | 783.67 | 783.67 | 18.17 | 97 | >sp\|P01833\|PIGR_HUMAN Polymeric immunoglobulin receptor OS=Homo sapiens OX=9606 GN=PIGR PE=1 SV=4 | 902 | 20201118_lss_colostrum_1ng_glyco_hcd203040_compound15.21850.21850.4 | controllerType=0 controllerNumber=1 scan=21850 | 45.4399 |
| 18254:3 | 5 | K.QIGLYPVLVIDSSGYVNPN[+1913.677]YTGR.I | HexNAc(4)Hex(5)NeuAc(1) | N[+1914] | 1480.3354 | 3 | 4438.9916 | 4438.9805 | 0 | 2.5 | 168 | Specific | 905.64 | 629.77 | 629.77 | 18.93 | 97 | >sp\|P01833\|PIGR_HUMAN Polymeric immunoglobulin receptor OS=Homo sapiens OX=9606 GN=PIGR PE=1 SV=4 | 902 | 20201118_lss_colostrum_1ng_glyco_hcd203040_compound15.21840.21840.3 | controllerType=0 controllerNumber=1 scan=21840 | 45.42 |
| 20192:4 | 5 | K.QIGLYPVLVIDSSGYVNPN[+2204.772]YTGR.I | HexNAc(4)Hex(5)NeuAc(2) | N[+2205] | 1183.2784 | 4 | 4730.0916 | 4730.0759 | 0 | 3.3 | 168 | Specific | 911.16 | 790.76 | 790.76 | 18.97 | 97 | >sp\|P01833\|PIGR_HUMAN Polymeric immunoglobulin receptor OS=Homo sapiens OX=9606 GN=PIGR PE=1 SV=4 | 902 | 20201118_lss_colostrum_1ng_glyco_hcd203040_compound15.23860.23860.4 | controllerType=0 controllerNumber=1 scan=23860 | 49.867 |
| 24620:3 | 5 | R.LSLLEEPGN[+3007.058]GTFTVILNQLTSR.D | HexNAc(5)Hex(6)Fuc(1)NeuAc(3) | N[+3007] | 1803.7994 | 3 | 5409.3837 | 5409.3505 | 0 | 6.1 | 413 | Specific | 239.04 | 203.86 | 203.86 | 5.66 | 97 | >sp\|P01833\|PIGR_HUMAN Polymeric immunoglobulin receptor OS=Homo sapiens OX=9606 GN=PIGR PE=1 SV=4 | 902 | 20201118_lss_colostrum_1ng_glyco_hcd203040_compound15.28475.28475.3 | controllerType=0 controllerNumber=1 scan=28475 | 59.526 |
| 27568:4 | 5 | R.[+42.011]LSLLEEPGN[+2351.851]GTFTVILNQLTSR.D | HexNAc(4)Hex(5)Fuc(3)NeuAc(1) | .[42], N[+2352] | 1199.7933 | 4 | 4796.1512 | 4796.1539 | 0 | -0.6 | 413 | Specific | 274.55 | 274.55 | 274.55 | 6.71 | 97 | >sp\|P01833\|PIGR_HUMAN Polymeric immunoglobulin receptor OS=Homo sapiens OX=9606 GN=PIGR PE=1 SV=4 | 902 | 20201118_lss_colostrum_1ng_glyco_hcd203040_compound15.31551.31551.2 | controllerType=0 controllerNumber=1 scan=31551 | 65.2728 |
| 27570:4 | 5 | R.[+42.011]LSLLEEPGN[+2351.851]GTFTVILNQLTSR.D | HexNAc(4)Hex(5)Fuc(3)NeuAc(1) | .[42], N[+2352] | 1199.7933 | 4 | 4796.1512 | 4796.1539 | 0 | -0.6 | 413 | Specific | 299.3 | 274.14 | 274.14 | 6.98 | 97 | >sp\|P01833\|PIGR_HUMAN Polymeric immunoglobulin receptor OS=Homo sapiens OX=9606 GN=PIGR PE=1 SV=4 | 902 | 20201118_lss_colostrum_1ng_glyco_hcd203040_compound15.31553.31553.2 | controllerType=0 controllerNumber=1 scan=31553 | 65.2776 |
| 25153:4 | 5 | R.LSLLEEPGN[+2642.946]GTFTVILNQLTSR.D | HexNAc(4)Hex(5)Fuc(3)NeuAc(2) | N[+2643] | 1262.0671 | 4 | 5045.2467 | 5045.2387 | 0 | 1.6 | 413 | Specific | 342.94 | 342.94 | 342.94 | 7.97 | 97 | >sp\|P01833\|PIGR_HUMAN Polymeric immunoglobulin receptor OS=Homo sapiens OX=9606 GN=PIGR PE=1 SV=4 | 902 | 20201118_lss_colostrum_1ng_glyco_hcd203040_compound15.29030.29030.4 | controllerType=0 controllerNumber=1 scan=29030 | 60.5809 |
| 27586:3 | 5 | R.[+42.011]LSLLEEPGN[+2351.851]GTFTVILNQLTSR.D | HexNAc(4)Hex(5)Fuc(3)NeuAc(1) | .[42], N[+2352] | 1599.3847 | 3 | 4796.1395 | 4796.1539 | 0 | -3 | 413 | Specific | 360.84 | 314.41 | 314.41 | 6.44 | 97 | >sp\|P01833\|PIGR_HUMAN Polymeric immunoglobulin receptor OS=Homo sapiens OX=9606 GN=PIGR PE=1 SV=4 | 902 | 20201118_lss_colostrum_1ng_glyco_hcd203040_compound15.31570.31570.3 | controllerType=0 controllerNumber=1 scan=31570 | 65.3114 |
| 23310:3 | 5 | R.[+42.011]LSLLEEPGN[+2205.793]GTFTVILNQLTSR.D | HexNAc(4)Hex(5)Fuc(2)NeuAc(1) | .[42], N[+2206] | 1550.7132 | 3 | 4650.125 | 4650.096 | 0 | 6.2 | 413 | Specific | 374.63 | 374.63 | 374.63 | 7.16 | 97 | >sp\|P01833\|PIGR_HUMAN Polymeric immunoglobulin receptor OS=Homo sapiens OX=9606 GN=PIGR PE=1 SV=4 | 902 | 20201118_lss_colostrum_1ng_glyco_hcd203040_compound15.27115.27115.3 | controllerType=0 controllerNumber=1 scan=27115 | 56.9504 |
| 24586:4 | 5 | R.LSLLEEPGN[+3007.058]GTFTVILNQLTSR.D | HexNAc(5)Hex(6)Fuc(1)NeuAc(3) | N[+3007] | 1353.0914 | 4 | 5409.3439 | 5409.3505 | 0 | -1.2 | 413 | Specific | 381.01 | 278.15 | 278.15 | 6.84 | 97 | >sp\|P01833\|PIGR_HUMAN Polymeric immunoglobulin receptor OS=Homo sapiens OX=9606 GN=PIGR PE=1 SV=4 | 902 | 20201118_lss_colostrum_1ng_glyco_hcd203040_compound15.28440.28440.4 | controllerType=0 controllerNumber=1 scan=28440 | 59.4608 |
| 23312:3 | 5 | R.[+42.011]LSLLEEPGN[+2205.793]GTFTVILNQLTSR.D | HexNAc(4)Hex(5)Fuc(2)NeuAc(1) | .[42], N[+2206] | 1550.7132 | 3 | 4650.125 | 4650.096 | 0 | 6.2 | 413 | Specific | 441.53 | 434.46 | 434.46 | 8.87 | 97 | >sp\|P01833\|PIGR_HUMAN Polymeric immunoglobulin receptor OS=Homo sapiens OX=9606 GN=PIGR PE=1 SV=4 | 902 | 20201118_lss_colostrum_1ng_glyco_hcd203040_compound15.27117.27117.3 | controllerType=0 controllerNumber=1 scan=27117 | 56.9548 |
| 22177:4 | 5 | R.LSLLEEPGN[+2351.851]GTFTVILNQLTSR.D | HexNAc(4)Hex(5)Fuc(3)NeuAc(1) | N[+2352] | 1189.2873 | 4 | 4754.1273 | 4754.1433 | 0 | -3.4 | 413 | Specific | 501.07 | 431.14 | 431.14 | 9.02 | 97 | >sp\|P01833\|PIGR_HUMAN Polymeric immunoglobulin receptor OS=Homo sapiens OX=9606 GN=PIGR PE=1 SV=4 | 902 | 20201118_lss_colostrum_1ng_glyco_hcd203040_compound15.25934.25934.4 | controllerType=0 controllerNumber=1 scan=25934 | 54.5035 |
| 25155:4 | 5 | R.LSLLEEPGN[+2642.946]GTFTVILNQLTSR.D | HexNAc(4)Hex(5)Fuc(3)NeuAc(2) | N[+2643] | 1262.0671 | 4 | 5045.2467 | 5045.2387 | 0 | 1.6 | 413 | Specific | 501.35 | 501.35 | 501.35 | 10.24 | 97 | >sp\|P01833\|PIGR_HUMAN Polymeric immunoglobulin receptor OS=Homo sapiens OX=9606 GN=PIGR PE=1 SV=4 | 902 | 20201118_lss_colostrum_1ng_glyco_hcd203040_compound15.29032.29032.4 | controllerType=0 controllerNumber=1 scan=29032 | 60.5884 |
| 22179:4 | 5 | R.LSLLEEPGN[+2351.851]GTFTVILNQLTSR.D | HexNAc(4)Hex(5)Fuc(3)NeuAc(1) | N[+2352] | 1189.2873 | 4 | 4754.1273 | 4754.1433 | 0 | -3.4 | 413 | Specific | 511.76 | 480.79 | 480.79 | 9.19 | 97 | >sp\|P01833\|PIGR_HUMAN Polymeric immunoglobulin receptor OS=Homo sapiens OX=9606 GN=PIGR PE=1 SV=4 | 902 | 20201118_lss_colostrum_1ng_glyco_hcd203040_compound15.25936.25936.4 | controllerType=0 controllerNumber=1 scan=25936 | 54.5091 |
| 24635:3 | 5 | R.LSLLEEPGN[+3008.078]GTFTVILNQLTSR.D | HexNAc(5)Hex(6)Fuc(3)NeuAc(2) | N[+3008] | 1804.1309 | 3 | 5410.3783 | 5410.3709 | 0 | 1.4 | 413 | Specific | 521.69 | 395.68 | 395.68 | 10.16 | 97 | >sp\|P01833\|PIGR_HUMAN Polymeric immunoglobulin receptor OS=Homo sapiens OX=9606 GN=PIGR PE=1 SV=4 | 902 | 20201118_lss_colostrum_1ng_glyco_hcd203040_compound15.28491.28491.3 | controllerType=0 controllerNumber=1 scan=28491 | 59.5575 |
| 22163:3 | 5 | R.LSLLEEPGN[+2350.830]GTFTVILNQLTSR.D | HexNAc(4)Hex(5)Fuc(1)NeuAc(2) | N[+2351] | 1585.0479 | 3 | 4753.1292 | 4753.1229 | 0 | 1.3 | 413 | Specific | 526.59 | 506.86 | 506.86 | 10.16 | 97 | >sp\|P01833\|PIGR_HUMAN Polymeric immunoglobulin receptor OS=Homo sapiens OX=9606 GN=PIGR PE=1 SV=4 | 902 | 20201118_lss_colostrum_1ng_glyco_hcd203040_compound15.25920.25920.3 | controllerType=0 controllerNumber=1 scan=25920 | 54.4758 |

Table S4. Cont.

| Query #:z | Protein Rank | Glycopeptide | Glycan | Modification Type(s) | Observed m/z | z | Observed (M+H) | Calc. mass (M+H) | Off-by-x error | Mass error (ppm) | StartPosition | Cleavage | Score | Delta | Delta Mod | \|Log Prob\| | # of unique peptides | Protein Name | Protein DB number | Comment | Scan # | Scan Time |
|---|---|---|---|---|---|---|---|---|---|---|---|---|---|---|---|---|---|---|---|---|---|---|
| 21941:3 | 5 | R.LSLLEEPGN[+2350.830]GTFTVILNQLTSR.D | HexNAc(4)Hex(5)Fuc(1)NeuAc(2) | N[+2351] | 1585.0588 | 3 | 4753.1618 | 4753.1229 | 0 | 8.2 | 413 | Specific | 589.45 | 488.76 | 488.76 | 10.66 | 97 | >sp\|P01833\|PIGR_HUMAN Polymeric immunoglobulin receptor OS=Homo sapiens OX=9606 GN=PIGR PE=1 SV=4 | 902 | 20201118_lss_colostrum_1ng_glyco_hcd203040_compound15.25687.25687.3 | controllerType=0 controllerNumber=1 scan=25687 | 53.9603 |
| 22161:3 | 5 | R.LSLLEEPGN[+2350.830]GTFTVILNQLTSR.D | HexNAc(4)Hex(5)Fuc(1)NeuAc(2) | N[+2351] | 1585.0479 | 3 | 4753.1292 | 4753.1229 | 0 | 1.3 | 413 | Specific | 612.21 | 513.39 | 513.39 | 12.77 | 97 | >sp\|P01833\|PIGR_HUMAN Polymeric immunoglobulin receptor OS=Homo sapiens OX=9606 GN=PIGR PE=1 SV=4 | 902 | 20201118_lss_colostrum_1ng_glyco_hcd203040_compound15.25918.25918.3 | controllerType=0 controllerNumber=1 scan=25918 | 54.471 |
| 24637:3 | 5 | R.LSLLEEPGN[+3008.078]GTFTVILNQLTSR.D | HexNAc(5)Hex(6)Fuc(3)NeuAc(2) | N[+3008] | 1804.1309 | 3 | 5410.3783 | 5410.3709 | 0 | 1.4 | 413 | Specific | 625.55 | 403.14 | 403.14 | 12.77 | 97 | >sp\|P01833\|PIGR_HUMAN Polymeric immunoglobulin receptor OS=Homo sapiens OX=9606 GN=PIGR PE=1 SV=4 | 902 | 20201118_lss_colostrum_1ng_glyco_hcd203040_compound15.28493.28493.3 | controllerType=0 controllerNumber=1 scan=28493 | 59.5616 |
| 19811:4 | 5 | R.LSLLEEPGN[+2076.750]GTFTVILNQLTSR.D | HexNAc(4)Hex(6)Fuc(2) | N[+2077] | 1120.5241 | 4 | 4479.0745 | 4479.0428 | 0 | 7.1 | 413 | Specific | 631.78 | 574.55 | 574.55 | 12.12 | 97 | >sp\|P01833\|PIGR_HUMAN Polymeric immunoglobulin receptor OS=Homo sapiens OX=9606 GN=PIGR PE=1 SV=4 | 902 | 20201118_lss_colostrum_1ng_glyco_hcd203040_compound15.23462.23462.4 | controllerType=0 controllerNumber=1 scan=23462 | 48.9417 |
| 19883:3 | 5 | R.LSLLEEPGN[+2059.735]GTFTVILNQLTSR.D | HexNAc(4)Hex(5)Fuc(1)NeuAc(1) | N[+2060] | 1488.0169 | 3 | 4462.0362 | 4462.0275 | 0 | 1.9 | 413 | Specific | 688.71 | 620.33 | 620.33 | 13.93 | 97 | >sp\|P01833\|PIGR_HUMAN Polymeric immunoglobulin receptor OS=Homo sapiens OX=9606 GN=PIGR PE=1 SV=4 | 902 | 20201118_lss_colostrum_1ng_glyco_hcd203040_compound15.23537.23537.3 | controllerType=0 controllerNumber=1 scan=23537 | 49.1295 |
| 21943:3 | 5 | R.LSLLEEPGN[+2350.830]GTFTVILNQLTSR.D | HexNAc(4)Hex(5)Fuc(1)NeuAc(2) | N[+2351] | 1585.0588 | 3 | 4753.1618 | 4753.1229 | 0 | 8.2 | 413 | Specific | 716.69 | 519.47 | 519.47 | 13.66 | 97 | >sp\|P01833\|PIGR_HUMAN Polymeric immunoglobulin receptor OS=Homo sapiens OX=9606 GN=PIGR PE=1 SV=4 | 902 | 20201118_lss_colostrum_1ng_glyco_hcd203040_compound15.25689.25689.3 | controllerType=0 controllerNumber=1 scan=25689 | 53.9652 |
| 21781:4 | 5 | R.LSLLEEPGN[+2496.888]GTFTVILNQLTSR.D | HexNAc(4)Hex(5)Fuc(2)NeuAc(2) | N[+2497] | 1225.5564 | 4 | 4899.2039 | 4899.1808 | 0 | 4.7 | 413 | Specific | 717.39 | 522.73 | 522.73 | 14.51 | 97 | >sp\|P01833\|PIGR_HUMAN Polymeric immunoglobulin receptor OS=Homo sapiens OX=9606 GN=PIGR PE=1 SV=4 | 902 | 20201118_lss_colostrum_1ng_glyco_hcd203040_compound15.25520.25520.4 | controllerType=0 controllerNumber=1 scan=25520 | 53.5686 |
| 21783:4 | 5 | R.LSLLEEPGN[+2496.888]GTFTVILNQLTSR.D | HexNAc(4)Hex(5)Fuc(2)NeuAc(2) | N[+2497] | 1225.5564 | 4 | 4899.2039 | 4899.1808 | 0 | 4.7 | 413 | Specific | 719.16 | 610.62 | 610.62 | 14.51 | 97 | >sp\|P01833\|PIGR_HUMAN Polymeric immunoglobulin receptor OS=Homo sapiens OX=9606 GN=PIGR PE=1 SV=4 | 902 | 20201118_lss_colostrum_1ng_glyco_hcd203040_compound15.25522.25522.4 | controllerType=0 controllerNumber=1 scan=25522 | 53.5738 |
| 19765:3 | 5 | R.LSLLEEPGN[+2351.851]GTFTVILNQLTSR.D | HexNAc(4)Hex(5)Fuc(3)NeuAc(1) | N[+2352] | 1585.3897 | 3 | 4754.1546 | 4754.1433 | 0 | 2.4 | 413 | Specific | 744.1 | 563.47 | 563.47 | 14.93 | 97 | >sp\|P01833\|PIGR_HUMAN Polymeric immunoglobulin receptor OS=Homo sapiens OX=9606 GN=PIGR PE=1 SV=4 | 902 | 20201118_lss_colostrum_1ng_glyco_hcd203040_compound15.23414.23414.3 | controllerType=0 controllerNumber=1 scan=23414 | 48.8259 |
| 19813:4 | 5 | R.LSLLEEPGN[+2076.750]GTFTVILNQLTSR.D | HexNAc(4)Hex(6)Fuc(2) | N[+2077] | 1120.5241 | 4 | 4479.0745 | 4479.0428 | 0 | 7.1 | 413 | Specific | 758.89 | 661.51 | 661.51 | 15.12 | 97 | >sp\|P01833\|PIGR_HUMAN Polymeric immunoglobulin receptor OS=Homo sapiens OX=9606 GN=PIGR PE=1 SV=4 | 902 | 20201118_lss_colostrum_1ng_glyco_hcd203040_compound15.23464.23464.4 | controllerType=0 controllerNumber=1 scan=23464 | 48.9461 |
| 19769:4 | 5 | R.LSLLEEPGN[+2351.851]GTFTVILNQLTSR.D | HexNAc(4)Hex(5)Fuc(3)NeuAc(1) | N[+2352] | 1189.294 | 4 | 4754.1541 | 4754.1433 | 0 | 2.3 | 413 | Specific | 778.22 | 647.32 | 647.32 | 16.17 | 97 | >sp\|P01833\|PIGR_HUMAN Polymeric immunoglobulin receptor OS=Homo sapiens OX=9606 GN=PIGR PE=1 SV=4 | 902 | 20201118_lss_colostrum_1ng_glyco_hcd203040_compound15.23418.23418.4 | controllerType=0 controllerNumber=1 scan=23418 | 48.8345 |
| 21776:4 | 5 | R.LSLLEEPGN[+2496.888]GTFTVILNQLTSR.D | HexNAc(4)Hex(5)Fuc(2)NeuAc(2) | N[+2497] | 1225.5523 | 4 | 4899.1873 | 4899.1808 | 0 | 1.3 | 413 | Specific | 860.07 | 675.57 | 675.57 | 18.01 | 97 | >sp\|P01833\|PIGR_HUMAN Polymeric immunoglobulin receptor OS=Homo sapiens OX=9606 GN=PIGR PE=1 SV=4 | 902 | 20201118_lss_colostrum_1ng_glyco_hcd203040_compound15.25514.25514.4 | controllerType=0 controllerNumber=1 scan=25514 | 53.5542 |
| 21777:3 | 5 | R.LSLLEEPGN[+2496.888]GTFTVILNQLTSR.D | HexNAc(4)Hex(5)Fuc(2)NeuAc(2) | N[+2497] | 1633.7388 | 3 | 4899.2019 | 4899.1808 | 0 | 4.3 | 413 | Specific | 911.73 | 727.1 | 727.1 | 18.73 | 97 | >sp\|P01833\|PIGR_HUMAN Polymeric immunoglobulin receptor OS=Homo sapiens OX=9606 GN=PIGR PE=1 SV=4 | 902 | 20201118_lss_colostrum_1ng_glyco_hcd203040_compound15.25516.25516.3 | controllerType=0 controllerNumber=1 scan=25516 | 53.5588 |
| 21774:4 | 5 | R.LSLLEEPGN[+2496.888]GTFTVILNQLTSR.D | HexNAc(4)Hex(5)Fuc(2)NeuAc(2) | N[+2497] | 1225.5523 | 4 | 4899.1873 | 4899.1808 | 0 | 1.3 | 413 | Specific | 912.05 | 701.67 | 701.67 | 19.01 | 97 | >sp\|P01833\|PIGR_HUMAN Polymeric immunoglobulin receptor OS=Homo sapiens OX=9606 GN=PIGR PE=1 SV=4 | 902 | 20201118_lss_colostrum_1ng_glyco_hcd203040_compound15.25512.25512.4 | controllerType=0 controllerNumber=1 scan=25512 | 53.547 |
| 19767:3 | 5 | R.LSLLEEPGN[+2351.851]GTFTVILNQLTSR.D | HexNAc(4)Hex(5)Fuc(3)NeuAc(1) | N[+2352] | 1585.3897 | 3 | 4754.1546 | 4754.1433 | 0 | 2.4 | 413 | Specific | 929.72 | 664.28 | 664.28 | 18.93 | 97 | >sp\|P01833\|PIGR_HUMAN Polymeric immunoglobulin receptor OS=Homo sapiens OX=9606 GN=PIGR PE=1 SV=4 | 902 | 20201118_lss_colostrum_1ng_glyco_hcd203040_compound15.23416.23416.3 | controllerType=0 controllerNumber=1 scan=23416 | 48.8299 |
| 19772:4 | 5 | R.LSLLEEPGN[+2351.851]GTFTVILNQLTSR.D | HexNAc(4)Hex(5)Fuc(3)NeuAc(1) | N[+2352] | 1189.294 | 4 | 4754.1541 | 4754.1433 | 0 | 2.3 | 413 | Specific | 931.9 | 867.64 | 867.64 | 19.17 | 97 | >sp\|P01833\|PIGR_HUMAN Polymeric immunoglobulin receptor OS=Homo sapiens OX=9606 GN=PIGR PE=1 SV=4 | 902 | 20201118_lss_colostrum_1ng_glyco_hcd203040_compound15.23421.23421.4 | controllerType=0 controllerNumber=1 scan=23421 | 48.8388 |
| 19885:3 | 5 | R.LSLLEEPGN[+2059.735]GTFTVILNQLTSR.D | HexNAc(4)Hex(5)Fuc(1)NeuAc(1) | N[+2060] | 1488.0169 | 3 | 4462.0362 | 4462.0275 | 0 | 1.9 | 413 | Specific | 943.33 | 639.13 | 639.13 | 18.93 | 97 | >sp\|P01833\|PIGR_HUMAN Polymeric immunoglobulin receptor OS=Homo sapiens OX=9606 GN=PIGR PE=1 SV=4 | 902 | 20201118_lss_colostrum_1ng_glyco_hcd203040_compound15.23539.23539.3 | controllerType=0 controllerNumber=1 scan=23539 | 49.1331 |
| 21626:3 | 5 | R.LSLLEEPGN[+2350.830]GTFTVILNQLTSR.D | HexNAc(4)Hex(5)Fuc(1)NeuAc(2) | N[+2351] | 1585.0517 | 3 | 4753.1405 | 4753.1229 | 0 | 3.7 | 413 | Specific | 950.04 | 668.21 | 668.21 | 19.73 | 97 | >sp\|P01833\|PIGR_HUMAN Polymeric immunoglobulin receptor OS=Homo sapiens OX=9606 GN=PIGR PE=1 SV=4 | 902 | 20201118_lss_colostrum_1ng_glyco_hcd203040_compound15.25358.25358.3 | controllerType=0 controllerNumber=1 scan=25358 | 53.2008 |
| 21779:3 | 5 | R.LSLLEEPGN[+2496.888]GTFTVILNQLTSR.D | HexNAc(4)Hex(5)Fuc(2)NeuAc(2) | N[+2497] | 1633.7388 | 3 | 4899.2019 | 4899.1808 | 0 | 4.3 | 413 | Specific | 960.81 | 706.77 | 706.77 | 19.73 | 97 | >sp\|P01833\|PIGR_HUMAN Polymeric immunoglobulin receptor OS=Homo sapiens OX=9606 GN=PIGR PE=1 SV=4 | 902 | 20201118_lss_colostrum_1ng_glyco_hcd203040_compound15.25518.25518.3 | controllerType=0 controllerNumber=1 scan=25518 | 53.5639 |
| 21631:4 | 5 | R.LSLLEEPGN[+2350.830]GTFTVILNQLTSR.D | HexNAc(4)Hex(5)Fuc(1)NeuAc(2) | N[+2351] | 1189.0402 | 4 | 4753.139 | 4753.1229 | 0 | 3.4 | 413 | Specific | 966.22 | 737.07 | 737.07 | 19.97 | 97 | >sp\|P01833\|PIGR_HUMAN Polymeric immunoglobulin receptor OS=Homo sapiens OX=9606 GN=PIGR PE=1 SV=4 | 902 | 20201118_lss_colostrum_1ng_glyco_hcd203040_compound15.25363.25363.4 | controllerType=0 controllerNumber=1 scan=25363 | 53.2145 |
| 21629:4 | 5 | R.LSLLEEPGN[+2350.830]GTFTVILNQLTSR.D | HexNAc(4)Hex(5)Fuc(1)NeuAc(2) | N[+2351] | 1189.0402 | 4 | 4753.139 | 4753.1229 | 0 | 3.4 | 413 | Specific | 1034 | 819.55 | 819.55 | 20.97 | 97 | >sp\|P01833\|PIGR_HUMAN Polymeric immunoglobulin receptor OS=Homo sapiens OX=9606 GN=PIGR PE=1 SV=4 | 902 | 20201118_lss_colostrum_1ng_glyco_hcd203040_compound15.25361.25361.4 | controllerType=0 controllerNumber=1 scan=25361 | 53.2055 |
| 21624:3 | 5 | R.LSLLEEPGN[+2350.830]GTFTVILNQLTSR.D | HexNAc(4)Hex(5)Fuc(1)NeuAc(2) | N[+2351] | 1585.0517 | 3 | 4753.1405 | 4753.1229 | 0 | 3.7 | 413 | Specific | 1149.5 | 904.66 | 904.66 | 22.73 | 97 | >sp\|P01833\|PIGR_HUMAN Polymeric immunoglobulin receptor OS=Homo sapiens OX=9606 GN=PIGR PE=1 SV=4 | 902 | 20201118_lss_colostrum_1ng_glyco_hcd203040_compound15.25356.25356.3 | controllerType=0 controllerNumber=1 scan=25356 | 53.1939 |
| 13724:3 | 5 | K.[+42.011]IIEGEPNLKVPGN[+2018.708]VTAVLGETLK.V | HexNAc(3)Hex(6)Fuc(1)NeuAc(1) | .[42], N[+2019] | 1484.7087 | 3 | 4452.1115 | 4452.0683 | 0 | 9.7 | 457 | Specific | 253.32 | 253.32 | 253.32 | 4.99 | 97 | >sp\|P01833\|PIGR_HUMAN Polymeric immunoglobulin receptor OS=Homo sapiens OX=9606 GN=PIGR PE=1 SV=4 | 902 | 20201118_lss_colostrum_1ng_glyco_hcd203040_compound15.17123.17123.3 | controllerType=0 controllerNumber=1 scan=17123 | 35.8316 |
| 17279:3 | 5 | K.IIEGEPNLKVPGN[+2350.830]VTAVLGETLK.V | HexNAc(4)Hex(5)Fuc(1)NeuAc(2) | N[+2351] | 1581.4048 | 3 | 4742.2 | 4742.1797 | 0 | 4.3 | 457 | Specific | 805.92 | 632.23 | 632.23 | 16.33 | 97 | >sp\|P01833\|PIGR_HUMAN Polymeric immunoglobulin receptor OS=Homo sapiens OX=9606 GN=PIGR PE=1 SV=4 | 902 | 20201118_lss_colostrum_1ng_glyco_hcd203040_compound15.20822.20822.3 | controllerType=0 controllerNumber=1 scan=20822 | 43.1944 |
| 15415:3 | 5 | K.IIEGEPNLKVPGN[+2059.735]VTAVLGETLK.V | HexNAc(4)Hex(5)Fuc(1)NeuAc(1) | N[+2060] | 1484.3726 | 3 | 4451.1033 | 4451.0843 | 0 | 4.3 | 457 | Specific | 809.98 | 726.79 | 726.79 | 16.33 | 97 | >sp\|P01833\|PIGR_HUMAN Polymeric immunoglobulin receptor OS=Homo sapiens OX=9606 GN=PIGR PE=1 SV=4 | 902 | 20201118_lss_colostrum_1ng_glyco_hcd203040_compound15.18882.18882.3 | controllerType=0 controllerNumber=1 scan=18882 | 39.215 |
| 15358:3 | 5 | K.IIEGEPNLKVPGN[+2351.851]VTAVLGETLK.V | HexNAc(4)Hex(5)Fuc(3)NeuAc(1) | N[+2352] | 1581.7427 | 3 | 4743.2136 | 4743.2001 | 0 | 2.9 | 457 | Specific | 837.65 | 799.36 | 799.36 | 16.52 | 97 | >sp\|P01833\|PIGR_HUMAN Polymeric immunoglobulin receptor OS=Homo sapiens OX=9606 GN=PIGR PE=1 SV=4 | 902 | 20201118_lss_colostrum_1ng_glyco_hcd203040_compound15.18823.18823.3 | controllerType=0 controllerNumber=1 scan=18823 | 39.0903 |
| 15380:3 | 5 | K.IIEGEPNLKVPGN[+2205.793]VTAVLGETLK.V | HexNAc(4)Hex(5)Fuc(2)NeuAc(1) | N[+2206] | 1533.0568 | 3 | 4597.1559 | 4597.1422 | 0 | 3 | 457 | Specific | 867.61 | 763.97 | 763.97 | 17.52 | 97 | >sp\|P01833\|PIGR_HUMAN Polymeric immunoglobulin receptor OS=Homo sapiens OX=9606 GN=PIGR PE=1 SV=4 | 902 | 20201118_lss_colostrum_1ng_glyco_hcd203040_compound15.18846.18846.3 | controllerType=0 controllerNumber=1 scan=18846 | 39.1396 |
| 15356:3 | 5 | K.IIEGEPNLKVPGN[+2351.851]VTAVLGETLK.V | HexNAc(4)Hex(5)Fuc(3)NeuAc(1) | N[+2352] | 1581.7427 | 3 | 4743.2136 | 4743.2001 | 0 | 2.9 | 457 | Specific | 965.4 | 932.73 | 932.73 | 19.52 | 97 | >sp\|P01833\|PIGR_HUMAN Polymeric immunoglobulin receptor OS=Homo sapiens OX=9606 GN=PIGR PE=1 SV=4 | 902 | 20201118_lss_colostrum_1ng_glyco_hcd203040_compound15.18821.18821.3 | controllerType=0 controllerNumber=1 scan=18821 | 39.0842 |
| 17277:3 | 5 | K.IIEGEPNLKVPGN[+2350.830]VTAVLGETLK.V | HexNAc(4)Hex(5)Fuc(1)NeuAc(2) | N[+2351] | 1581.4048 | 3 | 4742.2 | 4742.1797 | 0 | 4.3 | 457 | Specific | 976.27 | 883.59 | 883.59 | 19.33 | 97 | >sp\|P01833\|PIGR_HUMAN Polymeric immunoglobulin receptor OS=Homo sapiens OX=9606 GN=PIGR PE=1 SV=4 | 902 | 20201118_lss_colostrum_1ng_glyco_hcd203040_compound15.20820.20820.3 | controllerType=0 controllerNumber=1 scan=20820 | 43.189 |
| 15378:3 | 5 | K.IIEGEPNLKVPGN[+2205.793]VTAVLGETLK.V | HexNAc(4)Hex(5)Fuc(2)NeuAc(1) | N[+2206] | 1533.0568 | 3 | 4597.1559 | 4597.1422 | 0 | 3 | 457 | Specific | 1053.44 | 943.14 | 943.14 | 21.52 | 97 | >sp\|P01833\|PIGR_HUMAN Polymeric immunoglobulin receptor OS=Homo sapiens OX=9606 GN=PIGR PE=1 SV=4 | 902 | 20201118_lss_colostrum_1ng_glyco_hcd203040_compound15.18844.18844.3 | controllerType=0 controllerNumber=1 scan=18844 | 39.1319 |
| 15412:3 | 5 | K.IIEGEPNLKVPGN[+2059.735]VTAVLGETLK.V | HexNAc(4)Hex(5)Fuc(1)NeuAc(1) | N[+2060] | 1484.3726 | 3 | 4451.1033 | 4451.0843 | 0 | 4.3 | 457 | Specific | 1065.41 | 986.88 | 986.88 | 21.33 | 97 | >sp\|P01833\|PIGR_HUMAN Polymeric immunoglobulin receptor OS=Homo sapiens OX=9606 GN=PIGR PE=1 SV=4 | 902 | 20201118_lss_colostrum_1ng_glyco_hcd203040_compound15.18879.18879.3 | controllerType=0 controllerNumber=1 scan=18879 | 39.208 |
| 13912:5 | 5 | K.IIEGEPNLKVPGN[+1768.640]VTAVLGETLKVPC[+57.021]HFPC[+57.021]K.F | HexNAc(4)Hex(5)Fuc(1) | C[+57]*2, N[+1769] | 1037.8977 | 5 | 5185.4594 | 5185.4464 | 0 | 2.5 | 457 | Specific | 396.08 | 396.08 | 396.08 | 9.03 | 97 | >sp\|P01833\|PIGR_HUMAN Polymeric immunoglobulin receptor OS=Homo sapiens OX=9606 GN=PIGR PE=1 SV=4 | 902 | 20201118_lss_colostrum_1ng_glyco_hcd203040_compound15.17319.17319.5 | controllerType=0 controllerNumber=1 scan=17319 | 36.2544 |
| 13834:5 | 5 | K.IIEGEPNLKVPGN[+2059.735]VTAVLGETLKVPC[+57.021]HFPC[+57.021]K.F | HexNAc(4)Hex(5)Fuc(1)NeuAc(1) | C[+57]*2, N[+2060] | 1096.1211 | 5 | 5476.5764 | 5476.5418 | 0 | 6.3 | 457 | Specific | 398.32 | 398.32 | 398.32 | 7.01 | 97 | >sp\|P01833\|PIGR_HUMAN Polymeric immunoglobulin receptor OS=Homo sapiens OX=9606 GN=PIGR PE=1 SV=4 | 902 | 20201118_lss_colostrum_1ng_glyco_hcd203040_compound15.17238.17238.3 | controllerType=0 controllerNumber=1 scan=17238 | 36.0917 |
| 13831:5 | 5 | K.IIEGEPNLKVPGN[+2059.735]VTAVLGETLKVPC[+57.021]HFPC[+57.021]K.F | HexNAc(4)Hex(5)Fuc(1)NeuAc(1) | C[+57]*2, N[+2060] | 1096.1211 | 5 | 5476.5764 | 5476.5418 | 0 | 6.3 | 457 | Specific | 403.9 | 403.9 | 403.9 | 8.03 | 97 | >sp\|P01833\|PIGR_HUMAN Polymeric immunoglobulin receptor OS=Homo sapiens OX=9606 GN=PIGR PE=1 SV=4 | 902 | 20201118_lss_colostrum_1ng_glyco_hcd203040_compound15.17235.17235.3 | controllerType=0 controllerNumber=1 scan=17235 | 36.0793 |
| 14844:5 | 5 | K.IIEGEPNLKVPGN[+2570.925]VTAVLGETLKVPC[+57.021]HFPC[+57.021]K.F | HexNAc(5)Hex(6)Fuc(2)NeuAc(1) | C[+57]*2, N[+2571] | 1198.3543 | 5 | 5987.7421 | 5987.7319 | 0 | 1.7 | 457 | Specific | 495.59 | 311.61 | 311.61 | 7.32 | 97 | >sp\|P01833\|PIGR_HUMAN Polymeric immunoglobulin receptor OS=Homo sapiens OX=9606 GN=PIGR PE=1 SV=4 | 902 | 20201118_lss_colostrum_1ng_glyco_hcd203040_compound15.18286.18286.5 | controllerType=0 controllerNumber=1 scan=18286 | 38.1057 |
| 13854:4 | 5 | K.IIEGEPNLKVPGN[+1913.677]VTAVLGETLKVPC[+57.021]HFPC[+57.021]K.F | HexNAc(4)Hex(5)NeuAc(1) | C[+57]*2, N[+1914] | 1333.3824 | 4 | 5330.508 | 5330.4839 | 0 | 4.5 | 457 | Specific | 552.6 | 514.68 | 514.68 | 8.76 | 97 | >sp\|P01833\|PIGR_HUMAN Polymeric immunoglobulin receptor OS=Homo sapiens OX=9606 GN=PIGR PE=1 SV=4 | 902 | 20201118_lss_colostrum_1ng_glyco_hcd203040_compound15.17258.17258.4 | controllerType=0 controllerNumber=1 scan=17258 | 36.1298 |
| 13851:4 | 5 | K.IIEGEPNLKVPGN[+1913.677]VTAVLGETLKVPC[+57.021]HFPC[+57.021]K.F | HexNAc(4)Hex(5)NeuAc(1) | C[+57]*2, N[+1914] | 1333.3824 | 4 | 5330.508 | 5330.4839 | 0 | 4.5 | 457 | Specific | 626.83 | 626.83 | 626.83 | 11.35 | 97 | >sp\|P01833\|PIGR_HUMAN Polymeric immunoglobulin receptor OS=Homo sapiens OX=9606 GN=PIGR PE=1 SV=4 | 902 | 20201118_lss_colostrum_1ng_glyco_hcd203040_compound15.17255.17255.4 | controllerType=0 controllerNumber=1 scan=17255 | 36.1189 |
| 13914:5 | 5 | K.IIEGEPNLKVPGN[+1768.640]VTAVLGETLKVPC[+57.021]HFPC[+57.021]K.F | HexNAc(4)Hex(5)Fuc(1) | C[+57]*2, N[+1769] | 1037.8977 | 5 | 5185.4594 | 5185.4464 | 0 | 2.5 | 457 | Specific | 668.85 | 544.32 | 544.32 | 13.17 | 97 | >sp\|P01833\|PIGR_HUMAN Polymeric immunoglobulin receptor OS=Homo sapiens OX=9606 GN=PIGR PE=1 SV=4 | 902 | 20201118_lss_colostrum_1ng_glyco_hcd203040_compound15.17321.17321.5 | controllerType=0 controllerNumber=1 scan=17321 | 36.2578 |
| 14863:4 | 5 | K.IIEGEPNLKVPGN[+2059.735]VTAVLGETLKVPC[+57.021]HFPC[+57.021]K.F | HexNAc(4)Hex(5)Fuc(1)NeuAc(1) | C[+57]*2, N[+2060] | 1369.8964 | 4 | 5476.5636 | 5476.5418 | 0 | 4 | 457 | Specific | 678.69 | 678.69 | 678.69 | 12.35 | 97 | >sp\|P01833\|PIGR_HUMAN Polymeric immunoglobulin receptor OS=Homo sapiens OX=9606 GN=PIGR PE=1 SV=4 | 902 | 20201118_lss_colostrum_1ng_glyco_hcd203040_compound15.18306.18306.4 | controllerType=0 controllerNumber=1 scan=18306 | 38.1528 |
| 15003:4 | 5 | K.IIEGEPNLKVPGN[+2204.772]VTAVLGETLKVPC[+57.021]HFPC[+57.021]K.F | HexNAc(4)Hex(5)NeuAc(2) | C[+57]*2, N[+2205] | 1406.156 | 4 | 5621.6022 | 5621.5793 | 0 | 4.1 | 457 | Specific | 710.8 | 710.8 | 710.8 | 13.35 | 97 | >sp\|P01833\|PIGR_HUMAN Polymeric immunoglobulin receptor OS=Homo sapiens OX=9606 GN=PIGR PE=1 SV=4 | 902 | 20201118_lss_colostrum_1ng_glyco_hcd203040_compound15.18452.18452.2 | controllerType=0 controllerNumber=1 scan=18452 | 38.4361 |
| 14842:5 | 5 | K.IIEGEPNLKVPGN[+2570.925]VTAVLGETLKVPC[+57.021]HFPC[+57.021]K.F | HexNAc(5)Hex(6)Fuc(2)NeuAc(1) | C[+57]*2, N[+2571] | 1198.3543 | 5 | 5987.7421 | 5987.7319 | 0 | 1.7 | 457 | Specific | 714.52 | 497.84 | 497.84 | 14.01 | 97 | >sp\|P01833\|PIGR_HUMAN Polymeric immunoglobulin receptor OS=Homo sapiens OX=9606 GN=PIGR PE=1 SV=4 | 902 | 20201118_lss_colostrum_1ng_glyco_hcd203040_compound15.18284.18284.5 | controllerType=0 controllerNumber=1 scan=18284 | 38.0972 |
| 15006:4 | 5 | K.IIEGEPNLKVPGN[+2204.772]VTAVLGETLKVPC[+57.021]HFPC[+57.021]K.F | HexNAc(4)Hex(5)NeuAc(2) | C[+57]*2, N[+2205] | 1406.156 | 4 | 5621.6022 | 5621.5793 | 0 | 4.1 | 457 | Specific | 731.25 | 480.14 | 480.14 | 13.35 | 97 | >sp\|P01833\|PIGR_HUMAN Polymeric immunoglobulin receptor OS=Homo sapiens OX=9606 GN=PIGR PE=1 SV=4 | 902 | 20201118_lss_colostrum_1ng_glyco_hcd203040_compound15.18455.18455.2 | controllerType=0 controllerNumber=1 scan=18455 | 38.4406 |
| 14962:5 | 5 | K.IIEGEPNLKVPGN[+2351.851]VTAVLGETLKVPC[+57.021]HFPC[+57.021]K.F | HexNAc(4)Hex(5)Fuc(3)NeuAc(1) | C[+57]*2, N[+2352] | 1154.5416 | 5 | 5768.6789 | 5768.6576 | 0 | 3.7 | 457 | Specific | 732.67 | 732.67 | 732.67 | 13.98 | 97 | >sp\|P01833\|PIGR_HUMAN Polymeric immunoglobulin receptor OS=Homo sapiens OX=9606 GN=PIGR PE=1 SV=4 | 902 | 20201118_lss_colostrum_1ng_glyco_hcd203040_compound15.18409.18409.5 | controllerType=0 controllerNumber=1 scan=18409 | 38.3613 |
| 16465:4 | 5 | K.IIEGEPNLKVPGN[+2350.830]VTAVLGETLKVPC[+57.021]HFPC[+57.021]K.F | HexNAc(4)Hex(5)Fuc(1)NeuAc(2) | C[+57]*2, N[+2351] | 1442.6696 | 4 | 5767.6564 | 5767.6372 | 0 | 3.3 | 457 | Specific | 743.88 | 586.6 | 586.6 | 13.35 | 97 | >sp\|P01833\|PIGR_HUMAN Polymeric immunoglobulin receptor OS=Homo sapiens OX=9606 GN=PIGR PE=1 SV=4 | 902 | 20201118_lss_colostrum_1ng_glyco_hcd203040_compound15.19974.19974.4 | controllerType=0 controllerNumber=1 scan=19974 | 41.3547 |
| 16457:5 | 5 | K.IIEGEPNLKVPGN[+2350.830]VTAVLGETLKVPC[+57.021]HFPC[+57.021]K.F | HexNAc(4)Hex(5)Fuc(1)NeuAc(2) | C[+57]*2, N[+2351] | 1154.3362 | 5 | 5767.6518 | 5767.6372 | 0 | 2.5 | 457 | Specific | 771.98 | 545.2 | 545.2 | 15.17 | 97 | >sp\|P01833\|PIGR_HUMAN Polymeric immunoglobulin receptor OS=Homo sapiens OX=9606 GN=PIGR PE=1 SV=4 | 902 | 20201118_lss_colostrum_1ng_glyco_hcd203040_compound15.19966.19966.5 | controllerType=0 controllerNumber=1 scan=19966 | 41.3353 |
| 14975:4 | 5 | K.IIEGEPNLKVPGN[+2205.793]VTAVLGETLKVPC[+57.021]HFPC[+57.021]K.F | HexNAc(4)Hex(5)Fuc(2)NeuAc(1) | C[+57]*2, N[+2206] | 1406.4082 | 4 | 5622.611 | 5622.5997 | 0 | 2 | 457 | Specific | 774.15 | 746.28 | 746.28 | 14.55 | 97 | >sp\|P01833\|PIGR_HUMAN Polymeric immunoglobulin receptor OS=Homo sapiens OX=9606 GN=PIGR PE=1 SV=4 | 902 | 20201118_lss_colostrum_1ng_glyco_hcd203040_compound15.18423.18423.4 | controllerType=0 controllerNumber=1 scan=18423 | 38.3919 |
| 14955:4 | 5 | K.IIEGEPNLKVPGN[+2205.793]VTAVLGETLKVPC[+57.021]HFPC[+57.021]K.F | HexNAc(4)Hex(5)Fuc(2)NeuAc(1) | C[+57]*2, N[+2206] | 1406.4075 | 4 | 5622.6081 | 5622.5997 | 0 | 1.5 | 457 | Specific | 775.53 | 775.53 | 775.53 | 14.39 | 97 | >sp\|P01833\|PIGR_HUMAN Polymeric immunoglobulin receptor OS=Homo sapiens OX=9606 GN=PIGR PE=1 SV=4 | 902 | 20201118_lss_colostrum_1ng_glyco_hcd203040_compound15.18402.18402.4 | controllerType=0 controllerNumber=1 scan=18402 | 38.3456 |
| 16462:4 | 5 | K.IIEGEPNLKVPGN[+2350.830]VTAVLGETLKVPC[+57.021]HFPC[+57.021]K.F | HexNAc(4)Hex(5)Fuc(1)NeuAc(2) | C[+57]*2, N[+2351] | 1442.6696 | 4 | 5767.6564 | 5767.6372 | 0 | 3.3 | 457 | Specific | 796.14 | 656.85 | 656.85 | 14.35 | 97 | >sp\|P01833\|PIGR_HUMAN Polymeric immunoglobulin receptor OS=Homo sapiens OX=9606 GN=PIGR PE=1 SV=4 | 902 | 20201118_lss_colostrum_1ng_glyco_hcd203040_compound15.19971.19971.4 | controllerType=0 controllerNumber=1 scan=19971 | 41.3428 |
| 14959:5 | 5 | K.IIEGEPNLKVPGN[+2351.851]VTAVLGETLKVPC[+57.021]HFPC[+57.021]K.F | HexNAc(4)Hex(5)Fuc(3)NeuAc(1) | C[+57]*2, N[+2352] | 1154.5416 | 5 | 5768.6789 | 5768.6576 | 0 | 3.7 | 457 | Specific | 801.64 | 769.72 | 769.72 | 15.98 | 97 | >sp\|P01833\|PIGR_HUMAN Polymeric immunoglobulin receptor OS=Homo sapiens OX=9606 GN=PIGR PE=1 SV=4 | 902 | 20201118_lss_colostrum_1ng_glyco_hcd203040_compound15.18406.18406.5 | controllerType=0 controllerNumber=1 scan=18406 | 38.3506 |
| 14978:4 | 5 | K.IIEGEPNLKVPGN[+2205.793]VTAVLGETLKVPC[+57.021]HFPC[+57.021]K.F | HexNAc(4)Hex(5)Fuc(2)NeuAc(1) | C[+57]*2, N[+2206] | 1406.4082 | 4 | 5622.611 | 5622.5997 | 0 | 2 | 457 | Specific | 812.71 | 752.29 | 752.29 | 15.55 | 97 | >sp\|P01833\|PIGR_HUMAN Polymeric immunoglobulin receptor OS=Homo sapiens OX=9606 GN=PIGR PE=1 SV=4 | 902 | 20201118_lss_colostrum_1ng_glyco_hcd203040_compound15.18426.18426.4 | controllerType=0 controllerNumber=1 scan=18426 | 38.3958 |
| 14902:5 | 5 | K.IIEGEPNLKVPGN[+2059.735]VTAVLGETLKVPC[+57.021]HFPC[+57.021]K.F | HexNAc(4)Hex(5)Fuc(1)NeuAc(1) | C[+57]*2, N[+2060] | 1096.1165 | 5 | 5476.5532 | 5476.5418 | 0 | 2.1 | 457 | Specific | 836.43 | 721.03 | 721.03 | 16.17 | 97 | >sp\|P01833\|PIGR_HUMAN Polymeric immunoglobulin receptor OS=Homo sapiens OX=9606 GN=PIGR PE=1 SV=4 | 902 | 20201118_lss_colostrum_1ng_glyco_hcd203040_compound15.18347.18347.5 | controllerType=0 controllerNumber=1 scan=18347 | 38.2388 |
| 16460:5 | 5 | K.IIEGEPNLKVPGN[+2350.830]VTAVLGETLKVPC[+57.021]HFPC[+57.021]K.F | HexNAc(4)Hex(5)Fuc(1)NeuAc(2) | C[+57]*2, N[+2351] | 1154.3362 | 5 | 5767.6518 | 5767.6372 | 0 | 2.5 | 457 | Specific | 841.5 | 578.2 | 578.2 | 16.17 | 97 | >sp\|P01833\|PIGR_HUMAN Polymeric immunoglobulin receptor OS=Homo sapiens OX=9606 GN=PIGR PE=1 SV=4 | 902 | 20201118_lss_colostrum_1ng_glyco_hcd203040_compound15.19969.19969.5 | controllerType=0 controllerNumber=1 scan=19969 | 41.3398 |
| 14898:4 | 5 | K.IIEGEPNLKVPGN[+2059.735]VTAVLGETLKVPC[+57.021]HFPC[+57.021]K.F | HexNAc(4)Hex(5)Fuc(1)NeuAc(1) | C[+57]*2, N[+2060] | 1369.8933 | 4 | 5476.5514 | 5476.5418 | 0 | 1.8 | 457 | Specific | 879.4 | 706.1 | 706.1 | 16.39 | 97 | >sp\|P01833\|PIGR_HUMAN Polymeric immunoglobulin receptor OS=Homo sapiens OX=9606 GN=PIGR PE=1 SV=4 | 902 | 20201118_lss_colostrum_1ng_glyco_hcd203040_compound15.18343.18343.4 | controllerType=0 controllerNumber=1 scan=18343 | 38.232 |
| 13868:5 | 5 | K.IIEGEPNLKVPGN[+1768.640]VTAVLGETLKVPC[+57.021]HFPC[+57.021]K.F | HexNAc(4)Hex(5)Fuc(1) | C[+57]*2, N[+1769] | 1037.897 | 5 | 5185.4558 | 5185.4464 | 0 | 1.8 | 457 | Specific | 888.53 | 792.75 | 792.75 | 17.01 | 97 | >sp\|P01833\|PIGR_HUMAN Polymeric immunoglobulin receptor OS=Homo sapiens OX=9606 GN=PIGR PE=1 SV=4 | 902 | 20201118_lss_colostrum_1ng_glyco_hcd203040_compound15.17273.17273.5 | controllerType=0 controllerNumber=1 scan=17273 | 36.1637 |
| 14884:5 | 5 | K.IIEGEPNLKVPGN[+2351.851]VTAVLGETLKVPC[+57.021]HFPC[+57.021]K.F | HexNAc(4)Hex(5)Fuc(3)NeuAc(1) | C[+57]*2, N[+2352] | 1154.5388 | 5 | 5768.665 | 5768.6576 | 0 | 1.3 | 457 | Specific | 918.97 | 897.31 | 897.31 | 18.01 | 97 | >sp\|P01833\|PIGR_HUMAN Polymeric immunoglobulin receptor OS=Homo sapiens OX=9606 GN=PIGR PE=1 SV=4 | 902 | 20201118_lss_colostrum_1ng_glyco_hcd203040_compound15.18328.18328.5 | controllerType=0 controllerNumber=1 scan=18328 | 38.1985 |
| 13870:5 | 5 | K.IIEGEPNLKVPGN[+1768.640]VTAVLGETLKVPC[+57.021]HFPC[+57.021]K.F | HexNAc(4)Hex(5)Fuc(1) | C[+57]*2, N[+1769] | 1037.897 | 5 | 5185.4558 | 5185.4464 | 0 | 1.8 | 457 | Specific | 965.44 | 768.69 | 768.69 | 19.01 | 97 | >sp\|P01833\|PIGR_HUMAN Polymeric immunoglobulin receptor OS=Homo sapiens OX=9606 GN=PIGR PE=1 SV=4 | 902 | 20201118_lss_colostrum_1ng_glyco_hcd203040_compound15.17275.17275.5 | controllerType=0 controllerNumber=1 scan=17275 | 36.1697 |
| 14896:4 | 5 | K.IIEGEPNLKVPGN[+2059.735]VTAVLGETLKVPC[+57.021]HFPC[+57.021]K.F | HexNAc(4)Hex(5)Fuc(1)NeuAc(1) | C[+57]*2, N[+2060] | 1369.8933 | 4 | 5476.5514 | 5476.5418 | 0 | 1.8 | 457 | Specific | 988.47 | 929.65 | 929.65 | 18.39 | 97 | >sp\|P01833\|PIGR_HUMAN Polymeric immunoglobulin receptor OS=Homo sapiens OX=9606 GN=PIGR PE=1 SV=4 | 902 | 20201118_lss_colostrum_1ng_glyco_hcd203040_compound15.18341.18341.4 | controllerType=0 controllerNumber=1 scan=18341 | 38.2265 |
| 14881:5 | 5 | K.IIEGEPNLKVPGN[+2351.851]VTAVLGETLKVPC[+57.021]HFPC[+57.021]K.F | HexNAc(4)Hex(5)Fuc(3)NeuAc(1) | C[+57]*2, N[+2352] | 1154.5388 | 5 | 5768.665 | 5768.6576 | 0 | 1.3 | 457 | Specific | 1012.05 | 1012.05 | 1012.05 | 20.01 | 97 | >sp\|P01833\|PIGR_HUMAN Polymeric immunoglobulin receptor OS=Homo sapiens OX=9606 GN=PIGR PE=1 SV=4 | 902 | 20201118_lss_colostrum_1ng_glyco_hcd203040_compound15.18325.18325.5 | controllerType=0 controllerNumber=1 scan=18325 | 38.1907 |
| 14953:4 | 5 | K.IIEGEPNLKVPGN[+2205.793]VTAVLGETLKVPC[+57.021]HFPC[+57.021]K.F | HexNAc(4)Hex(5)Fuc(2)NeuAc(1) | C[+57]*2, N[+2206] | 1406.4075 | 4 | 5622.6081 | 5622.5997 | 0 | 1.5 | 457 | Specific | 1047.94 | 916.94 | 916.94 | 19.39 | 97 | >sp\|P01833\|PIGR_HUMAN Polymeric immunoglobulin receptor OS=Homo sapiens OX=9606 GN=PIGR PE=1 SV=4 | 902 | 20201118_lss_colostrum_1ng_glyco_hcd203040_compound15.18400.18400.4 | controllerType=0 controllerNumber=1 scan=18400 | 38.3396 |
| 14899:5 | 5 | K.IIEGEPNLKVPGN[+2059.735]VTAVLGETLKVPC[+57.021]HFPC[+57.021]K.F | HexNAc(4)Hex(5)Fuc(1)NeuAc(1) | C[+57]*2, N[+2060] | 1096.1165 | 5 | 5476.5532 | 5476.5418 | 0 | 2.1 | 457 | Specific | 1066.74 | 934.17 | 934.17 | 21.17 | 97 | >sp\|P01833\|PIGR_HUMAN Polymeric immunoglobulin receptor OS=Homo sapiens OX=9606 GN=PIGR PE=1 SV=4 | 902 | 20201118_lss_colostrum_1ng_glyco_hcd203040_compound15.18344.18344.5 | controllerType=0 controllerNumber=1 scan=18344 | 38.2332 |
| 14860:4 | 5 | K.IIEGEPNLKVPGN[+2059.735]VTAVLGETLKVPC[+57.021]HFPC[+57.021]K.F | HexNAc(4)Hex(5)Fuc(1)NeuAc(1) | C[+57]*2, N[+2060] | 1369.8964 | 4 | 5476.5636 | 5476.5418 | 0 | 4 | 457 | Specific | 1116.18 | 979.17 | 979.17 | 21.35 | 97 | >sp\|P01833\|PIGR_HUMAN Polymeric immunoglobulin receptor OS=Homo sapiens OX=9606 GN=PIGR PE=1 SV=4 | 902 | 20201118_lss_colostrum_1ng_glyco_hcd203040_compound15.18303.18303.4 | controllerType=0 controllerNumber=1 scan=18303 | 38.1372 |
| 16056:3 | 5 | K.VPGN[+2351.851]VTAVLGETLK.V | HexNAc(4)Hex(5)Fuc(3)NeuAc(1) | N[+2352] | 1250.5539 | 3 | 3749.6471 | 3749.6506 | 0 | -0.9 | 466 | Specific | 362.25 | 362.25 | 362.25 | 6.55 | 97 | >sp\|P01833\|PIGR_HUMAN Polymeric immunoglobulin receptor OS=Homo sapiens OX=9606 GN=PIGR PE=1 SV=4 | 902 | 20201118_lss_colostrum_1ng_glyco_hcd203040_compound15.19548.19548.3 | controllerType=0 controllerNumber=1 scan=19548 | 40.4643 |
| 16058:3 | 5 | K.VPGN[+2351.851]VTAVLGETLK.V | HexNAc(4)Hex(5)Fuc(3)NeuAc(1) | N[+2352] | 1250.5539 | 3 | 3749.6471 | 3749.6506 | 0 | -0.9 | 466 | Specific | 427.57 | 323.67 | 323.67 | 5.06 | 97 | >sp\|P01833\|PIGR_HUMAN Polymeric immunoglobulin receptor OS=Homo sapiens OX=9606 GN=PIGR PE=1 SV=4 | 902 | 20201118_lss_colostrum_1ng_glyco_hcd203040_compound15.19550.19550.3 | controllerType=0 controllerNumber=1 scan=19550 | 40.4677 |
| 13062:3 | 5 | K.VPGN[+2205.793]VTAVLGETLK.V | HexNAc(4)Hex(5)Fuc(2)NeuAc(1) | N[+2206] | 1201.8703 | 3 | 3603.5963 | 3603.5927 | 0 | 1 | 466 | Specific | 433.51 | 380.66 | 380.66 | 9.17 | 97 | >sp\|P01833\|PIGR_HUMAN Polymeric immunoglobulin receptor OS=Homo sapiens OX=9606 GN=PIGR PE=1 SV=4 | 902 | 20201118_lss_colostrum_1ng_glyco_hcd203040_compound15.16435.16435.3 | controllerType=0 controllerNumber=1 scan=16435 | [illegible] |
| 13065:3 | 5 | K.VPGN[+2205.793]VTAVLGETLK.V | HexNAc(4)Hex(5)Fuc(2)NeuAc(1) | N[+2206] | 1201.8703 | 3 | 3603.5963 | 3603.5927 | 0 | 1 | 466 | Specific | 658.47 | 579.35 | 579.35 | 13.18 | 97 | >sp\|P01833\|PIGR_HUMAN Polymeric immunoglobulin receptor OS=Homo sapiens OX=9606 GN=PIGR PE=1 SV=4 | 902 | 20201118_lss_colostrum_1ng_glyco_hcd203040_compound15.16438.16438.3 | controllerType=0 controllerNumber=1 scan=16438 | [illegible] |
| 13248:3 | 5 | K.VPGN[+2059.735]VTAVLGETLKVPC[+57.021]HFPC[+57.021]K.F | HexNAc(4)Hex(5)Fuc(1)NeuAc(1) | C[+57]*2, N[+2060] | 1495.0077 | 3 | 4483.0085 | 4482.9923 | 0 | 3.6 | 466 | Specific | 472.87 | 472.87 | 472.87 | 8.88 | 97 | >sp\|P01833\|PIGR_HUMAN Polymeric immunoglobulin receptor OS=Homo sapiens OX=9606 GN=PIGR PE=1 SV=4 | 902 | 20201118_lss_colostrum_1ng_glyco_hcd203040_compound15.16628.16628.3 | controllerType=0 controllerNumber=1 scan=16628 | [illegible] |
| 13251:3 | 5 | K.VPGN[+2059.735]VTAVLGETLKVPC[+57.021]HFPC[+57.021]K.F | HexNAc(4)Hex(5)Fuc(1)NeuAc(1) | C[+57]*2, N[+2060] | 1495.0077 | 3 | 4483.0085 | 4482.9923 | 0 | 3.6 | 466 | Specific | 473.81 | 438.2 | 438.2 | 8.88 | 97 | >sp\|P01833\|PIGR_HUMAN Polymeric immunoglobulin receptor OS=Homo sapiens OX=9606 GN=PIGR PE=1 SV=4 | 902 | 20201118_lss_colostrum_1ng_glyco_hcd203040_compound15.16631.16631.3 | controllerType=0 controllerNumber=1 scan=16631 | [illegible] |
| 13265:4 | 5 | K.VPGN[+2059.735]VTAVLGETLKVPC[+57.021]HFPC[+57.021]K.F | HexNAc(4)Hex(5)Fuc(1)NeuAc(1) | C[+57]*2, N[+2060] | 1121.506 | 4 | 4483.0023 | 4482.9923 | 0 | 2.2 | 466 | Specific | 541.07 | 533.69 | 533.69 | 9.93 | 97 | >sp\|P01833\|PIGR_HUMAN Polymeric immunoglobulin receptor OS=Homo sapiens OX=9606 GN=PIGR PE=1 SV=4 | 902 | 20201118_lss_colostrum_1ng_glyco_hcd203040_compound15.16646.16646.2 | controllerType=0 controllerNumber=1 scan=16646 | 34.8744 |
| 13268:4 | 5 | K.VPGN[+2059.735]VTAVLGETLKVPC[+57.021]HFPC[+57.021]K.F | HexNAc(4)Hex(5)Fuc(1)NeuAc(1) | C[+57]*2, N[+2060] | 1121.506 | 4 | 4483.0023 | 4482.9923 | 0 | 2.2 | 466 | Specific | 578.89 | 410.41 | 410.41 | 11.52 | 97 | >sp\|P01833\|PIGR_HUMAN Polymeric immunoglobulin receptor OS=Homo sapiens OX=9606 GN=PIGR PE=1 SV=4 | 902 | 20201118_lss_colostrum_1ng_glyco_hcd203040_compound15.16649.16649.2 | controllerType=0 controllerNumber=1 scan=16649 | 34.8811 |
| 13096:4 | 5 | K.VPGN[+2351.851]VTAVLGETLKVPC[+57.021]HFPC[+57.021]K.F | HexNAc(4)Hex(5)Fuc(3)NeuAc(1) | C[+57]*2, N[+2352] | 1194.5361 | 4 | 4775.1224 | 4775.1081 | 0 | 3 | 466 | Specific | 597.91 | 441.68 | 441.68 | 11.52 | 97 | >sp\|P01833\|PIGR_HUMAN Polymeric immunoglobulin receptor OS=Homo sapiens OX=9606 GN=PIGR PE=1 SV=4 | 902 | 20201118_lss_colostrum_1ng_glyco_hcd203040_compound15.16470.16470.4 | controllerType=0 controllerNumber=1 scan=16470 | 34.5078 |
| 13125:3 | 5 | K.VPGN[+2205.793]VTAVLGETLKVPC[+57.021]HFPC[+57.021]K.F | HexNAc(4)Hex(5)Fuc(2)NeuAc(1) | C[+57]*2, N[+2206] | 1543.6928 | 3 | 4629.0639 | 4629.0502 | 0 | 2.9 | 466 | Specific | 658.28 | 586.31 | 586.31 | 12.9 | 97 | >sp\|P01833\|PIGR_HUMAN Polymeric immunoglobulin receptor OS=Homo sapiens OX=9606 GN=PIGR PE=1 SV=4 | 902 | 20201118_lss_colostrum_1ng_glyco_hcd203040_compound15.16500.16500.3 | controllerType=0 controllerNumber=1 scan=16500 | 34.5837 |
| 13094:4 | 5 | K.VPGN[+2351.851]VTAVLGETLKVPC[+57.021]HFPC[+57.021]K.F | HexNAc(4)Hex(5)Fuc(3)NeuAc(1) | C[+57]*2, N[+2352] | 1194.5361 | 4 | 4775.1224 | 4775.1081 | 0 | 3 | 466 | Specific | 666.09 | 666.09 | 666.09 | 13.52 | 97 | >sp\|P01833\|PIGR_HUMAN Polymeric immunoglobulin receptor OS=Homo sapiens OX=9606 GN=PIGR PE=1 SV=4 | 902 | 20201118_lss_colostrum_1ng_glyco_hcd203040_compound15.16468.16468.4 | controllerType=0 controllerNumber=1 scan=16468 | 34.5037 |
| 13090:4 | 5 | K.VPGN[+2205.793]VTAVLGETLKVPC[+57.021]HFPC[+57.021]K.F | HexNAc(4)Hex(5)Fuc(2)NeuAc(1) | C[+57]*2, N[+2206] | 1158.0199 | 4 | 4629.0579 | 4629.0502 | 0 | 1.7 | 466 | Specific | 725.59 | 671.77 | 671.77 | 14.37 | 97 | >sp\|P01833\|PIGR_HUMAN Polymeric immunoglobulin receptor OS=Homo sapiens OX=9606 GN=PIGR PE=1 SV=4 | 902 | 20201118_lss_colostrum_1ng_glyco_hcd203040_compound15.16464.16464.4 | controllerType=0 controllerNumber=1 scan=16464 | 34.4978 |
| 13189:4 | 5 | K.VPGN[+2059.735]VTAVLGETLKVPC[+57.021]HFPC[+57.021]K.F | HexNAc(4)Hex(5)Fuc(1)NeuAc(1) | C[+57]*2, N[+2060] | 1121.5054 | 4 | 4482.9998 | 4482.9923 | 0 | 1.7 | 466 | Specific | 732.28 | 619.53 | 619.53 | 14.37 | 97 | >sp\|P01833\|PIGR_HUMAN Polymeric immunoglobulin receptor OS=Homo sapiens OX=9606 GN=PIGR PE=1 SV=4 | 902 | 20201118_lss_colostrum_1ng_glyco_hcd203040_compound15.16567.16567.4 | controllerType=0 controllerNumber=1 scan=16567 | 34.7082 |
| 13191:4 | 5 | K.VPGN[+2059.735]VTAVLGETLKVPC[+57.021]HFPC[+57.021]K.F | HexNAc(4)Hex(5)Fuc(1)NeuAc(1) | C[+57]*2, N[+2060] | 1121.5054 | 4 | 4482.9998 | 4482.9923 | 0 | 1.7 | 466 | Specific | 732.96 | 656.38 | 656.38 | 14.37 | 97 | >sp\|P01833\|PIGR_HUMAN Polymeric immunoglobulin receptor OS=Homo sapiens OX=9606 GN=PIGR PE=1 SV=4 | 902 | 20201118_lss_colostrum_1ng_glyco_hcd203040_compound15.16569.16569.4 | controllerType=0 controllerNumber=1 scan=16569 | 34.7135 |
| 13123:3 | 5 | K.VPGN[+2205.793]VTAVLGETLKVPC[+57.021]HFPC[+57.021]K.F | HexNAc(4)Hex(5)Fuc(2)NeuAc(1) | C[+57]*2, N[+2206] | 1543.6928 | 3 | 4629.0639 | 4629.0502 | 0 | 2.9 | 466 | Specific | 772.67 | 697.22 | 697.22 | 14.9 | 97 | >sp\|P01833\|PIGR_HUMAN Polymeric immunoglobulin receptor OS=Homo sapiens OX=9606 GN=PIGR PE=1 SV=4 | 902 | 20201118_lss_colostrum_1ng_glyco_hcd203040_compound15.16498.16498.3 | controllerType=0 controllerNumber=1 scan=16498 | 34.5691 |
| 13087:4 | 5 | K.VPGN[+2205.793]VTAVLGETLKVPC[+57.021]HFPC[+57.021]K.F | HexNAc(4)Hex(5)Fuc(2)NeuAc(1) | C[+57]*2, N[+2206] | 1158.0199 | 4 | 4629.0579 | 4629.0502 | 0 | 1.7 | 466 | Specific | 877.32 | 636.34 | 636.34 | 17.37 | 97 | >sp\|P01833\|PIGR_HUMAN Polymeric immunoglobulin receptor OS=Homo sapiens OX=9606 GN=PIGR PE=1 SV=4 | 902 | 20201118_lss_colostrum_1ng_glyco_hcd203040_compound15.16461.16461.4 | controllerType=0 controllerNumber=1 scan=16461 | 34.492 |
| 05531:3 | 5 | K.WN[+2059.735]NTGC[+57.021]QALPSQDEGPSK.A | HexNAc(4)Hex(5)Fuc(1)NeuAc(1) | C[+57], N[+2060] | 1350.2111 | 3 | 4048.6188 | 4048.6116 | 0 | 1.8 | 498 | Specific | 389.47 | 330.06 | 330.06 | 7.4 | 97 | >sp\|P01833\|PIGR_HUMAN Polymeric immunoglobulin receptor OS=Homo sapiens OX=9606 GN=PIGR PE=1 SV=4 | 902 | 20201118_lss_colostrum_1ng_glyco_hcd203040_compound15.8619.8619.3 | controllerType=0 controllerNumber=1 scan=8619 | 20.776 |
| 05529:3 | 5 | K.WN[+2059.735]NTGC[+57.021]QALPSQDEGPSK.A | HexNAc(4)Hex(5)Fuc(1)NeuAc(1) | C[+57], N[+2060] | 1350.2111 | 3 | 4048.6188 | 4048.6116 | 0 | 1.8 | 498 | Specific | 432.79 | 396.86 | 396.86 | 9.46 | 97 | >sp\|P01833\|PIGR_HUMAN Polymeric immunoglobulin receptor OS=Homo sapiens OX=9606 GN=PIGR PE=1 SV=4 | 902 | 20201118_lss_colostrum_1ng_glyco_hcd203040_compound15.8617.8617.3 | controllerType=0 controllerNumber=1 scan=8617 | 20.772 |
| 05430:3 | 5 | K.WN[+2205.793]NTGC[+57.021]QALPSQDEGPSK.A | HexNAc(4)Hex(5)Fuc(2)NeuAc(1) | C[+57], N[+2206] | 1398.8961 | 3 | 4194.6736 | 4194.6695 | 0 | 1 | 498 | Specific | 520.17 | 469.89 | 469.89 | 9.86 | 97 | >sp\|P01833\|PIGR_HUMAN Polymeric immunoglobulin receptor OS=Homo sapiens OX=9606 GN=PIGR PE=1 SV=4 | 902 | 20201118_lss_colostrum_1ng_glyco_hcd203040_compound15.8515.8515.3 | controllerType=0 controllerNumber=1 scan=8515 | 20.6041 |
| 05428:3 | 5 | K.WN[+2205.793]NTGC[+57.021]QALPSQDEGPSK.A | HexNAc(4)Hex(5)Fuc(2)NeuAc(1) | C[+57], N[+2206] | 1398.8961 | 3 | 4194.6736 | 4194.6695 | 0 | 1 | 498 | Specific | 520.63 | 479.3 | 479.3 | 9.86 | 97 | >sp\|P01833\|PIGR_HUMAN Polymeric immunoglobulin receptor OS=Homo sapiens OX=9606 GN=PIGR PE=1 SV=4 | 902 | 20201118_lss_colostrum_1ng_glyco_hcd203040_compound15.8513.8513.3 | controllerType=0 controllerNumber=1 scan=8513 | 20.6006 |
| 07626:3 | 5 | K.WN[+2204.772]NTGC[+57.021]QALPSQDEGPSK.A | HexNAc(4)Hex(5)NeuAc(2) | C[+57], N[+2205] | 1398.5585 | 3 | 4193.661 | 4193.6491 | 0 | 2.8 | 498 | Specific | 529.43 | 462.25 | 462.25 | 10.02 | 97 | >sp\|P01833\|PIGR_HUMAN Polymeric immunoglobulin receptor OS=Homo sapiens OX=9606 GN=PIGR PE=1 SV=4 | 902 | 20201118_lss_colostrum_1ng_glyco_hcd203040_compound15.10788.10788.3 | controllerType=0 controllerNumber=1 scan=10788 | 24.316 |
| 05704:3 | 5 | K.WN[+1548.545]NTGC[+57.021]QALPSQDEGPSK.A | HexNAc(3)Hex(4)NeuAc(1) | C[+57], N[+1549] | 1179.8134 | 3 | 3537.4257 | 3537.4215 | 0 | 1.2 | 498 | Specific | 609.99 | 548.08 | 548.08 | 12.47 | 97 | >sp\|P01833\|PIGR_HUMAN Polymeric immunoglobulin receptor OS=Homo sapiens OX=9606 GN=PIGR PE=1 SV=4 | 902 | 20201118_lss_colostrum_1ng_glyco_hcd203040_compound15.8798.8798.3 | controllerType=0 controllerNumber=1 scan=8798 | 21.0776 |
| 03910:2 | 5 | K.WN[+1216.423]NTGC[+57.021]QALPSQDEGPSK.A | HexNAc(2)Hex(5) | C[+57], N[+1216] | 1603.1578 | 2 | 3205.3082 | 3205.2995 | 0 | 2.7 | 498 | Specific | 629.79 | 557.72 | 557.72 | 12.7 | 97 | >sp\|P01833\|PIGR_HUMAN Polymeric immunoglobulin receptor OS=Homo sapiens OX=9606 GN=PIGR PE=1 SV=4 | 902 | 20201118_lss_colostrum_1ng_glyco_hcd203040_compound15.6942.6942.2 | controllerType=0 controllerNumber=1 scan=6942 | 18.0611 |
| 05706:3 | 5 | K.WN[+1548.545]NTGC[+57.021]QALPSQDEGPSK.A | HexNAc(3)Hex(4)NeuAc(1) | C[+57], N[+1549] | 1179.8134 | 3 | 3537.4257 | 3537.4215 | 0 | 1.2 | 498 | Specific | 675.67 | 576.66 | 576.66 | 13.47 | 97 | >sp\|P01833\|PIGR_HUMAN Polymeric immunoglobulin receptor OS=Homo sapiens OX=9606 GN=PIGR PE=1 SV=4 | 902 | 20201118_lss_colostrum_1ng_glyco_hcd203040_compound15.8800.8800.3 | controllerType=0 controllerNumber=1 scan=8800 | 21.0813 |
| 07628:3 | 5 | K.WN[+2204.772]NTGC[+57.021]QALPSQDEGPSK.A | HexNAc(4)Hex(5)NeuAc(2) | C[+57], N[+2205] | 1398.5585 | 3 | 4193.661 | 4193.6491 | 0 | 2.8 | 498 | Specific | 691.38 | 578.75 | 578.75 | 13.63 | 97 | >sp\|P01833\|PIGR_HUMAN Polymeric immunoglobulin receptor OS=Homo sapiens OX=9606 GN=PIGR PE=1 SV=4 | 902 | 20201118_lss_colostrum_1ng_glyco_hcd203040_compound15.10790.10790.3 | controllerType=0 controllerNumber=1 scan=10790 | 24.3198 |
| 03925:2 | 5 | K.WN[+1216.423]NTGC[+57.021]QALPSQDEGPSK.A | HexNAc(2)Hex(5) | C[+57], N[+1216] | 1603.1512 | 2 | 3205.2951 | 3205.2995 | 0 | -1.4 | 498 | Specific | 699.74 | 630.49 | 630.49 | 12.62 | 97 | >sp\|P01833\|PIGR_HUMAN Polymeric immunoglobulin receptor OS=Homo sapiens OX=9606 GN=PIGR PE=1 SV=4 | 902 | 20201118_lss_colostrum_1ng_glyco_hcd203040_compound15.6958.6958.2 | controllerType=0 controllerNumber=1 scan=6958 | 18.0885 |
| 03912:2 | 5 | K.WN[+1216.423]NTGC[+57.021]QALPSQDEGPSK.A | HexNAc(2)Hex(5) | C[+57], N[+1216] | 1603.1578 | 2 | 3205.3082 | 3205.2995 | 0 | 2.7 | 498 | Specific | 722.19 | 618.5 | 618.5 | 14.7 | 97 | >sp\|P01833\|PIGR_HUMAN Polymeric immunoglobulin receptor OS=Homo sapiens OX=9606 GN=PIGR PE=1 SV=4 | 902 | 20201118_lss_colostrum_1ng_glyco_hcd203040_compound15.6944.6944.2 | controllerType=0 controllerNumber=1 scan=6944 | 18.0649 |
| 03927:2 | 5 | K.WN[+1216.423]NTGC[+57.021]QALPSQDEGPSK.A | HexNAc(2)Hex(5) | C[+57], N[+1216] | 1603.1512 | 2 | 3205.2951 | 3205.2995 | 0 | -1.4 | 498 | Specific | 727.84 | 636.64 | 636.64 | 13.62 | 97 | >sp\|P01833\|PIGR_HUMAN Polymeric immunoglobulin receptor OS=Homo sapiens OX=9606 GN=PIGR PE=1 SV=4 | 902 | 20201118_lss_colostrum_1ng_glyco_hcd203040_compound15.6960.6960.2 | controllerType=0 controllerNumber=1 scan=6960 | 18.0922 |
| 05548:3 | 5 | K.WN[+1913.677]NTGC[+57.021]QALPSQDEGPSK.A | HexNAc(4)Hex(5)NeuAc(1) | C[+57], N[+1914] | 1301.525 | 3 | 3902.5603 | 3902.5537 | 0 | 1.7 | 498 | Specific | 809.6 | 739.75 | 739.75 | 16.47 | 97 | >sp\|P01833\|PIGR_HUMAN Polymeric immunoglobulin receptor OS=Homo sapiens OX=9606 GN=PIGR PE=1 SV=4 | 902 | 20201118_lss_colostrum_1ng_glyco_hcd203040_compound15.8637.8637.3 | controllerType=0 controllerNumber=1 scan=8637 | 20.808 |
| 05522:3 | 5 | K.WN[+1872.651]NTGC[+57.021]QALPSQDEGPSK.A | HexNAc(3)Hex(6)NeuAc(1) | C[+57], N[+1873] | 1287.8504 | 3 | 3861.5366 | 3861.5272 | 0 | 2.5 | 498 | Specific | 825.48 | 716.43 | 716.43 | 16.63 | 97 | >sp\|P01833\|PIGR_HUMAN Polymeric immunoglobulin receptor OS=Homo sapiens OX=9606 GN=PIGR PE=1 SV=4 | 902 | 20201118_lss_colostrum_1ng_glyco_hcd203040_compound15.8610.8610.3 | controllerType=0 controllerNumber=1 scan=8610 | 20.7598 |
| 05550:3 | 5 | K.WN[+1913.677]NTGC[+57.021]QALPSQDEGPSK.A | HexNAc(4)Hex(5)NeuAc(1) | C[+57], N[+1914] | 1301.525 | 3 | 3902.5603 | 3902.5537 | 0 | 1.7 | 498 | Specific | 830.66 | 743.13 | 743.13 | 16.47 | 97 | >sp\|P01833\|PIGR_HUMAN Polymeric immunoglobulin receptor OS=Homo sapiens OX=9606 GN=PIGR PE=1 SV=4 | 902 | 20201118_lss_colostrum_1ng_glyco_hcd203040_compound15.8639.8639.3 | controllerType=0 controllerNumber=1 scan=8639 | 20.812 |
| 05586:3 | 5 | K.WN[+1710.598]NTGC[+57.021]QALPSQDEGPSK.A | HexNAc(3)Hex(5)NeuAc(1) | C[+57], N[+1711] | 1233.8327 | 3 | 3699.4835 | 3699.4743 | 0 | 2.5 | 498 | Specific | 864.02 | 772.14 | 772.14 | 17.63 | 97 | >sp\|P01833\|PIGR_HUMAN Polymeric immunoglobulin receptor OS=Homo sapiens OX=9606 GN=PIGR PE=1 SV=4 | 902 | 20201118_lss_colostrum_1ng_glyco_hcd203040_compound15.8676.8676.3 | controllerType=0 controllerNumber=1 scan=8676 | 20.8775 |
| 05520:3 | 5 | K.WN[+1872.651]NTGC[+57.021]QALPSQDEGPSK.A | HexNAc(3)Hex(6)NeuAc(1) | C[+57], N[+1873] | 1287.8504 | 3 | 3861.5366 | 3861.5272 | 0 | 2.5 | 498 | Specific | 867.29 | 799.97 | 799.97 | 17.63 | 97 | >sp\|P01833\|PIGR_HUMAN Polymeric immunoglobulin receptor OS=Homo sapiens OX=9606 GN=PIGR PE=1 SV=4 | 902 | 20201118_lss_colostrum_1ng_glyco_hcd203040_compound15.8608.8608.3 | controllerType=0 controllerNumber=1 scan=8608 | 20.7547 |
| 05584:3 | 5 | K.WN[+1710.598]NTGC[+57.021]QALPSQDEGPSK.A | HexNAc(3)Hex(5)NeuAc(1) | C[+57], N[+1711] | 1233.8327 | 3 | 3699.4835 | 3699.4743 | 0 | 2.5 | 498 | Specific | 984.29 | 895.36 | 895.36 | 19.63 | 97 | >sp\|P01833\|PIGR_HUMAN Polymeric immunoglobulin receptor OS=Homo sapiens OX=9606 GN=PIGR PE=1 SV=4 | 902 | 20201118_lss_colostrum_1ng_glyco_hcd203040_compound15.8674.8674.3 | controllerType=0 controllerNumber=1 scan=8674 | 20.8727 |
| 03896:3 | 5 | K.WN[+1216.423]NTGC[+57.021]QALPSQDEGPSK.A | HexNAc(2)Hex(5) | C[+57], N[+1216] | 1069.1065 | 3 | 3205.3049 | 3205.2995 | 0 | 1.7 | 498 | Specific | 1010.19 | 982.49 | 982.49 | 20.47 | 97 | >sp\|P01833\|PIGR_HUMAN Polymeric immunoglobulin receptor OS=Homo sapiens OX=9606 GN=PIGR PE=1 SV=4 | 902 | 20201118_lss_colostrum_1ng_glyco_hcd203040_compound15.6928.6928.3 | controllerType=0 controllerNumber=1 scan=6928 | 18.0383 |
| 03898:3 | 5 | K.WN[+1216.423]NTGC[+57.021]QALPSQDEGPSK.A | HexNAc(2)Hex(5) | C[+57], N[+1216] | 1069.1065 | 3 | 3205.3049 | 3205.2995 | 0 | 1.7 | 498 | Specific | 1048.07 | 1021.15 | 1021.15 | 20.47 | 97 | >sp\|P01833\|PIGR_HUMAN Polymeric immunoglobulin receptor OS=Homo sapiens OX=9606 GN=PIGR PE=1 SV=4 | 902 | 20201118_lss_colostrum_1ng_glyco_hcd203040_compound15.6930.6930.3 | controllerType=0 controllerNumber=1 scan=6930 | 18.0431 |
| 19139:3 | 6 | R.LSLHRPALEDLLLGSEAN[+2204.772]LTC[+57.021]TLTGLR.D | HexNAc(4)Hex(5)NeuAc(2) | C[+57], N[+2205] | 1723.466 | 3 | 5168.3835 | 5168.3707 | 0 | 2.5 | 127 | Specific | 285.45 | 285.45 | 285.45 | 6.3 | 82 | >sp\|P01876\|IGHA1_HUMAN Immunoglobulin heavy constant alpha 1 OS=Homo sapiens OX=9606 GN=IGHA1 PE=1 SV=2 | 255 | 20201118_lss_colostrum_1ng_glyco_hcd203040_compound15.22760.22760.3 | controllerType=0 controllerNumber=1 scan=22760 | 47.3157 |
| 19128:4 | 6 | R.LSLHRPALEDLLLGSEAN[+2407.852]LTC[+57.021]TLTGLR.D | HexNAc(5)Hex(5)NeuAc(2) | C[+57], N[+2408] | 1343.6193 | 4 | 5371.4552 | 5371.4501 | 0 | 1 | 127 | Specific | 287.12 | 287.12 | 287.12 | 6.77 | 82 | >sp\|P01876\|IGHA1_HUMAN Immunoglobulin heavy constant alpha 1 OS=Homo sapiens OX=9606 GN=IGHA1 PE=1 SV=2 | 255 | 20201118_lss_colostrum_1ng_glyco_hcd203040_compound15.22749.22749.4 | controllerType=0 controllerNumber=1 scan=22749 | 47.2965 |
| 17173:3 | 6 | R.LSLHRPALEDLLLGSEAN[+1054.370]LTC[+57.021]TLTGLR.D | HexNAc(2)Hex(4) | C[+57], N[+1054] | 1339.9993 | 3 | 4017.9834 | 4017.9683 | 0 | 3.8 | 127 | Specific | 303.36 | 303.36 | 303.36 | 6.49 | 82 | >sp\|P01876\|IGHA1_HUMAN Immunoglobulin heavy constant alpha 1 OS=Homo sapiens OX=9606 GN=IGHA1 PE=1 SV=2 | 255 | 20201118_lss_colostrum_1ng_glyco_hcd203040_compound15.20711.20711.3 | controllerType=0 controllerNumber=1 scan=20711 | 42.9522 |
| 17993:4 | 6 | R.LSLHRPALEDLLLGSEAN[+2116.756]LTC[+57.021]TLTGLR.D | HexNAc(5)Hex(5)NeuAc(1) | C[+57], N[+2117] | 1270.8479 | 4 | 5080.3698 | 5080.3547 | 0 | 3 | 127 | Specific | 322.17 | 322.17 | 322.17 | 7.4 | 82 | >sp\|P01876\|IGHA1_HUMAN Immunoglobulin heavy constant alpha 1 OS=Homo sapiens OX=9606 GN=IGHA1 PE=1 SV=2 | 255 | 20201118_lss_colostrum_1ng_glyco_hcd203040_compound15.21568.21568.4 | controllerType=0 controllerNumber=1 scan=21568 | 44.7843 |
| 17175:3 | 6 | R.LSLHRPALEDLLLGSEAN[+1054.370]LTC[+57.021]TLTGLR.D | HexNAc(2)Hex(4) | C[+57], N[+1054] | 1339.9993 | 3 | 4017.9834 | 4017.9683 | 0 | 3.8 | 127 | Specific | 345.74 | 345.74 | 345.74 | 6.9 | 82 | >sp\|P01876\|IGHA1_HUMAN Immunoglobulin heavy constant alpha 1 OS=Homo sapiens OX=9606 GN=IGHA1 PE=1 SV=2 | 255 | 20201118_lss_colostrum_1ng_glyco_hcd203040_compound15.20713.20713.3 | controllerType=0 controllerNumber=1 scan=20713 | 42.9591 |
| 17034:4 | 6 | R.LSLHRPALEDLLLGSEAN[+1581.555]LTC[+57.021]TLTGLR.D | HexNAc(3)Hex(6) | C[+57], N[+1582] | 1137.0469 | 4 | 4545.1658 | 4545.1533 | 0 | 2.8 | 127 | Specific | 392.38 | 369.43 | 369.43 | 8.11 | 82 | >sp\|P01876\|IGHA1_HUMAN Immunoglobulin heavy constant alpha 1 OS=Homo sapiens OX=9606 GN=IGHA1 PE=1 SV=2 | 255 | 20201118_lss_colostrum_1ng_glyco_hcd203040_compound15.20566.20566.4 | controllerType=0 controllerNumber=1 scan=20566 | 42.6185 |
| 17071:4 | 6 | R.LSLHRPALEDLLLGSEAN[+1257.449]LTC[+57.021]TLTGLR.D | HexNAc(3)Hex(4) | C[+57], N[+1257] | 1056.0184 | 4 | 4221.0516 | 4221.0477 | 0 | 0.9 | 127 | Specific | 397.2 | 397.2 | 397.2 | 9.46 | 82 | >sp\|P01876\|IGHA1_HUMAN Immunoglobulin heavy constant alpha 1 OS=Homo sapiens OX=9606 GN=IGHA1 PE=1 SV=2 | 255 | 20201118_lss_colostrum_1ng_glyco_hcd203040_compound15.20605.20605.4 | controllerType=0 controllerNumber=1 scan=20605 | 42.7088 |
| 17027:4 | 6 | R.LSLHRPALEDLLLGSEAN[+1864.634]LTC[+57.021]TLTGLR.D | HexNAc(2)Hex(9) | C[+57], N[+1865] | 1207.8122 | 4 | 4828.2269 | 4828.2324 | 0 | -1.1 | 127 | Specific | 428.97 | 428.97 | 428.97 | 8.68 | 82 | >sp\|P01876\|IGHA1_HUMAN Immunoglobulin heavy constant alpha 1 OS=Homo sapiens OX=9606 GN=IGHA1 PE=1 SV=2 | 255 | 20201118_lss_colostrum_1ng_glyco_hcd203040_compound15.20559.20559.4 | controllerType=0 controllerNumber=1 scan=20559 | 42.6048 |
| 19127:4 | 6 | R.LSLHRPALEDLLLGSEAN[+2204.772]LTC[+57.021]TLTGLR.D | HexNAc(4)Hex(5)NeuAc(2) | C[+57], N[+2205] | 1292.8522 | 4 | 5168.3869 | 5168.3707 | 0 | 3.1 | 127 | Specific | 481.91 | 442.37 | 442.37 | 9.94 | 82 | >sp\|P01876\|IGHA1_HUMAN Immunoglobulin heavy constant alpha 1 OS=Homo sapiens OX=9606 GN=IGHA1 PE=1 SV=2 | 255 | 20201118_lss_colostrum_1ng_glyco_hcd203040_compound15.22748.22748.4 | controllerType=0 controllerNumber=1 scan=22748 | 47.2957 |
| 17138:4 | 6 | R.LSLHRPALEDLLLGSEAN[+1622.582]LTC[+57.021]TLTGLR.D | HexNAc(4)Hex(5) | C[+57], N[+1623] | 1147.3033 | 4 | 4586.1912 | 4586.1799 | 0 | 2.5 | 127 | Specific | 485.18 | 485.18 | 485.18 | 9.94 | 82 | >sp\|P01876\|IGHA1_HUMAN Immunoglobulin heavy constant alpha 1 OS=Homo sapiens OX=9606 GN=IGHA1 PE=1 SV=2 | 255 | 20201118_lss_colostrum_1ng_glyco_hcd203040_compound15.20675.20675.4 | controllerType=0 controllerNumber=1 scan=20675 | 42.8593 |
| 17090:3 | 6 | R.LSLHRPALEDLLLGSEAN[+1501.555]LTC[+57.021]TLTGLR.D | HexNAc(5)Hex(3) | C[+57], N[+1502] | 1489.0604 | 3 | 4465.1665 | 4465.1536 | 0 | 2.9 | 127 | Specific | 492.79 | 492.79 | 492.79 | 9.37 | 82 | >sp\|P01876\|IGHA1_HUMAN Immunoglobulin heavy constant alpha 1 OS=Homo sapiens OX=9606 GN=IGHA1 PE=1 SV=2 | 255 | 20201118_lss_colostrum_1ng_glyco_hcd203040_compound15.20625.20625.6 | controllerType=0 controllerNumber=1 scan=20625 | 42.7453 |
| 16985:4 | 6 | R.LSLHRPALEDLLLGSEAN[+1825.661]LTC[+57.021]TLTGLR.D | HexNAc(5)Hex(5) | C[+57], N[+1826] | 1198.0751 | 4 | 4789.2786 | 4789.2592 | 0 | 4 | 127 | Specific | 496.57 | 496.57 | 496.57 | 9.79 | 82 | >sp\|P01876\|IGHA1_HUMAN Immunoglobulin heavy constant alpha 1 OS=Homo sapiens OX=9606 GN=IGHA1 PE=1 SV=2 | 255 | 20201118_lss_colostrum_1ng_glyco_hcd203040_compound15.20515.20515.4 | controllerType=0 controllerNumber=1 scan=20515 | 42.4856 |
| 19142:3 | 6 | R.LSLHRPALEDLLLGSEAN[+2204.772]LTC[+57.021]TLTGLR.D | HexNAc(4)Hex(5)NeuAc(2) | C[+57], N[+2205] | 1723.466 | 3 | 5168.3835 | 5168.3707 | 0 | 2.5 | 127 | Specific | 501.64 | 454.85 | 454.85 | 9.37 | 82 | >sp\|P01876\|IGHA1_HUMAN Immunoglobulin heavy constant alpha 1 OS=Homo sapiens OX=9606 GN=IGHA1 PE=1 SV=2 | 255 | 20201118_lss_colostrum_1ng_glyco_hcd203040_compound15.22763.22763.3 | controllerType=0 controllerNumber=1 scan=22763 | 47.3204 |
| 17064:5 | 6 | R.LSLHRPALEDLLLGSEAN[+1501.555]LTC[+57.021]TLTGLR.D | HexNAc(5)Hex(3) | C[+57], N[+1502] | 893.8386 | 5 | 4465.1641 | 4465.1536 | 0 | 2.4 | 127 | Specific | 507.13 | 507.13 | 507.13 | 9.92 | 82 | >sp\|P01876\|IGHA1_HUMAN Immunoglobulin heavy constant alpha 1 OS=Homo sapiens OX=9606 GN=IGHA1 PE=1 SV=2 | 255 | 20201118_lss_colostrum_1ng_glyco_hcd203040_compound15.20598.20598.5 | controllerType=0 controllerNumber=1 scan=20598 | 42.6994 |
| 18072:3 | 6 | R.LSLHRPALEDLLLGSEAN[+1751.624]LTC[+57.021]TLTGLR.D | HexNAc(4)Hex(4)NeuAc(1) | C[+57], N[+1752] | 1572.4178 | 3 | 4715.2388 | 4715.2225 | 0 | 3.5 | 127 | Specific | 511.49 | 493.28 | 493.28 | 9.34 | 82 | >sp\|P01876\|IGHA1_HUMAN Immunoglobulin heavy constant alpha 1 OS=Homo sapiens OX=9606 GN=IGHA1 PE=1 SV=2 | 255 | 20201118_lss_colostrum_1ng_glyco_hcd203040_compound15.21650.21650.3 | controllerType=0 controllerNumber=1 scan=21650 | 44.9608 |
| 18093:3 | 6 | R.LSLHRPALEDLLLGSEAN[+1548.545]LTC[+57.021]TLTGLR.D | HexNAc(3)Hex(4)NeuAc(1) | C[+57], N[+1549] | 1504.7243 | 3 | 4512.1583 | 4512.1431 | 0 | 3.4 | 127 | Specific | 517.19 | 517.19 | 517.19 | 9.34 | 82 | >sp\|P01876\|IGHA1_HUMAN Immunoglobulin heavy constant alpha 1 OS=Homo sapiens OX=9606 GN=IGHA1 PE=1 SV=2 | 255 | 20201118_lss_colostrum_1ng_glyco_hcd203040_compound15.21672.21672.3 | controllerType=0 controllerNumber=1 scan=21672 | 45.0042 |
| 17037:4 | 6 | R.LSLHRPALEDLLLGSEAN[+1581.555]LTC[+57.021]TLTGLR.D | HexNAc(3)Hex(6) | C[+57], N[+1582] | 1137.0469 | 4 | 4545.1658 | 4545.1533 | 0 | 2.8 | 127 | Specific | 518.21 | 332.49 | 332.49 | 7.27 | 82 | >sp\|P01876\|IGHA1_HUMAN Immunoglobulin heavy constant alpha 1 OS=Homo sapiens OX=9606 GN=IGHA1 PE=1 SV=2 | 255 | 20201118_lss_colostrum_1ng_glyco_hcd203040_compound15.20569.20569.4 | controllerType=0 controllerNumber=1 scan=20569 | 42.6283 |
| 16978:3 | 6 | R.LSLHRPALEDLLLGSEAN[+1864.634]LTC[+57.021]TLTGLR.D | HexNAc(2)Hex(9) | C[+57], N[+1865] | 1610.0888 | 3 | 4828.2519 | 4828.2324 | 0 | 4 | 127 | Specific | 550.18 | 535.16 | 535.16 | 9.35 | 82 | >sp\|P01876\|IGHA1_HUMAN Immunoglobulin heavy constant alpha 1 OS=Homo sapiens OX=9606 GN=IGHA1 PE=1 SV=2 | 255 | 20201118_lss_colostrum_1ng_glyco_hcd203040_compound15.20508.20508.3 | controllerType=0 controllerNumber=1 scan=20508 | 42.4706 |
| 17031:5 | 6 | R.LSLHRPALEDLLLGSEAN[+1663.608]LTC[+57.021]TLTGLR.D | HexNAc(5)Hex(4) | C[+57], N[+1664] | 926.2487 | 5 | 4627.2146 | 4627.2064 | 0 | 1.8 | 127 | Specific | 553.26 | 553.26 | 553.26 | 9.94 | 82 | >sp\|P01876\|IGHA1_HUMAN Immunoglobulin heavy constant alpha 1 OS=Homo sapiens OX=9606 GN=IGHA1 PE=1 SV=2 | 255 | 20201118_lss_colostrum_1ng_glyco_hcd203040_compound15.20563.20563.5 | controllerType=0 controllerNumber=1 scan=20563 | 42.615 |
| 17028:5 | 6 | R.LSLHRPALEDLLLGSEAN[+1663.608]LTC[+57.021]TLTGLR.D | HexNAc(5)Hex(4) | C[+57], N[+1664] | 926.2487 | 5 | 4627.2146 | 4627.2064 | 0 | 1.8 | 127 | Specific | 566.36 | 566.36 | 566.36 | 10.2 | 82 | >sp\|P01876\|IGHA1_HUMAN Immunoglobulin heavy constant alpha 1 OS=Homo sapiens OX=9606 GN=IGHA1 PE=1 SV=2 | 255 | 20201118_lss_colostrum_1ng_glyco_hcd203040_compound15.20560.20560.5 | controllerType=0 controllerNumber=1 scan=20560 | 42.606 |
| 18042:3 | 6 | R.LSLHRPALEDLLLGSEAN[+1710.598]LTC[+57.021]TLTGLR.D | HexNAc(3)Hex(5)NeuAc(1) | C[+57], N[+1711] | 1558.7421 | 3 | 4674.2118 | 4674.1959 | 0 | 3.4 | 127 | Specific | 575.03 | 542.68 | 542.68 | 9.6 | 82 | >sp\|P01876\|IGHA1_HUMAN Immunoglobulin heavy constant alpha 1 OS=Homo sapiens OX=9606 GN=IGHA1 PE=1 SV=2 | 255 | 20201118_lss_colostrum_1ng_glyco_hcd203040_compound15.21618.21618.3 | controllerType=0 controllerNumber=1 scan=21618 | 44.8814 |
| 18096:3 | 6 | R.LSLHRPALEDLLLGSEAN[+1548.545]LTC[+57.021]TLTGLR.D | HexNAc(3)Hex(4)NeuAc(1) | C[+57], N[+1549] | 1504.7243 | 3 | 4512.1583 | 4512.1431 | 0 | 3.4 | 127 | Specific | 575.36 | 544.01 | 544.01 | 9.6 | 82 | >sp\|P01876\|IGHA1_HUMAN Immunoglobulin heavy constant alpha 1 OS=Homo sapiens OX=9606 GN=IGHA1 PE=1 SV=2 | 255 | 20201118_lss_colostrum_1ng_glyco_hcd203040_compound15.21675.21675.3 | controllerType=0 controllerNumber=1 scan=21675 | 45.0137 |
| 18039:3 | 6 | R.LSLHRPALEDLLLGSEAN[+1710.598]LTC[+57.021]TLTGLR.D | HexNAc(3)Hex(5)NeuAc(1) | C[+57], N[+1711] | 1558.7421 | 3 | 4674.2118 | 4674.1959 | 0 | 3.4 | 127 | Specific | 593.42 | 491.59 | 491.59 | 10.94 | 82 | >sp\|P01876\|IGHA1_HUMAN Immunoglobulin heavy constant alpha 1 OS=Homo sapiens OX=9606 GN=IGHA1 PE=1 SV=2 | 255 | 20201118_lss_colostrum_1ng_glyco_hcd203040_compound15.21615.21615.3 | controllerType=0 controllerNumber=1 scan=21615 | 44.8745 |
| 17134:4 | 6 | R.LSLHRPALEDLLLGSEAN[+1298.476]LTC[+57.021]TLTGLR.D | HexNAc(4)Hex(3) | C[+57], N[+1298] | 1066.2771 | 4 | 4262.0867 | 4262.0742 | 0 | 2.9 | 127 | Specific | 604.01 | 558.45 | 558.45 | 12.76 | 82 | >sp\|P01876\|IGHA1_HUMAN Immunoglobulin heavy constant alpha 1 OS=Homo sapiens OX=9606 GN=IGHA1 PE=1 SV=2 | 255 | 20201118_lss_colostrum_1ng_glyco_hcd203040_compound15.20671.20671.4 | controllerType=0 controllerNumber=1 scan=20671 | 42.8499 |
| 17018:3 | 6 | R.LSLHRPALEDLLLGSEAN[+1663.608]LTC[+57.021]TLTGLR.D | HexNAc(5)Hex(4) | C[+57], N[+1664] | 1543.0778 | 3 | 4627.2189 | 4627.2064 | 0 | 2.7 | 127 | Specific | 616.69 | 606.82 | 606.82 | 12.14 | 82 | >sp\|P01876\|IGHA1_HUMAN Immunoglobulin heavy constant alpha 1 OS=Homo sapiens OX=9606 GN=IGHA1 PE=1 SV=2 | 255 | 20201118_lss_colostrum_1ng_glyco_hcd203040_compound15.20550.20550.3 | controllerType=0 controllerNumber=1 scan=20550 | 42.5887 |
| 17092:3 | 6 | R.LSLHRPALEDLLLGSEAN[+1501.555]LTC[+57.021]TLTGLR.D | HexNAc(5)Hex(3) | C[+57], N[+1502] | 1489.0604 | 3 | 4465.1665 | 4465.1536 | 0 | 2.9 | 127 | Specific | 621.19 | 573.45 | 573.45 | 12.14 | 82 | >sp\|P01876\|IGHA1_HUMAN Immunoglobulin heavy constant alpha 1 OS=Homo sapiens OX=9606 GN=IGHA1 PE=1 SV=2 | 255 | 20201118_lss_colostrum_1ng_glyco_hcd203040_compound15.20627.20627.6 | controllerType=0 controllerNumber=1 scan=20627 | 42.7496 |
| 17006:3 | 6 | R.LSLHRPALEDLLLGSEAN[+1702.581]LTC[+57.021]TLTGLR.D | HexNAc(2)Hex(8) | C[+57], N[+1703] | 1556.0693 | 3 | 4666.1933 | 4666.1796 | 0 | 2.9 | 127 | Specific | 625.08 | 625.08 | 625.08 | 12.14 | 82 | >sp\|P01876\|IGHA1_HUMAN Immunoglobulin heavy constant alpha 1 OS=Homo sapiens OX=9606 GN=IGHA1 PE=1 SV=2 | 255 | 20201118_lss_colostrum_1ng_glyco_hcd203040_compound15.20537.20537.3 | controllerType=0 controllerNumber=1 scan=20537 | 42.5528 |
| 17063:3 | 6 | R.LSLHRPALEDLLLGSEAN[+1501.555]LTC[+57.021]TLTGLR.D | HexNAc(5)Hex(3) | C[+57], N[+1502] | 1489.0604 | 3 | 4465.1665 | 4465.1536 | 0 | 2.9 | 127 | Specific | 627.63 | 610.49 | 610.49 | 12.14 | 82 | >sp\|P01876\|IGHA1_HUMAN Immunoglobulin heavy constant alpha 1 OS=Homo sapiens OX=9606 GN=IGHA1 PE=1 SV=2 | 255 | 20201118_lss_colostrum_1ng_glyco_hcd203040_compound15.20597.20597.3 | controllerType=0 controllerNumber=1 scan=20597 | 42.6983 |
| 17019:3 | 6 | R.LSLHRPALEDLLLGSEAN[+1622.582]LTC[+57.021]TLTGLR.D | HexNAc(4)Hex(5) | C[+57], N[+1623] | 1529.402 | 3 | 4586.1916 | 4586.1799 | 0 | 2.5 | 127 | Specific | 644.39 | 562.55 | 562.55 | 12.14 | 82 | >sp\|P01876\|IGHA1_HUMAN Immunoglobulin heavy constant alpha 1 OS=Homo sapiens OX=9606 GN=IGHA1 PE=1 SV=2 | 255 | 20201118_lss_colostrum_1ng_glyco_hcd203040_compound15.20551.20551.3 | controllerType=0 controllerNumber=1 scan=20551 | 42.5912 |
| 17016:3 | 6 | R.LSLHRPALEDLLLGSEAN[+1663.608]LTC[+57.021]TLTGLR.D | HexNAc(5)Hex(4) | C[+57], N[+1664] | 1543.0778 | 3 | 4627.2189 | 4627.2064 | 0 | 2.7 | 127 | Specific | 648.28 | 629.45 | 629.45 | 12.14 | 82 | >sp\|P01876\|IGHA1_HUMAN Immunoglobulin heavy constant alpha 1 OS=Homo sapiens OX=9606 GN=IGHA1 PE=1 SV=2 | 255 | 20201118_lss_colostrum_1ng_glyco_hcd203040_compound15.20548.20548.3 | controllerType=0 controllerNumber=1 scan=20548 | 42.5837 |
| 17013:3 | 6 | R.LSLHRPALEDLLLGSEAN[+1540.529]LTC[+57.021]TLTGLR.D | HexNAc(2)Hex(7) | C[+57], N[+1541] | 1502.0511 | 3 | 4504.1387 | 4504.1268 | 0 | 2.6 | 127 | Specific | 661.46 | 631.32 | 631.32 | 13.14 | 82 | >sp\|P01876\|IGHA1_HUMAN Immunoglobulin heavy constant alpha 1 OS=Homo sapiens OX=9606 GN=IGHA1 PE=1 SV=2 | 255 | 20201118_lss_colostrum_1ng_glyco_hcd203040_compound15.20544.20544.3 | controllerType=0 controllerNumber=1 scan=20544 | 42.567 |
| 16980:3 | 6 | R.LSLHRPALEDLLLGSEAN[+1864.634]LTC[+57.021]TLTGLR.D | HexNAc(2)Hex(9) | C[+57], N[+1865] | 1610.0888 | 3 | 4828.2519 | 4828.2324 | 0 | 4 | 127 | Specific | 664.11 | 620.6 | 620.6 | 12.94 | 82 | >sp\|P01876\|IGHA1_HUMAN Immunoglobulin heavy constant alpha 1 OS=Homo sapiens OX=9606 GN=IGHA1 PE=1 SV=2 | 255 | 20201118_lss_colostrum_1ng_glyco_hcd203040_compound15.20510.20510.3 | controllerType=0 controllerNumber=1 scan=20510 | 42.4749 |
| 17044:3 | 6 | R.LSLHRPALEDLLLGSEAN[+1460.529]LTC[+57.021]TLTGLR.D | HexNAc(4)Hex(4) | C[+57], N[+1461] | 1475.3837 | 3 | 4424.1366 | 4424.1271 | 0 | 2.2 | 127 | Specific | 668.34 | 633.43 | 633.43 | 13.14 | 82 | >sp\|P01876\|IGHA1_HUMAN Immunoglobulin heavy constant alpha 1 OS=Homo sapiens OX=9606 GN=IGHA1 PE=1 SV=2 | 255 | 20201118_lss_colostrum_1ng_glyco_hcd203040_compound15.20577.20577.3 | controllerType=0 controllerNumber=1 scan=20577 | 42.6453 |
| 17017:3 | 6 | R.LSLHRPALEDLLLGSEAN[+1622.582]LTC[+57.021]TLTGLR.D | HexNAc(4)Hex(5) | C[+57], N[+1623] | 1529.402 | 3 | 4586.1916 | 4586.1799 | 0 | 2.5 | 127 | Specific | 671.19 | 589.63 | 589.63 | 13.14 | 82 | >sp\|P01876\|IGHA1_HUMAN Immunoglobulin heavy constant alpha 1 OS=Homo sapiens OX=9606 GN=IGHA1 PE=1 SV=2 | 255 | 20201118_lss_colostrum_1ng_glyco_hcd203040_compound15.20549.20549.3 | controllerType=0 controllerNumber=1 scan=20549 | 42.585 |
| 17047:3 | 6 | R.LSLHRPALEDLLLGSEAN[+1460.529]LTC[+57.021]TLTGLR.D | HexNAc(4)Hex(4) | C[+57], N[+1461] | 1475.3837 | 3 | 4424.1366 | 4424.1271 | 0 | 2.2 | 127 | Specific | 674.22 | 585.28 | 585.28 | 13.14 | 82 | >sp\|P01876\|IGHA1_HUMAN Immunoglobulin heavy constant alpha 1 OS=Homo sapiens OX=9606 GN=IGHA1 PE=1 SV=2 | 255 | 20201118_lss_colostrum_1ng_glyco_hcd203040_compound15.20580.20580.3 | controllerType=0 controllerNumber=1 scan=20580 | 42.6566 |
| 17181:4 | 6 | R.LSLHRPALEDLLLGSEAN[+1460.529]LTC[+57.021]TLTGLR.D | HexNAc(4)Hex(4) | C[+57], N[+1461] | 1106.7901 | 4 | 4424.1385 | 4424.1271 | 0 | 2.6 | 127 | Specific | 677.71 | 616.18 | 616.18 | 13.76 | 82 | >sp\|P01876\|IGHA1_HUMAN Immunoglobulin heavy constant alpha 1 OS=Homo sapiens OX=9606 GN=IGHA1 PE=1 SV=2 | 255 | 20201118_lss_colostrum_1ng_glyco_hcd203040_compound15.20720.20720.4 | controllerType=0 controllerNumber=1 scan=20720 | 42.9739 |
| 17061:3 | 6 | R.LSLHRPALEDLLLGSEAN[+1501.555]LTC[+57.021]TLTGLR.D | HexNAc(5)Hex(3) | C[+57], N[+1502] | 1489.0604 | 3 | 4465.1665 | 4465.1536 | 0 | 2.9 | 127 | Specific | 689.19 | 689.19 | 689.19 | 13.14 | 82 | >sp\|P01876\|IGHA1_HUMAN Immunoglobulin heavy constant alpha 1 OS=Homo sapiens OX=9606 GN=IGHA1 PE=1 SV=2 | 255 | 20201118_lss_colostrum_1ng_glyco_hcd203040_compound15.20595.20595.3 | controllerType=0 controllerNumber=1 scan=20595 | 42.6927 |
| 17009:3 | 6 | R.LSLHRPALEDLLLGSEAN[+1702.581]LTC[+57.021]TLTGLR.D | HexNAc(2)Hex(8) | C[+57], N[+1703] | 1556.0693 | 3 | 4666.1933 | 4666.1796 | 0 | 2.9 | 127 | Specific | 689.58 | 624.15 | 624.15 | 13.14 | 82 | >sp\|P01876\|IGHA1_HUMAN Immunoglobulin heavy constant alpha 1 OS=Homo sapiens OX=9606 GN=IGHA1 PE=1 SV=2 | 255 | 20201118_lss_colostrum_1ng_glyco_hcd203040_compound15.20540.20540.3 | controllerType=0 controllerNumber=1 scan=20540 | 42.5598 |

Table S4. Cont.

| Query #:z | Protein Rank | Glycopeptide | Glycan | Modification Type(s) | Observed m/z | z | Observed (M+H) | Calc. mass (M+H) | Off-by-x error | Mass error (ppm) | StartPosition | Cleavage | Score | Delta | Delta Mod | \|Log Prob\| | # of unique peptides | Protein Name | Protein DB number | Comment | Scan # | Scan Time |
|---|---|---|---|---|---|---|---|---|---|---|---|---|---|---|---|---|---|---|---|---|---|---|
| 17024:3 | 6 | R.LSLHRPALEDLLLGSEAN[+1378.476]LTC[+57.021]TLTGLR.D | HexNAc(2)Hex(6) | C[+57], N[+1378] | 1448.0335 | 3 | 4342.086 | 4342.074 | 0 | 2.8 | 127 | Specific | 697.93 | 610.03 | 610.03 | 13.14 | 82 | >sp\|P01876\|IGHA1_HUMAN Immunoglobulin heavy constant alpha 1 OS=Homo sapiens OX=9606 GN=IGHA1 PE=1 SV=2 | 255 | 20201118_lss_colostrum_1ng_glyco_hcd203040_compound15.20556.20556.3 | controllerType=0 controllerNumber=1 scan=20556 | 42.6009 |
| 17000:4 | 6 | R.LSLHRPALEDLLLGSEAN[+1663.608]LTC[+57.021]TLTGLR.D | HexNAc(5)Hex(4) | C[+57], N[+1664] | 1157.5601 | 4 | 4627.2186 | 4627.2064 | 0 | 2.6 | 127 | Specific | 699.86 | 698.42 | 698.42 | 13.76 | 82 | >sp\|P01876\|IGHA1_HUMAN Immunoglobulin heavy constant alpha 1 OS=Homo sapiens OX=9606 GN=IGHA1 PE=1 SV=2 | 255 | 20201118_lss_colostrum_1ng_glyco_hcd203040_compound15.20531.20531.4 | controllerType=0 controllerNumber=1 scan=20531 | 42.54 |
| 17011:4 | 6 | R.LSLHRPALEDLLLGSEAN[+1622.582]LTC[+57.021]TLTGLR.D | HexNAc(4)Hex(5) | C[+57], N[+1623] | 1147.3034 | 4 | 4586.1917 | 4586.1799 | 0 | 2.6 | 127 | Specific | 706.87 | 706.87 | 706.87 | 14.76 | 82 | >sp\|P01876\|IGHA1_HUMAN Immunoglobulin heavy constant alpha 1 OS=Homo sapiens OX=9606 GN=IGHA1 PE=1 SV=2 | 255 | 20201118_lss_colostrum_1ng_glyco_hcd203040_compound15.20542.20542.4 | controllerType=0 controllerNumber=1 scan=20542 | 42.562 |
| 18032:4 | 6 | R.LSLHRPALEDLLLGSEAN[+1751.624]LTC[+57.021]TLTGLR.D | HexNAc(4)Hex(4)NeuAc(1) | C[+57], N[+1752] | 1179.5649 | 4 | 4715.2376 | 4715.2225 | 0 | 3.2 | 127 | Specific | 710.41 | 710.41 | 710.41 | 14.57 | 82 | >sp\|P01876\|IGHA1_HUMAN Immunoglobulin heavy constant alpha 1 OS=Homo sapiens OX=9606 GN=IGHA1 PE=1 SV=2 | 255 | 20201118_lss_colostrum_1ng_glyco_hcd203040_compound15.21608.21608.4 | controllerType=0 controllerNumber=1 scan=21608 | 44.8593 |
| 17021:3 | 6 | R.LSLHRPALEDLLLGSEAN[+1378.476]LTC[+57.021]TLTGLR.D | HexNAc(2)Hex(6) | C[+57], N[+1378] | 1448.0335 | 3 | 4342.086 | 4342.074 | 0 | 2.8 | 127 | Specific | 719.69 | 693.91 | 693.91 | 14.14 | 82 | >sp\|P01876\|IGHA1_HUMAN Immunoglobulin heavy constant alpha 1 OS=Homo sapiens OX=9606 GN=IGHA1 PE=1 SV=2 | 255 | 20201118_lss_colostrum_1ng_glyco_hcd203040_compound15.20553.20553.3 | controllerType=0 controllerNumber=1 scan=20553 | 42.5935 |
| 17060:4 | 6 | R.LSLHRPALEDLLLGSEAN[+1501.555]LTC[+57.021]TLTGLR.D | HexNAc(5)Hex(3) | C[+57], N[+1502] | 1117.0477 | 4 | 4465.1688 | 4465.1536 | 0 | 3.4 | 127 | Specific | 722.61 | 722.61 | 722.61 | 14.57 | 82 | >sp\|P01876\|IGHA1_HUMAN Immunoglobulin heavy constant alpha 1 OS=Homo sapiens OX=9606 GN=IGHA1 PE=1 SV=2 | 255 | 20201118_lss_colostrum_1ng_glyco_hcd203040_compound15.20594.20594.4 | controllerType=0 controllerNumber=1 scan=20594 | 42.6913 |
| 19131:4 | 6 | R.LSLHRPALEDLLLGSEAN[+2407.852]LTC[+57.021]TLTGLR.D | HexNAc(5)Hex(5)NeuAc(2) | C[+57], N[+2408] | 1343.6193 | 4 | 5371.4552 | 5371.4501 | 0 | 1 | 127 | Specific | 723.69 | 575.97 | 575.97 | 14.6 | 82 | >sp\|P01876\|IGHA1_HUMAN Immunoglobulin heavy constant alpha 1 OS=Homo sapiens OX=9606 GN=IGHA1 PE=1 SV=2 | 255 | 20201118_lss_colostrum_1ng_glyco_hcd203040_compound15.22752.22752.4 | controllerType=0 controllerNumber=1 scan=22752 | 47.3009 |
| 18086:4 | 6 | R.LSLHRPALEDLLLGSEAN[+1548.545]LTC[+57.021]TLTGLR.D | HexNAc(3)Hex(4)NeuAc(1) | C[+57], N[+1549] | 1128.7947 | 4 | 4512.1571 | 4512.1431 | 0 | 3.1 | 127 | Specific | 731.86 | 731.86 | 731.86 | 14.76 | 82 | >sp\|P01876\|IGHA1_HUMAN Immunoglobulin heavy constant alpha 1 OS=Homo sapiens OX=9606 GN=IGHA1 PE=1 SV=2 | 255 | 20201118_lss_colostrum_1ng_glyco_hcd203040_compound15.21665.21665.4 | controllerType=0 controllerNumber=1 scan=21665 | 44.9927 |
| 17184:4 | 6 | R.LSLHRPALEDLLLGSEAN[+1460.529]LTC[+57.021]TLTGLR.D | HexNAc(4)Hex(4) | C[+57], N[+1461] | 1106.7901 | 4 | 4424.1385 | 4424.1271 | 0 | 2.6 | 127 | Specific | 736.96 | 612.22 | 612.22 | 14.76 | 82 | >sp\|P01876\|IGHA1_HUMAN Immunoglobulin heavy constant alpha 1 OS=Homo sapiens OX=9606 GN=IGHA1 PE=1 SV=2 | 255 | 20201118_lss_colostrum_1ng_glyco_hcd203040_compound15.20723.20723.4 | controllerType=0 controllerNumber=1 scan=20723 | 42.9794 |
| 17141:4 | 6 | R.LSLHRPALEDLLLGSEAN[+1622.582]LTC[+57.021]TLTGLR.D | HexNAc(4)Hex(5) | C[+57], N[+1623] | 1147.3033 | 4 | 4586.1912 | 4586.1799 | 0 | 2.5 | 127 | Specific | 747.16 | 665.96 | 665.96 | 14.76 | 82 | >sp\|P01876\|IGHA1_HUMAN Immunoglobulin heavy constant alpha 1 OS=Homo sapiens OX=9606 GN=IGHA1 PE=1 SV=2 | 255 | 20201118_lss_colostrum_1ng_glyco_hcd203040_compound15.20678.20678.4 | controllerType=0 controllerNumber=1 scan=20678 | 42.8758 |
| 17153:4 | 6 | R.LSLHRPALEDLLLGSEAN[+1054.370]LTC[+57.021]TLTGLR.D | HexNAc(2)Hex(4) | C[+57], N[+1054] | 1005.2483 | 4 | 4017.9715 | 4017.9683 | 0 | 0.8 | 127 | Specific | 748.44 | 748.44 | 748.44 | 14.6 | 82 | >sp\|P01876\|IGHA1_HUMAN Immunoglobulin heavy constant alpha 1 OS=Homo sapiens OX=9606 GN=IGHA1 PE=1 SV=2 | 255 | 20201118_lss_colostrum_1ng_glyco_hcd203040_compound15.20691.20691.4 | controllerType=0 controllerNumber=1 scan=20691 | 42.9116 |
| 17010:3 | 6 | R.LSLHRPALEDLLLGSEAN[+1540.529]LTC[+57.021]TLTGLR.D | HexNAc(2)Hex(7) | C[+57], N[+1541] | 1502.0511 | 3 | 4504.1387 | 4504.1268 | 0 | 2.6 | 127 | Specific | 748.58 | 678.74 | 678.74 | 14.14 | 82 | >sp\|P01876\|IGHA1_HUMAN Immunoglobulin heavy constant alpha 1 OS=Homo sapiens OX=9606 GN=IGHA1 PE=1 SV=2 | 255 | 20201118_lss_colostrum_1ng_glyco_hcd203040_compound15.20541.20541.3 | controllerType=0 controllerNumber=1 scan=20541 | 42.5609 |
| 16988:4 | 6 | R.LSLHRPALEDLLLGSEAN[+1825.661]LTC[+57.021]TLTGLR.D | HexNAc(5)Hex(5) | C[+57], N[+1826] | 1198.0751 | 4 | 4789.2786 | 4789.2592 | 0 | 4 | 127 | Specific | 749.27 | 639.85 | 639.85 | 14.57 | 82 | >sp\|P01876\|IGHA1_HUMAN Immunoglobulin heavy constant alpha 1 OS=Homo sapiens OX=9606 GN=IGHA1 PE=1 SV=2 | 255 | 20201118_lss_colostrum_1ng_glyco_hcd203040_compound15.20518.20518.4 | controllerType=0 controllerNumber=1 scan=20518 | 42.498 |
| 16995:4 | 6 | R.LSLHRPALEDLLLGSEAN[+1702.581]LTC[+57.021]TLTGLR.D | HexNAc(2)Hex(8) | C[+57], N[+1703] | 1167.3025 | 4 | 4666.1883 | 4666.1796 | 0 | 1.9 | 127 | Specific | 758.59 | 758.59 | 758.59 | 15.76 | 82 | >sp\|P01876\|IGHA1_HUMAN Immunoglobulin heavy constant alpha 1 OS=Homo sapiens OX=9606 GN=IGHA1 PE=1 SV=2 | 255 | 20201118_lss_colostrum_1ng_glyco_hcd203040_compound15.20526.20526.4 | controllerType=0 controllerNumber=1 scan=20526 | 42.5236 |
| 17995:4 | 6 | R.LSLHRPALEDLLLGSEAN[+2116.756]LTC[+57.021]TLTGLR.D | HexNAc(5)Hex(5)NeuAc(1) | C[+57], N[+2117] | 1270.8479 | 4 | 5080.3698 | 5080.3547 | 0 | 3 | 127 | Specific | 759.05 | 689.55 | 689.55 | 15.76 | 82 | >sp\|P01876\|IGHA1_HUMAN Immunoglobulin heavy constant alpha 1 OS=Homo sapiens OX=9606 GN=IGHA1 PE=1 SV=2 | 255 | 20201118_lss_colostrum_1ng_glyco_hcd203040_compound15.21570.21570.4 | controllerType=0 controllerNumber=1 scan=21570 | 44.7882 |
| 17048:4 | 6 | R.LSLHRPALEDLLLGSEAN[+1419.502]LTC[+57.021]TLTGLR.D | HexNAc(3)Hex(5) | C[+57], N[+1420] | 1096.5339 | 4 | 4383.1136 | 4383.1005 | 0 | 3 | 127 | Specific | 759.4 | 735.64 | 735.64 | 15.76 | 82 | >sp\|P01876\|IGHA1_HUMAN Immunoglobulin heavy constant alpha 1 OS=Homo sapiens OX=9606 GN=IGHA1 PE=1 SV=2 | 255 | 20201118_lss_colostrum_1ng_glyco_hcd203040_compound15.20581.20581.4 | controllerType=0 controllerNumber=1 scan=20581 | 42.6577 |
| 18035:4 | 6 | R.LSLHRPALEDLLLGSEAN[+1751.624]LTC[+57.021]TLTGLR.D | HexNAc(4)Hex(4)NeuAc(1) | C[+57], N[+1752] | 1179.5649 | 4 | 4715.2376 | 4715.2225 | 0 | 3.2 | 127 | Specific | 761.47 | 761.47 | 761.47 | 15.57 | 82 | >sp\|P01876\|IGHA1_HUMAN Immunoglobulin heavy constant alpha 1 OS=Homo sapiens OX=9606 GN=IGHA1 PE=1 SV=2 | 255 | 20201118_lss_colostrum_1ng_glyco_hcd203040_compound15.21611.21611.4 | controllerType=0 controllerNumber=1 scan=21611 | 44.8655 |
| 17042:3 | 6 | R.LSLHRPALEDLLLGSEAN[+1216.423]LTC[+57.021]TLTGLR.D | HexNAc(2)Hex(5) | C[+57], N[+1216] | 1394.0149 | 3 | 4180.0303 | 4180.0211 | 0 | 2.2 | 127 | Specific | 769.48 | 722.52 | 722.52 | 15.14 | 82 | >sp\|P01876\|IGHA1_HUMAN Immunoglobulin heavy constant alpha 1 OS=Homo sapiens OX=9606 GN=IGHA1 PE=1 SV=2 | 255 | 20201118_lss_colostrum_1ng_glyco_hcd203040_compound15.20575.20575.3 | controllerType=0 controllerNumber=1 scan=20575 | 42.6431 |
| 17045:3 | 6 | R.LSLHRPALEDLLLGSEAN[+1216.423]LTC[+57.021]TLTGLR.D | HexNAc(2)Hex(5) | C[+57], N[+1216] | 1394.0149 | 3 | 4180.0303 | 4180.0211 | 0 | 2.2 | 127 | Specific | 777.08 | 773.82 | 773.82 | 15.14 | 82 | >sp\|P01876\|IGHA1_HUMAN Immunoglobulin heavy constant alpha 1 OS=Homo sapiens OX=9606 GN=IGHA1 PE=1 SV=2 | 255 | 20201118_lss_colostrum_1ng_glyco_hcd203040_compound15.20578.20578.3 | controllerType=0 controllerNumber=1 scan=20578 | 42.6508 |
| 17002:4 | 6 | R.LSLHRPALEDLLLGSEAN[+1663.608]LTC[+57.021]TLTGLR.D | HexNAc(5)Hex(4) | C[+57], N[+1664] | 1157.5601 | 4 | 4627.2186 | 4627.2064 | 0 | 2.6 | 127 | Specific | 793.51 | 750.29 | 750.29 | 15.76 | 82 | >sp\|P01876\|IGHA1_HUMAN Immunoglobulin heavy constant alpha 1 OS=Homo sapiens OX=9606 GN=IGHA1 PE=1 SV=2 | 255 | 20201118_lss_colostrum_1ng_glyco_hcd203040_compound15.20533.20533.4 | controllerType=0 controllerNumber=1 scan=20533 | 42.5487 |
| 18026:4 | 6 | R.LSLHRPALEDLLLGSEAN[+1710.598]LTC[+57.021]TLTGLR.D | HexNAc(3)Hex(5)NeuAc(1) | C[+57], N[+1711] | 1169.3083 | 4 | 4674.2113 | 4674.1959 | 0 | 3.3 | 127 | Specific | 795.33 | 757.47 | 757.47 | 15.57 | 82 | >sp\|P01876\|IGHA1_HUMAN Immunoglobulin heavy constant alpha 1 OS=Homo sapiens OX=9606 GN=IGHA1 PE=1 SV=2 | 255 | 20201118_lss_colostrum_1ng_glyco_hcd203040_compound15.21602.21602.4 | controllerType=0 controllerNumber=1 scan=21602 | 44.8467 |
| 17043:4 | 6 | R.LSLHRPALEDLLLGSEAN[+1460.529]LTC[+57.021]TLTGLR.D | HexNAc(4)Hex(4) | C[+57], N[+1461] | 1106.7928 | 4 | 4424.1492 | 4424.1271 | 0 | 5 | 127 | Specific | 795.87 | 795.87 | 795.87 | 15.11 | 82 | >sp\|P01876\|IGHA1_HUMAN Immunoglobulin heavy constant alpha 1 OS=Homo sapiens OX=9606 GN=IGHA1 PE=1 SV=2 | 255 | 20201118_lss_colostrum_1ng_glyco_hcd203040_compound15.20576.20576.4 | controllerType=0 controllerNumber=1 scan=20576 | 42.6442 |
| 17062:4 | 6 | R.LSLHRPALEDLLLGSEAN[+1501.555]LTC[+57.021]TLTGLR.D | HexNAc(5)Hex(3) | C[+57], N[+1502] | 1117.0477 | 4 | 4465.1688 | 4465.1536 | 0 | 3.4 | 127 | Specific | 799.3 | 799.3 | 799.3 | 15.57 | 82 | >sp\|P01876\|IGHA1_HUMAN Immunoglobulin heavy constant alpha 1 OS=Homo sapiens OX=9606 GN=IGHA1 PE=1 SV=2 | 255 | 20201118_lss_colostrum_1ng_glyco_hcd203040_compound15.20596.20596.4 | controllerType=0 controllerNumber=1 scan=20596 | 42.696 |
| 19130:4 | 6 | R.LSLHRPALEDLLLGSEAN[+2204.772]LTC[+57.021]TLTGLR.D | HexNAc(4)Hex(5)NeuAc(2) | C[+57], N[+2205] | 1292.8522 | 4 | 5168.3869 | 5168.3707 | 0 | 3.1 | 127 | Specific | 799.34 | 691.21 | 691.21 | 15.76 | 82 | >sp\|P01876\|IGHA1_HUMAN Immunoglobulin heavy constant alpha 1 OS=Homo sapiens OX=9606 GN=IGHA1 PE=1 SV=2 | 255 | 20201118_lss_colostrum_1ng_glyco_hcd203040_compound15.22751.22751.4 | controllerType=0 controllerNumber=1 scan=22751 | 47.2996 |
| 17136:4 | 6 | R.LSLHRPALEDLLLGSEAN[+1298.476]LTC[+57.021]TLTGLR.D | HexNAc(4)Hex(3) | C[+57], N[+1298] | 1066.2771 | 4 | 4262.0867 | 4262.0742 | 0 | 2.9 | 127 | Specific | 802.2 | 775.03 | 775.03 | 16.76 | 82 | >sp\|P01876\|IGHA1_HUMAN Immunoglobulin heavy constant alpha 1 OS=Homo sapiens OX=9606 GN=IGHA1 PE=1 SV=2 | 255 | 20201118_lss_colostrum_1ng_glyco_hcd203040_compound15.20673.20673.4 | controllerType=0 controllerNumber=1 scan=20673 | 42.857 |
| 17014:4 | 6 | R.LSLHRPALEDLLLGSEAN[+1622.582]LTC[+57.021]TLTGLR.D | HexNAc(4)Hex(5) | C[+57], N[+1623] | 1147.3034 | 4 | 4586.1917 | 4586.1799 | 0 | 2.6 | 127 | Specific | 832.41 | 723.46 | 723.46 | 16.76 | 82 | >sp\|P01876\|IGHA1_HUMAN Immunoglobulin heavy constant alpha 1 OS=Homo sapiens OX=9606 GN=IGHA1 PE=1 SV=2 | 255 | 20201118_lss_colostrum_1ng_glyco_hcd203040_compound15.20545.20545.4 | controllerType=0 controllerNumber=1 scan=20545 | 42.5721 |
| 18089:4 | 6 | R.LSLHRPALEDLLLGSEAN[+1548.545]LTC[+57.021]TLTGLR.D | HexNAc(3)Hex(4)NeuAc(1) | C[+57], N[+1549] | 1128.7947 | 4 | 4512.1571 | 4512.1431 | 0 | 3.1 | 127 | Specific | 847.64 | 847.64 | 847.64 | 16.76 | 82 | >sp\|P01876\|IGHA1_HUMAN Immunoglobulin heavy constant alpha 1 OS=Homo sapiens OX=9606 GN=IGHA1 PE=1 SV=2 | 255 | 20201118_lss_colostrum_1ng_glyco_hcd203040_compound15.21668.21668.4 | controllerType=0 controllerNumber=1 scan=21668 | 44.9995 |
| 18029:4 | 6 | R.LSLHRPALEDLLLGSEAN[+1710.598]LTC[+57.021]TLTGLR.D | HexNAc(3)Hex(5)NeuAc(1) | C[+57], N[+1711] | 1169.3083 | 4 | 4674.2113 | 4674.1959 | 0 | 3.3 | 127 | Specific | 855.88 | 821.5 | 821.5 | 17.57 | 82 | >sp\|P01876\|IGHA1_HUMAN Immunoglobulin heavy constant alpha 1 OS=Homo sapiens OX=9606 GN=IGHA1 PE=1 SV=2 | 255 | 20201118_lss_colostrum_1ng_glyco_hcd203040_compound15.21605.21605.4 | controllerType=0 controllerNumber=1 scan=21605 | 44.8526 |
| 17046:4 | 6 | R.LSLHRPALEDLLLGSEAN[+1460.529]LTC[+57.021]TLTGLR.D | HexNAc(4)Hex(4) | C[+57], N[+1461] | 1106.7928 | 4 | 4424.1492 | 4424.1271 | 0 | 5 | 127 | Specific | 857.55 | 815.98 | 815.98 | 17.11 | 82 | >sp\|P01876\|IGHA1_HUMAN Immunoglobulin heavy constant alpha 1 OS=Homo sapiens OX=9606 GN=IGHA1 PE=1 SV=2 | 255 | 20201118_lss_colostrum_1ng_glyco_hcd203040_compound15.20579.20579.4 | controllerType=0 controllerNumber=1 scan=20579 | 42.6559 |
| 17051:4 | 6 | R.LSLHRPALEDLLLGSEAN[+1419.502]LTC[+57.021]TLTGLR.D | HexNAc(3)Hex(5) | C[+57], N[+1420] | 1096.5339 | 4 | 4383.1136 | 4383.1005 | 0 | 3 | 127 | Specific | 860.81 | 833.98 | 833.98 | 17.76 | 82 | >sp\|P01876\|IGHA1_HUMAN Immunoglobulin heavy constant alpha 1 OS=Homo sapiens OX=9606 GN=IGHA1 PE=1 SV=2 | 255 | 20201118_lss_colostrum_1ng_glyco_hcd203040_compound15.20584.20584.4 | controllerType=0 controllerNumber=1 scan=20584 | 42.6627 |
| 17030:4 | 6 | R.LSLHRPALEDLLLGSEAN[+1864.634]LTC[+57.021]TLTGLR.D | HexNAc(2)Hex(9) | C[+57], N[+1865] | 1207.8122 | 4 | 4828.2269 | 4828.2324 | 0 | -1.1 | 127 | Specific | 881.28 | 655.26 | 655.26 | 16.69 | 82 | >sp\|P01876\|IGHA1_HUMAN Immunoglobulin heavy constant alpha 1 OS=Homo sapiens OX=9606 GN=IGHA1 PE=1 SV=2 | 255 | 20201118_lss_colostrum_1ng_glyco_hcd203040_compound15.20562.20562.4 | controllerType=0 controllerNumber=1 scan=20562 | 42.6131 |
| 16955:4 | 6 | R.LSLHRPALEDLLLGSEAN[+1702.581]LTC[+57.021]TLTGLR.D | HexNAc(2)Hex(8) | C[+57], N[+1703] | 1167.302 | 4 | 4666.1863 | 4666.1796 | 0 | 1.4 | 127 | Specific | 892.33 | 892.33 | 892.33 | 17.6 | 82 | >sp\|P01876\|IGHA1_HUMAN Immunoglobulin heavy constant alpha 1 OS=Homo sapiens OX=9606 GN=IGHA1 PE=1 SV=2 | 255 | 20201118_lss_colostrum_1ng_glyco_hcd203040_compound15.20484.20484.4 | controllerType=0 controllerNumber=1 scan=20484 | 42.4108 |
| 16998:4 | 6 | R.LSLHRPALEDLLLGSEAN[+1540.529]LTC[+57.021]TLTGLR.D | HexNAc(2)Hex(7) | C[+57], N[+1541] | 1126.7897 | 4 | 4504.137 | 4504.1268 | 0 | 2.3 | 127 | Specific | 900.68 | 819.78 | 819.78 | 18.76 | 82 | >sp\|P01876\|IGHA1_HUMAN Immunoglobulin heavy constant alpha 1 OS=Homo sapiens OX=9606 GN=IGHA1 PE=1 SV=2 | 255 | 20201118_lss_colostrum_1ng_glyco_hcd203040_compound15.20529.20529.4 | controllerType=0 controllerNumber=1 scan=20529 | 42.5377 |
| 17039:4 | 6 | R.LSLHRPALEDLLLGSEAN[+1378.476]LTC[+57.021]TLTGLR.D | HexNAc(2)Hex(6) | C[+57], N[+1378] | 1086.2774 | 4 | 4342.0877 | 4342.074 | 0 | 3.2 | 127 | Specific | 908.24 | 866.56 | 866.56 | 18.57 | 82 | >sp\|P01876\|IGHA1_HUMAN Immunoglobulin heavy constant alpha 1 OS=Homo sapiens OX=9606 GN=IGHA1 PE=1 SV=2 | 255 | 20201118_lss_colostrum_1ng_glyco_hcd203040_compound15.20572.20572.4 | controllerType=0 controllerNumber=1 scan=20572 | 42.6338 |
| 17156:4 | 6 | R.LSLHRPALEDLLLGSEAN[+1054.370]LTC[+57.021]TLTGLR.D | HexNAc(2)Hex(4) | C[+57], N[+1054] | 1005.2483 | 4 | 4017.9715 | 4017.9683 | 0 | 0.8 | 127 | Specific | 930.61 | 912.54 | 912.54 | 18.6 | 82 | >sp\|P01876\|IGHA1_HUMAN Immunoglobulin heavy constant alpha 1 OS=Homo sapiens OX=9606 GN=IGHA1 PE=1 SV=2 | 255 | 20201118_lss_colostrum_1ng_glyco_hcd203040_compound15.20694.20694.4 | controllerType=0 controllerNumber=1 scan=20694 | 42.9163 |
| 17038:4 | 6 | R.LSLHRPALEDLLLGSEAN[+1216.423]LTC[+57.021]TLTGLR.D | HexNAc(2)Hex(5) | C[+57], N[+1216] | 1045.7641 | 4 | 4180.0345 | 4180.0211 | 0 | 3.2 | 127 | Specific | 943.76 | 867.39 | 867.39 | 18.57 | 82 | >sp\|P01876\|IGHA1_HUMAN Immunoglobulin heavy constant alpha 1 OS=Homo sapiens OX=9606 GN=IGHA1 PE=1 SV=2 | 255 | 20201118_lss_colostrum_1ng_glyco_hcd203040_compound15.20571.20571.4 | controllerType=0 controllerNumber=1 scan=20571 | 42.6325 |
| 17041:4 | 6 | R.LSLHRPALEDLLLGSEAN[+1378.476]LTC[+57.021]TLTGLR.D | HexNAc(2)Hex(6) | C[+57], N[+1378] | 1086.2774 | 4 | 4342.0877 | 4342.074 | 0 | 3.2 | 127 | Specific | 976.44 | 902.88 | 902.88 | 19.57 | 82 | >sp\|P01876\|IGHA1_HUMAN Immunoglobulin heavy constant alpha 1 OS=Homo sapiens OX=9606 GN=IGHA1 PE=1 SV=2 | 255 | 20201118_lss_colostrum_1ng_glyco_hcd203040_compound15.20574.20574.4 | controllerType=0 controllerNumber=1 scan=20574 | 42.6419 |
| 17040:4 | 6 | R.LSLHRPALEDLLLGSEAN[+1216.423]LTC[+57.021]TLTGLR.D | HexNAc(2)Hex(5) | C[+57], N[+1216] | 1045.7641 | 4 | 4180.0345 | 4180.0211 | 0 | 3.2 | 127 | Specific | 994.09 | 924.3 | 924.3 | 19.57 | 82 | >sp\|P01876\|IGHA1_HUMAN Immunoglobulin heavy constant alpha 1 OS=Homo sapiens OX=9606 GN=IGHA1 PE=1 SV=2 | 255 | 20201118_lss_colostrum_1ng_glyco_hcd203040_compound15.20573.20573.4 | controllerType=0 controllerNumber=1 scan=20573 | 42.6399 |
| 17001:4 | 6 | R.LSLHRPALEDLLLGSEAN[+1540.529]LTC[+57.021]TLTGLR.D | HexNAc(2)Hex(7) | C[+57], N[+1541] | 1126.7897 | 4 | 4504.137 | 4504.1268 | 0 | 2.3 | 127 | Specific | 1006.35 | 948.71 | 948.71 | 20.76 | 82 | >sp\|P01876\|IGHA1_HUMAN Immunoglobulin heavy constant alpha 1 OS=Homo sapiens OX=9606 GN=IGHA1 PE=1 SV=2 | 255 | 20201118_lss_colostrum_1ng_glyco_hcd203040_compound15.20532.20532.4 | controllerType=0 controllerNumber=1 scan=20532 | 42.5442 |
| 16997:4 | 6 | R.LSLHRPALEDLLLGSEAN[+1702.581]LTC[+57.021]TLTGLR.D | HexNAc(2)Hex(8) | C[+57], N[+1703] | 1167.3025 | 4 | 4666.1883 | 4666.1796 | 0 | 1.9 | 127 | Specific | 1041.91 | 982.57 | 982.57 | 20.76 | 82 | >sp\|P01876\|IGHA1_HUMAN Immunoglobulin heavy constant alpha 1 OS=Homo sapiens OX=9606 GN=IGHA1 PE=1 SV=2 | 255 | 20201118_lss_colostrum_1ng_glyco_hcd203040_compound15.20528.20528.4 | controllerType=0 controllerNumber=1 scan=20528 | 42.5366 |
| 16957:4 | 6 | R.LSLHRPALEDLLLGSEAN[+1702.581]LTC[+57.021]TLTGLR.D | HexNAc(2)Hex(8) | C[+57], N[+1703] | 1167.302 | 4 | 4666.1863 | 4666.1796 | 0 | 1.4 | 127 | Specific | 1062.57 | 1044.94 | 1044.94 | 21.6 | 82 | >sp\|P01876\|IGHA1_HUMAN Immunoglobulin heavy constant alpha 1 OS=Homo sapiens OX=9606 GN=IGHA1 PE=1 SV=2 | 255 | 20201118_lss_colostrum_1ng_glyco_hcd203040_compound15.20486.20486.4 | controllerType=0 controllerNumber=1 scan=20486 | 42.4177 |
| 17074:4 | 6 | R.LSLHRPALEDLLLGSEAN[+1257.449]LTC[+57.021]TLTGLR.D | HexNAc(3)Hex(4) | C[+57], N[+1257] | 1056.0184 | 4 | 4221.0516 | 4221.0477 | 0 | 0.9 | 127 | Specific | 1138.1 | 1080.17 | 1080.17 | 22.6 | 82 | >sp\|P01876\|IGHA1_HUMAN Immunoglobulin heavy constant alpha 1 OS=Homo sapiens OX=9606 GN=IGHA1 PE=1 SV=2 | 255 | 20201118_lss_colostrum_1ng_glyco_hcd203040_compound15.20608.20608.4 | controllerType=0 controllerNumber=1 scan=20608 | 42.7124 |
| 12820:3 | 6 | R.LAGKPTHVN[+2117.777]VSVVMAEVDGTC[+57.021].Y | HexNAc(5)Hex(5)Fuc(2) | C[+57], N[+2118] | 1434.626 | 3 | 4301.8635 | 4301.8555 | 0 | 1.9 | 332 | CRagged | 210.06 | 195.91 | 195.91 | 3.77 | 82 | >sp\|P01876\|IGHA1_HUMAN Immunoglobulin heavy constant alpha 1 OS=Homo sapiens OX=9606 GN=IGHA1 PE=1 SV=2 | 255 | 20201118_lss_colostrum_1ng_glyco_hcd203040_compound15.16183.16183.3 | controllerType=0 controllerNumber=1 scan=16183 | 33.9402 |
| 12693:3 | 6 | R.LAGKPTHVN[+2263.835]VSVVMAEVDGTC[+57.021].Y | HexNAc(5)Hex(5)Fuc(3) | C[+57], N[+2264] | 1483.3097 | 3 | 4447.9147 | 4447.9135 | 0 | 0.3 | 332 | CRagged | 235.02 | 210.12 | 210.12 | 4.5 | 82 | >sp\|P01876\|IGHA1_HUMAN Immunoglobulin heavy constant alpha 1 OS=Homo sapiens OX=9606 GN=IGHA1 PE=1 SV=2 | 255 | 20201118_lss_colostrum_1ng_glyco_hcd203040_compound15.16052.16052.3 | controllerType=0 controllerNumber=1 scan=16052 | 33.7047 |
| 12690:3 | 6 | R.LAGKPTHVN[+2263.835]VSVVMAEVDGTC[+57.021].Y | HexNAc(5)Hex(5)Fuc(3) | C[+57], N[+2264] | 1483.3097 | 3 | 4447.9147 | 4447.9135 | 0 | 0.3 | 332 | CRagged | 298.28 | 281.81 | 281.81 | 6.45 | 82 | >sp\|P01876\|IGHA1_HUMAN Immunoglobulin heavy constant alpha 1 OS=Homo sapiens OX=9606 GN=IGHA1 PE=1 SV=2 | 255 | 20201118_lss_colostrum_1ng_glyco_hcd203040_compound15.16049.16049.3 | controllerType=0 controllerNumber=1 scan=16049 | 33.6995 |
| 11018:3 | 6 | R.LAGKPTHVN[+2133.772]VSVVMAEVDGTC[+57.021].Y | HexNAc(5)Hex(6)Fuc(1) | C[+57], N[+2134] | 1439.9606 | 3 | 4317.8673 | 4317.8505 | 0 | 3.9 | 332 | CRagged | 323.76 | 205.73 | 84.43 | 6.62 | 82 | >sp\|P01876\|IGHA1_HUMAN Immunoglobulin heavy constant alpha 1 OS=Homo sapiens OX=9606 GN=IGHA1 PE=1 SV=2 | 255 | 20201118_lss_colostrum_1ng_glyco_hcd203040_compound15.14313.14313.3 | controllerType=0 controllerNumber=1 scan=14313 | 30.5221 |
| 11130:4 | 6 | R.LAGKPTHVN[+1809.666]VSVVMAEVDGTC[+57.021]Y.- | HexNAc(5)Hex(4)Fuc(1) | C[+57], N[+1810] | 1039.9589 | 4 | 4156.8138 | 4156.8081 | 0 | 1.4 | 332 | CRagged | 249.43 | 155.72 | 155.72 | 3.54 | 82 | >sp\|P01876\|IGHA1_HUMAN Immunoglobulin heavy constant alpha 1 OS=Homo sapiens OX=9606 GN=IGHA1 PE=1 SV=2 | 255 | 20201118_lss_colostrum_1ng_glyco_hcd203040_compound15.14430.14430.4 | controllerType=0 controllerNumber=1 scan=14430 | 30.7698 |
| 11224:3 | 6 | R.LAGKPTHVN[+1095.397]VSVVMAEVDGTC[+57.021]Y.- | HexNAc(3)Hex(3) | C[+57], N[+1095] | 1148.1887 | 3 | 3442.5517 | 3442.5387 | 0 | 3.8 | 332 | CRagged | 329.63 | 227.63 | 227.63 | 6.96 | 82 | >sp\|P01876\|IGHA1_HUMAN Immunoglobulin heavy constant alpha 1 OS=Homo sapiens OX=9606 GN=IGHA1 PE=1 SV=2 | 255 | 20201118_lss_colostrum_1ng_glyco_hcd203040_compound15.14528.14528.3 | controllerType=0 controllerNumber=1 scan=14528 | 30.9603 |
| 11164:4 | 6 | R.LAGKPTHVN[+1647.613]VSVVMAEVDGTC[+57.021]Y.- | HexNAc(5)Hex(3)Fuc(1) | C[+57], N[+1648] | 999.4456 | 4 | 3994.7606 | 3994.7553 | 0 | 1.3 | 332 | CRagged | 362.39 | 362.39 | 362.39 | 7.08 | 82 | >sp\|P01876\|IGHA1_HUMAN Immunoglobulin heavy constant alpha 1 OS=Homo sapiens OX=9606 GN=IGHA1 PE=1 SV=2 | 255 | 20201118_lss_colostrum_1ng_glyco_hcd203040_compound15.14465.14465.4 | controllerType=0 controllerNumber=1 scan=14465 | 30.841 |
| 11162:4 | 6 | R.LAGKPTHVN[+1647.613]VSVVMAEVDGTC[+57.021]Y.- | HexNAc(5)Hex(3)Fuc(1) | C[+57], N[+1648] | 999.4456 | 4 | 3994.7606 | 3994.7553 | 0 | 1.3 | 332 | CRagged | 426.3 | 426.3 | 426.3 | 9.76 | 82 | >sp\|P01876\|IGHA1_HUMAN Immunoglobulin heavy constant alpha 1 OS=Homo sapiens OX=9606 GN=IGHA1 PE=1 SV=2 | 255 | 20201118_lss_colostrum_1ng_glyco_hcd203040_compound15.14463.14463.4 | controllerType=0 controllerNumber=1 scan=14463 | 30.8366 |
| 12779:4 | 6 | R.LAGKPTHVN[+2262.814]VSVVMAEVDGTC[+57.021]Y.- | HexNAc(5)Hex(5)Fuc(1)NeuAc(1) | C[+57], N[+2263] | 1153.247 | 4 | 4609.9661 | 4609.9564 | 0 | 2.1 | 332 | CRagged | 428.96 | 428.96 | 428.96 | 9.92 | 82 | >sp\|P01876\|IGHA1_HUMAN Immunoglobulin heavy constant alpha 1 OS=Homo sapiens OX=9606 GN=IGHA1 PE=1 SV=2 | 255 | 20201118_lss_colostrum_1ng_glyco_hcd203040_compound15.16141.16141.4 | controllerType=0 controllerNumber=1 scan=16141 | 33.8587 |
| 11128:2 | 6 | R.LAGKPTHVN[+1216.423]VSVVMAEVDGTC[+57.021]Y.- | HexNAc(2)Hex(5) | C[+57], N[+1216] | 1782.2917 | 2 | 3563.5761 | 3563.5649 | 0 | 3.1 | 332 | CRagged | 431.52 | 431.52 | 214.25 | 9.36 | 82 | >sp\|P01876\|IGHA1_HUMAN Immunoglobulin heavy constant alpha 1 OS=Homo sapiens OX=9606 GN=IGHA1 PE=1 SV=2 | 255 | 20201118_lss_colostrum_1ng_glyco_hcd203040_compound15.14428.14428.2 | controllerType=0 controllerNumber=1 scan=14428 | 30.7668 |
| 09942:3 | 6 | R.LAGKPTHVN[+1809.666]VSVVMAEVDGTC[+57.021]Y.- | HexNAc(5)Hex(4)Fuc(1) | C[+57], N[+1810] | 1386.2752 | 3 | 4156.8111 | 4156.8081 | 0 | 0.7 | 332 | CRagged | 444.6 | 408.72 | 324.2 | 9.93 | 82 | >sp\|P01876\|IGHA1_HUMAN Immunoglobulin heavy constant alpha 1 OS=Homo sapiens OX=9606 GN=IGHA1 PE=1 SV=2 | 255 | 20201118_lss_colostrum_1ng_glyco_hcd203040_compound15.13197.13197.3 | controllerType=0 controllerNumber=1 scan=13197 | 28.4788 |
| 12777:3 | 6 | R.LAGKPTHVN[+2262.814]VSVVMAEVDGTC[+57.021]Y.- | HexNAc(5)Hex(5)Fuc(1)NeuAc(1) | C[+57], N[+2263] | 1537.3273 | 3 | 4609.9674 | 4609.9564 | 0 | 2.4 | 332 | CRagged | 448.2 | 448.2 | 448.2 | 10.09 | 82 | >sp\|P01876\|IGHA1_HUMAN Immunoglobulin heavy constant alpha 1 OS=Homo sapiens OX=9606 GN=IGHA1 PE=1 SV=2 | 255 | 20201118_lss_colostrum_1ng_glyco_hcd203040_compound15.16139.16139.3 | controllerType=0 controllerNumber=1 scan=16139 | 33.8568 |
| 11232:3 | 6 | R.LAGKPTHVN[+1444.534]VSVVMAEVDGTC[+57.021]Y.- | HexNAc(4)Hex(3)Fuc(1) | C[+57], N[+1445] | 1264.5671 | 3 | 3791.6867 | 3791.6759 | 0 | 2.8 | 332 | CRagged | 480.3 | 480.3 | 480.3 | 10.17 | 82 | >sp\|P01876\|IGHA1_HUMAN Immunoglobulin heavy constant alpha 1 OS=Homo sapiens OX=9606 GN=IGHA1 PE=1 SV=2 | 255 | 20201118_lss_colostrum_1ng_glyco_hcd203040_compound15.14536.14536.3 | controllerType=0 controllerNumber=1 scan=14536 | 30.977 |
| 11305:3 | 6 | R.LAGKPTHVN[+1647.613]VSVVMAEVDGTC[+57.021]Y.- | HexNAc(5)Hex(3)Fuc(1) | C[+57], N[+1648] | 1332.2582 | 3 | 3994.7602 | 3994.7553 | 0 | 1.2 | 332 | CRagged | 481.93 | 478 | 478 | 10.01 | 82 | >sp\|P01876\|IGHA1_HUMAN Immunoglobulin heavy constant alpha 1 OS=Homo sapiens OX=9606 GN=IGHA1 PE=1 SV=2 | 255 | 20201118_lss_colostrum_1ng_glyco_hcd203040_compound15.14612.14612.3 | controllerType=0 controllerNumber=1 scan=14612 | 31.1339 |
| 12869:3 | 6 | R.LAGKPTHVN[+1710.598]VSVVMAEVDGTC[+57.021]Y.- | HexNAc(3)Hex(5)NeuAc(1) | C[+57], N[+1711] | 1353.2516 | 3 | 4057.7404 | 4057.7397 | 0 | 0.2 | 332 | CRagged | 494.39 | 446.02 | 310.09 | 9.6 | 82 | >sp\|P01876\|IGHA1_HUMAN Immunoglobulin heavy constant alpha 1 OS=Homo sapiens OX=9606 GN=IGHA1 PE=1 SV=2 | 255 | 20201118_lss_colostrum_1ng_glyco_hcd203040_compound15.16234.16234.3 | controllerType=0 controllerNumber=1 scan=16234 | 34.0556 |
| 09940:3 | 6 | R.LAGKPTHVN[+1809.666]VSVVMAEVDGTC[+57.021]Y.- | HexNAc(5)Hex(4)Fuc(1) | C[+57], N[+1810] | 1386.2752 | 3 | 4156.8111 | 4156.8081 | 0 | 0.7 | 332 | CRagged | 515.56 | 515.56 | 369.01 | 10.23 | 82 | >sp\|P01876\|IGHA1_HUMAN Immunoglobulin heavy constant alpha 1 OS=Homo sapiens OX=9606 GN=IGHA1 PE=1 SV=2 | 255 | 20201118_lss_colostrum_1ng_glyco_hcd203040_compound15.13195.13195.3 | controllerType=0 controllerNumber=1 scan=13195 | 28.4723 |
| 12793:3 | 6 | R.LAGKPTHVN[+2059.735]VSVVMAEVDGTC[+57.021]Y.- | HexNAc(4)Hex(5)Fuc(1)NeuAc(1) | C[+57], N[+2060] | 1469.6337 | 3 | 4406.8866 | 4406.877 | 0 | 2.2 | 332 | CRagged | 535.05 | 323.72 | 323.72 | 8.47 | 82 | >sp\|P01876\|IGHA1_HUMAN Immunoglobulin heavy constant alpha 1 OS=Homo sapiens OX=9606 GN=IGHA1 PE=1 SV=2 | 255 | 20201118_lss_colostrum_1ng_glyco_hcd203040_compound15.16155.16155.3 | controllerType=0 controllerNumber=1 scan=16155 | 33.8844 |
| 11307:3 | 6 | R.LAGKPTHVN[+1647.613]VSVVMAEVDGTC[+57.021]Y.- | HexNAc(5)Hex(3)Fuc(1) | C[+57], N[+1648] | 1332.2582 | 3 | 3994.7602 | 3994.7553 | 0 | 1.2 | 332 | CRagged | 549.36 | 431.97 | 431.97 | 10.24 | 82 | >sp\|P01876\|IGHA1_HUMAN Immunoglobulin heavy constant alpha 1 OS=Homo sapiens OX=9606 GN=IGHA1 PE=1 SV=2 | 255 | 20201118_lss_colostrum_1ng_glyco_hcd203040_compound15.14614.14614.3 | controllerType=0 controllerNumber=1 scan=14614 | 31.1376 |
| 12781:4 | 6 | R.LAGKPTHVN[+2262.814]VSVVMAEVDGTC[+57.021]Y.- | HexNAc(5)Hex(5)Fuc(1)NeuAc(1) | C[+57], N[+2263] | 1153.247 | 4 | 4609.9661 | 4609.9564 | 0 | 2.1 | 332 | CRagged | 555.44 | 454.08 | 454.08 | 10.33 | 82 | >sp\|P01876\|IGHA1_HUMAN Immunoglobulin heavy constant alpha 1 OS=Homo sapiens OX=9606 GN=IGHA1 PE=1 SV=2 | 255 | 20201118_lss_colostrum_1ng_glyco_hcd203040_compound15.16143.16143.4 | controllerType=0 controllerNumber=1 scan=16143 | 33.8625 |
| 14598:3 | 6 | R.LAGKPTHVN[+2205.793]VSVVMAEVDGTC[+57.021]Y.- | HexNAc(4)Hex(5)Fuc(2)NeuAc(1) | C[+57], N[+2206] | 1518.3129 | 3 | 4552.9242 | 4552.9349 | 0 | -2.4 | 332 | CRagged | 560.58 | 519.21 | 519.21 | 9.3 | 82 | >sp\|P01876\|IGHA1_HUMAN Immunoglobulin heavy constant alpha 1 OS=Homo sapiens OX=9606 GN=IGHA1 PE=1 SV=2 | 255 | 20201118_lss_colostrum_1ng_glyco_hcd203040_compound15.18030.18030.3 | controllerType=0 controllerNumber=1 scan=18030 | 37.5705 |
| 14495:3 | 6 | R.LAGKPTHVN[+2350.830]VSVVMAEVDGTC[+57.021]Y.- | HexNAc(4)Hex(5)Fuc(1)NeuAc(2) | C[+57], N[+2351] | 1566.6723 | 3 | 4698.0023 | 4697.9724 | 0 | 6.4 | 332 | CRagged | 561.98 | 479.66 | 479.66 | 9.61 | 82 | >sp\|P01876\|IGHA1_HUMAN Immunoglobulin heavy constant alpha 1 OS=Homo sapiens OX=9606 GN=IGHA1 PE=1 SV=2 | 255 | 20201118_lss_colostrum_1ng_glyco_hcd203040_compound15.17923.17923.3 | controllerType=0 controllerNumber=1 scan=17923 | 37.3428 |
| 11000:3 | 6 | R.LAGKPTHVN[+1825.661]VSVVMAEVDGTC[+57.021]Y.- | HexNAc(5)Hex(5) | C[+57], N[+1826] | 1391.6071 | 3 | 4172.8068 | 4172.8031 | 0 | 0.9 | 332 | CRagged | 564.53 | 482.79 | 412.15 | 10.5 | 82 | >sp\|P01876\|IGHA1_HUMAN Immunoglobulin heavy constant alpha 1 OS=Homo sapiens OX=9606 GN=IGHA1 PE=1 SV=2 | 255 | 20201118_lss_colostrum_1ng_glyco_hcd203040_compound15.14295.14295.3 | controllerType=0 controllerNumber=1 scan=14295 | 30.4912 |
| 12866:3 | 6 | R.LAGKPTHVN[+1710.598]VSVVMAEVDGTC[+57.021]Y.- | HexNAc(3)Hex(5)NeuAc(1) | C[+57], N[+1711] | 1353.2516 | 3 | 4057.7404 | 4057.7397 | 0 | 0.2 | 332 | CRagged | 565.01 | 565.01 | 358.85 | 10.03 | 82 | >sp\|P01876\|IGHA1_HUMAN Immunoglobulin heavy constant alpha 1 OS=Homo sapiens OX=9606 GN=IGHA1 PE=1 SV=2 | 255 | 20201118_lss_colostrum_1ng_glyco_hcd203040_compound15.16231.16231.3 | controllerType=0 controllerNumber=1 scan=16231 | [illegible] |
| 14493:3 | 6 | R.LAGKPTHVN[+2350.830]VSVVMAEVDGTC[+57.021]Y.- | HexNAc(4)Hex(5)Fuc(1)NeuAc(2) | C[+57], N[+2351] | 1566.6723 | 3 | 4698.0023 | 4697.9724 | 0 | 6.4 | 332 | CRagged | 577.61 | 514.97 | 514.97 | 9.61 | 82 | >sp\|P01876\|IGHA1_HUMAN Immunoglobulin heavy constant alpha 1 OS=Homo sapiens OX=9606 GN=IGHA1 PE=1 SV=2 | 255 | 20201118_lss_colostrum_1ng_glyco_hcd203040_compound15.17921.17921.3 | controllerType=0 controllerNumber=1 scan=17921 | [illegible] |
| 14596:3 | 6 | R.LAGKPTHVN[+2205.793]VSVVMAEVDGTC[+57.021]Y.- | HexNAc(4)Hex(5)Fuc(2)NeuAc(1) | C[+57], N[+2206] | 1518.3129 | 3 | 4552.9242 | 4552.9349 | 0 | -2.4 | 332 | CRagged | 592.3 | 475.51 | 475.51 | 10.64 | 82 | >sp\|P01876\|IGHA1_HUMAN Immunoglobulin heavy constant alpha 1 OS=Homo sapiens OX=9606 GN=IGHA1 PE=1 SV=2 | 255 | 20201118_lss_colostrum_1ng_glyco_hcd203040_compound15.18028.18028.3 | controllerType=0 controllerNumber=1 scan=18028 | [illegible] |
| 12790:3 | 6 | R.LAGKPTHVN[+2059.735]VSVVMAEVDGTC[+57.021]Y.- | HexNAc(4)Hex(5)Fuc(1)NeuAc(1) | C[+57], N[+2060] | 1469.6337 | 3 | 4406.8866 | 4406.877 | 0 | 2.2 | 332 | CRagged | 595.88 | 580.68 | 580.68 | 11.99 | 82 | >sp\|P01876\|IGHA1_HUMAN Immunoglobulin heavy constant alpha 1 OS=Homo sapiens OX=9606 GN=IGHA1 PE=1 SV=2 | 255 | 20201118_lss_colostrum_1ng_glyco_hcd203040_compound15.16152.16152.3 | controllerType=0 controllerNumber=1 scan=16152 | 33.8795 |
| 12691:3 | 6 | R.LAGKPTHVN[+2116.756]VSVVMAEVDGTC[+57.021]Y.- | HexNAc(5)Hex(5)NeuAc(1) | C[+57], N[+2117] | 1488.6424 | 3 | 4463.9126 | 4463.8985 | 0 | 3.2 | 332 | CRagged | 596.64 | 480.18 | 480.18 | 11.99 | 82 | >sp\|P01876\|IGHA1_HUMAN Immunoglobulin heavy constant alpha 1 OS=Homo sapiens OX=9606 GN=IGHA1 PE=1 SV=2 | 255 | 20201118_lss_colostrum_1ng_glyco_hcd203040_compound15.16050.16050.3 | controllerType=0 controllerNumber=1 scan=16050 | 33.7006 |
| 12771:3 | 6 | R.LAGKPTHVN[+2059.735]VSVVMAEVDGTC[+57.021]Y.- | HexNAc(4)Hex(5)Fuc(1)NeuAc(1) | C[+57], N[+2060] | 1469.6345 | 3 | 4406.8888 | 4406.877 | 0 | 2.7 | 332 | CRagged | 605.23 | 501.07 | 501.07 | 12.99 | 82 | >sp\|P01876\|IGHA1_HUMAN Immunoglobulin heavy constant alpha 1 OS=Homo sapiens OX=9606 GN=IGHA1 PE=1 SV=2 | 255 | 20201118_lss_colostrum_1ng_glyco_hcd203040_compound15.16132.16132.3 | controllerType=0 controllerNumber=1 scan=16132 | 33.8423 |
| 12763:3 | 6 | R.LAGKPTHVN[+1872.651]VSVVMAEVDGTC[+57.021]Y.- | HexNAc(3)Hex(6)NeuAc(1) | C[+57], N[+1873] | 1407.2731 | 3 | 4219.8048 | 4219.7926 | 0 | 2.9 | 332 | CRagged | 614.65 | 614.65 | 418.03 | 12.99 | 82 | >sp\|P01876\|IGHA1_HUMAN Immunoglobulin heavy constant alpha 1 OS=Homo sapiens OX=9606 GN=IGHA1 PE=1 SV=2 | 255 | 20201118_lss_colostrum_1ng_glyco_hcd203040_compound15.16124.16124.3 | controllerType=0 controllerNumber=1 scan=16124 | 33.8286 |
| 11003:3 | 6 | R.LAGKPTHVN[+1825.661]VSVVMAEVDGTC[+57.021]Y.- | HexNAc(5)Hex(5) | C[+57], N[+1826] | 1391.6071 | 3 | 4172.8068 | 4172.8031 | 0 | 0.9 | 332 | CRagged | 621.7 | 544.16 | 433.66 | 12.84 | 82 | >sp\|P01876\|IGHA1_HUMAN Immunoglobulin heavy constant alpha 1 OS=Homo sapiens OX=9606 GN=IGHA1 PE=1 SV=2 | 255 | 20201118_lss_colostrum_1ng_glyco_hcd203040_compound15.14298.14298.3 | controllerType=0 controllerNumber=1 scan=14298 | 30.4958 |
| 11234:3 | 6 | R.LAGKPTHVN[+1444.534]VSVVMAEVDGTC[+57.021]Y.- | HexNAc(4)Hex(3)Fuc(1) | C[+57], N[+1445] | 1264.5671 | 3 | 3791.6867 | 3791.6759 | 0 | 2.8 | 332 | CRagged | 632.24 | 484.69 | 484.69 | 12.99 | 82 | >sp\|P01876\|IGHA1_HUMAN Immunoglobulin heavy constant alpha 1 OS=Homo sapiens OX=9606 GN=IGHA1 PE=1 SV=2 | 255 | 20201118_lss_colostrum_1ng_glyco_hcd203040_compound15.14538.14538.3 | controllerType=0 controllerNumber=1 scan=14538 | 30.9805 |
| 12685:4 | 6 | R.LAGKPTHVN[+2116.756]VSVVMAEVDGTC[+57.021]Y.- | HexNAc(5)Hex(5)NeuAc(1) | C[+57], N[+2117] | 1116.7344 | 4 | 4463.9158 | 4463.8985 | 0 | 3.9 | 332 | CRagged | 633.36 | 522.92 | 522.92 | 12.73 | 82 | >sp\|P01876\|IGHA1_HUMAN Immunoglobulin heavy constant alpha 1 OS=Homo sapiens OX=9606 GN=IGHA1 PE=1 SV=2 | 255 | 20201118_lss_colostrum_1ng_glyco_hcd203040_compound15.16043.16043.4 | controllerType=0 controllerNumber=1 scan=16043 | 33.6868 |
| 10954:3 | 6 | R.LAGKPTHVN[+1702.581]VSVVMAEVDGTC[+57.021]Y.- | HexNAc(2)Hex(8) | C[+57], N[+1703] | 1350.581 | 3 | 4049.7284 | 4049.7234 | 0 | 1.2 | 332 | CRagged | 633.87 | 633.87 | 401.47 | 12.84 | 82 | >sp\|P01876\|IGHA1_HUMAN Immunoglobulin heavy constant alpha 1 OS=Homo sapiens OX=9606 GN=IGHA1 PE=1 SV=2 | 255 | 20201118_lss_colostrum_1ng_glyco_hcd203040_compound15.14247.14247.3 | controllerType=0 controllerNumber=1 scan=14247 | 30.3869 |
| 14494:4 | 6 | R.LAGKPTHVN[+2553.910]VSVVMAEVDGTC[+57.021]Y.- | HexNAc(5)Hex(5)Fuc(1)NeuAc(2) | C[+57], N[+2554] | 1226.0246 | 4 | 4901.0765 | 4901.0518 | 0 | 5 | 332 | CRagged | 635.01 | 495.65 | 495.65 | 12.27 | 82 | >sp\|P01876\|IGHA1_HUMAN Immunoglobulin heavy constant alpha 1 OS=Homo sapiens OX=9606 GN=IGHA1 PE=1 SV=2 | 255 | 20201118_lss_colostrum_1ng_glyco_hcd203040_compound15.17922.17922.4 | controllerType=0 controllerNumber=1 scan=17922 | 37.3409 |
| 14489:3 | 6 | R.LAGKPTHVN[+2553.910]VSVVMAEVDGTC[+57.021]Y.- | HexNAc(5)Hex(5)Fuc(1)NeuAc(2) | C[+57], N[+2554] | 1634.3644 | 3 | 4901.0788 | 4901.0518 | 0 | 5.5 | 332 | CRagged | 647.8 | 463.14 | 463.14 | 12.34 | 82 | >sp\|P01876\|IGHA1_HUMAN Immunoglobulin heavy constant alpha 1 OS=Homo sapiens OX=9606 GN=IGHA1 PE=1 SV=2 | 255 | 20201118_lss_colostrum_1ng_glyco_hcd203040_compound15.17917.17917.3 | controllerType=0 controllerNumber=1 scan=17917 | 37.3311 |
| 12686:4 | 6 | R.LAGKPTHVN[+2116.756]VSVVMAEVDGTC[+57.021]Y.- | HexNAc(5)Hex(5)NeuAc(1) | C[+57], N[+2117] | 1116.7344 | 4 | 4463.9158 | 4463.8985 | 0 | 3.9 | 332 | CRagged | 649.7 | 489.44 | 489.44 | 12.73 | 82 | >sp\|P01876\|IGHA1_HUMAN Immunoglobulin heavy constant alpha 1 OS=Homo sapiens OX=9606 GN=IGHA1 PE=1 SV=2 | 255 | 20201118_lss_colostrum_1ng_glyco_hcd203040_compound15.16044.16044.4 | controllerType=0 controllerNumber=1 scan=16044 | 33.6911 |
| 09961:3 | 6 | R.LAGKPTHVN[+1216.423]VSVVMAEVDGTC[+57.021]Y.- | HexNAc(2)Hex(5) | C[+57], N[+1216] | 1188.5273 | 3 | 3563.5673 | 3563.5649 | 0 | 0.7 | 332 | CRagged | 671.68 | 606.89 | 375.43 | 13.84 | 82 | >sp\|P01876\|IGHA1_HUMAN Immunoglobulin heavy constant alpha 1 OS=Homo sapiens OX=9606 GN=IGHA1 PE=1 SV=2 | 255 | 20201118_lss_colostrum_1ng_glyco_hcd203040_compound15.13217.13217.3 | controllerType=0 controllerNumber=1 scan=13217 | 28.5141 |
| 11226:3 | 6 | R.LAGKPTHVN[+1095.397]VSVVMAEVDGTC[+57.021]Y.- | HexNAc(3)Hex(3) | C[+57], N[+1095] | 1148.1887 | 3 | 3442.5517 | 3442.5387 | 0 | 3.8 | 332 | CRagged | 674.17 | 527.47 | 527.47 | 13.8 | 82 | >sp\|P01876\|IGHA1_HUMAN Immunoglobulin heavy constant alpha 1 OS=Homo sapiens OX=9606 GN=IGHA1 PE=1 SV=2 | 255 | 20201118_lss_colostrum_1ng_glyco_hcd203040_compound15.14530.14530.3 | controllerType=0 controllerNumber=1 scan=14530 | 30.9659 |
| 14486:3 | 6 | R.LAGKPTHVN[+2553.910]VSVVMAEVDGTC[+57.021]Y.- | HexNAc(5)Hex(5)Fuc(1)NeuAc(2) | C[+57], N[+2554] | 1634.3644 | 3 | 4901.0788 | 4901.0518 | 0 | 5.5 | 332 | CRagged | 686.95 | 500.31 | 500.31 | 13.34 | 82 | >sp\|P01876\|IGHA1_HUMAN Immunoglobulin heavy constant alpha 1 OS=Homo sapiens OX=9606 GN=IGHA1 PE=1 SV=2 | 255 | 20201118_lss_colostrum_1ng_glyco_hcd203040_compound15.17914.17914.3 | controllerType=0 controllerNumber=1 scan=17914 | 37.3258 |
| 09908:3 | 6 | R.LAGKPTHVN[+1378.476]VSVVMAEVDGTC[+57.021]Y.- | HexNAc(2)Hex(6) | C[+57], N[+1378] | 1242.5472 | 3 | 3725.627 | 3725.6178 | 0 | 2.5 | 332 | CRagged | 687.06 | 687.06 | 414.86 | 13.99 | 82 | >sp\|P01876\|IGHA1_HUMAN Immunoglobulin heavy constant alpha 1 OS=Homo sapiens OX=9606 GN=IGHA1 PE=1 SV=2 | 255 | 20201118_lss_colostrum_1ng_glyco_hcd203040_compound15.13162.13162.3 | controllerType=0 controllerNumber=1 scan=13162 | 28.4153 |
| 12787:3 | 6 | R.LAGKPTHVN[+1872.651]VSVVMAEVDGTC[+57.021]Y.- | HexNAc(3)Hex(6)NeuAc(1) | C[+57], N[+1873] | 1407.2724 | 3 | 4219.8026 | 4219.7926 | 0 | 2.4 | 332 | CRagged | 708.18 | 613.37 | 426.28 | 14.99 | 82 | >sp\|P01876\|IGHA1_HUMAN Immunoglobulin heavy constant alpha 1 OS=Homo sapiens OX=9606 GN=IGHA1 PE=1 SV=2 | 255 | 20201118_lss_colostrum_1ng_glyco_hcd203040_compound15.16149.16149.3 | controllerType=0 controllerNumber=1 scan=16149 | 33.8746 |
| 11054:3 | 6 | R.LAGKPTHVN[+1663.608]VSVVMAEVDGTC[+57.021]Y.- | HexNAc(5)Hex(4) | C[+57], N[+1664] | 1337.5903 | 3 | 4010.7562 | 4010.7502 | 0 | 1.5 | 332 | CRagged | 718.47 | 718.47 | 447.79 | 14.84 | 82 | >sp\|P01876\|IGHA1_HUMAN Immunoglobulin heavy constant alpha 1 OS=Homo sapiens OX=9606 GN=IGHA1 PE=1 SV=2 | 255 | 20201118_lss_colostrum_1ng_glyco_hcd203040_compound15.14351.14351.3 | controllerType=0 controllerNumber=1 scan=14351 | 30.587 |
| 14492:4 | 6 | R.LAGKPTHVN[+2553.910]VSVVMAEVDGTC[+57.021]Y.- | HexNAc(5)Hex(5)Fuc(1)NeuAc(2) | C[+57], N[+2554] | 1226.0246 | 4 | 4901.0765 | 4901.0518 | 0 | 5 | 332 | CRagged | 725.69 | 528.9 | 528.9 | 14.27 | 82 | >sp\|P01876\|IGHA1_HUMAN Immunoglobulin heavy constant alpha 1 OS=Homo sapiens OX=9606 GN=IGHA1 PE=1 SV=2 | 255 | 20201118_lss_colostrum_1ng_glyco_hcd203040_compound15.17920.17920.4 | controllerType=0 controllerNumber=1 scan=17920 | 37.3357 |
| 12780:3 | 6 | R.LAGKPTHVN[+2262.814]VSVVMAEVDGTC[+57.021]Y.- | HexNAc(5)Hex(5)Fuc(1)NeuAc(1) | C[+57], N[+2263] | 1537.3273 | 3 | 4609.9674 | 4609.9564 | 0 | 2.4 | 332 | CRagged | 726.52 | 574.17 | 574.17 | 14.99 | 82 | >sp\|P01876\|IGHA1_HUMAN Immunoglobulin heavy constant alpha 1 OS=Homo sapiens OX=9606 GN=IGHA1 PE=1 SV=2 | 255 | 20201118_lss_colostrum_1ng_glyco_hcd203040_compound15.16142.16142.3 | controllerType=0 controllerNumber=1 scan=16142 | 33.8603 |
| 11131:2 | 6 | R.LAGKPTHVN[+1216.423]VSVVMAEVDGTC[+57.021]Y.- | HexNAc(2)Hex(5) | C[+57], N[+1216] | 1782.2917 | 2 | 3563.5761 | 3563.5649 | 0 | 3.1 | 332 | CRagged | 744.53 | 679.52 | 315.55 | 14.37 | 82 | >sp\|P01876\|IGHA1_HUMAN Immunoglobulin heavy constant alpha 1 OS=Homo sapiens OX=9606 GN=IGHA1 PE=1 SV=2 | 255 | 20201118_lss_colostrum_1ng_glyco_hcd203040_compound15.14431.14431.2 | controllerType=0 controllerNumber=1 scan=14431 | 30.7761 |
| 12784:3 | 6 | R.LAGKPTHVN[+1872.651]VSVVMAEVDGTC[+57.021]Y.- | HexNAc(3)Hex(6)NeuAc(1) | C[+57], N[+1873] | 1407.2724 | 3 | 4219.8026 | 4219.7926 | 0 | 2.4 | 332 | CRagged | 744.54 | 736.43 | 488.09 | 14.99 | 82 | >sp\|P01876\|IGHA1_HUMAN Immunoglobulin heavy constant alpha 1 OS=Homo sapiens OX=9606 GN=IGHA1 PE=1 SV=2 | 255 | 20201118_lss_colostrum_1ng_glyco_hcd203040_compound15.16146.16146.3 | controllerType=0 controllerNumber=1 scan=16146 | 33.8666 |
| 11037:3 | 6 | R.LAGKPTHVN[+1809.666]VSVVMAEVDGTC[+57.021]Y.- | HexNAc(5)Hex(4)Fuc(1) | C[+57], N[+1810] | 1386.2757 | 3 | 4156.8125 | 4156.8081 | 0 | 1.1 | 332 | CRagged | 746.62 | 746.62 | 489.87 | 14.84 | 82 | >sp\|P01876\|IGHA1_HUMAN Immunoglobulin heavy constant alpha 1 OS=Homo sapiens OX=9606 GN=IGHA1 PE=1 SV=2 | 255 | 20201118_lss_colostrum_1ng_glyco_hcd203040_compound15.14333.14333.3 | controllerType=0 controllerNumber=1 scan=14333 | 30.5548 |
| 11165:3 | 6 | R.LAGKPTHVN[+1054.370]VSVVMAEVDGTC[+57.021]Y.- | HexNAc(2)Hex(4) | C[+57], N[+1054] | 1134.5103 | 3 | 3401.5163 | 3401.5121 | 0 | 1.2 | 332 | CRagged | 747.69 | 747.69 | 394.99 | 14.84 | 82 | >sp\|P01876\|IGHA1_HUMAN Immunoglobulin heavy constant alpha 1 OS=Homo sapiens OX=9606 GN=IGHA1 PE=1 SV=2 | 255 | 20201118_lss_colostrum_1ng_glyco_hcd203040_compound15.14466.14466.3 | controllerType=0 controllerNumber=1 scan=14466 | 30.8428 |
| 12773:3 | 6 | R.LAGKPTHVN[+2059.735]VSVVMAEVDGTC[+57.021]Y.- | HexNAc(4)Hex(5)Fuc(1)NeuAc(1) | C[+57], N[+2060] | 1469.6345 | 3 | 4406.8888 | 4406.877 | 0 | 2.7 | 332 | CRagged | 748.33 | 655.76 | 655.76 | 14.99 | 82 | >sp\|P01876\|IGHA1_HUMAN Immunoglobulin heavy constant alpha 1 OS=Homo sapiens OX=9606 GN=IGHA1 PE=1 SV=2 | 255 | 20201118_lss_colostrum_1ng_glyco_hcd203040_compound15.16134.16134.3 | controllerType=0 controllerNumber=1 scan=16134 | 33.8467 |
| 11142:3 | 6 | R.LAGKPTHVN[+1647.613]VSVVMAEVDGTC[+57.021]Y.- | HexNAc(5)Hex(3)Fuc(1) | C[+57], N[+1648] | 1332.2576 | 3 | 3994.7583 | 3994.7553 | 0 | 0.8 | 332 | CRagged | 757.6 | 757.6 | 757.6 | 15.84 | 82 | >sp\|P01876\|IGHA1_HUMAN Immunoglobulin heavy constant alpha 1 OS=Homo sapiens OX=9606 GN=IGHA1 PE=1 SV=2 | 255 | 20201118_lss_colostrum_1ng_glyco_hcd203040_compound15.14443.14443.3 | controllerType=0 controllerNumber=1 scan=14443 | 30.8009 |
| 12694:3 | 6 | R.LAGKPTHVN[+2116.756]VSVVMAEVDGTC[+57.021]Y.- | HexNAc(5)Hex(5)NeuAc(1) | C[+57], N[+2117] | 1488.6424 | 3 | 4463.9126 | 4463.8985 | 0 | 3.2 | 332 | CRagged | 774.98 | 552.99 | 552.99 | 15.99 | 82 | >sp\|P01876\|IGHA1_HUMAN Immunoglobulin heavy constant alpha 1 OS=Homo sapiens OX=9606 GN=IGHA1 PE=1 SV=2 | 255 | 20201118_lss_colostrum_1ng_glyco_hcd203040_compound15.16053.16053.3 | controllerType=0 controllerNumber=1 scan=16053 | 33.706 |
| 09963:3 | 6 | R.LAGKPTHVN[+1216.423]VSVVMAEVDGTC[+57.021]Y.- | HexNAc(2)Hex(5) | C[+57], N[+1216] | 1188.5273 | 3 | 3563.5673 | 3563.5649 | 0 | 0.7 | 332 | CRagged | 777.38 | 688.19 | 476.35 | 15.84 | 82 | >sp\|P01876\|IGHA1_HUMAN Immunoglobulin heavy constant alpha 1 OS=Homo sapiens OX=9606 GN=IGHA1 PE=1 SV=2 | 255 | 20201118_lss_colostrum_1ng_glyco_hcd203040_compound15.13219.13219.3 | controllerType=0 controllerNumber=1 scan=13219 | 28.5195 |
| 11039:3 | 6 | R.LAGKPTHVN[+1702.581]VSVVMAEVDGTC[+57.021]Y.- | HexNAc(2)Hex(8) | C[+57], N[+1703] | 1350.5801 | 3 | 4049.7259 | 4049.7234 | 0 | 0.6 | 332 | CRagged | 790.84 | 774.71 | 446.55 | 15.84 | 82 | >sp\|P01876\|IGHA1_HUMAN Immunoglobulin heavy constant alpha 1 OS=Homo sapiens OX=9606 GN=IGHA1 PE=1 SV=2 | 255 | 20201118_lss_colostrum_1ng_glyco_hcd203040_compound15.14335.14335.3 | controllerType=0 controllerNumber=1 scan=14335 | 30.557 |
| 11023:3 | 6 | R.LAGKPTHVN[+1378.476]VSVVMAEVDGTC[+57.021]Y.- | HexNAc(2)Hex(6) | C[+57], N[+1378] | 1242.5458 | 3 | 3725.6229 | 3725.6178 | 0 | 1.4 | 332 | CRagged | 801.43 | 801.43 | 449.98 | 16.84 | 82 | >sp\|P01876\|IGHA1_HUMAN Immunoglobulin heavy constant alpha 1 OS=Homo sapiens OX=9606 GN=IGHA1 PE=1 SV=2 | 255 | 20201118_lss_colostrum_1ng_glyco_hcd203040_compound15.14319.14319.3 | controllerType=0 controllerNumber=1 scan=14319 | 30.536 |
| 11025:3 | 6 | R.LAGKPTHVN[+1378.476]VSVVMAEVDGTC[+57.021]Y.- | HexNAc(2)Hex(6) | C[+57], N[+1378] | 1242.5458 | 3 | 3725.6229 | 3725.6178 | 0 | 1.4 | 332 | CRagged | 815.45 | 760.17 | 526.22 | 16.84 | 82 | >sp\|P01876\|IGHA1_HUMAN Immunoglobulin heavy constant alpha 1 OS=Homo sapiens OX=9606 GN=IGHA1 PE=1 SV=2 | 255 | 20201118_lss_colostrum_1ng_glyco_hcd203040_compound15.14321.14321.3 | controllerType=0 controllerNumber=1 scan=14321 | 30.5414 |
| 11058:3 | 6 | R.LAGKPTHVN[+1809.666]VSVVMAEVDGTC[+57.021]Y.- | HexNAc(5)Hex(4)Fuc(1) | C[+57], N[+1810] | 1386.2763 | 3 | 4156.8144 | 4156.8081 | 0 | 1.5 | 332 | CRagged | 817.29 | 678.9 | 506.4 | 16.84 | 82 | >sp\|P01876\|IGHA1_HUMAN Immunoglobulin heavy constant alpha 1 OS=Homo sapiens OX=9606 GN=IGHA1 PE=1 SV=2 | 255 | 20201118_lss_colostrum_1ng_glyco_hcd203040_compound15.14355.14355.3 | controllerType=0 controllerNumber=1 scan=14355 | 30.5951 |
| 11057:3 | 6 | R.LAGKPTHVN[+1663.608]VSVVMAEVDGTC[+57.021]Y.- | HexNAc(5)Hex(4) | C[+57], N[+1664] | 1337.5903 | 3 | 4010.7562 | 4010.7502 | 0 | 1.5 | 332 | CRagged | 819.09 | 819.09 | 488.02 | 16.84 | 82 | >sp\|P01876\|IGHA1_HUMAN Immunoglobulin heavy constant alpha 1 OS=Homo sapiens OX=9606 GN=IGHA1 PE=1 SV=2 | 255 | 20201118_lss_colostrum_1ng_glyco_hcd203040_compound15.14354.14354.3 | controllerType=0 controllerNumber=1 scan=14354 | 30.5915 |
| 11144:3 | 6 | R.LAGKPTHVN[+1647.613]VSVVMAEVDGTC[+57.021]Y.- | HexNAc(5)Hex(3)Fuc(1) | C[+57], N[+1648] | 1332.2576 | 3 | 3994.7583 | 3994.7553 | 0 | 0.8 | 332 | CRagged | 819.15 | 693.06 | 693.06 | 16.84 | 82 | >sp\|P01876\|IGHA1_HUMAN Immunoglobulin heavy constant alpha 1 OS=Homo sapiens OX=9606 GN=IGHA1 PE=1 SV=2 | 255 | 20201118_lss_colostrum_1ng_glyco_hcd203040_compound15.14445.14445.3 | controllerType=0 controllerNumber=1 scan=14445 | 30.8066 |
| 11040:3 | 6 | R.LAGKPTHVN[+1809.666]VSVVMAEVDGTC[+57.021]Y.- | HexNAc(5)Hex(4)Fuc(1) | C[+57], N[+1810] | 1386.2757 | 3 | 4156.8125 | 4156.8081 | 0 | 1.1 | 332 | CRagged | 834.13 | 787.33 | 555.27 | 16.84 | 82 | >sp\|P01876\|IGHA1_HUMAN Immunoglobulin heavy constant alpha 1 OS=Homo sapiens OX=9606 GN=IGHA1 PE=1 SV=2 | 255 | 20201118_lss_colostrum_1ng_glyco_hcd203040_compound15.14336.14336.3 | controllerType=0 controllerNumber=1 scan=14336 | 30.5617 |
| 10958:3 | 6 | R.LAGKPTHVN[+1540.529]VSVVMAEVDGTC[+57.021]Y.- | HexNAc(2)Hex(7) | C[+57], N[+1541] | 1296.5632 | 3 | 3887.6749 | 3887.6706 | 0 | 1.1 | 332 | CRagged | 834.85 | 792.8 | 528.12 | 16.84 | 82 | >sp\|P01876\|IGHA1_HUMAN Immunoglobulin heavy constant alpha 1 OS=Homo sapiens OX=9606 GN=IGHA1 PE=1 SV=2 | 255 | 20201118_lss_colostrum_1ng_glyco_hcd203040_compound15.14251.14251.3 | controllerType=0 controllerNumber=1 scan=14251 | 30.3957 |
| 09910:3 | 6 | R.LAGKPTHVN[+1378.476]VSVVMAEVDGTC[+57.021]Y.- | HexNAc(2)Hex(6) | C[+57], N[+1378] | 1242.5472 | 3 | 3725.627 | 3725.6178 | 0 | 2.5 | 332 | CRagged | 835.9 | 815.28 | 500.55 | 16.99 | 82 | >sp\|P01876\|IGHA1_HUMAN Immunoglobulin heavy constant alpha 1 OS=Homo sapiens OX=9606 GN=IGHA1 PE=1 SV=2 | 255 | 20201118_lss_colostrum_1ng_glyco_hcd203040_compound15.13164.13164.3 | controllerType=0 controllerNumber=1 scan=13164 | 28.42 |
| 11167:3 | 6 | R.LAGKPTHVN[+1054.370]VSVVMAEVDGTC[+57.021]Y.- | HexNAc(2)Hex(4) | C[+57], N[+1054] | 1134.5103 | 3 | 3401.5163 | 3401.5121 | 0 | 1.2 | 332 | CRagged | 842.83 | 813.27 | 502.14 | 16.84 | 82 | >sp\|P01876\|IGHA1_HUMAN Immunoglobulin heavy constant alpha 1 OS=Homo sapiens OX=9606 GN=IGHA1 PE=1 SV=2 | 255 | 20201118_lss_colostrum_1ng_glyco_hcd203040_compound15.14468.14468.3 | controllerType=0 controllerNumber=1 scan=14468 | 30.8479 |
| 12765:3 | 6 | R.LAGKPTHVN[+1872.651]VSVVMAEVDGTC[+57.021]Y.- | HexNAc(3)Hex(6)NeuAc(1) | C[+57], N[+1873] | 1407.2731 | 3 | 4219.8048 | 4219.7926 | 0 | 2.9 | 332 | CRagged | 846.95 | 792.66 | 483.79 | 16.99 | 82 | >sp\|P01876\|IGHA1_HUMAN Immunoglobulin heavy constant alpha 1 OS=Homo sapiens OX=9606 GN=IGHA1 PE=1 SV=2 | 255 | 20201118_lss_colostrum_1ng_glyco_hcd203040_compound15.16126.16126.3 | controllerType=0 controllerNumber=1 scan=16126 | 33.8323 |
| 11056:3 | 6 | R.LAGKPTHVN[+1809.666]VSVVMAEVDGTC[+57.021]Y.- | HexNAc(5)Hex(4)Fuc(1) | C[+57], N[+1810] | 1386.2763 | 3 | 4156.8144 | 4156.8081 | 0 | 1.5 | 332 | CRagged | 855.01 | 855.01 | 562.95 | 17.84 | 82 | >sp\|P01876\|IGHA1_HUMAN Immunoglobulin heavy constant alpha 1 OS=Homo sapiens OX=9606 GN=IGHA1 PE=1 SV=2 | 255 | 20201118_lss_colostrum_1ng_glyco_hcd203040_compound15.14353.14353.3 | controllerType=0 controllerNumber=1 scan=14353 | 30.5893 |
| 10956:3 | 6 | R.LAGKPTHVN[+1540.529]VSVVMAEVDGTC[+57.021]Y.- | HexNAc(2)Hex(7) | C[+57], N[+1541] | 1296.5632 | 3 | 3887.6749 | 3887.6706 | 0 | 1.1 | 332 | CRagged | 855.98 | 710.96 | 537.75 | 17.84 | 82 | >sp\|P01876\|IGHA1_HUMAN Immunoglobulin heavy constant alpha 1 OS=Homo sapiens OX=9606 GN=IGHA1 PE=1 SV=2 | 255 | 20201118_lss_colostrum_1ng_glyco_hcd203040_compound15.14249.14249.3 | controllerType=0 controllerNumber=1 scan=14249 | 30.3895 |
| 11117:3 | 6 | R.LAGKPTHVN[+1501.555]VSVVMAEVDGTC[+57.021]Y.- | HexNAc(5)Hex(3) | C[+57], N[+1502] | 1283.5732 | 3 | 3848.705 | 3848.6974 | 0 | 2 | 332 | CRagged | 872.53 | 872.53 | 872.53 | 17.99 | 82 | >sp\|P01876\|IGHA1_HUMAN Immunoglobulin heavy constant alpha 1 OS=Homo sapiens OX=9606 GN=IGHA1 PE=1 SV=2 | 255 | 20201118_lss_colostrum_1ng_glyco_hcd203040_compound15.14417.14417.3 | controllerType=0 controllerNumber=1 scan=14417 | 30.7476 |
| 10957:3 | 6 | R.LAGKPTHVN[+1702.581]VSVVMAEVDGTC[+57.021]Y.- | HexNAc(2)Hex(8) | C[+57], N[+1703] | 1350.581 | 3 | 4049.7284 | 4049.7234 | 0 | 1.2 | 332 | CRagged | 880.68 | 838.8 | 526.72 | 17.84 | 82 | >sp\|P01876\|IGHA1_HUMAN Immunoglobulin heavy constant alpha 1 OS=Homo sapiens OX=9606 GN=IGHA1 PE=1 SV=2 | 255 | 20201118_lss_colostrum_1ng_glyco_hcd203040_compound15.14250.14250.3 | controllerType=0 controllerNumber=1 scan=14250 | 30.3914 |
| 11059:3 | 6 | R.LAGKPTHVN[+1216.423]VSVVMAEVDGTC[+57.021]Y.- | HexNAc(2)Hex(5) | C[+57], N[+1216] | 1188.5283 | 3 | 3563.5702 | 3563.5649 | 0 | 1.5 | 332 | CRagged | 885.25 | 885.25 | 533.5 | 17.84 | 82 | >sp\|P01876\|IGHA1_HUMAN Immunoglobulin heavy constant alpha 1 OS=Homo sapiens OX=9606 GN=IGHA1 PE=1 SV=2 | 255 | 20201118_lss_colostrum_1ng_glyco_hcd203040_compound15.14356.14356.3 | controllerType=0 controllerNumber=1 scan=14356 | 30.5959 |
| 11119:3 | 6 | R.LAGKPTHVN[+1501.555]VSVVMAEVDGTC[+57.021]Y.- | HexNAc(5)Hex(3) | C[+57], N[+1502] | 1283.5732 | 3 | 3848.705 | 3848.6974 | 0 | 2 | 332 | CRagged | 897.21 | 865.04 | 865.04 | 17.99 | 82 | >sp\|P01876\|IGHA1_HUMAN Immunoglobulin heavy constant alpha 1 OS=Homo sapiens OX=9606 GN=IGHA1 PE=1 SV=2 | 255 | 20201118_lss_colostrum_1ng_glyco_hcd203040_compound15.14419.14419.3 | controllerType=0 controllerNumber=1 scan=14419 | 30.7528 |
| 11063:3 | 6 | R.LAGKPTHVN[+1216.423]VSVVMAEVDGTC[+57.021]Y.- | HexNAc(2)Hex(5) | C[+57], N[+1216] | 1188.5283 | 3 | 3563.5702 | 3563.5649 | 0 | 1.5 | 332 | CRagged | 906.86 | 891.09 | 550.69 | 18.84 | 82 | >sp\|P01876\|IGHA1_HUMAN Immunoglobulin heavy constant alpha 1 OS=Homo sapiens OX=9606 GN=IGHA1 PE=1 SV=2 | 255 | 20201118_lss_colostrum_1ng_glyco_hcd203040_compound15.14360.14360.3 | controllerType=0 controllerNumber=1 scan=14360 | 30.609 |
| 09167:3 | 6 | R.LAGKPTHVN[+1216.423]VSVVM[+15.995]AEVDGTC[+57.021]Y.- | HexNAc(2)Hex(5) | C[+57], M[+16], N[+1216] | 1193.8606 | 3 | 3579.5672 | 3579.5599 | 0 | 2.1 | 332 | CRagged | 221.27 | 221.27 | 221.27 | 4.24 | 82 | >sp\|P01876\|IGHA1_HUMAN Immunoglobulin heavy constant alpha 1 OS=Homo sapiens OX=9606 GN=IGHA1 PE=1 SV=2 | 255 | 20201118_lss_colostrum_1ng_glyco_hcd203040_compound15.12391.12391.3 | controllerType=0 controllerNumber=1 scan=12391 | 27.025 |
| 09119:3 | 6 | R.LAGKPTHVN[+1647.613]VSVVM[+15.995]AEVDGTC[+57.021]Y.- | HexNAc(5)Hex(3)Fuc(1) | C[+57], M[+16], N[+1648] | 1337.593 | 3 | 4010.7643 | 4010.7502 | 0 | 3.5 | 332 | CRagged | 291.6 | 102.02 | 102.02 | 5.59 | 82 | >sp\|P01876\|IGHA1_HUMAN Immunoglobulin heavy constant alpha 1 OS=Homo sapiens OX=9606 GN=IGHA1 PE=1 SV=2 | 255 | 20201118_lss_colostrum_1ng_glyco_hcd203040_compound15.12341.12341.3 | controllerType=0 controllerNumber=1 scan=12341 | 26.9424 |
| 09165:3 | 6 | R.LAGKPTHVN[+1216.423]VSVVM[+15.995]AEVDGTC[+57.021]Y.- | HexNAc(2)Hex(5) | C[+57], M[+16], N[+1216] | 1193.8606 | 3 | 3579.5672 | 3579.5599 | 0 | 2.1 | 332 | CRagged | 293.47 | 293.47 | 293.47 | 7.55 | 82 | >sp\|P01876\|IGHA1_HUMAN Immunoglobulin heavy constant alpha 1 OS=Homo sapiens OX=9606 GN=IGHA1 PE=1 SV=2 | 255 | 20201118_lss_colostrum_1ng_glyco_hcd203040_compound15.12389.12389.3 | controllerType=0 controllerNumber=1 scan=12389 | 27.0206 |
| 09054:3 | 6 | R.LAGKPTHVN[+1378.476]VSVVM[+15.995]AEVDGTC[+57.021]Y.- | HexNAc(2)Hex(6) | C[+57], M[+16], N[+1378] | 1247.8768 | 3 | 3741.6157 | 3741.6127 | 0 | 0.8 | 332 | CRagged | 451.33 | 451.33 | 451.33 | 9.93 | 82 | >sp\|P01876\|IGHA1_HUMAN Immunoglobulin heavy constant alpha 1 OS=Homo sapiens OX=9606 GN=IGHA1 PE=1 SV=2 | 255 | 20201118_lss_colostrum_1ng_glyco_hcd203040_compound15.12274.12274.3 | controllerType=0 controllerNumber=1 scan=12274 | 26.8282 |
| 09052:3 | 6 | R.LAGKPTHVN[+1378.476]VSVVM[+15.995]AEVDGTC[+57.021]Y.- | HexNAc(2)Hex(6) | C[+57], M[+16], N[+1378] | 1247.8768 | 3 | 3741.6157 | 3741.6127 | 0 | 0.8 | 332 | CRagged | 468.17 | 468.17 | 468.17 | 10.01 | 82 | >sp\|P01876\|IGHA1_HUMAN Immunoglobulin heavy constant alpha 1 OS=Homo sapiens OX=9606 GN=IGHA1 PE=1 SV=2 | 255 | 20201118_lss_colostrum_1ng_glyco_hcd203040_compound15.12272.12272.3 | controllerType=0 controllerNumber=1 scan=12272 | 26.8245 |
| 08712:3 | 6 | R.LAGKPTHVN[+1216.423]VSVVM[+15.995]AEVDGTC[+57.021]Y.- | HexNAc(2)Hex(5) | C[+57], M[+16], N[+1216] | 1193.858 | 3 | 3579.5593 | 3579.5599 | 0 | -0.1 | 332 | CRagged | 621.84 | 621.84 | 621.84 | 12.37 | 82 | >sp\|P01876\|IGHA1_HUMAN Immunoglobulin heavy constant alpha 1 OS=Homo sapiens OX=9606 GN=IGHA1 PE=1 SV=2 | 255 | 20201118_lss_colostrum_1ng_glyco_hcd203040_compound15.11916.11916.3 | controllerType=0 controllerNumber=1 scan=11916 | 26.1938 |
| 08555:3 | 6 | R.LAGKPTHVN[+1216.423]VSVVM[+15.995]AEVDGTC[+57.021]Y.- | HexNAc(2)Hex(5) | C[+57], M[+16], N[+1216] | 1193.8589 | 3 | 3579.5623 | 3579.5599 | 0 | 0.7 | 332 | CRagged | 730.6 | 617.34 | 617.34 | 14.84 | 82 | >sp\|P01876\|IGHA1_HUMAN Immunoglobulin heavy constant alpha 1 OS=Homo sapiens OX=9606 GN=IGHA1 PE=1 SV=2 | 255 | 20201118_lss_colostrum_1ng_glyco_hcd203040_compound15.11753.11753.3 | controllerType=0 controllerNumber=1 scan=11753 | 25.9151 |
| 08714:3 | 6 | R.LAGKPTHVN[+1216.423]VSVVM[+15.995]AEVDGTC[+57.021]Y.- | HexNAc(2)Hex(5) | C[+57], M[+16], N[+1216] | 1193.858 | 3 | 3579.5593 | 3579.5599 | 0 | -0.1 | 332 | CRagged | 740.59 | 666.84 | 666.84 | 14.37 | 82 | >sp\|P01876\|IGHA1_HUMAN Immunoglobulin heavy constant alpha 1 OS=Homo sapiens OX=9606 GN=IGHA1 PE=1 SV=2 | 255 | 20201118_lss_colostrum_1ng_glyco_hcd203040_compound15.11918.11918.3 | controllerType=0 controllerNumber=1 scan=11918 | 26.1983 |
| 08557:3 | 6 | R.LAGKPTHVN[+1216.423]VSVVM[+15.995]AEVDGTC[+57.021]Y.- | HexNAc(2)Hex(5) | C[+57], M[+16], N[+1216] | 1193.8589 | 3 | 3579.5623 | 3579.5599 | 0 | 0.7 | 332 | CRagged | 742.1 | 742.1 | 742.1 | 14.84 | 82 | >sp\|P01876\|IGHA1_HUMAN Immunoglobulin heavy constant alpha 1 OS=Homo sapiens OX=9606 GN=IGHA1 PE=1 SV=2 | 255 | 20201118_lss_colostrum_1ng_glyco_hcd203040_compound15.11755.11755.3 | controllerType=0 controllerNumber=1 scan=11755 | 25.9203 |
| 00104:4 | 9 | K.KKEDALN[+2059.735]ETRESETK.L | HexNAc(4)Hex(5)Fuc(1)NeuAc(1) | N[+2060] | 960.1645 | 4 | 3837.6363 | 3837.6276 | 0 | 2.3 | 80 | Specific | 215.93 | 214.77 | 214.77 | 3.83 | 67 | >sp\|P10909\|CLUS_HUMAN Clusterin OS=Homo sapiens OX=9606 GN=CLU PE=1 SV=1 | 1467 | 20201118_lss_colostrum_1ng_glyco_hcd203040_compound15.2991.2991.4 | controllerType=0 controllerNumber=1 scan=2991 | 11.2618 |
| 00106:4 | 9 | K.KKEDALN[+2059.735]ETRESETK.L | HexNAc(4)Hex(5)Fuc(1)NeuAc(1) | N[+2060] | 960.1645 | 4 | 3837.6363 | 3837.6276 | 0 | 2.3 | 80 | Specific | 399.95 | 277.85 | 277.85 | 5.96 | 67 | >sp\|P10909\|CLUS_HUMAN Clusterin OS=Homo sapiens OX=9606 GN=CLU PE=1 SV=1 | 1467 | 20201118_lss_colostrum_1ng_glyco_hcd203040_compound15.2993.2993.4 | controllerType=0 controllerNumber=1 scan=2993 | 11.2674 |
| 13579:3 | 9 | K.LKELPGVC[+57.021]N[+2059.735]ETMMALWEEC[+57.021]KPC[+57.021]LK.Q | HexNAc(4)Hex(5)Fuc(1)NeuAc(1) | C[+57]*3, N[+2060] | 1666.0462 | 3 | 4996.1241 | 4996.1196 | 0 | 0.9 | 95 | Specific | 220.06 | 220.06 | 220.06 | 4.3 | 67 | >sp\|P10909\|CLUS_HUMAN Clusterin OS=Homo sapiens OX=9606 GN=CLU PE=1 SV=1 | 1467 | 20201118_lss_colostrum_1ng_glyco_hcd203040_compound15.16972.16972.3 | controllerType=0 controllerNumber=1 scan=16972 | 35.5072 |
| 13577:3 | 9 | K.LKELPGVC[+57.021]N[+2059.735]ETMMALWEEC[+57.021]KPC[+57.021]LK.Q | HexNAc(4)Hex(5)Fuc(1)NeuAc(1) | C[+57]*3, N[+2060] | 1666.0462 | 3 | 4996.1241 | 4996.1196 | 0 | 0.9 | 95 | Specific | 275.19 | 263.92 | 263.92 | 5.91 | 67 | >sp\|P10909\|CLUS_HUMAN Clusterin OS=Homo sapiens OX=9606 GN=CLU PE=1 SV=1 | 1467 | 20201118_lss_colostrum_1ng_glyco_hcd203040_compound15.16970.16970.3 | controllerType=0 controllerNumber=1 scan=16970 | 35.5034 |
| 13565:4 | 9 | K.LKELPGVC[+57.021]N[+2059.735]ETMMALWEEC[+57.021]KPC[+57.021]LK.Q | HexNAc(4)Hex(5)Fuc(1)NeuAc(1) | C[+57]*3, N[+2060] | 1249.7885 | 4 | 4996.132 | 4996.1196 | 0 | 2.5 | 95 | Specific | 287.24 | 287.24 | 287.24 | 6.69 | 67 | >sp\|P10909\|CLUS_HUMAN Clusterin OS=Homo sapiens OX=9606 GN=CLU PE=1 SV=1 | 1467 | 20201118_lss_colostrum_1ng_glyco_hcd203040_compound15.16958.16958.4 | controllerType=0 controllerNumber=1 scan=16958 | 35.4888 |
| 13568:4 | 9 | K.LKELPGVC[+57.021]N[+2059.735]ETMMALWEEC[+57.021]KPC[+57.021]LK.Q | HexNAc(4)Hex(5)Fuc(1)NeuAc(1) | C[+57]*3, N[+2060] | 1249.7885 | 4 | 4996.132 | 4996.1196 | 0 | 2.5 | 95 | Specific | 796.57 | 734.64 | 734.64 | 15.52 | 67 | >sp\|P10909\|CLUS_HUMAN Clusterin OS=Homo sapiens OX=9606 GN=CLU PE=1 SV=1 | 1467 | 20201118_lss_colostrum_1ng_glyco_hcd203040_compound15.16961.16961.4 | controllerType=0 controllerNumber=1 scan=16961 | 35.4924 |
| 13445:4 | 9 | K.LKELPGVC[+57.021]N[+2205.793]ETMMALWEEC[+57.021]KPC[+57.021]LK.Q | HexNAc(4)Hex(5)Fuc(2)NeuAc(1) | C[+57]*3, N[+2206] | 1286.304 | 4 | 5142.194 | 5142.1776 | 0 | 3.2 | 95 | Specific | 875.58 | 875.39 | 875.39 | 17.52 | 67 | >sp\|P10909\|CLUS_HUMAN Clusterin OS=Homo sapiens OX=9606 GN=CLU PE=1 SV=1 | 1467 | 20201118_lss_colostrum_1ng_glyco_hcd203040_compound15.16833.16833.4 | controllerType=0 controllerNumber=1 scan=16833 | 35.2614 |
| 13443:4 | 9 | K.LKELPGVC[+57.021]N[+2205.793]ETMMALWEEC[+57.021]KPC[+57.021]LK.Q | HexNAc(4)Hex(5)Fuc(2)NeuAc(1) | C[+57]*3, N[+2206] | 1286.304 | 4 | 5142.194 | 5142.1776 | 0 | 3.2 | 95 | Specific | 1080.95 | 1016.85 | 1016.85 | 21.52 | 67 | >sp\|P10909\|CLUS_HUMAN Clusterin OS=Homo sapiens OX=9606 GN=CLU PE=1 SV=1 | 1467 | 20201118_lss_colostrum_1ng_glyco_hcd203040_compound15.16831.16831.4 | controllerType=0 controllerNumber=1 scan=16831 | 35.2551 |
| 15216:4 | 9 | K.ELPGVC[+57.021]N[+2059.735]ETMMALWEEC[+57.021]KPC[+57.021]LK.Q | HexNAc(4)Hex(5)Fuc(1)NeuAc(1) | C[+57]*3, N[+2060] | 1189.4942 | 4 | 4754.9549 | 4754.9406 | 0 | 3 | 97 | Specific | 487.6 | 479.35 | 479.35 | 10.11 | 67 | >sp\|P10909\|CLUS_HUMAN Clusterin OS=Homo sapiens OX=9606 GN=CLU PE=1 SV=1 | 1467 | 20201118_lss_colostrum_1ng_glyco_hcd203040_compound15.18673.18673.4 | controllerType=0 controllerNumber=1 scan=18673 | 38.8188 |
| 15219:4 | 9 | K.ELPGVC[+57.021]N[+2059.735]ETMMALWEEC[+57.021]KPC[+57.021]LK.Q | HexNAc(4)Hex(5)Fuc(1)NeuAc(1) | C[+57]*3, N[+2060] | 1189.4942 | 4 | 4754.9549 | 4754.9406 | 0 | 3 | 97 | Specific | 737.13 | 600.27 | 600.27 | 14.93 | 67 | >sp\|P10909\|CLUS_HUMAN Clusterin OS=Homo sapiens OX=9606 GN=CLU PE=1 SV=1 | 1467 | 20201118_lss_colostrum_1ng_glyco_hcd203040_compound15.18676.18676.4 | controllerType=0 controllerNumber=1 scan=18676 | 38.8229 |
| 19472:3 | 9 | R.QLEEFLN[+2059.735]QSSPFYFWMNGDR.I | HexNAc(4)Hex(5)Fuc(1)NeuAc(1) | N[+2060] | 1523.2946 | 3 | 4567.8693 | 4567.8638 | 0 | 1.2 | 139 | Specific | 735.43 | 655.59 | 655.59 | 14.47 | 67 | >sp\|P10909\|CLUS_HUMAN Clusterin OS=Homo sapiens OX=9606 GN=CLU PE=1 SV=1 | 1467 | 20201118_lss_colostrum_1ng_glyco_hcd203040_compound15.23108.23108.3 | controllerType=0 controllerNumber=1 scan=23108 | 48.1329 |
| 19475:3 | 9 | R.QLEEFLN[+2059.735]QSSPFYFWMNGDR.I | HexNAc(4)Hex(5)Fuc(1)NeuAc(1) | N[+2060] | 1523.2946 | 3 | 4567.8693 | 4567.8638 | 0 | 1.2 | 139 | Specific | 764.71 | 544.27 | 544.27 | 15.47 | 67 | >sp\|P10909\|CLUS_HUMAN Clusterin OS=Homo sapiens OX=9606 GN=CLU PE=1 SV=1 | 1467 | 20201118_lss_colostrum_1ng_glyco_hcd203040_compound15.23111.23111.3 | controllerType=0 controllerNumber=1 scan=23111 | 48.1376 |
| 00098:3 | 9 | R.EIRHN[+2351.851]STGC[+57.021]LR.M | HexNAc(4)Hex(5)Fuc(3)NeuAc(1) | C[+57], N[+2352] | 1232.1791 | 3 | 3694.5228 | 3694.5152 | 0 | 2.1 | 287 | Specific | 238.99 | 149.76 | 149.76 | 4.3 | 67 | >sp\|P10909\|CLUS_HUMAN Clusterin OS=Homo sapiens OX=9606 GN=CLU PE=1 SV=1 | 1467 | 20201118_lss_colostrum_1ng_glyco_hcd203040_compound15.2984.2984.3 | controllerType=0 controllerNumber=1 scan=2984 | 11.2462 |
| 20173:4 | 9 | K.MLN[+2076.750]TSSLLEQLNEQFNWVSR.L | HexNAc(4)Hex(6)Fuc(2) | N[+2077] | 1122.246 | 4 | 4485.9622 | 4485.937 | 0 | 5.6 | 352 | Specific | 300.09 | 300.09 | 300.09 | 6.77 | 67 | >sp\|P10909\|CLUS_HUMAN Clusterin OS=Homo sapiens OX=9606 GN=CLU PE=1 SV=1 | 1467 | 20201118_lss_colostrum_1ng_glyco_hcd203040_compound15.23840.23840.4 | controllerType=0 controllerNumber=1 scan=23840 | 49.8131 |

Table S4. Cont.

| Query #:z | Protein Rank | Glycopeptide | Glycan | Modification Type(s) | Observed m/z | z | Observed (M+H) | Calc. mass (M+H) | Off-by-x error | Mass error (ppm) | StartPosition | Cleavage | Score | Delta | Delta Mod | \|Log Prob\| | # of unique peptides | Protein Name | Protein DB number | Comment | Scan # | Scan Time |
|---|---|---|---|---|---|---|---|---|---|---|---|---|---|---|---|---|---|---|---|---|---|---|
| 22347:3 | 9 | K.MLN[+2350.830]TSSLLEQLNEQFNWVSR.L | HexNAc(4)Hex(5)Fuc(1)NeuAc(2) | N[+2351] | 1587.351 | 3 | 4760.0385 | 4760.0171 | 0 | 4.5 | 352 | Specific | 315.01 | 315.01 | 315.01 | 6.61 | 67 | >sp\|P10909\|CLUS_HUMAN Clusterin OS=Homo sapiens OX=9606 GN=CLU PE=1 SV=1 | 1467 | 20201118_lss_colostrum_1ng_glyco_hcd203040_compound15.26111.26111.3 | controllerType=0 controllerNumber=1 scan=26111 | 54.8618 |
| 20170:3 | 9 | K.MLN[+2076.750]TSSLLEQLNEQFNWVSR.L | HexNAc(4)Hex(6)Fuc(2) | N[+2077] | 1495.9932 | 3 | 4485.9651 | 4485.937 | 0 | 6.3 | 352 | Specific | 358.02 | 321.3 | 321.3 | 6.45 | 67 | >sp\|P10909\|CLUS_HUMAN Clusterin OS=Homo sapiens OX=9606 GN=CLU PE=1 SV=1 | 1467 | 20201118_lss_colostrum_1ng_glyco_hcd203040_compound15.23837.23837.3 | controllerType=0 controllerNumber=1 scan=23837 | 49.8097 |
| 22149:4 | 9 | K.MLN[+2350.830]TSSLLEQLNEQFNWVSR.L | HexNAc(4)Hex(5)Fuc(1)NeuAc(2) | N[+2351] | 1190.7655 | 4 | 4760.0404 | 4760.0171 | 0 | 4.9 | 352 | Specific | 389.13 | 374.3 | 374.3 | 7.56 | 67 | >sp\|P10909\|CLUS_HUMAN Clusterin OS=Homo sapiens OX=9606 GN=CLU PE=1 SV=1 | 1467 | 20201118_lss_colostrum_1ng_glyco_hcd203040_compound15.25905.25905.3 | controllerType=0 controllerNumber=1 scan=25905 | 54.4446 |
| 22206:4 | 9 | K.MLN[+2350.830]TSSLLEQLNEQFNWVSR.L | HexNAc(4)Hex(5)Fuc(1)NeuAc(2) | N[+2351] | 1190.7652 | 4 | 4760.0389 | 4760.0171 | 0 | 4.6 | 352 | Specific | 416.53 | 354.14 | 354.14 | 7.42 | 67 | >sp\|P10909\|CLUS_HUMAN Clusterin OS=Homo sapiens OX=9606 GN=CLU PE=1 SV=1 | 1467 | 20201118_lss_colostrum_1ng_glyco_hcd203040_compound15.25965.25965.4 | controllerType=0 controllerNumber=1 scan=25965 | 54.564 |
| 22172:4 | 9 | K.MLN[+2350.830]TSSLLEQLNEQFNWVSR.L | HexNAc(4)Hex(5)Fuc(1)NeuAc(2) | N[+2351] | 1190.7638 | 4 | 4760.0335 | 4760.0171 | 0 | 3.5 | 352 | Specific | 441.9 | 383.61 | 383.61 | 9.66 | 67 | >sp\|P10909\|CLUS_HUMAN Clusterin OS=Homo sapiens OX=9606 GN=CLU PE=1 SV=1 | 1467 | 20201118_lss_colostrum_1ng_glyco_hcd203040_compound15.25929.25929.3 | controllerType=0 controllerNumber=1 scan=25929 | 54.4934 |
| 20176:4 | 9 | K.MLN[+2076.750]TSSLLEQLNEQFNWVSR.L | HexNAc(4)Hex(6)Fuc(2) | N[+2077] | 1122.246 | 4 | 4485.9622 | 4485.937 | 0 | 5.6 | 352 | Specific | 468.12 | 443.4 | 443.4 | 9.4 | 67 | >sp\|P10909\|CLUS_HUMAN Clusterin OS=Homo sapiens OX=9606 GN=CLU PE=1 SV=1 | 1467 | 20201118_lss_colostrum_1ng_glyco_hcd203040_compound15.23843.23843.4 | controllerType=0 controllerNumber=1 scan=23843 | 49.8199 |
| 22349:3 | 9 | K.MLN[+2350.830]TSSLLEQLNEQFNWVSR.L | HexNAc(4)Hex(5)Fuc(1)NeuAc(2) | N[+2351] | 1587.351 | 3 | 4760.0385 | 4760.0171 | 0 | 4.5 | 352 | Specific | 474.94 | 389.93 | 389.93 | 9.15 | 67 | >sp\|P10909\|CLUS_HUMAN Clusterin OS=Homo sapiens OX=9606 GN=CLU PE=1 SV=1 | 1467 | 20201118_lss_colostrum_1ng_glyco_hcd203040_compound15.26113.26113.3 | controllerType=0 controllerNumber=1 scan=26113 | 54.8657 |
| 22174:4 | 9 | K.MLN[+2350.830]TSSLLEQLNEQFNWVSR.L | HexNAc(4)Hex(5)Fuc(1)NeuAc(2) | N[+2351] | 1190.7638 | 4 | 4760.0335 | 4760.0171 | 0 | 3.5 | 352 | Specific | 546.69 | 488.04 | 488.04 | 10.08 | 67 | >sp\|P10909\|CLUS_HUMAN Clusterin OS=Homo sapiens OX=9606 GN=CLU PE=1 SV=1 | 1467 | 20201118_lss_colostrum_1ng_glyco_hcd203040_compound15.25931.25931.3 | controllerType=0 controllerNumber=1 scan=25931 | 54.4986 |
| 22207:3 | 9 | K.MLN[+2350.830]TSSLLEQLNEQFNWVSR.L | HexNAc(4)Hex(5)Fuc(1)NeuAc(2) | N[+2351] | 1587.3504 | 3 | 4760.0366 | 4760.0171 | 0 | 4.1 | 352 | Specific | 570.13 | 526.99 | 526.99 | 10.09 | 67 | >sp\|P10909\|CLUS_HUMAN Clusterin OS=Homo sapiens OX=9606 GN=CLU PE=1 SV=1 | 1467 | 20201118_lss_colostrum_1ng_glyco_hcd203040_compound15.25966.25966.3 | controllerType=0 controllerNumber=1 scan=25966 | 54.5655 |
| 20404:3 | 9 | K.MLN[+1913.677]TSSLLEQLNEQFNWVSR.L | HexNAc(4)Hex(5)NeuAc(1) | N[+1914] | 1441.6313 | 3 | 4322.8793 | 4322.8637 | 0 | 3.6 | 352 | Specific | 572.66 | 500.34 | 500.34 | 10.09 | 67 | >sp\|P10909\|CLUS_HUMAN Clusterin OS=Homo sapiens OX=9606 GN=CLU PE=1 SV=1 | 1467 | 20201118_lss_colostrum_1ng_glyco_hcd203040_compound15.24082.24082.3 | controllerType=0 controllerNumber=1 scan=24082 | 50.4017 |
| 22152:4 | 9 | K.MLN[+2350.830]TSSLLEQLNEQFNWVSR.L | HexNAc(4)Hex(5)Fuc(1)NeuAc(2) | N[+2351] | 1190.7655 | 4 | 4760.0404 | 4760.0171 | 0 | 4.9 | 352 | Specific | 611.08 | 484.04 | 484.04 | 12.22 | 67 | >sp\|P10909\|CLUS_HUMAN Clusterin OS=Homo sapiens OX=9606 GN=CLU PE=1 SV=1 | 1467 | 20201118_lss_colostrum_1ng_glyco_hcd203040_compound15.25908.25908.3 | controllerType=0 controllerNumber=1 scan=25908 | 54.4499 |
| 20479:3 | 9 | K.MLN[+1710.598]TSSLLEQLNEQFNWVSR.L | HexNAc(3)Hex(5)NeuAc(1) | N[+1711] | 1373.9385 | 3 | 4119.801 | 4119.7844 | 0 | 4 | 352 | Specific | 660.15 | 611.53 | 278.77 | 13.43 | 67 | >sp\|P10909\|CLUS_HUMAN Clusterin OS=Homo sapiens OX=9606 GN=CLU PE=1 SV=1 | 1467 | 20201118_lss_colostrum_1ng_glyco_hcd203040_compound15.24160.24160.3 | controllerType=0 controllerNumber=1 scan=24160 | 50.5706 |
| 20481:3 | 9 | K.MLN[+1710.598]TSSLLEQLNEQFNWVSR.L | HexNAc(3)Hex(5)NeuAc(1) | N[+1711] | 1373.9385 | 3 | 4119.801 | 4119.7844 | 0 | 4 | 352 | Specific | 716.59 | 478.54 | 342.79 | 14.43 | 67 | >sp\|P10909\|CLUS_HUMAN Clusterin OS=Homo sapiens OX=9606 GN=CLU PE=1 SV=1 | 1467 | 20201118_lss_colostrum_1ng_glyco_hcd203040_compound15.24162.24162.3 | controllerType=0 controllerNumber=1 scan=24162 | 50.5786 |
| 22493:3 | 9 | K.MLN[+2204.772]TSSLLEQLNEQFNWVSR.L | HexNAc(4)Hex(5)NeuAc(2) | N[+2205] | 1538.6663 | 3 | 4613.9844 | 4613.9592 | 0 | 5.5 | 352 | Specific | 736.28 | 596.66 | 596.66 | 13.97 | 67 | >sp\|P10909\|CLUS_HUMAN Clusterin OS=Homo sapiens OX=9606 GN=CLU PE=1 SV=1 | 1467 | 20201118_lss_colostrum_1ng_glyco_hcd203040_compound15.26263.26263.3 | controllerType=0 controllerNumber=1 scan=26263 | 55.1865 |
| 20430:3 | 9 | K.MLN[+1872.651]TSSLLEQLNEQFNWVSR.L | HexNAc(3)Hex(6)NeuAc(1) | N[+1873] | 1427.9573 | 3 | 4281.8574 | 4281.8372 | 0 | 4.7 | 352 | Specific | 760.1 | 663.4 | 279.08 | 14.97 | 67 | >sp\|P10909\|CLUS_HUMAN Clusterin OS=Homo sapiens OX=9606 GN=CLU PE=1 SV=1 | 1467 | 20201118_lss_colostrum_1ng_glyco_hcd203040_compound15.24109.24109.3 | controllerType=0 controllerNumber=1 scan=24109 | 50.4571 |
| 22495:3 | 9 | K.MLN[+2204.772]TSSLLEQLNEQFNWVSR.L | HexNAc(4)Hex(5)NeuAc(2) | N[+2205] | 1538.6663 | 3 | 4613.9844 | 4613.9592 | 0 | 5.5 | 352 | Specific | 763.83 | 586.8 | 586.8 | 14.97 | 67 | >sp\|P10909\|CLUS_HUMAN Clusterin OS=Homo sapiens OX=9606 GN=CLU PE=1 SV=1 | 1467 | 20201118_lss_colostrum_1ng_glyco_hcd203040_compound15.26265.26265.3 | controllerType=0 controllerNumber=1 scan=26265 | 55.191 |
| 22208:4 | 9 | K.MLN[+2350.830]TSSLLEQLNEQFNWVSR.L | HexNAc(4)Hex(5)Fuc(1)NeuAc(2) | N[+2351] | 1190.7652 | 4 | 4760.0389 | 4760.0171 | 0 | 4.6 | 352 | Specific | 850.12 | 603.37 | 603.37 | 17.22 | 67 | >sp\|P10909\|CLUS_HUMAN Clusterin OS=Homo sapiens OX=9606 GN=CLU PE=1 SV=1 | 1467 | 20201118_lss_colostrum_1ng_glyco_hcd203040_compound15.25967.25967.4 | controllerType=0 controllerNumber=1 scan=25967 | 54.5679 |
| 22209:3 | 9 | K.MLN[+2350.830]TSSLLEQLNEQFNWVSR.L | HexNAc(4)Hex(5)Fuc(1)NeuAc(2) | N[+2351] | 1587.3504 | 3 | 4760.0366 | 4760.0171 | 0 | 4.1 | 352 | Specific | 856.13 | 612.87 | 612.87 | 17.43 | 67 | >sp\|P10909\|CLUS_HUMAN Clusterin OS=Homo sapiens OX=9606 GN=CLU PE=1 SV=1 | 1467 | 20201118_lss_colostrum_1ng_glyco_hcd203040_compound15.25968.25968.3 | controllerType=0 controllerNumber=1 scan=25968 | 54.5698 |
| 20115:3 | 9 | K.MLN[+2351.851]TSSLLEQLNEQFNWVSR.L | HexNAc(4)Hex(5)Fuc(3)NeuAc(1) | N[+2352] | 1587.6883 | 3 | 4761.0503 | 4761.0375 | 0 | 2.7 | 352 | Specific | 880.93 | 739.17 | 739.17 | 17.63 | 67 | >sp\|P10909\|CLUS_HUMAN Clusterin OS=Homo sapiens OX=9606 GN=CLU PE=1 SV=1 | 1467 | 20201118_lss_colostrum_1ng_glyco_hcd203040_compound15.23779.23779.3 | controllerType=0 controllerNumber=1 scan=23779 | 49.6669 |
| 20118:3 | 9 | K.MLN[+2351.851]TSSLLEQLNEQFNWVSR.L | HexNAc(4)Hex(5)Fuc(3)NeuAc(1) | N[+2352] | 1587.6883 | 3 | 4761.0503 | 4761.0375 | 0 | 2.7 | 352 | Specific | 908.16 | 727.12 | 727.12 | 18.63 | 67 | >sp\|P10909\|CLUS_HUMAN Clusterin OS=Homo sapiens OX=9606 GN=CLU PE=1 SV=1 | 1467 | 20201118_lss_colostrum_1ng_glyco_hcd203040_compound15.23782.23782.3 | controllerType=0 controllerNumber=1 scan=23782 | 49.6722 |
| 20167:3 | 9 | K.MLN[+2059.735]TSSLLEQLNEQFNWVSR.L | HexNAc(4)Hex(5)Fuc(1)NeuAc(1) | N[+2060] | 1490.3185 | 3 | 4468.9411 | 4468.9217 | 0 | 4.3 | 352 | Specific | 910.92 | 697.69 | 697.69 | 18.43 | 67 | >sp\|P10909\|CLUS_HUMAN Clusterin OS=Homo sapiens OX=9606 GN=CLU PE=1 SV=1 | 1467 | 20201118_lss_colostrum_1ng_glyco_hcd203040_compound15.23834.23834.3 | controllerType=0 controllerNumber=1 scan=23834 | 49.8043 |
| 20099:3 | 9 | K.MLN[+2351.851]TSSLLEQLNEQFNWVSR.L | HexNAc(4)Hex(5)Fuc(3)NeuAc(1) | N[+2352] | 1587.6927 | 3 | 4761.0635 | 4761.0375 | 0 | 5.5 | 352 | Specific | 914.04 | 691.56 | 691.56 | 17.97 | 67 | >sp\|P10909\|CLUS_HUMAN Clusterin OS=Homo sapiens OX=9606 GN=CLU PE=1 SV=1 | 1467 | 20201118_lss_colostrum_1ng_glyco_hcd203040_compound15.23762.23762.3 | controllerType=0 controllerNumber=1 scan=23762 | 49.6202 |
| 20174:4 | 9 | K.MLN[+2059.735]TSSLLEQLNEQFNWVSR.L | HexNAc(4)Hex(5)Fuc(1)NeuAc(1) | N[+2060] | 1117.9903 | 4 | 4468.9393 | 4468.9217 | 0 | 3.9 | 352 | Specific | 945.36 | 693.51 | 693.51 | 18.67 | 67 | >sp\|P10909\|CLUS_HUMAN Clusterin OS=Homo sapiens OX=9606 GN=CLU PE=1 SV=1 | 1467 | 20201118_lss_colostrum_1ng_glyco_hcd203040_compound15.23841.23841.4 | controllerType=0 controllerNumber=1 scan=23841 | 49.8156 |
| 20169:3 | 9 | K.MLN[+2059.735]TSSLLEQLNEQFNWVSR.L | HexNAc(4)Hex(5)Fuc(1)NeuAc(1) | N[+2060] | 1490.3185 | 3 | 4468.9411 | 4468.9217 | 0 | 4.3 | 352 | Specific | 946.11 | 683.34 | 683.34 | 18.43 | 67 | >sp\|P10909\|CLUS_HUMAN Clusterin OS=Homo sapiens OX=9606 GN=CLU PE=1 SV=1 | 1467 | 20201118_lss_colostrum_1ng_glyco_hcd203040_compound15.23836.23836.3 | controllerType=0 controllerNumber=1 scan=23836 | 49.8089 |
| 20136:3 | 9 | K.MLN[+2205.793]TSSLLEQLNEQFNWVSR.L | HexNAc(4)Hex(5)Fuc(2)NeuAc(1) | N[+2206] | 1539.004 | 3 | 4614.9973 | 4614.9796 | 0 | 3.9 | 352 | Specific | 970.88 | 655.87 | 655.87 | 19.43 | 67 | >sp\|P10909\|CLUS_HUMAN Clusterin OS=Homo sapiens OX=9606 GN=CLU PE=1 SV=1 | 1467 | 20201118_lss_colostrum_1ng_glyco_hcd203040_compound15.23801.23801.3 | controllerType=0 controllerNumber=1 scan=23801 | 49.7304 |
| 20406:3 | 9 | K.MLN[+1913.677]TSSLLEQLNEQFNWVSR.L | HexNAc(4)Hex(5)NeuAc(1) | N[+1914] | 1441.6313 | 3 | 4322.8793 | 4322.8637 | 0 | 3.6 | 352 | Specific | 971.53 | 708.42 | 708.42 | 19.43 | 67 | >sp\|P10909\|CLUS_HUMAN Clusterin OS=Homo sapiens OX=9606 GN=CLU PE=1 SV=1 | 1467 | 20201118_lss_colostrum_1ng_glyco_hcd203040_compound15.24084.24084.3 | controllerType=0 controllerNumber=1 scan=24084 | 50.4056 |
| 20131:4 | 9 | K.MLN[+2205.793]TSSLLEQLNEQFNWVSR.L | HexNAc(4)Hex(5)Fuc(2)NeuAc(1) | N[+2206] | 1154.5039 | 4 | 4614.994 | 4614.9796 | 0 | 3.1 | 352 | Specific | 980.32 | 753.3 | 753.3 | 19.87 | 67 | >sp\|P10909\|CLUS_HUMAN Clusterin OS=Homo sapiens OX=9606 GN=CLU PE=1 SV=1 | 1467 | 20201118_lss_colostrum_1ng_glyco_hcd203040_compound15.23796.23796.4 | controllerType=0 controllerNumber=1 scan=23796 | 49.7164 |
| 20432:3 | 9 | K.MLN[+1872.651]TSSLLEQLNEQFNWVSR.L | HexNAc(3)Hex(6)NeuAc(1) | N[+1873] | 1427.9573 | 3 | 4281.8574 | 4281.8372 | 0 | 4.7 | 352 | Specific | 995.2 | 809.79 | 481.75 | 18.97 | 67 | >sp\|P10909\|CLUS_HUMAN Clusterin OS=Homo sapiens OX=9606 GN=CLU PE=1 SV=1 | 1467 | 20201118_lss_colostrum_1ng_glyco_hcd203040_compound15.24111.24111.3 | controllerType=0 controllerNumber=1 scan=24111 | 50.4631 |
| 20171:4 | 9 | K.MLN[+2059.735]TSSLLEQLNEQFNWVSR.L | HexNAc(4)Hex(5)Fuc(1)NeuAc(1) | N[+2060] | 1117.9903 | 4 | 4468.9393 | 4468.9217 | 0 | 3.9 | 352 | Specific | 1033.38 | 774.07 | 774.07 | 20.67 | 67 | >sp\|P10909\|CLUS_HUMAN Clusterin OS=Homo sapiens OX=9606 GN=CLU PE=1 SV=1 | 1467 | 20201118_lss_colostrum_1ng_glyco_hcd203040_compound15.23838.23838.4 | controllerType=0 controllerNumber=1 scan=23838 | 49.8108 |
| 20133:3 | 9 | K.MLN[+2205.793]TSSLLEQLNEQFNWVSR.L | HexNAc(4)Hex(5)Fuc(2)NeuAc(1) | N[+2206] | 1539.004 | 3 | 4614.9973 | 4614.9796 | 0 | 3.9 | 352 | Specific | 1048.91 | 761.71 | 761.71 | 20.43 | 67 | >sp\|P10909\|CLUS_HUMAN Clusterin OS=Homo sapiens OX=9606 GN=CLU PE=1 SV=1 | 1467 | 20201118_lss_colostrum_1ng_glyco_hcd203040_compound15.23798.23798.3 | controllerType=0 controllerNumber=1 scan=23798 | 49.7187 |
| 20129:4 | 9 | K.MLN[+2205.793]TSSLLEQLNEQFNWVSR.L | HexNAc(4)Hex(5)Fuc(2)NeuAc(1) | N[+2206] | 1154.5039 | 4 | 4614.994 | 4614.9796 | 0 | 3.1 | 352 | Specific | 1057.05 | 818.14 | 818.14 | 21.87 | 67 | >sp\|P10909\|CLUS_HUMAN Clusterin OS=Homo sapiens OX=9606 GN=CLU PE=1 SV=1 | 1467 | 20201118_lss_colostrum_1ng_glyco_hcd203040_compound15.23794.23794.4 | controllerType=0 controllerNumber=1 scan=23794 | 49.7053 |
| 20096:3 | 9 | K.MLN[+2351.851]TSSLLEQLNEQFNWVSR.L | HexNAc(4)Hex(5)Fuc(3)NeuAc(1) | N[+2352] | 1587.6927 | 3 | 4761.0635 | 4761.0375 | 0 | 5.5 | 352 | Specific | 1089.9 | 849.94 | 849.94 | 20.97 | 67 | >sp\|P10909\|CLUS_HUMAN Clusterin OS=Homo sapiens OX=9606 GN=CLU PE=1 SV=1 | 1467 | 20201118_lss_colostrum_1ng_glyco_hcd203040_compound15.23759.23759.3 | controllerType=0 controllerNumber=1 scan=23759 | 49.6118 |
| 19727:4 | 9 | K.M[+15.995]LN[+2059.735]TSSLLEQLNEQFNWVSR.L | HexNAc(4)Hex(5)Fuc(1)NeuAc(1) | M[+16], N[+2060] | 1121.9906 | 4 | 4484.9407 | 4484.9166 | 0 | 5.4 | 352 | Specific | 607.4 | 514.63 | 514.63 | 12.22 | 67 | >sp\|P10909\|CLUS_HUMAN Clusterin OS=Homo sapiens OX=9606 GN=CLU PE=1 SV=1 | 1467 | 20201118_lss_colostrum_1ng_glyco_hcd203040_compound15.23374.23374.4 | controllerType=0 controllerNumber=1 scan=23374 | 48.7362 |
| 19706:3 | 9 | K.M[+15.995]LN[+2059.735]TSSLLEQLNEQFNWVSR.L | HexNAc(4)Hex(5)Fuc(1)NeuAc(1) | M[+16], N[+2060] | 1495.648 | 3 | 4484.9295 | 4484.9166 | 0 | 2.9 | 352 | Specific | 798.97 | 614.22 | 614.22 | 15.63 | 67 | >sp\|P10909\|CLUS_HUMAN Clusterin OS=Homo sapiens OX=9606 GN=CLU PE=1 SV=1 | 1467 | 20201118_lss_colostrum_1ng_glyco_hcd203040_compound15.23352.23352.3 | controllerType=0 controllerNumber=1 scan=23352 | 48.692 |
| 19721:3 | 9 | K.M[+15.995]LN[+2059.735]TSSLLEQLNEQFNWVSR.L | HexNAc(4)Hex(5)Fuc(1)NeuAc(1) | M[+16], N[+2060] | 1495.6502 | 3 | 4484.9361 | 4484.9166 | 0 | 4.3 | 352 | Specific | 805.84 | 630.33 | 630.33 | 16.43 | 67 | >sp\|P10909\|CLUS_HUMAN Clusterin OS=Homo sapiens OX=9606 GN=CLU PE=1 SV=1 | 1467 | 20201118_lss_colostrum_1ng_glyco_hcd203040_compound15.23368.23368.3 | controllerType=0 controllerNumber=1 scan=23368 | 48.7247 |
| 19724:3 | 9 | K.M[+15.995]LN[+2059.735]TSSLLEQLNEQFNWVSR.L | HexNAc(4)Hex(5)Fuc(1)NeuAc(1) | M[+16], N[+2060] | 1495.6502 | 3 | 4484.9361 | 4484.9166 | 0 | 4.3 | 352 | Specific | 957.3 | 671.68 | 671.68 | 19.43 | 67 | >sp\|P10909\|CLUS_HUMAN Clusterin OS=Homo sapiens OX=9606 GN=CLU PE=1 SV=1 | 1467 | 20201118_lss_colostrum_1ng_glyco_hcd203040_compound15.23371.23371.3 | controllerType=0 controllerNumber=1 scan=23371 | 48.7312 |
| 19709:3 | 9 | K.M[+15.995]LN[+2059.735]TSSLLEQLNEQFNWVSR.L | HexNAc(4)Hex(5)Fuc(1)NeuAc(1) | M[+16], N[+2060] | 1495.648 | 3 | 4484.9295 | 4484.9166 | 0 | 2.9 | 352 | Specific | 959.05 | 654.92 | 654.92 | 19.63 | 67 | >sp\|P10909\|CLUS_HUMAN Clusterin OS=Homo sapiens OX=9606 GN=CLU PE=1 SV=1 | 1467 | 20201118_lss_colostrum_1ng_glyco_hcd203040_compound15.23355.23355.3 | controllerType=0 controllerNumber=1 scan=23355 | 48.6986 |
| 10056:3 | 9 | R.LAN[+2351.851]LTQGEDQYYLR.V | HexNAc(4)Hex(5)Fuc(3)NeuAc(1) | N[+2352] | 1345.9018 | 3 | 4035.6908 | 4035.6844 | 0 | 1.6 | 372 | Specific | 264.02 | 264.02 | 264.02 | 6.27 | 67 | >sp\|P10909\|CLUS_HUMAN Clusterin OS=Homo sapiens OX=9606 GN=CLU PE=1 SV=1 | 1467 | 20201118_lss_colostrum_1ng_glyco_hcd203040_compound15.13315.13315.3 | controllerType=0 controllerNumber=1 scan=13315 | 28.7018 |
| 09687:3 | 9 | R.LAN[+2862.021]LTQGEDQYYLR.V | HexNAc(5)Hex(6)Fuc(2)NeuAc(2) | N[+2862] | 1515.9705 | 3 | 4545.897 | 4545.8541 | 0 | 9.4 | 372 | Specific | 294.79 | 286.85 | 286.85 | 5.15 | 67 | >sp\|P10909\|CLUS_HUMAN Clusterin OS=Homo sapiens OX=9606 GN=CLU PE=1 SV=1 | 1467 | 20201118_lss_colostrum_1ng_glyco_hcd203040_compound15.12932.12932.3 | controllerType=0 controllerNumber=1 scan=12932 | 27.9825 |
| 09685:3 | 9 | R.LAN[+2862.021]LTQGEDQYYLR.V | HexNAc(5)Hex(6)Fuc(2)NeuAc(2) | N[+2862] | 1515.9705 | 3 | 4545.897 | 4545.8541 | 0 | 9.4 | 372 | Specific | 340.19 | 340.19 | 340.19 | 5.68 | 67 | >sp\|P10909\|CLUS_HUMAN Clusterin OS=Homo sapiens OX=9606 GN=CLU PE=1 SV=1 | 1467 | 20201118_lss_colostrum_1ng_glyco_hcd203040_compound15.12930.12930.3 | controllerType=0 controllerNumber=1 scan=12930 | 27.9785 |
| 10058:3 | 9 | R.LAN[+2351.851]LTQGEDQYYLR.V | HexNAc(4)Hex(5)Fuc(3)NeuAc(1) | N[+2352] | 1345.9018 | 3 | 4035.6908 | 4035.6844 | 0 | 1.6 | 372 | Specific | 362 | 322.37 | 322.37 | 7.05 | 67 | >sp\|P10909\|CLUS_HUMAN Clusterin OS=Homo sapiens OX=9606 GN=CLU PE=1 SV=1 | 1467 | 20201118_lss_colostrum_1ng_glyco_hcd203040_compound15.13317.13317.3 | controllerType=0 controllerNumber=1 scan=13317 | 28.7059 |
| 09913:3 | 9 | R.LAN[+2570.925]LTQGEDQYYLR.V | HexNAc(5)Hex(6)Fuc(2)NeuAc(1) | N[+2571] | 1418.9299 | 3 | 4254.775 | 4254.7587 | 0 | 3.8 | 372 | Specific | 390.72 | 360.92 | 360.92 | 7.49 | 67 | >sp\|P10909\|CLUS_HUMAN Clusterin OS=Homo sapiens OX=9606 GN=CLU PE=1 SV=1 | 1467 | 20201118_lss_colostrum_1ng_glyco_hcd203040_compound15.13167.13167.3 | controllerType=0 controllerNumber=1 scan=13167 | 28.4246 |
| 09848:3 | 9 | R.LAN[+2716.983]LTQGEDQYYLR.V | HexNAc(5)Hex(6)Fuc(3)NeuAc(1) | N[+2717] | 1467.6185 | 3 | 4400.8408 | 4400.8166 | 0 | 5.5 | 372 | Specific | 457.3 | 373.02 | 373.02 | 7.27 | 67 | >sp\|P10909\|CLUS_HUMAN Clusterin OS=Homo sapiens OX=9606 GN=CLU PE=1 SV=1 | 1467 | 20201118_lss_colostrum_1ng_glyco_hcd203040_compound15.13099.13099.3 | controllerType=0 controllerNumber=1 scan=13099 | 28.2845 |
| 10178:3 | 9 | R.LAN[+2059.735]LTQGEDQYYLR.V | HexNAc(4)Hex(5)Fuc(1)NeuAc(1) | N[+2060] | 1248.5284 | 3 | 3743.5706 | 3743.5686 | 0 | 0.5 | 372 | Specific | 514.42 | 514.42 | 514.42 | 9.11 | 67 | >sp\|P10909\|CLUS_HUMAN Clusterin OS=Homo sapiens OX=9606 GN=CLU PE=1 SV=1 | 1467 | 20201118_lss_colostrum_1ng_glyco_hcd203040_compound15.13442.13442.3 | controllerType=0 controllerNumber=1 scan=13442 | 28.9462 |
| 09916:3 | 9 | R.LAN[+2570.925]LTQGEDQYYLR.V | HexNAc(5)Hex(6)Fuc(2)NeuAc(1) | N[+2571] | 1418.9299 | 3 | 4254.775 | 4254.7587 | 0 | 3.8 | 372 | Specific | 542.24 | 458.29 | 458.29 | 9.55 | 67 | >sp\|P10909\|CLUS_HUMAN Clusterin OS=Homo sapiens OX=9606 GN=CLU PE=1 SV=1 | 1467 | 20201118_lss_colostrum_1ng_glyco_hcd203040_compound15.13170.13170.3 | controllerType=0 controllerNumber=1 scan=13170 | 28.4294 |
| 10180:3 | 9 | R.LAN[+2059.735]LTQGEDQYYLR.V | HexNAc(4)Hex(5)Fuc(1)NeuAc(1) | N[+2060] | 1248.5284 | 3 | 3743.5706 | 3743.5686 | 0 | 0.5 | 372 | Specific | 675.27 | 632.07 | 632.07 | 12.72 | 67 | >sp\|P10909\|CLUS_HUMAN Clusterin OS=Homo sapiens OX=9606 GN=CLU PE=1 SV=1 | 1467 | 20201118_lss_colostrum_1ng_glyco_hcd203040_compound15.13444.13444.3 | controllerType=0 controllerNumber=1 scan=13444 | 28.95 |
| 15172:3 | 10 | R.QQQHLFGSN[+2204.772]VTDC[+57.021]SGNFC[+57.021]LFR.S | HexNAc(4)Hex(5)NeuAc(2) | C[+57]*2, N[+2205] | 1573.978 | 3 | 4719.9193 | 4719.8966 | 0 | 4.8 | 622 | Specific | 412.1 | 412.1 | 412.1 | 8.95 | 48 | >sp\|P02787\|TRFE_HUMAN Serotransferrin OS=Homo sapiens OX=9606 GN=TF PE=1 SV=3 | 1111 | 20201118_lss_colostrum_1ng_glyco_hcd203040_compound15.18627.18627.3 | controllerType=0 controllerNumber=1 scan=18627 | 38.7298 |
| 15175:3 | 10 | R.QQQHLFGSN[+2204.772]VTDC[+57.021]SGNFC[+57.021]LFR.S | HexNAc(4)Hex(5)NeuAc(2) | C[+57]*2, N[+2205] | 1573.978 | 3 | 4719.9193 | 4719.8966 | 0 | 4.8 | 622 | Specific | 595.06 | 506.37 | 506.37 | 11.04 | 48 | >sp\|P02787\|TRFE_HUMAN Serotransferrin OS=Homo sapiens OX=9606 GN=TF PE=1 SV=3 | 1111 | 20201118_lss_colostrum_1ng_glyco_hcd203040_compound15.18630.18630.3 | controllerType=0 controllerNumber=1 scan=18630 | 38.7375 |
| 22728:4 | 11 | A.[+42.011]SPTSPKVFPLSLDSTPQDGNVVVAC[+57.021]LVQGFFPQEPLSVTWSESGQN | HexNAc(4)Hex(4) | .[42], C[+57], N[+1461] | 1719.7928 | 4 | 6876.1494 | 6876.213 | 0 | -9.2 | 2 | NRagged | 234.94 | 227.04 | 227.04 | 3.31 | 61 | >sp\|P01877\|IGHA2_HUMAN Immunoglobulin heavy constant alpha 2 OS=Homo sapiens OX=9606 GN=IGHA2 PE=1 SV=4 | 576 | 20201118_lss_colostrum_1ng_glyco_hcd203040_compound15.26509.26509.4 | controllerType=0 controllerNumber=1 scan=26509 | 55.7219 |
| 22534:4 | 11 | A.SPTSPKVFPLSLDSTPQDGNVVVAC[+57.021]LVQGFFPQEPLSVTWSESGQN[+1460.5 | HexNAc(4)Hex(4) | C[+57], N[+1461] | 1709.2916 | 4 | 6834.1445 | 6834.2024 | 0 | -8.5 | 2 | NRagged | 237.93 | 211.72 | 211.72 | 3.5 | 61 | >sp\|P01877\|IGHA2_HUMAN Immunoglobulin heavy constant alpha 2 OS=Homo sapiens OX=9606 GN=IGHA2 PE=1 SV=4 | 576 | 20201118_lss_colostrum_1ng_glyco_hcd203040_compound15.26306.26306.4 | controllerType=0 controllerNumber=1 scan=26306 | 55.2813 |
| 22536:4 | 11 | A.SPTSPKVFPLSLDSTPQDGNVVVAC[+57.021]LVQGFFPQEPLSVTWSESGQN[+1460.5 | HexNAc(4)Hex(4) | C[+57], N[+1461] | 1709.2916 | 4 | 6834.1445 | 6834.2024 | 0 | -8.5 | 2 | NRagged | 285.73 | 285.73 | 285.73 | 3.54 | 61 | >sp\|P01877\|IGHA2_HUMAN Immunoglobulin heavy constant alpha 2 OS=Homo sapiens OX=9606 GN=IGHA2 PE=1 SV=4 | 576 | 20201118_lss_colostrum_1ng_glyco_hcd203040_compound15.26308.26308.4 | controllerType=0 controllerNumber=1 scan=26308 | 55.2858 |
| 24822:4 | 11 | K.VFPLSLDSTPQDGNVVVAC[+57.021]LVQGFFPQEPLSVTWSESGQN[+2553.910]VTAI | HexNAc(5)Hex(5)Fuc(1)NeuAc(2) | C[+57], N[+2554] | 1833.3283 | 4 | 7330.2915 | 7330.2711 | 0 | 2.8 | 8 | Specific | 231.63 | 231.63 | 231.63 | 7.37 | 61 | >sp\|P01877\|IGHA2_HUMAN Immunoglobulin heavy constant alpha 2 OS=Homo sapiens OX=9606 GN=IGHA2 PE=1 SV=4 | 576 | 20201118_lss_colostrum_1ng_glyco_hcd203040_compound15.28686.28686.4 | controllerType=0 controllerNumber=1 scan=28686 | 59.9298 |
| 24828:5 | 11 | K.VFPLSLDSTPQDGNVVVAC[+57.021]LVQGFFPQEPLSVTWSESGQN[+2351.851]VTAI | HexNAc(4)Hex(5)Fuc(3)NeuAc(1) | C[+57], N[+2352] | 1426.4496 | 5 | 7128.219 | 7128.2121 | 0 | 1 | 8 | Specific | 268.6 | 268.6 | 268.6 | 7.61 | 61 | >sp\|P01877\|IGHA2_HUMAN Immunoglobulin heavy constant alpha 2 OS=Homo sapiens OX=9606 GN=IGHA2 PE=1 SV=4 | 576 | 20201118_lss_colostrum_1ng_glyco_hcd203040_compound15.28692.28692.5 | controllerType=0 controllerNumber=1 scan=28692 | 59.9419 |
| 22470:5 | 11 | K.[+42.011]VFPLSLDSTPQDGNVVVAC[+57.021]LVQGFFPQEPLSVTWSESGQN[+2059 | HexNAc(4)Hex(5)Fuc(1)NeuAc(1) | .[42], C[+57], N[+2060] | 1376.4296 | 5 | 6878.1189 | 6878.1069 | 0 | 1.7 | 8 | Specific | 274.62 | 274.62 | 274.62 | 6.33 | 61 | >sp\|P01877\|IGHA2_HUMAN Immunoglobulin heavy constant alpha 2 OS=Homo sapiens OX=9606 GN=IGHA2 PE=1 SV=4 | 576 | 20201118_lss_colostrum_1ng_glyco_hcd203040_compound15.26239.26239.5 | controllerType=0 controllerNumber=1 scan=26239 | 55.1359 |
| 21137:4 | 11 | K.VFPLSLDSTPQDGNVVVAC[+57.021]LVQGFFPQEPLSVTWSESGQN[+1971.719]VTAI | HexNAc(5)Hex(5)Fuc(1) | C[+57], N[+1972] | 1687.7804 | 4 | 6748.0996 | 6748.0803 | 0 | 2.9 | 8 | Specific | 284.86 | 284.86 | 284.86 | 7.77 | 61 | >sp\|P01877\|IGHA2_HUMAN Immunoglobulin heavy constant alpha 2 OS=Homo sapiens OX=9606 GN=IGHA2 PE=1 SV=4 | 576 | 20201118_lss_colostrum_1ng_glyco_hcd203040_compound15.24848.24848.2 | controllerType=0 controllerNumber=1 scan=24848 | 52.1324 |
| 22471:5 | 11 | K.[+42.011]VFPLSLDSTPQDGNVVVAC[+57.021]LVQGFFPQEPLSVTWSESGQN[+2059 | HexNAc(4)Hex(5)Fuc(1)NeuAc(1) | .[42], C[+57], N[+2060] | 1376.4296 | 5 | 6878.1189 | 6878.1069 | 0 | 1.7 | 8 | Specific | 306.33 | 306.33 | 306.33 | 6.58 | 61 | >sp\|P01877\|IGHA2_HUMAN Immunoglobulin heavy constant alpha 2 OS=Homo sapiens OX=9606 GN=IGHA2 PE=1 SV=4 | 576 | 20201118_lss_colostrum_1ng_glyco_hcd203040_compound15.26240.26240.5 | controllerType=0 controllerNumber=1 scan=26240 | 55.1406 |
| 22710:4 | 11 | K.VFPLSLDSTPQDGNVVVAC[+57.021]LVQGFFPQEPLSVTWSESGQN[+2262.814]VTAI | HexNAc(5)Hex(5)Fuc(1)NeuAc(1) | C[+57], N[+2263] | 1760.5556 | 4 | 7039.2007 | 7039.1757 | 0 | 3.5 | 8 | Specific | 314.99 | 314.99 | 314.99 | 8.04 | 61 | >sp\|P01877\|IGHA2_HUMAN Immunoglobulin heavy constant alpha 2 OS=Homo sapiens OX=9606 GN=IGHA2 PE=1 SV=4 | 576 | 20201118_lss_colostrum_1ng_glyco_hcd203040_compound15.26490.26490.4 | controllerType=0 controllerNumber=1 scan=26490 | 55.6848 |
| 22538:5 | 11 | K.VFPLSLDSTPQDGNVVVAC[+57.021]LVQGFFPQEPLSVTWSESGQN[+2059.735]VTAI | HexNAc(4)Hex(5)Fuc(1)NeuAc(1) | C[+57], N[+2060] | 1368.0331 | 5 | 6836.1365 | 6836.0963 | 0 | 5.9 | 8 | Specific | 326.67 | 326.67 | 326.67 | 7.19 | 61 | >sp\|P01877\|IGHA2_HUMAN Immunoglobulin heavy constant alpha 2 OS=Homo sapiens OX=9606 GN=IGHA2 PE=1 SV=4 | 576 | 20201118_lss_colostrum_1ng_glyco_hcd203040_compound15.26310.26310.5 | controllerType=0 controllerNumber=1 scan=26310 | 55.2892 |
| 22635:4 | 11 | K.VFPLSLDSTPQDGNVVVAC[+57.021]LVQGFFPQEPLSVTWSESGQN[+2279.830]VTAI | HexNAc(5)Hex(6)Fuc(2) | C[+57], N[+2280] | 1764.8137 | 4 | 7056.2329 | 7056.191 | 0 | 5.9 | 8 | Specific | 345.08 | 307.37 | 307.37 | 7.24 | 61 | >sp\|P01877\|IGHA2_HUMAN Immunoglobulin heavy constant alpha 2 OS=Homo sapiens OX=9606 GN=IGHA2 PE=1 SV=4 | 576 | 20201118_lss_colostrum_1ng_glyco_hcd203040_compound15.26412.26412.4 | controllerType=0 controllerNumber=1 scan=26412 | 55.5205 |
| 21087:6 | 11 | K.VFPLSLDSTPQDGNVVVAC[+57.021]LVQGFFPQEPLSVTWSESGQN[+1647.613]VTAI | HexNAc(5)Hex(3)Fuc(1) | C[+57], N[+1648] | 1071.5053 | 6 | 6423.9954 | 6423.9746 | 0 | 3.2 | 8 | Specific | 380.77 | 380.77 | 380.77 | 10.26 | 61 | >sp\|P01877\|IGHA2_HUMAN Immunoglobulin heavy constant alpha 2 OS=Homo sapiens OX=9606 GN=IGHA2 PE=1 SV=4 | 576 | 20201118_lss_colostrum_1ng_glyco_hcd203040_compound15.24796.24796.6 | controllerType=0 controllerNumber=1 scan=24796 | 52.0118 |
| 22546:5 | 11 | K.VFPLSLDSTPQDGNVVVAC[+57.021]LVQGFFPQEPLSVTWSESGQN[+2059.735]VTAI | HexNAc(4)Hex(5)Fuc(1)NeuAc(1) | C[+57], N[+2060] | 1368.027 | 5 | 6836.106 | 6836.0963 | 0 | 1.4 | 8 | Specific | 385.02 | 385.02 | 385.02 | 10.3 | 61 | >sp\|P01877\|IGHA2_HUMAN Immunoglobulin heavy constant alpha 2 OS=Homo sapiens OX=9606 GN=IGHA2 PE=1 SV=4 | 576 | 20201118_lss_colostrum_1ng_glyco_hcd203040_compound15.26319.26319.5 | controllerType=0 controllerNumber=1 scan=26319 | 55.3093 |
| 21052:5 | 11 | K.VFPLSLDSTPQDGNVVVAC[+57.021]LVQGFFPQEPLSVTWSESGQN[+1647.613]VTAI | HexNAc(5)Hex(3)Fuc(1) | C[+57], N[+1648] | 1285.6062 | 5 | 6424.0019 | 6423.9746 | 0 | 4.2 | 8 | Specific | 399.98 | 399.98 | 399.98 | 10.26 | 61 | >sp\|P01877\|IGHA2_HUMAN Immunoglobulin heavy constant alpha 2 OS=Homo sapiens OX=9606 GN=IGHA2 PE=1 SV=4 | 576 | 20201118_lss_colostrum_1ng_glyco_hcd203040_compound15.24760.24760.5 | controllerType=0 controllerNumber=1 scan=24760 | 51.9348 |
| 21139:4 | 11 | K.VFPLSLDSTPQDGNVVVAC[+57.021]LVQGFFPQEPLSVTWSESGQN[+1971.719]VTAI | HexNAc(5)Hex(5)Fuc(1) | C[+57], N[+1972] | 1687.7804 | 4 | 6748.0996 | 6748.0803 | 0 | 2.9 | 8 | Specific | 422.52 | 391.01 | 391.01 | 10.59 | 61 | >sp\|P01877\|IGHA2_HUMAN Immunoglobulin heavy constant alpha 2 OS=Homo sapiens OX=9606 GN=IGHA2 PE=1 SV=4 | 576 | 20201118_lss_colostrum_1ng_glyco_hcd203040_compound15.24850.24850.2 | controllerType=0 controllerNumber=1 scan=24850 | 52.1365 |
| 24824:4 | 11 | K.VFPLSLDSTPQDGNVVVAC[+57.021]LVQGFFPQEPLSVTWSESGQN[+2553.910]VTAI | HexNAc(5)Hex(5)Fuc(1)NeuAc(2) | C[+57], N[+2554] | 1833.3283 | 4 | 7330.2915 | 7330.2711 | 0 | 2.8 | 8 | Specific | 434.01 | 400.2 | 400.2 | 10.59 | 61 | >sp\|P01877\|IGHA2_HUMAN Immunoglobulin heavy constant alpha 2 OS=Homo sapiens OX=9606 GN=IGHA2 PE=1 SV=4 | 576 | 20201118_lss_colostrum_1ng_glyco_hcd203040_compound15.28688.28688.4 | controllerType=0 controllerNumber=1 scan=28688 | [illegible] |
| 24817:5 | 11 | K.VFPLSLDSTPQDGNVVVAC[+57.021]LVQGFFPQEPLSVTWSESGQN[+2553.910]VTAI | HexNAc(5)Hex(5)Fuc(1)NeuAc(2) | C[+57], N[+2554] | 1466.865 | 5 | 7330.2961 | 7330.2711 | 0 | 3.4 | 8 | Specific | 459.58 | 449.69 | 449.69 | 10.5 | 61 | >sp\|P01877\|IGHA2_HUMAN Immunoglobulin heavy constant alpha 2 OS=Homo sapiens OX=9606 GN=IGHA2 PE=1 SV=4 | 576 | 20201118_lss_colostrum_1ng_glyco_hcd203040_compound15.28681.28681.5 | controllerType=0 controllerNumber=1 scan=28681 | 59.9194 |
| 22568:4 | 11 | K.VFPLSLDSTPQDGNVVVAC[+57.021]LVQGFFPQEPLSVTWSESGQN[+2117.777]VTAI | HexNAc(5)Hex(5)Fuc(2) | C[+57], N[+2118] | 1724.3006 | 4 | 6894.1807 | 6894.1382 | 0 | 6.2 | 8 | Specific | 475.13 | 475.13 | 475.13 | 9.73 | 61 | >sp\|P01877\|IGHA2_HUMAN Immunoglobulin heavy constant alpha 2 OS=Homo sapiens OX=9606 GN=IGHA2 PE=1 SV=4 | 576 | 20201118_lss_colostrum_1ng_glyco_hcd203040_compound15.26342.26342.4 | controllerType=0 controllerNumber=1 scan=26342 | [illegible] |
| 22540:5 | 11 | K.VFPLSLDSTPQDGNVVVAC[+57.021]LVQGFFPQEPLSVTWSESGQN[+2059.735]VTAI | HexNAc(4)Hex(5)Fuc(1)NeuAc(1) | C[+57], N[+2060] | 1368.0331 | 5 | 6836.1365 | 6836.0963 | 0 | 5.9 | 8 | Specific | 483.1 | 483.1 | 483.1 | 9.73 | 61 | >sp\|P01877\|IGHA2_HUMAN Immunoglobulin heavy constant alpha 2 OS=Homo sapiens OX=9606 GN=IGHA2 PE=1 SV=4 | 576 | 20201118_lss_colostrum_1ng_glyco_hcd203040_compound15.26312.26312.5 | controllerType=0 controllerNumber=1 scan=26312 | 55.2948 |
| 22569:5 | 11 | K.VFPLSLDSTPQDGNVVVAC[+57.021]LVQGFFPQEPLSVTWSESGQN[+2117.777]VTAI | HexNAc(5)Hex(5)Fuc(2) | C[+57], N[+2118] | 1379.6409 | 5 | 6894.1755 | 6894.1382 | 0 | 5.4 | 8 | Specific | 483.3 | 483.3 | 483.3 | 10.13 | 61 | >sp\|P01877\|IGHA2_HUMAN Immunoglobulin heavy constant alpha 2 OS=Homo sapiens OX=9606 GN=IGHA2 PE=1 SV=4 | 576 | 20201118_lss_colostrum_1ng_glyco_hcd203040_compound15.26343.26343.5 | controllerType=0 controllerNumber=1 scan=26343 | 55.3594 |
| 21089:6 | 11 | K.VFPLSLDSTPQDGNVVVAC[+57.021]LVQGFFPQEPLSVTWSESGQN[+1647.613]VTAI | HexNAc(5)Hex(3)Fuc(1) | C[+57], N[+1648] | 1071.5053 | 6 | 6423.9954 | 6423.9746 | 0 | 3.2 | 8 | Specific | 483.4 | 483.4 | 483.4 | 10.59 | 61 | >sp\|P01877\|IGHA2_HUMAN Immunoglobulin heavy constant alpha 2 OS=Homo sapiens OX=9606 GN=IGHA2 PE=1 SV=4 | 576 | 20201118_lss_colostrum_1ng_glyco_hcd203040_compound15.24798.24798.6 | controllerType=0 controllerNumber=1 scan=24798 | 52.0167 |
| 22548:5 | 11 | K.VFPLSLDSTPQDGNVVVAC[+57.021]LVQGFFPQEPLSVTWSESGQN[+2059.735]VTAI | HexNAc(4)Hex(5)Fuc(1)NeuAc(1) | C[+57], N[+2060] | 1368.027 | 5 | 6836.106 | 6836.0963 | 0 | 1.4 | 8 | Specific | 533.57 | 533.57 | 533.57 | 10.85 | 61 | >sp\|P01877\|IGHA2_HUMAN Immunoglobulin heavy constant alpha 2 OS=Homo sapiens OX=9606 GN=IGHA2 PE=1 SV=4 | 576 | 20201118_lss_colostrum_1ng_glyco_hcd203040_compound15.26321.26321.5 | controllerType=0 controllerNumber=1 scan=26321 | 55.3136 |
| 22392:5 | 11 | K.VFPLSLDSTPQDGNVVVAC[+57.021]LVQGFFPQEPLSVTWSESGQN[+2262.814]VTAI | HexNAc(5)Hex(5)Fuc(1)NeuAc(1) | C[+57], N[+2263] | 1408.6474 | 5 | 7039.2078 | 7039.1757 | 0 | 4.6 | 8 | Specific | 556.65 | 544.85 | 544.85 | 10.61 | 61 | >sp\|P01877\|IGHA2_HUMAN Immunoglobulin heavy constant alpha 2 OS=Homo sapiens OX=9606 GN=IGHA2 PE=1 SV=4 | 576 | 20201118_lss_colostrum_1ng_glyco_hcd203040_compound15.26158.26158.4 | controllerType=0 controllerNumber=1 scan=26158 | 54.9728 |
| 24919:5 | 11 | K.VFPLSLDSTPQDGNVVVAC[+57.021]LVQGFFPQEPLSVTWSESGQN[+2553.910]VTAI | HexNAc(5)Hex(5)Fuc(1)NeuAc(2) | C[+57], N[+2554] | 1466.8661 | 5 | 7330.3016 | 7330.2711 | 0 | 4.2 | 8 | Specific | 586.17 | 586.17 | 586.17 | 12.41 | 61 | >sp\|P01877\|IGHA2_HUMAN Immunoglobulin heavy constant alpha 2 OS=Homo sapiens OX=9606 GN=IGHA2 PE=1 SV=4 | 576 | 20201118_lss_colostrum_1ng_glyco_hcd203040_compound15.28787.28787.5 | controllerType=0 controllerNumber=1 scan=28787 | 60.1211 |
| 20982:4 | 11 | K.VFPLSLDSTPQDGNVVVAC[+57.021]LVQGFFPQEPLSVTWSESGQN[+1809.666]VTAI | HexNAc(5)Hex(4)Fuc(1) | C[+57], N[+1810] | 1647.2675 | 4 | 6586.048 | 6586.0275 | 0 | 3.1 | 8 | Specific | 749.88 | 738.44 | 738.44 | 15.6 | 61 | >sp\|P01877\|IGHA2_HUMAN Immunoglobulin heavy constant alpha 2 OS=Homo sapiens OX=9606 GN=IGHA2 PE=1 SV=4 | 576 | 20201118_lss_colostrum_1ng_glyco_hcd203040_compound15.24687.24687.4 | controllerType=0 controllerNumber=1 scan=24687 | 51.7755 |
| 24921:5 | 11 | K.VFPLSLDSTPQDGNVVVAC[+57.021]LVQGFFPQEPLSVTWSESGQN[+2553.910]VTAI | HexNAc(5)Hex(5)Fuc(1)NeuAc(2) | C[+57], N[+2554] | 1466.8661 | 5 | 7330.3016 | 7330.2711 | 0 | 4.2 | 8 | Specific | 755.04 | 733.42 | 733.42 | 16.41 | 61 | >sp\|P01877\|IGHA2_HUMAN Immunoglobulin heavy constant alpha 2 OS=Homo sapiens OX=9606 GN=IGHA2 PE=1 SV=4 | 576 | 20201118_lss_colostrum_1ng_glyco_hcd203040_compound15.28789.28789.5 | controllerType=0 controllerNumber=1 scan=28789 | 60.1273 |
| 24819:5 | 11 | K.VFPLSLDSTPQDGNVVVAC[+57.021]LVQGFFPQEPLSVTWSESGQN[+2553.910]VTAI | HexNAc(5)Hex(5)Fuc(1)NeuAc(2) | C[+57], N[+2554] | 1466.865 | 5 | 7330.2961 | 7330.2711 | 0 | 3.4 | 8 | Specific | 769.37 | 769.37 | 769.37 | 16.41 | 61 | >sp\|P01877\|IGHA2_HUMAN Immunoglobulin heavy constant alpha 2 OS=Homo sapiens OX=9606 GN=IGHA2 PE=1 SV=4 | 576 | 20201118_lss_colostrum_1ng_glyco_hcd203040_compound15.28683.28683.5 | controllerType=0 controllerNumber=1 scan=28683 | 59.9243 |
| 22394:5 | 11 | K.VFPLSLDSTPQDGNVVVAC[+57.021]LVQGFFPQEPLSVTWSESGQN[+2262.814]VTAI | HexNAc(5)Hex(5)Fuc(1)NeuAc(1) | C[+57], N[+2263] | 1408.6474 | 5 | 7039.2078 | 7039.1757 | 0 | 4.6 | 8 | Specific | 785.01 | 784.22 | 784.22 | 15.95 | 61 | >sp\|P01877\|IGHA2_HUMAN Immunoglobulin heavy constant alpha 2 OS=Homo sapiens OX=9606 GN=IGHA2 PE=1 SV=4 | 576 | 20201118_lss_colostrum_1ng_glyco_hcd203040_compound15.26160.26160.4 | controllerType=0 controllerNumber=1 scan=26160 | 54.9783 |
| 21164:4 | 11 | K.VFPLSLDSTPQDGNVVVAC[+57.021]LVQGFFPQEPLSVTWSESGQN[+1809.666]VTAI | HexNAc(5)Hex(4)Fuc(1) | C[+57], N[+1810] | 1647.27 | 4 | 6586.0583 | 6586.0275 | 0 | 4.7 | 8 | Specific | 806.85 | 723.88 | 723.88 | 16.95 | 61 | >sp\|P01877\|IGHA2_HUMAN Immunoglobulin heavy constant alpha 2 OS=Homo sapiens OX=9606 GN=IGHA2 PE=1 SV=4 | 576 | 20201118_lss_colostrum_1ng_glyco_hcd203040_compound15.24877.24877.4 | controllerType=0 controllerNumber=1 scan=24877 | 52.1934 |
| 20985:4 | 11 | K.VFPLSLDSTPQDGNVVVAC[+57.021]LVQGFFPQEPLSVTWSESGQN[+1809.666]VTAI | HexNAc(5)Hex(4)Fuc(1) | C[+57], N[+1810] | 1647.2675 | 4 | 6586.048 | 6586.0275 | 0 | 3.1 | 8 | Specific | 818.35 | 818.35 | 818.35 | 17.6 | 61 | >sp\|P01877\|IGHA2_HUMAN Immunoglobulin heavy constant alpha 2 OS=Homo sapiens OX=9606 GN=IGHA2 PE=1 SV=4 | 576 | 20201118_lss_colostrum_1ng_glyco_hcd203040_compound15.24690.24690.4 | controllerType=0 controllerNumber=1 scan=24690 | 51.7827 |
| 21166:4 | 11 | K.VFPLSLDSTPQDGNVVVAC[+57.021]LVQGFFPQEPLSVTWSESGQN[+1809.666]VTAI | HexNAc(5)Hex(4)Fuc(1) | C[+57], N[+1810] | 1647.27 | 4 | 6586.0583 | 6586.0275 | 0 | 4.7 | 8 | Specific | 832.92 | 821.59 | 821.59 | 16.95 | 61 | >sp\|P01877\|IGHA2_HUMAN Immunoglobulin heavy constant alpha 2 OS=Homo sapiens OX=9606 GN=IGHA2 PE=1 SV=4 | 576 | 20201118_lss_colostrum_1ng_glyco_hcd203040_compound15.24879.24879.4 | controllerType=0 controllerNumber=1 scan=24879 | 52.2006 |
| 21032:4 | 11 | K.VFPLSLDSTPQDGNVVVAC[+57.021]LVQGFFPQEPLSVTWSESGQN[+1647.613]VTAI | HexNAc(5)Hex(3)Fuc(1) | C[+57], N[+1648] | 1606.7562 | 4 | 6424.0031 | 6423.9746 | 0 | 4.4 | 8 | Specific | 843.47 | 789.52 | 789.52 | 17.41 | 61 | >sp\|P01877\|IGHA2_HUMAN Immunoglobulin heavy constant alpha 2 OS=Homo sapiens OX=9606 GN=IGHA2 PE=1 SV=4 | 576 | 20201118_lss_colostrum_1ng_glyco_hcd203040_compound15.24739.24739.4 | controllerType=0 controllerNumber=1 scan=24739 | 51.8852 |
| 21053:4 | 11 | K.VFPLSLDSTPQDGNVVVAC[+57.021]LVQGFFPQEPLSVTWSESGQN[+1647.613]VTAI | HexNAc(5)Hex(3)Fuc(1) | C[+57], N[+1648] | 1606.7577 | 4 | 6424.0089 | 6423.9746 | 0 | 5.3 | 8 | Specific | 868.09 | 838.69 | 838.69 | 17.95 | 61 | >sp\|P01877\|IGHA2_HUMAN Immunoglobulin heavy constant alpha 2 OS=Homo sapiens OX=9606 GN=IGHA2 PE=1 SV=4 | 576 | 20201118_lss_colostrum_1ng_glyco_hcd203040_compound15.24761.24761.4 | controllerType=0 controllerNumber=1 scan=24761 | 51.9359 |
| 21034:4 | 11 | K.VFPLSLDSTPQDGNVVVAC[+57.021]LVQGFFPQEPLSVTWSESGQN[+1647.613]VTAI | HexNAc(5)Hex(3)Fuc(1) | C[+57], N[+1648] | 1606.7562 | 4 | 6424.0031 | 6423.9746 | 0 | 4.4 | 8 | Specific | 878.86 | 878.86 | 878.86 | 18.41 | 61 | >sp\|P01877\|IGHA2_HUMAN Immunoglobulin heavy constant alpha 2 OS=Homo sapiens OX=9606 GN=IGHA2 PE=1 SV=4 | 576 | 20201118_lss_colostrum_1ng_glyco_hcd203040_compound15.24741.24741.4 | controllerType=0 controllerNumber=1 scan=24741 | 51.8956 |
| 21055:4 | 11 | K.VFPLSLDSTPQDGNVVVAC[+57.021]LVQGFFPQEPLSVTWSESGQN[+1647.613]VTAI | HexNAc(5)Hex(3)Fuc(1) | C[+57], N[+1648] | 1606.7577 | 4 | 6424.0089 | 6423.9746 | 0 | 5.3 | 8 | Specific | 966.32 | 966.32 | 966.32 | 19.95 | 61 | >sp\|P01877\|IGHA2_HUMAN Immunoglobulin heavy constant alpha 2 OS=Homo sapiens OX=9606 GN=IGHA2 PE=1 SV=4 | 576 | 20201118_lss_colostrum_1ng_glyco_hcd203040_compound15.24763.24763.4 | controllerType=0 controllerNumber=1 scan=24763 | 51.9434 |
| 01818:3 | 11 | K.HYTN[+2059.735]SSQDVTVPC[+57.021]R.V | HexNAc(4)Hex(5)Fuc(1)NeuAc(1) | C[+57], N[+2060] | 1241.8377 | 3 | 3723.4986 | 3723.4842 | 0 | 3.8 | 89 | Specific | 212.34 | 140.08 | 140.08 | 3.98 | 61 | >sp\|P01877\|IGHA2_HUMAN Immunoglobulin heavy constant alpha 2 OS=Homo sapiens OX=9606 GN=IGHA2 PE=1 SV=4 | 576 | 20201118_lss_colostrum_1ng_glyco_hcd203040_compound15.4776.4776.3 | controllerType=0 controllerNumber=1 scan=4776 | 14.3582 |
| 01816:3 | 11 | K.HYTN[+2059.735]SSQDVTVPC[+57.021]R.V | HexNAc(4)Hex(5)Fuc(1)NeuAc(1) | C[+57], N[+2060] | 1241.8377 | 3 | 3723.4986 | 3723.4842 | 0 | 3.8 | 89 | Specific | 237.32 | 166.42 | 166.42 | 5.39 | 61 | >sp\|P01877\|IGHA2_HUMAN Immunoglobulin heavy constant alpha 2 OS=Homo sapiens OX=9606 GN=IGHA2 PE=1 SV=4 | 576 | 20201118_lss_colostrum_1ng_glyco_hcd203040_compound15.4774.4774.3 | controllerType=0 controllerNumber=1 scan=4774 | 14.3542 |
| 01042:2 | 11 | K.HYTN[+1647.613]SSQDVTVPC[+57.021]R.V | HexNAc(5)Hex(3)Fuc(1) | C[+57], N[+1648] | 1656.1922 | 2 | 3311.3771 | 3311.3625 | 0 | 4.4 | 89 | Specific | 276.28 | 193.12 | 193.12 | 4.03 | 61 | >sp\|P01877\|IGHA2_HUMAN Immunoglobulin heavy constant alpha 2 OS=Homo sapiens OX=9606 GN=IGHA2 PE=1 SV=4 | 576 | 20201118_lss_colostrum_1ng_glyco_hcd203040_compound15.3971.3971.2 | controllerType=0 controllerNumber=1 scan=3971 | 12.9956 |
| 00922:3 | 11 | K.HYTN[+1606.587]SSQDVTVPC[+57.021]R.V | HexNAc(4)Hex(4)Fuc(1) | C[+57], N[+1607] | 1090.786 | 3 | 3270.3436 | 3270.336 | 0 | 2.3 | 89 | Specific | 329.42 | 250.7 | 250.7 | 6.86 | 61 | >sp\|P01877\|IGHA2_HUMAN Immunoglobulin heavy constant alpha 2 OS=Homo sapiens OX=9606 GN=IGHA2 PE=1 SV=4 | 576 | 20201118_lss_colostrum_1ng_glyco_hcd203040_compound15.3847.3847.3 | controllerType=0 controllerNumber=1 scan=3847 | 12.781 |
| 00924:3 | 11 | K.HYTN[+1606.587]SSQDVTVPC[+57.021]R.V | HexNAc(4)Hex(4)Fuc(1) | C[+57], N[+1607] | 1090.786 | 3 | 3270.3436 | 3270.336 | 0 | 2.3 | 89 | Specific | 476.78 | 322.62 | 322.62 | 7.66 | 61 | >sp\|P01877\|IGHA2_HUMAN Immunoglobulin heavy constant alpha 2 OS=Homo sapiens OX=9606 GN=IGHA2 PE=1 SV=4 | 576 | 20201118_lss_colostrum_1ng_glyco_hcd203040_compound15.3849.3849.3 | controllerType=0 controllerNumber=1 scan=3849 | 12.7857 |
| 01850:3 | 11 | K.HYTN[+2262.814]SSQDVTVPC[+57.021]R.V | HexNAc(5)Hex(5)Fuc(1)NeuAc(1) | C[+57], N[+2263] | 1309.5295 | 3 | 3926.5739 | 3926.5636 | 0 | 2.6 | 89 | Specific | 549.44 | 312.48 | 312.48 | 7.89 | 61 | >sp\|P01877\|IGHA2_HUMAN Immunoglobulin heavy constant alpha 2 OS=Homo sapiens OX=9606 GN=IGHA2 PE=1 SV=4 | 576 | 20201118_lss_colostrum_1ng_glyco_hcd203040_compound15.4810.4810.3 | controllerType=0 controllerNumber=1 scan=4810 | 14.4194 |
| 00945:3 | 11 | K.HYTN[+1809.666]SSQDVTVPC[+57.021]R.V | HexNAc(5)Hex(4)Fuc(1) | C[+57], N[+1810] | 1158.4803 | 3 | 3473.4263 | 3473.4154 | 0 | 3.1 | 89 | Specific | 571.51 | 512.33 | 512.33 | 10.07 | 61 | >sp\|P01877\|IGHA2_HUMAN Immunoglobulin heavy constant alpha 2 OS=Homo sapiens OX=9606 GN=IGHA2 PE=1 SV=4 | 576 | 20201118_lss_colostrum_1ng_glyco_hcd203040_compound15.3871.3871.3 | controllerType=0 controllerNumber=1 scan=3871 | 12.8245 |
| 01852:3 | 11 | K.HYTN[+2262.814]SSQDVTVPC[+57.021]R.V | HexNAc(5)Hex(5)Fuc(1)NeuAc(1) | C[+57], N[+2263] | 1309.5295 | 3 | 3926.5739 | 3926.5636 | 0 | 2.6 | 89 | Specific | 632.43 | 420.72 | 420.72 | 12.41 | 61 | >sp\|P01877\|IGHA2_HUMAN Immunoglobulin heavy constant alpha 2 OS=Homo sapiens OX=9606 GN=IGHA2 PE=1 SV=4 | 576 | 20201118_lss_colostrum_1ng_glyco_hcd203040_compound15.4812.4812.3 | controllerType=0 controllerNumber=1 scan=4812 | 14.4235 |
| 00968:3 | 11 | K.HYTN[+1647.613]SSQDVTVPC[+57.021]R.V | HexNAc(5)Hex(3)Fuc(1) | C[+57], N[+1648] | 1104.4618 | 3 | 3311.3709 | 3311.3625 | 0 | 2.5 | 89 | Specific | 651.2 | 569.24 | 569.24 | 13.41 | 61 | >sp\|P01877\|IGHA2_HUMAN Immunoglobulin heavy constant alpha 2 OS=Homo sapiens OX=9606 GN=IGHA2 PE=1 SV=4 | 576 | 20201118_lss_colostrum_1ng_glyco_hcd203040_compound15.3895.3895.3 | controllerType=0 controllerNumber=1 scan=3895 | 12.8686 |
| 00947:3 | 11 | K.HYTN[+1809.666]SSQDVTVPC[+57.021]R.V | HexNAc(5)Hex(4)Fuc(1) | C[+57], N[+1810] | 1158.4803 | 3 | 3473.4263 | 3473.4154 | 0 | 3.1 | 89 | Specific | 651.63 | 640.58 | 640.58 | 13.41 | 61 | >sp\|P01877\|IGHA2_HUMAN Immunoglobulin heavy constant alpha 2 OS=Homo sapiens OX=9606 GN=IGHA2 PE=1 SV=4 | 576 | 20201118_lss_colostrum_1ng_glyco_hcd203040_compound15.3873.3873.3 | controllerType=0 controllerNumber=1 scan=3873 | 12.8294 |
| 00969:3 | 11 | K.HYTN[+1444.534]SSQDVTVPC[+57.021]R.V | HexNAc(4)Hex(3)Fuc(1) | C[+57], N[+1445] | 1036.768 | 3 | 3108.2894 | 3108.2832 | 0 | 2 | 89 | Specific | 657.38 | 520.95 | 520.95 | 13.41 | 61 | >sp\|P01877\|IGHA2_HUMAN Immunoglobulin heavy constant alpha 2 OS=Homo sapiens OX=9606 GN=IGHA2 PE=1 SV=4 | 576 | 20201118_lss_colostrum_1ng_glyco_hcd203040_compound15.3896.3896.3 | controllerType=0 controllerNumber=1 scan=3896 | 12.8705 |
| 00966:3 | 11 | K.HYTN[+1647.613]SSQDVTVPC[+57.021]R.V | HexNAc(5)Hex(3)Fuc(1) | C[+57], N[+1648] | 1104.4618 | 3 | 3311.3709 | 3311.3625 | 0 | 2.5 | 89 | Specific | 666.12 | 546.51 | 546.51 | 13.41 | 61 | >sp\|P01877\|IGHA2_HUMAN Immunoglobulin heavy constant alpha 2 OS=Homo sapiens OX=9606 GN=IGHA2 PE=1 SV=4 | 576 | 20201118_lss_colostrum_1ng_glyco_hcd203040_compound15.3893.3893.3 | controllerType=0 controllerNumber=1 scan=3893 | 12.8641 |
| 00967:3 | 11 | K.HYTN[+1444.534]SSQDVTVPC[+57.021]R.V | HexNAc(4)Hex(3)Fuc(1) | C[+57], N[+1445] | 1036.768 | 3 | 3108.2894 | 3108.2832 | 0 | 2 | 89 | Specific | 685.61 | 548.3 | 548.3 | 13.41 | 61 | >sp\|P01877\|IGHA2_HUMAN Immunoglobulin heavy constant alpha 2 OS=Homo sapiens OX=9606 GN=IGHA2 PE=1 SV=4 | 576 | 20201118_lss_colostrum_1ng_glyco_hcd203040_compound15.3894.3894.3 | controllerType=0 controllerNumber=1 scan=3894 | 12.8656 |
| 03757:5 | 11 | K.HYTN[+2263.835]SSQDVTVPC[+57.021]RVPPPPPC[+57.021]C[+57.021]HPR.L | HexNAc(5)Hex(5)Fuc(3) | C[+57]*3, N[+2264] | 1045.2443 | 5 | 5222.1922 | 5222.1903 | 0 | 0.4 | 89 | Specific | 308.05 | 308.05 | 308.05 | 6.22 | 61 | >sp\|P01877\|IGHA2_HUMAN Immunoglobulin heavy constant alpha 2 OS=Homo sapiens OX=9606 GN=IGHA2 PE=1 SV=4 | 576 | 20201118_lss_colostrum_1ng_glyco_hcd203040_compound15.6783.6783.5 | controllerType=0 controllerNumber=1 scan=6783 | 17.786 |
| 03755:5 | 11 | K.HYTN[+2263.835]SSQDVTVPC[+57.021]RVPPPPPC[+57.021]C[+57.021]HPR.L | HexNAc(5)Hex(5)Fuc(3) | C[+57]*3, N[+2264] | 1045.2443 | 5 | 5222.1922 | 5222.1903 | 0 | 0.4 | 89 | Specific | 325.15 | 325.15 | 325.15 | 6.31 | 61 | >sp\|P01877\|IGHA2_HUMAN Immunoglobulin heavy constant alpha 2 OS=Homo sapiens OX=9606 GN=IGHA2 PE=1 SV=4 | 576 | 20201118_lss_colostrum_1ng_glyco_hcd203040_compound15.6781.6781.5 | controllerType=0 controllerNumber=1 scan=6781 | 17.7825 |
| 03070:4 | 11 | K.HYTN[+1809.666]SSQDVTVPC[+57.021]RVPPPPPC[+57.021]C[+57.021]HPR.L | HexNAc(5)Hex(4)Fuc(1) | C[+57]*3, N[+1810] | 1192.7637 | 4 | 4768.033 | 4768.0217 | 0 | 2.4 | 89 | Specific | 356.44 | 279.69 | 279.69 | 6.36 | 61 | >sp\|P01877\|IGHA2_HUMAN Immunoglobulin heavy constant alpha 2 OS=Homo sapiens OX=9606 GN=IGHA2 PE=1 SV=4 | 576 | 20201118_lss_colostrum_1ng_glyco_hcd203040_compound15.6072.6072.4 | controllerType=0 controllerNumber=1 scan=6072 | 16.585 |
| 03068:4 | 11 | K.HYTN[+1809.666]SSQDVTVPC[+57.021]RVPPPPPC[+57.021]C[+57.021]HPR.L | HexNAc(5)Hex(4)Fuc(1) | C[+57]*3, N[+1810] | 1192.7637 | 4 | 4768.033 | 4768.0217 | 0 | 2.4 | 89 | Specific | 404.05 | 376.91 | 376.91 | 7.07 | 61 | >sp\|P01877\|IGHA2_HUMAN Immunoglobulin heavy constant alpha 2 OS=Homo sapiens OX=9606 GN=IGHA2 PE=1 SV=4 | 576 | 20201118_lss_colostrum_1ng_glyco_hcd203040_compound15.6070.6070.4 | controllerType=0 controllerNumber=1 scan=6070 | 16.5815 |
| 03005:5 | 11 | K.HYTN[+1647.613]SSQDVTVPC[+57.021]RVPPPPPC[+57.021]C[+57.021]HPR.L | HexNAc(5)Hex(3)Fuc(1) | C[+57]*3, N[+1648] | 922.001 | 5 | 4605.9759 | 4605.9689 | 0 | 1.5 | 89 | Specific | 663.38 | 576.44 | 576.44 | 13.14 | 61 | >sp\|P01877\|IGHA2_HUMAN Immunoglobulin heavy constant alpha 2 OS=Homo sapiens OX=9606 GN=IGHA2 PE=1 SV=4 | 576 | 20201118_lss_colostrum_1ng_glyco_hcd203040_compound15.6005.6005.5 | controllerType=0 controllerNumber=1 scan=6005 | 16.4712 |
| 03031:4 | 11 | K.HYTN[+1647.613]SSQDVTVPC[+57.021]RVPPPPPC[+57.021]C[+57.021]HPR.L | HexNAc(5)Hex(3)Fuc(1) | C[+57]*3, N[+1648] | 1152.2508 | 4 | 4605.9813 | 4605.9689 | 0 | 2.7 | 89 | Specific | 770.85 | 556.02 | 556.02 | 14.67 | 61 | >sp\|P01877\|IGHA2_HUMAN Immunoglobulin heavy constant alpha 2 OS=Homo sapiens OX=9606 GN=IGHA2 PE=1 SV=4 | 576 | 20201118_lss_colostrum_1ng_glyco_hcd203040_compound15.6032.6032.4 | controllerType=0 controllerNumber=1 scan=6032 | 16.5196 |
| 03033:4 | 11 | K.HYTN[+1647.613]SSQDVTVPC[+57.021]RVPPPPPC[+57.021]C[+57.021]HPR.L | HexNAc(5)Hex(3)Fuc(1) | C[+57]*3, N[+1648] | 1152.2508 | 4 | 4605.9813 | 4605.9689 | 0 | 2.7 | 89 | Specific | 776.59 | 526.17 | 526.17 | 14.67 | 61 | >sp\|P01877\|IGHA2_HUMAN Immunoglobulin heavy constant alpha 2 OS=Homo sapiens OX=9606 GN=IGHA2 PE=1 SV=4 | 576 | 20201118_lss_colostrum_1ng_glyco_hcd203040_compound15.6034.6034.4 | controllerType=0 controllerNumber=1 scan=6034 | 16.5242 |
| 03008:5 | 11 | K.HYTN[+1647.613]SSQDVTVPC[+57.021]RVPPPPPC[+57.021]C[+57.021]HPR.L | HexNAc(5)Hex(3)Fuc(1) | C[+57]*3, N[+1648] | 922.001 | 5 | 4605.9759 | 4605.9689 | 0 | 1.5 | 89 | Specific | 820.77 | 728.35 | 728.35 | 16.14 | 61 | >sp\|P01877\|IGHA2_HUMAN Immunoglobulin heavy constant alpha 2 OS=Homo sapiens OX=9606 GN=IGHA2 PE=1 SV=4 | 576 | 20201118_lss_colostrum_1ng_glyco_hcd203040_compound15.6008.6008.5 | controllerType=0 controllerNumber=1 scan=6008 | 16.4769 |
| 04508:2 | 11 | K.TPLTAN[+2059.735]ITK.S | HexNAc(4)Hex(5)Fuc(1)NeuAc(1) | N[+2060] | 1509.6518 | 2 | 3018.2963 | 3018.2917 | 0 | 1.5 | 200 | Specific | 336.03 | 316.91 | 316.91 | 6.84 | 61 | >sp\|P01877\|IGHA2_HUMAN Immunoglobulin heavy constant alpha 2 OS=Homo sapiens OX=9606 GN=IGHA2 PE=1 SV=4 | 576 | 20201118_lss_colostrum_1ng_glyco_hcd203040_compound15.7562.7562.2 | controllerType=0 controllerNumber=1 scan=7562 | 19.0436 |
| 04535:2 | 11 | K.TPLTAN[+2262.814]ITK.S | HexNAc(5)Hex(5)Fuc(1)NeuAc(1) | N[+2263] | 1611.1927 | 2 | 3221.3781 | 3221.3711 | 0 | 2.2 | 200 | Specific | 350.95 | 279.6 | 279.6 | 6.81 | 61 | >sp\|P01877\|IGHA2_HUMAN Immunoglobulin heavy constant alpha 2 OS=Homo sapiens OX=9606 GN=IGHA2 PE=1 SV=4 | 576 | 20201118_lss_colostrum_1ng_glyco_hcd203040_compound15.7590.7590.2 | controllerType=0 controllerNumber=1 scan=7590 | 19.0882 |
| 04533:2 | 11 | K.TPLTAN[+2262.814]ITK.S | HexNAc(5)Hex(5)Fuc(1)NeuAc(1) | N[+2263] | 1611.1927 | 2 | 3221.3781 | 3221.3711 | 0 | 2.2 | 200 | Specific | 367.8 | 238.5 | 238.5 | 6.65 | 61 | >sp\|P01877\|IGHA2_HUMAN Immunoglobulin heavy constant alpha 2 OS=Homo sapiens OX=9606 GN=IGHA2 PE=1 SV=4 | 576 | 20201118_lss_colostrum_1ng_glyco_hcd203040_compound15.7588.7588.2 | controllerType=0 controllerNumber=1 scan=7588 | 19.0845 |
| 04516:2 | 11 | K.TPLTAN[+2262.814]ITK.S | HexNAc(5)Hex(5)Fuc(1)NeuAc(1) | N[+2263] | 1611.1893 | 2 | 3221.3712 | 3221.3711 | 0 | 0 | 200 | Specific | 426.39 | 180.09 | 180.09 | 6.02 | 61 | >sp\|P01877\|IGHA2_HUMAN Immunoglobulin heavy constant alpha 2 OS=Homo sapiens OX=9606 GN=IGHA2 PE=1 SV=4 | 576 | 20201118_lss_colostrum_1ng_glyco_hcd203040_compound15.7570.7570.2 | controllerType=0 controllerNumber=1 scan=7570 | 19.0557 |
| 02628:3 | 11 | K.TPLTAN[+1971.719]ITK.S | HexNAc(5)Hex(5)Fuc(1) | N[+1972] | 977.4326 | 3 | 2930.2832 | 2930.2757 | 0 | 2.6 | 200 | Specific | 435.22 | 174.17 | 174.17 | 6.19 | 61 | >sp\|P01877\|IGHA2_HUMAN Immunoglobulin heavy constant alpha 2 OS=Homo sapiens OX=9606 GN=IGHA2 PE=1 SV=4 | 576 | 20201118_lss_colostrum_1ng_glyco_hcd203040_compound15.5614.5614.3 | controllerType=0 controllerNumber=1 scan=5614 | 15.7482 |
| 02643:2 | 11 | K.TPLTAN[+1971.719]ITK.S | HexNAc(5)Hex(5)Fuc(1) | N[+1972] | 1465.6459 | 2 | 2930.2846 | 2930.2757 | 0 | 3 | 200 | Specific | 452.04 | 228.05 | 228.05 | 6.95 | 61 | >sp\|P01877\|IGHA2_HUMAN Immunoglobulin heavy constant alpha 2 OS=Homo sapiens OX=9606 GN=IGHA2 PE=1 SV=4 | 576 | 20201118_lss_colostrum_1ng_glyco_hcd203040_compound15.5629.5629.2 | controllerType=0 controllerNumber=1 scan=5629 | 15.7728 |
| 04518:2 | 11 | K.TPLTAN[+2262.814]ITK.S | HexNAc(5)Hex(5)Fuc(1)NeuAc(1) | N[+2263] | 1611.1893 | 2 | 3221.3712 | 3221.3711 | 0 | 0 | 200 | Specific | 473.48 | 239.97 | 239.97 | 6.41 | 61 | >sp\|P01877\|IGHA2_HUMAN Immunoglobulin heavy constant alpha 2 OS=Homo sapiens OX=9606 GN=IGHA2 PE=1 SV=4 | 576 | 20201118_lss_colostrum_1ng_glyco_hcd203040_compound15.7572.7572.2 | controllerType=0 controllerNumber=1 scan=7572 | 19.0593 |
| 04487:3 | 11 | K.TPLTAN[+2262.814]ITK.S | HexNAc(5)Hex(5)Fuc(1)NeuAc(1) | N[+2263] | 1074.4637 | 3 | 3221.3764 | 3221.3711 | 0 | 1.7 | 200 | Specific | 474.34 | 229.86 | 229.86 | 6.81 | 61 | >sp\|P01877\|IGHA2_HUMAN Immunoglobulin heavy constant alpha 2 OS=Homo sapiens OX=9606 GN=IGHA2 PE=1 SV=4 | 576 | 20201118_lss_colostrum_1ng_glyco_hcd203040_compound15.7540.7540.3 | controllerType=0 controllerNumber=1 scan=7540 | 19.0061 |
| 02630:3 | 11 | K.TPLTAN[+1971.719]ITK.S | HexNAc(5)Hex(5)Fuc(1) | N[+1972] | 977.4326 | 3 | 2930.2832 | 2930.2757 | 0 | 2.6 | 200 | Specific | 482.17 | 275.33 | 275.33 | 7.25 | 61 | >sp\|P01877\|IGHA2_HUMAN Immunoglobulin heavy constant alpha 2 OS=Homo sapiens OX=9606 GN=IGHA2 PE=1 SV=4 | 576 | 20201118_lss_colostrum_1ng_glyco_hcd203040_compound15.5616.5616.3 | controllerType=0 controllerNumber=1 scan=5616 | 15.7528 |
| 02715:2 | 11 | K.TPLTAN[+1606.587]ITK.S | HexNAc(4)Hex(4)Fuc(1) | N[+1607] | 1283.0785 | 2 | 2565.1498 | 2565.1435 | 0 | 2.5 | 200 | Specific | 486.67 | 298.84 | 298.84 | 7.44 | 61 | >sp\|P01877\|IGHA2_HUMAN Immunoglobulin heavy constant alpha 2 OS=Homo sapiens OX=9606 GN=IGHA2 PE=1 SV=4 | 576 | 20201118_lss_colostrum_1ng_glyco_hcd203040_compound15.5704.5704.2 | controllerType=0 controllerNumber=1 scan=5704 | 15.9418 |
| 02645:2 | 11 | K.TPLTAN[+1971.719]ITK.S | HexNAc(5)Hex(5)Fuc(1) | N[+1972] | 1465.6459 | 2 | 2930.2846 | 2930.2757 | 0 | 3 | 200 | Specific | 505.25 | 316.02 | 316.02 | 7.55 | 61 | >sp\|P01877\|IGHA2_HUMAN Immunoglobulin heavy constant alpha 2 OS=Homo sapiens OX=9606 GN=IGHA2 PE=1 SV=4 | 576 | 20201118_lss_colostrum_1ng_glyco_hcd203040_compound15.5631.5631.2 | controllerType=0 controllerNumber=1 scan=5631 | 15.7768 |
| 02713:2 | 11 | K.TPLTAN[+1606.587]ITK.S | HexNAc(4)Hex(4)Fuc(1) | N[+1607] | 1283.0785 | 2 | 2565.1498 | 2565.1435 | 0 | 2.5 | 200 | Specific | 508.38 | 261.97 | 261.97 | 7.36 | 61 | >sp\|P01877\|IGHA2_HUMAN Immunoglobulin heavy constant alpha 2 OS=Homo sapiens OX=9606 GN=IGHA2 PE=1 SV=4 | 576 | 20201118_lss_colostrum_1ng_glyco_hcd203040_compound15.5702.5702.2 | controllerType=0 controllerNumber=1 scan=5702 | 15.9369 |
| 02762:3 | 11 | K.TPLTAN[+1647.613]ITK.S | HexNAc(5)Hex(3)Fuc(1) | N[+1648] | 869.3968 | 3 | 2606.1757 | 2606.17 | 0 | 2.2 | 200 | Specific | 517.97 | 229.12 | 229.12 | 7.18 | 61 | >sp\|P01877\|IGHA2_HUMAN Immunoglobulin heavy constant alpha 2 OS=Homo sapiens OX=9606 GN=IGHA2 PE=1 SV=4 | 576 | 20201118_lss_colostrum_1ng_glyco_hcd203040_compound15.5753.5753.3 | controllerType=0 controllerNumber=1 scan=5753 | 16.0323 |
| 02693:3 | 11 | K.TPLTAN[+1809.666]ITK.S | HexNAc(5)Hex(4)Fuc(1) | N[+1810] | 923.4144 | 3 | 2768.2286 | 2768.2228 | 0 | 2.1 | 200 | Specific | 532.79 | 285.16 | 285.16 | 7.47 | 61 | >sp\|P01877\|IGHA2_HUMAN Immunoglobulin heavy constant alpha 2 OS=Homo sapiens OX=9606 GN=IGHA2 PE=1 SV=4 | 576 | 20201118_lss_colostrum_1ng_glyco_hcd203040_compound15.5682.5682.3 | controllerType=0 controllerNumber=1 scan=5682 | 15.9035 |
| 04490:3 | 11 | K.TPLTAN[+2262.814]ITK.S | HexNAc(5)Hex(5)Fuc(1)NeuAc(1) | N[+2263] | 1074.4637 | 3 | 3221.3764 | 3221.3711 | 0 | 1.7 | 200 | Specific | 535.17 | 252.11 | 252.11 | 7.32 | 61 | >sp\|P01877\|IGHA2_HUMAN Immunoglobulin heavy constant alpha 2 OS=Homo sapiens OX=9606 GN=IGHA2 PE=1 SV=4 | 576 | 20201118_lss_colostrum_1ng_glyco_hcd203040_compound15.7543.7543.3 | controllerType=0 controllerNumber=1 scan=7543 | 19.0119 |
| 02751:2 | 11 | K.TPLTAN[+1501.555]ITK.S | HexNAc(5)Hex(3) | N[+1502] | 1230.5627 | 2 | 2460.1181 | 2460.1121 | 0 | 2.4 | 200 | Specific | 558.28 | 327.39 | 327.39 | 7.99 | 61 | >sp\|P01877\|IGHA2_HUMAN Immunoglobulin heavy constant alpha 2 OS=Homo sapiens OX=9606 GN=IGHA2 PE=1 SV=4 | 576 | 20201118_lss_colostrum_1ng_glyco_hcd203040_compound15.5742.5742.2 | controllerType=0 controllerNumber=1 scan=5742 | 16.0119 |
| 02691:3 | 11 | K.TPLTAN[+1809.666]ITK.S | HexNAc(5)Hex(4)Fuc(1) | N[+1810] | 923.4144 | 3 | 2768.2286 | 2768.2228 | 0 | 2.1 | 200 | Specific | 559.23 | 257.66 | 257.66 | 7.73 | 61 | >sp\|P01877\|IGHA2_HUMAN Immunoglobulin heavy constant alpha 2 OS=Homo sapiens OX=9606 GN=IGHA2 PE=1 SV=4 | 576 | 20201118_lss_colostrum_1ng_glyco_hcd203040_compound15.5680.5680.3 | controllerType=0 controllerNumber=1 scan=5680 | 15.8982 |
| 02753:2 | 11 | K.TPLTAN[+1501.555]ITK.S | HexNAc(5)Hex(3) | N[+1502] | 1230.5627 | 2 | 2460.1181 | 2460.1121 | 0 | 2.4 | 200 | Specific | 575.55 | 333.68 | 333.68 | 8.2 | 61 | >sp\|P01877\|IGHA2_HUMAN Immunoglobulin heavy constant alpha 2 OS=Homo sapiens OX=9606 GN=IGHA2 PE=1 SV=4 | 576 | 20201118_lss_colostrum_1ng_glyco_hcd203040_compound15.5744.5744.2 | controllerType=0 controllerNumber=1 scan=5744 | 16.0156 |
| 02764:3 | 11 | K.TPLTAN[+1647.613]ITK.S | HexNAc(5)Hex(3)Fuc(1) | N[+1648] | 869.3968 | 3 | 2606.1757 | 2606.17 | 0 | 2.2 | 200 | Specific | 588.74 | 289.6 | 289.6 | 9.19 | 61 | >sp\|P01877\|IGHA2_HUMAN Immunoglobulin heavy constant alpha 2 OS=Homo sapiens OX=9606 GN=IGHA2 PE=1 SV=4 | 576 | 20201118_lss_colostrum_1ng_glyco_hcd203040_compound15.5755.5755.3 | controllerType=0 controllerNumber=1 scan=5755 | 16.0384 |
| 02680:2 | 11 | K.TPLTAN[+1809.666]ITK.S | HexNAc(5)Hex(4)Fuc(1) | N[+1810] | 1384.6185 | 2 | 2768.2296 | 2768.2228 | 0 | 2.5 | 200 | Specific | 605.6 | 338.16 | 338.16 | 10.54 | 61 | >sp\|P01877\|IGHA2_HUMAN Immunoglobulin heavy constant alpha 2 OS=Homo sapiens OX=9606 GN=IGHA2 PE=1 SV=4 | 576 | 20201118_lss_colostrum_1ng_glyco_hcd203040_compound15.5668.5668.2 | controllerType=0 controllerNumber=1 scan=5668 | 15.868 |
| 02678:2 | 11 | K.TPLTAN[+1809.666]ITK.S | HexNAc(5)Hex(4)Fuc(1) | N[+1810] | 1384.6185 | 2 | 2768.2296 | 2768.2228 | 0 | 2.5 | 200 | Specific | 608.64 | 359.79 | 359.79 | 10.74 | 61 | >sp\|P01877\|IGHA2_HUMAN Immunoglobulin heavy constant alpha 2 OS=Homo sapiens OX=9606 GN=IGHA2 PE=1 SV=4 | 576 | 20201118_lss_colostrum_1ng_glyco_hcd203040_compound15.5666.5666.2 | controllerType=0 controllerNumber=1 scan=5666 | 15.8623 |
| 02747:2 | 11 | K.TPLTAN[+1647.613]ITK.S | HexNAc(5)Hex(3)Fuc(1) | N[+1648] | 1303.5912 | 2 | 2606.1752 | 2606.17 | 0 | 2 | 200 | Specific | 618.17 | 345.62 | 345.62 | 10.54 | 61 | >sp\|P01877\|IGHA2_HUMAN Immunoglobulin heavy constant alpha 2 OS=Homo sapiens OX=9606 GN=IGHA2 PE=1 SV=4 | 576 | 20201118_lss_colostrum_1ng_glyco_hcd203040_compound15.5738.5738.2 | controllerType=0 controllerNumber=1 scan=5738 | 16.0067 |
| 02769:2 | 11 | K.TPLTAN[+1444.534]ITK.S | HexNAc(4)Hex(3)Fuc(1) | N[+1445] | 1202.0513 | 2 | 2403.0953 | 2403.0906 | 0 | 2 | 200 | Specific | 624.62 | 315.34 | 315.34 | 10.32 | 61 | >sp\|P01877\|IGHA2_HUMAN Immunoglobulin heavy constant alpha 2 OS=Homo sapiens OX=9606 GN=IGHA2 PE=1 SV=4 | 576 | 20201118_lss_colostrum_1ng_glyco_hcd203040_compound15.5760.5760.2 | controllerType=0 controllerNumber=1 scan=5760 | 16.0508 |
| 02745:2 | 11 | K.TPLTAN[+1647.613]ITK.S | HexNAc(5)Hex(3)Fuc(1) | N[+1648] | 1303.5912 | 2 | 2606.1752 | 2606.17 | 0 | 2 | 200 | Specific | 640.45 | 368.27 | 368.27 | 10.74 | 61 | >sp\|P01877\|IGHA2_HUMAN Immunoglobulin heavy constant alpha 2 OS=Homo sapiens OX=9606 GN=IGHA2 PE=1 SV=4 | 576 | 20201118_lss_colostrum_1ng_glyco_hcd203040_compound15.5736.5736.2 | controllerType=0 controllerNumber=1 scan=5736 | 15.9999 |
| 02767:2 | 11 | K.TPLTAN[+1444.534]ITK.S | HexNAc(4)Hex(3)Fuc(1) | N[+1445] | 1202.0513 | 2 | 2403.0953 | 2403.0906 | 0 | 2 | 200 | Specific | 665.37 | 365.21 | 365.21 | 11.74 | 61 | >sp\|P01877\|IGHA2_HUMAN Immunoglobulin heavy constant alpha 2 OS=Homo sapiens OX=9606 GN=IGHA2 PE=1 SV=4 | 576 | 20201118_lss_colostrum_1ng_glyco_hcd203040_compound15.5758.5758.2 | controllerType=0 controllerNumber=1 scan=5758 | 16.0432 |
| 10200:3 | 11 | R.MAGKPTHIN[+1647.613]VSVVMAEADGTC[+57.021]Y.- | HexNAc(5)Hex(3)Fuc(1) | C[+57], N[+1648] | 1333.572 | 3 | 3998.7013 | 3998.6961 | 0 | 1.3 | 319 | CRagged | 277.35 | 277.35 | 277.35 | 7 | 61 | >sp\|P01877\|IGHA2_HUMAN Immunoglobulin heavy constant alpha 2 OS=Homo sapiens OX=9606 GN=IGHA2 PE=1 SV=4 | 576 | 20201118_lss_colostrum_1ng_glyco_hcd203040_compound15.13465.13465.3 | controllerType=0 controllerNumber=1 scan=13465 | 28.9881 |
| 11776:3 | 11 | R.MAGKPTHIN[+1710.598]VSVVMAEADGTC[+57.021]Y.- | HexNAc(3)Hex(5)NeuAc(1) | C[+57], N[+1711] | 1354.5676 | 3 | 4061.6881 | 4061.6805 | 0 | 1.9 | 319 | CRagged | 289.06 | 289.06 | 213.4 | 7.42 | 61 | >sp\|P01877\|IGHA2_HUMAN Immunoglobulin heavy constant alpha 2 OS=Homo sapiens OX=9606 GN=IGHA2 PE=1 SV=4 | 576 | 20201118_lss_colostrum_1ng_glyco_hcd203040_compound15.15100.15100.3 | controllerType=0 controllerNumber=1 scan=15100 | 31.9943 |
| 11779:3 | 11 | R.MAGKPTHIN[+1710.598]VSVVMAEADGTC[+57.021]Y.- | HexNAc(3)Hex(5)NeuAc(1) | C[+57], N[+1711] | 1354.5676 | 3 | 4061.6881 | 4061.6805 | 0 | 1.9 | 319 | CRagged | 313.05 | 313.05 | 259.95 | 7.63 | 61 | >sp\|P01877\|IGHA2_HUMAN Immunoglobulin heavy constant alpha 2 OS=Homo sapiens OX=9606 GN=IGHA2 PE=1 SV=4 | 576 | 20201118_lss_colostrum_1ng_glyco_hcd203040_compound15.15103.15103.3 | controllerType=0 controllerNumber=1 scan=15103 | 31.9992 |
| 10202:3 | 11 | R.MAGKPTHIN[+1647.613]VSVVMAEADGTC[+57.021]Y.- | HexNAc(5)Hex(3)Fuc(1) | C[+57], N[+1648] | 1333.572 | 3 | 3998.7013 | 3998.6961 | 0 | 1.3 | 319 | CRagged | 404.39 | 404.39 | 404.39 | 9.74 | 61 | >sp\|P01877\|IGHA2_HUMAN Immunoglobulin heavy constant alpha 2 OS=Homo sapiens OX=9606 GN=IGHA2 PE=1 SV=4 | 576 | 20201118_lss_colostrum_1ng_glyco_hcd203040_compound15.13467.13467.3 | controllerType=0 controllerNumber=1 scan=13467 | 28.9917 |
| 09969:3 | 11 | R.MAGKPTHIN[+1702.581]VSVVMAEADGTC[+57.021]Y.- | HexNAc(2)Hex(8) | C[+57], N[+1703] | 1351.8943 | 3 | 4053.6685 | 4053.6642 | 0 | 1.1 | 319 | CRagged | 468.1 | 468.1 | 345.86 | 10.01 | 61 | >sp\|P01877\|IGHA2_HUMAN Immunoglobulin heavy constant alpha 2 OS=Homo sapiens OX=9606 GN=IGHA2 PE=1 SV=4 | 576 | 20201118_lss_colostrum_1ng_glyco_hcd203040_compound15.13225.13225.3 | controllerType=0 controllerNumber=1 scan=13225 | 28.531 |
| 10184:3 | 11 | R.MAGKPTHIN[+1501.555]VSVVMAEADGTC[+57.021]Y.- | HexNAc(5)Hex(3) | C[+57], N[+1502] | 1284.8851 | 3 | 3852.6406 | 3852.6382 | 0 | 0.6 | 319 | CRagged | 553.93 | 343.78 | 343.78 | 8.53 | 61 | >sp\|P01877\|IGHA2_HUMAN Immunoglobulin heavy constant alpha 2 OS=Homo sapiens OX=9606 GN=IGHA2 PE=1 SV=4 | 576 | 20201118_lss_colostrum_1ng_glyco_hcd203040_compound15.13448.13448.3 | controllerType=0 controllerNumber=1 scan=13448 | 28.9569 |

Table S4. Cont.

| Query #:z | Protein Rank | Glycopeptide | Glycan | Modification Type(s) | Observed m/z | z | Observed (M+H) | Calc. mass (M+H) | Off-by-x error | Mass error (ppm) | StartPosition | Cleavage | Score | Delta | Delta Mod | \|Log Prob\| | # of unique peptides | Protein Name | Protein DB number | Comment | Scan # | Scan Time |
|---|---|---|---|---|---|---|---|---|---|---|---|---|---|---|---|---|---|---|---|---|---|---|
| 10075:3 | 11 | R.MAGKPTHIN[+1663.608]VSVVMAEADGTC[+57.021]Y.- | HexNAc(5)Hex(4) | C[+57], N[+1664] | 1338.9059 | 3 | 4014.7033 | 4014.691 | 0 | 3.1 | 319 | CRagged | 595.2 | 595.2 | 381.5 | 11.99 | 61 | >sp\|P01877\|IGHA2_HUMAN Immunoglobulin heavy constant alpha 2 OS=Homo sapiens OX=9606 GN=IGHA2 PE=1 SV=4 | 576 | 20201118_lss_colostrum_1ng_glyco_hcd203040_compound15.13335.13335.3 | controllerType=0 controllerNumber=1 scan=13335 | 28.7365 |
| 10181:3 | 11 | R.MAGKPTHIN[+1501.555]VSVVMAEADGTC[+57.021]Y.- | HexNAc(5)Hex(3) | C[+57], N[+1502] | 1284.8851 | 3 | 3852.6406 | 3852.6382 | 0 | 0.6 | 319 | CRagged | 604.23 | 604.23 | 604.23 | 12.84 | 61 | >sp\|P01877\|IGHA2_HUMAN Immunoglobulin heavy constant alpha 2 OS=Homo sapiens OX=9606 GN=IGHA2 PE=1 SV=4 | 576 | 20201118_lss_colostrum_1ng_glyco_hcd203040_compound15.13445.13445.3 | controllerType=0 controllerNumber=1 scan=13445 | 28.952 |
| 10077:3 | 11 | R.MAGKPTHIN[+1663.608]VSVVMAEADGTC[+57.021]Y.- | HexNAc(5)Hex(4) | C[+57], N[+1664] | 1338.9059 | 3 | 4014.7033 | 4014.691 | 0 | 3.1 | 319 | CRagged | 694.68 | 661.77 | 383.33 | 13.99 | 61 | >sp\|P01877\|IGHA2_HUMAN Immunoglobulin heavy constant alpha 2 OS=Homo sapiens OX=9606 GN=IGHA2 PE=1 SV=4 | 576 | 20201118_lss_colostrum_1ng_glyco_hcd203040_compound15.13337.13337.3 | controllerType=0 controllerNumber=1 scan=13337 | 28.7415 |
| 10044:3 | 11 | R.MAGKPTHIN[+1378.476]VSVVMAEADGTC[+57.021]Y.- | HexNAc(2)Hex(6) | C[+57], N[+1378] | 1243.8594 | 3 | 3729.5637 | 3729.5585 | 0 | 1.4 | 319 | CRagged | 872.75 | 749.68 | 493.87 | 17.84 | 61 | >sp\|P01877\|IGHA2_HUMAN Immunoglobulin heavy constant alpha 2 OS=Homo sapiens OX=9606 GN=IGHA2 PE=1 SV=4 | 576 | 20201118_lss_colostrum_1ng_glyco_hcd203040_compound15.13303.13303.3 | controllerType=0 controllerNumber=1 scan=13303 | 28.6765 |
| 09972:3 | 11 | R.MAGKPTHIN[+1702.581]VSVVMAEADGTC[+57.021]Y.- | HexNAc(2)Hex(8) | C[+57], N[+1703] | 1351.8943 | 3 | 4053.6685 | 4053.6642 | 0 | 1.1 | 319 | CRagged | 893.47 | 700.76 | 480.34 | 17.84 | 61 | >sp\|P01877\|IGHA2_HUMAN Immunoglobulin heavy constant alpha 2 OS=Homo sapiens OX=9606 GN=IGHA2 PE=1 SV=4 | 576 | 20201118_lss_colostrum_1ng_glyco_hcd203040_compound15.13228.13228.3 | controllerType=0 controllerNumber=1 scan=13228 | 28.5356 |
| 10094:3 | 11 | R.MAGKPTHIN[+1216.423]VSVVMAEADGTC[+57.021]Y.- | HexNAc(2)Hex(5) | C[+57], N[+1216] | 1189.8447 | 3 | 3567.5194 | 3567.5057 | 0 | 3.8 | 319 | CRagged | 932.21 | 932.21 | 545.68 | 18.8 | 61 | >sp\|P01877\|IGHA2_HUMAN Immunoglobulin heavy constant alpha 2 OS=Homo sapiens OX=9606 GN=IGHA2 PE=1 SV=4 | 576 | 20201118_lss_colostrum_1ng_glyco_hcd203040_compound15.13355.13355.3 | controllerType=0 controllerNumber=1 scan=13355 | 28.7816 |
| 10092:3 | 11 | R.MAGKPTHIN[+1216.423]VSVVMAEADGTC[+57.021]Y.- | HexNAc(2)Hex(5) | C[+57], N[+1216] | 1189.8447 | 3 | 3567.5194 | 3567.5057 | 0 | 3.8 | 319 | CRagged | 959.1 | 813.02 | 547.14 | 19.8 | 61 | >sp\|P01877\|IGHA2_HUMAN Immunoglobulin heavy constant alpha 2 OS=Homo sapiens OX=9606 GN=IGHA2 PE=1 SV=4 | 576 | 20201118_lss_colostrum_1ng_glyco_hcd203040_compound15.13353.13353.3 | controllerType=0 controllerNumber=1 scan=13353 | 28.7735 |
| 10042:3 | 11 | R.MAGKPTHIN[+1378.476]VSVVMAEADGTC[+57.021]Y.- | HexNAc(2)Hex(6) | C[+57], N[+1378] | 1243.8594 | 3 | 3729.5637 | 3729.5585 | 0 | 1.4 | 319 | CRagged | 967.89 | 946.99 | 586.63 | 19.84 | 61 | >sp\|P01877\|IGHA2_HUMAN Immunoglobulin heavy constant alpha 2 OS=Homo sapiens OX=9606 GN=IGHA2 PE=1 SV=4 | 576 | 20201118_lss_colostrum_1ng_glyco_hcd203040_compound15.13301.13301.3 | controllerType=0 controllerNumber=1 scan=13301 | 28.67 |
| 09671:3 | 11 | R.M[+15.995]AGKPTHIN[+1216.423]VSVVMAEADGTC[+57.021]Y.- | HexNAc(2)Hex(5) | C[+57], M[+16], N[+1216] | 1195.1774 | 3 | 3583.5177 | 3583.5006 | 0 | 4.8 | 319 | CRagged | 505.78 | 505.78 | 191.76 | 9.57 | 61 | >sp\|P01877\|IGHA2_HUMAN Immunoglobulin heavy constant alpha 2 OS=Homo sapiens OX=9606 GN=IGHA2 PE=1 SV=4 | 576 | 20201118_lss_colostrum_1ng_glyco_hcd203040_compound15.12915.12915.3 | controllerType=0 controllerNumber=1 scan=12915 | 27.9506 |
| 09669:3 | 11 | R.M[+15.995]AGKPTHIN[+1216.423]VSVVMAEADGTC[+57.021]Y.- | HexNAc(2)Hex(5) | C[+57], M[+16], N[+1216] | 1195.1774 | 3 | 3583.5177 | 3583.5006 | 0 | 4.8 | 319 | CRagged | 555.81 | 549.35 | 271.34 | 10 | 61 | >sp\|P01877\|IGHA2_HUMAN Immunoglobulin heavy constant alpha 2 OS=Homo sapiens OX=9606 GN=IGHA2 PE=1 SV=4 | 576 | 20201118_lss_colostrum_1ng_glyco_hcd203040_compound15.12913.12913.3 | controllerType=0 controllerNumber=1 scan=12913 | 27.946 |
| 09645:3 | 11 | R.M[+15.995]AGKPTHIN[+1378.476]VSVVMAEADGTC[+57.021]Y.- | HexNAc(2)Hex(6) | C[+57], M[+16], N[+1378] | 1249.1932 | 3 | 3745.565 | 3745.5535 | 0 | 3.1 | 319 | CRagged | 580.96 | 580.96 | 177.43 | 11.99 | 61 | >sp\|P01877\|IGHA2_HUMAN Immunoglobulin heavy constant alpha 2 OS=Homo sapiens OX=9606 GN=IGHA2 PE=1 SV=4 | 576 | 20201118_lss_colostrum_1ng_glyco_hcd203040_compound15.12888.12888.3 | controllerType=0 controllerNumber=1 scan=12888 | 27.9009 |
| 09643:3 | 11 | R.M[+15.995]AGKPTHIN[+1378.476]VSVVMAEADGTC[+57.021]Y.- | HexNAc(2)Hex(6) | C[+57], M[+16], N[+1378] | 1249.1932 | 3 | 3745.565 | 3745.5535 | 0 | 3.1 | 319 | CRagged | 626.65 | 626.65 | 282.65 | 12.99 | 61 | >sp\|P01877\|IGHA2_HUMAN Immunoglobulin heavy constant alpha 2 OS=Homo sapiens OX=9606 GN=IGHA2 PE=1 SV=4 | 576 | 20201118_lss_colostrum_1ng_glyco_hcd203040_compound15.12886.12886.3 | controllerType=0 controllerNumber=1 scan=12886 | 27.8966 |
| 25747:4 | 13 | R.QLAHQSN[+2861.000]STNIFFSPVSIATAFAMLSLGTK.A | HexNAc(5)Hex(6)NeuAc(3) | N[+2861] | 1511.416 | 4 | 6042.6422 | 6042.6351 | 0 | 1.2 | 64 | Specific | 201.84 | 201.84 | 201.84 | 5.2 | 50 | >sp\|P01009\|A1AT_HUMAN Alpha-1-antitrypsin OS=Homo sapiens OX=9606 GN=SERPINA1 PE=1 SV=3 | 642 | 20201118_lss_colostrum_1ng_glyco_hcd203040_compound15.29647.29647.4 | controllerType=0 controllerNumber=1 scan=29647 | 61.7413 |
| 22880:3 | 13 | R.[+42.011]QLAHQSN[+2190.793]STNIFFSPVSIATAFAMLSLGTK.A | HexNAc(6)Hex(6) | .[42], N[+2191] | 1805.4763 | 3 | 5414.4143 | 5414.4388 | 0 | -4.5 | 64 | Specific | 212.72 | 194.69 | 194.69 | 3.9 | 50 | >sp\|P01009\|A1AT_HUMAN Alpha-1-antitrypsin OS=Homo sapiens OX=9606 GN=SERPINA1 PE=1 SV=3 | 642 | 20201118_lss_colostrum_1ng_glyco_hcd203040_compound15.26667.26667.3 | controllerType=0 controllerNumber=1 scan=26667 | 56.0639 |
| 22850:3 | 13 | R.[+42.011]QLAHQSN[+2190.793]STNIFFSPVSIATAFAMLSLGTK.A | HexNAc(6)Hex(6) | .[42], N[+2191] | 1805.479 | 3 | 5414.4223 | 5414.4388 | 0 | -3 | 64 | Specific | 253.75 | 253.75 | 253.75 | 5.63 | 50 | >sp\|P01009\|A1AT_HUMAN Alpha-1-antitrypsin OS=Homo sapiens OX=9606 GN=SERPINA1 PE=1 SV=3 | 642 | 20201118_lss_colostrum_1ng_glyco_hcd203040_compound15.26636.26636.3 | controllerType=0 controllerNumber=1 scan=26636 | 55.9915 |
| 22877:3 | 13 | R.[+42.011]QLAHQSN[+2190.793]STNIFFSPVSIATAFAMLSLGTK.A | HexNAc(6)Hex(6) | .[42], N[+2191] | 1805.4763 | 3 | 5414.4143 | 5414.4388 | 0 | -4.5 | 64 | Specific | 263.06 | 263.06 | 263.06 | 5.61 | 50 | >sp\|P01009\|A1AT_HUMAN Alpha-1-antitrypsin OS=Homo sapiens OX=9606 GN=SERPINA1 PE=1 SV=3 | 642 | 20201118_lss_colostrum_1ng_glyco_hcd203040_compound15.26664.26664.3 | controllerType=0 controllerNumber=1 scan=26664 | 56.0509 |
| 22852:3 | 13 | R.[+42.011]QLAHQSN[+2190.793]STNIFFSPVSIATAFAMLSLGTK.A | HexNAc(6)Hex(6) | .[42], N[+2191] | 1805.479 | 3 | 5414.4223 | 5414.4388 | 0 | -3 | 64 | Specific | 270.74 | 264.81 | 264.81 | 5.7 | 50 | >sp\|P01009\|A1AT_HUMAN Alpha-1-antitrypsin OS=Homo sapiens OX=9606 GN=SERPINA1 PE=1 SV=3 | 642 | 20201118_lss_colostrum_1ng_glyco_hcd203040_compound15.26638.26638.3 | controllerType=0 controllerNumber=1 scan=26638 | 55.9955 |
| 22784:3 | 13 | R.QLAHQSN[+2350.830]STNIFFSPVSIATAFAMLSLGTK.A | HexNAc(4)Hex(5)Fuc(1)NeuAc(2) | N[+2351] | 1844.8307 | 3 | 5532.4774 | 5532.4654 | 0 | 2.2 | 64 | Specific | 286.6 | 274.49 | 274.49 | 7.06 | 50 | >sp\|P01009\|A1AT_HUMAN Alpha-1-antitrypsin OS=Homo sapiens OX=9606 GN=SERPINA1 PE=1 SV=3 | 642 | 20201118_lss_colostrum_1ng_glyco_hcd203040_compound15.26567.26567.3 | controllerType=0 controllerNumber=1 scan=26567 | 55.8518 |
| 22854:4 | 13 | R.QLAHQSN[+2350.830]STNIFFSPVSIATAFAMLSLGTK.A | HexNAc(4)Hex(5)Fuc(1)NeuAc(2) | N[+2351] | 1383.8694 | 4 | 5532.4557 | 5532.4654 | 0 | -1.7 | 64 | Specific | 319.55 | 318.02 | 318.02 | 6.39 | 50 | >sp\|P01009\|A1AT_HUMAN Alpha-1-antitrypsin OS=Homo sapiens OX=9606 GN=SERPINA1 PE=1 SV=3 | 642 | 20201118_lss_colostrum_1ng_glyco_hcd203040_compound15.26640.26640.4 | controllerType=0 controllerNumber=1 scan=26640 | 55.9996 |
| 25607:4 | 13 | R.QLAHQSN[+2861.000]STNIFFSPVSIATAFAMLSLGTK.A | HexNAc(5)Hex(6)NeuAc(3) | N[+2861] | 1511.4214 | 4 | 6042.6637 | 6042.6351 | 0 | 4.7 | 64 | Specific | 329.15 | 302.33 | 302.33 | 6.75 | 50 | >sp\|P01009\|A1AT_HUMAN Alpha-1-antitrypsin OS=Homo sapiens OX=9606 GN=SERPINA1 PE=1 SV=3 | 642 | 20201118_lss_colostrum_1ng_glyco_hcd203040_compound15.29502.29502.4 | controllerType=0 controllerNumber=1 scan=29502 | 61.4724 |
| 22782:3 | 13 | R.QLAHQSN[+2350.830]STNIFFSPVSIATAFAMLSLGTK.A | HexNAc(4)Hex(5)Fuc(1)NeuAc(2) | N[+2351] | 1844.8307 | 3 | 5532.4774 | 5532.4654 | 0 | 2.2 | 64 | Specific | 339.08 | 339.08 | 339.08 | 7.86 | 50 | >sp\|P01009\|A1AT_HUMAN Alpha-1-antitrypsin OS=Homo sapiens OX=9606 GN=SERPINA1 PE=1 SV=3 | 642 | 20201118_lss_colostrum_1ng_glyco_hcd203040_compound15.26565.26565.3 | controllerType=0 controllerNumber=1 scan=26565 | 55.8483 |
| 25392:4 | 13 | R.QLAHQSN[+2861.000]STNIFFSPVSIATAFAMLSLGTK.A | HexNAc(5)Hex(6)NeuAc(3) | N[+2861] | 1511.4248 | 4 | 6042.6774 | 6042.6351 | 0 | 7 | 64 | Specific | 342.19 | 342.19 | 342.19 | 6.74 | 50 | >sp\|P01009\|A1AT_HUMAN Alpha-1-antitrypsin OS=Homo sapiens OX=9606 GN=SERPINA1 PE=1 SV=3 | 642 | 20201118_lss_colostrum_1ng_glyco_hcd203040_compound15.29278.29278.4 | controllerType=0 controllerNumber=1 scan=29278 | 61.0526 |
| 22849:4 | 13 | R.[+42.011]QLAHQSN[+2190.793]STNIFFSPVSIATAFAMLSLGTK.A | HexNAc(6)Hex(6) | .[42], N[+2191] | 1354.3586 | 4 | 5414.4128 | 5414.4388 | 0 | -4.8 | 64 | Specific | 377.34 | 364.9 | 364.9 | 6.59 | 50 | >sp\|P01009\|A1AT_HUMAN Alpha-1-antitrypsin OS=Homo sapiens OX=9606 GN=SERPINA1 PE=1 SV=3 | 642 | 20201118_lss_colostrum_1ng_glyco_hcd203040_compound15.26635.26635.4 | controllerType=0 controllerNumber=1 scan=26635 | 55.9896 |
| 21073:4 | 13 | R.[+42.011]QLAHQSN[+1872.651]STNIFFSPVSIATAFAMLSLGTK.A | HexNAc(3)Hex(6)NeuAc(1) | .[42], N[+1873] | 1274.8401 | 4 | 5096.3385 | 5096.2961 | 0 | 8.3 | 64 | Specific | 380.74 | 360.82 | 360.82 | 5.59 | 50 | >sp\|P01009\|A1AT_HUMAN Alpha-1-antitrypsin OS=Homo sapiens OX=9606 GN=SERPINA1 PE=1 SV=3 | 642 | 20201118_lss_colostrum_1ng_glyco_hcd203040_compound15.24781.24781.4 | controllerType=0 controllerNumber=1 scan=24781 | 51.9783 |
| 25394:4 | 13 | R.QLAHQSN[+2861.000]STNIFFSPVSIATAFAMLSLGTK.A | HexNAc(5)Hex(6)NeuAc(3) | N[+2861] | 1511.4248 | 4 | 6042.6774 | 6042.6351 | 0 | 7 | 64 | Specific | 382.63 | 374.66 | 374.66 | 7.12 | 50 | >sp\|P01009\|A1AT_HUMAN Alpha-1-antitrypsin OS=Homo sapiens OX=9606 GN=SERPINA1 PE=1 SV=3 | 642 | 20201118_lss_colostrum_1ng_glyco_hcd203040_compound15.29280.29280.4 | controllerType=0 controllerNumber=1 scan=29280 | 61.0578 |
| 21071:4 | 13 | R.[+42.011]QLAHQSN[+1872.651]STNIFFSPVSIATAFAMLSLGTK.A | HexNAc(3)Hex(6)NeuAc(1) | .[42], N[+1873] | 1274.8401 | 4 | 5096.3385 | 5096.2961 | 0 | 8.3 | 64 | Specific | 385.41 | 373.89 | 373.89 | 6.91 | 50 | >sp\|P01009\|A1AT_HUMAN Alpha-1-antitrypsin OS=Homo sapiens OX=9606 GN=SERPINA1 PE=1 SV=3 | 642 | 20201118_lss_colostrum_1ng_glyco_hcd203040_compound15.24779.24779.4 | controllerType=0 controllerNumber=1 scan=24779 | 51.9728 |
| 25609:4 | 13 | R.QLAHQSN[+2861.000]STNIFFSPVSIATAFAMLSLGTK.A | HexNAc(5)Hex(6)NeuAc(3) | N[+2861] | 1511.4214 | 4 | 6042.6637 | 6042.6351 | 0 | 4.7 | 64 | Specific | 389.5 | 372.49 | 372.49 | 7.52 | 50 | >sp\|P01009\|A1AT_HUMAN Alpha-1-antitrypsin OS=Homo sapiens OX=9606 GN=SERPINA1 PE=1 SV=3 | 642 | 20201118_lss_colostrum_1ng_glyco_hcd203040_compound15.29504.29504.4 | controllerType=0 controllerNumber=1 scan=29504 | 61.4763 |
| 22847:4 | 13 | R.[+42.011]QLAHQSN[+2190.793]STNIFFSPVSIATAFAMLSLGTK.A | HexNAc(6)Hex(6) | .[42], N[+2191] | 1354.3586 | 4 | 5414.4128 | 5414.4388 | 0 | -4.8 | 64 | Specific | 440.39 | 440.39 | 440.39 | 8.3 | 50 | >sp\|P01009\|A1AT_HUMAN Alpha-1-antitrypsin OS=Homo sapiens OX=9606 GN=SERPINA1 PE=1 SV=3 | 642 | 20201118_lss_colostrum_1ng_glyco_hcd203040_compound15.26633.26633.4 | controllerType=0 controllerNumber=1 scan=26633 | 55.983 |
| 22856:4 | 13 | R.QLAHQSN[+2350.830]STNIFFSPVSIATAFAMLSLGTK.A | HexNAc(4)Hex(5)Fuc(1)NeuAc(2) | N[+2351] | 1383.8694 | 4 | 5532.4557 | 5532.4654 | 0 | -1.7 | 64 | Specific | 459.85 | 439.41 | 439.41 | 8.85 | 50 | >sp\|P01009\|A1AT_HUMAN Alpha-1-antitrypsin OS=Homo sapiens OX=9606 GN=SERPINA1 PE=1 SV=3 | 642 | 20201118_lss_colostrum_1ng_glyco_hcd203040_compound15.26642.26642.4 | controllerType=0 controllerNumber=1 scan=26642 | 56.0032 |
| 22756:4 | 13 | R.QLAHQSN[+2350.830]STNIFFSPVSIATAFAMLSLGTK.A | HexNAc(4)Hex(5)Fuc(1)NeuAc(2) | N[+2351] | 1383.8755 | 4 | 5532.4801 | 5532.4654 | 0 | 2.7 | 64 | Specific | 806.02 | 757.64 | 757.64 | 16.83 | 50 | >sp\|P01009\|A1AT_HUMAN Alpha-1-antitrypsin OS=Homo sapiens OX=9606 GN=SERPINA1 PE=1 SV=3 | 642 | 20201118_lss_colostrum_1ng_glyco_hcd203040_compound15.26538.26538.4 | controllerType=0 controllerNumber=1 scan=26538 | 55.7835 |
| 22775:4 | 13 | R.QLAHQSN[+2350.830]STNIFFSPVSIATAFAMLSLGTK.A | HexNAc(4)Hex(5)Fuc(1)NeuAc(2) | N[+2351] | 1383.8751 | 4 | 5532.4787 | 5532.4654 | 0 | 2.4 | 64 | Specific | 816.13 | 741.05 | 741.05 | 16.83 | 50 | >sp\|P01009\|A1AT_HUMAN Alpha-1-antitrypsin OS=Homo sapiens OX=9606 GN=SERPINA1 PE=1 SV=3 | 642 | 20201118_lss_colostrum_1ng_glyco_hcd203040_compound15.26558.26558.4 | controllerType=0 controllerNumber=1 scan=26558 | 55.8277 |
| 22758:4 | 13 | R.QLAHQSN[+2350.830]STNIFFSPVSIATAFAMLSLGTK.A | HexNAc(4)Hex(5)Fuc(1)NeuAc(2) | N[+2351] | 1383.8755 | 4 | 5532.4801 | 5532.4654 | 0 | 2.7 | 64 | Specific | 821.48 | 760.65 | 760.65 | 16.83 | 50 | >sp\|P01009\|A1AT_HUMAN Alpha-1-antitrypsin OS=Homo sapiens OX=9606 GN=SERPINA1 PE=1 SV=3 | 642 | 20201118_lss_colostrum_1ng_glyco_hcd203040_compound15.26540.26540.4 | controllerType=0 controllerNumber=1 scan=26540 | 55.7886 |
| 22781:3 | 13 | R.QLAHQSN[+2204.772]STNIFFSPVSIATAFAMLSLGTK.A | HexNAc(4)Hex(5)NeuAc(2) | N[+2205] | 1796.1453 | 3 | 5386.4215 | 5386.4075 | 0 | 2.6 | 64 | Specific | 828.84 | 809.03 | 524.38 | 16.9 | 50 | >sp\|P01009\|A1AT_HUMAN Alpha-1-antitrypsin OS=Homo sapiens OX=9606 GN=SERPINA1 PE=1 SV=3 | 642 | 20201118_lss_colostrum_1ng_glyco_hcd203040_compound15.26564.26564.3 | controllerType=0 controllerNumber=1 scan=26564 | 55.8464 |
| 22770:3 | 13 | R.QLAHQSN[+2204.772]STNIFFSPVSIATAFAMLSLGTK.A | HexNAc(4)Hex(5)NeuAc(2) | N[+2205] | 1796.1464 | 3 | 5386.4248 | 5386.4075 | 0 | 3.2 | 64 | Specific | 837.17 | 800.34 | 800.34 | 16.9 | 50 | >sp\|P01009\|A1AT_HUMAN Alpha-1-antitrypsin OS=Homo sapiens OX=9606 GN=SERPINA1 PE=1 SV=3 | 642 | 20201118_lss_colostrum_1ng_glyco_hcd203040_compound15.26552.26552.3 | controllerType=0 controllerNumber=1 scan=26552 | 55.8154 |
| 22772:3 | 13 | R.QLAHQSN[+2204.772]STNIFFSPVSIATAFAMLSLGTK.A | HexNAc(4)Hex(5)NeuAc(2) | N[+2205] | 1796.1464 | 3 | 5386.4248 | 5386.4075 | 0 | 3.2 | 64 | Specific | 843.59 | 754.56 | 754.56 | 16.9 | 50 | >sp\|P01009\|A1AT_HUMAN Alpha-1-antitrypsin OS=Homo sapiens OX=9606 GN=SERPINA1 PE=1 SV=3 | 642 | 20201118_lss_colostrum_1ng_glyco_hcd203040_compound15.26554.26554.3 | controllerType=0 controllerNumber=1 scan=26554 | 55.8198 |
| 22779:3 | 13 | R.QLAHQSN[+2204.772]STNIFFSPVSIATAFAMLSLGTK.A | HexNAc(4)Hex(5)NeuAc(2) | N[+2205] | 1796.1453 | 3 | 5386.4215 | 5386.4075 | 0 | 2.6 | 64 | Specific | 886.82 | 816.12 | 567.14 | 17.9 | 50 | >sp\|P01009\|A1AT_HUMAN Alpha-1-antitrypsin OS=Homo sapiens OX=9606 GN=SERPINA1 PE=1 SV=3 | 642 | 20201118_lss_colostrum_1ng_glyco_hcd203040_compound15.26562.26562.3 | controllerType=0 controllerNumber=1 scan=26562 | 55.8391 |
| 22776:4 | 13 | R.QLAHQSN[+2204.772]STNIFFSPVSIATAFAMLSLGTK.A | HexNAc(4)Hex(5)NeuAc(2) | N[+2205] | 1347.3607 | 4 | 5386.4211 | 5386.4075 | 0 | 2.5 | 64 | Specific | 897.3 | 848.43 | 597 | 17.83 | 50 | >sp\|P01009\|A1AT_HUMAN Alpha-1-antitrypsin OS=Homo sapiens OX=9606 GN=SERPINA1 PE=1 SV=3 | 642 | 20201118_lss_colostrum_1ng_glyco_hcd203040_compound15.26559.26559.4 | controllerType=0 controllerNumber=1 scan=26559 | 55.8336 |
| 22777:4 | 13 | R.QLAHQSN[+2350.830]STNIFFSPVSIATAFAMLSLGTK.A | HexNAc(4)Hex(5)Fuc(1)NeuAc(2) | N[+2351] | 1383.8751 | 4 | 5532.4787 | 5532.4654 | 0 | 2.4 | 64 | Specific | 899.26 | 813.42 | 813.42 | 17.83 | 50 | >sp\|P01009\|A1AT_HUMAN Alpha-1-antitrypsin OS=Homo sapiens OX=9606 GN=SERPINA1 PE=1 SV=3 | 642 | 20201118_lss_colostrum_1ng_glyco_hcd203040_compound15.26560.26560.4 | controllerType=0 controllerNumber=1 scan=26560 | 55.8358 |
| 22774:4 | 13 | R.QLAHQSN[+2204.772]STNIFFSPVSIATAFAMLSLGTK.A | HexNAc(4)Hex(5)NeuAc(2) | N[+2205] | 1347.3607 | 4 | 5386.4211 | 5386.4075 | 0 | 2.5 | 64 | Specific | 925.49 | 898.63 | 578.29 | 18.83 | 50 | >sp\|P01009\|A1AT_HUMAN Alpha-1-antitrypsin OS=Homo sapiens OX=9606 GN=SERPINA1 PE=1 SV=3 | 642 | 20201118_lss_colostrum_1ng_glyco_hcd203040_compound15.26557.26557.4 | controllerType=0 controllerNumber=1 scan=26557 | 55.8263 |
| 23888:4 | 13 | K.ADTHDEILEGLNFN[+3517.228]LTEIPEAQIHEGFQELLR.T | HexNAc(6)Hex(7)NeuAc(4) | N[+3517] | 1803.0239 | 4 | 7209.0737 | 7209.0515 | 0 | 3.1 | 94 | Specific | 330.4 | 276.91 | 276.91 | 7.63 | 50 | >sp\|P01009\|A1AT_HUMAN Alpha-1-antitrypsin OS=Homo sapiens OX=9606 GN=SERPINA1 PE=1 SV=3 | 642 | 20201118_lss_colostrum_1ng_glyco_hcd203040_compound15.27715.27715.4 | controllerType=0 controllerNumber=1 scan=27715 | 58.1026 |
| 23890:4 | 13 | K.ADTHDEILEGLNFN[+3517.228]LTEIPEAQIHEGFQELLR.T | HexNAc(6)Hex(7)NeuAc(4) | N[+3517] | 1803.0239 | 4 | 7209.0737 | 7209.0515 | 0 | 3.1 | 94 | Specific | 393.61 | 392.12 | 392.12 | 10.03 | 50 | >sp\|P01009\|A1AT_HUMAN Alpha-1-antitrypsin OS=Homo sapiens OX=9606 GN=SERPINA1 PE=1 SV=3 | 642 | 20201118_lss_colostrum_1ng_glyco_hcd203040_compound15.27717.27717.4 | controllerType=0 controllerNumber=1 scan=27717 | 58.106 |
| 23750:4 | 13 | K.ADTHDEILEGLNFN[+3517.228]LTEIPEAQIHEGFQELLR.T | HexNAc(6)Hex(7)NeuAc(4) | N[+3517] | 1803.0268 | 4 | 7209.0854 | 7209.0515 | 0 | 4.7 | 94 | Specific | 412.01 | 412.01 | 412.01 | 9.43 | 50 | >sp\|P01009\|A1AT_HUMAN Alpha-1-antitrypsin OS=Homo sapiens OX=9606 GN=SERPINA1 PE=1 SV=3 | 642 | 20201118_lss_colostrum_1ng_glyco_hcd203040_compound15.27572.27572.4 | controllerType=0 controllerNumber=1 scan=27572 | 57.8364 |
| 20305:4 | 13 | K.[+42.011]ADTHDEILEGLNFN[+2190.793]LTEIPEAQIHEGFQELLR.T | HexNAc(6)Hex(6) | .[42], N[+2191] | 1481.9106 | 4 | 5924.6208 | 5924.6276 | 0 | -1.2 | 94 | Specific | 419.46 | 411.23 | 411.23 | 9.01 | 50 | >sp\|P01009\|A1AT_HUMAN Alpha-1-antitrypsin OS=Homo sapiens OX=9606 GN=SERPINA1 PE=1 SV=3 | 642 | 20201118_lss_colostrum_1ng_glyco_hcd203040_compound15.23978.23978.4 | controllerType=0 controllerNumber=1 scan=23978 | 50.1495 |
| 20302:4 | 13 | K.[+42.011]ADTHDEILEGLNFN[+2190.793]LTEIPEAQIHEGFQELLR.T | HexNAc(6)Hex(6) | .[42], N[+2191] | 1481.9106 | 4 | 5924.6208 | 5924.6276 | 0 | -1.2 | 94 | Specific | 446.64 | 399.31 | 399.31 | 9.2 | 50 | >sp\|P01009\|A1AT_HUMAN Alpha-1-antitrypsin OS=Homo sapiens OX=9606 GN=SERPINA1 PE=1 SV=3 | 642 | 20201118_lss_colostrum_1ng_glyco_hcd203040_compound15.23975.23975.4 | controllerType=0 controllerNumber=1 scan=23975 | 50.1418 |
| 21600:4 | 13 | K.ADTHDEILEGLNFN[+3007.058]LTEIPEAQIHEGFQELLR.T | HexNAc(5)Hex(6)Fuc(1)NeuAc(3) | N[+3007] | 1675.4718 | 4 | 6698.8652 | 6698.8818 | 0 | -2.5 | 94 | Specific | 671.92 | 590.55 | 590.55 | 12.82 | 50 | >sp\|P01009\|A1AT_HUMAN Alpha-1-antitrypsin OS=Homo sapiens OX=9606 GN=SERPINA1 PE=1 SV=3 | 642 | 20201118_lss_colostrum_1ng_glyco_hcd203040_compound15.25331.25331.4 | controllerType=0 controllerNumber=1 scan=25331 | 53.1383 |
| 21646:4 | 13 | K.ADTHDEILEGLNFN[+2861.000]LTEIPEAQIHEGFQELLR.T | HexNAc(5)Hex(6)NeuAc(3) | N[+2861] | 1638.9683 | 4 | 6552.8512 | 6552.8239 | 0 | 4.2 | 94 | Specific | 821.19 | 711.94 | 711.94 | 16.98 | 50 | >sp\|P01009\|A1AT_HUMAN Alpha-1-antitrypsin OS=Homo sapiens OX=9606 GN=SERPINA1 PE=1 SV=3 | 642 | 20201118_lss_colostrum_1ng_glyco_hcd203040_compound15.25379.25379.4 | controllerType=0 controllerNumber=1 scan=25379 | 53.2468 |
| 21602:4 | 13 | K.ADTHDEILEGLNFN[+3007.058]LTEIPEAQIHEGFQELLR.T | HexNAc(5)Hex(6)Fuc(1)NeuAc(3) | N[+3007] | 1675.4718 | 4 | 6698.8652 | 6698.8818 | 0 | -2.5 | 94 | Specific | 865.79 | 550.36 | 550.36 | 16.82 | 50 | >sp\|P01009\|A1AT_HUMAN Alpha-1-antitrypsin OS=Homo sapiens OX=9606 GN=SERPINA1 PE=1 SV=3 | 642 | 20201118_lss_colostrum_1ng_glyco_hcd203040_compound15.25333.25333.4 | controllerType=0 controllerNumber=1 scan=25333 | 53.1427 |
| 21644:4 | 13 | K.ADTHDEILEGLNFN[+2861.000]LTEIPEAQIHEGFQELLR.T | HexNAc(5)Hex(6)NeuAc(3) | N[+2861] | 1638.9683 | 4 | 6552.8512 | 6552.8239 | 0 | 4.2 | 94 | Specific | 914 | 914 | 914 | 18.98 | 50 | >sp\|P01009\|A1AT_HUMAN Alpha-1-antitrypsin OS=Homo sapiens OX=9606 GN=SERPINA1 PE=1 SV=3 | 642 | 20201118_lss_colostrum_1ng_glyco_hcd203040_compound15.25377.25377.4 | controllerType=0 controllerNumber=1 scan=25377 | 53.2413 |
| 21617:5 | 13 | K.ADTHDEILEGLNFN[+2861.000]LTEIPEAQIHEGFQELLR.T | HexNAc(5)Hex(6)NeuAc(3) | N[+2861] | 1311.3728 | 5 | 6552.8349 | 6552.8239 | 0 | 1.7 | 94 | Specific | 1114.39 | 1073.26 | 1073.26 | 22.95 | 50 | >sp\|P01009\|A1AT_HUMAN Alpha-1-antitrypsin OS=Homo sapiens OX=9606 GN=SERPINA1 PE=1 SV=3 | 642 | 20201118_lss_colostrum_1ng_glyco_hcd203040_compound15.25348.25348.2 | controllerType=0 controllerNumber=1 scan=25348 | 53.1741 |
| 21619:5 | 13 | K.ADTHDEILEGLNFN[+2861.000]LTEIPEAQIHEGFQELLR.T | HexNAc(5)Hex(6)NeuAc(3) | N[+2861] | 1311.3728 | 5 | 6552.8349 | 6552.8239 | 0 | 1.7 | 94 | Specific | 1134.23 | 978.51 | 978.51 | 22.95 | 50 | >sp\|P01009\|A1AT_HUMAN Alpha-1-antitrypsin OS=Homo sapiens OX=9606 GN=SERPINA1 PE=1 SV=3 | 642 | 20201118_lss_colostrum_1ng_glyco_hcd203040_compound15.25350.25350.2 | controllerType=0 controllerNumber=1 scan=25350 | 53.1808 |
| 19338:5 | 13 | K.[+42.011]YLGN[+2190.793]ATAIFFLPDEGKLQHLENELTHDIITK.F | HexNAc(6)Hex(6) | .[42], N[+2191] | 1155.5305 | 5 | 5773.6235 | 5773.6411 | 0 | -3 | 268 | Specific | 334.74 | 281.65 | 281.65 | 5.63 | 50 | >sp\|P01009\|A1AT_HUMAN Alpha-1-antitrypsin OS=Homo sapiens OX=9606 GN=SERPINA1 PE=1 SV=3 | 642 | 20201118_lss_colostrum_1ng_glyco_hcd203040_compound15.22968.22968.5 | controllerType=0 controllerNumber=1 scan=22968 | 47.7972 |
| 19336:5 | 13 | K.[+42.011]YLGN[+2190.793]ATAIFFLPDEGKLQHLENELTHDIITK.F | HexNAc(6)Hex(6) | .[42], N[+2191] | 1155.5305 | 5 | 5773.6235 | 5773.6411 | 0 | -3 | 268 | Specific | 350.54 | 350.54 | 350.54 | 6.03 | 50 | >sp\|P01009\|A1AT_HUMAN Alpha-1-antitrypsin OS=Homo sapiens OX=9606 GN=SERPINA1 PE=1 SV=3 | 642 | 20201118_lss_colostrum_1ng_glyco_hcd203040_compound15.22966.22966.5 | controllerType=0 controllerNumber=1 scan=22966 | 47.7929 |
| 19339:4 | 13 | K.[+42.011]YLGN[+2190.793]ATAIFFLPDEGKLQHLENELTHDIITK.F | HexNAc(6)Hex(6) | .[42], N[+2191] | 1444.1616 | 4 | 5773.6247 | 5773.6411 | 0 | -2.8 | 268 | Specific | 393.4 | 393.4 | 393.4 | 7.3 | 50 | >sp\|P01009\|A1AT_HUMAN Alpha-1-antitrypsin OS=Homo sapiens OX=9606 GN=SERPINA1 PE=1 SV=3 | 642 | 20201118_lss_colostrum_1ng_glyco_hcd203040_compound15.22969.22969.4 | controllerType=0 controllerNumber=1 scan=22969 | 47.8021 |
| 19337:4 | 13 | K.[+42.011]YLGN[+2190.793]ATAIFFLPDEGKLQHLENELTHDIITK.F | HexNAc(6)Hex(6) | .[42], N[+2191] | 1444.1616 | 4 | 5773.6247 | 5773.6411 | 0 | -2.8 | 268 | Specific | 483.47 | 483.47 | 483.47 | 7.63 | 50 | >sp\|P01009\|A1AT_HUMAN Alpha-1-antitrypsin OS=Homo sapiens OX=9606 GN=SERPINA1 PE=1 SV=3 | 642 | 20201118_lss_colostrum_1ng_glyco_hcd203040_compound15.22967.22967.4 | controllerType=0 controllerNumber=1 scan=22967 | 47.7943 |
| 19275:5 | 13 | K.YLGN[+2204.772]ATAIFFLPDEGKLQHLENELTHDIITK.F | HexNAc(4)Hex(5)NeuAc(2) | N[+2205] | 1149.9305 | 5 | 5745.6236 | 5745.6098 | 0 | 2.4 | 268 | Specific | 892.98 | 844.26 | 844.26 | 17.43 | 50 | >sp\|P01009\|A1AT_HUMAN Alpha-1-antitrypsin OS=Homo sapiens OX=9606 GN=SERPINA1 PE=1 SV=3 | 642 | 20201118_lss_colostrum_1ng_glyco_hcd203040_compound15.22902.22902.5 | controllerType=0 controllerNumber=1 scan=22902 | 47.6519 |
| 19276:4 | 13 | K.YLGN[+2204.772]ATAIFFLPDEGKLQHLENELTHDIITK.F | HexNAc(4)Hex(5)NeuAc(2) | N[+2205] | 1437.162 | 4 | 5745.6261 | 5745.6098 | 0 | 2.9 | 268 | Specific | 979.38 | 916.56 | 916.56 | 18.8 | 50 | >sp\|P01009\|A1AT_HUMAN Alpha-1-antitrypsin OS=Homo sapiens OX=9606 GN=SERPINA1 PE=1 SV=3 | 642 | 20201118_lss_colostrum_1ng_glyco_hcd203040_compound15.22903.22903.4 | controllerType=0 controllerNumber=1 scan=22903 | 47.6567 |
| 19272:5 | 13 | K.YLGN[+2204.772]ATAIFFLPDEGKLQHLENELTHDIITK.F | HexNAc(4)Hex(5)NeuAc(2) | N[+2205] | 1149.9305 | 5 | 5745.6236 | 5745.6098 | 0 | 2.4 | 268 | Specific | 1120.94 | 1075.1 | 1075.1 | 22.43 | 50 | >sp\|P01009\|A1AT_HUMAN Alpha-1-antitrypsin OS=Homo sapiens OX=9606 GN=SERPINA1 PE=1 SV=3 | 642 | 20201118_lss_colostrum_1ng_glyco_hcd203040_compound15.22899.22899.5 | controllerType=0 controllerNumber=1 scan=22899 | 47.6445 |
| 19273:4 | 13 | K.YLGN[+2204.772]ATAIFFLPDEGKLQHLENELTHDIITK.F | HexNAc(4)Hex(5)NeuAc(2) | N[+2205] | 1437.162 | 4 | 5745.6261 | 5745.6098 | 0 | 2.9 | 268 | Specific | 1254.78 | 1178.18 | 1178.18 | 24.8 | 50 | >sp\|P01009\|A1AT_HUMAN Alpha-1-antitrypsin OS=Homo sapiens OX=9606 GN=SERPINA1 PE=1 SV=3 | 642 | 20201118_lss_colostrum_1ng_glyco_hcd203040_compound15.22900.22900.4 | controllerType=0 controllerNumber=1 scan=22900 | 47.6457 |
| 13405:3 | 14 | R.FELLGC[+57.021]ELNGC[+57.021]ANPLGLKN[+1216.423]NSIPDK.Q | HexNAc(2)Hex(5) | C[+57]*2, N[+1216] | 1330.6006 | 3 | 3989.7874 | 3989.7876 | 0 | -0.1 | 220 | Specific | 653.45 | 581.27 | 581.27 | 12.67 | 40 | >sp\|Q08431\|MFGM_HUMAN Lactadherin OS=Homo sapiens OX=9606 GN=MFGE8 PE=1 SV=3 | 1064 | 20201118_lss_colostrum_1ng_glyco_hcd203040_compound15.16791.16791.3 | controllerType=0 controllerNumber=1 scan=16791 | 35.1642 |
| 13403:3 | 14 | R.FELLGC[+57.021]ELNGC[+57.021]ANPLGLKN[+1216.423]NSIPDK.Q | HexNAc(2)Hex(5) | C[+57]*2, N[+1216] | 1330.6006 | 3 | 3989.7874 | 3989.7876 | 0 | -0.1 | 220 | Specific | 814.08 | 707.61 | 707.61 | 15.67 | 40 | >sp\|Q08431\|MFGM_HUMAN Lactadherin OS=Homo sapiens OX=9606 GN=MFGE8 PE=1 SV=3 | 1064 | 20201118_lss_colostrum_1ng_glyco_hcd203040_compound15.16789.16789.3 | controllerType=0 controllerNumber=1 scan=16789 | 35.1597 |
| 14300:4 | 14 | R.FELLGC[+57.021]ELNGC[+57.021]ANPLGLKN[+1872.651]NSIPDKQITASSSYK.T | HexNAc(3)Hex(6)NeuAc(1) | C[+57]*2, N[+1873] | 1403.6241 | 4 | 5611.4748 | 5611.497 | 0 | -4 | 220 | Specific | 514.98 | 514.98 | 514.98 | 8.47 | 40 | >sp\|Q08431\|MFGM_HUMAN Lactadherin OS=Homo sapiens OX=9606 GN=MFGE8 PE=1 SV=3 | 1064 | 20201118_lss_colostrum_1ng_glyco_hcd203040_compound15.17721.17721.4 | controllerType=0 controllerNumber=1 scan=17721 | 36.9909 |
| 14303:4 | 14 | R.FELLGC[+57.021]ELNGC[+57.021]ANPLGLKN[+1872.651]NSIPDKQITASSSYK.T | HexNAc(3)Hex(6)NeuAc(1) | C[+57]*2, N[+1873] | 1403.6241 | 4 | 5611.4748 | 5611.497 | 0 | -4 | 220 | Specific | 536.92 | 369.08 | 369.08 | 6.97 | 40 | >sp\|Q08431\|MFGM_HUMAN Lactadherin OS=Homo sapiens OX=9606 GN=MFGE8 PE=1 SV=3 | 1064 | 20201118_lss_colostrum_1ng_glyco_hcd203040_compound15.17724.17724.4 | controllerType=0 controllerNumber=1 scan=17724 | 36.9954 |
| 14246:4 | 14 | R.FELLGC[+57.021]ELNGC[+57.021]ANPLGLKN[+1872.651]NSIPDKQITASSSYK.T | HexNAc(3)Hex(6)NeuAc(1) | C[+57]*2, N[+1873] | 1403.6304 | 4 | 5611.4997 | 5611.497 | 0 | 0.5 | 220 | Specific | 648.21 | 481.59 | 481.59 | 11.83 | 40 | >sp\|Q08431\|MFGM_HUMAN Lactadherin OS=Homo sapiens OX=9606 GN=MFGE8 PE=1 SV=3 | 1064 | 20201118_lss_colostrum_1ng_glyco_hcd203040_compound15.17665.17665.4 | controllerType=0 controllerNumber=1 scan=17665 | 36.9045 |
| 12621:4 | 14 | R.FELLGC[+57.021]ELNGC[+57.021]ANPLGLKN[+1378.476]NSIPDKQITASSSYK.T | HexNAc(2)Hex(6) | C[+57]*2, N[+1378] | 1280.0867 | 4 | 5117.3249 | 5117.3223 | 0 | 0.5 | 220 | Specific | 690.07 | 606.8 | 606.8 | 12.83 | 40 | >sp\|Q08431\|MFGM_HUMAN Lactadherin OS=Homo sapiens OX=9606 GN=MFGE8 PE=1 SV=3 | 1064 | 20201118_lss_colostrum_1ng_glyco_hcd203040_compound15.15977.15977.4 | controllerType=0 controllerNumber=1 scan=15977 | [illegible] |
| 14249:4 | 14 | R.FELLGC[+57.021]ELNGC[+57.021]ANPLGLKN[+1872.651]NSIPDKQITASSSYK.T | HexNAc(3)Hex(6)NeuAc(1) | C[+57]*2, N[+1873] | 1403.6304 | 4 | 5611.4997 | 5611.497 | 0 | 0.5 | 220 | Specific | 772.46 | 618.45 | 618.45 | 14.83 | 40 | >sp\|Q08431\|MFGM_HUMAN Lactadherin OS=Homo sapiens OX=9606 GN=MFGE8 PE=1 SV=3 | 1064 | 20201118_lss_colostrum_1ng_glyco_hcd203040_compound15.17668.17668.4 | controllerType=0 controllerNumber=1 scan=17668 | [illegible] |
| 12623:4 | 14 | R.FELLGC[+57.021]ELNGC[+57.021]ANPLGLKN[+1378.476]NSIPDKQITASSSYK.T | HexNAc(2)Hex(6) | C[+57]*2, N[+1378] | 1280.0867 | 4 | 5117.3249 | 5117.3223 | 0 | 0.5 | 220 | Specific | 852.46 | 596.94 | 596.94 | 16.83 | 40 | >sp\|Q08431\|MFGM_HUMAN Lactadherin OS=Homo sapiens OX=9606 GN=MFGE8 PE=1 SV=3 | 1064 | 20201118_lss_colostrum_1ng_glyco_hcd203040_compound15.15979.15979.4 | controllerType=0 controllerNumber=1 scan=15979 | [illegible] |
| 12660:4 | 14 | R.FELLGC[+57.021]ELNGC[+57.021]ANPLGLKN[+1216.423]NSIPDKQITASSSYK.T | HexNAc(2)Hex(5) | C[+57]*2, N[+1216] | 1239.5762 | 4 | 4955.2829 | 4955.2694 | 0 | 2.7 | 220 | Specific | 1070.54 | 919.41 | 919.41 | 21.45 | 40 | >sp\|Q08431\|MFGM_HUMAN Lactadherin OS=Homo sapiens OX=9606 GN=MFGE8 PE=1 SV=3 | 1064 | 20201118_lss_colostrum_1ng_glyco_hcd203040_compound15.16018.16018.4 | controllerType=0 controllerNumber=1 scan=16018 | [illegible] |
| 12658:4 | 14 | R.FELLGC[+57.021]ELNGC[+57.021]ANPLGLKN[+1216.423]NSIPDKQITASSSYK.T | HexNAc(2)Hex(5) | C[+57]*2, N[+1216] | 1239.5762 | 4 | 4955.2829 | 4955.2694 | 0 | 2.7 | 220 | Specific | 1124.63 | 1029.83 | 1029.83 | 22.45 | 40 | >sp\|Q08431\|MFGM_HUMAN Lactadherin OS=Homo sapiens OX=9606 GN=MFGE8 PE=1 SV=3 | 1064 | 20201118_lss_colostrum_1ng_glyco_hcd203040_compound15.16016.16016.4 | controllerType=0 controllerNumber=1 scan=16016 | 33.6419 |
| 01830:3 | 14 | K.N[+1378.476]NSIPDKQITASSSYK.T | HexNAc(2)Hex(6) | N[+1378] | 1044.4583 | 3 | 3131.3603 | 3131.352 | 0 | 2.7 | 238 | Specific | 390.24 | 187.27 | 187.27 | 6.35 | 40 | >sp\|Q08431\|MFGM_HUMAN Lactadherin OS=Homo sapiens OX=9606 GN=MFGE8 PE=1 SV=3 | 1064 | 20201118_lss_colostrum_1ng_glyco_hcd203040_compound15.4789.4789.3 | controllerType=0 controllerNumber=1 scan=4789 | 14.3821 |
| 01832:3 | 14 | K.N[+1378.476]NSIPDKQITASSSYK.T | HexNAc(2)Hex(6) | N[+1378] | 1044.4583 | 3 | 3131.3603 | 3131.352 | 0 | 2.7 | 238 | Specific | 402.07 | 231.3 | 231.3 | 6.6 | 40 | >sp\|Q08431\|MFGM_HUMAN Lactadherin OS=Homo sapiens OX=9606 GN=MFGE8 PE=1 SV=3 | 1064 | 20201118_lss_colostrum_1ng_glyco_hcd203040_compound15.4791.4791.3 | controllerType=0 controllerNumber=1 scan=4791 | 14.3857 |
| 01741:3 | 14 | K.N[+1540.529]NSIPDKQITASSSYK.T | HexNAc(2)Hex(7) | N[+1541] | 1098.4759 | 3 | 3293.413 | 3293.4048 | 0 | 2.5 | 238 | Specific | 502.8 | 215.31 | 215.31 | 6.94 | 40 | >sp\|Q08431\|MFGM_HUMAN Lactadherin OS=Homo sapiens OX=9606 GN=MFGE8 PE=1 SV=3 | 1064 | 20201118_lss_colostrum_1ng_glyco_hcd203040_compound15.4697.4697.3 | controllerType=0 controllerNumber=1 scan=4697 | 14.2251 |
| 01739:3 | 14 | K.N[+1540.529]NSIPDKQITASSSYK.T | HexNAc(2)Hex(7) | N[+1541] | 1098.4759 | 3 | 3293.413 | 3293.4048 | 0 | 2.5 | 238 | Specific | 521.99 | 275.8 | 275.8 | 7.38 | 40 | >sp\|Q08431\|MFGM_HUMAN Lactadherin OS=Homo sapiens OX=9606 GN=MFGE8 PE=1 SV=3 | 1064 | 20201118_lss_colostrum_1ng_glyco_hcd203040_compound15.4695.4695.3 | controllerType=0 controllerNumber=1 scan=4695 | 14.2214 |
| 01874:3 | 14 | K.N[+1216.423]NSIPDKQITASSSYK.T | HexNAc(2)Hex(5) | N[+1216] | 990.4401 | 3 | 2969.3057 | 2969.2991 | 0 | 2.2 | 238 | Specific | 721.98 | 497.88 | 497.88 | 14.09 | 40 | >sp\|Q08431\|MFGM_HUMAN Lactadherin OS=Homo sapiens OX=9606 GN=MFGE8 PE=1 SV=3 | 1064 | 20201118_lss_colostrum_1ng_glyco_hcd203040_compound15.4835.4835.3 | controllerType=0 controllerNumber=1 scan=4835 | 14.4647 |
| 01876:3 | 14 | K.N[+1216.423]NSIPDKQITASSSYK.T | HexNAc(2)Hex(5) | N[+1216] | 990.4401 | 3 | 2969.3057 | 2969.2991 | 0 | 2.2 | 238 | Specific | 763.42 | 518.44 | 518.44 | 15.09 | 40 | >sp\|Q08431\|MFGM_HUMAN Lactadherin OS=Homo sapiens OX=9606 GN=MFGE8 PE=1 SV=3 | 1064 | 20201118_lss_colostrum_1ng_glyco_hcd203040_compound15.4837.4837.3 | controllerType=0 controllerNumber=1 scan=4837 | 14.4693 |
| 08430:3 | 14 | K.VAYSN[+2059.735]DSANWTEYQDPR.T | HexNAc(4)Hex(5)Fuc(1)NeuAc(1) | N[+2060] | 1359.2125 | 3 | 4075.6228 | 4075.6079 | 0 | 3.7 | 321 | Specific | 220.88 | 149.54 | 6.01 | 4.05 | 40 | >sp\|Q08431\|MFGM_HUMAN Lactadherin OS=Homo sapiens OX=9606 GN=MFGE8 PE=1 SV=3 | 1064 | 20201118_lss_colostrum_1ng_glyco_hcd203040_compound15.11623.11623.3 | controllerType=0 controllerNumber=1 scan=11623 | 25.6806 |
| 08433:3 | 14 | K.VAYSN[+2059.735]DSANWTEYQDPR.T | HexNAc(4)Hex(5)Fuc(1)NeuAc(1) | N[+2060] | 1359.2125 | 3 | 4075.6228 | 4075.6079 | 0 | 3.7 | 321 | Specific | 291.11 | 223.52 | 4.29 | 6.44 | 40 | >sp\|Q08431\|MFGM_HUMAN Lactadherin OS=Homo sapiens OX=9606 GN=MFGE8 PE=1 SV=3 | 1064 | 20201118_lss_colostrum_1ng_glyco_hcd203040_compound15.11626.11626.3 | controllerType=0 controllerNumber=1 scan=11626 | 25.6848 |
| 08353:3 | 14 | K.VAYSN[+2059.735]DSANWTEYQDPR.T | HexNAc(4)Hex(5)Fuc(1)NeuAc(1) | N[+2060] | 1359.2118 | 3 | 4075.621 | 4075.6079 | 0 | 3.2 | 321 | Specific | 395.44 | 341.29 | 6.6 | 7.44 | 40 | >sp\|Q08431\|MFGM_HUMAN Lactadherin OS=Homo sapiens OX=9606 GN=MFGE8 PE=1 SV=3 | 1064 | 20201118_lss_colostrum_1ng_glyco_hcd203040_compound15.11543.11543.3 | controllerType=0 controllerNumber=1 scan=11543 | 25.5514 |
| 08356:3 | 14 | K.VAYSN[+2059.735]DSANWTEYQDPR.T | HexNAc(4)Hex(5)Fuc(1)NeuAc(1) | N[+2060] | 1359.2118 | 3 | 4075.621 | 4075.6079 | 0 | 3.2 | 321 | Specific | 553.01 | 485.94 | 4.21 | 9.7 | 40 | >sp\|Q08431\|MFGM_HUMAN Lactadherin OS=Homo sapiens OX=9606 GN=MFGE8 PE=1 SV=3 | 1064 | 20201118_lss_colostrum_1ng_glyco_hcd203040_compound15.11546.11546.3 | controllerType=0 controllerNumber=1 scan=11546 | 25.5562 |
| 23317:4 | 15 | K.APDKNVIFSPLSISTALAFLSLGAHN[+2861.000]TTLTEILK.G | HexNAc(5)Hex(6)NeuAc(3) | N[+2861] | 1611.7557 | 4 | 6444.0011 | 6443.9782 | 0 | 3.6 | 68 | Specific | 206.18 | 206.18 | 206.18 | 5.04 | 40 | >sp\|P01011\|AACT_HUMAN Alpha-1-antichymotrypsin OS=Homo sapiens OX=9606 GN=SERPINA3 PE=1 SV=2 | 1314 | 20201118_lss_colostrum_1ng_glyco_hcd203040_compound15.27122.27122.4 | controllerType=0 controllerNumber=1 scan=27122 | 56.9646 |
| 23315:4 | 15 | K.APDKNVIFSPLSISTALAFLSLGAHN[+2861.000]TTLTEILK.G | HexNAc(5)Hex(6)NeuAc(3) | N[+2861] | 1611.7557 | 4 | 6444.0011 | 6443.9782 | 0 | 3.6 | 68 | Specific | 297.41 | 297.41 | 297.41 | 7.06 | 40 | >sp\|P01011\|AACT_HUMAN Alpha-1-antichymotrypsin OS=Homo sapiens OX=9606 GN=SERPINA3 PE=1 SV=2 | 1314 | 20201118_lss_colostrum_1ng_glyco_hcd203040_compound15.27120.27120.4 | controllerType=0 controllerNumber=1 scan=27120 | 56.9597 |
| 21769:5 | 15 | K.APDKNVIFSPLSISTALAFLSLGAHN[+2350.830]TTLTEILK.G | HexNAc(4)Hex(5)Fuc(1)NeuAc(2) | N[+2351] | 1187.5721 | 5 | 5933.8316 | 5933.8085 | 0 | 3.9 | 68 | Specific | 757.52 | 757.52 | 757.52 | 15.44 | 40 | >sp\|P01011\|AACT_HUMAN Alpha-1-antichymotrypsin OS=Homo sapiens OX=9606 GN=SERPINA3 PE=1 SV=2 | 1314 | 20201118_lss_colostrum_1ng_glyco_hcd203040_compound15.25507.25507.5 | controllerType=0 controllerNumber=1 scan=25507 | 53.5386 |
| 21767:5 | 15 | K.APDKNVIFSPLSISTALAFLSLGAHN[+2350.830]TTLTEILK.G | HexNAc(4)Hex(5)Fuc(1)NeuAc(2) | N[+2351] | 1187.5721 | 5 | 5933.8316 | 5933.8085 | 0 | 3.9 | 68 | Specific | 949.99 | 928.2 | 928.2 | 18.44 | 40 | >sp\|P01011\|AACT_HUMAN Alpha-1-antichymotrypsin OS=Homo sapiens OX=9606 GN=SERPINA3 PE=1 SV=2 | 1314 | 20201118_lss_colostrum_1ng_glyco_hcd203040_compound15.25505.25505.5 | controllerType=0 controllerNumber=1 scan=25505 | 53.5317 |
| 21282:4 | 15 | K.YTGN[+2861.000]ASALFILPDQDKMEEVEAMLLPETLKR.W | HexNAc(5)Hex(6)NeuAc(3) | N[+2861] | 1596.7133 | 4 | 6383.8312 | 6383.7859 | 0 | 7.1 | 268 | Specific | 513.94 | 239.47 | 239.47 | 6.12 | 40 | >sp\|P01011\|AACT_HUMAN Alpha-1-antichymotrypsin OS=Homo sapiens OX=9606 GN=SERPINA3 PE=1 SV=2 | 1314 | 20201118_lss_colostrum_1ng_glyco_hcd203040_compound15.25000.25000.4 | controllerType=0 controllerNumber=1 scan=25000 | 52.4604 |
| 21284:4 | 15 | K.YTGN[+2861.000]ASALFILPDQDKMEEVEAMLLPETLKR.W | HexNAc(5)Hex(6)NeuAc(3) | N[+2861] | 1596.7133 | 4 | 6383.8312 | 6383.7859 | 0 | 7.1 | 268 | Specific | 523.04 | 293.13 | 293.13 | 6.53 | 40 | >sp\|P01011\|AACT_HUMAN Alpha-1-antichymotrypsin OS=Homo sapiens OX=9606 GN=SERPINA3 PE=1 SV=2 | 1314 | 20201118_lss_colostrum_1ng_glyco_hcd203040_compound15.25002.25002.4 | controllerType=0 controllerNumber=1 scan=25002 | 52.4683 |
| 11312:3 | 16 | R.TVDKSTGKPTLYN[+1378.476]VSLVMSDTAGTC[+57.021]Y.- | HexNAc(2)Hex(6) | C[+57], N[+1378] | 1396.2761 | 3 | 4186.8137 | 4186.8187 | 0 | -1.2 | 428 | CRagged | 325.59 | 307.08 | 244.34 | 5.79 | 32 | >sp\|P01871\|IGHM_HUMAN Immunoglobulin heavy constant mu OS=Homo sapiens OX=9606 GN=IGHM PE=1 SV=4 | 328 | 20201118_lss_colostrum_1ng_glyco_hcd203040_compound15.14619.14619.3 | controllerType=0 controllerNumber=1 scan=14619 | 31.146 |
| 21561:4 | 18 | K.DIDGYGGIALPELIC[+57.021]TMFHTSGYDTQAIVENN[+1686.586]ESTEYGLFQISNK. | HexNAc(2)Hex(7)Fuc(1) | C[+57], N[+1687] | 1675.2302 | 4 | 6697.8989 | 6697.9153 | 0 | -2.4 | 33 | Specific | 529.39 | 529.39 | 529.39 | 9.4 | 25 | >sp\|P00709\|LALBA_HUMAN Alpha-lactalbumin OS=Homo sapiens OX=9606 GN=LALBA PE=1 SV=1 | 468 | 20201118_lss_colostrum_1ng_glyco_hcd203040_compound15.25290.25290.3 | controllerType=0 controllerNumber=1 scan=25290 | 53.0395 |
| 21563:4 | 18 | K.DIDGYGGIALPELIC[+57.021]TMFHTSGYDTQAIVENN[+1686.586]ESTEYGLFQISNK. | HexNAc(2)Hex(7)Fuc(1) | C[+57], N[+1687] | 1675.2302 | 4 | 6697.8989 | 6697.9153 | 0 | -2.4 | 33 | Specific | 648.61 | 639.38 | 639.38 | 12 | 25 | >sp\|P00709\|LALBA_HUMAN Alpha-lactalbumin OS=Homo sapiens OX=9606 GN=LALBA PE=1 SV=1 | 468 | 20201118_lss_colostrum_1ng_glyco_hcd203040_compound15.25292.25292.3 | controllerType=0 controllerNumber=1 scan=25292 | 53.0433 |
| 07511:3 | 19 | R.LSPN[+2350.830]ASAEHLELR.W | HexNAc(4)Hex(5)Fuc(1)NeuAc(2) | N[+2351] | 1263.2034 | 3 | 3787.5957 | 3787.5796 | 0 | 4.3 | 52 | Specific | 337.33 | 302.9 | 302.9 | 6.89 | 29 | >sp\|Q13410\|BT1A1_HUMAN Butyrophilin subfamily 1 member A1 OS=Homo sapiens OX=9606 GN=BTN1A1 PE=1 SV=3 | 179 | 20201118_lss_colostrum_1ng_glyco_hcd203040_compound15.10669.10669.3 | controllerType=0 controllerNumber=1 scan=10669 | 24.1153 |
| 07513:3 | 19 | R.LSPN[+2350.830]ASAEHLELR.W | HexNAc(4)Hex(5)Fuc(1)NeuAc(2) | N[+2351] | 1263.2034 | 3 | 3787.5957 | 3787.5796 | 0 | 4.3 | 52 | Specific | 356.71 | 262.55 | 262.55 | 6.89 | 29 | >sp\|Q13410\|BT1A1_HUMAN Butyrophilin subfamily 1 member A1 OS=Homo sapiens OX=9606 GN=BTN1A1 PE=1 SV=3 | 179 | 20201118_lss_colostrum_1ng_glyco_hcd203040_compound15.10671.10671.3 | controllerType=0 controllerNumber=1 scan=10671 | 24.1191 |
| 07483:3 | 19 | R.LSPN[+2350.830]ASAEHLELR.W | HexNAc(4)Hex(5)Fuc(1)NeuAc(2) | N[+2351] | 1263.2022 | 3 | 3787.5921 | 3787.5796 | 0 | 3.3 | 52 | Specific | 468.85 | 335.48 | 335.48 | 7.68 | 29 | >sp\|Q13410\|BT1A1_HUMAN Butyrophilin subfamily 1 member A1 OS=Homo sapiens OX=9606 GN=BTN1A1 PE=1 SV=3 | 179 | 20201118_lss_colostrum_1ng_glyco_hcd203040_compound15.10640.10640.3 | controllerType=0 controllerNumber=1 scan=10640 | 24.0693 |
| 07485:3 | 19 | R.LSPN[+2350.830]ASAEHLELR.W | HexNAc(4)Hex(5)Fuc(1)NeuAc(2) | N[+2351] | 1263.2022 | 3 | 3787.5921 | 3787.5796 | 0 | 3.3 | 52 | Specific | 484.55 | 385.72 | 385.72 | 9.39 | 29 | >sp\|Q13410\|BT1A1_HUMAN Butyrophilin subfamily 1 member A1 OS=Homo sapiens OX=9606 GN=BTN1A1 PE=1 SV=3 | 179 | 20201118_lss_colostrum_1ng_glyco_hcd203040_compound15.10642.10642.3 | controllerType=0 controllerNumber=1 scan=10642 | 24.0731 |
| 01796:3 | 22 | R.LRN[+1216.423]VSWATGR.S | HexNAc(2)Hex(5) | N[+1216] | 792.693 | 3 | 2376.0645 | 2376.0559 | 0 | 3.6 | 149 | Specific | 472.67 | 234.98 | 234.98 | 6.37 | 23 | >sp\|P08571\|CD14_HUMAN Monocyte differentiation antigen CD14 OS=Homo sapiens OX=9606 GN=CD14 PE=1 SV=2 | 1232 | 20201118_lss_colostrum_1ng_glyco_hcd203040_compound15.4754.4754.3 | controllerType=0 controllerNumber=1 scan=4754 | 14.3241 |
| 18011:3 | 22 | R.C[+57.021]MWSSALNSLN[+1864.634]LSFAGLEQVPK.G | HexNAc(2)Hex(9) | C[+57], N[+1865] | 1439.622 | 3 | 4316.8515 | 4316.8341 | 0 | 4 | 272 | Specific | 463.86 | 463.72 | 463.72 | 9.82 | 23 | >sp\|P08571\|CD14_HUMAN Monocyte differentiation antigen CD14 OS=Homo sapiens OX=9606 GN=CD14 PE=1 SV=2 | 1232 | 20201118_lss_colostrum_1ng_glyco_hcd203040_compound15.21586.21586.3 | controllerType=0 controllerNumber=1 scan=21586 | 44.8115 |
| 17988:3 | 22 | R.C[+57.021]MWSSALNSLN[+1864.634]LSFAGLEQVPK.G | HexNAc(2)Hex(9) | C[+57], N[+1865] | 1439.6192 | 3 | 4316.843 | 4316.8341 | 0 | 2.1 | 272 | Specific | 793.31 | 744.59 | 744.59 | 15.93 | 23 | >sp\|P08571\|CD14_HUMAN Monocyte differentiation antigen CD14 OS=Homo sapiens OX=9606 GN=CD14 PE=1 SV=2 | 1232 | 20201118_lss_colostrum_1ng_glyco_hcd203040_compound15.21562.21562.3 | controllerType=0 controllerNumber=1 scan=21562 | 44.7714 |
| 18014:3 | 22 | R.C[+57.021]MWSSALNSLN[+1864.634]LSFAGLEQVPK.G | HexNAc(2)Hex(9) | C[+57], N[+1865] | 1439.622 | 3 | 4316.8515 | 4316.8341 | 0 | 4 | 272 | Specific | 860.78 | 683.04 | 683.04 | 17.73 | 23 | >sp\|P08571\|CD14_HUMAN Monocyte differentiation antigen CD14 OS=Homo sapiens OX=9606 GN=CD14 PE=1 SV=2 | 1232 | 20201118_lss_colostrum_1ng_glyco_hcd203040_compound15.21589.21589.3 | controllerType=0 controllerNumber=1 scan=21589 | 44.8195 |
| 17991:3 | 22 | R.C[+57.021]MWSSALNSLN[+1864.634]LSFAGLEQVPK.G | HexNAc(2)Hex(9) | C[+57], N[+1865] | 1439.6192 | 3 | 4316.843 | 4316.8341 | 0 | 2.1 | 272 | Specific | 1304.54 | 1231.3 | 1231.3 | 26.93 | 23 | >sp\|P08571\|CD14_HUMAN Monocyte differentiation antigen CD14 OS=Homo sapiens OX=9606 GN=CD14 PE=1 SV=2 | 1232 | 20201118_lss_colostrum_1ng_glyco_hcd203040_compound15.21565.21565.3 | controllerType=0 controllerNumber=1 scan=21565 | 44.7757 |
| 11255:3 | 23 | R.VN[+1216.423]LTTNTIAVTQTLPNAAYNNR.F | HexNAc(2)Hex(5) | N[+1216] | 1202.5651 | 3 | 3605.6808 | 3605.6699 | 0 | 3 | 351 | Specific | 573.5 | 573.5 | 573.5 | 10.59 | 26 | >sp\|Q6UX06\|OLFM4_HUMAN Olfactomedin-4 OS=Homo sapiens OX=9606 GN=OLFM4 PE=1 SV=1 | 624 | 20201118_lss_colostrum_1ng_glyco_hcd203040_compound15.14560.14560.3 | controllerType=0 controllerNumber=1 scan=14560 | 31.0284 |
| 11257:3 | 23 | R.VN[+1216.423]LTTNTIAVTQTLPNAAYNNR.F | HexNAc(2)Hex(5) | N[+1216] | 1202.5651 | 3 | 3605.6808 | 3605.6699 | 0 | 3 | 351 | Specific | 760.55 | 700.8 | 700.8 | 15.93 | 26 | >sp\|Q6UX06\|OLFM4_HUMAN Olfactomedin-4 OS=Homo sapiens OX=9606 GN=OLFM4 PE=1 SV=1 | 624 | 20201118_lss_colostrum_1ng_glyco_hcd203040_compound15.14562.14562.3 | controllerType=0 controllerNumber=1 scan=14562 | 31.0323 |
| 11236:3 | 23 | R.VN[+1216.423]LTTNTIAVTQTLPNAAYNNR.F | HexNAc(2)Hex(5) | N[+1216] | 1202.5655 | 3 | 3605.6819 | 3605.6699 | 0 | 3.3 | 351 | Specific | 865.47 | 865.47 | 865.47 | 17.73 | 26 | >sp\|Q6UX06\|OLFM4_HUMAN Olfactomedin-4 OS=Homo sapiens OX=9606 GN=OLFM4 PE=1 SV=1 | 624 | 20201118_lss_colostrum_1ng_glyco_hcd203040_compound15.14540.14540.3 | controllerType=0 controllerNumber=1 scan=14540 | 30.987 |
| 11238:3 | 23 | R.VN[+1216.423]LTTNTIAVTQTLPNAAYNNR.F | HexNAc(2)Hex(5) | N[+1216] | 1202.5655 | 3 | 3605.6819 | 3605.6699 | 0 | 3.3 | 351 | Specific | 1109.26 | 1101.88 | 1101.88 | 22.73 | 26 | >sp\|Q6UX06\|OLFM4_HUMAN Olfactomedin-4 OS=Homo sapiens OX=9606 GN=OLFM4 PE=1 SV=1 | 624 | 20201118_lss_colostrum_1ng_glyco_hcd203040_compound15.14542.14542.3 | controllerType=0 controllerNumber=1 scan=14542 | 30.9924 |
| 01890:4 | 25 | R.EKQTDEIKDTRN[+2059.735]ESTQNC[+57.021]VVAEPEK.M | HexNAc(4)Hex(5)Fuc(1)NeuAc(1) | C[+57], N[+2060] | 1252.7891 | 4 | 5008.1344 | 5008.1251 | 0 | 1.9 | 58 | Specific | 213.38 | 213.38 | 213.38 | 3.58 | 24 | >sp\|P47710\|CASA1_HUMAN Alpha-S1-casein OS=Homo sapiens OX=9606 GN=CSN1S1 PE=1 SV=1 | 989 | 20201118_lss_colostrum_1ng_glyco_hcd203040_compound15.4851.4851.4 | controllerType=0 controllerNumber=1 scan=4851 | 14.4908 |
| 01892:4 | 25 | R.EKQTDEIKDTRN[+2059.735]ESTQNC[+57.021]VVAEPEK.M | HexNAc(4)Hex(5)Fuc(1)NeuAc(1) | C[+57], N[+2060] | 1252.7891 | 4 | 5008.1344 | 5008.1251 | 0 | 1.9 | 58 | Specific | 321.41 | 245.28 | 245.28 | 5.34 | 24 | >sp\|P47710\|CASA1_HUMAN Alpha-S1-casein OS=Homo sapiens OX=9606 GN=CSN1S1 PE=1 SV=1 | 989 | 20201118_lss_colostrum_1ng_glyco_hcd203040_compound15.4853.4853.4 | controllerType=0 controllerNumber=1 scan=4853 | 14.4944 |
| 03626:2 | 27 | K.AAIPSALDTN[+1216.423]SSK.S | HexNAc(2)Hex(5) | N[+1216] | 1246.0468 | 2 | 2491.0863 | 2491.0815 | 0 | 1.9 | 542 | Specific | 501.46 | 411.36 | 411.36 | 9.63 | 20 | >sp\|Q08380\|LG3BP_HUMAN Galectin-3-binding protein OS=Homo sapiens OX=9606 GN=LGALS3BP PE=1 SV=1 | 1562 | 20201118_lss_colostrum_1ng_glyco_hcd203040_compound15.6648.6648.2 | controllerType=0 controllerNumber=1 scan=6648 | 17.5571 |
| 03628:2 | 27 | K.AAIPSALDTN[+1216.423]SSK.S | HexNAc(2)Hex(5) | N[+1216] | 1246.0468 | 2 | 2491.0863 | 2491.0815 | 0 | 1.9 | 542 | Specific | 557.46 | 458.53 | 458.53 | 10.07 | 20 | >sp\|Q08380\|LG3BP_HUMAN Galectin-3-binding protein OS=Homo sapiens OX=9606 GN=LGALS3BP PE=1 SV=1 | 1562 | 20201118_lss_colostrum_1ng_glyco_hcd203040_compound15.6650.6650.2 | controllerType=0 controllerNumber=1 scan=6650 | 17.5608 |
| 12812:3 | 34 | R.IIVPLNNREN[+2350.830]ISDPTSPLR.T | HexNAc(4)Hex(5)Fuc(1)NeuAc(2) | N[+2351] | 1500.3441 | 3 | 4499.0176 | 4499.0075 | 0 | 2.2 | 62 | Specific | 385.59 | 365.73 | 365.73 | 7.57 | 20 | >sp\|P01591\|IGJ_HUMAN Immunoglobulin J chain OS=Homo sapiens OX=9606 GN=JCHAIN PE=1 SV=4 | 1392 | 20201118_lss_colostrum_1ng_glyco_hcd203040_compound15.16175.16175.3 | controllerType=0 controllerNumber=1 scan=16175 | 33.9273 |
| 09463:3 | 34 | R.IIVPLNNREN[+1768.640]ISDPTSPLR.T | HexNAc(4)Hex(5)Fuc(1) | N[+1769] | 1306.2812 | 3 | 3916.829 | 3916.8167 | 0 | 3.2 | 62 | Specific | 402.51 | 363.02 | 363.02 | 7.62 | 20 | >sp\|P01591\|IGJ_HUMAN Immunoglobulin J chain OS=Homo sapiens OX=9606 GN=JCHAIN PE=1 SV=4 | 1392 | 20201118_lss_colostrum_1ng_glyco_hcd203040_compound15.12700.12700.3 | controllerType=0 controllerNumber=1 scan=12700 | 27.5834 |
| 09465:3 | 34 | R.IIVPLNNREN[+1768.640]ISDPTSPLR.T | HexNAc(4)Hex(5)Fuc(1) | N[+1769] | 1306.2812 | 3 | 3916.829 | 3916.8167 | 0 | 3.2 | 62 | Specific | 403.05 | 361.43 | 361.43 | 7.62 | 20 | >sp\|P01591\|IGJ_HUMAN Immunoglobulin J chain OS=Homo sapiens OX=9606 GN=JCHAIN PE=1 SV=4 | 1392 | 20201118_lss_colostrum_1ng_glyco_hcd203040_compound15.12702.12702.3 | controllerType=0 controllerNumber=1 scan=12702 | 27.5874 |
| 12815:3 | 34 | R.IIVPLNNREN[+2350.830]ISDPTSPLR.T | HexNAc(4)Hex(5)Fuc(1)NeuAc(2) | N[+2351] | 1500.3441 | 3 | 4499.0176 | 4499.0075 | 0 | 2.2 | 62 | Specific | 497.38 | 480.94 | 480.94 | 9.45 | 20 | >sp\|P01591\|IGJ_HUMAN Immunoglobulin J chain OS=Homo sapiens OX=9606 GN=JCHAIN PE=1 SV=4 | 1392 | 20201118_lss_colostrum_1ng_glyco_hcd203040_compound15.16178.16178.3 | controllerType=0 controllerNumber=1 scan=16178 | 33.932 |
| 10992:3 | 34 | R.IIVPLNNREN[+1856.656]ISDPTSPLR.T | HexNAc(3)Hex(5)Fuc(1)NeuAc(1) | N[+1857] | 1335.6199 | 3 | 4004.8452 | 4004.8327 | 0 | 3.1 | 62 | Specific | 500.95 | 500.95 | 500.95 | 9.45 | 20 | >sp\|P01591\|IGJ_HUMAN Immunoglobulin J chain OS=Homo sapiens OX=9606 GN=JCHAIN PE=1 SV=4 | 1392 | 20201118_lss_colostrum_1ng_glyco_hcd203040_compound15.14286.14286.3 | controllerType=0 controllerNumber=1 scan=14286 | 30.4741 |
| 09381:3 | 34 | R.IIVPLNNREN[+1809.666]ISDPTSPLR.T | HexNAc(5)Hex(4)Fuc(1) | N[+1810] | 1319.9546 | 3 | 3957.8494 | 3957.8432 | 0 | 1.6 | 62 | Specific | 504.57 | 396.92 | 396.92 | 9.3 | 20 | >sp\|P01591\|IGJ_HUMAN Immunoglobulin J chain OS=Homo sapiens OX=9606 GN=JCHAIN PE=1 SV=4 | 1392 | 20201118_lss_colostrum_1ng_glyco_hcd203040_compound15.12615.12615.3 | controllerType=0 controllerNumber=1 scan=12615 | 27.4246 |
| 12875:3 | 34 | R.IIVPLNNREN[+2204.772]ISDPTSPLR.T | HexNAc(4)Hex(5)NeuAc(2) | N[+2205] | 1451.6569 | 3 | 4352.9562 | 4352.9496 | 0 | 1.5 | 62 | Specific | 517.39 | 496.04 | 496.04 | 9.46 | 20 | >sp\|P01591\|IGJ_HUMAN Immunoglobulin J chain OS=Homo sapiens OX=9606 GN=JCHAIN PE=1 SV=4 | 1392 | 20201118_lss_colostrum_1ng_glyco_hcd203040_compound15.16240.16240.3 | controllerType=0 controllerNumber=1 scan=16240 | 34.0676 |
| 09378:3 | 34 | R.IIVPLNNREN[+1809.666]ISDPTSPLR.T | HexNAc(5)Hex(4)Fuc(1) | N[+1810] | 1319.9546 | 3 | 3957.8494 | 3957.8432 | 0 | 1.6 | 62 | Specific | 530.35 | 488.06 | 488.06 | 9.46 | 20 | >sp\|P01591\|IGJ_HUMAN Immunoglobulin J chain OS=Homo sapiens OX=9606 GN=JCHAIN PE=1 SV=4 | 1392 | 20201118_lss_colostrum_1ng_glyco_hcd203040_compound15.12612.12612.3 | controllerType=0 controllerNumber=1 scan=12612 | 27.4198 |
| 10994:3 | 34 | R.IIVPLNNREN[+1856.656]ISDPTSPLR.T | HexNAc(3)Hex(5)Fuc(1)NeuAc(1) | N[+1857] | 1335.6199 | 3 | 4004.8452 | 4004.8327 | 0 | 3.1 | 62 | Specific | 548.21 | 548.21 | 548.21 | 9.63 | 20 | >sp\|P01591\|IGJ_HUMAN Immunoglobulin J chain OS=Homo sapiens OX=9606 GN=JCHAIN PE=1 SV=4 | 1392 | 20201118_lss_colostrum_1ng_glyco_hcd203040_compound15.14288.14288.3 | controllerType=0 controllerNumber=1 scan=14288 | 30.4796 |
| 12873:3 | 34 | R.IIVPLNNREN[+2204.772]ISDPTSPLR.T | HexNAc(4)Hex(5)NeuAc(2) | N[+2205] | 1451.6569 | 3 | 4352.9562 | 4352.9496 | 0 | 1.5 | 62 | Specific | 552.58 | 552.58 | 552.58 | 9.47 | 20 | >sp\|P01591\|IGJ_HUMAN Immunoglobulin J chain OS=Homo sapiens OX=9606 GN=JCHAIN PE=1 SV=4 | 1392 | 20201118_lss_colostrum_1ng_glyco_hcd203040_compound15.16238.16238.3 | controllerType=0 controllerNumber=1 scan=16238 | 34.0628 |
| 09364:3 | 34 | R.IIVPLNNREN[+1809.666]ISDPTSPLR.T | HexNAc(5)Hex(4)Fuc(1) | N[+1810] | 1319.9551 | 3 | 3957.8509 | 3957.8432 | 0 | 1.9 | 62 | Specific | 553.67 | 432.45 | 432.45 | 9.63 | 20 | >sp\|P01591\|IGJ_HUMAN Immunoglobulin J chain OS=Homo sapiens OX=9606 GN=JCHAIN PE=1 SV=4 | 1392 | 20201118_lss_colostrum_1ng_glyco_hcd203040_compound15.12597.12597.3 | controllerType=0 controllerNumber=1 scan=12597 | 27.3909 |
| 09362:3 | 34 | R.IIVPLNNREN[+1809.666]ISDPTSPLR.T | HexNAc(5)Hex(4)Fuc(1) | N[+1810] | 1319.9551 | 3 | 3957.8509 | 3957.8432 | 0 | 1.9 | 62 | Specific | 580.1 | 492.81 | 492.81 | 11.23 | 20 | >sp\|P01591\|IGJ_HUMAN Immunoglobulin J chain OS=Homo sapiens OX=9606 GN=JCHAIN PE=1 SV=4 | 1392 | 20201118_lss_colostrum_1ng_glyco_hcd203040_compound15.12595.12595.3 | controllerType=0 controllerNumber=1 scan=12595 | 27.3858 |
| 09388:3 | 34 | R.IIVPLNNREN[+1663.608]ISDPTSPLR.T | HexNAc(5)Hex(4) | N[+1664] | 1271.2674 | 3 | 3811.7876 | 3811.7853 | 0 | 0.6 | 62 | Specific | 624.15 | 581.99 | 581.99 | 12.07 | 20 | >sp\|P01591\|IGJ_HUMAN Immunoglobulin J chain OS=Homo sapiens OX=9606 GN=JCHAIN PE=1 SV=4 | 1392 | 20201118_lss_colostrum_1ng_glyco_hcd203040_compound15.12622.12622.3 | controllerType=0 controllerNumber=1 scan=12622 | 27.4371 |
| 09386:3 | 34 | R.IIVPLNNREN[+1663.608]ISDPTSPLR.T | HexNAc(5)Hex(4) | N[+1664] | 1271.2674 | 3 | 3811.7876 | 3811.7853 | 0 | 0.6 | 62 | Specific | 628.58 | 601.59 | 601.59 | 12.07 | 20 | >sp\|P01591\|IGJ_HUMAN Immunoglobulin J chain OS=Homo sapiens OX=9606 GN=JCHAIN PE=1 SV=4 | 1392 | 20201118_lss_colostrum_1ng_glyco_hcd203040_compound15.12620.12620.3 | controllerType=0 controllerNumber=1 scan=12620 | 27.4321 |
| 04779:3 | 34 | R.EN[+1663.608]ISDPTSPLR.T | HexNAc(5)Hex(4) | N[+1664] | 964.7498 | 3 | 2892.235 | 2892.225 | 0 | 3.5 | 70 | Specific | 247.5 | 105.1 | 105.1 | 3.5 | 20 | >sp\|P01591\|IGJ_HUMAN Immunoglobulin J chain OS=Homo sapiens OX=9606 GN=JCHAIN PE=1 SV=4 | 1392 | 20201118_lss_colostrum_1ng_glyco_hcd203040_compound15.7842.7842.3 | controllerType=0 controllerNumber=1 scan=7842 | 19.4986 |
| 21550:4 | 42 | R.QLDMLDLSN[+1913.677]NSLASVPEGLWASLGQPNWDMR.D | HexNAc(4)Hex(5)NeuAc(1) | N[+1914] | 1343.5914 | 4 | 5371.3439 | 5371.3285 | 0 | 2.9 | 261 | Specific | 1015.13 | 1013.78 | 1013.78 | 21.07 | 15 | >sp\|P02750\|A2GL_HUMAN Leucine-rich alpha-2-glycoprotein OS=Homo sapiens OX=9606 GN=LRG1 PE=1 SV=2 | 518 | 20201118_lss_colostrum_1ng_glyco_hcd203040_compound15.25278.25278.4 | controllerType=0 controllerNumber=1 scan=25278 | 53.013 |
| 21552:4 | 42 | R.QLDMLDLSN[+1913.677]NSLASVPEGLWASLGQPNWDMR.D | HexNAc(4)Hex(5)NeuAc(1) | N[+1914] | 1343.5914 | 4 | 5371.3439 | 5371.3285 | 0 | 2.9 | 261 | Specific | 1169.71 | 1148.93 | 1148.93 | 24.07 | 15 | >sp\|P02750\|A2GL_HUMAN Leucine-rich alpha-2-glycoprotein OS=Homo sapiens OX=9606 GN=LRG1 PE=1 SV=2 | 518 | 20201118_lss_colostrum_1ng_glyco_hcd203040_compound15.25280.25280.4 | controllerType=0 controllerNumber=1 scan=25280 | 53.0179 |
| 16923:3 | 44 | R.NYIVPILWLN[+1864.634]ETGTIGDEKANMFR.S | HexNAc(2)Hex(9) | N[+1865] | 1553.6951 | 3 | 4659.0708 | 4659.0574 | 0 | 2.9 | 408 | Specific | 897.89 | 897.89 | 897.89 | 17.52 | 14 | >sp\|P16671\|CD36_HUMAN Platelet glycoprotein 4 OS=Homo sapiens OX=9606 GN=CD36 PE=1 SV=2 | 812 | 20201118_lss_colostrum_1ng_glyco_hcd203040_compound15.20450.20450.3 | controllerType=0 controllerNumber=1 scan=20450 | 42.3403 |
| 16926:3 | 44 | R.NYIVPILWLN[+1864.634]ETGTIGDEKANMFR.S | HexNAc(2)Hex(9) | N[+1865] | 1553.6951 | 3 | 4659.0708 | 4659.0574 | 0 | 2.9 | 408 | Specific | 949.66 | 849.25 | 849.25 | 18.52 | 14 | >sp\|P16671\|CD36_HUMAN Platelet glycoprotein 4 OS=Homo sapiens OX=9606 GN=CD36 PE=1 SV=2 | 812 | 20201118_lss_colostrum_1ng_glyco_hcd203040_compound15.20453.20453.3 | controllerType=0 controllerNumber=1 scan=20453 | 42.3485 |
| 00741:2 | 53 | R.SYN[+1038.375]DSVDPR.I | HexNAc(2)Hex(3)Fuc(1) | N[+1038] | 1045.9237 | 2 | 2090.8402 | 2090.8395 | 0 | 0.4 | 481 | Specific | 242.59 | 204.42 | 204.42 | 5.41 | 14 | >sp\|P05164\|PERM_HUMAN Myeloperoxidase OS=Homo sapiens OX=9606 GN=MPO PE=1 SV=1 | 12 | 20201118_lss_colostrum_1ng_glyco_hcd203040_compound15.3659.3659.2 | controllerType=0 controllerNumber=1 scan=3659 | 12.4529 |
| 00742:2 | 53 | R.SYN[+1038.375]DSVDPR.I | HexNAc(2)Hex(3)Fuc(1) | N[+1038] | 1045.9237 | 2 | 2090.8402 | 2090.8395 | 0 | 0.4 | 481 | Specific | 255.15 | 222.88 | 222.88 | 5.43 | 14 | >sp\|P05164\|PERM_HUMAN Myeloperoxidase OS=Homo sapiens OX=9606 GN=MPO PE=1 SV=1 | 12 | 20201118_lss_colostrum_1ng_glyco_hcd203040_compound15.3660.3660.2 | controllerType=0 controllerNumber=1 scan=3660 | 12.4568 |
| 23418:3 | 77 | R.AVN[+2204.772]ITSENLIDDVVSLIR.G | HexNAc(4)Hex(5)NeuAc(2) | N[+2205] | 1392.6268 | 3 | 4175.8657 | 4175.8481 | 0 | 4.2 | 413 | Specific | 540.78 | 452.65 | 452.65 | 9.84 | 10 | >sp\|P02748\|CO9_HUMAN Complement component C9 OS=Homo sapiens OX=9606 GN=C9 PE=1 SV=2 | 1547 | 20201118_lss_colostrum_1ng_glyco_hcd203040_compound15.27227.27227.3 | controllerType=0 controllerNumber=1 scan=27227 | 57.1699 |
| 23420:3 | 77 | R.AVN[+2204.772]ITSENLIDDVVSLIR.G | HexNAc(4)Hex(5)NeuAc(2) | N[+2205] | 1392.6268 | 3 | 4175.8657 | 4175.8481 | 0 | 4.2 | 413 | Specific | 646 | 534.96 | 534.96 | 12.43 | 10 | >sp\|P02748\|CO9_HUMAN Complement component C9 OS=Homo sapiens OX=9606 GN=C9 PE=1 SV=2 | 1547 | 20201118_lss_colostrum_1ng_glyco_hcd203040_compound15.27229.27229.3 | controllerType=0 controllerNumber=1 scan=27229 | 57.1751 |
| 16345:5 | 397 | K.LIDEVIEDTRYTLPLTEGKAN[+2350.830]VTVLDTQIR.K | HexNAc(4)Hex(5)Fuc(1)NeuAc(2) | N[+2351] | 1154.1364 | 5 | 5766.6527 | 5766.6622 | 0 | -1.7 | 694 | Specific | 211.22 | 199.29 | 199.29 | 3.2 | 3 | >sp\|Q9NZM1\|MYOF_HUMAN Myoferlin OS=Homo sapiens OX=9606 GN=MYOF PE=1 SV=1 | 71 | 20201118_lss_colostrum_1ng_glyco_hcd203040_compound15.19849.19849.5 | controllerType=0 controllerNumber=1 scan=19849 | 41.0654 |
| 16347:5 | 397 | K.LIDEVIEDTRYTLPLTEGKAN[+2350.830]VTVLDTQIR.K | HexNAc(4)Hex(5)Fuc(1)NeuAc(2) | N[+2351] | 1154.1364 | 5 | 5766.6527 | 5766.6622 | 0 | -1.7 | 694 | Specific | 250.34 | 39.03 | 39.03 | 3.24 | 3 | >sp\|Q9NZM1\|MYOF_HUMAN Myoferlin OS=Homo sapiens OX=9606 GN=MYOF PE=1 SV=1 | 71 | 20201118_lss_colostrum_1ng_glyco_hcd203040_compound15.19851.19851.5 | controllerType=0 controllerNumber=1 scan=19851 | 41.0748 |
| 17781:4 | 1 | R.RTAGWNVPIGTLRPFLN[+2279.830]WTGPPEPIEAAVAR.F | HexNAc(5)Hex(6)Fuc(2) | N[+2280] | 1417.4213 | 4 | 5666.6632 | 5666.6417 | 0 | 3.8 | 140 | Specific | 211.05 | 142.31 | 142.31 | 3.15 | 146 | >sp\|P02788\|TRFL_HUMAN Lactotransferrin OS=Homo sapiens OX=9606 GN=LTF PE=1 SV=6 | 1270 | 20201118_lss_milk_1ng_glyco_hcd203040_compound15.20893.20893.2 | controllerType=0 controllerNumber=1 scan=20893 | 41.1824 |
| 19968:4 | 1 | R.RTAGWNVPIGTLRPFLN[+2350.830]WTGPPEPIEAAVAR.F | HexNAc(4)Hex(5)Fuc(1)NeuAc(2) | N[+2351] | 1435.1682 | 4 | 5737.651 | 5737.6424 | 0 | 1.5 | 140 | Specific | 376.07 | 370.61 | 370.61 | 7.71 | 146 | >sp\|P02788\|TRFL_HUMAN Lactotransferrin OS=Homo sapiens OX=9606 GN=LTF PE=1 SV=6 | 1270 | 20201118_lss_milk_1ng_glyco_hcd203040_compound15.23173.23173.4 | controllerType=0 controllerNumber=1 scan=23173 | 46.0132 |
| 19882:5 | 1 | R.RTAGWNVPIGTLRPFLN[+2350.830]WTGPPEPIEAAVAR.F | HexNAc(4)Hex(5)Fuc(1)NeuAc(2) | N[+2351] | 1148.337 | 5 | 5737.6561 | 5737.6424 | 0 | 2.4 | 140 | Specific | 437.61 | 420.19 | 420.19 | 9.55 | 146 | >sp\|P02788\|TRFL_HUMAN Lactotransferrin OS=Homo sapiens OX=9606 GN=LTF PE=1 SV=6 | 1270 | 20201118_lss_milk_1ng_glyco_hcd203040_compound15.23084.23084.5 | controllerType=0 controllerNumber=1 scan=23084 | 45.853 |
| 19862:4 | 1 | R.RTAGWNVPIGTLRPFLN[+2350.830]WTGPPEPIEAAVAR.F | HexNAc(4)Hex(5)Fuc(1)NeuAc(2) | N[+2351] | 1435.1714 | 4 | 5737.6637 | 5737.6424 | 0 | 3.7 | 140 | Specific | 481.98 | 455.91 | 455.91 | 9.73 | 146 | >sp\|P02788\|TRFL_HUMAN Lactotransferrin OS=Homo sapiens OX=9606 GN=LTF PE=1 SV=6 | 1270 | 20201118_lss_milk_1ng_glyco_hcd203040_compound15.23064.23064.4 | controllerType=0 controllerNumber=1 scan=23064 | 45.8248 |
| 19885:5 | 1 | R.RTAGWNVPIGTLRPFLN[+2350.830]WTGPPEPIEAAVAR.F | HexNAc(4)Hex(5)Fuc(1)NeuAc(2) | N[+2351] | 1148.337 | 5 | 5737.6561 | 5737.6424 | 0 | 2.4 | 140 | Specific | 485.68 | 443.75 | 443.75 | 9.69 | 146 | >sp\|P02788\|TRFL_HUMAN Lactotransferrin OS=Homo sapiens OX=9606 GN=LTF PE=1 SV=6 | 1270 | 20201118_lss_milk_1ng_glyco_hcd203040_compound15.23087.23087.5 | controllerType=0 controllerNumber=1 scan=23087 | 45.8585 |
| 18714:4 | 1 | R.RTAGWNVPIGTLRPFLN[+2570.925]WTGPPEPIEAAVAR.F | HexNAc(5)Hex(6)Fuc(2)NeuAc(1) | N[+2571] | 1490.1947 | 4 | 5957.757 | 5957.7371 | 0 | 3.3 | 140 | Specific | 492.53 | 453.49 | 453.49 | 9.8 | 146 | >sp\|P02788\|TRFL_HUMAN Lactotransferrin OS=Homo sapiens OX=9606 GN=LTF PE=1 SV=6 | 1270 | 20201118_lss_milk_1ng_glyco_hcd203040_compound15.21867.21867.4 | controllerType=0 controllerNumber=1 scan=21867 | 43.3345 |

Table S4. Cont.

| Query #:z | Protein Rank | Glycopeptide | Glycan | Modification Type(s) | Observed m/z | z | Observed (M+H) | Calc. mass (M+H) | Off-by-x error | Mass error (ppm) | StartPosition | Cleavage | Score | Delta | Delta Mod | \|Log Prob\| | # of unique peptides | Protein Name | Protein DB number | Comment | Scan # | Scan Time |
|---|---|---|---|---|---|---|---|---|---|---|---|---|---|---|---|---|---|---|---|---|---|---|
| 18824:5 | 1 | R.RTAGWNVPIGTLRPFLN[+2205.793]WTGPPEPIEAAVAR.F | HexNAc(4)Hex(5)Fuc(2)NeuAc(1) | N[+2206] | 1119.3313 | 5 | 5592.6274 | 5592.6049 | 0 | 4 | 140 | Specific | 522.43 | 522.43 | 522.43 | 9.85 | 146 | >sp\|P02788\|TRFL_HUMAN Lactotransferrin OS=Homo sapiens OX=9606 GN=LTF PE=1 SV=6 | 1270 | 20201118_lss_milk_1ng_glyco_hcd203040_compound15.21981.21981.5 | controllerType=0 controllerNumber=1 scan=21981 | 43.5531 |
| 17856:4 | 1 | R.RTAGWNVPIGTLRPFLN[+1768.640]WTGPPEPIEAAVAR.F | HexNAc(4)Hex(5)Fuc(1) | N[+1769] | 1289.622 | 4 | 5155.466 | 5155.4516 | 0 | 2.8 | 140 | Specific | 524.37 | 461.39 | 461.39 | 9.75 | 146 | >sp\|P02788\|TRFL_HUMAN Lactotransferrin OS=Homo sapiens OX=9606 GN=LTF PE=1 SV=6 | 1270 | 20201118_lss_milk_1ng_glyco_hcd203040_compound15.20972.20972.4 | controllerType=0 controllerNumber=1 scan=20972 | 41.3577 |
| 18717:4 | 1 | R.RTAGWNVPIGTLRPFLN[+2570.925]WTGPPEPIEAAVAR.F | HexNAc(5)Hex(6)Fuc(2)NeuAc(1) | N[+2571] | 1490.1947 | 4 | 5957.757 | 5957.7371 | 0 | 3.3 | 140 | Specific | 574.07 | 456.71 | 456.71 | 10.29 | 146 | >sp\|P02788\|TRFL_HUMAN Lactotransferrin OS=Homo sapiens OX=9606 GN=LTF PE=1 SV=6 | 1270 | 20201118_lss_milk_1ng_glyco_hcd203040_compound15.21870.21870.4 | controllerType=0 controllerNumber=1 scan=21870 | 43.3396 |
| 18816:4 | 1 | R.RTAGWNVPIGTLRPFLN[+2205.793]WTGPPEPIEAAVAR.F | HexNAc(4)Hex(5)Fuc(2)NeuAc(1) | N[+2206] | 1398.9109 | 4 | 5592.6217 | 5592.6049 | 0 | 3 | 140 | Specific | 575.18 | 538.92 | 538.92 | 10.18 | 146 | >sp\|P02788\|TRFL_HUMAN Lactotransferrin OS=Homo sapiens OX=9606 GN=LTF PE=1 SV=6 | 1270 | 20201118_lss_milk_1ng_glyco_hcd203040_compound15.21973.21973.4 | controllerType=0 controllerNumber=1 scan=21973 | 43.5314 |
| 19865:4 | 1 | R.RTAGWNVPIGTLRPFLN[+2350.830]WTGPPEPIEAAVAR.F | HexNAc(4)Hex(5)Fuc(1)NeuAc(2) | N[+2351] | 1435.1714 | 4 | 5737.6637 | 5737.6424 | 0 | 3.7 | 140 | Specific | 576.71 | 487.38 | 487.38 | 11.56 | 146 | >sp\|P02788\|TRFL_HUMAN Lactotransferrin OS=Homo sapiens OX=9606 GN=LTF PE=1 SV=6 | 1270 | 20201118_lss_milk_1ng_glyco_hcd203040_compound15.23067.23067.4 | controllerType=0 controllerNumber=1 scan=23067 | 45.8291 |
| 18825:4 | 1 | R.RTAGWNVPIGTLRPFLN[+2059.735]WTGPPEPIEAAVAR.F | HexNAc(4)Hex(5)Fuc(1)NeuAc(1) | N[+2060] | 1362.3964 | 4 | 5446.5636 | 5446.547 | 0 | 3.1 | 140 | Specific | 584.16 | 584.16 | 584.16 | 11.56 | 146 | >sp\|P02788\|TRFL_HUMAN Lactotransferrin OS=Homo sapiens OX=9606 GN=LTF PE=1 SV=6 | 1270 | 20201118_lss_milk_1ng_glyco_hcd203040_compound15.21982.21982.4 | controllerType=0 controllerNumber=1 scan=21982 | 43.5543 |
| 18828:4 | 1 | R.RTAGWNVPIGTLRPFLN[+2059.735]WTGPPEPIEAAVAR.F | HexNAc(4)Hex(5)Fuc(1)NeuAc(1) | N[+2060] | 1362.3964 | 4 | 5446.5636 | 5446.547 | 0 | 3.1 | 140 | Specific | 629.51 | 525.36 | 525.36 | 12.56 | 146 | >sp\|P02788\|TRFL_HUMAN Lactotransferrin OS=Homo sapiens OX=9606 GN=LTF PE=1 SV=6 | 1270 | 20201118_lss_milk_1ng_glyco_hcd203040_compound15.21985.21985.4 | controllerType=0 controllerNumber=1 scan=21985 | 43.5664 |
| 18827:5 | 1 | R.RTAGWNVPIGTLRPFLN[+2205.793]WTGPPEPIEAAVAR.F | HexNAc(4)Hex(5)Fuc(2)NeuAc(1) | N[+2206] | 1119.3313 | 5 | 5592.6274 | 5592.6049 | 0 | 4 | 140 | Specific | 636.21 | 585.86 | 585.86 | 12.55 | 146 | >sp\|P02788\|TRFL_HUMAN Lactotransferrin OS=Homo sapiens OX=9606 GN=LTF PE=1 SV=6 | 1270 | 20201118_lss_milk_1ng_glyco_hcd203040_compound15.21984.21984.5 | controllerType=0 controllerNumber=1 scan=21984 | 43.5653 |
| 17859:4 | 1 | R.RTAGWNVPIGTLRPFLN[+1768.640]WTGPPEPIEAAVAR.F | HexNAc(4)Hex(5)Fuc(1) | N[+1769] | 1289.622 | 4 | 5155.466 | 5155.4516 | 0 | 2.8 | 140 | Specific | 637.32 | 533.2 | 533.2 | 12.45 | 146 | >sp\|P02788\|TRFL_HUMAN Lactotransferrin OS=Homo sapiens OX=9606 GN=LTF PE=1 SV=6 | 1270 | 20201118_lss_milk_1ng_glyco_hcd203040_compound15.20975.20975.4 | controllerType=0 controllerNumber=1 scan=20975 | 41.3735 |
| 18819:4 | 1 | R.RTAGWNVPIGTLRPFLN[+2205.793]WTGPPEPIEAAVAR.F | HexNAc(4)Hex(5)Fuc(2)NeuAc(1) | N[+2206] | 1398.9109 | 4 | 5592.6217 | 5592.6049 | 0 | 3 | 140 | Specific | 649.62 | 523.82 | 523.82 | 12.45 | 146 | >sp\|P02788\|TRFL_HUMAN Lactotransferrin OS=Homo sapiens OX=9606 GN=LTF PE=1 SV=6 | 1270 | 20201118_lss_milk_1ng_glyco_hcd203040_compound15.21976.21976.4 | controllerType=0 controllerNumber=1 scan=21976 | 43.5376 |
| 19089:4 | 1 | R.[+42.011]TAGWNVPIGTLRPFLN[+1872.651]WTGPPEPIEAAVAR.F | HexNAc(3)Hex(6)NeuAc(1) | .[42], N[+1873] | 1287.1111 | 4 | 5145.4225 | 5145.372 | 0 | 9.8 | 141 | Specific | 246.58 | 115.89 | 115.89 | 2.14 | 146 | >sp\|P02788\|TRFL_HUMAN Lactotransferrin OS=Homo sapiens OX=9606 GN=LTF PE=1 SV=6 | 1270 | 20201118_lss_milk_1ng_glyco_hcd203040_compound15.22257.22257.4 | controllerType=0 controllerNumber=1 scan=22257 | 44.1042 |
| 20007:5 | 1 | R.TAGWNVPIGTLRPFLN[+2570.925]WTGPPEPIEAAVAR.F | HexNAc(5)Hex(6)Fuc(2)NeuAc(1) | N[+2571] | 1161.136 | 5 | 5801.6508 | 5801.636 | 0 | 2.6 | 141 | Specific | 265.56 | 265.56 | 265.56 | 7.19 | 146 | >sp\|P02788\|TRFL_HUMAN Lactotransferrin OS=Homo sapiens OX=9606 GN=LTF PE=1 SV=6 | 1270 | 20201118_lss_milk_1ng_glyco_hcd203040_compound15.23214.23214.5 | controllerType=0 controllerNumber=1 scan=23214 | 46.1179 |
| 19090:3 | 1 | R.[+42.011]TAGWNVPIGTLRPFLN[+1872.651]WTGPPEPIEAAVAR.F | HexNAc(3)Hex(6)NeuAc(1) | .[42], N[+1873] | 1715.8119 | 3 | 5145.421 | 5145.372 | 0 | 9.5 | 141 | Specific | 281.06 | 159.91 | 159.91 | 3.34 | 146 | >sp\|P02788\|TRFL_HUMAN Lactotransferrin OS=Homo sapiens OX=9606 GN=LTF PE=1 SV=6 | 1270 | 20201118_lss_milk_1ng_glyco_hcd203040_compound15.22258.22258.3 | controllerType=0 controllerNumber=1 scan=22258 | 44.1055 |
| 21352:3 | 1 | R.TAGWNVPIGTLRPFLN[+2350.830]WTGPPEPIEAAVAR.F | HexNAc(4)Hex(5)Fuc(1)NeuAc(2) | N[+2351] | 1861.1944 | 3 | 5581.5687 | 5581.5413 | 0 | 4.9 | 141 | Specific | 335.95 | 191.35 | 191.35 | 6.76 | 146 | >sp\|P02788\|TRFL_HUMAN Lactotransferrin OS=Homo sapiens OX=9606 GN=LTF PE=1 SV=6 | 1270 | 20201118_lss_milk_1ng_glyco_hcd203040_compound15.24622.24622.3 | controllerType=0 controllerNumber=1 scan=24622 | 49.26 |
| 20129:3 | 1 | R.[+42.011]TAGWNVPIGTLRPFLN[+1362.481]WTGPPEPIEAAVAR.F | HexNAc(2)Hex(5)Fuc(1) | .[42], N[+1362] | 1545.7473 | 3 | 4635.2273 | 4635.2023 | 0 | 5.4 | 141 | Specific | 355.83 | 319.25 | 319.25 | 6.84 | 146 | >sp\|P02788\|TRFL_HUMAN Lactotransferrin OS=Homo sapiens OX=9606 GN=LTF PE=1 SV=6 | 1270 | 20201118_lss_milk_1ng_glyco_hcd203040_compound15.23342.23342.3 | controllerType=0 controllerNumber=1 scan=23342 | 46.4362 |
| 20099:4 | 1 | R.[+42.011]TAGWNVPIGTLRPFLN[+2336.851]WTGPPEPIEAAVAR.F | HexNAc(6)Hex(6)Fuc(1) | .[42], N[+2337] | 1403.153 | 4 | 5609.59 | 5609.5726 | 0 | 3.1 | 141 | Specific | 371.59 | 368.1 | 368.1 | 8.08 | 146 | >sp\|P02788\|TRFL_HUMAN Lactotransferrin OS=Homo sapiens OX=9606 GN=LTF PE=1 SV=6 | 1270 | 20201118_lss_milk_1ng_glyco_hcd203040_compound15.23310.23310.4 | controllerType=0 controllerNumber=1 scan=23310 | 46.3345 |
| 20005:3 | 1 | R.TAGWNVPIGTLRPFLN[+2570.925]WTGPPEPIEAAVAR.F | HexNAc(5)Hex(6)Fuc(2)NeuAc(1) | N[+2571] | 1934.5568 | 3 | 5801.656 | 5801.636 | 0 | 3.5 | 141 | Specific | 383.54 | 260.51 | 260.51 | 7.6 | 146 | >sp\|P02788\|TRFL_HUMAN Lactotransferrin OS=Homo sapiens OX=9606 GN=LTF PE=1 SV=6 | 1270 | 20201118_lss_milk_1ng_glyco_hcd203040_compound15.23212.23212.3 | controllerType=0 controllerNumber=1 scan=23212 | 46.1155 |
| 20096:4 | 1 | R.[+42.011]TAGWNVPIGTLRPFLN[+2336.851]WTGPPEPIEAAVAR.F | HexNAc(6)Hex(6)Fuc(1) | .[42], N[+2337] | 1403.153 | 4 | 5609.59 | 5609.5726 | 0 | 3.1 | 141 | Specific | 404.4 | 404.4 | 404.4 | 9.37 | 146 | >sp\|P02788\|TRFL_HUMAN Lactotransferrin OS=Homo sapiens OX=9606 GN=LTF PE=1 SV=6 | 1270 | 20201118_lss_milk_1ng_glyco_hcd203040_compound15.23307.23307.4 | controllerType=0 controllerNumber=1 scan=23307 | 46.3181 |
| 19995:4 | 1 | R.TAGWNVPIGTLRPFLN[+2570.925]WTGPPEPIEAAVAR.F | HexNAc(5)Hex(6)Fuc(2)NeuAc(1) | N[+2571] | 1451.1704 | 4 | 5801.6598 | 5801.636 | 0 | 4.1 | 141 | Specific | 486.73 | 411.34 | 411.34 | 9.79 | 146 | >sp\|P02788\|TRFL_HUMAN Lactotransferrin OS=Homo sapiens OX=9606 GN=LTF PE=1 SV=6 | 1270 | 20201118_lss_milk_1ng_glyco_hcd203040_compound15.23202.23202.4 | controllerType=0 controllerNumber=1 scan=23202 | 46.0979 |
| 20132:3 | 1 | R.[+42.011]TAGWNVPIGTLRPFLN[+1362.481]WTGPPEPIEAAVAR.F | HexNAc(2)Hex(5)Fuc(1) | .[42], N[+1362] | 1545.7473 | 3 | 4635.2273 | 4635.2023 | 0 | 5.4 | 141 | Specific | 500.42 | 460.31 | 460.31 | 9.06 | 146 | >sp\|P02788\|TRFL_HUMAN Lactotransferrin OS=Homo sapiens OX=9606 GN=LTF PE=1 SV=6 | 1270 | 20201118_lss_milk_1ng_glyco_hcd203040_compound15.23345.23345.3 | controllerType=0 controllerNumber=1 scan=23345 | 46.4446 |
| 20063:3 | 1 | R.TAGWNVPIGTLRPFLN[+2351.851]WTGPPEPIEAAVAR.F | HexNAc(4)Hex(5)Fuc(3)NeuAc(1) | N[+2352] | 1861.5348 | 3 | 5582.5897 | 5582.5617 | 0 | 5 | 141 | Specific | 516.54 | 427.25 | 427.25 | 9.52 | 146 | >sp\|P02788\|TRFL_HUMAN Lactotransferrin OS=Homo sapiens OX=9606 GN=LTF PE=1 SV=6 | 1270 | 20201118_lss_milk_1ng_glyco_hcd203040_compound15.23272.23272.3 | controllerType=0 controllerNumber=1 scan=23272 | 46.2424 |
| 19072:4 | 1 | R.TAGWNVPIGTLRPFLN[+2279.830]WTGPPEPIEAAVAR.F | HexNAc(5)Hex(6)Fuc(2) | N[+2280] | 1378.396 | 4 | 5510.5622 | 5510.5406 | 0 | 3.9 | 141 | Specific | 556.47 | 474.96 | 474.96 | 10.28 | 146 | >sp\|P02788\|TRFL_HUMAN Lactotransferrin OS=Homo sapiens OX=9606 GN=LTF PE=1 SV=6 | 1270 | 20201118_lss_milk_1ng_glyco_hcd203040_compound15.22239.22239.4 | controllerType=0 controllerNumber=1 scan=22239 | 44.0646 |
| 20009:4 | 1 | R.TAGWNVPIGTLRPFLN[+2424.867]WTGPPEPIEAAVAR.F | HexNAc(5)Hex(5)Fuc(1)NeuAc(1) | N[+2425] | 1414.6558 | 4 | 5655.6012 | 5655.5781 | 0 | 4.1 | 141 | Specific | 564.67 | 454.03 | 454.03 | 10.28 | 146 | >sp\|P02788\|TRFL_HUMAN Lactotransferrin OS=Homo sapiens OX=9606 GN=LTF PE=1 SV=6 | 1270 | 20201118_lss_milk_1ng_glyco_hcd203040_compound15.23216.23216.4 | controllerType=0 controllerNumber=1 scan=23216 | 46.1201 |
| 21130:4 | 1 | R.TAGWNVPIGTLRPFLN[+2862.021]WTGPPEPIEAAVAR.F | HexNAc(5)Hex(6)Fuc(2)NeuAc(2) | N[+2862] | 1523.9419 | 4 | 6092.7458 | 6092.7314 | 0 | 2.4 | 141 | Specific | 575.82 | 533.6 | 533.6 | 10.17 | 146 | >sp\|P02788\|TRFL_HUMAN Lactotransferrin OS=Homo sapiens OX=9606 GN=LTF PE=1 SV=6 | 1270 | 20201118_lss_milk_1ng_glyco_hcd203040_compound15.24388.24388.4 | controllerType=0 controllerNumber=1 scan=24388 | 48.6836 |
| 20060:3 | 1 | R.TAGWNVPIGTLRPFLN[+2351.851]WTGPPEPIEAAVAR.F | HexNAc(4)Hex(5)Fuc(3)NeuAc(1) | N[+2352] | 1861.5348 | 3 | 5582.5897 | 5582.5617 | 0 | 5 | 141 | Specific | 621.25 | 549.95 | 549.95 | 12.22 | 146 | >sp\|P02788\|TRFL_HUMAN Lactotransferrin OS=Homo sapiens OX=9606 GN=LTF PE=1 SV=6 | 1270 | 20201118_lss_milk_1ng_glyco_hcd203040_compound15.23269.23269.3 | controllerType=0 controllerNumber=1 scan=23269 | 46.2313 |
| 20067:3 | 1 | R.TAGWNVPIGTLRPFLN[+2205.793]WTGPPEPIEAAVAR.F | HexNAc(4)Hex(5)Fuc(2)NeuAc(1) | N[+2206] | 1812.8474 | 3 | 5436.5276 | 5436.5038 | 0 | 4.4 | 141 | Specific | 623.58 | 509.5 | 509.5 | 12.22 | 146 | >sp\|P02788\|TRFL_HUMAN Lactotransferrin OS=Homo sapiens OX=9606 GN=LTF PE=1 SV=6 | 1270 | 20201118_lss_milk_1ng_glyco_hcd203040_compound15.23276.23276.3 | controllerType=0 controllerNumber=1 scan=23276 | 46.2514 |
| 21355:3 | 1 | R.TAGWNVPIGTLRPFLN[+2350.830]WTGPPEPIEAAVAR.F | HexNAc(4)Hex(5)Fuc(1)NeuAc(2) | N[+2351] | 1861.1944 | 3 | 5581.5687 | 5581.5413 | 0 | 4.9 | 141 | Specific | 628.75 | 462.4 | 462.4 | 12.22 | 146 | >sp\|P02788\|TRFL_HUMAN Lactotransferrin OS=Homo sapiens OX=9606 GN=LTF PE=1 SV=6 | 1270 | 20201118_lss_milk_1ng_glyco_hcd203040_compound15.24625.24625.3 | controllerType=0 controllerNumber=1 scan=24625 | 49.264 |
| 20052:4 | 1 | R.TAGWNVPIGTLRPFLN[+2205.793]WTGPPEPIEAAVAR.F | HexNAc(4)Hex(5)Fuc(2)NeuAc(1) | N[+2206] | 1359.8882 | 4 | 5436.5309 | 5436.5038 | 0 | 5 | 141 | Specific | 655.04 | 633.29 | 633.29 | 13.21 | 146 | >sp\|P02788\|TRFL_HUMAN Lactotransferrin OS=Homo sapiens OX=9606 GN=LTF PE=1 SV=6 | 1270 | 20201118_lss_milk_1ng_glyco_hcd203040_compound15.23261.23261.4 | controllerType=0 controllerNumber=1 scan=23261 | 46.2152 |
| 20053:4 | 1 | R.TAGWNVPIGTLRPFLN[+2351.851]WTGPPEPIEAAVAR.F | HexNAc(4)Hex(5)Fuc(3)NeuAc(1) | N[+2352] | 1396.4009 | 4 | 5582.5817 | 5582.5617 | 0 | 3.6 | 141 | Specific | 670.6 | 625.6 | 625.6 | 13.55 | 146 | >sp\|P02788\|TRFL_HUMAN Lactotransferrin OS=Homo sapiens OX=9606 GN=LTF PE=1 SV=6 | 1270 | 20201118_lss_milk_1ng_glyco_hcd203040_compound15.23262.23262.4 | controllerType=0 controllerNumber=1 scan=23262 | 46.2161 |
| 20011:4 | 1 | R.TAGWNVPIGTLRPFLN[+2424.867]WTGPPEPIEAAVAR.F | HexNAc(5)Hex(5)Fuc(1)NeuAc(1) | N[+2425] | 1414.6558 | 4 | 5655.6012 | 5655.5781 | 0 | 4.1 | 141 | Specific | 670.6 | 528.81 | 528.81 | 13.55 | 146 | >sp\|P02788\|TRFL_HUMAN Lactotransferrin OS=Homo sapiens OX=9606 GN=LTF PE=1 SV=6 | 1270 | 20201118_lss_milk_1ng_glyco_hcd203040_compound15.23218.23218.4 | controllerType=0 controllerNumber=1 scan=23218 | 46.1249 |
| 19065:4 | 1 | R.TAGWNVPIGTLRPFLN[+1768.640]WTGPPEPIEAAVAR.F | HexNAc(4)Hex(5)Fuc(1) | N[+1769] | 1250.5983 | 4 | 4999.3712 | 4999.3504 | 0 | 4.2 | 141 | Specific | 679.02 | 647.84 | 647.84 | 13.55 | 146 | >sp\|P02788\|TRFL_HUMAN Lactotransferrin OS=Homo sapiens OX=9606 GN=LTF PE=1 SV=6 | 1270 | 20201118_lss_milk_1ng_glyco_hcd203040_compound15.22232.22232.4 | controllerType=0 controllerNumber=1 scan=22232 | 44.0458 |
| 21132:4 | 1 | R.TAGWNVPIGTLRPFLN[+2862.021]WTGPPEPIEAAVAR.F | HexNAc(5)Hex(6)Fuc(2)NeuAc(2) | N[+2862] | 1523.9419 | 4 | 6092.7458 | 6092.7314 | 0 | 2.4 | 141 | Specific | 691.79 | 499.6 | 499.6 | 13.44 | 146 | >sp\|P02788\|TRFL_HUMAN Lactotransferrin OS=Homo sapiens OX=9606 GN=LTF PE=1 SV=6 | 1270 | 20201118_lss_milk_1ng_glyco_hcd203040_compound15.24390.24390.4 | controllerType=0 controllerNumber=1 scan=24390 | 48.6883 |
| 20079:3 | 1 | R.TAGWNVPIGTLRPFLN[+2059.735]WTGPPEPIEAAVAR.F | HexNAc(4)Hex(5)Fuc(1)NeuAc(1) | N[+2060] | 1764.1711 | 3 | 5290.4988 | 5290.4459 | 0 | 10 | 141 | Specific | 700.16 | 700.16 | 700.16 | 13.18 | 146 | >sp\|P02788\|TRFL_HUMAN Lactotransferrin OS=Homo sapiens OX=9606 GN=LTF PE=1 SV=6 | 1270 | 20201118_lss_milk_1ng_glyco_hcd203040_compound15.23289.23289.3 | controllerType=0 controllerNumber=1 scan=23289 | 46.2746 |
| 20064:3 | 1 | R.TAGWNVPIGTLRPFLN[+2205.793]WTGPPEPIEAAVAR.F | HexNAc(4)Hex(5)Fuc(2)NeuAc(1) | N[+2206] | 1812.8474 | 3 | 5436.5276 | 5436.5038 | 0 | 4.4 | 141 | Specific | 703.2 | 589.89 | 589.89 | 14.22 | 146 | >sp\|P02788\|TRFL_HUMAN Lactotransferrin OS=Homo sapiens OX=9606 GN=LTF PE=1 SV=6 | 1270 | 20201118_lss_milk_1ng_glyco_hcd203040_compound15.23273.23273.3 | controllerType=0 controllerNumber=1 scan=23273 | 46.2436 |
| 19071:3 | 1 | R.TAGWNVPIGTLRPFLN[+1768.640]WTGPPEPIEAAVAR.F | HexNAc(4)Hex(5)Fuc(1) | N[+1769] | 1667.1294 | 3 | 4999.3735 | 4999.3504 | 0 | 4.6 | 141 | Specific | 737.38 | 617.53 | 617.53 | 14.22 | 146 | >sp\|P02788\|TRFL_HUMAN Lactotransferrin OS=Homo sapiens OX=9606 GN=LTF PE=1 SV=6 | 1270 | 20201118_lss_milk_1ng_glyco_hcd203040_compound15.22238.22238.3 | controllerType=0 controllerNumber=1 scan=22238 | 44.059 |
| 20082:3 | 1 | R.TAGWNVPIGTLRPFLN[+2059.735]WTGPPEPIEAAVAR.F | HexNAc(4)Hex(5)Fuc(1)NeuAc(1) | N[+2060] | 1764.1711 | 3 | 5290.4988 | 5290.4459 | 0 | 10 | 141 | Specific | 737.44 | 650.31 | 650.31 | 13.18 | 146 | >sp\|P02788\|TRFL_HUMAN Lactotransferrin OS=Homo sapiens OX=9606 GN=LTF PE=1 SV=6 | 1270 | 20201118_lss_milk_1ng_glyco_hcd203040_compound15.23292.23292.3 | controllerType=0 controllerNumber=1 scan=23292 | 46.2848 |
| 20010:5 | 1 | R.TAGWNVPIGTLRPFLN[+2570.925]WTGPPEPIEAAVAR.F | HexNAc(5)Hex(6)Fuc(2)NeuAc(1) | N[+2571] | 1161.136 | 5 | 5801.6508 | 5801.636 | 0 | 2.6 | 141 | Specific | 752.53 | 621.99 | 621.99 | 15.67 | 146 | >sp\|P02788\|TRFL_HUMAN Lactotransferrin OS=Homo sapiens OX=9606 GN=LTF PE=1 SV=6 | 1270 | 20201118_lss_milk_1ng_glyco_hcd203040_compound15.23217.23217.5 | controllerType=0 controllerNumber=1 scan=23217 | 46.1218 |
| 19068:3 | 1 | R.TAGWNVPIGTLRPFLN[+1768.640]WTGPPEPIEAAVAR.F | HexNAc(4)Hex(5)Fuc(1) | N[+1769] | 1667.1294 | 3 | 4999.3735 | 4999.3504 | 0 | 4.6 | 141 | Specific | 756.52 | 657.86 | 657.86 | 15.22 | 146 | >sp\|P02788\|TRFL_HUMAN Lactotransferrin OS=Homo sapiens OX=9606 GN=LTF PE=1 SV=6 | 1270 | 20201118_lss_milk_1ng_glyco_hcd203040_compound15.22235.22235.3 | controllerType=0 controllerNumber=1 scan=22235 | 44.0534 |
| 19997:4 | 1 | R.TAGWNVPIGTLRPFLN[+2570.925]WTGPPEPIEAAVAR.F | HexNAc(5)Hex(6)Fuc(2)NeuAc(1) | N[+2571] | 1451.1704 | 4 | 5801.6598 | 5801.636 | 0 | 4.1 | 141 | Specific | 780.16 | 560.39 | 560.39 | 15.55 | 146 | >sp\|P02788\|TRFL_HUMAN Lactotransferrin OS=Homo sapiens OX=9606 GN=LTF PE=1 SV=6 | 1270 | 20201118_lss_milk_1ng_glyco_hcd203040_compound15.23204.23204.4 | controllerType=0 controllerNumber=1 scan=23204 | 46.1025 |
| 20056:4 | 1 | R.TAGWNVPIGTLRPFLN[+2351.851]WTGPPEPIEAAVAR.F | HexNAc(4)Hex(5)Fuc(3)NeuAc(1) | N[+2352] | 1396.4009 | 4 | 5582.5817 | 5582.5617 | 0 | 3.6 | 141 | Specific | 786.52 | 675.59 | 675.59 | 15.55 | 146 | >sp\|P02788\|TRFL_HUMAN Lactotransferrin OS=Homo sapiens OX=9606 GN=LTF PE=1 SV=6 | 1270 | 20201118_lss_milk_1ng_glyco_hcd203040_compound15.23265.23265.4 | controllerType=0 controllerNumber=1 scan=23265 | 46.2261 |
| 20055:4 | 1 | R.TAGWNVPIGTLRPFLN[+2205.793]WTGPPEPIEAAVAR.F | HexNAc(4)Hex(5)Fuc(2)NeuAc(1) | N[+2206] | 1359.8882 | 4 | 5436.5309 | 5436.5038 | 0 | 5 | 141 | Specific | 807.86 | 701.6 | 701.6 | 16.21 | 146 | >sp\|P02788\|TRFL_HUMAN Lactotransferrin OS=Homo sapiens OX=9606 GN=LTF PE=1 SV=6 | 1270 | 20201118_lss_milk_1ng_glyco_hcd203040_compound15.23264.23264.4 | controllerType=0 controllerNumber=1 scan=23264 | 46.2221 |
| 19067:4 | 1 | R.TAGWNVPIGTLRPFLN[+1768.640]WTGPPEPIEAAVAR.F | HexNAc(4)Hex(5)Fuc(1) | N[+1769] | 1250.5983 | 4 | 4999.3712 | 4999.3504 | 0 | 4.2 | 141 | Specific | 860.76 | 759.46 | 759.46 | 17.55 | 146 | >sp\|P02788\|TRFL_HUMAN Lactotransferrin OS=Homo sapiens OX=9606 GN=LTF PE=1 SV=6 | 1270 | 20201118_lss_milk_1ng_glyco_hcd203040_compound15.22234.22234.4 | controllerType=0 controllerNumber=1 scan=22234 | 44.0525 |
| 17366:3 | 1 | R.TAGWNIPMGLLFN[+2279.830]QTGSC[+57.021]K.F | HexNAc(5)Hex(6)Fuc(2) | C[+57], N[+2280] | 1458.9564 | 3 | 4374.8545 | 4374.8396 | 0 | 3.4 | 485 | Specific | 286.83 | 286.83 | 187.21 | 6.81 | 146 | >sp\|P02788\|TRFL_HUMAN Lactotransferrin OS=Homo sapiens OX=9606 GN=LTF PE=1 SV=6 | 1270 | 20201118_lss_milk_1ng_glyco_hcd203040_compound15.20461.20461.3 | controllerType=0 controllerNumber=1 scan=20461 | 40.3007 |
| 17368:3 | 1 | R.TAGWNIPMGLLFN[+2279.830]QTGSC[+57.021]K.F | HexNAc(5)Hex(6)Fuc(2) | C[+57], N[+2280] | 1458.9564 | 3 | 4374.8545 | 4374.8396 | 0 | 3.4 | 485 | Specific | 333.71 | 333.71 | 291.21 | 7.46 | 146 | >sp\|P02788\|TRFL_HUMAN Lactotransferrin OS=Homo sapiens OX=9606 GN=LTF PE=1 SV=6 | 1270 | 20201118_lss_milk_1ng_glyco_hcd203040_compound15.20463.20463.3 | controllerType=0 controllerNumber=1 scan=20463 | 40.3039 |
| 17930:2 | 1 | R.TAGWNIPMGLLFN[+1216.423]QTGSC[+57.021]K.F | HexNAc(2)Hex(5) | C[+57], N[+1216] | 1656.2255 | 2 | 3311.4437 | 3311.4328 | 0 | 3.3 | 485 | Specific | 418.82 | 418.82 | 189.35 | 9.18 | 146 | >sp\|P02788\|TRFL_HUMAN Lactotransferrin OS=Homo sapiens OX=9606 GN=LTF PE=1 SV=6 | 1270 | 20201118_lss_milk_1ng_glyco_hcd203040_compound15.21049.21049.2 | controllerType=0 controllerNumber=1 scan=21049 | 41.531 |
| 16638:3 | 1 | R.TAGWNIPMGLLFN[+1768.640]QTGSC[+57.021]K.F | HexNAc(4)Hex(5)Fuc(1) | C[+57], N[+1769] | 1288.56 | 3 | 3863.6654 | 3863.6495 | 0 | 4.1 | 485 | Specific | 429.43 | 429.43 | 429.43 | 9.17 | 146 | >sp\|P02788\|TRFL_HUMAN Lactotransferrin OS=Homo sapiens OX=9606 GN=LTF PE=1 SV=6 | 1270 | 20201118_lss_milk_1ng_glyco_hcd203040_compound15.20745.20745.3 | controllerType=0 controllerNumber=1 scan=20745 | 40.879 |
| 19492:4 | 1 | R.TAGWNIPMGLLFN[+2205.793]QTGSC[+57.021]K.F | HexNAc(4)Hex(5)Fuc(2)NeuAc(1) | C[+57], N[+2206] | 1075.9612 | 4 | 4300.8231 | 4300.8028 | 0 | 4.7 | 485 | Specific | 433 | 330.75 | 330.75 | 7.61 | 146 | >sp\|P02788\|TRFL_HUMAN Lactotransferrin OS=Homo sapiens OX=9606 GN=LTF PE=1 SV=6 | 1270 | 20201118_lss_milk_1ng_glyco_hcd203040_compound15.22678.22678.4 | controllerType=0 controllerNumber=1 scan=22678 | 44.9911 |
| 17586:2 | 1 | R.TAGWNIPMGLLFN[+1768.640]QTGSC[+57.021]K.F | HexNAc(4)Hex(5)Fuc(1) | C[+57], N[+1769] | 1932.3397 | 2 | 3863.6721 | 3863.6495 | 0 | 5.9 | 485 | Specific | 481.53 | 443.38 | 443.38 | 8.69 | 146 | >sp\|P02788\|TRFL_HUMAN Lactotransferrin OS=Homo sapiens OX=9606 GN=LTF PE=1 SV=6 | 1270 | 20201118_lss_milk_1ng_glyco_hcd203040_compound15.20690.20690.2 | controllerType=0 controllerNumber=1 scan=20690 | 40.7553 |
| 19494:4 | 1 | R.TAGWNIPMGLLFN[+2205.793]QTGSC[+57.021]K.F | HexNAc(4)Hex(5)Fuc(2)NeuAc(1) | C[+57], N[+2206] | 1075.9612 | 4 | 4300.8231 | 4300.8028 | 0 | 4.7 | 485 | Specific | 514.12 | 370.53 | 370.53 | 8.21 | 146 | >sp\|P02788\|TRFL_HUMAN Lactotransferrin OS=Homo sapiens OX=9606 GN=LTF PE=1 SV=6 | 1270 | 20201118_lss_milk_1ng_glyco_hcd203040_compound15.22680.22680.4 | controllerType=0 controllerNumber=1 scan=22680 | 44.9971 |
| 17588:2 | 1 | R.TAGWNIPMGLLFN[+1768.640]QTGSC[+57.021]K.F | HexNAc(4)Hex(5)Fuc(1) | C[+57], N[+1769] | 1932.3397 | 2 | 3863.6721 | 3863.6495 | 0 | 5.9 | 485 | Specific | 524.24 | 433.93 | 433.93 | 8.82 | 146 | >sp\|P02788\|TRFL_HUMAN Lactotransferrin OS=Homo sapiens OX=9606 GN=LTF PE=1 SV=6 | 1270 | 20201118_lss_milk_1ng_glyco_hcd203040_compound15.20692.20692.2 | controllerType=0 controllerNumber=1 scan=20692 | 40.7587 |
| 17761:3 | 1 | R.TAGWNIPMGLLFN[+1622.582]QTGSC[+57.021]K.F | HexNAc(4)Hex(5) | C[+57], N[+1623] | 1239.8732 | 3 | 3717.6051 | 3717.5916 | 0 | 3.7 | 485 | Specific | 578.66 | 578.66 | 225.14 | 11.25 | 146 | >sp\|P02788\|TRFL_HUMAN Lactotransferrin OS=Homo sapiens OX=9606 GN=LTF PE=1 SV=6 | 1270 | 20201118_lss_milk_1ng_glyco_hcd203040_compound15.20873.20873.3 | controllerType=0 controllerNumber=1 scan=20873 | 41.1383 |
| 19572:3 | 1 | R.TAGWNIPMGLLFN[+2205.793]QTGSC[+57.021]K.F | HexNAc(4)Hex(5)Fuc(2)NeuAc(1) | C[+57], N[+2206] | 1434.2803 | 3 | 4300.8264 | 4300.8028 | 0 | 5.5 | 485 | Specific | 591.64 | 586.53 | 586.53 | 10.51 | 146 | >sp\|P02788\|TRFL_HUMAN Lactotransferrin OS=Homo sapiens OX=9606 GN=LTF PE=1 SV=6 | 1270 | 20201118_lss_milk_1ng_glyco_hcd203040_compound15.22725.22725.3 | controllerType=0 controllerNumber=1 scan=22725 | 45.0983 |
| 17653:3 | 1 | R.TAGWNIPMGLLFN[+1606.587]QTGSC[+57.021]K.F | HexNAc(4)Hex(4)Fuc(1) | C[+57], N[+1607] | 1234.5406 | 3 | 3701.6072 | 3701.5966 | 0 | 2.9 | 485 | Specific | 668.89 | 609.49 | 609.49 | 13.14 | 146 | >sp\|P02788\|TRFL_HUMAN Lactotransferrin OS=Homo sapiens OX=9606 GN=LTF PE=1 SV=6 | 1270 | 20201118_lss_milk_1ng_glyco_hcd203040_compound15.20760.20760.3 | controllerType=0 controllerNumber=1 scan=20760 | 40.903 |
| 21438:3 | 1 | R.TAGWNIPMGLLFN[+2350.830]QTGSC[+57.021]K.F | HexNAc(4)Hex(5)Fuc(1)NeuAc(2) | C[+57], N[+2351] | 1482.6273 | 3 | 4445.8672 | 4445.8403 | 0 | 6.1 | 485 | Specific | 674.46 | 658.92 | 658.92 | 12.51 | 146 | >sp\|P02788\|TRFL_HUMAN Lactotransferrin OS=Homo sapiens OX=9606 GN=LTF PE=1 SV=6 | 1270 | 20201118_lss_milk_1ng_glyco_hcd203040_compound15.24712.24712.3 | controllerType=0 controllerNumber=1 scan=24712 | 49.4521 |
| 19540:3 | 1 | R.TAGWNIPMGLLFN[+2205.793]QTGSC[+57.021]K.F | HexNAc(4)Hex(5)Fuc(2)NeuAc(1) | C[+57], N[+2206] | 1434.2803 | 3 | 4300.8264 | 4300.8028 | 0 | 5.5 | 485 | Specific | 691.47 | 586.3 | 586.3 | 12.51 | 146 | >sp\|P02788\|TRFL_HUMAN Lactotransferrin OS=Homo sapiens OX=9606 GN=LTF PE=1 SV=6 | 1270 | 20201118_lss_milk_1ng_glyco_hcd203040_compound15.22728.22728.3 | controllerType=0 controllerNumber=1 scan=22728 | 45.1192 |
| 19404:3 | 1 | R.TAGWNIPMGLLFN[+2205.793]QTGSC[+57.021]K.F | HexNAc(4)Hex(5)Fuc(2)NeuAc(1) | C[+57], N[+2206] | 1434.2785 | 3 | 4300.8209 | 4300.8028 | 0 | 4.2 | 485 | Specific | 746.8 | 738.15 | 738.15 | 13.9 | 146 | >sp\|P02788\|TRFL_HUMAN Lactotransferrin OS=Homo sapiens OX=9606 GN=LTF PE=1 SV=6 | 1270 | 20201118_lss_milk_1ng_glyco_hcd203040_compound15.22587.22587.3 | controllerType=0 controllerNumber=1 scan=22587 | 44.8078 |
| 21418:4 | 1 | R.TAGWNIPMGLLFN[+2350.830]QTGSC[+57.021]K.F | HexNAc(4)Hex(5)Fuc(1)NeuAc(2) | C[+57], N[+2351] | 1112.2192 | 4 | 4445.8548 | 4445.8403 | 0 | 3.3 | 485 | Specific | 773.26 | 626.52 | 626.52 | 15.48 | 146 | >sp\|P02788\|TRFL_HUMAN Lactotransferrin OS=Homo sapiens OX=9606 GN=LTF PE=1 SV=6 | 1270 | 20201118_lss_milk_1ng_glyco_hcd203040_compound15.24691.24691.4 | controllerType=0 controllerNumber=1 scan=24691 | 49.4053 |
| 17932:2 | 1 | R.TAGWNIPMGLLFN[+1216.423]QTGSC[+57.021]K.F | HexNAc(2)Hex(5) | C[+57], N[+1216] | 1656.2255 | 2 | 3311.4437 | 3311.4328 | 0 | 3.3 | 485 | Specific | 816.96 | 735.67 | 262.67 | 16.26 | 146 | >sp\|P02788\|TRFL_HUMAN Lactotransferrin OS=Homo sapiens OX=9606 GN=LTF PE=1 SV=6 | 1270 | 20201118_lss_milk_1ng_glyco_hcd203040_compound15.21051.21051.2 | controllerType=0 controllerNumber=1 scan=21051 | 41.5359 |
| 17641:3 | 1 | R.TAGWNIPMGLLFN[+1768.640]QTGSC[+57.021]K.F | HexNAc(4)Hex(5)Fuc(1) | C[+57], N[+1769] | 1288.56 | 3 | 3863.6654 | 3863.6495 | 0 | 4.1 | 485 | Specific | 826.67 | 810.43 | 810.43 | 16.25 | 146 | >sp\|P02788\|TRFL_HUMAN Lactotransferrin OS=Homo sapiens OX=9606 GN=LTF PE=1 SV=6 | 1270 | 20201118_lss_milk_1ng_glyco_hcd203040_compound15.20748.20748.3 | controllerType=0 controllerNumber=1 scan=20748 | 40.8831 |
| 21421:4 | 1 | R.TAGWNIPMGLLFN[+2350.830]QTGSC[+57.021]K.F | HexNAc(4)Hex(5)Fuc(1)NeuAc(2) | C[+57], N[+2351] | 1112.2192 | 4 | 4445.8548 | 4445.8403 | 0 | 3.3 | 485 | Specific | 843.53 | 640.63 | 640.63 | 16.48 | 146 | >sp\|P02788\|TRFL_HUMAN Lactotransferrin OS=Homo sapiens OX=9606 GN=LTF PE=1 SV=6 | 1270 | 20201118_lss_milk_1ng_glyco_hcd203040_compound15.24694.24694.4 | controllerType=0 controllerNumber=1 scan=24694 | 49.4107 |
| 21588:3 | 1 | R.TAGWNIPMGLLFN[+2204.772]QTGSC[+57.021]K.F | HexNAc(4)Hex(4)NeuAc(2) | C[+57], N[+2205] | 1433.9383 | 3 | 4299.8003 | 4299.7824 | 0 | 4.2 | 485 | Specific | 862.9 | 818.25 | 818.25 | 16.9 | 146 | >sp\|P02788\|TRFL_HUMAN Lactotransferrin OS=Homo sapiens OX=9606 GN=LTF PE=1 SV=6 | 1270 | 20201118_lss_milk_1ng_glyco_hcd203040_compound15.24869.24869.3 | controllerType=0 controllerNumber=1 scan=24869 | 49.831 |
| 17655:3 | 1 | R.TAGWNIPMGLLFN[+1606.587]QTGSC[+57.021]K.F | HexNAc(4)Hex(4)Fuc(1) | C[+57], N[+1607] | 1234.5406 | 3 | 3701.6072 | 3701.5966 | 0 | 2.9 | 485 | Specific | 869.74 | 674.39 | 674.39 | 17.14 | 146 | >sp\|P02788\|TRFL_HUMAN Lactotransferrin OS=Homo sapiens OX=9606 GN=LTF PE=1 SV=6 | 1270 | 20201118_lss_milk_1ng_glyco_hcd203040_compound15.20762.20762.3 | controllerType=0 controllerNumber=1 scan=20762 | 40.9082 |
| 21590:3 | 1 | R.TAGWNIPMGLLFN[+2204.772]QTGSC[+57.021]K.F | HexNAc(4)Hex(4)NeuAc(2) | C[+57], N[+2205] | 1433.9383 | 3 | 4299.8003 | 4299.7824 | 0 | 4.2 | 485 | Specific | 885.75 | 828.37 | 828.37 | 16.9 | 146 | >sp\|P02788\|TRFL_HUMAN Lactotransferrin OS=Homo sapiens OX=9606 GN=LTF PE=1 SV=6 | 1270 | 20201118_lss_milk_1ng_glyco_hcd203040_compound15.24871.24871.3 | controllerType=0 controllerNumber=1 scan=24871 | 49.8387 |
| 21440:3 | 1 | R.TAGWNIPMGLLFN[+2350.830]QTGSC[+57.021]K.F | HexNAc(4)Hex(5)Fuc(1)NeuAc(2) | C[+57], N[+2351] | 1482.6273 | 3 | 4445.8672 | 4445.8403 | 0 | 6.1 | 485 | Specific | 905.88 | 864.17 | 864.17 | 17.51 | 146 | >sp\|P02788\|TRFL_HUMAN Lactotransferrin OS=Homo sapiens OX=9606 GN=LTF PE=1 SV=6 | 1270 | 20201118_lss_milk_1ng_glyco_hcd203040_compound15.24714.24714.3 | controllerType=0 controllerNumber=1 scan=24714 | [illegible] |
| 19570:3 | 1 | R.TAGWNIPMGLLFN[+1913.677]QTGSC[+57.021]K.F | HexNAc(4)Hex(5)NeuAc(1) | C[+57], N[+1914] | 1336.9056 | 3 | 4008.7022 | 4008.687 | 0 | 3.8 | 485 | Specific | 927.36 | 920.61 | 920.61 | 18.25 | 146 | >sp\|P02788\|TRFL_HUMAN Lactotransferrin OS=Homo sapiens OX=9606 GN=LTF PE=1 SV=6 | 1270 | 20201118_lss_milk_1ng_glyco_hcd203040_compound15.22760.22760.3 | controllerType=0 controllerNumber=1 scan=22760 | [illegible] |
| 17763:3 | 1 | R.TAGWNIPMGLLFN[+1622.582]QTGSC[+57.021]K.F | HexNAc(4)Hex(5) | C[+57], N[+1623] | 1239.8732 | 3 | 3717.6051 | 3717.5916 | 0 | 3.7 | 485 | Specific | 931.3 | 892.17 | 428.3 | 18.25 | 146 | >sp\|P02788\|TRFL_HUMAN Lactotransferrin OS=Homo sapiens OX=9606 GN=LTF PE=1 SV=6 | 1270 | 20201118_lss_milk_1ng_glyco_hcd203040_compound15.20875.20875.3 | controllerType=0 controllerNumber=1 scan=20875 | [illegible] |
| 19429:3 | 1 | R.TAGWNIPMGLLFN[+2059.735]QTGSC[+57.021]K.F | HexNAc(4)Hex(5)Fuc(1)NeuAc(1) | C[+57], N[+2060] | 1385.5909 | 3 | 4154.7581 | 4154.7449 | 0 | 3.2 | 485 | Specific | 931.52 | 898.66 | 898.66 | 18.25 | 146 | >sp\|P02788\|TRFL_HUMAN Lactotransferrin OS=Homo sapiens OX=9606 GN=LTF PE=1 SV=6 | 1270 | 20201118_lss_milk_1ng_glyco_hcd203040_compound15.22613.22613.3 | controllerType=0 controllerNumber=1 scan=22613 | [illegible] |
| 19406:3 | 1 | R.TAGWNIPMGLLFN[+2205.793]QTGSC[+57.021]K.F | HexNAc(4)Hex(5)Fuc(2)NeuAc(1) | C[+57], N[+2206] | 1434.2785 | 3 | 4300.8209 | 4300.8028 | 0 | 4.2 | 485 | Specific | 938.64 | 891.34 | 891.34 | 17.9 | 146 | >sp\|P02788\|TRFL_HUMAN Lactotransferrin OS=Homo sapiens OX=9606 GN=LTF PE=1 SV=6 | 1270 | 20201118_lss_milk_1ng_glyco_hcd203040_compound15.22589.22589.3 | controllerType=0 controllerNumber=1 scan=22589 | 44.8116 |
| 19567:3 | 1 | R.TAGWNIPMGLLFN[+1913.677]QTGSC[+57.021]K.F | HexNAc(4)Hex(5)NeuAc(1) | C[+57], N[+1914] | 1336.9056 | 3 | 4008.7022 | 4008.687 | 0 | 3.8 | 485 | Specific | 944.6 | 944.6 | 944.6 | 18.25 | 146 | >sp\|P02788\|TRFL_HUMAN Lactotransferrin OS=Homo sapiens OX=9606 GN=LTF PE=1 SV=6 | 1270 | 20201118_lss_milk_1ng_glyco_hcd203040_compound15.22757.22757.3 | controllerType=0 controllerNumber=1 scan=22757 | 45.1881 |
| 17890:3 | 1 | R.TAGWNIPMGLLFN[+1216.423]QTGSC[+57.021]K.F | HexNAc(2)Hex(5) | C[+57], N[+1216] | 1104.4875 | 3 | 3311.4478 | 3311.4328 | 0 | 4.5 | 485 | Specific | 987.09 | 883.36 | 437.78 | 18.9 | 146 | >sp\|P02788\|TRFL_HUMAN Lactotransferrin OS=Homo sapiens OX=9606 GN=LTF PE=1 SV=6 | 1270 | 20201118_lss_milk_1ng_glyco_hcd203040_compound15.21007.21007.3 | controllerType=0 controllerNumber=1 scan=21007 | 41.44 |
| 17892:3 | 1 | R.TAGWNIPMGLLFN[+1216.423]QTGSC[+57.021]K.F | HexNAc(2)Hex(5) | C[+57], N[+1216] | 1104.4875 | 3 | 3311.4478 | 3311.4328 | 0 | 4.5 | 485 | Specific | 1007.26 | 861.57 | 472.77 | 19.9 | 146 | >sp\|P02788\|TRFL_HUMAN Lactotransferrin OS=Homo sapiens OX=9606 GN=LTF PE=1 SV=6 | 1270 | 20201118_lss_milk_1ng_glyco_hcd203040_compound15.21009.21009.3 | controllerType=0 controllerNumber=1 scan=21009 | 41.449 |
| 19422:3 | 1 | R.TAGWNIPMGLLFN[+2059.735]QTGSC[+57.021]K.F | HexNAc(4)Hex(5)Fuc(1)NeuAc(1) | C[+57], N[+2060] | 1385.5909 | 3 | 4154.7581 | 4154.7449 | 0 | 3.2 | 485 | Specific | 1009.72 | 913.17 | 913.17 | 20.25 | 146 | >sp\|P02788\|TRFL_HUMAN Lactotransferrin OS=Homo sapiens OX=9606 GN=LTF PE=1 SV=6 | 1270 | 20201118_lss_milk_1ng_glyco_hcd203040_compound15.22611.22611.3 | controllerType=0 controllerNumber=1 scan=22611 | 44.8561 |
| 19807:3 | 1 | R.TAGWNIPMGLLFN[+1913.677]QTGSC[+57.021]KFDEYFSQSC[+57.021]APGSDPR.S | HexNAc(4)Hex(5)NeuAc(1) | C[+57]*2, N[+1914] | 1951.489 | 3 | 5852.4524 | 5852.4341 | 0 | 3.1 | 485 | Specific | 201.05 | 124 | 124 | 4.28 | 146 | >sp\|P02788\|TRFL_HUMAN Lactotransferrin OS=Homo sapiens OX=9606 GN=LTF PE=1 SV=6 | 1270 | 20201118_lss_milk_1ng_glyco_hcd203040_compound15.23007.23007.3 | controllerType=0 controllerNumber=1 scan=23007 | 45.7256 |
| 18688:5 | 1 | R.TAGWNIPMGLLFN[+1768.640]QTGSC[+57.021]KFDEYFSQSC[+57.021]APGSDPR.S | HexNAc(4)Hex(5)Fuc(1) | C[+57]*2, N[+1769] | 1142.2861 | 5 | 5707.4016 | 5707.3966 | 0 | 0.9 | 485 | Specific | 283.01 | 283.01 | 283.01 | 6.83 | 146 | >sp\|P02788\|TRFL_HUMAN Lactotransferrin OS=Homo sapiens OX=9606 GN=LTF PE=1 SV=6 | 1270 | 20201118_lss_milk_1ng_glyco_hcd203040_compound15.21840.21840.5 | controllerType=0 controllerNumber=1 scan=21840 | 43.2778 |
| 18656:4 | 1 | R.[+42.011]TAGWNIPMGLLFN[+1872.651]QTGSC[+57.021]KFDEYFSQSC[+57.021]APGSDPR.S | HexNAc(3)Hex(6)NeuAc(1) | .[42], C[+57]*2, N[+1873] | 1464.1229 | 4 | 5853.4699 | 5853.4181 | 0 | 8.8 | 485 | Specific | 321.36 | 307.43 | 307.43 | 4.61 | 146 | >sp\|P02788\|TRFL_HUMAN Lactotransferrin OS=Homo sapiens OX=9606 GN=LTF PE=1 SV=6 | 1270 | 20201118_lss_milk_1ng_glyco_hcd203040_compound15.21807.21807.4 | controllerType=0 controllerNumber=1 scan=21807 | 43.2095 |
| 18618:4 | 1 | R.TAGWNIPMGLLFN[+2076.750]QTGSC[+57.021]KFDEYFSQSC[+57.021]APGSDPR.S | HexNAc(4)Hex(6)Fuc(2) | C[+57]*2, N[+2077] | 1504.6422 | 4 | 6015.547 | 6015.5074 | 0 | 6.6 | 485 | Specific | 340.82 | 297.82 | 297.82 | 6.2 | 146 | >sp\|P02788\|TRFL_HUMAN Lactotransferrin OS=Homo sapiens OX=9606 GN=LTF PE=1 SV=6 | 1270 | 20201118_lss_milk_1ng_glyco_hcd203040_compound15.21767.21767.4 | controllerType=0 controllerNumber=1 scan=21767 | 43.1178 |
| 18616:4 | 1 | R.TAGWNIPMGLLFN[+2076.750]QTGSC[+57.021]KFDEYFSQSC[+57.021]APGSDPR.S | HexNAc(4)Hex(6)Fuc(2) | C[+57]*2, N[+2077] | 1504.6422 | 4 | 6015.547 | 6015.5074 | 0 | 6.6 | 485 | Specific | 365.45 | 302.4 | 302.4 | 6.26 | 146 | >sp\|P02788\|TRFL_HUMAN Lactotransferrin OS=Homo sapiens OX=9606 GN=LTF PE=1 SV=6 | 1270 | 20201118_lss_milk_1ng_glyco_hcd203040_compound15.21765.21765.4 | controllerType=0 controllerNumber=1 scan=21765 | 43.1134 |
| 18723:3 | 1 | R.TAGWNIPMGLLFN[+1622.582]QTGSC[+57.021]KFDEYFSQSC[+57.021]APGSDPR.S | HexNAc(4)Hex(5) | C[+57]*2, N[+1623] | 1854.4554 | 3 | 5561.3517 | 5561.3387 | 0 | 2.3 | 485 | Specific | 434.59 | 365.78 | 116.99 | 8.03 | 146 | >sp\|P02788\|TRFL_HUMAN Lactotransferrin OS=Homo sapiens OX=9606 GN=LTF PE=1 SV=6 | 1270 | 20201118_lss_milk_1ng_glyco_hcd203040_compound15.21876.21876.3 | controllerType=0 controllerNumber=1 scan=21876 | 43.3473 |
| 19831:4 | 1 | R.TAGWNIPMGLLFN[+1913.677]QTGSC[+57.021]KFDEYFSQSC[+57.021]APGSDPR.S | HexNAc(4)Hex(5)NeuAc(1) | C[+57]*2, N[+1914] | 1463.8684 | 4 | 5852.4517 | 5852.4341 | 0 | 3 | 485 | Specific | 459.2 | 410.69 | 410.69 | 9.49 | 146 | >sp\|P02788\|TRFL_HUMAN Lactotransferrin OS=Homo sapiens OX=9606 GN=LTF PE=1 SV=6 | 1270 | 20201118_lss_milk_1ng_glyco_hcd203040_compound15.23032.23032.4 | controllerType=0 controllerNumber=1 scan=23032 | 45.7705 |
| 19836:3 | 1 | R.TAGWNIPMGLLFN[+1913.677]QTGSC[+57.021]KFDEYFSQSC[+57.021]APGSDPR.S | HexNAc(4)Hex(5)NeuAc(1) | C[+57]*2, N[+1914] | 1951.4888 | 3 | 5852.4506 | 5852.4341 | 0 | 2.8 | 485 | Specific | 512 | 441.99 | 441.99 | 9.63 | 146 | >sp\|P02788\|TRFL_HUMAN Lactotransferrin OS=Homo sapiens OX=9606 GN=LTF PE=1 SV=6 | 1270 | 20201118_lss_milk_1ng_glyco_hcd203040_compound15.23037.23037.3 | controllerType=0 controllerNumber=1 scan=23037 | 45.7774 |
| 19839:3 | 1 | R.TAGWNIPMGLLFN[+1913.677]QTGSC[+57.021]KFDEYFSQSC[+57.021]APGSDPR.S | HexNAc(4)Hex(5)NeuAc(1) | C[+57]*2, N[+1914] | 1951.4888 | 3 | 5852.4506 | 5852.4341 | 0 | 2.8 | 485 | Specific | 531.06 | 413.91 | 413.91 | 9.85 | 146 | >sp\|P02788\|TRFL_HUMAN Lactotransferrin OS=Homo sapiens OX=9606 GN=LTF PE=1 SV=6 | 1270 | 20201118_lss_milk_1ng_glyco_hcd203040_compound15.23040.23040.3 | controllerType=0 controllerNumber=1 scan=23040 | 45.7818 |
| 19699:4 | 1 | R.TAGWNIPMGLLFN[+2205.793]QTGSC[+57.021]KFDEYFSQSC[+57.021]APGSDPR.S | HexNAc(4)Hex(5)Fuc(2)NeuAc(1) | C[+57]*2, N[+2206] | 1536.8975 | 4 | 6144.568 | 6144.5499 | 0 | 2.9 | 485 | Specific | 636.14 | 616.04 | 616.04 | 12.32 | 146 | >sp\|P02788\|TRFL_HUMAN Lactotransferrin OS=Homo sapiens OX=9606 GN=LTF PE=1 SV=6 | 1270 | 20201118_lss_milk_1ng_glyco_hcd203040_compound15.22895.22895.4 | controllerType=0 controllerNumber=1 scan=22895 | 45.5014 |
| 18640:3 | 1 | R.TAGWNIPMGLLFN[+1768.640]QTGSC[+57.021]KFDEYFSQSC[+57.021]APGSDPR.S | HexNAc(4)Hex(5)Fuc(1) | C[+57]*2, N[+1769] | 1903.1445 | 3 | 5707.4189 | 5707.3966 | 0 | 3.9 | 485 | Specific | 656.55 | 588.13 | 588.13 | 13.43 | 146 | >sp\|P02788\|TRFL_HUMAN Lactotransferrin OS=Homo sapiens OX=9606 GN=LTF PE=1 SV=6 | 1270 | 20201118_lss_milk_1ng_glyco_hcd203040_compound15.21790.21790.3 | controllerType=0 controllerNumber=1 scan=21790 | 43.1604 |
| 18707:4 | 1 | R.TAGWNIPMGLLFN[+1622.582]QTGSC[+57.021]KFDEYFSQSC[+57.021]APGSDPR.S | HexNAc(4)Hex(5) | C[+57]*2, N[+1623] | 1391.0942 | 4 | 5561.3551 | 5561.3387 | 0 | 3 | 485 | Specific | 659.98 | 618.23 | 186.65 | 13.32 | 146 | >sp\|P02788\|TRFL_HUMAN Lactotransferrin OS=Homo sapiens OX=9606 GN=LTF PE=1 SV=6 | 1270 | 20201118_lss_milk_1ng_glyco_hcd203040_compound15.21860.21860.4 | controllerType=0 controllerNumber=1 scan=21860 | 43.3202 |
| 18726:3 | 1 | R.TAGWNIPMGLLFN[+1622.582]QTGSC[+57.021]KFDEYFSQSC[+57.021]APGSDPR.S | HexNAc(4)Hex(5) | C[+57]*2, N[+1623] | 1854.4554 | 3 | 5561.3517 | 5561.3387 | 0 | 2.3 | 485 | Specific | 666.64 | 548.85 | 197.19 | 13.33 | 146 | >sp\|P02788\|TRFL_HUMAN Lactotransferrin OS=Homo sapiens OX=9606 GN=LTF PE=1 SV=6 | 1270 | 20201118_lss_milk_1ng_glyco_hcd203040_compound15.21879.21879.3 | controllerType=0 controllerNumber=1 scan=21879 | 43.3515 |
| 18690:5 | 1 | R.TAGWNIPMGLLFN[+1768.640]QTGSC[+57.021]KFDEYFSQSC[+57.021]APGSDPR.S | HexNAc(4)Hex(5)Fuc(1) | C[+57]*2, N[+1769] | 1142.2861 | 5 | 5707.4016 | 5707.3966 | 0 | 0.9 | 485 | Specific | 672.01 | 559 | 559 | 13.27 | 146 | >sp\|P02788\|TRFL_HUMAN Lactotransferrin OS=Homo sapiens OX=9606 GN=LTF PE=1 SV=6 | 1270 | 20201118_lss_milk_1ng_glyco_hcd203040_compound15.21842.21842.5 | controllerType=0 controllerNumber=1 scan=21842 | 43.281 |
| 19810:3 | 1 | R.TAGWNIPMGLLFN[+1913.677]QTGSC[+57.021]KFDEYFSQSC[+57.021]APGSDPR.S | HexNAc(4)Hex(5)NeuAc(1) | C[+57]*2, N[+1914] | 1951.489 | 3 | 5852.4524 | 5852.4341 | 0 | 3.1 | 485 | Specific | 742.62 | 634.29 | 634.29 | 14.43 | 146 | >sp\|P02788\|TRFL_HUMAN Lactotransferrin OS=Homo sapiens OX=9606 GN=LTF PE=1 SV=6 | 1270 | 20201118_lss_milk_1ng_glyco_hcd203040_compound15.23010.23010.3 | controllerType=0 controllerNumber=1 scan=23010 | 45.7291 |
| 20995:4 | 1 | R.TAGWNIPMGLLFN[+2350.830]QTGSC[+57.021]KFDEYFSQSC[+57.021]APGSDPR.S | HexNAc(4)Hex(5)Fuc(1)NeuAc(2) | C[+57]*2, N[+2351] | 1573.1619 | 4 | 6289.6258 | 6289.5875 | 0 | 6.1 | 485 | Specific | 765.59 | 744.84 | 744.84 | 14.68 | 146 | >sp\|P02788\|TRFL_HUMAN Lactotransferrin OS=Homo sapiens OX=9606 GN=LTF PE=1 SV=6 | 1270 | 20201118_lss_milk_1ng_glyco_hcd203040_compound15.24248.24248.4 | controllerType=0 controllerNumber=1 scan=24248 | 48.4086 |
| 18643:3 | 1 | R.TAGWNIPMGLLFN[+1768.640]QTGSC[+57.021]KFDEYFSQSC[+57.021]APGSDPR.S | HexNAc(4)Hex(5)Fuc(1) | C[+57]*2, N[+1769] | 1903.1445 | 3 | 5707.4189 | 5707.3966 | 0 | 3.9 | 485 | Specific | 786.41 | 652.53 | 652.53 | 15.43 | 146 | >sp\|P02788\|TRFL_HUMAN Lactotransferrin OS=Homo sapiens OX=9606 GN=LTF PE=1 SV=6 | 1270 | 20201118_lss_milk_1ng_glyco_hcd203040_compound15.21793.21793.3 | controllerType=0 controllerNumber=1 scan=21793 | 43.1666 |
| 19833:4 | 1 | R.TAGWNIPMGLLFN[+1913.677]QTGSC[+57.021]KFDEYFSQSC[+57.021]APGSDPR.S | HexNAc(4)Hex(5)NeuAc(1) | C[+57]*2, N[+1914] | 1463.8684 | 4 | 5852.4517 | 5852.4341 | 0 | 3 | 485 | Specific | 837.83 | 756.24 | 756.24 | 16.32 | 146 | >sp\|P02788\|TRFL_HUMAN Lactotransferrin OS=Homo sapiens OX=9606 GN=LTF PE=1 SV=6 | 1270 | 20201118_lss_milk_1ng_glyco_hcd203040_compound15.23034.23034.4 | controllerType=0 controllerNumber=1 scan=23034 | 45.774 |
| 18710:4 | 1 | R.TAGWNIPMGLLFN[+1622.582]QTGSC[+57.021]KFDEYFSQSC[+57.021]APGSDPR.S | HexNAc(4)Hex(5) | C[+57]*2, N[+1623] | 1391.0942 | 4 | 5561.3551 | 5561.3387 | 0 | 3 | 485 | Specific | 891.6 | 863.14 | 348.87 | 17.32 | 146 | >sp\|P02788\|TRFL_HUMAN Lactotransferrin OS=Homo sapiens OX=9606 GN=LTF PE=1 SV=6 | 1270 | 20201118_lss_milk_1ng_glyco_hcd203040_compound15.21863.21863.4 | controllerType=0 controllerNumber=1 scan=21863 | 43.3245 |
| 19700:5 | 1 | R.TAGWNIPMGLLFN[+2059.735]QTGSC[+57.021]KFDEYFSQSC[+57.021]APGSDPR.S | HexNAc(4)Hex(5)Fuc(1)NeuAc(1) | C[+57]*2, N[+2060] | 1200.5078 | 5 | 5998.51 | 5998.492 | 0 | 3 | 485 | Specific | 893.93 | 893.93 | 893.93 | 17.55 | 146 | >sp\|P02788\|TRFL_HUMAN Lactotransferrin OS=Homo sapiens OX=9606 GN=LTF PE=1 SV=6 | 1270 | 20201118_lss_milk_1ng_glyco_hcd203040_compound15.22896.22896.5 | controllerType=0 controllerNumber=1 scan=22896 | 45.5027 |
| 19701:4 | 1 | R.TAGWNIPMGLLFN[+2205.793]QTGSC[+57.021]KFDEYFSQSC[+57.021]APGSDPR.S | HexNAc(4)Hex(5)Fuc(2)NeuAc(1) | C[+57]*2, N[+2206] | 1536.8975 | 4 | 6144.568 | 6144.5499 | 0 | 2.9 | 485 | Specific | 918.49 | 776.96 | 776.96 | 18.32 | 146 | >sp\|P02788\|TRFL_HUMAN Lactotransferrin OS=Homo sapiens OX=9606 GN=LTF PE=1 SV=6 | 1270 | 20201118_lss_milk_1ng_glyco_hcd203040_compound15.22897.22897.4 | controllerType=0 controllerNumber=1 scan=22897 | 45.5076 |
| 20997:4 | 1 | R.TAGWNIPMGLLFN[+2350.830]QTGSC[+57.021]KFDEYFSQSC[+57.021]APGSDPR.S | HexNAc(4)Hex(5)Fuc(1)NeuAc(2) | C[+57]*2, N[+2351] | 1573.1619 | 4 | 6289.6256 | 6289.5875 | 0 | 6.1 | 485 | Specific | 919.38 | 896.59 | 896.59 | 17.68 | 146 | >sp\|P02788\|TRFL_HUMAN Lactotransferrin OS=Homo sapiens OX=9606 GN=LTF PE=1 SV=6 | 1270 | 20201118_lss_milk_1ng_glyco_hcd203040_compound15.24250.24250.4 | controllerType=0 controllerNumber=1 scan=24250 | 48.4135 |
| 19681:4 | 1 | R.TAGWNIPMGLLFN[+2059.735]QTGSC[+57.021]KFDEYFSQSC[+57.021]APGSDPR.S | HexNAc(4)Hex(5)Fuc(1)NeuAc(1) | C[+57]*2, N[+2060] | 1500.3848 | 4 | 5998.5172 | 5998.492 | 0 | 4.2 | 485 | Specific | 941.38 | 892.12 | 892.12 | 18.43 | 146 | >sp\|P02788\|TRFL_HUMAN Lactotransferrin OS=Homo sapiens OX=9606 GN=LTF PE=1 SV=6 | 1270 | 20201118_lss_milk_1ng_glyco_hcd203040_compound15.22876.22876.4 | controllerType=0 controllerNumber=1 scan=22876 | 45.4515 |
| 19702:5 | 1 | R.TAGWNIPMGLLFN[+2059.735]QTGSC[+57.021]KFDEYFSQSC[+57.021]APGSDPR.S | HexNAc(4)Hex(5)Fuc(1)NeuAc(1) | C[+57]*2, N[+2060] | 1200.5078 | 5 | 5998.51 | 5998.492 | 0 | 3 | 485 | Specific | 941.45 | 885.16 | 885.16 | 18.55 | 146 | >sp\|P02788\|TRFL_HUMAN Lactotransferrin OS=Homo sapiens OX=9606 GN=LTF PE=1 SV=6 | 1270 | 20201118_lss_milk_1ng_glyco_hcd203040_compound15.22898.22898.5 | controllerType=0 controllerNumber=1 scan=22898 | 45.5104 |
| 18635:4 | 1 | R.TAGWNIPMGLLFN[+1768.640]QTGSC[+57.021]KFDEYFSQSC[+57.021]APGSDPR.S | HexNAc(4)Hex(5)Fuc(1) | C[+57]*2, N[+1769] | 1427.6106 | 4 | 5707.4206 | 5707.3966 | 0 | 4.2 | 485 | Specific | 941.96 | 919.31 | 919.31 | 18.49 | 146 | >sp\|P02788\|TRFL_HUMAN Lactotransferrin OS=Homo sapiens OX=9606 GN=LTF PE=1 SV=6 | 1270 | 20201118_lss_milk_1ng_glyco_hcd203040_compound15.21785.21785.4 | controllerType=0 controllerNumber=1 scan=21785 | 43.155 |
| 19675:4 | 1 | R.TAGWNIPMGLLFN[+2351.851]QTGSC[+57.021]KFDEYFSQSC[+57.021]APGSDPR.S | HexNAc(4)Hex(5)Fuc(3)NeuAc(1) | C[+57]*2, N[+2352] | 1573.4111 | 4 | 6290.6227 | 6290.6079 | 0 | 2.4 | 485 | Specific | 943.29 | 914.38 | 914.38 | 18.32 | 146 | >sp\|P02788\|TRFL_HUMAN Lactotransferrin OS=Homo sapiens OX=9606 GN=LTF PE=1 SV=6 | 1270 | 20201118_lss_milk_1ng_glyco_hcd203040_compound15.22869.22869.4 | controllerType=0 controllerNumber=1 scan=22869 | 45.4374 |
| 19683:4 | 1 | R.TAGWNIPMGLLFN[+2059.735]QTGSC[+57.021]KFDEYFSQSC[+57.021]APGSDPR.S | HexNAc(4)Hex(5)Fuc(1)NeuAc(1) | C[+57]*2, N[+2060] | 1500.3848 | 4 | 5998.5172 | 5998.492 | 0 | 4.2 | 485 | Specific | 958.87 | 877.57 | 877.57 | 19.43 | 146 | >sp\|P02788\|TRFL_HUMAN Lactotransferrin OS=Homo sapiens OX=9606 GN=LTF PE=1 SV=6 | 1270 | 20201118_lss_milk_1ng_glyco_hcd203040_compound15.22878.22878.4 | controllerType=0 controllerNumber=1 scan=22878 | 45.4584 |
| 18633:4 | 1 | R.TAGWNIPMGLLFN[+1768.640]QTGSC[+57.021]KFDEYFSQSC[+57.021]APGSDPR.S | HexNAc(4)Hex(5)Fuc(1) | C[+57]*2, N[+1769] | 1427.6106 | 4 | 5707.4206 | 5707.3966 | 0 | 4.2 | 485 | Specific | 981.69 | 947.4 | 947.4 | 19.08 | 146 | >sp\|P02788\|TRFL_HUMAN Lactotransferrin OS=Homo sapiens OX=9606 GN=LTF PE=1 SV=6 | 1270 | 20201118_lss_milk_1ng_glyco_hcd203040_compound15.21783.21783.4 | controllerType=0 controllerNumber=1 scan=21783 | 43.1501 |
| 19672:4 | 1 | R.TAGWNIPMGLLFN[+2351.851]QTGSC[+57.021]KFDEYFSQSC[+57.021]APGSDPR.S | HexNAc(4)Hex(5)Fuc(3)NeuAc(1) | C[+57]*2, N[+2352] | 1573.4111 | 4 | 6290.6227 | 6290.6079 | 0 | 2.4 | 485 | Specific | 1057.93 | 1020.12 | 1020.12 | 21.32 | 146 | >sp\|P02788\|TRFL_HUMAN Lactotransferrin OS=Homo sapiens OX=9606 GN=LTF PE=1 SV=6 | 1270 | 20201118_lss_milk_1ng_glyco_hcd203040_compound15.22866.22866.4 | controllerType=0 controllerNumber=1 scan=22866 | 45.4296 |
| 18792:4 | 1 | R.TAGWNIPM[+15.995]GLLFN[+2059.735]QTGSC[+57.021]KFDEYFSQSC[+57.021]APGSDPR.S | HexNAc(4)Hex(5)Fuc(1)NeuAc(1) | C[+57]*2, M[+16], N[+2060] | 1504.3842 | 4 | 6014.5148 | 6014.487 | 0 | 4.6 | 485 | Specific | 422.4 | 431.08 | 431.08 | 9.38 | 146 | >sp\|P02788\|TRFL_HUMAN Lactotransferrin OS=Homo sapiens OX=9606 GN=LTF PE=1 SV=6 | 1270 | 20201118_lss_milk_1ng_glyco_hcd203040_compound15.21948.21948.4 | controllerType=0 controllerNumber=1 scan=21948 | 43.4775 |
| 18794:4 | 1 | R.TAGWNIPM[+15.995]GLLFN[+2059.735]QTGSC[+57.021]KFDEYFSQSC[+57.021]APGSDPR.S | HexNAc(4)Hex(5)Fuc(1)NeuAc(1) | C[+57]*2, M[+16], N[+2060] | 1504.3842 | 4 | 6014.5148 | 6014.487 | 0 | 4.6 | 485 | Specific | 704.72 | 589.32 | 589.32 | 14.08 | 146 | >sp\|P02788\|TRFL_HUMAN Lactotransferrin OS=Homo sapiens OX=9606 GN=LTF PE=1 SV=6 | 1270 | 20201118_lss_milk_1ng_glyco_hcd203040_compound15.21950.21950.4 | controllerType=0 controllerNumber=1 scan=21950 | 43.4813 |
| 10025:3 | 1 | R.N[+2205.793]GSDC[+57.021]PDKFC[+57.021]LFQSETK.N | HexNAc(4)Hex(5)Fuc(2)NeuAc(1) | C[+57]*2, N[+2206] | 1413.5668 | 3 | 4238.6859 | 4238.6667 | 0 | 4.5 | 642 | Specific | 338.43 | 281.13 | 281.13 | 6.28 | 146 | >sp\|P02788\|TRFL_HUMAN Lactotransferrin OS=Homo sapiens OX=9606 GN=LTF PE=1 SV=6 | 1270 | 20201118_lss_milk_1ng_glyco_hcd203040_compound15.12823.12823.3 | controllerType=0 controllerNumber=1 scan=12823 | 26.5976 |
| 10080:3 | 1 | R.N[+2059.735]GSDC[+57.021]PDKFC[+57.021]LFQSETK.N | HexNAc(4)Hex(5)Fuc(1)NeuAc(1) | C[+57]*2, N[+2060] | 1364.8793 | 3 | 4092.6234 | 4092.6088 | 0 | 3.6 | 642 | Specific | 353.03 | 353.03 | 353.03 | 6.99 | 146 | >sp\|P02788\|TRFL_HUMAN Lactotransferrin OS=Homo sapiens OX=9606 GN=LTF PE=1 SV=6 | 1270 | 20201118_lss_milk_1ng_glyco_hcd203040_compound15.12880.12880.3 | controllerType=0 controllerNumber=1 scan=12880 | 26.6993 |
| 10027:3 | 1 | R.N[+2205.793]GSDC[+57.021]PDKFC[+57.021]LFQSETK.N | HexNAc(4)Hex(5)Fuc(2)NeuAc(1) | C[+57]*2, N[+2206] | 1413.5668 | 3 | 4238.6859 | 4238.6667 | 0 | 4.5 | 642 | Specific | 357.74 | 274.57 | 274.57 | 6.77 | 146 | >sp\|P02788\|TRFL_HUMAN Lactotransferrin OS=Homo sapiens OX=9606 GN=LTF PE=1 SV=6 | 1270 | 20201118_lss_milk_1ng_glyco_hcd203040_compound15.12825.12825.3 | controllerType=0 controllerNumber=1 scan=12825 | 26.6011 |
| 09969:3 | 1 | R.N[+2205.793]GSDC[+57.021]PDKFC[+57.021]LFQSETK.N | HexNAc(4)Hex(5)Fuc(2)NeuAc(1) | C[+57]*2, N[+2206] | 1413.5632 | 3 | 4238.6749 | 4238.6667 | 0 | 1.9 | 642 | Specific | 601.37 | 456.17 | 456.17 | 11.58 | 146 | >sp\|P02788\|TRFL_HUMAN Lactotransferrin OS=Homo sapiens OX=9606 GN=LTF PE=1 SV=6 | 1270 | 20201118_lss_milk_1ng_glyco_hcd203040_compound15.12764.12764.3 | controllerType=0 controllerNumber=1 scan=12764 | 26.4857 |
| 10078:3 | 1 | R.N[+2059.735]GSDC[+57.021]PDKFC[+57.021]LFQSETK.N | HexNAc(4)Hex(5)Fuc(1)NeuAc(1) | C[+57]*2, N[+2060] | 1364.8793 | 3 | 4092.6234 | 4092.6088 | 0 | 3.6 | 642 | Specific | 619.81 | 470.38 | 470.38 | 11.68 | 146 | >sp\|P02788\|TRFL_HUMAN Lactotransferrin OS=Homo sapiens OX=9606 GN=LTF PE=1 SV=6 | 1270 | 20201118_lss_milk_1ng_glyco_hcd203040_compound15.12878.12878.3 | controllerType=0 controllerNumber=1 scan=12878 | 26.6947 |
| 09968:3 | 1 | R.N[+2205.793]GSDC[+57.021]PDKFC[+57.021]LFQSETK.N | HexNAc(4)Hex(5)Fuc(2)NeuAc(1) | C[+57]*2, N[+2206] | 1413.5632 | 3 | 4238.6749 | 4238.6667 | 0 | 1.9 | 642 | Specific | 691.65 | 528.87 | 528.87 | 12.58 | 146 | >sp\|P02788\|TRFL_HUMAN Lactotransferrin OS=Homo sapiens OX=9606 GN=LTF PE=1 SV=6 | 1270 | 20201118_lss_milk_1ng_glyco_hcd203040_compound15.12763.12763.3 | controllerType=0 controllerNumber=1 scan=12763 | 26.4815 |
| 05617:3 | 4 | R.FSDGLESN[+1864.634]SSTQFEVKK.Y | HexNAc(2)Hex(9) | N[+1865] | 1256.5223 | 3 | 3767.5523 | 3767.5421 | 0 | 2.7 | 219 | Specific | 747.75 | 586.28 | 586.28 | 13.58 | 90 | >sp\|P0C0L5\|CO4B_HUMAN Complement C4-B OS=Homo sapiens OX=9606 GN=C4B PE=1 SV=2 | 1424 | 20201118_lss_milk_1ng_glyco_hcd203040_compound15.8251.8251.3 | controllerType=0 controllerNumber=1 scan=8251 | 19.1095 |
| 05615:3 | 4 | R.FSDGLESN[+1864.634]SSTQFEVKK.Y | HexNAc(2)Hex(9) | N[+1865] | 1256.5223 | 3 | 3767.5523 | 3767.5421 | 0 | 2.7 | 219 | Specific | 919.09 | 666.43 | 666.43 | 17.58 | 90 | >sp\|P0C0L5\|CO4B_HUMAN Complement C4-B OS=Homo sapiens OX=9606 GN=C4B PE=1 SV=2 | 1424 | 20201118_lss_milk_1ng_glyco_hcd203040_compound15.8249.8249.3 | controllerType=0 controllerNumber=1 scan=8249 | 19.104 |
| 18140:3 | 7 | R.GLSFDVSLEVSQGPGLLN[+2716.983]DTK.V | HexNAc(5)Hex(6)Fuc(3)NeuAc(1) | N[+2717] | 1631.7076 | 3 | 4893.1082 | 4893.0962 | 0 | 2.5 | 118 | Specific | 666.71 | 607.49 | 607.49 | 13.14 | 62 | >sp\|P01833\|PIGR_HUMAN Polymeric immunoglobulin receptor OS=Homo sapiens OX=9606 GN=PIGR PE=1 SV=4 | 902 | 20201118_lss_milk_1ng_glyco_hcd203040_compound15.21267.21267.3 | controllerType=0 controllerNumber=1 scan=21267 | 41.9742 |
| 18143:3 | 7 | R.GLSFDVSLEVSQGPGLLN[+2716.983]DTK.V | HexNAc(5)Hex(6)Fuc(3)NeuAc(1) | N[+2717] | 1631.7076 | 3 | 4893.1082 | 4893.0962 | 0 | 2.5 | 118 | Specific | 764.64 | 719.37 | 719.37 | 15.14 | 62 | >sp\|P01833\|PIGR_HUMAN Polymeric immunoglobulin receptor OS=Homo sapiens OX=9606 GN=PIGR PE=1 SV=4 | 902 | 20201118_lss_milk_1ng_glyco_hcd203040_compound15.21270.21270.3 | controllerType=0 controllerNumber=1 scan=21270 | 41.9787 |
| 20260:4 | 7 | R.GLSFDVSLEVSQGPGLLN[+2350.830]DTK.V | HexNAc(4)Hex(5)Fuc(1)NeuAc(2) | N[+2351] | 1132.4971 | 4 | 4526.9666 | 4526.9436 | 0 | 5.1 | 118 | Specific | 858.82 | 695.06 | 695.06 | 17.13 | 62 | >sp\|P01833\|PIGR_HUMAN Polymeric immunoglobulin receptor OS=Homo sapiens OX=9606 GN=PIGR PE=1 SV=4 | 902 | 20201118_lss_milk_1ng_glyco_hcd203040_compound15.23480.23480.4 | controllerType=0 controllerNumber=1 scan=23480 | 46.7325 |
| 20262:4 | 7 | R.GLSFDVSLEVSQGPGLLN[+2350.830]DTK.V | HexNAc(4)Hex(5)Fuc(1)NeuAc(2) | N[+2351] | 1132.4971 | 4 | 4526.9666 | 4526.9436 | 0 | 5.1 | 118 | Specific | 945.92 | 625.02 | 625.02 | 18.13 | 62 | >sp\|P01833\|PIGR_HUMAN Polymeric immunoglobulin receptor OS=Homo sapiens OX=9606 GN=PIGR PE=1 SV=4 | 902 | 20201118_lss_milk_1ng_glyco_hcd203040_compound15.23482.23482.4 | controllerType=0 controllerNumber=1 scan=23482 | 46.739 |
| 19635:3 | 7 | K.QIGLYPVLVIDSSGYVNPN[+2059.735]YTGR.I | HexNAc(4)Hex(5)Fuc(1)NeuAc(1) | N[+2060] | 1529.0239 | 3 | 4585.057 | 4585.0384 | 0 | 4.1 | 168 | Specific | 594.48 | 487.94 | 487.94 | 11.48 | 62 | >sp\|P01833\|PIGR_HUMAN Polymeric immunoglobulin receptor OS=Homo sapiens OX=9606 GN=PIGR PE=1 SV=4 | 902 | 20201118_lss_milk_1ng_glyco_hcd203040_compound15.22828.22828.3 | controllerType=0 controllerNumber=1 scan=22828 | 45.3423 |
| 19602:4 | 7 | K.QIGLYPVLVIDSSGYVNPN[+2059.735]YTGR.I | HexNAc(4)Hex(5)Fuc(1)NeuAc(1) | N[+2060] | 1147.0192 | 4 | 4585.055 | 4585.0384 | 0 | 3.6 | 168 | Specific | 707.36 | 667.02 | 667.02 | 14.71 | 62 | >sp\|P01833\|PIGR_HUMAN Polymeric immunoglobulin receptor OS=Homo sapiens OX=9606 GN=PIGR PE=1 SV=4 | 902 | 20201118_lss_milk_1ng_glyco_hcd203040_compound15.22793.22793.4 | controllerType=0 controllerNumber=1 scan=22793 | 45.2642 |
| 19604:4 | 7 | K.QIGLYPVLVIDSSGYVNPN[+2059.735]YTGR.I | HexNAc(4)Hex(5)Fuc(1)NeuAc(1) | N[+2060] | 1147.0192 | 4 | 4585.055 | 4585.0384 | 0 | 3.6 | 168 | Specific | 871.85 | 626.3 | 626.3 | 17.71 | 62 | >sp\|P01833\|PIGR_HUMAN Polymeric immunoglobulin receptor OS=Homo sapiens OX=9606 GN=PIGR PE=1 SV=4 | 902 | 20201118_lss_milk_1ng_glyco_hcd203040_compound15.22795.22795.4 | controllerType=0 controllerNumber=1 scan=22795 | 45.2706 |
| 19638:3 | 7 | K.QIGLYPVLVIDSSGYVNPN[+2059.735]YTGR.I | HexNAc(4)Hex(5)Fuc(1)NeuAc(1) | N[+2060] | 1529.0239 | 3 | 4585.057 | 4585.0384 | 0 | 4.1 | 168 | Specific | 884.89 | 651.56 | 651.56 | 17.48 | 62 | >sp\|P01833\|PIGR_HUMAN Polymeric immunoglobulin receptor OS=Homo sapiens OX=9606 GN=PIGR PE=1 SV=4 | 902 | 20201118_lss_milk_1ng_glyco_hcd203040_compound15.22831.22831.3 | controllerType=0 controllerNumber=1 scan=22831 | 45.3513 |
| 29328:3 | 7 | R.[+42.011]LSLLEEPGN[+2351.851]GTFTVILNQLTSR.D | HexNAc(4)Hex(5)Fuc(3)NeuAc(1) | .[42], N[+2352] | 1599.3868 | 3 | 4796.1458 | 4796.1539 | 0 | -1.7 | 413 | Specific | 240.99 | 240.99 | 240.99 | 4.91 | 62 | >sp\|P01833\|PIGR_HUMAN Polymeric immunoglobulin receptor OS=Homo sapiens OX=9606 GN=PIGR PE=1 SV=4 | 902 | 20201118_lss_milk_1ng_glyco_hcd203040_compound15.32950.32950.3 | controllerType=0 controllerNumber=1 scan=32950 | 65.3664 |
| 23337:3 | 7 | R.[+42.011]LSLLEEPGN[+2336.851]GTFTVILNQLTSR.D | HexNAc(6)Hex(6)Fuc(1) | .[42], N[+2337] | 1594.3812 | 3 | 4781.1289 | 4781.1542 | 0 | -5.3 | 413 | Specific | 332.4 | 239.82 | 239.82 | 3.79 | 62 | >sp\|P01833\|PIGR_HUMAN Polymeric immunoglobulin receptor OS=Homo sapiens OX=9606 GN=PIGR PE=1 SV=4 | 902 | 20201118_lss_milk_1ng_glyco_hcd203040_compound15.26693.26693.3 | controllerType=0 controllerNumber=1 scan=26693 | 53.5902 |
| 23284:3 | 7 | R.[+42.011]LSLLEEPGN[+2336.851]GTFTVILNQLTSR.D | HexNAc(6)Hex(6)Fuc(1) | .[42], N[+2337] | 1594.3836 | 3 | 4781.1363 | 4781.1542 | 0 | -3.8 | 413 | Specific | 375.66 | 349.83 | 349.83 | 6 | 62 | >sp\|P01833\|PIGR_HUMAN Polymeric immunoglobulin receptor OS=Homo sapiens OX=9606 GN=PIGR PE=1 SV=4 | 902 | 20201118_lss_milk_1ng_glyco_hcd203040_compound15.26638.26638.3 | controllerType=0 controllerNumber=1 scan=26638 | 53.4826 |
| 23286:3 | 7 | R.[+42.011]LSLLEEPGN[+2336.851]GTFTVILNQLTSR.D | HexNAc(6)Hex(6)Fuc(1) | .[42], N[+2337] | 1594.3836 | 3 | 4781.1363 | 4781.1542 | 0 | -3.8 | 413 | Specific | 382.35 | 307.65 | 307.65 | 5.71 | 62 | >sp\|P01833\|PIGR_HUMAN Polymeric immunoglobulin receptor OS=Homo sapiens OX=9606 GN=PIGR PE=1 SV=4 | 902 | 20201118_lss_milk_1ng_glyco_hcd203040_compound15.26640.26640.3 | controllerType=0 controllerNumber=1 scan=26640 | 53.4884 |
| 23033:4 | 7 | R.LSLLEEPGN[+2861.000]GTFTVILNQLTSR.D | HexNAc(5)Hex(6)NeuAc(3) | N[+2861] | 1316.5883 | 4 | 5263.3263 | 5263.2926 | 0 | 6.4 | 413 | Specific | 394.74 | 394.74 | 394.74 | 8.07 | 62 | >sp\|P01833\|PIGR_HUMAN Polymeric immunoglobulin receptor OS=Homo sapiens OX=9606 GN=PIGR PE=1 SV=4 | 902 | 20201118_lss_milk_1ng_glyco_hcd203040_compound15.26376.26376.4 | controllerType=0 controllerNumber=1 scan=26376 | 52.9357 |
| 21257:3 | 7 | R.LSLLEEPGN[+2205.793]GTFTVILNQLTSR.D | HexNAc(4)Hex(5)Fuc(2)NeuAc(1) | N[+2206] | 1536.7083 | 3 | 4608.1103 | 4608.0854 | 0 | 5.4 | 413 | Specific | 401.18 | 373 | 373 | 7.1 | 62 | >sp\|P01833\|PIGR_HUMAN Polymeric immunoglobulin receptor OS=Homo sapiens OX=9606 GN=PIGR PE=1 SV=4 | 902 | 20201118_lss_milk_1ng_glyco_hcd203040_compound15.24521.24521.3 | controllerType=0 controllerNumber=1 scan=24521 | 49.012 |
| 23031:4 | 7 | R.LSLLEEPGN[+2861.000]GTFTVILNQLTSR.D | HexNAc(5)Hex(6)NeuAc(3) | N[+2861] | 1316.5883 | 4 | 5263.3263 | 5263.2926 | 0 | 6.4 | 413 | Specific | 485.55 | 453.53 | 453.53 | 8.5 | 62 | >sp\|P01833\|PIGR_HUMAN Polymeric immunoglobulin receptor OS=Homo sapiens OX=9606 GN=PIGR PE=1 SV=4 | 902 | 20201118_lss_milk_1ng_glyco_hcd203040_compound15.26374.26374.4 | controllerType=0 controllerNumber=1 scan=26374 | 52.9316 |
| 21259:3 | 7 | R.LSLLEEPGN[+2205.793]GTFTVILNQLTSR.D | HexNAc(4)Hex(5)Fuc(2)NeuAc(1) | N[+2206] | 1536.7083 | 3 | 4608.1103 | 4608.0854 | 0 | 5.4 | 413 | Specific | 526.2 | 393.08 | 393.08 | 8.81 | 62 | >sp\|P01833\|PIGR_HUMAN Polymeric immunoglobulin receptor OS=Homo sapiens OX=9606 GN=PIGR PE=1 SV=4 | 902 | 20201118_lss_milk_1ng_glyco_hcd203040_compound15.24523.24523.3 | controllerType=0 controllerNumber=1 scan=24523 | 49.0155 |
| 21215:4 | 7 | R.LSLLEEPGN[+2076.750]GTFTVILNQLTSR.D | HexNAc(4)Hex(6)Fuc(2) | N[+2077] | 1120.5225 | 4 | 4479.0682 | 4479.0428 | 0 | 5.7 | 413 | Specific | 544.84 | 428.52 | 428.52 | 9.26 | 62 | >sp\|P01833\|PIGR_HUMAN Polymeric immunoglobulin receptor OS=Homo sapiens OX=9606 GN=PIGR PE=1 SV=4 | 902 | 20201118_lss_milk_1ng_glyco_hcd203040_compound15.24477.24477.4 | controllerType=0 controllerNumber=1 scan=24477 | 48.9123 |
| 21213:4 | 7 | R.LSLLEEPGN[+2076.750]GTFTVILNQLTSR.D | HexNAc(4)Hex(6)Fuc(2) | N[+2077] | 1120.5225 | 4 | 4479.0682 | 4479.0428 | 0 | 5.7 | 413 | Specific | 582.87 | 570.3 | 570.3 | 10.73 | 62 | >sp\|P01833\|PIGR_HUMAN Polymeric immunoglobulin receptor OS=Homo sapiens OX=9606 GN=PIGR PE=1 SV=4 | 902 | 20201118_lss_milk_1ng_glyco_hcd203040_compound15.24475.24475.4 | controllerType=0 controllerNumber=1 scan=24475 | 48.9071 |
| 23233:3 | 7 | R.LSLLEEPGN[+2350.830]GTFTVILNQLTSR.D | HexNAc(4)Hex(5)Fuc(1)NeuAc(2) | N[+2351] | 1585.0519 | 3 | 4753.1413 | 4753.1229 | 0 | 3.9 | 413 | Specific | 752.46 | 574.81 | 574.81 | 15.25 | 62 | >sp\|P01833\|PIGR_HUMAN Polymeric immunoglobulin receptor OS=Homo sapiens OX=9606 GN=PIGR PE=1 SV=4 | 902 | 20201118_lss_milk_1ng_glyco_hcd203040_compound15.26585.26585.3 | controllerType=0 controllerNumber=1 scan=26585 | 53.3822 |
| 23236:3 | 7 | R.LSLLEEPGN[+2350.830]GTFTVILNQLTSR.D | HexNAc(4)Hex(5)Fuc(1)NeuAc(2) | N[+2351] | 1585.0519 | 3 | 4753.1413 | 4753.1229 | 0 | 3.9 | 413 | Specific | 939.53 | 681.93 | 681.93 | 18.25 | 62 | >sp\|P01833\|PIGR_HUMAN Polymeric immunoglobulin receptor OS=Homo sapiens OX=9606 GN=PIGR PE=1 SV=4 | 902 | 20201118_lss_milk_1ng_glyco_hcd203040_compound15.26588.26588.3 | controllerType=0 controllerNumber=1 scan=26588 | 53.3854 |
| 23214:4 | 7 | R.LSLLEEPGN[+2350.830]GTFTVILNQLTSR.D | HexNAc(4)Hex(5)Fuc(1)NeuAc(2) | N[+2351] | 1189.0418 | 4 | 4753.1453 | 4753.1229 | 0 | 4.7 | 413 | Specific | 987.41 | 745.57 | 745.57 | 19.13 | 62 | >sp\|P01833\|PIGR_HUMAN Polymeric immunoglobulin receptor OS=Homo sapiens OX=9606 GN=PIGR PE=1 SV=4 | 902 | 20201118_lss_milk_1ng_glyco_hcd203040_compound15.26565.26565.4 | controllerType=0 controllerNumber=1 scan=26565 | 53.3359 |
| 23212:4 | 7 | R.LSLLEEPGN[+2350.830]GTFTVILNQLTSR.D | HexNAc(4)Hex(5)Fuc(1)NeuAc(2) | N[+2351] | 1189.0418 | 4 | 4753.1453 | 4753.1229 | 0 | 4.7 | 413 | Specific | 1040.62 | 825.94 | 825.94 | 20.13 | 62 | >sp\|P01833\|PIGR_HUMAN Polymeric immunoglobulin receptor OS=Homo sapiens OX=9606 GN=PIGR PE=1 SV=4 | 902 | 20201118_lss_milk_1ng_glyco_hcd203040_compound15.26563.26563.4 | controllerType=0 controllerNumber=1 scan=26563 | 53.3311 |
| 16890:3 | 7 | K.IIEGEPNLKVPGN[+2205.793]YTVAVLGETLK.V | HexNAc(4)Hex(5)Fuc(2)NeuAc(1) | N[+2206] | 1533.0582 | 3 | 4597.16 | 4597.1422 | 0 | 3.9 | 457 | Specific | 393.01 | 393.01 | 393.01 | 8.91 | 62 | >sp\|P01833\|PIGR_HUMAN Polymeric immunoglobulin receptor OS=Homo sapiens OX=9606 GN=PIGR PE=1 SV=4 | 902 | 20201118_lss_milk_1ng_glyco_hcd203040_compound15.19963.19963.3 | controllerType=0 controllerNumber=1 scan=19963 | 39.4259 |
| 16887:3 | 7 | K.IIEGEPNLKVPGN[+2205.793]YTVAVLGETLK.V | HexNAc(4)Hex(5)Fuc(2)NeuAc(1) | N[+2206] | 1533.0582 | 3 | 4597.16 | 4597.1422 | 0 | 3.9 | 457 | Specific | 536.67 | 536.67 | 536.67 | 9.62 | 62 | >sp\|P01833\|PIGR_HUMAN Polymeric immunoglobulin receptor OS=Homo sapiens OX=9606 GN=PIGR PE=1 SV=4 | 902 | 20201118_lss_milk_1ng_glyco_hcd203040_compound15.19960.19960.3 | controllerType=0 controllerNumber=1 scan=19960 | 39.4214 |

Table S4. Cont.

| Query #:z | Protein Rank | Glycopeptide | Glycan | Modification Type(s) | Observed m/z | z | Observed (M+H) | Calc. mass (M+H) | Off-by-x error | Mass error (ppm) | StartPosition | Cleavage | Score | Delta | Delta Mod | \|Log Prob\| | # of unique peptides | Protein Name | Protein DB number | Comment | Scan # | Scan Time |
|---|---|---|---|---|---|---|---|---|---|---|---|---|---|---|---|---|---|---|---|---|---|---|
| 16832:4 | 7 | K.IIEGEPNLKVPGN[+2205.793]VTAVLGETLK.V | HexNAc(4)Hex(5)Fuc(2)NeuAc(1) | N[+2206] | 1150.047 | 4 | 4597.1663 | 4597.1422 | 0 | 5.2 | 457 | Specific | 667.84 | 654.61 | 654.61 | 12.74 | 62 | >sp\|P01833\|PIGR_HUMAN Polymeric immunoglobulin receptor OS=Homo sapiens OX=9606 GN=PIGR PE=1 SV=4 | 902 | 20201118_lss_milk_1ng_glyco_hcd203040_compound15.19903.19903.4 | controllerType=0 controllerNumber=1 scan=19903 | 39.2918 |
| 16835:4 | 7 | K.IIEGEPNLKVPGN[+2205.793]VTAVLGETLK.V | HexNAc(4)Hex(5)Fuc(2)NeuAc(1) | N[+2206] | 1150.047 | 4 | 4597.1663 | 4597.1422 | 0 | 5.2 | 457 | Specific | 699.01 | 629.02 | 629.02 | 12.74 | 62 | >sp\|P01833\|PIGR_HUMAN Polymeric immunoglobulin receptor OS=Homo sapiens OX=9606 GN=PIGR PE=1 SV=4 | 902 | 20201118_lss_milk_1ng_glyco_hcd203040_compound15.19906.19906.4 | controllerType=0 controllerNumber=1 scan=19906 | 39.2994 |
| 16775:3 | 7 | K.IIEGEPNLKVPGN[+2205.793]VTAVLGETLK.V | HexNAc(4)Hex(5)Fuc(2)NeuAc(1) | N[+2206] | 1533.0568 | 3 | 4597.1559 | 4597.1422 | 0 | 3 | 457 | Specific | 770.72 | 700.14 | 700.14 | 14.99 | 62 | >sp\|P01833\|PIGR_HUMAN Polymeric immunoglobulin receptor OS=Homo sapiens OX=9606 GN=PIGR PE=1 SV=4 | 902 | 20201118_lss_milk_1ng_glyco_hcd203040_compound15.19843.19843.3 | controllerType=0 controllerNumber=1 scan=19843 | 39.1691 |
| 18649:3 | 7 | K.IIEGEPNLKVPGN[+2350.830]VTAVLGETLK.V | HexNAc(4)Hex(5)Fuc(1)NeuAc(2) | N[+2351] | 1581.4046 | 3 | 4742.1992 | 4742.1797 | 0 | 4.1 | 457 | Specific | 773.15 | 713.07 | 713.07 | 15.09 | 62 | >sp\|P01833\|PIGR_HUMAN Polymeric immunoglobulin receptor OS=Homo sapiens OX=9606 GN=PIGR PE=1 SV=4 | 902 | 20201118_lss_milk_1ng_glyco_hcd203040_compound15.21799.21799.3 | controllerType=0 controllerNumber=1 scan=21799 | 43.1969 |
| 16830:3 | 7 | K.IIEGEPNLKVPGN[+2059.735]VTAVLGETLK.V | HexNAc(4)Hex(5)Fuc(1)NeuAc(1) | N[+2060] | 1484.3724 | 3 | 4451.1026 | 4451.0843 | 0 | 4.1 | 457 | Specific | 814.62 | 626.34 | 626.34 | 16.09 | 62 | >sp\|P01833\|PIGR_HUMAN Polymeric immunoglobulin receptor OS=Homo sapiens OX=9606 GN=PIGR PE=1 SV=4 | 902 | 20201118_lss_milk_1ng_glyco_hcd203040_compound15.19901.19901.3 | controllerType=0 controllerNumber=1 scan=19901 | 39.2887 |
| 16772:3 | 7 | K.IIEGEPNLKVPGN[+2205.793]VTAVLGETLK.V | HexNAc(4)Hex(5)Fuc(2)NeuAc(1) | N[+2206] | 1533.0568 | 3 | 4597.1559 | 4597.1422 | 0 | 3 | 457 | Specific | 950.6 | 950.6 | 950.6 | 18.99 | 62 | >sp\|P01833\|PIGR_HUMAN Polymeric immunoglobulin receptor OS=Homo sapiens OX=9606 GN=PIGR PE=1 SV=4 | 902 | 20201118_lss_milk_1ng_glyco_hcd203040_compound15.19840.19840.3 | controllerType=0 controllerNumber=1 scan=19840 | 39.1639 |
| 18647:3 | 7 | K.IIEGEPNLKVPGN[+2350.830]VTAVLGETLK.V | HexNAc(4)Hex(5)Fuc(1)NeuAc(2) | N[+2351] | 1581.4046 | 3 | 4742.1992 | 4742.1797 | 0 | 4.1 | 457 | Specific | 961.06 | 866.62 | 866.62 | 19.09 | 62 | >sp\|P01833\|PIGR_HUMAN Polymeric immunoglobulin receptor OS=Homo sapiens OX=9606 GN=PIGR PE=1 SV=4 | 902 | 20201118_lss_milk_1ng_glyco_hcd203040_compound15.21797.21797.3 | controllerType=0 controllerNumber=1 scan=21797 | 43.1797 |
| 16828:3 | 7 | K.IIEGEPNLKVPGN[+2059.735]VTAVLGETLK.V | HexNAc(4)Hex(5)Fuc(1)NeuAc(1) | N[+2060] | 1484.3724 | 3 | 4451.1026 | 4451.0843 | 0 | 4.1 | 457 | Specific | 1064.6 | 967.14 | 967.14 | 21.09 | 62 | >sp\|P01833\|PIGR_HUMAN Polymeric immunoglobulin receptor OS=Homo sapiens OX=9606 GN=PIGR PE=1 SV=4 | 902 | 20201118_lss_milk_1ng_glyco_hcd203040_compound15.19899.19899.3 | controllerType=0 controllerNumber=1 scan=19899 | 39.2833 |
| 16384:5 | 7 | K.IIEGEPNLKVPGN[+2059.735]VTAVLGETLKVPC[+57.021]HFPC[+57.021]K.F | HexNAc(4)Hex(5)Fuc(1)NeuAc(1) | C[+57]*2, N[+2060] | 1096.1163 | 5 | 5476.5526 | 5476.5418 | 0 | 2 | 457 | Specific | 280.25 | 259.49 | 259.49 | 4.93 | 62 | >sp\|P01833\|PIGR_HUMAN Polymeric immunoglobulin receptor OS=Homo sapiens OX=9606 GN=PIGR PE=1 SV=4 | 902 | 20201118_lss_milk_1ng_glyco_hcd203040_compound15.19421.19421.3 | controllerType=0 controllerNumber=1 scan=19421 | 38.4365 |
| 16387:5 | 7 | K.IIEGEPNLKVPGN[+2059.735]VTAVLGETLKVPC[+57.021]HFPC[+57.021]K.F | HexNAc(4)Hex(5)Fuc(1)NeuAc(1) | C[+57]*2, N[+2060] | 1096.1163 | 5 | 5476.5526 | 5476.5418 | 0 | 2 | 457 | Specific | 489.8 | 435.01 | 435.01 | 9.1 | 62 | >sp\|P01833\|PIGR_HUMAN Polymeric immunoglobulin receptor OS=Homo sapiens OX=9606 GN=PIGR PE=1 SV=4 | 902 | 20201118_lss_milk_1ng_glyco_hcd203040_compound15.19424.19424.3 | controllerType=0 controllerNumber=1 scan=19424 | 38.4422 |
| 16256:5 | 7 | K.IIEGEPNLKVPGN[+2059.735]VTAVLGETLKVPC[+57.021]HFPC[+57.021]K.F | HexNAc(4)Hex(5)Fuc(1)NeuAc(1) | C[+57]*2, N[+2060] | 1096.1158 | 5 | 5476.5501 | 5476.5418 | 0 | 1.5 | 457 | Specific | 554.53 | 513.55 | 513.55 | 9.31 | 62 | >sp\|P01833\|PIGR_HUMAN Polymeric immunoglobulin receptor OS=Homo sapiens OX=9606 GN=PIGR PE=1 SV=4 | 902 | 20201118_lss_milk_1ng_glyco_hcd203040_compound15.19289.19289.5 | controllerType=0 controllerNumber=1 scan=19289 | 38.2131 |
| 16320:4 | 7 | K.IIEGEPNLKVPGN[+2204.772]VTAVLGETLKVPC[+57.021]HFPC[+57.021]K.F | HexNAc(4)Hex(5)NeuAc(2) | C[+57]*2, N[+2205] | 1406.1615 | 4 | 5621.6242 | 5621.5793 | 0 | 8 | 457 | Specific | 635.28 | 615.24 | 615.24 | 9.96 | 62 | >sp\|P01833\|PIGR_HUMAN Polymeric immunoglobulin receptor OS=Homo sapiens OX=9606 GN=PIGR PE=1 SV=4 | 902 | 20201118_lss_milk_1ng_glyco_hcd203040_compound15.19355.19355.2 | controllerType=0 controllerNumber=1 scan=19355 | 38.3272 |
| 17766:4 | 7 | K.IIEGEPNLKVPGN[+2350.830]VTAVLGETLKVPC[+57.021]HFPC[+57.021]K.F | HexNAc(4)Hex(5)Fuc(1)NeuAc(2) | C[+57]*2, N[+2351] | 1442.6708 | 4 | 5767.6613 | 5767.6372 | 0 | 4.2 | 457 | Specific | 700.97 | 464.8 | 464.8 | 13.45 | 62 | >sp\|P01833\|PIGR_HUMAN Polymeric immunoglobulin receptor OS=Homo sapiens OX=9606 GN=PIGR PE=1 SV=4 | 902 | 20201118_lss_milk_1ng_glyco_hcd203040_compound15.20878.20878.4 | controllerType=0 controllerNumber=1 scan=20878 | 41.1495 |
| 17764:5 | 7 | K.IIEGEPNLKVPGN[+2350.830]VTAVLGETLKVPC[+57.021]HFPC[+57.021]K.F | HexNAc(4)Hex(5)Fuc(1)NeuAc(2) | C[+57]*2, N[+2351] | 1154.3366 | 5 | 5767.6536 | 5767.6372 | 0 | 2.8 | 457 | Specific | 729.09 | 634.24 | 634.24 | 13.86 | 62 | >sp\|P01833\|PIGR_HUMAN Polymeric immunoglobulin receptor OS=Homo sapiens OX=9606 GN=PIGR PE=1 SV=4 | 902 | 20201118_lss_milk_1ng_glyco_hcd203040_compound15.20876.20876.5 | controllerType=0 controllerNumber=1 scan=20876 | 41.1472 |
| 16323:4 | 7 | K.IIEGEPNLKVPGN[+2204.772]VTAVLGETLKVPC[+57.021]HFPC[+57.021]K.F | HexNAc(4)Hex(5)NeuAc(2) | C[+57]*2, N[+2205] | 1406.1615 | 4 | 5621.6242 | 5621.5793 | 0 | 8 | 457 | Specific | 736.58 | 736.58 | 736.58 | 11.96 | 62 | >sp\|P01833\|PIGR_HUMAN Polymeric immunoglobulin receptor OS=Homo sapiens OX=9606 GN=PIGR PE=1 SV=4 | 902 | 20201118_lss_milk_1ng_glyco_hcd203040_compound15.19358.19358.2 | controllerType=0 controllerNumber=1 scan=19358 | 38.3328 |
| 15167:5 | 7 | K.IIEGEPNLKVPGN[+1768.640]VTAVLGETLKVPC[+57.021]HFPC[+57.021]K.F | HexNAc(4)Hex(5)Fuc(1) | C[+57]*2, N[+1769] | 1037.8972 | 5 | 5185.457 | 5185.4464 | 0 | 2 | 457 | Specific | 743.82 | 642.84 | 642.84 | 13.86 | 62 | >sp\|P01833\|PIGR_HUMAN Polymeric immunoglobulin receptor OS=Homo sapiens OX=9606 GN=PIGR PE=1 SV=4 | 902 | 20201118_lss_milk_1ng_glyco_hcd203040_compound15.18159.18159.5 | controllerType=0 controllerNumber=1 scan=18159 | 36.129 |
| 17768:4 | 7 | K.IIEGEPNLKVPGN[+2350.830]VTAVLGETLKVPC[+57.021]HFPC[+57.021]K.F | HexNAc(4)Hex(5)Fuc(1)NeuAc(2) | C[+57]*2, N[+2351] | 1442.6708 | 4 | 5767.6613 | 5767.6372 | 0 | 4.2 | 457 | Specific | 755.97 | 607.19 | 607.19 | 14.45 | 62 | >sp\|P01833\|PIGR_HUMAN Polymeric immunoglobulin receptor OS=Homo sapiens OX=9606 GN=PIGR PE=1 SV=4 | 902 | 20201118_lss_milk_1ng_glyco_hcd203040_compound15.20880.20880.4 | controllerType=0 controllerNumber=1 scan=20880 | 41.1548 |
| 16259:5 | 7 | K.IIEGEPNLKVPGN[+2059.735]VTAVLGETLKVPC[+57.021]HFPC[+57.021]K.F | HexNAc(4)Hex(5)Fuc(1)NeuAc(1) | C[+57]*2, N[+2060] | 1096.1158 | 5 | 5476.5501 | 5476.5418 | 0 | 1.5 | 457 | Specific | 776.41 | 651.74 | 651.74 | 14.59 | 62 | >sp\|P01833\|PIGR_HUMAN Polymeric immunoglobulin receptor OS=Homo sapiens OX=9606 GN=PIGR PE=1 SV=4 | 902 | 20201118_lss_milk_1ng_glyco_hcd203040_compound15.19292.19292.5 | controllerType=0 controllerNumber=1 scan=19292 | 38.2174 |
| 16232:4 | 7 | K.IIEGEPNLKVPGN[+2205.793]VTAVLGETLKVPC[+57.021]HFPC[+57.021]K.F | HexNAc(4)Hex(5)Fuc(2)NeuAc(1) | C[+57]*2, N[+2206] | 1406.4108 | 4 | 5622.6212 | 5622.5997 | 0 | 3.8 | 457 | Specific | 798.68 | 798.68 | 798.68 | 14.45 | 62 | >sp\|P01833\|PIGR_HUMAN Polymeric immunoglobulin receptor OS=Homo sapiens OX=9606 GN=PIGR PE=1 SV=4 | 902 | 20201118_lss_milk_1ng_glyco_hcd203040_compound15.19264.19264.4 | controllerType=0 controllerNumber=1 scan=19264 | 38.1671 |
| 15169:5 | 7 | K.IIEGEPNLKVPGN[+1768.640]VTAVLGETLKVPC[+57.021]HFPC[+57.021]K.F | HexNAc(4)Hex(5)Fuc(1) | C[+57]*2, N[+1769] | 1037.8972 | 5 | 5185.457 | 5185.4464 | 0 | 2 | 457 | Specific | 804.53 | 694.73 | 694.73 | 15.86 | 62 | >sp\|P01833\|PIGR_HUMAN Polymeric immunoglobulin receptor OS=Homo sapiens OX=9606 GN=PIGR PE=1 SV=4 | 902 | 20201118_lss_milk_1ng_glyco_hcd203040_compound15.18161.18161.5 | controllerType=0 controllerNumber=1 scan=18161 | 36.1346 |
| 16240:4 | 7 | K.IIEGEPNLKVPGN[+2059.735]VTAVLGETLKVPC[+57.021]HFPC[+57.021]K.F | HexNAc(4)Hex(5)Fuc(1)NeuAc(1) | C[+57]*2, N[+2060] | 1369.8911 | 4 | 5476.5426 | 5476.5418 | 0 | 0.2 | 457 | Specific | 809.17 | 722.87 | 722.87 | 14.43 | 62 | >sp\|P01833\|PIGR_HUMAN Polymeric immunoglobulin receptor OS=Homo sapiens OX=9606 GN=PIGR PE=1 SV=4 | 902 | 20201118_lss_milk_1ng_glyco_hcd203040_compound15.19272.19272.2 | controllerType=0 controllerNumber=1 scan=19272 | 38.1851 |
| 16208:4 | 7 | K.IIEGEPNLKVPGN[+2205.793]VTAVLGETLKVPC[+57.021]HFPC[+57.021]K.F | HexNAc(4)Hex(5)Fuc(2)NeuAc(1) | C[+57]*2, N[+2206] | 1406.4104 | 4 | 5622.6198 | 5622.5997 | 0 | 3.6 | 457 | Specific | 809.55 | 809.55 | 809.55 | 15.45 | 62 | >sp\|P01833\|PIGR_HUMAN Polymeric immunoglobulin receptor OS=Homo sapiens OX=9606 GN=PIGR PE=1 SV=4 | 902 | 20201118_lss_milk_1ng_glyco_hcd203040_compound15.19239.19239.4 | controllerType=0 controllerNumber=1 scan=19239 | 38.118 |
| 16217:4 | 7 | K.IIEGEPNLKVPGN[+2059.735]VTAVLGETLKVPC[+57.021]HFPC[+57.021]K.F | HexNAc(4)Hex(5)Fuc(1)NeuAc(1) | C[+57]*2, N[+2060] | 1369.8933 | 4 | 5476.5514 | 5476.5418 | 0 | 1.8 | 457 | Specific | 832.65 | 731.23 | 731.23 | 15.07 | 62 | >sp\|P01833\|PIGR_HUMAN Polymeric immunoglobulin receptor OS=Homo sapiens OX=9606 GN=PIGR PE=1 SV=4 | 902 | 20201118_lss_milk_1ng_glyco_hcd203040_compound15.19248.19248.4 | controllerType=0 controllerNumber=1 scan=19248 | 38.1336 |
| 16193:5 | 7 | K.IIEGEPNLKVPGN[+2205.793]VTAVLGETLKVPC[+57.021]HFPC[+57.021]K.F | HexNAc(4)Hex(5)Fuc(2)NeuAc(1) | C[+57]*2, N[+2206] | 1125.3291 | 5 | 5622.6164 | 5622.5997 | 0 | 3 | 457 | Specific | 847.56 | 847.56 | 847.56 | 15.86 | 62 | >sp\|P01833\|PIGR_HUMAN Polymeric immunoglobulin receptor OS=Homo sapiens OX=9606 GN=PIGR PE=1 SV=4 | 902 | 20201118_lss_milk_1ng_glyco_hcd203040_compound15.19223.19223.5 | controllerType=0 controllerNumber=1 scan=19223 | 38.0863 |
| 17767:5 | 7 | K.IIEGEPNLKVPGN[+2350.830]VTAVLGETLKVPC[+57.021]HFPC[+57.021]K.F | HexNAc(4)Hex(5)Fuc(1)NeuAc(2) | C[+57]*2, N[+2351] | 1154.3366 | 5 | 5767.6536 | 5767.6372 | 0 | 2.8 | 457 | Specific | 877.97 | 681.16 | 681.16 | 16.86 | 62 | >sp\|P01833\|PIGR_HUMAN Polymeric immunoglobulin receptor OS=Homo sapiens OX=9606 GN=PIGR PE=1 SV=4 | 902 | 20201118_lss_milk_1ng_glyco_hcd203040_compound15.20879.20879.5 | controllerType=0 controllerNumber=1 scan=20879 | 41.152 |
| 16238:4 | 7 | K.IIEGEPNLKVPGN[+2059.735]VTAVLGETLKVPC[+57.021]HFPC[+57.021]K.F | HexNAc(4)Hex(5)Fuc(1)NeuAc(1) | C[+57]*2, N[+2060] | 1369.8911 | 4 | 5476.5426 | 5476.5418 | 0 | 0.2 | 457 | Specific | 914.76 | 884.87 | 884.87 | 16.43 | 62 | >sp\|P01833\|PIGR_HUMAN Polymeric immunoglobulin receptor OS=Homo sapiens OX=9606 GN=PIGR PE=1 SV=4 | 902 | 20201118_lss_milk_1ng_glyco_hcd203040_compound15.19270.19270.2 | controllerType=0 controllerNumber=1 scan=19270 | 38.1802 |
| 16214:4 | 7 | K.IIEGEPNLKVPGN[+2059.735]VTAVLGETLKVPC[+57.021]HFPC[+57.021]K.F | HexNAc(4)Hex(5)Fuc(1)NeuAc(1) | C[+57]*2, N[+2060] | 1369.8933 | 4 | 5476.5514 | 5476.5418 | 0 | 1.8 | 457 | Specific | 999.47 | 869.52 | 869.52 | 18.07 | 62 | >sp\|P01833\|PIGR_HUMAN Polymeric immunoglobulin receptor OS=Homo sapiens OX=9606 GN=PIGR PE=1 SV=4 | 902 | 20201118_lss_milk_1ng_glyco_hcd203040_compound15.19245.19245.4 | controllerType=0 controllerNumber=1 scan=19245 | 38.1285 |
| 16205:4 | 7 | K.IIEGEPNLKVPGN[+2205.793]VTAVLGETLKVPC[+57.021]HFPC[+57.021]K.F | HexNAc(4)Hex(5)Fuc(2)NeuAc(1) | C[+57]*2, N[+2206] | 1406.4104 | 4 | 5622.6198 | 5622.5997 | 0 | 3.6 | 457 | Specific | 1033.36 | 966.28 | 966.28 | 19.45 | 62 | >sp\|P01833\|PIGR_HUMAN Polymeric immunoglobulin receptor OS=Homo sapiens OX=9606 GN=PIGR PE=1 SV=4 | 902 | 20201118_lss_milk_1ng_glyco_hcd203040_compound15.19236.19236.4 | controllerType=0 controllerNumber=1 scan=19236 | 38.1111 |
| 16230:4 | 7 | K.IIEGEPNLKVPGN[+2205.793]VTAVLGETLKVPC[+57.021]HFPC[+57.021]K.F | HexNAc(4)Hex(5)Fuc(2)NeuAc(1) | C[+57]*2, N[+2206] | 1406.4108 | 4 | 5622.6212 | 5622.5997 | 0 | 3.8 | 457 | Specific | 1050.16 | 901.95 | 901.95 | 20.45 | 62 | >sp\|P01833\|PIGR_HUMAN Polymeric immunoglobulin receptor OS=Homo sapiens OX=9606 GN=PIGR PE=1 SV=4 | 902 | 20201118_lss_milk_1ng_glyco_hcd203040_compound15.19262.19262.4 | controllerType=0 controllerNumber=1 scan=19262 | 38.1613 |
| 16191:5 | 7 | K.IIEGEPNLKVPGN[+2205.793]VTAVLGETLKVPC[+57.021]HFPC[+57.021]K.F | HexNAc(4)Hex(5)Fuc(2)NeuAc(1) | C[+57]*2, N[+2206] | 1125.3291 | 5 | 5622.6164 | 5622.5997 | 0 | 3 | 457 | Specific | 1146.31 | 1070.37 | 1070.37 | 21.86 | 62 | >sp\|P01833\|PIGR_HUMAN Polymeric immunoglobulin receptor OS=Homo sapiens OX=9606 GN=PIGR PE=1 SV=4 | 902 | 20201118_lss_milk_1ng_glyco_hcd203040_compound15.19221.19221.5 | controllerType=0 controllerNumber=1 scan=19221 | 38.0776 |
| 16244:4 | 7 | K.VPGN[+2350.830]VTAVLGETLKVPC[+57.021]HFPC[+57.021]K.F | HexNAc(4)Hex(5)Fuc(1)NeuAc(2) | C[+57]*2, N[+2351] | 1194.2814 | 4 | 4774.1038 | 4774.0877 | 0 | 3.4 | 466 | Specific | 355.65 | 345.13 | 345.13 | 7.17 | 62 | >sp\|P01833\|PIGR_HUMAN Polymeric immunoglobulin receptor OS=Homo sapiens OX=9606 GN=PIGR PE=1 SV=4 | 902 | 20201118_lss_milk_1ng_glyco_hcd203040_compound15.19276.19276.2 | controllerType=0 controllerNumber=1 scan=19276 | 38.1907 |
| 16247:4 | 7 | K.VPGN[+2350.830]VTAVLGETLKVPC[+57.021]HFPC[+57.021]K.F | HexNAc(4)Hex(5)Fuc(1)NeuAc(2) | C[+57]*2, N[+2351] | 1194.2814 | 4 | 4774.1038 | 4774.0877 | 0 | 3.4 | 466 | Specific | 699.36 | 399.36 | 399.36 | 12.86 | 62 | >sp\|P01833\|PIGR_HUMAN Polymeric immunoglobulin receptor OS=Homo sapiens OX=9606 GN=PIGR PE=1 SV=4 | 902 | 20201118_lss_milk_1ng_glyco_hcd203040_compound15.19279.19279.2 | controllerType=0 controllerNumber=1 scan=19279 | 38.1944 |
| 06759:3 | 7 | K.WN[+1710.598]NTGC[+57.021]QALPSQDEGPSK.A | HexNAc(3)Hex(5)NeuAc(1) | C[+57], N[+1711] | 1233.8344 | 3 | 3699.4887 | 3699.4743 | 0 | 3.9 | 498 | Specific | 247.65 | 201.33 | 201.33 | 6.2 | 62 | >sp\|P01833\|PIGR_HUMAN Polymeric immunoglobulin receptor OS=Homo sapiens OX=9606 GN=PIGR PE=1 SV=4 | 902 | 20201118_lss_milk_1ng_glyco_hcd203040_compound15.9431.9431.3 | controllerType=0 controllerNumber=1 scan=9431 | 20.9809 |
| 06812:3 | 7 | K.WN[+1548.545]NTGC[+57.021]QALPSQDEGPSK.A | HexNAc(3)Hex(4)NeuAc(1) | C[+57], N[+1549] | 1179.8176 | 3 | 3537.4381 | 3537.4215 | 0 | 4.7 | 498 | Specific | 250.33 | 215.38 | 215.38 | 6.01 | 62 | >sp\|P01833\|PIGR_HUMAN Polymeric immunoglobulin receptor OS=Homo sapiens OX=9606 GN=PIGR PE=1 SV=4 | 902 | 20201118_lss_milk_1ng_glyco_hcd203040_compound15.9485.9485.3 | controllerType=0 controllerNumber=1 scan=9485 | 21.0667 |
| 06756:3 | 7 | K.WN[+1710.598]NTGC[+57.021]QALPSQDEGPSK.A | HexNAc(3)Hex(5)NeuAc(1) | C[+57], N[+1711] | 1233.8344 | 3 | 3699.4887 | 3699.4743 | 0 | 3.9 | 498 | Specific | 250.34 | 171.79 | 171.79 | 5.8 | 62 | >sp\|P01833\|PIGR_HUMAN Polymeric immunoglobulin receptor OS=Homo sapiens OX=9606 GN=PIGR PE=1 SV=4 | 902 | 20201118_lss_milk_1ng_glyco_hcd203040_compound15.9428.9428.3 | controllerType=0 controllerNumber=1 scan=9428 | 20.9766 |
| 06699:3 | 7 | K.WN[+1913.677]NTGC[+57.021]QALPSQDEGPSK.A | HexNAc(4)Hex(5)NeuAc(1) | C[+57], N[+1914] | 1301.528 | 3 | 3902.5695 | 3902.5537 | 0 | 4 | 498 | Specific | 267.57 | 233.29 | 233.29 | 6.35 | 62 | >sp\|P01833\|PIGR_HUMAN Polymeric immunoglobulin receptor OS=Homo sapiens OX=9606 GN=PIGR PE=1 SV=4 | 902 | 20201118_lss_milk_1ng_glyco_hcd203040_compound15.9369.9369.3 | controllerType=0 controllerNumber=1 scan=9369 | 20.8839 |
| 06814:3 | 7 | K.WN[+1548.545]NTGC[+57.021]QALPSQDEGPSK.A | HexNAc(3)Hex(4)NeuAc(1) | C[+57], N[+1549] | 1179.8176 | 3 | 3537.4381 | 3537.4215 | 0 | 4.7 | 498 | Specific | 404.78 | 327.86 | 327.86 | 6.81 | 62 | >sp\|P01833\|PIGR_HUMAN Polymeric immunoglobulin receptor OS=Homo sapiens OX=9606 GN=PIGR PE=1 SV=4 | 902 | 20201118_lss_milk_1ng_glyco_hcd203040_compound15.9487.9487.3 | controllerType=0 controllerNumber=1 scan=9487 | 21.0706 |
| 06514:3 | 7 | K.WN[+2059.735]NTGC[+57.021]QALPSQDEGPSK.A | HexNAc(4)Hex(5)Fuc(1)NeuAc(1) | C[+57], N[+2060] | 1350.212 | 3 | 4048.6214 | 4048.6116 | 0 | 2.4 | 498 | Specific | 682.57 | 612.2 | 612.2 | 12.96 | 62 | >sp\|P01833\|PIGR_HUMAN Polymeric immunoglobulin receptor OS=Homo sapiens OX=9606 GN=PIGR PE=1 SV=4 | 902 | 20201118_lss_milk_1ng_glyco_hcd203040_compound15.9178.9178.3 | controllerType=0 controllerNumber=1 scan=9178 | 20.5736 |
| 06516:3 | 7 | K.WN[+2059.735]NTGC[+57.021]QALPSQDEGPSK.A | HexNAc(4)Hex(5)Fuc(1)NeuAc(1) | C[+57], N[+2060] | 1350.212 | 3 | 4048.6214 | 4048.6116 | 0 | 2.4 | 498 | Specific | 725.28 | 601.39 | 601.39 | 13.96 | 62 | >sp\|P01833\|PIGR_HUMAN Polymeric immunoglobulin receptor OS=Homo sapiens OX=9606 GN=PIGR PE=1 SV=4 | 902 | 20201118_lss_milk_1ng_glyco_hcd203040_compound15.9180.9180.3 | controllerType=0 controllerNumber=1 scan=9180 | 20.5775 |
| 19420:3 | 8 | R.LSLHRPALEDLLLGSEAN[+1872.651]LTC[+57.021]TLTGLR.D | HexNAc(3)Hex(6)NeuAc(1) | C[+57], N[+1873] | 1612.7668 | 3 | 4836.2858 | 4836.2487 | 0 | 7.7 | 127 | Specific | 245.03 | 245.03 | 245.03 | 4.34 | 58 | >sp\|P01876\|IGHA1_HUMAN Immunoglobulin heavy constant alpha 1 OS=Homo sapiens OX=9606 GN=IGHA1 PE=1 SV=2 | 255 | 20201118_lss_milk_1ng_glyco_hcd203040_compound15.22603.22603.3 | controllerType=0 controllerNumber=1 scan=22603 | 44.84 |
| 19460:3 | 8 | R.LSLHRPALEDLLLGSEAN[+1751.624]LTC[+57.021]TLTGLR.D | HexNAc(4)Hex(4)NeuAc(1) | C[+57], N[+1752] | 1572.4188 | 3 | 4715.2417 | 4715.2225 | 0 | 4.1 | 127 | Specific | 285.17 | 285.17 | 285.17 | 6.18 | 58 | >sp\|P01876\|IGHA1_HUMAN Immunoglobulin heavy constant alpha 1 OS=Homo sapiens OX=9606 GN=IGHA1 PE=1 SV=2 | 255 | 20201118_lss_milk_1ng_glyco_hcd203040_compound15.22645.22645.3 | controllerType=0 controllerNumber=1 scan=22645 | 44.9224 |
| 19495:3 | 8 | R.LSLHRPALEDLLLGSEAN[+1548.545]LTC[+57.021]TLTGLR.D | HexNAc(3)Hex(4)NeuAc(1) | C[+57], N[+1549] | 1504.7227 | 3 | 4512.1536 | 4512.1431 | 0 | 2.3 | 127 | Specific | 339.9 | 339.9 | 339.9 | 6.73 | 58 | >sp\|P01876\|IGHA1_HUMAN Immunoglobulin heavy constant alpha 1 OS=Homo sapiens OX=9606 GN=IGHA1 PE=1 SV=2 | 255 | 20201118_lss_milk_1ng_glyco_hcd203040_compound15.22681.22681.3 | controllerType=0 controllerNumber=1 scan=22681 | 44.999 |
| 18440:3 | 8 | R.LSLHRPALEDLLLGSEAN[+1460.529]LTC[+57.021]TLTGLR.D | HexNAc(4)Hex(4) | C[+57], N[+1461] | 1475.3884 | 3 | 4424.1505 | 4424.1271 | 0 | 5.3 | 127 | Specific | 477.97 | 477.97 | 477.97 | 8.45 | 58 | >sp\|P01876\|IGHA1_HUMAN Immunoglobulin heavy constant alpha 1 OS=Homo sapiens OX=9606 GN=IGHA1 PE=1 SV=2 | 255 | 20201118_lss_milk_1ng_glyco_hcd203040_compound15.21582.21582.3 | controllerType=0 controllerNumber=1 scan=21582 | 42.7275 |
| 19454:4 | 8 | R.LSLHRPALEDLLLGSEAN[+1751.624]LTC[+57.021]TLTGLR.D | HexNAc(4)Hex(4)NeuAc(1) | C[+57], N[+1752] | 1179.5651 | 4 | 4715.2386 | 4715.2225 | 0 | 3.4 | 127 | Specific | 479.72 | 479.72 | 479.72 | 9.31 | 58 | >sp\|P01876\|IGHA1_HUMAN Immunoglobulin heavy constant alpha 1 OS=Homo sapiens OX=9606 GN=IGHA1 PE=1 SV=2 | 255 | 20201118_lss_milk_1ng_glyco_hcd203040_compound15.22639.22639.4 | controllerType=0 controllerNumber=1 scan=22639 | 44.9112 |
| 18420:4 | 8 | R.LSLHRPALEDLLLGSEAN[+1663.608]LTC[+57.021]TLTGLR.D | HexNAc(5)Hex(4) | C[+57], N[+1664] | 1157.5612 | 4 | 4627.223 | 4627.2064 | 0 | 3.6 | 127 | Specific | 504.38 | 504.38 | 504.38 | 9.38 | 58 | >sp\|P01876\|IGHA1_HUMAN Immunoglobulin heavy constant alpha 1 OS=Homo sapiens OX=9606 GN=IGHA1 PE=1 SV=2 | 255 | 20201118_lss_milk_1ng_glyco_hcd203040_compound15.21561.21561.4 | controllerType=0 controllerNumber=1 scan=21561 | 42.6676 |
| 18521:4 | 8 | R.LSLHRPALEDLLLGSEAN[+1054.370]LTC[+57.021]TLTGLR.D | HexNAc(2)Hex(4) | C[+57], N[+1054] | 1005.2521 | 4 | 4017.9866 | 4017.9683 | 0 | 4.6 | 127 | Specific | 514.07 | 510.78 | 510.78 | 9.1 | 58 | >sp\|P01876\|IGHA1_HUMAN Immunoglobulin heavy constant alpha 1 OS=Homo sapiens OX=9606 GN=IGHA1 PE=1 SV=2 | 255 | 20201118_lss_milk_1ng_glyco_hcd203040_compound15.21666.21666.4 | controllerType=0 controllerNumber=1 scan=21666 | 42.9112 |
| 19423:3 | 8 | R.LSLHRPALEDLLLGSEAN[+1872.651]LTC[+57.021]TLTGLR.D | HexNAc(3)Hex(6)NeuAc(1) | C[+57], N[+1873] | 1612.7668 | 3 | 4836.2858 | 4836.2487 | 0 | 7.7 | 127 | Specific | 517.93 | 471.47 | 471.47 | 7.44 | 58 | >sp\|P01876\|IGHA1_HUMAN Immunoglobulin heavy constant alpha 1 OS=Homo sapiens OX=9606 GN=IGHA1 PE=1 SV=2 | 255 | 20201118_lss_milk_1ng_glyco_hcd203040_compound15.22606.22606.3 | controllerType=0 controllerNumber=1 scan=22606 | 44.8439 |
| 18359:4 | 8 | R.LSLHRPALEDLLLGSEAN[+1825.661]LTC[+57.021]TLTGLR.D | HexNAc(5)Hex(5) | C[+57], N[+1826] | 1198.0755 | 4 | 4789.2801 | 4789.2592 | 0 | 4.4 | 127 | Specific | 553.24 | 519.21 | 519.21 | 9.52 | 58 | >sp\|P01876\|IGHA1_HUMAN Immunoglobulin heavy constant alpha 1 OS=Homo sapiens OX=9606 GN=IGHA1 PE=1 SV=2 | 255 | 20201118_lss_milk_1ng_glyco_hcd203040_compound15.21497.21497.4 | controllerType=0 controllerNumber=1 scan=21497 | 42.5072 |
| 18387:4 | 8 | R.LSLHRPALEDLLLGSEAN[+1864.634]LTC[+57.021]TLTGLR.D | HexNAc(2)Hex(9) | C[+57], N[+1865] | 1207.8174 | 4 | 4828.2479 | 4828.2324 | 0 | 3.2 | 127 | Specific | 567.88 | 567.88 | 567.88 | 9.87 | 58 | >sp\|P01876\|IGHA1_HUMAN Immunoglobulin heavy constant alpha 1 OS=Homo sapiens OX=9606 GN=IGHA1 PE=1 SV=2 | 255 | 20201118_lss_milk_1ng_glyco_hcd203040_compound15.21526.21526.4 | controllerType=0 controllerNumber=1 scan=21526 | 42.5847 |
| 19497:3 | 8 | R.LSLHRPALEDLLLGSEAN[+1548.545]LTC[+57.021]TLTGLR.D | HexNAc(3)Hex(4)NeuAc(1) | C[+57], N[+1549] | 1504.7227 | 3 | 4512.1536 | 4512.1431 | 0 | 2.3 | 127 | Specific | 573.99 | 564.91 | 564.91 | 9.24 | 58 | >sp\|P01876\|IGHA1_HUMAN Immunoglobulin heavy constant alpha 1 OS=Homo sapiens OX=9606 GN=IGHA1 PE=1 SV=2 | 255 | 20201118_lss_milk_1ng_glyco_hcd203040_compound15.22683.22683.3 | controllerType=0 controllerNumber=1 scan=22683 | 45.0024 |
| 19478:4 | 8 | R.LSLHRPALEDLLLGSEAN[+1548.545]LTC[+57.021]TLTGLR.D | HexNAc(3)Hex(4)NeuAc(1) | C[+57], N[+1549] | 1128.7948 | 4 | 4512.1575 | 4512.1431 | 0 | 3.2 | 127 | Specific | 585.3 | 585.3 | 585.3 | 11.14 | 58 | >sp\|P01876\|IGHA1_HUMAN Immunoglobulin heavy constant alpha 1 OS=Homo sapiens OX=9606 GN=IGHA1 PE=1 SV=2 | 255 | 20201118_lss_milk_1ng_glyco_hcd203040_compound15.22664.22664.4 | controllerType=0 controllerNumber=1 scan=22664 | 44.9645 |
| 19462:3 | 8 | R.LSLHRPALEDLLLGSEAN[+1751.624]LTC[+57.021]TLTGLR.D | HexNAc(4)Hex(4)NeuAc(1) | C[+57], N[+1752] | 1572.4188 | 3 | 4715.2417 | 4715.2225 | 0 | 4.1 | 127 | Specific | 586.78 | 552.16 | 552.16 | 10.63 | 58 | >sp\|P01876\|IGHA1_HUMAN Immunoglobulin heavy constant alpha 1 OS=Homo sapiens OX=9606 GN=IGHA1 PE=1 SV=2 | 255 | 20201118_lss_milk_1ng_glyco_hcd203040_compound15.22647.22647.3 | controllerType=0 controllerNumber=1 scan=22647 | 44.9276 |
| 18376:3 | 8 | R.LSLHRPALEDLLLGSEAN[+1540.529]LTC[+57.021]TLTGLR.D | HexNAc(2)Hex(7) | C[+57], N[+1541] | 1502.0525 | 3 | 4504.1431 | 4504.1268 | 0 | 3.6 | 127 | Specific | 587.07 | 568.95 | 568.95 | 10.63 | 58 | >sp\|P01876\|IGHA1_HUMAN Immunoglobulin heavy constant alpha 1 OS=Homo sapiens OX=9606 GN=IGHA1 PE=1 SV=2 | 255 | 20201118_lss_milk_1ng_glyco_hcd203040_compound15.21515.21515.3 | controllerType=0 controllerNumber=1 scan=21515 | 42.56 |
| 19408:4 | 8 | R.LSLHRPALEDLLLGSEAN[+1872.651]LTC[+57.021]TLTGLR.D | HexNAc(3)Hex(6)NeuAc(1) | C[+57], N[+1873] | 1209.823 | 4 | 4836.2703 | 4836.2487 | 0 | 4.5 | 127 | Specific | 620.59 | 615.2 | 615.2 | 11.8 | 58 | >sp\|P01876\|IGHA1_HUMAN Immunoglobulin heavy constant alpha 1 OS=Homo sapiens OX=9606 GN=IGHA1 PE=1 SV=2 | 255 | 20201118_lss_milk_1ng_glyco_hcd203040_compound15.22591.22591.4 | controllerType=0 controllerNumber=1 scan=22591 | 44.8156 |
| 18379:3 | 8 | R.LSLHRPALEDLLLGSEAN[+1540.529]LTC[+57.021]TLTGLR.D | HexNAc(2)Hex(7) | C[+57], N[+1541] | 1502.0525 | 3 | 4504.1431 | 4504.1268 | 0 | 3.6 | 127 | Specific | 630.57 | 598.2 | 598.2 | 11.63 | 58 | >sp\|P01876\|IGHA1_HUMAN Immunoglobulin heavy constant alpha 1 OS=Homo sapiens OX=9606 GN=IGHA1 PE=1 SV=2 | 255 | 20201118_lss_milk_1ng_glyco_hcd203040_compound15.21518.21518.3 | controllerType=0 controllerNumber=1 scan=21518 | 42.5646 |
| 18336:3 | 8 | R.LSLHRPALEDLLLGSEAN[+1702.581]LTC[+57.021]TLTGLR.D | HexNAc(2)Hex(8) | C[+57], N[+1703] | 1556.0705 | 3 | 4666.1969 | 4666.1796 | 0 | 3.7 | 127 | Specific | 636.47 | 583.77 | 583.77 | 11.63 | 58 | >sp\|P01876\|IGHA1_HUMAN Immunoglobulin heavy constant alpha 1 OS=Homo sapiens OX=9606 GN=IGHA1 PE=1 SV=2 | 255 | 20201118_lss_milk_1ng_glyco_hcd203040_compound15.21473.21473.3 | controllerType=0 controllerNumber=1 scan=21473 | 42.4505 |
| 18443:3 | 8 | R.LSLHRPALEDLLLGSEAN[+1460.529]LTC[+57.021]TLTGLR.D | HexNAc(4)Hex(4) | C[+57], N[+1461] | 1475.3884 | 3 | 4424.1505 | 4424.1271 | 0 | 5.3 | 127 | Specific | 638.29 | 579.46 | 579.46 | 11.28 | 58 | >sp\|P01876\|IGHA1_HUMAN Immunoglobulin heavy constant alpha 1 OS=Homo sapiens OX=9606 GN=IGHA1 PE=1 SV=2 | 255 | 20201118_lss_milk_1ng_glyco_hcd203040_compound15.21585.21585.3 | controllerType=0 controllerNumber=1 scan=21585 | 42.736 |
| 18397:3 | 8 | R.LSLHRPALEDLLLGSEAN[+1622.582]LTC[+57.021]TLTGLR.D | HexNAc(4)Hex(5) | C[+57], N[+1623] | 1529.4026 | 3 | 4586.1934 | 4586.1799 | 0 | 2.9 | 127 | Specific | 640.61 | 576.29 | 576.29 | 11.52 | 58 | >sp\|P01876\|IGHA1_HUMAN Immunoglobulin heavy constant alpha 1 OS=Homo sapiens OX=9606 GN=IGHA1 PE=1 SV=2 | 255 | 20201118_lss_milk_1ng_glyco_hcd203040_compound15.21537.21537.3 | controllerType=0 controllerNumber=1 scan=21537 | 42.6101 |
| 18399:3 | 8 | R.LSLHRPALEDLLLGSEAN[+1622.582]LTC[+57.021]TLTGLR.D | HexNAc(4)Hex(5) | C[+57], N[+1623] | 1529.4026 | 3 | 4586.1934 | 4586.1799 | 0 | 2.9 | 127 | Specific | 643.11 | 542.65 | 542.65 | 11.52 | 58 | >sp\|P01876\|IGHA1_HUMAN Immunoglobulin heavy constant alpha 1 OS=Homo sapiens OX=9606 GN=IGHA1 PE=1 SV=2 | 255 | 20201118_lss_milk_1ng_glyco_hcd203040_compound15.21539.21539.3 | controllerType=0 controllerNumber=1 scan=21539 | 42.6158 |
| 18427:3 | 8 | R.LSLHRPALEDLLLGSEAN[+1501.555]LTC[+57.021]TLTGLR.D | HexNAc(5)Hex(3) | C[+57], N[+1502] | 1489.0616 | 3 | 4465.1702 | 4465.1536 | 0 | 3.7 | 127 | Specific | 648.17 | 599.84 | 599.84 | 11.63 | 58 | >sp\|P01876\|IGHA1_HUMAN Immunoglobulin heavy constant alpha 1 OS=Homo sapiens OX=9606 GN=IGHA1 PE=1 SV=2 | 255 | 20201118_lss_milk_1ng_glyco_hcd203040_compound15.21568.21568.3 | controllerType=0 controllerNumber=1 scan=21568 | 42.6924 |
| 18424:3 | 8 | R.LSLHRPALEDLLLGSEAN[+1501.555]LTC[+57.021]TLTGLR.D | HexNAc(5)Hex(3) | C[+57], N[+1502] | 1489.0616 | 3 | 4465.1702 | 4465.1536 | 0 | 3.7 | 127 | Specific | 657.09 | 657.09 | 657.09 | 12.63 | 58 | >sp\|P01876\|IGHA1_HUMAN Immunoglobulin heavy constant alpha 1 OS=Homo sapiens OX=9606 GN=IGHA1 PE=1 SV=2 | 255 | 20201118_lss_milk_1ng_glyco_hcd203040_compound15.21565.21565.3 | controllerType=0 controllerNumber=1 scan=21565 | 42.6762 |
| 18435:4 | 8 | R.LSLHRPALEDLLLGSEAN[+1460.529]LTC[+57.021]TLTGLR.D | HexNAc(4)Hex(4) | C[+57], N[+1461] | 1106.7948 | 4 | 4424.1575 | 4424.1271 | 0 | 6.9 | 127 | Specific | 659.67 | 630.31 | 630.31 | 11.93 | 58 | >sp\|P01876\|IGHA1_HUMAN Immunoglobulin heavy constant alpha 1 OS=Homo sapiens OX=9606 GN=IGHA1 PE=1 SV=2 | 255 | 20201118_lss_milk_1ng_glyco_hcd203040_compound15.21577.21577.4 | controllerType=0 controllerNumber=1 scan=21577 | 42.7181 |
| 18419:4 | 8 | R.LSLHRPALEDLLLGSEAN[+1501.555]LTC[+57.021]TLTGLR.D | HexNAc(5)Hex(3) | C[+57], N[+1502] | 1117.0491 | 4 | 4465.1746 | 4465.1536 | 0 | 4.7 | 127 | Specific | 680.6 | 680.6 | 680.6 | 12.8 | 58 | >sp\|P01876\|IGHA1_HUMAN Immunoglobulin heavy constant alpha 1 OS=Homo sapiens OX=9606 GN=IGHA1 PE=1 SV=2 | 255 | 20201118_lss_milk_1ng_glyco_hcd203040_compound15.21560.21560.4 | controllerType=0 controllerNumber=1 scan=21560 | 42.6662 |
| 18373:4 | 8 | R.LSLHRPALEDLLLGSEAN[+1622.582]LTC[+57.021]TLTGLR.D | HexNAc(4)Hex(5) | C[+57], N[+1623] | 1147.305 | 4 | 4586.1981 | 4586.1799 | 0 | 4 | 127 | Specific | 712.95 | 712.95 | 712.95 | 14.14 | 58 | >sp\|P01876\|IGHA1_HUMAN Immunoglobulin heavy constant alpha 1 OS=Homo sapiens OX=9606 GN=IGHA1 PE=1 SV=2 | 255 | 20201118_lss_milk_1ng_glyco_hcd203040_compound15.21512.21512.4 | controllerType=0 controllerNumber=1 scan=21512 | [illegible] |
| 18428:3 | 8 | R.LSLHRPALEDLLLGSEAN[+1216.423]LTC[+57.021]TLTGLR.D | HexNAc(2)Hex(5) | C[+57], N[+1216] | 1394.0159 | 3 | 4180.0332 | 4180.0211 | 0 | 2.9 | 127 | Specific | 714.49 | 683.94 | 683.94 | 13.52 | 58 | >sp\|P01876\|IGHA1_HUMAN Immunoglobulin heavy constant alpha 1 OS=Homo sapiens OX=9606 GN=IGHA1 PE=1 SV=2 | 255 | 20201118_lss_milk_1ng_glyco_hcd203040_compound15.21569.21569.3 | controllerType=0 controllerNumber=1 scan=21569 | [illegible] |
| 18361:4 | 8 | R.LSLHRPALEDLLLGSEAN[+1825.661]LTC[+57.021]TLTGLR.D | HexNAc(5)Hex(5) | C[+57], N[+1826] | 1198.0755 | 4 | 4789.2801 | 4789.2592 | 0 | 4.4 | 127 | Specific | 727.5 | 643.21 | 643.21 | 13.8 | 58 | >sp\|P01876\|IGHA1_HUMAN Immunoglobulin heavy constant alpha 1 OS=Homo sapiens OX=9606 GN=IGHA1 PE=1 SV=2 | 255 | 20201118_lss_milk_1ng_glyco_hcd203040_compound15.21499.21499.4 | controllerType=0 controllerNumber=1 scan=21499 | [illegible] |
| 19456:4 | 8 | R.LSLHRPALEDLLLGSEAN[+1751.624]LTC[+57.021]TLTGLR.D | HexNAc(4)Hex(4)NeuAc(1) | C[+57], N[+1752] | 1179.5651 | 4 | 4715.2386 | 4715.2225 | 0 | 3.4 | 127 | Specific | 733.3 | 733.3 | 733.3 | 14.14 | 58 | >sp\|P01876\|IGHA1_HUMAN Immunoglobulin heavy constant alpha 1 OS=Homo sapiens OX=9606 GN=IGHA1 PE=1 SV=2 | 255 | 20201118_lss_milk_1ng_glyco_hcd203040_compound15.22641.22641.4 | controllerType=0 controllerNumber=1 scan=22641 | [illegible] |
| 18422:4 | 8 | R.LSLHRPALEDLLLGSEAN[+1663.608]LTC[+57.021]TLTGLR.D | HexNAc(5)Hex(4) | C[+57], N[+1664] | 1157.5612 | 4 | 4627.223 | 4627.2064 | 0 | 3.6 | 127 | Specific | 739.75 | 710.82 | 710.82 | 14.14 | 58 | >sp\|P01876\|IGHA1_HUMAN Immunoglobulin heavy constant alpha 1 OS=Homo sapiens OX=9606 GN=IGHA1 PE=1 SV=2 | 255 | 20201118_lss_milk_1ng_glyco_hcd203040_compound15.21563.21563.4 | controllerType=0 controllerNumber=1 scan=21563 | 42.6739 |
| 18446:4 | 8 | R.LSLHRPALEDLLLGSEAN[+1298.476]LTC[+57.021]TLTGLR.D | HexNAc(4)Hex(3) | C[+57], N[+1298] | 1066.277 | 4 | 4262.0863 | 4262.0742 | 0 | 2.8 | 127 | Specific | 744.86 | 744.86 | 744.86 | 14.04 | 58 | >sp\|P01876\|IGHA1_HUMAN Immunoglobulin heavy constant alpha 1 OS=Homo sapiens OX=9606 GN=IGHA1 PE=1 SV=2 | 255 | 20201118_lss_milk_1ng_glyco_hcd203040_compound15.21588.21588.4 | controllerType=0 controllerNumber=1 scan=21588 | 42.7403 |
| 18392:3 | 8 | R.LSLHRPALEDLLLGSEAN[+1378.476]LTC[+57.021]TLTGLR.D | HexNAc(2)Hex(6) | C[+57], N[+1378] | 1448.035 | 3 | 4342.0904 | 4342.074 | 0 | 3.8 | 127 | Specific | 745.63 | 711.13 | 711.13 | 13.63 | 58 | >sp\|P01876\|IGHA1_HUMAN Immunoglobulin heavy constant alpha 1 OS=Homo sapiens OX=9606 GN=IGHA1 PE=1 SV=2 | 255 | 20201118_lss_milk_1ng_glyco_hcd203040_compound15.21531.21531.3 | controllerType=0 controllerNumber=1 scan=21531 | 42.5918 |
| 18511:4 | 8 | R.LSLHRPALEDLLLGSEAN[+1095.397]LTC[+57.021]TLTGLR.D | HexNAc(3)Hex(3) | C[+57], N[+1095] | 1015.5076 | 4 | 4059.0086 | 4058.9949 | 0 | 3.4 | 127 | Specific | 750.73 | 750.73 | 750.73 | 15.14 | 58 | >sp\|P01876\|IGHA1_HUMAN Immunoglobulin heavy constant alpha 1 OS=Homo sapiens OX=9606 GN=IGHA1 PE=1 SV=2 | 255 | 20201118_lss_milk_1ng_glyco_hcd203040_compound15.21656.21656.4 | controllerType=0 controllerNumber=1 scan=21656 | 42.8906 |
| 18395:3 | 8 | R.LSLHRPALEDLLLGSEAN[+1378.476]LTC[+57.021]TLTGLR.D | HexNAc(2)Hex(6) | C[+57], N[+1378] | 1448.035 | 3 | 4342.0904 | 4342.074 | 0 | 3.8 | 127 | Specific | 753.13 | 668 | 668 | 14.63 | 58 | >sp\|P01876\|IGHA1_HUMAN Immunoglobulin heavy constant alpha 1 OS=Homo sapiens OX=9606 GN=IGHA1 PE=1 SV=2 | 255 | 20201118_lss_milk_1ng_glyco_hcd203040_compound15.21534.21534.3 | controllerType=0 controllerNumber=1 scan=21534 | 42.6044 |
| 18333:3 | 8 | R.LSLHRPALEDLLLGSEAN[+1702.581]LTC[+57.021]TLTGLR.D | HexNAc(2)Hex(8) | C[+57], N[+1703] | 1556.0705 | 3 | 4666.1969 | 4666.1796 | 0 | 3.7 | 127 | Specific | 757.41 | 748.94 | 748.94 | 14.63 | 58 | >sp\|P01876\|IGHA1_HUMAN Immunoglobulin heavy constant alpha 1 OS=Homo sapiens OX=9606 GN=IGHA1 PE=1 SV=2 | 255 | 20201118_lss_milk_1ng_glyco_hcd203040_compound15.21470.21470.3 | controllerType=0 controllerNumber=1 scan=21470 | 42.4446 |
| 18436:5 | 8 | R.LSLHRPALEDLLLGSEAN[+1501.555]LTC[+57.021]TLTGLR.D | HexNAc(5)Hex(3) | C[+57], N[+1502] | 893.84 | 5 | 4465.1708 | 4465.1536 | 0 | 3.9 | 127 | Specific | 758.2 | 758.2 | 758.2 | 15.13 | 58 | >sp\|P01876\|IGHA1_HUMAN Immunoglobulin heavy constant alpha 1 OS=Homo sapiens OX=9606 GN=IGHA1 PE=1 SV=2 | 255 | 20201118_lss_milk_1ng_glyco_hcd203040_compound15.21578.21578.5 | controllerType=0 controllerNumber=1 scan=21578 | 42.7196 |
| 18438:5 | 8 | R.LSLHRPALEDLLLGSEAN[+1501.555]LTC[+57.021]TLTGLR.D | HexNAc(5)Hex(3) | C[+57], N[+1502] | 893.84 | 5 | 4465.1708 | 4465.1536 | 0 | 3.9 | 127 | Specific | 766.26 | 766.26 | 766.26 | 15.13 | 58 | >sp\|P01876\|IGHA1_HUMAN Immunoglobulin heavy constant alpha 1 OS=Homo sapiens OX=9606 GN=IGHA1 PE=1 SV=2 | 255 | 20201118_lss_milk_1ng_glyco_hcd203040_compound15.21580.21580.5 | controllerType=0 controllerNumber=1 scan=21580 | 42.7252 |
| 18401:4 | 8 | R.LSLHRPALEDLLLGSEAN[+1419.502]LTC[+57.021]TLTGLR.D | HexNAc(3)Hex(5) | C[+57], N[+1420] | 1096.534 | 4 | 4383.1141 | 4383.1005 | 0 | 3.1 | 127 | Specific | 767.62 | 767.62 | 767.62 | 15.14 | 58 | >sp\|P01876\|IGHA1_HUMAN Immunoglobulin heavy constant alpha 1 OS=Homo sapiens OX=9606 GN=IGHA1 PE=1 SV=2 | 255 | 20201118_lss_milk_1ng_glyco_hcd203040_compound15.21541.21541.4 | controllerType=0 controllerNumber=1 scan=21541 | 42.6216 |
| 18425:3 | 8 | R.LSLHRPALEDLLLGSEAN[+1216.423]LTC[+57.021]TLTGLR.D | HexNAc(2)Hex(5) | C[+57], N[+1216] | 1394.0159 | 3 | 4180.0332 | 4180.0211 | 0 | 2.9 | 127 | Specific | 774.82 | 695.24 | 695.24 | 14.52 | 58 | >sp\|P01876\|IGHA1_HUMAN Immunoglobulin heavy constant alpha 1 OS=Homo sapiens OX=9606 GN=IGHA1 PE=1 SV=2 | 255 | 20201118_lss_milk_1ng_glyco_hcd203040_compound15.21566.21566.3 | controllerType=0 controllerNumber=1 scan=21566 | 42.6773 |
| 19480:4 | 8 | R.LSLHRPALEDLLLGSEAN[+1548.545]LTC[+57.021]TLTGLR.D | HexNAc(3)Hex(4)NeuAc(1) | C[+57], N[+1549] | 1128.7948 | 4 | 4512.1575 | 4512.1431 | 0 | 3.2 | 127 | Specific | 810.11 | 764.35 | 764.35 | 16.14 | 58 | >sp\|P01876\|IGHA1_HUMAN Immunoglobulin heavy constant alpha 1 OS=Homo sapiens OX=9606 GN=IGHA1 PE=1 SV=2 | 255 | 20201118_lss_milk_1ng_glyco_hcd203040_compound15.22666.22666.4 | controllerType=0 controllerNumber=1 scan=22666 | 44.9697 |
| 18421:4 | 8 | R.LSLHRPALEDLLLGSEAN[+1501.555]LTC[+57.021]TLTGLR.D | HexNAc(5)Hex(3) | C[+57], N[+1502] | 1117.0491 | 4 | 4465.1746 | 4465.1536 | 0 | 4.7 | 127 | Specific | 812.86 | 812.86 | 812.86 | 15.8 | 58 | >sp\|P01876\|IGHA1_HUMAN Immunoglobulin heavy constant alpha 1 OS=Homo sapiens OX=9606 GN=IGHA1 PE=1 SV=2 | 255 | 20201118_lss_milk_1ng_glyco_hcd203040_compound15.21562.21562.4 | controllerType=0 controllerNumber=1 scan=21562 | 42.6727 |
| 18375:4 | 8 | R.LSLHRPALEDLLLGSEAN[+1622.582]LTC[+57.021]TLTGLR.D | HexNAc(4)Hex(5) | C[+57], N[+1623] | 1147.305 | 4 | 4586.1981 | 4586.1799 | 0 | 4 | 127 | Specific | 827.94 | 720.91 | 720.91 | 16.14 | 58 | >sp\|P01876\|IGHA1_HUMAN Immunoglobulin heavy constant alpha 1 OS=Homo sapiens OX=9606 GN=IGHA1 PE=1 SV=2 | 255 | 20201118_lss_milk_1ng_glyco_hcd203040_compound15.21514.21514.4 | controllerType=0 controllerNumber=1 scan=21514 | 42.5589 |
| 18437:4 | 8 | R.LSLHRPALEDLLLGSEAN[+1460.529]LTC[+57.021]TLTGLR.D | HexNAc(4)Hex(4) | C[+57], N[+1461] | 1106.7948 | 4 | 4424.1575 | 4424.1271 | 0 | 6.9 | 127 | Specific | 831.82 | 717.27 | 717.27 | 14.93 | 58 | >sp\|P01876\|IGHA1_HUMAN Immunoglobulin heavy constant alpha 1 OS=Homo sapiens OX=9606 GN=IGHA1 PE=1 SV=2 | 255 | 20201118_lss_milk_1ng_glyco_hcd203040_compound15.21579.21579.4 | controllerType=0 controllerNumber=1 scan=21579 | 42.7229 |
| 18372:4 | 8 | R.LSLHRPALEDLLLGSEAN[+1540.529]LTC[+57.021]TLTGLR.D | HexNAc(2)Hex(7) | C[+57], N[+1541] | 1126.7909 | 4 | 4504.1419 | 4504.1268 | 0 | 3.4 | 127 | Specific | 851.44 | 803.46 | 803.46 | 17.14 | 58 | >sp\|P01876\|IGHA1_HUMAN Immunoglobulin heavy constant alpha 1 OS=Homo sapiens OX=9606 GN=IGHA1 PE=1 SV=2 | 255 | 20201118_lss_milk_1ng_glyco_hcd203040_compound15.21511.21511.4 | controllerType=0 controllerNumber=1 scan=21511 | 42.5511 |
| 18449:4 | 8 | R.LSLHRPALEDLLLGSEAN[+1298.476]LTC[+57.021]TLTGLR.D | HexNAc(4)Hex(3) | C[+57], N[+1298] | 1066.277 | 4 | 4262.0863 | 4262.0742 | 0 | 2.8 | 127 | Specific | 863.9 | 857.83 | 857.83 | 17.04 | 58 | >sp\|P01876\|IGHA1_HUMAN Immunoglobulin heavy constant alpha 1 OS=Homo sapiens OX=9606 GN=IGHA1 PE=1 SV=2 | 255 | 20201118_lss_milk_1ng_glyco_hcd203040_compound15.21591.21591.4 | controllerType=0 controllerNumber=1 scan=21591 | 42.753 |
| 18423:4 | 8 | R.LSLHRPALEDLLLGSEAN[+1216.423]LTC[+57.021]TLTGLR.D | HexNAc(2)Hex(5) | C[+57], N[+1216] | 1045.7641 | 4 | 4180.0345 | 4180.0211 | 0 | 3.2 | 127 | Specific | 864.12 | 814.35 | 814.35 | 17.14 | 58 | >sp\|P01876\|IGHA1_HUMAN Immunoglobulin heavy constant alpha 1 OS=Homo sapiens OX=9606 GN=IGHA1 PE=1 SV=2 | 255 | 20201118_lss_milk_1ng_glyco_hcd203040_compound15.21564.21564.4 | controllerType=0 controllerNumber=1 scan=21564 | 42.675 |
| 19411:4 | 8 | R.LSLHRPALEDLLLGSEAN[+1872.651]LTC[+57.021]TLTGLR.D | HexNAc(3)Hex(6)NeuAc(1) | C[+57], N[+1873] | 1209.823 | 4 | 4836.2703 | 4836.2487 | 0 | 4.5 | 127 | Specific | 875.93 | 867.44 | 867.44 | 16.8 | 58 | >sp\|P01876\|IGHA1_HUMAN Immunoglobulin heavy constant alpha 1 OS=Homo sapiens OX=9606 GN=IGHA1 PE=1 SV=2 | 255 | 20201118_lss_milk_1ng_glyco_hcd203040_compound15.22594.22594.4 | controllerType=0 controllerNumber=1 scan=22594 | 44.8203 |
| 18404:4 | 8 | R.LSLHRPALEDLLLGSEAN[+1419.502]LTC[+57.021]TLTGLR.D | HexNAc(3)Hex(5) | C[+57], N[+1420] | 1096.534 | 4 | 4383.1141 | 4383.1005 | 0 | 3.1 | 127 | Specific | 882.9 | 749.2 | 749.2 | 17.14 | 58 | >sp\|P01876\|IGHA1_HUMAN Immunoglobulin heavy constant alpha 1 OS=Homo sapiens OX=9606 GN=IGHA1 PE=1 SV=2 | 255 | 20201118_lss_milk_1ng_glyco_hcd203040_compound15.21544.21544.4 | controllerType=0 controllerNumber=1 scan=21544 | 42.6281 |
| 18377:4 | 8 | R.LSLHRPALEDLLLGSEAN[+1378.476]LTC[+57.021]TLTGLR.D | HexNAc(2)Hex(6) | C[+57], N[+1378] | 1086.278 | 4 | 4342.0902 | 4342.074 | 0 | 3.7 | 127 | Specific | 883.9 | 796.68 | 796.68 | 17.14 | 58 | >sp\|P01876\|IGHA1_HUMAN Immunoglobulin heavy constant alpha 1 OS=Homo sapiens OX=9606 GN=IGHA1 PE=1 SV=2 | 255 | 20201118_lss_milk_1ng_glyco_hcd203040_compound15.21516.21516.4 | controllerType=0 controllerNumber=1 scan=21516 | 42.5612 |
| 18524:4 | 8 | R.LSLHRPALEDLLLGSEAN[+1054.370]LTC[+57.021]TLTGLR.D | HexNAc(2)Hex(4) | C[+57], N[+1054] | 1005.2521 | 4 | 4017.9866 | 4017.9683 | 0 | 4.6 | 127 | Specific | 904.8 | 847.32 | 847.32 | 17.8 | 58 | >sp\|P01876\|IGHA1_HUMAN Immunoglobulin heavy constant alpha 1 OS=Homo sapiens OX=9606 GN=IGHA1 PE=1 SV=2 | 255 | 20201118_lss_milk_1ng_glyco_hcd203040_compound15.21669.21669.4 | controllerType=0 controllerNumber=1 scan=21669 | 42.9163 |
| 18329:4 | 8 | R.LSLHRPALEDLLLGSEAN[+1702.581]LTC[+57.021]TLTGLR.D | HexNAc(2)Hex(8) | C[+57], N[+1703] | 1167.3052 | 4 | 4666.199 | 4666.1796 | 0 | 4.2 | 127 | Specific | 914.54 | 914.54 | 914.54 | 18.14 | 58 | >sp\|P01876\|IGHA1_HUMAN Immunoglobulin heavy constant alpha 1 OS=Homo sapiens OX=9606 GN=IGHA1 PE=1 SV=2 | 255 | 20201118_lss_milk_1ng_glyco_hcd203040_compound15.21466.21466.4 | controllerType=0 controllerNumber=1 scan=21466 | 42.4369 |
| 18390:4 | 8 | R.LSLHRPALEDLLLGSEAN[+1864.634]LTC[+57.021]TLTGLR.D | HexNAc(2)Hex(9) | C[+57], N[+1865] | 1207.8174 | 4 | 4828.2479 | 4828.2324 | 0 | 3.2 | 127 | Specific | 957.51 | 909.42 | 909.42 | 19.14 | 58 | >sp\|P01876\|IGHA1_HUMAN Immunoglobulin heavy constant alpha 1 OS=Homo sapiens OX=9606 GN=IGHA1 PE=1 SV=2 | 255 | 20201118_lss_milk_1ng_glyco_hcd203040_compound15.21529.21529.4 | controllerType=0 controllerNumber=1 scan=21529 | 42.5888 |
| 18513:4 | 8 | R.LSLHRPALEDLLLGSEAN[+1095.397]LTC[+57.021]TLTGLR.D | HexNAc(3)Hex(3) | C[+57], N[+1095] | 1015.5076 | 4 | 4059.0086 | 4058.9949 | 0 | 3.4 | 127 | Specific | 970.46 | 939.61 | 939.61 | 19.14 | 58 | >sp\|P01876\|IGHA1_HUMAN Immunoglobulin heavy constant alpha 1 OS=Homo sapiens OX=9606 GN=IGHA1 PE=1 SV=2 | 255 | 20201118_lss_milk_1ng_glyco_hcd203040_compound15.21658.21658.4 | controllerType=0 controllerNumber=1 scan=21658 | 42.8949 |
| 18447:4 | 8 | R.LSLHRPALEDLLLGSEAN[+1257.449]LTC[+57.021]TLTGLR.D | HexNAc(3)Hex(4) | C[+57], N[+1257] | 1056.0197 | 4 | 4221.057 | 4221.0477 | 0 | 2.2 | 127 | Specific | 977.58 | 928.31 | 928.31 | 19.04 | 58 | >sp\|P01876\|IGHA1_HUMAN Immunoglobulin heavy constant alpha 1 OS=Homo sapiens OX=9606 GN=IGHA1 PE=1 SV=2 | 255 | 20201118_lss_milk_1ng_glyco_hcd203040_compound15.21589.21589.4 | controllerType=0 controllerNumber=1 scan=21589 | 42.7414 |
| 18426:4 | 8 | R.LSLHRPALEDLLLGSEAN[+1216.423]LTC[+57.021]TLTGLR.D | HexNAc(2)Hex(5) | C[+57], N[+1216] | 1045.7641 | 4 | 4180.0345 | 4180.0211 | 0 | 3.2 | 127 | Specific | 1002.22 | 1002.22 | 1002.22 | 20.14 | 58 | >sp\|P01876\|IGHA1_HUMAN Immunoglobulin heavy constant alpha 1 OS=Homo sapiens OX=9606 GN=IGHA1 PE=1 SV=2 | 255 | 20201118_lss_milk_1ng_glyco_hcd203040_compound15.21567.21567.4 | controllerType=0 controllerNumber=1 scan=21567 | 42.6818 |
| 18331:4 | 8 | R.LSLHRPALEDLLLGSEAN[+1702.581]LTC[+57.021]TLTGLR.D | HexNAc(2)Hex(8) | C[+57], N[+1703] | 1167.3052 | 4 | 4666.199 | 4666.1796 | 0 | 4.2 | 127 | Specific | 1010.94 | 961.3 | 961.3 | 20.14 | 58 | >sp\|P01876\|IGHA1_HUMAN Immunoglobulin heavy constant alpha 1 OS=Homo sapiens OX=9606 GN=IGHA1 PE=1 SV=2 | 255 | 20201118_lss_milk_1ng_glyco_hcd203040_compound15.21468.21468.4 | controllerType=0 controllerNumber=1 scan=21468 | 42.4427 |
| 18380:4 | 8 | R.LSLHRPALEDLLLGSEAN[+1378.476]LTC[+57.021]TLTGLR.D | HexNAc(2)Hex(6) | C[+57], N[+1378] | 1086.278 | 4 | 4342.0902 | 4342.074 | 0 | 3.7 | 127 | Specific | 1016.84 | 1016.84 | 1016.84 | 20.14 | 58 | >sp\|P01876\|IGHA1_HUMAN Immunoglobulin heavy constant alpha 1 OS=Homo sapiens OX=9606 GN=IGHA1 PE=1 SV=2 | 255 | 20201118_lss_milk_1ng_glyco_hcd203040_compound15.21519.21519.4 | controllerType=0 controllerNumber=1 scan=21519 | 42.5729 |
| 18374:4 | 8 | R.LSLHRPALEDLLLGSEAN[+1540.529]LTC[+57.021]TLTGLR.D | HexNAc(2)Hex(7) | C[+57], N[+1541] | 1126.7909 | 4 | 4504.1419 | 4504.1268 | 0 | 3.4 | 127 | Specific | 1045.21 | 1045.21 | 1045.21 | 20.14 | 58 | >sp\|P01876\|IGHA1_HUMAN Immunoglobulin heavy constant alpha 1 OS=Homo sapiens OX=9606 GN=IGHA1 PE=1 SV=2 | 255 | 20201118_lss_milk_1ng_glyco_hcd203040_compound15.21513.21513.4 | controllerType=0 controllerNumber=1 scan=21513 | 42.5561 |
| 18450:4 | 8 | R.LSLHRPALEDLLLGSEAN[+1257.449]LTC[+57.021]TLTGLR.D | HexNAc(3)Hex(4) | C[+57], N[+1257] | 1056.0197 | 4 | 4221.057 | 4221.0477 | 0 | 2.2 | 127 | Specific | 1101.34 | 1041.17 | 1041.17 | 22.04 | 58 | >sp\|P01876\|IGHA1_HUMAN Immunoglobulin heavy constant alpha 1 OS=Homo sapiens OX=9606 GN=IGHA1 PE=1 SV=2 | 255 | 20201118_lss_milk_1ng_glyco_hcd203040_compound15.21592.21592.4 | controllerType=0 controllerNumber=1 scan=21592 | 42.7541 |
| 12363:3 | 8 | R.LAGKPTHVN[+1095.397]VSVVMAEVDGTC[+57.021]Y.- | HexNAc(3)Hex(3) | C[+57], N[+1095] | 1148.1889 | 3 | 3442.5521 | 3442.5387 | 0 | 3.9 | 332 | CRagged | 211.75 | 211.75 | 211.75 | 3.89 | 58 | >sp\|P01876\|IGHA1_HUMAN Immunoglobulin heavy constant alpha 1 OS=Homo sapiens OX=9606 GN=IGHA1 PE=1 SV=2 | 255 | 20201118_lss_milk_1ng_glyco_hcd203040_compound15.15243.15243.3 | controllerType=0 controllerNumber=1 scan=15243 | 30.8353 |
| 15856:3 | 8 | R.LAGKPTHVN[+2350.830]VSVVMAEVDGTC[+57.021]Y.- | HexNAc(4)Hex(5)Fuc(1)NeuAc(2) | C[+57], N[+2351] | 1566.6686 | 3 | 4697.9914 | 4697.9724 | 0 | 4 | 332 | CRagged | 235.2 | 235.2 | 235.2 | 6.45 | 58 | >sp\|P01876\|IGHA1_HUMAN Immunoglobulin heavy constant alpha 1 OS=Homo sapiens OX=9606 GN=IGHA1 PE=1 SV=2 | 255 | 20201118_lss_milk_1ng_glyco_hcd203040_compound15.18873.18873.3 | controllerType=0 controllerNumber=1 scan=18873 | 37.3946 |
| 11097:3 | 8 | R.LAGKPTHVN[+1378.476]VSVVMAEVDGTC[+57.021]Y.- | HexNAc(2)Hex(6) | C[+57], N[+1378] | 1242.5494 | 3 | 3725.6336 | 3725.6178 | 0 | 4.2 | 332 | CRagged | 249.08 | 160.9 | 160.9 | 3.95 | 58 | >sp\|P01876\|IGHA1_HUMAN Immunoglobulin heavy constant alpha 1 OS=Homo sapiens OX=9606 GN=IGHA1 PE=1 SV=2 | 255 | 20201118_lss_milk_1ng_glyco_hcd203040_compound15.13931.13931.3 | controllerType=0 controllerNumber=1 scan=13931 | 28.4566 |
| 11101:3 | 8 | R.LAGKPTHVN[+1809.666]VSVVMAEVDGTC[+57.021]Y.- | HexNAc(5)Hex(4)Fuc(1) | C[+57], N[+1810] | 1386.2791 | 3 | 4156.8228 | 4156.8081 | 0 | 3.5 | 332 | CRagged | 272.43 | 272.43 | 251.86 | 6.8 | 58 | >sp\|P01876\|IGHA1_HUMAN Immunoglobulin heavy constant alpha 1 OS=Homo sapiens OX=9606 GN=IGHA1 PE=1 SV=2 | 255 | 20201118_lss_milk_1ng_glyco_hcd203040_compound15.13935.13935.3 | controllerType=0 controllerNumber=1 scan=13935 | 28.464 |
| 12307:3 | 8 | R.LAGKPTHVN[+1663.608]VSVVMAEVDGTC[+57.021]Y.- | HexNAc(5)Hex(4) | C[+57], N[+1664] | 1337.5897 | 3 | 4010.7544 | 4010.7502 | 0 | 1 | 332 | CRagged | 274.08 | 274.08 | 156.12 | 6.42 | 58 | >sp\|P01876\|IGHA1_HUMAN Immunoglobulin heavy constant alpha 1 OS=Homo sapiens OX=9606 GN=IGHA1 PE=1 SV=2 | 255 | 20201118_lss_milk_1ng_glyco_hcd203040_compound15.15185.15185.3 | controllerType=0 controllerNumber=1 scan=15185 | 30.733 |
| 11095:3 | 8 | R.LAGKPTHVN[+1378.476]VSVVMAEVDGTC[+57.021]Y.- | HexNAc(2)Hex(6) | C[+57], N[+1378] | 1242.5494 | 3 | 3725.6336 | 3725.6178 | 0 | 4.2 | 332 | CRagged | 288.03 | 288.03 | 221.69 | 6.47 | 58 | >sp\|P01876\|IGHA1_HUMAN Immunoglobulin heavy constant alpha 1 OS=Homo sapiens OX=9606 GN=IGHA1 PE=1 SV=2 | 255 | 20201118_lss_milk_1ng_glyco_hcd203040_compound15.13929.13929.3 | controllerType=0 controllerNumber=1 scan=13929 | 28.4526 |
| 14057:3 | 8 | R.LAGKPTHVN[+2059.735]VSVVMAEVDGTC[+57.021]Y.- | HexNAc(4)Hex(5)Fuc(1)NeuAc(1) | C[+57], N[+2060] | 1469.6343 | 3 | 4406.8885 | 4406.877 | 0 | 2.6 | 332 | CRagged | 350.18 | 289.53 | 289.53 | 5.84 | 58 | >sp\|P01876\|IGHA1_HUMAN Immunoglobulin heavy constant alpha 1 OS=Homo sapiens OX=9606 GN=IGHA1 PE=1 SV=2 | 255 | 20201118_lss_milk_1ng_glyco_hcd203040_compound15.17006.17006.3 | controllerType=0 controllerNumber=1 scan=17006 | 33.838 |
| 12309:3 | 8 | R.LAGKPTHVN[+1663.608]VSVVMAEVDGTC[+57.021]Y.- | HexNAc(5)Hex(4) | C[+57], N[+1664] | 1337.5897 | 3 | 4010.7544 | 4010.7502 | 0 | 1 | 332 | CRagged | 360.63 | 360.63 | 41.58 | 7.41 | 58 | >sp\|P01876\|IGHA1_HUMAN Immunoglobulin heavy constant alpha 1 OS=Homo sapiens OX=9606 GN=IGHA1 PE=1 SV=2 | 255 | 20201118_lss_milk_1ng_glyco_hcd203040_compound15.15187.15187.3 | controllerType=0 controllerNumber=1 scan=15187 | 30.7365 |
| 14055:3 | 8 | R.LAGKPTHVN[+2059.735]VSVVMAEVDGTC[+57.021]Y.- | HexNAc(4)Hex(5)Fuc(1)NeuAc(1) | C[+57], N[+2060] | 1469.6343 | 3 | 4406.8885 | 4406.877 | 0 | 2.6 | 332 | CRagged | 396.86 | 350.47 | 350.47 | 7.53 | 58 | >sp\|P01876\|IGHA1_HUMAN Immunoglobulin heavy constant alpha 1 OS=Homo sapiens OX=9606 GN=IGHA1 PE=1 SV=2 | 255 | 20201118_lss_milk_1ng_glyco_hcd203040_compound15.17004.17004.3 | controllerType=0 controllerNumber=1 scan=17004 | 33.8346 |
| 12318:3 | 8 | R.LAGKPTHVN[+1647.613]VSVVMAEVDGTC[+57.021]Y.- | HexNAc(5)Hex(3)Fuc(1) | C[+57], N[+1648] | 1332.2609 | 3 | 3994.7682 | 3994.7553 | 0 | 3.2 | 332 | CRagged | 446.82 | 197.17 | 197.17 | 7.26 | 58 | >sp\|P01876\|IGHA1_HUMAN Immunoglobulin heavy constant alpha 1 OS=Homo sapiens OX=9606 GN=IGHA1 PE=1 SV=2 | 255 | 20201118_lss_milk_1ng_glyco_hcd203040_compound15.15197.15197.3 | controllerType=0 controllerNumber=1 scan=15197 | 30.7588 |
| 15845:3 | 8 | R.LAGKPTHVN[+2553.910]VSVVMAEVDGTC[+57.021]Y.- | HexNAc(5)Hex(5)Fuc(1)NeuAc(2) | C[+57], N[+2554] | 1634.361 | 3 | 4901.0685 | 4901.0518 | 0 | 3.4 | 332 | CRagged | 512.45 | 372.3 | 372.3 | 8.33 | 58 | >sp\|P01876\|IGHA1_HUMAN Immunoglobulin heavy constant alpha 1 OS=Homo sapiens OX=9606 GN=IGHA1 PE=1 SV=2 | 255 | 20201118_lss_milk_1ng_glyco_hcd203040_compound15.18862.18862.3 | controllerType=0 controllerNumber=1 scan=18862 | 37.3773 |
| 14049:3 | 8 | R.LAGKPTHVN[+1872.651]VSVVMAEVDGTC[+57.021]Y.- | HexNAc(3)Hex(6)NeuAc(1) | C[+57], N[+1873] | 1407.274 | 3 | 4219.8074 | 4219.7926 | 0 | 3.5 | 332 | CRagged | 519.02 | 436.33 | 312.15 | 9.56 | 58 | >sp\|P01876\|IGHA1_HUMAN Immunoglobulin heavy constant alpha 1 OS=Homo sapiens OX=9606 GN=IGHA1 PE=1 SV=2 | 255 | 20201118_lss_milk_1ng_glyco_hcd203040_compound15.16998.16998.3 | controllerType=0 controllerNumber=1 scan=16998 | 33.8235 |
| 11125:3 | 8 | R.LAGKPTHVN[+1216.423]VSVVMAEVDGTC[+57.021]Y.- | HexNAc(2)Hex(5) | C[+57], N[+1216] | 1188.5305 | 3 | 3563.5768 | 3563.5649 | 0 | 3.3 | 332 | CRagged | 537.47 | 496.28 | 350.57 | 9.78 | 58 | >sp\|P01876\|IGHA1_HUMAN Immunoglobulin heavy constant alpha 1 OS=Homo sapiens OX=9606 GN=IGHA1 PE=1 SV=2 | 255 | 20201118_lss_milk_1ng_glyco_hcd203040_compound15.13960.13960.3 | controllerType=0 controllerNumber=1 scan=13960 | 28.5056 |
| 13980:4 | 8 | R.LAGKPTHVN[+2262.814]VSVVMAEVDGTC[+57.021]Y.- | HexNAc(5)Hex(5)Fuc(1)NeuAc(1) | C[+57], N[+2263] | 1153.2475 | 4 | 4609.9681 | 4609.9564 | 0 | 2.5 | 332 | CRagged | 541.25 | 279.25 | 279.25 | 7.96 | 58 | >sp\|P01876\|IGHA1_HUMAN Immunoglobulin heavy constant alpha 1 OS=Homo sapiens OX=9606 GN=IGHA1 PE=1 SV=2 | 255 | 20201118_lss_milk_1ng_glyco_hcd203040_compound15.16926.16926.4 | controllerType=0 controllerNumber=1 scan=16926 | 33.6781 |
| 14047:3 | 8 | R.LAGKPTHVN[+1872.651]VSVVMAEVDGTC[+57.021]Y.- | HexNAc(3)Hex(6)NeuAc(1) | C[+57], N[+1873] | 1407.274 | 3 | 4219.8074 | 4219.7926 | 0 | 3.5 | 332 | CRagged | 590.6 | 590.6 | 398.6 | 11.26 | 58 | >sp\|P01876\|IGHA1_HUMAN Immunoglobulin heavy constant alpha 1 OS=Homo sapiens OX=9606 GN=IGHA1 PE=1 SV=2 | 255 | 20201118_lss_milk_1ng_glyco_hcd203040_compound15.16996.16996.3 | controllerType=0 controllerNumber=1 scan=16996 | 33.8193 |
| 15810:4 | 8 | R.LAGKPTHVN[+2553.910]VSVVMAEVDGTC[+57.021]Y.- | HexNAc(5)Hex(5)Fuc(1)NeuAc(2) | C[+57], N[+2554] | 1226.0223 | 4 | 4901.0672 | 4901.0518 | 0 | 3.1 | 332 | CRagged | 593.4 | 473.46 | 473.46 | 11.25 | 58 | >sp\|P01876\|IGHA1_HUMAN Immunoglobulin heavy constant alpha 1 OS=Homo sapiens OX=9606 GN=IGHA1 PE=1 SV=2 | 255 | 20201118_lss_milk_1ng_glyco_hcd203040_compound15.18825.18825.4 | controllerType=0 controllerNumber=1 scan=18825 | 37.3111 |
| 14108:3 | 8 | R.LAGKPTHVN[+1751.624]VSVVMAEVDGTC[+57.021]Y.- | HexNAc(4)Hex(4)NeuAc(1) | C[+57], N[+1752] | 1366.9319 | 3 | 4098.7813 | 4098.7663 | 0 | 3.7 | 332 | CRagged | 597.32 | 585.9 | 585.9 | 11.26 | 58 | >sp\|P01876\|IGHA1_HUMAN Immunoglobulin heavy constant alpha 1 OS=Homo sapiens OX=9606 GN=IGHA1 PE=1 SV=2 | 255 | 20201118_lss_milk_1ng_glyco_hcd203040_compound15.17059.17059.3 | controllerType=0 controllerNumber=1 scan=17059 | 33.9269 |
| 12321:4 | 8 | R.LAGKPTHVN[+1647.613]VSVVMAEVDGTC[+57.021]Y.- | HexNAc(5)Hex(3)Fuc(1) | C[+57], N[+1648] | 999.4474 | 4 | 3994.7679 | 3994.7553 | 0 | 3.1 | 332 | CRagged | 604.85 | 503.19 | 503.19 | 12.25 | 58 | >sp\|P01876\|IGHA1_HUMAN Immunoglobulin heavy constant alpha 1 OS=Homo sapiens OX=9606 GN=IGHA1 PE=1 SV=2 | 255 | 20201118_lss_milk_1ng_glyco_hcd203040_compound15.15200.15200.4 | controllerType=0 controllerNumber=1 scan=15200 | 30.7624 |
| 11127:3 | 8 | R.LAGKPTHVN[+1216.423]VSVVMAEVDGTC[+57.021]Y.- | HexNAc(2)Hex(5) | C[+57], N[+1216] | 1188.5305 | 3 | 3563.5768 | 3563.5649 | 0 | 3.3 | 332 | CRagged | 619.01 | 571.01 | 397.16 | 12.26 | 58 | >sp\|P01876\|IGHA1_HUMAN Immunoglobulin heavy constant alpha 1 OS=Homo sapiens OX=9606 GN=IGHA1 PE=1 SV=2 | 255 | 20201118_lss_milk_1ng_glyco_hcd203040_compound15.13962.13962.3 | controllerType=0 controllerNumber=1 scan=13962 | 28.5118 |
| 12303:4 | 8 | R.LAGKPTHVN[+1501.555]VSVVMAEVDGTC[+57.021]Y.- | HexNAc(5)Hex(3) | C[+57], N[+1502] | 962.9329 | 4 | 3848.7098 | 3848.6974 | 0 | 3.2 | 332 | CRagged | 620.22 | 620.22 | 620.22 | 12.25 | 58 | >sp\|P01876\|IGHA1_HUMAN Immunoglobulin heavy constant alpha 1 OS=Homo sapiens OX=9606 GN=IGHA1 PE=1 SV=2 | 255 | 20201118_lss_milk_1ng_glyco_hcd203040_compound15.15181.15181.4 | controllerType=0 controllerNumber=1 scan=15181 | 30.725 |
| 12366:3 | 8 | R.LAGKPTHVN[+1095.397]VSVVMAEVDGTC[+57.021]Y.- | HexNAc(3)Hex(3) | C[+57], N[+1095] | 1148.1889 | 3 | 3442.5521 | 3442.5387 | 0 | 3.9 | 332 | CRagged | 623.61 | 499.82 | 499.82 | 12.26 | 58 | >sp\|P01876\|IGHA1_HUMAN Immunoglobulin heavy constant alpha 1 OS=Homo sapiens OX=9606 GN=IGHA1 PE=1 SV=2 | 255 | 20201118_lss_milk_1ng_glyco_hcd203040_compound15.15246.15246.3 | controllerType=0 controllerNumber=1 scan=15246 | 30.8393 |
| 12226:3 | 8 | R.LAGKPTHVN[+1663.608]VSVVMAEVDGTC[+57.021]Y.- | HexNAc(5)Hex(4) | C[+57], N[+1664] | 1337.5908 | 3 | 4010.7577 | 4010.7502 | 0 | 1.9 | 332 | CRagged | 628.96 | 628.62 | 397.82 | 12.15 | 58 | >sp\|P01876\|IGHA1_HUMAN Immunoglobulin heavy constant alpha 1 OS=Homo sapiens OX=9606 GN=IGHA1 PE=1 SV=2 | 255 | 20201118_lss_milk_1ng_glyco_hcd203040_compound15.15101.15101.3 | controllerType=0 controllerNumber=1 scan=15101 | 30.5468 |
| 12300:4 | 8 | R.LAGKPTHVN[+1501.555]VSVVMAEVDGTC[+57.021]Y.- | HexNAc(5)Hex(3) | C[+57], N[+1502] | 962.9329 | 4 | 3848.7098 | 3848.6974 | 0 | 3.2 | 332 | CRagged | 645.28 | 630.9 | 630.9 | 12.25 | 58 | >sp\|P01876\|IGHA1_HUMAN Immunoglobulin heavy constant alpha 1 OS=Homo sapiens OX=9606 GN=IGHA1 PE=1 SV=2 | 255 | 20201118_lss_milk_1ng_glyco_hcd203040_compound15.15178.15178.4 | controllerType=0 controllerNumber=1 scan=15178 | 30.7176 |
| 13982:4 | 8 | R.LAGKPTHVN[+2262.814]VSVVMAEVDGTC[+57.021]Y.- | HexNAc(5)Hex(5)Fuc(1)NeuAc(1) | C[+57], N[+2263] | 1153.2475 | 4 | 4609.9681 | 4609.9564 | 0 | 2.5 | 332 | CRagged | 686.94 | 595.27 | 595.27 | 13.14 | 58 | >sp\|P01876\|IGHA1_HUMAN Immunoglobulin heavy constant alpha 1 OS=Homo sapiens OX=9606 GN=IGHA1 PE=1 SV=2 | 255 | 20201118_lss_milk_1ng_glyco_hcd203040_compound15.16928.16928.4 | controllerType=0 controllerNumber=1 scan=16928 | 33.686 |
| 15808:4 | 8 | R.LAGKPTHVN[+2553.910]VSVVMAEVDGTC[+57.021]Y.- | HexNAc(5)Hex(5)Fuc(1)NeuAc(2) | C[+57], N[+2554] | 1226.0223 | 4 | 4901.0672 | 4901.0518 | 0 | 3.1 | 332 | CRagged | 697.09 | 523.06 | 523.06 | 13.25 | 58 | >sp\|P01876\|IGHA1_HUMAN Immunoglobulin heavy constant alpha 1 OS=Homo sapiens OX=9606 GN=IGHA1 PE=1 SV=2 | 255 | 20201118_lss_milk_1ng_glyco_hcd203040_compound15.18823.18823.4 | controllerType=0 controllerNumber=1 scan=18823 | 37.3052 |
| 13988:3 | 8 | R.LAGKPTHVN[+2262.814]VSVVMAEVDGTC[+57.021]Y.- | HexNAc(5)Hex(5)Fuc(1)NeuAc(1) | C[+57], N[+2263] | 1537.3295 | 3 | 4609.974 | 4609.9564 | 0 | 3.8 | 332 | CRagged | 699.87 | 548.18 | 548.18 | 13.26 | 58 | >sp\|P01876\|IGHA1_HUMAN Immunoglobulin heavy constant alpha 1 OS=Homo sapiens OX=9606 GN=IGHA1 PE=1 SV=2 | 255 | 20201118_lss_milk_1ng_glyco_hcd203040_compound15.16935.16935.3 | controllerType=0 controllerNumber=1 scan=16935 | 33.7034 |
| 13985:3 | 8 | R.LAGKPTHVN[+2262.814]VSVVMAEVDGTC[+57.021]Y.- | HexNAc(5)Hex(5)Fuc(1)NeuAc(1) | C[+57], N[+2263] | 1537.3295 | 3 | 4609.974 | 4609.9564 | 0 | 3.8 | 332 | CRagged | 707.98 | 494.15 | 494.15 | 14.26 | 58 | >sp\|P01876\|IGHA1_HUMAN Immunoglobulin heavy constant alpha 1 OS=Homo sapiens OX=9606 GN=IGHA1 PE=1 SV=2 | 255 | 20201118_lss_milk_1ng_glyco_hcd203040_compound15.16932.16932.3 | controllerType=0 controllerNumber=1 scan=16932 | 33.6947 |
| 14111:3 | 8 | R.LAGKPTHVN[+1751.624]VSVVMAEVDGTC[+57.021]Y.- | HexNAc(4)Hex(4)NeuAc(1) | C[+57], N[+1752] | 1366.9319 | 3 | 4098.7813 | 4098.7663 | 0 | 3.7 | 332 | CRagged | 717.79 | 717.79 | 717.79 | 14.26 | 58 | >sp\|P01876\|IGHA1_HUMAN Immunoglobulin heavy constant alpha 1 OS=Homo sapiens OX=9606 GN=IGHA1 PE=1 SV=2 | 255 | 20201118_lss_milk_1ng_glyco_hcd203040_compound15.17062.17062.3 | controllerType=0 controllerNumber=1 scan=17062 | 33.9318 |
| 12229:3 | 8 | R.LAGKPTHVN[+1663.608]VSVVMAEVDGTC[+57.021]Y.- | HexNAc(5)Hex(4) | C[+57], N[+1664] | 1337.5908 | 3 | 4010.7577 | 4010.7502 | 0 | 1.9 | 332 | CRagged | 783.27 | 783.27 | 475.79 | 15.15 | 58 | >sp\|P01876\|IGHA1_HUMAN Immunoglobulin heavy constant alpha 1 OS=Homo sapiens OX=9606 GN=IGHA1 PE=1 SV=2 | 255 | 20201118_lss_milk_1ng_glyco_hcd203040_compound15.15104.15104.3 | controllerType=0 controllerNumber=1 scan=15104 | 30.5553 |
| 12316:3 | 8 | R.LAGKPTHVN[+1647.613]VSVVMAEVDGTC[+57.021]Y.- | HexNAc(5)Hex(3)Fuc(1) | C[+57], N[+1648] | 1332.2609 | 3 | 3994.7682 | 3994.7553 | 0 | 3.2 | 332 | CRagged | 796.67 | 652.02 | 652.02 | 15.26 | 58 | >sp\|P01876\|IGHA1_HUMAN Immunoglobulin heavy constant alpha 1 OS=Homo sapiens OX=9606 GN=IGHA1 PE=1 SV=2 | 255 | 20201118_lss_milk_1ng_glyco_hcd203040_compound15.15195.15195.3 | controllerType=0 controllerNumber=1 scan=15195 | 30.7531 |
| 12209:3 | 8 | R.LAGKPTHVN[+1809.666]VSVVMAEVDGTC[+57.021]Y.- | HexNAc(5)Hex(4)Fuc(1) | C[+57], N[+1810] | 1386.2768 | 3 | 4156.8158 | 4156.8081 | 0 | 1.8 | 332 | CRagged | 830.59 | 778.28 | 538 | 16.15 | 58 | >sp\|P01876\|IGHA1_HUMAN Immunoglobulin heavy constant alpha 1 OS=Homo sapiens OX=9606 GN=IGHA1 PE=1 SV=2 | 255 | 20201118_lss_milk_1ng_glyco_hcd203040_compound15.15083.15083.3 | controllerType=0 controllerNumber=1 scan=15083 | 30.5105 |
| 12206:3 | 8 | R.LAGKPTHVN[+1809.666]VSVVMAEVDGTC[+57.021]Y.- | HexNAc(5)Hex(4)Fuc(1) | C[+57], N[+1810] | 1386.2768 | 3 | 4156.8158 | 4156.8081 | 0 | 1.8 | 332 | CRagged | 855.43 | 744.64 | 536.5 | 17.15 | 58 | >sp\|P01876\|IGHA1_HUMAN Immunoglobulin heavy constant alpha 1 OS=Homo sapiens OX=9606 GN=IGHA1 PE=1 SV=2 | 255 | 20201118_lss_milk_1ng_glyco_hcd203040_compound15.15080.15080.3 | controllerType=0 controllerNumber=1 scan=15080 | 30.5042 |
| 12274:3 | 8 | R.LAGKPTHVN[+1501.555]VSVVMAEVDGTC[+57.021]Y.- | HexNAc(5)Hex(3) | C[+57], N[+1502] | 1283.575 | 3 | 3848.7105 | 3848.6974 | 0 | 3.4 | 332 | CRagged | 863.84 | 863.84 | 863.84 | 17.26 | 58 | >sp\|P01876\|IGHA1_HUMAN Immunoglobulin heavy constant alpha 1 OS=Homo sapiens OX=9606 GN=IGHA1 PE=1 SV=2 | 255 | 20201118_lss_milk_1ng_glyco_hcd203040_compound15.15151.15151.3 | controllerType=0 controllerNumber=1 scan=15151 | 30.6703 |
| 12249:3 | 8 | R.LAGKPTHVN[+1216.423]VSVVMAEVDGTC[+57.021]Y.- | HexNAc(2)Hex(5) | C[+57], N[+1216] | 1188.5303 | 3 | 3563.5764 | 3563.5649 | 0 | 3.2 | 332 | CRagged | 873.34 | 860.89 | 510.57 | 17.26 | 58 | >sp\|P01876\|IGHA1_HUMAN Immunoglobulin heavy constant alpha 1 OS=Homo sapiens OX=9606 GN=IGHA1 PE=1 SV=2 | 255 | 20201118_lss_milk_1ng_glyco_hcd203040_compound15.15125.15125.3 | controllerType=0 controllerNumber=1 scan=15125 | 30.6045 |
| 12247:3 | 8 | R.LAGKPTHVN[+1216.423]VSVVMAEVDGTC[+57.021]Y.- | HexNAc(2)Hex(5) | C[+57], N[+1216] | 1188.5303 | 3 | 3563.5764 | 3563.5649 | 0 | 3.2 | 332 | CRagged | 890.57 | 890.57 | 521.54 | 17.26 | 58 | >sp\|P01876\|IGHA1_HUMAN Immunoglobulin heavy constant alpha 1 OS=Homo sapiens OX=9606 GN=IGHA1 PE=1 SV=2 | 255 | 20201118_lss_milk_1ng_glyco_hcd203040_compound15.15123.15123.3 | controllerType=0 controllerNumber=1 scan=15123 | 30.5965 |
| 12271:3 | 8 | R.LAGKPTHVN[+1501.555]VSVVMAEVDGTC[+57.021]Y.- | HexNAc(5)Hex(3) | C[+57], N[+1502] | 1283.575 | 3 | 3848.7105 | 3848.6974 | 0 | 3.4 | 332 | CRagged | 899.31 | 899.31 | 899.31 | 17.26 | 58 | >sp\|P01876\|IGHA1_HUMAN Immunoglobulin heavy constant alpha 1 OS=Homo sapiens OX=9606 GN=IGHA1 PE=1 SV=2 | 255 | 20201118_lss_milk_1ng_glyco_hcd203040_compound15.15148.15148.3 | controllerType=0 controllerNumber=1 scan=15148 | 30.6584 |
| 12139:3 | 8 | R.LAGKPTHVN[+1702.581]VSVVMAEVDGTC[+57.021]Y.- | HexNAc(2)Hex(8) | C[+57], N[+1703] | 1350.5838 | 3 | 4049.7368 | 4049.7234 | 0 | 3.3 | 332 | CRagged | 906.1 | 833.96 | 536.7 | 18.26 | 58 | >sp\|P01876\|IGHA1_HUMAN Immunoglobulin heavy constant alpha 1 OS=Homo sapiens OX=9606 GN=IGHA1 PE=1 SV=2 | 255 | 20201118_lss_milk_1ng_glyco_hcd203040_compound15.15010.15010.3 | controllerType=0 controllerNumber=1 scan=15010 | 30.368 |
| 12180:3 | 8 | R.LAGKPTHVN[+1378.476]VSVVMAEVDGTC[+57.021]Y.- | HexNAc(2)Hex(6) | C[+57], N[+1378] | 1242.5466 | 3 | 3725.6251 | 3725.6178 | 0 | 2 | 332 | CRagged | 907.52 | 907.52 | 550.77 | 18.15 | 58 | >sp\|P01876\|IGHA1_HUMAN Immunoglobulin heavy constant alpha 1 OS=Homo sapiens OX=9606 GN=IGHA1 PE=1 SV=2 | 255 | 20201118_lss_milk_1ng_glyco_hcd203040_compound15.15053.15053.3 | controllerType=0 controllerNumber=1 scan=15053 | 30.4532 |

Table S4. Cont.

| Query #:z | Protein Rank | Glycopeptide | Glycan | Modification Type(s) | Observed m/z | z | Observed (M+H) | Calc. mass (M+H) | Off-by-x error | Mass error (ppm) | StartPosition | Cleavage | Score | Delta | Delta Mod | \|Log Prob\| | # of unique peptides | Protein Name | Protein DB number | Comment | Scan # | Scan Time |
|---|---|---|---|---|---|---|---|---|---|---|---|---|---|---|---|---|---|---|---|---|---|---|
| 12183:3 | 8 | R.LAGKPTHVN[+1378.476]VSVVMAEVDGTC[+57.021]Y.- | HexNAc(2)Hex(6) | C[+57], N[+1378] | 1242.5466 | 3 | 3725.6251 | 3725.6178 | 0 | 2 | 332 | CRagged | 908.45 | 884.79 | 543.24 | 18.15 | 58 | >sp\|P01876\|IGHA1_HUMAN Immunoglobulin heavy constant alpha 1 OS=Homo sapiens OX=9606 GN=IGHA1 PE=1 SV=2 | 255 | 20201118_lss_milk_1ng_glyco_hcd203040_compound15.15056.15056.3 | controllerType=0 controllerNumber=1 scan=15056 | 30.4623 |
| 12136:3 | 8 | R.LAGKPTHVN[+1702.581]VSVVMAEVDGTC[+57.021]Y.- | HexNAc(2)Hex(8) | C[+57], N[+1703] | 1350.5838 | 3 | 4049.7368 | 4049.7234 | 0 | 3.3 | 332 | CRagged | 972.7 | 972.7 | 625.03 | 19.26 | 58 | >sp\|P01876\|IGHA1_HUMAN Immunoglobulin heavy constant alpha 1 OS=Homo sapiens OX=9606 GN=IGHA1 PE=1 SV=2 | 255 | 20201118_lss_milk_1ng_glyco_hcd203040_compound15.15007.15007.3 | controllerType=0 controllerNumber=1 scan=15007 | 30.3599 |
| 10167:3 | 8 | R.LAGKPTHVN[+1216.423]VSVVM[+15.995]AEVDGTC[+57.021]Y.- | HexNAc(2)Hex(5) | C[+57], M[+16], N[+1216] | 1193.8626 | 3 | 3579.5733 | 3579.5599 | 0 | 3.7 | 332 | CRagged | 205.62 | 205.62 | 205.62 | 5.07 | 58 | >sp\|P01876\|IGHA1_HUMAN Immunoglobulin heavy constant alpha 1 OS=Homo sapiens OX=9606 GN=IGHA1 PE=1 SV=2 | 255 | 20201118_lss_milk_1ng_glyco_hcd203040_compound15.12969.12969.3 | controllerType=0 controllerNumber=1 scan=12969 | 26.8416 |
| 10168:3 | 8 | R.LAGKPTHVN[+1216.423]VSVVM[+15.995]AEVDGTC[+57.021]Y.- | HexNAc(2)Hex(5) | C[+57], M[+16], N[+1216] | 1193.8626 | 3 | 3579.5733 | 3579.5599 | 0 | 3.7 | 332 | CRagged | 338.32 | 338.32 | 338.32 | 7.47 | 58 | >sp\|P01876\|IGHA1_HUMAN Immunoglobulin heavy constant alpha 1 OS=Homo sapiens OX=9606 GN=IGHA1 PE=1 SV=2 | 255 | 20201118_lss_milk_1ng_glyco_hcd203040_compound15.12970.12970.3 | controllerType=0 controllerNumber=1 scan=12970 | 26.8448 |
| 17888:4 | 9 | K.RNIAAFGGDPNN[+1913.677]ITLFGESAGGASVSLQTLSPYNK.G | HexNAc(4)Hex(5)NeuAc(1) | N[+1914] | 1370.8783 | 4 | 5480.4914 | 5480.4644 | 0 | 4.9 | 196 | Specific | 988.74 | 927.17 | 927.17 | 19.47 | 56 | >sp\|P19835\|CEL_HUMAN Bile salt-activated lipase OS=Homo sapiens OX=9606 GN=CEL PE=1 SV=3 | 476 | 20201118_lss_milk_1ng_glyco_hcd203040_compound15.21005.21005.4 | controllerType=0 controllerNumber=1 scan=21005 | 41.4374 |
| 17891:4 | 9 | K.RNIAAFGGDPNN[+1913.677]ITLFGESAGGASVSLQTLSPYNK.G | HexNAc(4)Hex(5)NeuAc(1) | N[+1914] | 1370.8783 | 4 | 5480.4914 | 5480.4644 | 0 | 4.9 | 196 | Specific | 993.81 | 913.11 | 913.11 | 19.47 | 56 | >sp\|P19835\|CEL_HUMAN Bile salt-activated lipase OS=Homo sapiens OX=9606 GN=CEL PE=1 SV=3 | 476 | 20201118_lss_milk_1ng_glyco_hcd203040_compound15.21008.21008.4 | controllerType=0 controllerNumber=1 scan=21008 | 41.4439 |
| 19684:3 | 9 | R.NIAAFGGDPNN[+1913.677]ITLFGESAGGASVSLQTLSPYNK.G | HexNAc(4)Hex(5)NeuAc(1) | N[+1914] | 1775.4682 | 3 | 5324.3901 | 5324.3633 | 0 | 5 | 197 | Specific | 532.83 | 532.83 | 532.83 | 10 | 56 | >sp\|P19835\|CEL_HUMAN Bile salt-activated lipase OS=Homo sapiens OX=9606 GN=CEL PE=1 SV=3 | 476 | 20201118_lss_milk_1ng_glyco_hcd203040_compound15.22879.22879.3 | controllerType=0 controllerNumber=1 scan=22879 | 45.4595 |
| 19687:3 | 9 | R.NIAAFGGDPNN[+1913.677]ITLFGESAGGASVSLQTLSPYNK.G | HexNAc(4)Hex(5)NeuAc(1) | N[+1914] | 1775.4682 | 3 | 5324.3901 | 5324.3633 | 0 | 5 | 197 | Specific | 927.54 | 759.92 | 759.92 | 18.47 | 56 | >sp\|P19835\|CEL_HUMAN Bile salt-activated lipase OS=Homo sapiens OX=9606 GN=CEL PE=1 SV=3 | 476 | 20201118_lss_milk_1ng_glyco_hcd203040_compound15.22882.22882.3 | controllerType=0 controllerNumber=1 scan=22882 | 45.4634 |
| 09337:3 | 10 | K.TAHC[+57.021]N[+2059.735]ESFYFLC[+57.021]K.R | HexNAc(4)Hex(5)Fuc(1)NeuAc(1) | C[+57]*2, N[+2060] | 1246.1584 | 3 | 3736.4606 | 3736.4545 | 0 | 1.6 | 1201 | Specific | 468.41 | 404.39 | 404.39 | 8.75 | 67 | >sp\|P22897\|MRC1_HUMAN Macrophage mannose receptor 1 OS=Homo sapiens OX=9606 GN=MRC1 PE=1 SV=1 | 1049 | 20201118_lss_milk_1ng_glyco_hcd203040_compound15.12101.12101.3 | controllerType=0 controllerNumber=1 scan=12101 | 25.372 |
| 09339:3 | 10 | K.TAHC[+57.021]N[+2059.735]ESFYFLC[+57.021]K.R | HexNAc(4)Hex(5)Fuc(1)NeuAc(1) | C[+57]*2, N[+2060] | 1246.1584 | 3 | 3736.4606 | 3736.4545 | 0 | 1.6 | 1201 | Specific | 479.82 | 400.25 | 400.25 | 8.75 | 67 | >sp\|P22897\|MRC1_HUMAN Macrophage mannose receptor 1 OS=Homo sapiens OX=9606 GN=MRC1 PE=1 SV=1 | 1049 | 20201118_lss_milk_1ng_glyco_hcd203040_compound15.12103.12103.3 | controllerType=0 controllerNumber=1 scan=12103 | 25.3755 |
| 23376:4 | 12 | K.APDKNVIFSPLSISTALAFLSLGAHN[+2204.772]TTLTEILK.G | HexNAc(4)Hex(5)NeuAc(2) | N[+2205] | 1447.6974 | 4 | 5787.7677 | 5787.7506 | 0 | 3 | 68 | Specific | 221.99 | 221.99 | 221.99 | 5.29 | 36 | >sp\|P01011\|AACT_HUMAN Alpha-1-antichymotrypsin OS=Homo sapiens OX=9606 GN=SERPINA3 PE=1 SV=2 | 1314 | 20201118_lss_milk_1ng_glyco_hcd203040_compound15.26733.26733.4 | controllerType=0 controllerNumber=1 scan=26733 | 53.6696 |
| 25148:4 | 12 | K.APDKNVIFSPLSISTALAFLSLGAHN[+2861.000]TTLTEILK.G | HexNAc(5)Hex(6)NeuAc(3) | N[+2861] | 1611.7625 | 4 | 6444.028 | 6443.9782 | 0 | 7.7 | 68 | Specific | 262.52 | 262.52 | 262.52 | 5.47 | 36 | >sp\|P01011\|AACT_HUMAN Alpha-1-antichymotrypsin OS=Homo sapiens OX=9606 GN=SERPINA3 PE=1 SV=2 | 1314 | 20201118_lss_milk_1ng_glyco_hcd203040_compound15.28576.28576.4 | controllerType=0 controllerNumber=1 scan=28576 | 57.3206 |
| 23374:4 | 12 | K.APDKNVIFSPLSISTALAFLSLGAHN[+2204.772]TTLTEILK.G | HexNAc(4)Hex(5)NeuAc(2) | N[+2205] | 1447.6974 | 4 | 5787.7677 | 5787.7506 | 0 | 3 | 68 | Specific | 268.14 | 268.14 | 268.14 | 6.85 | 36 | >sp\|P01011\|AACT_HUMAN Alpha-1-antichymotrypsin OS=Homo sapiens OX=9606 GN=SERPINA3 PE=1 SV=2 | 1314 | 20201118_lss_milk_1ng_glyco_hcd203040_compound15.26731.26731.4 | controllerType=0 controllerNumber=1 scan=26731 | 53.6662 |
| 23303:5 | 12 | K.APDKNVIFSPLSISTALAFLSLGAHN[+2350.830]TTLTEILK.G | HexNAc(4)Hex(5)Fuc(1)NeuAc(2) | N[+2351] | 1187.5708 | 5 | 5933.8249 | 5933.8085 | 0 | 2.8 | 68 | Specific | 380.68 | 307.61 | 307.61 | 7.34 | 36 | >sp\|P01011\|AACT_HUMAN Alpha-1-antichymotrypsin OS=Homo sapiens OX=9606 GN=SERPINA3 PE=1 SV=2 | 1314 | 20201118_lss_milk_1ng_glyco_hcd203040_compound15.26657.26657.5 | controllerType=0 controllerNumber=1 scan=26657 | 53.5229 |
| 23340:4 | 12 | K.APDKNVIFSPLSISTALAFLSLGAHN[+2350.830]TTLTEILK.G | HexNAc(4)Hex(5)Fuc(1)NeuAc(2) | N[+2351] | 1484.215 | 4 | 5933.838 | 5933.8085 | 0 | 5 | 68 | Specific | 489.03 | 489.03 | 489.03 | 9.33 | 36 | >sp\|P01011\|AACT_HUMAN Alpha-1-antichymotrypsin OS=Homo sapiens OX=9606 GN=SERPINA3 PE=1 SV=2 | 1314 | 20201118_lss_milk_1ng_glyco_hcd203040_compound15.26696.26696.4 | controllerType=0 controllerNumber=1 scan=26696 | 53.5976 |
| 23301:5 | 12 | K.APDKNVIFSPLSISTALAFLSLGAHN[+2350.830]TTLTEILK.G | HexNAc(4)Hex(5)Fuc(1)NeuAc(2) | N[+2351] | 1187.5708 | 5 | 5933.8249 | 5933.8085 | 0 | 2.8 | 68 | Specific | 531.13 | 531.13 | 531.13 | 9.84 | 36 | >sp\|P01011\|AACT_HUMAN Alpha-1-antichymotrypsin OS=Homo sapiens OX=9606 GN=SERPINA3 PE=1 SV=2 | 1314 | 20201118_lss_milk_1ng_glyco_hcd203040_compound15.26655.26655.5 | controllerType=0 controllerNumber=1 scan=26655 | 53.517 |
| 23338:4 | 12 | K.APDKNVIFSPLSISTALAFLSLGAHN[+2350.830]TTLTEILK.G | HexNAc(4)Hex(5)Fuc(1)NeuAc(2) | N[+2351] | 1484.215 | 4 | 5933.838 | 5933.8085 | 0 | 5 | 68 | Specific | 687.25 | 687.25 | 687.25 | 13.09 | 36 | >sp\|P01011\|AACT_HUMAN Alpha-1-antichymotrypsin OS=Homo sapiens OX=9606 GN=SERPINA3 PE=1 SV=2 | 1314 | 20201118_lss_milk_1ng_glyco_hcd203040_compound15.26694.26694.4 | controllerType=0 controllerNumber=1 scan=26694 | 53.5913 |
| 23298:4 | 12 | K.APDKNVIFSPLSISTALAFLSLGAHN[+2350.830]TTLTEILK.G | HexNAc(4)Hex(5)Fuc(1)NeuAc(2) | N[+2351] | 1484.2134 | 4 | 5933.8317 | 5933.8085 | 0 | 3.9 | 68 | Specific | 775.98 | 722.9 | 722.9 | 15.43 | 36 | >sp\|P01011\|AACT_HUMAN Alpha-1-antichymotrypsin OS=Homo sapiens OX=9606 GN=SERPINA3 PE=1 SV=2 | 1314 | 20201118_lss_milk_1ng_glyco_hcd203040_compound15.26652.26652.4 | controllerType=0 controllerNumber=1 scan=26652 | 53.512 |
| 23296:4 | 12 | K.APDKNVIFSPLSISTALAFLSLGAHN[+2350.830]TTLTEILK.G | HexNAc(4)Hex(5)Fuc(1)NeuAc(2) | N[+2351] | 1484.2134 | 4 | 5933.8317 | 5933.8085 | 0 | 3.9 | 68 | Specific | 1003.48 | 935.47 | 935.47 | 20.43 | 36 | >sp\|P01011\|AACT_HUMAN Alpha-1-antichymotrypsin OS=Homo sapiens OX=9606 GN=SERPINA3 PE=1 SV=2 | 1314 | 20201118_lss_milk_1ng_glyco_hcd203040_compound15.26650.26650.4 | controllerType=0 controllerNumber=1 scan=26650 | 53.5069 |
| 08582:3 | 13 | R.LSPN[+2350.830]ASAEHLELR.W | HexNAc(4)Hex(5)Fuc(1)NeuAc(2) | N[+2351] | 1263.2001 | 3 | 3787.5858 | 3787.5796 | 0 | 1.7 | 52 | Specific | 554.95 | 410.63 | 410.63 | 9.31 | 37 | >sp\|Q13410\|BT1A1_HUMAN Butyrophilin subfamily 1 member A1 OS=Homo sapiens OX=9606 GN=BTN1A1 PE=1 SV=3 | 179 | 20201118_lss_milk_1ng_glyco_hcd203040_compound15.11316.11316.3 | controllerType=0 controllerNumber=1 scan=11316 | 24.086 |
| 08579:3 | 13 | R.LSPN[+2350.830]ASAEHLELR.W | HexNAc(4)Hex(5)Fuc(1)NeuAc(2) | N[+2351] | 1263.2001 | 3 | 3787.5858 | 3787.5796 | 0 | 1.7 | 52 | Specific | 557.95 | 350.83 | 350.83 | 7.87 | 37 | >sp\|Q13410\|BT1A1_HUMAN Butyrophilin subfamily 1 member A1 OS=Homo sapiens OX=9606 GN=BTN1A1 PE=1 SV=3 | 179 | 20201118_lss_milk_1ng_glyco_hcd203040_compound15.11313.11313.3 | controllerType=0 controllerNumber=1 scan=11313 | 24.0817 |
| 06480:3 | 13 | R.LSPN[+2205.793]ASAEHLELR.W | HexNAc(4)Hex(5)Fuc(2)NeuAc(1) | N[+2206] | 1214.8547 | 3 | 3642.5495 | 3642.5421 | 0 | 2 | 52 | Specific | 577.66 | 389.5 | 389.5 | 10.85 | 37 | >sp\|Q13410\|BT1A1_HUMAN Butyrophilin subfamily 1 member A1 OS=Homo sapiens OX=9606 GN=BTN1A1 PE=1 SV=3 | 179 | 20201118_lss_milk_1ng_glyco_hcd203040_compound15.9143.9143.3 | controllerType=0 controllerNumber=1 scan=9143 | 20.5193 |
| 06478:3 | 13 | R.LSPN[+2205.793]ASAEHLELR.W | HexNAc(4)Hex(5)Fuc(2)NeuAc(1) | N[+2206] | 1214.8547 | 3 | 3642.5495 | 3642.5421 | 0 | 2 | 52 | Specific | 592.55 | 345.53 | 345.53 | 9.42 | 37 | >sp\|Q13410\|BT1A1_HUMAN Butyrophilin subfamily 1 member A1 OS=Homo sapiens OX=9606 GN=BTN1A1 PE=1 SV=3 | 179 | 20201118_lss_milk_1ng_glyco_hcd203040_compound15.9141.9141.3 | controllerType=0 controllerNumber=1 scan=9141 | 20.5145 |
| 24445:3 | 15 | R.[+42.011]QLAHQSN[+2190.793]STNIFFSPVSIATAFAMLSLGTK.A | HexNAc(6)Hex(6) | .[42], N[+2191] | 1805.4819 | 3 | 5414.4311 | 5414.4388 | 0 | -1.4 | 64 | Specific | 201.74 | 201.74 | 201.74 | 4.2 | 38 | >sp\|P01009\|A1AT_HUMAN Alpha-1-antitrypsin OS=Homo sapiens OX=9606 GN=SERPINA1 PE=1 SV=3 | 642 | 20201118_lss_milk_1ng_glyco_hcd203040_compound15.27846.27846.3 | controllerType=0 controllerNumber=1 scan=27846 | 55.9376 |
| 24512:4 | 15 | R.[+42.011]QLAHQSN[+2190.793]STNIFFSPVSIATAFAMLSLGTK.A | HexNAc(6)Hex(6) | .[42], N[+2191] | 1354.361 | 4 | 5414.422 | 5414.4388 | 0 | -3.1 | 64 | Specific | 278.86 | 278.86 | 278.86 | 5.53 | 38 | >sp\|P01009\|A1AT_HUMAN Alpha-1-antitrypsin OS=Homo sapiens OX=9606 GN=SERPINA1 PE=1 SV=3 | 642 | 20201118_lss_milk_1ng_glyco_hcd203040_compound15.27916.27916.4 | controllerType=0 controllerNumber=1 scan=27916 | 56.0759 |
| 24452:5 | 15 | R.QLAHQSN[+2204.772]STNIFFSPVSIATAFAMLSLGTK.A | HexNAc(4)Hex(5)NeuAc(2) | N[+2205] | 1078.0911 | 5 | 5386.4262 | 5386.4075 | 0 | 3.5 | 64 | Specific | 323 | 323 | 323 | 7.7 | 38 | >sp\|P01009\|A1AT_HUMAN Alpha-1-antitrypsin OS=Homo sapiens OX=9606 GN=SERPINA1 PE=1 SV=3 | 642 | 20201118_lss_milk_1ng_glyco_hcd203040_compound15.27853.27853.5 | controllerType=0 controllerNumber=1 scan=27853 | 55.9499 |
| 24450:5 | 15 | R.QLAHQSN[+2204.772]STNIFFSPVSIATAFAMLSLGTK.A | HexNAc(4)Hex(5)NeuAc(2) | N[+2205] | 1078.0911 | 5 | 5386.4262 | 5386.4075 | 0 | 3.5 | 64 | Specific | 344.66 | 344.66 | 344.66 | 8.09 | 38 | >sp\|P01009\|A1AT_HUMAN Alpha-1-antitrypsin OS=Homo sapiens OX=9606 GN=SERPINA1 PE=1 SV=3 | 642 | 20201118_lss_milk_1ng_glyco_hcd203040_compound15.27851.27851.5 | controllerType=0 controllerNumber=1 scan=27851 | 55.9453 |
| 24514:4 | 15 | R.[+42.011]QLAHQSN[+2190.793]STNIFFSPVSIATAFAMLSLGTK.A | HexNAc(6)Hex(6) | .[42], N[+2191] | 1354.361 | 4 | 5414.422 | 5414.4388 | 0 | -3.1 | 64 | Specific | 348.28 | 339.34 | 339.34 | 6.3 | 38 | >sp\|P01009\|A1AT_HUMAN Alpha-1-antitrypsin OS=Homo sapiens OX=9606 GN=SERPINA1 PE=1 SV=3 | 642 | 20201118_lss_milk_1ng_glyco_hcd203040_compound15.27918.27918.4 | controllerType=0 controllerNumber=1 scan=27918 | 56.0805 |
| 24438:4 | 15 | R.[+42.011]QLAHQSN[+2190.793]STNIFFSPVSIATAFAMLSLGTK.A | HexNAc(6)Hex(6) | .[42], N[+2191] | 1354.361 | 4 | 5414.422 | 5414.4388 | 0 | -3.1 | 64 | Specific | 362.45 | 362.45 | 362.45 | 6.52 | 38 | >sp\|P01009\|A1AT_HUMAN Alpha-1-antitrypsin OS=Homo sapiens OX=9606 GN=SERPINA1 PE=1 SV=3 | 642 | 20201118_lss_milk_1ng_glyco_hcd203040_compound15.27839.27839.4 | controllerType=0 controllerNumber=1 scan=27839 | 55.9238 |
| 24440:4 | 15 | R.[+42.011]QLAHQSN[+2190.793]STNIFFSPVSIATAFAMLSLGTK.A | HexNAc(6)Hex(6) | .[42], N[+2191] | 1354.361 | 4 | 5414.422 | 5414.4388 | 0 | -3.1 | 64 | Specific | 382.31 | 365.83 | 365.83 | 6.52 | 38 | >sp\|P01009\|A1AT_HUMAN Alpha-1-antitrypsin OS=Homo sapiens OX=9606 GN=SERPINA1 PE=1 SV=3 | 642 | 20201118_lss_milk_1ng_glyco_hcd203040_compound15.27841.27841.4 | controllerType=0 controllerNumber=1 scan=27841 | 55.9289 |
| 24419:3 | 15 | R.QLAHQSN[+2204.772]STNIFFSPVSIATAFAMLSLGTK.A | HexNAc(4)Hex(5)NeuAc(2) | N[+2205] | 1796.1483 | 3 | 5386.4303 | 5386.4075 | 0 | 4.2 | 64 | Specific | 840.39 | 785.08 | 785.08 | 16.56 | 38 | >sp\|P01009\|A1AT_HUMAN Alpha-1-antitrypsin OS=Homo sapiens OX=9606 GN=SERPINA1 PE=1 SV=3 | 642 | 20201118_lss_milk_1ng_glyco_hcd203040_compound15.27819.27819.3 | controllerType=0 controllerNumber=1 scan=27819 | 55.8827 |
| 24416:4 | 15 | R.QLAHQSN[+2204.772]STNIFFSPVSIATAFAMLSLGTK.A | HexNAc(4)Hex(5)NeuAc(2) | N[+2205] | 1347.3624 | 4 | 5386.4279 | 5386.4075 | 0 | 3.8 | 64 | Specific | 920.93 | 893.87 | 893.87 | 18.55 | 38 | >sp\|P01009\|A1AT_HUMAN Alpha-1-antitrypsin OS=Homo sapiens OX=9606 GN=SERPINA1 PE=1 SV=3 | 642 | 20201118_lss_milk_1ng_glyco_hcd203040_compound15.27816.27816.4 | controllerType=0 controllerNumber=1 scan=27816 | 55.8766 |
| 24414:4 | 15 | R.QLAHQSN[+2204.772]STNIFFSPVSIATAFAMLSLGTK.A | HexNAc(4)Hex(5)NeuAc(2) | N[+2205] | 1347.3624 | 4 | 5386.4279 | 5386.4075 | 0 | 3.8 | 64 | Specific | 935.5 | 896.46 | 896.46 | 18.55 | 38 | >sp\|P01009\|A1AT_HUMAN Alpha-1-antitrypsin OS=Homo sapiens OX=9606 GN=SERPINA1 PE=1 SV=3 | 642 | 20201118_lss_milk_1ng_glyco_hcd203040_compound15.27814.27814.4 | controllerType=0 controllerNumber=1 scan=27814 | 55.8698 |
| 24417:3 | 15 | R.QLAHQSN[+2204.772]STNIFFSPVSIATAFAMLSLGTK.A | HexNAc(4)Hex(5)NeuAc(2) | N[+2205] | 1796.1483 | 3 | 5386.4303 | 5386.4075 | 0 | 4.2 | 64 | Specific | 952.04 | 900.63 | 900.63 | 19.56 | 38 | >sp\|P01009\|A1AT_HUMAN Alpha-1-antitrypsin OS=Homo sapiens OX=9606 GN=SERPINA1 PE=1 SV=3 | 642 | 20201118_lss_milk_1ng_glyco_hcd203040_compound15.27817.27817.3 | controllerType=0 controllerNumber=1 scan=27817 | 55.8779 |
| 23133:5 | 15 | K.ADTHDEILEGLNFN[+2861.000]LTEIPEAQIHEGFQELLR.T | HexNAc(5)Hex(6)NeuAc(3) | N[+2861] | 1311.3754 | 5 | 6552.8477 | 6552.8239 | 0 | 3.6 | 94 | Specific | 693.26 | 693.26 | 693.26 | 13.55 | 38 | >sp\|P01009\|A1AT_HUMAN Alpha-1-antitrypsin OS=Homo sapiens OX=9606 GN=SERPINA1 PE=1 SV=3 | 642 | 20201118_lss_milk_1ng_glyco_hcd203040_compound15.26480.26480.5 | controllerType=0 controllerNumber=1 scan=26480 | 53.1521 |
| 23146:5 | 15 | K.ADTHDEILEGLNFN[+2861.000]LTEIPEAQIHEGFQELLR.T | HexNAc(5)Hex(6)NeuAc(3) | N[+2861] | 1311.3756 | 5 | 6552.849 | 6552.8239 | 0 | 3.8 | 94 | Specific | 864.48 | 712.11 | 712.11 | 17.55 | 38 | >sp\|P01009\|A1AT_HUMAN Alpha-1-antitrypsin OS=Homo sapiens OX=9606 GN=SERPINA1 PE=1 SV=3 | 642 | 20201118_lss_milk_1ng_glyco_hcd203040_compound15.26494.26494.5 | controllerType=0 controllerNumber=1 scan=26494 | 53.1837 |
| 23148:5 | 15 | K.ADTHDEILEGLNFN[+2861.000]LTEIPEAQIHEGFQELLR.T | HexNAc(5)Hex(6)NeuAc(3) | N[+2861] | 1311.3756 | 5 | 6552.849 | 6552.8239 | 0 | 3.8 | 94 | Specific | 895.89 | 723.43 | 723.43 | 17.55 | 38 | >sp\|P01009\|A1AT_HUMAN Alpha-1-antitrypsin OS=Homo sapiens OX=9606 GN=SERPINA1 PE=1 SV=3 | 642 | 20201118_lss_milk_1ng_glyco_hcd203040_compound15.26496.26496.5 | controllerType=0 controllerNumber=1 scan=26496 | 53.1883 |
| 23135:5 | 15 | K.ADTHDEILEGLNFN[+2861.000]LTEIPEAQIHEGFQELLR.T | HexNAc(5)Hex(6)NeuAc(3) | N[+2861] | 1311.3754 | 5 | 6552.8477 | 6552.8239 | 0 | 3.6 | 94 | Specific | 937.08 | 826.37 | 826.37 | 18.55 | 38 | >sp\|P01009\|A1AT_HUMAN Alpha-1-antitrypsin OS=Homo sapiens OX=9606 GN=SERPINA1 PE=1 SV=3 | 642 | 20201118_lss_milk_1ng_glyco_hcd203040_compound15.26482.26482.5 | controllerType=0 controllerNumber=1 scan=26482 | 53.1599 |
| 20688:5 | 15 | K.YLGN[+2204.772]ATAIFFLPDEGKLQHLENELTHDIITK.F | HexNAc(4)Hex(5)NeuAc(2) | N[+2205] | 1149.9318 | 5 | 5745.6297 | 5745.6098 | 0 | 3.5 | 268 | Specific | 897.18 | 760.5 | 760.5 | 17.17 | 38 | >sp\|P01009\|A1AT_HUMAN Alpha-1-antitrypsin OS=Homo sapiens OX=9606 GN=SERPINA1 PE=1 SV=3 | 642 | 20201118_lss_milk_1ng_glyco_hcd203040_compound15.23926.23926.5 | controllerType=0 controllerNumber=1 scan=23926 | 47.6802 |
| 20690:5 | 15 | K.YLGN[+2204.772]ATAIFFLPDEGKLQHLENELTHDIITK.F | HexNAc(4)Hex(5)NeuAc(2) | N[+2205] | 1149.9318 | 5 | 5745.6297 | 5745.6098 | 0 | 3.5 | 268 | Specific | 912.8 | 844.21 | 844.21 | 18.17 | 38 | >sp\|P01009\|A1AT_HUMAN Alpha-1-antitrypsin OS=Homo sapiens OX=9606 GN=SERPINA1 PE=1 SV=3 | 642 | 20201118_lss_milk_1ng_glyco_hcd203040_compound15.23928.23928.5 | controllerType=0 controllerNumber=1 scan=23928 | 47.685 |
| 20689:4 | 15 | K.YLGN[+2204.772]ATAIFFLPDEGKLQHLENELTHDIITK.F | HexNAc(4)Hex(5)NeuAc(2) | N[+2205] | 1437.1637 | 4 | 5745.633 | 5745.6098 | 0 | 4 | 268 | Specific | 931.75 | 885.53 | 885.53 | 17.65 | 38 | >sp\|P01009\|A1AT_HUMAN Alpha-1-antitrypsin OS=Homo sapiens OX=9606 GN=SERPINA1 PE=1 SV=3 | 642 | 20201118_lss_milk_1ng_glyco_hcd203040_compound15.23927.23927.4 | controllerType=0 controllerNumber=1 scan=23927 | 47.6832 |
| 20687:4 | 15 | K.YLGN[+2204.772]ATAIFFLPDEGKLQHLENELTHDIITK.F | HexNAc(4)Hex(5)NeuAc(2) | N[+2205] | 1437.1637 | 4 | 5745.633 | 5745.6098 | 0 | 4 | 268 | Specific | 1223.08 | 1097.37 | 1097.37 | 23.65 | 38 | >sp\|P01009\|A1AT_HUMAN Alpha-1-antitrypsin OS=Homo sapiens OX=9606 GN=SERPINA1 PE=1 SV=3 | 642 | 20201118_lss_milk_1ng_glyco_hcd203040_compound15.23925.23925.4 | controllerType=0 controllerNumber=1 scan=23925 | 47.6785 |
| 14832:3 | 16 | K.LKELPGVC[+57.021]N[+2059.735]ETMMALWEEC[+57.021]KPC[+57.021]LK.Q | HexNAc(4)Hex(5)Fuc(1)NeuAc(1) | C[+57]*3, N[+2060] | 1666.0459 | 3 | 4996.123 | 4996.1196 | 0 | 0.7 | 95 | Specific | 488.77 | 439.74 | 439.74 | 7.8 | 35 | >sp\|P10909\|CLUS_HUMAN Clusterin OS=Homo sapiens OX=9606 GN=CLU PE=1 SV=1 | 1467 | 20201118_lss_milk_1ng_glyco_hcd203040_compound15.17811.17811.3 | controllerType=0 controllerNumber=1 scan=17811 | 35.4123 |
| 14830:3 | 16 | K.LKELPGVC[+57.021]N[+2059.735]ETMMALWEEC[+57.021]KPC[+57.021]LK.Q | HexNAc(4)Hex(5)Fuc(1)NeuAc(1) | C[+57]*3, N[+2060] | 1666.0459 | 3 | 4996.123 | 4996.1196 | 0 | 0.7 | 95 | Specific | 520.95 | 407.41 | 407.41 | 7.86 | 35 | >sp\|P10909\|CLUS_HUMAN Clusterin OS=Homo sapiens OX=9606 GN=CLU PE=1 SV=1 | 1467 | 20201118_lss_milk_1ng_glyco_hcd203040_compound15.17809.17809.3 | controllerType=0 controllerNumber=1 scan=17809 | 35.4088 |
| 14777:4 | 16 | K.LKELPGVC[+57.021]N[+2205.793]ETMMALWEEC[+57.021]KPC[+57.021]LK.Q | HexNAc(4)Hex(5)Fuc(2)NeuAc(1) | C[+57]*3, N[+2206] | 1286.3055 | 4 | 5142.2004 | 5142.1776 | 0 | 4.4 | 95 | Specific | 642.96 | 642.96 | 642.96 | 11.75 | 35 | >sp\|P10909\|CLUS_HUMAN Clusterin OS=Homo sapiens OX=9606 GN=CLU PE=1 SV=1 | 1467 | 20201118_lss_milk_1ng_glyco_hcd203040_compound15.17754.17754.4 | controllerType=0 controllerNumber=1 scan=17754 | 35.2944 |
| 14799:4 | 16 | K.LKELPGVC[+57.021]N[+2059.735]ETMMALWEEC[+57.021]KPC[+57.021]LK.Q | HexNAc(4)Hex(5)Fuc(1)NeuAc(1) | C[+57]*3, N[+2060] | 1249.7894 | 4 | 4996.1359 | 4996.1196 | 0 | 3.3 | 95 | Specific | 887.06 | 766.64 | 766.64 | 17.09 | 35 | >sp\|P10909\|CLUS_HUMAN Clusterin OS=Homo sapiens OX=9606 GN=CLU PE=1 SV=1 | 1467 | 20201118_lss_milk_1ng_glyco_hcd203040_compound15.17777.17777.4 | controllerType=0 controllerNumber=1 scan=17777 | 35.3484 |
| 14780:4 | 16 | K.LKELPGVC[+57.021]N[+2205.793]ETMMALWEEC[+57.021]KPC[+57.021]LK.Q | HexNAc(4)Hex(5)Fuc(2)NeuAc(1) | C[+57]*3, N[+2206] | 1286.3055 | 4 | 5142.2004 | 5142.1776 | 0 | 4.4 | 95 | Specific | 893.1 | 826.12 | 826.12 | 16.75 | 35 | >sp\|P10909\|CLUS_HUMAN Clusterin OS=Homo sapiens OX=9606 GN=CLU PE=1 SV=1 | 1467 | 20201118_lss_milk_1ng_glyco_hcd203040_compound15.17757.17757.4 | controllerType=0 controllerNumber=1 scan=17757 | 35.299 |
| 14797:4 | 16 | K.LKELPGVC[+57.021]N[+2059.735]ETMMALWEEC[+57.021]KPC[+57.021]LK.Q | HexNAc(4)Hex(5)Fuc(1)NeuAc(1) | C[+57]*3, N[+2060] | 1249.7894 | 4 | 4996.1359 | 4996.1196 | 0 | 3.3 | 95 | Specific | 968.04 | 900.96 | 900.96 | 19.09 | 35 | >sp\|P10909\|CLUS_HUMAN Clusterin OS=Homo sapiens OX=9606 GN=CLU PE=1 SV=1 | 1467 | 20201118_lss_milk_1ng_glyco_hcd203040_compound15.17775.17775.4 | controllerType=0 controllerNumber=1 scan=17775 | 35.3433 |
| 23830:4 | 16 | K.[+42.011]MLN[+2336.851]TSSLLEQLNEQFNWVSR.L | HexNAc(6)Hex(6)Fuc(1) | .[42], N[+2337] | 1197.764 | 4 | 4788.034 | 4788.0484 | 0 | -3 | 352 | Specific | 276.65 | 264.36 | 264.36 | 5.46 | 35 | >sp\|P10909\|CLUS_HUMAN Clusterin OS=Homo sapiens OX=9606 GN=CLU PE=1 SV=1 | 1467 | 20201118_lss_milk_1ng_glyco_hcd203040_compound15.27206.27206.4 | controllerType=0 controllerNumber=1 scan=27206 | 54.6632 |
| 23689:4 | 16 | K.MLN[+2350.830]TSSLLEQLNEQFNWVSR.L | HexNAc(4)Hex(5)Fuc(1)NeuAc(2) | N[+2351] | 1190.7648 | 4 | 4760.0374 | 4760.0171 | 0 | 4.3 | 352 | Specific | 570.62 | 441.64 | 441.64 | 9.86 | 35 | >sp\|P10909\|CLUS_HUMAN Clusterin OS=Homo sapiens OX=9606 GN=CLU PE=1 SV=1 | 1467 | 20201118_lss_milk_1ng_glyco_hcd203040_compound15.27059.27059.4 | controllerType=0 controllerNumber=1 scan=27059 | 54.3385 |
| 21598:4 | 16 | K.MLN[+2059.735]TSSLLEQLNEQFNWVSR.L | HexNAc(4)Hex(5)Fuc(1)NeuAc(1) | N[+2060] | 1117.9877 | 4 | 4468.929 | 4468.9217 | 0 | 1.6 | 352 | Specific | 613.44 | 543.67 | 543.67 | 12.1 | 35 | >sp\|P10909\|CLUS_HUMAN Clusterin OS=Homo sapiens OX=9606 GN=CLU PE=1 SV=1 | 1467 | 20201118_lss_milk_1ng_glyco_hcd203040_compound15.24880.24880.4 | controllerType=0 controllerNumber=1 scan=24880 | 49.8598 |
| 23691:4 | 16 | K.MLN[+2350.830]TSSLLEQLNEQFNWVSR.L | HexNAc(4)Hex(5)Fuc(1)NeuAc(2) | N[+2351] | 1190.7648 | 4 | 4760.0374 | 4760.0171 | 0 | 4.3 | 352 | Specific | 718.19 | 634.49 | 634.49 | 14.13 | 35 | >sp\|P10909\|CLUS_HUMAN Clusterin OS=Homo sapiens OX=9606 GN=CLU PE=1 SV=1 | 1467 | 20201118_lss_milk_1ng_glyco_hcd203040_compound15.27061.27061.4 | controllerType=0 controllerNumber=1 scan=27061 | 54.3439 |
| 23770:4 | 16 | K.MLN[+2350.830]TSSLLEQLNEQFNWVSR.L | HexNAc(4)Hex(5)Fuc(1)NeuAc(2) | N[+2351] | 1190.7647 | 4 | 4760.0369 | 4760.0171 | 0 | 4.2 | 352 | Specific | 723.74 | 501.22 | 501.22 | 14.13 | 35 | >sp\|P10909\|CLUS_HUMAN Clusterin OS=Homo sapiens OX=9606 GN=CLU PE=1 SV=1 | 1467 | 20201118_lss_milk_1ng_glyco_hcd203040_compound15.27144.27144.4 | controllerType=0 controllerNumber=1 scan=27144 | 54.5235 |
| 23772:4 | 16 | K.MLN[+2350.830]TSSLLEQLNEQFNWVSR.L | HexNAc(4)Hex(5)Fuc(1)NeuAc(2) | N[+2351] | 1190.7647 | 4 | 4760.0369 | 4760.0171 | 0 | 4.2 | 352 | Specific | 757.26 | 509.25 | 509.25 | 15.13 | 35 | >sp\|P10909\|CLUS_HUMAN Clusterin OS=Homo sapiens OX=9606 GN=CLU PE=1 SV=1 | 1467 | 20201118_lss_milk_1ng_glyco_hcd203040_compound15.27146.27146.4 | controllerType=0 controllerNumber=1 scan=27146 | 54.5283 |
| 21600:4 | 16 | K.MLN[+2059.735]TSSLLEQLNEQFNWVSR.L | HexNAc(4)Hex(5)Fuc(1)NeuAc(1) | N[+2060] | 1117.9877 | 4 | 4468.929 | 4468.9217 | 0 | 1.6 | 352 | Specific | 793.42 | 617.3 | 617.3 | 15.1 | 35 | >sp\|P10909\|CLUS_HUMAN Clusterin OS=Homo sapiens OX=9606 GN=CLU PE=1 SV=1 | 1467 | 20201118_lss_milk_1ng_glyco_hcd203040_compound15.24882.24882.4 | controllerType=0 controllerNumber=1 scan=24882 | 49.8636 |
| 21816:3 | 16 | K.MLN[+1913.677]TSSLLEQLNEQFNWVSR.L | HexNAc(4)Hex(5)NeuAc(1) | N[+1914] | 1441.6301 | 3 | 4322.8756 | 4322.8637 | 0 | 2.7 | 352 | Specific | 848.03 | 675.69 | 675.69 | 16.14 | 35 | >sp\|P10909\|CLUS_HUMAN Clusterin OS=Homo sapiens OX=9606 GN=CLU PE=1 SV=1 | 1467 | 20201118_lss_milk_1ng_glyco_hcd203040_compound15.25107.25107.3 | controllerType=0 controllerNumber=1 scan=25107 | 50.342 |
| 23716:3 | 16 | K.MLN[+2350.830]TSSLLEQLNEQFNWVSR.L | HexNAc(4)Hex(5)Fuc(1)NeuAc(2) | N[+2351] | 1587.3525 | 3 | 4760.0429 | 4760.0171 | 0 | 5.4 | 352 | Specific | 918.55 | 658.63 | 658.63 | 17.51 | 35 | >sp\|P10909\|CLUS_HUMAN Clusterin OS=Homo sapiens OX=9606 GN=CLU PE=1 SV=1 | 1467 | 20201118_lss_milk_1ng_glyco_hcd203040_compound15.27087.27087.3 | controllerType=0 controllerNumber=1 scan=27087 | 54.4019 |
| 23714:3 | 16 | K.MLN[+2350.830]TSSLLEQLNEQFNWVSR.L | HexNAc(4)Hex(5)Fuc(1)NeuAc(2) | N[+2351] | 1587.3525 | 3 | 4760.0429 | 4760.0171 | 0 | 5.4 | 352 | Specific | 1038.94 | 803.35 | 803.35 | 19.51 | 35 | >sp\|P10909\|CLUS_HUMAN Clusterin OS=Homo sapiens OX=9606 GN=CLU PE=1 SV=1 | 1467 | 20201118_lss_milk_1ng_glyco_hcd203040_compound15.27085.27085.3 | controllerType=0 controllerNumber=1 scan=27085 | 54.397 |
| 11239:3 | 16 | R.LAN[+2205.793]LTQGEDQYYLR.V | HexNAc(4)Hex(5)Fuc(2)NeuAc(1) | N[+2206] | 1297.2145 | 3 | 3889.6291 | 3889.6265 | 0 | 0.7 | 372 | Specific | 246.9 | 186.99 | 186.99 | 4.57 | 35 | >sp\|P10909\|CLUS_HUMAN Clusterin OS=Homo sapiens OX=9606 GN=CLU PE=1 SV=1 | 1467 | 20201118_lss_milk_1ng_glyco_hcd203040_compound15.14079.14079.3 | controllerType=0 controllerNumber=1 scan=14079 | 28.76 |
| 11214:3 | 16 | R.LAN[+2205.793]LTQGEDQYYLR.V | HexNAc(4)Hex(5)Fuc(2)NeuAc(1) | N[+2206] | 1297.2162 | 3 | 3889.6342 | 3889.6265 | 0 | 2 | 372 | Specific | 337.18 | 295.53 | 295.53 | 6.8 | 35 | >sp\|P10909\|CLUS_HUMAN Clusterin OS=Homo sapiens OX=9606 GN=CLU PE=1 SV=1 | 1467 | 20201118_lss_milk_1ng_glyco_hcd203040_compound15.14053.14053.3 | controllerType=0 controllerNumber=1 scan=14053 | 28.7068 |
| 13888:3 | 16 | R.LAN[+2350.830]LTQGEDQYYLR.V | HexNAc(4)Hex(5)Fuc(1)NeuAc(2) | N[+2351] | 1345.5624 | 3 | 4034.6728 | 4034.664 | 0 | 2.2 | 372 | Specific | 455.56 | 436.27 | 436.27 | 8.95 | 35 | >sp\|P10909\|CLUS_HUMAN Clusterin OS=Homo sapiens OX=9606 GN=CLU PE=1 SV=1 | 1467 | 20201118_lss_milk_1ng_glyco_hcd203040_compound15.16831.16831.3 | controllerType=0 controllerNumber=1 scan=16831 | 33.5112 |
| 11217:3 | 16 | R.LAN[+2205.793]LTQGEDQYYLR.V | HexNAc(4)Hex(5)Fuc(2)NeuAc(1) | N[+2206] | 1297.2162 | 3 | 3889.6342 | 3889.6265 | 0 | 2 | 372 | Specific | 520.96 | 324.85 | 324.85 | 7.42 | 35 | >sp\|P10909\|CLUS_HUMAN Clusterin OS=Homo sapiens OX=9606 GN=CLU PE=1 SV=1 | 1467 | 20201118_lss_milk_1ng_glyco_hcd203040_compound15.14056.14056.3 | controllerType=0 controllerNumber=1 scan=14056 | 28.7105 |
| 13891:3 | 16 | R.LAN[+2350.830]LTQGEDQYYLR.V | HexNAc(4)Hex(5)Fuc(1)NeuAc(2) | N[+2351] | 1345.5624 | 3 | 4034.6728 | 4034.664 | 0 | 2.2 | 372 | Specific | 608 | 496.69 | 496.69 | 11.84 | 35 | >sp\|P10909\|CLUS_HUMAN Clusterin OS=Homo sapiens OX=9606 GN=CLU PE=1 SV=1 | 1467 | 20201118_lss_milk_1ng_glyco_hcd203040_compound15.16834.16834.3 | controllerType=0 controllerNumber=1 scan=16834 | 33.5156 |
| 24067:6 | 17 | K.VFPLSLDSTPQDGNVVVAC[+57.021]LVQGFFPQEPLSVTWSESGQN[+2263.835]VTAI | HexNAc(5)Hex(5)Fuc(3) | C[+57], N[+2264] | 1174.2044 | 6 | 7040.19 | 7040.1961 | 0 | -0.9 | 8 | Specific | 268.01 | 266.83 | 266.83 | 5.02 | 39 | >sp\|P01877\|IGHA2_HUMAN Immunoglobulin heavy constant alpha 2 OS=Homo sapiens OX=9606 GN=IGHA2 PE=1 SV=4 | 576 | 20201118_lss_milk_1ng_glyco_hcd203040_compound15.27452.27452.6 | controllerType=0 controllerNumber=1 scan=27452 | [illegible] |
| 22812:4 | 17 | K.[+42.011]VFPLSLDSTPQDGNVVVAC[+57.021]LVQGFFPQEPLSVTWSESGQN[+1606 | HexNAc(4)Hex(4)Fuc(1) | .[42], C[+57], N[+1607] | 1607.0022 | 4 | 6424.987 | 6424.9587 | 0 | 4.4 | 8 | Specific | 269.81 | 269.81 | 269.81 | 5.86 | 39 | >sp\|P01877\|IGHA2_HUMAN Immunoglobulin heavy constant alpha 2 OS=Homo sapiens OX=9606 GN=IGHA2 PE=1 SV=4 | 576 | 20201118_lss_milk_1ng_glyco_hcd203040_compound15.26147.26147.4 | controllerType=0 controllerNumber=1 scan=26147 | [illegible] |
| 22814:4 | 17 | K.[+42.011]VFPLSLDSTPQDGNVVVAC[+57.021]LVQGFFPQEPLSVTWSESGQN[+1606 | HexNAc(4)Hex(4)Fuc(1) | .[42], C[+57], N[+1607] | 1607.0022 | 4 | 6424.987 | 6424.9587 | 0 | 4.4 | 8 | Specific | 273.78 | 254.21 | 254.21 | 5.64 | 39 | >sp\|P01877\|IGHA2_HUMAN Immunoglobulin heavy constant alpha 2 OS=Homo sapiens OX=9606 GN=IGHA2 PE=1 SV=4 | 576 | 20201118_lss_milk_1ng_glyco_hcd203040_compound15.26149.26149.4 | controllerType=0 controllerNumber=1 scan=26149 | [illegible] |
| 24175:4 | 17 | K.VFPLSLDSTPQDGNVVVAC[+57.021]LVQGFFPQEPLSVTWSESGQN[+2262.814]VTAI | HexNAc(5)Hex(5)Fuc(1)NeuAc(1) | C[+57], N[+2263] | 1760.5532 | 4 | 7039.1909 | 7039.1757 | 0 | 2.2 | 8 | Specific | 399.08 | 356.72 | 356.72 | 8.63 | 39 | >sp\|P01877\|IGHA2_HUMAN Immunoglobulin heavy constant alpha 2 OS=Homo sapiens OX=9606 GN=IGHA2 PE=1 SV=4 | 576 | 20201118_lss_milk_1ng_glyco_hcd203040_compound15.27565.27565.4 | controllerType=0 controllerNumber=1 scan=27565 | [illegible] |
| 22492:4 | 17 | K.VFPLSLDSTPQDGNVVVAC[+57.021]LVQGFFPQEPLSVTWSESGQN[+1809.666]VTAI | HexNAc(5)Hex(4)Fuc(1) | C[+57], N[+1810] | 1647.2676 | 4 | 6586.0485 | 6586.0275 | 0 | 3.2 | 8 | Specific | 595.64 | 553.35 | 553.35 | 12.15 | 39 | >sp\|P01877\|IGHA2_HUMAN Immunoglobulin heavy constant alpha 2 OS=Homo sapiens OX=9606 GN=IGHA2 PE=1 SV=4 | 576 | 20201118_lss_milk_1ng_glyco_hcd203040_compound15.25812.25812.2 | controllerType=0 controllerNumber=1 scan=25812 | 51.839 |
| 22660:4 | 17 | K.VFPLSLDSTPQDGNVVVAC[+57.021]LVQGFFPQEPLSVTWSESGQN[+1647.613]VTAI | HexNAc(5)Hex(3)Fuc(1) | C[+57], N[+1648] | 1606.7557 | 4 | 6424.0011 | 6423.9746 | 0 | 4.1 | 8 | Specific | 818.66 | 743.92 | 743.92 | 17.15 | 39 | >sp\|P01877\|IGHA2_HUMAN Immunoglobulin heavy constant alpha 2 OS=Homo sapiens OX=9606 GN=IGHA2 PE=1 SV=4 | 576 | 20201118_lss_milk_1ng_glyco_hcd203040_compound15.25988.25988.4 | controllerType=0 controllerNumber=1 scan=25988 | 52.1684 |
| 22662:4 | 17 | K.VFPLSLDSTPQDGNVVVAC[+57.021]LVQGFFPQEPLSVTWSESGQN[+1647.613]VTAI | HexNAc(5)Hex(3)Fuc(1) | C[+57], N[+1648] | 1606.7557 | 4 | 6424.0011 | 6423.9746 | 0 | 4.1 | 8 | Specific | 830.24 | 802.24 | 802.24 | 17.15 | 39 | >sp\|P01877\|IGHA2_HUMAN Immunoglobulin heavy constant alpha 2 OS=Homo sapiens OX=9606 GN=IGHA2 PE=1 SV=4 | 576 | 20201118_lss_milk_1ng_glyco_hcd203040_compound15.25990.25990.4 | controllerType=0 controllerNumber=1 scan=25990 | 52.1754 |
| 22494:4 | 17 | K.VFPLSLDSTPQDGNVVVAC[+57.021]LVQGFFPQEPLSVTWSESGQN[+1809.666]VTAI | HexNAc(5)Hex(4)Fuc(1) | C[+57], N[+1810] | 1647.2676 | 4 | 6586.0485 | 6586.0275 | 0 | 3.2 | 8 | Specific | 880.61 | 769.37 | 769.37 | 18.15 | 39 | >sp\|P01877\|IGHA2_HUMAN Immunoglobulin heavy constant alpha 2 OS=Homo sapiens OX=9606 GN=IGHA2 PE=1 SV=4 | 576 | 20201118_lss_milk_1ng_glyco_hcd203040_compound15.25814.25814.2 | controllerType=0 controllerNumber=1 scan=25814 | 51.8464 |
| 02941:3 | 17 | K.HYTN[+2262.814]SSQDVTVPC[+57.021]R.V | HexNAc(5)Hex(5)Fuc(1)NeuAc(1) | C[+57], N[+2263] | 1309.5303 | 3 | 3926.5764 | 3926.5636 | 0 | 3.3 | 89 | Specific | 201.68 | 140.71 | 140.71 | 3.83 | 39 | >sp\|P01877\|IGHA2_HUMAN Immunoglobulin heavy constant alpha 2 OS=Homo sapiens OX=9606 GN=IGHA2 PE=1 SV=4 | 576 | 20201118_lss_milk_1ng_glyco_hcd203040_compound15.5483.5483.3 | controllerType=0 controllerNumber=1 scan=5483 | 14.5044 |
| 02212:3 | 17 | K.HYTN[+1647.613]SSQDVTVPC[+57.021]R.V | HexNAc(5)Hex(3)Fuc(1) | C[+57], N[+1648] | 1104.4617 | 3 | 3311.3705 | 3311.3625 | 0 | 2.4 | 89 | Specific | 221.42 | 169.56 | 169.56 | 4.28 | 39 | >sp\|P01877\|IGHA2_HUMAN Immunoglobulin heavy constant alpha 2 OS=Homo sapiens OX=9606 GN=IGHA2 PE=1 SV=4 | 576 | 20201118_lss_milk_1ng_glyco_hcd203040_compound15.4726.4726.3 | controllerType=0 controllerNumber=1 scan=4726 | 13.225 |
| 02210:3 | 17 | K.HYTN[+1647.613]SSQDVTVPC[+57.021]R.V | HexNAc(5)Hex(3)Fuc(1) | C[+57], N[+1648] | 1104.4617 | 3 | 3311.3705 | 3311.3625 | 0 | 2.4 | 89 | Specific | 247.62 | 118.62 | 118.62 | 5.01 | 39 | >sp\|P01877\|IGHA2_HUMAN Immunoglobulin heavy constant alpha 2 OS=Homo sapiens OX=9606 GN=IGHA2 PE=1 SV=4 | 576 | 20201118_lss_milk_1ng_glyco_hcd203040_compound15.4724.4724.3 | controllerType=0 controllerNumber=1 scan=4724 | 13.2217 |
| 02835:3 | 17 | K.HYTN[+2059.735]SSQDVTVPC[+57.021]R.V | HexNAc(4)Hex(5)Fuc(1)NeuAc(1) | C[+57], N[+2060] | 1241.839 | 3 | 3723.5026 | 3723.4842 | 0 | 4.9 | 89 | Specific | 283.03 | 195.2 | 195.2 | 5.77 | 39 | >sp\|P01877\|IGHA2_HUMAN Immunoglobulin heavy constant alpha 2 OS=Homo sapiens OX=9606 GN=IGHA2 PE=1 SV=4 | 576 | 20201118_lss_milk_1ng_glyco_hcd203040_compound15.5373.5373.3 | controllerType=0 controllerNumber=1 scan=5373 | 14.3148 |
| 01981:3 | 17 | K.HYTN[+1971.719]SSQDVTVPC[+57.021]R.V | HexNAc(5)Hex(5)Fuc(1) | C[+57], N[+1972] | 1212.4972 | 3 | 3635.4772 | 3635.4682 | 0 | 2.5 | 89 | Specific | 298.71 | 288.75 | 288.75 | 6.67 | 39 | >sp\|P01877\|IGHA2_HUMAN Immunoglobulin heavy constant alpha 2 OS=Homo sapiens OX=9606 GN=IGHA2 PE=1 SV=4 | 576 | 20201118_lss_milk_1ng_glyco_hcd203040_compound15.4487.4487.3 | controllerType=0 controllerNumber=1 scan=4487 | 12.8277 |
| 01983:3 | 17 | K.HYTN[+1971.719]SSQDVTVPC[+57.021]R.V | HexNAc(5)Hex(5)Fuc(1) | C[+57], N[+1972] | 1212.4972 | 3 | 3635.4772 | 3635.4682 | 0 | 2.5 | 89 | Specific | 366.09 | 300.53 | 300.53 | 6.91 | 39 | >sp\|P01877\|IGHA2_HUMAN Immunoglobulin heavy constant alpha 2 OS=Homo sapiens OX=9606 GN=IGHA2 PE=1 SV=4 | 576 | 20201118_lss_milk_1ng_glyco_hcd203040_compound15.4489.4489.3 | controllerType=0 controllerNumber=1 scan=4489 | 12.8317 |
| 02833:3 | 17 | K.HYTN[+2059.735]SSQDVTVPC[+57.021]R.V | HexNAc(4)Hex(5)Fuc(1)NeuAc(1) | C[+57], N[+2060] | 1241.839 | 3 | 3723.5026 | 3723.4842 | 0 | 4.9 | 89 | Specific | 374.59 | 254.79 | 254.79 | 6.42 | 39 | >sp\|P01877\|IGHA2_HUMAN Immunoglobulin heavy constant alpha 2 OS=Homo sapiens OX=9606 GN=IGHA2 PE=1 SV=4 | 576 | 20201118_lss_milk_1ng_glyco_hcd203040_compound15.5371.5371.3 | controllerType=0 controllerNumber=1 scan=5371 | 14.3101 |
| 02016:3 | 17 | K.HYTN[+1647.613]SSQDVTVPC[+57.021]R.V | HexNAc(5)Hex(3)Fuc(1) | C[+57], N[+1648] | 1104.4615 | 3 | 3311.3698 | 3311.3625 | 0 | 2.2 | 89 | Specific | 557.92 | 517.68 | 517.68 | 9.58 | 39 | >sp\|P01877\|IGHA2_HUMAN Immunoglobulin heavy constant alpha 2 OS=Homo sapiens OX=9606 GN=IGHA2 PE=1 SV=4 | 576 | 20201118_lss_milk_1ng_glyco_hcd203040_compound15.4523.4523.2 | controllerType=0 controllerNumber=1 scan=4523 | 12.891 |
| 01979:3 | 17 | K.HYTN[+1809.666]SSQDVTVPC[+57.021]R.V | HexNAc(5)Hex(4)Fuc(1) | C[+57], N[+1810] | 1158.4792 | 3 | 3473.423 | 3473.4154 | 0 | 2.2 | 89 | Specific | 581.58 | 456.55 | 456.55 | 10.85 | 39 | >sp\|P01877\|IGHA2_HUMAN Immunoglobulin heavy constant alpha 2 OS=Homo sapiens OX=9606 GN=IGHA2 PE=1 SV=4 | 576 | 20201118_lss_milk_1ng_glyco_hcd203040_compound15.4485.4485.3 | controllerType=0 controllerNumber=1 scan=4485 | 12.8244 |
| 02014:3 | 17 | K.HYTN[+1647.613]SSQDVTVPC[+57.021]R.V | HexNAc(5)Hex(3)Fuc(1) | C[+57], N[+1648] | 1104.4615 | 3 | 3311.3698 | 3311.3625 | 0 | 2.2 | 89 | Specific | 598.01 | 403.67 | 403.67 | 10.85 | 39 | >sp\|P01877\|IGHA2_HUMAN Immunoglobulin heavy constant alpha 2 OS=Homo sapiens OX=9606 GN=IGHA2 PE=1 SV=4 | 576 | 20201118_lss_milk_1ng_glyco_hcd203040_compound15.4521.4521.2 | controllerType=0 controllerNumber=1 scan=4521 | 12.8857 |
| 01977:3 | 17 | K.HYTN[+1809.666]SSQDVTVPC[+57.021]R.V | HexNAc(5)Hex(4)Fuc(1) | C[+57], N[+1810] | 1158.4792 | 3 | 3473.423 | 3473.4154 | 0 | 2.2 | 89 | Specific | 649.4 | 511.27 | 511.27 | 11.85 | 39 | >sp\|P01877\|IGHA2_HUMAN Immunoglobulin heavy constant alpha 2 OS=Homo sapiens OX=9606 GN=IGHA2 PE=1 SV=4 | 576 | 20201118_lss_milk_1ng_glyco_hcd203040_compound15.4483.4483.3 | controllerType=0 controllerNumber=1 scan=4483 | 12.8196 |
| 02000:3 | 17 | K.HYTN[+1647.613]SSQDVTVPC[+57.021]R.V | HexNAc(5)Hex(3)Fuc(1) | C[+57], N[+1648] | 1104.4615 | 3 | 3311.3698 | 3311.3625 | 0 | 2.2 | 89 | Specific | 704.3 | 584.42 | 584.42 | 13.85 | 39 | >sp\|P01877\|IGHA2_HUMAN Immunoglobulin heavy constant alpha 2 OS=Homo sapiens OX=9606 GN=IGHA2 PE=1 SV=4 | 576 | 20201118_lss_milk_1ng_glyco_hcd203040_compound15.4507.4507.3 | controllerType=0 controllerNumber=1 scan=4507 | 12.8626 |
| 02002:3 | 17 | K.HYTN[+1647.613]SSQDVTVPC[+57.021]R.V | HexNAc(5)Hex(3)Fuc(1) | C[+57], N[+1648] | 1104.4615 | 3 | 3311.3698 | 3311.3625 | 0 | 2.2 | 89 | Specific | 714.49 | 599.2 | 599.2 | 13.85 | 39 | >sp\|P01877\|IGHA2_HUMAN Immunoglobulin heavy constant alpha 2 OS=Homo sapiens OX=9606 GN=IGHA2 PE=1 SV=4 | 576 | 20201118_lss_milk_1ng_glyco_hcd203040_compound15.4509.4509.3 | controllerType=0 controllerNumber=1 scan=4509 | 12.8674 |
| 04113:4 | 17 | K.HYTN[+1809.666]SSQDVTVPC[+57.021]RVPPPPPC[+57.021]C[+57.021]HPR.L | HexNAc(5)Hex(4)Fuc(1) | C[+57]*3, N[+1810] | 1192.7642 | 4 | 4768.035 | 4768.0217 | 0 | 2.8 | 89 | Specific | 410.32 | 183.67 | 183.67 | 5.78 | 39 | >sp\|P01877\|IGHA2_HUMAN Immunoglobulin heavy constant alpha 2 OS=Homo sapiens OX=9606 GN=IGHA2 PE=1 SV=4 | 576 | 20201118_lss_milk_1ng_glyco_hcd203040_compound15.6696.6696.4 | controllerType=0 controllerNumber=1 scan=6696 | 16.5729 |
| 04115:4 | 17 | K.HYTN[+1809.666]SSQDVTVPC[+57.021]RVPPPPPC[+57.021]C[+57.021]HPR.L | HexNAc(5)Hex(4)Fuc(1) | C[+57]*3, N[+1810] | 1192.7642 | 4 | 4768.035 | 4768.0217 | 0 | 2.8 | 89 | Specific | 475.4 | 434.16 | 434.16 | 8.64 | 39 | >sp\|P01877\|IGHA2_HUMAN Immunoglobulin heavy constant alpha 2 OS=Homo sapiens OX=9606 GN=IGHA2 PE=1 SV=4 | 576 | 20201118_lss_milk_1ng_glyco_hcd203040_compound15.6698.6698.4 | controllerType=0 controllerNumber=1 scan=6698 | 16.5764 |
| 04510:5 | 17 | K.HYTN[+1647.613]SSQDVTVPC[+57.021]RVPPPPPC[+57.021]C[+57.021]HPR.L | HexNAc(5)Hex(3)Fuc(1) | C[+57]*3, N[+1648] | 922.0025 | 5 | 4605.9832 | 4605.9689 | 0 | 3.1 | 89 | Specific | 494.56 | 386.34 | 386.34 | 9.34 | 39 | >sp\|P01877\|IGHA2_HUMAN Immunoglobulin heavy constant alpha 2 OS=Homo sapiens OX=9606 GN=IGHA2 PE=1 SV=4 | 576 | 20201118_lss_milk_1ng_glyco_hcd203040_compound15.7106.7106.5 | controllerType=0 controllerNumber=1 scan=7106 | 17.2455 |
| 04053:5 | 17 | K.HYTN[+1647.613]SSQDVTVPC[+57.021]RVPPPPPC[+57.021]C[+57.021]HPR.L | HexNAc(5)Hex(3)Fuc(1) | C[+57]*3, N[+1648] | 922.0025 | 5 | 4605.9832 | 4605.9689 | 0 | 3.1 | 89 | Specific | 704.67 | 490.29 | 490.29 | 14.09 | 39 | >sp\|P01877\|IGHA2_HUMAN Immunoglobulin heavy constant alpha 2 OS=Homo sapiens OX=9606 GN=IGHA2 PE=1 SV=4 | 576 | 20201118_lss_milk_1ng_glyco_hcd203040_compound15.6634.6634.5 | controllerType=0 controllerNumber=1 scan=6634 | 16.4651 |
| 04055:5 | 17 | K.HYTN[+1647.613]SSQDVTVPC[+57.021]RVPPPPPC[+57.021]C[+57.021]HPR.L | HexNAc(5)Hex(3)Fuc(1) | C[+57]*3, N[+1648] | 922.0025 | 5 | 4605.9832 | 4605.9689 | 0 | 3.1 | 89 | Specific | 769.25 | 581.08 | 581.08 | 15.09 | 39 | >sp\|P01877\|IGHA2_HUMAN Immunoglobulin heavy constant alpha 2 OS=Homo sapiens OX=9606 GN=IGHA2 PE=1 SV=4 | 576 | 20201118_lss_milk_1ng_glyco_hcd203040_compound15.6636.6636.5 | controllerType=0 controllerNumber=1 scan=6636 | 16.4701 |
| 04082:4 | 17 | K.HYTN[+1647.613]SSQDVTVPC[+57.021]RVPPPPPC[+57.021]C[+57.021]HPR.L | HexNAc(5)Hex(3)Fuc(1) | C[+57]*3, N[+1648] | 1152.2516 | 4 | 4605.9847 | 4605.9689 | 0 | 3.4 | 89 | Specific | 840.92 | 603.22 | 603.22 | 15.58 | 39 | >sp\|P01877\|IGHA2_HUMAN Immunoglobulin heavy constant alpha 2 OS=Homo sapiens OX=9606 GN=IGHA2 PE=1 SV=4 | 576 | 20201118_lss_milk_1ng_glyco_hcd203040_compound15.6664.6664.4 | controllerType=0 controllerNumber=1 scan=6664 | 16.5182 |
| 04080:4 | 17 | K.HYTN[+1647.613]SSQDVTVPC[+57.021]RVPPPPPC[+57.021]C[+57.021]HPR.L | HexNAc(5)Hex(3)Fuc(1) | C[+57]*3, N[+1648] | 1152.2516 | 4 | 4605.9847 | 4605.9689 | 0 | 3.4 | 89 | Specific | 899.75 | 743.28 | 743.28 | 16.58 | 39 | >sp\|P01877\|IGHA2_HUMAN Immunoglobulin heavy constant alpha 2 OS=Homo sapiens OX=9606 GN=IGHA2 PE=1 SV=4 | 576 | 20201118_lss_milk_1ng_glyco_hcd203040_compound15.6662.6662.4 | controllerType=0 controllerNumber=1 scan=6662 | 16.5129 |
| 03718:2 | 17 | K.TPLTAN[+1971.719]ITK.S | HexNAc(5)Hex(5)Fuc(1) | N[+1972] | 1465.6484 | 2 | 2930.2894 | 2930.2757 | 0 | 4.7 | 200 | Specific | 207.18 | 71.36 | 71.36 | 2.2 | 39 | >sp\|P01877\|IGHA2_HUMAN Immunoglobulin heavy constant alpha 2 OS=Homo sapiens OX=9606 GN=IGHA2 PE=1 SV=4 | 576 | 20201118_lss_milk_1ng_glyco_hcd203040_compound15.6286.6286.2 | controllerType=0 controllerNumber=1 scan=6286 | 15.8386 |
| 03680:3 | 17 | K.TPLTAN[+1971.719]ITK.S | HexNAc(5)Hex(5)Fuc(1) | N[+1972] | 977.4326 | 3 | 2930.2834 | 2930.2757 | 0 | 2.6 | 200 | Specific | 237.97 | 161.88 | 161.88 | 5.01 | 39 | >sp\|P01877\|IGHA2_HUMAN Immunoglobulin heavy constant alpha 2 OS=Homo sapiens OX=9606 GN=IGHA2 PE=1 SV=4 | 576 | 20201118_lss_milk_1ng_glyco_hcd203040_compound15.6247.6247.3 | controllerType=0 controllerNumber=1 scan=6247 | 15.768 |
| 05614:3 | 17 | K.TPLTAN[+2262.814]ITK.S | HexNAc(5)Hex(5)Fuc(1)NeuAc(1) | N[+2263] | 1074.4634 | 3 | 3221.3757 | 3221.3711 | 0 | 1.4 | 200 | Specific | 263.41 | 145.57 | 145.57 | 4.83 | 39 | >sp\|P01877\|IGHA2_HUMAN Immunoglobulin heavy constant alpha 2 OS=Homo sapiens OX=9606 GN=IGHA2 PE=1 SV=4 | 576 | 20201118_lss_milk_1ng_glyco_hcd203040_compound15.8248.8248.3 | controllerType=0 controllerNumber=1 scan=8248 | 19.1025 |
| 05581:3 | 17 | K.TPLTAN[+2262.814]ITK.S | HexNAc(5)Hex(5)Fuc(1)NeuAc(1) | N[+2263] | 1074.4657 | 3 | 3221.3826 | 3221.3711 | 0 | 3.6 | 200 | Specific | 291.17 | 118.01 | 118.01 | 4.48 | 39 | >sp\|P01877\|IGHA2_HUMAN Immunoglobulin heavy constant alpha 2 OS=Homo sapiens OX=9606 GN=IGHA2 PE=1 SV=4 | 576 | 20201118_lss_milk_1ng_glyco_hcd203040_compound15.8214.8214.3 | controllerType=0 controllerNumber=1 scan=8214 | 19.048 |
| 05570:3 | 17 | K.TPLTAN[+2059.735]ITK.S | HexNAc(4)Hex(5)Fuc(1)NeuAc(1) | N[+2060] | 1006.7708 | 3 | 3018.298 | 3018.2917 | 0 | 2.1 | 200 | Specific | 298.83 | 273.94 | 273.94 | 6.42 | 39 | >sp\|P01877\|IGHA2_HUMAN Immunoglobulin heavy constant alpha 2 OS=Homo sapiens OX=9606 GN=IGHA2 PE=1 SV=4 | 576 | 20201118_lss_milk_1ng_glyco_hcd203040_compound15.8202.8202.3 | controllerType=0 controllerNumber=1 scan=8202 | 19.0277 |
| 03720:2 | 17 | K.TPLTAN[+1971.719]ITK.S | HexNAc(5)Hex(5)Fuc(1) | N[+1972] | 1465.6484 | 2 | 2930.2894 | 2930.2757 | 0 | 4.7 | 200 | Specific | 338.07 | 247.19 | 247.19 | 6.1 | 39 | >sp\|P01877\|IGHA2_HUMAN Immunoglobulin heavy constant alpha 2 OS=Homo sapiens OX=9606 GN=IGHA2 PE=1 SV=4 | 576 | 20201118_lss_milk_1ng_glyco_hcd203040_compound15.6288.6288.2 | controllerType=0 controllerNumber=1 scan=6288 | 15.8426 |
| 05616:3 | 17 | K.TPLTAN[+2262.814]ITK.S | HexNAc(5)Hex(5)Fuc(1)NeuAc(1) | N[+2263] | 1074.4634 | 3 | 3221.3757 | 3221.3711 | 0 | 1.4 | 200 | Specific | 361.85 | 172.99 | 172.99 | 5.6 | 39 | >sp\|P01877\|IGHA2_HUMAN Immunoglobulin heavy constant alpha 2 OS=Homo sapiens OX=9606 GN=IGHA2 PE=1 SV=4 | 576 | 20201118_lss_milk_1ng_glyco_hcd203040_compound15.8250.8250.3 | controllerType=0 controllerNumber=1 scan=8250 | 19.1063 |
| 03837:2 | 17 | K.TPLTAN[+1501.555]ITK.S | HexNAc(5)Hex(3) | N[+1502] | 1230.5649 | 2 | 2460.1224 | 2460.1121 | 0 | 4.2 | 200 | Specific | 368.22 | 176.68 | 176.68 | 5.65 | 39 | >sp\|P01877\|IGHA2_HUMAN Immunoglobulin heavy constant alpha 2 OS=Homo sapiens OX=9606 GN=IGHA2 PE=1 SV=4 | 576 | 20201118_lss_milk_1ng_glyco_hcd203040_compound15.6410.6410.2 | controllerType=0 controllerNumber=1 scan=6410 | 16.0864 |
| 05583:3 | 17 | K.TPLTAN[+2262.814]ITK.S | HexNAc(5)Hex(5)Fuc(1)NeuAc(1) | N[+2263] | 1074.4657 | 3 | 3221.3826 | 3221.3711 | 0 | 3.6 | 200 | Specific | 420.4 | 210.37 | 210.37 | 6.55 | 39 | >sp\|P01877\|IGHA2_HUMAN Immunoglobulin heavy constant alpha 2 OS=Homo sapiens OX=9606 GN=IGHA2 PE=1 SV=4 | 576 | 20201118_lss_milk_1ng_glyco_hcd203040_compound15.8216.8216.3 | controllerType=0 controllerNumber=1 scan=8216 | 19.052 |
| 03682:3 | 17 | K.TPLTAN[+1971.719]ITK.S | HexNAc(5)Hex(5)Fuc(1) | N[+1972] | 977.4326 | 3 | 2930.2834 | 2930.2757 | 0 | 2.6 | 200 | Specific | 435.7 | 216.49 | 216.49 | 6.78 | 39 | >sp\|P01877\|IGHA2_HUMAN Immunoglobulin heavy constant alpha 2 OS=Homo sapiens OX=9606 GN=IGHA2 PE=1 SV=4 | 576 | 20201118_lss_milk_1ng_glyco_hcd203040_compound15.6249.6249.3 | controllerType=0 controllerNumber=1 scan=6249 | 15.7718 |
| 03839:2 | 17 | K.TPLTAN[+1501.555]ITK.S | HexNAc(5)Hex(3) | N[+1502] | 1230.5649 | 2 | 2460.1224 | 2460.1121 | 0 | 4.2 | 200 | Specific | 458.05 | 256.54 | 256.54 | 6.55 | 39 | >sp\|P01877\|IGHA2_HUMAN Immunoglobulin heavy constant alpha 2 OS=Homo sapiens OX=9606 GN=IGHA2 PE=1 SV=4 | 576 | 20201118_lss_milk_1ng_glyco_hcd203040_compound15.6412.6412.2 | controllerType=0 controllerNumber=1 scan=6412 | 16.0906 |
| 03821:3 | 17 | K.TPLTAN[+1647.613]ITK.S | HexNAc(5)Hex(3)Fuc(1) | N[+1648] | 869.3976 | 3 | 2606.1781 | 2606.17 | 0 | 3.1 | 200 | Specific | 480.78 | 202.79 | 202.79 | 6.8 | 39 | >sp\|P01877\|IGHA2_HUMAN Immunoglobulin heavy constant alpha 2 OS=Homo sapiens OX=9606 GN=IGHA2 PE=1 SV=4 | 576 | 20201118_lss_milk_1ng_glyco_hcd203040_compound15.6393.6393.3 | controllerType=0 controllerNumber=1 scan=6393 | 16.0325 |
| 03741:3 | 17 | K.TPLTAN[+1809.666]ITK.S | HexNAc(5)Hex(4)Fuc(1) | N[+1810] | 923.4144 | 3 | 2768.2286 | 2768.2228 | 0 | 2.1 | 200 | Specific | 488.64 | 211.61 | 211.61 | 6.76 | 39 | >sp\|P01877\|IGHA2_HUMAN Immunoglobulin heavy constant alpha 2 OS=Homo sapiens OX=9606 GN=IGHA2 PE=1 SV=4 | 576 | 20201118_lss_milk_1ng_glyco_hcd203040_compound15.6310.6310.3 | controllerType=0 controllerNumber=1 scan=6310 | 15.8835 |
| 03743:3 | 17 | K.TPLTAN[+1809.666]ITK.S | HexNAc(5)Hex(4)Fuc(1) | N[+1810] | 923.4144 | 3 | 2768.2286 | 2768.2228 | 0 | 2.1 | 200 | Specific | 539.2 | 283.75 | 283.75 | 7.46 | 39 | >sp\|P01877\|IGHA2_HUMAN Immunoglobulin heavy constant alpha 2 OS=Homo sapiens OX=9606 GN=IGHA2 PE=1 SV=4 | 576 | 20201118_lss_milk_1ng_glyco_hcd203040_compound15.6312.6312.3 | controllerType=0 controllerNumber=1 scan=6312 | 15.8883 |
| 03823:3 | 17 | K.TPLTAN[+1647.613]ITK.S | HexNAc(5)Hex(3)Fuc(1) | N[+1648] | 869.3976 | 3 | 2606.1781 | 2606.17 | 0 | 3.1 | 200 | Specific | 581.62 | 279.94 | 279.94 | 9.02 | 39 | >sp\|P01877\|IGHA2_HUMAN Immunoglobulin heavy constant alpha 2 OS=Homo sapiens OX=9606 GN=IGHA2 PE=1 SV=4 | 576 | 20201118_lss_milk_1ng_glyco_hcd203040_compound15.6395.6395.3 | controllerType=0 controllerNumber=1 scan=6395 | 16.0524 |
| 03755:2 | 17 | K.TPLTAN[+1809.666]ITK.S | HexNAc(5)Hex(4)Fuc(1) | N[+1810] | 1384.6182 | 2 | 2768.2291 | 2768.2228 | 0 | 2.3 | 200 | Specific | 613.31 | 307.54 | 307.54 | 9.91 | 39 | >sp\|P01877\|IGHA2_HUMAN Immunoglobulin heavy constant alpha 2 OS=Homo sapiens OX=9606 GN=IGHA2 PE=1 SV=4 | 576 | 20201118_lss_milk_1ng_glyco_hcd203040_compound15.6325.6325.2 | controllerType=0 controllerNumber=1 scan=6325 | 15.9128 |
| 03805:2 | 17 | K.TPLTAN[+1647.613]ITK.S | HexNAc(5)Hex(3)Fuc(1) | N[+1648] | 1303.5927 | 2 | 2606.1781 | 2606.17 | 0 | 3.1 | 200 | Specific | 614.65 | 359.35 | 359.35 | 10.52 | 39 | >sp\|P01877\|IGHA2_HUMAN Immunoglobulin heavy constant alpha 2 OS=Homo sapiens OX=9606 GN=IGHA2 PE=1 SV=4 | 576 | 20201118_lss_milk_1ng_glyco_hcd203040_compound15.6377.6377.2 | controllerType=0 controllerNumber=1 scan=6377 | 16.0038 |
| 03803:2 | 17 | K.TPLTAN[+1647.613]ITK.S | HexNAc(5)Hex(3)Fuc(1) | N[+1648] | 1303.5927 | 2 | 2606.1781 | 2606.17 | 0 | 3.1 | 200 | Specific | 629.7 | 357.74 | 357.74 | 10.52 | 39 | >sp\|P01877\|IGHA2_HUMAN Immunoglobulin heavy constant alpha 2 OS=Homo sapiens OX=9606 GN=IGHA2 PE=1 SV=4 | 576 | 20201118_lss_milk_1ng_glyco_hcd203040_compound15.6375.6375.2 | controllerType=0 controllerNumber=1 scan=6375 | 15.9968 |
| 03753:2 | 17 | K.TPLTAN[+1809.666]ITK.S | HexNAc(5)Hex(4)Fuc(1) | N[+1810] | 1384.6182 | 2 | 2768.2291 | 2768.2228 | 0 | 2.3 | 200 | Specific | 646.89 | 364.74 | 364.74 | 10.41 | 39 | >sp\|P01877\|IGHA2_HUMAN Immunoglobulin heavy constant alpha 2 OS=Homo sapiens OX=9606 GN=IGHA2 PE=1 SV=4 | 576 | 20201118_lss_milk_1ng_glyco_hcd203040_compound15.6323.6323.2 | controllerType=0 controllerNumber=1 scan=6323 | 15.9079 |
| 03800:2 | 17 | K.TPLTAN[+1444.534]ITK.S | HexNAc(4)Hex(3)Fuc(1) | N[+1445] | 1202.0503 | 2 | 2403.0934 | 2403.0906 | 0 | 1.1 | 200 | Specific | 652.9 | 333.1 | 333.1 | 10.93 | 39 | >sp\|P01877\|IGHA2_HUMAN Immunoglobulin heavy constant alpha 2 OS=Homo sapiens OX=9606 GN=IGHA2 PE=1 SV=4 | 576 | 20201118_lss_milk_1ng_glyco_hcd203040_compound15.6371.6371.2 | controllerType=0 controllerNumber=1 scan=6371 | 15.9889 |
| 03798:2 | 17 | K.TPLTAN[+1444.534]ITK.S | HexNAc(4)Hex(3)Fuc(1) | N[+1445] | 1202.0503 | 2 | 2403.0934 | 2403.0906 | 0 | 1.1 | 200 | Specific | 653.82 | 374.93 | 374.93 | 11.14 | 39 | >sp\|P01877\|IGHA2_HUMAN Immunoglobulin heavy constant alpha 2 OS=Homo sapiens OX=9606 GN=IGHA2 PE=1 SV=4 | 576 | 20201118_lss_milk_1ng_glyco_hcd203040_compound15.6369.6369.2 | controllerType=0 controllerNumber=1 scan=6369 | 15.9842 |
| 11155:3 | 17 | R.MAGKPTHIN[+1702.581]VSVVMAEADGTC[+57.021]Y.- | HexNAc(2)Hex(8) | C[+57], N[+1703] | 1351.8964 | 3 | 4053.6747 | 4053.6642 | 0 | 2.6 | 319 | CRagged | 677.46 | 677.46 | 409.34 | 13.15 | 39 | >sp\|P01877\|IGHA2_HUMAN Immunoglobulin heavy constant alpha 2 OS=Homo sapiens OX=9606 GN=IGHA2 PE=1 SV=4 | 576 | 20201118_lss_milk_1ng_glyco_hcd203040_compound15.13992.13992.3 | controllerType=0 controllerNumber=1 scan=13992 | 28.5898 |
| 11161:3 | 17 | R.MAGKPTHIN[+1540.529]VSVVMAEADGTC[+57.021]Y.- | HexNAc(2)Hex(7) | C[+57], N[+1541] | 1297.8786 | 3 | 3891.6212 | 3891.6114 | 0 | 2.5 | 319 | CRagged | 696.61 | 647.54 | 406.81 | 13.15 | 39 | >sp\|P01877\|IGHA2_HUMAN Immunoglobulin heavy constant alpha 2 OS=Homo sapiens OX=9606 GN=IGHA2 PE=1 SV=4 | 576 | 20201118_lss_milk_1ng_glyco_hcd203040_compound15.13998.13998.3 | controllerType=0 controllerNumber=1 scan=13998 | 28.5998 |
| 11157:3 | 17 | R.MAGKPTHIN[+1702.581]VSVVMAEADGTC[+57.021]Y.- | HexNAc(2)Hex(8) | C[+57], N[+1703] | 1351.8964 | 3 | 4053.6747 | 4053.6642 | 0 | 2.6 | 319 | CRagged | 743.29 | 743.29 | 403.16 | 14.15 | 39 | >sp\|P01877\|IGHA2_HUMAN Immunoglobulin heavy constant alpha 2 OS=Homo sapiens OX=9606 GN=IGHA2 PE=1 SV=4 | 576 | 20201118_lss_milk_1ng_glyco_hcd203040_compound15.13994.13994.3 | controllerType=0 controllerNumber=1 scan=13994 | 28.5939 |
| 11163:3 | 17 | R.MAGKPTHIN[+1540.529]VSVVMAEADGTC[+57.021]Y.- | HexNAc(2)Hex(7) | C[+57], N[+1541] | 1297.8786 | 3 | 3891.6212 | 3891.6114 | 0 | 2.5 | 319 | CRagged | 820.02 | 807.45 | 464.56 | 16.15 | 39 | >sp\|P01877\|IGHA2_HUMAN Immunoglobulin heavy constant alpha 2 OS=Homo sapiens OX=9606 GN=IGHA2 PE=1 SV=4 | 576 | 20201118_lss_milk_1ng_glyco_hcd203040_compound15.14000.14000.3 | controllerType=0 controllerNumber=1 scan=14000 | 28.6044 |
| 11188:3 | 17 | R.MAGKPTHIN[+1378.476]VSVVMAEADGTC[+57.021]Y.- | HexNAc(2)Hex(6) | C[+57], N[+1378] | 1243.8604 | 3 | 3729.5667 | 3729.5585 | 0 | 2.2 | 319 | CRagged | 859.75 | 859.75 | 482.47 | 17.15 | 39 | >sp\|P01877\|IGHA2_HUMAN Immunoglobulin heavy constant alpha 2 OS=Homo sapiens OX=9606 GN=IGHA2 PE=1 SV=4 | 576 | 20201118_lss_milk_1ng_glyco_hcd203040_compound15.14026.14026.3 | controllerType=0 controllerNumber=1 scan=14026 | 28.6507 |
| 11235:3 | 17 | R.MAGKPTHIN[+1216.423]VSVVMAEADGTC[+57.021]Y.- | HexNAc(2)Hex(5) | C[+57], N[+1216] | 1189.8433 | 3 | 3567.5154 | 3567.5057 | 0 | 2.7 | 319 | CRagged | 883.13 | 883.13 | 511.57 | 17.15 | 39 | >sp\|P01877\|IGHA2_HUMAN Immunoglobulin heavy constant alpha 2 OS=Homo sapiens OX=9606 GN=IGHA2 PE=1 SV=4 | 576 | 20201118_lss_milk_1ng_glyco_hcd203040_compound15.14075.14075.3 | controllerType=0 controllerNumber=1 scan=14075 | 28.7542 |
| 11185:3 | 17 | R.MAGKPTHIN[+1378.476]VSVVMAEADGTC[+57.021]Y.- | HexNAc(2)Hex(6) | C[+57], N[+1378] | 1243.8604 | 3 | 3729.5667 | 3729.5585 | 0 | 2.2 | 319 | CRagged | 919.06 | 806.26 | 525.05 | 18.15 | 39 | >sp\|P01877\|IGHA2_HUMAN Immunoglobulin heavy constant alpha 2 OS=Homo sapiens OX=9606 GN=IGHA2 PE=1 SV=4 | 576 | 20201118_lss_milk_1ng_glyco_hcd203040_compound15.14023.14023.3 | controllerType=0 controllerNumber=1 scan=14023 | 28.6439 |
| 11233:3 | 17 | R.MAGKPTHIN[+1216.423]VSVVMAEADGTC[+57.021]Y.- | HexNAc(2)Hex(5) | C[+57], N[+1216] | 1189.8433 | 3 | 3567.5154 | 3567.5057 | 0 | 2.7 | 319 | CRagged | 945.33 | 799.05 | 517.52 | 18.15 | 39 | >sp\|P01877\|IGHA2_HUMAN Immunoglobulin heavy constant alpha 2 OS=Homo sapiens OX=9606 GN=IGHA2 PE=1 SV=4 | 576 | 20201118_lss_milk_1ng_glyco_hcd203040_compound15.14073.14073.3 | controllerType=0 controllerNumber=1 scan=14073 | 28.7484 |
| 10392:5 | 19 | K.[+42.011]QTDEIKDTRN[+2018.708]ESTQNC[+57.021]VVAEPEKMESSISSSSEEMSLS | HexNAc(3)Hex(6)Fuc(1)NeuAc(1) | .[42], C[+57], N[+2019] | 1291.1503 | 5 | 6451.7222 | 6451.6996 | 0 | 3.5 | 60 | Specific | 374.51 | 293.29 | 293.29 | 4.66 | 30 | >sp\|P47710\|CASA1_HUMAN Alpha-S1-casein OS=Homo sapiens OX=9606 GN=CSN1S1 PE=1 SV=1 | 989 | 20201118_lss_milk_1ng_glyco_hcd203040_compound15.13202.13202.5 | controllerType=0 controllerNumber=1 scan=13202 | 27.2256 |
| 10389:5 | 19 | K.[+42.011]QTDEIKDTRN[+2018.708]ESTQNC[+57.021]VVAEPEKMESSISSSSEEMSLS | HexNAc(3)Hex(6)Fuc(1)NeuAc(1) | .[42], C[+57], N[+2019] | 1291.1503 | 5 | 6451.7222 | 6451.6996 | 0 | 3.5 | 60 | Specific | 469.7 | 385.2 | 385.2 | 8.24 | 30 | >sp\|P47710\|CASA1_HUMAN Alpha-S1-casein OS=Homo sapiens OX=9606 GN=CSN1S1 PE=1 SV=1 | 989 | 20201118_lss_milk_1ng_glyco_hcd203040_compound15.13199.13199.5 | controllerType=0 controllerNumber=1 scan=13199 | 27.2201 |
| 02838:3 | 30 | R.LRN[+1216.423]VSWATGR.S | HexNAc(2)Hex(5) | N[+1216] | 792.6931 | 3 | 2376.0647 | 2376.0559 | 0 | 3.7 | 149 | Specific | 222.02 | 59.37 | 59.37 | 2.17 | 21 | >sp\|P08571\|CD14_HUMAN Monocyte differentiation antigen CD14 OS=Homo sapiens OX=9606 GN=CD14 PE=1 SV=2 | 1232 | 20201118_lss_milk_1ng_glyco_hcd203040_compound15.5376.5376.3 | controllerType=0 controllerNumber=1 scan=5376 | 14.3206 |
| 15076:3 | 31 | K.VVLHPN[+2204.772]YSQVDIGLIK.L | HexNAc(4)Hex(5)NeuAc(2) | N[+2205] | 1333.9445 | 3 | 3999.819 | 3999.7837 | 0 | 8.8 | 236 | Specific | 643.53 | 573.27 | 573.27 | 10.7 | 21 | >sp\|P00738\|HPT_HUMAN Haptoglobin OS=Homo sapiens OX=9606 GN=HP PE=1 SV=1 | 407 | 20201118_lss_milk_1ng_glyco_hcd203040_compound15.18064.18064.3 | controllerType=0 controllerNumber=1 scan=18064 | 35.9085 |
| 15074:3 | 31 | K.VVLHPN[+2204.772]YSQVDIGLIK.L | HexNAc(4)Hex(5)NeuAc(2) | N[+2205] | 1333.9445 | 3 | 3999.819 | 3999.7837 | 0 | 8.8 | 236 | Specific | 782.66 | 704.91 | 704.91 | 13.7 | 21 | >sp\|P00738\|HPT_HUMAN Haptoglobin OS=Homo sapiens OX=9606 GN=HP PE=1 SV=1 | 407 | 20201118_lss_milk_1ng_glyco_hcd203040_compound15.18062.18062.3 | controllerType=0 controllerNumber=1 scan=18062 | 35.904 |
| 12159:4 | 51 | R.IIVPLNNREN[+1913.677]ISDPTSPLR.T | HexNAc(4)Hex(5)NeuAc(1) | N[+1914] | 1016.2232 | 4 | 4061.8712 | 4061.8542 | 0 | 4.2 | 62 | Specific | 279.97 | 279.97 | 279.97 | 4.93 | 17 | >sp\|P01591\|IGJ_HUMAN Immunoglobulin J chain OS=Homo sapiens OX=9606 GN=JCHAIN PE=1 SV=4 | 1392 | 20201118_lss_milk_1ng_glyco_hcd203040_compound15.15031.15031.4 | controllerType=0 controllerNumber=1 scan=15031 | 30.4113 |
| 12161:4 | 51 | R.IIVPLNNREN[+1913.677]ISDPTSPLR.T | HexNAc(4)Hex(5)NeuAc(1) | N[+1914] | 1016.2232 | 4 | 4061.8712 | 4061.8542 | 0 | 4.2 | 62 | Specific | 328.97 | 323.01 | 323.01 | 5.01 | 17 | >sp\|P01591\|IGJ_HUMAN Immunoglobulin J chain OS=Homo sapiens OX=9606 GN=JCHAIN PE=1 SV=4 | 1392 | 20201118_lss_milk_1ng_glyco_hcd203040_compound15.15033.15033.4 | controllerType=0 controllerNumber=1 scan=15033 | 30.4157 |
| 10534:3 | 51 | R.IIVPLNNREN[+1809.666]ISDPTSPLR.T | HexNAc(5)Hex(4)Fuc(1) | N[+1810] | 1319.9568 | 3 | 3957.856 | 3957.8432 | 0 | 3.2 | 62 | Specific | 449.83 | 347.34 | 347.34 | 7.53 | 17 | >sp\|P01591\|IGJ_HUMAN Immunoglobulin J chain OS=Homo sapiens OX=9606 GN=JCHAIN PE=1 SV=4 | 1392 | 20201118_lss_milk_1ng_glyco_hcd203040_compound15.13349.13349.3 | controllerType=0 controllerNumber=1 scan=13349 | 27.477 |
| 10532:3 | 51 | R.IIVPLNNREN[+1809.666]ISDPTSPLR.T | HexNAc(5)Hex(4)Fuc(1) | N[+1810] | 1319.9568 | 3 | 3957.856 | 3957.8432 | 0 | 3.2 | 62 | Specific | 482.94 | 407.01 | 407.01 | 9.11 | 17 | >sp\|P01591\|IGJ_HUMAN Immunoglobulin J chain OS=Homo sapiens OX=9606 GN=JCHAIN PE=1 SV=4 | 1392 | 20201118_lss_milk_1ng_glyco_hcd203040_compound15.13347.13347.3 | controllerType=0 controllerNumber=1 scan=13347 | 27.4727 |
| 12128:3 | 51 | R.IIVPLNNREN[+1913.677]ISDPTSPLR.T | HexNAc(4)Hex(5)NeuAc(1) | N[+1914] | 1354.6277 | 3 | 4061.8687 | 4061.8542 | 0 | 3.6 | 62 | Specific | 566.34 | 510.33 | 510.33 | 9.59 | 17 | >sp\|P01591\|IGJ_HUMAN Immunoglobulin J chain OS=Homo sapiens OX=9606 GN=JCHAIN PE=1 SV=4 | 1392 | 20201118_lss_milk_1ng_glyco_hcd203040_compound15.14999.14999.3 | controllerType=0 controllerNumber=1 scan=14999 | 30.3484 |
| 12126:3 | 51 | R.IIVPLNNREN[+1913.677]ISDPTSPLR.T | HexNAc(4)Hex(5)NeuAc(1) | N[+1914] | 1354.6277 | 3 | 4061.8687 | 4061.8542 | 0 | 3.6 | 62 | Specific | 567.29 | 529.62 | 529.62 | 9.59 | 17 | >sp\|P01591\|IGJ_HUMAN Immunoglobulin J chain OS=Homo sapiens OX=9606 GN=JCHAIN PE=1 SV=4 | 1392 | 20201118_lss_milk_1ng_glyco_hcd203040_compound15.14997.14997.3 | controllerType=0 controllerNumber=1 scan=14997 | 30.3419 |
| 10552:3 | 51 | R.IIVPLNNREN[+1663.608]ISDPTSPLR.T | HexNAc(5)Hex(4) | N[+1664] | 1271.2704 | 3 | 3811.7968 | 3811.7853 | 0 | 3 | 62 | Specific | 579.15 | 551.8 | 551.8 | 10.76 | 17 | >sp\|P01591\|IGJ_HUMAN Immunoglobulin J chain OS=Homo sapiens OX=9606 GN=JCHAIN PE=1 SV=4 | 1392 | 20201118_lss_milk_1ng_glyco_hcd203040_compound15.13368.13368.3 | controllerType=0 controllerNumber=1 scan=13368 | 27.519 |
| 10550:3 | 51 | R.IIVPLNNREN[+1663.608]ISDPTSPLR.T | HexNAc(5)Hex(4) | N[+1664] | 1271.2704 | 3 | 3811.7968 | 3811.7853 | 0 | 3 | 62 | Specific | 623.4 | 582.58 | 582.58 | 11.76 | 17 | >sp\|P01591\|IGJ_HUMAN Immunoglobulin J chain OS=Homo sapiens OX=9606 GN=JCHAIN PE=1 SV=4 | 1392 | 20201118_lss_milk_1ng_glyco_hcd203040_compound15.13366.13366.3 | controllerType=0 controllerNumber=1 scan=13366 | 27.5138 |
| 25073:3 | 147 | R.AVN[+2204.772]ITSENLIDDVVSLIR.G | HexNAc(4)Hex(5)NeuAc(2) | N[+2205] | 1392.6249 | 3 | 4175.8603 | 4175.8481 | 0 | 2.9 | 413 | Specific | 405.15 | 394.57 | 394.57 | 8.77 | 6 | >sp\|P02748\|CO9_HUMAN Complement component C9 OS=Homo sapiens OX=9606 GN=C9 PE=1 SV=2 | 1547 | 20201118_lss_milk_1ng_glyco_hcd203040_compound15.28498.28498.3 | controllerType=0 controllerNumber=1 scan=28498 | 57.1805 |
| 25075:3 | 147 | R.AVN[+2204.772]ITSENLIDDVVSLIR.G | HexNAc(4)Hex(5)NeuAc(2) | N[+2205] | 1392.6249 | 3 | 4175.8603 | 4175.8481 | 0 | 2.9 | 413 | Specific | 544.72 | 482.13 | 482.13 | 9.49 | 6 | >sp\|P02748\|CO9_HUMAN Complement component C9 OS=Homo sapiens OX=9606 GN=C9 PE=1 SV=2 | 1547 | 20201118_lss_milk_1ng_glyco_hcd203040_compound15.28500.28500.3 | controllerType=0 controllerNumber=1 scan=28500 | 57.1846 |

Table S4. Cont.

| Query #:z | Protein Rank | Glycopeptide | Glycan | Modification Type(s) | Observed m/z | z | Observed (M+H) | Calc. mass (M+H) | Off-by-x error | Mass error (ppm) | StartPosition | Cleavage | Score | Delta | Delta Mod | \|Log Prob\| | # of unique peptides | Protein Name | Protein DB number | Comment | Scan # | Scan Time |
|---|---|---|---|---|---|---|---|---|---|---|---|---|---|---|---|---|---|---|---|---|---|---|
| 07527:3 | 471 | R.[+42.011]VEDAYILTC[+57.021]N[+730.264]VSLEYEK.T | HexNAc(2)Hex(2) | .[42], C[+57], N[+730] | 940.0959 | 3 | 2818.2732 | 2818.2486 | 0 | 8.7 | 225 | Specific | 314.94 | 216.7 | 216.7 | 2.56 | 4 | >sp\|P40227\|TCPZ_HUMAN T-complex protein 1 subunit zeta OS=Homo sapiens OX=9606 GN=CCT6A PE=1 SV=3 | 1115 | 20201118_lss_milk_1ng_glyco_hcd203040_compound15.10225.10225.3 | controllerType=0 controllerNumber=1 scan=10225 | 22.2829 |
| 09952:4 | 1 | K.TVLTPATNHMGN[+1378.476]VTFTIPANR.E | HexNAc(2)Hex(6) | N[+1378] | 909.166 | 4 | 3633.6422 | 3633.6358 | 0 | 1.8 | 74 | Specific | 503.88 | 503.88 | 284.26 | 10.13 | 145 | >sp\|P01024\|CO3_HUMAN Complement C3 OS=Homo sapiens OX=9606 GN=C3 PE=1 SV=2 | 492 | 20201116_xd_colostrum_1ng_glyco_hcd203040_compound15.12949.12949.4 | controllerType=0 controllerNumber=1 scan=12949 | 28.3209 |
| 09955:4 | 1 | K.TVLTPATNHMGN[+1378.476]VTFTIPANR.E | HexNAc(2)Hex(6) | N[+1378] | 909.166 | 4 | 3633.6422 | 3633.6358 | 0 | 1.8 | 74 | Specific | 550.73 | 392.04 | 307.13 | 9.61 | 145 | >sp\|P01024\|CO3_HUMAN Complement C3 OS=Homo sapiens OX=9606 GN=C3 PE=1 SV=2 | 492 | 20201116_xd_colostrum_1ng_glyco_hcd203040_compound15.12952.12952.4 | controllerType=0 controllerNumber=1 scan=12952 | 28.3281 |
| 09921:3 | 1 | K.TVLTPATNHMGN[+1540.529]VTFTIPANR.E | HexNAc(2)Hex(7) | N[+1541] | 1265.9053 | 3 | 3795.7014 | 3795.6886 | 0 | 3.4 | 74 | Specific | 635.72 | 635.72 | 329.07 | 12.77 | 145 | >sp\|P01024\|CO3_HUMAN Complement C3 OS=Homo sapiens OX=9606 GN=C3 PE=1 SV=2 | 492 | 20201116_xd_colostrum_1ng_glyco_hcd203040_compound15.12917.12917.3 | controllerType=0 controllerNumber=1 scan=12917 | 28.2362 |
| 09918:3 | 1 | K.TVLTPATNHMGN[+1540.529]VTFTIPANR.E | HexNAc(2)Hex(7) | N[+1541] | 1265.9053 | 3 | 3795.7014 | 3795.6886 | 0 | 3.4 | 74 | Specific | 683.49 | 683.49 | 356.23 | 13.77 | 145 | >sp\|P01024\|CO3_HUMAN Complement C3 OS=Homo sapiens OX=9606 GN=C3 PE=1 SV=2 | 492 | 20201116_xd_colostrum_1ng_glyco_hcd203040_compound15.12914.12914.3 | controllerType=0 controllerNumber=1 scan=12914 | 28.2321 |
| 09977:3 | 1 | K.TVLTPATNHMGN[+1216.423]VTFTIPANR.E | HexNAc(2)Hex(5) | N[+1216] | 1157.8677 | 3 | 3471.5886 | 3471.583 | 0 | 1.6 | 74 | Specific | 870.7 | 817.86 | 455.82 | 17.81 | 145 | >sp\|P01024\|CO3_HUMAN Complement C3 OS=Homo sapiens OX=9606 GN=C3 PE=1 SV=2 | 492 | 20201116_xd_colostrum_1ng_glyco_hcd203040_compound15.12975.12975.3 | controllerType=0 controllerNumber=1 scan=12975 | 28.3879 |
| 09917:3 | 1 | K.TVLTPATNHMGN[+1378.476]VTFTIPANR.E | HexNAc(2)Hex(6) | N[+1378] | 1211.887 | 3 | 3633.6465 | 3633.6358 | 0 | 2.9 | 74 | Specific | 993.01 | 886.3 | 578.97 | 19.96 | 145 | >sp\|P01024\|CO3_HUMAN Complement C3 OS=Homo sapiens OX=9606 GN=C3 PE=1 SV=2 | 492 | 20201116_xd_colostrum_1ng_glyco_hcd203040_compound15.12913.12913.3 | controllerType=0 controllerNumber=1 scan=12913 | 28.2308 |
| 09974:3 | 1 | K.TVLTPATNHMGN[+1216.423]VTFTIPANR.E | HexNAc(2)Hex(5) | N[+1216] | 1157.8677 | 3 | 3471.5886 | 3471.583 | 0 | 1.6 | 74 | Specific | 1109.79 | 1071.29 | 614.16 | 22.81 | 145 | >sp\|P01024\|CO3_HUMAN Complement C3 OS=Homo sapiens OX=9606 GN=C3 PE=1 SV=2 | 492 | 20201116_xd_colostrum_1ng_glyco_hcd203040_compound15.12972.12972.3 | controllerType=0 controllerNumber=1 scan=12972 | 28.3704 |
| 09915:3 | 1 | K.TVLTPATNHMGN[+1378.476]VTFTIPANR.E | HexNAc(2)Hex(6) | N[+1378] | 1211.887 | 3 | 3633.6465 | 3633.6358 | 0 | 2.9 | 74 | Specific | 1154.28 | 1137.64 | 632.68 | 23.96 | 145 | >sp\|P01024\|CO3_HUMAN Complement C3 OS=Homo sapiens OX=9606 GN=C3 PE=1 SV=2 | 492 | 20201116_xd_colostrum_1ng_glyco_hcd203040_compound15.12911.12911.3 | controllerType=0 controllerNumber=1 scan=12911 | 28.2251 |
| 16109:4 | 2 | R.RTAGWNVPIGTLRPFLN[+2424.867]WTGPPEPIEAAVAR.F | HexNAc(5)Hex(6)Fuc(1)NeuAc(1) | N[+2425] | 1453.6792 | 4 | 5811.695 | 5811.6792 | 0 | 2.7 | 140 | Specific | 325.19 | 207.54 | 207.54 | 6.56 | 134 | >sp\|P02788\|TRFL_HUMAN Lactotransferrin OS=Homo sapiens OX=9606 GN=LTF PE=1 SV=6 | 1270 | 20201116_xd_colostrum_1ng_glyco_hcd203040_compound15.19352.19352.4 | controllerType=0 controllerNumber=1 scan=19352 | 41.1437 |
| 16111:4 | 2 | R.RTAGWNVPIGTLRPFLN[+2424.867]WTGPPEPIEAAVAR.F | HexNAc(5)Hex(6)Fuc(1)NeuAc(1) | N[+2425] | 1453.6792 | 4 | 5811.695 | 5811.6792 | 0 | 2.7 | 140 | Specific | 392.21 | 383.7 | 383.7 | 9.35 | 134 | >sp\|P02788\|TRFL_HUMAN Lactotransferrin OS=Homo sapiens OX=9606 GN=LTF PE=1 SV=6 | 1270 | 20201116_xd_colostrum_1ng_glyco_hcd203040_compound15.19354.19354.4 | controllerType=0 controllerNumber=1 scan=19354 | 41.1502 |
| 17105:3 | 2 | R.RTAGWNVPIGTLRPFLN[+2350.830]WTGPPEPIEAAVAR.F | HexNAc(4)Hex(5)Fuc(1)NeuAc(2) | N[+2351] | 1913.2323 | 3 | 5737.6822 | 5737.6424 | 0 | 6.9 | 140 | Specific | 411.95 | 362.77 | 362.77 | 6.34 | 134 | >sp\|P02788\|TRFL_HUMAN Lactotransferrin OS=Homo sapiens OX=9606 GN=LTF PE=1 SV=6 | 1270 | 20201116_xd_colostrum_1ng_glyco_hcd203040_compound15.20393.20393.3 | controllerType=0 controllerNumber=1 scan=20393 | 43.5032 |
| 16162:4 | 2 | R.RTAGWNVPIGTLRPFLN[+2205.793]WTGPPEPIEAAVAR.F | HexNAc(4)Hex(5)Fuc(2)NeuAc(1) | N[+2206] | 1398.9149 | 4 | 5592.6379 | 5592.6049 | 0 | 5.9 | 140 | Specific | 427.78 | 391.86 | 391.86 | 9.44 | 134 | >sp\|P02788\|TRFL_HUMAN Lactotransferrin OS=Homo sapiens OX=9606 GN=LTF PE=1 SV=6 | 1270 | 20201116_xd_colostrum_1ng_glyco_hcd203040_compound15.19408.19408.4 | controllerType=0 controllerNumber=1 scan=19408 | 41.2678 |
| 16164:4 | 2 | R.RTAGWNVPIGTLRPFLN[+2205.793]WTGPPEPIEAAVAR.F | HexNAc(4)Hex(5)Fuc(2)NeuAc(1) | N[+2206] | 1398.9149 | 4 | 5592.6379 | 5592.6049 | 0 | 5.9 | 140 | Specific | 582.18 | 512.94 | 512.94 | 11.57 | 134 | >sp\|P02788\|TRFL_HUMAN Lactotransferrin OS=Homo sapiens OX=9606 GN=LTF PE=1 SV=6 | 1270 | 20201116_xd_colostrum_1ng_glyco_hcd203040_compound15.19410.19410.4 | controllerType=0 controllerNumber=1 scan=19410 | 41.2734 |
| 17291:4 | 2 | R.[+42.011]TAGWNVPIGTLRPFLN[+2018.708]WTGPPEPIEAAVAR.F | HexNAc(3)Hex(6)Fuc(1)NeuAc(1) | .[42], N[+2019] | 1323.6244 | 4 | 5291.4757 | 5291.4299 | 0 | 8.7 | 141 | Specific | 261.23 | 261.23 | 261.23 | 5.05 | 134 | >sp\|P02788\|TRFL_HUMAN Lactotransferrin OS=Homo sapiens OX=9606 GN=LTF PE=1 SV=6 | 1270 | 20201116_xd_colostrum_1ng_glyco_hcd203040_compound15.20590.20590.4 | controllerType=0 controllerNumber=1 scan=20590 | 43.9858 |
| 17289:4 | 2 | R.[+42.011]TAGWNVPIGTLRPFLN[+2018.708]WTGPPEPIEAAVAR.F | HexNAc(3)Hex(6)Fuc(1)NeuAc(1) | .[42], N[+2019] | 1323.6244 | 4 | 5291.4757 | 5291.4299 | 0 | 8.7 | 141 | Specific | 292.8 | 254.74 | 254.74 | 6.35 | 134 | >sp\|P02788\|TRFL_HUMAN Lactotransferrin OS=Homo sapiens OX=9606 GN=LTF PE=1 SV=6 | 1270 | 20201116_xd_colostrum_1ng_glyco_hcd203040_compound15.20588.20588.4 | controllerType=0 controllerNumber=1 scan=20588 | 43.9782 |
| 18363:5 | 2 | R.TAGWNVPIGTLRPFLN[+2076.750]WTGPPEPIEAAVAR.F | HexNAc(4)Hex(6)Fuc(2) | N[+2077] | 1062.3016 | 5 | 5307.4791 | 5307.4612 | 0 | 3.4 | 141 | Specific | 305.26 | 290.95 | 290.95 | 6.38 | 134 | >sp\|P02788\|TRFL_HUMAN Lactotransferrin OS=Homo sapiens OX=9606 GN=LTF PE=1 SV=6 | 1270 | 20201116_xd_colostrum_1ng_glyco_hcd203040_compound15.21709.21709.5 | controllerType=0 controllerNumber=1 scan=21709 | 46.3877 |
| 19575:4 | 2 | R.TAGWNVPIGTLRPFLN[+2571.946]WTGPPEPIEAAVAR.F | HexNAc(5)Hex(6)Fuc(4) | N[+2572] | 1451.4181 | 4 | 5802.6505 | 5802.6564 | 0 | -1 | 141 | Specific | 311.37 | 311.37 | 311.37 | 5.99 | 134 | >sp\|P02788\|TRFL_HUMAN Lactotransferrin OS=Homo sapiens OX=9606 GN=LTF PE=1 SV=6 | 1270 | 20201116_xd_colostrum_1ng_glyco_hcd203040_compound15.22975.22975.4 | controllerType=0 controllerNumber=1 scan=22975 | 49.2034 |
| 18239:4 | 2 | R.TAGWNVPIGTLRPFLN[+2716.983]WTGPPEPIEAAVAR.F | HexNAc(5)Hex(6)Fuc(3)NeuAc(1) | N[+2717] | 1487.6859 | 4 | 5947.7218 | 5947.6939 | 0 | 4.7 | 141 | Specific | 328.46 | 328.46 | 328.46 | 7.85 | 134 | >sp\|P02788\|TRFL_HUMAN Lactotransferrin OS=Homo sapiens OX=9606 GN=LTF PE=1 SV=6 | 1270 | 20201116_xd_colostrum_1ng_glyco_hcd203040_compound15.21579.21579.4 | controllerType=0 controllerNumber=1 scan=21579 | 46.0701 |
| 19560:3 | 2 | R.TAGWNVPIGTLRPFLN[+2496.888]WTGPPEPIEAAVAR.F | HexNAc(4)Hex(5)Fuc(2)NeuAc(2) | N[+2497] | 1909.8725 | 3 | 5727.603 | 5727.5992 | 0 | 0.7 | 141 | Specific | 331.52 | 271.91 | 271.91 | 7.31 | 134 | >sp\|P02788\|TRFL_HUMAN Lactotransferrin OS=Homo sapiens OX=9606 GN=LTF PE=1 SV=6 | 1270 | 20201116_xd_colostrum_1ng_glyco_hcd203040_compound15.22960.22960.3 | controllerType=0 controllerNumber=1 scan=22960 | 49.1675 |
| 18416:4 | 2 | R.TAGWNVPIGTLRPFLN[+2076.750]WTGPPEPIEAAVAR.F | HexNAc(4)Hex(6)Fuc(2) | N[+2077] | 1327.6267 | 4 | 5307.485 | 5307.4612 | 0 | 4.5 | 141 | Specific | 370.97 | 370.97 | 370.97 | 8.25 | 134 | >sp\|P02788\|TRFL_HUMAN Lactotransferrin OS=Homo sapiens OX=9606 GN=LTF PE=1 SV=6 | 1270 | 20201116_xd_colostrum_1ng_glyco_hcd203040_compound15.21764.21764.4 | controllerType=0 controllerNumber=1 scan=21764 | 46.4951 |
| 18353:4 | 2 | R.[+42.011]TAGWNVPIGTLRPFLN[+2336.851]WTGPPEPIEAAVAR.F | HexNAc(6)Hex(6)Fuc(1) | .[42], N[+2337] | 1403.1459 | 4 | 5609.5617 | 5609.5726 | 0 | -1.9 | 141 | Specific | 394.84 | 394.84 | 394.84 | 9.13 | 134 | >sp\|P02788\|TRFL_HUMAN Lactotransferrin OS=Homo sapiens OX=9606 GN=LTF PE=1 SV=6 | 1270 | 20201116_xd_colostrum_1ng_glyco_hcd203040_compound15.21699.21699.4 | controllerType=0 controllerNumber=1 scan=21699 | 46.3562 |
| 18346:3 | 2 | R.TAGWNVPIGTLRPFLN[+2059.735]WTGPPEPIEAAVAR.F | HexNAc(4)Hex(5)Fuc(1)NeuAc(1) | N[+2060] | 1764.1589 | 3 | 5290.4621 | 5290.4459 | 0 | 3.1 | 141 | Specific | 409.06 | 360.68 | 360.68 | 8.49 | 134 | >sp\|P02788\|TRFL_HUMAN Lactotransferrin OS=Homo sapiens OX=9606 GN=LTF PE=1 SV=6 | 1270 | 20201116_xd_colostrum_1ng_glyco_hcd203040_compound15.21692.21692.3 | controllerType=0 controllerNumber=1 scan=21692 | 46.3455 |
| 19556:3 | 2 | R.TAGWNVPIGTLRPFLN[+2350.830]WTGPPEPIEAAVAR.F | HexNAc(4)Hex(5)Fuc(1)NeuAc(2) | N[+2351] | 1861.1898 | 3 | 5581.5548 | 5581.5413 | 0 | 2.4 | 141 | Specific | 494.04 | 340.53 | 340.53 | 8.74 | 134 | >sp\|P02788\|TRFL_HUMAN Lactotransferrin OS=Homo sapiens OX=9606 GN=LTF PE=1 SV=6 | 1270 | 20201116_xd_colostrum_1ng_glyco_hcd203040_compound15.22956.22956.3 | controllerType=0 controllerNumber=1 scan=22956 | 49.1592 |
| 18323:3 | 2 | R.TAGWNVPIGTLRPFLN[+2205.793]WTGPPEPIEAAVAR.F | HexNAc(4)Hex(5)Fuc(2)NeuAc(1) | N[+2206] | 1812.8453 | 3 | 5436.5213 | 5436.5038 | 0 | 3.2 | 141 | Specific | 526.66 | 485.55 | 485.55 | 10.47 | 134 | >sp\|P02788\|TRFL_HUMAN Lactotransferrin OS=Homo sapiens OX=9606 GN=LTF PE=1 SV=6 | 1270 | 20201116_xd_colostrum_1ng_glyco_hcd203040_compound15.21668.21668.2 | controllerType=0 controllerNumber=1 scan=21668 | 46.2919 |
| 18301:4 | 2 | R.TAGWNVPIGTLRPFLN[+2570.925]WTGPPEPIEAAVAR.F | HexNAc(5)Hex(6)Fuc(2)NeuAc(1) | N[+2571] | 1451.1736 | 4 | 5801.6725 | 5801.636 | 0 | 6.3 | 141 | Specific | 528.23 | 423.04 | 423.04 | 9.77 | 134 | >sp\|P02788\|TRFL_HUMAN Lactotransferrin OS=Homo sapiens OX=9606 GN=LTF PE=1 SV=6 | 1270 | 20201116_xd_colostrum_1ng_glyco_hcd203040_compound15.21644.21644.4 | controllerType=0 controllerNumber=1 scan=21644 | 46.2221 |
| 18468:4 | 2 | R.TAGWNVPIGTLRPFLN[+2351.851]WTGPPEPIEAAVAR.F | HexNAc(4)Hex(5)Fuc(3)NeuAc(1) | N[+2352] | 1396.401 | 4 | 5582.5822 | 5582.5617 | 0 | 3.7 | 141 | Specific | 535.34 | 524.21 | 524.21 | 10.56 | 134 | >sp\|P02788\|TRFL_HUMAN Lactotransferrin OS=Homo sapiens OX=9606 GN=LTF PE=1 SV=6 | 1270 | 20201116_xd_colostrum_1ng_glyco_hcd203040_compound15.21818.21818.4 | controllerType=0 controllerNumber=1 scan=21818 | 46.5934 |
| 18303:4 | 2 | R.TAGWNVPIGTLRPFLN[+2570.925]WTGPPEPIEAAVAR.F | HexNAc(5)Hex(6)Fuc(2)NeuAc(1) | N[+2571] | 1451.1736 | 4 | 5801.6725 | 5801.636 | 0 | 6.3 | 141 | Specific | 547.62 | 529.63 | 529.63 | 10.01 | 134 | >sp\|P02788\|TRFL_HUMAN Lactotransferrin OS=Homo sapiens OX=9606 GN=LTF PE=1 SV=6 | 1270 | 20201116_xd_colostrum_1ng_glyco_hcd203040_compound15.21646.21646.4 | controllerType=0 controllerNumber=1 scan=21646 | 46.2312 |
| 18307:3 | 2 | R.TAGWNVPIGTLRPFLN[+2351.851]WTGPPEPIEAAVAR.F | HexNAc(4)Hex(5)Fuc(3)NeuAc(1) | N[+2352] | 1861.5308 | 3 | 5582.578 | 5582.5617 | 0 | 2.9 | 141 | Specific | 578.35 | 499.15 | 499.15 | 11.1 | 134 | >sp\|P02788\|TRFL_HUMAN Lactotransferrin OS=Homo sapiens OX=9606 GN=LTF PE=1 SV=6 | 1270 | 20201116_xd_colostrum_1ng_glyco_hcd203040_compound15.21651.21651.3 | controllerType=0 controllerNumber=1 scan=21651 | 46.2484 |
| 17298:3 | 2 | R.TAGWNVPIGTLRPFLN[+2059.735]WTGPPEPIEAAVAR.F | HexNAc(4)Hex(5)Fuc(1)NeuAc(1) | N[+2060] | 1764.1612 | 3 | 5290.4691 | 5290.4459 | 0 | 4.4 | 141 | Specific | 597.18 | 547.4 | 547.4 | 11.97 | 134 | >sp\|P02788\|TRFL_HUMAN Lactotransferrin OS=Homo sapiens OX=9606 GN=LTF PE=1 SV=6 | 1270 | 20201116_xd_colostrum_1ng_glyco_hcd203040_compound15.20597.20597.3 | controllerType=0 controllerNumber=1 scan=20597 | 44.0078 |
| 18326:3 | 2 | R.TAGWNVPIGTLRPFLN[+2205.793]WTGPPEPIEAAVAR.F | HexNAc(4)Hex(5)Fuc(2)NeuAc(1) | N[+2206] | 1812.8453 | 3 | 5436.5213 | 5436.5038 | 0 | 3.2 | 141 | Specific | 603.92 | 513.13 | 513.13 | 13.2 | 134 | >sp\|P02788\|TRFL_HUMAN Lactotransferrin OS=Homo sapiens OX=9606 GN=LTF PE=1 SV=6 | 1270 | 20201116_xd_colostrum_1ng_glyco_hcd203040_compound15.21671.21671.2 | controllerType=0 controllerNumber=1 scan=21671 | 46.3024 |
| 19562:3 | 2 | R.TAGWNVPIGTLRPFLN[+2496.888]WTGPPEPIEAAVAR.F | HexNAc(4)Hex(5)Fuc(2)NeuAc(2) | N[+2497] | 1909.8725 | 3 | 5727.603 | 5727.5992 | 0 | 0.7 | 141 | Specific | 632.81 | 565.48 | 565.48 | 12.97 | 134 | >sp\|P02788\|TRFL_HUMAN Lactotransferrin OS=Homo sapiens OX=9606 GN=LTF PE=1 SV=6 | 1270 | 20201116_xd_colostrum_1ng_glyco_hcd203040_compound15.22962.22962.3 | controllerType=0 controllerNumber=1 scan=22962 | 49.1714 |
| 18298:3 | 2 | R.TAGWNVPIGTLRPFLN[+2205.793]WTGPPEPIEAAVAR.F | HexNAc(4)Hex(5)Fuc(2)NeuAc(1) | N[+2206] | 1812.8514 | 3 | 5436.5397 | 5436.5038 | 0 | 6.6 | 141 | Specific | 634.63 | 530.32 | 530.32 | 12.41 | 134 | >sp\|P02788\|TRFL_HUMAN Lactotransferrin OS=Homo sapiens OX=9606 GN=LTF PE=1 SV=6 | 1270 | 20201116_xd_colostrum_1ng_glyco_hcd203040_compound15.21641.21641.3 | controllerType=0 controllerNumber=1 scan=21641 | 46.2125 |
| 19578:4 | 2 | R.TAGWNVPIGTLRPFLN[+2496.888]WTGPPEPIEAAVAR.F | HexNAc(4)Hex(5)Fuc(2)NeuAc(2) | N[+2497] | 1432.6586 | 4 | 5727.6125 | 5727.5992 | 0 | 2.3 | 141 | Specific | 635.66 | 578.27 | 578.27 | 13.49 | 134 | >sp\|P02788\|TRFL_HUMAN Lactotransferrin OS=Homo sapiens OX=9606 GN=LTF PE=1 SV=6 | 1270 | 20201116_xd_colostrum_1ng_glyco_hcd203040_compound15.22979.22979.4 | controllerType=0 controllerNumber=1 scan=22979 | 49.2135 |
| 19558:3 | 2 | R.TAGWNVPIGTLRPFLN[+2350.830]WTGPPEPIEAAVAR.F | HexNAc(4)Hex(5)Fuc(1)NeuAc(2) | N[+2351] | 1861.1898 | 3 | 5581.5548 | 5581.5413 | 0 | 2.4 | 141 | Specific | 645.96 | 449.77 | 449.77 | 13.4 | 134 | >sp\|P02788\|TRFL_HUMAN Lactotransferrin OS=Homo sapiens OX=9606 GN=LTF PE=1 SV=6 | 1270 | 20201116_xd_colostrum_1ng_glyco_hcd203040_compound15.22958.22958.3 | controllerType=0 controllerNumber=1 scan=22958 | 49.1636 |
| 17296:3 | 2 | R.TAGWNVPIGTLRPFLN[+2059.735]WTGPPEPIEAAVAR.F | HexNAc(4)Hex(5)Fuc(1)NeuAc(1) | N[+2060] | 1764.1612 | 3 | 5290.4691 | 5290.4459 | 0 | 4.4 | 141 | Specific | 646.12 | 613.13 | 613.13 | 12.97 | 134 | >sp\|P02788\|TRFL_HUMAN Lactotransferrin OS=Homo sapiens OX=9606 GN=LTF PE=1 SV=6 | 1270 | 20201116_xd_colostrum_1ng_glyco_hcd203040_compound15.20595.20595.3 | controllerType=0 controllerNumber=1 scan=20595 | 44.0025 |
| 18305:3 | 2 | R.TAGWNVPIGTLRPFLN[+2351.851]WTGPPEPIEAAVAR.F | HexNAc(4)Hex(5)Fuc(3)NeuAc(1) | N[+2352] | 1861.5308 | 3 | 5582.578 | 5582.5617 | 0 | 2.9 | 141 | Specific | 679.87 | 593.24 | 593.24 | 14.4 | 134 | >sp\|P02788\|TRFL_HUMAN Lactotransferrin OS=Homo sapiens OX=9606 GN=LTF PE=1 SV=6 | 1270 | 20201116_xd_colostrum_1ng_glyco_hcd203040_compound15.21649.21649.3 | controllerType=0 controllerNumber=1 scan=21649 | 46.2371 |
| 18290:4 | 2 | R.TAGWNVPIGTLRPFLN[+2205.793]WTGPPEPIEAAVAR.F | HexNAc(4)Hex(5)Fuc(2)NeuAc(1) | N[+2206] | 1359.8909 | 4 | 5436.5417 | 5436.5038 | 0 | 7 | 141 | Specific | 697.35 | 639.7 | 639.7 | 13.5 | 134 | >sp\|P02788\|TRFL_HUMAN Lactotransferrin OS=Homo sapiens OX=9606 GN=LTF PE=1 SV=6 | 1270 | 20201116_xd_colostrum_1ng_glyco_hcd203040_compound15.21633.21633.4 | controllerType=0 controllerNumber=1 scan=21633 | 46.1789 |
| 18304:4 | 2 | R.TAGWNVPIGTLRPFLN[+2351.851]WTGPPEPIEAAVAR.F | HexNAc(4)Hex(5)Fuc(3)NeuAc(1) | N[+2352] | 1396.3997 | 4 | 5582.5768 | 5582.5617 | 0 | 2.7 | 141 | Specific | 713.9 | 679.71 | 679.71 | 15.49 | 134 | >sp\|P02788\|TRFL_HUMAN Lactotransferrin OS=Homo sapiens OX=9606 GN=LTF PE=1 SV=6 | 1270 | 20201116_xd_colostrum_1ng_glyco_hcd203040_compound15.21648.21648.4 | controllerType=0 controllerNumber=1 scan=21648 | 46.2357 |
| 18241:4 | 2 | R.TAGWNVPIGTLRPFLN[+2716.983]WTGPPEPIEAAVAR.F | HexNAc(5)Hex(6)Fuc(3)NeuAc(1) | N[+2717] | 1487.6859 | 4 | 5947.7218 | 5947.6939 | 0 | 4.7 | 141 | Specific | 726.38 | 597.19 | 597.19 | 15.06 | 134 | >sp\|P02788\|TRFL_HUMAN Lactotransferrin OS=Homo sapiens OX=9606 GN=LTF PE=1 SV=6 | 1270 | 20201116_xd_colostrum_1ng_glyco_hcd203040_compound15.21581.21581.4 | controllerType=0 controllerNumber=1 scan=21581 | 46.0753 |
| 18296:3 | 2 | R.TAGWNVPIGTLRPFLN[+2205.793]WTGPPEPIEAAVAR.F | HexNAc(4)Hex(5)Fuc(2)NeuAc(1) | N[+2206] | 1812.8514 | 3 | 5436.5397 | 5436.5038 | 0 | 6.6 | 141 | Specific | 728.31 | 626.65 | 626.65 | 14.41 | 134 | >sp\|P02788\|TRFL_HUMAN Lactotransferrin OS=Homo sapiens OX=9606 GN=LTF PE=1 SV=6 | 1270 | 20201116_xd_colostrum_1ng_glyco_hcd203040_compound15.21639.21639.3 | controllerType=0 controllerNumber=1 scan=21639 | 46.2059 |
| 19531:4 | 2 | R.TAGWNVPIGTLRPFLN[+2350.830]WTGPPEPIEAAVAR.F | HexNAc(4)Hex(5)Fuc(1)NeuAc(2) | N[+2351] | 1396.1499 | 4 | 5581.5778 | 5581.5413 | 0 | 6.5 | 141 | Specific | 745.48 | 639.4 | 639.4 | 14.5 | 134 | >sp\|P02788\|TRFL_HUMAN Lactotransferrin OS=Homo sapiens OX=9606 GN=LTF PE=1 SV=6 | 1270 | 20201116_xd_colostrum_1ng_glyco_hcd203040_compound15.22929.22929.4 | controllerType=0 controllerNumber=1 scan=22929 | 49.0904 |
| 19580:4 | 2 | R.TAGWNVPIGTLRPFLN[+2496.888]WTGPPEPIEAAVAR.F | HexNAc(4)Hex(5)Fuc(2)NeuAc(2) | N[+2497] | 1432.6586 | 4 | 5727.6125 | 5727.5992 | 0 | 2.3 | 141 | Specific | 745.98 | 601.37 | 601.37 | 15.49 | 134 | >sp\|P02788\|TRFL_HUMAN Lactotransferrin OS=Homo sapiens OX=9606 GN=LTF PE=1 SV=6 | 1270 | 20201116_xd_colostrum_1ng_glyco_hcd203040_compound15.22981.22981.4 | controllerType=0 controllerNumber=1 scan=22981 | 49.2192 |
| 18348:3 | 2 | R.TAGWNVPIGTLRPFLN[+2059.735]WTGPPEPIEAAVAR.F | HexNAc(4)Hex(5)Fuc(1)NeuAc(1) | N[+2060] | 1764.1589 | 3 | 5290.4621 | 5290.4459 | 0 | 3.1 | 141 | Specific | 759.64 | 636.11 | 636.11 | 16.2 | 134 | >sp\|P02788\|TRFL_HUMAN Lactotransferrin OS=Homo sapiens OX=9606 GN=LTF PE=1 SV=6 | 1270 | 20201116_xd_colostrum_1ng_glyco_hcd203040_compound15.21694.21694.3 | controllerType=0 controllerNumber=1 scan=21694 | 46.3501 |
| 18354:5 | 2 | R.TAGWNVPIGTLRPFLN[+2059.735]WTGPPEPIEAAVAR.F | HexNAc(4)Hex(5)Fuc(1)NeuAc(1) | N[+2060] | 1058.8982 | 5 | 5290.4619 | 5290.4459 | 0 | 3 | 141 | Specific | 767.88 | 709.06 | 709.06 | 16.57 | 134 | >sp\|P02788\|TRFL_HUMAN Lactotransferrin OS=Homo sapiens OX=9606 GN=LTF PE=1 SV=6 | 1270 | 20201116_xd_colostrum_1ng_glyco_hcd203040_compound15.21700.21700.5 | controllerType=0 controllerNumber=1 scan=21700 | 46.3576 |
| 18345:4 | 2 | R.TAGWNVPIGTLRPFLN[+2059.735]WTGPPEPIEAAVAR.F | HexNAc(4)Hex(5)Fuc(1)NeuAc(1) | N[+2060] | 1323.3713 | 4 | 5290.4635 | 5290.4459 | 0 | 3.3 | 141 | Specific | 771.23 | 701.38 | 701.38 | 16.29 | 134 | >sp\|P02788\|TRFL_HUMAN Lactotransferrin OS=Homo sapiens OX=9606 GN=LTF PE=1 SV=6 | 1270 | 20201116_xd_colostrum_1ng_glyco_hcd203040_compound15.21691.21691.4 | controllerType=0 controllerNumber=1 scan=21691 | 46.3441 |
| 18311:5 | 2 | R.TAGWNVPIGTLRPFLN[+2205.793]WTGPPEPIEAAVAR.F | HexNAc(4)Hex(5)Fuc(2)NeuAc(1) | N[+2206] | 1088.1091 | 5 | 5436.5166 | 5436.5038 | 0 | 2.4 | 141 | Specific | 781.01 | 731.91 | 731.91 | 16.77 | 134 | >sp\|P02788\|TRFL_HUMAN Lactotransferrin OS=Homo sapiens OX=9606 GN=LTF PE=1 SV=6 | 1270 | 20201116_xd_colostrum_1ng_glyco_hcd203040_compound15.21655.21655.5 | controllerType=0 controllerNumber=1 scan=21655 | 46.2598 |
| 18306:4 | 2 | R.TAGWNVPIGTLRPFLN[+2351.851]WTGPPEPIEAAVAR.F | HexNAc(4)Hex(5)Fuc(3)NeuAc(1) | N[+2352] | 1396.3997 | 4 | 5582.5768 | 5582.5617 | 0 | 2.7 | 141 | Specific | 820.34 | 690.72 | 690.72 | 17.49 | 134 | >sp\|P02788\|TRFL_HUMAN Lactotransferrin OS=Homo sapiens OX=9606 GN=LTF PE=1 SV=6 | 1270 | 20201116_xd_colostrum_1ng_glyco_hcd203040_compound15.21650.21650.4 | controllerType=0 controllerNumber=1 scan=21650 | 46.2449 |
| 19557:5 | 2 | R.TAGWNVPIGTLRPFLN[+2350.830]WTGPPEPIEAAVAR.F | HexNAc(4)Hex(5)Fuc(1)NeuAc(2) | N[+2351] | 1117.1171 | 5 | 5581.5562 | 5581.5413 | 0 | 2.7 | 141 | Specific | 857.62 | 738.56 | 738.56 | 18.77 | 134 | >sp\|P02788\|TRFL_HUMAN Lactotransferrin OS=Homo sapiens OX=9606 GN=LTF PE=1 SV=6 | 1270 | 20201116_xd_colostrum_1ng_glyco_hcd203040_compound15.22957.22957.5 | controllerType=0 controllerNumber=1 scan=22957 | 49.1606 |
| 18293:4 | 2 | R.TAGWNVPIGTLRPFLN[+2205.793]WTGPPEPIEAAVAR.F | HexNAc(4)Hex(5)Fuc(2)NeuAc(1) | N[+2206] | 1359.8909 | 4 | 5436.5417 | 5436.5038 | 0 | 7 | 141 | Specific | 870.32 | 696.4 | 696.4 | 17.5 | 134 | >sp\|P02788\|TRFL_HUMAN Lactotransferrin OS=Homo sapiens OX=9606 GN=LTF PE=1 SV=6 | 1270 | 20201116_xd_colostrum_1ng_glyco_hcd203040_compound15.21636.21636.4 | controllerType=0 controllerNumber=1 scan=21636 | 46.1965 |
| 18308:5 | 2 | R.TAGWNVPIGTLRPFLN[+2351.851]WTGPPEPIEAAVAR.F | HexNAc(4)Hex(5)Fuc(3)NeuAc(1) | N[+2352] | 1117.3208 | 5 | 5582.5749 | 5582.5617 | 0 | 2.4 | 141 | Specific | 878.75 | 838.84 | 838.84 | 18.77 | 134 | >sp\|P02788\|TRFL_HUMAN Lactotransferrin OS=Homo sapiens OX=9606 GN=LTF PE=1 SV=6 | 1270 | 20201116_xd_colostrum_1ng_glyco_hcd203040_compound15.21652.21652.5 | controllerType=0 controllerNumber=1 scan=21652 | 46.2495 |
| 19533:4 | 2 | R.TAGWNVPIGTLRPFLN[+2350.830]WTGPPEPIEAAVAR.F | HexNAc(4)Hex(5)Fuc(1)NeuAc(2) | N[+2351] | 1396.1499 | 4 | 5581.5778 | 5581.5413 | 0 | 6.5 | 141 | Specific | 890.44 | 710.55 | 710.55 | 17.5 | 134 | >sp\|P02788\|TRFL_HUMAN Lactotransferrin OS=Homo sapiens OX=9606 GN=LTF PE=1 SV=6 | 1270 | 20201116_xd_colostrum_1ng_glyco_hcd203040_compound15.22931.22931.4 | controllerType=0 controllerNumber=1 scan=22931 | 49.0999 |
| 18471:4 | 2 | R.TAGWNVPIGTLRPFLN[+2351.851]WTGPPEPIEAAVAR.F | HexNAc(4)Hex(5)Fuc(3)NeuAc(1) | N[+2352] | 1396.401 | 4 | 5582.5822 | 5582.5617 | 0 | 3.7 | 141 | Specific | 894.43 | 753.04 | 753.04 | 18.29 | 134 | >sp\|P02788\|TRFL_HUMAN Lactotransferrin OS=Homo sapiens OX=9606 GN=LTF PE=1 SV=6 | 1270 | 20201116_xd_colostrum_1ng_glyco_hcd203040_compound15.21821.21821.4 | controllerType=0 controllerNumber=1 scan=21821 | 46.5976 |
| 18347:4 | 2 | R.TAGWNVPIGTLRPFLN[+2059.735]WTGPPEPIEAAVAR.F | HexNAc(4)Hex(5)Fuc(1)NeuAc(1) | N[+2060] | 1323.3713 | 4 | 5290.4635 | 5290.4459 | 0 | 3.3 | 141 | Specific | 894.5 | 740.44 | 740.44 | 18.29 | 134 | >sp\|P02788\|TRFL_HUMAN Lactotransferrin OS=Homo sapiens OX=9606 GN=LTF PE=1 SV=6 | 1270 | 20201116_xd_colostrum_1ng_glyco_hcd203040_compound15.21693.21693.4 | controllerType=0 controllerNumber=1 scan=21693 | 46.349 |
| 18310:5 | 2 | R.TAGWNVPIGTLRPFLN[+2351.851]WTGPPEPIEAAVAR.F | HexNAc(4)Hex(5)Fuc(3)NeuAc(1) | N[+2352] | 1117.3208 | 5 | 5582.5749 | 5582.5617 | 0 | 2.4 | 141 | Specific | 905.03 | 880.51 | 880.51 | 19.77 | 134 | >sp\|P02788\|TRFL_HUMAN Lactotransferrin OS=Homo sapiens OX=9606 GN=LTF PE=1 SV=6 | 1270 | 20201116_xd_colostrum_1ng_glyco_hcd203040_compound15.21654.21654.5 | controllerType=0 controllerNumber=1 scan=21654 | 46.2589 |
| 18313:5 | 2 | R.TAGWNVPIGTLRPFLN[+2205.793]WTGPPEPIEAAVAR.F | HexNAc(4)Hex(5)Fuc(2)NeuAc(1) | N[+2206] | 1088.1091 | 5 | 5436.5166 | 5436.5038 | 0 | 2.4 | 141 | Specific | 933.97 | 765.18 | 765.18 | 19.77 | 134 | >sp\|P02788\|TRFL_HUMAN Lactotransferrin OS=Homo sapiens OX=9606 GN=LTF PE=1 SV=6 | 1270 | 20201116_xd_colostrum_1ng_glyco_hcd203040_compound15.21657.21657.5 | controllerType=0 controllerNumber=1 scan=21657 | 46.2693 |
| 19559:5 | 2 | R.TAGWNVPIGTLRPFLN[+2350.830]WTGPPEPIEAAVAR.F | HexNAc(4)Hex(5)Fuc(1)NeuAc(2) | N[+2351] | 1117.1171 | 5 | 5581.5562 | 5581.5413 | 0 | 2.7 | 141 | Specific | 944.65 | 773.68 | 773.68 | 19.77 | 134 | >sp\|P02788\|TRFL_HUMAN Lactotransferrin OS=Homo sapiens OX=9606 GN=LTF PE=1 SV=6 | 1270 | 20201116_xd_colostrum_1ng_glyco_hcd203040_compound15.22959.22959.5 | controllerType=0 controllerNumber=1 scan=22959 | 49.1663 |
| 18357:5 | 2 | R.TAGWNVPIGTLRPFLN[+2059.735]WTGPPEPIEAAVAR.F | HexNAc(4)Hex(5)Fuc(1)NeuAc(1) | N[+2060] | 1058.8982 | 5 | 5290.4619 | 5290.4459 | 0 | 3 | 141 | Specific | 955.97 | 865.83 | 865.83 | 20.57 | 134 | >sp\|P02788\|TRFL_HUMAN Lactotransferrin OS=Homo sapiens OX=9606 GN=LTF PE=1 SV=6 | 1270 | 20201116_xd_colostrum_1ng_glyco_hcd203040_compound15.21703.21703.5 | controllerType=0 controllerNumber=1 scan=21703 | 46.3667 |
| 17848:3 | 2 | R.TAGWNIPMGLLFN[+1913.677]QTGSC[+57.021]K.F | HexNAc(4)Hex(5)NeuAc(1) | C[+57], N[+1914] | 1336.9056 | 3 | 4008.7022 | 4008.687 | 0 | 3.8 | 485 | Specific | 394.11 | 390.07 | 390.07 | 9.62 | 134 | >sp\|P02788\|TRFL_HUMAN Lactotransferrin OS=Homo sapiens OX=9606 GN=LTF PE=1 SV=6 | 1270 | 20201116_xd_colostrum_1ng_glyco_hcd203040_compound15.21171.21171.3 | controllerType=0 controllerNumber=1 scan=21171 | 45.2547 |
| 17821:3 | 2 | R.TAGWNIPMGLLFN[+2205.793]QTGSC[+57.021]K.F | HexNAc(4)Hex(5)Fuc(2)NeuAc(1) | C[+57], N[+2206] | 1434.2752 | 3 | 4300.8111 | 4300.8028 | 0 | 1.9 | 485 | Specific | 545.62 | 470.43 | 470.43 | 10.41 | 134 | >sp\|P02788\|TRFL_HUMAN Lactotransferrin OS=Homo sapiens OX=9606 GN=LTF PE=1 SV=6 | 1270 | 20201116_xd_colostrum_1ng_glyco_hcd203040_compound15.21142.21142.3 | controllerType=0 controllerNumber=1 scan=21142 | 45.1916 |
| 15877:3 | 2 | R.TAGWNIPMGLLFN[+1768.640]QTGSC[+57.021]K.F | HexNAc(4)Hex(5)Fuc(1) | C[+57], N[+1769] | 1288.5571 | 3 | 3863.6566 | 3863.6495 | 0 | 1.9 | 485 | Specific | 555.59 | 516.54 | 516.54 | 10.41 | 134 | >sp\|P02788\|TRFL_HUMAN Lactotransferrin OS=Homo sapiens OX=9606 GN=LTF PE=1 SV=6 | 1270 | 20201116_xd_colostrum_1ng_glyco_hcd203040_compound15.19111.19111.3 | controllerType=0 controllerNumber=1 scan=19111 | 40.6258 |
| 17649:3 | 2 | R.TAGWNIPMGLLFN[+2205.793]QTGSC[+57.021]K.F | HexNAc(4)Hex(5)Fuc(2)NeuAc(1) | C[+57], N[+2206] | 1434.278 | 3 | 4300.8195 | 4300.8028 | 0 | 3.9 | 485 | Specific | 610.31 | 585.81 | 585.81 | 12.86 | 134 | >sp\|P02788\|TRFL_HUMAN Lactotransferrin OS=Homo sapiens OX=9606 GN=LTF PE=1 SV=6 | 1270 | 20201116_xd_colostrum_1ng_glyco_hcd203040_compound15.20964.20964.3 | controllerType=0 controllerNumber=1 scan=20964 | 44.8116 |
| 19786:3 | 2 | R.TAGWNIPMGLLFN[+2204.772]QTGSC[+57.021]K.F | HexNAc(4)Hex(5)NeuAc(2) | C[+57], N[+2205] | 1433.9378 | 3 | 4299.7989 | 4299.7824 | 0 | 3.8 | 485 | Specific | 654.71 | 622.65 | 622.65 | 13.86 | 134 | >sp\|P02788\|TRFL_HUMAN Lactotransferrin OS=Homo sapiens OX=9606 GN=LTF PE=1 SV=6 | 1270 | 20201116_xd_colostrum_1ng_glyco_hcd203040_compound15.23197.23197.3 | controllerType=0 controllerNumber=1 scan=23197 | 49.7145 |
| 17672:4 | 2 | R.TAGWNIPMGLLFN[+2205.793]QTGSC[+57.021]K.F | HexNAc(4)Hex(5)Fuc(2)NeuAc(1) | C[+57], N[+2206] | 1075.9594 | 4 | 4300.8157 | 4300.8028 | 0 | 3 | 485 | Specific | 700.73 | 485.35 | 485.35 | 15.14 | 134 | >sp\|P02788\|TRFL_HUMAN Lactotransferrin OS=Homo sapiens OX=9606 GN=LTF PE=1 SV=6 | 1270 | 20201116_xd_colostrum_1ng_glyco_hcd203040_compound15.20987.20987.4 | controllerType=0 controllerNumber=1 scan=20987 | [illegible] |
| 16261:3 | 2 | R.TAGWNIPMGLLFN[+1216.423]QTGSC[+57.021]K.F | HexNAc(2)Hex(5) | C[+57], N[+1216] | 1104.4867 | 3 | 3311.4456 | 3311.4328 | 0 | 3.9 | 485 | Specific | 711.94 | 677.89 | 229.49 | 14.86 | 134 | >sp\|P02788\|TRFL_HUMAN Lactotransferrin OS=Homo sapiens OX=9606 GN=LTF PE=1 SV=6 | 1270 | 20201116_xd_colostrum_1ng_glyco_hcd203040_compound15.19511.19511.3 | controllerType=0 controllerNumber=1 scan=19511 | [illegible] |
| 17779:3 | 2 | R.TAGWNIPMGLLFN[+1856.656]QTGSC[+57.021]K.F | HexNAc(3)Hex(5)Fuc(1)NeuAc(1) | C[+57], N[+1857] | 1317.8996 | 3 | 3951.6842 | 3951.6655 | 0 | 4.7 | 485 | Specific | 719.85 | 674.04 | 285.18 | 14.62 | 134 | >sp\|P02788\|TRFL_HUMAN Lactotransferrin OS=Homo sapiens OX=9606 GN=LTF PE=1 SV=6 | 1270 | 20201116_xd_colostrum_1ng_glyco_hcd203040_compound15.21099.21099.3 | controllerType=0 controllerNumber=1 scan=21099 | [illegible] |
| 17604:3 | 2 | R.TAGWNIPMGLLFN[+2351.851]QTGSC[+57.021]K.F | HexNAc(4)Hex(5)Fuc(3)NeuAc(1) | C[+57], N[+2352] | 1482.9651 | 3 | 4446.8809 | 4446.8607 | 0 | 4.5 | 485 | Specific | 752.89 | 734.29 | 734.29 | 15.62 | 134 | >sp\|P02788\|TRFL_HUMAN Lactotransferrin OS=Homo sapiens OX=9606 GN=LTF PE=1 SV=6 | 1270 | 20201116_xd_colostrum_1ng_glyco_hcd203040_compound15.20917.20917.3 | controllerType=0 controllerNumber=1 scan=20917 | [illegible] |
| 17871:3 | 2 | R.TAGWNIPMGLLFN[+1872.651]QTGSC[+57.021]K.F | HexNAc(3)Hex(6)NeuAc(1) | C[+57], N[+1873] | 1323.2295 | 3 | 3967.6741 | 3967.6604 | 0 | 3.4 | 485 | Specific | 774.25 | 763.11 | 336.86 | 15.86 | 134 | >sp\|P02788\|TRFL_HUMAN Lactotransferrin OS=Homo sapiens OX=9606 GN=LTF PE=1 SV=6 | 1270 | 20201116_xd_colostrum_1ng_glyco_hcd203040_compound15.21195.21195.3 | controllerType=0 controllerNumber=1 scan=21195 | 45.3225 |
| 19788:3 | 2 | R.TAGWNIPMGLLFN[+2204.772]QTGSC[+57.021]K.F | HexNAc(4)Hex(5)NeuAc(2) | C[+57], N[+2205] | 1433.9378 | 3 | 4299.7989 | 4299.7824 | 0 | 3.8 | 485 | Specific | 778.85 | 697.86 | 697.86 | 15.86 | 134 | >sp\|P02788\|TRFL_HUMAN Lactotransferrin OS=Homo sapiens OX=9606 GN=LTF PE=1 SV=6 | 1270 | 20201116_xd_colostrum_1ng_glyco_hcd203040_compound15.23199.23199.3 | controllerType=0 controllerNumber=1 scan=23199 | 49.7188 |
| 16264:3 | 2 | R.TAGWNIPMGLLFN[+1216.423]QTGSC[+57.021]K.F | HexNAc(2)Hex(5) | C[+57], N[+1216] | 1104.4867 | 3 | 3311.4456 | 3311.4328 | 0 | 3.9 | 485 | Specific | 807.52 | 684.73 | 343.98 | 16.86 | 134 | >sp\|P02788\|TRFL_HUMAN Lactotransferrin OS=Homo sapiens OX=9606 GN=LTF PE=1 SV=6 | 1270 | 20201116_xd_colostrum_1ng_glyco_hcd203040_compound15.19514.19514.3 | controllerType=0 controllerNumber=1 scan=19514 | 41.5131 |
| 19628:4 | 2 | R.TAGWNIPMGLLFN[+2350.830]QTGSC[+57.021]K.F | HexNAc(4)Hex(5)Fuc(1)NeuAc(2) | C[+57], N[+2351] | 1112.2187 | 4 | 4445.8529 | 4445.8403 | 0 | 2.8 | 485 | Specific | 834.19 | 760.37 | 760.37 | 17.34 | 134 | >sp\|P02788\|TRFL_HUMAN Lactotransferrin OS=Homo sapiens OX=9606 GN=LTF PE=1 SV=6 | 1270 | 20201116_xd_colostrum_1ng_glyco_hcd203040_compound15.23031.23031.4 | controllerType=0 controllerNumber=1 scan=23031 | 49.3385 |
| 19626:4 | 2 | R.TAGWNIPMGLLFN[+2350.830]QTGSC[+57.021]K.F | HexNAc(4)Hex(5)Fuc(1)NeuAc(2) | C[+57], N[+2351] | 1112.2187 | 4 | 4445.8529 | 4445.8403 | 0 | 2.8 | 485 | Specific | 845.2 | 689.49 | 689.49 | 17.34 | 134 | >sp\|P02788\|TRFL_HUMAN Lactotransferrin OS=Homo sapiens OX=9606 GN=LTF PE=1 SV=6 | 1270 | 20201116_xd_colostrum_1ng_glyco_hcd203040_compound15.23029.23029.4 | controllerType=0 controllerNumber=1 scan=23029 | 49.3254 |
| 17675:3 | 2 | R.TAGWNIPMGLLFN[+2059.735]QTGSC[+57.021]K.F | HexNAc(4)Hex(5)Fuc(1)NeuAc(1) | C[+57], N[+2060] | 1385.59 | 3 | 4154.7555 | 4154.7449 | 0 | 2.6 | 485 | Specific | 893.9 | 804.03 | 804.03 | 18.05 | 134 | >sp\|P02788\|TRFL_HUMAN Lactotransferrin OS=Homo sapiens OX=9606 GN=LTF PE=1 SV=6 | 1270 | 20201116_xd_colostrum_1ng_glyco_hcd203040_compound15.20991.20991.3 | controllerType=0 controllerNumber=1 scan=20991 | 44.8684 |
| 17782:3 | 2 | R.TAGWNIPMGLLFN[+1856.656]QTGSC[+57.021]K.F | HexNAc(3)Hex(5)Fuc(1)NeuAc(1) | C[+57], N[+1857] | 1317.8996 | 3 | 3951.6842 | 3951.6655 | 0 | 4.7 | 485 | Specific | 899.5 | 649.94 | 446.26 | 17.62 | 134 | >sp\|P02788\|TRFL_HUMAN Lactotransferrin OS=Homo sapiens OX=9606 GN=LTF PE=1 SV=6 | 1270 | 20201116_xd_colostrum_1ng_glyco_hcd203040_compound15.21102.21102.3 | controllerType=0 controllerNumber=1 scan=21102 | 45.1075 |
| 15880:3 | 2 | R.TAGWNIPMGLLFN[+1768.640]QTGSC[+57.021]K.F | HexNAc(4)Hex(5)Fuc(1) | C[+57], N[+1769] | 1288.5571 | 3 | 3863.6566 | 3863.6495 | 0 | 1.9 | 485 | Specific | 900.7 | 597.55 | 597.55 | 18.9 | 134 | >sp\|P02788\|TRFL_HUMAN Lactotransferrin OS=Homo sapiens OX=9606 GN=LTF PE=1 SV=6 | 1270 | 20201116_xd_colostrum_1ng_glyco_hcd203040_compound15.19114.19114.3 | controllerType=0 controllerNumber=1 scan=19114 | 40.6308 |
| 17607:3 | 2 | R.TAGWNIPMGLLFN[+2351.851]QTGSC[+57.021]K.F | HexNAc(4)Hex(5)Fuc(3)NeuAc(1) | C[+57], N[+2352] | 1482.9651 | 3 | 4446.8809 | 4446.8607 | 0 | 4.5 | 485 | Specific | 900.9 | 789.02 | 789.02 | 18.62 | 134 | >sp\|P02788\|TRFL_HUMAN Lactotransferrin OS=Homo sapiens OX=9606 GN=LTF PE=1 SV=6 | 1270 | 20201116_xd_colostrum_1ng_glyco_hcd203040_compound15.20920.20920.3 | controllerType=0 controllerNumber=1 scan=20920 | 44.7139 |
| 17851:3 | 2 | R.TAGWNIPMGLLFN[+1913.677]QTGSC[+57.021]K.F | HexNAc(4)Hex(5)NeuAc(1) | C[+57], N[+1914] | 1336.9056 | 3 | 4008.7022 | 4008.687 | 0 | 3.8 | 485 | Specific | 904.18 | 638.42 | 638.42 | 18.86 | 134 | >sp\|P02788\|TRFL_HUMAN Lactotransferrin OS=Homo sapiens OX=9606 GN=LTF PE=1 SV=6 | 1270 | 20201116_xd_colostrum_1ng_glyco_hcd203040_compound15.21174.21174.3 | controllerType=0 controllerNumber=1 scan=21174 | 45.2589 |
| 19627:3 | 2 | R.TAGWNIPMGLLFN[+2350.830]QTGSC[+57.021]K.F | HexNAc(4)Hex(5)Fuc(1)NeuAc(2) | C[+57], N[+2351] | 1482.6236 | 3 | 4445.8562 | 4445.8403 | 0 | 3.6 | 485 | Specific | 914.31 | 890.9 | 890.9 | 18.86 | 134 | >sp\|P02788\|TRFL_HUMAN Lactotransferrin OS=Homo sapiens OX=9606 GN=LTF PE=1 SV=6 | 1270 | 20201116_xd_colostrum_1ng_glyco_hcd203040_compound15.23030.23030.3 | controllerType=0 controllerNumber=1 scan=23030 | 49.3352 |
| 17677:3 | 2 | R.TAGWNIPMGLLFN[+2059.735]QTGSC[+57.021]K.F | HexNAc(4)Hex(5)Fuc(1)NeuAc(1) | C[+57], N[+2060] | 1385.59 | 3 | 4154.7555 | 4154.7449 | 0 | 2.6 | 485 | Specific | 920.66 | 717.48 | 717.48 | 19.05 | 134 | >sp\|P02788\|TRFL_HUMAN Lactotransferrin OS=Homo sapiens OX=9606 GN=LTF PE=1 SV=6 | 1270 | 20201116_xd_colostrum_1ng_glyco_hcd203040_compound15.20993.20993.3 | controllerType=0 controllerNumber=1 scan=20993 | 44.8735 |
| 17651:3 | 2 | R.TAGWNIPMGLLFN[+2205.793]QTGSC[+57.021]K.F | HexNAc(4)Hex(5)Fuc(2)NeuAc(1) | C[+57], N[+2206] | 1434.278 | 3 | 4300.8195 | 4300.8028 | 0 | 3.9 | 485 | Specific | 940.7 | 724.78 | 724.78 | 18.86 | 134 | >sp\|P02788\|TRFL_HUMAN Lactotransferrin OS=Homo sapiens OX=9606 GN=LTF PE=1 SV=6 | 1270 | 20201116_xd_colostrum_1ng_glyco_hcd203040_compound15.20966.20966.3 | controllerType=0 controllerNumber=1 scan=20966 | 44.8158 |
| 17874:3 | 2 | R.TAGWNIPMGLLFN[+1872.651]QTGSC[+57.021]K.F | HexNAc(3)Hex(6)NeuAc(1) | C[+57], N[+1873] | 1323.2295 | 3 | 3967.6741 | 3967.6604 | 0 | 3.4 | 485 | Specific | 971.94 | 741.22 | 404.22 | 19.86 | 134 | >sp\|P02788\|TRFL_HUMAN Lactotransferrin OS=Homo sapiens OX=9606 GN=LTF PE=1 SV=6 | 1270 | 20201116_xd_colostrum_1ng_glyco_hcd203040_compound15.21198.21198.3 | controllerType=0 controllerNumber=1 scan=21198 | 45.3296 |
| 19625:3 | 2 | R.TAGWNIPMGLLFN[+2350.830]QTGSC[+57.021]K.F | HexNAc(4)Hex(5)Fuc(1)NeuAc(2) | C[+57], N[+2351] | 1482.6236 | 3 | 4445.8562 | 4445.8403 | 0 | 3.6 | 485 | Specific | 988.84 | 946.58 | 946.58 | 19.86 | 134 | >sp\|P02788\|TRFL_HUMAN Lactotransferrin OS=Homo sapiens OX=9606 GN=LTF PE=1 SV=6 | 1270 | 20201116_xd_colostrum_1ng_glyco_hcd203040_compound15.23028.23028.3 | controllerType=0 controllerNumber=1 scan=23028 | 49.324 |
| 17961:4 | 2 | R.TAGWNIPMGLLFN[+2205.793]QTGSC[+57.021]KFDEYFSQSC[+57.021]APGSDPR.S | HexNAc(4)Hex(5)Fuc(2)NeuAc(1) | C[+57]*2, N[+2206] | 1536.8983 | 4 | 6144.5714 | 6144.5499 | 0 | 3.5 | 485 | Specific | 254.29 | 227.61 | 227.61 | 6.45 | 134 | >sp\|P02788\|TRFL_HUMAN Lactotransferrin OS=Homo sapiens OX=9606 GN=LTF PE=1 SV=6 | 1270 | 20201116_xd_colostrum_1ng_glyco_hcd203040_compound15.21289.21289.3 | controllerType=0 controllerNumber=1 scan=21289 | 45.5143 |
| 16973:4 | 2 | R.TAGWNIPMGLLFN[+1930.692]QTGSC[+57.021]KFDEYFSQSC[+57.021]APGSDPR.S | HexNAc(4)Hex(6)Fuc(1) | C[+57]*2, N[+1931] | 1468.1218 | 4 | 5869.4655 | 5869.4494 | 0 | 2.7 | 485 | Specific | 271.64 | 257.43 | 257.43 | 6.91 | 134 | >sp\|P02788\|TRFL_HUMAN Lactotransferrin OS=Homo sapiens OX=9606 GN=LTF PE=1 SV=6 | 1270 | 20201116_xd_colostrum_1ng_glyco_hcd203040_compound15.20254.20254.4 | controllerType=0 controllerNumber=1 scan=20254 | 43.1659 |
| 18042:4 | 2 | R.[+42.011]TAGWNIPMGLLFN[+1872.651]QTGSC[+57.021]KFDEYFSQSC[+57.021]A | HexNAc(3)Hex(6)NeuAc(1) | .[42], C[+57]*2, N[+1873] | 1464.1176 | 4 | 5853.4484 | 5853.4181 | 0 | 5.2 | 485 | Specific | 286.29 | 256.04 | 256.04 | 6.25 | 134 | >sp\|P02788\|TRFL_HUMAN Lactotransferrin OS=Homo sapiens OX=9606 GN=LTF PE=1 SV=6 | 1270 | 20201116_xd_colostrum_1ng_glyco_hcd203040_compound15.21373.21373.2 | controllerType=0 controllerNumber=1 scan=21373 | 45.6482 |
| 18045:4 | 2 | R.[+42.011]TAGWNIPMGLLFN[+1872.651]QTGSC[+57.021]KFDEYFSQSC[+57.021]A | HexNAc(3)Hex(6)NeuAc(1) | .[42], C[+57]*2, N[+1873] | 1464.1176 | 4 | 5853.4484 | 5853.4181 | 0 | 5.2 | 485 | Specific | 288.22 | 183.12 | 183.12 | 5.66 | 134 | >sp\|P02788\|TRFL_HUMAN Lactotransferrin OS=Homo sapiens OX=9606 GN=LTF PE=1 SV=6 | 1270 | 20201116_xd_colostrum_1ng_glyco_hcd203040_compound15.21376.21376.2 | controllerType=0 controllerNumber=1 scan=21376 | 45.6521 |
| 17004:4 | 2 | R.TAGWNIPMGLLFN[+2076.750]QTGSC[+57.021]KFDEYFSQSC[+57.021]APGSDPR.S | HexNAc(4)Hex(6)Fuc(2) | C[+57]*2, N[+2077] | 1504.6378 | 4 | 6015.5295 | 6015.5074 | 0 | 3.7 | 485 | Specific | 335.84 | 335.84 | 335.84 | 6.62 | 134 | >sp\|P02788\|TRFL_HUMAN Lactotransferrin OS=Homo sapiens OX=9606 GN=LTF PE=1 SV=6 | 1270 | 20201116_xd_colostrum_1ng_glyco_hcd203040_compound15.20286.20286.4 | controllerType=0 controllerNumber=1 scan=20286 | 43.2361 |
| 16975:4 | 2 | R.TAGWNIPMGLLFN[+1930.692]QTGSC[+57.021]KFDEYFSQSC[+57.021]APGSDPR.S | HexNAc(4)Hex(6)Fuc(1) | C[+57]*2, N[+1931] | 1468.1218 | 4 | 5869.4655 | 5869.4494 | 0 | 2.7 | 485 | Specific | 394.96 | 363.32 | 363.32 | 8.19 | 134 | >sp\|P02788\|TRFL_HUMAN Lactotransferrin OS=Homo sapiens OX=9606 GN=LTF PE=1 SV=6 | 1270 | 20201116_xd_colostrum_1ng_glyco_hcd203040_compound15.20256.20256.4 | controllerType=0 controllerNumber=1 scan=20256 | 43.1715 |
| 18146:3 | 2 | R.TAGWNIPMGLLFN[+1710.598]QTGSC[+57.021]KFDEYFSQSC[+57.021]APGSDPR.S | HexNAc(3)Hex(5)NeuAc(1) | C[+57]*2, N[+1711] | 1883.7873 | 3 | 5649.3474 | 5649.3548 | 0 | -1.3 | 485 | Specific | 478.08 | 433.11 | 148.51 | 8.89 | 134 | >sp\|P02788\|TRFL_HUMAN Lactotransferrin OS=Homo sapiens OX=9606 GN=LTF PE=1 SV=6 | 1270 | 20201116_xd_colostrum_1ng_glyco_hcd203040_compound15.21481.21481.3 | controllerType=0 controllerNumber=1 scan=21481 | 45.8353 |
| 17758:4 | 2 | R.TAGWNIPMGLLFN[+2862.021]QTGSC[+57.021]KFDEYFSQSC[+57.021]APGSDPR.S | HexNAc(5)Hex(6)Fuc(2)NeuAc(2) | C[+57]*2, N[+2862] | 1700.9605 | 4 | 6800.8203 | 6800.7776 | 0 | 6.3 | 485 | Specific | 504.96 | 504.96 | 504.96 | 9.17 | 134 | >sp\|P02788\|TRFL_HUMAN Lactotransferrin OS=Homo sapiens OX=9606 GN=LTF PE=1 SV=6 | 1270 | 20201116_xd_colostrum_1ng_glyco_hcd203040_compound15.21077.21077.4 | controllerType=0 controllerNumber=1 scan=21077 | 45.0386 |
| 19228:5 | 2 | R.TAGWNIPMGLLFN[+2350.830]QTGSC[+57.021]KFDEYFSQSC[+57.021]APGSDPR.S | HexNAc(4)Hex(5)Fuc(1)NeuAc(2) | C[+57]*2, N[+2351] | 1258.7283 | 5 | 6289.6123 | 6289.5875 | 0 | 3.9 | 485 | Specific | 528.73 | 528.73 | 528.73 | 10.29 | 134 | >sp\|P02788\|TRFL_HUMAN Lactotransferrin OS=Homo sapiens OX=9606 GN=LTF PE=1 SV=6 | 1270 | 20201116_xd_colostrum_1ng_glyco_hcd203040_compound15.22614.22614.5 | controllerType=0 controllerNumber=1 scan=22614 | 48.418 |
| 18077:4 | 2 | R.TAGWNIPMGLLFN[+1872.651]QTGSC[+57.021]KFDEYFSQSC[+57.021]APGSDPR.S | HexNAc(3)Hex(6)NeuAc(1) | C[+57]*2, N[+1873] | 1453.6107 | 4 | 5811.4211 | 5811.4076 | 0 | 2.3 | 485 | Specific | 618.15 | 522.54 | 80.05 | 12.94 | 134 | >sp\|P02788\|TRFL_HUMAN Lactotransferrin OS=Homo sapiens OX=9606 GN=LTF PE=1 SV=6 | 1270 | 20201116_xd_colostrum_1ng_glyco_hcd203040_compound15.21410.21410.2 | controllerType=0 controllerNumber=1 scan=21410 | 45.7092 |
| 17884:4 | 2 | R.TAGWNIPMGLLFN[+2351.851]QTGSC[+57.021]KFDEYFSQSC[+57.021]APGSDPR.S | HexNAc(4)Hex(5)Fuc(3)NeuAc(1) | C[+57]*2, N[+2352] | 1573.4144 | 4 | 6290.6359 | 6290.6079 | 0 | 4.5 | 485 | Specific | 708.88 | 685.78 | 685.78 | 14.51 | 134 | >sp\|P02788\|TRFL_HUMAN Lactotransferrin OS=Homo sapiens OX=9606 GN=LTF PE=1 SV=6 | 1270 | 20201116_xd_colostrum_1ng_glyco_hcd203040_compound15.21209.21209.4 | controllerType=0 controllerNumber=1 scan=21209 | 45.3519 |
| 17760:4 | 2 | R.TAGWNIPMGLLFN[+2862.021]QTGSC[+57.021]KFDEYFSQSC[+57.021]APGSDPR.S | HexNAc(5)Hex(6)Fuc(2)NeuAc(2) | C[+57]*2, N[+2862] | 1700.9605 | 4 | 6800.8203 | 6800.7776 | 0 | 6.3 | 485 | Specific | 709.21 | 699.07 | 699.07 | 13.95 | 134 | >sp\|P02788\|TRFL_HUMAN Lactotransferrin OS=Homo sapiens OX=9606 GN=LTF PE=1 SV=6 | 1270 | 20201116_xd_colostrum_1ng_glyco_hcd203040_compound15.21079.21079.4 | controllerType=0 controllerNumber=1 scan=21079 | 45.0432 |
| 19227:4 | 2 | R.TAGWNIPMGLLFN[+2350.830]QTGSC[+57.021]KFDEYFSQSC[+57.021]APGSDPR.S | HexNAc(4)Hex(5)Fuc(1)NeuAc(2) | C[+57]*2, N[+2351] | 1573.1589 | 4 | 6289.6139 | 6289.5875 | 0 | 4.2 | 485 | Specific | 724.93 | 678.29 | 678.29 | 14.51 | 134 | >sp\|P02788\|TRFL_HUMAN Lactotransferrin OS=Homo sapiens OX=9606 GN=LTF PE=1 SV=6 | 1270 | 20201116_xd_colostrum_1ng_glyco_hcd203040_compound15.22613.22613.4 | controllerType=0 controllerNumber=1 scan=22613 | 48.4165 |
| 17100:3 | 2 | R.TAGWNIPMGLLFN[+1216.423]QTGSC[+57.021]KFDEYFSQSC[+57.021]APGSDPR.S | HexNAc(2)Hex(5) | C[+57]*2, N[+1216] | 1719.0706 | 3 | 5155.1974 | 5155.18 | 0 | 3.4 | 485 | Specific | 755.25 | 702.5 | 181.64 | 15.65 | 134 | >sp\|P02788\|TRFL_HUMAN Lactotransferrin OS=Homo sapiens OX=9606 GN=LTF PE=1 SV=6 | 1270 | 20201116_xd_colostrum_1ng_glyco_hcd203040_compound15.20388.20388.3 | controllerType=0 controllerNumber=1 scan=20388 | 43.4917 |
| 17885:5 | 2 | R.TAGWNIPMGLLFN[+2351.851]QTGSC[+57.021]KFDEYFSQSC[+57.021]APGSDPR.S | HexNAc(4)Hex(5)Fuc(3)NeuAc(1) | C[+57]*2, N[+2352] | 1258.9314 | 5 | 6290.6279 | 6290.6079 | 0 | 3.2 | 485 | Specific | 765.88 | 765.88 | 765.88 | 16.02 | 134 | >sp\|P02788\|TRFL_HUMAN Lactotransferrin OS=Homo sapiens OX=9606 GN=LTF PE=1 SV=6 | 1270 | 20201116_xd_colostrum_1ng_glyco_hcd203040_compound15.21210.21210.5 | controllerType=0 controllerNumber=1 scan=21210 | 45.3549 |
| 17103:3 | 2 | R.TAGWNIPMGLLFN[+1216.423]QTGSC[+57.021]KFDEYFSQSC[+57.021]APGSDPR.S | HexNAc(2)Hex(5) | C[+57]*2, N[+1216] | 1719.0706 | 3 | 5155.1974 | 5155.18 | 0 | 3.4 | 485 | Specific | 780.57 | 722.66 | 210.67 | 15.65 | 134 | >sp\|P02788\|TRFL_HUMAN Lactotransferrin OS=Homo sapiens OX=9606 GN=LTF PE=1 SV=6 | 1270 | 20201116_xd_colostrum_1ng_glyco_hcd203040_compound15.20391.20391.3 | controllerType=0 controllerNumber=1 scan=20391 | 43.4963 |
| 19229:4 | 2 | R.TAGWNIPMGLLFN[+2350.830]QTGSC[+57.021]KFDEYFSQSC[+57.021]APGSDPR.S | HexNAc(4)Hex(5)Fuc(1)NeuAc(2) | C[+57]*2, N[+2351] | 1573.1589 | 4 | 6289.6139 | 6289.5875 | 0 | 4.2 | 485 | Specific | 803.64 | 779.61 | 779.61 | 16.51 | 134 | >sp\|P02788\|TRFL_HUMAN Lactotransferrin OS=Homo sapiens OX=9606 GN=LTF PE=1 SV=6 | 1270 | 20201116_xd_colostrum_1ng_glyco_hcd203040_compound15.22615.22615.4 | controllerType=0 controllerNumber=1 scan=22615 | 48.4208 |
| 17887:4 | 2 | R.TAGWNIPMGLLFN[+2351.851]QTGSC[+57.021]KFDEYFSQSC[+57.021]APGSDPR.S | HexNAc(4)Hex(5)Fuc(3)NeuAc(1) | C[+57]*2, N[+2352] | 1573.4144 | 4 | 6290.6359 | 6290.6079 | 0 | 4.5 | 485 | Specific | 844.72 | 817.95 | 817.95 | 16.51 | 134 | >sp\|P02788\|TRFL_HUMAN Lactotransferrin OS=Homo sapiens OX=9606 GN=LTF PE=1 SV=6 | 1270 | 20201116_xd_colostrum_1ng_glyco_hcd203040_compound15.21212.21212.4 | controllerType=0 controllerNumber=1 scan=21212 | 45.3585 |
| 17964:4 | 2 | R.TAGWNIPMGLLFN[+2205.793]QTGSC[+57.021]KFDEYFSQSC[+57.021]APGSDPR.S | HexNAc(4)Hex(5)Fuc(2)NeuAc(1) | C[+57]*2, N[+2206] | 1536.8983 | 4 | 6144.5714 | 6144.5499 | 0 | 3.5 | 485 | Specific | 879.5 | 836.61 | 836.61 | 17.74 | 134 | >sp\|P02788\|TRFL_HUMAN Lactotransferrin OS=Homo sapiens OX=9606 GN=LTF PE=1 SV=6 | 1270 | 20201116_xd_colostrum_1ng_glyco_hcd203040_compound15.21292.21292.3 | controllerType=0 controllerNumber=1 scan=21292 | 45.5218 |
| 17919:4 | 2 | R.TAGWNIPMGLLFN[+2059.735]QTGSC[+57.021]KFDEYFSQSC[+57.021]APGSDPR.S | HexNAc(4)Hex(5)Fuc(1)NeuAc(1) | C[+57]*2, N[+2060] | 1500.3844 | 4 | 5998.5158 | 5998.492 | 0 | 4 | 485 | Specific | 900.67 | 865.75 | 865.75 | 18.74 | 134 | >sp\|P02788\|TRFL_HUMAN Lactotransferrin OS=Homo sapiens OX=9606 GN=LTF PE=1 SV=6 | 1270 | 20201116_xd_colostrum_1ng_glyco_hcd203040_compound15.21245.21245.4 | controllerType=0 controllerNumber=1 scan=21245 | 45.4319 |
| 17916:4 | 2 | R.TAGWNIPMGLLFN[+2059.735]QTGSC[+57.021]KFDEYFSQSC[+57.021]APGSDPR.S | HexNAc(4)Hex(5)Fuc(1)NeuAc(1) | C[+57]*2, N[+2060] | 1500.3844 | 4 | 5998.5158 | 5998.492 | 0 | 4 | 485 | Specific | 911.42 | 875.67 | 875.67 | 18.74 | 134 | >sp\|P02788\|TRFL_HUMAN Lactotransferrin OS=Homo sapiens OX=9606 GN=LTF PE=1 SV=6 | 1270 | 20201116_xd_colostrum_1ng_glyco_hcd203040_compound15.21242.21242.4 | controllerType=0 controllerNumber=1 scan=21242 | 45.4197 |
| 17888:5 | 2 | R.TAGWNIPMGLLFN[+2351.851]QTGSC[+57.021]KFDEYFSQSC[+57.021]APGSDPR.S | HexNAc(4)Hex(5)Fuc(3)NeuAc(1) | C[+57]*2, N[+2352] | 1258.9314 | 5 | 6290.6279 | 6290.6079 | 0 | 3.2 | 485 | Specific | 938.38 | 938.38 | 938.38 | 19.02 | 134 | >sp\|P02788\|TRFL_HUMAN Lactotransferrin OS=Homo sapiens OX=9606 GN=LTF PE=1 SV=6 | 1270 | 20201116_xd_colostrum_1ng_glyco_hcd203040_compound15.21213.21213.5 | controllerType=0 controllerNumber=1 scan=21213 | 45.3597 |
| 18080:4 | 2 | R.TAGWNIPMGLLFN[+1872.651]QTGSC[+57.021]KFDEYFSQSC[+57.021]APGSDPR.S | HexNAc(3)Hex(6)NeuAc(1) | C[+57]*2, N[+1873] | 1453.6107 | 4 | 5811.4211 | 5811.4076 | 0 | 2.3 | 485 | Specific | 944.81 | 848.49 | 332.69 | 18.94 | 134 | >sp\|P02788\|TRFL_HUMAN Lactotransferrin OS=Homo sapiens OX=9606 GN=LTF PE=1 SV=6 | 1270 | 20201116_xd_colostrum_1ng_glyco_hcd203040_compound15.21413.21413.2 | controllerType=0 controllerNumber=1 scan=21413 | 45.7128 |
| 09006:3 | 2 | R.N[+2351.851]GSDC[+57.021]PDKFC[+57.021]LFQSETK.N | HexNAc(4)Hex(5)Fuc(3)NeuAc(1) | C[+57]*2, N[+2352] | 1462.2516 | 3 | 4384.7404 | 4384.7247 | 0 | 3.6 | 642 | Specific | 244.07 | 206.44 | 206.44 | 5.78 | 134 | >sp\|P02788\|TRFL_HUMAN Lactotransferrin OS=Homo sapiens OX=9606 GN=LTF PE=1 SV=6 | 1270 | 20201116_xd_colostrum_1ng_glyco_hcd203040_compound15.11968.11968.3 | controllerType=0 controllerNumber=1 scan=11968 | 26.4997 |
| 09009:3 | 2 | R.N[+2351.851]GSDC[+57.021]PDKFC[+57.021]LFQSETK.N | HexNAc(4)Hex(5)Fuc(3)NeuAc(1) | C[+57]*2, N[+2352] | 1462.2516 | 3 | 4384.7404 | 4384.7247 | 0 | 3.6 | 642 | Specific | 384.03 | 277.87 | 277.87 | 6.71 | 134 | >sp\|P02788\|TRFL_HUMAN Lactotransferrin OS=Homo sapiens OX=9606 GN=LTF PE=1 SV=6 | 1270 | 20201116_xd_colostrum_1ng_glyco_hcd203040_compound15.11971.11971.3 | controllerType=0 controllerNumber=1 scan=11971 | 26.5036 |
| 04149:5 | 3 | R.C[+57.021]IN[+2351.851]GTC[+57.021]YC[+57.021]EEGFTGEDC[+57.021]GKPTC[ | HexNAc(4)Hex(5)Fuc(3)NeuAc(1) | C[+57]*6, N[+2352] | 1203.0653 | 5 | 6011.2974 | 6011.2791 | 0 | 3.1 | 325 | Specific | 295.79 | 295.79 | 295.79 | 7.61 | 118 | >sp\|P24821\|TENA_HUMAN Tenascin OS=Homo sapiens OX=9606 GN=TNC PE=1 SV=3 | 486 | 20201116_xd_colostrum_1ng_glyco_hcd203040_compound15.6932.6932.5 | controllerType=0 controllerNumber=1 scan=6932 | 17.6396 |
| 04093:5 | 3 | R.C[+57.021]IN[+2351.851]GTC[+57.021]YC[+57.021]EEGFTGEDC[+57.021]GKPTC[ | HexNAc(4)Hex(5)Fuc(3)NeuAc(1) | C[+57]*6, N[+2352] | 1203.0653 | 5 | 6011.2974 | 6011.2791 | 0 | 3.1 | 325 | Specific | 747.39 | 747.39 | 747.39 | 15.4 | 118 | >sp\|P24821\|TENA_HUMAN Tenascin OS=Homo sapiens OX=9606 GN=TNC PE=1 SV=3 | 486 | 20201116_xd_colostrum_1ng_glyco_hcd203040_compound15.6874.6874.5 | controllerType=0 controllerNumber=1 scan=6874 | 17.5426 |
| 04091:5 | 3 | R.C[+57.021]IN[+2351.851]GTC[+57.021]YC[+57.021]EEGFTGEDC[+57.021]GKPTC[ | HexNAc(4)Hex(5)Fuc(3)NeuAc(1) | C[+57]*6, N[+2352] | 1203.0653 | 5 | 6011.2974 | 6011.2791 | 0 | 3.1 | 325 | Specific | 855.35 | 855.35 | 855.35 | 18.4 | 118 | >sp\|P24821\|TENA_HUMAN Tenascin OS=Homo sapiens OX=9606 GN=TNC PE=1 SV=3 | 486 | 20201116_xd_colostrum_1ng_glyco_hcd203040_compound15.6872.6872.5 | controllerType=0 controllerNumber=1 scan=6872 | 17.5383 |
| 09026:4 | 3 | R.QTGLAPGQEYEISLHIVKN[+2059.735]NTR.G | HexNAc(4)Hex(5)Fuc(1)NeuAc(1) | N[+2060] | 1132.7652 | 4 | 4528.0389 | 4528.0241 | 0 | 3.3 | 770 | Specific | 423.09 | 247.63 | 247.63 | 7.81 | 118 | >sp\|P24821\|TENA_HUMAN Tenascin OS=Homo sapiens OX=9606 GN=TNC PE=1 SV=3 | 486 | 20201116_xd_colostrum_1ng_glyco_hcd203040_compound15.11988.11988.4 | controllerType=0 controllerNumber=1 scan=11988 | 26.5345 |
| 09028:4 | 3 | R.QTGLAPGQEYEISLHIVKN[+2059.735]NTR.G | HexNAc(4)Hex(5)Fuc(1)NeuAc(1) | N[+2060] | 1132.7652 | 4 | 4528.0389 | 4528.0241 | 0 | 3.3 | 770 | Specific | 478.08 | 331.84 | 331.84 | 7.51 | 118 | >sp\|P24821\|TENA_HUMAN Tenascin OS=Homo sapiens OX=9606 GN=TNC PE=1 SV=3 | 486 | 20201116_xd_colostrum_1ng_glyco_hcd203040_compound15.11990.11990.4 | controllerType=0 controllerNumber=1 scan=11990 | 26.5402 |
| 08931:4 | 3 | R.QTGLAPGQEYEISLHIVKN[+2351.851]NTR.G | HexNAc(4)Hex(5)Fuc(3)NeuAc(1) | N[+2352] | 1205.7929 | 4 | 4820.1497 | 4820.14 | 0 | 2 | 770 | Specific | 518.66 | 394.33 | 394.33 | 10.53 | 118 | >sp\|P24821\|TENA_HUMAN Tenascin OS=Homo sapiens OX=9606 GN=TNC PE=1 SV=3 | 486 | 20201116_xd_colostrum_1ng_glyco_hcd203040_compound15.11890.11890.4 | controllerType=0 controllerNumber=1 scan=11890 | 26.3611 |
| 08929:4 | 3 | R.QTGLAPGQEYEISLHIVKN[+2351.851]NTR.G | HexNAc(4)Hex(5)Fuc(3)NeuAc(1) | N[+2352] | 1205.7929 | 4 | 4820.1497 | 4820.14 | 0 | 2 | 770 | Specific | 534.67 | 500.58 | 500.58 | 10.58 | 118 | >sp\|P24821\|TENA_HUMAN Tenascin OS=Homo sapiens OX=9606 GN=TNC PE=1 SV=3 | 486 | 20201116_xd_colostrum_1ng_glyco_hcd203040_compound15.11888.11888.4 | controllerType=0 controllerNumber=1 scan=11888 | 26.3563 |
| 14672:2 | 3 | R.LN[+1216.423]YSLPTGQWVGVQLPR.N | HexNAc(2)Hex(5) | N[+1216] | 1572.7399 | 2 | 3144.4725 | 3144.4617 | 0 | 3.4 | 1017 | Specific | 835.28 | 680.28 | 680.28 | 16.46 | 118 | >sp\|P24821\|TENA_HUMAN Tenascin OS=Homo sapiens OX=9606 GN=TNC PE=1 SV=3 | 486 | 20201116_xd_colostrum_1ng_glyco_hcd203040_compound15.17856.17856.2 | controllerType=0 controllerNumber=1 scan=17856 | 38.0837 |
| 14669:2 | 3 | R.LN[+1216.423]YSLPTGQWVGVQLPR.N | HexNAc(2)Hex(5) | N[+1216] | 1572.7399 | 2 | 3144.4725 | 3144.4617 | 0 | 3.4 | 1017 | Specific | 934.51 | 807.38 | 807.38 | 18.46 | 118 | >sp\|P24821\|TENA_HUMAN Tenascin OS=Homo sapiens OX=9606 GN=TNC PE=1 SV=3 | 486 | 20201116_xd_colostrum_1ng_glyco_hcd203040_compound15.17853.17853.2 | controllerType=0 controllerNumber=1 scan=17853 | 38.0775 |
| 14666:3 | 3 | R.LN[+1216.423]YSLPTGQWVGVQLPR.N | HexNAc(2)Hex(5) | N[+1216] | 1048.8316 | 3 | 3144.4802 | 3144.4617 | 0 | 5.9 | 1017 | Specific | 1090.47 | 999.56 | 999.56 | 20.91 | 118 | >sp\|P24821\|TENA_HUMAN Tenascin OS=Homo sapiens OX=9606 GN=TNC PE=1 SV=3 | 486 | 20201116_xd_colostrum_1ng_glyco_hcd203040_compound15.17850.17850.3 | controllerType=0 controllerNumber=1 scan=17850 | 38.0709 |
| 14668:3 | 3 | R.LN[+1216.423]YSLPTGQWVGVQLPR.N | HexNAc(2)Hex(5) | N[+1216] | 1048.8316 | 3 | 3144.4802 | 3144.4617 | 0 | 5.9 | 1017 | Specific | 1153.85 | 987.69 | 987.69 | 22.91 | 118 | >sp\|P24821\|TENA_HUMAN Tenascin OS=Homo sapiens OX=9606 GN=TNC PE=1 SV=3 | 486 | 20201116_xd_colostrum_1ng_glyco_hcd203040_compound15.17852.17852.3 | controllerType=0 controllerNumber=1 scan=17852 | 38.0764 |
| 03993:3 | 3 | R.N[+2351.851]TTSYVLR.G | HexNAc(4)Hex(5)Fuc(3)NeuAc(1) | N[+2352] | 1102.4599 | 3 | 3305.365 | 3305.3558 | 0 | 2.8 | 1034 | Specific | 230.82 | 140.03 | 140.03 | 5.2 | 118 | >sp\|P24821\|TENA_HUMAN Tenascin OS=Homo sapiens OX=9606 GN=TNC PE=1 SV=3 | 486 | 20201116_xd_colostrum_1ng_glyco_hcd203040_compound15.6770.6770.3 | controllerType=0 controllerNumber=1 scan=6770 | 17.3603 |
| 04181:3 | 3 | R.N[+2059.735]TTSYVLR.G | HexNAc(4)Hex(5)Fuc(1)NeuAc(1) | N[+2060] | 1005.0869 | 3 | 3013.2463 | 3013.24 | 0 | 2.1 | 1034 | Specific | 233.52 | 138.28 | 138.28 | 4.3 | 118 | >sp\|P24821\|TENA_HUMAN Tenascin OS=Homo sapiens OX=9606 GN=TNC PE=1 SV=3 | 486 | 20201116_xd_colostrum_1ng_glyco_hcd203040_compound15.6965.6965.3 | controllerType=0 controllerNumber=1 scan=6965 | 17.696 |
| 22645:4 | 4 | R.LSLHRPALEDLLLGSEAN[+1501.555]LTC[+57.021]TLTGLR.D | HexNAc(5)Hex(3) | C[+57], N[+1502] | 1117.0463 | 4 | 4465.1634 | 4465.1536 | 0 | 2.2 | 127 | Specific | 212.79 | 212.79 | 212.79 | 4.36 | 109 | >sp\|P01876\|IGHA1_HUMAN Immunoglobulin heavy constant alpha 1 OS=Homo sapiens OX=9606 GN=IGHA1 PE=1 SV=2 | 255 | 20201116_xd_colostrum_1ng_glyco_hcd203040_compound15.26184.26184.2 | controllerType=0 controllerNumber=1 scan=26184 | 56.0728 |
| 24344:4 | 4 | R.LSLHRPALEDLLLGSEAN[+1501.555]LTC[+57.021]TLTGLR.D | HexNAc(5)Hex(3) | C[+57], N[+1502] | 1117.0434 | 4 | 4465.1517 | 4465.1536 | 0 | -0.4 | 127 | Specific | 226.48 | 226.48 | 226.48 | 5.66 | 109 | >sp\|P01876\|IGHA1_HUMAN Immunoglobulin heavy constant alpha 1 OS=Homo sapiens OX=9606 GN=IGHA1 PE=1 SV=2 | 255 | 20201116_xd_colostrum_1ng_glyco_hcd203040_compound15.27952.27952.3 | controllerType=0 controllerNumber=1 scan=27952 | 59.4056 |
| 19093:3 | 4 | R.[+42.011]LSLHRPALEDLLLGSEAN[+1955.724]LTC[+57.021]TLTGLR.D | HexNAc(5)Hex(4)Fuc(2) | .[42], C[+57], N[+1956] | 1654.4428 | 3 | 4961.314 | 4961.3328 | 0 | -3.8 | 127 | Specific | 253.29 | 171.53 | 171.53 | 3.84 | 109 | >sp\|P01876\|IGHA1_HUMAN Immunoglobulin heavy constant alpha 1 OS=Homo sapiens OX=9606 GN=IGHA1 PE=1 SV=2 | 255 | 20201116_xd_colostrum_1ng_glyco_hcd203040_compound15.22473.22473.3 | controllerType=0 controllerNumber=1 scan=22473 | 48.0892 |
| 33929:4 | 4 | R.LSLHRPALEDLLLGSEAN[+1216.423]LTC[+57.021]TLTGLR.D | HexNAc(2)Hex(5) | C[+57], N[+1216] | 1045.7625 | 4 | 4180.0281 | 4180.0211 | 0 | 1.7 | 127 | Specific | 253.41 | 253.41 | 253.41 | 6.95 | 109 | >sp\|P01876\|IGHA1_HUMAN Immunoglobulin heavy constant alpha 1 OS=Homo sapiens OX=9606 GN=IGHA1 PE=1 SV=2 | 255 | 20201116_xd_colostrum_1ng_glyco_hcd203040_compound15.40958.40958.4 | controllerType=0 controllerNumber=1 scan=40958 | 89.7439 |
| 22153:4 | 4 | R.LSLHRPALEDLLLGSEAN[+1501.555]LTC[+57.021]TLTGLR.D | HexNAc(5)Hex(3) | C[+57], N[+1502] | 1117.0461 | 4 | 4465.1624 | 4465.1536 | 0 | 2 | 127 | Specific | 254.07 | 141.7 | 141.7 | 5.06 | 109 | >sp\|P01876\|IGHA1_HUMAN Immunoglobulin heavy constant alpha 1 OS=Homo sapiens OX=9606 GN=IGHA1 PE=1 SV=2 | 255 | 20201116_xd_colostrum_1ng_glyco_hcd203040_compound15.25669.25669.4 | controllerType=0 controllerNumber=1 scan=25669 | 55.0223 |
| 22627:4 | 4 | R.LSLHRPALEDLLLGSEAN[+1501.555]LTC[+57.021]TLTGLR.D | HexNAc(5)Hex(3) | C[+57], N[+1502] | 1117.0404 | 4 | 4465.14 | 4465.1536 | 0 | -3.1 | 127 | Specific | 256.49 | 156.19 | 156.19 | 3.77 | 109 | >sp\|P01876\|IGHA1_HUMAN Immunoglobulin heavy constant alpha 1 OS=Homo sapiens OX=9606 GN=IGHA1 PE=1 SV=2 | 255 | 20201116_xd_colostrum_1ng_glyco_hcd203040_compound15.26165.26165.4 | controllerType=0 controllerNumber=1 scan=26165 | 56.0362 |
| 22647:4 | 4 | R.LSLHRPALEDLLLGSEAN[+1501.555]LTC[+57.021]TLTGLR.D | HexNAc(5)Hex(3) | C[+57], N[+1502] | 1117.0463 | 4 | 4465.1634 | 4465.1536 | 0 | 2.2 | 127 | Specific | 277.96 | 132.93 | 132.93 | 5.95 | 109 | >sp\|P01876\|IGHA1_HUMAN Immunoglobulin heavy constant alpha 1 OS=Homo sapiens OX=9606 GN=IGHA1 PE=1 SV=2 | 255 | 20201116_xd_colostrum_1ng_glyco_hcd203040_compound15.26186.26186.2 | controllerType=0 controllerNumber=1 scan=26186 | 56.0778 |
| 33764:4 | 4 | R.LSLHRPALEDLLLGSEAN[+1216.423]LTC[+57.021]TLTGLR.D | HexNAc(2)Hex(5) | C[+57], N[+1216] | 1045.7636 | 4 | 4180.0325 | 4180.0211 | 0 | 2.7 | 127 | Specific | 287.56 | 287.56 | 287.56 | 7.28 | 109 | >sp\|P01876\|IGHA1_HUMAN Immunoglobulin heavy constant alpha 1 OS=Homo sapiens OX=9606 GN=IGHA1 PE=1 SV=2 | 255 | 20201116_xd_colostrum_1ng_glyco_hcd203040_compound15.40769.40769.2 | controllerType=0 controllerNumber=1 scan=40769 | 89.3725 |
| 33460:4 | 4 | R.LSLHRPALEDLLLGSEAN[+1216.423]LTC[+57.021]TLTGLR.D | HexNAc(2)Hex(5) | C[+57], N[+1216] | 1045.7631 | 4 | 4180.0306 | 4180.0211 | 0 | 2.3 | 127 | Specific | 295.99 | 295.99 | 295.99 | 7.28 | 109 | >sp\|P01876\|IGHA1_HUMAN Immunoglobulin heavy constant alpha 1 OS=Homo sapiens OX=9606 GN=IGHA1 PE=1 SV=2 | 255 | 20201116_xd_colostrum_1ng_glyco_hcd203040_compound15.40431.40431.4 | controllerType=0 controllerNumber=1 scan=40431 | 88.713 |
| 16704:3 | 4 | R.LSLHRPALEDLLLGSEAN[+1825.661]LTC[+57.021]TLTGLR.D | HexNAc(5)Hex(5) | C[+57], N[+1826] | 1597.1005 | 3 | 4789.287 | 4789.2592 | 0 | 5.8 | 127 | Specific | 296.15 | 271.14 | 271.14 | 5.8 | 109 | >sp\|P01876\|IGHA1_HUMAN Immunoglobulin heavy constant alpha 1 OS=Homo sapiens OX=9606 GN=IGHA1 PE=1 SV=2 | 255 | 20201116_xd_colostrum_1ng_glyco_hcd203040_compound15.19972.19972.3 | controllerType=0 controllerNumber=1 scan=19972 | 42.455 |
| 33493:4 | 4 | R.LSLHRPALEDLLLGSEAN[+1501.555]LTC[+57.021]TLTGLR.D | HexNAc(5)Hex(3) | C[+57], N[+1502] | 1117.0433 | 4 | 4465.1512 | 4465.1536 | 0 | -0.5 | 127 | Specific | 317.5 | 317.5 | 317.5 | 6.54 | 109 | >sp\|P01876\|IGHA1_HUMAN Immunoglobulin heavy constant alpha 1 OS=Homo sapiens OX=9606 GN=IGHA1 PE=1 SV=2 | 255 | 20201116_xd_colostrum_1ng_glyco_hcd203040_compound15.40467.40467.4 | controllerType=0 controllerNumber=1 scan=40467 | 88.7827 |
| 20443:4 | 4 | R.LSLHRPALEDLLLGSEAN[+1501.555]LTC[+57.021]TLTGLR.D | HexNAc(5)Hex(3) | C[+57], N[+1502] | 1117.0463 | 4 | 4465.1634 | 4465.1536 | 0 | 2.2 | 127 | Specific | 318.98 | 216.93 | 216.93 | 6.72 | 109 | >sp\|P01876\|IGHA1_HUMAN Immunoglobulin heavy constant alpha 1 OS=Homo sapiens OX=9606 GN=IGHA1 PE=1 SV=2 | 255 | 20201116_xd_colostrum_1ng_glyco_hcd203040_compound15.23883.23883.3 | controllerType=0 controllerNumber=1 scan=23883 | 51.2285 |
| 33445:4 | 4 | R.LSLHRPALEDLLLGSEAN[+1216.423]LTC[+57.021]TLTGLR.D | HexNAc(2)Hex(5) | C[+57], N[+1216] | 1045.7627 | 4 | 4180.0291 | 4180.0211 | 0 | 1.9 | 127 | Specific | 324.07 | 324.07 | 324.07 | 7.71 | 109 | >sp\|P01876\|IGHA1_HUMAN Immunoglobulin heavy constant alpha 1 OS=Homo sapiens OX=9606 GN=IGHA1 PE=1 SV=2 | 255 | 20201116_xd_colostrum_1ng_glyco_hcd203040_compound15.40415.40415.2 | controllerType=0 controllerNumber=1 scan=40415 | 88.6833 |
| 17679:3 | 4 | R.LSLHRPALEDLLLGSEAN[+1751.624]LTC[+57.021]TLTGLR.D | HexNAc(4)Hex(4)NeuAc(1) | C[+57], N[+1752] | 1572.4166 | 3 | 4715.2351 | 4715.2225 | 0 | 2.7 | 127 | Specific | 328.3 | 293.79 | 293.79 | 6.95 | 109 | >sp\|P01876\|IGHA1_HUMAN Immunoglobulin heavy constant alpha 1 OS=Homo sapiens OX=9606 GN=IGHA1 PE=1 SV=2 | 255 | 20201116_xd_colostrum_1ng_glyco_hcd203040_compound15.20995.20995.3 | controllerType=0 controllerNumber=1 scan=20995 | 44.8757 |
| 33462:4 | 4 | R.LSLHRPALEDLLLGSEAN[+1216.423]LTC[+57.021]TLTGLR.D | HexNAc(2)Hex(5) | C[+57], N[+1216] | 1045.7631 | 4 | 4180.0306 | 4180.0211 | 0 | 2.3 | 127 | Specific | 330.02 | 330.02 | 330.02 | 7.86 | 109 | >sp\|P01876\|IGHA1_HUMAN Immunoglobulin heavy constant alpha 1 OS=Homo sapiens OX=9606 GN=IGHA1 PE=1 SV=2 | 255 | 20201116_xd_colostrum_1ng_glyco_hcd203040_compound15.40433.40433.4 | controllerType=0 controllerNumber=1 scan=40433 | 88.7178 |

Table S4. Cont.

| Query #:z | Protein Rank | Glycopeptide | Glycan | Modification Type(s) | Observed m/z | z | Observed (M+H) | Calc. mass (M+H) | Off-by-x error | Mass error (ppm) | StartPosition | Cleavage | Score | Delta | Delta Mod | \|Log Prob\| | # of unique peptides | Protein Name | Protein DB number | Comment | Scan # | Scan Time |
|---|---|---|---|---|---|---|---|---|---|---|---|---|---|---|---|---|---|---|---|---|---|---|
| 19042:4 | 4 | R.[+42.011]LSLHRPALEDLLLGSEAN[+2116.756]LTC[+57.021]TLTGLR.D | HexNAc(5)Hex(5)NeuAc(1) | .[42], C[+57], N[+2117] | 1281.3506 | 4 | 5122.3805 | 5122.3652 | 0 | 3 | 127 | Specific | 333.96 | 270.15 | 111.36 | 7.41 | 109 | >sp\|P01876\|IGHA1_HUMAN Immunoglobulin heavy constant alpha 1 OS=Homo sapiens OX=9606 GN=IGHA1 PE=1 SV=2 | 255 | 20201116_xd_colostrum_1ng_glyco_hcd203040_compound15.22420.22420.4 | controllerType=0 controllerNumber=1 scan=22420 | 47.9735 |
| 17577:4 | 4 | R.LSLHRPALEDLLLGSEAN[+2278.809]LTC[+57.021]TLTGLR.D | HexNAc(5)Hex(6)NeuAc(1) | C[+57], N[+2279] | 1311.3539 | 4 | 5242.3937 | 5242.4075 | 0 | -2.6 | 127 | Specific | 336.6 | 273.44 | 273.44 | 6.3 | 109 | >sp\|P01876\|IGHA1_HUMAN Immunoglobulin heavy constant alpha 1 OS=Homo sapiens OX=9606 GN=IGHA1 PE=1 SV=2 | 255 | 20201116_xd_colostrum_1ng_glyco_hcd203040_compound15.20889.20889.4 | controllerType=0 controllerNumber=1 scan=20889 | 44.6471 |
| 19102:5 | 4 | R.[+42.011]LSLHRPALEDLLLGSEAN[+1955.724]LTC[+57.021]TLTGLR.D | HexNAc(5)Hex(4)Fuc(2) | .[42], C[+57], N[+1956] | 993.071 | 5 | 4961.3261 | 4961.3328 | 0 | -1.3 | 127 | Specific | 336.75 | 336.75 | 336.75 | 6.92 | 109 | >sp\|P01876\|IGHA1_HUMAN Immunoglobulin heavy constant alpha 1 OS=Homo sapiens OX=9606 GN=IGHA1 PE=1 SV=2 | 255 | 20201116_xd_colostrum_1ng_glyco_hcd203040_compound15.22482.22482.5 | controllerType=0 controllerNumber=1 scan=22482 | 48.1071 |
| 18721:3 | 4 | R.LSLHRPALEDLLLGSEAN[+2407.852]LTC[+57.021]TLTGLR.D | HexNAc(5)Hex(5)NeuAc(2) | C[+57], N[+2408] | 1791.161 | 3 | 5371.4684 | 5371.4501 | 0 | 3.4 | 127 | Specific | 337.85 | 257.9 | 257.9 | 6.69 | 109 | >sp\|P01876\|IGHA1_HUMAN Immunoglobulin heavy constant alpha 1 OS=Homo sapiens OX=9606 GN=IGHA1 PE=1 SV=2 | 255 | 20201116_xd_colostrum_1ng_glyco_hcd203040_compound15.22084.22084.3 | controllerType=0 controllerNumber=1 scan=22084 | 47.2085 |
| 19090:4 | 4 | R.[+42.011]LSLHRPALEDLLLGSEAN[+1955.724]LTC[+57.021]TLTGLR.D | HexNAc(5)Hex(4)Fuc(2) | .[42], C[+57], N[+1956] | 1241.0862 | 4 | 4961.3229 | 4961.3328 | 0 | -2 | 127 | Specific | 340 | 328.73 | 328.73 | 6.83 | 109 | >sp\|P01876\|IGHA1_HUMAN Immunoglobulin heavy constant alpha 1 OS=Homo sapiens OX=9606 GN=IGHA1 PE=1 SV=2 | 255 | 20201116_xd_colostrum_1ng_glyco_hcd203040_compound15.22470.22470.4 | controllerType=0 controllerNumber=1 scan=22470 | 48.0818 |
| 17156:4 | 4 | R.LSLHRPALEDLLLGSEAN[+1501.555]LTC[+57.021]TLTGLR.D | HexNAc(5)Hex(3) | C[+57], N[+1502] | 1117.04 | 4 | 4465.1382 | 4465.1536 | 0 | -3.5 | 127 | Specific | 354.98 | 311.43 | 311.43 | 5.52 | 109 | >sp\|P01876\|IGHA1_HUMAN Immunoglobulin heavy constant alpha 1 OS=Homo sapiens OX=9606 GN=IGHA1 PE=1 SV=2 | 255 | 20201116_xd_colostrum_1ng_glyco_hcd203040_compound15.20446.20446.4 | controllerType=0 controllerNumber=1 scan=20446 | 43.6225 |
| 16792:5 | 4 | R.LSLHRPALEDLLLGSEAN[+1501.555]LTC[+57.021]TLTGLR.D | HexNAc(5)Hex(3) | C[+57], N[+1502] | 893.8383 | 5 | 4465.1626 | 4465.1536 | 0 | 2 | 127 | Specific | 355.02 | 355.02 | 355.02 | 7.96 | 109 | >sp\|P01876\|IGHA1_HUMAN Immunoglobulin heavy constant alpha 1 OS=Homo sapiens OX=9606 GN=IGHA1 PE=1 SV=2 | 255 | 20201116_xd_colostrum_1ng_glyco_hcd203040_compound15.20065.20065.5 | controllerType=0 controllerNumber=1 scan=20065 | 42.7268 |
| 33628:4 | 4 | R.LSLHRPALEDLLLGSEAN[+1216.423]LTC[+57.021]TLTGLR.D | HexNAc(2)Hex(5) | C[+57], N[+1216] | 1045.7632 | 4 | 4180.0311 | 4180.0211 | 0 | 2.4 | 127 | Specific | 373.86 | 373.86 | 373.86 | 8.26 | 109 | >sp\|P01876\|IGHA1_HUMAN Immunoglobulin heavy constant alpha 1 OS=Homo sapiens OX=9606 GN=IGHA1 PE=1 SV=2 | 255 | 20201116_xd_colostrum_1ng_glyco_hcd203040_compound15.40617.40617.3 | controllerType=0 controllerNumber=1 scan=40617 | 89.0783 |
| 20445:4 | 4 | R.LSLHRPALEDLLLGSEAN[+1501.555]LTC[+57.021]TLTGLR.D | HexNAc(5)Hex(3) | C[+57], N[+1502] | 1117.0463 | 4 | 4465.1634 | 4465.1536 | 0 | 2.2 | 127 | Specific | 404.5 | 308.16 | 308.16 | 7.95 | 109 | >sp\|P01876\|IGHA1_HUMAN Immunoglobulin heavy constant alpha 1 OS=Homo sapiens OX=9606 GN=IGHA1 PE=1 SV=2 | 255 | 20201116_xd_colostrum_1ng_glyco_hcd203040_compound15.23885.23885.3 | controllerType=0 controllerNumber=1 scan=23885 | 51.2336 |
| 19105:5 | 4 | R.[+42.011]LSLHRPALEDLLLGSEAN[+1955.724]LTC[+57.021]TLTGLR.D | HexNAc(5)Hex(4)Fuc(2) | .[42], C[+57], N[+1956] | 993.071 | 5 | 4961.3261 | 4961.3328 | 0 | -1.3 | 127 | Specific | 406.73 | 389.92 | 389.92 | 8.12 | 109 | >sp\|P01876\|IGHA1_HUMAN Immunoglobulin heavy constant alpha 1 OS=Homo sapiens OX=9606 GN=IGHA1 PE=1 SV=2 | 255 | 20201116_xd_colostrum_1ng_glyco_hcd203040_compound15.22485.22485.5 | controllerType=0 controllerNumber=1 scan=22485 | 48.1147 |
| 19044:4 | 4 | R.[+42.011]LSLHRPALEDLLLGSEAN[+2116.756]LTC[+57.021]TLTGLR.D | HexNAc(5)Hex(5)NeuAc(1) | .[42], C[+57], N[+2117] | 1281.3506 | 4 | 5122.3805 | 5122.3652 | 0 | 3 | 127 | Specific | 406.86 | 338.55 | 180.75 | 8.1 | 109 | >sp\|P01876\|IGHA1_HUMAN Immunoglobulin heavy constant alpha 1 OS=Homo sapiens OX=9606 GN=IGHA1 PE=1 SV=2 | 255 | 20201116_xd_colostrum_1ng_glyco_hcd203040_compound15.22422.22422.4 | controllerType=0 controllerNumber=1 scan=22422 | 47.9785 |
| 19108:4 | 4 | R.[+42.011]LSLHRPALEDLLLGSEAN[+1710.598]LTC[+57.021]TLTGLR.D | HexNAc(3)Hex(5)NeuAc(1) | .[42], C[+57], N[+1711] | 1179.8089 | 4 | 4716.2137 | 4716.2065 | 0 | 1.5 | 127 | Specific | 426.13 | 422.03 | 422.03 | 9.79 | 109 | >sp\|P01876\|IGHA1_HUMAN Immunoglobulin heavy constant alpha 1 OS=Homo sapiens OX=9606 GN=IGHA1 PE=1 SV=2 | 255 | 20201116_xd_colostrum_1ng_glyco_hcd203040_compound15.22488.22488.4 | controllerType=0 controllerNumber=1 scan=22488 | 48.12 |
| 16891:4 | 4 | R.LSLHRPALEDLLLGSEAN[+1622.582]LTC[+57.021]TLTGLR.D | HexNAc(4)Hex(5) | C[+57], N[+1623] | 1147.3031 | 4 | 4586.1907 | 4586.1799 | 0 | 2.4 | 127 | Specific | 431.91 | 431.91 | 431.91 | 9.94 | 109 | >sp\|P01876\|IGHA1_HUMAN Immunoglobulin heavy constant alpha 1 OS=Homo sapiens OX=9606 GN=IGHA1 PE=1 SV=2 | 255 | 20201116_xd_colostrum_1ng_glyco_hcd203040_compound15.20168.20168.4 | controllerType=0 controllerNumber=1 scan=20168 | 42.9413 |
| 17558:4 | 4 | R.LSLHRPALEDLLLGSEAN[+2262.814]LTC[+57.021]TLTGLR.D | HexNAc(5)Hex(5)Fuc(1)NeuAc(1) | C[+57], N[+2263] | 1307.3628 | 4 | 5226.4294 | 5226.4126 | 0 | 3.2 | 127 | Specific | 443.52 | 443.52 | 443.52 | 9.82 | 109 | >sp\|P01876\|IGHA1_HUMAN Immunoglobulin heavy constant alpha 1 OS=Homo sapiens OX=9606 GN=IGHA1 PE=1 SV=2 | 255 | 20201116_xd_colostrum_1ng_glyco_hcd203040_compound15.20869.20869.4 | controllerType=0 controllerNumber=1 scan=20869 | 44.5931 |
| 19092:4 | 4 | R.[+42.011]LSLHRPALEDLLLGSEAN[+1955.724]LTC[+57.021]TLTGLR.D | HexNAc(5)Hex(4)Fuc(2) | .[42], C[+57], N[+1956] | 1241.0862 | 4 | 4961.3229 | 4961.3328 | 0 | -2 | 127 | Specific | 476.02 | 446.84 | 446.84 | 9.02 | 109 | >sp\|P01876\|IGHA1_HUMAN Immunoglobulin heavy constant alpha 1 OS=Homo sapiens OX=9606 GN=IGHA1 PE=1 SV=2 | 255 | 20201116_xd_colostrum_1ng_glyco_hcd203040_compound15.22472.22472.4 | controllerType=0 controllerNumber=1 scan=22472 | 48.0874 |
| 16673:4 | 4 | R.LSLHRPALEDLLLGSEAN[+1825.661]LTC[+57.021]TLTGLR.D | HexNAc(5)Hex(5) | C[+57], N[+1826] | 1198.0728 | 4 | 4789.2694 | 4789.2592 | 0 | 2.1 | 127 | Specific | 484.29 | 484.29 | 484.29 | 10.17 | 109 | >sp\|P01876\|IGHA1_HUMAN Immunoglobulin heavy constant alpha 1 OS=Homo sapiens OX=9606 GN=IGHA1 PE=1 SV=2 | 255 | 20201116_xd_colostrum_1ng_glyco_hcd203040_compound15.19940.19940.4 | controllerType=0 controllerNumber=1 scan=19940 | 42.3804 |
| 16392:4 | 4 | R.LSLHRPALEDLLLGSEAN[+1501.555]LTC[+57.021]TLTGLR.D | HexNAc(5)Hex(3) | C[+57], N[+1502] | 1117.0458 | 4 | 4465.1614 | 4465.1536 | 0 | 1.8 | 127 | Specific | 484.39 | 419.62 | 419.62 | 10.02 | 109 | >sp\|P01876\|IGHA1_HUMAN Immunoglobulin heavy constant alpha 1 OS=Homo sapiens OX=9606 GN=IGHA1 PE=1 SV=2 | 255 | 20201116_xd_colostrum_1ng_glyco_hcd203040_compound15.19647.19647.4 | controllerType=0 controllerNumber=1 scan=19647 | 41.7564 |
| 16794:5 | 4 | R.LSLHRPALEDLLLGSEAN[+1501.555]LTC[+57.021]TLTGLR.D | HexNAc(5)Hex(3) | C[+57], N[+1502] | 893.8383 | 5 | 4465.1626 | 4465.1536 | 0 | 2 | 127 | Specific | 496.57 | 496.57 | 496.57 | 10.11 | 109 | >sp\|P01876\|IGHA1_HUMAN Immunoglobulin heavy constant alpha 1 OS=Homo sapiens OX=9606 GN=IGHA1 PE=1 SV=2 | 255 | 20201116_xd_colostrum_1ng_glyco_hcd203040_compound15.20067.20067.5 | controllerType=0 controllerNumber=1 scan=20067 | 42.7334 |
| 17707:3 | 4 | R.LSLHRPALEDLLLGSEAN[+1548.545]LTC[+57.021]TLTGLR.D | HexNAc(3)Hex(4)NeuAc(1) | C[+57], N[+1549] | 1504.7261 | 3 | 4512.1638 | 4512.1431 | 0 | 4.6 | 127 | Specific | 522.83 | 522.83 | 522.83 | 9.35 | 109 | >sp\|P01876\|IGHA1_HUMAN Immunoglobulin heavy constant alpha 1 OS=Homo sapiens OX=9606 GN=IGHA1 PE=1 SV=2 | 255 | 20201116_xd_colostrum_1ng_glyco_hcd203040_compound15.21024.21024.3 | controllerType=0 controllerNumber=1 scan=21024 | 44.9297 |
| 18689:4 | 4 | R.LSLHRPALEDLLLGSEAN[+2407.852]LTC[+57.021]TLTGLR.D | HexNAc(5)Hex(5)NeuAc(2) | C[+57], N[+2408] | 1343.6215 | 4 | 5371.464 | 5371.4501 | 0 | 2.6 | 127 | Specific | 532.15 | 443.84 | 443.84 | 10.34 | 109 | >sp\|P01876\|IGHA1_HUMAN Immunoglobulin heavy constant alpha 1 OS=Homo sapiens OX=9606 GN=IGHA1 PE=1 SV=2 | 255 | 20201116_xd_colostrum_1ng_glyco_hcd203040_compound15.22051.22051.4 | controllerType=0 controllerNumber=1 scan=22051 | 47.147 |
| 16781:3 | 4 | R.LSLHRPALEDLLLGSEAN[+1460.529]LTC[+57.021]TLTGLR.D | HexNAc(4)Hex(4) | C[+57], N[+1461] | 1475.3859 | 3 | 4424.1432 | 4424.1271 | 0 | 3.7 | 127 | Specific | 537.82 | 514.4 | 514.4 | 9.59 | 109 | >sp\|P01876\|IGHA1_HUMAN Immunoglobulin heavy constant alpha 1 OS=Homo sapiens OX=9606 GN=IGHA1 PE=1 SV=2 | 255 | 20201116_xd_colostrum_1ng_glyco_hcd203040_compound15.20053.20053.3 | controllerType=0 controllerNumber=1 scan=20053 | 42.7032 |
| 17614:3 | 4 | R.LSLHRPALEDLLLGSEAN[+2116.756]LTC[+57.021]TLTGLR.D | HexNAc(5)Hex(5)NeuAc(1) | C[+57], N[+2117] | 1694.13 | 3 | 5080.3754 | 5080.3547 | 0 | 4.1 | 127 | Specific | 538.21 | 504.23 | 504.23 | 9.35 | 109 | >sp\|P01876\|IGHA1_HUMAN Immunoglobulin heavy constant alpha 1 OS=Homo sapiens OX=9606 GN=IGHA1 PE=1 SV=2 | 255 | 20201116_xd_colostrum_1ng_glyco_hcd203040_compound15.20927.20927.2 | controllerType=0 controllerNumber=1 scan=20927 | 44.7301 |
| 19110:4 | 4 | R.[+42.011]LSLHRPALEDLLLGSEAN[+1710.598]LTC[+57.021]TLTGLR.D | HexNAc(3)Hex(5)NeuAc(1) | .[42], C[+57], N[+1711] | 1179.8089 | 4 | 4716.2137 | 4716.2065 | 0 | 1.5 | 127 | Specific | 547.9 | 486.74 | 486.74 | 10.42 | 109 | >sp\|P01876\|IGHA1_HUMAN Immunoglobulin heavy constant alpha 1 OS=Homo sapiens OX=9606 GN=IGHA1 PE=1 SV=2 | 255 | 20201116_xd_colostrum_1ng_glyco_hcd203040_compound15.22490.22490.4 | controllerType=0 controllerNumber=1 scan=22490 | 48.1252 |
| 17605:3 | 4 | R.LSLHRPALEDLLLGSEAN[+1872.651]LTC[+57.021]TLTGLR.D | HexNAc(3)Hex(6)NeuAc(1) | C[+57], N[+1873] | 1612.7617 | 3 | 4836.2704 | 4836.2487 | 0 | 4.5 | 127 | Specific | 574.67 | 574.67 | 574.67 | 9.78 | 109 | >sp\|P01876\|IGHA1_HUMAN Immunoglobulin heavy constant alpha 1 OS=Homo sapiens OX=9606 GN=IGHA1 PE=1 SV=2 | 255 | 20201116_xd_colostrum_1ng_glyco_hcd203040_compound15.20918.20918.3 | controllerType=0 controllerNumber=1 scan=20918 | 44.7107 |
| 16707:3 | 4 | R.LSLHRPALEDLLLGSEAN[+1825.661]LTC[+57.021]TLTGLR.D | HexNAc(5)Hex(5) | C[+57], N[+1826] | 1597.1005 | 3 | 4789.287 | 4789.2592 | 0 | 5.8 | 127 | Specific | 588.94 | 471.37 | 471.37 | 10.68 | 109 | >sp\|P01876\|IGHA1_HUMAN Immunoglobulin heavy constant alpha 1 OS=Homo sapiens OX=9606 GN=IGHA1 PE=1 SV=2 | 255 | 20201116_xd_colostrum_1ng_glyco_hcd203040_compound15.19975.19975.3 | controllerType=0 controllerNumber=1 scan=19975 | 42.4605 |
| 17682:3 | 4 | R.LSLHRPALEDLLLGSEAN[+1751.624]LTC[+57.021]TLTGLR.D | HexNAc(4)Hex(4)NeuAc(1) | C[+57], N[+1752] | 1572.4166 | 3 | 4715.2351 | 4715.2225 | 0 | 2.7 | 127 | Specific | 590.86 | 587.43 | 587.43 | 11.51 | 109 | >sp\|P01876\|IGHA1_HUMAN Immunoglobulin heavy constant alpha 1 OS=Homo sapiens OX=9606 GN=IGHA1 PE=1 SV=2 | 255 | 20201116_xd_colostrum_1ng_glyco_hcd203040_compound15.20998.20998.3 | controllerType=0 controllerNumber=1 scan=20998 | 44.8796 |
| 17534:4 | 4 | R.LSLHRPALEDLLLGSEAN[+1913.677]LTC[+57.021]TLTGLR.D | HexNAc(4)Hex(5)NeuAc(1) | C[+57], N[+1914] | 1220.0746 | 4 | 4877.2767 | 4877.2753 | 0 | 0.3 | 127 | Specific | 592.29 | 592.29 | 592.29 | 11.64 | 109 | >sp\|P01876\|IGHA1_HUMAN Immunoglobulin heavy constant alpha 1 OS=Homo sapiens OX=9606 GN=IGHA1 PE=1 SV=2 | 255 | 20201116_xd_colostrum_1ng_glyco_hcd203040_compound15.20844.20844.4 | controllerType=0 controllerNumber=1 scan=20844 | 44.5366 |
| 17600:4 | 4 | R.LSLHRPALEDLLLGSEAN[+1913.677]LTC[+57.021]TLTGLR.D | HexNAc(4)Hex(5)NeuAc(1) | C[+57], N[+1914] | 1220.0795 | 4 | 4877.2962 | 4877.2753 | 0 | 4.3 | 127 | Specific | 592.99 | 592.99 | 592.99 | 11.64 | 109 | >sp\|P01876\|IGHA1_HUMAN Immunoglobulin heavy constant alpha 1 OS=Homo sapiens OX=9606 GN=IGHA1 PE=1 SV=2 | 255 | 20201116_xd_colostrum_1ng_glyco_hcd203040_compound15.20913.20913.4 | controllerType=0 controllerNumber=1 scan=20913 | 44.7017 |
| 16783:3 | 4 | R.LSLHRPALEDLLLGSEAN[+1501.555]LTC[+57.021]TLTGLR.D | HexNAc(5)Hex(3) | C[+57], N[+1502] | 1489.0608 | 3 | 4465.168 | 4465.1536 | 0 | 3.2 | 127 | Specific | 595.68 | 595.68 | 595.68 | 11.32 | 109 | >sp\|P01876\|IGHA1_HUMAN Immunoglobulin heavy constant alpha 1 OS=Homo sapiens OX=9606 GN=IGHA1 PE=1 SV=2 | 255 | 20201116_xd_colostrum_1ng_glyco_hcd203040_compound15.20055.20055.2 | controllerType=0 controllerNumber=1 scan=20055 | 42.7055 |
| 17579:4 | 4 | R.LSLHRPALEDLLLGSEAN[+2278.809]LTC[+57.021]TLTGLR.D | HexNAc(5)Hex(6)NeuAc(1) | C[+57], N[+2279] | 1311.3539 | 4 | 5242.3937 | 5242.4075 | 0 | -2.6 | 127 | Specific | 598.96 | 537.49 | 537.49 | 10.95 | 109 | >sp\|P01876\|IGHA1_HUMAN Immunoglobulin heavy constant alpha 1 OS=Homo sapiens OX=9606 GN=IGHA1 PE=1 SV=2 | 255 | 20201116_xd_colostrum_1ng_glyco_hcd203040_compound15.20891.20891.4 | controllerType=0 controllerNumber=1 scan=20891 | 44.6594 |
| 16665:4 | 4 | R.LSLHRPALEDLLLGSEAN[+1864.634]LTC[+57.021]TLTGLR.D | HexNAc(2)Hex(9) | C[+57], N[+1865] | 1207.8144 | 4 | 4828.2357 | 4828.2324 | 0 | 0.7 | 127 | Specific | 604.03 | 604.03 | 604.03 | 12.64 | 109 | >sp\|P01876\|IGHA1_HUMAN Immunoglobulin heavy constant alpha 1 OS=Homo sapiens OX=9606 GN=IGHA1 PE=1 SV=2 | 255 | 20201116_xd_colostrum_1ng_glyco_hcd203040_compound15.19931.19931.4 | controllerType=0 controllerNumber=1 scan=19931 | 42.3445 |
| 17559:3 | 4 | R.LSLHRPALEDLLLGSEAN[+1913.677]LTC[+57.021]TLTGLR.D | HexNAc(4)Hex(5)NeuAc(1) | C[+57], N[+1914] | 1626.4355 | 3 | 4877.2919 | 4877.2753 | 0 | 3.4 | 127 | Specific | 604.88 | 604.88 | 604.88 | 12.32 | 109 | >sp\|P01876\|IGHA1_HUMAN Immunoglobulin heavy constant alpha 1 OS=Homo sapiens OX=9606 GN=IGHA1 PE=1 SV=2 | 255 | 20201116_xd_colostrum_1ng_glyco_hcd203040_compound15.20870.20870.3 | controllerType=0 controllerNumber=1 scan=20870 | 44.5939 |
| 17710:3 | 4 | R.LSLHRPALEDLLLGSEAN[+1548.545]LTC[+57.021]TLTGLR.D | HexNAc(3)Hex(4)NeuAc(1) | C[+57], N[+1549] | 1504.7261 | 3 | 4512.1638 | 4512.1431 | 0 | 4.6 | 127 | Specific | 605.6 | 595.05 | 595.05 | 12.08 | 109 | >sp\|P01876\|IGHA1_HUMAN Immunoglobulin heavy constant alpha 1 OS=Homo sapiens OX=9606 GN=IGHA1 PE=1 SV=2 | 255 | 20201116_xd_colostrum_1ng_glyco_hcd203040_compound15.21027.21027.3 | controllerType=0 controllerNumber=1 scan=21027 | 44.934 |
| 16683:3 | 4 | R.LSLHRPALEDLLLGSEAN[+1663.608]LTC[+57.021]TLTGLR.D | HexNAc(5)Hex(4) | C[+57], N[+1664] | 1543.0768 | 3 | 4627.216 | 4627.2064 | 0 | 2.1 | 127 | Specific | 616.14 | 568.37 | 568.37 | 12.36 | 109 | >sp\|P01876\|IGHA1_HUMAN Immunoglobulin heavy constant alpha 1 OS=Homo sapiens OX=9606 GN=IGHA1 PE=1 SV=2 | 255 | 20201116_xd_colostrum_1ng_glyco_hcd203040_compound15.19950.19950.3 | controllerType=0 controllerNumber=1 scan=19950 | 42.4052 |
| 16784:3 | 4 | R.LSLHRPALEDLLLGSEAN[+1460.529]LTC[+57.021]TLTGLR.D | HexNAc(4)Hex(4) | C[+57], N[+1461] | 1475.3859 | 3 | 4424.1432 | 4424.1271 | 0 | 3.7 | 127 | Specific | 619.51 | 537.07 | 537.07 | 12.32 | 109 | >sp\|P01876\|IGHA1_HUMAN Immunoglobulin heavy constant alpha 1 OS=Homo sapiens OX=9606 GN=IGHA1 PE=1 SV=2 | 255 | 20201116_xd_colostrum_1ng_glyco_hcd203040_compound15.20056.20056.3 | controllerType=0 controllerNumber=1 scan=20056 | 42.7073 |
| 17608:3 | 4 | R.LSLHRPALEDLLLGSEAN[+1872.651]LTC[+57.021]TLTGLR.D | HexNAc(3)Hex(6)NeuAc(1) | C[+57], N[+1873] | 1612.7617 | 3 | 4836.2704 | 4836.2487 | 0 | 4.5 | 127 | Specific | 623.34 | 563.56 | 563.56 | 12.08 | 109 | >sp\|P01876\|IGHA1_HUMAN Immunoglobulin heavy constant alpha 1 OS=Homo sapiens OX=9606 GN=IGHA1 PE=1 SV=2 | 255 | 20201116_xd_colostrum_1ng_glyco_hcd203040_compound15.20921.20921.3 | controllerType=0 controllerNumber=1 scan=20921 | 44.7182 |
| 16687:3 | 4 | R.LSLHRPALEDLLLGSEAN[+1702.581]LTC[+57.021]TLTGLR.D | HexNAc(2)Hex(8) | C[+57], N[+1703] | 1556.0681 | 3 | 4666.1896 | 4666.1796 | 0 | 2.1 | 127 | Specific | 631.12 | 585.76 | 585.76 | 12.51 | 109 | >sp\|P01876\|IGHA1_HUMAN Immunoglobulin heavy constant alpha 1 OS=Homo sapiens OX=9606 GN=IGHA1 PE=1 SV=2 | 255 | 20201116_xd_colostrum_1ng_glyco_hcd203040_compound15.19954.19954.3 | controllerType=0 controllerNumber=1 scan=19954 | 42.416 |
| 16686:3 | 4 | R.LSLHRPALEDLLLGSEAN[+1663.608]LTC[+57.021]TLTGLR.D | HexNAc(5)Hex(4) | C[+57], N[+1664] | 1543.0768 | 3 | 4627.216 | 4627.2064 | 0 | 2.1 | 127 | Specific | 632.95 | 558.66 | 558.66 | 12.36 | 109 | >sp\|P01876\|IGHA1_HUMAN Immunoglobulin heavy constant alpha 1 OS=Homo sapiens OX=9606 GN=IGHA1 PE=1 SV=2 | 255 | 20201116_xd_colostrum_1ng_glyco_hcd203040_compound15.19953.19953.3 | controllerType=0 controllerNumber=1 scan=19953 | 42.4141 |
| 18671:4 | 4 | R.LSLHRPALEDLLLGSEAN[+2407.852]LTC[+57.021]TLTGLR.D | HexNAc(5)Hex(5)NeuAc(2) | C[+57], N[+2408] | 1343.6201 | 4 | 5371.4587 | 5371.4501 | 0 | 1.6 | 127 | Specific | 637.36 | 636.97 | 636.97 | 12.91 | 109 | >sp\|P01876\|IGHA1_HUMAN Immunoglobulin heavy constant alpha 1 OS=Homo sapiens OX=9606 GN=IGHA1 PE=1 SV=2 | 255 | 20201116_xd_colostrum_1ng_glyco_hcd203040_compound15.22032.22032.4 | controllerType=0 controllerNumber=1 scan=22032 | 47.0934 |
| 16769:3 | 4 | R.LSLHRPALEDLLLGSEAN[+1501.555]LTC[+57.021]TLTGLR.D | HexNAc(5)Hex(3) | C[+57], N[+1502] | 1489.0602 | 3 | 4465.1662 | 4465.1536 | 0 | 2.8 | 127 | Specific | 641.01 | 594.51 | 594.51 | 12.51 | 109 | >sp\|P01876\|IGHA1_HUMAN Immunoglobulin heavy constant alpha 1 OS=Homo sapiens OX=9606 GN=IGHA1 PE=1 SV=2 | 255 | 20201116_xd_colostrum_1ng_glyco_hcd203040_compound15.20040.20040.3 | controllerType=0 controllerNumber=1 scan=20040 | 42.6624 |
| 16774:3 | 4 | R.LSLHRPALEDLLLGSEAN[+1298.476]LTC[+57.021]TLTGLR.D | HexNAc(4)Hex(3) | C[+57], N[+1298] | 1421.3669 | 3 | 4262.0861 | 4262.0742 | 0 | 2.8 | 127 | Specific | 645.09 | 604.25 | 604.25 | 12.51 | 109 | >sp\|P01876\|IGHA1_HUMAN Immunoglobulin heavy constant alpha 1 OS=Homo sapiens OX=9606 GN=IGHA1 PE=1 SV=2 | 255 | 20201116_xd_colostrum_1ng_glyco_hcd203040_compound15.20046.20046.3 | controllerType=0 controllerNumber=1 scan=20046 | 42.6857 |
| 17626:3 | 4 | R.LSLHRPALEDLLLGSEAN[+1710.598]LTC[+57.021]TLTGLR.D | HexNAc(3)Hex(5)NeuAc(1) | C[+57], N[+1711] | 1558.7416 | 3 | 4674.2103 | 4674.1959 | 0 | 3.1 | 127 | Specific | 648.47 | 560.74 | 560.74 | 12.32 | 109 | >sp\|P01876\|IGHA1_HUMAN Immunoglobulin heavy constant alpha 1 OS=Homo sapiens OX=9606 GN=IGHA1 PE=1 SV=2 | 255 | 20201116_xd_colostrum_1ng_glyco_hcd203040_compound15.20940.20940.3 | controllerType=0 controllerNumber=1 scan=20940 | 44.7582 |
| 16786:3 | 4 | R.LSLHRPALEDLLLGSEAN[+1501.555]LTC[+57.021]TLTGLR.D | HexNAc(5)Hex(3) | C[+57], N[+1502] | 1489.0608 | 3 | 4465.168 | 4465.1536 | 0 | 3.2 | 127 | Specific | 649.46 | 606.01 | 606.01 | 12.32 | 109 | >sp\|P01876\|IGHA1_HUMAN Immunoglobulin heavy constant alpha 1 OS=Homo sapiens OX=9606 GN=IGHA1 PE=1 SV=2 | 255 | 20201116_xd_colostrum_1ng_glyco_hcd203040_compound15.20058.20058.2 | controllerType=0 controllerNumber=1 scan=20058 | 42.7136 |
| 16684:3 | 4 | R.LSLHRPALEDLLLGSEAN[+1702.581]LTC[+57.021]TLTGLR.D | HexNAc(2)Hex(8) | C[+57], N[+1703] | 1556.0681 | 3 | 4666.1896 | 4666.1796 | 0 | 2.1 | 127 | Specific | 651.01 | 651.01 | 651.01 | 13.51 | 109 | >sp\|P01876\|IGHA1_HUMAN Immunoglobulin heavy constant alpha 1 OS=Homo sapiens OX=9606 GN=IGHA1 PE=1 SV=2 | 255 | 20201116_xd_colostrum_1ng_glyco_hcd203040_compound15.19951.19951.3 | controllerType=0 controllerNumber=1 scan=19951 | 42.4064 |
| 17628:3 | 4 | R.LSLHRPALEDLLLGSEAN[+1710.598]LTC[+57.021]TLTGLR.D | HexNAc(3)Hex(5)NeuAc(1) | C[+57], N[+1711] | 1558.7416 | 3 | 4674.2103 | 4674.1959 | 0 | 3.1 | 127 | Specific | 652.36 | 567 | 567 | 13.32 | 109 | >sp\|P01876\|IGHA1_HUMAN Immunoglobulin heavy constant alpha 1 OS=Homo sapiens OX=9606 GN=IGHA1 PE=1 SV=2 | 255 | 20201116_xd_colostrum_1ng_glyco_hcd203040_compound15.20942.20942.3 | controllerType=0 controllerNumber=1 scan=20942 | 44.7657 |
| 16766:3 | 4 | R.LSLHRPALEDLLLGSEAN[+1501.555]LTC[+57.021]TLTGLR.D | HexNAc(5)Hex(3) | C[+57], N[+1502] | 1489.0602 | 3 | 4465.1662 | 4465.1536 | 0 | 2.8 | 127 | Specific | 656.66 | 656.66 | 656.66 | 13.51 | 109 | >sp\|P01876\|IGHA1_HUMAN Immunoglobulin heavy constant alpha 1 OS=Homo sapiens OX=9606 GN=IGHA1 PE=1 SV=2 | 255 | 20201116_xd_colostrum_1ng_glyco_hcd203040_compound15.20037.20037.3 | controllerType=0 controllerNumber=1 scan=20037 | 42.6372 |
| 16772:3 | 4 | R.LSLHRPALEDLLLGSEAN[+1298.476]LTC[+57.021]TLTGLR.D | HexNAc(4)Hex(3) | C[+57], N[+1298] | 1421.3669 | 3 | 4262.0861 | 4262.0742 | 0 | 2.8 | 127 | Specific | 657.37 | 657.37 | 657.37 | 13.51 | 109 | >sp\|P01876\|IGHA1_HUMAN Immunoglobulin heavy constant alpha 1 OS=Homo sapiens OX=9606 GN=IGHA1 PE=1 SV=2 | 255 | 20201116_xd_colostrum_1ng_glyco_hcd203040_compound15.20044.20044.3 | controllerType=0 controllerNumber=1 scan=20044 | 42.676 |
| 17562:3 | 4 | R.LSLHRPALEDLLLGSEAN[+1913.677]LTC[+57.021]TLTGLR.D | HexNAc(4)Hex(5)NeuAc(1) | C[+57], N[+1914] | 1626.4355 | 3 | 4877.2919 | 4877.2753 | 0 | 3.4 | 127 | Specific | 661.54 | 586.56 | 586.56 | 13.32 | 109 | >sp\|P01876\|IGHA1_HUMAN Immunoglobulin heavy constant alpha 1 OS=Homo sapiens OX=9606 GN=IGHA1 PE=1 SV=2 | 255 | 20201116_xd_colostrum_1ng_glyco_hcd203040_compound15.20873.20873.3 | controllerType=0 controllerNumber=1 scan=20873 | 44.6029 |
| 18691:4 | 4 | R.LSLHRPALEDLLLGSEAN[+2204.772]LTC[+57.021]TLTGLR.D | HexNAc(4)Hex(5)NeuAc(2) | C[+57], N[+2205] | 1292.8518 | 4 | 5168.3854 | 5168.3707 | 0 | 2.8 | 127 | Specific | 666.16 | 643.02 | 643.02 | 14.07 | 109 | >sp\|P01876\|IGHA1_HUMAN Immunoglobulin heavy constant alpha 1 OS=Homo sapiens OX=9606 GN=IGHA1 PE=1 SV=2 | 255 | 20201116_xd_colostrum_1ng_glyco_hcd203040_compound15.22053.22053.4 | controllerType=0 controllerNumber=1 scan=22053 | 47.1494 |
| 16394:4 | 4 | R.LSLHRPALEDLLLGSEAN[+1501.555]LTC[+57.021]TLTGLR.D | HexNAc(5)Hex(3) | C[+57], N[+1502] | 1117.0458 | 4 | 4465.1614 | 4465.1536 | 0 | 1.8 | 127 | Specific | 676.59 | 400.38 | 400.38 | 13.91 | 109 | >sp\|P01876\|IGHA1_HUMAN Immunoglobulin heavy constant alpha 1 OS=Homo sapiens OX=9606 GN=IGHA1 PE=1 SV=2 | 255 | 20201116_xd_colostrum_1ng_glyco_hcd203040_compound15.19649.19649.4 | controllerType=0 controllerNumber=1 scan=19649 | 41.765 |
| 16779:3 | 4 | R.LSLHRPALEDLLLGSEAN[+1054.370]LTC[+57.021]TLTGLR.D | HexNAc(2)Hex(4) | C[+57], N[+1054] | 1339.9975 | 3 | 4017.9779 | 4017.9683 | 0 | 2.4 | 127 | Specific | 691.71 | 687.14 | 687.14 | 13.51 | 109 | >sp\|P01876\|IGHA1_HUMAN Immunoglobulin heavy constant alpha 1 OS=Homo sapiens OX=9606 GN=IGHA1 PE=1 SV=2 | 255 | 20201116_xd_colostrum_1ng_glyco_hcd203040_compound15.20051.20051.3 | controllerType=0 controllerNumber=1 scan=20051 | 42.7006 |
| 16697:4 | 4 | R.LSLHRPALEDLLLGSEAN[+1581.555]LTC[+57.021]TLTGLR.D | HexNAc(3)Hex(6) | C[+57], N[+1582] | 1137.0474 | 4 | 4545.1678 | 4545.1533 | 0 | 3.2 | 127 | Specific | 702.31 | 702.31 | 702.31 | 14.87 | 109 | >sp\|P01876\|IGHA1_HUMAN Immunoglobulin heavy constant alpha 1 OS=Homo sapiens OX=9606 GN=IGHA1 PE=1 SV=2 | 255 | 20201116_xd_colostrum_1ng_glyco_hcd203040_compound15.19965.19965.4 | controllerType=0 controllerNumber=1 scan=19965 | 42.4444 |
| 18673:4 | 4 | R.LSLHRPALEDLLLGSEAN[+2407.852]LTC[+57.021]TLTGLR.D | HexNAc(5)Hex(5)NeuAc(2) | C[+57], N[+2408] | 1343.6201 | 4 | 5371.4587 | 5371.4501 | 0 | 1.6 | 127 | Specific | 706.52 | 640.52 | 640.52 | 14.91 | 109 | >sp\|P01876\|IGHA1_HUMAN Immunoglobulin heavy constant alpha 1 OS=Homo sapiens OX=9606 GN=IGHA1 PE=1 SV=2 | 255 | 20201116_xd_colostrum_1ng_glyco_hcd203040_compound15.22034.22034.4 | controllerType=0 controllerNumber=1 scan=22034 | 47.1012 |
| 16674:4 | 4 | R.LSLHRPALEDLLLGSEAN[+1663.608]LTC[+57.021]TLTGLR.D | HexNAc(5)Hex(4) | C[+57], N[+1664] | 1157.5595 | 4 | 4627.2161 | 4627.2064 | 0 | 2.1 | 127 | Specific | 711.49 | 711.49 | 711.49 | 15.07 | 109 | >sp\|P01876\|IGHA1_HUMAN Immunoglobulin heavy constant alpha 1 OS=Homo sapiens OX=9606 GN=IGHA1 PE=1 SV=2 | 255 | 20201116_xd_colostrum_1ng_glyco_hcd203040_compound15.19941.19941.4 | controllerType=0 controllerNumber=1 scan=19941 | 42.3818 |
| 16658:4 | 4 | R.LSLHRPALEDLLLGSEAN[+1663.608]LTC[+57.021]TLTGLR.D | HexNAc(5)Hex(4) | C[+57], N[+1664] | 1157.5583 | 4 | 4627.2113 | 4627.2064 | 0 | 1 | 127 | Specific | 714.24 | 708.58 | 708.58 | 14.91 | 109 | >sp\|P01876\|IGHA1_HUMAN Immunoglobulin heavy constant alpha 1 OS=Homo sapiens OX=9606 GN=IGHA1 PE=1 SV=2 | 255 | 20201116_xd_colostrum_1ng_glyco_hcd203040_compound15.19924.19924.4 | controllerType=0 controllerNumber=1 scan=19924 | 42.3282 |
| 17601:4 | 4 | R.LSLHRPALEDLLLGSEAN[+1872.651]LTC[+57.021]TLTGLR.D | HexNAc(3)Hex(6)NeuAc(1) | C[+57], N[+1873] | 1209.824 | 4 | 4836.2742 | 4836.2487 | 0 | 5.3 | 127 | Specific | 714.74 | 714.74 | 714.74 | 14.24 | 109 | >sp\|P01876\|IGHA1_HUMAN Immunoglobulin heavy constant alpha 1 OS=Homo sapiens OX=9606 GN=IGHA1 PE=1 SV=2 | 255 | 20201116_xd_colostrum_1ng_glyco_hcd203040_compound15.20914.20914.4 | controllerType=0 controllerNumber=1 scan=20914 | 44.7032 |
| 16767:4 | 4 | R.LSLHRPALEDLLLGSEAN[+1460.529]LTC[+57.021]TLTGLR.D | HexNAc(4)Hex(4) | C[+57], N[+1461] | 1106.7896 | 4 | 4424.1365 | 4424.1271 | 0 | 2.1 | 127 | Specific | 718.97 | 691.06 | 691.06 | 15.07 | 109 | >sp\|P01876\|IGHA1_HUMAN Immunoglobulin heavy constant alpha 1 OS=Homo sapiens OX=9606 GN=IGHA1 PE=1 SV=2 | 255 | 20201116_xd_colostrum_1ng_glyco_hcd203040_compound15.20038.20038.4 | controllerType=0 controllerNumber=1 scan=20038 | 42.6384 |
| 16776:3 | 4 | R.LSLHRPALEDLLLGSEAN[+1054.370]LTC[+57.021]TLTGLR.D | HexNAc(2)Hex(4) | C[+57], N[+1054] | 1339.9975 | 3 | 4017.9779 | 4017.9683 | 0 | 2.4 | 127 | Specific | 721.05 | 692.34 | 692.34 | 14.51 | 109 | >sp\|P01876\|IGHA1_HUMAN Immunoglobulin heavy constant alpha 1 OS=Homo sapiens OX=9606 GN=IGHA1 PE=1 SV=2 | 255 | 20201116_xd_colostrum_1ng_glyco_hcd203040_compound15.20048.20048.3 | controllerType=0 controllerNumber=1 scan=20048 | 42.6879 |
| 17653:5 | 4 | R.LSLHRPALEDLLLGSEAN[+1710.598]LTC[+57.021]TLTGLR.D | HexNAc(3)Hex(5)NeuAc(1) | C[+57], N[+1711] | 935.6472 | 5 | 4674.2068 | 4674.1959 | 0 | 2.3 | 127 | Specific | 732.04 | 732.04 | 732.04 | 15.16 | 109 | >sp\|P01876\|IGHA1_HUMAN Immunoglobulin heavy constant alpha 1 OS=Homo sapiens OX=9606 GN=IGHA1 PE=1 SV=2 | 255 | 20201116_xd_colostrum_1ng_glyco_hcd203040_compound15.20968.20968.5 | controllerType=0 controllerNumber=1 scan=20968 | 44.8202 |
| 18694:4 | 4 | R.LSLHRPALEDLLLGSEAN[+2204.772]LTC[+57.021]TLTGLR.D | HexNAc(4)Hex(5)NeuAc(2) | C[+57], N[+2205] | 1292.8518 | 4 | 5168.3854 | 5168.3707 | 0 | 2.8 | 127 | Specific | 736.46 | 702.31 | 702.31 | 15.07 | 109 | >sp\|P01876\|IGHA1_HUMAN Immunoglobulin heavy constant alpha 1 OS=Homo sapiens OX=9606 GN=IGHA1 PE=1 SV=2 | 255 | 20201116_xd_colostrum_1ng_glyco_hcd203040_compound15.22056.22056.4 | controllerType=0 controllerNumber=1 scan=22056 | 47.1559 |
| 16693:4 | 4 | R.LSLHRPALEDLLLGSEAN[+1622.582]LTC[+57.021]TLTGLR.D | HexNAc(4)Hex(5) | C[+57], N[+1623] | 1147.3049 | 4 | 4586.1976 | 4586.1799 | 0 | 3.9 | 127 | Specific | 736.51 | 736.51 | 736.51 | 14.87 | 109 | >sp\|P01876\|IGHA1_HUMAN Immunoglobulin heavy constant alpha 1 OS=Homo sapiens OX=9606 GN=IGHA1 PE=1 SV=2 | 255 | 20201116_xd_colostrum_1ng_glyco_hcd203040_compound15.19961.19961.4 | controllerType=0 controllerNumber=1 scan=19961 | 42.435 |
| 17561:4 | 4 | R.LSLHRPALEDLLLGSEAN[+2262.814]LTC[+57.021]TLTGLR.D | HexNAc(5)Hex(5)Fuc(1)NeuAc(1) | C[+57], N[+2263] | 1307.3628 | 4 | 5226.4294 | 5226.4126 | 0 | 3.2 | 127 | Specific | 746.21 | 654.02 | 654.02 | 14.87 | 109 | >sp\|P01876\|IGHA1_HUMAN Immunoglobulin heavy constant alpha 1 OS=Homo sapiens OX=9606 GN=IGHA1 PE=1 SV=2 | 255 | 20201116_xd_colostrum_1ng_glyco_hcd203040_compound15.20872.20872.4 | controllerType=0 controllerNumber=1 scan=20872 | 44.597 |
| 17536:4 | 4 | R.LSLHRPALEDLLLGSEAN[+1913.677]LTC[+57.021]TLTGLR.D | HexNAc(4)Hex(5)NeuAc(1) | C[+57], N[+1914] | 1220.0746 | 4 | 4877.2767 | 4877.2753 | 0 | 0.3 | 127 | Specific | 754.33 | 747.71 | 747.71 | 15.64 | 109 | >sp\|P01876\|IGHA1_HUMAN Immunoglobulin heavy constant alpha 1 OS=Homo sapiens OX=9606 GN=IGHA1 PE=1 SV=2 | 255 | 20201116_xd_colostrum_1ng_glyco_hcd203040_compound15.20846.20846.4 | controllerType=0 controllerNumber=1 scan=20846 | 44.5463 |
| 17630:4 | 4 | R.LSLHRPALEDLLLGSEAN[+1751.624]LTC[+57.021]TLTGLR.D | HexNAc(4)Hex(4)NeuAc(1) | C[+57], N[+1752] | 1179.564 | 4 | 4715.2342 | 4715.2225 | 0 | 2.5 | 127 | Specific | 755.95 | 755.95 | 755.95 | 16.07 | 109 | >sp\|P01876\|IGHA1_HUMAN Immunoglobulin heavy constant alpha 1 OS=Homo sapiens OX=9606 GN=IGHA1 PE=1 SV=2 | 255 | 20201116_xd_colostrum_1ng_glyco_hcd203040_compound15.20944.20944.4 | controllerType=0 controllerNumber=1 scan=20944 | [illegible] |
| 18692:4 | 4 | R.LSLHRPALEDLLLGSEAN[+2407.852]LTC[+57.021]TLTGLR.D | HexNAc(5)Hex(5)NeuAc(2) | C[+57], N[+2408] | 1343.6215 | 4 | 5371.464 | 5371.4501 | 0 | 2.6 | 127 | Specific | 762.95 | 585.34 | 585.34 | 16.07 | 109 | >sp\|P01876\|IGHA1_HUMAN Immunoglobulin heavy constant alpha 1 OS=Homo sapiens OX=9606 GN=IGHA1 PE=1 SV=2 | 255 | 20201116_xd_colostrum_1ng_glyco_hcd203040_compound15.22054.22054.4 | controllerType=0 controllerNumber=1 scan=22054 | [illegible] |
| 16675:4 | 4 | R.LSLHRPALEDLLLGSEAN[+1825.661]LTC[+57.021]TLTGLR.D | HexNAc(5)Hex(5) | C[+57], N[+1826] | 1198.0728 | 4 | 4789.2694 | 4789.2592 | 0 | 2.1 | 127 | Specific | 767.29 | 611.65 | 611.65 | 16.07 | 109 | >sp\|P01876\|IGHA1_HUMAN Immunoglobulin heavy constant alpha 1 OS=Homo sapiens OX=9606 GN=IGHA1 PE=1 SV=2 | 255 | 20201116_xd_colostrum_1ng_glyco_hcd203040_compound15.19942.19942.4 | controllerType=0 controllerNumber=1 scan=19942 | [illegible] |
| 17602:4 | 4 | R.LSLHRPALEDLLLGSEAN[+1913.677]LTC[+57.021]TLTGLR.D | HexNAc(4)Hex(5)NeuAc(1) | C[+57], N[+1914] | 1220.0795 | 4 | 4877.2962 | 4877.2753 | 0 | 4.3 | 127 | Specific | 768.24 | 739.52 | 739.52 | 15.64 | 109 | >sp\|P01876\|IGHA1_HUMAN Immunoglobulin heavy constant alpha 1 OS=Homo sapiens OX=9606 GN=IGHA1 PE=1 SV=2 | 255 | 20201116_xd_colostrum_1ng_glyco_hcd203040_compound15.20915.20915.4 | controllerType=0 controllerNumber=1 scan=20915 | [illegible] |
| 16761:4 | 4 | R.LSLHRPALEDLLLGSEAN[+1501.555]LTC[+57.021]TLTGLR.D | HexNAc(5)Hex(3) | C[+57], N[+1502] | 1117.0466 | 4 | 4465.1644 | 4465.1536 | 0 | 2.4 | 127 | Specific | 769.46 | 769.46 | 769.46 | 16.07 | 109 | >sp\|P01876\|IGHA1_HUMAN Immunoglobulin heavy constant alpha 1 OS=Homo sapiens OX=9606 GN=IGHA1 PE=1 SV=2 | 255 | 20201116_xd_colostrum_1ng_glyco_hcd203040_compound15.20032.20032.5 | controllerType=0 controllerNumber=1 scan=20032 | 42.6175 |
| 16763:4 | 4 | R.LSLHRPALEDLLLGSEAN[+1501.555]LTC[+57.021]TLTGLR.D | HexNAc(5)Hex(3) | C[+57], N[+1502] | 1117.0466 | 4 | 4465.1644 | 4465.1536 | 0 | 2.4 | 127 | Specific | 770.88 | 723.2 | 723.2 | 16.07 | 109 | >sp\|P01876\|IGHA1_HUMAN Immunoglobulin heavy constant alpha 1 OS=Homo sapiens OX=9606 GN=IGHA1 PE=1 SV=2 | 255 | 20201116_xd_colostrum_1ng_glyco_hcd203040_compound15.20034.20034.5 | controllerType=0 controllerNumber=1 scan=20034 | 42.6265 |
| 17566:5 | 4 | R.LSLHRPALEDLLLGSEAN[+1913.677]LTC[+57.021]TLTGLR.D | HexNAc(4)Hex(5)NeuAc(1) | C[+57], N[+1914] | 976.2634 | 5 | 4877.2881 | 4877.2753 | 0 | 2.6 | 127 | Specific | 772.35 | 772.35 | 772.35 | 16.16 | 109 | >sp\|P01876\|IGHA1_HUMAN Immunoglobulin heavy constant alpha 1 OS=Homo sapiens OX=9606 GN=IGHA1 PE=1 SV=2 | 255 | 20201116_xd_colostrum_1ng_glyco_hcd203040_compound15.20877.20877.5 | controllerType=0 controllerNumber=1 scan=20877 | 44.6079 |
| 16661:4 | 4 | R.LSLHRPALEDLLLGSEAN[+1663.608]LTC[+57.021]TLTGLR.D | HexNAc(5)Hex(4) | C[+57], N[+1664] | 1157.5583 | 4 | 4627.2113 | 4627.2064 | 0 | 1 | 127 | Specific | 773.94 | 737.63 | 737.63 | 15.91 | 109 | >sp\|P01876\|IGHA1_HUMAN Immunoglobulin heavy constant alpha 1 OS=Homo sapiens OX=9606 GN=IGHA1 PE=1 SV=2 | 255 | 20201116_xd_colostrum_1ng_glyco_hcd203040_compound15.19927.19927.4 | controllerType=0 controllerNumber=1 scan=19927 | 42.3403 |
| 17569:5 | 4 | R.LSLHRPALEDLLLGSEAN[+1913.677]LTC[+57.021]TLTGLR.D | HexNAc(4)Hex(5)NeuAc(1) | C[+57], N[+1914] | 976.2634 | 5 | 4877.2881 | 4877.2753 | 0 | 2.6 | 127 | Specific | 795.86 | 795.86 | 795.86 | 16.16 | 109 | >sp\|P01876\|IGHA1_HUMAN Immunoglobulin heavy constant alpha 1 OS=Homo sapiens OX=9606 GN=IGHA1 PE=1 SV=2 | 255 | 20201116_xd_colostrum_1ng_glyco_hcd203040_compound15.20880.20880.5 | controllerType=0 controllerNumber=1 scan=20880 | 44.6202 |
| 17633:4 | 4 | R.LSLHRPALEDLLLGSEAN[+1751.624]LTC[+57.021]TLTGLR.D | HexNAc(4)Hex(4)NeuAc(1) | C[+57], N[+1752] | 1179.564 | 4 | 4715.2342 | 4715.2225 | 0 | 2.5 | 127 | Specific | 801.03 | 801.03 | 801.03 | 17.07 | 109 | >sp\|P01876\|IGHA1_HUMAN Immunoglobulin heavy constant alpha 1 OS=Homo sapiens OX=9606 GN=IGHA1 PE=1 SV=2 | 255 | 20201116_xd_colostrum_1ng_glyco_hcd203040_compound15.20947.20947.4 | controllerType=0 controllerNumber=1 scan=20947 | 44.7745 |
| 16856:3 | 4 | R.LSLHRPALEDLLLGSEAN[+892.317]LTC[+57.021]TLTGLR.D | HexNAc(2)Hex(3) | C[+57], N[+892] | 1285.9797 | 3 | 3855.9245 | 3855.9155 | 0 | 2.3 | 127 | Specific | 801.23 | 788.85 | 788.85 | 16.51 | 109 | >sp\|P01876\|IGHA1_HUMAN Immunoglobulin heavy constant alpha 1 OS=Homo sapiens OX=9606 GN=IGHA1 PE=1 SV=2 | 255 | 20201116_xd_colostrum_1ng_glyco_hcd203040_compound15.20131.20131.3 | controllerType=0 controllerNumber=1 scan=20131 | 42.8671 |
| 17625:4 | 4 | R.LSLHRPALEDLLLGSEAN[+1710.598]LTC[+57.021]TLTGLR.D | HexNAc(3)Hex(5)NeuAc(1) | C[+57], N[+1711] | 1169.3081 | 4 | 4674.2108 | 4674.1959 | 0 | 3.2 | 127 | Specific | 804.35 | 740.21 | 740.21 | 16.87 | 109 | >sp\|P01876\|IGHA1_HUMAN Immunoglobulin heavy constant alpha 1 OS=Homo sapiens OX=9606 GN=IGHA1 PE=1 SV=2 | 255 | 20201116_xd_colostrum_1ng_glyco_hcd203040_compound15.20939.20939.4 | controllerType=0 controllerNumber=1 scan=20939 | 44.7568 |
| 16676:4 | 4 | R.LSLHRPALEDLLLGSEAN[+1663.608]LTC[+57.021]TLTGLR.D | HexNAc(5)Hex(4) | C[+57], N[+1664] | 1157.5595 | 4 | 4627.2161 | 4627.2064 | 0 | 2.1 | 127 | Specific | 805.65 | 761.38 | 761.38 | 17.07 | 109 | >sp\|P01876\|IGHA1_HUMAN Immunoglobulin heavy constant alpha 1 OS=Homo sapiens OX=9606 GN=IGHA1 PE=1 SV=2 | 255 | 20201116_xd_colostrum_1ng_glyco_hcd203040_compound15.19943.19943.4 | controllerType=0 controllerNumber=1 scan=19943 | 42.3896 |
| 16853:3 | 4 | R.LSLHRPALEDLLLGSEAN[+892.317]LTC[+57.021]TLTGLR.D | HexNAc(2)Hex(3) | C[+57], N[+892] | 1285.9797 | 3 | 3855.9245 | 3855.9155 | 0 | 2.3 | 127 | Specific | 806.33 | 806.33 | 806.33 | 16.51 | 109 | >sp\|P01876\|IGHA1_HUMAN Immunoglobulin heavy constant alpha 1 OS=Homo sapiens OX=9606 GN=IGHA1 PE=1 SV=2 | 255 | 20201116_xd_colostrum_1ng_glyco_hcd203040_compound15.20128.20128.3 | controllerType=0 controllerNumber=1 scan=20128 | 42.8606 |
| 16894:4 | 4 | R.LSLHRPALEDLLLGSEAN[+1622.582]LTC[+57.021]TLTGLR.D | HexNAc(4)Hex(5) | C[+57], N[+1623] | 1147.3031 | 4 | 4586.1907 | 4586.1799 | 0 | 2.4 | 127 | Specific | 809.51 | 717.48 | 717.48 | 17.07 | 109 | >sp\|P01876\|IGHA1_HUMAN Immunoglobulin heavy constant alpha 1 OS=Homo sapiens OX=9606 GN=IGHA1 PE=1 SV=2 | 255 | 20201116_xd_colostrum_1ng_glyco_hcd203040_compound15.20171.20171.4 | controllerType=0 controllerNumber=1 scan=20171 | 42.9482 |
| 16695:4 | 4 | R.LSLHRPALEDLLLGSEAN[+1622.582]LTC[+57.021]TLTGLR.D | HexNAc(4)Hex(5) | C[+57], N[+1623] | 1147.3049 | 4 | 4586.1976 | 4586.1799 | 0 | 3.9 | 127 | Specific | 836.76 | 716.83 | 716.83 | 16.87 | 109 | >sp\|P01876\|IGHA1_HUMAN Immunoglobulin heavy constant alpha 1 OS=Homo sapiens OX=9606 GN=IGHA1 PE=1 SV=2 | 255 | 20201116_xd_colostrum_1ng_glyco_hcd203040_compound15.19963.19963.4 | controllerType=0 controllerNumber=1 scan=19963 | 42.4407 |
| 16771:4 | 4 | R.LSLHRPALEDLLLGSEAN[+1298.476]LTC[+57.021]TLTGLR.D | HexNAc(4)Hex(3) | C[+57], N[+1298] | 1066.2769 | 4 | 4262.0858 | 4262.0742 | 0 | 2.7 | 127 | Specific | 849.17 | 849.17 | 849.17 | 17.07 | 109 | >sp\|P01876\|IGHA1_HUMAN Immunoglobulin heavy constant alpha 1 OS=Homo sapiens OX=9606 GN=IGHA1 PE=1 SV=2 | 255 | 20201116_xd_colostrum_1ng_glyco_hcd203040_compound15.20043.20043.4 | controllerType=0 controllerNumber=1 scan=20043 | 42.6745 |
| 16820:4 | 4 | R.LSLHRPALEDLLLGSEAN[+892.317]LTC[+57.021]TLTGLR.D | HexNAc(2)Hex(3) | C[+57], N[+892] | 964.7366 | 4 | 3855.9246 | 3855.9155 | 0 | 2.4 | 127 | Specific | 850.84 | 850.84 | 850.84 | 18.07 | 109 | >sp\|P01876\|IGHA1_HUMAN Immunoglobulin heavy constant alpha 1 OS=Homo sapiens OX=9606 GN=IGHA1 PE=1 SV=2 | 255 | 20201116_xd_colostrum_1ng_glyco_hcd203040_compound15.20094.20094.4 | controllerType=0 controllerNumber=1 scan=20094 | 42.7861 |
| 17656:5 | 4 | R.LSLHRPALEDLLLGSEAN[+1710.598]LTC[+57.021]TLTGLR.D | HexNAc(3)Hex(5)NeuAc(1) | C[+57], N[+1711] | 935.6472 | 5 | 4674.2068 | 4674.1959 | 0 | 2.3 | 127 | Specific | 870.74 | 870.74 | 870.74 | 18.16 | 109 | >sp\|P01876\|IGHA1_HUMAN Immunoglobulin heavy constant alpha 1 OS=Homo sapiens OX=9606 GN=IGHA1 PE=1 SV=2 | 255 | 20201116_xd_colostrum_1ng_glyco_hcd203040_compound15.20971.20971.5 | controllerType=0 controllerNumber=1 scan=20971 | 44.8247 |
| 17603:4 | 4 | R.LSLHRPALEDLLLGSEAN[+1872.651]LTC[+57.021]TLTGLR.D | HexNAc(3)Hex(6)NeuAc(1) | C[+57], N[+1873] | 1209.824 | 4 | 4836.2742 | 4836.2487 | 0 | 5.3 | 127 | Specific | 870.81 | 870.81 | 870.81 | 17.24 | 109 | >sp\|P01876\|IGHA1_HUMAN Immunoglobulin heavy constant alpha 1 OS=Homo sapiens OX=9606 GN=IGHA1 PE=1 SV=2 | 255 | 20201116_xd_colostrum_1ng_glyco_hcd203040_compound15.20916.20916.4 | controllerType=0 controllerNumber=1 scan=20916 | 44.7084 |
| 16770:4 | 4 | R.LSLHRPALEDLLLGSEAN[+1460.529]LTC[+57.021]TLTGLR.D | HexNAc(4)Hex(4) | C[+57], N[+1461] | 1106.7896 | 4 | 4424.1365 | 4424.1271 | 0 | 2.1 | 127 | Specific | 885.3 | 775.74 | 775.74 | 18.07 | 109 | >sp\|P01876\|IGHA1_HUMAN Immunoglobulin heavy constant alpha 1 OS=Homo sapiens OX=9606 GN=IGHA1 PE=1 SV=2 | 255 | 20201116_xd_colostrum_1ng_glyco_hcd203040_compound15.20041.20041.4 | controllerType=0 controllerNumber=1 scan=20041 | 42.6708 |
| 16700:4 | 4 | R.LSLHRPALEDLLLGSEAN[+1581.555]LTC[+57.021]TLTGLR.D | HexNAc(3)Hex(6) | C[+57], N[+1582] | 1137.0474 | 4 | 4545.1678 | 4545.1533 | 0 | 3.2 | 127 | Specific | 889.18 | 876.34 | 876.34 | 17.87 | 109 | >sp\|P01876\|IGHA1_HUMAN Immunoglobulin heavy constant alpha 1 OS=Homo sapiens OX=9606 GN=IGHA1 PE=1 SV=2 | 255 | 20201116_xd_colostrum_1ng_glyco_hcd203040_compound15.19968.19968.4 | controllerType=0 controllerNumber=1 scan=19968 | 42.4497 |
| 17627:4 | 4 | R.LSLHRPALEDLLLGSEAN[+1710.598]LTC[+57.021]TLTGLR.D | HexNAc(3)Hex(5)NeuAc(1) | C[+57], N[+1711] | 1169.3081 | 4 | 4674.2108 | 4674.1959 | 0 | 3.2 | 127 | Specific | 891.18 | 857 | 857 | 17.87 | 109 | >sp\|P01876\|IGHA1_HUMAN Immunoglobulin heavy constant alpha 1 OS=Homo sapiens OX=9606 GN=IGHA1 PE=1 SV=2 | 255 | 20201116_xd_colostrum_1ng_glyco_hcd203040_compound15.20941.20941.4 | controllerType=0 controllerNumber=1 scan=20941 | 44.7628 |
| 16667:4 | 4 | R.LSLHRPALEDLLLGSEAN[+1864.634]LTC[+57.021]TLTGLR.D | HexNAc(2)Hex(9) | C[+57], N[+1865] | 1207.8144 | 4 | 4828.2357 | 4828.2324 | 0 | 0.7 | 127 | Specific | 894.09 | 894.09 | 894.09 | 17.64 | 109 | >sp\|P01876\|IGHA1_HUMAN Immunoglobulin heavy constant alpha 1 OS=Homo sapiens OX=9606 GN=IGHA1 PE=1 SV=2 | 255 | 20201116_xd_colostrum_1ng_glyco_hcd203040_compound15.19933.19933.4 | controllerType=0 controllerNumber=1 scan=19933 | 42.3545 |
| 17609:5 | 4 | R.LSLHRPALEDLLLGSEAN[+1872.651]LTC[+57.021]TLTGLR.D | HexNAc(3)Hex(6)NeuAc(1) | C[+57], N[+1873] | 968.0589 | 5 | 4836.2652 | 4836.2487 | 0 | 3.4 | 127 | Specific | 907.32 | 907.32 | 907.32 | 18.96 | 109 | >sp\|P01876\|IGHA1_HUMAN Immunoglobulin heavy constant alpha 1 OS=Homo sapiens OX=9606 GN=IGHA1 PE=1 SV=2 | 255 | 20201116_xd_colostrum_1ng_glyco_hcd203040_compound15.20922.20922.5 | controllerType=0 controllerNumber=1 scan=20922 | 44.7192 |
| 16796:4 | 4 | R.LSLHRPALEDLLLGSEAN[+1257.449]LTC[+57.021]TLTGLR.D | HexNAc(3)Hex(4) | C[+57], N[+1257] | 1056.0197 | 4 | 4221.057 | 4221.0477 | 0 | 2.2 | 127 | Specific | 912.39 | 912.39 | 912.39 | 19.07 | 109 | >sp\|P01876\|IGHA1_HUMAN Immunoglobulin heavy constant alpha 1 OS=Homo sapiens OX=9606 GN=IGHA1 PE=1 SV=2 | 255 | 20201116_xd_colostrum_1ng_glyco_hcd203040_compound15.20069.20069.4 | controllerType=0 controllerNumber=1 scan=20069 | 42.7356 |
| 16775:4 | 4 | R.LSLHRPALEDLLLGSEAN[+1054.370]LTC[+57.021]TLTGLR.D | HexNAc(2)Hex(4) | C[+57], N[+1054] | 1005.2504 | 4 | 4017.9798 | 4017.9683 | 0 | 2.9 | 127 | Specific | 912.59 | 912.59 | 912.59 | 19.07 | 109 | >sp\|P01876\|IGHA1_HUMAN Immunoglobulin heavy constant alpha 1 OS=Homo sapiens OX=9606 GN=IGHA1 PE=1 SV=2 | 255 | 20201116_xd_colostrum_1ng_glyco_hcd203040_compound15.20047.20047.4 | controllerType=0 controllerNumber=1 scan=20047 | 42.6868 |
| 16705:4 | 4 | R.LSLHRPALEDLLLGSEAN[+1216.423]LTC[+57.021]TLTGLR.D | HexNAc(2)Hex(5) | C[+57], N[+1216] | 1045.7629 | 4 | 4180.0296 | 4180.0211 | 0 | 2 | 127 | Specific | 915 | 901.4 | 901.4 | 19.07 | 109 | >sp\|P01876\|IGHA1_HUMAN Immunoglobulin heavy constant alpha 1 OS=Homo sapiens OX=9606 GN=IGHA1 PE=1 SV=2 | 255 | 20201116_xd_colostrum_1ng_glyco_hcd203040_compound15.19973.19973.4 | controllerType=0 controllerNumber=1 scan=19973 | 42.4561 |
| 16762:4 | 4 | R.LSLHRPALEDLLLGSEAN[+1216.423]LTC[+57.021]TLTGLR.D | HexNAc(2)Hex(5) | C[+57], N[+1216] | 1045.7633 | 4 | 4180.0316 | 4180.0211 | 0 | 2.5 | 127 | Specific | 921.34 | 884.7 | 884.7 | 19.07 | 109 | >sp\|P01876\|IGHA1_HUMAN Immunoglobulin heavy constant alpha 1 OS=Homo sapiens OX=9606 GN=IGHA1 PE=1 SV=2 | 255 | 20201116_xd_colostrum_1ng_glyco_hcd203040_compound15.20033.20033.4 | controllerType=0 controllerNumber=1 scan=20033 | 42.6189 |
| 16677:4 | 4 | R.LSLHRPALEDLLLGSEAN[+1540.529]LTC[+57.021]TLTGLR.D | HexNAc(2)Hex(7) | C[+57], N[+1541] | 1126.789 | 4 | 4504.1341 | 4504.1268 | 0 | 1.6 | 127 | Specific | 924.84 | 852.69 | 852.69 | 18.91 | 109 | >sp\|P01876\|IGHA1_HUMAN Immunoglobulin heavy constant alpha 1 OS=Homo sapiens OX=9606 GN=IGHA1 PE=1 SV=2 | 255 | 20201116_xd_colostrum_1ng_glyco_hcd203040_compound15.19944.19944.4 | controllerType=0 controllerNumber=1 scan=19944 | 42.3908 |
| 17606:5 | 4 | R.LSLHRPALEDLLLGSEAN[+1872.651]LTC[+57.021]TLTGLR.D | HexNAc(3)Hex(6)NeuAc(1) | C[+57], N[+1873] | 968.0589 | 5 | 4836.2652 | 4836.2487 | 0 | 3.4 | 127 | Specific | 934.2 | 934.2 | 934.2 | 18.96 | 109 | >sp\|P01876\|IGHA1_HUMAN Immunoglobulin heavy constant alpha 1 OS=Homo sapiens OX=9606 GN=IGHA1 PE=1 SV=2 | 255 | 20201116_xd_colostrum_1ng_glyco_hcd203040_compound15.20919.20919.5 | controllerType=0 controllerNumber=1 scan=20919 | 44.7118 |
| 16773:4 | 4 | R.LSLHRPALEDLLLGSEAN[+1298.476]LTC[+57.021]TLTGLR.D | HexNAc(4)Hex(3) | C[+57], N[+1298] | 1066.2769 | 4 | 4262.0858 | 4262.0742 | 0 | 2.7 | 127 | Specific | 946.96 | 905.38 | 905.38 | 19.07 | 109 | >sp\|P01876\|IGHA1_HUMAN Immunoglobulin heavy constant alpha 1 OS=Homo sapiens OX=9606 GN=IGHA1 PE=1 SV=2 | 255 | 20201116_xd_colostrum_1ng_glyco_hcd203040_compound15.20045.20045.4 | controllerType=0 controllerNumber=1 scan=20045 | 42.6811 |
| 16778:4 | 4 | R.LSLHRPALEDLLLGSEAN[+1054.370]LTC[+57.021]TLTGLR.D | HexNAc(2)Hex(4) | C[+57], N[+1054] | 1005.2504 | 4 | 4017.9798 | 4017.9683 | 0 | 2.9 | 127 | Specific | 958.5 | 958.5 | 958.5 | 20.07 | 109 | >sp\|P01876\|IGHA1_HUMAN Immunoglobulin heavy constant alpha 1 OS=Homo sapiens OX=9606 GN=IGHA1 PE=1 SV=2 | 255 | 20201116_xd_colostrum_1ng_glyco_hcd203040_compound15.20050.20050.4 | controllerType=0 controllerNumber=1 scan=20050 | 42.6946 |
| 16823:4 | 4 | R.LSLHRPALEDLLLGSEAN[+892.317]LTC[+57.021]TLTGLR.D | HexNAc(2)Hex(3) | C[+57], N[+892] | 964.7366 | 4 | 3855.9246 | 3855.9155 | 0 | 2.4 | 127 | Specific | 995.35 | 995.35 | 995.35 | 20.07 | 109 | >sp\|P01876\|IGHA1_HUMAN Immunoglobulin heavy constant alpha 1 OS=Homo sapiens OX=9606 GN=IGHA1 PE=1 SV=2 | 255 | 20201116_xd_colostrum_1ng_glyco_hcd203040_compound15.20097.20097.4 | controllerType=0 controllerNumber=1 scan=20097 | 42.7943 |
| 16708:4 | 4 | R.LSLHRPALEDLLLGSEAN[+1216.423]LTC[+57.021]TLTGLR.D | HexNAc(2)Hex(5) | C[+57], N[+1216] | 1045.7629 | 4 | 4180.0296 | 4180.0211 | 0 | 2 | 127 | Specific | 1000.63 | 1000.63 | 1000.63 | 21.07 | 109 | >sp\|P01876\|IGHA1_HUMAN Immunoglobulin heavy constant alpha 1 OS=Homo sapiens OX=9606 GN=IGHA1 PE=1 SV=2 | 255 | 20201116_xd_colostrum_1ng_glyco_hcd203040_compound15.19976.19976.4 | controllerType=0 controllerNumber=1 scan=19976 | 42.4676 |
| 16764:4 | 4 | R.LSLHRPALEDLLLGSEAN[+1216.423]LTC[+57.021]TLTGLR.D | HexNAc(2)Hex(5) | C[+57], N[+1216] | 1045.7633 | 4 | 4180.0316 | 4180.0211 | 0 | 2.5 | 127 | Specific | 1008.39 | 1008.39 | 1008.39 | 21.07 | 109 | >sp\|P01876\|IGHA1_HUMAN Immunoglobulin heavy constant alpha 1 OS=Homo sapiens OX=9606 GN=IGHA1 PE=1 SV=2 | 255 | 20201116_xd_colostrum_1ng_glyco_hcd203040_compound15.20035.20035.4 | controllerType=0 controllerNumber=1 scan=20035 | 42.6352 |
| 16680:4 | 4 | R.LSLHRPALEDLLLGSEAN[+1540.529]LTC[+57.021]TLTGLR.D | HexNAc(2)Hex(7) | C[+57], N[+1541] | 1126.789 | 4 | 4504.1341 | 4504.1268 | 0 | 1.6 | 127 | Specific | 1046.5 | 1004.75 | 1004.75 | 20.91 | 109 | >sp\|P01876\|IGHA1_HUMAN Immunoglobulin heavy constant alpha 1 OS=Homo sapiens OX=9606 GN=IGHA1 PE=1 SV=2 | 255 | 20201116_xd_colostrum_1ng_glyco_hcd203040_compound15.19947.19947.4 | controllerType=0 controllerNumber=1 scan=19947 | 42.3981 |
| 16799:4 | 4 | R.LSLHRPALEDLLLGSEAN[+1257.449]LTC[+57.021]TLTGLR.D | HexNAc(3)Hex(4) | C[+57], N[+1257] | 1056.0197 | 4 | 4221.057 | 4221.0477 | 0 | 2.2 | 127 | Specific | 1056.31 | 1049.2 | 1049.2 | 22.07 | 109 | >sp\|P01876\|IGHA1_HUMAN Immunoglobulin heavy constant alpha 1 OS=Homo sapiens OX=9606 GN=IGHA1 PE=1 SV=2 | 255 | 20201116_xd_colostrum_1ng_glyco_hcd203040_compound15.20072.20072.4 | controllerType=0 controllerNumber=1 scan=20072 | 42.7447 |
| 11285:4 | 4 | K.TIDRLAGKPTHVN[+1216.423]VSVVMAEVDGTC[+57.021]Y.- | HexNAc(2)Hex(5) | C[+57], N[+1216] | 1012.9664 | 4 | 4048.8438 | 4048.8247 | 0 | 4.7 | 328 | CRagged | 358.33 | 358.33 | 291.74 | 7.28 | 109 | >sp\|P01876\|IGHA1_HUMAN Immunoglobulin heavy constant alpha 1 OS=Homo sapiens OX=9606 GN=IGHA1 PE=1 SV=2 | 255 | 20201116_xd_colostrum_1ng_glyco_hcd203040_compound15.14335.14335.4 | controllerType=0 controllerNumber=1 scan=14335 | 31.1215 |
| 11288:4 | 4 | K.TIDRLAGKPTHVN[+1216.423]VSVVMAEVDGTC[+57.021]Y.- | HexNAc(2)Hex(5) | C[+57], N[+1216] | 1012.9664 | 4 | 4048.8438 | 4048.8247 | 0 | 4.7 | 328 | CRagged | 577.74 | 367.69 | 367.69 | 8.53 | 109 | >sp\|P01876\|IGHA1_HUMAN Immunoglobulin heavy constant alpha 1 OS=Homo sapiens OX=9606 GN=IGHA1 PE=1 SV=2 | 255 | 20201116_xd_colostrum_1ng_glyco_hcd203040_compound15.14338.14338.4 | controllerType=0 controllerNumber=1 scan=14338 | 31.1269 |
| 12547:3 | 4 | R.LAGKPTHVN[+2263.835]VSVVMAEVDGTC[+57.021].Y | HexNAc(5)Hex(5)Fuc(3) | C[+57], N[+2264] | 1483.3093 | 3 | 4447.9132 | 4447.9135 | 0 | -0.1 | 332 | CRagged | 210.4 | 190.76 | 190.76 | 4.28 | 109 | >sp\|P01876\|IGHA1_HUMAN Immunoglobulin heavy constant alpha 1 OS=Homo sapiens OX=9606 GN=IGHA1 PE=1 SV=2 | 255 | 20201116_xd_colostrum_1ng_glyco_hcd203040_compound15.15646.15646.3 | controllerType=0 controllerNumber=1 scan=15646 | 33.5913 |
| 10076:3 | 4 | R.LAGKPTHVN[+1809.666]VSVVMAEVDGTC[+57.021].Y | HexNAc(5)Hex(4)Fuc(1) | C[+57], N[+1810] | 1331.9236 | 3 | 3993.7564 | 3993.7448 | 0 | 2.9 | 332 | CRagged | 254.5 | 254.5 | 99.21 | 7.14 | 109 | >sp\|P01876\|IGHA1_HUMAN Immunoglobulin heavy constant alpha 1 OS=Homo sapiens OX=9606 GN=IGHA1 PE=1 SV=2 | 255 | 20201116_xd_colostrum_1ng_glyco_hcd203040_compound15.13079.13079.3 | controllerType=0 controllerNumber=1 scan=13079 | 28.6064 |
| 12550:3 | 4 | R.LAGKPTHVN[+2263.835]VSVVMAEVDGTC[+57.021].Y | HexNAc(5)Hex(5)Fuc(3) | C[+57], N[+2264] | 1483.3093 | 3 | 4447.9132 | 4447.9135 | 0 | -0.1 | 332 | CRagged | 264.18 | 264.18 | 264.18 | 4.65 | 109 | >sp\|P01876\|IGHA1_HUMAN Immunoglobulin heavy constant alpha 1 OS=Homo sapiens OX=9606 GN=IGHA1 PE=1 SV=2 | 255 | 20201116_xd_colostrum_1ng_glyco_hcd203040_compound15.15649.15649.3 | controllerType=0 controllerNumber=1 scan=15649 | 33.5987 |
| 11590:3 | 4 | R.LAGKPTHVN[+2262.814]VSVVMAEVDGTC[+57.021].Y | HexNAc(5)Hex(5)Fuc(1)NeuAc(1) | C[+57], N[+2263] | 1482.9739 | 3 | 4446.9072 | 4446.8931 | 0 | 3.2 | 332 | CRagged | 388.73 | 249.85 | 249.85 | 7.5 | 109 | >sp\|P01876\|IGHA1_HUMAN Immunoglobulin heavy constant alpha 1 OS=Homo sapiens OX=9606 GN=IGHA1 PE=1 SV=2 | 255 | 20201116_xd_colostrum_1ng_glyco_hcd203040_compound15.14652.14652.3 | controllerType=0 controllerNumber=1 scan=14652 | 31.7292 |
| 11588:3 | 4 | R.LAGKPTHVN[+2262.814]VSVVMAEVDGTC[+57.021].Y | HexNAc(5)Hex(5)Fuc(1)NeuAc(1) | C[+57], N[+2263] | 1482.9739 | 3 | 4446.9072 | 4446.8931 | 0 | 3.2 | 332 | CRagged | 411.05 | 395.25 | 395.25 | 9.7 | 109 | >sp\|P01876\|IGHA1_HUMAN Immunoglobulin heavy constant alpha 1 OS=Homo sapiens OX=9606 GN=IGHA1 PE=1 SV=2 | 255 | 20201116_xd_colostrum_1ng_glyco_hcd203040_compound15.14650.14650.3 | controllerType=0 controllerNumber=1 scan=14650 | 31.7247 |
| 10079:3 | 4 | R.LAGKPTHVN[+1809.666]VSVVMAEVDGTC[+57.021].Y | HexNAc(5)Hex(4)Fuc(1) | C[+57], N[+1810] | 1331.9236 | 3 | 3993.7564 | 3993.7448 | 0 | 2.9 | 332 | CRagged | 441.78 | 381.72 | 234.15 | 8.5 | 109 | >sp\|P01876\|IGHA1_HUMAN Immunoglobulin heavy constant alpha 1 OS=Homo sapiens OX=9606 GN=IGHA1 PE=1 SV=2 | 255 | 20201116_xd_colostrum_1ng_glyco_hcd203040_compound15.13082.13082.3 | controllerType=0 controllerNumber=1 scan=13082 | 28.6105 |
| 10148:3 | 4 | R.LAGKPTHVN[+1054.370]VSVVMAEVDGTC[+57.021].Y | HexNAc(2)Hex(4) | C[+57], N[+1054] | 1080.1586 | 3 | 3238.4612 | 3238.4488 | 0 | 3.8 | 332 | CRagged | 579.05 | 512.21 | 272.12 | 10.61 | 109 | >sp\|P01876\|IGHA1_HUMAN Immunoglobulin heavy constant alpha 1 OS=Homo sapiens OX=9606 GN=IGHA1 PE=1 SV=2 | 255 | 20201116_xd_colostrum_1ng_glyco_hcd203040_compound15.13154.13154.3 | controllerType=0 controllerNumber=1 scan=13154 | 28.7667 |
| 08841:3 | 4 | R.LAGKPTHVN[+1054.370]VSVVMAEVDGTC[+57.021].Y | HexNAc(2)Hex(4) | C[+57], N[+1054] | 1080.1563 | 3 | 3238.4543 | 3238.4488 | 0 | 1.7 | 332 | CRagged | 734.43 | 734.43 | 443.2 | 14.95 | 109 | >sp\|P01876\|IGHA1_HUMAN Immunoglobulin heavy constant alpha 1 OS=Homo sapiens OX=9606 GN=IGHA1 PE=1 SV=2 | 255 | 20201116_xd_colostrum_1ng_glyco_hcd203040_compound15.11797.11797.3 | controllerType=0 controllerNumber=1 scan=11797 | 26.1918 |
| 10151:3 | 4 | R.LAGKPTHVN[+1054.370]VSVVMAEVDGTC[+57.021].Y | HexNAc(2)Hex(4) | C[+57], N[+1054] | 1080.1586 | 3 | 3238.4612 | 3238.4488 | 0 | 3.8 | 332 | CRagged | 771.5 | 659.53 | 419.73 | 15.91 | 109 | >sp\|P01876\|IGHA1_HUMAN Immunoglobulin heavy constant alpha 1 OS=Homo sapiens OX=9606 GN=IGHA1 PE=1 SV=2 | 255 | 20201116_xd_colostrum_1ng_glyco_hcd203040_compound15.13157.13157.3 | controllerType=0 controllerNumber=1 scan=13157 | 28.7738 |
| 08843:3 | 4 | R.LAGKPTHVN[+1054.370]VSVVMAEVDGTC[+57.021].Y | HexNAc(2)Hex(4) | C[+57], N[+1054] | 1080.1563 | 3 | 3238.4543 | 3238.4488 | 0 | 1.7 | 332 | CRagged | 774.17 | 774.17 | 469.62 | 15.95 | 109 | >sp\|P01876\|IGHA1_HUMAN Immunoglobulin heavy constant alpha 1 OS=Homo sapiens OX=9606 GN=IGHA1 PE=1 SV=2 | 255 | 20201116_xd_colostrum_1ng_glyco_hcd203040_compound15.11799.11799.3 | controllerType=0 controllerNumber=1 scan=11799 | 26.2005 |
| 08793:3 | 4 | R.LAGKPTHVN[+1216.423]VSVVMAEVDGTC[+57.021].Y | HexNAc(2)Hex(5) | C[+57], N[+1216] | 1134.1746 | 3 | 3400.5092 | 3400.5016 | 0 | 2.2 | 332 | CRagged | 805.17 | 709.69 | 473 | 17.11 | 109 | >sp\|P01876\|IGHA1_HUMAN Immunoglobulin heavy constant alpha 1 OS=Homo sapiens OX=9606 GN=IGHA1 PE=1 SV=2 | 255 | 20201116_xd_colostrum_1ng_glyco_hcd203040_compound15.11747.11747.3 | controllerType=0 controllerNumber=1 scan=11747 | 26.0787 |
| 08795:3 | 4 | R.LAGKPTHVN[+1216.423]VSVVMAEVDGTC[+57.021].Y | HexNAc(2)Hex(5) | C[+57], N[+1216] | 1134.1746 | 3 | 3400.5092 | 3400.5016 | 0 | 2.2 | 332 | CRagged | 808.13 | 806.47 | 523.68 | 17.11 | 109 | >sp\|P01876\|IGHA1_HUMAN Immunoglobulin heavy constant alpha 1 OS=Homo sapiens OX=9606 GN=IGHA1 PE=1 SV=2 | 255 | 20201116_xd_colostrum_1ng_glyco_hcd203040_compound15.11749.11749.3 | controllerType=0 controllerNumber=1 scan=11749 | 26.0844 |
| 10978:4 | 4 | R.LAGKPTHVN[+1809.666]VSVVMAEVDGTC[+57.021]Y.- | HexNAc(5)Hex(4)Fuc(1) | C[+57], N[+1810] | 1039.9584 | 4 | 4156.8118 | 4156.8081 | 0 | 0.9 | 332 | CRagged | 216.45 | 166.73 | 166.73 | 3.99 | 109 | >sp\|P01876\|IGHA1_HUMAN Immunoglobulin heavy constant alpha 1 OS=Homo sapiens OX=9606 GN=IGHA1 PE=1 SV=2 | 255 | 20201116_xd_colostrum_1ng_glyco_hcd203040_compound15.14016.14016.4 | controllerType=0 controllerNumber=1 scan=14016 | 30.4733 |
| 12535:4 | 4 | R.LAGKPTHVN[+2424.867]VSVVMAEVDGTC[+57.021]Y.- | HexNAc(5)Hex(6)Fuc(1)NeuAc(1) | C[+57], N[+2425] | 1193.7597 | 4 | 4772.0169 | 4772.0092 | 0 | 1.6 | 332 | CRagged | 233.08 | 233.08 | 221.68 | 7.01 | 109 | >sp\|P01876\|IGHA1_HUMAN Immunoglobulin heavy constant alpha 1 OS=Homo sapiens OX=9606 GN=IGHA1 PE=1 SV=2 | 255 | 20201116_xd_colostrum_1ng_glyco_hcd203040_compound15.15633.15633.4 | controllerType=0 controllerNumber=1 scan=15633 | 33.5699 |
| 12548:4 | 4 | R.LAGKPTHVN[+1913.677]VSVVMAEVDGTC[+57.021]Y.- | HexNAc(4)Hex(5)NeuAc(1) | C[+57], N[+1914] | 1065.9622 | 4 | 4260.827 | 4260.8191 | 0 | 1.8 | 332 | CRagged | 233.2 | 200.58 | 200.58 | 5.82 | 109 | >sp\|P01876\|IGHA1_HUMAN Immunoglobulin heavy constant alpha 1 OS=Homo sapiens OX=9606 GN=IGHA1 PE=1 SV=2 | 255 | 20201116_xd_colostrum_1ng_glyco_hcd203040_compound15.15647.15647.4 | controllerType=0 controllerNumber=1 scan=15647 | 33.5924 |
| 11135:4 | 4 | R.LAGKPTHVN[+1647.613]VSVVMAEVDGTC[+57.021]Y.- | HexNAc(5)Hex(3)Fuc(1) | C[+57], N[+1648] | 999.4471 | 4 | 3994.7664 | 3994.7553 | 0 | 2.8 | 332 | CRagged | 244.81 | 124.33 | 124.33 | 4.11 | 109 | >sp\|P01876\|IGHA1_HUMAN Immunoglobulin heavy constant alpha 1 OS=Homo sapiens OX=9606 GN=IGHA1 PE=1 SV=2 | 255 | 20201116_xd_colostrum_1ng_glyco_hcd203040_compound15.14179.14179.4 | controllerType=0 controllerNumber=1 scan=14179 | 30.795 |
| 10602:3 | 4 | R.LAGKPTHVN[+1378.476]VSVVMAEVDGTC[+57.021]Y.- | HexNAc(2)Hex(6) | C[+57], N[+1378] | 1242.5422 | 3 | 3725.612 | 3725.6178 | 0 | -1.6 | 332 | CRagged | 245.05 | 245.05 | 213.62 | 5.49 | 109 | >sp\|P01876\|IGHA1_HUMAN Immunoglobulin heavy constant alpha 1 OS=Homo sapiens OX=9606 GN=IGHA1 PE=1 SV=2 | 255 | 20201116_xd_colostrum_1ng_glyco_hcd203040_compound15.13625.13625.3 | controllerType=0 controllerNumber=1 scan=13625 | 29.6983 |
| 10070:4 | 4 | R.LAGKPTHVN[+1647.613]VSVVMAEVDGTC[+57.021]Y.- | HexNAc(5)Hex(3)Fuc(1) | C[+57], N[+1648] | 999.4454 | 4 | 3994.7598 | 3994.7553 | 0 | 1.1 | 332 | CRagged | 258.55 | 216.88 | 216.88 | 5.62 | 109 | >sp\|P01876\|IGHA1_HUMAN Immunoglobulin heavy constant alpha 1 OS=Homo sapiens OX=9606 GN=IGHA1 PE=1 SV=2 | 255 | 20201116_xd_colostrum_1ng_glyco_hcd203040_compound15.13072.13072.4 | controllerType=0 controllerNumber=1 scan=13072 | 28.59 |
| 11386:3 | 4 | R.LAGKPTHVN[+2262.814]VSVVMAEVDGTC[+57.021]Y.- | HexNAc(5)Hex(5)Fuc(1)NeuAc(1) | C[+57], N[+2263] | 1537.3348 | 3 | 4609.9898 | 4609.9564 | 0 | 7.2 | 332 | CRagged | 262.49 | 262.49 | 262.49 | 6.42 | 109 | >sp\|P01876\|IGHA1_HUMAN Immunoglobulin heavy constant alpha 1 OS=Homo sapiens OX=9606 GN=IGHA1 PE=1 SV=2 | 255 | 20201116_xd_colostrum_1ng_glyco_hcd203040_compound15.14440.14440.3 | controllerType=0 controllerNumber=1 scan=14440 | 31.325 |
| 11327:4 | 4 | R.LAGKPTHVN[+2262.814]VSVVMAEVDGTC[+57.021]Y.- | HexNAc(5)Hex(5)Fuc(1)NeuAc(1) | C[+57], N[+2263] | 1153.2472 | 4 | 4609.9671 | 4609.9564 | 0 | 2.3 | 332 | CRagged | 307.76 | 307.76 | 307.76 | 8.04 | 109 | >sp\|P01876\|IGHA1_HUMAN Immunoglobulin heavy constant alpha 1 OS=Homo sapiens OX=9606 GN=IGHA1 PE=1 SV=2 | 255 | 20201116_xd_colostrum_1ng_glyco_hcd203040_compound15.14379.14379.4 | controllerType=0 controllerNumber=1 scan=14379 | 31.2105 |
| 12616:3 | 4 | R.LAGKPTHVN[+2059.735]VSVVMAEVDGTC[+57.021]Y.- | HexNAc(4)Hex(5)Fuc(1)NeuAc(1) | C[+57], N[+2060] | 1469.6335 | 3 | 4406.8859 | 4406.877 | 0 | 2 | 332 | CRagged | 329.78 | 312.1 | 312.1 | 8.11 | 109 | >sp\|P01876\|IGHA1_HUMAN Immunoglobulin heavy constant alpha 1 OS=Homo sapiens OX=9606 GN=IGHA1 PE=1 SV=2 | 255 | 20201116_xd_colostrum_1ng_glyco_hcd203040_compound15.15718.15718.3 | controllerType=0 controllerNumber=1 scan=15718 | 33.7484 |
| 11068:3 | 4 | R.LAGKPTHVN[+1419.502]VSVVMAEVDGTC[+57.021]Y.- | HexNAc(3)Hex(5) | C[+57], N[+1420] | 1256.2214 | 3 | 3766.6496 | 3766.6443 | 0 | 1.4 | 332 | CRagged | 363.7 | 258.13 | 258.13 | 7.07 | 109 | >sp\|P01876\|IGHA1_HUMAN Immunoglobulin heavy constant alpha 1 OS=Homo sapiens OX=9606 GN=IGHA1 PE=1 SV=2 | 255 | 20201116_xd_colostrum_1ng_glyco_hcd203040_compound15.14109.14109.3 | controllerType=0 controllerNumber=1 scan=14109 | 30.66 |
| 11389:3 | 4 | R.LAGKPTHVN[+2262.814]VSVVMAEVDGTC[+57.021]Y.- | HexNAc(5)Hex(5)Fuc(1)NeuAc(1) | C[+57], N[+2263] | 1537.3348 | 3 | 4609.9898 | 4609.9564 | 0 | 7.2 | 332 | CRagged | 395.8 | 395.8 | 395.8 | 9.24 | 109 | >sp\|P01876\|IGHA1_HUMAN Immunoglobulin heavy constant alpha 1 OS=Homo sapiens OX=9606 GN=IGHA1 PE=1 SV=2 | 255 | 20201116_xd_colostrum_1ng_glyco_hcd203040_compound15.14443.14443.3 | controllerType=0 controllerNumber=1 scan=14443 | 31.3287 |
| 10633:3 | 4 | R.LAGKPTHVN[+1216.423]VSVVMAEVDGTC[+57.021]Y.- | HexNAc(2)Hex(5) | C[+57], N[+1216] | 1188.5291 | 3 | 3563.5728 | 3563.5649 | 0 | 2.2 | 332 | CRagged | 432.61 | 432.61 | 241.92 | 10.49 | 109 | >sp\|P01876\|IGHA1_HUMAN Immunoglobulin heavy constant alpha 1 OS=Homo sapiens OX=9606 GN=IGHA1 PE=1 SV=2 | 255 | 20201116_xd_colostrum_1ng_glyco_hcd203040_compound15.13658.13658.3 | controllerType=0 controllerNumber=1 scan=13658 | 29.7641 |
| 09980:3 | 4 | R.LAGKPTHVN[+1581.555]VSVVMAEVDGTC[+57.021]Y.- | HexNAc(3)Hex(6) | C[+57], N[+1582] | 1310.2414 | 3 | 3928.7096 | 3928.6971 | 0 | 3.2 | 332 | CRagged | 437.97 | 437.97 | 344.03 | 10.3 | 109 | >sp\|P01876\|IGHA1_HUMAN Immunoglobulin heavy constant alpha 1 OS=Homo sapiens OX=9606 GN=IGHA1 PE=1 SV=2 | 255 | 20201116_xd_colostrum_1ng_glyco_hcd203040_compound15.12979.12979.3 | controllerType=0 controllerNumber=1 scan=12979 | 28.3957 |
| 11064:3 | 4 | R.LAGKPTHVN[+1622.582]VSVVMAEVDGTC[+57.021]Y.- | HexNAc(4)Hex(5) | C[+57], N[+1623] | 1323.9157 | 3 | 3969.7326 | 3969.7237 | 0 | 2.2 | 332 | CRagged | 440.28 | 398.95 | 331.01 | 10.56 | 109 | >sp\|P01876\|IGHA1_HUMAN Immunoglobulin heavy constant alpha 1 OS=Homo sapiens OX=9606 GN=IGHA1 PE=1 SV=2 | 255 | 20201116_xd_colostrum_1ng_glyco_hcd203040_compound15.14105.14105.3 | controllerType=0 controllerNumber=1 scan=14105 | 30.6536 |
| 14380:4 | 4 | R.LAGKPTHVN[+2204.772]VSVVMAEVDGTC[+57.021]Y.- | HexNAc(4)Hex(5)NeuAc(2) | C[+57], N[+2205] | 1138.736 | 4 | 4551.9222 | 4551.9145 | 0 | 1.7 | 332 | CRagged | 443.5 | 434.05 | 428.28 | 10.5 | 109 | >sp\|P01876\|IGHA1_HUMAN Immunoglobulin heavy constant alpha 1 OS=Homo sapiens OX=9606 GN=IGHA1 PE=1 SV=2 | 255 | 20201116_xd_colostrum_1ng_glyco_hcd203040_compound15.17552.17552.4 | controllerType=0 controllerNumber=1 scan=17552 | 37.4228 |
| 12538:4 | 4 | R.LAGKPTHVN[+2424.867]VSVVMAEVDGTC[+57.021]Y.- | HexNAc(5)Hex(6)Fuc(1)NeuAc(1) | C[+57], N[+2425] | 1193.7597 | 4 | 4772.0169 | 4772.0092 | 0 | 1.6 | 332 | CRagged | 444.25 | 384.55 | 323.03 | 9.58 | 109 | >sp\|P01876\|IGHA1_HUMAN Immunoglobulin heavy constant alpha 1 OS=Homo sapiens OX=9606 GN=IGHA1 PE=1 SV=2 | 255 | 20201116_xd_colostrum_1ng_glyco_hcd203040_compound15.15636.15636.4 | controllerType=0 controllerNumber=1 scan=15636 | 33.5734 |
| 11062:3 | 4 | R.LAGKPTHVN[+1971.719]VSVVMAEVDGTC[+57.021]Y.- | HexNAc(5)Hex(5)Fuc(1) | C[+57], N[+1972] | 1440.2989 | 3 | 4318.8821 | 4318.861 | 0 | 4.9 | 332 | CRagged | 461.59 | 416.06 | 331.19 | 8.71 | 109 | >sp\|P01876\|IGHA1_HUMAN Immunoglobulin heavy constant alpha 1 OS=Homo sapiens OX=9606 GN=IGHA1 PE=1 SV=2 | 255 | 20201116_xd_colostrum_1ng_glyco_hcd203040_compound15.14103.14103.3 | controllerType=0 controllerNumber=1 scan=14103 | 30.6505 |
| 14311:4 | 4 | R.LAGKPTHVN[+2350.830]VSVVMAEVDGTC[+57.021]Y.- | HexNAc(4)Hex(5)Fuc(1)NeuAc(2) | C[+57], N[+2351] | 1175.252 | 4 | 4697.9862 | 4697.9724 | 0 | 2.9 | 332 | CRagged | 463.38 | 454.43 | 454.43 | 10.78 | 109 | >sp\|P01876\|IGHA1_HUMAN Immunoglobulin heavy constant alpha 1 OS=Homo sapiens OX=9606 GN=IGHA1 PE=1 SV=2 | 255 | 20201116_xd_colostrum_1ng_glyco_hcd203040_compound15.17480.17480.4 | controllerType=0 controllerNumber=1 scan=17480 | 37.2607 |
| 12596:4 | 4 | R.LAGKPTHVN[+2059.735]VSVVMAEVDGTC[+57.021]Y.- | HexNAc(4)Hex(5)Fuc(1)NeuAc(1) | C[+57], N[+2060] | 1102.4775 | 4 | 4406.888 | 4406.877 | 0 | 2.5 | 332 | CRagged | 470.09 | 325.95 | 325.95 | 9.02 | 109 | >sp\|P01876\|IGHA1_HUMAN Immunoglobulin heavy constant alpha 1 OS=Homo sapiens OX=9606 GN=IGHA1 PE=1 SV=2 | 255 | 20201116_xd_colostrum_1ng_glyco_hcd203040_compound15.15697.15697.4 | controllerType=0 controllerNumber=1 scan=15697 | 33.7095 |
| 11060:3 | 4 | R.LAGKPTHVN[+1971.719]VSVVMAEVDGTC[+57.021]Y.- | HexNAc(5)Hex(5)Fuc(1) | C[+57], N[+1972] | 1440.2989 | 3 | 4318.8821 | 4318.861 | 0 | 4.9 | 332 | CRagged | 483.3 | 355.51 | 355.51 | 8.61 | 109 | >sp\|P01876\|IGHA1_HUMAN Immunoglobulin heavy constant alpha 1 OS=Homo sapiens OX=9606 GN=IGHA1 PE=1 SV=2 | 255 | 20201116_xd_colostrum_1ng_glyco_hcd203040_compound15.14101.14101.3 | controllerType=0 controllerNumber=1 scan=14101 | 30.6455 |

Table S4. Cont.

| Query #:z | Protein Rank | Glycopeptide | Glycan | Modification Type(s) | Observed m/z | z | Observed (M+H) | Calc. mass (M+H) | Off-by-x error | Mass error (ppm) | StartPosition | Cleavage | Score | Delta | Delta Mod | \|Log Prob\| | # of unique peptides | Protein Name | Protein DB number | Comment | Scan # | Scan Time |
|---|---|---|---|---|---|---|---|---|---|---|---|---|---|---|---|---|---|---|---|---|---|---|
| 12584:4 | 4 | R.LAGKPTHVN[+2059.735]VSVVMAEVDGTC[+57.021]Y.- | HexNAc(4)Hex(5)Fuc(1)NeuAc(1) | C[+57], N[+2060] | 1102.4754 | 4 | 4406.8797 | 4406.877 | 0 | 0.6 | 332 | CRagged | 487.41 | 352.91 | 352.91 | 8.7 | 109 | >sp\|P01876\|IGHA1_HUMAN Immunoglobulin heavy constant alpha 1 OS=Homo sapiens OX=9606 GN=IGHA1 PE=1 SV=2 | 255 | 20201116_xd_colostrum_1ng_glyco_hcd203040_compound15.15684.15684.4 | controllerType=0 controllerNumber=1 scan=15684 | 33.666 |
| 09964:4 | 4 | R.LAGKPTHVN[+1809.666]VSVVMAEVDGTC[+57.021]Y.- | HexNAc(5)Hex(4)Fuc(1) | C[+57], N[+1810] | 1039.9582 | 4 | 4156.8109 | 4156.8081 | 0 | 0.7 | 332 | CRagged | 499.88 | 499.88 | 407.21 | 10.38 | 109 | >sp\|P01876\|IGHA1_HUMAN Immunoglobulin heavy constant alpha 1 OS=Homo sapiens OX=9606 GN=IGHA1 PE=1 SV=2 | 255 | 20201116_xd_colostrum_1ng_glyco_hcd203040_compound15.12962.12962.4 | controllerType=0 controllerNumber=1 scan=12962 | 28.3475 |
| 10874:3 | 4 | R.LAGKPTHVN[+2174.798]VSVVMAEVDGTC[+57.021]Y.- | HexNAc(6)Hex(5)Fuc(1) | C[+57], N[+2175] | 1507.9911 | 3 | 4521.9589 | 4521.9403 | 0 | 4.1 | 332 | CRagged | 506.21 | 506.21 | 374.76 | 10.41 | 109 | >sp\|P01876\|IGHA1_HUMAN Immunoglobulin heavy constant alpha 1 OS=Homo sapiens OX=9606 GN=IGHA1 PE=1 SV=2 | 255 | 20201116_xd_colostrum_1ng_glyco_hcd203040_compound15.13907.13907.3 | controllerType=0 controllerNumber=1 scan=13907 | 30.2044 |
| 09983:3 | 4 | R.LAGKPTHVN[+1606.587]VSVVMAEVDGTC[+57.021]Y.- | HexNAc(4)Hex(4)Fuc(1) | C[+57], N[+1607] | 1318.584 | 3 | 3953.7376 | 3953.7288 | 0 | 2.2 | 332 | CRagged | 514.1 | 514.1 | 514.1 | 10.84 | 109 | >sp\|P01876\|IGHA1_HUMAN Immunoglobulin heavy constant alpha 1 OS=Homo sapiens OX=9606 GN=IGHA1 PE=1 SV=2 | 255 | 20201116_xd_colostrum_1ng_glyco_hcd203040_compound15.12982.12982.3 | controllerType=0 controllerNumber=1 scan=12982 | 28.403 |
| 10114:3 | 4 | R.LAGKPTHVN[+1095.397]VSVVMAEVDGTC[+57.021]Y.- | HexNAc(3)Hex(3) | C[+57], N[+1095] | 1148.1894 | 3 | 3442.5535 | 3442.5387 | 0 | 4.3 | 332 | CRagged | 530.69 | 524.01 | 524.01 | 10.46 | 109 | >sp\|P01876\|IGHA1_HUMAN Immunoglobulin heavy constant alpha 1 OS=Homo sapiens OX=9606 GN=IGHA1 PE=1 SV=2 | 255 | 20201116_xd_colostrum_1ng_glyco_hcd203040_compound15.13118.13118.3 | controllerType=0 controllerNumber=1 scan=13118 | 28.6799 |
| 10112:3 | 4 | R.LAGKPTHVN[+1095.397]VSVVMAEVDGTC[+57.021]Y.- | HexNAc(3)Hex(3) | C[+57], N[+1095] | 1148.1894 | 3 | 3442.5535 | 3442.5387 | 0 | 4.3 | 332 | CRagged | 531.61 | 505.61 | 505.61 | 10.46 | 109 | >sp\|P01876\|IGHA1_HUMAN Immunoglobulin heavy constant alpha 1 OS=Homo sapiens OX=9606 GN=IGHA1 PE=1 SV=2 | 255 | 20201116_xd_colostrum_1ng_glyco_hcd203040_compound15.13116.13116.3 | controllerType=0 controllerNumber=1 scan=13116 | 28.6762 |
| 11330:4 | 4 | R.LAGKPTHVN[+2262.814]VSVVMAEVDGTC[+57.021]Y.- | HexNAc(5)Hex(5)Fuc(1)NeuAc(1) | C[+57], N[+2263] | 1153.2472 | 4 | 4609.9671 | 4609.9564 | 0 | 2.3 | 332 | CRagged | 534.48 | 409.41 | 409.41 | 10.98 | 109 | >sp\|P01876\|IGHA1_HUMAN Immunoglobulin heavy constant alpha 1 OS=Homo sapiens OX=9606 GN=IGHA1 PE=1 SV=2 | 255 | 20201116_xd_colostrum_1ng_glyco_hcd203040_compound15.14382.14382.4 | controllerType=0 controllerNumber=1 scan=14382 | 31.2201 |
| 14312:3 | 4 | R.LAGKPTHVN[+2350.830]VSVVMAEVDGTC[+57.021]Y.- | HexNAc(4)Hex(5)Fuc(1)NeuAc(2) | C[+57], N[+2351] | 1566.6675 | 3 | 4697.9881 | 4697.9724 | 0 | 3.3 | 332 | CRagged | 546.98 | 546.98 | 546.98 | 10.93 | 109 | >sp\|P01876\|IGHA1_HUMAN Immunoglobulin heavy constant alpha 1 OS=Homo sapiens OX=9606 GN=IGHA1 PE=1 SV=2 | 255 | 20201116_xd_colostrum_1ng_glyco_hcd203040_compound15.17481.17481.3 | controllerType=0 controllerNumber=1 scan=17481 | 37.2618 |
| 09978:4 | 4 | R.LAGKPTHVN[+1663.608]VSVVMAEVDGTC[+57.021]Y.- | HexNAc(5)Hex(4) | C[+57], N[+1664] | 1003.4471 | 4 | 4010.7664 | 4010.7502 | 0 | 4 | 332 | CRagged | 553.21 | 313.79 | 313.79 | 8.88 | 109 | >sp\|P01876\|IGHA1_HUMAN Immunoglobulin heavy constant alpha 1 OS=Homo sapiens OX=9606 GN=IGHA1 PE=1 SV=2 | 255 | 20201116_xd_colostrum_1ng_glyco_hcd203040_compound15.12976.12976.4 | controllerType=0 controllerNumber=1 scan=12976 | 28.3898 |
| 10024:2 | 4 | R.LAGKPTHVN[+1216.423]VSVVMAEVDGTC[+57.021]Y.- | HexNAc(2)Hex(5) | C[+57], N[+1216] | 1782.2899 | 2 | 3563.5725 | 3563.5649 | 0 | 2.1 | 332 | CRagged | 553.6 | 553.6 | 289.93 | 10.57 | 109 | >sp\|P01876\|IGHA1_HUMAN Immunoglobulin heavy constant alpha 1 OS=Homo sapiens OX=9606 GN=IGHA1 PE=1 SV=2 | 255 | 20201116_xd_colostrum_1ng_glyco_hcd203040_compound15.13024.13024.2 | controllerType=0 controllerNumber=1 scan=13024 | 28.4893 |
| 14383:4 | 4 | R.LAGKPTHVN[+2204.772]VSVVMAEVDGTC[+57.021]Y.- | HexNAc(4)Hex(5)NeuAc(2) | C[+57], N[+2205] | 1138.736 | 4 | 4551.9222 | 4551.9145 | 0 | 1.7 | 332 | CRagged | 553.85 | 553.85 | 459.45 | 11.06 | 109 | >sp\|P01876\|IGHA1_HUMAN Immunoglobulin heavy constant alpha 1 OS=Homo sapiens OX=9606 GN=IGHA1 PE=1 SV=2 | 255 | 20201116_xd_colostrum_1ng_glyco_hcd203040_compound15.17555.17555.4 | controllerType=0 controllerNumber=1 scan=17555 | 37.4269 |
| 11248:3 | 4 | R.LAGKPTHVN[+1216.423]VSVVMAEVDGTC[+57.021]Y.- | HexNAc(2)Hex(5) | C[+57], N[+1216] | 1188.528 | 3 | 3563.5695 | 3563.5649 | 0 | 1.3 | 332 | CRagged | 555.06 | 555.06 | 337.96 | 10.97 | 109 | >sp\|P01876\|IGHA1_HUMAN Immunoglobulin heavy constant alpha 1 OS=Homo sapiens OX=9606 GN=IGHA1 PE=1 SV=2 | 255 | 20201116_xd_colostrum_1ng_glyco_hcd203040_compound15.14297.14297.3 | controllerType=0 controllerNumber=1 scan=14297 | 31.0416 |
| 11067:3 | 4 | R.LAGKPTHVN[+1622.582]VSVVMAEVDGTC[+57.021]Y.- | HexNAc(4)Hex(5) | C[+57], N[+1623] | 1323.9157 | 3 | 3969.7326 | 3969.7237 | 0 | 2.2 | 332 | CRagged | 561.96 | 432.28 | 348.77 | 11.32 | 109 | >sp\|P01876\|IGHA1_HUMAN Immunoglobulin heavy constant alpha 1 OS=Homo sapiens OX=9606 GN=IGHA1 PE=1 SV=2 | 255 | 20201116_xd_colostrum_1ng_glyco_hcd203040_compound15.14108.14108.3 | controllerType=0 controllerNumber=1 scan=14108 | 30.6574 |
| 11481:3 | 4 | R.LAGKPTHVN[+1710.598]VSVVMAEVDGTC[+57.021]Y.- | HexNAc(3)Hex(5)NeuAc(1) | C[+57], N[+1711] | 1353.2552 | 3 | 4057.751 | 4057.7397 | 0 | 2.8 | 332 | CRagged | 564.33 | 564.33 | 379.41 | 11.32 | 109 | >sp\|P01876\|IGHA1_HUMAN Immunoglobulin heavy constant alpha 1 OS=Homo sapiens OX=9606 GN=IGHA1 PE=1 SV=2 | 255 | 20201116_xd_colostrum_1ng_glyco_hcd203040_compound15.14539.14539.3 | controllerType=0 controllerNumber=1 scan=14539 | 31.4986 |
| 12522:4 | 4 | R.LAGKPTHVN[+2116.756]VSVVMAEVDGTC[+57.021]Y.- | HexNAc(5)Hex(5)NeuAc(1) | C[+57], N[+2117] | 1116.7325 | 4 | 4463.908 | 4463.8985 | 0 | 2.1 | 332 | CRagged | 570.78 | 499.46 | 499.46 | 11.41 | 109 | >sp\|P01876\|IGHA1_HUMAN Immunoglobulin heavy constant alpha 1 OS=Homo sapiens OX=9606 GN=IGHA1 PE=1 SV=2 | 255 | 20201116_xd_colostrum_1ng_glyco_hcd203040_compound15.15620.15620.4 | controllerType=0 controllerNumber=1 scan=15620 | 33.5428 |
| 09975:4 | 4 | R.LAGKPTHVN[+1663.608]VSVVMAEVDGTC[+57.021]Y.- | HexNAc(5)Hex(4) | C[+57], N[+1664] | 1003.4471 | 4 | 4010.7664 | 4010.7502 | 0 | 4 | 332 | CRagged | 576.61 | 460.56 | 460.56 | 10.98 | 109 | >sp\|P01876\|IGHA1_HUMAN Immunoglobulin heavy constant alpha 1 OS=Homo sapiens OX=9606 GN=IGHA1 PE=1 SV=2 | 255 | 20201116_xd_colostrum_1ng_glyco_hcd203040_compound15.12973.12973.4 | controllerType=0 controllerNumber=1 scan=12973 | 28.3719 |
| 12551:4 | 4 | R.LAGKPTHVN[+1913.677]VSVVMAEVDGTC[+57.021]Y.- | HexNAc(4)Hex(5)NeuAc(1) | C[+57], N[+1914] | 1065.9622 | 4 | 4260.827 | 4260.8191 | 0 | 1.8 | 332 | CRagged | 584.92 | 429.76 | 429.76 | 12.56 | 109 | >sp\|P01876\|IGHA1_HUMAN Immunoglobulin heavy constant alpha 1 OS=Homo sapiens OX=9606 GN=IGHA1 PE=1 SV=2 | 255 | 20201116_xd_colostrum_1ng_glyco_hcd203040_compound15.15650.15650.4 | controllerType=0 controllerNumber=1 scan=15650 | 33.5998 |
| 14307:4 | 4 | R.LAGKPTHVN[+2553.910]VSVVMAEVDGTC[+57.021]Y.- | HexNAc(5)Hex(5)Fuc(1)NeuAc(2) | C[+57], N[+2554] | 1226.0231 | 4 | 4901.0706 | 4901.0518 | 0 | 3.8 | 332 | CRagged | 587.1 | 444.49 | 444.49 | 12.51 | 109 | >sp\|P01876\|IGHA1_HUMAN Immunoglobulin heavy constant alpha 1 OS=Homo sapiens OX=9606 GN=IGHA1 PE=1 SV=2 | 255 | 20201116_xd_colostrum_1ng_glyco_hcd203040_compound15.17476.17476.4 | controllerType=0 controllerNumber=1 scan=17476 | 37.2539 |
| 10026:2 | 4 | R.LAGKPTHVN[+1216.423]VSVVMAEVDGTC[+57.021]Y.- | HexNAc(2)Hex(5) | C[+57], N[+1216] | 1782.2899 | 2 | 3563.5725 | 3563.5649 | 0 | 2.1 | 332 | CRagged | 590.41 | 428.93 | 217.95 | 12.06 | 109 | >sp\|P01876\|IGHA1_HUMAN Immunoglobulin heavy constant alpha 1 OS=Homo sapiens OX=9606 GN=IGHA1 PE=1 SV=2 | 255 | 20201116_xd_colostrum_1ng_glyco_hcd203040_compound15.13026.13026.2 | controllerType=0 controllerNumber=1 scan=13026 | 28.4953 |
| 10876:3 | 4 | R.LAGKPTHVN[+2174.798]VSVVMAEVDGTC[+57.021]Y.- | HexNAc(6)Hex(5)Fuc(1) | C[+57], N[+2175] | 1507.9911 | 3 | 4521.9589 | 4521.9403 | 0 | 4.1 | 332 | CRagged | 593.65 | 493.82 | 384.79 | 12.19 | 109 | >sp\|P01876\|IGHA1_HUMAN Immunoglobulin heavy constant alpha 1 OS=Homo sapiens OX=9606 GN=IGHA1 PE=1 SV=2 | 255 | 20201116_xd_colostrum_1ng_glyco_hcd203040_compound15.13909.13909.3 | controllerType=0 controllerNumber=1 scan=13909 | 30.208 |
| 09967:4 | 4 | R.LAGKPTHVN[+1809.666]VSVVMAEVDGTC[+57.021]Y.- | HexNAc(5)Hex(4)Fuc(1) | C[+57], N[+1810] | 1039.9582 | 4 | 4156.8109 | 4156.8081 | 0 | 0.7 | 332 | CRagged | 597.8 | 566.33 | 474.37 | 12.28 | 109 | >sp\|P01876\|IGHA1_HUMAN Immunoglobulin heavy constant alpha 1 OS=Homo sapiens OX=9606 GN=IGHA1 PE=1 SV=2 | 255 | 20201116_xd_colostrum_1ng_glyco_hcd203040_compound15.12965.12965.4 | controllerType=0 controllerNumber=1 scan=12965 | 28.3523 |
| 12671:3 | 4 | R.LAGKPTHVN[+1751.624]VSVVMAEVDGTC[+57.021]Y.- | HexNAc(4)Hex(4)NeuAc(1) | C[+57], N[+1752] | 1366.9283 | 3 | 4098.7703 | 4098.7663 | 0 | 1 | 332 | CRagged | 599.78 | 543.73 | 543.73 | 12.46 | 109 | >sp\|P01876\|IGHA1_HUMAN Immunoglobulin heavy constant alpha 1 OS=Homo sapiens OX=9606 GN=IGHA1 PE=1 SV=2 | 255 | 20201116_xd_colostrum_1ng_glyco_hcd203040_compound15.15775.15775.3 | controllerType=0 controllerNumber=1 scan=15775 | 33.8698 |
| 10954:4 | 4 | R.LAGKPTHVN[+1663.608]VSVVMAEVDGTC[+57.021]Y.- | HexNAc(5)Hex(4) | C[+57], N[+1664] | 1003.4448 | 4 | 4010.7574 | 4010.7502 | 0 | 1.8 | 332 | CRagged | 605.76 | 528.35 | 392.05 | 13.56 | 109 | >sp\|P01876\|IGHA1_HUMAN Immunoglobulin heavy constant alpha 1 OS=Homo sapiens OX=9606 GN=IGHA1 PE=1 SV=2 | 255 | 20201116_xd_colostrum_1ng_glyco_hcd203040_compound15.13991.13991.4 | controllerType=0 controllerNumber=1 scan=13991 | 30.4189 |
| 10951:4 | 4 | R.LAGKPTHVN[+1663.608]VSVVMAEVDGTC[+57.021]Y.- | HexNAc(5)Hex(4) | C[+57], N[+1664] | 1003.4448 | 4 | 4010.7574 | 4010.7502 | 0 | 1.8 | 332 | CRagged | 613.44 | 613.44 | 460.83 | 13.56 | 109 | >sp\|P01876\|IGHA1_HUMAN Immunoglobulin heavy constant alpha 1 OS=Homo sapiens OX=9606 GN=IGHA1 PE=1 SV=2 | 255 | 20201116_xd_colostrum_1ng_glyco_hcd203040_compound15.13988.13988.4 | controllerType=0 controllerNumber=1 scan=13988 | 30.414 |
| 10005:4 | 4 | R.LAGKPTHVN[+1216.423]VSVVMAEVDGTC[+57.021]Y.- | HexNAc(2)Hex(5) | C[+57], N[+1216] | 891.6484 | 4 | 3563.5716 | 3563.5649 | 0 | 1.9 | 332 | CRagged | 616.91 | 615.14 | 462.4 | 13.56 | 109 | >sp\|P01876\|IGHA1_HUMAN Immunoglobulin heavy constant alpha 1 OS=Homo sapiens OX=9606 GN=IGHA1 PE=1 SV=2 | 255 | 20201116_xd_colostrum_1ng_glyco_hcd203040_compound15.13005.13005.4 | controllerType=0 controllerNumber=1 scan=13005 | 28.4469 |
| 10976:3 | 4 | R.LAGKPTHVN[+1460.529]VSVVMAEVDGTC[+57.021]Y.- | HexNAc(4)Hex(4) | C[+57], N[+1461] | 1269.8985 | 3 | 3807.6809 | 3807.6709 | 0 | 2.6 | 332 | CRagged | 634.23 | 634.23 | 367.62 | 13.62 | 109 | >sp\|P01876\|IGHA1_HUMAN Immunoglobulin heavy constant alpha 1 OS=Homo sapiens OX=9606 GN=IGHA1 PE=1 SV=2 | 255 | 20201116_xd_colostrum_1ng_glyco_hcd203040_compound15.14014.14014.3 | controllerType=0 controllerNumber=1 scan=14014 | 30.4708 |
| 12595:3 | 4 | R.LAGKPTHVN[+1872.651]VSVVMAEVDGTC[+57.021]Y.- | HexNAc(3)Hex(6)NeuAc(1) | C[+57], N[+1873] | 1407.2718 | 3 | 4219.8008 | 4219.7926 | 0 | 2 | 332 | CRagged | 637.28 | 562.03 | 366.68 | 13.46 | 109 | >sp\|P01876\|IGHA1_HUMAN Immunoglobulin heavy constant alpha 1 OS=Homo sapiens OX=9606 GN=IGHA1 PE=1 SV=2 | 255 | 20201116_xd_colostrum_1ng_glyco_hcd203040_compound15.15696.15696.3 | controllerType=0 controllerNumber=1 scan=15696 | 33.7077 |
| 12587:4 | 4 | R.LAGKPTHVN[+2059.735]VSVVMAEVDGTC[+57.021]Y.- | HexNAc(4)Hex(5)Fuc(1)NeuAc(1) | C[+57], N[+2060] | 1102.4754 | 4 | 4406.8797 | 4406.877 | 0 | 0.6 | 332 | CRagged | 638.17 | 406.86 | 406.86 | 13.28 | 109 | >sp\|P01876\|IGHA1_HUMAN Immunoglobulin heavy constant alpha 1 OS=Homo sapiens OX=9606 GN=IGHA1 PE=1 SV=2 | 255 | 20201116_xd_colostrum_1ng_glyco_hcd203040_compound15.15687.15687.4 | controllerType=0 controllerNumber=1 scan=15687 | 33.6711 |
| 12599:4 | 4 | R.LAGKPTHVN[+2059.735]VSVVMAEVDGTC[+57.021]Y.- | HexNAc(4)Hex(5)Fuc(1)NeuAc(1) | C[+57], N[+2060] | 1102.4775 | 4 | 4406.888 | 4406.877 | 0 | 2.5 | 332 | CRagged | 638.66 | 363.04 | 363.04 | 12.2 | 109 | >sp\|P01876\|IGHA1_HUMAN Immunoglobulin heavy constant alpha 1 OS=Homo sapiens OX=9606 GN=IGHA1 PE=1 SV=2 | 255 | 20201116_xd_colostrum_1ng_glyco_hcd203040_compound15.15700.15700.4 | controllerType=0 controllerNumber=1 scan=15700 | 33.7131 |
| 12542:4 | 4 | R.LAGKPTHVN[+2262.814]VSVVMAEVDGTC[+57.021]Y.- | HexNAc(5)Hex(5)Fuc(1)NeuAc(1) | C[+57], N[+2263] | 1153.2466 | 4 | 4609.9647 | 4609.9564 | 0 | 1.8 | 332 | CRagged | 652.59 | 525.1 | 525.1 | 14.56 | 109 | >sp\|P01876\|IGHA1_HUMAN Immunoglobulin heavy constant alpha 1 OS=Homo sapiens OX=9606 GN=IGHA1 PE=1 SV=2 | 255 | 20201116_xd_colostrum_1ng_glyco_hcd203040_compound15.15641.15641.4 | controllerType=0 controllerNumber=1 scan=15641 | 33.5834 |
| 09935:3 | 4 | R.LAGKPTHVN[+1825.661]VSVVMAEVDGTC[+57.021]Y.- | HexNAc(5)Hex(5) | C[+57], N[+1826] | 1391.6089 | 3 | 4172.8123 | 4172.8031 | 0 | 2.2 | 332 | CRagged | 652.93 | 652.93 | 385.32 | 14.62 | 109 | >sp\|P01876\|IGHA1_HUMAN Immunoglobulin heavy constant alpha 1 OS=Homo sapiens OX=9606 GN=IGHA1 PE=1 SV=2 | 255 | 20201116_xd_colostrum_1ng_glyco_hcd203040_compound15.12931.12931.3 | controllerType=0 controllerNumber=1 scan=12931 | 28.2708 |
| 12544:4 | 4 | R.LAGKPTHVN[+2262.814]VSVVMAEVDGTC[+57.021]Y.- | HexNAc(5)Hex(5)Fuc(1)NeuAc(1) | C[+57], N[+2263] | 1153.2466 | 4 | 4609.9647 | 4609.9564 | 0 | 1.8 | 332 | CRagged | 654.88 | 547.23 | 547.23 | 14.56 | 109 | >sp\|P01876\|IGHA1_HUMAN Immunoglobulin heavy constant alpha 1 OS=Homo sapiens OX=9606 GN=IGHA1 PE=1 SV=2 | 255 | 20201116_xd_colostrum_1ng_glyco_hcd203040_compound15.15643.15643.4 | controllerType=0 controllerNumber=1 scan=15643 | 33.5878 |
| 10936:4 | 4 | R.LAGKPTHVN[+1825.661]VSVVMAEVDGTC[+57.021]Y.- | HexNAc(5)Hex(5) | C[+57], N[+1826] | 1043.9582 | 4 | 4172.8109 | 4172.8031 | 0 | 1.9 | 332 | CRagged | 660.68 | 660.68 | 482.09 | 14.56 | 109 | >sp\|P01876\|IGHA1_HUMAN Immunoglobulin heavy constant alpha 1 OS=Homo sapiens OX=9606 GN=IGHA1 PE=1 SV=2 | 255 | 20201116_xd_colostrum_1ng_glyco_hcd203040_compound15.13972.13972.4 | controllerType=0 controllerNumber=1 scan=13972 | 30.3655 |
| 09982:3 | 4 | R.LAGKPTHVN[+1581.555]VSVVMAEVDGTC[+57.021]Y.- | HexNAc(3)Hex(6) | C[+57], N[+1582] | 1310.2414 | 3 | 3928.7096 | 3928.6971 | 0 | 3.2 | 332 | CRagged | 661.44 | 639.03 | 414.79 | 14.42 | 109 | >sp\|P01876\|IGHA1_HUMAN Immunoglobulin heavy constant alpha 1 OS=Homo sapiens OX=9606 GN=IGHA1 PE=1 SV=2 | 255 | 20201116_xd_colostrum_1ng_glyco_hcd203040_compound15.12981.12981.3 | controllerType=0 controllerNumber=1 scan=12981 | 28.4011 |
| 11152:3 | 4 | R.LAGKPTHVN[+892.317]VSVVMAEVDGTC[+57.021]Y.- | HexNAc(2)Hex(3) | C[+57], N[+892] | 1080.495 | 3 | 3239.4705 | 3239.4593 | 0 | 3.5 | 332 | CRagged | 675.69 | 675.69 | 346.29 | 14.42 | 109 | >sp\|P01876\|IGHA1_HUMAN Immunoglobulin heavy constant alpha 1 OS=Homo sapiens OX=9606 GN=IGHA1 PE=1 SV=2 | 255 | 20201116_xd_colostrum_1ng_glyco_hcd203040_compound15.14197.14197.3 | controllerType=0 controllerNumber=1 scan=14197 | 30.8325 |
| 10007:4 | 4 | R.LAGKPTHVN[+1216.423]VSVVMAEVDGTC[+57.021]Y.- | HexNAc(2)Hex(5) | C[+57], N[+1216] | 891.6484 | 4 | 3563.5716 | 3563.5649 | 0 | 1.9 | 332 | CRagged | 679.58 | 671.44 | 483.38 | 14.56 | 109 | >sp\|P01876\|IGHA1_HUMAN Immunoglobulin heavy constant alpha 1 OS=Homo sapiens OX=9606 GN=IGHA1 PE=1 SV=2 | 255 | 20201116_xd_colostrum_1ng_glyco_hcd203040_compound15.13007.13007.4 | controllerType=0 controllerNumber=1 scan=13007 | 28.4523 |
| 10939:4 | 4 | R.LAGKPTHVN[+1825.661]VSVVMAEVDGTC[+57.021]Y.- | HexNAc(5)Hex(5) | C[+57], N[+1826] | 1043.9582 | 4 | 4172.8109 | 4172.8031 | 0 | 1.9 | 332 | CRagged | 684.05 | 649.45 | 482.37 | 14.56 | 109 | >sp\|P01876\|IGHA1_HUMAN Immunoglobulin heavy constant alpha 1 OS=Homo sapiens OX=9606 GN=IGHA1 PE=1 SV=2 | 255 | 20201116_xd_colostrum_1ng_glyco_hcd203040_compound15.13975.13975.4 | controllerType=0 controllerNumber=1 scan=13975 | 30.379 |
| 11022:2 | 4 | R.LAGKPTHVN[+1216.423]VSVVMAEVDGTC[+57.021]Y.- | HexNAc(2)Hex(5) | C[+57], N[+1216] | 1782.291 | 2 | 3563.5747 | 3563.5649 | 0 | 2.7 | 332 | CRagged | 684.11 | 684.11 | 335.49 | 14.06 | 109 | >sp\|P01876\|IGHA1_HUMAN Immunoglobulin heavy constant alpha 1 OS=Homo sapiens OX=9606 GN=IGHA1 PE=1 SV=2 | 255 | 20201116_xd_colostrum_1ng_glyco_hcd203040_compound15.14062.14062.2 | controllerType=0 controllerNumber=1 scan=14062 | 30.565 |
| 14298:3 | 4 | R.LAGKPTHVN[+2553.910]VSVVMAEVDGTC[+57.021]Y.- | HexNAc(5)Hex(5)Fuc(1)NeuAc(2) | C[+57], N[+2554] | 1634.3591 | 3 | 4901.0626 | 4901.0518 | 0 | 2.2 | 332 | CRagged | 689.7 | 484.56 | 484.56 | 14.62 | 109 | >sp\|P01876\|IGHA1_HUMAN Immunoglobulin heavy constant alpha 1 OS=Homo sapiens OX=9606 GN=IGHA1 PE=1 SV=2 | 255 | 20201116_xd_colostrum_1ng_glyco_hcd203040_compound15.17466.17466.3 | controllerType=0 controllerNumber=1 scan=17466 | 37.2335 |
| 14309:4 | 4 | R.LAGKPTHVN[+2553.910]VSVVMAEVDGTC[+57.021]Y.- | HexNAc(5)Hex(5)Fuc(1)NeuAc(2) | C[+57], N[+2554] | 1226.0231 | 4 | 4901.0706 | 4901.0518 | 0 | 3.8 | 332 | CRagged | 691.14 | 506.21 | 506.21 | 14.51 | 109 | >sp\|P01876\|IGHA1_HUMAN Immunoglobulin heavy constant alpha 1 OS=Homo sapiens OX=9606 GN=IGHA1 PE=1 SV=2 | 255 | 20201116_xd_colostrum_1ng_glyco_hcd203040_compound15.17478.17478.4 | controllerType=0 controllerNumber=1 scan=17478 | 37.2585 |
| 11070:3 | 4 | R.LAGKPTHVN[+1501.555]VSVVMAEVDGTC[+57.021]Y.- | HexNAc(5)Hex(3) | C[+57], N[+1502] | 1283.5727 | 3 | 3848.7035 | 3848.6974 | 0 | 1.6 | 332 | CRagged | 693.61 | 693.61 | 693.61 | 14.46 | 109 | >sp\|P01876\|IGHA1_HUMAN Immunoglobulin heavy constant alpha 1 OS=Homo sapiens OX=9606 GN=IGHA1 PE=1 SV=2 | 255 | 20201116_xd_colostrum_1ng_glyco_hcd203040_compound15.14112.14112.3 | controllerType=0 controllerNumber=1 scan=14112 | 30.6664 |
| 11484:3 | 4 | R.LAGKPTHVN[+1710.598]VSVVMAEVDGTC[+57.021]Y.- | HexNAc(3)Hex(5)NeuAc(1) | C[+57], N[+1711] | 1353.2552 | 3 | 4057.751 | 4057.7397 | 0 | 2.8 | 332 | CRagged | 700.82 | 695.91 | 434 | 15.62 | 109 | >sp\|P01876\|IGHA1_HUMAN Immunoglobulin heavy constant alpha 1 OS=Homo sapiens OX=9606 GN=IGHA1 PE=1 SV=2 | 255 | 20201116_xd_colostrum_1ng_glyco_hcd203040_compound15.14542.14542.3 | controllerType=0 controllerNumber=1 scan=14542 | 31.5057 |
| 10922:3 | 4 | R.LAGKPTHVN[+1825.661]VSVVMAEVDGTC[+57.021]Y.- | HexNAc(5)Hex(5) | C[+57], N[+1826] | 1391.6094 | 3 | 4172.8137 | 4172.8031 | 0 | 2.6 | 332 | CRagged | 702.45 | 685.7 | 435.16 | 15.62 | 109 | >sp\|P01876\|IGHA1_HUMAN Immunoglobulin heavy constant alpha 1 OS=Homo sapiens OX=9606 GN=IGHA1 PE=1 SV=2 | 255 | 20201116_xd_colostrum_1ng_glyco_hcd203040_compound15.13957.13957.3 | controllerType=0 controllerNumber=1 scan=13957 | 30.3285 |
| 14314:4 | 4 | R.LAGKPTHVN[+2350.830]VSVVMAEVDGTC[+57.021]Y.- | HexNAc(4)Hex(5)Fuc(1)NeuAc(2) | C[+57], N[+2351] | 1175.252 | 4 | 4697.9862 | 4697.9724 | 0 | 2.9 | 332 | CRagged | 705.67 | 639.73 | 639.73 | 15.71 | 109 | >sp\|P01876\|IGHA1_HUMAN Immunoglobulin heavy constant alpha 1 OS=Homo sapiens OX=9606 GN=IGHA1 PE=1 SV=2 | 255 | 20201116_xd_colostrum_1ng_glyco_hcd203040_compound15.17483.17483.4 | controllerType=0 controllerNumber=1 scan=17483 | 37.2648 |
| 10636:3 | 4 | R.LAGKPTHVN[+1216.423]VSVVMAEVDGTC[+57.021]Y.- | HexNAc(2)Hex(5) | C[+57], N[+1216] | 1188.5291 | 3 | 3563.5728 | 3563.5649 | 0 | 2.2 | 332 | CRagged | 708.36 | 679.74 | 370.39 | 15.62 | 109 | >sp\|P01876\|IGHA1_HUMAN Immunoglobulin heavy constant alpha 1 OS=Homo sapiens OX=9606 GN=IGHA1 PE=1 SV=2 | 255 | 20201116_xd_colostrum_1ng_glyco_hcd203040_compound15.13661.13661.3 | controllerType=0 controllerNumber=1 scan=13661 | 29.7683 |
| 11098:3 | 4 | R.LAGKPTHVN[+1444.534]VSVVMAEVDGTC[+57.021]Y.- | HexNAc(4)Hex(3)Fuc(1) | C[+57], N[+1445] | 1264.566 | 3 | 3791.6834 | 3791.6759 | 0 | 2 | 332 | CRagged | 708.39 | 629.07 | 629.07 | 15.46 | 109 | >sp\|P01876\|IGHA1_HUMAN Immunoglobulin heavy constant alpha 1 OS=Homo sapiens OX=9606 GN=IGHA1 PE=1 SV=2 | 255 | 20201116_xd_colostrum_1ng_glyco_hcd203040_compound15.14141.14141.3 | controllerType=0 controllerNumber=1 scan=14141 | 30.7199 |
| 14315:3 | 4 | R.LAGKPTHVN[+2350.830]VSVVMAEVDGTC[+57.021]Y.- | HexNAc(4)Hex(5)Fuc(1)NeuAc(2) | C[+57], N[+2351] | 1566.6675 | 3 | 4697.9881 | 4697.9724 | 0 | 3.3 | 332 | CRagged | 710.95 | 615.8 | 615.8 | 15.42 | 109 | >sp\|P01876\|IGHA1_HUMAN Immunoglobulin heavy constant alpha 1 OS=Homo sapiens OX=9606 GN=IGHA1 PE=1 SV=2 | 255 | 20201116_xd_colostrum_1ng_glyco_hcd203040_compound15.17484.17484.3 | controllerType=0 controllerNumber=1 scan=17484 | 37.2668 |
| 12541:3 | 4 | R.LAGKPTHVN[+2262.814]VSVVMAEVDGTC[+57.021]Y.- | HexNAc(5)Hex(5)Fuc(1)NeuAc(1) | C[+57], N[+2263] | 1537.3268 | 3 | 4609.966 | 4609.9564 | 0 | 2.1 | 332 | CRagged | 720.58 | 481.76 | 481.76 | 15.46 | 109 | >sp\|P01876\|IGHA1_HUMAN Immunoglobulin heavy constant alpha 1 OS=Homo sapiens OX=9606 GN=IGHA1 PE=1 SV=2 | 255 | 20201116_xd_colostrum_1ng_glyco_hcd203040_compound15.15640.15640.3 | controllerType=0 controllerNumber=1 scan=15640 | 33.5817 |
| 14377:3 | 4 | R.LAGKPTHVN[+2204.772]VSVVMAEVDGTC[+57.021]Y.- | HexNAc(4)Hex(5)NeuAc(2) | C[+57], N[+2205] | 1517.9802 | 3 | 4551.9259 | 4551.9145 | 0 | 2.5 | 332 | CRagged | 720.87 | 472.78 | 472.78 | 15.62 | 109 | >sp\|P01876\|IGHA1_HUMAN Immunoglobulin heavy constant alpha 1 OS=Homo sapiens OX=9606 GN=IGHA1 PE=1 SV=2 | 255 | 20201116_xd_colostrum_1ng_glyco_hcd203040_compound15.17549.17549.3 | controllerType=0 controllerNumber=1 scan=17549 | 37.4189 |
| 10930:3 | 4 | R.LAGKPTHVN[+1971.719]VSVVMAEVDGTC[+57.021]Y.- | HexNAc(5)Hex(5)Fuc(1) | C[+57], N[+1972] | 1440.2955 | 3 | 4318.8718 | 4318.861 | 0 | 2.5 | 332 | CRagged | 721.75 | 580.75 | 463.78 | 15.62 | 109 | >sp\|P01876\|IGHA1_HUMAN Immunoglobulin heavy constant alpha 1 OS=Homo sapiens OX=9606 GN=IGHA1 PE=1 SV=2 | 255 | 20201116_xd_colostrum_1ng_glyco_hcd203040_compound15.13966.13966.3 | controllerType=0 controllerNumber=1 scan=13966 | 30.352 |
| 12543:3 | 4 | R.LAGKPTHVN[+2262.814]VSVVMAEVDGTC[+57.021]Y.- | HexNAc(5)Hex(5)Fuc(1)NeuAc(1) | C[+57], N[+2263] | 1537.3268 | 3 | 4609.966 | 4609.9564 | 0 | 2.1 | 332 | CRagged | 723.62 | 587.39 | 587.39 | 15.46 | 109 | >sp\|P01876\|IGHA1_HUMAN Immunoglobulin heavy constant alpha 1 OS=Homo sapiens OX=9606 GN=IGHA1 PE=1 SV=2 | 255 | 20201116_xd_colostrum_1ng_glyco_hcd203040_compound15.15642.15642.3 | controllerType=0 controllerNumber=1 scan=15642 | 33.586 |
| 11329:3 | 4 | R.LAGKPTHVN[+2262.814]VSVVMAEVDGTC[+57.021]Y.- | HexNAc(5)Hex(5)Fuc(1)NeuAc(1) | C[+57], N[+2263] | 1537.3281 | 3 | 4609.9696 | 4609.9564 | 0 | 2.9 | 332 | CRagged | 725.38 | 569.21 | 569.21 | 15.62 | 109 | >sp\|P01876\|IGHA1_HUMAN Immunoglobulin heavy constant alpha 1 OS=Homo sapiens OX=9606 GN=IGHA1 PE=1 SV=2 | 255 | 20201116_xd_colostrum_1ng_glyco_hcd203040_compound15.14381.14381.3 | controllerType=0 controllerNumber=1 scan=14381 | 31.2185 |
| 12619:3 | 4 | R.LAGKPTHVN[+2059.735]VSVVMAEVDGTC[+57.021]Y.- | HexNAc(4)Hex(5)Fuc(1)NeuAc(1) | C[+57], N[+2060] | 1469.6335 | 3 | 4406.8859 | 4406.877 | 0 | 2 | 332 | CRagged | 726.61 | 706.13 | 706.13 | 15.46 | 109 | >sp\|P01876\|IGHA1_HUMAN Immunoglobulin heavy constant alpha 1 OS=Homo sapiens OX=9606 GN=IGHA1 PE=1 SV=2 | 255 | 20201116_xd_colostrum_1ng_glyco_hcd203040_compound15.15721.15721.3 | controllerType=0 controllerNumber=1 scan=15721 | 33.7518 |
| 09890:3 | 4 | R.LAGKPTHVN[+1702.581]VSVVMAEVDGTC[+57.021]Y.- | HexNAc(2)Hex(8) | C[+57], N[+1703] | 1350.5825 | 3 | 4049.7328 | 4049.7234 | 0 | 2.3 | 332 | CRagged | 730.84 | 730.84 | 481.65 | 15.62 | 109 | >sp\|P01876\|IGHA1_HUMAN Immunoglobulin heavy constant alpha 1 OS=Homo sapiens OX=9606 GN=IGHA1 PE=1 SV=2 | 255 | 20201116_xd_colostrum_1ng_glyco_hcd203040_compound15.12885.12885.3 | controllerType=0 controllerNumber=1 scan=12885 | 28.1742 |
| 14293:4 | 4 | R.LAGKPTHVN[+2350.830]VSVVMAEVDGTC[+57.021]Y.- | HexNAc(4)Hex(5)Fuc(1)NeuAc(2) | C[+57], N[+2351] | 1175.2505 | 4 | 4697.9803 | 4697.9724 | 0 | 1.7 | 332 | CRagged | 737.7 | 630.26 | 630.26 | 15.56 | 109 | >sp\|P01876\|IGHA1_HUMAN Immunoglobulin heavy constant alpha 1 OS=Homo sapiens OX=9606 GN=IGHA1 PE=1 SV=2 | 255 | 20201116_xd_colostrum_1ng_glyco_hcd203040_compound15.17461.17461.4 | controllerType=0 controllerNumber=1 scan=17461 | 37.2218 |
| 10965:3 | 4 | R.LAGKPTHVN[+1809.666]VSVVMAEVDGTC[+57.021]Y.- | HexNAc(5)Hex(4)Fuc(1) | C[+57], N[+1810] | 1386.2759 | 3 | 4156.8133 | 4156.8081 | 0 | 1.2 | 332 | CRagged | 740.27 | 737.94 | 424.21 | 15.46 | 109 | >sp\|P01876\|IGHA1_HUMAN Immunoglobulin heavy constant alpha 1 OS=Homo sapiens OX=9606 GN=IGHA1 PE=1 SV=2 | 255 | 20201116_xd_colostrum_1ng_glyco_hcd203040_compound15.14003.14003.3 | controllerType=0 controllerNumber=1 scan=14003 | 30.4499 |
| 09986:3 | 4 | R.LAGKPTHVN[+1606.587]VSVVMAEVDGTC[+57.021]Y.- | HexNAc(4)Hex(4)Fuc(1) | C[+57], N[+1607] | 1318.584 | 3 | 3953.7376 | 3953.7288 | 0 | 2.2 | 332 | CRagged | 742.68 | 640.71 | 640.71 | 15.62 | 109 | >sp\|P01876\|IGHA1_HUMAN Immunoglobulin heavy constant alpha 1 OS=Homo sapiens OX=9606 GN=IGHA1 PE=1 SV=2 | 255 | 20201116_xd_colostrum_1ng_glyco_hcd203040_compound15.12985.12985.3 | controllerType=0 controllerNumber=1 scan=12985 | 28.4077 |
| 14375:3 | 4 | R.LAGKPTHVN[+2204.772]VSVVMAEVDGTC[+57.021]Y.- | HexNAc(4)Hex(5)NeuAc(2) | C[+57], N[+2205] | 1517.9802 | 3 | 4551.9259 | 4551.9145 | 0 | 2.5 | 332 | CRagged | 750.58 | 506.38 | 506.38 | 16.62 | 109 | >sp\|P01876\|IGHA1_HUMAN Immunoglobulin heavy constant alpha 1 OS=Homo sapiens OX=9606 GN=IGHA1 PE=1 SV=2 | 255 | 20201116_xd_colostrum_1ng_glyco_hcd203040_compound15.17547.17547.3 | controllerType=0 controllerNumber=1 scan=17547 | 37.4146 |
| 09937:3 | 4 | R.LAGKPTHVN[+1825.661]VSVVMAEVDGTC[+57.021]Y.- | HexNAc(5)Hex(5) | C[+57], N[+1826] | 1391.6089 | 3 | 4172.8123 | 4172.8031 | 0 | 2.2 | 332 | CRagged | 753.25 | 664.18 | 388.89 | 16.62 | 109 | >sp\|P01876\|IGHA1_HUMAN Immunoglobulin heavy constant alpha 1 OS=Homo sapiens OX=9606 GN=IGHA1 PE=1 SV=2 | 255 | 20201116_xd_colostrum_1ng_glyco_hcd203040_compound15.12933.12933.3 | controllerType=0 controllerNumber=1 scan=12933 | 28.2789 |
| 12643:3 | 4 | R.LAGKPTHVN[+1710.598]VSVVMAEVDGTC[+57.021]Y.- | HexNAc(3)Hex(5)NeuAc(1) | C[+57], N[+1711] | 1353.2548 | 3 | 4057.7499 | 4057.7397 | 0 | 2.5 | 332 | CRagged | 753.59 | 753.59 | 456.07 | 16.62 | 109 | >sp\|P01876\|IGHA1_HUMAN Immunoglobulin heavy constant alpha 1 OS=Homo sapiens OX=9606 GN=IGHA1 PE=1 SV=2 | 255 | 20201116_xd_colostrum_1ng_glyco_hcd203040_compound15.15746.15746.3 | controllerType=0 controllerNumber=1 scan=15746 | 33.8057 |
| 09920:3 | 4 | R.LAGKPTHVN[+1971.719]VSVVMAEVDGTC[+57.021]Y.- | HexNAc(5)Hex(5)Fuc(1) | C[+57], N[+1972] | 1440.2957 | 3 | 4318.8726 | 4318.861 | 0 | 2.7 | 332 | CRagged | 758.71 | 754.84 | 484.84 | 16.62 | 109 | >sp\|P01876\|IGHA1_HUMAN Immunoglobulin heavy constant alpha 1 OS=Homo sapiens OX=9606 GN=IGHA1 PE=1 SV=2 | 255 | 20201116_xd_colostrum_1ng_glyco_hcd203040_compound15.12916.12916.3 | controllerType=0 controllerNumber=1 scan=12916 | 28.2344 |
| 12521:3 | 4 | R.LAGKPTHVN[+2116.756]VSVVMAEVDGTC[+57.021]Y.- | HexNAc(5)Hex(5)NeuAc(1) | C[+57], N[+2117] | 1488.6412 | 3 | 4463.909 | 4463.8985 | 0 | 2.3 | 332 | CRagged | 759.07 | 567.42 | 567.42 | 16.62 | 109 | >sp\|P01876\|IGHA1_HUMAN Immunoglobulin heavy constant alpha 1 OS=Homo sapiens OX=9606 GN=IGHA1 PE=1 SV=2 | 255 | 20201116_xd_colostrum_1ng_glyco_hcd203040_compound15.15619.15619.3 | controllerType=0 controllerNumber=1 scan=15619 | 33.5409 |
| 11326:3 | 4 | R.LAGKPTHVN[+2262.814]VSVVMAEVDGTC[+57.021]Y.- | HexNAc(5)Hex(5)Fuc(1)NeuAc(1) | C[+57], N[+2263] | 1537.3281 | 3 | 4609.9696 | 4609.9564 | 0 | 2.9 | 332 | CRagged | 759.08 | 474.68 | 474.68 | 16.62 | 109 | >sp\|P01876\|IGHA1_HUMAN Immunoglobulin heavy constant alpha 1 OS=Homo sapiens OX=9606 GN=IGHA1 PE=1 SV=2 | 255 | 20201116_xd_colostrum_1ng_glyco_hcd203040_compound15.14378.14378.3 | controllerType=0 controllerNumber=1 scan=14378 | 31.2093 |
| 09923:3 | 4 | R.LAGKPTHVN[+1971.719]VSVVMAEVDGTC[+57.021]Y.- | HexNAc(5)Hex(5)Fuc(1) | C[+57], N[+1972] | 1440.2957 | 3 | 4318.8726 | 4318.861 | 0 | 2.7 | 332 | CRagged | 759.6 | 721.94 | 513 | 16.62 | 109 | >sp\|P01876\|IGHA1_HUMAN Immunoglobulin heavy constant alpha 1 OS=Homo sapiens OX=9606 GN=IGHA1 PE=1 SV=2 | 255 | 20201116_xd_colostrum_1ng_glyco_hcd203040_compound15.12919.12919.3 | controllerType=0 controllerNumber=1 scan=12919 | [illegible] |
| 10057:3 | 4 | R.LAGKPTHVN[+1501.555]VSVVMAEVDGTC[+57.021]Y.- | HexNAc(5)Hex(3) | C[+57], N[+1502] | 1283.5739 | 3 | 3848.7072 | 3848.6974 | 0 | 2.5 | 332 | CRagged | 760.22 | 760.22 | 760.22 | 16.62 | 109 | >sp\|P01876\|IGHA1_HUMAN Immunoglobulin heavy constant alpha 1 OS=Homo sapiens OX=9606 GN=IGHA1 PE=1 SV=2 | 255 | 20201116_xd_colostrum_1ng_glyco_hcd203040_compound15.13059.13059.3 | controllerType=0 controllerNumber=1 scan=13059 | [illegible] |
| 12534:3 | 4 | R.LAGKPTHVN[+1913.677]VSVVMAEVDGTC[+57.021]Y.- | HexNAc(4)Hex(5)NeuAc(1) | C[+57], N[+1914] | 1420.9477 | 3 | 4260.8285 | 4260.8191 | 0 | 2.2 | 332 | CRagged | 761.23 | 544.11 | 544.11 | 16.62 | 109 | >sp\|P01876\|IGHA1_HUMAN Immunoglobulin heavy constant alpha 1 OS=Homo sapiens OX=9606 GN=IGHA1 PE=1 SV=2 | 255 | 20201116_xd_colostrum_1ng_glyco_hcd203040_compound15.15632.15632.3 | controllerType=0 controllerNumber=1 scan=15632 | [illegible] |
| 10955:3 | 4 | R.LAGKPTHVN[+1581.555]VSVVMAEVDGTC[+57.021]Y.- | HexNAc(3)Hex(6) | C[+57], N[+1582] | 1310.2407 | 3 | 3928.7074 | 3928.6971 | 0 | 2.6 | 332 | CRagged | 762.26 | 748.84 | 457.6 | 16.62 | 109 | >sp\|P01876\|IGHA1_HUMAN Immunoglobulin heavy constant alpha 1 OS=Homo sapiens OX=9606 GN=IGHA1 PE=1 SV=2 | 255 | 20201116_xd_colostrum_1ng_glyco_hcd203040_compound15.13992.13992.3 | controllerType=0 controllerNumber=1 scan=13992 | [illegible] |
| 11071:3 | 4 | R.LAGKPTHVN[+1647.613]VSVVMAEVDGTC[+57.021]Y.- | HexNAc(5)Hex(3)Fuc(1) | C[+57], N[+1648] | 1332.2593 | 3 | 3994.7635 | 3994.7553 | 0 | 2 | 332 | CRagged | 762.3 | 718.05 | 718.05 | 16.62 | 109 | >sp\|P01876\|IGHA1_HUMAN Immunoglobulin heavy constant alpha 1 OS=Homo sapiens OX=9606 GN=IGHA1 PE=1 SV=2 | 255 | 20201116_xd_colostrum_1ng_glyco_hcd203040_compound15.14113.14113.3 | controllerType=0 controllerNumber=1 scan=14113 | 30.6679 |
| 10971:3 | 4 | R.LAGKPTHVN[+1606.587]VSVVMAEVDGTC[+57.021]Y.- | HexNAc(4)Hex(4)Fuc(1) | C[+57], N[+1607] | 1318.5828 | 3 | 3953.7339 | 3953.7288 | 0 | 1.3 | 332 | CRagged | 780.16 | 780.16 | 780.16 | 16.46 | 109 | >sp\|P01876\|IGHA1_HUMAN Immunoglobulin heavy constant alpha 1 OS=Homo sapiens OX=9606 GN=IGHA1 PE=1 SV=2 | 255 | 20201116_xd_colostrum_1ng_glyco_hcd203040_compound15.14009.14009.3 | controllerType=0 controllerNumber=1 scan=14009 | 30.4591 |
| 11025:2 | 4 | R.LAGKPTHVN[+1216.423]VSVVMAEVDGTC[+57.021]Y.- | HexNAc(2)Hex(5) | C[+57], N[+1216] | 1782.291 | 2 | 3563.5747 | 3563.5649 | 0 | 2.7 | 332 | CRagged | 781.43 | 701.35 | 310.48 | 16.06 | 109 | >sp\|P01876\|IGHA1_HUMAN Immunoglobulin heavy constant alpha 1 OS=Homo sapiens OX=9606 GN=IGHA1 PE=1 SV=2 | 255 | 20201116_xd_colostrum_1ng_glyco_hcd203040_compound15.14065.14065.2 | controllerType=0 controllerNumber=1 scan=14065 | 30.5728 |
| 12641:3 | 4 | R.LAGKPTHVN[+1710.598]VSVVMAEVDGTC[+57.021]Y.- | HexNAc(3)Hex(5)NeuAc(1) | C[+57], N[+1711] | 1353.2548 | 3 | 4057.7499 | 4057.7397 | 0 | 2.5 | 332 | CRagged | 785.43 | 661.9 | 439.03 | 16.62 | 109 | >sp\|P01876\|IGHA1_HUMAN Immunoglobulin heavy constant alpha 1 OS=Homo sapiens OX=9606 GN=IGHA1 PE=1 SV=2 | 255 | 20201116_xd_colostrum_1ng_glyco_hcd203040_compound15.15744.15744.3 | controllerType=0 controllerNumber=1 scan=15744 | 33.8 |
| 11154:3 | 4 | R.LAGKPTHVN[+1378.476]VSVVMAEVDGTC[+57.021]Y.- | HexNAc(2)Hex(6) | C[+57], N[+1378] | 1242.5469 | 3 | 3725.6262 | 3725.6178 | 0 | 2.3 | 332 | CRagged | 786.66 | 656.42 | 462.7 | 16.62 | 109 | >sp\|P01876\|IGHA1_HUMAN Immunoglobulin heavy constant alpha 1 OS=Homo sapiens OX=9606 GN=IGHA1 PE=1 SV=2 | 255 | 20201116_xd_colostrum_1ng_glyco_hcd203040_compound15.14199.14199.3 | controllerType=0 controllerNumber=1 scan=14199 | 30.8347 |
| 11015:2 | 4 | R.LAGKPTHVN[+1216.423]VSVVMAEVDGTC[+57.021]Y.- | HexNAc(2)Hex(5) | C[+57], N[+1216] | 1782.2927 | 2 | 3563.5781 | 3563.5649 | 0 | 3.7 | 332 | CRagged | 790.15 | 698.97 | 304.12 | 15.87 | 109 | >sp\|P01876\|IGHA1_HUMAN Immunoglobulin heavy constant alpha 1 OS=Homo sapiens OX=9606 GN=IGHA1 PE=1 SV=2 | 255 | 20201116_xd_colostrum_1ng_glyco_hcd203040_compound15.14054.14054.2 | controllerType=0 controllerNumber=1 scan=14054 | 30.5533 |
| 12674:3 | 4 | R.LAGKPTHVN[+1751.624]VSVVMAEVDGTC[+57.021]Y.- | HexNAc(4)Hex(4)NeuAc(1) | C[+57], N[+1752] | 1366.9283 | 3 | 4098.7703 | 4098.7663 | 0 | 1 | 332 | CRagged | 793.89 | 665.44 | 665.44 | 16.46 | 109 | >sp\|P01876\|IGHA1_HUMAN Immunoglobulin heavy constant alpha 1 OS=Homo sapiens OX=9606 GN=IGHA1 PE=1 SV=2 | 255 | 20201116_xd_colostrum_1ng_glyco_hcd203040_compound15.15778.15778.3 | controllerType=0 controllerNumber=1 scan=15778 | 33.8781 |
| 11156:3 | 4 | R.LAGKPTHVN[+1378.476]VSVVMAEVDGTC[+57.021]Y.- | HexNAc(2)Hex(6) | C[+57], N[+1378] | 1242.5469 | 3 | 3725.6262 | 3725.6178 | 0 | 2.3 | 332 | CRagged | 795.04 | 545 | 479.32 | 16.62 | 109 | >sp\|P01876\|IGHA1_HUMAN Immunoglobulin heavy constant alpha 1 OS=Homo sapiens OX=9606 GN=IGHA1 PE=1 SV=2 | 255 | 20201116_xd_colostrum_1ng_glyco_hcd203040_compound15.14201.14201.3 | controllerType=0 controllerNumber=1 scan=14201 | 30.8418 |
| 10040:3 | 4 | R.LAGKPTHVN[+1054.370]VSVVMAEVDGTC[+57.021]Y.- | HexNAc(2)Hex(4) | C[+57], N[+1054] | 1134.5121 | 3 | 3401.5218 | 3401.5121 | 0 | 2.8 | 332 | CRagged | 806.33 | 764.22 | 492.73 | 17.62 | 109 | >sp\|P01876\|IGHA1_HUMAN Immunoglobulin heavy constant alpha 1 OS=Homo sapiens OX=9606 GN=IGHA1 PE=1 SV=2 | 255 | 20201116_xd_colostrum_1ng_glyco_hcd203040_compound15.13041.13041.3 | controllerType=0 controllerNumber=1 scan=13041 | 28.5342 |
| 09957:3 | 4 | R.LAGKPTHVN[+1663.608]VSVVMAEVDGTC[+57.021]Y.- | HexNAc(5)Hex(4) | C[+57], N[+1664] | 1337.5897 | 3 | 4010.7544 | 4010.7502 | 0 | 1 | 332 | CRagged | 808.93 | 808.93 | 496.92 | 17.46 | 109 | >sp\|P01876\|IGHA1_HUMAN Immunoglobulin heavy constant alpha 1 OS=Homo sapiens OX=9606 GN=IGHA1 PE=1 SV=2 | 255 | 20201116_xd_colostrum_1ng_glyco_hcd203040_compound15.12955.12955.3 | controllerType=0 controllerNumber=1 scan=12955 | 28.334 |
| 11100:3 | 4 | R.LAGKPTHVN[+1444.534]VSVVMAEVDGTC[+57.021]Y.- | HexNAc(4)Hex(3)Fuc(1) | C[+57], N[+1445] | 1264.566 | 3 | 3791.6834 | 3791.6759 | 0 | 2 | 332 | CRagged | 811.18 | 728.76 | 728.76 | 17.46 | 109 | >sp\|P01876\|IGHA1_HUMAN Immunoglobulin heavy constant alpha 1 OS=Homo sapiens OX=9606 GN=IGHA1 PE=1 SV=2 | 255 | 20201116_xd_colostrum_1ng_glyco_hcd203040_compound15.14143.14143.3 | controllerType=0 controllerNumber=1 scan=14143 | 30.7249 |
| 09959:3 | 4 | R.LAGKPTHVN[+1663.608]VSVVMAEVDGTC[+57.021]Y.- | HexNAc(5)Hex(4) | C[+57], N[+1664] | 1337.5897 | 3 | 4010.7544 | 4010.7502 | 0 | 1 | 332 | CRagged | 814.02 | 780.82 | 516.35 | 17.46 | 109 | >sp\|P01876\|IGHA1_HUMAN Immunoglobulin heavy constant alpha 1 OS=Homo sapiens OX=9606 GN=IGHA1 PE=1 SV=2 | 255 | 20201116_xd_colostrum_1ng_glyco_hcd203040_compound15.12957.12957.3 | controllerType=0 controllerNumber=1 scan=12957 | 28.3414 |
| 10906:3 | 4 | R.LAGKPTHVN[+1702.581]VSVVMAEVDGTC[+57.021]Y.- | HexNAc(2)Hex(8) | C[+57], N[+1703] | 1350.5827 | 3 | 4049.7335 | 4049.7234 | 0 | 2.5 | 332 | CRagged | 821.9 | 800.48 | 518.01 | 17.62 | 109 | >sp\|P01876\|IGHA1_HUMAN Immunoglobulin heavy constant alpha 1 OS=Homo sapiens OX=9606 GN=IGHA1 PE=1 SV=2 | 255 | 20201116_xd_colostrum_1ng_glyco_hcd203040_compound15.13941.13941.3 | controllerType=0 controllerNumber=1 scan=13941 | 30.2935 |
| 14291:4 | 4 | R.LAGKPTHVN[+2350.830]VSVVMAEVDGTC[+57.021]Y.- | HexNAc(4)Hex(5)Fuc(1)NeuAc(2) | C[+57], N[+2351] | 1175.2505 | 4 | 4697.9803 | 4697.9724 | 0 | 1.7 | 332 | CRagged | 824.26 | 808.38 | 808.38 | 17.56 | 109 | >sp\|P01876\|IGHA1_HUMAN Immunoglobulin heavy constant alpha 1 OS=Homo sapiens OX=9606 GN=IGHA1 PE=1 SV=2 | 255 | 20201116_xd_colostrum_1ng_glyco_hcd203040_compound15.17459.17459.4 | controllerType=0 controllerNumber=1 scan=17459 | 37.2166 |
| 12593:3 | 4 | R.LAGKPTHVN[+1872.651]VSVVMAEVDGTC[+57.021]Y.- | HexNAc(3)Hex(6)NeuAc(1) | C[+57], N[+1873] | 1407.2718 | 3 | 4219.8008 | 4219.7926 | 0 | 2 | 332 | CRagged | 826.01 | 686.84 | 538.6 | 17.46 | 109 | >sp\|P01876\|IGHA1_HUMAN Immunoglobulin heavy constant alpha 1 OS=Homo sapiens OX=9606 GN=IGHA1 PE=1 SV=2 | 255 | 20201116_xd_colostrum_1ng_glyco_hcd203040_compound15.15694.15694.3 | controllerType=0 controllerNumber=1 scan=15694 | 33.6942 |
| 10944:3 | 4 | R.LAGKPTHVN[+1663.608]VSVVMAEVDGTC[+57.021]Y.- | HexNAc(5)Hex(4) | C[+57], N[+1664] | 1337.5912 | 3 | 4010.7592 | 4010.7502 | 0 | 2.2 | 332 | CRagged | 829.04 | 769.16 | 524.96 | 17.62 | 109 | >sp\|P01876\|IGHA1_HUMAN Immunoglobulin heavy constant alpha 1 OS=Homo sapiens OX=9606 GN=IGHA1 PE=1 SV=2 | 255 | 20201116_xd_colostrum_1ng_glyco_hcd203040_compound15.13981.13981.3 | controllerType=0 controllerNumber=1 scan=13981 | 30.4036 |
| 12519:4 | 4 | R.LAGKPTHVN[+2116.756]VSVVMAEVDGTC[+57.021]Y.- | HexNAc(5)Hex(5)NeuAc(1) | C[+57], N[+2117] | 1116.7325 | 4 | 4463.908 | 4463.8985 | 0 | 2.1 | 332 | CRagged | 829.72 | 562.96 | 562.96 | 17.71 | 109 | >sp\|P01876\|IGHA1_HUMAN Immunoglobulin heavy constant alpha 1 OS=Homo sapiens OX=9606 GN=IGHA1 PE=1 SV=2 | 255 | 20201116_xd_colostrum_1ng_glyco_hcd203040_compound15.15617.15617.4 | controllerType=0 controllerNumber=1 scan=15617 | 33.5339 |
| 09949:3 | 4 | R.LAGKPTHVN[+1809.666]VSVVMAEVDGTC[+57.021]Y.- | HexNAc(5)Hex(4)Fuc(1) | C[+57], N[+1810] | 1386.277 | 3 | 4156.8166 | 4156.8081 | 0 | 2 | 332 | CRagged | 832.74 | 771.8 | 554.43 | 17.46 | 109 | >sp\|P01876\|IGHA1_HUMAN Immunoglobulin heavy constant alpha 1 OS=Homo sapiens OX=9606 GN=IGHA1 PE=1 SV=2 | 255 | 20201116_xd_colostrum_1ng_glyco_hcd203040_compound15.12946.12946.3 | controllerType=0 controllerNumber=1 scan=12946 | 28.3161 |
| 11013:2 | 4 | R.LAGKPTHVN[+1216.423]VSVVMAEVDGTC[+57.021]Y.- | HexNAc(2)Hex(5) | C[+57], N[+1216] | 1782.2927 | 2 | 3563.5781 | 3563.5649 | 0 | 3.7 | 332 | CRagged | 834.3 | 810.9 | 441.38 | 16.87 | 109 | >sp\|P01876\|IGHA1_HUMAN Immunoglobulin heavy constant alpha 1 OS=Homo sapiens OX=9606 GN=IGHA1 PE=1 SV=2 | 255 | 20201116_xd_colostrum_1ng_glyco_hcd203040_compound15.14052.14052.2 | controllerType=0 controllerNumber=1 scan=14052 | 30.5477 |
| 09933:4 | 4 | R.LAGKPTHVN[+1971.719]VSVVMAEVDGTC[+57.021]Y.- | HexNAc(5)Hex(5)Fuc(1) | C[+57], N[+1972] | 1080.4725 | 4 | 4318.868 | 4318.861 | 0 | 1.6 | 332 | CRagged | 837.31 | 770.06 | 508 | 17.56 | 109 | >sp\|P01876\|IGHA1_HUMAN Immunoglobulin heavy constant alpha 1 OS=Homo sapiens OX=9606 GN=IGHA1 PE=1 SV=2 | 255 | 20201116_xd_colostrum_1ng_glyco_hcd203040_compound15.12929.12929.4 | controllerType=0 controllerNumber=1 scan=12929 | 28.2675 |
| 10942:3 | 4 | R.LAGKPTHVN[+1663.608]VSVVMAEVDGTC[+57.021]Y.- | HexNAc(5)Hex(4) | C[+57], N[+1664] | 1337.5912 | 3 | 4010.7592 | 4010.7502 | 0 | 2.2 | 332 | CRagged | 840.48 | 741.35 | 525.15 | 17.62 | 109 | >sp\|P01876\|IGHA1_HUMAN Immunoglobulin heavy constant alpha 1 OS=Homo sapiens OX=9606 GN=IGHA1 PE=1 SV=2 | 255 | 20201116_xd_colostrum_1ng_glyco_hcd203040_compound15.13979.13979.3 | controllerType=0 controllerNumber=1 scan=13979 | 30.3968 |
| 10952:3 | 4 | R.LAGKPTHVN[+1581.555]VSVVMAEVDGTC[+57.021]Y.- | HexNAc(3)Hex(6) | C[+57], N[+1582] | 1310.2407 | 3 | 3928.7074 | 3928.6971 | 0 | 2.6 | 332 | CRagged | 841.05 | 841.05 | 511.03 | 17.62 | 109 | >sp\|P01876\|IGHA1_HUMAN Immunoglobulin heavy constant alpha 1 OS=Homo sapiens OX=9606 GN=IGHA1 PE=1 SV=2 | 255 | 20201116_xd_colostrum_1ng_glyco_hcd203040_compound15.13989.13989.3 | controllerType=0 controllerNumber=1 scan=13989 | 30.4151 |
| 09946:3 | 4 | R.LAGKPTHVN[+1809.666]VSVVMAEVDGTC[+57.021]Y.- | HexNAc(5)Hex(4)Fuc(1) | C[+57], N[+1810] | 1386.277 | 3 | 4156.8166 | 4156.8081 | 0 | 2 | 332 | CRagged | 844.65 | 844.65 | 563.12 | 17.46 | 109 | >sp\|P01876\|IGHA1_HUMAN Immunoglobulin heavy constant alpha 1 OS=Homo sapiens OX=9606 GN=IGHA1 PE=1 SV=2 | 255 | 20201116_xd_colostrum_1ng_glyco_hcd203040_compound15.12943.12943.3 | controllerType=0 controllerNumber=1 scan=12943 | 28.3087 |
| 09892:3 | 4 | R.LAGKPTHVN[+1702.581]VSVVMAEVDGTC[+57.021]Y.- | HexNAc(2)Hex(8) | C[+57], N[+1703] | 1350.5825 | 3 | 4049.7328 | 4049.7234 | 0 | 2.3 | 332 | CRagged | 844.84 | 826.19 | 499.68 | 17.62 | 109 | >sp\|P01876\|IGHA1_HUMAN Immunoglobulin heavy constant alpha 1 OS=Homo sapiens OX=9606 GN=IGHA1 PE=1 SV=2 | 255 | 20201116_xd_colostrum_1ng_glyco_hcd203040_compound15.12887.12887.3 | controllerType=0 controllerNumber=1 scan=12887 | 28.1808 |
| 11072:3 | 4 | R.LAGKPTHVN[+1501.555]VSVVMAEVDGTC[+57.021]Y.- | HexNAc(5)Hex(3) | C[+57], N[+1502] | 1283.5727 | 3 | 3848.7035 | 3848.6974 | 0 | 1.6 | 332 | CRagged | 846.38 | 825.71 | 825.71 | 17.46 | 109 | >sp\|P01876\|IGHA1_HUMAN Immunoglobulin heavy constant alpha 1 OS=Homo sapiens OX=9606 GN=IGHA1 PE=1 SV=2 | 255 | 20201116_xd_colostrum_1ng_glyco_hcd203040_compound15.14114.14114.3 | controllerType=0 controllerNumber=1 scan=14114 | 30.6708 |
| 09938:3 | 4 | R.LAGKPTHVN[+1378.476]VSVVMAEVDGTC[+57.021]Y.- | HexNAc(2)Hex(6) | C[+57], N[+1378] | 1242.5468 | 3 | 3725.6259 | 3725.6178 | 0 | 2.2 | 332 | CRagged | 846.8 | 759.22 | 510.36 | 17.62 | 109 | >sp\|P01876\|IGHA1_HUMAN Immunoglobulin heavy constant alpha 1 OS=Homo sapiens OX=9606 GN=IGHA1 PE=1 SV=2 | 255 | 20201116_xd_colostrum_1ng_glyco_hcd203040_compound15.12935.12935.3 | controllerType=0 controllerNumber=1 scan=12935 | 28.2838 |
| 11250:3 | 4 | R.LAGKPTHVN[+1216.423]VSVVMAEVDGTC[+57.021]Y.- | HexNAc(2)Hex(5) | C[+57], N[+1216] | 1188.528 | 3 | 3563.5695 | 3563.5649 | 0 | 1.3 | 332 | CRagged | 848.18 | 781.81 | 454.88 | 17.46 | 109 | >sp\|P01876\|IGHA1_HUMAN Immunoglobulin heavy constant alpha 1 OS=Homo sapiens OX=9606 GN=IGHA1 PE=1 SV=2 | 255 | 20201116_xd_colostrum_1ng_glyco_hcd203040_compound15.14299.14299.3 | controllerType=0 controllerNumber=1 scan=14299 | 31.0504 |
| 10037:3 | 4 | R.LAGKPTHVN[+1054.370]VSVVMAEVDGTC[+57.021]Y.- | HexNAc(2)Hex(4) | C[+57], N[+1054] | 1134.5121 | 3 | 3401.5218 | 3401.5121 | 0 | 2.8 | 332 | CRagged | 848.2 | 814.59 | 492.66 | 17.62 | 109 | >sp\|P01876\|IGHA1_HUMAN Immunoglobulin heavy constant alpha 1 OS=Homo sapiens OX=9606 GN=IGHA1 PE=1 SV=2 | 255 | 20201116_xd_colostrum_1ng_glyco_hcd203040_compound15.13038.13038.3 | controllerType=0 controllerNumber=1 scan=13038 | 28.522 |
| 09919:3 | 4 | R.LAGKPTHVN[+1540.529]VSVVMAEVDGTC[+57.021]Y.- | HexNAc(2)Hex(7) | C[+57], N[+1541] | 1296.5656 | 3 | 3887.6823 | 3887.6706 | 0 | 3 | 332 | CRagged | 849.89 | 736.78 | 526 | 17.42 | 109 | >sp\|P01876\|IGHA1_HUMAN Immunoglobulin heavy constant alpha 1 OS=Homo sapiens OX=9606 GN=IGHA1 PE=1 SV=2 | 255 | 20201116_xd_colostrum_1ng_glyco_hcd203040_compound15.12915.12915.3 | controllerType=0 controllerNumber=1 scan=12915 | 28.2332 |
| 09931:4 | 4 | R.LAGKPTHVN[+1971.719]VSVVMAEVDGTC[+57.021]Y.- | HexNAc(5)Hex(5)Fuc(1) | C[+57], N[+1972] | 1080.4725 | 4 | 4318.868 | 4318.861 | 0 | 1.6 | 332 | CRagged | 852.07 | 833.96 | 570.17 | 18.56 | 109 | >sp\|P01876\|IGHA1_HUMAN Immunoglobulin heavy constant alpha 1 OS=Homo sapiens OX=9606 GN=IGHA1 PE=1 SV=2 | 255 | 20201116_xd_colostrum_1ng_glyco_hcd203040_compound15.12927.12927.4 | controllerType=0 controllerNumber=1 scan=12927 | 28.2531 |
| 11073:3 | 4 | R.LAGKPTHVN[+1647.613]VSVVMAEVDGTC[+57.021]Y.- | HexNAc(5)Hex(3)Fuc(1) | C[+57], N[+1648] | 1332.2593 | 3 | 3994.7635 | 3994.7553 | 0 | 2 | 332 | CRagged | 852.14 | 774.46 | 774.46 | 18.62 | 109 | >sp\|P01876\|IGHA1_HUMAN Immunoglobulin heavy constant alpha 1 OS=Homo sapiens OX=9606 GN=IGHA1 PE=1 SV=2 | 255 | 20201116_xd_colostrum_1ng_glyco_hcd203040_compound15.14115.14115.3 | controllerType=0 controllerNumber=1 scan=14115 | 30.6737 |
| 10927:3 | 4 | R.LAGKPTHVN[+1971.719]VSVVMAEVDGTC[+57.021]Y.- | HexNAc(5)Hex(5)Fuc(1) | C[+57], N[+1972] | 1440.2955 | 3 | 4318.8718 | 4318.861 | 0 | 2.5 | 332 | CRagged | 853.63 | 853.63 | 567.76 | 18.62 | 109 | >sp\|P01876\|IGHA1_HUMAN Immunoglobulin heavy constant alpha 1 OS=Homo sapiens OX=9606 GN=IGHA1 PE=1 SV=2 | 255 | 20201116_xd_colostrum_1ng_glyco_hcd203040_compound15.13963.13963.3 | controllerType=0 controllerNumber=1 scan=13963 | 30.3444 |
| 09922:3 | 4 | R.LAGKPTHVN[+1540.529]VSVVMAEVDGTC[+57.021]Y.- | HexNAc(2)Hex(7) | C[+57], N[+1541] | 1296.5656 | 3 | 3887.6823 | 3887.6706 | 0 | 3 | 332 | CRagged | 854.24 | 838.78 | 487.78 | 18.42 | 109 | >sp\|P01876\|IGHA1_HUMAN Immunoglobulin heavy constant alpha 1 OS=Homo sapiens OX=9606 GN=IGHA1 PE=1 SV=2 | 255 | 20201116_xd_colostrum_1ng_glyco_hcd203040_compound15.12918.12918.3 | controllerType=0 controllerNumber=1 scan=12918 | 28.2405 |
| 14295:3 | 4 | R.LAGKPTHVN[+2553.910]VSVVMAEVDGTC[+57.021]Y.- | HexNAc(5)Hex(5)Fuc(1)NeuAc(2) | C[+57], N[+2554] | 1634.3591 | 3 | 4901.0626 | 4901.0518 | 0 | 2.2 | 332 | CRagged | 854.36 | 553.85 | 553.85 | 18.62 | 109 | >sp\|P01876\|IGHA1_HUMAN Immunoglobulin heavy constant alpha 1 OS=Homo sapiens OX=9606 GN=IGHA1 PE=1 SV=2 | 255 | 20201116_xd_colostrum_1ng_glyco_hcd203040_compound15.17463.17463.3 | controllerType=0 controllerNumber=1 scan=17463 | 37.2271 |
| 10967:3 | 4 | R.LAGKPTHVN[+1809.666]VSVVMAEVDGTC[+57.021]Y.- | HexNAc(5)Hex(4)Fuc(1) | C[+57], N[+1810] | 1386.2759 | 3 | 4156.8133 | 4156.8081 | 0 | 1.2 | 332 | CRagged | 855.84 | 803.96 | 534.03 | 18.46 | 109 | >sp\|P01876\|IGHA1_HUMAN Immunoglobulin heavy constant alpha 1 OS=Homo sapiens OX=9606 GN=IGHA1 PE=1 SV=2 | 255 | 20201116_xd_colostrum_1ng_glyco_hcd203040_compound15.14005.14005.3 | controllerType=0 controllerNumber=1 scan=14005 | 30.4549 |
| 10973:3 | 4 | R.LAGKPTHVN[+1606.587]VSVVMAEVDGTC[+57.021]Y.- | HexNAc(4)Hex(4)Fuc(1) | C[+57], N[+1607] | 1318.5828 | 3 | 3953.7339 | 3953.7288 | 0 | 1.3 | 332 | CRagged | 857.66 | 788.41 | 788.41 | 18.46 | 109 | >sp\|P01876\|IGHA1_HUMAN Immunoglobulin heavy constant alpha 1 OS=Homo sapiens OX=9606 GN=IGHA1 PE=1 SV=2 | 255 | 20201116_xd_colostrum_1ng_glyco_hcd203040_compound15.14011.14011.3 | controllerType=0 controllerNumber=1 scan=14011 | 30.4677 |
| 11155:3 | 4 | R.LAGKPTHVN[+892.317]VSVVMAEVDGTC[+57.021]Y.- | HexNAc(2)Hex(3) | C[+57], N[+892] | 1080.495 | 3 | 3239.4705 | 3239.4593 | 0 | 3.5 | 332 | CRagged | 875.21 | 875.21 | 529.96 | 18.42 | 109 | >sp\|P01876\|IGHA1_HUMAN Immunoglobulin heavy constant alpha 1 OS=Homo sapiens OX=9606 GN=IGHA1 PE=1 SV=2 | 255 | 20201116_xd_colostrum_1ng_glyco_hcd203040_compound15.14200.14200.3 | controllerType=0 controllerNumber=1 scan=14200 | 30.838 |
| 09972:3 | 4 | R.LAGKPTHVN[+1216.423]VSVVMAEVDGTC[+57.021]Y.- | HexNAc(2)Hex(5) | C[+57], N[+1216] | 1188.528 | 3 | 3563.5695 | 3563.5649 | 0 | 1.3 | 332 | CRagged | 876.27 | 876.27 | 499.34 | 18.46 | 109 | >sp\|P01876\|IGHA1_HUMAN Immunoglobulin heavy constant alpha 1 OS=Homo sapiens OX=9606 GN=IGHA1 PE=1 SV=2 | 255 | 20201116_xd_colostrum_1ng_glyco_hcd203040_compound15.12970.12970.3 | controllerType=0 controllerNumber=1 scan=12970 | 28.3681 |
| 11058:3 | 4 | R.LAGKPTHVN[+1647.613]VSVVMAEVDGTC[+57.021]Y.- | HexNAc(5)Hex(3)Fuc(1) | C[+57], N[+1648] | 1332.258 | 3 | 3994.7594 | 3994.7553 | 0 | 1 | 332 | CRagged | 878.28 | 725.5 | 725.5 | 18.46 | 109 | >sp\|P01876\|IGHA1_HUMAN Immunoglobulin heavy constant alpha 1 OS=Homo sapiens OX=9606 GN=IGHA1 PE=1 SV=2 | 255 | 20201116_xd_colostrum_1ng_glyco_hcd203040_compound15.14099.14099.3 | controllerType=0 controllerNumber=1 scan=14099 | 30.643 |
| 12518:3 | 4 | R.LAGKPTHVN[+2116.756]VSVVMAEVDGTC[+57.021]Y.- | HexNAc(5)Hex(5)NeuAc(1) | C[+57], N[+2117] | 1488.6412 | 3 | 4463.909 | 4463.8985 | 0 | 2.3 | 332 | CRagged | 882.02 | 578.43 | 578.43 | 18.62 | 109 | >sp\|P01876\|IGHA1_HUMAN Immunoglobulin heavy constant alpha 1 OS=Homo sapiens OX=9606 GN=IGHA1 PE=1 SV=2 | 255 | 20201116_xd_colostrum_1ng_glyco_hcd203040_compound15.15616.15616.3 | controllerType=0 controllerNumber=1 scan=15616 | 33.533 |
| 09940:3 | 4 | R.LAGKPTHVN[+1378.476]VSVVMAEVDGTC[+57.021]Y.- | HexNAc(2)Hex(6) | C[+57], N[+1378] | 1242.5468 | 3 | 3725.6259 | 3725.6178 | 0 | 2.2 | 332 | CRagged | 884.92 | 867.36 | 549.9 | 18.62 | 109 | >sp\|P01876\|IGHA1_HUMAN Immunoglobulin heavy constant alpha 1 OS=Homo sapiens OX=9606 GN=IGHA1 PE=1 SV=2 | 255 | 20201116_xd_colostrum_1ng_glyco_hcd203040_compound15.12937.12937.3 | controllerType=0 controllerNumber=1 scan=12937 | 28.2962 |
| 10924:3 | 4 | R.LAGKPTHVN[+1540.529]VSVVMAEVDGTC[+57.021]Y.- | HexNAc(2)Hex(7) | C[+57], N[+1541] | 1296.5646 | 3 | 3887.6793 | 3887.6706 | 0 | 2.3 | 332 | CRagged | 888.14 | 888.14 | 585.15 | 18.62 | 109 | >sp\|P01876\|IGHA1_HUMAN Immunoglobulin heavy constant alpha 1 OS=Homo sapiens OX=9606 GN=IGHA1 PE=1 SV=2 | 255 | 20201116_xd_colostrum_1ng_glyco_hcd203040_compound15.13960.13960.3 | controllerType=0 controllerNumber=1 scan=13960 | 30.3361 |
| 09976:3 | 4 | R.LAGKPTHVN[+1216.423]VSVVMAEVDGTC[+57.021]Y.- | HexNAc(2)Hex(5) | C[+57], N[+1216] | 1188.528 | 3 | 3563.5695 | 3563.5649 | 0 | 1.3 | 332 | CRagged | 893.39 | 884.09 | 521.65 | 18.46 | 109 | >sp\|P01876\|IGHA1_HUMAN Immunoglobulin heavy constant alpha 1 OS=Homo sapiens OX=9606 GN=IGHA1 PE=1 SV=2 | 255 | 20201116_xd_colostrum_1ng_glyco_hcd203040_compound15.12974.12974.3 | controllerType=0 controllerNumber=1 scan=12974 | 28.3861 |
| 10979:3 | 4 | R.LAGKPTHVN[+1460.529]VSVVMAEVDGTC[+57.021]Y.- | HexNAc(4)Hex(4) | C[+57], N[+1461] | 1269.8985 | 3 | 3807.6809 | 3807.6709 | 0 | 2.6 | 332 | CRagged | 902.02 | 875.42 | 540.81 | 19.62 | 109 | >sp\|P01876\|IGHA1_HUMAN Immunoglobulin heavy constant alpha 1 OS=Homo sapiens OX=9606 GN=IGHA1 PE=1 SV=2 | 255 | 20201116_xd_colostrum_1ng_glyco_hcd203040_compound15.14017.14017.3 | controllerType=0 controllerNumber=1 scan=14017 | 30.4789 |
| 10926:3 | 4 | R.LAGKPTHVN[+1540.529]VSVVMAEVDGTC[+57.021]Y.- | HexNAc(2)Hex(7) | C[+57], N[+1541] | 1296.5646 | 3 | 3887.6793 | 3887.6706 | 0 | 2.3 | 332 | CRagged | 907.94 | 821.83 | 534.4 | 19.62 | 109 | >sp\|P01876\|IGHA1_HUMAN Immunoglobulin heavy constant alpha 1 OS=Homo sapiens OX=9606 GN=IGHA1 PE=1 SV=2 | 255 | 20201116_xd_colostrum_1ng_glyco_hcd203040_compound15.13962.13962.3 | controllerType=0 controllerNumber=1 scan=13962 | 30.3435 |
| 10904:3 | 4 | R.LAGKPTHVN[+1702.581]VSVVMAEVDGTC[+57.021]Y.- | HexNAc(2)Hex(8) | C[+57], N[+1703] | 1350.5827 | 3 | 4049.7335 | 4049.7234 | 0 | 2.5 | 332 | CRagged | 925.67 | 925.67 | 597.09 | 19.62 | 109 | >sp\|P01876\|IGHA1_HUMAN Immunoglobulin heavy constant alpha 1 OS=Homo sapiens OX=9606 GN=IGHA1 PE=1 SV=2 | 255 | 20201116_xd_colostrum_1ng_glyco_hcd203040_compound15.13939.13939.3 | controllerType=0 controllerNumber=1 scan=13939 | 30.2867 |
| 10919:3 | 4 | R.LAGKPTHVN[+1825.661]VSVVMAEVDGTC[+57.021]Y.- | HexNAc(5)Hex(5) | C[+57], N[+1826] | 1391.6094 | 3 | 4172.8137 | 4172.8031 | 0 | 2.6 | 332 | CRagged | 925.99 | 925.99 | 618.38 | 19.62 | 109 | >sp\|P01876\|IGHA1_HUMAN Immunoglobulin heavy constant alpha 1 OS=Homo sapiens OX=9606 GN=IGHA1 PE=1 SV=2 | 255 | 20201116_xd_colostrum_1ng_glyco_hcd203040_compound15.13954.13954.3 | controllerType=0 controllerNumber=1 scan=13954 | 30.3192 |
| 11050:3 | 4 | R.LAGKPTHVN[+1054.370]VSVVMAEVDGTC[+57.021]Y.- | HexNAc(2)Hex(4) | C[+57], N[+1054] | 1134.5108 | 3 | 3401.5178 | 3401.5121 | 0 | 1.7 | 332 | CRagged | 930.33 | 930.33 | 550.77 | 19.46 | 109 | >sp\|P01876\|IGHA1_HUMAN Immunoglobulin heavy constant alpha 1 OS=Homo sapiens OX=9606 GN=IGHA1 PE=1 SV=2 | 255 | 20201116_xd_colostrum_1ng_glyco_hcd203040_compound15.14091.14091.3 | controllerType=0 controllerNumber=1 scan=14091 | 30.6162 |
| 11055:3 | 4 | R.LAGKPTHVN[+1647.613]VSVVMAEVDGTC[+57.021]Y.- | HexNAc(5)Hex(3)Fuc(1) | C[+57], N[+1648] | 1332.258 | 3 | 3994.7594 | 3994.7553 | 0 | 1 | 332 | CRagged | 933.88 | 930.17 | 930.17 | 19.46 | 109 | >sp\|P01876\|IGHA1_HUMAN Immunoglobulin heavy constant alpha 1 OS=Homo sapiens OX=9606 GN=IGHA1 PE=1 SV=2 | 255 | 20201116_xd_colostrum_1ng_glyco_hcd203040_compound15.14096.14096.3 | controllerType=0 controllerNumber=1 scan=14096 | 30.6358 |
| 10946:4 | 4 | R.LAGKPTHVN[+1971.719]VSVVMAEVDGTC[+57.021]Y.- | HexNAc(5)Hex(5)Fuc(1) | C[+57], N[+1972] | 1080.4732 | 4 | 4318.8709 | 4318.861 | 0 | 2.3 | 332 | CRagged | 940.45 | 940.45 | 590.86 | 19.71 | 109 | >sp\|P01876\|IGHA1_HUMAN Immunoglobulin heavy constant alpha 1 OS=Homo sapiens OX=9606 GN=IGHA1 PE=1 SV=2 | 255 | 20201116_xd_colostrum_1ng_glyco_hcd203040_compound15.13983.13983.4 | controllerType=0 controllerNumber=1 scan=13983 | 30.4054 |
| 10949:4 | 4 | R.LAGKPTHVN[+1971.719]VSVVMAEVDGTC[+57.021]Y.- | HexNAc(5)Hex(5)Fuc(1) | C[+57], N[+1972] | 1080.4732 | 4 | 4318.8709 | 4318.861 | 0 | 2.3 | 332 | CRagged | 968.28 | 889.63 | 606.34 | 20.71 | 109 | >sp\|P01876\|IGHA1_HUMAN Immunoglobulin heavy constant alpha 1 OS=Homo sapiens OX=9606 GN=IGHA1 PE=1 SV=2 | 255 | 20201116_xd_colostrum_1ng_glyco_hcd203040_compound15.13986.13986.4 | controllerType=0 controllerNumber=1 scan=13986 | 30.4119 |
| 12531:3 | 4 | R.LAGKPTHVN[+1913.677]VSVVMAEVDGTC[+57.021]Y.- | HexNAc(4)Hex(5)NeuAc(1) | C[+57], N[+1914] | 1420.9477 | 3 | 4260.8285 | 4260.8191 | 0 | 2.2 | 332 | CRagged | 988.48 | 607.24 | 607.24 | 20.62 | 109 | >sp\|P01876\|IGHA1_HUMAN Immunoglobulin heavy constant alpha 1 OS=Homo sapiens OX=9606 GN=IGHA1 PE=1 SV=2 | 255 | 20201116_xd_colostrum_1ng_glyco_hcd203040_compound15.15629.15629.3 | controllerType=0 controllerNumber=1 scan=15629 | 33.5593 |
| 08155:3 | 4 | R.LAGKPTHVN[+1054.370]VSVVM[+15.995]AEVDGTC[+57.021]Y.- | HexNAc(2)Hex(4) | C[+57], M[+16], N[+1054] | 1139.8428 | 3 | 3417.5139 | 3417.507 | 0 | 2 | 332 | CRagged | 255.75 | 251.94 | 251.94 | 5.88 | 109 | >sp\|P01876\|IGHA1_HUMAN Immunoglobulin heavy constant alpha 1 OS=Homo sapiens OX=9606 GN=IGHA1 PE=1 SV=2 | 255 | 20201116_xd_colostrum_1ng_glyco_hcd203040_compound15.11084.11084.3 | controllerType=0 controllerNumber=1 scan=11084 | 24.8467 |
| 08892:3 | 4 | R.LAGKPTHVN[+1216.423]VSVVM[+15.995]AEVDGTC[+57.021]Y.- | HexNAc(2)Hex(5) | C[+57], M[+16], N[+1216] | 1193.8596 | 3 | 3579.5641 | 3579.5599 | 0 | 1.2 | 332 | CRagged | 269.2 | 230.45 | 230.45 | 7.12 | 109 | >sp\|P01876\|IGHA1_HUMAN Immunoglobulin heavy constant alpha 1 OS=Homo sapiens OX=9606 GN=IGHA1 PE=1 SV=2 | 255 | 20201116_xd_colostrum_1ng_glyco_hcd203040_compound15.11850.11850.3 | controllerType=0 controllerNumber=1 scan=11850 | 26.2986 |
| 09154:3 | 4 | R.LAGKPTHVN[+1054.370]VSVVM[+15.995]AEVDGTC[+57.021]Y.- | HexNAc(2)Hex(4) | C[+57], M[+16], N[+1054] | 1139.8439 | 3 | 3417.5171 | 3417.507 | 0 | 3 | 332 | CRagged | 296.33 | 296.33 | 296.33 | 6.28 | 109 | >sp\|P01876\|IGHA1_HUMAN Immunoglobulin heavy constant alpha 1 OS=Homo sapiens OX=9606 GN=IGHA1 PE=1 SV=2 | 255 | 20201116_xd_colostrum_1ng_glyco_hcd203040_compound15.12121.12121.3 | controllerType=0 controllerNumber=1 scan=12121 | 26.7892 |
| 10857:3 | 4 | R.LAGKPTHVN[+1216.423]VSVVM[+15.995]AEVDGTC[+57.021]Y.- | HexNAc(2)Hex(5) | C[+57], M[+16], N[+1216] | 1193.866 | 3 | 3579.5835 | 3579.5599 | 0 | 6.6 | 332 | CRagged | 356.82 | 306.9 | 306.9 | 7.36 | 109 | >sp\|P01876\|IGHA1_HUMAN Immunoglobulin heavy constant alpha 1 OS=Homo sapiens OX=9606 GN=IGHA1 PE=1 SV=2 | 255 | 20201116_xd_colostrum_1ng_glyco_hcd203040_compound15.13889.13889.3 | controllerType=0 controllerNumber=1 scan=13889 | 30.1744 |
| 09156:3 | 4 | R.LAGKPTHVN[+1054.370]VSVVM[+15.995]AEVDGTC[+57.021]Y.- | HexNAc(2)Hex(4) | C[+57], M[+16], N[+1054] | 1139.8439 | 3 | 3417.5171 | 3417.507 | 0 | 3 | 332 | CRagged | 363.57 | 291.05 | 291.05 | 6.88 | 109 | >sp\|P01876\|IGHA1_HUMAN Immunoglobulin heavy constant alpha 1 OS=Homo sapiens OX=9606 GN=IGHA1 PE=1 SV=2 | 255 | 20201116_xd_colostrum_1ng_glyco_hcd203040_compound15.12123.12123.3 | controllerType=0 controllerNumber=1 scan=12123 | 26.7942 |
| 08573:3 | 4 | R.LAGKPTHVN[+1216.423]VSVVM[+15.995]AEVDGTC[+57.021]Y.- | HexNAc(2)Hex(5) | C[+57], M[+16], N[+1216] | 1193.8597 | 3 | 3579.5645 | 3579.5599 | 0 | 1.3 | 332 | CRagged | 454.15 | 409.74 | 409.74 | 10.41 | 109 | >sp\|P01876\|IGHA1_HUMAN Immunoglobulin heavy constant alpha 1 OS=Homo sapiens OX=9606 GN=IGHA1 PE=1 SV=2 | 255 | 20201116_xd_colostrum_1ng_glyco_hcd203040_compound15.11518.11518.3 | controllerType=0 controllerNumber=1 scan=11518 | 25.6583 |
| 08663:3 | 4 | R.LAGKPTHVN[+1378.476]VSVVM[+15.995]AEVDGTC[+57.021]Y.- | HexNAc(2)Hex(6) | C[+57], M[+16], N[+1378] | 1247.8785 | 3 | 3741.6209 | 3741.6127 | 0 | 2.2 | 332 | CRagged | 487.27 | 487.27 | 487.27 | 10.72 | 109 | >sp\|P01876\|IGHA1_HUMAN Immunoglobulin heavy constant alpha 1 OS=Homo sapiens OX=9606 GN=IGHA1 PE=1 SV=2 | 255 | 20201116_xd_colostrum_1ng_glyco_hcd203040_compound15.11612.11612.3 | controllerType=0 controllerNumber=1 scan=11612 | 25.8336 |
| 08575:3 | 4 | R.LAGKPTHVN[+1216.423]VSVVM[+15.995]AEVDGTC[+57.021]Y.- | HexNAc(2)Hex(5) | C[+57], M[+16], N[+1216] | 1193.8597 | 3 | 3579.5645 | 3579.5599 | 0 | 1.3 | 332 | CRagged | 508.21 | 447.71 | 447.71 | 10.69 | 109 | >sp\|P01876\|IGHA1_HUMAN Immunoglobulin heavy constant alpha 1 OS=Homo sapiens OX=9606 GN=IGHA1 PE=1 SV=2 | 255 | 20201116_xd_colostrum_1ng_glyco_hcd203040_compound15.11520.11520.3 | controllerType=0 controllerNumber=1 scan=11520 | 25.6622 |
| 08895:3 | 4 | R.LAGKPTHVN[+1216.423]VSVVM[+15.995]AEVDGTC[+57.021]Y.- | HexNAc(2)Hex(5) | C[+57], M[+16], N[+1216] | 1193.8596 | 3 | 3579.5641 | 3579.5599 | 0 | 1.2 | 332 | CRagged | 547.38 | 539.72 | 539.72 | 10.97 | 109 | >sp\|P01876\|IGHA1_HUMAN Immunoglobulin heavy constant alpha 1 OS=Homo sapiens OX=9606 GN=IGHA1 PE=1 SV=2 | 255 | 20201116_xd_colostrum_1ng_glyco_hcd203040_compound15.11853.11853.3 | controllerType=0 controllerNumber=1 scan=11853 | 26.3024 |
| 08801:3 | 4 | R.LAGKPTHVN[+1809.666]VSVVM[+15.995]AEVDGTC[+57.021]Y.- | HexNAc(5)Hex(4)Fuc(1) | C[+57], M[+16], N[+1810] | 1391.6087 | 3 | 4172.8115 | 4172.8031 | 0 | 2 | 332 | CRagged | 562.25 | 521.04 | 248.5 | 11.17 | 109 | >sp\|P01876\|IGHA1_HUMAN Immunoglobulin heavy constant alpha 1 OS=Homo sapiens OX=9606 GN=IGHA1 PE=1 SV=2 | 255 | 20201116_xd_colostrum_1ng_glyco_hcd203040_compound15.11755.11755.3 | controllerType=0 controllerNumber=1 scan=11755 | 26.1069 |
| 08783:3 | 4 | R.LAGKPTHVN[+1054.370]VSVVM[+15.995]AEVDGTC[+57.021]Y.- | HexNAc(2)Hex(4) | C[+57], M[+16], N[+1054] | 1139.8416 | 3 | 3417.5102 | 3417.507 | 0 | 0.9 | 332 | CRagged | 563.25 | 563.25 | 563.25 | 10.89 | 109 | >sp\|P01876\|IGHA1_HUMAN Immunoglobulin heavy constant alpha 1 OS=Homo sapiens OX=9606 GN=IGHA1 PE=1 SV=2 | 255 | 20201116_xd_colostrum_1ng_glyco_hcd203040_compound15.11736.11736.3 | controllerType=0 controllerNumber=1 scan=11736 | 26.0557 |
| 08786:3 | 4 | R.LAGKPTHVN[+1054.370]VSVVM[+15.995]AEVDGTC[+57.021]Y.- | HexNAc(2)Hex(4) | C[+57], M[+16], N[+1054] | 1139.8416 | 3 | 3417.5102 | 3417.507 | 0 | 0.9 | 332 | CRagged | 632.56 | 632.56 | 632.56 | 13.19 | 109 | >sp\|P01876\|IGHA1_HUMAN Immunoglobulin heavy constant alpha 1 OS=Homo sapiens OX=9606 GN=IGHA1 PE=1 SV=2 | 255 | 20201116_xd_colostrum_1ng_glyco_hcd203040_compound15.11739.11739.3 | controllerType=0 controllerNumber=1 scan=11739 | 26.0625 |
| 08800:3 | 4 | R.LAGKPTHVN[+1378.476]VSVVM[+15.995]AEVDGTC[+57.021]Y.- | HexNAc(2)Hex(6) | C[+57], M[+16], N[+1378] | 1247.8787 | 3 | 3741.6216 | 3741.6127 | 0 | 2.4 | 332 | CRagged | 643.79 | 643.79 | 643.79 | 13.62 | 109 | >sp\|P01876\|IGHA1_HUMAN Immunoglobulin heavy constant alpha 1 OS=Homo sapiens OX=9606 GN=IGHA1 PE=1 SV=2 | 255 | 20201116_xd_colostrum_1ng_glyco_hcd203040_compound15.11754.11754.3 | controllerType=0 controllerNumber=1 scan=11754 | 26.1058 |
| 08804:3 | 4 | R.LAGKPTHVN[+1809.666]VSVVM[+15.995]AEVDGTC[+57.021]Y.- | HexNAc(5)Hex(4)Fuc(1) | C[+57], M[+16], N[+1810] | 1391.6087 | 3 | 4172.8115 | 4172.8031 | 0 | 2 | 332 | CRagged | 653.46 | 615.62 | 360.62 | 14.46 | 109 | >sp\|P01876\|IGHA1_HUMAN Immunoglobulin heavy constant alpha 1 OS=Homo sapiens OX=9606 GN=IGHA1 PE=1 SV=2 | 255 | 20201116_xd_colostrum_1ng_glyco_hcd203040_compound15.11758.11758.3 | controllerType=0 controllerNumber=1 scan=11758 | 26.1176 |
| 08803:3 | 4 | R.LAGKPTHVN[+1378.476]VSVVM[+15.995]AEVDGTC[+57.021]Y.- | HexNAc(2)Hex(6) | C[+57], M[+16], N[+1378] | 1247.8787 | 3 | 3741.6216 | 3741.6127 | 0 | 2.4 | 332 | CRagged | 720.08 | 690.26 | 690.26 | 15.62 | 109 | >sp\|P01876\|IGHA1_HUMAN Immunoglobulin heavy constant alpha 1 OS=Homo sapiens OX=9606 GN=IGHA1 PE=1 SV=2 | 255 | 20201116_xd_colostrum_1ng_glyco_hcd203040_compound15.11757.11757.3 | controllerType=0 controllerNumber=1 scan=11757 | 26.1156 |
| 18489:4 | 5 | R.GLSFDVSLEVSQGPGLLN[+2350.830]DTK.V | HexNAc(4)Hex(5)Fuc(1)NeuAc(2) | N[+2351] | 1132.4963 | 4 | 4526.9632 | 4526.9436 | 0 | 4.3 | 118 | Specific | 535.52 | 522.69 | 522.69 | 10.18 | 117 | >sp\|P01833\|PIGR_HUMAN Polymeric immunoglobulin receptor OS=Homo sapiens OX=9606 GN=PIGR PE=1 SV=4 | 902 | 20201116_xd_colostrum_1ng_glyco_hcd203040_compound15.21840.21840.4 | controllerType=0 controllerNumber=1 scan=21840 | 46.6365 |

Table S4. Cont.

| Query #:z | Protein Rank | Glycopeptide | Glycan | Modification Type(s) | Observed m/z | z | Observed (M+H) | Calc. mass (M+H) | Off-by-x error | Mass error (ppm) | StartPosition | Cleavage | Score | Delta | Delta Mod | \|Log Prob\| | # of unique peptides | Protein Name | Protein DB number | Comment | Scan # | Scan Time |
|---|---|---|---|---|---|---|---|---|---|---|---|---|---|---|---|---|---|---|---|---|---|---|
| 16458:3 | 5 | R.GLSFDVSLEVSQGPGLLN[+2862.021]DTK.V | HexNAc(5)Hex(6)Fuc(2)NeuAc(2) | N[+2862] | 1680.0608 | 3 | 5038.1677 | 5038.1337 | 0 | 6.8 | 118 | Specific | 668.76 | 644.46 | 644.46 | 13.07 | 117 | >sp\|P01833\|PIGR_HUMAN Polymeric immunoglobulin receptor OS=Homo sapiens OX=9606 GN=PIGR PE=1 SV=4 | 902 | 20201116_xd_colostrum_1ng_glyco_hcd203040_compound15.19716.19716.3 | controllerType=0 controllerNumber=1 scan=19716 | 41.923 |
| 16455:3 | 5 | R.GLSFDVSLEVSQGPGLLN[+2862.021]DTK.V | HexNAc(5)Hex(6)Fuc(2)NeuAc(2) | N[+2862] | 1680.0608 | 3 | 5038.1677 | 5038.1337 | 0 | 6.8 | 118 | Specific | 742.82 | 690.87 | 690.87 | 14.07 | 117 | >sp\|P01833\|PIGR_HUMAN Polymeric immunoglobulin receptor OS=Homo sapiens OX=9606 GN=PIGR PE=1 SV=4 | 902 | 20201116_xd_colostrum_1ng_glyco_hcd203040_compound15.19713.19713.3 | controllerType=0 controllerNumber=1 scan=19713 | 41.909 |
| 16710:3 | 5 | R.GLSFDVSLEVSQGPGLLN[+2351.851]DTK.V | HexNAc(4)Hex(5)Fuc(3)NeuAc(1) | N[+2352] | 1509.9974 | 3 | 4527.9776 | 4527.964 | 0 | 3 | 118 | Specific | 829.72 | 761.65 | 761.65 | 16.86 | 117 | >sp\|P01833\|PIGR_HUMAN Polymeric immunoglobulin receptor OS=Homo sapiens OX=9606 GN=PIGR PE=1 SV=4 | 902 | 20201116_xd_colostrum_1ng_glyco_hcd203040_compound15.19978.19978.3 | controllerType=0 controllerNumber=1 scan=19978 | 42.4699 |
| 18492:4 | 5 | R.GLSFDVSLEVSQGPGLLN[+2350.830]DTK.V | HexNAc(4)Hex(5)Fuc(1)NeuAc(2) | N[+2351] | 1132.4963 | 4 | 4526.9632 | 4526.9436 | 0 | 4.3 | 118 | Specific | 877.4 | 823.43 | 823.43 | 17.91 | 117 | >sp\|P01833\|PIGR_HUMAN Polymeric immunoglobulin receptor OS=Homo sapiens OX=9606 GN=PIGR PE=1 SV=4 | 902 | 20201116_xd_colostrum_1ng_glyco_hcd203040_compound15.21843.21843.4 | controllerType=0 controllerNumber=1 scan=21843 | 46.641 |
| 16713:3 | 5 | R.GLSFDVSLEVSQGPGLLN[+2351.851]DTK.V | HexNAc(4)Hex(5)Fuc(3)NeuAc(1) | N[+2352] | 1509.9974 | 3 | 4527.9776 | 4527.964 | 0 | 3 | 118 | Specific | 878.78 | 795.36 | 795.36 | 17.86 | 117 | >sp\|P01833\|PIGR_HUMAN Polymeric immunoglobulin receptor OS=Homo sapiens OX=9606 GN=PIGR PE=1 SV=4 | 902 | 20201116_xd_colostrum_1ng_glyco_hcd203040_compound15.19981.19981.3 | controllerType=0 controllerNumber=1 scan=19981 | 42.4753 |
| 18478:3 | 5 | R.GLSFDVSLEVSQGPGLLN[+2350.830]DTK.V | HexNAc(4)Hex(5)Fuc(1)NeuAc(2) | N[+2351] | 1509.6586 | 3 | 4526.9613 | 4526.9436 | 0 | 3.9 | 118 | Specific | 930 | 886.14 | 886.14 | 18.86 | 117 | >sp\|P01833\|PIGR_HUMAN Polymeric immunoglobulin receptor OS=Homo sapiens OX=9606 GN=PIGR PE=1 SV=4 | 902 | 20201116_xd_colostrum_1ng_glyco_hcd203040_compound15.21829.21829.3 | controllerType=0 controllerNumber=1 scan=21829 | 46.6186 |
| 18476:3 | 5 | R.GLSFDVSLEVSQGPGLLN[+2350.830]DTK.V | HexNAc(4)Hex(5)Fuc(1)NeuAc(2) | N[+2351] | 1509.6586 | 3 | 4526.9613 | 4526.9436 | 0 | 3.9 | 118 | Specific | 1074.75 | 1023.99 | 1023.99 | 21.86 | 117 | >sp\|P01833\|PIGR_HUMAN Polymeric immunoglobulin receptor OS=Homo sapiens OX=9606 GN=PIGR PE=1 SV=4 | 902 | 20201116_xd_colostrum_1ng_glyco_hcd203040_compound15.21827.21827.3 | controllerType=0 controllerNumber=1 scan=21827 | 46.6106 |
| 18807:3 | 5 | R.GLSFDVSLEVSQGPGLLN[+2350.830]DTKVYTVDLGR.T | HexNAc(4)Hex(5)Fuc(1)NeuAc(2) | N[+2351] | 1810.8217 | 3 | 5430.4507 | 5430.425 | 0 | 4.7 | 118 | Specific | 275.62 | 275.62 | 275.62 | 6.61 | 117 | >sp\|P01833\|PIGR_HUMAN Polymeric immunoglobulin receptor OS=Homo sapiens OX=9606 GN=PIGR PE=1 SV=4 | 902 | 20201116_xd_colostrum_1ng_glyco_hcd203040_compound15.22173.22173.3 | controllerType=0 controllerNumber=1 scan=22173 | 47.4145 |
| 18805:3 | 5 | R.GLSFDVSLEVSQGPGLLN[+2350.830]DTKVYTVDLGR.T | HexNAc(4)Hex(5)Fuc(1)NeuAc(2) | N[+2351] | 1810.8217 | 3 | 5430.4507 | 5430.425 | 0 | 4.7 | 118 | Specific | 331.23 | 331.23 | 331.23 | 7.46 | 117 | >sp\|P01833\|PIGR_HUMAN Polymeric immunoglobulin receptor OS=Homo sapiens OX=9606 GN=PIGR PE=1 SV=4 | 902 | 20201116_xd_colostrum_1ng_glyco_hcd203040_compound15.22171.22171.3 | controllerType=0 controllerNumber=1 scan=22171 | 47.4104 |
| 17544:3 | 5 | R.GLSFDVSLEVSQGPGLLN[+2351.851]DTKVYTVDLGR.T | HexNAc(4)Hex(5)Fuc(3)NeuAc(1) | N[+2352] | 1811.1601 | 3 | 5431.4658 | 5431.4454 | 0 | 3.8 | 118 | Specific | 430.4 | 430.4 | 430.4 | 9.77 | 117 | >sp\|P01833\|PIGR_HUMAN Polymeric immunoglobulin receptor OS=Homo sapiens OX=9606 GN=PIGR PE=1 SV=4 | 902 | 20201116_xd_colostrum_1ng_glyco_hcd203040_compound15.20854.20854.3 | controllerType=0 controllerNumber=1 scan=20854 | 44.5627 |
| 17548:3 | 5 | R.GLSFDVSLEVSQGPGLLN[+2205.793]DTKVYTVDLGR.T | HexNAc(4)Hex(5)Fuc(2)NeuAc(1) | N[+2206] | 1762.4696 | 3 | 5285.3942 | 5285.3875 | 0 | 1.3 | 118 | Specific | 503.92 | 503.92 | 503.92 | 10.16 | 117 | >sp\|P01833\|PIGR_HUMAN Polymeric immunoglobulin receptor OS=Homo sapiens OX=9606 GN=PIGR PE=1 SV=4 | 902 | 20201116_xd_colostrum_1ng_glyco_hcd203040_compound15.20858.20858.3 | controllerType=0 controllerNumber=1 scan=20858 | 44.5695 |
| 17547:3 | 5 | R.GLSFDVSLEVSQGPGLLN[+2351.851]DTKVYTVDLGR.T | HexNAc(4)Hex(5)Fuc(3)NeuAc(1) | N[+2352] | 1811.1601 | 3 | 5431.4658 | 5431.4454 | 0 | 3.8 | 118 | Specific | 520.1 | 486.3 | 486.3 | 10.17 | 117 | >sp\|P01833\|PIGR_HUMAN Polymeric immunoglobulin receptor OS=Homo sapiens OX=9606 GN=PIGR PE=1 SV=4 | 902 | 20201116_xd_colostrum_1ng_glyco_hcd203040_compound15.20857.20857.3 | controllerType=0 controllerNumber=1 scan=20857 | 44.5664 |
| 17545:3 | 5 | R.GLSFDVSLEVSQGPGLLN[+2205.793]DTKVYTVDLGR.T | HexNAc(4)Hex(5)Fuc(2)NeuAc(1) | N[+2206] | 1762.4696 | 3 | 5285.3942 | 5285.3875 | 0 | 1.3 | 118 | Specific | 580.82 | 493.43 | 493.43 | 11.94 | 117 | >sp\|P01833\|PIGR_HUMAN Polymeric immunoglobulin receptor OS=Homo sapiens OX=9606 GN=PIGR PE=1 SV=4 | 902 | 20201116_xd_colostrum_1ng_glyco_hcd203040_compound15.20855.20855.3 | controllerType=0 controllerNumber=1 scan=20855 | 44.5638 |
| 18819:4 | 5 | R.GLSFDVSLEVSQGPGLLN[+2350.830]DTKVYTVDLGR.T | HexNAc(4)Hex(5)Fuc(1)NeuAc(2) | N[+2351] | 1358.3656 | 4 | 5430.4406 | 5430.425 | 0 | 2.9 | 118 | Specific | 645.15 | 583.55 | 583.55 | 13.18 | 117 | >sp\|P01833\|PIGR_HUMAN Polymeric immunoglobulin receptor OS=Homo sapiens OX=9606 GN=PIGR PE=1 SV=4 | 902 | 20201116_xd_colostrum_1ng_glyco_hcd203040_compound15.22186.22186.4 | controllerType=0 controllerNumber=1 scan=22186 | 47.4418 |
| 18821:4 | 5 | R.GLSFDVSLEVSQGPGLLN[+2350.830]DTKVYTVDLGR.T | HexNAc(4)Hex(5)Fuc(1)NeuAc(2) | N[+2351] | 1358.3656 | 4 | 5430.4406 | 5430.425 | 0 | 2.9 | 118 | Specific | 720.77 | 518.61 | 518.61 | 15.18 | 117 | >sp\|P01833\|PIGR_HUMAN Polymeric immunoglobulin receptor OS=Homo sapiens OX=9606 GN=PIGR PE=1 SV=4 | 902 | 20201116_xd_colostrum_1ng_glyco_hcd203040_compound15.22188.22188.4 | controllerType=0 controllerNumber=1 scan=22188 | 47.4463 |
| 21655:3 | 5 | K.QIGLYPVLVIDSSGYVNPN[+2205.793]YTGR.I | HexNAc(4)Hex(5)Fuc(2)NeuAc(1) | N[+2206] | 1577.7104 | 3 | 4731.1166 | 4731.0963 | 0 | 4.3 | 168 | Specific | 231.88 | 231.88 | 231.88 | 5.19 | 117 | >sp\|P01833\|PIGR_HUMAN Polymeric immunoglobulin receptor OS=Homo sapiens OX=9606 GN=PIGR PE=1 SV=4 | 902 | 20201116_xd_colostrum_1ng_glyco_hcd203040_compound15.25150.25150.3 | controllerType=0 controllerNumber=1 scan=25150 | 53.9586 |
| 21653:3 | 5 | K.QIGLYPVLVIDSSGYVNPN[+2205.793]YTGR.I | HexNAc(4)Hex(5)Fuc(2)NeuAc(1) | N[+2206] | 1577.7104 | 3 | 4731.1166 | 4731.0963 | 0 | 4.3 | 168 | Specific | 276.65 | 257.83 | 257.83 | 5.7 | 117 | >sp\|P01833\|PIGR_HUMAN Polymeric immunoglobulin receptor OS=Homo sapiens OX=9606 GN=PIGR PE=1 SV=4 | 902 | 20201116_xd_colostrum_1ng_glyco_hcd203040_compound15.25148.25148.3 | controllerType=0 controllerNumber=1 scan=25148 | 53.9524 |
| 21675:3 | 5 | K.QIGLYPVLVIDSSGYVNPN[+2188.740]YTGR.I | HexNAc(2)Hex(11) | N[+2189] | 1572.0325 | 3 | 4714.083 | 4714.0433 | 0 | 8.4 | 168 | Specific | 313.03 | 281.18 | 281.18 | 6.46 | 117 | >sp\|P01833\|PIGR_HUMAN Polymeric immunoglobulin receptor OS=Homo sapiens OX=9606 GN=PIGR PE=1 SV=4 | 902 | 20201116_xd_colostrum_1ng_glyco_hcd203040_compound15.25171.25171.3 | controllerType=0 controllerNumber=1 scan=25171 | 54.0035 |
| 21613:3 | 5 | K.QIGLYPVLVIDSSGYVNPN[+2205.793]YTGR.I | HexNAc(4)Hex(5)Fuc(2)NeuAc(1) | N[+2206] | 1577.709 | 3 | 4731.1126 | 4731.0963 | 0 | 3.4 | 168 | Specific | 313.47 | 300.76 | 300.76 | 7.64 | 117 | >sp\|P01833\|PIGR_HUMAN Polymeric immunoglobulin receptor OS=Homo sapiens OX=9606 GN=PIGR PE=1 SV=4 | 902 | 20201116_xd_colostrum_1ng_glyco_hcd203040_compound15.25106.25106.3 | controllerType=0 controllerNumber=1 scan=25106 | 53.8566 |
| 17932:3 | 5 | K.QIGLYPVLVIDSSGYVNPN[+1872.651]YTGR.I | HexNAc(3)Hex(6)NeuAc(1) | N[+1873] | 1466.6638 | 3 | 4397.9767 | 4397.9539 | 0 | 5.2 | 168 | Specific | 353.58 | 353.58 | 353.58 | 7.83 | 117 | >sp\|P01833\|PIGR_HUMAN Polymeric immunoglobulin receptor OS=Homo sapiens OX=9606 GN=PIGR PE=1 SV=4 | 902 | 20201116_xd_colostrum_1ng_glyco_hcd203040_compound15.21259.21259.3 | controllerType=0 controllerNumber=1 scan=21259 | 45.4645 |
| 16211:4 | 5 | K.QIGLYPVLVIDSSGYVNPN[+1913.677]YTGR.I | HexNAc(4)Hex(5)NeuAc(1) | N[+1914] | 1110.5091 | 4 | 4439.0145 | 4438.9805 | 0 | 7.7 | 168 | Specific | 369.95 | 336.49 | 336.49 | 6.66 | 117 | >sp\|P01833\|PIGR_HUMAN Polymeric immunoglobulin receptor OS=Homo sapiens OX=9606 GN=PIGR PE=1 SV=4 | 902 | 20201116_xd_colostrum_1ng_glyco_hcd203040_compound15.19459.19459.4 | controllerType=0 controllerNumber=1 scan=19459 | 41.3944 |
| 21677:3 | 5 | K.QIGLYPVLVIDSSGYVNPN[+2188.740]YTGR.I | HexNAc(2)Hex(11) | N[+2189] | 1572.0325 | 3 | 4714.083 | 4714.0433 | 0 | 8.4 | 168 | Specific | 373.26 | 255.2 | 255.2 | 6.94 | 117 | >sp\|P01833\|PIGR_HUMAN Polymeric immunoglobulin receptor OS=Homo sapiens OX=9606 GN=PIGR PE=1 SV=4 | 902 | 20201116_xd_colostrum_1ng_glyco_hcd203040_compound15.25173.25173.3 | controllerType=0 controllerNumber=1 scan=25173 | 54.0083 |
| 17941:3 | 5 | K.QIGLYPVLVIDSSGYVNPN[+1913.677]YTGR.I | HexNAc(4)Hex(5)NeuAc(1) | N[+1914] | 1480.3389 | 3 | 4439.0021 | 4438.9805 | 0 | 4.9 | 168 | Specific | 389.65 | 334.76 | 334.76 | 7.36 | 117 | >sp\|P01833\|PIGR_HUMAN Polymeric immunoglobulin receptor OS=Homo sapiens OX=9606 GN=PIGR PE=1 SV=4 | 902 | 20201116_xd_colostrum_1ng_glyco_hcd203040_compound15.21268.21268.3 | controllerType=0 controllerNumber=1 scan=21268 | 45.4779 |
| 17780:3 | 5 | K.QIGLYPVLVIDSSGYVNPN[+2351.851]YTGR.I | HexNAc(4)Hex(5)Fuc(3)NeuAc(1) | N[+2352] | 1626.4006 | 3 | 4877.1872 | 4877.1542 | 0 | 6.8 | 168 | Specific | 516.62 | 427.79 | 427.79 | 9.94 | 117 | >sp\|P01833\|PIGR_HUMAN Polymeric immunoglobulin receptor OS=Homo sapiens OX=9606 GN=PIGR PE=1 SV=4 | 902 | 20201116_xd_colostrum_1ng_glyco_hcd203040_compound15.21100.21100.3 | controllerType=0 controllerNumber=1 scan=21100 | 45.1042 |
| 17810:3 | 5 | K.QIGLYPVLVIDSSGYVNPN[+2351.851]YTGR.I | HexNAc(4)Hex(5)Fuc(3)NeuAc(1) | N[+2352] | 1626.3974 | 3 | 4877.1776 | 4877.1542 | 0 | 4.8 | 168 | Specific | 557.41 | 417.56 | 417.56 | 10.79 | 117 | >sp\|P01833\|PIGR_HUMAN Polymeric immunoglobulin receptor OS=Homo sapiens OX=9606 GN=PIGR PE=1 SV=4 | 902 | 20201116_xd_colostrum_1ng_glyco_hcd203040_compound15.21131.21131.3 | controllerType=0 controllerNumber=1 scan=21131 | 45.1655 |
| 17861:3 | 5 | K.QIGLYPVLVIDSSGYVNPN[+2059.735]YTGR.I | HexNAc(4)Hex(5)Fuc(1)NeuAc(1) | N[+2060] | 1529.0234 | 3 | 4585.0556 | 4585.0384 | 0 | 3.7 | 168 | Specific | 561.65 | 561.65 | 561.65 | 11.21 | 117 | >sp\|P01833\|PIGR_HUMAN Polymeric immunoglobulin receptor OS=Homo sapiens OX=9606 GN=PIGR PE=1 SV=4 | 902 | 20201116_xd_colostrum_1ng_glyco_hcd203040_compound15.21185.21185.3 | controllerType=0 controllerNumber=1 scan=21185 | 45.2994 |
| 17865:4 | 5 | K.QIGLYPVLVIDSSGYVNPN[+2059.735]YTGR.I | HexNAc(4)Hex(5)Fuc(1)NeuAc(1) | N[+2060] | 1147.0177 | 4 | 4585.0491 | 4585.0384 | 0 | 2.3 | 168 | Specific | 580.55 | 571.34 | 571.34 | 12.99 | 117 | >sp\|P01833\|PIGR_HUMAN Polymeric immunoglobulin receptor OS=Homo sapiens OX=9606 GN=PIGR PE=1 SV=4 | 902 | 20201116_xd_colostrum_1ng_glyco_hcd203040_compound15.21189.21189.4 | controllerType=0 controllerNumber=1 scan=21189 | 45.3118 |
| 16213:4 | 5 | K.QIGLYPVLVIDSSGYVNPN[+1913.677]YTGR.I | HexNAc(4)Hex(5)NeuAc(1) | N[+1914] | 1110.5091 | 4 | 4439.0145 | 4438.9805 | 0 | 7.7 | 168 | Specific | 671.45 | 556.17 | 556.17 | 13.85 | 117 | >sp\|P01833\|PIGR_HUMAN Polymeric immunoglobulin receptor OS=Homo sapiens OX=9606 GN=PIGR PE=1 SV=4 | 902 | 20201116_xd_colostrum_1ng_glyco_hcd203040_compound15.19461.19461.4 | controllerType=0 controllerNumber=1 scan=19461 | 41.3978 |
| 17935:3 | 5 | K.QIGLYPVLVIDSSGYVNPN[+1872.651]YTGR.I | HexNAc(3)Hex(6)NeuAc(1) | N[+1873] | 1466.6638 | 3 | 4397.9767 | 4397.9539 | 0 | 5.2 | 168 | Specific | 699.1 | 643.8 | 643.8 | 13.88 | 117 | >sp\|P01833\|PIGR_HUMAN Polymeric immunoglobulin receptor OS=Homo sapiens OX=9606 GN=PIGR PE=1 SV=4 | 902 | 20201116_xd_colostrum_1ng_glyco_hcd203040_compound15.21262.21262.3 | controllerType=0 controllerNumber=1 scan=21262 | 45.4687 |
| 17813:3 | 5 | K.QIGLYPVLVIDSSGYVNPN[+2351.851]YTGR.I | HexNAc(4)Hex(5)Fuc(3)NeuAc(1) | N[+2352] | 1626.3974 | 3 | 4877.1776 | 4877.1542 | 0 | 4.8 | 168 | Specific | 712.23 | 609.02 | 609.02 | 15.28 | 117 | >sp\|P01833\|PIGR_HUMAN Polymeric immunoglobulin receptor OS=Homo sapiens OX=9606 GN=PIGR PE=1 SV=4 | 902 | 20201116_xd_colostrum_1ng_glyco_hcd203040_compound15.21134.21134.3 | controllerType=0 controllerNumber=1 scan=21134 | 45.1755 |
| 19862:3 | 5 | K.QIGLYPVLVIDSSGYVNPN[+2350.830]YTGR.I | HexNAc(4)Hex(5)Fuc(1)NeuAc(2) | N[+2351] | 1626.0502 | 3 | 4876.1361 | 4876.1338 | 0 | 0.5 | 168 | Specific | 716.75 | 582.48 | 582.48 | 15.28 | 117 | >sp\|P01833\|PIGR_HUMAN Polymeric immunoglobulin receptor OS=Homo sapiens OX=9606 GN=PIGR PE=1 SV=4 | 902 | 20201116_xd_colostrum_1ng_glyco_hcd203040_compound15.23276.23276.3 | controllerType=0 controllerNumber=1 scan=23276 | 49.8983 |
| 19773:4 | 5 | K.QIGLYPVLVIDSSGYVNPN[+2350.830]YTGR.I | HexNAc(4)Hex(5)Fuc(1)NeuAc(2) | N[+2351] | 1219.797 | 4 | 4876.1663 | 4876.1338 | 0 | 6.7 | 168 | Specific | 748.71 | 661.59 | 661.59 | 15 | 117 | >sp\|P01833\|PIGR_HUMAN Polymeric immunoglobulin receptor OS=Homo sapiens OX=9606 GN=PIGR PE=1 SV=4 | 902 | 20201116_xd_colostrum_1ng_glyco_hcd203040_compound15.23183.23183.4 | controllerType=0 controllerNumber=1 scan=23183 | 49.6827 |
| 19772:3 | 5 | K.QIGLYPVLVIDSSGYVNPN[+2350.830]YTGR.I | HexNAc(4)Hex(5)Fuc(1)NeuAc(2) | N[+2351] | 1626.0626 | 3 | 4876.1731 | 4876.1338 | 0 | 8.1 | 168 | Specific | 793.71 | 727.2 | 727.2 | 15.42 | 117 | >sp\|P01833\|PIGR_HUMAN Polymeric immunoglobulin receptor OS=Homo sapiens OX=9606 GN=PIGR PE=1 SV=4 | 902 | 20201116_xd_colostrum_1ng_glyco_hcd203040_compound15.23182.23182.3 | controllerType=0 controllerNumber=1 scan=23182 | 49.6808 |
| 19770:3 | 5 | K.QIGLYPVLVIDSSGYVNPN[+2350.830]YTGR.I | HexNAc(4)Hex(5)Fuc(1)NeuAc(2) | N[+2351] | 1626.0626 | 3 | 4876.1731 | 4876.1338 | 0 | 8.1 | 168 | Specific | 803.79 | 656.34 | 656.34 | 16.42 | 117 | >sp\|P01833\|PIGR_HUMAN Polymeric immunoglobulin receptor OS=Homo sapiens OX=9606 GN=PIGR PE=1 SV=4 | 902 | 20201116_xd_colostrum_1ng_glyco_hcd203040_compound15.23180.23180.3 | controllerType=0 controllerNumber=1 scan=23180 | 49.6762 |
| 19864:3 | 5 | K.QIGLYPVLVIDSSGYVNPN[+2350.830]YTGR.I | HexNAc(4)Hex(5)Fuc(1)NeuAc(2) | N[+2351] | 1626.0502 | 3 | 4876.1361 | 4876.1338 | 0 | 0.5 | 168 | Specific | 810.04 | 574.26 | 574.26 | 17.28 | 117 | >sp\|P01833\|PIGR_HUMAN Polymeric immunoglobulin receptor OS=Homo sapiens OX=9606 GN=PIGR PE=1 SV=4 | 902 | 20201116_xd_colostrum_1ng_glyco_hcd203040_compound15.23278.23278.3 | controllerType=0 controllerNumber=1 scan=23278 | 49.9046 |
| 19768:3 | 5 | K.QIGLYPVLVIDSSGYVNPN[+2204.772]YTGR.I | HexNAc(4)Hex(5)NeuAc(2) | N[+2205] | 1577.3741 | 3 | 4730.1077 | 4730.0759 | 0 | 6.7 | 168 | Specific | 819.76 | 556.82 | 556.82 | 16.72 | 117 | >sp\|P01833\|PIGR_HUMAN Polymeric immunoglobulin receptor OS=Homo sapiens OX=9606 GN=PIGR PE=1 SV=4 | 902 | 20201116_xd_colostrum_1ng_glyco_hcd203040_compound15.23178.23178.3 | controllerType=0 controllerNumber=1 scan=23178 | 49.6692 |
| 19769:4 | 5 | K.QIGLYPVLVIDSSGYVNPN[+2204.772]YTGR.I | HexNAc(4)Hex(5)NeuAc(2) | N[+2205] | 1183.283 | 4 | 4730.1102 | 4730.0759 | 0 | 7.2 | 168 | Specific | 830.3 | 746.21 | 746.21 | 16.85 | 117 | >sp\|P01833\|PIGR_HUMAN Polymeric immunoglobulin receptor OS=Homo sapiens OX=9606 GN=PIGR PE=1 SV=4 | 902 | 20201116_xd_colostrum_1ng_glyco_hcd203040_compound15.23179.23179.4 | controllerType=0 controllerNumber=1 scan=23179 | 49.6743 |
| 17868:4 | 5 | K.QIGLYPVLVIDSSGYVNPN[+2059.735]YTGR.I | HexNAc(4)Hex(5)Fuc(1)NeuAc(1) | N[+2060] | 1147.0177 | 4 | 4585.0491 | 4585.0384 | 0 | 2.3 | 168 | Specific | 842.29 | 838.57 | 838.57 | 17.99 | 117 | >sp\|P01833\|PIGR_HUMAN Polymeric immunoglobulin receptor OS=Homo sapiens OX=9606 GN=PIGR PE=1 SV=4 | 902 | 20201116_xd_colostrum_1ng_glyco_hcd203040_compound15.21192.21192.4 | controllerType=0 controllerNumber=1 scan=21192 | 45.3189 |
| 19775:4 | 5 | K.QIGLYPVLVIDSSGYVNPN[+2350.830]YTGR.I | HexNAc(4)Hex(5)Fuc(1)NeuAc(2) | N[+2351] | 1219.797 | 4 | 4876.1663 | 4876.1338 | 0 | 6.7 | 168 | Specific | 853.56 | 659.66 | 659.66 | 18 | 117 | >sp\|P01833\|PIGR_HUMAN Polymeric immunoglobulin receptor OS=Homo sapiens OX=9606 GN=PIGR PE=1 SV=4 | 902 | 20201116_xd_colostrum_1ng_glyco_hcd203040_compound15.23185.23185.4 | controllerType=0 controllerNumber=1 scan=23185 | 49.688 |
| 17783:3 | 5 | K.QIGLYPVLVIDSSGYVNPN[+2351.851]YTGR.I | HexNAc(4)Hex(5)Fuc(3)NeuAc(1) | N[+2352] | 1626.4006 | 3 | 4877.1872 | 4877.1542 | 0 | 6.8 | 168 | Specific | 854.58 | 739.47 | 739.47 | 17.72 | 117 | >sp\|P01833\|PIGR_HUMAN Polymeric immunoglobulin receptor OS=Homo sapiens OX=9606 GN=PIGR PE=1 SV=4 | 902 | 20201116_xd_colostrum_1ng_glyco_hcd203040_compound15.21103.21103.3 | controllerType=0 controllerNumber=1 scan=21103 | 45.109 |
| 19766:3 | 5 | K.QIGLYPVLVIDSSGYVNPN[+2204.772]YTGR.I | HexNAc(4)Hex(5)NeuAc(2) | N[+2205] | 1577.3741 | 3 | 4730.1077 | 4730.0759 | 0 | 6.7 | 168 | Specific | 869.85 | 664.2 | 664.2 | 17.72 | 117 | >sp\|P01833\|PIGR_HUMAN Polymeric immunoglobulin receptor OS=Homo sapiens OX=9606 GN=PIGR PE=1 SV=4 | 902 | 20201116_xd_colostrum_1ng_glyco_hcd203040_compound15.23176.23176.3 | controllerType=0 controllerNumber=1 scan=23176 | 49.6619 |
| 19767:4 | 5 | K.QIGLYPVLVIDSSGYVNPN[+2204.772]YTGR.I | HexNAc(4)Hex(5)NeuAc(2) | N[+2205] | 1183.283 | 4 | 4730.1102 | 4730.0759 | 0 | 7.2 | 168 | Specific | 881.87 | 746.34 | 746.34 | 17.85 | 117 | >sp\|P01833\|PIGR_HUMAN Polymeric immunoglobulin receptor OS=Homo sapiens OX=9606 GN=PIGR PE=1 SV=4 | 902 | 20201116_xd_colostrum_1ng_glyco_hcd203040_compound15.23177.23177.4 | controllerType=0 controllerNumber=1 scan=23177 | 49.6633 |
| 17863:3 | 5 | K.QIGLYPVLVIDSSGYVNPN[+2059.735]YTGR.I | HexNAc(4)Hex(5)Fuc(1)NeuAc(1) | N[+2060] | 1529.0234 | 3 | 4585.0556 | 4585.0384 | 0 | 3.7 | 168 | Specific | 933.53 | 678.78 | 678.78 | 19.51 | 117 | >sp\|P01833\|PIGR_HUMAN Polymeric immunoglobulin receptor OS=Homo sapiens OX=9606 GN=PIGR PE=1 SV=4 | 902 | 20201116_xd_colostrum_1ng_glyco_hcd203040_compound15.21187.21187.3 | controllerType=0 controllerNumber=1 scan=21187 | 45.3095 |
| 19356:4 | 5 | K.[+42.011]AQYEGRLSLLEEPGN[+1622.582]GTFTVILNQLTSR.D | HexNAc(4)Hex(5) | .[42], N[+1623] | 1193.5534 | 4 | 4771.1917 | 4771.2089 | 0 | -3.6 | 407 | Specific | 265.45 | 265.45 | 265.45 | 5.44 | 117 | >sp\|P01833\|PIGR_HUMAN Polymeric immunoglobulin receptor OS=Homo sapiens OX=9606 GN=PIGR PE=1 SV=4 | 902 | 20201116_xd_colostrum_1ng_glyco_hcd203040_compound15.22747.22747.4 | controllerType=0 controllerNumber=1 scan=22747 | 48.6877 |
| 19381:3 | 5 | K.[+42.011]AQYEGRLSLLEEPGN[+1622.582]GTFTVILNQLTSR.D | HexNAc(4)Hex(5) | .[42], N[+1623] | 1591.065 | 3 | 4771.1804 | 4771.2089 | 0 | -6 | 407 | Specific | 304.29 | 286.36 | 286.36 | 4.87 | 117 | >sp\|P01833\|PIGR_HUMAN Polymeric immunoglobulin receptor OS=Homo sapiens OX=9606 GN=PIGR PE=1 SV=4 | 902 | 20201116_xd_colostrum_1ng_glyco_hcd203040_compound15.22773.22773.3 | controllerType=0 controllerNumber=1 scan=22773 | 48.7334 |
| 17816:3 | 5 | K.[+42.011]AQYEGRLSLLEEPGN[+2117.777]GTFTVILNQLTSR.D | HexNAc(5)Hex(5)Fuc(2) | .[42], N[+2118] | 1756.1319 | 3 | 5266.3812 | 5266.4041 | 0 | -4.3 | 407 | Specific | 308.89 | 155.16 | 155.16 | 4.49 | 117 | >sp\|P01833\|PIGR_HUMAN Polymeric immunoglobulin receptor OS=Homo sapiens OX=9606 GN=PIGR PE=1 SV=4 | 902 | 20201116_xd_colostrum_1ng_glyco_hcd203040_compound15.21137.21137.3 | controllerType=0 controllerNumber=1 scan=21137 | 45.1797 |
| 19383:3 | 5 | K.[+42.011]AQYEGRLSLLEEPGN[+1622.582]GTFTVILNQLTSR.D | HexNAc(4)Hex(5) | .[42], N[+1623] | 1591.065 | 3 | 4771.1804 | 4771.2089 | 0 | -6 | 407 | Specific | 324.27 | 320.41 | 320.41 | 5.38 | 117 | >sp\|P01833\|PIGR_HUMAN Polymeric immunoglobulin receptor OS=Homo sapiens OX=9606 GN=PIGR PE=1 SV=4 | 902 | 20201116_xd_colostrum_1ng_glyco_hcd203040_compound15.22775.22775.3 | controllerType=0 controllerNumber=1 scan=22775 | 48.7385 |
| 17819:3 | 5 | K.[+42.011]AQYEGRLSLLEEPGN[+2117.777]GTFTVILNQLTSR.D | HexNAc(5)Hex(5)Fuc(2) | .[42], N[+2118] | 1756.1319 | 3 | 5266.3812 | 5266.4041 | 0 | -4.3 | 407 | Specific | 353.51 | 229.15 | 229.15 | 5.48 | 117 | >sp\|P01833\|PIGR_HUMAN Polymeric immunoglobulin receptor OS=Homo sapiens OX=9606 GN=PIGR PE=1 SV=4 | 902 | 20201116_xd_colostrum_1ng_glyco_hcd203040_compound15.21140.21140.3 | controllerType=0 controllerNumber=1 scan=21140 | 45.1865 |
| 19803:3 | 5 | R.LSLLEEPGN[+2205.793]GTFTVILNQLTSR.D | HexNAc(4)Hex(5)Fuc(2)NeuAc(1) | N[+2206] | 1536.6981 | 3 | 4608.0796 | 4608.0854 | 0 | -1.3 | 413 | Specific | 222.03 | 101.68 | 101.68 | 4.71 | 117 | >sp\|P01833\|PIGR_HUMAN Polymeric immunoglobulin receptor OS=Homo sapiens OX=9606 GN=PIGR PE=1 SV=4 | 902 | 20201116_xd_colostrum_1ng_glyco_hcd203040_compound15.23214.23214.3 | controllerType=0 controllerNumber=1 scan=23214 | 49.7503 |
| 24562:4 | 5 | R.LSLLEEPGN[+3154.136]GTFTVILNQLTSR.D | HexNAc(5)Hex(6)Fuc(4)NeuAc(2) | N[+3154] | 1389.8651 | 4 | 5556.4386 | 5556.4288 | 0 | 1.8 | 413 | Specific | 251.06 | 202.41 | 202.41 | 7.28 | 117 | >sp\|P01833\|PIGR_HUMAN Polymeric immunoglobulin receptor OS=Homo sapiens OX=9606 GN=PIGR PE=1 SV=4 | 902 | 20201116_xd_colostrum_1ng_glyco_hcd203040_compound15.28178.28178.4 | controllerType=0 controllerNumber=1 scan=28178 | 59.8206 |
| 20063:4 | 5 | R.LSLLEEPGN[+2351.851]GTFTVILNQLTSR.D | HexNAc(4)Hex(5)Fuc(3)NeuAc(1) | N[+2352] | 1189.2901 | 4 | 4754.1385 | 4754.1433 | 0 | -1 | 413 | Specific | 296.89 | 209.81 | 209.81 | 5.78 | 117 | >sp\|P01833\|PIGR_HUMAN Polymeric immunoglobulin receptor OS=Homo sapiens OX=9606 GN=PIGR PE=1 SV=4 | 902 | 20201116_xd_colostrum_1ng_glyco_hcd203040_compound15.23486.23486.4 | controllerType=0 controllerNumber=1 scan=23486 | 50.3706 |
| 21134:3 | 5 | R.LSLLEEPGN[+2861.000]GTFTVILNQLTSR.D | HexNAc(5)Hex(6)NeuAc(3) | N[+2861] | 1755.1125 | 3 | 5263.323 | 5263.2926 | 0 | 5.8 | 413 | Specific | 312.37 | 259.2 | 259.2 | 6.94 | 117 | >sp\|P01833\|PIGR_HUMAN Polymeric immunoglobulin receptor OS=Homo sapiens OX=9606 GN=PIGR PE=1 SV=4 | 902 | 20201116_xd_colostrum_1ng_glyco_hcd203040_compound15.24606.24606.3 | controllerType=0 controllerNumber=1 scan=24606 | 52.78 |
| 19456:3 | 5 | R.[+42.011]LSLLEEPGN[+2336.851]GTFTVILNQLTSR.D | HexNAc(6)Hex(6)Fuc(1) | .[42], N[+2337] | 1594.3863 | 3 | 4781.1443 | 4781.1542 | 0 | -2.1 | 413 | Specific | 315.18 | 308.59 | 308.59 | 6.93 | 117 | >sp\|P01833\|PIGR_HUMAN Polymeric immunoglobulin receptor OS=Homo sapiens OX=9606 GN=PIGR PE=1 SV=4 | 902 | 20201116_xd_colostrum_1ng_glyco_hcd203040_compound15.22851.22851.3 | controllerType=0 controllerNumber=1 scan=22851 | 48.9103 |
| 21843:3 | 5 | R.LSLLEEPGN[+2351.851]GTFTVILNQLTSR.D | HexNAc(4)Hex(5)Fuc(3)NeuAc(1) | N[+2352] | 1585.3801 | 3 | 4754.1256 | 4754.1433 | 0 | -3.7 | 413 | Specific | 323.32 | 213.69 | 213.69 | 6.32 | 117 | >sp\|P01833\|PIGR_HUMAN Polymeric immunoglobulin receptor OS=Homo sapiens OX=9606 GN=PIGR PE=1 SV=4 | 902 | 20201116_xd_colostrum_1ng_glyco_hcd203040_compound15.25347.25347.3 | controllerType=0 controllerNumber=1 scan=25347 | 54.3673 |
| 19459:3 | 5 | R.[+42.011]LSLLEEPGN[+2336.851]GTFTVILNQLTSR.D | HexNAc(6)Hex(6)Fuc(1) | .[42], N[+2337] | 1594.3863 | 3 | 4781.1443 | 4781.1542 | 0 | -2.1 | 413 | Specific | 323.74 | 294.34 | 294.34 | 7.03 | 117 | >sp\|P01833\|PIGR_HUMAN Polymeric immunoglobulin receptor OS=Homo sapiens OX=9606 GN=PIGR PE=1 SV=4 | 902 | 20201116_xd_colostrum_1ng_glyco_hcd203040_compound15.22854.22854.3 | controllerType=0 controllerNumber=1 scan=22854 | 48.9142 |
| 24356:4 | 5 | R.LSLLEEPGN[+3007.058]GTFTVILNQLTSR.D | HexNAc(5)Hex(6)Fuc(1)NeuAc(3) | N[+3007] | 1353.0933 | 4 | 5409.3512 | 5409.3505 | 0 | 0.1 | 413 | Specific | 326.3 | 326.3 | 326.3 | 8.36 | 117 | >sp\|P01833\|PIGR_HUMAN Polymeric immunoglobulin receptor OS=Homo sapiens OX=9606 GN=PIGR PE=1 SV=4 | 902 | 20201116_xd_colostrum_1ng_glyco_hcd203040_compound15.27964.27964.3 | controllerType=0 controllerNumber=1 scan=27964 | 59.4274 |
| 19243:3 | 5 | R.LSLLEEPGN[+2571.946]GTFTVILNQLTSR.D | HexNAc(5)Hex(6)Fuc(4) | N[+2572] | 1658.7354 | 3 | 4974.1918 | 4974.238 | 0 | -9.3 | 413 | Specific | 382.62 | 180.56 | 180.56 | 6.11 | 117 | >sp\|P01833\|PIGR_HUMAN Polymeric immunoglobulin receptor OS=Homo sapiens OX=9606 GN=PIGR PE=1 SV=4 | 902 | 20201116_xd_colostrum_1ng_glyco_hcd203040_compound15.22629.22629.4 | controllerType=0 controllerNumber=1 scan=22629 | 48.4439 |
| 21132:3 | 5 | R.LSLLEEPGN[+2861.000]GTFTVILNQLTSR.D | HexNAc(5)Hex(6)NeuAc(3) | N[+2861] | 1755.1125 | 3 | 5263.323 | 5263.2926 | 0 | 5.8 | 413 | Specific | 384.57 | 354.92 | 354.92 | 7.95 | 117 | >sp\|P01833\|PIGR_HUMAN Polymeric immunoglobulin receptor OS=Homo sapiens OX=9606 GN=PIGR PE=1 SV=4 | 902 | 20201116_xd_colostrum_1ng_glyco_hcd203040_compound15.24604.24604.3 | controllerType=0 controllerNumber=1 scan=24604 | 52.7759 |
| 21383:5 | 5 | R.LSLLEEPGN[+2351.851]GTFTVILNQLTSR.D | HexNAc(4)Hex(5)Fuc(3)NeuAc(1) | N[+2352] | 951.6334 | 5 | 4754.1378 | 4754.1433 | 0 | -1.2 | 413 | Specific | 392.75 | 356.18 | 356.18 | 8.04 | 117 | >sp\|P01833\|PIGR_HUMAN Polymeric immunoglobulin receptor OS=Homo sapiens OX=9606 GN=PIGR PE=1 SV=4 | 902 | 20201116_xd_colostrum_1ng_glyco_hcd203040_compound15.24865.24865.5 | controllerType=0 controllerNumber=1 scan=24865 | 53.3263 |
| 19805:3 | 5 | R.LSLLEEPGN[+2205.793]GTFTVILNQLTSR.D | HexNAc(4)Hex(5)Fuc(2)NeuAc(1) | N[+2206] | 1536.6981 | 3 | 4608.0796 | 4608.0854 | 0 | -1.3 | 413 | Specific | 403.46 | 402.55 | 402.55 | 9.47 | 117 | >sp\|P01833\|PIGR_HUMAN Polymeric immunoglobulin receptor OS=Homo sapiens OX=9606 GN=PIGR PE=1 SV=4 | 902 | 20201116_xd_colostrum_1ng_glyco_hcd203040_compound15.23216.23216.3 | controllerType=0 controllerNumber=1 scan=23216 | 49.7538 |
| 21845:3 | 5 | R.LSLLEEPGN[+2351.851]GTFTVILNQLTSR.D | HexNAc(4)Hex(5)Fuc(3)NeuAc(1) | N[+2352] | 1585.3801 | 3 | 4754.1256 | 4754.1433 | 0 | -3.7 | 413 | Specific | 405.39 | 355.93 | 355.93 | 7.46 | 117 | >sp\|P01833\|PIGR_HUMAN Polymeric immunoglobulin receptor OS=Homo sapiens OX=9606 GN=PIGR PE=1 SV=4 | 902 | 20201116_xd_colostrum_1ng_glyco_hcd203040_compound15.25349.25349.3 | controllerType=0 controllerNumber=1 scan=25349 | 54.3716 |
| 20056:3 | 5 | R.LSLLEEPGN[+2351.851]GTFTVILNQLTSR.D | HexNAc(4)Hex(5)Fuc(3)NeuAc(1) | N[+2352] | 1585.3816 | 3 | 4754.1304 | 4754.1433 | 0 | -2.7 | 413 | Specific | 452.5 | 431.76 | 431.76 | 9.54 | 117 | >sp\|P01833\|PIGR_HUMAN Polymeric immunoglobulin receptor OS=Homo sapiens OX=9606 GN=PIGR PE=1 SV=4 | 902 | 20201116_xd_colostrum_1ng_glyco_hcd203040_compound15.23479.23479.3 | controllerType=0 controllerNumber=1 scan=23479 | 50.3567 |
| 20065:4 | 5 | R.LSLLEEPGN[+2351.851]GTFTVILNQLTSR.D | HexNAc(4)Hex(5)Fuc(3)NeuAc(1) | N[+2352] | 1189.2901 | 4 | 4754.1385 | 4754.1433 | 0 | -1 | 413 | Specific | 465.94 | 438.08 | 438.08 | 10.03 | 117 | >sp\|P01833\|PIGR_HUMAN Polymeric immunoglobulin receptor OS=Homo sapiens OX=9606 GN=PIGR PE=1 SV=4 | 902 | 20201116_xd_colostrum_1ng_glyco_hcd203040_compound15.23488.23488.4 | controllerType=0 controllerNumber=1 scan=23488 | 50.3793 |
| 19431:4 | 5 | R.LSLLEEPGN[+2076.750]GTFTVILNQLTSR.D | HexNAc(4)Hex(6)Fuc(2) | N[+2077] | 1120.5219 | 4 | 4479.0657 | 4479.0428 | 0 | 5.1 | 413 | Specific | 467.16 | 407.34 | 407.34 | 10.23 | 117 | >sp\|P01833\|PIGR_HUMAN Polymeric immunoglobulin receptor OS=Homo sapiens OX=9606 GN=PIGR PE=1 SV=4 | 902 | 20201116_xd_colostrum_1ng_glyco_hcd203040_compound15.22825.22825.4 | controllerType=0 controllerNumber=1 scan=22825 | [illegible] |
| 19267:3 | 5 | R.LSLLEEPGN[+2862.021]GTFTVILNQLTSR.D | HexNAc(5)Hex(6)Fuc(2)NeuAc(2) | N[+2862] | 1755.4544 | 3 | 5264.3487 | 5264.313 | 0 | 6.8 | 413 | Specific | 473.4 | 373.17 | 373.17 | 8.29 | 117 | >sp\|P01833\|PIGR_HUMAN Polymeric immunoglobulin receptor OS=Homo sapiens OX=9606 GN=PIGR PE=1 SV=4 | 902 | 20201116_xd_colostrum_1ng_glyco_hcd203040_compound15.22654.22654.3 | controllerType=0 controllerNumber=1 scan=22654 | [illegible] |
| 19265:3 | 5 | R.LSLLEEPGN[+2862.021]GTFTVILNQLTSR.D | HexNAc(5)Hex(6)Fuc(2)NeuAc(2) | N[+2862] | 1755.4544 | 3 | 5264.3487 | 5264.313 | 0 | 6.8 | 413 | Specific | 507.83 | 494.58 | 494.58 | 9.94 | 117 | >sp\|P01833\|PIGR_HUMAN Polymeric immunoglobulin receptor OS=Homo sapiens OX=9606 GN=PIGR PE=1 SV=4 | 902 | 20201116_xd_colostrum_1ng_glyco_hcd203040_compound15.22652.22652.3 | controllerType=0 controllerNumber=1 scan=22652 | [illegible] |
| 21385:5 | 5 | R.LSLLEEPGN[+2351.851]GTFTVILNQLTSR.D | HexNAc(4)Hex(5)Fuc(3)NeuAc(1) | N[+2352] | 951.6334 | 5 | 4754.1378 | 4754.1433 | 0 | -1.2 | 413 | Specific | 514.77 | 470.23 | 470.23 | 10.19 | 117 | >sp\|P01833\|PIGR_HUMAN Polymeric immunoglobulin receptor OS=Homo sapiens OX=9606 GN=PIGR PE=1 SV=4 | 902 | 20201116_xd_colostrum_1ng_glyco_hcd203040_compound15.24867.24867.5 | controllerType=0 controllerNumber=1 scan=24867 | [illegible] |
| 20058:3 | 5 | R.LSLLEEPGN[+2351.851]GTFTVILNQLTSR.D | HexNAc(4)Hex(5)Fuc(3)NeuAc(1) | N[+2352] | 1585.3816 | 3 | 4754.1304 | 4754.1433 | 0 | -2.7 | 413 | Specific | 538.67 | 315.74 | 315.74 | 7.96 | 117 | >sp\|P01833\|PIGR_HUMAN Polymeric immunoglobulin receptor OS=Homo sapiens OX=9606 GN=PIGR PE=1 SV=4 | 902 | 20201116_xd_colostrum_1ng_glyco_hcd203040_compound15.23481.23481.3 | controllerType=0 controllerNumber=1 scan=23481 | 50.3622 |
| 19434:4 | 5 | R.LSLLEEPGN[+2076.750]GTFTVILNQLTSR.D | HexNAc(4)Hex(6)Fuc(2) | N[+2077] | 1120.5219 | 4 | 4479.0657 | 4479.0428 | 0 | 5.1 | 413 | Specific | 561.51 | 489.38 | 489.38 | 10.86 | 117 | >sp\|P01833\|PIGR_HUMAN Polymeric immunoglobulin receptor OS=Homo sapiens OX=9606 GN=PIGR PE=1 SV=4 | 902 | 20201116_xd_colostrum_1ng_glyco_hcd203040_compound15.22828.22828.4 | controllerType=0 controllerNumber=1 scan=22828 | 48.8611 |
| 21448:3 | 5 | R.LSLLEEPGN[+2350.830]GTFTVILNQLTSR.D | HexNAc(4)Hex(5)Fuc(1)NeuAc(2) | N[+2351] | 1585.0523 | 3 | 4753.1424 | 4753.1229 | 0 | 4.1 | 413 | Specific | 562.28 | 536.67 | 536.67 | 10.98 | 117 | >sp\|P01833\|PIGR_HUMAN Polymeric immunoglobulin receptor OS=Homo sapiens OX=9606 GN=PIGR PE=1 SV=4 | 902 | 20201116_xd_colostrum_1ng_glyco_hcd203040_compound15.24934.24934.3 | controllerType=0 controllerNumber=1 scan=24934 | 53.4911 |
| 19245:3 | 5 | R.LSLLEEPGN[+2571.946]GTFTVILNQLTSR.D | HexNAc(5)Hex(6)Fuc(4) | N[+2572] | 1658.7354 | 3 | 4974.1918 | 4974.238 | 0 | -9.3 | 413 | Specific | 581.07 | 385.09 | 385.09 | 11.15 | 117 | >sp\|P01833\|PIGR_HUMAN Polymeric immunoglobulin receptor OS=Homo sapiens OX=9606 GN=PIGR PE=1 SV=4 | 902 | 20201116_xd_colostrum_1ng_glyco_hcd203040_compound15.22631.22631.4 | controllerType=0 controllerNumber=1 scan=22631 | 48.4476 |
| 21083:4 | 5 | R.LSLLEEPGN[+3007.058]GTFTVILNQLTSR.D | HexNAc(5)Hex(6)Fuc(1)NeuAc(3) | N[+3007] | 1353.1024 | 4 | 5409.3879 | 5409.3505 | 0 | 6.9 | 413 | Specific | 597.69 | 438.64 | 438.64 | 12 | 117 | >sp\|P01833\|PIGR_HUMAN Polymeric immunoglobulin receptor OS=Homo sapiens OX=9606 GN=PIGR PE=1 SV=4 | 902 | 20201116_xd_colostrum_1ng_glyco_hcd203040_compound15.24553.24553.4 | controllerType=0 controllerNumber=1 scan=24553 | 52.6769 |
| 19870:3 | 5 | R.LSLLEEPGN[+1458.442]GTFTVILNQLTSR.D | HexNAc(2)Hex(6)Phospho(1) | N[+1458] | 1287.5849 | 3 | 3860.7401 | 3860.7346 | 0 | 1.4 | 413 | Specific | 680.99 | 680.99 | 680.99 | 14.56 | 117 | >sp\|P01833\|PIGR_HUMAN Polymeric immunoglobulin receptor OS=Homo sapiens OX=9606 GN=PIGR PE=1 SV=4 | 902 | 20201116_xd_colostrum_1ng_glyco_hcd203040_compound15.23285.23285.3 | controllerType=0 controllerNumber=1 scan=23285 | 49.9194 |
| 19270:3 | 5 | R.LSLLEEPGN[+2715.963]GTFTVILNQLTSR.D | HexNAc(5)Hex(6)Fuc(1)NeuAc(2) | N[+2716] | 1706.7725 | 3 | 5118.3031 | 5118.2551 | 0 | 9.4 | 413 | Specific | 688.25 | 498.57 | 498.57 | 13.24 | 117 | >sp\|P01833\|PIGR_HUMAN Polymeric immunoglobulin receptor OS=Homo sapiens OX=9606 GN=PIGR PE=1 SV=4 | 902 | 20201116_xd_colostrum_1ng_glyco_hcd203040_compound15.22657.22657.3 | controllerType=0 controllerNumber=1 scan=22657 | 48.498 |
| 19240:4 | 5 | R.LSLLEEPGN[+2863.041]GTFTVILNQLTSR.D | HexNAc(5)Hex(6)Fuc(4)NeuAc(1) | N[+2863] | 1317.0929 | 4 | 5265.3498 | 5265.3334 | 0 | 3.1 | 413 | Specific | 716.02 | 294.25 | 294.25 | 13.68 | 117 | >sp\|P01833\|PIGR_HUMAN Polymeric immunoglobulin receptor OS=Homo sapiens OX=9606 GN=PIGR PE=1 SV=4 | 902 | 20201116_xd_colostrum_1ng_glyco_hcd203040_compound15.22626.22626.4 | controllerType=0 controllerNumber=1 scan=22626 | 48.4368 |
| 19866:3 | 5 | R.LSLLEEPGN[+1458.442]GTFTVILNQLTSR.D | HexNAc(2)Hex(6)Phospho(1) | N[+1458] | 1287.5826 | 3 | 3860.7332 | 3860.7346 | 0 | -0.4 | 413 | Specific | 725.46 | 725.46 | 725.46 | 14.85 | 117 | >sp\|P01833\|PIGR_HUMAN Polymeric immunoglobulin receptor OS=Homo sapiens OX=9606 GN=PIGR PE=1 SV=4 | 902 | 20201116_xd_colostrum_1ng_glyco_hcd203040_compound15.23280.23280.3 | controllerType=0 controllerNumber=1 scan=23280 | 49.9079 |
| 19242:4 | 5 | R.LSLLEEPGN[+2863.041]GTFTVILNQLTSR.D | HexNAc(5)Hex(6)Fuc(4)NeuAc(1) | N[+2863] | 1317.0929 | 4 | 5265.3498 | 5265.3334 | 0 | 3.1 | 413 | Specific | 745.99 | 352.14 | 352.14 | 14.11 | 117 | >sp\|P01833\|PIGR_HUMAN Polymeric immunoglobulin receptor OS=Homo sapiens OX=9606 GN=PIGR PE=1 SV=4 | 902 | 20201116_xd_colostrum_1ng_glyco_hcd203040_compound15.22628.22628.4 | controllerType=0 controllerNumber=1 scan=22628 | 48.442 |
| 21085:4 | 5 | R.LSLLEEPGN[+3007.058]GTFTVILNQLTSR.D | HexNAc(5)Hex(6)Fuc(1)NeuAc(3) | N[+3007] | 1353.1024 | 4 | 5409.3879 | 5409.3505 | 0 | 6.9 | 413 | Specific | 771.98 | 616.02 | 616.02 | 16 | 117 | >sp\|P01833\|PIGR_HUMAN Polymeric immunoglobulin receptor OS=Homo sapiens OX=9606 GN=PIGR PE=1 SV=4 | 902 | 20201116_xd_colostrum_1ng_glyco_hcd203040_compound15.24555.24555.4 | controllerType=0 controllerNumber=1 scan=24555 | 52.6805 |
| 19268:3 | 5 | R.LSLLEEPGN[+2715.963]GTFTVILNQLTSR.D | HexNAc(5)Hex(6)Fuc(1)NeuAc(2) | N[+2716] | 1706.7725 | 3 | 5118.3031 | 5118.2551 | 0 | 9.4 | 413 | Specific | 784.6 | 601.19 | 601.19 | 15.24 | 117 | >sp\|P01833\|PIGR_HUMAN Polymeric immunoglobulin receptor OS=Homo sapiens OX=9606 GN=PIGR PE=1 SV=4 | 902 | 20201116_xd_colostrum_1ng_glyco_hcd203040_compound15.22655.22655.3 | controllerType=0 controllerNumber=1 scan=22655 | 48.4938 |
| 19872:3 | 5 | R.LSLLEEPGN[+1458.442]GTFTVILNQLTSR.D | HexNAc(2)Hex(6)Phospho(1) | N[+1458] | 1287.5849 | 3 | 3860.7401 | 3860.7346 | 0 | 1.4 | 413 | Specific | 790.99 | 685.58 | 685.58 | 16.56 | 117 | >sp\|P01833\|PIGR_HUMAN Polymeric immunoglobulin receptor OS=Homo sapiens OX=9606 GN=PIGR PE=1 SV=4 | 902 | 20201116_xd_colostrum_1ng_glyco_hcd203040_compound15.23287.23287.3 | controllerType=0 controllerNumber=1 scan=23287 | 49.9243 |
| 21313:4 | 5 | R.LSLLEEPGN[+2496.888]GTFTVILNQLTSR.D | HexNAc(4)Hex(5)Fuc(2)NeuAc(2) | N[+2497] | 1225.5547 | 4 | 4899.1971 | 4899.1808 | 0 | 3.3 | 413 | Specific | 804.55 | 629.15 | 629.15 | 17.8 | 117 | >sp\|P01833\|PIGR_HUMAN Polymeric immunoglobulin receptor OS=Homo sapiens OX=9606 GN=PIGR PE=1 SV=4 | 902 | 20201116_xd_colostrum_1ng_glyco_hcd203040_compound15.24792.24792.4 | controllerType=0 controllerNumber=1 scan=24792 | 53.1687 |
| 19405:4 | 5 | R.LSLLEEPGN[+2059.735]GTFTVILNQLTSR.D | HexNAc(4)Hex(5)Fuc(1)NeuAc(1) | N[+2060] | 1116.2653 | 4 | 4462.0394 | 4462.0275 | 0 | 2.7 | 413 | Specific | 819.26 | 618.74 | 618.74 | 17.99 | 117 | >sp\|P01833\|PIGR_HUMAN Polymeric immunoglobulin receptor OS=Homo sapiens OX=9606 GN=PIGR PE=1 SV=4 | 902 | 20201116_xd_colostrum_1ng_glyco_hcd203040_compound15.22797.22797.4 | controllerType=0 controllerNumber=1 scan=22797 | 48.7824 |
| 19403:4 | 5 | R.LSLLEEPGN[+2059.735]GTFTVILNQLTSR.D | HexNAc(4)Hex(5)Fuc(1)NeuAc(1) | N[+2060] | 1116.2653 | 4 | 4462.0394 | 4462.0275 | 0 | 2.7 | 413 | Specific | 821.02 | 648.9 | 648.9 | 17.99 | 117 | >sp\|P01833\|PIGR_HUMAN Polymeric immunoglobulin receptor OS=Homo sapiens OX=9606 GN=PIGR PE=1 SV=4 | 902 | 20201116_xd_colostrum_1ng_glyco_hcd203040_compound15.22795.22795.4 | controllerType=0 controllerNumber=1 scan=22795 | 48.7755 |
| 19147:3 | 5 | R.LSLLEEPGN[+2863.041]GTFTVILNQLTSR.D | HexNAc(5)Hex(6)Fuc(4)NeuAc(1) | N[+2863] | 1755.7888 | 3 | 5265.3518 | 5265.3334 | 0 | 3.5 | 413 | Specific | 826.47 | 410.22 | 410.22 | 17.51 | 117 | >sp\|P01833\|PIGR_HUMAN Polymeric immunoglobulin receptor OS=Homo sapiens OX=9606 GN=PIGR PE=1 SV=4 | 902 | 20201116_xd_colostrum_1ng_glyco_hcd203040_compound15.22529.22529.3 | controllerType=0 controllerNumber=1 scan=22529 | 48.2103 |
| 19140:4 | 5 | R.LSLLEEPGN[+2863.041]GTFTVILNQLTSR.D | HexNAc(5)Hex(6)Fuc(4)NeuAc(1) | N[+2863] | 1317.093 | 4 | 5265.3503 | 5265.3334 | 0 | 3.2 | 413 | Specific | 836.04 | 437.86 | 437.86 | 17.8 | 117 | >sp\|P01833\|PIGR_HUMAN Polymeric immunoglobulin receptor OS=Homo sapiens OX=9606 GN=PIGR PE=1 SV=4 | 902 | 20201116_xd_colostrum_1ng_glyco_hcd203040_compound15.22522.22522.4 | controllerType=0 controllerNumber=1 scan=22522 | 48.1902 |
| 19868:3 | 5 | R.LSLLEEPGN[+1458.442]GTFTVILNQLTSR.D | HexNAc(2)Hex(6)Phospho(1) | N[+1458] | 1287.5826 | 3 | 3860.7332 | 3860.7346 | 0 | -0.4 | 413 | Specific | 862.73 | 747.43 | 747.43 | 17.85 | 117 | >sp\|P01833\|PIGR_HUMAN Polymeric immunoglobulin receptor OS=Homo sapiens OX=9606 GN=PIGR PE=1 SV=4 | 902 | 20201116_xd_colostrum_1ng_glyco_hcd203040_compound15.23282.23282.3 | controllerType=0 controllerNumber=1 scan=23282 | 49.9133 |
| 19144:3 | 5 | R.LSLLEEPGN[+2863.041]GTFTVILNQLTSR.D | HexNAc(5)Hex(6)Fuc(4)NeuAc(1) | N[+2863] | 1755.7888 | 3 | 5265.3518 | 5265.3334 | 0 | 3.5 | 413 | Specific | 863.67 | 433.04 | 433.04 | 18.51 | 117 | >sp\|P01833\|PIGR_HUMAN Polymeric immunoglobulin receptor OS=Homo sapiens OX=9606 GN=PIGR PE=1 SV=4 | 902 | 20201116_xd_colostrum_1ng_glyco_hcd203040_compound15.22526.22526.3 | controllerType=0 controllerNumber=1 scan=22526 | 48.2015 |
| 21315:4 | 5 | R.LSLLEEPGN[+2496.888]GTFTVILNQLTSR.D | HexNAc(4)Hex(5)Fuc(2)NeuAc(2) | N[+2497] | 1225.5547 | 4 | 4899.1971 | 4899.1808 | 0 | 3.3 | 413 | Specific | 878.91 | 780.46 | 780.46 | 18.8 | 117 | >sp\|P01833\|PIGR_HUMAN Polymeric immunoglobulin receptor OS=Homo sapiens OX=9606 GN=PIGR PE=1 SV=4 | 902 | 20201116_xd_colostrum_1ng_glyco_hcd203040_compound15.24794.24794.4 | controllerType=0 controllerNumber=1 scan=24794 | 53.1735 |
| 19142:4 | 5 | R.LSLLEEPGN[+2863.041]GTFTVILNQLTSR.D | HexNAc(5)Hex(6)Fuc(4)NeuAc(1) | N[+2863] | 1317.093 | 4 | 5265.3503 | 5265.3334 | 0 | 3.2 | 413 | Specific | 900.43 | 497.52 | 497.52 | 19.8 | 117 | >sp\|P01833\|PIGR_HUMAN Polymeric immunoglobulin receptor OS=Homo sapiens OX=9606 GN=PIGR PE=1 SV=4 | 902 | 20201116_xd_colostrum_1ng_glyco_hcd203040_compound15.22524.22524.4 | controllerType=0 controllerNumber=1 scan=22524 | 48.1984 |
| 19428:3 | 5 | R.LSLLEEPGN[+2059.735]GTFTVILNQLTSR.D | HexNAc(4)Hex(5)Fuc(1)NeuAc(1) | N[+2060] | 1488.0195 | 3 | 4462.0438 | 4462.0275 | 0 | 3.7 | 413 | Specific | 955.84 | 708.61 | 708.61 | 20.51 | 117 | >sp\|P01833\|PIGR_HUMAN Polymeric immunoglobulin receptor OS=Homo sapiens OX=9606 GN=PIGR PE=1 SV=4 | 902 | 20201116_xd_colostrum_1ng_glyco_hcd203040_compound15.22822.22822.3 | controllerType=0 controllerNumber=1 scan=22822 | 48.8506 |
| 19343:4 | 5 | R.LSLLEEPGN[+2351.851]GTFTVILNQLTSR.D | HexNAc(4)Hex(5)Fuc(3)NeuAc(1) | N[+2352] | 1189.295 | 4 | 4754.158 | 4754.1433 | 0 | 3.1 | 413 | Specific | 976.71 | 748.91 | 748.91 | 20.8 | 117 | >sp\|P01833\|PIGR_HUMAN Polymeric immunoglobulin receptor OS=Homo sapiens OX=9606 GN=PIGR PE=1 SV=4 | 902 | 20201116_xd_colostrum_1ng_glyco_hcd203040_compound15.22733.22733.4 | controllerType=0 controllerNumber=1 scan=22733 | 48.6592 |
| 21308:4 | 5 | R.LSLLEEPGN[+2350.830]GTFTVILNQLTSR.D | HexNAc(4)Hex(5)Fuc(1)NeuAc(2) | N[+2351] | 1189.0425 | 4 | 4753.1483 | 4753.1229 | 0 | 5.3 | 413 | Specific | 978.23 | 764 | 764 | 20.16 | 117 | >sp\|P01833\|PIGR_HUMAN Polymeric immunoglobulin receptor OS=Homo sapiens OX=9606 GN=PIGR PE=1 SV=4 | 902 | 20201116_xd_colostrum_1ng_glyco_hcd203040_compound15.24787.24787.4 | controllerType=0 controllerNumber=1 scan=24787 | 53.1583 |
| 19337:3 | 5 | R.LSLLEEPGN[+2351.851]GTFTVILNQLTSR.D | HexNAc(4)Hex(5)Fuc(3)NeuAc(1) | N[+2352] | 1585.3909 | 3 | 4754.1582 | 4754.1433 | 0 | 3.1 | 413 | Specific | 995.48 | 707.05 | 707.05 | 20.51 | 117 | >sp\|P01833\|PIGR_HUMAN Polymeric immunoglobulin receptor OS=Homo sapiens OX=9606 GN=PIGR PE=1 SV=4 | 902 | 20201116_xd_colostrum_1ng_glyco_hcd203040_compound15.22727.22727.3 | controllerType=0 controllerNumber=1 scan=22727 | 48.6429 |
| 21306:4 | 5 | R.LSLLEEPGN[+2350.830]GTFTVILNQLTSR.D | HexNAc(4)Hex(5)Fuc(1)NeuAc(2) | N[+2351] | 1189.0425 | 4 | 4753.1483 | 4753.1229 | 0 | 5.3 | 413 | Specific | 1026.37 | 817.72 | 817.72 | 21.16 | 117 | >sp\|P01833\|PIGR_HUMAN Polymeric immunoglobulin receptor OS=Homo sapiens OX=9606 GN=PIGR PE=1 SV=4 | 902 | 20201116_xd_colostrum_1ng_glyco_hcd203040_compound15.24785.24785.4 | controllerType=0 controllerNumber=1 scan=24785 | 53.1508 |
| 19426:3 | 5 | R.LSLLEEPGN[+2059.735]GTFTVILNQLTSR.D | HexNAc(4)Hex(5)Fuc(1)NeuAc(1) | N[+2060] | 1488.0195 | 3 | 4462.0438 | 4462.0275 | 0 | 3.7 | 413 | Specific | 1045.79 | 817.36 | 817.36 | 21.51 | 117 | >sp\|P01833\|PIGR_HUMAN Polymeric immunoglobulin receptor OS=Homo sapiens OX=9606 GN=PIGR PE=1 SV=4 | 902 | 20201116_xd_colostrum_1ng_glyco_hcd203040_compound15.22820.22820.3 | controllerType=0 controllerNumber=1 scan=22820 | 48.842 |
| 19341:4 | 5 | R.LSLLEEPGN[+2351.851]GTFTVILNQLTSR.D | HexNAc(4)Hex(5)Fuc(3)NeuAc(1) | N[+2352] | 1189.295 | 4 | 4754.158 | 4754.1433 | 0 | 3.1 | 413 | Specific | 1070.93 | 974.34 | 974.34 | 22.8 | 117 | >sp\|P01833\|PIGR_HUMAN Polymeric immunoglobulin receptor OS=Homo sapiens OX=9606 GN=PIGR PE=1 SV=4 | 902 | 20201116_xd_colostrum_1ng_glyco_hcd203040_compound15.22731.22731.4 | controllerType=0 controllerNumber=1 scan=22731 | 48.6498 |
| 19334:3 | 5 | R.LSLLEEPGN[+2351.851]GTFTVILNQLTSR.D | HexNAc(4)Hex(5)Fuc(3)NeuAc(1) | N[+2352] | 1585.3909 | 3 | 4754.1582 | 4754.1433 | 0 | 3.1 | 413 | Specific | 1089.15 | 913.01 | 913.01 | 22.51 | 117 | >sp\|P01833\|PIGR_HUMAN Polymeric immunoglobulin receptor OS=Homo sapiens OX=9606 GN=PIGR PE=1 SV=4 | 902 | 20201116_xd_colostrum_1ng_glyco_hcd203040_compound15.22724.22724.3 | controllerType=0 controllerNumber=1 scan=22724 | 48.6355 |
| 13609:3 | 5 | K.[+42.011]IIEGEPNLKVPGN[+2018.708]VTAVLGETLK.V | HexNAc(3)Hex(6)Fuc(1)NeuAc(1) | .[42], N[+2019] | 1484.7086 | 3 | 4452.1111 | 4452.0683 | 0 | 9.6 | 457 | Specific | 245.48 | 207.11 | 207.11 | 4.36 | 117 | >sp\|P01833\|PIGR_HUMAN Polymeric immunoglobulin receptor OS=Homo sapiens OX=9606 GN=PIGR PE=1 SV=4 | 902 | 20201116_xd_colostrum_1ng_glyco_hcd203040_compound15.16750.16750.3 | controllerType=0 controllerNumber=1 scan=16750 | 35.7794 |
| 13607:3 | 5 | K.[+42.011]IIEGEPNLKVPGN[+2018.708]VTAVLGETLK.V | HexNAc(3)Hex(6)Fuc(1)NeuAc(1) | .[42], N[+2019] | 1484.7086 | 3 | 4452.1111 | 4452.0683 | 0 | 9.6 | 457 | Specific | 340.95 | 322.25 | 322.25 | 6.72 | 117 | >sp\|P01833\|PIGR_HUMAN Polymeric immunoglobulin receptor OS=Homo sapiens OX=9606 GN=PIGR PE=1 SV=4 | 902 | 20201116_xd_colostrum_1ng_glyco_hcd203040_compound15.16748.16748.3 | controllerType=0 controllerNumber=1 scan=16748 | 35.7707 |
| 16986:4 | 5 | K.IIEGEPNLKVPGN[+2350.830]VTAVLGETLK.V | HexNAc(4)Hex(5)Fuc(1)NeuAc(2) | N[+2351] | 1186.3052 | 4 | 4742.199 | 4742.1797 | 0 | 4.1 | 457 | Specific | 484.24 | 484.24 | 484.24 | 10.08 | 117 | >sp\|P01833\|PIGR_HUMAN Polymeric immunoglobulin receptor OS=Homo sapiens OX=9606 GN=PIGR PE=1 SV=4 | 902 | 20201116_xd_colostrum_1ng_glyco_hcd203040_compound15.20268.20268.2 | controllerType=0 controllerNumber=1 scan=20268 | 43.2004 |
| 15294:3 | 5 | K.IIEGEPNLKVPGN[+2204.772]VTAVLGETLK.V | HexNAc(4)Hex(5)NeuAc(2) | N[+2205] | 1532.7189 | 3 | 4596.1422 | 4596.1218 | 0 | 4.4 | 457 | Specific | 583.94 | 583.94 | 583.94 | 11.88 | 117 | >sp\|P01833\|PIGR_HUMAN Polymeric immunoglobulin receptor OS=Homo sapiens OX=9606 GN=PIGR PE=1 SV=4 | 902 | 20201116_xd_colostrum_1ng_glyco_hcd203040_compound15.18504.18504.3 | controllerType=0 controllerNumber=1 scan=18504 | 39.4022 |
| 16988:4 | 5 | K.IIEGEPNLKVPGN[+2350.830]VTAVLGETLK.V | HexNAc(4)Hex(5)Fuc(1)NeuAc(2) | N[+2351] | 1186.3052 | 4 | 4742.199 | 4742.1797 | 0 | 4.1 | 457 | Specific | 594.32 | 419.28 | 419.28 | 11.97 | 117 | >sp\|P01833\|PIGR_HUMAN Polymeric immunoglobulin receptor OS=Homo sapiens OX=9606 GN=PIGR PE=1 SV=4 | 902 | 20201116_xd_colostrum_1ng_glyco_hcd203040_compound15.20270.20270.2 | controllerType=0 controllerNumber=1 scan=20270 | 43.2042 |
| 15202:4 | 5 | K.IIEGEPNLKVPGN[+2205.793]VTAVLGETLK.V | HexNAc(4)Hex(5)Fuc(2)NeuAc(1) | N[+2206] | 1150.043 | 4 | 4597.1502 | 4597.1422 | 0 | 1.7 | 457 | Specific | 690.82 | 639.25 | 639.25 | 14.25 | 117 | >sp\|P01833\|PIGR_HUMAN Polymeric immunoglobulin receptor OS=Homo sapiens OX=9606 GN=PIGR PE=1 SV=4 | 902 | 20201116_xd_colostrum_1ng_glyco_hcd203040_compound15.18408.18408.4 | controllerType=0 controllerNumber=1 scan=18408 | 39.1811 |
| 15291:3 | 5 | K.IIEGEPNLKVPGN[+2204.772]VTAVLGETLK.V | HexNAc(4)Hex(5)NeuAc(2) | N[+2205] | 1532.7189 | 3 | 4596.1422 | 4596.1218 | 0 | 4.4 | 457 | Specific | 714 | 654.47 | 654.47 | 14.88 | 117 | >sp\|P01833\|PIGR_HUMAN Polymeric immunoglobulin receptor OS=Homo sapiens OX=9606 GN=PIGR PE=1 SV=4 | 902 | 20201116_xd_colostrum_1ng_glyco_hcd203040_compound15.18501.18501.3 | controllerType=0 controllerNumber=1 scan=18501 | 39.3978 |
| 13596:3 | 5 | K.IIEGEPNLKVPGN[+2205.793]VTAVLGETLK.V | HexNAc(4)Hex(5)Fuc(2)NeuAc(1) | N[+2206] | 1533.0617 | 3 | 4597.1706 | 4597.1422 | 0 | 6.2 | 457 | Specific | 734.34 | 721.99 | 721.99 | 14.33 | 117 | >sp\|P01833\|PIGR_HUMAN Polymeric immunoglobulin receptor OS=Homo sapiens OX=9606 GN=PIGR PE=1 SV=4 | 902 | 20201116_xd_colostrum_1ng_glyco_hcd203040_compound15.16737.16737.3 | controllerType=0 controllerNumber=1 scan=16737 | 35.752 |
| 15199:4 | 5 | K.IIEGEPNLKVPGN[+2205.793]VTAVLGETLK.V | HexNAc(4)Hex(5)Fuc(2)NeuAc(1) | N[+2206] | 1150.043 | 4 | 4597.1502 | 4597.1422 | 0 | 1.7 | 457 | Specific | 740.51 | 740.51 | 740.51 | 15.25 | 117 | >sp\|P01833\|PIGR_HUMAN Polymeric immunoglobulin receptor OS=Homo sapiens OX=9606 GN=PIGR PE=1 SV=4 | 902 | 20201116_xd_colostrum_1ng_glyco_hcd203040_compound15.18405.18405.4 | controllerType=0 controllerNumber=1 scan=18405 | 39.1759 |
| 16993:3 | 5 | K.IIEGEPNLKVPGN[+2350.830]VTAVLGETLK.V | HexNAc(4)Hex(5)Fuc(1)NeuAc(2) | N[+2351] | 1581.4026 | 3 | 4742.1934 | 4742.1797 | 0 | 2.9 | 457 | Specific | 742.33 | 712.33 | 712.33 | 15.31 | 117 | >sp\|P01833\|PIGR_HUMAN Polymeric immunoglobulin receptor OS=Homo sapiens OX=9606 GN=PIGR PE=1 SV=4 | 902 | 20201116_xd_colostrum_1ng_glyco_hcd203040_compound15.20275.20275.3 | controllerType=0 controllerNumber=1 scan=20275 | 43.2139 |
| 16974:4 | 5 | K.IIEGEPNLKVPGN[+2350.830]VTAVLGETLK.V | HexNAc(4)Hex(5)Fuc(1)NeuAc(2) | N[+2351] | 1186.302 | 4 | 4742.1863 | 4742.1797 | 0 | 1.4 | 457 | Specific | 788.89 | 683.27 | 683.27 | 16.25 | 117 | >sp\|P01833\|PIGR_HUMAN Polymeric immunoglobulin receptor OS=Homo sapiens OX=9606 GN=PIGR PE=1 SV=4 | 902 | 20201116_xd_colostrum_1ng_glyco_hcd203040_compound15.20255.20255.4 | controllerType=0 controllerNumber=1 scan=20255 | 43.1696 |
| 13594:3 | 5 | K.IIEGEPNLKVPGN[+2205.793]VTAVLGETLK.V | HexNAc(4)Hex(5)Fuc(2)NeuAc(1) | N[+2206] | 1533.0617 | 3 | 4597.1706 | 4597.1422 | 0 | 6.2 | 457 | Specific | 817.82 | 812.78 | 812.78 | 16.33 | 117 | >sp\|P01833\|PIGR_HUMAN Polymeric immunoglobulin receptor OS=Homo sapiens OX=9606 GN=PIGR PE=1 SV=4 | 902 | 20201116_xd_colostrum_1ng_glyco_hcd203040_compound15.16735.16735.3 | controllerType=0 controllerNumber=1 scan=16735 | 35.7457 |
| 15147:4 | 5 | K.IIEGEPNLKVPGN[+2351.851]VTAVLGETLK.V | HexNAc(4)Hex(5)Fuc(3)NeuAc(1) | N[+2352] | 1186.5585 | 4 | 4743.2122 | 4743.2001 | 0 | 2.6 | 457 | Specific | 834.17 | 773.03 | 773.03 | 17.4 | 117 | >sp\|P01833\|PIGR_HUMAN Polymeric immunoglobulin receptor OS=Homo sapiens OX=9606 GN=PIGR PE=1 SV=4 | 902 | 20201116_xd_colostrum_1ng_glyco_hcd203040_compound15.18351.18351.4 | controllerType=0 controllerNumber=1 scan=18351 | 39.0607 |
| 15143:3 | 5 | K.IIEGEPNLKVPGN[+2351.851]VTAVLGETLK.V | HexNAc(4)Hex(5)Fuc(3)NeuAc(1) | N[+2352] | 1581.742 | 3 | 4743.2114 | 4743.2001 | 0 | 2.4 | 457 | Specific | 840.34 | 721.2 | 721.2 | 17.31 | 117 | >sp\|P01833\|PIGR_HUMAN Polymeric immunoglobulin receptor OS=Homo sapiens OX=9606 GN=PIGR PE=1 SV=4 | 902 | 20201116_xd_colostrum_1ng_glyco_hcd203040_compound15.18346.18346.3 | controllerType=0 controllerNumber=1 scan=18346 | 39.0492 |
| 16971:4 | 5 | K.IIEGEPNLKVPGN[+2350.830]VTAVLGETLK.V | HexNAc(4)Hex(5)Fuc(1)NeuAc(2) | N[+2351] | 1186.302 | 4 | 4742.1863 | 4742.1797 | 0 | 1.4 | 457 | Specific | 842.83 | 746.65 | 746.65 | 17.25 | 117 | >sp\|P01833\|PIGR_HUMAN Polymeric immunoglobulin receptor OS=Homo sapiens OX=9606 GN=PIGR PE=1 SV=4 | 902 | 20201116_xd_colostrum_1ng_glyco_hcd203040_compound15.20252.20252.4 | controllerType=0 controllerNumber=1 scan=20252 | 43.1573 |
| 16990:3 | 5 | K.IIEGEPNLKVPGN[+2350.830]VTAVLGETLK.V | HexNAc(4)Hex(5)Fuc(1)NeuAc(2) | N[+2351] | 1581.4026 | 3 | 4742.1934 | 4742.1797 | 0 | 2.9 | 457 | Specific | 901.55 | 838.72 | 838.72 | 19.31 | 117 | >sp\|P01833\|PIGR_HUMAN Polymeric immunoglobulin receptor OS=Homo sapiens OX=9606 GN=PIGR PE=1 SV=4 | 902 | 20201116_xd_colostrum_1ng_glyco_hcd203040_compound15.20272.20272.3 | controllerType=0 controllerNumber=1 scan=20272 | 43.2087 |
| 15145:4 | 5 | K.IIEGEPNLKVPGN[+2351.851]VTAVLGETLK.V | HexNAc(4)Hex(5)Fuc(3)NeuAc(1) | N[+2352] | 1186.5585 | 4 | 4743.2122 | 4743.2001 | 0 | 2.6 | 457 | Specific | 931.41 | 895.13 | 895.13 | 19.4 | 117 | >sp\|P01833\|PIGR_HUMAN Polymeric immunoglobulin receptor OS=Homo sapiens OX=9606 GN=PIGR PE=1 SV=4 | 902 | 20201116_xd_colostrum_1ng_glyco_hcd203040_compound15.18349.18349.4 | controllerType=0 controllerNumber=1 scan=18349 | 39.0552 |
| 15140:3 | 5 | K.IIEGEPNLKVPGN[+2351.851]VTAVLGETLK.V | HexNAc(4)Hex(5)Fuc(3)NeuAc(1) | N[+2352] | 1581.742 | 3 | 4743.2114 | 4743.2001 | 0 | 2.4 | 457 | Specific | 980.71 | 979.65 | 979.65 | 20.31 | 117 | >sp\|P01833\|PIGR_HUMAN Polymeric immunoglobulin receptor OS=Homo sapiens OX=9606 GN=PIGR PE=1 SV=4 | 902 | 20201116_xd_colostrum_1ng_glyco_hcd203040_compound15.18343.18343.3 | controllerType=0 controllerNumber=1 scan=18343 | 39.0403 |
| 13670:5 | 5 | K.[+42.011]IIEGEPNLKVPGN[+2018.708]VTAVLGETLKVPC[+57.021]HFPC[+57.021]K | HexNAc(3)Hex(6)Fuc(1)NeuAc(1) | .[42], C[+57]*2, N[+2019] | 1096.3201 | 5 | 5477.5712 | 5477.5258 | 0 | 8.3 | 457 | Specific | 275.85 | 221.02 | 221.02 | 5.02 | 117 | >sp\|P01833\|PIGR_HUMAN Polymeric immunoglobulin receptor OS=Homo sapiens OX=9606 GN=PIGR PE=1 SV=4 | 902 | 20201116_xd_colostrum_1ng_glyco_hcd203040_compound15.16815.16815.5 | controllerType=0 controllerNumber=1 scan=16815 | 35.9439 |
| 16108:6 | 5 | K.IIEGEPNLKVPGN[+2350.830]VTAVLGETLKVPC[+57.021]HFPC[+57.021]K.F | HexNAc(4)Hex(5)Fuc(1)NeuAc(2) | C[+57]*2, N[+2351] | 962.1155 | 6 | 5767.6564 | 5767.6372 | 0 | 3.3 | 457 | Specific | 293.02 | 293.02 | 293.02 | 6.04 | 117 | >sp\|P01833\|PIGR_HUMAN Polymeric immunoglobulin receptor OS=Homo sapiens OX=9606 GN=PIGR PE=1 SV=4 | 902 | 20201116_xd_colostrum_1ng_glyco_hcd203040_compound15.19351.19351.6 | controllerType=0 controllerNumber=1 scan=19351 | 41.1426 |
| 13766:5 | 5 | K.IIEGEPNLKVPGN[+2351.851]VTAVLGETLKVPC[+57.021]HFPC[+57.021]K.F | HexNAc(4)Hex(5)Fuc(3)NeuAc(1) | C[+57]*2, N[+2352] | 1154.5431 | 5 | 5768.6864 | 5768.6576 | 0 | 5 | 457 | Specific | 317.09 | 317.09 | 317.09 | 6.64 | 117 | >sp\|P01833\|PIGR_HUMAN Polymeric immunoglobulin receptor OS=Homo sapiens OX=9606 GN=PIGR PE=1 SV=4 | 902 | 20201116_xd_colostrum_1ng_glyco_hcd203040_compound15.16915.16915.5 | controllerType=0 controllerNumber=1 scan=16915 | 36.1847 |
| 16110:6 | 5 | K.IIEGEPNLKVPGN[+2350.830]VTAVLGETLKVPC[+57.021]HFPC[+57.021]K.F | HexNAc(4)Hex(5)Fuc(1)NeuAc(2) | C[+57]*2, N[+2351] | 962.1155 | 6 | 5767.6564 | 5767.6372 | 0 | 3.3 | 457 | Specific | 321.67 | 271.66 | 271.66 | 6.98 | 117 | >sp\|P01833\|PIGR_HUMAN Polymeric immunoglobulin receptor OS=Homo sapiens OX=9606 GN=PIGR PE=1 SV=4 | 902 | 20201116_xd_colostrum_1ng_glyco_hcd203040_compound15.19353.19353.6 | controllerType=0 controllerNumber=1 scan=19353 | 41.1483 |
| 13674:4 | 5 | K.[+42.011]IIEGEPNLKVPGN[+2018.708]VTAVLGETLKVPC[+57.021]HFPC[+57.021]K | HexNAc(3)Hex(6)Fuc(1)NeuAc(1) | .[42], C[+57]*2, N[+2019] | 1370.1494 | 4 | 5477.5758 | 5477.5258 | 0 | 9.1 | 457 | Specific | 344.36 | 232.24 | 232.24 | 4.86 | 117 | >sp\|P01833\|PIGR_HUMAN Polymeric immunoglobulin receptor OS=Homo sapiens OX=9606 GN=PIGR PE=1 SV=4 | 902 | 20201116_xd_colostrum_1ng_glyco_hcd203040_compound15.16819.16819.4 | controllerType=0 controllerNumber=1 scan=16819 | 35.9537 |
| 13718:5 | 5 | K.IIEGEPNLKVPGN[+2571.946]VTAVLGETLKVPC[+57.021]HFPC[+57.021]K.F | HexNAc(5)Hex(6)Fuc(4) | C[+57]*2, N[+2572] | 1198.5593 | 5 | 5988.7675 | 5988.7523 | 0 | 2.5 | 457 | Specific | 388.34 | 388.34 | 388.34 | 9.5 | 117 | >sp\|P01833\|PIGR_HUMAN Polymeric immunoglobulin receptor OS=Homo sapiens OX=9606 GN=PIGR PE=1 SV=4 | 902 | 20201116_xd_colostrum_1ng_glyco_hcd203040_compound15.16865.16865.4 | controllerType=0 controllerNumber=1 scan=16865 | 36.0721 |
| 14775:6 | 5 | K.IIEGEPNLKVPGN[+2205.793]VTAVLGETLKVPC[+57.021]HFPC[+57.021]K.F | HexNAc(4)Hex(5)Fuc(2)NeuAc(1) | C[+57]*2, N[+2206] | 937.9436 | 6 | 5622.6255 | 5622.5997 | 0 | 4.6 | 457 | Specific | 421.92 | 410.71 | 410.71 | 9.27 | 117 | >sp\|P01833\|PIGR_HUMAN Polymeric immunoglobulin receptor OS=Homo sapiens OX=9606 GN=PIGR PE=1 SV=4 | 902 | 20201116_xd_colostrum_1ng_glyco_hcd203040_compound15.17963.17963.6 | controllerType=0 controllerNumber=1 scan=17963 | 38.3191 |
| 13721:5 | 5 | K.IIEGEPNLKVPGN[+2571.946]VTAVLGETLKVPC[+57.021]HFPC[+57.021]K.F | HexNAc(5)Hex(6)Fuc(4) | C[+57]*2, N[+2572] | 1198.5593 | 5 | 5988.7675 | 5988.7523 | 0 | 2.5 | 457 | Specific | 429.84 | 417.33 | 417.33 | 8.81 | 117 | >sp\|P01833\|PIGR_HUMAN Polymeric immunoglobulin receptor OS=Homo sapiens OX=9606 GN=PIGR PE=1 SV=4 | 902 | 20201116_xd_colostrum_1ng_glyco_hcd203040_compound15.16868.16868.4 | controllerType=0 controllerNumber=1 scan=16868 | 36.0762 |
| 14656:4 | 5 | K.IIEGEPNLKVPGN[+2863.041]VTAVLGETLKVPC[+57.021]HFPC[+57.021]K.F | HexNAc(5)Hex(6)Fuc(4)NeuAc(1) | C[+57]*2, N[+2863] | 1570.7233 | 4 | 6279.8712 | 6279.8477 | 0 | 3.7 | 457 | Specific | 437.24 | 339.95 | 339.95 | 7.11 | 117 | >sp\|P01833\|PIGR_HUMAN Polymeric immunoglobulin receptor OS=Homo sapiens OX=9606 GN=PIGR PE=1 SV=4 | 902 | 20201116_xd_colostrum_1ng_glyco_hcd203040_compound15.17839.17839.4 | controllerType=0 controllerNumber=1 scan=17839 | 38.0452 |
| 14654:4 | 5 | K.IIEGEPNLKVPGN[+2863.041]VTAVLGETLKVPC[+57.021]HFPC[+57.021]K.F | HexNAc(5)Hex(6)Fuc(4)NeuAc(1) | C[+57]*2, N[+2863] | 1570.7233 | 4 | 6279.8712 | 6279.8477 | 0 | 3.7 | 457 | Specific | 495.08 | 495.08 | 495.08 | 9.09 | 117 | >sp\|P01833\|PIGR_HUMAN Polymeric immunoglobulin receptor OS=Homo sapiens OX=9606 GN=PIGR PE=1 SV=4 | 902 | 20201116_xd_colostrum_1ng_glyco_hcd203040_compound15.17837.17837.4 | controllerType=0 controllerNumber=1 scan=17837 | 38.0388 |
| 14778:6 | 5 | K.IIEGEPNLKVPGN[+2205.793]VTAVLGETLKVPC[+57.021]HFPC[+57.021]K.F | HexNAc(4)Hex(5)Fuc(2)NeuAc(1) | C[+57]*2, N[+2206] | 937.9436 | 6 | 5622.6255 | 5622.5997 | 0 | 4.6 | 457 | Specific | 514.91 | 477.42 | 477.42 | 9.62 | 117 | >sp\|P01833\|PIGR_HUMAN Polymeric immunoglobulin receptor OS=Homo sapiens OX=9606 GN=PIGR PE=1 SV=4 | 902 | 20201116_xd_colostrum_1ng_glyco_hcd203040_compound15.17966.17966.6 | controllerType=0 controllerNumber=1 scan=17966 | 38.324 |
| 14995:4 | 5 | K.IIEGEPNLKVPGN[+2205.793]VTAVLGETLKVPC[+57.021]HFPC[+57.021]K.F | HexNAc(4)Hex(5)Fuc(2)NeuAc(1) | C[+57]*2, N[+2206] | 1406.4119 | 4 | 5622.6256 | 5622.5997 | 0 | 4.6 | 457 | Specific | 516.43 | 516.43 | 516.43 | 8.98 | 117 | >sp\|P01833\|PIGR_HUMAN Polymeric immunoglobulin receptor OS=Homo sapiens OX=9606 GN=PIGR PE=1 SV=4 | 902 | 20201116_xd_colostrum_1ng_glyco_hcd203040_compound15.18191.18191.4 | controllerType=0 controllerNumber=1 scan=18191 | 38.7427 |
| 14997:4 | 5 | K.IIEGEPNLKVPGN[+2205.793]VTAVLGETLKVPC[+57.021]HFPC[+57.021]K.F | HexNAc(4)Hex(5)Fuc(2)NeuAc(1) | C[+57]*2, N[+2206] | 1406.4119 | 4 | 5622.6256 | 5622.5997 | 0 | 4.6 | 457 | Specific | 533.43 | 457.87 | 457.87 | 9.02 | 117 | >sp\|P01833\|PIGR_HUMAN Polymeric immunoglobulin receptor OS=Homo sapiens OX=9606 GN=PIGR PE=1 SV=4 | 902 | 20201116_xd_colostrum_1ng_glyco_hcd203040_compound15.18193.18193.4 | controllerType=0 controllerNumber=1 scan=18193 | 38.7465 |
| 13703:5 | 5 | K.IIEGEPNLKVPGN[+2263.835]VTAVLGETLKVPC[+57.021]HFPC[+57.021]K.F | HexNAc(5)Hex(5)Fuc(3) | C[+57]*2, N[+2264] | 1136.9351 | 5 | 5680.6462 | 5680.6416 | 0 | 0.8 | 457 | Specific | 569.33 | 569.33 | 569.33 | 10.01 | 117 | >sp\|P01833\|PIGR_HUMAN Polymeric immunoglobulin receptor OS=Homo sapiens OX=9606 GN=PIGR PE=1 SV=4 | 902 | 20201116_xd_colostrum_1ng_glyco_hcd203040_compound15.16849.16849.5 | controllerType=0 controllerNumber=1 scan=16849 | 36.0379 |
| 14694:4 | 5 | K.IIEGEPNLKVPGN[+2351.851]VTAVLGETLKVPC[+57.021]HFPC[+57.021]K.F | HexNAc(4)Hex(5)Fuc(3)NeuAc(1) | C[+57]*2, N[+2352] | 1442.9243 | 4 | 5768.6754 | 5768.6576 | 0 | 3.1 | 457 | Specific | 615.62 | 543.54 | 543.54 | 11.99 | 117 | >sp\|P01833\|PIGR_HUMAN Polymeric immunoglobulin receptor OS=Homo sapiens OX=9606 GN=PIGR PE=1 SV=4 | 902 | 20201116_xd_colostrum_1ng_glyco_hcd203040_compound15.17879.17879.4 | controllerType=0 controllerNumber=1 scan=17879 | 38.1392 |
| 13782:5 | 5 | K.IIEGEPNLKVPGN[+1768.640]VTAVLGETLKVPC[+57.021]HFPC[+57.021]K.F | HexNAc(4)Hex(5)Fuc(1) | C[+57]*2, N[+1769] | 1037.8916 | 5 | 5185.4289 | 5185.4464 | 0 | -3.4 | 457 | Specific | 649.43 | 605.51 | 605.51 | 11.38 | 117 | >sp\|P01833\|PIGR_HUMAN Polymeric immunoglobulin receptor OS=Homo sapiens OX=9606 GN=PIGR PE=1 SV=4 | 902 | 20201116_xd_colostrum_1ng_glyco_hcd203040_compound15.16931.16931.5 | controllerType=0 controllerNumber=1 scan=16931 | 36.2261 |
| 14580:5 | 5 | K.IIEGEPNLKVPGN[+2863.041]VTAVLGETLKVPC[+57.021]HFPC[+57.021]K.F | HexNAc(5)Hex(6)Fuc(4)NeuAc(1) | C[+57]*2, N[+2863] | 1256.7771 | 5 | 6279.8564 | 6279.8477 | 0 | 1.4 | 457 | Specific | 649.91 | 583.55 | 583.55 | 12.58 | 117 | >sp\|P01833\|PIGR_HUMAN Polymeric immunoglobulin receptor OS=Homo sapiens OX=9606 GN=PIGR PE=1 SV=4 | 902 | 20201116_xd_colostrum_1ng_glyco_hcd203040_compound15.17760.17760.5 | controllerType=0 controllerNumber=1 scan=17760 | 37.8639 |
| 13705:5 | 5 | K.IIEGEPNLKVPGN[+2263.835]VTAVLGETLKVPC[+57.021]HFPC[+57.021]K.F | HexNAc(5)Hex(5)Fuc(3) | C[+57]*2, N[+2264] | 1136.9351 | 5 | 5680.6462 | 5680.6416 | 0 | 0.8 | 457 | Specific | 652.08 | 652.08 | 652.08 | 13.31 | 117 | >sp\|P01833\|PIGR_HUMAN Polymeric immunoglobulin receptor OS=Homo sapiens OX=9606 GN=PIGR PE=1 SV=4 | 902 | 20201116_xd_colostrum_1ng_glyco_hcd203040_compound15.16851.16851.5 | controllerType=0 controllerNumber=1 scan=16851 | 36.0434 |
| 16166:5 | 5 | K.IIEGEPNLKVPGN[+2496.888]VTAVLGETLKVPC[+57.021]HFPC[+57.021]K.F | HexNAc(4)Hex(5)Fuc(2)NeuAc(2) | C[+57]*2, N[+2497] | 1183.5472 | 5 | 5913.7071 | 5913.6951 | 0 | 2 | 457 | Specific | 656.61 | 578.03 | 578.03 | 13.58 | 117 | >sp\|P01833\|PIGR_HUMAN Polymeric immunoglobulin receptor OS=Homo sapiens OX=9606 GN=PIGR PE=1 SV=4 | 902 | 20201116_xd_colostrum_1ng_glyco_hcd203040_compound15.19412.19412.5 | controllerType=0 controllerNumber=1 scan=19412 | 41.2762 |
| 14535:5 | 5 | K.IIEGEPNLKVPGN[+2863.041]VTAVLGETLKVPC[+57.021]HFPC[+57.021]K.F | HexNAc(5)Hex(6)Fuc(4)NeuAc(1) | C[+57]*2, N[+2863] | 1256.7784 | 5 | 6279.8631 | 6279.8477 | 0 | 2.4 | 457 | Specific | 669.25 | 622.04 | 622.04 | 13.74 | 117 | >sp\|P01833\|PIGR_HUMAN Polymeric immunoglobulin receptor OS=Homo sapiens OX=9606 GN=PIGR PE=1 SV=4 | 902 | 20201116_xd_colostrum_1ng_glyco_hcd203040_compound15.17713.17713.5 | controllerType=0 controllerNumber=1 scan=17713 | 37.7568 |

Table S4. Cont.

| Query #:z | Protein Rank | Glycopeptide | Glycan | Modification Type(s) | Observed m/z | z | Observed (M+H) | Calc. mass (M+H) | Off-by-x error | Mass error (ppm) | StartPosition | Cleavage | Score | Delta | Delta Mod | \|Log Prob\| | # of unique peptides | Protein Name | Protein DB number | Comment | Scan # | Scan Time |
|---|---|---|---|---|---|---|---|---|---|---|---|---|---|---|---|---|---|---|---|---|---|---|
| 13784:5 | 5 | K.IIEGEPNLKVPGN[+1768.640]VTAVLGETLKVPC[+57.021]HFPC[+57.021]K.F | HexNAc(4)Hex(5)Fuc(1) | C[+57]*2, N[+1769] | 1037.8916 | 5 | 5185.4289 | 5185.4464 | 0 | -3.4 | 457 | Specific | 682 | 572.44 | 572.44 | 12.38 | 117 | >sp\|P01833\|PIGR_HUMAN Polymeric immunoglobulin receptor OS=Homo sapiens OX=9606 GN=PIGR PE=1 SV=4 | 902 | 20201116_xd_colostrum_1ng_glyco_hcd203040_compound15.16933.16933.5 | controllerType=0 controllerNumber=1 scan=16933 | 36.2316 |
| 14577:5 | 5 | K.IIEGEPNLKVPGN[+2863.041]VTAVLGETLKVPC[+57.021]HFPC[+57.021]K.F | HexNAc(5)Hex(6)Fuc(4)NeuAc(1) | C[+57]*2, N[+2863] | 1256.7771 | 5 | 6279.8564 | 6279.8477 | 0 | 1.4 | 457 | Specific | 687.09 | 687.09 | 687.09 | 13.58 | 117 | >sp\|P01833\|PIGR_HUMAN Polymeric immunoglobulin receptor OS=Homo sapiens OX=9606 GN=PIGR PE=1 SV=4 | 902 | 20201116_xd_colostrum_1ng_glyco_hcd203040_compound15.17757.17757.5 | controllerType=0 controllerNumber=1 scan=17757 | 37.8591 |
| 13768:5 | 5 | K.IIEGEPNLKVPGN[+2351.851]VTAVLGETLKVPC[+57.021]HFPC[+57.021]K.F | HexNAc(4)Hex(5)Fuc(3)NeuAc(1) | C[+57]*2, N[+2352] | 1154.5431 | 5 | 5768.6864 | 5768.6576 | 0 | 5 | 457 | Specific | 744.5 | 675.74 | 675.74 | 14.31 | 117 | >sp\|P01833\|PIGR_HUMAN Polymeric immunoglobulin receptor OS=Homo sapiens OX=9606 GN=PIGR PE=1 SV=4 | 902 | 20201116_xd_colostrum_1ng_glyco_hcd203040_compound15.16917.16917.5 | controllerType=0 controllerNumber=1 scan=16917 | 36.1978 |
| 16069:4 | 5 | K.IIEGEPNLKVPGN[+2350.830]VTAVLGETLKVPC[+57.021]HFPC[+57.021]K.F | HexNAc(4)Hex(5)Fuc(1)NeuAc(2) | C[+57]*2, N[+2351] | 1442.6693 | 4 | 5767.6554 | 5767.6372 | 0 | 3.2 | 457 | Specific | 757.81 | 603.99 | 603.99 | 14.99 | 117 | >sp\|P01833\|PIGR_HUMAN Polymeric immunoglobulin receptor OS=Homo sapiens OX=9606 GN=PIGR PE=1 SV=4 | 902 | 20201116_xd_colostrum_1ng_glyco_hcd203040_compound15.19311.19311.4 | controllerType=0 controllerNumber=1 scan=19311 | 41.0503 |
| 14861:4 | 5 | K.IIEGEPNLKVPGN[+1856.656]VTAVLGETLKVPC[+57.021]HFPC[+57.021]K.F | HexNAc(3)Hex(5)Fuc(1)NeuAc(1) | C[+57]*2, N[+1857] | 1319.1263 | 4 | 5273.4836 | 5273.4624 | 0 | 4 | 457 | Specific | 765.25 | 672.29 | 672.29 | 14.99 | 117 | >sp\|P01833\|PIGR_HUMAN Polymeric immunoglobulin receptor OS=Homo sapiens OX=9606 GN=PIGR PE=1 SV=4 | 902 | 20201116_xd_colostrum_1ng_glyco_hcd203040_compound15.18052.18052.4 | controllerType=0 controllerNumber=1 scan=18052 | 38.4898 |
| 14729:4 | 5 | K.IIEGEPNLKVPGN[+2059.735]VTAVLGETLKVPC[+57.021]HFPC[+57.021]K.F | HexNAc(4)Hex(5)Fuc(1)NeuAc(1) | C[+57]*2, N[+2060] | 1369.8927 | 4 | 5476.549 | 5476.5418 | 0 | 1.3 | 457 | Specific | 782.52 | 748.83 | 748.83 | 15.03 | 117 | >sp\|P01833\|PIGR_HUMAN Polymeric immunoglobulin receptor OS=Homo sapiens OX=9606 GN=PIGR PE=1 SV=4 | 902 | 20201116_xd_colostrum_1ng_glyco_hcd203040_compound15.17915.17915.2 | controllerType=0 controllerNumber=1 scan=17915 | 38.2188 |
| 14783:5 | 5 | K.IIEGEPNLKVPGN[+2059.735]VTAVLGETLKVPC[+57.021]HFPC[+57.021]K.F | HexNAc(4)Hex(5)Fuc(1)NeuAc(1) | C[+57]*2, N[+2060] | 1096.1169 | 5 | 5476.5556 | 5476.5418 | 0 | 2.5 | 457 | Specific | 806.07 | 738.09 | 738.09 | 16.74 | 117 | >sp\|P01833\|PIGR_HUMAN Polymeric immunoglobulin receptor OS=Homo sapiens OX=9606 GN=PIGR PE=1 SV=4 | 902 | 20201116_xd_colostrum_1ng_glyco_hcd203040_compound15.17972.17972.5 | controllerType=0 controllerNumber=1 scan=17972 | 38.3401 |
| 14737:4 | 5 | K.IIEGEPNLKVPGN[+2205.793]VTAVLGETLKVPC[+57.021]HFPC[+57.021]K.F | HexNAc(4)Hex(5)Fuc(2)NeuAc(1) | C[+57]*2, N[+2206] | 1406.4094 | 4 | 5622.6159 | 5622.5997 | 0 | 2.9 | 457 | Specific | 808.08 | 808.08 | 808.08 | 16.18 | 117 | >sp\|P01833\|PIGR_HUMAN Polymeric immunoglobulin receptor OS=Homo sapiens OX=9606 GN=PIGR PE=1 SV=4 | 902 | 20201116_xd_colostrum_1ng_glyco_hcd203040_compound15.17924.17924.4 | controllerType=0 controllerNumber=1 scan=17924 | 38.2475 |
| 16169:5 | 5 | K.IIEGEPNLKVPGN[+2496.888]VTAVLGETLKVPC[+57.021]HFPC[+57.021]K.F | HexNAc(4)Hex(5)Fuc(2)NeuAc(2) | C[+57]*2, N[+2497] | 1183.5472 | 5 | 5913.7071 | 5913.6951 | 0 | 2 | 457 | Specific | 822.93 | 696.34 | 696.34 | 16.58 | 117 | >sp\|P01833\|PIGR_HUMAN Polymeric immunoglobulin receptor OS=Homo sapiens OX=9606 GN=PIGR PE=1 SV=4 | 902 | 20201116_xd_colostrum_1ng_glyco_hcd203040_compound15.19415.19415.5 | controllerType=0 controllerNumber=1 scan=19415 | 41.2854 |
| 14787:4 | 5 | K.IIEGEPNLKVPGN[+2059.735]VTAVLGETLKVPC[+57.021]HFPC[+57.021]K.F | HexNAc(4)Hex(5)Fuc(1)NeuAc(1) | C[+57]*2, N[+2060] | 1369.8936 | 4 | 5476.5524 | 5476.5418 | 0 | 1.9 | 457 | Specific | 837.36 | 662.57 | 662.57 | 16.03 | 117 | >sp\|P01833\|PIGR_HUMAN Polymeric immunoglobulin receptor OS=Homo sapiens OX=9606 GN=PIGR PE=1 SV=4 | 902 | 20201116_xd_colostrum_1ng_glyco_hcd203040_compound15.17976.17976.4 | controllerType=0 controllerNumber=1 scan=17976 | 38.347 |
| 14786:5 | 5 | K.IIEGEPNLKVPGN[+2059.735]VTAVLGETLKVPC[+57.021]HFPC[+57.021]K.F | HexNAc(4)Hex(5)Fuc(1)NeuAc(1) | C[+57]*2, N[+2060] | 1096.1169 | 5 | 5476.5556 | 5476.5418 | 0 | 2.5 | 457 | Specific | 868.49 | 868.49 | 868.49 | 17.74 | 117 | >sp\|P01833\|PIGR_HUMAN Polymeric immunoglobulin receptor OS=Homo sapiens OX=9606 GN=PIGR PE=1 SV=4 | 902 | 20201116_xd_colostrum_1ng_glyco_hcd203040_compound15.17975.17975.5 | controllerType=0 controllerNumber=1 scan=17975 | 38.3444 |
| 14689:5 | 5 | K.IIEGEPNLKVPGN[+2351.851]VTAVLGETLKVPC[+57.021]HFPC[+57.021]K.F | HexNAc(4)Hex(5)Fuc(3)NeuAc(1) | C[+57]*2, N[+2352] | 1154.5398 | 5 | 5768.6699 | 5768.6576 | 0 | 2.1 | 457 | Specific | 886.1 | 859.17 | 859.17 | 17.74 | 117 | >sp\|P01833\|PIGR_HUMAN Polymeric immunoglobulin receptor OS=Homo sapiens OX=9606 GN=PIGR PE=1 SV=4 | 902 | 20201116_xd_colostrum_1ng_glyco_hcd203040_compound15.17874.17874.5 | controllerType=0 controllerNumber=1 scan=17874 | 38.1276 |
| 16070:5 | 5 | K.IIEGEPNLKVPGN[+2350.830]VTAVLGETLKVPC[+57.021]HFPC[+57.021]K.F | HexNAc(4)Hex(5)Fuc(1)NeuAc(2) | C[+57]*2, N[+2351] | 1154.3366 | 5 | 5767.6536 | 5767.6372 | 0 | 2.8 | 457 | Specific | 891.23 | 614.2 | 614.2 | 17.74 | 117 | >sp\|P01833\|PIGR_HUMAN Polymeric immunoglobulin receptor OS=Homo sapiens OX=9606 GN=PIGR PE=1 SV=4 | 902 | 20201116_xd_colostrum_1ng_glyco_hcd203040_compound15.19312.19312.5 | controllerType=0 controllerNumber=1 scan=19312 | 41.0538 |
| 14532:5 | 5 | K.IIEGEPNLKVPGN[+2863.041]VTAVLGETLKVPC[+57.021]HFPC[+57.021]K.F | HexNAc(5)Hex(6)Fuc(4)NeuAc(1) | C[+57]*2, N[+2863] | 1256.7784 | 5 | 6279.8631 | 6279.8477 | 0 | 2.4 | 457 | Specific | 926.13 | 921.77 | 921.77 | 18.74 | 117 | >sp\|P01833\|PIGR_HUMAN Polymeric immunoglobulin receptor OS=Homo sapiens OX=9606 GN=PIGR PE=1 SV=4 | 902 | 20201116_xd_colostrum_1ng_glyco_hcd203040_compound15.17710.17710.5 | controllerType=0 controllerNumber=1 scan=17710 | 37.7472 |
| 14727:4 | 5 | K.IIEGEPNLKVPGN[+2059.735]VTAVLGETLKVPC[+57.021]HFPC[+57.021]K.F | HexNAc(4)Hex(5)Fuc(1)NeuAc(1) | C[+57]*2, N[+2060] | 1369.8927 | 4 | 5476.549 | 5476.5418 | 0 | 1.3 | 457 | Specific | 930.31 | 886.73 | 886.73 | 18.03 | 117 | >sp\|P01833\|PIGR_HUMAN Polymeric immunoglobulin receptor OS=Homo sapiens OX=9606 GN=PIGR PE=1 SV=4 | 902 | 20201116_xd_colostrum_1ng_glyco_hcd203040_compound15.17913.17913.2 | controllerType=0 controllerNumber=1 scan=17913 | 38.2071 |
| 14785:4 | 5 | K.IIEGEPNLKVPGN[+2059.735]VTAVLGETLKVPC[+57.021]HFPC[+57.021]K.F | HexNAc(4)Hex(5)Fuc(1)NeuAc(1) | C[+57]*2, N[+2060] | 1369.8936 | 4 | 5476.5524 | 5476.5418 | 0 | 1.9 | 457 | Specific | 935.5 | 842.7 | 842.7 | 18.03 | 117 | >sp\|P01833\|PIGR_HUMAN Polymeric immunoglobulin receptor OS=Homo sapiens OX=9606 GN=PIGR PE=1 SV=4 | 902 | 20201116_xd_colostrum_1ng_glyco_hcd203040_compound15.17974.17974.4 | controllerType=0 controllerNumber=1 scan=17974 | 38.3419 |
| 14736:5 | 5 | K.IIEGEPNLKVPGN[+2205.793]VTAVLGETLKVPC[+57.021]HFPC[+57.021]K.F | HexNAc(4)Hex(5)Fuc(2)NeuAc(1) | C[+57]*2, N[+2206] | 1125.3284 | 5 | 5622.6128 | 5622.5997 | 0 | 2.3 | 457 | Specific | 958.59 | 879.81 | 879.81 | 19.74 | 117 | >sp\|P01833\|PIGR_HUMAN Polymeric immunoglobulin receptor OS=Homo sapiens OX=9606 GN=PIGR PE=1 SV=4 | 902 | 20201116_xd_colostrum_1ng_glyco_hcd203040_compound15.17923.17923.5 | controllerType=0 controllerNumber=1 scan=17923 | 38.2433 |
| 14858:4 | 5 | K.IIEGEPNLKVPGN[+1856.656]VTAVLGETLKVPC[+57.021]HFPC[+57.021]K.F | HexNAc(3)Hex(5)Fuc(1)NeuAc(1) | C[+57]*2, N[+1857] | 1319.1263 | 4 | 5273.4836 | 5273.4624 | 0 | 4 | 457 | Specific | 997.14 | 864.9 | 864.9 | 18.99 | 117 | >sp\|P01833\|PIGR_HUMAN Polymeric immunoglobulin receptor OS=Homo sapiens OX=9606 GN=PIGR PE=1 SV=4 | 902 | 20201116_xd_colostrum_1ng_glyco_hcd203040_compound15.18049.18049.4 | controllerType=0 controllerNumber=1 scan=18049 | 38.4839 |
| 16068:5 | 5 | K.IIEGEPNLKVPGN[+2350.830]VTAVLGETLKVPC[+57.021]HFPC[+57.021]K.F | HexNAc(4)Hex(5)Fuc(1)NeuAc(2) | C[+57]*2, N[+2351] | 1154.3366 | 5 | 5767.6536 | 5767.6372 | 0 | 2.8 | 457 | Specific | 1002.9 | 802.32 | 802.32 | 20.74 | 117 | >sp\|P01833\|PIGR_HUMAN Polymeric immunoglobulin receptor OS=Homo sapiens OX=9606 GN=PIGR PE=1 SV=4 | 902 | 20201116_xd_colostrum_1ng_glyco_hcd203040_compound15.19310.19310.5 | controllerType=0 controllerNumber=1 scan=19310 | 41.0471 |
| 16067:4 | 5 | K.IIEGEPNLKVPGN[+2350.830]VTAVLGETLKVPC[+57.021]HFPC[+57.021]K.F | HexNAc(4)Hex(5)Fuc(1)NeuAc(2) | C[+57]*2, N[+2351] | 1442.6693 | 4 | 5767.6554 | 5767.6372 | 0 | 3.2 | 457 | Specific | 1003.7 | 766.59 | 766.59 | 19.99 | 117 | >sp\|P01833\|PIGR_HUMAN Polymeric immunoglobulin receptor OS=Homo sapiens OX=9606 GN=PIGR PE=1 SV=4 | 902 | 20201116_xd_colostrum_1ng_glyco_hcd203040_compound15.19309.19309.4 | controllerType=0 controllerNumber=1 scan=19309 | 41.0453 |
| 14687:5 | 5 | K.IIEGEPNLKVPGN[+2351.851]VTAVLGETLKVPC[+57.021]HFPC[+57.021]K.F | HexNAc(4)Hex(5)Fuc(3)NeuAc(1) | C[+57]*2, N[+2352] | 1154.5398 | 5 | 5768.6699 | 5768.6576 | 0 | 2.1 | 457 | Specific | 1028.87 | 1028.87 | 1028.87 | 20.74 | 117 | >sp\|P01833\|PIGR_HUMAN Polymeric immunoglobulin receptor OS=Homo sapiens OX=9606 GN=PIGR PE=1 SV=4 | 902 | 20201116_xd_colostrum_1ng_glyco_hcd203040_compound15.17872.17872.5 | controllerType=0 controllerNumber=1 scan=17872 | 38.1217 |
| 14734:5 | 5 | K.IIEGEPNLKVPGN[+2205.793]VTAVLGETLKVPC[+57.021]HFPC[+57.021]K.F | HexNAc(4)Hex(5)Fuc(2)NeuAc(1) | C[+57]*2, N[+2206] | 1125.3284 | 5 | 5622.6128 | 5622.5997 | 0 | 2.3 | 457 | Specific | 1052.46 | 968.86 | 968.86 | 21.74 | 117 | >sp\|P01833\|PIGR_HUMAN Polymeric immunoglobulin receptor OS=Homo sapiens OX=9606 GN=PIGR PE=1 SV=4 | 902 | 20201116_xd_colostrum_1ng_glyco_hcd203040_compound15.17921.17921.5 | controllerType=0 controllerNumber=1 scan=17921 | 38.2329 |
| 14735:4 | 5 | K.IIEGEPNLKVPGN[+2205.793]VTAVLGETLKVPC[+57.021]HFPC[+57.021]K.F | HexNAc(4)Hex(5)Fuc(2)NeuAc(1) | C[+57]*2, N[+2206] | 1406.4094 | 4 | 5622.6159 | 5622.5997 | 0 | 2.9 | 457 | Specific | 1092.15 | 1063.32 | 1063.32 | 21.18 | 117 | >sp\|P01833\|PIGR_HUMAN Polymeric immunoglobulin receptor OS=Homo sapiens OX=9606 GN=PIGR PE=1 SV=4 | 902 | 20201116_xd_colostrum_1ng_glyco_hcd203040_compound15.17922.17922.4 | controllerType=0 controllerNumber=1 scan=17922 | 38.2343 |
| 14691:4 | 5 | K.IIEGEPNLKVPGN[+2351.851]VTAVLGETLKVPC[+57.021]HFPC[+57.021]K.F | HexNAc(4)Hex(5)Fuc(3)NeuAc(1) | C[+57]*2, N[+2352] | 1442.9243 | 4 | 5768.6754 | 5768.6576 | 0 | 3.1 | 457 | Specific | 1111.91 | 1111.91 | 1111.91 | 21.99 | 117 | >sp\|P01833\|PIGR_HUMAN Polymeric immunoglobulin receptor OS=Homo sapiens OX=9606 GN=PIGR PE=1 SV=4 | 902 | 20201116_xd_colostrum_1ng_glyco_hcd203040_compound15.17876.17876.4 | controllerType=0 controllerNumber=1 scan=17876 | 38.1321 |
| 13036:4 | 5 | K.VPGN[+2205.793]VTAVLGETLKVPC[+57.021]HFPC[+57.021]K.F | HexNAc(4)Hex(5)Fuc(2)NeuAc(1) | C[+57]*2, N[+2206] | 1158.0251 | 4 | 4629.0784 | 4629.0502 | 0 | 6.1 | 466 | Specific | 452.34 | 452.34 | 452.34 | 9.27 | 117 | >sp\|P01833\|PIGR_HUMAN Polymeric immunoglobulin receptor OS=Homo sapiens OX=9606 GN=PIGR PE=1 SV=4 | 902 | 20201116_xd_colostrum_1ng_glyco_hcd203040_compound15.16155.16155.2 | controllerType=0 controllerNumber=1 scan=16155 | 34.6077 |
| 13039:4 | 5 | K.VPGN[+2205.793]VTAVLGETLKVPC[+57.021]HFPC[+57.021]K.F | HexNAc(4)Hex(5)Fuc(2)NeuAc(1) | C[+57]*2, N[+2206] | 1158.0251 | 4 | 4629.0784 | 4629.0502 | 0 | 6.1 | 466 | Specific | 653.17 | 575.53 | 575.53 | 13.33 | 117 | >sp\|P01833\|PIGR_HUMAN Polymeric immunoglobulin receptor OS=Homo sapiens OX=9606 GN=PIGR PE=1 SV=4 | 902 | 20201116_xd_colostrum_1ng_glyco_hcd203040_compound15.16158.16158.2 | controllerType=0 controllerNumber=1 scan=16158 | 34.6132 |
| 12925:4 | 5 | K.VPGN[+2351.851]VTAVLGETLKVPC[+57.021]HFPC[+57.021]K.F | HexNAc(4)Hex(5)Fuc(3)NeuAc(1) | C[+57]*2, N[+2352] | 1194.5353 | 4 | 4775.1195 | 4775.1081 | 0 | 2.4 | 466 | Specific | 833.23 | 512.46 | 512.46 | 17.31 | 117 | >sp\|P01833\|PIGR_HUMAN Polymeric immunoglobulin receptor OS=Homo sapiens OX=9606 GN=PIGR PE=1 SV=4 | 902 | 20201116_xd_colostrum_1ng_glyco_hcd203040_compound15.16040.16040.4 | controllerType=0 controllerNumber=1 scan=16040 | 34.386 |
| 12922:4 | 5 | K.VPGN[+2351.851]VTAVLGETLKVPC[+57.021]HFPC[+57.021]K.F | HexNAc(4)Hex(5)Fuc(3)NeuAc(1) | C[+57]*2, N[+2352] | 1194.5353 | 4 | 4775.1195 | 4775.1081 | 0 | 2.4 | 466 | Specific | 898.83 | 877.42 | 877.42 | 18.31 | 117 | >sp\|P01833\|PIGR_HUMAN Polymeric immunoglobulin receptor OS=Homo sapiens OX=9606 GN=PIGR PE=1 SV=4 | 902 | 20201116_xd_colostrum_1ng_glyco_hcd203040_compound15.16037.16037.4 | controllerType=0 controllerNumber=1 scan=16037 | 34.3802 |
| 07689:4 | 5 | K.YWC[+57.021]KWN[+2205.793]NTGC[+57.021]QALPSQDEGPSK.A | HexNAc(4)Hex(5)Fuc(2)NeuAc(1) | C[+57]*2, N[+2206] | 1208.7419 | 4 | 4831.9456 | 4831.9378 | 0 | 1.6 | 494 | Specific | 370.72 | 370.72 | 370.72 | 8.44 | 117 | >sp\|P01833\|PIGR_HUMAN Polymeric immunoglobulin receptor OS=Homo sapiens OX=9606 GN=PIGR PE=1 SV=4 | 902 | 20201116_xd_colostrum_1ng_glyco_hcd203040_compound15.10601.10601.4 | controllerType=0 controllerNumber=1 scan=10601 | 23.9594 |
| 07687:4 | 5 | K.YWC[+57.021]KWN[+2205.793]NTGC[+57.021]QALPSQDEGPSK.A | HexNAc(4)Hex(5)Fuc(2)NeuAc(1) | C[+57]*2, N[+2206] | 1208.7419 | 4 | 4831.9456 | 4831.9378 | 0 | 1.6 | 494 | Specific | 438.79 | 336.64 | 336.64 | 8.37 | 117 | >sp\|P01833\|PIGR_HUMAN Polymeric immunoglobulin receptor OS=Homo sapiens OX=9606 GN=PIGR PE=1 SV=4 | 902 | 20201116_xd_colostrum_1ng_glyco_hcd203040_compound15.10599.10599.4 | controllerType=0 controllerNumber=1 scan=10599 | 23.9558 |
| 07861:3 | 5 | K.YWC[+57.021]KWN[+1710.598]NTGC[+57.021]QALPSQDEGPSK.A | HexNAc(3)Hex(5)NeuAc(1) | C[+57]*2, N[+1711] | 1446.2543 | 3 | 4336.7484 | 4336.7426 | 0 | 1.3 | 494 | Specific | 640.81 | 614.66 | 614.66 | 13.16 | 117 | >sp\|P01833\|PIGR_HUMAN Polymeric immunoglobulin receptor OS=Homo sapiens OX=9606 GN=PIGR PE=1 SV=4 | 902 | 20201116_xd_colostrum_1ng_glyco_hcd203040_compound15.10779.10779.3 | controllerType=0 controllerNumber=1 scan=10779 | 24.2766 |
| 07863:3 | 5 | K.YWC[+57.021]KWN[+1710.598]NTGC[+57.021]QALPSQDEGPSK.A | HexNAc(3)Hex(5)NeuAc(1) | C[+57]*2, N[+1711] | 1446.2543 | 3 | 4336.7484 | 4336.7426 | 0 | 1.3 | 494 | Specific | 683.93 | 658.06 | 658.06 | 14.16 | 117 | >sp\|P01833\|PIGR_HUMAN Polymeric immunoglobulin receptor OS=Homo sapiens OX=9606 GN=PIGR PE=1 SV=4 | 902 | 20201116_xd_colostrum_1ng_glyco_hcd203040_compound15.10781.10781.3 | controllerType=0 controllerNumber=1 scan=10781 | 24.2804 |
| 07791:3 | 5 | K.YWC[+57.021]KWN[+1872.651]NTGC[+57.021]QALPSQDEGPSK.A | HexNAc(3)Hex(6)NeuAc(1) | C[+57]*2, N[+1873] | 1500.2757 | 3 | 4498.8125 | 4498.7954 | 0 | 3.8 | 494 | Specific | 739.98 | 727.04 | 727.04 | 15.12 | 117 | >sp\|P01833\|PIGR_HUMAN Polymeric immunoglobulin receptor OS=Homo sapiens OX=9606 GN=PIGR PE=1 SV=4 | 902 | 20201116_xd_colostrum_1ng_glyco_hcd203040_compound15.10707.10707.3 | controllerType=0 controllerNumber=1 scan=10707 | 24.1455 |
| 07794:3 | 5 | K.YWC[+57.021]KWN[+1872.651]NTGC[+57.021]QALPSQDEGPSK.A | HexNAc(3)Hex(6)NeuAc(1) | C[+57]*2, N[+1873] | 1500.2757 | 3 | 4498.8125 | 4498.7954 | 0 | 3.8 | 494 | Specific | 810.19 | 597.4 | 597.4 | 17.12 | 117 | >sp\|P01833\|PIGR_HUMAN Polymeric immunoglobulin receptor OS=Homo sapiens OX=9606 GN=PIGR PE=1 SV=4 | 902 | 20201116_xd_colostrum_1ng_glyco_hcd203040_compound15.10710.10710.3 | controllerType=0 controllerNumber=1 scan=10710 | 24.1493 |
| 06121:3 | 5 | K.WN[+1548.545]NTGC[+57.021]QALPSQDEGPSK.A | HexNAc(3)Hex(4)NeuAc(1) | C[+57], N[+1549] | 1179.8151 | 3 | 3537.4308 | 3537.4215 | 0 | 2.6 | 498 | Specific | 207.68 | 178.54 | 178.54 | 5.03 | 117 | >sp\|P01833\|PIGR_HUMAN Polymeric immunoglobulin receptor OS=Homo sapiens OX=9606 GN=PIGR PE=1 SV=4 | 902 | 20201116_xd_colostrum_1ng_glyco_hcd203040_compound15.8975.8975.3 | controllerType=0 controllerNumber=1 scan=8975 | 21.1034 |
| 06107:3 | 5 | K.WN[+1710.598]NTGC[+57.021]QALPSQDEGPSK.A | HexNAc(3)Hex(5)NeuAc(1) | C[+57], N[+1711] | 1233.8307 | 3 | 3699.4777 | 3699.4743 | 0 | 0.9 | 498 | Specific | 263.88 | 245.11 | 245.11 | 6.6 | 117 | >sp\|P01833\|PIGR_HUMAN Polymeric immunoglobulin receptor OS=Homo sapiens OX=9606 GN=PIGR PE=1 SV=4 | 902 | 20201116_xd_colostrum_1ng_glyco_hcd203040_compound15.8960.8960.3 | controllerType=0 controllerNumber=1 scan=8960 | 21.075 |
| 07911:3 | 5 | K.WN[+2204.772]NTGC[+57.021]QALPSQDEGPSK.A | HexNAc(4)Hex(5)NeuAc(2) | C[+57], N[+2205] | 1398.5592 | 3 | 4193.663 | 4193.6491 | 0 | 3.3 | 498 | Specific | 270.56 | 220.18 | 220.18 | 6.42 | 117 | >sp\|P01833\|PIGR_HUMAN Polymeric immunoglobulin receptor OS=Homo sapiens OX=9606 GN=PIGR PE=1 SV=4 | 902 | 20201116_xd_colostrum_1ng_glyco_hcd203040_compound15.10831.10831.3 | controllerType=0 controllerNumber=1 scan=10831 | 24.3791 |
| 06123:3 | 5 | K.WN[+1548.545]NTGC[+57.021]QALPSQDEGPSK.A | HexNAc(3)Hex(4)NeuAc(1) | C[+57], N[+1549] | 1179.8151 | 3 | 3537.4308 | 3537.4215 | 0 | 2.6 | 498 | Specific | 327.83 | 280.29 | 280.29 | 7.4 | 117 | >sp\|P01833\|PIGR_HUMAN Polymeric immunoglobulin receptor OS=Homo sapiens OX=9606 GN=PIGR PE=1 SV=4 | 902 | 20201116_xd_colostrum_1ng_glyco_hcd203040_compound15.8977.8977.3 | controllerType=0 controllerNumber=1 scan=8977 | 21.1074 |
| 06081:3 | 5 | K.WN[+1548.545]NTGC[+57.021]QALPSQDEGPSK.A | HexNAc(3)Hex(4)NeuAc(1) | C[+57], N[+1549] | 1179.8148 | 3 | 3537.4297 | 3537.4215 | 0 | 2.3 | 498 | Specific | 530.35 | 463.67 | 463.67 | 10.32 | 117 | >sp\|P01833\|PIGR_HUMAN Polymeric immunoglobulin receptor OS=Homo sapiens OX=9606 GN=PIGR PE=1 SV=4 | 902 | 20201116_xd_colostrum_1ng_glyco_hcd203040_compound15.8934.8934.3 | controllerType=0 controllerNumber=1 scan=8934 | 21.0332 |
| 05937:3 | 5 | K.WN[+1913.677]NTGC[+57.021]QALPSQDEGPSK.A | HexNAc(4)Hex(5)NeuAc(1) | C[+57], N[+1914] | 1301.5262 | 3 | 3902.564 | 3902.5537 | 0 | 2.6 | 498 | Specific | 589.2 | 516.24 | 516.24 | 12.05 | 117 | >sp\|P01833\|PIGR_HUMAN Polymeric immunoglobulin receptor OS=Homo sapiens OX=9606 GN=PIGR PE=1 SV=4 | 902 | 20201116_xd_colostrum_1ng_glyco_hcd203040_compound15.8785.8785.3 | controllerType=0 controllerNumber=1 scan=8785 | 20.7739 |
| 06084:3 | 5 | K.WN[+1548.545]NTGC[+57.021]QALPSQDEGPSK.A | HexNAc(3)Hex(4)NeuAc(1) | C[+57], N[+1549] | 1179.8148 | 3 | 3537.4297 | 3537.4215 | 0 | 2.3 | 498 | Specific | 591.26 | 536.08 | 536.08 | 12.05 | 117 | >sp\|P01833\|PIGR_HUMAN Polymeric immunoglobulin receptor OS=Homo sapiens OX=9606 GN=PIGR PE=1 SV=4 | 902 | 20201116_xd_colostrum_1ng_glyco_hcd203040_compound15.8937.8937.3 | controllerType=0 controllerNumber=1 scan=8937 | 21.0374 |
| 05939:3 | 5 | K.WN[+1913.677]NTGC[+57.021]QALPSQDEGPSK.A | HexNAc(4)Hex(5)NeuAc(1) | C[+57], N[+1914] | 1301.5262 | 3 | 3902.564 | 3902.5537 | 0 | 2.6 | 498 | Specific | 670.45 | 598.4 | 598.4 | 14.05 | 117 | >sp\|P01833\|PIGR_HUMAN Polymeric immunoglobulin receptor OS=Homo sapiens OX=9606 GN=PIGR PE=1 SV=4 | 902 | 20201116_xd_colostrum_1ng_glyco_hcd203040_compound15.8787.8787.3 | controllerType=0 controllerNumber=1 scan=8787 | 20.7775 |
| 05830:3 | 5 | K.WN[+2059.735]NTGC[+57.021]QALPSQDEGPSK.A | HexNAc(4)Hex(5)Fuc(1)NeuAc(1) | C[+57], N[+2060] | 1350.2115 | 3 | 4048.6199 | 4048.6116 | 0 | 2 | 498 | Specific | 737.74 | 660.72 | 660.72 | 15.05 | 117 | >sp\|P01833\|PIGR_HUMAN Polymeric immunoglobulin receptor OS=Homo sapiens OX=9606 GN=PIGR PE=1 SV=4 | 902 | 20201116_xd_colostrum_1ng_glyco_hcd203040_compound15.8674.8674.3 | controllerType=0 controllerNumber=1 scan=8674 | 20.5761 |
| 05832:3 | 5 | K.WN[+2059.735]NTGC[+57.021]QALPSQDEGPSK.A | HexNAc(4)Hex(5)Fuc(1)NeuAc(1) | C[+57], N[+2060] | 1350.2115 | 3 | 4048.6199 | 4048.6116 | 0 | 2 | 498 | Specific | 759.48 | 684.88 | 684.88 | 16.05 | 117 | >sp\|P01833\|PIGR_HUMAN Polymeric immunoglobulin receptor OS=Homo sapiens OX=9606 GN=PIGR PE=1 SV=4 | 902 | 20201116_xd_colostrum_1ng_glyco_hcd203040_compound15.8676.8676.3 | controllerType=0 controllerNumber=1 scan=8676 | 20.5802 |
| 05796:3 | 5 | K.WN[+2205.793]NTGC[+57.021]QALPSQDEGPSK.A | HexNAc(4)Hex(5)Fuc(2)NeuAc(1) | C[+57], N[+2206] | 1398.8965 | 3 | 4194.6751 | 4194.6695 | 0 | 1.3 | 498 | Specific | 801.81 | 757.89 | 757.89 | 16.9 | 117 | >sp\|P01833\|PIGR_HUMAN Polymeric immunoglobulin receptor OS=Homo sapiens OX=9606 GN=PIGR PE=1 SV=4 | 902 | 20201116_xd_colostrum_1ng_glyco_hcd203040_compound15.8639.8639.3 | controllerType=0 controllerNumber=1 scan=8639 | 20.519 |
| 05879:3 | 5 | K.WN[+1872.651]NTGC[+57.021]QALPSQDEGPSK.A | HexNAc(3)Hex(6)NeuAc(1) | C[+57], N[+1873] | 1287.8503 | 3 | 3861.5363 | 3861.5272 | 0 | 2.4 | 498 | Specific | 821.53 | 739.21 | 739.21 | 17.05 | 117 | >sp\|P01833\|PIGR_HUMAN Polymeric immunoglobulin receptor OS=Homo sapiens OX=9606 GN=PIGR PE=1 SV=4 | 902 | 20201116_xd_colostrum_1ng_glyco_hcd203040_compound15.8724.8724.3 | controllerType=0 controllerNumber=1 scan=8724 | 20.6639 |
| 05970:3 | 5 | K.WN[+1710.598]NTGC[+57.021]QALPSQDEGPSK.A | HexNAc(3)Hex(5)NeuAc(1) | C[+57], N[+1711] | 1233.8322 | 3 | 3699.4821 | 3699.4743 | 0 | 2.1 | 498 | Specific | 875.2 | 770.11 | 770.11 | 18.05 | 117 | >sp\|P01833\|PIGR_HUMAN Polymeric immunoglobulin receptor OS=Homo sapiens OX=9606 GN=PIGR PE=1 SV=4 | 902 | 20201116_xd_colostrum_1ng_glyco_hcd203040_compound15.8819.8819.3 | controllerType=0 controllerNumber=1 scan=8819 | 20.8319 |
| 05794:3 | 5 | K.WN[+2205.793]NTGC[+57.021]QALPSQDEGPSK.A | HexNAc(4)Hex(5)Fuc(2)NeuAc(1) | C[+57], N[+2206] | 1398.8965 | 3 | 4194.6751 | 4194.6695 | 0 | 1.3 | 498 | Specific | 894.67 | 843.33 | 843.33 | 17.9 | 117 | >sp\|P01833\|PIGR_HUMAN Polymeric immunoglobulin receptor OS=Homo sapiens OX=9606 GN=PIGR PE=1 SV=4 | 902 | 20201116_xd_colostrum_1ng_glyco_hcd203040_compound15.8637.8637.3 | controllerType=0 controllerNumber=1 scan=8637 | 20.5142 |
| 05967:3 | 5 | K.WN[+1710.598]NTGC[+57.021]QALPSQDEGPSK.A | HexNAc(3)Hex(5)NeuAc(1) | C[+57], N[+1711] | 1233.8322 | 3 | 3699.4821 | 3699.4743 | 0 | 2.1 | 498 | Specific | 914.53 | 838.37 | 838.37 | 19.05 | 117 | >sp\|P01833\|PIGR_HUMAN Polymeric immunoglobulin receptor OS=Homo sapiens OX=9606 GN=PIGR PE=1 SV=4 | 902 | 20201116_xd_colostrum_1ng_glyco_hcd203040_compound15.8816.8816.3 | controllerType=0 controllerNumber=1 scan=8816 | 20.8273 |
| 05877:3 | 5 | K.WN[+1872.651]NTGC[+57.021]QALPSQDEGPSK.A | HexNAc(3)Hex(6)NeuAc(1) | C[+57], N[+1873] | 1287.8503 | 3 | 3861.5363 | 3861.5272 | 0 | 2.4 | 498 | Specific | 999.62 | 950.21 | 950.21 | 20.05 | 117 | >sp\|P01833\|PIGR_HUMAN Polymeric immunoglobulin receptor OS=Homo sapiens OX=9606 GN=PIGR PE=1 SV=4 | 902 | 20201116_xd_colostrum_1ng_glyco_hcd203040_compound15.8722.8722.3 | controllerType=0 controllerNumber=1 scan=8722 | 20.659 |
| 22502:5 | 9 | R.QLAHQSN[+2350.830]STNIFFSPVSIATAFAMLSLGTK.A | HexNAc(4)Hex(5)Fuc(1)NeuAc(2) | N[+2351] | 1107.3022 | 5 | 5532.4821 | 5532.4654 | 0 | 3 | 64 | Specific | 325.16 | 325.16 | 325.16 | 8.56 | 48 | >sp\|P01009\|A1AT_HUMAN Alpha-1-antitrypsin OS=Homo sapiens OX=9606 GN=SERPINA1 PE=1 SV=3 | 642 | 20201116_xd_colostrum_1ng_glyco_hcd203040_compound15.26034.26034.5 | controllerType=0 controllerNumber=1 scan=26034 | 55.7702 |
| 25217:4 | 9 | R.QLAHQSN[+2861.000]STNIFFSPVSIATAFAMLSLGTK.A | HexNAc(5)Hex(6)NeuAc(3) | N[+2861] | 1511.4215 | 4 | 6042.6642 | 6042.6351 | 0 | 4.8 | 64 | Specific | 381.47 | 327.67 | 327.67 | 8.06 | 48 | >sp\|P01009\|A1AT_HUMAN Alpha-1-antitrypsin OS=Homo sapiens OX=9606 GN=SERPINA1 PE=1 SV=3 | 642 | 20201116_xd_colostrum_1ng_glyco_hcd203040_compound15.28858.28858.4 | controllerType=0 controllerNumber=1 scan=28858 | 61.0764 |
| 25219:4 | 9 | R.QLAHQSN[+2861.000]STNIFFSPVSIATAFAMLSLGTK.A | HexNAc(5)Hex(6)NeuAc(3) | N[+2861] | 1511.4215 | 4 | 6042.6642 | 6042.6351 | 0 | 4.8 | 64 | Specific | 422.22 | 392.77 | 392.77 | 9.93 | 48 | >sp\|P01009\|A1AT_HUMAN Alpha-1-antitrypsin OS=Homo sapiens OX=9606 GN=SERPINA1 PE=1 SV=3 | 642 | 20201116_xd_colostrum_1ng_glyco_hcd203040_compound15.28860.28860.4 | controllerType=0 controllerNumber=1 scan=28860 | 61.0805 |
| 22504:5 | 9 | R.QLAHQSN[+2350.830]STNIFFSPVSIATAFAMLSLGTK.A | HexNAc(4)Hex(5)Fuc(1)NeuAc(2) | N[+2351] | 1107.3022 | 5 | 5532.4821 | 5532.4654 | 0 | 3 | 64 | Specific | 458.84 | 425.77 | 425.77 | 10.72 | 48 | >sp\|P01009\|A1AT_HUMAN Alpha-1-antitrypsin OS=Homo sapiens OX=9606 GN=SERPINA1 PE=1 SV=3 | 642 | 20201116_xd_colostrum_1ng_glyco_hcd203040_compound15.26036.26036.5 | controllerType=0 controllerNumber=1 scan=26036 | 55.774 |
| 25245:4 | 9 | R.QLAHQSN[+2861.000]STNIFFSPVSIATAFAMLSLGTK.A | HexNAc(5)Hex(6)NeuAc(3) | N[+2861] | 1511.4213 | 4 | 6042.6632 | 6042.6351 | 0 | 4.7 | 64 | Specific | 483.99 | 443.94 | 443.94 | 10.16 | 48 | >sp\|P01009\|A1AT_HUMAN Alpha-1-antitrypsin OS=Homo sapiens OX=9606 GN=SERPINA1 PE=1 SV=3 | 642 | 20201116_xd_colostrum_1ng_glyco_hcd203040_compound15.28887.28887.4 | controllerType=0 controllerNumber=1 scan=28887 | 61.1307 |
| 25253:4 | 9 | R.QLAHQSN[+2861.000]STNIFFSPVSIATAFAMLSLGTK.A | HexNAc(5)Hex(6)NeuAc(3) | N[+2861] | 1511.418 | 4 | 6042.6501 | 6042.6351 | 0 | 2.5 | 64 | Specific | 493.62 | 471.02 | 471.02 | 10.59 | 48 | >sp\|P01009\|A1AT_HUMAN Alpha-1-antitrypsin OS=Homo sapiens OX=9606 GN=SERPINA1 PE=1 SV=3 | 642 | 20201116_xd_colostrum_1ng_glyco_hcd203040_compound15.28896.28896.4 | controllerType=0 controllerNumber=1 scan=28896 | 61.1479 |
| 20752:4 | 9 | R.QLAHQSN[+1913.677]STNIFFSPVSIATAFAMLSLGTK.A | HexNAc(4)Hex(5)NeuAc(1) | N[+1914] | 1274.5861 | 4 | 5095.3224 | 5095.3121 | 0 | 2 | 64 | Specific | 511.23 | 511.23 | 511.23 | 10.56 | 48 | >sp\|P01009\|A1AT_HUMAN Alpha-1-antitrypsin OS=Homo sapiens OX=9606 GN=SERPINA1 PE=1 SV=3 | 642 | 20201116_xd_colostrum_1ng_glyco_hcd203040_compound15.24206.24206.4 | controllerType=0 controllerNumber=1 scan=24206 | 51.9033 |
| 20754:4 | 9 | R.QLAHQSN[+1913.677]STNIFFSPVSIATAFAMLSLGTK.A | HexNAc(4)Hex(5)NeuAc(1) | N[+1914] | 1274.5861 | 4 | 5095.3224 | 5095.3121 | 0 | 2 | 64 | Specific | 533.95 | 487.79 | 487.79 | 10.61 | 48 | >sp\|P01009\|A1AT_HUMAN Alpha-1-antitrypsin OS=Homo sapiens OX=9606 GN=SERPINA1 PE=1 SV=3 | 642 | 20201116_xd_colostrum_1ng_glyco_hcd203040_compound15.24208.24208.4 | controllerType=0 controllerNumber=1 scan=24208 | 51.9071 |
| 22490:3 | 9 | R.QLAHQSN[+2350.830]STNIFFSPVSIATAFAMLSLGTK.A | HexNAc(4)Hex(5)Fuc(1)NeuAc(2) | N[+2351] | 1844.8342 | 3 | 5532.488 | 5532.4654 | 0 | 4.1 | 64 | Specific | 554.46 | 495.82 | 495.82 | 10.47 | 48 | >sp\|P01009\|A1AT_HUMAN Alpha-1-antitrypsin OS=Homo sapiens OX=9606 GN=SERPINA1 PE=1 SV=3 | 642 | 20201116_xd_colostrum_1ng_glyco_hcd203040_compound15.26022.26022.3 | controllerType=0 controllerNumber=1 scan=26022 | 55.7456 |
| 22492:3 | 9 | R.QLAHQSN[+2350.830]STNIFFSPVSIATAFAMLSLGTK.A | HexNAc(4)Hex(5)Fuc(1)NeuAc(2) | N[+2351] | 1844.8342 | 3 | 5532.488 | 5532.4654 | 0 | 4.1 | 64 | Specific | 584.24 | 534.71 | 534.71 | 11.97 | 48 | >sp\|P01009\|A1AT_HUMAN Alpha-1-antitrypsin OS=Homo sapiens OX=9606 GN=SERPINA1 PE=1 SV=3 | 642 | 20201116_xd_colostrum_1ng_glyco_hcd203040_compound15.26024.26024.3 | controllerType=0 controllerNumber=1 scan=26024 | 55.7494 |
| 25247:4 | 9 | R.QLAHQSN[+2861.000]STNIFFSPVSIATAFAMLSLGTK.A | HexNAc(5)Hex(6)NeuAc(3) | N[+2861] | 1511.4213 | 4 | 6042.6632 | 6042.6351 | 0 | 4.7 | 64 | Specific | 597.86 | 557.18 | 557.18 | 12.06 | 48 | >sp\|P01009\|A1AT_HUMAN Alpha-1-antitrypsin OS=Homo sapiens OX=9606 GN=SERPINA1 PE=1 SV=3 | 642 | 20201116_xd_colostrum_1ng_glyco_hcd203040_compound15.28889.28889.4 | controllerType=0 controllerNumber=1 scan=28889 | 61.1346 |
| 22506:5 | 9 | R.QLAHQSN[+2204.772]STNIFFSPVSIATAFAMLSLGTK.A | HexNAc(4)Hex(5)NeuAc(2) | N[+2205] | 1078.0929 | 5 | 5386.4354 | 5386.4075 | 0 | 5.2 | 64 | Specific | 607.37 | 576.58 | 576.58 | 12.94 | 48 | >sp\|P01009\|A1AT_HUMAN Alpha-1-antitrypsin OS=Homo sapiens OX=9606 GN=SERPINA1 PE=1 SV=3 | 642 | 20201116_xd_colostrum_1ng_glyco_hcd203040_compound15.26038.26038.5 | controllerType=0 controllerNumber=1 scan=26038 | 55.7774 |
| 25255:4 | 9 | R.QLAHQSN[+2861.000]STNIFFSPVSIATAFAMLSLGTK.A | HexNAc(5)Hex(6)NeuAc(3) | N[+2861] | 1511.418 | 4 | 6042.6501 | 6042.6351 | 0 | 2.5 | 64 | Specific | 642.45 | 567.21 | 567.21 | 13.49 | 48 | >sp\|P01009\|A1AT_HUMAN Alpha-1-antitrypsin OS=Homo sapiens OX=9606 GN=SERPINA1 PE=1 SV=3 | 642 | 20201116_xd_colostrum_1ng_glyco_hcd203040_compound15.28898.28898.4 | controllerType=0 controllerNumber=1 scan=28898 | 61.1525 |
| 22491:3 | 9 | R.QLAHQSN[+2204.772]STNIFFSPVSIATAFAMLSLGTK.A | HexNAc(4)Hex(5)NeuAc(2) | N[+2205] | 1796.1501 | 3 | 5386.4358 | 5386.4075 | 0 | 5.3 | 64 | Specific | 703.44 | 668.93 | 668.93 | 14.57 | 48 | >sp\|P01009\|A1AT_HUMAN Alpha-1-antitrypsin OS=Homo sapiens OX=9606 GN=SERPINA1 PE=1 SV=3 | 642 | 20201116_xd_colostrum_1ng_glyco_hcd203040_compound15.26023.26023.3 | controllerType=0 controllerNumber=1 scan=26023 | [illegible] |
| 22493:3 | 9 | R.QLAHQSN[+2204.772]STNIFFSPVSIATAFAMLSLGTK.A | HexNAc(4)Hex(5)NeuAc(2) | N[+2205] | 1796.1501 | 3 | 5386.4358 | 5386.4075 | 0 | 5.3 | 64 | Specific | 729.14 | 698.64 | 698.64 | 14.57 | 48 | >sp\|P01009\|A1AT_HUMAN Alpha-1-antitrypsin OS=Homo sapiens OX=9606 GN=SERPINA1 PE=1 SV=3 | 642 | 20201116_xd_colostrum_1ng_glyco_hcd203040_compound15.26025.26025.3 | controllerType=0 controllerNumber=1 scan=26025 | [illegible] |
| 22508:5 | 9 | R.QLAHQSN[+2204.772]STNIFFSPVSIATAFAMLSLGTK.A | HexNAc(4)Hex(5)NeuAc(2) | N[+2205] | 1078.0929 | 5 | 5386.4354 | 5386.4075 | 0 | 5.2 | 64 | Specific | 802.66 | 701.78 | 701.78 | 16.94 | 48 | >sp\|P01009\|A1AT_HUMAN Alpha-1-antitrypsin OS=Homo sapiens OX=9606 GN=SERPINA1 PE=1 SV=3 | 642 | 20201116_xd_colostrum_1ng_glyco_hcd203040_compound15.26040.26040.5 | controllerType=0 controllerNumber=1 scan=26040 | [illegible] |
| 22486:4 | 9 | R.QLAHQSN[+2350.830]STNIFFSPVSIATAFAMLSLGTK.A | HexNAc(4)Hex(5)Fuc(1)NeuAc(2) | N[+2351] | 1383.8787 | 4 | 5532.4928 | 5532.4654 | 0 | 5 | 64 | Specific | 880.51 | 826.66 | 826.66 | 18.06 | 48 | >sp\|P01009\|A1AT_HUMAN Alpha-1-antitrypsin OS=Homo sapiens OX=9606 GN=SERPINA1 PE=1 SV=3 | 642 | 20201116_xd_colostrum_1ng_glyco_hcd203040_compound15.26018.26018.4 | controllerType=0 controllerNumber=1 scan=26018 | [illegible] |
| 22488:4 | 9 | R.QLAHQSN[+2350.830]STNIFFSPVSIATAFAMLSLGTK.A | HexNAc(4)Hex(5)Fuc(1)NeuAc(2) | N[+2351] | 1383.8787 | 4 | 5532.4928 | 5532.4654 | 0 | 5 | 64 | Specific | 881.76 | 755.7 | 755.7 | 18.06 | 48 | >sp\|P01009\|A1AT_HUMAN Alpha-1-antitrypsin OS=Homo sapiens OX=9606 GN=SERPINA1 PE=1 SV=3 | 642 | 20201116_xd_colostrum_1ng_glyco_hcd203040_compound15.26020.26020.4 | controllerType=0 controllerNumber=1 scan=26020 | 55.7402 |
| 22489:4 | 9 | R.QLAHQSN[+2204.772]STNIFFSPVSIATAFAMLSLGTK.A | HexNAc(4)Hex(5)NeuAc(2) | N[+2205] | 1347.364 | 4 | 5386.4342 | 5386.4075 | 0 | 5 | 64 | Specific | 896.29 | 889.29 | 889.29 | 18.06 | 48 | >sp\|P01009\|A1AT_HUMAN Alpha-1-antitrypsin OS=Homo sapiens OX=9606 GN=SERPINA1 PE=1 SV=3 | 642 | 20201116_xd_colostrum_1ng_glyco_hcd203040_compound15.26021.26021.4 | controllerType=0 controllerNumber=1 scan=26021 | 55.7437 |
| 22487:4 | 9 | R.QLAHQSN[+2204.772]STNIFFSPVSIATAFAMLSLGTK.A | HexNAc(4)Hex(5)NeuAc(2) | N[+2205] | 1347.364 | 4 | 5386.4342 | 5386.4075 | 0 | 5 | 64 | Specific | 906.26 | 870.34 | 870.34 | 19.06 | 48 | >sp\|P01009\|A1AT_HUMAN Alpha-1-antitrypsin OS=Homo sapiens OX=9606 GN=SERPINA1 PE=1 SV=3 | 642 | 20201116_xd_colostrum_1ng_glyco_hcd203040_compound15.26019.26019.4 | controllerType=0 controllerNumber=1 scan=26019 | 55.735 |
| 21656:4 | 9 | R.QLAHQSN[+2204.772]STNIFFSPVSIATAFAM[+15.995]LSLGTK.A | HexNAc(4)Hex(5)NeuAc(2) | M[+16], N[+2205] | 1351.3615 | 4 | 5402.424 | 5402.4024 | 0 | 4 | 64 | Specific | 550.24 | 550.24 | 550.24 | 10.8 | 48 | >sp\|P01009\|A1AT_HUMAN Alpha-1-antitrypsin OS=Homo sapiens OX=9606 GN=SERPINA1 PE=1 SV=3 | 642 | 20201116_xd_colostrum_1ng_glyco_hcd203040_compound15.25151.25151.4 | controllerType=0 controllerNumber=1 scan=25151 | 53.9605 |
| 21658:4 | 9 | R.QLAHQSN[+2204.772]STNIFFSPVSIATAFAM[+15.995]LSLGTK.A | HexNAc(4)Hex(5)NeuAc(2) | M[+16], N[+2205] | 1351.3615 | 4 | 5402.424 | 5402.4024 | 0 | 4 | 64 | Specific | 698.42 | 597.49 | 597.49 | 14.29 | 48 | >sp\|P01009\|A1AT_HUMAN Alpha-1-antitrypsin OS=Homo sapiens OX=9606 GN=SERPINA1 PE=1 SV=3 | 642 | 20201116_xd_colostrum_1ng_glyco_hcd203040_compound15.25153.25153.4 | controllerType=0 controllerNumber=1 scan=25153 | 53.9655 |
| 19878:4 | 9 | K.ADTHDEILEGLNFN[+2569.905]LTEIPEAQIHEGFQELLR.T | HexNAc(5)Hex(6)NeuAc(2) | N[+2570] | 1566.1869 | 4 | 6261.7257 | 6261.7285 | 0 | -0.4 | 94 | Specific | 354.16 | 354.16 | 354.16 | 7.24 | 48 | >sp\|P01009\|A1AT_HUMAN Alpha-1-antitrypsin OS=Homo sapiens OX=9606 GN=SERPINA1 PE=1 SV=3 | 642 | 20201116_xd_colostrum_1ng_glyco_hcd203040_compound15.23293.23293.4 | controllerType=0 controllerNumber=1 scan=23293 | 49.9358 |
| 19883:4 | 9 | K.ADTHDEILEGLNFN[+2350.830]LTEIPEAQIHEGFQELLR.T | HexNAc(4)Hex(5)Fuc(1)NeuAc(2) | N[+2351] | 1511.4192 | 4 | 6042.6549 | 6042.6542 | 0 | 0.1 | 94 | Specific | 520.61 | 417.52 | 417.52 | 9.99 | 48 | >sp\|P01009\|A1AT_HUMAN Alpha-1-antitrypsin OS=Homo sapiens OX=9606 GN=SERPINA1 PE=1 SV=3 | 642 | 20201116_xd_colostrum_1ng_glyco_hcd203040_compound15.23298.23298.4 | controllerType=0 controllerNumber=1 scan=23298 | 49.9452 |
| 19885:4 | 9 | K.ADTHDEILEGLNFN[+2350.830]LTEIPEAQIHEGFQELLR.T | HexNAc(4)Hex(5)Fuc(1)NeuAc(2) | N[+2351] | 1511.4192 | 4 | 6042.6549 | 6042.6542 | 0 | 0.1 | 94 | Specific | 537.4 | 497.62 | 497.62 | 9.99 | 48 | >sp\|P01009\|A1AT_HUMAN Alpha-1-antitrypsin OS=Homo sapiens OX=9606 GN=SERPINA1 PE=1 SV=3 | 642 | 20201116_xd_colostrum_1ng_glyco_hcd203040_compound15.23300.23300.4 | controllerType=0 controllerNumber=1 scan=23300 | 49.9503 |
| 19880:4 | 9 | K.ADTHDEILEGLNFN[+2569.905]LTEIPEAQIHEGFQELLR.T | HexNAc(5)Hex(6)NeuAc(2) | N[+2570] | 1566.1869 | 4 | 6261.7257 | 6261.7285 | 0 | -0.4 | 94 | Specific | 611.74 | 577.28 | 577.28 | 12.29 | 48 | >sp\|P01009\|A1AT_HUMAN Alpha-1-antitrypsin OS=Homo sapiens OX=9606 GN=SERPINA1 PE=1 SV=3 | 642 | 20201116_xd_colostrum_1ng_glyco_hcd203040_compound15.23295.23295.4 | controllerType=0 controllerNumber=1 scan=23295 | 49.9399 |
| 21330:4 | 9 | K.ADTHDEILEGLNFN[+2861.000]LTEIPEAQIHEGFQELLR.T | HexNAc(5)Hex(6)NeuAc(3) | N[+2861] | 1638.9701 | 4 | 6552.8586 | 6552.8239 | 0 | 5.3 | 94 | Specific | 660.93 | 547.87 | 547.87 | 13.32 | 48 | >sp\|P01009\|A1AT_HUMAN Alpha-1-antitrypsin OS=Homo sapiens OX=9606 GN=SERPINA1 PE=1 SV=3 | 642 | 20201116_xd_colostrum_1ng_glyco_hcd203040_compound15.24810.24810.3 | controllerType=0 controllerNumber=1 scan=24810 | 53.2119 |
| 21300:4 | 9 | K.ADTHDEILEGLNFN[+2861.000]LTEIPEAQIHEGFQELLR.T | HexNAc(5)Hex(6)NeuAc(3) | N[+2861] | 1638.9702 | 4 | 6552.859 | 6552.8239 | 0 | 5.4 | 94 | Specific | 932.09 | 932.09 | 932.09 | 18.32 | 48 | >sp\|P01009\|A1AT_HUMAN Alpha-1-antitrypsin OS=Homo sapiens OX=9606 GN=SERPINA1 PE=1 SV=3 | 642 | 20201116_xd_colostrum_1ng_glyco_hcd203040_compound15.24779.24779.4 | controllerType=0 controllerNumber=1 scan=24779 | 53.1383 |
| 21275:4 | 9 | K.ADTHDEILEGLNFN[+2861.000]LTEIPEAQIHEGFQELLR.T | HexNAc(5)Hex(6)NeuAc(3) | N[+2861] | 1638.9691 | 4 | 6552.8546 | 6552.8239 | 0 | 4.7 | 94 | Specific | 1009.68 | 812.5 | 812.5 | 20.72 | 48 | >sp\|P01009\|A1AT_HUMAN Alpha-1-antitrypsin OS=Homo sapiens OX=9606 GN=SERPINA1 PE=1 SV=3 | 642 | 20201116_xd_colostrum_1ng_glyco_hcd203040_compound15.24753.24753.4 | controllerType=0 controllerNumber=1 scan=24753 | 53.0852 |
| 21273:4 | 9 | K.ADTHDEILEGLNFN[+2861.000]LTEIPEAQIHEGFQELLR.T | HexNAc(5)Hex(6)NeuAc(3) | N[+2861] | 1638.9691 | 4 | 6552.8546 | 6552.8239 | 0 | 4.7 | 94 | Specific | 1037.67 | 1037.67 | 1037.67 | 20.72 | 48 | >sp\|P01009\|A1AT_HUMAN Alpha-1-antitrypsin OS=Homo sapiens OX=9606 GN=SERPINA1 PE=1 SV=3 | 642 | 20201116_xd_colostrum_1ng_glyco_hcd203040_compound15.24751.24751.4 | controllerType=0 controllerNumber=1 scan=24751 | 53.0795 |
| 21298:4 | 9 | K.ADTHDEILEGLNFN[+2861.000]LTEIPEAQIHEGFQELLR.T | HexNAc(5)Hex(6)NeuAc(3) | N[+2861] | 1638.9702 | 4 | 6552.859 | 6552.8239 | 0 | 5.4 | 94 | Specific | 1131.73 | 910.27 | 910.27 | 22.32 | 48 | >sp\|P01009\|A1AT_HUMAN Alpha-1-antitrypsin OS=Homo sapiens OX=9606 GN=SERPINA1 PE=1 SV=3 | 642 | 20201116_xd_colostrum_1ng_glyco_hcd203040_compound15.24777.24777.4 | controllerType=0 controllerNumber=1 scan=24777 | 53.1336 |
| 18917:6 | 9 | K.YLGN[+2204.772]ATAIFFLPDEGKLQHLENELTHDIITK.F | HexNAc(4)Hex(5)NeuAc(2) | N[+2205] | 958.4437 | 6 | 5745.6259 | 5745.6098 | 0 | 2.8 | 268 | Specific | 808 | 696.15 | 696.15 | 17.18 | 48 | >sp\|P01009\|A1AT_HUMAN Alpha-1-antitrypsin OS=Homo sapiens OX=9606 GN=SERPINA1 PE=1 SV=3 | 642 | 20201116_xd_colostrum_1ng_glyco_hcd203040_compound15.22288.22288.6 | controllerType=0 controllerNumber=1 scan=22288 | 47.6589 |
| 18920:6 | 9 | K.YLGN[+2204.772]ATAIFFLPDEGKLQHLENELTHDIITK.F | HexNAc(4)Hex(5)NeuAc(2) | N[+2205] | 958.4437 | 6 | 5745.6259 | 5745.6098 | 0 | 2.8 | 268 | Specific | 825.45 | 648 | 648 | 17.18 | 48 | >sp\|P01009\|A1AT_HUMAN Alpha-1-antitrypsin OS=Homo sapiens OX=9606 GN=SERPINA1 PE=1 SV=3 | 642 | 20201116_xd_colostrum_1ng_glyco_hcd203040_compound15.22291.22291.6 | controllerType=0 controllerNumber=1 scan=22291 | 47.6663 |
| 20138:5 | 10 | K.APDKNVIFSPLSISTALAFLSLGAHN[+2351.851]TTLTEILK.G | HexNAc(4)Hex(5)Fuc(3)NeuAc(1) | N[+2352] | 1187.7745 | 5 | 5934.8436 | 5934.8289 | 0 | 2.5 | 68 | Specific | 729.58 | 662.55 | 662.55 | 14.94 | 42 | >sp\|P01011\|AACT_HUMAN Alpha-1-antichymotrypsin OS=Homo sapiens OX=9606 GN=SERPINA3 PE=1 SV=2 | 1314 | 20201116_xd_colostrum_1ng_glyco_hcd203040_compound15.23564.23564.3 | controllerType=0 controllerNumber=1 scan=23564 | 50.5442 |
| 20136:5 | 10 | K.APDKNVIFSPLSISTALAFLSLGAHN[+2351.851]TTLTEILK.G | HexNAc(4)Hex(5)Fuc(3)NeuAc(1) | N[+2352] | 1187.7745 | 5 | 5934.8436 | 5934.8289 | 0 | 2.5 | 68 | Specific | 846.85 | 833.86 | 833.86 | 16.94 | 42 | >sp\|P01011\|AACT_HUMAN Alpha-1-antichymotrypsin OS=Homo sapiens OX=9606 GN=SERPINA3 PE=1 SV=2 | 1314 | 20201116_xd_colostrum_1ng_glyco_hcd203040_compound15.23562.23562.3 | controllerType=0 controllerNumber=1 scan=23562 | 50.5394 |
| 24582:6 | 11 | K.VFPLSLDSTPQDGNVVVAC[+57.021]LVQGFFPQEPLSVTWSESGQN[+2553.910]VTAI | HexNAc(5)Hex(5)Fuc(1)NeuAc(2) | C[+57], N[+2554] | 1222.5552 | 6 | 7330.295 | 7330.2711 | 0 | 3.3 | 123 | Specific | 204.2 | 204.2 | 204.2 | 5.28 | 50 | >sp\|P0DOX2\|IGA2_HUMAN Immunoglobulin alpha-2 heavy chain OS=Homo sapiens OX=9606 PE=1 SV=2 | 631 | 20201116_xd_colostrum_1ng_glyco_hcd203040_compound15.28200.28200.6 | controllerType=0 controllerNumber=1 scan=28200 | 59.8644 |
| 24659:4 | 11 | K.VFPLSLDSTPQDGNVVVAC[+57.021]LVQGFFPQEPLSVTWSESGQN[+2553.910]VTAI | HexNAc(5)Hex(5)Fuc(1)NeuAc(2) | C[+57], N[+2554] | 1833.3382 | 4 | 7330.3311 | 7330.2711 | 0 | 8.2 | 123 | Specific | 240.78 | 240.78 | 240.78 | 6.64 | 50 | >sp\|P0DOX2\|IGA2_HUMAN Immunoglobulin alpha-2 heavy chain OS=Homo sapiens OX=9606 PE=1 SV=2 | 631 | 20201116_xd_colostrum_1ng_glyco_hcd203040_compound15.28280.28280.4 | controllerType=0 controllerNumber=1 scan=28280 | 60.0178 |
| 22359:4 | 11 | K.VFPLSLDSTPQDGNVVVAC[+57.021]LVQGFFPQEPLSVTWSESGQN[+2117.777]VTAI | HexNAc(5)Hex(5)Fuc(2) | C[+57], N[+2118] | 1724.3033 | 4 | 6894.1914 | 6894.1382 | 0 | 7.7 | 123 | Specific | 244.2 | 229.47 | 229.47 | 6.78 | 50 | >sp\|P0DOX2\|IGA2_HUMAN Immunoglobulin alpha-2 heavy chain OS=Homo sapiens OX=9606 PE=1 SV=2 | 631 | 20201116_xd_colostrum_1ng_glyco_hcd203040_compound15.25885.25885.4 | controllerType=0 controllerNumber=1 scan=25885 | 55.47 |
| 20808:5 | 11 | K.VFPLSLDSTPQDGNVVVAC[+57.021]LVQGFFPQEPLSVTWSESGQN[+1606.587]VTAI | HexNAc(4)Hex(4)Fuc(1) | C[+57], N[+1607] | 1277.398 | 5 | 6382.9607 | 6382.9481 | 0 | 2 | 123 | Specific | 252.06 | 252.06 | 252.06 | 6.53 | 50 | >sp\|P0DOX2\|IGA2_HUMAN Immunoglobulin alpha-2 heavy chain OS=Homo sapiens OX=9606 PE=1 SV=2 | 631 | 20201116_xd_colostrum_1ng_glyco_hcd203040_compound15.24265.24265.5 | controllerType=0 controllerNumber=1 scan=24265 | 52.0401 |
| 22182:4 | 11 | K.VFPLSLDSTPQDGNVVVAC[+57.021]LVQGFFPQEPLSVTWSESGQN[+2279.830]VTAI | HexNAc(5)Hex(6)Fuc(2) | C[+57], N[+2280] | 1764.8143 | 4 | 7056.2353 | 7056.191 | 0 | 6.3 | 123 | Specific | 261.19 | 261.19 | 261.19 | 7.19 | 50 | >sp\|P0DOX2\|IGA2_HUMAN Immunoglobulin alpha-2 heavy chain OS=Homo sapiens OX=9606 PE=1 SV=2 | 631 | 20201116_xd_colostrum_1ng_glyco_hcd203040_compound15.25700.25700.4 | controllerType=0 controllerNumber=1 scan=25700 | 55.0974 |
| 22258:5 | 11 | K.VFPLSLDSTPQDGNVVVAC[+57.021]LVQGFFPQEPLSVTWSESGQN[+2279.830]VTAI | HexNAc(5)Hex(6)Fuc(2) | C[+57], N[+2280] | 1412.0506 | 5 | 7056.2238 | 7056.191 | 0 | 4.6 | 123 | Specific | 264.28 | 264.28 | 264.28 | 6.4 | 50 | >sp\|P0DOX2\|IGA2_HUMAN Immunoglobulin alpha-2 heavy chain OS=Homo sapiens OX=9606 PE=1 SV=2 | 631 | 20201116_xd_colostrum_1ng_glyco_hcd203040_compound15.25779.25779.5 | controllerType=0 controllerNumber=1 scan=25779 | 55.2569 |
| 21299:5 | 11 | K.VFPLSLDSTPQDGNVVVAC[+57.021]LVQGFFPQEPLSVTWSESGQN[+1809.666]VTAI | HexNAc(5)Hex(4)Fuc(1) | C[+57], N[+1810] | 1318.0154 | 5 | 6586.0478 | 6586.0275 | 0 | 3.1 | 123 | Specific | 279.77 | 279.77 | 279.77 | 7.15 | 50 | >sp\|P0DOX2\|IGA2_HUMAN Immunoglobulin alpha-2 heavy chain OS=Homo sapiens OX=9606 PE=1 SV=2 | 631 | 20201116_xd_colostrum_1ng_glyco_hcd203040_compound15.24778.24778.5 | controllerType=0 controllerNumber=1 scan=24778 | 53.1363 |
| 24584:6 | 11 | K.VFPLSLDSTPQDGNVVVAC[+57.021]LVQGFFPQEPLSVTWSESGQN[+2553.910]VTAI | HexNAc(5)Hex(5)Fuc(1)NeuAc(2) | C[+57], N[+2554] | 1222.5552 | 6 | 7330.295 | 7330.2711 | 0 | 3.3 | 123 | Specific | 283.51 | 283.51 | 283.51 | 7.42 | 50 | >sp\|P0DOX2\|IGA2_HUMAN Immunoglobulin alpha-2 heavy chain OS=Homo sapiens OX=9606 PE=1 SV=2 | 631 | 20201116_xd_colostrum_1ng_glyco_hcd203040_compound15.28202.28202.6 | controllerType=0 controllerNumber=1 scan=28202 | 59.8696 |
| 22361:4 | 11 | K.VFPLSLDSTPQDGNVVVAC[+57.021]LVQGFFPQEPLSVTWSESGQN[+2117.777]VTAI | HexNAc(5)Hex(5)Fuc(2) | C[+57], N[+2118] | 1724.3033 | 4 | 6894.1914 | 6894.1382 | 0 | 7.7 | 123 | Specific | 284.57 | 283.64 | 283.64 | 7.28 | 50 | >sp\|P0DOX2\|IGA2_HUMAN Immunoglobulin alpha-2 heavy chain OS=Homo sapiens OX=9606 PE=1 SV=2 | 631 | 20201116_xd_colostrum_1ng_glyco_hcd203040_compound15.25887.25887.4 | controllerType=0 controllerNumber=1 scan=25887 | 55.4741 |
| 24608:5 | 11 | K.VFPLSLDSTPQDGNVVVAC[+57.021]LVQGFFPQEPLSVTWSESGQN[+2350.830]VTAI | HexNAc(4)Hex(5)Fuc(1)NeuAc(2) | C[+57], N[+2351] | 1426.2503 | 5 | 7127.2223 | 7127.1917 | 0 | 4.3 | 123 | Specific | 286.77 | 286.77 | 286.77 | 7.99 | 50 | >sp\|P0DOX2\|IGA2_HUMAN Immunoglobulin alpha-2 heavy chain OS=Homo sapiens OX=9606 PE=1 SV=2 | 631 | 20201116_xd_colostrum_1ng_glyco_hcd203040_compound15.28227.28227.5 | controllerType=0 controllerNumber=1 scan=28227 | 59.9175 |
| 21325:5 | 11 | K.VFPLSLDSTPQDGNVVVAC[+57.021]LVQGFFPQEPLSVTWSESGQN[+1647.613]VTAI | HexNAc(5)Hex(3)Fuc(1) | C[+57], N[+1648] | 1285.6074 | 5 | 6424.008 | 6423.9746 | 0 | 5.2 | 123 | Specific | 304.38 | 304.38 | 304.38 | 6.79 | 50 | >sp\|P0DOX2\|IGA2_HUMAN Immunoglobulin alpha-2 heavy chain OS=Homo sapiens OX=9606 PE=1 SV=2 | 631 | 20201116_xd_colostrum_1ng_glyco_hcd203040_compound15.24805.24805.5 | controllerType=0 controllerNumber=1 scan=24805 | 53.1974 |
| 22158:4 | 11 | K.VFPLSLDSTPQDGNVVVAC[+57.021]LVQGFFPQEPLSVTWSESGQN[+2059.735]VTAI | HexNAc(4)Hex(5)Fuc(1)NeuAc(1) | C[+57], N[+2060] | 1709.7828 | 4 | 6836.1094 | 6836.0963 | 0 | 1.9 | 123 | Specific | 305.17 | 305.17 | 305.17 | 8.39 | 50 | >sp\|P0DOX2\|IGA2_HUMAN Immunoglobulin alpha-2 heavy chain OS=Homo sapiens OX=9606 PE=1 SV=2 | 631 | 20201116_xd_colostrum_1ng_glyco_hcd203040_compound15.25674.25674.4 | controllerType=0 controllerNumber=1 scan=25674 | 55.0315 |
| 22365:5 | 11 | K.VFPLSLDSTPQDGNVVVAC[+57.021]LVQGFFPQEPLSVTWSESGQN[+2117.777]VTAI | HexNAc(5)Hex(5)Fuc(2) | C[+57], N[+2118] | 1379.6416 | 5 | 6894.1791 | 6894.1382 | 0 | 5.9 | 123 | Specific | 334.35 | 334.35 | 334.35 | 8.17 | 50 | >sp\|P0DOX2\|IGA2_HUMAN Immunoglobulin alpha-2 heavy chain OS=Homo sapiens OX=9606 PE=1 SV=2 | 631 | 20201116_xd_colostrum_1ng_glyco_hcd203040_compound15.25891.25891.5 | controllerType=0 controllerNumber=1 scan=25891 | 55.4844 |
| 24661:4 | 11 | K.VFPLSLDSTPQDGNVVVAC[+57.021]LVQGFFPQEPLSVTWSESGQN[+2553.910]VTAI | HexNAc(5)Hex(5)Fuc(1)NeuAc(2) | C[+57], N[+2554] | 1833.3382 | 4 | 7330.3311 | 7330.2711 | 0 | 8.2 | 123 | Specific | 344.02 | 296.52 | 296.52 | 7.45 | 50 | >sp\|P0DOX2\|IGA2_HUMAN Immunoglobulin alpha-2 heavy chain OS=Homo sapiens OX=9606 PE=1 SV=2 | 631 | 20201116_xd_colostrum_1ng_glyco_hcd203040_compound15.28282.28282.4 | controllerType=0 controllerNumber=1 scan=28282 | 60.0238 |
| 22232:4 | 11 | K.VFPLSLDSTPQDGNVVVAC[+57.021]LVQGFFPQEPLSVTWSESGQN[+2262.814]VTAI | HexNAc(5)Hex(5)Fuc(1)NeuAc(1) | C[+57], N[+2263] | 1760.5573 | 4 | 7039.2075 | 7039.1757 | 0 | 4.5 | 123 | Specific | 414.22 | 414.22 | 414.22 | 10.57 | 50 | >sp\|P0DOX2\|IGA2_HUMAN Immunoglobulin alpha-2 heavy chain OS=Homo sapiens OX=9606 PE=1 SV=2 | 631 | 20201116_xd_colostrum_1ng_glyco_hcd203040_compound15.25751.25751.4 | controllerType=0 controllerNumber=1 scan=25751 | 55.192 |
| 22160:4 | 11 | K.VFPLSLDSTPQDGNVVVAC[+57.021]LVQGFFPQEPLSVTWSESGQN[+2059.735]VTAI | HexNAc(4)Hex(5)Fuc(1)NeuAc(1) | C[+57], N[+2060] | 1709.7828 | 4 | 6836.1094 | 6836.0963 | 0 | 1.9 | 123 | Specific | 420.73 | 402.49 | 402.49 | 10.93 | 50 | >sp\|P0DOX2\|IGA2_HUMAN Immunoglobulin alpha-2 heavy chain OS=Homo sapiens OX=9606 PE=1 SV=2 | 631 | 20201116_xd_colostrum_1ng_glyco_hcd203040_compound15.25676.25676.4 | controllerType=0 controllerNumber=1 scan=25676 | 55.0354 |
| 20726:4 | 11 | K.VFPLSLDSTPQDGNVVVAC[+57.021]LVQGFFPQEPLSVTWSESGQN[+1663.608]VTAI | HexNAc(5)Hex(4) | C[+57], N[+1664] | 1610.7609 | 4 | 6440.0216 | 6439.9696 | 0 | 8.1 | 123 | Specific | 428.05 | 427.83 | 427.83 | 9.8 | 50 | >sp\|P0DOX2\|IGA2_HUMAN Immunoglobulin alpha-2 heavy chain OS=Homo sapiens OX=9606 PE=1 SV=2 | 631 | 20201116_xd_colostrum_1ng_glyco_hcd203040_compound15.24179.24179.4 | controllerType=0 controllerNumber=1 scan=24179 | 51.8453 |
| 20822:5 | 11 | K.VFPLSLDSTPQDGNVVVAC[+57.021]LVQGFFPQEPLSVTWSESGQN[+1444.534]VTAI | HexNAc(4)Hex(3)Fuc(1) | C[+57], N[+1445] | 1244.9893 | 5 | 6220.9172 | 6220.8953 | 0 | 3.5 | 123 | Specific | 434.55 | 418.12 | 418.12 | 9.75 | 50 | >sp\|P0DOX2\|IGA2_HUMAN Immunoglobulin alpha-2 heavy chain OS=Homo sapiens OX=9606 PE=1 SV=2 | 631 | 20201116_xd_colostrum_1ng_glyco_hcd203040_compound15.24280.24280.4 | controllerType=0 controllerNumber=1 scan=24280 | 52.076 |
| 20717:4 | 11 | K.VFPLSLDSTPQDGNVVVAC[+57.021]LVQGFFPQEPLSVTWSESGQN[+1809.666]VTAI | HexNAc(5)Hex(4)Fuc(1) | C[+57], N[+1810] | 1647.2693 | 4 | 6586.0553 | 6586.0275 | 0 | 4.2 | 123 | Specific | 440.04 | 437.19 | 437.19 | 10.72 | 50 | >sp\|P0DOX2\|IGA2_HUMAN Immunoglobulin alpha-2 heavy chain OS=Homo sapiens OX=9606 PE=1 SV=2 | 631 | 20201116_xd_colostrum_1ng_glyco_hcd203040_compound15.24170.24170.4 | controllerType=0 controllerNumber=1 scan=24170 | 51.8233 |
| 22244:6 | 11 | K.VFPLSLDSTPQDGNVVVAC[+57.021]LVQGFFPQEPLSVTWSESGQN[+2262.814]VTAI | HexNAc(5)Hex(5)Fuc(1)NeuAc(1) | C[+57], N[+2263] | 1174.0405 | 6 | 7039.2064 | 7039.1757 | 0 | 4.4 | 123 | Specific | 490.2 | 490.2 | 490.2 | 10.88 | 50 | >sp\|P0DOX2\|IGA2_HUMAN Immunoglobulin alpha-2 heavy chain OS=Homo sapiens OX=9606 PE=1 SV=2 | 631 | 20201116_xd_colostrum_1ng_glyco_hcd203040_compound15.25764.25764.6 | controllerType=0 controllerNumber=1 scan=25764 | 55.2209 |
| 22217:5 | 11 | K.VFPLSLDSTPQDGNVVVAC[+57.021]LVQGFFPQEPLSVTWSESGQN[+2117.777]VTAI | HexNAc(5)Hex(5)Fuc(2) | C[+57], N[+2118] | 1379.6409 | 5 | 6894.1755 | 6894.1382 | 0 | 5.4 | 123 | Specific | 493.41 | 493.41 | 493.41 | 10.48 | 50 | >sp\|P0DOX2\|IGA2_HUMAN Immunoglobulin alpha-2 heavy chain OS=Homo sapiens OX=9606 PE=1 SV=2 | 631 | 20201116_xd_colostrum_1ng_glyco_hcd203040_compound15.25736.25736.5 | controllerType=0 controllerNumber=1 scan=25736 | 55.162 |
| 22234:4 | 11 | K.VFPLSLDSTPQDGNVVVAC[+57.021]LVQGFFPQEPLSVTWSESGQN[+2262.814]VTAI | HexNAc(5)Hex(5)Fuc(1)NeuAc(1) | C[+57], N[+2263] | 1760.5573 | 4 | 7039.2075 | 7039.1757 | 0 | 4.5 | 123 | Specific | 503.13 | 496.6 | 496.6 | 11 | 50 | >sp\|P0DOX2\|IGA2_HUMAN Immunoglobulin alpha-2 heavy chain OS=Homo sapiens OX=9606 PE=1 SV=2 | 631 | 20201116_xd_colostrum_1ng_glyco_hcd203040_compound15.25753.25753.4 | controllerType=0 controllerNumber=1 scan=25753 | 55.1959 |
| 24632:5 | 11 | K.VFPLSLDSTPQDGNVVVAC[+57.021]LVQGFFPQEPLSVTWSESGQN[+2553.910]VTAI | HexNAc(5)Hex(5)Fuc(1)NeuAc(2) | C[+57], N[+2554] | 1466.8659 | 5 | 7330.3003 | 7330.2711 | 0 | 4 | 123 | Specific | 539.63 | 530.75 | 530.75 | 11.28 | 50 | >sp\|P0DOX2\|IGA2_HUMAN Immunoglobulin alpha-2 heavy chain OS=Homo sapiens OX=9606 PE=1 SV=2 | 631 | 20201116_xd_colostrum_1ng_glyco_hcd203040_compound15.28252.28252.5 | controllerType=0 controllerNumber=1 scan=28252 | 59.9642 |
| 22219:5 | 11 | K.VFPLSLDSTPQDGNVVVAC[+57.021]LVQGFFPQEPLSVTWSESGQN[+2117.777]VTAI | HexNAc(5)Hex(5)Fuc(2) | C[+57], N[+2118] | 1379.6409 | 5 | 6894.1755 | 6894.1382 | 0 | 5.4 | 123 | Specific | 585.51 | 585.51 | 585.51 | 12.38 | 50 | >sp\|P0DOX2\|IGA2_HUMAN Immunoglobulin alpha-2 heavy chain OS=Homo sapiens OX=9606 PE=1 SV=2 | 631 | 20201116_xd_colostrum_1ng_glyco_hcd203040_compound15.25738.25738.5 | controllerType=0 controllerNumber=1 scan=25738 | 55.1674 |
| 20824:5 | 11 | K.VFPLSLDSTPQDGNVVVAC[+57.021]LVQGFFPQEPLSVTWSESGQN[+1444.534]VTAI | HexNAc(4)Hex(3)Fuc(1) | C[+57], N[+1445] | 1244.9893 | 5 | 6220.9172 | 6220.8953 | 0 | 3.5 | 123 | Specific | 586.54 | 554.71 | 554.71 | 12.21 | 50 | >sp\|P0DOX2\|IGA2_HUMAN Immunoglobulin alpha-2 heavy chain OS=Homo sapiens OX=9606 PE=1 SV=2 | 631 | 20201116_xd_colostrum_1ng_glyco_hcd203040_compound15.24282.24282.4 | controllerType=0 controllerNumber=1 scan=24282 | 52.0827 |
| 24658:5 | 11 | K.VFPLSLDSTPQDGNVVVAC[+57.021]LVQGFFPQEPLSVTWSESGQN[+2553.910]VTAI | HexNAc(5)Hex(5)Fuc(1)NeuAc(2) | C[+57], N[+2554] | 1466.8667 | 5 | 7330.3046 | 7330.2711 | 0 | 4.6 | 123 | Specific | 590.15 | 590.15 | 590.15 | 12.78 | 50 | >sp\|P0DOX2\|IGA2_HUMAN Immunoglobulin alpha-2 heavy chain OS=Homo sapiens OX=9606 PE=1 SV=2 | 631 | 20201116_xd_colostrum_1ng_glyco_hcd203040_compound15.28279.28279.5 | controllerType=0 controllerNumber=1 scan=28279 | 60.0162 |
| 22874:5 | 11 | K.VFPLSLDSTPQDGNVVVAC[+57.021]LVQGFFPQEPLSVTWSESGQN[+2262.814]VTAI | HexNAc(5)Hex(5)Fuc(1)NeuAc(1) | C[+57], N[+2263] | 1408.6491 | 5 | 7039.2164 | 7039.1757 | 0 | 5.8 | 123 | Specific | 596.96 | 596.96 | 596.96 | 12.38 | 50 | >sp\|P0DOX2\|IGA2_HUMAN Immunoglobulin alpha-2 heavy chain OS=Homo sapiens OX=9606 PE=1 SV=2 | 631 | 20201116_xd_colostrum_1ng_glyco_hcd203040_compound15.26424.26424.5 | controllerType=0 controllerNumber=1 scan=26424 | 56.5505 |
| 22184:4 | 11 | K.VFPLSLDSTPQDGNVVVAC[+57.021]LVQGFFPQEPLSVTWSESGQN[+2279.830]VTAI | HexNAc(5)Hex(6)Fuc(2) | C[+57], N[+2280] | 1764.8143 | 4 | 7056.2353 | 7056.191 | 0 | 6.3 | 123 | Specific | 602.14 | 571.46 | 571.46 | 13.22 | 50 | >sp\|P0DOX2\|IGA2_HUMAN Immunoglobulin alpha-2 heavy chain OS=Homo sapiens OX=9606 PE=1 SV=2 | 631 | 20201116_xd_colostrum_1ng_glyco_hcd203040_compound15.25702.25702.4 | controllerType=0 controllerNumber=1 scan=25702 | 55.1019 |
| 22237:4 | 11 | K.VFPLSLDSTPQDGNVVVAC[+57.021]LVQGFFPQEPLSVTWSESGQN[+2262.814]VTAI | HexNAc(5)Hex(5)Fuc(1)NeuAc(1) | C[+57], N[+2263] | 1760.559 | 4 | 7039.2144 | 7039.1757 | 0 | 5.5 | 123 | Specific | 609.49 | 609.49 | 609.49 | 13.38 | 50 | >sp\|P0DOX2\|IGA2_HUMAN Immunoglobulin alpha-2 heavy chain OS=Homo sapiens OX=9606 PE=1 SV=2 | 631 | 20201116_xd_colostrum_1ng_glyco_hcd203040_compound15.25757.25757.4 | controllerType=0 controllerNumber=1 scan=25757 | 55.2038 |
| 21302:4 | 11 | K.VFPLSLDSTPQDGNVVVAC[+57.021]LVQGFFPQEPLSVTWSESGQN[+1809.666]VTAI | HexNAc(5)Hex(4)Fuc(1) | C[+57], N[+1810] | 1647.2692 | 4 | 6586.0548 | 6586.0275 | 0 | 4.2 | 123 | Specific | 612.58 | 612.58 | 612.58 | 13.78 | 50 | >sp\|P0DOX2\|IGA2_HUMAN Immunoglobulin alpha-2 heavy chain OS=Homo sapiens OX=9606 PE=1 SV=2 | 631 | 20201116_xd_colostrum_1ng_glyco_hcd203040_compound15.24781.24781.4 | controllerType=0 controllerNumber=1 scan=24781 | 53.1416 |
| 20947:4 | 11 | K.VFPLSLDSTPQDGNVVVAC[+57.021]LVQGFFPQEPLSVTWSESGQN[+1444.534]VTAI | HexNAc(4)Hex(3)Fuc(1) | C[+57], N[+1445] | 1555.9846 | 4 | 6220.9167 | 6220.8953 | 0 | 3.4 | 123 | Specific | 615.72 | 608.33 | 608.33 | 14.01 | 50 | >sp\|P0DOX2\|IGA2_HUMAN Immunoglobulin alpha-2 heavy chain OS=Homo sapiens OX=9606 PE=1 SV=2 | 631 | 20201116_xd_colostrum_1ng_glyco_hcd203040_compound15.24410.24410.4 | controllerType=0 controllerNumber=1 scan=24410 | 52.356 |
| 22890:5 | 11 | K.VFPLSLDSTPQDGNVVVAC[+57.021]LVQGFFPQEPLSVTWSESGQN[+2262.814]VTAI | HexNAc(5)Hex(5)Fuc(1)NeuAc(1) | C[+57], N[+2263] | 1408.6438 | 5 | 7039.1901 | 7039.1757 | 0 | 2 | 123 | Specific | 623.03 | 600.93 | 600.93 | 14.05 | 50 | >sp\|P0DOX2\|IGA2_HUMAN Immunoglobulin alpha-2 heavy chain OS=Homo sapiens OX=9606 PE=1 SV=2 | 631 | 20201116_xd_colostrum_1ng_glyco_hcd203040_compound15.26441.26441.5 | controllerType=0 controllerNumber=1 scan=26441 | 56.5849 |
| 20716:4 | 11 | K.VFPLSLDSTPQDGNVVVAC[+57.021]LVQGFFPQEPLSVTWSESGQN[+1971.719]VTAI | HexNAc(5)Hex(5)Fuc(1) | C[+57], N[+1972] | 1687.7801 | 4 | 6748.0986 | 6748.0803 | 0 | 2.7 | 123 | Specific | 673.27 | 646.11 | 646.11 | 15.21 | 50 | >sp\|P0DOX2\|IGA2_HUMAN Immunoglobulin alpha-2 heavy chain OS=Homo sapiens OX=9606 PE=1 SV=2 | 631 | 20201116_xd_colostrum_1ng_glyco_hcd203040_compound15.24169.24169.4 | controllerType=0 controllerNumber=1 scan=24169 | 51.8165 |
| 20771:4 | 11 | K.VFPLSLDSTPQDGNVVVAC[+57.021]LVQGFFPQEPLSVTWSESGQN[+1444.534]VTAI | HexNAc(4)Hex(3)Fuc(1) | C[+57], N[+1445] | 1555.9849 | 4 | 6220.9176 | 6220.8953 | 0 | 3.6 | 123 | Specific | 676.49 | 660.19 | 660.19 | 15.01 | 50 | >sp\|P0DOX2\|IGA2_HUMAN Immunoglobulin alpha-2 heavy chain OS=Homo sapiens OX=9606 PE=1 SV=2 | 631 | 20201116_xd_colostrum_1ng_glyco_hcd203040_compound15.24226.24226.4 | controllerType=0 controllerNumber=1 scan=24226 | 51.9482 |
| 21320:4 | 11 | K.VFPLSLDSTPQDGNVVVAC[+57.021]LVQGFFPQEPLSVTWSESGQN[+1647.613]VTAI | HexNAc(5)Hex(3)Fuc(1) | C[+57], N[+1648] | 1606.7537 | 4 | 6423.993 | 6423.9746 | 0 | 2.9 | 123 | Specific | 702.36 | 702.36 | 702.36 | 16.21 | 50 | >sp\|P0DOX2\|IGA2_HUMAN Immunoglobulin alpha-2 heavy chain OS=Homo sapiens OX=9606 PE=1 SV=2 | 631 | 20201116_xd_colostrum_1ng_glyco_hcd203040_compound15.24800.24800.4 | controllerType=0 controllerNumber=1 scan=24800 | 53.1862 |
| 20804:4 | 11 | K.VFPLSLDSTPQDGNVVVAC[+57.021]LVQGFFPQEPLSVTWSESGQN[+1606.587]VTAI | HexNAc(4)Hex(4)Fuc(1) | C[+57], N[+1607] | 1596.496 | 4 | 6382.9621 | 6382.9481 | 0 | 2.2 | 123 | Specific | 710.63 | 694.06 | 694.06 | 16.21 | 50 | >sp\|P0DOX2\|IGA2_HUMAN Immunoglobulin alpha-2 heavy chain OS=Homo sapiens OX=9606 PE=1 SV=2 | 631 | 20201116_xd_colostrum_1ng_glyco_hcd203040_compound15.24261.24261.4 | controllerType=0 controllerNumber=1 scan=24261 | 52.0311 |
| 24660:5 | 11 | K.VFPLSLDSTPQDGNVVVAC[+57.021]LVQGFFPQEPLSVTWSESGQN[+2553.910]VTAI | HexNAc(5)Hex(5)Fuc(1)NeuAc(2) | C[+57], N[+2554] | 1466.8667 | 5 | 7330.3046 | 7330.2711 | 0 | 4.6 | 123 | Specific | 723.37 | 723.37 | 723.37 | 15.78 | 50 | >sp\|P0DOX2\|IGA2_HUMAN Immunoglobulin alpha-2 heavy chain OS=Homo sapiens OX=9606 PE=1 SV=2 | 631 | 20201116_xd_colostrum_1ng_glyco_hcd203040_compound15.28281.28281.5 | controllerType=0 controllerNumber=1 scan=28281 | 60.0219 |
| 22236:5 | 11 | K.VFPLSLDSTPQDGNVVVAC[+57.021]LVQGFFPQEPLSVTWSESGQN[+2262.814]VTAI | HexNAc(5)Hex(5)Fuc(1)NeuAc(1) | C[+57], N[+2263] | 1408.6471 | 5 | 7039.2066 | 7039.1757 | 0 | 4.4 | 123 | Specific | 725.48 | 682.14 | 682.14 | 15.78 | 50 | >sp\|P0DOX2\|IGA2_HUMAN Immunoglobulin alpha-2 heavy chain OS=Homo sapiens OX=9606 PE=1 SV=2 | 631 | 20201116_xd_colostrum_1ng_glyco_hcd203040_compound15.25756.25756.5 | controllerType=0 controllerNumber=1 scan=25756 | 55.2024 |
| 22876:5 | 11 | K.VFPLSLDSTPQDGNVVVAC[+57.021]LVQGFFPQEPLSVTWSESGQN[+2262.814]VTAI | HexNAc(5)Hex(5)Fuc(1)NeuAc(1) | C[+57], N[+2263] | 1408.6491 | 5 | 7039.2164 | 7039.1757 | 0 | 5.8 | 123 | Specific | 725.69 | 717.81 | 717.81 | 15.38 | 50 | >sp\|P0DOX2\|IGA2_HUMAN Immunoglobulin alpha-2 heavy chain OS=Homo sapiens OX=9606 PE=1 SV=2 | 631 | 20201116_xd_colostrum_1ng_glyco_hcd203040_compound15.26426.26426.5 | controllerType=0 controllerNumber=1 scan=26426 | 56.5556 |
| 22892:5 | 11 | K.VFPLSLDSTPQDGNVVVAC[+57.021]LVQGFFPQEPLSVTWSESGQN[+2262.814]VTAI | HexNAc(5)Hex(5)Fuc(1)NeuAc(1) | C[+57], N[+2263] | 1408.6438 | 5 | 7039.1901 | 7039.1757 | 0 | 2 | 123 | Specific | 760.9 | 716.25 | 716.25 | 17.05 | 50 | >sp\|P0DOX2\|IGA2_HUMAN Immunoglobulin alpha-2 heavy chain OS=Homo sapiens OX=9606 PE=1 SV=2 | 631 | 20201116_xd_colostrum_1ng_glyco_hcd203040_compound15.26443.26443.5 | controllerType=0 controllerNumber=1 scan=26443 | 56.5909 |
| 20715:4 | 11 | K.VFPLSLDSTPQDGNVVVAC[+57.021]LVQGFFPQEPLSVTWSESGQN[+1809.666]VTAI | HexNAc(5)Hex(4)Fuc(1) | C[+57], N[+1810] | 1647.2693 | 4 | 6586.0553 | 6586.0275 | 0 | 4.2 | 123 | Specific | 762.42 | 731.47 | 731.47 | 16.78 | 50 | >sp\|P0DOX2\|IGA2_HUMAN Immunoglobulin alpha-2 heavy chain OS=Homo sapiens OX=9606 PE=1 SV=2 | 631 | 20201116_xd_colostrum_1ng_glyco_hcd203040_compound15.24168.24168.4 | controllerType=0 controllerNumber=1 scan=24168 | 51.8151 |
| 20756:4 | 11 | K.VFPLSLDSTPQDGNVVVAC[+57.021]LVQGFFPQEPLSVTWSESGQN[+1647.613]VTAI | HexNAc(5)Hex(3)Fuc(1) | C[+57], N[+1648] | 1606.7568 | 4 | 6424.0055 | 6423.9746 | 0 | 4.8 | 123 | Specific | 842.42 | 789.04 | 789.04 | 17.78 | 50 | >sp\|P0DOX2\|IGA2_HUMAN Immunoglobulin alpha-2 heavy chain OS=Homo sapiens OX=9606 PE=1 SV=2 | 631 | 20201116_xd_colostrum_1ng_glyco_hcd203040_compound15.24211.24211.4 | controllerType=0 controllerNumber=1 scan=24211 | 51.9139 |
| 20758:4 | 11 | K.VFPLSLDSTPQDGNVVVAC[+57.021]LVQGFFPQEPLSVTWSESGQN[+1647.613]VTAI | HexNAc(5)Hex(3)Fuc(1) | C[+57], N[+1648] | 1606.7568 | 4 | 6424.0055 | 6423.9746 | 0 | 4.8 | 123 | Specific | 846.54 | 841.96 | 841.96 | 17.78 | 50 | >sp\|P0DOX2\|IGA2_HUMAN Immunoglobulin alpha-2 heavy chain OS=Homo sapiens OX=9606 PE=1 SV=2 | 631 | 20201116_xd_colostrum_1ng_glyco_hcd203040_compound15.24213.24213.4 | controllerType=0 controllerNumber=1 scan=24213 | 51.9219 |
| 20807:4 | 11 | K.VFPLSLDSTPQDGNVVVAC[+57.021]LVQGFFPQEPLSVTWSESGQN[+1606.587]VTAI | HexNAc(4)Hex(4)Fuc(1) | C[+57], N[+1607] | 1596.496 | 4 | 6382.9621 | 6382.9481 | 0 | 2.2 | 123 | Specific | 846.61 | 759.48 | 759.48 | 18.21 | 50 | >sp\|P0DOX2\|IGA2_HUMAN Immunoglobulin alpha-2 heavy chain OS=Homo sapiens OX=9606 PE=1 SV=2 | 631 | 20201116_xd_colostrum_1ng_glyco_hcd203040_compound15.24264.24264.4 | controllerType=0 controllerNumber=1 scan=24264 | 52.0382 |
| 24634:5 | 11 | K.VFPLSLDSTPQDGNVVVAC[+57.021]LVQGFFPQEPLSVTWSESGQN[+2553.910]VTAI | HexNAc(5)Hex(5)Fuc(1)NeuAc(2) | C[+57], N[+2554] | 1466.8659 | 5 | 7330.3003 | 7330.2711 | 0 | 4 | 123 | Specific | 862.41 | 808.54 | 808.54 | 19.01 | 50 | >sp\|P0DOX2\|IGA2_HUMAN Immunoglobulin alpha-2 heavy chain OS=Homo sapiens OX=9606 PE=1 SV=2 | 631 | 20201116_xd_colostrum_1ng_glyco_hcd203040_compound15.28254.28254.5 | controllerType=0 controllerNumber=1 scan=28254 | 59.9713 |
| 22238:5 | 11 | K.VFPLSLDSTPQDGNVVVAC[+57.021]LVQGFFPQEPLSVTWSESGQN[+2262.814]VTAI | HexNAc(5)Hex(5)Fuc(1)NeuAc(1) | C[+57], N[+2263] | 1408.6471 | 5 | 7039.2066 | 7039.1757 | 0 | 4.4 | 123 | Specific | 862.7 | 849.2 | 849.2 | 18.78 | 50 | >sp\|P0DOX2\|IGA2_HUMAN Immunoglobulin alpha-2 heavy chain OS=Homo sapiens OX=9606 PE=1 SV=2 | 631 | 20201116_xd_colostrum_1ng_glyco_hcd203040_compound15.25758.25758.5 | controllerType=0 controllerNumber=1 scan=25758 | 55.2091 |
| 21304:4 | 11 | K.VFPLSLDSTPQDGNVVVAC[+57.021]LVQGFFPQEPLSVTWSESGQN[+1809.666]VTAI | HexNAc(5)Hex(4)Fuc(1) | C[+57], N[+1810] | 1647.2692 | 4 | 6586.0548 | 6586.0275 | 0 | 4.2 | 123 | Specific | 863.38 | 754.37 | 754.37 | 18.78 | 50 | >sp\|P0DOX2\|IGA2_HUMAN Immunoglobulin alpha-2 heavy chain OS=Homo sapiens OX=9606 PE=1 SV=2 | 631 | 20201116_xd_colostrum_1ng_glyco_hcd203040_compound15.24783.24783.4 | controllerType=0 controllerNumber=1 scan=24783 | 53.1474 |
| 21322:4 | 11 | K.VFPLSLDSTPQDGNVVVAC[+57.021]LVQGFFPQEPLSVTWSESGQN[+1647.613]VTAI | HexNAc(5)Hex(3)Fuc(1) | C[+57], N[+1648] | 1606.7537 | 4 | 6423.993 | 6423.9746 | 0 | 2.9 | 123 | Specific | 868 | 837.09 | 837.09 | 19.21 | 50 | >sp\|P0DOX2\|IGA2_HUMAN Immunoglobulin alpha-2 heavy chain OS=Homo sapiens OX=9606 PE=1 SV=2 | 631 | 20201116_xd_colostrum_1ng_glyco_hcd203040_compound15.24802.24802.4 | controllerType=0 controllerNumber=1 scan=24802 | 53.1917 |
| 22239:4 | 11 | K.VFPLSLDSTPQDGNVVVAC[+57.021]LVQGFFPQEPLSVTWSESGQN[+2262.814]VTAI | HexNAc(5)Hex(5)Fuc(1)NeuAc(1) | C[+57], N[+2263] | 1760.559 | 4 | 7039.2144 | 7039.1757 | 0 | 5.5 | 123 | Specific | 938.38 | 906.4 | 906.4 | 19.38 | 50 | >sp\|P0DOX2\|IGA2_HUMAN Immunoglobulin alpha-2 heavy chain OS=Homo sapiens OX=9606 PE=1 SV=2 | 631 | 20201116_xd_colostrum_1ng_glyco_hcd203040_compound15.25759.25759.4 | controllerType=0 controllerNumber=1 scan=25759 | 55.2117 |
| 20773:4 | 11 | K.VFPLSLDSTPQDGNVVVAC[+57.021]LVQGFFPQEPLSVTWSESGQN[+1444.534]VTAI | HexNAc(4)Hex(3)Fuc(1) | C[+57], N[+1445] | 1555.9849 | 4 | 6220.9176 | 6220.8953 | 0 | 3.6 | 123 | Specific | 948.62 | 821.7 | 821.7 | 20.01 | 50 | >sp\|P0DOX2\|IGA2_HUMAN Immunoglobulin alpha-2 heavy chain OS=Homo sapiens OX=9606 PE=1 SV=2 | 631 | 20201116_xd_colostrum_1ng_glyco_hcd203040_compound15.24228.24228.4 | controllerType=0 controllerNumber=1 scan=24228 | 51.9559 |

Table S4. Cont.

| Query #:z | Protein Rank | Glycopeptide | Glycan | Modification Type(s) | Observed m/z | z | Observed (M+H) | Calc. mass (M+H) | Off-by-x error | Mass error (ppm) | StartPosition | Cleavage | Score | Delta | Delta Mod | \|Log Prob\| | # of unique peptides | Protein Name | Protein DB number | Comment | Scan # | Scan Time |
|---|---|---|---|---|---|---|---|---|---|---|---|---|---|---|---|---|---|---|---|---|---|---|
| 20949:4 | 11 | K.VFPLSLDSTPQDGNVVVAC[+57.021]LVQGFFPQEPLSVTWSESGQN[+1444.534]VTAI | HexNAc(4)Hex(3)Fuc(1) | C[+57], N[+1445] | 1555.9846 | 4 | 6220.9167 | 6220.8953 | 0 | 3.4 | 123 | Specific | 990.29 | 826.91 | 826.91 | 21.01 | 50 | >sp\|P0DOX2\|IGA2_HUMAN Immunoglobulin alpha-2 heavy chain OS=Homo sapiens OX=9606 PE=1 SV=2 | 631 | 20201116_xd_colostrum_1ng_glyco_hcd203040_compound15.24412.24412.4 | controllerType=0 controllerNumber=1 scan=24412 | 52.3619 |
| 13347:5 | 11 | R.DLC[+57.021]GC[+57.021]YSVSSVLPGC[+57.021]AQPWNHGETFTC[+57.021]TAA | HexNAc(4)Hex(3)Fuc(1) | C[+57]*4, N[+1445] | 1267.567 | 5 | 6333.806 | 6333.8129 | 0 | -1.1 | 280 | Specific | 283.13 | 283.13 | 283.13 | 6.71 | 50 | >sp\|P0DOX2\|IGA2_HUMAN Immunoglobulin alpha-2 heavy chain OS=Homo sapiens OX=9606 PE=1 SV=2 | 631 | 20201116_xd_colostrum_1ng_glyco_hcd203040_compound15.16478.16478.5 | controllerType=0 controllerNumber=1 scan=16478 | 35.2074 |
| 13349:5 | 11 | R.DLC[+57.021]GC[+57.021]YSVSSVLPGC[+57.021]AQPWNHGETFTC[+57.021]TAA | HexNAc(4)Hex(3)Fuc(1) | C[+57]*4, N[+1445] | 1267.567 | 5 | 6333.806 | 6333.8129 | 0 | -1.1 | 280 | Specific | 360.77 | 266.98 | 266.98 | 5.68 | 50 | >sp\|P0DOX2\|IGA2_HUMAN Immunoglobulin alpha-2 heavy chain OS=Homo sapiens OX=9606 PE=1 SV=2 | 631 | 20201116_xd_colostrum_1ng_glyco_hcd203040_compound15.16480.16480.5 | controllerType=0 controllerNumber=1 scan=16480 | 35.2115 |
| 13225:4 | 11 | R.DLC[+57.021]GC[+57.021]YSVSSVLPGC[+57.021]AQPWNHGETFTC[+57.021]TAA | HexNAc(5)Hex(4)Fuc(1) | C[+57]*4, N[+1810] | 1675.4936 | 4 | 6698.9526 | 6698.9451 | 0 | 1.1 | 280 | Specific | 509.58 | 509.58 | 509.58 | 10.05 | 50 | >sp\|P0DOX2\|IGA2_HUMAN Immunoglobulin alpha-2 heavy chain OS=Homo sapiens OX=9606 PE=1 SV=2 | 631 | 20201116_xd_colostrum_1ng_glyco_hcd203040_compound15.16351.16351.4 | controllerType=0 controllerNumber=1 scan=16351 | 34.9756 |
| 13216:5 | 11 | R.DLC[+57.021]GC[+57.021]YSVSSVLPGC[+57.021]AQPWNHGETFTC[+57.021]TAA | HexNAc(5)Hex(4)Fuc(1) | C[+57]*4, N[+1810] | 1340.597 | 5 | 6698.9557 | 6698.9451 | 0 | 1.6 | 280 | Specific | 582.82 | 526.4 | 526.4 | 12.38 | 50 | >sp\|P0DOX2\|IGA2_HUMAN Immunoglobulin alpha-2 heavy chain OS=Homo sapiens OX=9606 PE=1 SV=2 | 631 | 20201116_xd_colostrum_1ng_glyco_hcd203040_compound15.16342.16342.5 | controllerType=0 controllerNumber=1 scan=16342 | 34.9566 |
| 13228:4 | 11 | R.DLC[+57.021]GC[+57.021]YSVSSVLPGC[+57.021]AQPWNHGETFTC[+57.021]TAA | HexNAc(5)Hex(4)Fuc(1) | C[+57]*4, N[+1810] | 1675.4936 | 4 | 6698.9526 | 6698.9451 | 0 | 1.1 | 280 | Specific | 681.02 | 543.45 | 543.45 | 13.82 | 50 | >sp\|P0DOX2\|IGA2_HUMAN Immunoglobulin alpha-2 heavy chain OS=Homo sapiens OX=9606 PE=1 SV=2 | 631 | 20201116_xd_colostrum_1ng_glyco_hcd203040_compound15.16354.16354.4 | controllerType=0 controllerNumber=1 scan=16354 | 34.9821 |
| 13213:5 | 11 | R.DLC[+57.021]GC[+57.021]YSVSSVLPGC[+57.021]AQPWNHGETFTC[+57.021]TAA | HexNAc(5)Hex(4)Fuc(1) | C[+57]*4, N[+1810] | 1340.597 | 5 | 6698.9557 | 6698.9451 | 0 | 1.6 | 280 | Specific | 769.67 | 769.67 | 769.67 | 16.38 | 50 | >sp\|P0DOX2\|IGA2_HUMAN Immunoglobulin alpha-2 heavy chain OS=Homo sapiens OX=9606 PE=1 SV=2 | 631 | 20201116_xd_colostrum_1ng_glyco_hcd203040_compound15.16339.16339.5 | controllerType=0 controllerNumber=1 scan=16339 | 34.9503 |
| 07516:3 | 11 | K.TPLTAN[+2553.910]ITK.S | HexNAc(5)Hex(5)Fuc(1)NeuAc(2) | N[+2554] | 1171.4962 | 3 | 3512.4739 | 3512.4665 | 0 | 2.1 | 315 | Specific | 250.07 | 71.58 | 71.58 | 4.71 | 50 | >sp\|P0DOX2\|IGA2_HUMAN Immunoglobulin alpha-2 heavy chain OS=Homo sapiens OX=9606 PE=1 SV=2 | 631 | 20201116_xd_colostrum_1ng_glyco_hcd203040_compound15.10422.10422.3 | controllerType=0 controllerNumber=1 scan=10422 | 23.6402 |
| 03344:2 | 11 | K.TPLTAN[+1647.613]ITK.S | HexNAc(5)Hex(3)Fuc(1) | N[+1648] | 1303.5927 | 2 | 2606.1781 | 2606.17 | 0 | 3.1 | 315 | Specific | 339.42 | 107.66 | 107.66 | 4.9 | 50 | >sp\|P0DOX2\|IGA2_HUMAN Immunoglobulin alpha-2 heavy chain OS=Homo sapiens OX=9606 PE=1 SV=2 | 631 | 20201116_xd_colostrum_1ng_glyco_hcd203040_compound15.6097.6097.2 | controllerType=0 controllerNumber=1 scan=6097 | 16.1789 |
| 03453:2 | 11 | K.TPLTAN[+1444.534]ITK.S | HexNAc(4)Hex(3)Fuc(1) | N[+1445] | 1202.049 | 2 | 2403.0907 | 2403.0906 | 0 | 0 | 315 | Specific | 358.29 | 315.89 | 315.89 | 6.72 | 50 | >sp\|P0DOX2\|IGA2_HUMAN Immunoglobulin alpha-2 heavy chain OS=Homo sapiens OX=9606 PE=1 SV=2 | 631 | 20201116_xd_colostrum_1ng_glyco_hcd203040_compound15.6210.6210.2 | controllerType=0 controllerNumber=1 scan=6210 | 16.3866 |
| 04998:2 | 11 | K.TPLTAN[+2059.735]ITK.S | HexNAc(4)Hex(5)Fuc(1)NeuAc(1) | N[+2060] | 1509.6525 | 2 | 3018.2977 | 3018.2917 | 0 | 2 | 315 | Specific | 382.97 | 262.3 | 262.3 | 6.97 | 50 | >sp\|P0DOX2\|IGA2_HUMAN Immunoglobulin alpha-2 heavy chain OS=Homo sapiens OX=9606 PE=1 SV=2 | 631 | 20201116_xd_colostrum_1ng_glyco_hcd203040_compound15.7812.7812.2 | controllerType=0 controllerNumber=1 scan=7812 | 19.0815 |
| 07518:3 | 11 | K.TPLTAN[+2553.910]ITK.S | HexNAc(5)Hex(5)Fuc(1)NeuAc(2) | N[+2554] | 1171.4962 | 3 | 3512.4739 | 3512.4665 | 0 | 2.1 | 315 | Specific | 385.53 | 212.89 | 212.89 | 6.68 | 50 | >sp\|P0DOX2\|IGA2_HUMAN Immunoglobulin alpha-2 heavy chain OS=Homo sapiens OX=9606 PE=1 SV=2 | 631 | 20201116_xd_colostrum_1ng_glyco_hcd203040_compound15.10424.10424.3 | controllerType=0 controllerNumber=1 scan=10424 | 23.6446 |
| 03016:3 | 11 | K.TPLTAN[+2117.777]ITK.S | HexNAc(5)Hex(5)Fuc(2) | N[+2118] | 1026.1186 | 3 | 3076.3411 | 3076.3336 | 0 | 2.5 | 315 | Specific | 398.2 | 248.19 | 248.19 | 7.09 | 50 | >sp\|P0DOX2\|IGA2_HUMAN Immunoglobulin alpha-2 heavy chain OS=Homo sapiens OX=9606 PE=1 SV=2 | 631 | 20201116_xd_colostrum_1ng_glyco_hcd203040_compound15.5756.5756.3 | controllerType=0 controllerNumber=1 scan=5756 | 15.4803 |
| 05000:2 | 11 | K.TPLTAN[+2059.735]ITK.S | HexNAc(4)Hex(5)Fuc(1)NeuAc(1) | N[+2060] | 1509.6525 | 2 | 3018.2977 | 3018.2917 | 0 | 2 | 315 | Specific | 457.23 | 224.37 | 224.37 | 6.86 | 50 | >sp\|P0DOX2\|IGA2_HUMAN Immunoglobulin alpha-2 heavy chain OS=Homo sapiens OX=9606 PE=1 SV=2 | 631 | 20201116_xd_colostrum_1ng_glyco_hcd203040_compound15.7814.7814.2 | controllerType=0 controllerNumber=1 scan=7814 | 19.0852 |
| 03223:3 | 11 | K.TPLTAN[+1809.666]ITK.S | HexNAc(5)Hex(4)Fuc(1) | N[+1810] | 923.4156 | 3 | 2768.2321 | 2768.2228 | 0 | 3.4 | 315 | Specific | 484.78 | 215.31 | 215.31 | 6.83 | 50 | >sp\|P0DOX2\|IGA2_HUMAN Immunoglobulin alpha-2 heavy chain OS=Homo sapiens OX=9606 PE=1 SV=2 | 631 | 20201116_xd_colostrum_1ng_glyco_hcd203040_compound15.5971.5971.3 | controllerType=0 controllerNumber=1 scan=5971 | 15.8725 |
| 03164:3 | 11 | K.TPLTAN[+1971.719]ITK.S | HexNAc(5)Hex(5)Fuc(1) | N[+1972] | 977.4315 | 3 | 2930.2801 | 2930.2757 | 0 | 1.5 | 315 | Specific | 491.2 | 210.8 | 210.8 | 6.87 | 50 | >sp\|P0DOX2\|IGA2_HUMAN Immunoglobulin alpha-2 heavy chain OS=Homo sapiens OX=9606 PE=1 SV=2 | 631 | 20201116_xd_colostrum_1ng_glyco_hcd203040_compound15.5909.5909.3 | controllerType=0 controllerNumber=1 scan=5909 | 15.7402 |
| 03284:3 | 11 | K.TPLTAN[+1647.613]ITK.S | HexNAc(5)Hex(3)Fuc(1) | N[+1648] | 869.3977 | 3 | 2606.1786 | 2606.17 | 0 | 3.3 | 315 | Specific | 502.53 | 222.58 | 222.58 | 7.04 | 50 | >sp\|P0DOX2\|IGA2_HUMAN Immunoglobulin alpha-2 heavy chain OS=Homo sapiens OX=9606 PE=1 SV=2 | 631 | 20201116_xd_colostrum_1ng_glyco_hcd203040_compound15.6035.6035.3 | controllerType=0 controllerNumber=1 scan=6035 | 16.0451 |
| 04966:3 | 11 | K.TPLTAN[+2262.814]ITK.S | HexNAc(5)Hex(5)Fuc(1)NeuAc(1) | N[+2263] | 1074.4641 | 3 | 3221.3779 | 3221.3711 | 0 | 2.1 | 315 | Specific | 511.91 | 254.7 | 254.7 | 7.55 | 50 | >sp\|P0DOX2\|IGA2_HUMAN Immunoglobulin alpha-2 heavy chain OS=Homo sapiens OX=9606 PE=1 SV=2 | 631 | 20201116_xd_colostrum_1ng_glyco_hcd203040_compound15.7779.7779.3 | controllerType=0 controllerNumber=1 scan=7779 | 19.0187 |
| 03213:2 | 11 | K.TPLTAN[+1403.507]ITK.S | HexNAc(3)Hex(4)Fuc(1) | N[+1404] | 1181.5396 | 2 | 2362.0719 | 2362.0641 | 0 | 3.3 | 315 | Specific | 529.17 | 269.35 | 269.35 | 7.28 | 50 | >sp\|P0DOX2\|IGA2_HUMAN Immunoglobulin alpha-2 heavy chain OS=Homo sapiens OX=9606 PE=1 SV=2 | 631 | 20201116_xd_colostrum_1ng_glyco_hcd203040_compound15.5960.5960.2 | controllerType=0 controllerNumber=1 scan=5960 | 15.847 |
| 03297:2 | 11 | K.TPLTAN[+1501.555]ITK.S | HexNAc(5)Hex(3) | N[+1502] | 1230.565 | 2 | 2460.1227 | 2460.1121 | 0 | 4.3 | 315 | Specific | 538.85 | 307.22 | 307.22 | 7.35 | 50 | >sp\|P0DOX2\|IGA2_HUMAN Immunoglobulin alpha-2 heavy chain OS=Homo sapiens OX=9606 PE=1 SV=2 | 631 | 20201116_xd_colostrum_1ng_glyco_hcd203040_compound15.6048.6048.2 | controllerType=0 controllerNumber=1 scan=6048 | 16.0728 |
| 03167:3 | 11 | K.TPLTAN[+1971.719]ITK.S | HexNAc(5)Hex(5)Fuc(1) | N[+1972] | 977.4315 | 3 | 2930.2801 | 2930.2757 | 0 | 1.5 | 315 | Specific | 539.18 | 288.11 | 288.11 | 7.51 | 50 | >sp\|P0DOX2\|IGA2_HUMAN Immunoglobulin alpha-2 heavy chain OS=Homo sapiens OX=9606 PE=1 SV=2 | 631 | 20201116_xd_colostrum_1ng_glyco_hcd203040_compound15.5912.5912.3 | controllerType=0 controllerNumber=1 scan=5912 | 15.7475 |
| 04960:3 | 11 | K.TPLTAN[+2059.735]ITK.S | HexNAc(4)Hex(5)Fuc(1)NeuAc(1) | N[+2060] | 1006.7703 | 3 | 3018.2963 | 3018.2917 | 0 | 1.5 | 315 | Specific | 543.89 | 247.63 | 247.63 | 7.69 | 50 | >sp\|P0DOX2\|IGA2_HUMAN Immunoglobulin alpha-2 heavy chain OS=Homo sapiens OX=9606 PE=1 SV=2 | 631 | 20201116_xd_colostrum_1ng_glyco_hcd203040_compound15.7772.7772.3 | controllerType=0 controllerNumber=1 scan=7772 | 19.0034 |
| 03295:2 | 11 | K.TPLTAN[+1501.555]ITK.S | HexNAc(5)Hex(3) | N[+1502] | 1230.565 | 2 | 2460.1227 | 2460.1121 | 0 | 4.3 | 315 | Specific | 548.8 | 322.76 | 322.76 | 7.74 | 50 | >sp\|P0DOX2\|IGA2_HUMAN Immunoglobulin alpha-2 heavy chain OS=Homo sapiens OX=9606 PE=1 SV=2 | 631 | 20201116_xd_colostrum_1ng_glyco_hcd203040_compound15.6046.6046.2 | controllerType=0 controllerNumber=1 scan=6046 | 16.0676 |
| 04969:3 | 11 | K.TPLTAN[+2262.814]ITK.S | HexNAc(5)Hex(5)Fuc(1)NeuAc(1) | N[+2263] | 1074.4641 | 3 | 3221.3779 | 3221.3711 | 0 | 2.1 | 315 | Specific | 553.05 | 292.13 | 292.13 | 7.9 | 50 | >sp\|P0DOX2\|IGA2_HUMAN Immunoglobulin alpha-2 heavy chain OS=Homo sapiens OX=9606 PE=1 SV=2 | 631 | 20201116_xd_colostrum_1ng_glyco_hcd203040_compound15.7782.7782.3 | controllerType=0 controllerNumber=1 scan=7782 | 19.0239 |
| 03225:3 | 11 | K.TPLTAN[+1809.666]ITK.S | HexNAc(5)Hex(4)Fuc(1) | N[+1810] | 923.4156 | 3 | 2768.2321 | 2768.2228 | 0 | 3.4 | 315 | Specific | 558.26 | 281.51 | 281.51 | 7.71 | 50 | >sp\|P0DOX2\|IGA2_HUMAN Immunoglobulin alpha-2 heavy chain OS=Homo sapiens OX=9606 PE=1 SV=2 | 631 | 20201116_xd_colostrum_1ng_glyco_hcd203040_compound15.5973.5973.3 | controllerType=0 controllerNumber=1 scan=5973 | 15.8796 |
| 03287:3 | 11 | K.TPLTAN[+1647.613]ITK.S | HexNAc(5)Hex(3)Fuc(1) | N[+1648] | 869.3977 | 3 | 2606.1786 | 2606.17 | 0 | 3.3 | 315 | Specific | 561.64 | 268.53 | 268.53 | 7.81 | 50 | >sp\|P0DOX2\|IGA2_HUMAN Immunoglobulin alpha-2 heavy chain OS=Homo sapiens OX=9606 PE=1 SV=2 | 631 | 20201116_xd_colostrum_1ng_glyco_hcd203040_compound15.6038.6038.3 | controllerType=0 controllerNumber=1 scan=6038 | 16.0522 |
| 03351:2 | 11 | K.TPLTAN[+1038.375]ITK.S | HexNAc(2)Hex(3)Fuc(1) | N[+1038] | 998.974 | 2 | 1996.9408 | 1996.9319 | 0 | 4.5 | 315 | Specific | 568.72 | 303.83 | 303.83 | 7.78 | 50 | >sp\|P0DOX2\|IGA2_HUMAN Immunoglobulin alpha-2 heavy chain OS=Homo sapiens OX=9606 PE=1 SV=2 | 631 | 20201116_xd_colostrum_1ng_glyco_hcd203040_compound15.6105.6105.2 | controllerType=0 controllerNumber=1 scan=6105 | 16.1939 |
| 03222:3 | 11 | K.TPLTAN[+1606.587]ITK.S | HexNAc(4)Hex(4)Fuc(1) | N[+1607] | 855.7222 | 3 | 2565.152 | 2565.1435 | 0 | 3.3 | 315 | Specific | 572.56 | 229.48 | 229.48 | 7.52 | 50 | >sp\|P0DOX2\|IGA2_HUMAN Immunoglobulin alpha-2 heavy chain OS=Homo sapiens OX=9606 PE=1 SV=2 | 631 | 20201116_xd_colostrum_1ng_glyco_hcd203040_compound15.5970.5970.3 | controllerType=0 controllerNumber=1 scan=5970 | 15.8711 |
| 03180:2 | 11 | K.TPLTAN[+1971.719]ITK.S | HexNAc(5)Hex(5)Fuc(1) | N[+1972] | 1465.6465 | 2 | 2930.2858 | 2930.2757 | 0 | 3.5 | 315 | Specific | 573.1 | 310.52 | 310.52 | 8.02 | 50 | >sp\|P0DOX2\|IGA2_HUMAN Immunoglobulin alpha-2 heavy chain OS=Homo sapiens OX=9606 PE=1 SV=2 | 631 | 20201116_xd_colostrum_1ng_glyco_hcd203040_compound15.5926.5926.2 | controllerType=0 controllerNumber=1 scan=5926 | 15.7766 |
| 03349:2 | 11 | K.TPLTAN[+1038.375]ITK.S | HexNAc(2)Hex(3)Fuc(1) | N[+1038] | 998.974 | 2 | 1996.9408 | 1996.9319 | 0 | 4.5 | 315 | Specific | 574.51 | 325.51 | 325.51 | 7.93 | 50 | >sp\|P0DOX2\|IGA2_HUMAN Immunoglobulin alpha-2 heavy chain OS=Homo sapiens OX=9606 PE=1 SV=2 | 631 | 20201116_xd_colostrum_1ng_glyco_hcd203040_compound15.6103.6103.2 | controllerType=0 controllerNumber=1 scan=6103 | 16.1901 |
| 03298:3 | 11 | K.TPLTAN[+1444.534]ITK.S | HexNAc(4)Hex(3)Fuc(1) | N[+1445] | 801.7048 | 3 | 2403.0999 | 2403.0906 | 0 | 3.8 | 315 | Specific | 576.62 | 272.11 | 272.11 | 7.81 | 50 | >sp\|P0DOX2\|IGA2_HUMAN Immunoglobulin alpha-2 heavy chain OS=Homo sapiens OX=9606 PE=1 SV=2 | 631 | 20201116_xd_colostrum_1ng_glyco_hcd203040_compound15.6050.6050.3 | controllerType=0 controllerNumber=1 scan=6050 | 16.0773 |
| 04962:3 | 11 | K.TPLTAN[+2059.735]ITK.S | HexNAc(4)Hex(5)Fuc(1)NeuAc(1) | N[+2060] | 1006.7703 | 3 | 3018.2963 | 3018.2917 | 0 | 1.5 | 315 | Specific | 578.62 | 278.52 | 278.52 | 7.85 | 50 | >sp\|P0DOX2\|IGA2_HUMAN Immunoglobulin alpha-2 heavy chain OS=Homo sapiens OX=9606 PE=1 SV=2 | 631 | 20201116_xd_colostrum_1ng_glyco_hcd203040_compound15.7774.7774.3 | controllerType=0 controllerNumber=1 scan=7774 | 19.0098 |
| 03206:2 | 11 | K.TPLTAN[+1606.587]ITK.S | HexNAc(4)Hex(4)Fuc(1) | N[+1607] | 1283.079 | 2 | 2565.1508 | 2565.1435 | 0 | 2.8 | 315 | Specific | 584.83 | 333.63 | 333.63 | 9.66 | 50 | >sp\|P0DOX2\|IGA2_HUMAN Immunoglobulin alpha-2 heavy chain OS=Homo sapiens OX=9606 PE=1 SV=2 | 631 | 20201116_xd_colostrum_1ng_glyco_hcd203040_compound15.5953.5953.2 | controllerType=0 controllerNumber=1 scan=5953 | 15.832 |
| 03345:2 | 11 | K.TPLTAN[+1647.613]ITK.S | HexNAc(5)Hex(3)Fuc(1) | N[+1648] | 1303.5927 | 2 | 2606.1781 | 2606.17 | 0 | 3.1 | 315 | Specific | 585.1 | 254.11 | 254.11 | 9.04 | 50 | >sp\|P0DOX2\|IGA2_HUMAN Immunoglobulin alpha-2 heavy chain OS=Homo sapiens OX=9606 PE=1 SV=2 | 631 | 20201116_xd_colostrum_1ng_glyco_hcd203040_compound15.6098.6098.2 | controllerType=0 controllerNumber=1 scan=6098 | 16.1818 |
| 03283:2 | 11 | K.TPLTAN[+1647.613]ITK.S | HexNAc(5)Hex(3)Fuc(1) | N[+1648] | 1303.5934 | 2 | 2606.1796 | 2606.17 | 0 | 3.7 | 315 | Specific | 597.59 | 308.74 | 308.74 | 9.31 | 50 | >sp\|P0DOX2\|IGA2_HUMAN Immunoglobulin alpha-2 heavy chain OS=Homo sapiens OX=9606 PE=1 SV=2 | 631 | 20201116_xd_colostrum_1ng_glyco_hcd203040_compound15.6034.6034.2 | controllerType=0 controllerNumber=1 scan=6034 | 16.044 |
| 03182:2 | 11 | K.TPLTAN[+1971.719]ITK.S | HexNAc(5)Hex(5)Fuc(1) | N[+1972] | 1465.6465 | 2 | 2930.2858 | 2930.2757 | 0 | 3.5 | 315 | Specific | 600.43 | 300.67 | 300.67 | 10.11 | 50 | >sp\|P0DOX2\|IGA2_HUMAN Immunoglobulin alpha-2 heavy chain OS=Homo sapiens OX=9606 PE=1 SV=2 | 631 | 20201116_xd_colostrum_1ng_glyco_hcd203040_compound15.5928.5928.2 | controllerType=0 controllerNumber=1 scan=5928 | 15.7819 |
| 03243:2 | 11 | K.TPLTAN[+1809.666]ITK.S | HexNAc(5)Hex(4)Fuc(1) | N[+1810] | 1384.6181 | 2 | 2768.2289 | 2768.2228 | 0 | 2.2 | 315 | Specific | 603.12 | 306.56 | 306.56 | 10.51 | 50 | >sp\|P0DOX2\|IGA2_HUMAN Immunoglobulin alpha-2 heavy chain OS=Homo sapiens OX=9606 PE=1 SV=2 | 631 | 20201116_xd_colostrum_1ng_glyco_hcd203040_compound15.5992.5992.2 | controllerType=0 controllerNumber=1 scan=5992 | 15.9272 |
| 03241:2 | 11 | K.TPLTAN[+1809.666]ITK.S | HexNAc(5)Hex(4)Fuc(1) | N[+1810] | 1384.6181 | 2 | 2768.2289 | 2768.2228 | 0 | 2.2 | 315 | Specific | 604.84 | 319.49 | 319.49 | 10.51 | 50 | >sp\|P0DOX2\|IGA2_HUMAN Immunoglobulin alpha-2 heavy chain OS=Homo sapiens OX=9606 PE=1 SV=2 | 631 | 20201116_xd_colostrum_1ng_glyco_hcd203040_compound15.5990.5990.2 | controllerType=0 controllerNumber=1 scan=5990 | 15.9204 |
| 04982:2 | 11 | K.TPLTAN[+2262.814]ITK.S | HexNAc(5)Hex(5)Fuc(1)NeuAc(1) | N[+2263] | 1611.193 | 2 | 3221.3788 | 3221.3711 | 0 | 2.4 | 315 | Specific | 618.46 | 297.75 | 297.75 | 10.3 | 50 | >sp\|P0DOX2\|IGA2_HUMAN Immunoglobulin alpha-2 heavy chain OS=Homo sapiens OX=9606 PE=1 SV=2 | 631 | 20201116_xd_colostrum_1ng_glyco_hcd203040_compound15.7795.7795.2 | controllerType=0 controllerNumber=1 scan=7795 | 19.0475 |
| 04985:2 | 11 | K.TPLTAN[+2262.814]ITK.S | HexNAc(5)Hex(5)Fuc(1)NeuAc(1) | N[+2263] | 1611.193 | 2 | 3221.3788 | 3221.3711 | 0 | 2.4 | 315 | Specific | 618.65 | 340.97 | 340.97 | 10.73 | 50 | >sp\|P0DOX2\|IGA2_HUMAN Immunoglobulin alpha-2 heavy chain OS=Homo sapiens OX=9606 PE=1 SV=2 | 631 | 20201116_xd_colostrum_1ng_glyco_hcd203040_compound15.7798.7798.2 | controllerType=0 controllerNumber=1 scan=7798 | 19.0539 |
| 03282:2 | 11 | K.TPLTAN[+1444.534]ITK.S | HexNAc(4)Hex(3)Fuc(1) | N[+1445] | 1202.0535 | 2 | 2403.0997 | 2403.0906 | 0 | 3.8 | 315 | Specific | 626.53 | 324.86 | 324.86 | 10.46 | 50 | >sp\|P0DOX2\|IGA2_HUMAN Immunoglobulin alpha-2 heavy chain OS=Homo sapiens OX=9606 PE=1 SV=2 | 631 | 20201116_xd_colostrum_1ng_glyco_hcd203040_compound15.6033.6033.2 | controllerType=0 controllerNumber=1 scan=6033 | 16.0387 |
| 03281:2 | 11 | K.TPLTAN[+1647.613]ITK.S | HexNAc(5)Hex(3)Fuc(1) | N[+1648] | 1303.5934 | 2 | 2606.1796 | 2606.17 | 0 | 3.7 | 315 | Specific | 627.92 | 354.77 | 354.77 | 10.54 | 50 | >sp\|P0DOX2\|IGA2_HUMAN Immunoglobulin alpha-2 heavy chain OS=Homo sapiens OX=9606 PE=1 SV=2 | 631 | 20201116_xd_colostrum_1ng_glyco_hcd203040_compound15.6032.6032.2 | controllerType=0 controllerNumber=1 scan=6032 | 16.0334 |
| 03224:3 | 11 | K.TPLTAN[+1606.587]ITK.S | HexNAc(4)Hex(4)Fuc(1) | N[+1607] | 855.7222 | 3 | 2565.152 | 2565.1435 | 0 | 3.3 | 315 | Specific | 629.58 | 288.88 | 288.88 | 10.2 | 50 | >sp\|P0DOX2\|IGA2_HUMAN Immunoglobulin alpha-2 heavy chain OS=Homo sapiens OX=9606 PE=1 SV=2 | 631 | 20201116_xd_colostrum_1ng_glyco_hcd203040_compound15.5972.5972.3 | controllerType=0 controllerNumber=1 scan=5972 | 15.8763 |
| 03204:2 | 11 | K.TPLTAN[+1606.587]ITK.S | HexNAc(4)Hex(4)Fuc(1) | N[+1607] | 1283.079 | 2 | 2565.1508 | 2565.1435 | 0 | 2.8 | 315 | Specific | 638.28 | 351.75 | 351.75 | 10.73 | 50 | >sp\|P0DOX2\|IGA2_HUMAN Immunoglobulin alpha-2 heavy chain OS=Homo sapiens OX=9606 PE=1 SV=2 | 631 | 20201116_xd_colostrum_1ng_glyco_hcd203040_compound15.5951.5951.2 | controllerType=0 controllerNumber=1 scan=5951 | 15.8235 |
| 03279:2 | 11 | K.TPLTAN[+1444.534]ITK.S | HexNAc(4)Hex(3)Fuc(1) | N[+1445] | 1202.0535 | 2 | 2403.0997 | 2403.0906 | 0 | 3.8 | 315 | Specific | 641.29 | 366.68 | 366.68 | 10.72 | 50 | >sp\|P0DOX2\|IGA2_HUMAN Immunoglobulin alpha-2 heavy chain OS=Homo sapiens OX=9606 PE=1 SV=2 | 631 | 20201116_xd_colostrum_1ng_glyco_hcd203040_compound15.6030.6030.2 | controllerType=0 controllerNumber=1 scan=6030 | 16.0313 |
| 03215:2 | 11 | K.TPLTAN[+1403.507]ITK.S | HexNAc(3)Hex(4)Fuc(1) | N[+1404] | 1181.5396 | 2 | 2362.0719 | 2362.0641 | 0 | 3.3 | 315 | Specific | 643.5 | 308.56 | 308.56 | 10.31 | 50 | >sp\|P0DOX2\|IGA2_HUMAN Immunoglobulin alpha-2 heavy chain OS=Homo sapiens OX=9606 PE=1 SV=2 | 631 | 20201116_xd_colostrum_1ng_glyco_hcd203040_compound15.5962.5962.2 | controllerType=0 controllerNumber=1 scan=5962 | 15.8525 |
| 03300:3 | 11 | K.TPLTAN[+1444.534]ITK.S | HexNAc(4)Hex(3)Fuc(1) | N[+1445] | 801.7048 | 3 | 2403.0999 | 2403.0906 | 0 | 3.8 | 315 | Specific | 655.52 | 293.49 | 293.49 | 11.2 | 50 | >sp\|P0DOX2\|IGA2_HUMAN Immunoglobulin alpha-2 heavy chain OS=Homo sapiens OX=9606 PE=1 SV=2 | 631 | 20201116_xd_colostrum_1ng_glyco_hcd203040_compound15.6052.6052.3 | controllerType=0 controllerNumber=1 scan=6052 | 16.0825 |
| 03306:2 | 11 | K.TPLTAN[+1241.454]ITK.S | HexNAc(3)Hex(3)Fuc(1) | N[+1241] | 1100.513 | 2 | 2200.0187 | 2200.0113 | 0 | 3.4 | 315 | Specific | 663.6 | 378.43 | 378.43 | 11.72 | 50 | >sp\|P0DOX2\|IGA2_HUMAN Immunoglobulin alpha-2 heavy chain OS=Homo sapiens OX=9606 PE=1 SV=2 | 631 | 20201116_xd_colostrum_1ng_glyco_hcd203040_compound15.6058.6058.2 | controllerType=0 controllerNumber=1 scan=6058 | 16.0974 |
| 03309:2 | 11 | K.TPLTAN[+1241.454]ITK.S | HexNAc(3)Hex(3)Fuc(1) | N[+1241] | 1100.513 | 2 | 2200.0187 | 2200.0113 | 0 | 3.4 | 315 | Specific | 670.16 | 314.81 | 314.81 | 11.31 | 50 | >sp\|P0DOX2\|IGA2_HUMAN Immunoglobulin alpha-2 heavy chain OS=Homo sapiens OX=9606 PE=1 SV=2 | 631 | 20201116_xd_colostrum_1ng_glyco_hcd203040_compound15.6061.6061.2 | controllerType=0 controllerNumber=1 scan=6061 | 16.1033 |
| 00640:3 | 12 | K.YKN[+2262.814]NSDISSTR.G | HexNAc(5)Hex(5)Fuc(1)NeuAc(1) | N[+2263] | 1183.1546 | 3 | 3547.4492 | 3547.4322 | 0 | 4.8 | 44 | Specific | 383.7 | 215.58 | 215.58 | 5.96 | 36 | >sp\|P01871\|IGHM_HUMAN Immunoglobulin heavy constant mu OS=Homo sapiens OX=9606 GN=IGHM PE=1 SV=4 | 328 | 20201116_xd_colostrum_1ng_glyco_hcd203040_compound15.3290.3290.3 | controllerType=0 controllerNumber=1 scan=3290 | 11.2527 |
| 00642:3 | 12 | K.YKN[+2262.814]NSDISSTR.G | HexNAc(5)Hex(5)Fuc(1)NeuAc(1) | N[+2263] | 1183.1546 | 3 | 3547.4492 | 3547.4322 | 0 | 4.8 | 44 | Specific | 448.65 | 358.21 | 358.21 | 7.05 | 36 | >sp\|P01871\|IGHM_HUMAN Immunoglobulin heavy constant mu OS=Homo sapiens OX=9606 GN=IGHM PE=1 SV=4 | 328 | 20201116_xd_colostrum_1ng_glyco_hcd203040_compound15.3292.3292.3 | controllerType=0 controllerNumber=1 scan=3292 | 11.2574 |
| 02443:3 | 13 | K.N[+1216.423]NSIPDKQITASSSYK.T | HexNAc(2)Hex(5) | N[+1216] | 990.44 | 3 | 2969.3054 | 2969.2991 | 0 | 2.1 | 238 | Specific | 565.54 | 306.08 | 306.08 | 8.14 | 27 | >sp\|Q08431\|MFGM_HUMAN Lactadherin OS=Homo sapiens OX=9606 GN=MFGE8 PE=1 SV=3 | 1064 | 20201116_xd_colostrum_1ng_glyco_hcd203040_compound15.5162.5162.3 | controllerType=0 controllerNumber=1 scan=5162 | 14.4908 |
| 02441:3 | 13 | K.N[+1216.423]NSIPDKQITASSSYK.T | HexNAc(2)Hex(5) | N[+1216] | 990.44 | 3 | 2969.3054 | 2969.2991 | 0 | 2.1 | 238 | Specific | 581.76 | 334.5 | 334.5 | 9.59 | 27 | >sp\|Q08431\|MFGM_HUMAN Lactadherin OS=Homo sapiens OX=9606 GN=MFGE8 PE=1 SV=3 | 1064 | 20201116_xd_colostrum_1ng_glyco_hcd203040_compound15.5160.5160.3 | controllerType=0 controllerNumber=1 scan=5160 | 14.4862 |
| 11231:3 | 18 | R.VN[+1216.423]LTTNTIAVTQTLPNAAYNNR.F | HexNAc(2)Hex(5) | N[+1216] | 1202.5643 | 3 | 3605.6782 | 3605.6699 | 0 | 2.3 | 351 | Specific | 751.77 | 751.77 | 751.77 | 16.71 | 24 | >sp\|Q6UX06\|OLFM4_HUMAN Olfactomedin-4 OS=Homo sapiens OX=9606 GN=OLFM4 PE=1 SV=1 | 624 | 20201116_xd_colostrum_1ng_glyco_hcd203040_compound15.14279.14279.3 | controllerType=0 controllerNumber=1 scan=14279 | 30.991 |
| 04111:2 | 19 | K.AAIPSALDTN[+1216.423]SSK.S | HexNAc(2)Hex(5) | N[+1216] | 1246.0483 | 2 | 2491.0892 | 2491.0815 | 0 | 3.1 | 542 | Specific | 604.15 | 445.4 | 445.4 | 12.32 | 24 | >sp\|Q08380\|LG3BP_HUMAN Galectin-3-binding protein OS=Homo sapiens OX=9606 GN=LGALS3BP PE=1 SV=1 | 1562 | 20201116_xd_colostrum_1ng_glyco_hcd203040_compound15.6892.6892.2 | controllerType=0 controllerNumber=1 scan=6892 | 17.5715 |
| 04109:2 | 19 | K.AAIPSALDTN[+1216.423]SSK.S | HexNAc(2)Hex(5) | N[+1216] | 1246.0483 | 2 | 2491.0892 | 2491.0815 | 0 | 3.1 | 542 | Specific | 616.42 | 532.66 | 532.66 | 12.32 | 24 | >sp\|Q08380\|LG3BP_HUMAN Galectin-3-binding protein OS=Homo sapiens OX=9606 GN=LGALS3BP PE=1 SV=1 | 1562 | 20201116_xd_colostrum_1ng_glyco_hcd203040_compound15.6890.6890.2 | controllerType=0 controllerNumber=1 scan=6890 | 17.567 |
| 13364:3 | 22 | K.LKELPGVC[+57.021]N[+2351.851]ETMMALWEEC[+57.021]KPC[+57.021]LK.Q | HexNAc(4)Hex(5)Fuc(3)NeuAc(1) | C[+57]*3, N[+2352] | 1763.4194 | 3 | 5288.2436 | 5288.2355 | 0 | 1.5 | 95 | Specific | 285.94 | 266.07 | 266.07 | 6.72 | 24 | >sp\|P10909\|CLUS_HUMAN Clusterin OS=Homo sapiens OX=9606 GN=CLU PE=1 SV=1 | 1467 | 20201116_xd_colostrum_1ng_glyco_hcd203040_compound15.16495.16495.3 | controllerType=0 controllerNumber=1 scan=16495 | 35.2407 |
| 13361:3 | 22 | K.LKELPGVC[+57.021]N[+2351.851]ETMMALWEEC[+57.021]KPC[+57.021]LK.Q | HexNAc(4)Hex(5)Fuc(3)NeuAc(1) | C[+57]*3, N[+2352] | 1763.4194 | 3 | 5288.2436 | 5288.2355 | 0 | 1.5 | 95 | Specific | 360.73 | 360.73 | 360.73 | 7.8 | 24 | >sp\|P10909\|CLUS_HUMAN Clusterin OS=Homo sapiens OX=9606 GN=CLU PE=1 SV=1 | 1467 | 20201116_xd_colostrum_1ng_glyco_hcd203040_compound15.16492.16492.3 | controllerType=0 controllerNumber=1 scan=16492 | 35.2361 |
| 13436:4 | 22 | K.LKELPGVC[+57.021]N[+2059.735]ETMMALWEEC[+57.021]KPC[+57.021]LK.Q | HexNAc(4)Hex(5)Fuc(1)NeuAc(1) | C[+57]*3, N[+2060] | 1249.7902 | 4 | 4996.139 | 4996.1196 | 0 | 3.9 | 95 | Specific | 699.25 | 509.78 | 509.78 | 14.12 | 24 | >sp\|P10909\|CLUS_HUMAN Clusterin OS=Homo sapiens OX=9606 GN=CLU PE=1 SV=1 | 1467 | 20201116_xd_colostrum_1ng_glyco_hcd203040_compound15.16570.16570.4 | controllerType=0 controllerNumber=1 scan=16570 | 35.4116 |
| 13434:4 | 22 | K.LKELPGVC[+57.021]N[+2059.735]ETMMALWEEC[+57.021]KPC[+57.021]LK.Q | HexNAc(4)Hex(5)Fuc(1)NeuAc(1) | C[+57]*3, N[+2060] | 1249.7902 | 4 | 4996.139 | 4996.1196 | 0 | 3.9 | 95 | Specific | 780.42 | 756.14 | 756.14 | 16.12 | 24 | >sp\|P10909\|CLUS_HUMAN Clusterin OS=Homo sapiens OX=9606 GN=CLU PE=1 SV=1 | 1467 | 20201116_xd_colostrum_1ng_glyco_hcd203040_compound15.16568.16568.4 | controllerType=0 controllerNumber=1 scan=16568 | 35.4071 |
| 19824:4 | 22 | K.MLN[+2059.735]TSSLLEQLNEQFNWVSR.L | HexNAc(4)Hex(5)Fuc(1)NeuAc(1) | N[+2060] | 1117.9847 | 4 | 4468.9168 | 4468.9217 | 0 | -1.1 | 352 | Specific | 610.63 | 513.09 | 513.09 | 12.31 | 24 | >sp\|P10909\|CLUS_HUMAN Clusterin OS=Homo sapiens OX=9606 GN=CLU PE=1 SV=1 | 1467 | 20201116_xd_colostrum_1ng_glyco_hcd203040_compound15.23237.23237.4 | controllerType=0 controllerNumber=1 scan=23237 | 49.8152 |
| 21847:4 | 22 | K.MLN[+2350.830]TSSLLEQLNEQFNWVSR.L | HexNAc(4)Hex(5)Fuc(1)NeuAc(2) | N[+2351] | 1190.7632 | 4 | 4760.0311 | 4760.0171 | 0 | 2.9 | 352 | Specific | 632.24 | 553.12 | 553.12 | 13.34 | 24 | >sp\|P10909\|CLUS_HUMAN Clusterin OS=Homo sapiens OX=9606 GN=CLU PE=1 SV=1 | 1467 | 20201116_xd_colostrum_1ng_glyco_hcd203040_compound15.25351.25351.4 | controllerType=0 controllerNumber=1 scan=25351 | 54.3754 |
| 19826:4 | 22 | K.MLN[+2059.735]TSSLLEQLNEQFNWVSR.L | HexNAc(4)Hex(5)Fuc(1)NeuAc(1) | N[+2060] | 1117.9847 | 4 | 4468.9168 | 4468.9217 | 0 | -1.1 | 352 | Specific | 693.64 | 426.1 | 426.1 | 13.31 | 24 | >sp\|P10909\|CLUS_HUMAN Clusterin OS=Homo sapiens OX=9606 GN=CLU PE=1 SV=1 | 1467 | 20201116_xd_colostrum_1ng_glyco_hcd203040_compound15.23239.23239.4 | controllerType=0 controllerNumber=1 scan=23239 | 49.8218 |
| 21849:4 | 22 | K.MLN[+2350.830]TSSLLEQLNEQFNWVSR.L | HexNAc(4)Hex(5)Fuc(1)NeuAc(2) | N[+2351] | 1190.7632 | 4 | 4760.0311 | 4760.0171 | 0 | 2.9 | 352 | Specific | 777.11 | 542.92 | 542.92 | 16.34 | 24 | >sp\|P10909\|CLUS_HUMAN Clusterin OS=Homo sapiens OX=9606 GN=CLU PE=1 SV=1 | 1467 | 20201116_xd_colostrum_1ng_glyco_hcd203040_compound15.25353.25353.4 | controllerType=0 controllerNumber=1 scan=25353 | [illegible] |
| 19729:3 | 22 | K.MLN[+2351.851]TSSLLEQLNEQFNWVSR.L | HexNAc(4)Hex(5)Fuc(3)NeuAc(1) | N[+2352] | 1587.6893 | 3 | 4761.0532 | 4761.0375 | 0 | 3.3 | 352 | Specific | 873.37 | 735.04 | 735.04 | 17.86 | 24 | >sp\|P10909\|CLUS_HUMAN Clusterin OS=Homo sapiens OX=9606 GN=CLU PE=1 SV=1 | 1467 | 20201116_xd_colostrum_1ng_glyco_hcd203040_compound15.23136.23136.2 | controllerType=0 controllerNumber=1 scan=23136 | [illegible] |
| 19728:3 | 22 | K.MLN[+2351.851]TSSLLEQLNEQFNWVSR.L | HexNAc(4)Hex(5)Fuc(3)NeuAc(1) | N[+2352] | 1587.6893 | 3 | 4761.0532 | 4761.0375 | 0 | 3.3 | 352 | Specific | 899.72 | 618.05 | 618.05 | 17.86 | 24 | >sp\|P10909\|CLUS_HUMAN Clusterin OS=Homo sapiens OX=9606 GN=CLU PE=1 SV=1 | 1467 | 20201116_xd_colostrum_1ng_glyco_hcd203040_compound15.23135.23135.3 | controllerType=0 controllerNumber=1 scan=23135 | [illegible] |
| 19726:3 | 22 | K.MLN[+2351.851]TSSLLEQLNEQFNWVSR.L | HexNAc(4)Hex(5)Fuc(3)NeuAc(1) | N[+2352] | 1587.6893 | 3 | 4761.0532 | 4761.0375 | 0 | 3.3 | 352 | Specific | 940.72 | 744.57 | 744.57 | 18.86 | 24 | >sp\|P10909\|CLUS_HUMAN Clusterin OS=Homo sapiens OX=9606 GN=CLU PE=1 SV=1 | 1467 | 20201116_xd_colostrum_1ng_glyco_hcd203040_compound15.23133.23133.3 | controllerType=0 controllerNumber=1 scan=23133 | [illegible] |
| 19731:3 | 22 | K.MLN[+2351.851]TSSLLEQLNEQFNWVSR.L | HexNAc(4)Hex(5)Fuc(3)NeuAc(1) | N[+2352] | 1587.6893 | 3 | 4761.0532 | 4761.0375 | 0 | 3.3 | 352 | Specific | 994.17 | 703.51 | 703.51 | 19.86 | 24 | >sp\|P10909\|CLUS_HUMAN Clusterin OS=Homo sapiens OX=9606 GN=CLU PE=1 SV=1 | 1467 | 20201116_xd_colostrum_1ng_glyco_hcd203040_compound15.23138.23138.2 | controllerType=0 controllerNumber=1 scan=23138 | 49.5763 |
| 10963:3 | 23 | R.IIVPLNNREN[+2116.756]ISDPTSPLR.T | HexNAc(5)Hex(5)NeuAc(1) | N[+2117] | 1422.3199 | 3 | 4264.9451 | 4264.9335 | 0 | 2.7 | 62 | Specific | 279.11 | 266.49 | 266.49 | 6.6 | 25 | >sp\|P01591\|IGJ_HUMAN Immunoglobulin J chain OS=Homo sapiens OX=9606 GN=JCHAIN PE=1 SV=4 | 1392 | 20201116_xd_colostrum_1ng_glyco_hcd203040_compound15.14000.14000.3 | controllerType=0 controllerNumber=1 scan=14000 | 30.4422 |
| 12771:4 | 23 | R.IIVPLNNREN[+2204.772]ISDPTSPLR.T | HexNAc(4)Hex(5)NeuAc(2) | N[+2205] | 1088.9938 | 4 | 4352.9534 | 4352.9496 | 0 | 0.9 | 62 | Specific | 289.29 | 289.29 | 289.29 | 6.53 | 25 | >sp\|P01591\|IGJ_HUMAN Immunoglobulin J chain OS=Homo sapiens OX=9606 GN=JCHAIN PE=1 SV=4 | 1392 | 20201116_xd_colostrum_1ng_glyco_hcd203040_compound15.15880.15880.4 | controllerType=0 controllerNumber=1 scan=15880 | 34.0702 |
| 11160:3 | 23 | R.IIVPLNNREN[+1548.545]ISDPTSPLR.T | HexNAc(3)Hex(4)NeuAc(1) | N[+1549] | 1232.9095 | 3 | 3696.7139 | 3696.722 | 0 | -2.2 | 62 | Specific | 294.35 | 282.98 | 282.98 | 4.23 | 25 | >sp\|P01591\|IGJ_HUMAN Immunoglobulin J chain OS=Homo sapiens OX=9606 GN=JCHAIN PE=1 SV=4 | 1392 | 20201116_xd_colostrum_1ng_glyco_hcd203040_compound15.14205.14205.3 | controllerType=0 controllerNumber=1 scan=14205 | 30.8475 |
| 09631:3 | 23 | R.IIVPLNNREN[+1501.555]ISDPTSPLR.T | HexNAc(5)Hex(3) | N[+1502] | 1217.2513 | 3 | 3649.7393 | 3649.7325 | 0 | 1.9 | 62 | Specific | 308.13 | 308.13 | 308.13 | 6.84 | 25 | >sp\|P01591\|IGJ_HUMAN Immunoglobulin J chain OS=Homo sapiens OX=9606 GN=JCHAIN PE=1 SV=4 | 1392 | 20201116_xd_colostrum_1ng_glyco_hcd203040_compound15.12617.12617.3 | controllerType=0 controllerNumber=1 scan=12617 | 27.7084 |
| 09634:3 | 23 | R.IIVPLNNREN[+1501.555]ISDPTSPLR.T | HexNAc(5)Hex(3) | N[+1502] | 1217.2513 | 3 | 3649.7393 | 3649.7325 | 0 | 1.9 | 62 | Specific | 420.12 | 420.12 | 420.12 | 9.38 | 25 | >sp\|P01591\|IGJ_HUMAN Immunoglobulin J chain OS=Homo sapiens OX=9606 GN=JCHAIN PE=1 SV=4 | 1392 | 20201116_xd_colostrum_1ng_glyco_hcd203040_compound15.12620.12620.3 | controllerType=0 controllerNumber=1 scan=12620 | 27.7142 |
| 10964:3 | 23 | R.IIVPLNNREN[+2116.756]ISDPTSPLR.T | HexNAc(5)Hex(5)NeuAc(1) | N[+2117] | 1422.3199 | 3 | 4264.9451 | 4264.9335 | 0 | 2.7 | 62 | Specific | 445.95 | 393.59 | 393.59 | 9.6 | 25 | >sp\|P01591\|IGJ_HUMAN Immunoglobulin J chain OS=Homo sapiens OX=9606 GN=JCHAIN PE=1 SV=4 | 1392 | 20201116_xd_colostrum_1ng_glyco_hcd203040_compound15.14001.14001.3 | controllerType=0 controllerNumber=1 scan=14001 | 30.4454 |
| 10899:4 | 23 | R.IIVPLNNREN[+2059.735]ISDPTSPLR.T | HexNAc(4)Hex(5)Fuc(1)NeuAc(1) | N[+2060] | 1052.7354 | 4 | 4207.9197 | 4207.9121 | 0 | 1.8 | 62 | Specific | 466.26 | 466.26 | 466.26 | 9.67 | 25 | >sp\|P01591\|IGJ_HUMAN Immunoglobulin J chain OS=Homo sapiens OX=9606 GN=JCHAIN PE=1 SV=4 | 1392 | 20201116_xd_colostrum_1ng_glyco_hcd203040_compound15.13933.13933.4 | controllerType=0 controllerNumber=1 scan=13933 | 30.2688 |
| 12774:4 | 23 | R.IIVPLNNREN[+2204.772]ISDPTSPLR.T | HexNAc(4)Hex(5)NeuAc(2) | N[+2205] | 1088.9938 | 4 | 4352.9534 | 4352.9496 | 0 | 0.9 | 62 | Specific | 478.54 | 421.4 | 421.4 | 9.39 | 25 | >sp\|P01591\|IGJ_HUMAN Immunoglobulin J chain OS=Homo sapiens OX=9606 GN=JCHAIN PE=1 SV=4 | 1392 | 20201116_xd_colostrum_1ng_glyco_hcd203040_compound15.15883.15883.4 | controllerType=0 controllerNumber=1 scan=15883 | 34.0739 |
| 10902:4 | 23 | R.IIVPLNNREN[+2059.735]ISDPTSPLR.T | HexNAc(4)Hex(5)Fuc(1)NeuAc(1) | N[+2060] | 1052.7354 | 4 | 4207.9197 | 4207.9121 | 0 | 1.8 | 62 | Specific | 519 | 467.07 | 467.07 | 9.82 | 25 | >sp\|P01591\|IGJ_HUMAN Immunoglobulin J chain OS=Homo sapiens OX=9606 GN=JCHAIN PE=1 SV=4 | 1392 | 20201116_xd_colostrum_1ng_glyco_hcd203040_compound15.13936.13936.4 | controllerType=0 controllerNumber=1 scan=13936 | 30.2798 |
| 12668:3 | 23 | R.IIVPLNNREN[+2350.830]ISDPTSPLR.T | HexNAc(4)Hex(5)Fuc(1)NeuAc(2) | N[+2351] | 1500.3422 | 3 | 4499.0121 | 4499.0075 | 0 | 1 | 62 | Specific | 524.14 | 455.44 | 455.44 | 9.78 | 25 | >sp\|P01591\|IGJ_HUMAN Immunoglobulin J chain OS=Homo sapiens OX=9606 GN=JCHAIN PE=1 SV=4 | 1392 | 20201116_xd_colostrum_1ng_glyco_hcd203040_compound15.15772.15772.3 | controllerType=0 controllerNumber=1 scan=15772 | 33.8656 |
| 09551:3 | 23 | R.IIVPLNNREN[+1768.640]ISDPTSPLR.T | HexNAc(4)Hex(5)Fuc(1) | N[+1769] | 1306.2798 | 3 | 3916.825 | 3916.8167 | 0 | 2.1 | 62 | Specific | 526.32 | 489.09 | 489.09 | 9.93 | 25 | >sp\|P01591\|IGJ_HUMAN Immunoglobulin J chain OS=Homo sapiens OX=9606 GN=JCHAIN PE=1 SV=4 | 1392 | 20201116_xd_colostrum_1ng_glyco_hcd203040_compound15.12534.12534.3 | controllerType=0 controllerNumber=1 scan=12534 | 27.5423 |
| 12665:3 | 23 | R.IIVPLNNREN[+2350.830]ISDPTSPLR.T | HexNAc(4)Hex(5)Fuc(1)NeuAc(2) | N[+2351] | 1500.3422 | 3 | 4499.0121 | 4499.0075 | 0 | 1 | 62 | Specific | 529.46 | 475.66 | 475.66 | 9.78 | 25 | >sp\|P01591\|IGJ_HUMAN Immunoglobulin J chain OS=Homo sapiens OX=9606 GN=JCHAIN PE=1 SV=4 | 1392 | 20201116_xd_colostrum_1ng_glyco_hcd203040_compound15.15769.15769.3 | controllerType=0 controllerNumber=1 scan=15769 | 33.8604 |
| 10886:3 | 23 | R.IIVPLNNREN[+2059.735]ISDPTSPLR.T | HexNAc(4)Hex(5)Fuc(1)NeuAc(1) | N[+2060] | 1403.3118 | 3 | 4207.9209 | 4207.9121 | 0 | 2.1 | 62 | Specific | 545.34 | 503.12 | 503.12 | 10.16 | 25 | >sp\|P01591\|IGJ_HUMAN Immunoglobulin J chain OS=Homo sapiens OX=9606 GN=JCHAIN PE=1 SV=4 | 1392 | 20201116_xd_colostrum_1ng_glyco_hcd203040_compound15.13920.13920.3 | controllerType=0 controllerNumber=1 scan=13920 | 30.2416 |
| 10884:3 | 23 | R.IIVPLNNREN[+2059.735]ISDPTSPLR.T | HexNAc(4)Hex(5)Fuc(1)NeuAc(1) | N[+2060] | 1403.3118 | 3 | 4207.9209 | 4207.9121 | 0 | 2.1 | 62 | Specific | 547.89 | 511.55 | 511.55 | 10.16 | 25 | >sp\|P01591\|IGJ_HUMAN Immunoglobulin J chain OS=Homo sapiens OX=9606 GN=JCHAIN PE=1 SV=4 | 1392 | 20201116_xd_colostrum_1ng_glyco_hcd203040_compound15.13918.13918.3 | controllerType=0 controllerNumber=1 scan=13918 | 30.2352 |
| 10994:3 | 23 | R.IIVPLNNREN[+1856.656]ISDPTSPLR.T | HexNAc(3)Hex(5)Fuc(1)NeuAc(1) | N[+1857] | 1335.6161 | 3 | 4004.8339 | 4004.8327 | 0 | 0.3 | 62 | Specific | 552.23 | 499.22 | 499.22 | 9.74 | 25 | >sp\|P01591\|IGJ_HUMAN Immunoglobulin J chain OS=Homo sapiens OX=9606 GN=JCHAIN PE=1 SV=4 | 1392 | 20201116_xd_colostrum_1ng_glyco_hcd203040_compound15.14033.14033.3 | controllerType=0 controllerNumber=1 scan=14033 | 30.5148 |
| 09553:3 | 23 | R.IIVPLNNREN[+1768.640]ISDPTSPLR.T | HexNAc(4)Hex(5)Fuc(1) | N[+1769] | 1306.2798 | 3 | 3916.825 | 3916.8167 | 0 | 2.1 | 62 | Specific | 560.45 | 498.16 | 498.16 | 10.36 | 25 | >sp\|P01591\|IGJ_HUMAN Immunoglobulin J chain OS=Homo sapiens OX=9606 GN=JCHAIN PE=1 SV=4 | 1392 | 20201116_xd_colostrum_1ng_glyco_hcd203040_compound15.12536.12536.3 | controllerType=0 controllerNumber=1 scan=12536 | 27.5532 |
| 10991:3 | 23 | R.IIVPLNNREN[+1856.656]ISDPTSPLR.T | HexNAc(3)Hex(5)Fuc(1)NeuAc(1) | N[+1857] | 1335.6161 | 3 | 4004.8339 | 4004.8327 | 0 | 0.3 | 62 | Specific | 598.26 | 580.81 | 580.81 | 11.23 | 25 | >sp\|P01591\|IGJ_HUMAN Immunoglobulin J chain OS=Homo sapiens OX=9606 GN=JCHAIN PE=1 SV=4 | 1392 | 20201116_xd_colostrum_1ng_glyco_hcd203040_compound15.14030.14030.3 | controllerType=0 controllerNumber=1 scan=14030 | 30.5067 |
| 14424:3 | 26 | K.TFYWDFYTN[+2351.851]R.T | HexNAc(4)Hex(5)Fuc(3)NeuAc(1) | N[+2352] | 1255.5015 | 3 | 3764.49 | 3764.4777 | 0 | 3.3 | 97 | Specific | 542.19 | 451.92 | 451.92 | 9.82 | 15 | >sp\|P12273\|PIP_HUMAN Prolactin-inducible protein OS=Homo sapiens OX=9606 GN=PIP PE=1 SV=1 | 647 | 20201116_xd_colostrum_1ng_glyco_hcd203040_compound15.17598.17598.3 | controllerType=0 controllerNumber=1 scan=17598 | 37.5205 |
| 10575:4 | 27 | R.ADGTVNQIEGEATPVN[+1054.370]LTEPAKLEVK.F | HexNAc(2)Hex(4) | N[+1054] | 945.2029 | 4 | 3777.7896 | 3777.7798 | 0 | 2.6 | 83 | Specific | 466.89 | 444.65 | 444.65 | 9.92 | 16 | >sp\|P05090\|APOD_HUMAN Apolipoprotein D OS=Homo sapiens OX=9606 GN=APOD PE=1 SV=1 | 594 | 20201116_xd_colostrum_1ng_glyco_hcd203040_compound15.13597.13597.2 | controllerType=0 controllerNumber=1 scan=13597 | 29.6393 |
| 10577:4 | 27 | R.ADGTVNQIEGEATPVN[+1054.370]LTEPAKLEVK.F | HexNAc(2)Hex(4) | N[+1054] | 945.2029 | 4 | 3777.7896 | 3777.7798 | 0 | 2.6 | 83 | Specific | 601.24 | 506.94 | 506.94 | 12.85 | 16 | >sp\|P05090\|APOD_HUMAN Apolipoprotein D OS=Homo sapiens OX=9606 GN=APOD PE=1 SV=1 | 594 | 20201116_xd_colostrum_1ng_glyco_hcd203040_compound15.13599.13599.2 | controllerType=0 controllerNumber=1 scan=13599 | 29.6441 |
| 10583:3 | 27 | R.ADGTVNQIEGEATPVN[+1054.370]LTEPAKLEVK.F | HexNAc(2)Hex(4) | N[+1054] | 1259.9328 | 3 | 3777.7838 | 3777.7798 | 0 | 1.1 | 83 | Specific | 666.37 | 643.88 | 643.88 | 13.61 | 16 | >sp\|P05090\|APOD_HUMAN Apolipoprotein D OS=Homo sapiens OX=9606 GN=APOD PE=1 SV=1 | 594 | 20201116_xd_colostrum_1ng_glyco_hcd203040_compound15.13606.13606.3 | controllerType=0 controllerNumber=1 scan=13606 | 29.6557 |
| 10586:3 | 27 | R.ADGTVNQIEGEATPVN[+1054.370]LTEPAKLEVK.F | HexNAc(2)Hex(4) | N[+1054] | 1259.9328 | 3 | 3777.7838 | 3777.7798 | 0 | 1.1 | 83 | Specific | 761.2 | 675.28 | 675.28 | 15.61 | 16 | >sp\|P05090\|APOD_HUMAN Apolipoprotein D OS=Homo sapiens OX=9606 GN=APOD PE=1 SV=1 | 594 | 20201116_xd_colostrum_1ng_glyco_hcd203040_compound15.13609.13609.3 | controllerType=0 controllerNumber=1 scan=13609 | 29.6599 |
| 19835:5 | 28 | R.SDHGSSISC[+57.021]QPPAEIPGYLPADTVHLAVEFFN[+2351.851]LTHLPANLLQGASI | HexNAc(4)Hex(5)Fuc(3)NeuAc(1) | C[+57], N[+2352] | 1451.0711 | 5 | 7251.3263 | 7251.2918 | 0 | 4.8 | 48 | Specific | 745.25 | 745.25 | 745.25 | 15.55 | 18 | >sp\|P02750\|A2GL_HUMAN Leucine-rich alpha-2-glycoprotein OS=Homo sapiens OX=9606 GN=LRG1 PE=1 SV=2 | 518 | 20201116_xd_colostrum_1ng_glyco_hcd203040_compound15.23248.23248.5 | controllerType=0 controllerNumber=1 scan=23248 | 49.8421 |
| 19837:5 | 28 | R.SDHGSSISC[+57.021]QPPAEIPGYLPADTVHLAVEFFN[+2351.851]LTHLPANLLQGASI | HexNAc(4)Hex(5)Fuc(3)NeuAc(1) | C[+57], N[+2352] | 1451.0711 | 5 | 7251.3263 | 7251.2918 | 0 | 4.8 | 48 | Specific | 797.7 | 781.87 | 781.87 | 16.55 | 18 | >sp\|P02750\|A2GL_HUMAN Leucine-rich alpha-2-glycoprotein OS=Homo sapiens OX=9606 GN=LRG1 PE=1 SV=2 | 518 | 20201116_xd_colostrum_1ng_glyco_hcd203040_compound15.23250.23250.5 | controllerType=0 controllerNumber=1 scan=23250 | 49.8477 |
| 02335:3 | 34 | R.LRN[+1216.423]VSWATGR.S | HexNAc(2)Hex(5) | N[+1216] | 792.6927 | 3 | 2376.0636 | 2376.0559 | 0 | 3.2 | 149 | Specific | 327.82 | 127.78 | 127.78 | 5.2 | 11 | >sp\|P08571\|CD14_HUMAN Monocyte differentiation antigen CD14 OS=Homo sapiens OX=9606 GN=CD14 PE=1 SV=2 | 1232 | 20201116_xd_colostrum_1ng_glyco_hcd203040_compound15.5050.5050.3 | controllerType=0 controllerNumber=1 scan=5050 | 14.2862 |
| 15666:3 | 45 | R.NYIVPILWLN[+1864.634]ETGTIGDEKANM[+15.995]FR.S | HexNAc(2)Hex(9) | M[+16], N[+1865] | 1559.028 | 3 | 4675.0695 | 4675.0523 | 0 | 3.7 | 408 | Specific | 435.8 | 419.49 | 419.49 | 9.99 | 11 | >sp\|P16671\|CD36_HUMAN Platelet glycoprotein 4 OS=Homo sapiens OX=9606 GN=CD36 PE=1 SV=2 | 812 | 20201116_xd_colostrum_1ng_glyco_hcd203040_compound15.18890.18890.3 | controllerType=0 controllerNumber=1 scan=18890 | 40.1527 |
| 12821:5 | 60 | K.SAVQGPPERDLC[+57.021]GC[+57.021]YSVSSVLPGC[+57.021]AQPWNHGETFTC[+ | HexNAc(5)Hex(5)Fuc(1)NeuAc(1) | C[+57]*4, N[+2263] | 1615.5225 | 5 | 8073.5834 | 8073.5601 | 0 | 2.9 | 156 | Specific | 277.7 | 202.8 | 202.8 | 4.15 | 6 | >sp\|P01877\|IGHA2_HUMAN Immunoglobulin heavy constant alpha 2 OS=Homo sapiens OX=9606 GN=IGHA2 PE=1 SV=4 | 576 | 20201116_xd_colostrum_1ng_glyco_hcd203040_compound15.15932.15932.5 | controllerType=0 controllerNumber=1 scan=15932 | 34.1649 |
| 12265:5 | 60 | K.SAVQGPPERDLC[+57.021]GC[+57.021]YSVSSVLPGC[+57.021]AQPWNHGETFTC[+ | HexNAc(5)Hex(4)Fuc(1) | C[+57]*4, N[+1810] | 1524.8913 | 5 | 7620.4273 | 7620.4119 | 0 | 2 | 156 | Specific | 416.28 | 400.21 | 400.21 | 6.78 | 6 | >sp\|P01877\|IGHA2_HUMAN Immunoglobulin heavy constant alpha 2 OS=Homo sapiens OX=9606 GN=IGHA2 PE=1 SV=4 | 576 | 20201116_xd_colostrum_1ng_glyco_hcd203040_compound15.15353.15353.5 | controllerType=0 controllerNumber=1 scan=15353 | 33.0126 |
| 12798:6 | 60 | K.SAVQGPPERDLC[+57.021]GC[+57.021]YSVSSVLPGC[+57.021]AQPWNHGETFTC[+ | HexNAc(5)Hex(5)Fuc(3) | C[+57]*4, N[+2264] | 1346.6017 | 6 | 8074.5741 | 8074.5805 | 0 | -0.8 | 156 | Specific | 432.97 | 175.91 | 175.91 | 5.45 | 6 | >sp\|P01877\|IGHA2_HUMAN Immunoglobulin heavy constant alpha 2 OS=Homo sapiens OX=9606 GN=IGHA2 PE=1 SV=4 | 576 | 20201116_xd_colostrum_1ng_glyco_hcd203040_compound15.15908.15908.2 | controllerType=0 controllerNumber=1 scan=15908 | 34.1154 |
| 12824:5 | 60 | K.SAVQGPPERDLC[+57.021]GC[+57.021]YSVSSVLPGC[+57.021]AQPWNHGETFTC[+ | HexNAc(5)Hex(5)Fuc(1)NeuAc(1) | C[+57]*4, N[+2263] | 1615.5225 | 5 | 8073.5834 | 8073.5601 | 0 | 2.9 | 156 | Specific | 448.11 | 448.11 | 448.11 | 7.31 | 6 | >sp\|P01877\|IGHA2_HUMAN Immunoglobulin heavy constant alpha 2 OS=Homo sapiens OX=9606 GN=IGHA2 PE=1 SV=4 | 576 | 20201116_xd_colostrum_1ng_glyco_hcd203040_compound15.15935.15935.5 | controllerType=0 controllerNumber=1 scan=15935 | 34.1696 |
| 12255:6 | 60 | K.SAVQGPPERDLC[+57.021]GC[+57.021]YSVSSVLPGC[+57.021]AQPWNHGETFTC[+ | HexNAc(5)Hex(4)Fuc(1) | C[+57]*4, N[+1810] | 1270.9122 | 6 | 7620.4366 | 7620.4119 | 0 | 3.2 | 156 | Specific | 503.04 | 503.04 | 503.04 | 8.13 | 6 | >sp\|P01877\|IGHA2_HUMAN Immunoglobulin heavy constant alpha 2 OS=Homo sapiens OX=9606 GN=IGHA2 PE=1 SV=4 | 576 | 20201116_xd_colostrum_1ng_glyco_hcd203040_compound15.15343.15343.6 | controllerType=0 controllerNumber=1 scan=15343 | 33.0002 |
| 12801:6 | 60 | K.SAVQGPPERDLC[+57.021]GC[+57.021]YSVSSVLPGC[+57.021]AQPWNHGETFTC[+ | HexNAc(5)Hex(5)Fuc(3) | C[+57]*4, N[+2264] | 1346.6017 | 6 | 8074.5741 | 8074.5805 | 0 | -0.8 | 156 | Specific | 638.24 | 262.85 | 262.85 | 9.17 | 6 | >sp\|P01877\|IGHA2_HUMAN Immunoglobulin heavy constant alpha 2 OS=Homo sapiens OX=9606 GN=IGHA2 PE=1 SV=4 | 576 | 20201116_xd_colostrum_1ng_glyco_hcd203040_compound15.15911.15911.2 | controllerType=0 controllerNumber=1 scan=15911 | 34.1208 |
| 17469:4 | 1 | R.[+42.011]RTAGWNVPIGTLRPFLN[+2018.708]WTGPPEPIEAAVAR.F | HexNAc(3)Hex(6)Fuc(1)NeuAc(1) | .[42], N[+2019] | 1362.6509 | 4 | 5447.5817 | 5447.531 | 0 | 9.3 | 140 | Specific | 221.04 | 187.35 | 187.35 | 3.63 | 158 | >sp\|P02788\|TRFL_HUMAN Lactotransferrin OS=Homo sapiens OX=9606 GN=LTF PE=1 SV=6 | 1270 | 20201113_xd_milk_1ng_glyco_hcd203040_compound15.20851.20851.4 | controllerType=0 controllerNumber=1 scan=20851 | 41.5016 |
| 18456:5 | 1 | R.RTAGWNVPIGTLRPFLN[+2205.793]WTGPPEPIEAAVAR.F | HexNAc(4)Hex(5)Fuc(2)NeuAc(1) | N[+2206] | 1119.3326 | 5 | 5592.6341 | 5592.6049 | 0 | 5.2 | 140 | Specific | 477.18 | 477.18 | 477.18 | 10.42 | 158 | >sp\|P02788\|TRFL_HUMAN Lactotransferrin OS=Homo sapiens OX=9606 GN=LTF PE=1 SV=6 | 1270 | 20201113_xd_milk_1ng_glyco_hcd203040_compound15.21877.21877.5 | controllerType=0 controllerNumber=1 scan=21877 | 43.6252 |
| 18295:4 | 1 | R.RTAGWNVPIGTLRPFLN[+2716.983]WTGPPEPIEAAVAR.F | HexNAc(5)Hex(6)Fuc(3)NeuAc(1) | N[+2717] | 1526.709 | 4 | 6103.8141 | 6103.795 | 0 | 3.1 | 140 | Specific | 489.02 | 370.26 | 370.26 | 9.74 | 158 | >sp\|P02788\|TRFL_HUMAN Lactotransferrin OS=Homo sapiens OX=9606 GN=LTF PE=1 SV=6 | 1270 | 20201113_xd_milk_1ng_glyco_hcd203040_compound15.21710.21710.4 | controllerType=0 controllerNumber=1 scan=21710 | 43.3086 |
| 19467:4 | 1 | R.RTAGWNVPIGTLRPFLN[+2350.830]WTGPPEPIEAAVAR.F | HexNAc(4)Hex(5)Fuc(1)NeuAc(2) | N[+2351] | 1435.1708 | 4 | 5737.6613 | 5737.6424 | 0 | 3.3 | 140 | Specific | 494.37 | 401.04 | 401.04 | 11.25 | 158 | >sp\|P02788\|TRFL_HUMAN Lactotransferrin OS=Homo sapiens OX=9606 GN=LTF PE=1 SV=6 | 1270 | 20201113_xd_milk_1ng_glyco_hcd203040_compound15.22930.22930.4 | controllerType=0 controllerNumber=1 scan=22930 | 45.8668 |
| 18297:4 | 1 | R.RTAGWNVPIGTLRPFLN[+2716.983]WTGPPEPIEAAVAR.F | HexNAc(5)Hex(6)Fuc(3)NeuAc(1) | N[+2717] | 1526.709 | 4 | 6103.8141 | 6103.795 | 0 | 3.1 | 140 | Specific | 499.64 | 421.59 | 421.59 | 11.25 | 158 | >sp\|P02788\|TRFL_HUMAN Lactotransferrin OS=Homo sapiens OX=9606 GN=LTF PE=1 SV=6 | 1270 | 20201113_xd_milk_1ng_glyco_hcd203040_compound15.21712.21712.4 | controllerType=0 controllerNumber=1 scan=21712 | 43.3133 |
| 19464:4 | 1 | R.RTAGWNVPIGTLRPFLN[+2350.830]WTGPPEPIEAAVAR.F | HexNAc(4)Hex(5)Fuc(1)NeuAc(2) | N[+2351] | 1435.1708 | 4 | 5737.6613 | 5737.6424 | 0 | 3.3 | 140 | Specific | 503.2 | 421.02 | 421.02 | 11.25 | 158 | >sp\|P02788\|TRFL_HUMAN Lactotransferrin OS=Homo sapiens OX=9606 GN=LTF PE=1 SV=6 | 1270 | 20201113_xd_milk_1ng_glyco_hcd203040_compound15.22927.22927.4 | controllerType=0 controllerNumber=1 scan=22927 | 45.8595 |
| 18450:4 | 1 | R.RTAGWNVPIGTLRPFLN[+2205.793]WTGPPEPIEAAVAR.F | HexNAc(4)Hex(5)Fuc(2)NeuAc(1) | N[+2206] | 1398.9139 | 4 | 5592.6339 | 5592.6049 | 0 | 5.2 | 140 | Specific | 509.67 | 509.67 | 509.67 | 10.6 | 158 | >sp\|P02788\|TRFL_HUMAN Lactotransferrin OS=Homo sapiens OX=9606 GN=LTF PE=1 SV=6 | 1270 | 20201113_xd_milk_1ng_glyco_hcd203040_compound15.21871.21871.4 | controllerType=0 controllerNumber=1 scan=21871 | 43.6133 |
| 18454:4 | 1 | R.RTAGWNVPIGTLRPFLN[+2351.851]WTGPPEPIEAAVAR.F | HexNAc(4)Hex(5)Fuc(3)NeuAc(1) | N[+2352] | 1435.4296 | 4 | 5738.6964 | 5738.6628 | 0 | 5.9 | 140 | Specific | 517.41 | 420.52 | 420.52 | 10.73 | 158 | >sp\|P02788\|TRFL_HUMAN Lactotransferrin OS=Homo sapiens OX=9606 GN=LTF PE=1 SV=6 | 1270 | 20201113_xd_milk_1ng_glyco_hcd203040_compound15.21875.21875.4 | controllerType=0 controllerNumber=1 scan=21875 | 43.6227 |
| 18459:5 | 1 | R.RTAGWNVPIGTLRPFLN[+2205.793]WTGPPEPIEAAVAR.F | HexNAc(4)Hex(5)Fuc(2)NeuAc(1) | N[+2206] | 1119.3326 | 5 | 5592.6341 | 5592.6049 | 0 | 5.2 | 140 | Specific | 534.99 | 511.98 | 511.98 | 10.84 | 158 | >sp\|P02788\|TRFL_HUMAN Lactotransferrin OS=Homo sapiens OX=9606 GN=LTF PE=1 SV=6 | 1270 | 20201113_xd_milk_1ng_glyco_hcd203040_compound15.21880.21880.5 | controllerType=0 controllerNumber=1 scan=21880 | 43.6384 |
| 18457:4 | 1 | R.RTAGWNVPIGTLRPFLN[+2351.851]WTGPPEPIEAAVAR.F | HexNAc(4)Hex(5)Fuc(3)NeuAc(1) | N[+2352] | 1435.4296 | 4 | 5738.6964 | 5738.6628 | 0 | 5.9 | 140 | Specific | 540.43 | 367.68 | 367.68 | 9.36 | 158 | >sp\|P02788\|TRFL_HUMAN Lactotransferrin OS=Homo sapiens OX=9606 GN=LTF PE=1 SV=6 | 1270 | 20201113_xd_milk_1ng_glyco_hcd203040_compound15.21878.21878.4 | controllerType=0 controllerNumber=1 scan=21878 | 43.6329 |
| 17545:4 | 1 | R.RTAGWNVPIGTLRPFLN[+1768.640]WTGPPEPIEAAVAR.F | HexNAc(4)Hex(5)Fuc(1) | N[+1769] | 1289.6216 | 4 | 5155.4646 | 5155.4516 | 0 | 2.5 | 140 | Specific | 547.28 | 467.86 | 467.86 | 11.71 | 158 | >sp\|P02788\|TRFL_HUMAN Lactotransferrin OS=Homo sapiens OX=9606 GN=LTF PE=1 SV=6 | 1270 | 20201113_xd_milk_1ng_glyco_hcd203040_compound15.20930.20930.4 | controllerType=0 controllerNumber=1 scan=20930 | 41.6646 |
| 18342:4 | 1 | R.RTAGWNVPIGTLRPFLN[+2570.925]WTGPPEPIEAAVAR.F | HexNAc(5)Hex(6)Fuc(2)NeuAc(1) | N[+2571] | 1490.1926 | 4 | 5957.7487 | 5957.7371 | 0 | 1.9 | 140 | Specific | 565.26 | 444.77 | 444.77 | 11.92 | 158 | >sp\|P02788\|TRFL_HUMAN Lactotransferrin OS=Homo sapiens OX=9606 GN=LTF PE=1 SV=6 | 1270 | 20201113_xd_milk_1ng_glyco_hcd203040_compound15.21759.21759.4 | controllerType=0 controllerNumber=1 scan=21759 | 43.4004 |
| 18340:4 | 1 | R.RTAGWNVPIGTLRPFLN[+2570.925]WTGPPEPIEAAVAR.F | HexNAc(5)Hex(6)Fuc(2)NeuAc(1) | N[+2571] | 1490.1926 | 4 | 5957.7487 | 5957.7371 | 0 | 1.9 | 140 | Specific | 578.06 | 399.88 | 399.88 | 13.19 | 158 | >sp\|P02788\|TRFL_HUMAN Lactotransferrin OS=Homo sapiens OX=9606 GN=LTF PE=1 SV=6 | 1270 | 20201113_xd_milk_1ng_glyco_hcd203040_compound15.21757.21757.4 | controllerType=0 controllerNumber=1 scan=21757 | 43.3952 |
| 18430:4 | 1 | R.RTAGWNVPIGTLRPFLN[+2351.851]WTGPPEPIEAAVAR.F | HexNAc(4)Hex(5)Fuc(3)NeuAc(1) | N[+2352] | 1435.4268 | 4 | 5738.6852 | 5738.6628 | 0 | 3.9 | 140 | Specific | 579.65 | 433.48 | 433.48 | 12.68 | 158 | >sp\|P02788\|TRFL_HUMAN Lactotransferrin OS=Homo sapiens OX=9606 GN=LTF PE=1 SV=6 | 1270 | 20201113_xd_milk_1ng_glyco_hcd203040_compound15.21850.21850.4 | controllerType=0 controllerNumber=1 scan=21850 | 43.5643 |
| 18452:4 | 1 | R.RTAGWNVPIGTLRPFLN[+2205.793]WTGPPEPIEAAVAR.F | HexNAc(4)Hex(5)Fuc(2)NeuAc(1) | N[+2206] | 1398.9139 | 4 | 5592.6339 | 5592.6049 | 0 | 5.2 | 140 | Specific | 582.96 | 468.85 | 468.85 | 12.34 | 158 | >sp\|P02788\|TRFL_HUMAN Lactotransferrin OS=Homo sapiens OX=9606 GN=LTF PE=1 SV=6 | 1270 | 20201113_xd_milk_1ng_glyco_hcd203040_compound15.21873.21873.4 | controllerType=0 controllerNumber=1 scan=21873 | 43.6205 |
| 18433:4 | 1 | R.RTAGWNVPIGTLRPFLN[+2351.851]WTGPPEPIEAAVAR.F | HexNAc(4)Hex(5)Fuc(3)NeuAc(1) | N[+2352] | 1435.4268 | 4 | 5738.6852 | 5738.6628 | 0 | 3.9 | 140 | Specific | 584.62 | 405.11 | 405.11 | 12.68 | 158 | >sp\|P02788\|TRFL_HUMAN Lactotransferrin OS=Homo sapiens OX=9606 GN=LTF PE=1 SV=6 | 1270 | 20201113_xd_milk_1ng_glyco_hcd203040_compound15.21853.21853.4 | controllerType=0 controllerNumber=1 scan=21853 | 43.5702 |
| 19607:3 | 1 | R.TAGWNVPIGTLRPFLN[+2570.925]WTGPPEPIEAAVAR.F | HexNAc(5)Hex(6)Fuc(2)NeuAc(1) | N[+2571] | 1934.556 | 3 | 5801.6534 | 5801.636 | 0 | 3 | 141 | Specific | 216.35 | 117.47 | 117.47 | 5.41 | 158 | >sp\|P02788\|TRFL_HUMAN Lactotransferrin OS=Homo sapiens OX=9606 GN=LTF PE=1 SV=6 | 1270 | 20201113_xd_milk_1ng_glyco_hcd203040_compound15.23075.23075.3 | controllerType=0 controllerNumber=1 scan=23075 | 46.1285 |
| 19769:4 | 1 | R.[+42.011]TAGWNVPIGTLRPFLN[+2336.851]WTGPPEPIEAAVAR.F | HexNAc(6)Hex(6)Fuc(1) | .[42], N[+2337] | 1403.1493 | 4 | 5609.5754 | 5609.5726 | 0 | 0.5 | 141 | Specific | 216.77 | 216.77 | 216.77 | 6.44 | 158 | >sp\|P02788\|TRFL_HUMAN Lactotransferrin OS=Homo sapiens OX=9606 GN=LTF PE=1 SV=6 | 1270 | 20201113_xd_milk_1ng_glyco_hcd203040_compound15.23243.23243.4 | controllerType=0 controllerNumber=1 scan=23243 | 46.4942 |
| 18675:3 | 1 | R.TAGWNVPIGTLRPFLN[+2279.830]WTGPPEPIEAAVAR.F | HexNAc(5)Hex(6)Fuc(2) | N[+2280] | 1837.5223 | 3 | 5510.5524 | 5510.5406 | 0 | 2.1 | 141 | Specific | 238.54 | 219.14 | 219.14 | 7.88 | 158 | >sp\|P02788\|TRFL_HUMAN Lactotransferrin OS=Homo sapiens OX=9606 GN=LTF PE=1 SV=6 | 1270 | 20201113_xd_milk_1ng_glyco_hcd203040_compound15.22105.22105.3 | controllerType=0 controllerNumber=1 scan=22105 | 44.0938 |
| 18690:4 | 1 | R.[+42.011]TAGWNVPIGTLRPFLN[+2018.708]WTGPPEPIEAAVAR.F | HexNAc(3)Hex(6)Fuc(1)NeuAc(1) | .[42], N[+2019] | 1323.6251 | 4 | 5291.4787 | 5291.4299 | 0 | 9.2 | 141 | Specific | 245.63 | 245.63 | 245.63 | 4.9 | 158 | >sp\|P02788\|TRFL_HUMAN Lactotransferrin OS=Homo sapiens OX=9606 GN=LTF PE=1 SV=6 | 1270 | 20201113_xd_milk_1ng_glyco_hcd203040_compound15.22120.22120.4 | controllerType=0 controllerNumber=1 scan=22120 | 44.1135 |
| 19772:5 | 1 | R.TAGWNVPIGTLRPFLN[+2076.750]WTGPPEPIEAAVAR.F | HexNAc(4)Hex(6)Fuc(2) | N[+2077] | 1062.3047 | 5 | 5307.4943 | 5307.4612 | 0 | 6.2 | 141 | Specific | 253.99 | 253.99 | 253.99 | 7.19 | 158 | >sp\|P02788\|TRFL_HUMAN Lactotransferrin OS=Homo sapiens OX=9606 GN=LTF PE=1 SV=6 | 1270 | 20201113_xd_milk_1ng_glyco_hcd203040_compound15.23246.23246.5 | controllerType=0 controllerNumber=1 scan=23246 | 46.4984 |
| 18703:4 | 1 | R.[+42.011]TAGWNVPIGTLRPFLN[+1872.651]WTGPPEPIEAAVAR.F | HexNAc(3)Hex(6)NeuAc(1) | .[42], N[+1873] | 1287.1082 | 4 | 5145.4108 | 5145.372 | 0 | 7.5 | 141 | Specific | 267.64 | 202.97 | 202.97 | 6.86 | 158 | >sp\|P02788\|TRFL_HUMAN Lactotransferrin OS=Homo sapiens OX=9606 GN=LTF PE=1 SV=6 | 1270 | 20201113_xd_milk_1ng_glyco_hcd203040_compound15.22134.22134.4 | controllerType=0 controllerNumber=1 scan=22134 | 44.1382 |
| 17444:4 | 1 | R.[+42.011]TAGWNVPIGTLRPFLN[+2028.740]WTGPPEPIEAAVAR.F | HexNAc(6)Hex(5) | .[42], N[+2029] | 1326.1342 | 4 | 5301.5148 | 5301.4618 | 0 | 10 | 141 | Specific | 269.64 | 115.09 | 115.09 | 6.01 | 158 | >sp\|P02788\|TRFL_HUMAN Lactotransferrin OS=Homo sapiens OX=9606 GN=LTF PE=1 SV=6 | 1270 | 20201113_xd_milk_1ng_glyco_hcd203040_compound15.20825.20825.4 | controllerType=0 controllerNumber=1 scan=20825 | 41.4505 |
| 18689:3 | 1 | R.[+42.011]TAGWNVPIGTLRPFLN[+2018.708]WTGPPEPIEAAVAR.F | HexNAc(3)Hex(6)Fuc(1)NeuAc(1) | .[42], N[+2019] | 1764.4984 | 3 | 5291.4806 | 5291.4299 | 0 | 9.6 | 141 | Specific | 279.29 | 216.02 | 216.02 | 5.36 | 158 | >sp\|P02788\|TRFL_HUMAN Lactotransferrin OS=Homo sapiens OX=9606 GN=LTF PE=1 SV=6 | 1270 | 20201113_xd_milk_1ng_glyco_hcd203040_compound15.22119.22119.3 | controllerType=0 controllerNumber=1 scan=22119 | 44.112 |
| 19760:5 | 1 | R.TAGWNVPIGTLRPFLN[+2204.772]WTGPPEPIEAAVAR.F | HexNAc(4)Hex(5)NeuAc(2) | N[+2205] | 1087.91 | 5 | 5435.5211 | 5435.4834 | 0 | 6.9 | 141 | Specific | 285.12 | 285.12 | 285.12 | 7.26 | 158 | >sp\|P02788\|TRFL_HUMAN Lactotransferrin OS=Homo sapiens OX=9606 GN=LTF PE=1 SV=6 | 1270 | 20201113_xd_milk_1ng_glyco_hcd203040_compound15.23234.23234.5 | controllerType=0 controllerNumber=1 scan=23234 | 46.4709 |
| 18705:4 | 1 | R.[+42.011]TAGWNVPIGTLRPFLN[+1872.651]WTGPPEPIEAAVAR.F | HexNAc(3)Hex(6)NeuAc(1) | .[42], N[+1873] | 1287.1082 | 4 | 5145.4108 | 5145.372 | 0 | 7.5 | 141 | Specific | 286.63 | 157.56 | 157.56 | 4.66 | 158 | >sp\|P02788\|TRFL_HUMAN Lactotransferrin OS=Homo sapiens OX=9606 GN=LTF PE=1 SV=6 | 1270 | 20201113_xd_milk_1ng_glyco_hcd203040_compound15.22136.22136.4 | controllerType=0 controllerNumber=1 scan=22136 | 44.1476 |
| 20176:4 | 1 | R.TAGWNVPIGTLRPFLN[+2059.735]WTGPPEPIEAAVAR.F | HexNAc(4)Hex(5)Fuc(1)NeuAc(1) | N[+2060] | 1323.3727 | 4 | 5290.4689 | 5290.4459 | 0 | 4.4 | 141 | Specific | 302.59 | 302.59 | 302.59 | 8.16 | 158 | >sp\|P02788\|TRFL_HUMAN Lactotransferrin OS=Homo sapiens OX=9606 GN=LTF PE=1 SV=6 | 1270 | 20201113_xd_milk_1ng_glyco_hcd203040_compound15.23668.23668.2 | controllerType=0 controllerNumber=1 scan=23668 | 47.2816 |
| 18701:5 | 1 | R.TAGWNVPIGTLRPFLN[+2425.887]WTGPPEPIEAAVAR.F | HexNAc(5)Hex(6)Fuc(3) | N[+2426] | 1132.1274 | 5 | 5656.6079 | 5656.5985 | 0 | 1.7 | 141 | Specific | 305.86 | 255.52 | 255.52 | 5.9 | 158 | >sp\|P02788\|TRFL_HUMAN Lactotransferrin OS=Homo sapiens OX=9606 GN=LTF PE=1 SV=6 | 1270 | 20201113_xd_milk_1ng_glyco_hcd203040_compound15.22131.22131.5 | controllerType=0 controllerNumber=1 scan=22131 | 44.1323 |
| 19682:3 | 1 | R.TAGWNVPIGTLRPFLN[+2424.867]WTGPPEPIEAAVAR.F | HexNAc(5)Hex(6)Fuc(1)NeuAc(1) | N[+2425] | 1885.8775 | 3 | 5655.618 | 5655.5781 | 0 | 7.1 | 141 | Specific | 306.12 | 181.79 | 181.79 | 7.02 | 158 | >sp\|P02788\|TRFL_HUMAN Lactotransferrin OS=Homo sapiens OX=9606 GN=LTF PE=1 SV=6 | 1270 | 20201113_xd_milk_1ng_glyco_hcd203040_compound15.23152.23152.3 | controllerType=0 controllerNumber=1 scan=23152 | 46.276 |
| 18704:3 | 1 | R.[+42.011]TAGWNVPIGTLRPFLN[+1872.651]WTGPPEPIEAAVAR.F | HexNAc(3)Hex(6)NeuAc(1) | .[42], N[+1873] | 1715.8064 | 3 | 5145.4045 | 5145.372 | 0 | 6.3 | 141 | Specific | 309.45 | 193.56 | 193.56 | 7.02 | 158 | >sp\|P02788\|TRFL_HUMAN Lactotransferrin OS=Homo sapiens OX=9606 GN=LTF PE=1 SV=6 | 1270 | 20201113_xd_milk_1ng_glyco_hcd203040_compound15.22135.22135.3 | controllerType=0 controllerNumber=1 scan=22135 | 44.1402 |
| 21074:5 | 1 | R.TAGWNVPIGTLRPFLN[+2350.830]WTGPPEPIEAAVAR.F | HexNAc(4)Hex(5)Fuc(1)NeuAc(2) | N[+2351] | 1117.1171 | 5 | 5581.5564 | 5581.5413 | 0 | 2.7 | 141 | Specific | 320.91 | 293.95 | 293.95 | 8.64 | 158 | >sp\|P02788\|TRFL_HUMAN Lactotransferrin OS=Homo sapiens OX=9606 GN=LTF PE=1 SV=6 | 1270 | 20201113_xd_milk_1ng_glyco_hcd203040_compound15.24606.24606.5 | controllerType=0 controllerNumber=1 scan=24606 | 49.3132 |
| 19685:3 | 1 | R.TAGWNVPIGTLRPFLN[+2424.867]WTGPPEPIEAAVAR.F | HexNAc(5)Hex(6)Fuc(1)NeuAc(1) | N[+2425] | 1885.8775 | 3 | 5655.618 | 5655.5781 | 0 | 7.1 | 141 | Specific | 324.68 | 232.55 | 232.55 | 7.36 | 158 | >sp\|P02788\|TRFL_HUMAN Lactotransferrin OS=Homo sapiens OX=9606 GN=LTF PE=1 SV=6 | 1270 | 20201113_xd_milk_1ng_glyco_hcd203040_compound15.23155.23155.3 | controllerType=0 controllerNumber=1 scan=23155 | 46.2849 |
| 19762:3 | 1 | R.[+42.011]TAGWNVPIGTLRPFLN[+2190.793]WTGPPEPIEAAVAR.F | HexNAc(6)Hex(6) | .[42], N[+2191] | 1821.8452 | 3 | 5463.521 | 5463.5147 | 0 | 1.2 | 141 | Specific | 333.9 | 48.79 | 48.79 | 6.64 | 158 | >sp\|P02788\|TRFL_HUMAN Lactotransferrin OS=Homo sapiens OX=9606 GN=LTF PE=1 SV=6 | 1270 | 20201113_xd_milk_1ng_glyco_hcd203040_compound15.23236.23236.3 | controllerType=0 controllerNumber=1 scan=23236 | 46.4838 |
| 20889:4 | 1 | R.TAGWNVPIGTLRPFLN[+2862.021]WTGPPEPIEAAVAR.F | HexNAc(5)Hex(6)Fuc(2)NeuAc(2) | N[+2862] | 1523.9385 | 4 | 6092.7321 | 6092.7314 | 0 | 0.1 | 141 | Specific | 353.58 | 212.81 | 212.81 | 7.86 | 158 | >sp\|P02788\|TRFL_HUMAN Lactotransferrin OS=Homo sapiens OX=9606 GN=LTF PE=1 SV=6 | 1270 | 20201113_xd_milk_1ng_glyco_hcd203040_compound15.24413.24413.3 | controllerType=0 controllerNumber=1 scan=24413 | 48.9042 |
| 18680:3 | 1 | R.TAGWNVPIGTLRPFLN[+2425.887]WTGPPEPIEAAVAR.F | HexNAc(5)Hex(6)Fuc(3) | N[+2426] | 1886.2086 | 3 | 5656.6112 | 5656.5985 | 0 | 2.3 | 141 | Specific | 382.75 | 247.9 | 247.9 | 8.84 | 158 | >sp\|P02788\|TRFL_HUMAN Lactotransferrin OS=Homo sapiens OX=9606 GN=LTF PE=1 SV=6 | 1270 | 20201113_xd_milk_1ng_glyco_hcd203040_compound15.22110.22110.3 | controllerType=0 controllerNumber=1 scan=22110 | 44.1006 |

Table S4. Cont.

| Query #:z | Protein Rank | Glycopeptide | Glycan | Modification Type(s) | Observed m/z | z | Observed (M+H) | Calc. mass (M+H) | Off-by-x error | Mass error (ppm) | StartPosition | Cleavage | Score | Delta | Delta Mod | \|Log Prob\| | # of unique peptides | Protein Name | Protein DB number | Comment | Scan # | Scan Time |
|---|---|---|---|---|---|---|---|---|---|---|---|---|---|---|---|---|---|---|---|---|---|---|
| 19610:5 | 1 | R.TAGWNVPIGTLRPFLN[+2570.925]WTGPPEPIEAAVAR.F | HexNAc(5)Hex(6)Fuc(2)NeuAc(1) | N[+2571] | 1161.1361 | 5 | 5801.6514 | 5801.636 | 0 | 2.7 | 141 | Specific | 395.82 | 369.59 | 369.59 | 9.41 | 158 | >sp\|P02788\|TRFL_HUMAN Lactotransferrin OS=Homo sapiens OX=9606 GN=LTF PE=1 SV=6 | 1270 | 20201113_xd_milk_1ng_glyco_hcd203040_compound15.23078.23078.5 | controllerType=0 controllerNumber=1 scan=23078 | 46.1338 |
| 19551:3 | 1 | R.TAGWNVPIGTLRPFLN[+2715.963]WTGPPEPIEAAVAR.F | HexNAc(5)Hex(6)Fuc(1)NeuAc(2) | N[+2716] | 1982.9063 | 3 | 5946.7045 | 5946.6735 | 0 | 5.2 | 141 | Specific | 395.85 | 225.55 | 225.55 | 7.85 | 158 | >sp\|P02788\|TRFL_HUMAN Lactotransferrin OS=Homo sapiens OX=9606 GN=LTF PE=1 SV=6 | 1270 | 20201113_xd_milk_1ng_glyco_hcd203040_compound15.23017.23017.3 | controllerType=0 controllerNumber=1 scan=23017 | 46.0257 |
| 21019:3 | 1 | R.TAGWNVPIGTLRPFLN[+2350.830]WTGPPEPIEAAVAR.F | HexNAc(4)Hex(5)Fuc(1)NeuAc(2) | N[+2351] | 1861.1848 | 3 | 5581.5398 | 5581.5413 | 0 | -0.3 | 141 | Specific | 396.59 | 213.66 | 213.66 | 7.96 | 158 | >sp\|P02788\|TRFL_HUMAN Lactotransferrin OS=Homo sapiens OX=9606 GN=LTF PE=1 SV=6 | 1270 | 20201113_xd_milk_1ng_glyco_hcd203040_compound15.24549.24549.3 | controllerType=0 controllerNumber=1 scan=24549 | 49.2036 |
| 18637:4 | 1 | R.TAGWNVPIGTLRPFLN[+2644.962]WTGPPEPIEAAVAR.F | HexNAc(6)Hex(7)Fuc(2) | N[+2645] | 1469.6753 | 4 | 5875.6794 | 5875.6727 | 0 | 1.1 | 141 | Specific | 408.02 | 355.84 | 355.84 | 9.22 | 158 | >sp\|P02788\|TRFL_HUMAN Lactotransferrin OS=Homo sapiens OX=9606 GN=LTF PE=1 SV=6 | 1270 | 20201113_xd_milk_1ng_glyco_hcd203040_compound15.22065.22065.4 | controllerType=0 controllerNumber=1 scan=22065 | 44.0272 |
| 19748:5 | 1 | R.TAGWNVPIGTLRPFLN[+2059.735]WTGPPEPIEAAVAR.F | HexNAc(4)Hex(5)Fuc(1)NeuAc(1) | N[+2060] | 1058.8989 | 5 | 5290.4655 | 5290.4459 | 0 | 3.7 | 141 | Specific | 415.07 | 415.07 | 415.07 | 10.8 | 158 | >sp\|P02788\|TRFL_HUMAN Lactotransferrin OS=Homo sapiens OX=9606 GN=LTF PE=1 SV=6 | 1270 | 20201113_xd_milk_1ng_glyco_hcd203040_compound15.23221.23221.5 | controllerType=0 controllerNumber=1 scan=23221 | 46.4373 |
| 18754:3 | 1 | R.TAGWNVPIGTLRPFLN[+1768.640]WTGPPEPIEAAVAR.F | HexNAc(4)Hex(5)Fuc(1) | N[+1769] | 1667.1374 | 3 | 4999.3977 | 4999.3504 | 0 | 9.5 | 141 | Specific | 417.2 | 417.2 | 417.2 | 9.81 | 158 | >sp\|P02788\|TRFL_HUMAN Lactotransferrin OS=Homo sapiens OX=9606 GN=LTF PE=1 SV=6 | 1270 | 20201113_xd_milk_1ng_glyco_hcd203040_compound15.22187.22187.3 | controllerType=0 controllerNumber=1 scan=22187 | 44.2487 |
| 19609:3 | 1 | R.TAGWNVPIGTLRPFLN[+2570.925]WTGPPEPIEAAVAR.F | HexNAc(5)Hex(6)Fuc(2)NeuAc(1) | N[+2571] | 1934.556 | 3 | 5801.6534 | 5801.636 | 0 | 3 | 141 | Specific | 458.45 | 402.35 | 402.35 | 11.02 | 158 | >sp\|P02788\|TRFL_HUMAN Lactotransferrin OS=Homo sapiens OX=9606 GN=LTF PE=1 SV=6 | 1270 | 20201113_xd_milk_1ng_glyco_hcd203040_compound15.23077.23077.3 | controllerType=0 controllerNumber=1 scan=23077 | 46.1327 |
| 21018:4 | 1 | R.TAGWNVPIGTLRPFLN[+2350.830]WTGPPEPIEAAVAR.F | HexNAc(4)Hex(5)Fuc(1)NeuAc(2) | N[+2351] | 1396.1395 | 4 | 5581.5363 | 5581.5413 | 0 | -0.9 | 141 | Specific | 482.75 | 296.19 | 296.19 | 8.4 | 158 | >sp\|P02788\|TRFL_HUMAN Lactotransferrin OS=Homo sapiens OX=9606 GN=LTF PE=1 SV=6 | 1270 | 20201113_xd_milk_1ng_glyco_hcd203040_compound15.24548.24548.4 | controllerType=0 controllerNumber=1 scan=24548 | 49.2017 |
| 19713:3 | 1 | R.TAGWNVPIGTLRPFLN[+2205.793]WTGPPEPIEAAVAR.F | HexNAc(4)Hex(5)Fuc(2)NeuAc(1) | N[+2206] | 1812.8457 | 3 | 5436.5224 | 5436.5038 | 0 | 3.4 | 141 | Specific | 490.74 | 454.22 | 454.22 | 11.25 | 158 | >sp\|P02788\|TRFL_HUMAN Lactotransferrin OS=Homo sapiens OX=9606 GN=LTF PE=1 SV=6 | 1270 | 20201113_xd_milk_1ng_glyco_hcd203040_compound15.23185.23185.3 | controllerType=0 controllerNumber=1 scan=23185 | 46.3618 |
| 18642:3 | 1 | R.TAGWNVPIGTLRPFLN[+2279.830]WTGPPEPIEAAVAR.F | HexNAc(5)Hex(6)Fuc(2) | N[+2280] | 1837.5199 | 3 | 5510.545 | 5510.5406 | 0 | 0.8 | 141 | Specific | 494.72 | 430.46 | 430.46 | 11.28 | 158 | >sp\|P02788\|TRFL_HUMAN Lactotransferrin OS=Homo sapiens OX=9606 GN=LTF PE=1 SV=6 | 1270 | 20201113_xd_milk_1ng_glyco_hcd203040_compound15.22071.22071.3 | controllerType=0 controllerNumber=1 scan=22071 | 44.0365 |
| 19675:4 | 1 | R.TAGWNVPIGTLRPFLN[+2424.867]WTGPPEPIEAAVAR.F | HexNAc(5)Hex(6)Fuc(1)NeuAc(1) | N[+2425] | 1414.6598 | 4 | 5655.6173 | 5655.5781 | 0 | 6.9 | 141 | Specific | 502.16 | 368.54 | 368.54 | 8.92 | 158 | >sp\|P02788\|TRFL_HUMAN Lactotransferrin OS=Homo sapiens OX=9606 GN=LTF PE=1 SV=6 | 1270 | 20201113_xd_milk_1ng_glyco_hcd203040_compound15.23145.23145.4 | controllerType=0 controllerNumber=1 scan=23145 | 46.2632 |
| 21185:4 | 1 | R.TAGWNVPIGTLRPFLN[+2351.851]WTGPPEPIEAAVAR.F | HexNAc(4)Hex(5)Fuc(3)NeuAc(1) | N[+2352] | 1396.3945 | 4 | 5582.5563 | 5582.5617 | 0 | -1 | 141 | Specific | 509.26 | 477.89 | 477.89 | 10.49 | 158 | >sp\|P02788\|TRFL_HUMAN Lactotransferrin OS=Homo sapiens OX=9606 GN=LTF PE=1 SV=6 | 1270 | 20201113_xd_milk_1ng_glyco_hcd203040_compound15.24721.24721.4 | controllerType=0 controllerNumber=1 scan=24721 | 49.5528 |
| 19749:3 | 1 | R.TAGWNVPIGTLRPFLN[+2205.793]WTGPPEPIEAAVAR.F | HexNAc(4)Hex(5)Fuc(2)NeuAc(1) | N[+2206] | 1812.8487 | 3 | 5436.5316 | 5436.5038 | 0 | 5.1 | 141 | Specific | 517.55 | 456.63 | 456.63 | 10.73 | 158 | >sp\|P02788\|TRFL_HUMAN Lactotransferrin OS=Homo sapiens OX=9606 GN=LTF PE=1 SV=6 | 1270 | 20201113_xd_milk_1ng_glyco_hcd203040_compound15.23222.23222.3 | controllerType=0 controllerNumber=1 scan=23222 | 46.4456 |
| 19553:3 | 1 | R.TAGWNVPIGTLRPFLN[+2715.963]WTGPPEPIEAAVAR.F | HexNAc(5)Hex(6)Fuc(1)NeuAc(2) | N[+2716] | 1982.9063 | 3 | 5946.7045 | 5946.6735 | 0 | 5.2 | 141 | Specific | 524.39 | 422.11 | 422.11 | 10.73 | 158 | >sp\|P02788\|TRFL_HUMAN Lactotransferrin OS=Homo sapiens OX=9606 GN=LTF PE=1 SV=6 | 1270 | 20201113_xd_milk_1ng_glyco_hcd203040_compound15.23019.23019.3 | controllerType=0 controllerNumber=1 scan=23019 | 46.0301 |
| 18645:3 | 1 | R.TAGWNVPIGTLRPFLN[+2279.830]WTGPPEPIEAAVAR.F | HexNAc(5)Hex(6)Fuc(2) | N[+2280] | 1837.5199 | 3 | 5510.545 | 5510.5406 | 0 | 0.8 | 141 | Specific | 534.03 | 425.11 | 425.11 | 11.55 | 158 | >sp\|P02788\|TRFL_HUMAN Lactotransferrin OS=Homo sapiens OX=9606 GN=LTF PE=1 SV=6 | 1270 | 20201113_xd_milk_1ng_glyco_hcd203040_compound15.22074.22074.3 | controllerType=0 controllerNumber=1 scan=22074 | 44.0407 |
| 21017:3 | 1 | R.TAGWNVPIGTLRPFLN[+2350.830]WTGPPEPIEAAVAR.F | HexNAc(4)Hex(5)Fuc(1)NeuAc(2) | N[+2351] | 1861.1848 | 3 | 5581.5398 | 5581.5413 | 0 | -0.3 | 141 | Specific | 543.44 | 373.15 | 373.15 | 9.63 | 158 | >sp\|P02788\|TRFL_HUMAN Lactotransferrin OS=Homo sapiens OX=9606 GN=LTF PE=1 SV=6 | 1270 | 20201113_xd_milk_1ng_glyco_hcd203040_compound15.24547.24547.3 | controllerType=0 controllerNumber=1 scan=24547 | 49.1972 |
| 18678:3 | 1 | R.TAGWNVPIGTLRPFLN[+2279.830]WTGPPEPIEAAVAR.F | HexNAc(5)Hex(6)Fuc(2) | N[+2280] | 1837.5223 | 3 | 5510.5524 | 5510.5406 | 0 | 2.1 | 141 | Specific | 544.63 | 423.02 | 423.02 | 11.71 | 158 | >sp\|P02788\|TRFL_HUMAN Lactotransferrin OS=Homo sapiens OX=9606 GN=LTF PE=1 SV=6 | 1270 | 20201113_xd_milk_1ng_glyco_hcd203040_compound15.22108.22108.3 | controllerType=0 controllerNumber=1 scan=22108 | 44.0975 |
| 19606:4 | 1 | R.TAGWNVPIGTLRPFLN[+2570.925]WTGPPEPIEAAVAR.F | HexNAc(5)Hex(6)Fuc(2)NeuAc(1) | N[+2571] | 1451.1703 | 4 | 5801.6593 | 5801.636 | 0 | 4 | 141 | Specific | 566.09 | 454.21 | 454.21 | 11.39 | 158 | >sp\|P02788\|TRFL_HUMAN Lactotransferrin OS=Homo sapiens OX=9606 GN=LTF PE=1 SV=6 | 1270 | 20201113_xd_milk_1ng_glyco_hcd203040_compound15.23074.23074.4 | controllerType=0 controllerNumber=1 scan=23074 | 46.1271 |
| 19543:4 | 1 | R.TAGWNVPIGTLRPFLN[+2716.983]WTGPPEPIEAAVAR.F | HexNAc(5)Hex(6)Fuc(3)NeuAc(1) | N[+2717] | 1487.6819 | 4 | 5947.7057 | 5947.6939 | 0 | 2 | 141 | Specific | 590.11 | 578.5 | 578.5 | 13.17 | 158 | >sp\|P02788\|TRFL_HUMAN Lactotransferrin OS=Homo sapiens OX=9606 GN=LTF PE=1 SV=6 | 1270 | 20201113_xd_milk_1ng_glyco_hcd203040_compound15.23009.23009.4 | controllerType=0 controllerNumber=1 scan=23009 | 46.0162 |
| 20891:4 | 1 | R.TAGWNVPIGTLRPFLN[+2862.021]WTGPPEPIEAAVAR.F | HexNAc(5)Hex(6)Fuc(2)NeuAc(2) | N[+2862] | 1523.9385 | 4 | 6092.7321 | 6092.7314 | 0 | 0.1 | 141 | Specific | 590.92 | 342.14 | 342.14 | 10.91 | 158 | >sp\|P02788\|TRFL_HUMAN Lactotransferrin OS=Homo sapiens OX=9606 GN=LTF PE=1 SV=6 | 1270 | 20201113_xd_milk_1ng_glyco_hcd203040_compound15.24415.24415.3 | controllerType=0 controllerNumber=1 scan=24415 | 48.9094 |
| 19926:4 | 1 | R.TAGWNVPIGTLRPFLN[+2059.735]WTGPPEPIEAAVAR.F | HexNAc(4)Hex(5)Fuc(1)NeuAc(1) | N[+2060] | 1323.3696 | 4 | 5290.4567 | 5290.4459 | 0 | 2 | 141 | Specific | 598.02 | 501.8 | 501.8 | 13.17 | 158 | >sp\|P02788\|TRFL_HUMAN Lactotransferrin OS=Homo sapiens OX=9606 GN=LTF PE=1 SV=6 | 1270 | 20201113_xd_milk_1ng_glyco_hcd203040_compound15.23409.23409.4 | controllerType=0 controllerNumber=1 scan=23409 | 46.7899 |
| 19695:3 | 1 | R.TAGWNVPIGTLRPFLN[+2205.793]WTGPPEPIEAAVAR.F | HexNAc(4)Hex(5)Fuc(2)NeuAc(1) | N[+2206] | 1812.8448 | 3 | 5436.5199 | 5436.5038 | 0 | 3 | 141 | Specific | 605.38 | 486.58 | 486.58 | 14 | 158 | >sp\|P02788\|TRFL_HUMAN Lactotransferrin OS=Homo sapiens OX=9606 GN=LTF PE=1 SV=6 | 1270 | 20201113_xd_milk_1ng_glyco_hcd203040_compound15.23166.23166.3 | controllerType=0 controllerNumber=1 scan=23166 | 46.3126 |
| 19670:5 | 1 | R.TAGWNVPIGTLRPFLN[+2351.851]WTGPPEPIEAAVAR.F | HexNAc(4)Hex(5)Fuc(3)NeuAc(1) | N[+2352] | 1117.3269 | 5 | 5582.6054 | 5582.5617 | 0 | 7.8 | 141 | Specific | 607.13 | 596.37 | 596.37 | 12.97 | 158 | >sp\|P02788\|TRFL_HUMAN Lactotransferrin OS=Homo sapiens OX=9606 GN=LTF PE=1 SV=6 | 1270 | 20201113_xd_milk_1ng_glyco_hcd203040_compound15.23140.23140.2 | controllerType=0 controllerNumber=1 scan=23140 | 46.2517 |
| 18644:4 | 1 | R.TAGWNVPIGTLRPFLN[+2279.830]WTGPPEPIEAAVAR.F | HexNAc(5)Hex(6)Fuc(2) | N[+2280] | 1378.392 | 4 | 5510.5461 | 5510.5406 | 0 | 1 | 141 | Specific | 632.67 | 580.22 | 580.22 | 14 | 158 | >sp\|P02788\|TRFL_HUMAN Lactotransferrin OS=Homo sapiens OX=9606 GN=LTF PE=1 SV=6 | 1270 | 20201113_xd_milk_1ng_glyco_hcd203040_compound15.22073.22073.4 | controllerType=0 controllerNumber=1 scan=22073 | 44.0388 |
| 19727:5 | 1 | R.TAGWNVPIGTLRPFLN[+2059.735]WTGPPEPIEAAVAR.F | HexNAc(4)Hex(5)Fuc(1)NeuAc(1) | N[+2060] | 1058.8978 | 5 | 5290.46 | 5290.4459 | 0 | 2.7 | 141 | Specific | 633.03 | 520.51 | 520.51 | 14.1 | 158 | >sp\|P02788\|TRFL_HUMAN Lactotransferrin OS=Homo sapiens OX=9606 GN=LTF PE=1 SV=6 | 1270 | 20201113_xd_milk_1ng_glyco_hcd203040_compound15.23199.23199.3 | controllerType=0 controllerNumber=1 scan=23199 | 46.3886 |
| 19736:3 | 1 | R.TAGWNVPIGTLRPFLN[+2059.735]WTGPPEPIEAAVAR.F | HexNAc(4)Hex(5)Fuc(1)NeuAc(1) | N[+2060] | 1764.1627 | 3 | 5290.4735 | 5290.4459 | 0 | 5.2 | 141 | Specific | 635.57 | 575.86 | 575.86 | 13.34 | 158 | >sp\|P02788\|TRFL_HUMAN Lactotransferrin OS=Homo sapiens OX=9606 GN=LTF PE=1 SV=6 | 1270 | 20201113_xd_milk_1ng_glyco_hcd203040_compound15.23209.23209.3 | controllerType=0 controllerNumber=1 scan=23209 | 46.4082 |
| 19692:3 | 1 | R.TAGWNVPIGTLRPFLN[+2205.793]WTGPPEPIEAAVAR.F | HexNAc(4)Hex(5)Fuc(2)NeuAc(1) | N[+2206] | 1812.8448 | 3 | 5436.5199 | 5436.5038 | 0 | 3 | 141 | Specific | 644.47 | 577.53 | 577.53 | 14 | 158 | >sp\|P02788\|TRFL_HUMAN Lactotransferrin OS=Homo sapiens OX=9606 GN=LTF PE=1 SV=6 | 1270 | 20201113_xd_milk_1ng_glyco_hcd203040_compound15.23163.23163.3 | controllerType=0 controllerNumber=1 scan=23163 | 46.3058 |
| 19613:5 | 1 | R.TAGWNVPIGTLRPFLN[+2570.925]WTGPPEPIEAAVAR.F | HexNAc(5)Hex(6)Fuc(2)NeuAc(1) | N[+2571] | 1161.1361 | 5 | 5801.6514 | 5801.636 | 0 | 2.7 | 141 | Specific | 646.25 | 530.86 | 530.86 | 14.1 | 158 | >sp\|P02788\|TRFL_HUMAN Lactotransferrin OS=Homo sapiens OX=9606 GN=LTF PE=1 SV=6 | 1270 | 20201113_xd_milk_1ng_glyco_hcd203040_compound15.23081.23081.5 | controllerType=0 controllerNumber=1 scan=23081 | 46.1381 |
| 18679:4 | 1 | R.TAGWNVPIGTLRPFLN[+2425.887]WTGPPEPIEAAVAR.F | HexNAc(5)Hex(6)Fuc(3) | N[+2426] | 1414.9103 | 4 | 5656.6193 | 5656.5985 | 0 | 3.7 | 141 | Specific | 647.47 | 360.36 | 360.36 | 12.47 | 158 | >sp\|P02788\|TRFL_HUMAN Lactotransferrin OS=Homo sapiens OX=9606 GN=LTF PE=1 SV=6 | 1270 | 20201113_xd_milk_1ng_glyco_hcd203040_compound15.22109.22109.4 | controllerType=0 controllerNumber=1 scan=22109 | 44.099 |
| 19578:4 | 1 | R.TAGWNVPIGTLRPFLN[+2863.041]WTGPPEPIEAAVAR.F | HexNAc(5)Hex(6)Fuc(4)NeuAc(1) | N[+2863] | 1524.1981 | 4 | 6093.7707 | 6093.7518 | 0 | 3.1 | 141 | Specific | 658.37 | 542.58 | 542.58 | 14.97 | 158 | >sp\|P02788\|TRFL_HUMAN Lactotransferrin OS=Homo sapiens OX=9606 GN=LTF PE=1 SV=6 | 1270 | 20201113_xd_milk_1ng_glyco_hcd203040_compound15.23045.23045.4 | controllerType=0 controllerNumber=1 scan=23045 | 46.0764 |
| 18756:3 | 1 | R.TAGWNVPIGTLRPFLN[+1768.640]WTGPPEPIEAAVAR.F | HexNAc(4)Hex(5)Fuc(1) | N[+1769] | 1667.1374 | 3 | 4999.3977 | 4999.3504 | 0 | 9.5 | 141 | Specific | 670.65 | 588.71 | 588.71 | 13.92 | 158 | >sp\|P02788\|TRFL_HUMAN Lactotransferrin OS=Homo sapiens OX=9606 GN=LTF PE=1 SV=6 | 1270 | 20201113_xd_milk_1ng_glyco_hcd203040_compound15.22189.22189.3 | controllerType=0 controllerNumber=1 scan=22189 | 44.2542 |
| 19687:4 | 1 | R.TAGWNVPIGTLRPFLN[+2205.793]WTGPPEPIEAAVAR.F | HexNAc(4)Hex(5)Fuc(2)NeuAc(1) | N[+2206] | 1359.8855 | 4 | 5436.5202 | 5436.5038 | 0 | 3 | 141 | Specific | 684.5 | 606.72 | 606.72 | 14.97 | 158 | >sp\|P02788\|TRFL_HUMAN Lactotransferrin OS=Homo sapiens OX=9606 GN=LTF PE=1 SV=6 | 1270 | 20201113_xd_milk_1ng_glyco_hcd203040_compound15.23158.23158.4 | controllerType=0 controllerNumber=1 scan=23158 | 46.2908 |
| 19715:3 | 1 | R.TAGWNVPIGTLRPFLN[+2205.793]WTGPPEPIEAAVAR.F | HexNAc(4)Hex(5)Fuc(2)NeuAc(1) | N[+2206] | 1812.8457 | 3 | 5436.5224 | 5436.5038 | 0 | 3.4 | 141 | Specific | 685.59 | 637.66 | 637.66 | 15 | 158 | >sp\|P02788\|TRFL_HUMAN Lactotransferrin OS=Homo sapiens OX=9606 GN=LTF PE=1 SV=6 | 1270 | 20201113_xd_milk_1ng_glyco_hcd203040_compound15.23187.23187.3 | controllerType=0 controllerNumber=1 scan=23187 | 46.3667 |
| 19735:4 | 1 | R.TAGWNVPIGTLRPFLN[+2059.735]WTGPPEPIEAAVAR.F | HexNAc(4)Hex(5)Fuc(1)NeuAc(1) | N[+2060] | 1323.3734 | 4 | 5290.4718 | 5290.4459 | 0 | 4.9 | 141 | Specific | 689.5 | 640.11 | 640.11 | 14.66 | 158 | >sp\|P02788\|TRFL_HUMAN Lactotransferrin OS=Homo sapiens OX=9606 GN=LTF PE=1 SV=6 | 1270 | 20201113_xd_milk_1ng_glyco_hcd203040_compound15.23208.23208.4 | controllerType=0 controllerNumber=1 scan=23208 | 46.4068 |
| 18646:4 | 1 | R.TAGWNVPIGTLRPFLN[+2279.830]WTGPPEPIEAAVAR.F | HexNAc(5)Hex(6)Fuc(2) | N[+2280] | 1378.392 | 4 | 5510.5461 | 5510.5406 | 0 | 1 | 141 | Specific | 696.3 | 622.09 | 622.09 | 15 | 158 | >sp\|P02788\|TRFL_HUMAN Lactotransferrin OS=Homo sapiens OX=9606 GN=LTF PE=1 SV=6 | 1270 | 20201113_xd_milk_1ng_glyco_hcd203040_compound15.22075.22075.4 | controllerType=0 controllerNumber=1 scan=22075 | 44.0432 |
| 21016:4 | 1 | R.TAGWNVPIGTLRPFLN[+2350.830]WTGPPEPIEAAVAR.F | HexNAc(4)Hex(5)Fuc(1)NeuAc(2) | N[+2351] | 1396.1395 | 4 | 5581.5363 | 5581.5413 | 0 | -0.9 | 141 | Specific | 696.54 | 607.38 | 607.38 | 14.24 | 158 | >sp\|P02788\|TRFL_HUMAN Lactotransferrin OS=Homo sapiens OX=9606 GN=LTF PE=1 SV=6 | 1270 | 20201113_xd_milk_1ng_glyco_hcd203040_compound15.24546.24546.4 | controllerType=0 controllerNumber=1 scan=24546 | 49.1958 |
| 21188:4 | 1 | R.TAGWNVPIGTLRPFLN[+2351.851]WTGPPEPIEAAVAR.F | HexNAc(4)Hex(5)Fuc(3)NeuAc(1) | N[+2352] | 1396.3945 | 4 | 5582.5563 | 5582.5617 | 0 | -1 | 141 | Specific | 703.56 | 622.65 | 622.65 | 15.24 | 158 | >sp\|P02788\|TRFL_HUMAN Lactotransferrin OS=Homo sapiens OX=9606 GN=LTF PE=1 SV=6 | 1270 | 20201113_xd_milk_1ng_glyco_hcd203040_compound15.24724.24724.4 | controllerType=0 controllerNumber=1 scan=24724 | 49.558 |
| 19608:4 | 1 | R.TAGWNVPIGTLRPFLN[+2570.925]WTGPPEPIEAAVAR.F | HexNAc(5)Hex(6)Fuc(2)NeuAc(1) | N[+2571] | 1451.1703 | 4 | 5801.6593 | 5801.636 | 0 | 4 | 141 | Specific | 717.24 | 493.85 | 493.85 | 15.66 | 158 | >sp\|P02788\|TRFL_HUMAN Lactotransferrin OS=Homo sapiens OX=9606 GN=LTF PE=1 SV=6 | 1270 | 20201113_xd_milk_1ng_glyco_hcd203040_compound15.23076.23076.4 | controllerType=0 controllerNumber=1 scan=23076 | 46.1319 |
| 19738:3 | 1 | R.TAGWNVPIGTLRPFLN[+2059.735]WTGPPEPIEAAVAR.F | HexNAc(4)Hex(5)Fuc(1)NeuAc(1) | N[+2060] | 1764.1627 | 3 | 5290.4735 | 5290.4459 | 0 | 5.2 | 141 | Specific | 720.78 | 635.76 | 635.76 | 15.34 | 158 | >sp\|P02788\|TRFL_HUMAN Lactotransferrin OS=Homo sapiens OX=9606 GN=LTF PE=1 SV=6 | 1270 | 20201113_xd_milk_1ng_glyco_hcd203040_compound15.23211.23211.3 | controllerType=0 controllerNumber=1 scan=23211 | 46.4144 |
| 19678:4 | 1 | R.TAGWNVPIGTLRPFLN[+2424.867]WTGPPEPIEAAVAR.F | HexNAc(5)Hex(6)Fuc(1)NeuAc(1) | N[+2425] | 1414.6598 | 4 | 5655.6173 | 5655.5781 | 0 | 6.9 | 141 | Specific | 725.66 | 599.94 | 599.94 | 15.17 | 158 | >sp\|P02788\|TRFL_HUMAN Lactotransferrin OS=Homo sapiens OX=9606 GN=LTF PE=1 SV=6 | 1270 | 20201113_xd_milk_1ng_glyco_hcd203040_compound15.23148.23148.4 | controllerType=0 controllerNumber=1 scan=23148 | 46.2696 |
| 19546:4 | 1 | R.TAGWNVPIGTLRPFLN[+2716.983]WTGPPEPIEAAVAR.F | HexNAc(5)Hex(6)Fuc(3)NeuAc(1) | N[+2717] | 1487.6819 | 4 | 5947.7057 | 5947.6939 | 0 | 2 | 141 | Specific | 726.12 | 634.63 | 634.63 | 16.17 | 158 | >sp\|P02788\|TRFL_HUMAN Lactotransferrin OS=Homo sapiens OX=9606 GN=LTF PE=1 SV=6 | 1270 | 20201113_xd_milk_1ng_glyco_hcd203040_compound15.23012.23012.4 | controllerType=0 controllerNumber=1 scan=23012 | 46.0204 |
| 18753:4 | 1 | R.TAGWNVPIGTLRPFLN[+1768.640]WTGPPEPIEAAVAR.F | HexNAc(4)Hex(5)Fuc(1) | N[+1769] | 1250.604 | 4 | 4999.3942 | 4999.3504 | 0 | 8.8 | 141 | Specific | 730.63 | 730.63 | 730.63 | 14.9 | 158 | >sp\|P02788\|TRFL_HUMAN Lactotransferrin OS=Homo sapiens OX=9606 GN=LTF PE=1 SV=6 | 1270 | 20201113_xd_milk_1ng_glyco_hcd203040_compound15.22186.22186.4 | controllerType=0 controllerNumber=1 scan=22186 | 44.2473 |
| 19763:5 | 1 | R.TAGWNVPIGTLRPFLN[+2204.772]WTGPPEPIEAAVAR.F | HexNAc(4)Hex(5)NeuAc(2) | N[+2205] | 1087.91 | 5 | 5435.5211 | 5435.4834 | 0 | 6.9 | 141 | Specific | 759.12 | 532.6 | 532.6 | 16.11 | 158 | >sp\|P02788\|TRFL_HUMAN Lactotransferrin OS=Homo sapiens OX=9606 GN=LTF PE=1 SV=6 | 1270 | 20201113_xd_milk_1ng_glyco_hcd203040_compound15.23237.23237.5 | controllerType=0 controllerNumber=1 scan=23237 | 46.4847 |
| 21077:5 | 1 | R.TAGWNVPIGTLRPFLN[+2350.830]WTGPPEPIEAAVAR.F | HexNAc(4)Hex(5)Fuc(1)NeuAc(2) | N[+2351] | 1117.1171 | 5 | 5581.5564 | 5581.5413 | 0 | 2.7 | 141 | Specific | 776.15 | 543.79 | 543.79 | 17.1 | 158 | >sp\|P02788\|TRFL_HUMAN Lactotransferrin OS=Homo sapiens OX=9606 GN=LTF PE=1 SV=6 | 1270 | 20201113_xd_milk_1ng_glyco_hcd203040_compound15.24609.24609.5 | controllerType=0 controllerNumber=1 scan=24609 | 49.3203 |
| 19737:4 | 1 | R.TAGWNVPIGTLRPFLN[+2059.735]WTGPPEPIEAAVAR.F | HexNAc(4)Hex(5)Fuc(1)NeuAc(1) | N[+2060] | 1323.3734 | 4 | 5290.4718 | 5290.4459 | 0 | 4.9 | 141 | Specific | 805.59 | 643.09 | 643.09 | 17.66 | 158 | >sp\|P02788\|TRFL_HUMAN Lactotransferrin OS=Homo sapiens OX=9606 GN=LTF PE=1 SV=6 | 1270 | 20201113_xd_milk_1ng_glyco_hcd203040_compound15.23210.23210.4 | controllerType=0 controllerNumber=1 scan=23210 | 46.4125 |
| 19673:5 | 1 | R.TAGWNVPIGTLRPFLN[+2351.851]WTGPPEPIEAAVAR.F | HexNAc(4)Hex(5)Fuc(3)NeuAc(1) | N[+2352] | 1117.3269 | 5 | 5582.6054 | 5582.5617 | 0 | 7.8 | 141 | Specific | 809.2 | 720.48 | 720.48 | 16.97 | 158 | >sp\|P02788\|TRFL_HUMAN Lactotransferrin OS=Homo sapiens OX=9606 GN=LTF PE=1 SV=6 | 1270 | 20201113_xd_milk_1ng_glyco_hcd203040_compound15.23143.23143.2 | controllerType=0 controllerNumber=1 scan=23143 | 46.2602 |
| 19775:5 | 1 | R.TAGWNVPIGTLRPFLN[+2076.750]WTGPPEPIEAAVAR.F | HexNAc(4)Hex(6)Fuc(2) | N[+2077] | 1062.3047 | 5 | 5307.4943 | 5307.4612 | 0 | 6.2 | 141 | Specific | 816.12 | 698.63 | 698.63 | 17.11 | 158 | >sp\|P02788\|TRFL_HUMAN Lactotransferrin OS=Homo sapiens OX=9606 GN=LTF PE=1 SV=6 | 1270 | 20201113_xd_milk_1ng_glyco_hcd203040_compound15.23249.23249.5 | controllerType=0 controllerNumber=1 scan=23249 | 46.5057 |
| 20179:4 | 1 | R.TAGWNVPIGTLRPFLN[+2059.735]WTGPPEPIEAAVAR.F | HexNAc(4)Hex(5)Fuc(1)NeuAc(1) | N[+2060] | 1323.3727 | 4 | 5290.4689 | 5290.4459 | 0 | 4.4 | 141 | Specific | 830.6 | 830.6 | 830.6 | 17.66 | 158 | >sp\|P02788\|TRFL_HUMAN Lactotransferrin OS=Homo sapiens OX=9606 GN=LTF PE=1 SV=6 | 1270 | 20201113_xd_milk_1ng_glyco_hcd203040_compound15.23671.23671.2 | controllerType=0 controllerNumber=1 scan=23671 | 47.2862 |
| 19689:4 | 1 | R.TAGWNVPIGTLRPFLN[+2205.793]WTGPPEPIEAAVAR.F | HexNAc(4)Hex(5)Fuc(2)NeuAc(1) | N[+2206] | 1359.8855 | 4 | 5436.5202 | 5436.5038 | 0 | 3 | 141 | Specific | 831.74 | 722.3 | 722.3 | 17.97 | 158 | >sp\|P02788\|TRFL_HUMAN Lactotransferrin OS=Homo sapiens OX=9606 GN=LTF PE=1 SV=6 | 1270 | 20201113_xd_milk_1ng_glyco_hcd203040_compound15.23160.23160.4 | controllerType=0 controllerNumber=1 scan=23160 | 46.2987 |
| 19724:5 | 1 | R.TAGWNVPIGTLRPFLN[+2059.735]WTGPPEPIEAAVAR.F | HexNAc(4)Hex(5)Fuc(1)NeuAc(1) | N[+2060] | 1058.8978 | 5 | 5290.46 | 5290.4459 | 0 | 2.7 | 141 | Specific | 835.12 | 759.79 | 759.79 | 18.1 | 158 | >sp\|P02788\|TRFL_HUMAN Lactotransferrin OS=Homo sapiens OX=9606 GN=LTF PE=1 SV=6 | 1270 | 20201113_xd_milk_1ng_glyco_hcd203040_compound15.23196.23196.3 | controllerType=0 controllerNumber=1 scan=23196 | 46.3805 |
| 19929:4 | 1 | R.TAGWNVPIGTLRPFLN[+2059.735]WTGPPEPIEAAVAR.F | HexNAc(4)Hex(5)Fuc(1)NeuAc(1) | N[+2060] | 1323.3696 | 4 | 5290.4567 | 5290.4459 | 0 | 2 | 141 | Specific | 857.94 | 709.44 | 709.44 | 19.17 | 158 | >sp\|P02788\|TRFL_HUMAN Lactotransferrin OS=Homo sapiens OX=9606 GN=LTF PE=1 SV=6 | 1270 | 20201113_xd_milk_1ng_glyco_hcd203040_compound15.23412.23412.4 | controllerType=0 controllerNumber=1 scan=23412 | 46.7946 |
| 18755:4 | 1 | R.TAGWNVPIGTLRPFLN[+1768.640]WTGPPEPIEAAVAR.F | HexNAc(4)Hex(5)Fuc(1) | N[+1769] | 1250.604 | 4 | 4999.3942 | 4999.3504 | 0 | 8.8 | 141 | Specific | 864.97 | 778.01 | 778.01 | 17.9 | 158 | >sp\|P02788\|TRFL_HUMAN Lactotransferrin OS=Homo sapiens OX=9606 GN=LTF PE=1 SV=6 | 1270 | 20201113_xd_milk_1ng_glyco_hcd203040_compound15.22188.22188.4 | controllerType=0 controllerNumber=1 scan=22188 | 44.2523 |
| 19745:5 | 1 | R.TAGWNVPIGTLRPFLN[+2059.735]WTGPPEPIEAAVAR.F | HexNAc(4)Hex(5)Fuc(1)NeuAc(1) | N[+2060] | 1058.8989 | 5 | 5290.4655 | 5290.4459 | 0 | 3.7 | 141 | Specific | 877.02 | 850.02 | 850.02 | 18.91 | 158 | >sp\|P02788\|TRFL_HUMAN Lactotransferrin OS=Homo sapiens OX=9606 GN=LTF PE=1 SV=6 | 1270 | 20201113_xd_milk_1ng_glyco_hcd203040_compound15.23218.23218.5 | controllerType=0 controllerNumber=1 scan=23218 | 46.4285 |
| 19001:3 | 1 | R.TAGWNIPMGLLFN[+2351.851]QTGSC[+57.021]K.F | HexNAc(4)Hex(5)Fuc(3)NeuAc(1) | C[+57], N[+2352] | 1482.9639 | 3 | 4446.8772 | 4446.8607 | 0 | 3.7 | 485 | Specific | 369.8 | 296.95 | 296.95 | 8.05 | 158 | >sp\|P02788\|TRFL_HUMAN Lactotransferrin OS=Homo sapiens OX=9606 GN=LTF PE=1 SV=6 | 1270 | 20201113_xd_milk_1ng_glyco_hcd203040_compound15.22445.22445.3 | controllerType=0 controllerNumber=1 scan=22445 | 44.8238 |
| 18968:3 | 1 | R.TAGWNIPMGLLFN[+2424.867]QTGSC[+57.021]K.F | HexNAc(5)Hex(6)Fuc(1)NeuAc(1) | C[+57], N[+2425] | 1507.3007 | 3 | 4519.8876 | 4519.8771 | 0 | 2.3 | 485 | Specific | 440.68 | 384.99 | 163.12 | 10.38 | 158 | >sp\|P02788\|TRFL_HUMAN Lactotransferrin OS=Homo sapiens OX=9606 GN=LTF PE=1 SV=6 | 1270 | 20201113_xd_milk_1ng_glyco_hcd203040_compound15.22410.22410.3 | controllerType=0 controllerNumber=1 scan=22410 | 44.7492 |
| 18970:3 | 1 | R.TAGWNIPMGLLFN[+2424.867]QTGSC[+57.021]K.F | HexNAc(5)Hex(6)Fuc(1)NeuAc(1) | C[+57], N[+2425] | 1507.3007 | 3 | 4519.8876 | 4519.8771 | 0 | 2.3 | 485 | Specific | 490.31 | 444.2 | 297.82 | 10.64 | 158 | >sp\|P02788\|TRFL_HUMAN Lactotransferrin OS=Homo sapiens OX=9606 GN=LTF PE=1 SV=6 | 1270 | 20201113_xd_milk_1ng_glyco_hcd203040_compound15.22412.22412.3 | controllerType=0 controllerNumber=1 scan=22412 | 44.7559 |
| 16998:3 | 1 | R.TAGWNIPMGLLFN[+2133.772]QTGSC[+57.021]K.F | HexNAc(5)Hex(6)Fuc(1) | C[+57], N[+2134] | 1410.2656 | 3 | 4228.7821 | 4228.7817 | 0 | 0.1 | 485 | Specific | 503.79 | 503.79 | 248.8 | 10.06 | 158 | >sp\|P02788\|TRFL_HUMAN Lactotransferrin OS=Homo sapiens OX=9606 GN=LTF PE=1 SV=6 | 1270 | 20201113_xd_milk_1ng_glyco_hcd203040_compound15.20361.20361.3 | controllerType=0 controllerNumber=1 scan=20361 | 40.5203 |
| 19274:3 | 1 | R.TAGWNIPMGLLFN[+1872.651]QTGSC[+57.021]K.F | HexNAc(3)Hex(6)NeuAc(1) | C[+57], N[+1873] | 1323.231 | 3 | 3967.6785 | 3967.6604 | 0 | 4.6 | 485 | Specific | 534.47 | 534.47 | 242.35 | 10.4 | 158 | >sp\|P02788\|TRFL_HUMAN Lactotransferrin OS=Homo sapiens OX=9606 GN=LTF PE=1 SV=6 | 1270 | 20201113_xd_milk_1ng_glyco_hcd203040_compound15.22729.22729.3 | controllerType=0 controllerNumber=1 scan=22729 | 45.4592 |
| 19370:3 | 1 | R.TAGWNIPMGLLFN[+1548.545]QTGSC[+57.021]K.F | HexNAc(3)Hex(4)NeuAc(1) | C[+57], N[+1549] | 1215.1943 | 3 | 3643.5683 | 3643.5548 | 0 | 3.7 | 485 | Specific | 613.46 | 530.74 | 530.74 | 13.19 | 158 | >sp\|P02788\|TRFL_HUMAN Lactotransferrin OS=Homo sapiens OX=9606 GN=LTF PE=1 SV=6 | 1270 | 20201113_xd_milk_1ng_glyco_hcd203040_compound15.22829.22829.3 | controllerType=0 controllerNumber=1 scan=22829 | 45.6708 |
| 17392:2 | 1 | R.TAGWNIPMGLLFN[+1622.582]QTGSC[+57.021]K.F | HexNAc(4)Hex(5) | C[+57], N[+1623] | 1859.3016 | 2 | 3717.5959 | 3717.5916 | 0 | 1.2 | 485 | Specific | 622.9 | 584.7 | 246.75 | 13.24 | 158 | >sp\|P02788\|TRFL_HUMAN Lactotransferrin OS=Homo sapiens OX=9606 GN=LTF PE=1 SV=6 | 1270 | 20201113_xd_milk_1ng_glyco_hcd203040_compound15.20771.20771.2 | controllerType=0 controllerNumber=1 scan=20771 | 41.35 |
| 16929:3 | 1 | R.TAGWNIPMGLLFN[+2279.830]QTGSC[+57.021]K.F | HexNAc(5)Hex(6)Fuc(2) | C[+57], N[+2280] | 1458.954 | 3 | 4374.8476 | 4374.8396 | 0 | 1.8 | 485 | Specific | 662.51 | 644.84 | 321.91 | 14.39 | 158 | >sp\|P02788\|TRFL_HUMAN Lactotransferrin OS=Homo sapiens OX=9606 GN=LTF PE=1 SV=6 | 1270 | 20201113_xd_milk_1ng_glyco_hcd203040_compound15.20289.20289.3 | controllerType=0 controllerNumber=1 scan=20289 | 40.3704 |
| 17395:2 | 1 | R.TAGWNIPMGLLFN[+1622.582]QTGSC[+57.021]K.F | HexNAc(4)Hex(5) | C[+57], N[+1623] | 1859.3016 | 2 | 3717.5959 | 3717.5916 | 0 | 1.2 | 485 | Specific | 663.74 | 546.34 | 227.01 | 14.24 | 158 | >sp\|P02788\|TRFL_HUMAN Lactotransferrin OS=Homo sapiens OX=9606 GN=LTF PE=1 SV=6 | 1270 | 20201113_xd_milk_1ng_glyco_hcd203040_compound15.20774.20774.2 | controllerType=0 controllerNumber=1 scan=20774 | 41.3576 |
| 19310:3 | 1 | R.TAGWNIPMGLLFN[+1710.598]QTGSC[+57.021]K.F | HexNAc(3)Hex(5)NeuAc(1) | C[+57], N[+1711] | 1269.2071 | 3 | 3805.6067 | 3805.6076 | 0 | -0.2 | 485 | Specific | 686.36 | 599.32 | 256.53 | 13.81 | 158 | >sp\|P02788\|TRFL_HUMAN Lactotransferrin OS=Homo sapiens OX=9606 GN=LTF PE=1 SV=6 | 1270 | 20201113_xd_milk_1ng_glyco_hcd203040_compound15.22767.22767.3 | controllerType=0 controllerNumber=1 scan=22767 | 45.5472 |
| 19373:3 | 1 | R.TAGWNIPMGLLFN[+1548.545]QTGSC[+57.021]K.F | HexNAc(3)Hex(4)NeuAc(1) | C[+57], N[+1549] | 1215.1943 | 3 | 3643.5683 | 3643.5548 | 0 | 3.7 | 485 | Specific | 754.82 | 616 | 616 | 16.19 | 158 | >sp\|P02788\|TRFL_HUMAN Lactotransferrin OS=Homo sapiens OX=9606 GN=LTF PE=1 SV=6 | 1270 | 20201113_xd_milk_1ng_glyco_hcd203040_compound15.22832.22832.3 | controllerType=0 controllerNumber=1 scan=22832 | [illegible] |
| 17172:2 | 1 | R.TAGWNIPMGLLFN[+1768.640]QTGSC[+57.021]K.F | HexNAc(4)Hex(5)Fuc(1) | C[+57], N[+1769] | 1932.327 | 2 | 3863.6467 | 3863.6495 | 0 | -0.7 | 485 | Specific | 768.13 | 654.67 | 654.67 | 15.83 | 158 | >sp\|P02788\|TRFL_HUMAN Lactotransferrin OS=Homo sapiens OX=9606 GN=LTF PE=1 SV=6 | 1270 | 20201113_xd_milk_1ng_glyco_hcd203040_compound15.20542.20542.2 | controllerType=0 controllerNumber=1 scan=20542 | [illegible] |
| 16956:3 | 1 | R.TAGWNIPMGLLFN[+2279.830]QTGSC[+57.021]K.F | HexNAc(5)Hex(6)Fuc(2) | C[+57], N[+2280] | 1458.9581 | 3 | 4374.8596 | 4374.8396 | 0 | 4.6 | 485 | Specific | 794.85 | 792.54 | 400.15 | 15.88 | 158 | >sp\|P02788\|TRFL_HUMAN Lactotransferrin OS=Homo sapiens OX=9606 GN=LTF PE=1 SV=6 | 1270 | 20201113_xd_milk_1ng_glyco_hcd203040_compound15.20317.20317.3 | controllerType=0 controllerNumber=1 scan=20317 | [illegible] |
| 19313:3 | 1 | R.TAGWNIPMGLLFN[+1710.598]QTGSC[+57.021]K.F | HexNAc(3)Hex(5)NeuAc(1) | C[+57], N[+1711] | 1269.2071 | 3 | 3805.6067 | 3805.6076 | 0 | -0.2 | 485 | Specific | 818.52 | 643.79 | 356.31 | 16.81 | 158 | >sp\|P02788\|TRFL_HUMAN Lactotransferrin OS=Homo sapiens OX=9606 GN=LTF PE=1 SV=6 | 1270 | 20201113_xd_milk_1ng_glyco_hcd203040_compound15.22770.22770.3 | controllerType=0 controllerNumber=1 scan=22770 | [illegible] |
| 19204:4 | 1 | R.TAGWNIPMGLLFN[+1913.677]QTGSC[+57.021]K.F | HexNAc(4)Hex(5)NeuAc(1) | C[+57], N[+1914] | 1002.928 | 4 | 4008.69 | 4008.687 | 0 | 0.8 | 485 | Specific | 818.94 | 656.77 | 656.77 | 17.16 | 158 | >sp\|P02788\|TRFL_HUMAN Lactotransferrin OS=Homo sapiens OX=9606 GN=LTF PE=1 SV=6 | 1270 | 20201113_xd_milk_1ng_glyco_hcd203040_compound15.22656.22656.4 | controllerType=0 controllerNumber=1 scan=22656 | 45.2714 |
| 19207:4 | 1 | R.TAGWNIPMGLLFN[+1913.677]QTGSC[+57.021]K.F | HexNAc(4)Hex(5)NeuAc(1) | C[+57], N[+1914] | 1002.928 | 4 | 4008.69 | 4008.687 | 0 | 0.8 | 485 | Specific | 820.67 | 626.99 | 626.99 | 17.16 | 158 | >sp\|P02788\|TRFL_HUMAN Lactotransferrin OS=Homo sapiens OX=9606 GN=LTF PE=1 SV=6 | 1270 | 20201113_xd_milk_1ng_glyco_hcd203040_compound15.22659.22659.4 | controllerType=0 controllerNumber=1 scan=22659 | 45.2818 |
| 17186:4 | 1 | R.TAGWNIPMGLLFN[+1768.640]QTGSC[+57.021]K.F | HexNAc(4)Hex(5)Fuc(1) | C[+57], N[+1769] | 966.6661 | 4 | 3863.6424 | 3863.6495 | 0 | -1.8 | 485 | Specific | 820.97 | 656.26 | 656.26 | 16.39 | 158 | >sp\|P02788\|TRFL_HUMAN Lactotransferrin OS=Homo sapiens OX=9606 GN=LTF PE=1 SV=6 | 1270 | 20201113_xd_milk_1ng_glyco_hcd203040_compound15.20556.20556.4 | controllerType=0 controllerNumber=1 scan=20556 | 40.8996 |
| 17169:2 | 1 | R.TAGWNIPMGLLFN[+1768.640]QTGSC[+57.021]K.F | HexNAc(4)Hex(5)Fuc(1) | C[+57], N[+1769] | 1932.327 | 2 | 3863.6467 | 3863.6495 | 0 | -0.7 | 485 | Specific | 829.44 | 822.65 | 822.65 | 16.83 | 158 | >sp\|P02788\|TRFL_HUMAN Lactotransferrin OS=Homo sapiens OX=9606 GN=LTF PE=1 SV=6 | 1270 | 20201113_xd_milk_1ng_glyco_hcd203040_compound15.20539.20539.2 | controllerType=0 controllerNumber=1 scan=20539 | 40.8671 |
| 19277:3 | 1 | R.TAGWNIPMGLLFN[+1872.651]QTGSC[+57.021]K.F | HexNAc(3)Hex(6)NeuAc(1) | C[+57], N[+1873] | 1323.231 | 3 | 3967.6785 | 3967.6604 | 0 | 4.6 | 485 | Specific | 831.63 | 768.14 | 386.02 | 16.88 | 158 | >sp\|P02788\|TRFL_HUMAN Lactotransferrin OS=Homo sapiens OX=9606 GN=LTF PE=1 SV=6 | 1270 | 20201113_xd_milk_1ng_glyco_hcd203040_compound15.22732.22732.3 | controllerType=0 controllerNumber=1 scan=22732 | 45.4631 |
| 16932:3 | 1 | R.TAGWNIPMGLLFN[+2279.830]QTGSC[+57.021]K.F | HexNAc(5)Hex(6)Fuc(2) | C[+57], N[+2280] | 1458.954 | 3 | 4374.8476 | 4374.8396 | 0 | 1.8 | 485 | Specific | 835.25 | 789.08 | 426.12 | 17.39 | 158 | >sp\|P02788\|TRFL_HUMAN Lactotransferrin OS=Homo sapiens OX=9606 GN=LTF PE=1 SV=6 | 1270 | 20201113_xd_milk_1ng_glyco_hcd203040_compound15.20292.20292.3 | controllerType=0 controllerNumber=1 scan=20292 | 40.3743 |
| 16953:3 | 1 | R.TAGWNIPMGLLFN[+2279.830]QTGSC[+57.021]K.F | HexNAc(5)Hex(6)Fuc(2) | C[+57], N[+2280] | 1458.9581 | 3 | 4374.8596 | 4374.8396 | 0 | 4.6 | 485 | Specific | 838.04 | 728.55 | 439.73 | 16.88 | 158 | >sp\|P02788\|TRFL_HUMAN Lactotransferrin OS=Homo sapiens OX=9606 GN=LTF PE=1 SV=6 | 1270 | 20201113_xd_milk_1ng_glyco_hcd203040_compound15.20314.20314.3 | controllerType=0 controllerNumber=1 scan=20314 | 40.4143 |
| 19188:4 | 1 | R.TAGWNIPMGLLFN[+1913.677]QTGSC[+57.021]K.F | HexNAc(4)Hex(5)NeuAc(1) | C[+57], N[+1914] | 1002.9282 | 4 | 4008.691 | 4008.687 | 0 | 1 | 485 | Specific | 849.32 | 760.55 | 760.55 | 17.16 | 158 | >sp\|P02788\|TRFL_HUMAN Lactotransferrin OS=Homo sapiens OX=9606 GN=LTF PE=1 SV=6 | 1270 | 20201113_xd_milk_1ng_glyco_hcd203040_compound15.22639.22639.2 | controllerType=0 controllerNumber=1 scan=22639 | 45.2234 |
| 17183:4 | 1 | R.TAGWNIPMGLLFN[+1768.640]QTGSC[+57.021]K.F | HexNAc(4)Hex(5)Fuc(1) | C[+57], N[+1769] | 966.6661 | 4 | 3863.6424 | 3863.6495 | 0 | -1.8 | 485 | Specific | 849.55 | 723.45 | 723.45 | 16.39 | 158 | >sp\|P02788\|TRFL_HUMAN Lactotransferrin OS=Homo sapiens OX=9606 GN=LTF PE=1 SV=6 | 1270 | 20201113_xd_milk_1ng_glyco_hcd203040_compound15.20553.20553.4 | controllerType=0 controllerNumber=1 scan=20553 | 40.8924 |
| 17001:3 | 1 | R.TAGWNIPMGLLFN[+2133.772]QTGSC[+57.021]K.F | HexNAc(5)Hex(6)Fuc(1) | C[+57], N[+2134] | 1410.2656 | 3 | 4228.7821 | 4228.7817 | 0 | 0.1 | 485 | Specific | 865.49 | 704.64 | 460.14 | 17.81 | 158 | >sp\|P02788\|TRFL_HUMAN Lactotransferrin OS=Homo sapiens OX=9606 GN=LTF PE=1 SV=6 | 1270 | 20201113_xd_milk_1ng_glyco_hcd203040_compound15.20364.20364.3 | controllerType=0 controllerNumber=1 scan=20364 | 40.5242 |
| 19004:3 | 1 | R.TAGWNIPMGLLFN[+2351.851]QTGSC[+57.021]K.F | HexNAc(4)Hex(5)Fuc(3)NeuAc(1) | C[+57], N[+2352] | 1482.9639 | 3 | 4446.8772 | 4446.8607 | 0 | 3.7 | 485 | Specific | 881.63 | 855.2 | 855.2 | 18.19 | 158 | >sp\|P02788\|TRFL_HUMAN Lactotransferrin OS=Homo sapiens OX=9606 GN=LTF PE=1 SV=6 | 1270 | 20201113_xd_milk_1ng_glyco_hcd203040_compound15.22448.22448.3 | controllerType=0 controllerNumber=1 scan=22448 | 44.8278 |
| 21096:3 | 1 | R.TAGWNIPMGLLFN[+2350.830]QTGSC[+57.021]K.F | HexNAc(4)Hex(5)Fuc(1)NeuAc(2) | C[+57], N[+2351] | 1482.6224 | 3 | 4445.8526 | 4445.8403 | 0 | 2.8 | 485 | Specific | 881.91 | 862.4 | 862.4 | 18.19 | 158 | >sp\|P02788\|TRFL_HUMAN Lactotransferrin OS=Homo sapiens OX=9606 GN=LTF PE=1 SV=6 | 1270 | 20201113_xd_milk_1ng_glyco_hcd203040_compound15.24629.24629.3 | controllerType=0 controllerNumber=1 scan=24629 | 49.3626 |
| 19175:3 | 1 | R.TAGWNIPMGLLFN[+1913.677]QTGSC[+57.021]K.F | HexNAc(4)Hex(5)NeuAc(1) | C[+57], N[+1914] | 1336.9023 | 3 | 4008.6923 | 4008.687 | 0 | 1.3 | 485 | Specific | 891.38 | 891.38 | 891.38 | 18.22 | 158 | >sp\|P02788\|TRFL_HUMAN Lactotransferrin OS=Homo sapiens OX=9606 GN=LTF PE=1 SV=6 | 1270 | 20201113_xd_milk_1ng_glyco_hcd203040_compound15.22626.22626.3 | controllerType=0 controllerNumber=1 scan=22626 | 45.204 |
| 17126:3 | 1 | R.TAGWNIPMGLLFN[+1768.640]QTGSC[+57.021]K.F | HexNAc(4)Hex(5)Fuc(1) | C[+57], N[+1769] | 1288.5619 | 3 | 3863.6713 | 3863.6495 | 0 | 5.6 | 485 | Specific | 907.37 | 907.37 | 907.37 | 18.54 | 158 | >sp\|P02788\|TRFL_HUMAN Lactotransferrin OS=Homo sapiens OX=9606 GN=LTF PE=1 SV=6 | 1270 | 20201113_xd_milk_1ng_glyco_hcd203040_compound15.20494.20494.3 | controllerType=0 controllerNumber=1 scan=20494 | 40.7729 |
| 21244:3 | 1 | R.TAGWNIPMGLLFN[+2204.772]QTGSC[+57.021]K.F | HexNAc(4)Hex(5)NeuAc(2) | C[+57], N[+2205] | 1433.9374 | 3 | 4299.7978 | 4299.7824 | 0 | 3.6 | 485 | Specific | 918.53 | 886.1 | 886.1 | 19.19 | 158 | >sp\|P02788\|TRFL_HUMAN Lactotransferrin OS=Homo sapiens OX=9606 GN=LTF PE=1 SV=6 | 1270 | 20201113_xd_milk_1ng_glyco_hcd203040_compound15.24783.24783.3 | controllerType=0 controllerNumber=1 scan=24783 | 49.7109 |
| 17344:3 | 1 | R.TAGWNIPMGLLFN[+1622.582]QTGSC[+57.021]K.F | HexNAc(4)Hex(5) | C[+57], N[+1623] | 1239.8747 | 3 | 3717.6095 | 3717.5916 | 0 | 4.8 | 485 | Specific | 922.65 | 902.51 | 466.52 | 18.88 | 158 | >sp\|P02788\|TRFL_HUMAN Lactotransferrin OS=Homo sapiens OX=9606 GN=LTF PE=1 SV=6 | 1270 | 20201113_xd_milk_1ng_glyco_hcd203040_compound15.20721.20721.3 | controllerType=0 controllerNumber=1 scan=20721 | 41.2365 |
| 17342:3 | 1 | R.TAGWNIPMGLLFN[+1622.582]QTGSC[+57.021]K.F | HexNAc(4)Hex(5) | C[+57], N[+1623] | 1239.8747 | 3 | 3717.6095 | 3717.5916 | 0 | 4.8 | 485 | Specific | 925.35 | 925.35 | 467.18 | 18.88 | 158 | >sp\|P02788\|TRFL_HUMAN Lactotransferrin OS=Homo sapiens OX=9606 GN=LTF PE=1 SV=6 | 1270 | 20201113_xd_milk_1ng_glyco_hcd203040_compound15.20719.20719.3 | controllerType=0 controllerNumber=1 scan=20719 | 41.2314 |
| 21242:3 | 1 | R.TAGWNIPMGLLFN[+2204.772]QTGSC[+57.021]K.F | HexNAc(4)Hex(5)NeuAc(2) | C[+57], N[+2205] | 1433.9374 | 3 | 4299.7978 | 4299.7824 | 0 | 3.6 | 485 | Specific | 930.62 | 815.31 | 815.31 | 19.19 | 158 | >sp\|P02788\|TRFL_HUMAN Lactotransferrin OS=Homo sapiens OX=9606 GN=LTF PE=1 SV=6 | 1270 | 20201113_xd_milk_1ng_glyco_hcd203040_compound15.24781.24781.3 | controllerType=0 controllerNumber=1 scan=24781 | 49.7025 |
| 18991:3 | 1 | R.TAGWNIPMGLLFN[+2205.793]QTGSC[+57.021]K.F | HexNAc(4)Hex(5)Fuc(2)NeuAc(1) | C[+57], N[+2206] | 1434.2752 | 3 | 4300.8111 | 4300.8028 | 0 | 1.9 | 485 | Specific | 931.87 | 921.51 | 921.51 | 19.39 | 158 | >sp\|P02788\|TRFL_HUMAN Lactotransferrin OS=Homo sapiens OX=9606 GN=LTF PE=1 SV=6 | 1270 | 20201113_xd_milk_1ng_glyco_hcd203040_compound15.22434.22434.3 | controllerType=0 controllerNumber=1 scan=22434 | 44.7996 |
| 17124:3 | 1 | R.TAGWNIPMGLLFN[+1768.640]QTGSC[+57.021]K.F | HexNAc(4)Hex(5)Fuc(1) | C[+57], N[+1769] | 1288.5619 | 3 | 3863.6713 | 3863.6495 | 0 | 5.6 | 485 | Specific | 941.79 | 941.79 | 941.79 | 18.54 | 158 | >sp\|P02788\|TRFL_HUMAN Lactotransferrin OS=Homo sapiens OX=9606 GN=LTF PE=1 SV=6 | 1270 | 20201113_xd_milk_1ng_glyco_hcd203040_compound15.20492.20492.3 | controllerType=0 controllerNumber=1 scan=20492 | 40.7673 |
| 19172:3 | 1 | R.TAGWNIPMGLLFN[+1913.677]QTGSC[+57.021]K.F | HexNAc(4)Hex(5)NeuAc(1) | C[+57], N[+1914] | 1336.9023 | 3 | 4008.6923 | 4008.687 | 0 | 1.3 | 485 | Specific | 959.42 | 901.5 | 901.5 | 20.22 | 158 | >sp\|P02788\|TRFL_HUMAN Lactotransferrin OS=Homo sapiens OX=9606 GN=LTF PE=1 SV=6 | 1270 | 20201113_xd_milk_1ng_glyco_hcd203040_compound15.22623.22623.3 | controllerType=0 controllerNumber=1 scan=22623 | 45.1979 |
| 19033:3 | 1 | R.TAGWNIPMGLLFN[+2059.735]QTGSC[+57.021]K.F | HexNAc(4)Hex(5)Fuc(1)NeuAc(1) | C[+57], N[+2060] | 1385.5905 | 3 | 4154.757 | 4154.7449 | 0 | 2.9 | 485 | Specific | 963.63 | 693.83 | 693.83 | 20.19 | 158 | >sp\|P02788\|TRFL_HUMAN Lactotransferrin OS=Homo sapiens OX=9606 GN=LTF PE=1 SV=6 | 1270 | 20201113_xd_milk_1ng_glyco_hcd203040_compound15.22478.22478.3 | controllerType=0 controllerNumber=1 scan=22478 | 44.8861 |
| 19191:4 | 1 | R.TAGWNIPMGLLFN[+1913.677]QTGSC[+57.021]K.F | HexNAc(4)Hex(5)NeuAc(1) | C[+57], N[+1914] | 1002.9282 | 4 | 4008.691 | 4008.687 | 0 | 1 | 485 | Specific | 973.7 | 894.1 | 894.1 | 20.16 | 158 | >sp\|P02788\|TRFL_HUMAN Lactotransferrin OS=Homo sapiens OX=9606 GN=LTF PE=1 SV=6 | 1270 | 20201113_xd_milk_1ng_glyco_hcd203040_compound15.22642.22642.2 | controllerType=0 controllerNumber=1 scan=22642 | 45.2317 |
| 21094:3 | 1 | R.TAGWNIPMGLLFN[+2350.830]QTGSC[+57.021]K.F | HexNAc(4)Hex(5)Fuc(1)NeuAc(2) | C[+57], N[+2351] | 1482.6224 | 3 | 4445.8526 | 4445.8403 | 0 | 2.8 | 485 | Specific | 990.28 | 947.81 | 947.81 | 20.19 | 158 | >sp\|P02788\|TRFL_HUMAN Lactotransferrin OS=Homo sapiens OX=9606 GN=LTF PE=1 SV=6 | 1270 | 20201113_xd_milk_1ng_glyco_hcd203040_compound15.24627.24627.3 | controllerType=0 controllerNumber=1 scan=24627 | 49.3568 |
| 18988:3 | 1 | R.TAGWNIPMGLLFN[+2205.793]QTGSC[+57.021]K.F | HexNAc(4)Hex(5)Fuc(2)NeuAc(1) | C[+57], N[+2206] | 1434.2752 | 3 | 4300.8111 | 4300.8028 | 0 | 1.9 | 485 | Specific | 991.94 | 867.01 | 867.01 | 20.39 | 158 | >sp\|P02788\|TRFL_HUMAN Lactotransferrin OS=Homo sapiens OX=9606 GN=LTF PE=1 SV=6 | 1270 | 20201113_xd_milk_1ng_glyco_hcd203040_compound15.22431.22431.3 | controllerType=0 controllerNumber=1 scan=22431 | 44.7941 |
| 19030:3 | 1 | R.TAGWNIPMGLLFN[+2059.735]QTGSC[+57.021]K.F | HexNAc(4)Hex(5)Fuc(1)NeuAc(1) | C[+57], N[+2060] | 1385.5905 | 3 | 4154.757 | 4154.7449 | 0 | 2.9 | 485 | Specific | 1003.22 | 859.73 | 859.73 | 21.19 | 158 | >sp\|P02788\|TRFL_HUMAN Lactotransferrin OS=Homo sapiens OX=9606 GN=LTF PE=1 SV=6 | 1270 | 20201113_xd_milk_1ng_glyco_hcd203040_compound15.22475.22475.3 | controllerType=0 controllerNumber=1 scan=22475 | 44.8784 |
| 16999:3 | 1 | R.TAGWNIPM[+15.995]GLLFN[+1913.677]QTGSC[+57.021]K.F | HexNAc(4)Hex(5)NeuAc(1) | C[+57], M[+16], N[+1914] | 1342.2353 | 3 | 4024.6913 | 4024.6819 | 0 | 2.3 | 485 | Specific | 237.79 | 198.44 | 198.44 | 6.93 | 158 | >sp\|P02788\|TRFL_HUMAN Lactotransferrin OS=Homo sapiens OX=9606 GN=LTF PE=1 SV=6 | 1270 | 20201113_xd_milk_1ng_glyco_hcd203040_compound15.20362.20362.3 | controllerType=0 controllerNumber=1 scan=20362 | 40.5214 |
| 14650:3 | 1 | R.TAGWNIPM[+15.995]GLLFN[+1622.582]QTGSC[+57.021]K.F | HexNAc(4)Hex(5) | C[+57], M[+16], N[+1623] | 1245.209 | 3 | 3733.6126 | 3733.5865 | 0 | 7 | 485 | Specific | 293.79 | 293.79 | 293.79 | 6.9 | 158 | >sp\|P02788\|TRFL_HUMAN Lactotransferrin OS=Homo sapiens OX=9606 GN=LTF PE=1 SV=6 | 1270 | 20201113_xd_milk_1ng_glyco_hcd203040_compound15.17904.17904.3 | controllerType=0 controllerNumber=1 scan=17904 | 35.9604 |
| 14623:3 | 1 | R.TAGWNIPM[+15.995]GLLFN[+1622.582]QTGSC[+57.021]K.F | HexNAc(4)Hex(5) | C[+57], M[+16], N[+1623] | 1245.2065 | 3 | 3733.6049 | 3733.5865 | 0 | 4.9 | 485 | Specific | 339.85 | 339.85 | 339.85 | 7.93 | 158 | >sp\|P02788\|TRFL_HUMAN Lactotransferrin OS=Homo sapiens OX=9606 GN=LTF PE=1 SV=6 | 1270 | 20201113_xd_milk_1ng_glyco_hcd203040_compound15.17876.17876.3 | controllerType=0 controllerNumber=1 scan=17876 | 35.908 |
| 17002:3 | 1 | R.TAGWNIPM[+15.995]GLLFN[+1913.677]QTGSC[+57.021]K.F | HexNAc(4)Hex(5)NeuAc(1) | C[+57], M[+16], N[+1914] | 1342.2353 | 3 | 4024.6913 | 4024.6819 | 0 | 2.3 | 485 | Specific | 510.62 | 430.39 | 430.39 | 10.64 | 158 | >sp\|P02788\|TRFL_HUMAN Lactotransferrin OS=Homo sapiens OX=9606 GN=LTF PE=1 SV=6 | 1270 | 20201113_xd_milk_1ng_glyco_hcd203040_compound15.20365.20365.3 | controllerType=0 controllerNumber=1 scan=20365 | 40.5262 |
| 14652:3 | 1 | R.TAGWNIPM[+15.995]GLLFN[+1622.582]QTGSC[+57.021]K.F | HexNAc(4)Hex(5) | C[+57], M[+16], N[+1623] | 1245.209 | 3 | 3733.6126 | 3733.5865 | 0 | 7 | 485 | Specific | 514.02 | 514.02 | 514.02 | 9.78 | 158 | >sp\|P02788\|TRFL_HUMAN Lactotransferrin OS=Homo sapiens OX=9606 GN=LTF PE=1 SV=6 | 1270 | 20201113_xd_milk_1ng_glyco_hcd203040_compound15.17906.17906.3 | controllerType=0 controllerNumber=1 scan=17906 | 35.9645 |
| 14625:3 | 1 | R.TAGWNIPM[+15.995]GLLFN[+1622.582]QTGSC[+57.021]K.F | HexNAc(4)Hex(5) | C[+57], M[+16], N[+1623] | 1245.2065 | 3 | 3733.6049 | 3733.5865 | 0 | 4.9 | 485 | Specific | 554.74 | 528.93 | 528.93 | 10.4 | 158 | >sp\|P02788\|TRFL_HUMAN Lactotransferrin OS=Homo sapiens OX=9606 GN=LTF PE=1 SV=6 | 1270 | 20201113_xd_milk_1ng_glyco_hcd203040_compound15.17878.17878.3 | controllerType=0 controllerNumber=1 scan=17878 | 35.9115 |
| 19470:4 | 1 | R.[+42.011]TAGWNIPMGLLFN[+1872.651]QTGSC[+57.021]KFDEYFSQSC[+57.021]A | HexNAc(3)Hex(6)NeuAc(1) | .[42], C[+57]*2, N[+1873] | 1464.1166 | 4 | 5853.4445 | 5853.4181 | 0 | 4.5 | 485 | Specific | 208.48 | 208.48 | 208.48 | 4.59 | 158 | >sp\|P02788\|TRFL_HUMAN Lactotransferrin OS=Homo sapiens OX=9606 GN=LTF PE=1 SV=6 | 1270 | 20201113_xd_milk_1ng_glyco_hcd203040_compound15.22933.22933.4 | controllerType=0 controllerNumber=1 scan=22933 | 45.8722 |
| 19329:4 | 1 | R.TAGWNIPMGLLFN[+2059.735]QTGSC[+57.021]KFDEYFSQSC[+57.021]APGSDPR.S | HexNAc(4)Hex(5)Fuc(1)NeuAc(1) | C[+57]*2, N[+2060] | 1500.382 | 4 | 5998.506 | 5998.492 | 0 | 2.3 | 485 | Specific | 213.02 | 213.02 | 213.02 | 5.09 | 158 | >sp\|P02788\|TRFL_HUMAN Lactotransferrin OS=Homo sapiens OX=9606 GN=LTF PE=1 SV=6 | 1270 | 20201113_xd_milk_1ng_glyco_hcd203040_compound15.22787.22787.4 | controllerType=0 controllerNumber=1 scan=22787 | 45.5889 |
| 18314:3 | 1 | R.TAGWNIPMGLLFN[+1768.640]QTGSC[+57.021]KFDEYFSQSC[+57.021]APGSDPR.S | HexNAc(4)Hex(5)Fuc(1) | C[+57]*2, N[+1769] | 1903.1417 | 3 | 5707.4105 | 5707.3966 | 0 | 2.4 | 485 | Specific | 248.21 | 207.16 | 207.16 | 6.46 | 158 | >sp\|P02788\|TRFL_HUMAN Lactotransferrin OS=Homo sapiens OX=9606 GN=LTF PE=1 SV=6 | 1270 | 20201113_xd_milk_1ng_glyco_hcd203040_compound15.21730.21730.3 | controllerType=0 controllerNumber=1 scan=21730 | 43.3468 |
| 19408:4 | 1 | R.TAGWNIPMGLLFN[+2076.750]QTGSC[+57.021]KFDEYFSQSC[+57.021]APGSDPR.S | HexNAc(4)Hex(6)Fuc(2) | C[+57]*2, N[+2077] | 1504.6383 | 4 | 6015.5314 | 6015.5074 | 0 | 4 | 485 | Specific | 269.18 | 256.62 | 256.62 | 6.26 | 158 | >sp\|P02788\|TRFL_HUMAN Lactotransferrin OS=Homo sapiens OX=9606 GN=LTF PE=1 SV=6 | 1270 | 20201113_xd_milk_1ng_glyco_hcd203040_compound15.22868.22868.2 | controllerType=0 controllerNumber=1 scan=22868 | 45.7407 |
| 19160:4 | 1 | R.TAGWNIPMGLLFN[+2862.021]QTGSC[+57.021]KFDEYFSQSC[+57.021]APGSDPR.S | HexNAc(5)Hex(6)Fuc(2)NeuAc(2) | C[+57]*2, N[+2862] | 1700.9616 | 4 | 6800.8247 | 6800.7776 | 0 | 6.9 | 485 | Specific | 333.22 | 333.22 | 333.22 | 6.21 | 158 | >sp\|P02788\|TRFL_HUMAN Lactotransferrin OS=Homo sapiens OX=9606 GN=LTF PE=1 SV=6 | 1270 | 20201113_xd_milk_1ng_glyco_hcd203040_compound15.22610.22610.4 | controllerType=0 controllerNumber=1 scan=22610 | 45.1703 |
| 19473:4 | 1 | R.[+42.011]TAGWNIPMGLLFN[+1872.651]QTGSC[+57.021]KFDEYFSQSC[+57.021]A | HexNAc(3)Hex(6)NeuAc(1) | .[42], C[+57]*2, N[+1873] | 1464.1166 | 4 | 5853.4445 | 5853.4181 | 0 | 4.5 | 485 | Specific | 334.13 | 145.74 | 145.74 | 4.07 | 158 | >sp\|P02788\|TRFL_HUMAN Lactotransferrin OS=Homo sapiens OX=9606 GN=LTF PE=1 SV=6 | 1270 | 20201113_xd_milk_1ng_glyco_hcd203040_compound15.22936.22936.4 | controllerType=0 controllerNumber=1 scan=22936 | 45.8758 |
| 19316:4 | 1 | R.TAGWNIPMGLLFN[+2205.793]QTGSC[+57.021]KFDEYFSQSC[+57.021]APGSDPR.S | HexNAc(4)Hex(5)Fuc(2)NeuAc(1) | C[+57]*2, N[+2206] | 1536.8972 | 4 | 6144.5671 | 6144.5499 | 0 | 2.8 | 485 | Specific | 357.12 | 350.6 | 350.6 | 7.56 | 158 | >sp\|P02788\|TRFL_HUMAN Lactotransferrin OS=Homo sapiens OX=9606 GN=LTF PE=1 SV=6 | 1270 | 20201113_xd_milk_1ng_glyco_hcd203040_compound15.22773.22773.3 | controllerType=0 controllerNumber=1 scan=22773 | 45.5582 |
| 19162:4 | 1 | R.TAGWNIPMGLLFN[+2862.021]QTGSC[+57.021]KFDEYFSQSC[+57.021]APGSDPR.S | HexNAc(5)Hex(6)Fuc(2)NeuAc(2) | C[+57]*2, N[+2862] | 1700.9616 | 4 | 6800.8247 | 6800.7776 | 0 | 6.9 | 485 | Specific | 444.71 | 407.14 | 407.14 | 8.67 | 158 | >sp\|P02788\|TRFL_HUMAN Lactotransferrin OS=Homo sapiens OX=9606 GN=LTF PE=1 SV=6 | 1270 | 20201113_xd_milk_1ng_glyco_hcd203040_compound15.22612.22612.4 | controllerType=0 controllerNumber=1 scan=22612 | 45.1738 |
| 19319:4 | 1 | R.TAGWNIPMGLLFN[+2205.793]QTGSC[+57.021]KFDEYFSQSC[+57.021]APGSDPR.S | HexNAc(4)Hex(5)Fuc(2)NeuAc(1) | C[+57]*2, N[+2206] | 1536.8972 | 4 | 6144.5671 | 6144.5499 | 0 | 2.8 | 485 | Specific | 511.25 | 461.33 | 461.33 | 9.83 | 158 | >sp\|P02788\|TRFL_HUMAN Lactotransferrin OS=Homo sapiens OX=9606 GN=LTF PE=1 SV=6 | 1270 | 20201113_xd_milk_1ng_glyco_hcd203040_compound15.22776.22776.3 | controllerType=0 controllerNumber=1 scan=22776 | 45.5653 |
| 19333:4 | 1 | R.TAGWNIPMGLLFN[+2205.793]QTGSC[+57.021]KFDEYFSQSC[+57.021]APGSDPR.S | HexNAc(4)Hex(5)Fuc(2)NeuAc(1) | C[+57]*2, N[+2206] | 1536.895 | 4 | 6144.5583 | 6144.5499 | 0 | 1.4 | 485 | Specific | 544.65 | 518.19 | 518.19 | 10 | 158 | >sp\|P02788\|TRFL_HUMAN Lactotransferrin OS=Homo sapiens OX=9606 GN=LTF PE=1 SV=6 | 1270 | 20201113_xd_milk_1ng_glyco_hcd203040_compound15.22791.22791.4 | controllerType=0 controllerNumber=1 scan=22791 | 45.5945 |
| 19395:3 | 1 | R.TAGWNIPMGLLFN[+1913.677]QTGSC[+57.021]KFDEYFSQSC[+57.021]APGSDPR.S | HexNAc(4)Hex(5)NeuAc(1) | C[+57]*2, N[+1914] | 1951.4881 | 3 | 5852.4498 | 5852.4341 | 0 | 2.7 | 485 | Specific | 559.38 | 458 | 458 | 10.4 | 158 | >sp\|P02788\|TRFL_HUMAN Lactotransferrin OS=Homo sapiens OX=9606 GN=LTF PE=1 SV=6 | 1270 | 20201113_xd_milk_1ng_glyco_hcd203040_compound15.22855.22855.3 | controllerType=0 controllerNumber=1 scan=22855 | 45.7203 |
| 18317:3 | 1 | R.TAGWNIPMGLLFN[+1768.640]QTGSC[+57.021]KFDEYFSQSC[+57.021]APGSDPR.S | HexNAc(4)Hex(5)Fuc(1) | C[+57]*2, N[+1769] | 1903.1417 | 3 | 5707.4105 | 5707.3966 | 0 | 2.4 | 485 | Specific | 616.26 | 510.36 | 510.36 | 12.67 | 158 | >sp\|P02788\|TRFL_HUMAN Lactotransferrin OS=Homo sapiens OX=9606 GN=LTF PE=1 SV=6 | 1270 | 20201113_xd_milk_1ng_glyco_hcd203040_compound15.21733.21733.3 | controllerType=0 controllerNumber=1 scan=21733 | 43.35 |
| 19309:4 | 1 | R.TAGWNIPMGLLFN[+2351.851]QTGSC[+57.021]KFDEYFSQSC[+57.021]APGSDPR.S | HexNAc(4)Hex(5)Fuc(3)NeuAc(1) | C[+57]*2, N[+2352] | 1573.4154 | 4 | 6290.6398 | 6290.6079 | 0 | 5.1 | 485 | Specific | 712.36 | 711.21 | 711.21 | 13.8 | 158 | >sp\|P02788\|TRFL_HUMAN Lactotransferrin OS=Homo sapiens OX=9606 GN=LTF PE=1 SV=6 | 1270 | 20201113_xd_milk_1ng_glyco_hcd203040_compound15.22766.22766.4 | controllerType=0 controllerNumber=1 scan=22766 | 45.5461 |
| 20680:4 | 1 | R.TAGWNIPMGLLFN[+2350.830]QTGSC[+57.021]KFDEYFSQSC[+57.021]APGSDPR.S | HexNAc(4)Hex(5)Fuc(1)NeuAc(2) | C[+57]*2, N[+2351] | 1573.1674 | 4 | 6289.6476 | 6289.5875 | 0 | 9.6 | 485 | Specific | 743.44 | 648.13 | 648.13 | 13.69 | 158 | >sp\|P02788\|TRFL_HUMAN Lactotransferrin OS=Homo sapiens OX=9606 GN=LTF PE=1 SV=6 | 1270 | 20201113_xd_milk_1ng_glyco_hcd203040_compound15.24194.24194.4 | controllerType=0 controllerNumber=1 scan=24194 | 48.4152 |
| 19397:3 | 1 | R.TAGWNIPMGLLFN[+1913.677]QTGSC[+57.021]KFDEYFSQSC[+57.021]APGSDPR.S | HexNAc(4)Hex(5)NeuAc(1) | C[+57]*2, N[+1914] | 1951.4881 | 3 | 5852.4498 | 5852.4341 | 0 | 2.7 | 485 | Specific | 758.27 | 629.91 | 629.91 | 15.67 | 158 | >sp\|P02788\|TRFL_HUMAN Lactotransferrin OS=Homo sapiens OX=9606 GN=LTF PE=1 SV=6 | 1270 | 20201113_xd_milk_1ng_glyco_hcd203040_compound15.22857.22857.3 | controllerType=0 controllerNumber=1 scan=22857 | 45.7238 |
| 18284:4 | 1 | R.TAGWNIPMGLLFN[+1768.640]QTGSC[+57.021]KFDEYFSQSC[+57.021]APGSDPR.S | HexNAc(4)Hex(5)Fuc(1) | C[+57]*2, N[+1769] | 1427.609 | 4 | 5707.4142 | 5707.3966 | 0 | 3.1 | 485 | Specific | 796.2 | 743.25 | 743.25 | 15.45 | 158 | >sp\|P02788\|TRFL_HUMAN Lactotransferrin OS=Homo sapiens OX=9606 GN=LTF PE=1 SV=6 | 1270 | 20201113_xd_milk_1ng_glyco_hcd203040_compound15.21699.21699.4 | controllerType=0 controllerNumber=1 scan=21699 | 43.286 |
| 19312:4 | 1 | R.TAGWNIPMGLLFN[+2351.851]QTGSC[+57.021]KFDEYFSQSC[+57.021]APGSDPR.S | HexNAc(4)Hex(5)Fuc(3)NeuAc(1) | C[+57]*2, N[+2352] | 1573.4154 | 4 | 6290.6398 | 6290.6079 | 0 | 5.1 | 485 | Specific | 891.15 | 846.76 | 846.76 | 16.8 | 158 | >sp\|P02788\|TRFL_HUMAN Lactotransferrin OS=Homo sapiens OX=9606 GN=LTF PE=1 SV=6 | 1270 | 20201113_xd_milk_1ng_glyco_hcd203040_compound15.22769.22769.4 | controllerType=0 controllerNumber=1 scan=22769 | 45.5525 |
| 18286:4 | 1 | R.TAGWNIPMGLLFN[+1768.640]QTGSC[+57.021]KFDEYFSQSC[+57.021]APGSDPR.S | HexNAc(4)Hex(5)Fuc(1) | C[+57]*2, N[+1769] | 1427.609 | 4 | 5707.4142 | 5707.3966 | 0 | 3.1 | 485 | Specific | 896.67 | 879.68 | 879.68 | 17.45 | 158 | >sp\|P02788\|TRFL_HUMAN Lactotransferrin OS=Homo sapiens OX=9606 GN=LTF PE=1 SV=6 | 1270 | 20201113_xd_milk_1ng_glyco_hcd203040_compound15.21701.21701.4 | controllerType=0 controllerNumber=1 scan=21701 | 43.2909 |
| 20683:4 | 1 | R.TAGWNIPMGLLFN[+2350.830]QTGSC[+57.021]KFDEYFSQSC[+57.021]APGSDPR.S | HexNAc(4)Hex(5)Fuc(1)NeuAc(2) | C[+57]*2, N[+2351] | 1573.1674 | 4 | 6289.6476 | 6289.5875 | 0 | 9.6 | 485 | Specific | 913.6 | 847.39 | 847.39 | 17.69 | 158 | >sp\|P02788\|TRFL_HUMAN Lactotransferrin OS=Homo sapiens OX=9606 GN=LTF PE=1 SV=6 | 1270 | 20201113_xd_milk_1ng_glyco_hcd203040_compound15.24197.24197.4 | controllerType=0 controllerNumber=1 scan=24197 | 48.4254 |
| 19331:4 | 1 | R.TAGWNIPMGLLFN[+2059.735]QTGSC[+57.021]KFDEYFSQSC[+57.021]APGSDPR.S | HexNAc(4)Hex(5)Fuc(1)NeuAc(1) | C[+57]*2, N[+2060] | 1500.382 | 4 | 5998.506 | 5998.492 | 0 | 2.3 | 485 | Specific | 915.73 | 865.16 | 865.16 | 18.65 | 158 | >sp\|P02788\|TRFL_HUMAN Lactotransferrin OS=Homo sapiens OX=9606 GN=LTF PE=1 SV=6 | 1270 | 20201113_xd_milk_1ng_glyco_hcd203040_compound15.22789.22789.4 | controllerType=0 controllerNumber=1 scan=22789 | 45.5924 |
| 19336:4 | 1 | R.TAGWNIPMGLLFN[+2205.793]QTGSC[+57.021]KFDEYFSQSC[+57.021]APGSDPR.S | HexNAc(4)Hex(5)Fuc(2)NeuAc(1) | C[+57]*2, N[+2206] | 1536.895 | 4 | 6144.5583 | 6144.5499 | 0 | 1.4 | 485 | Specific | 956.02 | 899.1 | 899.1 | 19.48 | 158 | >sp\|P02788\|TRFL_HUMAN Lactotransferrin OS=Homo sapiens OX=9606 GN=LTF PE=1 SV=6 | 1270 | 20201113_xd_milk_1ng_glyco_hcd203040_compound15.22794.22794.4 | controllerType=0 controllerNumber=1 scan=22794 | 45.5986 |
| 18281:4 | 1 | R.TAGWNIPMGLLFN[+1768.640]QTGSC[+57.021]KFDEYFSQSC[+57.021]APGSDPR.S | HexNAc(4)Hex(5)Fuc(1) | C[+57]*2, N[+1769] | 1427.61 | 4 | 5707.4181 | 5707.3966 | 0 | 3.8 | 485 | Specific | 999.12 | 955.79 | 955.79 | 19.45 | 158 | >sp\|P02788\|TRFL_HUMAN Lactotransferrin OS=Homo sapiens OX=9606 GN=LTF PE=1 SV=6 | 1270 | 20201113_xd_milk_1ng_glyco_hcd203040_compound15.21695.21695.4 | controllerType=0 controllerNumber=1 scan=21695 | 43.2779 |
| 18278:4 | 1 | R.TAGWNIPMGLLFN[+1768.640]QTGSC[+57.021]KFDEYFSQSC[+57.021]APGSDPR.S | HexNAc(4)Hex(5)Fuc(1) | C[+57]*2, N[+1769] | 1427.61 | 4 | 5707.4181 | 5707.3966 | 0 | 3.8 | 485 | Specific | 1037.42 | 978.92 | 978.92 | 20.45 | 158 | >sp\|P02788\|TRFL_HUMAN Lactotransferrin OS=Homo sapiens OX=9606 GN=LTF PE=1 SV=6 | 1270 | 20201113_xd_milk_1ng_glyco_hcd203040_compound15.21692.21692.4 | controllerType=0 controllerNumber=1 scan=21692 | 43.2706 |
| 09253:3 | 1 | R.N[+2571.946]GSDC[+57.021]PDKFC[+57.021]LFQSETK.N | HexNAc(5)Hex(6)Fuc(4) | C[+57]*2, N[+2572] | 1535.6042 | 3 | 4604.798 | 4604.8193 | 0 | -4.6 | 642 | Specific | 319.33 | 244.95 | 244.95 | 5.21 | 158 | >sp\|P02788\|TRFL_HUMAN Lactotransferrin OS=Homo sapiens OX=9606 GN=LTF PE=1 SV=6 | 1270 | 20201113_xd_milk_1ng_glyco_hcd203040_compound15.12295.12295.3 | controllerType=0 controllerNumber=1 scan=12295 | 26.2863 |
| 09251:3 | 1 | R.N[+2571.946]GSDC[+57.021]PDKFC[+57.021]LFQSETK.N | HexNAc(5)Hex(6)Fuc(4) | C[+57]*2, N[+2572] | 1535.6042 | 3 | 4604.798 | 4604.8193 | 0 | -4.6 | 642 | Specific | 319.81 | 191.7 | 191.7 | 4.91 | 158 | >sp\|P02788\|TRFL_HUMAN Lactotransferrin OS=Homo sapiens OX=9606 GN=LTF PE=1 SV=6 | 1270 | 20201113_xd_milk_1ng_glyco_hcd203040_compound15.12293.12293.3 | controllerType=0 controllerNumber=1 scan=12293 | 26.2825 |
| 09447:3 | 1 | R.N[+2351.851]GSDC[+57.021]PDKFC[+57.021]LFQSETK.N | HexNAc(4)Hex(5)Fuc(3)NeuAc(1) | C[+57]*2, N[+2352] | 1462.2493 | 3 | 4384.7334 | 4384.7247 | 0 | 2 | 642 | Specific | 324.06 | 273.65 | 273.65 | 7.03 | 158 | >sp\|P02788\|TRFL_HUMAN Lactotransferrin OS=Homo sapiens OX=9606 GN=LTF PE=1 SV=6 | 1270 | 20201113_xd_milk_1ng_glyco_hcd203040_compound15.12501.12501.3 | controllerType=0 controllerNumber=1 scan=12501 | 26.6193 |
| 09449:3 | 1 | R.N[+2351.851]GSDC[+57.021]PDKFC[+57.021]LFQSETK.N | HexNAc(4)Hex(5)Fuc(3)NeuAc(1) | C[+57]*2, N[+2352] | 1462.2493 | 3 | 4384.7334 | 4384.7247 | 0 | 2 | 642 | Specific | 340.85 | 212.81 | 212.81 | 6.97 | 158 | >sp\|P02788\|TRFL_HUMAN Lactotransferrin OS=Homo sapiens OX=9606 GN=LTF PE=1 SV=6 | 1270 | 20201113_xd_milk_1ng_glyco_hcd203040_compound15.12503.12503.3 | controllerType=0 controllerNumber=1 scan=12503 | 26.6231 |
| 09459:3 | 1 | R.N[+2351.851]GSDC[+57.021]PDKFC[+57.021]LFQSETK.N | HexNAc(4)Hex(5)Fuc(3)NeuAc(1) | C[+57]*2, N[+2352] | 1462.2501 | 3 | 4384.7356 | 4384.7247 | 0 | 2.5 | 642 | Specific | 371.47 | 239.74 | 239.74 | 7.18 | 158 | >sp\|P02788\|TRFL_HUMAN Lactotransferrin OS=Homo sapiens OX=9606 GN=LTF PE=1 SV=6 | 1270 | 20201113_xd_milk_1ng_glyco_hcd203040_compound15.12514.12514.3 | controllerType=0 controllerNumber=1 scan=12514 | 26.6436 |
| 11912:3 | 1 | R.N[+2351.851]GSDC[+57.021]PDKFC[+57.021]LFQSETK.N | HexNAc(4)Hex(5)Fuc(3)NeuAc(1) | C[+57]*2, N[+2352] | 1462.2459 | 3 | 4384.7232 | 4384.7247 | 0 | -0.3 | 642 | Specific | 391.34 | 335.78 | 335.78 | 7.26 | 158 | >sp\|P02788\|TRFL_HUMAN Lactotransferrin OS=Homo sapiens OX=9606 GN=LTF PE=1 SV=6 | 1270 | 20201113_xd_milk_1ng_glyco_hcd203040_compound15.15054.15054.3 | controllerType=0 controllerNumber=1 scan=15054 | 30.9198 |
| 11914:3 | 1 | R.N[+2351.851]GSDC[+57.021]PDKFC[+57.021]LFQSETK.N | HexNAc(4)Hex(5)Fuc(3)NeuAc(1) | C[+57]*2, N[+2352] | 1462.2459 | 3 | 4384.7232 | 4384.7247 | 0 | -0.3 | 642 | Specific | 410.59 | 317.69 | 317.69 | 7.1 | 158 | >sp\|P02788\|TRFL_HUMAN Lactotransferrin OS=Homo sapiens OX=9606 GN=LTF PE=1 SV=6 | 1270 | 20201113_xd_milk_1ng_glyco_hcd203040_compound15.15056.15056.3 | controllerType=0 controllerNumber=1 scan=15056 | 30.9235 |
| 09457:3 | 1 | R.N[+2351.851]GSDC[+57.021]PDKFC[+57.021]LFQSETK.N | HexNAc(4)Hex(5)Fuc(3)NeuAc(1) | C[+57]*2, N[+2352] | 1462.2501 | 3 | 4384.7356 | 4384.7247 | 0 | 2.5 | 642 | Specific | 429.65 | 338.17 | 338.17 | 8.02 | 158 | >sp\|P02788\|TRFL_HUMAN Lactotransferrin OS=Homo sapiens OX=9606 GN=LTF PE=1 SV=6 | 1270 | 20201113_xd_milk_1ng_glyco_hcd203040_compound15.12512.12512.3 | controllerType=0 controllerNumber=1 scan=12512 | 26.6394 |
| 09483:3 | 1 | R.N[+2205.793]GSDC[+57.021]PDKFC[+57.021]LFQSETK.N | HexNAc(4)Hex(5)Fuc(2)NeuAc(1) | C[+57]*2, N[+2206] | 1413.5637 | 3 | 4238.6764 | 4238.6667 | 0 | 2.3 | 642 | Specific | 515.11 | 374.07 | 374.07 | 8.58 | 158 | >sp\|P02788\|TRFL_HUMAN Lactotransferrin OS=Homo sapiens OX=9606 GN=LTF PE=1 SV=6 | 1270 | 20201113_xd_milk_1ng_glyco_hcd203040_compound15.12539.12539.3 | controllerType=0 controllerNumber=1 scan=12539 | 26.6866 |
| 09451:3 | 1 | R.N[+2205.793]GSDC[+57.021]PDKFC[+57.021]LFQSETK.N | HexNAc(4)Hex(5)Fuc(2)NeuAc(1) | C[+57]*2, N[+2206] | 1413.5623 | 3 | 4238.6724 | 4238.6667 | 0 | 1.3 | 642 | Specific | 515.66 | 381.34 | 381.34 | 9.91 | 158 | >sp\|P02788\|TRFL_HUMAN Lactotransferrin OS=Homo sapiens OX=9606 GN=LTF PE=1 SV=6 | 1270 | 20201113_xd_milk_1ng_glyco_hcd203040_compound15.12506.12506.3 | controllerType=0 controllerNumber=1 scan=12506 | 26.629 |
| 09577:3 | 1 | R.N[+2059.735]GSDC[+57.021]PDKFC[+57.021]LFQSETK.N | HexNAc(4)Hex(5)Fuc(1)NeuAc(1) | C[+57]*2, N[+2060] | 1364.8768 | 3 | 4092.6157 | 4092.6088 | 0 | 1.7 | 642 | Specific | 518.71 | 428.28 | 428.28 | 10.08 | 158 | >sp\|P02788\|TRFL_HUMAN Lactotransferrin OS=Homo sapiens OX=9606 GN=LTF PE=1 SV=6 | 1270 | 20201113_xd_milk_1ng_glyco_hcd203040_compound15.12636.12636.3 | controllerType=0 controllerNumber=1 scan=12636 | 26.8384 |
| 09480:3 | 1 | R.N[+2205.793]GSDC[+57.021]PDKFC[+57.021]LFQSETK.N | HexNAc(4)Hex(5)Fuc(2)NeuAc(1) | C[+57]*2, N[+2206] | 1413.5637 | 3 | 4238.6764 | 4238.6667 | 0 | 2.3 | 642 | Specific | 562.3 | 403.37 | 403.37 | 10.43 | 158 | >sp\|P02788\|TRFL_HUMAN Lactotransferrin OS=Homo sapiens OX=9606 GN=LTF PE=1 SV=6 | 1270 | 20201113_xd_milk_1ng_glyco_hcd203040_compound15.12536.12536.3 | controllerType=0 controllerNumber=1 scan=12536 | 26.6826 |
| 09579:3 | 1 | R.N[+2059.735]GSDC[+57.021]PDKFC[+57.021]LFQSETK.N | HexNAc(4)Hex(5)Fuc(1)NeuAc(1) | C[+57]*2, N[+2060] | 1364.8768 | 3 | 4092.6157 | 4092.6088 | 0 | 1.7 | 642 | Specific | 589.86 | 391.16 | 391.16 | 11.7 | 158 | >sp\|P02788\|TRFL_HUMAN Lactotransferrin OS=Homo sapiens OX=9606 GN=LTF PE=1 SV=6 | 1270 | 20201113_xd_milk_1ng_glyco_hcd203040_compound15.12638.12638.3 | controllerType=0 controllerNumber=1 scan=12638 | 26.8424 |

Table S4. Cont.

| Query #:z | Protein Rank | Glycopeptide | Glycan | Modification Type(s) | Observed m/z | z | Observed (M+H) | Calc. mass (M+H) | Off-by-x error | Mass error (ppm) | StartPosition | Cleavage | Score | Delta | Delta Mod | \|Log Prob\| | # of unique peptides | Protein Name | Protein DB number | Comment | Scan # | Scan Time |
|---|---|---|---|---|---|---|---|---|---|---|---|---|---|---|---|---|---|---|---|---|---|---|
| 09453:3 | 1 | R.N[+2205.793]GSDC[+57.021]PDKFC[+57.021]LFQSETK.N | HexNAc(4)Hex(5)Fuc(2)NeuAc(1) | C[+57]*2, N[+2206] | 1413.5623 | 3 | 4238.6724 | 4238.6667 | 0 | 1.3 | 642 | Specific | 617.45 | 408.51 | 408.51 | 12.53 | 158 | >sp\|P02788\|TRFL_HUMAN Lactotransferrin OS=Homo sapiens OX=9606 GN=LTF PE=1 SV=6 | 1270 | 20201113_xd_milk_1ng_glyco_hcd203040_compound15.12508.12508.3 | controllerType=0 controllerNumber=1 scan=12508 | 26.6331 |
| 10446:3 | 3 | K.TVLTPATNHMGN[+1704.635]VTFTIPANR.E | HexNAc(6)Hex(3) | N[+1705] | 1320.5903 | 3 | 3959.7563 | 3959.7948 | 0 | -9.7 | 74 | Specific | 701.05 | 701.05 | 701.05 | 13.52 | 114 | >sp\|P01024\|CO3_HUMAN Complement C3 OS=Homo sapiens OX=9606 GN=C3 PE=1 SV=2 | 492 | 20201113_xd_milk_1ng_glyco_hcd203040_compound15.13536.13536.3 | controllerType=0 controllerNumber=1 scan=13536 | 28.3037 |
| 10496:3 | 3 | K.TVLTPATNHMGN[+1540.529]VTFTIPANR.E | HexNAc(2)Hex(7) | N[+1541] | 1265.905 | 3 | 3795.7003 | 3795.6886 | 0 | 3.1 | 74 | Specific | 761.81 | 761.81 | 434.35 | 16.22 | 114 | >sp\|P01024\|CO3_HUMAN Complement C3 OS=Homo sapiens OX=9606 GN=C3 PE=1 SV=2 | 492 | 20201113_xd_milk_1ng_glyco_hcd203040_compound15.13588.13588.3 | controllerType=0 controllerNumber=1 scan=13588 | 28.3931 |
| 10523:3 | 3 | K.TVLTPATNHMGN[+1378.476]VTFTIPANR.E | HexNAc(2)Hex(6) | N[+1378] | 1211.8863 | 3 | 3633.6443 | 3633.6358 | 0 | 2.3 | 74 | Specific | 913.34 | 689.8 | 497.14 | 19.41 | 114 | >sp\|P01024\|CO3_HUMAN Complement C3 OS=Homo sapiens OX=9606 GN=C3 PE=1 SV=2 | 492 | 20201113_xd_milk_1ng_glyco_hcd203040_compound15.13616.13616.3 | controllerType=0 controllerNumber=1 scan=13616 | 28.4442 |
| 10444:3 | 3 | K.TVLTPATNHMGN[+1704.635]VTFTIPANR.E | HexNAc(6)Hex(3) | N[+1705] | 1320.5903 | 3 | 3959.7563 | 3959.7948 | 0 | -9.7 | 74 | Specific | 931.77 | 931.77 | 931.77 | 17.52 | 114 | >sp\|P01024\|CO3_HUMAN Complement C3 OS=Homo sapiens OX=9606 GN=C3 PE=1 SV=2 | 492 | 20201113_xd_milk_1ng_glyco_hcd203040_compound15.13534.13534.3 | controllerType=0 controllerNumber=1 scan=13534 | 28.2995 |
| 10499:3 | 3 | K.TVLTPATNHMGN[+1540.529]VTFTIPANR.E | HexNAc(2)Hex(7) | N[+1541] | 1265.905 | 3 | 3795.7003 | 3795.6886 | 0 | 3.1 | 74 | Specific | 969.48 | 871.19 | 498.4 | 20.22 | 114 | >sp\|P01024\|CO3_HUMAN Complement C3 OS=Homo sapiens OX=9606 GN=C3 PE=1 SV=2 | 492 | 20201113_xd_milk_1ng_glyco_hcd203040_compound15.13591.13591.3 | controllerType=0 controllerNumber=1 scan=13591 | 28.3965 |
| 10525:3 | 3 | K.TVLTPATNHMGN[+1378.476]VTFTIPANR.E | HexNAc(2)Hex(6) | N[+1378] | 1211.8863 | 3 | 3633.6443 | 3633.6358 | 0 | 2.3 | 74 | Specific | 986.65 | 914.36 | 535.93 | 20.41 | 114 | >sp\|P01024\|CO3_HUMAN Complement C3 OS=Homo sapiens OX=9606 GN=C3 PE=1 SV=2 | 492 | 20201113_xd_milk_1ng_glyco_hcd203040_compound15.13618.13618.3 | controllerType=0 controllerNumber=1 scan=13618 | 28.449 |
| 07674:3 | 4 | R.FSDGLESN[+1864.634]SSTQFEVK.K | HexNAc(2)Hex(9) | N[+1865] | 1213.8237 | 3 | 3639.4565 | 3639.4472 | 0 | 2.6 | 219 | Specific | 855.07 | 855.07 | 855.07 | 18.16 | 97 | >sp\|P0C0L5\|CO4B_HUMAN Complement C4-B OS=Homo sapiens OX=9606 GN=C4B PE=1 SV=2 | 1424 | 20201113_xd_milk_1ng_glyco_hcd203040_compound15.10653.10653.3 | controllerType=0 controllerNumber=1 scan=10653 | 23.5859 |
| 07677:3 | 4 | R.FSDGLESN[+1864.634]SSTQFEVK.K | HexNAc(2)Hex(9) | N[+1865] | 1213.8237 | 3 | 3639.4565 | 3639.4472 | 0 | 2.6 | 219 | Specific | 940.95 | 864.05 | 864.05 | 19.16 | 97 | >sp\|P0C0L5\|CO4B_HUMAN Complement C4-B OS=Homo sapiens OX=9606 GN=C4B PE=1 SV=2 | 1424 | 20201113_xd_milk_1ng_glyco_hcd203040_compound15.10656.10656.3 | controllerType=0 controllerNumber=1 scan=10656 | 23.5895 |
| 05085:3 | 4 | R.FSDGLESN[+1864.634]SSTQFEVKK.Y | HexNAc(2)Hex(9) | N[+1865] | 1256.5202 | 3 | 3767.546 | 3767.5421 | 0 | 1 | 219 | Specific | 765.01 | 597.54 | 597.54 | 15.53 | 97 | >sp\|P0C0L5\|CO4B_HUMAN Complement C4-B OS=Homo sapiens OX=9606 GN=C4B PE=1 SV=2 | 1424 | 20201113_xd_milk_1ng_glyco_hcd203040_compound15.7976.7976.3 | controllerType=0 controllerNumber=1 scan=7976 | 19.3316 |
| 05083:3 | 4 | R.FSDGLESN[+1864.634]SSTQFEVKK.Y | HexNAc(2)Hex(9) | N[+1865] | 1256.5202 | 3 | 3767.546 | 3767.5421 | 0 | 1 | 219 | Specific | 861.71 | 737.81 | 737.81 | 17.53 | 97 | >sp\|P0C0L5\|CO4B_HUMAN Complement C4-B OS=Homo sapiens OX=9606 GN=C4B PE=1 SV=2 | 1424 | 20201113_xd_milk_1ng_glyco_hcd203040_compound15.7974.7974.3 | controllerType=0 controllerNumber=1 scan=7974 | 19.3271 |
| 18077:3 | 7 | R.GLSFDVSLEVSQGPGLLN[+2059.735]DTK.V | HexNAc(4)Hex(5)Fuc(1)NeuAc(1) | N[+2060] | 1412.6276 | 3 | 4235.8683 | 4235.8481 | 0 | 4.8 | 118 | Specific | 551.05 | 551.05 | 551.05 | 10.4 | 71 | >sp\|P01833\|PIGR_HUMAN Polymeric immunoglobulin receptor OS=Homo sapiens OX=9606 GN=PIGR PE=1 SV=4 | 902 | 20201113_xd_milk_1ng_glyco_hcd203040_compound15.21483.21483.3 | controllerType=0 controllerNumber=1 scan=21483 | 42.8128 |
| 18080:3 | 7 | R.GLSFDVSLEVSQGPGLLN[+2059.735]DTK.V | HexNAc(4)Hex(5)Fuc(1)NeuAc(1) | N[+2060] | 1412.6276 | 3 | 4235.8683 | 4235.8481 | 0 | 4.8 | 118 | Specific | 969.13 | 884.18 | 884.18 | 19.88 | 71 | >sp\|P01833\|PIGR_HUMAN Polymeric immunoglobulin receptor OS=Homo sapiens OX=9606 GN=PIGR PE=1 SV=4 | 902 | 20201113_xd_milk_1ng_glyco_hcd203040_compound15.21486.21486.3 | controllerType=0 controllerNumber=1 scan=21486 | 42.8169 |
| 19223:4 | 7 | K.QIGLYPVLVIDSSGYVNPN[+2059.735]YTGR.I | HexNAc(4)Hex(5)Fuc(1)NeuAc(1) | N[+2060] | 1147.0157 | 4 | 4585.0408 | 4585.0384 | 0 | 0.5 | 168 | Specific | 283.59 | 236.95 | 236.95 | 5.44 | 71 | >sp\|P01833\|PIGR_HUMAN Polymeric immunoglobulin receptor OS=Homo sapiens OX=9606 GN=PIGR PE=1 SV=4 | 902 | 20201113_xd_milk_1ng_glyco_hcd203040_compound15.22675.22675.3 | controllerType=0 controllerNumber=1 scan=22675 | 45.3119 |
| 19285:3 | 7 | K.QIGLYPVLVIDSSGYVNPN[+1913.677]YTGR.I | HexNAc(4)Hex(5)NeuAc(1) | N[+1914] | 1480.3378 | 3 | 4438.9989 | 4438.9805 | 0 | 4.2 | 168 | Specific | 553.59 | 468.16 | 468.16 | 10.65 | 71 | >sp\|P01833\|PIGR_HUMAN Polymeric immunoglobulin receptor OS=Homo sapiens OX=9606 GN=PIGR PE=1 SV=4 | 902 | 20201113_xd_milk_1ng_glyco_hcd203040_compound15.22741.22741.3 | controllerType=0 controllerNumber=1 scan=22741 | 45.4921 |
| 19222:4 | 7 | K.QIGLYPVLVIDSSGYVNPN[+2059.735]YTGR.I | HexNAc(4)Hex(5)Fuc(1)NeuAc(1) | N[+2060] | 1147.0157 | 4 | 4585.0408 | 4585.0384 | 0 | 0.5 | 168 | Specific | 648.53 | 536.85 | 536.85 | 13.4 | 71 | >sp\|P01833\|PIGR_HUMAN Polymeric immunoglobulin receptor OS=Homo sapiens OX=9606 GN=PIGR PE=1 SV=4 | 902 | 20201113_xd_milk_1ng_glyco_hcd203040_compound15.22674.22674.3 | controllerType=0 controllerNumber=1 scan=22674 | 45.3058 |
| 21241:3 | 7 | K.QIGLYPVLVIDSSGYVNPN[+2204.772]YTGR.I | HexNAc(4)Hex(5)NeuAc(2) | N[+2205] | 1577.3597 | 3 | 4730.0645 | 4730.0759 | 0 | -2.4 | 168 | Specific | 725.87 | 544.37 | 544.37 | 14.18 | 71 | >sp\|P01833\|PIGR_HUMAN Polymeric immunoglobulin receptor OS=Homo sapiens OX=9606 GN=PIGR PE=1 SV=4 | 902 | 20201113_xd_milk_1ng_glyco_hcd203040_compound15.24780.24780.4 | controllerType=0 controllerNumber=1 scan=24780 | 49.7014 |
| 19269:3 | 7 | K.QIGLYPVLVIDSSGYVNPN[+1913.677]YTGR.I | HexNAc(4)Hex(5)NeuAc(1) | N[+1914] | 1480.3383 | 3 | 4439.0004 | 4438.9805 | 0 | 4.5 | 168 | Specific | 823.45 | 733.57 | 733.57 | 17.13 | 71 | >sp\|P01833\|PIGR_HUMAN Polymeric immunoglobulin receptor OS=Homo sapiens OX=9606 GN=PIGR PE=1 SV=4 | 902 | 20201113_xd_milk_1ng_glyco_hcd203040_compound15.22724.22724.3 | controllerType=0 controllerNumber=1 scan=22724 | 45.4299 |
| 21238:3 | 7 | K.QIGLYPVLVIDSSGYVNPN[+2204.772]YTGR.I | HexNAc(4)Hex(5)NeuAc(2) | N[+2205] | 1577.3597 | 3 | 4730.0645 | 4730.0759 | 0 | -2.4 | 168 | Specific | 855.65 | 824.34 | 824.34 | 17.18 | 71 | >sp\|P01833\|PIGR_HUMAN Polymeric immunoglobulin receptor OS=Homo sapiens OX=9606 GN=PIGR PE=1 SV=4 | 902 | 20201113_xd_milk_1ng_glyco_hcd203040_compound15.24777.24777.4 | controllerType=0 controllerNumber=1 scan=24777 | 49.6817 |
| 19288:3 | 7 | K.QIGLYPVLVIDSSGYVNPN[+1913.677]YTGR.I | HexNAc(4)Hex(5)NeuAc(1) | N[+1914] | 1480.3378 | 3 | 4438.9989 | 4438.9805 | 0 | 4.2 | 168 | Specific | 871.73 | 747.38 | 747.38 | 18.13 | 71 | >sp\|P01833\|PIGR_HUMAN Polymeric immunoglobulin receptor OS=Homo sapiens OX=9606 GN=PIGR PE=1 SV=4 | 902 | 20201113_xd_milk_1ng_glyco_hcd203040_compound15.22744.22744.3 | controllerType=0 controllerNumber=1 scan=22744 | 45.5028 |
| 19272:3 | 7 | K.QIGLYPVLVIDSSGYVNPN[+1913.677]YTGR.I | HexNAc(4)Hex(5)NeuAc(1) | N[+1914] | 1480.3383 | 3 | 4439.0004 | 4438.9805 | 0 | 4.5 | 168 | Specific | 884.52 | 628.52 | 628.52 | 18.13 | 71 | >sp\|P01833\|PIGR_HUMAN Polymeric immunoglobulin receptor OS=Homo sapiens OX=9606 GN=PIGR PE=1 SV=4 | 902 | 20201113_xd_milk_1ng_glyco_hcd203040_compound15.22727.22727.3 | controllerType=0 controllerNumber=1 scan=22727 | 45.4535 |
| 20826:4 | 7 | K.[+42.011]AQYEGRLSLLEEPGN[+1622.582]GTFTVILNQLTSR.D | HexNAc(4)Hex(5) | .[42], N[+1623] | 1193.5505 | 4 | 4771.18 | 4771.2089 | 0 | -6.1 | 407 | Specific | 287.52 | 284.45 | 284.45 | 5.59 | 71 | >sp\|P01833\|PIGR_HUMAN Polymeric immunoglobulin receptor OS=Homo sapiens OX=9606 GN=PIGR PE=1 SV=4 | 902 | 20201113_xd_milk_1ng_glyco_hcd203040_compound15.24347.24347.2 | controllerType=0 controllerNumber=1 scan=24347 | 48.7515 |
| 20828:4 | 7 | K.[+42.011]AQYEGRLSLLEEPGN[+1622.582]GTFTVILNQLTSR.D | HexNAc(4)Hex(5) | .[42], N[+1623] | 1193.5505 | 4 | 4771.18 | 4771.2089 | 0 | -6.1 | 407 | Specific | 308.69 | 308.69 | 308.69 | 4.43 | 71 | >sp\|P01833\|PIGR_HUMAN Polymeric immunoglobulin receptor OS=Homo sapiens OX=9606 GN=PIGR PE=1 SV=4 | 902 | 20201113_xd_milk_1ng_glyco_hcd203040_compound15.24349.24349.2 | controllerType=0 controllerNumber=1 scan=24349 | 48.7555 |
| 23003:4 | 7 | R.LSLLEEPGN[+2350.830]GTFTVILNQLTSR.D | HexNAc(4)Hex(5)Fuc(1)NeuAc(2) | N[+2351] | 1189.0424 | 4 | 4753.1478 | 4753.1229 | 0 | 5.2 | 413 | Specific | 201.68 | 201.68 | 201.68 | 3.25 | 71 | >sp\|P01833\|PIGR_HUMAN Polymeric immunoglobulin receptor OS=Homo sapiens OX=9606 GN=PIGR PE=1 SV=4 | 902 | 20201113_xd_milk_1ng_glyco_hcd203040_compound15.26616.26616.4 | controllerType=0 controllerNumber=1 scan=26616 | 53.4116 |
| 19384:3 | 7 | R.LSLLEEPGN[+1913.677]GTFTVILNQLTSR.D | HexNAc(4)Hex(5)NeuAc(1) | N[+1914] | 1439.3318 | 3 | 4315.981 | 4315.9696 | 0 | 2.6 | 413 | Specific | 339.51 | 339.51 | 339.51 | 8.44 | 71 | >sp\|P01833\|PIGR_HUMAN Polymeric immunoglobulin receptor OS=Homo sapiens OX=9606 GN=PIGR PE=1 SV=4 | 902 | 20201113_xd_milk_1ng_glyco_hcd203040_compound15.22844.22844.3 | controllerType=0 controllerNumber=1 scan=22844 | 45.7034 |
| 23015:3 | 7 | R.[+42.011]LSLLEEPGN[+2336.851]GTFTVILNQLTSR.D | HexNAc(6)Hex(6)Fuc(1) | .[42], N[+2337] | 1594.3831 | 3 | 4781.1348 | 4781.1542 | 0 | -4.1 | 413 | Specific | 354.46 | 339.27 | 339.27 | 7.14 | 71 | >sp\|P01833\|PIGR_HUMAN Polymeric immunoglobulin receptor OS=Homo sapiens OX=9606 GN=PIGR PE=1 SV=4 | 902 | 20201113_xd_milk_1ng_glyco_hcd203040_compound15.26629.26629.3 | controllerType=0 controllerNumber=1 scan=26629 | 53.4394 |
| 23013:3 | 7 | R.[+42.011]LSLLEEPGN[+2336.851]GTFTVILNQLTSR.D | HexNAc(6)Hex(6)Fuc(1) | .[42], N[+2337] | 1594.3831 | 3 | 4781.1348 | 4781.1542 | 0 | -4.1 | 413 | Specific | 399.66 | 334.85 | 334.85 | 7.22 | 71 | >sp\|P01833\|PIGR_HUMAN Polymeric immunoglobulin receptor OS=Homo sapiens OX=9606 GN=PIGR PE=1 SV=4 | 902 | 20201113_xd_milk_1ng_glyco_hcd203040_compound15.26627.26627.3 | controllerType=0 controllerNumber=1 scan=26627 | 53.4345 |
| 20709:3 | 7 | R.LSLLEEPGN[+2570.925]GTFTVILNQLTSR.D | HexNAc(5)Hex(6)Fuc(2)NeuAc(1) | N[+2571] | 1658.4176 | 3 | 4973.2381 | 4973.2176 | 0 | 4.1 | 413 | Specific | 625.71 | 454.49 | 454.49 | 12.88 | 71 | >sp\|P01833\|PIGR_HUMAN Polymeric immunoglobulin receptor OS=Homo sapiens OX=9606 GN=PIGR PE=1 SV=4 | 902 | 20201113_xd_milk_1ng_glyco_hcd203040_compound15.24224.24224.3 | controllerType=0 controllerNumber=1 scan=24224 | 48.4861 |
| 19387:3 | 7 | R.LSLLEEPGN[+1913.677]GTFTVILNQLTSR.D | HexNAc(4)Hex(5)NeuAc(1) | N[+1914] | 1439.3318 | 3 | 4315.981 | 4315.9696 | 0 | 2.6 | 413 | Specific | 778.32 | 737.61 | 737.61 | 16.39 | 71 | >sp\|P01833\|PIGR_HUMAN Polymeric immunoglobulin receptor OS=Homo sapiens OX=9606 GN=PIGR PE=1 SV=4 | 902 | 20201113_xd_milk_1ng_glyco_hcd203040_compound15.22847.22847.3 | controllerType=0 controllerNumber=1 scan=22847 | 45.7078 |
| 20711:3 | 7 | R.LSLLEEPGN[+2570.925]GTFTVILNQLTSR.D | HexNAc(5)Hex(6)Fuc(2)NeuAc(1) | N[+2571] | 1658.4176 | 3 | 4973.2381 | 4973.2176 | 0 | 4.1 | 413 | Specific | 857.17 | 546.68 | 546.68 | 17.88 | 71 | >sp\|P01833\|PIGR_HUMAN Polymeric immunoglobulin receptor OS=Homo sapiens OX=9606 GN=PIGR PE=1 SV=4 | 902 | 20201113_xd_milk_1ng_glyco_hcd203040_compound15.24226.24226.3 | controllerType=0 controllerNumber=1 scan=24226 | 48.4898 |
| 20837:3 | 7 | R.LSLLEEPGN[+2205.793]GTFTVILNQLTSR.D | HexNAc(4)Hex(5)Fuc(2)NeuAc(1) | N[+2206] | 1536.7078 | 3 | 4608.1089 | 4608.0854 | 0 | 5.1 | 413 | Specific | 926.93 | 658.33 | 658.33 | 18.54 | 71 | >sp\|P01833\|PIGR_HUMAN Polymeric immunoglobulin receptor OS=Homo sapiens OX=9606 GN=PIGR PE=1 SV=4 | 902 | 20201113_xd_milk_1ng_glyco_hcd203040_compound15.24359.24359.3 | controllerType=0 controllerNumber=1 scan=24359 | 48.7796 |
| 22874:3 | 7 | R.LSLLEEPGN[+2350.830]GTFTVILNQLTSR.D | HexNAc(4)Hex(5)Fuc(1)NeuAc(2) | N[+2351] | 1585.0551 | 3 | 4753.1508 | 4753.1229 | 0 | 5.9 | 413 | Specific | 964.55 | 696.46 | 696.46 | 19.54 | 71 | >sp\|P01833\|PIGR_HUMAN Polymeric immunoglobulin receptor OS=Homo sapiens OX=9606 GN=PIGR PE=1 SV=4 | 902 | 20201113_xd_milk_1ng_glyco_hcd203040_compound15.26482.26482.3 | controllerType=0 controllerNumber=1 scan=26482 | 53.1379 |
| 22872:3 | 7 | R.LSLLEEPGN[+2350.830]GTFTVILNQLTSR.D | HexNAc(4)Hex(5)Fuc(1)NeuAc(2) | N[+2351] | 1585.0551 | 3 | 4753.1508 | 4753.1229 | 0 | 5.9 | 413 | Specific | 1016.63 | 750.42 | 750.42 | 20.54 | 71 | >sp\|P01833\|PIGR_HUMAN Polymeric immunoglobulin receptor OS=Homo sapiens OX=9606 GN=PIGR PE=1 SV=4 | 902 | 20201113_xd_milk_1ng_glyco_hcd203040_compound15.26480.26480.3 | controllerType=0 controllerNumber=1 scan=26480 | 53.132 |
| 20835:3 | 7 | R.LSLLEEPGN[+2205.793]GTFTVILNQLTSR.D | HexNAc(4)Hex(5)Fuc(2)NeuAc(1) | N[+2206] | 1536.7078 | 3 | 4608.1089 | 4608.0854 | 0 | 5.1 | 413 | Specific | 1055.98 | 841.5 | 841.5 | 21.54 | 71 | >sp\|P01833\|PIGR_HUMAN Polymeric immunoglobulin receptor OS=Homo sapiens OX=9606 GN=PIGR PE=1 SV=4 | 902 | 20201113_xd_milk_1ng_glyco_hcd203040_compound15.24357.24357.3 | controllerType=0 controllerNumber=1 scan=24357 | 48.7743 |
| 16381:3 | 7 | K.[+42.011]IIEGEPNLKVPGN[+2018.708]VTAVLGETLK.V | HexNAc(3)Hex(6)Fuc(1)NeuAc(1) | .[42], N[+2019] | 1484.7026 | 3 | 4452.0932 | 4452.0683 | 0 | 5.6 | 457 | Specific | 296.7 | 283.18 | 283.18 | 6.63 | 71 | >sp\|P01833\|PIGR_HUMAN Polymeric immunoglobulin receptor OS=Homo sapiens OX=9606 GN=PIGR PE=1 SV=4 | 902 | 20201113_xd_milk_1ng_glyco_hcd203040_compound15.19717.19717.3 | controllerType=0 controllerNumber=1 scan=19717 | 39.2993 |
| 16416:3 | 7 | K.[+42.011]IIEGEPNLKVPGN[+1872.651]VTAVLGETLK.V | HexNAc(3)Hex(6)NeuAc(1) | .[42], N[+1873] | 1436.0137 | 3 | 4306.0266 | 4306.0104 | 0 | 3.8 | 457 | Specific | 325.12 | 310.14 | 310.14 | 7.21 | 71 | >sp\|P01833\|PIGR_HUMAN Polymeric immunoglobulin receptor OS=Homo sapiens OX=9606 GN=PIGR PE=1 SV=4 | 902 | 20201113_xd_milk_1ng_glyco_hcd203040_compound15.19754.19754.3 | controllerType=0 controllerNumber=1 scan=19754 | 39.372 |
| 16418:3 | 7 | K.[+42.011]IIEGEPNLKVPGN[+1872.651]VTAVLGETLK.V | HexNAc(3)Hex(6)NeuAc(1) | .[42], N[+1873] | 1436.0137 | 3 | 4306.0266 | 4306.0104 | 0 | 3.8 | 457 | Specific | 382.24 | 337.59 | 337.59 | 7.81 | 71 | >sp\|P01833\|PIGR_HUMAN Polymeric immunoglobulin receptor OS=Homo sapiens OX=9606 GN=PIGR PE=1 SV=4 | 902 | 20201113_xd_milk_1ng_glyco_hcd203040_compound15.19756.19756.3 | controllerType=0 controllerNumber=1 scan=19756 | 39.3764 |
| 18306:3 | 7 | K.IIEGEPNLKVPGN[+2350.830]VTAVLGETLK.V | HexNAc(4)Hex(5)Fuc(1)NeuAc(2) | N[+2351] | 1581.4033 | 3 | 4742.1952 | 4742.1797 | 0 | 3.3 | 457 | Specific | 463.42 | 463.42 | 463.42 | 10.01 | 71 | >sp\|P01833\|PIGR_HUMAN Polymeric immunoglobulin receptor OS=Homo sapiens OX=9606 GN=PIGR PE=1 SV=4 | 902 | 20201113_xd_milk_1ng_glyco_hcd203040_compound15.21721.21721.4 | controllerType=0 controllerNumber=1 scan=21721 | 43.3289 |
| 16398:3 | 7 | K.IIEGEPNLKVPGN[+2059.735]VTAVLGETLK.V | HexNAc(4)Hex(5)Fuc(1)NeuAc(1) | N[+2060] | 1484.3701 | 3 | 4451.0956 | 4451.0843 | 0 | 2.5 | 457 | Specific | 539.56 | 454.19 | 454.19 | 10.7 | 71 | >sp\|P01833\|PIGR_HUMAN Polymeric immunoglobulin receptor OS=Homo sapiens OX=9606 GN=PIGR PE=1 SV=4 | 902 | 20201113_xd_milk_1ng_glyco_hcd203040_compound15.19735.19735.3 | controllerType=0 controllerNumber=1 scan=19735 | 39.3337 |
| 18309:3 | 7 | K.IIEGEPNLKVPGN[+2350.830]VTAVLGETLK.V | HexNAc(4)Hex(5)Fuc(1)NeuAc(2) | N[+2351] | 1581.4033 | 3 | 4742.1952 | 4742.1797 | 0 | 3.3 | 457 | Specific | 597.95 | 552.29 | 552.29 | 11.98 | 71 | >sp\|P01833\|PIGR_HUMAN Polymeric immunoglobulin receptor OS=Homo sapiens OX=9606 GN=PIGR PE=1 SV=4 | 902 | 20201113_xd_milk_1ng_glyco_hcd203040_compound15.21724.21724.4 | controllerType=0 controllerNumber=1 scan=21724 | 43.3337 |
| 16427:4 | 7 | K.IIEGEPNLKVPGN[+2205.793]VTAVLGETLK.V | HexNAc(4)Hex(5)Fuc(2)NeuAc(1) | N[+2206] | 1150.0463 | 4 | 4597.1634 | 4597.1422 | 0 | 4.6 | 457 | Specific | 647.97 | 647.97 | 647.97 | 12.65 | 71 | >sp\|P01833\|PIGR_HUMAN Polymeric immunoglobulin receptor OS=Homo sapiens OX=9606 GN=PIGR PE=1 SV=4 | 902 | 20201113_xd_milk_1ng_glyco_hcd203040_compound15.19766.19766.4 | controllerType=0 controllerNumber=1 scan=19766 | 39.3984 |
| 16425:4 | 7 | K.IIEGEPNLKVPGN[+2205.793]VTAVLGETLK.V | HexNAc(4)Hex(5)Fuc(2)NeuAc(1) | N[+2206] | 1150.0463 | 4 | 4597.1634 | 4597.1422 | 0 | 4.6 | 457 | Specific | 686.17 | 686.17 | 686.17 | 13.65 | 71 | >sp\|P01833\|PIGR_HUMAN Polymeric immunoglobulin receptor OS=Homo sapiens OX=9606 GN=PIGR PE=1 SV=4 | 902 | 20201113_xd_milk_1ng_glyco_hcd203040_compound15.19764.19764.4 | controllerType=0 controllerNumber=1 scan=19764 | 39.3907 |
| 16392:3 | 7 | K.IIEGEPNLKVPGN[+2204.772]VTAVLGETLK.V | HexNAc(4)Hex(5)NeuAc(2) | N[+2205] | 1532.7206 | 3 | 4596.1474 | 4596.1218 | 0 | 5.6 | 457 | Specific | 717.39 | 700.72 | 700.72 | 14.33 | 71 | >sp\|P01833\|PIGR_HUMAN Polymeric immunoglobulin receptor OS=Homo sapiens OX=9606 GN=PIGR PE=1 SV=4 | 902 | 20201113_xd_milk_1ng_glyco_hcd203040_compound15.19729.19729.3 | controllerType=0 controllerNumber=1 scan=19729 | 39.3226 |
| 16389:3 | 7 | K.IIEGEPNLKVPGN[+2204.772]VTAVLGETLK.V | HexNAc(4)Hex(5)NeuAc(2) | N[+2205] | 1532.7206 | 3 | 4596.1474 | 4596.1218 | 0 | 5.6 | 457 | Specific | 1010.29 | 987.07 | 987.07 | 20.33 | 71 | >sp\|P01833\|PIGR_HUMAN Polymeric immunoglobulin receptor OS=Homo sapiens OX=9606 GN=PIGR PE=1 SV=4 | 902 | 20201113_xd_milk_1ng_glyco_hcd203040_compound15.19726.19726.3 | controllerType=0 controllerNumber=1 scan=19726 | 39.3181 |
| 14818:4 | 7 | K.[+42.011]IIEGEPNLKVPGN[+1872.651]VTAVLGETLKVPC[+57.021]HFPC[+57.021] | HexNAc(3)Hex(6)NeuAc(1) | .[42], C[+57]*2, N[+1873] | 1333.632 | 4 | 5331.506 | 5331.4679 | 0 | 7.1 | 457 | Specific | 227.7 | 213.52 | 213.52 | 5.66 | 71 | >sp\|P01833\|PIGR_HUMAN Polymeric immunoglobulin receptor OS=Homo sapiens OX=9606 GN=PIGR PE=1 SV=4 | 902 | 20201113_xd_milk_1ng_glyco_hcd203040_compound15.18079.18079.4 | controllerType=0 controllerNumber=1 scan=18079 | 36.3072 |
| 14820:4 | 7 | K.[+42.011]IIEGEPNLKVPGN[+1872.651]VTAVLGETLKVPC[+57.021]HFPC[+57.021] | HexNAc(3)Hex(6)NeuAc(1) | .[42], C[+57]*2, N[+1873] | 1333.632 | 4 | 5331.506 | 5331.4679 | 0 | 7.1 | 457 | Specific | 270.84 | 202.54 | 202.54 | 3.95 | 71 | >sp\|P01833\|PIGR_HUMAN Polymeric immunoglobulin receptor OS=Homo sapiens OX=9606 GN=PIGR PE=1 SV=4 | 902 | 20201113_xd_milk_1ng_glyco_hcd203040_compound15.18081.18081.4 | controllerType=0 controllerNumber=1 scan=18081 | 36.311 |
| 17434:4 | 7 | K.IIEGEPNLKVPGN[+2350.830]VTAVLGETLKVPC[+57.021]HFPC[+57.021]K.F | HexNAc(4)Hex(5)Fuc(1)NeuAc(2) | C[+57]*2, N[+2351] | 1442.6721 | 4 | 5767.6667 | 5767.6372 | 0 | 5.1 | 457 | Specific | 584.23 | 420.36 | 420.36 | 11.11 | 71 | >sp\|P01833\|PIGR_HUMAN Polymeric immunoglobulin receptor OS=Homo sapiens OX=9606 GN=PIGR PE=1 SV=4 | 902 | 20201113_xd_milk_1ng_glyco_hcd203040_compound15.20814.20814.4 | controllerType=0 controllerNumber=1 scan=20814 | 41.4297 |
| 15958:4 | 7 | K.IIEGEPNLKVPGN[+2205.793]VTAVLGETLKVPC[+57.021]HFPC[+57.021]K.F | HexNAc(4)Hex(5)Fuc(2)NeuAc(1) | C[+57]*2, N[+2206] | 1406.4077 | 4 | 5622.609 | 5622.5997 | 0 | 1.6 | 457 | Specific | 607.12 | 607.12 | 607.12 | 12.96 | 71 | >sp\|P01833\|PIGR_HUMAN Polymeric immunoglobulin receptor OS=Homo sapiens OX=9606 GN=PIGR PE=1 SV=4 | 902 | 20201113_xd_milk_1ng_glyco_hcd203040_compound15.19263.19263.4 | controllerType=0 controllerNumber=1 scan=19263 | 38.4502 |
| 16052:4 | 7 | K.IIEGEPNLKVPGN[+2059.735]VTAVLGETLKVPC[+57.021]HFPC[+57.021]K.F | HexNAc(4)Hex(5)Fuc(1)NeuAc(1) | C[+57]*2, N[+2060] | 1369.89 | 4 | 5476.5382 | 5476.5418 | 0 | -0.7 | 457 | Specific | 647.31 | 517.49 | 517.49 | 12.03 | 71 | >sp\|P01833\|PIGR_HUMAN Polymeric immunoglobulin receptor OS=Homo sapiens OX=9606 GN=PIGR PE=1 SV=4 | 902 | 20201113_xd_milk_1ng_glyco_hcd203040_compound15.19361.19361.2 | controllerType=0 controllerNumber=1 scan=19361 | 38.6257 |
| 17431:4 | 7 | K.IIEGEPNLKVPGN[+2350.830]VTAVLGETLKVPC[+57.021]HFPC[+57.021]K.F | HexNAc(4)Hex(5)Fuc(1)NeuAc(2) | C[+57]*2, N[+2351] | 1442.6721 | 4 | 5767.6667 | 5767.6372 | 0 | 5.1 | 457 | Specific | 656.92 | 510.33 | 510.33 | 13.11 | 71 | >sp\|P01833\|PIGR_HUMAN Polymeric immunoglobulin receptor OS=Homo sapiens OX=9606 GN=PIGR PE=1 SV=4 | 902 | 20201113_xd_milk_1ng_glyco_hcd203040_compound15.20811.20811.4 | controllerType=0 controllerNumber=1 scan=20811 | 41.4242 |
| 15994:4 | 7 | K.IIEGEPNLKVPGN[+2205.793]VTAVLGETLKVPC[+57.021]HFPC[+57.021]K.F | HexNAc(4)Hex(5)Fuc(2)NeuAc(1) | C[+57]*2, N[+2206] | 1406.4119 | 4 | 5622.6256 | 5622.5997 | 0 | 4.6 | 457 | Specific | 711.38 | 685.27 | 685.27 | 14.45 | 71 | >sp\|P01833\|PIGR_HUMAN Polymeric immunoglobulin receptor OS=Homo sapiens OX=9606 GN=PIGR PE=1 SV=4 | 902 | 20201113_xd_milk_1ng_glyco_hcd203040_compound15.19301.19301.2 | controllerType=0 controllerNumber=1 scan=19301 | 38.5158 |
| 16050:4 | 7 | K.IIEGEPNLKVPGN[+2059.735]VTAVLGETLKVPC[+57.021]HFPC[+57.021]K.F | HexNAc(4)Hex(5)Fuc(1)NeuAc(1) | C[+57]*2, N[+2060] | 1369.89 | 4 | 5476.5382 | 5476.5418 | 0 | -0.7 | 457 | Specific | 717.98 | 694.78 | 694.78 | 14.03 | 71 | >sp\|P01833\|PIGR_HUMAN Polymeric immunoglobulin receptor OS=Homo sapiens OX=9606 GN=PIGR PE=1 SV=4 | 902 | 20201113_xd_milk_1ng_glyco_hcd203040_compound15.19359.19359.2 | controllerType=0 controllerNumber=1 scan=19359 | 38.6211 |
| 15987:4 | 7 | K.IIEGEPNLKVPGN[+2059.735]VTAVLGETLKVPC[+57.021]HFPC[+57.021]K.F | HexNAc(4)Hex(5)Fuc(1)NeuAc(1) | C[+57]*2, N[+2060] | 1369.8938 | 4 | 5476.5534 | 5476.5418 | 0 | 2.1 | 457 | Specific | 729.44 | 717.92 | 717.92 | 14.96 | 71 | >sp\|P01833\|PIGR_HUMAN Polymeric immunoglobulin receptor OS=Homo sapiens OX=9606 GN=PIGR PE=1 SV=4 | 902 | 20201113_xd_milk_1ng_glyco_hcd203040_compound15.19294.19294.2 | controllerType=0 controllerNumber=1 scan=19294 | 38.5048 |
| 15991:4 | 7 | K.IIEGEPNLKVPGN[+2205.793]VTAVLGETLKVPC[+57.021]HFPC[+57.021]K.F | HexNAc(4)Hex(5)Fuc(2)NeuAc(1) | C[+57]*2, N[+2206] | 1406.4119 | 4 | 5622.6256 | 5622.5997 | 0 | 4.6 | 457 | Specific | 750.48 | 750.48 | 750.48 | 15.45 | 71 | >sp\|P01833\|PIGR_HUMAN Polymeric immunoglobulin receptor OS=Homo sapiens OX=9606 GN=PIGR PE=1 SV=4 | 902 | 20201113_xd_milk_1ng_glyco_hcd203040_compound15.19298.19298.2 | controllerType=0 controllerNumber=1 scan=19298 | 38.5109 |
| 15990:4 | 7 | K.IIEGEPNLKVPGN[+2059.735]VTAVLGETLKVPC[+57.021]HFPC[+57.021]K.F | HexNAc(4)Hex(5)Fuc(1)NeuAc(1) | C[+57]*2, N[+2060] | 1369.8938 | 4 | 5476.5534 | 5476.5418 | 0 | 2.1 | 457 | Specific | 837.8 | 772.2 | 772.2 | 16.96 | 71 | >sp\|P01833\|PIGR_HUMAN Polymeric immunoglobulin receptor OS=Homo sapiens OX=9606 GN=PIGR PE=1 SV=4 | 902 | 20201113_xd_milk_1ng_glyco_hcd203040_compound15.19297.19297.2 | controllerType=0 controllerNumber=1 scan=19297 | 38.509 |
| 15954:5 | 7 | K.IIEGEPNLKVPGN[+2205.793]VTAVLGETLKVPC[+57.021]HFPC[+57.021]K.F | HexNAc(4)Hex(5)Fuc(2)NeuAc(1) | C[+57]*2, N[+2206] | 1125.3276 | 5 | 5622.6089 | 5622.5997 | 0 | 1.6 | 457 | Specific | 859.34 | 819.35 | 819.35 | 18.51 | 71 | >sp\|P01833\|PIGR_HUMAN Polymeric immunoglobulin receptor OS=Homo sapiens OX=9606 GN=PIGR PE=1 SV=4 | 902 | 20201113_xd_milk_1ng_glyco_hcd203040_compound15.19259.19259.5 | controllerType=0 controllerNumber=1 scan=19259 | 38.4413 |
| 15956:4 | 7 | K.IIEGEPNLKVPGN[+2205.793]VTAVLGETLKVPC[+57.021]HFPC[+57.021]K.F | HexNAc(4)Hex(5)Fuc(2)NeuAc(1) | C[+57]*2, N[+2206] | 1406.4077 | 4 | 5622.609 | 5622.5997 | 0 | 1.6 | 457 | Specific | 1158.56 | 1044.54 | 1044.54 | 23.96 | 71 | >sp\|P01833\|PIGR_HUMAN Polymeric immunoglobulin receptor OS=Homo sapiens OX=9606 GN=PIGR PE=1 SV=4 | 902 | 20201113_xd_milk_1ng_glyco_hcd203040_compound15.19261.19261.4 | controllerType=0 controllerNumber=1 scan=19261 | 38.4445 |
| 15952:5 | 7 | K.IIEGEPNLKVPGN[+2205.793]VTAVLGETLKVPC[+57.021]HFPC[+57.021]K.F | HexNAc(4)Hex(5)Fuc(2)NeuAc(1) | C[+57]*2, N[+2206] | 1125.3276 | 5 | 5622.6089 | 5622.5997 | 0 | 1.6 | 457 | Specific | 1168.36 | 1099.31 | 1099.31 | 24.51 | 71 | >sp\|P01833\|PIGR_HUMAN Polymeric immunoglobulin receptor OS=Homo sapiens OX=9606 GN=PIGR PE=1 SV=4 | 902 | 20201113_xd_milk_1ng_glyco_hcd203040_compound15.19257.19257.5 | controllerType=0 controllerNumber=1 scan=19257 | 38.4354 |
| 14060:3 | 7 | K.VPGN[+2059.735]VTAVLGETLK.V | HexNAc(4)Hex(5)Fuc(1)NeuAc(1) | N[+2060] | 1153.1833 | 3 | 3457.5353 | 3457.5348 | 0 | 0.1 | 466 | Specific | 314.02 | 92.06 | 92.06 | 3.82 | 71 | >sp\|P01833\|PIGR_HUMAN Polymeric immunoglobulin receptor OS=Homo sapiens OX=9606 GN=PIGR PE=1 SV=4 | 902 | 20201113_xd_milk_1ng_glyco_hcd203040_compound15.17292.17292.3 | controllerType=0 controllerNumber=1 scan=17292 | 34.829 |
| 13974:3 | 7 | K.VPGN[+2205.793]VTAVLGETLK.V | HexNAc(4)Hex(5)Fuc(2)NeuAc(1) | N[+2206] | 1201.8731 | 3 | 3603.6048 | 3603.5927 | 0 | 3.3 | 466 | Specific | 389.21 | 338.19 | 338.19 | 8.11 | 71 | >sp\|P01833\|PIGR_HUMAN Polymeric immunoglobulin receptor OS=Homo sapiens OX=9606 GN=PIGR PE=1 SV=4 | 902 | 20201113_xd_milk_1ng_glyco_hcd203040_compound15.17203.17203.3 | controllerType=0 controllerNumber=1 scan=17203 | 34.6277 |
| 14057:3 | 7 | K.VPGN[+2059.735]VTAVLGETLK.V | HexNAc(4)Hex(5)Fuc(1)NeuAc(1) | N[+2060] | 1153.1833 | 3 | 3457.5353 | 3457.5348 | 0 | 0.1 | 466 | Specific | 412.75 | 261.14 | 261.14 | 7.31 | 71 | >sp\|P01833\|PIGR_HUMAN Polymeric immunoglobulin receptor OS=Homo sapiens OX=9606 GN=PIGR PE=1 SV=4 | 902 | 20201113_xd_milk_1ng_glyco_hcd203040_compound15.17289.17289.3 | controllerType=0 controllerNumber=1 scan=17289 | 34.8201 |
| 13976:3 | 7 | K.VPGN[+2205.793]VTAVLGETLK.V | HexNAc(4)Hex(5)Fuc(2)NeuAc(1) | N[+2206] | 1201.8731 | 3 | 3603.6048 | 3603.5927 | 0 | 3.3 | 466 | Specific | 596.88 | 596.88 | 596.88 | 11.97 | 71 | >sp\|P01833\|PIGR_HUMAN Polymeric immunoglobulin receptor OS=Homo sapiens OX=9606 GN=PIGR PE=1 SV=4 | 902 | 20201113_xd_milk_1ng_glyco_hcd203040_compound15.17205.17205.3 | controllerType=0 controllerNumber=1 scan=17205 | 34.6324 |
| 16955:3 | 7 | K.VPGN[+2350.830]VTAVLGETLK.V | HexNAc(4)Hex(5)Fuc(1)NeuAc(2) | N[+2351] | 1250.2226 | 3 | 3748.6532 | 3748.6302 | 0 | 6.1 | 466 | Specific | 615.87 | 435.33 | 435.33 | 12.17 | 71 | >sp\|P01833\|PIGR_HUMAN Polymeric immunoglobulin receptor OS=Homo sapiens OX=9606 GN=PIGR PE=1 SV=4 | 902 | 20201113_xd_milk_1ng_glyco_hcd203040_compound15.20316.20316.3 | controllerType=0 controllerNumber=1 scan=20316 | 40.4168 |
| 16957:3 | 7 | K.VPGN[+2350.830]VTAVLGETLK.V | HexNAc(4)Hex(5)Fuc(1)NeuAc(2) | N[+2351] | 1250.2226 | 3 | 3748.6532 | 3748.6302 | 0 | 6.1 | 466 | Specific | 635.59 | 469.79 | 469.79 | 12.17 | 71 | >sp\|P01833\|PIGR_HUMAN Polymeric immunoglobulin receptor OS=Homo sapiens OX=9606 GN=PIGR PE=1 SV=4 | 902 | 20201113_xd_milk_1ng_glyco_hcd203040_compound15.20318.20318.3 | controllerType=0 controllerNumber=1 scan=20318 | 40.4[illegible]28 |
| 12598:4 | 7 | K.VPGN[+2059.735]VTAVLGETLKVPC[+57.021]HFPC[+57.021]K.F | HexNAc(4)Hex(5)Fuc(1)NeuAc(1) | C[+57]*2, N[+2060] | 1121.5097 | 4 | 4483.0169 | 4482.9923 | 0 | 5.5 | 466 | Specific | 278.86 | 278.86 | 278.86 | 6.23 | 71 | >sp\|P01833\|PIGR_HUMAN Polymeric immunoglobulin receptor OS=Homo sapiens OX=9606 GN=PIGR PE=1 SV=4 | 902 | 20201113_xd_milk_1ng_glyco_hcd203040_compound15.15763.15763.3 | controllerType=0 controllerNumber=1 scan=15763 | 3[illegible]08 |
| 12468:4 | 7 | K.VPGN[+2059.735]VTAVLGETLKVPC[+57.021]HFPC[+57.021]K.F | HexNAc(4)Hex(5)Fuc(1)NeuAc(1) | C[+57]*2, N[+2060] | 1121.5113 | 4 | 4483.0233 | 4482.9923 | 0 | 6.9 | 466 | Specific | 579.09 | 537.66 | 537.66 | 10.94 | 71 | >sp\|P01833\|PIGR_HUMAN Polymeric immunoglobulin receptor OS=Homo sapiens OX=9606 GN=PIGR PE=1 SV=4 | 902 | 20201113_xd_milk_1ng_glyco_hcd203040_compound15.15629.15629.2 | controllerType=0 controllerNumber=1 scan=15629 | 31.[illegible]46 |
| 12578:4 | 7 | K.VPGN[+1913.677]VTAVLGETLKVPC[+57.021]HFPC[+57.021]K.F | HexNAc(4)Hex(5)NeuAc(1) | C[+57]*2, N[+1914] | 1084.995 | 4 | 4336.9583 | 4336.9344 | 0 | 5.5 | 466 | Specific | 589.02 | 589 | 589 | 11.08 | 71 | >sp\|P01833\|PIGR_HUMAN Polymeric immunoglobulin receptor OS=Homo sapiens OX=9606 GN=PIGR PE=1 SV=4 | 902 | 20201113_xd_milk_1ng_glyco_hcd203040_compound15.15743.15743.2 | controllerType=0 controllerNumber=1 scan=15743 | 3[illegible]92 |
| 12470:4 | 7 | K.VPGN[+2059.735]VTAVLGETLKVPC[+57.021]HFPC[+57.021]K.F | HexNAc(4)Hex(5)Fuc(1)NeuAc(1) | C[+57]*2, N[+2060] | 1121.5113 | 4 | 4483.0233 | 4482.9923 | 0 | 6.9 | 466 | Specific | 624.88 | 531.74 | 531.74 | 11.94 | 71 | >sp\|P01833\|PIGR_HUMAN Polymeric immunoglobulin receptor OS=Homo sapiens OX=9606 GN=PIGR PE=1 SV=4 | 902 | 20201113_xd_milk_1ng_glyco_hcd203040_compound15.15631.15631.2 | controllerType=0 controllerNumber=1 scan=15631 | 31.8784 |
| 12581:4 | 7 | K.VPGN[+1913.677]VTAVLGETLKVPC[+57.021]HFPC[+57.021]K.F | HexNAc(4)Hex(5)NeuAc(1) | C[+57]*2, N[+1914] | 1084.995 | 4 | 4336.9583 | 4336.9344 | 0 | 5.5 | 466 | Specific | 651.07 | 497.11 | 497.11 | 13.08 | 71 | >sp\|P01833\|PIGR_HUMAN Polymeric immunoglobulin receptor OS=Homo sapiens OX=9606 GN=PIGR PE=1 SV=4 | 902 | 20201113_xd_milk_1ng_glyco_hcd203040_compound15.15746.15746.2 | controllerType=0 controllerNumber=1 scan=15746 | 32.0845 |
| 14096:4 | 7 | K.VPGN[+2059.735]VTAVLGETLKVPC[+57.021]HFPC[+57.021]K.F | HexNAc(4)Hex(5)Fuc(1)NeuAc(1) | C[+57]*2, N[+2060] | 1121.5066 | 4 | 4483.0047 | 4482.9923 | 0 | 2.8 | 466 | Specific | 708.51 | 334.14 | 334.14 | 13.06 | 71 | >sp\|P01833\|PIGR_HUMAN Polymeric immunoglobulin receptor OS=Homo sapiens OX=9606 GN=PIGR PE=1 SV=4 | 902 | 20201113_xd_milk_1ng_glyco_hcd203040_compound15.17330.17330.4 | controllerType=0 controllerNumber=1 scan=17330 | 34.9096 |
| 14094:4 | 7 | K.VPGN[+2059.735]VTAVLGETLKVPC[+57.021]HFPC[+57.021]K.F | HexNAc(4)Hex(5)Fuc(1)NeuAc(1) | C[+57]*2, N[+2060] | 1121.5066 | 4 | 4483.0047 | 4482.9923 | 0 | 2.8 | 466 | Specific | 926.02 | 785.73 | 785.73 | 18.74 | 71 | >sp\|P01833\|PIGR_HUMAN Polymeric immunoglobulin receptor OS=Homo sapiens OX=9606 GN=PIGR PE=1 SV=4 | 902 | 20201113_xd_milk_1ng_glyco_hcd203040_compound15.17328.17328.4 | controllerType=0 controllerNumber=1 scan=17328 | 34.9012 |
| 04272:3 | 7 | K.WN[+1768.640]NTGC[+57.021]QALPSQDEGPSK.A | HexNAc(4)Hex(5)Fuc(1) | C[+57], N[+1769] | 1253.1804 | 3 | 3757.5265 | 3757.5162 | 0 | 2.7 | 498 | Specific | 237.02 | 237.02 | 237.02 | 6.81 | 71 | >sp\|P01833\|PIGR_HUMAN Polymeric immunoglobulin receptor OS=Homo sapiens OX=9606 GN=PIGR PE=1 SV=4 | 902 | 20201113_xd_milk_1ng_glyco_hcd203040_compound15.7134.7134.3 | controllerType=0 controllerNumber=1 scan=7134 | 17.983 |
| 08091:3 | 7 | K.WN[+2204.772]NTGC[+57.021]QALPSQDEGPSK.A | HexNAc(4)Hex(5)NeuAc(2) | C[+57], N[+2205] | 1398.5534 | 3 | 4193.6457 | 4193.6491 | 0 | -0.8 | 498 | Specific | 266.18 | 217.71 | 217.71 | 6.02 | 71 | >sp\|P01833\|PIGR_HUMAN Polymeric immunoglobulin receptor OS=Homo sapiens OX=9606 GN=PIGR PE=1 SV=4 | 902 | 20201113_xd_milk_1ng_glyco_hcd203040_compound15.11087.11087.3 | controllerType=0 controllerNumber=1 scan=11087 | 24.3303 |
| 04274:3 | 7 | K.WN[+1768.640]NTGC[+57.021]QALPSQDEGPSK.A | HexNAc(4)Hex(5)Fuc(1) | C[+57], N[+1769] | 1253.1804 | 3 | 3757.5265 | 3757.5162 | 0 | 2.7 | 498 | Specific | 301.4 | 287.9 | 287.9 | 7.26 | 71 | >sp\|P01833\|PIGR_HUMAN Polymeric immunoglobulin receptor OS=Homo sapiens OX=9606 GN=PIGR PE=1 SV=4 | 902 | 20201113_xd_milk_1ng_glyco_hcd203040_compound15.7136.7136.3 | controllerType=0 controllerNumber=1 scan=7136 | 17.9869 |
| 08093:3 | 7 | K.WN[+2204.772]NTGC[+57.021]QALPSQDEGPSK.A | HexNAc(4)Hex(5)NeuAc(2) | C[+57], N[+2205] | 1398.5534 | 3 | 4193.6457 | 4193.6491 | 0 | -0.8 | 498 | Specific | 347.3 | 281.7 | 281.7 | 6.82 | 71 | >sp\|P01833\|PIGR_HUMAN Polymeric immunoglobulin receptor OS=Homo sapiens OX=9606 GN=PIGR PE=1 SV=4 | 902 | 20201113_xd_milk_1ng_glyco_hcd203040_compound15.11089.11089.3 | controllerType=0 controllerNumber=1 scan=11089 | 24.3343 |
| 06203:3 | 7 | K.WN[+1548.545]NTGC[+57.021]QALPSQDEGPSK.A | HexNAc(3)Hex(4)NeuAc(1) | C[+57], N[+1549] | 1179.8138 | 3 | 3537.4268 | 3537.4215 | 0 | 1.5 | 498 | Specific | 348.38 | 329.17 | 329.17 | 7.81 | 71 | >sp\|P01833\|PIGR_HUMAN Polymeric immunoglobulin receptor OS=Homo sapiens OX=9606 GN=PIGR PE=1 SV=4 | 902 | 20201113_xd_milk_1ng_glyco_hcd203040_compound15.9130.9130.3 | controllerType=0 controllerNumber=1 scan=9130 | 21.1751 |
| 06205:3 | 7 | K.WN[+1548.545]NTGC[+57.021]QALPSQDEGPSK.A | HexNAc(3)Hex(4)NeuAc(1) | C[+57], N[+1549] | 1179.8138 | 3 | 3537.4268 | 3537.4215 | 0 | 1.5 | 498 | Specific | 375.91 | 306.7 | 306.7 | 7.84 | 71 | >sp\|P01833\|PIGR_HUMAN Polymeric immunoglobulin receptor OS=Homo sapiens OX=9606 GN=PIGR PE=1 SV=4 | 902 | 20201113_xd_milk_1ng_glyco_hcd203040_compound15.9132.9132.3 | controllerType=0 controllerNumber=1 scan=9132 | 21.1805 |
| 06179:3 | 7 | K.WN[+1548.545]NTGC[+57.021]QALPSQDEGPSK.A | HexNAc(3)Hex(4)NeuAc(1) | C[+57], N[+1549] | 1179.8139 | 3 | 3537.4271 | 3537.4215 | 0 | 1.6 | 498 | Specific | 406.41 | 359.89 | 359.89 | 8.55 | 71 | >sp\|P01833\|PIGR_HUMAN Polymeric immunoglobulin receptor OS=Homo sapiens OX=9606 GN=PIGR PE=1 SV=4 | 902 | 20201113_xd_milk_1ng_glyco_hcd203040_compound15.9105.9105.3 | controllerType=0 controllerNumber=1 scan=9105 | 21.1356 |
| 06011:3 | 7 | K.WN[+1872.651]NTGC[+57.021]QALPSQDEGPSK.A | HexNAc(3)Hex(6)NeuAc(1) | C[+57], N[+1873] | 1287.8499 | 3 | 3861.5352 | 3861.5272 | 0 | 2.1 | 498 | Specific | 573.25 | 514.4 | 514.4 | 10.89 | 71 | >sp\|P01833\|PIGR_HUMAN Polymeric immunoglobulin receptor OS=Homo sapiens OX=9606 GN=PIGR PE=1 SV=4 | 902 | 20201113_xd_milk_1ng_glyco_hcd203040_compound15.8932.8932.3 | controllerType=0 controllerNumber=1 scan=8932 | 20.8561 |
| 06181:3 | 7 | K.WN[+1548.545]NTGC[+57.021]QALPSQDEGPSK.A | HexNAc(3)Hex(4)NeuAc(1) | C[+57], N[+1549] | 1179.8139 | 3 | 3537.4271 | 3537.4215 | 0 | 1.6 | 498 | Specific | 591.85 | 521.18 | 521.18 | 12.16 | 71 | >sp\|P01833\|PIGR_HUMAN Polymeric immunoglobulin receptor OS=Homo sapiens OX=9606 GN=PIGR PE=1 SV=4 | 902 | 20201113_xd_milk_1ng_glyco_hcd203040_compound15.9107.9107.3 | controllerType=0 controllerNumber=1 scan=9107 | 21.1391 |
| 05990:3 | 7 | K.WN[+1872.651]NTGC[+57.021]QALPSQDEGPSK.A | HexNAc(3)Hex(6)NeuAc(1) | C[+57], N[+1873] | 1287.8515 | 3 | 3861.5399 | 3861.5272 | 0 | 3.3 | 498 | Specific | 652.21 | 618.15 | 618.15 | 13.96 | 71 | >sp\|P01833\|PIGR_HUMAN Polymeric immunoglobulin receptor OS=Homo sapiens OX=9606 GN=PIGR PE=1 SV=4 | 902 | 20201113_xd_milk_1ng_glyco_hcd203040_compound15.8910.8910.3 | controllerType=0 controllerNumber=1 scan=8910 | 20.8171 |
| 06095:3 | 7 | K.WN[+1710.598]NTGC[+57.021]QALPSQDEGPSK.A | HexNAc(3)Hex(5)NeuAc(1) | C[+57], N[+1711] | 1233.8323 | 3 | 3699.4824 | 3699.4743 | 0 | 2.2 | 498 | Specific | 709.24 | 620.13 | 620.13 | 15.16 | 71 | >sp\|P01833\|PIGR_HUMAN Polymeric immunoglobulin receptor OS=Homo sapiens OX=9606 GN=PIGR PE=1 SV=4 | 902 | 20201113_xd_milk_1ng_glyco_hcd203040_compound15.9019.9019.3 | controllerType=0 controllerNumber=1 scan=9019 | 21.0038 |
| 05992:3 | 7 | K.WN[+1872.651]NTGC[+57.021]QALPSQDEGPSK.A | HexNAc(3)Hex(6)NeuAc(1) | C[+57], N[+1873] | 1287.8515 | 3 | 3861.5399 | 3861.5272 | 0 | 3.3 | 498 | Specific | 722.39 | 671.11 | 671.11 | 14.96 | 71 | >sp\|P01833\|PIGR_HUMAN Polymeric immunoglobulin receptor OS=Homo sapiens OX=9606 GN=PIGR PE=1 SV=4 | 902 | 20201113_xd_milk_1ng_glyco_hcd203040_compound15.8912.8912.3 | controllerType=0 controllerNumber=1 scan=8912 | 20.8209 |
| 06005:3 | 7 | K.WN[+1913.677]NTGC[+57.021]QALPSQDEGPSK.A | HexNAc(4)Hex(5)NeuAc(1) | C[+57], N[+1914] | 1301.5254 | 3 | 3902.5618 | 3902.5537 | 0 | 2.1 | 498 | Specific | 738.9 | 636.48 | 636.48 | 15.16 | 71 | >sp\|P01833\|PIGR_HUMAN Polymeric immunoglobulin receptor OS=Homo sapiens OX=9606 GN=PIGR PE=1 SV=4 | 902 | 20201113_xd_milk_1ng_glyco_hcd203040_compound15.8926.8926.3 | controllerType=0 controllerNumber=1 scan=8926 | 20.8453 |
| 06093:3 | 7 | K.WN[+1710.598]NTGC[+57.021]QALPSQDEGPSK.A | HexNAc(3)Hex(5)NeuAc(1) | C[+57], N[+1711] | 1233.8323 | 3 | 3699.4824 | 3699.4743 | 0 | 2.2 | 498 | Specific | 739.14 | 677.38 | 677.38 | 15.16 | 71 | >sp\|P01833\|PIGR_HUMAN Polymeric immunoglobulin receptor OS=Homo sapiens OX=9606 GN=PIGR PE=1 SV=4 | 902 | 20201113_xd_milk_1ng_glyco_hcd203040_compound15.9017.9017.3 | controllerType=0 controllerNumber=1 scan=9017 | 20.9991 |
| 06009:3 | 7 | K.WN[+1872.651]NTGC[+57.021]QALPSQDEGPSK.A | HexNAc(3)Hex(6)NeuAc(1) | C[+57], N[+1873] | 1287.8499 | 3 | 3861.5352 | 3861.5272 | 0 | 2.1 | 498 | Specific | 741.88 | 693.32 | 693.32 | 15.16 | 71 | >sp\|P01833\|PIGR_HUMAN Polymeric immunoglobulin receptor OS=Homo sapiens OX=9606 GN=PIGR PE=1 SV=4 | 902 | 20201113_xd_milk_1ng_glyco_hcd203040_compound15.8930.8930.3 | controllerType=0 controllerNumber=1 scan=8930 | 20.8523 |
| 05898:3 | 7 | K.WN[+2059.735]NTGC[+57.021]QALPSQDEGPSK.A | HexNAc(4)Hex(5)Fuc(1)NeuAc(1) | C[+57], N[+2060] | 1350.2109 | 3 | 4048.6181 | 4048.6116 | 0 | 1.6 | 498 | Specific | 783.84 | 700.42 | 700.42 | 16.16 | 71 | >sp\|P01833\|PIGR_HUMAN Polymeric immunoglobulin receptor OS=Homo sapiens OX=9606 GN=PIGR PE=1 SV=4 | 902 | 20201113_xd_milk_1ng_glyco_hcd203040_compound15.8815.8815.3 | controllerType=0 controllerNumber=1 scan=8815 | 20.6628 |
| 05896:3 | 7 | K.WN[+2059.735]NTGC[+57.021]QALPSQDEGPSK.A | HexNAc(4)Hex(5)Fuc(1)NeuAc(1) | C[+57], N[+2060] | 1350.2109 | 3 | 4048.6181 | 4048.6116 | 0 | 1.6 | 498 | Specific | 785.89 | 717.22 | 717.22 | 16.16 | 71 | >sp\|P01833\|PIGR_HUMAN Polymeric immunoglobulin receptor OS=Homo sapiens OX=9606 GN=PIGR PE=1 SV=4 | 902 | 20201113_xd_milk_1ng_glyco_hcd203040_compound15.8813.8813.3 | controllerType=0 controllerNumber=1 scan=8813 | 20.6585 |
| 06007:3 | 7 | K.WN[+1913.677]NTGC[+57.021]QALPSQDEGPSK.A | HexNAc(4)Hex(5)NeuAc(1) | C[+57], N[+1914] | 1301.5254 | 3 | 3902.5618 | 3902.5537 | 0 | 2.1 | 498 | Specific | 786.5 | 692.31 | 692.31 | 16.16 | 71 | >sp\|P01833\|PIGR_HUMAN Polymeric immunoglobulin receptor OS=Homo sapiens OX=9606 GN=PIGR PE=1 SV=4 | 902 | 20201113_xd_milk_1ng_glyco_hcd203040_compound15.8928.8928.3 | controllerType=0 controllerNumber=1 scan=8928 | 20.8492 |
| 13633:4 | 8 | K.KGN[+2059.735]TTLNSFVIPSESDVPTHC[+57.021]PSQWWPYAGHC[+57.021]YK.I | HexNAc(4)Hex(5)Fuc(1)NeuAc(1) | C[+57]*2, N[+2060] | 1506.6466 | 4 | 6023.5646 | 6023.5567 | 0 | 1.3 | 342 | Specific | 249.64 | 249.64 | 249.64 | 6.51 | 59 | >sp\|P22897\|MRC1_HUMAN Macrophage mannose receptor 1 OS=Homo sapiens OX=9606 GN=MRC1 PE=1 SV=1 | 1049 | 20201113_xd_milk_1ng_glyco_hcd203040_compound15.16848.16848.2 | controllerType=0 controllerNumber=1 scan=16848 | 33.9154 |
| 13636:4 | 8 | K.KGN[+2059.735]TTLNSFVIPSESDVPTHC[+57.021]PSQWWPYAGHC[+57.021]YK.I | HexNAc(4)Hex(5)Fuc(1)NeuAc(1) | C[+57]*2, N[+2060] | 1506.6466 | 4 | 6023.5646 | 6023.5567 | 0 | 1.3 | 342 | Specific | 370.42 | 370.42 | 370.42 | 7.71 | 59 | >sp\|P22897\|MRC1_HUMAN Macrophage mannose receptor 1 OS=Homo sapiens OX=9606 GN=MRC1 PE=1 SV=1 | 1049 | 20201113_xd_milk_1ng_glyco_hcd203040_compound15.16851.16851.2 | controllerType=0 controllerNumber=1 scan=16851 | 33.9197 |
| 17936:4 | 9 | R.LSLHRPALEDLLLGSEAN[+1825.661]LTC[+57.021]TLTGLR.D | HexNAc(5)Hex(5) | C[+57], N[+1826] | 1198.0722 | 4 | 4789.2669 | 4789.2592 | 0 | 1.6 | 127 | Specific | 216.08 | 216.08 | 216.08 | 4.06 | 55 | >sp\|P01876\|IGHA1_HUMAN Immunoglobulin heavy constant alpha 1 OS=Homo sapiens OX=9606 GN=IGHA1 PE=1 SV=2 | 255 | 20201113_xd_milk_1ng_glyco_hcd203040_compound15.21336.21336.3 | controllerType=0 controllerNumber=1 scan=21336 | 42.4607 |
| 17903:4 | 9 | R.LSLHRPALEDLLLGSEAN[+1864.634]LTC[+57.021]TLTGLR.D | HexNAc(2)Hex(9) | C[+57], N[+1865] | 1207.8154 | 4 | 4828.2396 | 4828.2324 | 0 | 1.5 | 127 | Specific | 340.37 | 338.53 | 338.53 | 8.42 | 55 | >sp\|P01876\|IGHA1_HUMAN Immunoglobulin heavy constant alpha 1 OS=Homo sapiens OX=9606 GN=IGHA1 PE=1 SV=2 | 255 | 20201113_xd_milk_1ng_glyco_hcd203040_compound15.21302.21302.4 | controllerType=0 controllerNumber=1 scan=21302 | 42.3965 |
| 19002:4 | 9 | R.LSLHRPALEDLLLGSEAN[+1872.651]LTC[+57.021]TLTGLR.D | HexNAc(3)Hex(6)NeuAc(1) | C[+57], N[+1873] | 1209.8222 | 4 | 4836.2669 | 4836.2487 | 0 | 3.8 | 127 | Specific | 384.52 | 384.52 | 384.52 | 10.16 | 55 | >sp\|P01876\|IGHA1_HUMAN Immunoglobulin heavy constant alpha 1 OS=Homo sapiens OX=9606 GN=IGHA1 PE=1 SV=2 | 255 | 20201113_xd_milk_1ng_glyco_hcd203040_compound15.22446.22446.4 | controllerType=0 controllerNumber=1 scan=22446 | 44.8249 |
| 19041:3 | 9 | R.LSLHRPALEDLLLGSEAN[+1710.598]LTC[+57.021]TLTGLR.D | HexNAc(3)Hex(5)NeuAc(1) | C[+57], N[+1711] | 1558.7398 | 3 | 4674.2049 | 4674.1959 | 0 | 1.9 | 127 | Specific | 416.31 | 416.31 | 416.31 | 9.87 | 55 | >sp\|P01876\|IGHA1_HUMAN Immunoglobulin heavy constant alpha 1 OS=Homo sapiens OX=9606 GN=IGHA1 PE=1 SV=2 | 255 | 20201113_xd_milk_1ng_glyco_hcd203040_compound15.22486.22486.3 | controllerType=0 controllerNumber=1 scan=22486 | 44.9054 |
| 18359:4 | 9 | R.LSLHRPALEDLLLGSEAN[+1501.555]LTC[+57.021]TLTGLR.D | HexNAc(5)Hex(3) | C[+57], N[+1502] | 1117.0414 | 4 | 4465.1439 | 4465.1536 | 0 | -2.2 | 127 | Specific | 430.38 | 430.38 | 430.38 | 9.06 | 55 | >sp\|P01876\|IGHA1_HUMAN Immunoglobulin heavy constant alpha 1 OS=Homo sapiens OX=9606 GN=IGHA1 PE=1 SV=2 | 255 | 20201113_xd_milk_1ng_glyco_hcd203040_compound15.21776.21776.4 | controllerType=0 controllerNumber=1 scan=21776 | 43.4284 |
| 17961:3 | 9 | R.LSLHRPALEDLLLGSEAN[+1702.581]LTC[+57.021]TLTGLR.D | HexNAc(2)Hex(8) | C[+57], N[+1703] | 1556.0666 | 3 | 4666.1852 | 4666.1796 | 0 | 1.2 | 127 | Specific | 453.69 | 439.38 | 439.38 | 9.83 | 55 | >sp\|P01876\|IGHA1_HUMAN Immunoglobulin heavy constant alpha 1 OS=Homo sapiens OX=9606 GN=IGHA1 PE=1 SV=2 | 255 | 20201113_xd_milk_1ng_glyco_hcd203040_compound15.21362.21362.3 | controllerType=0 controllerNumber=1 scan=21362 | 42.5106 |
| 19044:3 | 9 | R.LSLHRPALEDLLLGSEAN[+1710.598]LTC[+57.021]TLTGLR.D | HexNAc(3)Hex(5)NeuAc(1) | C[+57], N[+1711] | 1558.7398 | 3 | 4674.2049 | 4674.1959 | 0 | 1.9 | 127 | Specific | 463.74 | 412.47 | 412.47 | 10 | 55 | >sp\|P01876\|IGHA1_HUMAN Immunoglobulin heavy constant alpha 1 OS=Homo sapiens OX=9606 GN=IGHA1 PE=1 SV=2 | 255 | 20201113_xd_milk_1ng_glyco_hcd203040_compound15.22489.22489.3 | controllerType=0 controllerNumber=1 scan=22489 | 44.9105 |
| 20114:4 | 9 | R.LSLHRPALEDLLLGSEAN[+2407.852]LTC[+57.021]TLTGLR.D | HexNAc(5)Hex(5)NeuAc(2) | C[+57], N[+2408] | 1343.6211 | 4 | 5371.4626 | 5371.4501 | 0 | 2.3 | 127 | Specific | 526.01 | 352.62 | 352.62 | 9.24 | 55 | >sp\|P01876\|IGHA1_HUMAN Immunoglobulin heavy constant alpha 1 OS=Homo sapiens OX=9606 GN=IGHA1 PE=1 SV=2 | 255 | 20201113_xd_milk_1ng_glyco_hcd203040_compound15.23604.23604.2 | controllerType=0 controllerNumber=1 scan=23604 | 47.1659 |
| 18042:5 | 9 | R.LSLHRPALEDLLLGSEAN[+1501.555]LTC[+57.021]TLTGLR.D | HexNAc(5)Hex(3) | C[+57], N[+1502] | 893.8438 | 5 | 4465.1898 | 4465.1536 | 0 | 8.1 | 127 | Specific | 540.75 | 540.75 | 540.75 | 9.9 | 55 | >sp\|P01876\|IGHA1_HUMAN Immunoglobulin heavy constant alpha 1 OS=Homo sapiens OX=9606 GN=IGHA1 PE=1 SV=2 | 255 | 20201113_xd_milk_1ng_glyco_hcd203040_compound15.21446.21446.5 | controllerType=0 controllerNumber=1 scan=21446 | 42.6971 |
| 18111:3 | 9 | R.LSLHRPALEDLLLGSEAN[+1095.397]LTC[+57.021]TLTGLR.D | HexNAc(3)Hex(3) | C[+57], N[+1095] | 1353.6746 | 3 | 4059.0093 | 4058.9949 | 0 | 3.6 | 127 | Specific | 543.36 | 540.59 | 540.59 | 10.31 | 55 | >sp\|P01876\|IGHA1_HUMAN Immunoglobulin heavy constant alpha 1 OS=Homo sapiens OX=9606 GN=IGHA1 PE=1 SV=2 | 255 | 20201113_xd_milk_1ng_glyco_hcd203040_compound15.21518.21518.3 | controllerType=0 controllerNumber=1 scan=21518 | 42.8866 |
| 18044:5 | 9 | R.LSLHRPALEDLLLGSEAN[+1501.555]LTC[+57.021]TLTGLR.D | HexNAc(5)Hex(3) | C[+57], N[+1502] | 893.8438 | 5 | 4465.1898 | 4465.1536 | 0 | 8.1 | 127 | Specific | 551.16 | 551.16 | 551.16 | 9.9 | 55 | >sp\|P01876\|IGHA1_HUMAN Immunoglobulin heavy constant alpha 1 OS=Homo sapiens OX=9606 GN=IGHA1 PE=1 SV=2 | 255 | 20201113_xd_milk_1ng_glyco_hcd203040_compound15.21448.21448.5 | controllerType=0 controllerNumber=1 scan=21448 | 42.7051 |
| 17915:3 | 9 | R.LSLHRPALEDLLLGSEAN[+1864.634]LTC[+57.021]TLTGLR.D | HexNAc(2)Hex(9) | C[+57], N[+1865] | 1610.0851 | 3 | 4828.2409 | 4828.2324 | 0 | 1.7 | 127 | Specific | 552.32 | 552.32 | 552.32 | 10.5 | 55 | >sp\|P01876\|IGHA1_HUMAN Immunoglobulin heavy constant alpha 1 OS=Homo sapiens OX=9606 GN=IGHA1 PE=1 SV=2 | 255 | 20201113_xd_milk_1ng_glyco_hcd203040_compound15.21314.21314.3 | controllerType=0 controllerNumber=1 scan=21314 | 42.4158 |
| 19040:3 | 9 | R.LSLHRPALEDLLLGSEAN[+1751.624]LTC[+57.021]TLTGLR.D | HexNAc(4)Hex(4)NeuAc(1) | C[+57], N[+1752] | 1572.4145 | 3 | 4715.2289 | 4715.2225 | 0 | 1.4 | 127 | Specific | 554.78 | 552.31 | 552.31 | 10.33 | 55 | >sp\|P01876\|IGHA1_HUMAN Immunoglobulin heavy constant alpha 1 OS=Homo sapiens OX=9606 GN=IGHA1 PE=1 SV=2 | 255 | 20201113_xd_milk_1ng_glyco_hcd203040_compound15.22485.22485.3 | controllerType=0 controllerNumber=1 scan=22485 | 44.9035 |
| 18041:3 | 9 | R.LSLHRPALEDLLLGSEAN[+1622.582]LTC[+57.021]TLTGLR.D | HexNAc(4)Hex(5) | C[+57], N[+1623] | 1529.4086 | 3 | 4586.2113 | 4586.1799 | 0 | 6.9 | 127 | Specific | 557.89 | 512.54 | 512.54 | 9.71 | 55 | >sp\|P01876\|IGHA1_HUMAN Immunoglobulin heavy constant alpha 1 OS=Homo sapiens OX=9606 GN=IGHA1 PE=1 SV=2 | 255 | 20201113_xd_milk_1ng_glyco_hcd203040_compound15.21445.21445.3 | controllerType=0 controllerNumber=1 scan=21445 | 42.6953 |
| 17918:3 | 9 | R.LSLHRPALEDLLLGSEAN[+1864.634]LTC[+57.021]TLTGLR.D | HexNAc(2)Hex(9) | C[+57], N[+1865] | 1610.0851 | 3 | 4828.2409 | 4828.2324 | 0 | 1.7 | 127 | Specific | 568.96 | 514.66 | 514.66 | 10.71 | 55 | >sp\|P01876\|IGHA1_HUMAN Immunoglobulin heavy constant alpha 1 OS=Homo sapiens OX=9606 GN=IGHA1 PE=1 SV=2 | 255 | 20201113_xd_milk_1ng_glyco_hcd203040_compound15.21317.21317.3 | controllerType=0 controllerNumber=1 scan=21317 | 42.4224 |
| 18039:3 | 9 | R.LSLHRPALEDLLLGSEAN[+1622.582]LTC[+57.021]TLTGLR.D | HexNAc(4)Hex(5) | C[+57], N[+1623] | 1529.4086 | 3 | 4586.2113 | 4586.1799 | 0 | 6.9 | 127 | Specific | 583.88 | 583.88 | 583.88 | 10.98 | 55 | >sp\|P01876\|IGHA1_HUMAN Immunoglobulin heavy constant alpha 1 OS=Homo sapiens OX=9606 GN=IGHA1 PE=1 SV=2 | 255 | 20201113_xd_milk_1ng_glyco_hcd203040_compound15.21443.21443.3 | controllerType=0 controllerNumber=1 scan=21443 | 42.6834 |
| 19037:3 | 9 | R.LSLHRPALEDLLLGSEAN[+1751.624]LTC[+57.021]TLTGLR.D | HexNAc(4)Hex(4)NeuAc(1) | C[+57], N[+1752] | 1572.4145 | 3 | 4715.2289 | 4715.2225 | 0 | 1.4 | 127 | Specific | 588.51 | 588.51 | 588.51 | 11.81 | 55 | >sp\|P01876\|IGHA1_HUMAN Immunoglobulin heavy constant alpha 1 OS=Homo sapiens OX=9606 GN=IGHA1 PE=1 SV=2 | 255 | 20201113_xd_milk_1ng_glyco_hcd203040_compound15.22482.22482.3 | controllerType=0 controllerNumber=1 scan=22482 | 44.895 |
| 17992:3 | 9 | R.LSLHRPALEDLLLGSEAN[+1501.555]LTC[+57.021]TLTGLR.D | HexNAc(5)Hex(3) | C[+57], N[+1502] | 1489.0587 | 3 | 4465.1614 | 4465.1536 | 0 | 1.7 | 127 | Specific | 603.28 | 544.71 | 544.71 | 12.98 | 55 | >sp\|P01876\|IGHA1_HUMAN Immunoglobulin heavy constant alpha 1 OS=Homo sapiens OX=9606 GN=IGHA1 PE=1 SV=2 | 255 | 20201113_xd_milk_1ng_glyco_hcd203040_compound15.21394.21394.3 | controllerType=0 controllerNumber=1 scan=21394 | 42.5716 |
| 17964:3 | 9 | R.LSLHRPALEDLLLGSEAN[+1702.581]LTC[+57.021]TLTGLR.D | HexNAc(2)Hex(8) | C[+57], N[+1703] | 1556.0666 | 3 | 4666.1852 | 4666.1796 | 0 | 1.2 | 127 | Specific | 606.76 | 539.44 | 539.44 | 12.81 | 55 | >sp\|P01876\|IGHA1_HUMAN Immunoglobulin heavy constant alpha 1 OS=Homo sapiens OX=9606 GN=IGHA1 PE=1 SV=2 | 255 | 20201113_xd_milk_1ng_glyco_hcd203040_compound15.21365.21365.3 | controllerType=0 controllerNumber=1 scan=21365 | 42.5146 |
| 19035:4 | 9 | R.LSLHRPALEDLLLGSEAN[+1751.624]LTC[+57.021]TLTGLR.D | HexNAc(4)Hex(4)NeuAc(1) | C[+57], N[+1752] | 1179.5636 | 4 | 4715.2327 | 4715.2225 | 0 | 2.2 | 127 | Specific | 623.43 | 582.71 | 582.71 | 13.53 | 55 | >sp\|P01876\|IGHA1_HUMAN Immunoglobulin heavy constant alpha 1 OS=Homo sapiens OX=9606 GN=IGHA1 PE=1 SV=2 | 255 | 20201113_xd_milk_1ng_glyco_hcd203040_compound15.22480.22480.4 | controllerType=0 controllerNumber=1 scan=22480 | 44.8928 |
| 18095:4 | 9 | R.LSLHRPALEDLLLGSEAN[+1663.608]LTC[+57.021]TLTGLR.D | HexNAc(5)Hex(4) | C[+57], N[+1664] | 1157.5614 | 4 | 4627.2239 | 4627.2064 | 0 | 3.8 | 127 | Specific | 624.44 | 411.2 | 411.2 | 13.34 | 55 | >sp\|P01876\|IGHA1_HUMAN Immunoglobulin heavy constant alpha 1 OS=Homo sapiens OX=9606 GN=IGHA1 PE=1 SV=2 | 255 | 20201113_xd_milk_1ng_glyco_hcd203040_compound15.21501.21501.4 | controllerType=0 controllerNumber=1 scan=21501 | 42.8499 |
| 17954:3 | 9 | R.LSLHRPALEDLLLGSEAN[+1540.529]LTC[+57.021]TLTGLR.D | HexNAc(2)Hex(7) | C[+57], N[+1541] | 1502.0488 | 3 | 4504.1317 | 4504.1268 | 0 | 1.1 | 127 | Specific | 626.18 | 604.15 | 604.15 | 12.81 | 55 | >sp\|P01876\|IGHA1_HUMAN Immunoglobulin heavy constant alpha 1 OS=Homo sapiens OX=9606 GN=IGHA1 PE=1 SV=2 | 255 | 20201113_xd_milk_1ng_glyco_hcd203040_compound15.21354.21354.3 | controllerType=0 controllerNumber=1 scan=21354 | 42.4914 |
| 18113:3 | 9 | R.LSLHRPALEDLLLGSEAN[+1095.397]LTC[+57.021]TLTGLR.D | HexNAc(3)Hex(3) | C[+57], N[+1095] | 1353.6746 | 3 | 4059.0093 | 4058.9949 | 0 | 3.6 | 127 | Specific | 626.62 | 485.34 | 485.34 | 12.78 | 55 | >sp\|P01876\|IGHA1_HUMAN Immunoglobulin heavy constant alpha 1 OS=Homo sapiens OX=9606 GN=IGHA1 PE=1 SV=2 | 255 | 20201113_xd_milk_1ng_glyco_hcd203040_compound15.21520.21520.3 | controllerType=0 controllerNumber=1 scan=21520 | 42.8924 |
| 17995:3 | 9 | R.LSLHRPALEDLLLGSEAN[+1501.555]LTC[+57.021]TLTGLR.D | HexNAc(5)Hex(3) | C[+57], N[+1502] | 1489.0587 | 3 | 4465.1614 | 4465.1536 | 0 | 1.7 | 127 | Specific | 633.72 | 599.29 | 599.29 | 12.98 | 55 | >sp\|P01876\|IGHA1_HUMAN Immunoglobulin heavy constant alpha 1 OS=Homo sapiens OX=9606 GN=IGHA1 PE=1 SV=2 | 255 | 20201113_xd_milk_1ng_glyco_hcd203040_compound15.21397.21397.3 | controllerType=0 controllerNumber=1 scan=21397 | 42.5778 |
| 18361:4 | 9 | R.LSLHRPALEDLLLGSEAN[+1501.555]LTC[+57.021]TLTGLR.D | HexNAc(5)Hex(3) | C[+57], N[+1502] | 1117.0414 | 4 | 4465.1439 | 4465.1536 | 0 | -2.2 | 127 | Specific | 650.22 | 584.1 | 584.1 | 13.07 | 55 | >sp\|P01876\|IGHA1_HUMAN Immunoglobulin heavy constant alpha 1 OS=Homo sapiens OX=9606 GN=IGHA1 PE=1 SV=2 | 255 | 20201113_xd_milk_1ng_glyco_hcd203040_compound15.21778.21778.4 | controllerType=0 controllerNumber=1 scan=21778 | 43.4336 |
| 18067:3 | 9 | R.LSLHRPALEDLLLGSEAN[+1501.555]LTC[+57.021]TLTGLR.D | HexNAc(5)Hex(3) | C[+57], N[+1502] | 1489.0596 | 3 | 4465.1643 | 4465.1536 | 0 | 2.4 | 127 | Specific | 654 | 599.43 | 599.43 | 13.98 | 55 | >sp\|P01876\|IGHA1_HUMAN Immunoglobulin heavy constant alpha 1 OS=Homo sapiens OX=9606 GN=IGHA1 PE=1 SV=2 | 255 | 20201113_xd_milk_1ng_glyco_hcd203040_compound15.21472.21472.2 | controllerType=0 controllerNumber=1 scan=21472 | 42.7814 |
| 18094:3 | 9 | R.LSLHRPALEDLLLGSEAN[+1095.397]LTC[+57.021]TLTGLR.D | HexNAc(3)Hex(3) | C[+57], N[+1095] | 1353.676 | 3 | 4059.0133 | 4058.9949 | 0 | 4.6 | 127 | Specific | 657.98 | 656.7 | 656.7 | 13.47 | 55 | >sp\|P01876\|IGHA1_HUMAN Immunoglobulin heavy constant alpha 1 OS=Homo sapiens OX=9606 GN=IGHA1 PE=1 SV=2 | 255 | 20201113_xd_milk_1ng_glyco_hcd203040_compound15.21500.21500.3 | controllerType=0 controllerNumber=1 scan=21500 | 42.8478 |
| 18054:4 | 9 | R.LSLHRPALEDLLLGSEAN[+1501.555]LTC[+57.021]TLTGLR.D | HexNAc(5)Hex(3) | C[+57], N[+1502] | 1117.0467 | 4 | 4465.1649 | 4465.1536 | 0 | 2.5 | 127 | Specific | 666.66 | 666.66 | 666.66 | 14.53 | 55 | >sp\|P01876\|IGHA1_HUMAN Immunoglobulin heavy constant alpha 1 OS=Homo sapiens OX=9606 GN=IGHA1 PE=1 SV=2 | 255 | 20201113_xd_milk_1ng_glyco_hcd203040_compound15.21459.21459.4 | controllerType=0 controllerNumber=1 scan=21459 | 42.7433 |
| 18065:3 | 9 | R.LSLHRPALEDLLLGSEAN[+1501.555]LTC[+57.021]TLTGLR.D | HexNAc(5)Hex(3) | C[+57], N[+1502] | 1489.0596 | 3 | 4465.1643 | 4465.1536 | 0 | 2.4 | 127 | Specific | 677.88 | 677.88 | 677.88 | 13.98 | 55 | >sp\|P01876\|IGHA1_HUMAN Immunoglobulin heavy constant alpha 1 OS=Homo sapiens OX=9606 GN=IGHA1 PE=1 SV=2 | 255 | 20201113_xd_milk_1ng_glyco_hcd203040_compound15.21470.21470.2 | controllerType=0 controllerNumber=1 scan=21470 | 42.7749 |
| 17957:4 | 9 | R.LSLHRPALEDLLLGSEAN[+1702.581]LTC[+57.021]TLTGLR.D | HexNAc(2)Hex(8) | C[+57], N[+1703] | 1167.3018 | 4 | 4666.1854 | 4666.1796 | 0 | 1.2 | 127 | Specific | 678.11 | 678.11 | 678.11 | 14.36 | 55 | >sp\|P01876\|IGHA1_HUMAN Immunoglobulin heavy constant alpha 1 OS=Homo sapiens OX=9606 GN=IGHA1 PE=1 SV=2 | 255 | 20201113_xd_milk_1ng_glyco_hcd203040_compound15.21358.21358.4 | controllerType=0 controllerNumber=1 scan=21358 | 42.5025 |
| 17986:4 | 9 | R.LSLHRPALEDLLLGSEAN[+1501.555]LTC[+57.021]TLTGLR.D | HexNAc(5)Hex(3) | C[+57], N[+1502] | 1117.0459 | 4 | 4465.1619 | 4465.1536 | 0 | 1.9 | 127 | Specific | 683.85 | 683.85 | 683.85 | 14.53 | 55 | >sp\|P01876\|IGHA1_HUMAN Immunoglobulin heavy constant alpha 1 OS=Homo sapiens OX=9606 GN=IGHA1 PE=1 SV=2 | 255 | 20201113_xd_milk_1ng_glyco_hcd203040_compound15.21388.21388.4 | controllerType=0 controllerNumber=1 scan=21388 | 42.5612 |
| 18040:3 | 9 | R.LSLHRPALEDLLLGSEAN[+1216.423]LTC[+57.021]TLTGLR.D | HexNAc(2)Hex(5) | C[+57], N[+1216] | 1394.023 | 3 | 4180.0545 | 4180.0211 | 0 | 8 | 127 | Specific | 683.86 | 653.64 | 653.64 | 12.85 | 55 | >sp\|P01876\|IGHA1_HUMAN Immunoglobulin heavy constant alpha 1 OS=Homo sapiens OX=9606 GN=IGHA1 PE=1 SV=2 | 255 | 20201113_xd_milk_1ng_glyco_hcd203040_compound15.21444.21444.3 | controllerType=0 controllerNumber=1 scan=21444 | 42.6903 |
| 18096:3 | 9 | R.LSLHRPALEDLLLGSEAN[+1095.397]LTC[+57.021]TLTGLR.D | HexNAc(3)Hex(3) | C[+57], N[+1095] | 1353.676 | 3 | 4059.0133 | 4058.9949 | 0 | 4.6 | 127 | Specific | 684.38 | 654.36 | 654.36 | 13.47 | 55 | >sp\|P01876\|IGHA1_HUMAN Immunoglobulin heavy constant alpha 1 OS=Homo sapiens OX=9606 GN=IGHA1 PE=1 SV=2 | 255 | 20201113_xd_milk_1ng_glyco_hcd203040_compound15.21502.21502.3 | controllerType=0 controllerNumber=1 scan=21502 | 42.8541 |
| 19036:4 | 9 | R.LSLHRPALEDLLLGSEAN[+1710.598]LTC[+57.021]TLTGLR.D | HexNAc(3)Hex(5)NeuAc(1) | C[+57], N[+1711] | 1169.3075 | 4 | 4674.2083 | 4674.1959 | 0 | 2.7 | 127 | Specific | 708.8 | 681.69 | 681.69 | 15.34 | 55 | >sp\|P01876\|IGHA1_HUMAN Immunoglobulin heavy constant alpha 1 OS=Homo sapiens OX=9606 GN=IGHA1 PE=1 SV=2 | 255 | 20201113_xd_milk_1ng_glyco_hcd203040_compound15.22481.22481.4 | controllerType=0 controllerNumber=1 scan=22481 | 44.8939 |
| 18037:3 | 9 | R.LSLHRPALEDLLLGSEAN[+1216.423]LTC[+57.021]TLTGLR.D | HexNAc(2)Hex(5) | C[+57], N[+1216] | 1394.023 | 3 | 4180.0545 | 4180.0211 | 0 | 8 | 127 | Specific | 729.99 | 729.99 | 729.99 | 13.85 | 55 | >sp\|P01876\|IGHA1_HUMAN Immunoglobulin heavy constant alpha 1 OS=Homo sapiens OX=9606 GN=IGHA1 PE=1 SV=2 | 255 | 20201113_xd_milk_1ng_glyco_hcd203040_compound15.21441.21441.3 | controllerType=0 controllerNumber=1 scan=21441 | 42.679 |
| 17951:3 | 9 | R.LSLHRPALEDLLLGSEAN[+1540.529]LTC[+57.021]TLTGLR.D | HexNAc(2)Hex(7) | C[+57], N[+1541] | 1502.0488 | 3 | 4504.1317 | 4504.1268 | 0 | 1.1 | 127 | Specific | 730.06 | 694.94 | 694.94 | 14.81 | 55 | >sp\|P01876\|IGHA1_HUMAN Immunoglobulin heavy constant alpha 1 OS=Homo sapiens OX=9606 GN=IGHA1 PE=1 SV=2 | 255 | 20201113_xd_milk_1ng_glyco_hcd203040_compound15.21351.21351.3 | controllerType=0 controllerNumber=1 scan=21351 | 42.485 |
| 19005:4 | 9 | R.LSLHRPALEDLLLGSEAN[+1872.651]LTC[+57.021]TLTGLR.D | HexNAc(3)Hex(6)NeuAc(1) | C[+57], N[+1873] | 1209.8222 | 4 | 4836.2669 | 4836.2487 | 0 | 3.8 | 127 | Specific | 740.31 | 652.11 | 652.11 | 15.34 | 55 | >sp\|P01876\|IGHA1_HUMAN Immunoglobulin heavy constant alpha 1 OS=Homo sapiens OX=9606 GN=IGHA1 PE=1 SV=2 | 255 | 20201113_xd_milk_1ng_glyco_hcd203040_compound15.22449.22449.4 | controllerType=0 controllerNumber=1 scan=22449 | 44.8292 |
| 18056:4 | 9 | R.LSLHRPALEDLLLGSEAN[+1501.555]LTC[+57.021]TLTGLR.D | HexNAc(5)Hex(3) | C[+57], N[+1502] | 1117.0467 | 4 | 4465.1649 | 4465.1536 | 0 | 2.5 | 127 | Specific | 777.16 | 676.64 | 676.64 | 16.53 | 55 | >sp\|P01876\|IGHA1_HUMAN Immunoglobulin heavy constant alpha 1 OS=Homo sapiens OX=9606 GN=IGHA1 PE=1 SV=2 | 255 | 20201113_xd_milk_1ng_glyco_hcd203040_compound15.21461.21461.4 | controllerType=0 controllerNumber=1 scan=21461 | 42.753 |
| 17989:4 | 9 | R.LSLHRPALEDLLLGSEAN[+1501.555]LTC[+57.021]TLTGLR.D | HexNAc(5)Hex(3) | C[+57], N[+1502] | 1117.0459 | 4 | 4465.1619 | 4465.1536 | 0 | 1.9 | 127 | Specific | 777.71 | 719.26 | 719.26 | 16.53 | 55 | >sp\|P01876\|IGHA1_HUMAN Immunoglobulin heavy constant alpha 1 OS=Homo sapiens OX=9606 GN=IGHA1 PE=1 SV=2 | 255 | 20201113_xd_milk_1ng_glyco_hcd203040_compound15.21391.21391.4 | controllerType=0 controllerNumber=1 scan=21391 | 42.5678 |
| 19038:4 | 9 | R.LSLHRPALEDLLLGSEAN[+1751.624]LTC[+57.021]TLTGLR.D | HexNAc(4)Hex(4)NeuAc(1) | C[+57], N[+1752] | 1179.5636 | 4 | 4715.2327 | 4715.2225 | 0 | 2.2 | 127 | Specific | 802.83 | 742.74 | 742.74 | 17.53 | 55 | >sp\|P01876\|IGHA1_HUMAN Immunoglobulin heavy constant alpha 1 OS=Homo sapiens OX=9606 GN=IGHA1 PE=1 SV=2 | 255 | 20201113_xd_milk_1ng_glyco_hcd203040_compound15.22483.22483.4 | controllerType=0 controllerNumber=1 scan=22483 | 44.8983 |
| 19039:4 | 9 | R.LSLHRPALEDLLLGSEAN[+1710.598]LTC[+57.021]TLTGLR.D | HexNAc(3)Hex(5)NeuAc(1) | C[+57], N[+1711] | 1169.3075 | 4 | 4674.2083 | 4674.1959 | 0 | 2.7 | 127 | Specific | 871.46 | 829.85 | 829.85 | 18.34 | 55 | >sp\|P01876\|IGHA1_HUMAN Immunoglobulin heavy constant alpha 1 OS=Homo sapiens OX=9606 GN=IGHA1 PE=1 SV=2 | 255 | 20201113_xd_milk_1ng_glyco_hcd203040_compound15.22484.22484.4 | controllerType=0 controllerNumber=1 scan=22484 | 44.9021 |
| 17958:4 | 9 | R.LSLHRPALEDLLLGSEAN[+1540.529]LTC[+57.021]TLTGLR.D | HexNAc(2)Hex(7) | C[+57], N[+1541] | 1126.7878 | 4 | 4504.1292 | 4504.1268 | 0 | 0.5 | 127 | Specific | 924.97 | 830.88 | 830.88 | 19.36 | 55 | >sp\|P01876\|IGHA1_HUMAN Immunoglobulin heavy constant alpha 1 OS=Homo sapiens OX=9606 GN=IGHA1 PE=1 SV=2 | 255 | 20201113_xd_milk_1ng_glyco_hcd203040_compound15.21359.21359.4 | controllerType=0 controllerNumber=1 scan=21359 | 42.5039 |
| 18017:4 | 9 | R.LSLHRPALEDLLLGSEAN[+1216.423]LTC[+57.021]TLTGLR.D | HexNAc(2)Hex(5) | C[+57], N[+1216] | 1045.7699 | 4 | 4180.0579 | 4180.0211 | 0 | 8.8 | 127 | Specific | 932.04 | 932.04 | 932.04 | 18.26 | 55 | >sp\|P01876\|IGHA1_HUMAN Immunoglobulin heavy constant alpha 1 OS=Homo sapiens OX=9606 GN=IGHA1 PE=1 SV=2 | 255 | 20201113_xd_milk_1ng_glyco_hcd203040_compound15.21420.21420.4 | controllerType=0 controllerNumber=1 scan=21420 | 42.6254 |
| 17985:4 | 9 | R.LSLHRPALEDLLLGSEAN[+1378.476]LTC[+57.021]TLTGLR.D | HexNAc(2)Hex(6) | C[+57], N[+1378] | 1086.2758 | 4 | 4342.0814 | 4342.074 | 0 | 1.7 | 127 | Specific | 947.5 | 856.2 | 856.2 | 19.53 | 55 | >sp\|P01876\|IGHA1_HUMAN Immunoglobulin heavy constant alpha 1 OS=Homo sapiens OX=9606 GN=IGHA1 PE=1 SV=2 | 255 | 20201113_xd_milk_1ng_glyco_hcd203040_compound15.21387.21387.4 | controllerType=0 controllerNumber=1 scan=21387 | 42.5605 |

Table S4. Cont.

| Query #:z | Protein Rank | Glycopeptide | Glycan | Modification Type(s) | Observed m/z | z | Observed (M+H) | Calc. mass (M+H) | Off-by-x error | Mass error (ppm) | StartPosition | Cleavage | Score | Delta | Delta Mod | \|Log Prob\| | # of unique peptides | Protein Name | Protein DB number | Comment | Scan # | Scan Time |
|---|---|---|---|---|---|---|---|---|---|---|---|---|---|---|---|---|---|---|---|---|---|---|
| 17988:4 | 9 | R.LSLHRPALEDLLLGSEAN[+1378.476]LTC[+57.021]TLTGLR.D | HexNAc(2)Hex(6) | C[+57], N[+1378] | 1086.2758 | 4 | 4342.0814 | 4342.074 | 0 | 1.7 | 127 | Specific | 958 | 958 | 958 | 20.53 | 55 | >sp\|P01876\|IGHA1_HUMAN Immunoglobulin heavy constant alpha 1 OS=Homo sapiens OX=9606 GN=IGHA1 PE=1 SV=2 | 255 | 20201113_xd_milk_1ng_glyco_hcd203040_compound15.21390.21390.4 | controllerType=0 controllerNumber=1 scan=21390 | 42.5651 |
| 17905:4 | 9 | R.LSLHRPALEDLLLGSEAN[+1864.634]LTC[+57.021]TLTGLR.D | HexNAc(2)Hex(9) | C[+57], N[+1865] | 1207.8154 | 4 | 4828.2396 | 4828.2324 | 0 | 1.5 | 127 | Specific | 986.37 | 986.37 | 986.37 | 20.36 | 55 | >sp\|P01876\|IGHA1_HUMAN Immunoglobulin heavy constant alpha 1 OS=Homo sapiens OX=9606 GN=IGHA1 PE=1 SV=2 | 255 | 20201113_xd_milk_1ng_glyco_hcd203040_compound15.21304.21304.4 | controllerType=0 controllerNumber=1 scan=21304 | 42.4002 |
| 17959:4 | 9 | R.LSLHRPALEDLLLGSEAN[+1702.581]LTC[+57.021]TLTGLR.D | HexNAc(2)Hex(8) | C[+57], N[+1703] | 1167.3018 | 4 | 4666.1854 | 4666.1796 | 0 | 1.2 | 127 | Specific | 1018.22 | 1009.27 | 1009.27 | 21.36 | 55 | >sp\|P01876\|IGHA1_HUMAN Immunoglobulin heavy constant alpha 1 OS=Homo sapiens OX=9606 GN=IGHA1 PE=1 SV=2 | 255 | 20201113_xd_milk_1ng_glyco_hcd203040_compound15.21360.21360.4 | controllerType=0 controllerNumber=1 scan=21360 | 42.5067 |
| 17960:4 | 9 | R.LSLHRPALEDLLLGSEAN[+1540.529]LTC[+57.021]TLTGLR.D | HexNAc(2)Hex(7) | C[+57], N[+1541] | 1126.7878 | 4 | 4504.1292 | 4504.1268 | 0 | 0.5 | 127 | Specific | 1038.02 | 959.01 | 959.01 | 21.36 | 55 | >sp\|P01876\|IGHA1_HUMAN Immunoglobulin heavy constant alpha 1 OS=Homo sapiens OX=9606 GN=IGHA1 PE=1 SV=2 | 255 | 20201113_xd_milk_1ng_glyco_hcd203040_compound15.21361.21361.4 | controllerType=0 controllerNumber=1 scan=21361 | 42.5094 |
| 20024:3 | 9 | R.[+42.011]PALEDLLLGSEAN[+1460.529]LTC[+57.021]TLTGLR.D | HexNAc(4)Hex(4) | .[42], C[+57], N[+1461] | 1287.2719 | 3 | 3859.8012 | 3859.7774 | 0 | 6.1 | 132 | Specific | 281.16 | 281.16 | 281.16 | 6.54 | 55 | >sp\|P01876\|IGHA1_HUMAN Immunoglobulin heavy constant alpha 1 OS=Homo sapiens OX=9606 GN=IGHA1 PE=1 SV=2 | 255 | 20201113_xd_milk_1ng_glyco_hcd203040_compound15.23511.23511.3 | controllerType=0 controllerNumber=1 scan=23511 | 46.9896 |
| 20027:3 | 9 | R.[+42.011]PALEDLLLGSEAN[+1460.529]LTC[+57.021]TLTGLR.D | HexNAc(4)Hex(4) | .[42], C[+57], N[+1461] | 1287.2719 | 3 | 3859.8012 | 3859.7774 | 0 | 6.1 | 132 | Specific | 351.81 | 343.09 | 343.09 | 7.45 | 55 | >sp\|P01876\|IGHA1_HUMAN Immunoglobulin heavy constant alpha 1 OS=Homo sapiens OX=9606 GN=IGHA1 PE=1 SV=2 | 255 | 20201113_xd_milk_1ng_glyco_hcd203040_compound15.23514.23514.3 | controllerType=0 controllerNumber=1 scan=23514 | 46.9939 |
| 13614:3 | 9 | R.LAGKPTHVN[+1872.651]VSVVMAEVDGTC[+57.021].Y | HexNAc(3)Hex(6)NeuAc(1) | C[+57], N[+1873] | 1352.9224 | 3 | 4056.7527 | 4056.7292 | 0 | 5.8 | 332 | CRagged | 223.45 | 196.48 | 43.75 | 3.93 | 55 | >sp\|P01876\|IGHA1_HUMAN Immunoglobulin heavy constant alpha 1 OS=Homo sapiens OX=9606 GN=IGHA1 PE=1 SV=2 | 255 | 20201113_xd_milk_1ng_glyco_hcd203040_compound15.16828.16828.3 | controllerType=0 controllerNumber=1 scan=16828 | 33.883 |
| 13593:3 | 9 | R.LAGKPTHVN[+2018.708]VSVVM[+15.995]AEVDGTC[+57.021].Y | HexNAc(3)Hex(6)Fuc(1)NeuAc(1) | C[+57], M[+16], N[+2019] | 1406.9365 | 3 | 4218.7948 | 4218.7821 | 0 | 3 | 332 | CRagged | 206.37 | 178.02 | 178.02 | 3.16 | 55 | >sp\|P01876\|IGHA1_HUMAN Immunoglobulin heavy constant alpha 1 OS=Homo sapiens OX=9606 GN=IGHA1 PE=1 SV=2 | 255 | 20201113_xd_milk_1ng_glyco_hcd203040_compound15.16807.16807.3 | controllerType=0 controllerNumber=1 scan=16807 | 33.8476 |
| 15433:3 | 9 | R.LAGKPTHVN[+2553.910]VSVVMAEVDGTC[+57.021]Y.- | HexNAc(5)Hex(5)Fuc(1)NeuAc(2) | C[+57], N[+2554] | 1634.3597 | 3 | 4901.0645 | 4901.0518 | 0 | 2.6 | 332 | CRagged | 242.37 | 242.37 | 242.37 | 7.26 | 55 | >sp\|P01876\|IGHA1_HUMAN Immunoglobulin heavy constant alpha 1 OS=Homo sapiens OX=9606 GN=IGHA1 PE=1 SV=2 | 255 | 20201113_xd_milk_1ng_glyco_hcd203040_compound15.18718.18718.3 | controllerType=0 controllerNumber=1 scan=18718 | 37.416 |
| 11832:3 | 9 | R.LAGKPTHVN[+1378.476]VSVVMAEVDGTC[+57.021]Y.- | HexNAc(2)Hex(6) | C[+57], N[+1378] | 1242.5475 | 3 | 3725.6281 | 3725.6178 | 0 | 2.8 | 332 | CRagged | 243.51 | 204.86 | 168.58 | 5.38 | 55 | >sp\|P01876\|IGHA1_HUMAN Immunoglobulin heavy constant alpha 1 OS=Homo sapiens OX=9606 GN=IGHA1 PE=1 SV=2 | 255 | 20201113_xd_milk_1ng_glyco_hcd203040_compound15.14971.14971.3 | controllerType=0 controllerNumber=1 scan=14971 | 30.7735 |
| 15431:3 | 9 | R.LAGKPTHVN[+2553.910]VSVVMAEVDGTC[+57.021]Y.- | HexNAc(5)Hex(5)Fuc(1)NeuAc(2) | C[+57], N[+2554] | 1634.3597 | 3 | 4901.0645 | 4901.0518 | 0 | 2.6 | 332 | CRagged | 258.81 | 258.81 | 258.81 | 7.5 | 55 | >sp\|P01876\|IGHA1_HUMAN Immunoglobulin heavy constant alpha 1 OS=Homo sapiens OX=9606 GN=IGHA1 PE=1 SV=2 | 255 | 20201113_xd_milk_1ng_glyco_hcd203040_compound15.18716.18716.3 | controllerType=0 controllerNumber=1 scan=18716 | 37.4126 |
| 15405:4 | 9 | R.LAGKPTHVN[+2553.910]VSVVMAEVDGTC[+57.021]Y.- | HexNAc(5)Hex(5)Fuc(1)NeuAc(2) | C[+57], N[+2554] | 1226.0177 | 4 | 4901.0491 | 4901.0518 | 0 | -0.5 | 332 | CRagged | 315.5 | 210.02 | 210.02 | 6.82 | 55 | >sp\|P01876\|IGHA1_HUMAN Immunoglobulin heavy constant alpha 1 OS=Homo sapiens OX=9606 GN=IGHA1 PE=1 SV=2 | 255 | 20201113_xd_milk_1ng_glyco_hcd203040_compound15.18688.18688.2 | controllerType=0 controllerNumber=1 scan=18688 | 37.3633 |
| 11705:3 | 9 | R.LAGKPTHVN[+1825.661]VSVVMAEVDGTC[+57.021]Y.- | HexNAc(5)Hex(5) | C[+57], N[+1826] | 1391.6107 | 3 | 4172.8174 | 4172.8031 | 0 | 3.4 | 332 | CRagged | 399.08 | 249.84 | 249.84 | 6.16 | 55 | >sp\|P01876\|IGHA1_HUMAN Immunoglobulin heavy constant alpha 1 OS=Homo sapiens OX=9606 GN=IGHA1 PE=1 SV=2 | 255 | 20201113_xd_milk_1ng_glyco_hcd203040_compound15.14839.14839.3 | controllerType=0 controllerNumber=1 scan=14839 | 30.5312 |
| 13509:4 | 9 | R.LAGKPTHVN[+2262.814]VSVVMAEVDGTC[+57.021]Y.- | HexNAc(5)Hex(5)Fuc(1)NeuAc(1) | C[+57], N[+2263] | 1153.2482 | 4 | 4609.971 | 4609.9564 | 0 | 3.2 | 332 | CRagged | 405.37 | 405.37 | 405.37 | 10.09 | 55 | >sp\|P01876\|IGHA1_HUMAN Immunoglobulin heavy constant alpha 1 OS=Homo sapiens OX=9606 GN=IGHA1 PE=1 SV=2 | 255 | 20201113_xd_milk_1ng_glyco_hcd203040_compound15.16719.16719.4 | controllerType=0 controllerNumber=1 scan=16719 | 33.6891 |
| 11707:3 | 9 | R.LAGKPTHVN[+1825.661]VSVVMAEVDGTC[+57.021]Y.- | HexNAc(5)Hex(5) | C[+57], N[+1826] | 1391.6107 | 3 | 4172.8174 | 4172.8031 | 0 | 3.4 | 332 | CRagged | 438.71 | 438.71 | 365.43 | 10.21 | 55 | >sp\|P01876\|IGHA1_HUMAN Immunoglobulin heavy constant alpha 1 OS=Homo sapiens OX=9606 GN=IGHA1 PE=1 SV=2 | 255 | 20201113_xd_milk_1ng_glyco_hcd203040_compound15.14841.14841.3 | controllerType=0 controllerNumber=1 scan=14841 | 30.5367 |
| 10611:3 | 9 | R.LAGKPTHVN[+1216.423]VSVVMAEVDGTC[+57.021]Y.- | HexNAc(2)Hex(5) | C[+57], N[+1216] | 1188.5296 | 3 | 3563.5742 | 3563.5649 | 0 | 2.6 | 332 | CRagged | 475.23 | 475.23 | 289.3 | 10.51 | 55 | >sp\|P01876\|IGHA1_HUMAN Immunoglobulin heavy constant alpha 1 OS=Homo sapiens OX=9606 GN=IGHA1 PE=1 SV=2 | 255 | 20201113_xd_milk_1ng_glyco_hcd203040_compound15.13707.13707.3 | controllerType=0 controllerNumber=1 scan=13707 | 28.603 |
| 10614:3 | 9 | R.LAGKPTHVN[+1216.423]VSVVMAEVDGTC[+57.021]Y.- | HexNAc(2)Hex(5) | C[+57], N[+1216] | 1188.5296 | 3 | 3563.5742 | 3563.5649 | 0 | 2.6 | 332 | CRagged | 576.26 | 501.08 | 366.87 | 11.14 | 55 | >sp\|P01876\|IGHA1_HUMAN Immunoglobulin heavy constant alpha 1 OS=Homo sapiens OX=9606 GN=IGHA1 PE=1 SV=2 | 255 | 20201113_xd_milk_1ng_glyco_hcd203040_compound15.13710.13710.3 | controllerType=0 controllerNumber=1 scan=13710 | 28.6116 |
| 11715:3 | 9 | R.LAGKPTHVN[+1809.666]VSVVMAEVDGTC[+57.021]Y.- | HexNAc(5)Hex(4)Fuc(1) | C[+57], N[+1810] | 1386.2762 | 3 | 4156.814 | 4156.8081 | 0 | 1.4 | 332 | CRagged | 585.35 | 585.35 | 403.14 | 12.24 | 55 | >sp\|P01876\|IGHA1_HUMAN Immunoglobulin heavy constant alpha 1 OS=Homo sapiens OX=9606 GN=IGHA1 PE=1 SV=2 | 255 | 20201113_xd_milk_1ng_glyco_hcd203040_compound15.14850.14850.3 | controllerType=0 controllerNumber=1 scan=14850 | 30.5619 |
| 13503:3 | 9 | R.LAGKPTHVN[+2262.814]VSVVMAEVDGTC[+57.021]Y.- | HexNAc(5)Hex(5)Fuc(1)NeuAc(1) | C[+57], N[+2263] | 1537.3271 | 3 | 4609.9667 | 4609.9564 | 0 | 2.2 | 332 | CRagged | 633.31 | 436.11 | 436.11 | 13.41 | 55 | >sp\|P01876\|IGHA1_HUMAN Immunoglobulin heavy constant alpha 1 OS=Homo sapiens OX=9606 GN=IGHA1 PE=1 SV=2 | 255 | 20201113_xd_milk_1ng_glyco_hcd203040_compound15.16713.16713.3 | controllerType=0 controllerNumber=1 scan=16713 | 33.68 |
| 11834:3 | 9 | R.LAGKPTHVN[+1378.476]VSVVMAEVDGTC[+57.021]Y.- | HexNAc(2)Hex(6) | C[+57], N[+1378] | 1242.5475 | 3 | 3725.6281 | 3725.6178 | 0 | 2.8 | 332 | CRagged | 633.71 | 526.35 | 418.32 | 13.22 | 55 | >sp\|P01876\|IGHA1_HUMAN Immunoglobulin heavy constant alpha 1 OS=Homo sapiens OX=9606 GN=IGHA1 PE=1 SV=2 | 255 | 20201113_xd_milk_1ng_glyco_hcd203040_compound15.14973.14973.3 | controllerType=0 controllerNumber=1 scan=14973 | 30.7775 |
| 11818:3 | 9 | R.LAGKPTHVN[+1809.666]VSVVMAEVDGTC[+57.021]Y.- | HexNAc(5)Hex(4)Fuc(1) | C[+57], N[+1810] | 1386.2773 | 3 | 4156.8173 | 4156.8081 | 0 | 2.2 | 332 | CRagged | 647.16 | 647.16 | 427.58 | 13.41 | 55 | >sp\|P01876\|IGHA1_HUMAN Immunoglobulin heavy constant alpha 1 OS=Homo sapiens OX=9606 GN=IGHA1 PE=1 SV=2 | 255 | 20201113_xd_milk_1ng_glyco_hcd203040_compound15.14957.14957.3 | controllerType=0 controllerNumber=1 scan=14957 | 30.7463 |
| 13504:3 | 9 | R.LAGKPTHVN[+1913.677]VSVVMAEVDGTC[+57.021]Y.- | HexNAc(4)Hex(5)NeuAc(1) | C[+57], N[+1914] | 1420.9479 | 3 | 4260.8292 | 4260.8191 | 0 | 2.4 | 332 | CRagged | 697.66 | 488.53 | 488.53 | 14.41 | 55 | >sp\|P01876\|IGHA1_HUMAN Immunoglobulin heavy constant alpha 1 OS=Homo sapiens OX=9606 GN=IGHA1 PE=1 SV=2 | 255 | 20201113_xd_milk_1ng_glyco_hcd203040_compound15.16714.16714.3 | controllerType=0 controllerNumber=1 scan=16714 | 33.6811 |
| 11786:3 | 9 | R.LAGKPTHVN[+1501.555]VSVVMAEVDGTC[+57.021]Y.- | HexNAc(5)Hex(3) | C[+57], N[+1502] | 1283.572 | 3 | 3848.7013 | 3848.6974 | 0 | 1 | 332 | CRagged | 727.99 | 727.99 | 727.99 | 15.24 | 55 | >sp\|P01876\|IGHA1_HUMAN Immunoglobulin heavy constant alpha 1 OS=Homo sapiens OX=9606 GN=IGHA1 PE=1 SV=2 | 255 | 20201113_xd_milk_1ng_glyco_hcd203040_compound15.14924.14924.3 | controllerType=0 controllerNumber=1 scan=14924 | 30.6871 |
| 11788:3 | 9 | R.LAGKPTHVN[+1501.555]VSVVMAEVDGTC[+57.021]Y.- | HexNAc(5)Hex(3) | C[+57], N[+1502] | 1283.572 | 3 | 3848.7013 | 3848.6974 | 0 | 1 | 332 | CRagged | 734.07 | 732.48 | 732.48 | 15.24 | 55 | >sp\|P01876\|IGHA1_HUMAN Immunoglobulin heavy constant alpha 1 OS=Homo sapiens OX=9606 GN=IGHA1 PE=1 SV=2 | 255 | 20201113_xd_milk_1ng_glyco_hcd203040_compound15.14926.14926.3 | controllerType=0 controllerNumber=1 scan=14926 | 30.6933 |
| 11816:3 | 9 | R.LAGKPTHVN[+1501.555]VSVVMAEVDGTC[+57.021]Y.- | HexNAc(5)Hex(3) | C[+57], N[+1502] | 1283.5727 | 3 | 3848.7035 | 3848.6974 | 0 | 1.6 | 332 | CRagged | 736.55 | 736.55 | 736.55 | 15.41 | 55 | >sp\|P01876\|IGHA1_HUMAN Immunoglobulin heavy constant alpha 1 OS=Homo sapiens OX=9606 GN=IGHA1 PE=1 SV=2 | 255 | 20201113_xd_milk_1ng_glyco_hcd203040_compound15.14955.14955.3 | controllerType=0 controllerNumber=1 scan=14955 | 30.7441 |
| 13507:3 | 9 | R.LAGKPTHVN[+1913.677]VSVVMAEVDGTC[+57.021]Y.- | HexNAc(4)Hex(5)NeuAc(1) | C[+57], N[+1914] | 1420.9479 | 3 | 4260.8292 | 4260.8191 | 0 | 2.4 | 332 | CRagged | 765.83 | 516.9 | 516.9 | 16.41 | 55 | >sp\|P01876\|IGHA1_HUMAN Immunoglobulin heavy constant alpha 1 OS=Homo sapiens OX=9606 GN=IGHA1 PE=1 SV=2 | 255 | 20201113_xd_milk_1ng_glyco_hcd203040_compound15.16717.16717.3 | controllerType=0 controllerNumber=1 scan=16717 | 33.6864 |
| 11820:3 | 9 | R.LAGKPTHVN[+1809.666]VSVVMAEVDGTC[+57.021]Y.- | HexNAc(5)Hex(4)Fuc(1) | C[+57], N[+1810] | 1386.2773 | 3 | 4156.8173 | 4156.8081 | 0 | 2.2 | 332 | CRagged | 770.8 | 715.42 | 496.87 | 16.41 | 55 | >sp\|P01876\|IGHA1_HUMAN Immunoglobulin heavy constant alpha 1 OS=Homo sapiens OX=9606 GN=IGHA1 PE=1 SV=2 | 255 | 20201113_xd_milk_1ng_glyco_hcd203040_compound15.14959.14959.3 | controllerType=0 controllerNumber=1 scan=14959 | 30.7518 |
| 13506:3 | 9 | R.LAGKPTHVN[+2262.814]VSVVMAEVDGTC[+57.021]Y.- | HexNAc(5)Hex(5)Fuc(1)NeuAc(1) | C[+57], N[+2263] | 1537.3271 | 3 | 4609.9667 | 4609.9564 | 0 | 2.2 | 332 | CRagged | 784.77 | 599.37 | 599.37 | 16.41 | 55 | >sp\|P01876\|IGHA1_HUMAN Immunoglobulin heavy constant alpha 1 OS=Homo sapiens OX=9606 GN=IGHA1 PE=1 SV=2 | 255 | 20201113_xd_milk_1ng_glyco_hcd203040_compound15.16716.16716.3 | controllerType=0 controllerNumber=1 scan=16716 | 33.6847 |
| 11839:3 | 9 | R.LAGKPTHVN[+1647.613]VSVVMAEVDGTC[+57.021]Y.- | HexNAc(5)Hex(3)Fuc(1) | C[+57], N[+1648] | 1332.2586 | 3 | 3994.7613 | 3994.7553 | 0 | 1.5 | 332 | CRagged | 784.83 | 640.86 | 640.86 | 16.24 | 55 | >sp\|P01876\|IGHA1_HUMAN Immunoglobulin heavy constant alpha 1 OS=Homo sapiens OX=9606 GN=IGHA1 PE=1 SV=2 | 255 | 20201113_xd_milk_1ng_glyco_hcd203040_compound15.14979.14979.3 | controllerType=0 controllerNumber=1 scan=14979 | 30.8009 |
| 11819:3 | 9 | R.LAGKPTHVN[+1501.555]VSVVMAEVDGTC[+57.021]Y.- | HexNAc(5)Hex(3) | C[+57], N[+1502] | 1283.5727 | 3 | 3848.7035 | 3848.6974 | 0 | 1.6 | 332 | CRagged | 790.22 | 790.22 | 790.22 | 16.41 | 55 | >sp\|P01876\|IGHA1_HUMAN Immunoglobulin heavy constant alpha 1 OS=Homo sapiens OX=9606 GN=IGHA1 PE=1 SV=2 | 255 | 20201113_xd_milk_1ng_glyco_hcd203040_compound15.14958.14958.3 | controllerType=0 controllerNumber=1 scan=14958 | 30.7491 |
| 11718:3 | 9 | R.LAGKPTHVN[+1809.666]VSVVMAEVDGTC[+57.021]Y.- | HexNAc(5)Hex(4)Fuc(1) | C[+57], N[+1810] | 1386.2762 | 3 | 4156.814 | 4156.8081 | 0 | 1.4 | 332 | CRagged | 791.76 | 791.76 | 510.63 | 16.24 | 55 | >sp\|P01876\|IGHA1_HUMAN Immunoglobulin heavy constant alpha 1 OS=Homo sapiens OX=9606 GN=IGHA1 PE=1 SV=2 | 255 | 20201113_xd_milk_1ng_glyco_hcd203040_compound15.14853.14853.3 | controllerType=0 controllerNumber=1 scan=14853 | 30.5656 |
| 11841:3 | 9 | R.LAGKPTHVN[+1647.613]VSVVMAEVDGTC[+57.021]Y.- | HexNAc(5)Hex(3)Fuc(1) | C[+57], N[+1648] | 1332.2586 | 3 | 3994.7613 | 3994.7553 | 0 | 1.5 | 332 | CRagged | 808.39 | 738.22 | 738.22 | 17.24 | 55 | >sp\|P01876\|IGHA1_HUMAN Immunoglobulin heavy constant alpha 1 OS=Homo sapiens OX=9606 GN=IGHA1 PE=1 SV=2 | 255 | 20201113_xd_milk_1ng_glyco_hcd203040_compound15.14981.14981.3 | controllerType=0 controllerNumber=1 scan=14981 | 30.8058 |
| 11667:3 | 9 | R.LAGKPTHVN[+1702.581]VSVVMAEVDGTC[+57.021]Y.- | HexNAc(2)Hex(8) | C[+57], N[+1703] | 1350.5828 | 3 | 4049.7339 | 4049.7234 | 0 | 2.6 | 332 | CRagged | 821.32 | 821.32 | 519.14 | 17.41 | 55 | >sp\|P01876\|IGHA1_HUMAN Immunoglobulin heavy constant alpha 1 OS=Homo sapiens OX=9606 GN=IGHA1 PE=1 SV=2 | 255 | 20201113_xd_milk_1ng_glyco_hcd203040_compound15.14799.14799.3 | controllerType=0 controllerNumber=1 scan=14799 | 30.4067 |
| 11703:3 | 9 | R.LAGKPTHVN[+1378.476]VSVVMAEVDGTC[+57.021]Y.- | HexNAc(2)Hex(6) | C[+57], N[+1378] | 1242.5477 | 3 | 3725.6284 | 3725.6178 | 0 | 2.9 | 332 | CRagged | 845.42 | 784.1 | 508.67 | 17.22 | 55 | >sp\|P01876\|IGHA1_HUMAN Immunoglobulin heavy constant alpha 1 OS=Homo sapiens OX=9606 GN=IGHA1 PE=1 SV=2 | 255 | 20201113_xd_milk_1ng_glyco_hcd203040_compound15.14837.14837.3 | controllerType=0 controllerNumber=1 scan=14837 | 30.5214 |
| 11669:3 | 9 | R.LAGKPTHVN[+1702.581]VSVVMAEVDGTC[+57.021]Y.- | HexNAc(2)Hex(8) | C[+57], N[+1703] | 1350.5828 | 3 | 4049.7339 | 4049.7234 | 0 | 2.6 | 332 | CRagged | 848.36 | 811.82 | 547.97 | 17.41 | 55 | >sp\|P01876\|IGHA1_HUMAN Immunoglobulin heavy constant alpha 1 OS=Homo sapiens OX=9606 GN=IGHA1 PE=1 SV=2 | 255 | 20201113_xd_milk_1ng_glyco_hcd203040_compound15.14801.14801.3 | controllerType=0 controllerNumber=1 scan=14801 | 30.4124 |
| 11701:3 | 9 | R.LAGKPTHVN[+1663.608]VSVVMAEVDGTC[+57.021]Y.- | HexNAc(5)Hex(4) | C[+57], N[+1664] | 1337.5917 | 3 | 4010.7606 | 4010.7502 | 0 | 2.6 | 332 | CRagged | 896.98 | 896.98 | 587.37 | 18.41 | 55 | >sp\|P01876\|IGHA1_HUMAN Immunoglobulin heavy constant alpha 1 OS=Homo sapiens OX=9606 GN=IGHA1 PE=1 SV=2 | 255 | 20201113_xd_milk_1ng_glyco_hcd203040_compound15.14835.14835.3 | controllerType=0 controllerNumber=1 scan=14835 | 30.5037 |
| 11719:3 | 9 | R.LAGKPTHVN[+1216.423]VSVVMAEVDGTC[+57.021]Y.- | HexNAc(2)Hex(5) | C[+57], N[+1216] | 1188.5286 | 3 | 3563.5713 | 3563.5649 | 0 | 1.8 | 332 | CRagged | 911.98 | 857.76 | 571.72 | 19.41 | 55 | >sp\|P01876\|IGHA1_HUMAN Immunoglobulin heavy constant alpha 1 OS=Homo sapiens OX=9606 GN=IGHA1 PE=1 SV=2 | 255 | 20201113_xd_milk_1ng_glyco_hcd203040_compound15.14854.14854.3 | controllerType=0 controllerNumber=1 scan=14854 | 30.5707 |
| 11717:3 | 9 | R.LAGKPTHVN[+1216.423]VSVVMAEVDGTC[+57.021]Y.- | HexNAc(2)Hex(5) | C[+57], N[+1216] | 1188.5286 | 3 | 3563.5713 | 3563.5649 | 0 | 1.8 | 332 | CRagged | 924.31 | 893.45 | 558.84 | 19.41 | 55 | >sp\|P01876\|IGHA1_HUMAN Immunoglobulin heavy constant alpha 1 OS=Homo sapiens OX=9606 GN=IGHA1 PE=1 SV=2 | 255 | 20201113_xd_milk_1ng_glyco_hcd203040_compound15.14852.14852.3 | controllerType=0 controllerNumber=1 scan=14852 | 30.5637 |
| 11700:3 | 9 | R.LAGKPTHVN[+1378.476]VSVVMAEVDGTC[+57.021]Y.- | HexNAc(2)Hex(6) | C[+57], N[+1378] | 1242.5477 | 3 | 3725.6284 | 3725.6178 | 0 | 2.9 | 332 | CRagged | 936.58 | 936.58 | 579.67 | 19.22 | 55 | >sp\|P01876\|IGHA1_HUMAN Immunoglobulin heavy constant alpha 1 OS=Homo sapiens OX=9606 GN=IGHA1 PE=1 SV=2 | 255 | 20201113_xd_milk_1ng_glyco_hcd203040_compound15.14834.14834.3 | controllerType=0 controllerNumber=1 scan=14834 | 30.5022 |
| 19265:4 | 10 | R.NIAAFGGDPNN[+1913.677]ITLFGESAGGASVSLQTLSPYNK.G | HexNAc(4)Hex(5)NeuAc(1) | N[+1914] | 1331.8497 | 4 | 5324.3771 | 5324.3633 | 0 | 2.6 | 197 | Specific | 837.92 | 835.18 | 835.18 | 17.06 | 47 | >sp\|P19835\|CEL_HUMAN Bile salt-activated lipase OS=Homo sapiens OX=9606 GN=CEL PE=1 SV=3 | 476 | 20201113_xd_milk_1ng_glyco_hcd203040_compound15.22719.22719.4 | controllerType=0 controllerNumber=1 scan=22719 | 45.422 |
| 19264:4 | 10 | R.NIAAFGGDPNN[+1913.677]ITLFGESAGGASVSLQTLSPYNK.G | HexNAc(4)Hex(5)NeuAc(1) | N[+1914] | 1331.8497 | 4 | 5324.3771 | 5324.3633 | 0 | 2.6 | 197 | Specific | 970.35 | 949.96 | 949.96 | 20.06 | 47 | >sp\|P19835\|CEL_HUMAN Bile salt-activated lipase OS=Homo sapiens OX=9606 GN=CEL PE=1 SV=3 | 476 | 20201113_xd_milk_1ng_glyco_hcd203040_compound15.22718.22718.4 | controllerType=0 controllerNumber=1 scan=22718 | 45.4154 |
| 23135:4 | 11 | K.APDKNVIFSPLSISTALAFLSLGAHN[+2204.772]TTLTEILK.G | HexNAc(4)Hex(5)NeuAc(2) | N[+2205] | 1447.6974 | 4 | 5787.7677 | 5787.7506 | 0 | 3 | 68 | Specific | 290.12 | 290.12 | 290.12 | 6.98 | 41 | >sp\|P01011\|AACT_HUMAN Alpha-1-antichymotrypsin OS=Homo sapiens OX=9606 GN=SERPINA3 PE=1 SV=2 | 1314 | 20201113_xd_milk_1ng_glyco_hcd203040_compound15.26754.26754.4 | controllerType=0 controllerNumber=1 scan=26754 | 53.6889 |
| 23137:4 | 11 | K.APDKNVIFSPLSISTALAFLSLGAHN[+2204.772]TTLTEILK.G | HexNAc(4)Hex(5)NeuAc(2) | N[+2205] | 1447.6974 | 4 | 5787.7677 | 5787.7506 | 0 | 3 | 68 | Specific | 294.93 | 294.93 | 294.93 | 6.98 | 41 | >sp\|P01011\|AACT_HUMAN Alpha-1-antichymotrypsin OS=Homo sapiens OX=9606 GN=SERPINA3 PE=1 SV=2 | 1314 | 20201113_xd_milk_1ng_glyco_hcd203040_compound15.26756.26756.4 | controllerType=0 controllerNumber=1 scan=26756 | 53.693 |
| 23166:4 | 11 | K.APDKNVIFSPLSISTALAFLSLGAHN[+2204.772]TTLTEILK.G | HexNAc(4)Hex(5)NeuAc(2) | N[+2205] | 1447.6978 | 4 | 5787.7692 | 5787.7506 | 0 | 3.2 | 68 | Specific | 299.99 | 299.99 | 299.99 | 6.98 | 41 | >sp\|P01011\|AACT_HUMAN Alpha-1-antichymotrypsin OS=Homo sapiens OX=9606 GN=SERPINA3 PE=1 SV=2 | 1314 | 20201113_xd_milk_1ng_glyco_hcd203040_compound15.26786.26786.4 | controllerType=0 controllerNumber=1 scan=26786 | 53.7684 |
| 21704:4 | 11 | K.APDKNVIFSPLSISTALAFLSLGAHN[+2059.735]TTLTEILK.G | HexNAc(4)Hex(5)Fuc(1)NeuAc(1) | N[+2060] | 1411.4454 | 4 | 5642.7599 | 5642.7131 | 0 | 8.3 | 68 | Specific | 931.94 | 931.94 | 931.94 | 17.54 | 41 | >sp\|P01011\|AACT_HUMAN Alpha-1-antichymotrypsin OS=Homo sapiens OX=9606 GN=SERPINA3 PE=1 SV=2 | 1314 | 20201113_xd_milk_1ng_glyco_hcd203040_compound15.25262.25262.4 | controllerType=0 controllerNumber=1 scan=25262 | 50.7286 |
| 21702:4 | 11 | K.APDKNVIFSPLSISTALAFLSLGAHN[+2059.735]TTLTEILK.G | HexNAc(4)Hex(5)Fuc(1)NeuAc(1) | N[+2060] | 1411.4454 | 4 | 5642.7599 | 5642.7131 | 0 | 8.3 | 68 | Specific | 1153 | 1144.11 | 1144.11 | 22.54 | 41 | >sp\|P01011\|AACT_HUMAN Alpha-1-antichymotrypsin OS=Homo sapiens OX=9606 GN=SERPINA3 PE=1 SV=2 | 1314 | 20201113_xd_milk_1ng_glyco_hcd203040_compound15.25260.25260.4 | controllerType=0 controllerNumber=1 scan=25260 | 50.7207 |
| 22941:4 | 11 | K.NVIFSPLSISTALAFLSLGAHN[+2205.793]TTLTEILK.G | HexNAc(4)Hex(5)Fuc(2)NeuAc(1) | N[+2206] | 1345.1471 | 4 | 5377.5666 | 5377.5592 | 0 | 1.4 | 72 | Specific | 222.31 | 217.66 | 217.66 | 7.69 | 41 | >sp\|P01011\|AACT_HUMAN Alpha-1-antichymotrypsin OS=Homo sapiens OX=9606 GN=SERPINA3 PE=1 SV=2 | 1314 | 20201113_xd_milk_1ng_glyco_hcd203040_compound15.26552.26552.4 | controllerType=0 controllerNumber=1 scan=26552 | 53.2851 |
| 24771:4 | 11 | K.NVIFSPLSISTALAFLSLGAHN[+2350.830]TTLTEILK.G | HexNAc(4)Hex(5)Fuc(1)NeuAc(2) | N[+2351] | 1381.4092 | 4 | 5522.6149 | 5522.5967 | 0 | 3.3 | 72 | Specific | 286.9 | 286.9 | 286.9 | 8.12 | 41 | >sp\|P01011\|AACT_HUMAN Alpha-1-antichymotrypsin OS=Homo sapiens OX=9606 GN=SERPINA3 PE=1 SV=2 | 1314 | 20201113_xd_milk_1ng_glyco_hcd203040_compound15.28457.28457.4 | controllerType=0 controllerNumber=1 scan=28457 | 57.0387 |
| 16089:3 | 11 | R.[+42.011]TLN[+2018.708]QSSDELQLSMGNAMFVK.E | HexNAc(3)Hex(6)Fuc(1)NeuAc(1) | .[42], N[+2019] | 1425.2708 | 3 | 4273.7978 | 4273.7766 | 0 | 5 | 125 | Specific | 299.09 | 122.01 | 122.01 | 5.83 | 41 | >sp\|P01011\|AACT_HUMAN Alpha-1-antichymotrypsin OS=Homo sapiens OX=9606 GN=SERPINA3 PE=1 SV=2 | 1314 | 20201113_xd_milk_1ng_glyco_hcd203040_compound15.19399.19399.3 | controllerType=0 controllerNumber=1 scan=19399 | 38.6881 |
| 16086:3 | 11 | R.[+42.011]TLN[+2018.708]QSSDELQLSMGNAMFVK.E | HexNAc(3)Hex(6)Fuc(1)NeuAc(1) | .[42], N[+2019] | 1425.2708 | 3 | 4273.7978 | 4273.7766 | 0 | 5 | 125 | Specific | 310.95 | 202.54 | 202.54 | 6.85 | 41 | >sp\|P01011\|AACT_HUMAN Alpha-1-antichymotrypsin OS=Homo sapiens OX=9606 GN=SERPINA3 PE=1 SV=2 | 1314 | 20201113_xd_milk_1ng_glyco_hcd203040_compound15.19396.19396.3 | controllerType=0 controllerNumber=1 scan=19396 | 38.6836 |
| 08026:3 | 12 | R.LSPN[+2350.830]ASAEHLELR.W | HexNAc(4)Hex(5)Fuc(1)NeuAc(2) | N[+2351] | 1263.2011 | 3 | 3787.5888 | 3787.5796 | 0 | 2.4 | 52 | Specific | 206.89 | 130.4 | 130.4 | 4.62 | 34 | >sp\|Q13410\|BT1A1_HUMAN Butyrophilin subfamily 1 member A1 OS=Homo sapiens OX=9606 GN=BTN1A1 PE=1 SV=3 | 179 | 20201113_xd_milk_1ng_glyco_hcd203040_compound15.11020.11020.3 | controllerType=0 controllerNumber=1 scan=11020 | 24.2283 |
| 08024:3 | 12 | R.LSPN[+2350.830]ASAEHLELR.W | HexNAc(4)Hex(5)Fuc(1)NeuAc(2) | N[+2351] | 1263.2011 | 3 | 3787.5888 | 3787.5796 | 0 | 2.4 | 52 | Specific | 327.85 | 206.99 | 206.99 | 7.2 | 34 | >sp\|Q13410\|BT1A1_HUMAN Butyrophilin subfamily 1 member A1 OS=Homo sapiens OX=9606 GN=BTN1A1 PE=1 SV=3 | 179 | 20201113_xd_milk_1ng_glyco_hcd203040_compound15.11018.11018.3 | controllerType=0 controllerNumber=1 scan=11018 | 24.2225 |
| 07994:3 | 12 | R.LSPN[+2350.830]ASAEHLELR.W | HexNAc(4)Hex(5)Fuc(1)NeuAc(2) | N[+2351] | 1263.1987 | 3 | 3787.5814 | 3787.5796 | 0 | 0.5 | 52 | Specific | 540.89 | 499.34 | 499.34 | 10.54 | 34 | >sp\|Q13410\|BT1A1_HUMAN Butyrophilin subfamily 1 member A1 OS=Homo sapiens OX=9606 GN=BTN1A1 PE=1 SV=3 | 179 | 20201113_xd_milk_1ng_glyco_hcd203040_compound15.10987.10987.3 | controllerType=0 controllerNumber=1 scan=10987 | 24.1683 |
| 05913:3 | 12 | R.LSPN[+2205.793]ASAEHLELR.W | HexNAc(4)Hex(5)Fuc(2)NeuAc(1) | N[+2206] | 1214.8554 | 3 | 3642.5517 | 3642.5421 | 0 | 2.6 | 52 | Specific | 553.13 | 341.03 | 341.03 | 9.03 | 34 | >sp\|Q13410\|BT1A1_HUMAN Butyrophilin subfamily 1 member A1 OS=Homo sapiens OX=9606 GN=BTN1A1 PE=1 SV=3 | 179 | 20201113_xd_milk_1ng_glyco_hcd203040_compound15.8831.8831.3 | controllerType=0 controllerNumber=1 scan=8831 | 20.6899 |
| 07997:3 | 12 | R.LSPN[+2350.830]ASAEHLELR.W | HexNAc(4)Hex(5)Fuc(1)NeuAc(2) | N[+2351] | 1263.1987 | 3 | 3787.5814 | 3787.5796 | 0 | 0.5 | 52 | Specific | 564.11 | 523.56 | 523.56 | 10.75 | 34 | >sp\|Q13410\|BT1A1_HUMAN Butyrophilin subfamily 1 member A1 OS=Homo sapiens OX=9606 GN=BTN1A1 PE=1 SV=3 | 179 | 20201113_xd_milk_1ng_glyco_hcd203040_compound15.10990.10990.3 | controllerType=0 controllerNumber=1 scan=10990 | 24.1734 |
| 05915:3 | 12 | R.LSPN[+2205.793]ASAEHLELR.W | HexNAc(4)Hex(5)Fuc(2)NeuAc(1) | N[+2206] | 1214.8554 | 3 | 3642.5517 | 3642.5421 | 0 | 2.6 | 52 | Specific | 584.09 | 393.52 | 393.52 | 12.19 | 34 | >sp\|Q13410\|BT1A1_HUMAN Butyrophilin subfamily 1 member A1 OS=Homo sapiens OX=9606 GN=BTN1A1 PE=1 SV=3 | 179 | 20201113_xd_milk_1ng_glyco_hcd203040_compound15.8833.8833.3 | controllerType=0 controllerNumber=1 scan=8833 | 20.6947 |
| 02316:3 | 13 | K.N[+1378.476]NSIPDKQITASSSYK.T | HexNAc(2)Hex(6) | N[+1378] | 1044.4575 | 3 | 3131.3581 | 3131.352 | 0 | 2 | 238 | Specific | 264.39 | 132.21 | 132.21 | 5.47 | 29 | >sp\|Q08431\|MFGM_HUMAN Lactadherin OS=Homo sapiens OX=9606 GN=MFGE8 PE=1 SV=3 | 1064 | 20201113_xd_milk_1ng_glyco_hcd203040_compound15.5105.5105.3 | controllerType=0 controllerNumber=1 scan=5105 | 14.571 |
| 02230:3 | 13 | K.N[+1540.529]NSIPDKQITASSSYK.T | HexNAc(2)Hex(7) | N[+1541] | 1098.4746 | 3 | 3293.4094 | 3293.4048 | 0 | 1.4 | 238 | Specific | 314.3 | 153.98 | 153.98 | 5.86 | 29 | >sp\|Q08431\|MFGM_HUMAN Lactadherin OS=Homo sapiens OX=9606 GN=MFGE8 PE=1 SV=3 | 1064 | 20201113_xd_milk_1ng_glyco_hcd203040_compound15.5016.5016.3 | controllerType=0 controllerNumber=1 scan=5016 | 14.421 |
| 02367:3 | 13 | K.N[+1216.423]NSIPDKQITASSSYK.T | HexNAc(2)Hex(5) | N[+1216] | 990.4404 | 3 | 2969.3066 | 2969.2991 | 0 | 2.5 | 238 | Specific | 393.6 | 369.16 | 369.16 | 8.01 | 29 | >sp\|Q08431\|MFGM_HUMAN Lactadherin OS=Homo sapiens OX=9606 GN=MFGE8 PE=1 SV=3 | 1064 | 20201113_xd_milk_1ng_glyco_hcd203040_compound15.5158.5158.3 | controllerType=0 controllerNumber=1 scan=5158 | 14.6624 |
| 02228:3 | 13 | K.N[+1540.529]NSIPDKQITASSSYK.T | HexNAc(2)Hex(7) | N[+1541] | 1098.4746 | 3 | 3293.4094 | 3293.4048 | 0 | 1.4 | 238 | Specific | 403.04 | 223.68 | 223.68 | 6.97 | 29 | >sp\|Q08431\|MFGM_HUMAN Lactadherin OS=Homo sapiens OX=9606 GN=MFGE8 PE=1 SV=3 | 1064 | 20201113_xd_milk_1ng_glyco_hcd203040_compound15.5014.5014.3 | controllerType=0 controllerNumber=1 scan=5014 | 14.4175 |
| 02369:3 | 13 | K.N[+1216.423]NSIPDKQITASSSYK.T | HexNAc(2)Hex(5) | N[+1216] | 990.4404 | 3 | 2969.3066 | 2969.2991 | 0 | 2.5 | 238 | Specific | 469.95 | 328.28 | 328.28 | 7.88 | 29 | >sp\|Q08431\|MFGM_HUMAN Lactadherin OS=Homo sapiens OX=9606 GN=MFGE8 PE=1 SV=3 | 1064 | 20201113_xd_milk_1ng_glyco_hcd203040_compound15.5160.5160.3 | controllerType=0 controllerNumber=1 scan=5160 | 14.6659 |
| 24193:4 | 14 | R.QLAHQSN[+2204.772]STNIFFSPVSIATAFAMLSLGTK.A | HexNAc(4)Hex(5)NeuAc(2) | N[+2205] | 1347.3566 | 4 | 5386.4045 | 5386.4075 | 0 | -0.6 | 64 | Specific | 791.79 | 727.95 | 653.1 | 16.59 | 28 | >sp\|P01009\|A1AT_HUMAN Alpha-1-antitrypsin OS=Homo sapiens OX=9606 GN=SERPINA1 PE=1 SV=3 | 642 | 20201113_xd_milk_1ng_glyco_hcd203040_compound15.27856.27856.4 | controllerType=0 controllerNumber=1 scan=27856 | 55.8965 |
| 24195:3 | 14 | R.QLAHQSN[+2204.772]STNIFFSPVSIATAFAMLSLGTK.A | HexNAc(4)Hex(5)NeuAc(2) | N[+2205] | 1796.1445 | 3 | 5386.4189 | 5386.4075 | 0 | 2.1 | 64 | Specific | 843.94 | 813.65 | 560.8 | 18.19 | 28 | >sp\|P01009\|A1AT_HUMAN Alpha-1-antitrypsin OS=Homo sapiens OX=9606 GN=SERPINA1 PE=1 SV=3 | 642 | 20201113_xd_milk_1ng_glyco_hcd203040_compound15.27858.27858.3 | controllerType=0 controllerNumber=1 scan=27858 | 55.9011 |
| 24197:3 | 14 | R.QLAHQSN[+2204.772]STNIFFSPVSIATAFAMLSLGTK.A | HexNAc(4)Hex(5)NeuAc(2) | N[+2205] | 1796.1445 | 3 | 5386.4189 | 5386.4075 | 0 | 2.1 | 64 | Specific | 852.72 | 786.92 | 519.3 | 19.19 | 28 | >sp\|P01009\|A1AT_HUMAN Alpha-1-antitrypsin OS=Homo sapiens OX=9606 GN=SERPINA1 PE=1 SV=3 | 642 | 20201113_xd_milk_1ng_glyco_hcd203040_compound15.27860.27860.3 | controllerType=0 controllerNumber=1 scan=27860 | 55.9054 |
| 24167:4 | 14 | R.QLAHQSN[+2204.772]STNIFFSPVSIATAFAMLSLGTK.A | HexNAc(4)Hex(5)NeuAc(2) | N[+2205] | 1347.3702 | 4 | 5386.4591 | 5386.4075 | 0 | 9.6 | 64 | Specific | 867.47 | 810.12 | 810.12 | 18.21 | 28 | >sp\|P01009\|A1AT_HUMAN Alpha-1-antitrypsin OS=Homo sapiens OX=9606 GN=SERPINA1 PE=1 SV=3 | 642 | 20201113_xd_milk_1ng_glyco_hcd203040_compound15.27829.27829.4 | controllerType=0 controllerNumber=1 scan=27829 | 55.[illegible]48 |
| 24191:4 | 14 | R.QLAHQSN[+2204.772]STNIFFSPVSIATAFAMLSLGTK.A | HexNAc(4)Hex(5)NeuAc(2) | N[+2205] | 1347.3566 | 4 | 5386.4045 | 5386.4075 | 0 | -0.6 | 64 | Specific | 927.07 | 875.22 | 560.3 | 19.59 | 28 | >sp\|P01009\|A1AT_HUMAN Alpha-1-antitrypsin OS=Homo sapiens OX=9606 GN=SERPINA1 PE=1 SV=3 | 642 | 20201113_xd_milk_1ng_glyco_hcd203040_compound15.27854.27854.4 | controllerType=0 controllerNumber=1 scan=27854 | 55.8[illegible]02 |
| 20523:3 | 14 | K.YLGN[+2204.772]ATAIFFLPDEGK.L | HexNAc(4)Hex(5)NeuAc(2) | N[+2205] | 1320.9046 | 3 | 3960.6992 | 3960.6677 | 0 | 8 | 268 | Specific | 457.23 | 363.84 | 363.84 | 7.54 | 28 | >sp\|P01009\|A1AT_HUMAN Alpha-1-antitrypsin OS=Homo sapiens OX=9606 GN=SERPINA1 PE=1 SV=3 | 642 | 20201113_xd_milk_1ng_glyco_hcd203040_compound15.24030.24030.2 | controllerType=0 controllerNumber=1 scan=24030 | 48.[illegible]972 |
| 20561:3 | 14 | K.YLGN[+2205.793]ATAIFFLPDEGK.L | HexNAc(4)Hex(5)Fuc(2)NeuAc(1) | N[+2206] | 1321.2314 | 3 | 3961.6796 | 3961.6881 | 0 | -2.1 | 268 | Specific | 496.76 | 371.1 | 371.1 | 7.45 | 28 | >sp\|P01009\|A1AT_HUMAN Alpha-1-antitrypsin OS=Homo sapiens OX=9606 GN=SERPINA1 PE=1 SV=3 | 642 | 20201113_xd_milk_1ng_glyco_hcd203040_compound15.24070.24070.3 | controllerType=0 controllerNumber=1 scan=24070 | 48.1357 |
| 20525:3 | 14 | K.YLGN[+2204.772]ATAIFFLPDEGK.L | HexNAc(4)Hex(5)NeuAc(2) | N[+2205] | 1320.9046 | 3 | 3960.6992 | 3960.6677 | 0 | 8 | 268 | Specific | 594.98 | 471.33 | 471.33 | 11.03 | 28 | >sp\|P01009\|A1AT_HUMAN Alpha-1-antitrypsin OS=Homo sapiens OX=9606 GN=SERPINA1 PE=1 SV=3 | 642 | 20201113_xd_milk_1ng_glyco_hcd203040_compound15.24032.24032.2 | controllerType=0 controllerNumber=1 scan=24032 | 48.0517 |
| 20564:3 | 14 | K.YLGN[+2205.793]ATAIFFLPDEGK.L | HexNAc(4)Hex(5)Fuc(2)NeuAc(1) | N[+2206] | 1321.2314 | 3 | 3961.6796 | 3961.6881 | 0 | -2.1 | 268 | Specific | 719.99 | 492.9 | 492.9 | 13.7 | 28 | >sp\|P01009\|A1AT_HUMAN Alpha-1-antitrypsin OS=Homo sapiens OX=9606 GN=SERPINA1 PE=1 SV=3 | 642 | 20201113_xd_milk_1ng_glyco_hcd203040_compound15.24073.24073.3 | controllerType=0 controllerNumber=1 scan=24073 | 48.1401 |
| 14427:3 | 16 | K.LKELPGVC[+57.021]N[+2059.735]ETMMALWEEC[+57.021]KPC[+57.021]LK.Q | HexNAc(4)Hex(5)Fuc(1)NeuAc(1) | C[+57]*3, N[+2060] | 1666.0542 | 3 | 4996.1479 | 4996.1196 | 0 | 5.7 | 95 | Specific | 296.66 | 282.85 | 282.85 | 6.08 | 27 | >sp\|P10909\|CLUS_HUMAN Clusterin OS=Homo sapiens OX=9606 GN=CLU PE=1 SV=1 | 1467 | 20201113_xd_milk_1ng_glyco_hcd203040_compound15.17673.17673.3 | controllerType=0 controllerNumber=1 scan=17673 | 35.5492 |
| 14425:3 | 16 | K.LKELPGVC[+57.021]N[+2059.735]ETMMALWEEC[+57.021]KPC[+57.021]LK.Q | HexNAc(4)Hex(5)Fuc(1)NeuAc(1) | C[+57]*3, N[+2060] | 1666.0542 | 3 | 4996.1479 | 4996.1196 | 0 | 5.7 | 95 | Specific | 362.75 | 362.75 | 362.75 | 7.06 | 27 | >sp\|P10909\|CLUS_HUMAN Clusterin OS=Homo sapiens OX=9606 GN=CLU PE=1 SV=1 | 1467 | 20201113_xd_milk_1ng_glyco_hcd203040_compound15.17671.17671.3 | controllerType=0 controllerNumber=1 scan=17671 | 35.5451 |
| 14435:4 | 16 | K.LKELPGVC[+57.021]N[+2059.735]ETMMALWEEC[+57.021]KPC[+57.021]LK.Q | HexNAc(4)Hex(5)Fuc(1)NeuAc(1) | C[+57]*3, N[+2060] | 1249.7881 | 4 | 4996.1305 | 4996.1196 | 0 | 2.2 | 95 | Specific | 688.01 | 688.01 | 688.01 | 14.18 | 27 | >sp\|P10909\|CLUS_HUMAN Clusterin OS=Homo sapiens OX=9606 GN=CLU PE=1 SV=1 | 1467 | 20201113_xd_milk_1ng_glyco_hcd203040_compound15.17682.17682.4 | controllerType=0 controllerNumber=1 scan=17682 | 35.5656 |
| 14438:4 | 16 | K.LKELPGVC[+57.021]N[+2059.735]ETMMALWEEC[+57.021]KPC[+57.021]LK.Q | HexNAc(4)Hex(5)Fuc(1)NeuAc(1) | C[+57]*3, N[+2060] | 1249.7881 | 4 | 4996.1305 | 4996.1196 | 0 | 2.2 | 95 | Specific | 831.43 | 828.12 | 828.12 | 17.18 | 27 | >sp\|P10909\|CLUS_HUMAN Clusterin OS=Homo sapiens OX=9606 GN=CLU PE=1 SV=1 | 1467 | 20201113_xd_milk_1ng_glyco_hcd203040_compound15.17685.17685.4 | controllerType=0 controllerNumber=1 scan=17685 | 35.5702 |
| 23786:3 | 16 | K.MLN[+2204.772]TSSLLEQLNEQFNWVSR.L | HexNAc(4)Hex(5)NeuAc(2) | N[+2205] | 1538.6627 | 3 | 4613.9734 | 4613.9592 | 0 | 3.1 | 352 | Specific | 421.76 | 301.11 | 301.11 | 8.16 | 27 | >sp\|P10909\|CLUS_HUMAN Clusterin OS=Homo sapiens OX=9606 GN=CLU PE=1 SV=1 | 1467 | 20201113_xd_milk_1ng_glyco_hcd203040_compound15.27431.27431.3 | controllerType=0 controllerNumber=1 scan=27431 | 55.0565 |
| 23784:3 | 16 | K.MLN[+2204.772]TSSLLEQLNEQFNWVSR.L | HexNAc(4)Hex(5)NeuAc(2) | N[+2205] | 1538.6627 | 3 | 4613.9734 | 4613.9592 | 0 | 3.1 | 352 | Specific | 579.26 | 459.96 | 459.96 | 12.19 | 27 | >sp\|P10909\|CLUS_HUMAN Clusterin OS=Homo sapiens OX=9606 GN=CLU PE=1 SV=1 | 1467 | 20201113_xd_milk_1ng_glyco_hcd203040_compound15.27429.27429.3 | controllerType=0 controllerNumber=1 scan=27429 | 55.0516 |
| 21577:3 | 16 | K.MLN[+1913.677]TSSLLEQLNEQFNWVSR.L | HexNAc(4)Hex(5)NeuAc(1) | N[+1914] | 1441.6375 | 3 | 4322.898 | 4322.8637 | 0 | 7.9 | 352 | Specific | 648.96 | 566.29 | 566.29 | 12.26 | 27 | >sp\|P10909\|CLUS_HUMAN Clusterin OS=Homo sapiens OX=9606 GN=CLU PE=1 SV=1 | 1467 | 20201113_xd_milk_1ng_glyco_hcd203040_compound15.25129.25129.3 | controllerType=0 controllerNumber=1 scan=25129 | 50.3952 |
| 21580:3 | 16 | K.MLN[+1913.677]TSSLLEQLNEQFNWVSR.L | HexNAc(4)Hex(5)NeuAc(1) | N[+1914] | 1441.6375 | 3 | 4322.898 | 4322.8637 | 0 | 7.9 | 352 | Specific | 738.88 | 507.46 | 507.46 | 14.26 | 27 | >sp\|P10909\|CLUS_HUMAN Clusterin OS=Homo sapiens OX=9606 GN=CLU PE=1 SV=1 | 1467 | 20201113_xd_milk_1ng_glyco_hcd203040_compound15.25132.25132.3 | controllerType=0 controllerNumber=1 scan=25132 | 50.4018 |
| 21231:3 | 16 | K.MLN[+2350.830]TSSLLEQLNEQFNWVSR.L | HexNAc(4)Hex(5)Fuc(1)NeuAc(2) | N[+2351] | 1587.357 | 3 | 4760.0564 | 4760.0171 | 0 | 8.3 | 352 | Specific | 787.44 | 560.57 | 560.57 | 15.26 | 27 | >sp\|P10909\|CLUS_HUMAN Clusterin OS=Homo sapiens OX=9606 GN=CLU PE=1 SV=1 | 1467 | 20201113_xd_milk_1ng_glyco_hcd203040_compound15.24769.24769.3 | controllerType=0 controllerNumber=1 scan=24769 | 49.6557 |
| 21233:3 | 16 | K.MLN[+2350.830]TSSLLEQLNEQFNWVSR.L | HexNAc(4)Hex(5)Fuc(1)NeuAc(2) | N[+2351] | 1587.357 | 3 | 4760.0564 | 4760.0171 | 0 | 8.3 | 352 | Specific | 908.94 | 650.42 | 650.42 | 18.26 | 27 | >sp\|P10909\|CLUS_HUMAN Clusterin OS=Homo sapiens OX=9606 GN=CLU PE=1 SV=1 | 1467 | 20201113_xd_milk_1ng_glyco_hcd203040_compound15.24771.24771.3 | controllerType=0 controllerNumber=1 scan=24771 | 49.6629 |
| 02205:3 | 20 | R.LRN[+1216.423]VSWATGR.S | HexNAc(2)Hex(5) | N[+1216] | 792.6919 | 3 | 2376.061 | 2376.0559 | 0 | 2.1 | 149 | Specific | 286.22 | 127.69 | 127.69 | 5.27 | 20 | >sp\|P08571\|CD14_HUMAN Monocyte differentiation antigen CD14 OS=Homo sapiens OX=9606 GN=CD14 PE=1 SV=2 | 1232 | 20201113_xd_milk_1ng_glyco_hcd203040_compound15.4990.4990.3 | controllerType=0 controllerNumber=1 scan=4990 | 14.3743 |
| 23925:4 | 41 | A.[+42.011]SPTSPKVFPLSLDSTPQDGNVVVAC[+57.021]LVQGFFPQEPLSVTWSESGQN | HexNAc(4)Hex(4) | .[42], C[+57], N[+1461] | 1719.792 | 4 | 6876.146 | 6876.213 | 0 | -9.7 | 2 | NRagged | 274.16 | 235.53 | 235.53 | 3.25 | 17 | >sp\|P01877\|IGHA2_HUMAN Immunoglobulin heavy constant alpha 2 OS=Homo sapiens OX=9606 GN=IGHA2 PE=1 SV=4 | 576 | 20201113_xd_milk_1ng_glyco_hcd203040_compound15.27576.27576.4 | controllerType=0 controllerNumber=1 scan=27576 | 55.366 |
| 23927:4 | 41 | A.[+42.011]SPTSPKVFPLSLDSTPQDGNVVVAC[+57.021]LVQGFFPQEPLSVTWSESGQN | HexNAc(4)Hex(4) | .[42], C[+57], N[+1461] | 1719.792 | 4 | 6876.146 | 6876.213 | 0 | -9.7 | 2 | NRagged | 320.85 | 306.51 | 306.51 | 3.58 | 17 | >sp\|P01877\|IGHA2_HUMAN Immunoglobulin heavy constant alpha 2 OS=Homo sapiens OX=9606 GN=IGHA2 PE=1 SV=4 | 576 | 20201113_xd_milk_1ng_glyco_hcd203040_compound15.27578.27578.4 | controllerType=0 controllerNumber=1 scan=27578 | 55.3711 |
| 22251:4 | 41 | K.VFPLSLDSTPQDGNVVVAC[+57.021]LVQGFFPQEPLSVTWSESGQN[+1971.719]VTAI | HexNAc(5)Hex(5)Fuc(1) | C[+57], N[+1972] | 1687.781 | 4 | 6748.102 | 6748.0803 | 0 | 3.2 | 8 | Specific | 415.88 | 405.88 | 405.88 | 10.76 | 17 | >sp\|P01877\|IGHA2_HUMAN Immunoglobulin heavy constant alpha 2 OS=Homo sapiens OX=9606 GN=IGHA2 PE=1 SV=4 | 576 | 20201113_xd_milk_1ng_glyco_hcd203040_compound15.25831.25831.4 | controllerType=0 controllerNumber=1 scan=25831 | 51.8511 |
| 22253:4 | 41 | K.VFPLSLDSTPQDGNVVVAC[+57.021]LVQGFFPQEPLSVTWSESGQN[+1971.719]VTAI | HexNAc(5)Hex(5)Fuc(1) | C[+57], N[+1972] | 1687.781 | 4 | 6748.102 | 6748.0803 | 0 | 3.2 | 8 | Specific | 468.6 | 433.95 | 433.95 | 10.96 | 17 | >sp\|P01877\|IGHA2_HUMAN Immunoglobulin heavy constant alpha 2 OS=Homo sapiens OX=9606 GN=IGHA2 PE=1 SV=4 | 576 | 20201113_xd_milk_1ng_glyco_hcd203040_compound15.25833.25833.4 | controllerType=0 controllerNumber=1 scan=25833 | 51.8567 |
| 23857:5 | 41 | K.VFPLSLDSTPQDGNVVVAC[+57.021]LVQGFFPQEPLSVTWSESGQN[+2262.814]VTAI | HexNAc(5)Hex(5)Fuc(1)NeuAc(1) | C[+57], N[+2263] | 1408.6414 | 5 | 7039.1779 | 7039.1757 | 0 | 0.3 | 8 | Specific | 476.8 | 476.8 | 476.8 | 10.57 | 17 | >sp\|P01877\|IGHA2_HUMAN Immunoglobulin heavy constant alpha 2 OS=Homo sapiens OX=9606 GN=IGHA2 PE=1 SV=4 | 576 | 20201113_xd_milk_1ng_glyco_hcd203040_compound15.27504.27504.5 | controllerType=0 controllerNumber=1 scan=27504 | 55.2002 |
| 23907:5 | 41 | K.VFPLSLDSTPQDGNVVVAC[+57.021]LVQGFFPQEPLSVTWSESGQN[+2262.814]VTAI | HexNAc(5)Hex(5)Fuc(1)NeuAc(1) | C[+57], N[+2263] | 1408.6408 | 5 | 7039.1748 | 7039.1757 | 0 | -0.1 | 8 | Specific | 503.7 | 503.7 | 503.7 | 10.73 | 17 | >sp\|P01877\|IGHA2_HUMAN Immunoglobulin heavy constant alpha 2 OS=Homo sapiens OX=9606 GN=IGHA2 PE=1 SV=4 | 576 | 20201113_xd_milk_1ng_glyco_hcd203040_compound15.27557.27557.5 | controllerType=0 controllerNumber=1 scan=27557 | 55.3318 |
| 23905:5 | 41 | K.VFPLSLDSTPQDGNVVVAC[+57.021]LVQGFFPQEPLSVTWSESGQN[+2262.814]VTAI | HexNAc(5)Hex(5)Fuc(1)NeuAc(1) | C[+57], N[+2263] | 1408.6408 | 5 | 7039.1748 | 7039.1757 | 0 | -0.1 | 8 | Specific | 525.87 | 525.87 | 525.87 | 10.86 | 17 | >sp\|P01877\|IGHA2_HUMAN Immunoglobulin heavy constant alpha 2 OS=Homo sapiens OX=9606 GN=IGHA2 PE=1 SV=4 | 576 | 20201113_xd_milk_1ng_glyco_hcd203040_compound15.27555.27555.5 | controllerType=0 controllerNumber=1 scan=27555 | 55.3267 |
| 22336:4 | 41 | K.VFPLSLDSTPQDGNVVVAC[+57.021]LVQGFFPQEPLSVTWSESGQN[+1647.613]VTAI | HexNAc(5)Hex(3)Fuc(1) | C[+57], N[+1648] | 1606.7538 | 4 | 6423.9933 | 6423.9746 | 0 | 2.9 | 8 | Specific | 779.15 | 709.52 | 709.52 | 16.86 | 17 | >sp\|P01877\|IGHA2_HUMAN Immunoglobulin heavy constant alpha 2 OS=Homo sapiens OX=9606 GN=IGHA2 PE=1 SV=4 | 576 | 20201113_xd_milk_1ng_glyco_hcd203040_compound15.25920.25920.4 | controllerType=0 controllerNumber=1 scan=25920 | 52.0237 |
| 22339:4 | 41 | K.VFPLSLDSTPQDGNVVVAC[+57.021]LVQGFFPQEPLSVTWSESGQN[+1647.613]VTAI | HexNAc(5)Hex(3)Fuc(1) | C[+57], N[+1648] | 1606.7538 | 4 | 6423.9933 | 6423.9746 | 0 | 2.9 | 8 | Specific | 943.09 | 930.09 | 930.09 | 19.86 | 17 | >sp\|P01877\|IGHA2_HUMAN Immunoglobulin heavy constant alpha 2 OS=Homo sapiens OX=9606 GN=IGHA2 PE=1 SV=4 | 576 | 20201113_xd_milk_1ng_glyco_hcd203040_compound15.25923.25923.4 | controllerType=0 controllerNumber=1 scan=25923 | 52.0317 |
| 22280:4 | 41 | K.VFPLSLDSTPQDGNVVVAC[+57.021]LVQGFFPQEPLSVTWSESGQN[+1809.666]VTAI | HexNAc(5)Hex(4)Fuc(1) | C[+57], N[+1810] | 1647.2662 | 4 | 6586.0431 | 6586.0275 | 0 | 2.4 | 8 | Specific | 946.56 | 924.05 | 924.05 | 20.06 | 17 | >sp\|P01877\|IGHA2_HUMAN Immunoglobulin heavy constant alpha 2 OS=Homo sapiens OX=9606 GN=IGHA2 PE=1 SV=4 | 576 | 20201113_xd_milk_1ng_glyco_hcd203040_compound15.25862.25862.4 | controllerType=0 controllerNumber=1 scan=25862 | 51.9145 |
| 05021:2 | 41 | K.TPLTAN[+2263.835]ITK.S | HexNAc(5)Hex(5)Fuc(3) | N[+2264] | 1611.6906 | 2 | 3222.3739 | 3222.3915 | 0 | -5.4 | 200 | Specific | 202.52 | 180.75 | 180.75 | 3.74 | 17 | >sp\|P01877\|IGHA2_HUMAN Immunoglobulin heavy constant alpha 2 OS=Homo sapiens OX=9606 GN=IGHA2 PE=1 SV=4 | 576 | 20201113_xd_milk_1ng_glyco_hcd203040_compound15.7909.7909.2 | controllerType=0 controllerNumber=1 scan=7909 | 19.2259 |
| 04998:3 | 41 | K.TPLTAN[+2262.814]ITK.S | HexNAc(5)Hex(5)Fuc(1)NeuAc(1) | N[+2263] | 1074.4627 | 3 | 3221.3735 | 3221.3711 | 0 | 0.7 | 200 | Specific | 215.38 | 64.79 | 64.79 | 3.65 | 17 | >sp\|P01877\|IGHA2_HUMAN Immunoglobulin heavy constant alpha 2 OS=Homo sapiens OX=9606 GN=IGHA2 PE=1 SV=4 | 576 | 20201113_xd_milk_1ng_glyco_hcd203040_compound15.7886.7886.3 | controllerType=0 controllerNumber=1 scan=7886 | 19.1883 |
| 05023:2 | 41 | K.TPLTAN[+2263.835]ITK.S | HexNAc(5)Hex(5)Fuc(3) | N[+2264] | 1611.6906 | 2 | 3222.3739 | 3222.3915 | 0 | -5.4 | 200 | Specific | 220.68 | 179.88 | 179.88 | 3.74 | 17 | >sp\|P01877\|IGHA2_HUMAN Immunoglobulin heavy constant alpha 2 OS=Homo sapiens OX=9606 GN=IGHA2 PE=1 SV=4 | 576 | 20201113_xd_milk_1ng_glyco_hcd203040_compound15.7911.7911.2 | controllerType=0 controllerNumber=1 scan=7911 | 19.2299 |
| 05001:3 | 41 | K.TPLTAN[+2262.814]ITK.S | HexNAc(5)Hex(5)Fuc(1)NeuAc(1) | N[+2263] | 1074.4627 | 3 | 3221.3735 | 3221.3711 | 0 | 0.7 | 200 | Specific | 282.36 | 133.97 | 133.97 | 5.56 | 17 | >sp\|P01877\|IGHA2_HUMAN Immunoglobulin heavy constant alpha 2 OS=Homo sapiens OX=9606 GN=IGHA2 PE=1 SV=4 | 576 | 20201113_xd_milk_1ng_glyco_hcd203040_compound15.7889.7889.3 | controllerType=0 controllerNumber=1 scan=7889 | 19.1969 |
| 04970:3 | 41 | K.TPLTAN[+2262.814]ITK.S | HexNAc(5)Hex(5)Fuc(1)NeuAc(1) | N[+2263] | 1074.4627 | 3 | 3221.3735 | 3221.3711 | 0 | 0.7 | 200 | Specific | 342.42 | 101.52 | 101.52 | 5.71 | 17 | >sp\|P01877\|IGHA2_HUMAN Immunoglobulin heavy constant alpha 2 OS=Homo sapiens OX=9606 GN=IGHA2 PE=1 SV=4 | 576 | 20201113_xd_milk_1ng_glyco_hcd203040_compound15.7857.7857.3 | controllerType=0 controllerNumber=1 scan=7857 | 19.1424 |
| 03130:3 | 41 | K.TPLTAN[+1971.719]ITK.S | HexNAc(5)Hex(5)Fuc(1) | N[+1972] | 977.4337 | 3 | 2930.2865 | 2930.2757 | 0 | 3.7 | 200 | Specific | 418.18 | 154.86 | 154.86 | 6.48 | 17 | >sp\|P01877\|IGHA2_HUMAN Immunoglobulin heavy constant alpha 2 OS=Homo sapiens OX=9606 GN=IGHA2 PE=1 SV=4 | 576 | 20201113_xd_milk_1ng_glyco_hcd203040_compound15.5948.5948.3 | controllerType=0 controllerNumber=1 scan=5948 | 15.9771 |
| 03214:3 | 41 | K.TPLTAN[+1809.666]ITK.S | HexNAc(5)Hex(4)Fuc(1) | N[+1810] | 923.4143 | 3 | 2768.2283 | 2768.2228 | 0 | 2 | 200 | Specific | 453.33 | 174.85 | 174.85 | 7.13 | 17 | >sp\|P01877\|IGHA2_HUMAN Immunoglobulin heavy constant alpha 2 OS=Homo sapiens OX=9606 GN=IGHA2 PE=1 SV=4 | 576 | 20201113_xd_milk_1ng_glyco_hcd203040_compound15.6035.6035.3 | controllerType=0 controllerNumber=1 scan=6035 | 16.1303 |
| 03145:2 | 41 | K.TPLTAN[+1971.719]ITK.S | HexNAc(5)Hex(5)Fuc(1) | N[+1972] | 1465.6475 | 2 | 2930.2877 | 2930.2757 | 0 | 4.1 | 200 | Specific | 457.79 | 256.27 | 256.27 | 7.29 | 17 | >sp\|P01877\|IGHA2_HUMAN Immunoglobulin heavy constant alpha 2 OS=Homo sapiens OX=9606 GN=IGHA2 PE=1 SV=4 | 576 | 20201113_xd_milk_1ng_glyco_hcd203040_compound15.5963.5963.2 | controllerType=0 controllerNumber=1 scan=5963 | 16.0032 |
| 03147:2 | 41 | K.TPLTAN[+1971.719]ITK.S | HexNAc(5)Hex(5)Fuc(1) | N[+1972] | 1465.6475 | 2 | 2930.2877 | 2930.2757 | 0 | 4.1 | 200 | Specific | 465.11 | 266.13 | 266.13 | 7.29 | 17 | >sp\|P01877\|IGHA2_HUMAN Immunoglobulin heavy constant alpha 2 OS=Homo sapiens OX=9606 GN=IGHA2 PE=1 SV=4 | 576 | 20201113_xd_milk_1ng_glyco_hcd203040_compound15.5965.5965.2 | controllerType=0 controllerNumber=1 scan=5965 | 16.007 |
| 04973:3 | 41 | K.TPLTAN[+2262.814]ITK.S | HexNAc(5)Hex(5)Fuc(1)NeuAc(1) | N[+2263] | 1074.4627 | 3 | 3221.3735 | 3221.3711 | 0 | 0.7 | 200 | Specific | 474.61 | 244.22 | 244.22 | 7.54 | 17 | >sp\|P01877\|IGHA2_HUMAN Immunoglobulin heavy constant alpha 2 OS=Homo sapiens OX=9606 GN=IGHA2 PE=1 SV=4 | 576 | 20201113_xd_milk_1ng_glyco_hcd203040_compound15.7860.7860.3 | controllerType=0 controllerNumber=1 scan=7860 | 19.1478 |
| 03132:3 | 41 | K.TPLTAN[+1971.719]ITK.S | HexNAc(5)Hex(5)Fuc(1) | N[+1972] | 977.4337 | 3 | 2930.2865 | 2930.2757 | 0 | 3.7 | 200 | Specific | 474.97 | 275.75 | 275.75 | 7.68 | 17 | >sp\|P01877\|IGHA2_HUMAN Immunoglobulin heavy constant alpha 2 OS=Homo sapiens OX=9606 GN=IGHA2 PE=1 SV=4 | 576 | 20201113_xd_milk_1ng_glyco_hcd203040_compound15.5950.5950.3 | controllerType=0 controllerNumber=1 scan=5950 | 15.9816 |
| 03266:3 | 41 | K.TPLTAN[+1647.613]ITK.S | HexNAc(5)Hex(3)Fuc(1) | N[+1648] | 869.3967 | 3 | 2606.1755 | 2606.17 | 0 | 2.1 | 200 | Specific | 481.19 | 178.96 | 178.96 | 7.21 | 17 | >sp\|P01877\|IGHA2_HUMAN Immunoglobulin heavy constant alpha 2 OS=Homo sapiens OX=9606 GN=IGHA2 PE=1 SV=4 | 576 | 20201113_xd_milk_1ng_glyco_hcd203040_compound15.6089.6089.3 | controllerType=0 controllerNumber=1 scan=6089 | 16.2322 |
| 03272:2 | 41 | K.TPLTAN[+1501.555]ITK.S | HexNAc(5)Hex(3) | N[+1502] | 1230.5617 | 2 | 2460.1161 | 2460.1121 | 0 | 1.6 | 200 | Specific | 494.31 | 279.42 | 279.42 | 8.06 | 17 | >sp\|P01877\|IGHA2_HUMAN Immunoglobulin heavy constant alpha 2 OS=Homo sapiens OX=9606 GN=IGHA2 PE=1 SV=4 | 576 | 20201113_xd_milk_1ng_glyco_hcd203040_compound15.6095.6095.2 | controllerType=0 controllerNumber=1 scan=6095 | 16.2564 |
| 03268:3 | 41 | K.TPLTAN[+1647.613]ITK.S | HexNAc(5)Hex(3)Fuc(1) | N[+1648] | 869.3967 | 3 | 2606.1755 | 2606.17 | 0 | 2.1 | 200 | Specific | 546.59 | 250.49 | 250.49 | 8.28 | 17 | >sp\|P01877\|IGHA2_HUMAN Immunoglobulin heavy constant alpha 2 OS=Homo sapiens OX=9606 GN=IGHA2 PE=1 SV=4 | 576 | 20201113_xd_milk_1ng_glyco_hcd203040_compound15.6091.6091.3 | controllerType=0 controllerNumber=1 scan=6091 | 16.249 |
| 03217:3 | 41 | K.TPLTAN[+1809.666]ITK.S | HexNAc(5)Hex(4)Fuc(1) | N[+1810] | 923.4143 | 3 | 2768.2283 | 2768.2228 | 0 | 2 | 200 | Specific | 551.95 | 269.41 | 269.41 | 8.31 | 17 | >sp\|P01877\|IGHA2_HUMAN Immunoglobulin heavy constant alpha 2 OS=Homo sapiens OX=9606 GN=IGHA2 PE=1 SV=4 | 576 | 20201113_xd_milk_1ng_glyco_hcd203040_compound15.6038.6038.3 | controllerType=0 controllerNumber=1 scan=6038 | 16.1352 |
| 03270:2 | 41 | K.TPLTAN[+1501.555]ITK.S | HexNAc(5)Hex(3) | N[+1502] | 1230.5617 | 2 | 2460.1161 | 2460.1121 | 0 | 1.6 | 200 | Specific | 554.65 | 341.15 | 341.15 | 8.79 | 17 | >sp\|P01877\|IGHA2_HUMAN Immunoglobulin heavy constant alpha 2 OS=Homo sapiens OX=9606 GN=IGHA2 PE=1 SV=4 | 576 | 20201113_xd_milk_1ng_glyco_hcd203040_compound15.6093.6093.2 | controllerType=0 controllerNumber=1 scan=6093 | 16.2515 |
| 03247:2 | 41 | K.TPLTAN[+1647.613]ITK.S | HexNAc(5)Hex(3)Fuc(1) | N[+1648] | 1303.5918 | 2 | 2606.1764 | 2606.17 | 0 | 2.5 | 200 | Specific | 596.47 | 324.61 | 324.61 | 10.03 | 17 | >sp\|P01877\|IGHA2_HUMAN Immunoglobulin heavy constant alpha 2 OS=Homo sapiens OX=9606 GN=IGHA2 PE=1 SV=4 | 576 | 20201113_xd_milk_1ng_glyco_hcd203040_compound15.6069.6069.2 | controllerType=0 controllerNumber=1 scan=6069 | 16.1972 |
| 03245:2 | 41 | K.TPLTAN[+1647.613]ITK.S | HexNAc(5)Hex(3)Fuc(1) | N[+1648] | 1303.5918 | 2 | 2606.1764 | 2606.17 | 0 | 2.5 | 200 | Specific | 619.83 | 352.93 | 352.93 | 11.27 | 17 | >sp\|P01877\|IGHA2_HUMAN Immunoglobulin heavy constant alpha 2 OS=Homo sapiens OX=9606 GN=IGHA2 PE=1 SV=4 | 576 | 20201113_xd_milk_1ng_glyco_hcd203040_compound15.6067.6067.2 | controllerType=0 controllerNumber=1 scan=6067 | 16.1898 |
| 03292:2 | 41 | K.TPLTAN[+1444.534]ITK.S | HexNAc(4)Hex(3)Fuc(1) | N[+1445] | 1202.053 | 2 | 2403.0988 | 2403.0906 | 0 | 3.4 | 200 | Specific | 630.39 | 309.14 | 309.14 | 10.82 | 17 | >sp\|P01877\|IGHA2_HUMAN Immunoglobulin heavy constant alpha 2 OS=Homo sapiens OX=9606 GN=IGHA2 PE=1 SV=4 | 576 | 20201113_xd_milk_1ng_glyco_hcd203040_compound15.6116.6116.2 | controllerType=0 controllerNumber=1 scan=6116 | 16.3058 |
| 03186:2 | 41 | K.TPLTAN[+1809.666]ITK.S | HexNAc(5)Hex(4)Fuc(1) | N[+1810] | 1384.6172 | 2 | 2768.2272 | 2768.2228 | 0 | 1.6 | 200 | Specific | 640.21 | 334.47 | 334.47 | 11.27 | 17 | >sp\|P01877\|IGHA2_HUMAN Immunoglobulin heavy constant alpha 2 OS=Homo sapiens OX=9606 GN=IGHA2 PE=1 SV=4 | 576 | 20201113_xd_milk_1ng_glyco_hcd203040_compound15.6006.6006.2 | controllerType=0 controllerNumber=1 scan=6006 | 16.0835 |
| 03290:2 | 41 | K.TPLTAN[+1444.534]ITK.S | HexNAc(4)Hex(3)Fuc(1) | N[+1445] | 1202.053 | 2 | 2403.0988 | 2403.0906 | 0 | 3.4 | 200 | Specific | 650.89 | 357.34 | 357.34 | 12.24 | 17 | >sp\|P01877\|IGHA2_HUMAN Immunoglobulin heavy constant alpha 2 OS=Homo sapiens OX=9606 GN=IGHA2 PE=1 SV=4 | 576 | 20201113_xd_milk_1ng_glyco_hcd203040_compound15.6114.6114.2 | controllerType=0 controllerNumber=1 scan=6114 | 16.3004 |
| 03184:2 | 41 | K.TPLTAN[+1809.666]ITK.S | HexNAc(5)Hex(4)Fuc(1) | N[+1810] | 1384.6172 | 2 | 2768.2272 | 2768.2228 | 0 | 1.6 | 200 | Specific | 654.85 | 355.47 | 355.47 | 12.27 | 17 | >sp\|P01877\|IGHA2_HUMAN Immunoglobulin heavy constant alpha 2 OS=Homo sapiens OX=9606 GN=IGHA2 PE=1 SV=4 | 576 | 20201113_xd_milk_1ng_glyco_hcd203040_compound15.6004.6004.2 | controllerType=0 controllerNumber=1 scan=6004 | 16.0785 |
| 10125:3 | 53 | R.IIVPLNNREN[+1663.608]ISDPTSPLR.T | HexNAc(5)Hex(4) | N[+1664] | 1271.2631 | 3 | 3811.7748 | 3811.7853 | 0 | -2.8 | 62 | Specific | 240.18 | 240.18 | 240.18 | 5.32 | 14 | >sp\|P01591\|IGJ_HUMAN Immunoglobulin J chain OS=Homo sapiens OX=9606 GN=JCHAIN PE=1 SV=4 | 1392 | 20201113_xd_milk_1ng_glyco_hcd203040_compound15.13204.13204.3 | controllerType=0 controllerNumber=1 scan=13204 | 27.7764 |
| 10127:3 | 53 | R.IIVPLNNREN[+1663.608]ISDPTSPLR.T | HexNAc(5)Hex(4) | N[+1664] | 1271.2631 | 3 | 3811.7748 | 3811.7853 | 0 | -2.8 | 62 | Specific | 241.29 | 174.77 | 174.77 | 3.47 | 14 | >sp\|P01591\|IGJ_HUMAN Immunoglobulin J chain OS=Homo sapiens OX=9606 GN=JCHAIN PE=1 SV=4 | 1392 | 20201113_xd_milk_1ng_glyco_hcd203040_compound15.13206.13206.3 | controllerType=0 controllerNumber=1 scan=13206 | 27.7831 |
| 11733:3 | 53 | R.IIVPLNNREN[+1913.677]ISDPTSPLR.T | HexNAc(4)Hex(5)NeuAc(1) | N[+1914] | 1354.6255 | 3 | 4061.8621 | 4061.8542 | 0 | 1.9 | 62 | Specific | 339.57 | 307.78 | 307.78 | 7.73 | 14 | >sp\|P01591\|IGJ_HUMAN Immunoglobulin J chain OS=Homo sapiens OX=9606 GN=JCHAIN PE=1 SV=4 | 1392 | 20201113_xd_milk_1ng_glyco_hcd203040_compound15.14868.14868.3 | controllerType=0 controllerNumber=1 scan=14868 | 30.5892 |
| 13720:3 | 53 | R.IIVPLNNREN[+2204.772]ISDPTSPLR.T | HexNAc(4)Hex(5)NeuAc(2) | N[+2205] | 1451.6607 | 3 | 4352.9676 | 4352.9496 | 0 | 4.1 | 62 | Specific | 454.25 | 437.73 | 437.73 | 9.44 | 14 | >sp\|P01591\|IGJ_HUMAN Immunoglobulin J chain OS=Homo sapiens OX=9606 GN=JCHAIN PE=1 SV=4 | 1392 | 20201113_xd_milk_1ng_glyco_hcd203040_compound15.16938.16938.3 | controllerType=0 controllerNumber=1 scan=16938 | 34.0957 |
| 11666:3 | 53 | R.IIVPLNNREN[+2059.735]ISDPTSPLR.T | HexNAc(4)Hex(5)Fuc(1)NeuAc(1) | N[+2060] | 1403.3122 | 3 | 4207.922 | 4207.9121 | 0 | 2.4 | 62 | Specific | 541.65 | 522.44 | 522.44 | 10.45 | 14 | >sp\|P01591\|IGJ_HUMAN Immunoglobulin J chain OS=Homo sapiens OX=9606 GN=JCHAIN PE=1 SV=4 | 1392 | 20201113_xd_milk_1ng_glyco_hcd203040_compound15.14798.14798.3 | controllerType=0 controllerNumber=1 scan=14798 | 30.4049 |
| 11664:3 | 53 | R.IIVPLNNREN[+2059.735]ISDPTSPLR.T | HexNAc(4)Hex(5)Fuc(1)NeuAc(1) | N[+2060] | 1403.3122 | 3 | 4207.922 | 4207.9121 | 0 | 2.4 | 62 | Specific | 544.41 | 544.41 | 544.41 | 10.45 | 14 | >sp\|P01591\|IGJ_HUMAN Immunoglobulin J chain OS=Homo sapiens OX=9606 GN=JCHAIN PE=1 SV=4 | 1392 | 20201113_xd_milk_1ng_glyco_hcd203040_compound15.14796.14796.3 | controllerType=0 controllerNumber=1 scan=14796 | 30.3947 |
| 13718:3 | 53 | R.IIVPLNNREN[+2204.772]ISDPTSPLR.T | HexNAc(4)Hex(5)NeuAc(2) | N[+2205] | 1451.6607 | 3 | 4352.9676 | 4352.9496 | 0 | 4.1 | 62 | Specific | 545.82 | 532.37 | 532.37 | 9.94 | 14 | >sp\|P01591\|IGJ_HUMAN Immunoglobulin J chain OS=Homo sapiens OX=9606 GN=JCHAIN PE=1 SV=4 | 1392 | 20201113_xd_milk_1ng_glyco_hcd203040_compound15.16936.16936.3 | controllerType=0 controllerNumber=1 scan=16936 | 34.0914 |
| 10068:3 | 53 | R.IIVPLNNREN[+1663.608]ISDPTSPLR.T | HexNAc(5)Hex(4) | N[+1664] | 1271.2671 | 3 | 3811.7869 | 3811.7853 | 0 | 0.4 | 62 | Specific | 614.75 | 602.5 | 602.5 | 12.35 | 14 | >sp\|P01591\|IGJ_HUMAN Immunoglobulin J chain OS=Homo sapiens OX=9606 GN=JCHAIN PE=1 SV=4 | 1392 | 20201113_xd_milk_1ng_glyco_hcd203040_compound15.13144.13144.3 | controllerType=0 controllerNumber=1 scan=13144 | 27.641 |
| 10066:3 | 53 | R.IIVPLNNREN[+1663.608]ISDPTSPLR.T | HexNAc(5)Hex(4) | N[+1664] | 1271.2671 | 3 | 3811.7869 | 3811.7853 | 0 | 0.4 | 62 | Specific | 625.6 | 593.94 | 593.94 | 12.35 | 14 | >sp\|P01591\|IGJ_HUMAN Immunoglobulin J chain OS=Homo sapiens OX=9606 GN=JCHAIN PE=1 SV=4 | 1392 | 20201113_xd_milk_1ng_glyco_hcd203040_compound15.13142.13142.3 | controllerType=0 controllerNumber=1 scan=13142 | 27.6363 |
| 06917:3 | 216 | R.[+42.011]VEDAYILTC[+57.021]N[+730.264]VSLEYEK.T | HexNAc(2)Hex(2) | .[42], C[+57], N[+730] | 940.0947 | 3 | 2818.2695 | 2818.2486 | 0 | 7.4 | 225 | Specific | 268.43 | 209.99 | 209.99 | 3.18 | 8 | >sp\|P40227\|TCPZ_HUMAN T-complex protein 1 subunit zeta OS=Homo sapiens OX=9606 GN=CCT6A PE=1 SV=3 | 1115 | 20201113_xd_milk_1ng_glyco_hcd203040_compound15.9868.9868.3 | controllerType=0 controllerNumber=1 scan=9868 | 22.3519 |
| 03686:4 | 1 | R.C[+57.021]IN[+2059.735]GTC[+57.021]YC[+57.021]EEGFTGEDC[+57.021]GKPTC[ | HexNAc(4)Hex(5)Fuc(1)NeuAc(1) | C[+57]*6, N[+2060] | 1430.5519 | 4 | 5719.1857 | 5719.1633 | 0 | 3.9 | 325 | Specific | 290.88 | 290.88 | 290.88 | 7.4 | 171 | >sp\|P24821\|TENA_HUMAN Tenascin OS=Homo sapiens OX=9606 GN=TNC PE=1 SV=3 | 486 | 20201118_ly_colostrum_1ng_glyco_hcd203040_compound15.6702.6702.4 | controllerType=0 controllerNumber=1 scan=6702 | 17.7877 |
| 03668:4 | 1 | R.C[+57.021]IN[+2059.735]GTC[+57.021]YC[+57.021]EEGFTGEDC[+57.021]GKPTC[ | HexNAc(4)Hex(5)Fuc(1)NeuAc(1) | C[+57]*6, N[+2060] | 1430.5516 | 4 | 5719.1847 | 5719.1633 | 0 | 3.8 | 325 | Specific | 313.35 | 313.35 | 313.35 | 7.7 | 171 | >sp\|P24821\|TENA_HUMAN Tenascin OS=Homo sapiens OX=9606 GN=TNC PE=1 SV=3 | 486 | 20201118_ly_colostrum_1ng_glyco_hcd203040_compound15.6683.6683.4 | controllerType=0 controllerNumber=1 scan=6683 | 17.7541 |
| 03670:4 | 1 | R.C[+57.021]IN[+2059.735]GTC[+57.021]YC[+57.021]EEGFTGEDC[+57.021]GKPTC[ | HexNAc(4)Hex(5)Fuc(1)NeuAc(1) | C[+57]*6, N[+2060] | 1430.5516 | 4 | 5719.1847 | 5719.1633 | 0 | 3.8 | 325 | Specific | 435.49 | 435.49 | 435.49 | 9.94 | 171 | >sp\|P24821\|TENA_HUMAN Tenascin OS=Homo sapiens OX=9606 GN=TNC PE=1 SV=3 | 486 | 20201118_ly_colostrum_1ng_glyco_hcd203040_compound15.6685.6685.4 | controllerType=0 controllerNumber=1 scan=6685 | 17.7577 |
| 04565:4 | 1 | R.C[+57.021]IN[+2350.830]GTC[+57.021]YC[+57.021]EEGFTGEDC[+57.021]GKPTC[ | HexNAc(4)Hex(5)Fuc(1)NeuAc(2) | C[+57]*6, N[+2351] | 1503.3243 | 4 | 6010.2755 | 6010.2587 | 0 | 2.8 | 325 | Specific | 445.14 | 445.14 | 445.14 | 10.1 | 171 | >sp\|P24821\|TENA_HUMAN Tenascin OS=Homo sapiens OX=9606 GN=TNC PE=1 SV=3 | 486 | 20201118_ly_colostrum_1ng_glyco_hcd203040_compound15.7614.7614.4 | controllerType=0 controllerNumber=1 scan=7614 | 19.3764 |
| 04567:4 | 1 | R.C[+57.021]IN[+2350.830]GTC[+57.021]YC[+57.021]EEGFTGEDC[+57.021]GKPTC[ | HexNAc(4)Hex(5)Fuc(1)NeuAc(2) | C[+57]*6, N[+2351] | 1503.3243 | 4 | 6010.2755 | 6010.2587 | 0 | 2.8 | 325 | Specific | 473.38 | 473.38 | 473.38 | 10.17 | 171 | >sp\|P24821\|TENA_HUMAN Tenascin OS=Homo sapiens OX=9606 GN=TNC PE=1 SV=3 | 486 | 20201118_ly_colostrum_1ng_glyco_hcd203040_compound15.7616.7616.4 | controllerType=0 controllerNumber=1 scan=7616 | 19.3802 |
| 03671:5 | 1 | R.C[+57.021]IN[+2059.735]GTC[+57.021]YC[+57.021]EEGFTGEDC[+57.021]GKPTC[ | HexNAc(4)Hex(5)Fuc(1)NeuAc(1) | C[+57]*6, N[+2060] | 1144.6428 | 5 | 5719.185 | 5719.1633 | 0 | 3.8 | 325 | Specific | 494.69 | 494.69 | 494.69 | 10.69 | 171 | >sp\|P24821\|TENA_HUMAN Tenascin OS=Homo sapiens OX=9606 GN=TNC PE=1 SV=3 | 486 | 20201118_ly_colostrum_1ng_glyco_hcd203040_compound15.6686.6686.5 | controllerType=0 controllerNumber=1 scan=6686 | 17.7605 |
| 03669:5 | 1 | R.C[+57.021]IN[+2059.735]GTC[+57.021]YC[+57.021]EEGFTGEDC[+57.021]GKPTC[ | HexNAc(4)Hex(5)Fuc(1)NeuAc(1) | C[+57]*6, N[+2060] | 1144.6428 | 5 | 5719.185 | 5719.1633 | 0 | 3.8 | 325 | Specific | 563.97 | 563.97 | 563.97 | 11.22 | 171 | >sp\|P24821\|TENA_HUMAN Tenascin OS=Homo sapiens OX=9606 GN=TNC PE=1 SV=3 | 486 | 20201118_ly_colostrum_1ng_glyco_hcd203040_compound15.6684.6684.5 | controllerType=0 controllerNumber=1 scan=6684 | 17.7557 |
| 03600:4 | 1 | R.C[+57.021]IN[+2351.851]GTC[+57.021]YC[+57.021]EEGFTGEDC[+57.021]GKPTC[ | HexNAc(4)Hex(5)Fuc(3)NeuAc(1) | C[+57]*6, N[+2352] | 1503.5793 | 4 | 6011.2956 | 6011.2791 | 0 | 2.7 | 325 | Specific | 777.16 | 777.16 | 777.16 | 16.1 | 171 | >sp\|P24821\|TENA_HUMAN Tenascin OS=Homo sapiens OX=9606 GN=TNC PE=1 SV=3 | 486 | 20201118_ly_colostrum_1ng_glyco_hcd203040_compound15.6613.6613.4 | controllerType=0 controllerNumber=1 scan=6613 | 17.6276 |
| 03598:4 | 1 | R.C[+57.021]IN[+2351.851]GTC[+57.021]YC[+57.021]EEGFTGEDC[+57.021]GKPTC[ | HexNAc(4)Hex(5)Fuc(3)NeuAc(1) | C[+57]*6, N[+2352] | 1503.5793 | 4 | 6011.2956 | 6011.2791 | 0 | 2.7 | 325 | Specific | 895.08 | 895.08 | 895.08 | 18.1 | 171 | >sp\|P24821\|TENA_HUMAN Tenascin OS=Homo sapiens OX=9606 GN=TNC PE=1 SV=3 | 486 | 20201118_ly_colostrum_1ng_glyco_hcd203040_compound15.6611.6611.4 | controllerType=0 controllerNumber=1 scan=6611 | 17.6226 |
| 03646:5 | 1 | R.C[+57.021]IN[+2059.735]GTC[+57.021]YC[+57.021]EEGFTGEDC[+57.021]GKPTC[ | HexNAc(4)Hex(5)Fuc(1)NeuAc(1) | C[+57]*6, N[+2060] | 1144.6404 | 5 | 5719.1728 | 5719.1633 | 0 | 1.7 | 325 | Specific | 906.71 | 906.71 | 906.71 | 19.28 | 171 | >sp\|P24821\|TENA_HUMAN Tenascin OS=Homo sapiens OX=9606 GN=TNC PE=1 SV=3 | 486 | 20201118_ly_colostrum_1ng_glyco_hcd203040_compound15.6660.6660.5 | controllerType=0 controllerNumber=1 scan=6660 | 17.7087 |
| 03648:5 | 1 | R.C[+57.021]IN[+2059.735]GTC[+57.021]YC[+57.021]EEGFTGEDC[+57.021]GKPTC[ | HexNAc(4)Hex(5)Fuc(1)NeuAc(1) | C[+57]*6, N[+2060] | 1144.6404 | 5 | 5719.1728 | 5719.1633 | 0 | 1.7 | 325 | Specific | 956.36 | 956.36 | 956.36 | 20.28 | 171 | >sp\|P24821\|TENA_HUMAN Tenascin OS=Homo sapiens OX=9606 GN=TNC PE=1 SV=3 | 486 | 20201118_ly_colostrum_1ng_glyco_hcd203040_compound15.6662.6662.5 | controllerType=0 controllerNumber=1 scan=6662 | 17.713 |
| 03579:5 | 1 | R.C[+57.021]IN[+2351.851]GTC[+57.021]YC[+57.021]EEGFTGEDC[+57.021]GKPTC[ | HexNAc(4)Hex(5)Fuc(3)NeuAc(1) | C[+57]*6, N[+2352] | 1203.0642 | 5 | 6011.292 | 6011.2791 | 0 | 2.1 | 325 | Specific | 1014.84 | 1014.84 | 1014.84 | 21.52 | 171 | >sp\|P24821\|TENA_HUMAN Tenascin OS=Homo sapiens OX=9606 GN=TNC PE=1 SV=3 | 486 | 20201118_ly_colostrum_1ng_glyco_hcd203040_compound15.6591.6591.5 | controllerType=0 controllerNumber=1 scan=6591 | 17.5843 |

Table S4. Cont.

| Query #:z | Protein Rank | Glycopeptide | Glycan | Modification Type(s) | Observed m/z | z | Observed (M+H) | Calc. mass (M+H) | Off-by-x error | Mass error (ppm) | StartPosition | Cleavage | Score | Delta | Delta Mod | \|Log Prob\| | # of unique peptides | Protein Name | Protein DB number | Comment | Scan # | Scan Time |
|---|---|---|---|---|---|---|---|---|---|---|---|---|---|---|---|---|---|---|---|---|---|---|
| 03577:5 | 1 | R.C[+57.021]IN[+2351.851]GTC[+57.021]YC[+57.021]EEGFTGEDC[+57.021]GKPTC[ | HexNAc(4)Hex(5)Fuc(3)NeuAc(1) | C[+57]*6, N[+2352] | 1203.0642 | 5 | 6011.292 | 6011.2791 | 0 | 2.1 | 325 | Specific | 1081.47 | 1081.47 | 1081.47 | 22.52 | 171 | >sp\|P24821\|TENA_HUMAN Tenascin OS=Homo sapiens OX=9606 GN=TNC PE=1 SV=3 | 486 | 20201118_ly_colostrum_1ng_glyco_hcd203040_compound15.6589.6589.5 | controllerType=0 controllerNumber=1 scan=6589 | 17.5795 |
| 09764:4 | 1 | R.QTGLAPGQEYEISLHIVKN[+2350.830]NTR.G | HexNAc(4)Hex(5)Fuc(1)NeuAc(2) | N[+2351] | 1205.5376 | 4 | 4819.1287 | 4819.1196 | 0 | 1.9 | 770 | Specific | 405.06 | 229.39 | 229.39 | 7.4 | 171 | >sp\|P24821\|TENA_HUMAN Tenascin OS=Homo sapiens OX=9606 GN=TNC PE=1 SV=3 | 486 | 20201118_ly_colostrum_1ng_glyco_hcd203040_compound15.13006.13006.4 | controllerType=0 controllerNumber=1 scan=13006 | 29.3645 |
| 09766:4 | 1 | R.QTGLAPGQEYEISLHIVKN[+2350.830]NTR.G | HexNAc(4)Hex(5)Fuc(1)NeuAc(2) | N[+2351] | 1205.5376 | 4 | 4819.1287 | 4819.1196 | 0 | 1.9 | 770 | Specific | 445.6 | 273.47 | 273.47 | 7.8 | 171 | >sp\|P24821\|TENA_HUMAN Tenascin OS=Homo sapiens OX=9606 GN=TNC PE=1 SV=3 | 486 | 20201118_ly_colostrum_1ng_glyco_hcd203040_compound15.13008.13008.4 | controllerType=0 controllerNumber=1 scan=13008 | 29.3694 |
| 08346:4 | 1 | R.QTGLAPGQEYEISLHIVKN[+2205.793]NTR.G | HexNAc(4)Hex(5)Fuc(2)NeuAc(1) | N[+2206] | 1169.2791 | 4 | 4674.0946 | 4674.0821 | 0 | 2.7 | 770 | Specific | 506.55 | 405.69 | 405.69 | 10.43 | 171 | >sp\|P24821\|TENA_HUMAN Tenascin OS=Homo sapiens OX=9606 GN=TNC PE=1 SV=3 | 486 | 20201118_ly_colostrum_1ng_glyco_hcd203040_compound15.11531.11531.4 | controllerType=0 controllerNumber=1 scan=11531 | 26.4084 |
| 08344:4 | 1 | R.QTGLAPGQEYEISLHIVKN[+2205.793]NTR.G | HexNAc(4)Hex(5)Fuc(2)NeuAc(1) | N[+2206] | 1169.2791 | 4 | 4674.0946 | 4674.0821 | 0 | 2.7 | 770 | Specific | 624.73 | 452.85 | 452.85 | 13.15 | 171 | >sp\|P24821\|TENA_HUMAN Tenascin OS=Homo sapiens OX=9606 GN=TNC PE=1 SV=3 | 486 | 20201118_ly_colostrum_1ng_glyco_hcd203040_compound15.11529.11529.4 | controllerType=0 controllerNumber=1 scan=11529 | 26.4028 |
| 12447:4 | 1 | K.FDRYRLN[+1216.423]YSLPTGQWVGVQLPR.N | HexNAc(2)Hex(5) | N[+1216] | 971.2116 | 4 | 3881.8245 | 3881.8226 | 0 | 0.5 | 1012 | Specific | 258.87 | 174.4 | 174.4 | 3.51 | 171 | >sp\|P24821\|TENA_HUMAN Tenascin OS=Homo sapiens OX=9606 GN=TNC PE=1 SV=3 | 486 | 20201118_ly_colostrum_1ng_glyco_hcd203040_compound15.15794.15794.4 | controllerType=0 controllerNumber=1 scan=15794 | 34.8857 |
| 12444:4 | 1 | K.FDRYRLN[+1216.423]YSLPTGQWVGVQLPR.N | HexNAc(2)Hex(5) | N[+1216] | 971.2116 | 4 | 3881.8245 | 3881.8226 | 0 | 0.5 | 1012 | Specific | 291.5 | 200.91 | 200.91 | 4.25 | 171 | >sp\|P24821\|TENA_HUMAN Tenascin OS=Homo sapiens OX=9606 GN=TNC PE=1 SV=3 | 486 | 20201118_ly_colostrum_1ng_glyco_hcd203040_compound15.15791.15791.4 | controllerType=0 controllerNumber=1 scan=15791 | 34.8746 |
| 12426:3 | 1 | K.FDRYRLN[+1216.423]YSLPTGQWVGVQLPR.N | HexNAc(2)Hex(5) | N[+1216] | 1294.616 | 3 | 3881.8335 | 3881.8226 | 0 | 2.8 | 1012 | Specific | 451.91 | 338.34 | 338.34 | 5.25 | 171 | >sp\|P24821\|TENA_HUMAN Tenascin OS=Homo sapiens OX=9606 GN=TNC PE=1 SV=3 | 486 | 20201118_ly_colostrum_1ng_glyco_hcd203040_compound15.15772.15772.3 | controllerType=0 controllerNumber=1 scan=15772 | 34.8359 |
| 13960:3 | 1 | R.LN[+1216.423]YSLPTGQWVGVQLPR.N | HexNAc(2)Hex(5) | N[+1216] | 1048.8311 | 3 | 3144.4787 | 3144.4617 | 0 | 5.4 | 1017 | Specific | 270.25 | 252.81 | 252.81 | 4.17 | 171 | >sp\|P24821\|TENA_HUMAN Tenascin OS=Homo sapiens OX=9606 GN=TNC PE=1 SV=3 | 486 | 20201118_ly_colostrum_1ng_glyco_hcd203040_compound15.17367.17367.3 | controllerType=0 controllerNumber=1 scan=17367 | 38.1405 |
| 13957:3 | 1 | R.LN[+1216.423]YSLPTGQWVGVQLPR.N | HexNAc(2)Hex(5) | N[+1216] | 1048.8311 | 3 | 3144.4787 | 3144.4617 | 0 | 5.4 | 1017 | Specific | 295.13 | 295.13 | 295.13 | 4.88 | 171 | >sp\|P24821\|TENA_HUMAN Tenascin OS=Homo sapiens OX=9606 GN=TNC PE=1 SV=3 | 486 | 20201118_ly_colostrum_1ng_glyco_hcd203040_compound15.17364.17364.3 | controllerType=0 controllerNumber=1 scan=17364 | 38.1308 |
| 13949:2 | 1 | R.LN[+1216.423]YSLPTGQWVGVQLPR.N | HexNAc(2)Hex(5) | N[+1216] | 1572.7393 | 2 | 3144.4713 | 3144.4617 | 0 | 3.1 | 1017 | Specific | 839.13 | 791.22 | 791.22 | 16.97 | 171 | >sp\|P24821\|TENA_HUMAN Tenascin OS=Homo sapiens OX=9606 GN=TNC PE=1 SV=3 | 486 | 20201118_ly_colostrum_1ng_glyco_hcd203040_compound15.17356.17356.2 | controllerType=0 controllerNumber=1 scan=17356 | 38.1194 |
| 13946:2 | 1 | R.LN[+1216.423]YSLPTGQWVGVQLPR.N | HexNAc(2)Hex(5) | N[+1216] | 1572.7393 | 2 | 3144.4713 | 3144.4617 | 0 | 3.1 | 1017 | Specific | 892.53 | 776.53 | 776.53 | 17.97 | 171 | >sp\|P24821\|TENA_HUMAN Tenascin OS=Homo sapiens OX=9606 GN=TNC PE=1 SV=3 | 486 | 20201118_ly_colostrum_1ng_glyco_hcd203040_compound15.17353.17353.2 | controllerType=0 controllerNumber=1 scan=17353 | 38.1065 |
| 13942:3 | 1 | R.LN[+1216.423]YSLPTGQWVGVQLPR.N | HexNAc(2)Hex(5) | N[+1216] | 1048.8311 | 3 | 3144.4787 | 3144.4617 | 0 | 5.4 | 1017 | Specific | 1082.68 | 1011.68 | 1011.68 | 21.57 | 171 | >sp\|P24821\|TENA_HUMAN Tenascin OS=Homo sapiens OX=9606 GN=TNC PE=1 SV=3 | 486 | 20201118_ly_colostrum_1ng_glyco_hcd203040_compound15.17349.17349.3 | controllerType=0 controllerNumber=1 scan=17349 | 38.0994 |
| 13944:3 | 1 | R.LN[+1216.423]YSLPTGQWVGVQLPR.N | HexNAc(2)Hex(5) | N[+1216] | 1048.8311 | 3 | 3144.4787 | 3144.4617 | 0 | 5.4 | 1017 | Specific | 1098.22 | 944.36 | 944.36 | 21.57 | 171 | >sp\|P24821\|TENA_HUMAN Tenascin OS=Homo sapiens OX=9606 GN=TNC PE=1 SV=3 | 486 | 20201118_ly_colostrum_1ng_glyco_hcd203040_compound15.17351.17351.3 | controllerType=0 controllerNumber=1 scan=17351 | 38.1045 |
| 01848:2 | 1 | R.N[+1768.640]TTSYVLR.G | HexNAc(4)Hex(5)Fuc(1) | N[+1769] | 1361.5798 | 2 | 2722.1522 | 2722.1446 | 0 | 2.8 | 1034 | Specific | 215.79 | 136.35 | 136.35 | 3.89 | 171 | >sp\|P24821\|TENA_HUMAN Tenascin OS=Homo sapiens OX=9606 GN=TNC PE=1 SV=3 | 486 | 20201118_ly_colostrum_1ng_glyco_hcd203040_compound15.4798.4798.2 | controllerType=0 controllerNumber=1 scan=4798 | 14.3577 |
| 03459:3 | 1 | R.N[+2351.851]TTSYVLR.G | HexNAc(4)Hex(5)Fuc(3)NeuAc(1) | N[+2352] | 1102.4597 | 3 | 3305.3647 | 3305.3558 | 0 | 2.7 | 1034 | Specific | 216.12 | 130.2 | 130.2 | 3.39 | 171 | >sp\|P24821\|TENA_HUMAN Tenascin OS=Homo sapiens OX=9606 GN=TNC PE=1 SV=3 | 486 | 20201118_ly_colostrum_1ng_glyco_hcd203040_compound15.6466.6466.3 | controllerType=0 controllerNumber=1 scan=6466 | 17.344 |
| 01850:2 | 1 | R.N[+1768.640]TTSYVLR.G | HexNAc(4)Hex(5)Fuc(1) | N[+1769] | 1361.5798 | 2 | 2722.1522 | 2722.1446 | 0 | 2.8 | 1034 | Specific | 328.69 | 228.69 | 228.69 | 6.69 | 171 | >sp\|P24821\|TENA_HUMAN Tenascin OS=Homo sapiens OX=9606 GN=TNC PE=1 SV=3 | 486 | 20201118_ly_colostrum_1ng_glyco_hcd203040_compound15.4800.4800.2 | controllerType=0 controllerNumber=1 scan=4800 | 14.362 |
| 03633:2 | 1 | R.N[+2059.735]TTSYVLR.G | HexNAc(4)Hex(5)Fuc(1)NeuAc(1) | N[+2060] | 1507.1266 | 2 | 3013.246 | 3013.24 | 0 | 2 | 1034 | Specific | 343.83 | 196.2 | 196.2 | 6.45 | 171 | >sp\|P24821\|TENA_HUMAN Tenascin OS=Homo sapiens OX=9606 GN=TNC PE=1 SV=3 | 486 | 20201118_ly_colostrum_1ng_glyco_hcd203040_compound15.6647.6647.2 | controllerType=0 controllerNumber=1 scan=6647 | 17.6857 |
| 15395:4 | 2 | R.RTAGWNVPIGTLRPFLN[+2279.830]WTGPPEPIEAAVAR.F | HexNAc(5)Hex(6)Fuc(2) | N[+2280] | 1417.4165 | 4 | 5666.6442 | 5666.6417 | 0 | 0.4 | 140 | Specific | 208.87 | 177.91 | 177.91 | 3.35 | 126 | >sp\|P02788\|TRFL_HUMAN Lactotransferrin OS=Homo sapiens OX=9606 GN=LTF PE=1 SV=6 | 1270 | 20201118_ly_colostrum_1ng_glyco_hcd203040_compound15.18861.18861.4 | controllerType=0 controllerNumber=1 scan=18861 | 41.2386 |
| 17412:5 | 2 | R.RTAGWNVPIGTLRPFLN[+2351.851]WTGPPEPIEAAVAR.F | HexNAc(4)Hex(5)Fuc(3)NeuAc(1) | N[+2352] | 1148.5372 | 5 | 5738.6571 | 5738.6628 | 0 | -1 | 140 | Specific | 236.59 | 236.59 | 236.59 | 6.13 | 126 | >sp\|P02788\|TRFL_HUMAN Lactotransferrin OS=Homo sapiens OX=9606 GN=LTF PE=1 SV=6 | 1270 | 20201118_ly_colostrum_1ng_glyco_hcd203040_compound15.20968.20968.5 | controllerType=0 controllerNumber=1 scan=20968 | 45.9319 |
| 17599:4 | 2 | R.[+42.011]RTAGWNVPIGTLRPFLN[+2026.687]WTGPPEPIEAAVAR.F | HexNAc(2)Hex(10) | .[42], N[+2027] | 1364.6426 | 4 | 5455.5485 | 5455.5096 | 0 | 7.1 | 140 | Specific | 311.31 | 311.31 | 311.31 | 6.26 | 126 | >sp\|P02788\|TRFL_HUMAN Lactotransferrin OS=Homo sapiens OX=9606 GN=LTF PE=1 SV=6 | 1270 | 20201118_ly_colostrum_1ng_glyco_hcd203040_compound15.21164.21164.4 | controllerType=0 controllerNumber=1 scan=21164 | 46.393 |
| 17597:4 | 2 | R.[+42.011]RTAGWNVPIGTLRPFLN[+2026.687]WTGPPEPIEAAVAR.F | HexNAc(2)Hex(10) | .[42], N[+2027] | 1364.6426 | 4 | 5455.5485 | 5455.5096 | 0 | 7.1 | 140 | Specific | 321.51 | 321.51 | 321.51 | 7.5 | 126 | >sp\|P02788\|TRFL_HUMAN Lactotransferrin OS=Homo sapiens OX=9606 GN=LTF PE=1 SV=6 | 1270 | 20201118_ly_colostrum_1ng_glyco_hcd203040_compound15.21162.21162.4 | controllerType=0 controllerNumber=1 scan=21162 | 46.3782 |
| 16247:4 | 2 | R.RTAGWNVPIGTLRPFLN[+2863.041]WTGPPEPIEAAVAR.F | HexNAc(5)Hex(6)Fuc(4)NeuAc(1) | N[+2863] | 1563.2236 | 4 | 6249.8727 | 6249.8529 | 0 | 3.2 | 140 | Specific | 373.63 | 373.63 | 373.63 | 7.81 | 126 | >sp\|P02788\|TRFL_HUMAN Lactotransferrin OS=Homo sapiens OX=9606 GN=LTF PE=1 SV=6 | 1270 | 20201118_ly_colostrum_1ng_glyco_hcd203040_compound15.19751.19751.4 | controllerType=0 controllerNumber=1 scan=19751 | 43.2531 |
| 17415:5 | 2 | R.RTAGWNVPIGTLRPFLN[+2351.851]WTGPPEPIEAAVAR.F | HexNAc(4)Hex(5)Fuc(3)NeuAc(1) | N[+2352] | 1148.5372 | 5 | 5738.6571 | 5738.6628 | 0 | -1 | 140 | Specific | 482.99 | 357.05 | 357.05 | 8.02 | 126 | >sp\|P02788\|TRFL_HUMAN Lactotransferrin OS=Homo sapiens OX=9606 GN=LTF PE=1 SV=6 | 1270 | 20201118_ly_colostrum_1ng_glyco_hcd203040_compound15.20971.20971.5 | controllerType=0 controllerNumber=1 scan=20971 | 45.9374 |
| 16249:4 | 2 | R.RTAGWNVPIGTLRPFLN[+2863.041]WTGPPEPIEAAVAR.F | HexNAc(5)Hex(6)Fuc(4)NeuAc(1) | N[+2863] | 1563.2236 | 4 | 6249.8727 | 6249.8529 | 0 | 3.2 | 140 | Specific | 518.86 | 442.45 | 442.45 | 10.81 | 126 | >sp\|P02788\|TRFL_HUMAN Lactotransferrin OS=Homo sapiens OX=9606 GN=LTF PE=1 SV=6 | 1270 | 20201118_ly_colostrum_1ng_glyco_hcd203040_compound15.19753.19753.4 | controllerType=0 controllerNumber=1 scan=19753 | 43.2579 |
| 16341:4 | 2 | R.RTAGWNVPIGTLRPFLN[+2351.851]WTGPPEPIEAAVAR.F | HexNAc(4)Hex(5)Fuc(3)NeuAc(1) | N[+2352] | 1435.4258 | 4 | 5738.6813 | 5738.6628 | 0 | 3.2 | 140 | Specific | 557.24 | 389.08 | 389.08 | 11.04 | 126 | >sp\|P02788\|TRFL_HUMAN Lactotransferrin OS=Homo sapiens OX=9606 GN=LTF PE=1 SV=6 | 1270 | 20201118_ly_colostrum_1ng_glyco_hcd203040_compound15.19849.19849.4 | controllerType=0 controllerNumber=1 scan=19849 | 43.4652 |
| 16339:4 | 2 | R.RTAGWNVPIGTLRPFLN[+2351.851]WTGPPEPIEAAVAR.F | HexNAc(4)Hex(5)Fuc(3)NeuAc(1) | N[+2352] | 1435.4258 | 4 | 5738.6813 | 5738.6628 | 0 | 3.2 | 140 | Specific | 576.27 | 378.28 | 378.28 | 9.75 | 126 | >sp\|P02788\|TRFL_HUMAN Lactotransferrin OS=Homo sapiens OX=9606 GN=LTF PE=1 SV=6 | 1270 | 20201118_ly_colostrum_1ng_glyco_hcd203040_compound15.19847.19847.4 | controllerType=0 controllerNumber=1 scan=19847 | 43.4593 |
| 17626:3 | 2 | R.TAGWNVPIGTLRPFLN[+2350.830]WTGPPEPIEAAVAR.F | HexNAc(4)Hex(5)Fuc(1)NeuAc(2) | N[+2351] | 1861.1993 | 3 | 5581.5834 | 5581.5413 | 0 | 7.5 | 141 | Specific | 238.95 | 107.02 | 107.02 | 4.75 | 126 | >sp\|P02788\|TRFL_HUMAN Lactotransferrin OS=Homo sapiens OX=9606 GN=LTF PE=1 SV=6 | 1270 | 20201118_ly_colostrum_1ng_glyco_hcd203040_compound15.21193.21193.3 | controllerType=0 controllerNumber=1 scan=21193 | 46.4862 |
| 18787:4 | 2 | R.TAGWNVPIGTLRPFLN[+2569.905]WTGPPEPIEAAVAR.F | HexNAc(5)Hex(6)NeuAc(2) | N[+2570] | 1450.9135 | 4 | 5800.632 | 5800.6156 | 0 | 2.8 | 141 | Specific | 282.39 | 265.33 | 265.33 | 7.91 | 126 | >sp\|P02788\|TRFL_HUMAN Lactotransferrin OS=Homo sapiens OX=9606 GN=LTF PE=1 SV=6 | 1270 | 20201118_ly_colostrum_1ng_glyco_hcd203040_compound15.22406.22406.4 | controllerType=0 controllerNumber=1 scan=22406 | 49.2563 |
| 16630:3 | 2 | R.TAGWNVPIGTLRPFLN[+1913.677]WTGPPEPIEAAVAR.F | HexNAc(4)Hex(5)NeuAc(1) | N[+1914] | 1715.4801 | 3 | 5144.4256 | 5144.388 | 0 | 7.3 | 141 | Specific | 302.88 | 302.88 | 302.88 | 7.23 | 126 | >sp\|P02788\|TRFL_HUMAN Lactotransferrin OS=Homo sapiens OX=9606 GN=LTF PE=1 SV=6 | 1270 | 20201118_ly_colostrum_1ng_glyco_hcd203040_compound15.20150.20150.3 | controllerType=0 controllerNumber=1 scan=20150 | 44.1177 |
| 17629:3 | 2 | R.TAGWNVPIGTLRPFLN[+2350.830]WTGPPEPIEAAVAR.F | HexNAc(4)Hex(5)Fuc(1)NeuAc(2) | N[+2351] | 1861.1993 | 3 | 5581.5834 | 5581.5413 | 0 | 7.5 | 141 | Specific | 308.85 | 308.85 | 308.85 | 7.23 | 126 | >sp\|P02788\|TRFL_HUMAN Lactotransferrin OS=Homo sapiens OX=9606 GN=LTF PE=1 SV=6 | 1270 | 20201118_ly_colostrum_1ng_glyco_hcd203040_compound15.21196.21196.3 | controllerType=0 controllerNumber=1 scan=21196 | 46.4931 |
| 17624:5 | 2 | R.TAGWNVPIGTLRPFLN[+2059.735]WTGPPEPIEAAVAR.F | HexNAc(4)Hex(5)Fuc(1)NeuAc(1) | N[+2060] | 1058.8994 | 5 | 5290.468 | 5290.4459 | 0 | 4.2 | 141 | Specific | 309.29 | 309.29 | 309.29 | 6.87 | 126 | >sp\|P02788\|TRFL_HUMAN Lactotransferrin OS=Homo sapiens OX=9606 GN=LTF PE=1 SV=6 | 1270 | 20201118_ly_colostrum_1ng_glyco_hcd203040_compound15.21191.21191.5 | controllerType=0 controllerNumber=1 scan=21191 | 46.4839 |
| 16626:4 | 2 | R.TAGWNVPIGTLRPFLN[+1913.677]WTGPPEPIEAAVAR.F | HexNAc(4)Hex(5)NeuAc(1) | N[+1914] | 1286.8621 | 4 | 5144.4264 | 5144.388 | 0 | 7.5 | 141 | Specific | 326.3 | 326.3 | 326.3 | 7.66 | 126 | >sp\|P02788\|TRFL_HUMAN Lactotransferrin OS=Homo sapiens OX=9606 GN=LTF PE=1 SV=6 | 1270 | 20201118_ly_colostrum_1ng_glyco_hcd203040_compound15.20146.20146.4 | controllerType=0 controllerNumber=1 scan=20146 | 44.111 |
| 17421:3 | 2 | R.TAGWNVPIGTLRPFLN[+2716.983]WTGPPEPIEAAVAR.F | HexNAc(5)Hex(6)Fuc(3)NeuAc(1) | N[+2717] | 1983.2429 | 3 | 5947.7141 | 5947.6939 | 0 | 3.4 | 141 | Specific | 339.56 | 339.56 | 339.56 | 8.39 | 126 | >sp\|P02788\|TRFL_HUMAN Lactotransferrin OS=Homo sapiens OX=9606 GN=LTF PE=1 SV=6 | 1270 | 20201118_ly_colostrum_1ng_glyco_hcd203040_compound15.20978.20978.3 | controllerType=0 controllerNumber=1 scan=20978 | 45.9542 |
| 18792:3 | 2 | R.TAGWNVPIGTLRPFLN[+2496.888]WTGPPEPIEAAVAR.F | HexNAc(4)Hex(5)Fuc(2)NeuAc(2) | N[+2497] | 1909.8762 | 3 | 5727.614 | 5727.5992 | 0 | 2.6 | 141 | Specific | 340.69 | 316.13 | 316.13 | 8.32 | 126 | >sp\|P02788\|TRFL_HUMAN Lactotransferrin OS=Homo sapiens OX=9606 GN=LTF PE=1 SV=6 | 1270 | 20201118_ly_colostrum_1ng_glyco_hcd203040_compound15.22411.22411.3 | controllerType=0 controllerNumber=1 scan=22411 | 49.2673 |
| 16633:3 | 2 | R.TAGWNVPIGTLRPFLN[+1913.677]WTGPPEPIEAAVAR.F | HexNAc(4)Hex(5)NeuAc(1) | N[+1914] | 1715.4801 | 3 | 5144.4256 | 5144.388 | 0 | 7.3 | 141 | Specific | 352.44 | 352.44 | 352.44 | 7.7 | 126 | >sp\|P02788\|TRFL_HUMAN Lactotransferrin OS=Homo sapiens OX=9606 GN=LTF PE=1 SV=6 | 1270 | 20201118_ly_colostrum_1ng_glyco_hcd203040_compound15.20153.20153.3 | controllerType=0 controllerNumber=1 scan=20153 | 44.1221 |
| 18790:3 | 2 | R.TAGWNVPIGTLRPFLN[+2496.888]WTGPPEPIEAAVAR.F | HexNAc(4)Hex(5)Fuc(2)NeuAc(2) | N[+2497] | 1909.8762 | 3 | 5727.614 | 5727.5992 | 0 | 2.6 | 141 | Specific | 354.63 | 277.06 | 277.06 | 8.1 | 126 | >sp\|P02788\|TRFL_HUMAN Lactotransferrin OS=Homo sapiens OX=9606 GN=LTF PE=1 SV=6 | 1270 | 20201118_ly_colostrum_1ng_glyco_hcd203040_compound15.22409.22409.3 | controllerType=0 controllerNumber=1 scan=22409 | 49.2622 |
| 17463:3 | 2 | R.TAGWNVPIGTLRPFLN[+2569.905]WTGPPEPIEAAVAR.F | HexNAc(5)Hex(6)NeuAc(2) | N[+2570] | 1934.2258 | 3 | 5800.6628 | 5800.6156 | 0 | 8.1 | 141 | Specific | 386.69 | 248.55 | 248.55 | 6.8 | 126 | >sp\|P02788\|TRFL_HUMAN Lactotransferrin OS=Homo sapiens OX=9606 GN=LTF PE=1 SV=6 | 1270 | 20201118_ly_colostrum_1ng_glyco_hcd203040_compound15.21021.21021.3 | controllerType=0 controllerNumber=1 scan=21021 | 46.0513 |
| 17612:4 | 2 | R.TAGWNVPIGTLRPFLN[+2205.793]WTGPPEPIEAAVAR.F | HexNAc(4)Hex(5)Fuc(2)NeuAc(1) | N[+2206] | 1359.8848 | 4 | 5436.5172 | 5436.5038 | 0 | 2.5 | 141 | Specific | 388.89 | 388.89 | 388.89 | 10.37 | 126 | >sp\|P02788\|TRFL_HUMAN Lactotransferrin OS=Homo sapiens OX=9606 GN=LTF PE=1 SV=6 | 1270 | 20201118_ly_colostrum_1ng_glyco_hcd203040_compound15.21179.21179.4 | controllerType=0 controllerNumber=1 scan=21179 | 46.4594 |
| 16578:4 | 2 | R.TAGWNVPIGTLRPFLN[+2351.851]WTGPPEPIEAAVAR.F | HexNAc(4)Hex(5)Fuc(3)NeuAc(1) | N[+2352] | 1396.4037 | 4 | 5582.5929 | 5582.5617 | 0 | 5.6 | 141 | Specific | 394.4 | 394.4 | 394.4 | 9.82 | 126 | >sp\|P02788\|TRFL_HUMAN Lactotransferrin OS=Homo sapiens OX=9606 GN=LTF PE=1 SV=6 | 1270 | 20201118_ly_colostrum_1ng_glyco_hcd203040_compound15.20095.20095.4 | controllerType=0 controllerNumber=1 scan=20095 | 43.975 |
| 18804:4 | 2 | R.TAGWNVPIGTLRPFLN[+2350.830]WTGPPEPIEAAVAR.F | HexNAc(4)Hex(5)Fuc(1)NeuAc(2) | N[+2351] | 1396.1472 | 4 | 5581.5671 | 5581.5413 | 0 | 4.6 | 141 | Specific | 432.63 | 300.5 | 300.5 | 8.14 | 126 | >sp\|P02788\|TRFL_HUMAN Lactotransferrin OS=Homo sapiens OX=9606 GN=LTF PE=1 SV=6 | 1270 | 20201118_ly_colostrum_1ng_glyco_hcd203040_compound15.22424.22424.4 | controllerType=0 controllerNumber=1 scan=22424 | 49.3023 |
| 17522:4 | 2 | R.TAGWNVPIGTLRPFLN[+2716.983]WTGPPEPIEAAVAR.F | HexNAc(5)Hex(6)Fuc(3)NeuAc(1) | N[+2717] | 1487.6899 | 4 | 5947.7379 | 5947.6939 | 0 | 7.4 | 141 | Specific | 449.59 | 439.6 | 439.6 | 9.82 | 126 | >sp\|P02788\|TRFL_HUMAN Lactotransferrin OS=Homo sapiens OX=9606 GN=LTF PE=1 SV=6 | 1270 | 20201118_ly_colostrum_1ng_glyco_hcd203040_compound15.21083.21083.4 | controllerType=0 controllerNumber=1 scan=21083 | 46.197 |
| 16593:4 | 2 | R.TAGWNVPIGTLRPFLN[+2059.735]WTGPPEPIEAAVAR.F | HexNAc(4)Hex(5)Fuc(1)NeuAc(1) | N[+2060] | 1323.3765 | 4 | 5290.484 | 5290.4459 | 0 | 7.2 | 141 | Specific | 455.45 | 455.45 | 455.45 | 9.82 | 126 | >sp\|P02788\|TRFL_HUMAN Lactotransferrin OS=Homo sapiens OX=9606 GN=LTF PE=1 SV=6 | 1270 | 20201118_ly_colostrum_1ng_glyco_hcd203040_compound15.20111.20111.4 | controllerType=0 controllerNumber=1 scan=20111 | 44.0305 |
| 17424:3 | 2 | R.TAGWNVPIGTLRPFLN[+2716.983]WTGPPEPIEAAVAR.F | HexNAc(5)Hex(6)Fuc(3)NeuAc(1) | N[+2717] | 1983.2429 | 3 | 5947.7141 | 5947.6939 | 0 | 3.4 | 141 | Specific | 471.37 | 423.57 | 423.57 | 10.61 | 126 | >sp\|P02788\|TRFL_HUMAN Lactotransferrin OS=Homo sapiens OX=9606 GN=LTF PE=1 SV=6 | 1270 | 20201118_ly_colostrum_1ng_glyco_hcd203040_compound15.20981.20981.3 | controllerType=0 controllerNumber=1 scan=20981 | 45.9581 |
| 17510:4 | 2 | R.TAGWNVPIGTLRPFLN[+2571.946]WTGPPEPIEAAVAR.F | HexNAc(5)Hex(6)Fuc(4) | N[+2572] | 1451.4213 | 4 | 5802.6632 | 5802.6564 | 0 | 1.2 | 141 | Specific | 472.1 | 464.04 | 464.04 | 10.51 | 126 | >sp\|P02788\|TRFL_HUMAN Lactotransferrin OS=Homo sapiens OX=9606 GN=LTF PE=1 SV=6 | 1270 | 20201118_ly_colostrum_1ng_glyco_hcd203040_compound15.21071.21071.4 | controllerType=0 controllerNumber=1 scan=21071 | 46.1706 |
| 17613:4 | 2 | R.TAGWNVPIGTLRPFLN[+2351.851]WTGPPEPIEAAVAR.F | HexNAc(4)Hex(5)Fuc(3)NeuAc(1) | N[+2352] | 1396.4006 | 4 | 5582.5807 | 5582.5617 | 0 | 3.4 | 141 | Specific | 512.93 | 496.55 | 496.55 | 10.98 | 126 | >sp\|P02788\|TRFL_HUMAN Lactotransferrin OS=Homo sapiens OX=9606 GN=LTF PE=1 SV=6 | 1270 | 20201118_ly_colostrum_1ng_glyco_hcd203040_compound15.21180.21180.4 | controllerType=0 controllerNumber=1 scan=21180 | 46.4612 |
| 17627:5 | 2 | R.TAGWNVPIGTLRPFLN[+2059.735]WTGPPEPIEAAVAR.F | HexNAc(4)Hex(5)Fuc(1)NeuAc(1) | N[+2060] | 1058.8994 | 5 | 5290.468 | 5290.4459 | 0 | 4.2 | 141 | Specific | 527.13 | 527.13 | 527.13 | 10.58 | 126 | >sp\|P02788\|TRFL_HUMAN Lactotransferrin OS=Homo sapiens OX=9606 GN=LTF PE=1 SV=6 | 1270 | 20201118_ly_colostrum_1ng_glyco_hcd203040_compound15.21194.21194.5 | controllerType=0 controllerNumber=1 scan=21194 | 46.4886 |
| 17525:4 | 2 | R.TAGWNVPIGTLRPFLN[+2716.983]WTGPPEPIEAAVAR.F | HexNAc(5)Hex(6)Fuc(3)NeuAc(1) | N[+2717] | 1487.6899 | 4 | 5947.7379 | 5947.6939 | 0 | 7.4 | 141 | Specific | 534.3 | 480.35 | 480.35 | 10.13 | 126 | >sp\|P02788\|TRFL_HUMAN Lactotransferrin OS=Homo sapiens OX=9606 GN=LTF PE=1 SV=6 | 1270 | 20201118_ly_colostrum_1ng_glyco_hcd203040_compound15.21086.21086.4 | controllerType=0 controllerNumber=1 scan=21086 | 46.2042 |
| 18807:3 | 2 | R.TAGWNVPIGTLRPFLN[+2350.830]WTGPPEPIEAAVAR.F | HexNAc(4)Hex(5)Fuc(1)NeuAc(2) | N[+2351] | 1861.1942 | 3 | 5581.568 | 5581.5413 | 0 | 4.8 | 141 | Specific | 543.55 | 344 | 344 | 9.08 | 126 | >sp\|P02788\|TRFL_HUMAN Lactotransferrin OS=Homo sapiens OX=9606 GN=LTF PE=1 SV=6 | 1270 | 20201118_ly_colostrum_1ng_glyco_hcd203040_compound15.22427.22427.3 | controllerType=0 controllerNumber=1 scan=22427 | 49.3074 |
| 18777:3 | 2 | R.TAGWNVPIGTLRPFLN[+2350.830]WTGPPEPIEAAVAR.F | HexNAc(4)Hex(5)Fuc(1)NeuAc(2) | N[+2351] | 1861.1927 | 3 | 5581.5636 | 5581.5413 | 0 | 4 | 141 | Specific | 555.24 | 380.93 | 380.93 | 11.04 | 126 | >sp\|P02788\|TRFL_HUMAN Lactotransferrin OS=Homo sapiens OX=9606 GN=LTF PE=1 SV=6 | 1270 | 20201118_ly_colostrum_1ng_glyco_hcd203040_compound15.22395.22395.2 | controllerType=0 controllerNumber=1 scan=22395 | 49.2311 |
| 17692:4 | 2 | R.TAGWNVPIGTLRPFLN[+2059.735]WTGPPEPIEAAVAR.F | HexNAc(4)Hex(5)Fuc(1)NeuAc(1) | N[+2060] | 1323.3737 | 4 | 5290.4728 | 5290.4459 | 0 | 5.1 | 141 | Specific | 568.88 | 470.87 | 470.87 | 10.81 | 126 | >sp\|P02788\|TRFL_HUMAN Lactotransferrin OS=Homo sapiens OX=9606 GN=LTF PE=1 SV=6 | 1270 | 20201118_ly_colostrum_1ng_glyco_hcd203040_compound15.21262.21262.4 | controllerType=0 controllerNumber=1 scan=21262 | 46.6338 |
| 17539:3 | 2 | R.TAGWNVPIGTLRPFLN[+2205.793]WTGPPEPIEAAVAR.F | HexNAc(4)Hex(5)Fuc(2)NeuAc(1) | N[+2206] | 1812.8444 | 3 | 5436.5188 | 5436.5038 | 0 | 2.8 | 141 | Specific | 572.6 | 475.39 | 475.39 | 11.19 | 126 | >sp\|P02788\|TRFL_HUMAN Lactotransferrin OS=Homo sapiens OX=9606 GN=LTF PE=1 SV=6 | 1270 | 20201118_ly_colostrum_1ng_glyco_hcd203040_compound15.21101.21101.3 | controllerType=0 controllerNumber=1 scan=21101 | 46.2384 |
| 17445:4 | 2 | R.TAGWNVPIGTLRPFLN[+2569.905]WTGPPEPIEAAVAR.F | HexNAc(5)Hex(6)NeuAc(2) | N[+2570] | 1450.9191 | 4 | 5800.6545 | 5800.6156 | 0 | 6.7 | 141 | Specific | 605.37 | 470.2 | 470.2 | 12.84 | 126 | >sp\|P02788\|TRFL_HUMAN Lactotransferrin OS=Homo sapiens OX=9606 GN=LTF PE=1 SV=6 | 1270 | 20201118_ly_colostrum_1ng_glyco_hcd203040_compound15.21003.21003.4 | controllerType=0 controllerNumber=1 scan=21003 | 46.0035 |
| 17377:4 | 2 | R.TAGWNVPIGTLRPFLN[+2863.041]WTGPPEPIEAAVAR.F | HexNAc(5)Hex(6)Fuc(4)NeuAc(1) | N[+2863] | 1524.1963 | 4 | 6093.7633 | 6093.7518 | 0 | 1.9 | 141 | Specific | 624.38 | 545.85 | 545.85 | 13.44 | 126 | >sp\|P02788\|TRFL_HUMAN Lactotransferrin OS=Homo sapiens OX=9606 GN=LTF PE=1 SV=6 | 1270 | 20201118_ly_colostrum_1ng_glyco_hcd203040_compound15.20932.20932.4 | controllerType=0 controllerNumber=1 scan=20932 | 45.844 |
| 16629:4 | 2 | R.TAGWNVPIGTLRPFLN[+1913.677]WTGPPEPIEAAVAR.F | HexNAc(4)Hex(5)NeuAc(1) | N[+1914] | 1286.8621 | 4 | 5144.4264 | 5144.388 | 0 | 7.5 | 141 | Specific | 637.74 | 616.13 | 616.13 | 12.82 | 126 | >sp\|P02788\|TRFL_HUMAN Lactotransferrin OS=Homo sapiens OX=9606 GN=LTF PE=1 SV=6 | 1270 | 20201118_ly_colostrum_1ng_glyco_hcd203040_compound15.20149.20149.4 | controllerType=0 controllerNumber=1 scan=20149 | 44.1158 |
| 17512:4 | 2 | R.TAGWNVPIGTLRPFLN[+2571.946]WTGPPEPIEAAVAR.F | HexNAc(5)Hex(6)Fuc(4) | N[+2572] | 1451.4213 | 4 | 5802.6632 | 5802.6564 | 0 | 1.2 | 141 | Specific | 657.3 | 616.17 | 616.17 | 14.44 | 126 | >sp\|P02788\|TRFL_HUMAN Lactotransferrin OS=Homo sapiens OX=9606 GN=LTF PE=1 SV=6 | 1270 | 20201118_ly_colostrum_1ng_glyco_hcd203040_compound15.21073.21073.4 | controllerType=0 controllerNumber=1 scan=21073 | 46.1768 |
| 17586:3 | 2 | R.TAGWNVPIGTLRPFLN[+2059.735]WTGPPEPIEAAVAR.F | HexNAc(4)Hex(5)Fuc(1)NeuAc(1) | N[+2060] | 1764.1598 | 3 | 5290.4647 | 5290.4459 | 0 | 3.6 | 141 | Specific | 659.81 | 577.15 | 577.15 | 14.54 | 126 | >sp\|P02788\|TRFL_HUMAN Lactotransferrin OS=Homo sapiens OX=9606 GN=LTF PE=1 SV=6 | 1270 | 20201118_ly_colostrum_1ng_glyco_hcd203040_compound15.21151.21151.3 | controllerType=0 controllerNumber=1 scan=21151 | 46.336 |
| 18779:3 | 2 | R.TAGWNVPIGTLRPFLN[+2350.830]WTGPPEPIEAAVAR.F | HexNAc(4)Hex(5)Fuc(1)NeuAc(2) | N[+2351] | 1861.1927 | 3 | 5581.5636 | 5581.5413 | 0 | 4 | 141 | Specific | 661.72 | 497.1 | 497.1 | 14.54 | 126 | >sp\|P02788\|TRFL_HUMAN Lactotransferrin OS=Homo sapiens OX=9606 GN=LTF PE=1 SV=6 | 1270 | 20201118_ly_colostrum_1ng_glyco_hcd203040_compound15.22397.22397.2 | controllerType=0 controllerNumber=1 scan=22397 | 49.2354 |
| 17537:3 | 2 | R.TAGWNVPIGTLRPFLN[+2205.793]WTGPPEPIEAAVAR.F | HexNAc(4)Hex(5)Fuc(2)NeuAc(1) | N[+2206] | 1812.8444 | 3 | 5436.5188 | 5436.5038 | 0 | 2.8 | 141 | Specific | 664.49 | 555.06 | 555.06 | 14.52 | 126 | >sp\|P02788\|TRFL_HUMAN Lactotransferrin OS=Homo sapiens OX=9606 GN=LTF PE=1 SV=6 | 1270 | 20201118_ly_colostrum_1ng_glyco_hcd203040_compound15.21099.21099.3 | controllerType=0 controllerNumber=1 scan=21099 | 46.2315 |
| 17447:4 | 2 | R.TAGWNVPIGTLRPFLN[+2569.905]WTGPPEPIEAAVAR.F | HexNAc(5)Hex(6)NeuAc(2) | N[+2570] | 1450.9191 | 4 | 5800.6545 | 5800.6156 | 0 | 6.7 | 141 | Specific | 681.64 | 448.49 | 448.49 | 13.84 | 126 | >sp\|P02788\|TRFL_HUMAN Lactotransferrin OS=Homo sapiens OX=9606 GN=LTF PE=1 SV=6 | 1270 | 20201118_ly_colostrum_1ng_glyco_hcd203040_compound15.21005.21005.4 | controllerType=0 controllerNumber=1 scan=21005 | 46.0098 |
| 17589:3 | 2 | R.TAGWNVPIGTLRPFLN[+2059.735]WTGPPEPIEAAVAR.F | HexNAc(4)Hex(5)Fuc(1)NeuAc(1) | N[+2060] | 1764.1598 | 3 | 5290.4647 | 5290.4459 | 0 | 3.6 | 141 | Specific | 689.91 | 598.1 | 598.1 | 14.54 | 126 | >sp\|P02788\|TRFL_HUMAN Lactotransferrin OS=Homo sapiens OX=9606 GN=LTF PE=1 SV=6 | 1270 | 20201118_ly_colostrum_1ng_glyco_hcd203040_compound15.21154.21154.3 | controllerType=0 controllerNumber=1 scan=21154 | 46.3542 |
| 17379:4 | 2 | R.TAGWNVPIGTLRPFLN[+2863.041]WTGPPEPIEAAVAR.F | HexNAc(5)Hex(6)Fuc(4)NeuAc(1) | N[+2863] | 1524.1963 | 4 | 6093.7633 | 6093.7518 | 0 | 1.9 | 141 | Specific | 693.27 | 539.06 | 539.06 | 14.44 | 126 | >sp\|P02788\|TRFL_HUMAN Lactotransferrin OS=Homo sapiens OX=9606 GN=LTF PE=1 SV=6 | 1270 | 20201118_ly_colostrum_1ng_glyco_hcd203040_compound15.20934.20934.4 | controllerType=0 controllerNumber=1 scan=20934 | 45.8533 |
| 16596:4 | 2 | R.TAGWNVPIGTLRPFLN[+2059.735]WTGPPEPIEAAVAR.F | HexNAc(4)Hex(5)Fuc(1)NeuAc(1) | N[+2060] | 1323.3765 | 4 | 5290.484 | 5290.4459 | 0 | 7.2 | 141 | Specific | 697.54 | 697.54 | 697.54 | 13.82 | 126 | >sp\|P02788\|TRFL_HUMAN Lactotransferrin OS=Homo sapiens OX=9606 GN=LTF PE=1 SV=6 | 1270 | 20201118_ly_colostrum_1ng_glyco_hcd203040_compound15.20114.20114.4 | controllerType=0 controllerNumber=1 scan=20114 | 44.035 |
| 18782:4 | 2 | R.TAGWNVPIGTLRPFLN[+2496.888]WTGPPEPIEAAVAR.F | HexNAc(4)Hex(5)Fuc(2)NeuAc(2) | N[+2497] | 1432.6597 | 4 | 5727.6169 | 5727.5992 | 0 | 3.1 | 141 | Specific | 709.02 | 620.8 | 620.8 | 15.7 | 126 | >sp\|P02788\|TRFL_HUMAN Lactotransferrin OS=Homo sapiens OX=9606 GN=LTF PE=1 SV=6 | 1270 | 20201118_ly_colostrum_1ng_glyco_hcd203040_compound15.22401.22401.4 | controllerType=0 controllerNumber=1 scan=22401 | [illegible] |
| 17585:4 | 2 | R.TAGWNVPIGTLRPFLN[+2059.735]WTGPPEPIEAAVAR.F | HexNAc(4)Hex(5)Fuc(1)NeuAc(1) | N[+2060] | 1323.3716 | 4 | 5290.4645 | 5290.4459 | 0 | 3.5 | 141 | Specific | 734.06 | 663.9 | 663.9 | 15.7 | 126 | >sp\|P02788\|TRFL_HUMAN Lactotransferrin OS=Homo sapiens OX=9606 GN=LTF PE=1 SV=6 | 1270 | 20201118_ly_colostrum_1ng_glyco_hcd203040_compound15.21150.21150.4 | controllerType=0 controllerNumber=1 scan=21150 | [illegible] |
| 16581:4 | 2 | R.TAGWNVPIGTLRPFLN[+2351.851]WTGPPEPIEAAVAR.F | HexNAc(4)Hex(5)Fuc(3)NeuAc(1) | N[+2352] | 1396.4037 | 4 | 5582.5929 | 5582.5617 | 0 | 5.6 | 141 | Specific | 747.96 | 614.07 | 614.07 | 15.14 | 126 | >sp\|P02788\|TRFL_HUMAN Lactotransferrin OS=Homo sapiens OX=9606 GN=LTF PE=1 SV=6 | 1270 | 20201118_ly_colostrum_1ng_glyco_hcd203040_compound15.20098.20098.4 | controllerType=0 controllerNumber=1 scan=20098 | [illegible] |
| 17695:4 | 2 | R.TAGWNVPIGTLRPFLN[+2059.735]WTGPPEPIEAAVAR.F | HexNAc(4)Hex(5)Fuc(1)NeuAc(1) | N[+2060] | 1323.3737 | 4 | 5290.4728 | 5290.4459 | 0 | 5.1 | 141 | Specific | 793.47 | 602.45 | 602.45 | 16.14 | 126 | >sp\|P02788\|TRFL_HUMAN Lactotransferrin OS=Homo sapiens OX=9606 GN=LTF PE=1 SV=6 | 1270 | 20201118_ly_colostrum_1ng_glyco_hcd203040_compound15.21265.21265.4 | controllerType=0 controllerNumber=1 scan=21265 | [illegible] |
| 18784:4 | 2 | R.TAGWNVPIGTLRPFLN[+2496.888]WTGPPEPIEAAVAR.F | HexNAc(4)Hex(5)Fuc(2)NeuAc(2) | N[+2497] | 1432.6597 | 4 | 5727.6169 | 5727.5992 | 0 | 3.1 | 141 | Specific | 794.52 | 671.4 | 671.4 | 16.7 | 126 | >sp\|P02788\|TRFL_HUMAN Lactotransferrin OS=Homo sapiens OX=9606 GN=LTF PE=1 SV=6 | 1270 | 20201118_ly_colostrum_1ng_glyco_hcd203040_compound15.22403.22403.4 | controllerType=0 controllerNumber=1 scan=22403 | 49.2504 |
| 18806:4 | 2 | R.TAGWNVPIGTLRPFLN[+2350.830]WTGPPEPIEAAVAR.F | HexNAc(4)Hex(5)Fuc(1)NeuAc(2) | N[+2351] | 1396.1472 | 4 | 5581.5671 | 5581.5413 | 0 | 4.6 | 141 | Specific | 801.06 | 637.68 | 637.68 | 17.41 | 126 | >sp\|P02788\|TRFL_HUMAN Lactotransferrin OS=Homo sapiens OX=9606 GN=LTF PE=1 SV=6 | 1270 | 20201118_ly_colostrum_1ng_glyco_hcd203040_compound15.22426.22426.4 | controllerType=0 controllerNumber=1 scan=22426 | 49.3065 |
| 17614:4 | 2 | R.TAGWNVPIGTLRPFLN[+2205.793]WTGPPEPIEAAVAR.F | HexNAc(4)Hex(5)Fuc(2)NeuAc(1) | N[+2206] | 1359.8848 | 4 | 5436.5172 | 5436.5038 | 0 | 2.5 | 141 | Specific | 839.68 | 717.95 | 717.95 | 17.68 | 126 | >sp\|P02788\|TRFL_HUMAN Lactotransferrin OS=Homo sapiens OX=9606 GN=LTF PE=1 SV=6 | 1270 | 20201118_ly_colostrum_1ng_glyco_hcd203040_compound15.21181.21181.4 | controllerType=0 controllerNumber=1 scan=21181 | 46.4627 |
| 17615:4 | 2 | R.TAGWNVPIGTLRPFLN[+2351.851]WTGPPEPIEAAVAR.F | HexNAc(4)Hex(5)Fuc(3)NeuAc(1) | N[+2352] | 1396.4006 | 4 | 5582.5807 | 5582.5617 | 0 | 3.4 | 141 | Specific | 848.47 | 684.98 | 684.98 | 17.7 | 126 | >sp\|P02788\|TRFL_HUMAN Lactotransferrin OS=Homo sapiens OX=9606 GN=LTF PE=1 SV=6 | 1270 | 20201118_ly_colostrum_1ng_glyco_hcd203040_compound15.21182.21182.4 | controllerType=0 controllerNumber=1 scan=21182 | 46.4657 |
| 17588:4 | 2 | R.TAGWNVPIGTLRPFLN[+2059.735]WTGPPEPIEAAVAR.F | HexNAc(4)Hex(5)Fuc(1)NeuAc(1) | N[+2060] | 1323.3716 | 4 | 5290.4645 | 5290.4459 | 0 | 3.5 | 141 | Specific | 861.69 | 709.45 | 709.45 | 18.7 | 126 | >sp\|P02788\|TRFL_HUMAN Lactotransferrin OS=Homo sapiens OX=9606 GN=LTF PE=1 SV=6 | 1270 | 20201118_ly_colostrum_1ng_glyco_hcd203040_compound15.21153.21153.4 | controllerType=0 controllerNumber=1 scan=21153 | 46.3523 |
| 16928:3 | 2 | R.TAGWNIPMGLLFN[+2205.793]QTGSC[+57.021]K.F | HexNAc(4)Hex(5)Fuc(2)NeuAc(1) | C[+57], N[+2206] | 1434.2831 | 3 | 4300.8349 | 4300.8028 | 0 | 7.5 | 485 | Specific | 703.59 | 614.6 | 614.6 | 14.25 | 126 | >sp\|P02788\|TRFL_HUMAN Lactotransferrin OS=Homo sapiens OX=9606 GN=LTF PE=1 SV=6 | 1270 | 20201118_ly_colostrum_1ng_glyco_hcd203040_compound15.20463.20463.3 | controllerType=0 controllerNumber=1 scan=20463 | 44.8438 |
| 18856:3 | 2 | R.TAGWNIPMGLLFN[+2350.830]QTGSC[+57.021]K.F | HexNAc(4)Hex(5)Fuc(1)NeuAc(2) | C[+57], N[+2351] | 1482.6225 | 3 | 4445.8529 | 4445.8403 | 0 | 2.8 | 485 | Specific | 740.39 | 721.36 | 721.36 | 15.11 | 126 | >sp\|P02788\|TRFL_HUMAN Lactotransferrin OS=Homo sapiens OX=9606 GN=LTF PE=1 SV=6 | 1270 | 20201118_ly_colostrum_1ng_glyco_hcd203040_compound15.22478.22478.3 | controllerType=0 controllerNumber=1 scan=22478 | 49.421 |
| 16931:3 | 2 | R.TAGWNIPMGLLFN[+2205.793]QTGSC[+57.021]K.F | HexNAc(4)Hex(5)Fuc(2)NeuAc(1) | C[+57], N[+2206] | 1434.2831 | 3 | 4300.8349 | 4300.8028 | 0 | 7.5 | 485 | Specific | 860.51 | 782.33 | 782.33 | 17.25 | 126 | >sp\|P02788\|TRFL_HUMAN Lactotransferrin OS=Homo sapiens OX=9606 GN=LTF PE=1 SV=6 | 1270 | 20201118_ly_colostrum_1ng_glyco_hcd203040_compound15.20466.20466.3 | controllerType=0 controllerNumber=1 scan=20466 | 44.8496 |
| 16885:3 | 2 | R.TAGWNIPMGLLFN[+2350.830]QTGSC[+57.021]K.F | HexNAc(4)Hex(5)Fuc(1)NeuAc(2) | C[+57], N[+2351] | 1482.6304 | 3 | 4445.8767 | 4445.8403 | 0 | 8.2 | 485 | Specific | 903.24 | 889.55 | 889.55 | 17.97 | 126 | >sp\|P02788\|TRFL_HUMAN Lactotransferrin OS=Homo sapiens OX=9606 GN=LTF PE=1 SV=6 | 1270 | 20201118_ly_colostrum_1ng_glyco_hcd203040_compound15.20417.20417.3 | controllerType=0 controllerNumber=1 scan=20417 | 44.7089 |
| 18859:3 | 2 | R.TAGWNIPMGLLFN[+2350.830]QTGSC[+57.021]K.F | HexNAc(4)Hex(5)Fuc(1)NeuAc(2) | C[+57], N[+2351] | 1482.6225 | 3 | 4445.8529 | 4445.8403 | 0 | 2.8 | 485 | Specific | 932.88 | 894.92 | 894.92 | 19.11 | 126 | >sp\|P02788\|TRFL_HUMAN Lactotransferrin OS=Homo sapiens OX=9606 GN=LTF PE=1 SV=6 | 1270 | 20201118_ly_colostrum_1ng_glyco_hcd203040_compound15.22481.22481.3 | controllerType=0 controllerNumber=1 scan=22481 | 49.4259 |
| 16883:3 | 2 | R.TAGWNIPMGLLFN[+2350.830]QTGSC[+57.021]K.F | HexNAc(4)Hex(5)Fuc(1)NeuAc(2) | C[+57], N[+2351] | 1482.6304 | 3 | 4445.8767 | 4445.8403 | 0 | 8.2 | 485 | Specific | 1007.58 | 857.91 | 857.91 | 19.97 | 126 | >sp\|P02788\|TRFL_HUMAN Lactotransferrin OS=Homo sapiens OX=9606 GN=LTF PE=1 SV=6 | 1270 | 20201118_ly_colostrum_1ng_glyco_hcd203040_compound15.20415.20415.3 | controllerType=0 controllerNumber=1 scan=20415 | 44.7038 |
| 16269:3 | 2 | R.TAGWNIPMGLLFN[+1768.640]QTGSC[+57.021]KFDEYFSQSC[+57.021]APGSDPR.S | HexNAc(4)Hex(5)Fuc(1) | C[+57]*2, N[+1769] | 1903.1425 | 3 | 5707.4131 | 5707.3966 | 0 | 2.9 | 485 | Specific | 260.28 | 193.05 | 193.05 | 7.24 | 126 | >sp\|P02788\|TRFL_HUMAN Lactotransferrin OS=Homo sapiens OX=9606 GN=LTF PE=1 SV=6 | 1270 | 20201118_ly_colostrum_1ng_glyco_hcd203040_compound15.19773.19773.3 | controllerType=0 controllerNumber=1 scan=19773 | 43.3022 |
| 17195:4 | 2 | R.TAGWNIPMGLLFN[+2205.793]QTGSC[+57.021]KFDEYFSQSC[+57.021]APGSDPR.S | HexNAc(4)Hex(5)Fuc(2)NeuAc(1) | C[+57]*2, N[+2206] | 1536.8986 | 4 | 6144.5724 | 6144.5499 | 0 | 3.7 | 485 | Specific | 294.13 | 272.45 | 272.45 | 8.01 | 126 | >sp\|P02788\|TRFL_HUMAN Lactotransferrin OS=Homo sapiens OX=9606 GN=LTF PE=1 SV=6 | 1270 | 20201118_ly_colostrum_1ng_glyco_hcd203040_compound15.20742.20742.3 | controllerType=0 controllerNumber=1 scan=20742 | 45.4881 |
| 17199:5 | 2 | R.TAGWNIPMGLLFN[+2351.851]QTGSC[+57.021]KFDEYFSQSC[+57.021]APGSDPR.S | HexNAc(4)Hex(5)Fuc(3)NeuAc(1) | C[+57]*2, N[+2352] | 1258.9296 | 5 | 6290.6189 | 6290.6079 | 0 | 1.8 | 485 | Specific | 336.8 | 336.8 | 336.8 | 7.26 | 126 | >sp\|P02788\|TRFL_HUMAN Lactotransferrin OS=Homo sapiens OX=9606 GN=LTF PE=1 SV=6 | 1270 | 20201118_ly_colostrum_1ng_glyco_hcd203040_compound15.20746.20746.5 | controllerType=0 controllerNumber=1 scan=20746 | 45.4933 |
| 17194:4 | 2 | R.TAGWNIPMGLLFN[+2350.830]QTGSC[+57.021]KFDEYFSQSC[+57.021]APGSDPR.S | HexNAc(4)Hex(5)Fuc(1)NeuAc(2) | C[+57]*2, N[+2351] | 1573.1583 | 4 | 6289.6115 | 6289.5875 | 0 | 3.8 | 485 | Specific | 349.71 | 340.25 | 340.25 | 8.77 | 126 | >sp\|P02788\|TRFL_HUMAN Lactotransferrin OS=Homo sapiens OX=9606 GN=LTF PE=1 SV=6 | 1270 | 20201118_ly_colostrum_1ng_glyco_hcd203040_compound15.20741.20741.4 | controllerType=0 controllerNumber=1 scan=20741 | 45.4873 |
| 16248:4 | 2 | R.TAGWNIPMGLLFN[+1768.640]QTGSC[+57.021]KFDEYFSQSC[+57.021]APGSDPR.S | HexNAc(4)Hex(5)Fuc(1) | C[+57]*2, N[+1769] | 1427.6101 | 4 | 5707.4186 | 5707.3966 | 0 | 3.9 | 485 | Specific | 405.9 | 384.4 | 384.4 | 10.67 | 126 | >sp\|P02788\|TRFL_HUMAN Lactotransferrin OS=Homo sapiens OX=9606 GN=LTF PE=1 SV=6 | 1270 | 20201118_ly_colostrum_1ng_glyco_hcd203040_compound15.19752.19752.4 | controllerType=0 controllerNumber=1 scan=19752 | 43.2545 |
| 16169:4 | 2 | R.TAGWNIPMGLLFN[+2351.851]QTGSC[+57.021]KFDEYFSQSC[+57.021]APGSDPR.S | HexNAc(4)Hex(5)Fuc(3)NeuAc(1) | C[+57]*2, N[+2352] | 1573.4186 | 4 | 6290.6525 | 6290.6079 | 0 | 7.1 | 485 | Specific | 560 | 560 | 560 | 10.4 | 126 | >sp\|P02788\|TRFL_HUMAN Lactotransferrin OS=Homo sapiens OX=9606 GN=LTF PE=1 SV=6 | 1270 | 20201118_ly_colostrum_1ng_glyco_hcd203040_compound15.19669.19669.4 | controllerType=0 controllerNumber=1 scan=19669 | 43.0441 |
| 16378:3 | 2 | R.TAGWNIPMGLLFN[+1216.423]QTGSC[+57.021]KFDEYFSQSC[+57.021]APGSDPR.S | HexNAc(2)Hex(5) | C[+57]*2, N[+1216] | 1719.0698 | 3 | 5155.1948 | 5155.18 | 0 | 2.9 | 485 | Specific | 670.18 | 633.7 | 167.7 | 14.6 | 126 | >sp\|P02788\|TRFL_HUMAN Lactotransferrin OS=Homo sapiens OX=9606 GN=LTF PE=1 SV=6 | 1270 | 20201118_ly_colostrum_1ng_glyco_hcd203040_compound15.19887.19887.3 | controllerType=0 controllerNumber=1 scan=19887 | 43.5401 |
| 17198:4 | 2 | R.TAGWNIPMGLLFN[+2205.793]QTGSC[+57.021]KFDEYFSQSC[+57.021]APGSDPR.S | HexNAc(4)Hex(5)Fuc(2)NeuAc(1) | C[+57]*2, N[+2206] | 1536.8986 | 4 | 6144.5724 | 6144.5499 | 0 | 3.7 | 485 | Specific | 774.79 | 751.66 | 751.66 | 16.78 | 126 | >sp\|P02788\|TRFL_HUMAN Lactotransferrin OS=Homo sapiens OX=9606 GN=LTF PE=1 SV=6 | 1270 | 20201118_ly_colostrum_1ng_glyco_hcd203040_compound15.20745.20745.3 | controllerType=0 controllerNumber=1 scan=20745 | 45.4923 |
| 18503:4 | 2 | R.TAGWNIPMGLLFN[+2204.772]QTGSC[+57.021]KFDEYFSQSC[+57.021]APGSDPR.S | HexNAc(4)Hex(5)NeuAc(2) | C[+57]*2, N[+2205] | 1536.6423 | 4 | 6143.5475 | 6143.5295 | 0 | 2.9 | 485 | Specific | 798.51 | 774.3 | 304.56 | 16.76 | 126 | >sp\|P02788\|TRFL_HUMAN Lactotransferrin OS=Homo sapiens OX=9606 GN=LTF PE=1 SV=6 | 1270 | 20201118_ly_colostrum_1ng_glyco_hcd203040_compound15.22109.22109.4 | controllerType=0 controllerNumber=1 scan=22109 | 48.5707 |
| 16171:4 | 2 | R.TAGWNIPMGLLFN[+2351.851]QTGSC[+57.021]KFDEYFSQSC[+57.021]APGSDPR.S | HexNAc(4)Hex(5)Fuc(3)NeuAc(1) | C[+57]*2, N[+2352] | 1573.4186 | 4 | 6290.6525 | 6290.6079 | 0 | 7.1 | 485 | Specific | 817.94 | 800.8 | 800.8 | 16.9 | 126 | >sp\|P02788\|TRFL_HUMAN Lactotransferrin OS=Homo sapiens OX=9606 GN=LTF PE=1 SV=6 | 1270 | 20201118_ly_colostrum_1ng_glyco_hcd203040_compound15.19671.19671.4 | controllerType=0 controllerNumber=1 scan=19671 | 43.0542 |
| 18506:4 | 2 | R.TAGWNIPMGLLFN[+2204.772]QTGSC[+57.021]KFDEYFSQSC[+57.021]APGSDPR.S | HexNAc(4)Hex(5)NeuAc(2) | C[+57]*2, N[+2205] | 1536.6423 | 4 | 6143.5475 | 6143.5295 | 0 | 2.9 | 485 | Specific | 831.57 | 799.06 | 442.56 | 17.76 | 126 | >sp\|P02788\|TRFL_HUMAN Lactotransferrin OS=Homo sapiens OX=9606 GN=LTF PE=1 SV=6 | 1270 | 20201118_ly_colostrum_1ng_glyco_hcd203040_compound15.22112.22112.4 | controllerType=0 controllerNumber=1 scan=22112 | 48.576 |
| 17191:4 | 2 | R.TAGWNIPMGLLFN[+2059.735]QTGSC[+57.021]KFDEYFSQSC[+57.021]APGSDPR.S | HexNAc(4)Hex(5)Fuc(1)NeuAc(1) | C[+57]*2, N[+2060] | 1500.3851 | 4 | 5998.5187 | 5998.492 | 0 | 4.4 | 485 | Specific | 881.84 | 845 | 845 | 18.49 | 126 | >sp\|P02788\|TRFL_HUMAN Lactotransferrin OS=Homo sapiens OX=9606 GN=LTF PE=1 SV=6 | 1270 | 20201118_ly_colostrum_1ng_glyco_hcd203040_compound15.20737.20737.4 | controllerType=0 controllerNumber=1 scan=20737 | 45.4806 |
| 17197:4 | 2 | R.TAGWNIPMGLLFN[+2350.830]QTGSC[+57.021]KFDEYFSQSC[+57.021]APGSDPR.S | HexNAc(4)Hex(5)Fuc(1)NeuAc(2) | C[+57]*2, N[+2351] | 1573.1583 | 4 | 6289.6115 | 6289.5875 | 0 | 3.8 | 485 | Specific | 889.93 | 834.34 | 834.34 | 18.78 | 126 | >sp\|P02788\|TRFL_HUMAN Lactotransferrin OS=Homo sapiens OX=9606 GN=LTF PE=1 SV=6 | 1270 | 20201118_ly_colostrum_1ng_glyco_hcd203040_compound15.20744.20744.4 | controllerType=0 controllerNumber=1 scan=20744 | 45.4905 |
| 18407:4 | 2 | R.TAGWNIPMGLLFN[+2350.830]QTGSC[+57.021]KFDEYFSQSC[+57.021]APGSDPR.S | HexNAc(4)Hex(5)Fuc(1)NeuAc(2) | C[+57]*2, N[+2351] | 1573.1576 | 4 | 6289.6086 | 6289.5875 | 0 | 3.4 | 485 | Specific | 899.75 | 871.74 | 871.74 | 18.78 | 126 | >sp\|P02788\|TRFL_HUMAN Lactotransferrin OS=Homo sapiens OX=9606 GN=LTF PE=1 SV=6 | 1270 | 20201118_ly_colostrum_1ng_glyco_hcd203040_compound15.22009.22009.4 | controllerType=0 controllerNumber=1 scan=22009 | 48.3597 |
| 18404:4 | 2 | R.TAGWNIPMGLLFN[+2350.830]QTGSC[+57.021]KFDEYFSQSC[+57.021]APGSDPR.S | HexNAc(4)Hex(5)Fuc(1)NeuAc(2) | C[+57]*2, N[+2351] | 1573.1576 | 4 | 6289.6086 | 6289.5875 | 0 | 3.4 | 485 | Specific | 937.36 | 898.48 | 898.48 | 19.78 | 126 | >sp\|P02788\|TRFL_HUMAN Lactotransferrin OS=Homo sapiens OX=9606 GN=LTF PE=1 SV=6 | 1270 | 20201118_ly_colostrum_1ng_glyco_hcd203040_compound15.22006.22006.4 | controllerType=0 controllerNumber=1 scan=22006 | 48.3487 |
| 17188:4 | 2 | R.TAGWNIPMGLLFN[+2059.735]QTGSC[+57.021]KFDEYFSQSC[+57.021]APGSDPR.S | HexNAc(4)Hex(5)Fuc(1)NeuAc(1) | C[+57]*2, N[+2060] | 1500.3851 | 4 | 5998.5187 | 5998.492 | 0 | 4.4 | 485 | Specific | 955.47 | 926.99 | 926.99 | 20.49 | 126 | >sp\|P02788\|TRFL_HUMAN Lactotransferrin OS=Homo sapiens OX=9606 GN=LTF PE=1 SV=6 | 1270 | 20201118_ly_colostrum_1ng_glyco_hcd203040_compound15.20734.20734.4 | controllerType=0 controllerNumber=1 scan=20734 | 45.4705 |
| 16250:4 | 2 | R.TAGWNIPMGLLFN[+1768.640]QTGSC[+57.021]KFDEYFSQSC[+57.021]APGSDPR.S | HexNAc(4)Hex(5)Fuc(1) | C[+57]*2, N[+1769] | 1427.6101 | 4 | 5707.4186 | 5707.3966 | 0 | 3.9 | 485 | Specific | 1009.32 | 897.97 | 897.97 | 21.78 | 126 | >sp\|P02788\|TRFL_HUMAN Lactotransferrin OS=Homo sapiens OX=9606 GN=LTF PE=1 SV=6 | 1270 | 20201118_ly_colostrum_1ng_glyco_hcd203040_compound15.19754.19754.4 | controllerType=0 controllerNumber=1 scan=19754 | 43.2598 |
| 15682:3 | 3 | R.GLSFDVSLEVSQGPGLLN[+2863.041]DTK.V | HexNAc(5)Hex(6)Fuc(4)NeuAc(1) | N[+2863] | 1680.3946 | 3 | 5039.1693 | 5039.1541 | 0 | 3 | 118 | Specific | 487.62 | 451.69 | 451.69 | 10.69 | 136 | >sp\|P01833\|PIGR_HUMAN Polymeric immunoglobulin receptor OS=Homo sapiens OX=9606 GN=PIGR PE=1 SV=4 | 902 | 20201118_ly_colostrum_1ng_glyco_hcd203040_compound15.19161.19161.3 | controllerType=0 controllerNumber=1 scan=19161 | 41.9018 |
| 15684:3 | 3 | R.GLSFDVSLEVSQGPGLLN[+2863.041]DTK.V | HexNAc(5)Hex(6)Fuc(4)NeuAc(1) | N[+2863] | 1680.3946 | 3 | 5039.1693 | 5039.1541 | 0 | 3 | 118 | Specific | 596.2 | 528.16 | 528.16 | 12.54 | 136 | >sp\|P01833\|PIGR_HUMAN Polymeric immunoglobulin receptor OS=Homo sapiens OX=9606 GN=PIGR PE=1 SV=4 | 902 | 20201118_ly_colostrum_1ng_glyco_hcd203040_compound15.19163.19163.3 | controllerType=0 controllerNumber=1 scan=19163 | 41.9063 |
| 17737:3 | 3 | R.GLSFDVSLEVSQGPGLLN[+2350.830]DTK.V | HexNAc(4)Hex(5)Fuc(1)NeuAc(2) | N[+2351] | 1509.6607 | 3 | 4526.9676 | 4526.9436 | 0 | 5.3 | 118 | Specific | 738.3 | 730.02 | 730.02 | 15 | 136 | >sp\|P01833\|PIGR_HUMAN Polymeric immunoglobulin receptor OS=Homo sapiens OX=9606 GN=PIGR PE=1 SV=4 | 902 | 20201118_ly_colostrum_1ng_glyco_hcd203040_compound15.21309.21309.3 | controllerType=0 controllerNumber=1 scan=21309 | 46.742 |
| 17730:4 | 3 | R.GLSFDVSLEVSQGPGLLN[+2350.830]DTK.V | HexNAc(4)Hex(5)Fuc(1)NeuAc(2) | N[+2351] | 1132.4967 | 4 | 4526.9652 | 4526.9436 | 0 | 4.8 | 118 | Specific | 808.32 | 680.14 | 680.14 | 17.13 | 136 | >sp\|P01833\|PIGR_HUMAN Polymeric immunoglobulin receptor OS=Homo sapiens OX=9606 GN=PIGR PE=1 SV=4 | 902 | 20201118_ly_colostrum_1ng_glyco_hcd203040_compound15.21301.21301.4 | controllerType=0 controllerNumber=1 scan=21301 | 46.7241 |
| 17733:4 | 3 | R.GLSFDVSLEVSQGPGLLN[+2350.830]DTK.V | HexNAc(4)Hex(5)Fuc(1)NeuAc(2) | N[+2351] | 1132.4967 | 4 | 4526.9652 | 4526.9436 | 0 | 4.8 | 118 | Specific | 921.57 | 863.89 | 863.89 | 19.13 | 136 | >sp\|P01833\|PIGR_HUMAN Polymeric immunoglobulin receptor OS=Homo sapiens OX=9606 GN=PIGR PE=1 SV=4 | 902 | 20201118_ly_colostrum_1ng_glyco_hcd203040_compound15.21304.21304.4 | controllerType=0 controllerNumber=1 scan=21304 | 46.7324 |
| 17740:3 | 3 | R.GLSFDVSLEVSQGPGLLN[+2350.830]DTK.V | HexNAc(4)Hex(5)Fuc(1)NeuAc(2) | N[+2351] | 1509.6607 | 3 | 4526.9676 | 4526.9436 | 0 | 5.3 | 118 | Specific | 1020.13 | 963.1 | 963.1 | 21 | 136 | >sp\|P01833\|PIGR_HUMAN Polymeric immunoglobulin receptor OS=Homo sapiens OX=9606 GN=PIGR PE=1 SV=4 | 902 | 20201118_ly_colostrum_1ng_glyco_hcd203040_compound15.21312.21312.3 | controllerType=0 controllerNumber=1 scan=21312 | 46.7458 |
| 18192:3 | 3 | R.GLSFDVSLEVSQGPGLLN[+2351.851]DTKVYTVDLGR.T | HexNAc(4)Hex(5)Fuc(3)NeuAc(1) | N[+2352] | 1811.1472 | 3 | 5431.427 | 5431.4454 | 0 | -3.4 | 118 | Specific | 280.67 | 249.38 | 249.38 | 6.22 | 136 | >sp\|P01833\|PIGR_HUMAN Polymeric immunoglobulin receptor OS=Homo sapiens OX=9606 GN=PIGR PE=1 SV=4 | 902 | 20201118_ly_colostrum_1ng_glyco_hcd203040_compound15.21784.21784.3 | controllerType=0 controllerNumber=1 scan=21784 | 47.836 |
| 18190:3 | 3 | R.GLSFDVSLEVSQGPGLLN[+2351.851]DTKVYTVDLGR.T | HexNAc(4)Hex(5)Fuc(3)NeuAc(1) | N[+2352] | 1811.1472 | 3 | 5431.427 | 5431.4454 | 0 | -3.4 | 118 | Specific | 349.61 | 334.81 | 334.81 | 7.11 | 136 | >sp\|P01833\|PIGR_HUMAN Polymeric immunoglobulin receptor OS=Homo sapiens OX=9606 GN=PIGR PE=1 SV=4 | 902 | 20201118_ly_colostrum_1ng_glyco_hcd203040_compound15.21782.21782.3 | controllerType=0 controllerNumber=1 scan=21782 | 47.8323 |
| 18142:4 | 3 | R.GLSFDVSLEVSQGPGLLN[+2350.830]DTKVYTVDLGR.T | HexNAc(4)Hex(5)Fuc(1)NeuAc(2) | N[+2351] | 1358.3673 | 4 | 5430.4474 | 5430.425 | 0 | 4.1 | 118 | Specific | 692.85 | 692.85 | 692.85 | 14.18 | 136 | >sp\|P01833\|PIGR_HUMAN Polymeric immunoglobulin receptor OS=Homo sapiens OX=9606 GN=PIGR PE=1 SV=4 | 902 | 20201118_ly_colostrum_1ng_glyco_hcd203040_compound15.21732.21732.4 | controllerType=0 controllerNumber=1 scan=21732 | 47.7241 |
| 16771:3 | 3 | R.GLSFDVSLEVSQGPGLLN[+2351.851]DTKVYTVDLGR.T | HexNAc(4)Hex(5)Fuc(3)NeuAc(1) | N[+2352] | 1811.1555 | 3 | 5431.4519 | 5431.4454 | 0 | 1.2 | 118 | Specific | 764.69 | 663.43 | 663.43 | 16.06 | 136 | >sp\|P01833\|PIGR_HUMAN Polymeric immunoglobulin receptor OS=Homo sapiens OX=9606 GN=PIGR PE=1 SV=4 | 902 | 20201118_ly_colostrum_1ng_glyco_hcd203040_compound15.20297.20297.3 | controllerType=0 controllerNumber=1 scan=20297 | 44.443 |
| 16768:3 | 3 | R.GLSFDVSLEVSQGPGLLN[+2351.851]DTKVYTVDLGR.T | HexNAc(4)Hex(5)Fuc(3)NeuAc(1) | N[+2352] | 1811.1555 | 3 | 5431.4519 | 5431.4454 | 0 | 1.2 | 118 | Specific | 793.48 | 658.27 | 658.27 | 16.06 | 136 | >sp\|P01833\|PIGR_HUMAN Polymeric immunoglobulin receptor OS=Homo sapiens OX=9606 GN=PIGR PE=1 SV=4 | 902 | 20201118_ly_colostrum_1ng_glyco_hcd203040_compound15.20294.20294.3 | controllerType=0 controllerNumber=1 scan=20294 | 44.4382 |
| 18144:4 | 3 | R.GLSFDVSLEVSQGPGLLN[+2350.830]DTKVYTVDLGR.T | HexNAc(4)Hex(5)Fuc(1)NeuAc(2) | N[+2351] | 1358.3673 | 4 | 5430.4474 | 5430.425 | 0 | 4.1 | 118 | Specific | 807.07 | 703.44 | 703.44 | 17.18 | 136 | >sp\|P01833\|PIGR_HUMAN Polymeric immunoglobulin receptor OS=Homo sapiens OX=9606 GN=PIGR PE=1 SV=4 | 902 | 20201118_ly_colostrum_1ng_glyco_hcd203040_compound15.21734.21734.4 | controllerType=0 controllerNumber=1 scan=21734 | 47.7294 |
| 20877:3 | 3 | K.QIGLYPVLVIDSSGYVNPN[+2205.793]YTGR.I | HexNAc(4)Hex(5)Fuc(2)NeuAc(1) | N[+2206] | 1577.7098 | 3 | 4731.1148 | 4731.0963 | 0 | 3.9 | 168 | Specific | 221.21 | 221.21 | 221.21 | 5.02 | 136 | >sp\|P01833\|PIGR_HUMAN Polymeric immunoglobulin receptor OS=Homo sapiens OX=9606 GN=PIGR PE=1 SV=4 | 902 | 20201118_ly_colostrum_1ng_glyco_hcd203040_compound15.24588.24588.3 | controllerType=0 controllerNumber=1 scan=24588 | 54.0195 |
| 20880:3 | 3 | K.QIGLYPVLVIDSSGYVNPN[+2205.793]YTGR.I | HexNAc(4)Hex(5)Fuc(2)NeuAc(1) | N[+2206] | 1577.7098 | 3 | 4731.1148 | 4731.0963 | 0 | 3.9 | 168 | Specific | 231.59 | 231.59 | 231.59 | 5.02 | 136 | >sp\|P01833\|PIGR_HUMAN Polymeric immunoglobulin receptor OS=Homo sapiens OX=9606 GN=PIGR PE=1 SV=4 | 902 | 20201118_ly_colostrum_1ng_glyco_hcd203040_compound15.24591.24591.3 | controllerType=0 controllerNumber=1 scan=24591 | 54.0269 |
| 17109:3 | 3 | K.QIGLYPVLVIDSSGYVNPN[+2076.750]YTGR.I | HexNAc(4)Hex(6)Fuc(2) | N[+2077] | 1534.6977 | 3 | 4602.0785 | 4602.0537 | 0 | 5.4 | 168 | Specific | 257.04 | 257.04 | 257.04 | 5.21 | 136 | >sp\|P01833\|PIGR_HUMAN Polymeric immunoglobulin receptor OS=Homo sapiens OX=9606 GN=PIGR PE=1 SV=4 | 902 | 20201118_ly_colostrum_1ng_glyco_hcd203040_compound15.20652.20652.3 | controllerType=0 controllerNumber=1 scan=20652 | 45.2789 |
| 17153:4 | 3 | K.QIGLYPVLVIDSSGYVNPN[+2059.735]YTGR.I | HexNAc(4)Hex(5)Fuc(1)NeuAc(1) | N[+2060] | 1147.0159 | 4 | 4585.0418 | 4585.0384 | 0 | 0.7 | 168 | Specific | 303.49 | 303.49 | 303.49 | 7.35 | 136 | >sp\|P01833\|PIGR_HUMAN Polymeric immunoglobulin receptor OS=Homo sapiens OX=9606 GN=PIGR PE=1 SV=4 | 902 | 20201118_ly_colostrum_1ng_glyco_hcd203040_compound15.20698.20698.4 | controllerType=0 controllerNumber=1 scan=20698 | 45.3864 |
| 17047:4 | 3 | K.QIGLYPVLVIDSSGYVNPN[+2351.851]YTGR.I | HexNAc(4)Hex(5)Fuc(3)NeuAc(1) | N[+2352] | 1220.054 | 4 | 4877.1942 | 4877.1542 | 0 | 8.2 | 168 | Specific | 334.26 | 334.26 | 334.26 | 6.91 | 136 | >sp\|P01833\|PIGR_HUMAN Polymeric immunoglobulin receptor OS=Homo sapiens OX=9606 GN=PIGR PE=1 SV=4 | 902 | 20201118_ly_colostrum_1ng_glyco_hcd203040_compound15.20587.20587.4 | controllerType=0 controllerNumber=1 scan=20587 | 45.1232 |
| 17110:4 | 3 | K.QIGLYPVLVIDSSGYVNPN[+2076.750]YTGR.I | HexNAc(4)Hex(6)Fuc(2) | N[+2077] | 1151.2753 | 4 | 4602.0794 | 4602.0537 | 0 | 5.6 | 168 | Specific | 338.68 | 275.08 | 275.08 | 5.16 | 136 | >sp\|P01833\|PIGR_HUMAN Polymeric immunoglobulin receptor OS=Homo sapiens OX=9606 GN=PIGR PE=1 SV=4 | 902 | 20201118_ly_colostrum_1ng_glyco_hcd203040_compound15.20653.20653.4 | controllerType=0 controllerNumber=1 scan=20653 | 45.2799 |
| 18943:4 | 3 | K.QIGLYPVLVIDSSGYVNPN[+2350.830]YTGR.I | HexNAc(4)Hex(5)Fuc(1)NeuAc(2) | N[+2351] | 1219.7919 | 4 | 4876.1458 | 4876.1338 | 0 | 2.5 | 168 | Specific | 371.01 | 335.02 | 335.02 | 8.18 | 136 | >sp\|P01833\|PIGR_HUMAN Polymeric immunoglobulin receptor OS=Homo sapiens OX=9606 GN=PIGR PE=1 SV=4 | 902 | 20201118_ly_colostrum_1ng_glyco_hcd203040_compound15.22569.22569.4 | controllerType=0 controllerNumber=1 scan=22569 | 49.6446 |
| 19049:3 | 3 | K.QIGLYPVLVIDSSGYVNPN[+2350.830]YTGR.I | HexNAc(4)Hex(5)Fuc(1)NeuAc(2) | N[+2351] | 1626.0572 | 3 | 4876.157 | 4876.1338 | 0 | 4.8 | 168 | Specific | 430.55 | 322.92 | 322.92 | 8.17 | 136 | >sp\|P01833\|PIGR_HUMAN Polymeric immunoglobulin receptor OS=Homo sapiens OX=9606 GN=PIGR PE=1 SV=4 | 902 | 20201118_ly_colostrum_1ng_glyco_hcd203040_compound15.22680.22680.3 | controllerType=0 controllerNumber=1 scan=22680 | 49.8883 |
| 19169:3 | 3 | K.QIGLYPVLVIDSSGYVNPN[+2205.793]YTGR.I | HexNAc(4)Hex(5)Fuc(2)NeuAc(1) | N[+2206] | 1577.7042 | 3 | 4731.098 | 4731.0963 | 0 | 0.4 | 168 | Specific | 508.51 | 419.5 | 419.5 | 10.19 | 136 | >sp\|P01833\|PIGR_HUMAN Polymeric immunoglobulin receptor OS=Homo sapiens OX=9606 GN=PIGR PE=1 SV=4 | 902 | 20201118_ly_colostrum_1ng_glyco_hcd203040_compound15.22806.22806.3 | controllerType=0 controllerNumber=1 scan=22806 | 50.219 |
| 19051:3 | 3 | K.QIGLYPVLVIDSSGYVNPN[+2350.830]YTGR.I | HexNAc(4)Hex(5)Fuc(1)NeuAc(2) | N[+2351] | 1626.0572 | 3 | 4876.157 | 4876.1338 | 0 | 4.8 | 168 | Specific | 580.18 | 353.94 | 353.94 | 10.41 | 136 | >sp\|P01833\|PIGR_HUMAN Polymeric immunoglobulin receptor OS=Homo sapiens OX=9606 GN=PIGR PE=1 SV=4 | 902 | 20201118_ly_colostrum_1ng_glyco_hcd203040_compound15.22682.22682.3 | controllerType=0 controllerNumber=1 scan=22682 | 49.8956 |
| 19127:3 | 3 | K.QIGLYPVLVIDSSGYVNPN[+2204.772]YTGR.I | HexNAc(4)Hex(5)NeuAc(2) | N[+2205] | 1577.3671 | 3 | 4730.0868 | 4730.0759 | 0 | 2.3 | 168 | Specific | 649.06 | 501.73 | 501.73 | 13.35 | 136 | >sp\|P01833\|PIGR_HUMAN Polymeric immunoglobulin receptor OS=Homo sapiens OX=9606 GN=PIGR PE=1 SV=4 | 902 | 20201118_ly_colostrum_1ng_glyco_hcd203040_compound15.22762.22762.3 | controllerType=0 controllerNumber=1 scan=22762 | 50.115 |
| 17050:4 | 3 | K.QIGLYPVLVIDSSGYVNPN[+2351.851]YTGR.I | HexNAc(4)Hex(5)Fuc(3)NeuAc(1) | N[+2352] | 1220.054 | 4 | 4877.1942 | 4877.1542 | 0 | 8.2 | 168 | Specific | 655.86 | 580.77 | 580.77 | 13.07 | 136 | >sp\|P01833\|PIGR_HUMAN Polymeric immunoglobulin receptor OS=Homo sapiens OX=9606 GN=PIGR PE=1 SV=4 | 902 | 20201118_ly_colostrum_1ng_glyco_hcd203040_compound15.20590.20590.4 | controllerType=0 controllerNumber=1 scan=20590 | 45.1381 |
| 17156:4 | 3 | K.QIGLYPVLVIDSSGYVNPN[+2059.735]YTGR.I | HexNAc(4)Hex(5)Fuc(1)NeuAc(1) | N[+2060] | 1147.0159 | 4 | 4585.0418 | 4585.0384 | 0 | 0.7 | 168 | Specific | 694.22 | 598.6 | 598.6 | 13.78 | 136 | >sp\|P01833\|PIGR_HUMAN Polymeric immunoglobulin receptor OS=Homo sapiens OX=9606 GN=PIGR PE=1 SV=4 | 902 | 20201118_ly_colostrum_1ng_glyco_hcd203040_compound15.20701.20701.4 | controllerType=0 controllerNumber=1 scan=20701 | 45.3903 |
| 19034:5 | 3 | K.QIGLYPVLVIDSSGYVNPN[+2204.772]YTGR.I | HexNAc(4)Hex(5)NeuAc(2) | N[+2205] | 946.8236 | 5 | 4730.0887 | 4730.0759 | 0 | 2.7 | 168 | Specific | 718.06 | 567.55 | 567.55 | 15.21 | 136 | >sp\|P01833\|PIGR_HUMAN Polymeric immunoglobulin receptor OS=Homo sapiens OX=9606 GN=PIGR PE=1 SV=4 | 902 | 20201118_ly_colostrum_1ng_glyco_hcd203040_compound15.22664.22664.5 | controllerType=0 controllerNumber=1 scan=22664 | 49.8492 |
| 17042:3 | 3 | K.QIGLYPVLVIDSSGYVNPN[+2351.851]YTGR.I | HexNAc(4)Hex(5)Fuc(3)NeuAc(1) | N[+2352] | 1626.3972 | 3 | 4877.1769 | 4877.1542 | 0 | 4.7 | 168 | Specific | 740.33 | 663.02 | 663.02 | 15.08 | 136 | >sp\|P01833\|PIGR_HUMAN Polymeric immunoglobulin receptor OS=Homo sapiens OX=9606 GN=PIGR PE=1 SV=4 | 902 | 20201118_ly_colostrum_1ng_glyco_hcd203040_compound15.20582.20582.3 | controllerType=0 controllerNumber=1 scan=20582 | 45.1123 |
| 19130:3 | 3 | K.QIGLYPVLVIDSSGYVNPN[+2204.772]YTGR.I | HexNAc(4)Hex(5)NeuAc(2) | N[+2205] | 1577.3671 | 3 | 4730.0868 | 4730.0759 | 0 | 2.3 | 168 | Specific | 766.07 | 672.46 | 672.46 | 16.35 | 136 | >sp\|P01833\|PIGR_HUMAN Polymeric immunoglobulin receptor OS=Homo sapiens OX=9606 GN=PIGR PE=1 SV=4 | 902 | 20201118_ly_colostrum_1ng_glyco_hcd203040_compound15.22765.22765.3 | controllerType=0 controllerNumber=1 scan=22765 | 50.1203 |
| 18935:3 | 3 | K.QIGLYPVLVIDSSGYVNPN[+2350.830]YTGR.I | HexNAc(4)Hex(5)Fuc(1)NeuAc(2) | N[+2351] | 1626.054 | 3 | 4876.1475 | 4876.1338 | 0 | 2.8 | 168 | Specific | 767.81 | 613.53 | 613.53 | 16.35 | 136 | >sp\|P01833\|PIGR_HUMAN Polymeric immunoglobulin receptor OS=Homo sapiens OX=9606 GN=PIGR PE=1 SV=4 | 902 | 20201118_ly_colostrum_1ng_glyco_hcd203040_compound15.22561.22561.3 | controllerType=0 controllerNumber=1 scan=22561 | 49.6286 |
| 19032:5 | 3 | K.QIGLYPVLVIDSSGYVNPN[+2204.772]YTGR.I | HexNAc(4)Hex(5)NeuAc(2) | N[+2205] | 946.8236 | 5 | 4730.0887 | 4730.0759 | 0 | 2.7 | 168 | Specific | 787.91 | 720.8 | 720.8 | 16.21 | 136 | >sp\|P01833\|PIGR_HUMAN Polymeric immunoglobulin receptor OS=Homo sapiens OX=9606 GN=PIGR PE=1 SV=4 | 902 | 20201118_ly_colostrum_1ng_glyco_hcd203040_compound15.22662.22662.5 | controllerType=0 controllerNumber=1 scan=22662 | 49.8426 |
| 17066:4 | 3 | K.QIGLYPVLVIDSSGYVNPN[+2205.793]YTGR.I | HexNAc(4)Hex(5)Fuc(2)NeuAc(1) | N[+2206] | 1183.5339 | 4 | 4731.1136 | 4731.0963 | 0 | 3.7 | 168 | Specific | 820.37 | 658.76 | 658.76 | 17.24 | 136 | >sp\|P01833\|PIGR_HUMAN Polymeric immunoglobulin receptor OS=Homo sapiens OX=9606 GN=PIGR PE=1 SV=4 | 902 | 20201118_ly_colostrum_1ng_glyco_hcd203040_compound15.20607.20607.4 | controllerType=0 controllerNumber=1 scan=20607 | 45.1842 |
| 17045:3 | 3 | K.QIGLYPVLVIDSSGYVNPN[+2351.851]YTGR.I | HexNAc(4)Hex(5)Fuc(3)NeuAc(1) | N[+2352] | 1626.3972 | 3 | 4877.1769 | 4877.1542 | 0 | 4.7 | 168 | Specific | 842.89 | 762.31 | 762.31 | 17.08 | 136 | >sp\|P01833\|PIGR_HUMAN Polymeric immunoglobulin receptor OS=Homo sapiens OX=9606 GN=PIGR PE=1 SV=4 | 902 | 20201118_ly_colostrum_1ng_glyco_hcd203040_compound15.20585.20585.3 | controllerType=0 controllerNumber=1 scan=20585 | 45.1204 |
| 17163:3 | 3 | K.QIGLYPVLVIDSSGYVNPN[+1913.677]YTGR.I | HexNAc(4)Hex(5)NeuAc(1) | N[+1914] | 1480.3344 | 3 | 4438.9887 | 4438.9805 | 0 | 1.8 | 168 | Specific | 849.82 | 734.47 | 734.47 | 17.12 | 136 | >sp\|P01833\|PIGR_HUMAN Polymeric immunoglobulin receptor OS=Homo sapiens OX=9606 GN=PIGR PE=1 SV=4 | 902 | 20201118_ly_colostrum_1ng_glyco_hcd203040_compound15.20708.20708.3 | controllerType=0 controllerNumber=1 scan=20708 | 45.4047 |
| 17166:3 | 3 | K.QIGLYPVLVIDSSGYVNPN[+1913.677]YTGR.I | HexNAc(4)Hex(5)NeuAc(1) | N[+1914] | 1480.3344 | 3 | 4438.9887 | 4438.9805 | 0 | 1.8 | 168 | Specific | 885.46 | 582.04 | 582.04 | 18.12 | 136 | >sp\|P01833\|PIGR_HUMAN Polymeric immunoglobulin receptor OS=Homo sapiens OX=9606 GN=PIGR PE=1 SV=4 | 902 | 20201118_ly_colostrum_1ng_glyco_hcd203040_compound15.20711.20711.3 | controllerType=0 controllerNumber=1 scan=20711 | 45.4114 |
| 18945:4 | 3 | K.QIGLYPVLVIDSSGYVNPN[+2350.830]YTGR.I | HexNAc(4)Hex(5)Fuc(1)NeuAc(2) | N[+2351] | 1219.7919 | 4 | 4876.1458 | 4876.1338 | 0 | 2.5 | 168 | Specific | 886.84 | 652.04 | 652.04 | 18.21 | 136 | >sp\|P01833\|PIGR_HUMAN Polymeric immunoglobulin receptor OS=Homo sapiens OX=9606 GN=PIGR PE=1 SV=4 | 902 | 20201118_ly_colostrum_1ng_glyco_hcd203040_compound15.22571.22571.4 | controllerType=0 controllerNumber=1 scan=22571 | 49.6492 |
| 17088:3 | 3 | K.QIGLYPVLVIDSSGYVNPN[+2059.735]YTGR.I | HexNAc(4)Hex(5)Fuc(1)NeuAc(1) | N[+2060] | 1529.0247 | 3 | 4585.0596 | 4585.0384 | 0 | 4.6 | 168 | Specific | 893.69 | 625.45 | 625.45 | 18.08 | 136 | >sp\|P01833\|PIGR_HUMAN Polymeric immunoglobulin receptor OS=Homo sapiens OX=9606 GN=PIGR PE=1 SV=4 | 902 | 20201118_ly_colostrum_1ng_glyco_hcd203040_compound15.20630.20630.3 | controllerType=0 controllerNumber=1 scan=20630 | 45.232 |
| 17058:3 | 3 | K.QIGLYPVLVIDSSGYVNPN[+2205.793]YTGR.I | HexNAc(4)Hex(5)Fuc(2)NeuAc(1) | N[+2206] | 1577.71 | 3 | 4731.1155 | 4731.0963 | 0 | 4.1 | 168 | Specific | 899.72 | 849.77 | 849.77 | 18.08 | 136 | >sp\|P01833\|PIGR_HUMAN Polymeric immunoglobulin receptor OS=Homo sapiens OX=9606 GN=PIGR PE=1 SV=4 | 902 | 20201118_ly_colostrum_1ng_glyco_hcd203040_compound15.20599.20599.3 | controllerType=0 controllerNumber=1 scan=20599 | 45.161 |
| 17060:3 | 3 | K.QIGLYPVLVIDSSGYVNPN[+2205.793]YTGR.I | HexNAc(4)Hex(5)Fuc(2)NeuAc(1) | N[+2206] | 1577.71 | 3 | 4731.1155 | 4731.0963 | 0 | 4.1 | 168 | Specific | 912.95 | 842.37 | 842.37 | 19.08 | 136 | >sp\|P01833\|PIGR_HUMAN Polymeric immunoglobulin receptor OS=Homo sapiens OX=9606 GN=PIGR PE=1 SV=4 | 902 | 20201118_ly_colostrum_1ng_glyco_hcd203040_compound15.20601.20601.3 | controllerType=0 controllerNumber=1 scan=20601 | 45.168 |
| 17085:3 | 3 | K.QIGLYPVLVIDSSGYVNPN[+2059.735]YTGR.I | HexNAc(4)Hex(5)Fuc(1)NeuAc(1) | N[+2060] | 1529.0247 | 3 | 4585.0596 | 4585.0384 | 0 | 4.6 | 168 | Specific | 919.23 | 760.43 | 760.43 | 19.08 | 136 | >sp\|P01833\|PIGR_HUMAN Polymeric immunoglobulin receptor OS=Homo sapiens OX=9606 GN=PIGR PE=1 SV=4 | 902 | 20201118_ly_colostrum_1ng_glyco_hcd203040_compound15.20627.20627.3 | controllerType=0 controllerNumber=1 scan=20627 | 45.2232 |
| 18761:3 | 3 | K.AQYEGRLSLLEEPGN[+1647.613]GTFTVILNQLTSR.D | HexNAc(5)Hex(3)Fuc(1) | N[+1648] | 1585.4233 | 3 | 4754.2553 | 4754.23 | 0 | 5.3 | 407 | Specific | 215.8 | 215.8 | 215.8 | 5.08 | 136 | >sp\|P01833\|PIGR_HUMAN Polymeric immunoglobulin receptor OS=Homo sapiens OX=9606 GN=PIGR PE=1 SV=4 | 902 | 20201118_ly_colostrum_1ng_glyco_hcd203040_compound15.22378.22378.3 | controllerType=0 controllerNumber=1 scan=22378 | 49.1859 |

Table S4. Cont.

| Query #:z | Protein Rank | Glycopeptide | Glycan | Modification Type(s) | Observed m/z | z | Observed (M+H) | Calc. mass (M+H) | Off-by-x error | Mass error (ppm) | StartPosition | Cleavage | Score | Delta | Delta Mod | \|Log Prob\| | # of unique peptides | Protein Name | Protein DB number | Comment | Scan # | Scan Time |
|---|---|---|---|---|---|---|---|---|---|---|---|---|---|---|---|---|---|---|---|---|---|---|
| 18759:3 | 3 | K.AQYEGRLSLLEEPGN[+1647.613]GTFTVILNQLTSR.D | HexNAc(5)Hex(3)Fuc(1) | N[+1648] | 1585.4233 | 3 | 4754.2553 | 4754.23 | 0 | 5.3 | 407 | Specific | 284.9 | 284.9 | 284.9 | 7.1 | 136 | >sp\|P01833\|PIGR_HUMAN Polymeric immunoglobulin receptor OS=Homo sapiens OX=9606 GN=PIGR PE=1 SV=4 | 902 | 20201118_ly_colostrum_1ng_glyco_hcd203040_compound15.22376.22376.3 | controllerType=0 controllerNumber=1 scan=22376 | 49.1806 |
| 18553:4 | 3 | K.[+42.011]AQYEGRLSLLEEPGN[+1622.582]GTFTVILNQLTSR.D | HexNAc(4)Hex(5) | .[42], N[+1623] | 1193.5513 | 4 | 4771.1834 | 4771.2089 | 0 | -5.3 | 407 | Specific | 327.61 | 301.42 | 301.42 | 6.89 | 136 | >sp\|P01833\|PIGR_HUMAN Polymeric immunoglobulin receptor OS=Homo sapiens OX=9606 GN=PIGR PE=1 SV=4 | 902 | 20201118_ly_colostrum_1ng_glyco_hcd203040_compound15.22161.22161.5 | controllerType=0 controllerNumber=1 scan=22161 | 48.6951 |
| 18551:4 | 3 | K.[+42.011]AQYEGRLSLLEEPGN[+1622.582]GTFTVILNQLTSR.D | HexNAc(4)Hex(5) | .[42], N[+1623] | 1193.5513 | 4 | 4771.1834 | 4771.2089 | 0 | -5.3 | 407 | Specific | 366.08 | 220.74 | 220.74 | 6.36 | 136 | >sp\|P01833\|PIGR_HUMAN Polymeric immunoglobulin receptor OS=Homo sapiens OX=9606 GN=PIGR PE=1 SV=4 | 902 | 20201118_ly_colostrum_1ng_glyco_hcd203040_compound15.22159.22159.5 | controllerType=0 controllerNumber=1 scan=22159 | 48.688 |
| 18684:4 | 3 | R.LSLLEEPGN[+2076.750]GTFTVILNQLTSR.D | HexNAc(4)Hex(6)Fuc(2) | N[+2077] | 1120.5231 | 4 | 4479.0706 | 4479.0428 | 0 | 6.2 | 413 | Specific | 241.33 | 181.07 | 181.07 | 3.65 | 136 | >sp\|P01833\|PIGR_HUMAN Polymeric immunoglobulin receptor OS=Homo sapiens OX=9606 GN=PIGR PE=1 SV=4 | 902 | 20201118_ly_colostrum_1ng_glyco_hcd203040_compound15.22298.22298.4 | controllerType=0 controllerNumber=1 scan=22298 | 48.9919 |
| 26667:4 | 3 | R.[+42.011]LSLLEEPGN[+2351.851]GTFTVILNQLTSR.D | HexNAc(4)Hex(5)Fuc(3)NeuAc(1) | .[42], N[+2352] | 1199.7909 | 4 | 4796.1419 | 4796.1539 | 0 | -2.5 | 413 | Specific | 245.34 | 244.35 | 244.35 | 5.99 | 136 | >sp\|P01833\|PIGR_HUMAN Polymeric immunoglobulin receptor OS=Homo sapiens OX=9606 GN=PIGR PE=1 SV=4 | 902 | 20201118_ly_colostrum_1ng_glyco_hcd203040_compound15.30619.30619.3 | controllerType=0 controllerNumber=1 scan=30619 | 65.4088 |
| 18599:3 | 3 | R.LSLLEEPGN[+2351.851]GTFTVILNQLTSR.D | HexNAc(4)Hex(5)Fuc(3)NeuAc(1) | N[+2352] | 1585.3926 | 3 | 4754.1634 | 4754.1433 | 0 | 4.2 | 413 | Specific | 255.13 | 215.06 | 215.06 | 6.84 | 136 | >sp\|P01833\|PIGR_HUMAN Polymeric immunoglobulin receptor OS=Homo sapiens OX=9606 GN=PIGR PE=1 SV=4 | 902 | 20201118_ly_colostrum_1ng_glyco_hcd203040_compound15.22209.22209.2 | controllerType=0 controllerNumber=1 scan=22209 | 48.8008 |
| 20740:3 | 3 | R.[+42.011]LSLLEEPGN[+2482.909]GTFTVILNQLTSR.D | HexNAc(6)Hex(6)Fuc(2) | .[42], N[+2483] | 1643.0728 | 3 | 4927.2039 | 4927.2121 | 0 | -1.7 | 413 | Specific | 294.34 | 273.19 | 273.19 | 6.64 | 136 | >sp\|P01833\|PIGR_HUMAN Polymeric immunoglobulin receptor OS=Homo sapiens OX=9606 GN=PIGR PE=1 SV=4 | 902 | 20201118_ly_colostrum_1ng_glyco_hcd203040_compound15.24445.24445.3 | controllerType=0 controllerNumber=1 scan=24445 | 53.6978 |
| 26616:3 | 3 | R.[+42.011]LSLLEEPGN[+2351.851]GTFTVILNQLTSR.D | HexNAc(4)Hex(5)Fuc(3)NeuAc(1) | .[42], N[+2352] | 1599.3841 | 3 | 4796.1377 | 4796.1539 | 0 | -3.4 | 413 | Specific | 314.18 | 314.18 | 314.18 | 6.95 | 136 | >sp\|P01833\|PIGR_HUMAN Polymeric immunoglobulin receptor OS=Homo sapiens OX=9606 GN=PIGR PE=1 SV=4 | 902 | 20201118_ly_colostrum_1ng_glyco_hcd203040_compound15.30566.30566.3 | controllerType=0 controllerNumber=1 scan=30566 | 65.3103 |
| 22290:4 | 3 | R.[+42.011]LSLLEEPGN[+2205.793]GTFTVILNQLTSR.D | HexNAc(4)Hex(5)Fuc(2)NeuAc(1) | .[42], N[+2206] | 1163.2825 | 4 | 4650.1082 | 4650.096 | 0 | 2.6 | 413 | Specific | 320.58 | 320.58 | 320.58 | 8.24 | 136 | >sp\|P01833\|PIGR_HUMAN Polymeric immunoglobulin receptor OS=Homo sapiens OX=9606 GN=PIGR PE=1 SV=4 | 902 | 20201118_ly_colostrum_1ng_glyco_hcd203040_compound15.26060.26060.4 | controllerType=0 controllerNumber=1 scan=26060 | 56.9946 |
| 23506:3 | 3 | R.LSLLEEPGN[+3153.116]GTFTVILNQLTSR.D | HexNAc(5)Hex(6)Fuc(2)NeuAc(3) | N[+3153] | 1852.4817 | 3 | 5555.4304 | 5555.4084 | 0 | 4 | 413 | Specific | 324.44 | 319.55 | 319.55 | 8.23 | 136 | >sp\|P01833\|PIGR_HUMAN Polymeric immunoglobulin receptor OS=Homo sapiens OX=9606 GN=PIGR PE=1 SV=4 | 902 | 20201118_ly_colostrum_1ng_glyco_hcd203040_compound15.27321.27321.3 | controllerType=0 controllerNumber=1 scan=27321 | 59.3233 |
| 26618:3 | 3 | R.[+42.011]LSLLEEPGN[+2351.851]GTFTVILNQLTSR.D | HexNAc(4)Hex(5)Fuc(3)NeuAc(1) | .[42], N[+2352] | 1599.3841 | 3 | 4796.1377 | 4796.1539 | 0 | -3.4 | 413 | Specific | 326.94 | 274.14 | 274.14 | 6.82 | 136 | >sp\|P01833\|PIGR_HUMAN Polymeric immunoglobulin receptor OS=Homo sapiens OX=9606 GN=PIGR PE=1 SV=4 | 902 | 20201118_ly_colostrum_1ng_glyco_hcd203040_compound15.30568.30568.3 | controllerType=0 controllerNumber=1 scan=30568 | 65.3146 |
| 20738:3 | 3 | R.[+42.011]LSLLEEPGN[+2482.909]GTFTVILNQLTSR.D | HexNAc(6)Hex(6)Fuc(2) | .[42], N[+2483] | 1643.0728 | 3 | 4927.2039 | 4927.2121 | 0 | -1.7 | 413 | Specific | 363.65 | 349.37 | 349.37 | 7.43 | 136 | >sp\|P01833\|PIGR_HUMAN Polymeric immunoglobulin receptor OS=Homo sapiens OX=9606 GN=PIGR PE=1 SV=4 | 902 | 20201118_ly_colostrum_1ng_glyco_hcd203040_compound15.24443.24443.3 | controllerType=0 controllerNumber=1 scan=24443 | 53.6921 |
| 19083:3 | 3 | R.LSLLEEPGN[+1200.428]GTFTVILNQLTSR.D | HexNAc(2)Hex(4)Fuc(1) | N[+1200] | 1201.5862 | 3 | 3602.7442 | 3602.7205 | 0 | 6.6 | 413 | Specific | 383.27 | 295.53 | 295.53 | 7.32 | 136 | >sp\|P01833\|PIGR_HUMAN Polymeric immunoglobulin receptor OS=Homo sapiens OX=9606 GN=PIGR PE=1 SV=4 | 902 | 20201118_ly_colostrum_1ng_glyco_hcd203040_compound15.22715.22715.2 | controllerType=0 controllerNumber=1 scan=22715 | 49.9793 |
| 21114:3 | 3 | R.LSLLEEPGN[+2350.830]GTFTVILNQLTSR.D | HexNAc(4)Hex(5)Fuc(1)NeuAc(2) | N[+2351] | 1585.0462 | 3 | 4753.124 | 4753.1229 | 0 | 0.2 | 413 | Specific | 388.85 | 312.85 | 312.85 | 7.9 | 136 | >sp\|P01833\|PIGR_HUMAN Polymeric immunoglobulin receptor OS=Homo sapiens OX=9606 GN=PIGR PE=1 SV=4 | 902 | 20201118_ly_colostrum_1ng_glyco_hcd203040_compound15.24835.24835.3 | controllerType=0 controllerNumber=1 scan=24835 | 54.537 |
| 26642:3 | 3 | R.[+42.011]LSLLEEPGN[+2351.851]GTFTVILNQLTSR.D | HexNAc(4)Hex(5)Fuc(3)NeuAc(1) | .[42], N[+2352] | 1599.3865 | 3 | 4796.145 | 4796.1539 | 0 | -1.8 | 413 | Specific | 403.17 | 355.98 | 355.98 | 7.63 | 136 | >sp\|P01833\|PIGR_HUMAN Polymeric immunoglobulin receptor OS=Homo sapiens OX=9606 GN=PIGR PE=1 SV=4 | 902 | 20201118_ly_colostrum_1ng_glyco_hcd203040_compound15.30593.30593.4 | controllerType=0 controllerNumber=1 scan=30593 | 65.3613 |
| 26644:3 | 3 | R.[+42.011]LSLLEEPGN[+2351.851]GTFTVILNQLTSR.D | HexNAc(4)Hex(5)Fuc(3)NeuAc(1) | .[42], N[+2352] | 1599.3865 | 3 | 4796.145 | 4796.1539 | 0 | -1.8 | 413 | Specific | 451.97 | 382.57 | 382.57 | 9.41 | 136 | >sp\|P01833\|PIGR_HUMAN Polymeric immunoglobulin receptor OS=Homo sapiens OX=9606 GN=PIGR PE=1 SV=4 | 902 | 20201118_ly_colostrum_1ng_glyco_hcd203040_compound15.30595.30595.4 | controllerType=0 controllerNumber=1 scan=30595 | 65.365 |
| 19085:3 | 3 | R.LSLLEEPGN[+1200.428]GTFTVILNQLTSR.D | HexNAc(2)Hex(4)Fuc(1) | N[+1200] | 1201.5862 | 3 | 3602.7442 | 3602.7205 | 0 | 6.6 | 413 | Specific | 452.17 | 385.02 | 385.02 | 9.71 | 136 | >sp\|P01833\|PIGR_HUMAN Polymeric immunoglobulin receptor OS=Homo sapiens OX=9606 GN=PIGR PE=1 SV=4 | 902 | 20201118_ly_colostrum_1ng_glyco_hcd203040_compound15.22717.22717.2 | controllerType=0 controllerNumber=1 scan=22717 | 49.9855 |
| 20254:3 | 3 | R.LSLLEEPGN[+2861.000]GTFTVILNQLTSR.D | HexNAc(5)Hex(6)NeuAc(3) | N[+2861] | 1755.1158 | 3 | 5263.3329 | 5263.2926 | 0 | 7.6 | 413 | Specific | 457.51 | 457.51 | 457.51 | 9.68 | 136 | >sp\|P01833\|PIGR_HUMAN Polymeric immunoglobulin receptor OS=Homo sapiens OX=9606 GN=PIGR PE=1 SV=4 | 902 | 20201118_ly_colostrum_1ng_glyco_hcd203040_compound15.23937.23937.3 | controllerType=0 controllerNumber=1 scan=23937 | 52.5877 |
| 21116:3 | 3 | R.LSLLEEPGN[+2350.830]GTFTVILNQLTSR.D | HexNAc(4)Hex(5)Fuc(1)NeuAc(2) | N[+2351] | 1585.0462 | 3 | 4753.124 | 4753.1229 | 0 | 0.2 | 413 | Specific | 480.32 | 350.59 | 350.59 | 8.59 | 136 | >sp\|P01833\|PIGR_HUMAN Polymeric immunoglobulin receptor OS=Homo sapiens OX=9606 GN=PIGR PE=1 SV=4 | 902 | 20201118_ly_colostrum_1ng_glyco_hcd203040_compound15.24837.24837.3 | controllerType=0 controllerNumber=1 scan=24837 | 54.5413 |
| 18695:4 | 3 | R.LSLLEEPGN[+2059.735]GTFTVILNQLTSR.D | HexNAc(4)Hex(5)Fuc(1)NeuAc(1) | N[+2060] | 1116.2638 | 4 | 4462.0335 | 4462.0275 | 0 | 1.3 | 413 | Specific | 498.99 | 497.62 | 497.62 | 10.32 | 136 | >sp\|P01833\|PIGR_HUMAN Polymeric immunoglobulin receptor OS=Homo sapiens OX=9606 GN=PIGR PE=1 SV=4 | 902 | 20201118_ly_colostrum_1ng_glyco_hcd203040_compound15.22310.22310.4 | controllerType=0 controllerNumber=1 scan=22310 | 49.0474 |
| 18631:4 | 3 | R.LSLLEEPGN[+2076.750]GTFTVILNQLTSR.D | HexNAc(4)Hex(6)Fuc(2) | N[+2077] | 1120.5219 | 4 | 4479.0657 | 4479.0428 | 0 | 5.1 | 413 | Specific | 505.98 | 431.13 | 431.13 | 10.13 | 136 | >sp\|P01833\|PIGR_HUMAN Polymeric immunoglobulin receptor OS=Homo sapiens OX=9606 GN=PIGR PE=1 SV=4 | 902 | 20201118_ly_colostrum_1ng_glyco_hcd203040_compound15.22243.22243.4 | controllerType=0 controllerNumber=1 scan=22243 | 48.8784 |
| 18748:3 | 3 | R.LSLLEEPGN[+2205.793]GTFTVILNQLTSR.D | HexNAc(4)Hex(5)Fuc(2)NeuAc(1) | N[+2206] | 1536.7146 | 3 | 4608.1292 | 4608.0854 | 0 | 9.5 | 413 | Specific | 510.85 | 188.88 | 188.88 | 6.94 | 136 | >sp\|P01833\|PIGR_HUMAN Polymeric immunoglobulin receptor OS=Homo sapiens OX=9606 GN=PIGR PE=1 SV=4 | 902 | 20201118_ly_colostrum_1ng_glyco_hcd203040_compound15.22365.22365.3 | controllerType=0 controllerNumber=1 scan=22365 | 49.1571 |
| 20375:4 | 3 | R.LSLLEEPGN[+3008.078]GTFTVILNQLTSR.D | HexNAc(5)Hex(6)Fuc(3)NeuAc(2) | N[+3008] | 1353.3579 | 4 | 5410.4098 | 5410.3709 | 0 | 7.2 | 413 | Specific | 511.91 | 427.43 | 427.43 | 9.81 | 136 | >sp\|P01833\|PIGR_HUMAN Polymeric immunoglobulin receptor OS=Homo sapiens OX=9606 GN=PIGR PE=1 SV=4 | 902 | 20201118_ly_colostrum_1ng_glyco_hcd203040_compound15.24063.24063.4 | controllerType=0 controllerNumber=1 scan=24063 | 52.8528 |
| 20257:3 | 3 | R.LSLLEEPGN[+2861.000]GTFTVILNQLTSR.D | HexNAc(5)Hex(6)NeuAc(3) | N[+2861] | 1755.1158 | 3 | 5263.3329 | 5263.2926 | 0 | 7.6 | 413 | Specific | 529.51 | 374.22 | 374.22 | 8.53 | 136 | >sp\|P01833\|PIGR_HUMAN Polymeric immunoglobulin receptor OS=Homo sapiens OX=9606 GN=PIGR PE=1 SV=4 | 902 | 20201118_ly_colostrum_1ng_glyco_hcd203040_compound15.23940.23940.3 | controllerType=0 controllerNumber=1 scan=23940 | 52.5924 |
| 20811:3 | 3 | R.LSLLEEPGN[+2350.830]GTFTVILNQLTSR.D | HexNAc(4)Hex(5)Fuc(1)NeuAc(2) | N[+2351] | 1585.049 | 3 | 4753.1325 | 4753.1229 | 0 | 2 | 413 | Specific | 539.66 | 482.36 | 482.36 | 10.63 | 136 | >sp\|P01833\|PIGR_HUMAN Polymeric immunoglobulin receptor OS=Homo sapiens OX=9606 GN=PIGR PE=1 SV=4 | 902 | 20201118_ly_colostrum_1ng_glyco_hcd203040_compound15.24519.24519.3 | controllerType=0 controllerNumber=1 scan=24519 | 53.8642 |
| 18634:3 | 3 | R.LSLLEEPGN[+2076.750]GTFTVILNQLTSR.D | HexNAc(4)Hex(6)Fuc(2) | N[+2077] | 1493.6939 | 3 | 4479.0672 | 4479.0428 | 0 | 5.4 | 413 | Specific | 570.54 | 498.88 | 498.88 | 10.68 | 136 | >sp\|P01833\|PIGR_HUMAN Polymeric immunoglobulin receptor OS=Homo sapiens OX=9606 GN=PIGR PE=1 SV=4 | 902 | 20201118_ly_colostrum_1ng_glyco_hcd203040_compound15.22246.22246.3 | controllerType=0 controllerNumber=1 scan=22246 | 48.8869 |
| 20377:4 | 3 | R.LSLLEEPGN[+3008.078]GTFTVILNQLTSR.D | HexNAc(5)Hex(6)Fuc(3)NeuAc(2) | N[+3008] | 1353.3579 | 4 | 5410.4098 | 5410.3709 | 0 | 7.2 | 413 | Specific | 578.99 | 422.27 | 422.27 | 10.22 | 136 | >sp\|P01833\|PIGR_HUMAN Polymeric immunoglobulin receptor OS=Homo sapiens OX=9606 GN=PIGR PE=1 SV=4 | 902 | 20201118_ly_colostrum_1ng_glyco_hcd203040_compound15.24065.24065.4 | controllerType=0 controllerNumber=1 scan=24065 | 52.857 |
| 18750:3 | 3 | R.LSLLEEPGN[+2205.793]GTFTVILNQLTSR.D | HexNAc(4)Hex(5)Fuc(2)NeuAc(1) | N[+2206] | 1536.7146 | 3 | 4608.1292 | 4608.0854 | 0 | 9.5 | 413 | Specific | 594.09 | 356.44 | 356.44 | 9.55 | 136 | >sp\|P01833\|PIGR_HUMAN Polymeric immunoglobulin receptor OS=Homo sapiens OX=9606 GN=PIGR PE=1 SV=4 | 902 | 20201118_ly_colostrum_1ng_glyco_hcd203040_compound15.22367.22367.3 | controllerType=0 controllerNumber=1 scan=22367 | 49.1638 |
| 20584:4 | 3 | R.LSLLEEPGN[+2496.888]GTFTVILNQLTSR.D | HexNAc(4)Hex(5)Fuc(2)NeuAc(2) | N[+2497] | 1225.5544 | 4 | 4899.1956 | 4899.1808 | 0 | 3 | 413 | Specific | 632.14 | 551.42 | 551.42 | 13.43 | 136 | >sp\|P01833\|PIGR_HUMAN Polymeric immunoglobulin receptor OS=Homo sapiens OX=9606 GN=PIGR PE=1 SV=4 | 902 | 20201118_ly_colostrum_1ng_glyco_hcd203040_compound15.24282.24282.4 | controllerType=0 controllerNumber=1 scan=24282 | 53.3281 |
| 18633:4 | 3 | R.LSLLEEPGN[+2076.750]GTFTVILNQLTSR.D | HexNAc(4)Hex(6)Fuc(2) | N[+2077] | 1120.5219 | 4 | 4479.0657 | 4479.0428 | 0 | 5.1 | 413 | Specific | 657.35 | 607.34 | 607.34 | 13.86 | 136 | >sp\|P01833\|PIGR_HUMAN Polymeric immunoglobulin receptor OS=Homo sapiens OX=9606 GN=PIGR PE=1 SV=4 | 902 | 20201118_ly_colostrum_1ng_glyco_hcd203040_compound15.22245.22245.4 | controllerType=0 controllerNumber=1 scan=22245 | 48.885 |
| 18636:3 | 3 | R.LSLLEEPGN[+2076.750]GTFTVILNQLTSR.D | HexNAc(4)Hex(6)Fuc(2) | N[+2077] | 1493.6939 | 3 | 4479.0672 | 4479.0428 | 0 | 5.4 | 413 | Specific | 667.35 | 598.7 | 598.7 | 14 | 136 | >sp\|P01833\|PIGR_HUMAN Polymeric immunoglobulin receptor OS=Homo sapiens OX=9606 GN=PIGR PE=1 SV=4 | 902 | 20201118_ly_colostrum_1ng_glyco_hcd203040_compound15.22248.22248.3 | controllerType=0 controllerNumber=1 scan=22248 | 48.8923 |
| 18690:4 | 3 | R.LSLLEEPGN[+2204.772]GTFTVILNQLTSR.D | HexNAc(4)Hex(5)NeuAc(2) | N[+2205] | 1152.5303 | 4 | 4607.0994 | 4607.065 | 0 | 7.5 | 413 | Specific | 669.61 | 350.3 | 350.3 | 11.87 | 136 | >sp\|P01833\|PIGR_HUMAN Polymeric immunoglobulin receptor OS=Homo sapiens OX=9606 GN=PIGR PE=1 SV=4 | 902 | 20201118_ly_colostrum_1ng_glyco_hcd203040_compound15.22304.22304.4 | controllerType=0 controllerNumber=1 scan=22304 | 49.0299 |
| 20813:3 | 3 | R.LSLLEEPGN[+2350.830]GTFTVILNQLTSR.D | HexNAc(4)Hex(5)Fuc(1)NeuAc(2) | N[+2351] | 1585.049 | 3 | 4753.1325 | 4753.1229 | 0 | 2 | 413 | Specific | 673.26 | 626.38 | 626.38 | 14.31 | 136 | >sp\|P01833\|PIGR_HUMAN Polymeric immunoglobulin receptor OS=Homo sapiens OX=9606 GN=PIGR PE=1 SV=4 | 902 | 20201118_ly_colostrum_1ng_glyco_hcd203040_compound15.24521.24521.3 | controllerType=0 controllerNumber=1 scan=24521 | 53.8679 |
| 20582:4 | 3 | R.LSLLEEPGN[+2496.888]GTFTVILNQLTSR.D | HexNAc(4)Hex(5)Fuc(2)NeuAc(2) | N[+2497] | 1225.5544 | 4 | 4899.1956 | 4899.1808 | 0 | 3 | 413 | Specific | 682.42 | 615.04 | 615.04 | 14.43 | 136 | >sp\|P01833\|PIGR_HUMAN Polymeric immunoglobulin receptor OS=Homo sapiens OX=9606 GN=PIGR PE=1 SV=4 | 902 | 20201118_ly_colostrum_1ng_glyco_hcd203040_compound15.24280.24280.4 | controllerType=0 controllerNumber=1 scan=24280 | 53.3237 |
| 18687:4 | 3 | R.LSLLEEPGN[+2204.772]GTFTVILNQLTSR.D | HexNAc(4)Hex(5)NeuAc(2) | N[+2205] | 1152.5303 | 4 | 4607.0994 | 4607.065 | 0 | 7.5 | 413 | Specific | 686.55 | 343.63 | 343.63 | 11.87 | 136 | >sp\|P01833\|PIGR_HUMAN Polymeric immunoglobulin receptor OS=Homo sapiens OX=9606 GN=PIGR PE=1 SV=4 | 902 | 20201118_ly_colostrum_1ng_glyco_hcd203040_compound15.22301.22301.4 | controllerType=0 controllerNumber=1 scan=22301 | 49.0158 |
| 20613:4 | 3 | R.LSLLEEPGN[+2553.910]GTFTVILNQLTSR.D | HexNAc(5)Hex(5)Fuc(1)NeuAc(2) | N[+2554] | 1239.809 | 4 | 4956.214 | 4956.2023 | 0 | 2.4 | 413 | Specific | 709.41 | 709.41 | 709.41 | 15.4 | 136 | >sp\|P01833\|PIGR_HUMAN Polymeric immunoglobulin receptor OS=Homo sapiens OX=9606 GN=PIGR PE=1 SV=4 | 902 | 20201118_ly_colostrum_1ng_glyco_hcd203040_compound15.24312.24312.4 | controllerType=0 controllerNumber=1 scan=24312 | 53.3914 |
| 20741:3 | 3 | R.LSLLEEPGN[+2350.830]GTFTVILNQLTSR.D | HexNAc(4)Hex(5)Fuc(1)NeuAc(2) | N[+2351] | 1585.0556 | 3 | 4753.1522 | 4753.1229 | 0 | 6.2 | 413 | Specific | 729.61 | 516.33 | 516.33 | 14.71 | 136 | >sp\|P01833\|PIGR_HUMAN Polymeric immunoglobulin receptor OS=Homo sapiens OX=9606 GN=PIGR PE=1 SV=4 | 902 | 20201118_ly_colostrum_1ng_glyco_hcd203040_compound15.24446.24446.3 | controllerType=0 controllerNumber=1 scan=24446 | 53.6997 |
| 18414:4 | 3 | R.LSLLEEPGN[+2863.041]GTFTVILNQLTSR.D | HexNAc(5)Hex(6)Fuc(4)NeuAc(1) | N[+2863] | 1317.0955 | 4 | 5265.36 | 5265.3334 | 0 | 5.1 | 413 | Specific | 757.74 | 391.34 | 391.34 | 16.13 | 136 | >sp\|P01833\|PIGR_HUMAN Polymeric immunoglobulin receptor OS=Homo sapiens OX=9606 GN=PIGR PE=1 SV=4 | 902 | 20201118_ly_colostrum_1ng_glyco_hcd203040_compound15.22016.22016.4 | controllerType=0 controllerNumber=1 scan=22016 | 48.3704 |
| 18698:4 | 3 | R.LSLLEEPGN[+2059.735]GTFTVILNQLTSR.D | HexNAc(4)Hex(5)Fuc(1)NeuAc(1) | N[+2060] | 1116.2638 | 4 | 4462.0335 | 4462.0275 | 0 | 1.3 | 413 | Specific | 766.64 | 576.79 | 576.79 | 16.17 | 136 | >sp\|P01833\|PIGR_HUMAN Polymeric immunoglobulin receptor OS=Homo sapiens OX=9606 GN=PIGR PE=1 SV=4 | 902 | 20201118_ly_colostrum_1ng_glyco_hcd203040_compound15.22313.22313.4 | controllerType=0 controllerNumber=1 scan=22313 | 49.0561 |
| 20616:4 | 3 | R.LSLLEEPGN[+2553.910]GTFTVILNQLTSR.D | HexNAc(5)Hex(5)Fuc(1)NeuAc(2) | N[+2554] | 1239.809 | 4 | 4956.214 | 4956.2023 | 0 | 2.4 | 413 | Specific | 767.18 | 619.73 | 619.73 | 16.4 | 136 | >sp\|P01833\|PIGR_HUMAN Polymeric immunoglobulin receptor OS=Homo sapiens OX=9606 GN=PIGR PE=1 SV=4 | 902 | 20201118_ly_colostrum_1ng_glyco_hcd203040_compound15.24315.24315.4 | controllerType=0 controllerNumber=1 scan=24315 | 53.4016 |
| 18332:3 | 3 | R.LSLLEEPGN[+2863.041]GTFTVILNQLTSR.D | HexNAc(5)Hex(6)Fuc(4)NeuAc(1) | N[+2863] | 1755.7887 | 3 | 5265.3514 | 5265.3334 | 0 | 3.4 | 413 | Specific | 861.39 | 438.48 | 438.48 | 18.57 | 136 | >sp\|P01833\|PIGR_HUMAN Polymeric immunoglobulin receptor OS=Homo sapiens OX=9606 GN=PIGR PE=1 SV=4 | 902 | 20201118_ly_colostrum_1ng_glyco_hcd203040_compound15.21930.21930.3 | controllerType=0 controllerNumber=1 scan=21930 | 48.1713 |
| 20743:3 | 3 | R.LSLLEEPGN[+2350.830]GTFTVILNQLTSR.D | HexNAc(4)Hex(5)Fuc(1)NeuAc(2) | N[+2351] | 1585.0556 | 3 | 4753.1522 | 4753.1229 | 0 | 6.2 | 413 | Specific | 876.14 | 612.71 | 612.71 | 17.71 | 136 | >sp\|P01833\|PIGR_HUMAN Polymeric immunoglobulin receptor OS=Homo sapiens OX=9606 GN=PIGR PE=1 SV=4 | 902 | 20201118_ly_colostrum_1ng_glyco_hcd203040_compound15.24448.24448.3 | controllerType=0 controllerNumber=1 scan=24448 | 53.7036 |
| 18416:4 | 3 | R.LSLLEEPGN[+2863.041]GTFTVILNQLTSR.D | HexNAc(5)Hex(6)Fuc(4)NeuAc(1) | N[+2863] | 1317.0955 | 4 | 5265.36 | 5265.3334 | 0 | 5.1 | 413 | Specific | 876.9 | 416.53 | 416.53 | 18.13 | 136 | >sp\|P01833\|PIGR_HUMAN Polymeric immunoglobulin receptor OS=Homo sapiens OX=9606 GN=PIGR PE=1 SV=4 | 902 | 20201118_ly_colostrum_1ng_glyco_hcd203040_compound15.22018.22018.4 | controllerType=0 controllerNumber=1 scan=22018 | 48.3742 |
| 18563:3 | 3 | R.LSLLEEPGN[+2351.851]GTFTVILNQLTSR.D | HexNAc(4)Hex(5)Fuc(3)NeuAc(1) | N[+2352] | 1585.392 | 3 | 4754.1615 | 4754.1433 | 0 | 3.8 | 413 | Specific | 915.71 | 658.44 | 658.44 | 19.57 | 136 | >sp\|P01833\|PIGR_HUMAN Polymeric immunoglobulin receptor OS=Homo sapiens OX=9606 GN=PIGR PE=1 SV=4 | 902 | 20201118_ly_colostrum_1ng_glyco_hcd203040_compound15.22172.22172.3 | controllerType=0 controllerNumber=1 scan=22172 | 48.7201 |
| 20641:3 | 3 | R.LSLLEEPGN[+2496.888]GTFTVILNQLTSR.D | HexNAc(4)Hex(5)Fuc(2)NeuAc(2) | N[+2497] | 1633.7348 | 3 | 4899.1898 | 4899.1808 | 0 | 1.8 | 413 | Specific | 931.49 | 638.46 | 638.46 | 19.31 | 136 | >sp\|P01833\|PIGR_HUMAN Polymeric immunoglobulin receptor OS=Homo sapiens OX=9606 GN=PIGR PE=1 SV=4 | 902 | 20201118_ly_colostrum_1ng_glyco_hcd203040_compound15.24341.24341.3 | controllerType=0 controllerNumber=1 scan=24341 | 53.4615 |
| 18612:3 | 3 | R.LSLLEEPGN[+2059.735]GTFTVILNQLTSR.D | HexNAc(4)Hex(5)Fuc(1)NeuAc(1) | N[+2060] | 1488.0197 | 3 | 4462.0446 | 4462.0275 | 0 | 3.8 | 413 | Specific | 957.83 | 713.11 | 713.11 | 20.57 | 136 | >sp\|P01833\|PIGR_HUMAN Polymeric immunoglobulin receptor OS=Homo sapiens OX=9606 GN=PIGR PE=1 SV=4 | 902 | 20201118_ly_colostrum_1ng_glyco_hcd203040_compound15.22223.22223.3 | controllerType=0 controllerNumber=1 scan=22223 | 48.8375 |
| 18569:3 | 3 | R.LSLLEEPGN[+2205.793]GTFTVILNQLTSR.D | HexNAc(4)Hex(5)Fuc(2)NeuAc(1) | N[+2206] | 1536.706 | 3 | 4608.1034 | 4608.0854 | 0 | 3.9 | 413 | Specific | 971.63 | 732.38 | 732.38 | 20.57 | 136 | >sp\|P01833\|PIGR_HUMAN Polymeric immunoglobulin receptor OS=Homo sapiens OX=9606 GN=PIGR PE=1 SV=4 | 902 | 20201118_ly_colostrum_1ng_glyco_hcd203040_compound15.22178.22178.3 | controllerType=0 controllerNumber=1 scan=22178 | 48.7359 |
| 18330:3 | 3 | R.LSLLEEPGN[+2863.041]GTFTVILNQLTSR.D | HexNAc(5)Hex(6)Fuc(4)NeuAc(1) | N[+2863] | 1755.7887 | 3 | 5265.3514 | 5265.3334 | 0 | 3.4 | 413 | Specific | 1004.48 | 529.15 | 529.15 | 21.57 | 136 | >sp\|P01833\|PIGR_HUMAN Polymeric immunoglobulin receptor OS=Homo sapiens OX=9606 GN=PIGR PE=1 SV=4 | 902 | 20201118_ly_colostrum_1ng_glyco_hcd203040_compound15.21928.21928.3 | controllerType=0 controllerNumber=1 scan=21928 | 48.1614 |
| 18564:4 | 3 | R.LSLLEEPGN[+2351.851]GTFTVILNQLTSR.D | HexNAc(4)Hex(5)Fuc(3)NeuAc(1) | N[+2352] | 1189.2961 | 4 | 4754.1624 | 4754.1433 | 0 | 4 | 413 | Specific | 1009.66 | 752.12 | 752.12 | 21.43 | 136 | >sp\|P01833\|PIGR_HUMAN Polymeric immunoglobulin receptor OS=Homo sapiens OX=9606 GN=PIGR PE=1 SV=4 | 902 | 20201118_ly_colostrum_1ng_glyco_hcd203040_compound15.22173.22173.4 | controllerType=0 controllerNumber=1 scan=22173 | 48.7236 |
| 18577:4 | 3 | R.LSLLEEPGN[+2205.793]GTFTVILNQLTSR.D | HexNAc(4)Hex(5)Fuc(2)NeuAc(1) | N[+2206] | 1152.7802 | 4 | 4608.0989 | 4608.0854 | 0 | 2.9 | 413 | Specific | 1015.52 | 757.7 | 757.7 | 21.4 | 136 | >sp\|P01833\|PIGR_HUMAN Polymeric immunoglobulin receptor OS=Homo sapiens OX=9606 GN=PIGR PE=1 SV=4 | 902 | 20201118_ly_colostrum_1ng_glyco_hcd203040_compound15.22186.22186.4 | controllerType=0 controllerNumber=1 scan=22186 | 48.7552 |
| 20639:3 | 3 | R.LSLLEEPGN[+2496.888]GTFTVILNQLTSR.D | HexNAc(4)Hex(5)Fuc(2)NeuAc(2) | N[+2497] | 1633.7348 | 3 | 4899.1898 | 4899.1808 | 0 | 1.8 | 413 | Specific | 1070.45 | 811.46 | 811.46 | 22.31 | 136 | >sp\|P01833\|PIGR_HUMAN Polymeric immunoglobulin receptor OS=Homo sapiens OX=9606 GN=PIGR PE=1 SV=4 | 902 | 20201118_ly_colostrum_1ng_glyco_hcd203040_compound15.24339.24339.3 | controllerType=0 controllerNumber=1 scan=24339 | 53.4552 |
| 18567:3 | 3 | R.LSLLEEPGN[+2205.793]GTFTVILNQLTSR.D | HexNAc(4)Hex(5)Fuc(2)NeuAc(1) | N[+2206] | 1536.706 | 3 | 4608.1034 | 4608.0854 | 0 | 3.9 | 413 | Specific | 1073.21 | 822.6 | 822.6 | 22.57 | 136 | >sp\|P01833\|PIGR_HUMAN Polymeric immunoglobulin receptor OS=Homo sapiens OX=9606 GN=PIGR PE=1 SV=4 | 902 | 20201118_ly_colostrum_1ng_glyco_hcd203040_compound15.22176.22176.3 | controllerType=0 controllerNumber=1 scan=22176 | 48.727 |
| 18561:3 | 3 | R.LSLLEEPGN[+2351.851]GTFTVILNQLTSR.D | HexNAc(4)Hex(5)Fuc(3)NeuAc(1) | N[+2352] | 1585.392 | 3 | 4754.1615 | 4754.1433 | 0 | 3.8 | 413 | Specific | 1081.27 | 1080.69 | 1080.69 | 22.57 | 136 | >sp\|P01833\|PIGR_HUMAN Polymeric immunoglobulin receptor OS=Homo sapiens OX=9606 GN=PIGR PE=1 SV=4 | 902 | 20201118_ly_colostrum_1ng_glyco_hcd203040_compound15.22170.22170.3 | controllerType=0 controllerNumber=1 scan=22170 | 48.7146 |
| 18610:3 | 3 | R.LSLLEEPGN[+2059.735]GTFTVILNQLTSR.D | HexNAc(4)Hex(5)Fuc(1)NeuAc(1) | N[+2060] | 1488.0197 | 3 | 4462.0446 | 4462.0275 | 0 | 3.8 | 413 | Specific | 1081.36 | 847.55 | 847.55 | 22.57 | 136 | >sp\|P01833\|PIGR_HUMAN Polymeric immunoglobulin receptor OS=Homo sapiens OX=9606 GN=PIGR PE=1 SV=4 | 902 | 20201118_ly_colostrum_1ng_glyco_hcd203040_compound15.22221.22221.3 | controllerType=0 controllerNumber=1 scan=22221 | 48.8263 |
| 18562:4 | 3 | R.LSLLEEPGN[+2351.851]GTFTVILNQLTSR.D | HexNAc(4)Hex(5)Fuc(3)NeuAc(1) | N[+2352] | 1189.2961 | 4 | 4754.1624 | 4754.1433 | 0 | 4 | 413 | Specific | 1138.05 | 1130.12 | 1130.12 | 23.43 | 136 | >sp\|P01833\|PIGR_HUMAN Polymeric immunoglobulin receptor OS=Homo sapiens OX=9606 GN=PIGR PE=1 SV=4 | 902 | 20201118_ly_colostrum_1ng_glyco_hcd203040_compound15.22171.22171.4 | controllerType=0 controllerNumber=1 scan=22171 | 48.7163 |
| 18574:4 | 3 | R.LSLLEEPGN[+2205.793]GTFTVILNQLTSR.D | HexNAc(4)Hex(5)Fuc(2)NeuAc(1) | N[+2206] | 1152.7802 | 4 | 4608.0989 | 4608.0854 | 0 | 2.9 | 413 | Specific | 1176.1 | 947.58 | 947.58 | 24.4 | 136 | >sp\|P01833\|PIGR_HUMAN Polymeric immunoglobulin receptor OS=Homo sapiens OX=9606 GN=PIGR PE=1 SV=4 | 902 | 20201118_ly_colostrum_1ng_glyco_hcd203040_compound15.22183.22183.4 | controllerType=0 controllerNumber=1 scan=22183 | 48.7424 |
| 15824:6 | 3 | R.TTVEIKIIEGEPNLKVPGN[+2351.851]VTAVLGETLKVPC[+57.021]HFPC[+57.021]K.F | HexNAc(4)Hex(5)Fuc(3)NeuAc(1) | C[+57]*2, N[+2352] | 1074.1772 | 6 | 6440.0266 | 6440.043 | 0 | -2.5 | 451 | Specific | 728.24 | 701.22 | 701.22 | 14.18 | 136 | >sp\|P01833\|PIGR_HUMAN Polymeric immunoglobulin receptor OS=Homo sapiens OX=9606 GN=PIGR PE=1 SV=4 | 902 | 20201118_ly_colostrum_1ng_glyco_hcd203040_compound15.19309.19309.3 | controllerType=0 controllerNumber=1 scan=19309 | 42.2226 |
| 15865:5 | 3 | R.TTVEIKIIEGEPNLKVPGN[+2350.830]VTAVLGETLKVPC[+57.021]HFPC[+57.021]K.F | HexNAc(4)Hex(5)Fuc(1)NeuAc(2) | C[+57]*2, N[+2351] | 1288.6102 | 5 | 6439.0221 | 6439.0226 | 0 | -0.1 | 451 | Specific | 733.5 | 672.43 | 672.43 | 14.2 | 136 | >sp\|P01833\|PIGR_HUMAN Polymeric immunoglobulin receptor OS=Homo sapiens OX=9606 GN=PIGR PE=1 SV=4 | 902 | 20201118_ly_colostrum_1ng_glyco_hcd203040_compound15.19352.19352.5 | controllerType=0 controllerNumber=1 scan=19352 | 42.3148 |
| 14822:6 | 3 | R.TTVEIKIIEGEPNLKVPGN[+2351.851]VTAVLGETLKVPC[+57.021]HFPC[+57.021]K.F | HexNAc(4)Hex(5)Fuc(3)NeuAc(1) | C[+57]*2, N[+2352] | 1074.1836 | 6 | 6440.0654 | 6440.043 | 0 | 3.5 | 451 | Specific | 773.48 | 702.29 | 702.29 | 16.48 | 136 | >sp\|P01833\|PIGR_HUMAN Polymeric immunoglobulin receptor OS=Homo sapiens OX=9606 GN=PIGR PE=1 SV=4 | 902 | 20201118_ly_colostrum_1ng_glyco_hcd203040_compound15.18265.18265.6 | controllerType=0 controllerNumber=1 scan=18265 | 39.9696 |
| 14820:6 | 3 | R.TTVEIKIIEGEPNLKVPGN[+2351.851]VTAVLGETLKVPC[+57.021]HFPC[+57.021]K.F | HexNAc(4)Hex(5)Fuc(3)NeuAc(1) | C[+57]*2, N[+2352] | 1074.1836 | 6 | 6440.0654 | 6440.043 | 0 | 3.5 | 451 | Specific | 773.6 | 773.6 | 773.6 | 16.48 | 136 | >sp\|P01833\|PIGR_HUMAN Polymeric immunoglobulin receptor OS=Homo sapiens OX=9606 GN=PIGR PE=1 SV=4 | 902 | 20201118_ly_colostrum_1ng_glyco_hcd203040_compound15.18263.18263.6 | controllerType=0 controllerNumber=1 scan=18263 | 39.9645 |
| 15863:5 | 3 | R.TTVEIKIIEGEPNLKVPGN[+2350.830]VTAVLGETLKVPC[+57.021]HFPC[+57.021]K.F | HexNAc(4)Hex(5)Fuc(1)NeuAc(2) | C[+57]*2, N[+2351] | 1288.6102 | 5 | 6439.0221 | 6439.0226 | 0 | -0.1 | 451 | Specific | 799.09 | 798.95 | 798.95 | 15.2 | 136 | >sp\|P01833\|PIGR_HUMAN Polymeric immunoglobulin receptor OS=Homo sapiens OX=9606 GN=PIGR PE=1 SV=4 | 902 | 20201118_ly_colostrum_1ng_glyco_hcd203040_compound15.19350.19350.5 | controllerType=0 controllerNumber=1 scan=19350 | 42.3099 |
| 14827:6 | 3 | R.TTVEIKIIEGEPNLKVPGN[+2205.793]VTAVLGETLKVPC[+57.021]HFPC[+57.021]K.F | HexNAc(4)Hex(5)Fuc(2)NeuAc(1) | C[+57]*2, N[+2206] | 1049.8415 | 6 | 6294.0125 | 6293.9851 | 0 | 4.3 | 451 | Specific | 887.85 | 887.85 | 887.85 | 18.18 | 136 | >sp\|P01833\|PIGR_HUMAN Polymeric immunoglobulin receptor OS=Homo sapiens OX=9606 GN=PIGR PE=1 SV=4 | 902 | 20201118_ly_colostrum_1ng_glyco_hcd203040_compound15.18270.18270.3 | controllerType=0 controllerNumber=1 scan=18270 | 39.9806 |
| 14830:6 | 3 | R.TTVEIKIIEGEPNLKVPGN[+2205.793]VTAVLGETLKVPC[+57.021]HFPC[+57.021]K.F | HexNAc(4)Hex(5)Fuc(2)NeuAc(1) | C[+57]*2, N[+2206] | 1049.8415 | 6 | 6294.0125 | 6293.9851 | 0 | 4.3 | 451 | Specific | 905.88 | 779.99 | 779.99 | 19.18 | 136 | >sp\|P01833\|PIGR_HUMAN Polymeric immunoglobulin receptor OS=Homo sapiens OX=9606 GN=PIGR PE=1 SV=4 | 902 | 20201118_ly_colostrum_1ng_glyco_hcd203040_compound15.18273.18273.3 | controllerType=0 controllerNumber=1 scan=18273 | 39.9891 |
| 15826:6 | 3 | R.TTVEIKIIEGEPNLKVPGN[+2351.851]VTAVLGETLKVPC[+57.021]HFPC[+57.021]K.F | HexNAc(4)Hex(5)Fuc(3)NeuAc(1) | C[+57]*2, N[+2352] | 1074.1772 | 6 | 6440.0266 | 6440.043 | 0 | -2.5 | 451 | Specific | 932.34 | 740.91 | 740.91 | 18.18 | 136 | >sp\|P01833\|PIGR_HUMAN Polymeric immunoglobulin receptor OS=Homo sapiens OX=9606 GN=PIGR PE=1 SV=4 | 902 | 20201118_ly_colostrum_1ng_glyco_hcd203040_compound15.19311.19311.3 | controllerType=0 controllerNumber=1 scan=19311 | [illegible] |
| 16228:3 | 3 | K.IIEGEPNLKVPGN[+2350.830]VTAVLGETLK.V | HexNAc(4)Hex(5)Fuc(1)NeuAc(2) | N[+2351] | 1581.4025 | 3 | 4742.193 | 4742.1797 | 0 | 2.8 | 457 | Specific | 760.72 | 565.21 | 565.21 | 15.96 | 136 | >sp\|P01833\|PIGR_HUMAN Polymeric immunoglobulin receptor OS=Homo sapiens OX=9606 GN=PIGR PE=1 SV=4 | 902 | 20201118_ly_colostrum_1ng_glyco_hcd203040_compound15.19731.19731.3 | controllerType=0 controllerNumber=1 scan=19731 | [illegible] |
| 16226:3 | 3 | K.IIEGEPNLKVPGN[+2350.830]VTAVLGETLK.V | HexNAc(4)Hex(5)Fuc(1)NeuAc(2) | N[+2351] | 1581.4025 | 3 | 4742.193 | 4742.1797 | 0 | 2.8 | 457 | Specific | 827.14 | 827.14 | 827.14 | 16.96 | 136 | >sp\|P01833\|PIGR_HUMAN Polymeric immunoglobulin receptor OS=Homo sapiens OX=9606 GN=PIGR PE=1 SV=4 | 902 | 20201118_ly_colostrum_1ng_glyco_hcd203040_compound15.19729.19729.3 | controllerType=0 controllerNumber=1 scan=19729 | [illegible] |
| 12988:5 | 3 | K.[+42.011]IIEGEPNLKVPGN[+2018.708]VTAVLGETLKVPC[+57.021]HFPC[+57.021] | HexNAc(3)Hex(6)Fuc(1)NeuAc(1) | .[42], C[+57]*2, N[+2019] | 1096.3206 | 5 | 5477.5737 | 5477.5258 | 0 | 8.7 | 457 | Specific | 202.17 | 141.86 | 141.86 | 2.38 | 136 | >sp\|P01833\|PIGR_HUMAN Polymeric immunoglobulin receptor OS=Homo sapiens OX=9606 GN=PIGR PE=1 SV=4 | 902 | 20201118_ly_colostrum_1ng_glyco_hcd203040_compound15.16357.16357.5 | controllerType=0 controllerNumber=1 scan=16357 | [illegible] |
| 15438:6 | 3 | K.IIEGEPNLKVPGN[+2350.830]VTAVLGETLKVPC[+57.021]HFPC[+57.021]K.F | HexNAc(4)Hex(5)Fuc(1)NeuAc(2) | C[+57]*2, N[+2351] | 962.1172 | 6 | 5767.667 | 5767.6372 | 0 | 5.2 | 457 | Specific | 217.85 | 174.12 | 174.12 | 3.53 | 136 | >sp\|P01833\|PIGR_HUMAN Polymeric immunoglobulin receptor OS=Homo sapiens OX=9606 GN=PIGR PE=1 SV=4 | 902 | 20201118_ly_colostrum_1ng_glyco_hcd203040_compound15.18906.18906.6 | controllerType=0 controllerNumber=1 scan=18906 | 41.3353 |
| 16388:4 | 3 | K.[+42.011]IIEGEPNLKVPGN[+2351.851]VTAVLGETLKVPC[+57.021]HFPC[+57.021] | HexNAc(4)Hex(5)Fuc(3)NeuAc(1) | .[42], C[+57]*2, N[+2352] | 1453.4227 | 4 | 5810.6691 | 5810.6682 | 0 | 0.2 | 457 | Specific | 294.84 | 239.47 | 239.47 | 6.23 | 136 | >sp\|P01833\|PIGR_HUMAN Polymeric immunoglobulin receptor OS=Homo sapiens OX=9606 GN=PIGR PE=1 SV=4 | 902 | 20201118_ly_colostrum_1ng_glyco_hcd203040_compound15.19897.19897.4 | controllerType=0 controllerNumber=1 scan=19897 | 43.556 |
| 12999:5 | 3 | K.IIEGEPNLKVPGN[+2263.835]VTAVLGETLKVPC[+57.021]HFPC[+57.021]K.F | HexNAc(5)Hex(5)Fuc(3) | C[+57]*2, N[+2264] | 1136.936 | 5 | 5680.6511 | 5680.6416 | 0 | 1.7 | 457 | Specific | 305.16 | 305.16 | 305.16 | 6.01 | 136 | >sp\|P01833\|PIGR_HUMAN Polymeric immunoglobulin receptor OS=Homo sapiens OX=9606 GN=PIGR PE=1 SV=4 | 902 | 20201118_ly_colostrum_1ng_glyco_hcd203040_compound15.16368.16368.3 | controllerType=0 controllerNumber=1 scan=16368 | 36.0765 |
| 15031:4 | 3 | K.IIEGEPNLKVPGN[+3008.078]VTAVLGETLKVPC[+57.021]HFPC[+57.021]K.F | HexNAc(5)Hex(6)Fuc(3)NeuAc(2) | C[+57]*2, N[+3008] | 1606.9835 | 4 | 6424.9123 | 6424.8852 | 0 | 4.2 | 457 | Specific | 374.97 | 374.97 | 374.97 | 7.68 | 136 | >sp\|P01833\|PIGR_HUMAN Polymeric immunoglobulin receptor OS=Homo sapiens OX=9606 GN=PIGR PE=1 SV=4 | 902 | 20201118_ly_colostrum_1ng_glyco_hcd203040_compound15.18482.18482.4 | controllerType=0 controllerNumber=1 scan=18482 | 40.4452 |
| 15441:6 | 3 | K.IIEGEPNLKVPGN[+2350.830]VTAVLGETLKVPC[+57.021]HFPC[+57.021]K.F | HexNAc(4)Hex(5)Fuc(1)NeuAc(2) | C[+57]*2, N[+2351] | 962.1172 | 6 | 5767.667 | 5767.6372 | 0 | 5.2 | 457 | Specific | 378.58 | 364.23 | 364.23 | 6.51 | 136 | >sp\|P01833\|PIGR_HUMAN Polymeric immunoglobulin receptor OS=Homo sapiens OX=9606 GN=PIGR PE=1 SV=4 | 902 | 20201118_ly_colostrum_1ng_glyco_hcd203040_compound15.18909.18909.6 | controllerType=0 controllerNumber=1 scan=18909 | 41.3404 |
| 16391:4 | 3 | K.[+42.011]IIEGEPNLKVPGN[+2351.851]VTAVLGETLKVPC[+57.021]HFPC[+57.021] | HexNAc(4)Hex(5)Fuc(3)NeuAc(1) | .[42], C[+57]*2, N[+2352] | 1453.4227 | 4 | 5810.6691 | 5810.6682 | 0 | 0.2 | 457 | Specific | 423.46 | 352.86 | 352.86 | 6.47 | 136 | >sp\|P01833\|PIGR_HUMAN Polymeric immunoglobulin receptor OS=Homo sapiens OX=9606 GN=PIGR PE=1 SV=4 | 902 | 20201118_ly_colostrum_1ng_glyco_hcd203040_compound15.19900.19900.4 | controllerType=0 controllerNumber=1 scan=19900 | 43.5684 |
| 13006:5 | 3 | K.IIEGEPNLKVPGN[+2279.830]VTAVLGETLKVPC[+57.021]HFPC[+57.021]K.F | HexNAc(5)Hex(6)Fuc(2) | C[+57]*2, N[+2280] | 1140.1355 | 5 | 5696.6484 | 5696.6365 | 0 | 2.1 | 457 | Specific | 478.92 | 470.69 | 470.69 | 9.97 | 136 | >sp\|P01833\|PIGR_HUMAN Polymeric immunoglobulin receptor OS=Homo sapiens OX=9606 GN=PIGR PE=1 SV=4 | 902 | 20201118_ly_colostrum_1ng_glyco_hcd203040_compound15.16375.16375.5 | controllerType=0 controllerNumber=1 scan=16375 | 36.0907 |
| 14148:4 | 3 | K.IIEGEPNLKVPGN[+2351.851]VTAVLGETLKVPC[+57.021]HFPC[+57.021]K.F | HexNAc(4)Hex(5)Fuc(3)NeuAc(1) | C[+57]*2, N[+2352] | 1442.9229 | 4 | 5768.6696 | 5768.6576 | 0 | 2.1 | 457 | Specific | 498.56 | 498.56 | 498.56 | 9.64 | 136 | >sp\|P01833\|PIGR_HUMAN Polymeric immunoglobulin receptor OS=Homo sapiens OX=9606 GN=PIGR PE=1 SV=4 | 902 | 20201118_ly_colostrum_1ng_glyco_hcd203040_compound15.17563.17563.4 | controllerType=0 controllerNumber=1 scan=17563 | 38.5718 |
| 14981:5 | 3 | K.IIEGEPNLKVPGN[+3154.136]VTAVLGETLKVPC[+57.021]HFPC[+57.021]K.F | HexNAc(5)Hex(6)Fuc(4)NeuAc(2) | C[+57]*2, N[+3154] | 1315.0018 | 5 | 6570.9801 | 6570.9432 | 0 | 5.6 | 457 | Specific | 510.77 | 461.98 | 461.98 | 9.87 | 136 | >sp\|P01833\|PIGR_HUMAN Polymeric immunoglobulin receptor OS=Homo sapiens OX=9606 GN=PIGR PE=1 SV=4 | 902 | 20201118_ly_colostrum_1ng_glyco_hcd203040_compound15.18430.18430.5 | controllerType=0 controllerNumber=1 scan=18430 | 40.3396 |
| 14007:5 | 3 | K.IIEGEPNLKVPGN[+2862.021]VTAVLGETLKVPC[+57.021]HFPC[+57.021]K.F | HexNAc(5)Hex(6)Fuc(2)NeuAc(2) | C[+57]*2, N[+2862] | 1256.5797 | 5 | 6278.8695 | 6278.8273 | 0 | 6.7 | 457 | Specific | 512.24 | 487.96 | 487.96 | 9.57 | 136 | >sp\|P01833\|PIGR_HUMAN Polymeric immunoglobulin receptor OS=Homo sapiens OX=9606 GN=PIGR PE=1 SV=4 | 902 | 20201118_ly_colostrum_1ng_glyco_hcd203040_compound15.17416.17416.5 | controllerType=0 controllerNumber=1 scan=17416 | 38.2619 |
| 15029:4 | 3 | K.IIEGEPNLKVPGN[+3008.078]VTAVLGETLKVPC[+57.021]HFPC[+57.021]K.F | HexNAc(5)Hex(6)Fuc(3)NeuAc(2) | C[+57]*2, N[+3008] | 1606.9835 | 4 | 6424.9123 | 6424.8852 | 0 | 4.2 | 457 | Specific | 522.83 | 522.83 | 522.83 | 9.77 | 136 | >sp\|P01833\|PIGR_HUMAN Polymeric immunoglobulin receptor OS=Homo sapiens OX=9606 GN=PIGR PE=1 SV=4 | 902 | 20201118_ly_colostrum_1ng_glyco_hcd203040_compound15.18480.18480.4 | controllerType=0 controllerNumber=1 scan=18480 | 40.4403 |
| 13969:5 | 3 | K.IIEGEPNLKVPGN[+2862.021]VTAVLGETLKVPC[+57.021]HFPC[+57.021]K.F | HexNAc(5)Hex(6)Fuc(2)NeuAc(2) | C[+57]*2, N[+2862] | 1256.5757 | 5 | 6278.8493 | 6278.8273 | 0 | 3.5 | 457 | Specific | 541.54 | 541.54 | 541.54 | 10.66 | 136 | >sp\|P01833\|PIGR_HUMAN Polymeric immunoglobulin receptor OS=Homo sapiens OX=9606 GN=PIGR PE=1 SV=4 | 902 | 20201118_ly_colostrum_1ng_glyco_hcd203040_compound15.17377.17377.5 | controllerType=0 controllerNumber=1 scan=17377 | 38.164 |
| 13972:5 | 3 | K.IIEGEPNLKVPGN[+2862.021]VTAVLGETLKVPC[+57.021]HFPC[+57.021]K.F | HexNAc(5)Hex(6)Fuc(2)NeuAc(2) | C[+57]*2, N[+2862] | 1256.5757 | 5 | 6278.8493 | 6278.8273 | 0 | 3.5 | 457 | Specific | 565.63 | 526.2 | 526.2 | 10.84 | 136 | >sp\|P01833\|PIGR_HUMAN Polymeric immunoglobulin receptor OS=Homo sapiens OX=9606 GN=PIGR PE=1 SV=4 | 902 | 20201118_ly_colostrum_1ng_glyco_hcd203040_compound15.17380.17380.5 | controllerType=0 controllerNumber=1 scan=17380 | 38.1687 |
| 14004:5 | 3 | K.IIEGEPNLKVPGN[+2862.021]VTAVLGETLKVPC[+57.021]HFPC[+57.021]K.F | HexNAc(5)Hex(6)Fuc(2)NeuAc(2) | C[+57]*2, N[+2862] | 1256.5797 | 5 | 6278.8695 | 6278.8273 | 0 | 6.7 | 457 | Specific | 569.23 | 502.28 | 502.28 | 9.98 | 136 | >sp\|P01833\|PIGR_HUMAN Polymeric immunoglobulin receptor OS=Homo sapiens OX=9606 GN=PIGR PE=1 SV=4 | 902 | 20201118_ly_colostrum_1ng_glyco_hcd203040_compound15.17413.17413.5 | controllerType=0 controllerNumber=1 scan=17413 | 38.2509 |
| 13915:4 | 3 | K.IIEGEPNLKVPGN[+2716.983]VTAVLGETLKVPC[+57.021]HFPC[+57.021]K.F | HexNAc(5)Hex(6)Fuc(3)NeuAc(1) | C[+57]*2, N[+2717] | 1534.2081 | 4 | 6133.8107 | 6133.7898 | 0 | 3.4 | 457 | Specific | 573.33 | 573.33 | 573.33 | 10.42 | 136 | >sp\|P01833\|PIGR_HUMAN Polymeric immunoglobulin receptor OS=Homo sapiens OX=9606 GN=PIGR PE=1 SV=4 | 902 | 20201118_ly_colostrum_1ng_glyco_hcd203040_compound15.17320.17320.4 | controllerType=0 controllerNumber=1 scan=17320 | 38.0302 |
| 13008:5 | 3 | K.IIEGEPNLKVPGN[+2279.830]VTAVLGETLKVPC[+57.021]HFPC[+57.021]K.F | HexNAc(5)Hex(6)Fuc(2) | C[+57]*2, N[+2280] | 1140.1355 | 5 | 5696.6484 | 5696.6365 | 0 | 2.1 | 457 | Specific | 576.71 | 451.67 | 451.67 | 10.58 | 136 | >sp\|P01833\|PIGR_HUMAN Polymeric immunoglobulin receptor OS=Homo sapiens OX=9606 GN=PIGR PE=1 SV=4 | 902 | 20201118_ly_colostrum_1ng_glyco_hcd203040_compound15.16377.16377.5 | controllerType=0 controllerNumber=1 scan=16377 | 36.0952 |
| 13002:5 | 3 | K.IIEGEPNLKVPGN[+2263.835]VTAVLGETLKVPC[+57.021]HFPC[+57.021]K.F | HexNAc(5)Hex(5)Fuc(3) | C[+57]*2, N[+2264] | 1136.936 | 5 | 5680.6511 | 5680.6416 | 0 | 1.7 | 457 | Specific | 576.94 | 554.02 | 554.02 | 10.58 | 136 | >sp\|P01833\|PIGR_HUMAN Polymeric immunoglobulin receptor OS=Homo sapiens OX=9606 GN=PIGR PE=1 SV=4 | 902 | 20201118_ly_colostrum_1ng_glyco_hcd203040_compound15.16371.16371.3 | controllerType=0 controllerNumber=1 scan=16371 | 36.0843 |
| 13918:4 | 3 | K.IIEGEPNLKVPGN[+2716.983]VTAVLGETLKVPC[+57.021]HFPC[+57.021]K.F | HexNAc(5)Hex(6)Fuc(3)NeuAc(1) | C[+57]*2, N[+2717] | 1534.2081 | 4 | 6133.8107 | 6133.7898 | 0 | 3.4 | 457 | Specific | 608.6 | 494.45 | 494.45 | 12.75 | 136 | >sp\|P01833\|PIGR_HUMAN Polymeric immunoglobulin receptor OS=Homo sapiens OX=9606 GN=PIGR PE=1 SV=4 | 902 | 20201118_ly_colostrum_1ng_glyco_hcd203040_compound15.17323.17323.4 | controllerType=0 controllerNumber=1 scan=17323 | 38.0389 |
| 13852:5 | 3 | K.IIEGEPNLKVPGN[+2716.983]VTAVLGETLKVPC[+57.021]HFPC[+57.021]K.F | HexNAc(5)Hex(6)Fuc(3)NeuAc(1) | C[+57]*2, N[+2717] | 1227.5678 | 5 | 6133.8097 | 6133.7898 | 0 | 3.2 | 457 | Specific | 637.48 | 534.56 | 534.56 | 13.16 | 136 | >sp\|P01833\|PIGR_HUMAN Polymeric immunoglobulin receptor OS=Homo sapiens OX=9606 GN=PIGR PE=1 SV=4 | 902 | 20201118_ly_colostrum_1ng_glyco_hcd203040_compound15.17255.17255.5 | controllerType=0 controllerNumber=1 scan=17255 | 37.8909 |
| 14174:4 | 3 | K.IIEGEPNLKVPGN[+2350.830]VTAVLGETLKVPC[+57.021]HFPC[+57.021]K.F | HexNAc(4)Hex(5)Fuc(1)NeuAc(2) | C[+57]*2, N[+2351] | 1442.673 | 4 | 5767.6701 | 5767.6372 | 0 | 5.7 | 457 | Specific | 660.95 | 568.66 | 568.66 | 13.18 | 136 | >sp\|P01833\|PIGR_HUMAN Polymeric immunoglobulin receptor OS=Homo sapiens OX=9606 GN=PIGR PE=1 SV=4 | 902 | 20201118_ly_colostrum_1ng_glyco_hcd203040_compound15.17590.17590.4 | controllerType=0 controllerNumber=1 scan=17590 | 38.6325 |
| 15361:5 | 3 | K.IIEGEPNLKVPGN[+2496.888]VTAVLGETLKVPC[+57.021]HFPC[+57.021]K.F | HexNAc(4)Hex(5)Fuc(2)NeuAc(2) | C[+57]*2, N[+2497] | 1183.5469 | 5 | 5913.7053 | 5913.6951 | 0 | 1.7 | 457 | Specific | 663.27 | 594.84 | 594.84 | 13.9 | 136 | >sp\|P01833\|PIGR_HUMAN Polymeric immunoglobulin receptor OS=Homo sapiens OX=9606 GN=PIGR PE=1 SV=4 | 902 | 20201118_ly_colostrum_1ng_glyco_hcd203040_compound15.18826.18826.5 | controllerType=0 controllerNumber=1 scan=18826 | 41.1428 |
| 14151:4 | 3 | K.IIEGEPNLKVPGN[+2351.851]VTAVLGETLKVPC[+57.021]HFPC[+57.021]K.F | HexNAc(4)Hex(5)Fuc(3)NeuAc(1) | C[+57]*2, N[+2352] | 1442.9229 | 4 | 5768.6696 | 5768.6576 | 0 | 2.1 | 457 | Specific | 673.97 | 642.9 | 642.9 | 13.49 | 136 | >sp\|P01833\|PIGR_HUMAN Polymeric immunoglobulin receptor OS=Homo sapiens OX=9606 GN=PIGR PE=1 SV=4 | 902 | 20201118_ly_colostrum_1ng_glyco_hcd203040_compound15.17566.17566.4 | controllerType=0 controllerNumber=1 scan=17566 | 38.5767 |
| 12978:5 | 3 | K.IIEGEPNLKVPGN[+2425.887]VTAVLGETLKVPC[+57.021]HFPC[+57.021]K.F | HexNAc(5)Hex(6)Fuc(3) | C[+57]*2, N[+2426] | 1169.3478 | 5 | 5842.7098 | 5842.6944 | 0 | 2.6 | 457 | Specific | 677.95 | 677.95 | 677.95 | 14.13 | 136 | >sp\|P01833\|PIGR_HUMAN Polymeric immunoglobulin receptor OS=Homo sapiens OX=9606 GN=PIGR PE=1 SV=4 | 902 | 20201118_ly_colostrum_1ng_glyco_hcd203040_compound15.16346.16346.5 | controllerType=0 controllerNumber=1 scan=16346 | 36.0205 |
| 12981:5 | 3 | K.IIEGEPNLKVPGN[+2425.887]VTAVLGETLKVPC[+57.021]HFPC[+57.021]K.F | HexNAc(5)Hex(6)Fuc(3) | C[+57]*2, N[+2426] | 1169.3478 | 5 | 5842.7098 | 5842.6944 | 0 | 2.6 | 457 | Specific | 681.72 | 681.72 | 681.72 | 14.13 | 136 | >sp\|P01833\|PIGR_HUMAN Polymeric immunoglobulin receptor OS=Homo sapiens OX=9606 GN=PIGR PE=1 SV=4 | 902 | 20201118_ly_colostrum_1ng_glyco_hcd203040_compound15.16349.16349.5 | controllerType=0 controllerNumber=1 scan=16349 | 36.0256 |
| 15437:5 | 3 | K.IIEGEPNLKVPGN[+2553.910]VTAVLGETLKVPC[+57.021]HFPC[+57.021]K.F | HexNAc(5)Hex(5)Fuc(1)NeuAc(2) | C[+57]*2, N[+2554] | 1194.9573 | 5 | 5970.7573 | 5970.7166 | 0 | 6.8 | 457 | Specific | 693.13 | 682 | 682 | 13.3 | 136 | >sp\|P01833\|PIGR_HUMAN Polymeric immunoglobulin receptor OS=Homo sapiens OX=9606 GN=PIGR PE=1 SV=4 | 902 | 20201118_ly_colostrum_1ng_glyco_hcd203040_compound15.18905.18905.5 | controllerType=0 controllerNumber=1 scan=18905 | 41.3335 |
| 13854:5 | 3 | K.IIEGEPNLKVPGN[+2716.983]VTAVLGETLKVPC[+57.021]HFPC[+57.021]K.F | HexNAc(5)Hex(6)Fuc(3)NeuAc(1) | C[+57]*2, N[+2717] | 1227.5678 | 5 | 6133.8097 | 6133.7898 | 0 | 3.2 | 457 | Specific | 699.32 | 692.5 | 692.5 | 14.16 | 136 | >sp\|P01833\|PIGR_HUMAN Polymeric immunoglobulin receptor OS=Homo sapiens OX=9606 GN=PIGR PE=1 SV=4 | 902 | 20201118_ly_colostrum_1ng_glyco_hcd203040_compound15.17257.17257.5 | controllerType=0 controllerNumber=1 scan=17257 | 37.8957 |
| 14171:4 | 3 | K.IIEGEPNLKVPGN[+2350.830]VTAVLGETLKVPC[+57.021]HFPC[+57.021]K.F | HexNAc(4)Hex(5)Fuc(1)NeuAc(2) | C[+57]*2, N[+2351] | 1442.673 | 4 | 5767.6701 | 5767.6372 | 0 | 5.7 | 457 | Specific | 703.59 | 554.62 | 554.62 | 14.18 | 136 | >sp\|P01833\|PIGR_HUMAN Polymeric immunoglobulin receptor OS=Homo sapiens OX=9606 GN=PIGR PE=1 SV=4 | 902 | 20201118_ly_colostrum_1ng_glyco_hcd203040_compound15.17587.17587.4 | controllerType=0 controllerNumber=1 scan=17587 | 38.6243 |
| 15312:4 | 3 | K.IIEGEPNLKVPGN[+2350.830]VTAVLGETLKVPC[+57.021]HFPC[+57.021]K.F | HexNAc(4)Hex(5)Fuc(1)NeuAc(2) | C[+57]*2, N[+2351] | 1442.6692 | 4 | 5767.6549 | 5767.6372 | 0 | 3.1 | 457 | Specific | 707.33 | 707.33 | 707.33 | 14.72 | 136 | >sp\|P01833\|PIGR_HUMAN Polymeric immunoglobulin receptor OS=Homo sapiens OX=9606 GN=PIGR PE=1 SV=4 | 902 | 20201118_ly_colostrum_1ng_glyco_hcd203040_compound15.18775.18775.4 | controllerType=0 controllerNumber=1 scan=18775 | 41.0443 |
| 15369:4 | 3 | K.IIEGEPNLKVPGN[+2496.888]VTAVLGETLKVPC[+57.021]HFPC[+57.021]K.F | HexNAc(4)Hex(5)Fuc(2)NeuAc(2) | C[+57]*2, N[+2497] | 1479.1838 | 4 | 5913.7135 | 5913.6951 | 0 | 3.1 | 457 | Specific | 717.17 | 642.22 | 642.22 | 14.75 | 136 | >sp\|P01833\|PIGR_HUMAN Polymeric immunoglobulin receptor OS=Homo sapiens OX=9606 GN=PIGR PE=1 SV=4 | 902 | 20201118_ly_colostrum_1ng_glyco_hcd203040_compound15.18834.18834.4 | controllerType=0 controllerNumber=1 scan=18834 | 41.1619 |
| 13877:5 | 3 | K.IIEGEPNLKVPGN[+2716.983]VTAVLGETLKVPC[+57.021]HFPC[+57.021]K.F | HexNAc(5)Hex(6)Fuc(3)NeuAc(1) | C[+57]*2, N[+2717] | 1227.5669 | 5 | 6133.8054 | 6133.7898 | 0 | 2.5 | 457 | Specific | 724.75 | 683.17 | 683.17 | 15.13 | 136 | >sp\|P01833\|PIGR_HUMAN Polymeric immunoglobulin receptor OS=Homo sapiens OX=9606 GN=PIGR PE=1 SV=4 | 902 | 20201118_ly_colostrum_1ng_glyco_hcd203040_compound15.17281.17281.5 | controllerType=0 controllerNumber=1 scan=17281 | 37.9472 |
| 14979:5 | 3 | K.IIEGEPNLKVPGN[+3154.136]VTAVLGETLKVPC[+57.021]HFPC[+57.021]K.F | HexNAc(5)Hex(6)Fuc(4)NeuAc(2) | C[+57]*2, N[+3154] | 1315.0018 | 5 | 6570.9801 | 6570.9432 | 0 | 5.6 | 457 | Specific | 740.33 | 641.75 | 641.75 | 14.59 | 136 | >sp\|P01833\|PIGR_HUMAN Polymeric immunoglobulin receptor OS=Homo sapiens OX=9606 GN=PIGR PE=1 SV=4 | 902 | 20201118_ly_colostrum_1ng_glyco_hcd203040_compound15.18428.18428.5 | controllerType=0 controllerNumber=1 scan=18428 | 40.3345 |
| 14017:4 | 3 | K.IIEGEPNLKVPGN[+2059.735]VTAVLGETLKVPC[+57.021]HFPC[+57.021]K.F | HexNAc(4)Hex(5)Fuc(1)NeuAc(1) | C[+57]*2, N[+2060] | 1369.894 | 4 | 5476.5544 | 5476.5418 | 0 | 2.3 | 457 | Specific | 745.64 | 745.64 | 745.64 | 14.72 | 136 | >sp\|P01833\|PIGR_HUMAN Polymeric immunoglobulin receptor OS=Homo sapiens OX=9606 GN=PIGR PE=1 SV=4 | 902 | 20201118_ly_colostrum_1ng_glyco_hcd203040_compound15.17427.17427.4 | controllerType=0 controllerNumber=1 scan=17427 | 38.2852 |
| 15383:5 | 3 | K.IIEGEPNLKVPGN[+2553.910]VTAVLGETLKVPC[+57.021]HFPC[+57.021]K.F | HexNAc(5)Hex(5)Fuc(1)NeuAc(2) | C[+57]*2, N[+2554] | 1194.9507 | 5 | 5970.7243 | 5970.7166 | 0 | 1.3 | 457 | Specific | 754.91 | 663.08 | 663.08 | 15.9 | 136 | >sp\|P01833\|PIGR_HUMAN Polymeric immunoglobulin receptor OS=Homo sapiens OX=9606 GN=PIGR PE=1 SV=4 | 902 | 20201118_ly_colostrum_1ng_glyco_hcd203040_compound15.18849.18849.5 | controllerType=0 controllerNumber=1 scan=18849 | 41.2101 |
| 13948:4 | 3 | K.IIEGEPNLKVPGN[+2351.851]VTAVLGETLKVPC[+57.021]HFPC[+57.021]K.F | HexNAc(4)Hex(5)Fuc(3)NeuAc(1) | C[+57]*2, N[+2352] | 1442.924 | 4 | 5768.674 | 5768.6576 | 0 | 2.8 | 457 | Specific | 757.33 | 747.76 | 747.76 | 15.72 | 136 | >sp\|P01833\|PIGR_HUMAN Polymeric immunoglobulin receptor OS=Homo sapiens OX=9606 GN=PIGR PE=1 SV=4 | 902 | 20201118_ly_colostrum_1ng_glyco_hcd203040_compound15.17355.17355.4 | controllerType=0 controllerNumber=1 scan=17355 | 38.1119 |
| 15435:5 | 3 | K.IIEGEPNLKVPGN[+2553.910]VTAVLGETLKVPC[+57.021]HFPC[+57.021]K.F | HexNAc(5)Hex(5)Fuc(1)NeuAc(2) | C[+57]*2, N[+2554] | 1194.9573 | 5 | 5970.7573 | 5970.7166 | 0 | 6.8 | 457 | Specific | 764.65 | 758.96 | 758.96 | 15.3 | 136 | >sp\|P01833\|PIGR_HUMAN Polymeric immunoglobulin receptor OS=Homo sapiens OX=9606 GN=PIGR PE=1 SV=4 | 902 | 20201118_ly_colostrum_1ng_glyco_hcd203040_compound15.18903.18903.5 | controllerType=0 controllerNumber=1 scan=18903 | 41.3269 |
| 14135:4 | 3 | K.IIEGEPNLKVPGN[+2205.793]VTAVLGETLKVPC[+57.021]HFPC[+57.021]K.F | HexNAc(4)Hex(5)Fuc(2)NeuAc(1) | C[+57]*2, N[+2206] | 1406.415 | 4 | 5622.6383 | 5622.5997 | 0 | 6.9 | 457 | Specific | 764.73 | 764.73 | 764.73 | 14.89 | 136 | >sp\|P01833\|PIGR_HUMAN Polymeric immunoglobulin receptor OS=Homo sapiens OX=9606 GN=PIGR PE=1 SV=4 | 902 | 20201118_ly_colostrum_1ng_glyco_hcd203040_compound15.17550.17550.4 | controllerType=0 controllerNumber=1 scan=17550 | 38.5467 |
| 15290:5 | 3 | K.IIEGEPNLKVPGN[+2350.830]VTAVLGETLKVPC[+57.021]HFPC[+57.021]K.F | HexNAc(4)Hex(5)Fuc(1)NeuAc(2) | C[+57]*2, N[+2351] | 1154.3362 | 5 | 5767.6518 | 5767.6372 | 0 | 2.5 | 457 | Specific | 783.53 | 680.66 | 680.66 | 16.13 | 136 | >sp\|P01833\|PIGR_HUMAN Polymeric immunoglobulin receptor OS=Homo sapiens OX=9606 GN=PIGR PE=1 SV=4 | 902 | 20201118_ly_colostrum_1ng_glyco_hcd203040_compound15.18752.18752.5 | controllerType=0 controllerNumber=1 scan=18752 | 40.9916 |
| 14035:4 | 3 | K.IIEGEPNLKVPGN[+2205.793]VTAVLGETLKVPC[+57.021]HFPC[+57.021]K.F | HexNAc(4)Hex(5)Fuc(2)NeuAc(1) | C[+57]*2, N[+2206] | 1406.4104 | 4 | 5622.6198 | 5622.5997 | 0 | 3.6 | 457 | Specific | 791.8 | 791.8 | 791.8 | 15.75 | 136 | >sp\|P01833\|PIGR_HUMAN Polymeric immunoglobulin receptor OS=Homo sapiens OX=9606 GN=PIGR PE=1 SV=4 | 902 | 20201118_ly_colostrum_1ng_glyco_hcd203040_compound15.17446.17446.4 | controllerType=0 controllerNumber=1 scan=17446 | 38.3277 |
| 13874:5 | 3 | K.IIEGEPNLKVPGN[+2716.983]VTAVLGETLKVPC[+57.021]HFPC[+57.021]K.F | HexNAc(5)Hex(6)Fuc(3)NeuAc(1) | C[+57]*2, N[+2717] | 1227.5669 | 5 | 6133.8054 | 6133.7898 | 0 | 2.5 | 457 | Specific | 797.74 | 693.67 | 693.67 | 16.13 | 136 | >sp\|P01833\|PIGR_HUMAN Polymeric immunoglobulin receptor OS=Homo sapiens OX=9606 GN=PIGR PE=1 SV=4 | 902 | 20201118_ly_colostrum_1ng_glyco_hcd203040_compound15.17278.17278.5 | controllerType=0 controllerNumber=1 scan=17278 | 37.9396 |
| 15386:5 | 3 | K.IIEGEPNLKVPGN[+2553.910]VTAVLGETLKVPC[+57.021]HFPC[+57.021]K.F | HexNAc(5)Hex(5)Fuc(1)NeuAc(2) | C[+57]*2, N[+2554] | 1194.9507 | 5 | 5970.7243 | 5970.7166 | 0 | 1.3 | 457 | Specific | 802.22 | 627.08 | 627.08 | 16.9 | 136 | >sp\|P01833\|PIGR_HUMAN Polymeric immunoglobulin receptor OS=Homo sapiens OX=9606 GN=PIGR PE=1 SV=4 | 902 | 20201118_ly_colostrum_1ng_glyco_hcd203040_compound15.18852.18852.5 | controllerType=0 controllerNumber=1 scan=18852 | 41.2205 |
| 14133:4 | 3 | K.IIEGEPNLKVPGN[+2205.793]VTAVLGETLKVPC[+57.021]HFPC[+57.021]K.F | HexNAc(4)Hex(5)Fuc(2)NeuAc(1) | C[+57]*2, N[+2206] | 1406.415 | 4 | 5622.6383 | 5622.5997 | 0 | 6.9 | 457 | Specific | 804.06 | 804.06 | 804.06 | 15.89 | 136 | >sp\|P01833\|PIGR_HUMAN Polymeric immunoglobulin receptor OS=Homo sapiens OX=9606 GN=PIGR PE=1 SV=4 | 902 | 20201118_ly_colostrum_1ng_glyco_hcd203040_compound15.17548.17548.4 | controllerType=0 controllerNumber=1 scan=17548 | 38.5419 |
| 15311:5 | 3 | K.IIEGEPNLKVPGN[+2350.830]VTAVLGETLKVPC[+57.021]HFPC[+57.021]K.F | HexNAc(4)Hex(5)Fuc(1)NeuAc(2) | C[+57]*2, N[+2351] | 1154.3362 | 5 | 5767.6518 | 5767.6372 | 0 | 2.5 | 457 | Specific | 824.13 | 625.88 | 625.88 | 17.13 | 136 | >sp\|P01833\|PIGR_HUMAN Polymeric immunoglobulin receptor OS=Homo sapiens OX=9606 GN=PIGR PE=1 SV=4 | 902 | 20201118_ly_colostrum_1ng_glyco_hcd203040_compound15.18774.18774.5 | controllerType=0 controllerNumber=1 scan=18774 | 41.0407 |
| 14034:5 | 3 | K.IIEGEPNLKVPGN[+2205.793]VTAVLGETLKVPC[+57.021]HFPC[+57.021]K.F | HexNAc(4)Hex(5)Fuc(2)NeuAc(1) | C[+57]*2, N[+2206] | 1125.3295 | 5 | 5622.6182 | 5622.5997 | 0 | 3.3 | 457 | Specific | 854.08 | 854.08 | 854.08 | 18.16 | 136 | >sp\|P01833\|PIGR_HUMAN Polymeric immunoglobulin receptor OS=Homo sapiens OX=9606 GN=PIGR PE=1 SV=4 | 902 | 20201118_ly_colostrum_1ng_glyco_hcd203040_compound15.17445.17445.5 | controllerType=0 controllerNumber=1 scan=17445 | 38.3242 |
| 15364:5 | 3 | K.IIEGEPNLKVPGN[+2496.888]VTAVLGETLKVPC[+57.021]HFPC[+57.021]K.F | HexNAc(4)Hex(5)Fuc(2)NeuAc(2) | C[+57]*2, N[+2497] | 1183.5469 | 5 | 5913.7053 | 5913.6951 | 0 | 1.7 | 457 | Specific | 861.98 | 820.94 | 820.94 | 17.9 | 136 | >sp\|P01833\|PIGR_HUMAN Polymeric immunoglobulin receptor OS=Homo sapiens OX=9606 GN=PIGR PE=1 SV=4 | 902 | 20201118_ly_colostrum_1ng_glyco_hcd203040_compound15.18829.18829.5 | controllerType=0 controllerNumber=1 scan=18829 | 41.1528 |
| 14032:4 | 3 | K.IIEGEPNLKVPGN[+2205.793]VTAVLGETLKVPC[+57.021]HFPC[+57.021]K.F | HexNAc(4)Hex(5)Fuc(2)NeuAc(1) | C[+57]*2, N[+2206] | 1406.4104 | 4 | 5622.6198 | 5622.5997 | 0 | 3.6 | 457 | Specific | 911.86 | 882.18 | 882.18 | 18.75 | 136 | >sp\|P01833\|PIGR_HUMAN Polymeric immunoglobulin receptor OS=Homo sapiens OX=9606 GN=PIGR PE=1 SV=4 | 902 | 20201118_ly_colostrum_1ng_glyco_hcd203040_compound15.17443.17443.4 | controllerType=0 controllerNumber=1 scan=17443 | 38.3197 |
| 14011:5 | 3 | K.IIEGEPNLKVPGN[+2059.735]VTAVLGETLKVPC[+57.021]HFPC[+57.021]K.F | HexNAc(4)Hex(5)Fuc(1)NeuAc(1) | C[+57]*2, N[+2060] | 1096.1172 | 5 | 5476.5568 | 5476.5418 | 0 | 2.7 | 457 | Specific | 958.2 | 877.68 | 877.68 | 20.13 | 136 | >sp\|P01833\|PIGR_HUMAN Polymeric immunoglobulin receptor OS=Homo sapiens OX=9606 GN=PIGR PE=1 SV=4 | 902 | 20201118_ly_colostrum_1ng_glyco_hcd203040_compound15.17421.17421.5 | controllerType=0 controllerNumber=1 scan=17421 | 38.2746 |
| 14014:4 | 3 | K.IIEGEPNLKVPGN[+2059.735]VTAVLGETLKVPC[+57.021]HFPC[+57.021]K.F | HexNAc(4)Hex(5)Fuc(1)NeuAc(1) | C[+57]*2, N[+2060] | 1369.894 | 4 | 5476.5544 | 5476.5418 | 0 | 2.3 | 457 | Specific | 962.41 | 903.33 | 903.33 | 19.72 | 136 | >sp\|P01833\|PIGR_HUMAN Polymeric immunoglobulin receptor OS=Homo sapiens OX=9606 GN=PIGR PE=1 SV=4 | 902 | 20201118_ly_colostrum_1ng_glyco_hcd203040_compound15.17424.17424.4 | controllerType=0 controllerNumber=1 scan=17424 | 38.2787 |
| 15310:4 | 3 | K.IIEGEPNLKVPGN[+2350.830]VTAVLGETLKVPC[+57.021]HFPC[+57.021]K.F | HexNAc(4)Hex(5)Fuc(1)NeuAc(2) | C[+57]*2, N[+2351] | 1442.6692 | 4 | 5767.6549 | 5767.6372 | 0 | 3.1 | 457 | Specific | 984.52 | 731.4 | 731.4 | 19.72 | 136 | >sp\|P01833\|PIGR_HUMAN Polymeric immunoglobulin receptor OS=Homo sapiens OX=9606 GN=PIGR PE=1 SV=4 | 902 | 20201118_ly_colostrum_1ng_glyco_hcd203040_compound15.18773.18773.4 | controllerType=0 controllerNumber=1 scan=18773 | 41.0362 |
| 13981:5 | 3 | K.IIEGEPNLKVPGN[+2351.851]VTAVLGETLKVPC[+57.021]HFPC[+57.021]K.F | HexNAc(4)Hex(5)Fuc(3)NeuAc(1) | C[+57]*2, N[+2352] | 1154.5404 | 5 | 5768.6729 | 5768.6576 | 0 | 2.7 | 457 | Specific | 986.26 | 986.26 | 986.26 | 20.13 | 136 | >sp\|P01833\|PIGR_HUMAN Polymeric immunoglobulin receptor OS=Homo sapiens OX=9606 GN=PIGR PE=1 SV=4 | 902 | 20201118_ly_colostrum_1ng_glyco_hcd203040_compound15.17390.17390.5 | controllerType=0 controllerNumber=1 scan=17390 | 38.2082 |
| 15366:4 | 3 | K.IIEGEPNLKVPGN[+2496.888]VTAVLGETLKVPC[+57.021]HFPC[+57.021]K.F | HexNAc(4)Hex(5)Fuc(2)NeuAc(2) | C[+57]*2, N[+2497] | 1479.1838 | 4 | 5913.7135 | 5913.6951 | 0 | 3.1 | 457 | Specific | 991.53 | 865.37 | 865.37 | 19.75 | 136 | >sp\|P01833\|PIGR_HUMAN Polymeric immunoglobulin receptor OS=Homo sapiens OX=9606 GN=PIGR PE=1 SV=4 | 902 | 20201118_ly_colostrum_1ng_glyco_hcd203040_compound15.18831.18831.4 | controllerType=0 controllerNumber=1 scan=18831 | 41.1554 |
| 14031:5 | 3 | K.IIEGEPNLKVPGN[+2205.793]VTAVLGETLKVPC[+57.021]HFPC[+57.021]K.F | HexNAc(4)Hex(5)Fuc(2)NeuAc(1) | C[+57]*2, N[+2206] | 1125.3295 | 5 | 5622.6182 | 5622.5997 | 0 | 3.3 | 457 | Specific | 1004.52 | 940.02 | 940.02 | 21.16 | 136 | >sp\|P01833\|PIGR_HUMAN Polymeric immunoglobulin receptor OS=Homo sapiens OX=9606 GN=PIGR PE=1 SV=4 | 902 | 20201118_ly_colostrum_1ng_glyco_hcd203040_compound15.17442.17442.5 | controllerType=0 controllerNumber=1 scan=17442 | 38.3179 |
| 15309:5 | 3 | K.IIEGEPNLKVPGN[+2350.830]VTAVLGETLKVPC[+57.021]HFPC[+57.021]K.F | HexNAc(4)Hex(5)Fuc(1)NeuAc(2) | C[+57]*2, N[+2351] | 1154.3362 | 5 | 5767.6518 | 5767.6372 | 0 | 2.5 | 457 | Specific | 1012.89 | 792.65 | 792.65 | 21.13 | 136 | >sp\|P01833\|PIGR_HUMAN Polymeric immunoglobulin receptor OS=Homo sapiens OX=9606 GN=PIGR PE=1 SV=4 | 902 | 20201118_ly_colostrum_1ng_glyco_hcd203040_compound15.18772.18772.5 | controllerType=0 controllerNumber=1 scan=18772 | 41.0348 |
| 13945:4 | 3 | K.IIEGEPNLKVPGN[+2351.851]VTAVLGETLKVPC[+57.021]HFPC[+57.021]K.F | HexNAc(4)Hex(5)Fuc(3)NeuAc(1) | C[+57]*2, N[+2352] | 1442.924 | 4 | 5768.674 | 5768.6576 | 0 | 2.8 | 457 | Specific | 1050.57 | 1050.57 | 1050.57 | 21.72 | 136 | >sp\|P01833\|PIGR_HUMAN Polymeric immunoglobulin receptor OS=Homo sapiens OX=9606 GN=PIGR PE=1 SV=4 | 902 | 20201118_ly_colostrum_1ng_glyco_hcd203040_compound15.17352.17352.4 | controllerType=0 controllerNumber=1 scan=17352 | 38.1053 |
| 15288:5 | 3 | K.IIEGEPNLKVPGN[+2350.830]VTAVLGETLKVPC[+57.021]HFPC[+57.021]K.F | HexNAc(4)Hex(5)Fuc(1)NeuAc(2) | C[+57]*2, N[+2351] | 1154.3362 | 5 | 5767.6518 | 5767.6372 | 0 | 2.5 | 457 | Specific | 1077 | 857.72 | 857.72 | 22.13 | 136 | >sp\|P01833\|PIGR_HUMAN Polymeric immunoglobulin receptor OS=Homo sapiens OX=9606 GN=PIGR PE=1 SV=4 | 902 | 20201118_ly_colostrum_1ng_glyco_hcd203040_compound15.18750.18750.5 | controllerType=0 controllerNumber=1 scan=18750 | 40.985 |
| 14009:5 | 3 | K.IIEGEPNLKVPGN[+2059.735]VTAVLGETLKVPC[+57.021]HFPC[+57.021]K.F | HexNAc(4)Hex(5)Fuc(1)NeuAc(1) | C[+57]*2, N[+2060] | 1096.1172 | 5 | 5476.5568 | 5476.5418 | 0 | 2.7 | 457 | Specific | 1128.87 | 1047.75 | 1047.75 | 23.13 | 136 | >sp\|P01833\|PIGR_HUMAN Polymeric immunoglobulin receptor OS=Homo sapiens OX=9606 GN=PIGR PE=1 SV=4 | 902 | 20201118_ly_colostrum_1ng_glyco_hcd203040_compound15.17419.17419.5 | controllerType=0 controllerNumber=1 scan=17419 | 38.2679 |
| 12297:4 | 3 | K.VPGN[+2205.793]VTAVLGETLKVPC[+57.021]HFPC[+57.021]K.F | HexNAc(4)Hex(5)Fuc(2)NeuAc(1) | C[+57]*2, N[+2206] | 1158.0248 | 4 | 4629.0775 | 4629.0502 | 0 | 5.9 | 466 | Specific | 405.62 | 256.07 | 256.07 | 7.23 | 136 | >sp\|P01833\|PIGR_HUMAN Polymeric immunoglobulin receptor OS=Homo sapiens OX=9606 GN=PIGR PE=1 SV=4 | 902 | 20201118_ly_colostrum_1ng_glyco_hcd203040_compound15.15638.15638.4 | controllerType=0 controllerNumber=1 scan=15638 | 34.5792 |
| 12300:4 | 3 | K.VPGN[+2205.793]VTAVLGETLKVPC[+57.021]HFPC[+57.021]K.F | HexNAc(4)Hex(5)Fuc(2)NeuAc(1) | C[+57]*2, N[+2206] | 1158.0248 | 4 | 4629.0775 | 4629.0502 | 0 | 5.9 | 466 | Specific | 562.35 | 400.48 | 400.48 | 10.29 | 136 | >sp\|P01833\|PIGR_HUMAN Polymeric immunoglobulin receptor OS=Homo sapiens OX=9606 GN=PIGR PE=1 SV=4 | 902 | 20201118_ly_colostrum_1ng_glyco_hcd203040_compound15.15641.15641.4 | controllerType=0 controllerNumber=1 scan=15641 | 34.5836 |
| 09190:4 | 3 | K.FSSYEKYWC[+57.021]KWN[+1872.651]NTGC[+57.021]QALPSQDEGPSK.A | HexNAc(3)Hex(6)NeuAc(1) | C[+57]*2, N[+1873] | 1310.7899 | 4 | 5240.1379 | 5240.1288 | 0 | 1.7 | 488 | Specific | 565.79 | 336.01 | 336.01 | 8.93 | 136 | >sp\|P01833\|PIGR_HUMAN Polymeric immunoglobulin receptor OS=Homo sapiens OX=9606 GN=PIGR PE=1 SV=4 | 902 | 20201118_ly_colostrum_1ng_glyco_hcd203040_compound15.12409.12409.4 | controllerType=0 controllerNumber=1 scan=12409 | 28.1509 |
| 09192:4 | 3 | K.FSSYEKYWC[+57.021]KWN[+1872.651]NTGC[+57.021]QALPSQDEGPSK.A | HexNAc(3)Hex(6)NeuAc(1) | C[+57]*2, N[+1873] | 1310.7899 | 4 | 5240.1379 | 5240.1288 | 0 | 1.7 | 488 | Specific | 706.89 | 468.74 | 468.74 | 14.98 | 136 | >sp\|P01833\|PIGR_HUMAN Polymeric immunoglobulin receptor OS=Homo sapiens OX=9606 GN=PIGR PE=1 SV=4 | 902 | 20201118_ly_colostrum_1ng_glyco_hcd203040_compound15.12411.12411.4 | controllerType=0 controllerNumber=1 scan=12411 | 28.1543 |
| 09080:4 | 3 | K.FSSYEKYWC[+57.021]KWN[+2205.793]NTGC[+57.021]QALPSQDEGPSK.A | HexNAc(4)Hex(5)Fuc(2)NeuAc(1) | C[+57]*2, N[+2206] | 1394.0761 | 4 | 5573.2824 | 5573.2711 | 0 | 2 | 488 | Specific | 774.48 | 774.48 | 774.48 | 15.98 | 136 | >sp\|P01833\|PIGR_HUMAN Polymeric immunoglobulin receptor OS=Homo sapiens OX=9606 GN=PIGR PE=1 SV=4 | 902 | 20201118_ly_colostrum_1ng_glyco_hcd203040_compound15.12294.12294.4 | controllerType=0 controllerNumber=1 scan=12294 | 27.912 |
| 09078:4 | 3 | K.FSSYEKYWC[+57.021]KWN[+2205.793]NTGC[+57.021]QALPSQDEGPSK.A | HexNAc(4)Hex(5)Fuc(2)NeuAc(1) | C[+57]*2, N[+2206] | 1394.0761 | 4 | 5573.2824 | 5573.2711 | 0 | 2 | 488 | Specific | 827.26 | 680.79 | 680.79 | 16.98 | 136 | >sp\|P01833\|PIGR_HUMAN Polymeric immunoglobulin receptor OS=Homo sapiens OX=9606 GN=PIGR PE=1 SV=4 | 902 | 20201118_ly_colostrum_1ng_glyco_hcd203040_compound15.12292.12292.4 | controllerType=0 controllerNumber=1 scan=12292 | 27.9078 |
| 09147:4 | 3 | K.FSSYEKYWC[+57.021]KWN[+1872.651]NTGC[+57.021]QALPSQDEGPSK.A | HexNAc(3)Hex(6)NeuAc(1) | C[+57]*2, N[+1873] | 1310.7907 | 4 | 5240.1408 | 5240.1288 | 0 | 2.3 | 488 | Specific | 928.11 | 607.35 | 607.35 | 19.21 | 136 | >sp\|P01833\|PIGR_HUMAN Polymeric immunoglobulin receptor OS=Homo sapiens OX=9606 GN=PIGR PE=1 SV=4 | 902 | 20201118_ly_colostrum_1ng_glyco_hcd203040_compound15.12364.12364.4 | controllerType=0 controllerNumber=1 scan=12364 | 28.0521 |
| 09150:4 | 3 | K.FSSYEKYWC[+57.021]KWN[+1872.651]NTGC[+57.021]QALPSQDEGPSK.A | HexNAc(3)Hex(6)NeuAc(1) | C[+57]*2, N[+1873] | 1310.7907 | 4 | 5240.1408 | 5240.1288 | 0 | 2.3 | 488 | Specific | 999.64 | 658.94 | 658.94 | 20.21 | 136 | >sp\|P01833\|PIGR_HUMAN Polymeric immunoglobulin receptor OS=Homo sapiens OX=9606 GN=PIGR PE=1 SV=4 | 902 | 20201118_ly_colostrum_1ng_glyco_hcd203040_compound15.12367.12367.4 | controllerType=0 controllerNumber=1 scan=12367 | 28.0604 |
| 09879:5 | 3 | K.FSSYEKYWC[+57.021]KWN[+2204.772]NTGC[+57.021]QALPSQDEGPSKAFVNC[+5 | HexNAc(4)Hex(5)NeuAc(2) | C[+57]*3, N[+2205] | 1353.7607 | 5 | 6764.7742 | 6764.7439 | 0 | 4.5 | 488 | Specific | 369.66 | 369.66 | 369.66 | 7.71 | 136 | >sp\|P01833\|PIGR_HUMAN Polymeric immunoglobulin receptor OS=Homo sapiens OX=9606 GN=PIGR PE=1 SV=4 | 902 | 20201118_ly_colostrum_1ng_glyco_hcd203040_compound15.13125.13125.3 | controllerType=0 controllerNumber=1 scan=13125 | 29.621 |
| 09880:5 | 3 | K.FSSYEKYWC[+57.021]KWN[+2204.772]NTGC[+57.021]QALPSQDEGPSKAFVNC[+5 | HexNAc(4)Hex(5)NeuAc(2) | C[+57]*3, N[+2205] | 1353.7607 | 5 | 6764.7742 | 6764.7439 | 0 | 4.5 | 488 | Specific | 481.17 | 481.17 | 481.17 | 9.63 | 136 | >sp\|P01833\|PIGR_HUMAN Polymeric immunoglobulin receptor OS=Homo sapiens OX=9606 GN=PIGR PE=1 SV=4 | 902 | 20201118_ly_colostrum_1ng_glyco_hcd203040_compound15.13126.13126.3 | controllerType=0 controllerNumber=1 scan=13126 | 29.6253 |
| 09952:5 | 3 | K.FSSYEKYWC[+57.021]KWN[+1710.598]NTGC[+57.021]QALPSQDEGPSKAFVNC[+5 | HexNAc(3)Hex(5)NeuAc(1) | C[+57]*3, N[+1711] | 1254.9207 | 5 | 6270.5742 | 6270.5691 | 0 | 0.8 | 488 | Specific | 1134.18 | 1108.66 | 1108.66 | 22.32 | 136 | >sp\|P01833\|PIGR_HUMAN Polymeric immunoglobulin receptor OS=Homo sapiens OX=9606 GN=PIGR PE=1 SV=4 | 902 | 20201118_ly_colostrum_1ng_glyco_hcd203040_compound15.13201.13201.5 | controllerType=0 controllerNumber=1 scan=13201 | 29.7688 |
| 09950:5 | 3 | K.FSSYEKYWC[+57.021]KWN[+1710.598]NTGC[+57.021]QALPSQDEGPSKAFVNC[+5 | HexNAc(3)Hex(5)NeuAc(1) | C[+57]*3, N[+1711] | 1254.9207 | 5 | 6270.5742 | 6270.5691 | 0 | 0.8 | 488 | Specific | 1294.75 | 1272.08 | 1272.08 | 25.32 | 136 | >sp\|P01833\|PIGR_HUMAN Polymeric immunoglobulin receptor OS=Homo sapiens OX=9606 GN=PIGR PE=1 SV=4 | 902 | 20201118_ly_colostrum_1ng_glyco_hcd203040_compound15.13199.13199.5 | controllerType=0 controllerNumber=1 scan=13199 | 29.7636 |
| 07347:3 | 3 | K.YWC[+57.021]KWN[+1710.598]NTGC[+57.021]QALPSQDEGPSK.A | HexNAc(3)Hex(5)NeuAc(1) | C[+57]*2, N[+1711] | 1446.2621 | 3 | 4336.7719 | 4336.7426 | 0 | 6.8 | 494 | Specific | 224.25 | 203.51 | 203.51 | 5.76 | 136 | >sp\|P01833\|PIGR_HUMAN Polymeric immunoglobulin receptor OS=Homo sapiens OX=9606 GN=PIGR PE=1 SV=4 | 902 | 20201118_ly_colostrum_1ng_glyco_hcd203040_compound15.10494.10494.3 | controllerType=0 controllerNumber=1 scan=10494 | 24.4081 |

Table S4. Cont.

| Query #:z | Protein Rank | Glycopeptide | Glycan | Modification Type(s) | Observed m/z | z | Observed (M+H) | Calc. mass (M+H) | Off-by-x error | Mass error (ppm) | StartPosition | Cleavage | Score | Delta | Delta Mod | \|Log Prob\| | # of unique peptides | Protein Name | Protein DB number | Comment | Scan # | Scan Time |
|---|---|---|---|---|---|---|---|---|---|---|---|---|---|---|---|---|---|---|---|---|---|---|
| 07345:3 | 3 | K.YWC[+57.021]KWN[+1710.598]NTGC[+57.021]QALPSQDEGPSK.A | HexNAc(3)Hex(5)NeuAc(1) | C[+57]*2, N[+1711] | 1446.2621 | 3 | 4336.7719 | 4336.7426 | 0 | 6.8 | 494 | Specific | 241.07 | 241.07 | 241.07 | 6.18 | 136 | >sp\|P01833\|PIGR_HUMAN Polymeric immunoglobulin receptor OS=Homo sapiens OX=9606 GN=PIGR PE=1 SV=4 | 902 | 20201118_ly_colostrum_1ng_glyco_hcd203040_compound15.10492.10492.3 | controllerType=0 controllerNumber=1 scan=10492 | 24.4042 |
| 07229:3 | 3 | K.YWC[+57.021]KWN[+1913.677]NTGC[+57.021]QALPSQDEGPSK.A | HexNAc(4)Hex(5)NeuAc(1) | C[+57]*2, N[+1914] | 1513.9496 | 3 | 4539.8344 | 4539.822 | 0 | 2.7 | 494 | Specific | 356.44 | 345.47 | 345.47 | 8.2 | 136 | >sp\|P01833\|PIGR_HUMAN Polymeric immunoglobulin receptor OS=Homo sapiens OX=9606 GN=PIGR PE=1 SV=4 | 902 | 20201118_ly_colostrum_1ng_glyco_hcd203040_compound15.10372.10372.3 | controllerType=0 controllerNumber=1 scan=10372 | 24.1792 |
| 07231:3 | 3 | K.YWC[+57.021]KWN[+1913.677]NTGC[+57.021]QALPSQDEGPSK.A | HexNAc(4)Hex(5)NeuAc(1) | C[+57]*2, N[+1914] | 1513.9496 | 3 | 4539.8344 | 4539.822 | 0 | 2.7 | 494 | Specific | 396.98 | 382.82 | 382.82 | 9.84 | 136 | >sp\|P01833\|PIGR_HUMAN Polymeric immunoglobulin receptor OS=Homo sapiens OX=9606 GN=PIGR PE=1 SV=4 | 902 | 20201118_ly_colostrum_1ng_glyco_hcd203040_compound15.10374.10374.3 | controllerType=0 controllerNumber=1 scan=10374 | 24.1827 |
| 07222:3 | 3 | K.YWC[+57.021]KWN[+1872.651]NTGC[+57.021]QALPSQDEGPSK.A | HexNAc(3)Hex(6)NeuAc(1) | C[+57]*2, N[+1873] | 1500.2752 | 3 | 4498.8111 | 4498.7954 | 0 | 3.5 | 494 | Specific | 839.24 | 829.3 | 829.3 | 17.18 | 136 | >sp\|P01833\|PIGR_HUMAN Polymeric immunoglobulin receptor OS=Homo sapiens OX=9606 GN=PIGR PE=1 SV=4 | 902 | 20201118_ly_colostrum_1ng_glyco_hcd203040_compound15.10365.10365.3 | controllerType=0 controllerNumber=1 scan=10365 | 24.1663 |
| 07224:3 | 3 | K.YWC[+57.021]KWN[+1872.651]NTGC[+57.021]QALPSQDEGPSK.A | HexNAc(3)Hex(6)NeuAc(1) | C[+57]*2, N[+1873] | 1500.2752 | 3 | 4498.8111 | 4498.7954 | 0 | 3.5 | 494 | Specific | 841.25 | 806.19 | 806.19 | 17.18 | 136 | >sp\|P01833\|PIGR_HUMAN Polymeric immunoglobulin receptor OS=Homo sapiens OX=9606 GN=PIGR PE=1 SV=4 | 902 | 20201118_ly_colostrum_1ng_glyco_hcd203040_compound15.10367.10367.3 | controllerType=0 controllerNumber=1 scan=10367 | 24.1723 |
| 08618:4 | 3 | K.YWC[+57.021]KWN[+1872.651]NTGC[+57.021]QALPSQDEGPSKAFVNC[+57.021]I | HexNAc(3)Hex(6)NeuAc(1) | C[+57]*3, N[+1873] | 1423.5814 | 4 | 5691.3039 | 5691.2885 | 0 | 2.7 | 494 | Specific | 1100.07 | 1046.3 | 1046.3 | 23.49 | 136 | >sp\|P01833\|PIGR_HUMAN Polymeric immunoglobulin receptor OS=Homo sapiens OX=9606 GN=PIGR PE=1 SV=4 | 902 | 20201118_ly_colostrum_1ng_glyco_hcd203040_compound15.11814.11814.4 | controllerType=0 controllerNumber=1 scan=11814 | 26.9627 |
| 08675:4 | 3 | K.YWC[+57.021]KWN[+1710.598]NTGC[+57.021]QALPSQDEGPSKAFVNC[+57.021]I | HexNAc(3)Hex(5)NeuAc(1) | C[+57]*3, N[+1711] | 1383.0673 | 4 | 5529.2472 | 5529.2357 | 0 | 2.1 | 494 | Specific | 1107.28 | 1038.67 | 1038.67 | 23.26 | 136 | >sp\|P01833\|PIGR_HUMAN Polymeric immunoglobulin receptor OS=Homo sapiens OX=9606 GN=PIGR PE=1 SV=4 | 902 | 20201118_ly_colostrum_1ng_glyco_hcd203040_compound15.11873.11873.4 | controllerType=0 controllerNumber=1 scan=11873 | 27.0882 |
| 08616:4 | 3 | K.YWC[+57.021]KWN[+1872.651]NTGC[+57.021]QALPSQDEGPSKAFVNC[+57.021]I | HexNAc(3)Hex(6)NeuAc(1) | C[+57]*3, N[+1873] | 1423.5814 | 4 | 5691.3039 | 5691.2885 | 0 | 2.7 | 494 | Specific | 1242.07 | 1216.8 | 1216.8 | 25.49 | 136 | >sp\|P01833\|PIGR_HUMAN Polymeric immunoglobulin receptor OS=Homo sapiens OX=9606 GN=PIGR PE=1 SV=4 | 902 | 20201118_ly_colostrum_1ng_glyco_hcd203040_compound15.11812.11812.4 | controllerType=0 controllerNumber=1 scan=11812 | 26.9555 |
| 08673:4 | 3 | K.YWC[+57.021]KWN[+1710.598]NTGC[+57.021]QALPSQDEGPSKAFVNC[+57.021]I | HexNAc(3)Hex(5)NeuAc(1) | C[+57]*3, N[+1711] | 1383.0673 | 4 | 5529.2472 | 5529.2357 | 0 | 2.1 | 494 | Specific | 1257.69 | 1253.33 | 1253.33 | 26.26 | 136 | >sp\|P01833\|PIGR_HUMAN Polymeric immunoglobulin receptor OS=Homo sapiens OX=9606 GN=PIGR PE=1 SV=4 | 902 | 20201118_ly_colostrum_1ng_glyco_hcd203040_compound15.11871.11871.4 | controllerType=0 controllerNumber=1 scan=11871 | 27.0818 |
| 05277:3 | 3 | K.WN[+2059.735]NTGC[+57.021]QALPSQDEGPSK.A | HexNAc(4)Hex(5)Fuc(1)NeuAc(1) | C[+57], N[+2060] | 1350.2116 | 3 | 4048.6203 | 4048.6116 | 0 | 2.1 | 498 | Specific | 535.92 | 470.39 | 470.39 | 10.42 | 136 | >sp\|P01833\|PIGR_HUMAN Polymeric immunoglobulin receptor OS=Homo sapiens OX=9606 GN=PIGR PE=1 SV=4 | 902 | 20201118_ly_colostrum_1ng_glyco_hcd203040_compound15.8351.8351.3 | controllerType=0 controllerNumber=1 scan=8351 | 20.6401 |
| 05279:3 | 3 | K.WN[+2059.735]NTGC[+57.021]QALPSQDEGPSK.A | HexNAc(4)Hex(5)Fuc(1)NeuAc(1) | C[+57], N[+2060] | 1350.2116 | 3 | 4048.6203 | 4048.6116 | 0 | 2.1 | 498 | Specific | 537.01 | 472.04 | 472.04 | 10.42 | 136 | >sp\|P01833\|PIGR_HUMAN Polymeric immunoglobulin receptor OS=Homo sapiens OX=9606 GN=PIGR PE=1 SV=4 | 902 | 20201118_ly_colostrum_1ng_glyco_hcd203040_compound15.8353.8353.3 | controllerType=0 controllerNumber=1 scan=8353 | 20.6442 |
| 05312:3 | 3 | K.WN[+1872.651]NTGC[+57.021]QALPSQDEGPSK.A | HexNAc(3)Hex(6)NeuAc(1) | C[+57], N[+1873] | 1287.8498 | 3 | 3861.5348 | 3861.5272 | 0 | 2 | 498 | Specific | 789.12 | 693.82 | 693.82 | 15.87 | 136 | >sp\|P01833\|PIGR_HUMAN Polymeric immunoglobulin receptor OS=Homo sapiens OX=9606 GN=PIGR PE=1 SV=4 | 902 | 20201118_ly_colostrum_1ng_glyco_hcd203040_compound15.8387.8387.3 | controllerType=0 controllerNumber=1 scan=8387 | 20.699 |
| 05405:3 | 3 | K.WN[+1710.598]NTGC[+57.021]QALPSQDEGPSK.A | HexNAc(3)Hex(5)NeuAc(1) | C[+57], N[+1711] | 1233.8325 | 3 | 3699.4828 | 3699.4743 | 0 | 2.3 | 498 | Specific | 852.1 | 756.91 | 756.91 | 18.11 | 136 | >sp\|P01833\|PIGR_HUMAN Polymeric immunoglobulin receptor OS=Homo sapiens OX=9606 GN=PIGR PE=1 SV=4 | 902 | 20201118_ly_colostrum_1ng_glyco_hcd203040_compound15.8483.8483.3 | controllerType=0 controllerNumber=1 scan=8483 | 20.8559 |
| 05402:3 | 3 | K.WN[+1710.598]NTGC[+57.021]QALPSQDEGPSK.A | HexNAc(3)Hex(5)NeuAc(1) | C[+57], N[+1711] | 1233.8325 | 3 | 3699.4828 | 3699.4743 | 0 | 2.3 | 498 | Specific | 875.06 | 810.18 | 810.18 | 18.11 | 136 | >sp\|P01833\|PIGR_HUMAN Polymeric immunoglobulin receptor OS=Homo sapiens OX=9606 GN=PIGR PE=1 SV=4 | 902 | 20201118_ly_colostrum_1ng_glyco_hcd203040_compound15.8480.8480.3 | controllerType=0 controllerNumber=1 scan=8480 | 20.8502 |
| 05310:3 | 3 | K.WN[+1872.651]NTGC[+57.021]QALPSQDEGPSK.A | HexNAc(3)Hex(6)NeuAc(1) | C[+57], N[+1873] | 1287.8498 | 3 | 3861.5348 | 3861.5272 | 0 | 2 | 498 | Specific | 924.25 | 861.57 | 861.57 | 18.87 | 136 | >sp\|P01833\|PIGR_HUMAN Polymeric immunoglobulin receptor OS=Homo sapiens OX=9606 GN=PIGR PE=1 SV=4 | 902 | 20201118_ly_colostrum_1ng_glyco_hcd203040_compound15.8385.8385.3 | controllerType=0 controllerNumber=1 scan=8385 | 20.6946 |
| 07792:4 | 3 | K.WN[+2059.735]NTGC[+57.021]QALPSQDEGPSKAFVNC[+57.021]DENSR.L | HexNAc(4)Hex(5)Fuc(1)NeuAc(1) | C[+57]*2, N[+2060] | 1311.0382 | 4 | 5241.131 | 5241.1047 | 0 | 5 | 498 | Specific | 228.08 | 228.08 | 228.08 | 6.57 | 136 | >sp\|P01833\|PIGR_HUMAN Polymeric immunoglobulin receptor OS=Homo sapiens OX=9606 GN=PIGR PE=1 SV=4 | 902 | 20201118_ly_colostrum_1ng_glyco_hcd203040_compound15.10956.10956.4 | controllerType=0 controllerNumber=1 scan=10956 | 25.2753 |
| 09244:4 | 3 | K.WN[+2204.772]NTGC[+57.021]QALPSQDEGPSKAFVNC[+57.021]DENSR.L | HexNAc(4)Hex(5)NeuAc(2) | C[+57]*2, N[+2205] | 1347.2944 | 4 | 5386.1559 | 5386.1422 | 0 | 2.5 | 498 | Specific | 228.47 | 228.47 | 228.47 | 7.11 | 136 | >sp\|P01833\|PIGR_HUMAN Polymeric immunoglobulin receptor OS=Homo sapiens OX=9606 GN=PIGR PE=1 SV=4 | 902 | 20201118_ly_colostrum_1ng_glyco_hcd203040_compound15.12464.12464.4 | controllerType=0 controllerNumber=1 scan=12464 | 28.2495 |
| 07794:4 | 3 | K.WN[+2059.735]NTGC[+57.021]QALPSQDEGPSKAFVNC[+57.021]DENSR.L | HexNAc(4)Hex(5)Fuc(1)NeuAc(1) | C[+57]*2, N[+2060] | 1311.0382 | 4 | 5241.131 | 5241.1047 | 0 | 5 | 498 | Specific | 293.84 | 293.84 | 293.84 | 7.26 | 136 | >sp\|P01833\|PIGR_HUMAN Polymeric immunoglobulin receptor OS=Homo sapiens OX=9606 GN=PIGR PE=1 SV=4 | 902 | 20201118_ly_colostrum_1ng_glyco_hcd203040_compound15.10958.10958.4 | controllerType=0 controllerNumber=1 scan=10958 | 25.2801 |
| 09245:4 | 3 | K.WN[+2204.772]NTGC[+57.021]QALPSQDEGPSKAFVNC[+57.021]DENSR.L | HexNAc(4)Hex(5)NeuAc(2) | C[+57]*2, N[+2205] | 1347.2944 | 4 | 5386.1559 | 5386.1422 | 0 | 2.5 | 498 | Specific | 319.44 | 319.44 | 319.44 | 8.1 | 136 | >sp\|P01833\|PIGR_HUMAN Polymeric immunoglobulin receptor OS=Homo sapiens OX=9606 GN=PIGR PE=1 SV=4 | 902 | 20201118_ly_colostrum_1ng_glyco_hcd203040_compound15.12465.12465.4 | controllerType=0 controllerNumber=1 scan=12465 | 28.2529 |
| 07814:3 | 3 | K.WN[+1872.651]NTGC[+57.021]QALPSQDEGPSKAFVNC[+57.021]DENSR.L | HexNAc(3)Hex(6)NeuAc(1) | C[+57]*2, N[+1873] | 1685.3476 | 3 | 5054.0283 | 5054.0203 | 0 | 1.6 | 498 | Specific | 766.88 | 727.71 | 727.71 | 16.14 | 136 | >sp\|P01833\|PIGR_HUMAN Polymeric immunoglobulin receptor OS=Homo sapiens OX=9606 GN=PIGR PE=1 SV=4 | 902 | 20201118_ly_colostrum_1ng_glyco_hcd203040_compound15.10979.10979.3 | controllerType=0 controllerNumber=1 scan=10979 | 25.3236 |
| 07812:3 | 3 | K.WN[+1872.651]NTGC[+57.021]QALPSQDEGPSKAFVNC[+57.021]DENSR.L | HexNAc(3)Hex(6)NeuAc(1) | C[+57]*2, N[+1873] | 1685.3476 | 3 | 5054.0283 | 5054.0203 | 0 | 1.6 | 498 | Specific | 783.25 | 742.02 | 742.02 | 16.14 | 136 | >sp\|P01833\|PIGR_HUMAN Polymeric immunoglobulin receptor OS=Homo sapiens OX=9606 GN=PIGR PE=1 SV=4 | 902 | 20201118_ly_colostrum_1ng_glyco_hcd203040_compound15.10977.10977.3 | controllerType=0 controllerNumber=1 scan=10977 | 25.3197 |
| 07807:4 | 3 | K.WN[+1872.651]NTGC[+57.021]QALPSQDEGPSKAFVNC[+57.021]DENSR.L | HexNAc(3)Hex(6)NeuAc(1) | C[+57]*2, N[+1873] | 1264.2634 | 4 | 5054.0319 | 5054.0203 | 0 | 2.3 | 498 | Specific | 855.36 | 855.36 | 855.36 | 18.53 | 136 | >sp\|P01833\|PIGR_HUMAN Polymeric immunoglobulin receptor OS=Homo sapiens OX=9606 GN=PIGR PE=1 SV=4 | 902 | 20201118_ly_colostrum_1ng_glyco_hcd203040_compound15.10972.10972.4 | controllerType=0 controllerNumber=1 scan=10972 | 25.3105 |
| 07809:4 | 3 | K.WN[+1872.651]NTGC[+57.021]QALPSQDEGPSKAFVNC[+57.021]DENSR.L | HexNAc(3)Hex(6)NeuAc(1) | C[+57]*2, N[+1873] | 1264.2634 | 4 | 5054.0319 | 5054.0203 | 0 | 2.3 | 498 | Specific | 974.87 | 929.79 | 929.79 | 20.53 | 136 | >sp\|P01833\|PIGR_HUMAN Polymeric immunoglobulin receptor OS=Homo sapiens OX=9606 GN=PIGR PE=1 SV=4 | 902 | 20201118_ly_colostrum_1ng_glyco_hcd203040_compound15.10974.10974.4 | controllerType=0 controllerNumber=1 scan=10974 | 25.3152 |
| 22665:4 | 5 | R.LSLHRPALEDLLLGSEAN[+1501.555]LTC[+57.021]TLTGLR.D | HexNAc(5)Hex(3) | C[+57], N[+1502] | 1117.0457 | 4 | 4465.161 | 4465.1536 | 0 | 1.6 | 127 | Specific | 201.6 | 201.6 | 201.6 | 5.61 | 123 | >sp\|P01876\|IGHA1_HUMAN Immunoglobulin heavy constant alpha 1 OS=Homo sapiens OX=9606 GN=IGHA1 PE=1 SV=2 | 255 | 20201118_ly_colostrum_1ng_glyco_hcd203040_compound15.26449.26449.4 | controllerType=0 controllerNumber=1 scan=26449 | 57.7226 |
| 16954:5 | 5 | R.[+42.011]LSLHRPALEDLLLGSEAN[+1710.598]LTC[+57.021]TLTGLR.D | HexNAc(3)Hex(5)NeuAc(1) | .[42], C[+57], N[+1711] | 944.0527 | 5 | 4716.2344 | 4716.2065 | 0 | 5.9 | 127 | Specific | 211.34 | 211.34 | 211.34 | 3.85 | 123 | >sp\|P01876\|IGHA1_HUMAN Immunoglobulin heavy constant alpha 1 OS=Homo sapiens OX=9606 GN=IGHA1 PE=1 SV=2 | 255 | 20201118_ly_colostrum_1ng_glyco_hcd203040_compound15.20490.20490.5 | controllerType=0 controllerNumber=1 scan=20490 | 44.9065 |
| 16063:3 | 5 | R.[+42.011]LSLHRPALEDLLLGSEAN[+1216.423]LTC[+57.021]TLTGLR.D | HexNAc(2)Hex(5) | .[42], C[+57], N[+1216] | 1408.0264 | 3 | 4222.0647 | 4222.0317 | 0 | 7.8 | 127 | Specific | 217.99 | 217.99 | 217.99 | 3.44 | 123 | >sp\|P01876\|IGHA1_HUMAN Immunoglobulin heavy constant alpha 1 OS=Homo sapiens OX=9606 GN=IGHA1 PE=1 SV=2 | 255 | 20201118_ly_colostrum_1ng_glyco_hcd203040_compound15.19558.19558.3 | controllerType=0 controllerNumber=1 scan=19558 | 42.7905 |
| 16075:3 | 5 | R.LSLHRPALEDLLLGSEAN[+1501.555]LTC[+57.021]TLTGLR.D | HexNAc(5)Hex(3) | C[+57], N[+1502] | 1489.0607 | 3 | 4465.1676 | 4465.1536 | 0 | 3.1 | 127 | Specific | 233.89 | 233.89 | 233.89 | 6.79 | 123 | >sp\|P01876\|IGHA1_HUMAN Immunoglobulin heavy constant alpha 1 OS=Homo sapiens OX=9606 GN=IGHA1 PE=1 SV=2 | 255 | 20201118_ly_colostrum_1ng_glyco_hcd203040_compound15.19571.19571.6 | controllerType=0 controllerNumber=1 scan=19571 | 42.8281 |
| 21394:4 | 5 | R.LSLHRPALEDLLLGSEAN[+1501.555]LTC[+57.021]TLTGLR.D | HexNAc(5)Hex(3) | C[+57], N[+1502] | 1117.0459 | 4 | 4465.1619 | 4465.1536 | 0 | 1.9 | 127 | Specific | 234.87 | 234.87 | 234.87 | 6.95 | 123 | >sp\|P01876\|IGHA1_HUMAN Immunoglobulin heavy constant alpha 1 OS=Homo sapiens OX=9606 GN=IGHA1 PE=1 SV=2 | 255 | 20201118_ly_colostrum_1ng_glyco_hcd203040_compound15.25126.25126.4 | controllerType=0 controllerNumber=1 scan=25126 | 55.1233 |
| 17971:3 | 5 | R.LSLHRPALEDLLLGSEAN[+2204.772]LTC[+57.021]TLTGLR.D | HexNAc(4)Hex(5)NeuAc(2) | C[+57], N[+2205] | 1723.469 | 3 | 5168.3923 | 5168.3707 | 0 | 4.2 | 127 | Specific | 235.4 | 235.4 | 235.4 | 6.5 | 123 | >sp\|P01876\|IGHA1_HUMAN Immunoglobulin heavy constant alpha 1 OS=Homo sapiens OX=9606 GN=IGHA1 PE=1 SV=2 | 255 | 20201118_ly_colostrum_1ng_glyco_hcd203040_compound15.21553.21553.3 | controllerType=0 controllerNumber=1 scan=21553 | 47.2877 |
| 16061:3 | 5 | R.[+42.011]LSLHRPALEDLLLGSEAN[+1216.423]LTC[+57.021]TLTGLR.D | HexNAc(2)Hex(5) | .[42], C[+57], N[+1216] | 1408.0264 | 3 | 4222.0647 | 4222.0317 | 0 | 7.8 | 127 | Specific | 252.47 | 252.47 | 252.47 | 6.18 | 123 | >sp\|P01876\|IGHA1_HUMAN Immunoglobulin heavy constant alpha 1 OS=Homo sapiens OX=9606 GN=IGHA1 PE=1 SV=2 | 255 | 20201118_ly_colostrum_1ng_glyco_hcd203040_compound15.19556.19556.3 | controllerType=0 controllerNumber=1 scan=19556 | 42.779 |
| 16113:3 | 5 | R.LSLHRPALEDLLLGSEAN[+1501.555]LTC[+57.021]TLTGLR.D | HexNAc(5)Hex(3) | C[+57], N[+1502] | 1489.0602 | 3 | 4465.1662 | 4465.1536 | 0 | 2.8 | 127 | Specific | 265.28 | 265.28 | 265.28 | 7.25 | 123 | >sp\|P01876\|IGHA1_HUMAN Immunoglobulin heavy constant alpha 1 OS=Homo sapiens OX=9606 GN=IGHA1 PE=1 SV=2 | 255 | 20201118_ly_colostrum_1ng_glyco_hcd203040_compound15.19610.19610.2 | controllerType=0 controllerNumber=1 scan=19610 | 42.9191 |
| 16064:5 | 5 | R.LSLHRPALEDLLLGSEAN[+1524.534]LTC[+57.021]TLTGLR.D | HexNAc(2)Hex(6)Fuc(1) | C[+57], N[+1525] | 898.4298 | 5 | 4488.1197 | 4488.1319 | 0 | -2.7 | 127 | Specific | 278.26 | 278.26 | 278.26 | 5.1 | 123 | >sp\|P01876\|IGHA1_HUMAN Immunoglobulin heavy constant alpha 1 OS=Homo sapiens OX=9606 GN=IGHA1 PE=1 SV=2 | 255 | 20201118_ly_colostrum_1ng_glyco_hcd203040_compound15.19559.19559.2 | controllerType=0 controllerNumber=1 scan=19559 | 42.7924 |
| 22063:4 | 5 | R.LSLHRPALEDLLLGSEAN[+1501.555]LTC[+57.021]TLTGLR.D | HexNAc(5)Hex(3) | C[+57], N[+1502] | 1117.0437 | 4 | 4465.1531 | 4465.1536 | 0 | -0.1 | 127 | Specific | 292.73 | 283.3 | 283.3 | 7.03 | 123 | >sp\|P01876\|IGHA1_HUMAN Immunoglobulin heavy constant alpha 1 OS=Homo sapiens OX=9606 GN=IGHA1 PE=1 SV=2 | 255 | 20201118_ly_colostrum_1ng_glyco_hcd203040_compound15.25823.25823.4 | controllerType=0 controllerNumber=1 scan=25823 | 56.5302 |
| 26649:4 | 5 | R.LSLHRPALEDLLLGSEAN[+1216.423]LTC[+57.021]TLTGLR.D | HexNAc(2)Hex(5) | C[+57], N[+1216] | 1045.7638 | 4 | 4180.0335 | 4180.0211 | 0 | 3 | 127 | Specific | 303.56 | 303.56 | 303.56 | 8.16 | 123 | >sp\|P01876\|IGHA1_HUMAN Immunoglobulin heavy constant alpha 1 OS=Homo sapiens OX=9606 GN=IGHA1 PE=1 SV=2 | 255 | 20201118_ly_colostrum_1ng_glyco_hcd203040_compound15.30600.30600.3 | controllerType=0 controllerNumber=1 scan=30600 | 65.3745 |
| 22370:4 | 5 | R.LSLHRPALEDLLLGSEAN[+1501.555]LTC[+57.021]TLTGLR.D | HexNAc(5)Hex(3) | C[+57], N[+1502] | 1117.0463 | 4 | 4465.1634 | 4465.1536 | 0 | 2.2 | 127 | Specific | 314.65 | 314.65 | 314.65 | 8.16 | 123 | >sp\|P01876\|IGHA1_HUMAN Immunoglobulin heavy constant alpha 1 OS=Homo sapiens OX=9606 GN=IGHA1 PE=1 SV=2 | 255 | 20201118_ly_colostrum_1ng_glyco_hcd203040_compound15.26143.26143.3 | controllerType=0 controllerNumber=1 scan=26143 | 57.1512 |
| 18967:4 | 5 | R.LSLHRPALEDLLLGSEAN[+1501.555]LTC[+57.021]TLTGLR.D | HexNAc(5)Hex(3) | C[+57], N[+1502] | 1117.0436 | 4 | 4465.1527 | 4465.1536 | 0 | -0.2 | 127 | Specific | 317.18 | 304.05 | 304.05 | 6.37 | 123 | >sp\|P01876\|IGHA1_HUMAN Immunoglobulin heavy constant alpha 1 OS=Homo sapiens OX=9606 GN=IGHA1 PE=1 SV=2 | 255 | 20201118_ly_colostrum_1ng_glyco_hcd203040_compound15.22594.22594.4 | controllerType=0 controllerNumber=1 scan=22594 | 49.6951 |
| 34986:4 | 5 | R.LSLHRPALEDLLLGSEAN[+1501.555]LTC[+57.021]TLTGLR.D | HexNAc(5)Hex(3) | C[+57], N[+1502] | 1117.0455 | 4 | 4465.16 | 4465.1536 | 0 | 1.4 | 127 | Specific | 329.05 | 329.05 | 329.05 | 8.21 | 123 | >sp\|P01876\|IGHA1_HUMAN Immunoglobulin heavy constant alpha 1 OS=Homo sapiens OX=9606 GN=IGHA1 PE=1 SV=2 | 255 | 20201118_ly_colostrum_1ng_glyco_hcd203040_compound15.41323.41323.4 | controllerType=0 controllerNumber=1 scan=41323 | 88.7959 |
| 34894:4 | 5 | R.LSLHRPALEDLLLGSEAN[+1216.423]LTC[+57.021]TLTGLR.D | HexNAc(2)Hex(5) | C[+57], N[+1216] | 1045.7641 | 4 | 4180.0345 | 4180.0211 | 0 | 3.2 | 127 | Specific | 352.2 | 352.2 | 352.2 | 8.66 | 123 | >sp\|P01876\|IGHA1_HUMAN Immunoglobulin heavy constant alpha 1 OS=Homo sapiens OX=9606 GN=IGHA1 PE=1 SV=2 | 255 | 20201118_ly_colostrum_1ng_glyco_hcd203040_compound15.41221.41221.2 | controllerType=0 controllerNumber=1 scan=41221 | 88.6014 |
| 23747:4 | 5 | R.LSLHRPALEDLLLGSEAN[+1216.423]LTC[+57.021]TLTGLR.D | HexNAc(2)Hex(5) | C[+57], N[+1216] | 1045.7662 | 4 | 4180.0428 | 4180.0211 | 0 | 5.2 | 127 | Specific | 358.32 | 358.32 | 358.32 | 8.09 | 123 | >sp\|P01876\|IGHA1_HUMAN Immunoglobulin heavy constant alpha 1 OS=Homo sapiens OX=9606 GN=IGHA1 PE=1 SV=2 | 255 | 20201118_ly_colostrum_1ng_glyco_hcd203040_compound15.27571.27571.4 | controllerType=0 controllerNumber=1 scan=27571 | 59.7936 |
| 21396:4 | 5 | R.LSLHRPALEDLLLGSEAN[+1501.555]LTC[+57.021]TLTGLR.D | HexNAc(5)Hex(3) | C[+57], N[+1502] | 1117.0459 | 4 | 4465.1619 | 4465.1536 | 0 | 1.9 | 127 | Specific | 361.05 | 328.36 | 328.36 | 8.33 | 123 | >sp\|P01876\|IGHA1_HUMAN Immunoglobulin heavy constant alpha 1 OS=Homo sapiens OX=9606 GN=IGHA1 PE=1 SV=2 | 255 | 20201118_ly_colostrum_1ng_glyco_hcd203040_compound15.25128.25128.4 | controllerType=0 controllerNumber=1 scan=25128 | 55.1275 |
| 22667:4 | 5 | R.LSLHRPALEDLLLGSEAN[+1501.555]LTC[+57.021]TLTGLR.D | HexNAc(5)Hex(3) | C[+57], N[+1502] | 1117.0457 | 4 | 4465.161 | 4465.1536 | 0 | 1.6 | 127 | Specific | 361.29 | 244.71 | 244.71 | 7.79 | 123 | >sp\|P01876\|IGHA1_HUMAN Immunoglobulin heavy constant alpha 1 OS=Homo sapiens OX=9606 GN=IGHA1 PE=1 SV=2 | 255 | 20201118_ly_colostrum_1ng_glyco_hcd203040_compound15.26451.26451.4 | controllerType=0 controllerNumber=1 scan=26451 | 57.7267 |
| 19548:4 | 5 | R.LSLHRPALEDLLLGSEAN[+1501.555]LTC[+57.021]TLTGLR.D | HexNAc(5)Hex(3) | C[+57], N[+1502] | 1117.0478 | 4 | 4465.1693 | 4465.1536 | 0 | 3.5 | 127 | Specific | 365.11 | 365.11 | 365.11 | 8.85 | 123 | >sp\|P01876\|IGHA1_HUMAN Immunoglobulin heavy constant alpha 1 OS=Homo sapiens OX=9606 GN=IGHA1 PE=1 SV=2 | 255 | 20201118_ly_colostrum_1ng_glyco_hcd203040_compound15.23202.23202.4 | controllerType=0 controllerNumber=1 scan=23202 | 51.0727 |
| 35598:4 | 5 | R.LSLHRPALEDLLLGSEAN[+1216.423]LTC[+57.021]TLTGLR.D | HexNAc(2)Hex(5) | C[+57], N[+1216] | 1045.7607 | 4 | 4180.0208 | 4180.0211 | 0 | -0.1 | 127 | Specific | 381.91 | 381.91 | 381.91 | 9.45 | 123 | >sp\|P01876\|IGHA1_HUMAN Immunoglobulin heavy constant alpha 1 OS=Homo sapiens OX=9606 GN=IGHA1 PE=1 SV=2 | 255 | 20201118_ly_colostrum_1ng_glyco_hcd203040_compound15.41984.41984.4 | controllerType=0 controllerNumber=1 scan=41984 | 90.0342 |
| 15944:5 | 5 | R.LSLHRPALEDLLLGSEAN[+1702.581]LTC[+57.021]TLTGLR.D | HexNAc(2)Hex(8) | C[+57], N[+1703] | 934.0354 | 5 | 4666.148 | 4666.1796 | 0 | -6.8 | 127 | Specific | 382.43 | 382.43 | 382.43 | 7.6 | 123 | >sp\|P01876\|IGHA1_HUMAN Immunoglobulin heavy constant alpha 1 OS=Homo sapiens OX=9606 GN=IGHA1 PE=1 SV=2 | 255 | 20201118_ly_colostrum_1ng_glyco_hcd203040_compound15.19434.19434.5 | controllerType=0 controllerNumber=1 scan=19434 | 42.5232 |
| 16067:5 | 5 | R.LSLHRPALEDLLLGSEAN[+1524.534]LTC[+57.021]TLTGLR.D | HexNAc(2)Hex(6)Fuc(1) | C[+57], N[+1525] | 898.4298 | 5 | 4488.1197 | 4488.1319 | 0 | -2.7 | 127 | Specific | 394.34 | 394.34 | 394.34 | 9.17 | 123 | >sp\|P01876\|IGHA1_HUMAN Immunoglobulin heavy constant alpha 1 OS=Homo sapiens OX=9606 GN=IGHA1 PE=1 SV=2 | 255 | 20201118_ly_colostrum_1ng_glyco_hcd203040_compound15.19562.19562.2 | controllerType=0 controllerNumber=1 scan=19562 | 42.7998 |
| 16118:3 | 5 | R.LSLHRPALEDLLLGSEAN[+1095.397]LTC[+57.021]TLTGLR.D | HexNAc(3)Hex(3) | C[+57], N[+1095] | 1353.6743 | 3 | 4059.0082 | 4058.9949 | 0 | 3.3 | 127 | Specific | 398.69 | 398.69 | 398.69 | 9.89 | 123 | >sp\|P01876\|IGHA1_HUMAN Immunoglobulin heavy constant alpha 1 OS=Homo sapiens OX=9606 GN=IGHA1 PE=1 SV=2 | 255 | 20201118_ly_colostrum_1ng_glyco_hcd203040_compound15.19616.19616.3 | controllerType=0 controllerNumber=1 scan=19616 | 42.9297 |
| 16973:4 | 5 | R.LSLHRPALEDLLLGSEAN[+1913.677]LTC[+57.021]TLTGLR.D | HexNAc(4)Hex(5)NeuAc(1) | C[+57], N[+1914] | 1220.0769 | 4 | 4877.286 | 4877.2753 | 0 | 2.2 | 127 | Specific | 415.53 | 410.14 | 410.14 | 10.48 | 123 | >sp\|P01876\|IGHA1_HUMAN Immunoglobulin heavy constant alpha 1 OS=Homo sapiens OX=9606 GN=IGHA1 PE=1 SV=2 | 255 | 20201118_ly_colostrum_1ng_glyco_hcd203040_compound15.20510.20510.4 | controllerType=0 controllerNumber=1 scan=20510 | 44.9447 |
| 21926:4 | 5 | R.LSLHRPALEDLLLGSEAN[+1501.555]LTC[+57.021]TLTGLR.D | HexNAc(5)Hex(3) | C[+57], N[+1502] | 1117.0467 | 4 | 4465.1649 | 4465.1536 | 0 | 2.5 | 127 | Specific | 427.07 | 373.08 | 373.08 | 8.94 | 123 | >sp\|P01876\|IGHA1_HUMAN Immunoglobulin heavy constant alpha 1 OS=Homo sapiens OX=9606 GN=IGHA1 PE=1 SV=2 | 255 | 20201118_ly_colostrum_1ng_glyco_hcd203040_compound15.25681.25681.4 | controllerType=0 controllerNumber=1 scan=25681 | 56.2646 |
| 34987:4 | 5 | R.LSLHRPALEDLLLGSEAN[+1216.423]LTC[+57.021]TLTGLR.D | HexNAc(2)Hex(5) | C[+57], N[+1216] | 1045.7629 | 4 | 4180.0296 | 4180.0211 | 0 | 2 | 127 | Specific | 433.13 | 433.13 | 433.13 | 10.41 | 123 | >sp\|P01876\|IGHA1_HUMAN Immunoglobulin heavy constant alpha 1 OS=Homo sapiens OX=9606 GN=IGHA1 PE=1 SV=2 | 255 | 20201118_ly_colostrum_1ng_glyco_hcd203040_compound15.41324.41324.2 | controllerType=0 controllerNumber=1 scan=41324 | 88.7976 |
| 34950:4 | 5 | R.LSLHRPALEDLLLGSEAN[+1216.423]LTC[+57.021]TLTGLR.D | HexNAc(2)Hex(5) | C[+57], N[+1216] | 1045.7637 | 4 | 4180.033 | 4180.0211 | 0 | 2.8 | 127 | Specific | 435.7 | 435.7 | 435.7 | 10.41 | 123 | >sp\|P01876\|IGHA1_HUMAN Immunoglobulin heavy constant alpha 1 OS=Homo sapiens OX=9606 GN=IGHA1 PE=1 SV=2 | 255 | 20201118_ly_colostrum_1ng_glyco_hcd203040_compound15.41284.41284.4 | controllerType=0 controllerNumber=1 scan=41284 | 88.7226 |
| 18969:4 | 5 | R.LSLHRPALEDLLLGSEAN[+1501.555]LTC[+57.021]TLTGLR.D | HexNAc(5)Hex(3) | C[+57], N[+1502] | 1117.0436 | 4 | 4465.1527 | 4465.1536 | 0 | -0.2 | 127 | Specific | 450.04 | 368.69 | 368.69 | 8.29 | 123 | >sp\|P01876\|IGHA1_HUMAN Immunoglobulin heavy constant alpha 1 OS=Homo sapiens OX=9606 GN=IGHA1 PE=1 SV=2 | 255 | 20201118_ly_colostrum_1ng_glyco_hcd203040_compound15.22596.22596.4 | controllerType=0 controllerNumber=1 scan=22596 | 49.7002 |
| 16947:3 | 5 | R.LSLHRPALEDLLLGSEAN[+1751.624]LTC[+57.021]TLTGLR.D | HexNAc(4)Hex(4)NeuAc(1) | C[+57], N[+1752] | 1572.4177 | 3 | 4715.2384 | 4715.2225 | 0 | 3.4 | 127 | Specific | 450.71 | 450.71 | 450.71 | 10.21 | 123 | >sp\|P01876\|IGHA1_HUMAN Immunoglobulin heavy constant alpha 1 OS=Homo sapiens OX=9606 GN=IGHA1 PE=1 SV=2 | 255 | 20201118_ly_colostrum_1ng_glyco_hcd203040_compound15.20483.20483.3 | controllerType=0 controllerNumber=1 scan=20483 | 44.8934 |
| 16943:3 | 5 | R.LSLHRPALEDLLLGSEAN[+1710.598]LTC[+57.021]TLTGLR.D | HexNAc(3)Hex(5)NeuAc(1) | C[+57], N[+1711] | 1558.7419 | 3 | 4674.2111 | 4674.1959 | 0 | 3.2 | 127 | Specific | 461.81 | 440.7 | 440.7 | 10.27 | 123 | >sp\|P01876\|IGHA1_HUMAN Immunoglobulin heavy constant alpha 1 OS=Homo sapiens OX=9606 GN=IGHA1 PE=1 SV=2 | 255 | 20201118_ly_colostrum_1ng_glyco_hcd203040_compound15.20479.20479.3 | controllerType=0 controllerNumber=1 scan=20479 | 44.8868 |
| 15971:4 | 5 | R.LSLHRPALEDLLLGSEAN[+1622.582]LTC[+57.021]TLTGLR.D | HexNAc(4)Hex(5) | C[+57], N[+1623] | 1147.3024 | 4 | 4586.1878 | 4586.1799 | 0 | 1.7 | 127 | Specific | 463.35 | 463.35 | 463.35 | 10.43 | 123 | >sp\|P01876\|IGHA1_HUMAN Immunoglobulin heavy constant alpha 1 OS=Homo sapiens OX=9606 GN=IGHA1 PE=1 SV=2 | 255 | 20201118_ly_colostrum_1ng_glyco_hcd203040_compound15.19463.19463.4 | controllerType=0 controllerNumber=1 scan=19463 | 42.5942 |
| 16081:3 | 5 | R.LSLHRPALEDLLLGSEAN[+1663.608]LTC[+57.021]TLTGLR.D | HexNAc(5)Hex(4) | C[+57], N[+1664] | 1543.0787 | 3 | 4627.2215 | 4627.2064 | 0 | 3.2 | 127 | Specific | 464.19 | 446.61 | 446.61 | 10.27 | 123 | >sp\|P01876\|IGHA1_HUMAN Immunoglobulin heavy constant alpha 1 OS=Homo sapiens OX=9606 GN=IGHA1 PE=1 SV=2 | 255 | 20201118_ly_colostrum_1ng_glyco_hcd203040_compound15.19577.19577.3 | controllerType=0 controllerNumber=1 scan=19577 | 42.8382 |
| 34988:4 | 5 | R.LSLHRPALEDLLLGSEAN[+1501.555]LTC[+57.021]TLTGLR.D | HexNAc(5)Hex(3) | C[+57], N[+1502] | 1117.0455 | 4 | 4465.16 | 4465.1536 | 0 | 1.4 | 127 | Specific | 471.35 | 471.35 | 471.35 | 10.43 | 123 | >sp\|P01876\|IGHA1_HUMAN Immunoglobulin heavy constant alpha 1 OS=Homo sapiens OX=9606 GN=IGHA1 PE=1 SV=2 | 255 | 20201118_ly_colostrum_1ng_glyco_hcd203040_compound15.41325.41325.4 | controllerType=0 controllerNumber=1 scan=41325 | 88.7999 |
| 15911:3 | 5 | R.LSLHRPALEDLLLGSEAN[+1864.634]LTC[+57.021]TLTGLR.D | HexNAc(2)Hex(9) | C[+57], N[+1865] | 1610.0894 | 3 | 4828.2537 | 4828.2324 | 0 | 4.4 | 127 | Specific | 482.37 | 477.06 | 477.06 | 10.06 | 123 | >sp\|P01876\|IGHA1_HUMAN Immunoglobulin heavy constant alpha 1 OS=Homo sapiens OX=9606 GN=IGHA1 PE=1 SV=2 | 255 | 20201118_ly_colostrum_1ng_glyco_hcd203040_compound15.19400.19400.3 | controllerType=0 controllerNumber=1 scan=19400 | 42.4335 |
| 16293:4 | 5 | R.LSLHRPALEDLLLGSEAN[+1501.555]LTC[+57.021]TLTGLR.D | HexNAc(5)Hex(3) | C[+57], N[+1502] | 1117.0496 | 4 | 4465.1766 | 4465.1536 | 0 | 5.1 | 127 | Specific | 493.08 | 493.08 | 493.08 | 10.2 | 123 | >sp\|P01876\|IGHA1_HUMAN Immunoglobulin heavy constant alpha 1 OS=Homo sapiens OX=9606 GN=IGHA1 PE=1 SV=2 | 255 | 20201118_ly_colostrum_1ng_glyco_hcd203040_compound15.19799.19799.4 | controllerType=0 controllerNumber=1 scan=19799 | 43.3609 |
| 16071:4 | 5 | R.LSLHRPALEDLLLGSEAN[+1647.613]LTC[+57.021]TLTGLR.D | HexNAc(5)Hex(3)Fuc(1) | C[+57], N[+1648] | 1153.5591 | 4 | 4611.2147 | 4611.2115 | 0 | 0.7 | 127 | Specific | 499.88 | 499.88 | 499.88 | 10.31 | 123 | >sp\|P01876\|IGHA1_HUMAN Immunoglobulin heavy constant alpha 1 OS=Homo sapiens OX=9606 GN=IGHA1 PE=1 SV=2 | 255 | 20201118_ly_colostrum_1ng_glyco_hcd203040_compound15.19567.19567.4 | controllerType=0 controllerNumber=1 scan=19567 | 42.8199 |
| 15977:3 | 5 | R.LSLHRPALEDLLLGSEAN[+1622.582]LTC[+57.021]TLTGLR.D | HexNAc(4)Hex(5) | C[+57], N[+1623] | 1529.4028 | 3 | 4586.1937 | 4586.1799 | 0 | 3 | 127 | Specific | 512.21 | 466.4 | 466.4 | 10.48 | 123 | >sp\|P01876\|IGHA1_HUMAN Immunoglobulin heavy constant alpha 1 OS=Homo sapiens OX=9606 GN=IGHA1 PE=1 SV=2 | 255 | 20201118_ly_colostrum_1ng_glyco_hcd203040_compound15.19469.19469.3 | controllerType=0 controllerNumber=1 scan=19469 | 42.6033 |
| 16108:3 | 5 | R.LSLHRPALEDLLLGSEAN[+1622.582]LTC[+57.021]TLTGLR.D | HexNAc(4)Hex(5) | C[+57], N[+1623] | 1529.3993 | 3 | 4586.1835 | 4586.1799 | 0 | 0.8 | 127 | Specific | 515.58 | 515.58 | 515.58 | 10.02 | 123 | >sp\|P01876\|IGHA1_HUMAN Immunoglobulin heavy constant alpha 1 OS=Homo sapiens OX=9606 GN=IGHA1 PE=1 SV=2 | 255 | 20201118_ly_colostrum_1ng_glyco_hcd203040_compound15.19605.19605.3 | controllerType=0 controllerNumber=1 scan=19605 | 42.909 |
| 17974:3 | 5 | R.LSLHRPALEDLLLGSEAN[+2204.772]LTC[+57.021]TLTGLR.D | HexNAc(4)Hex(5)NeuAc(2) | C[+57], N[+2205] | 1723.469 | 3 | 5168.3923 | 5168.3707 | 0 | 4.2 | 127 | Specific | 521.77 | 472.39 | 472.39 | 10.23 | 123 | >sp\|P01876\|IGHA1_HUMAN Immunoglobulin heavy constant alpha 1 OS=Homo sapiens OX=9606 GN=IGHA1 PE=1 SV=2 | 255 | 20201118_ly_colostrum_1ng_glyco_hcd203040_compound15.21556.21556.3 | controllerType=0 controllerNumber=1 scan=21556 | 47.2935 |
| 16033:3 | 5 | R.LSLHRPALEDLLLGSEAN[+1216.423]LTC[+57.021]TLTGLR.D | HexNAc(2)Hex(5) | C[+57], N[+1216] | 1394.0146 | 3 | 4180.0292 | 4180.0211 | 0 | 1.9 | 127 | Specific | 530.49 | 530.49 | 530.49 | 10.26 | 123 | >sp\|P01876\|IGHA1_HUMAN Immunoglobulin heavy constant alpha 1 OS=Homo sapiens OX=9606 GN=IGHA1 PE=1 SV=2 | 255 | 20201118_ly_colostrum_1ng_glyco_hcd203040_compound15.19527.19527.6 | controllerType=0 controllerNumber=1 scan=19527 | 42.7173 |
| 15569:4 | 5 | R.LSLHRPALEDLLLGSEAN[+1501.555]LTC[+57.021]TLTGLR.D | HexNAc(5)Hex(3) | C[+57], N[+1502] | 1117.044 | 4 | 4465.1541 | 4465.1536 | 0 | 0.1 | 127 | Specific | 546.12 | 546.12 | 546.12 | 10.66 | 123 | >sp\|P01876\|IGHA1_HUMAN Immunoglobulin heavy constant alpha 1 OS=Homo sapiens OX=9606 GN=IGHA1 PE=1 SV=2 | 255 | 20201118_ly_colostrum_1ng_glyco_hcd203040_compound15.19043.19043.4 | controllerType=0 controllerNumber=1 scan=19043 | [illegible] |
| 17731:4 | 5 | R.LSLHRPALEDLLLGSEAN[+1501.555]LTC[+57.021]TLTGLR.D | HexNAc(5)Hex(3) | C[+57], N[+1502] | 1117.0473 | 4 | 4465.1673 | 4465.1536 | 0 | 3.1 | 127 | Specific | 551.83 | 469.79 | 469.79 | 11.12 | 123 | >sp\|P01876\|IGHA1_HUMAN Immunoglobulin heavy constant alpha 1 OS=Homo sapiens OX=9606 GN=IGHA1 PE=1 SV=2 | 255 | 20201118_ly_colostrum_1ng_glyco_hcd203040_compound15.21302.21302.2 | controllerType=0 controllerNumber=1 scan=21302 | [illegible] |
| 15912:4 | 5 | R.LSLHRPALEDLLLGSEAN[+1825.661]LTC[+57.021]TLTGLR.D | HexNAc(5)Hex(5) | C[+57], N[+1826] | 1198.0746 | 4 | 4789.2767 | 4789.2592 | 0 | 3.6 | 127 | Specific | 554.69 | 554.69 | 554.69 | 11.12 | 123 | >sp\|P01876\|IGHA1_HUMAN Immunoglobulin heavy constant alpha 1 OS=Homo sapiens OX=9606 GN=IGHA1 PE=1 SV=2 | 255 | 20201118_ly_colostrum_1ng_glyco_hcd203040_compound15.19401.19401.4 | controllerType=0 controllerNumber=1 scan=19401 | [illegible] |
| 19550:4 | 5 | R.LSLHRPALEDLLLGSEAN[+1501.555]LTC[+57.021]TLTGLR.D | HexNAc(5)Hex(3) | C[+57], N[+1502] | 1117.0478 | 4 | 4465.1693 | 4465.1536 | 0 | 3.5 | 127 | Specific | 555.7 | 248.5 | 248.5 | 8.86 | 123 | >sp\|P01876\|IGHA1_HUMAN Immunoglobulin heavy constant alpha 1 OS=Homo sapiens OX=9606 GN=IGHA1 PE=1 SV=2 | 255 | 20201118_ly_colostrum_1ng_glyco_hcd203040_compound15.23204.23204.4 | controllerType=0 controllerNumber=1 scan=23204 | [illegible] |
| 15964:5 | 5 | R.LSLHRPALEDLLLGSEAN[+1663.608]LTC[+57.021]TLTGLR.D | HexNAc(5)Hex(4) | C[+57], N[+1664] | 926.2486 | 5 | 4627.214 | 4627.2064 | 0 | 1.6 | 127 | Specific | 557.59 | 557.59 | 557.59 | 11.03 | 123 | >sp\|P01876\|IGHA1_HUMAN Immunoglobulin heavy constant alpha 1 OS=Homo sapiens OX=9606 GN=IGHA1 PE=1 SV=2 | 255 | 20201118_ly_colostrum_1ng_glyco_hcd203040_compound15.19455.19455.5 | controllerType=0 controllerNumber=1 scan=19455 | 42.5784 |
| 16084:3 | 5 | R.LSLHRPALEDLLLGSEAN[+1663.608]LTC[+57.021]TLTGLR.D | HexNAc(5)Hex(4) | C[+57], N[+1664] | 1543.0787 | 3 | 4627.2215 | 4627.2064 | 0 | 3.2 | 127 | Specific | 559.99 | 507.04 | 507.04 | 10.71 | 123 | >sp\|P01876\|IGHA1_HUMAN Immunoglobulin heavy constant alpha 1 OS=Homo sapiens OX=9606 GN=IGHA1 PE=1 SV=2 | 255 | 20201118_ly_colostrum_1ng_glyco_hcd203040_compound15.19580.19580.3 | controllerType=0 controllerNumber=1 scan=19580 | 42.8465 |
| 15923:3 | 5 | R.LSLHRPALEDLLLGSEAN[+1702.581]LTC[+57.021]TLTGLR.D | HexNAc(2)Hex(8) | C[+57], N[+1703] | 1556.0689 | 3 | 4666.1922 | 4666.1796 | 0 | 2.7 | 127 | Specific | 561.72 | 561.72 | 561.72 | 10.86 | 123 | >sp\|P01876\|IGHA1_HUMAN Immunoglobulin heavy constant alpha 1 OS=Homo sapiens OX=9606 GN=IGHA1 PE=1 SV=2 | 255 | 20201118_ly_colostrum_1ng_glyco_hcd203040_compound15.19413.19413.3 | controllerType=0 controllerNumber=1 scan=19413 | 42.4802 |
| 17966:4 | 5 | R.LSLHRPALEDLLLGSEAN[+2204.772]LTC[+57.021]TLTGLR.D | HexNAc(4)Hex(5)NeuAc(2) | C[+57], N[+2205] | 1292.8521 | 4 | 5168.3864 | 5168.3707 | 0 | 3 | 127 | Specific | 562.43 | 562.43 | 562.43 | 11.27 | 123 | >sp\|P01876\|IGHA1_HUMAN Immunoglobulin heavy constant alpha 1 OS=Homo sapiens OX=9606 GN=IGHA1 PE=1 SV=2 | 255 | 20201118_ly_colostrum_1ng_glyco_hcd203040_compound15.21548.21548.4 | controllerType=0 controllerNumber=1 scan=21548 | 47.2759 |
| 16050:4 | 5 | R.LSLHRPALEDLLLGSEAN[+1501.555]LTC[+57.021]TLTGLR.D | HexNAc(5)Hex(3) | C[+57], N[+1502] | 1117.0494 | 4 | 4465.1756 | 4465.1536 | 0 | 4.9 | 127 | Specific | 563 | 563 | 563 | 11 | 123 | >sp\|P01876\|IGHA1_HUMAN Immunoglobulin heavy constant alpha 1 OS=Homo sapiens OX=9606 GN=IGHA1 PE=1 SV=2 | 255 | 20201118_ly_colostrum_1ng_glyco_hcd203040_compound15.19545.19545.4 | controllerType=0 controllerNumber=1 scan=19545 | 42.7531 |
| 15949:4 | 5 | R.LSLHRPALEDLLLGSEAN[+1663.608]LTC[+57.021]TLTGLR.D | HexNAc(5)Hex(4) | C[+57], N[+1664] | 1157.559 | 4 | 4627.2142 | 4627.2064 | 0 | 1.7 | 127 | Specific | 566.86 | 566.86 | 566.86 | 11.04 | 123 | >sp\|P01876\|IGHA1_HUMAN Immunoglobulin heavy constant alpha 1 OS=Homo sapiens OX=9606 GN=IGHA1 PE=1 SV=2 | 255 | 20201118_ly_colostrum_1ng_glyco_hcd203040_compound15.19440.19440.4 | controllerType=0 controllerNumber=1 scan=19440 | 42.5408 |
| 34952:4 | 5 | R.LSLHRPALEDLLLGSEAN[+1216.423]LTC[+57.021]TLTGLR.D | HexNAc(2)Hex(5) | C[+57], N[+1216] | 1045.7637 | 4 | 4180.033 | 4180.0211 | 0 | 2.8 | 127 | Specific | 573.92 | 573.92 | 573.92 | 11.27 | 123 | >sp\|P01876\|IGHA1_HUMAN Immunoglobulin heavy constant alpha 1 OS=Homo sapiens OX=9606 GN=IGHA1 PE=1 SV=2 | 255 | 20201118_ly_colostrum_1ng_glyco_hcd203040_compound15.41286.41286.4 | controllerType=0 controllerNumber=1 scan=41286 | 88.7261 |
| 16111:3 | 5 | R.LSLHRPALEDLLLGSEAN[+1622.582]LTC[+57.021]TLTGLR.D | HexNAc(4)Hex(5) | C[+57], N[+1623] | 1529.3993 | 3 | 4586.1835 | 4586.1799 | 0 | 0.8 | 127 | Specific | 584.33 | 472.61 | 472.61 | 11.75 | 123 | >sp\|P01876\|IGHA1_HUMAN Immunoglobulin heavy constant alpha 1 OS=Homo sapiens OX=9606 GN=IGHA1 PE=1 SV=2 | 255 | 20201118_ly_colostrum_1ng_glyco_hcd203040_compound15.19608.19608.3 | controllerType=0 controllerNumber=1 scan=19608 | 42.9161 |
| 16210:3 | 5 | R.LSLHRPALEDLLLGSEAN[+1501.555]LTC[+57.021]TLTGLR.D | HexNAc(5)Hex(3) | C[+57], N[+1502] | 1489.0587 | 3 | 4465.1614 | 4465.1536 | 0 | 1.7 | 127 | Specific | 586.9 | 586.9 | 586.9 | 11.95 | 123 | >sp\|P01876\|IGHA1_HUMAN Immunoglobulin heavy constant alpha 1 OS=Homo sapiens OX=9606 GN=IGHA1 PE=1 SV=2 | 255 | 20201118_ly_colostrum_1ng_glyco_hcd203040_compound15.19712.19712.3 | controllerType=0 controllerNumber=1 scan=19712 | 43.1692 |
| 15961:5 | 5 | R.LSLHRPALEDLLLGSEAN[+1663.608]LTC[+57.021]TLTGLR.D | HexNAc(5)Hex(4) | C[+57], N[+1664] | 926.2486 | 5 | 4627.214 | 4627.2064 | 0 | 1.6 | 127 | Specific | 587.78 | 587.78 | 587.78 | 12.52 | 123 | >sp\|P01876\|IGHA1_HUMAN Immunoglobulin heavy constant alpha 1 OS=Homo sapiens OX=9606 GN=IGHA1 PE=1 SV=2 | 255 | 20201118_ly_colostrum_1ng_glyco_hcd203040_compound15.19452.19452.5 | controllerType=0 controllerNumber=1 scan=19452 | 42.5718 |
| 16295:4 | 5 | R.LSLHRPALEDLLLGSEAN[+1501.555]LTC[+57.021]TLTGLR.D | HexNAc(5)Hex(3) | C[+57], N[+1502] | 1117.0496 | 4 | 4465.1766 | 4465.1536 | 0 | 5.1 | 127 | Specific | 590.07 | 497.83 | 497.83 | 12.05 | 123 | >sp\|P01876\|IGHA1_HUMAN Immunoglobulin heavy constant alpha 1 OS=Homo sapiens OX=9606 GN=IGHA1 PE=1 SV=2 | 255 | 20201118_ly_colostrum_1ng_glyco_hcd203040_compound15.19801.19801.4 | controllerType=0 controllerNumber=1 scan=19801 | 43.366 |
| 15947:5 | 5 | R.LSLHRPALEDLLLGSEAN[+1702.581]LTC[+57.021]TLTGLR.D | HexNAc(2)Hex(8) | C[+57], N[+1703] | 934.0354 | 5 | 4666.148 | 4666.1796 | 0 | -6.8 | 127 | Specific | 591.51 | 586.41 | 586.41 | 11.37 | 123 | >sp\|P01876\|IGHA1_HUMAN Immunoglobulin heavy constant alpha 1 OS=Homo sapiens OX=9606 GN=IGHA1 PE=1 SV=2 | 255 | 20201118_ly_colostrum_1ng_glyco_hcd203040_compound15.19437.19437.5 | controllerType=0 controllerNumber=1 scan=19437 | 42.5348 |
| 16213:3 | 5 | R.LSLHRPALEDLLLGSEAN[+1501.555]LTC[+57.021]TLTGLR.D | HexNAc(5)Hex(3) | C[+57], N[+1502] | 1489.0587 | 3 | 4465.1614 | 4465.1536 | 0 | 1.7 | 127 | Specific | 596.33 | 565.28 | 565.28 | 11.95 | 123 | >sp\|P01876\|IGHA1_HUMAN Immunoglobulin heavy constant alpha 1 OS=Homo sapiens OX=9606 GN=IGHA1 PE=1 SV=2 | 255 | 20201118_ly_colostrum_1ng_glyco_hcd203040_compound15.19715.19715.3 | controllerType=0 controllerNumber=1 scan=19715 | 43.175 |
| 15972:3 | 5 | R.LSLHRPALEDLLLGSEAN[+1378.476]LTC[+57.021]TLTGLR.D | HexNAc(2)Hex(6) | C[+57], N[+1378] | 1448.034 | 3 | 4342.0874 | 4342.074 | 0 | 3.1 | 127 | Specific | 603.71 | 561.54 | 561.54 | 13.21 | 123 | >sp\|P01876\|IGHA1_HUMAN Immunoglobulin heavy constant alpha 1 OS=Homo sapiens OX=9606 GN=IGHA1 PE=1 SV=2 | 255 | 20201118_ly_colostrum_1ng_glyco_hcd203040_compound15.19464.19464.3 | controllerType=0 controllerNumber=1 scan=19464 | 42.5957 |
| 16078:3 | 5 | R.LSLHRPALEDLLLGSEAN[+1501.555]LTC[+57.021]TLTGLR.D | HexNAc(5)Hex(3) | C[+57], N[+1502] | 1489.0607 | 3 | 4465.1676 | 4465.1536 | 0 | 3.1 | 127 | Specific | 606.42 | 567.41 | 567.41 | 13.21 | 123 | >sp\|P01876\|IGHA1_HUMAN Immunoglobulin heavy constant alpha 1 OS=Homo sapiens OX=9606 GN=IGHA1 PE=1 SV=2 | 255 | 20201118_ly_colostrum_1ng_glyco_hcd203040_compound15.19574.19574.6 | controllerType=0 controllerNumber=1 scan=19574 | 42.8351 |
| 16115:3 | 5 | R.LSLHRPALEDLLLGSEAN[+1501.555]LTC[+57.021]TLTGLR.D | HexNAc(5)Hex(3) | C[+57], N[+1502] | 1489.0602 | 3 | 4465.1662 | 4465.1536 | 0 | 2.8 | 127 | Specific | 607.59 | 607.59 | 607.59 | 13.18 | 123 | >sp\|P01876\|IGHA1_HUMAN Immunoglobulin heavy constant alpha 1 OS=Homo sapiens OX=9606 GN=IGHA1 PE=1 SV=2 | 255 | 20201118_ly_colostrum_1ng_glyco_hcd203040_compound15.19612.19612.2 | controllerType=0 controllerNumber=1 scan=19612 | 42.9221 |
| 15914:3 | 5 | R.LSLHRPALEDLLLGSEAN[+1864.634]LTC[+57.021]TLTGLR.D | HexNAc(2)Hex(9) | C[+57], N[+1865] | 1610.0894 | 3 | 4828.2537 | 4828.2324 | 0 | 4.4 | 127 | Specific | 607.86 | 607.86 | 607.86 | 12.91 | 123 | >sp\|P01876\|IGHA1_HUMAN Immunoglobulin heavy constant alpha 1 OS=Homo sapiens OX=9606 GN=IGHA1 PE=1 SV=2 | 255 | 20201118_ly_colostrum_1ng_glyco_hcd203040_compound15.19403.19403.3 | controllerType=0 controllerNumber=1 scan=19403 | 42.44 |
| 16974:3 | 5 | R.LSLHRPALEDLLLGSEAN[+1548.545]LTC[+57.021]TLTGLR.D | HexNAc(3)Hex(4)NeuAc(1) | C[+57], N[+1549] | 1504.7233 | 3 | 4512.1554 | 4512.1431 | 0 | 2.7 | 127 | Specific | 613.37 | 613.37 | 613.37 | 13.18 | 123 | >sp\|P01876\|IGHA1_HUMAN Immunoglobulin heavy constant alpha 1 OS=Homo sapiens OX=9606 GN=IGHA1 PE=1 SV=2 | 255 | 20201118_ly_colostrum_1ng_glyco_hcd203040_compound15.20511.20511.3 | controllerType=0 controllerNumber=1 scan=20511 | 44.9458 |
| 16895:3 | 5 | R.LSLHRPALEDLLLGSEAN[+1872.651]LTC[+57.021]TLTGLR.D | HexNAc(3)Hex(6)NeuAc(1) | C[+57], N[+1873] | 1612.7596 | 3 | 4836.2642 | 4836.2487 | 0 | 3.2 | 127 | Specific | 617.48 | 544.07 | 544.07 | 13.21 | 123 | >sp\|P01876\|IGHA1_HUMAN Immunoglobulin heavy constant alpha 1 OS=Homo sapiens OX=9606 GN=IGHA1 PE=1 SV=2 | 255 | 20201118_ly_colostrum_1ng_glyco_hcd203040_compound15.20428.20428.3 | controllerType=0 controllerNumber=1 scan=20428 | 44.7474 |
| 15948:3 | 5 | R.LSLHRPALEDLLLGSEAN[+1581.555]LTC[+57.021]TLTGLR.D | HexNAc(3)Hex(6) | C[+57], N[+1582] | 1515.7256 | 3 | 4545.1624 | 4545.1533 | 0 | 2 | 127 | Specific | 618.85 | 612.71 | 612.71 | 12.95 | 123 | >sp\|P01876\|IGHA1_HUMAN Immunoglobulin heavy constant alpha 1 OS=Homo sapiens OX=9606 GN=IGHA1 PE=1 SV=2 | 255 | 20201118_ly_colostrum_1ng_glyco_hcd203040_compound15.19438.19438.3 | controllerType=0 controllerNumber=1 scan=19438 | 42.5367 |
| 15970:4 | 5 | R.LSLHRPALEDLLLGSEAN[+1809.666]LTC[+57.021]TLTGLR.D | HexNAc(5)Hex(4)Fuc(1) | C[+57], N[+1810] | 1194.074 | 4 | 4773.2742 | 4773.2643 | 0 | 2.1 | 127 | Specific | 624.99 | 535.56 | 535.56 | 13.59 | 123 | >sp\|P01876\|IGHA1_HUMAN Immunoglobulin heavy constant alpha 1 OS=Homo sapiens OX=9606 GN=IGHA1 PE=1 SV=2 | 255 | 20201118_ly_colostrum_1ng_glyco_hcd203040_compound15.19461.19461.4 | controllerType=0 controllerNumber=1 scan=19461 | 42.5905 |
| 16942:4 | 5 | R.LSLHRPALEDLLLGSEAN[+1751.624]LTC[+57.021]TLTGLR.D | HexNAc(4)Hex(4)NeuAc(1) | C[+57], N[+1752] | 1179.564 | 4 | 4715.2342 | 4715.2225 | 0 | 2.5 | 127 | Specific | 625.73 | 625.73 | 625.73 | 13.59 | 123 | >sp\|P01876\|IGHA1_HUMAN Immunoglobulin heavy constant alpha 1 OS=Homo sapiens OX=9606 GN=IGHA1 PE=1 SV=2 | 255 | 20201118_ly_colostrum_1ng_glyco_hcd203040_compound15.20478.20478.4 | controllerType=0 controllerNumber=1 scan=20478 | 44.8854 |
| 16192:4 | 5 | R.LSLHRPALEDLLLGSEAN[+1622.582]LTC[+57.021]TLTGLR.D | HexNAc(4)Hex(5) | C[+57], N[+1623] | 1147.2989 | 4 | 4586.1737 | 4586.1799 | 0 | -1.4 | 127 | Specific | 626.05 | 526.58 | 526.58 | 12.46 | 123 | >sp\|P01876\|IGHA1_HUMAN Immunoglobulin heavy constant alpha 1 OS=Homo sapiens OX=9606 GN=IGHA1 PE=1 SV=2 | 255 | 20201118_ly_colostrum_1ng_glyco_hcd203040_compound15.19693.19693.4 | controllerType=0 controllerNumber=1 scan=19693 | 43.1098 |
| 15953:3 | 5 | R.LSLHRPALEDLLLGSEAN[+1663.608]LTC[+57.021]TLTGLR.D | HexNAc(5)Hex(4) | C[+57], N[+1664] | 1543.0776 | 3 | 4627.2182 | 4627.2064 | 0 | 2.5 | 127 | Specific | 626.24 | 605.93 | 605.93 | 13.18 | 123 | >sp\|P01876\|IGHA1_HUMAN Immunoglobulin heavy constant alpha 1 OS=Homo sapiens OX=9606 GN=IGHA1 PE=1 SV=2 | 255 | 20201118_ly_colostrum_1ng_glyco_hcd203040_compound15.19444.19444.3 | controllerType=0 controllerNumber=1 scan=19444 | 42.5525 |
| 16977:3 | 5 | R.LSLHRPALEDLLLGSEAN[+1548.545]LTC[+57.021]TLTGLR.D | HexNAc(3)Hex(4)NeuAc(1) | C[+57], N[+1549] | 1504.7233 | 3 | 4512.1554 | 4512.1431 | 0 | 2.7 | 127 | Specific | 634.4 | 633.95 | 633.95 | 13.18 | 123 | >sp\|P01876\|IGHA1_HUMAN Immunoglobulin heavy constant alpha 1 OS=Homo sapiens OX=9606 GN=IGHA1 PE=1 SV=2 | 255 | 20201118_ly_colostrum_1ng_glyco_hcd203040_compound15.20514.20514.3 | controllerType=0 controllerNumber=1 scan=20514 | 44.9509 |
| 16950:3 | 5 | R.LSLHRPALEDLLLGSEAN[+1751.624]LTC[+57.021]TLTGLR.D | HexNAc(4)Hex(4)NeuAc(1) | C[+57], N[+1752] | 1572.4177 | 3 | 4715.2384 | 4715.2225 | 0 | 3.4 | 127 | Specific | 639.53 | 639.53 | 639.53 | 13.21 | 123 | >sp\|P01876\|IGHA1_HUMAN Immunoglobulin heavy constant alpha 1 OS=Homo sapiens OX=9606 GN=IGHA1 PE=1 SV=2 | 255 | 20201118_ly_colostrum_1ng_glyco_hcd203040_compound15.20486.20486.3 | controllerType=0 controllerNumber=1 scan=20486 | 44.8974 |
| 15919:3 | 5 | R.LSLHRPALEDLLLGSEAN[+1540.529]LTC[+57.021]TLTGLR.D | HexNAc(2)Hex(7) | C[+57], N[+1541] | 1502.0534 | 3 | 4504.1457 | 4504.1268 | 0 | 4.2 | 127 | Specific | 639.94 | 586 | 586 | 12.91 | 123 | >sp\|P01876\|IGHA1_HUMAN Immunoglobulin heavy constant alpha 1 OS=Homo sapiens OX=9606 GN=IGHA1 PE=1 SV=2 | 255 | 20201118_ly_colostrum_1ng_glyco_hcd203040_compound15.19408.19408.3 | controllerType=0 controllerNumber=1 scan=19408 | 42.4555 |
| 15979:3 | 5 | R.LSLHRPALEDLLLGSEAN[+1501.555]LTC[+57.021]TLTGLR.D | HexNAc(5)Hex(3) | C[+57], N[+1502] | 1489.0608 | 3 | 4465.168 | 4465.1536 | 0 | 3.2 | 127 | Specific | 642.02 | 594.07 | 594.07 | 13.21 | 123 | >sp\|P01876\|IGHA1_HUMAN Immunoglobulin heavy constant alpha 1 OS=Homo sapiens OX=9606 GN=IGHA1 PE=1 SV=2 | 255 | 20201118_ly_colostrum_1ng_glyco_hcd203040_compound15.19471.19471.3 | controllerType=0 controllerNumber=1 scan=19471 | 42.6092 |
| 15945:3 | 5 | R.LSLHRPALEDLLLGSEAN[+1581.555]LTC[+57.021]TLTGLR.D | HexNAc(3)Hex(6) | C[+57], N[+1582] | 1515.7256 | 3 | 4545.1624 | 4545.1533 | 0 | 2 | 127 | Specific | 649.33 | 563.59 | 563.59 | 12.95 | 123 | >sp\|P01876\|IGHA1_HUMAN Immunoglobulin heavy constant alpha 1 OS=Homo sapiens OX=9606 GN=IGHA1 PE=1 SV=2 | 255 | 20201118_ly_colostrum_1ng_glyco_hcd203040_compound15.19435.19435.3 | controllerType=0 controllerNumber=1 scan=19435 | 42.5243 |
| 15956:3 | 5 | R.LSLHRPALEDLLLGSEAN[+1663.608]LTC[+57.021]TLTGLR.D | HexNAc(5)Hex(4) | C[+57], N[+1664] | 1543.0776 | 3 | 4627.2182 | 4627.2064 | 0 | 2.5 | 127 | Specific | 650.91 | 554.26 | 554.26 | 14.18 | 123 | >sp\|P01876\|IGHA1_HUMAN Immunoglobulin heavy constant alpha 1 OS=Homo sapiens OX=9606 GN=IGHA1 PE=1 SV=2 | 255 | 20201118_ly_colostrum_1ng_glyco_hcd203040_compound15.19447.19447.3 | controllerType=0 controllerNumber=1 scan=19447 | 42.5572 |
| 16945:3 | 5 | R.LSLHRPALEDLLLGSEAN[+1710.598]LTC[+57.021]TLTGLR.D | HexNAc(3)Hex(5)NeuAc(1) | C[+57], N[+1711] | 1558.7419 | 3 | 4674.2111 | 4674.1959 | 0 | 3.2 | 127 | Specific | 659.85 | 577.34 | 577.34 | 14.21 | 123 | >sp\|P01876\|IGHA1_HUMAN Immunoglobulin heavy constant alpha 1 OS=Homo sapiens OX=9606 GN=IGHA1 PE=1 SV=2 | 255 | 20201118_ly_colostrum_1ng_glyco_hcd203040_compound15.20481.20481.3 | controllerType=0 controllerNumber=1 scan=20481 | 44.8911 |
| 16897:3 | 5 | R.LSLHRPALEDLLLGSEAN[+1872.651]LTC[+57.021]TLTGLR.D | HexNAc(3)Hex(6)NeuAc(1) | C[+57], N[+1873] | 1612.7596 | 3 | 4836.2642 | 4836.2487 | 0 | 3.2 | 127 | Specific | 662.22 | 618.4 | 618.4 | 14.21 | 123 | >sp\|P01876\|IGHA1_HUMAN Immunoglobulin heavy constant alpha 1 OS=Homo sapiens OX=9606 GN=IGHA1 PE=1 SV=2 | 255 | 20201118_ly_colostrum_1ng_glyco_hcd203040_compound15.20430.20430.3 | controllerType=0 controllerNumber=1 scan=20430 | 44.7582 |
| 15980:3 | 5 | R.LSLHRPALEDLLLGSEAN[+1622.582]LTC[+57.021]TLTGLR.D | HexNAc(4)Hex(5) | C[+57], N[+1623] | 1529.4028 | 3 | 4586.1937 | 4586.1799 | 0 | 3 | 127 | Specific | 662.55 | 573.06 | 573.06 | 14.21 | 123 | >sp\|P01876\|IGHA1_HUMAN Immunoglobulin heavy constant alpha 1 OS=Homo sapiens OX=9606 GN=IGHA1 PE=1 SV=2 | 255 | 20201118_ly_colostrum_1ng_glyco_hcd203040_compound15.19472.19472.3 | controllerType=0 controllerNumber=1 scan=19472 | 42.6099 |
| 15975:4 | 5 | R.LSLHRPALEDLLLGSEAN[+1378.476]LTC[+57.021]TLTGLR.D | HexNAc(2)Hex(6) | C[+57], N[+1378] | 1086.2769 | 4 | 4342.0858 | 4342.074 | 0 | 2.7 | 127 | Specific | 663.11 | 663.11 | 663.11 | 14.59 | 123 | >sp\|P01876\|IGHA1_HUMAN Immunoglobulin heavy constant alpha 1 OS=Homo sapiens OX=9606 GN=IGHA1 PE=1 SV=2 | 255 | 20201118_ly_colostrum_1ng_glyco_hcd203040_compound15.19467.19467.4 | controllerType=0 controllerNumber=1 scan=19467 | 42.601 |
| 16029:4 | 5 | R.LSLHRPALEDLLLGSEAN[+1647.613]LTC[+57.021]TLTGLR.D | HexNAc(5)Hex(3)Fuc(1) | C[+57], N[+1648] | 1153.561 | 4 | 4611.222 | 4611.2115 | 0 | 2.3 | 127 | Specific | 663.43 | 663.43 | 663.43 | 14.59 | 123 | >sp\|P01876\|IGHA1_HUMAN Immunoglobulin heavy constant alpha 1 OS=Homo sapiens OX=9606 GN=IGHA1 PE=1 SV=2 | 255 | 20201118_ly_colostrum_1ng_glyco_hcd203040_compound15.19523.19523.4 | controllerType=0 controllerNumber=1 scan=19523 | 42.7072 |
| 15906:4 | 5 | R.LSLHRPALEDLLLGSEAN[+1864.634]LTC[+57.021]TLTGLR.D | HexNAc(2)Hex(9) | C[+57], N[+1865] | 1207.8169 | 4 | 4828.2459 | 4828.2324 | 0 | 2.8 | 127 | Specific | 666.74 | 666.74 | 666.74 | 14.59 | 123 | >sp\|P01876\|IGHA1_HUMAN Immunoglobulin heavy constant alpha 1 OS=Homo sapiens OX=9606 GN=IGHA1 PE=1 SV=2 | 255 | 20201118_ly_colostrum_1ng_glyco_hcd203040_compound15.19395.19395.4 | controllerType=0 controllerNumber=1 scan=19395 | 42.4211 |
| 15866:3 | 5 | R.LSLHRPALEDLLLGSEAN[+1864.634]LTC[+57.021]TLTGLR.D | HexNAc(2)Hex(9) | C[+57], N[+1865] | 1610.0881 | 3 | 4828.2497 | 4828.2324 | 0 | 3.6 | 127 | Specific | 669.17 | 669.17 | 669.17 | 14.21 | 123 | >sp\|P01876\|IGHA1_HUMAN Immunoglobulin heavy constant alpha 1 OS=Homo sapiens OX=9606 GN=IGHA1 PE=1 SV=2 | 255 | 20201118_ly_colostrum_1ng_glyco_hcd203040_compound15.19353.19353.3 | controllerType=0 controllerNumber=1 scan=19353 | 42.3182 |
| 15965:3 | 5 | R.LSLHRPALEDLLLGSEAN[+1460.529]LTC[+57.021]TLTGLR.D | HexNAc(4)Hex(4) | C[+57], N[+1461] | 1475.3834 | 3 | 4424.1355 | 4424.1271 | 0 | 1.9 | 127 | Specific | 670.61 | 640.31 | 640.31 | 13.95 | 123 | >sp\|P01876\|IGHA1_HUMAN Immunoglobulin heavy constant alpha 1 OS=Homo sapiens OX=9606 GN=IGHA1 PE=1 SV=2 | 255 | 20201118_ly_colostrum_1ng_glyco_hcd203040_compound15.19456.19456.3 | controllerType=0 controllerNumber=1 scan=19456 | 42.5796 |
| 16917:3 | 5 | R.LSLHRPALEDLLLGSEAN[+1710.598]LTC[+57.021]TLTGLR.D | HexNAc(3)Hex(5)NeuAc(1) | C[+57], N[+1711] | 1558.7436 | 3 | 4674.2162 | 4674.1959 | 0 | 4.3 | 127 | Specific | 672.34 | 602.94 | 602.94 | 13.91 | 123 | >sp\|P01876\|IGHA1_HUMAN Immunoglobulin heavy constant alpha 1 OS=Homo sapiens OX=9606 GN=IGHA1 PE=1 SV=2 | 255 | 20201118_ly_colostrum_1ng_glyco_hcd203040_compound15.20451.20451.3 | controllerType=0 controllerNumber=1 scan=20451 | 44.8194 |
| 15968:3 | 5 | R.LSLHRPALEDLLLGSEAN[+1460.529]LTC[+57.021]TLTGLR.D | HexNAc(4)Hex(4) | C[+57], N[+1461] | 1475.3834 | 3 | 4424.1355 | 4424.1271 | 0 | 1.9 | 127 | Specific | 672.88 | 555.76 | 555.76 | 13.95 | 123 | >sp\|P01876\|IGHA1_HUMAN Immunoglobulin heavy constant alpha 1 OS=Homo sapiens OX=9606 GN=IGHA1 PE=1 SV=2 | 255 | 20201118_ly_colostrum_1ng_glyco_hcd203040_compound15.19459.19459.3 | controllerType=0 controllerNumber=1 scan=19459 | 42.5865 |
| 16914:3 | 5 | R.LSLHRPALEDLLLGSEAN[+1710.598]LTC[+57.021]TLTGLR.D | HexNAc(3)Hex(5)NeuAc(1) | C[+57], N[+1711] | 1558.7436 | 3 | 4674.2162 | 4674.1959 | 0 | 4.3 | 127 | Specific | 672.96 | 620.44 | 620.44 | 13.91 | 123 | >sp\|P01876\|IGHA1_HUMAN Immunoglobulin heavy constant alpha 1 OS=Homo sapiens OX=9606 GN=IGHA1 PE=1 SV=2 | 255 | 20201118_ly_colostrum_1ng_glyco_hcd203040_compound15.20448.20448.3 | controllerType=0 controllerNumber=1 scan=20448 | 44.8085 |
| 15922:4 | 5 | R.LSLHRPALEDLLLGSEAN[+1702.581]LTC[+57.021]TLTGLR.D | HexNAc(2)Hex(8) | C[+57], N[+1703] | 1167.3013 | 4 | 4666.1834 | 4666.1796 | 0 | 0.8 | 127 | Specific | 678.27 | 678.27 | 678.27 | 14.16 | 123 | >sp\|P01876\|IGHA1_HUMAN Immunoglobulin heavy constant alpha 1 OS=Homo sapiens OX=9606 GN=IGHA1 PE=1 SV=2 | 255 | 20201118_ly_colostrum_1ng_glyco_hcd203040_compound15.19412.19412.4 | controllerType=0 controllerNumber=1 scan=19412 | 42.4788 |
| 15976:3 | 5 | R.LSLHRPALEDLLLGSEAN[+1501.555]LTC[+57.021]TLTGLR.D | HexNAc(5)Hex(3) | C[+57], N[+1502] | 1489.0608 | 3 | 4465.168 | 4465.1536 | 0 | 3.2 | 127 | Specific | 678.88 | 582.05 | 582.05 | 14.21 | 123 | >sp\|P01876\|IGHA1_HUMAN Immunoglobulin heavy constant alpha 1 OS=Homo sapiens OX=9606 GN=IGHA1 PE=1 SV=2 | 255 | 20201118_ly_colostrum_1ng_glyco_hcd203040_compound15.19468.19468.3 | controllerType=0 controllerNumber=1 scan=19468 | 42.6021 |
| 15925:3 | 5 | R.LSLHRPALEDLLLGSEAN[+1702.581]LTC[+57.021]TLTGLR.D | HexNAc(2)Hex(8) | C[+57], N[+1703] | 1556.0689 | 3 | 4666.1922 | 4666.1796 | 0 | 2.7 | 127 | Specific | 683.29 | 671 | 671 | 14.18 | 123 | >sp\|P01876\|IGHA1_HUMAN Immunoglobulin heavy constant alpha 1 OS=Homo sapiens OX=9606 GN=IGHA1 PE=1 SV=2 | 255 | 20201118_ly_colostrum_1ng_glyco_hcd203040_compound15.19415.19415.3 | controllerType=0 controllerNumber=1 scan=19415 | 42.4845 |
| 16036:3 | 5 | R.LSLHRPALEDLLLGSEAN[+1216.423]LTC[+57.021]TLTGLR.D | HexNAc(2)Hex(5) | C[+57], N[+1216] | 1394.0146 | 3 | 4180.0292 | 4180.0211 | 0 | 1.9 | 127 | Specific | 684.85 | 577.47 | 577.47 | 13.95 | 123 | >sp\|P01876\|IGHA1_HUMAN Immunoglobulin heavy constant alpha 1 OS=Homo sapiens OX=9606 GN=IGHA1 PE=1 SV=2 | 255 | 20201118_ly_colostrum_1ng_glyco_hcd203040_compound15.19530.19530.6 | controllerType=0 controllerNumber=1 scan=19530 | 42.7213 |
| 34989:4 | 5 | R.LSLHRPALEDLLLGSEAN[+1216.423]LTC[+57.021]TLTGLR.D | HexNAc(2)Hex(5) | C[+57], N[+1216] | 1045.7629 | 4 | 4180.0296 | 4180.0211 | 0 | 2 | 127 | Specific | 692.04 | 692.04 | 692.04 | 14.59 | 123 | >sp\|P01876\|IGHA1_HUMAN Immunoglobulin heavy constant alpha 1 OS=Homo sapiens OX=9606 GN=IGHA1 PE=1 SV=2 | 255 | 20201118_ly_colostrum_1ng_glyco_hcd203040_compound15.41326.41326.2 | controllerType=0 controllerNumber=1 scan=41326 | 88.8024 |
| 15916:3 | 5 | R.LSLHRPALEDLLLGSEAN[+1540.529]LTC[+57.021]TLTGLR.D | HexNAc(2)Hex(7) | C[+57], N[+1541] | 1502.0534 | 3 | 4504.1457 | 4504.1268 | 0 | 4.2 | 127 | Specific | 701.79 | 631.9 | 631.9 | 14.91 | 123 | >sp\|P01876\|IGHA1_HUMAN Immunoglobulin heavy constant alpha 1 OS=Homo sapiens OX=9606 GN=IGHA1 PE=1 SV=2 | 255 | 20201118_ly_colostrum_1ng_glyco_hcd203040_compound15.19405.19405.3 | controllerType=0 controllerNumber=1 scan=19405 | 42.4431 |
| 15967:4 | 5 | R.LSLHRPALEDLLLGSEAN[+1809.666]LTC[+57.021]TLTGLR.D | HexNAc(5)Hex(4)Fuc(1) | C[+57], N[+1810] | 1194.074 | 4 | 4773.2742 | 4773.2643 | 0 | 2.1 | 127 | Specific | 702.1 | 685.76 | 685.76 | 15.59 | 123 | >sp\|P01876\|IGHA1_HUMAN Immunoglobulin heavy constant alpha 1 OS=Homo sapiens OX=9606 GN=IGHA1 PE=1 SV=2 | 255 | 20201118_ly_colostrum_1ng_glyco_hcd203040_compound15.19458.19458.4 | controllerType=0 controllerNumber=1 scan=19458 | 42.5818 |
| 15864:3 | 5 | R.LSLHRPALEDLLLGSEAN[+1864.634]LTC[+57.021]TLTGLR.D | HexNAc(2)Hex(9) | C[+57], N[+1865] | 1610.0881 | 3 | 4828.2497 | 4828.2324 | 0 | 3.6 | 127 | Specific | 704.36 | 701.11 | 701.11 | 15.21 | 123 | >sp\|P01876\|IGHA1_HUMAN Immunoglobulin heavy constant alpha 1 OS=Homo sapiens OX=9606 GN=IGHA1 PE=1 SV=2 | 255 | 20201118_ly_colostrum_1ng_glyco_hcd203040_compound15.19351.19351.3 | controllerType=0 controllerNumber=1 scan=19351 | 42.3114 |
| 15974:3 | 5 | R.LSLHRPALEDLLLGSEAN[+1378.476]LTC[+57.021]TLTGLR.D | HexNAc(2)Hex(6) | C[+57], N[+1378] | 1448.034 | 3 | 4342.0874 | 4342.074 | 0 | 3.1 | 127 | Specific | 705.32 | 657.54 | 657.54 | 15.21 | 123 | >sp\|P01876\|IGHA1_HUMAN Immunoglobulin heavy constant alpha 1 OS=Homo sapiens OX=9606 GN=IGHA1 PE=1 SV=2 | 255 | 20201118_ly_colostrum_1ng_glyco_hcd203040_compound15.19466.19466.3 | controllerType=0 controllerNumber=1 scan=19466 | 42.5999 |
| 17734:4 | 5 | R.LSLHRPALEDLLLGSEAN[+1501.555]LTC[+57.021]TLTGLR.D | HexNAc(5)Hex(3) | C[+57], N[+1502] | 1117.0473 | 4 | 4465.1673 | 4465.1536 | 0 | 3.1 | 127 | Specific | 708.93 | 603.16 | 603.16 | 15.62 | 123 | >sp\|P01876\|IGHA1_HUMAN Immunoglobulin heavy constant alpha 1 OS=Homo sapiens OX=9606 GN=IGHA1 PE=1 SV=2 | 255 | 20201118_ly_colostrum_1ng_glyco_hcd203040_compound15.21305.21305.2 | controllerType=0 controllerNumber=1 scan=21305 | 46.7343 |
| 16131:3 | 5 | R.LSLHRPALEDLLLGSEAN[+892.317]LTC[+57.021]TLTGLR.D | HexNAc(2)Hex(3) | C[+57], N[+892] | 1285.9782 | 3 | 3855.9201 | 3855.9155 | 0 | 1.2 | 127 | Specific | 712.96 | 712.96 | 712.96 | 14.95 | 123 | >sp\|P01876\|IGHA1_HUMAN Immunoglobulin heavy constant alpha 1 OS=Homo sapiens OX=9606 GN=IGHA1 PE=1 SV=2 | 255 | 20201118_ly_colostrum_1ng_glyco_hcd203040_compound15.19629.19629.3 | controllerType=0 controllerNumber=1 scan=19629 | 42.9556 |
| 15958:3 | 5 | R.LSLHRPALEDLLLGSEAN[+1216.423]LTC[+57.021]TLTGLR.D | HexNAc(2)Hex(5) | C[+57], N[+1216] | 1394.0141 | 3 | 4180.0277 | 4180.0211 | 0 | 1.6 | 127 | Specific | 733.47 | 709.2 | 709.2 | 14.95 | 123 | >sp\|P01876\|IGHA1_HUMAN Immunoglobulin heavy constant alpha 1 OS=Homo sapiens OX=9606 GN=IGHA1 PE=1 SV=2 | 255 | 20201118_ly_colostrum_1ng_glyco_hcd203040_compound15.19449.19449.3 | controllerType=0 controllerNumber=1 scan=19449 | 42.5684 |
| 16056:3 | 5 | R.LSLHRPALEDLLLGSEAN[+1054.370]LTC[+57.021]TLTGLR.D | HexNAc(2)Hex(4) | C[+57], N[+1054] | 1339.9987 | 3 | 4017.9816 | 4017.9683 | 0 | 3.3 | 127 | Specific | 734.89 | 682.78 | 682.78 | 15.21 | 123 | >sp\|P01876\|IGHA1_HUMAN Immunoglobulin heavy constant alpha 1 OS=Homo sapiens OX=9606 GN=IGHA1 PE=1 SV=2 | 255 | 20201118_ly_colostrum_1ng_glyco_hcd203040_compound15.19551.19551.3 | controllerType=0 controllerNumber=1 scan=19551 | 42.7651 |
| 16926:4 | 5 | R.LSLHRPALEDLLLGSEAN[+2116.756]LTC[+57.021]TLTGLR.D | HexNAc(5)Hex(5)NeuAc(1) | C[+57], N[+2117] | 1270.8484 | 4 | 5080.3717 | 5080.3547 | 0 | 3.4 | 127 | Specific | 748.93 | 653.03 | 653.03 | 15.62 | 123 | >sp\|P01876\|IGHA1_HUMAN Immunoglobulin heavy constant alpha 1 OS=Homo sapiens OX=9606 GN=IGHA1 PE=1 SV=2 | 255 | 20201118_ly_colostrum_1ng_glyco_hcd203040_compound15.20461.20461.4 | controllerType=0 controllerNumber=1 scan=20461 | 44.8402 |
| 16120:3 | 5 | R.LSLHRPALEDLLLGSEAN[+1095.397]LTC[+57.021]TLTGLR.D | HexNAc(3)Hex(3) | C[+57], N[+1095] | 1353.6743 | 3 | 4059.0082 | 4058.9949 | 0 | 3.3 | 127 | Specific | 749.44 | 704.58 | 704.58 | 15.21 | 123 | >sp\|P01876\|IGHA1_HUMAN Immunoglobulin heavy constant alpha 1 OS=Homo sapiens OX=9606 GN=IGHA1 PE=1 SV=2 | 255 | 20201118_ly_colostrum_1ng_glyco_hcd203040_compound15.19618.19618.3 | controllerType=0 controllerNumber=1 scan=19618 | 42.9332 |
| 16059:3 | 5 | R.LSLHRPALEDLLLGSEAN[+1054.370]LTC[+57.021]TLTGLR.D | HexNAc(2)Hex(4) | C[+57], N[+1054] | 1339.9987 | 3 | 4017.9816 | 4017.9683 | 0 | 3.3 | 127 | Specific | 749.88 | 737.17 | 737.17 | 15.21 | 123 | >sp\|P01876\|IGHA1_HUMAN Immunoglobulin heavy constant alpha 1 OS=Homo sapiens OX=9606 GN=IGHA1 PE=1 SV=2 | 255 | 20201118_ly_colostrum_1ng_glyco_hcd203040_compound15.19554.19554.3 | controllerType=0 controllerNumber=1 scan=19554 | 42.7763 |
| 17969:4 | 5 | R.LSLHRPALEDLLLGSEAN[+2204.772]LTC[+57.021]TLTGLR.D | HexNAc(4)Hex(5)NeuAc(2) | C[+57], N[+2205] | 1292.8521 | 4 | 5168.3864 | 5168.3707 | 0 | 3 | 127 | Specific | 752.03 | 578.39 | 578.39 | 16.59 | 123 | >sp\|P01876\|IGHA1_HUMAN Immunoglobulin heavy constant alpha 1 OS=Homo sapiens OX=9606 GN=IGHA1 PE=1 SV=2 | 255 | 20201118_ly_colostrum_1ng_glyco_hcd203040_compound15.21551.21551.4 | controllerType=0 controllerNumber=1 scan=21551 | 47.2823 |
| 15984:5 | 5 | R.LSLHRPALEDLLLGSEAN[+1501.555]LTC[+57.021]TLTGLR.D | HexNAc(5)Hex(3) | C[+57], N[+1502] | 893.8383 | 5 | 4465.1623 | 4465.1536 | 0 | 1.9 | 127 | Specific | 752.73 | 752.73 | 752.73 | 16.52 | 123 | >sp\|P01876\|IGHA1_HUMAN Immunoglobulin heavy constant alpha 1 OS=Homo sapiens OX=9606 GN=IGHA1 PE=1 SV=2 | 255 | 20201118_ly_colostrum_1ng_glyco_hcd203040_compound15.19476.19476.5 | controllerType=0 controllerNumber=1 scan=19476 | 42.6169 |
| 16968:4 | 5 | R.LSLHRPALEDLLLGSEAN[+1548.545]LTC[+57.021]TLTGLR.D | HexNAc(3)Hex(4)NeuAc(1) | C[+57], N[+1549] | 1128.7946 | 4 | 4512.1566 | 4512.1431 | 0 | 3 | 127 | Specific | 754.93 | 754.93 | 754.93 | 16.59 | 123 | >sp\|P01876\|IGHA1_HUMAN Immunoglobulin heavy constant alpha 1 OS=Homo sapiens OX=9606 GN=IGHA1 PE=1 SV=2 | 255 | 20201118_ly_colostrum_1ng_glyco_hcd203040_compound15.20505.20505.4 | controllerType=0 controllerNumber=1 scan=20505 | 44.9354 |
| 15998:5 | 5 | R.LSLHRPALEDLLLGSEAN[+1460.529]LTC[+57.021]TLTGLR.D | HexNAc(4)Hex(4) | C[+57], N[+1461] | 885.6333 | 5 | 4424.1372 | 4424.1271 | 0 | 2.3 | 127 | Specific | 760.41 | 760.41 | 760.41 | 16.76 | 123 | >sp\|P01876\|IGHA1_HUMAN Immunoglobulin heavy constant alpha 1 OS=Homo sapiens OX=9606 GN=IGHA1 PE=1 SV=2 | 255 | 20201118_ly_colostrum_1ng_glyco_hcd203040_compound15.19490.19490.5 | controllerType=0 controllerNumber=1 scan=19490 | 42.6396 |
| 15981:5 | 5 | R.LSLHRPALEDLLLGSEAN[+1501.555]LTC[+57.021]TLTGLR.D | HexNAc(5)Hex(3) | C[+57], N[+1502] | 893.8383 | 5 | 4465.1623 | 4465.1536 | 0 | 1.9 | 127 | Specific | 763.04 | 763.04 | 763.04 | 16.52 | 123 | >sp\|P01876\|IGHA1_HUMAN Immunoglobulin heavy constant alpha 1 OS=Homo sapiens OX=9606 GN=IGHA1 PE=1 SV=2 | 255 | 20201118_ly_colostrum_1ng_glyco_hcd203040_compound15.19473.19473.5 | controllerType=0 controllerNumber=1 scan=19473 | 42.611 |
| 15955:3 | 5 | R.LSLHRPALEDLLLGSEAN[+1216.423]LTC[+57.021]TLTGLR.D | HexNAc(2)Hex(5) | C[+57], N[+1216] | 1394.0141 | 3 | 4180.0277 | 4180.0211 | 0 | 1.6 | 127 | Specific | 763.51 | 718.65 | 718.65 | 15.95 | 123 | >sp\|P01876\|IGHA1_HUMAN Immunoglobulin heavy constant alpha 1 OS=Homo sapiens OX=9606 GN=IGHA1 PE=1 SV=2 | 255 | 20201118_ly_colostrum_1ng_glyco_hcd203040_compound15.19446.19446.3 | controllerType=0 controllerNumber=1 scan=19446 | 42.5548 |
| 15954:4 | 5 | R.LSLHRPALEDLLLGSEAN[+1460.529]LTC[+57.021]TLTGLR.D | HexNAc(4)Hex(4) | C[+57], N[+1461] | 1106.7902 | 4 | 4424.139 | 4424.1271 | 0 | 2.7 | 127 | Specific | 772.34 | 772.34 | 772.34 | 16.59 | 123 | >sp\|P01876\|IGHA1_HUMAN Immunoglobulin heavy constant alpha 1 OS=Homo sapiens OX=9606 GN=IGHA1 PE=1 SV=2 | 255 | 20201118_ly_colostrum_1ng_glyco_hcd203040_compound15.19445.19445.4 | controllerType=0 controllerNumber=1 scan=19445 | 42.5537 |
| 16101:4 | 5 | R.LSLHRPALEDLLLGSEAN[+892.317]LTC[+57.021]TLTGLR.D | HexNAc(2)Hex(3) | C[+57], N[+892] | 964.7352 | 4 | 3855.919 | 3855.9155 | 0 | 0.9 | 127 | Specific | 772.95 | 772.95 | 772.95 | 16.16 | 123 | >sp\|P01876\|IGHA1_HUMAN Immunoglobulin heavy constant alpha 1 OS=Homo sapiens OX=9606 GN=IGHA1 PE=1 SV=2 | 255 | 20201118_ly_colostrum_1ng_glyco_hcd203040_compound15.19598.19598.4 | controllerType=0 controllerNumber=1 scan=19598 | 42.887 |
| 16944:4 | 5 | R.LSLHRPALEDLLLGSEAN[+1751.624]LTC[+57.021]TLTGLR.D | HexNAc(4)Hex(4)NeuAc(1) | C[+57], N[+1752] | 1179.564 | 4 | 4715.2342 | 4715.2225 | 0 | 2.5 | 127 | Specific | 779.22 | 779.22 | 779.22 | 16.59 | 123 | >sp\|P01876\|IGHA1_HUMAN Immunoglobulin heavy constant alpha 1 OS=Homo sapiens OX=9606 GN=IGHA1 PE=1 SV=2 | 255 | 20201118_ly_colostrum_1ng_glyco_hcd203040_compound15.20480.20480.4 | controllerType=0 controllerNumber=1 scan=20480 | 44.89 |
| 15999:5 | 5 | R.LSLHRPALEDLLLGSEAN[+1460.529]LTC[+57.021]TLTGLR.D | HexNAc(4)Hex(4) | C[+57], N[+1461] | 885.6333 | 5 | 4424.1372 | 4424.1271 | 0 | 2.3 | 127 | Specific | 781.4 | 781.4 | 781.4 | 16.76 | 123 | >sp\|P01876\|IGHA1_HUMAN Immunoglobulin heavy constant alpha 1 OS=Homo sapiens OX=9606 GN=IGHA1 PE=1 SV=2 | 255 | 20201118_ly_colostrum_1ng_glyco_hcd203040_compound15.19491.19491.5 | controllerType=0 controllerNumber=1 scan=19491 | 42.6449 |
| 16194:4 | 5 | R.LSLHRPALEDLLLGSEAN[+1622.582]LTC[+57.021]TLTGLR.D | HexNAc(4)Hex(5) | C[+57], N[+1623] | 1147.2989 | 4 | 4586.1737 | 4586.1799 | 0 | -1.4 | 127 | Specific | 783.34 | 615.13 | 615.13 | 15.46 | 123 | >sp\|P01876\|IGHA1_HUMAN Immunoglobulin heavy constant alpha 1 OS=Homo sapiens OX=9606 GN=IGHA1 PE=1 SV=2 | 255 | 20201118_ly_colostrum_1ng_glyco_hcd203040_compound15.19695.19695.4 | controllerType=0 controllerNumber=1 scan=19695 | 43.1181 |

Table S4. Cont.

| Query #:z | Protein Rank | Glycopeptide | Glycan | Modification Type(s) | Observed m/z | z | Observed (M+H) | Calc. mass (M+H) | Off-by-x error | Mass error (ppm) | StartPosition | Cleavage | Score | Delta | Delta Mod | \|Log Prob\| | # of unique peptides | Protein Name | Protein DB number | Comment | Scan # | Scan Time |
|---|---|---|---|---|---|---|---|---|---|---|---|---|---|---|---|---|---|---|---|---|---|---|
| 15951:4 | 5 | R.LSLHRPALEDLLLGSEAN[+1663.608]LTC[+57.021]TLTGLR.D | HexNAc(5)Hex(4) | C[+57], N[+1664] | 1157.559 | 4 | 4627.2142 | 4627.2064 | 0 | 1.7 | 127 | Specific | 784 | 710.76 | 710.76 | 16.36 | 123 | >sp\|P01876\|IGHA1_HUMAN Immunoglobulin heavy constant alpha 1 OS=Homo sapiens OX=9606 GN=IGHA1 PE=1 SV=2 | 255 | 20201118_ly_colostrum_1ng_glyco_hcd203040_compound15.19442.19442.4 | controllerType=0 controllerNumber=1 scan=19442 | 42.5451 |
| 15915:4 | 5 | R.LSLHRPALEDLLLGSEAN[+1825.661]LTC[+57.021]TLTGLR.D | HexNAc(5)Hex(5) | C[+57], N[+1826] | 1198.0746 | 4 | 4789.2767 | 4789.2592 | 0 | 3.6 | 127 | Specific | 793.56 | 698.16 | 698.16 | 16.62 | 123 | >sp\|P01876\|IGHA1_HUMAN Immunoglobulin heavy constant alpha 1 OS=Homo sapiens OX=9606 GN=IGHA1 PE=1 SV=2 | 255 | 20201118_ly_colostrum_1ng_glyco_hcd203040_compound15.19404.19404.4 | controllerType=0 controllerNumber=1 scan=19404 | 42.4419 |
| 16052:4 | 5 | R.LSLHRPALEDLLLGSEAN[+1501.555]LTC[+57.021]TLTGLR.D | HexNAc(5)Hex(3) | C[+57], N[+1502] | 1117.0494 | 4 | 4465.1756 | 4465.1536 | 0 | 4.9 | 127 | Specific | 794.91 | 794.91 | 794.91 | 16.32 | 123 | >sp\|P01876\|IGHA1_HUMAN Immunoglobulin heavy constant alpha 1 OS=Homo sapiens OX=9606 GN=IGHA1 PE=1 SV=2 | 255 | 20201118_ly_colostrum_1ng_glyco_hcd203040_compound15.19547.19547.4 | controllerType=0 controllerNumber=1 scan=19547 | 42.7574 |
| 16976:4 | 5 | R.LSLHRPALEDLLLGSEAN[+1913.677]LTC[+57.021]TLTGLR.D | HexNAc(4)Hex(5)NeuAc(1) | C[+57], N[+1914] | 1220.0769 | 4 | 4877.286 | 4877.2753 | 0 | 2.2 | 127 | Specific | 798.63 | 638.73 | 638.73 | 16.59 | 123 | >sp\|P01876\|IGHA1_HUMAN Immunoglobulin heavy constant alpha 1 OS=Homo sapiens OX=9606 GN=IGHA1 PE=1 SV=2 | 255 | 20201118_ly_colostrum_1ng_glyco_hcd203040_compound15.20513.20513.4 | controllerType=0 controllerNumber=1 scan=20513 | 44.949 |
| 15572:4 | 5 | R.LSLHRPALEDLLLGSEAN[+1501.555]LTC[+57.021]TLTGLR.D | HexNAc(5)Hex(3) | C[+57], N[+1502] | 1117.044 | 4 | 4465.1541 | 4465.1536 | 0 | 0.1 | 127 | Specific | 799.01 | 685.62 | 685.62 | 16.16 | 123 | >sp\|P01876\|IGHA1_HUMAN Immunoglobulin heavy constant alpha 1 OS=Homo sapiens OX=9606 GN=IGHA1 PE=1 SV=2 | 255 | 20201118_ly_colostrum_1ng_glyco_hcd203040_compound15.19046.19046.4 | controllerType=0 controllerNumber=1 scan=19046 | 41.6574 |
| 15973:4 | 5 | R.LSLHRPALEDLLLGSEAN[+1622.582]LTC[+57.021]TLTGLR.D | HexNAc(4)Hex(5) | C[+57], N[+1623] | 1147.3024 | 4 | 4586.1878 | 4586.1799 | 0 | 1.7 | 127 | Specific | 815.59 | 707.24 | 707.24 | 17.36 | 123 | >sp\|P01876\|IGHA1_HUMAN Immunoglobulin heavy constant alpha 1 OS=Homo sapiens OX=9606 GN=IGHA1 PE=1 SV=2 | 255 | 20201118_ly_colostrum_1ng_glyco_hcd203040_compound15.19465.19465.4 | controllerType=0 controllerNumber=1 scan=19465 | 42.5983 |
| 16073:4 | 5 | R.LSLHRPALEDLLLGSEAN[+1647.613]LTC[+57.021]TLTGLR.D | HexNAc(5)Hex(3)Fuc(1) | C[+57], N[+1648] | 1153.5591 | 4 | 4611.2147 | 4611.2115 | 0 | 0.7 | 127 | Specific | 824.09 | 742.1 | 742.1 | 17.16 | 123 | >sp\|P01876\|IGHA1_HUMAN Immunoglobulin heavy constant alpha 1 OS=Homo sapiens OX=9606 GN=IGHA1 PE=1 SV=2 | 255 | 20201118_ly_colostrum_1ng_glyco_hcd203040_compound15.19569.19569.4 | controllerType=0 controllerNumber=1 scan=19569 | 42.8258 |
| 16902:4 | 5 | R.LSLHRPALEDLLLGSEAN[+1872.651]LTC[+57.021]TLTGLR.D | HexNAc(3)Hex(6)NeuAc(1) | C[+57], N[+1873] | 1209.8241 | 4 | 4836.2747 | 4836.2487 | 0 | 5.4 | 127 | Specific | 840.42 | 840.42 | 840.42 | 17.05 | 123 | >sp\|P01876\|IGHA1_HUMAN Immunoglobulin heavy constant alpha 1 OS=Homo sapiens OX=9606 GN=IGHA1 PE=1 SV=2 | 255 | 20201118_ly_colostrum_1ng_glyco_hcd203040_compound15.20436.20436.4 | controllerType=0 controllerNumber=1 scan=20436 | 44.7838 |
| 16128:3 | 5 | R.LSLHRPALEDLLLGSEAN[+892.317]LTC[+57.021]TLTGLR.D | HexNAc(2)Hex(3) | C[+57], N[+892] | 1285.9782 | 3 | 3855.9201 | 3855.9155 | 0 | 1.2 | 127 | Specific | 840.95 | 840.95 | 840.95 | 16.95 | 123 | >sp\|P01876\|IGHA1_HUMAN Immunoglobulin heavy constant alpha 1 OS=Homo sapiens OX=9606 GN=IGHA1 PE=1 SV=2 | 255 | 20201118_ly_colostrum_1ng_glyco_hcd203040_compound15.19626.19626.3 | controllerType=0 controllerNumber=1 scan=19626 | 42.9502 |
| 16070:4 | 5 | R.LSLHRPALEDLLLGSEAN[+1054.370]LTC[+57.021]TLTGLR.D | HexNAc(2)Hex(4) | C[+57], N[+1054] | 1005.2506 | 4 | 4017.9805 | 4017.9683 | 0 | 3 | 127 | Specific | 844.5 | 836.49 | 836.49 | 17.59 | 123 | >sp\|P01876\|IGHA1_HUMAN Immunoglobulin heavy constant alpha 1 OS=Homo sapiens OX=9606 GN=IGHA1 PE=1 SV=2 | 255 | 20201118_ly_colostrum_1ng_glyco_hcd203040_compound15.19566.19566.4 | controllerType=0 controllerNumber=1 scan=19566 | 42.8185 |
| 16906:4 | 5 | R.LSLHRPALEDLLLGSEAN[+1710.598]LTC[+57.021]TLTGLR.D | HexNAc(3)Hex(5)NeuAc(1) | C[+57], N[+1711] | 1169.3092 | 4 | 4674.2152 | 4674.1959 | 0 | 4.1 | 127 | Specific | 854.8 | 854.8 | 854.8 | 18.32 | 123 | >sp\|P01876\|IGHA1_HUMAN Immunoglobulin heavy constant alpha 1 OS=Homo sapiens OX=9606 GN=IGHA1 PE=1 SV=2 | 255 | 20201118_ly_colostrum_1ng_glyco_hcd203040_compound15.20440.20440.4 | controllerType=0 controllerNumber=1 scan=20440 | 44.7908 |
| 16032:4 | 5 | R.LSLHRPALEDLLLGSEAN[+1647.613]LTC[+57.021]TLTGLR.D | HexNAc(5)Hex(3)Fuc(1) | C[+57], N[+1648] | 1153.561 | 4 | 4611.222 | 4611.2115 | 0 | 2.3 | 127 | Specific | 860.48 | 743.82 | 743.82 | 18.59 | 123 | >sp\|P01876\|IGHA1_HUMAN Immunoglobulin heavy constant alpha 1 OS=Homo sapiens OX=9606 GN=IGHA1 PE=1 SV=2 | 255 | 20201118_ly_colostrum_1ng_glyco_hcd203040_compound15.19526.19526.4 | controllerType=0 controllerNumber=1 scan=19526 | 42.7161 |
| 16905:5 | 5 | R.LSLHRPALEDLLLGSEAN[+1872.651]LTC[+57.021]TLTGLR.D | HexNAc(3)Hex(6)NeuAc(1) | C[+57], N[+1873] | 968.0587 | 5 | 4836.2643 | 4836.2487 | 0 | 3.2 | 127 | Specific | 864.49 | 864.49 | 864.49 | 18.78 | 123 | >sp\|P01876\|IGHA1_HUMAN Immunoglobulin heavy constant alpha 1 OS=Homo sapiens OX=9606 GN=IGHA1 PE=1 SV=2 | 255 | 20201118_ly_colostrum_1ng_glyco_hcd203040_compound15.20439.20439.5 | controllerType=0 controllerNumber=1 scan=20439 | 44.7896 |
| 15987:5 | 5 | R.LSLHRPALEDLLLGSEAN[+1216.423]LTC[+57.021]TLTGLR.D | HexNAc(2)Hex(5) | C[+57], N[+1216] | 836.8116 | 5 | 4180.0289 | 4180.0211 | 0 | 1.9 | 127 | Specific | 870.32 | 870.32 | 870.32 | 18.52 | 123 | >sp\|P01876\|IGHA1_HUMAN Immunoglobulin heavy constant alpha 1 OS=Homo sapiens OX=9606 GN=IGHA1 PE=1 SV=2 | 255 | 20201118_ly_colostrum_1ng_glyco_hcd203040_compound15.19479.19479.5 | controllerType=0 controllerNumber=1 scan=19479 | 42.6201 |
| 16904:4 | 5 | R.LSLHRPALEDLLLGSEAN[+1872.651]LTC[+57.021]TLTGLR.D | HexNAc(3)Hex(6)NeuAc(1) | C[+57], N[+1873] | 1209.8241 | 4 | 4836.2747 | 4836.2487 | 0 | 5.4 | 127 | Specific | 874.07 | 874.07 | 874.07 | 18.05 | 123 | >sp\|P01876\|IGHA1_HUMAN Immunoglobulin heavy constant alpha 1 OS=Homo sapiens OX=9606 GN=IGHA1 PE=1 SV=2 | 255 | 20201118_ly_colostrum_1ng_glyco_hcd203040_compound15.20438.20438.4 | controllerType=0 controllerNumber=1 scan=20438 | 44.7888 |
| 16908:5 | 5 | R.LSLHRPALEDLLLGSEAN[+1872.651]LTC[+57.021]TLTGLR.D | HexNAc(3)Hex(6)NeuAc(1) | C[+57], N[+1873] | 968.0587 | 5 | 4836.2643 | 4836.2487 | 0 | 3.2 | 127 | Specific | 878.72 | 844.83 | 844.83 | 18.78 | 123 | >sp\|P01876\|IGHA1_HUMAN Immunoglobulin heavy constant alpha 1 OS=Homo sapiens OX=9606 GN=IGHA1 PE=1 SV=2 | 255 | 20201118_ly_colostrum_1ng_glyco_hcd203040_compound15.20442.20442.5 | controllerType=0 controllerNumber=1 scan=20442 | 44.7949 |
| 15907:4 | 5 | R.LSLHRPALEDLLLGSEAN[+1540.529]LTC[+57.021]TLTGLR.D | HexNAc(2)Hex(7) | C[+57], N[+1541] | 1126.7913 | 4 | 4504.1434 | 4504.1268 | 0 | 3.7 | 127 | Specific | 891.4 | 802.87 | 802.87 | 18.62 | 123 | >sp\|P01876\|IGHA1_HUMAN Immunoglobulin heavy constant alpha 1 OS=Homo sapiens OX=9606 GN=IGHA1 PE=1 SV=2 | 255 | 20201118_ly_colostrum_1ng_glyco_hcd203040_compound15.19396.19396.4 | controllerType=0 controllerNumber=1 scan=19396 | 42.4225 |
| 15957:4 | 5 | R.LSLHRPALEDLLLGSEAN[+1460.529]LTC[+57.021]TLTGLR.D | HexNAc(4)Hex(4) | C[+57], N[+1461] | 1106.7902 | 4 | 4424.139 | 4424.1271 | 0 | 2.7 | 127 | Specific | 900.5 | 768.56 | 768.56 | 19.59 | 123 | >sp\|P01876\|IGHA1_HUMAN Immunoglobulin heavy constant alpha 1 OS=Homo sapiens OX=9606 GN=IGHA1 PE=1 SV=2 | 255 | 20201118_ly_colostrum_1ng_glyco_hcd203040_compound15.19448.19448.4 | controllerType=0 controllerNumber=1 scan=19448 | 42.5677 |
| 16909:4 | 5 | R.LSLHRPALEDLLLGSEAN[+1710.598]LTC[+57.021]TLTGLR.D | HexNAc(3)Hex(5)NeuAc(1) | C[+57], N[+1711] | 1169.3092 | 4 | 4674.2152 | 4674.1959 | 0 | 4.1 | 127 | Specific | 903.62 | 868.1 | 868.1 | 19.32 | 123 | >sp\|P01876\|IGHA1_HUMAN Immunoglobulin heavy constant alpha 1 OS=Homo sapiens OX=9606 GN=IGHA1 PE=1 SV=2 | 255 | 20201118_ly_colostrum_1ng_glyco_hcd203040_compound15.20443.20443.4 | controllerType=0 controllerNumber=1 scan=20443 | 44.803 |
| 16028:4 | 5 | R.LSLHRPALEDLLLGSEAN[+1257.449]LTC[+57.021]TLTGLR.D | HexNAc(3)Hex(4) | C[+57], N[+1257] | 1056.0185 | 4 | 4221.0521 | 4221.0477 | 0 | 1 | 127 | Specific | 906.73 | 883.78 | 883.78 | 19.16 | 123 | >sp\|P01876\|IGHA1_HUMAN Immunoglobulin heavy constant alpha 1 OS=Homo sapiens OX=9606 GN=IGHA1 PE=1 SV=2 | 255 | 20201118_ly_colostrum_1ng_glyco_hcd203040_compound15.19522.19522.4 | controllerType=0 controllerNumber=1 scan=19522 | 42.7066 |
| 16970:4 | 5 | R.LSLHRPALEDLLLGSEAN[+1548.545]LTC[+57.021]TLTGLR.D | HexNAc(3)Hex(4)NeuAc(1) | C[+57], N[+1549] | 1128.7946 | 4 | 4512.1566 | 4512.1431 | 0 | 3 | 127 | Specific | 926.88 | 926.88 | 926.88 | 19.59 | 123 | >sp\|P01876\|IGHA1_HUMAN Immunoglobulin heavy constant alpha 1 OS=Homo sapiens OX=9606 GN=IGHA1 PE=1 SV=2 | 255 | 20201118_ly_colostrum_1ng_glyco_hcd203040_compound15.20507.20507.4 | controllerType=0 controllerNumber=1 scan=20507 | 44.9414 |
| 15950:4 | 5 | R.LSLHRPALEDLLLGSEAN[+1216.423]LTC[+57.021]TLTGLR.D | HexNAc(2)Hex(5) | C[+57], N[+1216] | 1045.7635 | 4 | 4180.0321 | 4180.0211 | 0 | 2.6 | 127 | Specific | 953.03 | 911.93 | 911.93 | 20.59 | 123 | >sp\|P01876\|IGHA1_HUMAN Immunoglobulin heavy constant alpha 1 OS=Homo sapiens OX=9606 GN=IGHA1 PE=1 SV=2 | 255 | 20201118_ly_colostrum_1ng_glyco_hcd203040_compound15.19441.19441.4 | controllerType=0 controllerNumber=1 scan=19441 | 42.5422 |
| 16072:4 | 5 | R.LSLHRPALEDLLLGSEAN[+1054.370]LTC[+57.021]TLTGLR.D | HexNAc(2)Hex(4) | C[+57], N[+1054] | 1005.2506 | 4 | 4017.9805 | 4017.9683 | 0 | 3 | 127 | Specific | 968.7 | 968.7 | 968.7 | 20.59 | 123 | >sp\|P01876\|IGHA1_HUMAN Immunoglobulin heavy constant alpha 1 OS=Homo sapiens OX=9606 GN=IGHA1 PE=1 SV=2 | 255 | 20201118_ly_colostrum_1ng_glyco_hcd203040_compound15.19568.19568.4 | controllerType=0 controllerNumber=1 scan=19568 | 42.8245 |
| 16104:4 | 5 | R.LSLHRPALEDLLLGSEAN[+892.317]LTC[+57.021]TLTGLR.D | HexNAc(2)Hex(3) | C[+57], N[+892] | 964.7352 | 4 | 3855.919 | 3855.9155 | 0 | 0.9 | 127 | Specific | 991.86 | 991.86 | 991.86 | 20.16 | 123 | >sp\|P01876\|IGHA1_HUMAN Immunoglobulin heavy constant alpha 1 OS=Homo sapiens OX=9606 GN=IGHA1 PE=1 SV=2 | 255 | 20201118_ly_colostrum_1ng_glyco_hcd203040_compound15.19601.19601.4 | controllerType=0 controllerNumber=1 scan=19601 | 42.9046 |
| 15924:4 | 5 | R.LSLHRPALEDLLLGSEAN[+1702.581]LTC[+57.021]TLTGLR.D | HexNAc(2)Hex(8) | C[+57], N[+1703] | 1167.3013 | 4 | 4666.1834 | 4666.1796 | 0 | 0.8 | 127 | Specific | 1005.48 | 1005.48 | 1005.48 | 21.16 | 123 | >sp\|P01876\|IGHA1_HUMAN Immunoglobulin heavy constant alpha 1 OS=Homo sapiens OX=9606 GN=IGHA1 PE=1 SV=2 | 255 | 20201118_ly_colostrum_1ng_glyco_hcd203040_compound15.19414.19414.4 | controllerType=0 controllerNumber=1 scan=19414 | 42.483 |
| 15909:4 | 5 | R.LSLHRPALEDLLLGSEAN[+1540.529]LTC[+57.021]TLTGLR.D | HexNAc(2)Hex(7) | C[+57], N[+1541] | 1126.7913 | 4 | 4504.1434 | 4504.1268 | 0 | 3.7 | 127 | Specific | 1010.21 | 942.44 | 942.44 | 21.62 | 123 | >sp\|P01876\|IGHA1_HUMAN Immunoglobulin heavy constant alpha 1 OS=Homo sapiens OX=9606 GN=IGHA1 PE=1 SV=2 | 255 | 20201118_ly_colostrum_1ng_glyco_hcd203040_compound15.19398.19398.4 | controllerType=0 controllerNumber=1 scan=19398 | 42.4312 |
| 15952:4 | 5 | R.LSLHRPALEDLLLGSEAN[+1216.423]LTC[+57.021]TLTGLR.D | HexNAc(2)Hex(5) | C[+57], N[+1216] | 1045.7635 | 4 | 4180.0321 | 4180.0211 | 0 | 2.6 | 127 | Specific | 1010.84 | 957.81 | 957.81 | 21.59 | 123 | >sp\|P01876\|IGHA1_HUMAN Immunoglobulin heavy constant alpha 1 OS=Homo sapiens OX=9606 GN=IGHA1 PE=1 SV=2 | 255 | 20201118_ly_colostrum_1ng_glyco_hcd203040_compound15.19443.19443.4 | controllerType=0 controllerNumber=1 scan=19443 | 42.5514 |
| 15978:4 | 5 | R.LSLHRPALEDLLLGSEAN[+1378.476]LTC[+57.021]TLTGLR.D | HexNAc(2)Hex(6) | C[+57], N[+1378] | 1086.2769 | 4 | 4342.0858 | 4342.074 | 0 | 2.7 | 127 | Specific | 1013.82 | 972.89 | 972.89 | 21.59 | 123 | >sp\|P01876\|IGHA1_HUMAN Immunoglobulin heavy constant alpha 1 OS=Homo sapiens OX=9606 GN=IGHA1 PE=1 SV=2 | 255 | 20201118_ly_colostrum_1ng_glyco_hcd203040_compound15.19470.19470.4 | controllerType=0 controllerNumber=1 scan=19470 | 42.6051 |
| 15908:4 | 5 | R.LSLHRPALEDLLLGSEAN[+1864.634]LTC[+57.021]TLTGLR.D | HexNAc(2)Hex(9) | C[+57], N[+1865] | 1207.8169 | 4 | 4828.2459 | 4828.2324 | 0 | 2.8 | 127 | Specific | 1037.75 | 1037.75 | 1037.75 | 21.59 | 123 | >sp\|P01876\|IGHA1_HUMAN Immunoglobulin heavy constant alpha 1 OS=Homo sapiens OX=9606 GN=IGHA1 PE=1 SV=2 | 255 | 20201118_ly_colostrum_1ng_glyco_hcd203040_compound15.19397.19397.4 | controllerType=0 controllerNumber=1 scan=19397 | 42.4255 |
| 16031:4 | 5 | R.LSLHRPALEDLLLGSEAN[+1257.449]LTC[+57.021]TLTGLR.D | HexNAc(3)Hex(4) | C[+57], N[+1257] | 1056.0185 | 4 | 4221.0521 | 4221.0477 | 0 | 1 | 127 | Specific | 1067.25 | 1067.25 | 1067.25 | 22.16 | 123 | >sp\|P01876\|IGHA1_HUMAN Immunoglobulin heavy constant alpha 1 OS=Homo sapiens OX=9606 GN=IGHA1 PE=1 SV=2 | 255 | 20201118_ly_colostrum_1ng_glyco_hcd203040_compound15.19525.19525.4 | controllerType=0 controllerNumber=1 scan=19525 | 42.7152 |
| 15990:5 | 5 | R.LSLHRPALEDLLLGSEAN[+1216.423]LTC[+57.021]TLTGLR.D | HexNAc(2)Hex(5) | C[+57], N[+1216] | 836.8116 | 5 | 4180.0289 | 4180.0211 | 0 | 1.9 | 127 | Specific | 1076.15 | 1076.15 | 1076.15 | 22.52 | 123 | >sp\|P01876\|IGHA1_HUMAN Immunoglobulin heavy constant alpha 1 OS=Homo sapiens OX=9606 GN=IGHA1 PE=1 SV=2 | 255 | 20201118_ly_colostrum_1ng_glyco_hcd203040_compound15.19482.19482.5 | controllerType=0 controllerNumber=1 scan=19482 | 42.6262 |
| 17135:5 | 5 | R.LSLHRPALEDLLLGSEAN[+1460.529]LTC[+57.021]TLTGLRDASGVTFTWTPSSGK.S | HexNAc(4)Hex(4) | C[+57], N[+1461] | 1189.9788 | 5 | 5945.8647 | 5945.837 | 0 | 4.7 | 127 | Specific | 330.21 | 330.21 | 330.21 | 7.23 | 123 | >sp\|P01876\|IGHA1_HUMAN Immunoglobulin heavy constant alpha 1 OS=Homo sapiens OX=9606 GN=IGHA1 PE=1 SV=2 | 255 | 20201118_ly_colostrum_1ng_glyco_hcd203040_compound15.20679.20679.5 | controllerType=0 controllerNumber=1 scan=20679 | 45.3333 |
| 17098:5 | 5 | R.LSLHRPALEDLLLGSEAN[+1378.476]LTC[+57.021]TLTGLRDASGVTFTWTPSSGK.S | HexNAc(2)Hex(6) | C[+57], N[+1378] | 1173.5663 | 5 | 5863.8023 | 5863.7839 | 0 | 3.1 | 127 | Specific | 647.45 | 542.2 | 542.2 | 12.69 | 123 | >sp\|P01876\|IGHA1_HUMAN Immunoglobulin heavy constant alpha 1 OS=Homo sapiens OX=9606 GN=IGHA1 PE=1 SV=2 | 255 | 20201118_ly_colostrum_1ng_glyco_hcd203040_compound15.20640.20640.5 | controllerType=0 controllerNumber=1 scan=20640 | 45.2512 |
| 17101:5 | 5 | R.LSLHRPALEDLLLGSEAN[+1378.476]LTC[+57.021]TLTGLRDASGVTFTWTPSSGK.S | HexNAc(2)Hex(6) | C[+57], N[+1378] | 1173.5663 | 5 | 5863.8023 | 5863.7839 | 0 | 3.1 | 127 | Specific | 679.49 | 679.49 | 679.49 | 13.69 | 123 | >sp\|P01876\|IGHA1_HUMAN Immunoglobulin heavy constant alpha 1 OS=Homo sapiens OX=9606 GN=IGHA1 PE=1 SV=2 | 255 | 20201118_ly_colostrum_1ng_glyco_hcd203040_compound15.20643.20643.5 | controllerType=0 controllerNumber=1 scan=20643 | 45.2605 |
| 17138:5 | 5 | R.LSLHRPALEDLLLGSEAN[+1460.529]LTC[+57.021]TLTGLRDASGVTFTWTPSSGK.S | HexNAc(4)Hex(4) | C[+57], N[+1461] | 1189.9788 | 5 | 5945.8647 | 5945.837 | 0 | 4.7 | 127 | Specific | 785.52 | 567.81 | 567.81 | 15.39 | 123 | >sp\|P01876\|IGHA1_HUMAN Immunoglobulin heavy constant alpha 1 OS=Homo sapiens OX=9606 GN=IGHA1 PE=1 SV=2 | 255 | 20201118_ly_colostrum_1ng_glyco_hcd203040_compound15.20682.20682.5 | controllerType=0 controllerNumber=1 scan=20682 | 45.3383 |
| 17108:6 | 5 | R.LSLHRPALEDLLLGSEAN[+1501.555]LTC[+57.021]TLTGLRDASGVTFTWTPSSGK.S | HexNAc(5)Hex(3) | C[+57], N[+1502] | 998.6535 | 6 | 5986.8845 | 5986.8636 | 0 | 3.5 | 127 | Specific | 889.01 | 889.01 | 889.01 | 17.85 | 123 | >sp\|P01876\|IGHA1_HUMAN Immunoglobulin heavy constant alpha 1 OS=Homo sapiens OX=9606 GN=IGHA1 PE=1 SV=2 | 255 | 20201118_ly_colostrum_1ng_glyco_hcd203040_compound15.20651.20651.6 | controllerType=0 controllerNumber=1 scan=20651 | 45.2776 |
| 17105:4 | 5 | R.LSLHRPALEDLLLGSEAN[+1501.555]LTC[+57.021]TLTGLRDASGVTFTWTPSSGK.S | HexNAc(5)Hex(3) | C[+57], N[+1502] | 1497.4767 | 4 | 5986.8849 | 5986.8636 | 0 | 3.6 | 127 | Specific | 907.19 | 793.59 | 793.59 | 18.27 | 123 | >sp\|P01876\|IGHA1_HUMAN Immunoglobulin heavy constant alpha 1 OS=Homo sapiens OX=9606 GN=IGHA1 PE=1 SV=2 | 255 | 20201118_ly_colostrum_1ng_glyco_hcd203040_compound15.20648.20648.4 | controllerType=0 controllerNumber=1 scan=20648 | 45.2742 |
| 17104:5 | 5 | R.LSLHRPALEDLLLGSEAN[+1501.555]LTC[+57.021]TLTGLRDASGVTFTWTPSSGK.S | HexNAc(5)Hex(3) | C[+57], N[+1502] | 1198.1821 | 5 | 5986.8816 | 5986.8636 | 0 | 3 | 127 | Specific | 1066.1 | 1066.1 | 1066.1 | 21.66 | 123 | >sp\|P01876\|IGHA1_HUMAN Immunoglobulin heavy constant alpha 1 OS=Homo sapiens OX=9606 GN=IGHA1 PE=1 SV=2 | 255 | 20201118_ly_colostrum_1ng_glyco_hcd203040_compound15.20647.20647.5 | controllerType=0 controllerNumber=1 scan=20647 | 45.271 |
| 17102:5 | 5 | R.LSLHRPALEDLLLGSEAN[+1501.555]LTC[+57.021]TLTGLRDASGVTFTWTPSSGK.S | HexNAc(5)Hex(3) | C[+57], N[+1502] | 1198.1821 | 5 | 5986.8816 | 5986.8636 | 0 | 3 | 127 | Specific | 1071.77 | 907.04 | 907.04 | 21.66 | 123 | >sp\|P01876\|IGHA1_HUMAN Immunoglobulin heavy constant alpha 1 OS=Homo sapiens OX=9606 GN=IGHA1 PE=1 SV=2 | 255 | 20201118_ly_colostrum_1ng_glyco_hcd203040_compound15.20645.20645.5 | controllerType=0 controllerNumber=1 scan=20645 | 45.2651 |
| 17103:4 | 5 | R.LSLHRPALEDLLLGSEAN[+1501.555]LTC[+57.021]TLTGLRDASGVTFTWTPSSGK.S | HexNAc(5)Hex(3) | C[+57], N[+1502] | 1497.4767 | 4 | 5986.8849 | 5986.8636 | 0 | 3.6 | 127 | Specific | 1093.34 | 1028.11 | 1028.11 | 21.27 | 123 | >sp\|P01876\|IGHA1_HUMAN Immunoglobulin heavy constant alpha 1 OS=Homo sapiens OX=9606 GN=IGHA1 PE=1 SV=2 | 255 | 20201118_ly_colostrum_1ng_glyco_hcd203040_compound15.20646.20646.4 | controllerType=0 controllerNumber=1 scan=20646 | 45.2665 |
| 17111:6 | 5 | R.LSLHRPALEDLLLGSEAN[+1501.555]LTC[+57.021]TLTGLRDASGVTFTWTPSSGK.S | HexNAc(5)Hex(3) | C[+57], N[+1502] | 998.6535 | 6 | 5986.8845 | 5986.8636 | 0 | 3.5 | 127 | Specific | 1124.77 | 976.85 | 976.85 | 22.85 | 123 | >sp\|P01876\|IGHA1_HUMAN Immunoglobulin heavy constant alpha 1 OS=Homo sapiens OX=9606 GN=IGHA1 PE=1 SV=2 | 255 | 20201118_ly_colostrum_1ng_glyco_hcd203040_compound15.20654.20654.6 | controllerType=0 controllerNumber=1 scan=20654 | 45.2846 |
| 10725:3 | 5 | K.TIDRLAGKPTHVN[+1647.613]VSVVMAEVDGTC[+57.021]Y.- | HexNAc(5)Hex(3)Fuc(1) | C[+57], N[+1648] | 1494.0126 | 3 | 4480.0233 | 4480.0151 | 0 | 1.8 | 328 | CRagged | 265.11 | 265.11 | 265.11 | 6.8 | 123 | >sp\|P01876\|IGHA1_HUMAN Immunoglobulin heavy constant alpha 1 OS=Homo sapiens OX=9606 GN=IGHA1 PE=1 SV=2 | 255 | 20201118_ly_colostrum_1ng_glyco_hcd203040_compound15.14005.14005.3 | controllerType=0 controllerNumber=1 scan=14005 | 31.3699 |
| 09719:4 | 5 | K.TIDRLAGKPTHVN[+1216.423]VSVVMAEVDGTC[+57.021]Y.- | HexNAc(2)Hex(5) | C[+57], N[+1216] | 1012.9673 | 4 | 4048.8475 | 4048.8247 | 0 | 5.6 | 328 | CRagged | 321.57 | 321.57 | 275.31 | 6.2 | 123 | >sp\|P01876\|IGHA1_HUMAN Immunoglobulin heavy constant alpha 1 OS=Homo sapiens OX=9606 GN=IGHA1 PE=1 SV=2 | 255 | 20201118_ly_colostrum_1ng_glyco_hcd203040_compound15.12959.12959.4 | controllerType=0 controllerNumber=1 scan=12959 | 29.2608 |
| 10590:4 | 5 | K.TIDRLAGKPTHVN[+1825.661]VSVVMAEVDGTC[+57.021]Y.- | HexNAc(5)Hex(5) | C[+57], N[+1826] | 1165.2751 | 4 | 4658.0784 | 4658.0629 | 0 | 3.3 | 328 | CRagged | 328.19 | 316.38 | 299.55 | 8.06 | 123 | >sp\|P01876\|IGHA1_HUMAN Immunoglobulin heavy constant alpha 1 OS=Homo sapiens OX=9606 GN=IGHA1 PE=1 SV=2 | 255 | 20201118_ly_colostrum_1ng_glyco_hcd203040_compound15.13864.13864.4 | controllerType=0 controllerNumber=1 scan=13864 | 31.0656 |
| 09713:4 | 5 | K.TIDRLAGKPTHVN[+1216.423]VSVVMAEVDGTC[+57.021]Y.- | HexNAc(2)Hex(5) | C[+57], N[+1216] | 1012.9664 | 4 | 4048.8438 | 4048.8247 | 0 | 4.7 | 328 | CRagged | 341.38 | 321.18 | 187.21 | 6.62 | 123 | >sp\|P01876\|IGHA1_HUMAN Immunoglobulin heavy constant alpha 1 OS=Homo sapiens OX=9606 GN=IGHA1 PE=1 SV=2 | 255 | 20201118_ly_colostrum_1ng_glyco_hcd203040_compound15.12952.12952.2 | controllerType=0 controllerNumber=1 scan=12952 | 29.244 |
| 13207:4 | 5 | K.TIDRLAGKPTHVN[+2553.910]VSVVMAEVDGTC[+57.021]Y.- | HexNAc(5)Hex(5)Fuc(1)NeuAc(2) | C[+57], N[+2554] | 1347.3376 | 4 | 5386.3288 | 5386.3116 | 0 | 3.2 | 328 | CRagged | 343.55 | 286.23 | 286.23 | 8.02 | 123 | >sp\|P01876\|IGHA1_HUMAN Immunoglobulin heavy constant alpha 1 OS=Homo sapiens OX=9606 GN=IGHA1 PE=1 SV=2 | 255 | 20201118_ly_colostrum_1ng_glyco_hcd203040_compound15.16584.16584.4 | controllerType=0 controllerNumber=1 scan=16584 | 36.5245 |
| 10606:4 | 5 | K.TIDRLAGKPTHVN[+1540.529]VSVVMAEVDGTC[+57.021]Y.- | HexNAc(2)Hex(7) | C[+57], N[+1541] | 1093.9889 | 4 | 4372.9339 | 4372.9304 | 0 | 0.8 | 328 | CRagged | 371.22 | 371.22 | 300.09 | 8.16 | 123 | >sp\|P01876\|IGHA1_HUMAN Immunoglobulin heavy constant alpha 1 OS=Homo sapiens OX=9606 GN=IGHA1 PE=1 SV=2 | 255 | 20201118_ly_colostrum_1ng_glyco_hcd203040_compound15.13881.13881.4 | controllerType=0 controllerNumber=1 scan=13881 | 31.1095 |
| 10593:4 | 5 | K.TIDRLAGKPTHVN[+1825.661]VSVVMAEVDGTC[+57.021]Y.- | HexNAc(5)Hex(5) | C[+57], N[+1826] | 1165.2751 | 4 | 4658.0784 | 4658.0629 | 0 | 3.3 | 328 | CRagged | 374.45 | 328.78 | 254.19 | 7.4 | 123 | >sp\|P01876\|IGHA1_HUMAN Immunoglobulin heavy constant alpha 1 OS=Homo sapiens OX=9606 GN=IGHA1 PE=1 SV=2 | 255 | 20201118_ly_colostrum_1ng_glyco_hcd203040_compound15.13867.13867.4 | controllerType=0 controllerNumber=1 scan=13867 | 31.0701 |
| 10721:4 | 5 | K.TIDRLAGKPTHVN[+1647.613]VSVVMAEVDGTC[+57.021]Y.- | HexNAc(5)Hex(3)Fuc(1) | C[+57], N[+1648] | 1120.761 | 4 | 4480.0223 | 4480.0151 | 0 | 1.6 | 328 | CRagged | 430.07 | 331.95 | 331.95 | 6.76 | 123 | >sp\|P01876\|IGHA1_HUMAN Immunoglobulin heavy constant alpha 1 OS=Homo sapiens OX=9606 GN=IGHA1 PE=1 SV=2 | 255 | 20201118_ly_colostrum_1ng_glyco_hcd203040_compound15.14001.14001.4 | controllerType=0 controllerNumber=1 scan=14001 | 31.365 |
| 13210:4 | 5 | K.TIDRLAGKPTHVN[+2553.910]VSVVMAEVDGTC[+57.021]Y.- | HexNAc(5)Hex(5)Fuc(1)NeuAc(2) | C[+57], N[+2554] | 1347.3376 | 4 | 5386.3288 | 5386.3116 | 0 | 3.2 | 328 | CRagged | 441.23 | 376.32 | 376.32 | 8.93 | 123 | >sp\|P01876\|IGHA1_HUMAN Immunoglobulin heavy constant alpha 1 OS=Homo sapiens OX=9606 GN=IGHA1 PE=1 SV=2 | 255 | 20201118_ly_colostrum_1ng_glyco_hcd203040_compound15.16587.16587.4 | controllerType=0 controllerNumber=1 scan=16587 | 36.5291 |
| 10601:4 | 5 | K.TIDRLAGKPTHVN[+1663.608]VSVVMAEVDGTC[+57.021]Y.- | HexNAc(5)Hex(4) | C[+57], N[+1664] | 1124.7596 | 4 | 4496.0164 | 4496.01 | 0 | 1.4 | 328 | CRagged | 513.53 | 472.82 | 346.21 | 10.41 | 123 | >sp\|P01876\|IGHA1_HUMAN Immunoglobulin heavy constant alpha 1 OS=Homo sapiens OX=9606 GN=IGHA1 PE=1 SV=2 | 255 | 20201118_ly_colostrum_1ng_glyco_hcd203040_compound15.13876.13876.4 | controllerType=0 controllerNumber=1 scan=13876 | 31.0997 |
| 10611:4 | 5 | K.TIDRLAGKPTHVN[+1971.719]VSVVMAEVDGTC[+57.021]Y.- | HexNAc(5)Hex(5)Fuc(1) | C[+57], N[+1972] | 1201.7886 | 4 | 4804.1326 | 4804.1208 | 0 | 2.5 | 328 | CRagged | 519.03 | 519.03 | 423.37 | 10.64 | 123 | >sp\|P01876\|IGHA1_HUMAN Immunoglobulin heavy constant alpha 1 OS=Homo sapiens OX=9606 GN=IGHA1 PE=1 SV=2 | 255 | 20201118_ly_colostrum_1ng_glyco_hcd203040_compound15.13886.13886.4 | controllerType=0 controllerNumber=1 scan=13886 | 31.1267 |
| 09716:4 | 5 | K.TIDRLAGKPTHVN[+1216.423]VSVVMAEVDGTC[+57.021]Y.- | HexNAc(2)Hex(5) | C[+57], N[+1216] | 1012.9664 | 4 | 4048.8438 | 4048.8247 | 0 | 4.7 | 328 | CRagged | 521.66 | 512.11 | 337.51 | 10.41 | 123 | >sp\|P01876\|IGHA1_HUMAN Immunoglobulin heavy constant alpha 1 OS=Homo sapiens OX=9606 GN=IGHA1 PE=1 SV=2 | 255 | 20201118_ly_colostrum_1ng_glyco_hcd203040_compound15.12955.12955.2 | controllerType=0 controllerNumber=1 scan=12955 | 29.2535 |
| 11859:4 | 5 | K.TIDRLAGKPTHVN[+2262.814]VSVVMAEVDGTC[+57.021]Y.- | HexNAc(5)Hex(5)Fuc(1)NeuAc(1) | C[+57], N[+2263] | 1274.5618 | 4 | 5095.2253 | 5095.2162 | 0 | 1.8 | 328 | CRagged | 525.85 | 478.07 | 478.07 | 10.45 | 123 | >sp\|P01876\|IGHA1_HUMAN Immunoglobulin heavy constant alpha 1 OS=Homo sapiens OX=9606 GN=IGHA1 PE=1 SV=2 | 255 | 20201118_ly_colostrum_1ng_glyco_hcd203040_compound15.15183.15183.4 | controllerType=0 controllerNumber=1 scan=15183 | 33.646 |
| 10623:4 | 5 | K.TIDRLAGKPTHVN[+1809.666]VSVVMAEVDGTC[+57.021]Y.- | HexNAc(5)Hex(4)Fuc(1) | C[+57], N[+1810] | 1161.2757 | 4 | 4642.0809 | 4642.0679 | 0 | 2.8 | 328 | CRagged | 527.37 | 483.24 | 395.9 | 10.68 | 123 | >sp\|P01876\|IGHA1_HUMAN Immunoglobulin heavy constant alpha 1 OS=Homo sapiens OX=9606 GN=IGHA1 PE=1 SV=2 | 255 | 20201118_ly_colostrum_1ng_glyco_hcd203040_compound15.13899.13899.4 | controllerType=0 controllerNumber=1 scan=13899 | 31.1624 |
| 10620:4 | 5 | K.TIDRLAGKPTHVN[+1809.666]VSVVMAEVDGTC[+57.021]Y.- | HexNAc(5)Hex(4)Fuc(1) | C[+57], N[+1810] | 1161.2757 | 4 | 4642.0809 | 4642.0679 | 0 | 2.8 | 328 | CRagged | 532.89 | 522.43 | 424.45 | 10.68 | 123 | >sp\|P01876\|IGHA1_HUMAN Immunoglobulin heavy constant alpha 1 OS=Homo sapiens OX=9606 GN=IGHA1 PE=1 SV=2 | 255 | 20201118_ly_colostrum_1ng_glyco_hcd203040_compound15.13896.13896.4 | controllerType=0 controllerNumber=1 scan=13896 | 31.1548 |
| 09722:4 | 5 | K.TIDRLAGKPTHVN[+1216.423]VSVVMAEVDGTC[+57.021]Y.- | HexNAc(2)Hex(5) | C[+57], N[+1216] | 1012.9673 | 4 | 4048.8475 | 4048.8247 | 0 | 5.6 | 328 | CRagged | 564.82 | 554.76 | 405.53 | 10.5 | 123 | >sp\|P01876\|IGHA1_HUMAN Immunoglobulin heavy constant alpha 1 OS=Homo sapiens OX=9606 GN=IGHA1 PE=1 SV=2 | 255 | 20201118_ly_colostrum_1ng_glyco_hcd203040_compound15.12962.12962.4 | controllerType=0 controllerNumber=1 scan=12962 | 29.2654 |
| 10609:4 | 5 | K.TIDRLAGKPTHVN[+1540.529]VSVVMAEVDGTC[+57.021]Y.- | HexNAc(2)Hex(7) | C[+57], N[+1541] | 1093.9889 | 4 | 4372.9339 | 4372.9304 | 0 | 0.8 | 328 | CRagged | 572.76 | 571.14 | 415.85 | 10.61 | 123 | >sp\|P01876\|IGHA1_HUMAN Immunoglobulin heavy constant alpha 1 OS=Homo sapiens OX=9606 GN=IGHA1 PE=1 SV=2 | 255 | 20201118_ly_colostrum_1ng_glyco_hcd203040_compound15.13884.13884.4 | controllerType=0 controllerNumber=1 scan=13884 | 31.1235 |
| 11862:4 | 5 | K.TIDRLAGKPTHVN[+2262.814]VSVVMAEVDGTC[+57.021]Y.- | HexNAc(5)Hex(5)Fuc(1)NeuAc(1) | C[+57], N[+2263] | 1274.5618 | 4 | 5095.2253 | 5095.2162 | 0 | 1.8 | 328 | CRagged | 579.04 | 488.26 | 488.26 | 10.81 | 123 | >sp\|P01876\|IGHA1_HUMAN Immunoglobulin heavy constant alpha 1 OS=Homo sapiens OX=9606 GN=IGHA1 PE=1 SV=2 | 255 | 20201118_ly_colostrum_1ng_glyco_hcd203040_compound15.15186.15186.4 | controllerType=0 controllerNumber=1 scan=15186 | 33.6558 |
| 10628:3 | 5 | K.TIDRLAGKPTHVN[+1216.423]VSVVMAEVDGTC[+57.021]Y.- | HexNAc(2)Hex(5) | C[+57], N[+1216] | 1350.2833 | 3 | 4048.8352 | 4048.8247 | 0 | 2.6 | 328 | CRagged | 590.1 | 444.07 | 279.48 | 11.95 | 123 | >sp\|P01876\|IGHA1_HUMAN Immunoglobulin heavy constant alpha 1 OS=Homo sapiens OX=9606 GN=IGHA1 PE=1 SV=2 | 255 | 20201118_ly_colostrum_1ng_glyco_hcd203040_compound15.13904.13904.3 | controllerType=0 controllerNumber=1 scan=13904 | 31.1836 |
| 10613:3 | 5 | K.TIDRLAGKPTHVN[+1663.608]VSVVMAEVDGTC[+57.021]Y.- | HexNAc(5)Hex(4) | C[+57], N[+1664] | 1499.3447 | 3 | 4496.0194 | 4496.01 | 0 | 2.1 | 328 | CRagged | 595.41 | 525.55 | 322.67 | 11.95 | 123 | >sp\|P01876\|IGHA1_HUMAN Immunoglobulin heavy constant alpha 1 OS=Homo sapiens OX=9606 GN=IGHA1 PE=1 SV=2 | 255 | 20201118_ly_colostrum_1ng_glyco_hcd203040_compound15.13888.13888.3 | controllerType=0 controllerNumber=1 scan=13888 | 31.1313 |
| 10608:3 | 5 | K.TIDRLAGKPTHVN[+1378.476]VSVVMAEVDGTC[+57.021]Y.- | HexNAc(2)Hex(6) | C[+57], N[+1378] | 1404.2997 | 3 | 4210.8847 | 4210.8776 | 0 | 1.7 | 328 | CRagged | 596.03 | 512.73 | 315.76 | 11.72 | 123 | >sp\|P01876\|IGHA1_HUMAN Immunoglobulin heavy constant alpha 1 OS=Homo sapiens OX=9606 GN=IGHA1 PE=1 SV=2 | 255 | 20201118_ly_colostrum_1ng_glyco_hcd203040_compound15.13883.13883.3 | controllerType=0 controllerNumber=1 scan=13883 | 31.1216 |
| 10636:3 | 5 | K.TIDRLAGKPTHVN[+1809.666]VSVVMAEVDGTC[+57.021]Y.- | HexNAc(5)Hex(4)Fuc(1) | C[+57], N[+1810] | 1548.0328 | 3 | 4642.0838 | 4642.0679 | 0 | 3.4 | 328 | CRagged | 602.63 | 471.99 | 362.54 | 12.98 | 123 | >sp\|P01876\|IGHA1_HUMAN Immunoglobulin heavy constant alpha 1 OS=Homo sapiens OX=9606 GN=IGHA1 PE=1 SV=2 | 255 | 20201118_ly_colostrum_1ng_glyco_hcd203040_compound15.13912.13912.3 | controllerType=0 controllerNumber=1 scan=13912 | 31.2008 |
| 10614:4 | 5 | K.TIDRLAGKPTHVN[+1971.719]VSVVMAEVDGTC[+57.021]Y.- | HexNAc(5)Hex(5)Fuc(1) | C[+57], N[+1972] | 1201.7886 | 4 | 4804.1326 | 4804.1208 | 0 | 2.5 | 328 | CRagged | 617.74 | 519.63 | 450.56 | 13.37 | 123 | >sp\|P01876\|IGHA1_HUMAN Immunoglobulin heavy constant alpha 1 OS=Homo sapiens OX=9606 GN=IGHA1 PE=1 SV=2 | 255 | 20201118_ly_colostrum_1ng_glyco_hcd203040_compound15.13889.13889.4 | controllerType=0 controllerNumber=1 scan=13889 | 31.[illegible]66 |
| 10603:4 | 5 | K.TIDRLAGKPTHVN[+1663.608]VSVVMAEVDGTC[+57.021]Y.- | HexNAc(5)Hex(4) | C[+57], N[+1664] | 1124.7596 | 4 | 4496.0164 | 4496.01 | 0 | 1.4 | 328 | CRagged | 623.18 | 599.32 | 441.24 | 13.14 | 123 | >sp\|P01876\|IGHA1_HUMAN Immunoglobulin heavy constant alpha 1 OS=Homo sapiens OX=9606 GN=IGHA1 PE=1 SV=2 | 255 | 20201118_ly_colostrum_1ng_glyco_hcd203040_compound15.13878.13878.4 | controllerType=0 controllerNumber=1 scan=13878 | 31.[illegible]61 |
| 10728:3 | 5 | K.TIDRLAGKPTHVN[+1647.613]VSVVMAEVDGTC[+57.021]Y.- | HexNAc(5)Hex(3)Fuc(1) | C[+57], N[+1648] | 1494.0126 | 3 | 4480.0233 | 4480.0151 | 0 | 1.8 | 328 | CRagged | 649.01 | 525.75 | 525.75 | 12.72 | 123 | >sp\|P01876\|IGHA1_HUMAN Immunoglobulin heavy constant alpha 1 OS=Homo sapiens OX=9606 GN=IGHA1 PE=1 SV=2 | 255 | 20201118_ly_colostrum_1ng_glyco_hcd203040_compound15.14008.14008.3 | controllerType=0 controllerNumber=1 scan=14008 | 31.[illegible]36 |
| 10600:4 | 5 | K.TIDRLAGKPTHVN[+1378.476]VSVVMAEVDGTC[+57.021]Y.- | HexNAc(2)Hex(6) | C[+57], N[+1378] | 1053.4766 | 4 | 4210.8846 | 4210.8776 | 0 | 1.7 | 328 | CRagged | 655.83 | 595.21 | 400.69 | 14.14 | 123 | >sp\|P01876\|IGHA1_HUMAN Immunoglobulin heavy constant alpha 1 OS=Homo sapiens OX=9606 GN=IGHA1 PE=1 SV=2 | 255 | 20201118_ly_colostrum_1ng_glyco_hcd203040_compound15.13875.13875.4 | controllerType=0 controllerNumber=1 scan=13875 | 31.[illegible]83 |
| 10604:4 | 5 | K.TIDRLAGKPTHVN[+1216.423]VSVVMAEVDGTC[+57.021]Y.- | HexNAc(2)Hex(5) | C[+57], N[+1216] | 1012.9636 | 4 | 4048.8326 | 4048.8247 | 0 | 1.9 | 328 | CRagged | 656.32 | 497.99 | 443.18 | 14.14 | 123 | >sp\|P01876\|IGHA1_HUMAN Immunoglobulin heavy constant alpha 1 OS=Homo sapiens OX=9606 GN=IGHA1 PE=1 SV=2 | 255 | 20201118_ly_colostrum_1ng_glyco_hcd203040_compound15.13879.13879.4 | controllerType=0 controllerNumber=1 scan=13879 | 31.1072 |
| 10610:3 | 5 | K.TIDRLAGKPTHVN[+1663.608]VSVVMAEVDGTC[+57.021]Y.- | HexNAc(5)Hex(4) | C[+57], N[+1664] | 1499.3447 | 3 | 4496.0194 | 4496.01 | 0 | 2.1 | 328 | CRagged | 656.36 | 616.02 | 421.09 | 13.95 | 123 | >sp\|P01876\|IGHA1_HUMAN Immunoglobulin heavy constant alpha 1 OS=Homo sapiens OX=9606 GN=IGHA1 PE=1 SV=2 | 255 | 20201118_ly_colostrum_1ng_glyco_hcd203040_compound15.13885.13885.3 | controllerType=0 controllerNumber=1 scan=13885 | 31.1254 |
| 10625:3 | 5 | K.TIDRLAGKPTHVN[+1216.423]VSVVMAEVDGTC[+57.021]Y.- | HexNAc(2)Hex(5) | C[+57], N[+1216] | 1350.2833 | 3 | 4048.8352 | 4048.8247 | 0 | 2.6 | 328 | CRagged | 666.91 | 502.52 | 361.19 | 13.95 | 123 | >sp\|P01876\|IGHA1_HUMAN Immunoglobulin heavy constant alpha 1 OS=Homo sapiens OX=9606 GN=IGHA1 PE=1 SV=2 | 255 | 20201118_ly_colostrum_1ng_glyco_hcd203040_compound15.13901.13901.3 | controllerType=0 controllerNumber=1 scan=13901 | 31.1757 |
| 10605:3 | 5 | K.TIDRLAGKPTHVN[+1378.476]VSVVMAEVDGTC[+57.021]Y.- | HexNAc(2)Hex(6) | C[+57], N[+1378] | 1404.2997 | 3 | 4210.8847 | 4210.8776 | 0 | 1.7 | 328 | CRagged | 668.6 | 668.6 | 344.48 | 13.72 | 123 | >sp\|P01876\|IGHA1_HUMAN Immunoglobulin heavy constant alpha 1 OS=Homo sapiens OX=9606 GN=IGHA1 PE=1 SV=2 | 255 | 20201118_ly_colostrum_1ng_glyco_hcd203040_compound15.13880.13880.3 | controllerType=0 controllerNumber=1 scan=13880 | 31.1083 |
| 10633:3 | 5 | K.TIDRLAGKPTHVN[+1809.666]VSVVMAEVDGTC[+57.021]Y.- | HexNAc(5)Hex(4)Fuc(1) | C[+57], N[+1810] | 1548.0328 | 3 | 4642.0838 | 4642.0679 | 0 | 3.4 | 328 | CRagged | 696.33 | 542.54 | 484.94 | 13.98 | 123 | >sp\|P01876\|IGHA1_HUMAN Immunoglobulin heavy constant alpha 1 OS=Homo sapiens OX=9606 GN=IGHA1 PE=1 SV=2 | 255 | 20201118_ly_colostrum_1ng_glyco_hcd203040_compound15.13909.13909.3 | controllerType=0 controllerNumber=1 scan=13909 | 31.1917 |
| 10607:4 | 5 | K.TIDRLAGKPTHVN[+1216.423]VSVVMAEVDGTC[+57.021]Y.- | HexNAc(2)Hex(5) | C[+57], N[+1216] | 1012.9636 | 4 | 4048.8326 | 4048.8247 | 0 | 1.9 | 328 | CRagged | 723.71 | 723.71 | 499.8 | 15.14 | 123 | >sp\|P01876\|IGHA1_HUMAN Immunoglobulin heavy constant alpha 1 OS=Homo sapiens OX=9606 GN=IGHA1 PE=1 SV=2 | 255 | 20201118_ly_colostrum_1ng_glyco_hcd203040_compound15.13882.13882.4 | controllerType=0 controllerNumber=1 scan=13882 | 31.1159 |
| 10602:4 | 5 | K.TIDRLAGKPTHVN[+1378.476]VSVVMAEVDGTC[+57.021]Y.- | HexNAc(2)Hex(6) | C[+57], N[+1378] | 1053.4766 | 4 | 4210.8846 | 4210.8776 | 0 | 1.7 | 328 | CRagged | 739.86 | 672.63 | 470.15 | 15.14 | 123 | >sp\|P01876\|IGHA1_HUMAN Immunoglobulin heavy constant alpha 1 OS=Homo sapiens OX=9606 GN=IGHA1 PE=1 SV=2 | 255 | 20201118_ly_colostrum_1ng_glyco_hcd203040_compound15.13877.13877.4 | controllerType=0 controllerNumber=1 scan=13877 | 31.1041 |
| 11965:3 | 5 | R.LAGKPTHVN[+2117.777]VSVVMAEVDGTC[+57.021].Y | HexNAc(5)Hex(5)Fuc(2) | C[+57], N[+2118] | 1434.6244 | 3 | 4301.8588 | 4301.8555 | 0 | 0.8 | 332 | CRagged | 211.44 | 156.8 | 156.8 | 3.3 | 123 | >sp\|P01876\|IGHA1_HUMAN Immunoglobulin heavy constant alpha 1 OS=Homo sapiens OX=9606 GN=IGHA1 PE=1 SV=2 | 255 | 20201118_ly_colostrum_1ng_glyco_hcd203040_compound15.15293.15293.3 | controllerType=0 controllerNumber=1 scan=15293 | 33.8807 |
| 11962:3 | 5 | R.LAGKPTHVN[+2117.777]VSVVMAEVDGTC[+57.021].Y | HexNAc(5)Hex(5)Fuc(2) | C[+57], N[+2118] | 1434.6244 | 3 | 4301.8588 | 4301.8555 | 0 | 0.8 | 332 | CRagged | 309.85 | 251.12 | 251.12 | 6.38 | 123 | >sp\|P01876\|IGHA1_HUMAN Immunoglobulin heavy constant alpha 1 OS=Homo sapiens OX=9606 GN=IGHA1 PE=1 SV=2 | 255 | 20201118_ly_colostrum_1ng_glyco_hcd203040_compound15.15290.15290.3 | controllerType=0 controllerNumber=1 scan=15290 | 33.8755 |
| 11858:3 | 5 | R.LAGKPTHVN[+2263.835]VSVVMAEVDGTC[+57.021].Y | HexNAc(5)Hex(5)Fuc(3) | C[+57], N[+2264] | 1483.3104 | 3 | 4447.9165 | 4447.9135 | 0 | 0.7 | 332 | CRagged | 316.97 | 209.03 | 209.03 | 7.06 | 123 | >sp\|P01876\|IGHA1_HUMAN Immunoglobulin heavy constant alpha 1 OS=Homo sapiens OX=9606 GN=IGHA1 PE=1 SV=2 | 255 | 20201118_ly_colostrum_1ng_glyco_hcd203040_compound15.15182.15182.3 | controllerType=0 controllerNumber=1 scan=15182 | 33.6451 |
| 13715:4 | 5 | R.LAGKPTHVN[+2204.772]VSVVMAEVDGTC[+57.021]Y.- | HexNAc(4)Hex(5)NeuAc(2) | C[+57], N[+2205] | 1138.737 | 4 | 4551.9261 | 4551.9145 | 0 | 2.5 | 332 | CRagged | 203.84 | 39.72 | 39.72 | 2.26 | 123 | >sp\|P01876\|IGHA1_HUMAN Immunoglobulin heavy constant alpha 1 OS=Homo sapiens OX=9606 GN=IGHA1 PE=1 SV=2 | 255 | 20201118_ly_colostrum_1ng_glyco_hcd203040_compound15.17112.17112.4 | controllerType=0 controllerNumber=1 scan=17112 | 37.574 |
| 11914:4 | 5 | R.LAGKPTHVN[+1913.677]VSVVMAEVDGTC[+57.021]Y.- | HexNAc(4)Hex(5)NeuAc(1) | C[+57], N[+1914] | 1065.9629 | 4 | 4260.8299 | 4260.8191 | 0 | 2.5 | 332 | CRagged | 234.68 | 35.42 | 35.42 | 3.82 | 123 | >sp\|P01876\|IGHA1_HUMAN Immunoglobulin heavy constant alpha 1 OS=Homo sapiens OX=9606 GN=IGHA1 PE=1 SV=2 | 255 | 20201118_ly_colostrum_1ng_glyco_hcd203040_compound15.15240.15240.4 | controllerType=0 controllerNumber=1 scan=15240 | 33.7607 |
| 10389:4 | 5 | R.LAGKPTHVN[+1663.608]VSVVMAEVDGTC[+57.021]Y.- | HexNAc(5)Hex(4) | C[+57], N[+1664] | 1003.4425 | 4 | 4010.7484 | 4010.7502 | 0 | -0.5 | 332 | CRagged | 241.41 | 161.84 | 161.84 | 3.75 | 123 | >sp\|P01876\|IGHA1_HUMAN Immunoglobulin heavy constant alpha 1 OS=Homo sapiens OX=9606 GN=IGHA1 PE=1 SV=2 | 255 | 20201118_ly_colostrum_1ng_glyco_hcd203040_compound15.13654.13654.4 | controllerType=0 controllerNumber=1 scan=13654 | 30.6191 |
| 13616:4 | 5 | R.LAGKPTHVN[+2351.851]VSVVMAEVDGTC[+57.021]Y.- | HexNAc(4)Hex(5)Fuc(3)NeuAc(1) | C[+57], N[+2352] | 1175.5021 | 4 | 4698.9866 | 4698.9928 | 0 | -1.3 | 332 | CRagged | 262.13 | 119.42 | 119.42 | 3.3 | 123 | >sp\|P01876\|IGHA1_HUMAN Immunoglobulin heavy constant alpha 1 OS=Homo sapiens OX=9606 GN=IGHA1 PE=1 SV=2 | 255 | 20201118_ly_colostrum_1ng_glyco_hcd203040_compound15.17009.17009.4 | controllerType=0 controllerNumber=1 scan=17009 | 37.3676 |
| 13613:4 | 5 | R.LAGKPTHVN[+2351.851]VSVVMAEVDGTC[+57.021]Y.- | HexNAc(4)Hex(5)Fuc(3)NeuAc(1) | C[+57], N[+2352] | 1175.5021 | 4 | 4698.9866 | 4698.9928 | 0 | -1.3 | 332 | CRagged | 293.37 | 216.21 | 216.21 | 5.25 | 123 | >sp\|P01876\|IGHA1_HUMAN Immunoglobulin heavy constant alpha 1 OS=Homo sapiens OX=9606 GN=IGHA1 PE=1 SV=2 | 255 | 20201118_ly_colostrum_1ng_glyco_hcd203040_compound15.17006.17006.4 | controllerType=0 controllerNumber=1 scan=17006 | 37.3629 |
| 13718:4 | 5 | R.LAGKPTHVN[+2204.772]VSVVMAEVDGTC[+57.021]Y.- | HexNAc(4)Hex(5)NeuAc(2) | C[+57], N[+2205] | 1138.737 | 4 | 4551.9261 | 4551.9145 | 0 | 2.5 | 332 | CRagged | 307.2 | 212.63 | 212.63 | 5.3 | 123 | >sp\|P01876\|IGHA1_HUMAN Immunoglobulin heavy constant alpha 1 OS=Homo sapiens OX=9606 GN=IGHA1 PE=1 SV=2 | 255 | 20201118_ly_colostrum_1ng_glyco_hcd203040_compound15.17115.17115.4 | controllerType=0 controllerNumber=1 scan=17115 | 37.5782 |
| 09284:3 | 5 | R.LAGKPTHVN[+1702.581]VSVVMAEVDGTC[+57.021]Y.- | HexNAc(2)Hex(8) | C[+57], N[+1703] | 1350.5811 | 3 | 4049.7288 | 4049.7234 | 0 | 1.3 | 332 | CRagged | 313.51 | 313.51 | 228.08 | 7.72 | 123 | >sp\|P01876\|IGHA1_HUMAN Immunoglobulin heavy constant alpha 1 OS=Homo sapiens OX=9606 GN=IGHA1 PE=1 SV=2 | 255 | 20201118_ly_colostrum_1ng_glyco_hcd203040_compound15.12506.12506.3 | controllerType=0 controllerNumber=1 scan=12506 | 28.3278 |
| 11917:4 | 5 | R.LAGKPTHVN[+1913.677]VSVVMAEVDGTC[+57.021]Y.- | HexNAc(4)Hex(5)NeuAc(1) | C[+57], N[+1914] | 1065.9629 | 4 | 4260.8299 | 4260.8191 | 0 | 2.5 | 332 | CRagged | 326.52 | 45.14 | 45.14 | 3.68 | 123 | >sp\|P01876\|IGHA1_HUMAN Immunoglobulin heavy constant alpha 1 OS=Homo sapiens OX=9606 GN=IGHA1 PE=1 SV=2 | 255 | 20201118_ly_colostrum_1ng_glyco_hcd203040_compound15.15243.15243.4 | controllerType=0 controllerNumber=1 scan=15243 | 33.7653 |
| 09286:3 | 5 | R.LAGKPTHVN[+1702.581]VSVVMAEVDGTC[+57.021]Y.- | HexNAc(2)Hex(8) | C[+57], N[+1703] | 1350.5811 | 3 | 4049.7288 | 4049.7234 | 0 | 1.3 | 332 | CRagged | 336.62 | 336.62 | 316.45 | 7.99 | 123 | >sp\|P01876\|IGHA1_HUMAN Immunoglobulin heavy constant alpha 1 OS=Homo sapiens OX=9606 GN=IGHA1 PE=1 SV=2 | 255 | 20201118_ly_colostrum_1ng_glyco_hcd203040_compound15.12508.12508.3 | controllerType=0 controllerNumber=1 scan=12508 | 28.3326 |
| 13737:3 | 5 | R.LAGKPTHVN[+2204.772]VSVVMAEVDGTC[+57.021]Y.- | HexNAc(4)Hex(5)NeuAc(2) | C[+57], N[+2205] | 1517.9839 | 3 | 4551.9373 | 4551.9145 | 0 | 5 | 332 | CRagged | 388.31 | 264 | 264 | 7.4 | 123 | >sp\|P01876\|IGHA1_HUMAN Immunoglobulin heavy constant alpha 1 OS=Homo sapiens OX=9606 GN=IGHA1 PE=1 SV=2 | 255 | 20201118_ly_colostrum_1ng_glyco_hcd203040_compound15.17135.17135.3 | controllerType=0 controllerNumber=1 scan=17135 | 37.6176 |
| 10377:4 | 5 | R.LAGKPTHVN[+1809.666]VSVVMAEVDGTC[+57.021]Y.- | HexNAc(5)Hex(4)Fuc(1) | C[+57], N[+1810] | 1039.9594 | 4 | 4156.8157 | 4156.8081 | 0 | 1.8 | 332 | CRagged | 439.44 | 439.44 | 381.42 | 10.12 | 123 | >sp\|P01876\|IGHA1_HUMAN Immunoglobulin heavy constant alpha 1 OS=Homo sapiens OX=9606 GN=IGHA1 PE=1 SV=2 | 255 | 20201118_ly_colostrum_1ng_glyco_hcd203040_compound15.13642.13642.4 | controllerType=0 controllerNumber=1 scan=13642 | 30.5939 |
| 13739:3 | 5 | R.LAGKPTHVN[+2204.772]VSVVMAEVDGTC[+57.021]Y.- | HexNAc(4)Hex(5)NeuAc(2) | C[+57], N[+2205] | 1517.9839 | 3 | 4551.9373 | 4551.9145 | 0 | 5 | 332 | CRagged | 440.13 | 415.18 | 415.18 | 9.84 | 123 | >sp\|P01876\|IGHA1_HUMAN Immunoglobulin heavy constant alpha 1 OS=Homo sapiens OX=9606 GN=IGHA1 PE=1 SV=2 | 255 | 20201118_ly_colostrum_1ng_glyco_hcd203040_compound15.17137.17137.3 | controllerType=0 controllerNumber=1 scan=17137 | 37.6246 |
| 10374:4 | 5 | R.LAGKPTHVN[+1809.666]VSVVMAEVDGTC[+57.021]Y.- | HexNAc(5)Hex(4)Fuc(1) | C[+57], N[+1810] | 1039.9594 | 4 | 4156.8157 | 4156.8081 | 0 | 1.8 | 332 | CRagged | 451.3 | 424.49 | 350.95 | 10.31 | 123 | >sp\|P01876\|IGHA1_HUMAN Immunoglobulin heavy constant alpha 1 OS=Homo sapiens OX=9606 GN=IGHA1 PE=1 SV=2 | 255 | 20201118_ly_colostrum_1ng_glyco_hcd203040_compound15.13639.13639.4 | controllerType=0 controllerNumber=1 scan=13639 | 30.5897 |
| 10457:4 | 5 | R.LAGKPTHVN[+1647.613]VSVVMAEVDGTC[+57.021]Y.- | HexNAc(5)Hex(3)Fuc(1) | C[+57], N[+1648] | 999.4454 | 4 | 3994.7598 | 3994.7553 | 0 | 1.1 | 332 | CRagged | 492.17 | 436.18 | 436.18 | 10.46 | 123 | >sp\|P01876\|IGHA1_HUMAN Immunoglobulin heavy constant alpha 1 OS=Homo sapiens OX=9606 GN=IGHA1 PE=1 SV=2 | 255 | 20201118_ly_colostrum_1ng_glyco_hcd203040_compound15.13725.13725.4 | controllerType=0 controllerNumber=1 scan=13725 | 30.7842 |
| 10455:4 | 5 | R.LAGKPTHVN[+1647.613]VSVVMAEVDGTC[+57.021]Y.- | HexNAc(5)Hex(3)Fuc(1) | C[+57], N[+1648] | 999.4454 | 4 | 3994.7598 | 3994.7553 | 0 | 1.1 | 332 | CRagged | 509.22 | 464.14 | 464.14 | 10.58 | 123 | >sp\|P01876\|IGHA1_HUMAN Immunoglobulin heavy constant alpha 1 OS=Homo sapiens OX=9606 GN=IGHA1 PE=1 SV=2 | 255 | 20201118_ly_colostrum_1ng_glyco_hcd203040_compound15.13723.13723.4 | controllerType=0 controllerNumber=1 scan=13723 | 30.7779 |
| 09288:3 | 5 | R.LAGKPTHVN[+1971.719]VSVVMAEVDGTC[+57.021]Y.- | HexNAc(5)Hex(5)Fuc(1) | C[+57], N[+1972] | 1440.2941 | 3 | 4318.8678 | 4318.861 | 0 | 1.6 | 332 | CRagged | 515.95 | 507.21 | 370.02 | 10.42 | 123 | >sp\|P01876\|IGHA1_HUMAN Immunoglobulin heavy constant alpha 1 OS=Homo sapiens OX=9606 GN=IGHA1 PE=1 SV=2 | 255 | 20201118_ly_colostrum_1ng_glyco_hcd203040_compound15.12510.12510.3 | controllerType=0 controllerNumber=1 scan=12510 | 28.3358 |
| 10363:3 | 5 | R.LAGKPTHVN[+1581.555]VSVVMAEVDGTC[+57.021]Y.- | HexNAc(3)Hex(6) | C[+57], N[+1582] | 1310.2394 | 3 | 3928.7036 | 3928.6971 | 0 | 1.7 | 332 | CRagged | 523.13 | 465.34 | 413.7 | 10.47 | 123 | >sp\|P01876\|IGHA1_HUMAN Immunoglobulin heavy constant alpha 1 OS=Homo sapiens OX=9606 GN=IGHA1 PE=1 SV=2 | 255 | 20201118_ly_colostrum_1ng_glyco_hcd203040_compound15.13627.13627.3 | controllerType=0 controllerNumber=1 scan=13627 | 30.5665 |
| 10387:2 | 5 | R.LAGKPTHVN[+1216.423]VSVVMAEVDGTC[+57.021]Y.- | HexNAc(2)Hex(5) | C[+57], N[+1216] | 1782.2882 | 2 | 3563.5691 | 3563.5649 | 0 | 1.2 | 332 | CRagged | 549.22 | 549.22 | 233 | 10.24 | 123 | >sp\|P01876\|IGHA1_HUMAN Immunoglobulin heavy constant alpha 1 OS=Homo sapiens OX=9606 GN=IGHA1 PE=1 SV=2 | 255 | 20201118_ly_colostrum_1ng_glyco_hcd203040_compound15.13652.13652.2 | controllerType=0 controllerNumber=1 scan=13652 | 30.6158 |
| 09290:3 | 5 | R.LAGKPTHVN[+1971.719]VSVVMAEVDGTC[+57.021]Y.- | HexNAc(5)Hex(5)Fuc(1) | C[+57], N[+1972] | 1440.2941 | 3 | 4318.8678 | 4318.861 | 0 | 1.6 | 332 | CRagged | 557.04 | 557.04 | 355.85 | 10.65 | 123 | >sp\|P01876\|IGHA1_HUMAN Immunoglobulin heavy constant alpha 1 OS=Homo sapiens OX=9606 GN=IGHA1 PE=1 SV=2 | 255 | 20201118_ly_colostrum_1ng_glyco_hcd203040_compound15.12512.12512.3 | controllerType=0 controllerNumber=1 scan=12512 | 28.3394 |
| 11911:3 | 5 | R.LAGKPTHVN[+1913.677]VSVVMAEVDGTC[+57.021]Y.- | HexNAc(4)Hex(5)NeuAc(1) | C[+57], N[+1914] | 1420.9501 | 3 | 4260.8358 | 4260.8191 | 0 | 3.9 | 332 | CRagged | 567.38 | 433.81 | 433.81 | 11.08 | 123 | >sp\|P01876\|IGHA1_HUMAN Immunoglobulin heavy constant alpha 1 OS=Homo sapiens OX=9606 GN=IGHA1 PE=1 SV=2 | 255 | 20201118_ly_colostrum_1ng_glyco_hcd203040_compound15.15237.15237.3 | controllerType=0 controllerNumber=1 scan=15237 | 33.7553 |
| 09325:3 | 5 | R.LAGKPTHVN[+1663.608]VSVVMAEVDGTC[+57.021]Y.- | HexNAc(5)Hex(4) | C[+57], N[+1664] | 1337.5922 | 3 | 4010.7621 | 4010.7502 | 0 | 3 | 332 | CRagged | 568.33 | 568.33 | 392.04 | 11.06 | 123 | >sp\|P01876\|IGHA1_HUMAN Immunoglobulin heavy constant alpha 1 OS=Homo sapiens OX=9606 GN=IGHA1 PE=1 SV=2 | 255 | 20201118_ly_colostrum_1ng_glyco_hcd203040_compound15.12549.12549.3 | controllerType=0 controllerNumber=1 scan=12549 | 28.4134 |
| 10390:2 | 5 | R.LAGKPTHVN[+1216.423]VSVVMAEVDGTC[+57.021]Y.- | HexNAc(2)Hex(5) | C[+57], N[+1216] | 1782.2882 | 2 | 3563.5691 | 3563.5649 | 0 | 1.2 | 332 | CRagged | 583.25 | 535.57 | 212.93 | 11.73 | 123 | >sp\|P01876\|IGHA1_HUMAN Immunoglobulin heavy constant alpha 1 OS=Homo sapiens OX=9606 GN=IGHA1 PE=1 SV=2 | 255 | 20201118_ly_colostrum_1ng_glyco_hcd203040_compound15.13655.13655.2 | controllerType=0 controllerNumber=1 scan=13655 | 30.6267 |
| 11942:3 | 5 | R.LAGKPTHVN[+1872.651]VSVVMAEVDGTC[+57.021]Y.- | HexNAc(3)Hex(6)NeuAc(1) | C[+57], N[+1873] | 1407.2751 | 3 | 4219.8107 | 4219.7926 | 0 | 4.3 | 332 | CRagged | 606.08 | 469.68 | 381.49 | 13.11 | 123 | >sp\|P01876\|IGHA1_HUMAN Immunoglobulin heavy constant alpha 1 OS=Homo sapiens OX=9606 GN=IGHA1 PE=1 SV=2 | 255 | 20201118_ly_colostrum_1ng_glyco_hcd203040_compound15.15269.15269.3 | controllerType=0 controllerNumber=1 scan=15269 | 33.8285 |
| 13593:4 | 5 | R.LAGKPTHVN[+2553.910]VSVVMAEVDGTC[+57.021]Y.- | HexNAc(5)Hex(5)Fuc(1)NeuAc(2) | C[+57], N[+2554] | 1226.0235 | 4 | 4901.0721 | 4901.0518 | 0 | 4.1 | 332 | CRagged | 611.35 | 481.49 | 481.49 | 13.27 | 123 | >sp\|P01876\|IGHA1_HUMAN Immunoglobulin heavy constant alpha 1 OS=Homo sapiens OX=9606 GN=IGHA1 PE=1 SV=2 | 255 | 20201118_ly_colostrum_1ng_glyco_hcd203040_compound15.16985.16985.4 | controllerType=0 controllerNumber=1 scan=16985 | 37.3014 |
| 10485:3 | 5 | R.LAGKPTHVN[+1095.397]VSVVMAEVDGTC[+57.021]Y.- | HexNAc(3)Hex(3) | C[+57], N[+1095] | 1148.1877 | 3 | 3442.5484 | 3442.5387 | 0 | 2.8 | 332 | CRagged | 612.66 | 527.25 | 527.25 | 13.38 | 123 | >sp\|P01876\|IGHA1_HUMAN Immunoglobulin heavy constant alpha 1 OS=Homo sapiens OX=9606 GN=IGHA1 PE=1 SV=2 | 255 | 20201118_ly_colostrum_1ng_glyco_hcd203040_compound15.13755.13755.3 | controllerType=0 controllerNumber=1 scan=13755 | 30.8429 |
| 10484:3 | 5 | R.LAGKPTHVN[+1054.370]VSVVMAEVDGTC[+57.021]Y.- | HexNAc(2)Hex(4) | C[+57], N[+1054] | 1134.5113 | 3 | 3401.5192 | 3401.5121 | 0 | 2.1 | 332 | CRagged | 635.09 | 635.09 | 317.11 | 13.38 | 123 | >sp\|P01876\|IGHA1_HUMAN Immunoglobulin heavy constant alpha 1 OS=Homo sapiens OX=9606 GN=IGHA1 PE=1 SV=2 | 255 | 20201118_ly_colostrum_1ng_glyco_hcd203040_compound15.13754.13754.3 | controllerType=0 controllerNumber=1 scan=13754 | 30.8418 |
| 09283:3 | 5 | R.LAGKPTHVN[+1540.529]VSVVMAEVDGTC[+57.021]Y.- | HexNAc(2)Hex(7) | C[+57], N[+1541] | 1296.5643 | 3 | 3887.6782 | 3887.6706 | 0 | 2 | 332 | CRagged | 637.9 | 637.9 | 395.54 | 13.15 | 123 | >sp\|P01876\|IGHA1_HUMAN Immunoglobulin heavy constant alpha 1 OS=Homo sapiens OX=9606 GN=IGHA1 PE=1 SV=2 | 255 | 20201118_ly_colostrum_1ng_glyco_hcd203040_compound15.12505.12505.3 | controllerType=0 controllerNumber=1 scan=12505 | 28.3263 |
| 10483:3 | 5 | R.LAGKPTHVN[+1444.534]VSVVMAEVDGTC[+57.021]Y.- | HexNAc(4)Hex(3)Fuc(1) | C[+57], N[+1445] | 1264.5674 | 3 | 3791.6878 | 3791.6759 | 0 | 3.1 | 332 | CRagged | 640.15 | 640.15 | 640.15 | 13.41 | 123 | >sp\|P01876\|IGHA1_HUMAN Immunoglobulin heavy constant alpha 1 OS=Homo sapiens OX=9606 GN=IGHA1 PE=1 SV=2 | 255 | 20201118_ly_colostrum_1ng_glyco_hcd203040_compound15.13753.13753.3 | controllerType=0 controllerNumber=1 scan=13753 | 30.841 |
| 11910:3 | 5 | R.LAGKPTHVN[+2059.735]VSVVMAEVDGTC[+57.021]Y.- | HexNAc(4)Hex(5)Fuc(1)NeuAc(1) | C[+57], N[+2060] | 1469.636 | 3 | 4406.8936 | 4406.877 | 0 | 3.8 | 332 | CRagged | 645.01 | 602.14 | 602.14 | 13.41 | 123 | >sp\|P01876\|IGHA1_HUMAN Immunoglobulin heavy constant alpha 1 OS=Homo sapiens OX=9606 GN=IGHA1 PE=1 SV=2 | 255 | 20201118_ly_colostrum_1ng_glyco_hcd203040_compound15.15236.15236.3 | controllerType=0 controllerNumber=1 scan=15236 | 33.7535 |
| 11912:3 | 5 | R.LAGKPTHVN[+2059.735]VSVVMAEVDGTC[+57.021]Y.- | HexNAc(4)Hex(5)Fuc(1)NeuAc(1) | C[+57], N[+2060] | 1469.636 | 3 | 4406.8936 | 4406.877 | 0 | 3.8 | 332 | CRagged | 651.85 | 557.87 | 557.87 | 14.41 | 123 | >sp\|P01876\|IGHA1_HUMAN Immunoglobulin heavy constant alpha 1 OS=Homo sapiens OX=9606 GN=IGHA1 PE=1 SV=2 | 255 | 20201118_ly_colostrum_1ng_glyco_hcd203040_compound15.15238.15238.3 | controllerType=0 controllerNumber=1 scan=15238 | 33.7573 |
| 10407:2 | 5 | R.LAGKPTHVN[+1216.423]VSVVMAEVDGTC[+57.021]Y.- | HexNAc(2)Hex(5) | C[+57], N[+1216] | 1782.288 | 2 | 3563.5688 | 3563.5649 | 0 | 1.1 | 332 | CRagged | 659.35 | 646.71 | 287.46 | 13.73 | 123 | >sp\|P01876\|IGHA1_HUMAN Immunoglobulin heavy constant alpha 1 OS=Homo sapiens OX=9606 GN=IGHA1 PE=1 SV=2 | 255 | 20201118_ly_colostrum_1ng_glyco_hcd203040_compound15.13673.13673.2 | controllerType=0 controllerNumber=1 scan=13673 | 30.6628 |
| 11870:4 | 5 | R.LAGKPTHVN[+2116.756]VSVVMAEVDGTC[+57.021]Y.- | HexNAc(5)Hex(5)NeuAc(1) | C[+57], N[+2117] | 1116.7322 | 4 | 4463.9071 | 4463.8985 | 0 | 1.9 | 332 | CRagged | 676.21 | 525.51 | 525.51 | 14.31 | 123 | >sp\|P01876\|IGHA1_HUMAN Immunoglobulin heavy constant alpha 1 OS=Homo sapiens OX=9606 GN=IGHA1 PE=1 SV=2 | 255 | 20201118_ly_colostrum_1ng_glyco_hcd203040_compound15.15194.15194.4 | controllerType=0 controllerNumber=1 scan=15194 | 33.667 |
| 10296:3 | 5 | R.LAGKPTHVN[+1825.661]VSVVMAEVDGTC[+57.021]Y.- | HexNAc(5)Hex(5) | C[+57], N[+1826] | 1391.6093 | 3 | 4172.8134 | 4172.8031 | 0 | 2.5 | 332 | CRagged | 684.09 | 555.32 | 419.02 | 14.38 | 123 | >sp\|P01876\|IGHA1_HUMAN Immunoglobulin heavy constant alpha 1 OS=Homo sapiens OX=9606 GN=IGHA1 PE=1 SV=2 | 255 | 20201118_ly_colostrum_1ng_glyco_hcd203040_compound15.13558.13558.3 | controllerType=0 controllerNumber=1 scan=13558 | 30.4288 |
| 13604:3 | 5 | R.LAGKPTHVN[+2553.910]VSVVMAEVDGTC[+57.021]Y.- | HexNAc(5)Hex(5)Fuc(1)NeuAc(2) | C[+57], N[+2554] | 1634.3622 | 3 | 4901.0722 | 4901.0518 | 0 | 4.2 | 332 | CRagged | 684.13 | 493.71 | 493.71 | 14.11 | 123 | >sp\|P01876\|IGHA1_HUMAN Immunoglobulin heavy constant alpha 1 OS=Homo sapiens OX=9606 GN=IGHA1 PE=1 SV=2 | 255 | 20201118_ly_colostrum_1ng_glyco_hcd203040_compound15.16996.16996.3 | controllerType=0 controllerNumber=1 scan=16996 | 37.3402 |
| 13610:3 | 5 | R.LAGKPTHVN[+2350.830]VSVVMAEVDGTC[+57.021]Y.- | HexNAc(4)Hex(5)Fuc(1)NeuAc(2) | C[+57], N[+2351] | 1566.6677 | 3 | 4697.9884 | 4697.9724 | 0 | 3.4 | 332 | CRagged | 699.85 | 659.65 | 659.65 | 14.41 | 123 | >sp\|P01876\|IGHA1_HUMAN Immunoglobulin heavy constant alpha 1 OS=Homo sapiens OX=9606 GN=IGHA1 PE=1 SV=2 | 255 | 20201118_ly_colostrum_1ng_glyco_hcd203040_compound15.17003.17003.3 | controllerType=0 controllerNumber=1 scan=17003 | 37.3587 |
| 11867:4 | 5 | R.LAGKPTHVN[+2116.756]VSVVMAEVDGTC[+57.021]Y.- | HexNAc(5)Hex(5)NeuAc(1) | C[+57], N[+2117] | 1116.7322 | 4 | 4463.9071 | 4463.8985 | 0 | 1.9 | 332 | CRagged | 702.26 | 490.68 | 490.68 | 15.31 | 123 | >sp\|P01876\|IGHA1_HUMAN Immunoglobulin heavy constant alpha 1 OS=Homo sapiens OX=9606 GN=IGHA1 PE=1 SV=2 | 255 | 20201118_ly_colostrum_1ng_glyco_hcd203040_compound15.15191.15191.4 | controllerType=0 controllerNumber=1 scan=15191 | 33.6619 |
| 11855:4 | 5 | R.LAGKPTHVN[+2262.814]VSVVMAEVDGTC[+57.021]Y.- | HexNAc(5)Hex(5)Fuc(1)NeuAc(1) | C[+57], N[+2263] | 1153.2482 | 4 | 4609.971 | 4609.9564 | 0 | 3.2 | 332 | CRagged | 705.47 | 599.48 | 599.48 | 15.57 | 123 | >sp\|P01876\|IGHA1_HUMAN Immunoglobulin heavy constant alpha 1 OS=Homo sapiens OX=9606 GN=IGHA1 PE=1 SV=2 | 255 | 20201118_ly_colostrum_1ng_glyco_hcd203040_compound15.15178.15178.4 | controllerType=0 controllerNumber=1 scan=15178 | 33.6378 |
| 13608:3 | 5 | R.LAGKPTHVN[+2350.830]VSVVMAEVDGTC[+57.021]Y.- | HexNAc(4)Hex(5)Fuc(1)NeuAc(2) | C[+57], N[+2351] | 1566.6677 | 3 | 4697.9884 | 4697.9724 | 0 | 3.4 | 332 | CRagged | 705.7 | 684.93 | 684.93 | 15.41 | 123 | >sp\|P01876\|IGHA1_HUMAN Immunoglobulin heavy constant alpha 1 OS=Homo sapiens OX=9606 GN=IGHA1 PE=1 SV=2 | 255 | 20201118_ly_colostrum_1ng_glyco_hcd203040_compound15.17001.17001.3 | controllerType=0 controllerNumber=1 scan=17001 | 37.3512 |
| 10410:2 | 5 | R.LAGKPTHVN[+1216.423]VSVVMAEVDGTC[+57.021]Y.- | HexNAc(2)Hex(5) | C[+57], N[+1216] | 1782.288 | 2 | 3563.5688 | 3563.5649 | 0 | 1.1 | 332 | CRagged | 728.6 | 609.51 | 298.45 | 14.73 | 123 | >sp\|P01876\|IGHA1_HUMAN Immunoglobulin heavy constant alpha 1 OS=Homo sapiens OX=9606 GN=IGHA1 PE=1 SV=2 | 255 | 20201118_ly_colostrum_1ng_glyco_hcd203040_compound15.13676.13676.2 | controllerType=0 controllerNumber=1 scan=13676 | 30.6702 |
| 10393:3 | 5 | R.LAGKPTHVN[+1216.423]VSVVMAEVDGTC[+57.021]Y.- | HexNAc(2)Hex(5) | C[+57], N[+1216] | 1188.5275 | 3 | 3563.568 | 3563.5649 | 0 | 0.9 | 332 | CRagged | 730 | 730 | 396.81 | 14.95 | 123 | >sp\|P01876\|IGHA1_HUMAN Immunoglobulin heavy constant alpha 1 OS=Homo sapiens OX=9606 GN=IGHA1 PE=1 SV=2 | 255 | 20201118_ly_colostrum_1ng_glyco_hcd203040_compound15.13659.13659.3 | controllerType=0 controllerNumber=1 scan=13659 | 30.6345 |
| 09322:3 | 5 | R.LAGKPTHVN[+1809.666]VSVVMAEVDGTC[+57.021]Y.- | HexNAc(5)Hex(4)Fuc(1) | C[+57], N[+1810] | 1386.2783 | 3 | 4156.8202 | 4156.8081 | 0 | 2.9 | 332 | CRagged | 732.57 | 511.65 | 472.16 | 15.38 | 123 | >sp\|P01876\|IGHA1_HUMAN Immunoglobulin heavy constant alpha 1 OS=Homo sapiens OX=9606 GN=IGHA1 PE=1 SV=2 | 255 | 20201118_ly_colostrum_1ng_glyco_hcd203040_compound15.12546.12546.3 | controllerType=0 controllerNumber=1 scan=12546 | 28.4067 |
| 11863:3 | 5 | R.LAGKPTHVN[+2116.756]VSVVMAEVDGTC[+57.021]Y.- | HexNAc(5)Hex(5)NeuAc(1) | C[+57], N[+2117] | 1488.6414 | 3 | 4463.9097 | 4463.8985 | 0 | 2.5 | 332 | CRagged | 740.24 | 518.97 | 518.97 | 15.38 | 123 | >sp\|P01876\|IGHA1_HUMAN Immunoglobulin heavy constant alpha 1 OS=Homo sapiens OX=9606 GN=IGHA1 PE=1 SV=2 | 255 | 20201118_ly_colostrum_1ng_glyco_hcd203040_compound15.15187.15187.3 | controllerType=0 controllerNumber=1 scan=15187 | 33.657 |
| 09285:3 | 5 | R.LAGKPTHVN[+1540.529]VSVVMAEVDGTC[+57.021]Y.- | HexNAc(2)Hex(7) | C[+57], N[+1541] | 1296.5643 | 3 | 3887.6782 | 3887.6706 | 0 | 2 | 332 | CRagged | 743.8 | 743.8 | 441.24 | 15.15 | 123 | >sp\|P01876\|IGHA1_HUMAN Immunoglobulin heavy constant alpha 1 OS=Homo sapiens OX=9606 GN=IGHA1 PE=1 SV=2 | 255 | 20201118_ly_colostrum_1ng_glyco_hcd203040_compound15.12507.12507.3 | controllerType=0 controllerNumber=1 scan=12507 | 28.3307 |
| 10299:3 | 5 | R.LAGKPTHVN[+1825.661]VSVVMAEVDGTC[+57.021]Y.- | HexNAc(5)Hex(5) | C[+57], N[+1826] | 1391.6093 | 3 | 4172.8134 | 4172.8031 | 0 | 2.5 | 332 | CRagged | 744.67 | 744.67 | 470.05 | 15.38 | 123 | >sp\|P01876\|IGHA1_HUMAN Immunoglobulin heavy constant alpha 1 OS=Homo sapiens OX=9606 GN=IGHA1 PE=1 SV=2 | 255 | 20201118_ly_colostrum_1ng_glyco_hcd203040_compound15.13561.13561.3 | controllerType=0 controllerNumber=1 scan=13561 | 30.4409 |
| 09328:3 | 5 | R.LAGKPTHVN[+1663.608]VSVVMAEVDGTC[+57.021]Y.- | HexNAc(5)Hex(4) | C[+57], N[+1664] | 1337.5922 | 3 | 4010.7621 | 4010.7502 | 0 | 3 | 332 | CRagged | 749.78 | 749.78 | 391.17 | 15.38 | 123 | >sp\|P01876\|IGHA1_HUMAN Immunoglobulin heavy constant alpha 1 OS=Homo sapiens OX=9606 GN=IGHA1 PE=1 SV=2 | 255 | 20201118_ly_colostrum_1ng_glyco_hcd203040_compound15.12552.12552.3 | controllerType=0 controllerNumber=1 scan=12552 | 28.4179 |
| 11939:3 | 5 | R.LAGKPTHVN[+1872.651]VSVVMAEVDGTC[+57.021]Y.- | HexNAc(3)Hex(6)NeuAc(1) | C[+57], N[+1873] | 1407.2751 | 3 | 4219.8107 | 4219.7926 | 0 | 4.3 | 332 | CRagged | 751.54 | 700.37 | 474.33 | 16.11 | 123 | >sp\|P01876\|IGHA1_HUMAN Immunoglobulin heavy constant alpha 1 OS=Homo sapiens OX=9606 GN=IGHA1 PE=1 SV=2 | 255 | 20201118_ly_colostrum_1ng_glyco_hcd203040_compound15.15266.15266.3 | controllerType=0 controllerNumber=1 scan=15266 | 33.8184 |
| 10417:3 | 5 | R.LAGKPTHVN[+1501.555]VSVVMAEVDGTC[+57.021]Y.- | HexNAc(5)Hex(3) | C[+57], N[+1502] | 1283.5727 | 3 | 3848.7035 | 3848.6974 | 0 | 1.6 | 332 | CRagged | 752.53 | 752.53 | 752.53 | 16.15 | 123 | >sp\|P01876\|IGHA1_HUMAN Immunoglobulin heavy constant alpha 1 OS=Homo sapiens OX=9606 GN=IGHA1 PE=1 SV=2 | 255 | 20201118_ly_colostrum_1ng_glyco_hcd203040_compound15.13684.13684.3 | controllerType=0 controllerNumber=1 scan=13684 | 30.6855 |
| 10357:4 | 5 | R.LAGKPTHVN[+1971.719]VSVVMAEVDGTC[+57.021]Y.- | HexNAc(5)Hex(5)Fuc(1) | C[+57], N[+1972] | 1080.4729 | 4 | 4318.8698 | 4318.861 | 0 | 2 | 332 | CRagged | 755.9 | 755.9 | 436.98 | 16.31 | 123 | >sp\|P01876\|IGHA1_HUMAN Immunoglobulin heavy constant alpha 1 OS=Homo sapiens OX=9606 GN=IGHA1 PE=1 SV=2 | 255 | 20201118_ly_colostrum_1ng_glyco_hcd203040_compound15.13621.13621.4 | controllerType=0 controllerNumber=1 scan=13621 | 30.553 |
| 11913:3 | 5 | R.LAGKPTHVN[+1913.677]VSVVMAEVDGTC[+57.021]Y.- | HexNAc(4)Hex(5)NeuAc(1) | C[+57], N[+1914] | 1420.9501 | 3 | 4260.8358 | 4260.8191 | 0 | 3.9 | 332 | CRagged | 756.66 | 519.28 | 519.28 | 16.41 | 123 | >sp\|P01876\|IGHA1_HUMAN Immunoglobulin heavy constant alpha 1 OS=Homo sapiens OX=9606 GN=IGHA1 PE=1 SV=2 | 255 | 20201118_ly_colostrum_1ng_glyco_hcd203040_compound15.15239.15239.3 | controllerType=0 controllerNumber=1 scan=15239 | 33.7592 |
| 10366:3 | 5 | R.LAGKPTHVN[+1581.555]VSVVMAEVDGTC[+57.021]Y.- | HexNAc(3)Hex(6) | C[+57], N[+1582] | 1310.2394 | 3 | 3928.7036 | 3928.6971 | 0 | 1.7 | 332 | CRagged | 767.62 | 729.08 | 482.39 | 16.15 | 123 | >sp\|P01876\|IGHA1_HUMAN Immunoglobulin heavy constant alpha 1 OS=Homo sapiens OX=9606 GN=IGHA1 PE=1 SV=2 | 255 | 20201118_ly_colostrum_1ng_glyco_hcd203040_compound15.13630.13630.3 | controllerType=0 controllerNumber=1 scan=13630 | 30.573 |
| 11843:3 | 5 | R.LAGKPTHVN[+2262.814]VSVVMAEVDGTC[+57.021]Y.- | HexNAc(5)Hex(5)Fuc(1)NeuAc(1) | C[+57], N[+2263] | 1537.329 | 3 | 4609.9726 | 4609.9564 | 0 | 3.5 | 332 | CRagged | 771.56 | 629.07 | 629.07 | 16.41 | 123 | >sp\|P01876\|IGHA1_HUMAN Immunoglobulin heavy constant alpha 1 OS=Homo sapiens OX=9606 GN=IGHA1 PE=1 SV=2 | 255 | 20201118_ly_colostrum_1ng_glyco_hcd203040_compound15.15166.15166.3 | controllerType=0 controllerNumber=1 scan=15166 | 33.6087 |
| 10320:3 | 5 | R.LAGKPTHVN[+1663.608]VSVVMAEVDGTC[+57.021]Y.- | HexNAc(5)Hex(4) | C[+57], N[+1664] | 1337.5914 | 3 | 4010.7595 | 4010.7502 | 0 | 2.3 | 332 | CRagged | 773.33 | 773.33 | 501.53 | 16.38 | 123 | >sp\|P01876\|IGHA1_HUMAN Immunoglobulin heavy constant alpha 1 OS=Homo sapiens OX=9606 GN=IGHA1 PE=1 SV=2 | 255 | 20201118_ly_colostrum_1ng_glyco_hcd203040_compound15.13583.13583.3 | controllerType=0 controllerNumber=1 scan=13583 | 30.485 |
| 09394:3 | 5 | R.LAGKPTHVN[+1647.613]VSVVMAEVDGTC[+57.021]Y.- | HexNAc(5)Hex(3)Fuc(1) | C[+57], N[+1648] | 1332.2593 | 3 | 3994.7635 | 3994.7553 | 0 | 2 | 332 | CRagged | 773.81 | 582.19 | 582.19 | 16.15 | 123 | >sp\|P01876\|IGHA1_HUMAN Immunoglobulin heavy constant alpha 1 OS=Homo sapiens OX=9606 GN=IGHA1 PE=1 SV=2 | 255 | 20201118_ly_colostrum_1ng_glyco_hcd203040_compound15.12621.12621.3 | controllerType=0 controllerNumber=1 scan=12621 | 28.5735 |
| 13601:3 | 5 | R.LAGKPTHVN[+2553.910]VSVVMAEVDGTC[+57.021]Y.- | HexNAc(5)Hex(5)Fuc(1)NeuAc(2) | C[+57], N[+2554] | 1634.3622 | 3 | 4901.0722 | 4901.0518 | 0 | 4.2 | 332 | CRagged | 782.21 | 500.58 | 500.58 | 16.11 | 123 | >sp\|P01876\|IGHA1_HUMAN Immunoglobulin heavy constant alpha 1 OS=Homo sapiens OX=9606 GN=IGHA1 PE=1 SV=2 | 255 | 20201118_ly_colostrum_1ng_glyco_hcd203040_compound15.16993.16993.3 | controllerType=0 controllerNumber=1 scan=16993 | 37.3177 |
| 09324:3 | 5 | R.LAGKPTHVN[+1809.666]VSVVMAEVDGTC[+57.021]Y.- | HexNAc(5)Hex(4)Fuc(1) | C[+57], N[+1810] | 1386.2783 | 3 | 4156.8202 | 4156.8081 | 0 | 2.9 | 332 | CRagged | 787.03 | 787.03 | 527.31 | 16.38 | 123 | >sp\|P01876\|IGHA1_HUMAN Immunoglobulin heavy constant alpha 1 OS=Homo sapiens OX=9606 GN=IGHA1 PE=1 SV=2 | 255 | 20201118_ly_colostrum_1ng_glyco_hcd203040_compound15.12548.12548.3 | controllerType=0 controllerNumber=1 scan=12548 | 28.4115 |
| 09299:3 | 5 | R.LAGKPTHVN[+1378.476]VSVVMAEVDGTC[+57.021]Y.- | HexNAc(2)Hex(6) | C[+57], N[+1378] | 1242.5473 | 3 | 3725.6273 | 3725.6178 | 0 | 2.6 | 332 | CRagged | 792.06 | 792.06 | 453.12 | 16.38 | 123 | >sp\|P01876\|IGHA1_HUMAN Immunoglobulin heavy constant alpha 1 OS=Homo sapiens OX=9606 GN=IGHA1 PE=1 SV=2 | 255 | 20201118_ly_colostrum_1ng_glyco_hcd203040_compound15.12522.12522.3 | controllerType=0 controllerNumber=1 scan=12522 | 28.3568 |
| 10332:3 | 5 | R.LAGKPTHVN[+1809.666]VSVVMAEVDGTC[+57.021]Y.- | HexNAc(5)Hex(4)Fuc(1) | C[+57], N[+1810] | 1386.2767 | 3 | 4156.8155 | 4156.8081 | 0 | 1.8 | 332 | CRagged | 792.76 | 728.03 | 509.99 | 16.15 | 123 | >sp\|P01876\|IGHA1_HUMAN Immunoglobulin heavy constant alpha 1 OS=Homo sapiens OX=9606 GN=IGHA1 PE=1 SV=2 | 255 | 20201118_ly_colostrum_1ng_glyco_hcd203040_compound15.13595.13595.3 | controllerType=0 controllerNumber=1 scan=13595 | 30.5105 |
| 13591:4 | 5 | R.LAGKPTHVN[+2553.910]VSVVMAEVDGTC[+57.021]Y.- | HexNAc(5)Hex(5)Fuc(1)NeuAc(2) | C[+57], N[+2554] | 1226.0235 | 4 | 4901.0721 | 4901.0518 | 0 | 4.1 | 332 | CRagged | 796.69 | 565.07 | 565.07 | 16.27 | 123 | >sp\|P01876\|IGHA1_HUMAN Immunoglobulin heavy constant alpha 1 OS=Homo sapiens OX=9606 GN=IGHA1 PE=1 SV=2 | 255 | 20201118_ly_colostrum_1ng_glyco_hcd203040_compound15.16983.16983.4 | controllerType=0 controllerNumber=1 scan=16983 | 37.2957 |

Table S4. Cont.

| Query #:z | Protein Rank | Glycopeptide | Glycan | Modification Type(s) | Observed m/z | z | Observed (M+H) | Calc. mass (M+H) | Off-by-x error | Mass error (ppm) | StartPosition | Cleavage | Score | Delta | Delta Mod | \|Log Prob\| | # of unique peptides | Protein Name | Protein DB number | Comment | Scan # | Scan Time |
|---|---|---|---|---|---|---|---|---|---|---|---|---|---|---|---|---|---|---|---|---|---|---|
| 10419:3 | 5 | R.LAGKPTHVN[+1501.555]VSVVMAEVDGTC[+57.021]Y.- | HexNAc(5)Hex(3) | C[+57], N[+1502] | 1283.5727 | 3 | 3848.7035 | 3848.6974 | 0 | 1.6 | 332 | CRagged | 799.71 | 678.38 | 678.38 | 16.15 | 123 | >sp\|P01876\|IGHA1_HUMAN Immunoglobulin heavy constant alpha 1 OS=Homo sapiens OX=9606 GN=IGHA1 PE=1 SV=2 | 255 | 20201118_ly_colostrum_1ng_glyco_hcd203040_compound15.13686.13686.3 | controllerType=0 controllerNumber=1 scan=13686 | 30.691 |
| 11853:4 | 5 | R.LAGKPTHVN[+2262.814]VSVVMAEVDGTC[+57.021]Y.- | HexNAc(5)Hex(5)Fuc(1)NeuAc(1) | C[+57], N[+2263] | 1153.2482 | 4 | 4609.971 | 4609.9564 | 0 | 3.2 | 332 | CRagged | 807.16 | 593.63 | 593.63 | 17.57 | 123 | >sp\|P01876\|IGHA1_HUMAN Immunoglobulin heavy constant alpha 1 OS=Homo sapiens OX=9606 GN=IGHA1 PE=1 SV=2 | 255 | 20201118_ly_colostrum_1ng_glyco_hcd203040_compound15.15176.15176.4 | controllerType=0 controllerNumber=1 scan=15176 | 33.6286 |
| 09348:3 | 5 | R.LAGKPTHVN[+1216.423]VSVVMAEVDGTC[+57.021]Y.- | HexNAc(2)Hex(5) | C[+57], N[+1216] | 1188.5286 | 3 | 3563.5713 | 3563.5649 | 0 | 1.8 | 332 | CRagged | 815.09 | 815.09 | 464.68 | 17.15 | 123 | >sp\|P01876\|IGHA1_HUMAN Immunoglobulin heavy constant alpha 1 OS=Homo sapiens OX=9606 GN=IGHA1 PE=1 SV=2 | 255 | 20201118_ly_colostrum_1ng_glyco_hcd203040_compound15.12573.12573.3 | controllerType=0 controllerNumber=1 scan=12573 | 28.4615 |
| 10486:3 | 5 | R.LAGKPTHVN[+1444.534]VSVVMAEVDGTC[+57.021]Y.- | HexNAc(4)Hex(3)Fuc(1) | C[+57], N[+1445] | 1264.5674 | 3 | 3791.6878 | 3791.6759 | 0 | 3.1 | 332 | CRagged | 825.25 | 699.33 | 699.33 | 17.41 | 123 | >sp\|P01876\|IGHA1_HUMAN Immunoglobulin heavy constant alpha 1 OS=Homo sapiens OX=9606 GN=IGHA1 PE=1 SV=2 | 255 | 20201118_ly_colostrum_1ng_glyco_hcd203040_compound15.13756.13756.3 | controllerType=0 controllerNumber=1 scan=13756 | 30.8448 |
| 10291:3 | 5 | R.LAGKPTHVN[+1540.529]VSVVMAEVDGTC[+57.021]Y.- | HexNAc(2)Hex(7) | C[+57], N[+1541] | 1296.5649 | 3 | 3887.6801 | 3887.6706 | 0 | 2.4 | 332 | CRagged | 840.8 | 840.8 | 493.19 | 17.38 | 123 | >sp\|P01876\|IGHA1_HUMAN Immunoglobulin heavy constant alpha 1 OS=Homo sapiens OX=9606 GN=IGHA1 PE=1 SV=2 | 255 | 20201118_ly_colostrum_1ng_glyco_hcd203040_compound15.13553.13553.3 | controllerType=0 controllerNumber=1 scan=13553 | 30.4193 |
| 09329:3 | 5 | R.LAGKPTHVN[+1216.423]VSVVMAEVDGTC[+57.021]Y.- | HexNAc(2)Hex(5) | C[+57], N[+1216] | 1188.5308 | 3 | 3563.5779 | 3563.5649 | 0 | 3.6 | 332 | CRagged | 846.17 | 846.17 | 520.78 | 17.41 | 123 | >sp\|P01876\|IGHA1_HUMAN Immunoglobulin heavy constant alpha 1 OS=Homo sapiens OX=9606 GN=IGHA1 PE=1 SV=2 | 255 | 20201118_ly_colostrum_1ng_glyco_hcd203040_compound15.12553.12553.3 | controllerType=0 controllerNumber=1 scan=12553 | 28.4199 |
| 10425:3 | 5 | R.LAGKPTHVN[+1647.613]VSVVMAEVDGTC[+57.021]Y.- | HexNAc(5)Hex(3)Fuc(1) | C[+57], N[+1648] | 1332.2593 | 3 | 3994.7635 | 3994.7553 | 0 | 2 | 332 | CRagged | 848.26 | 848.26 | 848.26 | 17.15 | 123 | >sp\|P01876\|IGHA1_HUMAN Immunoglobulin heavy constant alpha 1 OS=Homo sapiens OX=9606 GN=IGHA1 PE=1 SV=2 | 255 | 20201118_ly_colostrum_1ng_glyco_hcd203040_compound15.13692.13692.3 | controllerType=0 controllerNumber=1 scan=13692 | 30.7024 |
| 10487:3 | 5 | R.LAGKPTHVN[+1054.370]VSVVMAEVDGTC[+57.021]Y.- | HexNAc(2)Hex(4) | C[+57], N[+1054] | 1134.5113 | 3 | 3401.5192 | 3401.5121 | 0 | 2.1 | 332 | CRagged | 851.98 | 851.98 | 517.03 | 18.38 | 123 | >sp\|P01876\|IGHA1_HUMAN Immunoglobulin heavy constant alpha 1 OS=Homo sapiens OX=9606 GN=IGHA1 PE=1 SV=2 | 255 | 20201118_ly_colostrum_1ng_glyco_hcd203040_compound15.13757.13757.3 | controllerType=0 controllerNumber=1 scan=13757 | 30.848 |
| 09301:3 | 5 | R.LAGKPTHVN[+1378.476]VSVVMAEVDGTC[+57.021]Y.- | HexNAc(2)Hex(6) | C[+57], N[+1378] | 1242.5473 | 3 | 3725.6273 | 3725.6178 | 0 | 2.6 | 332 | CRagged | 856.29 | 841.51 | 504.54 | 18.38 | 123 | >sp\|P01876\|IGHA1_HUMAN Immunoglobulin heavy constant alpha 1 OS=Homo sapiens OX=9606 GN=IGHA1 PE=1 SV=2 | 255 | 20201118_ly_colostrum_1ng_glyco_hcd203040_compound15.12524.12524.3 | controllerType=0 controllerNumber=1 scan=12524 | 28.3624 |
| 10293:3 | 5 | R.LAGKPTHVN[+1540.529]VSVVMAEVDGTC[+57.021]Y.- | HexNAc(2)Hex(7) | C[+57], N[+1541] | 1296.5649 | 3 | 3887.6801 | 3887.6706 | 0 | 2.4 | 332 | CRagged | 856.9 | 807.78 | 503.15 | 18.38 | 123 | >sp\|P01876\|IGHA1_HUMAN Immunoglobulin heavy constant alpha 1 OS=Homo sapiens OX=9606 GN=IGHA1 PE=1 SV=2 | 255 | 20201118_ly_colostrum_1ng_glyco_hcd203040_compound15.13555.13555.3 | controllerType=0 controllerNumber=1 scan=13555 | 30.4253 |
| 10270:3 | 5 | R.LAGKPTHVN[+1702.581]VSVVMAEVDGTC[+57.021]Y.- | HexNAc(2)Hex(8) | C[+57], N[+1703] | 1350.5829 | 3 | 4049.7343 | 4049.7234 | 0 | 2.7 | 332 | CRagged | 858.04 | 829.2 | 544.82 | 18.38 | 123 | >sp\|P01876\|IGHA1_HUMAN Immunoglobulin heavy constant alpha 1 OS=Homo sapiens OX=9606 GN=IGHA1 PE=1 SV=2 | 255 | 20201118_ly_colostrum_1ng_glyco_hcd203040_compound15.13531.13531.3 | controllerType=0 controllerNumber=1 scan=13531 | 30.3754 |
| 09332:3 | 5 | R.LAGKPTHVN[+1216.423]VSVVMAEVDGTC[+57.021]Y.- | HexNAc(2)Hex(5) | C[+57], N[+1216] | 1188.5308 | 3 | 3563.5779 | 3563.5649 | 0 | 3.6 | 332 | CRagged | 864.27 | 825.62 | 517.4 | 18.41 | 123 | >sp\|P01876\|IGHA1_HUMAN Immunoglobulin heavy constant alpha 1 OS=Homo sapiens OX=9606 GN=IGHA1 PE=1 SV=2 | 255 | 20201118_ly_colostrum_1ng_glyco_hcd203040_compound15.12556.12556.3 | controllerType=0 controllerNumber=1 scan=12556 | 28.4283 |
| 11860:3 | 5 | R.LAGKPTHVN[+2116.756]VSVVMAEVDGTC[+57.021]Y.- | HexNAc(5)Hex(5)NeuAc(1) | C[+57], N[+2117] | 1488.6414 | 3 | 4463.9097 | 4463.8985 | 0 | 2.5 | 332 | CRagged | 864.58 | 554.85 | 554.85 | 18.38 | 123 | >sp\|P01876\|IGHA1_HUMAN Immunoglobulin heavy constant alpha 1 OS=Homo sapiens OX=9606 GN=IGHA1 PE=1 SV=2 | 255 | 20201118_ly_colostrum_1ng_glyco_hcd203040_compound15.15184.15184.3 | controllerType=0 controllerNumber=1 scan=15184 | 33.6471 |
| 11840:3 | 5 | R.LAGKPTHVN[+2262.814]VSVVMAEVDGTC[+57.021]Y.- | HexNAc(5)Hex(5)Fuc(1)NeuAc(1) | C[+57], N[+2263] | 1537.329 | 3 | 4609.9726 | 4609.9564 | 0 | 3.5 | 332 | CRagged | 868.23 | 590.33 | 590.33 | 18.41 | 123 | >sp\|P01876\|IGHA1_HUMAN Immunoglobulin heavy constant alpha 1 OS=Homo sapiens OX=9606 GN=IGHA1 PE=1 SV=2 | 255 | 20201118_ly_colostrum_1ng_glyco_hcd203040_compound15.15163.15163.3 | controllerType=0 controllerNumber=1 scan=15163 | 33.5972 |
| 10488:3 | 5 | R.LAGKPTHVN[+1095.397]VSVVMAEVDGTC[+57.021]Y.- | HexNAc(3)Hex(3) | C[+57], N[+1095] | 1148.1877 | 3 | 3442.5484 | 3442.5387 | 0 | 2.8 | 332 | CRagged | 869.88 | 825.85 | 825.85 | 18.38 | 123 | >sp\|P01876\|IGHA1_HUMAN Immunoglobulin heavy constant alpha 1 OS=Homo sapiens OX=9606 GN=IGHA1 PE=1 SV=2 | 255 | 20201118_ly_colostrum_1ng_glyco_hcd203040_compound15.13758.13758.3 | controllerType=0 controllerNumber=1 scan=13758 | 30.8499 |
| 10319:3 | 5 | R.LAGKPTHVN[+1378.476]VSVVMAEVDGTC[+57.021]Y.- | HexNAc(2)Hex(6) | C[+57], N[+1378] | 1242.5469 | 3 | 3725.6262 | 3725.6178 | 0 | 2.3 | 332 | CRagged | 870.82 | 850.36 | 554.09 | 18.38 | 123 | >sp\|P01876\|IGHA1_HUMAN Immunoglobulin heavy constant alpha 1 OS=Homo sapiens OX=9606 GN=IGHA1 PE=1 SV=2 | 255 | 20201118_ly_colostrum_1ng_glyco_hcd203040_compound15.13582.13582.3 | controllerType=0 controllerNumber=1 scan=13582 | 30.4814 |
| 10422:3 | 5 | R.LAGKPTHVN[+1647.613]VSVVMAEVDGTC[+57.021]Y.- | HexNAc(5)Hex(3)Fuc(1) | C[+57], N[+1648] | 1332.2593 | 3 | 3994.7635 | 3994.7553 | 0 | 2 | 332 | CRagged | 874.69 | 827.85 | 827.85 | 18.15 | 123 | >sp\|P01876\|IGHA1_HUMAN Immunoglobulin heavy constant alpha 1 OS=Homo sapiens OX=9606 GN=IGHA1 PE=1 SV=2 | 255 | 20201118_ly_colostrum_1ng_glyco_hcd203040_compound15.13689.13689.3 | controllerType=0 controllerNumber=1 scan=13689 | 30.6953 |
| 10317:3 | 5 | R.LAGKPTHVN[+1663.608]VSVVMAEVDGTC[+57.021]Y.- | HexNAc(5)Hex(4) | C[+57], N[+1664] | 1337.5914 | 3 | 4010.7595 | 4010.7502 | 0 | 2.3 | 332 | CRagged | 878.44 | 738.45 | 570.13 | 18.38 | 123 | >sp\|P01876\|IGHA1_HUMAN Immunoglobulin heavy constant alpha 1 OS=Homo sapiens OX=9606 GN=IGHA1 PE=1 SV=2 | 255 | 20201118_ly_colostrum_1ng_glyco_hcd203040_compound15.13580.13580.3 | controllerType=0 controllerNumber=1 scan=13580 | 30.4754 |
| 09396:3 | 5 | R.LAGKPTHVN[+1647.613]VSVVMAEVDGTC[+57.021]Y.- | HexNAc(5)Hex(3)Fuc(1) | C[+57], N[+1648] | 1332.2593 | 3 | 3994.7635 | 3994.7553 | 0 | 2 | 332 | CRagged | 878.98 | 750.83 | 750.83 | 18.15 | 123 | >sp\|P01876\|IGHA1_HUMAN Immunoglobulin heavy constant alpha 1 OS=Homo sapiens OX=9606 GN=IGHA1 PE=1 SV=2 | 255 | 20201118_ly_colostrum_1ng_glyco_hcd203040_compound15.12623.12623.3 | controllerType=0 controllerNumber=1 scan=12623 | 28.5847 |
| 10268:3 | 5 | R.LAGKPTHVN[+1702.581]VSVVMAEVDGTC[+57.021]Y.- | HexNAc(2)Hex(8) | C[+57], N[+1703] | 1350.5829 | 3 | 4049.7343 | 4049.7234 | 0 | 2.7 | 332 | CRagged | 905.92 | 905.92 | 603.3 | 19.38 | 123 | >sp\|P01876\|IGHA1_HUMAN Immunoglobulin heavy constant alpha 1 OS=Homo sapiens OX=9606 GN=IGHA1 PE=1 SV=2 | 255 | 20201118_ly_colostrum_1ng_glyco_hcd203040_compound15.13529.13529.3 | controllerType=0 controllerNumber=1 scan=13529 | 30.3697 |
| 09350:3 | 5 | R.LAGKPTHVN[+1216.423]VSVVMAEVDGTC[+57.021]Y.- | HexNAc(2)Hex(5) | C[+57], N[+1216] | 1188.5286 | 3 | 3563.5713 | 3563.5649 | 0 | 1.8 | 332 | CRagged | 911.41 | 860.61 | 525.7 | 19.15 | 123 | >sp\|P01876\|IGHA1_HUMAN Immunoglobulin heavy constant alpha 1 OS=Homo sapiens OX=9606 GN=IGHA1 PE=1 SV=2 | 255 | 20201118_ly_colostrum_1ng_glyco_hcd203040_compound15.12575.12575.3 | controllerType=0 controllerNumber=1 scan=12575 | 28.468 |
| 10316:3 | 5 | R.LAGKPTHVN[+1378.476]VSVVMAEVDGTC[+57.021]Y.- | HexNAc(2)Hex(6) | C[+57], N[+1378] | 1242.5469 | 3 | 3725.6262 | 3725.6178 | 0 | 2.3 | 332 | CRagged | 911.69 | 789.49 | 554.67 | 19.38 | 123 | >sp\|P01876\|IGHA1_HUMAN Immunoglobulin heavy constant alpha 1 OS=Homo sapiens OX=9606 GN=IGHA1 PE=1 SV=2 | 255 | 20201118_ly_colostrum_1ng_glyco_hcd203040_compound15.13579.13579.3 | controllerType=0 controllerNumber=1 scan=13579 | 30.4743 |
| 10396:3 | 5 | R.LAGKPTHVN[+1216.423]VSVVMAEVDGTC[+57.021]Y.- | HexNAc(2)Hex(5) | C[+57], N[+1216] | 1188.5275 | 3 | 3563.568 | 3563.5649 | 0 | 0.9 | 332 | CRagged | 912.44 | 869.79 | 561.38 | 18.95 | 123 | >sp\|P01876\|IGHA1_HUMAN Immunoglobulin heavy constant alpha 1 OS=Homo sapiens OX=9606 GN=IGHA1 PE=1 SV=2 | 255 | 20201118_ly_colostrum_1ng_glyco_hcd203040_compound15.13662.13662.3 | controllerType=0 controllerNumber=1 scan=13662 | 30.6398 |
| 10329:3 | 5 | R.LAGKPTHVN[+1809.666]VSVVMAEVDGTC[+57.021]Y.- | HexNAc(5)Hex(4)Fuc(1) | C[+57], N[+1810] | 1386.2767 | 3 | 4156.8155 | 4156.8081 | 0 | 1.8 | 332 | CRagged | 917.72 | 917.72 | 608.22 | 19.15 | 123 | >sp\|P01876\|IGHA1_HUMAN Immunoglobulin heavy constant alpha 1 OS=Homo sapiens OX=9606 GN=IGHA1 PE=1 SV=2 | 255 | 20201118_ly_colostrum_1ng_glyco_hcd203040_compound15.13592.13592.3 | controllerType=0 controllerNumber=1 scan=13592 | 30.4998 |
| 10360:4 | 5 | R.LAGKPTHVN[+1971.719]VSVVMAEVDGTC[+57.021]Y.- | HexNAc(5)Hex(5)Fuc(1) | C[+57], N[+1972] | 1080.4729 | 4 | 4318.8698 | 4318.861 | 0 | 2 | 332 | CRagged | 953.04 | 947.27 | 586 | 20.31 | 123 | >sp\|P01876\|IGHA1_HUMAN Immunoglobulin heavy constant alpha 1 OS=Homo sapiens OX=9606 GN=IGHA1 PE=1 SV=2 | 255 | 20201118_ly_colostrum_1ng_glyco_hcd203040_compound15.13624.13624.4 | controllerType=0 controllerNumber=1 scan=13624 | 30.5622 |
| 08236:3 | 5 | R.LAGKPTHVN[+1663.608]VSVVM[+15.995]AEVDGTC[+57.021]Y.- | HexNAc(5)Hex(4) | C[+57], M[+16], N[+1664] | 1342.925 | 3 | 4026.7604 | 4026.7452 | 0 | 3.8 | 332 | CRagged | 201.54 | 191.93 | 191.93 | 5.66 | 123 | >sp\|P01876\|IGHA1_HUMAN Immunoglobulin heavy constant alpha 1 OS=Homo sapiens OX=9606 GN=IGHA1 PE=1 SV=2 | 255 | 20201118_ly_colostrum_1ng_glyco_hcd203040_compound15.11417.11417.3 | controllerType=0 controllerNumber=1 scan=11417 | 26.1841 |
| 09367:3 | 5 | R.LAGKPTHVN[+2262.814]VSVVM[+15.995]AEVDGTC[+57.021]Y.- | HexNAc(5)Hex(5)Fuc(1)NeuAc(1) | C[+57], M[+16], N[+2263] | 1542.6581 | 3 | 4625.9599 | 4625.9513 | 0 | 1.9 | 332 | CRagged | 212.49 | 212.49 | 120.04 | 5.4 | 123 | >sp\|P01876\|IGHA1_HUMAN Immunoglobulin heavy constant alpha 1 OS=Homo sapiens OX=9606 GN=IGHA1 PE=1 SV=2 | 255 | 20201118_ly_colostrum_1ng_glyco_hcd203040_compound15.12592.12592.3 | controllerType=0 controllerNumber=1 scan=12592 | 28.5041 |
| 09369:3 | 5 | R.LAGKPTHVN[+2262.814]VSVVM[+15.995]AEVDGTC[+57.021]Y.- | HexNAc(5)Hex(5)Fuc(1)NeuAc(1) | C[+57], M[+16], N[+2263] | 1542.6581 | 3 | 4625.9599 | 4625.9513 | 0 | 1.9 | 332 | CRagged | 242.5 | 242.5 | 180.98 | 5.54 | 123 | >sp\|P01876\|IGHA1_HUMAN Immunoglobulin heavy constant alpha 1 OS=Homo sapiens OX=9606 GN=IGHA1 PE=1 SV=2 | 255 | 20201118_ly_colostrum_1ng_glyco_hcd203040_compound15.12594.12594.3 | controllerType=0 controllerNumber=1 scan=12594 | 28.5081 |
| 08499:3 | 5 | R.LAGKPTHVN[+1663.608]VSVVM[+15.995]AEVDGTC[+57.021]Y.- | HexNAc(5)Hex(4) | C[+57], M[+16], N[+1664] | 1342.9233 | 3 | 4026.7553 | 4026.7452 | 0 | 2.5 | 332 | CRagged | 246.08 | 246.08 | 246.08 | 7.24 | 123 | >sp\|P01876\|IGHA1_HUMAN Immunoglobulin heavy constant alpha 1 OS=Homo sapiens OX=9606 GN=IGHA1 PE=1 SV=2 | 255 | 20201118_ly_colostrum_1ng_glyco_hcd203040_compound15.11690.11690.3 | controllerType=0 controllerNumber=1 scan=11690 | 26.7185 |
| 08094:3 | 5 | R.LAGKPTHVN[+1809.666]VSVVM[+15.995]AEVDGTC[+57.021]Y.- | HexNAc(5)Hex(4)Fuc(1) | C[+57], M[+16], N[+1810] | 1391.6086 | 3 | 4172.8112 | 4172.8031 | 0 | 1.9 | 332 | CRagged | 248.73 | 241.14 | 66.76 | 7.01 | 123 | >sp\|P01876\|IGHA1_HUMAN Immunoglobulin heavy constant alpha 1 OS=Homo sapiens OX=9606 GN=IGHA1 PE=1 SV=2 | 255 | 20201118_ly_colostrum_1ng_glyco_hcd203040_compound15.11270.11270.3 | controllerType=0 controllerNumber=1 scan=11270 | 25.8914 |
| 08238:3 | 5 | R.LAGKPTHVN[+1663.608]VSVVM[+15.995]AEVDGTC[+57.021]Y.- | HexNAc(5)Hex(4) | C[+57], M[+16], N[+1664] | 1342.925 | 3 | 4026.7604 | 4026.7452 | 0 | 3.8 | 332 | CRagged | 277.15 | 277.15 | 277.15 | 7.48 | 123 | >sp\|P01876\|IGHA1_HUMAN Immunoglobulin heavy constant alpha 1 OS=Homo sapiens OX=9606 GN=IGHA1 PE=1 SV=2 | 255 | 20201118_ly_colostrum_1ng_glyco_hcd203040_compound15.11419.11419.3 | controllerType=0 controllerNumber=1 scan=11419 | 26.188 |
| 08654:3 | 5 | R.LAGKPTHVN[+1809.666]VSVVM[+15.995]AEVDGTC[+57.021]Y.- | HexNAc(5)Hex(4)Fuc(1) | C[+57], M[+16], N[+1810] | 1391.6104 | 3 | 4172.8167 | 4172.8031 | 0 | 3.3 | 332 | CRagged | 346.65 | 268.7 | 226.12 | 7.99 | 123 | >sp\|P01876\|IGHA1_HUMAN Immunoglobulin heavy constant alpha 1 OS=Homo sapiens OX=9606 GN=IGHA1 PE=1 SV=2 | 255 | 20201118_ly_colostrum_1ng_glyco_hcd203040_compound15.11851.11851.3 | controllerType=0 controllerNumber=1 scan=11851 | 27.0369 |
| 08173:3 | 5 | R.LAGKPTHVN[+1647.613]VSVVM[+15.995]AEVDGTC[+57.021]Y.- | HexNAc(5)Hex(3)Fuc(1) | C[+57], M[+16], N[+1648] | 1337.5914 | 3 | 4010.7595 | 4010.7502 | 0 | 2.3 | 332 | CRagged | 360.53 | 360.53 | 262.72 | 8.61 | 123 | >sp\|P01876\|IGHA1_HUMAN Immunoglobulin heavy constant alpha 1 OS=Homo sapiens OX=9606 GN=IGHA1 PE=1 SV=2 | 255 | 20201118_ly_colostrum_1ng_glyco_hcd203040_compound15.11352.11352.3 | controllerType=0 controllerNumber=1 scan=11352 | 26.0579 |
| 08171:3 | 5 | R.LAGKPTHVN[+1647.613]VSVVM[+15.995]AEVDGTC[+57.021]Y.- | HexNAc(5)Hex(3)Fuc(1) | C[+57], M[+16], N[+1648] | 1337.5914 | 3 | 4010.7595 | 4010.7502 | 0 | 2.3 | 332 | CRagged | 385.38 | 296.15 | 217.32 | 7.99 | 123 | >sp\|P01876\|IGHA1_HUMAN Immunoglobulin heavy constant alpha 1 OS=Homo sapiens OX=9606 GN=IGHA1 PE=1 SV=2 | 255 | 20201118_ly_colostrum_1ng_glyco_hcd203040_compound15.11350.11350.3 | controllerType=0 controllerNumber=1 scan=11350 | 26.0528 |
| 08827:3 | 5 | R.LAGKPTHVN[+1216.423]VSVVM[+15.995]AEVDGTC[+57.021]Y.- | HexNAc(2)Hex(5) | C[+57], M[+16], N[+1216] | 1193.86 | 3 | 3579.5656 | 3579.5599 | 0 | 1.6 | 332 | CRagged | 409.84 | 409.84 | 409.84 | 10.03 | 123 | >sp\|P01876\|IGHA1_HUMAN Immunoglobulin heavy constant alpha 1 OS=Homo sapiens OX=9606 GN=IGHA1 PE=1 SV=2 | 255 | 20201118_ly_colostrum_1ng_glyco_hcd203040_compound15.12031.12031.3 | controllerType=0 controllerNumber=1 scan=12031 | 27.3931 |
| 08656:3 | 5 | R.LAGKPTHVN[+1809.666]VSVVM[+15.995]AEVDGTC[+57.021]Y.- | HexNAc(5)Hex(4)Fuc(1) | C[+57], M[+16], N[+1810] | 1391.6104 | 3 | 4172.8167 | 4172.8031 | 0 | 3.3 | 332 | CRagged | 441.68 | 321.44 | 298.94 | 8.68 | 123 | >sp\|P01876\|IGHA1_HUMAN Immunoglobulin heavy constant alpha 1 OS=Homo sapiens OX=9606 GN=IGHA1 PE=1 SV=2 | 255 | 20201118_ly_colostrum_1ng_glyco_hcd203040_compound15.11853.11853.3 | controllerType=0 controllerNumber=1 scan=11853 | 27.0407 |
| 08671:3 | 5 | R.LAGKPTHVN[+1216.423]VSVVM[+15.995]AEVDGTC[+57.021]Y.- | HexNAc(2)Hex(5) | C[+57], M[+16], N[+1216] | 1193.8621 | 3 | 3579.5718 | 3579.5599 | 0 | 3.3 | 332 | CRagged | 479.42 | 227.37 | 227.37 | 8.06 | 123 | >sp\|P01876\|IGHA1_HUMAN Immunoglobulin heavy constant alpha 1 OS=Homo sapiens OX=9606 GN=IGHA1 PE=1 SV=2 | 255 | 20201118_ly_colostrum_1ng_glyco_hcd203040_compound15.11869.11869.3 | controllerType=0 controllerNumber=1 scan=11869 | 27.0785 |
| 08829:3 | 5 | R.LAGKPTHVN[+1216.423]VSVVM[+15.995]AEVDGTC[+57.021]Y.- | HexNAc(2)Hex(5) | C[+57], M[+16], N[+1216] | 1193.86 | 3 | 3579.5656 | 3579.5599 | 0 | 1.6 | 332 | CRagged | 495.51 | 470.74 | 470.74 | 10.3 | 123 | >sp\|P01876\|IGHA1_HUMAN Immunoglobulin heavy constant alpha 1 OS=Homo sapiens OX=9606 GN=IGHA1 PE=1 SV=2 | 255 | 20201118_ly_colostrum_1ng_glyco_hcd203040_compound15.12033.12033.3 | controllerType=0 controllerNumber=1 scan=12033 | 27.397 |
| 08669:3 | 5 | R.LAGKPTHVN[+1216.423]VSVVM[+15.995]AEVDGTC[+57.021]Y.- | HexNAc(2)Hex(5) | C[+57], M[+16], N[+1216] | 1193.8621 | 3 | 3579.5718 | 3579.5599 | 0 | 3.3 | 332 | CRagged | 523.42 | 402.86 | 402.86 | 10.72 | 123 | >sp\|P01876\|IGHA1_HUMAN Immunoglobulin heavy constant alpha 1 OS=Homo sapiens OX=9606 GN=IGHA1 PE=1 SV=2 | 255 | 20201118_ly_colostrum_1ng_glyco_hcd203040_compound15.11867.11867.3 | controllerType=0 controllerNumber=1 scan=11867 | 27.0719 |
| 08097:3 | 5 | R.LAGKPTHVN[+1809.666]VSVVM[+15.995]AEVDGTC[+57.021]Y.- | HexNAc(5)Hex(4)Fuc(1) | C[+57], M[+16], N[+1810] | 1391.6086 | 3 | 4172.8112 | 4172.8031 | 0 | 1.9 | 332 | CRagged | 596.05 | 394.13 | 295.64 | 12.15 | 123 | >sp\|P01876\|IGHA1_HUMAN Immunoglobulin heavy constant alpha 1 OS=Homo sapiens OX=9606 GN=IGHA1 PE=1 SV=2 | 255 | 20201118_ly_colostrum_1ng_glyco_hcd203040_compound15.11273.11273.3 | controllerType=0 controllerNumber=1 scan=11273 | 25.8957 |
| 08120:3 | 5 | R.LAGKPTHVN[+1216.423]VSVVM[+15.995]AEVDGTC[+57.021]Y.- | HexNAc(2)Hex(5) | C[+57], M[+16], N[+1216] | 1193.8594 | 3 | 3579.5637 | 3579.5599 | 0 | 1.1 | 332 | CRagged | 726.4 | 726.4 | 726.4 | 15.15 | 123 | >sp\|P01876\|IGHA1_HUMAN Immunoglobulin heavy constant alpha 1 OS=Homo sapiens OX=9606 GN=IGHA1 PE=1 SV=2 | 255 | 20201118_ly_colostrum_1ng_glyco_hcd203040_compound15.11297.11297.3 | controllerType=0 controllerNumber=1 scan=11297 | 25.947 |
| 08123:3 | 5 | R.LAGKPTHVN[+1216.423]VSVVM[+15.995]AEVDGTC[+57.021]Y.- | HexNAc(2)Hex(5) | C[+57], M[+16], N[+1216] | 1193.8594 | 3 | 3579.5637 | 3579.5599 | 0 | 1.1 | 332 | CRagged | 844.45 | 844.45 | 844.45 | 17.15 | 123 | >sp\|P01876\|IGHA1_HUMAN Immunoglobulin heavy constant alpha 1 OS=Homo sapiens OX=9606 GN=IGHA1 PE=1 SV=2 | 255 | 20201118_ly_colostrum_1ng_glyco_hcd203040_compound15.11300.11300.3 | controllerType=0 controllerNumber=1 scan=11300 | 25.952 |
| 21472:4 | 6 | A.[+42.011]SPTSPKVFPLSLDSTPQDGNVVVAC[+57.021]LVQGFFPQEPLSVTWSESGQN | HexNAc(3)Hex(5) | .[42], C[+57], N[+1420] | 1709.5382 | 4 | 6835.1309 | 6835.1864 | 0 | -8.1 | 2 | NRagged | 286.47 | 263.65 | 263.65 | 3.84 | 97 | >sp\|P01877\|IGHA2_HUMAN Immunoglobulin heavy constant alpha 2 OS=Homo sapiens OX=9606 GN=IGHA2 PE=1 SV=4 | 576 | 20201118_ly_colostrum_1ng_glyco_hcd203040_compound15.25208.25208.4 | controllerType=0 controllerNumber=1 scan=25208 | 55.3169 |
| 21474:4 | 6 | A.[+42.011]SPTSPKVFPLSLDSTPQDGNVVVAC[+57.021]LVQGFFPQEPLSVTWSESGQN | HexNAc(3)Hex(5) | .[42], C[+57], N[+1420] | 1709.5382 | 4 | 6835.1309 | 6835.1864 | 0 | -8.1 | 2 | NRagged | 303.26 | 251.92 | 251.92 | 3.45 | 97 | >sp\|P01877\|IGHA2_HUMAN Immunoglobulin heavy constant alpha 2 OS=Homo sapiens OX=9606 GN=IGHA2 PE=1 SV=4 | 576 | 20201118_ly_colostrum_1ng_glyco_hcd203040_compound15.25210.25210.4 | controllerType=0 controllerNumber=1 scan=25210 | 55.3236 |
| 22197:4 | 6 | K.VFPLSLDSTPQDGNVVVAC[+57.021]LVQGFFPQEPLSVTWSESGQN[+2262.814]VTAI | HexNAc(5)Hex(5)Fuc(1)NeuAc(1) | C[+57], N[+2263] | 1760.5437 | 4 | 7039.1528 | 7039.1757 | 0 | -3.2 | 8 | Specific | 203.94 | 165.54 | 165.54 | 4.27 | 97 | >sp\|P01877\|IGHA2_HUMAN Immunoglobulin heavy constant alpha 2 OS=Homo sapiens OX=9606 GN=IGHA2 PE=1 SV=4 | 576 | 20201118_ly_colostrum_1ng_glyco_hcd203040_compound15.25962.25962.4 | controllerType=0 controllerNumber=1 scan=25962 | 56.8032 |
| 21705:5 | 6 | K.VFPLSLDSTPQDGNVVVAC[+57.021]LVQGFFPQEPLSVTWSESGQN[+2262.814]VTAI | HexNAc(5)Hex(5)Fuc(1)NeuAc(1) | C[+57], N[+2263] | 1408.6482 | 5 | 7039.2121 | 7039.1757 | 0 | 5.2 | 8 | Specific | 206.15 | 206.15 | 206.15 | 6.1 | 97 | >sp\|P01877\|IGHA2_HUMAN Immunoglobulin heavy constant alpha 2 OS=Homo sapiens OX=9606 GN=IGHA2 PE=1 SV=4 | 576 | 20201118_ly_colostrum_1ng_glyco_hcd203040_compound15.25450.25450.5 | controllerType=0 controllerNumber=1 scan=25450 | 55.8084 |
| 20048:5 | 6 | K.VFPLSLDSTPQDGNVVVAC[+57.021]LVQGFFPQEPLSVTWSESGQN[+1647.613]VTAI | HexNAc(5)Hex(3)Fuc(1) | C[+57], N[+1648] | 1285.6086 | 5 | 6424.0141 | 6423.9746 | 0 | 6.1 | 8 | Specific | 222.7 | 222.7 | 222.7 | 6.18 | 97 | >sp\|P01877\|IGHA2_HUMAN Immunoglobulin heavy constant alpha 2 OS=Homo sapiens OX=9606 GN=IGHA2 PE=1 SV=4 | 576 | 20201118_ly_colostrum_1ng_glyco_hcd203040_compound15.23723.23723.5 | controllerType=0 controllerNumber=1 scan=23723 | 52.1385 |
| 21495:5 | 6 | K.VFPLSLDSTPQDGNVVVAC[+57.021]LVQGFFPQEPLSVTWSESGQN[+2117.777]VTAI | HexNAc(5)Hex(5)Fuc(2) | C[+57], N[+2118] | 1379.6426 | 5 | 6894.184 | 6894.1382 | 0 | 6.6 | 8 | Specific | 238.41 | 238.41 | 238.41 | 7.14 | 97 | >sp\|P01877\|IGHA2_HUMAN Immunoglobulin heavy constant alpha 2 OS=Homo sapiens OX=9606 GN=IGHA2 PE=1 SV=4 | 576 | 20201118_ly_colostrum_1ng_glyco_hcd203040_compound15.25232.25232.5 | controllerType=0 controllerNumber=1 scan=25232 | 55.3666 |
| 21421:7 | 6 | K.VFPLSLDSTPQDGNVVVAC[+57.021]LVQGFFPQEPLSVTWSESGQN[+2262.814]VTAI | HexNAc(5)Hex(5)Fuc(1)NeuAc(1) | C[+57], N[+2263] | 1006.4651 | 7 | 7039.2121 | 7039.1757 | 0 | 5.2 | 8 | Specific | 253.77 | 253.77 | 253.77 | 5.75 | 97 | >sp\|P01877\|IGHA2_HUMAN Immunoglobulin heavy constant alpha 2 OS=Homo sapiens OX=9606 GN=IGHA2 PE=1 SV=4 | 576 | 20201118_ly_colostrum_1ng_glyco_hcd203040_compound15.25154.25154.7 | controllerType=0 controllerNumber=1 scan=25154 | 55.1902 |
| 21405:7 | 6 | K.VFPLSLDSTPQDGNVVVAC[+57.021]LVQGFFPQEPLSVTWSESGQN[+2262.814]VTAI | HexNAc(5)Hex(5)Fuc(1)NeuAc(1) | C[+57], N[+2263] | 1006.4649 | 7 | 7039.2104 | 7039.1757 | 0 | 4.9 | 8 | Specific | 255 | 255 | 255 | 6.51 | 97 | >sp\|P01877\|IGHA2_HUMAN Immunoglobulin heavy constant alpha 2 OS=Homo sapiens OX=9606 GN=IGHA2 PE=1 SV=4 | 576 | 20201118_ly_colostrum_1ng_glyco_hcd203040_compound15.25137.25137.7 | controllerType=0 controllerNumber=1 scan=25137 | 55.1467 |
| 20049:6 | 6 | K.VFPLSLDSTPQDGNVVVAC[+57.021]LVQGFFPQEPLSVTWSESGQN[+1647.613]VTAI | HexNAc(5)Hex(3)Fuc(1) | C[+57], N[+1648] | 1071.505 | 6 | 6423.9939 | 6423.9746 | 0 | 3 | 8 | Specific | 259.09 | 259.09 | 259.09 | 8.25 | 97 | >sp\|P01877\|IGHA2_HUMAN Immunoglobulin heavy constant alpha 2 OS=Homo sapiens OX=9606 GN=IGHA2 PE=1 SV=4 | 576 | 20201118_ly_colostrum_1ng_glyco_hcd203040_compound15.23724.23724.6 | controllerType=0 controllerNumber=1 scan=23724 | 52.1399 |
| 20302:4 | 6 | K.[+42.011]VFPLSLDSTPQDGNVVVAC[+57.021]LVQGFFPQEPLSVTWSESGQN[+1606 | HexNAc(4)Hex(4)Fuc(1) | .[42], C[+57], N[+1607] | 1606.9921 | 4 | 6424.9464 | 6424.9587 | 0 | -1.9 | 8 | Specific | 272.58 | 272.58 | 272.58 | 5.38 | 97 | >sp\|P01877\|IGHA2_HUMAN Immunoglobulin heavy constant alpha 2 OS=Homo sapiens OX=9606 GN=IGHA2 PE=1 SV=4 | 576 | 20201118_ly_colostrum_1ng_glyco_hcd203040_compound15.23987.23987.4 | controllerType=0 controllerNumber=1 scan=23987 | 52.6946 |
| 20300:4 | 6 | K.[+42.011]VFPLSLDSTPQDGNVVVAC[+57.021]LVQGFFPQEPLSVTWSESGQN[+1606 | HexNAc(4)Hex(4)Fuc(1) | .[42], C[+57], N[+1607] | 1606.9921 | 4 | 6424.9464 | 6424.9587 | 0 | -1.9 | 8 | Specific | 277.03 | 275.62 | 275.62 | 7.33 | 97 | >sp\|P01877\|IGHA2_HUMAN Immunoglobulin heavy constant alpha 2 OS=Homo sapiens OX=9606 GN=IGHA2 PE=1 SV=4 | 576 | 20201118_ly_colostrum_1ng_glyco_hcd203040_compound15.23985.23985.4 | controllerType=0 controllerNumber=1 scan=23985 | 52.6891 |
| 23929:4 | 6 | K.VFPLSLDSTPQDGNVVVAC[+57.021]LVQGFFPQEPLSVTWSESGQN[+2570.925]VTAI | HexNAc(5)Hex(6)Fuc(2)NeuAc(1) | C[+57], N[+2571] | 1837.5883 | 4 | 7347.3315 | 7347.2864 | 0 | 6.1 | 8 | Specific | 279.54 | 178.93 | 178.93 | 6.92 | 97 | >sp\|P01877\|IGHA2_HUMAN Immunoglobulin heavy constant alpha 2 OS=Homo sapiens OX=9606 GN=IGHA2 PE=1 SV=4 | 576 | 20201118_ly_colostrum_1ng_glyco_hcd203040_compound15.27760.27760.4 | controllerType=0 controllerNumber=1 scan=27760 | 60.1457 |
| 23847:4 | 6 | K.VFPLSLDSTPQDGNVVVAC[+57.021]LVQGFFPQEPLSVTWSESGQN[+2553.910]VTAI | HexNAc(5)Hex(5)Fuc(1)NeuAc(2) | C[+57], N[+2554] | 1833.3339 | 4 | 7330.314 | 7330.2711 | 0 | 5.8 | 8 | Specific | 296.31 | 296.31 | 296.31 | 8.12 | 97 | >sp\|P01877\|IGHA2_HUMAN Immunoglobulin heavy constant alpha 2 OS=Homo sapiens OX=9606 GN=IGHA2 PE=1 SV=4 | 576 | 20201118_ly_colostrum_1ng_glyco_hcd203040_compound15.27675.27675.4 | controllerType=0 controllerNumber=1 scan=27675 | 59.9849 |
| 19846:4 | 6 | K.[+42.011]VFPLSLDSTPQDGNVVVAC[+57.021]LVQGFFPQEPLSVTWSESGQN[+1768 | HexNAc(4)Hex(5)Fuc(1) | .[42], C[+57], N[+1769] | 1647.5183 | 4 | 6587.0514 | 6587.0115 | 0 | 6.1 | 8 | Specific | 324.57 | 300.39 | 300.39 | 6.38 | 97 | >sp\|P01877\|IGHA2_HUMAN Immunoglobulin heavy constant alpha 2 OS=Homo sapiens OX=9606 GN=IGHA2 PE=1 SV=4 | 576 | 20201118_ly_colostrum_1ng_glyco_hcd203040_compound15.23512.23512.4 | controllerType=0 controllerNumber=1 scan=23512 | 51.7117 |
| 19844:4 | 6 | K.[+42.011]VFPLSLDSTPQDGNVVVAC[+57.021]LVQGFFPQEPLSVTWSESGQN[+1768 | HexNAc(4)Hex(5)Fuc(1) | .[42], C[+57], N[+1769] | 1647.5183 | 4 | 6587.0514 | 6587.0115 | 0 | 6.1 | 8 | Specific | 378.79 | 368.26 | 368.26 | 7.61 | 97 | >sp\|P01877\|IGHA2_HUMAN Immunoglobulin heavy constant alpha 2 OS=Homo sapiens OX=9606 GN=IGHA2 PE=1 SV=4 | 576 | 20201118_ly_colostrum_1ng_glyco_hcd203040_compound15.23510.23510.4 | controllerType=0 controllerNumber=1 scan=23510 | 51.7064 |
| 21473:4 | 6 | K.VFPLSLDSTPQDGNVVVAC[+57.021]LVQGFFPQEPLSVTWSESGQN[+2117.777]VTAI | HexNAc(5)Hex(5)Fuc(2) | C[+57], N[+2118] | 1724.301 | 4 | 6894.1821 | 6894.1382 | 0 | 6.4 | 8 | Specific | 386.02 | 374.41 | 374.41 | 7.82 | 97 | >sp\|P01877\|IGHA2_HUMAN Immunoglobulin heavy constant alpha 2 OS=Homo sapiens OX=9606 GN=IGHA2 PE=1 SV=4 | 576 | 20201118_ly_colostrum_1ng_glyco_hcd203040_compound15.25209.25209.4 | controllerType=0 controllerNumber=1 scan=25209 | 55.3195 |
| 21407:7 | 6 | K.VFPLSLDSTPQDGNVVVAC[+57.021]LVQGFFPQEPLSVTWSESGQN[+2262.814]VTAI | HexNAc(5)Hex(5)Fuc(1)NeuAc(1) | C[+57], N[+2263] | 1006.4649 | 7 | 7039.2104 | 7039.1757 | 0 | 4.9 | 8 | Specific | 402.65 | 402.65 | 402.65 | 11.01 | 97 | >sp\|P01877\|IGHA2_HUMAN Immunoglobulin heavy constant alpha 2 OS=Homo sapiens OX=9606 GN=IGHA2 PE=1 SV=4 | 576 | 20201118_ly_colostrum_1ng_glyco_hcd203040_compound15.25139.25139.7 | controllerType=0 controllerNumber=1 scan=25139 | 55.1524 |
| 20057:4 | 6 | K.VFPLSLDSTPQDGNVVVAC[+57.021]LVQGFFPQEPLSVTWSESGQN[+1606.587]VTAI | HexNAc(4)Hex(4)Fuc(1) | C[+57], N[+1607] | 1596.4944 | 4 | 6382.9557 | 6382.9481 | 0 | 1.2 | 8 | Specific | 421.07 | 415.86 | 415.86 | 10.97 | 97 | >sp\|P01877\|IGHA2_HUMAN Immunoglobulin heavy constant alpha 2 OS=Homo sapiens OX=9606 GN=IGHA2 PE=1 SV=4 | 576 | 20201118_ly_colostrum_1ng_glyco_hcd203040_compound15.23732.23732.2 | controllerType=0 controllerNumber=1 scan=23732 | 52.1709 |
| 21496:4 | 6 | K.VFPLSLDSTPQDGNVVVAC[+57.021]LVQGFFPQEPLSVTWSESGQN[+2117.777]VTAI | HexNAc(5)Hex(5)Fuc(2) | C[+57], N[+2118] | 1724.3029 | 4 | 6894.1899 | 6894.1382 | 0 | 7.5 | 8 | Specific | 433.8 | 433.8 | 433.8 | 10.34 | 97 | >sp\|P01877\|IGHA2_HUMAN Immunoglobulin heavy constant alpha 2 OS=Homo sapiens OX=9606 GN=IGHA2 PE=1 SV=4 | 576 | 20201118_ly_colostrum_1ng_glyco_hcd203040_compound15.25233.25233.4 | controllerType=0 controllerNumber=1 scan=25233 | 55.3687 |
| 21521:5 | 6 | K.VFPLSLDSTPQDGNVVVAC[+57.021]LVQGFFPQEPLSVTWSESGQN[+2059.735]VTAI | HexNAc(4)Hex(5)Fuc(1)NeuAc(1) | C[+57], N[+2060] | 1368.0309 | 5 | 6836.1255 | 6836.0963 | 0 | 4.3 | 8 | Specific | 438.66 | 438.66 | 438.66 | 10.93 | 97 | >sp\|P01877\|IGHA2_HUMAN Immunoglobulin heavy constant alpha 2 OS=Homo sapiens OX=9606 GN=IGHA2 PE=1 SV=4 | 576 | 20201118_ly_colostrum_1ng_glyco_hcd203040_compound15.25259.25259.5 | controllerType=0 controllerNumber=1 scan=25259 | 55.4184 |
| 23849:4 | 6 | K.VFPLSLDSTPQDGNVVVAC[+57.021]LVQGFFPQEPLSVTWSESGQN[+2553.910]VTAI | HexNAc(5)Hex(5)Fuc(1)NeuAc(2) | C[+57], N[+2554] | 1833.3339 | 4 | 7330.314 | 7330.2711 | 0 | 5.8 | 8 | Specific | 487.4 | 482.04 | 482.04 | 11 | 97 | >sp\|P01877\|IGHA2_HUMAN Immunoglobulin heavy constant alpha 2 OS=Homo sapiens OX=9606 GN=IGHA2 PE=1 SV=4 | 576 | 20201118_ly_colostrum_1ng_glyco_hcd203040_compound15.27677.27677.4 | controllerType=0 controllerNumber=1 scan=27677 | 59.9911 |
| 21497:5 | 6 | K.VFPLSLDSTPQDGNVVVAC[+57.021]LVQGFFPQEPLSVTWSESGQN[+2117.777]VTAI | HexNAc(5)Hex(5)Fuc(2) | C[+57], N[+2118] | 1379.6426 | 5 | 6894.184 | 6894.1382 | 0 | 6.6 | 8 | Specific | 517.44 | 517.44 | 517.44 | 10.83 | 97 | >sp\|P01877\|IGHA2_HUMAN Immunoglobulin heavy constant alpha 2 OS=Homo sapiens OX=9606 GN=IGHA2 PE=1 SV=4 | 576 | 20201118_ly_colostrum_1ng_glyco_hcd203040_compound15.25234.25234.5 | controllerType=0 controllerNumber=1 scan=25234 | [illegible] |
| 21412:6 | 6 | K.VFPLSLDSTPQDGNVVVAC[+57.021]LVQGFFPQEPLSVTWSESGQN[+2262.814]VTAI | HexNAc(5)Hex(5)Fuc(1)NeuAc(1) | C[+57], N[+2263] | 1174.043 | 6 | 7039.2218 | 7039.1757 | 0 | 6.5 | 8 | Specific | 565.72 | 565.72 | 565.72 | 11.23 | 97 | >sp\|P01877\|IGHA2_HUMAN Immunoglobulin heavy constant alpha 2 OS=Homo sapiens OX=9606 GN=IGHA2 PE=1 SV=4 | 576 | 20201118_ly_colostrum_1ng_glyco_hcd203040_compound15.25145.25145.6 | controllerType=0 controllerNumber=1 scan=25145 | [illegible] |
| 20059:4 | 6 | K.VFPLSLDSTPQDGNVVVAC[+57.021]LVQGFFPQEPLSVTWSESGQN[+1606.587]VTAI | HexNAc(4)Hex(4)Fuc(1) | C[+57], N[+1607] | 1596.4944 | 4 | 6382.9557 | 6382.9481 | 0 | 1.2 | 8 | Specific | 574.28 | 547.57 | 547.57 | 11.83 | 97 | >sp\|P01877\|IGHA2_HUMAN Immunoglobulin heavy constant alpha 2 OS=Homo sapiens OX=9606 GN=IGHA2 PE=1 SV=4 | 576 | 20201118_ly_colostrum_1ng_glyco_hcd203040_compound15.23734.23734.2 | controllerType=0 controllerNumber=1 scan=23734 | [illegible] |
| 23846:5 | 6 | K.VFPLSLDSTPQDGNVVVAC[+57.021]LVQGFFPQEPLSVTWSESGQN[+2553.910]VTAI | HexNAc(5)Hex(5)Fuc(1)NeuAc(2) | C[+57], N[+2554] | 1466.8677 | 5 | 7330.3095 | 7330.2711 | 0 | 5.2 | 8 | Specific | 575.95 | 532.73 | 532.73 | 11.53 | 97 | >sp\|P01877\|IGHA2_HUMAN Immunoglobulin heavy constant alpha 2 OS=Homo sapiens OX=9606 GN=IGHA2 PE=1 SV=4 | 576 | 20201118_ly_colostrum_1ng_glyco_hcd203040_compound15.27674.27674.5 | controllerType=0 controllerNumber=1 scan=27674 | [illegible] |
| 20028:4 | 6 | K.VFPLSLDSTPQDGNVVVAC[+57.021]LVQGFFPQEPLSVTWSESGQN[+1971.719]VTAI | HexNAc(5)Hex(5)Fuc(1) | C[+57], N[+1972] | 1687.7833 | 4 | 6748.1113 | 6748.0803 | 0 | 4.6 | 8 | Specific | 595.54 | 582.97 | 582.97 | 13.12 | 97 | >sp\|P01877\|IGHA2_HUMAN Immunoglobulin heavy constant alpha 2 OS=Homo sapiens OX=9606 GN=IGHA2 PE=1 SV=4 | 576 | 20201118_ly_colostrum_1ng_glyco_hcd203040_compound15.23702.23702.4 | controllerType=0 controllerNumber=1 scan=23702 | 52.0889 |
| 21414:6 | 6 | K.VFPLSLDSTPQDGNVVVAC[+57.021]LVQGFFPQEPLSVTWSESGQN[+2262.814]VTAI | HexNAc(5)Hex(5)Fuc(1)NeuAc(1) | C[+57], N[+2263] | 1174.043 | 6 | 7039.2218 | 7039.1757 | 0 | 6.5 | 8 | Specific | 619.99 | 619.99 | 619.99 | 13.56 | 97 | >sp\|P01877\|IGHA2_HUMAN Immunoglobulin heavy constant alpha 2 OS=Homo sapiens OX=9606 GN=IGHA2 PE=1 SV=4 | 576 | 20201118_ly_colostrum_1ng_glyco_hcd203040_compound15.25147.25147.6 | controllerType=0 controllerNumber=1 scan=25147 | 55.1758 |
| 20070:5 | 6 | K.VFPLSLDSTPQDGNVVVAC[+57.021]LVQGFFPQEPLSVTWSESGQN[+1444.534]VTAI | HexNAc(4)Hex(3)Fuc(1) | C[+57], N[+1445] | 1244.9882 | 5 | 6220.9117 | 6220.8953 | 0 | 2.6 | 8 | Specific | 629.45 | 629.45 | 629.45 | 14.39 | 97 | >sp\|P01877\|IGHA2_HUMAN Immunoglobulin heavy constant alpha 2 OS=Homo sapiens OX=9606 GN=IGHA2 PE=1 SV=4 | 576 | 20201118_ly_colostrum_1ng_glyco_hcd203040_compound15.23746.23746.5 | controllerType=0 controllerNumber=1 scan=23746 | 52.2048 |
| 21409:4 | 6 | K.VFPLSLDSTPQDGNVVVAC[+57.021]LVQGFFPQEPLSVTWSESGQN[+2262.814]VTAI | HexNAc(5)Hex(5)Fuc(1)NeuAc(1) | C[+57], N[+2263] | 1760.5638 | 4 | 7039.2334 | 7039.1757 | 0 | 8.2 | 8 | Specific | 632.88 | 554.68 | 554.68 | 13.25 | 97 | >sp\|P01877\|IGHA2_HUMAN Immunoglobulin heavy constant alpha 2 OS=Homo sapiens OX=9606 GN=IGHA2 PE=1 SV=4 | 576 | 20201118_ly_colostrum_1ng_glyco_hcd203040_compound15.25142.25142.4 | controllerType=0 controllerNumber=1 scan=25142 | 55.1584 |
| 20103:4 | 6 | K.VFPLSLDSTPQDGNVVVAC[+57.021]LVQGFFPQEPLSVTWSESGQN[+1809.666]VTAI | HexNAc(5)Hex(4)Fuc(1) | C[+57], N[+1810] | 1647.2686 | 4 | 6586.0524 | 6586.0275 | 0 | 3.8 | 8 | Specific | 642.5 | 635.41 | 635.41 | 14.42 | 97 | >sp\|P01877\|IGHA2_HUMAN Immunoglobulin heavy constant alpha 2 OS=Homo sapiens OX=9606 GN=IGHA2 PE=1 SV=4 | 576 | 20201118_ly_colostrum_1ng_glyco_hcd203040_compound15.23780.23780.4 | controllerType=0 controllerNumber=1 scan=23780 | 52.2805 |
| 20051:6 | 6 | K.VFPLSLDSTPQDGNVVVAC[+57.021]LVQGFFPQEPLSVTWSESGQN[+1647.613]VTAI | HexNAc(5)Hex(3)Fuc(1) | C[+57], N[+1648] | 1071.505 | 6 | 6423.9939 | 6423.9746 | 0 | 3 | 8 | Specific | 681.84 | 657.93 | 657.93 | 15.39 | 97 | >sp\|P01877\|IGHA2_HUMAN Immunoglobulin heavy constant alpha 2 OS=Homo sapiens OX=9606 GN=IGHA2 PE=1 SV=4 | 576 | 20201118_ly_colostrum_1ng_glyco_hcd203040_compound15.23726.23726.6 | controllerType=0 controllerNumber=1 scan=23726 | 52.144 |
| 23848:5 | 6 | K.VFPLSLDSTPQDGNVVVAC[+57.021]LVQGFFPQEPLSVTWSESGQN[+2553.910]VTAI | HexNAc(5)Hex(5)Fuc(1)NeuAc(2) | C[+57], N[+2554] | 1466.8677 | 5 | 7330.3095 | 7330.2711 | 0 | 5.2 | 8 | Specific | 708.76 | 690.39 | 690.39 | 15.85 | 97 | >sp\|P01877\|IGHA2_HUMAN Immunoglobulin heavy constant alpha 2 OS=Homo sapiens OX=9606 GN=IGHA2 PE=1 SV=4 | 576 | 20201118_ly_colostrum_1ng_glyco_hcd203040_compound15.27676.27676.5 | controllerType=0 controllerNumber=1 scan=27676 | 59.9892 |
| 20029:4 | 6 | K.VFPLSLDSTPQDGNVVVAC[+57.021]LVQGFFPQEPLSVTWSESGQN[+1444.534]VTAI | HexNAc(4)Hex(3)Fuc(1) | C[+57], N[+1445] | 1555.9906 | 4 | 6220.9406 | 6220.8953 | 0 | 7.3 | 8 | Specific | 733.29 | 682.47 | 682.47 | 15.53 | 97 | >sp\|P01877\|IGHA2_HUMAN Immunoglobulin heavy constant alpha 2 OS=Homo sapiens OX=9606 GN=IGHA2 PE=1 SV=4 | 576 | 20201118_ly_colostrum_1ng_glyco_hcd203040_compound15.23703.23703.4 | controllerType=0 controllerNumber=1 scan=23703 | 52.0907 |
| 20023:4 | 6 | K.VFPLSLDSTPQDGNVVVAC[+57.021]LVQGFFPQEPLSVTWSESGQN[+1809.666]VTAI | HexNAc(5)Hex(4)Fuc(1) | C[+57], N[+1810] | 1647.2679 | 4 | 6586.05 | 6586.0275 | 0 | 3.4 | 8 | Specific | 738.4 | 636.88 | 636.88 | 16.42 | 97 | >sp\|P01877\|IGHA2_HUMAN Immunoglobulin heavy constant alpha 2 OS=Homo sapiens OX=9606 GN=IGHA2 PE=1 SV=4 | 576 | 20201118_ly_colostrum_1ng_glyco_hcd203040_compound15.23696.23696.4 | controllerType=0 controllerNumber=1 scan=23696 | 52.0761 |
| 21523:5 | 6 | K.VFPLSLDSTPQDGNVVVAC[+57.021]LVQGFFPQEPLSVTWSESGQN[+2059.735]VTAI | HexNAc(4)Hex(5)Fuc(1)NeuAc(1) | C[+57], N[+2060] | 1368.0309 | 5 | 6836.1255 | 6836.0963 | 0 | 4.3 | 8 | Specific | 743.33 | 743.33 | 743.33 | 16.12 | 97 | >sp\|P01877\|IGHA2_HUMAN Immunoglobulin heavy constant alpha 2 OS=Homo sapiens OX=9606 GN=IGHA2 PE=1 SV=4 | 576 | 20201118_ly_colostrum_1ng_glyco_hcd203040_compound15.25261.25261.5 | controllerType=0 controllerNumber=1 scan=25261 | 55.4236 |
| 20101:4 | 6 | K.VFPLSLDSTPQDGNVVVAC[+57.021]LVQGFFPQEPLSVTWSESGQN[+1809.666]VTAI | HexNAc(5)Hex(4)Fuc(1) | C[+57], N[+1810] | 1647.2686 | 4 | 6586.0524 | 6586.0275 | 0 | 3.8 | 8 | Specific | 746.51 | 686.38 | 686.38 | 16.42 | 97 | >sp\|P01877\|IGHA2_HUMAN Immunoglobulin heavy constant alpha 2 OS=Homo sapiens OX=9606 GN=IGHA2 PE=1 SV=4 | 576 | 20201118_ly_colostrum_1ng_glyco_hcd203040_compound15.23778.23778.4 | controllerType=0 controllerNumber=1 scan=23778 | 52.2753 |
| 21408:5 | 6 | K.VFPLSLDSTPQDGNVVVAC[+57.021]LVQGFFPQEPLSVTWSESGQN[+2262.814]VTAI | HexNAc(5)Hex(5)Fuc(1)NeuAc(1) | C[+57], N[+2263] | 1408.653 | 5 | 7039.2359 | 7039.1757 | 0 | 8.5 | 8 | Specific | 754.06 | 754.06 | 754.06 | 16.25 | 97 | >sp\|P01877\|IGHA2_HUMAN Immunoglobulin heavy constant alpha 2 OS=Homo sapiens OX=9606 GN=IGHA2 PE=1 SV=4 | 576 | 20201118_ly_colostrum_1ng_glyco_hcd203040_compound15.25141.25141.5 | controllerType=0 controllerNumber=1 scan=25141 | 55.1571 |
| 20025:4 | 6 | K.VFPLSLDSTPQDGNVVVAC[+57.021]LVQGFFPQEPLSVTWSESGQN[+1809.666]VTAI | HexNAc(5)Hex(4)Fuc(1) | C[+57], N[+1810] | 1647.2679 | 4 | 6586.05 | 6586.0275 | 0 | 3.4 | 8 | Specific | 774.24 | 774.24 | 774.24 | 17.42 | 97 | >sp\|P01877\|IGHA2_HUMAN Immunoglobulin heavy constant alpha 2 OS=Homo sapiens OX=9606 GN=IGHA2 PE=1 SV=4 | 576 | 20201118_ly_colostrum_1ng_glyco_hcd203040_compound15.23698.23698.4 | controllerType=0 controllerNumber=1 scan=23698 | 52.0813 |
| 19901:4 | 6 | K.VFPLSLDSTPQDGNVVVAC[+57.021]LVQGFFPQEPLSVTWSESGQN[+1809.666]VTAI | HexNAc(5)Hex(4)Fuc(1) | C[+57], N[+1810] | 1647.2706 | 4 | 6586.0607 | 6586.0275 | 0 | 5 | 8 | Specific | 798.41 | 713.57 | 713.57 | 16.85 | 97 | >sp\|P01877\|IGHA2_HUMAN Immunoglobulin heavy constant alpha 2 OS=Homo sapiens OX=9606 GN=IGHA2 PE=1 SV=4 | 576 | 20201118_ly_colostrum_1ng_glyco_hcd203040_compound15.23570.23570.4 | controllerType=0 controllerNumber=1 scan=23570 | 51.8242 |
| 19950:4 | 6 | K.VFPLSLDSTPQDGNVVVAC[+57.021]LVQGFFPQEPLSVTWSESGQN[+1647.613]VTAI | HexNAc(5)Hex(3)Fuc(1) | C[+57], N[+1648] | 1606.7589 | 4 | 6424.0138 | 6423.9746 | 0 | 6.1 | 8 | Specific | 813.02 | 789.78 | 789.78 | 17.56 | 97 | >sp\|P01877\|IGHA2_HUMAN Immunoglobulin heavy constant alpha 2 OS=Homo sapiens OX=9606 GN=IGHA2 PE=1 SV=4 | 576 | 20201118_ly_colostrum_1ng_glyco_hcd203040_compound15.23621.23621.4 | controllerType=0 controllerNumber=1 scan=23621 | 51.9279 |
| 20050:5 | 6 | K.VFPLSLDSTPQDGNVVVAC[+57.021]LVQGFFPQEPLSVTWSESGQN[+1647.613]VTAI | HexNAc(5)Hex(3)Fuc(1) | C[+57], N[+1648] | 1285.6086 | 5 | 6424.0141 | 6423.9746 | 0 | 6.1 | 8 | Specific | 835.57 | 835.57 | 835.57 | 17.56 | 97 | >sp\|P01877\|IGHA2_HUMAN Immunoglobulin heavy constant alpha 2 OS=Homo sapiens OX=9606 GN=IGHA2 PE=1 SV=4 | 576 | 20201118_ly_colostrum_1ng_glyco_hcd203040_compound15.23725.23725.5 | controllerType=0 controllerNumber=1 scan=23725 | 52.1426 |
| 19903:4 | 6 | K.VFPLSLDSTPQDGNVVVAC[+57.021]LVQGFFPQEPLSVTWSESGQN[+1809.666]VTAI | HexNAc(5)Hex(4)Fuc(1) | C[+57], N[+1810] | 1647.2706 | 4 | 6586.0607 | 6586.0275 | 0 | 5 | 8 | Specific | 835.77 | 820.06 | 820.06 | 17.85 | 97 | >sp\|P01877\|IGHA2_HUMAN Immunoglobulin heavy constant alpha 2 OS=Homo sapiens OX=9606 GN=IGHA2 PE=1 SV=4 | 576 | 20201118_ly_colostrum_1ng_glyco_hcd203040_compound15.23572.23572.4 | controllerType=0 controllerNumber=1 scan=23572 | 51.8318 |
| 20030:4 | 6 | K.VFPLSLDSTPQDGNVVVAC[+57.021]LVQGFFPQEPLSVTWSESGQN[+1971.719]VTAI | HexNAc(5)Hex(5)Fuc(1) | C[+57], N[+1972] | 1687.7833 | 4 | 6748.1113 | 6748.0803 | 0 | 4.6 | 8 | Specific | 837.97 | 798.17 | 798.17 | 18.12 | 97 | >sp\|P01877\|IGHA2_HUMAN Immunoglobulin heavy constant alpha 2 OS=Homo sapiens OX=9606 GN=IGHA2 PE=1 SV=4 | 576 | 20201118_ly_colostrum_1ng_glyco_hcd203040_compound15.23704.23704.4 | controllerType=0 controllerNumber=1 scan=23704 | 52.0942 |
| 21410:5 | 6 | K.VFPLSLDSTPQDGNVVVAC[+57.021]LVQGFFPQEPLSVTWSESGQN[+2262.814]VTAI | HexNAc(5)Hex(5)Fuc(1)NeuAc(1) | C[+57], N[+2263] | 1408.653 | 5 | 7039.2359 | 7039.1757 | 0 | 8.5 | 8 | Specific | 838.22 | 838.22 | 838.22 | 17.25 | 97 | >sp\|P01877\|IGHA2_HUMAN Immunoglobulin heavy constant alpha 2 OS=Homo sapiens OX=9606 GN=IGHA2 PE=1 SV=4 | 576 | 20201118_ly_colostrum_1ng_glyco_hcd203040_compound15.25143.25143.5 | controllerType=0 controllerNumber=1 scan=25143 | 55.1646 |
| 20072:5 | 6 | K.VFPLSLDSTPQDGNVVVAC[+57.021]LVQGFFPQEPLSVTWSESGQN[+1444.534]VTAI | HexNAc(4)Hex(3)Fuc(1) | C[+57], N[+1445] | 1244.9882 | 5 | 6220.9117 | 6220.8953 | 0 | 2.6 | 8 | Specific | 842.4 | 828.35 | 828.35 | 18.39 | 97 | >sp\|P01877\|IGHA2_HUMAN Immunoglobulin heavy constant alpha 2 OS=Homo sapiens OX=9606 GN=IGHA2 PE=1 SV=4 | 576 | 20201118_ly_colostrum_1ng_glyco_hcd203040_compound15.23748.23748.5 | controllerType=0 controllerNumber=1 scan=23748 | 52.2116 |
| 21411:4 | 6 | K.VFPLSLDSTPQDGNVVVAC[+57.021]LVQGFFPQEPLSVTWSESGQN[+2262.814]VTAI | HexNAc(5)Hex(5)Fuc(1)NeuAc(1) | C[+57], N[+2263] | 1760.5638 | 4 | 7039.2334 | 7039.1757 | 0 | 8.2 | 8 | Specific | 868.38 | 836.47 | 836.47 | 18.25 | 97 | >sp\|P01877\|IGHA2_HUMAN Immunoglobulin heavy constant alpha 2 OS=Homo sapiens OX=9606 GN=IGHA2 PE=1 SV=4 | 576 | 20201118_ly_colostrum_1ng_glyco_hcd203040_compound15.25144.25144.4 | controllerType=0 controllerNumber=1 scan=25144 | 55.1688 |
| 20031:4 | 6 | K.VFPLSLDSTPQDGNVVVAC[+57.021]LVQGFFPQEPLSVTWSESGQN[+1444.534]VTAI | HexNAc(4)Hex(3)Fuc(1) | C[+57], N[+1445] | 1555.9906 | 4 | 6220.9406 | 6220.8953 | 0 | 7.3 | 8 | Specific | 909.49 | 854.03 | 854.03 | 19.53 | 97 | >sp\|P01877\|IGHA2_HUMAN Immunoglobulin heavy constant alpha 2 OS=Homo sapiens OX=9606 GN=IGHA2 PE=1 SV=4 | 576 | 20201118_ly_colostrum_1ng_glyco_hcd203040_compound15.23705.23705.4 | controllerType=0 controllerNumber=1 scan=23705 | 52.0983 |
| 02962:7 | 6 | K.SVTC[+57.021]HVKHYTN[+1647.613]SSQDVTVPC[+57.021]RVPPPPPC[+57.021]C[ | HexNAc(5)Hex(3)Fuc(1) | C[+57]*4, N[+1648] | 774.7747 | 7 | 5417.379 | 5417.3699 | 0 | 1.7 | 82 | Specific | 427.99 | 427.99 | 427.99 | 10.07 | 97 | >sp\|P01877\|IGHA2_HUMAN Immunoglobulin heavy constant alpha 2 OS=Homo sapiens OX=9606 GN=IGHA2 PE=1 SV=4 | 576 | 20201118_ly_colostrum_1ng_glyco_hcd203040_compound15.5951.5951.2 | controllerType=0 controllerNumber=1 scan=5951 | 16.4198 |
| 03070:6 | 6 | K.SVTC[+57.021]HVKHYTN[+1647.613]SSQDVTVPC[+57.021]RVPPPPPC[+57.021]C[ | HexNAc(5)Hex(3)Fuc(1) | C[+57]*4, N[+1648] | 903.7366 | 6 | 5417.3829 | 5417.3699 | 0 | 2.4 | 82 | Specific | 460.38 | 332.23 | 332.23 | 8.42 | 97 | >sp\|P01877\|IGHA2_HUMAN Immunoglobulin heavy constant alpha 2 OS=Homo sapiens OX=9606 GN=IGHA2 PE=1 SV=4 | 576 | 20201118_ly_colostrum_1ng_glyco_hcd203040_compound15.6063.6063.6 | controllerType=0 controllerNumber=1 scan=6063 | 16.628 |
| 03071:6 | 6 | K.SVTC[+57.021]HVKHYTN[+1647.613]SSQDVTVPC[+57.021]RVPPPPPC[+57.021]C[ | HexNAc(5)Hex(3)Fuc(1) | C[+57]*4, N[+1648] | 903.7366 | 6 | 5417.3829 | 5417.3699 | 0 | 2.4 | 82 | Specific | 547.27 | 547.27 | 547.27 | 10.58 | 97 | >sp\|P01877\|IGHA2_HUMAN Immunoglobulin heavy constant alpha 2 OS=Homo sapiens OX=9606 GN=IGHA2 PE=1 SV=4 | 576 | 20201118_ly_colostrum_1ng_glyco_hcd203040_compound15.6064.6064.6 | controllerType=0 controllerNumber=1 scan=6064 | 16.6317 |
| 02963:7 | 6 | K.SVTC[+57.021]HVKHYTN[+1647.613]SSQDVTVPC[+57.021]RVPPPPPC[+57.021]C[ | HexNAc(5)Hex(3)Fuc(1) | C[+57]*4, N[+1648] | 774.7747 | 7 | 5417.379 | 5417.3699 | 0 | 1.7 | 82 | Specific | 612.54 | 612.54 | 612.54 | 13.26 | 97 | >sp\|P01877\|IGHA2_HUMAN Immunoglobulin heavy constant alpha 2 OS=Homo sapiens OX=9606 GN=IGHA2 PE=1 SV=4 | 576 | 20201118_ly_colostrum_1ng_glyco_hcd203040_compound15.5952.5952.2 | controllerType=0 controllerNumber=1 scan=5952 | 16.4239 |
| 02981:6 | 6 | K.SVTC[+57.021]HVKHYTN[+1647.613]SSQDVTVPC[+57.021]RVPPPPPC[+57.021]C[ | HexNAc(5)Hex(3)Fuc(1) | C[+57]*4, N[+1648] | 903.7369 | 6 | 5417.3848 | 5417.3699 | 0 | 2.7 | 82 | Specific | 686.31 | 686.31 | 686.31 | 14.08 | 97 | >sp\|P01877\|IGHA2_HUMAN Immunoglobulin heavy constant alpha 2 OS=Homo sapiens OX=9606 GN=IGHA2 PE=1 SV=4 | 576 | 20201118_ly_colostrum_1ng_glyco_hcd203040_compound15.5971.5971.6 | controllerType=0 controllerNumber=1 scan=5971 | 16.4667 |
| 02983:6 | 6 | K.SVTC[+57.021]HVKHYTN[+1647.613]SSQDVTVPC[+57.021]RVPPPPPC[+57.021]C[ | HexNAc(5)Hex(3)Fuc(1) | C[+57]*4, N[+1648] | 903.7369 | 6 | 5417.3848 | 5417.3699 | 0 | 2.7 | 82 | Specific | 873.24 | 873.24 | 873.24 | 18.08 | 97 | >sp\|P01877\|IGHA2_HUMAN Immunoglobulin heavy constant alpha 2 OS=Homo sapiens OX=9606 GN=IGHA2 PE=1 SV=4 | 576 | 20201118_ly_colostrum_1ng_glyco_hcd203040_compound15.5973.5973.6 | controllerType=0 controllerNumber=1 scan=5973 | 16.4703 |
| 02900:6 | 6 | K.SVTC[+57.021]HVKHYTN[+1647.613]SSQDVTVPC[+57.021]RVPPPPPC[+57.021]C[ | HexNAc(5)Hex(3)Fuc(1) | C[+57]*4, N[+1648] | 903.7362 | 6 | 5417.3807 | 5417.3699 | 0 | 2 | 82 | Specific | 1101.2 | 1004.52 | 1004.52 | 22.84 | 97 | >sp\|P01877\|IGHA2_HUMAN Immunoglobulin heavy constant alpha 2 OS=Homo sapiens OX=9606 GN=IGHA2 PE=1 SV=4 | 576 | 20201118_ly_colostrum_1ng_glyco_hcd203040_compound15.5887.5887.3 | controllerType=0 controllerNumber=1 scan=5887 | 16.2842 |
| 02898:6 | 6 | K.SVTC[+57.021]HVKHYTN[+1647.613]SSQDVTVPC[+57.021]RVPPPPPC[+57.021]C[ | HexNAc(5)Hex(3)Fuc(1) | C[+57]*4, N[+1648] | 903.7362 | 6 | 5417.3807 | 5417.3699 | 0 | 2 | 82 | Specific | 1154.68 | 1154.68 | 1154.68 | 23.84 | 97 | >sp\|P01877\|IGHA2_HUMAN Immunoglobulin heavy constant alpha 2 OS=Homo sapiens OX=9606 GN=IGHA2 PE=1 SV=4 | 576 | 20201118_ly_colostrum_1ng_glyco_hcd203040_compound15.5885.5885.3 | controllerType=0 controllerNumber=1 scan=5885 | 16.2788 |
| 01924:4 | 6 | K.HYTN[+2262.814]SSQDVTVPC[+57.021]R.V | HexNAc(5)Hex(5)Fuc(1)NeuAc(1) | C[+57], N[+2263] | 982.3992 | 4 | 3926.575 | 3926.5636 | 0 | 2.9 | 89 | Specific | 213.47 | 163.17 | 163.17 | 4.78 | 97 | >sp\|P01877\|IGHA2_HUMAN Immunoglobulin heavy constant alpha 2 OS=Homo sapiens OX=9606 GN=IGHA2 PE=1 SV=4 | 576 | 20201118_ly_colostrum_1ng_glyco_hcd203040_compound15.4876.4876.4 | controllerType=0 controllerNumber=1 scan=4876 | 14.4958 |
| 01042:2 | 6 | K.HYTN[+1647.613]SSQDVTVPC[+57.021]R.V | HexNAc(5)Hex(3)Fuc(1) | C[+57], N[+1648] | 1656.1917 | 2 | 3311.3761 | 3311.3625 | 0 | 4.1 | 89 | Specific | 258.39 | 258.39 | 258.39 | 5.81 | 97 | >sp\|P01877\|IGHA2_HUMAN Immunoglobulin heavy constant alpha 2 OS=Homo sapiens OX=9606 GN=IGHA2 PE=1 SV=4 | 576 | 20201118_ly_colostrum_1ng_glyco_hcd203040_compound15.3963.3963.2 | controllerType=0 controllerNumber=1 scan=3963 | 12.9623 |
| 01044:2 | 6 | K.HYTN[+1647.613]SSQDVTVPC[+57.021]R.V | HexNAc(5)Hex(3)Fuc(1) | C[+57], N[+1648] | 1656.1917 | 2 | 3311.3761 | 3311.3625 | 0 | 4.1 | 89 | Specific | 280.07 | 201.76 | 201.76 | 4.8 | 97 | >sp\|P01877\|IGHA2_HUMAN Immunoglobulin heavy constant alpha 2 OS=Homo sapiens OX=9606 GN=IGHA2 PE=1 SV=4 | 576 | 20201118_ly_colostrum_1ng_glyco_hcd203040_compound15.3965.3965.2 | controllerType=0 controllerNumber=1 scan=3965 | 12.9663 |
| 00944:3 | 6 | K.HYTN[+1971.719]SSQDVTVPC[+57.021]R.V | HexNAc(5)Hex(5)Fuc(1) | C[+57], N[+1972] | 1212.4982 | 3 | 3635.4801 | 3635.4682 | 0 | 3.3 | 89 | Specific | 587.86 | 482 | 482 | 11.66 | 97 | >sp\|P01877\|IGHA2_HUMAN Immunoglobulin heavy constant alpha 2 OS=Homo sapiens OX=9606 GN=IGHA2 PE=1 SV=4 | 576 | 20201118_ly_colostrum_1ng_glyco_hcd203040_compound15.3862.3862.3 | controllerType=0 controllerNumber=1 scan=3862 | 12.7814 |
| 00937:3 | 6 | K.HYTN[+1606.587]SSQDVTVPC[+57.021]R.V | HexNAc(4)Hex(4)Fuc(1) | C[+57], N[+1607] | 1090.7865 | 3 | 3270.345 | 3270.336 | 0 | 2.8 | 89 | Specific | 595.26 | 458.28 | 458.28 | 11.64 | 97 | >sp\|P01877\|IGHA2_HUMAN Immunoglobulin heavy constant alpha 2 OS=Homo sapiens OX=9606 GN=IGHA2 PE=1 SV=4 | 576 | 20201118_ly_colostrum_1ng_glyco_hcd203040_compound15.3855.3855.3 | controllerType=0 controllerNumber=1 scan=3855 | 12.7678 |
| 00939:3 | 6 | K.HYTN[+1606.587]SSQDVTVPC[+57.021]R.V | HexNAc(4)Hex(4)Fuc(1) | C[+57], N[+1607] | 1090.7865 | 3 | 3270.345 | 3270.336 | 0 | 2.8 | 89 | Specific | 610.65 | 500.74 | 500.74 | 12.64 | 97 | >sp\|P01877\|IGHA2_HUMAN Immunoglobulin heavy constant alpha 2 OS=Homo sapiens OX=9606 GN=IGHA2 PE=1 SV=4 | 576 | 20201118_ly_colostrum_1ng_glyco_hcd203040_compound15.3857.3857.3 | controllerType=0 controllerNumber=1 scan=3857 | 12.7718 |
| 00931:3 | 6 | K.HYTN[+1606.587]SSQDVTVPC[+57.021]R.V | HexNAc(4)Hex(4)Fuc(1) | C[+57], N[+1607] | 1090.7874 | 3 | 3270.3476 | 3270.336 | 0 | 3.5 | 89 | Specific | 620.44 | 433.73 | 433.73 | 12.66 | 97 | >sp\|P01877\|IGHA2_HUMAN Immunoglobulin heavy constant alpha 2 OS=Homo sapiens OX=9606 GN=IGHA2 PE=1 SV=4 | 576 | 20201118_ly_colostrum_1ng_glyco_hcd203040_compound15.3848.3848.3 | controllerType=0 controllerNumber=1 scan=3848 | 12.7544 |
| 00980:3 | 6 | K.HYTN[+1647.613]SSQDVTVPC[+57.021]R.V | HexNAc(5)Hex(3)Fuc(1) | C[+57], N[+1648] | 1104.4613 | 3 | 3311.3694 | 3311.3625 | 0 | 2.1 | 89 | Specific | 638.73 | 592.65 | 592.65 | 12.64 | 97 | >sp\|P01877\|IGHA2_HUMAN Immunoglobulin heavy constant alpha 2 OS=Homo sapiens OX=9606 GN=IGHA2 PE=1 SV=4 | 576 | 20201118_ly_colostrum_1ng_glyco_hcd203040_compound15.3899.3899.3 | controllerType=0 controllerNumber=1 scan=3899 | 12.8497 |
| 01852:3 | 6 | K.HYTN[+2262.814]SSQDVTVPC[+57.021]R.V | HexNAc(5)Hex(5)Fuc(1)NeuAc(1) | C[+57], N[+2263] | 1309.5291 | 3 | 3926.5728 | 3926.5636 | 0 | 2.3 | 89 | Specific | 648.83 | 423.91 | 423.91 | 12.64 | 97 | >sp\|P01877\|IGHA2_HUMAN Immunoglobulin heavy constant alpha 2 OS=Homo sapiens OX=9606 GN=IGHA2 PE=1 SV=4 | 576 | 20201118_ly_colostrum_1ng_glyco_hcd203040_compound15.4802.4802.3 | controllerType=0 controllerNumber=1 scan=4802 | 14.3655 |
| 01019:3 | 6 | K.HYTN[+1647.613]SSQDVTVPC[+57.021]R.V | HexNAc(5)Hex(3)Fuc(1) | C[+57], N[+1648] | 1104.4633 | 3 | 3311.3753 | 3311.3625 | 0 | 3.9 | 89 | Specific | 652.67 | 577.68 | 577.68 | 13.66 | 97 | >sp\|P01877\|IGHA2_HUMAN Immunoglobulin heavy constant alpha 2 OS=Homo sapiens OX=9606 GN=IGHA2 PE=1 SV=4 | 576 | 20201118_ly_colostrum_1ng_glyco_hcd203040_compound15.3940.3940.3 | controllerType=0 controllerNumber=1 scan=3940 | 12.925 |
| 01017:3 | 6 | K.HYTN[+1647.613]SSQDVTVPC[+57.021]R.V | HexNAc(5)Hex(3)Fuc(1) | C[+57], N[+1648] | 1104.4633 | 3 | 3311.3753 | 3311.3625 | 0 | 3.9 | 89 | Specific | 652.73 | 586.95 | 586.95 | 13.66 | 97 | >sp\|P01877\|IGHA2_HUMAN Immunoglobulin heavy constant alpha 2 OS=Homo sapiens OX=9606 GN=IGHA2 PE=1 SV=4 | 576 | 20201118_ly_colostrum_1ng_glyco_hcd203040_compound15.3938.3938.3 | controllerType=0 controllerNumber=1 scan=3938 | 12.9194 |
| 00965:3 | 6 | K.HYTN[+1809.666]SSQDVTVPC[+57.021]R.V | HexNAc(5)Hex(4)Fuc(1) | C[+57], N[+1810] | 1158.4794 | 3 | 3473.4237 | 3473.4154 | 0 | 2.4 | 89 | Specific | 665.92 | 639.75 | 639.75 | 13.64 | 97 | >sp\|P01877\|IGHA2_HUMAN Immunoglobulin heavy constant alpha 2 OS=Homo sapiens OX=9606 GN=IGHA2 PE=1 SV=4 | 576 | 20201118_ly_colostrum_1ng_glyco_hcd203040_compound15.3884.3884.3 | controllerType=0 controllerNumber=1 scan=3884 | 12.8213 |
| 00929:3 | 6 | K.HYTN[+1606.587]SSQDVTVPC[+57.021]R.V | HexNAc(4)Hex(4)Fuc(1) | C[+57], N[+1607] | 1090.7874 | 3 | 3270.3476 | 3270.336 | 0 | 3.5 | 89 | Specific | 669.28 | 591.81 | 591.81 | 13.66 | 97 | >sp\|P01877\|IGHA2_HUMAN Immunoglobulin heavy constant alpha 2 OS=Homo sapiens OX=9606 GN=IGHA2 PE=1 SV=4 | 576 | 20201118_ly_colostrum_1ng_glyco_hcd203040_compound15.3846.3846.3 | controllerType=0 controllerNumber=1 scan=3846 | 12.7504 |
| 01854:3 | 6 | K.HYTN[+2262.814]SSQDVTVPC[+57.021]R.V | HexNAc(5)Hex(5)Fuc(1)NeuAc(1) | C[+57], N[+2263] | 1309.5291 | 3 | 3926.5728 | 3926.5636 | 0 | 2.3 | 89 | Specific | 688.7 | 474.97 | 474.97 | 13.64 | 97 | >sp\|P01877\|IGHA2_HUMAN Immunoglobulin heavy constant alpha 2 OS=Homo sapiens OX=9606 GN=IGHA2 PE=1 SV=4 | 576 | 20201118_ly_colostrum_1ng_glyco_hcd203040_compound15.4804.4804.3 | controllerType=0 controllerNumber=1 scan=4804 | 14.3703 |
| 00967:3 | 6 | K.HYTN[+1809.666]SSQDVTVPC[+57.021]R.V | HexNAc(5)Hex(4)Fuc(1) | C[+57], N[+1810] | 1158.4794 | 3 | 3473.4237 | 3473.4154 | 0 | 2.4 | 89 | Specific | 689.3 | 597.93 | 597.93 | 13.64 | 97 | >sp\|P01877\|IGHA2_HUMAN Immunoglobulin heavy constant alpha 2 OS=Homo sapiens OX=9606 GN=IGHA2 PE=1 SV=4 | 576 | 20201118_ly_colostrum_1ng_glyco_hcd203040_compound15.3886.3886.3 | controllerType=0 controllerNumber=1 scan=3886 | 12.8265 |
| 00978:3 | 6 | K.HYTN[+1647.613]SSQDVTVPC[+57.021]R.V | HexNAc(5)Hex(3)Fuc(1) | C[+57], N[+1648] | 1104.4613 | 3 | 3311.3694 | 3311.3625 | 0 | 2.1 | 89 | Specific | 691.47 | 683.99 | 683.99 | 13.64 | 97 | >sp\|P01877\|IGHA2_HUMAN Immunoglobulin heavy constant alpha 2 OS=Homo sapiens OX=9606 GN=IGHA2 PE=1 SV=4 | 576 | 20201118_ly_colostrum_1ng_glyco_hcd203040_compound15.3897.3897.3 | controllerType=0 controllerNumber=1 scan=3897 | 12.844 |
| 00942:3 | 6 | K.HYTN[+1971.719]SSQDVTVPC[+57.021]R.V | HexNAc(5)Hex(5)Fuc(1) | C[+57], N[+1972] | 1212.4982 | 3 | 3635.4801 | 3635.4682 | 0 | 3.3 | 89 | Specific | 696.99 | 539.6 | 539.6 | 13.66 | 97 | >sp\|P01877\|IGHA2_HUMAN Immunoglobulin heavy constant alpha 2 OS=Homo sapiens OX=9606 GN=IGHA2 PE=1 SV=4 | 576 | 20201118_ly_colostrum_1ng_glyco_hcd203040_compound15.3860.3860.3 | controllerType=0 controllerNumber=1 scan=3860 | 12.7774 |
| 00988:3 | 6 | K.HYTN[+1444.534]SSQDVTVPC[+57.021]R.V | HexNAc(4)Hex(3)Fuc(1) | C[+57], N[+1445] | 1036.7692 | 3 | 3108.293 | 3108.2832 | 0 | 3.2 | 89 | Specific | 722.76 | 635.37 | 635.37 | 14.66 | 97 | >sp\|P01877\|IGHA2_HUMAN Immunoglobulin heavy constant alpha 2 OS=Homo sapiens OX=9606 GN=IGHA2 PE=1 SV=4 | 576 | 20201118_ly_colostrum_1ng_glyco_hcd203040_compound15.3908.3908.3 | controllerType=0 controllerNumber=1 scan=3908 | 12.8674 |
| 00960:3 | 6 | K.HYTN[+1444.534]SSQDVTVPC[+57.021]R.V | HexNAc(4)Hex(3)Fuc(1) | C[+57], N[+1445] | 1036.7688 | 3 | 3108.2919 | 3108.2832 | 0 | 2.8 | 89 | Specific | 726.24 | 637.55 | 637.55 | 14.64 | 97 | >sp\|P01877\|IGHA2_HUMAN Immunoglobulin heavy constant alpha 2 OS=Homo sapiens OX=9606 GN=IGHA2 PE=1 SV=4 | 576 | 20201118_ly_colostrum_1ng_glyco_hcd203040_compound15.3878.3878.3 | controllerType=0 controllerNumber=1 scan=3878 | 12.8078 |
| 00961:3 | 6 | K.HYTN[+1444.534]SSQDVTVPC[+57.021]R.V | HexNAc(4)Hex(3)Fuc(1) | C[+57], N[+1445] | 1036.7688 | 3 | 3108.2919 | 3108.2832 | 0 | 2.8 | 89 | Specific | 759.71 | 755.25 | 755.25 | 15.64 | 97 | >sp\|P01877\|IGHA2_HUMAN Immunoglobulin heavy constant alpha 2 OS=Homo sapiens OX=9606 GN=IGHA2 PE=1 SV=4 | 576 | 20201118_ly_colostrum_1ng_glyco_hcd203040_compound15.3879.3879.3 | controllerType=0 controllerNumber=1 scan=3879 | 12.8123 |
| 00990:3 | 6 | K.HYTN[+1444.534]SSQDVTVPC[+57.021]R.V | HexNAc(4)Hex(3)Fuc(1) | C[+57], N[+1445] | 1036.7692 | 3 | 3108.293 | 3108.2832 | 0 | 3.2 | 89 | Specific | 770.21 | 652.37 | 652.37 | 15.66 | 97 | >sp\|P01877\|IGHA2_HUMAN Immunoglobulin heavy constant alpha 2 OS=Homo sapiens OX=9606 GN=IGHA2 PE=1 SV=4 | 576 | 20201118_ly_colostrum_1ng_glyco_hcd203040_compound15.3910.3910.3 | controllerType=0 controllerNumber=1 scan=3910 | 12.8729 |
| 03000:3 | 6 | K.HYTN[+1809.666]SSQDVTVPC[+57.021]RVPPPPPC[+57.021]C[+57.021]HPR.L | HexNAc(5)Hex(4)Fuc(1) | C[+57]*3, N[+1810] | 1590.0167 | 3 | 4768.0354 | 4768.0217 | 0 | 2.9 | 89 | Specific | 296.26 | 239.18 | 239.18 | 5.92 | 97 | >sp\|P01877\|IGHA2_HUMAN Immunoglobulin heavy constant alpha 2 OS=Homo sapiens OX=9606 GN=IGHA2 PE=1 SV=4 | 576 | 20201118_ly_colostrum_1ng_glyco_hcd203040_compound15.5991.5991.3 | controllerType=0 controllerNumber=1 scan=5991 | 16.5035 |
| 03002:3 | 6 | K.HYTN[+1809.666]SSQDVTVPC[+57.021]RVPPPPPC[+57.021]C[+57.021]HPR.L | HexNAc(5)Hex(4)Fuc(1) | C[+57]*3, N[+1810] | 1590.0167 | 3 | 4768.0354 | 4768.0217 | 0 | 2.9 | 89 | Specific | 443.33 | 282.04 | 282.04 | 6.9 | 97 | >sp\|P01877\|IGHA2_HUMAN Immunoglobulin heavy constant alpha 2 OS=Homo sapiens OX=9606 GN=IGHA2 PE=1 SV=4 | 576 | 20201118_ly_colostrum_1ng_glyco_hcd203040_compound15.5993.5993.3 | controllerType=0 controllerNumber=1 scan=5993 | 16.5072 |
| 03591:4 | 6 | K.HYTN[+1809.666]SSQDVTVPC[+57.021]RVPPPPPC[+57.021]C[+57.021]HPR.L | HexNAc(5)Hex(4)Fuc(1) | C[+57]*3, N[+1810] | 1192.7633 | 4 | 4768.0316 | 4768.0217 | 0 | 2.1 | 89 | Specific | 449.65 | 381.24 | 381.24 | 9.55 | 97 | >sp\|P01877\|IGHA2_HUMAN Immunoglobulin heavy constant alpha 2 OS=Homo sapiens OX=9606 GN=IGHA2 PE=1 SV=4 | 576 | 20201118_ly_colostrum_1ng_glyco_hcd203040_compound15.6603.6603.4 | controllerType=0 controllerNumber=1 scan=6603 | 17.607 |
| 03058:3 | 6 | K.HYTN[+1444.534]SSQDVTVPC[+57.021]RVPPPPPC[+57.021]C[+57.021]HPR.L | HexNAc(4)Hex(3)Fuc(1) | C[+57]*3, N[+1445] | 1468.3025 | 3 | 4402.8931 | 4402.8895 | 0 | 0.8 | 89 | Specific | 451.18 | 451.18 | 451.18 | 8.55 | 97 | >sp\|P01877\|IGHA2_HUMAN Immunoglobulin heavy constant alpha 2 OS=Homo sapiens OX=9606 GN=IGHA2 PE=1 SV=4 | 576 | 20201118_ly_colostrum_1ng_glyco_hcd203040_compound15.6051.6051.3 | controllerType=0 controllerNumber=1 scan=6051 | 16.6072 |
| 03060:3 | 6 | K.HYTN[+1444.534]SSQDVTVPC[+57.021]RVPPPPPC[+57.021]C[+57.021]HPR.L | HexNAc(4)Hex(3)Fuc(1) | C[+57]*3, N[+1445] | 1468.3025 | 3 | 4402.8931 | 4402.8895 | 0 | 0.8 | 89 | Specific | 452.1 | 299.75 | 299.75 | 5.09 | 97 | >sp\|P01877\|IGHA2_HUMAN Immunoglobulin heavy constant alpha 2 OS=Homo sapiens OX=9606 GN=IGHA2 PE=1 SV=4 | 576 | 20201118_ly_colostrum_1ng_glyco_hcd203040_compound15.6053.6053.3 | controllerType=0 controllerNumber=1 scan=6053 | 16.6119 |
| 03662:5 | 6 | K.HYTN[+2059.735]SSQDVTVPC[+57.021]RVPPPPPC[+57.021]C[+57.021]HPR.L | HexNAc(4)Hex(5)Fuc(1)NeuAc(1) | C[+57]*3, N[+2060] | 1004.4271 | 5 | 5018.1062 | 5018.0905 | 0 | 3.1 | 89 | Specific | 466.36 | 367.57 | 367.57 | 8.59 | 97 | >sp\|P01877\|IGHA2_HUMAN Immunoglobulin heavy constant alpha 2 OS=Homo sapiens OX=9606 GN=IGHA2 PE=1 SV=4 | 576 | 20201118_ly_colostrum_1ng_glyco_hcd203040_compound15.6677.6677.5 | controllerType=0 controllerNumber=1 scan=6677 | 17.7435 |
| 04588:4 | 6 | K.HYTN[+2553.910]SSQDVTVPC[+57.021]RVPPPPPC[+57.021]C[+57.021]HPR.L | HexNAc(5)Hex(5)Fuc(1)NeuAc(2) | C[+57]*3, N[+2554] | 1378.8258 | 4 | 5512.2814 | 5512.2653 | 0 | 2.9 | 89 | Specific | 485.8 | 403.95 | 403.95 | 9.7 | 97 | >sp\|P01877\|IGHA2_HUMAN Immunoglobulin heavy constant alpha 2 OS=Homo sapiens OX=9606 GN=IGHA2 PE=1 SV=4 | 576 | 20201118_ly_colostrum_1ng_glyco_hcd203040_compound15.7638.7638.4 | controllerType=0 controllerNumber=1 scan=7638 | 19.421 |
| 03261:5 | 6 | K.HYTN[+1606.587]SSQDVTVPC[+57.021]RVPPPPPC[+57.021]C[+57.021]HPR.L | HexNAc(4)Hex(4)Fuc(1) | C[+57]*3, N[+1607] | 913.7957 | 5 | 4564.9496 | 4564.9423 | 0 | 1.6 | 89 | Specific | 512 | 512 | 512 | 10.01 | 97 | >sp\|P01877\|IGHA2_HUMAN Immunoglobulin heavy constant alpha 2 OS=Homo sapiens OX=9606 GN=IGHA2 PE=1 SV=4 | 576 | 20201118_ly_colostrum_1ng_glyco_hcd203040_compound15.6261.6261.2 | controllerType=0 controllerNumber=1 scan=6261 | 16.986 |
| 03141:4 | 6 | K.HYTN[+1606.587]SSQDVTVPC[+57.021]RVPPPPPC[+57.021]C[+57.021]HPR.L | HexNAc(4)Hex(4)Fuc(1) | C[+57]*3, N[+1607] | 1141.9926 | 4 | 4564.9486 | 4564.9423 | 0 | 1.4 | 89 | Specific | 520.75 | 463.55 | 463.55 | 9.63 | 97 | >sp\|P01877\|IGHA2_HUMAN Immunoglobulin heavy constant alpha 2 OS=Homo sapiens OX=9606 GN=IGHA2 PE=1 SV=4 | 576 | 20201118_ly_colostrum_1ng_glyco_hcd203040_compound15.6137.6137.4 | controllerType=0 controllerNumber=1 scan=6137 | 16.7646 |
| 03263:5 | 6 | K.HYTN[+1606.587]SSQDVTVPC[+57.021]RVPPPPPC[+57.021]C[+57.021]HPR.L | HexNAc(4)Hex(4)Fuc(1) | C[+57]*3, N[+1607] | 913.7957 | 5 | 4564.9496 | 4564.9423 | 0 | 1.6 | 89 | Specific | 532.51 | 385.19 | 385.19 | 10.05 | 97 | >sp\|P01877\|IGHA2_HUMAN Immunoglobulin heavy constant alpha 2 OS=Homo sapiens OX=9606 GN=IGHA2 PE=1 SV=4 | 576 | 20201118_ly_colostrum_1ng_glyco_hcd203040_compound15.6263.6263.2 | controllerType=0 controllerNumber=1 scan=6263 | 16.9902 |
| 03593:4 | 6 | K.HYTN[+1809.666]SSQDVTVPC[+57.021]RVPPPPPC[+57.021]C[+57.021]HPR.L | HexNAc(5)Hex(4)Fuc(1) | C[+57]*3, N[+1810] | 1192.7633 | 4 | 4768.0316 | 4768.0217 | 0 | 2.1 | 89 | Specific | 543.77 | 332.97 | 332.97 | 8.33 | 97 | >sp\|P01877\|IGHA2_HUMAN Immunoglobulin heavy constant alpha 2 OS=Homo sapiens OX=9606 GN=IGHA2 PE=1 SV=4 | 576 | 20201118_ly_colostrum_1ng_glyco_hcd203040_compound15.6605.6605.4 | controllerType=0 controllerNumber=1 scan=6605 | 17.6107 |
| 03663:4 | 6 | K.HYTN[+2059.735]SSQDVTVPC[+57.021]RVPPPPPC[+57.021]C[+57.021]HPR.L | HexNAc(4)Hex(5)Fuc(1)NeuAc(1) | C[+57]*3, N[+2060] | 1255.2824 | 4 | 5018.1076 | 5018.0905 | 0 | 3.4 | 89 | Specific | 544.07 | 484.56 | 484.56 | 10.08 | 97 | >sp\|P01877\|IGHA2_HUMAN Immunoglobulin heavy constant alpha 2 OS=Homo sapiens OX=9606 GN=IGHA2 PE=1 SV=4 | 576 | 20201118_ly_colostrum_1ng_glyco_hcd203040_compound15.6678.6678.4 | controllerType=0 controllerNumber=1 scan=6678 | 17.7445 |
| 03553:4 | 6 | K.HYTN[+1647.613]SSQDVTVPC[+57.021]RVPPPPPC[+57.021]C[+57.021]HPR.L | HexNAc(5)Hex(3)Fuc(1) | C[+57]*3, N[+1648] | 1152.2498 | 4 | 4605.9774 | 4605.9689 | 0 | 1.8 | 89 | Specific | 577.63 | 556.47 | 556.47 | 9.99 | 97 | >sp\|P01877\|IGHA2_HUMAN Immunoglobulin heavy constant alpha 2 OS=Homo sapiens OX=9606 GN=IGHA2 PE=1 SV=4 | 576 | 20201118_ly_colostrum_1ng_glyco_hcd203040_compound15.6564.6564.4 | controllerType=0 controllerNumber=1 scan=6564 | 17.533 |
| 03666:4 | 6 | K.HYTN[+2059.735]SSQDVTVPC[+57.021]RVPPPPPC[+57.021]C[+57.021]HPR.L | HexNAc(4)Hex(5)Fuc(1)NeuAc(1) | C[+57]*3, N[+2060] | 1255.2824 | 4 | 5018.1076 | 5018.0905 | 0 | 3.4 | 89 | Specific | 580.17 | 443.68 | 443.68 | 11.58 | 97 | >sp\|P01877\|IGHA2_HUMAN Immunoglobulin heavy constant alpha 2 OS=Homo sapiens OX=9606 GN=IGHA2 PE=1 SV=4 | 576 | 20201118_ly_colostrum_1ng_glyco_hcd203040_compound15.6681.6681.4 | controllerType=0 controllerNumber=1 scan=6681 | 17.7511 |
| 03555:4 | 6 | K.HYTN[+1647.613]SSQDVTVPC[+57.021]RVPPPPPC[+57.021]C[+57.021]HPR.L | HexNAc(5)Hex(3)Fuc(1) | C[+57]*3, N[+1648] | 1152.2498 | 4 | 4605.9774 | 4605.9689 | 0 | 1.8 | 89 | Specific | 586.49 | 377.57 | 377.57 | 9.85 | 97 | >sp\|P01877\|IGHA2_HUMAN Immunoglobulin heavy constant alpha 2 OS=Homo sapiens OX=9606 GN=IGHA2 PE=1 SV=4 | 576 | 20201118_ly_colostrum_1ng_glyco_hcd203040_compound15.6566.6566.4 | controllerType=0 controllerNumber=1 scan=6566 | 17.5371 |
| 03143:4 | 6 | K.HYTN[+1606.587]SSQDVTVPC[+57.021]RVPPPPPC[+57.021]C[+57.021]HPR.L | HexNAc(4)Hex(4)Fuc(1) | C[+57]*3, N[+1607] | 1141.9926 | 4 | 4564.9486 | 4564.9423 | 0 | 1.4 | 89 | Specific | 620.99 | 446.59 | 446.59 | 12.32 | 97 | >sp\|P01877\|IGHA2_HUMAN Immunoglobulin heavy constant alpha 2 OS=Homo sapiens OX=9606 GN=IGHA2 PE=1 SV=4 | 576 | 20201118_ly_colostrum_1ng_glyco_hcd203040_compound15.6139.6139.4 | controllerType=0 controllerNumber=1 scan=6139 | 16.7684 |
| 03549:5 | 6 | K.HYTN[+1647.613]SSQDVTVPC[+57.021]RVPPPPPC[+57.021]C[+57.021]HPR.L | HexNAc(5)Hex(3)Fuc(1) | C[+57]*3, N[+1648] | 922.0015 | 5 | 4605.9786 | 4605.9689 | 0 | 2.1 | 89 | Specific | 686.88 | 526.29 | 526.29 | 13.96 | 97 | >sp\|P01877\|IGHA2_HUMAN Immunoglobulin heavy constant alpha 2 OS=Homo sapiens OX=9606 GN=IGHA2 PE=1 SV=4 | 576 | 20201118_ly_colostrum_1ng_glyco_hcd203040_compound15.6560.6560.5 | controllerType=0 controllerNumber=1 scan=6560 | 17.5252 |

Table S4. Cont.

| Query #:z | Protein Rank | Glycopeptide | Glycan | Modification Type(s) | Observed m/z | z | Observed (M+H) | Calc. mass (M+H) | Off-by-x error | Mass error (ppm) | StartPosition | Cleavage | Score | Delta | Delta Mod | \|Log Prob\| | # of unique peptides | Protein Name | Protein DB number | Comment | Scan # | Scan Time |
|---|---|---|---|---|---|---|---|---|---|---|---|---|---|---|---|---|---|---|---|---|---|---|
| 02990:3 | 6 | K.HYTN[+1647.613]SSQDVTVPC[+57.021]RVPPPPPC[+57.021]C[+57.021]HPR.L | HexNAc(5)Hex(3)Fuc(1) | C[+57]*3, N[+1648] | 1535.9986 | 3 | 4605.9812 | 4605.9689 | 0 | 2.7 | 89 | Specific | 707.46 | 621.71 | 621.71 | 13.98 | 97 | >sp\|P01877\|IGHA2_HUMAN Immunoglobulin heavy constant alpha 2 OS=Homo sapiens OX=9606 GN=IGHA2 PE=1 SV=4 | 576 | 20201118_ly_colostrum_1ng_glyco_hcd203040_compound15.5981.5981.3 | controllerType=0 controllerNumber=1 scan=5981 | 16.4851 |
| 03360:5 | 6 | K.HYTN[+1444.534]SSQDVTVPC[+57.021]RVPPPPPC[+57.021]C[+57.021]HPR.L | HexNAc(4)Hex(3)Fuc(1) | C[+57]*3, N[+1445] | 881.386 | 5 | 4402.9009 | 4402.8895 | 0 | 2.6 | 89 | Specific | 709.2 | 467.26 | 467.26 | 14.96 | 97 | >sp\|P01877\|IGHA2_HUMAN Immunoglobulin heavy constant alpha 2 OS=Homo sapiens OX=9606 GN=IGHA2 PE=1 SV=4 | 576 | 20201118_ly_colostrum_1ng_glyco_hcd203040_compound15.6364.6364.5 | controllerType=0 controllerNumber=1 scan=6364 | 17.1688 |
| 04591:4 | 6 | K.HYTN[+2553.910]SSQDVTVPC[+57.021]RVPPPPPC[+57.021]C[+57.021]HPR.L | HexNAc(5)Hex(5)Fuc(1)NeuAc(2) | C[+57]*3, N[+2554] | 1378.8258 | 4 | 5512.2814 | 5512.2653 | 0 | 2.9 | 89 | Specific | 711.7 | 466.57 | 466.57 | 14.55 | 97 | >sp\|P01877\|IGHA2_HUMAN Immunoglobulin heavy constant alpha 2 OS=Homo sapiens OX=9606 GN=IGHA2 PE=1 SV=4 | 576 | 20201118_ly_colostrum_1ng_glyco_hcd203040_compound15.7641.7641.4 | controllerType=0 controllerNumber=1 scan=7641 | 19.4252 |
| 03576:5 | 6 | K.HYTN[+2262.814]SSQDVTVPC[+57.021]RVPPPPPC[+57.021]C[+57.021]HPR.L | HexNAc(5)Hex(5)Fuc(1)NeuAc(1) | C[+57]*3, N[+2263] | 1045.0422 | 5 | 5221.1821 | 5221.1699 | 0 | 2.3 | 89 | Specific | 715 | 544.42 | 544.42 | 14.96 | 97 | >sp\|P01877\|IGHA2_HUMAN Immunoglobulin heavy constant alpha 2 OS=Homo sapiens OX=9606 GN=IGHA2 PE=1 SV=4 | 576 | 20201118_ly_colostrum_1ng_glyco_hcd203040_compound15.6588.6588.5 | controllerType=0 controllerNumber=1 scan=6588 | 17.5776 |
| 03551:5 | 6 | K.HYTN[+1647.613]SSQDVTVPC[+57.021]RVPPPPPC[+57.021]C[+57.021]HPR.L | HexNAc(5)Hex(3)Fuc(1) | C[+57]*3, N[+1648] | 922.0015 | 5 | 4605.9786 | 4605.9689 | 0 | 2.1 | 89 | Specific | 719.79 | 587.06 | 587.06 | 14.96 | 97 | >sp\|P01877\|IGHA2_HUMAN Immunoglobulin heavy constant alpha 2 OS=Homo sapiens OX=9606 GN=IGHA2 PE=1 SV=4 | 576 | 20201118_ly_colostrum_1ng_glyco_hcd203040_compound15.6562.6562.5 | controllerType=0 controllerNumber=1 scan=6562 | 17.5298 |
| 03740:4 | 6 | K.HYTN[+2262.814]SSQDVTVPC[+57.021]RVPPPPPC[+57.021]C[+57.021]HPR.L | HexNAc(5)Hex(5)Fuc(1)NeuAc(1) | C[+57]*3, N[+2263] | 1306.0474 | 4 | 5221.1676 | 5221.1699 | 0 | -0.4 | 89 | Specific | 727.6 | 715.04 | 715.04 | 13.71 | 97 | >sp\|P01877\|IGHA2_HUMAN Immunoglobulin heavy constant alpha 2 OS=Homo sapiens OX=9606 GN=IGHA2 PE=1 SV=4 | 576 | 20201118_ly_colostrum_1ng_glyco_hcd203040_compound15.6758.6758.4 | controllerType=0 controllerNumber=1 scan=6758 | 17.8846 |
| 02964:5 | 6 | K.HYTN[+1809.666]SSQDVTVPC[+57.021]RVPPPPPC[+57.021]C[+57.021]HPR.L | HexNAc(5)Hex(4)Fuc(1) | C[+57]*3, N[+1810] | 954.4115 | 5 | 4768.0285 | 4768.0217 | 0 | 1.4 | 89 | Specific | 733.35 | 612.13 | 612.13 | 14.73 | 97 | >sp\|P01877\|IGHA2_HUMAN Immunoglobulin heavy constant alpha 2 OS=Homo sapiens OX=9606 GN=IGHA2 PE=1 SV=4 | 576 | 20201118_ly_colostrum_1ng_glyco_hcd203040_compound15.5954.5954.5 | controllerType=0 controllerNumber=1 scan=5954 | 16.4284 |
| 02971:5 | 6 | K.HYTN[+1647.613]SSQDVTVPC[+57.021]RVPPPPPC[+57.021]C[+57.021]HPR.L | HexNAc(5)Hex(3)Fuc(1) | C[+57]*3, N[+1648] | 922.001 | 5 | 4605.9759 | 4605.9689 | 0 | 1.5 | 89 | Specific | 734.06 | 547.02 | 547.02 | 14.73 | 97 | >sp\|P01877\|IGHA2_HUMAN Immunoglobulin heavy constant alpha 2 OS=Homo sapiens OX=9606 GN=IGHA2 PE=1 SV=4 | 576 | 20201118_ly_colostrum_1ng_glyco_hcd203040_compound15.5961.5961.5 | controllerType=0 controllerNumber=1 scan=5961 | 16.443 |
| 02966:5 | 6 | K.HYTN[+1809.666]SSQDVTVPC[+57.021]RVPPPPPC[+57.021]C[+57.021]HPR.L | HexNAc(5)Hex(4)Fuc(1) | C[+57]*3, N[+1810] | 954.4115 | 5 | 4768.0285 | 4768.0217 | 0 | 1.4 | 89 | Specific | 743 | 618.62 | 618.62 | 14.73 | 97 | >sp\|P01877\|IGHA2_HUMAN Immunoglobulin heavy constant alpha 2 OS=Homo sapiens OX=9606 GN=IGHA2 PE=1 SV=4 | 576 | 20201118_ly_colostrum_1ng_glyco_hcd203040_compound15.5956.5956.5 | controllerType=0 controllerNumber=1 scan=5956 | 16.4337 |
| 02968:5 | 6 | K.HYTN[+1647.613]SSQDVTVPC[+57.021]RVPPPPPC[+57.021]C[+57.021]HPR.L | HexNAc(5)Hex(3)Fuc(1) | C[+57]*3, N[+1648] | 922.001 | 5 | 4605.9759 | 4605.9689 | 0 | 1.5 | 89 | Specific | 748.1 | 529.15 | 529.15 | 14.73 | 97 | >sp\|P01877\|IGHA2_HUMAN Immunoglobulin heavy constant alpha 2 OS=Homo sapiens OX=9606 GN=IGHA2 PE=1 SV=4 | 576 | 20201118_ly_colostrum_1ng_glyco_hcd203040_compound15.5958.5958.5 | controllerType=0 controllerNumber=1 scan=5958 | 16.4369 |
| 03665:5 | 6 | K.HYTN[+2059.735]SSQDVTVPC[+57.021]RVPPPPPC[+57.021]C[+57.021]HPR.L | HexNAc(4)Hex(5)Fuc(1)NeuAc(1) | C[+57]*3, N[+2060] | 1004.4271 | 5 | 5018.1062 | 5018.0905 | 0 | 3.1 | 89 | Specific | 751.37 | 485.31 | 485.31 | 15.99 | 97 | >sp\|P01877\|IGHA2_HUMAN Immunoglobulin heavy constant alpha 2 OS=Homo sapiens OX=9606 GN=IGHA2 PE=1 SV=4 | 576 | 20201118_ly_colostrum_1ng_glyco_hcd203040_compound15.6680.6680.5 | controllerType=0 controllerNumber=1 scan=6680 | 17.7492 |
| 03574:5 | 6 | K.HYTN[+2262.814]SSQDVTVPC[+57.021]RVPPPPPC[+57.021]C[+57.021]HPR.L | HexNAc(5)Hex(5)Fuc(1)NeuAc(1) | C[+57]*3, N[+2263] | 1045.0422 | 5 | 5221.1821 | 5221.1699 | 0 | 2.3 | 89 | Specific | 757.59 | 491.05 | 491.05 | 15.96 | 97 | >sp\|P01877\|IGHA2_HUMAN Immunoglobulin heavy constant alpha 2 OS=Homo sapiens OX=9606 GN=IGHA2 PE=1 SV=4 | 576 | 20201118_ly_colostrum_1ng_glyco_hcd203040_compound15.6586.6586.5 | controllerType=0 controllerNumber=1 scan=6586 | 17.5713 |
| 02943:5 | 6 | K.HYTN[+1971.719]SSQDVTVPC[+57.021]RVPPPPPC[+57.021]C[+57.021]HPR.L | HexNAc(5)Hex(5)Fuc(1) | C[+57]*3, N[+1972] | 986.8235 | 5 | 4930.0884 | 4930.0745 | 0 | 2.8 | 89 | Specific | 785.91 | 657.67 | 657.67 | 15.96 | 97 | >sp\|P01877\|IGHA2_HUMAN Immunoglobulin heavy constant alpha 2 OS=Homo sapiens OX=9606 GN=IGHA2 PE=1 SV=4 | 576 | 20201118_ly_colostrum_1ng_glyco_hcd203040_compound15.5932.5932.5 | controllerType=0 controllerNumber=1 scan=5932 | 16.3851 |
| 03583:4 | 6 | K.HYTN[+2262.814]SSQDVTVPC[+57.021]RVPPPPPC[+57.021]C[+57.021]HPR.L | HexNAc(5)Hex(5)Fuc(1)NeuAc(1) | C[+57]*3, N[+2263] | 1306.0507 | 4 | 5221.1808 | 5221.1699 | 0 | 2.1 | 89 | Specific | 793.12 | 494.26 | 494.26 | 15.55 | 97 | >sp\|P01877\|IGHA2_HUMAN Immunoglobulin heavy constant alpha 2 OS=Homo sapiens OX=9606 GN=IGHA2 PE=1 SV=4 | 576 | 20201118_ly_colostrum_1ng_glyco_hcd203040_compound15.6595.6595.4 | controllerType=0 controllerNumber=1 scan=6595 | 17.5936 |
| 02992:3 | 6 | K.HYTN[+1647.613]SSQDVTVPC[+57.021]RVPPPPPC[+57.021]C[+57.021]HPR.L | HexNAc(5)Hex(3)Fuc(1) | C[+57]*3, N[+1648] | 1535.9986 | 3 | 4605.9812 | 4605.9689 | 0 | 2.7 | 89 | Specific | 793.69 | 629.73 | 629.73 | 14.98 | 97 | >sp\|P01877\|IGHA2_HUMAN Immunoglobulin heavy constant alpha 2 OS=Homo sapiens OX=9606 GN=IGHA2 PE=1 SV=4 | 576 | 20201118_ly_colostrum_1ng_glyco_hcd203040_compound15.5983.5983.3 | controllerType=0 controllerNumber=1 scan=5983 | 16.4896 |
| 02924:4 | 6 | K.HYTN[+1971.719]SSQDVTVPC[+57.021]RVPPPPPC[+57.021]C[+57.021]HPR.L | HexNAc(5)Hex(5)Fuc(1) | C[+57]*3, N[+1972] | 1233.2771 | 4 | 4930.0867 | 4930.0745 | 0 | 2.5 | 89 | Specific | 814.76 | 565.55 | 565.55 | 16.55 | 97 | >sp\|P01877\|IGHA2_HUMAN Immunoglobulin heavy constant alpha 2 OS=Homo sapiens OX=9606 GN=IGHA2 PE=1 SV=4 | 576 | 20201118_ly_colostrum_1ng_glyco_hcd203040_compound15.5912.5912.4 | controllerType=0 controllerNumber=1 scan=5912 | 16.3372 |
| 04585:5 | 6 | K.HYTN[+2553.910]SSQDVTVPC[+57.021]RVPPPPPC[+57.021]C[+57.021]HPR.L | HexNAc(5)Hex(5)Fuc(1)NeuAc(2) | C[+57]*3, N[+2554] | 1103.2627 | 5 | 5512.2844 | 5512.2653 | 0 | 3.5 | 89 | Specific | 815.76 | 647.18 | 647.18 | 16.99 | 97 | >sp\|P01877\|IGHA2_HUMAN Immunoglobulin heavy constant alpha 2 OS=Homo sapiens OX=9606 GN=IGHA2 PE=1 SV=4 | 576 | 20201118_ly_colostrum_1ng_glyco_hcd203040_compound15.7635.7635.5 | controllerType=0 controllerNumber=1 scan=7635 | 19.4166 |
| 02975:5 | 6 | K.HYTN[+1606.587]SSQDVTVPC[+57.021]RVPPPPPC[+57.021]C[+57.021]HPR.L | HexNAc(4)Hex(4)Fuc(1) | C[+57]*3, N[+1607] | 913.796 | 5 | 4564.9511 | 4564.9423 | 0 | 1.9 | 89 | Specific | 825.49 | 716.77 | 716.77 | 16.73 | 97 | >sp\|P01877\|IGHA2_HUMAN Immunoglobulin heavy constant alpha 2 OS=Homo sapiens OX=9606 GN=IGHA2 PE=1 SV=4 | 576 | 20201118_ly_colostrum_1ng_glyco_hcd203040_compound15.5965.5965.5 | controllerType=0 controllerNumber=1 scan=5965 | 16.4534 |
| 04583:5 | 6 | K.HYTN[+2553.910]SSQDVTVPC[+57.021]RVPPPPPC[+57.021]C[+57.021]HPR.L | HexNAc(5)Hex(5)Fuc(1)NeuAc(2) | C[+57]*3, N[+2554] | 1103.2627 | 5 | 5512.2844 | 5512.2653 | 0 | 3.5 | 89 | Specific | 841.84 | 814.82 | 814.82 | 16.99 | 97 | >sp\|P01877\|IGHA2_HUMAN Immunoglobulin heavy constant alpha 2 OS=Homo sapiens OX=9606 GN=IGHA2 PE=1 SV=4 | 576 | 20201118_ly_colostrum_1ng_glyco_hcd203040_compound15.7633.7633.5 | controllerType=0 controllerNumber=1 scan=7633 | 19.412 |
| 03743:4 | 6 | K.HYTN[+2262.814]SSQDVTVPC[+57.021]RVPPPPPC[+57.021]C[+57.021]HPR.L | HexNAc(5)Hex(5)Fuc(1)NeuAc(1) | C[+57]*3, N[+2263] | 1306.0474 | 4 | 5221.1676 | 5221.1699 | 0 | -0.4 | 89 | Specific | 847.94 | 549.09 | 549.09 | 15.71 | 97 | >sp\|P01877\|IGHA2_HUMAN Immunoglobulin heavy constant alpha 2 OS=Homo sapiens OX=9606 GN=IGHA2 PE=1 SV=4 | 576 | 20201118_ly_colostrum_1ng_glyco_hcd203040_compound15.6761.6761.4 | controllerType=0 controllerNumber=1 scan=6761 | 17.8888 |
| 02922:4 | 6 | K.HYTN[+1971.719]SSQDVTVPC[+57.021]RVPPPPPC[+57.021]C[+57.021]HPR.L | HexNAc(5)Hex(5)Fuc(1) | C[+57]*3, N[+1972] | 1233.2771 | 4 | 4930.0867 | 4930.0745 | 0 | 2.5 | 89 | Specific | 856.01 | 641.02 | 641.02 | 17.55 | 97 | >sp\|P01877\|IGHA2_HUMAN Immunoglobulin heavy constant alpha 2 OS=Homo sapiens OX=9606 GN=IGHA2 PE=1 SV=4 | 576 | 20201118_ly_colostrum_1ng_glyco_hcd203040_compound15.5910.5910.4 | controllerType=0 controllerNumber=1 scan=5910 | 16.328 |
| 02941:5 | 6 | K.HYTN[+1971.719]SSQDVTVPC[+57.021]RVPPPPPC[+57.021]C[+57.021]HPR.L | HexNAc(5)Hex(5)Fuc(1) | C[+57]*3, N[+1972] | 986.8235 | 5 | 4930.0884 | 4930.0745 | 0 | 2.8 | 89 | Specific | 864 | 710.46 | 710.46 | 17.96 | 97 | >sp\|P01877\|IGHA2_HUMAN Immunoglobulin heavy constant alpha 2 OS=Homo sapiens OX=9606 GN=IGHA2 PE=1 SV=4 | 576 | 20201118_ly_colostrum_1ng_glyco_hcd203040_compound15.5930.5930.5 | controllerType=0 controllerNumber=1 scan=5930 | 16.3783 |
| 03241:4 | 6 | K.HYTN[+1444.534]SSQDVTVPC[+57.021]RVPPPPPC[+57.021]C[+57.021]HPR.L | HexNAc(4)Hex(3)Fuc(1) | C[+57]*3, N[+1445] | 1101.4815 | 4 | 4402.9041 | 4402.8895 | 0 | 3.3 | 89 | Specific | 866.95 | 725.78 | 725.78 | 17.58 | 97 | >sp\|P01877\|IGHA2_HUMAN Immunoglobulin heavy constant alpha 2 OS=Homo sapiens OX=9606 GN=IGHA2 PE=1 SV=4 | 576 | 20201118_ly_colostrum_1ng_glyco_hcd203040_compound15.6241.6241.4 | controllerType=0 controllerNumber=1 scan=6241 | 16.9513 |
| 02910:5 | 6 | K.HYTN[+1971.719]SSQDVTVPC[+57.021]RVPPPPPC[+57.021]C[+57.021]HPR.L | HexNAc(5)Hex(5)Fuc(1) | C[+57]*3, N[+1972] | 986.8217 | 5 | 4930.0796 | 4930.0745 | 0 | 1 | 89 | Specific | 873.93 | 750.45 | 750.45 | 17.53 | 97 | >sp\|P01877\|IGHA2_HUMAN Immunoglobulin heavy constant alpha 2 OS=Homo sapiens OX=9606 GN=IGHA2 PE=1 SV=4 | 576 | 20201118_ly_colostrum_1ng_glyco_hcd203040_compound15.5897.5897.2 | controllerType=0 controllerNumber=1 scan=5897 | 16.3022 |
| 03581:4 | 6 | K.HYTN[+2262.814]SSQDVTVPC[+57.021]RVPPPPPC[+57.021]C[+57.021]HPR.L | HexNAc(5)Hex(5)Fuc(1)NeuAc(1) | C[+57]*3, N[+2263] | 1306.0507 | 4 | 5221.1808 | 5221.1699 | 0 | 2.1 | 89 | Specific | 877.49 | 615.1 | 615.1 | 17.55 | 97 | >sp\|P01877\|IGHA2_HUMAN Immunoglobulin heavy constant alpha 2 OS=Homo sapiens OX=9606 GN=IGHA2 PE=1 SV=4 | 576 | 20201118_ly_colostrum_1ng_glyco_hcd203040_compound15.6593.6593.4 | controllerType=0 controllerNumber=1 scan=6593 | 17.5881 |
| 02967:4 | 6 | K.HYTN[+1809.666]SSQDVTVPC[+57.021]RVPPPPPC[+57.021]C[+57.021]HPR.L | HexNAc(5)Hex(4)Fuc(1) | C[+57]*3, N[+1810] | 1192.7629 | 4 | 4768.0296 | 4768.0217 | 0 | 1.7 | 89 | Specific | 888.4 | 622.89 | 622.89 | 17.32 | 97 | >sp\|P01877\|IGHA2_HUMAN Immunoglobulin heavy constant alpha 2 OS=Homo sapiens OX=9606 GN=IGHA2 PE=1 SV=4 | 576 | 20201118_ly_colostrum_1ng_glyco_hcd203040_compound15.5957.5957.4 | controllerType=0 controllerNumber=1 scan=5957 | 16.4358 |
| 02965:4 | 6 | K.HYTN[+1809.666]SSQDVTVPC[+57.021]RVPPPPPC[+57.021]C[+57.021]HPR.L | HexNAc(5)Hex(4)Fuc(1) | C[+57]*3, N[+1810] | 1192.7629 | 4 | 4768.0296 | 4768.0217 | 0 | 1.7 | 89 | Specific | 908.16 | 700.08 | 700.08 | 18.32 | 97 | >sp\|P01877\|IGHA2_HUMAN Immunoglobulin heavy constant alpha 2 OS=Homo sapiens OX=9606 GN=IGHA2 PE=1 SV=4 | 576 | 20201118_ly_colostrum_1ng_glyco_hcd203040_compound15.5955.5955.4 | controllerType=0 controllerNumber=1 scan=5955 | 16.4299 |
| 02940:4 | 6 | K.HYTN[+1971.719]SSQDVTVPC[+57.021]RVPPPPPC[+57.021]C[+57.021]HPR.L | HexNAc(5)Hex(5)Fuc(1) | C[+57]*3, N[+1972] | 1233.278 | 4 | 4930.0902 | 4930.0745 | 0 | 3.2 | 89 | Specific | 909.82 | 707.43 | 707.43 | 18.58 | 97 | >sp\|P01877\|IGHA2_HUMAN Immunoglobulin heavy constant alpha 2 OS=Homo sapiens OX=9606 GN=IGHA2 PE=1 SV=4 | 576 | 20201118_ly_colostrum_1ng_glyco_hcd203040_compound15.5929.5929.4 | controllerType=0 controllerNumber=1 scan=5929 | 16.3759 |
| 03362:5 | 6 | K.HYTN[+1444.534]SSQDVTVPC[+57.021]RVPPPPPC[+57.021]C[+57.021]HPR.L | HexNAc(4)Hex(3)Fuc(1) | C[+57]*3, N[+1445] | 881.386 | 5 | 4402.9009 | 4402.8895 | 0 | 2.6 | 89 | Specific | 914.87 | 729.73 | 729.73 | 18.96 | 97 | >sp\|P01877\|IGHA2_HUMAN Immunoglobulin heavy constant alpha 2 OS=Homo sapiens OX=9606 GN=IGHA2 PE=1 SV=4 | 576 | 20201118_ly_colostrum_1ng_glyco_hcd203040_compound15.6366.6366.5 | controllerType=0 controllerNumber=1 scan=6366 | 17.173 |
| 03243:4 | 6 | K.HYTN[+1444.534]SSQDVTVPC[+57.021]RVPPPPPC[+57.021]C[+57.021]HPR.L | HexNAc(4)Hex(3)Fuc(1) | C[+57]*3, N[+1445] | 1101.4815 | 4 | 4402.9041 | 4402.8895 | 0 | 3.3 | 89 | Specific | 921.98 | 714.25 | 714.25 | 18.58 | 97 | >sp\|P01877\|IGHA2_HUMAN Immunoglobulin heavy constant alpha 2 OS=Homo sapiens OX=9606 GN=IGHA2 PE=1 SV=4 | 576 | 20201118_ly_colostrum_1ng_glyco_hcd203040_compound15.6243.6243.4 | controllerType=0 controllerNumber=1 scan=6243 | 16.9559 |
| 02976:5 | 6 | K.HYTN[+1444.534]SSQDVTVPC[+57.021]RVPPPPPC[+57.021]C[+57.021]HPR.L | HexNAc(4)Hex(3)Fuc(1) | C[+57]*3, N[+1445] | 881.3852 | 5 | 4402.8967 | 4402.8895 | 0 | 1.6 | 89 | Specific | 926.88 | 691.02 | 691.02 | 18.73 | 97 | >sp\|P01877\|IGHA2_HUMAN Immunoglobulin heavy constant alpha 2 OS=Homo sapiens OX=9606 GN=IGHA2 PE=1 SV=4 | 576 | 20201118_ly_colostrum_1ng_glyco_hcd203040_compound15.5966.5966.5 | controllerType=0 controllerNumber=1 scan=5966 | 16.4545 |
| 02942:4 | 6 | K.HYTN[+1971.719]SSQDVTVPC[+57.021]RVPPPPPC[+57.021]C[+57.021]HPR.L | HexNAc(5)Hex(5)Fuc(1) | C[+57]*3, N[+1972] | 1233.278 | 4 | 4930.0902 | 4930.0745 | 0 | 3.2 | 89 | Specific | 931.7 | 681.81 | 681.81 | 18.58 | 97 | >sp\|P01877\|IGHA2_HUMAN Immunoglobulin heavy constant alpha 2 OS=Homo sapiens OX=9606 GN=IGHA2 PE=1 SV=4 | 576 | 20201118_ly_colostrum_1ng_glyco_hcd203040_compound15.5931.5931.4 | controllerType=0 controllerNumber=1 scan=5931 | 16.3806 |
| 02980:4 | 6 | K.HYTN[+1606.587]SSQDVTVPC[+57.021]RVPPPPPC[+57.021]C[+57.021]HPR.L | HexNAc(4)Hex(4)Fuc(1) | C[+57]*3, N[+1607] | 1141.9937 | 4 | 4564.953 | 4564.9423 | 0 | 2.3 | 89 | Specific | 951.23 | 655.75 | 655.75 | 19.55 | 97 | >sp\|P01877\|IGHA2_HUMAN Immunoglobulin heavy constant alpha 2 OS=Homo sapiens OX=9606 GN=IGHA2 PE=1 SV=4 | 576 | 20201118_ly_colostrum_1ng_glyco_hcd203040_compound15.5970.5970.4 | controllerType=0 controllerNumber=1 scan=5970 | 16.4648 |
| 02908:5 | 6 | K.HYTN[+1971.719]SSQDVTVPC[+57.021]RVPPPPPC[+57.021]C[+57.021]HPR.L | HexNAc(5)Hex(5)Fuc(1) | C[+57]*3, N[+1972] | 986.8217 | 5 | 4930.0796 | 4930.0745 | 0 | 1 | 89 | Specific | 999.06 | 925.87 | 925.87 | 19.53 | 97 | >sp\|P01877\|IGHA2_HUMAN Immunoglobulin heavy constant alpha 2 OS=Homo sapiens OX=9606 GN=IGHA2 PE=1 SV=4 | 576 | 20201118_ly_colostrum_1ng_glyco_hcd203040_compound15.5895.5895.2 | controllerType=0 controllerNumber=1 scan=5895 | 16.2975 |
| 02969:4 | 6 | K.HYTN[+1647.613]SSQDVTVPC[+57.021]RVPPPPPC[+57.021]C[+57.021]HPR.L | HexNAc(5)Hex(3)Fuc(1) | C[+57]*3, N[+1648] | 1152.2492 | 4 | 4605.9749 | 4605.9689 | 0 | 1.3 | 89 | Specific | 1001.65 | 787.41 | 787.41 | 20.32 | 97 | >sp\|P01877\|IGHA2_HUMAN Immunoglobulin heavy constant alpha 2 OS=Homo sapiens OX=9606 GN=IGHA2 PE=1 SV=4 | 576 | 20201118_ly_colostrum_1ng_glyco_hcd203040_compound15.5959.5959.4 | controllerType=0 controllerNumber=1 scan=5959 | 16.4381 |
| 02977:4 | 6 | K.HYTN[+1606.587]SSQDVTVPC[+57.021]RVPPPPPC[+57.021]C[+57.021]HPR.L | HexNAc(4)Hex(4)Fuc(1) | C[+57]*3, N[+1607] | 1141.9937 | 4 | 4564.953 | 4564.9423 | 0 | 2.3 | 89 | Specific | 1003.1 | 786.08 | 786.08 | 20.55 | 97 | >sp\|P01877\|IGHA2_HUMAN Immunoglobulin heavy constant alpha 2 OS=Homo sapiens OX=9606 GN=IGHA2 PE=1 SV=4 | 576 | 20201118_ly_colostrum_1ng_glyco_hcd203040_compound15.5967.5967.4 | controllerType=0 controllerNumber=1 scan=5967 | 16.4558 |
| 02972:4 | 6 | K.HYTN[+1647.613]SSQDVTVPC[+57.021]RVPPPPPC[+57.021]C[+57.021]HPR.L | HexNAc(5)Hex(3)Fuc(1) | C[+57]*3, N[+1648] | 1152.2492 | 4 | 4605.9749 | 4605.9689 | 0 | 1.3 | 89 | Specific | 1006.03 | 728.29 | 728.29 | 20.32 | 97 | >sp\|P01877\|IGHA2_HUMAN Immunoglobulin heavy constant alpha 2 OS=Homo sapiens OX=9606 GN=IGHA2 PE=1 SV=4 | 576 | 20201118_ly_colostrum_1ng_glyco_hcd203040_compound15.5962.5962.4 | controllerType=0 controllerNumber=1 scan=5962 | 16.4493 |
| 02979:5 | 6 | K.HYTN[+1444.534]SSQDVTVPC[+57.021]RVPPPPPC[+57.021]C[+57.021]HPR.L | HexNAc(4)Hex(3)Fuc(1) | C[+57]*3, N[+1445] | 881.3852 | 5 | 4402.8967 | 4402.8895 | 0 | 1.6 | 89 | Specific | 1050 | 823.1 | 823.1 | 20.73 | 97 | >sp\|P01877\|IGHA2_HUMAN Immunoglobulin heavy constant alpha 2 OS=Homo sapiens OX=9606 GN=IGHA2 PE=1 SV=4 | 576 | 20201118_ly_colostrum_1ng_glyco_hcd203040_compound15.5969.5969.5 | controllerType=0 controllerNumber=1 scan=5969 | 16.4631 |
| 02970:4 | 6 | K.HYTN[+1444.534]SSQDVTVPC[+57.021]RVPPPPPC[+57.021]C[+57.021]HPR.L | HexNAc(4)Hex(3)Fuc(1) | C[+57]*3, N[+1445] | 1101.4799 | 4 | 4402.8978 | 4402.8895 | 0 | 1.9 | 89 | Specific | 1052.31 | 827.59 | 827.59 | 21.32 | 97 | >sp\|P01877\|IGHA2_HUMAN Immunoglobulin heavy constant alpha 2 OS=Homo sapiens OX=9606 GN=IGHA2 PE=1 SV=4 | 576 | 20201118_ly_colostrum_1ng_glyco_hcd203040_compound15.5960.5960.4 | controllerType=0 controllerNumber=1 scan=5960 | 16.4392 |
| 02973:4 | 6 | K.HYTN[+1444.534]SSQDVTVPC[+57.021]RVPPPPPC[+57.021]C[+57.021]HPR.L | HexNAc(4)Hex(3)Fuc(1) | C[+57]*3, N[+1445] | 1101.4799 | 4 | 4402.8978 | 4402.8895 | 0 | 1.9 | 89 | Specific | 1068.42 | 768.74 | 768.74 | 21.32 | 97 | >sp\|P01877\|IGHA2_HUMAN Immunoglobulin heavy constant alpha 2 OS=Homo sapiens OX=9606 GN=IGHA2 PE=1 SV=4 | 576 | 20201118_ly_colostrum_1ng_glyco_hcd203040_compound15.5963.5963.4 | controllerType=0 controllerNumber=1 scan=5963 | 16.4502 |
| 02978:5 | 6 | K.HYTN[+1606.587]SSQDVTVPC[+57.021]RVPPPPPC[+57.021]C[+57.021]HPR.L | HexNAc(4)Hex(4)Fuc(1) | C[+57]*3, N[+1607] | 913.796 | 5 | 4564.9511 | 4564.9423 | 0 | 1.9 | 89 | Specific | 1081.3 | 729.54 | 729.54 | 21.73 | 97 | >sp\|P01877\|IGHA2_HUMAN Immunoglobulin heavy constant alpha 2 OS=Homo sapiens OX=9606 GN=IGHA2 PE=1 SV=4 | 576 | 20201118_ly_colostrum_1ng_glyco_hcd203040_compound15.5968.5968.5 | controllerType=0 controllerNumber=1 scan=5968 | 16.4609 |
| 17542:5 | 6 | R.LSLHRPALEDLLLGSEAN[+1872.651]LTC[+57.021]TLTGLRDASGATFTWTPSSGK.S | HexNAc(3)Hex(6)NeuAc(1) | C[+57], N[+1873] | 1266.7952 | 5 | 6329.9467 | 6329.9274 | 0 | 3.1 | 114 | Specific | 237.9 | 237.9 | 237.9 | 6.27 | 97 | >sp\|P01877\|IGHA2_HUMAN Immunoglobulin heavy constant alpha 2 OS=Homo sapiens OX=9606 GN=IGHA2 PE=1 SV=4 | 576 | 20201118_ly_colostrum_1ng_glyco_hcd203040_compound15.21104.21104.5 | controllerType=0 controllerNumber=1 scan=21104 | 46.2418 |
| 17545:5 | 6 | R.LSLHRPALEDLLLGSEAN[+1872.651]LTC[+57.021]TLTGLRDASGATFTWTPSSGK.S | HexNAc(3)Hex(6)NeuAc(1) | C[+57], N[+1873] | 1266.7952 | 5 | 6329.9467 | 6329.9274 | 0 | 3.1 | 114 | Specific | 587.28 | 501.79 | 501.79 | 11.69 | 97 | >sp\|P01877\|IGHA2_HUMAN Immunoglobulin heavy constant alpha 2 OS=Homo sapiens OX=9606 GN=IGHA2 PE=1 SV=4 | 576 | 20201118_ly_colostrum_1ng_glyco_hcd203040_compound15.21107.21107.5 | controllerType=0 controllerNumber=1 scan=21107 | 46.2482 |
| 16877:4 | 6 | R.LSLHRPALEDLLLGSEAN[+1501.555]LTC[+57.021]TLTGLRDASGATFTWTPSSGK.S | HexNAc(5)Hex(3) | C[+57], N[+1502] | 1490.4711 | 4 | 5958.8625 | 5958.8323 | 0 | 5.1 | 114 | Specific | 908.76 | 718.57 | 718.57 | 17.7 | 97 | >sp\|P01877\|IGHA2_HUMAN Immunoglobulin heavy constant alpha 2 OS=Homo sapiens OX=9606 GN=IGHA2 PE=1 SV=4 | 576 | 20201118_ly_colostrum_1ng_glyco_hcd203040_compound15.20409.20409.4 | controllerType=0 controllerNumber=1 scan=20409 | 44.6833 |
| 16894:4 | 6 | R.LSLHRPALEDLLLGSEAN[+1216.423]LTC[+57.021]TLTGLRDASGATFTWTPSSGK.S | HexNAc(2)Hex(5) | C[+57], N[+1216] | 1419.1847 | 4 | 5673.717 | 5673.6998 | 0 | 3 | 114 | Specific | 932.27 | 817.47 | 817.47 | 18.24 | 97 | >sp\|P01877\|IGHA2_HUMAN Immunoglobulin heavy constant alpha 2 OS=Homo sapiens OX=9606 GN=IGHA2 PE=1 SV=4 | 576 | 20201118_ly_colostrum_1ng_glyco_hcd203040_compound15.20427.20427.4 | controllerType=0 controllerNumber=1 scan=20427 | 44.7459 |
| 16875:4 | 6 | R.LSLHRPALEDLLLGSEAN[+1501.555]LTC[+57.021]TLTGLRDASGATFTWTPSSGK.S | HexNAc(5)Hex(3) | C[+57], N[+1502] | 1490.4711 | 4 | 5958.8625 | 5958.8323 | 0 | 5.1 | 114 | Specific | 1010.51 | 1010.51 | 1010.51 | 19.7 | 97 | >sp\|P01877\|IGHA2_HUMAN Immunoglobulin heavy constant alpha 2 OS=Homo sapiens OX=9606 GN=IGHA2 PE=1 SV=4 | 576 | 20201118_ly_colostrum_1ng_glyco_hcd203040_compound15.20407.20407.4 | controllerType=0 controllerNumber=1 scan=20407 | 44.6744 |
| 16891:5 | 6 | R.LSLHRPALEDLLLGSEAN[+1216.423]LTC[+57.021]TLTGLRDASGATFTWTPSSGK.S | HexNAc(2)Hex(5) | C[+57], N[+1216] | 1135.5496 | 5 | 5673.7187 | 5673.6998 | 0 | 3.3 | 114 | Specific | 1064.86 | 1064.86 | 1064.86 | 21.69 | 97 | >sp\|P01877\|IGHA2_HUMAN Immunoglobulin heavy constant alpha 2 OS=Homo sapiens OX=9606 GN=IGHA2 PE=1 SV=4 | 576 | 20201118_ly_colostrum_1ng_glyco_hcd203040_compound15.20424.20424.5 | controllerType=0 controllerNumber=1 scan=20424 | 44.7304 |
| 16893:5 | 6 | R.LSLHRPALEDLLLGSEAN[+1216.423]LTC[+57.021]TLTGLRDASGATFTWTPSSGK.S | HexNAc(2)Hex(5) | C[+57], N[+1216] | 1135.5496 | 5 | 5673.7187 | 5673.6998 | 0 | 3.3 | 114 | Specific | 1158.95 | 1082.81 | 1082.81 | 23.69 | 97 | >sp\|P01877\|IGHA2_HUMAN Immunoglobulin heavy constant alpha 2 OS=Homo sapiens OX=9606 GN=IGHA2 PE=1 SV=4 | 576 | 20201118_ly_colostrum_1ng_glyco_hcd203040_compound15.20426.20426.5 | controllerType=0 controllerNumber=1 scan=20426 | 44.7405 |
| 16892:4 | 6 | R.LSLHRPALEDLLLGSEAN[+1216.423]LTC[+57.021]TLTGLRDASGATFTWTPSSGK.S | HexNAc(2)Hex(5) | C[+57], N[+1216] | 1419.1847 | 4 | 5673.717 | 5673.6998 | 0 | 3 | 114 | Specific | 1275.84 | 1275.84 | 1275.84 | 25.24 | 97 | >sp\|P01877\|IGHA2_HUMAN Immunoglobulin heavy constant alpha 2 OS=Homo sapiens OX=9606 GN=IGHA2 PE=1 SV=4 | 576 | 20201118_ly_colostrum_1ng_glyco_hcd203040_compound15.20425.20425.4 | controllerType=0 controllerNumber=1 scan=20425 | 44.7318 |
| 04391:2 | 6 | K.TPLTAN[+2059.735]ITK.S | HexNAc(4)Hex(5)Fuc(1)NeuAc(1) | N[+2060] | 1509.6546 | 2 | 3018.3019 | 3018.2917 | 0 | 3.4 | 200 | Specific | 397.24 | 180.59 | 180.59 | 6.41 | 97 | >sp\|P01877\|IGHA2_HUMAN Immunoglobulin heavy constant alpha 2 OS=Homo sapiens OX=9606 GN=IGHA2 PE=1 SV=4 | 576 | 20201118_ly_colostrum_1ng_glyco_hcd203040_compound15.7433.7433.2 | controllerType=0 controllerNumber=1 scan=7433 | 19.0343 |
| 04373:3 | 6 | K.TPLTAN[+2059.735]ITK.S | HexNAc(4)Hex(5)Fuc(1)NeuAc(1) | N[+2060] | 1006.7717 | 3 | 3018.3005 | 3018.2917 | 0 | 2.9 | 200 | Specific | 441.65 | 151.72 | 151.72 | 6.43 | 97 | >sp\|P01877\|IGHA2_HUMAN Immunoglobulin heavy constant alpha 2 OS=Homo sapiens OX=9606 GN=IGHA2 PE=1 SV=4 | 576 | 20201118_ly_colostrum_1ng_glyco_hcd203040_compound15.7415.7415.3 | controllerType=0 controllerNumber=1 scan=7415 | 19.0045 |
| 02658:3 | 6 | K.TPLTAN[+1971.719]ITK.S | HexNAc(5)Hex(5)Fuc(1) | N[+1972] | 977.4317 | 3 | 2930.2805 | 2930.2757 | 0 | 1.6 | 200 | Specific | 441.97 | 178.44 | 178.44 | 6.63 | 97 | >sp\|P01877\|IGHA2_HUMAN Immunoglobulin heavy constant alpha 2 OS=Homo sapiens OX=9606 GN=IGHA2 PE=1 SV=4 | 576 | 20201118_ly_colostrum_1ng_glyco_hcd203040_compound15.5635.5635.3 | controllerType=0 controllerNumber=1 scan=5635 | 15.7422 |
| 04376:3 | 6 | K.TPLTAN[+2059.735]ITK.S | HexNAc(4)Hex(5)Fuc(1)NeuAc(1) | N[+2060] | 1006.7717 | 3 | 3018.3005 | 3018.2917 | 0 | 2.9 | 200 | Specific | 462.34 | 230.24 | 230.24 | 7.35 | 97 | >sp\|P01877\|IGHA2_HUMAN Immunoglobulin heavy constant alpha 2 OS=Homo sapiens OX=9606 GN=IGHA2 PE=1 SV=4 | 576 | 20201118_ly_colostrum_1ng_glyco_hcd203040_compound15.7418.7418.3 | controllerType=0 controllerNumber=1 scan=7418 | 19.0094 |
| 04404:2 | 6 | K.TPLTAN[+2262.814]ITK.S | HexNAc(5)Hex(5)Fuc(1)NeuAc(1) | N[+2263] | 1611.1929 | 2 | 3221.3786 | 3221.3711 | 0 | 2.3 | 200 | Specific | 472.65 | 204.57 | 204.57 | 7.04 | 97 | >sp\|P01877\|IGHA2_HUMAN Immunoglobulin heavy constant alpha 2 OS=Homo sapiens OX=9606 GN=IGHA2 PE=1 SV=4 | 576 | 20201118_ly_colostrum_1ng_glyco_hcd203040_compound15.7447.7447.2 | controllerType=0 controllerNumber=1 scan=7447 | 19.0611 |
| 02777:3 | 6 | K.TPLTAN[+1647.613]ITK.S | HexNAc(5)Hex(3)Fuc(1) | N[+1648] | 869.3969 | 3 | 2606.1763 | 2606.17 | 0 | 2.4 | 200 | Specific | 481.41 | 195.88 | 195.88 | 7.29 | 97 | >sp\|P01877\|IGHA2_HUMAN Immunoglobulin heavy constant alpha 2 OS=Homo sapiens OX=9606 GN=IGHA2 PE=1 SV=4 | 576 | 20201118_ly_colostrum_1ng_glyco_hcd203040_compound15.5759.5759.3 | controllerType=0 controllerNumber=1 scan=5759 | 16.0172 |
| 02707:3 | 6 | K.TPLTAN[+1809.666]ITK.S | HexNAc(5)Hex(4)Fuc(1) | N[+1810] | 923.4144 | 3 | 2768.2286 | 2768.2228 | 0 | 2.1 | 200 | Specific | 484.68 | 204.81 | 204.81 | 7.05 | 97 | >sp\|P01877\|IGHA2_HUMAN Immunoglobulin heavy constant alpha 2 OS=Homo sapiens OX=9606 GN=IGHA2 PE=1 SV=4 | 576 | 20201118_ly_colostrum_1ng_glyco_hcd203040_compound15.5686.5686.3 | controllerType=0 controllerNumber=1 scan=5686 | 15.8583 |
| 04393:2 | 6 | K.TPLTAN[+2059.735]ITK.S | HexNAc(4)Hex(5)Fuc(1)NeuAc(1) | N[+2060] | 1509.6546 | 2 | 3018.3019 | 3018.2917 | 0 | 3.4 | 200 | Specific | 514.38 | 289.52 | 289.52 | 7.75 | 97 | >sp\|P01877\|IGHA2_HUMAN Immunoglobulin heavy constant alpha 2 OS=Homo sapiens OX=9606 GN=IGHA2 PE=1 SV=4 | 576 | 20201118_ly_colostrum_1ng_glyco_hcd203040_compound15.7435.7435.2 | controllerType=0 controllerNumber=1 scan=7435 | 19.038 |
| 02660:3 | 6 | K.TPLTAN[+1971.719]ITK.S | HexNAc(5)Hex(5)Fuc(1) | N[+1972] | 977.4317 | 3 | 2930.2805 | 2930.2757 | 0 | 1.6 | 200 | Specific | 515.39 | 275.62 | 275.62 | 7.61 | 97 | >sp\|P01877\|IGHA2_HUMAN Immunoglobulin heavy constant alpha 2 OS=Homo sapiens OX=9606 GN=IGHA2 PE=1 SV=4 | 576 | 20201118_ly_colostrum_1ng_glyco_hcd203040_compound15.5637.5637.3 | controllerType=0 controllerNumber=1 scan=5637 | 15.7474 |
| 02779:3 | 6 | K.TPLTAN[+1647.613]ITK.S | HexNAc(5)Hex(3)Fuc(1) | N[+1648] | 869.3969 | 3 | 2606.1763 | 2606.17 | 0 | 2.4 | 200 | Specific | 529.23 | 257.71 | 257.71 | 7.74 | 97 | >sp\|P01877\|IGHA2_HUMAN Immunoglobulin heavy constant alpha 2 OS=Homo sapiens OX=9606 GN=IGHA2 PE=1 SV=4 | 576 | 20201118_ly_colostrum_1ng_glyco_hcd203040_compound15.5761.5761.3 | controllerType=0 controllerNumber=1 scan=5761 | 16.0238 |
| 02709:3 | 6 | K.TPLTAN[+1809.666]ITK.S | HexNAc(5)Hex(4)Fuc(1) | N[+1810] | 923.4144 | 3 | 2768.2286 | 2768.2228 | 0 | 2.1 | 200 | Specific | 534.23 | 292.73 | 292.73 | 7.69 | 97 | >sp\|P01877\|IGHA2_HUMAN Immunoglobulin heavy constant alpha 2 OS=Homo sapiens OX=9606 GN=IGHA2 PE=1 SV=4 | 576 | 20201118_ly_colostrum_1ng_glyco_hcd203040_compound15.5688.5688.3 | controllerType=0 controllerNumber=1 scan=5688 | 15.8664 |
| 02823:2 | 6 | K.TPLTAN[+1647.613]ITK.S | HexNAc(5)Hex(3)Fuc(1) | N[+1648] | 1303.5923 | 2 | 2606.1774 | 2606.17 | 0 | 2.8 | 200 | Specific | 534.63 | 306.17 | 306.17 | 7.95 | 97 | >sp\|P01877\|IGHA2_HUMAN Immunoglobulin heavy constant alpha 2 OS=Homo sapiens OX=9606 GN=IGHA2 PE=1 SV=4 | 576 | 20201118_ly_colostrum_1ng_glyco_hcd203040_compound15.5807.5807.2 | controllerType=0 controllerNumber=1 scan=5807 | 16.1181 |
| 04406:2 | 6 | K.TPLTAN[+2262.814]ITK.S | HexNAc(5)Hex(5)Fuc(1)NeuAc(1) | N[+2263] | 1611.1929 | 2 | 3221.3786 | 3221.3711 | 0 | 2.3 | 200 | Specific | 543.32 | 279.73 | 279.73 | 7.91 | 97 | >sp\|P01877\|IGHA2_HUMAN Immunoglobulin heavy constant alpha 2 OS=Homo sapiens OX=9606 GN=IGHA2 PE=1 SV=4 | 576 | 20201118_ly_colostrum_1ng_glyco_hcd203040_compound15.7449.7449.2 | controllerType=0 controllerNumber=1 scan=7449 | 19.0652 |
| 04394:3 | 6 | K.TPLTAN[+2262.814]ITK.S | HexNAc(5)Hex(5)Fuc(1)NeuAc(1) | N[+2263] | 1074.4651 | 3 | 3221.3808 | 3221.3711 | 0 | 3 | 200 | Specific | 546.44 | 242.4 | 242.4 | 7.93 | 97 | >sp\|P01877\|IGHA2_HUMAN Immunoglobulin heavy constant alpha 2 OS=Homo sapiens OX=9606 GN=IGHA2 PE=1 SV=4 | 576 | 20201118_ly_colostrum_1ng_glyco_hcd203040_compound15.7437.7437.3 | controllerType=0 controllerNumber=1 scan=7437 | 19.0425 |
| 02681:2 | 6 | K.TPLTAN[+1971.719]ITK.S | HexNAc(5)Hex(5)Fuc(1) | N[+1972] | 1465.6469 | 2 | 2930.2865 | 2930.2757 | 0 | 3.7 | 200 | Specific | 566.44 | 332.7 | 332.7 | 8.51 | 97 | >sp\|P01877\|IGHA2_HUMAN Immunoglobulin heavy constant alpha 2 OS=Homo sapiens OX=9606 GN=IGHA2 PE=1 SV=4 | 576 | 20201118_ly_colostrum_1ng_glyco_hcd203040_compound15.5659.5659.2 | controllerType=0 controllerNumber=1 scan=5659 | 15.7963 |
| 02683:2 | 6 | K.TPLTAN[+1971.719]ITK.S | HexNAc(5)Hex(5)Fuc(1) | N[+1972] | 1465.6469 | 2 | 2930.2865 | 2930.2757 | 0 | 3.7 | 200 | Specific | 583.81 | 316.26 | 316.26 | 9.66 | 97 | >sp\|P01877\|IGHA2_HUMAN Immunoglobulin heavy constant alpha 2 OS=Homo sapiens OX=9606 GN=IGHA2 PE=1 SV=4 | 576 | 20201118_ly_colostrum_1ng_glyco_hcd203040_compound15.5661.5661.2 | controllerType=0 controllerNumber=1 scan=5661 | 15.8032 |
| 04396:3 | 6 | K.TPLTAN[+2262.814]ITK.S | HexNAc(5)Hex(5)Fuc(1)NeuAc(1) | N[+2263] | 1074.4651 | 3 | 3221.3808 | 3221.3711 | 0 | 3 | 200 | Specific | 586.26 | 265.74 | 265.74 | 9.57 | 97 | >sp\|P01877\|IGHA2_HUMAN Immunoglobulin heavy constant alpha 2 OS=Homo sapiens OX=9606 GN=IGHA2 PE=1 SV=4 | 576 | 20201118_ly_colostrum_1ng_glyco_hcd203040_compound15.7439.7439.3 | controllerType=0 controllerNumber=1 scan=7439 | 19.0[illegible]48 |
| 02774:2 | 6 | K.TPLTAN[+1647.613]ITK.S | HexNAc(5)Hex(3)Fuc(1) | N[+1648] | 1303.592 | 2 | 2606.1766 | 2606.17 | 0 | 2.5 | 200 | Specific | 594.02 | 325.13 | 325.13 | 9.81 | 97 | >sp\|P01877\|IGHA2_HUMAN Immunoglobulin heavy constant alpha 2 OS=Homo sapiens OX=9606 GN=IGHA2 PE=1 SV=4 | 576 | 20201118_ly_colostrum_1ng_glyco_hcd203040_compound15.5756.5756.2 | controllerType=0 controllerNumber=1 scan=5756 | 16.[illegible]21 |
| 02782:2 | 6 | K.TPLTAN[+1501.555]ITK.S | HexNAc(5)Hex(3) | N[+1502] | 1230.5638 | 2 | 2460.1203 | 2460.1121 | 0 | 3.3 | 200 | Specific | 599.26 | 318.67 | 318.67 | 9.66 | 97 | >sp\|P01877\|IGHA2_HUMAN Immunoglobulin heavy constant alpha 2 OS=Homo sapiens OX=9606 GN=IGHA2 PE=1 SV=4 | 576 | 20201118_ly_colostrum_1ng_glyco_hcd203040_compound15.5764.5764.2 | controllerType=0 controllerNumber=1 scan=5764 | 16.0291 |
| 02826:2 | 6 | K.TPLTAN[+1647.613]ITK.S | HexNAc(5)Hex(3)Fuc(1) | N[+1648] | 1303.5923 | 2 | 2606.1774 | 2606.17 | 0 | 2.8 | 200 | Specific | 609.58 | 320.12 | 320.12 | 10.63 | 97 | >sp\|P01877\|IGHA2_HUMAN Immunoglobulin heavy constant alpha 2 OS=Homo sapiens OX=9606 GN=IGHA2 PE=1 SV=4 | 576 | 20201118_ly_colostrum_1ng_glyco_hcd203040_compound15.5810.5810.2 | controllerType=0 controllerNumber=1 scan=5810 | 16.1222 |
| 02708:2 | 6 | K.TPLTAN[+1809.666]ITK.S | HexNAc(5)Hex(4)Fuc(1) | N[+1810] | 1384.6177 | 2 | 2768.2282 | 2768.2228 | 0 | 1.9 | 200 | Specific | 609.69 | 329.72 | 329.72 | 10.57 | 97 | >sp\|P01877\|IGHA2_HUMAN Immunoglobulin heavy constant alpha 2 OS=Homo sapiens OX=9606 GN=IGHA2 PE=1 SV=4 | 576 | 20201118_ly_colostrum_1ng_glyco_hcd203040_compound15.5687.5687.2 | controllerType=0 controllerNumber=1 scan=5687 | 15.863 |
| 02706:2 | 6 | K.TPLTAN[+1809.666]ITK.S | HexNAc(5)Hex(4)Fuc(1) | N[+1810] | 1384.6177 | 2 | 2768.2282 | 2768.2228 | 0 | 1.9 | 200 | Specific | 622.03 | 364.84 | 364.84 | 10.83 | 97 | >sp\|P01877\|IGHA2_HUMAN Immunoglobulin heavy constant alpha 2 OS=Homo sapiens OX=9606 GN=IGHA2 PE=1 SV=4 | 576 | 20201118_ly_colostrum_1ng_glyco_hcd203040_compound15.5685.5685.2 | controllerType=0 controllerNumber=1 scan=5685 | 15.8568 |
| 02712:2 | 6 | K.TPLTAN[+1606.587]ITK.S | HexNAc(4)Hex(4)Fuc(1) | N[+1607] | 1283.0777 | 2 | 2565.1481 | 2565.1435 | 0 | 1.8 | 200 | Specific | 624.04 | 301.25 | 301.25 | 10.4 | 97 | >sp\|P01877\|IGHA2_HUMAN Immunoglobulin heavy constant alpha 2 OS=Homo sapiens OX=9606 GN=IGHA2 PE=1 SV=4 | 576 | 20201118_ly_colostrum_1ng_glyco_hcd203040_compound15.5691.5691.2 | controllerType=0 controllerNumber=1 scan=5691 | 15.8725 |
| 02780:2 | 6 | K.TPLTAN[+1501.555]ITK.S | HexNAc(5)Hex(3) | N[+1502] | 1230.5638 | 2 | 2460.1203 | 2460.1121 | 0 | 3.3 | 200 | Specific | 626.59 | 312.96 | 312.96 | 10.66 | 97 | >sp\|P01877\|IGHA2_HUMAN Immunoglobulin heavy constant alpha 2 OS=Homo sapiens OX=9606 GN=IGHA2 PE=1 SV=4 | 576 | 20201118_ly_colostrum_1ng_glyco_hcd203040_compound15.5762.5762.2 | controllerType=0 controllerNumber=1 scan=5762 | 16.025 |
| 02772:2 | 6 | K.TPLTAN[+1647.613]ITK.S | HexNAc(5)Hex(3)Fuc(1) | N[+1648] | 1303.592 | 2 | 2606.1766 | 2606.17 | 0 | 2.5 | 200 | Specific | 637.23 | 363.55 | 363.55 | 11.06 | 97 | >sp\|P01877\|IGHA2_HUMAN Immunoglobulin heavy constant alpha 2 OS=Homo sapiens OX=9606 GN=IGHA2 PE=1 SV=4 | 576 | 20201118_ly_colostrum_1ng_glyco_hcd203040_compound15.5754.5754.2 | controllerType=0 controllerNumber=1 scan=5754 | 16.0038 |
| 02771:2 | 6 | K.TPLTAN[+1444.534]ITK.S | HexNAc(4)Hex(3)Fuc(1) | N[+1445] | 1202.0525 | 2 | 2403.0978 | 2403.0906 | 0 | 3 | 200 | Specific | 645.91 | 314.86 | 314.86 | 10.66 | 97 | >sp\|P01877\|IGHA2_HUMAN Immunoglobulin heavy constant alpha 2 OS=Homo sapiens OX=9606 GN=IGHA2 PE=1 SV=4 | 576 | 20201118_ly_colostrum_1ng_glyco_hcd203040_compound15.5752.5752.2 | controllerType=0 controllerNumber=1 scan=5752 | 15.9992 |
| 02770:2 | 6 | K.TPLTAN[+1444.534]ITK.S | HexNAc(4)Hex(3)Fuc(1) | N[+1445] | 1202.0525 | 2 | 2403.0978 | 2403.0906 | 0 | 3 | 200 | Specific | 650.11 | 371.34 | 371.34 | 12.09 | 97 | >sp\|P01877\|IGHA2_HUMAN Immunoglobulin heavy constant alpha 2 OS=Homo sapiens OX=9606 GN=IGHA2 PE=1 SV=4 | 576 | 20201118_ly_colostrum_1ng_glyco_hcd203040_compound15.5751.5751.2 | controllerType=0 controllerNumber=1 scan=5751 | 15.9926 |
| 02710:2 | 6 | K.TPLTAN[+1606.587]ITK.S | HexNAc(4)Hex(4)Fuc(1) | N[+1607] | 1283.0777 | 2 | 2565.1481 | 2565.1435 | 0 | 1.8 | 200 | Specific | 665.53 | 328.44 | 328.44 | 11.57 | 97 | >sp\|P01877\|IGHA2_HUMAN Immunoglobulin heavy constant alpha 2 OS=Homo sapiens OX=9606 GN=IGHA2 PE=1 SV=4 | 576 | 20201118_ly_colostrum_1ng_glyco_hcd203040_compound15.5689.5689.2 | controllerType=0 controllerNumber=1 scan=5689 | 15.8679 |
| 08746:3 | 6 | R.MAGKPTHIN[+1809.666]VSVVMAEADGTC[+57.021]Y.- | HexNAc(5)Hex(4)Fuc(1) | C[+57], N[+1810] | 1387.5835 | 3 | 4160.7359 | 4160.7489 | 0 | -3.1 | 319 | CRagged | 226.73 | 226.73 | 150.86 | 5.8 | 97 | >sp\|P01877\|IGHA2_HUMAN Immunoglobulin heavy constant alpha 2 OS=Homo sapiens OX=9606 GN=IGHA2 PE=1 SV=4 | 576 | 20201118_ly_colostrum_1ng_glyco_hcd203040_compound15.11947.11947.3 | controllerType=0 controllerNumber=1 scan=11947 | 27.2312 |
| 08748:3 | 6 | R.MAGKPTHIN[+1809.666]VSVVMAEADGTC[+57.021]Y.- | HexNAc(5)Hex(4)Fuc(1) | C[+57], N[+1810] | 1387.5835 | 3 | 4160.7359 | 4160.7489 | 0 | -3.1 | 319 | CRagged | 251.96 | 175.7 | 175.7 | 5.51 | 97 | >sp\|P01877\|IGHA2_HUMAN Immunoglobulin heavy constant alpha 2 OS=Homo sapiens OX=9606 GN=IGHA2 PE=1 SV=4 | 576 | 20201118_ly_colostrum_1ng_glyco_hcd203040_compound15.11949.11949.3 | controllerType=0 controllerNumber=1 scan=11949 | 27.2348 |
| 09515:3 | 6 | R.MAGKPTHIN[+1419.502]VSVVMAEADGTC[+57.021]Y.- | HexNAc(3)Hex(5) | C[+57], N[+1420] | 1257.5367 | 3 | 3770.5955 | 3770.5851 | 0 | 2.8 | 319 | CRagged | 257.87 | 231.35 | 175.4 | 7.1 | 97 | >sp\|P01877\|IGHA2_HUMAN Immunoglobulin heavy constant alpha 2 OS=Homo sapiens OX=9606 GN=IGHA2 PE=1 SV=4 | 576 | 20201118_ly_colostrum_1ng_glyco_hcd203040_compound15.12747.12747.3 | controllerType=0 controllerNumber=1 scan=12747 | 28.8422 |
| 09516:3 | 6 | R.MAGKPTHIN[+1622.582]VSVVMAEADGTC[+57.021]Y.- | HexNAc(4)Hex(5) | C[+57], N[+1623] | 1325.2316 | 3 | 3973.6803 | 3973.6645 | 0 | 4 | 319 | CRagged | 320.27 | 306.62 | 238.07 | 8.07 | 97 | >sp\|P01877\|IGHA2_HUMAN Immunoglobulin heavy constant alpha 2 OS=Homo sapiens OX=9606 GN=IGHA2 PE=1 SV=4 | 576 | 20201118_ly_colostrum_1ng_glyco_hcd203040_compound15.12748.12748.3 | controllerType=0 controllerNumber=1 scan=12748 | 28.8434 |
| 09467:3 | 6 | R.MAGKPTHIN[+1581.555]VSVVMAEADGTC[+57.021]Y.- | HexNAc(3)Hex(6) | C[+57], N[+1582] | 1311.5536 | 3 | 3932.6464 | 3932.6379 | 0 | 2.2 | 319 | CRagged | 339.11 | 339.11 | 244.14 | 8.22 | 97 | >sp\|P01877\|IGHA2_HUMAN Immunoglobulin heavy constant alpha 2 OS=Homo sapiens OX=9606 GN=IGHA2 PE=1 SV=4 | 576 | 20201118_ly_colostrum_1ng_glyco_hcd203040_compound15.12697.12697.3 | controllerType=0 controllerNumber=1 scan=12697 | 28.7435 |
| 09518:3 | 6 | R.MAGKPTHIN[+1622.582]VSVVMAEADGTC[+57.021]Y.- | HexNAc(4)Hex(5) | C[+57], N[+1623] | 1325.2316 | 3 | 3973.6803 | 3973.6645 | 0 | 4 | 319 | CRagged | 361.57 | 361.57 | 299.04 | 8.63 | 97 | >sp\|P01877\|IGHA2_HUMAN Immunoglobulin heavy constant alpha 2 OS=Homo sapiens OX=9606 GN=IGHA2 PE=1 SV=4 | 576 | 20201118_ly_colostrum_1ng_glyco_hcd203040_compound15.12750.12750.3 | controllerType=0 controllerNumber=1 scan=12750 | 28.8484 |
| 09555:3 | 6 | R.MAGKPTHIN[+1257.449]VSVVMAEADGTC[+57.021]Y.- | HexNAc(3)Hex(4) | C[+57], N[+1257] | 1203.5189 | 3 | 3608.542 | 3608.5323 | 0 | 2.7 | 319 | CRagged | 363.81 | 363.81 | 204.11 | 8.61 | 97 | >sp\|P01877\|IGHA2_HUMAN Immunoglobulin heavy constant alpha 2 OS=Homo sapiens OX=9606 GN=IGHA2 PE=1 SV=4 | 576 | 20201118_ly_colostrum_1ng_glyco_hcd203040_compound15.12788.12788.3 | controllerType=0 controllerNumber=1 scan=12788 | 28.9214 |
| 09517:3 | 6 | R.MAGKPTHIN[+1419.502]VSVVMAEADGTC[+57.021]Y.- | HexNAc(3)Hex(5) | C[+57], N[+1420] | 1257.5367 | 3 | 3770.5955 | 3770.5851 | 0 | 2.8 | 319 | CRagged | 364.46 | 364.46 | 299.35 | 8.61 | 97 | >sp\|P01877\|IGHA2_HUMAN Immunoglobulin heavy constant alpha 2 OS=Homo sapiens OX=9606 GN=IGHA2 PE=1 SV=4 | 576 | 20201118_ly_colostrum_1ng_glyco_hcd203040_compound15.12749.12749.3 | controllerType=0 controllerNumber=1 scan=12749 | 28.8459 |
| 09415:3 | 6 | R.MAGKPTHIN[+1825.661]VSVVMAEADGTC[+57.021]Y.- | HexNAc(5)Hex(5) | C[+57], N[+1826] | 1392.9236 | 3 | 4176.7564 | 4176.7438 | 0 | 3 | 319 | CRagged | 366.11 | 328.54 | 307.78 | 8.35 | 97 | >sp\|P01877\|IGHA2_HUMAN Immunoglobulin heavy constant alpha 2 OS=Homo sapiens OX=9606 GN=IGHA2 PE=1 SV=4 | 576 | 20201118_ly_colostrum_1ng_glyco_hcd203040_compound15.12643.12643.3 | controllerType=0 controllerNumber=1 scan=12643 | 28.6345 |
| 09417:3 | 6 | R.MAGKPTHIN[+1825.661]VSVVMAEADGTC[+57.021]Y.- | HexNAc(5)Hex(5) | C[+57], N[+1826] | 1392.9236 | 3 | 4176.7564 | 4176.7438 | 0 | 3 | 319 | CRagged | 412.35 | 412.35 | 226.41 | 10.27 | 97 | >sp\|P01877\|IGHA2_HUMAN Immunoglobulin heavy constant alpha 2 OS=Homo sapiens OX=9606 GN=IGHA2 PE=1 SV=4 | 576 | 20201118_ly_colostrum_1ng_glyco_hcd203040_compound15.12645.12645.3 | controllerType=0 controllerNumber=1 scan=12645 | 28.6377 |
| 10947:3 | 6 | R.MAGKPTHIN[+1872.651]VSVVMAEADGTC[+57.021]Y.- | HexNAc(3)Hex(6)NeuAc(1) | C[+57], N[+1873] | 1408.5851 | 3 | 4223.7409 | 4223.7333 | 0 | 1.8 | 319 | CRagged | 477.79 | 399.1 | 304.96 | 10.21 | 97 | >sp\|P01877\|IGHA2_HUMAN Immunoglobulin heavy constant alpha 2 OS=Homo sapiens OX=9606 GN=IGHA2 PE=1 SV=4 | 576 | 20201118_ly_colostrum_1ng_glyco_hcd203040_compound15.14236.14236.3 | controllerType=0 controllerNumber=1 scan=14236 | 31.8352 |
| 10925:3 | 6 | R.MAGKPTHIN[+1913.677]VSVVMAEADGTC[+57.021]Y.- | HexNAc(4)Hex(5)NeuAc(1) | C[+57], N[+1914] | 1422.2598 | 3 | 4264.7649 | 4264.7599 | 0 | 1.2 | 319 | CRagged | 498.58 | 439.95 | 439.95 | 10.3 | 97 | >sp\|P01877\|IGHA2_HUMAN Immunoglobulin heavy constant alpha 2 OS=Homo sapiens OX=9606 GN=IGHA2 PE=1 SV=4 | 576 | 20201118_ly_colostrum_1ng_glyco_hcd203040_compound15.14213.14213.3 | controllerType=0 controllerNumber=1 scan=14213 | 31.7942 |
| 09470:3 | 6 | R.MAGKPTHIN[+1581.555]VSVVMAEADGTC[+57.021]Y.- | HexNAc(3)Hex(6) | C[+57], N[+1582] | 1311.5536 | 3 | 3932.6464 | 3932.6379 | 0 | 2.2 | 319 | CRagged | 509.32 | 490.09 | 328.81 | 10.65 | 97 | >sp\|P01877\|IGHA2_HUMAN Immunoglobulin heavy constant alpha 2 OS=Homo sapiens OX=9606 GN=IGHA2 PE=1 SV=4 | 576 | 20201118_ly_colostrum_1ng_glyco_hcd203040_compound15.12700.12700.3 | controllerType=0 controllerNumber=1 scan=12700 | 28.7477 |
| 10922:3 | 6 | R.MAGKPTHIN[+1913.677]VSVVMAEADGTC[+57.021]Y.- | HexNAc(4)Hex(5)NeuAc(1) | C[+57], N[+1914] | 1422.2598 | 3 | 4264.7649 | 4264.7599 | 0 | 1.2 | 319 | CRagged | 546.48 | 431.98 | 431.98 | 10.65 | 97 | >sp\|P01877\|IGHA2_HUMAN Immunoglobulin heavy constant alpha 2 OS=Homo sapiens OX=9606 GN=IGHA2 PE=1 SV=4 | 576 | 20201118_ly_colostrum_1ng_glyco_hcd203040_compound15.14210.14210.3 | controllerType=0 controllerNumber=1 scan=14210 | 31.7898 |
| 09557:3 | 6 | R.MAGKPTHIN[+1257.449]VSVVMAEADGTC[+57.021]Y.- | HexNAc(3)Hex(4) | C[+57], N[+1257] | 1203.5189 | 3 | 3608.542 | 3608.5323 | 0 | 2.7 | 319 | CRagged | 547.47 | 523.16 | 350.25 | 10.88 | 97 | >sp\|P01877\|IGHA2_HUMAN Immunoglobulin heavy constant alpha 2 OS=Homo sapiens OX=9606 GN=IGHA2 PE=1 SV=4 | 576 | 20201118_ly_colostrum_1ng_glyco_hcd203040_compound15.12790.12790.3 | controllerType=0 controllerNumber=1 scan=12790 | 28.9249 |
| 11018:3 | 6 | R.MAGKPTHIN[+1548.545]VSVVMAEADGTC[+57.021]Y.- | HexNAc(3)Hex(4)NeuAc(1) | C[+57], N[+1549] | 1300.5511 | 3 | 3899.6387 | 3899.6277 | 0 | 2.8 | 319 | CRagged | 588.01 | 495.26 | 495.26 | 12.38 | 97 | >sp\|P01877\|IGHA2_HUMAN Immunoglobulin heavy constant alpha 2 OS=Homo sapiens OX=9606 GN=IGHA2 PE=1 SV=4 | 576 | 20201118_ly_colostrum_1ng_glyco_hcd203040_compound15.14310.14310.3 | controllerType=0 controllerNumber=1 scan=14310 | 31.9732 |
| 09590:3 | 6 | R.MAGKPTHIN[+1095.397]VSVVMAEADGTC[+57.021]Y.- | HexNAc(3)Hex(3) | C[+57], N[+1095] | 1149.5018 | 3 | 3446.4907 | 3446.4794 | 0 | 3.3 | 319 | CRagged | 590.57 | 590.57 | 590.57 | 12.41 | 97 | >sp\|P01877\|IGHA2_HUMAN Immunoglobulin heavy constant alpha 2 OS=Homo sapiens OX=9606 GN=IGHA2 PE=1 SV=4 | 576 | 20201118_ly_colostrum_1ng_glyco_hcd203040_compound15.12825.12825.3 | controllerType=0 controllerNumber=1 scan=12825 | 28.9906 |
| 09531:3 | 6 | R.MAGKPTHIN[+1501.555]VSVVMAEADGTC[+57.021]Y.- | HexNAc(5)Hex(3) | C[+57], N[+1502] | 1284.8882 | 3 | 3852.6502 | 3852.6382 | 0 | 3.1 | 319 | CRagged | 605.29 | 556.73 | 556.73 | 13.41 | 97 | >sp\|P01877\|IGHA2_HUMAN Immunoglobulin heavy constant alpha 2 OS=Homo sapiens OX=9606 GN=IGHA2 PE=1 SV=4 | 576 | 20201118_ly_colostrum_1ng_glyco_hcd203040_compound15.12764.12764.3 | controllerType=0 controllerNumber=1 scan=12764 | 28.8797 |
| 09461:3 | 6 | R.MAGKPTHIN[+1809.666]VSVVMAEADGTC[+57.021]Y.- | HexNAc(5)Hex(4)Fuc(1) | C[+57], N[+1810] | 1387.591 | 3 | 4160.7584 | 4160.7489 | 0 | 2.3 | 319 | CRagged | 610.53 | 556.44 | 364.78 | 13.38 | 97 | >sp\|P01877\|IGHA2_HUMAN Immunoglobulin heavy constant alpha 2 OS=Homo sapiens OX=9606 GN=IGHA2 PE=1 SV=4 | 576 | 20201118_ly_colostrum_1ng_glyco_hcd203040_compound15.12691.12691.3 | controllerType=0 controllerNumber=1 scan=12691 | 28.7312 |
| 10950:3 | 6 | R.MAGKPTHIN[+1872.651]VSVVMAEADGTC[+57.021]Y.- | HexNAc(3)Hex(6)NeuAc(1) | C[+57], N[+1873] | 1408.5851 | 3 | 4223.7409 | 4223.7333 | 0 | 1.8 | 319 | CRagged | 719.23 | 637.77 | 335.87 | 15.15 | 97 | >sp\|P01877\|IGHA2_HUMAN Immunoglobulin heavy constant alpha 2 OS=Homo sapiens OX=9606 GN=IGHA2 PE=1 SV=4 | 576 | 20201118_ly_colostrum_1ng_glyco_hcd203040_compound15.14239.14239.3 | controllerType=0 controllerNumber=1 scan=14239 | 31.8391 |
| 08686:3 | 6 | R.MAGKPTHIN[+1702.581]VSVVMAEADGTC[+57.021]Y.- | HexNAc(2)Hex(8) | C[+57], N[+1703] | 1351.8969 | 3 | 4053.6762 | 4053.6642 | 0 | 3 | 319 | CRagged | 741.94 | 741.94 | 422.19 | 15.38 | 97 | >sp\|P01877\|IGHA2_HUMAN Immunoglobulin heavy constant alpha 2 OS=Homo sapiens OX=9606 GN=IGHA2 PE=1 SV=4 | 576 | 20201118_ly_colostrum_1ng_glyco_hcd203040_compound15.11885.11885.3 | controllerType=0 controllerNumber=1 scan=11885 | 27.1142 |
| 08810:3 | 6 | R.MAGKPTHIN[+1216.423]VSVVMAEADGTC[+57.021]Y.- | HexNAc(2)Hex(5) | C[+57], N[+1216] | 1189.8431 | 3 | 3567.5147 | 3567.5057 | 0 | 2.5 | 319 | CRagged | 754.02 | 754.02 | 399.94 | 16.38 | 97 | >sp\|P01877\|IGHA2_HUMAN Immunoglobulin heavy constant alpha 2 OS=Homo sapiens OX=9606 GN=IGHA2 PE=1 SV=4 | 576 | 20201118_ly_colostrum_1ng_glyco_hcd203040_compound15.12014.12014.3 | controllerType=0 controllerNumber=1 scan=12014 | 27.3585 |
| 10878:3 | 6 | R.MAGKPTHIN[+2262.814]VSVVMAEADGTC[+57.021]Y.- | HexNAc(5)Hex(5)Fuc(1)NeuAc(1) | C[+57], N[+2263] | 1538.6408 | 3 | 4613.9079 | 4613.8972 | 0 | 2.3 | 319 | CRagged | 754.17 | 524.73 | 524.73 | 16.38 | 97 | >sp\|P01877\|IGHA2_HUMAN Immunoglobulin heavy constant alpha 2 OS=Homo sapiens OX=9606 GN=IGHA2 PE=1 SV=4 | 576 | 20201118_ly_colostrum_1ng_glyco_hcd203040_compound15.14164.14164.3 | controllerType=0 controllerNumber=1 scan=14164 | 31.6982 |
| 09432:3 | 6 | R.MAGKPTHIN[+1663.608]VSVVMAEADGTC[+57.021]Y.- | HexNAc(5)Hex(4) | C[+57], N[+1664] | 1338.9062 | 3 | 4014.704 | 4014.691 | 0 | 3.2 | 319 | CRagged | 766.49 | 752.54 | 450.6 | 16.41 | 97 | >sp\|P01877\|IGHA2_HUMAN Immunoglobulin heavy constant alpha 2 OS=Homo sapiens OX=9606 GN=IGHA2 PE=1 SV=4 | 576 | 20201118_ly_colostrum_1ng_glyco_hcd203040_compound15.12661.12661.3 | controllerType=0 controllerNumber=1 scan=12661 | 28.6721 |
| 08709:3 | 6 | R.MAGKPTHIN[+1540.529]VSVVMAEADGTC[+57.021]Y.- | HexNAc(2)Hex(7) | C[+57], N[+1541] | 1297.8785 | 3 | 3891.6209 | 3891.6114 | 0 | 2.4 | 319 | CRagged | 767.45 | 767.45 | 414.27 | 16.38 | 97 | >sp\|P01877\|IGHA2_HUMAN Immunoglobulin heavy constant alpha 2 OS=Homo sapiens OX=9606 GN=IGHA2 PE=1 SV=4 | 576 | 20201118_ly_colostrum_1ng_glyco_hcd203040_compound15.11909.11909.3 | controllerType=0 controllerNumber=1 scan=11909 | 27.1583 |
| 08689:3 | 6 | R.MAGKPTHIN[+1702.581]VSVVMAEADGTC[+57.021]Y.- | HexNAc(2)Hex(8) | C[+57], N[+1703] | 1351.8969 | 3 | 4053.6762 | 4053.6642 | 0 | 3 | 319 | CRagged | 782.71 | 715.17 | 426.94 | 16.38 | 97 | >sp\|P01877\|IGHA2_HUMAN Immunoglobulin heavy constant alpha 2 OS=Homo sapiens OX=9606 GN=IGHA2 PE=1 SV=4 | 576 | 20201118_ly_colostrum_1ng_glyco_hcd203040_compound15.11888.11888.3 | controllerType=0 controllerNumber=1 scan=11888 | 27.1187 |
| 09534:3 | 6 | R.MAGKPTHIN[+1647.613]VSVVMAEADGTC[+57.021]Y.- | HexNAc(5)Hex(3)Fuc(1) | C[+57], N[+1648] | 1333.5745 | 3 | 3998.709 | 3998.6961 | 0 | 3.2 | 319 | CRagged | 786.95 | 786.95 | 786.95 | 16.41 | 97 | >sp\|P01877\|IGHA2_HUMAN Immunoglobulin heavy constant alpha 2 OS=Homo sapiens OX=9606 GN=IGHA2 PE=1 SV=4 | 576 | 20201118_ly_colostrum_1ng_glyco_hcd203040_compound15.12767.12767.3 | controllerType=0 controllerNumber=1 scan=12767 | 28.8873 |
| 09371:3 | 6 | R.MAGKPTHIN[+1702.581]VSVVMAEADGTC[+57.021]Y.- | HexNAc(2)Hex(8) | C[+57], N[+1703] | 1351.8961 | 3 | 4053.6736 | 4053.6642 | 0 | 2.3 | 319 | CRagged | 788.05 | 626.3 | 484.96 | 16.38 | 97 | >sp\|P01877\|IGHA2_HUMAN Immunoglobulin heavy constant alpha 2 OS=Homo sapiens OX=9606 GN=IGHA2 PE=1 SV=4 | 576 | 20201118_ly_colostrum_1ng_glyco_hcd203040_compound15.12597.12597.3 | controllerType=0 controllerNumber=1 scan=12597 | 28.5163 |
| 10881:3 | 6 | R.MAGKPTHIN[+2262.814]VSVVMAEADGTC[+57.021]Y.- | HexNAc(5)Hex(5)Fuc(1)NeuAc(1) | C[+57], N[+2263] | 1538.6408 | 3 | 4613.9079 | 4613.8972 | 0 | 2.3 | 319 | CRagged | 788.25 | 574.76 | 574.76 | 16.38 | 97 | >sp\|P01877\|IGHA2_HUMAN Immunoglobulin heavy constant alpha 2 OS=Homo sapiens OX=9606 GN=IGHA2 PE=1 SV=4 | 576 | 20201118_ly_colostrum_1ng_glyco_hcd203040_compound15.14167.14167.3 | controllerType=0 controllerNumber=1 scan=14167 | 31.7033 |
| 09593:3 | 6 | R.MAGKPTHIN[+1095.397]VSVVMAEADGTC[+57.021]Y.- | HexNAc(3)Hex(3) | C[+57], N[+1095] | 1149.5018 | 3 | 3446.4907 | 3446.4794 | 0 | 3.3 | 319 | CRagged | 789.54 | 789.54 | 789.54 | 16.41 | 97 | >sp\|P01877\|IGHA2_HUMAN Immunoglobulin heavy constant alpha 2 OS=Homo sapiens OX=9606 GN=IGHA2 PE=1 SV=4 | 576 | 20201118_ly_colostrum_1ng_glyco_hcd203040_compound15.12828.12828.3 | controllerType=0 controllerNumber=1 scan=12828 | 29.0014 |
| 11021:3 | 6 | R.MAGKPTHIN[+1548.545]VSVVMAEADGTC[+57.021]Y.- | HexNAc(3)Hex(4)NeuAc(1) | C[+57], N[+1549] | 1300.5511 | 3 | 3899.6387 | 3899.6277 | 0 | 2.8 | 319 | CRagged | 790.34 | 708.95 | 708.95 | 16.38 | 97 | >sp\|P01877\|IGHA2_HUMAN Immunoglobulin heavy constant alpha 2 OS=Homo sapiens OX=9606 GN=IGHA2 PE=1 SV=4 | 576 | 20201118_ly_colostrum_1ng_glyco_hcd203040_compound15.14313.14313.3 | controllerType=0 controllerNumber=1 scan=14313 | 31.9779 |
| 09464:3 | 6 | R.MAGKPTHIN[+1809.666]VSVVMAEADGTC[+57.021]Y.- | HexNAc(5)Hex(4)Fuc(1) | C[+57], N[+1810] | 1387.591 | 3 | 4160.7584 | 4160.7489 | 0 | 2.3 | 319 | CRagged | 807.71 | 755.47 | 514.56 | 17.38 | 97 | >sp\|P01877\|IGHA2_HUMAN Immunoglobulin heavy constant alpha 2 OS=Homo sapiens OX=9606 GN=IGHA2 PE=1 SV=4 | 576 | 20201118_ly_colostrum_1ng_glyco_hcd203040_compound15.12694.12694.3 | controllerType=0 controllerNumber=1 scan=12694 | 28.7354 |
| 09434:3 | 6 | R.MAGKPTHIN[+1663.608]VSVVMAEADGTC[+57.021]Y.- | HexNAc(5)Hex(4) | C[+57], N[+1664] | 1338.9062 | 3 | 4014.704 | 4014.691 | 0 | 3.2 | 319 | CRagged | 809.67 | 615.95 | 458.22 | 17.41 | 97 | >sp\|P01877\|IGHA2_HUMAN Immunoglobulin heavy constant alpha 2 OS=Homo sapiens OX=9606 GN=IGHA2 PE=1 SV=4 | 576 | 20201118_ly_colostrum_1ng_glyco_hcd203040_compound15.12663.12663.3 | controllerType=0 controllerNumber=1 scan=12663 | 28.6767 |
| 09354:3 | 6 | R.MAGKPTHIN[+1864.634]VSVVMAEADGTC[+57.021]Y.- | HexNAc(2)Hex(9) | C[+57], N[+1865] | 1405.9139 | 3 | 4215.7271 | 4215.717 | 0 | 2.4 | 319 | CRagged | 824.29 | 824.29 | 824.29 | 17.38 | 97 | >sp\|P01877\|IGHA2_HUMAN Immunoglobulin heavy constant alpha 2 OS=Homo sapiens OX=9606 GN=IGHA2 PE=1 SV=4 | 576 | 20201118_ly_colostrum_1ng_glyco_hcd203040_compound15.12579.12579.3 | controllerType=0 controllerNumber=1 scan=12579 | 28.4756 |
| 09533:3 | 6 | R.MAGKPTHIN[+1501.555]VSVVMAEADGTC[+57.021]Y.- | HexNAc(5)Hex(3) | C[+57], N[+1502] | 1284.8882 | 3 | 3852.6502 | 3852.6382 | 0 | 3.1 | 319 | CRagged | 857.83 | 844.63 | 844.63 | 18.41 | 97 | >sp\|P01877\|IGHA2_HUMAN Immunoglobulin heavy constant alpha 2 OS=Homo sapiens OX=9606 GN=IGHA2 PE=1 SV=4 | 576 | 20201118_ly_colostrum_1ng_glyco_hcd203040_compound15.12766.12766.3 | controllerType=0 controllerNumber=1 scan=12766 | 28.8843 |
| 08712:3 | 6 | R.MAGKPTHIN[+1540.529]VSVVMAEADGTC[+57.021]Y.- | HexNAc(2)Hex(7) | C[+57], N[+1541] | 1297.8785 | 3 | 3891.6209 | 3891.6114 | 0 | 2.4 | 319 | CRagged | 868.29 | 868.29 | 469.86 | 18.38 | 97 | >sp\|P01877\|IGHA2_HUMAN Immunoglobulin heavy constant alpha 2 OS=Homo sapiens OX=9606 GN=IGHA2 PE=1 SV=4 | 576 | 20201118_ly_colostrum_1ng_glyco_hcd203040_compound15.11912.11912.3 | controllerType=0 controllerNumber=1 scan=11912 | 27.1639 |
| 09532:3 | 6 | R.MAGKPTHIN[+1647.613]VSVVMAEADGTC[+57.021]Y.- | HexNAc(5)Hex(3)Fuc(1) | C[+57], N[+1648] | 1333.5745 | 3 | 3998.709 | 3998.6961 | 0 | 3.2 | 319 | CRagged | 873.62 | 847.44 | 847.44 | 18.41 | 97 | >sp\|P01877\|IGHA2_HUMAN Immunoglobulin heavy constant alpha 2 OS=Homo sapiens OX=9606 GN=IGHA2 PE=1 SV=4 | 576 | 20201118_ly_colostrum_1ng_glyco_hcd203040_compound15.12765.12765.3 | controllerType=0 controllerNumber=1 scan=12765 | 28.8815 |
| 09373:3 | 6 | R.MAGKPTHIN[+1702.581]VSVVMAEADGTC[+57.021]Y.- | HexNAc(2)Hex(8) | C[+57], N[+1703] | 1351.8961 | 3 | 4053.6736 | 4053.6642 | 0 | 2.3 | 319 | CRagged | 879.86 | 714.79 | 505.62 | 18.38 | 97 | >sp\|P01877\|IGHA2_HUMAN Immunoglobulin heavy constant alpha 2 OS=Homo sapiens OX=9606 GN=IGHA2 PE=1 SV=4 | 576 | 20201118_ly_colostrum_1ng_glyco_hcd203040_compound15.12599.12599.3 | controllerType=0 controllerNumber=1 scan=12599 | 28.5216 |
| 09380:3 | 6 | R.MAGKPTHIN[+1540.529]VSVVMAEADGTC[+57.021]Y.- | HexNAc(2)Hex(7) | C[+57], N[+1541] | 1297.8782 | 3 | 3891.6201 | 3891.6114 | 0 | 2.3 | 319 | CRagged | 890.07 | 736.58 | 498.92 | 18.38 | 97 | >sp\|P01877\|IGHA2_HUMAN Immunoglobulin heavy constant alpha 2 OS=Homo sapiens OX=9606 GN=IGHA2 PE=1 SV=4 | 576 | 20201118_ly_colostrum_1ng_glyco_hcd203040_compound15.12606.12606.3 | controllerType=0 controllerNumber=1 scan=12606 | 28.5443 |
| 09401:3 | 6 | R.MAGKPTHIN[+1378.476]VSVVMAEADGTC[+57.021]Y.- | HexNAc(2)Hex(6) | C[+57], N[+1378] | 1243.8607 | 3 | 3729.5674 | 3729.5585 | 0 | 2.4 | 319 | CRagged | 891.85 | 742.81 | 505.09 | 18.38 | 97 | >sp\|P01877\|IGHA2_HUMAN Immunoglobulin heavy constant alpha 2 OS=Homo sapiens OX=9606 GN=IGHA2 PE=1 SV=4 | 576 | 20201118_ly_colostrum_1ng_glyco_hcd203040_compound15.12628.12628.3 | controllerType=0 controllerNumber=1 scan=12628 | 28.61 |
| 08813:3 | 6 | R.MAGKPTHIN[+1216.423]VSVVMAEADGTC[+57.021]Y.- | HexNAc(2)Hex(5) | C[+57], N[+1216] | 1189.8431 | 3 | 3567.5147 | 3567.5057 | 0 | 2.5 | 319 | CRagged | 897.35 | 749.53 | 497.01 | 18.38 | 97 | >sp\|P01877\|IGHA2_HUMAN Immunoglobulin heavy constant alpha 2 OS=Homo sapiens OX=9606 GN=IGHA2 PE=1 SV=4 | 576 | 20201118_ly_colostrum_1ng_glyco_hcd203040_compound15.12017.12017.3 | controllerType=0 controllerNumber=1 scan=12017 | 27.363 |
| 09377:3 | 6 | R.MAGKPTHIN[+1540.529]VSVVMAEADGTC[+57.021]Y.- | HexNAc(2)Hex(7) | C[+57], N[+1541] | 1297.8782 | 3 | 3891.6201 | 3891.6114 | 0 | 2.3 | 319 | CRagged | 898.39 | 898.39 | 502.63 | 18.38 | 97 | >sp\|P01877\|IGHA2_HUMAN Immunoglobulin heavy constant alpha 2 OS=Homo sapiens OX=9606 GN=IGHA2 PE=1 SV=4 | 576 | 20201118_ly_colostrum_1ng_glyco_hcd203040_compound15.12603.12603.3 | controllerType=0 controllerNumber=1 scan=12603 | 28.5324 |
| 09460:3 | 6 | R.MAGKPTHIN[+1216.423]VSVVMAEADGTC[+57.021]Y.- | HexNAc(2)Hex(5) | C[+57], N[+1216] | 1189.8433 | 3 | 3567.5154 | 3567.5057 | 0 | 2.7 | 319 | CRagged | 922.73 | 751.95 | 526.77 | 19.38 | 97 | >sp\|P01877\|IGHA2_HUMAN Immunoglobulin heavy constant alpha 2 OS=Homo sapiens OX=9606 GN=IGHA2 PE=1 SV=4 | 576 | 20201118_ly_colostrum_1ng_glyco_hcd203040_compound15.12690.12690.3 | controllerType=0 controllerNumber=1 scan=12690 | 28.7303 |
| 09351:3 | 6 | R.MAGKPTHIN[+1864.634]VSVVMAEADGTC[+57.021]Y.- | HexNAc(2)Hex(9) | C[+57], N[+1865] | 1405.9139 | 3 | 4215.7271 | 4215.717 | 0 | 2.4 | 319 | CRagged | 929.23 | 929.23 | 929.23 | 19.38 | 97 | >sp\|P01877\|IGHA2_HUMAN Immunoglobulin heavy constant alpha 2 OS=Homo sapiens OX=9606 GN=IGHA2 PE=1 SV=4 | 576 | 20201118_ly_colostrum_1ng_glyco_hcd203040_compound15.12576.12576.3 | controllerType=0 controllerNumber=1 scan=12576 | 28.4692 |
| 08740:3 | 6 | R.MAGKPTHIN[+1378.476]VSVVMAEADGTC[+57.021]Y.- | HexNAc(2)Hex(6) | C[+57], N[+1378] | 1243.8593 | 3 | 3729.5634 | 3729.5585 | 0 | 1.3 | 319 | CRagged | 930.53 | 930.53 | 516.91 | 19.15 | 97 | >sp\|P01877\|IGHA2_HUMAN Immunoglobulin heavy constant alpha 2 OS=Homo sapiens OX=9606 GN=IGHA2 PE=1 SV=4 | 576 | 20201118_ly_colostrum_1ng_glyco_hcd203040_compound15.11941.11941.3 | controllerType=0 controllerNumber=1 scan=11941 | 27.2227 |
| 08737:3 | 6 | R.MAGKPTHIN[+1378.476]VSVVMAEADGTC[+57.021]Y.- | HexNAc(2)Hex(6) | C[+57], N[+1378] | 1243.8593 | 3 | 3729.5634 | 3729.5585 | 0 | 1.3 | 319 | CRagged | 949.29 | 832.64 | 510.96 | 19.15 | 97 | >sp\|P01877\|IGHA2_HUMAN Immunoglobulin heavy constant alpha 2 OS=Homo sapiens OX=9606 GN=IGHA2 PE=1 SV=4 | 576 | 20201118_ly_colostrum_1ng_glyco_hcd203040_compound15.11938.11938.3 | controllerType=0 controllerNumber=1 scan=11938 | 27.215 |
| 09398:3 | 6 | R.MAGKPTHIN[+1378.476]VSVVMAEADGTC[+57.021]Y.- | HexNAc(2)Hex(6) | C[+57], N[+1378] | 1243.8607 | 3 | 3729.5674 | 3729.5585 | 0 | 2.4 | 319 | CRagged | 971.16 | 971.16 | 572.53 | 20.38 | 97 | >sp\|P01877\|IGHA2_HUMAN Immunoglobulin heavy constant alpha 2 OS=Homo sapiens OX=9606 GN=IGHA2 PE=1 SV=4 | 576 | 20201118_ly_colostrum_1ng_glyco_hcd203040_compound15.12625.12625.3 | controllerType=0 controllerNumber=1 scan=12625 | 28.5878 |
| 09458:3 | 6 | R.MAGKPTHIN[+1216.423]VSVVMAEADGTC[+57.021]Y.- | HexNAc(2)Hex(5) | C[+57], N[+1216] | 1189.8433 | 3 | 3567.5154 | 3567.5057 | 0 | 2.7 | 319 | CRagged | 978.47 | 817.78 | 581.81 | 20.38 | 97 | >sp\|P01877\|IGHA2_HUMAN Immunoglobulin heavy constant alpha 2 OS=Homo sapiens OX=9606 GN=IGHA2 PE=1 SV=4 | 576 | 20201118_ly_colostrum_1ng_glyco_hcd203040_compound15.12688.12688.3 | controllerType=0 controllerNumber=1 scan=12688 | 28.7225 |
| 09186:3 | 6 | R.M[+15.995]AGKPTHIN[+1378.476]VSVVMAEADGTC[+57.021]Y.- | HexNAc(2)Hex(6) | C[+57], M[+16], N[+1378] | 1249.1932 | 3 | 3745.565 | 3745.5535 | 0 | 3.1 | 319 | CRagged | 318.81 | 318.81 | 100.86 | 6.51 | 97 | >sp\|P01877\|IGHA2_HUMAN Immunoglobulin heavy constant alpha 2 OS=Homo sapiens OX=9606 GN=IGHA2 PE=1 SV=4 | 576 | 20201118_ly_colostrum_1ng_glyco_hcd203040_compound15.12404.12404.3 | controllerType=0 controllerNumber=1 scan=12404 | 28.1395 |
| 09039:3 | 6 | R.M[+15.995]AGKPTHIN[+1540.529]VSVVMAEADGTC[+57.021]Y.- | HexNAc(2)Hex(7) | C[+57], M[+16], N[+1541] | 1303.211 | 3 | 3907.6184 | 3907.6063 | 0 | 3.1 | 319 | CRagged | 327.75 | 327.75 | 133.39 | 8.25 | 97 | >sp\|P01877\|IGHA2_HUMAN Immunoglobulin heavy constant alpha 2 OS=Homo sapiens OX=9606 GN=IGHA2 PE=1 SV=4 | 576 | 20201118_ly_colostrum_1ng_glyco_hcd203040_compound15.12251.12251.3 | controllerType=0 controllerNumber=1 scan=12251 | 27.8267 |
| 09188:3 | 6 | R.M[+15.995]AGKPTHIN[+1378.476]VSVVMAEADGTC[+57.021]Y.- | HexNAc(2)Hex(6) | C[+57], M[+16], N[+1378] | 1249.1932 | 3 | 3745.565 | 3745.5535 | 0 | 3.1 | 319 | CRagged | 389.35 | 389.35 | 60.1 | 8.65 | 97 | >sp\|P01877\|IGHA2_HUMAN Immunoglobulin heavy constant alpha 2 OS=Homo sapiens OX=9606 GN=IGHA2 PE=1 SV=4 | 576 | 20201118_ly_colostrum_1ng_glyco_hcd203040_compound15.12406.12406.3 | controllerType=0 controllerNumber=1 scan=12406 | 28.145 |
| 09106:3 | 6 | R.M[+15.995]AGKPTHIN[+1501.555]VSVVMAEADGTC[+57.021]Y.- | HexNAc(5)Hex(3) | C[+57], M[+16], N[+1502] | 1290.2191 | 3 | 3868.6426 | 3868.6331 | 0 | 2.5 | 319 | CRagged | 410.81 | 410.81 | 175.46 | 10.27 | 97 | >sp\|P01877\|IGHA2_HUMAN Immunoglobulin heavy constant alpha 2 OS=Homo sapiens OX=9606 GN=IGHA2 PE=1 SV=4 | 576 | 20201118_ly_colostrum_1ng_glyco_hcd203040_compound15.12321.12321.3 | controllerType=0 controllerNumber=1 scan=12321 | 27.9655 |
| 09199:3 | 6 | R.M[+15.995]AGKPTHIN[+1216.423]VSVVMAEADGTC[+57.021]Y.- | HexNAc(2)Hex(5) | C[+57], M[+16], N[+1216] | 1195.1762 | 3 | 3583.5141 | 3583.5006 | 0 | 3.7 | 319 | CRagged | 414.52 | 414.52 | 178.89 | 10.29 | 97 | >sp\|P01877\|IGHA2_HUMAN Immunoglobulin heavy constant alpha 2 OS=Homo sapiens OX=9606 GN=IGHA2 PE=1 SV=4 | 576 | 20201118_ly_colostrum_1ng_glyco_hcd203040_compound15.12418.12418.3 | controllerType=0 controllerNumber=1 scan=12418 | 28.163 |
| 09202:3 | 6 | R.M[+15.995]AGKPTHIN[+1216.423]VSVVMAEADGTC[+57.021]Y.- | HexNAc(2)Hex(5) | C[+57], M[+16], N[+1216] | 1195.1762 | 3 | 3583.5141 | 3583.5006 | 0 | 3.7 | 319 | CRagged | 443.55 | 371.64 | 143.07 | 8.94 | 97 | >sp\|P01877\|IGHA2_HUMAN Immunoglobulin heavy constant alpha 2 OS=Homo sapiens OX=9606 GN=IGHA2 PE=1 SV=4 | 576 | 20201118_ly_colostrum_1ng_glyco_hcd203040_compound15.12421.12421.3 | controllerType=0 controllerNumber=1 scan=12421 | 28.1672 |
| 09041:3 | 6 | R.M[+15.995]AGKPTHIN[+1540.529]VSVVMAEADGTC[+57.021]Y.- | HexNAc(2)Hex(7) | C[+57], M[+16], N[+1541] | 1303.211 | 3 | 3907.6184 | 3907.6063 | 0 | 3.1 | 319 | CRagged | 453.81 | 453.81 | 154.79 | 10.41 | 97 | >sp\|P01877\|IGHA2_HUMAN Immunoglobulin heavy constant alpha 2 OS=Homo sapiens OX=9606 GN=IGHA2 PE=1 SV=4 | 576 | 20201118_ly_colostrum_1ng_glyco_hcd203040_compound15.12253.12253.3 | controllerType=0 controllerNumber=1 scan=12253 | 27.8318 |
| 08963:3 | 6 | R.M[+15.995]AGKPTHIN[+1702.581]VSVVMAEADGTC[+57.021]Y.- | HexNAc(2)Hex(8) | C[+57], M[+16], N[+1703] | 1357.2252 | 3 | 4069.6609 | 4069.6591 | 0 | 0.4 | 319 | CRagged | 469.82 | 469.82 | 180.75 | 10.01 | 97 | >sp\|P01877\|IGHA2_HUMAN Immunoglobulin heavy constant alpha 2 OS=Homo sapiens OX=9606 GN=IGHA2 PE=1 SV=4 | 576 | 20201118_ly_colostrum_1ng_glyco_hcd203040_compound15.12173.12173.3 | controllerType=0 controllerNumber=1 scan=12173 | 27.6926 |
| 08965:3 | 6 | R.M[+15.995]AGKPTHIN[+1702.581]VSVVMAEADGTC[+57.021]Y.- | HexNAc(2)Hex(8) | C[+57], M[+16], N[+1703] | 1357.2252 | 3 | 4069.6609 | 4069.6591 | 0 | 0.4 | 319 | CRagged | 516.94 | 516.94 | 198.56 | 10.22 | 97 | >sp\|P01877\|IGHA2_HUMAN Immunoglobulin heavy constant alpha 2 OS=Homo sapiens OX=9606 GN=IGHA2 PE=1 SV=4 | 576 | 20201118_ly_colostrum_1ng_glyco_hcd203040_compound15.12175.12175.3 | controllerType=0 controllerNumber=1 scan=12175 | 27.6997 |
| 07340:3 | 6 | R.MAGKPTHIN[+1216.423]VSVVM[+15.995]AEADGTC[+57.021]Y.- | HexNAc(2)Hex(5) | C[+57], M[+16], N[+1216] | 1195.1749 | 3 | 3583.51 | 3583.5006 | 0 | 2.6 | 319 | CRagged | 548.04 | 548.04 | 313.04 | 10.88 | 97 | >sp\|P01877\|IGHA2_HUMAN Immunoglobulin heavy constant alpha 2 OS=Homo sapiens OX=9606 GN=IGHA2 PE=1 SV=4 | 576 | 20201118_ly_colostrum_1ng_glyco_hcd203040_compound15.10487.10487.3 | controllerType=0 controllerNumber=1 scan=10487 | 24.3921 |

Table S4. Cont.

| Query #:z | Protein Rank | Glycopeptide | Glycan | Modification Type(s) | Observed m/z | z | Observed (M+H) | Calc. mass (M+H) | Off-by-x error | Mass error (ppm) | StartPosition | Cleavage | Score | Delta | Delta Mod | \|Log Prob\| | # of unique peptides | Protein Name | Protein DB number | Comment | Scan # | Scan Time |
|---|---|---|---|---|---|---|---|---|---|---|---|---|---|---|---|---|---|---|---|---|---|---|
| 08992:3 | 6 | R.M[+15.995]AGKPTHIN[+1378.476]VSVVMAEADGTC[+57.021]Y.- | HexNAc(2)Hex(6) | C[+57], M[+16], N[+1378] | 1249.1922 | 3 | 3745.562 | 3745.5535 | 0 | 2.3 | 319 | CRagged | 571.96 | 564.18 | 252.79 | 11.06 | 97 | >sp\|P01877\|IGHA2_HUMAN Immunoglobulin heavy constant alpha 2 OS=Homo sapiens OX=9606 GN=IGHA2 PE=1 SV=4 | 576 | 20201118_ly_colostrum_1ng_glyco_hcd203040_compound15.12203.12203.3 | controllerType=0 controllerNumber=1 scan=12203 | 27.745 |
| 09109:3 | 6 | R.M[+15.995]AGKPTHIN[+1501.555]VSVVMAEADGTC[+57.021]Y.- | HexNAc(5)Hex(3) | C[+57], M[+16], N[+1502] | 1290.2191 | 3 | 3868.6426 | 3868.6331 | 0 | 2.5 | 319 | CRagged | 577.36 | 577.36 | 219.21 | 11.06 | 97 | >sp\|P01877\|IGHA2_HUMAN Immunoglobulin heavy constant alpha 2 OS=Homo sapiens OX=9606 GN=IGHA2 PE=1 SV=4 | 576 | 20201118_ly_colostrum_1ng_glyco_hcd203040_compound15.12324.12324.3 | controllerType=0 controllerNumber=1 scan=12324 | 27.9759 |
| 09043:3 | 6 | R.M[+15.995]AGKPTHIN[+1216.423]VSVVMAEADGTC[+57.021]Y.- | HexNAc(2)Hex(5) | C[+57], M[+16], N[+1216] | 1195.175 | 3 | 3583.5104 | 3583.5006 | 0 | 2.7 | 319 | CRagged | 592.62 | 592.62 | 268.26 | 12.38 | 97 | >sp\|P01877\|IGHA2_HUMAN Immunoglobulin heavy constant alpha 2 OS=Homo sapiens OX=9606 GN=IGHA2 PE=1 SV=4 | 576 | 20201118_ly_colostrum_1ng_glyco_hcd203040_compound15.12256.12256.3 | controllerType=0 controllerNumber=1 scan=12256 | 27.8388 |
| 08977:3 | 6 | R.M[+15.995]AGKPTHIN[+1663.608]VSVVMAEADGTC[+57.021]Y.- | HexNAc(5)Hex(4) | C[+57], M[+16], N[+1664] | 1344.2369 | 3 | 4030.6961 | 4030.6859 | 0 | 2.5 | 319 | CRagged | 614.86 | 578.28 | 246.19 | 13.38 | 97 | >sp\|P01877\|IGHA2_HUMAN Immunoglobulin heavy constant alpha 2 OS=Homo sapiens OX=9606 GN=IGHA2 PE=1 SV=4 | 576 | 20201118_ly_colostrum_1ng_glyco_hcd203040_compound15.12187.12187.3 | controllerType=0 controllerNumber=1 scan=12187 | 27.7181 |
| 08975:3 | 6 | R.M[+15.995]AGKPTHIN[+1663.608]VSVVMAEADGTC[+57.021]Y.- | HexNAc(5)Hex(4) | C[+57], M[+16], N[+1664] | 1344.2369 | 3 | 4030.6961 | 4030.6859 | 0 | 2.5 | 319 | CRagged | 615.32 | 515.04 | 246.16 | 13.38 | 97 | >sp\|P01877\|IGHA2_HUMAN Immunoglobulin heavy constant alpha 2 OS=Homo sapiens OX=9606 GN=IGHA2 PE=1 SV=4 | 576 | 20201118_ly_colostrum_1ng_glyco_hcd203040_compound15.12185.12185.3 | controllerType=0 controllerNumber=1 scan=12185 | 27.7141 |
| 07343:3 | 6 | R.MAGKPTHIN[+1216.423]VSVVM[+15.995]AEADGTC[+57.021]Y.- | HexNAc(2)Hex(5) | C[+57], M[+16], N[+1216] | 1195.1749 | 3 | 3583.51 | 3583.5006 | 0 | 2.6 | 319 | CRagged | 636.16 | 554.37 | 266.86 | 13.38 | 97 | >sp\|P01877\|IGHA2_HUMAN Immunoglobulin heavy constant alpha 2 OS=Homo sapiens OX=9606 GN=IGHA2 PE=1 SV=4 | 576 | 20201118_ly_colostrum_1ng_glyco_hcd203040_compound15.10490.10490.3 | controllerType=0 controllerNumber=1 scan=10490 | 24.3974 |
| 08995:3 | 6 | R.M[+15.995]AGKPTHIN[+1378.476]VSVVMAEADGTC[+57.021]Y.- | HexNAc(2)Hex(6) | C[+57], M[+16], N[+1378] | 1249.1922 | 3 | 3745.562 | 3745.5535 | 0 | 2.3 | 319 | CRagged | 713.82 | 713.82 | 285.7 | 15.38 | 97 | >sp\|P01877\|IGHA2_HUMAN Immunoglobulin heavy constant alpha 2 OS=Homo sapiens OX=9606 GN=IGHA2 PE=1 SV=4 | 576 | 20201118_ly_colostrum_1ng_glyco_hcd203040_compound15.12206.12206.3 | controllerType=0 controllerNumber=1 scan=12206 | 27.7506 |
| 08944:3 | 6 | R.M[+15.995]AGKPTHIN[+1540.529]VSVVMAEADGTC[+57.021]Y.- | HexNAc(2)Hex(7) | C[+57], M[+16], N[+1541] | 1303.2098 | 3 | 3907.6148 | 3907.6063 | 0 | 2.2 | 319 | CRagged | 742.84 | 742.84 | 304.25 | 15.38 | 97 | >sp\|P01877\|IGHA2_HUMAN Immunoglobulin heavy constant alpha 2 OS=Homo sapiens OX=9606 GN=IGHA2 PE=1 SV=4 | 576 | 20201118_ly_colostrum_1ng_glyco_hcd203040_compound15.12153.12153.3 | controllerType=0 controllerNumber=1 scan=12153 | 27.6579 |
| 08947:3 | 6 | R.M[+15.995]AGKPTHIN[+1540.529]VSVVMAEADGTC[+57.021]Y.- | HexNAc(2)Hex(7) | C[+57], M[+16], N[+1541] | 1303.2098 | 3 | 3907.6148 | 3907.6063 | 0 | 2.2 | 319 | CRagged | 748.91 | 748.91 | 238.01 | 15.38 | 97 | >sp\|P01877\|IGHA2_HUMAN Immunoglobulin heavy constant alpha 2 OS=Homo sapiens OX=9606 GN=IGHA2 PE=1 SV=4 | 576 | 20201118_ly_colostrum_1ng_glyco_hcd203040_compound15.12156.12156.3 | controllerType=0 controllerNumber=1 scan=12156 | 27.6633 |
| 09045:3 | 6 | R.M[+15.995]AGKPTHIN[+1216.423]VSVVMAEADGTC[+57.021]Y.- | HexNAc(2)Hex(5) | C[+57], M[+16], N[+1216] | 1195.175 | 3 | 3583.5104 | 3583.5006 | 0 | 2.7 | 319 | CRagged | 790.33 | 790.33 | 288.16 | 16.38 | 97 | >sp\|P01877\|IGHA2_HUMAN Immunoglobulin heavy constant alpha 2 OS=Homo sapiens OX=9606 GN=IGHA2 PE=1 SV=4 | 576 | 20201118_ly_colostrum_1ng_glyco_hcd203040_compound15.12258.12258.3 | controllerType=0 controllerNumber=1 scan=12258 | 27.8435 |
| 09265:3 | 7 | K.TVLTPATNHMGN[+1378.476]VTFTIPANR.E | HexNAc(2)Hex(6) | N[+1378] | 1211.8878 | 3 | 3633.6487 | 3633.6358 | 0 | 3.6 | 74 | Specific | 517.64 | 517.64 | 252.89 | 10.68 | 93 | >sp\|P01024\|CO3_HUMAN Complement C3 OS=Homo sapiens OX=9606 GN=C3 PE=1 SV=2 | 492 | 20201118_ly_colostrum_1ng_glyco_hcd203040_compound15.12486.12486.3 | controllerType=0 controllerNumber=1 scan=12486 | 28.2917 |
| 09262:3 | 7 | K.TVLTPATNHMGN[+1378.476]VTFTIPANR.E | HexNAc(2)Hex(6) | N[+1378] | 1211.8878 | 3 | 3633.6487 | 3633.6358 | 0 | 3.6 | 74 | Specific | 597.53 | 597.53 | 342.45 | 12.41 | 93 | >sp\|P01024\|CO3_HUMAN Complement C3 OS=Homo sapiens OX=9606 GN=C3 PE=1 SV=2 | 492 | 20201118_ly_colostrum_1ng_glyco_hcd203040_compound15.12483.12483.3 | controllerType=0 controllerNumber=1 scan=12483 | 28.2866 |
| 04486:3 | 8 | R.FSDGLESN[+1702.581]SSTQFEVKK.Y | HexNAc(2)Hex(8) | N[+1703] | 1202.5032 | 3 | 3605.4951 | 3605.4893 | 0 | 1.6 | 219 | Specific | 313.35 | 122.4 | 122.4 | 5.43 | 74 | >sp\|P0C0L4\|CO4A_HUMAN Complement C4-A OS=Homo sapiens OX=9606 GN=C4A PE=1 SV=2 | 1235 | 20201118_ly_colostrum_1ng_glyco_hcd203040_compound15.7532.7532.3 | controllerType=0 controllerNumber=1 scan=7532 | 19.2225 |
| 04475:3 | 8 | R.FSDGLESN[+1702.581]SSTQFEVKK.Y | HexNAc(2)Hex(8) | N[+1703] | 1202.5036 | 3 | 3605.4962 | 3605.4893 | 0 | 1.9 | 219 | Specific | 396.61 | 361.7 | 361.7 | 7.71 | 74 | >sp\|P0C0L4\|CO4A_HUMAN Complement C4-A OS=Homo sapiens OX=9606 GN=C4A PE=1 SV=2 | 1235 | 20201118_ly_colostrum_1ng_glyco_hcd203040_compound15.7520.7520.3 | controllerType=0 controllerNumber=1 scan=7520 | 19.1937 |
| 04484:3 | 8 | R.FSDGLESN[+1702.581]SSTQFEVKK.Y | HexNAc(2)Hex(8) | N[+1703] | 1202.5032 | 3 | 3605.4951 | 3605.4893 | 0 | 1.6 | 219 | Specific | 408.86 | 365 | 365 | 7.91 | 74 | >sp\|P0C0L4\|CO4A_HUMAN Complement C4-A OS=Homo sapiens OX=9606 GN=C4A PE=1 SV=2 | 1235 | 20201118_ly_colostrum_1ng_glyco_hcd203040_compound15.7530.7530.3 | controllerType=0 controllerNumber=1 scan=7530 | 19.219 |
| 04457:3 | 8 | R.FSDGLESN[+1864.634]SSTQFEVKK.Y | HexNAc(2)Hex(9) | N[+1865] | 1256.5222 | 3 | 3767.5519 | 3767.5421 | 0 | 2.6 | 219 | Specific | 442.99 | 442.99 | 442.99 | 9.72 | 74 | >sp\|P0C0L4\|CO4A_HUMAN Complement C4-A OS=Homo sapiens OX=9606 GN=C4A PE=1 SV=2 | 1235 | 20201118_ly_colostrum_1ng_glyco_hcd203040_compound15.7502.7502.3 | controllerType=0 controllerNumber=1 scan=7502 | 19.1624 |
| 04455:3 | 8 | R.FSDGLESN[+1864.634]SSTQFEVKK.Y | HexNAc(2)Hex(9) | N[+1865] | 1256.5222 | 3 | 3767.5519 | 3767.5421 | 0 | 2.6 | 219 | Specific | 476.22 | 476.22 | 476.22 | 9.79 | 74 | >sp\|P0C0L4\|CO4A_HUMAN Complement C4-A OS=Homo sapiens OX=9606 GN=C4A PE=1 SV=2 | 1235 | 20201118_ly_colostrum_1ng_glyco_hcd203040_compound15.7500.7500.3 | controllerType=0 controllerNumber=1 scan=7500 | 19.1587 |
| 04473:3 | 8 | R.FSDGLESN[+1702.581]SSTQFEVKK.Y | HexNAc(2)Hex(8) | N[+1703] | 1202.5036 | 3 | 3605.4962 | 3605.4893 | 0 | 1.9 | 219 | Specific | 631.69 | 376.97 | 376.97 | 11.02 | 74 | >sp\|P0C0L4\|CO4A_HUMAN Complement C4-A OS=Homo sapiens OX=9606 GN=C4A PE=1 SV=2 | 1235 | 20201118_ly_colostrum_1ng_glyco_hcd203040_compound15.7518.7518.3 | controllerType=0 controllerNumber=1 scan=7518 | 19.19 |
| 14230:3 | 9 | R.QQQHLFGSN[+2204.772]VTDC[+57.021]SGNFC[+57.021]LFR.S | HexNAc(4)Hex(5)NeuAc(2) | C[+57]*2, N[+2205] | 1573.9846 | 3 | 4719.9392 | 4719.8966 | 0 | 9 | 622 | Specific | 266.83 | 241.32 | 241.32 | 5.99 | 50 | >sp\|P02787\|TRFE_HUMAN Serotransferrin OS=Homo sapiens OX=9606 GN=TF PE=1 SV=3 | 1111 | 20201118_ly_colostrum_1ng_glyco_hcd203040_compound15.17649.17649.3 | controllerType=0 controllerNumber=1 scan=17649 | 38.7584 |
| 00123:3 | 10 | K.YKN[+2262.814]NSDISSTR.G | HexNAc(5)Hex(5)Fuc(1)NeuAc(1) | N[+2263] | 1183.1585 | 3 | 3547.461 | 3547.4322 | 0 | 8.1 | 44 | Specific | 303.71 | 207.32 | 207.32 | 5.23 | 38 | >sp\|P01871\|IGHM_HUMAN Immunoglobulin heavy constant mu OS=Homo sapiens OX=9606 GN=IGHM PE=1 SV=4 | 328 | 20201118_ly_colostrum_1ng_glyco_hcd203040_compound15.3003.3003.3 | controllerType=0 controllerNumber=1 scan=3003 | 11.2585 |
| 00125:3 | 10 | K.YKN[+2262.814]NSDISSTR.G | HexNAc(5)Hex(5)Fuc(1)NeuAc(1) | N[+2263] | 1183.1585 | 3 | 3547.461 | 3547.4322 | 0 | 8.1 | 44 | Specific | 353.93 | 216.47 | 216.47 | 5.47 | 38 | >sp\|P01871\|IGHM_HUMAN Immunoglobulin heavy constant mu OS=Homo sapiens OX=9606 GN=IGHM PE=1 SV=4 | 328 | 20201118_ly_colostrum_1ng_glyco_hcd203040_compound15.3005.3005.3 | controllerType=0 controllerNumber=1 scan=3005 | 11.2634 |
| 10571:3 | 10 | R.TVDKSTGKPTLYN[+1378.476]VSLVMSDTAGTC[+57.021]Y.- | HexNAc(2)Hex(6) | C[+57], N[+1378] | 1396.282 | 3 | 4186.8316 | 4186.8187 | 0 | 3.1 | 428 | CRagged | 805.67 | 660.18 | 370.62 | 17.37 | 38 | >sp\|P01871\|IGHM_HUMAN Immunoglobulin heavy constant mu OS=Homo sapiens OX=9606 GN=IGHM PE=1 SV=4 | 328 | 20201118_ly_colostrum_1ng_glyco_hcd203040_compound15.13844.13844.3 | controllerType=0 controllerNumber=1 scan=13844 | 31.0261 |
| 10569:3 | 10 | R.TVDKSTGKPTLYN[+1378.476]VSLVMSDTAGTC[+57.021]Y.- | HexNAc(2)Hex(6) | C[+57], N[+1378] | 1396.282 | 3 | 4186.8316 | 4186.8187 | 0 | 3.1 | 428 | CRagged | 901.89 | 808.81 | 421.27 | 19.37 | 38 | >sp\|P01871\|IGHM_HUMAN Immunoglobulin heavy constant mu OS=Homo sapiens OX=9606 GN=IGHM PE=1 SV=4 | 328 | 20201118_ly_colostrum_1ng_glyco_hcd203040_compound15.13842.13842.3 | controllerType=0 controllerNumber=1 scan=13842 | 31.0189 |
| 21719:3 | 11 | R.QLAHQSN[+2350.830]STNIFFSPVSIATAFAMLSLGTK.A | HexNAc(4)Hex(5)Fuc(1)NeuAc(2) | N[+2351] | 1844.8336 | 3 | 5532.4862 | 5532.4654 | 0 | 3.8 | 64 | Specific | 269.44 | 206.3 | 206.3 | 7.18 | 46 | >sp\|P01009\|A1AT_HUMAN Alpha-1-antitrypsin OS=Homo sapiens OX=9606 GN=SERPINA1 PE=1 SV=3 | 642 | 20201118_ly_colostrum_1ng_glyco_hcd203040_compound15.25465.25465.3 | controllerType=0 controllerNumber=1 scan=25465 | 55.8363 |
| 24561:4 | 11 | R.QLAHQSN[+2861.000]STNIFFSPVSIATAFAMLSLGTK.A | HexNAc(5)Hex(6)NeuAc(3) | N[+2861] | 1511.4218 | 4 | 6042.6652 | 6042.6351 | 0 | 5 | 64 | Specific | 320.46 | 262.33 | 262.33 | 7.85 | 46 | >sp\|P01009\|A1AT_HUMAN Alpha-1-antitrypsin OS=Homo sapiens OX=9606 GN=SERPINA1 PE=1 SV=3 | 642 | 20201118_ly_colostrum_1ng_glyco_hcd203040_compound15.28416.28416.4 | controllerType=0 controllerNumber=1 scan=28416 | 61.3513 |
| 24563:4 | 11 | R.QLAHQSN[+2861.000]STNIFFSPVSIATAFAMLSLGTK.A | HexNAc(5)Hex(6)NeuAc(3) | N[+2861] | 1511.4218 | 4 | 6042.6652 | 6042.6351 | 0 | 5 | 64 | Specific | 336.96 | 336.96 | 336.96 | 8.25 | 46 | >sp\|P01009\|A1AT_HUMAN Alpha-1-antitrypsin OS=Homo sapiens OX=9606 GN=SERPINA1 PE=1 SV=3 | 642 | 20201118_ly_colostrum_1ng_glyco_hcd203040_compound15.28418.28418.4 | controllerType=0 controllerNumber=1 scan=28418 | 61.3552 |
| 24503:4 | 11 | R.QLAHQSN[+3007.058]STNIFFSPVSIATAFAMLSLGTK.A | HexNAc(5)Hex(6)Fuc(1)NeuAc(3) | N[+3007] | 1547.9346 | 4 | 6188.7165 | 6188.693 | 0 | 3.8 | 64 | Specific | 397.87 | 389.06 | 389.06 | 10.39 | 46 | >sp\|P01009\|A1AT_HUMAN Alpha-1-antitrypsin OS=Homo sapiens OX=9606 GN=SERPINA1 PE=1 SV=3 | 642 | 20201118_ly_colostrum_1ng_glyco_hcd203040_compound15.28356.28356.4 | controllerType=0 controllerNumber=1 scan=28356 | 61.2407 |
| 21880:3 | 11 | R.QLAHQSN[+2204.772]STNIFFSPVSIATAFAMLSLGTK.A | HexNAc(4)Hex(5)NeuAc(2) | N[+2205] | 1796.1511 | 3 | 5386.4387 | 5386.4075 | 0 | 5.8 | 64 | Specific | 405.94 | 350.43 | 350.43 | 8.19 | 46 | >sp\|P01009\|A1AT_HUMAN Alpha-1-antitrypsin OS=Homo sapiens OX=9606 GN=SERPINA1 PE=1 SV=3 | 642 | 20201118_ly_colostrum_1ng_glyco_hcd203040_compound15.25633.25633.3 | controllerType=0 controllerNumber=1 scan=25633 | 56.1722 |
| 24501:4 | 11 | R.QLAHQSN[+3007.058]STNIFFSPVSIATAFAMLSLGTK.A | HexNAc(5)Hex(6)Fuc(1)NeuAc(3) | N[+3007] | 1547.9346 | 4 | 6188.7165 | 6188.693 | 0 | 3.8 | 64 | Specific | 427.65 | 427.65 | 427.65 | 10.52 | 46 | >sp\|P01009\|A1AT_HUMAN Alpha-1-antitrypsin OS=Homo sapiens OX=9606 GN=SERPINA1 PE=1 SV=3 | 642 | 20201118_ly_colostrum_1ng_glyco_hcd203040_compound15.28354.28354.4 | controllerType=0 controllerNumber=1 scan=28354 | 61.2359 |
| 21878:3 | 11 | R.QLAHQSN[+2204.772]STNIFFSPVSIATAFAMLSLGTK.A | HexNAc(4)Hex(5)NeuAc(2) | N[+2205] | 1796.1511 | 3 | 5386.4387 | 5386.4075 | 0 | 5.8 | 64 | Specific | 442.26 | 416.72 | 416.72 | 9.98 | 46 | >sp\|P01009\|A1AT_HUMAN Alpha-1-antitrypsin OS=Homo sapiens OX=9606 GN=SERPINA1 PE=1 SV=3 | 642 | 20201118_ly_colostrum_1ng_glyco_hcd203040_compound15.25631.25631.3 | controllerType=0 controllerNumber=1 scan=25631 | 56.1672 |
| 21721:3 | 11 | R.QLAHQSN[+2350.830]STNIFFSPVSIATAFAMLSLGTK.A | HexNAc(4)Hex(5)Fuc(1)NeuAc(2) | N[+2351] | 1844.8336 | 3 | 5532.4862 | 5532.4654 | 0 | 3.8 | 64 | Specific | 486.3 | 395.37 | 395.37 | 10.69 | 46 | >sp\|P01009\|A1AT_HUMAN Alpha-1-antitrypsin OS=Homo sapiens OX=9606 GN=SERPINA1 PE=1 SV=3 | 642 | 20201118_ly_colostrum_1ng_glyco_hcd203040_compound15.25467.25467.3 | controllerType=0 controllerNumber=1 scan=25467 | 55.8402 |
| 21722:3 | 11 | R.QLAHQSN[+2204.772]STNIFFSPVSIATAFAMLSLGTK.A | HexNAc(4)Hex(5)NeuAc(2) | N[+2205] | 1796.1476 | 3 | 5386.4282 | 5386.4075 | 0 | 3.9 | 64 | Specific | 707.49 | 703.34 | 703.34 | 15.54 | 46 | >sp\|P01009\|A1AT_HUMAN Alpha-1-antitrypsin OS=Homo sapiens OX=9606 GN=SERPINA1 PE=1 SV=3 | 642 | 20201118_ly_colostrum_1ng_glyco_hcd203040_compound15.25468.25468.3 | controllerType=0 controllerNumber=1 scan=25468 | 55.8421 |
| 21724:3 | 11 | R.QLAHQSN[+2204.772]STNIFFSPVSIATAFAMLSLGTK.A | HexNAc(4)Hex(5)NeuAc(2) | N[+2205] | 1796.1476 | 3 | 5386.4282 | 5386.4075 | 0 | 3.9 | 64 | Specific | 729.97 | 664.4 | 664.4 | 15.54 | 46 | >sp\|P01009\|A1AT_HUMAN Alpha-1-antitrypsin OS=Homo sapiens OX=9606 GN=SERPINA1 PE=1 SV=3 | 642 | 20201118_ly_colostrum_1ng_glyco_hcd203040_compound15.25470.25470.3 | controllerType=0 controllerNumber=1 scan=25470 | 55.846 |
| 21002:4 | 11 | R.QLAHQSN[+2204.772]STNIFFSPVSIATAFAM[+15.995]LSLGTK.A | HexNAc(4)Hex(5)NeuAc(2) | M[+16], N[+2205] | 1351.361 | 4 | 5402.422 | 5402.4024 | 0 | 3.6 | 64 | Specific | 269.32 | 211.38 | 211.38 | 7.34 | 46 | >sp\|P01009\|A1AT_HUMAN Alpha-1-antitrypsin OS=Homo sapiens OX=9606 GN=SERPINA1 PE=1 SV=3 | 642 | 20201118_ly_colostrum_1ng_glyco_hcd203040_compound15.24718.24718.4 | controllerType=0 controllerNumber=1 scan=24718 | 54.2988 |
| 20528:5 | 11 | K.ADTHDEILEGLNFN[+2861.000]LTEIPEAQIHEGFQELLR.T | HexNAc(5)Hex(6)NeuAc(3) | N[+2861] | 1311.3752 | 5 | 6552.8471 | 6552.8239 | 0 | 3.5 | 94 | Specific | 886.11 | 758.77 | 758.77 | 19.06 | 46 | >sp\|P01009\|A1AT_HUMAN Alpha-1-antitrypsin OS=Homo sapiens OX=9606 GN=SERPINA1 PE=1 SV=3 | 642 | 20201118_ly_colostrum_1ng_glyco_hcd203040_compound15.24223.24223.4 | controllerType=0 controllerNumber=1 scan=24223 | 53.19 |
| 20530:5 | 11 | K.ADTHDEILEGLNFN[+2861.000]LTEIPEAQIHEGFQELLR.T | HexNAc(5)Hex(6)NeuAc(3) | N[+2861] | 1311.3752 | 5 | 6552.8471 | 6552.8239 | 0 | 3.5 | 94 | Specific | 960.89 | 942.86 | 942.86 | 21.06 | 46 | >sp\|P01009\|A1AT_HUMAN Alpha-1-antitrypsin OS=Homo sapiens OX=9606 GN=SERPINA1 PE=1 SV=3 | 642 | 20201118_ly_colostrum_1ng_glyco_hcd203040_compound15.24225.24225.4 | controllerType=0 controllerNumber=1 scan=24225 | 53.1958 |
| 18141:3 | 11 | K.YLGN[+2204.772]ATAIFFLPDEGKLQHLENELTHDIITK.F | HexNAc(4)Hex(5)NeuAc(2) | N[+2205] | 1915.8859 | 3 | 5745.6433 | 5745.6098 | 0 | 5.8 | 268 | Specific | 328.03 | 328.03 | 328.03 | 6.61 | 46 | >sp\|P01009\|A1AT_HUMAN Alpha-1-antitrypsin OS=Homo sapiens OX=9606 GN=SERPINA1 PE=1 SV=3 | 642 | 20201118_ly_colostrum_1ng_glyco_hcd203040_compound15.21731.21731.3 | controllerType=0 controllerNumber=1 scan=21731 | 47.7227 |
| 18163:4 | 11 | K.YLGN[+2351.851]ATAIFFLPDEGKLQHLENELTHDIITK.F | HexNAc(4)Hex(5)Fuc(3)NeuAc(1) | N[+2352] | 1473.9271 | 4 | 5892.6867 | 5892.6881 | 0 | -0.2 | 268 | Specific | 407.82 | 170.92 | 170.92 | 6.09 | 46 | >sp\|P01009\|A1AT_HUMAN Alpha-1-antitrypsin OS=Homo sapiens OX=9606 GN=SERPINA1 PE=1 SV=3 | 642 | 20201118_ly_colostrum_1ng_glyco_hcd203040_compound15.21754.21754.4 | controllerType=0 controllerNumber=1 scan=21754 | 47.769 |
| 18161:4 | 11 | K.YLGN[+2351.851]ATAIFFLPDEGKLQHLENELTHDIITK.F | HexNAc(4)Hex(5)Fuc(3)NeuAc(1) | N[+2352] | 1473.9271 | 4 | 5892.6867 | 5892.6881 | 0 | -0.2 | 268 | Specific | 429.48 | 429.48 | 429.48 | 8.85 | 46 | >sp\|P01009\|A1AT_HUMAN Alpha-1-antitrypsin OS=Homo sapiens OX=9606 GN=SERPINA1 PE=1 SV=3 | 642 | 20201118_ly_colostrum_1ng_glyco_hcd203040_compound15.21752.21752.4 | controllerType=0 controllerNumber=1 scan=21752 | 47.7652 |
| 18103:4 | 11 | K.YLGN[+2204.772]ATAIFFLPDEGKLQHLENELTHDIITK.F | HexNAc(4)Hex(5)NeuAc(2) | N[+2205] | 1437.1646 | 4 | 5745.6364 | 5745.6098 | 0 | 4.6 | 268 | Specific | 877.25 | 830.91 | 830.91 | 17.61 | 46 | >sp\|P01009\|A1AT_HUMAN Alpha-1-antitrypsin OS=Homo sapiens OX=9606 GN=SERPINA1 PE=1 SV=3 | 642 | 20201118_ly_colostrum_1ng_glyco_hcd203040_compound15.21691.21691.4 | controllerType=0 controllerNumber=1 scan=21691 | 47.6247 |
| 18098:5 | 11 | K.YLGN[+2204.772]ATAIFFLPDEGKLQHLENELTHDIITK.F | HexNAc(4)Hex(5)NeuAc(2) | N[+2205] | 1149.9325 | 5 | 5745.6334 | 5745.6098 | 0 | 4.1 | 268 | Specific | 997.24 | 997.24 | 997.24 | 20.02 | 46 | >sp\|P01009\|A1AT_HUMAN Alpha-1-antitrypsin OS=Homo sapiens OX=9606 GN=SERPINA1 PE=1 SV=3 | 642 | 20201118_ly_colostrum_1ng_glyco_hcd203040_compound15.21686.21686.5 | controllerType=0 controllerNumber=1 scan=21686 | 47.6117 |
| 18096:5 | 11 | K.YLGN[+2204.772]ATAIFFLPDEGKLQHLENELTHDIITK.F | HexNAc(4)Hex(5)NeuAc(2) | N[+2205] | 1149.9325 | 5 | 5745.6334 | 5745.6098 | 0 | 4.1 | 268 | Specific | 1158.89 | 1068.01 | 1068.01 | 24.02 | 46 | >sp\|P01009\|A1AT_HUMAN Alpha-1-antitrypsin OS=Homo sapiens OX=9606 GN=SERPINA1 PE=1 SV=3 | 642 | 20201118_ly_colostrum_1ng_glyco_hcd203040_compound15.21684.21684.5 | controllerType=0 controllerNumber=1 scan=21684 | 47.6023 |
| 18101:4 | 11 | K.YLGN[+2204.772]ATAIFFLPDEGKLQHLENELTHDIITK.F | HexNAc(4)Hex(5)NeuAc(2) | N[+2205] | 1437.1646 | 4 | 5745.6364 | 5745.6098 | 0 | 4.6 | 268 | Specific | 1310.73 | 1199.18 | 1199.18 | 26.61 | 46 | >sp\|P01009\|A1AT_HUMAN Alpha-1-antitrypsin OS=Homo sapiens OX=9606 GN=SERPINA1 PE=1 SV=3 | 642 | 20201118_ly_colostrum_1ng_glyco_hcd203040_compound15.21689.21689.4 | controllerType=0 controllerNumber=1 scan=21689 | 47.6157 |
| 19348:5 | 12 | K.APDKNVIFSPLSISTALAFLSLGAHN[+2350.830]TTLTEILK.G | HexNAc(4)Hex(5)Fuc(1)NeuAc(2) | N[+2351] | 1187.5739 | 5 | 5933.8402 | 5933.8085 | 0 | 5.3 | 68 | Specific | 339.33 | 339.33 | 339.33 | 8.11 | 37 | >sp\|P01011\|AACT_HUMAN Alpha-1-antichymotrypsin OS=Homo sapiens OX=9606 GN=SERPINA3 PE=1 SV=2 | 1314 | 20201118_ly_colostrum_1ng_glyco_hcd203040_compound15.22994.22994.5 | controllerType=0 controllerNumber=1 scan=22994 | 50.6424 |
| 19350:5 | 12 | K.APDKNVIFSPLSISTALAFLSLGAHN[+2350.830]TTLTEILK.G | HexNAc(4)Hex(5)Fuc(1)NeuAc(2) | N[+2351] | 1187.5739 | 5 | 5933.8402 | 5933.8085 | 0 | 5.3 | 68 | Specific | 346.39 | 346.39 | 346.39 | 8.31 | 37 | >sp\|P01011\|AACT_HUMAN Alpha-1-antichymotrypsin OS=Homo sapiens OX=9606 GN=SERPINA3 PE=1 SV=2 | 1314 | 20201118_ly_colostrum_1ng_glyco_hcd203040_compound15.22996.22996.5 | controllerType=0 controllerNumber=1 scan=22996 | 50.6464 |
| 19326:5 | 12 | K.APDKNVIFSPLSISTALAFLSLGAHN[+2351.851]TTLTEILK.G | HexNAc(4)Hex(5)Fuc(3)NeuAc(1) | N[+2352] | 1187.7754 | 5 | 5934.8479 | 5934.8289 | 0 | 3.2 | 68 | Specific | 423.02 | 423.02 | 423.02 | 10.65 | 37 | >sp\|P01011\|AACT_HUMAN Alpha-1-antichymotrypsin OS=Homo sapiens OX=9606 GN=SERPINA3 PE=1 SV=2 | 1314 | 20201118_ly_colostrum_1ng_glyco_hcd203040_compound15.22971.22971.5 | controllerType=0 controllerNumber=1 scan=22971 | 50.5916 |
| 19324:5 | 12 | K.APDKNVIFSPLSISTALAFLSLGAHN[+2351.851]TTLTEILK.G | HexNAc(4)Hex(5)Fuc(3)NeuAc(1) | N[+2352] | 1187.7754 | 5 | 5934.8479 | 5934.8289 | 0 | 3.2 | 68 | Specific | 486.91 | 486.91 | 486.91 | 10.98 | 37 | >sp\|P01011\|AACT_HUMAN Alpha-1-antichymotrypsin OS=Homo sapiens OX=9606 GN=SERPINA3 PE=1 SV=2 | 1314 | 20201118_ly_colostrum_1ng_glyco_hcd203040_compound15.22969.22969.5 | controllerType=0 controllerNumber=1 scan=22969 | 50.5874 |
| 19347:4 | 12 | K.APDKNVIFSPLSISTALAFLSLGAHN[+2351.851]TTLTEILK.G | HexNAc(4)Hex(5)Fuc(3)NeuAc(1) | N[+2352] | 1484.4674 | 4 | 5934.8478 | 5934.8289 | 0 | 3.2 | 68 | Specific | 542.78 | 542.78 | 542.78 | 11.17 | 37 | >sp\|P01011\|AACT_HUMAN Alpha-1-antichymotrypsin OS=Homo sapiens OX=9606 GN=SERPINA3 PE=1 SV=2 | 1314 | 20201118_ly_colostrum_1ng_glyco_hcd203040_compound15.22993.22993.4 | controllerType=0 controllerNumber=1 scan=22993 | 50.6406 |
| 19345:4 | 12 | K.APDKNVIFSPLSISTALAFLSLGAHN[+2351.851]TTLTEILK.G | HexNAc(4)Hex(5)Fuc(3)NeuAc(1) | N[+2352] | 1484.4674 | 4 | 5934.8478 | 5934.8289 | 0 | 3.2 | 68 | Specific | 688.45 | 688.45 | 688.45 | 14.67 | 37 | >sp\|P01011\|AACT_HUMAN Alpha-1-antichymotrypsin OS=Homo sapiens OX=9606 GN=SERPINA3 PE=1 SV=2 | 1314 | 20201118_ly_colostrum_1ng_glyco_hcd203040_compound15.22991.22991.4 | controllerType=0 controllerNumber=1 scan=22991 | 50.6364 |
| 11970:3 | 13 | R.IIVPLNNREN[+1856.656]ISDPTSPLR.T | HexNAc(3)Hex(5)Fuc(1)NeuAc(1) | N[+1857] | 1335.6168 | 3 | 4004.8357 | 4004.8327 | 0 | 0.8 | 62 | Specific | 228.03 | 214.75 | 214.75 | 4.06 | 34 | >sp\|P01591\|IGJ_HUMAN Immunoglobulin J chain OS=Homo sapiens OX=9606 GN=JCHAIN PE=1 SV=4 | 1392 | 20201118_ly_colostrum_1ng_glyco_hcd203040_compound15.15298.15298.3 | controllerType=0 controllerNumber=1 scan=15298 | 33.8885 |
| 10305:4 | 13 | R.IIVPLNNREN[+1913.677]ISDPTSPLR.T | HexNAc(4)Hex(5)NeuAc(1) | N[+1914] | 1016.2214 | 4 | 4061.8636 | 4061.8542 | 0 | 2.3 | 62 | Specific | 229.85 | 196.33 | 196.33 | 4.38 | 34 | >sp\|P01591\|IGJ_HUMAN Immunoglobulin J chain OS=Homo sapiens OX=9606 GN=JCHAIN PE=1 SV=4 | 1392 | 20201118_ly_colostrum_1ng_glyco_hcd203040_compound15.13567.13567.4 | controllerType=0 controllerNumber=1 scan=13567 | 30.4511 |
| 11968:3 | 13 | R.IIVPLNNREN[+1856.656]ISDPTSPLR.T | HexNAc(3)Hex(5)Fuc(1)NeuAc(1) | N[+1857] | 1335.6168 | 3 | 4004.8357 | 4004.8327 | 0 | 0.8 | 62 | Specific | 241.25 | 226.92 | 226.92 | 6 | 34 | >sp\|P01591\|IGJ_HUMAN Immunoglobulin J chain OS=Homo sapiens OX=9606 GN=JCHAIN PE=1 SV=4 | 1392 | 20201118_ly_colostrum_1ng_glyco_hcd203040_compound15.15296.15296.3 | controllerType=0 controllerNumber=1 scan=15296 | 33.8848 |
| 08905:3 | 13 | R.IIVPLNNREN[+1809.666]ISDPTSPLR.T | HexNAc(5)Hex(4)Fuc(1) | N[+1810] | 1319.9576 | 3 | 3957.8582 | 3957.8432 | 0 | 3.8 | 62 | Specific | 258.48 | 142.16 | 142.16 | 5.61 | 34 | >sp\|P01591\|IGJ_HUMAN Immunoglobulin J chain OS=Homo sapiens OX=9606 GN=JCHAIN PE=1 SV=4 | 1392 | 20201118_ly_colostrum_1ng_glyco_hcd203040_compound15.12113.12113.3 | controllerType=0 controllerNumber=1 scan=12113 | 27.5734 |
| 12051:3 | 13 | R.IIVPLNNREN[+2351.851]ISDPTSPLR.T | HexNAc(4)Hex(5)Fuc(3)NeuAc(1) | N[+2352] | 1500.6761 | 3 | 4500.0137 | 4500.0279 | 0 | -3.1 | 62 | Specific | 272.54 | 272.54 | 272.54 | 5.63 | 34 | >sp\|P01591\|IGJ_HUMAN Immunoglobulin J chain OS=Homo sapiens OX=9606 GN=JCHAIN PE=1 SV=4 | 1392 | 20201118_ly_colostrum_1ng_glyco_hcd203040_compound15.15383.15383.3 | controllerType=0 controllerNumber=1 scan=15383 | 34.063 |
| 10494:3 | 13 | R.IIVPLNNREN[+1694.603]ISDPTSPLR.T | HexNAc(3)Hex(4)Fuc(1)NeuAc(1) | N[+1695] | 1281.5988 | 3 | 3842.7819 | 3842.7799 | 0 | 0.5 | 62 | Specific | 278.95 | 278.64 | 278.64 | 4.99 | 34 | >sp\|P01591\|IGJ_HUMAN Immunoglobulin J chain OS=Homo sapiens OX=9606 GN=JCHAIN PE=1 SV=4 | 1392 | 20201118_ly_colostrum_1ng_glyco_hcd203040_compound15.13764.13764.3 | controllerType=0 controllerNumber=1 scan=13764 | 30.8598 |
| 10491:3 | 13 | R.IIVPLNNREN[+1694.603]ISDPTSPLR.T | HexNAc(3)Hex(4)Fuc(1)NeuAc(1) | N[+1695] | 1281.5988 | 3 | 3842.7819 | 3842.7799 | 0 | 0.5 | 62 | Specific | 281.3 | 281.3 | 281.3 | 6.56 | 34 | >sp\|P01591\|IGJ_HUMAN Immunoglobulin J chain OS=Homo sapiens OX=9606 GN=JCHAIN PE=1 SV=4 | 1392 | 20201118_ly_colostrum_1ng_glyco_hcd203040_compound15.13761.13761.3 | controllerType=0 controllerNumber=1 scan=13761 | 30.8541 |
| 08929:3 | 13 | R.IIVPLNNREN[+1622.582]ISDPTSPLR.T | HexNAc(4)Hex(5) | N[+1623] | 1257.5944 | 3 | 3770.7687 | 3770.7588 | 0 | 2.6 | 62 | Specific | 297.93 | 257.91 | 257.91 | 6.81 | 34 | >sp\|P01591\|IGJ_HUMAN Immunoglobulin J chain OS=Homo sapiens OX=9606 GN=JCHAIN PE=1 SV=4 | 1392 | 20201118_ly_colostrum_1ng_glyco_hcd203040_compound15.12137.12137.3 | controllerType=0 controllerNumber=1 scan=12137 | 27.6213 |
| 08819:3 | 13 | R.IIVPLNNREN[+1622.582]ISDPTSPLR.T | HexNAc(4)Hex(5) | N[+1623] | 1257.5932 | 3 | 3770.765 | 3770.7588 | 0 | 1.7 | 62 | Specific | 313.55 | 261.39 | 261.39 | 6.83 | 34 | >sp\|P01591\|IGJ_HUMAN Immunoglobulin J chain OS=Homo sapiens OX=9606 GN=JCHAIN PE=1 SV=4 | 1392 | 20201118_ly_colostrum_1ng_glyco_hcd203040_compound15.12023.12023.3 | controllerType=0 controllerNumber=1 scan=12023 | 27.3709 |
| 08927:3 | 13 | R.IIVPLNNREN[+1622.582]ISDPTSPLR.T | HexNAc(4)Hex(5) | N[+1623] | 1257.5944 | 3 | 3770.7687 | 3770.7588 | 0 | 2.6 | 62 | Specific | 326.09 | 289.91 | 289.91 | 7.21 | 34 | >sp\|P01591\|IGJ_HUMAN Immunoglobulin J chain OS=Homo sapiens OX=9606 GN=JCHAIN PE=1 SV=4 | 1392 | 20201118_ly_colostrum_1ng_glyco_hcd203040_compound15.12135.12135.3 | controllerType=0 controllerNumber=1 scan=12135 | 27.6155 |
| 12052:4 | 13 | R.IIVPLNNREN[+2204.772]ISDPTSPLR.T | HexNAc(4)Hex(5)NeuAc(2) | N[+2205] | 1088.9955 | 4 | 4352.9603 | 4352.9496 | 0 | 2.5 | 62 | Specific | 378.98 | 378.98 | 378.98 | 8.11 | 34 | >sp\|P01591\|IGJ_HUMAN Immunoglobulin J chain OS=Homo sapiens OX=9606 GN=JCHAIN PE=1 SV=4 | 1392 | 20201118_ly_colostrum_1ng_glyco_hcd203040_compound15.15384.15384.4 | controllerType=0 controllerNumber=1 scan=15384 | 34.064 |
| 08795:3 | 13 | R.IIVPLNNREN[+1768.640]ISDPTSPLR.T | HexNAc(4)Hex(5)Fuc(1) | N[+1769] | 1306.2794 | 3 | 3916.8235 | 3916.8167 | 0 | 1.7 | 62 | Specific | 393.78 | 333.15 | 333.15 | 7.45 | 34 | >sp\|P01591\|IGJ_HUMAN Immunoglobulin J chain OS=Homo sapiens OX=9606 GN=JCHAIN PE=1 SV=4 | 1392 | 20201118_ly_colostrum_1ng_glyco_hcd203040_compound15.11998.11998.3 | controllerType=0 controllerNumber=1 scan=11998 | 27.3219 |
| 12099:3 | 13 | R.IIVPLNNREN[+2204.772]ISDPTSPLR.T | HexNAc(4)Hex(5)NeuAc(2) | N[+2205] | 1451.6576 | 3 | 4352.9584 | 4352.9496 | 0 | 2 | 62 | Specific | 399.78 | 399.78 | 399.78 | 9.18 | 34 | >sp\|P01591\|IGJ_HUMAN Immunoglobulin J chain OS=Homo sapiens OX=9606 GN=JCHAIN PE=1 SV=4 | 1392 | 20201118_ly_colostrum_1ng_glyco_hcd203040_compound15.15433.15433.3 | controllerType=0 controllerNumber=1 scan=15433 | 34.1657 |
| 12028:3 | 13 | R.IIVPLNNREN[+2350.830]ISDPTSPLR.T | HexNAc(4)Hex(5)Fuc(1)NeuAc(2) | N[+2351] | 1500.3436 | 3 | 4499.0161 | 4499.0075 | 0 | 1.9 | 62 | Specific | 408.19 | 405.74 | 405.74 | 9.37 | 34 | >sp\|P01591\|IGJ_HUMAN Immunoglobulin J chain OS=Homo sapiens OX=9606 GN=JCHAIN PE=1 SV=4 | 1392 | 20201118_ly_colostrum_1ng_glyco_hcd203040_compound15.15359.15359.3 | controllerType=0 controllerNumber=1 scan=15359 | 34.0111 |
| 12054:3 | 13 | R.IIVPLNNREN[+2351.851]ISDPTSPLR.T | HexNAc(4)Hex(5)Fuc(3)NeuAc(1) | N[+2352] | 1500.6761 | 3 | 4500.0137 | 4500.0279 | 0 | -3.1 | 62 | Specific | 408.33 | 408.33 | 408.33 | 8.44 | 34 | >sp\|P01591\|IGJ_HUMAN Immunoglobulin J chain OS=Homo sapiens OX=9606 GN=JCHAIN PE=1 SV=4 | 1392 | 20201118_ly_colostrum_1ng_glyco_hcd203040_compound15.15386.15386.3 | controllerType=0 controllerNumber=1 scan=15386 | 34.0669 |
| 08821:3 | 13 | R.IIVPLNNREN[+1622.582]ISDPTSPLR.T | HexNAc(4)Hex(5) | N[+1623] | 1257.5932 | 3 | 3770.765 | 3770.7588 | 0 | 1.7 | 62 | Specific | 410.19 | 346.09 | 346.09 | 7.71 | 34 | >sp\|P01591\|IGJ_HUMAN Immunoglobulin J chain OS=Homo sapiens OX=9606 GN=JCHAIN PE=1 SV=4 | 1392 | 20201118_ly_colostrum_1ng_glyco_hcd203040_compound15.12025.12025.3 | controllerType=0 controllerNumber=1 scan=12025 | 27.3778 |
| 08886:3 | 13 | R.IIVPLNNREN[+1565.560]ISDPTSPLR.T | HexNAc(3)Hex(5)Fuc(1) | N[+1566] | 1238.5883 | 3 | 3713.7504 | 3713.7373 | 0 | 3.5 | 62 | Specific | 414.88 | 347.46 | 347.46 | 7.97 | 34 | >sp\|P01591\|IGJ_HUMAN Immunoglobulin J chain OS=Homo sapiens OX=9606 GN=JCHAIN PE=1 SV=4 | 1392 | 20201118_ly_colostrum_1ng_glyco_hcd203040_compound15.12093.12093.3 | controllerType=0 controllerNumber=1 scan=12093 | 27.5147 |
| 12055:4 | 13 | R.IIVPLNNREN[+2204.772]ISDPTSPLR.T | HexNAc(4)Hex(5)NeuAc(2) | N[+2205] | 1088.9955 | 4 | 4352.9603 | 4352.9496 | 0 | 2.5 | 62 | Specific | 429.94 | 387.13 | 387.13 | 9.69 | 34 | >sp\|P01591\|IGJ_HUMAN Immunoglobulin J chain OS=Homo sapiens OX=9606 GN=JCHAIN PE=1 SV=4 | 1392 | 20201118_ly_colostrum_1ng_glyco_hcd203040_compound15.15387.15387.4 | controllerType=0 controllerNumber=1 scan=15387 | 34.0695 |
| 08888:3 | 13 | R.IIVPLNNREN[+1565.560]ISDPTSPLR.T | HexNAc(3)Hex(5)Fuc(1) | N[+1566] | 1238.5883 | 3 | 3713.7504 | 3713.7373 | 0 | 3.5 | 62 | Specific | 450.42 | 366.01 | 366.01 | 8.28 | 34 | >sp\|P01591\|IGJ_HUMAN Immunoglobulin J chain OS=Homo sapiens OX=9606 GN=JCHAIN PE=1 SV=4 | 1392 | 20201118_ly_colostrum_1ng_glyco_hcd203040_compound15.12095.12095.3 | controllerType=0 controllerNumber=1 scan=12095 | 27.5391 |
| 08798:3 | 13 | R.IIVPLNNREN[+1768.640]ISDPTSPLR.T | HexNAc(4)Hex(5)Fuc(1) | N[+1769] | 1306.2794 | 3 | 3916.8235 | 3916.8167 | 0 | 1.7 | 62 | Specific | 458.72 | 363.79 | 363.79 | 8.02 | 34 | >sp\|P01591\|IGJ_HUMAN Immunoglobulin J chain OS=Homo sapiens OX=9606 GN=JCHAIN PE=1 SV=4 | 1392 | 20201118_ly_colostrum_1ng_glyco_hcd203040_compound15.12001.12001.3 | controllerType=0 controllerNumber=1 scan=12001 | 27.3259 |
| 08867:3 | 13 | R.IIVPLNNREN[+1768.640]ISDPTSPLR.T | HexNAc(4)Hex(5)Fuc(1) | N[+1769] | 1306.2809 | 3 | 3916.8283 | 3916.8167 | 0 | 3 | 62 | Specific | 460.69 | 356.01 | 356.01 | 8.12 | 34 | >sp\|P01591\|IGJ_HUMAN Immunoglobulin J chain OS=Homo sapiens OX=9606 GN=JCHAIN PE=1 SV=4 | 1392 | 20201118_ly_colostrum_1ng_glyco_hcd203040_compound15.12073.12073.3 | controllerType=0 controllerNumber=1 scan=12073 | 27.479 |
| 08842:3 | 13 | R.IIVPLNNREN[+1971.719]ISDPTSPLR.T | HexNAc(5)Hex(5)Fuc(1) | N[+1972] | 1373.9732 | 3 | 4119.9051 | 4119.896 | 0 | 2.2 | 62 | Specific | 470.05 | 348.4 | 348.4 | 8.12 | 34 | >sp\|P01591\|IGJ_HUMAN Immunoglobulin J chain OS=Homo sapiens OX=9606 GN=JCHAIN PE=1 SV=4 | 1392 | 20201118_ly_colostrum_1ng_glyco_hcd203040_compound15.12047.12047.3 | controllerType=0 controllerNumber=1 scan=12047 | 27.423 |
| 08908:3 | 13 | R.IIVPLNNREN[+1809.666]ISDPTSPLR.T | HexNAc(5)Hex(4)Fuc(1) | N[+1810] | 1319.9576 | 3 | 3957.8582 | 3957.8432 | 0 | 3.8 | 62 | Specific | 472.63 | 319.58 | 319.58 | 7.91 | 34 | >sp\|P01591\|IGJ_HUMAN Immunoglobulin J chain OS=Homo sapiens OX=9606 GN=JCHAIN PE=1 SV=4 | 1392 | 20201118_ly_colostrum_1ng_glyco_hcd203040_compound15.12116.12116.3 | controllerType=0 controllerNumber=1 scan=12116 | 27.5769 |
| 08922:3 | 13 | R.IIVPLNNREN[+1647.613]ISDPTSPLR.T | HexNAc(5)Hex(3)Fuc(1) | N[+1648] | 1265.9389 | 3 | 3795.8021 | 3795.7904 | 0 | 3.1 | 62 | Specific | 481.74 | 436.29 | 436.29 | 9.9 | 34 | >sp\|P01591\|IGJ_HUMAN Immunoglobulin J chain OS=Homo sapiens OX=9606 GN=JCHAIN PE=1 SV=4 | 1392 | 20201118_ly_colostrum_1ng_glyco_hcd203040_compound15.12130.12130.3 | controllerType=0 controllerNumber=1 scan=12130 | 27.6024 |
| 08839:3 | 13 | R.IIVPLNNREN[+1971.719]ISDPTSPLR.T | HexNAc(5)Hex(5)Fuc(1) | N[+1972] | 1373.9732 | 3 | 4119.9051 | 4119.896 | 0 | 2.2 | 62 | Specific | 489.13 | 393.88 | 393.88 | 9.87 | 34 | >sp\|P01591\|IGJ_HUMAN Immunoglobulin J chain OS=Homo sapiens OX=9606 GN=JCHAIN PE=1 SV=4 | 1392 | 20201118_ly_colostrum_1ng_glyco_hcd203040_compound15.12044.12044.3 | controllerType=0 controllerNumber=1 scan=12044 | 27.4184 |
| 12033:3 | 13 | R.IIVPLNNREN[+2204.772]ISDPTSPLR.T | HexNAc(4)Hex(5)NeuAc(2) | N[+2205] | 1451.6581 | 3 | 4352.9599 | 4352.9496 | 0 | 2.4 | 62 | Specific | 496.31 | 446.42 | 446.42 | 9.87 | 34 | >sp\|P01591\|IGJ_HUMAN Immunoglobulin J chain OS=Homo sapiens OX=9606 GN=JCHAIN PE=1 SV=4 | 1392 | 20201118_ly_colostrum_1ng_glyco_hcd203040_compound15.15364.15364.3 | controllerType=0 controllerNumber=1 scan=15364 | 34.0198 |
| 10221:3 | 13 | R.IIVPLNNREN[+2059.735]ISDPTSPLR.T | HexNAc(4)Hex(5)Fuc(1)NeuAc(1) | N[+2060] | 1403.3126 | 3 | 4207.9231 | 4207.9121 | 0 | 2.6 | 62 | Specific | 529.79 | 499.11 | 499.11 | 10.04 | 34 | >sp\|P01591\|IGJ_HUMAN Immunoglobulin J chain OS=Homo sapiens OX=9606 GN=JCHAIN PE=1 SV=4 | 1392 | 20201118_ly_colostrum_1ng_glyco_hcd203040_compound15.13480.13480.3 | controllerType=0 controllerNumber=1 scan=13480 | 30.2539 |
| 08924:3 | 13 | R.IIVPLNNREN[+1647.613]ISDPTSPLR.T | HexNAc(5)Hex(3)Fuc(1) | N[+1648] | 1265.9389 | 3 | 3795.8021 | 3795.7904 | 0 | 3.1 | 62 | Specific | 532.49 | 487.54 | 487.54 | 10.06 | 34 | >sp\|P01591\|IGJ_HUMAN Immunoglobulin J chain OS=Homo sapiens OX=9606 GN=JCHAIN PE=1 SV=4 | 1392 | 20201118_ly_colostrum_1ng_glyco_hcd203040_compound15.12132.12132.3 | controllerType=0 controllerNumber=1 scan=12132 | 27.6109 |
| 12031:3 | 13 | R.IIVPLNNREN[+2350.830]ISDPTSPLR.T | HexNAc(4)Hex(5)Fuc(1)NeuAc(2) | N[+2351] | 1500.3436 | 3 | 4499.0161 | 4499.0075 | 0 | 1.9 | 62 | Specific | 534.01 | 452.14 | 452.14 | 9.8 | 34 | >sp\|P01591\|IGJ_HUMAN Immunoglobulin J chain OS=Homo sapiens OX=9606 GN=JCHAIN PE=1 SV=4 | 1392 | 20201118_ly_colostrum_1ng_glyco_hcd203040_compound15.15362.15362.3 | controllerType=0 controllerNumber=1 scan=15362 | 34.0151 |
| 10322:3 | 13 | R.IIVPLNNREN[+1856.656]ISDPTSPLR.T | HexNAc(3)Hex(5)Fuc(1)NeuAc(1) | N[+1857] | 1335.6187 | 3 | 4004.8416 | 4004.8327 | 0 | 2.2 | 62 | Specific | 540.29 | 540.29 | 540.29 | 10.22 | 34 | >sp\|P01591\|IGJ_HUMAN Immunoglobulin J chain OS=Homo sapiens OX=9606 GN=JCHAIN PE=1 SV=4 | 1392 | 20201118_ly_colostrum_1ng_glyco_hcd203040_compound15.13585.13585.3 | controllerType=0 controllerNumber=1 scan=13585 | 30.4872 |
| 10223:3 | 13 | R.IIVPLNNREN[+2059.735]ISDPTSPLR.T | HexNAc(4)Hex(5)Fuc(1)NeuAc(1) | N[+2060] | 1403.3126 | 3 | 4207.9231 | 4207.9121 | 0 | 2.6 | 62 | Specific | 544.51 | 496.77 | 496.77 | 10.22 | 34 | >sp\|P01591\|IGJ_HUMAN Immunoglobulin J chain OS=Homo sapiens OX=9606 GN=JCHAIN PE=1 SV=4 | 1392 | 20201118_ly_colostrum_1ng_glyco_hcd203040_compound15.13482.13482.3 | controllerType=0 controllerNumber=1 scan=13482 | 30.2595 |
| 08837:3 | 13 | R.IIVPLNNREN[+1663.608]ISDPTSPLR.T | HexNAc(5)Hex(4) | N[+1664] | 1271.2698 | 3 | 3811.7949 | 3811.7853 | 0 | 2.5 | 62 | Specific | 544.85 | 528 | 528 | 10.22 | 34 | >sp\|P01591\|IGJ_HUMAN Immunoglobulin J chain OS=Homo sapiens OX=9606 GN=JCHAIN PE=1 SV=4 | 1392 | 20201118_ly_colostrum_1ng_glyco_hcd203040_compound15.12042.12042.3 | controllerType=0 controllerNumber=1 scan=12042 | 27.4135 |
| 08870:3 | 13 | R.IIVPLNNREN[+1768.640]ISDPTSPLR.T | HexNAc(4)Hex(5)Fuc(1) | N[+1769] | 1306.2809 | 3 | 3916.8283 | 3916.8167 | 0 | 3 | 62 | Specific | 546.99 | 458.45 | 458.45 | 10.22 | 34 | >sp\|P01591\|IGJ_HUMAN Immunoglobulin J chain OS=Homo sapiens OX=9606 GN=JCHAIN PE=1 SV=4 | 1392 | 20201118_ly_colostrum_1ng_glyco_hcd203040_compound15.12076.12076.3 | controllerType=0 controllerNumber=1 scan=12076 | 27.4851 |
| 10418:3 | 13 | R.IIVPLNNREN[+1694.603]ISDPTSPLR.T | HexNAc(3)Hex(4)Fuc(1)NeuAc(1) | N[+1695] | 1281.6013 | 3 | 3842.7892 | 3842.7799 | 0 | 2.4 | 62 | Specific | 547.86 | 547.86 | 547.86 | 10.22 | 34 | >sp\|P01591\|IGJ_HUMAN Immunoglobulin J chain OS=Homo sapiens OX=9606 GN=JCHAIN PE=1 SV=4 | 1392 | 20201118_ly_colostrum_1ng_glyco_hcd203040_compound15.13685.13685.3 | controllerType=0 controllerNumber=1 scan=13685 | 30.6872 |
| 10247:3 | 13 | R.IIVPLNNREN[+1913.677]ISDPTSPLR.T | HexNAc(4)Hex(5)NeuAc(1) | N[+1914] | 1354.6258 | 3 | 4061.8628 | 4061.8542 | 0 | 2.1 | 62 | Specific | 556.08 | 526.86 | 526.86 | 10.22 | 34 | >sp\|P01591\|IGJ_HUMAN Immunoglobulin J chain OS=Homo sapiens OX=9606 GN=JCHAIN PE=1 SV=4 | 1392 | 20201118_ly_colostrum_1ng_glyco_hcd203040_compound15.13507.13507.3 | controllerType=0 controllerNumber=1 scan=13507 | 30.3101 |
| 10420:3 | 13 | R.IIVPLNNREN[+1694.603]ISDPTSPLR.T | HexNAc(3)Hex(4)Fuc(1)NeuAc(1) | N[+1695] | 1281.6013 | 3 | 3842.7892 | 3842.7799 | 0 | 2.4 | 62 | Specific | 558.87 | 558.87 | 558.87 | 10.22 | 34 | >sp\|P01591\|IGJ_HUMAN Immunoglobulin J chain OS=Homo sapiens OX=9606 GN=JCHAIN PE=1 SV=4 | 1392 | 20201118_ly_colostrum_1ng_glyco_hcd203040_compound15.13687.13687.3 | controllerType=0 controllerNumber=1 scan=13687 | 30.6926 |
| 08834:3 | 13 | R.IIVPLNNREN[+1663.608]ISDPTSPLR.T | HexNAc(5)Hex(4) | N[+1664] | 1271.2698 | 3 | 3811.7949 | 3811.7853 | 0 | 2.5 | 62 | Specific | 563.24 | 544.79 | 544.79 | 10.4 | 34 | >sp\|P01591\|IGJ_HUMAN Immunoglobulin J chain OS=Homo sapiens OX=9606 GN=JCHAIN PE=1 SV=4 | 1392 | 20201118_ly_colostrum_1ng_glyco_hcd203040_compound15.12039.12039.3 | controllerType=0 controllerNumber=1 scan=12039 | 27.4083 |
| 12036:3 | 13 | R.IIVPLNNREN[+2204.772]ISDPTSPLR.T | HexNAc(4)Hex(5)NeuAc(2) | N[+2205] | 1451.6581 | 3 | 4352.9599 | 4352.9496 | 0 | 2.4 | 62 | Specific | 563.38 | 526.94 | 526.94 | 10.4 | 34 | >sp\|P01591\|IGJ_HUMAN Immunoglobulin J chain OS=Homo sapiens OX=9606 GN=JCHAIN PE=1 SV=4 | 1392 | 20201118_ly_colostrum_1ng_glyco_hcd203040_compound15.15367.15367.3 | controllerType=0 controllerNumber=1 scan=15367 | 34.0249 |
| 12101:3 | 13 | R.IIVPLNNREN[+2204.772]ISDPTSPLR.T | HexNAc(4)Hex(5)NeuAc(2) | N[+2205] | 1451.6576 | 3 | 4352.9584 | 4352.9496 | 0 | 2 | 62 | Specific | 574.04 | 545.52 | 545.52 | 10.16 | 34 | >sp\|P01591\|IGJ_HUMAN Immunoglobulin J chain OS=Homo sapiens OX=9606 GN=JCHAIN PE=1 SV=4 | 1392 | 20201118_ly_colostrum_1ng_glyco_hcd203040_compound15.15435.15435.3 | controllerType=0 controllerNumber=1 scan=15435 | 34.1695 |
| 10325:3 | 13 | R.IIVPLNNREN[+1856.656]ISDPTSPLR.T | HexNAc(3)Hex(5)Fuc(1)NeuAc(1) | N[+1857] | 1335.6187 | 3 | 4004.8416 | 4004.8327 | 0 | 2.2 | 62 | Specific | 578.77 | 578.77 | 578.77 | 10.4 | 34 | >sp\|P01591\|IGJ_HUMAN Immunoglobulin J chain OS=Homo sapiens OX=9606 GN=JCHAIN PE=1 SV=4 | 1392 | 20201118_ly_colostrum_1ng_glyco_hcd203040_compound15.13588.13588.3 | controllerType=0 controllerNumber=1 scan=13588 | 30.4953 |
| 10250:3 | 13 | R.IIVPLNNREN[+1913.677]ISDPTSPLR.T | HexNAc(4)Hex(5)NeuAc(1) | N[+1914] | 1354.6258 | 3 | 4061.8628 | 4061.8542 | 0 | 2.1 | 62 | Specific | 626.82 | 626.82 | 626.82 | 12.72 | 34 | >sp\|P01591\|IGJ_HUMAN Immunoglobulin J chain OS=Homo sapiens OX=9606 GN=JCHAIN PE=1 SV=4 | 1392 | 20201118_ly_colostrum_1ng_glyco_hcd203040_compound15.13510.13510.3 | controllerType=0 controllerNumber=1 scan=13510 | 30.3183 |
| 12601:3 | 14 | K.LKELPGVC[+57.021]N[+2351.851]ETMMALWEEC[+57.021]KPC[+57.021]LK.Q | HexNAc(4)Hex(5)Fuc(3)NeuAc(1) | C[+57]*3, N[+2352] | 1763.4226 | 3 | 5288.2532 | 5288.2355 | 0 | 3.3 | 95 | Specific | 677.47 | 624.9 | 624.9 | 13.58 | 33 | >sp\|P10909\|CLUS_HUMAN Clusterin OS=Homo sapiens OX=9606 GN=CLU PE=1 SV=1 | 1467 | 20201118_ly_colostrum_1ng_glyco_hcd203040_compound15.15954.15954.3 | controllerType=0 controllerNumber=1 scan=15954 | 35.1985 |
| 12599:3 | 14 | K.LKELPGVC[+57.021]N[+2351.851]ETMMALWEEC[+57.021]KPC[+57.021]LK.Q | HexNAc(4)Hex(5)Fuc(3)NeuAc(1) | C[+57]*3, N[+2352] | 1763.4226 | 3 | 5288.2532 | 5288.2355 | 0 | 3.3 | 95 | Specific | 762.05 | 744.61 | 744.61 | 15.58 | 33 | >sp\|P10909\|CLUS_HUMAN Clusterin OS=Homo sapiens OX=9606 GN=CLU PE=1 SV=1 | 1467 | 20201118_ly_colostrum_1ng_glyco_hcd203040_compound15.15952.15952.3 | controllerType=0 controllerNumber=1 scan=15952 | 35.1929 |
| 14128:4 | 14 | K.LKELPGVC[+57.021]N[+2350.830]ETMMALWEEC[+57.021]KPC[+57.021]LK.Q | HexNAc(4)Hex(5)Fuc(1)NeuAc(2) | C[+57]*3, N[+2351] | 1322.5641 | 4 | 5287.2345 | 5287.2151 | 0 | 3.7 | 95 | Specific | 773.36 | 609.46 | 609.46 | 15.99 | 33 | >sp\|P10909\|CLUS_HUMAN Clusterin OS=Homo sapiens OX=9606 GN=CLU PE=1 SV=1 | 1467 | 20201118_ly_colostrum_1ng_glyco_hcd203040_compound15.17542.17542.4 | controllerType=0 controllerNumber=1 scan=17542 | 38.532 |
| 12589:4 | 14 | K.LKELPGVC[+57.021]N[+2351.851]ETMMALWEEC[+57.021]KPC[+57.021]LK.Q | HexNAc(4)Hex(5)Fuc(3)NeuAc(1) | C[+57]*3, N[+2352] | 1322.8179 | 4 | 5288.2497 | 5288.2355 | 0 | 2.7 | 95 | Specific | 851.93 | 842.89 | 842.89 | 17.96 | 33 | >sp\|P10909\|CLUS_HUMAN Clusterin OS=Homo sapiens OX=9606 GN=CLU PE=1 SV=1 | 1467 | 20201118_ly_colostrum_1ng_glyco_hcd203040_compound15.15942.15942.4 | controllerType=0 controllerNumber=1 scan=15942 | 35.1795 |
| 12586:4 | 14 | K.LKELPGVC[+57.021]N[+2351.851]ETMMALWEEC[+57.021]KPC[+57.021]LK.Q | HexNAc(4)Hex(5)Fuc(3)NeuAc(1) | C[+57]*3, N[+2352] | 1322.8179 | 4 | 5288.2497 | 5288.2355 | 0 | 2.7 | 95 | Specific | 999.23 | 999.23 | 999.23 | 19.96 | 33 | >sp\|P10909\|CLUS_HUMAN Clusterin OS=Homo sapiens OX=9606 GN=CLU PE=1 SV=1 | 1467 | 20201118_ly_colostrum_1ng_glyco_hcd203040_compound15.15939.15939.4 | controllerType=0 controllerNumber=1 scan=15939 | 35.1739 |
| 14125:4 | 14 | K.LKELPGVC[+57.021]N[+2350.830]ETMMALWEEC[+57.021]KPC[+57.021]LK.Q | HexNAc(4)Hex(5)Fuc(1)NeuAc(2) | C[+57]*3, N[+2351] | 1322.5641 | 4 | 5287.2345 | 5287.2151 | 0 | 3.7 | 95 | Specific | 1034.46 | 826.03 | 826.03 | 20.99 | 33 | >sp\|P10909\|CLUS_HUMAN Clusterin OS=Homo sapiens OX=9606 GN=CLU PE=1 SV=1 | 1467 | 20201118_ly_colostrum_1ng_glyco_hcd203040_compound15.17539.17539.4 | controllerType=0 controllerNumber=1 scan=17539 | 38.5262 |
| 00129:4 | 14 | R.EIRHN[+2205.793]STGC[+57.021]LR.M | HexNAc(4)Hex(5)Fuc(2)NeuAc(1) | C[+57], N[+2206] | 887.8727 | 4 | 3548.4688 | 3548.4573 | 0 | 3.3 | 287 | Specific | 325.85 | 238.12 | 238.12 | 5.18 | 33 | >sp\|P10909\|CLUS_HUMAN Clusterin OS=Homo sapiens OX=9606 GN=CLU PE=1 SV=1 | 1467 | 20201118_ly_colostrum_1ng_glyco_hcd203040_compound15.3009.3009.4 | controllerType=0 controllerNumber=1 scan=3009 | 11.2714 |
| 21130:3 | 14 | K.[+42.011]MLN[+2336.851]TSSLLEQLNEQFNWVSR.L | HexNAc(6)Hex(6)Fuc(1) | .[42], N[+2337] | 1596.6778 | 3 | 4788.0188 | 4788.0484 | 0 | -6.2 | 352 | Specific | 252.91 | 193.59 | 193.59 | 5.73 | 33 | >sp\|P10909\|CLUS_HUMAN Clusterin OS=Homo sapiens OX=9606 GN=CLU PE=1 SV=1 | 1467 | 20201118_ly_colostrum_1ng_glyco_hcd203040_compound15.24851.24851.3 | controllerType=0 controllerNumber=1 scan=24851 | 54.5698 |
| 21128:3 | 14 | K.[+42.011]MLN[+2336.851]TSSLLEQLNEQFNWVSR.L | HexNAc(6)Hex(6)Fuc(1) | .[42], N[+2337] | 1596.6778 | 3 | 4788.0188 | 4788.0484 | 0 | -6.2 | 352 | Specific | 253.5 | 201.58 | 201.58 | 5.73 | 33 | >sp\|P10909\|CLUS_HUMAN Clusterin OS=Homo sapiens OX=9606 GN=CLU PE=1 SV=1 | 1467 | 20201118_ly_colostrum_1ng_glyco_hcd203040_compound15.24849.24849.3 | controllerType=0 controllerNumber=1 scan=24849 | 54.5655 |
| 19030:3 | 14 | K.MLN[+2205.793]TSSLLEQLNEQFNWVSR.L | HexNAc(4)Hex(5)Fuc(2)NeuAc(1) | N[+2206] | 1539.0124 | 3 | 4615.0226 | 4614.9796 | 0 | 9.3 | 352 | Specific | 411.22 | 346.46 | 346.46 | 7.44 | 33 | >sp\|P10909\|CLUS_HUMAN Clusterin OS=Homo sapiens OX=9606 GN=CLU PE=1 SV=1 | 1467 | 20201118_ly_colostrum_1ng_glyco_hcd203040_compound15.22660.22660.3 | controllerType=0 controllerNumber=1 scan=22660 | 49.8392 |
| 19028:3 | 14 | K.MLN[+2205.793]TSSLLEQLNEQFNWVSR.L | HexNAc(4)Hex(5)Fuc(2)NeuAc(1) | N[+2206] | 1539.0124 | 3 | 4615.0226 | 4614.9796 | 0 | 9.3 | 352 | Specific | 457.94 | 457.94 | 457.94 | 9.22 | 33 | >sp\|P10909\|CLUS_HUMAN Clusterin OS=Homo sapiens OX=9606 GN=CLU PE=1 SV=1 | 1467 | 20201118_ly_colostrum_1ng_glyco_hcd203040_compound15.22658.22658.3 | controllerType=0 controllerNumber=1 scan=22658 | 49.8317 |
| 18959:4 | 14 | K.MLN[+2351.851]TSSLLEQLNEQFNWVSR.L | HexNAc(4)Hex(5)Fuc(3)NeuAc(1) | N[+2352] | 1191.0197 | 4 | 4761.057 | 4761.0375 | 0 | 4.1 | 352 | Specific | 527.55 | 473.96 | 473.96 | 10.45 | 33 | >sp\|P10909\|CLUS_HUMAN Clusterin OS=Homo sapiens OX=9606 GN=CLU PE=1 SV=1 | 1467 | 20201118_ly_colostrum_1ng_glyco_hcd203040_compound15.22586.22586.4 | controllerType=0 controllerNumber=1 scan=22586 | 49.6788 |
| 21017:3 | 14 | K.MLN[+2350.830]TSSLLEQLNEQFNWVSR.L | HexNAc(4)Hex(5)Fuc(1)NeuAc(2) | N[+2351] | 1587.3567 | 3 | 4760.0557 | 4760.0171 | 0 | 8.1 | 352 | Specific | 706.81 | 567.45 | 567.45 | 14.4 | 33 | >sp\|P10909\|CLUS_HUMAN Clusterin OS=Homo sapiens OX=9606 GN=CLU PE=1 SV=1 | 1467 | 20201118_ly_colostrum_1ng_glyco_hcd203040_compound15.24734.24734.3 | controllerType=0 controllerNumber=1 scan=24734 | 54.3323 |
| 18983:4 | 14 | K.MLN[+2205.793]TSSLLEQLNEQFNWVSR.L | HexNAc(4)Hex(5)Fuc(2)NeuAc(1) | N[+2206] | 1154.5052 | 4 | 4614.9988 | 4614.9796 | 0 | 4.2 | 352 | Specific | 786.01 | 610.73 | 610.73 | 16.13 | 33 | >sp\|P10909\|CLUS_HUMAN Clusterin OS=Homo sapiens OX=9606 GN=CLU PE=1 SV=1 | 1467 | 20201118_ly_colostrum_1ng_glyco_hcd203040_compound15.22611.22611.4 | controllerType=0 controllerNumber=1 scan=22611 | 49.7287 |
| 21064:3 | 14 | K.MLN[+2350.830]TSSLLEQLNEQFNWVSR.L | HexNAc(4)Hex(5)Fuc(1)NeuAc(2) | N[+2351] | 1587.3503 | 3 | 4760.0363 | 4760.0171 | 0 | 4 | 352 | Specific | 808.21 | 586.27 | 586.27 | 17.57 | 33 | >sp\|P10909\|CLUS_HUMAN Clusterin OS=Homo sapiens OX=9606 GN=CLU PE=1 SV=1 | 1467 | 20201118_ly_colostrum_1ng_glyco_hcd203040_compound15.24783.24783.3 | controllerType=0 controllerNumber=1 scan=24783 | 54.4302 |
| 21065:4 | 14 | K.MLN[+2350.830]TSSLLEQLNEQFNWVSR.L | HexNAc(4)Hex(5)Fuc(1)NeuAc(2) | N[+2351] | 1190.7637 | 4 | 4760.033 | 4760.0171 | 0 | 3.4 | 352 | Specific | 815.31 | 599.54 | 599.54 | 17.43 | 33 | >sp\|P10909\|CLUS_HUMAN Clusterin OS=Homo sapiens OX=9606 GN=CLU PE=1 SV=1 | 1467 | 20201118_ly_colostrum_1ng_glyco_hcd203040_compound15.24784.24784.4 | controllerType=0 controllerNumber=1 scan=24784 | 54.4316 |
| 21019:3 | 14 | K.MLN[+2350.830]TSSLLEQLNEQFNWVSR.L | HexNAc(4)Hex(5)Fuc(1)NeuAc(2) | N[+2351] | 1587.3567 | 3 | 4760.0557 | 4760.0171 | 0 | 8.1 | 352 | Specific | 823.76 | 569.57 | 569.57 | 16.4 | 33 | >sp\|P10909\|CLUS_HUMAN Clusterin OS=Homo sapiens OX=9606 GN=CLU PE=1 SV=1 | 1467 | 20201118_ly_colostrum_1ng_glyco_hcd203040_compound15.24736.24736.3 | controllerType=0 controllerNumber=1 scan=24736 | 54.3363 |
| 19256:3 | 14 | K.MLN[+1872.651]TSSLLEQLNEQFNWVSR.L | HexNAc(3)Hex(6)NeuAc(1) | N[+1873] | 1427.956 | 3 | 4281.8534 | 4281.8372 | 0 | 3.8 | 352 | Specific | 829.43 | 798.6 | 352.58 | 17.57 | 33 | >sp\|P10909\|CLUS_HUMAN Clusterin OS=Homo sapiens OX=9606 GN=CLU PE=1 SV=1 | 1467 | 20201118_ly_colostrum_1ng_glyco_hcd203040_compound15.22898.22898.3 | controllerType=0 controllerNumber=1 scan=22898 | 50.4188 |
| 19258:3 | 14 | K.MLN[+1872.651]TSSLLEQLNEQFNWVSR.L | HexNAc(3)Hex(6)NeuAc(1) | N[+1873] | 1427.956 | 3 | 4281.8534 | 4281.8372 | 0 | 3.8 | 352 | Specific | 832.83 | 616.52 | 369.61 | 17.57 | 33 | >sp\|P10909\|CLUS_HUMAN Clusterin OS=Homo sapiens OX=9606 GN=CLU PE=1 SV=1 | 1467 | 20201118_ly_colostrum_1ng_glyco_hcd203040_compound15.22900.22900.3 | controllerType=0 controllerNumber=1 scan=22900 | 50.4273 |
| 18961:4 | 14 | K.MLN[+2351.851]TSSLLEQLNEQFNWVSR.L | HexNAc(4)Hex(5)Fuc(3)NeuAc(1) | N[+2352] | 1191.0197 | 4 | 4761.057 | 4761.0375 | 0 | 4.1 | 352 | Specific | 838.21 | 661.58 | 661.58 | 17.13 | 33 | >sp\|P10909\|CLUS_HUMAN Clusterin OS=Homo sapiens OX=9606 GN=CLU PE=1 SV=1 | 1467 | 20201118_ly_colostrum_1ng_glyco_hcd203040_compound15.22588.22588.4 | controllerType=0 controllerNumber=1 scan=22588 | 49.6833 |
| 21067:4 | 14 | K.MLN[+2350.830]TSSLLEQLNEQFNWVSR.L | HexNAc(4)Hex(5)Fuc(1)NeuAc(2) | N[+2351] | 1190.7637 | 4 | 4760.033 | 4760.0171 | 0 | 3.4 | 352 | Specific | 859.4 | 567.66 | 567.66 | 18.43 | 33 | >sp\|P10909\|CLUS_HUMAN Clusterin OS=Homo sapiens OX=9606 GN=CLU PE=1 SV=1 | 1467 | 20201118_ly_colostrum_1ng_glyco_hcd203040_compound15.24786.24786.4 | controllerType=0 controllerNumber=1 scan=24786 | 54.4372 |
| 21066:3 | 14 | K.MLN[+2350.830]TSSLLEQLNEQFNWVSR.L | HexNAc(4)Hex(5)Fuc(1)NeuAc(2) | N[+2351] | 1587.3503 | 3 | 4760.0363 | 4760.0171 | 0 | 4 | 352 | Specific | 914.3 | 641.88 | 641.88 | 19.57 | 33 | >sp\|P10909\|CLUS_HUMAN Clusterin OS=Homo sapiens OX=9606 GN=CLU PE=1 SV=1 | 1467 | 20201118_ly_colostrum_1ng_glyco_hcd203040_compound15.24785.24785.3 | controllerType=0 controllerNumber=1 scan=24785 | 54.4348 |
| 18985:4 | 14 | K.MLN[+2205.793]TSSLLEQLNEQFNWVSR.L | HexNAc(4)Hex(5)Fuc(2)NeuAc(1) | N[+2206] | 1154.5052 | 4 | 4614.9988 | 4614.9796 | 0 | 4.2 | 352 | Specific | 930.48 | 692.8 | 692.8 | 19.13 | 33 | >sp\|P10909\|CLUS_HUMAN Clusterin OS=Homo sapiens OX=9606 GN=CLU PE=1 SV=1 | 1467 | 20201118_ly_colostrum_1ng_glyco_hcd203040_compound15.22613.22613.4 | controllerType=0 controllerNumber=1 scan=22613 | 49.7332 |
| 18932:3 | 14 | K.MLN[+2351.851]TSSLLEQLNEQFNWVSR.L | HexNAc(4)Hex(5)Fuc(3)NeuAc(1) | N[+2352] | 1587.6915 | 3 | 4761.0598 | 4761.0375 | 0 | 4.7 | 352 | Specific | 970.57 | 666.22 | 666.22 | 20.27 | 33 | >sp\|P10909\|CLUS_HUMAN Clusterin OS=Homo sapiens OX=9606 GN=CLU PE=1 SV=1 | 1467 | 20201118_ly_colostrum_1ng_glyco_hcd203040_compound15.22557.22557.3 | controllerType=0 controllerNumber=1 scan=22557 | 49.6209 |
| 18930:3 | 14 | K.MLN[+2351.851]TSSLLEQLNEQFNWVSR.L | HexNAc(4)Hex(5)Fuc(3)NeuAc(1) | N[+2352] | 1587.6915 | 3 | 4761.0598 | 4761.0375 | 0 | 4.7 | 352 | Specific | 1091.35 | 854.98 | 854.98 | 22.27 | 33 | >sp\|P10909\|CLUS_HUMAN Clusterin OS=Homo sapiens OX=9606 GN=CLU PE=1 SV=1 | 1467 | 20201118_ly_colostrum_1ng_glyco_hcd203040_compound15.22555.22555.3 | controllerType=0 controllerNumber=1 scan=22555 | 49.6154 |
| 20508:3 | 14 | K.M[+15.995]LN[+2350.830]TSSLLEQLNEQFNWVSR.L | HexNAc(4)Hex(5)Fuc(1)NeuAc(2) | M[+16], N[+2351] | 1592.6826 | 3 | 4776.0331 | 4776.012 | 0 | 4.4 | 352 | Specific | 418.68 | 363.78 | 363.78 | 8.69 | 33 | >sp\|P10909\|CLUS_HUMAN Clusterin OS=Homo sapiens OX=9606 GN=CLU PE=1 SV=1 | 1467 | 20201118_ly_colostrum_1ng_glyco_hcd203040_compound15.24202.24202.3 | controllerType=0 controllerNumber=1 scan=24202 | 53.1481 |
| 20510:3 | 14 | K.M[+15.995]LN[+2350.830]TSSLLEQLNEQFNWVSR.L | HexNAc(4)Hex(5)Fuc(1)NeuAc(2) | M[+16], N[+2351] | 1592.6826 | 3 | 4776.0331 | 4776.012 | 0 | 4.4 | 352 | Specific | 514.78 | 237.59 | 237.59 | 8.14 | 33 | >sp\|P10909\|CLUS_HUMAN Clusterin OS=Homo sapiens OX=9606 GN=CLU PE=1 SV=1 | 1467 | 20201118_ly_colostrum_1ng_glyco_hcd203040_compound15.24204.24204.3 | controllerType=0 controllerNumber=1 scan=24204 | 53.1531 |
| 10568:2 | 17 | R.VN[+1216.423]LTTNTIAVTQTLPNAAYNNR.F | HexNAc(2)Hex(5) | N[+1216] | 1803.342 | 2 | 3605.6767 | 3605.6699 | 0 | 1.9 | 351 | Specific | 577.57 | 577.57 | 577.57 | 10.82 | 26 | >sp\|Q6UX06\|OLFM4_HUMAN Olfactomedin-4 OS=Homo sapiens OX=9606 GN=OLFM4 PE=1 SV=1 | 624 | 20201118_ly_colostrum_1ng_glyco_hcd203040_compound15.13841.13841.2 | controllerType=0 controllerNumber=1 scan=13841 | 31.0175 |
| 10570:2 | 17 | R.VN[+1216.423]LTTNTIAVTQTLPNAAYNNR.F | HexNAc(2)Hex(5) | N[+1216] | 1803.342 | 2 | 3605.6767 | 3605.6699 | 0 | 1.9 | 351 | Specific | 598.29 | 598.29 | 598.29 | 12.15 | 26 | >sp\|Q6UX06\|OLFM4_HUMAN Olfactomedin-4 OS=Homo sapiens OX=9606 GN=OLFM4 PE=1 SV=1 | 624 | 20201118_ly_colostrum_1ng_glyco_hcd203040_compound15.13843.13843.2 | controllerType=0 controllerNumber=1 scan=13843 | 31.0209 |
| 10537:3 | 17 | R.VN[+1216.423]LTTNTIAVTQTLPNAAYNNR.F | HexNAc(2)Hex(5) | N[+1216] | 1202.5648 | 3 | 3605.6797 | 3605.6699 | 0 | 2.7 | 351 | Specific | 926.11 | 926.11 | 926.11 | 19.54 | 26 | >sp\|Q6UX06\|OLFM4_HUMAN Olfactomedin-4 OS=Homo sapiens OX=9606 GN=OLFM4 PE=1 SV=1 | 624 | 20201118_ly_colostrum_1ng_glyco_hcd203040_compound15.13809.13809.3 | controllerType=0 controllerNumber=1 scan=13809 | 30.9539 |
| 10540:3 | 17 | R.VN[+1216.423]LTTNTIAVTQTLPNAAYNNR.F | HexNAc(2)Hex(5) | N[+1216] | 1202.5648 | 3 | 3605.6797 | 3605.6699 | 0 | 2.7 | 351 | Specific | 1200.5 | 1200.5 | 1200.5 | 25.54 | 26 | >sp\|Q6UX06\|OLFM4_HUMAN Olfactomedin-4 OS=Homo sapiens OX=9606 GN=OLFM4 PE=1 SV=1 | 624 | 20201118_ly_colostrum_1ng_glyco_hcd203040_compound15.13812.13812.3 | controllerType=0 controllerNumber=1 scan=13812 | 30.9619 |
| 18303:4 | 20 | K.AREDIFMETLKDIVEYYNDSN[+2350.830]GSHVLQGR.F | HexNAc(4)Hex(5)Fuc(1)NeuAc(2) | N[+2351] | 1438.3785 | 4 | 5750.4923 | 5750.4577 | 0 | 6 | 92 | Specific | 693.45 | 580.03 | 10.63 | 12.89 | 25 | >sp\|P25311\|ZA2G_HUMAN Zinc-alpha-2-glycoprotein OS=Homo sapiens OX=9606 GN=AZGP1 PE=1 SV=2 | 744 | 20201118_ly_colostrum_1ng_glyco_hcd203040_compound15.21900.21900.4 | controllerType=0 controllerNumber=1 scan=21900 | 48.0871 |
| 18306:4 | 20 | K.AREDIFMETLKDIVEYYNDSN[+2350.830]GSHVLQGR.F | HexNAc(4)Hex(5)Fuc(1)NeuAc(2) | N[+2351] | 1438.3785 | 4 | 5750.4923 | 5750.4577 | 0 | 6 | 92 | Specific | 702.04 | 514.27 | 9.54 | 13.89 | 25 | >sp\|P25311\|ZA2G_HUMAN Zinc-alpha-2-glycoprotein OS=Homo sapiens OX=9606 GN=AZGP1 PE=1 SV=2 | 744 | 20201118_ly_colostrum_1ng_glyco_hcd203040_compound15.21903.21903.4 | controllerType=0 controllerNumber=1 scan=21903 | 48.0926 |
| 03588:2 | 23 | K.AAIPSALDTN[+1216.423]SSK.S | HexNAc(2)Hex(5) | N[+1216] | 1246.0475 | 2 | 2491.0878 | 2491.0815 | 0 | 2.5 | 542 | Specific | 485.33 | 394.93 | 394.93 | 9.83 | 22 | >sp\|Q08380\|LG3BP_HUMAN Galectin-3-binding protein OS=Homo sapiens OX=9606 GN=LGALS3BP PE=1 SV=1 | 1562 | 20201118_ly_colostrum_1ng_glyco_hcd203040_compound15.6600.6600.2 | controllerType=0 controllerNumber=1 scan=6600 | 17.6014 |
| 03590:2 | 23 | K.AAIPSALDTN[+1216.423]SSK.S | HexNAc(2)Hex(5) | N[+1216] | 1246.0475 | 2 | 2491.0878 | 2491.0815 | 0 | 2.5 | 542 | Specific | 491.94 | 378.3 | 378.3 | 8.37 | 22 | >sp\|Q08380\|LG3BP_HUMAN Galectin-3-binding protein OS=Homo sapiens OX=9606 GN=LGALS3BP PE=1 SV=1 | 1562 | 20201118_ly_colostrum_1ng_glyco_hcd203040_compound15.6602.6602.2 | controllerType=0 controllerNumber=1 scan=6602 | 17.6051 |
| 00757:2 | 32 | R.SYN[+1038.375]DSVDPR.I | HexNAc(2)Hex(3)Fuc(1) | N[+1038] | 1045.9277 | 2 | 2090.848 | 2090.8395 | 0 | 4.1 | 481 | Specific | 248.55 | 105.38 | 105.38 | 4.49 | 18 | >sp\|P05164\|PERM_HUMAN Myeloperoxidase OS=Homo sapiens OX=9606 GN=MPO PE=1 SV=1 | 12 | 20201118_ly_colostrum_1ng_glyco_hcd203040_compound15.3668.3668.2 | controllerType=0 controllerNumber=1 scan=3668 | 12.44 |
| 00755:2 | 32 | R.SYN[+1038.375]DSVDPR.I | HexNAc(2)Hex(3)Fuc(1) | N[+1038] | 1045.9277 | 2 | 2090.848 | 2090.8395 | 0 | 4.1 | 481 | Specific | 272 | 232.98 | 232.98 | 6.05 | 18 | >sp\|P05164\|PERM_HUMAN Myeloperoxidase OS=Homo sapiens OX=9606 GN=MPO PE=1 SV=1 | 12 | 20201118_ly_colostrum_1ng_glyco_hcd203040_compound15.3666.3666.2 | controllerType=0 controllerNumber=1 scan=3666 | 12.4359 |
| 15028:3 | 36 | K.AALAAFNAQNN[+2204.772]GSNFQLEEISR.A | HexNAc(4)Hex(5)NeuAc(2) | N[+2205] | 1523.9894 | 3 | 4569.9538 | 4569.9256 | 0 | 6.2 | 166 | Specific | 368.42 | 298.18 | 298.18 | 7.32 | 11 | >sp\|P02765\|FETUA_HUMAN Alpha-2-HS-glycoprotein OS=Homo sapiens OX=9606 GN=AHSG PE=1 SV=2 | 135 | 20201118_ly_colostrum_1ng_glyco_hcd203040_compound15.18479.18479.2 | controllerType=0 controllerNumber=1 scan=18479 | 40.4388 |
| 15030:3 | 36 | K.AALAAFNAQNN[+2204.772]GSNFQLEEISR.A | HexNAc(4)Hex(5)NeuAc(2) | N[+2205] | 1523.9894 | 3 | 4569.9538 | 4569.9256 | 0 | 6.2 | 166 | Specific | 522.49 | 327.12 | 327.12 | 7.34 | 11 | >sp\|P02765\|FETUA_HUMAN Alpha-2-HS-glycoprotein OS=Homo sapiens OX=9606 GN=AHSG PE=1 SV=2 | 135 | 20201118_ly_colostrum_1ng_glyco_hcd203040_compound15.18481.18481.2 | controllerType=0 controllerNumber=1 scan=18481 | 40.4423 |
| 08283:3 | 74 | R.FVN[+876.322]VTVTPEDQC[+57.021]RPNNVC[+57.021]TGVLTR.R | HexNAc(2)Hex(2)Fuc(1) | C[+57]*2, N[+876] | 1218.2311 | 3 | 3652.6789 | 3652.6728 | 0 | 1.7 | 169 | Specific | 891.59 | 891.59 | 891.59 | 17.96 | 5 | >sp\|P20160\|CAP7_HUMAN Azurocidin OS=Homo sapiens OX=9606 GN=AZU1 PE=1 SV=3 | 850 | 20201118_ly_colostrum_1ng_glyco_hcd203040_compound15.11466.11466.3 | controllerType=0 controllerNumber=1 scan=11466 | 26.2869 |
| 08286:3 | 74 | R.FVN[+876.322]VTVTPEDQC[+57.021]RPNNVC[+57.021]TGVLTR.R | HexNAc(2)Hex(2)Fuc(1) | C[+57]*2, N[+876] | 1218.2311 | 3 | 3652.6789 | 3652.6728 | 0 | 1.7 | 169 | Specific | 935.78 | 931.28 | 931.28 | 18.96 | 5 | >sp\|P20160\|CAP7_HUMAN Azurocidin OS=Homo sapiens OX=9606 GN=AZU1 PE=1 SV=3 | 850 | 20201118_ly_colostrum_1ng_glyco_hcd203040_compound15.11469.11469.3 | controllerType=0 controllerNumber=1 scan=11469 | 26.2929 |
| 19637:3 | 2 | R.[+42.011]RTAGWNVPIGTLRPFLN[+1864.634]WTGPPEPIEAAVAR.F | HexNAc(2)Hex(9) | .[42], N[+1865] | 1765.1541 | 3 | 5293.4477 | 5293.4568 | 0 | -1.7 | 140 | Specific | 328.57 | 328.57 | 328.57 | 6.36 | 115 | >sp\|P02788\|TRFL_HUMAN Lactotransferrin OS=Homo sapiens OX=9606 GN=LTF PE=1 SV=6 | 1270 | 20201117_ly_milk_1ng_glyco_hcd203040_compound15.23090.23090.3 | controllerType=0 controllerNumber=1 scan=23090 | 46.5002 |
| 18253:4 | 2 | R.RTAGWNVPIGTLRPFLN[+2569.905]WTGPPEPIEAAVAR.F | HexNAc(5)Hex(6)NeuAc(2) | N[+2570] | 1489.943 | 4 | 5956.7502 | 5956.7167 | 0 | 5.6 | 140 | Specific | 453.52 | 453.52 | 453.52 | 9.23 | 115 | >sp\|P02788\|TRFL_HUMAN Lactotransferrin OS=Homo sapiens OX=9606 GN=LTF PE=1 SV=6 | 1270 | 20201117_ly_milk_1ng_glyco_hcd203040_compound15.21646.21646.4 | controllerType=0 controllerNumber=1 scan=21646 | 43.3824 |
| 17400:4 | 2 | R.RTAGWNVPIGTLRPFLN[+1768.640]WTGPPEPIEAAVAR.F | HexNAc(4)Hex(5)Fuc(1) | N[+1769] | 1289.6221 | 4 | 5155.4665 | 5155.4516 | 0 | 2.9 | 140 | Specific | 492.12 | 429.95 | 429.95 | 10.23 | 115 | >sp\|P02788\|TRFL_HUMAN Lactotransferrin OS=Homo sapiens OX=9606 GN=LTF PE=1 SV=6 | 1270 | 20201117_ly_milk_1ng_glyco_hcd203040_compound15.20756.20756.4 | controllerType=0 controllerNumber=1 scan=20756 | 41.4009 |
| 17397:4 | 2 | R.RTAGWNVPIGTLRPFLN[+1768.640]WTGPPEPIEAAVAR.F | HexNAc(4)Hex(5)Fuc(1) | N[+1769] | 1289.6234 | 4 | 5155.4718 | 5155.4516 | 0 | 3.9 | 140 | Specific | 538.33 | 484.65 | 484.65 | 10.16 | 115 | >sp\|P02788\|TRFL_HUMAN Lactotransferrin OS=Homo sapiens OX=9606 GN=LTF PE=1 SV=6 | 1270 | 20201117_ly_milk_1ng_glyco_hcd203040_compound15.20752.20752.4 | controllerType=0 controllerNumber=1 scan=20752 | 41.3905 |

Table S4. Cont.

| Query #:z | Protein Rank | Glycopeptide | Glycan | Modification Type(s) | Observed m/z | z | Observed (M+H) | Calc. mass (M+H) | Off-by-x error | Mass error (ppm) | StartPosition | Cleavage | Score | Delta | Delta Mod | \|Log Prob\| | # of unique peptides | Protein Name | Protein DB number | Comment | Scan # | Scan Time |
|---|---|---|---|---|---|---|---|---|---|---|---|---|---|---|---|---|---|---|---|---|---|---|
| 18330:4 | 2 | R.RTAGWNVPIGTLRPFLN[+2059.735]WTGPPEPIEAAVAR.F | HexNAc(4)Hex(5)Fuc(1)NeuAc(1) | N[+2060] | 1362.3967 | 4 | 5446.5651 | 5446.547 | 0 | 3.3 | 140 | Specific | 572.75 | 572.75 | 572.75 | 10.67 | 115 | >sp\|P02788\|TRFL_HUMAN Lactotransferrin OS=Homo sapiens OX=9606 GN=LTF PE=1 SV=6 | 1270 | 20201117_ly_milk_1ng_glyco_hcd203040_compound15.21726.21726.4 | controllerType=0 controllerNumber=1 scan=21726 | 43.5471 |
| 17398:4 | 2 | R.RTAGWNVPIGTLRPFLN[+1768.640]WTGPPEPIEAAVAR.F | HexNAc(4)Hex(5)Fuc(1) | N[+1769] | 1289.6234 | 4 | 5155.4718 | 5155.4516 | 0 | 3.9 | 140 | Specific | 577.81 | 442.89 | 442.89 | 11.57 | 115 | >sp\|P02788\|TRFL_HUMAN Lactotransferrin OS=Homo sapiens OX=9606 GN=LTF PE=1 SV=6 | 1270 | 20201117_ly_milk_1ng_glyco_hcd203040_compound15.20753.20753.4 | controllerType=0 controllerNumber=1 scan=20753 | 41.3949 |
| 18347:5 | 2 | R.RTAGWNVPIGTLRPFLN[+2059.735]WTGPPEPIEAAVAR.F | HexNAc(4)Hex(5)Fuc(1)NeuAc(1) | N[+2060] | 1090.118 | 5 | 5446.5611 | 5446.547 | 0 | 2.6 | 140 | Specific | 600.99 | 580.45 | 580.45 | 12.74 | 115 | >sp\|P02788\|TRFL_HUMAN Lactotransferrin OS=Homo sapiens OX=9606 GN=LTF PE=1 SV=6 | 1270 | 20201117_ly_milk_1ng_glyco_hcd203040_compound15.21743.21743.5 | controllerType=0 controllerNumber=1 scan=21743 | 43.5916 |
| 17402:4 | 2 | R.RTAGWNVPIGTLRPFLN[+1768.640]WTGPPEPIEAAVAR.F | HexNAc(4)Hex(5)Fuc(1) | N[+1769] | 1289.6221 | 4 | 5155.4665 | 5155.4516 | 0 | 2.9 | 140 | Specific | 621.05 | 495 | 495 | 12.84 | 115 | >sp\|P02788\|TRFL_HUMAN Lactotransferrin OS=Homo sapiens OX=9606 GN=LTF PE=1 SV=6 | 1270 | 20201117_ly_milk_1ng_glyco_hcd203040_compound15.20758.20758.4 | controllerType=0 controllerNumber=1 scan=20758 | 41.4054 |
| 18332:4 | 2 | R.RTAGWNVPIGTLRPFLN[+2059.735]WTGPPEPIEAAVAR.F | HexNAc(4)Hex(5)Fuc(1)NeuAc(1) | N[+2060] | 1362.3967 | 4 | 5446.5651 | 5446.547 | 0 | 3.3 | 140 | Specific | 634.04 | 509.69 | 509.69 | 12.84 | 115 | >sp\|P02788\|TRFL_HUMAN Lactotransferrin OS=Homo sapiens OX=9606 GN=LTF PE=1 SV=6 | 1270 | 20201117_ly_milk_1ng_glyco_hcd203040_compound15.21728.21728.4 | controllerType=0 controllerNumber=1 scan=21728 | 43.5543 |
| 18344:5 | 2 | R.RTAGWNVPIGTLRPFLN[+2059.735]WTGPPEPIEAAVAR.F | HexNAc(4)Hex(5)Fuc(1)NeuAc(1) | N[+2060] | 1090.118 | 5 | 5446.5611 | 5446.547 | 0 | 2.6 | 140 | Specific | 654.15 | 571.22 | 571.22 | 13.74 | 115 | >sp\|P02788\|TRFL_HUMAN Lactotransferrin OS=Homo sapiens OX=9606 GN=LTF PE=1 SV=6 | 1270 | 20201117_ly_milk_1ng_glyco_hcd203040_compound15.21740.21740.5 | controllerType=0 controllerNumber=1 scan=21740 | 43.5774 |
| 18622:4 | 2 | R.[+42.011]TAGWNVPIGTLRPFLN[+1872.651]WTGPPEPIEAAVAR.F | HexNAc(3)Hex(6)NeuAc(1) | .[42], N[+1873] | 1287.1112 | 4 | 5145.423 | 5145.372 | 0 | 9.9 | 141 | Specific | 239.28 | 135.99 | 135.99 | 3.33 | 115 | >sp\|P02788\|TRFL_HUMAN Lactotransferrin OS=Homo sapiens OX=9606 GN=LTF PE=1 SV=6 | 1270 | 20201117_ly_milk_1ng_glyco_hcd203040_compound15.22030.22030.4 | controllerType=0 controllerNumber=1 scan=22030 | 44.1821 |
| 19518:3 | 2 | R.TAGWNVPIGTLRPFLN[+2351.851]WTGPPEPIEAAVAR.F | HexNAc(4)Hex(5)Fuc(3)NeuAc(1) | N[+2352] | 1861.5301 | 3 | 5582.5758 | 5582.5617 | 0 | 2.5 | 141 | Specific | 265.35 | 265.35 | 265.35 | 7.18 | 115 | >sp\|P02788\|TRFL_HUMAN Lactotransferrin OS=Homo sapiens OX=9606 GN=LTF PE=1 SV=6 | 1270 | 20201117_ly_milk_1ng_glyco_hcd203040_compound15.22966.22966.3 | controllerType=0 controllerNumber=1 scan=22966 | 46.2808 |
| 18613:4 | 2 | R.TAGWNVPIGTLRPFLN[+2059.735]WTGPPEPIEAAVAR.F | HexNAc(4)Hex(5)Fuc(1)NeuAc(1) | N[+2060] | 1323.3738 | 4 | 5290.4733 | 5290.4459 | 0 | 5.2 | 141 | Specific | 308.79 | 308.79 | 308.79 | 6.74 | 115 | >sp\|P02788\|TRFL_HUMAN Lactotransferrin OS=Homo sapiens OX=9606 GN=LTF PE=1 SV=6 | 1270 | 20201117_ly_milk_1ng_glyco_hcd203040_compound15.22020.22020.4 | controllerType=0 controllerNumber=1 scan=22020 | 44.1604 |
| 19326:4 | 2 | R.TAGWNVPIGTLRPFLN[+2936.057]WTGPPEPIEAAVAR.F | HexNAc(6)Hex(7)Fuc(2)NeuAc(1) | N[+2936] | 1542.4513 | 4 | 6166.7834 | 6166.7682 | 0 | 2.5 | 141 | Specific | 390.03 | 349.96 | 349.96 | 8.38 | 115 | >sp\|P02788\|TRFL_HUMAN Lactotransferrin OS=Homo sapiens OX=9606 GN=LTF PE=1 SV=6 | 1270 | 20201117_ly_milk_1ng_glyco_hcd203040_compound15.22766.22766.4 | controllerType=0 controllerNumber=1 scan=22766 | 45.8621 |
| 19947:4 | 2 | R.TAGWNVPIGTLRPFLN[+2059.735]WTGPPEPIEAAVAR.F | HexNAc(4)Hex(5)Fuc(1)NeuAc(1) | N[+2060] | 1323.3661 | 4 | 5290.4425 | 5290.4459 | 0 | -0.6 | 141 | Specific | 438.49 | 423.49 | 423.49 | 8.9 | 115 | >sp\|P02788\|TRFL_HUMAN Lactotransferrin OS=Homo sapiens OX=9606 GN=LTF PE=1 SV=6 | 1270 | 20201117_ly_milk_1ng_glyco_hcd203040_compound15.23418.23418.4 | controllerType=0 controllerNumber=1 scan=23418 | 47.1853 |
| 19948:4 | 2 | R.TAGWNVPIGTLRPFLN[+2059.735]WTGPPEPIEAAVAR.F | HexNAc(4)Hex(5)Fuc(1)NeuAc(1) | N[+2060] | 1323.3661 | 4 | 5290.4425 | 5290.4459 | 0 | -0.6 | 141 | Specific | 564.61 | 502.3 | 502.3 | 9.58 | 115 | >sp\|P02788\|TRFL_HUMAN Lactotransferrin OS=Homo sapiens OX=9606 GN=LTF PE=1 SV=6 | 1270 | 20201117_ly_milk_1ng_glyco_hcd203040_compound15.23419.23419.4 | controllerType=0 controllerNumber=1 scan=23419 | 47.189 |
| 19538:4 | 2 | R.TAGWNVPIGTLRPFLN[+2351.851]WTGPPEPIEAAVAR.F | HexNAc(4)Hex(5)Fuc(3)NeuAc(1) | N[+2352] | 1396.3999 | 4 | 5582.5778 | 5582.5617 | 0 | 2.9 | 141 | Specific | 590.88 | 590.88 | 590.88 | 11.74 | 115 | >sp\|P02788\|TRFL_HUMAN Lactotransferrin OS=Homo sapiens OX=9606 GN=LTF PE=1 SV=6 | 1270 | 20201117_ly_milk_1ng_glyco_hcd203040_compound15.22987.22987.4 | controllerType=0 controllerNumber=1 scan=22987 | 46.3394 |
| 19520:3 | 2 | R.TAGWNVPIGTLRPFLN[+2351.851]WTGPPEPIEAAVAR.F | HexNAc(4)Hex(5)Fuc(3)NeuAc(1) | N[+2352] | 1861.5301 | 3 | 5582.5758 | 5582.5617 | 0 | 2.5 | 141 | Specific | 593.13 | 550.74 | 550.74 | 11.84 | 115 | >sp\|P02788\|TRFL_HUMAN Lactotransferrin OS=Homo sapiens OX=9606 GN=LTF PE=1 SV=6 | 1270 | 20201117_ly_milk_1ng_glyco_hcd203040_compound15.22968.22968.3 | controllerType=0 controllerNumber=1 scan=22968 | 46.2876 |
| 19329:4 | 2 | R.TAGWNVPIGTLRPFLN[+2936.057]WTGPPEPIEAAVAR.F | HexNAc(6)Hex(7)Fuc(2)NeuAc(1) | N[+2936] | 1542.4513 | 4 | 6166.7834 | 6166.7682 | 0 | 2.5 | 141 | Specific | 601.55 | 492.4 | 492.4 | 12.74 | 115 | >sp\|P02788\|TRFL_HUMAN Lactotransferrin OS=Homo sapiens OX=9606 GN=LTF PE=1 SV=6 | 1270 | 20201117_ly_milk_1ng_glyco_hcd203040_compound15.22769.22769.4 | controllerType=0 controllerNumber=1 scan=22769 | 45.8719 |
| 19483:5 | 2 | R.TAGWNVPIGTLRPFLN[+2351.851]WTGPPEPIEAAVAR.F | HexNAc(4)Hex(5)Fuc(3)NeuAc(1) | N[+2352] | 1117.3206 | 5 | 5582.5737 | 5582.5617 | 0 | 2.2 | 141 | Specific | 614.13 | 574.79 | 574.79 | 12.79 | 115 | >sp\|P02788\|TRFL_HUMAN Lactotransferrin OS=Homo sapiens OX=9606 GN=LTF PE=1 SV=6 | 1270 | 20201117_ly_milk_1ng_glyco_hcd203040_compound15.22929.22929.2 | controllerType=0 controllerNumber=1 scan=22929 | 46.2018 |
| 19480:3 | 2 | R.TAGWNVPIGTLRPFLN[+2205.793]WTGPPEPIEAAVAR.F | HexNAc(4)Hex(5)Fuc(2)NeuAc(1) | N[+2206] | 1812.8442 | 3 | 5436.5181 | 5436.5038 | 0 | 2.6 | 141 | Specific | 628.65 | 553.57 | 553.57 | 12.84 | 115 | >sp\|P02788\|TRFL_HUMAN Lactotransferrin OS=Homo sapiens OX=9606 GN=LTF PE=1 SV=6 | 1270 | 20201117_ly_milk_1ng_glyco_hcd203040_compound15.22926.22926.3 | controllerType=0 controllerNumber=1 scan=22926 | 46.1981 |
| 19468:4 | 2 | R.TAGWNVPIGTLRPFLN[+2205.793]WTGPPEPIEAAVAR.F | HexNAc(4)Hex(5)Fuc(2)NeuAc(1) | N[+2206] | 1359.8853 | 4 | 5436.5192 | 5436.5038 | 0 | 2.8 | 141 | Specific | 690.19 | 652.6 | 652.6 | 13.74 | 115 | >sp\|P02788\|TRFL_HUMAN Lactotransferrin OS=Homo sapiens OX=9606 GN=LTF PE=1 SV=6 | 1270 | 20201117_ly_milk_1ng_glyco_hcd203040_compound15.22914.22914.4 | controllerType=0 controllerNumber=1 scan=22914 | 46.1752 |
| 18580:3 | 2 | R.TAGWNVPIGTLRPFLN[+1768.640]WTGPPEPIEAAVAR.F | HexNAc(4)Hex(5)Fuc(1) | N[+1769] | 1667.1274 | 3 | 4999.3677 | 4999.3504 | 0 | 3.4 | 141 | Specific | 692.85 | 588.61 | 588.61 | 13.57 | 115 | >sp\|P02788\|TRFL_HUMAN Lactotransferrin OS=Homo sapiens OX=9606 GN=LTF PE=1 SV=6 | 1270 | 20201117_ly_milk_1ng_glyco_hcd203040_compound15.21986.21986.3 | controllerType=0 controllerNumber=1 scan=21986 | 44.0877 |
| 18578:3 | 2 | R.TAGWNVPIGTLRPFLN[+1768.640]WTGPPEPIEAAVAR.F | HexNAc(4)Hex(5)Fuc(1) | N[+1769] | 1667.1274 | 3 | 4999.3677 | 4999.3504 | 0 | 3.4 | 141 | Specific | 708.01 | 609.89 | 609.89 | 14.57 | 115 | >sp\|P02788\|TRFL_HUMAN Lactotransferrin OS=Homo sapiens OX=9606 GN=LTF PE=1 SV=6 | 1270 | 20201117_ly_milk_1ng_glyco_hcd203040_compound15.21984.21984.3 | controllerType=0 controllerNumber=1 scan=21984 | 44.0791 |
| 19478:3 | 2 | R.TAGWNVPIGTLRPFLN[+2205.793]WTGPPEPIEAAVAR.F | HexNAc(4)Hex(5)Fuc(2)NeuAc(1) | N[+2206] | 1812.8442 | 3 | 5436.5181 | 5436.5038 | 0 | 2.6 | 141 | Specific | 710.29 | 624.71 | 624.71 | 14.84 | 115 | >sp\|P02788\|TRFL_HUMAN Lactotransferrin OS=Homo sapiens OX=9606 GN=LTF PE=1 SV=6 | 1270 | 20201117_ly_milk_1ng_glyco_hcd203040_compound15.22924.22924.3 | controllerType=0 controllerNumber=1 scan=22924 | 46.1918 |
| 19503:3 | 2 | R.TAGWNVPIGTLRPFLN[+2059.735]WTGPPEPIEAAVAR.F | HexNAc(4)Hex(5)Fuc(1)NeuAc(1) | N[+2060] | 1764.1601 | 3 | 5290.4658 | 5290.4459 | 0 | 3.8 | 141 | Specific | 711.13 | 643.89 | 643.89 | 14.57 | 115 | >sp\|P02788\|TRFL_HUMAN Lactotransferrin OS=Homo sapiens OX=9606 GN=LTF PE=1 SV=6 | 1270 | 20201117_ly_milk_1ng_glyco_hcd203040_compound15.22950.22950.3 | controllerType=0 controllerNumber=1 scan=22950 | 46.2491 |
| 19506:3 | 2 | R.TAGWNVPIGTLRPFLN[+2059.735]WTGPPEPIEAAVAR.F | HexNAc(4)Hex(5)Fuc(1)NeuAc(1) | N[+2060] | 1764.1601 | 3 | 5290.4658 | 5290.4459 | 0 | 3.8 | 141 | Specific | 711.92 | 646.39 | 646.39 | 14.57 | 115 | >sp\|P02788\|TRFL_HUMAN Lactotransferrin OS=Homo sapiens OX=9606 GN=LTF PE=1 SV=6 | 1270 | 20201117_ly_milk_1ng_glyco_hcd203040_compound15.22953.22953.3 | controllerType=0 controllerNumber=1 scan=22953 | 46.2566 |
| 20816:4 | 2 | R.TAGWNVPIGTLRPFLN[+2350.830]WTGPPEPIEAAVAR.F | HexNAc(4)Hex(5)Fuc(1)NeuAc(2) | N[+2351] | 1396.1411 | 4 | 5581.5426 | 5581.5413 | 0 | 0.2 | 141 | Specific | 727.74 | 594.14 | 594.14 | 14.18 | 115 | >sp\|P02788\|TRFL_HUMAN Lactotransferrin OS=Homo sapiens OX=9606 GN=LTF PE=1 SV=6 | 1270 | 20201117_ly_milk_1ng_glyco_hcd203040_compound15.24328.24328.4 | controllerType=0 controllerNumber=1 scan=24328 | 49.1547 |
| 18574:4 | 2 | R.TAGWNVPIGTLRPFLN[+1768.640]WTGPPEPIEAAVAR.F | HexNAc(4)Hex(5)Fuc(1) | N[+1769] | 1250.5975 | 4 | 4999.3683 | 4999.3504 | 0 | 3.6 | 141 | Specific | 746.39 | 688.37 | 688.37 | 14.47 | 115 | >sp\|P02788\|TRFL_HUMAN Lactotransferrin OS=Homo sapiens OX=9606 GN=LTF PE=1 SV=6 | 1270 | 20201117_ly_milk_1ng_glyco_hcd203040_compound15.21980.21980.4 | controllerType=0 controllerNumber=1 scan=21980 | 44.0694 |
| 19470:4 | 2 | R.TAGWNVPIGTLRPFLN[+2205.793]WTGPPEPIEAAVAR.F | HexNAc(4)Hex(5)Fuc(2)NeuAc(1) | N[+2206] | 1359.8853 | 4 | 5436.5192 | 5436.5038 | 0 | 2.8 | 141 | Specific | 782.12 | 678.69 | 678.69 | 15.74 | 115 | >sp\|P02788\|TRFL_HUMAN Lactotransferrin OS=Homo sapiens OX=9606 GN=LTF PE=1 SV=6 | 1270 | 20201117_ly_milk_1ng_glyco_hcd203040_compound15.22916.22916.4 | controllerType=0 controllerNumber=1 scan=22916 | 46.1825 |
| 20818:4 | 2 | R.TAGWNVPIGTLRPFLN[+2350.830]WTGPPEPIEAAVAR.F | HexNAc(4)Hex(5)Fuc(1)NeuAc(2) | N[+2351] | 1396.1411 | 4 | 5581.5426 | 5581.5413 | 0 | 0.2 | 141 | Specific | 805.64 | 627.06 | 627.06 | 16.18 | 115 | >sp\|P02788\|TRFL_HUMAN Lactotransferrin OS=Homo sapiens OX=9606 GN=LTF PE=1 SV=6 | 1270 | 20201117_ly_milk_1ng_glyco_hcd203040_compound15.24330.24330.4 | controllerType=0 controllerNumber=1 scan=24330 | 49.164 |
| 19494:5 | 2 | R.TAGWNVPIGTLRPFLN[+2205.793]WTGPPEPIEAAVAR.F | HexNAc(4)Hex(5)Fuc(2)NeuAc(1) | N[+2206] | 1088.1101 | 5 | 5436.5214 | 5436.5038 | 0 | 3.3 | 141 | Specific | 826.56 | 705.64 | 705.64 | 16.79 | 115 | >sp\|P02788\|TRFL_HUMAN Lactotransferrin OS=Homo sapiens OX=9606 GN=LTF PE=1 SV=6 | 1270 | 20201117_ly_milk_1ng_glyco_hcd203040_compound15.22941.22941.5 | controllerType=0 controllerNumber=1 scan=22941 | 46.2252 |
| 18576:4 | 2 | R.TAGWNVPIGTLRPFLN[+1768.640]WTGPPEPIEAAVAR.F | HexNAc(4)Hex(5)Fuc(1) | N[+1769] | 1250.5975 | 4 | 4999.3683 | 4999.3504 | 0 | 3.6 | 141 | Specific | 829.17 | 744.2 | 744.2 | 16.47 | 115 | >sp\|P02788\|TRFL_HUMAN Lactotransferrin OS=Homo sapiens OX=9606 GN=LTF PE=1 SV=6 | 1270 | 20201117_ly_milk_1ng_glyco_hcd203040_compound15.21982.21982.4 | controllerType=0 controllerNumber=1 scan=21982 | 44.0769 |
| 19485:5 | 2 | R.TAGWNVPIGTLRPFLN[+2351.851]WTGPPEPIEAAVAR.F | HexNAc(4)Hex(5)Fuc(3)NeuAc(1) | N[+2352] | 1117.3206 | 5 | 5582.5737 | 5582.5617 | 0 | 2.2 | 141 | Specific | 876.02 | 751.82 | 751.82 | 17.79 | 115 | >sp\|P02788\|TRFL_HUMAN Lactotransferrin OS=Homo sapiens OX=9606 GN=LTF PE=1 SV=6 | 1270 | 20201117_ly_milk_1ng_glyco_hcd203040_compound15.22931.22931.2 | controllerType=0 controllerNumber=1 scan=22931 | 46.2073 |
| 19496:5 | 2 | R.TAGWNVPIGTLRPFLN[+2205.793]WTGPPEPIEAAVAR.F | HexNAc(4)Hex(5)Fuc(2)NeuAc(1) | N[+2206] | 1088.1101 | 5 | 5436.5214 | 5436.5038 | 0 | 3.3 | 141 | Specific | 968.49 | 744.87 | 744.87 | 19.79 | 115 | >sp\|P02788\|TRFL_HUMAN Lactotransferrin OS=Homo sapiens OX=9606 GN=LTF PE=1 SV=6 | 1270 | 20201117_ly_milk_1ng_glyco_hcd203040_compound15.22943.22943.5 | controllerType=0 controllerNumber=1 scan=22943 | 46.2333 |
| 17119:2 | 2 | R.TAGWNIPMGLLFN[+1768.640]QTGSC[+57.021]K.F | HexNAc(4)Hex(5)Fuc(1) | C[+57], N[+1769] | 1932.3347 | 2 | 3863.6621 | 3863.6495 | 0 | 3.3 | 485 | Specific | 381.62 | 306.37 | 306.37 | 7.26 | 115 | >sp\|P02788\|TRFL_HUMAN Lactotransferrin OS=Homo sapiens OX=9606 GN=LTF PE=1 SV=6 | 1270 | 20201117_ly_milk_1ng_glyco_hcd203040_compound15.20462.20462.2 | controllerType=0 controllerNumber=1 scan=20462 | 40.7742 |
| 20957:4 | 2 | R.TAGWNIPMGLLFN[+2350.830]QTGSC[+57.021]K.F | HexNAc(4)Hex(5)Fuc(1)NeuAc(2) | C[+57], N[+2351] | 1112.2177 | 4 | 4445.8489 | 4445.8403 | 0 | 1.9 | 485 | Specific | 414.83 | 364.7 | 364.7 | 7.81 | 115 | >sp\|P02788\|TRFL_HUMAN Lactotransferrin OS=Homo sapiens OX=9606 GN=LTF PE=1 SV=6 | 1270 | 20201117_ly_milk_1ng_glyco_hcd203040_compound15.24474.24474.4 | controllerType=0 controllerNumber=1 scan=24474 | 49.4762 |
| 18926:4 | 2 | R.TAGWNIPMGLLFN[+2059.735]QTGSC[+57.021]K.F | HexNAc(4)Hex(5)Fuc(1)NeuAc(1) | C[+57], N[+2060] | 1039.4452 | 4 | 4154.7591 | 4154.7449 | 0 | 3.4 | 485 | Specific | 493.02 | 273.9 | 273.9 | 7.46 | 115 | >sp\|P02788\|TRFL_HUMAN Lactotransferrin OS=Homo sapiens OX=9606 GN=LTF PE=1 SV=6 | 1270 | 20201117_ly_milk_1ng_glyco_hcd203040_compound15.22346.22346.4 | controllerType=0 controllerNumber=1 scan=22346 | 44.8702 |
| 18929:4 | 2 | R.TAGWNIPMGLLFN[+2059.735]QTGSC[+57.021]K.F | HexNAc(4)Hex(5)Fuc(1)NeuAc(1) | C[+57], N[+2060] | 1039.4452 | 4 | 4154.7591 | 4154.7449 | 0 | 3.4 | 485 | Specific | 498.88 | 227.56 | 227.56 | 6.98 | 115 | >sp\|P02788\|TRFL_HUMAN Lactotransferrin OS=Homo sapiens OX=9606 GN=LTF PE=1 SV=6 | 1270 | 20201117_ly_milk_1ng_glyco_hcd203040_compound15.22349.22349.4 | controllerType=0 controllerNumber=1 scan=22349 | 44.8874 |
| 20960:4 | 2 | R.TAGWNIPMGLLFN[+2350.830]QTGSC[+57.021]K.F | HexNAc(4)Hex(5)Fuc(1)NeuAc(2) | C[+57], N[+2351] | 1112.2177 | 4 | 4445.8489 | 4445.8403 | 0 | 1.9 | 485 | Specific | 506.92 | 361.42 | 361.42 | 8.17 | 115 | >sp\|P02788\|TRFL_HUMAN Lactotransferrin OS=Homo sapiens OX=9606 GN=LTF PE=1 SV=6 | 1270 | 20201117_ly_milk_1ng_glyco_hcd203040_compound15.24477.24477.4 | controllerType=0 controllerNumber=1 scan=24477 | 49.4818 |
| 17392:3 | 2 | R.TAGWNIPMGLLFN[+1460.529]QTGSC[+57.021]K.F | HexNAc(4)Hex(4) | C[+57], N[+1461] | 1185.8544 | 3 | 3555.5487 | 3555.5387 | 0 | 2.8 | 485 | Specific | 566.64 | 539.22 | 326.83 | 9.97 | 115 | >sp\|P02788\|TRFL_HUMAN Lactotransferrin OS=Homo sapiens OX=9606 GN=LTF PE=1 SV=6 | 1270 | 20201117_ly_milk_1ng_glyco_hcd203040_compound15.20747.20747.3 | controllerType=0 controllerNumber=1 scan=20747 | 41.3811 |
| 20933:3 | 2 | R.TAGWNIPMGLLFN[+2350.830]QTGSC[+57.021]K.F | HexNAc(4)Hex(5)Fuc(1)NeuAc(2) | C[+57], N[+2351] | 1482.6241 | 3 | 4445.8577 | 4445.8403 | 0 | 3.9 | 485 | Specific | 587.23 | 570.93 | 570.93 | 10.87 | 115 | >sp\|P02788\|TRFL_HUMAN Lactotransferrin OS=Homo sapiens OX=9606 GN=LTF PE=1 SV=6 | 1270 | 20201117_ly_milk_1ng_glyco_hcd203040_compound15.24450.24450.3 | controllerType=0 controllerNumber=1 scan=24450 | 49.4358 |
| 17464:2 | 2 | R.TAGWNIPMGLLFN[+1216.423]QTGSC[+57.021]K.F | HexNAc(2)Hex(5) | C[+57], N[+1216] | 1656.2247 | 2 | 3311.442 | 3311.4328 | 0 | 2.8 | 485 | Specific | 691.91 | 653.42 | 215.01 | 13.24 | 115 | >sp\|P02788\|TRFL_HUMAN Lactotransferrin OS=Homo sapiens OX=9606 GN=LTF PE=1 SV=6 | 1270 | 20201117_ly_milk_1ng_glyco_hcd203040_compound15.20822.20822.2 | controllerType=0 controllerNumber=1 scan=20822 | 41.5367 |
| 18928:4 | 2 | R.TAGWNIPMGLLFN[+2205.793]QTGSC[+57.021]K.F | HexNAc(4)Hex(5)Fuc(2)NeuAc(1) | C[+57], N[+2206] | 1075.9577 | 4 | 4300.8089 | 4300.8028 | 0 | 1.4 | 485 | Specific | 694.97 | 654.35 | 654.35 | 13.14 | 115 | >sp\|P02788\|TRFL_HUMAN Lactotransferrin OS=Homo sapiens OX=9606 GN=LTF PE=1 SV=6 | 1270 | 20201117_ly_milk_1ng_glyco_hcd203040_compound15.22348.22348.4 | controllerType=0 controllerNumber=1 scan=22348 | 44.8726 |
| 17260:3 | 2 | R.TAGWNIPMGLLFN[+1622.582]QTGSC[+57.021]K.F | HexNAc(4)Hex(5) | C[+57], N[+1623] | 1239.8735 | 3 | 3717.6059 | 3717.5916 | 0 | 3.8 | 485 | Specific | 761.04 | 761.04 | 342.59 | 14.87 | 115 | >sp\|P02788\|TRFL_HUMAN Lactotransferrin OS=Homo sapiens OX=9606 GN=LTF PE=1 SV=6 | 1270 | 20201117_ly_milk_1ng_glyco_hcd203040_compound15.20610.20610.3 | controllerType=0 controllerNumber=1 scan=20610 | 41.0899 |
| 18930:4 | 2 | R.TAGWNIPMGLLFN[+2205.793]QTGSC[+57.021]K.F | HexNAc(4)Hex(5)Fuc(2)NeuAc(1) | C[+57], N[+2206] | 1075.9577 | 4 | 4300.8089 | 4300.8028 | 0 | 1.4 | 485 | Specific | 761.06 | 670.21 | 670.21 | 15.14 | 115 | >sp\|P02788\|TRFL_HUMAN Lactotransferrin OS=Homo sapiens OX=9606 GN=LTF PE=1 SV=6 | 1270 | 20201117_ly_milk_1ng_glyco_hcd203040_compound15.22350.22350.4 | controllerType=0 controllerNumber=1 scan=22350 | 44.8889 |
| 17193:3 | 2 | R.TAGWNIPMGLLFN[+1606.587]QTGSC[+57.021]K.F | HexNAc(4)Hex(4)Fuc(1) | C[+57], N[+1607] | 1234.5401 | 3 | 3701.6057 | 3701.5966 | 0 | 2.5 | 485 | Specific | 761.34 | 761.34 | 761.34 | 15.14 | 115 | >sp\|P02788\|TRFL_HUMAN Lactotransferrin OS=Homo sapiens OX=9606 GN=LTF PE=1 SV=6 | 1270 | 20201117_ly_milk_1ng_glyco_hcd203040_compound15.20540.20540.3 | controllerType=0 controllerNumber=1 scan=20540 | 40.9372 |
| 19131:3 | 2 | R.TAGWNIPMGLLFN[+1751.624]QTGSC[+57.021]K.F | HexNAc(4)Hex(4)NeuAc(1) | C[+57], N[+1752] | 1282.8862 | 3 | 3846.6439 | 3846.6341 | 0 | 2.5 | 485 | Specific | 780.56 | 780.4 | 780.4 | 15.14 | 115 | >sp\|P02788\|TRFL_HUMAN Lactotransferrin OS=Homo sapiens OX=9606 GN=LTF PE=1 SV=6 | 1270 | 20201117_ly_milk_1ng_glyco_hcd203040_compound15.22563.22563.3 | controllerType=0 controllerNumber=1 scan=22563 | 45.4385 |
| 17100:4 | 2 | R.TAGWNIPMGLLFN[+1768.640]QTGSC[+57.021]K.F | HexNAc(4)Hex(5)Fuc(1) | C[+57], N[+1769] | 966.6711 | 4 | 3863.6624 | 3863.6495 | 0 | 3.4 | 485 | Specific | 811.55 | 657.14 | 657.14 | 15.93 | 115 | >sp\|P02788\|TRFL_HUMAN Lactotransferrin OS=Homo sapiens OX=9606 GN=LTF PE=1 SV=6 | 1270 | 20201117_ly_milk_1ng_glyco_hcd203040_compound15.20442.20442.4 | controllerType=0 controllerNumber=1 scan=20442 | 40.7297 |
| 17098:4 | 2 | R.TAGWNIPMGLLFN[+1768.640]QTGSC[+57.021]K.F | HexNAc(4)Hex(5)Fuc(1) | C[+57], N[+1769] | 966.6711 | 4 | 3863.6624 | 3863.6495 | 0 | 3.4 | 485 | Specific | 824.86 | 775.4 | 775.4 | 15.93 | 115 | >sp\|P02788\|TRFL_HUMAN Lactotransferrin OS=Homo sapiens OX=9606 GN=LTF PE=1 SV=6 | 1270 | 20201117_ly_milk_1ng_glyco_hcd203040_compound15.20440.20440.4 | controllerType=0 controllerNumber=1 scan=20440 | 40.7238 |
| 19060:4 | 2 | R.TAGWNIPMGLLFN[+1913.677]QTGSC[+57.021]K.F | HexNAc(4)Hex(5)NeuAc(1) | C[+57], N[+1914] | 1002.9281 | 4 | 4008.6907 | 4008.687 | 0 | 0.9 | 485 | Specific | 839.78 | 795.58 | 795.58 | 16.14 | 115 | >sp\|P02788\|TRFL_HUMAN Lactotransferrin OS=Homo sapiens OX=9606 GN=LTF PE=1 SV=6 | 1270 | 20201117_ly_milk_1ng_glyco_hcd203040_compound15.22487.22487.4 | controllerType=0 controllerNumber=1 scan=22487 | 45.2411 |
| 17031:3 | 2 | R.TAGWNIPMGLLFN[+1768.640]QTGSC[+57.021]K.F | HexNAc(4)Hex(5)Fuc(1) | C[+57], N[+1769] | 1288.5582 | 3 | 3863.6599 | 3863.6495 | 0 | 2.7 | 485 | Specific | 868.88 | 802.39 | 802.39 | 17.14 | 115 | >sp\|P02788\|TRFL_HUMAN Lactotransferrin OS=Homo sapiens OX=9606 GN=LTF PE=1 SV=6 | 1270 | 20201117_ly_milk_1ng_glyco_hcd203040_compound15.20371.20371.3 | controllerType=0 controllerNumber=1 scan=20371 | 40.5669 |
| 17262:3 | 2 | R.TAGWNIPMGLLFN[+1622.582]QTGSC[+57.021]K.F | HexNAc(4)Hex(5) | C[+57], N[+1623] | 1239.8735 | 3 | 3717.6059 | 3717.5916 | 0 | 3.8 | 485 | Specific | 887.33 | 872.53 | 429.11 | 16.87 | 115 | >sp\|P02788\|TRFL_HUMAN Lactotransferrin OS=Homo sapiens OX=9606 GN=LTF PE=1 SV=6 | 1270 | 20201117_ly_milk_1ng_glyco_hcd203040_compound15.20612.20612.3 | controllerType=0 controllerNumber=1 scan=20612 | 41.0948 |
| 19062:4 | 2 | R.TAGWNIPMGLLFN[+1913.677]QTGSC[+57.021]K.F | HexNAc(4)Hex(5)NeuAc(1) | C[+57], N[+1914] | 1002.9281 | 4 | 4008.6907 | 4008.687 | 0 | 0.9 | 485 | Specific | 888.2 | 856.15 | 856.15 | 17.14 | 115 | >sp\|P02788\|TRFL_HUMAN Lactotransferrin OS=Homo sapiens OX=9606 GN=LTF PE=1 SV=6 | 1270 | 20201117_ly_milk_1ng_glyco_hcd203040_compound15.22489.22489.4 | controllerType=0 controllerNumber=1 scan=22489 | 45.2481 |
| 18883:3 | 2 | R.TAGWNIPMGLLFN[+2205.793]QTGSC[+57.021]K.F | HexNAc(4)Hex(5)Fuc(2)NeuAc(1) | C[+57], N[+2206] | 1434.277 | 3 | 4300.8166 | 4300.8028 | 0 | 3.2 | 485 | Specific | 889.04 | 741.21 | 741.21 | 17.14 | 115 | >sp\|P02788\|TRFL_HUMAN Lactotransferrin OS=Homo sapiens OX=9606 GN=LTF PE=1 SV=6 | 1270 | 20201117_ly_milk_1ng_glyco_hcd203040_compound15.22302.22302.3 | controllerType=0 controllerNumber=1 scan=22302 | 44.7635 |
| 17195:3 | 2 | R.TAGWNIPMGLLFN[+1606.587]QTGSC[+57.021]K.F | HexNAc(4)Hex(4)Fuc(1) | C[+57], N[+1607] | 1234.5401 | 3 | 3701.6057 | 3701.5966 | 0 | 2.5 | 485 | Specific | 908.91 | 900.88 | 900.88 | 18.14 | 115 | >sp\|P02788\|TRFL_HUMAN Lactotransferrin OS=Homo sapiens OX=9606 GN=LTF PE=1 SV=6 | 1270 | 20201117_ly_milk_1ng_glyco_hcd203040_compound15.20542.20542.3 | controllerType=0 controllerNumber=1 scan=20542 | 40.9416 |
| 18885:3 | 2 | R.TAGWNIPMGLLFN[+2205.793]QTGSC[+57.021]K.F | HexNAc(4)Hex(5)Fuc(2)NeuAc(1) | C[+57], N[+2206] | 1434.277 | 3 | 4300.8166 | 4300.8028 | 0 | 3.2 | 485 | Specific | 910.87 | 910.37 | 910.37 | 18.14 | 115 | >sp\|P02788\|TRFL_HUMAN Lactotransferrin OS=Homo sapiens OX=9606 GN=LTF PE=1 SV=6 | 1270 | 20201117_ly_milk_1ng_glyco_hcd203040_compound15.22304.22304.3 | controllerType=0 controllerNumber=1 scan=22304 | 44.7686 |
| 19133:3 | 2 | R.TAGWNIPMGLLFN[+1751.624]QTGSC[+57.021]K.F | HexNAc(4)Hex(4)NeuAc(1) | C[+57], N[+1752] | 1282.8862 | 3 | 3846.6439 | 3846.6341 | 0 | 2.5 | 485 | Specific | 911.2 | 687.87 | 687.87 | 18.14 | 115 | >sp\|P02788\|TRFL_HUMAN Lactotransferrin OS=Homo sapiens OX=9606 GN=LTF PE=1 SV=6 | 1270 | 20201117_ly_milk_1ng_glyco_hcd203040_compound15.22565.22565.3 | controllerType=0 controllerNumber=1 scan=22565 | 45.4435 |
| 20935:3 | 2 | R.TAGWNIPMGLLFN[+2350.830]QTGSC[+57.021]K.F | HexNAc(4)Hex(5)Fuc(1)NeuAc(2) | C[+57], N[+2351] | 1482.6241 | 3 | 4445.8577 | 4445.8403 | 0 | 3.9 | 485 | Specific | 922.83 | 896.51 | 896.51 | 17.87 | 115 | >sp\|P02788\|TRFL_HUMAN Lactotransferrin OS=Homo sapiens OX=9606 GN=LTF PE=1 SV=6 | 1270 | 20201117_ly_milk_1ng_glyco_hcd203040_compound15.24452.24452.3 | controllerType=0 controllerNumber=1 scan=24452 | 49.4401 |
| 19031:3 | 2 | R.TAGWNIPMGLLFN[+1913.677]QTGSC[+57.021]K.F | HexNAc(4)Hex(5)NeuAc(1) | C[+57], N[+1914] | 1336.9039 | 3 | 4008.697 | 4008.687 | 0 | 2.5 | 485 | Specific | 930.91 | 914.81 | 914.81 | 18.14 | 115 | >sp\|P02788\|TRFL_HUMAN Lactotransferrin OS=Homo sapiens OX=9606 GN=LTF PE=1 SV=6 | 1270 | 20201117_ly_milk_1ng_glyco_hcd203040_compound15.22456.22456.3 | controllerType=0 controllerNumber=1 scan=22456 | 45.1508 |
| 17033:3 | 2 | R.TAGWNIPMGLLFN[+1768.640]QTGSC[+57.021]K.F | HexNAc(4)Hex(5)Fuc(1) | C[+57], N[+1769] | 1288.5582 | 3 | 3863.6599 | 3863.6495 | 0 | 2.7 | 485 | Specific | 939.56 | 939.56 | 939.56 | 18.14 | 115 | >sp\|P02788\|TRFL_HUMAN Lactotransferrin OS=Homo sapiens OX=9606 GN=LTF PE=1 SV=6 | 1270 | 20201117_ly_milk_1ng_glyco_hcd203040_compound15.20373.20373.3 | controllerType=0 controllerNumber=1 scan=20373 | 40.5715 |
| 18903:3 | 2 | R.TAGWNIPMGLLFN[+2059.735]QTGSC[+57.021]K.F | HexNAc(4)Hex(5)Fuc(1)NeuAc(1) | C[+57], N[+2060] | 1385.5909 | 3 | 4154.7581 | 4154.7449 | 0 | 3.2 | 485 | Specific | 946.68 | 943.04 | 943.04 | 18.14 | 115 | >sp\|P02788\|TRFL_HUMAN Lactotransferrin OS=Homo sapiens OX=9606 GN=LTF PE=1 SV=6 | 1270 | 20201117_ly_milk_1ng_glyco_hcd203040_compound15.22322.22322.3 | controllerType=0 controllerNumber=1 scan=22322 | 44.8262 |
| 18900:3 | 2 | R.TAGWNIPMGLLFN[+2059.735]QTGSC[+57.021]K.F | HexNAc(4)Hex(5)Fuc(1)NeuAc(1) | C[+57], N[+2060] | 1385.5909 | 3 | 4154.7581 | 4154.7449 | 0 | 3.2 | 485 | Specific | 961 | 943.49 | 943.49 | 19.14 | 115 | >sp\|P02788\|TRFL_HUMAN Lactotransferrin OS=Homo sapiens OX=9606 GN=LTF PE=1 SV=6 | 1270 | 20201117_ly_milk_1ng_glyco_hcd203040_compound15.22319.22319.3 | controllerType=0 controllerNumber=1 scan=22319 | 44.8064 |
| 17423:3 | 2 | R.TAGWNIPMGLLFN[+1216.423]QTGSC[+57.021]K.F | HexNAc(2)Hex(5) | C[+57], N[+1216] | 1104.4871 | 3 | 3311.4467 | 3311.4328 | 0 | 4.2 | 485 | Specific | 968.52 | 968.52 | 422.67 | 18.87 | 115 | >sp\|P02788\|TRFL_HUMAN Lactotransferrin OS=Homo sapiens OX=9606 GN=LTF PE=1 SV=6 | 1270 | 20201117_ly_milk_1ng_glyco_hcd203040_compound15.20780.20780.3 | controllerType=0 controllerNumber=1 scan=20780 | 41.4509 |
| 19028:3 | 2 | R.TAGWNIPMGLLFN[+1913.677]QTGSC[+57.021]K.F | HexNAc(4)Hex(5)NeuAc(1) | C[+57], N[+1914] | 1336.9039 | 3 | 4008.697 | 4008.687 | 0 | 2.5 | 485 | Specific | 982.97 | 977.29 | 977.29 | 19.14 | 115 | >sp\|P02788\|TRFL_HUMAN Lactotransferrin OS=Homo sapiens OX=9606 GN=LTF PE=1 SV=6 | 1270 | 20201117_ly_milk_1ng_glyco_hcd203040_compound15.22453.22453.3 | controllerType=0 controllerNumber=1 scan=22453 | 45.1296 |
| 17425:3 | 2 | R.TAGWNIPMGLLFN[+1216.423]QTGSC[+57.021]K.F | HexNAc(2)Hex(5) | C[+57], N[+1216] | 1104.4871 | 3 | 3311.4467 | 3311.4328 | 0 | 4.2 | 485 | Specific | 1028.07 | 880.12 | 459.55 | 19.87 | 115 | >sp\|P02788\|TRFL_HUMAN Lactotransferrin OS=Homo sapiens OX=9606 GN=LTF PE=1 SV=6 | 1270 | 20201117_ly_milk_1ng_glyco_hcd203040_compound15.20782.20782.3 | controllerType=0 controllerNumber=1 scan=20782 | 41.4564 |
| 17598:3 | 2 | R.TAGWNIPM[+15.995]GLLFN[+1913.677]QTGSC[+57.021]K.F | HexNAc(4)Hex(5)NeuAc(1) | C[+57], M[+16], N[+1914] | 1342.2342 | 3 | 4024.688 | 4024.6819 | 0 | 1.5 | 485 | Specific | 437.39 | 405.65 | 405.65 | 9.24 | 115 | >sp\|P02788\|TRFL_HUMAN Lactotransferrin OS=Homo sapiens OX=9606 GN=LTF PE=1 SV=6 | 1270 | 20201117_ly_milk_1ng_glyco_hcd203040_compound15.20962.20962.3 | controllerType=0 controllerNumber=1 scan=20962 | 41.8245 |
| 17600:3 | 2 | R.TAGWNIPM[+15.995]GLLFN[+1913.677]QTGSC[+57.021]K.F | HexNAc(4)Hex(5)NeuAc(1) | C[+57], M[+16], N[+1914] | 1342.2342 | 3 | 4024.688 | 4024.6819 | 0 | 1.5 | 485 | Specific | 659.72 | 558.43 | 558.43 | 13.09 | 115 | >sp\|P02788\|TRFL_HUMAN Lactotransferrin OS=Homo sapiens OX=9606 GN=LTF PE=1 SV=6 | 1270 | 20201117_ly_milk_1ng_glyco_hcd203040_compound15.20964.20964.3 | controllerType=0 controllerNumber=1 scan=20964 | 41.8283 |
| 17138:3 | 2 | R.TAGWNIPM[+15.995]GLLFN[+2059.735]QTGSC[+57.021]K.F | HexNAc(4)Hex(5)Fuc(1)NeuAc(1) | C[+57], M[+16], N[+2060] | 1390.9235 | 3 | 4170.756 | 4170.7398 | 0 | 3.9 | 485 | Specific | 713.71 | 571.07 | 571.07 | 13.87 | 115 | >sp\|P02788\|TRFL_HUMAN Lactotransferrin OS=Homo sapiens OX=9606 GN=LTF PE=1 SV=6 | 1270 | 20201117_ly_milk_1ng_glyco_hcd203040_compound15.20482.20482.3 | controllerType=0 controllerNumber=1 scan=20482 | 40.8128 |
| 17140:3 | 2 | R.TAGWNIPM[+15.995]GLLFN[+2059.735]QTGSC[+57.021]K.F | HexNAc(4)Hex(5)Fuc(1)NeuAc(1) | C[+57], M[+16], N[+2060] | 1390.9235 | 3 | 4170.756 | 4170.7398 | 0 | 3.9 | 485 | Specific | 802.55 | 694.76 | 694.76 | 15.87 | 115 | >sp\|P02788\|TRFL_HUMAN Lactotransferrin OS=Homo sapiens OX=9606 GN=LTF PE=1 SV=6 | 1270 | 20201117_ly_milk_1ng_glyco_hcd203040_compound15.20484.20484.3 | controllerType=0 controllerNumber=1 scan=20484 | 40.8173 |
| 19173:4 | 2 | R.TAGWNIPMGLLFN[+2205.793]QTGSC[+57.021]KFDEYFSQSC[+57.021]APGSDPR.S | HexNAc(4)Hex(5)Fuc(2)NeuAc(1) | C[+57]*2, N[+2206] | 1536.8939 | 4 | 6144.5539 | 6144.5499 | 0 | 0.6 | 485 | Specific | 270.37 | 246.69 | 246.69 | 6.22 | 115 | >sp\|P02788\|TRFL_HUMAN Lactotransferrin OS=Homo sapiens OX=9606 GN=LTF PE=1 SV=6 | 1270 | 20201117_ly_milk_1ng_glyco_hcd203040_compound15.22607.22607.4 | controllerType=0 controllerNumber=1 scan=22607 | 45.5442 |
| 18182:3 | 2 | R.TAGWNIPMGLLFN[+1768.640]QTGSC[+57.021]KFDEYFSQSC[+57.021]APGSDPR.S | HexNAc(4)Hex(5)Fuc(1) | C[+57]*2, N[+1769] | 1903.1441 | 3 | 5707.4178 | 5707.3966 | 0 | 3.7 | 485 | Specific | 557.69 | 426.45 | 426.45 | 10.1 | 115 | >sp\|P02788\|TRFL_HUMAN Lactotransferrin OS=Homo sapiens OX=9606 GN=LTF PE=1 SV=6 | 1270 | 20201117_ly_milk_1ng_glyco_hcd203040_compound15.21572.21572.3 | controllerType=0 controllerNumber=1 scan=21572 | 43.2239 |
| 19222:4 | 2 | R.TAGWNIPMGLLFN[+1913.677]QTGSC[+57.021]KFDEYFSQSC[+57.021]APGSDPR.S | HexNAc(4)Hex(5)NeuAc(1) | C[+57]*2, N[+1914] | 1463.8702 | 4 | 5852.4591 | 5852.4341 | 0 | 4.3 | 485 | Specific | 698.13 | 639.07 | 639.07 | 13.17 | 115 | >sp\|P02788\|TRFL_HUMAN Lactotransferrin OS=Homo sapiens OX=9606 GN=LTF PE=1 SV=6 | 1270 | 20201117_ly_milk_1ng_glyco_hcd203040_compound15.22658.22658.4 | controllerType=0 controllerNumber=1 scan=22658 | 45.66 |
| 19176:4 | 2 | R.TAGWNIPMGLLFN[+2205.793]QTGSC[+57.021]KFDEYFSQSC[+57.021]APGSDPR.S | HexNAc(4)Hex(5)Fuc(2)NeuAc(1) | C[+57]*2, N[+2206] | 1536.8939 | 4 | 6144.5539 | 6144.5499 | 0 | 0.6 | 485 | Specific | 702.56 | 702.56 | 702.56 | 13.88 | 115 | >sp\|P02788\|TRFL_HUMAN Lactotransferrin OS=Homo sapiens OX=9606 GN=LTF PE=1 SV=6 | 1270 | 20201117_ly_milk_1ng_glyco_hcd203040_compound15.22610.22610.4 | controllerType=0 controllerNumber=1 scan=22610 | 45.5482 |
| 19225:4 | 2 | R.TAGWNIPMGLLFN[+1913.677]QTGSC[+57.021]KFDEYFSQSC[+57.021]APGSDPR.S | HexNAc(4)Hex(5)NeuAc(1) | C[+57]*2, N[+1914] | 1463.8702 | 4 | 5852.4591 | 5852.4341 | 0 | 4.3 | 485 | Specific | 897.06 | 829.61 | 829.61 | 17.17 | 115 | >sp\|P02788\|TRFL_HUMAN Lactotransferrin OS=Homo sapiens OX=9606 GN=LTF PE=1 SV=6 | 1270 | 20201117_ly_milk_1ng_glyco_hcd203040_compound15.22661.22661.4 | controllerType=0 controllerNumber=1 scan=22661 | 45.6689 |
| 19150:4 | 2 | R.TAGWNIPMGLLFN[+2059.735]QTGSC[+57.021]KFDEYFSQSC[+57.021]APGSDPR.S | HexNAc(4)Hex(5)Fuc(1)NeuAc(1) | C[+57]*2, N[+2060] | 1500.3854 | 4 | 5998.5197 | 5998.492 | 0 | 4.6 | 485 | Specific | 944.23 | 916.92 | 916.92 | 18.17 | 115 | >sp\|P02788\|TRFL_HUMAN Lactotransferrin OS=Homo sapiens OX=9606 GN=LTF PE=1 SV=6 | 1270 | 20201117_ly_milk_1ng_glyco_hcd203040_compound15.22583.22583.4 | controllerType=0 controllerNumber=1 scan=22583 | 45.4889 |
| 19148:4 | 2 | R.TAGWNIPMGLLFN[+2059.735]QTGSC[+57.021]KFDEYFSQSC[+57.021]APGSDPR.S | HexNAc(4)Hex(5)Fuc(1)NeuAc(1) | C[+57]*2, N[+2060] | 1500.3854 | 4 | 5998.5197 | 5998.492 | 0 | 4.6 | 485 | Specific | 1041.8 | 986.73 | 986.73 | 20.17 | 115 | >sp\|P02788\|TRFL_HUMAN Lactotransferrin OS=Homo sapiens OX=9606 GN=LTF PE=1 SV=6 | 1270 | 20201117_ly_milk_1ng_glyco_hcd203040_compound15.22581.22581.4 | controllerType=0 controllerNumber=1 scan=22581 | 45.4828 |
| 11759:3 | 2 | R.N[+2350.830]GSDC[+57.021]PDKFC[+57.021]LFQSETK.N | HexNAc(4)Hex(5)Fuc(1)NeuAc(2) | C[+57]*2, N[+2351] | 1461.9117 | 3 | 4383.7205 | 4383.7043 | 0 | 3.7 | 642 | Specific | 375.31 | 298.33 | 298.33 | 6.39 | 115 | >sp\|P02788\|TRFL_HUMAN Lactotransferrin OS=Homo sapiens OX=9606 GN=LTF PE=1 SV=6 | 1270 | 20201117_ly_milk_1ng_glyco_hcd203040_compound15.14845.14845.3 | controllerType=0 controllerNumber=1 scan=14845 | 30.8911 |
| 11757:3 | 2 | R.N[+2350.830]GSDC[+57.021]PDKFC[+57.021]LFQSETK.N | HexNAc(4)Hex(5)Fuc(1)NeuAc(2) | C[+57]*2, N[+2351] | 1461.9117 | 3 | 4383.7205 | 4383.7043 | 0 | 3.7 | 642 | Specific | 381.11 | 334.03 | 334.03 | 6.68 | 115 | >sp\|P02788\|TRFL_HUMAN Lactotransferrin OS=Homo sapiens OX=9606 GN=LTF PE=1 SV=6 | 1270 | 20201117_ly_milk_1ng_glyco_hcd203040_compound15.14843.14843.3 | controllerType=0 controllerNumber=1 scan=14843 | 30.8869 |
| 09390:3 | 2 | R.N[+2059.735]GSDC[+57.021]PDKFC[+57.021]LFQSETK.N | HexNAc(4)Hex(5)Fuc(1)NeuAc(1) | C[+57]*2, N[+2060] | 1364.8779 | 3 | 4092.619 | 4092.6088 | 0 | 2.5 | 642 | Specific | 467.15 | 330.97 | 330.97 | 7.49 | 115 | >sp\|P02788\|TRFL_HUMAN Lactotransferrin OS=Homo sapiens OX=9606 GN=LTF PE=1 SV=6 | 1270 | 20201117_ly_milk_1ng_glyco_hcd203040_compound15.12393.12393.3 | controllerType=0 controllerNumber=1 scan=12393 | 26.6672 |
| 09278:3 | 2 | R.N[+2205.793]GSDC[+57.021]PDKFC[+57.021]LFQSETK.N | HexNAc(4)Hex(5)Fuc(2)NeuAc(1) | C[+57]*2, N[+2206] | 1413.5634 | 3 | 4238.6757 | 4238.6667 | 0 | 2.1 | 642 | Specific | 484.17 | 331.23 | 331.23 | 7.49 | 115 | >sp\|P02788\|TRFL_HUMAN Lactotransferrin OS=Homo sapiens OX=9606 GN=LTF PE=1 SV=6 | 1270 | 20201117_ly_milk_1ng_glyco_hcd203040_compound15.12277.12277.3 | controllerType=0 controllerNumber=1 scan=12277 | 26.4696 |
| 09274:3 | 2 | R.N[+2205.793]GSDC[+57.021]PDKFC[+57.021]LFQSETK.N | HexNAc(4)Hex(5)Fuc(2)NeuAc(1) | C[+57]*2, N[+2206] | 1413.5615 | 3 | 4238.6698 | 4238.6667 | 0 | 0.7 | 642 | Specific | 495.63 | 305.82 | 305.82 | 7.26 | 115 | >sp\|P02788\|TRFL_HUMAN Lactotransferrin OS=Homo sapiens OX=9606 GN=LTF PE=1 SV=6 | 1270 | 20201117_ly_milk_1ng_glyco_hcd203040_compound15.12272.12272.2 | controllerType=0 controllerNumber=1 scan=12272 | 26.4593 |
| 09392:3 | 2 | R.N[+2059.735]GSDC[+57.021]PDKFC[+57.021]LFQSETK.N | HexNAc(4)Hex(5)Fuc(1)NeuAc(1) | C[+57]*2, N[+2060] | 1364.8779 | 3 | 4092.619 | 4092.6088 | 0 | 2.5 | 642 | Specific | 500.74 | 332.98 | 332.98 | 7.61 | 115 | >sp\|P02788\|TRFL_HUMAN Lactotransferrin OS=Homo sapiens OX=9606 GN=LTF PE=1 SV=6 | 1270 | 20201117_ly_milk_1ng_glyco_hcd203040_compound15.12395.12395.3 | controllerType=0 controllerNumber=1 scan=12395 | 26.6713 |
| 09272:3 | 2 | R.N[+2205.793]GSDC[+57.021]PDKFC[+57.021]LFQSETK.N | HexNAc(4)Hex(5)Fuc(2)NeuAc(1) | C[+57]*2, N[+2206] | 1413.5615 | 3 | 4238.6698 | 4238.6667 | 0 | 0.7 | 642 | Specific | 514.44 | 416.1 | 416.1 | 9.1 | 115 | >sp\|P02788\|TRFL_HUMAN Lactotransferrin OS=Homo sapiens OX=9606 GN=LTF PE=1 SV=6 | 1270 | 20201117_ly_milk_1ng_glyco_hcd203040_compound15.12270.12270.2 | controllerType=0 controllerNumber=1 scan=12270 | 26.4554 |
| 09280:3 | 2 | R.N[+2205.793]GSDC[+57.021]PDKFC[+57.021]LFQSETK.N | HexNAc(4)Hex(5)Fuc(2)NeuAc(1) | C[+57]*2, N[+2206] | 1413.5634 | 3 | 4238.6757 | 4238.6667 | 0 | 2.1 | 642 | Specific | 530.62 | 368.82 | 368.82 | 7.75 | 115 | >sp\|P02788\|TRFL_HUMAN Lactotransferrin OS=Homo sapiens OX=9606 GN=LTF PE=1 SV=6 | 1270 | 20201117_ly_milk_1ng_glyco_hcd203040_compound15.12279.12279.3 | controllerType=0 controllerNumber=1 scan=12279 | 26.4741 |
| 15727:3 | 4 | R.LN[+1378.476]YSLPTGQWVGVQLPR.N | HexNAc(2)Hex(6) | N[+1378] | 1102.8449 | 3 | 3306.5201 | 3306.5145 | 0 | 1.7 | 1017 | Specific | 603.38 | 590.51 | 590.51 | 12.09 | 79 | >sp\|P24821\|TENA_HUMAN Tenascin OS=Homo sapiens OX=9606 GN=TNC PE=1 SV=3 | 486 | 20201117_ly_milk_1ng_glyco_hcd203040_compound15.18980.18980.3 | controllerType=0 controllerNumber=1 scan=18980 | 38.048 |
| 15730:3 | 4 | R.LN[+1378.476]YSLPTGQWVGVQLPR.N | HexNAc(2)Hex(6) | N[+1378] | 1102.8449 | 3 | 3306.5201 | 3306.5145 | 0 | 1.7 | 1017 | Specific | 646.83 | 545.13 | 545.13 | 12.09 | 79 | >sp\|P24821\|TENA_HUMAN Tenascin OS=Homo sapiens OX=9606 GN=TNC PE=1 SV=3 | 486 | 20201117_ly_milk_1ng_glyco_hcd203040_compound15.18983.18983.3 | controllerType=0 controllerNumber=1 scan=18983 | 38.0583 |
| 15646:3 | 4 | R.LN[+1540.529]YSLPTGQWVGVQLPR.N | HexNAc(2)Hex(7) | N[+1541] | 1156.8658 | 3 | 3468.5828 | 3468.5674 | 0 | 4.4 | 1017 | Specific | 646.9 | 646.9 | 646.9 | 11.87 | 79 | >sp\|P24821\|TENA_HUMAN Tenascin OS=Homo sapiens OX=9606 GN=TNC PE=1 SV=3 | 486 | 20201117_ly_milk_1ng_glyco_hcd203040_compound15.18896.18896.3 | controllerType=0 controllerNumber=1 scan=18896 | 37.8894 |
| 07853:3 | 5 | K.WEC[+57.021]KN[+2205.793]DTLLGIK.G | HexNAc(4)Hex(5)Fuc(2)NeuAc(1) | C[+57], N[+2206] | 1228.1913 | 3 | 3682.5595 | 3682.5444 | 0 | 4.1 | 100 | Specific | 231.2 | 67.5 | 67.5 | 3.5 | 78 | >sp\|P22897\|MRC1_HUMAN Macrophage mannose receptor 1 OS=Homo sapiens OX=9606 GN=MRC1 PE=1 SV=1 | 1049 | 20201117_ly_milk_1ng_glyco_hcd203040_compound15.10784.10784.3 | controllerType=0 controllerNumber=1 scan=10784 | 23.9708 |
| 07828:3 | 5 | K.WEC[+57.021]KN[+2205.793]DTLLGIK.G | HexNAc(4)Hex(5)Fuc(2)NeuAc(1) | C[+57], N[+2206] | 1228.1906 | 3 | 3682.5573 | 3682.5444 | 0 | 3.5 | 100 | Specific | 313.42 | 313.42 | 313.42 | 5.82 | 78 | >sp\|P22897\|MRC1_HUMAN Macrophage mannose receptor 1 OS=Homo sapiens OX=9606 GN=MRC1 PE=1 SV=1 | 1049 | 20201117_ly_milk_1ng_glyco_hcd203040_compound15.10758.10758.3 | controllerType=0 controllerNumber=1 scan=10758 | 23.9265 |
| 07952:3 | 5 | K.WEC[+57.021]KN[+2059.735]DTLLGIK.G | HexNAc(4)Hex(5)Fuc(1)NeuAc(1) | C[+57], N[+2060] | 1179.5029 | 3 | 3536.494 | 3536.4865 | 0 | 2.1 | 100 | Specific | 388.28 | 328.31 | 328.31 | 6.81 | 78 | >sp\|P22897\|MRC1_HUMAN Macrophage mannose receptor 1 OS=Homo sapiens OX=9606 GN=MRC1 PE=1 SV=1 | 1049 | 20201117_ly_milk_1ng_glyco_hcd203040_compound15.10887.10887.3 | controllerType=0 controllerNumber=1 scan=10887 | 24.15 |
| 07954:3 | 5 | K.WEC[+57.021]KN[+2059.735]DTLLGIK.G | HexNAc(4)Hex(5)Fuc(1)NeuAc(1) | C[+57], N[+2060] | 1179.5029 | 3 | 3536.494 | 3536.4865 | 0 | 2.1 | 100 | Specific | 531.45 | 275.19 | 275.19 | 6.97 | 78 | >sp\|P22897\|MRC1_HUMAN Macrophage mannose receptor 1 OS=Homo sapiens OX=9606 GN=MRC1 PE=1 SV=1 | 1049 | 20201117_ly_milk_1ng_glyco_hcd203040_compound15.10889.10889.3 | controllerType=0 controllerNumber=1 scan=10889 | 24.1542 |
| 07943:3 | 5 | K.WEC[+57.021]KN[+2059.735]DTLLGIK.G | HexNAc(4)Hex(5)Fuc(1)NeuAc(1) | C[+57], N[+2060] | 1179.5026 | 3 | 3536.4933 | 3536.4865 | 0 | 1.9 | 100 | Specific | 533.27 | 190.85 | 190.85 | 6.37 | 78 | >sp\|P22897\|MRC1_HUMAN Macrophage mannose receptor 1 OS=Homo sapiens OX=9606 GN=MRC1 PE=1 SV=1 | 1049 | 20201117_ly_milk_1ng_glyco_hcd203040_compound15.10877.10877.3 | controllerType=0 controllerNumber=1 scan=10877 | 24.1294 |
| 07945:3 | 5 | K.WEC[+57.021]KN[+2059.735]DTLLGIK.G | HexNAc(4)Hex(5)Fuc(1)NeuAc(1) | C[+57], N[+2060] | 1179.5026 | 3 | 3536.4933 | 3536.4865 | 0 | 1.9 | 100 | Specific | 558.44 | 444.32 | 444.32 | 9.01 | 78 | >sp\|P22897\|MRC1_HUMAN Macrophage mannose receptor 1 OS=Homo sapiens OX=9606 GN=MRC1 PE=1 SV=1 | 1049 | 20201117_ly_milk_1ng_glyco_hcd203040_compound15.10879.10879.3 | controllerType=0 controllerNumber=1 scan=10879 | 24.1334 |
| 13377:4 | 5 | K.KGN[+2059.735]TTLNSFVIPSESDVPTHC[+57.021]PSQWWPYAGHC[+57.021]YK.I | HexNAc(4)Hex(5)Fuc(1)NeuAc(1) | C[+57]*2, N[+2060] | 1506.6415 | 4 | 6023.5441 | 6023.5567 | 0 | -2.1 | 342 | Specific | 661.08 | 651.09 | 651.09 | 12.27 | 78 | >sp\|P22897\|MRC1_HUMAN Macrophage mannose receptor 1 OS=Homo sapiens OX=9606 GN=MRC1 PE=1 SV=1 | 1049 | 20201117_ly_milk_1ng_glyco_hcd203040_compound15.16543.16543.4 | controllerType=0 controllerNumber=1 scan=16543 | 33.7104 |
| 13375:4 | 5 | K.KGN[+2059.735]TTLNSFVIPSESDVPTHC[+57.021]PSQWWPYAGHC[+57.021]YK.I | HexNAc(4)Hex(5)Fuc(1)NeuAc(1) | C[+57]*2, N[+2060] | 1506.6415 | 4 | 6023.5441 | 6023.5567 | 0 | -2.1 | 342 | Specific | 699.29 | 699.29 | 699.29 | 12.27 | 78 | >sp\|P22897\|MRC1_HUMAN Macrophage mannose receptor 1 OS=Homo sapiens OX=9606 GN=MRC1 PE=1 SV=1 | 1049 | 20201117_ly_milk_1ng_glyco_hcd203040_compound15.16541.16541.4 | controllerType=0 controllerNumber=1 scan=16541 | 33.7067 |
| 16720:3 | 5 | R.VWIALNSN[+2205.793]LTDNQYTWTDKWR.V | HexNAc(4)Hex(5)Fuc(2)NeuAc(1) | N[+2206] | 1610.6998 | 3 | 4830.0847 | 4830.0821 | 0 | 0.6 | 1153 | Specific | 608.31 | 365.94 | 365.94 | 9.61 | 78 | >sp\|P22897\|MRC1_HUMAN Macrophage mannose receptor 1 OS=Homo sapiens OX=9606 GN=MRC1 PE=1 SV=1 | 1049 | 20201117_ly_milk_1ng_glyco_hcd203040_compound15.20048.20048.3 | controllerType=0 controllerNumber=1 scan=20048 | 39.9816 |
| 16718:3 | 5 | R.VWIALNSN[+2205.793]LTDNQYTWTDKWR.V | HexNAc(4)Hex(5)Fuc(2)NeuAc(1) | N[+2206] | 1610.6998 | 3 | 4830.0847 | 4830.0821 | 0 | 0.6 | 1153 | Specific | 653.88 | 494.73 | 494.73 | 12.09 | 78 | >sp\|P22897\|MRC1_HUMAN Macrophage mannose receptor 1 OS=Homo sapiens OX=9606 GN=MRC1 PE=1 SV=1 | 1049 | 20201117_ly_milk_1ng_glyco_hcd203040_compound15.20046.20046.3 | controllerType=0 controllerNumber=1 scan=20046 | 39.9779 |
| 08559:3 | 5 | K.TAHC[+57.021]N[+2205.793]ESFYFLC[+57.021]K.R | HexNAc(4)Hex(5)Fuc(2)NeuAc(1) | C[+57]*2, N[+2206] | 1294.8444 | 3 | 3882.5187 | 3882.5124 | 0 | 1.6 | 1201 | Specific | 302.16 | 201.28 | 201.28 | 6.02 | 78 | >sp\|P22897\|MRC1_HUMAN Macrophage mannose receptor 1 OS=Homo sapiens OX=9606 GN=MRC1 PE=1 SV=1 | 1049 | 20201117_ly_milk_1ng_glyco_hcd203040_compound15.11519.11519.3 | controllerType=0 controllerNumber=1 scan=11519 | 25.2026 |
| 08557:3 | 5 | K.TAHC[+57.021]N[+2205.793]ESFYFLC[+57.021]K.R | HexNAc(4)Hex(5)Fuc(2)NeuAc(1) | C[+57]*2, N[+2206] | 1294.8444 | 3 | 3882.5187 | 3882.5124 | 0 | 1.6 | 1201 | Specific | 372.25 | 318.06 | 318.06 | 7.03 | 78 | >sp\|P22897\|MRC1_HUMAN Macrophage mannose receptor 1 OS=Homo sapiens OX=9606 GN=MRC1 PE=1 SV=1 | 1049 | 20201117_ly_milk_1ng_glyco_hcd203040_compound15.11517.11517.3 | controllerType=0 controllerNumber=1 scan=11517 | 25.1992 |
| 08664:3 | 5 | K.TAHC[+57.021]N[+2059.735]ESFYFLC[+57.021]K.R | HexNAc(4)Hex(5)Fuc(1)NeuAc(1) | C[+57]*2, N[+2060] | 1246.1573 | 3 | 3736.4573 | 3736.4545 | 0 | 0.7 | 1201 | Specific | 667.12 | 561.03 | 561.03 | 13.02 | 78 | >sp\|P22897\|MRC1_HUMAN Macrophage mannose receptor 1 OS=Homo sapiens OX=9606 GN=MRC1 PE=1 SV=1 | 1049 | 20201117_ly_milk_1ng_glyco_hcd203040_compound15.11628.11628.3 | controllerType=0 controllerNumber=1 scan=11628 | 25.3802 |
| 08662:3 | 5 | K.TAHC[+57.021]N[+2059.735]ESFYFLC[+57.021]K.R | HexNAc(4)Hex(5)Fuc(1)NeuAc(1) | C[+57]*2, N[+2060] | 1246.1573 | 3 | 3736.4573 | 3736.4545 | 0 | 0.7 | 1201 | Specific | 834.15 | 656.73 | 656.73 | 16.02 | 78 | >sp\|P22897\|MRC1_HUMAN Macrophage mannose receptor 1 OS=Homo sapiens OX=9606 GN=MRC1 PE=1 SV=1 | 1049 | 20201117_ly_milk_1ng_glyco_hcd203040_compound15.11626.11626.3 | controllerType=0 controllerNumber=1 scan=11626 | 25.3761 |
| 17970:3 | 8 | R.GLSFDVSLEVSQGPGLLN[+2059.735]DTK.V | HexNAc(4)Hex(5)Fuc(1)NeuAc(1) | N[+2060] | 1412.6238 | 3 | 4235.857 | 4235.8481 | 0 | 2.1 | 118 | Specific | 530.61 | 342.61 | 342.61 | 8.19 | 63 | >sp\|P01833\|PIGR_HUMAN Polymeric immunoglobulin receptor OS=Homo sapiens OX=9606 GN=PIGR PE=1 SV=4 | 902 | 20201117_ly_milk_1ng_glyco_hcd203040_compound15.21351.21351.3 | controllerType=0 controllerNumber=1 scan=21351 | 42.7294 |
| 17792:3 | 8 | R.GLSFDVSLEVSQGPGLLN[+2569.905]DTK.V | HexNAc(5)Hex(6)NeuAc(2) | N[+2570] | 1582.6893 | 3 | 4746.0532 | 4746.0178 | 0 | 7.5 | 118 | Specific | 552.4 | 461.8 | 461.8 | 8.5 | 63 | >sp\|P01833\|PIGR_HUMAN Polymeric immunoglobulin receptor OS=Homo sapiens OX=9606 GN=PIGR PE=1 SV=4 | 902 | 20201117_ly_milk_1ng_glyco_hcd203040_compound15.21164.21164.3 | controllerType=0 controllerNumber=1 scan=21164 | 42.266 |
| 17790:3 | 8 | R.GLSFDVSLEVSQGPGLLN[+2569.905]DTK.V | HexNAc(5)Hex(6)NeuAc(2) | N[+2570] | 1582.6893 | 3 | 4746.0532 | 4746.0178 | 0 | 7.5 | 118 | Specific | 576.69 | 576.69 | 576.69 | 9.67 | 63 | >sp\|P01833\|PIGR_HUMAN Polymeric immunoglobulin receptor OS=Homo sapiens OX=9606 GN=PIGR PE=1 SV=4 | 902 | 20201117_ly_milk_1ng_glyco_hcd203040_compound15.21162.21162.3 | controllerType=0 controllerNumber=1 scan=21162 | 42.2622 |
| 19730:4 | 8 | R.GLSFDVSLEVSQGPGLLN[+2350.830]DTK.V | HexNAc(4)Hex(5)Fuc(1)NeuAc(2) | N[+2351] | 1132.4945 | 4 | 4526.9564 | 4526.9436 | 0 | 2.8 | 118 | Specific | 577.09 | 473.07 | 473.07 | 11.2 | 63 | >sp\|P01833\|PIGR_HUMAN Polymeric immunoglobulin receptor OS=Homo sapiens OX=9606 GN=PIGR PE=1 SV=4 | 902 | 20201117_ly_milk_1ng_glyco_hcd203040_compound15.23191.23191.4 | controllerType=0 controllerNumber=1 scan=23191 | 46.667 |
| 19725:3 | 8 | R.GLSFDVSLEVSQGPGLLN[+2350.830]DTK.V | HexNAc(4)Hex(5)Fuc(1)NeuAc(2) | N[+2351] | 1509.6569 | 3 | 4526.9562 | 4526.9436 | 0 | 2.8 | 118 | Specific | 712.39 | 701.66 | 701.66 | 14.14 | 63 | >sp\|P01833\|PIGR_HUMAN Polymeric immunoglobulin receptor OS=Homo sapiens OX=9606 GN=PIGR PE=1 SV=4 | 902 | 20201117_ly_milk_1ng_glyco_hcd203040_compound15.23186.23186.3 | controllerType=0 controllerNumber=1 scan=23186 | 46.6603 |
| 19760:4 | 8 | R.GLSFDVSLEVSQGPGLLN[+2350.830]DTK.V | HexNAc(4)Hex(5)Fuc(1)NeuAc(2) | N[+2351] | 1132.4932 | 4 | 4526.951 | 4526.9436 | 0 | 1.6 | 118 | Specific | 713.9 | 542.55 | 542.55 | 14.14 | 63 | >sp\|P01833\|PIGR_HUMAN Polymeric immunoglobulin receptor OS=Homo sapiens OX=9606 GN=PIGR PE=1 SV=4 | 902 | 20201117_ly_milk_1ng_glyco_hcd203040_compound15.23222.23222.4 | controllerType=0 controllerNumber=1 scan=23222 | 46.752 |
| 19752:3 | 8 | R.GLSFDVSLEVSQGPGLLN[+2350.830]DTK.V | HexNAc(4)Hex(5)Fuc(1)NeuAc(2) | N[+2351] | 1509.6566 | 3 | 4526.9551 | 4526.9436 | 0 | 2.6 | 118 | Specific | 748.16 | 732.07 | 732.07 | 14.14 | 63 | >sp\|P01833\|PIGR_HUMAN Polymeric immunoglobulin receptor OS=Homo sapiens OX=9606 GN=PIGR PE=1 SV=4 | 902 | 20201117_ly_milk_1ng_glyco_hcd203040_compound15.23214.23214.3 | controllerType=0 controllerNumber=1 scan=23214 | 46.7193 |
| 19733:4 | 8 | R.GLSFDVSLEVSQGPGLLN[+2350.830]DTK.V | HexNAc(4)Hex(5)Fuc(1)NeuAc(2) | N[+2351] | 1132.4945 | 4 | 4526.9564 | 4526.9436 | 0 | 2.8 | 118 | Specific | 896.23 | 835.1 | 835.1 | 17.2 | 63 | >sp\|P01833\|PIGR_HUMAN Polymeric immunoglobulin receptor OS=Homo sapiens OX=9606 GN=PIGR PE=1 SV=4 | 902 | 20201117_ly_milk_1ng_glyco_hcd203040_compound15.23194.23194.4 | controllerType=0 controllerNumber=1 scan=23194 | 46.677 |
| 17973:3 | 8 | R.GLSFDVSLEVSQGPGLLN[+2059.735]DTK.V | HexNAc(4)Hex(5)Fuc(1)NeuAc(1) | N[+2060] | 1412.6238 | 3 | 4235.857 | 4235.8481 | 0 | 2.1 | 118 | Specific | 961 | 888.86 | 888.86 | 19.14 | 63 | >sp\|P01833\|PIGR_HUMAN Polymeric immunoglobulin receptor OS=Homo sapiens OX=9606 GN=PIGR PE=1 SV=4 | 902 | 20201117_ly_milk_1ng_glyco_hcd203040_compound15.21354.21354.3 | controllerType=0 controllerNumber=1 scan=21354 | 42.7329 |
| 19755:3 | 8 | R.GLSFDVSLEVSQGPGLLN[+2350.830]DTK.V | HexNAc(4)Hex(5)Fuc(1)NeuAc(2) | N[+2351] | 1509.6566 | 3 | 4526.9551 | 4526.9436 | 0 | 2.6 | 118 | Specific | 983.84 | 941.12 | 941.12 | 19.14 | 63 | >sp\|P01833\|PIGR_HUMAN Polymeric immunoglobulin receptor OS=Homo sapiens OX=9606 GN=PIGR PE=1 SV=4 | 902 | 20201117_ly_milk_1ng_glyco_hcd203040_compound15.23217.23217.3 | controllerType=0 controllerNumber=1 scan=23217 | 46.7239 |
| 19728:3 | 8 | R.GLSFDVSLEVSQGPGLLN[+2350.830]DTK.V | HexNAc(4)Hex(5)Fuc(1)NeuAc(2) | N[+2351] | 1509.6569 | 3 | 4526.9562 | 4526.9436 | 0 | 2.8 | 118 | Specific | 1031.64 | 984.06 | 984.06 | 20.14 | 63 | >sp\|P01833\|PIGR_HUMAN Polymeric immunoglobulin receptor OS=Homo sapiens OX=9606 GN=PIGR PE=1 SV=4 | 902 | 20201117_ly_milk_1ng_glyco_hcd203040_compound15.23189.23189.3 | controllerType=0 controllerNumber=1 scan=23189 | 46.6645 |
| 19079:4 | 8 | K.QIGLYPVLVIDSSGYVNPN[+2059.735]YTGR.I | HexNAc(4)Hex(5)Fuc(1)NeuAc(1) | N[+2060] | 1147.0178 | 4 | 4585.0494 | 4585.0384 | 0 | 2.4 | 168 | Specific | 745.2 | 648.93 | 648.93 | 14.37 | 63 | >sp\|P01833\|PIGR_HUMAN Polymeric immunoglobulin receptor OS=Homo sapiens OX=9606 GN=PIGR PE=1 SV=4 | 902 | 20201117_ly_milk_1ng_glyco_hcd203040_compound15.22507.22507.4 | controllerType=0 controllerNumber=1 scan=22507 | 45.2864 |
| 19082:4 | 8 | K.QIGLYPVLVIDSSGYVNPN[+2059.735]YTGR.I | HexNAc(4)Hex(5)Fuc(1)NeuAc(1) | N[+2060] | 1147.0178 | 4 | 4585.0494 | 4585.0384 | 0 | 2.4 | 168 | Specific | 841.53 | 807.84 | 807.84 | 16.37 | 63 | >sp\|P01833\|PIGR_HUMAN Polymeric immunoglobulin receptor OS=Homo sapiens OX=9606 GN=PIGR PE=1 SV=4 | 902 | 20201117_ly_milk_1ng_glyco_hcd203040_compound15.22510.22510.4 | controllerType=0 controllerNumber=1 scan=22510 | 45.2925 |
| 20720:4 | 8 | R.LSLLEEPGN[+2059.735]GTFTVILNQLTSR.D | HexNAc(4)Hex(5)Fuc(1)NeuAc(1) | N[+2060] | 1116.2659 | 4 | 4462.0418 | 4462.0275 | 0 | 3.2 | 413 | Specific | 768.97 | 630.49 | 630.49 | 15.37 | 63 | >sp\|P01833\|PIGR_HUMAN Polymeric immunoglobulin receptor OS=Homo sapiens OX=9606 GN=PIGR PE=1 SV=4 | 902 | 20201117_ly_milk_1ng_glyco_hcd203040_compound15.24227.24227.4 | controllerType=0 controllerNumber=1 scan=24227 | 48.9328 |
| 20722:4 | 8 | R.LSLLEEPGN[+2059.735]GTFTVILNQLTSR.D | HexNAc(4)Hex(5)Fuc(1)NeuAc(1) | N[+2060] | 1116.2659 | 4 | 4462.0418 | 4462.0275 | 0 | 3.2 | 413 | Specific | 937.46 | 691.08 | 691.08 | 18.37 | 63 | >sp\|P01833\|PIGR_HUMAN Polymeric immunoglobulin receptor OS=Homo sapiens OX=9606 GN=PIGR PE=1 SV=4 | 902 | 20201117_ly_milk_1ng_glyco_hcd203040_compound15.24229.24229.4 | controllerType=0 controllerNumber=1 scan=24229 | 48.9389 |
| 20636:3 | 8 | R.LSLLEEPGN[+2205.793]GTFTVILNQLTSR.D | HexNAc(4)Hex(5)Fuc(2)NeuAc(1) | N[+2206] | 1536.7116 | 3 | 4608.1203 | 4608.0854 | 0 | 7.6 | 413 | Specific | 971.33 | 737.07 | 737.07 | 17.84 | 63 | >sp\|P01833\|PIGR_HUMAN Polymeric immunoglobulin receptor OS=Homo sapiens OX=9606 GN=PIGR PE=1 SV=4 | 902 | 20201117_ly_milk_1ng_glyco_hcd203040_compound15.24138.24138.3 | controllerType=0 controllerNumber=1 scan=24138 | 48.7106 |
| 22939:4 | 8 | R.LSLLEEPGN[+2350.830]GTFTVILNQLTSR.D | HexNAc(4)Hex(5)Fuc(1)NeuAc(2) | N[+2351] | 1189.0422 | 4 | 4753.1468 | 4753.1229 | 0 | 5 | 413 | Specific | 998.55 | 778.15 | 778.15 | 18.51 | 63 | >sp\|P01833\|PIGR_HUMAN Polymeric immunoglobulin receptor OS=Homo sapiens OX=9606 GN=PIGR PE=1 SV=4 | 902 | 20201117_ly_milk_1ng_glyco_hcd203040_compound15.26541.26541.4 | controllerType=0 controllerNumber=1 scan=26541 | 53.2409 |
| 20634:3 | 8 | R.LSLLEEPGN[+2205.793]GTFTVILNQLTSR.D | HexNAc(4)Hex(5)Fuc(2)NeuAc(1) | N[+2206] | 1536.7116 | 3 | 4608.1203 | 4608.0854 | 0 | 7.6 | 413 | Specific | 1079.64 | 845.03 | 845.03 | 19.84 | 63 | >sp\|P01833\|PIGR_HUMAN Polymeric immunoglobulin receptor OS=Homo sapiens OX=9606 GN=PIGR PE=1 SV=4 | 902 | 20201117_ly_milk_1ng_glyco_hcd203040_compound15.24136.24136.3 | controllerType=0 controllerNumber=1 scan=24136 | 48.7042 |
| 22937:4 | 8 | R.LSLLEEPGN[+2350.830]GTFTVILNQLTSR.D | HexNAc(4)Hex(5)Fuc(1)NeuAc(2) | N[+2351] | 1189.0422 | 4 | 4753.1468 | 4753.1229 | 0 | 5 | 413 | Specific | 1103.97 | 899.59 | 899.59 | 21.51 | 63 | >sp\|P01833\|PIGR_HUMAN Polymeric immunoglobulin receptor OS=Homo sapiens OX=9606 GN=PIGR PE=1 SV=4 | 902 | 20201117_ly_milk_1ng_glyco_hcd203040_compound15.26539.26539.4 | controllerType=0 controllerNumber=1 scan=26539 | 53.2329 |
| 18220:3 | 8 | K.IIEGEPNLKVPGN[+2350.830]VTAVLGETLK.V | HexNAc(4)Hex(5)Fuc(1)NeuAc(2) | N[+2351] | 1581.402 | 3 | 4742.1916 | 4742.1797 | 0 | 2.5 | 457 | Specific | 293.31 | 235.39 | 235.39 | 5.75 | 63 | >sp\|P01833\|PIGR_HUMAN Polymeric immunoglobulin receptor OS=Homo sapiens OX=9606 GN=PIGR PE=1 SV=4 | 902 | 20201117_ly_milk_1ng_glyco_hcd203040_compound15.21611.21611.3 | controllerType=0 controllerNumber=1 scan=21611 | 43.3148 |
| 18218:3 | 8 | K.IIEGEPNLKVPGN[+2350.830]VTAVLGETLK.V | HexNAc(4)Hex(5)Fuc(1)NeuAc(2) | N[+2351] | 1581.402 | 3 | 4742.1916 | 4742.1797 | 0 | 2.5 | 457 | Specific | 307.55 | 307.55 | 307.55 | 6.67 | 63 | >sp\|P01833\|PIGR_HUMAN Polymeric immunoglobulin receptor OS=Homo sapiens OX=9606 GN=PIGR PE=1 SV=4 | 902 | 20201117_ly_milk_1ng_glyco_hcd203040_compound15.21609.21609.3 | controllerType=0 controllerNumber=1 scan=21609 | 43.3116 |
| 16321:4 | 8 | K.IIEGEPNLKVPGN[+2204.772]VTAVLGETLK.V | HexNAc(4)Hex(5)NeuAc(2) | N[+2205] | 1149.7936 | 4 | 4596.1527 | 4596.1218 | 0 | 6.7 | 457 | Specific | 751.31 | 679.75 | 679.75 | 13.49 | 63 | >sp\|P01833\|PIGR_HUMAN Polymeric immunoglobulin receptor OS=Homo sapiens OX=9606 GN=PIGR PE=1 SV=4 | 902 | 20201117_ly_milk_1ng_glyco_hcd203040_compound15.19628.19628.2 | controllerType=0 controllerNumber=1 scan=19628 | 39.221 |
| 16296:3 | 8 | K.IIEGEPNLKVPGN[+2205.793]VTAVLGETLK.V | HexNAc(4)Hex(5)Fuc(2)NeuAc(1) | N[+2206] | 1533.0532 | 3 | 4597.1449 | 4597.1422 | 0 | 0.6 | 457 | Specific | 751.87 | 652.79 | 652.79 | 14.26 | 63 | >sp\|P01833\|PIGR_HUMAN Polymeric immunoglobulin receptor OS=Homo sapiens OX=9606 GN=PIGR PE=1 SV=4 | 902 | 20201117_ly_milk_1ng_glyco_hcd203040_compound15.19602.19602.3 | controllerType=0 controllerNumber=1 scan=19602 | 39.1778 |
| 16293:3 | 8 | K.IIEGEPNLKVPGN[+2205.793]VTAVLGETLK.V | HexNAc(4)Hex(5)Fuc(2)NeuAc(1) | N[+2206] | 1533.0532 | 3 | 4597.1449 | 4597.1422 | 0 | 0.6 | 457 | Specific | 790.96 | 708.45 | 708.45 | 14.26 | 63 | >sp\|P01833\|PIGR_HUMAN Polymeric immunoglobulin receptor OS=Homo sapiens OX=9606 GN=PIGR PE=1 SV=4 | 902 | 20201117_ly_milk_1ng_glyco_hcd203040_compound15.19599.19599.3 | controllerType=0 controllerNumber=1 scan=19599 | 39.1731 |
| 16371:4 | 8 | K.IIEGEPNLKVPGN[+2059.735]VTAVLGETLK.V | HexNAc(4)Hex(5)Fuc(1)NeuAc(1) | N[+2060] | 1113.528 | 4 | 4451.0902 | 4451.0843 | 0 | 1.3 | 457 | Specific | 795.84 | 592.09 | 592.09 | 14.66 | 63 | >sp\|P01833\|PIGR_HUMAN Polymeric immunoglobulin receptor OS=Homo sapiens OX=9606 GN=PIGR PE=1 SV=4 | 902 | 20201117_ly_milk_1ng_glyco_hcd203040_compound15.19681.19681.4 | controllerType=0 controllerNumber=1 scan=19681 | 39.3202 |
| 16319:4 | 8 | K.IIEGEPNLKVPGN[+2204.772]VTAVLGETLK.V | HexNAc(4)Hex(5)NeuAc(2) | N[+2205] | 1149.7936 | 4 | 4596.1527 | 4596.1218 | 0 | 6.7 | 457 | Specific | 798.4 | 798.4 | 798.4 | 13.49 | 63 | >sp\|P01833\|PIGR_HUMAN Polymeric immunoglobulin receptor OS=Homo sapiens OX=9606 GN=PIGR PE=1 SV=4 | 902 | 20201117_ly_milk_1ng_glyco_hcd203040_compound15.19626.19626.2 | controllerType=0 controllerNumber=1 scan=19626 | 39.2164 |
| 16372:3 | 8 | K.IIEGEPNLKVPGN[+2059.735]VTAVLGETLK.V | HexNAc(4)Hex(5)Fuc(1)NeuAc(1) | N[+2060] | 1484.3708 | 3 | 4451.0978 | 4451.0843 | 0 | 3 | 457 | Specific | 802.35 | 794.23 | 794.23 | 15.82 | 63 | >sp\|P01833\|PIGR_HUMAN Polymeric immunoglobulin receptor OS=Homo sapiens OX=9606 GN=PIGR PE=1 SV=4 | 902 | 20201117_ly_milk_1ng_glyco_hcd203040_compound15.19682.19682.3 | controllerType=0 controllerNumber=1 scan=19682 | 39.3223 |
| 16336:3 | 8 | K.IIEGEPNLKVPGN[+2059.735]VTAVLGETLK.V | HexNAc(4)Hex(5)Fuc(1)NeuAc(1) | N[+2060] | 1484.3697 | 3 | 4451.0945 | 4451.0843 | 0 | 2.3 | 457 | Specific | 816.89 | 628.71 | 628.71 | 15.82 | 63 | >sp\|P01833\|PIGR_HUMAN Polymeric immunoglobulin receptor OS=Homo sapiens OX=9606 GN=PIGR PE=1 SV=4 | 902 | 20201117_ly_milk_1ng_glyco_hcd203040_compound15.19644.19644.3 | controllerType=0 controllerNumber=1 scan=19644 | 39.2501 |
| 16373:4 | 8 | K.IIEGEPNLKVPGN[+2059.735]VTAVLGETLK.V | HexNAc(4)Hex(5)Fuc(1)NeuAc(1) | N[+2060] | 1113.528 | 4 | 4451.0902 | 4451.0843 | 0 | 1.3 | 457 | Specific | 821.01 | 738.54 | 738.54 | 15.66 | 63 | >sp\|P01833\|PIGR_HUMAN Polymeric immunoglobulin receptor OS=Homo sapiens OX=9606 GN=PIGR PE=1 SV=4 | 902 | 20201117_ly_milk_1ng_glyco_hcd203040_compound15.19683.19683.4 | controllerType=0 controllerNumber=1 scan=19683 | 39.3259 |
| 16327:3 | 8 | K.IIEGEPNLKVPGN[+2205.793]VTAVLGETLK.V | HexNAc(4)Hex(5)Fuc(2)NeuAc(1) | N[+2206] | 1533.0546 | 3 | 4597.1493 | 4597.1422 | 0 | 1.5 | 457 | Specific | 830.65 | 737.1 | 737.1 | 15.76 | 63 | >sp\|P01833\|PIGR_HUMAN Polymeric immunoglobulin receptor OS=Homo sapiens OX=9606 GN=PIGR PE=1 SV=4 | 902 | 20201117_ly_milk_1ng_glyco_hcd203040_compound15.19635.19635.3 | controllerType=0 controllerNumber=1 scan=19635 | 39.2348 |
| 16347:3 | 8 | K.IIEGEPNLKVPGN[+2059.735]VTAVLGETLK.V | HexNAc(4)Hex(5)Fuc(1)NeuAc(1) | N[+2060] | 1484.3698 | 3 | 4451.0949 | 4451.0843 | 0 | 2.4 | 457 | Specific | 834.64 | 760.62 | 760.62 | 15.82 | 63 | >sp\|P01833\|PIGR_HUMAN Polymeric immunoglobulin receptor OS=Homo sapiens OX=9606 GN=PIGR PE=1 SV=4 | 902 | 20201117_ly_milk_1ng_glyco_hcd203040_compound15.19656.19656.3 | controllerType=0 controllerNumber=1 scan=19656 | 39.2719 |
| 16325:3 | 8 | K.IIEGEPNLKVPGN[+2205.793]VTAVLGETLK.V | HexNAc(4)Hex(5)Fuc(2)NeuAc(1) | N[+2206] | 1533.0546 | 3 | 4597.1493 | 4597.1422 | 0 | 1.5 | 457 | Specific | 837.7 | 727.08 | 727.08 | 15.76 | 63 | >sp\|P01833\|PIGR_HUMAN Polymeric immunoglobulin receptor OS=Homo sapiens OX=9606 GN=PIGR PE=1 SV=4 | 902 | 20201117_ly_milk_1ng_glyco_hcd203040_compound15.19633.19633.3 | controllerType=0 controllerNumber=1 scan=19633 | 39.2312 |
| 16369:3 | 8 | K.IIEGEPNLKVPGN[+2059.735]VTAVLGETLK.V | HexNAc(4)Hex(5)Fuc(1)NeuAc(1) | N[+2060] | 1484.3708 | 3 | 4451.0978 | 4451.0843 | 0 | 3 | 457 | Specific | 916.43 | 899.82 | 899.82 | 17.82 | 63 | >sp\|P01833\|PIGR_HUMAN Polymeric immunoglobulin receptor OS=Homo sapiens OX=9606 GN=PIGR PE=1 SV=4 | 902 | 20201117_ly_milk_1ng_glyco_hcd203040_compound15.19679.19679.3 | controllerType=0 controllerNumber=1 scan=19679 | 39.3179 |
| 16345:3 | 8 | K.IIEGEPNLKVPGN[+2059.735]VTAVLGETLK.V | HexNAc(4)Hex(5)Fuc(1)NeuAc(1) | N[+2060] | 1484.3698 | 3 | 4451.0949 | 4451.0843 | 0 | 2.4 | 457 | Specific | 951.89 | 934.18 | 934.18 | 18.82 | 63 | >sp\|P01833\|PIGR_HUMAN Polymeric immunoglobulin receptor OS=Homo sapiens OX=9606 GN=PIGR PE=1 SV=4 | 902 | 20201117_ly_milk_1ng_glyco_hcd203040_compound15.19654.19654.3 | controllerType=0 controllerNumber=1 scan=19654 | 39.2669 |
| 16334:3 | 8 | K.IIEGEPNLKVPGN[+2059.735]VTAVLGETLK.V | HexNAc(4)Hex(5)Fuc(1)NeuAc(1) | N[+2060] | 1484.3697 | 3 | 4451.0945 | 4451.0843 | 0 | 2.3 | 457 | Specific | 983.91 | 933.7 | 933.7 | 18.82 | 63 | >sp\|P01833\|PIGR_HUMAN Polymeric immunoglobulin receptor OS=Homo sapiens OX=9606 GN=PIGR PE=1 SV=4 | 902 | 20201117_ly_milk_1ng_glyco_hcd203040_compound15.19642.19642.3 | controllerType=0 controllerNumber=1 scan=19642 | 39.2451 |
| 15903:5 | 8 | K.IIEGEPNLKVPGN[+2059.735]VTAVLGETLKVPC[+57.021]HFPC[+57.021]K.F | HexNAc(4)Hex(5)Fuc(1)NeuAc(1) | C[+57]*2, N[+2060] | 1096.1172 | 5 | 5476.5568 | 5476.5418 | 0 | 2.7 | 457 | Specific | 650.06 | 577.54 | 577.54 | 13.03 | 63 | >sp\|P01833\|PIGR_HUMAN Polymeric immunoglobulin receptor OS=Homo sapiens OX=9606 GN=PIGR PE=1 SV=4 | 902 | 20201117_ly_milk_1ng_glyco_hcd203040_compound15.19163.19163.5 | controllerType=0 controllerNumber=1 scan=19163 | 38.3947 |
| 15901:5 | 8 | K.IIEGEPNLKVPGN[+2059.735]VTAVLGETLKVPC[+57.021]HFPC[+57.021]K.F | HexNAc(4)Hex(5)Fuc(1)NeuAc(1) | C[+57]*2, N[+2060] | 1096.1172 | 5 | 5476.5568 | 5476.5418 | 0 | 2.7 | 457 | Specific | 708.67 | 553.63 | 553.63 | 14.03 | 63 | >sp\|P01833\|PIGR_HUMAN Polymeric immunoglobulin receptor OS=Homo sapiens OX=9606 GN=PIGR PE=1 SV=4 | 902 | 20201117_ly_milk_1ng_glyco_hcd203040_compound15.19161.19161.5 | controllerType=0 controllerNumber=1 scan=19161 | 38.3895 |

Table S4. Cont.

| Query #:z | Protein Rank | Glycopeptide | Glycan | Modification Type(s) | Observed m/z | z | Observed (M+H) | Calc. mass (M+H) | Off-by-x error | Mass error (ppm) | StartPosition | Cleavage | Score | Delta | Delta Mod | \|Log Prob\| | # of unique peptides | Protein Name | Protein DB number | Comment | Scan # | Scan Time |
|---|---|---|---|---|---|---|---|---|---|---|---|---|---|---|---|---|---|---|---|---|---|---|
| 15895:5 | 8 | K.IIEGEPNLKVPGN[+2205.793]VTAVLGETLKVPC[+57.021]HFPC[+57.021]K.F | HexNAc(4)Hex(5)Fuc(2)NeuAc(1) | C[+57]*2, N[+2206] | 1125.3275 | 5 | 5622.6085 | 5622.5997 | 0 | 1.6 | 457 | Specific | 724.27 | 637.81 | 637.81 | 13.97 | 63 | >sp\|P01833\|PIGR_HUMAN Polymeric immunoglobulin receptor OS=Homo sapiens OX=9606 GN=PIGR PE=1 SV=4 | 902 | 20201117_ly_milk_1ng_glyco_hcd203040_compound15.19155.19155.5 | controllerType=0 controllerNumber=1 scan=19155 | 38.3796 |
| 15898:5 | 8 | K.IIEGEPNLKVPGN[+2205.793]VTAVLGETLKVPC[+57.021]HFPC[+57.021]K.F | HexNAc(4)Hex(5)Fuc(2)NeuAc(1) | C[+57]*2, N[+2206] | 1125.3275 | 5 | 5622.6085 | 5622.5997 | 0 | 1.6 | 457 | Specific | 750.35 | 645.76 | 645.76 | 14.97 | 63 | >sp\|P01833\|PIGR_HUMAN Polymeric immunoglobulin receptor OS=Homo sapiens OX=9606 GN=PIGR PE=1 SV=4 | 902 | 20201117_ly_milk_1ng_glyco_hcd203040_compound15.19158.19158.5 | controllerType=0 controllerNumber=1 scan=19158 | 38.3848 |
| 15820:5 | 8 | K.IIEGEPNLKVPGN[+2205.793]VTAVLGETLKVPC[+57.021]HFPC[+57.021]K.F | HexNAc(4)Hex(5)Fuc(2)NeuAc(1) | C[+57]*2, N[+2206] | 1125.328 | 5 | 5622.6109 | 5622.5997 | 0 | 2 | 457 | Specific | 824.26 | 738.15 | 738.15 | 15.97 | 63 | >sp\|P01833\|PIGR_HUMAN Polymeric immunoglobulin receptor OS=Homo sapiens OX=9606 GN=PIGR PE=1 SV=4 | 902 | 20201117_ly_milk_1ng_glyco_hcd203040_compound15.19077.19077.5 | controllerType=0 controllerNumber=1 scan=19077 | 38.2392 |
| 15823:5 | 8 | K.IIEGEPNLKVPGN[+2205.793]VTAVLGETLKVPC[+57.021]HFPC[+57.021]K.F | HexNAc(4)Hex(5)Fuc(2)NeuAc(1) | C[+57]*2, N[+2206] | 1125.328 | 5 | 5622.6109 | 5622.5997 | 0 | 2 | 457 | Specific | 869.5 | 869.5 | 869.5 | 16.97 | 63 | >sp\|P01833\|PIGR_HUMAN Polymeric immunoglobulin receptor OS=Homo sapiens OX=9606 GN=PIGR PE=1 SV=4 | 902 | 20201117_ly_milk_1ng_glyco_hcd203040_compound15.19080.19080.5 | controllerType=0 controllerNumber=1 scan=19080 | 38.245 |
| 16993:3 | 8 | K.VPGN[+2350.830]VTAVLGETLK.V | HexNAc(4)Hex(5)Fuc(1)NeuAc(2) | N[+2351] | 1250.2211 | 3 | 3748.6488 | 3748.6302 | 0 | 5 | 466 | Specific | 286.57 | 234.76 | 234.76 | 5.06 | 63 | >sp\|P01833\|PIGR_HUMAN Polymeric immunoglobulin receptor OS=Homo sapiens OX=9606 GN=PIGR PE=1 SV=4 | 902 | 20201117_ly_milk_1ng_glyco_hcd203040_compound15.20331.20331.3 | controllerType=0 controllerNumber=1 scan=20331 | 40.4951 |
| 13916:3 | 8 | K.VPGN[+2059.735]VTAVLGETLK.V | HexNAc(4)Hex(5)Fuc(1)NeuAc(1) | N[+2060] | 1153.186 | 3 | 3457.5434 | 3457.5348 | 0 | 2.5 | 466 | Specific | 548.39 | 367.68 | 367.68 | 8.33 | 63 | >sp\|P01833\|PIGR_HUMAN Polymeric immunoglobulin receptor OS=Homo sapiens OX=9606 GN=PIGR PE=1 SV=4 | 902 | 20201117_ly_milk_1ng_glyco_hcd203040_compound15.17102.17102.3 | controllerType=0 controllerNumber=1 scan=17102 | 34.7249 |
| 13914:3 | 8 | K.VPGN[+2059.735]VTAVLGETLK.V | HexNAc(4)Hex(5)Fuc(1)NeuAc(1) | N[+2060] | 1153.186 | 3 | 3457.5434 | 3457.5348 | 0 | 2.5 | 466 | Specific | 581.03 | 559.17 | 559.17 | 10.98 | 63 | >sp\|P01833\|PIGR_HUMAN Polymeric immunoglobulin receptor OS=Homo sapiens OX=9606 GN=PIGR PE=1 SV=4 | 902 | 20201117_ly_milk_1ng_glyco_hcd203040_compound15.17100.17100.3 | controllerType=0 controllerNumber=1 scan=17100 | 34.7206 |
| 13760:3 | 8 | K.VPGN[+2205.793]VTAVLGETLK.V | HexNAc(4)Hex(5)Fuc(2)NeuAc(1) | N[+2206] | 1201.8735 | 3 | 3603.6059 | 3603.5927 | 0 | 3.7 | 466 | Specific | 714.48 | 431.62 | 431.62 | 13.71 | 63 | >sp\|P01833\|PIGR_HUMAN Polymeric immunoglobulin receptor OS=Homo sapiens OX=9606 GN=PIGR PE=1 SV=4 | 902 | 20201117_ly_milk_1ng_glyco_hcd203040_compound15.16940.16940.3 | controllerType=0 controllerNumber=1 scan=16940 | 34.4161 |
| 13758:3 | 8 | K.VPGN[+2205.793]VTAVLGETLK.V | HexNAc(4)Hex(5)Fuc(2)NeuAc(1) | N[+2206] | 1201.8735 | 3 | 3603.6059 | 3603.5927 | 0 | 3.7 | 466 | Specific | 773.6 | 501.26 | 501.26 | 14.71 | 63 | >sp\|P01833\|PIGR_HUMAN Polymeric immunoglobulin receptor OS=Homo sapiens OX=9606 GN=PIGR PE=1 SV=4 | 902 | 20201117_ly_milk_1ng_glyco_hcd203040_compound15.16938.16938.3 | controllerType=0 controllerNumber=1 scan=16938 | 34.4115 |
| 13820:4 | 8 | K.VPGN[+2204.772]VTAVLGETLKVPC[+57.021]HFPC[+57.021]K.F | HexNAc(4)Hex(5)NeuAc(2) | C[+57]*2, N[+2205] | 1157.7527 | 4 | 4627.9891 | 4628.0298 | 0 | -8.8 | 466 | Specific | 446.79 | 256.68 | 256.68 | 4.91 | 63 | >sp\|P01833\|PIGR_HUMAN Polymeric immunoglobulin receptor OS=Homo sapiens OX=9606 GN=PIGR PE=1 SV=4 | 902 | 20201117_ly_milk_1ng_glyco_hcd203040_compound15.17002.17002.3 | controllerType=0 controllerNumber=1 scan=17002 | 34.5287 |
| 08026:3 | 8 | K.WN[+2204.772]NTGC[+57.021]QALPSQDEGPSK.A | HexNAc(4)Hex(5)NeuAc(2) | C[+57], N[+2205] | 1398.5577 | 3 | 4193.6585 | 4193.6491 | 0 | 2.2 | 498 | Specific | 286.52 | 238.53 | 238.53 | 6.08 | 63 | >sp\|P01833\|PIGR_HUMAN Polymeric immunoglobulin receptor OS=Homo sapiens OX=9606 GN=PIGR PE=1 SV=4 | 902 | 20201117_ly_milk_1ng_glyco_hcd203040_compound15.10963.10963.3 | controllerType=0 controllerNumber=1 scan=10963 | 24.27 |
| 08028:3 | 8 | K.WN[+2204.772]NTGC[+57.021]QALPSQDEGPSK.A | HexNAc(4)Hex(5)NeuAc(2) | C[+57], N[+2205] | 1398.5577 | 3 | 4193.6585 | 4193.6491 | 0 | 2.2 | 498 | Specific | 353.35 | 282.94 | 282.94 | 7.13 | 63 | >sp\|P01833\|PIGR_HUMAN Polymeric immunoglobulin receptor OS=Homo sapiens OX=9606 GN=PIGR PE=1 SV=4 | 902 | 20201117_ly_milk_1ng_glyco_hcd203040_compound15.10965.10965.3 | controllerType=0 controllerNumber=1 scan=10965 | 24.274 |
| 05824:3 | 8 | K.WN[+2059.735]NTGC[+57.021]QALPSQDEGPSK.A | HexNAc(4)Hex(5)Fuc(1)NeuAc(1) | C[+57], N[+2060] | 1350.2128 | 3 | 4048.6239 | 4048.6116 | 0 | 3 | 498 | Specific | 629.22 | 561.29 | 561.29 | 12.14 | 63 | >sp\|P01833\|PIGR_HUMAN Polymeric immunoglobulin receptor OS=Homo sapiens OX=9606 GN=PIGR PE=1 SV=4 | 902 | 20201117_ly_milk_1ng_glyco_hcd203040_compound15.8685.8685.3 | controllerType=0 controllerNumber=1 scan=8685 | 20.5563 |
| 05822:3 | 8 | K.WN[+2059.735]NTGC[+57.021]QALPSQDEGPSK.A | HexNAc(4)Hex(5)Fuc(1)NeuAc(1) | C[+57], N[+2060] | 1350.2128 | 3 | 4048.6239 | 4048.6116 | 0 | 3 | 498 | Specific | 659.33 | 578.83 | 578.83 | 13.14 | 63 | >sp\|P01833\|PIGR_HUMAN Polymeric immunoglobulin receptor OS=Homo sapiens OX=9606 GN=PIGR PE=1 SV=4 | 902 | 20201117_ly_milk_1ng_glyco_hcd203040_compound15.8683.8683.3 | controllerType=0 controllerNumber=1 scan=8683 | 20.5527 |
| 05931:3 | 8 | K.WN[+1913.677]NTGC[+57.021]QALPSQDEGPSK.A | HexNAc(4)Hex(5)NeuAc(1) | C[+57], N[+1914] | 1301.5256 | 3 | 3902.5622 | 3902.5537 | 0 | 2.2 | 498 | Specific | 676.95 | 607.24 | 607.24 | 13.14 | 63 | >sp\|P01833\|PIGR_HUMAN Polymeric immunoglobulin receptor OS=Homo sapiens OX=9606 GN=PIGR PE=1 SV=4 | 902 | 20201117_ly_milk_1ng_glyco_hcd203040_compound15.8796.8796.3 | controllerType=0 controllerNumber=1 scan=8796 | 20.7543 |
| 05933:3 | 8 | K.WN[+1913.677]NTGC[+57.021]QALPSQDEGPSK.A | HexNAc(4)Hex(5)NeuAc(1) | C[+57], N[+1914] | 1301.5256 | 3 | 3902.5622 | 3902.5537 | 0 | 2.2 | 498 | Specific | 731.75 | 648.8 | 648.8 | 14.14 | 63 | >sp\|P01833\|PIGR_HUMAN Polymeric immunoglobulin receptor OS=Homo sapiens OX=9606 GN=PIGR PE=1 SV=4 | 902 | 20201117_ly_milk_1ng_glyco_hcd203040_compound15.8798.8798.3 | controllerType=0 controllerNumber=1 scan=8798 | 20.7604 |
| 17410:4 | 10 | K.RNIAAFGGDPNN[+1913.677]ITLFGESAGGASVSLQTLSPYNK.G | HexNAc(4)Hex(5)NeuAc(1) | N[+1914] | 1370.8744 | 4 | 5480.4757 | 5480.4644 | 0 | 2.1 | 196 | Specific | 1000.91 | 911.74 | 911.74 | 21.04 | 48 | >sp\|P19835\|CEL_HUMAN Bile salt-activated lipase OS=Homo sapiens OX=9606 GN=CEL PE=1 SV=3 | 476 | 20201117_ly_milk_1ng_glyco_hcd203040_compound15.20766.20766.4 | controllerType=0 controllerNumber=1 scan=20766 | 41.4219 |
| 17407:4 | 10 | K.RNIAAFGGDPNN[+1913.677]ITLFGESAGGASVSLQTLSPYNK.G | HexNAc(4)Hex(5)NeuAc(1) | N[+1914] | 1370.8744 | 4 | 5480.4757 | 5480.4644 | 0 | 2.1 | 196 | Specific | 1008.69 | 971.75 | 971.75 | 21.04 | 48 | >sp\|P19835\|CEL_HUMAN Bile salt-activated lipase OS=Homo sapiens OX=9606 GN=CEL PE=1 SV=3 | 476 | 20201117_ly_milk_1ng_glyco_hcd203040_compound15.20763.20763.4 | controllerType=0 controllerNumber=1 scan=20763 | 41.4128 |
| 18004:3 | 11 | R.LSLHRPALEDLLLGSEAN[+1501.555]LTC[+57.021]TLTGLR.D | HexNAc(5)Hex(3) | C[+57], N[+1502] | 1489.0618 | 3 | 4465.1709 | 4465.1536 | 0 | 3.9 | 127 | Specific | 260.92 | 260.92 | 260.92 | 5.67 | 51 | >sp\|P01876\|IGHA1_HUMAN Immunoglobulin heavy constant alpha 1 OS=Homo sapiens OX=9606 GN=IGHA1 PE=1 SV=2 | 255 | 20201117_ly_milk_1ng_glyco_hcd203040_compound15.21386.21386.2 | controllerType=0 controllerNumber=1 scan=21386 | 42.8208 |
| 18905:4 | 11 | R.LSLHRPALEDLLLGSEAN[+1913.677]LTC[+57.021]TLTGLR.D | HexNAc(4)Hex(5)NeuAc(1) | C[+57], N[+1914] | 1220.0787 | 4 | 4877.2928 | 4877.2753 | 0 | 3.6 | 127 | Specific | 265.43 | 265.43 | 265.43 | 6.3 | 51 | >sp\|P01876\|IGHA1_HUMAN Immunoglobulin heavy constant alpha 1 OS=Homo sapiens OX=9606 GN=IGHA1 PE=1 SV=2 | 255 | 20201117_ly_milk_1ng_glyco_hcd203040_compound15.22325.22325.4 | controllerType=0 controllerNumber=1 scan=22325 | 44.8314 |
| 19968:4 | 11 | R.LSLHRPALEDLLLGSEAN[+2204.772]LTC[+57.021]TLTGLR.D | HexNAc(4)Hex(5)NeuAc(2) | C[+57], N[+2205] | 1292.8499 | 4 | 5168.3776 | 5168.3707 | 0 | 1.3 | 127 | Specific | 292.25 | 273.17 | 273.17 | 6.71 | 51 | >sp\|P01876\|IGHA1_HUMAN Immunoglobulin heavy constant alpha 1 OS=Homo sapiens OX=9606 GN=IGHA1 PE=1 SV=2 | 255 | 20201117_ly_milk_1ng_glyco_hcd203040_compound15.23441.23441.4 | controllerType=0 controllerNumber=1 scan=23441 | 47.245 |
| 18018:4 | 11 | R.LSLHRPALEDLLLGSEAN[+1647.613]LTC[+57.021]TLTGLR.D | HexNAc(5)Hex(3)Fuc(1) | C[+57], N[+1648] | 1153.5622 | 4 | 4611.2269 | 4611.2115 | 0 | 3.3 | 127 | Specific | 332.17 | 332.17 | 332.17 | 7.1 | 51 | >sp\|P01876\|IGHA1_HUMAN Immunoglobulin heavy constant alpha 1 OS=Homo sapiens OX=9606 GN=IGHA1 PE=1 SV=2 | 255 | 20201117_ly_milk_1ng_glyco_hcd203040_compound15.21401.21401.4 | controllerType=0 controllerNumber=1 scan=21401 | 42.8525 |
| 18020:4 | 11 | R.LSLHRPALEDLLLGSEAN[+1647.613]LTC[+57.021]TLTGLR.D | HexNAc(5)Hex(3)Fuc(1) | C[+57], N[+1648] | 1153.5622 | 4 | 4611.2269 | 4611.2115 | 0 | 3.3 | 127 | Specific | 476.35 | 476.35 | 476.35 | 9.35 | 51 | >sp\|P01876\|IGHA1_HUMAN Immunoglobulin heavy constant alpha 1 OS=Homo sapiens OX=9606 GN=IGHA1 PE=1 SV=2 | 255 | 20201117_ly_milk_1ng_glyco_hcd203040_compound15.21403.21403.4 | controllerType=0 controllerNumber=1 scan=21403 | 42.8559 |
| 18937:4 | 11 | R.LSLHRPALEDLLLGSEAN[+1710.598]LTC[+57.021]TLTGLR.D | HexNAc(3)Hex(5)NeuAc(1) | C[+57], N[+1711] | 1169.3078 | 4 | 4674.2093 | 4674.1959 | 0 | 2.9 | 127 | Specific | 483.41 | 483.41 | 483.41 | 9.63 | 51 | >sp\|P01876\|IGHA1_HUMAN Immunoglobulin heavy constant alpha 1 OS=Homo sapiens OX=9606 GN=IGHA1 PE=1 SV=2 | 255 | 20201117_ly_milk_1ng_glyco_hcd203040_compound15.22358.22358.4 | controllerType=0 controllerNumber=1 scan=22358 | 44.9023 |
| 18907:4 | 11 | R.LSLHRPALEDLLLGSEAN[+1913.677]LTC[+57.021]TLTGLR.D | HexNAc(4)Hex(5)NeuAc(1) | C[+57], N[+1914] | 1220.0787 | 4 | 4877.2928 | 4877.2753 | 0 | 3.6 | 127 | Specific | 504.73 | 420.92 | 420.92 | 9.48 | 51 | >sp\|P01876\|IGHA1_HUMAN Immunoglobulin heavy constant alpha 1 OS=Homo sapiens OX=9606 GN=IGHA1 PE=1 SV=2 | 255 | 20201117_ly_milk_1ng_glyco_hcd203040_compound15.22327.22327.4 | controllerType=0 controllerNumber=1 scan=22327 | 44.8383 |
| 17900:3 | 11 | R.LSLHRPALEDLLLGSEAN[+1663.608]LTC[+57.021]TLTGLR.D | HexNAc(5)Hex(4) | C[+57], N[+1664] | 1543.0759 | 3 | 4627.213 | 4627.2064 | 0 | 1.4 | 127 | Specific | 579.2 | 525.48 | 525.48 | 10.55 | 51 | >sp\|P01876\|IGHA1_HUMAN Immunoglobulin heavy constant alpha 1 OS=Homo sapiens OX=9606 GN=IGHA1 PE=1 SV=2 | 255 | 20201117_ly_milk_1ng_glyco_hcd203040_compound15.21278.21278.3 | controllerType=0 controllerNumber=1 scan=21278 | 42.5534 |
| 17878:3 | 11 | R.LSLHRPALEDLLLGSEAN[+1540.529]LTC[+57.021]TLTGLR.D | HexNAc(2)Hex(7) | C[+57], N[+1541] | 1502.0512 | 3 | 4504.1391 | 4504.1268 | 0 | 2.7 | 127 | Specific | 580.32 | 542.82 | 542.82 | 10.61 | 51 | >sp\|P01876\|IGHA1_HUMAN Immunoglobulin heavy constant alpha 1 OS=Homo sapiens OX=9606 GN=IGHA1 PE=1 SV=2 | 255 | 20201117_ly_milk_1ng_glyco_hcd203040_compound15.21254.21254.3 | controllerType=0 controllerNumber=1 scan=21254 | 42.4884 |
| 17962:3 | 11 | R.LSLHRPALEDLLLGSEAN[+1460.529]LTC[+57.021]TLTGLR.D | HexNAc(4)Hex(4) | C[+57], N[+1461] | 1475.3857 | 3 | 4424.1425 | 4424.1271 | 0 | 3.5 | 127 | Specific | 601.92 | 530.61 | 530.61 | 11.34 | 51 | >sp\|P01876\|IGHA1_HUMAN Immunoglobulin heavy constant alpha 1 OS=Homo sapiens OX=9606 GN=IGHA1 PE=1 SV=2 | 255 | 20201117_ly_milk_1ng_glyco_hcd203040_compound15.21342.21342.3 | controllerType=0 controllerNumber=1 scan=21342 | 42.7032 |
| 17983:4 | 11 | R.LSLHRPALEDLLLGSEAN[+1524.534]LTC[+57.021]TLTGLR.D | HexNAc(2)Hex(6)Fuc(1) | C[+57], N[+1525] | 1122.7814 | 4 | 4488.1038 | 4488.1319 | 0 | -6.2 | 127 | Specific | 604.74 | 565.88 | 565.88 | 10.43 | 51 | >sp\|P01876\|IGHA1_HUMAN Immunoglobulin heavy constant alpha 1 OS=Homo sapiens OX=9606 GN=IGHA1 PE=1 SV=2 | 255 | 20201117_ly_milk_1ng_glyco_hcd203040_compound15.21365.21365.4 | controllerType=0 controllerNumber=1 scan=21365 | 42.7795 |
| 17965:3 | 11 | R.LSLHRPALEDLLLGSEAN[+1460.529]LTC[+57.021]TLTGLR.D | HexNAc(4)Hex(4) | C[+57], N[+1461] | 1475.3857 | 3 | 4424.1425 | 4424.1271 | 0 | 3.5 | 127 | Specific | 611.23 | 505.28 | 505.28 | 11.34 | 51 | >sp\|P01876\|IGHA1_HUMAN Immunoglobulin heavy constant alpha 1 OS=Homo sapiens OX=9606 GN=IGHA1 PE=1 SV=2 | 255 | 20201117_ly_milk_1ng_glyco_hcd203040_compound15.21345.21345.3 | controllerType=0 controllerNumber=1 scan=21345 | 42.7161 |
| 19970:4 | 11 | R.LSLHRPALEDLLLGSEAN[+2204.772]LTC[+57.021]TLTGLR.D | HexNAc(4)Hex(5)NeuAc(2) | C[+57], N[+2205] | 1292.8499 | 4 | 5168.3776 | 5168.3707 | 0 | 1.3 | 127 | Specific | 614.77 | 562.12 | 562.12 | 12.17 | 51 | >sp\|P01876\|IGHA1_HUMAN Immunoglobulin heavy constant alpha 1 OS=Homo sapiens OX=9606 GN=IGHA1 PE=1 SV=2 | 255 | 20201117_ly_milk_1ng_glyco_hcd203040_compound15.23443.23443.4 | controllerType=0 controllerNumber=1 scan=23443 | 47.2498 |
| 17910:3 | 11 | R.LSLHRPALEDLLLGSEAN[+1622.582]LTC[+57.021]TLTGLR.D | HexNAc(4)Hex(5) | C[+57], N[+1623] | 1529.4 | 3 | 4586.1853 | 4586.1799 | 0 | 1.2 | 127 | Specific | 623.19 | 522.02 | 522.02 | 11.55 | 51 | >sp\|P01876\|IGHA1_HUMAN Immunoglobulin heavy constant alpha 1 OS=Homo sapiens OX=9606 GN=IGHA1 PE=1 SV=2 | 255 | 20201117_ly_milk_1ng_glyco_hcd203040_compound15.21288.21288.3 | controllerType=0 controllerNumber=1 scan=21288 | 42.5764 |
| 17868:3 | 11 | R.LSLHRPALEDLLLGSEAN[+1702.581]LTC[+57.021]TLTGLR.D | HexNAc(2)Hex(8) | C[+57], N[+1703] | 1556.0699 | 3 | 4666.1951 | 4666.1796 | 0 | 3.3 | 127 | Specific | 627.19 | 589.15 | 589.15 | 11.34 | 51 | >sp\|P01876\|IGHA1_HUMAN Immunoglobulin heavy constant alpha 1 OS=Homo sapiens OX=9606 GN=IGHA1 PE=1 SV=2 | 255 | 20201117_ly_milk_1ng_glyco_hcd203040_compound15.21244.21244.3 | controllerType=0 controllerNumber=1 scan=21244 | 42.4606 |
| 18007:3 | 11 | R.LSLHRPALEDLLLGSEAN[+1501.555]LTC[+57.021]TLTGLR.D | HexNAc(5)Hex(3) | C[+57], N[+1502] | 1489.0618 | 3 | 4465.1709 | 4465.1536 | 0 | 3.9 | 127 | Specific | 628.87 | 610.59 | 610.59 | 11.34 | 51 | >sp\|P01876\|IGHA1_HUMAN Immunoglobulin heavy constant alpha 1 OS=Homo sapiens OX=9606 GN=IGHA1 PE=1 SV=2 | 255 | 20201117_ly_milk_1ng_glyco_hcd203040_compound15.21389.21389.2 | controllerType=0 controllerNumber=1 scan=21389 | 42.8264 |
| 17902:3 | 11 | R.LSLHRPALEDLLLGSEAN[+1663.608]LTC[+57.021]TLTGLR.D | HexNAc(5)Hex(4) | C[+57], N[+1664] | 1543.0759 | 3 | 4627.213 | 4627.2064 | 0 | 1.4 | 127 | Specific | 635.55 | 576.1 | 576.1 | 11.55 | 51 | >sp\|P01876\|IGHA1_HUMAN Immunoglobulin heavy constant alpha 1 OS=Homo sapiens OX=9606 GN=IGHA1 PE=1 SV=2 | 255 | 20201117_ly_milk_1ng_glyco_hcd203040_compound15.21280.21280.3 | controllerType=0 controllerNumber=1 scan=21280 | 42.5584 |
| 17908:3 | 11 | R.LSLHRPALEDLLLGSEAN[+1622.582]LTC[+57.021]TLTGLR.D | HexNAc(4)Hex(5) | C[+57], N[+1623] | 1529.4 | 3 | 4586.1853 | 4586.1799 | 0 | 1.2 | 127 | Specific | 639.92 | 586.83 | 586.83 | 11.55 | 51 | >sp\|P01876\|IGHA1_HUMAN Immunoglobulin heavy constant alpha 1 OS=Homo sapiens OX=9606 GN=IGHA1 PE=1 SV=2 | 255 | 20201117_ly_milk_1ng_glyco_hcd203040_compound15.21286.21286.3 | controllerType=0 controllerNumber=1 scan=21286 | 42.5717 |
| 17836:3 | 11 | R.LSLHRPALEDLLLGSEAN[+1864.634]LTC[+57.021]TLTGLR.D | HexNAc(2)Hex(9) | C[+57], N[+1865] | 1610.0871 | 3 | 4828.2467 | 4828.2324 | 0 | 3 | 127 | Specific | 641.58 | 620.67 | 620.67 | 11.61 | 51 | >sp\|P01876\|IGHA1_HUMAN Immunoglobulin heavy constant alpha 1 OS=Homo sapiens OX=9606 GN=IGHA1 PE=1 SV=2 | 255 | 20201117_ly_milk_1ng_glyco_hcd203040_compound15.21210.21210.3 | controllerType=0 controllerNumber=1 scan=21210 | 42.3696 |
| 17947:3 | 11 | R.LSLHRPALEDLLLGSEAN[+1501.555]LTC[+57.021]TLTGLR.D | HexNAc(5)Hex(3) | C[+57], N[+1502] | 1489.0612 | 3 | 4465.1691 | 4465.1536 | 0 | 3.5 | 127 | Specific | 655 | 655 | 655 | 12.34 | 51 | >sp\|P01876\|IGHA1_HUMAN Immunoglobulin heavy constant alpha 1 OS=Homo sapiens OX=9606 GN=IGHA1 PE=1 SV=2 | 255 | 20201117_ly_milk_1ng_glyco_hcd203040_compound15.21327.21327.3 | controllerType=0 controllerNumber=1 scan=21327 | 42.6657 |
| 17923:4 | 11 | R.LSLHRPALEDLLLGSEAN[+1419.502]LTC[+57.021]TLTGLR.D | HexNAc(3)Hex(5) | C[+57], N[+1420] | 1096.5339 | 4 | 4383.1136 | 4383.1005 | 0 | 3 | 127 | Specific | 662.17 | 661.82 | 661.82 | 13.23 | 51 | >sp\|P01876\|IGHA1_HUMAN Immunoglobulin heavy constant alpha 1 OS=Homo sapiens OX=9606 GN=IGHA1 PE=1 SV=2 | 255 | 20201117_ly_milk_1ng_glyco_hcd203040_compound15.21302.21302.4 | controllerType=0 controllerNumber=1 scan=21302 | 42.609 |
| 17949:3 | 11 | R.LSLHRPALEDLLLGSEAN[+1501.555]LTC[+57.021]TLTGLR.D | HexNAc(5)Hex(3) | C[+57], N[+1502] | 1489.0612 | 3 | 4465.1691 | 4465.1536 | 0 | 3.5 | 127 | Specific | 664.14 | 612.67 | 612.67 | 12.34 | 51 | >sp\|P01876\|IGHA1_HUMAN Immunoglobulin heavy constant alpha 1 OS=Homo sapiens OX=9606 GN=IGHA1 PE=1 SV=2 | 255 | 20201117_ly_milk_1ng_glyco_hcd203040_compound15.21329.21329.3 | controllerType=0 controllerNumber=1 scan=21329 | 42.6748 |
| 17890:4 | 11 | R.LSLHRPALEDLLLGSEAN[+1663.608]LTC[+57.021]TLTGLR.D | HexNAc(5)Hex(4) | C[+57], N[+1664] | 1157.5601 | 4 | 4627.2186 | 4627.2064 | 0 | 2.6 | 127 | Specific | 665.48 | 665.48 | 665.48 | 13.23 | 51 | >sp\|P01876\|IGHA1_HUMAN Immunoglobulin heavy constant alpha 1 OS=Homo sapiens OX=9606 GN=IGHA1 PE=1 SV=2 | 255 | 20201117_ly_milk_1ng_glyco_hcd203040_compound15.21267.21267.4 | controllerType=0 controllerNumber=1 scan=21267 | 42.5276 |
| 17980:5 | 11 | R.LSLHRPALEDLLLGSEAN[+1501.555]LTC[+57.021]TLTGLR.D | HexNAc(5)Hex(3) | C[+57], N[+1502] | 893.8373 | 5 | 4465.1574 | 4465.1536 | 0 | 0.9 | 127 | Specific | 673.45 | 673.45 | 673.45 | 13.07 | 51 | >sp\|P01876\|IGHA1_HUMAN Immunoglobulin heavy constant alpha 1 OS=Homo sapiens OX=9606 GN=IGHA1 PE=1 SV=2 | 255 | 20201117_ly_milk_1ng_glyco_hcd203040_compound15.21361.21361.5 | controllerType=0 controllerNumber=1 scan=21361 | 42.7721 |
| 17876:3 | 11 | R.LSLHRPALEDLLLGSEAN[+1540.529]LTC[+57.021]TLTGLR.D | HexNAc(2)Hex(7) | C[+57], N[+1541] | 1502.0512 | 3 | 4504.1391 | 4504.1268 | 0 | 2.7 | 127 | Specific | 688.36 | 648.09 | 648.09 | 12.61 | 51 | >sp\|P01876\|IGHA1_HUMAN Immunoglobulin heavy constant alpha 1 OS=Homo sapiens OX=9606 GN=IGHA1 PE=1 SV=2 | 255 | 20201117_ly_milk_1ng_glyco_hcd203040_compound15.21252.21252.3 | controllerType=0 controllerNumber=1 scan=21252 | 42.4816 |
| 17946:4 | 11 | R.LSLHRPALEDLLLGSEAN[+1501.555]LTC[+57.021]TLTGLR.D | HexNAc(5)Hex(3) | C[+57], N[+1502] | 1117.0509 | 4 | 4465.182 | 4465.1536 | 0 | 6.4 | 127 | Specific | 692.7 | 692.7 | 692.7 | 12 | 51 | >sp\|P01876\|IGHA1_HUMAN Immunoglobulin heavy constant alpha 1 OS=Homo sapiens OX=9606 GN=IGHA1 PE=1 SV=2 | 255 | 20201117_ly_milk_1ng_glyco_hcd203040_compound15.21326.21326.4 | controllerType=0 controllerNumber=1 scan=21326 | 42.6642 |
| 17834:3 | 11 | R.LSLHRPALEDLLLGSEAN[+1864.634]LTC[+57.021]TLTGLR.D | HexNAc(2)Hex(9) | C[+57], N[+1865] | 1610.0871 | 3 | 4828.2467 | 4828.2324 | 0 | 3 | 127 | Specific | 703.27 | 703.27 | 703.27 | 13.61 | 51 | >sp\|P01876\|IGHA1_HUMAN Immunoglobulin heavy constant alpha 1 OS=Homo sapiens OX=9606 GN=IGHA1 PE=1 SV=2 | 255 | 20201117_ly_milk_1ng_glyco_hcd203040_compound15.21208.21208.3 | controllerType=0 controllerNumber=1 scan=21208 | 42.364 |
| 17977:5 | 11 | R.LSLHRPALEDLLLGSEAN[+1501.555]LTC[+57.021]TLTGLR.D | HexNAc(5)Hex(3) | C[+57], N[+1502] | 893.8373 | 5 | 4465.1574 | 4465.1536 | 0 | 0.9 | 127 | Specific | 706.51 | 706.51 | 706.51 | 14.07 | 51 | >sp\|P01876\|IGHA1_HUMAN Immunoglobulin heavy constant alpha 1 OS=Homo sapiens OX=9606 GN=IGHA1 PE=1 SV=2 | 255 | 20201117_ly_milk_1ng_glyco_hcd203040_compound15.21358.21358.5 | controllerType=0 controllerNumber=1 scan=21358 | 42.7473 |
| 17865:3 | 11 | R.LSLHRPALEDLLLGSEAN[+1702.581]LTC[+57.021]TLTGLR.D | HexNAc(2)Hex(8) | C[+57], N[+1703] | 1556.0699 | 3 | 4666.1951 | 4666.1796 | 0 | 3.3 | 127 | Specific | 710.42 | 699.41 | 699.41 | 13.34 | 51 | >sp\|P01876\|IGHA1_HUMAN Immunoglobulin heavy constant alpha 1 OS=Homo sapiens OX=9606 GN=IGHA1 PE=1 SV=2 | 255 | 20201117_ly_milk_1ng_glyco_hcd203040_compound15.21241.21241.3 | controllerType=0 controllerNumber=1 scan=21241 | 42.4538 |
| 17985:4 | 11 | R.LSLHRPALEDLLLGSEAN[+1524.534]LTC[+57.021]TLTGLR.D | HexNAc(2)Hex(6)Fuc(1) | C[+57], N[+1525] | 1122.7814 | 4 | 4488.1038 | 4488.1319 | 0 | -6.2 | 127 | Specific | 721.35 | 623.83 | 623.83 | 12.43 | 51 | >sp\|P01876\|IGHA1_HUMAN Immunoglobulin heavy constant alpha 1 OS=Homo sapiens OX=9606 GN=IGHA1 PE=1 SV=2 | 255 | 20201117_ly_milk_1ng_glyco_hcd203040_compound15.21367.21367.4 | controllerType=0 controllerNumber=1 scan=21367 | 42.7864 |
| 17948:4 | 11 | R.LSLHRPALEDLLLGSEAN[+1501.555]LTC[+57.021]TLTGLR.D | HexNAc(5)Hex(3) | C[+57], N[+1502] | 1117.0509 | 4 | 4465.182 | 4465.1536 | 0 | 6.4 | 127 | Specific | 740.06 | 734.59 | 734.59 | 13 | 51 | >sp\|P01876\|IGHA1_HUMAN Immunoglobulin heavy constant alpha 1 OS=Homo sapiens OX=9606 GN=IGHA1 PE=1 SV=2 | 255 | 20201117_ly_milk_1ng_glyco_hcd203040_compound15.21328.21328.4 | controllerType=0 controllerNumber=1 scan=21328 | 42.6706 |
| 17950:4 | 11 | R.LSLHRPALEDLLLGSEAN[+1460.529]LTC[+57.021]TLTGLR.D | HexNAc(4)Hex(4) | C[+57], N[+1461] | 1106.7945 | 4 | 4424.1561 | 4424.1271 | 0 | 6.6 | 127 | Specific | 740.36 | 740.36 | 740.36 | 13 | 51 | >sp\|P01876\|IGHA1_HUMAN Immunoglobulin heavy constant alpha 1 OS=Homo sapiens OX=9606 GN=IGHA1 PE=1 SV=2 | 255 | 20201117_ly_milk_1ng_glyco_hcd203040_compound15.21330.21330.4 | controllerType=0 controllerNumber=1 scan=21330 | 42.676 |
| 17893:4 | 11 | R.LSLHRPALEDLLLGSEAN[+1663.608]LTC[+57.021]TLTGLR.D | HexNAc(5)Hex(4) | C[+57], N[+1664] | 1157.5601 | 4 | 4627.2186 | 4627.2064 | 0 | 2.6 | 127 | Specific | 741.7 | 677.75 | 677.75 | 14.23 | 51 | >sp\|P01876\|IGHA1_HUMAN Immunoglobulin heavy constant alpha 1 OS=Homo sapiens OX=9606 GN=IGHA1 PE=1 SV=2 | 255 | 20201117_ly_milk_1ng_glyco_hcd203040_compound15.21270.21270.4 | controllerType=0 controllerNumber=1 scan=21270 | 42.5345 |
| 18889:4 | 11 | R.LSLHRPALEDLLLGSEAN[+1872.651]LTC[+57.021]TLTGLR.D | HexNAc(3)Hex(6)NeuAc(1) | C[+57], N[+1873] | 1209.8212 | 4 | 4836.263 | 4836.2487 | 0 | 2.9 | 127 | Specific | 743.53 | 692.63 | 692.63 | 14.23 | 51 | >sp\|P01876\|IGHA1_HUMAN Immunoglobulin heavy constant alpha 1 OS=Homo sapiens OX=9606 GN=IGHA1 PE=1 SV=2 | 255 | 20201117_ly_milk_1ng_glyco_hcd203040_compound15.22308.22308.4 | controllerType=0 controllerNumber=1 scan=22308 | 44.7735 |
| 18011:4 | 11 | R.LSLHRPALEDLLLGSEAN[+1095.397]LTC[+57.021]TLTGLR.D | HexNAc(3)Hex(3) | C[+57], N[+1095] | 1015.5081 | 4 | 4059.0106 | 4058.9949 | 0 | 3.9 | 127 | Specific | 759.2 | 759.2 | 759.2 | 14.96 | 51 | >sp\|P01876\|IGHA1_HUMAN Immunoglobulin heavy constant alpha 1 OS=Homo sapiens OX=9606 GN=IGHA1 PE=1 SV=2 | 255 | 20201117_ly_milk_1ng_glyco_hcd203040_compound15.21394.21394.4 | controllerType=0 controllerNumber=1 scan=21394 | 42.8347 |
| 18940:4 | 11 | R.LSLHRPALEDLLLGSEAN[+1710.598]LTC[+57.021]TLTGLR.D | HexNAc(3)Hex(5)NeuAc(1) | C[+57], N[+1711] | 1169.3078 | 4 | 4674.2093 | 4674.1959 | 0 | 2.9 | 127 | Specific | 801.04 | 757.01 | 757.01 | 16.23 | 51 | >sp\|P01876\|IGHA1_HUMAN Immunoglobulin heavy constant alpha 1 OS=Homo sapiens OX=9606 GN=IGHA1 PE=1 SV=2 | 255 | 20201117_ly_milk_1ng_glyco_hcd203040_compound15.22361.22361.4 | controllerType=0 controllerNumber=1 scan=22361 | 44.9073 |
| 17966:4 | 11 | R.LSLHRPALEDLLLGSEAN[+1216.423]LTC[+57.021]TLTGLR.D | HexNAc(2)Hex(5) | C[+57], N[+1216] | 1045.762 | 4 | 4180.0262 | 4180.0211 | 0 | 1.2 | 127 | Specific | 802.21 | 768.67 | 768.67 | 16.17 | 51 | >sp\|P01876\|IGHA1_HUMAN Immunoglobulin heavy constant alpha 1 OS=Homo sapiens OX=9606 GN=IGHA1 PE=1 SV=2 | 255 | 20201117_ly_milk_1ng_glyco_hcd203040_compound15.21347.21347.4 | controllerType=0 controllerNumber=1 scan=21347 | 42.7206 |
| 18892:4 | 11 | R.LSLHRPALEDLLLGSEAN[+1872.651]LTC[+57.021]TLTGLR.D | HexNAc(3)Hex(6)NeuAc(1) | C[+57], N[+1873] | 1209.8212 | 4 | 4836.263 | 4836.2487 | 0 | 2.9 | 127 | Specific | 803.73 | 739.52 | 739.52 | 16.23 | 51 | >sp\|P01876\|IGHA1_HUMAN Immunoglobulin heavy constant alpha 1 OS=Homo sapiens OX=9606 GN=IGHA1 PE=1 SV=2 | 255 | 20201117_ly_milk_1ng_glyco_hcd203040_compound15.22311.22311.4 | controllerType=0 controllerNumber=1 scan=22311 | 44.7844 |
| 17953:4 | 11 | R.LSLHRPALEDLLLGSEAN[+1460.529]LTC[+57.021]TLTGLR.D | HexNAc(4)Hex(4) | C[+57], N[+1461] | 1106.7945 | 4 | 4424.1561 | 4424.1271 | 0 | 6.6 | 127 | Specific | 873.41 | 772.62 | 772.62 | 16 | 51 | >sp\|P01876\|IGHA1_HUMAN Immunoglobulin heavy constant alpha 1 OS=Homo sapiens OX=9606 GN=IGHA1 PE=1 SV=2 | 255 | 20201117_ly_milk_1ng_glyco_hcd203040_compound15.21333.21333.4 | controllerType=0 controllerNumber=1 scan=21333 | 42.6819 |
| 17925:4 | 11 | R.LSLHRPALEDLLLGSEAN[+1419.502]LTC[+57.021]TLTGLR.D | HexNAc(3)Hex(5) | C[+57], N[+1420] | 1096.5339 | 4 | 4383.1136 | 4383.1005 | 0 | 3 | 127 | Specific | 891.4 | 855.92 | 855.92 | 17.23 | 51 | >sp\|P01876\|IGHA1_HUMAN Immunoglobulin heavy constant alpha 1 OS=Homo sapiens OX=9606 GN=IGHA1 PE=1 SV=2 | 255 | 20201117_ly_milk_1ng_glyco_hcd203040_compound15.21304.21304.4 | controllerType=0 controllerNumber=1 scan=21304 | 42.614 |
| 17862:4 | 11 | R.LSLHRPALEDLLLGSEAN[+1702.581]LTC[+57.021]TLTGLR.D | HexNAc(2)Hex(8) | C[+57], N[+1703] | 1167.3036 | 4 | 4666.1927 | 4666.1796 | 0 | 2.8 | 127 | Specific | 899.35 | 899.35 | 899.35 | 17.23 | 51 | >sp\|P01876\|IGHA1_HUMAN Immunoglobulin heavy constant alpha 1 OS=Homo sapiens OX=9606 GN=IGHA1 PE=1 SV=2 | 255 | 20201117_ly_milk_1ng_glyco_hcd203040_compound15.21238.21238.4 | controllerType=0 controllerNumber=1 scan=21238 | 42.4464 |
| 17812:4 | 11 | R.LSLHRPALEDLLLGSEAN[+1864.634]LTC[+57.021]TLTGLR.D | HexNAc(2)Hex(9) | C[+57], N[+1865] | 1207.8166 | 4 | 4828.2445 | 4828.2324 | 0 | 2.5 | 127 | Specific | 904.29 | 904.29 | 904.29 | 18.23 | 51 | >sp\|P01876\|IGHA1_HUMAN Immunoglobulin heavy constant alpha 1 OS=Homo sapiens OX=9606 GN=IGHA1 PE=1 SV=2 | 255 | 20201117_ly_milk_1ng_glyco_hcd203040_compound15.21185.21185.4 | controllerType=0 controllerNumber=1 scan=21185 | 42.3109 |
| 17871:4 | 11 | R.LSLHRPALEDLLLGSEAN[+1540.529]LTC[+57.021]TLTGLR.D | HexNAc(2)Hex(7) | C[+57], N[+1541] | 1126.7904 | 4 | 4504.14 | 4504.1268 | 0 | 2.9 | 127 | Specific | 906.19 | 823.47 | 823.47 | 18.23 | 51 | >sp\|P01876\|IGHA1_HUMAN Immunoglobulin heavy constant alpha 1 OS=Homo sapiens OX=9606 GN=IGHA1 PE=1 SV=2 | 255 | 20201117_ly_milk_1ng_glyco_hcd203040_compound15.21247.21247.4 | controllerType=0 controllerNumber=1 scan=21247 | 42.4684 |
| 17978:4 | 11 | R.LSLHRPALEDLLLGSEAN[+1257.449]LTC[+57.021]TLTGLR.D | HexNAc(3)Hex(4) | C[+57], N[+1257] | 1056.0181 | 4 | 4221.0506 | 4221.0477 | 0 | 0.7 | 127 | Specific | 931.09 | 862.96 | 862.96 | 17.68 | 51 | >sp\|P01876\|IGHA1_HUMAN Immunoglobulin heavy constant alpha 1 OS=Homo sapiens OX=9606 GN=IGHA1 PE=1 SV=2 | 255 | 20201117_ly_milk_1ng_glyco_hcd203040_compound15.21359.21359.4 | controllerType=0 controllerNumber=1 scan=21359 | 42.7484 |
| 17825:4 | 11 | R.LSLHRPALEDLLLGSEAN[+1864.634]LTC[+57.021]TLTGLR.D | HexNAc(2)Hex(9) | C[+57], N[+1865] | 1207.8166 | 4 | 4828.2445 | 4828.2324 | 0 | 2.5 | 127 | Specific | 936.98 | 936.98 | 936.98 | 18.23 | 51 | >sp\|P01876\|IGHA1_HUMAN Immunoglobulin heavy constant alpha 1 OS=Homo sapiens OX=9606 GN=IGHA1 PE=1 SV=2 | 255 | 20201117_ly_milk_1ng_glyco_hcd203040_compound15.21199.21199.4 | controllerType=0 controllerNumber=1 scan=21199 | 42.3427 |
| 18013:4 | 11 | R.LSLHRPALEDLLLGSEAN[+1095.397]LTC[+57.021]TLTGLR.D | HexNAc(3)Hex(3) | C[+57], N[+1095] | 1015.5081 | 4 | 4059.0106 | 4058.9949 | 0 | 3.9 | 127 | Specific | 945.6 | 945.6 | 945.6 | 17.96 | 51 | >sp\|P01876\|IGHA1_HUMAN Immunoglobulin heavy constant alpha 1 OS=Homo sapiens OX=9606 GN=IGHA1 PE=1 SV=2 | 255 | 20201117_ly_milk_1ng_glyco_hcd203040_compound15.21396.21396.4 | controllerType=0 controllerNumber=1 scan=21396 | 42.8418 |
| 17873:4 | 11 | R.LSLHRPALEDLLLGSEAN[+1540.529]LTC[+57.021]TLTGLR.D | HexNAc(2)Hex(7) | C[+57], N[+1541] | 1126.7904 | 4 | 4504.14 | 4504.1268 | 0 | 2.9 | 127 | Specific | 990.04 | 926.18 | 926.18 | 19.23 | 51 | >sp\|P01876\|IGHA1_HUMAN Immunoglobulin heavy constant alpha 1 OS=Homo sapiens OX=9606 GN=IGHA1 PE=1 SV=2 | 255 | 20201117_ly_milk_1ng_glyco_hcd203040_compound15.21249.21249.4 | controllerType=0 controllerNumber=1 scan=21249 | 42.4779 |
| 17827:4 | 11 | R.LSLHRPALEDLLLGSEAN[+1864.634]LTC[+57.021]TLTGLR.D | HexNAc(2)Hex(9) | C[+57], N[+1865] | 1207.8166 | 4 | 4828.2445 | 4828.2324 | 0 | 2.5 | 127 | Specific | 992.63 | 992.63 | 992.63 | 19.23 | 51 | >sp\|P01876\|IGHA1_HUMAN Immunoglobulin heavy constant alpha 1 OS=Homo sapiens OX=9606 GN=IGHA1 PE=1 SV=2 | 255 | 20201117_ly_milk_1ng_glyco_hcd203040_compound15.21201.21201.4 | controllerType=0 controllerNumber=1 scan=21201 | 42.3492 |
| 17968:4 | 11 | R.LSLHRPALEDLLLGSEAN[+1216.423]LTC[+57.021]TLTGLR.D | HexNAc(2)Hex(5) | C[+57], N[+1216] | 1045.762 | 4 | 4180.0262 | 4180.0211 | 0 | 1.2 | 127 | Specific | 1000.41 | 827.03 | 827.03 | 20.17 | 51 | >sp\|P01876\|IGHA1_HUMAN Immunoglobulin heavy constant alpha 1 OS=Homo sapiens OX=9606 GN=IGHA1 PE=1 SV=2 | 255 | 20201117_ly_milk_1ng_glyco_hcd203040_compound15.21349.21349.4 | controllerType=0 controllerNumber=1 scan=21349 | [illegible] |
| 17864:4 | 11 | R.LSLHRPALEDLLLGSEAN[+1702.581]LTC[+57.021]TLTGLR.D | HexNAc(2)Hex(8) | C[+57], N[+1703] | 1167.3036 | 4 | 4666.1927 | 4666.1796 | 0 | 2.8 | 127 | Specific | 1023.2 | 1003.59 | 1003.59 | 20.23 | 51 | >sp\|P01876\|IGHA1_HUMAN Immunoglobulin heavy constant alpha 1 OS=Homo sapiens OX=9606 GN=IGHA1 PE=1 SV=2 | 255 | 20201117_ly_milk_1ng_glyco_hcd203040_compound15.21240.21240.4 | controllerType=0 controllerNumber=1 scan=21240 | 42.453 |
| 17814:4 | 11 | R.LSLHRPALEDLLLGSEAN[+1864.634]LTC[+57.021]TLTGLR.D | HexNAc(2)Hex(9) | C[+57], N[+1865] | 1207.8166 | 4 | 4828.2445 | 4828.2324 | 0 | 2.5 | 127 | Specific | 1038.58 | 977.61 | 977.61 | 20.23 | 51 | >sp\|P01876\|IGHA1_HUMAN Immunoglobulin heavy constant alpha 1 OS=Homo sapiens OX=9606 GN=IGHA1 PE=1 SV=2 | 255 | 20201117_ly_milk_1ng_glyco_hcd203040_compound15.21187.21187.4 | controllerType=0 controllerNumber=1 scan=21187 | 42.3191 |
| 17981:4 | 11 | R.LSLHRPALEDLLLGSEAN[+1257.449]LTC[+57.021]TLTGLR.D | HexNAc(3)Hex(4) | C[+57], N[+1257] | 1056.0181 | 4 | 4221.0506 | 4221.0477 | 0 | 0.7 | 127 | Specific | 1048.4 | 1048.4 | 1048.4 | 19.68 | 51 | >sp\|P01876\|IGHA1_HUMAN Immunoglobulin heavy constant alpha 1 OS=Homo sapiens OX=9606 GN=IGHA1 PE=1 SV=2 | 255 | 20201117_ly_milk_1ng_glyco_hcd203040_compound15.21362.21362.4 | controllerType=0 controllerNumber=1 scan=21362 | 42.7735 |
| 11588:3 | 11 | R.LAGKPTHVN[+1825.661]VSVVMAEVDGTC[+57.021].Y | HexNAc(5)Hex(5) | C[+57], N[+1826] | 1337.2552 | 3 | 4009.751 | 4009.7397 | 0 | 2.8 | 332 | CRagged | 254.21 | 211.22 | 5.14 | 4.74 | 51 | >sp\|P01876\|IGHA1_HUMAN Immunoglobulin heavy constant alpha 1 OS=Homo sapiens OX=9606 GN=IGHA1 PE=1 SV=2 | 255 | 20201117_ly_milk_1ng_glyco_hcd203040_compound15.14667.14667.3 | controllerType=0 controllerNumber=1 scan=14667 | 30.529 |
| 11585:3 | 11 | R.LAGKPTHVN[+1825.661]VSVVMAEVDGTC[+57.021].Y | HexNAc(5)Hex(5) | C[+57], N[+1826] | 1337.2552 | 3 | 4009.751 | 4009.7397 | 0 | 2.8 | 332 | CRagged | 362.16 | 246.97 | 95.26 | 6.71 | 51 | >sp\|P01876\|IGHA1_HUMAN Immunoglobulin heavy constant alpha 1 OS=Homo sapiens OX=9606 GN=IGHA1 PE=1 SV=2 | 255 | 20201117_ly_milk_1ng_glyco_hcd203040_compound15.14664.14664.3 | controllerType=0 controllerNumber=1 scan=14664 | 30.5244 |
| 13352:3 | 11 | R.LAGKPTHVN[+2117.777]VSVVMAEVDGTC[+57.021]Y.- | HexNAc(5)Hex(5)Fuc(2) | C[+57], N[+2118] | 1488.9734 | 3 | 4464.9058 | 4464.9189 | 0 | -2.9 | 332 | CRagged | 219.04 | 217.74 | 217.74 | 3.87 | 51 | >sp\|P01876\|IGHA1_HUMAN Immunoglobulin heavy constant alpha 1 OS=Homo sapiens OX=9606 GN=IGHA1 PE=1 SV=2 | 255 | 20201117_ly_milk_1ng_glyco_hcd203040_compound15.16517.16517.3 | controllerType=0 controllerNumber=1 scan=16517 | 33.6625 |
| 13354:3 | 11 | R.LAGKPTHVN[+2117.777]VSVVMAEVDGTC[+57.021]Y.- | HexNAc(5)Hex(5)Fuc(2) | C[+57], N[+2118] | 1488.9734 | 3 | 4464.9058 | 4464.9189 | 0 | -2.9 | 332 | CRagged | 248.87 | 194.48 | 194.48 | 3.99 | 51 | >sp\|P01876\|IGHA1_HUMAN Immunoglobulin heavy constant alpha 1 OS=Homo sapiens OX=9606 GN=IGHA1 PE=1 SV=2 | 255 | 20201117_ly_milk_1ng_glyco_hcd203040_compound15.16519.16519.3 | controllerType=0 controllerNumber=1 scan=16519 | 33.6659 |
| 15380:3 | 11 | R.LAGKPTHVN[+2553.910]VSVVMAEVDGTC[+57.021]Y.- | HexNAc(5)Hex(5)Fuc(1)NeuAc(2) | C[+57], N[+2554] | 1634.363 | 3 | 4901.0744 | 4901.0518 | 0 | 4.6 | 332 | CRagged | 273.06 | 264.78 | 264.78 | 6.48 | 51 | >sp\|P01876\|IGHA1_HUMAN Immunoglobulin heavy constant alpha 1 OS=Homo sapiens OX=9606 GN=IGHA1 PE=1 SV=2 | 255 | 20201117_ly_milk_1ng_glyco_hcd203040_compound15.18619.18619.3 | controllerType=0 controllerNumber=1 scan=18619 | 37.3746 |
| 11638:4 | 11 | R.LAGKPTHVN[+1809.666]VSVVMAEVDGTC[+57.021]Y.- | HexNAc(5)Hex(4)Fuc(1) | C[+57], N[+1810] | 1039.9587 | 4 | 4156.8128 | 4156.8081 | 0 | 1.1 | 332 | CRagged | 312.29 | 215.74 | 215.74 | 6.48 | 51 | >sp\|P01876\|IGHA1_HUMAN Immunoglobulin heavy constant alpha 1 OS=Homo sapiens OX=9606 GN=IGHA1 PE=1 SV=2 | 255 | 20201117_ly_milk_1ng_glyco_hcd203040_compound15.14719.14719.4 | controllerType=0 controllerNumber=1 scan=14719 | 30.6328 |
| 15378:3 | 11 | R.LAGKPTHVN[+2553.910]VSVVMAEVDGTC[+57.021]Y.- | HexNAc(5)Hex(5)Fuc(1)NeuAc(2) | C[+57], N[+2554] | 1634.363 | 3 | 4901.0744 | 4901.0518 | 0 | 4.6 | 332 | CRagged | 328.73 | 257.98 | 257.98 | 6.72 | 51 | >sp\|P01876\|IGHA1_HUMAN Immunoglobulin heavy constant alpha 1 OS=Homo sapiens OX=9606 GN=IGHA1 PE=1 SV=2 | 255 | 20201117_ly_milk_1ng_glyco_hcd203040_compound15.18617.18617.3 | controllerType=0 controllerNumber=1 scan=18617 | 37.3705 |
| 15375:4 | 11 | R.LAGKPTHVN[+2553.910]VSVVMAEVDGTC[+57.021]Y.- | HexNAc(5)Hex(5)Fuc(1)NeuAc(2) | C[+57], N[+2554] | 1226.0241 | 4 | 4901.0745 | 4901.0518 | 0 | 4.6 | 332 | CRagged | 370.52 | 324.52 | 324.52 | 6.77 | 51 | >sp\|P01876\|IGHA1_HUMAN Immunoglobulin heavy constant alpha 1 OS=Homo sapiens OX=9606 GN=IGHA1 PE=1 SV=2 | 255 | 20201117_ly_milk_1ng_glyco_hcd203040_compound15.18614.18614.4 | controllerType=0 controllerNumber=1 scan=18614 | 37.3656 |
| 11610:3 | 11 | R.LAGKPTHVN[+1663.608]VSVVMAEVDGTC[+57.021]Y.- | HexNAc(5)Hex(4) | C[+57], N[+1664] | 1337.5914 | 3 | 4010.7595 | 4010.7502 | 0 | 2.3 | 332 | CRagged | 411.73 | 322.81 | 304.96 | 7.99 | 51 | >sp\|P01876\|IGHA1_HUMAN Immunoglobulin heavy constant alpha 1 OS=Homo sapiens OX=9606 GN=IGHA1 PE=1 SV=2 | 255 | 20201117_ly_milk_1ng_glyco_hcd203040_compound15.14690.14690.3 | controllerType=0 controllerNumber=1 scan=14690 | 30.5734 |
| 15373:4 | 11 | R.LAGKPTHVN[+2553.910]VSVVMAEVDGTC[+57.021]Y.- | HexNAc(5)Hex(5)Fuc(1)NeuAc(2) | C[+57], N[+2554] | 1226.0241 | 4 | 4901.0745 | 4901.0518 | 0 | 4.6 | 332 | CRagged | 417.28 | 309.32 | 309.32 | 6.78 | 51 | >sp\|P01876\|IGHA1_HUMAN Immunoglobulin heavy constant alpha 1 OS=Homo sapiens OX=9606 GN=IGHA1 PE=1 SV=2 | 255 | 20201117_ly_milk_1ng_glyco_hcd203040_compound15.18612.18612.4 | controllerType=0 controllerNumber=1 scan=18612 | 37.3618 |
| 11641:4 | 11 | R.LAGKPTHVN[+1809.666]VSVVMAEVDGTC[+57.021]Y.- | HexNAc(5)Hex(4)Fuc(1) | C[+57], N[+1810] | 1039.9587 | 4 | 4156.8128 | 4156.8081 | 0 | 1.1 | 332 | CRagged | 465.81 | 400.17 | 354.57 | 9.5 | 51 | >sp\|P01876\|IGHA1_HUMAN Immunoglobulin heavy constant alpha 1 OS=Homo sapiens OX=9606 GN=IGHA1 PE=1 SV=2 | 255 | 20201117_ly_milk_1ng_glyco_hcd203040_compound15.14722.14722.4 | controllerType=0 controllerNumber=1 scan=14722 | 30.6371 |
| 13373:3 | 11 | R.LAGKPTHVN[+2262.814]VSVVMAEVDGTC[+57.021]Y.- | HexNAc(5)Hex(5)Fuc(1)NeuAc(1) | C[+57], N[+2263] | 1537.3261 | 3 | 4609.9638 | 4609.9564 | 0 | 1.6 | 332 | CRagged | 579.58 | 507.41 | 507.41 | 11.36 | 51 | >sp\|P01876\|IGHA1_HUMAN Immunoglobulin heavy constant alpha 1 OS=Homo sapiens OX=9606 GN=IGHA1 PE=1 SV=2 | 255 | 20201117_ly_milk_1ng_glyco_hcd203040_compound15.16539.16539.3 | controllerType=0 controllerNumber=1 scan=16539 | 33.7001 |
| 11704:4 | 11 | R.LAGKPTHVN[+1647.613]VSVVMAEVDGTC[+57.021]Y.- | HexNAc(5)Hex(3)Fuc(1) | C[+57], N[+1648] | 999.4463 | 4 | 3994.7633 | 3994.7553 | 0 | 2 | 332 | CRagged | 624.35 | 566.79 | 566.79 | 12.26 | 51 | >sp\|P01876\|IGHA1_HUMAN Immunoglobulin heavy constant alpha 1 OS=Homo sapiens OX=9606 GN=IGHA1 PE=1 SV=2 | 255 | 20201117_ly_milk_1ng_glyco_hcd203040_compound15.14788.14788.4 | controllerType=0 controllerNumber=1 scan=14788 | 30.7824 |
| 13370:3 | 11 | R.LAGKPTHVN[+2262.814]VSVVMAEVDGTC[+57.021]Y.- | HexNAc(5)Hex(5)Fuc(1)NeuAc(1) | C[+57], N[+2263] | 1537.3261 | 3 | 4609.9638 | 4609.9564 | 0 | 1.6 | 332 | CRagged | 667.71 | 494.55 | 494.55 | 13.36 | 51 | >sp\|P01876\|IGHA1_HUMAN Immunoglobulin heavy constant alpha 1 OS=Homo sapiens OX=9606 GN=IGHA1 PE=1 SV=2 | 255 | 20201117_ly_milk_1ng_glyco_hcd203040_compound15.16536.16536.3 | controllerType=0 controllerNumber=1 scan=16536 | 33.6945 |
| 11613:3 | 11 | R.LAGKPTHVN[+1663.608]VSVVMAEVDGTC[+57.021]Y.- | HexNAc(5)Hex(4) | C[+57], N[+1664] | 1337.5914 | 3 | 4010.7595 | 4010.7502 | 0 | 2.3 | 332 | CRagged | 706.76 | 706.76 | 441.48 | 14.42 | 51 | >sp\|P01876\|IGHA1_HUMAN Immunoglobulin heavy constant alpha 1 OS=Homo sapiens OX=9606 GN=IGHA1 PE=1 SV=2 | 255 | 20201117_ly_milk_1ng_glyco_hcd203040_compound15.14693.14693.3 | controllerType=0 controllerNumber=1 scan=14693 | 30.5776 |
| 11702:4 | 11 | R.LAGKPTHVN[+1647.613]VSVVMAEVDGTC[+57.021]Y.- | HexNAc(5)Hex(3)Fuc(1) | C[+57], N[+1648] | 999.4463 | 4 | 3994.7633 | 3994.7553 | 0 | 2 | 332 | CRagged | 713.18 | 632.62 | 632.62 | 14.26 | 51 | >sp\|P01876\|IGHA1_HUMAN Immunoglobulin heavy constant alpha 1 OS=Homo sapiens OX=9606 GN=IGHA1 PE=1 SV=2 | 255 | 20201117_ly_milk_1ng_glyco_hcd203040_compound15.14786.14786.4 | controllerType=0 controllerNumber=1 scan=14786 | 30.7772 |
| 11602:3 | 11 | R.LAGKPTHVN[+1378.476]VSVVMAEVDGTC[+57.021]Y.- | HexNAc(2)Hex(6) | C[+57], N[+1378] | 1242.5472 | 3 | 3725.627 | 3725.6178 | 0 | 2.5 | 332 | CRagged | 735.85 | 735.85 | 424.74 | 14.42 | 51 | >sp\|P01876\|IGHA1_HUMAN Immunoglobulin heavy constant alpha 1 OS=Homo sapiens OX=9606 GN=IGHA1 PE=1 SV=2 | 255 | 20201117_ly_milk_1ng_glyco_hcd203040_compound15.14682.14682.3 | controllerType=0 controllerNumber=1 scan=14682 | 30.5591 |
| 11699:3 | 11 | R.LAGKPTHVN[+1647.613]VSVVMAEVDGTC[+57.021]Y.- | HexNAc(5)Hex(3)Fuc(1) | C[+57], N[+1648] | 1332.2595 | 3 | 3994.7638 | 3994.7553 | 0 | 2.1 | 332 | CRagged | 769.2 | 609.85 | 609.85 | 15.42 | 51 | >sp\|P01876\|IGHA1_HUMAN Immunoglobulin heavy constant alpha 1 OS=Homo sapiens OX=9606 GN=IGHA1 PE=1 SV=2 | 255 | 20201117_ly_milk_1ng_glyco_hcd203040_compound15.14783.14783.3 | controllerType=0 controllerNumber=1 scan=14783 | 30.7736 |
| 11697:3 | 11 | R.LAGKPTHVN[+1647.613]VSVVMAEVDGTC[+57.021]Y.- | HexNAc(5)Hex(3)Fuc(1) | C[+57], N[+1648] | 1332.2595 | 3 | 3994.7638 | 3994.7553 | 0 | 2.1 | 332 | CRagged | 771.04 | 771.04 | 771.04 | 15.42 | 51 | >sp\|P01876\|IGHA1_HUMAN Immunoglobulin heavy constant alpha 1 OS=Homo sapiens OX=9606 GN=IGHA1 PE=1 SV=2 | 255 | 20201117_ly_milk_1ng_glyco_hcd203040_compound15.14781.14781.3 | controllerType=0 controllerNumber=1 scan=14781 | 30.768 |
| 11667:3 | 11 | R.LAGKPTHVN[+1501.555]VSVVMAEVDGTC[+57.021]Y.- | HexNAc(5)Hex(3) | C[+57], N[+1502] | 1283.5734 | 3 | 3848.7057 | 3848.6974 | 0 | 2.2 | 332 | CRagged | 792.89 | 691.7 | 691.7 | 15.42 | 51 | >sp\|P01876\|IGHA1_HUMAN Immunoglobulin heavy constant alpha 1 OS=Homo sapiens OX=9606 GN=IGHA1 PE=1 SV=2 | 255 | 20201117_ly_milk_1ng_glyco_hcd203040_compound15.14749.14749.3 | controllerType=0 controllerNumber=1 scan=14749 | 30.6856 |
| 11598:3 | 11 | R.LAGKPTHVN[+1809.666]VSVVMAEVDGTC[+57.021]Y.- | HexNAc(5)Hex(4)Fuc(1) | C[+57], N[+1810] | 1386.2762 | 3 | 4156.814 | 4156.8081 | 0 | 1.4 | 332 | CRagged | 795.3 | 640.26 | 514.98 | 15.36 | 51 | >sp\|P01876\|IGHA1_HUMAN Immunoglobulin heavy constant alpha 1 OS=Homo sapiens OX=9606 GN=IGHA1 PE=1 SV=2 | 255 | 20201117_ly_milk_1ng_glyco_hcd203040_compound15.14677.14677.3 | controllerType=0 controllerNumber=1 scan=14677 | 30.5501 |
| 11525:3 | 11 | R.LAGKPTHVN[+1702.581]VSVVMAEVDGTC[+57.021]Y.- | HexNAc(2)Hex(8) | C[+57], N[+1703] | 1350.5825 | 3 | 4049.7328 | 4049.7234 | 0 | 2.3 | 332 | CRagged | 803.64 | 803.64 | 483.92 | 16.42 | 51 | >sp\|P01876\|IGHA1_HUMAN Immunoglobulin heavy constant alpha 1 OS=Homo sapiens OX=9606 GN=IGHA1 PE=1 SV=2 | 255 | 20201117_ly_milk_1ng_glyco_hcd203040_compound15.14602.14602.3 | controllerType=0 controllerNumber=1 scan=14602 | 30.4039 |
| 11527:3 | 11 | R.LAGKPTHVN[+1702.581]VSVVMAEVDGTC[+57.021]Y.- | HexNAc(2)Hex(8) | C[+57], N[+1703] | 1350.5825 | 3 | 4049.7328 | 4049.7234 | 0 | 2.3 | 332 | CRagged | 806.97 | 781.82 | 512.27 | 16.42 | 51 | >sp\|P01876\|IGHA1_HUMAN Immunoglobulin heavy constant alpha 1 OS=Homo sapiens OX=9606 GN=IGHA1 PE=1 SV=2 | 255 | 20201117_ly_milk_1ng_glyco_hcd203040_compound15.14604.14604.3 | controllerType=0 controllerNumber=1 scan=14604 | 30.4091 |
| 11669:3 | 11 | R.LAGKPTHVN[+1501.555]VSVVMAEVDGTC[+57.021]Y.- | HexNAc(5)Hex(3) | C[+57], N[+1502] | 1283.5734 | 3 | 3848.7057 | 3848.6974 | 0 | 2.2 | 332 | CRagged | 860.87 | 737.49 | 737.49 | 17.42 | 51 | >sp\|P01876\|IGHA1_HUMAN Immunoglobulin heavy constant alpha 1 OS=Homo sapiens OX=9606 GN=IGHA1 PE=1 SV=2 | 255 | 20201117_ly_milk_1ng_glyco_hcd203040_compound15.14751.14751.3 | controllerType=0 controllerNumber=1 scan=14751 | 30.7099 |
| 11605:3 | 11 | R.LAGKPTHVN[+1378.476]VSVVMAEVDGTC[+57.021]Y.- | HexNAc(2)Hex(6) | C[+57], N[+1378] | 1242.5472 | 3 | 3725.627 | 3725.6178 | 0 | 2.5 | 332 | CRagged | 861.45 | 816.56 | 525.56 | 17.42 | 51 | >sp\|P01876\|IGHA1_HUMAN Immunoglobulin heavy constant alpha 1 OS=Homo sapiens OX=9606 GN=IGHA1 PE=1 SV=2 | 255 | 20201117_ly_milk_1ng_glyco_hcd203040_compound15.14685.14685.3 | controllerType=0 controllerNumber=1 scan=14685 | 30.5677 |
| 11629:3 | 11 | R.LAGKPTHVN[+1216.423]VSVVMAEVDGTC[+57.021]Y.- | HexNAc(2)Hex(5) | C[+57], N[+1216] | 1188.5286 | 3 | 3563.5713 | 3563.5649 | 0 | 1.8 | 332 | CRagged | 882.99 | 882.99 | 542.39 | 17.36 | 51 | >sp\|P01876\|IGHA1_HUMAN Immunoglobulin heavy constant alpha 1 OS=Homo sapiens OX=9606 GN=IGHA1 PE=1 SV=2 | 255 | 20201117_ly_milk_1ng_glyco_hcd203040_compound15.14710.14710.3 | controllerType=0 controllerNumber=1 scan=14710 | 30.6121 |
| 11631:3 | 11 | R.LAGKPTHVN[+1216.423]VSVVMAEVDGTC[+57.021]Y.- | HexNAc(2)Hex(5) | C[+57], N[+1216] | 1188.5286 | 3 | 3563.5713 | 3563.5649 | 0 | 1.8 | 332 | CRagged | 898.98 | 839.41 | 548.19 | 17.36 | 51 | >sp\|P01876\|IGHA1_HUMAN Immunoglobulin heavy constant alpha 1 OS=Homo sapiens OX=9606 GN=IGHA1 PE=1 SV=2 | 255 | 20201117_ly_milk_1ng_glyco_hcd203040_compound15.14712.14712.3 | controllerType=0 controllerNumber=1 scan=14712 | 30.6199 |
| 11597:3 | 11 | R.LAGKPTHVN[+1809.666]VSVVMAEVDGTC[+57.021]Y.- | HexNAc(5)Hex(4)Fuc(1) | C[+57], N[+1810] | 1386.2762 | 3 | 4156.814 | 4156.8081 | 0 | 1.4 | 332 | CRagged | 913.03 | 913.03 | 606.69 | 18.36 | 51 | >sp\|P01876\|IGHA1_HUMAN Immunoglobulin heavy constant alpha 1 OS=Homo sapiens OX=9606 GN=IGHA1 PE=1 SV=2 | 255 | 20201117_ly_milk_1ng_glyco_hcd203040_compound15.14676.14676.3 | controllerType=0 controllerNumber=1 scan=14676 | 30.5428 |
| 11615:3 | 11 | R.LAGKPTHVN[+1809.666]VSVVMAEVDGTC[+57.021]Y.- | HexNAc(5)Hex(4)Fuc(1) | C[+57], N[+1810] | 1386.2767 | 3 | 4156.8155 | 4156.8081 | 0 | 1.8 | 332 | CRagged | 929.56 | 929.56 | 606.86 | 18.36 | 51 | >sp\|P01876\|IGHA1_HUMAN Immunoglobulin heavy constant alpha 1 OS=Homo sapiens OX=9606 GN=IGHA1 PE=1 SV=2 | 255 | 20201117_ly_milk_1ng_glyco_hcd203040_compound15.14695.14695.3 | controllerType=0 controllerNumber=1 scan=14695 | 30.5816 |
| 07927:3 | 12 | R.LSPN[+2350.830]ASAEHLELR.W | HexNAc(4)Hex(5)Fuc(1)NeuAc(2) | N[+2351] | 1263.2007 | 3 | 3787.5877 | 3787.5796 | 0 | 2.1 | 52 | Specific | 598.28 | 538.39 | 538.39 | 11.08 | 32 | >sp\|Q13410\|BT1A1_HUMAN Butyrophilin subfamily 1 member A1 OS=Homo sapiens OX=9606 GN=BTN1A1 PE=1 SV=3 | 179 | 20201117_ly_milk_1ng_glyco_hcd203040_compound15.10861.10861.3 | controllerType=0 controllerNumber=1 scan=10861 | 24.0987 |
| 07929:3 | 12 | R.LSPN[+2350.830]ASAEHLELR.W | HexNAc(4)Hex(5)Fuc(1)NeuAc(2) | N[+2351] | 1263.2007 | 3 | 3787.5877 | 3787.5796 | 0 | 2.1 | 52 | Specific | 652.09 | 580.23 | 580.23 | 13.08 | 32 | >sp\|Q13410\|BT1A1_HUMAN Butyrophilin subfamily 1 member A1 OS=Homo sapiens OX=9606 GN=BTN1A1 PE=1 SV=3 | 179 | 20201117_ly_milk_1ng_glyco_hcd203040_compound15.10863.10863.3 | controllerType=0 controllerNumber=1 scan=10863 | 24.103 |
| 17835:3 | 12 | K.N[+2059.735]VSC[+57.021]YIQNLLLGQEK.K | HexNAc(4)Hex(5)Fuc(1)NeuAc(1) | C[+57], N[+2060] | 1280.2244 | 3 | 3838.6587 | 3838.6455 | 0 | 3.4 | 215 | Specific | 237.61 | 171.08 | 171.08 | 5.16 | 32 | >sp\|Q13410\|BT1A1_HUMAN Butyrophilin subfamily 1 member A1 OS=Homo sapiens OX=9606 GN=BTN1A1 PE=1 SV=3 | 179 | 20201117_ly_milk_1ng_glyco_hcd203040_compound15.21209.21209.3 | controllerType=0 controllerNumber=1 scan=21209 | 42.3654 |
| 17837:3 | 12 | K.N[+2059.735]VSC[+57.021]YIQNLLLGQEK.K | HexNAc(4)Hex(5)Fuc(1)NeuAc(1) | C[+57], N[+2060] | 1280.2244 | 3 | 3838.6587 | 3838.6455 | 0 | 3.4 | 215 | Specific | 284.07 | 215.3 | 215.3 | 4.64 | 32 | >sp\|Q13410\|BT1A1_HUMAN Butyrophilin subfamily 1 member A1 OS=Homo sapiens OX=9606 GN=BTN1A1 PE=1 SV=3 | 179 | 20201117_ly_milk_1ng_glyco_hcd203040_compound15.21211.21211.3 | controllerType=0 controllerNumber=1 scan=21211 | 42.3715 |
| 17889:3 | 12 | K.N[+1913.677]VSC[+57.021]YIQNLLLGQEK.K | HexNAc(4)Hex(5)NeuAc(1) | C[+57], N[+1914] | 1231.5368 | 3 | 3692.5958 | 3692.5876 | 0 | 2.2 | 215 | Specific | 596.35 | 575.58 | 575.58 | 10.98 | 32 | >sp\|Q13410\|BT1A1_HUMAN Butyrophilin subfamily 1 member A1 OS=Homo sapiens OX=9606 GN=BTN1A1 PE=1 SV=3 | 179 | 20201117_ly_milk_1ng_glyco_hcd203040_compound15.21266.21266.3 | controllerType=0 controllerNumber=1 scan=21266 | 42.5264 |
| 17892:3 | 12 | K.N[+1913.677]VSC[+57.021]YIQNLLLGQEK.K | HexNAc(4)Hex(5)NeuAc(1) | C[+57], N[+1914] | 1231.5368 | 3 | 3692.5958 | 3692.5876 | 0 | 2.2 | 215 | Specific | 651.86 | 462.43 | 462.43 | 12.98 | 32 | >sp\|Q13410\|BT1A1_HUMAN Butyrophilin subfamily 1 member A1 OS=Homo sapiens OX=9606 GN=BTN1A1 PE=1 SV=3 | 179 | 20201117_ly_milk_1ng_glyco_hcd203040_compound15.21269.21269.3 | controllerType=0 controllerNumber=1 scan=21269 | 42.5311 |
| 24339:4 | 13 | R.[+42.011]QLAHQSN[+2190.793]STNIFFSPVSIATAFAMLSLGTK.A | HexNAc(6)Hex(6) | .[42], N[+2191] | 1354.3597 | 4 | 5414.4171 | 5414.4388 | 0 | -4 | 64 | Specific | 210.49 | 210.49 | 210.49 | 3.84 | 33 | >sp\|P01009\|A1AT_HUMAN Alpha-1-antitrypsin OS=Homo sapiens OX=9606 GN=SERPINA1 PE=1 SV=3 | 642 | 20201117_ly_milk_1ng_glyco_hcd203040_compound15.27998.27998.4 | controllerType=0 controllerNumber=1 scan=27998 | 56.0369 |
| 27196:4 | 13 | R.QLAHQSN[+2861.000]STNIFFSPVSIATAFAMLSLGTK.A | HexNAc(5)Hex(6)NeuAc(3) | N[+2861] | 1511.4199 | 4 | 6042.6579 | 6042.6351 | 0 | 3.8 | 64 | Specific | 230.49 | 230.49 | 230.49 | 6.24 | 33 | >sp\|P01009\|A1AT_HUMAN Alpha-1-antitrypsin OS=Homo sapiens OX=9606 GN=SERPINA1 PE=1 SV=3 | 642 | 20201117_ly_milk_1ng_glyco_hcd203040_compound15.30966.30966.4 | controllerType=0 controllerNumber=1 scan=30966 | 61.3868 |
| 24285:4 | 13 | R.QLAHQSN[+2350.830]STNIFFSPVSIATAFAMLSLGTK.A | HexNAc(4)Hex(5)Fuc(1)NeuAc(2) | N[+2351] | 1383.8765 | 4 | 5532.484 | 5532.4654 | 0 | 3.4 | 64 | Specific | 234.63 | 169.44 | 169.44 | 5.92 | 33 | >sp\|P01009\|A1AT_HUMAN Alpha-1-antitrypsin OS=Homo sapiens OX=9606 GN=SERPINA1 PE=1 SV=3 | 642 | 20201117_ly_milk_1ng_glyco_hcd203040_compound15.27941.27941.2 | controllerType=0 controllerNumber=1 scan=27941 | 55.9219 |
| 24341:4 | 13 | R.[+42.011]QLAHQSN[+2190.793]STNIFFSPVSIATAFAMLSLGTK.A | HexNAc(6)Hex(6) | .[42], N[+2191] | 1354.3597 | 4 | 5414.4171 | 5414.4388 | 0 | -4 | 64 | Specific | 298.72 | 298.72 | 298.72 | 5.62 | 33 | >sp\|P01009\|A1AT_HUMAN Alpha-1-antitrypsin OS=Homo sapiens OX=9606 GN=SERPINA1 PE=1 SV=3 | 642 | 20201117_ly_milk_1ng_glyco_hcd203040_compound15.28000.28000.4 | controllerType=0 controllerNumber=1 scan=28000 | 56.0405 |
| 24240:3 | 13 | R.QLAHQSN[+2204.772]STNIFFSPVSIATAFAMLSLGTK.A | HexNAc(4)Hex(5)NeuAc(2) | N[+2205] | 1796.144 | 3 | 5386.4175 | 5386.4075 | 0 | 1.9 | 64 | Specific | 606.98 | 571.04 | 447.32 | 12.78 | 33 | >sp\|P01009\|A1AT_HUMAN Alpha-1-antitrypsin OS=Homo sapiens OX=9606 GN=SERPINA1 PE=1 SV=3 | 642 | 20201117_ly_milk_1ng_glyco_hcd203040_compound15.27894.27894.3 | controllerType=0 controllerNumber=1 scan=27894 | 55.8304 |
| 24242:3 | 13 | R.QLAHQSN[+2204.772]STNIFFSPVSIATAFAMLSLGTK.A | HexNAc(4)Hex(5)NeuAc(2) | N[+2205] | 1796.144 | 3 | 5386.4175 | 5386.4075 | 0 | 1.9 | 64 | Specific | 704.29 | 623.79 | 519.2 | 14.78 | 33 | >sp\|P01009\|A1AT_HUMAN Alpha-1-antitrypsin OS=Homo sapiens OX=9606 GN=SERPINA1 PE=1 SV=3 | 642 | 20201117_ly_milk_1ng_glyco_hcd203040_compound15.27896.27896.3 | controllerType=0 controllerNumber=1 scan=27896 | 55.8359 |
| 22901:4 | 13 | K.ADTHDEILEGLNFN[+2861.000]LTEIPEAQIHEGFQELLR.T | HexNAc(5)Hex(6)NeuAc(3) | N[+2861] | 1638.9688 | 4 | 6552.8532 | 6552.8239 | 0 | 4.5 | 94 | Specific | 966.04 | 876.23 | 876.23 | 19.57 | 33 | >sp\|P01009\|A1AT_HUMAN Alpha-1-antitrypsin OS=Homo sapiens OX=9606 GN=SERPINA1 PE=1 SV=3 | 642 | 20201117_ly_milk_1ng_glyco_hcd203040_compound15.26502.26502.4 | controllerType=0 controllerNumber=1 scan=26502 | 53.1582 |
| 22903:4 | 13 | K.ADTHDEILEGLNFN[+2861.000]LTEIPEAQIHEGFQELLR.T | HexNAc(5)Hex(6)NeuAc(3) | N[+2861] | 1638.9688 | 4 | 6552.8532 | 6552.8239 | 0 | 4.5 | 94 | Specific | 1005.41 | 844.53 | 844.53 | 20.57 | 33 | >sp\|P01009\|A1AT_HUMAN Alpha-1-antitrypsin OS=Homo sapiens OX=9606 GN=SERPINA1 PE=1 SV=3 | 642 | 20201117_ly_milk_1ng_glyco_hcd203040_compound15.26504.26504.4 | controllerType=0 controllerNumber=1 scan=26504 | 53.1626 |
| 22198:4 | 18 | K.VFPLSLDSTPQDGNVVVAC[+57.021]LVQGFFPQEPLSVTWSESGQN[+1647.613]VTAI | HexNAc(5)Hex(3)Fuc(1) | C[+57], N[+1648] | 1606.7579 | 4 | 6424.0099 | 6423.9746 | 0 | 5.5 | 8 | Specific | 209.56 | 209.56 | 209.56 | 3.85 | 31 | >sp\|P01877\|IGHA2_HUMAN Immunoglobulin heavy constant alpha 2 OS=Homo sapiens OX=9606 GN=IGHA2 PE=1 SV=4 | 576 | 20201117_ly_milk_1ng_glyco_hcd203040_compound15.25762.25762.4 | controllerType=0 controllerNumber=1 scan=25762 | 51.8603 |
| 22200:4 | 18 | K.VFPLSLDSTPQDGNVVVAC[+57.021]LVQGFFPQEPLSVTWSESGQN[+1647.613]VTAI | HexNAc(5)Hex(3)Fuc(1) | C[+57], N[+1648] | 1606.7579 | 4 | 6424.0099 | 6423.9746 | 0 | 5.5 | 8 | Specific | 356.92 | 356.92 | 356.92 | 7.5 | 31 | >sp\|P01877\|IGHA2_HUMAN Immunoglobulin heavy constant alpha 2 OS=Homo sapiens OX=9606 GN=IGHA2 PE=1 SV=4 | 576 | 20201117_ly_milk_1ng_glyco_hcd203040_compound15.25764.25764.4 | controllerType=0 controllerNumber=1 scan=25764 | 51.8644 |
| 22257:4 | 18 | K.VFPLSLDSTPQDGNVVVAC[+57.021]LVQGFFPQEPLSVTWSESGQN[+1647.613]VTAI | HexNAc(5)Hex(3)Fuc(1) | C[+57], N[+1648] | 1606.7533 | 4 | 6423.9914 | 6423.9746 | 0 | 2.6 | 8 | Specific | 578.89 | 578.89 | 578.89 | 11.99 | 31 | >sp\|P01877\|IGHA2_HUMAN Immunoglobulin heavy constant alpha 2 OS=Homo sapiens OX=9606 GN=IGHA2 PE=1 SV=4 | 576 | 20201117_ly_milk_1ng_glyco_hcd203040_compound15.25824.25824.4 | controllerType=0 controllerNumber=1 scan=25824 | 51.988 |
| 22260:4 | 18 | K.VFPLSLDSTPQDGNVVVAC[+57.021]LVQGFFPQEPLSVTWSESGQN[+1647.613]VTAI | HexNAc(5)Hex(3)Fuc(1) | C[+57], N[+1648] | 1606.7533 | 4 | 6423.9914 | 6423.9746 | 0 | 2.6 | 8 | Specific | 951.39 | 864.1 | 864.1 | 19.99 | 31 | >sp\|P01877\|IGHA2_HUMAN Immunoglobulin heavy constant alpha 2 OS=Homo sapiens OX=9606 GN=IGHA2 PE=1 SV=4 | 576 | 20201117_ly_milk_1ng_glyco_hcd203040_compound15.25827.25827.4 | controllerType=0 controllerNumber=1 scan=25827 | 51.9928 |
| 02337:3 | 18 | K.HYTN[+2262.814]SSQDVTVPC[+57.021]R.V | HexNAc(5)Hex(5)Fuc(1)NeuAc(1) | C[+57], N[+2263] | 1309.5298 | 3 | 3926.575 | 3926.5636 | 0 | 2.9 | 89 | Specific | 216.06 | 147.31 | 147.31 | 4.26 | 31 | >sp\|P01877\|IGHA2_HUMAN Immunoglobulin heavy constant alpha 2 OS=Homo sapiens OX=9606 GN=IGHA2 PE=1 SV=4 | 576 | 20201117_ly_milk_1ng_glyco_hcd203040_compound15.5072.5072.3 | controllerType=0 controllerNumber=1 scan=5072 | 14.4832 |
| 01458:3 | 18 | K.HYTN[+1971.719]SSQDVTVPC[+57.021]R.V | HexNAc(5)Hex(5)Fuc(1) | C[+57], N[+1972] | 1212.494 | 3 | 3635.4673 | 3635.4682 | 0 | -0.2 | 89 | Specific | 264.28 | 226.54 | 226.54 | 5.51 | 31 | >sp\|P01877\|IGHA2_HUMAN Immunoglobulin heavy constant alpha 2 OS=Homo sapiens OX=9606 GN=IGHA2 PE=1 SV=4 | 576 | 20201117_ly_milk_1ng_glyco_hcd203040_compound15.4158.4158.3 | controllerType=0 controllerNumber=1 scan=4158 | 12.8576 |
| 01456:3 | 18 | K.HYTN[+1971.719]SSQDVTVPC[+57.021]R.V | HexNAc(5)Hex(5)Fuc(1) | C[+57], N[+1972] | 1212.494 | 3 | 3635.4673 | 3635.4682 | 0 | -0.2 | 89 | Specific | 322.78 | 251.28 | 251.28 | 6.1 | 31 | >sp\|P01877\|IGHA2_HUMAN Immunoglobulin heavy constant alpha 2 OS=Homo sapiens OX=9606 GN=IGHA2 PE=1 SV=4 | 576 | 20201117_ly_milk_1ng_glyco_hcd203040_compound15.4156.4156.3 | controllerType=0 controllerNumber=1 scan=4156 | 12.8405 |
| 01491:3 | 18 | K.HYTN[+1647.613]SSQDVTVPC[+57.021]R.V | HexNAc(5)Hex(3)Fuc(1) | C[+57], N[+1648] | 1104.4618 | 3 | 3311.3709 | 3311.3625 | 0 | 2.5 | 89 | Specific | 630.29 | 405.54 | 405.54 | 12.08 | 31 | >sp\|P01877\|IGHA2_HUMAN Immunoglobulin heavy constant alpha 2 OS=Homo sapiens OX=9606 GN=IGHA2 PE=1 SV=4 | 576 | 20201117_ly_milk_1ng_glyco_hcd203040_compound15.4192.4192.2 | controllerType=0 controllerNumber=1 scan=4192 | 12.9262 |
| 01463:3 | 18 | K.HYTN[+1647.613]SSQDVTVPC[+57.021]R.V | HexNAc(5)Hex(3)Fuc(1) | C[+57], N[+1648] | 1104.4601 | 3 | 3311.3658 | 3311.3625 | 0 | 1 | 89 | Specific | 673.18 | 590.71 | 590.71 | 13.02 | 31 | >sp\|P01877\|IGHA2_HUMAN Immunoglobulin heavy constant alpha 2 OS=Homo sapiens OX=9606 GN=IGHA2 PE=1 SV=4 | 576 | 20201117_ly_milk_1ng_glyco_hcd203040_compound15.4163.4163.3 | controllerType=0 controllerNumber=1 scan=4163 | 12.8719 |
| 01461:3 | 18 | K.HYTN[+1647.613]SSQDVTVPC[+57.021]R.V | HexNAc(5)Hex(3)Fuc(1) | C[+57], N[+1648] | 1104.4601 | 3 | 3311.3658 | 3311.3625 | 0 | 1 | 89 | Specific | 690.49 | 609.69 | 609.69 | 13.02 | 31 | >sp\|P01877\|IGHA2_HUMAN Immunoglobulin heavy constant alpha 2 OS=Homo sapiens OX=9606 GN=IGHA2 PE=1 SV=4 | 576 | 20201117_ly_milk_1ng_glyco_hcd203040_compound15.4161.4161.3 | controllerType=0 controllerNumber=1 scan=4161 | 12.8625 |
| 01448:3 | 18 | K.HYTN[+1809.666]SSQDVTVPC[+57.021]R.V | HexNAc(5)Hex(4)Fuc(1) | C[+57], N[+1810] | 1158.4782 | 3 | 3473.4201 | 3473.4154 | 0 | 1.4 | 89 | Specific | 692.08 | 467.97 | 467.97 | 13.02 | 31 | >sp\|P01877\|IGHA2_HUMAN Immunoglobulin heavy constant alpha 2 OS=Homo sapiens OX=9606 GN=IGHA2 PE=1 SV=4 | 576 | 20201117_ly_milk_1ng_glyco_hcd203040_compound15.4148.4148.3 | controllerType=0 controllerNumber=1 scan=4148 | 12.8268 |
| 01450:3 | 18 | K.HYTN[+1809.666]SSQDVTVPC[+57.021]R.V | HexNAc(5)Hex(4)Fuc(1) | C[+57], N[+1810] | 1158.4782 | 3 | 3473.4201 | 3473.4154 | 0 | 1.4 | 89 | Specific | 708.02 | 476.36 | 476.36 | 14.02 | 31 | >sp\|P01877\|IGHA2_HUMAN Immunoglobulin heavy constant alpha 2 OS=Homo sapiens OX=9606 GN=IGHA2 PE=1 SV=4 | 576 | 20201117_ly_milk_1ng_glyco_hcd203040_compound15.4150.4150.3 | controllerType=0 controllerNumber=1 scan=4150 | 12.8312 |
| 01489:3 | 18 | K.HYTN[+1647.613]SSQDVTVPC[+57.021]R.V | HexNAc(5)Hex(3)Fuc(1) | C[+57], N[+1648] | 1104.4618 | 3 | 3311.3709 | 3311.3625 | 0 | 2.5 | 89 | Specific | 720.92 | 530.44 | 530.44 | 14.08 | 31 | >sp\|P01877\|IGHA2_HUMAN Immunoglobulin heavy constant alpha 2 OS=Homo sapiens OX=9606 GN=IGHA2 PE=1 SV=4 | 576 | 20201117_ly_milk_1ng_glyco_hcd203040_compound15.4190.4190.2 | controllerType=0 controllerNumber=1 scan=4190 | 12.917 |
| 03463:4 | 18 | K.HYTN[+1647.613]SSQDVTVPC[+57.021]RVPPPPPC[+57.021]C[+57.021]HPR.L | HexNAc(5)Hex(3)Fuc(1) | C[+57]*3, N[+1648] | 1152.2486 | 4 | 4605.9725 | 4605.9689 | 0 | 0.8 | 89 | Specific | 356.38 | 293.34 | 293.34 | 6.14 | 31 | >sp\|P01877\|IGHA2_HUMAN Immunoglobulin heavy constant alpha 2 OS=Homo sapiens OX=9606 GN=IGHA2 PE=1 SV=4 | 576 | 20201117_ly_milk_1ng_glyco_hcd203040_compound15.6240.6240.4 | controllerType=0 controllerNumber=1 scan=6240 | 16.532 |
| 03461:4 | 18 | K.HYTN[+1647.613]SSQDVTVPC[+57.021]RVPPPPPC[+57.021]C[+57.021]HPR.L | HexNAc(5)Hex(3)Fuc(1) | C[+57]*3, N[+1648] | 1152.2486 | 4 | 4605.9725 | 4605.9689 | 0 | 0.8 | 89 | Specific | 358.22 | 357.47 | 357.47 | 6.5 | 31 | >sp\|P01877\|IGHA2_HUMAN Immunoglobulin heavy constant alpha 2 OS=Homo sapiens OX=9606 GN=IGHA2 PE=1 SV=4 | 576 | 20201117_ly_milk_1ng_glyco_hcd203040_compound15.6238.6238.4 | controllerType=0 controllerNumber=1 scan=6238 | 16.5273 |
| 03444:5 | 18 | K.HYTN[+1647.613]SSQDVTVPC[+57.021]RVPPPPPC[+57.021]C[+57.021]HPR.L | HexNAc(5)Hex(3)Fuc(1) | C[+57]*3, N[+1648] | 922.0015 | 5 | 4605.9786 | 4605.9689 | 0 | 2.1 | 89 | Specific | 447.19 | 223.03 | 223.03 | 6.59 | 31 | >sp\|P01877\|IGHA2_HUMAN Immunoglobulin heavy constant alpha 2 OS=Homo sapiens OX=9606 GN=IGHA2 PE=1 SV=4 | 576 | 20201117_ly_milk_1ng_glyco_hcd203040_compound15.6220.6220.5 | controllerType=0 controllerNumber=1 scan=6220 | 16.4901 |
| 03446:5 | 18 | K.HYTN[+1647.613]SSQDVTVPC[+57.021]RVPPPPPC[+57.021]C[+57.021]HPR.L | HexNAc(5)Hex(3)Fuc(1) | C[+57]*3, N[+1648] | 922.0015 | 5 | 4605.9786 | 4605.9689 | 0 | 2.1 | 89 | Specific | 510.4 | 441.01 | 441.01 | 9.33 | 31 | >sp\|P01877\|IGHA2_HUMAN Immunoglobulin heavy constant alpha 2 OS=Homo sapiens OX=9606 GN=IGHA2 PE=1 SV=4 | 576 | 20201117_ly_milk_1ng_glyco_hcd203040_compound15.6222.6222.5 | controllerType=0 controllerNumber=1 scan=6222 | 16.4967 |
| 04964:3 | 18 | K.TPLTAN[+2262.814]ITK.S | HexNAc(5)Hex(5)Fuc(1)NeuAc(1) | N[+2263] | 1074.4649 | 3 | 3221.3801 | 3221.3711 | 0 | 2.8 | 200 | Specific | 218.67 | 216.01 | 216.01 | 4.39 | 31 | >sp\|P01877\|IGHA2_HUMAN Immunoglobulin heavy constant alpha 2 OS=Homo sapiens OX=9606 GN=IGHA2 PE=1 SV=4 | 576 | 20201117_ly_milk_1ng_glyco_hcd203040_compound15.7795.7795.3 | controllerType=0 controllerNumber=1 scan=7795 | 19.0991 |
| 03057:3 | 18 | K.TPLTAN[+1971.719]ITK.S | HexNAc(5)Hex(5)Fuc(1) | N[+1972] | 977.4333 | 3 | 2930.2852 | 2930.2757 | 0 | 3.3 | 200 | Specific | 252.88 | 252.82 | 252.82 | 5.86 | 31 | >sp\|P01877\|IGHA2_HUMAN Immunoglobulin heavy constant alpha 2 OS=Homo sapiens OX=9606 GN=IGHA2 PE=1 SV=4 | 576 | 20201117_ly_milk_1ng_glyco_hcd203040_compound15.5819.5819.3 | controllerType=0 controllerNumber=1 scan=5819 | 15.7744 |
| 03147:2 | 18 | K.TPLTAN[+1444.534]ITK.S | HexNAc(4)Hex(3)Fuc(1) | N[+1445] | 1202.053 | 2 | 2403.0988 | 2403.0906 | 0 | 3.4 | 200 | Specific | 367.38 | 201.31 | 201.31 | 6.02 | 31 | >sp\|P01877\|IGHA2_HUMAN Immunoglobulin heavy constant alpha 2 OS=Homo sapiens OX=9606 GN=IGHA2 PE=1 SV=4 | 576 | 20201117_ly_milk_1ng_glyco_hcd203040_compound15.5912.5912.2 | controllerType=0 controllerNumber=1 scan=5912 | 15.9451 |
| 03132:3 | 18 | K.TPLTAN[+1809.666]ITK.S | HexNAc(5)Hex(4)Fuc(1) | N[+1810] | 923.4165 | 3 | 2768.235 | 2768.2228 | 0 | 4.4 | 200 | Specific | 405.61 | 148.96 | 148.96 | 5.64 | 31 | >sp\|P01877\|IGHA2_HUMAN Immunoglobulin heavy constant alpha 2 OS=Homo sapiens OX=9606 GN=IGHA2 PE=1 SV=4 | 576 | 20201117_ly_milk_1ng_glyco_hcd203040_compound15.5897.5897.3 | controllerType=0 controllerNumber=1 scan=5897 | 15.9176 |
| 03149:2 | 18 | K.TPLTAN[+1444.534]ITK.S | HexNAc(4)Hex(3)Fuc(1) | N[+1445] | 1202.053 | 2 | 2403.0988 | 2403.0906 | 0 | 3.4 | 200 | Specific | 437.7 | 246.03 | 246.03 | 6.61 | 31 | >sp\|P01877\|IGHA2_HUMAN Immunoglobulin heavy constant alpha 2 OS=Homo sapiens OX=9606 GN=IGHA2 PE=1 SV=4 | 576 | 20201117_ly_milk_1ng_glyco_hcd203040_compound15.5914.5914.2 | controllerType=0 controllerNumber=1 scan=5914 | 15.9492 |

Table S4. Cont.

| Query #:z | Protein Rank | Glycopeptide | Glycan | Modification Type(s) | Observed m/z | z | Observed (M+H) | Calc. mass (M+H) | Off-by-x error | Mass error (ppm) | StartPosition | Cleavage | Score | Delta | Delta Mod | \|Log Prob\| | # of unique peptides | Protein Name | Protein DB number | Comment | Scan # | Scan Time |
|---|---|---|---|---|---|---|---|---|---|---|---|---|---|---|---|---|---|---|---|---|---|---|
| 03195:3 | 18 | K.TPLTAN[+1647.613]ITK.S | HexNAc(5)Hex(3)Fuc(1) | N[+1648] | 869.3974 | 3 | 2606.1777 | 2606.17 | 0 | 3 | 200 | Specific | 461 | 173.46 | 173.46 | 6.2 | 31 | >sp\|P01877\|IGHA2_HUMAN Immunoglobulin heavy constant alpha 2 OS=Homo sapiens OX=9606 GN=IGHA2 PE=1 SV=4 | 576 | 20201117_ly_milk_1ng_glyco_hcd203040_compound15.5962.5962.3 | controllerType=0 controllerNumber=1 scan=5962 | 16.0381 |
| 03134:3 | 18 | K.TPLTAN[+1809.666]ITK.S | HexNAc(5)Hex(4)Fuc(1) | N[+1810] | 923.4165 | 3 | 2768.235 | 2768.2228 | 0 | 4.4 | 200 | Specific | 503.65 | 262.45 | 262.45 | 6.87 | 31 | >sp\|P01877\|IGHA2_HUMAN Immunoglobulin heavy constant alpha 2 OS=Homo sapiens OX=9606 GN=IGHA2 PE=1 SV=4 | 576 | 20201117_ly_milk_1ng_glyco_hcd203040_compound15.5899.5899.3 | controllerType=0 controllerNumber=1 scan=5899 | 15.9219 |
| 03138:2 | 18 | K.TPLTAN[+1809.666]ITK.S | HexNAc(5)Hex(4)Fuc(1) | N[+1810] | 1384.6209 | 2 | 2768.2345 | 2768.2228 | 0 | 4.2 | 200 | Specific | 525.05 | 257.48 | 257.48 | 7.04 | 31 | >sp\|P01877\|IGHA2_HUMAN Immunoglobulin heavy constant alpha 2 OS=Homo sapiens OX=9606 GN=IGHA2 PE=1 SV=4 | 576 | 20201117_ly_milk_1ng_glyco_hcd203040_compound15.5903.5903.2 | controllerType=0 controllerNumber=1 scan=5903 | 15.9293 |
| 03197:3 | 18 | K.TPLTAN[+1647.613]ITK.S | HexNAc(5)Hex(3)Fuc(1) | N[+1648] | 869.3974 | 3 | 2606.1777 | 2606.17 | 0 | 3 | 200 | Specific | 544.89 | 262.16 | 262.16 | 7.21 | 31 | >sp\|P01877\|IGHA2_HUMAN Immunoglobulin heavy constant alpha 2 OS=Homo sapiens OX=9606 GN=IGHA2 PE=1 SV=4 | 576 | 20201117_ly_milk_1ng_glyco_hcd203040_compound15.5964.5964.3 | controllerType=0 controllerNumber=1 scan=5964 | 16.0432 |
| 03136:2 | 18 | K.TPLTAN[+1809.666]ITK.S | HexNAc(5)Hex(4)Fuc(1) | N[+1810] | 1384.6209 | 2 | 2768.2345 | 2768.2228 | 0 | 4.2 | 200 | Specific | 573.48 | 256.62 | 256.62 | 8.46 | 31 | >sp\|P01877\|IGHA2_HUMAN Immunoglobulin heavy constant alpha 2 OS=Homo sapiens OX=9606 GN=IGHA2 PE=1 SV=4 | 576 | 20201117_ly_milk_1ng_glyco_hcd203040_compound15.5901.5901.2 | controllerType=0 controllerNumber=1 scan=5901 | 15.9252 |
| 03183:2 | 18 | K.TPLTAN[+1647.613]ITK.S | HexNAc(5)Hex(3)Fuc(1) | N[+1648] | 1303.5931 | 2 | 2606.1788 | 2606.17 | 0 | 3.4 | 200 | Specific | 580.98 | 317.68 | 317.68 | 8.73 | 31 | >sp\|P01877\|IGHA2_HUMAN Immunoglobulin heavy constant alpha 2 OS=Homo sapiens OX=9606 GN=IGHA2 PE=1 SV=4 | 576 | 20201117_ly_milk_1ng_glyco_hcd203040_compound15.5950.5950.2 | controllerType=0 controllerNumber=1 scan=5950 | 16.0181 |
| 03181:2 | 18 | K.TPLTAN[+1647.613]ITK.S | HexNAc(5)Hex(3)Fuc(1) | N[+1648] | 1303.5931 | 2 | 2606.1788 | 2606.17 | 0 | 3.4 | 200 | Specific | 617.46 | 344.82 | 344.82 | 10.02 | 31 | >sp\|P01877\|IGHA2_HUMAN Immunoglobulin heavy constant alpha 2 OS=Homo sapiens OX=9606 GN=IGHA2 PE=1 SV=4 | 576 | 20201117_ly_milk_1ng_glyco_hcd203040_compound15.5948.5948.2 | controllerType=0 controllerNumber=1 scan=5948 | 16.0119 |
| 10699:3 | 18 | R.MAGKPTHIN[+1647.613]VSVVMAEADGTC[+57.021]Y.- | HexNAc(5)Hex(3)Fuc(1) | C[+57], N[+1648] | 1333.571 | 3 | 3998.6984 | 3998.6961 | 0 | 0.6 | 319 | CRagged | 212.53 | 188.6 | 188.6 | 4.57 | 31 | >sp\|P01877\|IGHA2_HUMAN Immunoglobulin heavy constant alpha 2 OS=Homo sapiens OX=9606 GN=IGHA2 PE=1 SV=4 | 576 | 20201117_ly_milk_1ng_glyco_hcd203040_compound15.13748.13748.3 | controllerType=0 controllerNumber=1 scan=13748 | 28.9738 |
| 10551:3 | 18 | R.MAGKPTHIN[+1663.608]VSVVMAEADGTC[+57.021]Y.- | HexNAc(5)Hex(4) | C[+57], N[+1664] | 1338.905 | 3 | 4014.7003 | 4014.691 | 0 | 2.3 | 319 | CRagged | 330.26 | 330.26 | 260.96 | 7.56 | 31 | >sp\|P01877\|IGHA2_HUMAN Immunoglobulin heavy constant alpha 2 OS=Homo sapiens OX=9606 GN=IGHA2 PE=1 SV=4 | 576 | 20201117_ly_milk_1ng_glyco_hcd203040_compound15.13594.13594.3 | controllerType=0 controllerNumber=1 scan=13594 | 28.6941 |
| 10701:3 | 18 | R.MAGKPTHIN[+1647.613]VSVVMAEADGTC[+57.021]Y.- | HexNAc(5)Hex(3)Fuc(1) | C[+57], N[+1648] | 1333.571 | 3 | 3998.6984 | 3998.6961 | 0 | 0.6 | 319 | CRagged | 335.7 | 335.7 | 335.7 | 7.16 | 31 | >sp\|P01877\|IGHA2_HUMAN Immunoglobulin heavy constant alpha 2 OS=Homo sapiens OX=9606 GN=IGHA2 PE=1 SV=4 | 576 | 20201117_ly_milk_1ng_glyco_hcd203040_compound15.13750.13750.3 | controllerType=0 controllerNumber=1 scan=13750 | 28.9779 |
| 10553:3 | 18 | R.MAGKPTHIN[+1663.608]VSVVMAEADGTC[+57.021]Y.- | HexNAc(5)Hex(4) | C[+57], N[+1664] | 1338.905 | 3 | 4014.7003 | 4014.691 | 0 | 2.3 | 319 | CRagged | 351.02 | 351.02 | 325.97 | 7.78 | 31 | >sp\|P01877\|IGHA2_HUMAN Immunoglobulin heavy constant alpha 2 OS=Homo sapiens OX=9606 GN=IGHA2 PE=1 SV=4 | 576 | 20201117_ly_milk_1ng_glyco_hcd203040_compound15.13596.13596.3 | controllerType=0 controllerNumber=1 scan=13596 | 28.6981 |
| 10644:3 | 18 | R.MAGKPTHIN[+1501.555]VSVVMAEADGTC[+57.021]Y.- | HexNAc(5)Hex(3) | C[+57], N[+1502] | 1284.8881 | 3 | 3852.6498 | 3852.6382 | 0 | 3 | 319 | CRagged | 570.14 | 442.88 | 442.88 | 10.25 | 31 | >sp\|P01877\|IGHA2_HUMAN Immunoglobulin heavy constant alpha 2 OS=Homo sapiens OX=9606 GN=IGHA2 PE=1 SV=4 | 576 | 20201117_ly_milk_1ng_glyco_hcd203040_compound15.13691.13691.3 | controllerType=0 controllerNumber=1 scan=13691 | 28.8816 |
| 10646:3 | 18 | R.MAGKPTHIN[+1501.555]VSVVMAEADGTC[+57.021]Y.- | HexNAc(5)Hex(3) | C[+57], N[+1502] | 1284.8881 | 3 | 3852.6498 | 3852.6382 | 0 | 3 | 319 | CRagged | 607.29 | 592.95 | 592.95 | 12.42 | 31 | >sp\|P01877\|IGHA2_HUMAN Immunoglobulin heavy constant alpha 2 OS=Homo sapiens OX=9606 GN=IGHA2 PE=1 SV=4 | 576 | 20201117_ly_milk_1ng_glyco_hcd203040_compound15.13693.13693.3 | controllerType=0 controllerNumber=1 scan=13693 | 28.8855 |
| 10486:3 | 18 | R.MAGKPTHIN[+1540.529]VSVVMAEADGTC[+57.021]Y.- | HexNAc(2)Hex(7) | C[+57], N[+1541] | 1297.8788 | 3 | 3891.622 | 3891.6114 | 0 | 2.7 | 319 | CRagged | 760.63 | 659.05 | 440.13 | 15.42 | 31 | >sp\|P01877\|IGHA2_HUMAN Immunoglobulin heavy constant alpha 2 OS=Homo sapiens OX=9606 GN=IGHA2 PE=1 SV=4 | 576 | 20201117_ly_milk_1ng_glyco_hcd203040_compound15.13527.13527.3 | controllerType=0 controllerNumber=1 scan=13527 | 28.564 |
| 10589:3 | 18 | R.MAGKPTHIN[+1216.423]VSVVMAEADGTC[+57.021]Y.- | HexNAc(2)Hex(5) | C[+57], N[+1216] | 1189.8426 | 3 | 3567.5132 | 3567.5057 | 0 | 2.1 | 319 | CRagged | 804.69 | 799.93 | 415.93 | 16.42 | 31 | >sp\|P01877\|IGHA2_HUMAN Immunoglobulin heavy constant alpha 2 OS=Homo sapiens OX=9606 GN=IGHA2 PE=1 SV=4 | 576 | 20201117_ly_milk_1ng_glyco_hcd203040_compound15.13634.13634.3 | controllerType=0 controllerNumber=1 scan=13634 | 28.7731 |
| 10460:3 | 18 | R.MAGKPTHIN[+1702.581]VSVVMAEADGTC[+57.021]Y.- | HexNAc(2)Hex(8) | C[+57], N[+1703] | 1351.8968 | 3 | 4053.6758 | 4053.6642 | 0 | 2.9 | 319 | CRagged | 823.92 | 823.92 | 498.63 | 16.42 | 31 | >sp\|P01877\|IGHA2_HUMAN Immunoglobulin heavy constant alpha 2 OS=Homo sapiens OX=9606 GN=IGHA2 PE=1 SV=4 | 576 | 20201117_ly_milk_1ng_glyco_hcd203040_compound15.13500.13500.3 | controllerType=0 controllerNumber=1 scan=13500 | 28.5111 |
| 10462:3 | 18 | R.MAGKPTHIN[+1702.581]VSVVMAEADGTC[+57.021]Y.- | HexNAc(2)Hex(8) | C[+57], N[+1703] | 1351.8968 | 3 | 4053.6758 | 4053.6642 | 0 | 2.9 | 319 | CRagged | 831.15 | 684.58 | 460.8 | 16.42 | 31 | >sp\|P01877\|IGHA2_HUMAN Immunoglobulin heavy constant alpha 2 OS=Homo sapiens OX=9606 GN=IGHA2 PE=1 SV=4 | 576 | 20201117_ly_milk_1ng_glyco_hcd203040_compound15.13502.13502.3 | controllerType=0 controllerNumber=1 scan=13502 | 28.516 |
| 10488:3 | 18 | R.MAGKPTHIN[+1540.529]VSVVMAEADGTC[+57.021]Y.- | HexNAc(2)Hex(7) | C[+57], N[+1541] | 1297.8788 | 3 | 3891.622 | 3891.6114 | 0 | 2.7 | 319 | CRagged | 846.25 | 846.25 | 459.94 | 16.42 | 31 | >sp\|P01877\|IGHA2_HUMAN Immunoglobulin heavy constant alpha 2 OS=Homo sapiens OX=9606 GN=IGHA2 PE=1 SV=4 | 576 | 20201117_ly_milk_1ng_glyco_hcd203040_compound15.13529.13529.3 | controllerType=0 controllerNumber=1 scan=13529 | 28.569 |
| 10536:3 | 18 | R.MAGKPTHIN[+1378.476]VSVVMAEADGTC[+57.021]Y.- | HexNAc(2)Hex(6) | C[+57], N[+1378] | 1243.8602 | 3 | 3729.5659 | 3729.5585 | 0 | 2 | 319 | CRagged | 851.95 | 677.64 | 510.83 | 17.36 | 31 | >sp\|P01877\|IGHA2_HUMAN Immunoglobulin heavy constant alpha 2 OS=Homo sapiens OX=9606 GN=IGHA2 PE=1 SV=4 | 576 | 20201117_ly_milk_1ng_glyco_hcd203040_compound15.13579.13579.3 | controllerType=0 controllerNumber=1 scan=13579 | 28.6668 |
| 10591:3 | 18 | R.MAGKPTHIN[+1216.423]VSVVMAEADGTC[+57.021]Y.- | HexNAc(2)Hex(5) | C[+57], N[+1216] | 1189.8426 | 3 | 3567.5132 | 3567.5057 | 0 | 2.1 | 319 | CRagged | 902.47 | 902.47 | 502.69 | 18.42 | 31 | >sp\|P01877\|IGHA2_HUMAN Immunoglobulin heavy constant alpha 2 OS=Homo sapiens OX=9606 GN=IGHA2 PE=1 SV=4 | 576 | 20201117_ly_milk_1ng_glyco_hcd203040_compound15.13636.13636.3 | controllerType=0 controllerNumber=1 scan=13636 | 28.7789 |
| 10534:3 | 18 | R.MAGKPTHIN[+1378.476]VSVVMAEADGTC[+57.021]Y.- | HexNAc(2)Hex(6) | C[+57], N[+1378] | 1243.8602 | 3 | 3729.5659 | 3729.5585 | 0 | 2 | 319 | CRagged | 950.53 | 846.26 | 560.9 | 19.36 | 31 | >sp\|P01877\|IGHA2_HUMAN Immunoglobulin heavy constant alpha 2 OS=Homo sapiens OX=9606 GN=IGHA2 PE=1 SV=4 | 576 | 20201117_ly_milk_1ng_glyco_hcd203040_compound15.13577.13577.3 | controllerType=0 controllerNumber=1 scan=13577 | 28.6595 |
| 25270:4 | 19 | K.DIDGYGGIALPELIC[+57.021]TMFHTSGYDTQAIVENN[+2350.830]ESTEYGLFQISNK. | HexNAc(4)Hex(5)Fuc(1)NeuAc(2) | C[+57], N[+2351] | 1841.307 | 4 | 7362.2061 | 7362.1592 | 0 | 6.4 | 33 | Specific | 264.18 | 264.18 | 264.18 | 6.04 | 24 | >sp\|P00709\|LALBA_HUMAN Alpha-lactalbumin OS=Homo sapiens OX=9606 GN=LALBA PE=1 SV=1 | 468 | 20201117_ly_milk_1ng_glyco_hcd203040_compound15.28965.28965.4 | controllerType=0 controllerNumber=1 scan=28965 | 57.7763 |
| 22896:4 | 19 | K.DIDGYGGIALPELIC[+57.021]TMFHTSGYDTQAIVENN[+1686.586]ESTEYGLFQISNK. | HexNAc(2)Hex(7)Fuc(1) | C[+57], N[+1687] | 1675.2308 | 4 | 6697.9014 | 6697.9153 | 0 | -2.1 | 33 | Specific | 334.96 | 214.7 | 214.7 | 6.07 | 24 | >sp\|P00709\|LALBA_HUMAN Alpha-lactalbumin OS=Homo sapiens OX=9606 GN=LALBA PE=1 SV=1 | 468 | 20201117_ly_milk_1ng_glyco_hcd203040_compound15.26497.26497.4 | controllerType=0 controllerNumber=1 scan=26497 | 53.1481 |
| 25272:4 | 19 | K.DIDGYGGIALPELIC[+57.021]TMFHTSGYDTQAIVENN[+2350.830]ESTEYGLFQISNK. | HexNAc(4)Hex(5)Fuc(1)NeuAc(2) | C[+57], N[+2351] | 1841.307 | 4 | 7362.2061 | 7362.1592 | 0 | 6.4 | 33 | Specific | 381.64 | 381.64 | 381.64 | 8.56 | 24 | >sp\|P00709\|LALBA_HUMAN Alpha-lactalbumin OS=Homo sapiens OX=9606 GN=LALBA PE=1 SV=1 | 468 | 20201117_ly_milk_1ng_glyco_hcd203040_compound15.28967.28967.4 | controllerType=0 controllerNumber=1 scan=28967 | 57.7808 |
| 22898:4 | 19 | K.DIDGYGGIALPELIC[+57.021]TMFHTSGYDTQAIVENN[+1686.586]ESTEYGLFQISNK. | HexNAc(2)Hex(7)Fuc(1) | C[+57], N[+1687] | 1675.2308 | 4 | 6697.9014 | 6697.9153 | 0 | -2.1 | 33 | Specific | 415.83 | 332.07 | 332.07 | 6.64 | 24 | >sp\|P00709\|LALBA_HUMAN Alpha-lactalbumin OS=Homo sapiens OX=9606 GN=LALBA PE=1 SV=1 | 468 | 20201117_ly_milk_1ng_glyco_hcd203040_compound15.26499.26499.4 | controllerType=0 controllerNumber=1 scan=26499 | 53.1532 |
| 22854:4 | 19 | K.DIDGYGGIALPELIC[+57.021]TMFHTSGYDTQAIVENN[+1686.586]ESTEYGLFQISNK. | HexNAc(2)Hex(7)Fuc(1) | C[+57], N[+1687] | 1675.2337 | 4 | 6697.9131 | 6697.9153 | 0 | -0.3 | 33 | Specific | 501.53 | 470.22 | 470.22 | 9.89 | 24 | >sp\|P00709\|LALBA_HUMAN Alpha-lactalbumin OS=Homo sapiens OX=9606 GN=LALBA PE=1 SV=1 | 468 | 20201117_ly_milk_1ng_glyco_hcd203040_compound15.26453.26453.2 | controllerType=0 controllerNumber=1 scan=26453 | 53.0602 |
| 22856:4 | 19 | K.DIDGYGGIALPELIC[+57.021]TMFHTSGYDTQAIVENN[+1686.586]ESTEYGLFQISNK. | HexNAc(2)Hex(7)Fuc(1) | C[+57], N[+1687] | 1675.2337 | 4 | 6697.9131 | 6697.9153 | 0 | -0.3 | 33 | Specific | 556.67 | 501.87 | 501.87 | 10.2 | 24 | >sp\|P00709\|LALBA_HUMAN Alpha-lactalbumin OS=Homo sapiens OX=9606 GN=LALBA PE=1 SV=1 | 468 | 20201117_ly_milk_1ng_glyco_hcd203040_compound15.26455.26455.2 | controllerType=0 controllerNumber=1 scan=26455 | 53.0648 |
| 02307:5 | 24 | R.EKQTDEIKDTRN[+2059.735]ESTQNC[+57.021]VVAEPEK.M | HexNAc(4)Hex(5)Fuc(1)NeuAc(1) | C[+57], N[+2060] | 1002.434 | 5 | 5008.141 | 5008.1251 | 0 | 3.2 | 58 | Specific | 272.43 | 272.43 | 272.43 | 5.29 | 26 | >sp\|P47710\|CASA1_HUMAN Alpha-S1-casein OS=Homo sapiens OX=9606 GN=CSN1S1 PE=1 SV=1 | 989 | 20201117_ly_milk_1ng_glyco_hcd203040_compound15.5040.5040.5 | controllerType=0 controllerNumber=1 scan=5040 | 14.4236 |
| 02306:5 | 24 | R.EKQTDEIKDTRN[+2059.735]ESTQNC[+57.021]VVAEPEK.M | HexNAc(4)Hex(5)Fuc(1)NeuAc(1) | C[+57], N[+2060] | 1002.434 | 5 | 5008.141 | 5008.1251 | 0 | 3.2 | 58 | Specific | 392.34 | 392.34 | 392.34 | 8.47 | 26 | >sp\|P47710\|CASA1_HUMAN Alpha-S1-casein OS=Homo sapiens OX=9606 GN=CSN1S1 PE=1 SV=1 | 989 | 20201117_ly_milk_1ng_glyco_hcd203040_compound15.5039.5039.5 | controllerType=0 controllerNumber=1 scan=5039 | 14.4194 |
| 09678:4 | 24 | K.QTDEIKDTRN[+2059.735]ESTQNC[+57.021]VVAEPEKMESSISSSSEEMSLSK.C | HexNAc(4)Hex(5)Fuc(1)NeuAc(1) | C[+57], N[+2060] | 1613.4408 | 4 | 6450.7414 | 6450.7156 | 0 | 4 | 60 | Specific | 418.35 | 318.81 | 318.81 | 5.37 | 26 | >sp\|P47710\|CASA1_HUMAN Alpha-S1-casein OS=Homo sapiens OX=9606 GN=CSN1S1 PE=1 SV=1 | 989 | 20201117_ly_milk_1ng_glyco_hcd203040_compound15.12691.12691.4 | controllerType=0 controllerNumber=1 scan=12691 | 27.1586 |
| 09680:4 | 24 | K.QTDEIKDTRN[+2059.735]ESTQNC[+57.021]VVAEPEKMESSISSSSEEMSLSK.C | HexNAc(4)Hex(5)Fuc(1)NeuAc(1) | C[+57], N[+2060] | 1613.4408 | 4 | 6450.7414 | 6450.7156 | 0 | 4 | 60 | Specific | 522.59 | 313.72 | 313.72 | 5.8 | 26 | >sp\|P47710\|CASA1_HUMAN Alpha-S1-casein OS=Homo sapiens OX=9606 GN=CSN1S1 PE=1 SV=1 | 989 | 20201117_ly_milk_1ng_glyco_hcd203040_compound15.12693.12693.4 | controllerType=0 controllerNumber=1 scan=12693 | 27.1625 |
| 14580:3 | 25 | K.VVLHPN[+2204.772]YSQVDIGLIK.L | HexNAc(4)Hex(5)NeuAc(2) | N[+2205] | 1333.9354 | 3 | 3999.7915 | 3999.7837 | 0 | 2 | 236 | Specific | 557.53 | 504.73 | 504.73 | 9.85 | 24 | >sp\|P00738\|HPT_HUMAN Haptoglobin OS=Homo sapiens OX=9606 GN=HP PE=1 SV=1 | 407 | 20201117_ly_milk_1ng_glyco_hcd203040_compound15.17789.17789.3 | controllerType=0 controllerNumber=1 scan=17789 | 35.9238 |
| 14577:3 | 25 | K.VVLHPN[+2204.772]YSQVDIGLIK.L | HexNAc(4)Hex(5)NeuAc(2) | N[+2205] | 1333.9354 | 3 | 3999.7915 | 3999.7837 | 0 | 2 | 236 | Specific | 621.99 | 580.21 | 580.21 | 12.02 | 24 | >sp\|P00738\|HPT_HUMAN Haptoglobin OS=Homo sapiens OX=9606 GN=HP PE=1 SV=1 | 407 | 20201117_ly_milk_1ng_glyco_hcd203040_compound15.17786.17786.3 | controllerType=0 controllerNumber=1 scan=17786 | 35.9187 |
| 14312:4 | 26 | K.LKELPGVC[+57.021]N[+2059.735]ETMMALWEEC[+57.021]KPC[+57.021]LK.Q | HexNAc(4)Hex(5)Fuc(1)NeuAc(1) | C[+57]*3, N[+2060] | 1249.7885 | 4 | 4996.132 | 4996.1196 | 0 | 2.5 | 95 | Specific | 587.13 | 587.13 | 587.13 | 10.82 | 26 | >sp\|P10909\|CLUS_HUMAN Clusterin OS=Homo sapiens OX=9606 GN=CLU PE=1 SV=1 | 1467 | 20201117_ly_milk_1ng_glyco_hcd203040_compound15.17512.17512.4 | controllerType=0 controllerNumber=1 scan=17512 | 35.4531 |
| 14314:4 | 26 | K.LKELPGVC[+57.021]N[+2059.735]ETMMALWEEC[+57.021]KPC[+57.021]LK.Q | HexNAc(4)Hex(5)Fuc(1)NeuAc(1) | C[+57]*3, N[+2060] | 1249.7885 | 4 | 4996.132 | 4996.1196 | 0 | 2.5 | 95 | Specific | 627.34 | 586.45 | 586.45 | 11.82 | 26 | >sp\|P10909\|CLUS_HUMAN Clusterin OS=Homo sapiens OX=9606 GN=CLU PE=1 SV=1 | 1467 | 20201117_ly_milk_1ng_glyco_hcd203040_compound15.17514.17514.4 | controllerType=0 controllerNumber=1 scan=17514 | 35.457 |
| 21112:4 | 26 | K.MLN[+2059.735]TSSLLEQLNEQFNWVSR.L | HexNAc(4)Hex(5)Fuc(1)NeuAc(1) | N[+2060] | 1117.9888 | 4 | 4468.9334 | 4468.9217 | 0 | 2.6 | 352 | Specific | 601.74 | 601.74 | 601.74 | 12.2 | 26 | >sp\|P10909\|CLUS_HUMAN Clusterin OS=Homo sapiens OX=9606 GN=CLU PE=1 SV=1 | 1467 | 20201117_ly_milk_1ng_glyco_hcd203040_compound15.24636.24636.4 | controllerType=0 controllerNumber=1 scan=24636 | 49.8247 |
| 23530:4 | 26 | K.MLN[+2350.830]TSSLLEQLNEQFNWVSR.L | HexNAc(4)Hex(5)Fuc(1)NeuAc(2) | N[+2351] | 1190.7624 | 4 | 4760.0277 | 4760.0171 | 0 | 2.2 | 352 | Specific | 620.9 | 551.46 | 551.46 | 12.2 | 26 | >sp\|P10909\|CLUS_HUMAN Clusterin OS=Homo sapiens OX=9606 GN=CLU PE=1 SV=1 | 1467 | 20201117_ly_milk_1ng_glyco_hcd203040_compound15.27157.27157.4 | controllerType=0 controllerNumber=1 scan=27157 | 54.4481 |
| 21114:4 | 26 | K.MLN[+2059.735]TSSLLEQLNEQFNWVSR.L | HexNAc(4)Hex(5)Fuc(1)NeuAc(1) | N[+2060] | 1117.9888 | 4 | 4468.9334 | 4468.9217 | 0 | 2.6 | 352 | Specific | 680.42 | 432.93 | 432.93 | 13.2 | 26 | >sp\|P10909\|CLUS_HUMAN Clusterin OS=Homo sapiens OX=9606 GN=CLU PE=1 SV=1 | 1467 | 20201117_ly_milk_1ng_glyco_hcd203040_compound15.24638.24638.4 | controllerType=0 controllerNumber=1 scan=24638 | 49.8376 |
| 23533:4 | 26 | K.MLN[+2350.830]TSSLLEQLNEQFNWVSR.L | HexNAc(4)Hex(5)Fuc(1)NeuAc(2) | N[+2351] | 1190.7624 | 4 | 4760.0277 | 4760.0171 | 0 | 2.2 | 352 | Specific | 720.5 | 489.38 | 489.38 | 14.2 | 26 | >sp\|P10909\|CLUS_HUMAN Clusterin OS=Homo sapiens OX=9606 GN=CLU PE=1 SV=1 | 1467 | 20201117_ly_milk_1ng_glyco_hcd203040_compound15.27160.27160.4 | controllerType=0 controllerNumber=1 scan=27160 | 54.4541 |
| 23483:3 | 26 | K.MLN[+2350.830]TSSLLEQLNEQFNWVSR.L | HexNAc(4)Hex(5)Fuc(1)NeuAc(2) | N[+2351] | 1587.3482 | 3 | 4760.0301 | 4760.0171 | 0 | 2.7 | 352 | Specific | 804.69 | 602.66 | 602.66 | 16.14 | 26 | >sp\|P10909\|CLUS_HUMAN Clusterin OS=Homo sapiens OX=9606 GN=CLU PE=1 SV=1 | 1467 | 20201117_ly_milk_1ng_glyco_hcd203040_compound15.27108.27108.3 | controllerType=0 controllerNumber=1 scan=27108 | 54.3549 |
| 23527:3 | 26 | K.MLN[+2350.830]TSSLLEQLNEQFNWVSR.L | HexNAc(4)Hex(5)Fuc(1)NeuAc(2) | N[+2351] | 1587.3473 | 3 | 4760.0275 | 4760.0171 | 0 | 2.2 | 352 | Specific | 821.09 | 599.95 | 599.95 | 16.14 | 26 | >sp\|P10909\|CLUS_HUMAN Clusterin OS=Homo sapiens OX=9606 GN=CLU PE=1 SV=1 | 1467 | 20201117_ly_milk_1ng_glyco_hcd203040_compound15.27154.27154.3 | controllerType=0 controllerNumber=1 scan=27154 | 54.4418 |
| 23529:3 | 26 | K.MLN[+2350.830]TSSLLEQLNEQFNWVSR.L | HexNAc(4)Hex(5)Fuc(1)NeuAc(2) | N[+2351] | 1587.3473 | 3 | 4760.0275 | 4760.0171 | 0 | 2.2 | 352 | Specific | 868.97 | 609.34 | 609.34 | 17.14 | 26 | >sp\|P10909\|CLUS_HUMAN Clusterin OS=Homo sapiens OX=9606 GN=CLU PE=1 SV=1 | 1467 | 20201117_ly_milk_1ng_glyco_hcd203040_compound15.27156.27156.3 | controllerType=0 controllerNumber=1 scan=27156 | 54.4463 |
| 23485:3 | 26 | K.MLN[+2350.830]TSSLLEQLNEQFNWVSR.L | HexNAc(4)Hex(5)Fuc(1)NeuAc(2) | N[+2351] | 1587.3482 | 3 | 4760.0301 | 4760.0171 | 0 | 2.7 | 352 | Specific | 917.49 | 641.49 | 641.49 | 18.14 | 26 | >sp\|P10909\|CLUS_HUMAN Clusterin OS=Homo sapiens OX=9606 GN=CLU PE=1 SV=1 | 1467 | 20201117_ly_milk_1ng_glyco_hcd203040_compound15.27110.27110.3 | controllerType=0 controllerNumber=1 scan=27110 | 54.3597 |
| 13257:3 | 26 | R.LAN[+2351.851]LTQGEDQYYLR.V | HexNAc(4)Hex(5)Fuc(3)NeuAc(1) | N[+2352] | 1345.8938 | 3 | 4035.6668 | 4035.6844 | 0 | -4.4 | 372 | Specific | 382.27 | 347.39 | 347.39 | 5.68 | 26 | >sp\|P10909\|CLUS_HUMAN Clusterin OS=Homo sapiens OX=9606 GN=CLU PE=1 SV=1 | 1467 | 20201117_ly_milk_1ng_glyco_hcd203040_compound15.16419.16419.3 | controllerType=0 controllerNumber=1 scan=16419 | 33.4888 |
| 13259:3 | 26 | R.LAN[+2351.851]LTQGEDQYYLR.V | HexNAc(4)Hex(5)Fuc(3)NeuAc(1) | N[+2352] | 1345.8938 | 3 | 4035.6668 | 4035.6844 | 0 | -4.4 | 372 | Specific | 507.55 | 446.83 | 446.83 | 7.82 | 26 | >sp\|P10909\|CLUS_HUMAN Clusterin OS=Homo sapiens OX=9606 GN=CLU PE=1 SV=1 | 1467 | 20201117_ly_milk_1ng_glyco_hcd203040_compound15.16421.16421.3 | controllerType=0 controllerNumber=1 scan=16421 | 33.4925 |
| 16947:3 | 54 | K.AALAAFNAQNN[+2204.772]GSNFQLEEISR.A | HexNAc(4)Hex(5)NeuAc(2) | N[+2205] | 1523.9905 | 3 | 4569.9569 | 4569.9256 | 0 | 6.9 | 166 | Specific | 303.3 | 239.62 | 239.62 | 5.02 | 9 | >sp\|P02765\|FETUA_HUMAN Alpha-2-HS-glycoprotein OS=Homo sapiens OX=9606 GN=AHSG PE=1 SV=2 | 135 | 20201117_ly_milk_1ng_glyco_hcd203040_compound15.20283.20283.3 | controllerType=0 controllerNumber=1 scan=20283 | 40.405 |
| 16945:3 | 54 | K.AALAAFNAQNN[+2204.772]GSNFQLEEISR.A | HexNAc(4)Hex(5)NeuAc(2) | N[+2205] | 1523.9905 | 3 | 4569.9569 | 4569.9256 | 0 | 6.9 | 166 | Specific | 408.47 | 318.9 | 318.9 | 6.36 | 9 | >sp\|P02765\|FETUA_HUMAN Alpha-2-HS-glycoprotein OS=Homo sapiens OX=9606 GN=AHSG PE=1 SV=2 | 135 | 20201117_ly_milk_1ng_glyco_hcd203040_compound15.20281.20281.3 | controllerType=0 controllerNumber=1 scan=20281 | 40.4011 |
| 09877:3 | 58 | R.IIVPLNNREN[+1809.666]ISDPTSPLR.T | HexNAc(5)Hex(4)Fuc(1) | N[+1810] | 1319.9587 | 3 | 3957.8615 | 3957.8432 | 0 | 4.6 | 62 | Specific | 281.04 | 218.41 | 218.41 | 5.31 | 13 | >sp\|P01591\|IGJ_HUMAN Immunoglobulin J chain OS=Homo sapiens OX=9606 GN=JCHAIN PE=1 SV=4 | 1392 | 20201117_ly_milk_1ng_glyco_hcd203040_compound15.12897.12897.3 | controllerType=0 controllerNumber=1 scan=12897 | 27.5125 |
| 09879:3 | 58 | R.IIVPLNNREN[+1809.666]ISDPTSPLR.T | HexNAc(5)Hex(4)Fuc(1) | N[+1810] | 1319.9587 | 3 | 3957.8615 | 3957.8432 | 0 | 4.6 | 62 | Specific | 306.52 | 203.18 | 203.18 | 5.37 | 13 | >sp\|P01591\|IGJ_HUMAN Immunoglobulin J chain OS=Homo sapiens OX=9606 GN=JCHAIN PE=1 SV=4 | 1392 | 20201117_ly_milk_1ng_glyco_hcd203040_compound15.12899.12899.3 | controllerType=0 controllerNumber=1 scan=12899 | 27.519 |
| 11622:3 | 58 | R.IIVPLNNREN[+1856.656]ISDPTSPLR.T | HexNAc(3)Hex(5)Fuc(1)NeuAc(1) | N[+1857] | 1335.6142 | 3 | 4004.828 | 4004.8327 | 0 | -1.2 | 62 | Specific | 318.21 | 318.21 | 318.21 | 5.5 | 13 | >sp\|P01591\|IGJ_HUMAN Immunoglobulin J chain OS=Homo sapiens OX=9606 GN=JCHAIN PE=1 SV=4 | 1392 | 20201117_ly_milk_1ng_glyco_hcd203040_compound15.14702.14702.3 | controllerType=0 controllerNumber=1 scan=14702 | 30.5938 |
| 11624:3 | 58 | R.IIVPLNNREN[+1856.656]ISDPTSPLR.T | HexNAc(3)Hex(5)Fuc(1)NeuAc(1) | N[+1857] | 1335.6142 | 3 | 4004.828 | 4004.8327 | 0 | -1.2 | 62 | Specific | 461.62 | 341.95 | 341.95 | 6.34 | 13 | >sp\|P01591\|IGJ_HUMAN Immunoglobulin J chain OS=Homo sapiens OX=9606 GN=JCHAIN PE=1 SV=4 | 1392 | 20201117_ly_milk_1ng_glyco_hcd203040_compound15.14704.14704.3 | controllerType=0 controllerNumber=1 scan=14704 | 30.5978 |
| 24927:3 | 164 | R.AVN[+2204.772]ITSENLIDDVVSLIR.G | HexNAc(4)Hex(5)NeuAc(2) | N[+2205] | 1392.6192 | 3 | 4175.843 | 4175.8481 | 0 | -1.2 | 413 | Specific | 288.56 | 288.56 | 288.56 | 5.69 | 6 | >sp\|P02748\|CO9_HUMAN Complement component C9 OS=Homo sapiens OX=9606 GN=C9 PE=1 SV=2 | 1547 | 20201117_ly_milk_1ng_glyco_hcd203040_compound15.28609.28609.3 | controllerType=0 controllerNumber=1 scan=28609 | 57.1622 |
| 24929:3 | 164 | R.AVN[+2204.772]ITSENLIDDVVSLIR.G | HexNAc(4)Hex(5)NeuAc(2) | N[+2205] | 1392.6192 | 3 | 4175.843 | 4175.8481 | 0 | -1.2 | 413 | Specific | 423 | 290.96 | 290.96 | 6.45 | 6 | >sp\|P02748\|CO9_HUMAN Complement component C9 OS=Homo sapiens OX=9606 GN=C9 PE=1 SV=2 | 1547 | 20201117_ly_milk_1ng_glyco_hcd203040_compound15.28611.28611.3 | controllerType=0 controllerNumber=1 scan=28611 | 57.1658 |
| 08368:3 | 1 | R.GN[+2059.735]FSTEGC[+57.021]GC[+57.021]VC[+57.021]EPGWK.G | HexNAc(4)Hex(5)Fuc(1)NeuAc(1) | C[+57]*3, N[+2060] | 1335.5107 | 3 | 4004.5175 | 4004.5023 | 0 | 3.8 | 165 | Specific | 720.26 | 601.6 | 601.6 | 14.88 | 152 | >sp\|P24821\|TENA_HUMAN Tenascin OS=Homo sapiens OX=9606 GN=TNC PE=1 SV=3 | 486 | 20201113_gy_colo_milk_1ng_glyco_hcd203040_compound15.11305.11305.3 | controllerType=0 controllerNumber=1 scan=11305 | 24.9892 |
| 08370:3 | 1 | R.GN[+2059.735]FSTEGC[+57.021]GC[+57.021]VC[+57.021]EPGWK.G | HexNAc(4)Hex(5)Fuc(1)NeuAc(1) | C[+57]*3, N[+2060] | 1335.5107 | 3 | 4004.5175 | 4004.5023 | 0 | 3.8 | 165 | Specific | 745.42 | 636.36 | 636.36 | 14.88 | 152 | >sp\|P24821\|TENA_HUMAN Tenascin OS=Homo sapiens OX=9606 GN=TNC PE=1 SV=3 | 486 | 20201113_gy_colo_milk_1ng_glyco_hcd203040_compound15.11307.11307.3 | controllerType=0 controllerNumber=1 scan=11307 | 24.9932 |
| 01910:3 | 1 | K.GPN[+2424.867]C[+57.021]SEPEC[+57.021]PGNC[+57.021]HLR.G | HexNAc(5)Hex(6)Fuc(1)NeuAc(1) | C[+57]*3, N[+2425] | 1436.8868 | 3 | 4308.6458 | 4308.6253 | 0 | 4.8 | 182 | Specific | 216.76 | 163.82 | 163.82 | 4.48 | 152 | >sp\|P24821\|TENA_HUMAN Tenascin OS=Homo sapiens OX=9606 GN=TNC PE=1 SV=3 | 486 | 20201113_gy_colo_milk_1ng_glyco_hcd203040_compound15.4617.4617.3 | controllerType=0 controllerNumber=1 scan=4617 | 13.715 |
| 01908:3 | 1 | K.GPN[+2424.867]C[+57.021]SEPEC[+57.021]PGNC[+57.021]HLR.G | HexNAc(5)Hex(6)Fuc(1)NeuAc(1) | C[+57]*3, N[+2425] | 1436.8868 | 3 | 4308.6458 | 4308.6253 | 0 | 4.8 | 182 | Specific | 237.35 | 224.83 | 224.83 | 6.25 | 152 | >sp\|P24821\|TENA_HUMAN Tenascin OS=Homo sapiens OX=9606 GN=TNC PE=1 SV=3 | 486 | 20201113_gy_colo_milk_1ng_glyco_hcd203040_compound15.4615.4615.3 | controllerType=0 controllerNumber=1 scan=4615 | 13.7103 |
| 01984:3 | 1 | K.GPN[+1913.677]C[+57.021]SEPEC[+57.021]PGNC[+57.021]HLR.G | HexNAc(4)Hex(5)NeuAc(1) | C[+57]*3, N[+1914] | 1266.4871 | 3 | 3797.4468 | 3797.4352 | 0 | 3.1 | 182 | Specific | 332.97 | 298.77 | 298.77 | 7.45 | 152 | >sp\|P24821\|TENA_HUMAN Tenascin OS=Homo sapiens OX=9606 GN=TNC PE=1 SV=3 | 486 | 20201113_gy_colo_milk_1ng_glyco_hcd203040_compound15.4694.4694.3 | controllerType=0 controllerNumber=1 scan=4694 | 13.8504 |
| 01137:3 | 1 | K.GPN[+1216.423]C[+57.021]SEPEC[+57.021]PGNC[+57.021]HLR.G | HexNAc(2)Hex(5) | C[+57]*3, N[+1216] | 1034.0732 | 3 | 3100.205 | 3100.181 | 0 | 7.7 | 182 | Specific | 338.13 | 338.13 | 338.13 | 6.87 | 152 | >sp\|P24821\|TENA_HUMAN Tenascin OS=Homo sapiens OX=9606 GN=TNC PE=1 SV=3 | 486 | 20201113_gy_colo_milk_1ng_glyco_hcd203040_compound15.3814.3814.3 | controllerType=0 controllerNumber=1 scan=3814 | 12.3068 |
| 01135:3 | 1 | K.GPN[+1216.423]C[+57.021]SEPEC[+57.021]PGNC[+57.021]HLR.G | HexNAc(2)Hex(5) | C[+57]*3, N[+1216] | 1034.0732 | 3 | 3100.205 | 3100.181 | 0 | 7.7 | 182 | Specific | 356.98 | 356.98 | 356.98 | 6.99 | 152 | >sp\|P24821\|TENA_HUMAN Tenascin OS=Homo sapiens OX=9606 GN=TNC PE=1 SV=3 | 486 | 20201113_gy_colo_milk_1ng_glyco_hcd203040_compound15.3812.3812.3 | controllerType=0 controllerNumber=1 scan=3812 | 12.3028 |
| 01986:3 | 1 | K.GPN[+1913.677]C[+57.021]SEPEC[+57.021]PGNC[+57.021]HLR.G | HexNAc(4)Hex(5)NeuAc(1) | C[+57]*3, N[+1914] | 1266.4871 | 3 | 3797.4468 | 3797.4352 | 0 | 3.1 | 182 | Specific | 364.72 | 283.13 | 283.13 | 7.51 | 152 | >sp\|P24821\|TENA_HUMAN Tenascin OS=Homo sapiens OX=9606 GN=TNC PE=1 SV=3 | 486 | 20201113_gy_colo_milk_1ng_glyco_hcd203040_compound15.4696.4696.3 | controllerType=0 controllerNumber=1 scan=4696 | 13.8543 |
| 01967:3 | 1 | K.GPN[+2059.735]C[+57.021]SEPEC[+57.021]PGNC[+57.021]HLR.G | HexNAc(4)Hex(5)Fuc(1)NeuAc(1) | C[+57]*3, N[+2060] | 1315.1735 | 3 | 3943.506 | 3943.4931 | 0 | 3.3 | 182 | Specific | 523.67 | 416.77 | 416.77 | 10.17 | 152 | >sp\|P24821\|TENA_HUMAN Tenascin OS=Homo sapiens OX=9606 GN=TNC PE=1 SV=3 | 486 | 20201113_gy_colo_milk_1ng_glyco_hcd203040_compound15.4676.4676.3 | controllerType=0 controllerNumber=1 scan=4676 | 13.8178 |
| 01965:3 | 1 | K.GPN[+2059.735]C[+57.021]SEPEC[+57.021]PGNC[+57.021]HLR.G | HexNAc(4)Hex(5)Fuc(1)NeuAc(1) | C[+57]*3, N[+2060] | 1315.1735 | 3 | 3943.506 | 3943.4931 | 0 | 3.3 | 182 | Specific | 548.13 | 510.22 | 510.22 | 10.32 | 152 | >sp\|P24821\|TENA_HUMAN Tenascin OS=Homo sapiens OX=9606 GN=TNC PE=1 SV=3 | 486 | 20201113_gy_colo_milk_1ng_glyco_hcd203040_compound15.4674.4674.3 | controllerType=0 controllerNumber=1 scan=4674 | 13.814 |
| 04223:4 | 1 | R.C[+57.021]IN[+2059.735]GTC[+57.021]YC[+57.021]EEGFTGEDC[+57.021]GKPTC[ | HexNAc(4)Hex(5)Fuc(1)NeuAc(1) | C[+57]*6, N[+2060] | 1430.5525 | 4 | 5719.1881 | 5719.1633 | 0 | 4.3 | 325 | Specific | 798.15 | 798.15 | 798.15 | 15.83 | 152 | >sp\|P24821\|TENA_HUMAN Tenascin OS=Homo sapiens OX=9606 GN=TNC PE=1 SV=3 | 486 | 20201113_gy_colo_milk_1ng_glyco_hcd203040_compound15.7013.7013.4 | controllerType=0 controllerNumber=1 scan=7013 | 17.8[illegible]24 |
| 04225:4 | 1 | R.C[+57.021]IN[+2059.735]GTC[+57.021]YC[+57.021]EEGFTGEDC[+57.021]GKPTC[ | HexNAc(4)Hex(5)Fuc(1)NeuAc(1) | C[+57]*6, N[+2060] | 1430.5525 | 4 | 5719.1881 | 5719.1633 | 0 | 4.3 | 325 | Specific | 824.31 | 824.31 | 824.31 | 16.83 | 152 | >sp\|P24821\|TENA_HUMAN Tenascin OS=Homo sapiens OX=9606 GN=TNC PE=1 SV=3 | 486 | 20201113_gy_colo_milk_1ng_glyco_hcd203040_compound15.7015.7015.4 | controllerType=0 controllerNumber=1 scan=7015 | 17.8268 |
| 04236:5 | 1 | R.C[+57.021]IN[+2059.735]GTC[+57.021]YC[+57.021]EEGFTGEDC[+57.021]GKPTC[ | HexNAc(4)Hex(5)Fuc(1)NeuAc(1) | C[+57]*6, N[+2060] | 1144.6409 | 5 | 5719.1753 | 5719.1633 | 0 | 2.1 | 325 | Specific | 947.53 | 947.53 | 947.53 | 19.75 | 152 | >sp\|P24821\|TENA_HUMAN Tenascin OS=Homo sapiens OX=9606 GN=TNC PE=1 SV=3 | 486 | 20201113_gy_colo_milk_1ng_glyco_hcd203040_compound15.7027.7027.5 | controllerType=0 controllerNumber=1 scan=7027 | 17.8495 |
| 04234:5 | 1 | R.C[+57.021]IN[+2059.735]GTC[+57.021]YC[+57.021]EEGFTGEDC[+57.021]GKPTC[ | HexNAc(4)Hex(5)Fuc(1)NeuAc(1) | C[+57]*6, N[+2060] | 1144.6409 | 5 | 5719.1753 | 5719.1633 | 0 | 2.1 | 325 | Specific | 1023.75 | 1023.75 | 1023.75 | 21.75 | 152 | >sp\|P24821\|TENA_HUMAN Tenascin OS=Homo sapiens OX=9606 GN=TNC PE=1 SV=3 | 486 | 20201113_gy_colo_milk_1ng_glyco_hcd203040_compound15.7025.7025.5 | controllerType=0 controllerNumber=1 scan=7025 | 17.845 |
| 09340:3 | 1 | R.QTGLAPGQEYEISLHIVKN[+2059.735]NTR.G | HexNAc(4)Hex(5)Fuc(1)NeuAc(1) | N[+2060] | 1510.0143 | 3 | 4528.0285 | 4528.0241 | 0 | 1 | 770 | Specific | 334.33 | 174.55 | 174.55 | 5.72 | 152 | >sp\|P24821\|TENA_HUMAN Tenascin OS=Homo sapiens OX=9606 GN=TNC PE=1 SV=3 | 486 | 20201113_gy_colo_milk_1ng_glyco_hcd203040_compound15.12312.12312.3 | controllerType=0 controllerNumber=1 scan=12312 | 26.7879 |
| 09262:4 | 1 | R.QTGLAPGQEYEISLHIVKN[+2059.735]NTR.G | HexNAc(4)Hex(5)Fuc(1)NeuAc(1) | N[+2060] | 1132.7637 | 4 | 4528.033 | 4528.0241 | 0 | 2 | 770 | Specific | 517.01 | 374.4 | 374.4 | 8.43 | 152 | >sp\|P24821\|TENA_HUMAN Tenascin OS=Homo sapiens OX=9606 GN=TNC PE=1 SV=3 | 486 | 20201113_gy_colo_milk_1ng_glyco_hcd203040_compound15.12231.12231.4 | controllerType=0 controllerNumber=1 scan=12231 | 26.6424 |
| 09261:3 | 1 | R.QTGLAPGQEYEISLHIVKN[+2059.735]NTR.G | HexNAc(4)Hex(5)Fuc(1)NeuAc(1) | N[+2060] | 1510.0169 | 3 | 4528.0362 | 4528.0241 | 0 | 2.7 | 770 | Specific | 585.81 | 383.43 | 383.43 | 11.03 | 152 | >sp\|P24821\|TENA_HUMAN Tenascin OS=Homo sapiens OX=9606 GN=TNC PE=1 SV=3 | 486 | 20201113_gy_colo_milk_1ng_glyco_hcd203040_compound15.12230.12230.3 | controllerType=0 controllerNumber=1 scan=12230 | 26.6388 |
| 09260:4 | 1 | R.QTGLAPGQEYEISLHIVKN[+2059.735]NTR.G | HexNAc(4)Hex(5)Fuc(1)NeuAc(1) | N[+2060] | 1132.7637 | 4 | 4528.033 | 4528.0241 | 0 | 2 | 770 | Specific | 694.93 | 453.72 | 453.72 | 13.6 | 152 | >sp\|P24821\|TENA_HUMAN Tenascin OS=Homo sapiens OX=9606 GN=TNC PE=1 SV=3 | 486 | 20201113_gy_colo_milk_1ng_glyco_hcd203040_compound15.12229.12229.4 | controllerType=0 controllerNumber=1 scan=12229 | 26.6349 |
| 09259:3 | 1 | R.QTGLAPGQEYEISLHIVKN[+2059.735]NTR.G | HexNAc(4)Hex(5)Fuc(1)NeuAc(1) | N[+2060] | 1510.0169 | 3 | 4528.0362 | 4528.0241 | 0 | 2.7 | 770 | Specific | 781.3 | 459.63 | 459.63 | 15.03 | 152 | >sp\|P24821\|TENA_HUMAN Tenascin OS=Homo sapiens OX=9606 GN=TNC PE=1 SV=3 | 486 | 20201113_gy_colo_milk_1ng_glyco_hcd203040_compound15.12228.12228.3 | controllerType=0 controllerNumber=1 scan=12228 | 26.6333 |
| 15244:2 | 1 | R.LN[+1378.476]YSLPTGQWVGVQLPR.N | HexNAc(2)Hex(6) | N[+1378] | 1653.767 | 2 | 3306.5267 | 3306.5145 | 0 | 3.7 | 1017 | Specific | 793.39 | 793.39 | 793.39 | 16.04 | 152 | >sp\|P24821\|TENA_HUMAN Tenascin OS=Homo sapiens OX=9606 GN=TNC PE=1 SV=3 | 486 | 20201113_gy_colo_milk_1ng_glyco_hcd203040_compound15.18437.18437.2 | controllerType=0 controllerNumber=1 scan=18437 | 38.2131 |
| 15267:2 | 1 | R.LN[+1216.423]YSLPTGQWVGVQLPR.N | HexNAc(2)Hex(5) | N[+1216] | 1572.7386 | 2 | 3144.4699 | 3144.4617 | 0 | 2.6 | 1017 | Specific | 808.94 | 731.04 | 731.04 | 17.04 | 152 | >sp\|P24821\|TENA_HUMAN Tenascin OS=Homo sapiens OX=9606 GN=TNC PE=1 SV=3 | 486 | 20201113_gy_colo_milk_1ng_glyco_hcd203040_compound15.18461.18461.2 | controllerType=0 controllerNumber=1 scan=18461 | 38.2613 |
| 15246:2 | 1 | R.LN[+1378.476]YSLPTGQWVGVQLPR.N | HexNAc(2)Hex(6) | N[+1378] | 1653.767 | 2 | 3306.5267 | 3306.5145 | 0 | 3.7 | 1017 | Specific | 812.72 | 647.39 | 647.39 | 17.04 | 152 | >sp\|P24821\|TENA_HUMAN Tenascin OS=Homo sapiens OX=9606 GN=TNC PE=1 SV=3 | 486 | 20201113_gy_colo_milk_1ng_glyco_hcd203040_compound15.18439.18439.2 | controllerType=0 controllerNumber=1 scan=18439 | 38.2171 |
| 15269:2 | 1 | R.LN[+1216.423]YSLPTGQWVGVQLPR.N | HexNAc(2)Hex(5) | N[+1216] | 1572.7386 | 2 | 3144.4699 | 3144.4617 | 0 | 2.6 | 1017 | Specific | 849.31 | 814.7 | 814.7 | 17.04 | 152 | >sp\|P24821\|TENA_HUMAN Tenascin OS=Homo sapiens OX=9606 GN=TNC PE=1 SV=3 | 486 | 20201113_gy_colo_milk_1ng_glyco_hcd203040_compound15.18463.18463.2 | controllerType=0 controllerNumber=1 scan=18463 | 38.2654 |
| 15203:3 | 1 | R.LN[+1540.529]YSLPTGQWVGVQLPR.N | HexNAc(2)Hex(7) | N[+1541] | 1156.8628 | 3 | 3468.574 | 3468.5674 | 0 | 1.9 | 1017 | Specific | 875.57 | 844.15 | 844.15 | 18.22 | 152 | >sp\|P24821\|TENA_HUMAN Tenascin OS=Homo sapiens OX=9606 GN=TNC PE=1 SV=3 | 486 | 20201113_gy_colo_milk_1ng_glyco_hcd203040_compound15.18395.18395.3 | controllerType=0 controllerNumber=1 scan=18395 | 38.1214 |
| 15181:3 | 1 | R.LN[+1540.529]YSLPTGQWVGVQLPR.N | HexNAc(2)Hex(7) | N[+1541] | 1156.8654 | 3 | 3468.5817 | 3468.5674 | 0 | 4.1 | 1017 | Specific | 887.97 | 802.87 | 802.87 | 17.88 | 152 | >sp\|P24821\|TENA_HUMAN Tenascin OS=Homo sapiens OX=9606 GN=TNC PE=1 SV=3 | 486 | 20201113_gy_colo_milk_1ng_glyco_hcd203040_compound15.18372.18372.3 | controllerType=0 controllerNumber=1 scan=18372 | 38.076 |
| 15205:3 | 1 | R.LN[+1540.529]YSLPTGQWVGVQLPR.N | HexNAc(2)Hex(7) | N[+1541] | 1156.8628 | 3 | 3468.574 | 3468.5674 | 0 | 1.9 | 1017 | Specific | 899.44 | 894.13 | 894.13 | 18.22 | 152 | >sp\|P24821\|TENA_HUMAN Tenascin OS=Homo sapiens OX=9606 GN=TNC PE=1 SV=3 | 486 | 20201113_gy_colo_milk_1ng_glyco_hcd203040_compound15.18397.18397.3 | controllerType=0 controllerNumber=1 scan=18397 | 38.1253 |
| 15140:3 | 1 | R.LN[+1702.581]YSLPTGQWVGVQLPR.N | HexNAc(2)Hex(8) | N[+1703] | 1210.8781 | 3 | 3630.6198 | 3630.6202 | 0 | -0.1 | 1017 | Specific | 963.93 | 865.45 | 865.45 | 19.66 | 152 | >sp\|P24821\|TENA_HUMAN Tenascin OS=Homo sapiens OX=9606 GN=TNC PE=1 SV=3 | 486 | 20201113_gy_colo_milk_1ng_glyco_hcd203040_compound15.18329.18329.3 | controllerType=0 controllerNumber=1 scan=18329 | 37.9872 |
| 15184:3 | 1 | R.LN[+1540.529]YSLPTGQWVGVQLPR.N | HexNAc(2)Hex(7) | N[+1541] | 1156.8654 | 3 | 3468.5817 | 3468.5674 | 0 | 4.1 | 1017 | Specific | 1035.48 | 1035.48 | 1035.48 | 20.88 | 152 | >sp\|P24821\|TENA_HUMAN Tenascin OS=Homo sapiens OX=9606 GN=TNC PE=1 SV=3 | 486 | 20201113_gy_colo_milk_1ng_glyco_hcd203040_compound15.18375.18375.3 | controllerType=0 controllerNumber=1 scan=18375 | 38.0803 |
| 15142:3 | 1 | R.LN[+1702.581]YSLPTGQWVGVQLPR.N | HexNAc(2)Hex(8) | N[+1703] | 1210.8781 | 3 | 3630.6198 | 3630.6202 | 0 | -0.1 | 1017 | Specific | 1054.81 | 851.49 | 851.49 | 21.66 | 152 | >sp\|P24821\|TENA_HUMAN Tenascin OS=Homo sapiens OX=9606 GN=TNC PE=1 SV=3 | 486 | 20201113_gy_colo_milk_1ng_glyco_hcd203040_compound15.18331.18331.3 | controllerType=0 controllerNumber=1 scan=18331 | 37.9916 |
| 15253:3 | 1 | R.LN[+1216.423]YSLPTGQWVGVQLPR.N | HexNAc(2)Hex(5) | N[+1216] | 1048.8296 | 3 | 3144.4743 | 3144.4617 | 0 | 4 | 1017 | Specific | 1058.18 | 1058.18 | 1058.18 | 21.88 | 152 | >sp\|P24821\|TENA_HUMAN Tenascin OS=Homo sapiens OX=9606 GN=TNC PE=1 SV=3 | 486 | 20201113_gy_colo_milk_1ng_glyco_hcd203040_compound15.18447.18447.3 | controllerType=0 controllerNumber=1 scan=18447 | 38.2319 |
| 15256:3 | 1 | R.LN[+1216.423]YSLPTGQWVGVQLPR.N | HexNAc(2)Hex(5) | N[+1216] | 1048.8296 | 3 | 3144.4743 | 3144.4617 | 0 | 4 | 1017 | Specific | 1088.72 | 926.95 | 926.95 | 21.88 | 152 | >sp\|P24821\|TENA_HUMAN Tenascin OS=Homo sapiens OX=9606 GN=TNC PE=1 SV=3 | 486 | 20201113_gy_colo_milk_1ng_glyco_hcd203040_compound15.18450.18450.3 | controllerType=0 controllerNumber=1 scan=18450 | 38.2401 |
| 02408:2 | 1 | R.N[+1216.423]TTSYVLR.G | HexNAc(2)Hex(5) | N[+1216] | 1085.47 | 2 | 2169.9328 | 2169.9279 | 0 | 2.2 | 1034 | Specific | 226.56 | 141.02 | 141.02 | 5.52 | 152 | >sp\|P24821\|TENA_HUMAN Tenascin OS=Homo sapiens OX=9606 GN=TNC PE=1 SV=3 | 486 | 20201113_gy_colo_milk_1ng_glyco_hcd203040_compound15.5135.5135.2 | controllerType=0 controllerNumber=1 scan=5135 | 14.6215 |
| 04204:3 | 1 | R.N[+2059.735]TTSYVLR.G | HexNAc(4)Hex(5)Fuc(1)NeuAc(1) | N[+2060] | 1005.0884 | 3 | 3013.2507 | 3013.24 | 0 | 3.5 | 1034 | Specific | 251.37 | 158.03 | 158.03 | 6.15 | 152 | >sp\|P24821\|TENA_HUMAN Tenascin OS=Homo sapiens OX=9606 GN=TNC PE=1 SV=3 | 486 | 20201113_gy_colo_milk_1ng_glyco_hcd203040_compound15.6994.6994.3 | controllerType=0 controllerNumber=1 scan=6994 | 17.7875 |
| 04207:2 | 1 | R.N[+2059.735]TTSYVLR.G | HexNAc(4)Hex(5)Fuc(1)NeuAc(1) | N[+2060] | 1507.1288 | 2 | 3013.2504 | 3013.24 | 0 | 3.4 | 1034 | Specific | 262.91 | 153.28 | 153.28 | 5.98 | 152 | >sp\|P24821\|TENA_HUMAN Tenascin OS=Homo sapiens OX=9606 GN=TNC PE=1 SV=3 | 486 | 20201113_gy_colo_milk_1ng_glyco_hcd203040_compound15.6997.6997.2 | controllerType=0 controllerNumber=1 scan=6997 | 17.7941 |
| 04206:3 | 1 | R.N[+2059.735]TTSYVLR.G | HexNAc(4)Hex(5)Fuc(1)NeuAc(1) | N[+2060] | 1005.0884 | 3 | 3013.2507 | 3013.24 | 0 | 3.5 | 1034 | Specific | 314.36 | 212.61 | 212.61 | 6.82 | 152 | >sp\|P24821\|TENA_HUMAN Tenascin OS=Homo sapiens OX=9606 GN=TNC PE=1 SV=3 | 486 | 20201113_gy_colo_milk_1ng_glyco_hcd203040_compound15.6996.6996.3 | controllerType=0 controllerNumber=1 scan=6996 | 17.7922 |
| 04209:2 | 1 | R.N[+2059.735]TTSYVLR.G | HexNAc(4)Hex(5)Fuc(1)NeuAc(1) | N[+2060] | 1507.1288 | 2 | 3013.2504 | 3013.24 | 0 | 3.4 | 1034 | Specific | 389.3 | 252.29 | 252.29 | 7.28 | 152 | >sp\|P24821\|TENA_HUMAN Tenascin OS=Homo sapiens OX=9606 GN=TNC PE=1 SV=3 | 486 | 20201113_gy_colo_milk_1ng_glyco_hcd203040_compound15.6999.6999.2 | controllerType=0 controllerNumber=1 scan=6999 | 17.798 |
| 18413:4 | 1 | R.LN[+2350.830]WTAADQAYEHFIIQVQEANKVEAAR.N | HexNAc(4)Hex(5)Fuc(1)NeuAc(2) | N[+2351] | 1367.358 | 4 | 5466.4103 | 5466.3899 | 0 | 3.7 | 1092 | Specific | 291.07 | 223.74 | 223.74 | 4.87 | 152 | >sp\|P24821\|TENA_HUMAN Tenascin OS=Homo sapiens OX=9606 GN=TNC PE=1 SV=3 | 486 | 20201113_gy_colo_milk_1ng_glyco_hcd203040_compound15.21741.21741.4 | controllerType=0 controllerNumber=1 scan=21741 | 45.0968 |
| 18410:4 | 1 | R.LN[+2350.830]WTAADQAYEHFIIQVQEANKVEAAR.N | HexNAc(4)Hex(5)Fuc(1)NeuAc(2) | N[+2351] | 1367.358 | 4 | 5466.4103 | 5466.3899 | 0 | 3.7 | 1092 | Specific | 427.06 | 388.91 | 388.91 | 9.52 | 152 | >sp\|P24821\|TENA_HUMAN Tenascin OS=Homo sapiens OX=9606 GN=TNC PE=1 SV=3 | 486 | 20201113_gy_colo_milk_1ng_glyco_hcd203040_compound15.21738.21738.4 | controllerType=0 controllerNumber=1 scan=21738 | 45.0898 |
| 17310:4 | 1 | R.LN[+2059.735]WTAADQAYEHFIIQVQEANKVEAAR.N | HexNAc(4)Hex(5)Fuc(1)NeuAc(1) | N[+2060] | 1294.5817 | 4 | 5175.3048 | 5175.2945 | 0 | 2 | 1092 | Specific | 636.91 | 612.84 | 612.84 | 12.64 | 152 | >sp\|P24821\|TENA_HUMAN Tenascin OS=Homo sapiens OX=9606 GN=TNC PE=1 SV=3 | 486 | 20201113_gy_colo_milk_1ng_glyco_hcd203040_compound15.20590.20590.4 | controllerType=0 controllerNumber=1 scan=20590 | 42.5893 |
| 17313:4 | 1 | R.LN[+2059.735]WTAADQAYEHFIIQVQEANKVEAAR.N | HexNAc(4)Hex(5)Fuc(1)NeuAc(1) | N[+2060] | 1294.5817 | 4 | 5175.3048 | 5175.2945 | 0 | 2 | 1092 | Specific | 1045.68 | 828.21 | 828.21 | 20.64 | 152 | >sp\|P24821\|TENA_HUMAN Tenascin OS=Homo sapiens OX=9606 GN=TNC PE=1 SV=3 | 486 | 20201113_gy_colo_milk_1ng_glyco_hcd203040_compound15.20593.20593.4 | controllerType=0 controllerNumber=1 scan=20593 | 42.6001 |
| 08026:2 | 1 | R.N[+2059.735]LTVPGSLR.A | HexNAc(4)Hex(5)Fuc(1)NeuAc(1) | N[+2060] | 1508.6506 | 2 | 3016.2938 | 3016.2873 | 0 | 2.2 | 1119 | Specific | 227.19 | 75.69 | 75.69 | 4.73 | 152 | >sp\|P24821\|TENA_HUMAN Tenascin OS=Homo sapiens OX=9606 GN=TNC PE=1 SV=3 | 486 | 20201113_gy_colo_milk_1ng_glyco_hcd203040_compound15.10950.10950.2 | controllerType=0 controllerNumber=1 scan=10950 | 24.385 |
| 08030:2 | 1 | R.N[+2059.735]LTVPGSLR.A | HexNAc(4)Hex(5)Fuc(1)NeuAc(1) | N[+2060] | 1508.6506 | 2 | 3016.2938 | 3016.2873 | 0 | 2.2 | 1119 | Specific | 228.67 | 74.91 | 74.91 | 4.73 | 152 | >sp\|P24821\|TENA_HUMAN Tenascin OS=Homo sapiens OX=9606 GN=TNC PE=1 SV=3 | 486 | 20201113_gy_colo_milk_1ng_glyco_hcd203040_compound15.10954.10954.3 | controllerType=0 controllerNumber=1 scan=10954 | 24.3925 |
| 08019:3 | 1 | R.N[+2059.735]LTVPGSLR.A | HexNAc(4)Hex(5)Fuc(1)NeuAc(1) | N[+2060] | 1006.1025 | 3 | 3016.293 | 3016.2873 | 0 | 1.9 | 1119 | Specific | 305.64 | 268.77 | 268.77 | 6.96 | 152 | >sp\|P24821\|TENA_HUMAN Tenascin OS=Homo sapiens OX=9606 GN=TNC PE=1 SV=3 | 486 | 20201113_gy_colo_milk_1ng_glyco_hcd203040_compound15.10943.10943.3 | controllerType=0 controllerNumber=1 scan=10943 | 24.3719 |
| 08028:2 | 1 | R.N[+2059.735]LTVPGSLR.A | HexNAc(4)Hex(5)Fuc(1)NeuAc(1) | N[+2060] | 1508.6506 | 2 | 3016.2938 | 3016.2873 | 0 | 2.2 | 1119 | Specific | 321.75 | 178.49 | 178.49 | 6.29 | 152 | >sp\|P24821\|TENA_HUMAN Tenascin OS=Homo sapiens OX=9606 GN=TNC PE=1 SV=3 | 486 | 20201113_gy_colo_milk_1ng_glyco_hcd203040_compound15.10952.10952.2 | controllerType=0 controllerNumber=1 scan=10952 | 24.3891 |
| 08032:2 | 1 | R.N[+2059.735]LTVPGSLR.A | HexNAc(4)Hex(5)Fuc(1)NeuAc(1) | N[+2060] | 1508.6506 | 2 | 3016.2938 | 3016.2873 | 0 | 2.2 | 1119 | Specific | 333.89 | 254.34 | 254.34 | 6.99 | 152 | >sp\|P24821\|TENA_HUMAN Tenascin OS=Homo sapiens OX=9606 GN=TNC PE=1 SV=3 | 486 | 20201113_gy_colo_milk_1ng_glyco_hcd203040_compound15.10956.10956.3 | controllerType=0 controllerNumber=1 scan=10956 | 24.3962 |
| 08021:3 | 1 | R.N[+2059.735]LTVPGSLR.A | HexNAc(4)Hex(5)Fuc(1)NeuAc(1) | N[+2060] | 1006.1025 | 3 | 3016.293 | 3016.2873 | 0 | 1.9 | 1119 | Specific | 347.9 | 168.87 | 168.87 | 6.64 | 152 | >sp\|P24821\|TENA_HUMAN Tenascin OS=Homo sapiens OX=9606 GN=TNC PE=1 SV=3 | 486 | 20201113_gy_colo_milk_1ng_glyco_hcd203040_compound15.10945.10945.3 | controllerType=0 controllerNumber=1 scan=10945 | 24.3768 |
| 11014:3 | 1 | K.VEAAQN[+2059.735]LTLPGSLR.A | HexNAc(4)Hex(5)Fuc(1)NeuAc(1) | N[+2060] | 1176.8564 | 3 | 3528.5546 | 3528.5468 | 0 | 2.2 | 1387 | Specific | 302.4 | 302.4 | 302.4 | 7.42 | 152 | >sp\|P24821\|TENA_HUMAN Tenascin OS=Homo sapiens OX=9606 GN=TNC PE=1 SV=3 | 486 | 20201113_gy_colo_milk_1ng_glyco_hcd203040_compound15.14046.14046.3 | controllerType=0 controllerNumber=1 scan=14046 | 29.9106 |
| 11016:3 | 1 | K.VEAAQN[+2059.735]LTLPGSLR.A | HexNAc(4)Hex(5)Fuc(1)NeuAc(1) | N[+2060] | 1176.8564 | 3 | 3528.5546 | 3528.5468 | 0 | 2.2 | 1387 | Specific | 447.33 | 322.48 | 322.48 | 8.19 | 152 | >sp\|P24821\|TENA_HUMAN Tenascin OS=Homo sapiens OX=9606 GN=TNC PE=1 SV=3 | 486 | 20201113_gy_colo_milk_1ng_glyco_hcd203040_compound15.14048.14048.3 | controllerType=0 controllerNumber=1 scan=14048 | 29.915 |
| 11158:3 | 1 | R.LLETVEYN[+2059.735]ISGAER.T | HexNAc(4)Hex(5)Fuc(1)NeuAc(1) | N[+2060] | 1218.5208 | 3 | 3653.5479 | 3653.5468 | 0 | 0.3 | 1478 | Specific | 258.82 | 165.46 | 165.46 | 5.8 | 152 | >sp\|P24821\|TENA_HUMAN Tenascin OS=Homo sapiens OX=9606 GN=TNC PE=1 SV=3 | 486 | 20201113_gy_colo_milk_1ng_glyco_hcd203040_compound15.14195.14195.3 | controllerType=0 controllerNumber=1 scan=14195 | 30.1715 |
| 11001:3 | 1 | R.LLETVEYN[+2424.867]ISGAER.T | HexNAc(5)Hex(6)Fuc(1)NeuAc(1) | N[+2425] | 1340.2381 | 3 | 4018.6997 | 4018.679 | 0 | 5.2 | 1478 | Specific | 377.08 | 377.08 | 377.08 | 7.7 | 152 | >sp\|P24821\|TENA_HUMAN Tenascin OS=Homo sapiens OX=9606 GN=TNC PE=1 SV=3 | 486 | 20201113_gy_colo_milk_1ng_glyco_hcd203040_compound15.14032.14032.3 | controllerType=0 controllerNumber=1 scan=14032 | 29.8875 |
| 11160:3 | 1 | R.LLETVEYN[+2059.735]ISGAER.T | HexNAc(4)Hex(5)Fuc(1)NeuAc(1) | N[+2060] | 1218.5208 | 3 | 3653.5479 | 3653.5468 | 0 | 0.3 | 1478 | Specific | 393.79 | 387.26 | 387.26 | 9.29 | 152 | >sp\|P24821\|TENA_HUMAN Tenascin OS=Homo sapiens OX=9606 GN=TNC PE=1 SV=3 | 486 | 20201113_gy_colo_milk_1ng_glyco_hcd203040_compound15.14197.14197.3 | controllerType=0 controllerNumber=1 scan=14197 | 30.1773 |
| 11003:3 | 1 | R.LLETVEYN[+2424.867]ISGAER.T | HexNAc(5)Hex(6)Fuc(1)NeuAc(1) | N[+2425] | 1340.2381 | 3 | 4018.6997 | 4018.679 | 0 | 5.2 | 1478 | Specific | 541.06 | 516.85 | 516.85 | 9.8 | 152 | >sp\|P24821\|TENA_HUMAN Tenascin OS=Homo sapiens OX=9606 GN=TNC PE=1 SV=3 | 486 | 20201113_gy_colo_milk_1ng_glyco_hcd203040_compound15.14034.14034.3 | controllerType=0 controllerNumber=1 scan=14034 | 29.8909 |
| 16880:4 | 2 | R.RTAGWNVPIGTLRPFLN[+1768.640]WTGPPEPIEAAVAR.F | HexNAc(4)Hex(5)Fuc(1) | N[+1769] | 1289.621 | 4 | 5155.4622 | 5155.4516 | 0 | 2.1 | 140 | Specific | 205.48 | 205.48 | 205.48 | 5.93 | 112 | >sp\|P02788\|TRFL_HUMAN Lactotransferrin OS=Homo sapiens OX=9606 GN=LTF PE=1 SV=6 | 1270 | 20201113_gy_colo_milk_1ng_glyco_hcd203040_compound15.20141.20141.4 | controllerType=0 controllerNumber=1 scan=20141 | 41.6108 |
| 17776:4 | 2 | R.RTAGWNVPIGTLRPFLN[+2571.946]WTGPPEPIEAAVAR.F | HexNAc(5)Hex(6)Fuc(4) | N[+2572] | 1490.4454 | 4 | 5958.7599 | 5958.7575 | 0 | 0.4 | 140 | Specific | 222.83 | 222.83 | 222.83 | 5.16 | 112 | >sp\|P02788\|TRFL_HUMAN Lactotransferrin OS=Homo sapiens OX=9606 GN=LTF PE=1 SV=6 | 1270 | 20201113_gy_colo_milk_1ng_glyco_hcd203040_compound15.21076.21076.4 | controllerType=0 controllerNumber=1 scan=21076 | 43.6157 |
| 17676:4 | 2 | R.RTAGWNVPIGTLRPFLN[+2424.867]WTGPPEPIEAAVAR.F | HexNAc(5)Hex(6)Fuc(1)NeuAc(1) | N[+2425] | 1453.6781 | 4 | 5811.6906 | 5811.6792 | 0 | 2 | 140 | Specific | 317.98 | 317.98 | 317.98 | 8.35 | 112 | >sp\|P02788\|TRFL_HUMAN Lactotransferrin OS=Homo sapiens OX=9606 GN=LTF PE=1 SV=6 | 1270 | 20201113_gy_colo_milk_1ng_glyco_hcd203040_compound15.20973.20973.4 | controllerType=0 controllerNumber=1 scan=20973 | 43.4204 |
| 18763:4 | 2 | R.RTAGWNVPIGTLRPFLN[+2350.830]WTGPPEPIEAAVAR.F | HexNAc(4)Hex(5)Fuc(1)NeuAc(2) | N[+2351] | 1435.1705 | 4 | 5737.6603 | 5737.6424 | 0 | 3.1 | 140 | Specific | 402.05 | 327.84 | 327.84 | 8.75 | 112 | >sp\|P02788\|TRFL_HUMAN Lactotransferrin OS=Homo sapiens OX=9606 GN=LTF PE=1 SV=6 | 1270 | 20201113_gy_colo_milk_1ng_glyco_hcd203040_compound15.22107.22107.4 | controllerType=0 controllerNumber=1 scan=22107 | 45.8751 |
| 18729:4 | 2 | R.RTAGWNVPIGTLRPFLN[+2350.830]WTGPPEPIEAAVAR.F | HexNAc(4)Hex(5)Fuc(1)NeuAc(2) | N[+2351] | 1435.1801 | 4 | 5737.6984 | 5737.6424 | 0 | 9.8 | 140 | Specific | 405.26 | 405.26 | 405.26 | 9.48 | 112 | >sp\|P02788\|TRFL_HUMAN Lactotransferrin OS=Homo sapiens OX=9606 GN=LTF PE=1 SV=6 | 1270 | 20201113_gy_colo_milk_1ng_glyco_hcd203040_compound15.22071.22071.3 | controllerType=0 controllerNumber=1 scan=22071 | 45.8085 |
| 16797:4 | 2 | R.RTAGWNVPIGTLRPFLN[+2279.830]WTGPPEPIEAAVAR.F | HexNAc(5)Hex(6)Fuc(2) | N[+2280] | 1417.4199 | 4 | 5666.6579 | 5666.6417 | 0 | 2.9 | 140 | Specific | 435.08 | 323.98 | 323.98 | 8.9 | 112 | >sp\|P02788\|TRFL_HUMAN Lactotransferrin OS=Homo sapiens OX=9606 GN=LTF PE=1 SV=6 | 1270 | 20201113_gy_colo_milk_1ng_glyco_hcd203040_compound15.20054.20054.4 | controllerType=0 controllerNumber=1 scan=20054 | 41.4261 |
| 16795:4 | 2 | R.RTAGWNVPIGTLRPFLN[+2279.830]WTGPPEPIEAAVAR.F | HexNAc(5)Hex(6)Fuc(2) | N[+2280] | 1417.4199 | 4 | 5666.6579 | 5666.6417 | 0 | 2.9 | 140 | Specific | 459.52 | 274.8 | 274.8 | 8.76 | 112 | >sp\|P02788\|TRFL_HUMAN Lactotransferrin OS=Homo sapiens OX=9606 GN=LTF PE=1 SV=6 | 1270 | 20201113_gy_colo_milk_1ng_glyco_hcd203040_compound15.20052.20052.4 | controllerType=0 controllerNumber=1 scan=20052 | 41.4207 |
| 17805:5 | 2 | R.RTAGWNVPIGTLRPFLN[+2204.772]WTGPPEPIEAAVAR.F | HexNAc(4)Hex(5)NeuAc(2) | N[+2205] | 1119.1282 | 5 | 5591.6118 | 5591.5845 | 0 | 4.9 | 140 | Specific | 461.76 | 414.63 | 414.63 | 10.34 | 112 | >sp\|P02788\|TRFL_HUMAN Lactotransferrin OS=Homo sapiens OX=9606 GN=LTF PE=1 SV=6 | 1270 | 20201113_gy_colo_milk_1ng_glyco_hcd203040_compound15.21106.21106.5 | controllerType=0 controllerNumber=1 scan=21106 | 43.6824 |
| 18732:4 | 2 | R.RTAGWNVPIGTLRPFLN[+2350.830]WTGPPEPIEAAVAR.F | HexNAc(4)Hex(5)Fuc(1)NeuAc(2) | N[+2351] | 1435.1801 | 4 | 5737.6984 | 5737.6424 | 0 | 9.8 | 140 | Specific | 464.47 | 464.47 | 464.47 | 9.78 | 112 | >sp\|P02788\|TRFL_HUMAN Lactotransferrin OS=Homo sapiens OX=9606 GN=LTF PE=1 SV=6 | 1270 | 20201113_gy_colo_milk_1ng_glyco_hcd203040_compound15.22074.22074.3 | controllerType=0 controllerNumber=1 scan=22074 | 45.8136 |
| 18766:4 | 2 | R.RTAGWNVPIGTLRPFLN[+2350.830]WTGPPEPIEAAVAR.F | HexNAc(4)Hex(5)Fuc(1)NeuAc(2) | N[+2351] | 1435.1705 | 4 | 5737.6603 | 5737.6424 | 0 | 3.1 | 140 | Specific | 528.17 | 455.01 | 455.01 | 11.05 | 112 | >sp\|P02788\|TRFL_HUMAN Lactotransferrin OS=Homo sapiens OX=9606 GN=LTF PE=1 SV=6 | 1270 | 20201113_gy_colo_milk_1ng_glyco_hcd203040_compound15.22110.22110.4 | controllerType=0 controllerNumber=1 scan=22110 | 45.8798 |
| 17782:4 | 2 | R.RTAGWNVPIGTLRPFLN[+2205.793]WTGPPEPIEAAVAR.F | HexNAc(4)Hex(5)Fuc(2)NeuAc(1) | N[+2206] | 1398.9085 | 4 | 5592.612 | 5592.6049 | 0 | 1.3 | 140 | Specific | 539.18 | 467.35 | 467.35 | 11.07 | 112 | >sp\|P02788\|TRFL_HUMAN Lactotransferrin OS=Homo sapiens OX=9606 GN=LTF PE=1 SV=6 | 1270 | 20201113_gy_colo_milk_1ng_glyco_hcd203040_compound15.21083.21083.4 | controllerType=0 controllerNumber=1 scan=21083 | 43.635 |
| 16883:4 | 2 | R.RTAGWNVPIGTLRPFLN[+1768.640]WTGPPEPIEAAVAR.F | HexNAc(4)Hex(5)Fuc(1) | N[+1769] | 1289.621 | 4 | 5155.4622 | 5155.4516 | 0 | 2.1 | 140 | Specific | 590.27 | 550.37 | 550.37 | 12.75 | 112 | >sp\|P02788\|TRFL_HUMAN Lactotransferrin OS=Homo sapiens OX=9606 GN=LTF PE=1 SV=6 | 1270 | 20201113_gy_colo_milk_1ng_glyco_hcd203040_compound15.20144.20144.4 | controllerType=0 controllerNumber=1 scan=20144 | 41.615 |
| 17678:4 | 2 | R.RTAGWNVPIGTLRPFLN[+2424.867]WTGPPEPIEAAVAR.F | HexNAc(5)Hex(6)Fuc(1)NeuAc(1) | N[+2425] | 1453.6781 | 4 | 5811.6906 | 5811.6792 | 0 | 2 | 140 | Specific | 613.08 | 526.28 | 526.28 | 13.75 | 112 | >sp\|P02788\|TRFL_HUMAN Lactotransferrin OS=Homo sapiens OX=9606 GN=LTF PE=1 SV=6 | 1270 | 20201113_gy_colo_milk_1ng_glyco_hcd203040_compound15.20975.20975.4 | controllerType=0 controllerNumber=1 scan=20975 | 43.4253 |
| 18012:3 | 2 | R.[+42.011]TAGWNVPIGTLRPFLN[+1872.651]WTGPPEPIEAAVAR.F | HexNAc(3)Hex(6)NeuAc(1) | .[42], N[+1873] | 1715.8081 | 3 | 5145.4097 | 5145.372 | 0 | 7.3 | 141 | Specific | 224.73 | 108.07 | 108.07 | 3.6 | 112 | >sp\|P02788\|TRFL_HUMAN Lactotransferrin OS=Homo sapiens OX=9606 GN=LTF PE=1 SV=6 | 1270 | 20201113_gy_colo_milk_1ng_glyco_hcd203040_compound15.21322.21322.3 | controllerType=0 controllerNumber=1 scan=21322 | 44.1304 |
| 18917:4 | 2 | R.TAGWNVPIGTLRPFLN[+2424.867]WTGPPEPIEAAVAR.F | HexNAc(5)Hex(6)Fuc(1)NeuAc(1) | N[+2425] | 1414.6553 | 4 | 5655.5993 | 5655.5781 | 0 | 3.8 | 141 | Specific | 225.47 | 225.47 | 225.47 | 7.65 | 112 | >sp\|P02788\|TRFL_HUMAN Lactotransferrin OS=Homo sapiens OX=9606 GN=LTF PE=1 SV=6 | 1270 | 20201113_gy_colo_milk_1ng_glyco_hcd203040_compound15.22267.22267.4 | controllerType=0 controllerNumber=1 scan=22267 | 46.202 |
| 21644:4 | 2 | R.TAGWNVPIGTLRPFLN[+3153.116]WTGPPEPIEAAVAR.F | HexNAc(5)Hex(6)Fuc(2)NeuAc(3) | N[+3153] | 1596.7214 | 4 | 6383.8639 | 6383.8268 | 0 | 5.8 | 141 | Specific | 233.05 | 152.69 | 152.69 | 6.45 | 112 | >sp\|P02788\|TRFL_HUMAN Lactotransferrin OS=Homo sapiens OX=9606 GN=LTF PE=1 SV=6 | 1270 | 20201113_gy_colo_milk_1ng_glyco_hcd203040_compound15.25111.25111.4 | controllerType=0 controllerNumber=1 scan=25111 | 52.359 |
| 19102:4 | 2 | R.TAGWNVPIGTLRPFLN[+2076.750]WTGPPEPIEAAVAR.F | HexNAc(4)Hex(6)Fuc(2) | N[+2077] | 1327.6323 | 4 | 5307.5075 | 5307.4612 | 0 | 8.7 | 141 | Specific | 246.59 | 246.59 | 246.59 | 4.98 | 112 | >sp\|P02788\|TRFL_HUMAN Lactotransferrin OS=Homo sapiens OX=9606 GN=LTF PE=1 SV=6 | 1270 | 20201113_gy_colo_milk_1ng_glyco_hcd203040_compound15.22459.22459.4 | controllerType=0 controllerNumber=1 scan=22459 | 46.6179 |
| 18018:4 | 2 | R.[+42.011]TAGWNVPIGTLRPFLN[+1872.651]WTGPPEPIEAAVAR.F | HexNAc(3)Hex(6)NeuAc(1) | .[42], N[+1873] | 1287.109 | 4 | 5145.4142 | 5145.372 | 0 | 8.2 | 141 | Specific | 277.12 | 238.5 | 238.5 | 6.85 | 112 | >sp\|P02788\|TRFL_HUMAN Lactotransferrin OS=Homo sapiens OX=9606 GN=LTF PE=1 SV=6 | 1270 | 20201113_gy_colo_milk_1ng_glyco_hcd203040_compound15.21328.21328.4 | controllerType=0 controllerNumber=1 scan=21328 | 44.1461 |
| 20051:4 | 2 | R.TAGWNVPIGTLRPFLN[+2863.041]WTGPPEPIEAAVAR.F | HexNAc(5)Hex(6)Fuc(4)NeuAc(1) | N[+2863] | 1524.192 | 4 | 6093.7462 | 6093.7518 | 0 | -0.9 | 141 | Specific | 280.71 | 280.71 | 280.71 | 7.15 | 112 | >sp\|P02788\|TRFL_HUMAN Lactotransferrin OS=Homo sapiens OX=9606 GN=LTF PE=1 SV=6 | 1270 | 20201113_gy_colo_milk_1ng_glyco_hcd203040_compound15.23448.23448.4 | controllerType=0 controllerNumber=1 scan=23448 | 48.7569 |
| 18020:4 | 2 | R.[+42.011]TAGWNVPIGTLRPFLN[+1872.651]WTGPPEPIEAAVAR.F | HexNAc(3)Hex(6)NeuAc(1) | .[42], N[+1873] | 1287.109 | 4 | 5145.4142 | 5145.372 | 0 | 8.2 | 141 | Specific | 288.76 | 135.99 | 135.99 | 4.72 | 112 | >sp\|P02788\|TRFL_HUMAN Lactotransferrin OS=Homo sapiens OX=9606 GN=LTF PE=1 SV=6 | 1270 | 20201113_gy_colo_milk_1ng_glyco_hcd203040_compound15.21330.21330.4 | controllerType=0 controllerNumber=1 scan=21330 | 44.1518 |
| 20048:4 | 2 | R.TAGWNVPIGTLRPFLN[+2863.041]WTGPPEPIEAAVAR.F | HexNAc(5)Hex(6)Fuc(4)NeuAc(1) | N[+2863] | 1524.192 | 4 | 6093.7462 | 6093.7518 | 0 | -0.9 | 141 | Specific | 298.71 | 298.71 | 298.71 | 7.46 | 112 | >sp\|P02788\|TRFL_HUMAN Lactotransferrin OS=Homo sapiens OX=9606 GN=LTF PE=1 SV=6 | 1270 | 20201113_gy_colo_milk_1ng_glyco_hcd203040_compound15.23445.23445.3 | controllerType=0 controllerNumber=1 scan=23445 | 48.7496 |
| 18039:4 | 2 | R.TAGWNVPIGTLRPFLN[+2059.735]WTGPPEPIEAAVAR.F | HexNAc(4)Hex(5)Fuc(1)NeuAc(1) | N[+2060] | 1323.3757 | 4 | 5290.4811 | 5290.4459 | 0 | 6.7 | 141 | Specific | 300.61 | 300.61 | 300.61 | 7.56 | 112 | >sp\|P02788\|TRFL_HUMAN Lactotransferrin OS=Homo sapiens OX=9606 GN=LTF PE=1 SV=6 | 1270 | 20201113_gy_colo_milk_1ng_glyco_hcd203040_compound15.21350.21350.2 | controllerType=0 controllerNumber=1 scan=21350 | 44.2041 |
| 21646:4 | 2 | R.TAGWNVPIGTLRPFLN[+3153.116]WTGPPEPIEAAVAR.F | HexNAc(5)Hex(6)Fuc(2)NeuAc(3) | N[+3153] | 1596.7214 | 4 | 6383.8639 | 6383.8268 | 0 | 5.8 | 141 | Specific | 311.59 | 227.5 | 227.5 | 5.47 | 112 | >sp\|P02788\|TRFL_HUMAN Lactotransferrin OS=Homo sapiens OX=9606 GN=LTF PE=1 SV=6 | 1270 | 20201113_gy_colo_milk_1ng_glyco_hcd203040_compound15.25113.25113.4 | controllerType=0 controllerNumber=1 scan=25113 | 52.3631 |
| 18974:3 | 2 | R.TAGWNVPIGTLRPFLN[+2351.851]WTGPPEPIEAAVAR.F | HexNAc(4)Hex(5)Fuc(3)NeuAc(1) | N[+2352] | 1861.5359 | 3 | 5582.593 | 5582.5617 | 0 | 5.6 | 141 | Specific | 334.55 | 334.55 | 334.55 | 7.86 | 112 | >sp\|P02788\|TRFL_HUMAN Lactotransferrin OS=Homo sapiens OX=9606 GN=LTF PE=1 SV=6 | 1270 | 20201113_gy_colo_milk_1ng_glyco_hcd203040_compound15.22326.22326.3 | controllerType=0 controllerNumber=1 scan=22326 | 46.319 |
| 18010:3 | 2 | R.[+42.011]TAGWNVPIGTLRPFLN[+1872.651]WTGPPEPIEAAVAR.F | HexNAc(3)Hex(6)NeuAc(1) | .[42], N[+1873] | 1715.8081 | 3 | 5145.4097 | 5145.372 | 0 | 7.3 | 141 | Specific | 335.48 | 243.09 | 243.09 | 7.21 | 112 | >sp\|P02788\|TRFL_HUMAN Lactotransferrin OS=Homo sapiens OX=9606 GN=LTF PE=1 SV=6 | 1270 | 20201113_gy_colo_milk_1ng_glyco_hcd203040_compound15.21320.21320.3 | controllerType=0 controllerNumber=1 scan=21320 | 44.1234 |
| 19061:4 | 2 | R.TAGWNVPIGTLRPFLN[+2059.735]WTGPPEPIEAAVAR.F | HexNAc(4)Hex(5)Fuc(1)NeuAc(1) | N[+2060] | 1323.369 | 4 | 5290.4543 | 5290.4459 | 0 | 1.6 | 141 | Specific | 359.95 | 359.95 | 359.95 | 9.05 | 112 | >sp\|P02788\|TRFL_HUMAN Lactotransferrin OS=Homo sapiens OX=9606 GN=LTF PE=1 SV=6 | 1270 | 20201113_gy_colo_milk_1ng_glyco_hcd203040_compound15.22417.22417.4 | controllerType=0 controllerNumber=1 scan=22417 | 46.5203 |
| 20050:4 | 2 | R.TAGWNVPIGTLRPFLN[+2863.041]WTGPPEPIEAAVAR.F | HexNAc(5)Hex(6)Fuc(4)NeuAc(1) | N[+2863] | 1524.192 | 4 | 6093.7462 | 6093.7518 | 0 | -0.9 | 141 | Specific | 370.16 | 348.45 | 348.45 | 8.23 | 112 | >sp\|P02788\|TRFL_HUMAN Lactotransferrin OS=Homo sapiens OX=9606 GN=LTF PE=1 SV=6 | 1270 | 20201113_gy_colo_milk_1ng_glyco_hcd203040_compound15.23447.23447.3 | controllerType=0 controllerNumber=1 scan=23447 | 48.755 |
| 18042:4 | 2 | R.TAGWNVPIGTLRPFLN[+2059.735]WTGPPEPIEAAVAR.F | HexNAc(4)Hex(5)Fuc(1)NeuAc(1) | N[+2060] | 1323.3757 | 4 | 5290.4811 | 5290.4459 | 0 | 6.7 | 141 | Specific | 412.4 | 412.4 | 412.4 | 10.06 | 112 | >sp\|P02788\|TRFL_HUMAN Lactotransferrin OS=Homo sapiens OX=9606 GN=LTF PE=1 SV=6 | 1270 | 20201113_gy_colo_milk_1ng_glyco_hcd203040_compound15.21353.21353.2 | controllerType=0 controllerNumber=1 scan=21353 | 44.2146 |
| 18891:5 | 2 | R.TAGWNVPIGTLRPFLN[+2570.925]WTGPPEPIEAAVAR.F | HexNAc(5)Hex(6)Fuc(2)NeuAc(1) | N[+2571] | 1161.1371 | 5 | 5801.6563 | 5801.636 | 0 | 3.5 | 141 | Specific | 433.85 | 351.57 | 351.57 | 9.36 | 112 | >sp\|P02788\|TRFL_HUMAN Lactotransferrin OS=Homo sapiens OX=9606 GN=LTF PE=1 SV=6 | 1270 | 20201113_gy_colo_milk_1ng_glyco_hcd203040_compound15.22240.22240.5 | controllerType=0 controllerNumber=1 scan=22240 | 46.1525 |
| 20318:3 | 2 | R.TAGWNVPIGTLRPFLN[+2350.830]WTGPPEPIEAAVAR.F | HexNAc(4)Hex(5)Fuc(1)NeuAc(2) | N[+2351] | 1861.1864 | 3 | 5581.5445 | 5581.5413 | 0 | 0.6 | 141 | Specific | 448.32 | 298.19 | 298.19 | 8.14 | 112 | >sp\|P02788\|TRFL_HUMAN Lactotransferrin OS=Homo sapiens OX=9606 GN=LTF PE=1 SV=6 | 1270 | 20201113_gy_colo_milk_1ng_glyco_hcd203040_compound15.23727.23727.2 | controllerType=0 controllerNumber=1 scan=23727 | 49.3785 |

Table S4. Cont.

| Query #:z | Protein Rank | Glycopeptide | Glycan | Modification Type(s) | Observed m/z | z | Observed (M+H) | Calc. mass (M+H) | Off-by-x error | Mass error (ppm) | StartPosition | Cleavage | Score | Delta | Delta Mod | \|Log Prob\| | # of unique peptides | Protein Name | Protein DB number | Comment | Scan # | Scan Time |
|---|---|---|---|---|---|---|---|---|---|---|---|---|---|---|---|---|---|---|---|---|---|---|
| 18848:3 | 2 | R.TAGWNVPIGTLRPFLN[+2570.925]WTGPPEPIEAAVAR.F | HexNAc(5)Hex(6)Fuc(2)NeuAc(1) | N[+2571] | 1934.5554 | 3 | 5801.6516 | 5801.636 | 0 | 2.7 | 141 | Specific | 495.75 | 361.07 | 361.07 | 9.53 | 112 | >sp\|P02788\|TRFL_HUMAN Lactotransferrin OS=Homo sapiens OX=9606 GN=LTF PE=1 SV=6 | 1270 | 20201113_gy_colo_milk_1ng_glyco_hcd203040_compound15.22195.22195.3 | controllerType=0 controllerNumber=1 scan=22195 | 46.0546 |
| 18851:3 | 2 | R.TAGWNVPIGTLRPFLN[+2570.925]WTGPPEPIEAAVAR.F | HexNAc(5)Hex(6)Fuc(2)NeuAc(1) | N[+2571] | 1934.5554 | 3 | 5801.6516 | 5801.636 | 0 | 2.7 | 141 | Specific | 563.12 | 445.66 | 445.66 | 11.39 | 112 | >sp\|P02788\|TRFL_HUMAN Lactotransferrin OS=Homo sapiens OX=9606 GN=LTF PE=1 SV=6 | 1270 | 20201113_gy_colo_milk_1ng_glyco_hcd203040_compound15.22198.22198.3 | controllerType=0 controllerNumber=1 scan=22198 | 46.0602 |
| 19033:3 | 2 | R.TAGWNVPIGTLRPFLN[+2205.793]WTGPPEPIEAAVAR.F | HexNAc(4)Hex(5)Fuc(2)NeuAc(1) | N[+2206] | 1812.843 | 3 | 5436.5144 | 5436.5038 | 0 | 2 | 141 | Specific | 582.92 | 489.95 | 489.95 | 12.75 | 112 | >sp\|P02788\|TRFL_HUMAN Lactotransferrin OS=Homo sapiens OX=9606 GN=LTF PE=1 SV=6 | 1270 | 20201113_gy_colo_milk_1ng_glyco_hcd203040_compound15.22387.22387.2 | controllerType=0 controllerNumber=1 scan=22387 | 46.4543 |
| 18977:3 | 2 | R.TAGWNVPIGTLRPFLN[+2351.851]WTGPPEPIEAAVAR.F | HexNAc(4)Hex(5)Fuc(3)NeuAc(1) | N[+2352] | 1861.5359 | 3 | 5582.593 | 5582.5617 | 0 | 5.6 | 141 | Specific | 604.37 | 511.8 | 511.8 | 13.06 | 112 | >sp\|P02788\|TRFL_HUMAN Lactotransferrin OS=Homo sapiens OX=9606 GN=LTF PE=1 SV=6 | 1270 | 20201113_gy_colo_milk_1ng_glyco_hcd203040_compound15.22329.22329.3 | controllerType=0 controllerNumber=1 scan=22329 | 46.3241 |
| 20053:4 | 2 | R.TAGWNVPIGTLRPFLN[+2863.041]WTGPPEPIEAAVAR.F | HexNAc(5)Hex(6)Fuc(4)NeuAc(1) | N[+2863] | 1524.192 | 4 | 6093.7462 | 6093.7518 | 0 | -0.9 | 141 | Specific | 606.51 | 513.95 | 513.95 | 13.07 | 112 | >sp\|P02788\|TRFL_HUMAN Lactotransferrin OS=Homo sapiens OX=9606 GN=LTF PE=1 SV=6 | 1270 | 20201113_gy_colo_milk_1ng_glyco_hcd203040_compound15.23450.23450.4 | controllerType=0 controllerNumber=1 scan=23450 | 48.7606 |
| 18823:4 | 2 | R.TAGWNVPIGTLRPFLN[+2716.983]WTGPPEPIEAAVAR.F | HexNAc(5)Hex(6)Fuc(3)NeuAc(1) | N[+2717] | 1487.6815 | 4 | 5947.7043 | 5947.6939 | 0 | 1.7 | 141 | Specific | 620.33 | 565.55 | 565.55 | 13.92 | 112 | >sp\|P02788\|TRFL_HUMAN Lactotransferrin OS=Homo sapiens OX=9606 GN=LTF PE=1 SV=6 | 1270 | 20201113_gy_colo_milk_1ng_glyco_hcd203040_compound15.22169.22169.4 | controllerType=0 controllerNumber=1 scan=22169 | 45.9931 |
| 18842:4 | 2 | R.TAGWNVPIGTLRPFLN[+2570.925]WTGPPEPIEAAVAR.F | HexNAc(5)Hex(6)Fuc(2)NeuAc(1) | N[+2571] | 1451.1702 | 4 | 5801.6588 | 5801.636 | 0 | 3.9 | 141 | Specific | 629.27 | 548.8 | 548.8 | 13.58 | 112 | >sp\|P02788\|TRFL_HUMAN Lactotransferrin OS=Homo sapiens OX=9606 GN=LTF PE=1 SV=6 | 1270 | 20201113_gy_colo_milk_1ng_glyco_hcd203040_compound15.22189.22189.4 | controllerType=0 controllerNumber=1 scan=22189 | 46.0376 |
| 18826:4 | 2 | R.TAGWNVPIGTLRPFLN[+2716.983]WTGPPEPIEAAVAR.F | HexNAc(5)Hex(6)Fuc(3)NeuAc(1) | N[+2717] | 1487.6815 | 4 | 5947.7043 | 5947.6939 | 0 | 1.7 | 141 | Specific | 660.36 | 532.54 | 532.54 | 14.92 | 112 | >sp\|P02788\|TRFL_HUMAN Lactotransferrin OS=Homo sapiens OX=9606 GN=LTF PE=1 SV=6 | 1270 | 20201113_gy_colo_milk_1ng_glyco_hcd203040_compound15.22172.22172.4 | controllerType=0 controllerNumber=1 scan=22172 | 45.9991 |
| 18969:4 | 2 | R.TAGWNVPIGTLRPFLN[+2205.793]WTGPPEPIEAAVAR.F | HexNAc(4)Hex(5)Fuc(2)NeuAc(1) | N[+2206] | 1359.8895 | 4 | 5436.5363 | 5436.5038 | 0 | 6 | 141 | Specific | 661.02 | 606.38 | 606.38 | 14.18 | 112 | >sp\|P02788\|TRFL_HUMAN Lactotransferrin OS=Homo sapiens OX=9606 GN=LTF PE=1 SV=6 | 1270 | 20201113_gy_colo_milk_1ng_glyco_hcd203040_compound15.22321.22321.4 | controllerType=0 controllerNumber=1 scan=22321 | 46.3086 |
| 18976:3 | 2 | R.TAGWNVPIGTLRPFLN[+2205.793]WTGPPEPIEAAVAR.F | HexNAc(4)Hex(5)Fuc(2)NeuAc(1) | N[+2206] | 1812.8501 | 3 | 5436.5356 | 5436.5038 | 0 | 5.9 | 141 | Specific | 670.58 | 611.12 | 611.12 | 14.01 | 112 | >sp\|P02788\|TRFL_HUMAN Lactotransferrin OS=Homo sapiens OX=9606 GN=LTF PE=1 SV=6 | 1270 | 20201113_gy_colo_milk_1ng_glyco_hcd203040_compound15.22328.22328.3 | controllerType=0 controllerNumber=1 scan=22328 | 46.323 |
| 18845:4 | 2 | R.TAGWNVPIGTLRPFLN[+2570.925]WTGPPEPIEAAVAR.F | HexNAc(5)Hex(6)Fuc(2)NeuAc(1) | N[+2571] | 1451.1702 | 4 | 5801.6588 | 5801.636 | 0 | 3.9 | 141 | Specific | 681.1 | 473.73 | 473.73 | 14.58 | 112 | >sp\|P02788\|TRFL_HUMAN Lactotransferrin OS=Homo sapiens OX=9606 GN=LTF PE=1 SV=6 | 1270 | 20201113_gy_colo_milk_1ng_glyco_hcd203040_compound15.22192.22192.4 | controllerType=0 controllerNumber=1 scan=22192 | 46.05 |
| 19064:3 | 2 | R.TAGWNVPIGTLRPFLN[+2059.735]WTGPPEPIEAAVAR.F | HexNAc(4)Hex(5)Fuc(1)NeuAc(1) | N[+2060] | 1764.156 | 3 | 5290.4534 | 5290.4459 | 0 | 1.4 | 141 | Specific | 684.28 | 632.98 | 632.98 | 14.61 | 112 | >sp\|P02788\|TRFL_HUMAN Lactotransferrin OS=Homo sapiens OX=9606 GN=LTF PE=1 SV=6 | 1270 | 20201113_gy_colo_milk_1ng_glyco_hcd203040_compound15.22420.22420.3 | controllerType=0 controllerNumber=1 scan=22420 | 46.526 |
| 18973:3 | 2 | R.TAGWNVPIGTLRPFLN[+2205.793]WTGPPEPIEAAVAR.F | HexNAc(4)Hex(5)Fuc(2)NeuAc(1) | N[+2206] | 1812.8501 | 3 | 5436.5356 | 5436.5038 | 0 | 5.9 | 141 | Specific | 699.7 | 614.49 | 614.49 | 14.01 | 112 | >sp\|P02788\|TRFL_HUMAN Lactotransferrin OS=Homo sapiens OX=9606 GN=LTF PE=1 SV=6 | 1270 | 20201113_gy_colo_milk_1ng_glyco_hcd203040_compound15.22325.22325.3 | controllerType=0 controllerNumber=1 scan=22325 | 46.3179 |
| 19079:5 | 2 | R.TAGWNVPIGTLRPFLN[+2059.735]WTGPPEPIEAAVAR.F | HexNAc(4)Hex(5)Fuc(1)NeuAc(1) | N[+2060] | 1058.8922 | 5 | 5290.432 | 5290.4459 | 0 | -2.6 | 141 | Specific | 707.5 | 600.22 | 600.22 | 15.01 | 112 | >sp\|P02788\|TRFL_HUMAN Lactotransferrin OS=Homo sapiens OX=9606 GN=LTF PE=1 SV=6 | 1270 | 20201113_gy_colo_milk_1ng_glyco_hcd203040_compound15.22435.22435.5 | controllerType=0 controllerNumber=1 scan=22435 | 46.5612 |
| 18870:4 | 2 | R.TAGWNVPIGTLRPFLN[+2862.021]WTGPPEPIEAAVAR.F | HexNAc(5)Hex(6)Fuc(2)NeuAc(2) | N[+2862] | 1523.9446 | 4 | 6092.7565 | 6092.7314 | 0 | 4.1 | 141 | Specific | 717.33 | 583.67 | 583.67 | 15.58 | 112 | >sp\|P02788\|TRFL_HUMAN Lactotransferrin OS=Homo sapiens OX=9606 GN=LTF PE=1 SV=6 | 1270 | 20201113_gy_colo_milk_1ng_glyco_hcd203040_compound15.22218.22218.4 | controllerType=0 controllerNumber=1 scan=22218 | 46.1072 |
| 18919:4 | 2 | R.TAGWNVPIGTLRPFLN[+2424.867]WTGPPEPIEAAVAR.F | HexNAc(5)Hex(6)Fuc(1)NeuAc(1) | N[+2425] | 1414.6553 | 4 | 5655.5993 | 5655.5781 | 0 | 3.8 | 141 | Specific | 738.11 | 595.43 | 595.43 | 15.92 | 112 | >sp\|P02788\|TRFL_HUMAN Lactotransferrin OS=Homo sapiens OX=9606 GN=LTF PE=1 SV=6 | 1270 | 20201113_gy_colo_milk_1ng_glyco_hcd203040_compound15.22269.22269.4 | controllerType=0 controllerNumber=1 scan=22269 | 46.2064 |
| 20254:5 | 2 | R.TAGWNVPIGTLRPFLN[+2350.830]WTGPPEPIEAAVAR.F | HexNAc(4)Hex(5)Fuc(1)NeuAc(2) | N[+2351] | 1117.1194 | 5 | 5581.5678 | 5581.5413 | 0 | 4.8 | 141 | Specific | 816.16 | 649.04 | 649.04 | 17.72 | 112 | >sp\|P02788\|TRFL_HUMAN Lactotransferrin OS=Homo sapiens OX=9606 GN=LTF PE=1 SV=6 | 1270 | 20201113_gy_colo_milk_1ng_glyco_hcd203040_compound15.23660.23660.5 | controllerType=0 controllerNumber=1 scan=23660 | 49.2222 |
| 20252:5 | 2 | R.TAGWNVPIGTLRPFLN[+2350.830]WTGPPEPIEAAVAR.F | HexNAc(4)Hex(5)Fuc(1)NeuAc(2) | N[+2351] | 1117.1194 | 5 | 5581.5678 | 5581.5413 | 0 | 4.8 | 141 | Specific | 844.24 | 761.11 | 761.11 | 17.72 | 112 | >sp\|P02788\|TRFL_HUMAN Lactotransferrin OS=Homo sapiens OX=9606 GN=LTF PE=1 SV=6 | 1270 | 20201113_gy_colo_milk_1ng_glyco_hcd203040_compound15.23658.23658.5 | controllerType=0 controllerNumber=1 scan=23658 | 49.2127 |
| 18971:4 | 2 | R.TAGWNVPIGTLRPFLN[+2205.793]WTGPPEPIEAAVAR.F | HexNAc(4)Hex(5)Fuc(2)NeuAc(1) | N[+2206] | 1359.8895 | 4 | 5436.5363 | 5436.5038 | 0 | 6 | 141 | Specific | 872.52 | 731.06 | 731.06 | 18.18 | 112 | >sp\|P02788\|TRFL_HUMAN Lactotransferrin OS=Homo sapiens OX=9606 GN=LTF PE=1 SV=6 | 1270 | 20201113_gy_colo_milk_1ng_glyco_hcd203040_compound15.22323.22323.4 | controllerType=0 controllerNumber=1 scan=22323 | 46.3152 |
| 19063:4 | 2 | R.TAGWNVPIGTLRPFLN[+2059.735]WTGPPEPIEAAVAR.F | HexNAc(4)Hex(5)Fuc(1)NeuAc(1) | N[+2060] | 1323.369 | 4 | 5290.4543 | 5290.4459 | 0 | 1.6 | 141 | Specific | 888.49 | 743.96 | 743.96 | 18.78 | 112 | >sp\|P02788\|TRFL_HUMAN Lactotransferrin OS=Homo sapiens OX=9606 GN=LTF PE=1 SV=6 | 1270 | 20201113_gy_colo_milk_1ng_glyco_hcd203040_compound15.22419.22419.4 | controllerType=0 controllerNumber=1 scan=22419 | 46.5251 |
| 16586:2 | 2 | R.TAGWNIPMGLLFN[+203.079]QTGSC[+57.021]K.F | HexNAc(1) | C[+57], N[+203] | 1149.5423 | 2 | 2298.0773 | 2298.0893 | 0 | -5.2 | 485 | Specific | 448.71 | 448.71 | 448.71 | 7.42 | 112 | >sp\|P02788\|TRFL_HUMAN Lactotransferrin OS=Homo sapiens OX=9606 GN=LTF PE=1 SV=6 | 1270 | 20201113_gy_colo_milk_1ng_glyco_hcd203040_compound15.19834.19834.2 | controllerType=0 controllerNumber=1 scan=19834 | 40.9503 |
| 18461:3 | 2 | R.TAGWNIPMGLLFN[+1856.656]QTGSC[+57.021]K.F | HexNAc(3)Hex(5)Fuc(1)NeuAc(1) | C[+57], N[+1857] | 1317.9057 | 3 | 3951.7025 | 3951.6655 | 0 | 9.4 | 485 | Specific | 489.41 | 441.71 | 203.47 | 9.29 | 112 | >sp\|P02788\|TRFL_HUMAN Lactotransferrin OS=Homo sapiens OX=9606 GN=LTF PE=1 SV=6 | 1270 | 20201113_gy_colo_milk_1ng_glyco_hcd203040_compound15.21791.21791.3 | controllerType=0 controllerNumber=1 scan=21791 | 45.2089 |
| 18340:4 | 2 | R.TAGWNIPMGLLFN[+2059.735]QTGSC[+57.021]K.F | HexNAc(4)Hex(5)Fuc(1)NeuAc(1) | C[+57], N[+2060] | 1039.4447 | 4 | 4154.7572 | 4154.7449 | 0 | 3 | 485 | Specific | 511.93 | 378.01 | 378.01 | 9.13 | 112 | >sp\|P02788\|TRFL_HUMAN Lactotransferrin OS=Homo sapiens OX=9606 GN=LTF PE=1 SV=6 | 1270 | 20201113_gy_colo_milk_1ng_glyco_hcd203040_compound15.21665.21665.2 | controllerType=0 controllerNumber=1 scan=21665 | 44.9232 |
| 20299:3 | 2 | R.TAGWNIPMGLLFN[+2350.830]QTGSC[+57.021]K.F | HexNAc(4)Hex(5)Fuc(1)NeuAc(2) | C[+57], N[+2351] | 1482.6188 | 3 | 4445.8419 | 4445.8403 | 0 | 0.4 | 485 | Specific | 536.05 | 536.05 | 536.05 | 9.96 | 112 | >sp\|P02788\|TRFL_HUMAN Lactotransferrin OS=Homo sapiens OX=9606 GN=LTF PE=1 SV=6 | 1270 | 20201113_gy_colo_milk_1ng_glyco_hcd203040_compound15.23708.23708.3 | controllerType=0 controllerNumber=1 scan=23708 | 49.3428 |
| 18336:3 | 2 | R.TAGWNIPMGLLFN[+2351.851]QTGSC[+57.021]K.F | HexNAc(4)Hex(5)Fuc(3)NeuAc(1) | C[+57], N[+2352] | 1482.9717 | 3 | 4446.9007 | 4446.8607 | 0 | 9 | 485 | Specific | 581.24 | 557.84 | 557.84 | 11.14 | 112 | >sp\|P02788\|TRFL_HUMAN Lactotransferrin OS=Homo sapiens OX=9606 GN=LTF PE=1 SV=6 | 1270 | 20201113_gy_colo_milk_1ng_glyco_hcd203040_compound15.21661.21661.3 | controllerType=0 controllerNumber=1 scan=21661 | 44.9148 |
| 16402:3 | 2 | R.TAGWNIPMGLLFN[+2133.772]QTGSC[+57.021]K.F | HexNAc(5)Hex(6)Fuc(1) | C[+57], N[+2134] | 1410.2718 | 3 | 4228.8008 | 4228.7817 | 0 | 4.5 | 485 | Specific | 605.31 | 563.29 | 309.92 | 12.88 | 112 | >sp\|P02788\|TRFL_HUMAN Lactotransferrin OS=Homo sapiens OX=9606 GN=LTF PE=1 SV=6 | 1270 | 20201113_gy_colo_milk_1ng_glyco_hcd203040_compound15.19643.19643.3 | controllerType=0 controllerNumber=1 scan=19643 | 40.5715 |
| 16589:2 | 2 | R.TAGWNIPMGLLFN[+203.079]QTGSC[+57.021]K.F | HexNAc(1) | C[+57], N[+203] | 1149.5423 | 2 | 2298.0773 | 2298.0893 | 0 | -5.2 | 485 | Specific | 622.62 | 574.6 | 574.6 | 9.97 | 112 | >sp\|P02788\|TRFL_HUMAN Lactotransferrin OS=Homo sapiens OX=9606 GN=LTF PE=1 SV=6 | 1270 | 20201113_gy_colo_milk_1ng_glyco_hcd203040_compound15.19837.19837.2 | controllerType=0 controllerNumber=1 scan=19837 | 40.9556 |
| 16751:3 | 2 | R.TAGWNIPMGLLFN[+1622.582]QTGSC[+57.021]K.F | HexNAc(4)Hex(5) | C[+57], N[+1623] | 1239.8769 | 3 | 3717.6161 | 3717.5916 | 0 | 6.6 | 485 | Specific | 674.95 | 674.95 | 256.53 | 13.48 | 112 | >sp\|P02788\|TRFL_HUMAN Lactotransferrin OS=Homo sapiens OX=9606 GN=LTF PE=1 SV=6 | 1270 | 20201113_gy_colo_milk_1ng_glyco_hcd203040_compound15.20006.20006.3 | controllerType=0 controllerNumber=1 scan=20006 | 41.2985 |
| 16404:3 | 2 | R.TAGWNIPMGLLFN[+2133.772]QTGSC[+57.021]K.F | HexNAc(5)Hex(6)Fuc(1) | C[+57], N[+2134] | 1410.2718 | 3 | 4228.8008 | 4228.7817 | 0 | 4.5 | 485 | Specific | 701.95 | 701.95 | 368.27 | 14.88 | 112 | >sp\|P02788\|TRFL_HUMAN Lactotransferrin OS=Homo sapiens OX=9606 GN=LTF PE=1 SV=6 | 1270 | 20201113_gy_colo_milk_1ng_glyco_hcd203040_compound15.19645.19645.3 | controllerType=0 controllerNumber=1 scan=19645 | 40.5755 |
| 16591:4 | 2 | R.TAGWNIPMGLLFN[+1768.640]QTGSC[+57.021]K.F | HexNAc(4)Hex(5)Fuc(1) | C[+57], N[+1769] | 966.6708 | 4 | 3863.6612 | 3863.6495 | 0 | 3 | 485 | Specific | 749.07 | 572.55 | 572.55 | 15.35 | 112 | >sp\|P02788\|TRFL_HUMAN Lactotransferrin OS=Homo sapiens OX=9606 GN=LTF PE=1 SV=6 | 1270 | 20201113_gy_colo_milk_1ng_glyco_hcd203040_compound15.19839.19839.4 | controllerType=0 controllerNumber=1 scan=19839 | 40.96 |
| 18472:4 | 2 | R.TAGWNIPMGLLFN[+1913.677]QTGSC[+57.021]K.F | HexNAc(4)Hex(5)NeuAc(1) | C[+57], N[+1914] | 1002.936 | 4 | 4008.722 | 4008.687 | 0 | 8.7 | 485 | Specific | 778.78 | 573.87 | 573.87 | 15.31 | 112 | >sp\|P02788\|TRFL_HUMAN Lactotransferrin OS=Homo sapiens OX=9606 GN=LTF PE=1 SV=6 | 1270 | 20201113_gy_colo_milk_1ng_glyco_hcd203040_compound15.21802.21802.2 | controllerType=0 controllerNumber=1 scan=21802 | 45.2302 |
| 18339:3 | 2 | R.TAGWNIPMGLLFN[+2351.851]QTGSC[+57.021]K.F | HexNAc(4)Hex(5)Fuc(3)NeuAc(1) | C[+57], N[+2352] | 1482.9717 | 3 | 4446.9007 | 4446.8607 | 0 | 9 | 485 | Specific | 779.2 | 693.74 | 693.74 | 15.14 | 112 | >sp\|P02788\|TRFL_HUMAN Lactotransferrin OS=Homo sapiens OX=9606 GN=LTF PE=1 SV=6 | 1270 | 20201113_gy_colo_milk_1ng_glyco_hcd203040_compound15.21664.21664.3 | controllerType=0 controllerNumber=1 scan=21664 | 44.9213 |
| 18464:3 | 2 | R.TAGWNIPMGLLFN[+1856.656]QTGSC[+57.021]K.F | HexNAc(3)Hex(5)Fuc(1)NeuAc(1) | C[+57], N[+1857] | 1317.9057 | 3 | 3951.7025 | 3951.6655 | 0 | 9.4 | 485 | Specific | 787.8 | 676.26 | 331.01 | 15.14 | 112 | >sp\|P02788\|TRFL_HUMAN Lactotransferrin OS=Homo sapiens OX=9606 GN=LTF PE=1 SV=6 | 1270 | 20201113_gy_colo_milk_1ng_glyco_hcd203040_compound15.21794.21794.3 | controllerType=0 controllerNumber=1 scan=21794 | 45.2134 |
| 16545:3 | 2 | R.TAGWNIPMGLLFN[+1768.640]QTGSC[+57.021]K.F | HexNAc(4)Hex(5)Fuc(1) | C[+57], N[+1769] | 1288.558 | 3 | 3863.6596 | 3863.6495 | 0 | 2.6 | 485 | Specific | 818.08 | 818.08 | 818.08 | 17.22 | 112 | >sp\|P02788\|TRFL_HUMAN Lactotransferrin OS=Homo sapiens OX=9606 GN=LTF PE=1 SV=6 | 1270 | 20201113_gy_colo_milk_1ng_glyco_hcd203040_compound15.19792.19792.3 | controllerType=0 controllerNumber=1 scan=19792 | 40.8747 |
| 20291:4 | 2 | R.TAGWNIPMGLLFN[+2350.830]QTGSC[+57.021]K.F | HexNAc(4)Hex(5)Fuc(1)NeuAc(2) | C[+57], N[+2351] | 1112.2172 | 4 | 4445.847 | 4445.8403 | 0 | 1.5 | 485 | Specific | 831 | 673.61 | 673.61 | 17.22 | 112 | >sp\|P02788\|TRFL_HUMAN Lactotransferrin OS=Homo sapiens OX=9606 GN=LTF PE=1 SV=6 | 1270 | 20201113_gy_colo_milk_1ng_glyco_hcd203040_compound15.23699.23699.4 | controllerType=0 controllerNumber=1 scan=23699 | 49.3182 |
| 16922:2 | 2 | R.TAGWNIPMGLLFN[+1216.423]QTGSC[+57.021]K.F | HexNAc(2)Hex(5) | C[+57], N[+1216] | 1656.2262 | 2 | 3311.4451 | 3311.4328 | 0 | 3.7 | 485 | Specific | 833.43 | 821.44 | 275.36 | 17.04 | 112 | >sp\|P02788\|TRFL_HUMAN Lactotransferrin OS=Homo sapiens OX=9606 GN=LTF PE=1 SV=6 | 1270 | 20201113_gy_colo_milk_1ng_glyco_hcd203040_compound15.20186.20186.2 | controllerType=0 controllerNumber=1 scan=20186 | 41.7377 |
| 18488:4 | 2 | R.TAGWNIPMGLLFN[+1913.677]QTGSC[+57.021]K.F | HexNAc(4)Hex(5)NeuAc(1) | C[+57], N[+1914] | 1002.9292 | 4 | 4008.6949 | 4008.687 | 0 | 2 | 485 | Specific | 836.7 | 616.07 | 616.07 | 17.35 | 112 | >sp\|P02788\|TRFL_HUMAN Lactotransferrin OS=Homo sapiens OX=9606 GN=LTF PE=1 SV=6 | 1270 | 20201113_gy_colo_milk_1ng_glyco_hcd203040_compound15.21819.21819.4 | controllerType=0 controllerNumber=1 scan=21819 | 45.2783 |
| 16753:3 | 2 | R.TAGWNIPMGLLFN[+1622.582]QTGSC[+57.021]K.F | HexNAc(4)Hex(5) | C[+57], N[+1623] | 1239.8769 | 3 | 3717.6161 | 3717.5916 | 0 | 6.6 | 485 | Specific | 839.61 | 707.33 | 369.77 | 16.48 | 112 | >sp\|P02788\|TRFL_HUMAN Lactotransferrin OS=Homo sapiens OX=9606 GN=LTF PE=1 SV=6 | 1270 | 20201113_gy_colo_milk_1ng_glyco_hcd203040_compound15.20008.20008.3 | controllerType=0 controllerNumber=1 scan=20008 | 41.3048 |
| 18342:4 | 2 | R.TAGWNIPMGLLFN[+2059.735]QTGSC[+57.021]K.F | HexNAc(4)Hex(5)Fuc(1)NeuAc(1) | C[+57], N[+2060] | 1039.4447 | 4 | 4154.7572 | 4154.7449 | 0 | 3 | 485 | Specific | 852.22 | 658.27 | 658.27 | 18.35 | 112 | >sp\|P02788\|TRFL_HUMAN Lactotransferrin OS=Homo sapiens OX=9606 GN=LTF PE=1 SV=6 | 1270 | 20201113_gy_colo_milk_1ng_glyco_hcd203040_compound15.21667.21667.2 | controllerType=0 controllerNumber=1 scan=21667 | 44.9289 |
| 16547:3 | 2 | R.TAGWNIPMGLLFN[+1768.640]QTGSC[+57.021]K.F | HexNAc(4)Hex(5)Fuc(1) | C[+57], N[+1769] | 1288.558 | 3 | 3863.6596 | 3863.6495 | 0 | 2.6 | 485 | Specific | 863.43 | 863.43 | 863.43 | 18.22 | 112 | >sp\|P02788\|TRFL_HUMAN Lactotransferrin OS=Homo sapiens OX=9606 GN=LTF PE=1 SV=6 | 1270 | 20201113_gy_colo_milk_1ng_glyco_hcd203040_compound15.19794.19794.3 | controllerType=0 controllerNumber=1 scan=19794 | 40.8793 |
| 20460:3 | 2 | R.TAGWNIPMGLLFN[+2204.772]QTGSC[+57.021]K.F | HexNAc(4)Hex(5)NeuAc(2) | C[+57], N[+2205] | 1433.9368 | 3 | 4299.7959 | 4299.7824 | 0 | 3.1 | 485 | Specific | 883.59 | 808.62 | 808.62 | 18.21 | 112 | >sp\|P02788\|TRFL_HUMAN Lactotransferrin OS=Homo sapiens OX=9606 GN=LTF PE=1 SV=6 | 1270 | 20201113_gy_colo_milk_1ng_glyco_hcd203040_compound15.23876.23876.3 | controllerType=0 controllerNumber=1 scan=23876 | 49.7143 |
| 20301:3 | 2 | R.TAGWNIPMGLLFN[+2350.830]QTGSC[+57.021]K.F | HexNAc(4)Hex(5)Fuc(1)NeuAc(2) | C[+57], N[+2351] | 1482.6188 | 3 | 4445.8419 | 4445.8403 | 0 | 0.4 | 485 | Specific | 893.38 | 751.5 | 751.5 | 17.66 | 112 | >sp\|P02788\|TRFL_HUMAN Lactotransferrin OS=Homo sapiens OX=9606 GN=LTF PE=1 SV=6 | 1270 | 20201113_gy_colo_milk_1ng_glyco_hcd203040_compound15.23710.23710.3 | controllerType=0 controllerNumber=1 scan=23710 | 49.3473 |
| 16920:2 | 2 | R.TAGWNIPMGLLFN[+1216.423]QTGSC[+57.021]K.F | HexNAc(2)Hex(5) | C[+57], N[+1216] | 1656.2262 | 2 | 3311.4451 | 3311.4328 | 0 | 3.7 | 485 | Specific | 902.23 | 850.45 | 335.92 | 19.04 | 112 | >sp\|P02788\|TRFL_HUMAN Lactotransferrin OS=Homo sapiens OX=9606 GN=LTF PE=1 SV=6 | 1270 | 20201113_gy_colo_milk_1ng_glyco_hcd203040_compound15.20184.20184.2 | controllerType=0 controllerNumber=1 scan=20184 | 41.7273 |
| 18460:3 | 2 | R.TAGWNIPMGLLFN[+1913.677]QTGSC[+57.021]K.F | HexNAc(4)Hex(5)NeuAc(1) | C[+57], N[+1914] | 1336.9134 | 3 | 4008.7256 | 4008.687 | 0 | 9.6 | 485 | Specific | 905.64 | 905.64 | 905.64 | 18.14 | 112 | >sp\|P02788\|TRFL_HUMAN Lactotransferrin OS=Homo sapiens OX=9606 GN=LTF PE=1 SV=6 | 1270 | 20201113_gy_colo_milk_1ng_glyco_hcd203040_compound15.21790.21790.3 | controllerType=0 controllerNumber=1 scan=21790 | 45.2076 |
| 18333:3 | 2 | R.TAGWNIPMGLLFN[+2059.735]QTGSC[+57.021]K.F | HexNAc(4)Hex(5)Fuc(1)NeuAc(1) | C[+57], N[+2060] | 1385.5917 | 3 | 4154.7606 | 4154.7449 | 0 | 3.8 | 485 | Specific | 913.08 | 913.08 | 913.08 | 18.88 | 112 | >sp\|P02788\|TRFL_HUMAN Lactotransferrin OS=Homo sapiens OX=9606 GN=LTF PE=1 SV=6 | 1270 | 20201113_gy_colo_milk_1ng_glyco_hcd203040_compound15.21658.21658.3 | controllerType=0 controllerNumber=1 scan=21658 | 44.9119 |
| 18293:3 | 2 | R.TAGWNIPMGLLFN[+2205.793]QTGSC[+57.021]K.F | HexNAc(4)Hex(5)Fuc(2)NeuAc(1) | C[+57], N[+2206] | 1434.2767 | 3 | 4300.8155 | 4300.8028 | 0 | 2.9 | 485 | Specific | 913.12 | 900.51 | 900.51 | 19.21 | 112 | >sp\|P02788\|TRFL_HUMAN Lactotransferrin OS=Homo sapiens OX=9606 GN=LTF PE=1 SV=6 | 1270 | 20201113_gy_colo_milk_1ng_glyco_hcd203040_compound15.21616.21616.3 | controllerType=0 controllerNumber=1 scan=21616 | 44.8165 |
| 18457:3 | 2 | R.TAGWNIPMGLLFN[+1913.677]QTGSC[+57.021]K.F | HexNAc(4)Hex(5)NeuAc(1) | C[+57], N[+1914] | 1336.9134 | 3 | 4008.7256 | 4008.687 | 0 | 9.6 | 485 | Specific | 941.46 | 932.96 | 932.96 | 18.14 | 112 | >sp\|P02788\|TRFL_HUMAN Lactotransferrin OS=Homo sapiens OX=9606 GN=LTF PE=1 SV=6 | 1270 | 20201113_gy_colo_milk_1ng_glyco_hcd203040_compound15.21787.21787.3 | controllerType=0 controllerNumber=1 scan=21787 | 45.1963 |
| 20293:4 | 2 | R.TAGWNIPMGLLFN[+2350.830]QTGSC[+57.021]K.F | HexNAc(4)Hex(5)Fuc(1)NeuAc(2) | C[+57], N[+2351] | 1112.2172 | 4 | 4445.847 | 4445.8403 | 0 | 1.5 | 485 | Specific | 950.06 | 798.36 | 798.36 | 20.22 | 112 | >sp\|P02788\|TRFL_HUMAN Lactotransferrin OS=Homo sapiens OX=9606 GN=LTF PE=1 SV=6 | 1270 | 20201113_gy_colo_milk_1ng_glyco_hcd203040_compound15.23701.23701.4 | controllerType=0 controllerNumber=1 scan=23701 | 49.3245 |
| 18331:3 | 2 | R.TAGWNIPMGLLFN[+2059.735]QTGSC[+57.021]K.F | HexNAc(4)Hex(5)Fuc(1)NeuAc(1) | C[+57], N[+2060] | 1385.5917 | 3 | 4154.7606 | 4154.7449 | 0 | 3.8 | 485 | Specific | 954.96 | 930.42 | 930.42 | 19.88 | 112 | >sp\|P02788\|TRFL_HUMAN Lactotransferrin OS=Homo sapiens OX=9606 GN=LTF PE=1 SV=6 | 1270 | 20201113_gy_colo_milk_1ng_glyco_hcd203040_compound15.21656.21656.3 | controllerType=0 controllerNumber=1 scan=21656 | 44.906 |
| 18558:3 | 2 | R.TAGWNIPMGLLFN[+1710.598]QTGSC[+57.021]K.F | HexNAc(3)Hex(5)NeuAc(1) | C[+57], N[+1711] | 1269.2099 | 3 | 3805.6151 | 3805.6076 | 0 | 2 | 485 | Specific | 957.23 | 791.41 | 398.87 | 20.22 | 112 | >sp\|P02788\|TRFL_HUMAN Lactotransferrin OS=Homo sapiens OX=9606 GN=LTF PE=1 SV=6 | 1270 | 20201113_gy_colo_milk_1ng_glyco_hcd203040_compound15.21894.21894.3 | controllerType=0 controllerNumber=1 scan=21894 | 45.4825 |
| 20458:3 | 2 | R.TAGWNIPMGLLFN[+2204.772]QTGSC[+57.021]K.F | HexNAc(4)Hex(5)NeuAc(2) | C[+57], N[+2205] | 1433.9368 | 3 | 4299.7959 | 4299.7824 | 0 | 3.1 | 485 | Specific | 962.49 | 951 | 951 | 20.21 | 112 | >sp\|P02788\|TRFL_HUMAN Lactotransferrin OS=Homo sapiens OX=9606 GN=LTF PE=1 SV=6 | 1270 | 20201113_gy_colo_milk_1ng_glyco_hcd203040_compound15.23874.23874.3 | controllerType=0 controllerNumber=1 scan=23874 | 49.7077 |
| 18290:3 | 2 | R.TAGWNIPMGLLFN[+2205.793]QTGSC[+57.021]K.F | HexNAc(4)Hex(5)Fuc(2)NeuAc(1) | C[+57], N[+2206] | 1434.2767 | 3 | 4300.8155 | 4300.8028 | 0 | 2.9 | 485 | Specific | 970.4 | 844.44 | 844.44 | 20.21 | 112 | >sp\|P02788\|TRFL_HUMAN Lactotransferrin OS=Homo sapiens OX=9606 GN=LTF PE=1 SV=6 | 1270 | 20201113_gy_colo_milk_1ng_glyco_hcd203040_compound15.21613.21613.3 | controllerType=0 controllerNumber=1 scan=21613 | 44.8064 |
| 18486:4 | 2 | R.TAGWNIPMGLLFN[+1913.677]QTGSC[+57.021]K.F | HexNAc(4)Hex(5)NeuAc(1) | C[+57], N[+1914] | 1002.9292 | 4 | 4008.6949 | 4008.687 | 0 | 2 | 485 | Specific | 978.39 | 931.16 | 931.16 | 20.35 | 112 | >sp\|P02788\|TRFL_HUMAN Lactotransferrin OS=Homo sapiens OX=9606 GN=LTF PE=1 SV=6 | 1270 | 20201113_gy_colo_milk_1ng_glyco_hcd203040_compound15.21817.21817.4 | controllerType=0 controllerNumber=1 scan=21817 | 45.2681 |
| 18556:3 | 2 | R.TAGWNIPMGLLFN[+1710.598]QTGSC[+57.021]K.F | HexNAc(3)Hex(5)NeuAc(1) | C[+57], N[+1711] | 1269.2099 | 3 | 3805.6151 | 3805.6076 | 0 | 2 | 485 | Specific | 983.38 | 849.95 | 433.95 | 20.22 | 112 | >sp\|P02788\|TRFL_HUMAN Lactotransferrin OS=Homo sapiens OX=9606 GN=LTF PE=1 SV=6 | 1270 | 20201113_gy_colo_milk_1ng_glyco_hcd203040_compound15.21892.21892.3 | controllerType=0 controllerNumber=1 scan=21892 | 45.4727 |
| 16906:3 | 2 | R.TAGWNIPMGLLFN[+1216.423]QTGSC[+57.021]K.F | HexNAc(2)Hex(5) | C[+57], N[+1216] | 1104.485 | 3 | 3311.4405 | 3311.4328 | 0 | 2.3 | 485 | Specific | 993.02 | 993.02 | 453.7 | 20.22 | 112 | >sp\|P02788\|TRFL_HUMAN Lactotransferrin OS=Homo sapiens OX=9606 GN=LTF PE=1 SV=6 | 1270 | 20201113_gy_colo_milk_1ng_glyco_hcd203040_compound15.20169.20169.3 | controllerType=0 controllerNumber=1 scan=20169 | 41.6741 |
| 16909:3 | 2 | R.TAGWNIPMGLLFN[+1216.423]QTGSC[+57.021]K.F | HexNAc(2)Hex(5) | C[+57], N[+1216] | 1104.485 | 3 | 3311.4405 | 3311.4328 | 0 | 2.3 | 485 | Specific | 1058.8 | 1058.8 | 485.83 | 22.22 | 112 | >sp\|P02788\|TRFL_HUMAN Lactotransferrin OS=Homo sapiens OX=9606 GN=LTF PE=1 SV=6 | 1270 | 20201113_gy_colo_milk_1ng_glyco_hcd203040_compound15.20172.20172.3 | controllerType=0 controllerNumber=1 scan=20172 | 41.6962 |
| 16149:3 | 2 | R.TAGWNIPM[+15.995]GLLFN[+2059.735]QTGSC[+57.021]K.F | HexNAc(4)Hex(5)Fuc(1)NeuAc(1) | C[+57], M[+16], N[+2060] | 1390.9233 | 3 | 4170.7553 | 4170.7398 | 0 | 3.7 | 485 | Specific | 333.15 | 283.89 | 283.89 | 6.12 | 112 | >sp\|P02788\|TRFL_HUMAN Lactotransferrin OS=Homo sapiens OX=9606 GN=LTF PE=1 SV=6 | 1270 | 20201113_gy_colo_milk_1ng_glyco_hcd203040_compound15.19380.19380.3 | controllerType=0 controllerNumber=1 scan=19380 | 40.0375 |
| 16152:3 | 2 | R.TAGWNIPM[+15.995]GLLFN[+2059.735]QTGSC[+57.021]K.F | HexNAc(4)Hex(5)Fuc(1)NeuAc(1) | C[+57], M[+16], N[+2060] | 1390.9233 | 3 | 4170.7553 | 4170.7398 | 0 | 3.7 | 485 | Specific | 629.87 | 569.25 | 569.25 | 12.88 | 112 | >sp\|P02788\|TRFL_HUMAN Lactotransferrin OS=Homo sapiens OX=9606 GN=LTF PE=1 SV=6 | 1270 | 20201113_gy_colo_milk_1ng_glyco_hcd203040_compound15.19383.19383.3 | controllerType=0 controllerNumber=1 scan=19383 | 40.042 |
| 16724:3 | 2 | R.TAGWNIPM[+15.995]GLLFN[+2059.735]QTGSC[+57.021]K.F | HexNAc(4)Hex(5)Fuc(1)NeuAc(1) | C[+57], M[+16], N[+2060] | 1390.9219 | 3 | 4170.7512 | 4170.7398 | 0 | 2.7 | 485 | Specific | 660.46 | 596.09 | 596.09 | 14.21 | 112 | >sp\|P02788\|TRFL_HUMAN Lactotransferrin OS=Homo sapiens OX=9606 GN=LTF PE=1 SV=6 | 1270 | 20201113_gy_colo_milk_1ng_glyco_hcd203040_compound15.19977.19977.3 | controllerType=0 controllerNumber=1 scan=19977 | 41.2347 |
| 16725:3 | 2 | R.TAGWNIPM[+15.995]GLLFN[+2059.735]QTGSC[+57.021]K.F | HexNAc(4)Hex(5)Fuc(1)NeuAc(1) | C[+57], M[+16], N[+2060] | 1390.9219 | 3 | 4170.7512 | 4170.7398 | 0 | 2.7 | 485 | Specific | 817.08 | 613.86 | 613.86 | 17.21 | 112 | >sp\|P02788\|TRFL_HUMAN Lactotransferrin OS=Homo sapiens OX=9606 GN=LTF PE=1 SV=6 | 1270 | 20201113_gy_colo_milk_1ng_glyco_hcd203040_compound15.19978.19978.3 | controllerType=0 controllerNumber=1 scan=19978 | 41.2386 |
| 09293:3 | 2 | R.N[+2205.793]GSDC[+57.021]PDKFC[+57.021]LFQSETK.N | HexNAc(4)Hex(5)Fuc(2)NeuAc(1) | C[+57]*2, N[+2206] | 1413.5624 | 3 | 4238.6728 | 4238.6667 | 0 | 1.4 | 642 | Specific | 345.68 | 242.43 | 242.43 | 6.74 | 112 | >sp\|P02788\|TRFL_HUMAN Lactotransferrin OS=Homo sapiens OX=9606 GN=LTF PE=1 SV=6 | 1270 | 20201113_gy_colo_milk_1ng_glyco_hcd203040_compound15.12263.12263.3 | controllerType=0 controllerNumber=1 scan=12263 | 26.6986 |
| 09120:3 | 2 | R.N[+2570.925]GSDC[+57.021]PDKFC[+57.021]LFQSETK.N | HexNAc(5)Hex(6)Fuc(2)NeuAc(1) | C[+57]*2, N[+2571] | 1535.2745 | 3 | 4603.8089 | 4603.7989 | 0 | 2.2 | 642 | Specific | 361.25 | 324.65 | 324.65 | 7.39 | 112 | >sp\|P02788\|TRFL_HUMAN Lactotransferrin OS=Homo sapiens OX=9606 GN=LTF PE=1 SV=6 | 1270 | 20201113_gy_colo_milk_1ng_glyco_hcd203040_compound15.12084.12084.3 | controllerType=0 controllerNumber=1 scan=12084 | 26.3831 |
| 09122:3 | 2 | R.N[+2570.925]GSDC[+57.021]PDKFC[+57.021]LFQSETK.N | HexNAc(5)Hex(6)Fuc(2)NeuAc(1) | C[+57]*2, N[+2571] | 1535.2745 | 3 | 4603.8089 | 4603.7989 | 0 | 2.2 | 642 | Specific | 372.78 | 336.04 | 336.04 | 7.47 | 112 | >sp\|P02788\|TRFL_HUMAN Lactotransferrin OS=Homo sapiens OX=9606 GN=LTF PE=1 SV=6 | 1270 | 20201113_gy_colo_milk_1ng_glyco_hcd203040_compound15.12086.12086.3 | controllerType=0 controllerNumber=1 scan=12086 | 26.3869 |
| 09291:3 | 2 | R.N[+2205.793]GSDC[+57.021]PDKFC[+57.021]LFQSETK.N | HexNAc(4)Hex(5)Fuc(2)NeuAc(1) | C[+57]*2, N[+2206] | 1413.5624 | 3 | 4238.6728 | 4238.6667 | 0 | 1.4 | 642 | Specific | 416.37 | 323.64 | 323.64 | 7.4 | 112 | >sp\|P02788\|TRFL_HUMAN Lactotransferrin OS=Homo sapiens OX=9606 GN=LTF PE=1 SV=6 | 1270 | 20201113_gy_colo_milk_1ng_glyco_hcd203040_compound15.12261.12261.3 | controllerType=0 controllerNumber=1 scan=12261 | [illegible] |
| 07627:3 | 3 | R.FSDGLESN[+1864.634]SSTQFEVK.K | HexNAc(2)Hex(9) | N[+1865] | 1213.8243 | 3 | 3639.4583 | 3639.4472 | 0 | 3.1 | 219 | Specific | 431.86 | 431.86 | 431.86 | 10 | 96 | >sp\|P0C0L4\|CO4A_HUMAN Complement C4-A OS=Homo sapiens OX=9606 GN=C4A PE=1 SV=2 | 1235 | 20201113_gy_colo_milk_1ng_glyco_hcd203040_compound15.10537.10537.3 | controllerType=0 controllerNumber=1 scan=10537 | [illegible] |
| 07629:3 | 3 | R.FSDGLESN[+1864.634]SSTQFEVK.K | HexNAc(2)Hex(9) | N[+1865] | 1213.8243 | 3 | 3639.4583 | 3639.4472 | 0 | 3.1 | 219 | Specific | 889.77 | 863.5 | 863.5 | 18.03 | 96 | >sp\|P0C0L4\|CO4A_HUMAN Complement C4-A OS=Homo sapiens OX=9606 GN=C4A PE=1 SV=2 | 1235 | 20201113_gy_colo_milk_1ng_glyco_hcd203040_compound15.10539.10539.3 | controllerType=0 controllerNumber=1 scan=10539 | [illegible] |
| 05156:3 | 3 | R.FSDGLESN[+1864.634]SSTQFEVKK.Y | HexNAc(2)Hex(9) | N[+1865] | 1256.5219 | 3 | 3767.5512 | 3767.5421 | 0 | 2.4 | 219 | Specific | 275.53 | 275.53 | 275.53 | 6.55 | 96 | >sp\|P0C0L4\|CO4A_HUMAN Complement C4-A OS=Homo sapiens OX=9606 GN=C4A PE=1 SV=2 | 1235 | 20201113_gy_colo_milk_1ng_glyco_hcd203040_compound15.7979.7979.3 | controllerType=0 controllerNumber=1 scan=7979 | [illegible] |
| 05154:3 | 3 | R.FSDGLESN[+1864.634]SSTQFEVKK.Y | HexNAc(2)Hex(9) | N[+1865] | 1256.5219 | 3 | 3767.5512 | 3767.5421 | 0 | 2.4 | 219 | Specific | 291.14 | 291.14 | 291.14 | 6.78 | 96 | >sp\|P0C0L4\|CO4A_HUMAN Complement C4-A OS=Homo sapiens OX=9606 GN=C4A PE=1 SV=2 | 1235 | 20201113_gy_colo_milk_1ng_glyco_hcd203040_compound15.7977.7977.3 | controllerType=0 controllerNumber=1 scan=7977 | 19.4104 |
| 05112:3 | 3 | R.FSDGLESN[+1864.634]SSTQFEVKK.Y | HexNAc(2)Hex(9) | N[+1865] | 1256.522 | 3 | 3767.5515 | 3767.5421 | 0 | 2.5 | 219 | Specific | 574.22 | 574.22 | 574.22 | 10.11 | 96 | >sp\|P0C0L4\|CO4A_HUMAN Complement C4-A OS=Homo sapiens OX=9606 GN=C4A PE=1 SV=2 | 1235 | 20201113_gy_colo_milk_1ng_glyco_hcd203040_compound15.7933.7933.3 | controllerType=0 controllerNumber=1 scan=7933 | 19.3281 |
| 05109:3 | 3 | R.FSDGLESN[+1864.634]SSTQFEVKK.Y | HexNAc(2)Hex(9) | N[+1865] | 1256.522 | 3 | 3767.5515 | 3767.5421 | 0 | 2.5 | 219 | Specific | 748.25 | 582.45 | 582.45 | 14.47 | 96 | >sp\|P0C0L4\|CO4A_HUMAN Complement C4-A OS=Homo sapiens OX=9606 GN=C4A PE=1 SV=2 | 1235 | 20201113_gy_colo_milk_1ng_glyco_hcd203040_compound15.7930.7930.3 | controllerType=0 controllerNumber=1 scan=7930 | 19.323 |
| 18241:5 | 4 | R.LSLHRPALEDLLLGSEAN[+2116.756]LTC[+57.021]TLTGLR.D | HexNAc(5)Hex(5)NeuAc(1) | C[+57], N[+2117] | 1016.8767 | 5 | 5080.3545 | 5080.3547 | 0 | 0 | 127 | Specific | 244.52 | 244.52 | 244.52 | 5.59 | 99 | >sp\|P01876\|IGHA1_HUMAN Immunoglobulin heavy constant alpha 1 OS=Homo sapiens OX=9606 GN=IGHA1 PE=1 SV=2 | 255 | 20201113_gy_colo_milk_1ng_glyco_hcd203040_compound15.21562.21562.5 | controllerType=0 controllerNumber=1 scan=21562 | 44.6909 |
| 19370:4 | 4 | R.LSLHRPALEDLLLGSEAN[+2204.772]LTC[+57.021]TLTGLR.D | HexNAc(4)Hex(5)NeuAc(2) | C[+57], N[+2205] | 1292.8516 | 4 | 5168.3844 | 5168.3707 | 0 | 2.7 | 127 | Specific | 274.1 | 256.24 | 256.24 | 7.11 | 99 | >sp\|P01876\|IGHA1_HUMAN Immunoglobulin heavy constant alpha 1 OS=Homo sapiens OX=9606 GN=IGHA1 PE=1 SV=2 | 255 | 20201113_gy_colo_milk_1ng_glyco_hcd203040_compound15.22738.22738.4 | controllerType=0 controllerNumber=1 scan=22738 | 47.1935 |
| 17376:3 | 4 | R.LSLHRPALEDLLLGSEAN[+1501.555]LTC[+57.021]TLTGLR.D | HexNAc(5)Hex(3) | C[+57], N[+1502] | 1489.0617 | 3 | 4465.1706 | 4465.1536 | 0 | 3.8 | 127 | Specific | 300.98 | 300.98 | 300.98 | 7.03 | 99 | >sp\|P01876\|IGHA1_HUMAN Immunoglobulin heavy constant alpha 1 OS=Homo sapiens OX=9606 GN=IGHA1 PE=1 SV=2 | 255 | 20201113_gy_colo_milk_1ng_glyco_hcd203040_compound15.20659.20659.6 | controllerType=0 controllerNumber=1 scan=20659 | 42.7228 |
| 34298:4 | 4 | R.LSLHRPALEDLLLGSEAN[+1501.555]LTC[+57.021]TLTGLR.D | HexNAc(5)Hex(3) | C[+57], N[+1502] | 1117.0448 | 4 | 4465.1575 | 4465.1536 | 0 | 0.9 | 127 | Specific | 312.15 | 312.15 | 312.15 | 7.48 | 99 | >sp\|P01876\|IGHA1_HUMAN Immunoglobulin heavy constant alpha 1 OS=Homo sapiens OX=9606 GN=IGHA1 PE=1 SV=2 | 255 | 20201113_gy_colo_milk_1ng_glyco_hcd203040_compound15.41324.41324.4 | controllerType=0 controllerNumber=1 scan=41324 | 88.863 |
| 18236:3 | 4 | R.LSLHRPALEDLLLGSEAN[+2116.756]LTC[+57.021]TLTGLR.D | HexNAc(5)Hex(5)NeuAc(1) | C[+57], N[+2117] | 1694.1263 | 3 | 5080.3644 | 5080.3547 | 0 | 1.9 | 127 | Specific | 315.95 | 308.46 | 308.46 | 7.04 | 99 | >sp\|P01876\|IGHA1_HUMAN Immunoglobulin heavy constant alpha 1 OS=Homo sapiens OX=9606 GN=IGHA1 PE=1 SV=2 | 255 | 20201113_gy_colo_milk_1ng_glyco_hcd203040_compound15.21557.21557.3 | controllerType=0 controllerNumber=1 scan=21557 | 44.6832 |
| 18263:4 | 4 | R.LSLHRPALEDLLLGSEAN[+1872.651]LTC[+57.021]TLTGLR.D | HexNAc(3)Hex(6)NeuAc(1) | C[+57], N[+1873] | 1209.8227 | 4 | 4836.2689 | 4836.2487 | 0 | 4.2 | 127 | Specific | 337.11 | 329.79 | 329.79 | 7.67 | 99 | >sp\|P01876\|IGHA1_HUMAN Immunoglobulin heavy constant alpha 1 OS=Homo sapiens OX=9606 GN=IGHA1 PE=1 SV=2 | 255 | 20201113_gy_colo_milk_1ng_glyco_hcd203040_compound15.21585.21585.3 | controllerType=0 controllerNumber=1 scan=21585 | 44.7462 |
| 16874:4 | 4 | R.LSLHRPALEDLLLGSEAN[+1622.582]LTC[+57.021]TLTGLR.D | HexNAc(4)Hex(5) | C[+57], N[+1623] | 1147.3044 | 4 | 4586.1956 | 4586.1799 | 0 | 3.4 | 127 | Specific | 339.74 | 263.88 | 263.88 | 7.54 | 99 | >sp\|P01876\|IGHA1_HUMAN Immunoglobulin heavy constant alpha 1 OS=Homo sapiens OX=9606 GN=IGHA1 PE=1 SV=2 | 255 | 20201113_gy_colo_milk_1ng_glyco_hcd203040_compound15.20134.20134.4 | controllerType=0 controllerNumber=1 scan=20134 | 41.5977 |
| 18266:4 | 4 | R.LSLHRPALEDLLLGSEAN[+1872.651]LTC[+57.021]TLTGLR.D | HexNAc(3)Hex(6)NeuAc(1) | C[+57], N[+1873] | 1209.8227 | 4 | 4836.2689 | 4836.2487 | 0 | 4.2 | 127 | Specific | 423.38 | 423.38 | 423.38 | 9.76 | 99 | >sp\|P01876\|IGHA1_HUMAN Immunoglobulin heavy constant alpha 1 OS=Homo sapiens OX=9606 GN=IGHA1 PE=1 SV=2 | 255 | 20201113_gy_colo_milk_1ng_glyco_hcd203040_compound15.21588.21588.3 | controllerType=0 controllerNumber=1 scan=21588 | 44.7515 |
| 18287:4 | 4 | R.LSLHRPALEDLLLGSEAN[+1751.624]LTC[+57.021]TLTGLR.D | HexNAc(4)Hex(4)NeuAc(1) | C[+57], N[+1752] | 1179.5636 | 4 | 4715.2327 | 4715.2225 | 0 | 2.2 | 127 | Specific | 440.45 | 440.45 | 440.45 | 10.19 | 99 | >sp\|P01876\|IGHA1_HUMAN Immunoglobulin heavy constant alpha 1 OS=Homo sapiens OX=9606 GN=IGHA1 PE=1 SV=2 | 255 | 20201113_gy_colo_milk_1ng_glyco_hcd203040_compound15.21610.21610.4 | controllerType=0 controllerNumber=1 scan=21610 | 44.8017 |
| 18258:4 | 4 | R.LSLHRPALEDLLLGSEAN[+1913.677]LTC[+57.021]TLTGLR.D | HexNAc(4)Hex(5)NeuAc(1) | C[+57], N[+1914] | 1220.0788 | 4 | 4877.2933 | 4877.2753 | 0 | 3.7 | 127 | Specific | 451.36 | 451.36 | 451.36 | 10.19 | 99 | >sp\|P01876\|IGHA1_HUMAN Immunoglobulin heavy constant alpha 1 OS=Homo sapiens OX=9606 GN=IGHA1 PE=1 SV=2 | 255 | 20201113_gy_colo_milk_1ng_glyco_hcd203040_compound15.21580.21580.4 | controllerType=0 controllerNumber=1 scan=21580 | 44.7376 |
| 17429:3 | 4 | R.LSLHRPALEDLLLGSEAN[+1298.476]LTC[+57.021]TLTGLR.D | HexNAc(4)Hex(3) | C[+57], N[+1298] | 1421.3672 | 3 | 4262.0872 | 4262.0742 | 0 | 3 | 127 | Specific | 452.82 | 447.21 | 447.21 | 9.75 | 99 | >sp\|P01876\|IGHA1_HUMAN Immunoglobulin heavy constant alpha 1 OS=Homo sapiens OX=9606 GN=IGHA1 PE=1 SV=2 | 255 | 20201113_gy_colo_milk_1ng_glyco_hcd203040_compound15.20714.20714.2 | controllerType=0 controllerNumber=1 scan=20714 | 42.8637 |
| 17426:3 | 4 | R.LSLHRPALEDLLLGSEAN[+1298.476]LTC[+57.021]TLTGLR.D | HexNAc(4)Hex(3) | C[+57], N[+1298] | 1421.3672 | 3 | 4262.0872 | 4262.0742 | 0 | 3 | 127 | Specific | 463.93 | 362.52 | 362.52 | 8.28 | 99 | >sp\|P01876\|IGHA1_HUMAN Immunoglobulin heavy constant alpha 1 OS=Homo sapiens OX=9606 GN=IGHA1 PE=1 SV=2 | 255 | 20201113_gy_colo_milk_1ng_glyco_hcd203040_compound15.20711.20711.2 | controllerType=0 controllerNumber=1 scan=20711 | 42.8576 |
| 18268:5 | 4 | R.LSLHRPALEDLLLGSEAN[+1913.677]LTC[+57.021]TLTGLR.D | HexNAc(4)Hex(5)NeuAc(1) | C[+57], N[+1914] | 976.2641 | 5 | 4877.2912 | 4877.2753 | 0 | 3.3 | 127 | Specific | 485.53 | 485.53 | 485.53 | 10.54 | 99 | >sp\|P01876\|IGHA1_HUMAN Immunoglobulin heavy constant alpha 1 OS=Homo sapiens OX=9606 GN=IGHA1 PE=1 SV=2 | 255 | 20201113_gy_colo_milk_1ng_glyco_hcd203040_compound15.21590.21590.5 | controllerType=0 controllerNumber=1 scan=21590 | 44.7542 |
| 18334:4 | 4 | R.LSLHRPALEDLLLGSEAN[+1710.598]LTC[+57.021]TLTGLR.D | HexNAc(3)Hex(5)NeuAc(1) | C[+57], N[+1711] | 1169.3089 | 4 | 4674.2137 | 4674.1959 | 0 | 3.8 | 127 | Specific | 488.57 | 488.57 | 488.57 | 10.03 | 99 | >sp\|P01876\|IGHA1_HUMAN Immunoglobulin heavy constant alpha 1 OS=Homo sapiens OX=9606 GN=IGHA1 PE=1 SV=2 | 255 | 20201113_gy_colo_milk_1ng_glyco_hcd203040_compound15.21659.21659.4 | controllerType=0 controllerNumber=1 scan=21659 | 44.9125 |
| 17301:3 | 4 | R.LSLHRPALEDLLLGSEAN[+1663.608]LTC[+57.021]TLTGLR.D | HexNAc(5)Hex(4) | C[+57], N[+1664] | 1543.0779 | 3 | 4627.2193 | 4627.2064 | 0 | 2.8 | 127 | Specific | 520.69 | 456.49 | 456.49 | 9.95 | 99 | >sp\|P01876\|IGHA1_HUMAN Immunoglobulin heavy constant alpha 1 OS=Homo sapiens OX=9606 GN=IGHA1 PE=1 SV=2 | 255 | 20201113_gy_colo_milk_1ng_glyco_hcd203040_compound15.20581.20581.3 | controllerType=0 controllerNumber=1 scan=20581 | 42.5646 |
| 17245:3 | 4 | R.LSLHRPALEDLLLGSEAN[+1864.634]LTC[+57.021]TLTGLR.D | HexNAc(2)Hex(9) | C[+57], N[+1865] | 1610.0851 | 3 | 4828.2409 | 4828.2324 | 0 | 1.7 | 127 | Specific | 523.58 | 523.58 | 523.58 | 9.95 | 99 | >sp\|P01876\|IGHA1_HUMAN Immunoglobulin heavy constant alpha 1 OS=Homo sapiens OX=9606 GN=IGHA1 PE=1 SV=2 | 255 | 20201113_gy_colo_milk_1ng_glyco_hcd203040_compound15.20523.20523.3 | controllerType=0 controllerNumber=1 scan=20523 | 42.4457 |
| 17344:5 | 4 | R.LSLHRPALEDLLLGSEAN[+1501.555]LTC[+57.021]TLTGLR.D | HexNAc(5)Hex(3) | C[+57], N[+1502] | 893.8395 | 5 | 4465.1684 | 4465.1536 | 0 | 3.3 | 127 | Specific | 527.57 | 527.57 | 527.57 | 10.69 | 99 | >sp\|P01876\|IGHA1_HUMAN Immunoglobulin heavy constant alpha 1 OS=Homo sapiens OX=9606 GN=IGHA1 PE=1 SV=2 | 255 | 20201113_gy_colo_milk_1ng_glyco_hcd203040_compound15.20626.20626.5 | controllerType=0 controllerNumber=1 scan=20626 | 42.6648 |
| 17395:3 | 4 | R.LSLHRPALEDLLLGSEAN[+1298.476]LTC[+57.021]TLTGLR.D | HexNAc(4)Hex(3) | C[+57], N[+1298] | 1421.3681 | 3 | 4262.0897 | 4262.0742 | 0 | 3.6 | 127 | Specific | 536.23 | 536.23 | 536.23 | 9.95 | 99 | >sp\|P01876\|IGHA1_HUMAN Immunoglobulin heavy constant alpha 1 OS=Homo sapiens OX=9606 GN=IGHA1 PE=1 SV=2 | 255 | 20201113_gy_colo_milk_1ng_glyco_hcd203040_compound15.20679.20679.3 | controllerType=0 controllerNumber=1 scan=20679 | 42.7697 |
| 17432:4 | 4 | R.LSLHRPALEDLLLGSEAN[+1524.534]LTC[+57.021]TLTGLR.D | HexNAc(2)Hex(6)Fuc(1) | C[+57], N[+1525] | 1122.7846 | 4 | 4488.1165 | 4488.1319 | 0 | -3.4 | 127 | Specific | 537.54 | 537.54 | 537.54 | 9.03 | 99 | >sp\|P01876\|IGHA1_HUMAN Immunoglobulin heavy constant alpha 1 OS=Homo sapiens OX=9606 GN=IGHA1 PE=1 SV=2 | 255 | 20201113_gy_colo_milk_1ng_glyco_hcd203040_compound15.20717.20717.4 | controllerType=0 controllerNumber=1 scan=20717 | 42.8679 |
| 17375:4 | 4 | R.LSLHRPALEDLLLGSEAN[+1524.534]LTC[+57.021]TLTGLR.D | HexNAc(2)Hex(6)Fuc(1) | C[+57], N[+1525] | 1122.7846 | 4 | 4488.1165 | 4488.1319 | 0 | -3.4 | 127 | Specific | 542.8 | 529.55 | 529.55 | 9.03 | 99 | >sp\|P01876\|IGHA1_HUMAN Immunoglobulin heavy constant alpha 1 OS=Homo sapiens OX=9606 GN=IGHA1 PE=1 SV=2 | 255 | 20201113_gy_colo_milk_1ng_glyco_hcd203040_compound15.20658.20658.4 | controllerType=0 controllerNumber=1 scan=20658 | 42.7217 |
| 18235:4 | 4 | R.LSLHRPALEDLLLGSEAN[+2116.756]LTC[+57.021]TLTGLR.D | HexNAc(5)Hex(5)NeuAc(1) | C[+57], N[+2117] | 1270.8461 | 4 | 5080.3625 | 5080.3547 | 0 | 1.5 | 127 | Specific | 567.09 | 567.09 | 567.09 | 10.73 | 99 | >sp\|P01876\|IGHA1_HUMAN Immunoglobulin heavy constant alpha 1 OS=Homo sapiens OX=9606 GN=IGHA1 PE=1 SV=2 | 255 | 20201113_gy_colo_milk_1ng_glyco_hcd203040_compound15.21556.21556.4 | controllerType=0 controllerNumber=1 scan=21556 | 44.6818 |
| 18374:5 | 4 | R.LSLHRPALEDLLLGSEAN[+1955.724]LTC[+57.021]TLTGLR.D | HexNAc(5)Hex(4)Fuc(2) | C[+57], N[+1956] | 984.6713 | 5 | 4919.3273 | 4919.3222 | 0 | 1 | 127 | Specific | 573.9 | 516.36 | 441.74 | 10.9 | 99 | >sp\|P01876\|IGHA1_HUMAN Immunoglobulin heavy constant alpha 1 OS=Homo sapiens OX=9606 GN=IGHA1 PE=1 SV=2 | 255 | 20201113_gy_colo_milk_1ng_glyco_hcd203040_compound15.21700.21700.5 | controllerType=0 controllerNumber=1 scan=21700 | 44.9994 |
| 17397:3 | 4 | R.LSLHRPALEDLLLGSEAN[+1298.476]LTC[+57.021]TLTGLR.D | HexNAc(4)Hex(3) | C[+57], N[+1298] | 1421.3681 | 3 | 4262.0897 | 4262.0742 | 0 | 3.6 | 127 | Specific | 575.34 | 537.26 | 537.26 | 10.29 | 99 | >sp\|P01876\|IGHA1_HUMAN Immunoglobulin heavy constant alpha 1 OS=Homo sapiens OX=9606 GN=IGHA1 PE=1 SV=2 | 255 | 20201113_gy_colo_milk_1ng_glyco_hcd203040_compound15.20681.20681.3 | controllerType=0 controllerNumber=1 scan=20681 | 42.7775 |
| 17271:3 | 4 | R.LSLHRPALEDLLLGSEAN[+1622.582]LTC[+57.021]TLTGLR.D | HexNAc(4)Hex(5) | C[+57], N[+1623] | 1529.4029 | 3 | 4586.1941 | 4586.1799 | 0 | 3.1 | 127 | Specific | 590.14 | 506.3 | 506.3 | 11.64 | 99 | >sp\|P01876\|IGHA1_HUMAN Immunoglobulin heavy constant alpha 1 OS=Homo sapiens OX=9606 GN=IGHA1 PE=1 SV=2 | 255 | 20201113_gy_colo_milk_1ng_glyco_hcd203040_compound15.20549.20549.3 | controllerType=0 controllerNumber=1 scan=20549 | 42.4998 |
| 17359:3 | 4 | R.LSLHRPALEDLLLGSEAN[+1257.449]LTC[+57.021]TLTGLR.D | HexNAc(3)Hex(4) | C[+57], N[+1257] | 1407.6885 | 3 | 4221.051 | 4221.0477 | 0 | 0.8 | 127 | Specific | 592.11 | 592.11 | 592.11 | 11.51 | 99 | >sp\|P01876\|IGHA1_HUMAN Immunoglobulin heavy constant alpha 1 OS=Homo sapiens OX=9606 GN=IGHA1 PE=1 SV=2 | 255 | 20201113_gy_colo_milk_1ng_glyco_hcd203040_compound15.20641.20641.3 | controllerType=0 controllerNumber=1 scan=20641 | 42.6932 |
| 19373:4 | 4 | R.LSLHRPALEDLLLGSEAN[+2204.772]LTC[+57.021]TLTGLR.D | HexNAc(4)Hex(5)NeuAc(2) | C[+57], N[+2205] | 1292.8516 | 4 | 5168.3844 | 5168.3707 | 0 | 2.7 | 127 | Specific | 593.41 | 592.9 | 592.9 | 12.22 | 99 | >sp\|P01876\|IGHA1_HUMAN Immunoglobulin heavy constant alpha 1 OS=Homo sapiens OX=9606 GN=IGHA1 PE=1 SV=2 | 255 | 20201113_gy_colo_milk_1ng_glyco_hcd203040_compound15.22741.22741.4 | controllerType=0 controllerNumber=1 scan=22741 | 47.1974 |
| 18338:3 | 4 | R.LSLHRPALEDLLLGSEAN[+1710.598]LTC[+57.021]TLTGLR.D | HexNAc(3)Hex(5)NeuAc(1) | C[+57], N[+1711] | 1558.743 | 3 | 4674.2144 | 4674.1959 | 0 | 3.9 | 127 | Specific | 602.87 | 564.53 | 564.53 | 12.3 | 99 | >sp\|P01876\|IGHA1_HUMAN Immunoglobulin heavy constant alpha 1 OS=Homo sapiens OX=9606 GN=IGHA1 PE=1 SV=2 | 255 | 20201113_gy_colo_milk_1ng_glyco_hcd203040_compound15.21663.21663.3 | controllerType=0 controllerNumber=1 scan=21663 | 44.9194 |
| 17305:3 | 4 | R.LSLHRPALEDLLLGSEAN[+1663.608]LTC[+57.021]TLTGLR.D | HexNAc(5)Hex(4) | C[+57], N[+1664] | 1543.0779 | 3 | 4627.2193 | 4627.2064 | 0 | 2.8 | 127 | Specific | 604.67 | 542.46 | 542.46 | 12.64 | 99 | >sp\|P01876\|IGHA1_HUMAN Immunoglobulin heavy constant alpha 1 OS=Homo sapiens OX=9606 GN=IGHA1 PE=1 SV=2 | 255 | 20201113_gy_colo_milk_1ng_glyco_hcd203040_compound15.20585.20585.3 | controllerType=0 controllerNumber=1 scan=20585 | 42.5839 |
| 18288:3 | 4 | R.LSLHRPALEDLLLGSEAN[+1751.624]LTC[+57.021]TLTGLR.D | HexNAc(4)Hex(4)NeuAc(1) | C[+57], N[+1752] | 1572.4162 | 3 | 4715.234 | 4715.2225 | 0 | 2.5 | 127 | Specific | 609.52 | 609.52 | 609.52 | 12.64 | 99 | >sp\|P01876\|IGHA1_HUMAN Immunoglobulin heavy constant alpha 1 OS=Homo sapiens OX=9606 GN=IGHA1 PE=1 SV=2 | 255 | 20201113_gy_colo_milk_1ng_glyco_hcd203040_compound15.21611.21611.3 | controllerType=0 controllerNumber=1 scan=21611 | 44.8041 |
| 18286:3 | 4 | R.LSLHRPALEDLLLGSEAN[+1751.624]LTC[+57.021]TLTGLR.D | HexNAc(4)Hex(4)NeuAc(1) | C[+57], N[+1752] | 1572.4162 | 3 | 4715.234 | 4715.2225 | 0 | 2.5 | 127 | Specific | 609.88 | 609.88 | 609.88 | 12.64 | 99 | >sp\|P01876\|IGHA1_HUMAN Immunoglobulin heavy constant alpha 1 OS=Homo sapiens OX=9606 GN=IGHA1 PE=1 SV=2 | 255 | 20201113_gy_colo_milk_1ng_glyco_hcd203040_compound15.21609.21609.3 | controllerType=0 controllerNumber=1 scan=21609 | 44.797 |
| 17247:3 | 4 | R.LSLHRPALEDLLLGSEAN[+1864.634]LTC[+57.021]TLTGLR.D | HexNAc(2)Hex(9) | C[+57], N[+1865] | 1610.0851 | 3 | 4828.2409 | 4828.2324 | 0 | 1.7 | 127 | Specific | 614.31 | 614.31 | 614.31 | 12.64 | 99 | >sp\|P01876\|IGHA1_HUMAN Immunoglobulin heavy constant alpha 1 OS=Homo sapiens OX=9606 GN=IGHA1 PE=1 SV=2 | 255 | 20201113_gy_colo_milk_1ng_glyco_hcd203040_compound15.20525.20525.3 | controllerType=0 controllerNumber=1 scan=20525 | 42.4509 |
| 17378:4 | 4 | R.LSLHRPALEDLLLGSEAN[+1524.534]LTC[+57.021]TLTGLR.D | HexNAc(2)Hex(6)Fuc(1) | C[+57], N[+1525] | 1122.7846 | 4 | 4488.1165 | 4488.1319 | 0 | -3.4 | 127 | Specific | 622.14 | 533.67 | 533.67 | 11.57 | 99 | >sp\|P01876\|IGHA1_HUMAN Immunoglobulin heavy constant alpha 1 OS=Homo sapiens OX=9606 GN=IGHA1 PE=1 SV=2 | 255 | 20201113_gy_colo_milk_1ng_glyco_hcd203040_compound15.20661.20661.4 | controllerType=0 controllerNumber=1 scan=20661 | 42.7274 |
| 17379:3 | 4 | R.LSLHRPALEDLLLGSEAN[+1501.555]LTC[+57.021]TLTGLR.D | HexNAc(5)Hex(3) | C[+57], N[+1502] | 1489.0617 | 3 | 4465.1706 | 4465.1536 | 0 | 3.8 | 127 | Specific | 622.4 | 583.74 | 583.74 | 12.64 | 99 | >sp\|P01876\|IGHA1_HUMAN Immunoglobulin heavy constant alpha 1 OS=Homo sapiens OX=9606 GN=IGHA1 PE=1 SV=2 | 255 | 20201113_gy_colo_milk_1ng_glyco_hcd203040_compound15.20662.20662.6 | controllerType=0 controllerNumber=1 scan=20662 | 42.7284 |
| 17320:3 | 4 | R.LSLHRPALEDLLLGSEAN[+1378.476]LTC[+57.021]TLTGLR.D | HexNAc(2)Hex(6) | C[+57], N[+1378] | 1448.0344 | 3 | 4342.0885 | 4342.074 | 0 | 3.4 | 127 | Specific | 629.58 | 598.48 | 598.48 | 12.64 | 99 | >sp\|P01876\|IGHA1_HUMAN Immunoglobulin heavy constant alpha 1 OS=Homo sapiens OX=9606 GN=IGHA1 PE=1 SV=2 | 255 | 20201113_gy_colo_milk_1ng_glyco_hcd203040_compound15.20601.20601.3 | controllerType=0 controllerNumber=1 scan=20601 | 42.6158 |
| 18238:3 | 4 | R.LSLHRPALEDLLLGSEAN[+2116.756]LTC[+57.021]TLTGLR.D | HexNAc(5)Hex(5)NeuAc(1) | C[+57], N[+2117] | 1694.1263 | 3 | 5080.3644 | 5080.3547 | 0 | 1.9 | 127 | Specific | 631.01 | 526.94 | 526.94 | 12.64 | 99 | >sp\|P01876\|IGHA1_HUMAN Immunoglobulin heavy constant alpha 1 OS=Homo sapiens OX=9606 GN=IGHA1 PE=1 SV=2 | 255 | 20201113_gy_colo_milk_1ng_glyco_hcd203040_compound15.21559.21559.3 | controllerType=0 controllerNumber=1 scan=21559 | 44.6872 |
| 18243:5 | 4 | R.LSLHRPALEDLLLGSEAN[+2116.756]LTC[+57.021]TLTGLR.D | HexNAc(5)Hex(5)NeuAc(1) | C[+57], N[+2117] | 1016.8767 | 5 | 5080.3545 | 5080.3547 | 0 | 0 | 127 | Specific | 634.62 | 577.87 | 577.87 | 12.83 | 99 | >sp\|P01876\|IGHA1_HUMAN Immunoglobulin heavy constant alpha 1 OS=Homo sapiens OX=9606 GN=IGHA1 PE=1 SV=2 | 255 | 20201113_gy_colo_milk_1ng_glyco_hcd203040_compound15.21564.21564.5 | controllerType=0 controllerNumber=1 scan=21564 | 44.6977 |
| 18265:3 | 4 | R.LSLHRPALEDLLLGSEAN[+1913.677]LTC[+57.021]TLTGLR.D | HexNAc(4)Hex(5)NeuAc(1) | C[+57], N[+1914] | 1626.4356 | 3 | 4877.2923 | 4877.2753 | 0 | 3.5 | 127 | Specific | 639.7 | 572.66 | 572.66 | 12.64 | 99 | >sp\|P01876\|IGHA1_HUMAN Immunoglobulin heavy constant alpha 1 OS=Homo sapiens OX=9606 GN=IGHA1 PE=1 SV=2 | 255 | 20201113_gy_colo_milk_1ng_glyco_hcd203040_compound15.21587.21587.3 | controllerType=0 controllerNumber=1 scan=21587 | 44.7485 |
| 17546:4 | 4 | R.LSLHRPALEDLLLGSEAN[+1501.555]LTC[+57.021]TLTGLR.D | HexNAc(5)Hex(3) | C[+57], N[+1502] | 1117.0458 | 4 | 4465.1614 | 4465.1536 | 0 | 1.8 | 127 | Specific | 640.04 | 640.04 | 640.04 | 13.22 | 99 | >sp\|P01876\|IGHA1_HUMAN Immunoglobulin heavy constant alpha 1 OS=Homo sapiens OX=9606 GN=IGHA1 PE=1 SV=2 | 255 | 20201113_gy_colo_milk_1ng_glyco_hcd203040_compound15.20836.20836.3 | controllerType=0 controllerNumber=1 scan=20836 | 43.1065 |
| 17362:3 | 4 | R.LSLHRPALEDLLLGSEAN[+1257.449]LTC[+57.021]TLTGLR.D | HexNAc(3)Hex(4) | C[+57], N[+1257] | 1407.6885 | 3 | 4221.051 | 4221.0477 | 0 | 0.8 | 127 | Specific | 649.1 | 631.31 | 631.31 | 12.51 | 99 | >sp\|P01876\|IGHA1_HUMAN Immunoglobulin heavy constant alpha 1 OS=Homo sapiens OX=9606 GN=IGHA1 PE=1 SV=2 | 255 | 20201113_gy_colo_milk_1ng_glyco_hcd203040_compound15.20644.20644.3 | controllerType=0 controllerNumber=1 scan=20644 | 42.698 |
| 17268:3 | 4 | R.LSLHRPALEDLLLGSEAN[+1622.582]LTC[+57.021]TLTGLR.D | HexNAc(4)Hex(5) | C[+57], N[+1623] | 1529.4029 | 3 | 4586.1941 | 4586.1799 | 0 | 3.1 | 127 | Specific | 651.53 | 597.99 | 597.99 | 13.64 | 99 | >sp\|P01876\|IGHA1_HUMAN Immunoglobulin heavy constant alpha 1 OS=Homo sapiens OX=9606 GN=IGHA1 PE=1 SV=2 | 255 | 20201113_gy_colo_milk_1ng_glyco_hcd203040_compound15.20546.20546.3 | controllerType=0 controllerNumber=1 scan=20546 | 42.4905 |
| 18195:4 | 4 | R.LSLHRPALEDLLLGSEAN[+2116.756]LTC[+57.021]TLTGLR.D | HexNAc(5)Hex(5)NeuAc(1) | C[+57], N[+2117] | 1270.8467 | 4 | 5080.3649 | 5080.3547 | 0 | 2 | 127 | Specific | 652.84 | 652.84 | 652.84 | 14.22 | 99 | >sp\|P01876\|IGHA1_HUMAN Immunoglobulin heavy constant alpha 1 OS=Homo sapiens OX=9606 GN=IGHA1 PE=1 SV=2 | 255 | 20201113_gy_colo_milk_1ng_glyco_hcd203040_compound15.21514.21514.2 | controllerType=0 controllerNumber=1 scan=21514 | 44.6027 |
| 17256:3 | 4 | R.LSLHRPALEDLLLGSEAN[+1540.529]LTC[+57.021]TLTGLR.D | HexNAc(2)Hex(7) | C[+57], N[+1541] | 1502.0501 | 3 | 4504.1358 | 4504.1268 | 0 | 2 | 127 | Specific | 659.42 | 659.42 | 659.42 | 13.64 | 99 | >sp\|P01876\|IGHA1_HUMAN Immunoglobulin heavy constant alpha 1 OS=Homo sapiens OX=9606 GN=IGHA1 PE=1 SV=2 | 255 | 20201113_gy_colo_milk_1ng_glyco_hcd203040_compound15.20534.20534.3 | controllerType=0 controllerNumber=1 scan=20534 | 42.4715 |
| 17308:3 | 4 | R.LSLHRPALEDLLLGSEAN[+1501.555]LTC[+57.021]TLTGLR.D | HexNAc(5)Hex(3) | C[+57], N[+1502] | 1489.0608 | 3 | 4465.168 | 4465.1536 | 0 | 3.2 | 127 | Specific | 661.39 | 661.39 | 661.39 | 13.64 | 99 | >sp\|P01876\|IGHA1_HUMAN Immunoglobulin heavy constant alpha 1 OS=Homo sapiens OX=9606 GN=IGHA1 PE=1 SV=2 | 255 | 20201113_gy_colo_milk_1ng_glyco_hcd203040_compound15.20588.20588.3 | controllerType=0 controllerNumber=1 scan=20588 | 42.587 |
| 17248:3 | 4 | R.LSLHRPALEDLLLGSEAN[+1702.581]LTC[+57.021]TLTGLR.D | HexNAc(2)Hex(8) | C[+57], N[+1703] | 1556.0668 | 3 | 4666.1859 | 4666.1796 | 0 | 1.4 | 127 | Specific | 661.49 | 640.27 | 640.27 | 13.51 | 99 | >sp\|P01876\|IGHA1_HUMAN Immunoglobulin heavy constant alpha 1 OS=Homo sapiens OX=9606 GN=IGHA1 PE=1 SV=2 | 255 | 20201113_gy_colo_milk_1ng_glyco_hcd203040_compound15.20526.20526.3 | controllerType=0 controllerNumber=1 scan=20526 | 42.4521 |
| 17221:4 | 4 | R.LSLHRPALEDLLLGSEAN[+1702.581]LTC[+57.021]TLTGLR.D | HexNAc(2)Hex(8) | C[+57], N[+1703] | 1167.3039 | 4 | 4666.1937 | 4666.1796 | 0 | 3 | 127 | Specific | 662.28 | 662.28 | 662.28 | 14.22 | 99 | >sp\|P01876\|IGHA1_HUMAN Immunoglobulin heavy constant alpha 1 OS=Homo sapiens OX=9606 GN=IGHA1 PE=1 SV=2 | 255 | 20201113_gy_colo_milk_1ng_glyco_hcd203040_compound15.20498.20498.4 | controllerType=0 controllerNumber=1 scan=20498 | 42.3984 |
| 17234:4 | 4 | R.LSLHRPALEDLLLGSEAN[+1663.608]LTC[+57.021]TLTGLR.D | HexNAc(5)Hex(4) | C[+57], N[+1664] | 1157.5602 | 4 | 4627.2191 | 4627.2064 | 0 | 2.7 | 127 | Specific | 666.08 | 656.77 | 656.77 | 14.22 | 99 | >sp\|P01876\|IGHA1_HUMAN Immunoglobulin heavy constant alpha 1 OS=Homo sapiens OX=9606 GN=IGHA1 PE=1 SV=2 | 255 | 20201113_gy_colo_milk_1ng_glyco_hcd203040_compound15.20511.20511.4 | controllerType=0 controllerNumber=1 scan=20511 | 42.4206 |
| 17326:3 | 4 | R.LSLHRPALEDLLLGSEAN[+1460.529]LTC[+57.021]TLTGLR.D | HexNAc(4)Hex(4) | C[+57], N[+1461] | 1475.3857 | 3 | 4424.1425 | 4424.1271 | 0 | 3.5 | 127 | Specific | 666.23 | 582.29 | 582.29 | 13.64 | 99 | >sp\|P01876\|IGHA1_HUMAN Immunoglobulin heavy constant alpha 1 OS=Homo sapiens OX=9606 GN=IGHA1 PE=1 SV=2 | 255 | 20201113_gy_colo_milk_1ng_glyco_hcd203040_compound15.20607.20607.3 | controllerType=0 controllerNumber=1 scan=20607 | 42.6274 |
| 17280:4 | 4 | R.LSLHRPALEDLLLGSEAN[+1825.661]LTC[+57.021]TLTGLR.D | HexNAc(5)Hex(5) | C[+57], N[+1826] | 1198.0752 | 4 | 4789.2791 | 4789.2592 | 0 | 4.1 | 127 | Specific | 668.49 | 617.09 | 617.09 | 13.88 | 99 | >sp\|P01876\|IGHA1_HUMAN Immunoglobulin heavy constant alpha 1 OS=Homo sapiens OX=9606 GN=IGHA1 PE=1 SV=2 | 255 | 20201113_gy_colo_milk_1ng_glyco_hcd203040_compound15.20559.20559.4 | controllerType=0 controllerNumber=1 scan=20559 | 42.519 |
| 17311:3 | 4 | R.LSLHRPALEDLLLGSEAN[+1501.555]LTC[+57.021]TLTGLR.D | HexNAc(5)Hex(3) | C[+57], N[+1502] | 1489.0608 | 3 | 4465.168 | 4465.1536 | 0 | 3.2 | 127 | Specific | 671.58 | 638.06 | 638.06 | 13.64 | 99 | >sp\|P01876\|IGHA1_HUMAN Immunoglobulin heavy constant alpha 1 OS=Homo sapiens OX=9606 GN=IGHA1 PE=1 SV=2 | 255 | 20201113_gy_colo_milk_1ng_glyco_hcd203040_compound15.20591.20591.3 | controllerType=0 controllerNumber=1 scan=20591 | 42.5953 |
| 17282:3 | 4 | R.LSLHRPALEDLLLGSEAN[+1378.476]LTC[+57.021]TLTGLR.D | HexNAc(2)Hex(6) | C[+57], N[+1378] | 1448.0348 | 3 | 4342.09 | 4342.074 | 0 | 3.7 | 127 | Specific | 675.24 | 633.94 | 633.94 | 13.64 | 99 | >sp\|P01876\|IGHA1_HUMAN Immunoglobulin heavy constant alpha 1 OS=Homo sapiens OX=9606 GN=IGHA1 PE=1 SV=2 | 255 | 20201113_gy_colo_milk_1ng_glyco_hcd203040_compound15.20561.20561.3 | controllerType=0 controllerNumber=1 scan=20561 | 42.5242 |
| 17548:4 | 4 | R.LSLHRPALEDLLLGSEAN[+1501.555]LTC[+57.021]TLTGLR.D | HexNAc(5)Hex(3) | C[+57], N[+1502] | 1117.0458 | 4 | 4465.1614 | 4465.1536 | 0 | 1.8 | 127 | Specific | 675.63 | 665.97 | 665.97 | 14.22 | 99 | >sp\|P01876\|IGHA1_HUMAN Immunoglobulin heavy constant alpha 1 OS=Homo sapiens OX=9606 GN=IGHA1 PE=1 SV=2 | 255 | 20201113_gy_colo_milk_1ng_glyco_hcd203040_compound15.20838.20838.3 | controllerType=0 controllerNumber=1 scan=20838 | 43.1126 |
| 17329:3 | 4 | R.LSLHRPALEDLLLGSEAN[+1460.529]LTC[+57.021]TLTGLR.D | HexNAc(4)Hex(4) | C[+57], N[+1461] | 1475.3857 | 3 | 4424.1425 | 4424.1271 | 0 | 3.5 | 127 | Specific | 678.77 | 570.86 | 570.86 | 13.64 | 99 | >sp\|P01876\|IGHA1_HUMAN Immunoglobulin heavy constant alpha 1 OS=Homo sapiens OX=9606 GN=IGHA1 PE=1 SV=2 | 255 | 20201113_gy_colo_milk_1ng_glyco_hcd203040_compound15.20610.20610.3 | controllerType=0 controllerNumber=1 scan=20610 | 42.6333 |
| 17403:3 | 4 | R.LSLHRPALEDLLLGSEAN[+1054.370]LTC[+57.021]TLTGLR.D | HexNAc(2)Hex(4) | C[+57], N[+1054] | 1339.9982 | 3 | 4017.9801 | 4017.9683 | 0 | 2.9 | 127 | Specific | 695.35 | 693.45 | 693.45 | 13.64 | 99 | >sp\|P01876\|IGHA1_HUMAN Immunoglobulin heavy constant alpha 1 OS=Homo sapiens OX=9606 GN=IGHA1 PE=1 SV=2 | 255 | 20201113_gy_colo_milk_1ng_glyco_hcd203040_compound15.20687.20687.3 | controllerType=0 controllerNumber=1 scan=20687 | 42.7912 |
| 17279:3 | 4 | R.LSLHRPALEDLLLGSEAN[+1378.476]LTC[+57.021]TLTGLR.D | HexNAc(2)Hex(6) | C[+57], N[+1378] | 1448.0348 | 3 | 4342.09 | 4342.074 | 0 | 3.7 | 127 | Specific | 697.02 | 697.02 | 697.02 | 13.64 | 99 | >sp\|P01876\|IGHA1_HUMAN Immunoglobulin heavy constant alpha 1 OS=Homo sapiens OX=9606 GN=IGHA1 PE=1 SV=2 | 255 | 20201113_gy_colo_milk_1ng_glyco_hcd203040_compound15.20558.20558.3 | controllerType=0 controllerNumber=1 scan=20558 | 42.5177 |
| 17251:3 | 4 | R.LSLHRPALEDLLLGSEAN[+1702.581]LTC[+57.021]TLTGLR.D | HexNAc(2)Hex(8) | C[+57], N[+1703] | 1556.0668 | 3 | 4666.1859 | 4666.1796 | 0 | 1.4 | 127 | Specific | 697.05 | 647.88 | 647.88 | 13.51 | 99 | >sp\|P01876\|IGHA1_HUMAN Immunoglobulin heavy constant alpha 1 OS=Homo sapiens OX=9606 GN=IGHA1 PE=1 SV=2 | 255 | 20201113_gy_colo_milk_1ng_glyco_hcd203040_compound15.20529.20529.3 | controllerType=0 controllerNumber=1 scan=20529 | 42.4577 |
| 18285:4 | 4 | R.LSLHRPALEDLLLGSEAN[+1751.624]LTC[+57.021]TLTGLR.D | HexNAc(4)Hex(4)NeuAc(1) | C[+57], N[+1752] | 1179.5636 | 4 | 4715.2327 | 4715.2225 | 0 | 2.2 | 127 | Specific | 697.39 | 686.45 | 686.45 | 14.22 | 99 | >sp\|P01876\|IGHA1_HUMAN Immunoglobulin heavy constant alpha 1 OS=Homo sapiens OX=9606 GN=IGHA1 PE=1 SV=2 | 255 | 20201113_gy_colo_milk_1ng_glyco_hcd203040_compound15.21608.21608.4 | controllerType=0 controllerNumber=1 scan=21608 | 44.7956 |
| 17316:3 | 4 | R.LSLHRPALEDLLLGSEAN[+1216.423]LTC[+57.021]TLTGLR.D | HexNAc(2)Hex(5) | C[+57], N[+1216] | 1394.0148 | 3 | 4180.0299 | 4180.0211 | 0 | 2.1 | 127 | Specific | 698.95 | 669 | 669 | 13.64 | 99 | >sp\|P01876\|IGHA1_HUMAN Immunoglobulin heavy constant alpha 1 OS=Homo sapiens OX=9606 GN=IGHA1 PE=1 SV=2 | 255 | 20201113_gy_colo_milk_1ng_glyco_hcd203040_compound15.20597.20597.3 | controllerType=0 controllerNumber=1 scan=20597 | 42.6105 |
| 17430:4 | 4 | R.LSLHRPALEDLLLGSEAN[+1647.613]LTC[+57.021]TLTGLR.D | HexNAc(5)Hex(3)Fuc(1) | C[+57], N[+1648] | 1153.5589 | 4 | 4611.2137 | 4611.2115 | 0 | 0.5 | 127 | Specific | 701.42 | 661.64 | 661.64 | 14.66 | 99 | >sp\|P01876\|IGHA1_HUMAN Immunoglobulin heavy constant alpha 1 OS=Homo sapiens OX=9606 GN=IGHA1 PE=1 SV=2 | 255 | 20201113_gy_colo_milk_1ng_glyco_hcd203040_compound15.20715.20715.4 | controllerType=0 controllerNumber=1 scan=20715 | 42.8647 |
| 17347:5 | 4 | R.LSLHRPALEDLLLGSEAN[+1501.555]LTC[+57.021]TLTGLR.D | HexNAc(5)Hex(3) | C[+57], N[+1502] | 893.8395 | 5 | 4465.1684 | 4465.1536 | 0 | 3.3 | 127 | Specific | 714.11 | 714.11 | 714.11 | 15.39 | 99 | >sp\|P01876\|IGHA1_HUMAN Immunoglobulin heavy constant alpha 1 OS=Homo sapiens OX=9606 GN=IGHA1 PE=1 SV=2 | 255 | 20201113_gy_colo_milk_1ng_glyco_hcd203040_compound15.20629.20629.5 | controllerType=0 controllerNumber=1 scan=20629 | 42.6689 |
| 17323:3 | 4 | R.LSLHRPALEDLLLGSEAN[+1378.476]LTC[+57.021]TLTGLR.D | HexNAc(2)Hex(6) | C[+57], N[+1378] | 1448.0344 | 3 | 4342.0885 | 4342.074 | 0 | 3.4 | 127 | Specific | 714.8 | 659.75 | 659.75 | 14.64 | 99 | >sp\|P01876\|IGHA1_HUMAN Immunoglobulin heavy constant alpha 1 OS=Homo sapiens OX=9606 GN=IGHA1 PE=1 SV=2 | 255 | 20201113_gy_colo_milk_1ng_glyco_hcd203040_compound15.20604.20604.3 | controllerType=0 controllerNumber=1 scan=20604 | 42.624 |
| 17319:4 | 4 | R.LSLHRPALEDLLLGSEAN[+1298.476]LTC[+57.021]TLTGLR.D | HexNAc(4)Hex(3) | C[+57], N[+1298] | 1066.2762 | 4 | 4262.0828 | 4262.0742 | 0 | 2 | 127 | Specific | 722.36 | 722.36 | 722.36 | 15.22 | 99 | >sp\|P01876\|IGHA1_HUMAN Immunoglobulin heavy constant alpha 1 OS=Homo sapiens OX=9606 GN=IGHA1 PE=1 SV=2 | 255 | 20201113_gy_colo_milk_1ng_glyco_hcd203040_compound15.20600.20600.4 | controllerType=0 controllerNumber=1 scan=20600 | 42.6147 |
| 17253:3 | 4 | R.LSLHRPALEDLLLGSEAN[+1540.529]LTC[+57.021]TLTGLR.D | HexNAc(2)Hex(7) | C[+57], N[+1541] | 1502.0501 | 3 | 4504.1358 | 4504.1268 | 0 | 2 | 127 | Specific | 722.83 | 657.77 | 657.77 | 14.64 | 99 | >sp\|P01876\|IGHA1_HUMAN Immunoglobulin heavy constant alpha 1 OS=Homo sapiens OX=9606 GN=IGHA1 PE=1 SV=2 | 255 | 20201113_gy_colo_milk_1ng_glyco_hcd203040_compound15.20531.20531.3 | controllerType=0 controllerNumber=1 scan=20531 | 42.4626 |
| 17296:4 | 4 | R.LSLHRPALEDLLLGSEAN[+1501.555]LTC[+57.021]TLTGLR.D | HexNAc(5)Hex(3) | C[+57], N[+1502] | 1117.0474 | 4 | 4465.1678 | 4465.1536 | 0 | 3.2 | 127 | Specific | 728.68 | 728.68 | 728.68 | 15.22 | 99 | >sp\|P01876\|IGHA1_HUMAN Immunoglobulin heavy constant alpha 1 OS=Homo sapiens OX=9606 GN=IGHA1 PE=1 SV=2 | 255 | 20201113_gy_colo_milk_1ng_glyco_hcd203040_compound15.20576.20576.4 | controllerType=0 controllerNumber=1 scan=20576 | 42.5527 |
| 17441:4 | 4 | R.LSLHRPALEDLLLGSEAN[+892.317]LTC[+57.021]TLTGLR.D | HexNAc(2)Hex(3) | C[+57], N[+892] | 964.7367 | 4 | 3855.9251 | 3855.9155 | 0 | 2.5 | 127 | Specific | 739.22 | 739.22 | 739.22 | 15.22 | 99 | >sp\|P01876\|IGHA1_HUMAN Immunoglobulin heavy constant alpha 1 OS=Homo sapiens OX=9606 GN=IGHA1 PE=1 SV=2 | 255 | 20201113_gy_colo_milk_1ng_glyco_hcd203040_compound15.20727.20727.4 | controllerType=0 controllerNumber=1 scan=20727 | 42.8878 |
| 17314:3 | 4 | R.LSLHRPALEDLLLGSEAN[+1216.423]LTC[+57.021]TLTGLR.D | HexNAc(2)Hex(5) | C[+57], N[+1216] | 1394.0148 | 3 | 4180.0299 | 4180.0211 | 0 | 2.1 | 127 | Specific | 739.39 | 694.79 | 694.79 | 14.64 | 99 | >sp\|P01876\|IGHA1_HUMAN Immunoglobulin heavy constant alpha 1 OS=Homo sapiens OX=9606 GN=IGHA1 PE=1 SV=2 | 255 | 20201113_gy_colo_milk_1ng_glyco_hcd203040_compound15.20595.20595.3 | controllerType=0 controllerNumber=1 scan=20595 | 42.6041 |

Table S4. Cont.

| Query #:z | Protein Rank | Glycopeptide | Glycan | Modification Type(s) | Observed m/z | z | Observed (M+H) | Calc. mass (M+H) | Off-by-x error | Mass error (ppm) | StartPosition | Cleavage | Score | Delta | Delta Mod | \|Log Prob\| | # of unique peptides | Protein Name | Protein DB number | Comment | Scan # | Scan Time |
|---|---|---|---|---|---|---|---|---|---|---|---|---|---|---|---|---|---|---|---|---|---|---|
| 17400:3 | 4 | R.LSLHRPALEDLLLGSEAN[+1054.370]LTC[+57.021]TLTGLR.D | HexNAc(2)Hex(4) | C[+57], N[+1054] | 1339.9982 | 3 | 4017.9801 | 4017.9683 | 0 | 2.9 | 127 | Specific | 743.19 | 708.67 | 708.67 | 14.64 | 99 | >sp\|P01876\|IGHA1_HUMAN Immunoglobulin heavy constant alpha 1 OS=Homo sapiens OX=9606 GN=IGHA1 PE=1 SV=2 | 255 | 20201113_gy_colo_milk_1ng_glyco_hcd203040_compound15.20684.20684.3 | controllerType=0 controllerNumber=1 scan=20684 | 42.7838 |
| 18271:5 | 4 | R.LSLHRPALEDLLLGSEAN[+1913.677]LTC[+57.021]TLTGLR.D | HexNAc(4)Hex(5)NeuAc(1) | C[+57], N[+1914] | 976.2641 | 5 | 4877.2912 | 4877.2753 | 0 | 3.3 | 127 | Specific | 746.92 | 746.92 | 746.92 | 15.39 | 99 | >sp\|P01876\|IGHA1_HUMAN Immunoglobulin heavy constant alpha 1 OS=Homo sapiens OX=9606 GN=IGHA1 PE=1 SV=2 | 255 | 20201113_gy_colo_milk_1ng_glyco_hcd203040_compound15.21593.21593.5 | controllerType=0 controllerNumber=1 scan=21593 | 44.7583 |
| 17446:3 | 4 | R.LSLHRPALEDLLLGSEAN[+892.317]LTC[+57.021]TLTGLR.D | HexNAc(2)Hex(3) | C[+57], N[+892] | 1285.9811 | 3 | 3855.9289 | 3855.9155 | 0 | 3.5 | 127 | Specific | 768.36 | 754.26 | 754.26 | 15.64 | 99 | >sp\|P01876\|IGHA1_HUMAN Immunoglobulin heavy constant alpha 1 OS=Homo sapiens OX=9606 GN=IGHA1 PE=1 SV=2 | 255 | 20201113_gy_colo_milk_1ng_glyco_hcd203040_compound15.20732.20732.3 | controllerType=0 controllerNumber=1 scan=20732 | 42.9 |
| 17303:4 | 4 | R.LSLHRPALEDLLLGSEAN[+1460.529]LTC[+57.021]TLTGLR.D | HexNAc(4)Hex(4) | C[+57], N[+1461] | 1106.7908 | 4 | 4424.1414 | 4424.1271 | 0 | 3.2 | 127 | Specific | 770.34 | 770.28 | 770.28 | 16.22 | 99 | >sp\|P01876\|IGHA1_HUMAN Immunoglobulin heavy constant alpha 1 OS=Homo sapiens OX=9606 GN=IGHA1 PE=1 SV=2 | 255 | 20201113_gy_colo_milk_1ng_glyco_hcd203040_compound15.20583.20583.4 | controllerType=0 controllerNumber=1 scan=20583 | 42.5669 |
| 18237:4 | 4 | R.LSLHRPALEDLLLGSEAN[+2116.756]LTC[+57.021]TLTGLR.D | HexNAc(5)Hex(5)NeuAc(1) | C[+57], N[+2117] | 1270.8461 | 4 | 5080.3625 | 5080.3547 | 0 | 1.5 | 127 | Specific | 771.56 | 668.07 | 668.07 | 16.08 | 99 | >sp\|P01876\|IGHA1_HUMAN Immunoglobulin heavy constant alpha 1 OS=Homo sapiens OX=9606 GN=IGHA1 PE=1 SV=2 | 255 | 20201113_gy_colo_milk_1ng_glyco_hcd203040_compound15.21558.21558.4 | controllerType=0 controllerNumber=1 scan=21558 | 44.6862 |
| 17443:3 | 4 | R.LSLHRPALEDLLLGSEAN[+892.317]LTC[+57.021]TLTGLR.D | HexNAc(2)Hex(3) | C[+57], N[+892] | 1285.9811 | 3 | 3855.9289 | 3855.9155 | 0 | 3.5 | 127 | Specific | 773.95 | 773.95 | 773.95 | 15.64 | 99 | >sp\|P01876\|IGHA1_HUMAN Immunoglobulin heavy constant alpha 1 OS=Homo sapiens OX=9606 GN=IGHA1 PE=1 SV=2 | 255 | 20201113_gy_colo_milk_1ng_glyco_hcd203040_compound15.20729.20729.3 | controllerType=0 controllerNumber=1 scan=20729 | 42.8896 |
| 17298:4 | 4 | R.LSLHRPALEDLLLGSEAN[+1501.555]LTC[+57.021]TLTGLR.D | HexNAc(5)Hex(3) | C[+57], N[+1502] | 1117.0474 | 4 | 4465.1678 | 4465.1536 | 0 | 3.2 | 127 | Specific | 779.78 | 779.78 | 779.78 | 16.22 | 99 | >sp\|P01876\|IGHA1_HUMAN Immunoglobulin heavy constant alpha 1 OS=Homo sapiens OX=9606 GN=IGHA1 PE=1 SV=2 | 255 | 20201113_gy_colo_milk_1ng_glyco_hcd203040_compound15.20578.20578.4 | controllerType=0 controllerNumber=1 scan=20578 | 42.5599 |
| 17434:4 | 4 | R.LSLHRPALEDLLLGSEAN[+1524.534]LTC[+57.021]TLTGLR.D | HexNAc(2)Hex(6)Fuc(1) | C[+57], N[+1525] | 1122.7846 | 4 | 4488.1165 | 4488.1319 | 0 | -3.4 | 127 | Specific | 781.77 | 781.77 | 781.77 | 14.57 | 99 | >sp\|P01876\|IGHA1_HUMAN Immunoglobulin heavy constant alpha 1 OS=Homo sapiens OX=9606 GN=IGHA1 PE=1 SV=2 | 255 | 20201113_gy_colo_milk_1ng_glyco_hcd203040_compound15.20719.20719.4 | controllerType=0 controllerNumber=1 scan=20719 | 42.8734 |
| 17321:4 | 4 | R.LSLHRPALEDLLLGSEAN[+1419.502]LTC[+57.021]TLTGLR.D | HexNAc(3)Hex(5) | C[+57], N[+1420] | 1096.5339 | 4 | 4383.1136 | 4383.1005 | 0 | 3 | 127 | Specific | 784.82 | 773.89 | 773.89 | 16.22 | 99 | >sp\|P01876\|IGHA1_HUMAN Immunoglobulin heavy constant alpha 1 OS=Homo sapiens OX=9606 GN=IGHA1 PE=1 SV=2 | 255 | 20201113_gy_colo_milk_1ng_glyco_hcd203040_compound15.20602.20602.4 | controllerType=0 controllerNumber=1 scan=20602 | 42.6172 |
| 18305:5 | 4 | R.LSLHRPALEDLLLGSEAN[+1751.624]LTC[+57.021]TLTGLR.D | HexNAc(4)Hex(4)NeuAc(1) | C[+57], N[+1752] | 943.8513 | 5 | 4715.2273 | 4715.2225 | 0 | 1 | 127 | Specific | 791.85 | 791.85 | 791.85 | 16.25 | 99 | >sp\|P01876\|IGHA1_HUMAN Immunoglobulin heavy constant alpha 1 OS=Homo sapiens OX=9606 GN=IGHA1 PE=1 SV=2 | 255 | 20201113_gy_colo_milk_1ng_glyco_hcd203040_compound15.21628.21628.5 | controllerType=0 controllerNumber=1 scan=21628 | 44.8406 |
| 17237:4 | 4 | R.LSLHRPALEDLLLGSEAN[+1663.608]LTC[+57.021]TLTGLR.D | HexNAc(5)Hex(4) | C[+57], N[+1664] | 1157.5602 | 4 | 4627.2191 | 4627.2064 | 0 | 2.7 | 127 | Specific | 796.22 | 659.22 | 659.22 | 16.22 | 99 | >sp\|P01876\|IGHA1_HUMAN Immunoglobulin heavy constant alpha 1 OS=Homo sapiens OX=9606 GN=IGHA1 PE=1 SV=2 | 255 | 20201113_gy_colo_milk_1ng_glyco_hcd203040_compound15.20514.20514.4 | controllerType=0 controllerNumber=1 scan=20514 | 42.4284 |
| 18198:4 | 4 | R.LSLHRPALEDLLLGSEAN[+2116.756]LTC[+57.021]TLTGLR.D | HexNAc(5)Hex(5)NeuAc(1) | C[+57], N[+2117] | 1270.8467 | 4 | 5080.3649 | 5080.3547 | 0 | 2 | 127 | Specific | 800.04 | 746.15 | 746.15 | 17.22 | 99 | >sp\|P01876\|IGHA1_HUMAN Immunoglobulin heavy constant alpha 1 OS=Homo sapiens OX=9606 GN=IGHA1 PE=1 SV=2 | 255 | 20201113_gy_colo_milk_1ng_glyco_hcd203040_compound15.21517.21517.2 | controllerType=0 controllerNumber=1 scan=21517 | 44.6077 |
| 18260:4 | 4 | R.LSLHRPALEDLLLGSEAN[+1913.677]LTC[+57.021]TLTGLR.D | HexNAc(4)Hex(5)NeuAc(1) | C[+57], N[+1914] | 1220.0788 | 4 | 4877.2933 | 4877.2753 | 0 | 3.7 | 127 | Specific | 805.05 | 771.39 | 771.39 | 17.22 | 99 | >sp\|P01876\|IGHA1_HUMAN Immunoglobulin heavy constant alpha 1 OS=Homo sapiens OX=9606 GN=IGHA1 PE=1 SV=2 | 255 | 20201113_gy_colo_milk_1ng_glyco_hcd203040_compound15.21582.21582.4 | controllerType=0 controllerNumber=1 scan=21582 | 44.7417 |
| 18307:5 | 4 | R.LSLHRPALEDLLLGSEAN[+1751.624]LTC[+57.021]TLTGLR.D | HexNAc(4)Hex(4)NeuAc(1) | C[+57], N[+1752] | 943.8513 | 5 | 4715.2273 | 4715.2225 | 0 | 1 | 127 | Specific | 808.3 | 808.3 | 808.3 | 17.25 | 99 | >sp\|P01876\|IGHA1_HUMAN Immunoglobulin heavy constant alpha 1 OS=Homo sapiens OX=9606 GN=IGHA1 PE=1 SV=2 | 255 | 20201113_gy_colo_milk_1ng_glyco_hcd203040_compound15.21630.21630.5 | controllerType=0 controllerNumber=1 scan=21630 | 44.8496 |
| 17244:4 | 4 | R.LSLHRPALEDLLLGSEAN[+1864.634]LTC[+57.021]TLTGLR.D | HexNAc(2)Hex(9) | C[+57], N[+1865] | 1207.8154 | 4 | 4828.2396 | 4828.2324 | 0 | 1.5 | 127 | Specific | 867.34 | 867.34 | 867.34 | 18.08 | 99 | >sp\|P01876\|IGHA1_HUMAN Immunoglobulin heavy constant alpha 1 OS=Homo sapiens OX=9606 GN=IGHA1 PE=1 SV=2 | 255 | 20201113_gy_colo_milk_1ng_glyco_hcd203040_compound15.20522.20522.4 | controllerType=0 controllerNumber=1 scan=20522 | 42.4443 |
| 18337:4 | 4 | R.LSLHRPALEDLLLGSEAN[+1710.598]LTC[+57.021]TLTGLR.D | HexNAc(3)Hex(5)NeuAc(1) | C[+57], N[+1711] | 1169.3089 | 4 | 4674.2137 | 4674.1959 | 0 | 3.8 | 127 | Specific | 869.01 | 826.56 | 826.56 | 17.88 | 99 | >sp\|P01876\|IGHA1_HUMAN Immunoglobulin heavy constant alpha 1 OS=Homo sapiens OX=9606 GN=IGHA1 PE=1 SV=2 | 255 | 20201113_gy_colo_milk_1ng_glyco_hcd203040_compound15.21662.21662.4 | controllerType=0 controllerNumber=1 scan=21662 | 44.9172 |
| 17307:4 | 4 | R.LSLHRPALEDLLLGSEAN[+1460.529]LTC[+57.021]TLTGLR.D | HexNAc(4)Hex(4) | C[+57], N[+1461] | 1106.7908 | 4 | 4424.1414 | 4424.1271 | 0 | 3.2 | 127 | Specific | 869.53 | 791.1 | 791.1 | 18.22 | 99 | >sp\|P01876\|IGHA1_HUMAN Immunoglobulin heavy constant alpha 1 OS=Homo sapiens OX=9606 GN=IGHA1 PE=1 SV=2 | 255 | 20201113_gy_colo_milk_1ng_glyco_hcd203040_compound15.20587.20587.4 | controllerType=0 controllerNumber=1 scan=20587 | 42.5859 |
| 17322:4 | 4 | R.LSLHRPALEDLLLGSEAN[+1298.476]LTC[+57.021]TLTGLR.D | HexNAc(4)Hex(3) | C[+57], N[+1298] | 1066.2762 | 4 | 4262.0828 | 4262.0742 | 0 | 2 | 127 | Specific | 880.72 | 880.72 | 880.72 | 18.22 | 99 | >sp\|P01876\|IGHA1_HUMAN Immunoglobulin heavy constant alpha 1 OS=Homo sapiens OX=9606 GN=IGHA1 PE=1 SV=2 | 255 | 20201113_gy_colo_milk_1ng_glyco_hcd203040_compound15.20603.20603.4 | controllerType=0 controllerNumber=1 scan=20603 | 42.6218 |
| 17346:4 | 4 | R.LSLHRPALEDLLLGSEAN[+1257.449]LTC[+57.021]TLTGLR.D | HexNAc(3)Hex(4) | C[+57], N[+1257] | 1056.016 | 4 | 4221.0423 | 4221.0477 | 0 | -1.3 | 127 | Specific | 900.67 | 854.42 | 854.42 | 18.37 | 99 | >sp\|P01876\|IGHA1_HUMAN Immunoglobulin heavy constant alpha 1 OS=Homo sapiens OX=9606 GN=IGHA1 PE=1 SV=2 | 255 | 20201113_gy_colo_milk_1ng_glyco_hcd203040_compound15.20628.20628.4 | controllerType=0 controllerNumber=1 scan=20628 | 42.6671 |
| 17324:4 | 4 | R.LSLHRPALEDLLLGSEAN[+1419.502]LTC[+57.021]TLTGLR.D | HexNAc(3)Hex(5) | C[+57], N[+1420] | 1096.5339 | 4 | 4383.1136 | 4383.1005 | 0 | 3 | 127 | Specific | 907.66 | 820.58 | 820.58 | 19.22 | 99 | >sp\|P01876\|IGHA1_HUMAN Immunoglobulin heavy constant alpha 1 OS=Homo sapiens OX=9606 GN=IGHA1 PE=1 SV=2 | 255 | 20201113_gy_colo_milk_1ng_glyco_hcd203040_compound15.20605.20605.4 | controllerType=0 controllerNumber=1 scan=20605 | 42.6251 |
| 17246:4 | 4 | R.LSLHRPALEDLLLGSEAN[+1864.634]LTC[+57.021]TLTGLR.D | HexNAc(2)Hex(9) | C[+57], N[+1865] | 1207.8154 | 4 | 4828.2396 | 4828.2324 | 0 | 1.5 | 127 | Specific | 924.05 | 924.05 | 924.05 | 19.08 | 99 | >sp\|P01876\|IGHA1_HUMAN Immunoglobulin heavy constant alpha 1 OS=Homo sapiens OX=9606 GN=IGHA1 PE=1 SV=2 | 255 | 20201113_gy_colo_milk_1ng_glyco_hcd203040_compound15.20524.20524.4 | controllerType=0 controllerNumber=1 scan=20524 | 42.4493 |
| 17300:4 | 4 | R.LSLHRPALEDLLLGSEAN[+1216.423]LTC[+57.021]TLTGLR.D | HexNAc(2)Hex(5) | C[+57], N[+1216] | 1045.7638 | 4 | 4180.0335 | 4180.0211 | 0 | 3 | 127 | Specific | 936.95 | 936.95 | 936.95 | 19.22 | 99 | >sp\|P01876\|IGHA1_HUMAN Immunoglobulin heavy constant alpha 1 OS=Homo sapiens OX=9606 GN=IGHA1 PE=1 SV=2 | 255 | 20201113_gy_colo_milk_1ng_glyco_hcd203040_compound15.20580.20580.4 | controllerType=0 controllerNumber=1 scan=20580 | 42.5635 |
| 17396:4 | 4 | R.LSLHRPALEDLLLGSEAN[+1054.370]LTC[+57.021]TLTGLR.D | HexNAc(2)Hex(4) | C[+57], N[+1054] | 1005.2509 | 4 | 4017.9818 | 4017.9683 | 0 | 3.3 | 127 | Specific | 937.67 | 918.89 | 918.89 | 19.22 | 99 | >sp\|P01876\|IGHA1_HUMAN Immunoglobulin heavy constant alpha 1 OS=Homo sapiens OX=9606 GN=IGHA1 PE=1 SV=2 | 255 | 20201113_gy_colo_milk_1ng_glyco_hcd203040_compound15.20680.20680.4 | controllerType=0 controllerNumber=1 scan=20680 | 42.771 |
| 17302:4 | 4 | R.LSLHRPALEDLLLGSEAN[+1378.476]LTC[+57.021]TLTGLR.D | HexNAc(2)Hex(6) | C[+57], N[+1378] | 1086.2769 | 4 | 4342.0858 | 4342.074 | 0 | 2.7 | 127 | Specific | 958.74 | 876.12 | 876.12 | 20.22 | 99 | >sp\|P01876\|IGHA1_HUMAN Immunoglobulin heavy constant alpha 1 OS=Homo sapiens OX=9606 GN=IGHA1 PE=1 SV=2 | 255 | 20201113_gy_colo_milk_1ng_glyco_hcd203040_compound15.20582.20582.4 | controllerType=0 controllerNumber=1 scan=20582 | 42.5658 |
| 17297:4 | 4 | R.LSLHRPALEDLLLGSEAN[+1540.529]LTC[+57.021]TLTGLR.D | HexNAc(2)Hex(7) | C[+57], N[+1541] | 1126.7891 | 4 | 4504.1346 | 4504.1268 | 0 | 1.7 | 127 | Specific | 962.74 | 892.79 | 892.79 | 20.22 | 99 | >sp\|P01876\|IGHA1_HUMAN Immunoglobulin heavy constant alpha 1 OS=Homo sapiens OX=9606 GN=IGHA1 PE=1 SV=2 | 255 | 20201113_gy_colo_milk_1ng_glyco_hcd203040_compound15.20577.20577.4 | controllerType=0 controllerNumber=1 scan=20577 | 42.5542 |
| 17444:4 | 4 | R.LSLHRPALEDLLLGSEAN[+892.317]LTC[+57.021]TLTGLR.D | HexNAc(2)Hex(3) | C[+57], N[+892] | 964.7367 | 4 | 3855.9251 | 3855.9155 | 0 | 2.5 | 127 | Specific | 979.41 | 979.41 | 979.41 | 20.22 | 99 | >sp\|P01876\|IGHA1_HUMAN Immunoglobulin heavy constant alpha 1 OS=Homo sapiens OX=9606 GN=IGHA1 PE=1 SV=2 | 255 | 20201113_gy_colo_milk_1ng_glyco_hcd203040_compound15.20730.20730.4 | controllerType=0 controllerNumber=1 scan=20730 | 42.8918 |
| 17306:4 | 4 | R.LSLHRPALEDLLLGSEAN[+1378.476]LTC[+57.021]TLTGLR.D | HexNAc(2)Hex(6) | C[+57], N[+1378] | 1086.2769 | 4 | 4342.0858 | 4342.074 | 0 | 2.7 | 127 | Specific | 1008.01 | 1008.01 | 1008.01 | 21.22 | 99 | >sp\|P01876\|IGHA1_HUMAN Immunoglobulin heavy constant alpha 1 OS=Homo sapiens OX=9606 GN=IGHA1 PE=1 SV=2 | 255 | 20201113_gy_colo_milk_1ng_glyco_hcd203040_compound15.20586.20586.4 | controllerType=0 controllerNumber=1 scan=20586 | 42.5848 |
| 17398:4 | 4 | R.LSLHRPALEDLLLGSEAN[+1054.370]LTC[+57.021]TLTGLR.D | HexNAc(2)Hex(4) | C[+57], N[+1054] | 1005.2509 | 4 | 4017.9818 | 4017.9683 | 0 | 3.3 | 127 | Specific | 1014.67 | 977.87 | 977.87 | 21.22 | 99 | >sp\|P01876\|IGHA1_HUMAN Immunoglobulin heavy constant alpha 1 OS=Homo sapiens OX=9606 GN=IGHA1 PE=1 SV=2 | 255 | 20201113_gy_colo_milk_1ng_glyco_hcd203040_compound15.20682.20682.4 | controllerType=0 controllerNumber=1 scan=20682 | 42.7815 |
| 17304:4 | 4 | R.LSLHRPALEDLLLGSEAN[+1216.423]LTC[+57.021]TLTGLR.D | HexNAc(2)Hex(5) | C[+57], N[+1216] | 1045.7638 | 4 | 4180.0335 | 4180.0211 | 0 | 3 | 127 | Specific | 1021.95 | 1021.95 | 1021.95 | 21.22 | 99 | >sp\|P01876\|IGHA1_HUMAN Immunoglobulin heavy constant alpha 1 OS=Homo sapiens OX=9606 GN=IGHA1 PE=1 SV=2 | 255 | 20201113_gy_colo_milk_1ng_glyco_hcd203040_compound15.20584.20584.4 | controllerType=0 controllerNumber=1 scan=20584 | 42.5751 |
| 17224:4 | 4 | R.LSLHRPALEDLLLGSEAN[+1702.581]LTC[+57.021]TLTGLR.D | HexNAc(2)Hex(8) | C[+57], N[+1703] | 1167.3039 | 4 | 4666.1937 | 4666.1796 | 0 | 3 | 127 | Specific | 1026.68 | 951.72 | 951.72 | 21.22 | 99 | >sp\|P01876\|IGHA1_HUMAN Immunoglobulin heavy constant alpha 1 OS=Homo sapiens OX=9606 GN=IGHA1 PE=1 SV=2 | 255 | 20201113_gy_colo_milk_1ng_glyco_hcd203040_compound15.20501.20501.4 | controllerType=0 controllerNumber=1 scan=20501 | 42.4039 |
| 17299:4 | 4 | R.LSLHRPALEDLLLGSEAN[+1540.529]LTC[+57.021]TLTGLR.D | HexNAc(2)Hex(7) | C[+57], N[+1541] | 1126.7891 | 4 | 4504.1346 | 4504.1268 | 0 | 1.7 | 127 | Specific | 1027.93 | 996.22 | 996.22 | 21.22 | 99 | >sp\|P01876\|IGHA1_HUMAN Immunoglobulin heavy constant alpha 1 OS=Homo sapiens OX=9606 GN=IGHA1 PE=1 SV=2 | 255 | 20201113_gy_colo_milk_1ng_glyco_hcd203040_compound15.20579.20579.4 | controllerType=0 controllerNumber=1 scan=20579 | 42.5624 |
| 17349:4 | 4 | R.LSLHRPALEDLLLGSEAN[+1257.449]LTC[+57.021]TLTGLR.D | HexNAc(3)Hex(4) | C[+57], N[+1257] | 1056.016 | 4 | 4221.0423 | 4221.0477 | 0 | -1.3 | 127 | Specific | 1095.13 | 1011.43 | 1011.43 | 21.37 | 99 | >sp\|P01876\|IGHA1_HUMAN Immunoglobulin heavy constant alpha 1 OS=Homo sapiens OX=9606 GN=IGHA1 PE=1 SV=2 | 255 | 20201113_gy_colo_milk_1ng_glyco_hcd203040_compound15.20631.20631.4 | controllerType=0 controllerNumber=1 scan=20631 | 42.6744 |
| 09041:3 | 4 | R.LAGKPTHVN[+1216.423]VSVVMAEVDGTC[+57.021].Y | HexNAc(2)Hex(5) | C[+57], N[+1216] | 1134.174 | 3 | 3400.5074 | 3400.5016 | 0 | 1.7 | 332 | CRagged | 207.28 | 163.33 | 163.33 | 3.36 | 99 | >sp\|P01876\|IGHA1_HUMAN Immunoglobulin heavy constant alpha 1 OS=Homo sapiens OX=9606 GN=IGHA1 PE=1 SV=2 | 255 | 20201113_gy_colo_milk_1ng_glyco_hcd203040_compound15.12001.12001.3 | controllerType=0 controllerNumber=1 scan=12001 | 26.2116 |
| 13000:3 | 4 | R.LAGKPTHVN[+2263.835]VSVVMAEVDGTC[+57.021].Y | HexNAc(5)Hex(5)Fuc(3) | C[+57], N[+2264] | 1483.3093 | 3 | 4447.9132 | 4447.9135 | 0 | -0.1 | 332 | CRagged | 208.53 | 205.08 | 205.08 | 3.2 | 99 | >sp\|P01876\|IGHA1_HUMAN Immunoglobulin heavy constant alpha 1 OS=Homo sapiens OX=9606 GN=IGHA1 PE=1 SV=2 | 255 | 20201113_gy_colo_milk_1ng_glyco_hcd203040_compound15.16107.16107.3 | controllerType=0 controllerNumber=1 scan=16107 | 33.7081 |
| 11680:3 | 4 | R.LAGKPTHVN[+1378.476]VSVVMAEVDGTC[+57.021].Y | HexNAc(2)Hex(6) | C[+57], N[+1378] | 1188.1912 | 3 | 3562.5591 | 3562.5544 | 0 | 1.3 | 332 | CRagged | 219.13 | 112.8 | 86.84 | 3.33 | 99 | >sp\|P01876\|IGHA1_HUMAN Immunoglobulin heavy constant alpha 1 OS=Homo sapiens OX=9606 GN=IGHA1 PE=1 SV=2 | 255 | 20201113_gy_colo_milk_1ng_glyco_hcd203040_compound15.14738.14738.3 | controllerType=0 controllerNumber=1 scan=14738 | 31.2864 |
| 09039:3 | 4 | R.LAGKPTHVN[+1216.423]VSVVMAEVDGTC[+57.021].Y | HexNAc(2)Hex(5) | C[+57], N[+1216] | 1134.174 | 3 | 3400.5074 | 3400.5016 | 0 | 1.7 | 332 | CRagged | 288.93 | 238.93 | 221.7 | 5.79 | 99 | >sp\|P01876\|IGHA1_HUMAN Immunoglobulin heavy constant alpha 1 OS=Homo sapiens OX=9606 GN=IGHA1 PE=1 SV=2 | 255 | 20201113_gy_colo_milk_1ng_glyco_hcd203040_compound15.11999.11999.3 | controllerType=0 controllerNumber=1 scan=11999 | 26.2053 |
| 12983:3 | 4 | R.LAGKPTHVN[+2263.835]VSVVMAEVDGTC[+57.021].Y | HexNAc(5)Hex(5)Fuc(3) | C[+57], N[+2264] | 1483.3071 | 3 | 4447.9066 | 4447.9135 | 0 | -1.5 | 332 | CRagged | 342.12 | 280.75 | 280.75 | 7.02 | 99 | >sp\|P01876\|IGHA1_HUMAN Immunoglobulin heavy constant alpha 1 OS=Homo sapiens OX=9606 GN=IGHA1 PE=1 SV=2 | 255 | 20201113_gy_colo_milk_1ng_glyco_hcd203040_compound15.16089.16089.3 | controllerType=0 controllerNumber=1 scan=16089 | 33.6752 |
| 13146:4 | 4 | R.LAGKPTHVN[+1751.624]VSVVMAEVDGTC[+57.021]Y.- | HexNAc(4)Hex(4)NeuAc(1) | C[+57], N[+1752] | 1025.4482 | 4 | 4098.7708 | 4098.7663 | 0 | 1.1 | 332 | CRagged | 252.76 | 143.18 | 143.18 | 4.03 | 99 | >sp\|P01876\|IGHA1_HUMAN Immunoglobulin heavy constant alpha 1 OS=Homo sapiens OX=9606 GN=IGHA1 PE=1 SV=2 | 255 | 20201113_gy_colo_milk_1ng_glyco_hcd203040_compound15.16259.16259.4 | controllerType=0 controllerNumber=1 scan=16259 | 34.0018 |
| 10438:3 | 4 | R.LAGKPTHVN[+1095.397]VSVVMAEVDGTC[+57.021]Y.- | HexNAc(3)Hex(3) | C[+57], N[+1095] | 1148.1881 | 3 | 3442.5499 | 3442.5387 | 0 | 3.3 | 332 | CRagged | 287.02 | 230.25 | 230.25 | 5.01 | 99 | >sp\|P01876\|IGHA1_HUMAN Immunoglobulin heavy constant alpha 1 OS=Homo sapiens OX=9606 GN=IGHA1 PE=1 SV=2 | 255 | 20201113_gy_colo_milk_1ng_glyco_hcd203040_compound15.13449.13449.3 | controllerType=0 controllerNumber=1 scan=13449 | 28.8423 |
| 13080:3 | 4 | R.LAGKPTHVN[+2059.735]VSVVMAEVDGTC[+57.021]Y.- | HexNAc(4)Hex(5)Fuc(1)NeuAc(1) | C[+57], N[+2060] | 1469.6318 | 3 | 4406.8808 | 4406.877 | 0 | 0.9 | 332 | CRagged | 287.45 | 259.49 | 259.49 | 7.16 | 99 | >sp\|P01876\|IGHA1_HUMAN Immunoglobulin heavy constant alpha 1 OS=Homo sapiens OX=9606 GN=IGHA1 PE=1 SV=2 | 255 | 20201113_gy_colo_milk_1ng_glyco_hcd203040_compound15.16191.16191.3 | controllerType=0 controllerNumber=1 scan=16191 | 33.8755 |
| 13025:4 | 4 | R.LAGKPTHVN[+2116.756]VSVVMAEVDGTC[+57.021]Y.- | HexNAc(5)Hex(5)NeuAc(1) | C[+57], N[+2117] | 1116.7321 | 4 | 4463.9066 | 4463.8985 | 0 | 1.8 | 332 | CRagged | 299.41 | 205.15 | 205.15 | 7.07 | 99 | >sp\|P01876\|IGHA1_HUMAN Immunoglobulin heavy constant alpha 1 OS=Homo sapiens OX=9606 GN=IGHA1 PE=1 SV=2 | 255 | 20201113_gy_colo_milk_1ng_glyco_hcd203040_compound15.16133.16133.4 | controllerType=0 controllerNumber=1 scan=16133 | 33.7542 |
| 12987:3 | 4 | R.LAGKPTHVN[+2262.814]VSVVMAEVDGTC[+57.021]Y.- | HexNAc(5)Hex(5)Fuc(1)NeuAc(1) | C[+57], N[+2263] | 1537.3267 | 3 | 4609.9656 | 4609.9564 | 0 | 2 | 332 | CRagged | 306.4 | 306.4 | 306.4 | 7.57 | 99 | >sp\|P01876\|IGHA1_HUMAN Immunoglobulin heavy constant alpha 1 OS=Homo sapiens OX=9606 GN=IGHA1 PE=1 SV=2 | 255 | 20201113_gy_colo_milk_1ng_glyco_hcd203040_compound15.16094.16094.3 | controllerType=0 controllerNumber=1 scan=16094 | 33.6898 |
| 12031:3 | 4 | R.LAGKPTHVN[+1216.423]VSVVMAEVDGTC[+57.021]Y.- | HexNAc(2)Hex(5) | C[+57], N[+1216] | 1188.5301 | 3 | 3563.5757 | 3563.5649 | 0 | 3 | 332 | CRagged | 317.48 | 317.48 | 219.19 | 7.78 | 99 | >sp\|P01876\|IGHA1_HUMAN Immunoglobulin heavy constant alpha 1 OS=Homo sapiens OX=9606 GN=IGHA1 PE=1 SV=2 | 255 | 20201113_gy_colo_milk_1ng_glyco_hcd203040_compound15.15102.15102.3 | controllerType=0 controllerNumber=1 scan=15102 | 31.925 |
| 11376:4 | 4 | R.LAGKPTHVN[+1809.666]VSVVMAEVDGTC[+57.021]Y.- | HexNAc(5)Hex(4)Fuc(1) | C[+57], N[+1810] | 1039.9591 | 4 | 4156.8148 | 4156.8081 | 0 | 1.6 | 332 | CRagged | 338.75 | 312.93 | 301.19 | 6.19 | 99 | >sp\|P01876\|IGHA1_HUMAN Immunoglobulin heavy constant alpha 1 OS=Homo sapiens OX=9606 GN=IGHA1 PE=1 SV=2 | 255 | 20201113_gy_colo_milk_1ng_glyco_hcd203040_compound15.14422.14422.4 | controllerType=0 controllerNumber=1 scan=14422 | 30.6653 |
| 15023:3 | 4 | R.LAGKPTHVN[+2205.793]VSVVMAEVDGTC[+57.021]Y.- | HexNAc(4)Hex(5)Fuc(2)NeuAc(1) | C[+57], N[+2206] | 1518.3167 | 3 | 4552.9356 | 4552.9349 | 0 | 0.1 | 332 | CRagged | 351.75 | 316.98 | 316.98 | 7.54 | 99 | >sp\|P01876\|IGHA1_HUMAN Immunoglobulin heavy constant alpha 1 OS=Homo sapiens OX=9606 GN=IGHA1 PE=1 SV=2 | 255 | 20201113_gy_colo_milk_1ng_glyco_hcd203040_compound15.18207.18207.3 | controllerType=0 controllerNumber=1 scan=18207 | 37.7222 |
| 13027:4 | 4 | R.LAGKPTHVN[+2116.756]VSVVMAEVDGTC[+57.021]Y.- | HexNAc(5)Hex(5)NeuAc(1) | C[+57], N[+2117] | 1116.7321 | 4 | 4463.9066 | 4463.8985 | 0 | 1.8 | 332 | CRagged | 353.01 | 227.1 | 227.1 | 6.14 | 99 | >sp\|P01876\|IGHA1_HUMAN Immunoglobulin heavy constant alpha 1 OS=Homo sapiens OX=9606 GN=IGHA1 PE=1 SV=2 | 255 | 20201113_gy_colo_milk_1ng_glyco_hcd203040_compound15.16135.16135.4 | controllerType=0 controllerNumber=1 scan=16135 | 33.7599 |
| 13014:3 | 4 | R.LAGKPTHVN[+2116.756]VSVVMAEVDGTC[+57.021]Y.- | HexNAc(5)Hex(5)NeuAc(1) | C[+57], N[+2117] | 1488.6399 | 3 | 4463.9053 | 4463.8985 | 0 | 1.5 | 332 | CRagged | 383.68 | 383.68 | 383.68 | 9.86 | 99 | >sp\|P01876\|IGHA1_HUMAN Immunoglobulin heavy constant alpha 1 OS=Homo sapiens OX=9606 GN=IGHA1 PE=1 SV=2 | 255 | 20201113_gy_colo_milk_1ng_glyco_hcd203040_compound15.16122.16122.3 | controllerType=0 controllerNumber=1 scan=16122 | 33.738 |
| 12033:3 | 4 | R.LAGKPTHVN[+1216.423]VSVVMAEVDGTC[+57.021]Y.- | HexNAc(2)Hex(5) | C[+57], N[+1216] | 1188.5301 | 3 | 3563.5757 | 3563.5649 | 0 | 3 | 332 | CRagged | 385.83 | 385.83 | 250.17 | 10 | 99 | >sp\|P01876\|IGHA1_HUMAN Immunoglobulin heavy constant alpha 1 OS=Homo sapiens OX=9606 GN=IGHA1 PE=1 SV=2 | 255 | 20201113_gy_colo_milk_1ng_glyco_hcd203040_compound15.15104.15104.3 | controllerType=0 controllerNumber=1 scan=15104 | 31.929 |
| 15021:3 | 4 | R.LAGKPTHVN[+2205.793]VSVVMAEVDGTC[+57.021]Y.- | HexNAc(4)Hex(5)Fuc(2)NeuAc(1) | C[+57], N[+2206] | 1518.3167 | 3 | 4552.9356 | 4552.9349 | 0 | 0.1 | 332 | CRagged | 409.44 | 314.64 | 314.64 | 7.41 | 99 | >sp\|P01876\|IGHA1_HUMAN Immunoglobulin heavy constant alpha 1 OS=Homo sapiens OX=9606 GN=IGHA1 PE=1 SV=2 | 255 | 20201113_gy_colo_milk_1ng_glyco_hcd203040_compound15.18205.18205.3 | controllerType=0 controllerNumber=1 scan=18205 | 37.7186 |
| 11538:3 | 4 | R.LAGKPTHVN[+1241.454]VSVVMAEVDGTC[+57.021]Y.- | HexNAc(3)Hex(3)Fuc(1) | C[+57], N[+1241] | 1196.8725 | 3 | 3588.6029 | 3588.5966 | 0 | 1.8 | 332 | CRagged | 484.91 | 484.91 | 484.91 | 10.33 | 99 | >sp\|P01876\|IGHA1_HUMAN Immunoglobulin heavy constant alpha 1 OS=Homo sapiens OX=9606 GN=IGHA1 PE=1 SV=2 | 255 | 20201113_gy_colo_milk_1ng_glyco_hcd203040_compound15.14591.14591.3 | controllerType=0 controllerNumber=1 scan=14591 | 31.0158 |
| 13144:4 | 4 | R.LAGKPTHVN[+1751.624]VSVVMAEVDGTC[+57.021]Y.- | HexNAc(4)Hex(4)NeuAc(1) | C[+57], N[+1752] | 1025.4482 | 4 | 4098.7708 | 4098.7663 | 0 | 1.1 | 332 | CRagged | 486.02 | 378.51 | 378.51 | 8.9 | 99 | >sp\|P01876\|IGHA1_HUMAN Immunoglobulin heavy constant alpha 1 OS=Homo sapiens OX=9606 GN=IGHA1 PE=1 SV=2 | 255 | 20201113_gy_colo_milk_1ng_glyco_hcd203040_compound15.16257.16257.4 | controllerType=0 controllerNumber=1 scan=16257 | 33.9951 |
| 11373:4 | 4 | R.LAGKPTHVN[+1809.666]VSVVMAEVDGTC[+57.021]Y.- | HexNAc(5)Hex(4)Fuc(1) | C[+57], N[+1810] | 1039.9591 | 4 | 4156.8148 | 4156.8081 | 0 | 1.6 | 332 | CRagged | 491.43 | 487.22 | 354.67 | 10.51 | 99 | >sp\|P01876\|IGHA1_HUMAN Immunoglobulin heavy constant alpha 1 OS=Homo sapiens OX=9606 GN=IGHA1 PE=1 SV=2 | 255 | 20201113_gy_colo_milk_1ng_glyco_hcd203040_compound15.14419.14419.4 | controllerType=0 controllerNumber=1 scan=14419 | 30.6572 |
| 14923:4 | 4 | R.LAGKPTHVN[+2204.772]VSVVMAEVDGTC[+57.021]Y.- | HexNAc(4)Hex(5)NeuAc(2) | C[+57], N[+2205] | 1138.7339 | 4 | 4551.9139 | 4551.9145 | 0 | -0.1 | 332 | CRagged | 492.7 | 484.46 | 432.06 | 9.95 | 99 | >sp\|P01876\|IGHA1_HUMAN Immunoglobulin heavy constant alpha 1 OS=Homo sapiens OX=9606 GN=IGHA1 PE=1 SV=2 | 255 | 20201113_gy_colo_milk_1ng_glyco_hcd203040_compound15.18103.18103.4 | controllerType=0 controllerNumber=1 scan=18103 | 37.5158 |
| 13161:3 | 4 | R.LAGKPTHVN[+1710.598]VSVVMAEVDGTC[+57.021]Y.- | HexNAc(3)Hex(5)NeuAc(1) | C[+57], N[+1711] | 1353.2553 | 3 | 4057.7514 | 4057.7397 | 0 | 2.9 | 332 | CRagged | 497.23 | 472.39 | 356.03 | 10.43 | 99 | >sp\|P01876\|IGHA1_HUMAN Immunoglobulin heavy constant alpha 1 OS=Homo sapiens OX=9606 GN=IGHA1 PE=1 SV=2 | 255 | 20201113_gy_colo_milk_1ng_glyco_hcd203040_compound15.16275.16275.3 | controllerType=0 controllerNumber=1 scan=16275 | 34.0489 |
| 10282:3 | 4 | R.LAGKPTHVN[+1663.608]VSVVMAEVDGTC[+57.021]Y.- | HexNAc(5)Hex(4) | C[+57], N[+1664] | 1337.5921 | 3 | 4010.7617 | 4010.7502 | 0 | 2.9 | 332 | CRagged | 507.97 | 507.97 | 201.32 | 10.43 | 99 | >sp\|P01876\|IGHA1_HUMAN Immunoglobulin heavy constant alpha 1 OS=Homo sapiens OX=9606 GN=IGHA1 PE=1 SV=2 | 255 | 20201113_gy_colo_milk_1ng_glyco_hcd203040_compound15.13288.13288.3 | controllerType=0 controllerNumber=1 scan=13288 | 28.5422 |
| 10288:3 | 4 | R.LAGKPTHVN[+1606.587]VSVVMAEVDGTC[+57.021]Y.- | HexNAc(4)Hex(4)Fuc(1) | C[+57], N[+1607] | 1318.584 | 3 | 3953.7376 | 3953.7288 | 0 | 2.2 | 332 | CRagged | 515.87 | 515.87 | 515.87 | 10.48 | 99 | >sp\|P01876\|IGHA1_HUMAN Immunoglobulin heavy constant alpha 1 OS=Homo sapiens OX=9606 GN=IGHA1 PE=1 SV=2 | 255 | 20201113_gy_colo_milk_1ng_glyco_hcd203040_compound15.13294.13294.3 | controllerType=0 controllerNumber=1 scan=13294 | 28.5511 |
| 13035:4 | 4 | R.LAGKPTHVN[+1913.677]VSVVMAEVDGTC[+57.021]Y.- | HexNAc(4)Hex(5)NeuAc(1) | C[+57], N[+1914] | 1065.9642 | 4 | 4260.8348 | 4260.8191 | 0 | 3.7 | 332 | CRagged | 517.78 | 475.58 | 475.58 | 10.65 | 99 | >sp\|P01876\|IGHA1_HUMAN Immunoglobulin heavy constant alpha 1 OS=Homo sapiens OX=9606 GN=IGHA1 PE=1 SV=2 | 255 | 20201113_gy_colo_milk_1ng_glyco_hcd203040_compound15.16144.16144.4 | controllerType=0 controllerNumber=1 scan=16144 | 33.7768 |
| 14804:4 | 4 | R.LAGKPTHVN[+2350.830]VSVVMAEVDGTC[+57.021]Y.- | HexNAc(4)Hex(5)Fuc(1)NeuAc(2) | C[+57], N[+2351] | 1175.2543 | 4 | 4697.9954 | 4697.9724 | 0 | 4.9 | 332 | CRagged | 520.85 | 520.85 | 520.85 | 9.96 | 99 | >sp\|P01876\|IGHA1_HUMAN Immunoglobulin heavy constant alpha 1 OS=Homo sapiens OX=9606 GN=IGHA1 PE=1 SV=2 | 255 | 20201113_gy_colo_milk_1ng_glyco_hcd203040_compound15.17980.17980.4 | controllerType=0 controllerNumber=1 scan=17980 | 37.2814 |
| 14830:3 | 4 | R.LAGKPTHVN[+2553.910]VSVVMAEVDGTC[+57.021]Y.- | HexNAc(5)Hex(5)Fuc(1)NeuAc(2) | C[+57], N[+2554] | 1634.3577 | 3 | 4901.0586 | 4901.0518 | 0 | 1.4 | 332 | CRagged | 554.56 | 404.25 | 404.25 | 10.5 | 99 | >sp\|P01876\|IGHA1_HUMAN Immunoglobulin heavy constant alpha 1 OS=Homo sapiens OX=9606 GN=IGHA1 PE=1 SV=2 | 255 | 20201113_gy_colo_milk_1ng_glyco_hcd203040_compound15.18007.18007.3 | controllerType=0 controllerNumber=1 scan=18007 | 37.3287 |
| 13032:4 | 4 | R.LAGKPTHVN[+1913.677]VSVVMAEVDGTC[+57.021]Y.- | HexNAc(4)Hex(5)NeuAc(1) | C[+57], N[+1914] | 1065.9642 | 4 | 4260.8348 | 4260.8191 | 0 | 3.7 | 332 | CRagged | 555.01 | 505.13 | 505.13 | 10.81 | 99 | >sp\|P01876\|IGHA1_HUMAN Immunoglobulin heavy constant alpha 1 OS=Homo sapiens OX=9606 GN=IGHA1 PE=1 SV=2 | 255 | 20201113_gy_colo_milk_1ng_glyco_hcd203040_compound15.16141.16141.4 | controllerType=0 controllerNumber=1 scan=16141 | 33.7716 |
| 11468:3 | 4 | R.LAGKPTHVN[+1298.476]VSVVMAEVDGTC[+57.021]Y.- | HexNAc(4)Hex(3) | C[+57], N[+1298] | 1215.8871 | 3 | 3645.6469 | 3645.618 | 0 | 7.9 | 332 | CRagged | 563.81 | 557.74 | 557.74 | 9.78 | 99 | >sp\|P01876\|IGHA1_HUMAN Immunoglobulin heavy constant alpha 1 OS=Homo sapiens OX=9606 GN=IGHA1 PE=1 SV=2 | 255 | 20201113_gy_colo_milk_1ng_glyco_hcd203040_compound15.14518.14518.3 | controllerType=0 controllerNumber=1 scan=14518 | 30.8743 |
| 10242:3 | 4 | R.LAGKPTHVN[+1540.529]VSVVMAEVDGTC[+57.021]Y.- | HexNAc(2)Hex(7) | C[+57], N[+1541] | 1296.5648 | 3 | 3887.6797 | 3887.6706 | 0 | 2.3 | 332 | CRagged | 575.05 | 575.05 | 368.69 | 10.82 | 99 | >sp\|P01876\|IGHA1_HUMAN Immunoglobulin heavy constant alpha 1 OS=Homo sapiens OX=9606 GN=IGHA1 PE=1 SV=2 | 255 | 20201113_gy_colo_milk_1ng_glyco_hcd203040_compound15.13246.13246.3 | controllerType=0 controllerNumber=1 scan=13246 | 28.4532 |
| 11611:3 | 4 | R.LAGKPTHVN[+1378.476]VSVVMAEVDGTC[+57.021]Y.- | HexNAc(2)Hex(6) | C[+57], N[+1378] | 1242.5475 | 3 | 3725.6281 | 3725.6178 | 0 | 2.8 | 332 | CRagged | 576.58 | 576.58 | 346.79 | 10.82 | 99 | >sp\|P01876\|IGHA1_HUMAN Immunoglobulin heavy constant alpha 1 OS=Homo sapiens OX=9606 GN=IGHA1 PE=1 SV=2 | 255 | 20201113_gy_colo_milk_1ng_glyco_hcd203040_compound15.14666.14666.3 | controllerType=0 controllerNumber=1 scan=14666 | 31.1451 |
| 10290:3 | 4 | R.LAGKPTHVN[+1606.587]VSVVMAEVDGTC[+57.021]Y.- | HexNAc(4)Hex(4)Fuc(1) | C[+57], N[+1607] | 1318.584 | 3 | 3953.7376 | 3953.7288 | 0 | 2.2 | 332 | CRagged | 578.06 | 479.28 | 479.28 | 10.82 | 99 | >sp\|P01876\|IGHA1_HUMAN Immunoglobulin heavy constant alpha 1 OS=Homo sapiens OX=9606 GN=IGHA1 PE=1 SV=2 | 255 | 20201113_gy_colo_milk_1ng_glyco_hcd203040_compound15.13296.13296.3 | controllerType=0 controllerNumber=1 scan=13296 | [illegible] |
| 14925:4 | 4 | R.LAGKPTHVN[+2204.772]VSVVMAEVDGTC[+57.021]Y.- | HexNAc(4)Hex(5)NeuAc(2) | C[+57], N[+2205] | 1138.7339 | 4 | 4551.9139 | 4551.9145 | 0 | -0.1 | 332 | CRagged | 598.53 | 409.77 | 409.77 | 11.79 | 99 | >sp\|P01876\|IGHA1_HUMAN Immunoglobulin heavy constant alpha 1 OS=Homo sapiens OX=9606 GN=IGHA1 PE=1 SV=2 | 255 | 20201113_gy_colo_milk_1ng_glyco_hcd203040_compound15.18105.18105.4 | controllerType=0 controllerNumber=1 scan=18105 | [illegible] |
| 11533:3 | 4 | R.LAGKPTHVN[+892.317]VSVVMAEVDGTC[+57.021]Y.- | HexNAc(2)Hex(3) | C[+57], N[+892] | 1080.494 | 3 | 3239.4676 | 3239.4593 | 0 | 2.6 | 332 | CRagged | 606.56 | 606.56 | 326.9 | 13.18 | 99 | >sp\|P01876\|IGHA1_HUMAN Immunoglobulin heavy constant alpha 1 OS=Homo sapiens OX=9606 GN=IGHA1 PE=1 SV=2 | 255 | 20201113_gy_colo_milk_1ng_glyco_hcd203040_compound15.14586.14586.3 | controllerType=0 controllerNumber=1 scan=14586 | [illegible] |
| 10280:3 | 4 | R.LAGKPTHVN[+1663.608]VSVVMAEVDGTC[+57.021]Y.- | HexNAc(5)Hex(4) | C[+57], N[+1664] | 1337.5921 | 3 | 4010.7617 | 4010.7502 | 0 | 2.9 | 332 | CRagged | 614.05 | 614.05 | 371.71 | 13.18 | 99 | >sp\|P01876\|IGHA1_HUMAN Immunoglobulin heavy constant alpha 1 OS=Homo sapiens OX=9606 GN=IGHA1 PE=1 SV=2 | 255 | 20201113_gy_colo_milk_1ng_glyco_hcd203040_compound15.13286.13286.3 | controllerType=0 controllerNumber=1 scan=13286 | [illegible] |
| 13194:3 | 4 | R.LAGKPTHVN[+1694.603]VSVVMAEVDGTC[+57.021]Y.- | HexNAc(3)Hex(4)Fuc(1)NeuAc(1) | C[+57], N[+1695] | 1347.924 | 3 | 4041.7575 | 4041.7448 | 0 | 3.1 | 332 | CRagged | 621.63 | 379.22 | 379.22 | 13.18 | 99 | >sp\|P01876\|IGHA1_HUMAN Immunoglobulin heavy constant alpha 1 OS=Homo sapiens OX=9606 GN=IGHA1 PE=1 SV=2 | 255 | 20201113_gy_colo_milk_1ng_glyco_hcd203040_compound15.16309.16309.3 | controllerType=0 controllerNumber=1 scan=16309 | 34.1105 |
| 11609:3 | 4 | R.LAGKPTHVN[+1378.476]VSVVMAEVDGTC[+57.021]Y.- | HexNAc(2)Hex(6) | C[+57], N[+1378] | 1242.5475 | 3 | 3725.6281 | 3725.6178 | 0 | 2.8 | 332 | CRagged | 626.07 | 626.07 | 383.93 | 13.18 | 99 | >sp\|P01876\|IGHA1_HUMAN Immunoglobulin heavy constant alpha 1 OS=Homo sapiens OX=9606 GN=IGHA1 PE=1 SV=2 | 255 | 20201113_gy_colo_milk_1ng_glyco_hcd203040_compound15.14664.14664.3 | controllerType=0 controllerNumber=1 scan=14664 | 31.1407 |
| 11443:4 | 4 | R.LAGKPTHVN[+1647.613]VSVVMAEVDGTC[+57.021]Y.- | HexNAc(5)Hex(3)Fuc(1) | C[+57], N[+1648] | 999.4461 | 4 | 3994.7628 | 3994.7553 | 0 | 1.9 | 332 | CRagged | 635.37 | 564.08 | 564.08 | 13.35 | 99 | >sp\|P01876\|IGHA1_HUMAN Immunoglobulin heavy constant alpha 1 OS=Homo sapiens OX=9606 GN=IGHA1 PE=1 SV=2 | 255 | 20201113_gy_colo_milk_1ng_glyco_hcd203040_compound15.14492.14492.4 | controllerType=0 controllerNumber=1 scan=14492 | 30.8277 |
| 11377:3 | 4 | R.LAGKPTHVN[+1460.529]VSVVMAEVDGTC[+57.021]Y.- | HexNAc(4)Hex(4) | C[+57], N[+1461] | 1269.8956 | 3 | 3807.6721 | 3807.6709 | 0 | 0.3 | 332 | CRagged | 636.17 | 538.8 | 375.82 | 12.62 | 99 | >sp\|P01876\|IGHA1_HUMAN Immunoglobulin heavy constant alpha 1 OS=Homo sapiens OX=9606 GN=IGHA1 PE=1 SV=2 | 255 | 20201113_gy_colo_milk_1ng_glyco_hcd203040_compound15.14423.14423.3 | controllerType=0 controllerNumber=1 scan=14423 | 30.6664 |
| 10255:3 | 4 | R.LAGKPTHVN[+1809.666]VSVVMAEVDGTC[+57.021]Y.- | HexNAc(5)Hex(4)Fuc(1) | C[+57], N[+1810] | 1386.2776 | 3 | 4156.8184 | 4156.8081 | 0 | 2.5 | 332 | CRagged | 644.88 | 566.99 | 464.28 | 13.18 | 99 | >sp\|P01876\|IGHA1_HUMAN Immunoglobulin heavy constant alpha 1 OS=Homo sapiens OX=9606 GN=IGHA1 PE=1 SV=2 | 255 | 20201113_gy_colo_milk_1ng_glyco_hcd203040_compound15.13260.13260.3 | controllerType=0 controllerNumber=1 scan=13260 | 28.4805 |
| 14837:4 | 4 | R.LAGKPTHVN[+2553.910]VSVVMAEVDGTC[+57.021]Y.- | HexNAc(5)Hex(5)Fuc(1)NeuAc(2) | C[+57], N[+2554] | 1226.0224 | 4 | 4901.0677 | 4901.0518 | 0 | 3.2 | 332 | CRagged | 645.49 | 491.11 | 491.11 | 13.35 | 99 | >sp\|P01876\|IGHA1_HUMAN Immunoglobulin heavy constant alpha 1 OS=Homo sapiens OX=9606 GN=IGHA1 PE=1 SV=2 | 255 | 20201113_gy_colo_milk_1ng_glyco_hcd203040_compound15.18014.18014.4 | controllerType=0 controllerNumber=1 scan=18014 | 37.3435 |
| 10244:3 | 4 | R.LAGKPTHVN[+1540.529]VSVVMAEVDGTC[+57.021]Y.- | HexNAc(2)Hex(7) | C[+57], N[+1541] | 1296.5648 | 3 | 3887.6797 | 3887.6706 | 0 | 2.3 | 332 | CRagged | 658.89 | 605.79 | 445.42 | 14.18 | 99 | >sp\|P01876\|IGHA1_HUMAN Immunoglobulin heavy constant alpha 1 OS=Homo sapiens OX=9606 GN=IGHA1 PE=1 SV=2 | 255 | 20201113_gy_colo_milk_1ng_glyco_hcd203040_compound15.13248.13248.3 | controllerType=0 controllerNumber=1 scan=13248 | 28.4575 |
| 14833:3 | 4 | R.LAGKPTHVN[+2553.910]VSVVMAEVDGTC[+57.021]Y.- | HexNAc(5)Hex(5)Fuc(1)NeuAc(2) | C[+57], N[+2554] | 1634.3577 | 3 | 4901.0586 | 4901.0518 | 0 | 1.4 | 332 | CRagged | 662.13 | 514.24 | 514.24 | 14.04 | 99 | >sp\|P01876\|IGHA1_HUMAN Immunoglobulin heavy constant alpha 1 OS=Homo sapiens OX=9606 GN=IGHA1 PE=1 SV=2 | 255 | 20201113_gy_colo_milk_1ng_glyco_hcd203040_compound15.18010.18010.3 | controllerType=0 controllerNumber=1 scan=18010 | 37.3326 |
| 13164:3 | 4 | R.LAGKPTHVN[+1710.598]VSVVMAEVDGTC[+57.021]Y.- | HexNAc(3)Hex(5)NeuAc(1) | C[+57], N[+1711] | 1353.2553 | 3 | 4057.7514 | 4057.7397 | 0 | 2.9 | 332 | CRagged | 663.29 | 578.4 | 386.62 | 14.18 | 99 | >sp\|P01876\|IGHA1_HUMAN Immunoglobulin heavy constant alpha 1 OS=Homo sapiens OX=9606 GN=IGHA1 PE=1 SV=2 | 255 | 20201113_gy_colo_milk_1ng_glyco_hcd203040_compound15.16278.16278.3 | controllerType=0 controllerNumber=1 scan=16278 | 34.0563 |
| 10350:3 | 4 | R.LAGKPTHVN[+1444.534]VSVVMAEVDGTC[+57.021]Y.- | HexNAc(4)Hex(3)Fuc(1) | C[+57], N[+1445] | 1264.5674 | 3 | 3791.6878 | 3791.6759 | 0 | 3.1 | 332 | CRagged | 665.98 | 665.98 | 665.98 | 14.18 | 99 | >sp\|P01876\|IGHA1_HUMAN Immunoglobulin heavy constant alpha 1 OS=Homo sapiens OX=9606 GN=IGHA1 PE=1 SV=2 | 255 | 20201113_gy_colo_milk_1ng_glyco_hcd203040_compound15.13358.13358.3 | controllerType=0 controllerNumber=1 scan=13358 | 28.6771 |
| 13197:3 | 4 | R.LAGKPTHVN[+1694.603]VSVVMAEVDGTC[+57.021]Y.- | HexNAc(3)Hex(4)Fuc(1)NeuAc(1) | C[+57], N[+1695] | 1347.924 | 3 | 4041.7575 | 4041.7448 | 0 | 3.1 | 332 | CRagged | 669.28 | 489.35 | 489.35 | 14.18 | 99 | >sp\|P01876\|IGHA1_HUMAN Immunoglobulin heavy constant alpha 1 OS=Homo sapiens OX=9606 GN=IGHA1 PE=1 SV=2 | 255 | 20201113_gy_colo_milk_1ng_glyco_hcd203040_compound15.16312.16312.3 | controllerType=0 controllerNumber=1 scan=16312 | 34.1162 |
| 11457:3 | 4 | R.LAGKPTHVN[+1054.370]VSVVMAEVDGTC[+57.021]Y.- | HexNAc(2)Hex(4) | C[+57], N[+1054] | 1134.5111 | 3 | 3401.5189 | 3401.5121 | 0 | 2 | 332 | CRagged | 672.73 | 672.73 | 361.87 | 14.18 | 99 | >sp\|P01876\|IGHA1_HUMAN Immunoglobulin heavy constant alpha 1 OS=Homo sapiens OX=9606 GN=IGHA1 PE=1 SV=2 | 255 | 20201113_gy_colo_milk_1ng_glyco_hcd203040_compound15.14507.14507.3 | controllerType=0 controllerNumber=1 scan=14507 | 30.8554 |
| 13028:3 | 4 | R.LAGKPTHVN[+1913.677]VSVVMAEVDGTC[+57.021]Y.- | HexNAc(4)Hex(5)NeuAc(1) | C[+57], N[+1914] | 1420.9499 | 3 | 4260.8351 | 4260.8191 | 0 | 3.8 | 332 | CRagged | 679.04 | 452.2 | 452.2 | 14.18 | 99 | >sp\|P01876\|IGHA1_HUMAN Immunoglobulin heavy constant alpha 1 OS=Homo sapiens OX=9606 GN=IGHA1 PE=1 SV=2 | 255 | 20201113_gy_colo_milk_1ng_glyco_hcd203040_compound15.16137.16137.3 | controllerType=0 controllerNumber=1 scan=16137 | 33.7646 |
| 10352:3 | 4 | R.LAGKPTHVN[+1444.534]VSVVMAEVDGTC[+57.021]Y.- | HexNAc(4)Hex(3)Fuc(1) | C[+57], N[+1445] | 1264.5674 | 3 | 3791.6878 | 3791.6759 | 0 | 3.1 | 332 | CRagged | 681.17 | 666.48 | 666.48 | 14.18 | 99 | >sp\|P01876\|IGHA1_HUMAN Immunoglobulin heavy constant alpha 1 OS=Homo sapiens OX=9606 GN=IGHA1 PE=1 SV=2 | 255 | 20201113_gy_colo_milk_1ng_glyco_hcd203040_compound15.13360.13360.3 | controllerType=0 controllerNumber=1 scan=13360 | 28.6817 |
| 14781:3 | 4 | R.LAGKPTHVN[+2350.830]VSVVMAEVDGTC[+57.021]Y.- | HexNAc(4)Hex(5)Fuc(1)NeuAc(2) | C[+57], N[+2351] | 1566.6663 | 3 | 4697.9844 | 4697.9724 | 0 | 2.5 | 332 | CRagged | 681.3 | 664.66 | 664.66 | 14.18 | 99 | >sp\|P01876\|IGHA1_HUMAN Immunoglobulin heavy constant alpha 1 OS=Homo sapiens OX=9606 GN=IGHA1 PE=1 SV=2 | 255 | 20201113_gy_colo_milk_1ng_glyco_hcd203040_compound15.17956.17956.3 | controllerType=0 controllerNumber=1 scan=17956 | 37.2418 |
| 14807:4 | 4 | R.LAGKPTHVN[+2350.830]VSVVMAEVDGTC[+57.021]Y.- | HexNAc(4)Hex(5)Fuc(1)NeuAc(2) | C[+57], N[+2351] | 1175.2543 | 4 | 4697.9954 | 4697.9724 | 0 | 4.9 | 332 | CRagged | 685.95 | 685.95 | 685.95 | 13.66 | 99 | >sp\|P01876\|IGHA1_HUMAN Immunoglobulin heavy constant alpha 1 OS=Homo sapiens OX=9606 GN=IGHA1 PE=1 SV=2 | 255 | 20201113_gy_colo_milk_1ng_glyco_hcd203040_compound15.17983.17983.4 | controllerType=0 controllerNumber=1 scan=17983 | 37.288 |
| 10353:3 | 4 | R.LAGKPTHVN[+1501.555]VSVVMAEVDGTC[+57.021]Y.- | HexNAc(5)Hex(3) | C[+57], N[+1502] | 1283.5745 | 3 | 3848.709 | 3848.6974 | 0 | 3 | 332 | CRagged | 694.2 | 680.63 | 680.63 | 14.18 | 99 | >sp\|P01876\|IGHA1_HUMAN Immunoglobulin heavy constant alpha 1 OS=Homo sapiens OX=9606 GN=IGHA1 PE=1 SV=2 | 255 | 20201113_gy_colo_milk_1ng_glyco_hcd203040_compound15.13361.13361.3 | controllerType=0 controllerNumber=1 scan=13361 | 28.6836 |
| 11395:2 | 4 | R.LAGKPTHVN[+1216.423]VSVVMAEVDGTC[+57.021]Y.- | HexNAc(2)Hex(5) | C[+57], N[+1216] | 1782.2882 | 2 | 3563.5691 | 3563.5649 | 0 | 1.2 | 332 | CRagged | 694.45 | 628.75 | 280.26 | 13.47 | 99 | >sp\|P01876\|IGHA1_HUMAN Immunoglobulin heavy constant alpha 1 OS=Homo sapiens OX=9606 GN=IGHA1 PE=1 SV=2 | 255 | 20201113_gy_colo_milk_1ng_glyco_hcd203040_compound15.14442.14442.2 | controllerType=0 controllerNumber=1 scan=14442 | 30.7204 |
| 13083:3 | 4 | R.LAGKPTHVN[+2059.735]VSVVMAEVDGTC[+57.021]Y.- | HexNAc(4)Hex(5)Fuc(1)NeuAc(1) | C[+57], N[+2060] | 1469.6318 | 3 | 4406.8808 | 4406.877 | 0 | 0.9 | 332 | CRagged | 699.2 | 598.95 | 598.95 | 14.04 | 99 | >sp\|P01876\|IGHA1_HUMAN Immunoglobulin heavy constant alpha 1 OS=Homo sapiens OX=9606 GN=IGHA1 PE=1 SV=2 | 255 | 20201113_gy_colo_milk_1ng_glyco_hcd203040_compound15.16194.16194.3 | controllerType=0 controllerNumber=1 scan=16194 | 33.8791 |
| 13017:3 | 4 | R.LAGKPTHVN[+2116.756]VSVVMAEVDGTC[+57.021]Y.- | HexNAc(5)Hex(5)NeuAc(1) | C[+57], N[+2117] | 1488.6399 | 3 | 4463.9053 | 4463.8985 | 0 | 1.5 | 332 | CRagged | 700.96 | 503.64 | 503.64 | 15.04 | 99 | >sp\|P01876\|IGHA1_HUMAN Immunoglobulin heavy constant alpha 1 OS=Homo sapiens OX=9606 GN=IGHA1 PE=1 SV=2 | 255 | 20201113_gy_colo_milk_1ng_glyco_hcd203040_compound15.16125.16125.3 | controllerType=0 controllerNumber=1 scan=16125 | 33.742 |
| 14913:3 | 4 | R.LAGKPTHVN[+2204.772]VSVVMAEVDGTC[+57.021]Y.- | HexNAc(4)Hex(5)NeuAc(2) | C[+57], N[+2205] | 1517.9767 | 3 | 4551.9157 | 4551.9145 | 0 | 0.3 | 332 | CRagged | 701.3 | 470.75 | 470.75 | 14.62 | 99 | >sp\|P01876\|IGHA1_HUMAN Immunoglobulin heavy constant alpha 1 OS=Homo sapiens OX=9606 GN=IGHA1 PE=1 SV=2 | 255 | 20201113_gy_colo_milk_1ng_glyco_hcd203040_compound15.18093.18093.3 | controllerType=0 controllerNumber=1 scan=18093 | 37.5003 |
| 14835:4 | 4 | R.LAGKPTHVN[+2553.910]VSVVMAEVDGTC[+57.021]Y.- | HexNAc(5)Hex(5)Fuc(1)NeuAc(2) | C[+57], N[+2554] | 1226.0224 | 4 | 4901.0677 | 4901.0518 | 0 | 3.2 | 332 | CRagged | 707.33 | 458.13 | 458.13 | 15.35 | 99 | >sp\|P01876\|IGHA1_HUMAN Immunoglobulin heavy constant alpha 1 OS=Homo sapiens OX=9606 GN=IGHA1 PE=1 SV=2 | 255 | 20201113_gy_colo_milk_1ng_glyco_hcd203040_compound15.18012.18012.4 | controllerType=0 controllerNumber=1 scan=18012 | 37.3392 |
| 11392:2 | 4 | R.LAGKPTHVN[+1216.423]VSVVMAEVDGTC[+57.021]Y.- | HexNAc(2)Hex(5) | C[+57], N[+1216] | 1782.2882 | 2 | 3563.5691 | 3563.5649 | 0 | 1.2 | 332 | CRagged | 715.11 | 686.68 | 348.78 | 14.47 | 99 | >sp\|P01876\|IGHA1_HUMAN Immunoglobulin heavy constant alpha 1 OS=Homo sapiens OX=9606 GN=IGHA1 PE=1 SV=2 | 255 | 20201113_gy_colo_milk_1ng_glyco_hcd203040_compound15.14439.14439.2 | controllerType=0 controllerNumber=1 scan=14439 | 30.7131 |
| 10351:3 | 4 | R.LAGKPTHVN[+1501.555]VSVVMAEVDGTC[+57.021]Y.- | HexNAc(5)Hex(3) | C[+57], N[+1502] | 1283.5745 | 3 | 3848.709 | 3848.6974 | 0 | 3 | 332 | CRagged | 715.38 | 595.48 | 595.48 | 15.18 | 99 | >sp\|P01876\|IGHA1_HUMAN Immunoglobulin heavy constant alpha 1 OS=Homo sapiens OX=9606 GN=IGHA1 PE=1 SV=2 | 255 | 20201113_gy_colo_milk_1ng_glyco_hcd203040_compound15.13359.13359.3 | controllerType=0 controllerNumber=1 scan=13359 | 28.6786 |
| 13030:3 | 4 | R.LAGKPTHVN[+1913.677]VSVVMAEVDGTC[+57.021]Y.- | HexNAc(4)Hex(5)NeuAc(1) | C[+57], N[+1914] | 1420.9499 | 3 | 4260.8351 | 4260.8191 | 0 | 3.8 | 332 | CRagged | 719.4 | 500.88 | 500.88 | 15.18 | 99 | >sp\|P01876\|IGHA1_HUMAN Immunoglobulin heavy constant alpha 1 OS=Homo sapiens OX=9606 GN=IGHA1 PE=1 SV=2 | 255 | 20201113_gy_colo_milk_1ng_glyco_hcd203040_compound15.16139.16139.3 | controllerType=0 controllerNumber=1 scan=16139 | 33.7694 |
| 10257:3 | 4 | R.LAGKPTHVN[+1809.666]VSVVMAEVDGTC[+57.021]Y.- | HexNAc(5)Hex(4)Fuc(1) | C[+57], N[+1810] | 1386.2776 | 3 | 4156.8184 | 4156.8081 | 0 | 2.5 | 332 | CRagged | 735.39 | 735.39 | 506.3 | 15.18 | 99 | >sp\|P01876\|IGHA1_HUMAN Immunoglobulin heavy constant alpha 1 OS=Homo sapiens OX=9606 GN=IGHA1 PE=1 SV=2 | 255 | 20201113_gy_colo_milk_1ng_glyco_hcd203040_compound15.13262.13262.3 | controllerType=0 controllerNumber=1 scan=13262 | 28.4867 |
| 11416:3 | 4 | R.LAGKPTHVN[+1647.613]VSVVMAEVDGTC[+57.021]Y.- | HexNAc(5)Hex(3)Fuc(1) | C[+57], N[+1648] | 1332.2587 | 3 | 3994.7616 | 3994.7553 | 0 | 1.6 | 332 | CRagged | 740.77 | 458.53 | 458.53 | 15.04 | 99 | >sp\|P01876\|IGHA1_HUMAN Immunoglobulin heavy constant alpha 1 OS=Homo sapiens OX=9606 GN=IGHA1 PE=1 SV=2 | 255 | 20201113_gy_colo_milk_1ng_glyco_hcd203040_compound15.14464.14464.3 | controllerType=0 controllerNumber=1 scan=14464 | 30.7718 |
| 11441:3 | 4 | R.LAGKPTHVN[+1298.476]VSVVMAEVDGTC[+57.021]Y.- | HexNAc(4)Hex(3) | C[+57], N[+1298] | 1215.8806 | 3 | 3645.6271 | 3645.618 | 0 | 2.5 | 332 | CRagged | 743.04 | 645.92 | 645.92 | 15.18 | 99 | >sp\|P01876\|IGHA1_HUMAN Immunoglobulin heavy constant alpha 1 OS=Homo sapiens OX=9606 GN=IGHA1 PE=1 SV=2 | 255 | 20201113_gy_colo_milk_1ng_glyco_hcd203040_compound15.14490.14490.3 | controllerType=0 controllerNumber=1 scan=14490 | 30.8171 |
| 11440:4 | 4 | R.LAGKPTHVN[+1647.613]VSVVMAEVDGTC[+57.021]Y.- | HexNAc(5)Hex(3)Fuc(1) | C[+57], N[+1648] | 999.4461 | 4 | 3994.7628 | 3994.7553 | 0 | 1.9 | 332 | CRagged | 747.34 | 738.66 | 738.66 | 15.35 | 99 | >sp\|P01876\|IGHA1_HUMAN Immunoglobulin heavy constant alpha 1 OS=Homo sapiens OX=9606 GN=IGHA1 PE=1 SV=2 | 255 | 20201113_gy_colo_milk_1ng_glyco_hcd203040_compound15.14489.14489.4 | controllerType=0 controllerNumber=1 scan=14489 | 30.816 |
| 11408:3 | 4 | R.LAGKPTHVN[+1501.555]VSVVMAEVDGTC[+57.021]Y.- | HexNAc(5)Hex(3) | C[+57], N[+1502] | 1283.5731 | 3 | 3848.7046 | 3848.6974 | 0 | 1.9 | 332 | CRagged | 750.8 | 652.61 | 652.61 | 16.18 | 99 | >sp\|P01876\|IGHA1_HUMAN Immunoglobulin heavy constant alpha 1 OS=Homo sapiens OX=9606 GN=IGHA1 PE=1 SV=2 | 255 | 20201113_gy_colo_milk_1ng_glyco_hcd203040_compound15.14456.14456.3 | controllerType=0 controllerNumber=1 scan=14456 | 30.7484 |
| 11341:3 | 4 | R.LAGKPTHVN[+1809.666]VSVVMAEVDGTC[+57.021]Y.- | HexNAc(5)Hex(4)Fuc(1) | C[+57], N[+1810] | 1386.2784 | 3 | 4156.8206 | 4156.8081 | 0 | 3 | 332 | CRagged | 752.64 | 714.13 | 479.15 | 16.18 | 99 | >sp\|P01876\|IGHA1_HUMAN Immunoglobulin heavy constant alpha 1 OS=Homo sapiens OX=9606 GN=IGHA1 PE=1 SV=2 | 255 | 20201113_gy_colo_milk_1ng_glyco_hcd203040_compound15.14386.14386.3 | controllerType=0 controllerNumber=1 scan=14386 | 30.5781 |
| 14779:3 | 4 | R.LAGKPTHVN[+2350.830]VSVVMAEVDGTC[+57.021]Y.- | HexNAc(4)Hex(5)Fuc(1)NeuAc(2) | C[+57], N[+2351] | 1566.6663 | 3 | 4697.9844 | 4697.9724 | 0 | 2.5 | 332 | CRagged | 758.76 | 758.76 | 758.76 | 16.18 | 99 | >sp\|P01876\|IGHA1_HUMAN Immunoglobulin heavy constant alpha 1 OS=Homo sapiens OX=9606 GN=IGHA1 PE=1 SV=2 | 255 | 20201113_gy_colo_milk_1ng_glyco_hcd203040_compound15.17954.17954.3 | controllerType=0 controllerNumber=1 scan=17954 | 37.2363 |
| 11337:3 | 4 | R.LAGKPTHVN[+1971.719]VSVVMAEVDGTC[+57.021]Y.- | HexNAc(5)Hex(5)Fuc(1) | C[+57], N[+1972] | 1440.2966 | 3 | 4318.8751 | 4318.861 | 0 | 3.3 | 332 | CRagged | 766.9 | 738.61 | 465.61 | 16.18 | 99 | >sp\|P01876\|IGHA1_HUMAN Immunoglobulin heavy constant alpha 1 OS=Homo sapiens OX=9606 GN=IGHA1 PE=1 SV=2 | 255 | 20201113_gy_colo_milk_1ng_glyco_hcd203040_compound15.14381.14381.3 | controllerType=0 controllerNumber=1 scan=14381 | 30.5646 |
| 12977:3 | 4 | R.LAGKPTHVN[+2262.814]VSVVMAEVDGTC[+57.021]Y.- | HexNAc(5)Hex(5)Fuc(1)NeuAc(1) | C[+57], N[+2263] | 1537.3244 | 3 | 4609.9586 | 4609.9564 | 0 | 0.5 | 332 | CRagged | 772.13 | 605.98 | 605.98 | 15.62 | 99 | >sp\|P01876\|IGHA1_HUMAN Immunoglobulin heavy constant alpha 1 OS=Homo sapiens OX=9606 GN=IGHA1 PE=1 SV=2 | 255 | 20201113_gy_colo_milk_1ng_glyco_hcd203040_compound15.16083.16083.3 | controllerType=0 controllerNumber=1 scan=16083 | 33.6641 |
| 13064:3 | 4 | R.LAGKPTHVN[+1872.651]VSVVMAEVDGTC[+57.021]Y.- | HexNAc(3)Hex(6)NeuAc(1) | C[+57], N[+1873] | 1407.272 | 3 | 4219.8015 | 4219.7926 | 0 | 2.1 | 332 | CRagged | 775.03 | 751.11 | 441.24 | 16.18 | 99 | >sp\|P01876\|IGHA1_HUMAN Immunoglobulin heavy constant alpha 1 OS=Homo sapiens OX=9606 GN=IGHA1 PE=1 SV=2 | 255 | 20201113_gy_colo_milk_1ng_glyco_hcd203040_compound15.16174.16174.3 | controllerType=0 controllerNumber=1 scan=16174 | 33.84 |
| 13062:3 | 4 | R.LAGKPTHVN[+1872.651]VSVVMAEVDGTC[+57.021]Y.- | HexNAc(3)Hex(6)NeuAc(1) | C[+57], N[+1873] | 1407.272 | 3 | 4219.8015 | 4219.7926 | 0 | 2.1 | 332 | CRagged | 775.96 | 715.08 | 465.73 | 16.18 | 99 | >sp\|P01876\|IGHA1_HUMAN Immunoglobulin heavy constant alpha 1 OS=Homo sapiens OX=9606 GN=IGHA1 PE=1 SV=2 | 255 | 20201113_gy_colo_milk_1ng_glyco_hcd203040_compound15.16172.16172.3 | controllerType=0 controllerNumber=1 scan=16172 | 33.8345 |
| 11460:3 | 4 | R.LAGKPTHVN[+1054.370]VSVVMAEVDGTC[+57.021]Y.- | HexNAc(2)Hex(4) | C[+57], N[+1054] | 1134.5111 | 3 | 3401.5189 | 3401.5121 | 0 | 2 | 332 | CRagged | 780.71 | 771.65 | 470.83 | 16.18 | 99 | >sp\|P01876\|IGHA1_HUMAN Immunoglobulin heavy constant alpha 1 OS=Homo sapiens OX=9606 GN=IGHA1 PE=1 SV=2 | 255 | 20201113_gy_colo_milk_1ng_glyco_hcd203040_compound15.14510.14510.3 | controllerType=0 controllerNumber=1 scan=14510 | 30.8634 |
| 11410:3 | 4 | R.LAGKPTHVN[+1501.555]VSVVMAEVDGTC[+57.021]Y.- | HexNAc(5)Hex(3) | C[+57], N[+1502] | 1283.5731 | 3 | 3848.7046 | 3848.6974 | 0 | 1.9 | 332 | CRagged | 796.52 | 796.52 | 796.52 | 16.18 | 99 | >sp\|P01876\|IGHA1_HUMAN Immunoglobulin heavy constant alpha 1 OS=Homo sapiens OX=9606 GN=IGHA1 PE=1 SV=2 | 255 | 20201113_gy_colo_milk_1ng_glyco_hcd203040_compound15.14458.14458.3 | controllerType=0 controllerNumber=1 scan=14458 | 30.7538 |
| 11375:3 | 4 | R.LAGKPTHVN[+1606.587]VSVVMAEVDGTC[+57.021]Y.- | HexNAc(4)Hex(4)Fuc(1) | C[+57], N[+1607] | 1318.5832 | 3 | 3953.735 | 3953.7288 | 0 | 1.6 | 332 | CRagged | 799.61 | 771.83 | 771.83 | 16.04 | 99 | >sp\|P01876\|IGHA1_HUMAN Immunoglobulin heavy constant alpha 1 OS=Homo sapiens OX=9606 GN=IGHA1 PE=1 SV=2 | 255 | 20201113_gy_colo_milk_1ng_glyco_hcd203040_compound15.14421.14421.3 | controllerType=0 controllerNumber=1 scan=14421 | 30.663 |
| 10330:3 | 4 | R.LAGKPTHVN[+1647.613]VSVVMAEVDGTC[+57.021]Y.- | HexNAc(5)Hex(3)Fuc(1) | C[+57], N[+1648] | 1332.2598 | 3 | 3994.7649 | 3994.7553 | 0 | 2.4 | 332 | CRagged | 812.32 | 680.61 | 680.61 | 17.18 | 99 | >sp\|P01876\|IGHA1_HUMAN Immunoglobulin heavy constant alpha 1 OS=Homo sapiens OX=9606 GN=IGHA1 PE=1 SV=2 | 255 | 20201113_gy_colo_milk_1ng_glyco_hcd203040_compound15.13338.13338.3 | controllerType=0 controllerNumber=1 scan=13338 | 28.6425 |
| 11442:3 | 4 | R.LAGKPTHVN[+1444.534]VSVVMAEVDGTC[+57.021]Y.- | HexNAc(4)Hex(3)Fuc(1) | C[+57], N[+1445] | 1264.5662 | 3 | 3791.6841 | 3791.6759 | 0 | 2.2 | 332 | CRagged | 823.31 | 823.31 | 823.31 | 17.18 | 99 | >sp\|P01876\|IGHA1_HUMAN Immunoglobulin heavy constant alpha 1 OS=Homo sapiens OX=9606 GN=IGHA1 PE=1 SV=2 | 255 | 20201113_gy_colo_milk_1ng_glyco_hcd203040_compound15.14491.14491.3 | controllerType=0 controllerNumber=1 scan=14491 | 30.8259 |
| 10256:3 | 4 | R.LAGKPTHVN[+1378.476]VSVVMAEVDGTC[+57.021]Y.- | HexNAc(2)Hex(6) | C[+57], N[+1378] | 1242.5469 | 3 | 3725.6262 | 3725.6178 | 0 | 2.3 | 332 | CRagged | 824.72 | 798.63 | 496.37 | 17.18 | 99 | >sp\|P01876\|IGHA1_HUMAN Immunoglobulin heavy constant alpha 1 OS=Homo sapiens OX=9606 GN=IGHA1 PE=1 SV=2 | 255 | 20201113_gy_colo_milk_1ng_glyco_hcd203040_compound15.13261.13261.3 | controllerType=0 controllerNumber=1 scan=13261 | 28.4852 |
| 11380:3 | 4 | R.LAGKPTHVN[+1460.529]VSVVMAEVDGTC[+57.021]Y.- | HexNAc(4)Hex(4) | C[+57], N[+1461] | 1269.8956 | 3 | 3807.6721 | 3807.6709 | 0 | 0.3 | 332 | CRagged | 827.2 | 801.89 | 482.02 | 16.62 | 99 | >sp\|P01876\|IGHA1_HUMAN Immunoglobulin heavy constant alpha 1 OS=Homo sapiens OX=9606 GN=IGHA1 PE=1 SV=2 | 255 | 20201113_gy_colo_milk_1ng_glyco_hcd203040_compound15.14426.14426.3 | controllerType=0 controllerNumber=1 scan=14426 | 30.6715 |
| 13165:3 | 4 | R.LAGKPTHVN[+1548.545]VSVVMAEVDGTC[+57.021]Y.- | HexNAc(3)Hex(4)NeuAc(1) | C[+57], N[+1549] | 1299.238 | 3 | 3895.6994 | 3895.6869 | 0 | 3.2 | 332 | CRagged | 828.45 | 828.45 | 828.45 | 17.18 | 99 | >sp\|P01876\|IGHA1_HUMAN Immunoglobulin heavy constant alpha 1 OS=Homo sapiens OX=9606 GN=IGHA1 PE=1 SV=2 | 255 | 20201113_gy_colo_milk_1ng_glyco_hcd203040_compound15.16279.16279.3 | controllerType=0 controllerNumber=1 scan=16279 | 34.0582 |
| 14911:3 | 4 | R.LAGKPTHVN[+2204.772]VSVVMAEVDGTC[+57.021]Y.- | HexNAc(4)Hex(5)NeuAc(2) | C[+57], N[+2205] | 1517.9767 | 3 | 4551.9157 | 4551.9145 | 0 | 0.3 | 332 | CRagged | 830.36 | 585.6 | 585.6 | 16.62 | 99 | >sp\|P01876\|IGHA1_HUMAN Immunoglobulin heavy constant alpha 1 OS=Homo sapiens OX=9606 GN=IGHA1 PE=1 SV=2 | 255 | 20201113_gy_colo_milk_1ng_glyco_hcd203040_compound15.18091.18091.3 | controllerType=0 controllerNumber=1 scan=18091 | 37.4945 |
| 11444:3 | 4 | R.LAGKPTHVN[+1298.476]VSVVMAEVDGTC[+57.021]Y.- | HexNAc(4)Hex(3) | C[+57], N[+1298] | 1215.8806 | 3 | 3645.6271 | 3645.618 | 0 | 2.5 | 332 | CRagged | 831.21 | 664.73 | 664.73 | 17.18 | 99 | >sp\|P01876\|IGHA1_HUMAN Immunoglobulin heavy constant alpha 1 OS=Homo sapiens OX=9606 GN=IGHA1 PE=1 SV=2 | 255 | 20201113_gy_colo_milk_1ng_glyco_hcd203040_compound15.14493.14493.3 | controllerType=0 controllerNumber=1 scan=14493 | 30.829 |
| 10328:3 | 4 | R.LAGKPTHVN[+1647.613]VSVVMAEVDGTC[+57.021]Y.- | HexNAc(5)Hex(3)Fuc(1) | C[+57], N[+1648] | 1332.2598 | 3 | 3994.7649 | 3994.7553 | 0 | 2.4 | 332 | CRagged | 838.63 | 765.77 | 765.77 | 17.18 | 99 | >sp\|P01876\|IGHA1_HUMAN Immunoglobulin heavy constant alpha 1 OS=Homo sapiens OX=9606 GN=IGHA1 PE=1 SV=2 | 255 | 20201113_gy_colo_milk_1ng_glyco_hcd203040_compound15.13336.13336.3 | controllerType=0 controllerNumber=1 scan=13336 | 28.6369 |
| 11323:3 | 4 | R.LAGKPTHVN[+1663.608]VSVVMAEVDGTC[+57.021]Y.- | HexNAc(5)Hex(4) | C[+57], N[+1664] | 1337.5925 | 3 | 4010.7628 | 4010.7502 | 0 | 3.1 | 332 | CRagged | 839.36 | 707.15 | 506.75 | 17.18 | 99 | >sp\|P01876\|IGHA1_HUMAN Immunoglobulin heavy constant alpha 1 OS=Homo sapiens OX=9606 GN=IGHA1 PE=1 SV=2 | 255 | 20201113_gy_colo_milk_1ng_glyco_hcd203040_compound15.14367.14367.3 | controllerType=0 controllerNumber=1 scan=14367 | 30.5282 |
| 11372:3 | 4 | R.LAGKPTHVN[+1606.587]VSVVMAEVDGTC[+57.021]Y.- | HexNAc(4)Hex(4)Fuc(1) | C[+57], N[+1607] | 1318.5832 | 3 | 3953.735 | 3953.7288 | 0 | 1.6 | 332 | CRagged | 844.42 | 844.42 | 844.42 | 17.04 | 99 | >sp\|P01876\|IGHA1_HUMAN Immunoglobulin heavy constant alpha 1 OS=Homo sapiens OX=9606 GN=IGHA1 PE=1 SV=2 | 255 | 20201113_gy_colo_milk_1ng_glyco_hcd203040_compound15.14418.14418.3 | controllerType=0 controllerNumber=1 scan=14418 | 30.656 |
| 10281:3 | 4 | R.LAGKPTHVN[+1216.423]VSVVMAEVDGTC[+57.021]Y.- | HexNAc(2)Hex(5) | C[+57], N[+1216] | 1188.53 | 3 | 3563.5753 | 3563.5649 | 0 | 2.9 | 332 | CRagged | 847.1 | 830.46 | 505.45 | 17.18 | 99 | >sp\|P01876\|IGHA1_HUMAN Immunoglobulin heavy constant alpha 1 OS=Homo sapiens OX=9606 GN=IGHA1 PE=1 SV=2 | 255 | 20201113_gy_colo_milk_1ng_glyco_hcd203040_compound15.13287.13287.3 | controllerType=0 controllerNumber=1 scan=13287 | 28.5408 |
| 11364:3 | 4 | R.LAGKPTHVN[+1216.423]VSVVMAEVDGTC[+57.021]Y.- | HexNAc(2)Hex(5) | C[+57], N[+1216] | 1188.529 | 3 | 3563.5724 | 3563.5649 | 0 | 2.1 | 332 | CRagged | 849.07 | 832.32 | 535.96 | 17.18 | 99 | >sp\|P01876\|IGHA1_HUMAN Immunoglobulin heavy constant alpha 1 OS=Homo sapiens OX=9606 GN=IGHA1 PE=1 SV=2 | 255 | 20201113_gy_colo_milk_1ng_glyco_hcd203040_compound15.14410.14410.3 | controllerType=0 controllerNumber=1 scan=14410 | 30.6382 |
| 13089:3 | 4 | R.LAGKPTHVN[+1751.624]VSVVMAEVDGTC[+57.021]Y.- | HexNAc(4)Hex(4)NeuAc(1) | C[+57], N[+1752] | 1366.9295 | 3 | 4098.7739 | 4098.7663 | 0 | 1.9 | 332 | CRagged | 850.82 | 812.67 | 812.67 | 18.18 | 99 | >sp\|P01876\|IGHA1_HUMAN Immunoglobulin heavy constant alpha 1 OS=Homo sapiens OX=9606 GN=IGHA1 PE=1 SV=2 | 255 | 20201113_gy_colo_milk_1ng_glyco_hcd203040_compound15.16200.16200.3 | controllerType=0 controllerNumber=1 scan=16200 | 33.891 |
| 10254:3 | 4 | R.LAGKPTHVN[+1378.476]VSVVMAEVDGTC[+57.021]Y.- | HexNAc(2)Hex(6) | C[+57], N[+1378] | 1242.5469 | 3 | 3725.6262 | 3725.6178 | 0 | 2.3 | 332 | CRagged | 852.89 | 852.89 | 522.71 | 18.18 | 99 | >sp\|P01876\|IGHA1_HUMAN Immunoglobulin heavy constant alpha 1 OS=Homo sapiens OX=9606 GN=IGHA1 PE=1 SV=2 | 255 | 20201113_gy_colo_milk_1ng_glyco_hcd203040_compound15.13259.13259.3 | controllerType=0 controllerNumber=1 scan=13259 | 28.4791 |
| 11299:3 | 4 | R.LAGKPTHVN[+1702.581]VSVVMAEVDGTC[+57.021]Y.- | HexNAc(2)Hex(8) | C[+57], N[+1703] | 1350.5855 | 3 | 4049.742 | 4049.7234 | 0 | 4.6 | 332 | CRagged | 861.35 | 830.09 | 502.68 | 17.84 | 99 | >sp\|P01876\|IGHA1_HUMAN Immunoglobulin heavy constant alpha 1 OS=Homo sapiens OX=9606 GN=IGHA1 PE=1 SV=2 | 255 | 20201113_gy_colo_milk_1ng_glyco_hcd203040_compound15.14342.14342.3 | controllerType=0 controllerNumber=1 scan=14342 | 30.4695 |
| 11471:3 | 4 | R.LAGKPTHVN[+1298.476]VSVVMAEVDGTC[+57.021]Y.- | HexNAc(4)Hex(3) | C[+57], N[+1298] | 1215.8871 | 3 | 3645.6469 | 3645.618 | 0 | 7.9 | 332 | CRagged | 861.61 | 730.71 | 730.71 | 17.14 | 99 | >sp\|P01876\|IGHA1_HUMAN Immunoglobulin heavy constant alpha 1 OS=Homo sapiens OX=9606 GN=IGHA1 PE=1 SV=2 | 255 | 20201113_gy_colo_milk_1ng_glyco_hcd203040_compound15.14521.14521.3 | controllerType=0 controllerNumber=1 scan=14521 | 30.8789 |
| 11540:3 | 4 | R.LAGKPTHVN[+1241.454]VSVVMAEVDGTC[+57.021]Y.- | HexNAc(3)Hex(3)Fuc(1) | C[+57], N[+1241] | 1196.8725 | 3 | 3588.6029 | 3588.5966 | 0 | 1.8 | 332 | CRagged | 861.75 | 713.97 | 713.97 | 18.18 | 99 | >sp\|P01876\|IGHA1_HUMAN Immunoglobulin heavy constant alpha 1 OS=Homo sapiens OX=9606 GN=IGHA1 PE=1 SV=2 | 255 | 20201113_gy_colo_milk_1ng_glyco_hcd203040_compound15.14593.14593.3 | controllerType=0 controllerNumber=1 scan=14593 | 31.0203 |
| 13162:3 | 4 | R.LAGKPTHVN[+1548.545]VSVVMAEVDGTC[+57.021]Y.- | HexNAc(3)Hex(4)NeuAc(1) | C[+57], N[+1549] | 1299.238 | 3 | 3895.6994 | 3895.6869 | 0 | 3.2 | 332 | CRagged | 867.72 | 653.71 | 653.71 | 18.18 | 99 | >sp\|P01876\|IGHA1_HUMAN Immunoglobulin heavy constant alpha 1 OS=Homo sapiens OX=9606 GN=IGHA1 PE=1 SV=2 | 255 | 20201113_gy_colo_milk_1ng_glyco_hcd203040_compound15.16276.16276.3 | controllerType=0 controllerNumber=1 scan=16276 | 34.05 |
| 11310:3 | 4 | R.LAGKPTHVN[+1540.529]VSVVMAEVDGTC[+57.021]Y.- | HexNAc(2)Hex(7) | C[+57], N[+1541] | 1296.5674 | 3 | 3887.6878 | 3887.6706 | 0 | 4.4 | 332 | CRagged | 870.15 | 812.05 | 517.12 | 17.84 | 99 | >sp\|P01876\|IGHA1_HUMAN Immunoglobulin heavy constant alpha 1 OS=Homo sapiens OX=9606 GN=IGHA1 PE=1 SV=2 | 255 | 20201113_gy_colo_milk_1ng_glyco_hcd203040_compound15.14353.14353.3 | controllerType=0 controllerNumber=1 scan=14353 | 30.4903 |
| 11535:3 | 4 | R.LAGKPTHVN[+892.317]VSVVMAEVDGTC[+57.021]Y.- | HexNAc(2)Hex(3) | C[+57], N[+892] | 1080.494 | 3 | 3239.4676 | 3239.4593 | 0 | 2.6 | 332 | CRagged | 882.01 | 864.9 | 537.91 | 18.18 | 99 | >sp\|P01876\|IGHA1_HUMAN Immunoglobulin heavy constant alpha 1 OS=Homo sapiens OX=9606 GN=IGHA1 PE=1 SV=2 | 255 | 20201113_gy_colo_milk_1ng_glyco_hcd203040_compound15.14588.14588.3 | controllerType=0 controllerNumber=1 scan=14588 | 31.0115 |
| 10279:3 | 4 | R.LAGKPTHVN[+1216.423]VSVVMAEVDGTC[+57.021]Y.- | HexNAc(2)Hex(5) | C[+57], N[+1216] | 1188.53 | 3 | 3563.5753 | 3563.5649 | 0 | 2.9 | 332 | CRagged | 883.42 | 883.42 | 521.55 | 18.18 | 99 | >sp\|P01876\|IGHA1_HUMAN Immunoglobulin heavy constant alpha 1 OS=Homo sapiens OX=9606 GN=IGHA1 PE=1 SV=2 | 255 | 20201113_gy_colo_milk_1ng_glyco_hcd203040_compound15.13285.13285.3 | controllerType=0 controllerNumber=1 scan=13285 | 28.5335 |
| 11439:3 | 4 | R.LAGKPTHVN[+1444.534]VSVVMAEVDGTC[+57.021]Y.- | HexNAc(4)Hex(3)Fuc(1) | C[+57], N[+1445] | 1264.5662 | 3 | 3791.6841 | 3791.6759 | 0 | 2.2 | 332 | CRagged | 884.72 | 884.72 | 884.72 | 18.18 | 99 | >sp\|P01876\|IGHA1_HUMAN Immunoglobulin heavy constant alpha 1 OS=Homo sapiens OX=9606 GN=IGHA1 PE=1 SV=2 | 255 | 20201113_gy_colo_milk_1ng_glyco_hcd203040_compound15.14488.14488.3 | controllerType=0 controllerNumber=1 scan=14488 | 30.8144 |
| 11339:3 | 4 | R.LAGKPTHVN[+1809.666]VSVVMAEVDGTC[+57.021]Y.- | HexNAc(5)Hex(4)Fuc(1) | C[+57], N[+1810] | 1386.2784 | 3 | 4156.8206 | 4156.8081 | 0 | 3 | 332 | CRagged | 891.79 | 891.79 | 600.99 | 18.18 | 99 | >sp\|P01876\|IGHA1_HUMAN Immunoglobulin heavy constant alpha 1 OS=Homo sapiens OX=9606 GN=IGHA1 PE=1 SV=2 | 255 | 20201113_gy_colo_milk_1ng_glyco_hcd203040_compound15.14384.14384.3 | controllerType=0 controllerNumber=1 scan=14384 | 30.5705 |
| 11413:3 | 4 | R.LAGKPTHVN[+1647.613]VSVVMAEVDGTC[+57.021]Y.- | HexNAc(5)Hex(3)Fuc(1) | C[+57], N[+1648] | 1332.2587 | 3 | 3994.7616 | 3994.7553 | 0 | 1.6 | 332 | CRagged | 894.74 | 679.25 | 679.25 | 18.04 | 99 | >sp\|P01876\|IGHA1_HUMAN Immunoglobulin heavy constant alpha 1 OS=Homo sapiens OX=9606 GN=IGHA1 PE=1 SV=2 | 255 | 20201113_gy_colo_milk_1ng_glyco_hcd203040_compound15.14461.14461.3 | controllerType=0 controllerNumber=1 scan=14461 | 30.7569 |
| 11327:3 | 4 | R.LAGKPTHVN[+1378.476]VSVVMAEVDGTC[+57.021]Y.- | HexNAc(2)Hex(6) | C[+57], N[+1378] | 1242.5479 | 3 | 3725.6292 | 3725.6178 | 0 | 3.1 | 332 | CRagged | 901.53 | 889.83 | 543.77 | 19.18 | 99 | >sp\|P01876\|IGHA1_HUMAN Immunoglobulin heavy constant alpha 1 OS=Homo sapiens OX=9606 GN=IGHA1 PE=1 SV=2 | 255 | 20201113_gy_colo_milk_1ng_glyco_hcd203040_compound15.14371.14371.3 | controllerType=0 controllerNumber=1 scan=14371 | 30.5445 |
| 11321:3 | 4 | R.LAGKPTHVN[+1663.608]VSVVMAEVDGTC[+57.021]Y.- | HexNAc(5)Hex(4) | C[+57], N[+1664] | 1337.5925 | 3 | 4010.7628 | 4010.7502 | 0 | 3.1 | 332 | CRagged | 904.12 | 735.8 | 602.4 | 19.18 | 99 | >sp\|P01876\|IGHA1_HUMAN Immunoglobulin heavy constant alpha 1 OS=Homo sapiens OX=9606 GN=IGHA1 PE=1 SV=2 | 255 | 20201113_gy_colo_milk_1ng_glyco_hcd203040_compound15.14365.14365.3 | controllerType=0 controllerNumber=1 scan=14365 | 30.5214 |
| 11335:3 | 4 | R.LAGKPTHVN[+1971.719]VSVVMAEVDGTC[+57.021]Y.- | HexNAc(5)Hex(5)Fuc(1) | C[+57], N[+1972] | 1440.2966 | 3 | 4318.8751 | 4318.861 | 0 | 3.3 | 332 | CRagged | 907.12 | 906.07 | 597.5 | 19.18 | 99 | >sp\|P01876\|IGHA1_HUMAN Immunoglobulin heavy constant alpha 1 OS=Homo sapiens OX=9606 GN=IGHA1 PE=1 SV=2 | 255 | 20201113_gy_colo_milk_1ng_glyco_hcd203040_compound15.14379.14379.3 | controllerType=0 controllerNumber=1 scan=14379 | 30.5582 |
| 11297:3 | 4 | R.LAGKPTHVN[+1702.581]VSVVMAEVDGTC[+57.021]Y.- | HexNAc(2)Hex(8) | C[+57], N[+1703] | 1350.5855 | 3 | 4049.742 | 4049.7234 | 0 | 4.6 | 332 | CRagged | 907.73 | 907.73 | 592.31 | 18.84 | 99 | >sp\|P01876\|IGHA1_HUMAN Immunoglobulin heavy constant alpha 1 OS=Homo sapiens OX=9606 GN=IGHA1 PE=1 SV=2 | 255 | 20201113_gy_colo_milk_1ng_glyco_hcd203040_compound15.14340.14340.3 | controllerType=0 controllerNumber=1 scan=14340 | 30.4633 |
| 11307:3 | 4 | R.LAGKPTHVN[+1540.529]VSVVMAEVDGTC[+57.021]Y.- | HexNAc(2)Hex(7) | C[+57], N[+1541] | 1296.5674 | 3 | 3887.6878 | 3887.6706 | 0 | 4.4 | 332 | CRagged | 908.36 | 908.36 | 609.53 | 18.84 | 99 | >sp\|P01876\|IGHA1_HUMAN Immunoglobulin heavy constant alpha 1 OS=Homo sapiens OX=9606 GN=IGHA1 PE=1 SV=2 | 255 | 20201113_gy_colo_milk_1ng_glyco_hcd203040_compound15.14350.14350.3 | controllerType=0 controllerNumber=1 scan=14350 | 30.4807 |

Table S4. Cont.

| Query #:z | Protein Rank | Glycopeptide | Glycan | Modification Type(s) | Observed m/z | z | Observed (M+H) | Calc. mass (M+H) | Off-by-x error | Mass error (ppm) | StartPosition | Cleavage | Score | Delta | Delta Mod | \|Log Prob\| | # of unique peptides | Protein Name | Protein DB number | Comment | Scan # | Scan Time |
|---|---|---|---|---|---|---|---|---|---|---|---|---|---|---|---|---|---|---|---|---|---|---|
| 11361:3 | 4 | R.LAGKPTHVN[+1216.423]VSVVMAEVDGTC[+57.021]Y.- | HexNAc(2)Hex(5) | C[+57], N[+1216] | 1188.529 | 3 | 3563.5724 | 3563.5649 | 0 | 2.1 | 332 | CRagged | 919.43 | 919.43 | 562.7 | 19.18 | 99 | >sp\|P01876\|IGHA1_HUMAN Immunoglobulin heavy constant alpha 1 OS=Homo sapiens OX=9606 GN=IGHA1 PE=1 SV=2 | 255 | 20201113_gy_colo_milk_1ng_glyco_hcd203040_compound15.14407.14407.3 | controllerType=0 controllerNumber=1 scan=14407 | 30.629 |
| 11325:3 | 4 | R.LAGKPTHVN[+1378.476]VSVVMAEVDGTC[+57.021]Y.- | HexNAc(2)Hex(6) | C[+57], N[+1378] | 1242.5479 | 3 | 3725.6292 | 3725.6178 | 0 | 3.1 | 332 | CRagged | 921.14 | 921.14 | 590.16 | 19.18 | 99 | >sp\|P01876\|IGHA1_HUMAN Immunoglobulin heavy constant alpha 1 OS=Homo sapiens OX=9606 GN=IGHA1 PE=1 SV=2 | 255 | 20201113_gy_colo_milk_1ng_glyco_hcd203040_compound15.14369.14369.3 | controllerType=0 controllerNumber=1 scan=14369 | 30.5306 |
| 12975:3 | 4 | R.LAGKPTHVN[+2262.814]VSVVMAEVDGTC[+57.021]Y.- | HexNAc(5)Hex(5)Fuc(1)NeuAc(1) | C[+57], N[+2263] | 1537.3244 | 3 | 4609.9586 | 4609.9564 | 0 | 0.5 | 332 | CRagged | 929.38 | 616.56 | 616.56 | 18.62 | 99 | >sp\|P01876\|IGHA1_HUMAN Immunoglobulin heavy constant alpha 1 OS=Homo sapiens OX=9606 GN=IGHA1 PE=1 SV=2 | 255 | 20201113_gy_colo_milk_1ng_glyco_hcd203040_compound15.16081.16081.3 | controllerType=0 controllerNumber=1 scan=16081 | 33.6553 |
| 13087:3 | 4 | R.LAGKPTHVN[+1751.624]VSVVMAEVDGTC[+57.021]Y.- | HexNAc(4)Hex(4)NeuAc(1) | C[+57], N[+1752] | 1366.9295 | 3 | 4098.7739 | 4098.7663 | 0 | 1.9 | 332 | CRagged | 943.99 | 812.8 | 812.8 | 19.18 | 99 | >sp\|P01876\|IGHA1_HUMAN Immunoglobulin heavy constant alpha 1 OS=Homo sapiens OX=9606 GN=IGHA1 PE=1 SV=2 | 255 | 20201113_gy_colo_milk_1ng_glyco_hcd203040_compound15.16198.16198.3 | controllerType=0 controllerNumber=1 scan=16198 | 33.8844 |
| 09604:3 | 4 | R.LAGKPTHVN[+1647.613]VSVVM[+15.995]AEVDGTC[+57.021]Y.- | HexNAc(5)Hex(3)Fuc(1) | C[+57], M[+16], N[+1648] | 1337.591 | 3 | 4010.7584 | 4010.7502 | 0 | 2 | 332 | CRagged | 202.9 | 202.9 | 138.54 | 3.7 | 99 | >sp\|P01876\|IGHA1_HUMAN Immunoglobulin heavy constant alpha 1 OS=Homo sapiens OX=9606 GN=IGHA1 PE=1 SV=2 | 255 | 20201113_gy_colo_milk_1ng_glyco_hcd203040_compound15.12585.12585.3 | controllerType=0 controllerNumber=1 scan=12585 | 27.2835 |
| 09519:3 | 4 | R.LAGKPTHVN[+1809.666]VSVVM[+15.995]AEVDGTC[+57.021]Y.- | HexNAc(5)Hex(4)Fuc(1) | C[+57], M[+16], N[+1810] | 1391.6076 | 3 | 4172.8082 | 4172.8031 | 0 | 1.2 | 332 | CRagged | 229.56 | 229.56 | 194.68 | 6.77 | 99 | >sp\|P01876\|IGHA1_HUMAN Immunoglobulin heavy constant alpha 1 OS=Homo sapiens OX=9606 GN=IGHA1 PE=1 SV=2 | 255 | 20201113_gy_colo_milk_1ng_glyco_hcd203040_compound15.12497.12497.3 | controllerType=0 controllerNumber=1 scan=12497 | 27.1266 |
| 09408:3 | 4 | R.LAGKPTHVN[+1809.666]VSVVM[+15.995]AEVDGTC[+57.021]Y.- | HexNAc(5)Hex(4)Fuc(1) | C[+57], M[+16], N[+1810] | 1391.6071 | 3 | 4172.8068 | 4172.8031 | 0 | 0.9 | 332 | CRagged | 251.78 | 251.78 | 233.94 | 5.81 | 99 | >sp\|P01876\|IGHA1_HUMAN Immunoglobulin heavy constant alpha 1 OS=Homo sapiens OX=9606 GN=IGHA1 PE=1 SV=2 | 255 | 20201113_gy_colo_milk_1ng_glyco_hcd203040_compound15.12382.12382.3 | controllerType=0 controllerNumber=1 scan=12382 | 26.912 |
| 09602:3 | 4 | R.LAGKPTHVN[+1647.613]VSVVM[+15.995]AEVDGTC[+57.021]Y.- | HexNAc(5)Hex(3)Fuc(1) | C[+57], M[+16], N[+1648] | 1337.591 | 3 | 4010.7584 | 4010.7502 | 0 | 2 | 332 | CRagged | 262.62 | 262.62 | 146.98 | 7.14 | 99 | >sp\|P01876\|IGHA1_HUMAN Immunoglobulin heavy constant alpha 1 OS=Homo sapiens OX=9606 GN=IGHA1 PE=1 SV=2 | 255 | 20201113_gy_colo_milk_1ng_glyco_hcd203040_compound15.12583.12583.3 | controllerType=0 controllerNumber=1 scan=12583 | 27.2791 |
| 09276:3 | 4 | R.LAGKPTHVN[+1216.423]VSVVM[+15.995]AEVDGTC[+57.021]Y.- | HexNAc(2)Hex(5) | C[+57], M[+16], N[+1216] | 1193.8614 | 3 | 3579.5696 | 3579.5599 | 0 | 2.7 | 332 | CRagged | 315.32 | 315.32 | 315.32 | 7.57 | 99 | >sp\|P01876\|IGHA1_HUMAN Immunoglobulin heavy constant alpha 1 OS=Homo sapiens OX=9606 GN=IGHA1 PE=1 SV=2 | 255 | 20201113_gy_colo_milk_1ng_glyco_hcd203040_compound15.12245.12245.3 | controllerType=0 controllerNumber=1 scan=12245 | 26.6642 |
| 09278:3 | 4 | R.LAGKPTHVN[+1216.423]VSVVM[+15.995]AEVDGTC[+57.021]Y.- | HexNAc(2)Hex(5) | C[+57], M[+16], N[+1216] | 1193.8614 | 3 | 3579.5696 | 3579.5599 | 0 | 2.7 | 332 | CRagged | 382.31 | 382.31 | 382.31 | 10 | 99 | >sp\|P01876\|IGHA1_HUMAN Immunoglobulin heavy constant alpha 1 OS=Homo sapiens OX=9606 GN=IGHA1 PE=1 SV=2 | 255 | 20201113_gy_colo_milk_1ng_glyco_hcd203040_compound15.12247.12247.3 | controllerType=0 controllerNumber=1 scan=12247 | 26.6685 |
| 08266:3 | 4 | R.LAGKPTHVN[+1216.423]VSVVM[+15.995]AEVDGTC[+57.021]Y.- | HexNAc(2)Hex(5) | C[+57], M[+16], N[+1216] | 1193.8585 | 3 | 3579.5608 | 3579.5599 | 0 | 0.3 | 332 | CRagged | 417.44 | 417.44 | 417.44 | 9.5 | 99 | >sp\|P01876\|IGHA1_HUMAN Immunoglobulin heavy constant alpha 1 OS=Homo sapiens OX=9606 GN=IGHA1 PE=1 SV=2 | 255 | 20201113_gy_colo_milk_1ng_glyco_hcd203040_compound15.11199.11199.3 | controllerType=0 controllerNumber=1 scan=11199 | 24.801 |
| 09058:3 | 4 | R.LAGKPTHVN[+1378.476]VSVVM[+15.995]AEVDGTC[+57.021]Y.- | HexNAc(2)Hex(6) | C[+57], M[+16], N[+1378] | 1247.8785 | 3 | 3741.6209 | 3741.6127 | 0 | 2.2 | 332 | CRagged | 428.53 | 428.53 | 428.53 | 10.06 | 99 | >sp\|P01876\|IGHA1_HUMAN Immunoglobulin heavy constant alpha 1 OS=Homo sapiens OX=9606 GN=IGHA1 PE=1 SV=2 | 255 | 20201113_gy_colo_milk_1ng_glyco_hcd203040_compound15.12019.12019.3 | controllerType=0 controllerNumber=1 scan=12019 | 26.2473 |
| 09056:3 | 4 | R.LAGKPTHVN[+1378.476]VSVVM[+15.995]AEVDGTC[+57.021]Y.- | HexNAc(2)Hex(6) | C[+57], M[+16], N[+1378] | 1247.8785 | 3 | 3741.6209 | 3741.6127 | 0 | 2.2 | 332 | CRagged | 456.69 | 397.19 | 397.19 | 10.28 | 99 | >sp\|P01876\|IGHA1_HUMAN Immunoglobulin heavy constant alpha 1 OS=Homo sapiens OX=9606 GN=IGHA1 PE=1 SV=2 | 255 | 20201113_gy_colo_milk_1ng_glyco_hcd203040_compound15.12017.12017.3 | controllerType=0 controllerNumber=1 scan=12017 | 26.2433 |
| 08268:3 | 4 | R.LAGKPTHVN[+1216.423]VSVVM[+15.995]AEVDGTC[+57.021]Y.- | HexNAc(2)Hex(5) | C[+57], M[+16], N[+1216] | 1193.8585 | 3 | 3579.5608 | 3579.5599 | 0 | 0.3 | 332 | CRagged | 548.95 | 548.95 | 548.95 | 10.07 | 99 | >sp\|P01876\|IGHA1_HUMAN Immunoglobulin heavy constant alpha 1 OS=Homo sapiens OX=9606 GN=IGHA1 PE=1 SV=2 | 255 | 20201113_gy_colo_milk_1ng_glyco_hcd203040_compound15.11201.11201.3 | controllerType=0 controllerNumber=1 scan=11201 | 24.8048 |
| 09558:3 | 4 | R.LAGKPTHVN[+1216.423]VSVVM[+15.995]AEVDGTC[+57.021]Y.- | HexNAc(2)Hex(5) | C[+57], M[+16], N[+1216] | 1193.8628 | 3 | 3579.574 | 3579.5599 | 0 | 4 | 332 | CRagged | 550.49 | 550.49 | 550.49 | 10.29 | 99 | >sp\|P01876\|IGHA1_HUMAN Immunoglobulin heavy constant alpha 1 OS=Homo sapiens OX=9606 GN=IGHA1 PE=1 SV=2 | 255 | 20201113_gy_colo_milk_1ng_glyco_hcd203040_compound15.12538.12538.3 | controllerType=0 controllerNumber=1 scan=12538 | 27.2036 |
| 09296:3 | 4 | R.LAGKPTHVN[+1216.423]VSVVM[+15.995]AEVDGTC[+57.021]Y.- | HexNAc(2)Hex(5) | C[+57], M[+16], N[+1216] | 1193.8616 | 3 | 3579.5703 | 3579.5599 | 0 | 2.9 | 332 | CRagged | 605.41 | 579.93 | 579.93 | 13.18 | 99 | >sp\|P01876\|IGHA1_HUMAN Immunoglobulin heavy constant alpha 1 OS=Homo sapiens OX=9606 GN=IGHA1 PE=1 SV=2 | 255 | 20201113_gy_colo_milk_1ng_glyco_hcd203040_compound15.12266.12266.3 | controllerType=0 controllerNumber=1 scan=12266 | 26.7058 |
| 08956:3 | 4 | R.LAGKPTHVN[+1216.423]VSVVM[+15.995]AEVDGTC[+57.021]Y.- | HexNAc(2)Hex(5) | C[+57], M[+16], N[+1216] | 1193.8618 | 3 | 3579.5707 | 3579.5599 | 0 | 3 | 332 | CRagged | 649.71 | 649.71 | 649.71 | 13.18 | 99 | >sp\|P01876\|IGHA1_HUMAN Immunoglobulin heavy constant alpha 1 OS=Homo sapiens OX=9606 GN=IGHA1 PE=1 SV=2 | 255 | 20201113_gy_colo_milk_1ng_glyco_hcd203040_compound15.11914.11914.3 | controllerType=0 controllerNumber=1 scan=11914 | 26.0651 |
| 09561:3 | 4 | R.LAGKPTHVN[+1216.423]VSVVM[+15.995]AEVDGTC[+57.021]Y.- | HexNAc(2)Hex(5) | C[+57], M[+16], N[+1216] | 1193.8628 | 3 | 3579.574 | 3579.5599 | 0 | 4 | 332 | CRagged | 662.31 | 660 | 660 | 13.84 | 99 | >sp\|P01876\|IGHA1_HUMAN Immunoglobulin heavy constant alpha 1 OS=Homo sapiens OX=9606 GN=IGHA1 PE=1 SV=2 | 255 | 20201113_gy_colo_milk_1ng_glyco_hcd203040_compound15.12541.12541.3 | controllerType=0 controllerNumber=1 scan=12541 | 27.2089 |
| 09298:3 | 4 | R.LAGKPTHVN[+1216.423]VSVVM[+15.995]AEVDGTC[+57.021]Y.- | HexNAc(2)Hex(5) | C[+57], M[+16], N[+1216] | 1193.8616 | 3 | 3579.5703 | 3579.5599 | 0 | 2.9 | 332 | CRagged | 687.05 | 687.05 | 687.05 | 14.18 | 99 | >sp\|P01876\|IGHA1_HUMAN Immunoglobulin heavy constant alpha 1 OS=Homo sapiens OX=9606 GN=IGHA1 PE=1 SV=2 | 255 | 20201113_gy_colo_milk_1ng_glyco_hcd203040_compound15.12268.12268.3 | controllerType=0 controllerNumber=1 scan=12268 | 26.7101 |
| 08964:3 | 4 | R.LAGKPTHVN[+1647.613]VSVVM[+15.995]AEVDGTC[+57.021]Y.- | HexNAc(5)Hex(3)Fuc(1) | C[+57], M[+16], N[+1648] | 1337.5912 | 3 | 4010.7592 | 4010.7502 | 0 | 2.2 | 332 | CRagged | 701.74 | 701.74 | 372.18 | 15.18 | 99 | >sp\|P01876\|IGHA1_HUMAN Immunoglobulin heavy constant alpha 1 OS=Homo sapiens OX=9606 GN=IGHA1 PE=1 SV=2 | 255 | 20201113_gy_colo_milk_1ng_glyco_hcd203040_compound15.11922.11922.3 | controllerType=0 controllerNumber=1 scan=11922 | 26.078 |
| 08966:3 | 4 | R.LAGKPTHVN[+1647.613]VSVVM[+15.995]AEVDGTC[+57.021]Y.- | HexNAc(5)Hex(3)Fuc(1) | C[+57], M[+16], N[+1648] | 1337.5912 | 3 | 4010.7592 | 4010.7502 | 0 | 2.2 | 332 | CRagged | 705.49 | 705.49 | 398 | 15.18 | 99 | >sp\|P01876\|IGHA1_HUMAN Immunoglobulin heavy constant alpha 1 OS=Homo sapiens OX=9606 GN=IGHA1 PE=1 SV=2 | 255 | 20201113_gy_colo_milk_1ng_glyco_hcd203040_compound15.11924.11924.3 | controllerType=0 controllerNumber=1 scan=11924 | 26.0825 |
| 08958:3 | 4 | R.LAGKPTHVN[+1216.423]VSVVM[+15.995]AEVDGTC[+57.021]Y.- | HexNAc(2)Hex(5) | C[+57], M[+16], N[+1216] | 1193.8618 | 3 | 3579.5707 | 3579.5599 | 0 | 3 | 332 | CRagged | 808.86 | 808.86 | 808.86 | 17.18 | 99 | >sp\|P01876\|IGHA1_HUMAN Immunoglobulin heavy constant alpha 1 OS=Homo sapiens OX=9606 GN=IGHA1 PE=1 SV=2 | 255 | 20201113_gy_colo_milk_1ng_glyco_hcd203040_compound15.11916.11916.3 | controllerType=0 controllerNumber=1 scan=11916 | 26.0693 |
| 10194:3 | 5 | K.TVLTPATNHMGN[+1540.529]VTFTIPANR.E | HexNAc(2)Hex(7) | N[+1541] | 1265.9033 | 3 | 3795.6952 | 3795.6886 | 0 | 1.7 | 74 | Specific | 777.64 | 777.64 | 362.92 | 16.18 | 110 | >sp\|P01024\|CO3_HUMAN Complement C3 OS=Homo sapiens OX=9606 GN=C3 PE=1 SV=2 | 492 | 20201113_gy_colo_milk_1ng_glyco_hcd203040_compound15.13196.13196.3 | controllerType=0 controllerNumber=1 scan=13196 | 28.3356 |
| 10220:3 | 5 | K.TVLTPATNHMGN[+1540.529]VTFTIPANR.E | HexNAc(2)Hex(7) | N[+1541] | 1265.9045 | 3 | 3795.6989 | 3795.6886 | 0 | 2.7 | 74 | Specific | 822.25 | 778.24 | 409.65 | 17.18 | 110 | >sp\|P01024\|CO3_HUMAN Complement C3 OS=Homo sapiens OX=9606 GN=C3 PE=1 SV=2 | 492 | 20201113_gy_colo_milk_1ng_glyco_hcd203040_compound15.13223.13223.3 | controllerType=0 controllerNumber=1 scan=13223 | 28.3882 |
| 10192:3 | 5 | K.TVLTPATNHMGN[+1540.529]VTFTIPANR.E | HexNAc(2)Hex(7) | N[+1541] | 1265.9033 | 3 | 3795.6952 | 3795.6886 | 0 | 1.7 | 74 | Specific | 862.89 | 851.83 | 442.27 | 18.18 | 110 | >sp\|P01024\|CO3_HUMAN Complement C3 OS=Homo sapiens OX=9606 GN=C3 PE=1 SV=2 | 492 | 20201113_gy_colo_milk_1ng_glyco_hcd203040_compound15.13194.13194.3 | controllerType=0 controllerNumber=1 scan=13194 | 28.3297 |
| 10218:3 | 5 | K.TVLTPATNHMGN[+1540.529]VTFTIPANR.E | HexNAc(2)Hex(7) | N[+1541] | 1265.9045 | 3 | 3795.6989 | 3795.6886 | 0 | 2.7 | 74 | Specific | 991.53 | 731.56 | 547.11 | 20.18 | 110 | >sp\|P01024\|CO3_HUMAN Complement C3 OS=Homo sapiens OX=9606 GN=C3 PE=1 SV=2 | 492 | 20201113_gy_colo_milk_1ng_glyco_hcd203040_compound15.13221.13221.3 | controllerType=0 controllerNumber=1 scan=13221 | 28.3834 |
| 17161:4 | 6 | R.GLSFDVSLEVSQGPGLLN[+2569.905]DTK.V | HexNAc(5)Hex(6)NeuAc(2) | N[+2570] | 1187.2646 | 4 | 4746.0364 | 4746.0178 | 0 | 3.9 | 118 | Specific | 266.42 | 214.34 | 214.34 | 5.15 | 82 | >sp\|P01833\|PIGR_HUMAN Polymeric immunoglobulin receptor OS=Homo sapiens OX=9606 GN=PIGR PE=1 SV=4 | 902 | 20201113_gy_colo_milk_1ng_glyco_hcd203040_compound15.20435.20435.2 | controllerType=0 controllerNumber=1 scan=20435 | 42.2635 |
| 17286:3 | 6 | R.GLSFDVSLEVSQGPGLLN[+2351.851]DTK.V | HexNAc(4)Hex(5)Fuc(3)NeuAc(1) | N[+2352] | 1510.0018 | 3 | 4527.9907 | 4527.964 | 0 | 5.9 | 118 | Specific | 694.82 | 577.97 | 577.97 | 13.61 | 82 | >sp\|P01833\|PIGR_HUMAN Polymeric immunoglobulin receptor OS=Homo sapiens OX=9606 GN=PIGR PE=1 SV=4 | 902 | 20201113_gy_colo_milk_1ng_glyco_hcd203040_compound15.20565.20565.3 | controllerType=0 controllerNumber=1 scan=20565 | 42.5288 |
| 17035:3 | 6 | R.GLSFDVSLEVSQGPGLLN[+2716.983]DTK.V | HexNAc(5)Hex(6)Fuc(3)NeuAc(1) | N[+2717] | 1631.7109 | 3 | 4893.1181 | 4893.0962 | 0 | 4.5 | 118 | Specific | 799.96 | 767.05 | 767.05 | 16.01 | 82 | >sp\|P01833\|PIGR_HUMAN Polymeric immunoglobulin receptor OS=Homo sapiens OX=9606 GN=PIGR PE=1 SV=4 | 902 | 20201113_gy_colo_milk_1ng_glyco_hcd203040_compound15.20304.20304.3 | controllerType=0 controllerNumber=1 scan=20304 | 41.9918 |
| 17089:3 | 6 | R.GLSFDVSLEVSQGPGLLN[+2569.905]DTK.V | HexNAc(5)Hex(6)NeuAc(2) | N[+2570] | 1582.6856 | 3 | 4746.0423 | 4746.0178 | 0 | 5.1 | 118 | Specific | 802.49 | 723.9 | 723.9 | 16.66 | 82 | >sp\|P01833\|PIGR_HUMAN Polymeric immunoglobulin receptor OS=Homo sapiens OX=9606 GN=PIGR PE=1 SV=4 | 902 | 20201113_gy_colo_milk_1ng_glyco_hcd203040_compound15.20361.20361.3 | controllerType=0 controllerNumber=1 scan=20361 | 42.1205 |
| 17091:3 | 6 | R.GLSFDVSLEVSQGPGLLN[+2569.905]DTK.V | HexNAc(5)Hex(6)NeuAc(2) | N[+2570] | 1582.6856 | 3 | 4746.0423 | 4746.0178 | 0 | 5.1 | 118 | Specific | 831.62 | 775.82 | 775.82 | 16.66 | 82 | >sp\|P01833\|PIGR_HUMAN Polymeric immunoglobulin receptor OS=Homo sapiens OX=9606 GN=PIGR PE=1 SV=4 | 902 | 20201113_gy_colo_milk_1ng_glyco_hcd203040_compound15.20363.20363.3 | controllerType=0 controllerNumber=1 scan=20363 | 42.1257 |
| 17144:3 | 6 | R.GLSFDVSLEVSQGPGLLN[+2570.925]DTK.V | HexNAc(5)Hex(6)Fuc(2)NeuAc(1) | N[+2571] | 1583.0192 | 3 | 4747.0431 | 4747.0382 | 0 | 1 | 118 | Specific | 839.7 | 838.03 | 838.03 | 17.21 | 82 | >sp\|P01833\|PIGR_HUMAN Polymeric immunoglobulin receptor OS=Homo sapiens OX=9606 GN=PIGR PE=1 SV=4 | 902 | 20201113_gy_colo_milk_1ng_glyco_hcd203040_compound15.20418.20418.3 | controllerType=0 controllerNumber=1 scan=20418 | 42.2316 |
| 17147:3 | 6 | R.GLSFDVSLEVSQGPGLLN[+2570.925]DTK.V | HexNAc(5)Hex(6)Fuc(2)NeuAc(1) | N[+2571] | 1583.0192 | 3 | 4747.0431 | 4747.0382 | 0 | 1 | 118 | Specific | 906.1 | 824.4 | 824.4 | 19.21 | 82 | >sp\|P01833\|PIGR_HUMAN Polymeric immunoglobulin receptor OS=Homo sapiens OX=9606 GN=PIGR PE=1 SV=4 | 902 | 20201113_gy_colo_milk_1ng_glyco_hcd203040_compound15.20421.20421.3 | controllerType=0 controllerNumber=1 scan=20421 | 42.2359 |
| 17289:3 | 6 | R.GLSFDVSLEVSQGPGLLN[+2351.851]DTK.V | HexNAc(4)Hex(5)Fuc(3)NeuAc(1) | N[+2352] | 1510.0018 | 3 | 4527.9907 | 4527.964 | 0 | 5.9 | 118 | Specific | 947.84 | 850.45 | 850.45 | 18.61 | 82 | >sp\|P01833\|PIGR_HUMAN Polymeric immunoglobulin receptor OS=Homo sapiens OX=9606 GN=PIGR PE=1 SV=4 | 902 | 20201113_gy_colo_milk_1ng_glyco_hcd203040_compound15.20568.20568.3 | controllerType=0 controllerNumber=1 scan=20568 | 42.5372 |
| 17032:3 | 6 | R.GLSFDVSLEVSQGPGLLN[+2716.983]DTK.V | HexNAc(5)Hex(6)Fuc(3)NeuAc(1) | N[+2717] | 1631.7109 | 3 | 4893.1181 | 4893.0962 | 0 | 4.5 | 118 | Specific | 965.82 | 936.87 | 936.87 | 20.01 | 82 | >sp\|P01833\|PIGR_HUMAN Polymeric immunoglobulin receptor OS=Homo sapiens OX=9606 GN=PIGR PE=1 SV=4 | 902 | 20201113_gy_colo_milk_1ng_glyco_hcd203040_compound15.20301.20301.3 | controllerType=0 controllerNumber=1 scan=20301 | 41.9865 |
| 22350:3 | 6 | K.QIGLYPVLVIDSSGYVNPN[+2188.740]YTGR.I | HexNAc(2)Hex(11) | N[+2189] | 1572.034 | 3 | 4714.0874 | 4714.0433 | 0 | 9.4 | 168 | Specific | 291.03 | 291.03 | 291.03 | 6.58 | 82 | >sp\|P01833\|PIGR_HUMAN Polymeric immunoglobulin receptor OS=Homo sapiens OX=9606 GN=PIGR PE=1 SV=4 | 902 | 20201113_gy_colo_milk_1ng_glyco_hcd203040_compound15.25847.25847.3 | controllerType=0 controllerNumber=1 scan=25847 | 53.8834 |
| 18378:3 | 6 | K.QIGLYPVLVIDSSGYVNPN[+2569.905]YTGR.I | HexNAc(5)Hex(6)NeuAc(2) | N[+2570] | 1699.088 | 3 | 5095.2494 | 5095.2081 | 0 | 8.1 | 168 | Specific | 315.45 | 222.87 | 222.87 | 6.3 | 82 | >sp\|P01833\|PIGR_HUMAN Polymeric immunoglobulin receptor OS=Homo sapiens OX=9606 GN=PIGR PE=1 SV=4 | 902 | 20201113_gy_colo_milk_1ng_glyco_hcd203040_compound15.21704.21704.3 | controllerType=0 controllerNumber=1 scan=21704 | 45.0055 |
| 22373:3 | 6 | K.QIGLYPVLVIDSSGYVNPN[+2188.740]YTGR.I | HexNAc(2)Hex(11) | N[+2189] | 1572.0261 | 3 | 4714.0636 | 4714.0433 | 0 | 4.3 | 168 | Specific | 317.12 | 279.32 | 279.32 | 7.33 | 82 | >sp\|P01833\|PIGR_HUMAN Polymeric immunoglobulin receptor OS=Homo sapiens OX=9606 GN=PIGR PE=1 SV=4 | 902 | 20201113_gy_colo_milk_1ng_glyco_hcd203040_compound15.25871.25871.3 | controllerType=0 controllerNumber=1 scan=25871 | 53.9352 |
| 22371:3 | 6 | K.QIGLYPVLVIDSSGYVNPN[+2188.740]YTGR.I | HexNAc(2)Hex(11) | N[+2189] | 1572.0261 | 3 | 4714.0636 | 4714.0433 | 0 | 4.3 | 168 | Specific | 321.12 | 313.49 | 313.49 | 7.41 | 82 | >sp\|P01833\|PIGR_HUMAN Polymeric immunoglobulin receptor OS=Homo sapiens OX=9606 GN=PIGR PE=1 SV=4 | 902 | 20201113_gy_colo_milk_1ng_glyco_hcd203040_compound15.25869.25869.3 | controllerType=0 controllerNumber=1 scan=25869 | 53.9296 |
| 22352:3 | 6 | K.QIGLYPVLVIDSSGYVNPN[+2188.740]YTGR.I | HexNAc(2)Hex(11) | N[+2189] | 1572.034 | 3 | 4714.0874 | 4714.0433 | 0 | 9.4 | 168 | Specific | 400.52 | 319.31 | 319.31 | 7.27 | 82 | >sp\|P01833\|PIGR_HUMAN Polymeric immunoglobulin receptor OS=Homo sapiens OX=9606 GN=PIGR PE=1 SV=4 | 902 | 20201113_gy_colo_milk_1ng_glyco_hcd203040_compound15.25849.25849.3 | controllerType=0 controllerNumber=1 scan=25849 | 53.8882 |
| 18380:3 | 6 | K.QIGLYPVLVIDSSGYVNPN[+2569.905]YTGR.I | HexNAc(5)Hex(6)NeuAc(2) | N[+2570] | 1699.088 | 3 | 5095.2494 | 5095.2081 | 0 | 8.1 | 168 | Specific | 635.93 | 565.15 | 565.15 | 12.31 | 82 | >sp\|P01833\|PIGR_HUMAN Polymeric immunoglobulin receptor OS=Homo sapiens OX=9606 GN=PIGR PE=1 SV=4 | 902 | 20201113_gy_colo_milk_1ng_glyco_hcd203040_compound15.21706.21706.3 | controllerType=0 controllerNumber=1 scan=21706 | 45.0144 |
| 18479:4 | 6 | K.QIGLYPVLVIDSSGYVNPN[+2204.772]YTGR.I | HexNAc(4)Hex(5)NeuAc(2) | N[+2205] | 1183.2826 | 4 | 4730.1087 | 4730.0759 | 0 | 6.9 | 168 | Specific | 742.9 | 677.7 | 677.7 | 14.54 | 82 | >sp\|P01833\|PIGR_HUMAN Polymeric immunoglobulin receptor OS=Homo sapiens OX=9606 GN=PIGR PE=1 SV=4 | 902 | 20201113_gy_colo_milk_1ng_glyco_hcd203040_compound15.21810.21810.4 | controllerType=0 controllerNumber=1 scan=21810 | 45.2443 |
| 20407:3 | 6 | K.QIGLYPVLVIDSSGYVNPN[+2350.830]YTGR.I | HexNAc(4)Hex(5)Fuc(1)NeuAc(2) | N[+2351] | 1626.0578 | 3 | 4876.1588 | 4876.1338 | 0 | 5.1 | 168 | Specific | 753.98 | 728.99 | 728.99 | 15.66 | 82 | >sp\|P01833\|PIGR_HUMAN Polymeric immunoglobulin receptor OS=Homo sapiens OX=9606 GN=PIGR PE=1 SV=4 | 902 | 20201113_gy_colo_milk_1ng_glyco_hcd203040_compound15.23820.23820.3 | controllerType=0 controllerNumber=1 scan=23820 | 49.5825 |
| 18481:4 | 6 | K.QIGLYPVLVIDSSGYVNPN[+2204.772]YTGR.I | HexNAc(4)Hex(5)NeuAc(2) | N[+2205] | 1183.2826 | 4 | 4730.1087 | 4730.0759 | 0 | 6.9 | 168 | Specific | 799.91 | 762.16 | 762.16 | 15.54 | 82 | >sp\|P01833\|PIGR_HUMAN Polymeric immunoglobulin receptor OS=Homo sapiens OX=9606 GN=PIGR PE=1 SV=4 | 902 | 20201113_gy_colo_milk_1ng_glyco_hcd203040_compound15.21812.21812.4 | controllerType=0 controllerNumber=1 scan=21812 | 45.255 |
| 20410:3 | 6 | K.QIGLYPVLVIDSSGYVNPN[+2350.830]YTGR.I | HexNAc(4)Hex(5)Fuc(1)NeuAc(2) | N[+2351] | 1626.0578 | 3 | 4876.1588 | 4876.1338 | 0 | 5.1 | 168 | Specific | 822.23 | 732.77 | 732.77 | 16.66 | 82 | >sp\|P01833\|PIGR_HUMAN Polymeric immunoglobulin receptor OS=Homo sapiens OX=9606 GN=PIGR PE=1 SV=4 | 902 | 20201113_gy_colo_milk_1ng_glyco_hcd203040_compound15.23823.23823.3 | controllerType=0 controllerNumber=1 scan=23823 | 49.5974 |
| 18502:4 | 6 | K.QIGLYPVLVIDSSGYVNPN[+2059.735]YTGR.I | HexNAc(4)Hex(5)Fuc(1)NeuAc(1) | N[+2060] | 1147.0186 | 4 | 4585.0526 | 4585.0384 | 0 | 3.1 | 168 | Specific | 831.05 | 796.25 | 796.25 | 17.49 | 82 | >sp\|P01833\|PIGR_HUMAN Polymeric immunoglobulin receptor OS=Homo sapiens OX=9606 GN=PIGR PE=1 SV=4 | 902 | 20201113_gy_colo_milk_1ng_glyco_hcd203040_compound15.21834.21834.4 | controllerType=0 controllerNumber=1 scan=21834 | 45.3136 |
| 18505:4 | 6 | K.QIGLYPVLVIDSSGYVNPN[+2059.735]YTGR.I | HexNAc(4)Hex(5)Fuc(1)NeuAc(1) | N[+2060] | 1147.0186 | 4 | 4585.0526 | 4585.0384 | 0 | 3.1 | 168 | Specific | 857.52 | 805.55 | 805.55 | 18.49 | 82 | >sp\|P01833\|PIGR_HUMAN Polymeric immunoglobulin receptor OS=Homo sapiens OX=9606 GN=PIGR PE=1 SV=4 | 902 | 20201113_gy_colo_milk_1ng_glyco_hcd203040_compound15.21837.21837.4 | controllerType=0 controllerNumber=1 scan=21837 | 45.3198 |
| 24910:4 | 6 | R.LSLLEEPGN[+3154.136]GTFTVILNQLTSR.D | HexNAc(5)Hex(6)Fuc(4)NeuAc(2) | N[+3154] | 1389.864 | 4 | 5556.4342 | 5556.4288 | 0 | 1 | 413 | Specific | 207.81 | 67.05 | 67.05 | 4.17 | 82 | >sp\|P01833\|PIGR_HUMAN Polymeric immunoglobulin receptor OS=Homo sapiens OX=9606 GN=PIGR PE=1 SV=4 | 902 | 20201113_gy_colo_milk_1ng_glyco_hcd203040_compound15.28513.28513.4 | controllerType=0 controllerNumber=1 scan=28513 | 59.0761 |
| 19959:4 | 6 | R.LSLLEEPGN[+2715.963]GTFTVILNQLTSR.D | HexNAc(5)Hex(6)Fuc(1)NeuAc(2) | N[+2716] | 1280.3248 | 4 | 5118.2775 | 5118.2551 | 0 | 4.4 | 413 | Specific | 225.8 | 225.8 | 225.8 | 6.87 | 82 | >sp\|P01833\|PIGR_HUMAN Polymeric immunoglobulin receptor OS=Homo sapiens OX=9606 GN=PIGR PE=1 SV=4 | 902 | 20201113_gy_colo_milk_1ng_glyco_hcd203040_compound15.23352.23352.4 | controllerType=0 controllerNumber=1 scan=23352 | 48.5391 |
| 27950:3 | 6 | R.[+42.011]LSLLEEPGN[+2350.830]GTFTVILNQLTSR.D | HexNAc(4)Hex(5)Fuc(1)NeuAc(2) | .[42], N[+2351] | 1599.053 | 3 | 4795.1446 | 4795.1335 | 0 | 2.3 | 413 | Specific | 233.23 | 233.23 | 233.23 | 7.3 | 82 | >sp\|P01833\|PIGR_HUMAN Polymeric immunoglobulin receptor OS=Homo sapiens OX=9606 GN=PIGR PE=1 SV=4 | 902 | 20201113_gy_colo_milk_1ng_glyco_hcd203040_compound15.31704.31704.3 | controllerType=0 controllerNumber=1 scan=31704 | 64.8831 |
| 24912:4 | 6 | R.LSLLEEPGN[+3154.136]GTFTVILNQLTSR.D | HexNAc(5)Hex(6)Fuc(4)NeuAc(2) | N[+3154] | 1389.864 | 4 | 5556.4342 | 5556.4288 | 0 | 1 | 413 | Specific | 290 | 249.75 | 249.75 | 7.55 | 82 | >sp\|P01833\|PIGR_HUMAN Polymeric immunoglobulin receptor OS=Homo sapiens OX=9606 GN=PIGR PE=1 SV=4 | 902 | 20201113_gy_colo_milk_1ng_glyco_hcd203040_compound15.28515.28515.4 | controllerType=0 controllerNumber=1 scan=28515 | 59.0808 |
| 22573:3 | 6 | R.LSLLEEPGN[+2351.851]GTFTVILNQLTSR.D | HexNAc(4)Hex(5)Fuc(3)NeuAc(1) | N[+2352] | 1585.3802 | 3 | 4754.126 | 4754.1433 | 0 | -3.6 | 413 | Specific | 320.76 | 228.46 | 228.46 | 6.04 | 82 | >sp\|P01833\|PIGR_HUMAN Polymeric immunoglobulin receptor OS=Homo sapiens OX=9606 GN=PIGR PE=1 SV=4 | 902 | 20201113_gy_colo_milk_1ng_glyco_hcd203040_compound15.26081.26081.3 | controllerType=0 controllerNumber=1 scan=26081 | 54.3881 |
| 22575:3 | 6 | R.LSLLEEPGN[+2351.851]GTFTVILNQLTSR.D | HexNAc(4)Hex(5)Fuc(3)NeuAc(1) | N[+2352] | 1585.3802 | 3 | 4754.126 | 4754.1433 | 0 | -3.6 | 413 | Specific | 355.32 | 271.55 | 271.55 | 6.5 | 82 | >sp\|P01833\|PIGR_HUMAN Polymeric immunoglobulin receptor OS=Homo sapiens OX=9606 GN=PIGR PE=1 SV=4 | 902 | 20201113_gy_colo_milk_1ng_glyco_hcd203040_compound15.26083.26083.3 | controllerType=0 controllerNumber=1 scan=26083 | 54.3925 |
| 28036:3 | 6 | R.[+42.011]LSLLEEPGN[+2351.851]GTFTVILNQLTSR.D | HexNAc(4)Hex(5)Fuc(3)NeuAc(1) | .[42], N[+2352] | 1599.3884 | 3 | 4796.1505 | 4796.1539 | 0 | -0.7 | 413 | Specific | 368.91 | 357.92 | 357.92 | 7.66 | 82 | >sp\|P01833\|PIGR_HUMAN Polymeric immunoglobulin receptor OS=Homo sapiens OX=9606 GN=PIGR PE=1 SV=4 | 902 | 20201113_gy_colo_milk_1ng_glyco_hcd203040_compound15.31795.31795.3 | controllerType=0 controllerNumber=1 scan=31795 | 65.0512 |
| 27952:3 | 6 | R.[+42.011]LSLLEEPGN[+2350.830]GTFTVILNQLTSR.D | HexNAc(4)Hex(5)Fuc(1)NeuAc(2) | .[42], N[+2351] | 1599.053 | 3 | 4795.1446 | 4795.1335 | 0 | 2.3 | 413 | Specific | 438.62 | 412.86 | 412.86 | 10.32 | 82 | >sp\|P01833\|PIGR_HUMAN Polymeric immunoglobulin receptor OS=Homo sapiens OX=9606 GN=PIGR PE=1 SV=4 | 902 | 20201113_gy_colo_milk_1ng_glyco_hcd203040_compound15.31706.31706.3 | controllerType=0 controllerNumber=1 scan=31706 | 64.8878 |
| 20116:4 | 6 | R.LSLLEEPGN[+2076.750]GTFTVILNQLTSR.D | HexNAc(4)Hex(6)Fuc(2) | N[+2077] | 1120.5193 | 4 | 4479.0555 | 4479.0428 | 0 | 2.8 | 413 | Specific | 468.93 | 409.94 | 409.94 | 10.59 | 82 | >sp\|P01833\|PIGR_HUMAN Polymeric immunoglobulin receptor OS=Homo sapiens OX=9606 GN=PIGR PE=1 SV=4 | 902 | 20201113_gy_colo_milk_1ng_glyco_hcd203040_compound15.23516.23516.4 | controllerType=0 controllerNumber=1 scan=23516 | 48.8968 |
| 27981:3 | 6 | R.[+42.011]LSLLEEPGN[+2351.851]GTFTVILNQLTSR.D | HexNAc(4)Hex(5)Fuc(3)NeuAc(1) | .[42], N[+2352] | 1599.3853 | 3 | 4796.1414 | 4796.1539 | 0 | -2.6 | 413 | Specific | 487.73 | 316.08 | 316.08 | 7.44 | 82 | >sp\|P01833\|PIGR_HUMAN Polymeric immunoglobulin receptor OS=Homo sapiens OX=9606 GN=PIGR PE=1 SV=4 | 902 | 20201113_gy_colo_milk_1ng_glyco_hcd203040_compound15.31736.31736.4 | controllerType=0 controllerNumber=1 scan=31736 | 64.9422 |
| 28038:3 | 6 | R.[+42.011]LSLLEEPGN[+2351.851]GTFTVILNQLTSR.D | HexNAc(4)Hex(5)Fuc(3)NeuAc(1) | .[42], N[+2352] | 1599.3884 | 3 | 4796.1505 | 4796.1539 | 0 | -0.7 | 413 | Specific | 514.29 | 427.25 | 427.25 | 9.75 | 82 | >sp\|P01833\|PIGR_HUMAN Polymeric immunoglobulin receptor OS=Homo sapiens OX=9606 GN=PIGR PE=1 SV=4 | 902 | 20201113_gy_colo_milk_1ng_glyco_hcd203040_compound15.31797.31797.3 | controllerType=0 controllerNumber=1 scan=31797 | 65.0551 |
| 20144:3 | 6 | R.LSLLEEPGN[+2351.851]GTFTVILNQLTSR.D | HexNAc(4)Hex(5)Fuc(3)NeuAc(1) | N[+2352] | 1585.3896 | 3 | 4754.1542 | 4754.1433 | 0 | 2.3 | 413 | Specific | 515.26 | 503.02 | 503.02 | 10.6 | 82 | >sp\|P01833\|PIGR_HUMAN Polymeric immunoglobulin receptor OS=Homo sapiens OX=9606 GN=PIGR PE=1 SV=4 | 902 | 20201113_gy_colo_milk_1ng_glyco_hcd203040_compound15.23545.23545.3 | controllerType=0 controllerNumber=1 scan=23545 | 48.9647 |
| 19875:3 | 6 | R.LSLLEEPGN[+2863.041]GTFTVILNQLTSR.D | HexNAc(5)Hex(6)Fuc(4)NeuAc(1) | N[+2863] | 1755.7885 | 3 | 5265.3511 | 5265.3334 | 0 | 3.3 | 413 | Specific | 526.01 | 271.81 | 271.81 | 8.37 | 82 | >sp\|P01833\|PIGR_HUMAN Polymeric immunoglobulin receptor OS=Homo sapiens OX=9606 GN=PIGR PE=1 SV=4 | 902 | 20201113_gy_colo_milk_1ng_glyco_hcd203040_compound15.23265.23265.3 | controllerType=0 controllerNumber=1 scan=23265 | 48.3474 |
| 19913:3 | 6 | R.LSLLEEPGN[+2862.021]GTFTVILNQLTSR.D | HexNAc(5)Hex(6)Fuc(2)NeuAc(2) | N[+2862] | 1755.453 | 3 | 5264.3443 | 5264.313 | 0 | 6 | 413 | Specific | 534.33 | 383.25 | 383.25 | 9.91 | 82 | >sp\|P01833\|PIGR_HUMAN Polymeric immunoglobulin receptor OS=Homo sapiens OX=9606 GN=PIGR PE=1 SV=4 | 902 | 20201113_gy_colo_milk_1ng_glyco_hcd203040_compound15.23304.23304.3 | controllerType=0 controllerNumber=1 scan=23304 | 48.4393 |
| 20002:3 | 6 | R.LSLLEEPGN[+2570.925]GTFTVILNQLTSR.D | HexNAc(5)Hex(6)Fuc(2)NeuAc(1) | N[+2571] | 1658.4192 | 3 | 4973.2429 | 4973.2176 | 0 | 5.1 | 413 | Specific | 541.65 | 411.96 | 411.96 | 10.12 | 82 | >sp\|P01833\|PIGR_HUMAN Polymeric immunoglobulin receptor OS=Homo sapiens OX=9606 GN=PIGR PE=1 SV=4 | 902 | 20201113_gy_colo_milk_1ng_glyco_hcd203040_compound15.23397.23397.3 | controllerType=0 controllerNumber=1 scan=23397 | 48.6343 |
| 19911:3 | 6 | R.LSLLEEPGN[+2862.021]GTFTVILNQLTSR.D | HexNAc(5)Hex(6)Fuc(2)NeuAc(2) | N[+2862] | 1755.453 | 3 | 5264.3443 | 5264.313 | 0 | 6 | 413 | Specific | 557.12 | 386.29 | 386.29 | 10.07 | 82 | >sp\|P01833\|PIGR_HUMAN Polymeric immunoglobulin receptor OS=Homo sapiens OX=9606 GN=PIGR PE=1 SV=4 | 902 | 20201113_gy_colo_milk_1ng_glyco_hcd203040_compound15.23302.23302.3 | controllerType=0 controllerNumber=1 scan=23302 | 48.4306 |
| 19961:4 | 6 | R.LSLLEEPGN[+2715.963]GTFTVILNQLTSR.D | HexNAc(5)Hex(6)Fuc(1)NeuAc(2) | N[+2716] | 1280.3248 | 4 | 5118.2775 | 5118.2551 | 0 | 4.4 | 413 | Specific | 632.42 | 526.01 | 526.01 | 13.15 | 82 | >sp\|P01833\|PIGR_HUMAN Polymeric immunoglobulin receptor OS=Homo sapiens OX=9606 GN=PIGR PE=1 SV=4 | 902 | 20201113_gy_colo_milk_1ng_glyco_hcd203040_compound15.23354.23354.4 | controllerType=0 controllerNumber=1 scan=23354 | [illegible] |
| 20000:3 | 6 | R.LSLLEEPGN[+2570.925]GTFTVILNQLTSR.D | HexNAc(5)Hex(6)Fuc(2)NeuAc(1) | N[+2571] | 1658.4192 | 3 | 4973.2429 | 4973.2176 | 0 | 5.1 | 413 | Specific | 638.32 | 553.42 | 553.42 | 12.66 | 82 | >sp\|P01833\|PIGR_HUMAN Polymeric immunoglobulin receptor OS=Homo sapiens OX=9606 GN=PIGR PE=1 SV=4 | 902 | 20201113_gy_colo_milk_1ng_glyco_hcd203040_compound15.23395.23395.3 | controllerType=0 controllerNumber=1 scan=23395 | [illegible] |
| 20119:4 | 6 | R.LSLLEEPGN[+2076.750]GTFTVILNQLTSR.D | HexNAc(4)Hex(6)Fuc(2) | N[+2077] | 1120.5193 | 4 | 4479.0555 | 4479.0428 | 0 | 2.8 | 413 | Specific | 688.21 | 559.51 | 559.51 | 14.49 | 82 | >sp\|P01833\|PIGR_HUMAN Polymeric immunoglobulin receptor OS=Homo sapiens OX=9606 GN=PIGR PE=1 SV=4 | 902 | 20201113_gy_colo_milk_1ng_glyco_hcd203040_compound15.23519.23519.4 | controllerType=0 controllerNumber=1 scan=23519 | [illegible] |
| 20185:3 | 6 | R.LSLLEEPGN[+2205.793]GTFTVILNQLTSR.D | HexNAc(4)Hex(5)Fuc(2)NeuAc(1) | N[+2206] | 1536.7115 | 3 | 4608.1199 | 4608.0854 | 0 | 7.5 | 413 | Specific | 762.87 | 529.29 | 529.29 | 15.41 | 82 | >sp\|P01833\|PIGR_HUMAN Polymeric immunoglobulin receptor OS=Homo sapiens OX=9606 GN=PIGR PE=1 SV=4 | 902 | 20201113_gy_colo_milk_1ng_glyco_hcd203040_compound15.23588.23588.3 | controllerType=0 controllerNumber=1 scan=23588 | [illegible] |
| 20187:3 | 6 | R.LSLLEEPGN[+2205.793]GTFTVILNQLTSR.D | HexNAc(4)Hex(5)Fuc(2)NeuAc(1) | N[+2206] | 1536.7115 | 3 | 4608.1199 | 4608.0854 | 0 | 7.5 | 413 | Specific | 767.07 | 481.71 | 481.71 | 15.41 | 82 | >sp\|P01833\|PIGR_HUMAN Polymeric immunoglobulin receptor OS=Homo sapiens OX=9606 GN=PIGR PE=1 SV=4 | 902 | 20201113_gy_colo_milk_1ng_glyco_hcd203040_compound15.23590.23590.3 | controllerType=0 controllerNumber=1 scan=23590 | 49.0652 |
| 20138:4 | 6 | R.LSLLEEPGN[+2205.793]GTFTVILNQLTSR.D | HexNAc(4)Hex(5)Fuc(2)NeuAc(1) | N[+2206] | 1152.7768 | 4 | 4608.0853 | 4608.0854 | 0 | 0 | 413 | Specific | 781.92 | 561.37 | 561.37 | 15.93 | 82 | >sp\|P01833\|PIGR_HUMAN Polymeric immunoglobulin receptor OS=Homo sapiens OX=9606 GN=PIGR PE=1 SV=4 | 902 | 20201113_gy_colo_milk_1ng_glyco_hcd203040_compound15.23539.23539.4 | controllerType=0 controllerNumber=1 scan=23539 | 48.9458 |
| 22145:3 | 6 | R.LSLLEEPGN[+2496.888]GTFTVILNQLTSR.D | HexNAc(4)Hex(5)Fuc(2)NeuAc(2) | N[+2497] | 1633.738 | 3 | 4899.1994 | 4899.1808 | 0 | 3.8 | 413 | Specific | 878.15 | 742 | 742 | 18.01 | 82 | >sp\|P01833\|PIGR_HUMAN Polymeric immunoglobulin receptor OS=Homo sapiens OX=9606 GN=PIGR PE=1 SV=4 | 902 | 20201113_gy_colo_milk_1ng_glyco_hcd203040_compound15.25633.25633.3 | controllerType=0 controllerNumber=1 scan=25633 | 53.4061 |
| 22147:3 | 6 | R.LSLLEEPGN[+2496.888]GTFTVILNQLTSR.D | HexNAc(4)Hex(5)Fuc(2)NeuAc(2) | N[+2497] | 1633.738 | 3 | 4899.1994 | 4899.1808 | 0 | 3.8 | 413 | Specific | 908.91 | 632.27 | 632.27 | 19.01 | 82 | >sp\|P01833\|PIGR_HUMAN Polymeric immunoglobulin receptor OS=Homo sapiens OX=9606 GN=PIGR PE=1 SV=4 | 902 | 20201113_gy_colo_milk_1ng_glyco_hcd203040_compound15.25635.25635.3 | controllerType=0 controllerNumber=1 scan=25635 | 53.4119 |
| 19912:3 | 6 | R.LSLLEEPGN[+2570.925]GTFTVILNQLTSR.D | HexNAc(5)Hex(6)Fuc(2)NeuAc(1) | N[+2571] | 1658.4176 | 3 | 4973.2381 | 4973.2176 | 0 | 4.1 | 413 | Specific | 918.71 | 648.85 | 648.85 | 19.01 | 82 | >sp\|P01833\|PIGR_HUMAN Polymeric immunoglobulin receptor OS=Homo sapiens OX=9606 GN=PIGR PE=1 SV=4 | 902 | 20201113_gy_colo_milk_1ng_glyco_hcd203040_compound15.23303.23303.3 | controllerType=0 controllerNumber=1 scan=23303 | 48.4369 |
| 19910:3 | 6 | R.LSLLEEPGN[+2570.925]GTFTVILNQLTSR.D | HexNAc(5)Hex(6)Fuc(2)NeuAc(1) | N[+2571] | 1658.4176 | 3 | 4973.2381 | 4973.2176 | 0 | 4.1 | 413 | Specific | 939.95 | 697.13 | 697.13 | 19.01 | 82 | >sp\|P01833\|PIGR_HUMAN Polymeric immunoglobulin receptor OS=Homo sapiens OX=9606 GN=PIGR PE=1 SV=4 | 902 | 20201113_gy_colo_milk_1ng_glyco_hcd203040_compound15.23301.23301.3 | controllerType=0 controllerNumber=1 scan=23301 | 48.4291 |
| 15810:4 | 6 | K.IIEGEPNLKVPGN[+2350.830]VTAVLGETLK.V | HexNAc(4)Hex(5)Fuc(1)NeuAc(2) | N[+2351] | 1186.3051 | 4 | 4742.1986 | 4742.1797 | 0 | 4 | 457 | Specific | 243.79 | 243.79 | 243.79 | 6.48 | 82 | >sp\|P01833\|PIGR_HUMAN Polymeric immunoglobulin receptor OS=Homo sapiens OX=9606 GN=PIGR PE=1 SV=4 | 902 | 20201113_gy_colo_milk_1ng_glyco_hcd203040_compound15.19027.19027.2 | controllerType=0 controllerNumber=1 scan=19027 | 39.3526 |
| 15812:4 | 6 | K.IIEGEPNLKVPGN[+2350.830]VTAVLGETLK.V | HexNAc(4)Hex(5)Fuc(1)NeuAc(2) | N[+2351] | 1186.3051 | 4 | 4742.1986 | 4742.1797 | 0 | 4 | 457 | Specific | 318.05 | 226.53 | 226.53 | 5.15 | 82 | >sp\|P01833\|PIGR_HUMAN Polymeric immunoglobulin receptor OS=Homo sapiens OX=9606 GN=PIGR PE=1 SV=4 | 902 | 20201113_gy_colo_milk_1ng_glyco_hcd203040_compound15.19029.19029.2 | controllerType=0 controllerNumber=1 scan=19029 | 39.3561 |
| 15815:4 | 6 | K.IIEGEPNLKVPGN[+2350.830]VTAVLGETLK.V | HexNAc(4)Hex(5)Fuc(1)NeuAc(2) | N[+2351] | 1186.3051 | 4 | 4742.1986 | 4742.1797 | 0 | 4 | 457 | Specific | 356.56 | 206.08 | 206.08 | 6.49 | 82 | >sp\|P01833\|PIGR_HUMAN Polymeric immunoglobulin receptor OS=Homo sapiens OX=9606 GN=PIGR PE=1 SV=4 | 902 | 20201113_gy_colo_milk_1ng_glyco_hcd203040_compound15.19032.19032.3 | controllerType=0 controllerNumber=1 scan=19032 | 39.3626 |
| 15818:4 | 6 | K.IIEGEPNLKVPGN[+2350.830]VTAVLGETLK.V | HexNAc(4)Hex(5)Fuc(1)NeuAc(2) | N[+2351] | 1186.3051 | 4 | 4742.1986 | 4742.1797 | 0 | 4 | 457 | Specific | 386.91 | 308.07 | 308.07 | 5.94 | 82 | >sp\|P01833\|PIGR_HUMAN Polymeric immunoglobulin receptor OS=Homo sapiens OX=9606 GN=PIGR PE=1 SV=4 | 902 | 20201113_gy_colo_milk_1ng_glyco_hcd203040_compound15.19035.19035.3 | controllerType=0 controllerNumber=1 scan=19035 | 39.3676 |
| 15893:4 | 6 | K.IIEGEPNLKVPGN[+2205.793]VTAVLGETLK.V | HexNAc(4)Hex(5)Fuc(2)NeuAc(1) | N[+2206] | 1150.0494 | 4 | 4597.1756 | 4597.1422 | 0 | 7.3 | 457 | Specific | 406.94 | 381.06 | 381.06 | 6.46 | 82 | >sp\|P01833\|PIGR_HUMAN Polymeric immunoglobulin receptor OS=Homo sapiens OX=9606 GN=PIGR PE=1 SV=4 | 902 | 20201113_gy_colo_milk_1ng_glyco_hcd203040_compound15.19113.19113.4 | controllerType=0 controllerNumber=1 scan=19113 | 39.5326 |
| 17630:4 | 6 | K.IIEGEPNLKVPGN[+2351.851]VTAVLGETLK.V | HexNAc(4)Hex(5)Fuc(3)NeuAc(1) | N[+2352] | 1186.5546 | 4 | 4743.1966 | 4743.2001 | 0 | -0.7 | 457 | Specific | 526.27 | 526.27 | 526.27 | 9.23 | 82 | >sp\|P01833\|PIGR_HUMAN Polymeric immunoglobulin receptor OS=Homo sapiens OX=9606 GN=PIGR PE=1 SV=4 | 902 | 20201113_gy_colo_milk_1ng_glyco_hcd203040_compound15.20924.20924.4 | controllerType=0 controllerNumber=1 scan=20924 | 43.3261 |
| 17633:4 | 6 | K.IIEGEPNLKVPGN[+2351.851]VTAVLGETLK.V | HexNAc(4)Hex(5)Fuc(3)NeuAc(1) | N[+2352] | 1186.5546 | 4 | 4743.1966 | 4743.2001 | 0 | -0.7 | 457 | Specific | 643.69 | 571.05 | 571.05 | 11.93 | 82 | >sp\|P01833\|PIGR_HUMAN Polymeric immunoglobulin receptor OS=Homo sapiens OX=9606 GN=PIGR PE=1 SV=4 | 902 | 20201113_gy_colo_milk_1ng_glyco_hcd203040_compound15.20927.20927.4 | controllerType=0 controllerNumber=1 scan=20927 | 43.3321 |
| 15803:4 | 6 | K.IIEGEPNLKVPGN[+2205.793]VTAVLGETLK.V | HexNAc(4)Hex(5)Fuc(2)NeuAc(1) | N[+2206] | 1150.0456 | 4 | 4597.1605 | 4597.1422 | 0 | 4 | 457 | Specific | 722.99 | 658.9 | 658.9 | 14.43 | 82 | >sp\|P01833\|PIGR_HUMAN Polymeric immunoglobulin receptor OS=Homo sapiens OX=9606 GN=PIGR PE=1 SV=4 | 902 | 20201113_gy_colo_milk_1ng_glyco_hcd203040_compound15.19020.19020.4 | controllerType=0 controllerNumber=1 scan=19020 | 39.3418 |
| 15877:4 | 6 | K.IIEGEPNLKVPGN[+2059.735]VTAVLGETLK.V | HexNAc(4)Hex(5)Fuc(1)NeuAc(1) | N[+2060] | 1113.5293 | 4 | 4451.0955 | 4451.0843 | 0 | 2.5 | 457 | Specific | 729.43 | 729.43 | 729.43 | 14.77 | 82 | >sp\|P01833\|PIGR_HUMAN Polymeric immunoglobulin receptor OS=Homo sapiens OX=9606 GN=PIGR PE=1 SV=4 | 902 | 20201113_gy_colo_milk_1ng_glyco_hcd203040_compound15.19097.19097.4 | controllerType=0 controllerNumber=1 scan=19097 | 39.4982 |
| 15871:3 | 6 | K.IIEGEPNLKVPGN[+2205.793]VTAVLGETLK.V | HexNAc(4)Hex(5)Fuc(2)NeuAc(1) | N[+2206] | 1533.0579 | 3 | 4597.1592 | 4597.1422 | 0 | 3.7 | 457 | Specific | 792.62 | 792.62 | 792.62 | 15.6 | 82 | >sp\|P01833\|PIGR_HUMAN Polymeric immunoglobulin receptor OS=Homo sapiens OX=9606 GN=PIGR PE=1 SV=4 | 902 | 20201113_gy_colo_milk_1ng_glyco_hcd203040_compound15.19091.19091.3 | controllerType=0 controllerNumber=1 scan=19091 | 39.4882 |
| 15873:3 | 6 | K.IIEGEPNLKVPGN[+2205.793]VTAVLGETLK.V | HexNAc(4)Hex(5)Fuc(2)NeuAc(1) | N[+2206] | 1533.0579 | 3 | 4597.1592 | 4597.1422 | 0 | 3.7 | 457 | Specific | 805.67 | 805.67 | 805.67 | 16.6 | 82 | >sp\|P01833\|PIGR_HUMAN Polymeric immunoglobulin receptor OS=Homo sapiens OX=9606 GN=PIGR PE=1 SV=4 | 902 | 20201113_gy_colo_milk_1ng_glyco_hcd203040_compound15.19093.19093.3 | controllerType=0 controllerNumber=1 scan=19093 | 39.492 |
| 17588:3 | 6 | K.IIEGEPNLKVPGN[+2350.830]VTAVLGETLK.V | HexNAc(4)Hex(5)Fuc(1)NeuAc(2) | N[+2351] | 1581.4072 | 3 | 4742.2069 | 4742.1797 | 0 | 5.7 | 457 | Specific | 808.45 | 745.67 | 745.67 | 15.91 | 82 | >sp\|P01833\|PIGR_HUMAN Polymeric immunoglobulin receptor OS=Homo sapiens OX=9606 GN=PIGR PE=1 SV=4 | 902 | 20201113_gy_colo_milk_1ng_glyco_hcd203040_compound15.20880.20880.4 | controllerType=0 controllerNumber=1 scan=20880 | 43.2189 |
| 15880:4 | 6 | K.IIEGEPNLKVPGN[+2059.735]VTAVLGETLK.V | HexNAc(4)Hex(5)Fuc(1)NeuAc(1) | N[+2060] | 1113.5293 | 4 | 4451.0955 | 4451.0843 | 0 | 2.5 | 457 | Specific | 817.2 | 717.88 | 717.88 | 16.77 | 82 | >sp\|P01833\|PIGR_HUMAN Polymeric immunoglobulin receptor OS=Homo sapiens OX=9606 GN=PIGR PE=1 SV=4 | 902 | 20201113_gy_colo_milk_1ng_glyco_hcd203040_compound15.19100.19100.4 | controllerType=0 controllerNumber=1 scan=19100 | 39.5034 |
| 15852:3 | 6 | K.IIEGEPNLKVPGN[+2059.735]VTAVLGETLK.V | HexNAc(4)Hex(5)Fuc(1)NeuAc(1) | N[+2060] | 1484.3708 | 3 | 4451.0978 | 4451.0843 | 0 | 3 | 457 | Specific | 830.8 | 722.46 | 722.46 | 16.6 | 82 | >sp\|P01833\|PIGR_HUMAN Polymeric immunoglobulin receptor OS=Homo sapiens OX=9606 GN=PIGR PE=1 SV=4 | 902 | 20201113_gy_colo_milk_1ng_glyco_hcd203040_compound15.19071.19071.3 | controllerType=0 controllerNumber=1 scan=19071 | 39.4427 |
| 15801:4 | 6 | K.IIEGEPNLKVPGN[+2205.793]VTAVLGETLK.V | HexNAc(4)Hex(5)Fuc(2)NeuAc(1) | N[+2206] | 1150.0456 | 4 | 4597.1605 | 4597.1422 | 0 | 4 | 457 | Specific | 893.77 | 813.59 | 813.59 | 17.43 | 82 | >sp\|P01833\|PIGR_HUMAN Polymeric immunoglobulin receptor OS=Homo sapiens OX=9606 GN=PIGR PE=1 SV=4 | 902 | 20201113_gy_colo_milk_1ng_glyco_hcd203040_compound15.19018.19018.4 | controllerType=0 controllerNumber=1 scan=19018 | 39.3365 |
| 15849:3 | 6 | K.IIEGEPNLKVPGN[+2059.735]VTAVLGETLK.V | HexNAc(4)Hex(5)Fuc(1)NeuAc(1) | N[+2060] | 1484.3708 | 3 | 4451.0978 | 4451.0843 | 0 | 3 | 457 | Specific | 1033.21 | 942.66 | 942.66 | 20.6 | 82 | >sp\|P01833\|PIGR_HUMAN Polymeric immunoglobulin receptor OS=Homo sapiens OX=9606 GN=PIGR PE=1 SV=4 | 902 | 20201113_gy_colo_milk_1ng_glyco_hcd203040_compound15.19068.19068.3 | controllerType=0 controllerNumber=1 scan=19068 | 39.4369 |
| 15444:5 | 6 | K.IIEGEPNLKVPGN[+2059.735]VTAVLGETLKVPC[+57.021]HFPC[+57.021]K.F | HexNAc(4)Hex(5)Fuc(1)NeuAc(1) | C[+57]*2, N[+2060] | 1096.1163 | 5 | 5476.5526 | 5476.5418 | 0 | 2 | 457 | Specific | 434.17 | 434.17 | 434.17 | 9.56 | 82 | >sp\|P01833\|PIGR_HUMAN Polymeric immunoglobulin receptor OS=Homo sapiens OX=9606 GN=PIGR PE=1 SV=4 | 902 | 20201113_gy_colo_milk_1ng_glyco_hcd203040_compound15.18645.18645.2 | controllerType=0 controllerNumber=1 scan=18645 | 38.6306 |
| 16811:5 | 6 | K.IIEGEPNLKVPGN[+2350.830]VTAVLGETLKVPC[+57.021]HFPC[+57.021]K.F | HexNAc(4)Hex(5)Fuc(1)NeuAc(2) | C[+57]*2, N[+2351] | 1154.3405 | 5 | 5767.6732 | 5767.6372 | 0 | 6.2 | 457 | Specific | 621.1 | 541.9 | 541.9 | 11.85 | 82 | >sp\|P01833\|PIGR_HUMAN Polymeric immunoglobulin receptor OS=Homo sapiens OX=9606 GN=PIGR PE=1 SV=4 | 902 | 20201113_gy_colo_milk_1ng_glyco_hcd203040_compound15.20069.20069.5 | controllerType=0 controllerNumber=1 scan=20069 | 41.4562 |
| 16790:5 | 6 | K.IIEGEPNLKVPGN[+2351.851]VTAVLGETLKVPC[+57.021]HFPC[+57.021]K.F | HexNAc(4)Hex(5)Fuc(3)NeuAc(1) | C[+57]*2, N[+2352] | 1154.5355 | 5 | 5768.6485 | 5768.6576 | 0 | -1.6 | 457 | Specific | 623.3 | 611.88 | 611.88 | 11.54 | 82 | >sp\|P01833\|PIGR_HUMAN Polymeric immunoglobulin receptor OS=Homo sapiens OX=9606 GN=PIGR PE=1 SV=4 | 902 | 20201113_gy_colo_milk_1ng_glyco_hcd203040_compound15.20047.20047.5 | controllerType=0 controllerNumber=1 scan=20047 | 41.4007 |
| 15402:4 | 6 | K.IIEGEPNLKVPGN[+2205.793]VTAVLGETLKVPC[+57.021]HFPC[+57.021]K.F | HexNAc(4)Hex(5)Fuc(2)NeuAc(1) | C[+57]*2, N[+2206] | 1406.4085 | 4 | 5622.612 | 5622.5997 | 0 | 2.2 | 457 | Specific | 700.89 | 662.14 | 662.14 | 14.02 | 82 | >sp\|P01833\|PIGR_HUMAN Polymeric immunoglobulin receptor OS=Homo sapiens OX=9606 GN=PIGR PE=1 SV=4 | 902 | 20201113_gy_colo_milk_1ng_glyco_hcd203040_compound15.18601.18601.4 | controllerType=0 controllerNumber=1 scan=18601 | 38.5461 |
| 15404:4 | 6 | K.IIEGEPNLKVPGN[+2205.793]VTAVLGETLKVPC[+57.021]HFPC[+57.021]K.F | HexNAc(4)Hex(5)Fuc(2)NeuAc(1) | C[+57]*2, N[+2206] | 1406.4085 | 4 | 5622.612 | 5622.5997 | 0 | 2.2 | 457 | Specific | 742.53 | 742.53 | 742.53 | 14.02 | 82 | >sp\|P01833\|PIGR_HUMAN Polymeric immunoglobulin receptor OS=Homo sapiens OX=9606 GN=PIGR PE=1 SV=4 | 902 | 20201113_gy_colo_milk_1ng_glyco_hcd203040_compound15.18603.18603.4 | controllerType=0 controllerNumber=1 scan=18603 | 38.5518 |
| 15447:5 | 6 | K.IIEGEPNLKVPGN[+2059.735]VTAVLGETLKVPC[+57.021]HFPC[+57.021]K.F | HexNAc(4)Hex(5)Fuc(1)NeuAc(1) | C[+57]*2, N[+2060] | 1096.1163 | 5 | 5476.5526 | 5476.5418 | 0 | 2 | 457 | Specific | 773.53 | 618.09 | 618.09 | 15.59 | 82 | >sp\|P01833\|PIGR_HUMAN Polymeric immunoglobulin receptor OS=Homo sapiens OX=9606 GN=PIGR PE=1 SV=4 | 902 | 20201113_gy_colo_milk_1ng_glyco_hcd203040_compound15.18648.18648.2 | controllerType=0 controllerNumber=1 scan=18648 | 38.6353 |
| 16813:5 | 6 | K.IIEGEPNLKVPGN[+2350.830]VTAVLGETLKVPC[+57.021]HFPC[+57.021]K.F | HexNAc(4)Hex(5)Fuc(1)NeuAc(2) | C[+57]*2, N[+2351] | 1154.3405 | 5 | 5767.6732 | 5767.6372 | 0 | 6.2 | 457 | Specific | 798.9 | 751.29 | 751.29 | 14.85 | 82 | >sp\|P01833\|PIGR_HUMAN Polymeric immunoglobulin receptor OS=Homo sapiens OX=9606 GN=PIGR PE=1 SV=4 | 902 | 20201113_gy_colo_milk_1ng_glyco_hcd203040_compound15.20071.20071.5 | controllerType=0 controllerNumber=1 scan=20071 | 41.46 |
| 16792:5 | 6 | K.IIEGEPNLKVPGN[+2351.851]VTAVLGETLKVPC[+57.021]HFPC[+57.021]K.F | HexNAc(4)Hex(5)Fuc(3)NeuAc(1) | C[+57]*2, N[+2352] | 1154.5355 | 5 | 5768.6485 | 5768.6576 | 0 | -1.6 | 457 | Specific | 805.27 | 768.04 | 768.04 | 15.54 | 82 | >sp\|P01833\|PIGR_HUMAN Polymeric immunoglobulin receptor OS=Homo sapiens OX=9606 GN=PIGR PE=1 SV=4 | 902 | 20201113_gy_colo_milk_1ng_glyco_hcd203040_compound15.20049.20049.5 | controllerType=0 controllerNumber=1 scan=20049 | 41.4067 |
| 13583:3 | 6 | K.VPGN[+2059.735]VTAVLGETLK.V | HexNAc(4)Hex(5)Fuc(1)NeuAc(1) | N[+2060] | 1153.1938 | 3 | 3457.5668 | 3457.5348 | 0 | 9.3 | 466 | Specific | 421.82 | 175.56 | 175.56 | 6.05 | 82 | >sp\|P01833\|PIGR_HUMAN Polymeric immunoglobulin receptor OS=Homo sapiens OX=9606 GN=PIGR PE=1 SV=4 | 902 | 20201113_gy_colo_milk_1ng_glyco_hcd203040_compound15.16713.16713.3 | controllerType=0 controllerNumber=1 scan=16713 | 34.8757 |
| 13586:3 | 6 | K.VPGN[+2059.735]VTAVLGETLK.V | HexNAc(4)Hex(5)Fuc(1)NeuAc(1) | N[+2060] | 1153.1938 | 3 | 3457.5668 | 3457.5348 | 0 | 9.3 | 466 | Specific | 498.35 | 419.17 | 419.17 | 7.45 | 82 | >sp\|P01833\|PIGR_HUMAN Polymeric immunoglobulin receptor OS=Homo sapiens OX=9606 GN=PIGR PE=1 SV=4 | 902 | 20201113_gy_colo_milk_1ng_glyco_hcd203040_compound15.16716.16716.3 | controllerType=0 controllerNumber=1 scan=16716 | 34.8798 |
| 13395:3 | 6 | K.VPGN[+2351.851]VTAVLGETLK.V | HexNAc(4)Hex(5)Fuc(3)NeuAc(1) | N[+2352] | 1250.5615 | 3 | 3749.6698 | 3749.6506 | 0 | 5.1 | 466 | Specific | 644.43 | 348.28 | 348.28 | 10.68 | 82 | >sp\|P01833\|PIGR_HUMAN Polymeric immunoglobulin receptor OS=Homo sapiens OX=9606 GN=PIGR PE=1 SV=4 | 902 | 20201113_gy_colo_milk_1ng_glyco_hcd203040_compound15.16517.16517.3 | controllerType=0 controllerNumber=1 scan=16517 | 34.4631 |
| 13397:3 | 6 | K.VPGN[+2351.851]VTAVLGETLK.V | HexNAc(4)Hex(5)Fuc(3)NeuAc(1) | N[+2352] | 1250.5615 | 3 | 3749.6698 | 3749.6506 | 0 | 5.1 | 466 | Specific | 661.33 | 518.12 | 518.12 | 13.35 | 82 | >sp\|P01833\|PIGR_HUMAN Polymeric immunoglobulin receptor OS=Homo sapiens OX=9606 GN=PIGR PE=1 SV=4 | 902 | 20201113_gy_colo_milk_1ng_glyco_hcd203040_compound15.16519.16519.3 | controllerType=0 controllerNumber=1 scan=16519 | 34.4735 |
| 13421:3 | 6 | K.VPGN[+2205.793]VTAVLGETLK.V | HexNAc(4)Hex(5)Fuc(2)NeuAc(1) | N[+2206] | 1201.8722 | 3 | 3603.6022 | 3603.5927 | 0 | 2.6 | 466 | Specific | 740.22 | 661.31 | 661.31 | 15.03 | 82 | >sp\|P01833\|PIGR_HUMAN Polymeric immunoglobulin receptor OS=Homo sapiens OX=9606 GN=PIGR PE=1 SV=4 | 902 | 20201113_gy_colo_milk_1ng_glyco_hcd203040_compound15.16544.16544.3 | controllerType=0 controllerNumber=1 scan=16544 | 34.5223 |
| 13419:3 | 6 | K.VPGN[+2205.793]VTAVLGETLK.V | HexNAc(4)Hex(5)Fuc(2)NeuAc(1) | N[+2206] | 1201.8722 | 3 | 3603.6022 | 3603.5927 | 0 | 2.6 | 466 | Specific | 756.25 | 749.81 | 749.81 | 16.03 | 82 | >sp\|P01833\|PIGR_HUMAN Polymeric immunoglobulin receptor OS=Homo sapiens OX=9606 GN=PIGR PE=1 SV=4 | 902 | 20201113_gy_colo_milk_1ng_glyco_hcd203040_compound15.16542.16542.3 | controllerType=0 controllerNumber=1 scan=16542 | 34.5137 |
| 16329:3 | 6 | K.VPGN[+2350.830]VTAVLGETLK.V | HexNAc(4)Hex(5)Fuc(1)NeuAc(2) | N[+2351] | 1250.222 | 3 | 3748.6514 | 3748.6302 | 0 | 5.7 | 466 | Specific | 766.45 | 626.15 | 626.15 | 15.35 | 82 | >sp\|P01833\|PIGR_HUMAN Polymeric immunoglobulin receptor OS=Homo sapiens OX=9606 GN=PIGR PE=1 SV=4 | 902 | 20201113_gy_colo_milk_1ng_glyco_hcd203040_compound15.19567.19567.3 | controllerType=0 controllerNumber=1 scan=19567 | 40.4092 |
| 16326:3 | 6 | K.VPGN[+2350.830]VTAVLGETLK.V | HexNAc(4)Hex(5)Fuc(1)NeuAc(2) | N[+2351] | 1250.222 | 3 | 3748.6514 | 3748.6302 | 0 | 5.7 | 466 | Specific | 772.93 | 554.24 | 554.24 | 15.35 | 82 | >sp\|P01833\|PIGR_HUMAN Polymeric immunoglobulin receptor OS=Homo sapiens OX=9606 GN=PIGR PE=1 SV=4 | 902 | 20201113_gy_colo_milk_1ng_glyco_hcd203040_compound15.19564.19564.3 | controllerType=0 controllerNumber=1 scan=19564 | 40.4043 |
| 13559:4 | 6 | K.VPGN[+2205.793]VTAVLGETLKVPC[+57.021]HFPC[+57.021]K.F | HexNAc(4)Hex(5)Fuc(2)NeuAc(1) | C[+57]*2, N[+2206] | 1158.0208 | 4 | 4629.0613 | 4629.0502 | 0 | 2.4 | 466 | Specific | 640.89 | 603.45 | 603.45 | 12.6 | 82 | >sp\|P01833\|PIGR_HUMAN Polymeric immunoglobulin receptor OS=Homo sapiens OX=9606 GN=PIGR PE=1 SV=4 | 902 | 20201113_gy_colo_milk_1ng_glyco_hcd203040_compound15.16688.16688.2 | controllerType=0 controllerNumber=1 scan=16688 | 34.8247 |
| 15300:4 | 6 | K.VPGN[+2350.830]VTAVLGETLKVPC[+57.021]HFPC[+57.021]K.F | HexNAc(4)Hex(5)Fuc(1)NeuAc(2) | C[+57]*2, N[+2351] | 1194.2835 | 4 | 4774.1121 | 4774.0877 | 0 | 5.1 | 466 | Specific | 757.48 | 515.02 | 515.02 | 14.91 | 82 | >sp\|P01833\|PIGR_HUMAN Polymeric immunoglobulin receptor OS=Homo sapiens OX=9606 GN=PIGR PE=1 SV=4 | 902 | 20201113_gy_colo_milk_1ng_glyco_hcd203040_compound15.18495.18495.4 | controllerType=0 controllerNumber=1 scan=18495 | 38.327 |
| 13562:4 | 6 | K.VPGN[+2205.793]VTAVLGETLKVPC[+57.021]HFPC[+57.021]K.F | HexNAc(4)Hex(5)Fuc(2)NeuAc(1) | C[+57]*2, N[+2206] | 1158.0208 | 4 | 4629.0613 | 4629.0502 | 0 | 2.4 | 466 | Specific | 785.75 | 776.6 | 776.6 | 15.6 | 82 | >sp\|P01833\|PIGR_HUMAN Polymeric immunoglobulin receptor OS=Homo sapiens OX=9606 GN=PIGR PE=1 SV=4 | 902 | 20201113_gy_colo_milk_1ng_glyco_hcd203040_compound15.16691.16691.2 | controllerType=0 controllerNumber=1 scan=16691 | 34.8326 |
| 13600:4 | 6 | K.VPGN[+2059.735]VTAVLGETLKVPC[+57.021]HFPC[+57.021]K.F | HexNAc(4)Hex(5)Fuc(1)NeuAc(1) | C[+57]*2, N[+2060] | 1121.505 | 4 | 4482.9984 | 4482.9923 | 0 | 1.3 | 466 | Specific | 808.42 | 487.73 | 487.73 | 16.46 | 82 | >sp\|P01833\|PIGR_HUMAN Polymeric immunoglobulin receptor OS=Homo sapiens OX=9606 GN=PIGR PE=1 SV=4 | 902 | 20201113_gy_colo_milk_1ng_glyco_hcd203040_compound15.16730.16730.2 | controllerType=0 controllerNumber=1 scan=16730 | 34.9055 |
| 15317:4 | 6 | K.VPGN[+2350.830]VTAVLGETLKVPC[+57.021]HFPC[+57.021]K.F | HexNAc(4)Hex(5)Fuc(1)NeuAc(2) | C[+57]*2, N[+2351] | 1194.2837 | 4 | 4774.1131 | 4774.0877 | 0 | 5.3 | 466 | Specific | 809.45 | 741.03 | 741.03 | 15.91 | 82 | >sp\|P01833\|PIGR_HUMAN Polymeric immunoglobulin receptor OS=Homo sapiens OX=9606 GN=PIGR PE=1 SV=4 | 902 | 20201113_gy_colo_milk_1ng_glyco_hcd203040_compound15.18513.18513.4 | controllerType=0 controllerNumber=1 scan=18513 | 38.37 |
| 15298:4 | 6 | K.VPGN[+2350.830]VTAVLGETLKVPC[+57.021]HFPC[+57.021]K.F | HexNAc(4)Hex(5)Fuc(1)NeuAc(2) | C[+57]*2, N[+2351] | 1194.2835 | 4 | 4774.1121 | 4774.0877 | 0 | 5.1 | 466 | Specific | 871.89 | 652.51 | 652.51 | 16.91 | 82 | >sp\|P01833\|PIGR_HUMAN Polymeric immunoglobulin receptor OS=Homo sapiens OX=9606 GN=PIGR PE=1 SV=4 | 902 | 20201113_gy_colo_milk_1ng_glyco_hcd203040_compound15.18493.18493.4 | controllerType=0 controllerNumber=1 scan=18493 | 38.3205 |
| 13597:4 | 6 | K.VPGN[+2059.735]VTAVLGETLKVPC[+57.021]HFPC[+57.021]K.F | HexNAc(4)Hex(5)Fuc(1)NeuAc(1) | C[+57]*2, N[+2060] | 1121.505 | 4 | 4482.9984 | 4482.9923 | 0 | 1.3 | 466 | Specific | 886.61 | 806.01 | 806.01 | 17.46 | 82 | >sp\|P01833\|PIGR_HUMAN Polymeric immunoglobulin receptor OS=Homo sapiens OX=9606 GN=PIGR PE=1 SV=4 | 902 | 20201113_gy_colo_milk_1ng_glyco_hcd203040_compound15.16727.16727.2 | controllerType=0 controllerNumber=1 scan=16727 | 34.8989 |
| 15315:4 | 6 | K.VPGN[+2350.830]VTAVLGETLKVPC[+57.021]HFPC[+57.021]K.F | HexNAc(4)Hex(5)Fuc(1)NeuAc(2) | C[+57]*2, N[+2351] | 1194.2837 | 4 | 4774.1131 | 4774.0877 | 0 | 5.3 | 466 | Specific | 892.61 | 777.29 | 777.29 | 16.91 | 82 | >sp\|P01833\|PIGR_HUMAN Polymeric immunoglobulin receptor OS=Homo sapiens OX=9606 GN=PIGR PE=1 SV=4 | 902 | 20201113_gy_colo_milk_1ng_glyco_hcd203040_compound15.18511.18511.4 | controllerType=0 controllerNumber=1 scan=18511 | 38.3649 |
| 06199:3 | 6 | K.WN[+1548.545]NTGC[+57.021]QALPSQDEGPSK.A | HexNAc(3)Hex(4)NeuAc(1) | C[+57], N[+1549] | 1179.8123 | 3 | 3537.4224 | 3537.4215 | 0 | 0.2 | 498 | Specific | 490.08 | 413.39 | 413.39 | 9.81 | 82 | >sp\|P01833\|PIGR_HUMAN Polymeric immunoglobulin receptor OS=Homo sapiens OX=9606 GN=PIGR PE=1 SV=4 | 902 | 20201113_gy_colo_milk_1ng_glyco_hcd203040_compound15.9059.9059.3 | controllerType=0 controllerNumber=1 scan=9059 | 21.1777 |
| 05895:3 | 6 | K.WN[+2205.793]NTGC[+57.021]QALPSQDEGPSK.A | HexNAc(4)Hex(5)Fuc(2)NeuAc(1) | C[+57], N[+2206] | 1398.8961 | 3 | 4194.6736 | 4194.6695 | 0 | 1 | 498 | Specific | 589.16 | 488.83 | 488.83 | 12.08 | 82 | >sp\|P01833\|PIGR_HUMAN Polymeric immunoglobulin receptor OS=Homo sapiens OX=9606 GN=PIGR PE=1 SV=4 | 902 | 20201113_gy_colo_milk_1ng_glyco_hcd203040_compound15.8745.8745.3 | controllerType=0 controllerNumber=1 scan=8745 | 20.6653 |
| 05897:3 | 6 | K.WN[+2205.793]NTGC[+57.021]QALPSQDEGPSK.A | HexNAc(4)Hex(5)Fuc(2)NeuAc(1) | C[+57], N[+2206] | 1398.8961 | 3 | 4194.6736 | 4194.6695 | 0 | 1 | 498 | Specific | 627.04 | 533.97 | 533.97 | 13.08 | 82 | >sp\|P01833\|PIGR_HUMAN Polymeric immunoglobulin receptor OS=Homo sapiens OX=9606 GN=PIGR PE=1 SV=4 | 902 | 20201113_gy_colo_milk_1ng_glyco_hcd203040_compound15.8747.8747.3 | controllerType=0 controllerNumber=1 scan=8747 | 20.6697 |
| 05926:3 | 6 | K.WN[+2059.735]NTGC[+57.021]QALPSQDEGPSK.A | HexNAc(4)Hex(5)Fuc(1)NeuAc(1) | C[+57], N[+2060] | 1350.2118 | 3 | 4048.621 | 4048.6116 | 0 | 2.3 | 498 | Specific | 649.89 | 557.73 | 557.73 | 13.22 | 82 | >sp\|P01833\|PIGR_HUMAN Polymeric immunoglobulin receptor OS=Homo sapiens OX=9606 GN=PIGR PE=1 SV=4 | 902 | 20201113_gy_colo_milk_1ng_glyco_hcd203040_compound15.8777.8777.3 | controllerType=0 controllerNumber=1 scan=8777 | 20.7167 |
| 04445:2 | 6 | K.WN[+1216.423]NTGC[+57.021]QALPSQDEGPSK.A | HexNAc(2)Hex(5) | C[+57], N[+1216] | 1603.1526 | 2 | 3205.298 | 3205.2995 | 0 | -0.5 | 498 | Specific | 653.07 | 553.43 | 553.43 | 13.48 | 82 | >sp\|P01833\|PIGR_HUMAN Polymeric immunoglobulin receptor OS=Homo sapiens OX=9606 GN=PIGR PE=1 SV=4 | 902 | 20201113_gy_colo_milk_1ng_glyco_hcd203040_compound15.7243.7243.2 | controllerType=0 controllerNumber=1 scan=7243 | 18.2043 |
| 06201:3 | 6 | K.WN[+1548.545]NTGC[+57.021]QALPSQDEGPSK.A | HexNAc(3)Hex(4)NeuAc(1) | C[+57], N[+1549] | 1179.8123 | 3 | 3537.4224 | 3537.4215 | 0 | 0.2 | 498 | Specific | 665.22 | 545.78 | 545.78 | 13.66 | 82 | >sp\|P01833\|PIGR_HUMAN Polymeric immunoglobulin receptor OS=Homo sapiens OX=9606 GN=PIGR PE=1 SV=4 | 902 | 20201113_gy_colo_milk_1ng_glyco_hcd203040_compound15.9061.9061.3 | controllerType=0 controllerNumber=1 scan=9061 | 21.1813 |
| 04443:2 | 6 | K.WN[+1216.423]NTGC[+57.021]QALPSQDEGPSK.A | HexNAc(2)Hex(5) | C[+57], N[+1216] | 1603.1526 | 2 | 3205.298 | 3205.2995 | 0 | -0.5 | 498 | Specific | 693.04 | 656.06 | 656.06 | 13.48 | 82 | >sp\|P01833\|PIGR_HUMAN Polymeric immunoglobulin receptor OS=Homo sapiens OX=9606 GN=PIGR PE=1 SV=4 | 902 | 20201113_gy_colo_milk_1ng_glyco_hcd203040_compound15.7241.7241.2 | controllerType=0 controllerNumber=1 scan=7241 | 18.1999 |
| 04477:3 | 6 | K.WN[+1216.423]NTGC[+57.021]QALPSQDEGPSK.A | HexNAc(2)Hex(5) | C[+57], N[+1216] | 1069.1076 | 3 | 3205.3082 | 3205.2995 | 0 | 2.7 | 498 | Specific | 731.77 | 707.84 | 707.84 | 15.22 | 82 | >sp\|P01833\|PIGR_HUMAN Polymeric immunoglobulin receptor OS=Homo sapiens OX=9606 GN=PIGR PE=1 SV=4 | 902 | 20201113_gy_colo_milk_1ng_glyco_hcd203040_compound15.7276.7276.3 | controllerType=0 controllerNumber=1 scan=7276 | 18.2584 |
| 07973:3 | 6 | K.WN[+2204.772]NTGC[+57.021]QALPSQDEGPSK.A | HexNAc(4)Hex(5)NeuAc(2) | C[+57], N[+2205] | 1398.5591 | 3 | 4193.6629 | 4193.6491 | 0 | 3.3 | 498 | Specific | 754.98 | 635.27 | 635.27 | 16.21 | 82 | >sp\|P01833\|PIGR_HUMAN Polymeric immunoglobulin receptor OS=Homo sapiens OX=9606 GN=PIGR PE=1 SV=4 | 902 | 20201113_gy_colo_milk_1ng_glyco_hcd203040_compound15.10895.10895.3 | controllerType=0 controllerNumber=1 scan=10895 | 24.2835 |
| 04475:3 | 6 | K.WN[+1216.423]NTGC[+57.021]QALPSQDEGPSK.A | HexNAc(2)Hex(5) | C[+57], N[+1216] | 1069.1076 | 3 | 3205.3082 | 3205.2995 | 0 | 2.7 | 498 | Specific | 766.05 | 723.07 | 723.07 | 16.22 | 82 | >sp\|P01833\|PIGR_HUMAN Polymeric immunoglobulin receptor OS=Homo sapiens OX=9606 GN=PIGR PE=1 SV=4 | 902 | 20201113_gy_colo_milk_1ng_glyco_hcd203040_compound15.7274.7274.3 | controllerType=0 controllerNumber=1 scan=7274 | 18.2549 |
| 05928:3 | 6 | K.WN[+2059.735]NTGC[+57.021]QALPSQDEGPSK.A | HexNAc(4)Hex(5)Fuc(1)NeuAc(1) | C[+57], N[+2060] | 1350.2118 | 3 | 4048.621 | 4048.6116 | 0 | 2.3 | 498 | Specific | 801.87 | 699.08 | 699.08 | 17.22 | 82 | >sp\|P01833\|PIGR_HUMAN Polymeric immunoglobulin receptor OS=Homo sapiens OX=9606 GN=PIGR PE=1 SV=4 | 902 | 20201113_gy_colo_milk_1ng_glyco_hcd203040_compound15.8779.8779.3 | controllerType=0 controllerNumber=1 scan=8779 | 20.7211 |
| 05997:3 | 6 | K.WN[+1913.677]NTGC[+57.021]QALPSQDEGPSK.A | HexNAc(4)Hex(5)NeuAc(1) | C[+57], N[+1914] | 1301.5252 | 3 | 3902.5611 | 3902.5537 | 0 | 1.9 | 498 | Specific | 807.72 | 720.33 | 720.33 | 17.22 | 82 | >sp\|P01833\|PIGR_HUMAN Polymeric immunoglobulin receptor OS=Homo sapiens OX=9606 GN=PIGR PE=1 SV=4 | 902 | 20201113_gy_colo_milk_1ng_glyco_hcd203040_compound15.8850.8850.3 | controllerType=0 controllerNumber=1 scan=8850 | 20.8374 |
| 07971:3 | 6 | K.WN[+2204.772]NTGC[+57.021]QALPSQDEGPSK.A | HexNAc(4)Hex(5)NeuAc(2) | C[+57], N[+2205] | 1398.5591 | 3 | 4193.6629 | 4193.6491 | 0 | 3.3 | 498 | Specific | 814.51 | 729.94 | 729.94 | 17.21 | 82 | >sp\|P01833\|PIGR_HUMAN Polymeric immunoglobulin receptor OS=Homo sapiens OX=9606 GN=PIGR PE=1 SV=4 | 902 | 20201113_gy_colo_milk_1ng_glyco_hcd203040_compound15.10893.10893.3 | controllerType=0 controllerNumber=1 scan=10893 | 24.2793 |
| 06078:3 | 6 | K.WN[+1710.598]NTGC[+57.021]QALPSQDEGPSK.A | HexNAc(3)Hex(5)NeuAc(1) | C[+57], N[+1711] | 1233.8327 | 3 | 3699.4835 | 3699.4743 | 0 | 2.5 | 498 | Specific | 862.54 | 755.08 | 755.08 | 18.22 | 82 | >sp\|P01833\|PIGR_HUMAN Polymeric immunoglobulin receptor OS=Homo sapiens OX=9606 GN=PIGR PE=1 SV=4 | 902 | 20201113_gy_colo_milk_1ng_glyco_hcd203040_compound15.8934.8934.3 | controllerType=0 controllerNumber=1 scan=8934 | 20.9747 |
| 05989:3 | 6 | K.WN[+1872.651]NTGC[+57.021]QALPSQDEGPSK.A | HexNAc(3)Hex(6)NeuAc(1) | C[+57], N[+1873] | 1287.8497 | 3 | 3861.5344 | 3861.5272 | 0 | 1.9 | 498 | Specific | 876.8 | 798.31 | 798.31 | 18.22 | 82 | >sp\|P01833\|PIGR_HUMAN Polymeric immunoglobulin receptor OS=Homo sapiens OX=9606 GN=PIGR PE=1 SV=4 | 902 | 20201113_gy_colo_milk_1ng_glyco_hcd203040_compound15.8842.8842.3 | controllerType=0 controllerNumber=1 scan=8842 | 20.8208 |
| 05991:3 | 6 | K.WN[+1872.651]NTGC[+57.021]QALPSQDEGPSK.A | HexNAc(3)Hex(6)NeuAc(1) | C[+57], N[+1873] | 1287.8497 | 3 | 3861.5344 | 3861.5272 | 0 | 1.9 | 498 | Specific | 889.46 | 794.95 | 794.95 | 18.22 | 82 | >sp\|P01833\|PIGR_HUMAN Polymeric immunoglobulin receptor OS=Homo sapiens OX=9606 GN=PIGR PE=1 SV=4 | 902 | 20201113_gy_colo_milk_1ng_glyco_hcd203040_compound15.8844.8844.3 | controllerType=0 controllerNumber=1 scan=8844 | 20.826 |
| 05995:3 | 6 | K.WN[+1913.677]NTGC[+57.021]QALPSQDEGPSK.A | HexNAc(4)Hex(5)NeuAc(1) | C[+57], N[+1914] | 1301.5252 | 3 | 3902.5611 | 3902.5537 | 0 | 1.9 | 498 | Specific | 907.28 | 824 | 824 | 19.22 | 82 | >sp\|P01833\|PIGR_HUMAN Polymeric immunoglobulin receptor OS=Homo sapiens OX=9606 GN=PIGR PE=1 SV=4 | 902 | 20201113_gy_colo_milk_1ng_glyco_hcd203040_compound15.8848.8848.3 | controllerType=0 controllerNumber=1 scan=8848 | 20.8328 |
| 06076:3 | 6 | K.WN[+1710.598]NTGC[+57.021]QALPSQDEGPSK.A | HexNAc(3)Hex(5)NeuAc(1) | C[+57], N[+1711] | 1233.8327 | 3 | 3699.4835 | 3699.4743 | 0 | 2.5 | 498 | Specific | 908.89 | 830.47 | 830.47 | 19.22 | 82 | >sp\|P01833\|PIGR_HUMAN Polymeric immunoglobulin receptor OS=Homo sapiens OX=9606 GN=PIGR PE=1 SV=4 | 902 | 20201113_gy_colo_milk_1ng_glyco_hcd203040_compound15.8932.8932.3 | controllerType=0 controllerNumber=1 scan=8932 | 20.9698 |
| 23346:3 | 11 | R.[+42.011]QLAHQSN[+2190.793]STNIFFSPVSIATAFAMLSLGTK.A | HexNAc(6)Hex(6) | .[42], N[+2191] | 1805.4759 | 3 | 5414.4132 | 5414.4388 | 0 | -4.7 | 64 | Specific | 211.13 | 211.13 | 211.13 | 4.63 | 42 | >sp\|P01009\|A1AT_HUMAN Alpha-1-antitrypsin OS=Homo sapiens OX=9606 GN=SERPINA1 PE=1 SV=3 | 642 | 20201113_gy_colo_milk_1ng_glyco_hcd203040_compound15.26885.26885.3 | controllerType=0 controllerNumber=1 scan=26885 | 56.0431 |
| 23406:4 | 11 | R.[+42.011]QLAHQSN[+2190.793]STNIFFSPVSIATAFAMLSLGTK.A | HexNAc(6)Hex(6) | .[42], N[+2191] | 1354.3579 | 4 | 5414.4098 | 5414.4388 | 0 | -5.3 | 64 | Specific | 223.92 | 223.92 | 223.92 | 6.33 | 42 | >sp\|P01009\|A1AT_HUMAN Alpha-1-antitrypsin OS=Homo sapiens OX=9606 GN=SERPINA1 PE=1 SV=3 | 642 | 20201113_gy_colo_milk_1ng_glyco_hcd203040_compound15.26948.26948.4 | controllerType=0 controllerNumber=1 scan=26948 | 56.164 |
| 23408:4 | 11 | R.[+42.011]QLAHQSN[+2190.793]STNIFFSPVSIATAFAMLSLGTK.A | HexNAc(6)Hex(6) | .[42], N[+2191] | 1354.3579 | 4 | 5414.4098 | 5414.4388 | 0 | -5.3 | 64 | Specific | 226.71 | 226.71 | 226.71 | 6.33 | 42 | >sp\|P01009\|A1AT_HUMAN Alpha-1-antitrypsin OS=Homo sapiens OX=9606 GN=SERPINA1 PE=1 SV=3 | 642 | 20201113_gy_colo_milk_1ng_glyco_hcd203040_compound15.26950.26950.4 | controllerType=0 controllerNumber=1 scan=26950 | 56.1682 |
| 26042:4 | 11 | R.QLAHQSN[+3007.058]STNIFFSPVSIATAFAMLSLGTK.A | HexNAc(5)Hex(6)Fuc(1)NeuAc(3) | N[+3007] | 1547.9352 | 4 | 6188.7189 | 6188.693 | 0 | 4.2 | 64 | Specific | 255.45 | 253.86 | 253.86 | 7.55 | 42 | >sp\|P01009\|A1AT_HUMAN Alpha-1-antitrypsin OS=Homo sapiens OX=9606 GN=SERPINA1 PE=1 SV=3 | 642 | 20201113_gy_colo_milk_1ng_glyco_hcd203040_compound15.29695.29695.4 | controllerType=0 controllerNumber=1 scan=29695 | 61.2194 |

Table S4. Cont.

| Query #:z | Protein Rank | Glycopeptide | Glycan | Modification Type(s) | Observed m/z | z | Observed (M+H) | Calc. mass (M+H) | Off-by-x error | Mass error (ppm) | StartPosition | Cleavage | Score | Delta | Delta Mod | \|Log Prob\| | # of unique peptides | Protein Name | Protein DB number | Comment | Scan # | Scan Time |
|---|---|---|---|---|---|---|---|---|---|---|---|---|---|---|---|---|---|---|---|---|---|---|
| 23252:4 | 11 | R.QLAHQSN[+2350.830]STNIFFSPVSIATAFAMLSLGTK.A | HexNAc(4)Hex(5)Fuc(1)NeuAc(2) | N[+2351] | 1383.8751 | 4 | 5532.4787 | 5532.4654 | 0 | 2.4 | 64 | Specific | 311.68 | 311.68 | 311.68 | 8.32 | 42 | >sp\|P01009\|A1AT_HUMAN Alpha-1-antitrypsin OS=Homo sapiens OX=9606 GN=SERPINA1 PE=1 SV=3 | 642 | 20201113_gy_colo_milk_1ng_glyco_hcd203040_compound15.26787.26787.4 | controllerType=0 controllerNumber=1 scan=26787 | 55.8164 |
| 26065:4 | 11 | R.QLAHQSN[+2861.000]STNIFFSPVSIATAFAMLSLGTK.A | HexNAc(5)Hex(6)NeuAc(3) | N[+2861] | 1511.4279 | 4 | 6042.6898 | 6042.6351 | 0 | 9 | 64 | Specific | 369.47 | 354.39 | 354.39 | 8 | 42 | >sp\|P01009\|A1AT_HUMAN Alpha-1-antitrypsin OS=Homo sapiens OX=9606 GN=SERPINA1 PE=1 SV=3 | 642 | 20201113_gy_colo_milk_1ng_glyco_hcd203040_compound15.29719.29719.4 | controllerType=0 controllerNumber=1 scan=29719 | 61.2684 |
| 23259:3 | 11 | R.QLAHQSN[+2350.830]STNIFFSPVSIATAFAMLSLGTK.A | HexNAc(4)Hex(5)Fuc(1)NeuAc(2) | N[+2351] | 1844.8316 | 3 | 5532.4803 | 5532.4654 | 0 | 2.7 | 64 | Specific | 390.86 | 344.13 | 344.13 | 8.9 | 42 | >sp\|P01009\|A1AT_HUMAN Alpha-1-antitrypsin OS=Homo sapiens OX=9606 GN=SERPINA1 PE=1 SV=3 | 642 | 20201113_gy_colo_milk_1ng_glyco_hcd203040_compound15.26794.26794.3 | controllerType=0 controllerNumber=1 scan=26794 | 55.8297 |
| 26063:4 | 11 | R.QLAHQSN[+2861.000]STNIFFSPVSIATAFAMLSLGTK.A | HexNAc(5)Hex(6)NeuAc(3) | N[+2861] | 1511.4279 | 4 | 6042.6898 | 6042.6351 | 0 | 9 | 64 | Specific | 400.18 | 386.26 | 386.26 | 9.66 | 42 | >sp\|P01009\|A1AT_HUMAN Alpha-1-antitrypsin OS=Homo sapiens OX=9606 GN=SERPINA1 PE=1 SV=3 | 642 | 20201113_gy_colo_milk_1ng_glyco_hcd203040_compound15.29717.29717.4 | controllerType=0 controllerNumber=1 scan=29717 | 61.2627 |
| 23274:3 | 11 | R.QLAHQSN[+2204.772]STNIFFSPVSIATAFAMLSLGTK.A | HexNAc(4)Hex(5)NeuAc(2) | N[+2205] | 1796.1401 | 3 | 5386.4057 | 5386.4075 | 0 | -0.3 | 64 | Specific | 474.45 | 461.58 | 407.78 | 10.34 | 42 | >sp\|P01009\|A1AT_HUMAN Alpha-1-antitrypsin OS=Homo sapiens OX=9606 GN=SERPINA1 PE=1 SV=3 | 642 | 20201113_gy_colo_milk_1ng_glyco_hcd203040_compound15.26810.26810.3 | controllerType=0 controllerNumber=1 scan=26810 | 55.8723 |
| 23286:4 | 11 | R.QLAHQSN[+2350.830]STNIFFSPVSIATAFAMLSLGTK.A | HexNAc(4)Hex(5)Fuc(1)NeuAc(2) | N[+2351] | 1383.875 | 4 | 5532.4782 | 5532.4654 | 0 | 2.3 | 64 | Specific | 633.66 | 574.13 | 574.13 | 13.92 | 42 | >sp\|P01009\|A1AT_HUMAN Alpha-1-antitrypsin OS=Homo sapiens OX=9606 GN=SERPINA1 PE=1 SV=3 | 642 | 20201113_gy_colo_milk_1ng_glyco_hcd203040_compound15.26823.26823.4 | controllerType=0 controllerNumber=1 scan=26823 | 55.8979 |
| 23267:4 | 11 | R.QLAHQSN[+2204.772]STNIFFSPVSIATAFAMLSLGTK.A | HexNAc(4)Hex(5)NeuAc(2) | N[+2205] | 1347.3573 | 4 | 5386.4074 | 5386.4075 | 0 | 0 | 64 | Specific | 711.83 | 704.99 | 507.8 | 15.36 | 42 | >sp\|P01009\|A1AT_HUMAN Alpha-1-antitrypsin OS=Homo sapiens OX=9606 GN=SERPINA1 PE=1 SV=3 | 642 | 20201113_gy_colo_milk_1ng_glyco_hcd203040_compound15.26803.26803.4 | controllerType=0 controllerNumber=1 scan=26803 | 55.8544 |
| 23288:4 | 11 | R.QLAHQSN[+2350.830]STNIFFSPVSIATAFAMLSLGTK.A | HexNAc(4)Hex(5)Fuc(1)NeuAc(2) | N[+2351] | 1383.875 | 4 | 5532.4782 | 5532.4654 | 0 | 2.3 | 64 | Specific | 745.56 | 593.78 | 593.78 | 15.92 | 42 | >sp\|P01009\|A1AT_HUMAN Alpha-1-antitrypsin OS=Homo sapiens OX=9606 GN=SERPINA1 PE=1 SV=3 | 642 | 20201113_gy_colo_milk_1ng_glyco_hcd203040_compound15.26825.26825.4 | controllerType=0 controllerNumber=1 scan=26825 | 55.9026 |
| 23250:4 | 11 | R.QLAHQSN[+2350.830]STNIFFSPVSIATAFAMLSLGTK.A | HexNAc(4)Hex(5)Fuc(1)NeuAc(2) | N[+2351] | 1383.8751 | 4 | 5532.4787 | 5532.4654 | 0 | 2.4 | 64 | Specific | 780.46 | 737.52 | 737.52 | 16.92 | 42 | >sp\|P01009\|A1AT_HUMAN Alpha-1-antitrypsin OS=Homo sapiens OX=9606 GN=SERPINA1 PE=1 SV=3 | 642 | 20201113_gy_colo_milk_1ng_glyco_hcd203040_compound15.26785.26785.4 | controllerType=0 controllerNumber=1 scan=26785 | 55.8116 |
| 23272:3 | 11 | R.QLAHQSN[+2204.772]STNIFFSPVSIATAFAMLSLGTK.A | HexNAc(4)Hex(5)NeuAc(2) | N[+2205] | 1796.1401 | 3 | 5386.4057 | 5386.4075 | 0 | -0.3 | 64 | Specific | 914.22 | 852.77 | 597.64 | 19.19 | 42 | >sp\|P01009\|A1AT_HUMAN Alpha-1-antitrypsin OS=Homo sapiens OX=9606 GN=SERPINA1 PE=1 SV=3 | 642 | 20201113_gy_colo_milk_1ng_glyco_hcd203040_compound15.26808.26808.3 | controllerType=0 controllerNumber=1 scan=26808 | 55.8674 |
| 23265:4 | 11 | R.QLAHQSN[+2204.772]STNIFFSPVSIATAFAMLSLGTK.A | HexNAc(4)Hex(5)NeuAc(2) | N[+2205] | 1347.3573 | 4 | 5386.4074 | 5386.4075 | 0 | 0 | 64 | Specific | 964.77 | 923.1 | 610.03 | 20.36 | 42 | >sp\|P01009\|A1AT_HUMAN Alpha-1-antitrypsin OS=Homo sapiens OX=9606 GN=SERPINA1 PE=1 SV=3 | 642 | 20201113_gy_colo_milk_1ng_glyco_hcd203040_compound15.26801.26801.4 | controllerType=0 controllerNumber=1 scan=26801 | 55.846 |
| 20614:3 | 11 | K.ADTHDEILEGLNFN[+2205.793]LTEIPEAQIHEGFQELLR.T | HexNAc(4)Hex(5)Fuc(2)NeuAc(1) | N[+2206] | 1966.5441 | 3 | 5897.6177 | 5897.6167 | 0 | 0.2 | 94 | Specific | 379.57 | 379.57 | 379.57 | 9.43 | 42 | >sp\|P01009\|A1AT_HUMAN Alpha-1-antitrypsin OS=Homo sapiens OX=9606 GN=SERPINA1 PE=1 SV=3 | 642 | 20201113_gy_colo_milk_1ng_glyco_hcd203040_compound15.24037.24037.3 | controllerType=0 controllerNumber=1 scan=24037 | 50.0925 |
| 20616:3 | 11 | K.ADTHDEILEGLNFN[+2205.793]LTEIPEAQIHEGFQELLR.T | HexNAc(4)Hex(5)Fuc(2)NeuAc(1) | N[+2206] | 1966.5441 | 3 | 5897.6177 | 5897.6167 | 0 | 0.2 | 94 | Specific | 391.09 | 334.44 | 334.44 | 7.61 | 42 | >sp\|P01009\|A1AT_HUMAN Alpha-1-antitrypsin OS=Homo sapiens OX=9606 GN=SERPINA1 PE=1 SV=3 | 642 | 20201113_gy_colo_milk_1ng_glyco_hcd203040_compound15.24039.24039.3 | controllerType=0 controllerNumber=1 scan=24039 | 50.097 |
| 20583:4 | 11 | K.ADTHDEILEGLNFN[+2204.772]LTEIPEAQIHEGFQELLR.T | HexNAc(4)Hex(5)NeuAc(2) | N[+2205] | 1474.9086 | 4 | 5896.6125 | 5896.5963 | 0 | 2.7 | 94 | Specific | 1122.04 | 836.03 | 836.03 | 23.75 | 42 | >sp\|P01009\|A1AT_HUMAN Alpha-1-antitrypsin OS=Homo sapiens OX=9606 GN=SERPINA1 PE=1 SV=3 | 642 | 20201113_gy_colo_milk_1ng_glyco_hcd203040_compound15.24005.24005.4 | controllerType=0 controllerNumber=1 scan=24005 | 50.0316 |
| 20581:4 | 11 | K.ADTHDEILEGLNFN[+2204.772]LTEIPEAQIHEGFQELLR.T | HexNAc(4)Hex(5)NeuAc(2) | N[+2205] | 1474.9086 | 4 | 5896.6125 | 5896.5963 | 0 | 2.7 | 94 | Specific | 1277.92 | 1035.19 | 1035.19 | 26.75 | 42 | >sp\|P01009\|A1AT_HUMAN Alpha-1-antitrypsin OS=Homo sapiens OX=9606 GN=SERPINA1 PE=1 SV=3 | 642 | 20201113_gy_colo_milk_1ng_glyco_hcd203040_compound15.24003.24003.4 | controllerType=0 controllerNumber=1 scan=24003 | 50.0263 |
| 19658:4 | 11 | K.YLGN[+2204.772]ATAIFFLPDEGKLQHLENELTHDIITK.F | HexNAc(4)Hex(5)NeuAc(2) | N[+2205] | 1437.1628 | 4 | 5745.6296 | 5745.6098 | 0 | 3.4 | 268 | Specific | 313.98 | 290.15 | 290.15 | 6.92 | 42 | >sp\|P01009\|A1AT_HUMAN Alpha-1-antitrypsin OS=Homo sapiens OX=9606 GN=SERPINA1 PE=1 SV=3 | 642 | 20201113_gy_colo_milk_1ng_glyco_hcd203040_compound15.23037.23037.4 | controllerType=0 controllerNumber=1 scan=23037 | 47.7929 |
| 19659:5 | 11 | K.YLGN[+2204.772]ATAIFFLPDEGKLQHLENELTHDIITK.F | HexNAc(4)Hex(5)NeuAc(2) | N[+2205] | 1149.9304 | 5 | 5745.623 | 5745.6098 | 0 | 2.3 | 268 | Specific | 500.29 | 445.47 | 445.47 | 10.42 | 42 | >sp\|P01009\|A1AT_HUMAN Alpha-1-antitrypsin OS=Homo sapiens OX=9606 GN=SERPINA1 PE=1 SV=3 | 642 | 20201113_gy_colo_milk_1ng_glyco_hcd203040_compound15.23038.23038.5 | controllerType=0 controllerNumber=1 scan=23038 | 47.7984 |
| 19657:5 | 11 | K.YLGN[+2204.772]ATAIFFLPDEGKLQHLENELTHDIITK.F | HexNAc(4)Hex(5)NeuAc(2) | N[+2205] | 1149.9304 | 5 | 5745.623 | 5745.6098 | 0 | 2.3 | 268 | Specific | 811.4 | 688.35 | 688.35 | 17.17 | 42 | >sp\|P01009\|A1AT_HUMAN Alpha-1-antitrypsin OS=Homo sapiens OX=9606 GN=SERPINA1 PE=1 SV=3 | 642 | 20201113_gy_colo_milk_1ng_glyco_hcd203040_compound15.23036.23036.5 | controllerType=0 controllerNumber=1 scan=23036 | 47.7875 |
| 19656:4 | 11 | K.YLGN[+2204.772]ATAIFFLPDEGKLQHLENELTHDIITK.F | HexNAc(4)Hex(5)NeuAc(2) | N[+2205] | 1437.1628 | 4 | 5745.6296 | 5745.6098 | 0 | 3.4 | 268 | Specific | 1321 | 1230.84 | 1230.84 | 26.6 | 42 | >sp\|P01009\|A1AT_HUMAN Alpha-1-antitrypsin OS=Homo sapiens OX=9606 GN=SERPINA1 PE=1 SV=3 | 642 | 20201113_gy_colo_milk_1ng_glyco_hcd203040_compound15.23035.23035.4 | controllerType=0 controllerNumber=1 scan=23035 | 47.7861 |
| 23918:4 | 14 | K.APDKNVIFSPLSISTALAFLSLGAHN[+2861.000]TTLTEILK.G | HexNAc(5)Hex(6)NeuAc(3) | N[+2861] | 1611.7607 | 4 | 6444.0212 | 6443.9782 | 0 | 6.7 | 68 | Specific | 283.72 | 283.72 | 283.72 | 5.72 | 36 | >sp\|P01011\|AACT_HUMAN Alpha-1-antichymotrypsin OS=Homo sapiens OX=9606 GN=SERPINA3 PE=1 SV=2 | 1314 | 20201113_gy_colo_milk_1ng_glyco_hcd203040_compound15.27480.27480.4 | controllerType=0 controllerNumber=1 scan=27480 | 57.1813 |
| 20828:4 | 14 | K.APDKNVIFSPLSISTALAFLSLGAHN[+2570.925]TTLTEILK.G | HexNAc(5)Hex(6)Fuc(2)NeuAc(1) | N[+2571] | 1539.238 | 4 | 6153.9303 | 6153.9032 | 0 | 4.4 | 68 | Specific | 354.66 | 354.66 | 354.66 | 7.11 | 36 | >sp\|P01011\|AACT_HUMAN Alpha-1-antichymotrypsin OS=Homo sapiens OX=9606 GN=SERPINA3 PE=1 SV=2 | 1314 | 20201113_gy_colo_milk_1ng_glyco_hcd203040_compound15.24261.24261.4 | controllerType=0 controllerNumber=1 scan=24261 | 50.576 |
| 20831:4 | 14 | K.APDKNVIFSPLSISTALAFLSLGAHN[+2570.925]TTLTEILK.G | HexNAc(5)Hex(6)Fuc(2)NeuAc(1) | N[+2571] | 1539.238 | 4 | 6153.9303 | 6153.9032 | 0 | 4.4 | 68 | Specific | 470.92 | 470.92 | 470.92 | 9.15 | 36 | >sp\|P01011\|AACT_HUMAN Alpha-1-antichymotrypsin OS=Homo sapiens OX=9606 GN=SERPINA3 PE=1 SV=2 | 1314 | 20201113_gy_colo_milk_1ng_glyco_hcd203040_compound15.24264.24264.4 | controllerType=0 controllerNumber=1 scan=24264 | 50.5804 |
| 23799:4 | 14 | K.NVIFSPLSISTALAFLSLGAHN[+2350.830]TTLTEILK.G | HexNAc(4)Hex(5)Fuc(1)NeuAc(2) | N[+2351] | 1381.4121 | 4 | 5522.6266 | 5522.5967 | 0 | 5.4 | 72 | Specific | 496.68 | 496.68 | 496.68 | 10.48 | 36 | >sp\|P01011\|AACT_HUMAN Alpha-1-antichymotrypsin OS=Homo sapiens OX=9606 GN=SERPINA3 PE=1 SV=2 | 1314 | 20201113_gy_colo_milk_1ng_glyco_hcd203040_compound15.27357.27357.4 | controllerType=0 controllerNumber=1 scan=27357 | 56.9463 |
| 23801:4 | 14 | K.NVIFSPLSISTALAFLSLGAHN[+2350.830]TTLTEILK.G | HexNAc(4)Hex(5)Fuc(1)NeuAc(2) | N[+2351] | 1381.4121 | 4 | 5522.6266 | 5522.5967 | 0 | 5.4 | 72 | Specific | 635.26 | 635.26 | 635.26 | 13.23 | 36 | >sp\|P01011\|AACT_HUMAN Alpha-1-antichymotrypsin OS=Homo sapiens OX=9606 GN=SERPINA3 PE=1 SV=2 | 1314 | 20201113_gy_colo_milk_1ng_glyco_hcd203040_compound15.27359.27359.4 | controllerType=0 controllerNumber=1 scan=27359 | 56.9523 |
| 02305:3 | 15 | K.N[+1540.529]NSIPDKQITASSSYK.T | HexNAc(2)Hex(7) | N[+1541] | 1098.476 | 3 | 3293.4134 | 3293.4048 | 0 | 2.6 | 238 | Specific | 387.66 | 168.82 | 168.82 | 6.18 | 29 | >sp\|Q08431\|MFGM_HUMAN Lactadherin OS=Homo sapiens OX=9606 GN=MFGE8 PE=1 SV=3 | 1064 | 20201113_gy_colo_milk_1ng_glyco_hcd203040_compound15.5028.5028.3 | controllerType=0 controllerNumber=1 scan=5028 | 14.4257 |
| 02307:3 | 15 | K.N[+1540.529]NSIPDKQITASSSYK.T | HexNAc(2)Hex(7) | N[+1541] | 1098.476 | 3 | 3293.4134 | 3293.4048 | 0 | 2.6 | 238 | Specific | 390.54 | 156.35 | 156.35 | 6.17 | 29 | >sp\|Q08431\|MFGM_HUMAN Lactadherin OS=Homo sapiens OX=9606 GN=MFGE8 PE=1 SV=3 | 1064 | 20201113_gy_colo_milk_1ng_glyco_hcd203040_compound15.5030.5030.3 | controllerType=0 controllerNumber=1 scan=5030 | 14.43 |
| 02423:3 | 15 | K.N[+1216.423]NSIPDKQITASSSYK.T | HexNAc(2)Hex(5) | N[+1216] | 990.4408 | 3 | 2969.3077 | 2969.2991 | 0 | 2.9 | 238 | Specific | 509.77 | 282.96 | 282.96 | 7.44 | 29 | >sp\|Q08431\|MFGM_HUMAN Lactadherin OS=Homo sapiens OX=9606 GN=MFGE8 PE=1 SV=3 | 1064 | 20201113_gy_colo_milk_1ng_glyco_hcd203040_compound15.5150.5150.3 | controllerType=0 controllerNumber=1 scan=5150 | 14.6478 |
| 02421:3 | 15 | K.N[+1216.423]NSIPDKQITASSSYK.T | HexNAc(2)Hex(5) | N[+1216] | 990.4408 | 3 | 2969.3077 | 2969.2991 | 0 | 2.9 | 238 | Specific | 661.08 | 424.12 | 424.12 | 13.28 | 29 | >sp\|Q08431\|MFGM_HUMAN Lactadherin OS=Homo sapiens OX=9606 GN=MFGE8 PE=1 SV=3 | 1064 | 20201113_gy_colo_milk_1ng_glyco_hcd203040_compound15.5148.5148.3 | controllerType=0 controllerNumber=1 scan=5148 | 14.6439 |
| 08654:3 | 15 | K.VAYSN[+2205.793]DSANWTEYQDPR.T | HexNAc(4)Hex(5)Fuc(2)NeuAc(1) | N[+2206] | 1407.8985 | 3 | 4221.6809 | 4221.6658 | 0 | 3.6 | 321 | Specific | 384.19 | 337.34 | 6.04 | 8.22 | 29 | >sp\|Q08431\|MFGM_HUMAN Lactadherin OS=Homo sapiens OX=9606 GN=MFGE8 PE=1 SV=3 | 1064 | 20201113_gy_colo_milk_1ng_glyco_hcd203040_compound15.11601.11601.3 | controllerType=0 controllerNumber=1 scan=11601 | 25.5129 |
| 08643:3 | 15 | K.VAYSN[+2205.793]DSANWTEYQDPR.T | HexNAc(4)Hex(5)Fuc(2)NeuAc(1) | N[+2206] | 1407.8969 | 3 | 4221.6762 | 4221.6658 | 0 | 2.5 | 321 | Specific | 526.42 | 444.14 | 7.9 | 10.52 | 29 | >sp\|Q08431\|MFGM_HUMAN Lactadherin OS=Homo sapiens OX=9606 GN=MFGE8 PE=1 SV=3 | 1064 | 20201113_gy_colo_milk_1ng_glyco_hcd203040_compound15.11589.11589.3 | controllerType=0 controllerNumber=1 scan=11589 | 25.4923 |
| 08656:3 | 15 | K.VAYSN[+2205.793]DSANWTEYQDPR.T | HexNAc(4)Hex(5)Fuc(2)NeuAc(1) | N[+2206] | 1407.8985 | 3 | 4221.6809 | 4221.6658 | 0 | 3.6 | 321 | Specific | 552.88 | 444.97 | 3.49 | 10.67 | 29 | >sp\|Q08431\|MFGM_HUMAN Lactadherin OS=Homo sapiens OX=9606 GN=MFGE8 PE=1 SV=3 | 1064 | 20201113_gy_colo_milk_1ng_glyco_hcd203040_compound15.11603.11603.3 | controllerType=0 controllerNumber=1 scan=11603 | 25.5163 |
| 08645:3 | 15 | K.VAYSN[+2205.793]DSANWTEYQDPR.T | HexNAc(4)Hex(5)Fuc(2)NeuAc(1) | N[+2206] | 1407.8969 | 3 | 4221.6762 | 4221.6658 | 0 | 2.5 | 321 | Specific | 594.58 | 520.08 | 3.22 | 12.22 | 29 | >sp\|Q08431\|MFGM_HUMAN Lactadherin OS=Homo sapiens OX=9606 GN=MFGE8 PE=1 SV=3 | 1064 | 20201113_gy_colo_milk_1ng_glyco_hcd203040_compound15.11591.11591.3 | controllerType=0 controllerNumber=1 scan=11591 | 25.496 |
| 13756:4 | 16 | K.LKELPGVC[+57.021]N[+2571.946]ETMMALWEEC[+57.021]KPC[+57.021]LK.Q | HexNAc(5)Hex(6)Fuc(4) | C[+57]*3, N[+2572] | 1377.8345 | 4 | 5508.3161 | 5508.3302 | 0 | -2.6 | 95 | Specific | 365.77 | 365.77 | 365.77 | 6.68 | 30 | >sp\|P10909\|CLUS_HUMAN Clusterin OS=Homo sapiens OX=9606 GN=CLU PE=1 SV=1 | 1467 | 20201113_gy_colo_milk_1ng_glyco_hcd203040_compound15.16892.16892.4 | controllerType=0 controllerNumber=1 scan=16892 | 35.2039 |
| 13758:4 | 16 | K.LKELPGVC[+57.021]N[+2571.946]ETMMALWEEC[+57.021]KPC[+57.021]LK.Q | HexNAc(5)Hex(6)Fuc(4) | C[+57]*3, N[+2572] | 1377.8345 | 4 | 5508.3161 | 5508.3302 | 0 | -2.6 | 95 | Specific | 396.04 | 396.04 | 396.04 | 8.23 | 30 | >sp\|P10909\|CLUS_HUMAN Clusterin OS=Homo sapiens OX=9606 GN=CLU PE=1 SV=1 | 1467 | 20201113_gy_colo_milk_1ng_glyco_hcd203040_compound15.16894.16894.4 | controllerType=0 controllerNumber=1 scan=16894 | 35.2075 |
| 13887:4 | 16 | K.LKELPGVC[+57.021]N[+2205.793]ETMMALWEEC[+57.021]KPC[+57.021]LK.Q | HexNAc(4)Hex(5)Fuc(2)NeuAc(1) | C[+57]*3, N[+2206] | 1286.304 | 4 | 5142.194 | 5142.1776 | 0 | 3.2 | 95 | Specific | 411.4 | 411.4 | 411.4 | 9.35 | 30 | >sp\|P10909\|CLUS_HUMAN Clusterin OS=Homo sapiens OX=9606 GN=CLU PE=1 SV=1 | 1467 | 20201113_gy_colo_milk_1ng_glyco_hcd203040_compound15.17028.17028.4 | controllerType=0 controllerNumber=1 scan=17028 | 35.4521 |
| 13889:4 | 16 | K.LKELPGVC[+57.021]N[+2205.793]ETMMALWEEC[+57.021]KPC[+57.021]LK.Q | HexNAc(4)Hex(5)Fuc(2)NeuAc(1) | C[+57]*3, N[+2206] | 1286.304 | 4 | 5142.194 | 5142.1776 | 0 | 3.2 | 95 | Specific | 654.69 | 528.08 | 528.08 | 13.47 | 30 | >sp\|P10909\|CLUS_HUMAN Clusterin OS=Homo sapiens OX=9606 GN=CLU PE=1 SV=1 | 1467 | 20201113_gy_colo_milk_1ng_glyco_hcd203040_compound15.17030.17030.4 | controllerType=0 controllerNumber=1 scan=17030 | 35.456 |
| 13924:4 | 16 | K.LKELPGVC[+57.021]N[+2059.735]ETMMALWEEC[+57.021]KPC[+57.021]LK.Q | HexNAc(4)Hex(5)Fuc(1)NeuAc(1) | C[+57]*3, N[+2060] | 1249.7889 | 4 | 4996.1339 | 4996.1196 | 0 | 2.9 | 95 | Specific | 867.37 | 729.96 | 729.96 | 17.47 | 30 | >sp\|P10909\|CLUS_HUMAN Clusterin OS=Homo sapiens OX=9606 GN=CLU PE=1 SV=1 | 1467 | 20201113_gy_colo_milk_1ng_glyco_hcd203040_compound15.17066.17066.4 | controllerType=0 controllerNumber=1 scan=17066 | 35.5377 |
| 13922:4 | 16 | K.LKELPGVC[+57.021]N[+2059.735]ETMMALWEEC[+57.021]KPC[+57.021]LK.Q | HexNAc(4)Hex(5)Fuc(1)NeuAc(1) | C[+57]*3, N[+2060] | 1249.7889 | 4 | 4996.1339 | 4996.1196 | 0 | 2.9 | 95 | Specific | 1011.08 | 985.25 | 985.25 | 20.47 | 30 | >sp\|P10909\|CLUS_HUMAN Clusterin OS=Homo sapiens OX=9606 GN=CLU PE=1 SV=1 | 1467 | 20201113_gy_colo_milk_1ng_glyco_hcd203040_compound15.17064.17064.4 | controllerType=0 controllerNumber=1 scan=17064 | 35.5326 |
| 21127:3 | 16 | K.MLN[+1458.442]TSSLLEQLNEQFNWVSR.L | HexNAc(2)Hex(6)Phospho(1) | N[+1458] | 1289.8832 | 3 | 3867.6352 | 3867.6287 | 0 | 1.7 | 352 | Specific | 308.79 | 217.81 | 217.81 | 7.21 | 30 | >sp\|P10909\|CLUS_HUMAN Clusterin OS=Homo sapiens OX=9606 GN=CLU PE=1 SV=1 | 1467 | 20201113_gy_colo_milk_1ng_glyco_hcd203040_compound15.24573.24573.3 | controllerType=0 controllerNumber=1 scan=24573 | 51.2656 |
| 21129:3 | 16 | K.MLN[+1458.442]TSSLLEQLNEQFNWVSR.L | HexNAc(2)Hex(6)Phospho(1) | N[+1458] | 1289.8832 | 3 | 3867.6352 | 3867.6287 | 0 | 1.7 | 352 | Specific | 617.21 | 522.62 | 522.62 | 13.22 | 30 | >sp\|P10909\|CLUS_HUMAN Clusterin OS=Homo sapiens OX=9606 GN=CLU PE=1 SV=1 | 1467 | 20201113_gy_colo_milk_1ng_glyco_hcd203040_compound15.24575.24575.3 | controllerType=0 controllerNumber=1 scan=24575 | 51.2691 |
| 20490:3 | 16 | K.MLN[+2205.793]TSSLLEQLNEQFNWVSR.L | HexNAc(4)Hex(5)Fuc(2)NeuAc(1) | N[+2206] | 1539.0027 | 3 | 4614.9937 | 4614.9796 | 0 | 3.1 | 352 | Specific | 671.35 | 460.38 | 460.38 | 14.21 | 30 | >sp\|P10909\|CLUS_HUMAN Clusterin OS=Homo sapiens OX=9606 GN=CLU PE=1 SV=1 | 1467 | 20201113_gy_colo_milk_1ng_glyco_hcd203040_compound15.23907.23907.3 | controllerType=0 controllerNumber=1 scan=23907 | 49.7813 |
| 22529:4 | 16 | K.MLN[+2350.830]TSSLLEQLNEQFNWVSR.L | HexNAc(4)Hex(5)Fuc(1)NeuAc(2) | N[+2351] | 1190.7654 | 4 | 4760.0399 | 4760.0171 | 0 | 4.8 | 352 | Specific | 825.3 | 594.18 | 594.18 | 16.66 | 30 | >sp\|P10909\|CLUS_HUMAN Clusterin OS=Homo sapiens OX=9606 GN=CLU PE=1 SV=1 | 1467 | 20201113_gy_colo_milk_1ng_glyco_hcd203040_compound15.26035.26035.4 | controllerType=0 controllerNumber=1 scan=26035 | 54.2955 |
| 20493:3 | 16 | K.MLN[+2205.793]TSSLLEQLNEQFNWVSR.L | HexNAc(4)Hex(5)Fuc(2)NeuAc(1) | N[+2206] | 1539.0027 | 3 | 4614.9937 | 4614.9796 | 0 | 3.1 | 352 | Specific | 840.85 | 564.5 | 564.5 | 17.21 | 30 | >sp\|P10909\|CLUS_HUMAN Clusterin OS=Homo sapiens OX=9606 GN=CLU PE=1 SV=1 | 1467 | 20201113_gy_colo_milk_1ng_glyco_hcd203040_compound15.23910.23910.3 | controllerType=0 controllerNumber=1 scan=23910 | 49.7867 |
| 22523:3 | 16 | K.MLN[+2350.830]TSSLLEQLNEQFNWVSR.L | HexNAc(4)Hex(5)Fuc(1)NeuAc(2) | N[+2351] | 1587.35 | 3 | 4760.0355 | 4760.0171 | 0 | 3.9 | 352 | Specific | 844.48 | 477.44 | 477.44 | 16.88 | 30 | >sp\|P10909\|CLUS_HUMAN Clusterin OS=Homo sapiens OX=9606 GN=CLU PE=1 SV=1 | 1467 | 20201113_gy_colo_milk_1ng_glyco_hcd203040_compound15.26029.26029.3 | controllerType=0 controllerNumber=1 scan=26029 | 54.2857 |
| 20526:4 | 16 | K.MLN[+2059.735]TSSLLEQLNEQFNWVSR.L | HexNAc(4)Hex(5)Fuc(1)NeuAc(1) | N[+2060] | 1117.9911 | 4 | 4468.9427 | 4468.9217 | 0 | 4.7 | 352 | Specific | 869.99 | 647.21 | 647.21 | 18.01 | 30 | >sp\|P10909\|CLUS_HUMAN Clusterin OS=Homo sapiens OX=9606 GN=CLU PE=1 SV=1 | 1467 | 20201113_gy_colo_milk_1ng_glyco_hcd203040_compound15.23945.23945.4 | controllerType=0 controllerNumber=1 scan=23945 | 49.8706 |
| 22532:4 | 16 | K.MLN[+2350.830]TSSLLEQLNEQFNWVSR.L | HexNAc(4)Hex(5)Fuc(1)NeuAc(2) | N[+2351] | 1190.7654 | 4 | 4760.0399 | 4760.0171 | 0 | 4.8 | 352 | Specific | 881.23 | 630.09 | 630.09 | 17.66 | 30 | >sp\|P10909\|CLUS_HUMAN Clusterin OS=Homo sapiens OX=9606 GN=CLU PE=1 SV=1 | 1467 | 20201113_gy_colo_milk_1ng_glyco_hcd203040_compound15.26038.26038.4 | controllerType=0 controllerNumber=1 scan=26038 | 54.3025 |
| 20495:3 | 16 | K.MLN[+2059.735]TSSLLEQLNEQFNWVSR.L | HexNAc(4)Hex(5)Fuc(1)NeuAc(1) | N[+2060] | 1490.3167 | 3 | 4468.9356 | 4468.9217 | 0 | 3.1 | 352 | Specific | 987.33 | 728.81 | 728.81 | 20.21 | 30 | >sp\|P10909\|CLUS_HUMAN Clusterin OS=Homo sapiens OX=9606 GN=CLU PE=1 SV=1 | 1467 | 20201113_gy_colo_milk_1ng_glyco_hcd203040_compound15.23912.23912.3 | controllerType=0 controllerNumber=1 scan=23912 | 49.7966 |
| 20529:4 | 16 | K.MLN[+2059.735]TSSLLEQLNEQFNWVSR.L | HexNAc(4)Hex(5)Fuc(1)NeuAc(1) | N[+2060] | 1117.9911 | 4 | 4468.9427 | 4468.9217 | 0 | 4.7 | 352 | Specific | 1009.7 | 756.55 | 756.55 | 21.01 | 30 | >sp\|P10909\|CLUS_HUMAN Clusterin OS=Homo sapiens OX=9606 GN=CLU PE=1 SV=1 | 1467 | 20201113_gy_colo_milk_1ng_glyco_hcd203040_compound15.23948.23948.4 | controllerType=0 controllerNumber=1 scan=23948 | 49.8755 |
| 20492:3 | 16 | K.MLN[+2059.735]TSSLLEQLNEQFNWVSR.L | HexNAc(4)Hex(5)Fuc(1)NeuAc(1) | N[+2060] | 1490.3167 | 3 | 4468.9356 | 4468.9217 | 0 | 3.1 | 352 | Specific | 1071.99 | 822.46 | 822.46 | 22.21 | 30 | >sp\|P10909\|CLUS_HUMAN Clusterin OS=Homo sapiens OX=9606 GN=CLU PE=1 SV=1 | 1467 | 20201113_gy_colo_milk_1ng_glyco_hcd203040_compound15.23909.23909.3 | controllerType=0 controllerNumber=1 scan=23909 | 49.7837 |
| 22521:3 | 16 | K.MLN[+2350.830]TSSLLEQLNEQFNWVSR.L | HexNAc(4)Hex(5)Fuc(1)NeuAc(2) | N[+2351] | 1587.35 | 3 | 4760.0355 | 4760.0171 | 0 | 3.9 | 352 | Specific | 1079.35 | 757.98 | 757.98 | 21.88 | 30 | >sp\|P10909\|CLUS_HUMAN Clusterin OS=Homo sapiens OX=9606 GN=CLU PE=1 SV=1 | 1467 | 20201113_gy_colo_milk_1ng_glyco_hcd203040_compound15.26027.26027.3 | controllerType=0 controllerNumber=1 scan=26027 | 54.2805 |
| 07941:3 | 17 | R.LSPN[+2350.830]ASAEHLELR.W | HexNAc(4)Hex(5)Fuc(1)NeuAc(2) | N[+2351] | 1263.2014 | 3 | 3787.5895 | 3787.5796 | 0 | 2.6 | 52 | Specific | 397.31 | 397.31 | 397.31 | 9.59 | 30 | >sp\|Q13410\|BT1A1_HUMAN Butyrophilin subfamily 1 member A1 OS=Homo sapiens OX=9606 GN=BTN1A1 PE=1 SV=3 | 179 | 20201113_gy_colo_milk_1ng_glyco_hcd203040_compound15.10862.10862.3 | controllerType=0 controllerNumber=1 scan=10862 | 24.2253 |
| 07943:3 | 17 | R.LSPN[+2350.830]ASAEHLELR.W | HexNAc(4)Hex(5)Fuc(1)NeuAc(2) | N[+2351] | 1263.2014 | 3 | 3787.5895 | 3787.5796 | 0 | 2.6 | 52 | Specific | 469.19 | 469.19 | 469.19 | 9.88 | 30 | >sp\|Q13410\|BT1A1_HUMAN Butyrophilin subfamily 1 member A1 OS=Homo sapiens OX=9606 GN=BTN1A1 PE=1 SV=3 | 179 | 20201113_gy_colo_milk_1ng_glyco_hcd203040_compound15.10864.10864.3 | controllerType=0 controllerNumber=1 scan=10864 | 24.2291 |
| 21660:5 | 19 | K.VFPLSLDSTPQDGNVVVAC[+57.021]LVQGFFPQEPLSVTWSESGQN[+1444.534]VTAI | HexNAc(4)Hex(3)Fuc(1) | C[+57], N[+1445] | 1244.9867 | 5 | 6220.9044 | 6220.8953 | 0 | 1.5 | 123 | Specific | 234.11 | 234.11 | 234.11 | 5.75 | 44 | >sp\|P0DOX2\|IGA2_HUMAN Immunoglobulin alpha-2 heavy chain OS=Homo sapiens OX=9606 PE=1 SV=2 | 631 | 20201113_gy_colo_milk_1ng_glyco_hcd203040_compound15.25127.25127.2 | controllerType=0 controllerNumber=1 scan=25127 | 52.3899 |
| 21679:6 | 19 | K.VFPLSLDSTPQDGNVVVAC[+57.021]LVQGFFPQEPLSVTWSESGQN[+1647.613]VTAI | HexNAc(5)Hex(3)Fuc(1) | C[+57], N[+1648] | 1071.5125 | 6 | 6424.0386 | 6423.9746 | 0 | 10 | 123 | Specific | 239.58 | 239.58 | 239.58 | 4.47 | 44 | >sp\|P0DOX2\|IGA2_HUMAN Immunoglobulin alpha-2 heavy chain OS=Homo sapiens OX=9606 PE=1 SV=2 | 631 | 20201113_gy_colo_milk_1ng_glyco_hcd203040_compound15.25147.25147.6 | controllerType=0 controllerNumber=1 scan=25147 | 52.4299 |
| 22974:6 | 19 | K.VFPLSLDSTPQDGNVVVAC[+57.021]LVQGFFPQEPLSVTWSESGQN[+2262.814]VTAI | HexNAc(5)Hex(5)Fuc(1)NeuAc(1) | C[+57], N[+2263] | 1174.0439 | 6 | 7039.2269 | 7039.1757 | 0 | 7.3 | 123 | Specific | 274.65 | 274.65 | 274.65 | 6.84 | 44 | >sp\|P0DOX2\|IGA2_HUMAN Immunoglobulin alpha-2 heavy chain OS=Homo sapiens OX=9606 PE=1 SV=2 | 631 | 20201113_gy_colo_milk_1ng_glyco_hcd203040_compound15.26498.26498.6 | controllerType=0 controllerNumber=1 scan=26498 | 55.2351 |
| 22903:4 | 19 | K.VFPLSLDSTPQDGNVVVAC[+57.021]LVQGFFPQEPLSVTWSESGQN[+2059.735]VTAI | HexNAc(4)Hex(5)Fuc(1)NeuAc(1) | C[+57], N[+2060] | 1709.786 | 4 | 6836.1221 | 6836.0963 | 0 | 3.8 | 123 | Specific | 320.08 | 282.54 | 282.54 | 7.88 | 44 | >sp\|P0DOX2\|IGA2_HUMAN Immunoglobulin alpha-2 heavy chain OS=Homo sapiens OX=9606 PE=1 SV=2 | 631 | 20201113_gy_colo_milk_1ng_glyco_hcd203040_compound15.26424.26424.4 | controllerType=0 controllerNumber=1 scan=26424 | 55.0776 |
| 21658:5 | 19 | K.VFPLSLDSTPQDGNVVVAC[+57.021]LVQGFFPQEPLSVTWSESGQN[+1444.534]VTAI | HexNAc(4)Hex(3)Fuc(1) | C[+57], N[+1445] | 1244.9867 | 5 | 6220.9044 | 6220.8953 | 0 | 1.5 | 123 | Specific | 337.86 | 337.86 | 337.86 | 7.41 | 44 | >sp\|P0DOX2\|IGA2_HUMAN Immunoglobulin alpha-2 heavy chain OS=Homo sapiens OX=9606 PE=1 SV=2 | 631 | 20201113_gy_colo_milk_1ng_glyco_hcd203040_compound15.25125.25125.2 | controllerType=0 controllerNumber=1 scan=25125 | 52.3844 |
| 21441:4 | 19 | K.VFPLSLDSTPQDGNVVVAC[+57.021]LVQGFFPQEPLSVTWSESGQN[+1971.719]VTAI | HexNAc(5)Hex(5)Fuc(1) | C[+57], N[+1972] | 1687.788 | 4 | 6748.1304 | 6748.0803 | 0 | 7.4 | 123 | Specific | 355.33 | 355.33 | 355.33 | 8.02 | 44 | >sp\|P0DOX2\|IGA2_HUMAN Immunoglobulin alpha-2 heavy chain OS=Homo sapiens OX=9606 PE=1 SV=2 | 631 | 20201113_gy_colo_milk_1ng_glyco_hcd203040_compound15.24900.24900.4 | controllerType=0 controllerNumber=1 scan=24900 | 51.9256 |
| 21651:5 | 19 | K.VFPLSLDSTPQDGNVVVAC[+57.021]LVQGFFPQEPLSVTWSESGQN[+1444.534]VTAI | HexNAc(4)Hex(3)Fuc(1) | C[+57], N[+1445] | 1244.9867 | 5 | 6220.9044 | 6220.8953 | 0 | 1.5 | 123 | Specific | 378.01 | 378.01 | 378.01 | 7.83 | 44 | >sp\|P0DOX2\|IGA2_HUMAN Immunoglobulin alpha-2 heavy chain OS=Homo sapiens OX=9606 PE=1 SV=2 | 631 | 20201113_gy_colo_milk_1ng_glyco_hcd203040_compound15.25118.25118.5 | controllerType=0 controllerNumber=1 scan=25118 | 52.3716 |
| 21593:5 | 19 | K.VFPLSLDSTPQDGNVVVAC[+57.021]LVQGFFPQEPLSVTWSESGQN[+1444.534]VTAI | HexNAc(4)Hex(3)Fuc(1) | C[+57], N[+1445] | 1244.9866 | 5 | 6220.9038 | 6220.8953 | 0 | 1.4 | 123 | Specific | 414.15 | 414.15 | 414.15 | 10.64 | 44 | >sp\|P0DOX2\|IGA2_HUMAN Immunoglobulin alpha-2 heavy chain OS=Homo sapiens OX=9606 PE=1 SV=2 | 631 | 20201113_gy_colo_milk_1ng_glyco_hcd203040_compound15.25058.25058.5 | controllerType=0 controllerNumber=1 scan=25058 | 52.2556 |
| 21590:5 | 19 | K.VFPLSLDSTPQDGNVVVAC[+57.021]LVQGFFPQEPLSVTWSESGQN[+1444.534]VTAI | HexNAc(4)Hex(3)Fuc(1) | C[+57], N[+1445] | 1244.9866 | 5 | 6220.9038 | 6220.8953 | 0 | 1.4 | 123 | Specific | 474.85 | 474.85 | 474.85 | 10.91 | 44 | >sp\|P0DOX2\|IGA2_HUMAN Immunoglobulin alpha-2 heavy chain OS=Homo sapiens OX=9606 PE=1 SV=2 | 631 | 20201113_gy_colo_milk_1ng_glyco_hcd203040_compound15.25055.25055.5 | controllerType=0 controllerNumber=1 scan=25055 | 52.2495 |
| 21443:4 | 19 | K.VFPLSLDSTPQDGNVVVAC[+57.021]LVQGFFPQEPLSVTWSESGQN[+1971.719]VTAI | HexNAc(5)Hex(5)Fuc(1) | C[+57], N[+1972] | 1687.788 | 4 | 6748.1304 | 6748.0803 | 0 | 7.4 | 123 | Specific | 485.84 | 484.4 | 484.4 | 10.11 | 44 | >sp\|P0DOX2\|IGA2_HUMAN Immunoglobulin alpha-2 heavy chain OS=Homo sapiens OX=9606 PE=1 SV=2 | 631 | 20201113_gy_colo_milk_1ng_glyco_hcd203040_compound15.24902.24902.4 | controllerType=0 controllerNumber=1 scan=24902 | 51.933 |
| 21624:4 | 19 | K.VFPLSLDSTPQDGNVVVAC[+57.021]LVQGFFPQEPLSVTWSESGQN[+1444.534]VTAI | HexNAc(4)Hex(3)Fuc(1) | C[+57], N[+1445] | 1555.9825 | 4 | 6220.9084 | 6220.8953 | 0 | 2.1 | 123 | Specific | 550.18 | 550.18 | 550.18 | 11.35 | 44 | >sp\|P0DOX2\|IGA2_HUMAN Immunoglobulin alpha-2 heavy chain OS=Homo sapiens OX=9606 PE=1 SV=2 | 631 | 20201113_gy_colo_milk_1ng_glyco_hcd203040_compound15.25090.25090.4 | controllerType=0 controllerNumber=1 scan=25090 | 52.318 |
| 22900:4 | 19 | K.VFPLSLDSTPQDGNVVVAC[+57.021]LVQGFFPQEPLSVTWSESGQN[+2262.814]VTAI | HexNAc(5)Hex(5)Fuc(1)NeuAc(1) | C[+57], N[+2263] | 1760.5542 | 4 | 7039.1948 | 7039.1757 | 0 | 2.7 | 123 | Specific | 562.84 | 518.94 | 518.94 | 11.54 | 44 | >sp\|P0DOX2\|IGA2_HUMAN Immunoglobulin alpha-2 heavy chain OS=Homo sapiens OX=9606 PE=1 SV=2 | 631 | 20201113_gy_colo_milk_1ng_glyco_hcd203040_compound15.26421.26421.4 | controllerType=0 controllerNumber=1 scan=26421 | [illegible] |
| 22905:4 | 19 | K.VFPLSLDSTPQDGNVVVAC[+57.021]LVQGFFPQEPLSVTWSESGQN[+2059.735]VTAI | HexNAc(4)Hex(5)Fuc(1)NeuAc(1) | C[+57], N[+2060] | 1709.786 | 4 | 6836.1221 | 6836.0963 | 0 | 3.8 | 123 | Specific | 660.77 | 609.22 | 609.22 | 14.55 | 44 | >sp\|P0DOX2\|IGA2_HUMAN Immunoglobulin alpha-2 heavy chain OS=Homo sapiens OX=9606 PE=1 SV=2 | 631 | 20201113_gy_colo_milk_1ng_glyco_hcd203040_compound15.26426.26426.4 | controllerType=0 controllerNumber=1 scan=26426 | [illegible] |
| 21392:4 | 19 | K.VFPLSLDSTPQDGNVVVAC[+57.021]LVQGFFPQEPLSVTWSESGQN[+1809.666]VTAI | HexNAc(5)Hex(4)Fuc(1) | C[+57], N[+1810] | 1647.2671 | 4 | 6586.0465 | 6586.0275 | 0 | 2.9 | 123 | Specific | 677.89 | 666.62 | 666.62 | 14.89 | 44 | >sp\|P0DOX2\|IGA2_HUMAN Immunoglobulin alpha-2 heavy chain OS=Homo sapiens OX=9606 PE=1 SV=2 | 631 | 20201113_gy_colo_milk_1ng_glyco_hcd203040_compound15.24849.24849.4 | controllerType=0 controllerNumber=1 scan=24849 | [illegible] |
| 21375:4 | 19 | K.VFPLSLDSTPQDGNVVVAC[+57.021]LVQGFFPQEPLSVTWSESGQN[+1971.719]VTAI | HexNAc(5)Hex(5)Fuc(1) | C[+57], N[+1972] | 1687.783 | 4 | 6748.1103 | 6748.0803 | 0 | 4.5 | 123 | Specific | 690.82 | 625.93 | 625.93 | 14.55 | 44 | >sp\|P0DOX2\|IGA2_HUMAN Immunoglobulin alpha-2 heavy chain OS=Homo sapiens OX=9606 PE=1 SV=2 | 631 | 20201113_gy_colo_milk_1ng_glyco_hcd203040_compound15.24831.24831.4 | controllerType=0 controllerNumber=1 scan=24831 | [illegible] |
| 21510:4 | 19 | K.VFPLSLDSTPQDGNVVVAC[+57.021]LVQGFFPQEPLSVTWSESGQN[+1647.613]VTAI | HexNAc(5)Hex(3)Fuc(1) | C[+57], N[+1648] | 1606.7561 | 4 | 6424.0026 | 6423.9746 | 0 | 4.4 | 123 | Specific | 720.11 | 672.89 | 672.89 | 15.55 | 44 | >sp\|P0DOX2\|IGA2_HUMAN Immunoglobulin alpha-2 heavy chain OS=Homo sapiens OX=9606 PE=1 SV=2 | 631 | 20201113_gy_colo_milk_1ng_glyco_hcd203040_compound15.24972.24972.4 | controllerType=0 controllerNumber=1 scan=24972 | 52.0716 |
| 22902:4 | 19 | K.VFPLSLDSTPQDGNVVVAC[+57.021]LVQGFFPQEPLSVTWSESGQN[+2262.814]VTAI | HexNAc(5)Hex(5)Fuc(1)NeuAc(1) | C[+57], N[+2263] | 1760.5542 | 4 | 7039.1948 | 7039.1757 | 0 | 2.7 | 123 | Specific | 791.29 | 776.92 | 776.92 | 16.89 | 44 | >sp\|P0DOX2\|IGA2_HUMAN Immunoglobulin alpha-2 heavy chain OS=Homo sapiens OX=9606 PE=1 SV=2 | 631 | 20201113_gy_colo_milk_1ng_glyco_hcd203040_compound15.26423.26423.4 | controllerType=0 controllerNumber=1 scan=26423 | 55.0757 |
| 21394:4 | 19 | K.VFPLSLDSTPQDGNVVVAC[+57.021]LVQGFFPQEPLSVTWSESGQN[+1809.666]VTAI | HexNAc(5)Hex(4)Fuc(1) | C[+57], N[+1810] | 1647.2671 | 4 | 6586.0465 | 6586.0275 | 0 | 2.9 | 123 | Specific | 817.05 | 777.62 | 777.62 | 17.89 | 44 | >sp\|P0DOX2\|IGA2_HUMAN Immunoglobulin alpha-2 heavy chain OS=Homo sapiens OX=9606 PE=1 SV=2 | 631 | 20201113_gy_colo_milk_1ng_glyco_hcd203040_compound15.24851.24851.4 | controllerType=0 controllerNumber=1 scan=24851 | 51.8255 |
| 21512:4 | 19 | K.VFPLSLDSTPQDGNVVVAC[+57.021]LVQGFFPQEPLSVTWSESGQN[+1647.613]VTAI | HexNAc(5)Hex(3)Fuc(1) | C[+57], N[+1648] | 1606.7561 | 4 | 6424.0026 | 6423.9746 | 0 | 4.4 | 123 | Specific | 930.73 | 923.85 | 923.85 | 19.55 | 44 | >sp\|P0DOX2\|IGA2_HUMAN Immunoglobulin alpha-2 heavy chain OS=Homo sapiens OX=9606 PE=1 SV=2 | 631 | 20201113_gy_colo_milk_1ng_glyco_hcd203040_compound15.24974.24974.4 | controllerType=0 controllerNumber=1 scan=24974 | 52.0783 |
| 21627:4 | 19 | K.VFPLSLDSTPQDGNVVVAC[+57.021]LVQGFFPQEPLSVTWSESGQN[+1444.534]VTAI | HexNAc(4)Hex(3)Fuc(1) | C[+57], N[+1445] | 1555.9825 | 4 | 6220.9084 | 6220.8953 | 0 | 2.1 | 123 | Specific | 962.5 | 939.65 | 939.65 | 20.89 | 44 | >sp\|P0DOX2\|IGA2_HUMAN Immunoglobulin alpha-2 heavy chain OS=Homo sapiens OX=9606 PE=1 SV=2 | 631 | 20201113_gy_colo_milk_1ng_glyco_hcd203040_compound15.25093.25093.4 | controllerType=0 controllerNumber=1 scan=25093 | 52.3229 |
| 03239:2 | 19 | K.TPLTAN[+1403.507]ITK.S | HexNAc(3)Hex(4)Fuc(1) | N[+1404] | 1181.5387 | 2 | 2362.0702 | 2362.0641 | 0 | 2.6 | 315 | Specific | 269.43 | 238.45 | 238.45 | 6.63 | 44 | >sp\|P0DOX2\|IGA2_HUMAN Immunoglobulin alpha-2 heavy chain OS=Homo sapiens OX=9606 PE=1 SV=2 | 631 | 20201113_gy_colo_milk_1ng_glyco_hcd203040_compound15.5994.5994.2 | controllerType=0 controllerNumber=1 scan=5994 | 16.0673 |
| 05034:3 | 19 | K.TPLTAN[+2262.814]ITK.S | HexNAc(5)Hex(5)Fuc(1)NeuAc(1) | N[+2263] | 1074.4649 | 3 | 3221.3801 | 3221.3711 | 0 | 2.8 | 315 | Specific | 274.33 | 103.11 | 103.11 | 5.05 | 44 | >sp\|P0DOX2\|IGA2_HUMAN Immunoglobulin alpha-2 heavy chain OS=Homo sapiens OX=9606 PE=1 SV=2 | 631 | 20201113_gy_colo_milk_1ng_glyco_hcd203040_compound15.7852.7852.3 | controllerType=0 controllerNumber=1 scan=7852 | 19.1863 |
| 03367:3 | 19 | K.TPLTAN[+1444.534]ITK.S | HexNAc(4)Hex(3)Fuc(1) | N[+1445] | 801.7044 | 3 | 2403.0986 | 2403.0906 | 0 | 3.3 | 315 | Specific | 286.27 | 134.33 | 134.33 | 5.95 | 44 | >sp\|P0DOX2\|IGA2_HUMAN Immunoglobulin alpha-2 heavy chain OS=Homo sapiens OX=9606 PE=1 SV=2 | 631 | 20201113_gy_colo_milk_1ng_glyco_hcd203040_compound15.6127.6127.3 | controllerType=0 controllerNumber=1 scan=6127 | 16.3267 |
| 03212:2 | 19 | K.TPLTAN[+1971.719]ITK.S | HexNAc(5)Hex(5)Fuc(1) | N[+1972] | 1465.6468 | 2 | 2930.2863 | 2930.2757 | 0 | 3.6 | 315 | Specific | 313.29 | 150.1 | 150.1 | 6.15 | 44 | >sp\|P0DOX2\|IGA2_HUMAN Immunoglobulin alpha-2 heavy chain OS=Homo sapiens OX=9606 PE=1 SV=2 | 631 | 20201113_gy_colo_milk_1ng_glyco_hcd203040_compound15.5966.5966.2 | controllerType=0 controllerNumber=1 scan=5966 | 16.0128 |
| 03149:2 | 19 | K.TPLTAN[+1606.587]ITK.S | HexNAc(4)Hex(4)Fuc(1) | N[+1607] | 1283.0782 | 2 | 2565.1491 | 2565.1435 | 0 | 2.2 | 315 | Specific | 325.76 | 247.02 | 247.02 | 6.87 | 44 | >sp\|P0DOX2\|IGA2_HUMAN Immunoglobulin alpha-2 heavy chain OS=Homo sapiens OX=9606 PE=1 SV=2 | 631 | 20201113_gy_colo_milk_1ng_glyco_hcd203040_compound15.5901.5901.2 | controllerType=0 controllerNumber=1 scan=5901 | 15.8983 |
| 03214:2 | 19 | K.TPLTAN[+1971.719]ITK.S | HexNAc(5)Hex(5)Fuc(1) | N[+1972] | 1465.6468 | 2 | 2930.2863 | 2930.2757 | 0 | 3.6 | 315 | Specific | 340.56 | 187.55 | 187.55 | 6.48 | 44 | >sp\|P0DOX2\|IGA2_HUMAN Immunoglobulin alpha-2 heavy chain OS=Homo sapiens OX=9606 PE=1 SV=2 | 631 | 20201113_gy_colo_milk_1ng_glyco_hcd203040_compound15.5968.5968.2 | controllerType=0 controllerNumber=1 scan=5968 | 16.0169 |
| 05042:2 | 19 | K.TPLTAN[+2059.735]ITK.S | HexNAc(4)Hex(5)Fuc(1)NeuAc(1) | N[+2060] | 1509.6532 | 2 | 3018.2992 | 3018.2917 | 0 | 2.5 | 315 | Specific | 345.9 | 120.77 | 120.77 | 5.56 | 44 | >sp\|P0DOX2\|IGA2_HUMAN Immunoglobulin alpha-2 heavy chain OS=Homo sapiens OX=9606 PE=1 SV=2 | 631 | 20201113_gy_colo_milk_1ng_glyco_hcd203040_compound15.7861.7861.2 | controllerType=0 controllerNumber=1 scan=7861 | 19.2059 |
| 03151:2 | 19 | K.TPLTAN[+1606.587]ITK.S | HexNAc(4)Hex(4)Fuc(1) | N[+1607] | 1283.0782 | 2 | 2565.1491 | 2565.1435 | 0 | 2.2 | 315 | Specific | 347.52 | 167.41 | 167.41 | 6.47 | 44 | >sp\|P0DOX2\|IGA2_HUMAN Immunoglobulin alpha-2 heavy chain OS=Homo sapiens OX=9606 PE=1 SV=2 | 631 | 20201113_gy_colo_milk_1ng_glyco_hcd203040_compound15.5903.5903.2 | controllerType=0 controllerNumber=1 scan=5903 | 15.9025 |
| 03331:3 | 19 | K.TPLTAN[+1809.666]ITK.S | HexNAc(5)Hex(4)Fuc(1) | N[+1810] | 923.4146 | 3 | 2768.2294 | 2768.2228 | 0 | 2.4 | 315 | Specific | 353.93 | 251.99 | 251.99 | 7.36 | 44 | >sp\|P0DOX2\|IGA2_HUMAN Immunoglobulin alpha-2 heavy chain OS=Homo sapiens OX=9606 PE=1 SV=2 | 631 | 20201113_gy_colo_milk_1ng_glyco_hcd203040_compound15.6090.6090.3 | controllerType=0 controllerNumber=1 scan=6090 | 16.2456 |
| 05022:3 | 19 | K.TPLTAN[+2059.735]ITK.S | HexNAc(4)Hex(5)Fuc(1)NeuAc(1) | N[+2060] | 1006.7718 | 3 | 3018.3009 | 3018.2917 | 0 | 3 | 315 | Specific | 380.61 | 139.6 | 139.6 | 6.37 | 44 | >sp\|P0DOX2\|IGA2_HUMAN Immunoglobulin alpha-2 heavy chain OS=Homo sapiens OX=9606 PE=1 SV=2 | 631 | 20201113_gy_colo_milk_1ng_glyco_hcd203040_compound15.7840.7840.3 | controllerType=0 controllerNumber=1 scan=7840 | 19.1638 |
| 03369:3 | 19 | K.TPLTAN[+1444.534]ITK.S | HexNAc(4)Hex(3)Fuc(1) | N[+1445] | 801.7044 | 3 | 2403.0986 | 2403.0906 | 0 | 3.3 | 315 | Specific | 382.14 | 348.53 | 348.53 | 7.98 | 44 | >sp\|P0DOX2\|IGA2_HUMAN Immunoglobulin alpha-2 heavy chain OS=Homo sapiens OX=9606 PE=1 SV=2 | 631 | 20201113_gy_colo_milk_1ng_glyco_hcd203040_compound15.6129.6129.3 | controllerType=0 controllerNumber=1 scan=6129 | 16.3316 |
| 03195:3 | 19 | K.TPLTAN[+1971.719]ITK.S | HexNAc(5)Hex(5)Fuc(1) | N[+1972] | 977.4324 | 3 | 2930.2827 | 2930.2757 | 0 | 2.4 | 315 | Specific | 383.29 | 98.84 | 98.84 | 5.69 | 44 | >sp\|P0DOX2\|IGA2_HUMAN Immunoglobulin alpha-2 heavy chain OS=Homo sapiens OX=9606 PE=1 SV=2 | 631 | 20201113_gy_colo_milk_1ng_glyco_hcd203040_compound15.5949.5949.3 | controllerType=0 controllerNumber=1 scan=5949 | 15.9851 |
| 03241:2 | 19 | K.TPLTAN[+1403.507]ITK.S | HexNAc(3)Hex(4)Fuc(1) | N[+1404] | 1181.5387 | 2 | 2362.0702 | 2362.0641 | 0 | 2.6 | 315 | Specific | 398.93 | 336.91 | 336.91 | 7.66 | 44 | >sp\|P0DOX2\|IGA2_HUMAN Immunoglobulin alpha-2 heavy chain OS=Homo sapiens OX=9606 PE=1 SV=2 | 631 | 20201113_gy_colo_milk_1ng_glyco_hcd203040_compound15.5996.5996.2 | controllerType=0 controllerNumber=1 scan=5996 | 16.0713 |
| 03333:3 | 19 | K.TPLTAN[+1809.666]ITK.S | HexNAc(5)Hex(4)Fuc(1) | N[+1810] | 923.4146 | 3 | 2768.2294 | 2768.2228 | 0 | 2.4 | 315 | Specific | 422.97 | 191.84 | 191.84 | 7.03 | 44 | >sp\|P0DOX2\|IGA2_HUMAN Immunoglobulin alpha-2 heavy chain OS=Homo sapiens OX=9606 PE=1 SV=2 | 631 | 20201113_gy_colo_milk_1ng_glyco_hcd203040_compound15.6092.6092.3 | controllerType=0 controllerNumber=1 scan=6092 | 16.25 |
| 05044:2 | 19 | K.TPLTAN[+2059.735]ITK.S | HexNAc(4)Hex(5)Fuc(1)NeuAc(1) | N[+2060] | 1509.6532 | 2 | 3018.2992 | 3018.2917 | 0 | 2.5 | 315 | Specific | 453.02 | 269.36 | 269.36 | 7.48 | 44 | >sp\|P0DOX2\|IGA2_HUMAN Immunoglobulin alpha-2 heavy chain OS=Homo sapiens OX=9606 PE=1 SV=2 | 631 | 20201113_gy_colo_milk_1ng_glyco_hcd203040_compound15.7863.7863.2 | controllerType=0 controllerNumber=1 scan=7863 | 19.2115 |
| 03197:3 | 19 | K.TPLTAN[+1971.719]ITK.S | HexNAc(5)Hex(5)Fuc(1) | N[+1972] | 977.4324 | 3 | 2930.2827 | 2930.2757 | 0 | 2.4 | 315 | Specific | 454.06 | 247.68 | 247.68 | 7.73 | 44 | >sp\|P0DOX2\|IGA2_HUMAN Immunoglobulin alpha-2 heavy chain OS=Homo sapiens OX=9606 PE=1 SV=2 | 631 | 20201113_gy_colo_milk_1ng_glyco_hcd203040_compound15.5951.5951.3 | controllerType=0 controllerNumber=1 scan=5951 | 15.9897 |
| 03330:3 | 19 | K.TPLTAN[+1647.613]ITK.S | HexNAc(5)Hex(3)Fuc(1) | N[+1648] | 869.3969 | 3 | 2606.1761 | 2606.17 | 0 | 2.3 | 315 | Specific | 459.14 | 183.31 | 183.31 | 7.15 | 44 | >sp\|P0DOX2\|IGA2_HUMAN Immunoglobulin alpha-2 heavy chain OS=Homo sapiens OX=9606 PE=1 SV=2 | 631 | 20201113_gy_colo_milk_1ng_glyco_hcd203040_compound15.6089.6089.3 | controllerType=0 controllerNumber=1 scan=6089 | 16.2439 |
| 05036:3 | 19 | K.TPLTAN[+2262.814]ITK.S | HexNAc(5)Hex(5)Fuc(1)NeuAc(1) | N[+2263] | 1074.4649 | 3 | 3221.3801 | 3221.3711 | 0 | 2.8 | 315 | Specific | 487.62 | 250.75 | 250.75 | 7.78 | 44 | >sp\|P0DOX2\|IGA2_HUMAN Immunoglobulin alpha-2 heavy chain OS=Homo sapiens OX=9606 PE=1 SV=2 | 631 | 20201113_gy_colo_milk_1ng_glyco_hcd203040_compound15.7854.7854.3 | controllerType=0 controllerNumber=1 scan=7854 | 19.1926 |
| 05024:3 | 19 | K.TPLTAN[+2059.735]ITK.S | HexNAc(4)Hex(5)Fuc(1)NeuAc(1) | N[+2060] | 1006.7718 | 3 | 3018.3009 | 3018.2917 | 0 | 3 | 315 | Specific | 508.68 | 259.57 | 259.57 | 7.8 | 44 | >sp\|P0DOX2\|IGA2_HUMAN Immunoglobulin alpha-2 heavy chain OS=Homo sapiens OX=9606 PE=1 SV=2 | 631 | 20201113_gy_colo_milk_1ng_glyco_hcd203040_compound15.7842.7842.3 | controllerType=0 controllerNumber=1 scan=7842 | 19.1693 |
| 03332:3 | 19 | K.TPLTAN[+1647.613]ITK.S | HexNAc(5)Hex(3)Fuc(1) | N[+1648] | 869.3969 | 3 | 2606.1761 | 2606.17 | 0 | 2.3 | 315 | Specific | 562.88 | 260.59 | 260.59 | 8.19 | 44 | >sp\|P0DOX2\|IGA2_HUMAN Immunoglobulin alpha-2 heavy chain OS=Homo sapiens OX=9606 PE=1 SV=2 | 631 | 20201113_gy_colo_milk_1ng_glyco_hcd203040_compound15.6091.6091.3 | controllerType=0 controllerNumber=1 scan=6091 | 16.2487 |
| 03317:2 | 19 | K.TPLTAN[+1501.555]ITK.S | HexNAc(5)Hex(3) | N[+1502] | 1230.563 | 2 | 2460.1188 | 2460.1121 | 0 | 2.7 | 315 | Specific | 569.61 | 339.79 | 339.79 | 8.64 | 44 | >sp\|P0DOX2\|IGA2_HUMAN Immunoglobulin alpha-2 heavy chain OS=Homo sapiens OX=9606 PE=1 SV=2 | 631 | 20201113_gy_colo_milk_1ng_glyco_hcd203040_compound15.6075.6075.2 | controllerType=0 controllerNumber=1 scan=6075 | 16.2177 |
| 03319:2 | 19 | K.TPLTAN[+1501.555]ITK.S | HexNAc(5)Hex(3) | N[+1502] | 1230.563 | 2 | 2460.1188 | 2460.1121 | 0 | 2.7 | 315 | Specific | 575.22 | 286.91 | 286.91 | 8.21 | 44 | >sp\|P0DOX2\|IGA2_HUMAN Immunoglobulin alpha-2 heavy chain OS=Homo sapiens OX=9606 PE=1 SV=2 | 631 | 20201113_gy_colo_milk_1ng_glyco_hcd203040_compound15.6077.6077.2 | controllerType=0 controllerNumber=1 scan=6077 | 16.2223 |
| 03308:2 | 19 | K.TPLTAN[+1647.613]ITK.S | HexNAc(5)Hex(3)Fuc(1) | N[+1648] | 1303.5923 | 2 | 2606.1774 | 2606.17 | 0 | 2.8 | 315 | Specific | 599.56 | 318.19 | 318.19 | 9.84 | 44 | >sp\|P0DOX2\|IGA2_HUMAN Immunoglobulin alpha-2 heavy chain OS=Homo sapiens OX=9606 PE=1 SV=2 | 631 | 20201113_gy_colo_milk_1ng_glyco_hcd203040_compound15.6066.6066.2 | controllerType=0 controllerNumber=1 scan=6066 | 16.204 |
| 03306:2 | 19 | K.TPLTAN[+1647.613]ITK.S | HexNAc(5)Hex(3)Fuc(1) | N[+1648] | 1303.5923 | 2 | 2606.1774 | 2606.17 | 0 | 2.8 | 315 | Specific | 607.58 | 352.71 | 352.71 | 10.99 | 44 | >sp\|P0DOX2\|IGA2_HUMAN Immunoglobulin alpha-2 heavy chain OS=Homo sapiens OX=9606 PE=1 SV=2 | 631 | 20201113_gy_colo_milk_1ng_glyco_hcd203040_compound15.6064.6064.2 | controllerType=0 controllerNumber=1 scan=6064 | 16.197 |
| 03252:2 | 19 | K.TPLTAN[+1809.666]ITK.S | HexNAc(5)Hex(4)Fuc(1) | N[+1810] | 1384.6176 | 2 | 2768.2279 | 2768.2228 | 0 | 1.8 | 315 | Specific | 608.01 | 352.78 | 352.78 | 10.99 | 44 | >sp\|P0DOX2\|IGA2_HUMAN Immunoglobulin alpha-2 heavy chain OS=Homo sapiens OX=9606 PE=1 SV=2 | 631 | 20201113_gy_colo_milk_1ng_glyco_hcd203040_compound15.6008.6008.2 | controllerType=0 controllerNumber=1 scan=6008 | 16.0948 |
| 03254:2 | 19 | K.TPLTAN[+1809.666]ITK.S | HexNAc(5)Hex(4)Fuc(1) | N[+1810] | 1384.6176 | 2 | 2768.2279 | 2768.2228 | 0 | 1.8 | 315 | Specific | 608.32 | 309.58 | 309.58 | 10.64 | 44 | >sp\|P0DOX2\|IGA2_HUMAN Immunoglobulin alpha-2 heavy chain OS=Homo sapiens OX=9606 PE=1 SV=2 | 631 | 20201113_gy_colo_milk_1ng_glyco_hcd203040_compound15.6010.6010.2 | controllerType=0 controllerNumber=1 scan=6010 | 16.1004 |
| 03258:2 | 19 | K.TPLTAN[+1444.534]ITK.S | HexNAc(4)Hex(3)Fuc(1) | N[+1445] | 1202.0513 | 2 | 2403.0953 | 2403.0906 | 0 | 2 | 315 | Specific | 612.22 | 327.27 | 327.27 | 10.84 | 44 | >sp\|P0DOX2\|IGA2_HUMAN Immunoglobulin alpha-2 heavy chain OS=Homo sapiens OX=9606 PE=1 SV=2 | 631 | 20201113_gy_colo_milk_1ng_glyco_hcd203040_compound15.6014.6014.2 | controllerType=0 controllerNumber=1 scan=6014 | 16.1067 |
| 03368:2 | 19 | K.TPLTAN[+1038.375]ITK.S | HexNAc(2)Hex(3)Fuc(1) | N[+1038] | 998.9722 | 2 | 1996.9371 | 1996.9319 | 0 | 2.6 | 315 | Specific | 614.19 | 357.62 | 357.62 | 10.99 | 44 | >sp\|P0DOX2\|IGA2_HUMAN Immunoglobulin alpha-2 heavy chain OS=Homo sapiens OX=9606 PE=1 SV=2 | 631 | 20201113_gy_colo_milk_1ng_glyco_hcd203040_compound15.6128.6128.2 | controllerType=0 controllerNumber=1 scan=6128 | 16.3282 |
| 03370:2 | 19 | K.TPLTAN[+1038.375]ITK.S | HexNAc(2)Hex(3)Fuc(1) | N[+1038] | 998.9722 | 2 | 1996.9371 | 1996.9319 | 0 | 2.6 | 315 | Specific | 629.92 | 307.55 | 307.55 | 10.64 | 44 | >sp\|P0DOX2\|IGA2_HUMAN Immunoglobulin alpha-2 heavy chain OS=Homo sapiens OX=9606 PE=1 SV=2 | 631 | 20201113_gy_colo_milk_1ng_glyco_hcd203040_compound15.6130.6130.2 | controllerType=0 controllerNumber=1 scan=6130 | 16.3342 |
| 03260:2 | 19 | K.TPLTAN[+1444.534]ITK.S | HexNAc(4)Hex(3)Fuc(1) | N[+1445] | 1202.0513 | 2 | 2403.0953 | 2403.0906 | 0 | 2 | 315 | Specific | 646.46 | 292.54 | 292.54 | 10.57 | 44 | >sp\|P0DOX2\|IGA2_HUMAN Immunoglobulin alpha-2 heavy chain OS=Homo sapiens OX=9606 PE=1 SV=2 | 631 | 20201113_gy_colo_milk_1ng_glyco_hcd203040_compound15.6016.6016.2 | controllerType=0 controllerNumber=1 scan=6016 | 16.1115 |
| 03339:2 | 19 | K.TPLTAN[+1241.454]ITK.S | HexNAc(3)Hex(3)Fuc(1) | N[+1241] | 1100.5116 | 2 | 2200.016 | 2200.0113 | 0 | 2.2 | 315 | Specific | 665.66 | 352.53 | 352.53 | 11.99 | 44 | >sp\|P0DOX2\|IGA2_HUMAN Immunoglobulin alpha-2 heavy chain OS=Homo sapiens OX=9606 PE=1 SV=2 | 631 | 20201113_gy_colo_milk_1ng_glyco_hcd203040_compound15.6098.6098.2 | controllerType=0 controllerNumber=1 scan=6098 | 16.2591 |
| 03341:2 | 19 | K.TPLTAN[+1241.454]ITK.S | HexNAc(3)Hex(3)Fuc(1) | N[+1241] | 1100.5116 | 2 | 2200.016 | 2200.0113 | 0 | 2.2 | 315 | Specific | 669.78 | 325 | 325 | 11.84 | 44 | >sp\|P0DOX2\|IGA2_HUMAN Immunoglobulin alpha-2 heavy chain OS=Homo sapiens OX=9606 PE=1 SV=2 | 631 | 20201113_gy_colo_milk_1ng_glyco_hcd203040_compound15.6100.6100.2 | controllerType=0 controllerNumber=1 scan=6100 | 16.264 |
| 14158:2 | 22 | K.LC[+57.021]DN[+1866.688]LSTK.N | HexNAc(6)Hex(4) | C[+57], N[+1867] | 1409.0765 | 2 | 2817.1456 | 2817.1487 | 0 | -1.1 | 285 | Specific | 293.25 | 204.37 | 204.37 | 2.52 | 27 | >sp\|P02774\|VTDB_HUMAN Vitamin D-binding protein OS=Homo sapiens OX=9606 GN=GC PE=1 SV=2 | 1103 | 20201113_gy_colo_milk_1ng_glyco_hcd203040_compound15.17309.17309.2 | controllerType=0 controllerNumber=1 scan=17309 | 35.986 |
| 01260:2 | 25 | R.SYN[+1038.375]DSVDPR.I | HexNAc(2)Hex(3)Fuc(1) | N[+1038] | 1045.9277 | 2 | 2090.848 | 2090.8395 | 0 | 4.1 | 481 | Specific | 248.2 | 205.66 | 205.66 | 5.89 | 28 | >sp\|P05164\|PERM_HUMAN Myeloperoxidase OS=Homo sapiens OX=9606 GN=MPO PE=1 SV=1 | 12 | 20201113_gy_colo_milk_1ng_glyco_hcd203040_compound15.3944.3944.2 | controllerType=0 controllerNumber=1 scan=3944 | 12.5392 |
| 01258:2 | 25 | R.SYN[+1038.375]DSVDPR.I | HexNAc(2)Hex(3)Fuc(1) | N[+1038] | 1045.9261 | 2 | 2090.8449 | 2090.8395 | 0 | 2.6 | 481 | Specific | 251.09 | 189.83 | 189.83 | 6.12 | 28 | >sp\|P05164\|PERM_HUMAN Myeloperoxidase OS=Homo sapiens OX=9606 GN=MPO PE=1 SV=1 | 12 | 20201113_gy_colo_milk_1ng_glyco_hcd203040_compound15.3941.3941.2 | controllerType=0 controllerNumber=1 scan=3941 | 12.5319 |
| 01262:2 | 25 | R.SYN[+1038.375]DSVDPR.I | HexNAc(2)Hex(3)Fuc(1) | N[+1038] | 1045.9277 | 2 | 2090.848 | 2090.8395 | 0 | 4.1 | 481 | Specific | 251.36 | 208.71 | 208.71 | 5.89 | 28 | >sp\|P05164\|PERM_HUMAN Myeloperoxidase OS=Homo sapiens OX=9606 GN=MPO PE=1 SV=1 | 12 | 20201113_gy_colo_milk_1ng_glyco_hcd203040_compound15.3946.3946.2 | controllerType=0 controllerNumber=1 scan=3946 | 12.5437 |
| 01256:2 | 25 | R.SYN[+1038.375]DSVDPR.I | HexNAc(2)Hex(3)Fuc(1) | N[+1038] | 1045.9261 | 2 | 2090.8449 | 2090.8395 | 0 | 2.6 | 481 | Specific | 275.79 | 238.11 | 238.11 | 6.63 | 28 | >sp\|P05164\|PERM_HUMAN Myeloperoxidase OS=Homo sapiens OX=9606 GN=MPO PE=1 SV=1 | 12 | 20201113_gy_colo_milk_1ng_glyco_hcd203040_compound15.3939.3939.2 | controllerType=0 controllerNumber=1 scan=3939 | 12.5278 |
| 02367:4 | 30 | R.EKQTDEIKDTRN[+2059.735]ESTQNC[+57.021]VVAEPEK.M | HexNAc(4)Hex(5)Fuc(1)NeuAc(1) | C[+57], N[+2060] | 1252.791 | 4 | 5008.1422 | 5008.1251 | 0 | 3.4 | 58 | Specific | 289.92 | 289.92 | 289.92 | 5.49 | 19 | >sp\|P47710\|CASA1_HUMAN Alpha-S1-casein OS=Homo sapiens OX=9606 GN=CSN1S1 PE=1 SV=1 | 989 | 20201113_gy_colo_milk_1ng_glyco_hcd203040_compound15.5092.5092.4 | controllerType=0 controllerNumber=1 scan=5092 | 14.5385 |
| 02407:4 | 30 | R.EKQTDEIKDTRN[+2059.735]ESTQNC[+57.021]VVAEPEK.M | HexNAc(4)Hex(5)Fuc(1)NeuAc(1) | C[+57], N[+2060] | 1252.7898 | 4 | 5008.1374 | 5008.1251 | 0 | 2.4 | 58 | Specific | 322.97 | 228.99 | 228.99 | 5.17 | 19 | >sp\|P47710\|CASA1_HUMAN Alpha-S1-casein OS=Homo sapiens OX=9606 GN=CSN1S1 PE=1 SV=1 | 989 | 20201113_gy_colo_milk_1ng_glyco_hcd203040_compound15.5134.5134.4 | controllerType=0 controllerNumber=1 scan=5134 | 14.6192 |
| 02400:5 | 30 | R.EKQTDEIKDTRN[+2059.735]ESTQNC[+57.021]VVAEPEK.M | HexNAc(4)Hex(5)Fuc(1)NeuAc(1) | C[+57], N[+2060] | 1002.435 | 5 | 5008.1459 | 5008.1251 | 0 | 4.1 | 58 | Specific | 327.14 | 327.14 | 327.14 | 6.2 | 19 | >sp\|P47710\|CASA1_HUMAN Alpha-S1-casein OS=Homo sapiens OX=9606 GN=CSN1S1 PE=1 SV=1 | 989 | 20201113_gy_colo_milk_1ng_glyco_hcd203040_compound15.5126.5126.5 | controllerType=0 controllerNumber=1 scan=5126 | 14.6026 |
| 02369:4 | 30 | R.EKQTDEIKDTRN[+2059.735]ESTQNC[+57.021]VVAEPEK.M | HexNAc(4)Hex(5)Fuc(1)NeuAc(1) | C[+57], N[+2060] | 1252.791 | 4 | 5008.1422 | 5008.1251 | 0 | 3.4 | 58 | Specific | 339.91 | 308.54 | 308.54 | 5.76 | 19 | >sp\|P47710\|CASA1_HUMAN Alpha-S1-casein OS=Homo sapiens OX=9606 GN=CSN1S1 PE=1 SV=1 | 989 | 20201113_gy_colo_milk_1ng_glyco_hcd203040_compound15.5094.5094.4 | controllerType=0 controllerNumber=1 scan=5094 | 14.5428 |
| 02405:4 | 30 | R.EKQTDEIKDTRN[+2059.735]ESTQNC[+57.021]VVAEPEK.M | HexNAc(4)Hex(5)Fuc(1)NeuAc(1) | C[+57], N[+2060] | 1252.7898 | 4 | 5008.1374 | 5008.1251 | 0 | 2.4 | 58 | Specific | 398.64 | 314.8 | 314.8 | 5.97 | 19 | >sp\|P47710\|CASA1_HUMAN Alpha-S1-casein OS=Homo sapiens OX=9606 GN=CSN1S1 PE=1 SV=1 | 989 | 20201113_gy_colo_milk_1ng_glyco_hcd203040_compound15.5132.5132.4 | controllerType=0 controllerNumber=1 scan=5132 | 14.6149 |
| 08833:5 | 31 | K.SC[+57.021]QHN[+2059.735]GTMYQHGEIFSAHELFPSR.L | HexNAc(4)Hex(5)Fuc(1)NeuAc(1) | C[+57], N[+2060] | 979.4015 | 5 | 4892.9785 | 4892.9919 | 0 | -2.7 | 110 | Specific | 312.31 | 312.31 | 312.31 | 5.79 | 19 | >sp\|Q6WN34\|CRDL2_HUMAN Chordin-like protein 2 OS=Homo sapiens OX=9606 GN=CHRDL2 PE=1 SV=1 | 1189 | 20201113_gy_colo_milk_1ng_glyco_hcd203040_compound15.11786.11786.5 | controllerType=0 controllerNumber=1 scan=11786 | 25.8355 |
| 08835:5 | 31 | K.SC[+57.021]QHN[+2059.735]GTMYQHGEIFSAHELFPSR.L | HexNAc(4)Hex(5)Fuc(1)NeuAc(1) | C[+57], N[+2060] | 979.4015 | 5 | 4892.9785 | 4892.9919 | 0 | -2.7 | 110 | Specific | 421.03 | 413.86 | 413.86 | 8.28 | 19 | >sp\|Q6WN34\|CRDL2_HUMAN Chordin-like protein 2 OS=Homo sapiens OX=9606 GN=CHRDL2 PE=1 SV=1 | 1189 | 20201113_gy_colo_milk_1ng_glyco_hcd203040_compound15.11788.11788.5 | controllerType=0 controllerNumber=1 scan=11788 | 25.8391 |
| 08775:5 | 31 | K.SC[+57.021]QHN[+2059.735]GTMYQHGEIFSAHELFPSR.L | HexNAc(4)Hex(5)Fuc(1)NeuAc(1) | C[+57], N[+2060] | 979.4086 | 5 | 4893.0139 | 4892.9919 | 0 | 4.5 | 110 | Specific | 698.65 | 697.1 | 697.1 | 13.7 | 19 | >sp\|Q6WN34\|CRDL2_HUMAN Chordin-like protein 2 OS=Homo sapiens OX=9606 GN=CHRDL2 PE=1 SV=1 | 1189 | 20201113_gy_colo_milk_1ng_glyco_hcd203040_compound15.11726.11726.3 | controllerType=0 controllerNumber=1 scan=11726 | 25.7238 |
| 08777:5 | 31 | K.SC[+57.021]QHN[+2059.735]GTMYQHGEIFSAHELFPSR.L | HexNAc(4)Hex(5)Fuc(1)NeuAc(1) | C[+57], N[+2060] | 979.4086 | 5 | 4893.0139 | 4892.9919 | 0 | 4.5 | 110 | Specific | 699.42 | 681.46 | 681.46 | 13.7 | 19 | >sp\|Q6WN34\|CRDL2_HUMAN Chordin-like protein 2 OS=Homo sapiens OX=9606 GN=CHRDL2 PE=1 SV=1 | 1189 | 20201113_gy_colo_milk_1ng_glyco_hcd203040_compound15.11728.11728.3 | controllerType=0 controllerNumber=1 scan=11728 | 25.7285 |
| 02272:3 | 33 | R.LRN[+1216.423]VSWATGR.S | HexNAc(2)Hex(5) | N[+1216] | 792.6923 | 3 | 2376.0623 | 2376.0559 | 0 | 2.7 | 149 | Specific | 298.09 | 109.27 | 109.27 | 4.78 | 16 | >sp\|P08571\|CD14_HUMAN Monocyte differentiation antigen CD14 OS=Homo sapiens OX=9606 GN=CD14 PE=1 SV=2 | 1232 | 20201113_gy_colo_milk_1ng_glyco_hcd203040_compound15.4994.4994.3 | controllerType=0 controllerNumber=1 scan=4994 | 14.3656 |
| 13248:3 | 36 | R.IIVPLNNREN[+2205.793]ISDPTSPLR.T | HexNAc(4)Hex(5)Fuc(2)NeuAc(1) | N[+2206] | 1451.9908 | 3 | 4353.9578 | 4353.97 | 0 | -2.8 | 62 | Specific | 253.98 | 253.98 | 253.98 | 4.79 | 25 | >sp\|P01591\|IGJ_HUMAN Immunoglobulin J chain OS=Homo sapiens OX=9606 GN=JCHAIN PE=1 SV=4 | 1392 | 20201113_gy_colo_milk_1ng_glyco_hcd203040_compound15.16365.16365.3 | controllerType=0 controllerNumber=1 scan=16365 | 34.2057 |
| 09834:3 | 36 | R.IIVPLNNREN[+1809.666]ISDPTSPLR.T | HexNAc(5)Hex(4)Fuc(1) | N[+1810] | 1319.9567 | 3 | 3957.8556 | 3957.8432 | 0 | 3.1 | 62 | Specific | 264.29 | 231.99 | 231.99 | 6.22 | 25 | >sp\|P01591\|IGJ_HUMAN Immunoglobulin J chain OS=Homo sapiens OX=9606 GN=JCHAIN PE=1 SV=4 | 1392 | 20201113_gy_colo_milk_1ng_glyco_hcd203040_compound15.12824.12824.3 | controllerType=0 controllerNumber=1 scan=12824 | 27.7059 |
| 09847:3 | 36 | R.IIVPLNNREN[+1460.529]ISDPTSPLR.T | HexNAc(4)Hex(4) | N[+1461] | 1203.5773 | 3 | 3608.7174 | 3608.7059 | 0 | 3.2 | 62 | Specific | 311.67 | 290.54 | 290.54 | 6.79 | 25 | >sp\|P01591\|IGJ_HUMAN Immunoglobulin J chain OS=Homo sapiens OX=9606 GN=JCHAIN PE=1 SV=4 | 1392 | 20201113_gy_colo_milk_1ng_glyco_hcd203040_compound15.12837.12837.3 | controllerType=0 controllerNumber=1 scan=12837 | 27.7278 |
| 09836:3 | 36 | R.IIVPLNNREN[+1809.666]ISDPTSPLR.T | HexNAc(5)Hex(4)Fuc(1) | N[+1810] | 1319.9567 | 3 | 3957.8556 | 3957.8432 | 0 | 3.1 | 62 | Specific | 335.42 | 261.43 | 261.43 | 6.79 | 25 | >sp\|P01591\|IGJ_HUMAN Immunoglobulin J chain OS=Homo sapiens OX=9606 GN=JCHAIN PE=1 SV=4 | 1392 | 20201113_gy_colo_milk_1ng_glyco_hcd203040_compound15.12826.12826.3 | controllerType=0 controllerNumber=1 scan=12826 | 27.7095 |
| 09849:3 | 36 | R.IIVPLNNREN[+1460.529]ISDPTSPLR.T | HexNAc(4)Hex(4) | N[+1461] | 1203.5773 | 3 | 3608.7174 | 3608.7059 | 0 | 3.2 | 62 | Specific | 341.67 | 341.67 | 341.67 | 7.41 | 25 | >sp\|P01591\|IGJ_HUMAN Immunoglobulin J chain OS=Homo sapiens OX=9606 GN=JCHAIN PE=1 SV=4 | 1392 | 20201113_gy_colo_milk_1ng_glyco_hcd203040_compound15.12839.12839.3 | controllerType=0 controllerNumber=1 scan=12839 | 27.7331 |
| 13251:3 | 36 | R.IIVPLNNREN[+2205.793]ISDPTSPLR.T | HexNAc(4)Hex(5)Fuc(2)NeuAc(1) | N[+2206] | 1451.9908 | 3 | 4353.9578 | 4353.97 | 0 | -2.8 | 62 | Specific | 357.79 | 357.79 | 357.79 | 5.89 | 25 | >sp\|P01591\|IGJ_HUMAN Immunoglobulin J chain OS=Homo sapiens OX=9606 GN=JCHAIN PE=1 SV=4 | 1392 | 20201113_gy_colo_milk_1ng_glyco_hcd203040_compound15.16368.16368.3 | controllerType=0 controllerNumber=1 scan=16368 | 34.2101 |
| 09916:3 | 36 | R.IIVPLNNREN[+1403.507]ISDPTSPLR.T | HexNAc(3)Hex(4)Fuc(1) | N[+1404] | 1184.569 | 3 | 3551.6925 | 3551.6845 | 0 | 2.3 | 62 | Specific | 368.42 | 341.35 | 341.35 | 7.54 | 25 | >sp\|P01591\|IGJ_HUMAN Immunoglobulin J chain OS=Homo sapiens OX=9606 GN=JCHAIN PE=1 SV=4 | 1392 | 20201113_gy_colo_milk_1ng_glyco_hcd203040_compound15.12908.12908.3 | controllerType=0 controllerNumber=1 scan=12908 | 27.8485 |
| 09813:3 | 36 | R.IIVPLNNREN[+1606.587]ISDPTSPLR.T | HexNAc(4)Hex(4)Fuc(1) | N[+1607] | 1252.2624 | 3 | 3754.7726 | 3754.7638 | 0 | 2.3 | 62 | Specific | 372.41 | 355.1 | 355.1 | 7.62 | 25 | >sp\|P01591\|IGJ_HUMAN Immunoglobulin J chain OS=Homo sapiens OX=9606 GN=JCHAIN PE=1 SV=4 | 1392 | 20201113_gy_colo_milk_1ng_glyco_hcd203040_compound15.12802.12802.3 | controllerType=0 controllerNumber=1 scan=12802 | 27.664 |
| 09918:3 | 36 | R.IIVPLNNREN[+1403.507]ISDPTSPLR.T | HexNAc(3)Hex(4)Fuc(1) | N[+1404] | 1184.569 | 3 | 3551.6925 | 3551.6845 | 0 | 2.3 | 62 | Specific | 389.09 | 338 | 338 | 7.62 | 25 | >sp\|P01591\|IGJ_HUMAN Immunoglobulin J chain OS=Homo sapiens OX=9606 GN=JCHAIN PE=1 SV=4 | 1392 | 20201113_gy_colo_milk_1ng_glyco_hcd203040_compound15.12910.12910.3 | controllerType=0 controllerNumber=1 scan=12910 | 27.854 |
| 09917:3 | 36 | R.IIVPLNNREN[+1647.613]ISDPTSPLR.T | HexNAc(5)Hex(3)Fuc(1) | N[+1648] | 1265.9362 | 3 | 3795.7941 | 3795.7904 | 0 | 1 | 62 | Specific | 395.16 | 335 | 335 | 7.33 | 25 | >sp\|P01591\|IGJ_HUMAN Immunoglobulin J chain OS=Homo sapiens OX=9606 GN=JCHAIN PE=1 SV=4 | 1392 | 20201113_gy_colo_milk_1ng_glyco_hcd203040_compound15.12909.12909.3 | controllerType=0 controllerNumber=1 scan=12909 | 27.8508 |
| 09792:3 | 36 | R.IIVPLNNREN[+1809.666]ISDPTSPLR.T | HexNAc(5)Hex(4)Fuc(1) | N[+1810] | 1319.9565 | 3 | 3957.8549 | 3957.8432 | 0 | 2.9 | 62 | Specific | 409.34 | 376.55 | 376.55 | 7.81 | 25 | >sp\|P01591\|IGJ_HUMAN Immunoglobulin J chain OS=Homo sapiens OX=9606 GN=JCHAIN PE=1 SV=4 | 1392 | 20201113_gy_colo_milk_1ng_glyco_hcd203040_compound15.12780.12780.3 | controllerType=0 controllerNumber=1 scan=12780 | 27.6249 |
| 09815:3 | 36 | R.IIVPLNNREN[+1606.587]ISDPTSPLR.T | HexNAc(4)Hex(4)Fuc(1) | N[+1607] | 1252.2624 | 3 | 3754.7726 | 3754.7638 | 0 | 2.3 | 62 | Specific | 462.3 | 389.39 | 389.39 | 9.57 | 25 | >sp\|P01591\|IGJ_HUMAN Immunoglobulin J chain OS=Homo sapiens OX=9606 GN=JCHAIN PE=1 SV=4 | 1392 | 20201113_gy_colo_milk_1ng_glyco_hcd203040_compound15.12804.12804.3 | controllerType=0 controllerNumber=1 scan=12804 | 27.6679 |
| 09914:3 | 36 | R.IIVPLNNREN[+1647.613]ISDPTSPLR.T | HexNAc(5)Hex(3)Fuc(1) | N[+1648] | 1265.9362 | 3 | 3795.7941 | 3795.7904 | 0 | 1 | 62 | Specific | 481.7 | 428.44 | 428.44 | 9.49 | 25 | >sp\|P01591\|IGJ_HUMAN Immunoglobulin J chain OS=Homo sapiens OX=9606 GN=JCHAIN PE=1 SV=4 | 1392 | 20201113_gy_colo_milk_1ng_glyco_hcd203040_compound15.12906.12906.3 | controllerType=0 controllerNumber=1 scan=12906 | 27.8459 |
| 11456:3 | 36 | R.IIVPLNNREN[+1694.603]ISDPTSPLR.T | HexNAc(3)Hex(4)Fuc(1)NeuAc(1) | N[+1695] | 1281.6005 | 3 | 3842.787 | 3842.7799 | 0 | 1.9 | 62 | Specific | 488.41 | 488.41 | 488.41 | 9.62 | 25 | >sp\|P01591\|IGJ_HUMAN Immunoglobulin J chain OS=Homo sapiens OX=9606 GN=JCHAIN PE=1 SV=4 | 1392 | 20201113_gy_colo_milk_1ng_glyco_hcd203040_compound15.14506.14506.3 | controllerType=0 controllerNumber=1 scan=14506 | 30.8543 |
| 13107:3 | 36 | R.IIVPLNNREN[+2350.830]ISDPTSPLR.T | HexNAc(4)Hex(5)Fuc(1)NeuAc(2) | N[+2351] | 1500.3439 | 3 | 4499.0172 | 4499.0075 | 0 | 2.2 | 62 | Specific | 502.25 | 466.32 | 466.32 | 9.72 | 25 | >sp\|P01591\|IGJ_HUMAN Immunoglobulin J chain OS=Homo sapiens OX=9606 GN=JCHAIN PE=1 SV=4 | 1392 | 20201113_gy_colo_milk_1ng_glyco_hcd203040_compound15.16219.16219.3 | controllerType=0 controllerNumber=1 scan=16219 | 33.9324 |
| 13104:3 | 36 | R.IIVPLNNREN[+2350.830]ISDPTSPLR.T | HexNAc(4)Hex(5)Fuc(1)NeuAc(2) | N[+2351] | 1500.3439 | 3 | 4499.0172 | 4499.0075 | 0 | 2.2 | 62 | Specific | 520.42 | 488.87 | 488.87 | 9.77 | 25 | >sp\|P01591\|IGJ_HUMAN Immunoglobulin J chain OS=Homo sapiens OX=9606 GN=JCHAIN PE=1 SV=4 | 1392 | 20201113_gy_colo_milk_1ng_glyco_hcd203040_compound15.16216.16216.3 | controllerType=0 controllerNumber=1 scan=16216 | 33.9268 |

Table S4. Cont.

| Query #:z | Protein Rank | Glycopeptide | Glycan | Modification Type(s) | Observed m/z | z | Observed (M+H) | Calc. mass (M+H) | Off-by-x error | Mass error (ppm) | StartPosition | Cleavage | Score | Delta | Delta Mod | \|Log Prob\| | # of unique peptides | Protein Name | Protein DB number | Comment | Scan # | Scan Time |
|---|---|---|---|---|---|---|---|---|---|---|---|---|---|---|---|---|---|---|---|---|---|---|
| 11459:3 | 36 | R.IIVPLNNREN[+1694.603]ISDPTSPLR.T | HexNAc(3)Hex(4)Fuc(1)NeuAc(1) | N[+1695] | 1281.6005 | 3 | 3842.787 | 3842.7799 | 0 | 1.9 | 62 | Specific | 532.56 | 486.34 | 486.34 | 9.77 | 25 | >sp\|P01591\|IGJ_HUMAN Immunoglobulin J chain OS=Homo sapiens OX=9606 GN=JCHAIN PE=1 SV=4 | 1392 | 20201113_gy_colo_milk_1ng_glyco_hcd203040_compound15.14509.14509.3 | controllerType=0 controllerNumber=1 scan=14509 | 30.8592 |
| 11367:3 | 36 | R.IIVPLNNREN[+1856.656]ISDPTSPLR.T | HexNAc(3)Hex(5)Fuc(1)NeuAc(1) | N[+1857] | 1335.6181 | 3 | 4004.8397 | 4004.8327 | 0 | 1.8 | 62 | Specific | 547.53 | 547.53 | 547.53 | 9.92 | 25 | >sp\|P01591\|IGJ_HUMAN Immunoglobulin J chain OS=Homo sapiens OX=9606 GN=JCHAIN PE=1 SV=4 | 1392 | 20201113_gy_colo_milk_1ng_glyco_hcd203040_compound15.14413.14413.3 | controllerType=0 controllerNumber=1 scan=14413 | 30.6478 |
| 09794:3 | 36 | R.IIVPLNNREN[+1809.666]ISDPTSPLR.T | HexNAc(5)Hex(4)Fuc(1) | N[+1810] | 1319.9565 | 3 | 3957.8549 | 3957.8432 | 0 | 2.9 | 62 | Specific | 548.84 | 471.22 | 471.22 | 9.92 | 25 | >sp\|P01591\|IGJ_HUMAN Immunoglobulin J chain OS=Homo sapiens OX=9606 GN=JCHAIN PE=1 SV=4 | 1392 | 20201113_gy_colo_milk_1ng_glyco_hcd203040_compound15.12782.12782.3 | controllerType=0 controllerNumber=1 scan=12782 | 27.6296 |
| 09837:3 | 36 | R.IIVPLNNREN[+1663.608]ISDPTSPLR.T | HexNAc(5)Hex(4) | N[+1664] | 1271.2693 | 3 | 3811.7935 | 3811.7853 | 0 | 2.1 | 62 | Specific | 557.12 | 543.4 | 543.4 | 9.92 | 25 | >sp\|P01591\|IGJ_HUMAN Immunoglobulin J chain OS=Homo sapiens OX=9606 GN=JCHAIN PE=1 SV=4 | 1392 | 20201113_gy_colo_milk_1ng_glyco_hcd203040_compound15.12827.12827.3 | controllerType=0 controllerNumber=1 scan=12827 | 27.7128 |
| 09835:3 | 36 | R.IIVPLNNREN[+1663.608]ISDPTSPLR.T | HexNAc(5)Hex(4) | N[+1664] | 1271.2693 | 3 | 3811.7935 | 3811.7853 | 0 | 2.1 | 62 | Specific | 561.47 | 536.24 | 536.24 | 10.11 | 25 | >sp\|P01591\|IGJ_HUMAN Immunoglobulin J chain OS=Homo sapiens OX=9606 GN=JCHAIN PE=1 SV=4 | 1392 | 20201113_gy_colo_milk_1ng_glyco_hcd203040_compound15.12825.12825.3 | controllerType=0 controllerNumber=1 scan=12825 | 27.7069 |
| 11370:3 | 36 | R.IIVPLNNREN[+1856.656]ISDPTSPLR.T | HexNAc(3)Hex(5)Fuc(1)NeuAc(1) | N[+1857] | 1335.6181 | 3 | 4004.8397 | 4004.8327 | 0 | 1.8 | 62 | Specific | 572.45 | 572.45 | 572.45 | 10.11 | 25 | >sp\|P01591\|IGJ_HUMAN Immunoglobulin J chain OS=Homo sapiens OX=9606 GN=JCHAIN PE=1 SV=4 | 1392 | 20201113_gy_colo_milk_1ng_glyco_hcd203040_compound15.14416.14416.3 | controllerType=0 controllerNumber=1 scan=14416 | 30.6539 |
| 07216:3 | 36 | R.EN[+1913.677]ISDPTSPLR.T | HexNAc(4)Hex(5)NeuAc(1) | N[+1914] | 1048.1074 | 3 | 3142.3076 | 3142.2938 | 0 | 4.4 | 70 | Specific | 237.02 | 163.83 | 163.83 | 3.75 | 25 | >sp\|P01591\|IGJ_HUMAN Immunoglobulin J chain OS=Homo sapiens OX=9606 GN=JCHAIN PE=1 SV=4 | 1392 | 20201113_gy_colo_milk_1ng_glyco_hcd203040_compound15.10111.10111.3 | controllerType=0 controllerNumber=1 scan=10111 | 22.9297 |
| 07214:3 | 36 | R.EN[+1913.677]ISDPTSPLR.T | HexNAc(4)Hex(5)NeuAc(1) | N[+1914] | 1048.1074 | 3 | 3142.3076 | 3142.2938 | 0 | 4.4 | 70 | Specific | 237.07 | 128.74 | 128.74 | 5.47 | 25 | >sp\|P01591\|IGJ_HUMAN Immunoglobulin J chain OS=Homo sapiens OX=9606 GN=JCHAIN PE=1 SV=4 | 1392 | 20201113_gy_colo_milk_1ng_glyco_hcd203040_compound15.10109.10109.3 | controllerType=0 controllerNumber=1 scan=10109 | 22.9254 |
| 09560:3 | 36 | R.EN[+2204.772]ISDPTSPLR.T | HexNAc(4)Hex(5)NeuAc(2) | N[+2205] | 1145.1392 | 3 | 3433.403 | 3433.3892 | 0 | 4 | 70 | Specific | 255.86 | 148.52 | 148.52 | 5.57 | 25 | >sp\|P01591\|IGJ_HUMAN Immunoglobulin J chain OS=Homo sapiens OX=9606 GN=JCHAIN PE=1 SV=4 | 1392 | 20201113_gy_colo_milk_1ng_glyco_hcd203040_compound15.12540.12540.3 | controllerType=0 controllerNumber=1 scan=12540 | 27.2061 |
| 07198:2 | 36 | R.EN[+1913.677]ISDPTSPLR.T | HexNAc(4)Hex(5)NeuAc(1) | N[+1914] | 1571.6574 | 2 | 3142.3075 | 3142.2938 | 0 | 4.4 | 70 | Specific | 265.59 | 129.63 | 129.63 | 5.32 | 25 | >sp\|P01591\|IGJ_HUMAN Immunoglobulin J chain OS=Homo sapiens OX=9606 GN=JCHAIN PE=1 SV=4 | 1392 | 20201113_gy_colo_milk_1ng_glyco_hcd203040_compound15.10092.10092.2 | controllerType=0 controllerNumber=1 scan=10092 | 22.8976 |
| 07196:2 | 36 | R.EN[+1913.677]ISDPTSPLR.T | HexNAc(4)Hex(5)NeuAc(1) | N[+1914] | 1571.6574 | 2 | 3142.3075 | 3142.2938 | 0 | 4.4 | 70 | Specific | 267.33 | 206.91 | 206.91 | 5.92 | 25 | >sp\|P01591\|IGJ_HUMAN Immunoglobulin J chain OS=Homo sapiens OX=9606 GN=JCHAIN PE=1 SV=4 | 1392 | 20201113_gy_colo_milk_1ng_glyco_hcd203040_compound15.10090.10090.2 | controllerType=0 controllerNumber=1 scan=10090 | 22.894 |
| 09562:3 | 36 | R.EN[+2204.772]ISDPTSPLR.T | HexNAc(4)Hex(5)NeuAc(2) | N[+2205] | 1145.1392 | 3 | 3433.403 | 3433.3892 | 0 | 4 | 70 | Specific | 383.18 | 192.66 | 192.66 | 6.74 | 25 | >sp\|P01591\|IGJ_HUMAN Immunoglobulin J chain OS=Homo sapiens OX=9606 GN=JCHAIN PE=1 SV=4 | 1392 | 20201113_gy_colo_milk_1ng_glyco_hcd203040_compound15.12542.12542.3 | controllerType=0 controllerNumber=1 scan=12542 | 27.2117 |
| 15558:3 | 46 | R.KLPPGLLAN[+2059.735]FTLLR.T | HexNAc(4)Hex(5)Fuc(1)NeuAc(1) | N[+2060] | 1204.9031 | 3 | 3612.6948 | 3612.6923 | 0 | 0.7 | 178 | Specific | 368.42 | 368.42 | 368.42 | 7.99 | 15 | >sp\|P02750\|A2GL_HUMAN Leucine-rich alpha-2-glycoprotein OS=Homo sapiens OX=9606 GN=LRG1 PE=1 SV=2 | 518 | 20201113_gy_colo_milk_1ng_glyco_hcd203040_compound15.18765.18765.3 | controllerType=0 controllerNumber=1 scan=18765 | 38.8717 |
| 15561:3 | 46 | R.KLPPGLLAN[+2059.735]FTLLR.T | HexNAc(4)Hex(5)Fuc(1)NeuAc(1) | N[+2060] | 1204.9031 | 3 | 3612.6948 | 3612.6923 | 0 | 0.7 | 178 | Specific | 490.41 | 452.01 | 452.01 | 8.61 | 15 | >sp\|P02750\|A2GL_HUMAN Leucine-rich alpha-2-glycoprotein OS=Homo sapiens OX=9606 GN=LRG1 PE=1 SV=2 | 518 | 20201113_gy_colo_milk_1ng_glyco_hcd203040_compound15.18768.18768.3 | controllerType=0 controllerNumber=1 scan=18768 | 38.8767 |
| 21950:4 | 46 | R.QLDMLDLSN[+1913.677]NSLASVPEGLWASLGQPNWDMR.D | HexNAc(4)Hex(5)NeuAc(1) | N[+1914] | 1343.5885 | 4 | 5371.3322 | 5371.3285 | 0 | 0.7 | 261 | Specific | 742.41 | 740.73 | 740.73 | 15.92 | 15 | >sp\|P02750\|A2GL_HUMAN Leucine-rich alpha-2-glycoprotein OS=Homo sapiens OX=9606 GN=LRG1 PE=1 SV=2 | 518 | 20201113_gy_colo_milk_1ng_glyco_hcd203040_compound15.25430.25430.2 | controllerType=0 controllerNumber=1 scan=25430 | 52.9914 |
| 21952:4 | 46 | R.QLDMLDLSN[+1913.677]NSLASVPEGLWASLGQPNWDMR.D | HexNAc(4)Hex(5)NeuAc(1) | N[+1914] | 1343.5885 | 4 | 5371.3322 | 5371.3285 | 0 | 0.7 | 261 | Specific | 1034.74 | 998.6 | 998.6 | 21.92 | 15 | >sp\|P02750\|A2GL_HUMAN Leucine-rich alpha-2-glycoprotein OS=Homo sapiens OX=9606 GN=LRG1 PE=1 SV=2 | 518 | 20201113_gy_colo_milk_1ng_glyco_hcd203040_compound15.25432.25432.2 | controllerType=0 controllerNumber=1 scan=25432 | 52.9971 |
| 10958:3 | 75 | R.ADGTVNQIEGEATPVN[+1054.370]LTEPAKLEVK.F | HexNAc(2)Hex(4) | N[+1054] | 1259.9355 | 3 | 3777.7919 | 3777.7798 | 0 | 3.2 | 83 | Specific | 622.43 | 622.43 | 622.43 | 12.47 | 10 | >sp\|P05090\|APOD_HUMAN Apolipoprotein D OS=Homo sapiens OX=9606 GN=APOD PE=1 SV=1 | 594 | 20201113_gy_colo_milk_1ng_glyco_hcd203040_compound15.13988.13988.3 | controllerType=0 controllerNumber=1 scan=13988 | 29.8058 |
| 10956:3 | 75 | R.ADGTVNQIEGEATPVN[+1054.370]LTEPAKLEVK.F | HexNAc(2)Hex(4) | N[+1054] | 1259.9355 | 3 | 3777.7919 | 3777.7798 | 0 | 3.2 | 83 | Specific | 764.44 | 764.44 | 764.44 | 15.47 | 10 | >sp\|P05090\|APOD_HUMAN Apolipoprotein D OS=Homo sapiens OX=9606 GN=APOD PE=1 SV=1 | 594 | 20201113_gy_colo_milk_1ng_glyco_hcd203040_compound15.13986.13986.3 | controllerType=0 controllerNumber=1 scan=13986 | 29.8018 |
| 18964:3 | 94 | R.GIASVLQELN[+876.322]VTVVTSLC[+57.021]R.R | HexNAc(2)Hex(2)Fuc(1) | C[+57], N[+876] | 979.1552 | 3 | 2935.4512 | 2935.4439 | 0 | 2.5 | 164 | Specific | 330.8 | 330.8 | 330.8 | 5.86 | 8 | >sp\|P08246\|ELNE_HUMAN Neutrophil elastase OS=Homo sapiens OX=9606 GN=ELANE PE=1 SV=1 | 1269 | 20201113_gy_colo_milk_1ng_glyco_hcd203040_compound15.22315.22315.2 | controllerType=0 controllerNumber=1 scan=22315 | 46.2943 |
| 18928:3 | 94 | R.GIASVLQELN[+876.322]VTVVTSLC[+57.021]R.R | HexNAc(2)Hex(2)Fuc(1) | C[+57], N[+876] | 979.1549 | 3 | 2935.4503 | 2935.4439 | 0 | 2.2 | 164 | Specific | 1025.01 | 1025.01 | 1025.01 | 21.22 | 8 | >sp\|P08246\|ELNE_HUMAN Neutrophil elastase OS=Homo sapiens OX=9606 GN=ELANE PE=1 SV=1 | 1269 | 20201113_gy_colo_milk_1ng_glyco_hcd203040_compound15.22278.22278.3 | controllerType=0 controllerNumber=1 scan=22278 | 46.2233 |
| 18931:3 | 94 | R.GIASVLQELN[+876.322]VTVVTSLC[+57.021]R.R | HexNAc(2)Hex(2)Fuc(1) | C[+57], N[+876] | 979.1549 | 3 | 2935.4503 | 2935.4439 | 0 | 2.2 | 164 | Specific | 1084.64 | 1062 | 1062 | 22.22 | 8 | >sp\|P08246\|ELNE_HUMAN Neutrophil elastase OS=Homo sapiens OX=9606 GN=ELANE PE=1 SV=1 | 1269 | 20201113_gy_colo_milk_1ng_glyco_hcd203040_compound15.22281.22281.3 | controllerType=0 controllerNumber=1 scan=22281 | 46.2297 |
| 23875:3 | 99 | R.AVN[+2204.772]ITSENLIDDVVSLIR.G | HexNAc(4)Hex(5)NeuAc(2) | N[+2205] | 1392.6314 | 3 | 4175.8797 | 4175.8481 | 0 | 7.5 | 413 | Specific | 449.59 | 449.59 | 449.59 | 9.24 | 10 | >sp\|P02748\|CO9_HUMAN Complement component C9 OS=Homo sapiens OX=9606 GN=C9 PE=1 SV=2 | 1547 | 20201113_gy_colo_milk_1ng_glyco_hcd203040_compound15.27436.27436.3 | controllerType=0 controllerNumber=1 scan=27436 | 57.1001 |
| 23877:3 | 99 | R.AVN[+2204.772]ITSENLIDDVVSLIR.G | HexNAc(4)Hex(5)NeuAc(2) | N[+2205] | 1392.6314 | 3 | 4175.8797 | 4175.8481 | 0 | 7.5 | 413 | Specific | 658.06 | 641.91 | 641.91 | 13.27 | 10 | >sp\|P02748\|CO9_HUMAN Complement component C9 OS=Homo sapiens OX=9606 GN=C9 PE=1 SV=2 | 1547 | 20201113_gy_colo_milk_1ng_glyco_hcd203040_compound15.27438.27438.3 | controllerType=0 controllerNumber=1 scan=27438 | 57.1043 |
| 14156:3 | 102 | R.EDFVLTAAHC[+57.021]WGSNIN[+203.079]VTLGAHNIQR.R | HexNAc(1) | C[+57], N[+203] | 1042.8415 | 3 | 3126.5098 | 3126.5062 | 0 | 1.2 | 56 | Specific | 408.97 | 408.97 | 408.97 | 7.63 | 7 | >sp\|P08311\|CATG_HUMAN Cathepsin G OS=Homo sapiens OX=9606 GN=CTSG PE=1 SV=2 | 1574 | 20201113_gy_colo_milk_1ng_glyco_hcd203040_compound15.17307.17307.3 | controllerType=0 controllerNumber=1 scan=17307 | 35.9828 |
| 14159:3 | 102 | R.EDFVLTAAHC[+57.021]WGSNIN[+203.079]VTLGAHNIQR.R | HexNAc(1) | C[+57], N[+203] | 1042.8415 | 3 | 3126.5098 | 3126.5062 | 0 | 1.2 | 56 | Specific | 794.99 | 697.57 | 697.57 | 13.81 | 7 | >sp\|P08311\|CATG_HUMAN Cathepsin G OS=Homo sapiens OX=9606 GN=CTSG PE=1 SV=2 | 1574 | 20201113_gy_colo_milk_1ng_glyco_hcd203040_compound15.17310.17310.3 | controllerType=0 controllerNumber=1 scan=17310 | 35.9879 |
| 14111:3 | 102 | R.EDFVLTAAHC[+57.021]WGSNIN[+203.079]VTLGAHNIQR.R | HexNAc(1) | C[+57], N[+203] | 1042.8399 | 3 | 3126.5051 | 3126.5062 | 0 | -0.3 | 56 | Specific | 981.24 | 895.96 | 895.96 | 18.91 | 7 | >sp\|P08311\|CATG_HUMAN Cathepsin G OS=Homo sapiens OX=9606 GN=CTSG PE=1 SV=2 | 1574 | 20201113_gy_colo_milk_1ng_glyco_hcd203040_compound15.17260.17260.3 | controllerType=0 controllerNumber=1 scan=17260 | 35.8952 |
| 14109:3 | 102 | R.EDFVLTAAHC[+57.021]WGSNIN[+203.079]VTLGAHNIQR.R | HexNAc(1) | C[+57], N[+203] | 1042.8399 | 3 | 3126.5051 | 3126.5062 | 0 | -0.3 | 56 | Specific | 1105.44 | 1009.05 | 1009.05 | 21.91 | 7 | >sp\|P08311\|CATG_HUMAN Cathepsin G OS=Homo sapiens OX=9606 GN=CTSG PE=1 SV=2 | 1574 | 20201113_gy_colo_milk_1ng_glyco_hcd203040_compound15.17258.17258.3 | controllerType=0 controllerNumber=1 scan=17258 | 35.8893 |
| 17185:4 | 1 | R.RTAGWNVPIGTLRPFLN[+1768.640]WTGPPEPIEAAVAR.F | HexNAc(4)Hex(5)Fuc(1) | N[+1769] | 1289.6196 | 4 | 5155.4566 | 5155.4516 | 0 | 1 | 140 | Specific | 485.32 | 405.01 | 405.01 | 10.12 | 130 | >sp\|P02788\|TRFL_HUMAN Lactotransferrin OS=Homo sapiens OX=9606 GN=LTF PE=1 SV=6 | 1270 | 20201113_gy_milk_1ng_glyco_hcd203040_compound15.20606.20606.4 | controllerType=0 controllerNumber=1 scan=20606 | 41.3921 |
| 18066:4 | 1 | R.RTAGWNVPIGTLRPFLN[+2570.925]WTGPPEPIEAAVAR.F | HexNAc(5)Hex(6)Fuc(2)NeuAc(1) | N[+2571] | 1490.1937 | 4 | 5957.7531 | 5957.7371 | 0 | 2.7 | 140 | Specific | 547.34 | 424.56 | 424.56 | 10.42 | 130 | >sp\|P02788\|TRFL_HUMAN Lactotransferrin OS=Homo sapiens OX=9606 GN=LTF PE=1 SV=6 | 1270 | 20201113_gy_milk_1ng_glyco_hcd203040_compound15.21529.21529.4 | controllerType=0 controllerNumber=1 scan=21529 | 43.3027 |
| 19106:4 | 1 | R.RTAGWNVPIGTLRPFLN[+2350.830]WTGPPEPIEAAVAR.F | HexNAc(4)Hex(5)Fuc(1)NeuAc(2) | N[+2351] | 1435.1685 | 4 | 5737.652 | 5737.6424 | 0 | 1.7 | 140 | Specific | 560.38 | 467.73 | 467.73 | 10.47 | 130 | >sp\|P02788\|TRFL_HUMAN Lactotransferrin OS=Homo sapiens OX=9606 GN=LTF PE=1 SV=6 | 1270 | 20201113_gy_milk_1ng_glyco_hcd203040_compound15.22622.22622.2 | controllerType=0 controllerNumber=1 scan=22622 | 45.7601 |
| 19103:4 | 1 | R.RTAGWNVPIGTLRPFLN[+2350.830]WTGPPEPIEAAVAR.F | HexNAc(4)Hex(5)Fuc(1)NeuAc(2) | N[+2351] | 1435.1685 | 4 | 5737.652 | 5737.6424 | 0 | 1.7 | 140 | Specific | 619 | 476.98 | 476.98 | 12.69 | 130 | >sp\|P02788\|TRFL_HUMAN Lactotransferrin OS=Homo sapiens OX=9606 GN=LTF PE=1 SV=6 | 1270 | 20201113_gy_milk_1ng_glyco_hcd203040_compound15.22619.22619.2 | controllerType=0 controllerNumber=1 scan=22619 | 45.7526 |
| 19614:3 | 1 | R.TAGWNVPIGTLRPFLN[+2059.735]WTGPPEPIEAAVAR.F | HexNAc(4)Hex(5)Fuc(1)NeuAc(1) | N[+2060] | 1764.1605 | 3 | 5290.4669 | 5290.4459 | 0 | 4 | 141 | Specific | 231.76 | 231.76 | 231.76 | 6.34 | 130 | >sp\|P02788\|TRFL_HUMAN Lactotransferrin OS=Homo sapiens OX=9606 GN=LTF PE=1 SV=6 | 1270 | 20201113_gy_milk_1ng_glyco_hcd203040_compound15.23155.23155.3 | controllerType=0 controllerNumber=1 scan=23155 | 46.8209 |
| 18364:3 | 1 | R.TAGWNVPIGTLRPFLN[+2425.887]WTGPPEPIEAAVAR.F | HexNAc(5)Hex(6)Fuc(3) | N[+2426] | 1886.2111 | 3 | 5656.6189 | 5656.5985 | 0 | 3.6 | 141 | Specific | 232.18 | 116.83 | 116.83 | 5.14 | 130 | >sp\|P02788\|TRFL_HUMAN Lactotransferrin OS=Homo sapiens OX=9606 GN=LTF PE=1 SV=6 | 1270 | 20201113_gy_milk_1ng_glyco_hcd203040_compound15.21840.21840.3 | controllerType=0 controllerNumber=1 scan=21840 | 43.9458 |
| 19347:4 | 1 | R.TAGWNVPIGTLRPFLN[+2076.750]WTGPPEPIEAAVAR.F | HexNAc(4)Hex(6)Fuc(2) | N[+2077] | 1327.6296 | 4 | 5307.4967 | 5307.4612 | 0 | 6.7 | 141 | Specific | 235.51 | 235.51 | 235.51 | 6.03 | 130 | >sp\|P02788\|TRFL_HUMAN Lactotransferrin OS=Homo sapiens OX=9606 GN=LTF PE=1 SV=6 | 1270 | 20201113_gy_milk_1ng_glyco_hcd203040_compound15.22874.22874.4 | controllerType=0 controllerNumber=1 scan=22874 | 46.2831 |
| 19344:4 | 1 | R.TAGWNVPIGTLRPFLN[+2076.750]WTGPPEPIEAAVAR.F | HexNAc(4)Hex(6)Fuc(2) | N[+2077] | 1327.6296 | 4 | 5307.4967 | 5307.4612 | 0 | 6.7 | 141 | Specific | 254.32 | 254.32 | 254.32 | 6.36 | 130 | >sp\|P02788\|TRFL_HUMAN Lactotransferrin OS=Homo sapiens OX=9606 GN=LTF PE=1 SV=6 | 1270 | 20201113_gy_milk_1ng_glyco_hcd203040_compound15.22871.22871.4 | controllerType=0 controllerNumber=1 scan=22871 | 46.2787 |
| 19617:3 | 1 | R.TAGWNVPIGTLRPFLN[+2059.735]WTGPPEPIEAAVAR.F | HexNAc(4)Hex(5)Fuc(1)NeuAc(1) | N[+2060] | 1764.1605 | 3 | 5290.4669 | 5290.4459 | 0 | 4 | 141 | Specific | 292.14 | 284.18 | 284.18 | 6.92 | 130 | >sp\|P02788\|TRFL_HUMAN Lactotransferrin OS=Homo sapiens OX=9606 GN=LTF PE=1 SV=6 | 1270 | 20201113_gy_milk_1ng_glyco_hcd203040_compound15.23158.23158.3 | controllerType=0 controllerNumber=1 scan=23158 | 46.8252 |
| 19281:3 | 1 | R.TAGWNVPIGTLRPFLN[+2570.925]WTGPPEPIEAAVAR.F | HexNAc(5)Hex(6)Fuc(2)NeuAc(1) | N[+2571] | 1934.564 | 3 | 5801.6774 | 5801.636 | 0 | 7.1 | 141 | Specific | 295.83 | 188.81 | 188.81 | 5.87 | 130 | >sp\|P02788\|TRFL_HUMAN Lactotransferrin OS=Homo sapiens OX=9606 GN=LTF PE=1 SV=6 | 1270 | 20201113_gy_milk_1ng_glyco_hcd203040_compound15.22805.22805.3 | controllerType=0 controllerNumber=1 scan=22805 | 46.1162 |
| 19284:3 | 1 | R.TAGWNVPIGTLRPFLN[+2570.925]WTGPPEPIEAAVAR.F | HexNAc(5)Hex(6)Fuc(2)NeuAc(1) | N[+2571] | 1934.564 | 3 | 5801.6774 | 5801.636 | 0 | 7.1 | 141 | Specific | 297.09 | 204.11 | 204.11 | 5.99 | 130 | >sp\|P02788\|TRFL_HUMAN Lactotransferrin OS=Homo sapiens OX=9606 GN=LTF PE=1 SV=6 | 1270 | 20201113_gy_milk_1ng_glyco_hcd203040_compound15.22808.22808.3 | controllerType=0 controllerNumber=1 scan=22808 | 46.1211 |
| 18366:3 | 1 | R.TAGWNVPIGTLRPFLN[+2425.887]WTGPPEPIEAAVAR.F | HexNAc(5)Hex(6)Fuc(3) | N[+2426] | 1886.2111 | 3 | 5656.6189 | 5656.5985 | 0 | 3.6 | 141 | Specific | 311.53 | 212.24 | 212.24 | 6.44 | 130 | >sp\|P02788\|TRFL_HUMAN Lactotransferrin OS=Homo sapiens OX=9606 GN=LTF PE=1 SV=6 | 1270 | 20201113_gy_milk_1ng_glyco_hcd203040_compound15.21842.21842.3 | controllerType=0 controllerNumber=1 scan=21842 | 43.9509 |
| 19539:3 | 1 | R.TAGWNVPIGTLRPFLN[+2059.735]WTGPPEPIEAAVAR.F | HexNAc(4)Hex(5)Fuc(1)NeuAc(1) | N[+2060] | 1764.1524 | 3 | 5290.4427 | 5290.4459 | 0 | -0.6 | 141 | Specific | 332.56 | 286.54 | 286.54 | 7.04 | 130 | >sp\|P02788\|TRFL_HUMAN Lactotransferrin OS=Homo sapiens OX=9606 GN=LTF PE=1 SV=6 | 1270 | 20201113_gy_milk_1ng_glyco_hcd203040_compound15.23077.23077.3 | controllerType=0 controllerNumber=1 scan=23077 | 46.6451 |
| 19369:4 | 1 | R.TAGWNVPIGTLRPFLN[+2059.735]WTGPPEPIEAAVAR.F | HexNAc(4)Hex(5)Fuc(1)NeuAc(1) | N[+2060] | 1323.3687 | 4 | 5290.4528 | 5290.4459 | 0 | 1.3 | 141 | Specific | 335.59 | 335.59 | 335.59 | 8.09 | 130 | >sp\|P02788\|TRFL_HUMAN Lactotransferrin OS=Homo sapiens OX=9606 GN=LTF PE=1 SV=6 | 1270 | 20201113_gy_milk_1ng_glyco_hcd203040_compound15.22897.22897.4 | controllerType=0 controllerNumber=1 scan=22897 | 46.3319 |
| 20836:3 | 1 | R.TAGWNVPIGTLRPFLN[+2351.851]WTGPPEPIEAAVAR.F | HexNAc(4)Hex(5)Fuc(3)NeuAc(1) | N[+2352] | 1861.5353 | 3 | 5582.5913 | 5582.5617 | 0 | 5.3 | 141 | Specific | 338.39 | 338.39 | 338.39 | 7.39 | 130 | >sp\|P02788\|TRFL_HUMAN Lactotransferrin OS=Homo sapiens OX=9606 GN=LTF PE=1 SV=6 | 1270 | 20201113_gy_milk_1ng_glyco_hcd203040_compound15.24437.24437.3 | controllerType=0 controllerNumber=1 scan=24437 | 49.3915 |
| 19188:4 | 1 | R.TAGWNVPIGTLRPFLN[+2716.983]WTGPPEPIEAAVAR.F | HexNAc(5)Hex(6)Fuc(3)NeuAc(1) | N[+2717] | 1487.6854 | 4 | 5947.7199 | 5947.6939 | 0 | 4.4 | 141 | Specific | 344.49 | 344.49 | 344.49 | 7.73 | 130 | >sp\|P02788\|TRFL_HUMAN Lactotransferrin OS=Homo sapiens OX=9606 GN=LTF PE=1 SV=6 | 1270 | 20201113_gy_milk_1ng_glyco_hcd203040_compound15.22708.22708.4 | controllerType=0 controllerNumber=1 scan=22708 | 45.8983 |
| 19274:4 | 1 | R.TAGWNVPIGTLRPFLN[+2863.041]WTGPPEPIEAAVAR.F | HexNAc(5)Hex(6)Fuc(4)NeuAc(1) | N[+2863] | 1524.2074 | 4 | 6093.8078 | 6093.7518 | 0 | 9.2 | 141 | Specific | 346.54 | 346.54 | 346.54 | 7.31 | 130 | >sp\|P02788\|TRFL_HUMAN Lactotransferrin OS=Homo sapiens OX=9606 GN=LTF PE=1 SV=6 | 1270 | 20201113_gy_milk_1ng_glyco_hcd203040_compound15.22797.22797.4 | controllerType=0 controllerNumber=1 scan=22797 | 46.1015 |
| 19371:4 | 1 | R.TAGWNVPIGTLRPFLN[+2059.735]WTGPPEPIEAAVAR.F | HexNAc(4)Hex(5)Fuc(1)NeuAc(1) | N[+2060] | 1323.3687 | 4 | 5290.4528 | 5290.4459 | 0 | 1.3 | 141 | Specific | 365.96 | 324.93 | 324.93 | 8.27 | 130 | >sp\|P02788\|TRFL_HUMAN Lactotransferrin OS=Homo sapiens OX=9606 GN=LTF PE=1 SV=6 | 1270 | 20201113_gy_milk_1ng_glyco_hcd203040_compound15.22899.22899.4 | controllerType=0 controllerNumber=1 scan=22899 | 46.3383 |
| 19200:3 | 1 | R.TAGWNVPIGTLRPFLN[+2716.983]WTGPPEPIEAAVAR.F | HexNAc(5)Hex(6)Fuc(3)NeuAc(1) | N[+2717] | 1983.2426 | 3 | 5947.7134 | 5947.6939 | 0 | 3.3 | 141 | Specific | 371.01 | 319.16 | 319.16 | 7.97 | 130 | >sp\|P02788\|TRFL_HUMAN Lactotransferrin OS=Homo sapiens OX=9606 GN=LTF PE=1 SV=6 | 1270 | 20201113_gy_milk_1ng_glyco_hcd203040_compound15.22720.22720.3 | controllerType=0 controllerNumber=1 scan=22720 | 45.9299 |
| 20805:4 | 1 | R.[+42.011]TAGWNVPIGTLRPFLN[+2336.851]WTGPPEPIEAAVAR.F | HexNAc(6)Hex(6)Fuc(1) | .[42], N[+2337] | 1403.1454 | 4 | 5609.5597 | 5609.5726 | 0 | -2.3 | 141 | Specific | 373.66 | 373.66 | 373.66 | 8.53 | 130 | >sp\|P02788\|TRFL_HUMAN Lactotransferrin OS=Homo sapiens OX=9606 GN=LTF PE=1 SV=6 | 1270 | 20201113_gy_milk_1ng_glyco_hcd203040_compound15.24405.24405.4 | controllerType=0 controllerNumber=1 scan=24405 | 49.3304 |
| 19425:3 | 1 | R.TAGWNVPIGTLRPFLN[+2205.793]WTGPPEPIEAAVAR.F | HexNAc(4)Hex(5)Fuc(2)NeuAc(1) | N[+2206] | 1812.8414 | 3 | 5436.5096 | 5436.5038 | 0 | 1.1 | 141 | Specific | 389.79 | 345.02 | 345.02 | 8.24 | 130 | >sp\|P02788\|TRFL_HUMAN Lactotransferrin OS=Homo sapiens OX=9606 GN=LTF PE=1 SV=6 | 1270 | 20201113_gy_milk_1ng_glyco_hcd203040_compound15.22955.22955.3 | controllerType=0 controllerNumber=1 scan=22955 | 46.4481 |
| 20796:4 | 1 | R.TAGWNVPIGTLRPFLN[+2350.830]WTGPPEPIEAAVAR.F | HexNAc(4)Hex(5)Fuc(1)NeuAc(2) | N[+2351] | 1396.1466 | 4 | 5581.5646 | 5581.5413 | 0 | 4.2 | 141 | Specific | 456.15 | 315.58 | 315.58 | 8.03 | 130 | >sp\|P02788\|TRFL_HUMAN Lactotransferrin OS=Homo sapiens OX=9606 GN=LTF PE=1 SV=6 | 1270 | 20201113_gy_milk_1ng_glyco_hcd203040_compound15.24396.24396.4 | controllerType=0 controllerNumber=1 scan=24396 | 49.3147 |
| 19197:3 | 1 | R.TAGWNVPIGTLRPFLN[+2716.983]WTGPPEPIEAAVAR.F | HexNAc(5)Hex(6)Fuc(3)NeuAc(1) | N[+2717] | 1983.2426 | 3 | 5947.7134 | 5947.6939 | 0 | 3.3 | 141 | Specific | 470.98 | 410.04 | 410.04 | 10.06 | 130 | >sp\|P02788\|TRFL_HUMAN Lactotransferrin OS=Homo sapiens OX=9606 GN=LTF PE=1 SV=6 | 1270 | 20201113_gy_milk_1ng_glyco_hcd203040_compound15.22717.22717.3 | controllerType=0 controllerNumber=1 scan=22717 | 45.9178 |
| 19427:3 | 1 | R.TAGWNVPIGTLRPFLN[+2205.793]WTGPPEPIEAAVAR.F | HexNAc(4)Hex(5)Fuc(2)NeuAc(1) | N[+2206] | 1812.8414 | 3 | 5436.5096 | 5436.5038 | 0 | 1.1 | 141 | Specific | 581.51 | 527.72 | 527.72 | 11.69 | 130 | >sp\|P02788\|TRFL_HUMAN Lactotransferrin OS=Homo sapiens OX=9606 GN=LTF PE=1 SV=6 | 1270 | 20201113_gy_milk_1ng_glyco_hcd203040_compound15.22957.22957.3 | controllerType=0 controllerNumber=1 scan=22957 | 46.4518 |
| 19276:4 | 1 | R.TAGWNVPIGTLRPFLN[+2570.925]WTGPPEPIEAAVAR.F | HexNAc(5)Hex(6)Fuc(2)NeuAc(1) | N[+2571] | 1451.1748 | 4 | 5801.6774 | 5801.636 | 0 | 7.1 | 141 | Specific | 607.23 | 522.52 | 522.52 | 12.02 | 130 | >sp\|P02788\|TRFL_HUMAN Lactotransferrin OS=Homo sapiens OX=9606 GN=LTF PE=1 SV=6 | 1270 | 20201113_gy_milk_1ng_glyco_hcd203040_compound15.22800.22800.4 | controllerType=0 controllerNumber=1 scan=22800 | 46.1075 |
| 19279:4 | 1 | R.TAGWNVPIGTLRPFLN[+2570.925]WTGPPEPIEAAVAR.F | HexNAc(5)Hex(6)Fuc(2)NeuAc(1) | N[+2571] | 1451.1748 | 4 | 5801.6774 | 5801.636 | 0 | 7.1 | 141 | Specific | 612.88 | 493.74 | 493.74 | 12.02 | 130 | >sp\|P02788\|TRFL_HUMAN Lactotransferrin OS=Homo sapiens OX=9606 GN=LTF PE=1 SV=6 | 1270 | 20201113_gy_milk_1ng_glyco_hcd203040_compound15.22803.22803.4 | controllerType=0 controllerNumber=1 scan=22803 | 46.113 |
| 19330:4 | 1 | R.TAGWNVPIGTLRPFLN[+2205.793]WTGPPEPIEAAVAR.F | HexNAc(4)Hex(5)Fuc(2)NeuAc(1) | N[+2206] | 1359.8828 | 4 | 5436.5094 | 5436.5038 | 0 | 1 | 141 | Specific | 706.94 | 665.02 | 665.02 | 14.78 | 130 | >sp\|P02788\|TRFL_HUMAN Lactotransferrin OS=Homo sapiens OX=9606 GN=LTF PE=1 SV=6 | 1270 | 20201113_gy_milk_1ng_glyco_hcd203040_compound15.22856.22856.4 | controllerType=0 controllerNumber=1 scan=22856 | 46.2462 |
| 19541:3 | 1 | R.TAGWNVPIGTLRPFLN[+2059.735]WTGPPEPIEAAVAR.F | HexNAc(4)Hex(5)Fuc(1)NeuAc(1) | N[+2060] | 1764.1524 | 3 | 5290.4427 | 5290.4459 | 0 | -0.6 | 141 | Specific | 723.48 | 690.22 | 690.22 | 14.42 | 130 | >sp\|P02788\|TRFL_HUMAN Lactotransferrin OS=Homo sapiens OX=9606 GN=LTF PE=1 SV=6 | 1270 | 20201113_gy_milk_1ng_glyco_hcd203040_compound15.23079.23079.3 | controllerType=0 controllerNumber=1 scan=23079 | 46.6493 |
| 20799:4 | 1 | R.TAGWNVPIGTLRPFLN[+2350.830]WTGPPEPIEAAVAR.F | HexNAc(4)Hex(5)Fuc(1)NeuAc(2) | N[+2351] | 1396.1466 | 4 | 5581.5646 | 5581.5413 | 0 | 4.2 | 141 | Specific | 769.71 | 553.36 | 553.36 | 15.43 | 130 | >sp\|P02788\|TRFL_HUMAN Lactotransferrin OS=Homo sapiens OX=9606 GN=LTF PE=1 SV=6 | 1270 | 20201113_gy_milk_1ng_glyco_hcd203040_compound15.24399.24399.4 | controllerType=0 controllerNumber=1 scan=24399 | 49.3188 |
| 19333:4 | 1 | R.TAGWNVPIGTLRPFLN[+2205.793]WTGPPEPIEAAVAR.F | HexNAc(4)Hex(5)Fuc(2)NeuAc(1) | N[+2206] | 1359.8828 | 4 | 5436.5094 | 5436.5038 | 0 | 1 | 141 | Specific | 774.02 | 647.49 | 647.49 | 15.78 | 130 | >sp\|P02788\|TRFL_HUMAN Lactotransferrin OS=Homo sapiens OX=9606 GN=LTF PE=1 SV=6 | 1270 | 20201113_gy_milk_1ng_glyco_hcd203040_compound15.22859.22859.4 | controllerType=0 controllerNumber=1 scan=22859 | 46.2558 |
| 16798:3 | 1 | R.[+42.011]TAGWNIPMGLLFN[+1872.651]QTGSC[+57.021]K.F | HexNAc(3)Hex(6)NeuAc(1) | .[42], C[+57], N[+1873] | 1337.2377 | 3 | 4009.6986 | 4009.671 | 0 | 6.9 | 485 | Specific | 220.05 | 220.05 | 220.05 | 4.97 | 130 | >sp\|P02788\|TRFL_HUMAN Lactotransferrin OS=Homo sapiens OX=9606 GN=LTF PE=1 SV=6 | 1270 | 20201113_gy_milk_1ng_glyco_hcd203040_compound15.20200.20200.3 | controllerType=0 controllerNumber=1 scan=20200 | 40.593 |
| 16957:3 | 1 | R.TAGWNIPMGLLFN[+1606.587]QTGSC[+57.021]K.F | HexNAc(4)Hex(4)Fuc(1) | C[+57], N[+1607] | 1234.5366 | 3 | 3701.5951 | 3701.5966 | 0 | -0.4 | 485 | Specific | 284.47 | 284.47 | 284.47 | 7.06 | 130 | >sp\|P02788\|TRFL_HUMAN Lactotransferrin OS=Homo sapiens OX=9606 GN=LTF PE=1 SV=6 | 1270 | 20201113_gy_milk_1ng_glyco_hcd203040_compound15.20366.20366.3 | controllerType=0 controllerNumber=1 scan=20366 | 40.9071 |
| 17406:3 | 1 | R.TAGWNIPMGLLFN[+1095.397]QTGSC[+57.021]K.F | HexNAc(3)Hex(3) | C[+57], N[+1095] | 1064.1499 | 3 | 3190.4352 | 3190.4065 | 0 | 9 | 485 | Specific | 338.47 | 283.18 | 283.18 | 5.26 | 130 | >sp\|P02788\|TRFL_HUMAN Lactotransferrin OS=Homo sapiens OX=9606 GN=LTF PE=1 SV=6 | 1270 | 20201113_gy_milk_1ng_glyco_hcd203040_compound15.20838.20838.3 | controllerType=0 controllerNumber=1 scan=20838 | 41.8434 |
| 17027:3 | 1 | R.TAGWNIPMGLLFN[+1606.587]QTGSC[+57.021]K.F | HexNAc(4)Hex(4)Fuc(1) | C[+57], N[+1607] | 1234.5438 | 3 | 3701.6167 | 3701.5966 | 0 | 5.4 | 485 | Specific | 400.78 | 400.78 | 400.78 | 9.25 | 130 | >sp\|P02788\|TRFL_HUMAN Lactotransferrin OS=Homo sapiens OX=9606 GN=LTF PE=1 SV=6 | 1270 | 20201113_gy_milk_1ng_glyco_hcd203040_compound15.20438.20438.3 | controllerType=0 controllerNumber=1 scan=20438 | 41.0434 |
| 17184:3 | 1 | R.TAGWNIPMGLLFN[+1378.476]QTGSC[+57.021]K.F | HexNAc(2)Hex(6) | C[+57], N[+1378] | 1158.5056 | 3 | 3473.5022 | 3473.4856 | 0 | 4.8 | 485 | Specific | 429.89 | 429.89 | 155.18 | 9.29 | 130 | >sp\|P02788\|TRFL_HUMAN Lactotransferrin OS=Homo sapiens OX=9606 GN=LTF PE=1 SV=6 | 1270 | 20201113_gy_milk_1ng_glyco_hcd203040_compound15.20605.20605.3 | controllerType=0 controllerNumber=1 scan=20605 | 41.3896 |
| 18715:3 | 1 | R.TAGWNIPMGLLFN[+2205.793]QTGSC[+57.021]K.F | HexNAc(4)Hex(5)Fuc(2)NeuAc(1) | C[+57], N[+2206] | 1434.2773 | 3 | 4300.8173 | 4300.8028 | 0 | 3.4 | 485 | Specific | 443.51 | 313.76 | 313.76 | 7.99 | 130 | >sp\|P02788\|TRFL_HUMAN Lactotransferrin OS=Homo sapiens OX=9606 GN=LTF PE=1 SV=6 | 1270 | 20201113_gy_milk_1ng_glyco_hcd203040_compound15.22212.22212.3 | controllerType=0 controllerNumber=1 scan=22212 | 44.8195 |
| 18673:3 | 1 | R.TAGWNIPMGLLFN[+2204.772]QTGSC[+57.021]K.F | HexNAc(4)Hex(5)NeuAc(2) | C[+57], N[+2205] | 1433.9367 | 3 | 4299.7956 | 4299.7824 | 0 | 3.1 | 485 | Specific | 462.99 | 376.78 | 376.78 | 10 | 130 | >sp\|P02788\|TRFL_HUMAN Lactotransferrin OS=Homo sapiens OX=9606 GN=LTF PE=1 SV=6 | 1270 | 20201113_gy_milk_1ng_glyco_hcd203040_compound15.22166.22166.3 | controllerType=0 controllerNumber=1 scan=22166 | [illegible] |
| 16879:3 | 1 | R.TAGWNIPMGLLFN[+1768.640]QTGSC[+57.021]K.F | HexNAc(4)Hex(5)Fuc(1) | C[+57], N[+1769] | 1288.5571 | 3 | 3863.6566 | 3863.6495 | 0 | 1.9 | 485 | Specific | 526.49 | 526.49 | 526.49 | 10.36 | 130 | >sp\|P02788\|TRFL_HUMAN Lactotransferrin OS=Homo sapiens OX=9606 GN=LTF PE=1 SV=6 | 1270 | 20201113_gy_milk_1ng_glyco_hcd203040_compound15.20285.20285.3 | controllerType=0 controllerNumber=1 scan=20285 | [illegible] |
| 17408:3 | 1 | R.TAGWNIPMGLLFN[+1095.397]QTGSC[+57.021]K.F | HexNAc(3)Hex(3) | C[+57], N[+1095] | 1064.1499 | 3 | 3190.4352 | 3190.4065 | 0 | 9 | 485 | Specific | 542.91 | 506.85 | 506.85 | 7.67 | 130 | >sp\|P02788\|TRFL_HUMAN Lactotransferrin OS=Homo sapiens OX=9606 GN=LTF PE=1 SV=6 | 1270 | 20201113_gy_milk_1ng_glyco_hcd203040_compound15.20840.20840.3 | controllerType=0 controllerNumber=1 scan=20840 | [illegible] |
| 17166:3 | 1 | R.TAGWNIPMGLLFN[+1378.476]QTGSC[+57.021]K.F | HexNAc(2)Hex(6) | C[+57], N[+1378] | 1158.5054 | 3 | 3473.5017 | 3473.4856 | 0 | 4.6 | 485 | Specific | 597.27 | 597.27 | 277.46 | 11.4 | 130 | >sp\|P02788\|TRFL_HUMAN Lactotransferrin OS=Homo sapiens OX=9606 GN=LTF PE=1 SV=6 | 1270 | 20201113_gy_milk_1ng_glyco_hcd203040_compound15.20586.20586.3 | controllerType=0 controllerNumber=1 scan=20586 | [illegible] |
| 17186:3 | 1 | R.TAGWNIPMGLLFN[+1378.476]QTGSC[+57.021]K.F | HexNAc(2)Hex(6) | C[+57], N[+1378] | 1158.5056 | 3 | 3473.5022 | 3473.4856 | 0 | 4.8 | 485 | Specific | 604.61 | 578.56 | 316.99 | 12.15 | 130 | >sp\|P02788\|TRFL_HUMAN Lactotransferrin OS=Homo sapiens OX=9606 GN=LTF PE=1 SV=6 | 1270 | 20201113_gy_milk_1ng_glyco_hcd203040_compound15.20607.20607.3 | controllerType=0 controllerNumber=1 scan=20607 | 41.395 |
| 18821:3 | 1 | R.TAGWNIPMGLLFN[+1913.677]QTGSC[+57.021]K.F | HexNAc(4)Hex(5)NeuAc(1) | C[+57], N[+1914] | 1336.905 | 3 | 4008.7003 | 4008.687 | 0 | 3.3 | 485 | Specific | 607.15 | 568.19 | 568.19 | 12.69 | 130 | >sp\|P02788\|TRFL_HUMAN Lactotransferrin OS=Homo sapiens OX=9606 GN=LTF PE=1 SV=6 | 1270 | 20201113_gy_milk_1ng_glyco_hcd203040_compound15.22323.22323.2 | controllerType=0 controllerNumber=1 scan=22323 | 45.048 |
| 16959:3 | 1 | R.TAGWNIPMGLLFN[+1606.587]QTGSC[+57.021]K.F | HexNAc(4)Hex(4)Fuc(1) | C[+57], N[+1607] | 1234.5366 | 3 | 3701.5951 | 3701.5966 | 0 | -0.4 | 485 | Specific | 633.28 | 599.61 | 599.61 | 12.48 | 130 | >sp\|P02788\|TRFL_HUMAN Lactotransferrin OS=Homo sapiens OX=9606 GN=LTF PE=1 SV=6 | 1270 | 20201113_gy_milk_1ng_glyco_hcd203040_compound15.20368.20368.3 | controllerType=0 controllerNumber=1 scan=20368 | 40.9107 |
| 18691:3 | 1 | R.TAGWNIPMGLLFN[+2205.793]QTGSC[+57.021]K.F | HexNAc(4)Hex(5)Fuc(2)NeuAc(1) | C[+57], N[+2206] | 1434.2792 | 3 | 4300.8231 | 4300.8028 | 0 | 4.7 | 485 | Specific | 664.24 | 534.19 | 534.19 | 13.15 | 130 | >sp\|P02788\|TRFL_HUMAN Lactotransferrin OS=Homo sapiens OX=9606 GN=LTF PE=1 SV=6 | 1270 | 20201113_gy_milk_1ng_glyco_hcd203040_compound15.22185.22185.3 | controllerType=0 controllerNumber=1 scan=22185 | 44.7652 |
| 18708:3 | 1 | R.TAGWNIPMGLLFN[+2205.793]QTGSC[+57.021]K.F | HexNAc(4)Hex(5)Fuc(2)NeuAc(1) | C[+57], N[+2206] | 1434.268 | 3 | 4300.7895 | 4300.8028 | 0 | -3.1 | 485 | Specific | 689.8 | 676.19 | 676.19 | 12.45 | 130 | >sp\|P02788\|TRFL_HUMAN Lactotransferrin OS=Homo sapiens OX=9606 GN=LTF PE=1 SV=6 | 1270 | 20201113_gy_milk_1ng_glyco_hcd203040_compound15.22204.22204.3 | controllerType=0 controllerNumber=1 scan=22204 | 44.8054 |
| 16848:3 | 1 | R.TAGWNIPMGLLFN[+1768.640]QTGSC[+57.021]K.F | HexNAc(4)Hex(5)Fuc(1) | C[+57], N[+1769] | 1288.5521 | 3 | 3863.6416 | 3863.6495 | 0 | -2 | 485 | Specific | 737.88 | 737.88 | 737.88 | 13.45 | 130 | >sp\|P02788\|TRFL_HUMAN Lactotransferrin OS=Homo sapiens OX=9606 GN=LTF PE=1 SV=6 | 1270 | 20201113_gy_milk_1ng_glyco_hcd203040_compound15.20252.20252.3 | controllerType=0 controllerNumber=1 scan=20252 | 40.6904 |
| 20763:3 | 1 | R.TAGWNIPMGLLFN[+2350.830]QTGSC[+57.021]K.F | HexNAc(4)Hex(5)Fuc(1)NeuAc(2) | C[+57], N[+2351] | 1482.623 | 3 | 4445.8544 | 4445.8403 | 0 | 3.2 | 485 | Specific | 761.86 | 761.86 | 761.86 | 15.69 | 130 | >sp\|P02788\|TRFL_HUMAN Lactotransferrin OS=Homo sapiens OX=9606 GN=LTF PE=1 SV=6 | 1270 | 20201113_gy_milk_1ng_glyco_hcd203040_compound15.24361.24361.3 | controllerType=0 controllerNumber=1 scan=24361 | 49.235 |
| 18711:3 | 1 | R.TAGWNIPMGLLFN[+2205.793]QTGSC[+57.021]K.F | HexNAc(4)Hex(5)Fuc(2)NeuAc(1) | C[+57], N[+2206] | 1434.268 | 3 | 4300.7895 | 4300.8028 | 0 | -3.1 | 485 | Specific | 787.41 | 626.14 | 626.14 | 14.45 | 130 | >sp\|P02788\|TRFL_HUMAN Lactotransferrin OS=Homo sapiens OX=9606 GN=LTF PE=1 SV=6 | 1270 | 20201113_gy_milk_1ng_glyco_hcd203040_compound15.22207.22207.3 | controllerType=0 controllerNumber=1 scan=22207 | 44.8102 |
| 18717:3 | 1 | R.TAGWNIPMGLLFN[+2205.793]QTGSC[+57.021]K.F | HexNAc(4)Hex(5)Fuc(2)NeuAc(1) | C[+57], N[+2206] | 1434.2773 | 3 | 4300.8173 | 4300.8028 | 0 | 3.4 | 485 | Specific | 797.61 | 796.66 | 796.66 | 15.4 | 130 | >sp\|P02788\|TRFL_HUMAN Lactotransferrin OS=Homo sapiens OX=9606 GN=LTF PE=1 SV=6 | 1270 | 20201113_gy_milk_1ng_glyco_hcd203040_compound15.22214.22214.3 | controllerType=0 controllerNumber=1 scan=22214 | 44.8232 |
| 17253:2 | 1 | R.TAGWNIPMGLLFN[+1216.423]QTGSC[+57.021]K.F | HexNAc(2)Hex(5) | C[+57], N[+1216] | 1656.2238 | 2 | 3311.4403 | 3311.4328 | 0 | 2.3 | 485 | Specific | 805.4 | 766.35 | 250.73 | 16.6 | 130 | >sp\|P02788\|TRFL_HUMAN Lactotransferrin OS=Homo sapiens OX=9606 GN=LTF PE=1 SV=6 | 1270 | 20201113_gy_milk_1ng_glyco_hcd203040_compound15.20678.20678.2 | controllerType=0 controllerNumber=1 scan=20678 | 41.5418 |
| 18695:3 | 1 | R.TAGWNIPMGLLFN[+2059.735]QTGSC[+57.021]K.F | HexNAc(4)Hex(5)Fuc(1)NeuAc(1) | C[+57], N[+2060] | 1385.5837 | 3 | 4154.7365 | 4154.7449 | 0 | -2 | 485 | Specific | 832.91 | 779.35 | 779.35 | 15.45 | 130 | >sp\|P02788\|TRFL_HUMAN Lactotransferrin OS=Homo sapiens OX=9606 GN=LTF PE=1 SV=6 | 1270 | 20201113_gy_milk_1ng_glyco_hcd203040_compound15.22190.22190.3 | controllerType=0 controllerNumber=1 scan=22190 | 44.774 |
| 18675:3 | 1 | R.TAGWNIPMGLLFN[+2204.772]QTGSC[+57.021]K.F | HexNAc(4)Hex(5)NeuAc(2) | C[+57], N[+2205] | 1433.9367 | 3 | 4299.7956 | 4299.7824 | 0 | 3.1 | 485 | Specific | 834.94 | 825.88 | 825.88 | 16.69 | 130 | >sp\|P02788\|TRFL_HUMAN Lactotransferrin OS=Homo sapiens OX=9606 GN=LTF PE=1 SV=6 | 1270 | 20201113_gy_milk_1ng_glyco_hcd203040_compound15.22168.22168.3 | controllerType=0 controllerNumber=1 scan=22168 | 44.7329 |
| 16881:3 | 1 | R.TAGWNIPMGLLFN[+1768.640]QTGSC[+57.021]K.F | HexNAc(4)Hex(5)Fuc(1) | C[+57], N[+1769] | 1288.5571 | 3 | 3863.6566 | 3863.6495 | 0 | 1.9 | 485 | Specific | 837.9 | 837.9 | 837.9 | 16.75 | 130 | >sp\|P02788\|TRFL_HUMAN Lactotransferrin OS=Homo sapiens OX=9606 GN=LTF PE=1 SV=6 | 1270 | 20201113_gy_milk_1ng_glyco_hcd203040_compound15.20287.20287.3 | controllerType=0 controllerNumber=1 scan=20287 | 40.752 |
| 17212:3 | 1 | R.TAGWNIPMGLLFN[+1216.423]QTGSC[+57.021]K.F | HexNAc(2)Hex(5) | C[+57], N[+1216] | 1104.4831 | 3 | 3311.4346 | 3311.4328 | 0 | 0.6 | 485 | Specific | 838.15 | 646.02 | 292.95 | 16.48 | 130 | >sp\|P02788\|TRFL_HUMAN Lactotransferrin OS=Homo sapiens OX=9606 GN=LTF PE=1 SV=6 | 1270 | 20201113_gy_milk_1ng_glyco_hcd203040_compound15.20635.20635.3 | controllerType=0 controllerNumber=1 scan=20635 | 41.4537 |
| 17082:3 | 1 | R.TAGWNIPMGLLFN[+1622.582]QTGSC[+57.021]K.F | HexNAc(4)Hex(5) | C[+57], N[+1623] | 1239.8697 | 3 | 3717.5945 | 3717.5916 | 0 | 0.8 | 485 | Specific | 844.87 | 728.68 | 337.26 | 16.75 | 130 | >sp\|P02788\|TRFL_HUMAN Lactotransferrin OS=Homo sapiens OX=9606 GN=LTF PE=1 SV=6 | 1270 | 20201113_gy_milk_1ng_glyco_hcd203040_compound15.20498.20498.3 | controllerType=0 controllerNumber=1 scan=20498 | 41.1792 |
| 17084:3 | 1 | R.TAGWNIPMGLLFN[+1622.582]QTGSC[+57.021]K.F | HexNAc(4)Hex(5) | C[+57], N[+1623] | 1239.8697 | 3 | 3717.5945 | 3717.5916 | 0 | 0.8 | 485 | Specific | 873.06 | 800.03 | 381.37 | 17.75 | 130 | >sp\|P02788\|TRFL_HUMAN Lactotransferrin OS=Homo sapiens OX=9606 GN=LTF PE=1 SV=6 | 1270 | 20201113_gy_milk_1ng_glyco_hcd203040_compound15.20500.20500.3 | controllerType=0 controllerNumber=1 scan=20500 | 41.1835 |
| 18693:3 | 1 | R.TAGWNIPMGLLFN[+2205.793]QTGSC[+57.021]K.F | HexNAc(4)Hex(5)Fuc(2)NeuAc(1) | C[+57], N[+2206] | 1434.2792 | 3 | 4300.8231 | 4300.8028 | 0 | 4.7 | 485 | Specific | 884.63 | 854.2 | 854.2 | 17.15 | 130 | >sp\|P02788\|TRFL_HUMAN Lactotransferrin OS=Homo sapiens OX=9606 GN=LTF PE=1 SV=6 | 1270 | 20201113_gy_milk_1ng_glyco_hcd203040_compound15.22187.22187.3 | controllerType=0 controllerNumber=1 scan=22187 | 44.7679 |
| 17068:3 | 1 | R.TAGWNIPMGLLFN[+1622.582]QTGSC[+57.021]K.F | HexNAc(4)Hex(5) | C[+57], N[+1623] | 1239.868 | 3 | 3717.5894 | 3717.5916 | 0 | -0.6 | 485 | Specific | 886.5 | 886.5 | 409.91 | 17.48 | 130 | >sp\|P02788\|TRFL_HUMAN Lactotransferrin OS=Homo sapiens OX=9606 GN=LTF PE=1 SV=6 | 1270 | 20201113_gy_milk_1ng_glyco_hcd203040_compound15.20481.20481.3 | controllerType=0 controllerNumber=1 scan=20481 | 41.1346 |
| 16846:3 | 1 | R.TAGWNIPMGLLFN[+1768.640]QTGSC[+57.021]K.F | HexNAc(4)Hex(5)Fuc(1) | C[+57], N[+1769] | 1288.5521 | 3 | 3863.6416 | 3863.6495 | 0 | -2 | 485 | Specific | 886.95 | 884.76 | 884.76 | 16.45 | 130 | >sp\|P02788\|TRFL_HUMAN Lactotransferrin OS=Homo sapiens OX=9606 GN=LTF PE=1 SV=6 | 1270 | 20201113_gy_milk_1ng_glyco_hcd203040_compound15.20250.20250.3 | controllerType=0 controllerNumber=1 scan=20250 | 40.686 |
| 18697:3 | 1 | R.TAGWNIPMGLLFN[+2059.735]QTGSC[+57.021]K.F | HexNAc(4)Hex(5)Fuc(1)NeuAc(1) | C[+57], N[+2060] | 1385.5837 | 3 | 4154.7365 | 4154.7449 | 0 | -2 | 485 | Specific | 888.37 | 728.45 | 728.45 | 16.45 | 130 | >sp\|P02788\|TRFL_HUMAN Lactotransferrin OS=Homo sapiens OX=9606 GN=LTF PE=1 SV=6 | 1270 | 20201113_gy_milk_1ng_glyco_hcd203040_compound15.22192.22192.3 | controllerType=0 controllerNumber=1 scan=22192 | 44.7806 |
| 17210:3 | 1 | R.TAGWNIPMGLLFN[+1216.423]QTGSC[+57.021]K.F | HexNAc(2)Hex(5) | C[+57], N[+1216] | 1104.4831 | 3 | 3311.4346 | 3311.4328 | 0 | 0.6 | 485 | Specific | 906.47 | 896.67 | 363.37 | 18.48 | 130 | >sp\|P02788\|TRFL_HUMAN Lactotransferrin OS=Homo sapiens OX=9606 GN=LTF PE=1 SV=6 | 1270 | 20201113_gy_milk_1ng_glyco_hcd203040_compound15.20633.20633.3 | controllerType=0 controllerNumber=1 scan=20633 | 41.4486 |
| 17251:2 | 1 | R.TAGWNIPMGLLFN[+1216.423]QTGSC[+57.021]K.F | HexNAc(2)Hex(5) | C[+57], N[+1216] | 1656.2238 | 2 | 3311.4403 | 3311.4328 | 0 | 2.3 | 485 | Specific | 914.32 | 878.08 | 339.42 | 18.6 | 130 | >sp\|P02788\|TRFL_HUMAN Lactotransferrin OS=Homo sapiens OX=9606 GN=LTF PE=1 SV=6 | 1270 | 20201113_gy_milk_1ng_glyco_hcd203040_compound15.20676.20676.2 | controllerType=0 controllerNumber=1 scan=20676 | 41.5359 |
| 17066:3 | 1 | R.TAGWNIPMGLLFN[+1622.582]QTGSC[+57.021]K.F | HexNAc(4)Hex(5) | C[+57], N[+1623] | 1239.868 | 3 | 3717.5894 | 3717.5916 | 0 | -0.6 | 485 | Specific | 919.89 | 782.08 | 481.22 | 18.48 | 130 | >sp\|P02788\|TRFL_HUMAN Lactotransferrin OS=Homo sapiens OX=9606 GN=LTF PE=1 SV=6 | 1270 | 20201113_gy_milk_1ng_glyco_hcd203040_compound15.20479.20479.3 | controllerType=0 controllerNumber=1 scan=20479 | 41.1233 |
| 20761:3 | 1 | R.TAGWNIPMGLLFN[+2350.830]QTGSC[+57.021]K.F | HexNAc(4)Hex(5)Fuc(1)NeuAc(2) | C[+57], N[+2351] | 1482.623 | 3 | 4445.8544 | 4445.8403 | 0 | 3.2 | 485 | Specific | 976.01 | 917.04 | 917.04 | 19.69 | 130 | >sp\|P02788\|TRFL_HUMAN Lactotransferrin OS=Homo sapiens OX=9606 GN=LTF PE=1 SV=6 | 1270 | 20201113_gy_milk_1ng_glyco_hcd203040_compound15.24359.24359.3 | controllerType=0 controllerNumber=1 scan=24359 | 49.2294 |
| 17450:3 | 1 | R.TAGWNIPM[+15.995]GLLFN[+2059.735]QTGSC[+57.021]K.F | HexNAc(4)Hex(5)Fuc(1)NeuAc(1) | C[+57], M[+16], N[+2060] | 1390.9201 | 3 | 4170.7457 | 4170.7398 | 0 | 1.4 | 485 | Specific | 253.81 | 199.36 | 199.36 | 4.34 | 130 | >sp\|P02788\|TRFL_HUMAN Lactotransferrin OS=Homo sapiens OX=9606 GN=LTF PE=1 SV=6 | 1270 | 20201113_gy_milk_1ng_glyco_hcd203040_compound15.20884.20884.3 | controllerType=0 controllerNumber=1 scan=20884 | 41.9313 |
| 16643:3 | 1 | R.TAGWNIPM[+15.995]GLLFN[+1913.677]QTGSC[+57.021]K.F | HexNAc(4)Hex(5)NeuAc(1) | C[+57], M[+16], N[+1914] | 1342.2361 | 3 | 4024.6939 | 4024.6819 | 0 | 3 | 485 | Specific | 321.88 | 321.88 | 321.88 | 7.79 | 130 | >sp\|P02788\|TRFL_HUMAN Lactotransferrin OS=Homo sapiens OX=9606 GN=LTF PE=1 SV=6 | 1270 | 20201113_gy_milk_1ng_glyco_hcd203040_compound15.20038.20038.3 | controllerType=0 controllerNumber=1 scan=20038 | 40.3092 |
| 17412:3 | 1 | R.TAGWNIPM[+15.995]GLLFN[+1913.677]QTGSC[+57.021]K.F | HexNAc(4)Hex(5)NeuAc(1) | C[+57], M[+16], N[+1914] | 1342.2408 | 3 | 4024.7078 | 4024.6819 | 0 | 6.4 | 485 | Specific | 505.93 | 468.71 | 468.71 | 9.46 | 130 | >sp\|P02788\|TRFL_HUMAN Lactotransferrin OS=Homo sapiens OX=9606 GN=LTF PE=1 SV=6 | 1270 | 20201113_gy_milk_1ng_glyco_hcd203040_compound15.20844.20844.3 | controllerType=0 controllerNumber=1 scan=20844 | 41.8552 |
| 18063:3 | 1 | R.TAGWNIPMGLLFN[+1768.640]QTGSC[+57.021]KFDEYFSQSC[+57.021]APGSDPR.S | HexNAc(4)Hex(5)Fuc(1) | C[+57]*2, N[+1769] | 1903.1408 | 3 | 5707.4079 | 5707.3966 | 0 | 2 | 485 | Specific | 296.21 | 235.62 | 235.62 | 6.87 | 130 | >sp\|P02788\|TRFL_HUMAN Lactotransferrin OS=Homo sapiens OX=9606 GN=LTF PE=1 SV=6 | 1270 | 20201113_gy_milk_1ng_glyco_hcd203040_compound15.21526.21526.3 | controllerType=0 controllerNumber=1 scan=21526 | 43.2971 |
| 20415:4 | 1 | R.TAGWNIPMGLLFN[+2204.772]QTGSC[+57.021]KFDEYFSQSC[+57.021]APGSDPR.S | HexNAc(4)Hex(5)NeuAc(2) | C[+57]*2, N[+2205] | 1536.6384 | 4 | 6143.5319 | 6143.5295 | 0 | 0.4 | 485 | Specific | 580.4 | 561.43 | 231.37 | 11.26 | 130 | >sp\|P02788\|TRFL_HUMAN Lactotransferrin OS=Homo sapiens OX=9606 GN=LTF PE=1 SV=6 | 1270 | 20201113_gy_milk_1ng_glyco_hcd203040_compound15.23998.23998.4 | controllerType=0 controllerNumber=1 scan=23998 | 48.4849 |
| 20417:4 | 1 | R.TAGWNIPMGLLFN[+2204.772]QTGSC[+57.021]KFDEYFSQSC[+57.021]APGSDPR.S | HexNAc(4)Hex(5)NeuAc(2) | C[+57]*2, N[+2205] | 1536.6384 | 4 | 6143.5319 | 6143.5295 | 0 | 0.4 | 485 | Specific | 603.58 | 595.01 | 338.45 | 12.26 | 130 | >sp\|P02788\|TRFL_HUMAN Lactotransferrin OS=Homo sapiens OX=9606 GN=LTF PE=1 SV=6 | 1270 | 20201113_gy_milk_1ng_glyco_hcd203040_compound15.24000.24000.4 | controllerType=0 controllerNumber=1 scan=24000 | 48.4889 |
| 19039:3 | 1 | R.TAGWNIPMGLLFN[+1913.677]QTGSC[+57.021]KFDEYFSQSC[+57.021]APGSDPR.S | HexNAc(4)Hex(5)NeuAc(1) | C[+57]*2, N[+1914] | 1951.4848 | 3 | 5852.4399 | 5852.4341 | 0 | 1 | 485 | Specific | 668.62 | 562.22 | 562.22 | 13.45 | 130 | >sp\|P02788\|TRFL_HUMAN Lactotransferrin OS=Homo sapiens OX=9606 GN=LTF PE=1 SV=6 | 1270 | 20201113_gy_milk_1ng_glyco_hcd203040_compound15.22552.22552.3 | controllerType=0 controllerNumber=1 scan=22552 | 45.6164 |
| 19048:3 | 1 | R.TAGWNIPMGLLFN[+1913.677]QTGSC[+57.021]KFDEYFSQSC[+57.021]APGSDPR.S | HexNAc(4)Hex(5)NeuAc(1) | C[+57]*2, N[+1914] | 1951.4896 | 3 | 5852.4542 | 5852.4341 | 0 | 3.4 | 485 | Specific | 740.09 | 593.24 | 593.24 | 14.09 | 130 | >sp\|P02788\|TRFL_HUMAN Lactotransferrin OS=Homo sapiens OX=9606 GN=LTF PE=1 SV=6 | 1270 | 20201113_gy_milk_1ng_glyco_hcd203040_compound15.22562.22562.3 | controllerType=0 controllerNumber=1 scan=22562 | 45.6415 |
| 18045:4 | 1 | R.TAGWNIPMGLLFN[+1622.582]QTGSC[+57.021]KFDEYFSQSC[+57.021]APGSDPR.S | HexNAc(4)Hex(5) | C[+57]*2, N[+1623] | 1391.0925 | 4 | 5561.3483 | 5561.3387 | 0 | 1.7 | 485 | Specific | 740.3 | 693.3 | 221.12 | 14.54 | 130 | >sp\|P02788\|TRFL_HUMAN Lactotransferrin OS=Homo sapiens OX=9606 GN=LTF PE=1 SV=6 | 1270 | 20201113_gy_milk_1ng_glyco_hcd203040_compound15.21507.21507.4 | controllerType=0 controllerNumber=1 scan=21507 | 43.2543 |
| 19046:3 | 1 | R.TAGWNIPMGLLFN[+1913.677]QTGSC[+57.021]KFDEYFSQSC[+57.021]APGSDPR.S | HexNAc(4)Hex(5)NeuAc(1) | C[+57]*2, N[+1914] | 1951.4896 | 3 | 5852.4542 | 5852.4341 | 0 | 3.4 | 485 | Specific | 746.7 | 616.97 | 616.97 | 14.09 | 130 | >sp\|P02788\|TRFL_HUMAN Lactotransferrin OS=Homo sapiens OX=9606 GN=LTF PE=1 SV=6 | 1270 | 20201113_gy_milk_1ng_glyco_hcd203040_compound15.22560.22560.3 | controllerType=0 controllerNumber=1 scan=22560 | 45.6372 |
| 18058:4 | 1 | R.TAGWNIPMGLLFN[+1768.640]QTGSC[+57.021]KFDEYFSQSC[+57.021]APGSDPR.S | HexNAc(4)Hex(5)Fuc(1) | C[+57]*2, N[+1769] | 1427.6129 | 4 | 5707.4298 | 5707.3966 | 0 | 5.8 | 485 | Specific | 756.55 | 680.9 | 680.9 | 14.94 | 130 | >sp\|P02788\|TRFL_HUMAN Lactotransferrin OS=Homo sapiens OX=9606 GN=LTF PE=1 SV=6 | 1270 | 20201113_gy_milk_1ng_glyco_hcd203040_compound15.21521.21521.4 | controllerType=0 controllerNumber=1 scan=21521 | 43.2879 |
| 18986:4 | 1 | R.TAGWNIPMGLLFN[+2059.735]QTGSC[+57.021]KFDEYFSQSC[+57.021]APGSDPR.S | HexNAc(4)Hex(5)Fuc(1)NeuAc(1) | C[+57]*2, N[+2060] | 1500.3864 | 4 | 5998.5236 | 5998.492 | 0 | 5.3 | 485 | Specific | 805.35 | 774.21 | 774.21 | 15.94 | 130 | >sp\|P02788\|TRFL_HUMAN Lactotransferrin OS=Homo sapiens OX=9606 GN=LTF PE=1 SV=6 | 1270 | 20201113_gy_milk_1ng_glyco_hcd203040_compound15.22497.22497.4 | controllerType=0 controllerNumber=1 scan=22497 | 45.4902 |
| 19041:3 | 1 | R.TAGWNIPMGLLFN[+1913.677]QTGSC[+57.021]KFDEYFSQSC[+57.021]APGSDPR.S | HexNAc(4)Hex(5)NeuAc(1) | C[+57]*2, N[+1914] | 1951.4848 | 3 | 5852.4399 | 5852.4341 | 0 | 1 | 485 | Specific | 816.27 | 683.32 | 683.32 | 16.45 | 130 | >sp\|P02788\|TRFL_HUMAN Lactotransferrin OS=Homo sapiens OX=9606 GN=LTF PE=1 SV=6 | 1270 | 20201113_gy_milk_1ng_glyco_hcd203040_compound15.22554.22554.3 | controllerType=0 controllerNumber=1 scan=22554 | 45.6195 |
| 19042:4 | 1 | R.TAGWNIPMGLLFN[+1913.677]QTGSC[+57.021]KFDEYFSQSC[+57.021]APGSDPR.S | HexNAc(4)Hex(5)NeuAc(1) | C[+57]*2, N[+1914] | 1463.8682 | 4 | 5852.4508 | 5852.4341 | 0 | 2.9 | 485 | Specific | 909.18 | 839.56 | 839.56 | 18.48 | 130 | >sp\|P02788\|TRFL_HUMAN Lactotransferrin OS=Homo sapiens OX=9606 GN=LTF PE=1 SV=6 | 1270 | 20201113_gy_milk_1ng_glyco_hcd203040_compound15.22556.22556.4 | controllerType=0 controllerNumber=1 scan=22556 | 45.624 |
| 19044:4 | 1 | R.TAGWNIPMGLLFN[+1913.677]QTGSC[+57.021]KFDEYFSQSC[+57.021]APGSDPR.S | HexNAc(4)Hex(5)NeuAc(1) | C[+57]*2, N[+1914] | 1463.8682 | 4 | 5852.4508 | 5852.4341 | 0 | 2.9 | 485 | Specific | 912.72 | 858.51 | 858.51 | 18.48 | 130 | >sp\|P02788\|TRFL_HUMAN Lactotransferrin OS=Homo sapiens OX=9606 GN=LTF PE=1 SV=6 | 1270 | 20201113_gy_milk_1ng_glyco_hcd203040_compound15.22558.22558.4 | controllerType=0 controllerNumber=1 scan=22558 | 45.6309 |
| 18047:4 | 1 | R.TAGWNIPMGLLFN[+1622.582]QTGSC[+57.021]KFDEYFSQSC[+57.021]APGSDPR.S | HexNAc(4)Hex(5) | C[+57]*2, N[+1623] | 1391.0925 | 4 | 5561.3483 | 5561.3387 | 0 | 1.7 | 485 | Specific | 919.97 | 883.04 | 338.43 | 18.54 | 130 | >sp\|P02788\|TRFL_HUMAN Lactotransferrin OS=Homo sapiens OX=9606 GN=LTF PE=1 SV=6 | 1270 | 20201113_gy_milk_1ng_glyco_hcd203040_compound15.21509.21509.4 | controllerType=0 controllerNumber=1 scan=21509 | 43.2591 |
| 19029:4 | 1 | R.TAGWNIPMGLLFN[+1913.677]QTGSC[+57.021]KFDEYFSQSC[+57.021]APGSDPR.S | HexNAc(4)Hex(5)NeuAc(1) | C[+57]*2, N[+1914] | 1463.8654 | 4 | 5852.4396 | 5852.4341 | 0 | 0.9 | 485 | Specific | 943.08 | 891.9 | 891.9 | 18.54 | 130 | >sp\|P02788\|TRFL_HUMAN Lactotransferrin OS=Homo sapiens OX=9606 GN=LTF PE=1 SV=6 | 1270 | 20201113_gy_milk_1ng_glyco_hcd203040_compound15.22542.22542.4 | controllerType=0 controllerNumber=1 scan=22542 | 45.5991 |
| 19027:4 | 1 | R.TAGWNIPMGLLFN[+1913.677]QTGSC[+57.021]KFDEYFSQSC[+57.021]APGSDPR.S | HexNAc(4)Hex(5)NeuAc(1) | C[+57]*2, N[+1914] | 1463.8654 | 4 | 5852.4396 | 5852.4341 | 0 | 0.9 | 485 | Specific | 991.42 | 907.66 | 907.66 | 19.54 | 130 | >sp\|P02788\|TRFL_HUMAN Lactotransferrin OS=Homo sapiens OX=9606 GN=LTF PE=1 SV=6 | 1270 | 20201113_gy_milk_1ng_glyco_hcd203040_compound15.22540.22540.4 | controllerType=0 controllerNumber=1 scan=22540 | 45.5925 |
| 11530:3 | 1 | R.N[+2350.830]GSDC[+57.021]PDKFC[+57.021]LFQSETK.N | HexNAc(4)Hex(5)Fuc(1)NeuAc(2) | C[+57]*2, N[+2351] | 1461.9108 | 3 | 4383.7179 | 4383.7043 | 0 | 3.1 | 642 | Specific | 255.14 | 210.18 | 210.18 | 5.96 | 130 | >sp\|P02788\|TRFL_HUMAN Lactotransferrin OS=Homo sapiens OX=9606 GN=LTF PE=1 SV=6 | 1270 | 20201113_gy_milk_1ng_glyco_hcd203040_compound15.14668.14668.3 | controllerType=0 controllerNumber=1 scan=14668 | 30.8525 |
| 11532:3 | 1 | R.N[+2350.830]GSDC[+57.021]PDKFC[+57.021]LFQSETK.N | HexNAc(4)Hex(5)Fuc(1)NeuAc(2) | C[+57]*2, N[+2351] | 1461.9108 | 3 | 4383.7179 | 4383.7043 | 0 | 3.1 | 642 | Specific | 264.77 | 176.83 | 176.83 | 5.84 | 130 | >sp\|P02788\|TRFL_HUMAN Lactotransferrin OS=Homo sapiens OX=9606 GN=LTF PE=1 SV=6 | 1270 | 20201113_gy_milk_1ng_glyco_hcd203040_compound15.14670.14670.3 | controllerType=0 controllerNumber=1 scan=14670 | 30.8561 |
| 09114:3 | 1 | R.N[+2059.735]GSDC[+57.021]PDKFC[+57.021]LFQSETK.N | HexNAc(4)Hex(5)Fuc(1)NeuAc(1) | C[+57]*2, N[+2060] | 1364.8779 | 3 | 4092.619 | 4092.6088 | 0 | 2.5 | 642 | Specific | 267.47 | 154.7 | 154.7 | 5.44 | 130 | >sp\|P02788\|TRFL_HUMAN Lactotransferrin OS=Homo sapiens OX=9606 GN=LTF PE=1 SV=6 | 1270 | 20201113_gy_milk_1ng_glyco_hcd203040_compound15.12163.12163.3 | controllerType=0 controllerNumber=1 scan=12163 | 26.6254 |
| 09112:3 | 1 | R.N[+2059.735]GSDC[+57.021]PDKFC[+57.021]LFQSETK.N | HexNAc(4)Hex(5)Fuc(1)NeuAc(1) | C[+57]*2, N[+2060] | 1364.8779 | 3 | 4092.619 | 4092.6088 | 0 | 2.5 | 642 | Specific | 288.99 | 217.67 | 217.67 | 6.53 | 130 | >sp\|P02788\|TRFL_HUMAN Lactotransferrin OS=Homo sapiens OX=9606 GN=LTF PE=1 SV=6 | 1270 | 20201113_gy_milk_1ng_glyco_hcd203040_compound15.12161.12161.3 | controllerType=0 controllerNumber=1 scan=12161 | 26.6218 |
| 09012:3 | 1 | R.N[+2205.793]GSDC[+57.021]PDKFC[+57.021]LFQSETK.N | HexNAc(4)Hex(5)Fuc(2)NeuAc(1) | C[+57]*2, N[+2206] | 1413.5607 | 3 | 4238.6676 | 4238.6667 | 0 | 0.2 | 642 | Specific | 387.49 | 247.05 | 247.05 | 6.93 | 130 | >sp\|P02788\|TRFL_HUMAN Lactotransferrin OS=Homo sapiens OX=9606 GN=LTF PE=1 SV=6 | 1270 | 20201113_gy_milk_1ng_glyco_hcd203040_compound15.12057.12057.3 | controllerType=0 controllerNumber=1 scan=12057 | 26.4433 |
| 09010:3 | 1 | R.N[+2205.793]GSDC[+57.021]PDKFC[+57.021]LFQSETK.N | HexNAc(4)Hex(5)Fuc(2)NeuAc(1) | C[+57]*2, N[+2206] | 1413.5607 | 3 | 4238.6676 | 4238.6667 | 0 | 0.2 | 642 | Specific | 394.58 | 271.49 | 271.49 | 6.96 | 130 | >sp\|P02788\|TRFL_HUMAN Lactotransferrin OS=Homo sapiens OX=9606 GN=LTF PE=1 SV=6 | 1270 | 20201113_gy_milk_1ng_glyco_hcd203040_compound15.12055.12055.3 | controllerType=0 controllerNumber=1 scan=12055 | 26.4394 |
| 07310:3 | 3 | R.FSDGLESN[+1864.634]SSTQFEVK.K | HexNAc(2)Hex(9) | N[+1865] | 1213.8211 | 3 | 3639.4488 | 3639.4472 | 0 | 0.4 | 219 | Specific | 627.88 | 627.71 | 627.71 | 12.17 | 89 | >sp\|P0C0L5\|CO4B_HUMAN Complement C4-B OS=Homo sapiens OX=9606 GN=C4B PE=1 SV=2 | 1424 | 20201113_gy_milk_1ng_glyco_hcd203040_compound15.10278.10278.3 | controllerType=0 controllerNumber=1 scan=10278 | 23.4421 |
| 07312:3 | 3 | R.FSDGLESN[+1864.634]SSTQFEVK.K | HexNAc(2)Hex(9) | N[+1865] | 1213.8211 | 3 | 3639.4488 | 3639.4472 | 0 | 0.4 | 219 | Specific | 636.25 | 636.25 | 636.25 | 12.17 | 89 | >sp\|P0C0L5\|CO4B_HUMAN Complement C4-B OS=Homo sapiens OX=9606 GN=C4B PE=1 SV=2 | 1424 | 20201113_gy_milk_1ng_glyco_hcd203040_compound15.10280.10280.3 | controllerType=0 controllerNumber=1 scan=10280 | 23.4465 |
| 04835:3 | 3 | R.FSDGLESN[+1864.634]SSTQFEVKK.Y | HexNAc(2)Hex(9) | N[+1865] | 1256.5191 | 3 | 3767.5427 | 3767.5421 | 0 | 0.2 | 219 | Specific | 314.45 | 314.45 | 314.45 | 6.83 | 89 | >sp\|P0C0L5\|CO4B_HUMAN Complement C4-B OS=Homo sapiens OX=9606 GN=C4B PE=1 SV=2 | 1424 | 20201113_gy_milk_1ng_glyco_hcd203040_compound15.7716.7716.3 | controllerType=0 controllerNumber=1 scan=7716 | 19.1987 |
| 04833:3 | 3 | R.FSDGLESN[+1864.634]SSTQFEVKK.Y | HexNAc(2)Hex(9) | N[+1865] | 1256.5191 | 3 | 3767.5427 | 3767.5421 | 0 | 0.2 | 219 | Specific | 376.34 | 376.34 | 376.34 | 8.93 | 89 | >sp\|P0C0L5\|CO4B_HUMAN Complement C4-B OS=Homo sapiens OX=9606 GN=C4B PE=1 SV=2 | 1424 | 20201113_gy_milk_1ng_glyco_hcd203040_compound15.7714.7714.3 | controllerType=0 controllerNumber=1 scan=7714 | 19.1946 |
| 08442:3 | 6 | K.TAHC[+57.021]N[+2059.735]ESFYFLC[+57.021]K.R | HexNAc(4)Hex(5)Fuc(1)NeuAc(1) | C[+57]*2, N[+2060] | 1246.1586 | 3 | 3736.4613 | 3736.4545 | 0 | 1.8 | 1201 | Specific | 396.21 | 315.33 | 315.33 | 7.75 | 77 | >sp\|P22897\|MRC1_HUMAN Macrophage mannose receptor 1 OS=Homo sapiens OX=9606 GN=MRC1 PE=1 SV=1 | 1049 | 20201113_gy_milk_1ng_glyco_hcd203040_compound15.11454.11454.3 | controllerType=0 controllerNumber=1 scan=11454 | 25.3854 |
| 08310:3 | 6 | K.TAHC[+57.021]N[+2205.793]ESFYFLC[+57.021]K.R | HexNAc(4)Hex(5)Fuc(2)NeuAc(1) | C[+57]*2, N[+2206] | 1294.8461 | 3 | 3882.5238 | 3882.5124 | 0 | 2.9 | 1201 | Specific | 401.07 | 336.91 | 336.91 | 7.84 | 77 | >sp\|P22897\|MRC1_HUMAN Macrophage mannose receptor 1 OS=Homo sapiens OX=9606 GN=MRC1 PE=1 SV=1 | 1049 | 20201113_gy_milk_1ng_glyco_hcd203040_compound15.11318.11318.3 | controllerType=0 controllerNumber=1 scan=11318 | 25.1602 |
| 08440:3 | 6 | K.TAHC[+57.021]N[+2059.735]ESFYFLC[+57.021]K.R | HexNAc(4)Hex(5)Fuc(1)NeuAc(1) | C[+57]*2, N[+2060] | 1246.1586 | 3 | 3736.4613 | 3736.4545 | 0 | 1.8 | 1201 | Specific | 417.89 | 381.47 | 381.47 | 9.49 | 77 | >sp\|P22897\|MRC1_HUMAN Macrophage mannose receptor 1 OS=Homo sapiens OX=9606 GN=MRC1 PE=1 SV=1 | 1049 | 20201113_gy_milk_1ng_glyco_hcd203040_compound15.11452.11452.3 | controllerType=0 controllerNumber=1 scan=11452 | 25.3813 |
| 22891:3 | 7 | R.[+42.011]LSLLEEPGN[+2336.851]GTFTVILNQLTSR.D | HexNAc(6)Hex(6)Fuc(1) | .[42], N[+2337] | 1594.3778 | 3 | 4781.1188 | 4781.1542 | 0 | -7.4 | 413 | Specific | 264.13 | 249.45 | 249.45 | 5.37 | 56 | >sp\|P01833\|PIGR_HUMAN Polymeric immunoglobulin receptor OS=Homo sapiens OX=9606 GN=PIGR PE=1 SV=4 | 902 | 20201113_gy_milk_1ng_glyco_hcd203040_compound15.26597.26597.3 | controllerType=0 controllerNumber=1 scan=26597 | 53.3346 |
| 20568:3 | 7 | R.LSLLEEPGN[+2076.750]GTFTVILNQLTSR.D | HexNAc(4)Hex(6)Fuc(2) | N[+2077] | 1493.692 | 3 | 4479.0613 | 4479.0428 | 0 | 4.1 | 413 | Specific | 329.55 | 327.99 | 327.99 | 7.61 | 56 | >sp\|P01833\|PIGR_HUMAN Polymeric immunoglobulin receptor OS=Homo sapiens OX=9606 GN=PIGR PE=1 SV=4 | 902 | 20201113_gy_milk_1ng_glyco_hcd203040_compound15.24158.24158.3 | controllerType=0 controllerNumber=1 scan=24158 | 48.8325 |
| 22893:3 | 7 | R.[+42.011]LSLLEEPGN[+2336.851]GTFTVILNQLTSR.D | HexNAc(6)Hex(6)Fuc(1) | .[42], N[+2337] | 1594.3778 | 3 | 4781.1188 | 4781.1542 | 0 | -7.4 | 413 | Specific | 363.58 | 238.54 | 238.54 | 5.88 | 56 | >sp\|P01833\|PIGR_HUMAN Polymeric immunoglobulin receptor OS=Homo sapiens OX=9606 GN=PIGR PE=1 SV=4 | 902 | 20201113_gy_milk_1ng_glyco_hcd203040_compound15.26599.26599.3 | controllerType=0 controllerNumber=1 scan=26599 | 53.3387 |
| 20566:3 | 7 | R.LSLLEEPGN[+2076.750]GTFTVILNQLTSR.D | HexNAc(4)Hex(6)Fuc(2) | N[+2077] | 1493.692 | 3 | 4479.0613 | 4479.0428 | 0 | 4.1 | 413 | Specific | 387.3 | 330.36 | 330.36 | 8.06 | 56 | >sp\|P01833\|PIGR_HUMAN Polymeric immunoglobulin receptor OS=Homo sapiens OX=9606 GN=PIGR PE=1 SV=4 | 902 | 20201113_gy_milk_1ng_glyco_hcd203040_compound15.24156.24156.3 | controllerType=0 controllerNumber=1 scan=24156 | 48.8288 |
| 22885:4 | 7 | R.LSLLEEPGN[+2350.830]GTFTVILNQLTSR.D | HexNAc(4)Hex(5)Fuc(1)NeuAc(2) | N[+2351] | 1189.0395 | 4 | 4753.1361 | 4753.1229 | 0 | 2.8 | 413 | Specific | 443.08 | 413.91 | 413.91 | 10.08 | 56 | >sp\|P01833\|PIGR_HUMAN Polymeric immunoglobulin receptor OS=Homo sapiens OX=9606 GN=PIGR PE=1 SV=4 | 902 | 20201113_gy_milk_1ng_glyco_hcd203040_compound15.26590.26590.4 | controllerType=0 controllerNumber=1 scan=26590 | 53.3225 |
| 22941:3 | 7 | R.LSLLEEPGN[+2350.830]GTFTVILNQLTSR.D | HexNAc(4)Hex(5)Fuc(1)NeuAc(2) | N[+2351] | 1585.0549 | 3 | 4753.15 | 4753.1229 | 0 | 5.7 | 413 | Specific | 546.71 | 446.58 | 446.58 | 10.05 | 56 | >sp\|P01833\|PIGR_HUMAN Polymeric immunoglobulin receptor OS=Homo sapiens OX=9606 GN=PIGR PE=1 SV=4 | 902 | 20201113_gy_milk_1ng_glyco_hcd203040_compound15.26650.26650.3 | controllerType=0 controllerNumber=1 scan=26650 | 53.4495 |
| 22968:3 | 7 | R.LSLLEEPGN[+2350.830]GTFTVILNQLTSR.D | HexNAc(4)Hex(5)Fuc(1)NeuAc(2) | N[+2351] | 1585.0551 | 3 | 4753.1508 | 4753.1229 | 0 | 5.9 | 413 | Specific | 583.91 | 479.13 | 479.13 | 11.27 | 56 | >sp\|P01833\|PIGR_HUMAN Polymeric immunoglobulin receptor OS=Homo sapiens OX=9606 GN=PIGR PE=1 SV=4 | 902 | 20201113_gy_milk_1ng_glyco_hcd203040_compound15.26678.26678.3 | controllerType=0 controllerNumber=1 scan=26678 | 53.4999 |
| 22970:3 | 7 | R.LSLLEEPGN[+2350.830]GTFTVILNQLTSR.D | HexNAc(4)Hex(5)Fuc(1)NeuAc(2) | N[+2351] | 1585.0551 | 3 | 4753.1508 | 4753.1229 | 0 | 5.9 | 413 | Specific | 589.19 | 529.11 | 529.11 | 11.27 | 56 | >sp\|P01833\|PIGR_HUMAN Polymeric immunoglobulin receptor OS=Homo sapiens OX=9606 GN=PIGR PE=1 SV=4 | 902 | 20201113_gy_milk_1ng_glyco_hcd203040_compound15.26680.26680.3 | controllerType=0 controllerNumber=1 scan=26680 | 53.5049 |
| 22943:3 | 7 | R.LSLLEEPGN[+2350.830]GTFTVILNQLTSR.D | HexNAc(4)Hex(5)Fuc(1)NeuAc(2) | N[+2351] | 1585.0549 | 3 | 4753.15 | 4753.1229 | 0 | 5.7 | 413 | Specific | 782.29 | 565.83 | 565.83 | 15.27 | 56 | >sp\|P01833\|PIGR_HUMAN Polymeric immunoglobulin receptor OS=Homo sapiens OX=9606 GN=PIGR PE=1 SV=4 | 902 | 20201113_gy_milk_1ng_glyco_hcd203040_compound15.26652.26652.3 | controllerType=0 controllerNumber=1 scan=26652 | 53.4535 |
| 18042:3 | 7 | K.IIEGEPNLKVPGN[+2204.772]VTAVLGETLK.V | HexNAc(4)Hex(5)NeuAc(2) | N[+2205] | 1532.7183 | 3 | 4596.1404 | 4596.1218 | 0 | 4 | 457 | Specific | 291.82 | 291.82 | 291.82 | 6.51 | 56 | >sp\|P01833\|PIGR_HUMAN Polymeric immunoglobulin receptor OS=Homo sapiens OX=9606 GN=PIGR PE=1 SV=4 | 902 | 20201113_gy_milk_1ng_glyco_hcd203040_compound15.21504.21504.3 | controllerType=0 controllerNumber=1 scan=21504 | 43.2485 |
| 16135:3 | 7 | K.IIEGEPNLKVPGN[+1913.677]VTAVLGETLK.V | HexNAc(4)Hex(5)NeuAc(1) | N[+1914] | 1435.6872 | 3 | 4305.047 | 4305.0264 | 0 | 4.8 | 457 | Specific | 451.99 | 446.61 | 446.61 | 8.98 | 56 | >sp\|P01833\|PIGR_HUMAN Polymeric immunoglobulin receptor OS=Homo sapiens OX=9606 GN=PIGR PE=1 SV=4 | 902 | 20201113_gy_milk_1ng_glyco_hcd203040_compound15.19506.19506.3 | controllerType=0 controllerNumber=1 scan=19506 | 39.3481 |
| 18044:3 | 7 | K.IIEGEPNLKVPGN[+2204.772]VTAVLGETLK.V | HexNAc(4)Hex(5)NeuAc(2) | N[+2205] | 1532.7183 | 3 | 4596.1404 | 4596.1218 | 0 | 4 | 457 | Specific | 493.04 | 465.37 | 465.37 | 9.35 | 56 | >sp\|P01833\|PIGR_HUMAN Polymeric immunoglobulin receptor OS=Homo sapiens OX=9606 GN=PIGR PE=1 SV=4 | 902 | 20201113_gy_milk_1ng_glyco_hcd203040_compound15.21506.21506.3 | controllerType=0 controllerNumber=1 scan=21506 | 43.2524 |
| 16137:3 | 7 | K.IIEGEPNLKVPGN[+1913.677]VTAVLGETLK.V | HexNAc(4)Hex(5)NeuAc(1) | N[+1914] | 1435.6872 | 3 | 4305.047 | 4305.0264 | 0 | 4.8 | 457 | Specific | 544.4 | 431.72 | 431.72 | 9.29 | 56 | >sp\|P01833\|PIGR_HUMAN Polymeric immunoglobulin receptor OS=Homo sapiens OX=9606 GN=PIGR PE=1 SV=4 | 902 | 20201113_gy_milk_1ng_glyco_hcd203040_compound15.19508.19508.3 | controllerType=0 controllerNumber=1 scan=19508 | 39.3516 |

Table S4. Cont.

| Query #:z | Protein Rank | Glycopeptide | Glycan | Modification Type(s) | Observed m/z | z | Observed (M+H) | Calc. mass (M+H) | Off-by-x error | Mass error (ppm) | StartPosition | Cleavage | Score | Delta | Delta Mod | \|Log Prob\| | # of unique peptides | Protein Name | Protein DB number | Comment | Scan # | Scan Time |
|---|---|---|---|---|---|---|---|---|---|---|---|---|---|---|---|---|---|---|---|---|---|---|
| 18030:3 | 7 | K.IIEGEPNLKVPGN[+2350.830]VTAVLGETLK.V | HexNAc(4)Hex(5)Fuc(1)NeuAc(2) | N[+2351] | 1581.4013 | 3 | 4742.1894 | 4742.1797 | 0 | 2 | 457 | Specific | 861.8 | 788.49 | 788.49 | 17.22 | 56 | >sp\|P01833\|PIGR_HUMAN Polymeric immunoglobulin receptor OS=Homo sapiens OX=9606 GN=PIGR PE=1 SV=4 | 902 | 20201113_gy_milk_1ng_glyco_hcd203040_compound15.21492.21492.3 | controllerType=0 controllerNumber=1 scan=21492 | 43.2254 |
| 18032:3 | 7 | K.IIEGEPNLKVPGN[+2350.830]VTAVLGETLK.V | HexNAc(4)Hex(5)Fuc(1)NeuAc(2) | N[+2351] | 1581.4013 | 3 | 4742.1894 | 4742.1797 | 0 | 2 | 457 | Specific | 879.04 | 771.34 | 771.34 | 17.22 | 56 | >sp\|P01833\|PIGR_HUMAN Polymeric immunoglobulin receptor OS=Homo sapiens OX=9606 GN=PIGR PE=1 SV=4 | 902 | 20201113_gy_milk_1ng_glyco_hcd203040_compound15.21494.21494.3 | controllerType=0 controllerNumber=1 scan=21494 | 43.2309 |
| 16122:3 | 7 | K.IIEGEPNLKVPGN[+2059.735]VTAVLGETLK.V | HexNAc(4)Hex(5)Fuc(1)NeuAc(1) | N[+2060] | 1484.376 | 3 | 4451.1135 | 4451.0843 | 0 | 6.6 | 457 | Specific | 880.41 | 854.93 | 854.93 | 16.52 | 56 | >sp\|P01833\|PIGR_HUMAN Polymeric immunoglobulin receptor OS=Homo sapiens OX=9606 GN=PIGR PE=1 SV=4 | 902 | 20201113_gy_milk_1ng_glyco_hcd203040_compound15.19493.19493.3 | controllerType=0 controllerNumber=1 scan=19493 | 39.3249 |
| 16120:3 | 7 | K.IIEGEPNLKVPGN[+2059.735]VTAVLGETLK.V | HexNAc(4)Hex(5)Fuc(1)NeuAc(1) | N[+2060] | 1484.376 | 3 | 4451.1135 | 4451.0843 | 0 | 6.6 | 457 | Specific | 1115.82 | 849.32 | 849.32 | 21.52 | 56 | >sp\|P01833\|PIGR_HUMAN Polymeric immunoglobulin receptor OS=Homo sapiens OX=9606 GN=PIGR PE=1 SV=4 | 902 | 20201113_gy_milk_1ng_glyco_hcd203040_compound15.19491.19491.3 | controllerType=0 controllerNumber=1 scan=19491 | 39.3185 |
| 15641:4 | 7 | K.IIEGEPNLKVPGN[+2205.793]VTAVLGETLKVPC[+57.021]HFPC[+57.021]K.F | HexNAc(4)Hex(5)Fuc(2)NeuAc(1) | C[+57]*2, N[+2206] | 1406.4065 | 4 | 5622.6042 | 5622.5997 | 0 | 0.8 | 457 | Specific | 757.38 | 757.38 | 757.38 | 14.42 | 56 | >sp\|P01833\|PIGR_HUMAN Polymeric immunoglobulin receptor OS=Homo sapiens OX=9606 GN=PIGR PE=1 SV=4 | 902 | 20201113_gy_milk_1ng_glyco_hcd203040_compound15.18951.18951.4 | controllerType=0 controllerNumber=1 scan=18951 | 38.2637 |
| 15647:5 | 7 | K.IIEGEPNLKVPGN[+2205.793]VTAVLGETLKVPC[+57.021]HFPC[+57.021]K.F | HexNAc(4)Hex(5)Fuc(2)NeuAc(1) | C[+57]*2, N[+2206] | 1125.328 | 5 | 5622.6109 | 5622.5997 | 0 | 2 | 457 | Specific | 963.5 | 873.4 | 873.4 | 19.01 | 56 | >sp\|P01833\|PIGR_HUMAN Polymeric immunoglobulin receptor OS=Homo sapiens OX=9606 GN=PIGR PE=1 SV=4 | 902 | 20201113_gy_milk_1ng_glyco_hcd203040_compound15.18957.18957.5 | controllerType=0 controllerNumber=1 scan=18957 | 38.2759 |
| 15639:4 | 7 | K.IIEGEPNLKVPGN[+2205.793]VTAVLGETLKVPC[+57.021]HFPC[+57.021]K.F | HexNAc(4)Hex(5)Fuc(2)NeuAc(1) | C[+57]*2, N[+2206] | 1406.4065 | 4 | 5622.6042 | 5622.5997 | 0 | 0.8 | 457 | Specific | 1059.74 | 911.16 | 911.16 | 20.42 | 56 | >sp\|P01833\|PIGR_HUMAN Polymeric immunoglobulin receptor OS=Homo sapiens OX=9606 GN=PIGR PE=1 SV=4 | 902 | 20201113_gy_milk_1ng_glyco_hcd203040_compound15.18949.18949.4 | controllerType=0 controllerNumber=1 scan=18949 | 38.2588 |
| 15645:5 | 7 | K.IIEGEPNLKVPGN[+2205.793]VTAVLGETLKVPC[+57.021]HFPC[+57.021]K.F | HexNAc(4)Hex(5)Fuc(2)NeuAc(1) | C[+57]*2, N[+2206] | 1125.328 | 5 | 5622.6109 | 5622.5997 | 0 | 2 | 457 | Specific | 1080.67 | 960.24 | 960.24 | 21.01 | 56 | >sp\|P01833\|PIGR_HUMAN Polymeric immunoglobulin receptor OS=Homo sapiens OX=9606 GN=PIGR PE=1 SV=4 | 902 | 20201113_gy_milk_1ng_glyco_hcd203040_compound15.18955.18955.5 | controllerType=0 controllerNumber=1 scan=18955 | 38.2696 |
| 13596:3 | 7 | K.VPGN[+2205.793]VTAVLGETLK.V | HexNAc(4)Hex(5)Fuc(2)NeuAc(1) | N[+2206] | 1201.8735 | 3 | 3603.6059 | 3603.5927 | 0 | 3.7 | 466 | Specific | 383.92 | 337.26 | 337.26 | 7.57 | 56 | >sp\|P01833\|PIGR_HUMAN Polymeric immunoglobulin receptor OS=Homo sapiens OX=9606 GN=PIGR PE=1 SV=4 | 902 | 20201113_gy_milk_1ng_glyco_hcd203040_compound15.16824.16824.3 | controllerType=0 controllerNumber=1 scan=16824 | 34.4703 |
| 13598:3 | 7 | K.VPGN[+2205.793]VTAVLGETLK.V | HexNAc(4)Hex(5)Fuc(2)NeuAc(1) | N[+2206] | 1201.8735 | 3 | 3603.6059 | 3603.5927 | 0 | 3.7 | 466 | Specific | 439.61 | 408.66 | 408.66 | 9.23 | 56 | >sp\|P01833\|PIGR_HUMAN Polymeric immunoglobulin receptor OS=Homo sapiens OX=9606 GN=PIGR PE=1 SV=4 | 902 | 20201113_gy_milk_1ng_glyco_hcd203040_compound15.16826.16826.3 | controllerType=0 controllerNumber=1 scan=16826 | 34.4741 |
| 13766:4 | 7 | K.VPGN[+2059.735]VTAVLGETLKVPC[+57.021]HFPC[+57.021]K.F | HexNAc(4)Hex(5)Fuc(1)NeuAc(1) | C[+57]*2, N[+2060] | 1121.5061 | 4 | 4483.0028 | 4482.9923 | 0 | 2.3 | 466 | Specific | 304.05 | 272.62 | 272.62 | 6.81 | 56 | >sp\|P01833\|PIGR_HUMAN Polymeric immunoglobulin receptor OS=Homo sapiens OX=9606 GN=PIGR PE=1 SV=4 | 902 | 20201113_gy_milk_1ng_glyco_hcd203040_compound15.17000.17000.4 | controllerType=0 controllerNumber=1 scan=17000 | 34.7846 |
| 13770:4 | 7 | K.VPGN[+2205.793]VTAVLGETLKVPC[+57.021]HFPC[+57.021]K.F | HexNAc(4)Hex(5)Fuc(2)NeuAc(1) | C[+57]*2, N[+2206] | 1158.0214 | 4 | 4629.0638 | 4629.0502 | 0 | 2.9 | 466 | Specific | 368.53 | 368.53 | 368.53 | 7.91 | 56 | >sp\|P01833\|PIGR_HUMAN Polymeric immunoglobulin receptor OS=Homo sapiens OX=9606 GN=PIGR PE=1 SV=4 | 902 | 20201113_gy_milk_1ng_glyco_hcd203040_compound15.17004.17004.4 | controllerType=0 controllerNumber=1 scan=17004 | 34.7917 |
| 13768:4 | 7 | K.VPGN[+2059.735]VTAVLGETLKVPC[+57.021]HFPC[+57.021]K.F | HexNAc(4)Hex(5)Fuc(1)NeuAc(1) | C[+57]*2, N[+2060] | 1121.5061 | 4 | 4483.0028 | 4482.9923 | 0 | 2.3 | 466 | Specific | 600.1 | 296.02 | 296.02 | 10.51 | 56 | >sp\|P01833\|PIGR_HUMAN Polymeric immunoglobulin receptor OS=Homo sapiens OX=9606 GN=PIGR PE=1 SV=4 | 902 | 20201113_gy_milk_1ng_glyco_hcd203040_compound15.17002.17002.4 | controllerType=0 controllerNumber=1 scan=17002 | 34.7883 |
| 05936:3 | 7 | K.WN[+1548.545]NTGC[+57.021]QALPSQDEGPSK.A | HexNAc(3)Hex(4)NeuAc(1) | C[+57], N[+1549] | 1179.8143 | 3 | 3537.4283 | 3537.4215 | 0 | 1.9 | 498 | Specific | 284.07 | 216.79 | 216.79 | 7.18 | 56 | >sp\|P01833\|PIGR_HUMAN Polymeric immunoglobulin receptor OS=Homo sapiens OX=9606 GN=PIGR PE=1 SV=4 | 902 | 20201113_gy_milk_1ng_glyco_hcd203040_compound15.8856.8856.3 | controllerType=0 controllerNumber=1 scan=8856 | 21.0761 |
| 05791:3 | 7 | K.WN[+1913.677]NTGC[+57.021]QALPSQDEGPSK.A | HexNAc(4)Hex(5)NeuAc(1) | C[+57], N[+1914] | 1301.5229 | 3 | 3902.5541 | 3902.5537 | 0 | 0.1 | 498 | Specific | 320.05 | 282.23 | 282.23 | 7.1 | 56 | >sp\|P01833\|PIGR_HUMAN Polymeric immunoglobulin receptor OS=Homo sapiens OX=9606 GN=PIGR PE=1 SV=4 | 902 | 20201113_gy_milk_1ng_glyco_hcd203040_compound15.8706.8706.3 | controllerType=0 controllerNumber=1 scan=8706 | 20.8238 |
| 05841:3 | 7 | K.WN[+1710.598]NTGC[+57.021]QALPSQDEGPSK.A | HexNAc(3)Hex(5)NeuAc(1) | C[+57], N[+1711] | 1233.832 | 3 | 3699.4813 | 3699.4743 | 0 | 1.9 | 498 | Specific | 369.97 | 309.17 | 309.17 | 8.09 | 56 | >sp\|P01833\|PIGR_HUMAN Polymeric immunoglobulin receptor OS=Homo sapiens OX=9606 GN=PIGR PE=1 SV=4 | 902 | 20201113_gy_milk_1ng_glyco_hcd203040_compound15.8758.8758.3 | controllerType=0 controllerNumber=1 scan=8758 | 20.9124 |
| 05719:3 | 7 | K.WN[+1872.651]NTGC[+57.021]QALPSQDEGPSK.A | HexNAc(3)Hex(6)NeuAc(1) | C[+57], N[+1873] | 1287.8513 | 3 | 3861.5392 | 3861.5272 | 0 | 3.1 | 498 | Specific | 473.15 | 444.96 | 444.96 | 10.12 | 56 | >sp\|P01833\|PIGR_HUMAN Polymeric immunoglobulin receptor OS=Homo sapiens OX=9606 GN=PIGR PE=1 SV=4 | 902 | 20201113_gy_milk_1ng_glyco_hcd203040_compound15.8631.8631.3 | controllerType=0 controllerNumber=1 scan=8631 | 20.6954 |
| 05843:3 | 7 | K.WN[+1710.598]NTGC[+57.021]QALPSQDEGPSK.A | HexNAc(3)Hex(5)NeuAc(1) | C[+57], N[+1711] | 1233.832 | 3 | 3699.4813 | 3699.4743 | 0 | 1.9 | 498 | Specific | 486.59 | 424.6 | 424.6 | 10.18 | 56 | >sp\|P01833\|PIGR_HUMAN Polymeric immunoglobulin receptor OS=Homo sapiens OX=9606 GN=PIGR PE=1 SV=4 | 902 | 20201113_gy_milk_1ng_glyco_hcd203040_compound15.8760.8760.3 | controllerType=0 controllerNumber=1 scan=8760 | 20.9165 |
| 05717:3 | 7 | K.WN[+1872.651]NTGC[+57.021]QALPSQDEGPSK.A | HexNAc(3)Hex(6)NeuAc(1) | C[+57], N[+1873] | 1287.8513 | 3 | 3861.5392 | 3861.5272 | 0 | 3.1 | 498 | Specific | 501.97 | 466.9 | 466.9 | 10.16 | 56 | >sp\|P01833\|PIGR_HUMAN Polymeric immunoglobulin receptor OS=Homo sapiens OX=9606 GN=PIGR PE=1 SV=4 | 902 | 20201113_gy_milk_1ng_glyco_hcd203040_compound15.8629.8629.3 | controllerType=0 controllerNumber=1 scan=8629 | 20.6914 |
| 05738:3 | 7 | K.WN[+1913.677]NTGC[+57.021]QALPSQDEGPSK.A | HexNAc(4)Hex(5)NeuAc(1) | C[+57], N[+1914] | 1301.5336 | 3 | 3902.5863 | 3902.5537 | 0 | 8.4 | 498 | Specific | 671.33 | 656.37 | 656.37 | 13.02 | 56 | >sp\|P01833\|PIGR_HUMAN Polymeric immunoglobulin receptor OS=Homo sapiens OX=9606 GN=PIGR PE=1 SV=4 | 902 | 20201113_gy_milk_1ng_glyco_hcd203040_compound15.8651.8651.3 | controllerType=0 controllerNumber=1 scan=8651 | 20.7311 |
| 05740:3 | 7 | K.WN[+1913.677]NTGC[+57.021]QALPSQDEGPSK.A | HexNAc(4)Hex(5)NeuAc(1) | C[+57], N[+1914] | 1301.5336 | 3 | 3902.5863 | 3902.5537 | 0 | 8.4 | 498 | Specific | 699.85 | 624.74 | 624.74 | 13.02 | 56 | >sp\|P01833\|PIGR_HUMAN Polymeric immunoglobulin receptor OS=Homo sapiens OX=9606 GN=PIGR PE=1 SV=4 | 902 | 20201113_gy_milk_1ng_glyco_hcd203040_compound15.8653.8653.3 | controllerType=0 controllerNumber=1 scan=8653 | 20.7347 |
| 10019:3 | 8 | K.TVLTPATNHMGN[+1540.529]VTFTIPANR.E | HexNAc(2)Hex(7) | N[+1541] | 1265.9041 | 3 | 3795.6978 | 3795.6886 | 0 | 2.4 | 74 | Specific | 466.78 | 466.78 | 228.12 | 10.15 | 70 | >sp\|P01024\|CO3_HUMAN Complement C3 OS=Homo sapiens OX=9606 GN=C3 PE=1 SV=2 | 492 | 20201113_gy_milk_1ng_glyco_hcd203040_compound15.13101.13101.3 | controllerType=0 controllerNumber=1 scan=13101 | 28.176 |
| 10021:3 | 8 | K.TVLTPATNHMGN[+1540.529]VTFTIPANR.E | HexNAc(2)Hex(7) | N[+1541] | 1265.9041 | 3 | 3795.6978 | 3795.6886 | 0 | 2.4 | 74 | Specific | 480.82 | 480.82 | 201.78 | 10.15 | 70 | >sp\|P01024\|CO3_HUMAN Complement C3 OS=Homo sapiens OX=9606 GN=C3 PE=1 SV=2 | 492 | 20201113_gy_milk_1ng_glyco_hcd203040_compound15.13103.13103.3 | controllerType=0 controllerNumber=1 scan=13103 | 28.1801 |
| 10115:3 | 8 | K.TVLTPATNHMGN[+1378.476]VTFTIPANR.E | HexNAc(2)Hex(6) | N[+1378] | 1211.8836 | 3 | 3633.6363 | 3633.6358 | 0 | 0.1 | 74 | Specific | 482.03 | 482.03 | 208.11 | 9.93 | 70 | >sp\|P01024\|CO3_HUMAN Complement C3 OS=Homo sapiens OX=9606 GN=C3 PE=1 SV=2 | 492 | 20201113_gy_milk_1ng_glyco_hcd203040_compound15.13200.13200.3 | controllerType=0 controllerNumber=1 scan=13200 | 28.3421 |
| 10113:3 | 8 | K.TVLTPATNHMGN[+1378.476]VTFTIPANR.E | HexNAc(2)Hex(6) | N[+1378] | 1211.8836 | 3 | 3633.6363 | 3633.6358 | 0 | 0.1 | 74 | Specific | 549.93 | 549.93 | 286.68 | 10.28 | 70 | >sp\|P01024\|CO3_HUMAN Complement C3 OS=Homo sapiens OX=9606 GN=C3 PE=1 SV=2 | 492 | 20201113_gy_milk_1ng_glyco_hcd203040_compound15.13198.13198.3 | controllerType=0 controllerNumber=1 scan=13198 | 28.337 |
| 17197:3 | 10 | K.RNIAAFGGDPNN[+1913.677]ITLFGESAGGASVSLQTLSPYNK.G | HexNAc(4)Hex(5)NeuAc(1) | N[+1914] | 1827.5028 | 3 | 5480.4938 | 5480.4644 | 0 | 5.4 | 196 | Specific | 239.67 | 131.95 | 131.95 | 5.14 | 41 | >sp\|P19835\|CEL_HUMAN Bile salt-activated lipase OS=Homo sapiens OX=9606 GN=CEL PE=1 SV=3 | 476 | 20201113_gy_milk_1ng_glyco_hcd203040_compound15.20618.20618.3 | controllerType=0 controllerNumber=1 scan=20618 | 41.4146 |
| 17195:3 | 10 | K.RNIAAFGGDPNN[+1913.677]ITLFGESAGGASVSLQTLSPYNK.G | HexNAc(4)Hex(5)NeuAc(1) | N[+1914] | 1827.5028 | 3 | 5480.4938 | 5480.4644 | 0 | 5.4 | 196 | Specific | 335.98 | 243.57 | 243.57 | 7.14 | 41 | >sp\|P19835\|CEL_HUMAN Bile salt-activated lipase OS=Homo sapiens OX=9606 GN=CEL PE=1 SV=3 | 476 | 20201113_gy_milk_1ng_glyco_hcd203040_compound15.20616.20616.3 | controllerType=0 controllerNumber=1 scan=20616 | 41.4106 |
| 17608:3 | 11 | R.LSLHRPALEDLLLGSEAN[+1702.581]LTC[+57.021]TLTGLR.D | HexNAc(2)Hex(8) | C[+57], N[+1703] | 1556.0786 | 3 | 4666.2212 | 4666.1796 | 0 | 8.9 | 127 | Specific | 206.92 | 206.92 | 206.92 | 4.58 | 42 | >sp\|P01876\|IGHA1_HUMAN Immunoglobulin heavy constant alpha 1 OS=Homo sapiens OX=9606 GN=IGHA1 PE=1 SV=2 | 255 | 20201113_gy_milk_1ng_glyco_hcd203040_compound15.21049.21049.3 | controllerType=0 controllerNumber=1 scan=21049 | 42.2837 |
| 17760:4 | 11 | R.LSLHRPALEDLLLGSEAN[+1298.476]LTC[+57.021]TLTGLR.D | HexNAc(4)Hex(3) | C[+57], N[+1298] | 1066.2736 | 4 | 4262.0726 | 4262.0742 | 0 | -0.4 | 127 | Specific | 274.56 | 274.56 | 274.56 | 7.44 | 42 | >sp\|P01876\|IGHA1_HUMAN Immunoglobulin heavy constant alpha 1 OS=Homo sapiens OX=9606 GN=IGHA1 PE=1 SV=2 | 255 | 20201113_gy_milk_1ng_glyco_hcd203040_compound15.21210.21210.4 | controllerType=0 controllerNumber=1 scan=21210 | 42.6226 |
| 17758:3 | 11 | R.LSLHRPALEDLLLGSEAN[+1501.555]LTC[+57.021]TLTGLR.D | HexNAc(5)Hex(3) | C[+57], N[+1502] | 1489.0562 | 3 | 4465.1541 | 4465.1536 | 0 | 0.1 | 127 | Specific | 301.49 | 230.66 | 230.66 | 5.62 | 42 | >sp\|P01876\|IGHA1_HUMAN Immunoglobulin heavy constant alpha 1 OS=Homo sapiens OX=9606 GN=IGHA1 PE=1 SV=2 | 255 | 20201113_gy_milk_1ng_glyco_hcd203040_compound15.21208.21208.3 | controllerType=0 controllerNumber=1 scan=21208 | 42.6194 |
| 17811:4 | 11 | R.LSLHRPALEDLLLGSEAN[+1054.370]LTC[+57.021]TLTGLR.D | HexNAc(2)Hex(4) | C[+57], N[+1054] | 1005.2543 | 4 | 4017.9952 | 4017.9683 | 0 | 6.7 | 127 | Specific | 394.1 | 394.1 | 394.1 | 9.68 | 42 | >sp\|P01876\|IGHA1_HUMAN Immunoglobulin heavy constant alpha 1 OS=Homo sapiens OX=9606 GN=IGHA1 PE=1 SV=2 | 255 | 20201113_gy_milk_1ng_glyco_hcd203040_compound15.21263.21263.4 | controllerType=0 controllerNumber=1 scan=21263 | 42.7412 |
| 17729:3 | 11 | R.LSLHRPALEDLLLGSEAN[+1501.555]LTC[+57.021]TLTGLR.D | HexNAc(5)Hex(3) | C[+57], N[+1502] | 1489.0569 | 3 | 4465.1563 | 4465.1536 | 0 | 0.6 | 127 | Specific | 492.88 | 452.92 | 452.92 | 9.89 | 42 | >sp\|P01876\|IGHA1_HUMAN Immunoglobulin heavy constant alpha 1 OS=Homo sapiens OX=9606 GN=IGHA1 PE=1 SV=2 | 255 | 20201113_gy_milk_1ng_glyco_hcd203040_compound15.21177.21177.3 | controllerType=0 controllerNumber=1 scan=21177 | 42.5549 |
| 17605:4 | 11 | R.LSLHRPALEDLLLGSEAN[+1702.581]LTC[+57.021]TLTGLR.D | HexNAc(2)Hex(8) | C[+57], N[+1703] | 1167.3108 | 4 | 4666.2214 | 4666.1796 | 0 | 9 | 127 | Specific | 521.07 | 521.07 | 521.07 | 10.17 | 42 | >sp\|P01876\|IGHA1_HUMAN Immunoglobulin heavy constant alpha 1 OS=Homo sapiens OX=9606 GN=IGHA1 PE=1 SV=2 | 255 | 20201113_gy_milk_1ng_glyco_hcd203040_compound15.21046.21046.4 | controllerType=0 controllerNumber=1 scan=21046 | 42.2747 |
| 17602:3 | 11 | R.LSLHRPALEDLLLGSEAN[+1864.634]LTC[+57.021]TLTGLR.D | HexNAc(2)Hex(9) | C[+57], N[+1865] | 1610.0969 | 3 | 4828.276 | 4828.2324 | 0 | 9 | 127 | Specific | 529.18 | 529.18 | 529.18 | 9.57 | 42 | >sp\|P01876\|IGHA1_HUMAN Immunoglobulin heavy constant alpha 1 OS=Homo sapiens OX=9606 GN=IGHA1 PE=1 SV=2 | 255 | 20201113_gy_milk_1ng_glyco_hcd203040_compound15.21042.21042.3 | controllerType=0 controllerNumber=1 scan=21042 | 42.2661 |
| 17679:3 | 11 | R.LSLHRPALEDLLLGSEAN[+1663.608]LTC[+57.021]TLTGLR.D | HexNAc(5)Hex(4) | C[+57], N[+1664] | 1543.0788 | 3 | 4627.2218 | 4627.2064 | 0 | 3.3 | 127 | Specific | 548.89 | 485.5 | 485.5 | 10.47 | 42 | >sp\|P01876\|IGHA1_HUMAN Immunoglobulin heavy constant alpha 1 OS=Homo sapiens OX=9606 GN=IGHA1 PE=1 SV=2 | 255 | 20201113_gy_milk_1ng_glyco_hcd203040_compound15.21124.21124.3 | controllerType=0 controllerNumber=1 scan=21124 | 42.4393 |
| 17756:3 | 11 | R.LSLHRPALEDLLLGSEAN[+1501.555]LTC[+57.021]TLTGLR.D | HexNAc(5)Hex(3) | C[+57], N[+1502] | 1489.0562 | 3 | 4465.1541 | 4465.1536 | 0 | 0.1 | 127 | Specific | 549.2 | 528.42 | 528.42 | 10.25 | 42 | >sp\|P01876\|IGHA1_HUMAN Immunoglobulin heavy constant alpha 1 OS=Homo sapiens OX=9606 GN=IGHA1 PE=1 SV=2 | 255 | 20201113_gy_milk_1ng_glyco_hcd203040_compound15.21206.21206.3 | controllerType=0 controllerNumber=1 scan=21206 | 42.6148 |
| 17654:3 | 11 | R.LSLHRPALEDLLLGSEAN[+1663.608]LTC[+57.021]TLTGLR.D | HexNAc(5)Hex(4) | C[+57], N[+1664] | 1543.0789 | 3 | 4627.2221 | 4627.2064 | 0 | 3.4 | 127 | Specific | 564.22 | 504.61 | 504.61 | 10.17 | 42 | >sp\|P01876\|IGHA1_HUMAN Immunoglobulin heavy constant alpha 1 OS=Homo sapiens OX=9606 GN=IGHA1 PE=1 SV=2 | 255 | 20201113_gy_milk_1ng_glyco_hcd203040_compound15.21098.21098.3 | controllerType=0 controllerNumber=1 scan=21098 | 42.3818 |
| 17808:3 | 11 | R.LSLHRPALEDLLLGSEAN[+1095.397]LTC[+57.021]TLTGLR.D | HexNAc(3)Hex(3) | C[+57], N[+1095] | 1353.6796 | 3 | 4059.0243 | 4058.9949 | 0 | 7.3 | 127 | Specific | 567.21 | 532.58 | 532.58 | 9.76 | 42 | >sp\|P01876\|IGHA1_HUMAN Immunoglobulin heavy constant alpha 1 OS=Homo sapiens OX=9606 GN=IGHA1 PE=1 SV=2 | 255 | 20201113_gy_milk_1ng_glyco_hcd203040_compound15.21260.21260.3 | controllerType=0 controllerNumber=1 scan=21260 | 42.7352 |
| 17677:3 | 11 | R.LSLHRPALEDLLLGSEAN[+1663.608]LTC[+57.021]TLTGLR.D | HexNAc(5)Hex(4) | C[+57], N[+1664] | 1543.0788 | 3 | 4627.2218 | 4627.2064 | 0 | 3.3 | 127 | Specific | 591.12 | 528.81 | 528.81 | 11.68 | 42 | >sp\|P01876\|IGHA1_HUMAN Immunoglobulin heavy constant alpha 1 OS=Homo sapiens OX=9606 GN=IGHA1 PE=1 SV=2 | 255 | 20201113_gy_milk_1ng_glyco_hcd203040_compound15.21122.21122.3 | controllerType=0 controllerNumber=1 scan=21122 | 42.4342 |
| 17607:4 | 11 | R.LSLHRPALEDLLLGSEAN[+1702.581]LTC[+57.021]TLTGLR.D | HexNAc(2)Hex(8) | C[+57], N[+1703] | 1167.3108 | 4 | 4666.2214 | 4666.1796 | 0 | 9 | 127 | Specific | 595.73 | 595.73 | 595.73 | 11.56 | 42 | >sp\|P01876\|IGHA1_HUMAN Immunoglobulin heavy constant alpha 1 OS=Homo sapiens OX=9606 GN=IGHA1 PE=1 SV=2 | 255 | 20201113_gy_milk_1ng_glyco_hcd203040_compound15.21048.21048.4 | controllerType=0 controllerNumber=1 scan=21048 | 42.2784 |
| 17639:3 | 11 | R.LSLHRPALEDLLLGSEAN[+1540.529]LTC[+57.021]TLTGLR.D | HexNAc(2)Hex(7) | C[+57], N[+1541] | 1502.0505 | 3 | 4504.1369 | 4504.1268 | 0 | 2.2 | 127 | Specific | 649.66 | 564.36 | 564.36 | 12.68 | 42 | >sp\|P01876\|IGHA1_HUMAN Immunoglobulin heavy constant alpha 1 OS=Homo sapiens OX=9606 GN=IGHA1 PE=1 SV=2 | 255 | 20201113_gy_milk_1ng_glyco_hcd203040_compound15.21083.21083.3 | controllerType=0 controllerNumber=1 scan=21083 | 42.3482 |
| 17600:3 | 11 | R.LSLHRPALEDLLLGSEAN[+1864.634]LTC[+57.021]TLTGLR.D | HexNAc(2)Hex(9) | C[+57], N[+1865] | 1610.0969 | 3 | 4828.276 | 4828.2324 | 0 | 9 | 127 | Specific | 664.44 | 664.44 | 664.44 | 12.96 | 42 | >sp\|P01876\|IGHA1_HUMAN Immunoglobulin heavy constant alpha 1 OS=Homo sapiens OX=9606 GN=IGHA1 PE=1 SV=2 | 255 | 20201113_gy_milk_1ng_glyco_hcd203040_compound15.21040.21040.3 | controllerType=0 controllerNumber=1 scan=21040 | 42.2613 |
| 17634:4 | 11 | R.LSLHRPALEDLLLGSEAN[+1540.529]LTC[+57.021]TLTGLR.D | HexNAc(2)Hex(7) | C[+57], N[+1541] | 1126.7892 | 4 | 4504.1351 | 4504.1268 | 0 | 1.8 | 127 | Specific | 679.22 | 658.45 | 658.45 | 14.34 | 42 | >sp\|P01876\|IGHA1_HUMAN Immunoglobulin heavy constant alpha 1 OS=Homo sapiens OX=9606 GN=IGHA1 PE=1 SV=2 | 255 | 20201113_gy_milk_1ng_glyco_hcd203040_compound15.21078.21078.4 | controllerType=0 controllerNumber=1 scan=21078 | 42.3371 |
| 17606:3 | 11 | R.LSLHRPALEDLLLGSEAN[+1702.581]LTC[+57.021]TLTGLR.D | HexNAc(2)Hex(8) | C[+57], N[+1703] | 1556.0786 | 3 | 4666.2212 | 4666.1796 | 0 | 8.9 | 127 | Specific | 679.85 | 675.12 | 675.12 | 12.96 | 42 | >sp\|P01876\|IGHA1_HUMAN Immunoglobulin heavy constant alpha 1 OS=Homo sapiens OX=9606 GN=IGHA1 PE=1 SV=2 | 255 | 20201113_gy_milk_1ng_glyco_hcd203040_compound15.21047.21047.3 | controllerType=0 controllerNumber=1 scan=21047 | 42.2762 |
| 17761:4 | 11 | R.LSLHRPALEDLLLGSEAN[+1501.555]LTC[+57.021]TLTGLR.D | HexNAc(5)Hex(3) | C[+57], N[+1502] | 1117.0441 | 4 | 4465.1546 | 4465.1536 | 0 | 0.2 | 127 | Specific | 686.06 | 686.06 | 686.06 | 14.06 | 42 | >sp\|P01876\|IGHA1_HUMAN Immunoglobulin heavy constant alpha 1 OS=Homo sapiens OX=9606 GN=IGHA1 PE=1 SV=2 | 255 | 20201113_gy_milk_1ng_glyco_hcd203040_compound15.21211.21211.4 | controllerType=0 controllerNumber=1 scan=21211 | 42.6237 |
| 17809:4 | 11 | R.LSLHRPALEDLLLGSEAN[+1054.370]LTC[+57.021]TLTGLR.D | HexNAc(2)Hex(4) | C[+57], N[+1054] | 1005.2543 | 4 | 4017.9952 | 4017.9683 | 0 | 6.7 | 127 | Specific | 705.88 | 705.88 | 705.88 | 14.58 | 42 | >sp\|P01876\|IGHA1_HUMAN Immunoglobulin heavy constant alpha 1 OS=Homo sapiens OX=9606 GN=IGHA1 PE=1 SV=2 | 255 | 20201113_gy_milk_1ng_glyco_hcd203040_compound15.21261.21261.4 | controllerType=0 controllerNumber=1 scan=21261 | 42.737 |
| 17637:3 | 11 | R.LSLHRPALEDLLLGSEAN[+1540.529]LTC[+57.021]TLTGLR.D | HexNAc(2)Hex(7) | C[+57], N[+1541] | 1502.0505 | 3 | 4504.1369 | 4504.1268 | 0 | 2.2 | 127 | Specific | 713.46 | 683.89 | 683.89 | 14.68 | 42 | >sp\|P01876\|IGHA1_HUMAN Immunoglobulin heavy constant alpha 1 OS=Homo sapiens OX=9606 GN=IGHA1 PE=1 SV=2 | 255 | 20201113_gy_milk_1ng_glyco_hcd203040_compound15.21081.21081.3 | controllerType=0 controllerNumber=1 scan=21081 | 42.3431 |
| 17807:4 | 11 | R.LSLHRPALEDLLLGSEAN[+1501.555]LTC[+57.021]TLTGLR.D | HexNAc(5)Hex(3) | C[+57], N[+1502] | 1117.0494 | 4 | 4465.1756 | 4465.1536 | 0 | 4.9 | 127 | Specific | 731.44 | 720.12 | 720.12 | 14.74 | 42 | >sp\|P01876\|IGHA1_HUMAN Immunoglobulin heavy constant alpha 1 OS=Homo sapiens OX=9606 GN=IGHA1 PE=1 SV=2 | 255 | 20201113_gy_milk_1ng_glyco_hcd203040_compound15.21259.21259.4 | controllerType=0 controllerNumber=1 scan=21259 | 42.7333 |
| 17762:3 | 11 | R.LSLHRPALEDLLLGSEAN[+1216.423]LTC[+57.021]TLTGLR.D | HexNAc(2)Hex(5) | C[+57], N[+1216] | 1394.0107 | 3 | 4180.0175 | 4180.0211 | 0 | -0.9 | 127 | Specific | 744.7 | 654.15 | 654.15 | 14.02 | 42 | >sp\|P01876\|IGHA1_HUMAN Immunoglobulin heavy constant alpha 1 OS=Homo sapiens OX=9606 GN=IGHA1 PE=1 SV=2 | 255 | 20201113_gy_milk_1ng_glyco_hcd203040_compound15.21212.21212.3 | controllerType=0 controllerNumber=1 scan=21212 | 42.6249 |
| 17704:4 | 11 | R.LSLHRPALEDLLLGSEAN[+1663.608]LTC[+57.021]TLTGLR.D | HexNAc(5)Hex(4) | C[+57], N[+1664] | 1157.557 | 4 | 4627.2064 | 4627.2064 | 0 | 0 | 127 | Specific | 750.11 | 661.61 | 661.61 | 16.06 | 42 | >sp\|P01876\|IGHA1_HUMAN Immunoglobulin heavy constant alpha 1 OS=Homo sapiens OX=9606 GN=IGHA1 PE=1 SV=2 | 255 | 20201113_gy_milk_1ng_glyco_hcd203040_compound15.21151.21151.4 | controllerType=0 controllerNumber=1 scan=21151 | 42.5027 |
| 17764:4 | 11 | R.LSLHRPALEDLLLGSEAN[+1501.555]LTC[+57.021]TLTGLR.D | HexNAc(5)Hex(3) | C[+57], N[+1502] | 1117.0441 | 4 | 4465.1546 | 4465.1536 | 0 | 0.2 | 127 | Specific | 751.15 | 746.79 | 746.79 | 16.06 | 42 | >sp\|P01876\|IGHA1_HUMAN Immunoglobulin heavy constant alpha 1 OS=Homo sapiens OX=9606 GN=IGHA1 PE=1 SV=2 | 255 | 20201113_gy_milk_1ng_glyco_hcd203040_compound15.21214.21214.4 | controllerType=0 controllerNumber=1 scan=21214 | 42.633 |
| 17636:4 | 11 | R.LSLHRPALEDLLLGSEAN[+1540.529]LTC[+57.021]TLTGLR.D | HexNAc(2)Hex(7) | C[+57], N[+1541] | 1126.7892 | 4 | 4504.1351 | 4504.1268 | 0 | 1.8 | 127 | Specific | 792.49 | 792.49 | 792.49 | 16.34 | 42 | >sp\|P01876\|IGHA1_HUMAN Immunoglobulin heavy constant alpha 1 OS=Homo sapiens OX=9606 GN=IGHA1 PE=1 SV=2 | 255 | 20201113_gy_milk_1ng_glyco_hcd203040_compound15.21080.21080.4 | controllerType=0 controllerNumber=1 scan=21080 | 42.3414 |
| 17765:3 | 11 | R.LSLHRPALEDLLLGSEAN[+1216.423]LTC[+57.021]TLTGLR.D | HexNAc(2)Hex(5) | C[+57], N[+1216] | 1394.0107 | 3 | 4180.0175 | 4180.0211 | 0 | -0.9 | 127 | Specific | 819.4 | 785.97 | 785.97 | 16.02 | 42 | >sp\|P01876\|IGHA1_HUMAN Immunoglobulin heavy constant alpha 1 OS=Homo sapiens OX=9606 GN=IGHA1 PE=1 SV=2 | 255 | 20201113_gy_milk_1ng_glyco_hcd203040_compound15.21215.21215.3 | controllerType=0 controllerNumber=1 scan=21215 | 42.6337 |
| 17593:4 | 11 | R.LSLHRPALEDLLLGSEAN[+1864.634]LTC[+57.021]TLTGLR.D | HexNAc(2)Hex(9) | C[+57], N[+1865] | 1207.8237 | 4 | 4828.2728 | 4828.2324 | 0 | 8.4 | 127 | Specific | 852.7 | 852.7 | 852.7 | 17.61 | 42 | >sp\|P01876\|IGHA1_HUMAN Immunoglobulin heavy constant alpha 1 OS=Homo sapiens OX=9606 GN=IGHA1 PE=1 SV=2 | 255 | 20201113_gy_milk_1ng_glyco_hcd203040_compound15.21033.21033.4 | controllerType=0 controllerNumber=1 scan=21033 | 42.2268 |
| 17763:4 | 11 | R.LSLHRPALEDLLLGSEAN[+1298.476]LTC[+57.021]TLTGLR.D | HexNAc(4)Hex(3) | C[+57], N[+1298] | 1066.2736 | 4 | 4262.0726 | 4262.0742 | 0 | -0.4 | 127 | Specific | 871.56 | 838.89 | 838.89 | 18.06 | 42 | >sp\|P01876\|IGHA1_HUMAN Immunoglobulin heavy constant alpha 1 OS=Homo sapiens OX=9606 GN=IGHA1 PE=1 SV=2 | 255 | 20201113_gy_milk_1ng_glyco_hcd203040_compound15.21213.21213.4 | controllerType=0 controllerNumber=1 scan=21213 | 42.6268 |
| 17803:4 | 11 | R.LSLHRPALEDLLLGSEAN[+1095.397]LTC[+57.021]TLTGLR.D | HexNAc(3)Hex(3) | C[+57], N[+1095] | 1015.5121 | 4 | 4059.0267 | 4058.9949 | 0 | 7.8 | 127 | Specific | 910.08 | 910.08 | 910.08 | 18.61 | 42 | >sp\|P01876\|IGHA1_HUMAN Immunoglobulin heavy constant alpha 1 OS=Homo sapiens OX=9606 GN=IGHA1 PE=1 SV=2 | 255 | 20201113_gy_milk_1ng_glyco_hcd203040_compound15.21255.21255.4 | controllerType=0 controllerNumber=1 scan=21255 | 42.7209 |
| 17806:4 | 11 | R.LSLHRPALEDLLLGSEAN[+1095.397]LTC[+57.021]TLTGLR.D | HexNAc(3)Hex(3) | C[+57], N[+1095] | 1015.5121 | 4 | 4059.0267 | 4058.9949 | 0 | 7.8 | 127 | Specific | 967.19 | 955.77 | 955.77 | 19.61 | 42 | >sp\|P01876\|IGHA1_HUMAN Immunoglobulin heavy constant alpha 1 OS=Homo sapiens OX=9606 GN=IGHA1 PE=1 SV=2 | 255 | 20201113_gy_milk_1ng_glyco_hcd203040_compound15.21258.21258.4 | controllerType=0 controllerNumber=1 scan=21258 | 42.731 |
| 17595:4 | 11 | R.LSLHRPALEDLLLGSEAN[+1864.634]LTC[+57.021]TLTGLR.D | HexNAc(2)Hex(9) | C[+57], N[+1865] | 1207.8237 | 4 | 4828.2728 | 4828.2324 | 0 | 8.4 | 127 | Specific | 1039.49 | 995.8 | 995.8 | 20.61 | 42 | >sp\|P01876\|IGHA1_HUMAN Immunoglobulin heavy constant alpha 1 OS=Homo sapiens OX=9606 GN=IGHA1 PE=1 SV=2 | 255 | 20201113_gy_milk_1ng_glyco_hcd203040_compound15.21035.21035.4 | controllerType=0 controllerNumber=1 scan=21035 | 42.2359 |
| 11533:3 | 11 | R.LAGKPTHVN[+892.317]VSVVMAEVDGTC[+57.021]Y.- | HexNAc(2)Hex(3) | C[+57], N[+892] | 1080.491 | 3 | 3239.4584 | 3239.4593 | 0 | -0.3 | 332 | CRagged | 246.57 | 246.57 | 203.19 | 6.63 | 42 | >sp\|P01876\|IGHA1_HUMAN Immunoglobulin heavy constant alpha 1 OS=Homo sapiens OX=9606 GN=IGHA1 PE=1 SV=2 | 255 | 20201113_gy_milk_1ng_glyco_hcd203040_compound15.14671.14671.3 | controllerType=0 controllerNumber=1 scan=14671 | 30.8581 |
| 13200:3 | 11 | R.LAGKPTHVN[+2059.735]VSVVMAEVDGTC[+57.021]Y.- | HexNAc(4)Hex(5)Fuc(1)NeuAc(1) | C[+57], N[+2060] | 1469.6325 | 3 | 4406.883 | 4406.877 | 0 | 1.3 | 332 | CRagged | 295.82 | 274.26 | 274.26 | 7.36 | 42 | >sp\|P01876\|IGHA1_HUMAN Immunoglobulin heavy constant alpha 1 OS=Homo sapiens OX=9606 GN=IGHA1 PE=1 SV=2 | 255 | 20201113_gy_milk_1ng_glyco_hcd203040_compound15.16412.16412.3 | controllerType=0 controllerNumber=1 scan=16412 | 33.7651 |
| 10191:3 | 11 | R.LAGKPTHVN[+1216.423]VSVVMAEVDGTC[+57.021]Y.- | HexNAc(2)Hex(5) | C[+57], N[+1216] | 1188.5281 | 3 | 3563.5698 | 3563.5649 | 0 | 1.4 | 332 | CRagged | 298.9 | 244.96 | 244.96 | 7.33 | 42 | >sp\|P01876\|IGHA1_HUMAN Immunoglobulin heavy constant alpha 1 OS=Homo sapiens OX=9606 GN=IGHA1 PE=1 SV=2 | 255 | 20201113_gy_milk_1ng_glyco_hcd203040_compound15.13279.13279.3 | controllerType=0 controllerNumber=1 scan=13279 | 28.4762 |
| 11535:3 | 11 | R.LAGKPTHVN[+892.317]VSVVMAEVDGTC[+57.021]Y.- | HexNAc(2)Hex(3) | C[+57], N[+892] | 1080.491 | 3 | 3239.4584 | 3239.4593 | 0 | -0.3 | 332 | CRagged | 324.06 | 324.06 | 258.03 | 7.59 | 42 | >sp\|P01876\|IGHA1_HUMAN Immunoglobulin heavy constant alpha 1 OS=Homo sapiens OX=9606 GN=IGHA1 PE=1 SV=2 | 255 | 20201113_gy_milk_1ng_glyco_hcd203040_compound15.14673.14673.3 | controllerType=0 controllerNumber=1 scan=14673 | 30.8615 |
| 11392:3 | 11 | R.LAGKPTHVN[+1501.555]VSVVMAEVDGTC[+57.021]Y.- | HexNAc(5)Hex(3) | C[+57], N[+1502] | 1283.5731 | 3 | 3848.7046 | 3848.6974 | 0 | 1.9 | 332 | CRagged | 332.02 | 243.02 | 243.02 | 7.37 | 42 | >sp\|P01876\|IGHA1_HUMAN Immunoglobulin heavy constant alpha 1 OS=Homo sapiens OX=9606 GN=IGHA1 PE=1 SV=2 | 255 | 20201113_gy_milk_1ng_glyco_hcd203040_compound15.14525.14525.3 | controllerType=0 controllerNumber=1 scan=14525 | 30.5885 |
| 10237:3 | 11 | R.LAGKPTHVN[+1647.613]VSVVMAEVDGTC[+57.021]Y.- | HexNAc(5)Hex(3)Fuc(1) | C[+57], N[+1648] | 1332.2598 | 3 | 3994.7649 | 3994.7553 | 0 | 2.4 | 332 | CRagged | 340.92 | 284.6 | 284.6 | 7.55 | 42 | >sp\|P01876\|IGHA1_HUMAN Immunoglobulin heavy constant alpha 1 OS=Homo sapiens OX=9606 GN=IGHA1 PE=1 SV=2 | 255 | 20201113_gy_milk_1ng_glyco_hcd203040_compound15.13326.13326.3 | controllerType=0 controllerNumber=1 scan=13326 | 28.5533 |
| 10195:3 | 11 | R.LAGKPTHVN[+1647.613]VSVVMAEVDGTC[+57.021]Y.- | HexNAc(5)Hex(3)Fuc(1) | C[+57], N[+1648] | 1332.2566 | 3 | 3994.7554 | 3994.7553 | 0 | 0 | 332 | CRagged | 398.5 | 398.5 | 398.5 | 9.6 | 42 | >sp\|P01876\|IGHA1_HUMAN Immunoglobulin heavy constant alpha 1 OS=Homo sapiens OX=9606 GN=IGHA1 PE=1 SV=2 | 255 | 20201113_gy_milk_1ng_glyco_hcd203040_compound15.13283.13283.3 | controllerType=0 controllerNumber=1 scan=13283 | 28.4826 |
| 10197:3 | 11 | R.LAGKPTHVN[+1647.613]VSVVMAEVDGTC[+57.021]Y.- | HexNAc(5)Hex(3)Fuc(1) | C[+57], N[+1648] | 1332.2566 | 3 | 3994.7554 | 3994.7553 | 0 | 0 | 332 | CRagged | 425.11 | 425.11 | 425.11 | 9.64 | 42 | >sp\|P01876\|IGHA1_HUMAN Immunoglobulin heavy constant alpha 1 OS=Homo sapiens OX=9606 GN=IGHA1 PE=1 SV=2 | 255 | 20201113_gy_milk_1ng_glyco_hcd203040_compound15.13285.13285.3 | controllerType=0 controllerNumber=1 scan=13285 | 28.4867 |
| 10235:3 | 11 | R.LAGKPTHVN[+1647.613]VSVVMAEVDGTC[+57.021]Y.- | HexNAc(5)Hex(3)Fuc(1) | C[+57], N[+1648] | 1332.2598 | 3 | 3994.7649 | 3994.7553 | 0 | 2.4 | 332 | CRagged | 433.41 | 309.98 | 309.98 | 8.15 | 42 | >sp\|P01876\|IGHA1_HUMAN Immunoglobulin heavy constant alpha 1 OS=Homo sapiens OX=9606 GN=IGHA1 PE=1 SV=2 | 255 | 20201113_gy_milk_1ng_glyco_hcd203040_compound15.13324.13324.3 | controllerType=0 controllerNumber=1 scan=13324 | [illegible] |
| 11426:3 | 11 | R.LAGKPTHVN[+1298.476]VSVVMAEVDGTC[+57.021]Y.- | HexNAc(4)Hex(3) | C[+57], N[+1298] | 1215.8882 | 3 | 3645.6502 | 3645.618 | 0 | 8.8 | 332 | CRagged | 499.92 | 385.18 | 385.18 | 9.46 | 42 | >sp\|P01876\|IGHA1_HUMAN Immunoglobulin heavy constant alpha 1 OS=Homo sapiens OX=9606 GN=IGHA1 PE=1 SV=2 | 255 | 20201113_gy_milk_1ng_glyco_hcd203040_compound15.14560.14560.3 | controllerType=0 controllerNumber=1 scan=14560 | 30.6538 |
| 13243:3 | 11 | R.LAGKPTHVN[+1751.624]VSVVMAEVDGTC[+57.021]Y.- | HexNAc(4)Hex(4)NeuAc(1) | C[+57], N[+1752] | 1366.9277 | 3 | 4098.7685 | 4098.7663 | 0 | 0.5 | 332 | CRagged | 586.23 | 586.23 | 586.23 | 11.5 | 42 | >sp\|P01876\|IGHA1_HUMAN Immunoglobulin heavy constant alpha 1 OS=Homo sapiens OX=9606 GN=IGHA1 PE=1 SV=2 | 255 | 20201113_gy_milk_1ng_glyco_hcd203040_compound15.16457.16457.3 | controllerType=0 controllerNumber=1 scan=16457 | 33.8426 |
| 11192:3 | 11 | R.LAGKPTHVN[+1702.581]VSVVMAEVDGTC[+57.021]Y.- | HexNAc(2)Hex(8) | C[+57], N[+1703] | 1350.5855 | 3 | 4049.742 | 4049.7234 | 0 | 4.6 | 332 | CRagged | 611.75 | 611.75 | 467.07 | 12.42 | 42 | >sp\|P01876\|IGHA1_HUMAN Immunoglobulin heavy constant alpha 1 OS=Homo sapiens OX=9606 GN=IGHA1 PE=1 SV=2 | 255 | 20201113_gy_milk_1ng_glyco_hcd203040_compound15.14317.14317.3 | controllerType=0 controllerNumber=1 scan=14317 | 30.2187 |
| 13245:3 | 11 | R.LAGKPTHVN[+1751.624]VSVVMAEVDGTC[+57.021]Y.- | HexNAc(4)Hex(4)NeuAc(1) | C[+57], N[+1752] | 1366.9277 | 3 | 4098.7685 | 4098.7663 | 0 | 0.5 | 332 | CRagged | 653.84 | 637.21 | 637.21 | 13.5 | 42 | >sp\|P01876\|IGHA1_HUMAN Immunoglobulin heavy constant alpha 1 OS=Homo sapiens OX=9606 GN=IGHA1 PE=1 SV=2 | 255 | 20201113_gy_milk_1ng_glyco_hcd203040_compound15.16459.16459.3 | controllerType=0 controllerNumber=1 scan=16459 | 33.8466 |
| 11428:3 | 11 | R.LAGKPTHVN[+1298.476]VSVVMAEVDGTC[+57.021]Y.- | HexNAc(4)Hex(3) | C[+57], N[+1298] | 1215.8882 | 3 | 3645.6502 | 3645.618 | 0 | 8.8 | 332 | CRagged | 656.83 | 463.38 | 463.38 | 12.99 | 42 | >sp\|P01876\|IGHA1_HUMAN Immunoglobulin heavy constant alpha 1 OS=Homo sapiens OX=9606 GN=IGHA1 PE=1 SV=2 | 255 | 20201113_gy_milk_1ng_glyco_hcd203040_compound15.14562.14562.3 | controllerType=0 controllerNumber=1 scan=14562 | 30.6586 |
| 11323:3 | 11 | R.LAGKPTHVN[+1378.476]VSVVMAEVDGTC[+57.021]Y.- | HexNAc(2)Hex(6) | C[+57], N[+1378] | 1242.5436 | 3 | 3725.6164 | 3725.6178 | 0 | -0.4 | 332 | CRagged | 678.21 | 678.21 | 408.69 | 13.5 | 42 | >sp\|P01876\|IGHA1_HUMAN Immunoglobulin heavy constant alpha 1 OS=Homo sapiens OX=9606 GN=IGHA1 PE=1 SV=2 | 255 | 20201113_gy_milk_1ng_glyco_hcd203040_compound15.14453.14453.3 | controllerType=0 controllerNumber=1 scan=14453 | 30.4648 |
| 11391:3 | 11 | R.LAGKPTHVN[+1647.613]VSVVMAEVDGTC[+57.021]Y.- | HexNAc(5)Hex(3)Fuc(1) | C[+57], N[+1648] | 1332.259 | 3 | 3994.7624 | 3994.7553 | 0 | 1.8 | 332 | CRagged | 734.19 | 644.12 | 644.12 | 14.78 | 42 | >sp\|P01876\|IGHA1_HUMAN Immunoglobulin heavy constant alpha 1 OS=Homo sapiens OX=9606 GN=IGHA1 PE=1 SV=2 | 255 | 20201113_gy_milk_1ng_glyco_hcd203040_compound15.14524.14524.3 | controllerType=0 controllerNumber=1 scan=14524 | 30.5868 |
| 11268:3 | 11 | R.LAGKPTHVN[+1540.529]VSVVMAEVDGTC[+57.021]Y.- | HexNAc(2)Hex(7) | C[+57], N[+1541] | 1296.5637 | 3 | 3887.6765 | 3887.6706 | 0 | 1.5 | 332 | CRagged | 777.52 | 717.28 | 479.27 | 15.78 | 42 | >sp\|P01876\|IGHA1_HUMAN Immunoglobulin heavy constant alpha 1 OS=Homo sapiens OX=9606 GN=IGHA1 PE=1 SV=2 | 255 | 20201113_gy_milk_1ng_glyco_hcd203040_compound15.14396.14396.3 | controllerType=0 controllerNumber=1 scan=14396 | 30.3604 |
| 11245:3 | 11 | R.LAGKPTHVN[+1702.581]VSVVMAEVDGTC[+57.021]Y.- | HexNAc(2)Hex(8) | C[+57], N[+1703] | 1350.5817 | 3 | 4049.7306 | 4049.7234 | 0 | 1.8 | 332 | CRagged | 783.06 | 746.12 | 474.94 | 15.78 | 42 | >sp\|P01876\|IGHA1_HUMAN Immunoglobulin heavy constant alpha 1 OS=Homo sapiens OX=9606 GN=IGHA1 PE=1 SV=2 | 255 | 20201113_gy_milk_1ng_glyco_hcd203040_compound15.14372.14372.3 | controllerType=0 controllerNumber=1 scan=14372 | 30.3161 |
| 11393:3 | 11 | R.LAGKPTHVN[+1647.613]VSVVMAEVDGTC[+57.021]Y.- | HexNAc(5)Hex(3)Fuc(1) | C[+57], N[+1648] | 1332.259 | 3 | 3994.7624 | 3994.7553 | 0 | 1.8 | 332 | CRagged | 815.82 | 635.08 | 635.08 | 16.78 | 42 | >sp\|P01876\|IGHA1_HUMAN Immunoglobulin heavy constant alpha 1 OS=Homo sapiens OX=9606 GN=IGHA1 PE=1 SV=2 | 255 | 20201113_gy_milk_1ng_glyco_hcd203040_compound15.14526.14526.3 | controllerType=0 controllerNumber=1 scan=14526 | 30.5915 |
| 11341:3 | 11 | R.LAGKPTHVN[+1216.423]VSVVMAEVDGTC[+57.021]Y.- | HexNAc(2)Hex(5) | C[+57], N[+1216] | 1188.5267 | 3 | 3563.5655 | 3563.5649 | 0 | 0.1 | 332 | CRagged | 852.06 | 852.06 | 508.88 | 17.5 | 42 | >sp\|P01876\|IGHA1_HUMAN Immunoglobulin heavy constant alpha 1 OS=Homo sapiens OX=9606 GN=IGHA1 PE=1 SV=2 | 255 | 20201113_gy_milk_1ng_glyco_hcd203040_compound15.14471.14471.3 | controllerType=0 controllerNumber=1 scan=14471 | 30.4916 |
| 11326:3 | 11 | R.LAGKPTHVN[+1378.476]VSVVMAEVDGTC[+57.021]Y.- | HexNAc(2)Hex(6) | C[+57], N[+1378] | 1242.5436 | 3 | 3725.6164 | 3725.6178 | 0 | -0.4 | 332 | CRagged | 869.07 | 822.74 | 547.88 | 17.5 | 42 | >sp\|P01876\|IGHA1_HUMAN Immunoglobulin heavy constant alpha 1 OS=Homo sapiens OX=9606 GN=IGHA1 PE=1 SV=2 | 255 | 20201113_gy_milk_1ng_glyco_hcd203040_compound15.14456.14456.3 | controllerType=0 controllerNumber=1 scan=14456 | 30.4688 |
| 11344:3 | 11 | R.LAGKPTHVN[+1216.423]VSVVMAEVDGTC[+57.021]Y.- | HexNAc(2)Hex(5) | C[+57], N[+1216] | 1188.5267 | 3 | 3563.5655 | 3563.5649 | 0 | 0.1 | 332 | CRagged | 887.99 | 870.07 | 531.59 | 17.5 | 42 | >sp\|P01876\|IGHA1_HUMAN Immunoglobulin heavy constant alpha 1 OS=Homo sapiens OX=9606 GN=IGHA1 PE=1 SV=2 | 255 | 20201113_gy_milk_1ng_glyco_hcd203040_compound15.14474.14474.3 | controllerType=0 controllerNumber=1 scan=14474 | 30.4994 |
| 11394:3 | 11 | R.LAGKPTHVN[+1501.555]VSVVMAEVDGTC[+57.021]Y.- | HexNAc(5)Hex(3) | C[+57], N[+1502] | 1283.5731 | 3 | 3848.7046 | 3848.6974 | 0 | 1.9 | 332 | CRagged | 894.12 | 894.12 | 894.12 | 17.78 | 42 | >sp\|P01876\|IGHA1_HUMAN Immunoglobulin heavy constant alpha 1 OS=Homo sapiens OX=9606 GN=IGHA1 PE=1 SV=2 | 255 | 20201113_gy_milk_1ng_glyco_hcd203040_compound15.14527.14527.3 | controllerType=0 controllerNumber=1 scan=14527 | 30.5925 |
| 11243:3 | 11 | R.LAGKPTHVN[+1702.581]VSVVMAEVDGTC[+57.021]Y.- | HexNAc(2)Hex(8) | C[+57], N[+1703] | 1350.5817 | 3 | 4049.7306 | 4049.7234 | 0 | 1.8 | 332 | CRagged | 905.37 | 905.37 | 593.53 | 18.78 | 42 | >sp\|P01876\|IGHA1_HUMAN Immunoglobulin heavy constant alpha 1 OS=Homo sapiens OX=9606 GN=IGHA1 PE=1 SV=2 | 255 | 20201113_gy_milk_1ng_glyco_hcd203040_compound15.14370.14370.3 | controllerType=0 controllerNumber=1 scan=14370 | 30.3106 |
| 11354:3 | 11 | R.LAGKPTHVN[+1809.666]VSVVMAEVDGTC[+57.021]Y.- | HexNAc(5)Hex(4)Fuc(1) | C[+57], N[+1810] | 1386.275 | 3 | 4156.8103 | 4156.8081 | 0 | 0.5 | 332 | CRagged | 914.01 | 780.52 | 587.41 | 18.5 | 42 | >sp\|P01876\|IGHA1_HUMAN Immunoglobulin heavy constant alpha 1 OS=Homo sapiens OX=9606 GN=IGHA1 PE=1 SV=2 | 255 | 20201113_gy_milk_1ng_glyco_hcd203040_compound15.14485.14485.3 | controllerType=0 controllerNumber=1 scan=14485 | 30.5174 |
| 07677:3 | 12 | R.LSPN[+2350.830]ASAEHLELR.W | HexNAc(4)Hex(5)Fuc(1)NeuAc(2) | N[+2351] | 1263.1995 | 3 | 3787.584 | 3787.5796 | 0 | 1.2 | 52 | Specific | 495.43 | 455.83 | 455.83 | 9.82 | 37 | >sp\|Q13410\|BT1A1_HUMAN Butyrophilin subfamily 1 member A1 OS=Homo sapiens OX=9606 GN=BTN1A1 PE=1 SV=3 | 179 | 20201113_gy_milk_1ng_glyco_hcd203040_compound15.10658.10658.3 | controllerType=0 controllerNumber=1 scan=10658 | 24.074 |
| 07679:3 | 12 | R.LSPN[+2350.830]ASAEHLELR.W | HexNAc(4)Hex(5)Fuc(1)NeuAc(2) | N[+2351] | 1263.1995 | 3 | 3787.584 | 3787.5796 | 0 | 1.2 | 52 | Specific | 526.31 | 444.2 | 444.2 | 9.96 | 37 | >sp\|Q13410\|BT1A1_HUMAN Butyrophilin subfamily 1 member A1 OS=Homo sapiens OX=9606 GN=BTN1A1 PE=1 SV=3 | 179 | 20201113_gy_milk_1ng_glyco_hcd203040_compound15.10660.10660.3 | controllerType=0 controllerNumber=1 scan=10660 | 24.0786 |
| 05628:3 | 12 | R.LSPN[+2205.793]ASAEHLELR.W | HexNAc(4)Hex(5)Fuc(2)NeuAc(1) | N[+2206] | 1214.8535 | 3 | 3642.5458 | 3642.5421 | 0 | 1 | 52 | Specific | 594.27 | 443.45 | 443.45 | 11.35 | 37 | >sp\|Q13410\|BT1A1_HUMAN Butyrophilin subfamily 1 member A1 OS=Homo sapiens OX=9606 GN=BTN1A1 PE=1 SV=3 | 179 | 20201113_gy_milk_1ng_glyco_hcd203040_compound15.8536.8536.3 | controllerType=0 controllerNumber=1 scan=8536 | 20.5337 |
| 05644:3 | 12 | R.LSPN[+2205.793]ASAEHLELR.W | HexNAc(4)Hex(5)Fuc(2)NeuAc(1) | N[+2206] | 1214.855 | 3 | 3642.5506 | 3642.5421 | 0 | 2.3 | 52 | Specific | 595.03 | 402.42 | 402.42 | 11.3 | 37 | >sp\|Q13410\|BT1A1_HUMAN Butyrophilin subfamily 1 member A1 OS=Homo sapiens OX=9606 GN=BTN1A1 PE=1 SV=3 | 179 | 20201113_gy_milk_1ng_glyco_hcd203040_compound15.8553.8553.3 | controllerType=0 controllerNumber=1 scan=8553 | 20.5665 |
| 05643:3 | 12 | R.LSPN[+2205.793]ASAEHLELR.W | HexNAc(4)Hex(5)Fuc(2)NeuAc(1) | N[+2206] | 1214.855 | 3 | 3642.5506 | 3642.5421 | 0 | 2.3 | 52 | Specific | 616.58 | 549.79 | 549.79 | 12.3 | 37 | >sp\|Q13410\|BT1A1_HUMAN Butyrophilin subfamily 1 member A1 OS=Homo sapiens OX=9606 GN=BTN1A1 PE=1 SV=3 | 179 | 20201113_gy_milk_1ng_glyco_hcd203040_compound15.8552.8552.3 | controllerType=0 controllerNumber=1 scan=8552 | 20.5621 |
| 05626:3 | 12 | R.LSPN[+2205.793]ASAEHLELR.W | HexNAc(4)Hex(5)Fuc(2)NeuAc(1) | N[+2206] | 1214.8535 | 3 | 3642.5458 | 3642.5421 | 0 | 1 | 52 | Specific | 650.47 | 543.12 | 543.12 | 13.35 | 37 | >sp\|Q13410\|BT1A1_HUMAN Butyrophilin subfamily 1 member A1 OS=Homo sapiens OX=9606 GN=BTN1A1 PE=1 SV=3 | 179 | 20201113_gy_milk_1ng_glyco_hcd203040_compound15.8534.8534.3 | controllerType=0 controllerNumber=1 scan=8534 | 20.5275 |
| 17887:3 | 12 | R.DTSAKN[+2059.735]VSC[+57.021]YIQNLLLGQEK.K | HexNAc(4)Hex(5)Fuc(1)NeuAc(1) | C[+57], N[+2060] | 1447.6318 | 3 | 4340.8808 | 4340.8842 | 0 | -0.8 | 210 | Specific | 413.95 | 335.63 | 335.63 | 5.48 | 37 | >sp\|Q13410\|BT1A1_HUMAN Butyrophilin subfamily 1 member A1 OS=Homo sapiens OX=9606 GN=BTN1A1 PE=1 SV=3 | 179 | 20201113_gy_milk_1ng_glyco_hcd203040_compound15.21343.21343.3 | controllerType=0 controllerNumber=1 scan=21343 | 42.9236 |
| 17885:3 | 12 | R.DTSAKN[+2059.735]VSC[+57.021]YIQNLLLGQEK.K | HexNAc(4)Hex(5)Fuc(1)NeuAc(1) | C[+57], N[+2060] | 1447.6318 | 3 | 4340.8808 | 4340.8842 | 0 | -0.8 | 210 | Specific | 548.86 | 354.5 | 354.5 | 7.87 | 37 | >sp\|Q13410\|BT1A1_HUMAN Butyrophilin subfamily 1 member A1 OS=Homo sapiens OX=9606 GN=BTN1A1 PE=1 SV=3 | 179 | 20201113_gy_milk_1ng_glyco_hcd203040_compound15.21341.21341.3 | controllerType=0 controllerNumber=1 scan=21341 | 42.9187 |
| 17666:3 | 12 | K.N[+1913.677]VSC[+57.021]YIQNLLLGQEK.K | HexNAc(4)Hex(5)NeuAc(1) | C[+57], N[+1914] | 1231.5378 | 3 | 3692.5988 | 3692.5876 | 0 | 3 | 215 | Specific | 245.37 | 245.37 | 245.37 | 6.52 | 37 | >sp\|Q13410\|BT1A1_HUMAN Butyrophilin subfamily 1 member A1 OS=Homo sapiens OX=9606 GN=BTN1A1 PE=1 SV=3 | 179 | 20201113_gy_milk_1ng_glyco_hcd203040_compound15.21111.21111.3 | controllerType=0 controllerNumber=1 scan=21111 | 42.4135 |
| 17697:3 | 12 | K.N[+1913.677]VSC[+57.021]YIQNLLLGQEK.K | HexNAc(4)Hex(5)NeuAc(1) | C[+57], N[+1914] | 1231.5428 | 3 | 3692.6138 | 3692.5876 | 0 | 7.1 | 215 | Specific | 325.97 | 275.76 | 275.76 | 6.31 | 37 | >sp\|Q13410\|BT1A1_HUMAN Butyrophilin subfamily 1 member A1 OS=Homo sapiens OX=9606 GN=BTN1A1 PE=1 SV=3 | 179 | 20201113_gy_milk_1ng_glyco_hcd203040_compound15.21143.21143.3 | controllerType=0 controllerNumber=1 scan=21143 | 42.4849 |
| 17668:3 | 12 | K.N[+1913.677]VSC[+57.021]YIQNLLLGQEK.K | HexNAc(4)Hex(5)NeuAc(1) | C[+57], N[+1914] | 1231.5378 | 3 | 3692.5988 | 3692.5876 | 0 | 3 | 215 | Specific | 638.15 | 455.02 | 455.02 | 12.39 | 37 | >sp\|Q13410\|BT1A1_HUMAN Butyrophilin subfamily 1 member A1 OS=Homo sapiens OX=9606 GN=BTN1A1 PE=1 SV=3 | 179 | 20201113_gy_milk_1ng_glyco_hcd203040_compound15.21113.21113.3 | controllerType=0 controllerNumber=1 scan=21113 | 42.4172 |
| 24287:3 | 13 | R.[+42.011]QLAHQSN[+2190.793]STNIFFSPVSIATAFAMLSLGTK.A | HexNAc(6)Hex(6) | .[42], N[+2191] | 1805.4818 | 3 | 5414.4308 | 5414.4388 | 0 | -1.5 | 64 | Specific | 267.38 | 267.38 | 267.38 | 6.31 | 33 | >sp\|P01009\|A1AT_HUMAN Alpha-1-antitrypsin OS=Homo sapiens OX=9606 GN=SERPINA1 PE=1 SV=3 | 642 | 20201113_gy_milk_1ng_glyco_hcd203040_compound15.28061.28061.3 | controllerType=0 controllerNumber=1 scan=28061 | 55.9849 |
| 20199:3 | 13 | K.YLGN[+2204.772]ATAIFFLPDEGK.L | HexNAc(4)Hex(5)NeuAc(2) | N[+2205] | 1320.9009 | 3 | 3960.6883 | 3960.6677 | 0 | 5.2 | 268 | Specific | 397.04 | 276.15 | 276.15 | 6.92 | 33 | >sp\|P01009\|A1AT_HUMAN Alpha-1-antitrypsin OS=Homo sapiens OX=9606 GN=SERPINA1 PE=1 SV=3 | 642 | 20201113_gy_milk_1ng_glyco_hcd203040_compound15.23772.23772.3 | controllerType=0 controllerNumber=1 scan=23772 | 48.0561 |
| 20201:3 | 13 | K.YLGN[+2204.772]ATAIFFLPDEGK.L | HexNAc(4)Hex(5)NeuAc(2) | N[+2205] | 1320.9009 | 3 | 3960.6883 | 3960.6677 | 0 | 5.2 | 268 | Specific | 616.65 | 569.97 | 569.97 | 11.84 | 33 | >sp\|P01009\|A1AT_HUMAN Alpha-1-antitrypsin OS=Homo sapiens OX=9606 GN=SERPINA1 PE=1 SV=3 | 642 | 20201113_gy_milk_1ng_glyco_hcd203040_compound15.23774.23774.3 | controllerType=0 controllerNumber=1 scan=23774 | 48.06 |
| 20011:4 | 13 | K.YLGN[+2204.772]ATAIFFLPDEGKLQHLENELTHDIITK.F | HexNAc(4)Hex(5)NeuAc(2) | N[+2205] | 1437.1628 | 4 | 5745.6296 | 5745.6098 | 0 | 3.4 | 268 | Specific | 985.63 | 906.59 | 906.59 | 18.24 | 33 | >sp\|P01009\|A1AT_HUMAN Alpha-1-antitrypsin OS=Homo sapiens OX=9606 GN=SERPINA1 PE=1 SV=3 | 642 | 20201113_gy_milk_1ng_glyco_hcd203040_compound15.23574.23574.4 | controllerType=0 controllerNumber=1 scan=23574 | 47.6578 |
| 20055:4 | 13 | K.YLGN[+2204.772]ATAIFFLPDEGKLQHLENELTHDIITK.F | HexNAc(4)Hex(5)NeuAc(2) | N[+2205] | 1437.1658 | 4 | 5745.6413 | 5745.6098 | 0 | 5.5 | 268 | Specific | 1007.98 | 952.43 | 952.43 | 18.99 | 33 | >sp\|P01009\|A1AT_HUMAN Alpha-1-antitrypsin OS=Homo sapiens OX=9606 GN=SERPINA1 PE=1 SV=3 | 642 | 20201113_gy_milk_1ng_glyco_hcd203040_compound15.23620.23620.4 | controllerType=0 controllerNumber=1 scan=23620 | 47.7458 |
| 09417:4 | 17 | K.QTDEIKDTRN[+2059.735]ESTQNC[+57.021]VVAEPEKMESSISSSSEEMSLSK.C | HexNAc(4)Hex(5)Fuc(1)NeuAc(1) | C[+57], N[+2060] | 1613.4332 | 4 | 6450.7111 | 6450.7156 | 0 | -0.7 | 60 | Specific | 235.6 | 153.79 | 153.79 | 3.24 | 28 | >sp\|P47710\|CASA1_HUMAN Alpha-S1-casein OS=Homo sapiens OX=9606 GN=CSN1S1 PE=1 SV=1 | 989 | 20201113_gy_milk_1ng_glyco_hcd203040_compound15.12476.12476.4 | controllerType=0 controllerNumber=1 scan=12476 | 27.1521 |
| 09415:4 | 17 | K.QTDEIKDTRN[+2059.735]ESTQNC[+57.021]VVAEPEKMESSISSSSEEMSLSK.C | HexNAc(4)Hex(5)Fuc(1)NeuAc(1) | C[+57], N[+2060] | 1613.4332 | 4 | 6450.7111 | 6450.7156 | 0 | -0.7 | 60 | Specific | 255.87 | 243.88 | 243.88 | 5.31 | 28 | >sp\|P47710\|CASA1_HUMAN Alpha-S1-casein OS=Homo sapiens OX=9606 GN=CSN1S1 PE=1 SV=1 | 989 | 20201113_gy_milk_1ng_glyco_hcd203040_compound15.12474.12474.4 | controllerType=0 controllerNumber=1 scan=12474 | 27.1485 |
| 09435:5 | 17 | K.QTDEIKDTRN[+2059.735]ESTQNC[+57.021]VVAEPEKMESSISSSSEEMSLSK.C | HexNAc(4)Hex(5)Fuc(1)NeuAc(1) | C[+57], N[+2060] | 1290.9501 | 5 | 6450.7213 | 6450.7156 | 0 | 0.9 | 60 | Specific | 319.1 | 295.49 | 295.49 | 7.23 | 28 | >sp\|P47710\|CASA1_HUMAN Alpha-S1-casein OS=Homo sapiens OX=9606 GN=CSN1S1 PE=1 SV=1 | 989 | 20201113_gy_milk_1ng_glyco_hcd203040_compound15.12495.12495.3 | controllerType=0 controllerNumber=1 scan=12495 | 27.1836 |
| 09459:4 | 17 | K.QTDEIKDTRN[+2059.735]ESTQNC[+57.021]VVAEPEKMESSISSSSEEMSLSK.C | HexNAc(4)Hex(5)Fuc(1)NeuAc(1) | C[+57], N[+2060] | 1613.4487 | 4 | 6450.7731 | 6450.7156 | 0 | 8.9 | 60 | Specific | 356.25 | 288 | 288 | 5.74 | 28 | >sp\|P47710\|CASA1_HUMAN Alpha-S1-casein OS=Homo sapiens OX=9606 GN=CSN1S1 PE=1 SV=1 | 989 | 20201113_gy_milk_1ng_glyco_hcd203040_compound15.12520.12520.4 | controllerType=0 controllerNumber=1 scan=12520 | 27.2284 |
| 09461:4 | 17 | K.QTDEIKDTRN[+2059.735]ESTQNC[+57.021]VVAEPEKMESSISSSSEEMSLSK.C | HexNAc(4)Hex(5)Fuc(1)NeuAc(1) | C[+57], N[+2060] | 1613.4487 | 4 | 6450.7731 | 6450.7156 | 0 | 8.9 | 60 | Specific | 386.13 | 241.98 | 241.98 | 5.89 | 28 | >sp\|P47710\|CASA1_HUMAN Alpha-S1-casein OS=Homo sapiens OX=9606 GN=CSN1S1 PE=1 SV=1 | 989 | 20201113_gy_milk_1ng_glyco_hcd203040_compound15.12522.12522.4 | controllerType=0 controllerNumber=1 scan=12522 | 27.2321 |
| 09437:5 | 17 | K.QTDEIKDTRN[+2059.735]ESTQNC[+57.021]VVAEPEKMESSISSSSEEMSLSK.C | HexNAc(4)Hex(5)Fuc(1)NeuAc(1) | C[+57], N[+2060] | 1290.9501 | 5 | 6450.7213 | 6450.7156 | 0 | 0.9 | 60 | Specific | 784.28 | 546.29 | 546.29 | 15.29 | 28 | >sp\|P47710\|CASA1_HUMAN Alpha-S1-casein OS=Homo sapiens OX=9606 GN=CSN1S1 PE=1 SV=1 | 989 | 20201113_gy_milk_1ng_glyco_hcd203040_compound15.12497.12497.3 | controllerType=0 controllerNumber=1 scan=12497 | 27.1873 |
| 23025:4 | 19 | K.APDKNVIFSPLSISTALAFLSLGAHN[+2350.830]TTLTEILK.G | HexNAc(4)Hex(5)Fuc(1)NeuAc(2) | N[+2351] | 1484.2079 | 4 | 5933.8097 | 5933.8085 | 0 | 0.2 | 68 | Specific | 259.79 | 259.79 | 259.79 | 6.51 | 25 | >sp\|P01011\|AACT_HUMAN Alpha-1-antichymotrypsin OS=Homo sapiens OX=9606 GN=SERPINA3 PE=1 SV=2 | 1314 | 20201113_gy_milk_1ng_glyco_hcd203040_compound15.26737.26737.2 | controllerType=0 controllerNumber=1 scan=26737 | 53.6089 |
| 22969:4 | 19 | K.APDKNVIFSPLSISTALAFLSLGAHN[+2350.830]TTLTEILK.G | HexNAc(4)Hex(5)Fuc(1)NeuAc(2) | N[+2351] | 1484.215 | 4 | 5933.838 | 5933.8085 | 0 | 5 | 68 | Specific | 317.88 | 317.88 | 317.88 | 6.79 | 25 | >sp\|P01011\|AACT_HUMAN Alpha-1-antichymotrypsin OS=Homo sapiens OX=9606 GN=SERPINA3 PE=1 SV=2 | 1314 | 20201113_gy_milk_1ng_glyco_hcd203040_compound15.26679.26679.4 | controllerType=0 controllerNumber=1 scan=26679 | 53.503 |
| 22967:4 | 19 | K.APDKNVIFSPLSISTALAFLSLGAHN[+2350.830]TTLTEILK.G | HexNAc(4)Hex(5)Fuc(1)NeuAc(2) | N[+2351] | 1484.215 | 4 | 5933.838 | 5933.8085 | 0 | 5 | 68 | Specific | 702.05 | 702.05 | 702.05 | 13.85 | 25 | >sp\|P01011\|AACT_HUMAN Alpha-1-antichymotrypsin OS=Homo sapiens OX=9606 GN=SERPINA3 PE=1 SV=2 | 1314 | 20201113_gy_milk_1ng_glyco_hcd203040_compound15.26677.26677.4 | controllerType=0 controllerNumber=1 scan=26677 | 53.4984 |
| 14075:4 | 22 | K.LKELPGVC[+57.021]N[+2059.735]ETMMALWEEC[+57.021]KPC[+57.021]LK.Q | HexNAc(4)Hex(5)Fuc(1)NeuAc(1) | C[+57]*3, N[+2060] | 1249.7876 | 4 | 4996.1286 | 4996.1196 | 0 | 1.8 | 95 | Specific | 289.38 | 289.38 | 289.38 | 5.39 | 22 | >sp\|P10909\|CLUS_HUMAN Clusterin OS=Homo sapiens OX=9606 GN=CLU PE=1 SV=1 | 1467 | 20201113_gy_milk_1ng_glyco_hcd203040_compound15.17320.17320.4 | controllerType=0 controllerNumber=1 scan=17320 | 35.362 |
| 14098:4 | 22 | K.LKELPGVC[+57.021]N[+2059.735]ETMMALWEEC[+57.021]KPC[+57.021]LK.Q | HexNAc(4)Hex(5)Fuc(1)NeuAc(1) | C[+57]*3, N[+2060] | 1249.7878 | 4 | 4996.1296 | 4996.1196 | 0 | 2 | 95 | Specific | 325.25 | 311.55 | 311.55 | 7.22 | 22 | >sp\|P10909\|CLUS_HUMAN Clusterin OS=Homo sapiens OX=9606 GN=CLU PE=1 SV=1 | 1467 | 20201113_gy_milk_1ng_glyco_hcd203040_compound15.17344.17344.4 | controllerType=0 controllerNumber=1 scan=17344 | 35.4003 |
| 14100:4 | 22 | K.LKELPGVC[+57.021]N[+2059.735]ETMMALWEEC[+57.021]KPC[+57.021]LK.Q | HexNAc(4)Hex(5)Fuc(1)NeuAc(1) | C[+57]*3, N[+2060] | 1249.7878 | 4 | 4996.1296 | 4996.1196 | 0 | 2 | 95 | Specific | 583.22 | 559.57 | 559.57 | 11.28 | 22 | >sp\|P10909\|CLUS_HUMAN Clusterin OS=Homo sapiens OX=9606 GN=CLU PE=1 SV=1 | 1467 | 20201113_gy_milk_1ng_glyco_hcd203040_compound15.17346.17346.4 | controllerType=0 controllerNumber=1 scan=17346 | 35.4038 |
| 14073:4 | 22 | K.LKELPGVC[+57.021]N[+2059.735]ETMMALWEEC[+57.021]KPC[+57.021]LK.Q | HexNAc(4)Hex(5)Fuc(1)NeuAc(1) | C[+57]*3, N[+2060] | 1249.7876 | 4 | 4996.1286 | 4996.1196 | 0 | 1.8 | 95 | Specific | 849.81 | 837.66 | 837.66 | 16.28 | 22 | >sp\|P10909\|CLUS_HUMAN Clusterin OS=Homo sapiens OX=9606 GN=CLU PE=1 SV=1 | 1467 | 20201113_gy_milk_1ng_glyco_hcd203040_compound15.17318.17318.4 | controllerType=0 controllerNumber=1 scan=17318 | 35.3574 |
| 23438:3 | 22 | K.MLN[+2350.830]TSSLLEQLNEQFNWVSR.L | HexNAc(4)Hex(5)Fuc(1)NeuAc(2) | N[+2351] | 1587.3556 | 3 | 4760.0524 | 4760.0171 | 0 | 7.4 | 352 | Specific | 473.24 | 460.62 | 460.62 | 9.45 | 22 | >sp\|P10909\|CLUS_HUMAN Clusterin OS=Homo sapiens OX=9606 GN=CLU PE=1 SV=1 | 1467 | 20201113_gy_milk_1ng_glyco_hcd203040_compound15.27168.27168.3 | controllerType=0 controllerNumber=1 scan=27168 | 54.3879 |
| 20972:3 | 22 | K.MLN[+2204.772]TSSLLEQLNEQFNWVSR.L | HexNAc(4)Hex(5)NeuAc(2) | N[+2205] | 1538.6561 | 3 | 4613.9536 | 4613.9592 | 0 | -1.2 | 352 | Specific | 562.99 | 473.62 | 228.18 | 9.82 | 22 | >sp\|P10909\|CLUS_HUMAN Clusterin OS=Homo sapiens OX=9606 GN=CLU PE=1 SV=1 | 1467 | 20201113_gy_milk_1ng_glyco_hcd203040_compound15.24579.24579.3 | controllerType=0 controllerNumber=1 scan=24579 | 49.7034 |
| 23436:3 | 22 | K.MLN[+2350.830]TSSLLEQLNEQFNWVSR.L | HexNAc(4)Hex(5)Fuc(1)NeuAc(2) | N[+2351] | 1587.3556 | 3 | 4760.0524 | 4760.0171 | 0 | 7.4 | 352 | Specific | 736.87 | 624.32 | 624.32 | 14.02 | 22 | >sp\|P10909\|CLUS_HUMAN Clusterin OS=Homo sapiens OX=9606 GN=CLU PE=1 SV=1 | 1467 | 20201113_gy_milk_1ng_glyco_hcd203040_compound15.27166.27166.3 | controllerType=0 controllerNumber=1 scan=27166 | 54.384 |
| 10324:3 | 22 | R.LAN[+2205.793]LTQGEDQYYLR.V | HexNAc(4)Hex(5)Fuc(2)NeuAc(1) | N[+2206] | 1297.215 | 3 | 3889.6305 | 3889.6265 | 0 | 1 | 372 | Specific | 202.35 | 202.35 | 202.35 | 5.06 | 22 | >sp\|P10909\|CLUS_HUMAN Clusterin OS=Homo sapiens OX=9606 GN=CLU PE=1 SV=1 | 1467 | 20201113_gy_milk_1ng_glyco_hcd203040_compound15.13416.13416.3 | controllerType=0 controllerNumber=1 scan=13416 | 28.7099 |
| 10325:3 | 22 | R.LAN[+2205.793]LTQGEDQYYLR.V | HexNAc(4)Hex(5)Fuc(2)NeuAc(1) | N[+2206] | 1297.215 | 3 | 3889.6305 | 3889.6265 | 0 | 1 | 372 | Specific | 268.95 | 138.6 | 138.6 | 5.78 | 22 | >sp\|P10909\|CLUS_HUMAN Clusterin OS=Homo sapiens OX=9606 GN=CLU PE=1 SV=1 | 1467 | 20201113_gy_milk_1ng_glyco_hcd203040_compound15.13417.13417.3 | controllerType=0 controllerNumber=1 scan=13417 | 28.7136 |
| 08452:4 | 26 | K.SC[+57.021]QHN[+2204.772]GTMYQHGEIFSAHELFPSR.L | HexNAc(4)Hex(5)NeuAc(2) | C[+57], N[+2205] | 1260.2692 | 4 | 5038.0548 | 5038.0294 | 0 | 5.1 | 110 | Specific | 344.97 | 344.97 | 344.97 | 6.88 | 19 | >sp\|Q6WN34\|CRDL2_HUMAN Chordin-like protein 2 OS=Homo sapiens OX=9606 GN=CHRDL2 PE=1 SV=1 | 1189 | 20201113_gy_milk_1ng_glyco_hcd203040_compound15.11465.11465.2 | controllerType=0 controllerNumber=1 scan=11465 | 25.4058 |
| 08454:4 | 26 | K.SC[+57.021]QHN[+2204.772]GTMYQHGEIFSAHELFPSR.L | HexNAc(4)Hex(5)NeuAc(2) | C[+57], N[+2205] | 1260.2692 | 4 | 5038.0548 | 5038.0294 | 0 | 5.1 | 110 | Specific | 366.01 | 366.01 | 366.01 | 7.27 | 19 | >sp\|Q6WN34\|CRDL2_HUMAN Chordin-like protein 2 OS=Homo sapiens OX=9606 GN=CHRDL2 PE=1 SV=1 | 1189 | 20201113_gy_milk_1ng_glyco_hcd203040_compound15.11467.11467.2 | controllerType=0 controllerNumber=1 scan=11467 | 25.4094 |
| 09610:3 | 39 | R.IIVPLNNREN[+1460.529]ISDPTSPLR.T | HexNAc(4)Hex(4) | N[+1461] | 1203.5818 | 3 | 3608.731 | 3608.7059 | 0 | 6.9 | 62 | Specific | 208.14 | 208.14 | 208.14 | 4.06 | 18 | >sp\|P01591\|IGJ_HUMAN Immunoglobulin J chain OS=Homo sapiens OX=9606 GN=JCHAIN PE=1 SV=4 | 1392 | 20201113_gy_milk_1ng_glyco_hcd203040_compound15.12678.12678.2 | controllerType=0 controllerNumber=1 scan=12678 | 27.4823 |
| 13275:3 | 39 | R.IIVPLNNREN[+2350.830]ISDPTSPLR.T | HexNAc(4)Hex(5)Fuc(1)NeuAc(2) | N[+2351] | 1500.3437 | 3 | 4499.0165 | 4499.0075 | 0 | 2 | 62 | Specific | 215.79 | 188.39 | 188.39 | 4.61 | 18 | >sp\|P01591\|IGJ_HUMAN Immunoglobulin J chain OS=Homo sapiens OX=9606 GN=JCHAIN PE=1 SV=4 | 1392 | 20201113_gy_milk_1ng_glyco_hcd203040_compound15.16490.16490.3 | controllerType=0 controllerNumber=1 scan=16490 | 33.9 |
| 13277:3 | 39 | R.IIVPLNNREN[+2350.830]ISDPTSPLR.T | HexNAc(4)Hex(5)Fuc(1)NeuAc(2) | N[+2351] | 1500.3437 | 3 | 4499.0165 | 4499.0075 | 0 | 2 | 62 | Specific | 307.15 | 307.15 | 307.15 | 7.05 | 18 | >sp\|P01591\|IGJ_HUMAN Immunoglobulin J chain OS=Homo sapiens OX=9606 GN=JCHAIN PE=1 SV=4 | 1392 | 20201113_gy_milk_1ng_glyco_hcd203040_compound15.16492.16492.3 | controllerType=0 controllerNumber=1 scan=16492 | 33.9037 |
| 13359:3 | 39 | R.IIVPLNNREN[+2204.772]ISDPTSPLR.T | HexNAc(4)Hex(5)NeuAc(2) | N[+2205] | 1451.6539 | 3 | 4352.947 | 4352.9496 | 0 | -0.6 | 62 | Specific | 314.82 | 314.82 | 314.82 | 6.83 | 18 | >sp\|P01591\|IGJ_HUMAN Immunoglobulin J chain OS=Homo sapiens OX=9606 GN=JCHAIN PE=1 SV=4 | 1392 | 20201113_gy_milk_1ng_glyco_hcd203040_compound15.16577.16577.3 | controllerType=0 controllerNumber=1 scan=16577 | 34.0462 |
| 11422:3 | 39 | R.IIVPLNNREN[+1856.656]ISDPTSPLR.T | HexNAc(3)Hex(5)Fuc(1)NeuAc(1) | N[+1857] | 1335.6174 | 3 | 4004.8375 | 4004.8327 | 0 | 1.2 | 62 | Specific | 321.03 | 261.1 | 261.1 | 6.75 | 18 | >sp\|P01591\|IGJ_HUMAN Immunoglobulin J chain OS=Homo sapiens OX=9606 GN=JCHAIN PE=1 SV=4 | 1392 | 20201113_gy_milk_1ng_glyco_hcd203040_compound15.14556.14556.3 | controllerType=0 controllerNumber=1 scan=14556 | 30.6471 |
| 11424:3 | 39 | R.IIVPLNNREN[+1856.656]ISDPTSPLR.T | HexNAc(3)Hex(5)Fuc(1)NeuAc(1) | N[+1857] | 1335.6174 | 3 | 4004.8375 | 4004.8327 | 0 | 1.2 | 62 | Specific | 322.88 | 322.88 | 322.88 | 7.25 | 18 | >sp\|P01591\|IGJ_HUMAN Immunoglobulin J chain OS=Homo sapiens OX=9606 GN=JCHAIN PE=1 SV=4 | 1392 | 20201113_gy_milk_1ng_glyco_hcd203040_compound15.14558.14558.3 | controllerType=0 controllerNumber=1 scan=14558 | 30.6507 |
| 09760:3 | 39 | R.IIVPLNNREN[+1403.507]ISDPTSPLR.T | HexNAc(3)Hex(4)Fuc(1) | N[+1404] | 1184.5688 | 3 | 3551.6918 | 3551.6845 | 0 | 2.1 | 62 | Specific | 342.17 | 317.07 | 317.07 | 7.26 | 18 | >sp\|P01591\|IGJ_HUMAN Immunoglobulin J chain OS=Homo sapiens OX=9606 GN=JCHAIN PE=1 SV=4 | 1392 | 20201113_gy_milk_1ng_glyco_hcd203040_compound15.12833.12833.3 | controllerType=0 controllerNumber=1 scan=12833 | 27.7399 |
| 09627:3 | 39 | R.IIVPLNNREN[+1606.587]ISDPTSPLR.T | HexNAc(4)Hex(4)Fuc(1) | N[+1607] | 1252.2577 | 3 | 3754.7587 | 3754.7638 | 0 | -1.4 | 62 | Specific | 352.43 | 280.21 | 280.21 | 6.28 | 18 | >sp\|P01591\|IGJ_HUMAN Immunoglobulin J chain OS=Homo sapiens OX=9606 GN=JCHAIN PE=1 SV=4 | 1392 | 20201113_gy_milk_1ng_glyco_hcd203040_compound15.12696.12696.3 | controllerType=0 controllerNumber=1 scan=12696 | 27.514 |
| 11482:3 | 39 | R.IIVPLNNREN[+1694.603]ISDPTSPLR.T | HexNAc(3)Hex(4)Fuc(1)NeuAc(1) | N[+1695] | 1281.6 | 3 | 3842.7855 | 3842.7799 | 0 | 1.5 | 62 | Specific | 352.57 | 301.3 | 301.3 | 7.32 | 18 | >sp\|P01591\|IGJ_HUMAN Immunoglobulin J chain OS=Homo sapiens OX=9606 GN=JCHAIN PE=1 SV=4 | 1392 | 20201113_gy_milk_1ng_glyco_hcd203040_compound15.14618.14618.3 | controllerType=0 controllerNumber=1 scan=14618 | 30.7614 |
| 09758:3 | 39 | R.IIVPLNNREN[+1403.507]ISDPTSPLR.T | HexNAc(3)Hex(4)Fuc(1) | N[+1404] | 1184.5688 | 3 | 3551.6918 | 3551.6845 | 0 | 2.1 | 62 | Specific | 370.85 | 370.85 | 370.85 | 7.8 | 18 | >sp\|P01591\|IGJ_HUMAN Immunoglobulin J chain OS=Homo sapiens OX=9606 GN=JCHAIN PE=1 SV=4 | 1392 | 20201113_gy_milk_1ng_glyco_hcd203040_compound15.12831.12831.3 | controllerType=0 controllerNumber=1 scan=12831 | 27.736 |
| 13361:3 | 39 | R.IIVPLNNREN[+2204.772]ISDPTSPLR.T | HexNAc(4)Hex(5)NeuAc(2) | N[+2205] | 1451.6539 | 3 | 4352.947 | 4352.9496 | 0 | -0.6 | 62 | Specific | 380.14 | 374.49 | 374.49 | 8.93 | 18 | >sp\|P01591\|IGJ_HUMAN Immunoglobulin J chain OS=Homo sapiens OX=9606 GN=JCHAIN PE=1 SV=4 | 1392 | 20201113_gy_milk_1ng_glyco_hcd203040_compound15.16579.16579.3 | controllerType=0 controllerNumber=1 scan=16579 | 34.0501 |
| 11480:3 | 39 | R.IIVPLNNREN[+1694.603]ISDPTSPLR.T | HexNAc(3)Hex(4)Fuc(1)NeuAc(1) | N[+1695] | 1281.6 | 3 | 3842.7855 | 3842.7799 | 0 | 1.5 | 62 | Specific | 408.15 | 408.15 | 408.15 | 9.27 | 18 | >sp\|P01591\|IGJ_HUMAN Immunoglobulin J chain OS=Homo sapiens OX=9606 GN=JCHAIN PE=1 SV=4 | 1392 | 20201113_gy_milk_1ng_glyco_hcd203040_compound15.14616.14616.3 | controllerType=0 controllerNumber=1 scan=14616 | 30.7575 |
| 09629:3 | 39 | R.IIVPLNNREN[+1606.587]ISDPTSPLR.T | HexNAc(4)Hex(4)Fuc(1) | N[+1607] | 1252.2577 | 3 | 3754.7587 | 3754.7638 | 0 | -1.4 | 62 | Specific | 410.02 | 321.41 | 321.41 | 6.99 | 18 | >sp\|P01591\|IGJ_HUMAN Immunoglobulin J chain OS=Homo sapiens OX=9606 GN=JCHAIN PE=1 SV=4 | 1392 | 20201113_gy_milk_1ng_glyco_hcd203040_compound15.12698.12698.3 | controllerType=0 controllerNumber=1 scan=12698 | 27.5178 |
| 11383:3 | 39 | R.IIVPLNNREN[+1856.656]ISDPTSPLR.T | HexNAc(3)Hex(5)Fuc(1)NeuAc(1) | N[+1857] | 1335.6174 | 3 | 4004.8375 | 4004.8327 | 0 | 1.2 | 62 | Specific | 496.8 | 496.8 | 496.8 | 9.63 | 18 | >sp\|P01591\|IGJ_HUMAN Immunoglobulin J chain OS=Homo sapiens OX=9606 GN=JCHAIN PE=1 SV=4 | 1392 | 20201113_gy_milk_1ng_glyco_hcd203040_compound15.14515.14515.3 | controllerType=0 controllerNumber=1 scan=14515 | 30.5701 |
| 11386:3 | 39 | R.IIVPLNNREN[+1856.656]ISDPTSPLR.T | HexNAc(3)Hex(5)Fuc(1)NeuAc(1) | N[+1857] | 1335.6174 | 3 | 4004.8375 | 4004.8327 | 0 | 1.2 | 62 | Specific | 510.45 | 510.45 | 510.45 | 9.63 | 18 | >sp\|P01591\|IGJ_HUMAN Immunoglobulin J chain OS=Homo sapiens OX=9606 GN=JCHAIN PE=1 SV=4 | 1392 | 20201113_gy_milk_1ng_glyco_hcd203040_compound15.14518.14518.3 | controllerType=0 controllerNumber=1 scan=14518 | 30.5752 |
| 09612:3 | 39 | R.IIVPLNNREN[+1460.529]ISDPTSPLR.T | HexNAc(4)Hex(4) | N[+1461] | 1203.5818 | 3 | 3608.731 | 3608.7059 | 0 | 6.9 | 62 | Specific | 526.67 | 506.34 | 506.34 | 9.01 | 18 | >sp\|P01591\|IGJ_HUMAN Immunoglobulin J chain OS=Homo sapiens OX=9606 GN=JCHAIN PE=1 SV=4 | 1392 | 20201113_gy_milk_1ng_glyco_hcd203040_compound15.12680.12680.2 | controllerType=0 controllerNumber=1 scan=12680 | 27.4859 |

Table S4. Cont.

| Query #:z | Protein Rank | Glycopeptide | Glycan | Modification Type(s) | Observed m/z | z | Observed (M+H) | Calc. mass (M+H) | Off-by-x error | Mass error (ppm) | StartPosition | Cleavage | Score | Delta | Delta Mod | \|Log Prob\| | # of unique peptides | Protein Name | Protein DB number | Comment | Scan # | Scan Time |
|---|---|---|---|---|---|---|---|---|---|---|---|---|---|---|---|---|---|---|---|---|---|---|
| 16614:3 | 42 | K.AALAAFNAQNN[+2204.772]GSNFQLEEISR.A | HexNAc(4)Hex(5)NeuAc(2) | N[+2205] | 1523.9828 | 3 | 4569.934 | 4569.9256 | 0 | 1.8 | 166 | Specific | 630.23 | 575.62 | 575.62 | 12.87 | 10 | >sp\|P02765\|FETUA_HUMAN Alpha-2-HS-glycoprotein OS=Homo sapiens OX=9606 GN=AHSG PE=1 SV=2 | 135 | 20201113_gy_milk_1ng_glyco_hcd203040_compound15.20008.20008.3 | controllerType=0 controllerNumber=1 scan=20008 | 40.252 |
| 15169:3 | 49 | K.RNYIVPILWLN[+1864.634]ETGTIGDEK.A | HexNAc(2)Hex(9) | N[+1865] | 1399.2975 | 3 | 4195.8781 | 4195.8685 | 0 | 2.3 | 407 | Specific | 451.28 | 451.28 | 451.28 | 9.9 | 14 | >sp\|P16671\|CD36_HUMAN Platelet glycoprotein 4 OS=Homo sapiens OX=9606 GN=CD36 PE=1 SV=2 | 812 | 20201113_gy_milk_1ng_glyco_hcd203040_compound15.18460.18460.3 | controllerType=0 controllerNumber=1 scan=18460 | 37.3976 |
| 15167:3 | 49 | K.RNYIVPILWLN[+1864.634]ETGTIGDEK.A | HexNAc(2)Hex(9) | N[+1865] | 1399.2975 | 3 | 4195.8781 | 4195.8685 | 0 | 2.3 | 407 | Specific | 636.23 | 636.23 | 636.23 | 12.6 | 14 | >sp\|P16671\|CD36_HUMAN Platelet glycoprotein 4 OS=Homo sapiens OX=9606 GN=CD36 PE=1 SV=2 | 812 | 20201113_gy_milk_1ng_glyco_hcd203040_compound15.18458.18458.3 | controllerType=0 controllerNumber=1 scan=18458 | 37.3937 |
| 22092:4 | 54 | K.VFPLSLDSTPQDGNVVVAC[+57.021]LVQGFFPQEPLSVTWSESGQN[+1809.666]VTAI | HexNAc(5)Hex(4)Fuc(1) | C[+57], N[+1810] | 1647.2749 | 4 | 6586.0778 | 6586.0275 | 0 | 7.6 | 123 | Specific | 469.48 | 469.48 | 469.48 | 10.07 | 15 | >sp\|P0DOX2\|IGA2_HUMAN Immunoglobulin alpha-2 heavy chain OS=Homo sapiens OX=9606 PE=1 SV=2 | 631 | 20201113_gy_milk_1ng_glyco_hcd203040_compound15.25749.25749.4 | controllerType=0 controllerNumber=1 scan=25749 | 51.8821 |
| 03050:2 | 54 | K.TPLTAN[+1501.555]ITK.S | HexNAc(5)Hex(3) | N[+1502] | 1230.5627 | 2 | 2460.1181 | 2460.1121 | 0 | 2.4 | 315 | Specific | 285.22 | 103.85 | 103.85 | 4.87 | 15 | >sp\|P0DOX2\|IGA2_HUMAN Immunoglobulin alpha-2 heavy chain OS=Homo sapiens OX=9606 PE=1 SV=2 | 631 | 20201113_gy_milk_1ng_glyco_hcd203040_compound15.5858.5858.2 | controllerType=0 controllerNumber=1 scan=5858 | 16.0916 |
| 02992:3 | 54 | K.TPLTAN[+1809.666]ITK.S | HexNAc(5)Hex(4)Fuc(1) | N[+1810] | 923.414 | 3 | 2768.2274 | 2768.2228 | 0 | 1.6 | 315 | Specific | 315.42 | 187.66 | 187.66 | 6.19 | 15 | >sp\|P0DOX2\|IGA2_HUMAN Immunoglobulin alpha-2 heavy chain OS=Homo sapiens OX=9606 PE=1 SV=2 | 631 | 20201113_gy_milk_1ng_glyco_hcd203040_compound15.5798.5798.3 | controllerType=0 controllerNumber=1 scan=5798 | 15.986 |
| 03057:3 | 54 | K.TPLTAN[+1647.613]ITK.S | HexNAc(5)Hex(3)Fuc(1) | N[+1648] | 869.3974 | 3 | 2606.1775 | 2606.17 | 0 | 2.9 | 315 | Specific | 324.18 | 52.97 | 52.97 | 4.61 | 15 | >sp\|P0DOX2\|IGA2_HUMAN Immunoglobulin alpha-2 heavy chain OS=Homo sapiens OX=9606 PE=1 SV=2 | 631 | 20201113_gy_milk_1ng_glyco_hcd203040_compound15.5866.5866.3 | controllerType=0 controllerNumber=1 scan=5866 | 16.1079 |
| 02974:2 | 54 | K.TPLTAN[+1444.534]ITK.S | HexNAc(4)Hex(3)Fuc(1) | N[+1445] | 1202.0512 | 2 | 2403.0951 | 2403.0906 | 0 | 1.9 | 315 | Specific | 401.66 | 194.88 | 194.88 | 6.67 | 15 | >sp\|P0DOX2\|IGA2_HUMAN Immunoglobulin alpha-2 heavy chain OS=Homo sapiens OX=9606 PE=1 SV=2 | 631 | 20201113_gy_milk_1ng_glyco_hcd203040_compound15.5780.5780.2 | controllerType=0 controllerNumber=1 scan=5780 | 15.9544 |
| 02983:2 | 54 | K.TPLTAN[+1809.666]ITK.S | HexNAc(5)Hex(4)Fuc(1) | N[+1810] | 1384.6172 | 2 | 2768.2272 | 2768.2228 | 0 | 1.6 | 315 | Specific | 413.58 | 196.67 | 196.67 | 6.7 | 15 | >sp\|P0DOX2\|IGA2_HUMAN Immunoglobulin alpha-2 heavy chain OS=Homo sapiens OX=9606 PE=1 SV=2 | 631 | 20201113_gy_milk_1ng_glyco_hcd203040_compound15.5789.5789.2 | controllerType=0 controllerNumber=1 scan=5789 | 15.969 |
| 02985:2 | 54 | K.TPLTAN[+1809.666]ITK.S | HexNAc(5)Hex(4)Fuc(1) | N[+1810] | 1384.6172 | 2 | 2768.2272 | 2768.2228 | 0 | 1.6 | 315 | Specific | 495.2 | 299.15 | 299.15 | 7.87 | 15 | >sp\|P0DOX2\|IGA2_HUMAN Immunoglobulin alpha-2 heavy chain OS=Homo sapiens OX=9606 PE=1 SV=2 | 631 | 20201113_gy_milk_1ng_glyco_hcd203040_compound15.5791.5791.2 | controllerType=0 controllerNumber=1 scan=5791 | 15.9725 |
| 03059:3 | 54 | K.TPLTAN[+1647.613]ITK.S | HexNAc(5)Hex(3)Fuc(1) | N[+1648] | 869.3974 | 3 | 2606.1775 | 2606.17 | 0 | 2.9 | 315 | Specific | 525.09 | 227.17 | 227.17 | 7.57 | 15 | >sp\|P0DOX2\|IGA2_HUMAN Immunoglobulin alpha-2 heavy chain OS=Homo sapiens OX=9606 PE=1 SV=2 | 631 | 20201113_gy_milk_1ng_glyco_hcd203040_compound15.5868.5868.3 | controllerType=0 controllerNumber=1 scan=5868 | 16.1119 |
| 02976:2 | 54 | K.TPLTAN[+1444.534]ITK.S | HexNAc(4)Hex(3)Fuc(1) | N[+1445] | 1202.0512 | 2 | 2403.0951 | 2403.0906 | 0 | 1.9 | 315 | Specific | 584.8 | 320.26 | 320.26 | 9.41 | 15 | >sp\|P0DOX2\|IGA2_HUMAN Immunoglobulin alpha-2 heavy chain OS=Homo sapiens OX=9606 PE=1 SV=2 | 631 | 20201113_gy_milk_1ng_glyco_hcd203040_compound15.5782.5782.2 | controllerType=0 controllerNumber=1 scan=5782 | 15.9581 |
| 03039:2 | 54 | K.TPLTAN[+1647.613]ITK.S | HexNAc(5)Hex(3)Fuc(1) | N[+1648] | 1303.5912 | 2 | 2606.1752 | 2606.17 | 0 | 2 | 315 | Specific | 591.47 | 308.72 | 308.72 | 9.35 | 15 | >sp\|P0DOX2\|IGA2_HUMAN Immunoglobulin alpha-2 heavy chain OS=Homo sapiens OX=9606 PE=1 SV=2 | 631 | 20201113_gy_milk_1ng_glyco_hcd203040_compound15.5847.5847.2 | controllerType=0 controllerNumber=1 scan=5847 | 16.0712 |
| 03041:2 | 54 | K.TPLTAN[+1647.613]ITK.S | HexNAc(5)Hex(3)Fuc(1) | N[+1648] | 1303.5912 | 2 | 2606.1752 | 2606.17 | 0 | 2 | 315 | Specific | 607.06 | 346.86 | 346.86 | 10.5 | 15 | >sp\|P0DOX2\|IGA2_HUMAN Immunoglobulin alpha-2 heavy chain OS=Homo sapiens OX=9606 PE=1 SV=2 | 631 | 20201113_gy_milk_1ng_glyco_hcd203040_compound15.5849.5849.2 | controllerType=0 controllerNumber=1 scan=5849 | 16.0767 |
| 14388:3 | 61 | K.VVLHPN[+2204.772]YSQVDIGLIK.L | HexNAc(4)Hex(5)NeuAc(2) | N[+2205] | 1333.9367 | 3 | 3999.7956 | 3999.7837 | 0 | 3 | 236 | Specific | 387.86 | 385.34 | 385.34 | 9.4 | 13 | >sp\|P00738\|HPT_HUMAN Haptoglobin OS=Homo sapiens OX=9606 GN=HP PE=1 SV=1 | 407 | 20201113_gy_milk_1ng_glyco_hcd203040_compound15.17647.17647.3 | controllerType=0 controllerNumber=1 scan=17647 | 35.9436 |
| 14386:3 | 61 | K.VVLHPN[+2204.772]YSQVDIGLIK.L | HexNAc(4)Hex(5)NeuAc(2) | N[+2205] | 1333.9367 | 3 | 3999.7956 | 3999.7837 | 0 | 3 | 236 | Specific | 524.52 | 524.52 | 524.52 | 9.91 | 13 | >sp\|P00738\|HPT_HUMAN Haptoglobin OS=Homo sapiens OX=9606 GN=HP PE=1 SV=1 | 407 | 20201113_gy_milk_1ng_glyco_hcd203040_compound15.17645.17645.3 | controllerType=0 controllerNumber=1 scan=17645 | 35.9397 |
| 06571:3 | 398 | R.[+42.011]VEDAYILTC[+57.021]N[+730.264]VSLEYEK.T | HexNAc(2)Hex(2) | .[42], C[+57], N[+730] | 940.0949 | 3 | 2818.2703 | 2818.2486 | 0 | 7.7 | 225 | Specific | 238.62 | 143.49 | 143.49 | 3.73 | 4 | >sp\|P40227\|TCPZ_HUMAN T-complex protein 1 subunit zeta OS=Homo sapiens OX=9606 GN=CCT6A PE=1 SV=3 | 1115 | 20201113_gy_milk_1ng_glyco_hcd203040_compound15.9512.9512.3 | controllerType=0 controllerNumber=1 scan=9512 | 22.1688 |
| 06573:3 | 398 | R.[+42.011]VEDAYILTC[+57.021]N[+730.264]VSLEYEK.T | HexNAc(2)Hex(2) | .[42], C[+57], N[+730] | 940.0949 | 3 | 2818.2703 | 2818.2486 | 0 | 7.7 | 225 | Specific | 283.56 | 203.87 | 203.87 | 4.55 | 4 | >sp\|P40227\|TCPZ_HUMAN T-complex protein 1 subunit zeta OS=Homo sapiens OX=9606 GN=CCT6A PE=1 SV=3 | 1115 | 20201113_gy_milk_1ng_glyco_hcd203040_compound15.9514.9514.3 | controllerType=0 controllerNumber=1 scan=9514 | 22.1729 |
| 17484:4 | 1 | R.RTAGWNVPIGTLRPFLN[+1930.692]WTGPPEPIEAAVAR.F | HexNAc(4)Hex(6)Fuc(1) | N[+1931] | 1330.1226 | 4 | 5317.4684 | 5317.5044 | 0 | -6.8 | 140 | Specific | 307.33 | 307.33 | 307.33 | 5.93 | 122 | >sp\|P02788\|TRFL_HUMAN Lactotransferrin OS=Homo sapiens OX=9606 GN=LTF PE=1 SV=6 | 1270 | 20201114_qwl_colo_milk_1ng_glyco_hcd203040_compound15.20952.20952.2 | controllerType=0 controllerNumber=1 scan=20952 | 46.3945 |
| 15346:4 | 1 | R.RTAGWNVPIGTLRPFLN[+2278.809]WTGPPEPIEAAVAR.F | HexNAc(5)Hex(6)NeuAc(1) | N[+2279] | 1417.1694 | 4 | 5665.6559 | 5665.6213 | 0 | 6.1 | 140 | Specific | 341.35 | 320.02 | 320.02 | 7.48 | 122 | >sp\|P02788\|TRFL_HUMAN Lactotransferrin OS=Homo sapiens OX=9606 GN=LTF PE=1 SV=6 | 1270 | 20201114_qwl_colo_milk_1ng_glyco_hcd203040_compound15.18703.18703.4 | controllerType=0 controllerNumber=1 scan=18703 | 41.1404 |
| 17481:4 | 1 | R.RTAGWNVPIGTLRPFLN[+1930.692]WTGPPEPIEAAVAR.F | HexNAc(4)Hex(6)Fuc(1) | N[+1931] | 1330.1226 | 4 | 5317.4684 | 5317.5044 | 0 | -6.8 | 140 | Specific | 342.42 | 342.42 | 342.42 | 6.51 | 122 | >sp\|P02788\|TRFL_HUMAN Lactotransferrin OS=Homo sapiens OX=9606 GN=LTF PE=1 SV=6 | 1270 | 20201114_qwl_colo_milk_1ng_glyco_hcd203040_compound15.20949.20949.2 | controllerType=0 controllerNumber=1 scan=20949 | 46.3877 |
| 15348:4 | 1 | R.RTAGWNVPIGTLRPFLN[+2278.809]WTGPPEPIEAAVAR.F | HexNAc(5)Hex(6)NeuAc(1) | N[+2279] | 1417.1694 | 4 | 5665.6559 | 5665.6213 | 0 | 6.1 | 140 | Specific | 433.64 | 422.56 | 422.56 | 9.41 | 122 | >sp\|P02788\|TRFL_HUMAN Lactotransferrin OS=Homo sapiens OX=9606 GN=LTF PE=1 SV=6 | 1270 | 20201114_qwl_colo_milk_1ng_glyco_hcd203040_compound15.18705.18705.4 | controllerType=0 controllerNumber=1 scan=18705 | 41.147 |
| 16249:4 | 1 | R.RTAGWNVPIGTLRPFLN[+2205.793]WTGPPEPIEAAVAR.F | HexNAc(4)Hex(5)Fuc(2)NeuAc(1) | N[+2206] | 1398.9104 | 4 | 5592.6198 | 5592.6049 | 0 | 2.7 | 140 | Specific | 519.96 | 519.96 | 519.96 | 10.24 | 122 | >sp\|P02788\|TRFL_HUMAN Lactotransferrin OS=Homo sapiens OX=9606 GN=LTF PE=1 SV=6 | 1270 | 20201114_qwl_colo_milk_1ng_glyco_hcd203040_compound15.19654.19654.4 | controllerType=0 controllerNumber=1 scan=19654 | 43.4837 |
| 15419:4 | 1 | R.RTAGWNVPIGTLRPFLN[+1768.640]WTGPPEPIEAAVAR.F | HexNAc(4)Hex(5)Fuc(1) | N[+1769] | 1289.6261 | 4 | 5155.4826 | 5155.4516 | 0 | 6 | 140 | Specific | 606.39 | 513.28 | 513.28 | 12.34 | 122 | >sp\|P02788\|TRFL_HUMAN Lactotransferrin OS=Homo sapiens OX=9606 GN=LTF PE=1 SV=6 | 1270 | 20201114_qwl_colo_milk_1ng_glyco_hcd203040_compound15.18780.18780.4 | controllerType=0 controllerNumber=1 scan=18780 | 41.3516 |
| 15421:4 | 1 | R.RTAGWNVPIGTLRPFLN[+1768.640]WTGPPEPIEAAVAR.F | HexNAc(4)Hex(5)Fuc(1) | N[+1769] | 1289.6261 | 4 | 5155.4826 | 5155.4516 | 0 | 6 | 140 | Specific | 633.18 | 633.18 | 633.18 | 12.34 | 122 | >sp\|P02788\|TRFL_HUMAN Lactotransferrin OS=Homo sapiens OX=9606 GN=LTF PE=1 SV=6 | 1270 | 20201114_qwl_colo_milk_1ng_glyco_hcd203040_compound15.18782.18782.4 | controllerType=0 controllerNumber=1 scan=18782 | 41.3601 |
| 16252:4 | 1 | R.RTAGWNVPIGTLRPFLN[+2205.793]WTGPPEPIEAAVAR.F | HexNAc(4)Hex(5)Fuc(2)NeuAc(1) | N[+2206] | 1398.9104 | 4 | 5592.6198 | 5592.6049 | 0 | 2.7 | 140 | Specific | 659.31 | 532.79 | 532.79 | 13.86 | 122 | >sp\|P02788\|TRFL_HUMAN Lactotransferrin OS=Homo sapiens OX=9606 GN=LTF PE=1 SV=6 | 1270 | 20201114_qwl_colo_milk_1ng_glyco_hcd203040_compound15.19657.19657.4 | controllerType=0 controllerNumber=1 scan=19657 | 43.4936 |
| 17430:3 | 1 | R.TAGWNVPIGTLRPFLN[+2205.793]WTGPPEPIEAAVAR.F | HexNAc(4)Hex(5)Fuc(2)NeuAc(1) | N[+2206] | 1812.8407 | 3 | 5436.5074 | 5436.5038 | 0 | 0.7 | 141 | Specific | 200.23 | 200.23 | 200.23 | 5.15 | 122 | >sp\|P02788\|TRFL_HUMAN Lactotransferrin OS=Homo sapiens OX=9606 GN=LTF PE=1 SV=6 | 1270 | 20201114_qwl_colo_milk_1ng_glyco_hcd203040_compound15.20896.20896.2 | controllerType=0 controllerNumber=1 scan=20896 | 46.2897 |
| 16503:3 | 1 | R.[+42.011]TAGWNVPIGTLRPFLN[+2018.708]WTGPPEPIEAAVAR.F | HexNAc(3)Hex(6)Fuc(1)NeuAc(1) | .[42], N[+2019] | 1764.4897 | 3 | 5291.4546 | 5291.4299 | 0 | 4.7 | 141 | Specific | 217.85 | 176.2 | 176.2 | 4.77 | 122 | >sp\|P02788\|TRFL_HUMAN Lactotransferrin OS=Homo sapiens OX=9606 GN=LTF PE=1 SV=6 | 1270 | 20201114_qwl_colo_milk_1ng_glyco_hcd203040_compound15.19920.19920.3 | controllerType=0 controllerNumber=1 scan=19920 | 44.0695 |
| 17419:4 | 1 | R.TAGWNVPIGTLRPFLN[+2351.851]WTGPPEPIEAAVAR.F | HexNAc(4)Hex(5)Fuc(3)NeuAc(1) | N[+2352] | 1396.3971 | 4 | 5582.5666 | 5582.5617 | 0 | 0.9 | 141 | Specific | 251.53 | 251.53 | 251.53 | 6.9 | 122 | >sp\|P02788\|TRFL_HUMAN Lactotransferrin OS=Homo sapiens OX=9606 GN=LTF PE=1 SV=6 | 1270 | 20201114_qwl_colo_milk_1ng_glyco_hcd203040_compound15.20885.20885.4 | controllerType=0 controllerNumber=1 scan=20885 | 46.2693 |
| 16463:3 | 1 | R.TAGWNVPIGTLRPFLN[+2425.887]WTGPPEPIEAAVAR.F | HexNAc(5)Hex(6)Fuc(3) | N[+2426] | 1886.2077 | 3 | 5656.6086 | 5656.5985 | 0 | 1.8 | 141 | Specific | 252.03 | 186.99 | 186.99 | 6.45 | 122 | >sp\|P02788\|TRFL_HUMAN Lactotransferrin OS=Homo sapiens OX=9606 GN=LTF PE=1 SV=6 | 1270 | 20201114_qwl_colo_milk_1ng_glyco_hcd203040_compound15.19878.19878.3 | controllerType=0 controllerNumber=1 scan=19878 | 43.9877 |
| 17426:3 | 1 | R.TAGWNVPIGTLRPFLN[+2351.851]WTGPPEPIEAAVAR.F | HexNAc(4)Hex(5)Fuc(3)NeuAc(1) | N[+2352] | 1861.5271 | 3 | 5582.5666 | 5582.5617 | 0 | 0.9 | 141 | Specific | 259.85 | 259.85 | 259.85 | 6.91 | 122 | >sp\|P02788\|TRFL_HUMAN Lactotransferrin OS=Homo sapiens OX=9606 GN=LTF PE=1 SV=6 | 1270 | 20201114_qwl_colo_milk_1ng_glyco_hcd203040_compound15.20892.20892.3 | controllerType=0 controllerNumber=1 scan=20892 | 46.2814 |
| 16523:3 | 1 | R.TAGWNVPIGTLRPFLN[+2279.830]WTGPPEPIEAAVAR.F | HexNAc(5)Hex(6)Fuc(2) | N[+2280] | 1837.5272 | 3 | 5510.567 | 5510.5406 | 0 | 4.8 | 141 | Specific | 269.05 | 253.9 | 253.9 | 6.85 | 122 | >sp\|P02788\|TRFL_HUMAN Lactotransferrin OS=Homo sapiens OX=9606 GN=LTF PE=1 SV=6 | 1270 | 20201114_qwl_colo_milk_1ng_glyco_hcd203040_compound15.19940.19940.3 | controllerType=0 controllerNumber=1 scan=19940 | 44.1287 |
| 16465:4 | 1 | R.TAGWNVPIGTLRPFLN[+2425.887]WTGPPEPIEAAVAR.F | HexNAc(5)Hex(6)Fuc(3) | N[+2426] | 1414.9094 | 4 | 5656.6159 | 5656.5985 | 0 | 3.1 | 141 | Specific | 274.04 | 211.62 | 211.62 | 6.9 | 122 | >sp\|P02788\|TRFL_HUMAN Lactotransferrin OS=Homo sapiens OX=9606 GN=LTF PE=1 SV=6 | 1270 | 20201114_qwl_colo_milk_1ng_glyco_hcd203040_compound15.19880.19880.4 | controllerType=0 controllerNumber=1 scan=19880 | 43.9908 |
| 17421:3 | 1 | R.TAGWNVPIGTLRPFLN[+2059.735]WTGPPEPIEAAVAR.F | HexNAc(4)Hex(5)Fuc(1)NeuAc(1) | N[+2060] | 1764.1554 | 3 | 5290.4515 | 5290.4459 | 0 | 1.1 | 141 | Specific | 289.07 | 289.07 | 289.07 | 7.2 | 122 | >sp\|P02788\|TRFL_HUMAN Lactotransferrin OS=Homo sapiens OX=9606 GN=LTF PE=1 SV=6 | 1270 | 20201114_qwl_colo_milk_1ng_glyco_hcd203040_compound15.20887.20887.3 | controllerType=0 controllerNumber=1 scan=20887 | 46.2762 |
| 17433:3 | 1 | R.TAGWNVPIGTLRPFLN[+2205.793]WTGPPEPIEAAVAR.F | HexNAc(4)Hex(5)Fuc(2)NeuAc(1) | N[+2206] | 1812.8407 | 3 | 5436.5074 | 5436.5038 | 0 | 0.7 | 141 | Specific | 289.14 | 289.14 | 289.14 | 7.2 | 122 | >sp\|P02788\|TRFL_HUMAN Lactotransferrin OS=Homo sapiens OX=9606 GN=LTF PE=1 SV=6 | 1270 | 20201114_qwl_colo_milk_1ng_glyco_hcd203040_compound15.20899.20899.2 | controllerType=0 controllerNumber=1 scan=20899 | 46.2961 |
| 18732:3 | 1 | R.[+42.011]TAGWNVPIGTLRPFLN[+2336.851]WTGPPEPIEAAVAR.F | HexNAc(6)Hex(6)Fuc(1) | .[42], N[+2337] | 1870.5354 | 3 | 5609.5915 | 5609.5726 | 0 | 3.4 | 141 | Specific | 311.35 | 311.35 | 311.35 | 6.77 | 122 | >sp\|P02788\|TRFL_HUMAN Lactotransferrin OS=Homo sapiens OX=9606 GN=LTF PE=1 SV=6 | 1270 | 20201114_qwl_colo_milk_1ng_glyco_hcd203040_compound15.22258.22258.3 | controllerType=0 controllerNumber=1 scan=22258 | 49.2297 |
| 16521:3 | 1 | R.TAGWNVPIGTLRPFLN[+2279.830]WTGPPEPIEAAVAR.F | HexNAc(5)Hex(6)Fuc(2) | N[+2280] | 1837.5272 | 3 | 5510.567 | 5510.5406 | 0 | 4.8 | 141 | Specific | 315.13 | 232.23 | 232.23 | 6.92 | 122 | >sp\|P02788\|TRFL_HUMAN Lactotransferrin OS=Homo sapiens OX=9606 GN=LTF PE=1 SV=6 | 1270 | 20201114_qwl_colo_milk_1ng_glyco_hcd203040_compound15.19938.19938.3 | controllerType=0 controllerNumber=1 scan=19938 | 44.1143 |
| 16462:3 | 1 | R.[+42.011]TAGWNVPIGTLRPFLN[+1872.651]WTGPPEPIEAAVAR.F | HexNAc(3)Hex(6)NeuAc(1) | .[42], N[+1873] | 1715.8115 | 3 | 5145.4199 | 5145.372 | 0 | 9.3 | 141 | Specific | 317.73 | 206.29 | 206.29 | 6.57 | 122 | >sp\|P02788\|TRFL_HUMAN Lactotransferrin OS=Homo sapiens OX=9606 GN=LTF PE=1 SV=6 | 1270 | 20201114_qwl_colo_milk_1ng_glyco_hcd203040_compound15.19877.19877.3 | controllerType=0 controllerNumber=1 scan=19877 | 43.9864 |
| 18726:4 | 1 | R.[+42.011]TAGWNVPIGTLRPFLN[+2336.851]WTGPPEPIEAAVAR.F | HexNAc(6)Hex(6)Fuc(1) | .[42], N[+2337] | 1403.1519 | 4 | 5609.5856 | 5609.5726 | 0 | 2.3 | 141 | Specific | 321.53 | 321.53 | 321.53 | 7.79 | 122 | >sp\|P02788\|TRFL_HUMAN Lactotransferrin OS=Homo sapiens OX=9606 GN=LTF PE=1 SV=6 | 1270 | 20201114_qwl_colo_milk_1ng_glyco_hcd203040_compound15.22252.22252.4 | controllerType=0 controllerNumber=1 scan=22252 | 49.2172 |
| 18728:4 | 1 | R.[+42.011]TAGWNVPIGTLRPFLN[+2336.851]WTGPPEPIEAAVAR.F | HexNAc(6)Hex(6)Fuc(1) | .[42], N[+2337] | 1403.1519 | 4 | 5609.5856 | 5609.5726 | 0 | 2.3 | 141 | Specific | 322.62 | 322.62 | 322.62 | 7.79 | 122 | >sp\|P02788\|TRFL_HUMAN Lactotransferrin OS=Homo sapiens OX=9606 GN=LTF PE=1 SV=6 | 1270 | 20201114_qwl_colo_milk_1ng_glyco_hcd203040_compound15.22254.22254.4 | controllerType=0 controllerNumber=1 scan=22254 | 49.2225 |
| 17432:3 | 1 | R.[+42.011]TAGWNVPIGTLRPFLN[+2190.793]WTGPPEPIEAAVAR.F | HexNAc(6)Hex(6) | .[42], N[+2191] | 1821.8379 | 3 | 5463.499 | 5463.5147 | 0 | -2.9 | 141 | Specific | 334.63 | 334.63 | 334.63 | 6.85 | 122 | >sp\|P02788\|TRFL_HUMAN Lactotransferrin OS=Homo sapiens OX=9606 GN=LTF PE=1 SV=6 | 1270 | 20201114_qwl_colo_milk_1ng_glyco_hcd203040_compound15.20898.20898.3 | controllerType=0 controllerNumber=1 scan=20898 | 46.2952 |
| 16460:3 | 1 | R.[+42.011]TAGWNVPIGTLRPFLN[+1872.651]WTGPPEPIEAAVAR.F | HexNAc(3)Hex(6)NeuAc(1) | .[42], N[+1873] | 1715.8115 | 3 | 5145.4199 | 5145.372 | 0 | 9.3 | 141 | Specific | 351.14 | 244.42 | 244.42 | 6.76 | 122 | >sp\|P02788\|TRFL_HUMAN Lactotransferrin OS=Homo sapiens OX=9606 GN=LTF PE=1 SV=6 | 1270 | 20201114_qwl_colo_milk_1ng_glyco_hcd203040_compound15.19875.19875.3 | controllerType=0 controllerNumber=1 scan=19875 | 43.971 |
| 17429:3 | 1 | R.[+42.011]TAGWNVPIGTLRPFLN[+2190.793]WTGPPEPIEAAVAR.F | HexNAc(6)Hex(6) | .[42], N[+2191] | 1821.8379 | 3 | 5463.499 | 5463.5147 | 0 | -2.9 | 141 | Specific | 362.86 | 351.52 | 351.52 | 6.8 | 122 | >sp\|P02788\|TRFL_HUMAN Lactotransferrin OS=Homo sapiens OX=9606 GN=LTF PE=1 SV=6 | 1270 | 20201114_qwl_colo_milk_1ng_glyco_hcd203040_compound15.20895.20895.3 | controllerType=0 controllerNumber=1 scan=20895 | 46.2885 |
| 18753:4 | 1 | R.TAGWNVPIGTLRPFLN[+2496.888]WTGPPEPIEAAVAR.F | HexNAc(4)Hex(5)Fuc(2)NeuAc(2) | N[+2497] | 1432.6628 | 4 | 5727.6296 | 5727.5992 | 0 | 5.3 | 141 | Specific | 364.12 | 329.55 | 329.55 | 7.61 | 122 | >sp\|P02788\|TRFL_HUMAN Lactotransferrin OS=Homo sapiens OX=9606 GN=LTF PE=1 SV=6 | 1270 | 20201114_qwl_colo_milk_1ng_glyco_hcd203040_compound15.22280.22280.4 | controllerType=0 controllerNumber=1 scan=22280 | 49.2758 |
| 16461:3 | 1 | R.TAGWNVPIGTLRPFLN[+2425.887]WTGPPEPIEAAVAR.F | HexNAc(5)Hex(6)Fuc(3) | N[+2426] | 1886.2077 | 3 | 5656.6086 | 5656.5985 | 0 | 1.8 | 141 | Specific | 405.11 | 209.03 | 209.03 | 7.29 | 122 | >sp\|P02788\|TRFL_HUMAN Lactotransferrin OS=Homo sapiens OX=9606 GN=LTF PE=1 SV=6 | 1270 | 20201114_qwl_colo_milk_1ng_glyco_hcd203040_compound15.19876.19876.3 | controllerType=0 controllerNumber=1 scan=19876 | 43.9726 |
| 17583:3 | 1 | R.TAGWNVPIGTLRPFLN[+2059.735]WTGPPEPIEAAVAR.F | HexNAc(4)Hex(5)Fuc(1)NeuAc(1) | N[+2060] | 1764.1589 | 3 | 5290.4621 | 5290.4459 | 0 | 3.1 | 141 | Specific | 440.77 | 436.73 | 436.73 | 9.93 | 122 | >sp\|P02788\|TRFL_HUMAN Lactotransferrin OS=Homo sapiens OX=9606 GN=LTF PE=1 SV=6 | 1270 | 20201114_qwl_colo_milk_1ng_glyco_hcd203040_compound15.21057.21057.3 | controllerType=0 controllerNumber=1 scan=21057 | 46.6664 |
| 16468:4 | 1 | R.TAGWNVPIGTLRPFLN[+2425.887]WTGPPEPIEAAVAR.F | HexNAc(5)Hex(6)Fuc(3) | N[+2426] | 1414.9094 | 4 | 5656.6159 | 5656.5985 | 0 | 3.1 | 141 | Specific | 499.92 | 341.43 | 341.43 | 8.52 | 122 | >sp\|P02788\|TRFL_HUMAN Lactotransferrin OS=Homo sapiens OX=9606 GN=LTF PE=1 SV=6 | 1270 | 20201114_qwl_colo_milk_1ng_glyco_hcd203040_compound15.19883.19883.4 | controllerType=0 controllerNumber=1 scan=19883 | 43.9948 |
| 17581:3 | 1 | R.TAGWNVPIGTLRPFLN[+2059.735]WTGPPEPIEAAVAR.F | HexNAc(4)Hex(5)Fuc(1)NeuAc(1) | N[+2060] | 1764.1589 | 3 | 5290.4621 | 5290.4459 | 0 | 3.1 | 141 | Specific | 529.16 | 477.61 | 477.61 | 10.24 | 122 | >sp\|P02788\|TRFL_HUMAN Lactotransferrin OS=Homo sapiens OX=9606 GN=LTF PE=1 SV=6 | 1270 | 20201114_qwl_colo_milk_1ng_glyco_hcd203040_compound15.21055.21055.3 | controllerType=0 controllerNumber=1 scan=21055 | 46.6624 |
| 16434:4 | 1 | R.TAGWNVPIGTLRPFLN[+2571.946]WTGPPEPIEAAVAR.F | HexNAc(5)Hex(6)Fuc(4) | N[+2572] | 1451.4338 | 4 | 5802.7135 | 5802.6564 | 0 | 9.9 | 141 | Specific | 546.63 | 500.44 | 500.44 | 9.85 | 122 | >sp\|P02788\|TRFL_HUMAN Lactotransferrin OS=Homo sapiens OX=9606 GN=LTF PE=1 SV=6 | 1270 | 20201114_qwl_colo_milk_1ng_glyco_hcd203040_compound15.19848.19848.4 | controllerType=0 controllerNumber=1 scan=19848 | 43.9258 |
| 16471:4 | 1 | R.TAGWNVPIGTLRPFLN[+1913.677]WTGPPEPIEAAVAR.F | HexNAc(4)Hex(5)NeuAc(1) | N[+1914] | 1286.8604 | 4 | 5144.4196 | 5144.388 | 0 | 6.1 | 141 | Specific | 560.53 | 560.53 | 560.53 | 10.01 | 122 | >sp\|P02788\|TRFL_HUMAN Lactotransferrin OS=Homo sapiens OX=9606 GN=LTF PE=1 SV=6 | 1270 | 20201114_qwl_colo_milk_1ng_glyco_hcd203040_compound15.19886.19886.4 | controllerType=0 controllerNumber=1 scan=19886 | 44.0044 |
| 16474:4 | 1 | R.TAGWNVPIGTLRPFLN[+1913.677]WTGPPEPIEAAVAR.F | HexNAc(4)Hex(5)NeuAc(1) | N[+1914] | 1286.8604 | 4 | 5144.4196 | 5144.388 | 0 | 6.1 | 141 | Specific | 659.2 | 639.83 | 639.83 | 13.33 | 122 | >sp\|P02788\|TRFL_HUMAN Lactotransferrin OS=Homo sapiens OX=9606 GN=LTF PE=1 SV=6 | 1270 | 20201114_qwl_colo_milk_1ng_glyco_hcd203040_compound15.19889.19889.4 | controllerType=0 controllerNumber=1 scan=19889 | 44.0123 |
| 18755:4 | 1 | R.TAGWNVPIGTLRPFLN[+2496.888]WTGPPEPIEAAVAR.F | HexNAc(4)Hex(5)Fuc(2)NeuAc(2) | N[+2497] | 1432.6628 | 4 | 5727.6296 | 5727.5992 | 0 | 5.3 | 141 | Specific | 685.8 | 569.72 | 569.72 | 13.51 | 122 | >sp\|P02788\|TRFL_HUMAN Lactotransferrin OS=Homo sapiens OX=9606 GN=LTF PE=1 SV=6 | 1270 | 20201114_qwl_colo_milk_1ng_glyco_hcd203040_compound15.22282.22282.4 | controllerType=0 controllerNumber=1 scan=22282 | 49.2804 |
| 17417:4 | 1 | R.TAGWNVPIGTLRPFLN[+2059.735]WTGPPEPIEAAVAR.F | HexNAc(4)Hex(5)Fuc(1)NeuAc(1) | N[+2060] | 1323.368 | 4 | 5290.4504 | 5290.4459 | 0 | 0.8 | 141 | Specific | 696.52 | 661.11 | 661.11 | 13.69 | 122 | >sp\|P02788\|TRFL_HUMAN Lactotransferrin OS=Homo sapiens OX=9606 GN=LTF PE=1 SV=6 | 1270 | 20201114_qwl_colo_milk_1ng_glyco_hcd203040_compound15.20883.20883.4 | controllerType=0 controllerNumber=1 scan=20883 | 46.267 |
| 17418:3 | 1 | R.TAGWNVPIGTLRPFLN[+2059.735]WTGPPEPIEAAVAR.F | HexNAc(4)Hex(5)Fuc(1)NeuAc(1) | N[+2060] | 1764.1554 | 3 | 5290.4515 | 5290.4459 | 0 | 1.1 | 141 | Specific | 722.42 | 618.06 | 618.06 | 14.7 | 122 | >sp\|P02788\|TRFL_HUMAN Lactotransferrin OS=Homo sapiens OX=9606 GN=LTF PE=1 SV=6 | 1270 | 20201114_qwl_colo_milk_1ng_glyco_hcd203040_compound15.20884.20884.3 | controllerType=0 controllerNumber=1 scan=20884 | 46.2685 |
| 17422:4 | 1 | R.TAGWNVPIGTLRPFLN[+2351.851]WTGPPEPIEAAVAR.F | HexNAc(4)Hex(5)Fuc(3)NeuAc(1) | N[+2352] | 1396.3971 | 4 | 5582.5666 | 5582.5617 | 0 | 0.9 | 141 | Specific | 804.1 | 666.61 | 666.61 | 16.69 | 122 | >sp\|P02788\|TRFL_HUMAN Lactotransferrin OS=Homo sapiens OX=9606 GN=LTF PE=1 SV=6 | 1270 | 20201114_qwl_colo_milk_1ng_glyco_hcd203040_compound15.20888.20888.4 | controllerType=0 controllerNumber=1 scan=20888 | 46.2769 |
| 17420:4 | 1 | R.TAGWNVPIGTLRPFLN[+2059.735]WTGPPEPIEAAVAR.F | HexNAc(4)Hex(5)Fuc(1)NeuAc(1) | N[+2060] | 1323.368 | 4 | 5290.4504 | 5290.4459 | 0 | 0.8 | 141 | Specific | 832.18 | 709.4 | 709.4 | 16.69 | 122 | >sp\|P02788\|TRFL_HUMAN Lactotransferrin OS=Homo sapiens OX=9606 GN=LTF PE=1 SV=6 | 1270 | 20201114_qwl_colo_milk_1ng_glyco_hcd203040_compound15.20886.20886.4 | controllerType=0 controllerNumber=1 scan=20886 | 46.2727 |
| 14891:3 | 1 | R.TAGWNIPMGLLFN[+2424.867]QTGSC[+57.021]K.F | HexNAc(5)Hex(6)Fuc(1)NeuAc(1) | C[+57], N[+2425] | 1507.3107 | 3 | 4519.9176 | 4519.8771 | 0 | 9 | 485 | Specific | 271.24 | 271.24 | 262.97 | 5.47 | 122 | >sp\|P02788\|TRFL_HUMAN Lactotransferrin OS=Homo sapiens OX=9606 GN=LTF PE=1 SV=6 | 1270 | 20201114_qwl_colo_milk_1ng_glyco_hcd203040_compound15.18229.18229.3 | controllerType=0 controllerNumber=1 scan=18229 | 40.1804 |
| 19045:3 | 1 | R.TAGWNIPMGLLFN[+2204.772]QTGSC[+57.021]K.F | HexNAc(4)Hex(5)NeuAc(2) | C[+57], N[+2205] | 1433.9437 | 3 | 4299.8164 | 4299.7824 | 0 | 7.9 | 485 | Specific | 325.26 | 287.88 | 287.88 | 5.18 | 122 | >sp\|P02788\|TRFL_HUMAN Lactotransferrin OS=Homo sapiens OX=9606 GN=LTF PE=1 SV=6 | 1270 | 20201114_qwl_colo_milk_1ng_glyco_hcd203040_compound15.22584.22584.3 | controllerType=0 controllerNumber=1 scan=22584 | 49.9171 |
| 17046:3 | 1 | R.TAGWNIPMGLLFN[+1694.603]QTGSC[+57.021]K.F | HexNAc(3)Hex(4)Fuc(1)NeuAc(1) | C[+57], N[+1695] | 1263.8716 | 3 | 3789.6004 | 3789.6127 | 0 | -3.2 | 485 | Specific | 387.27 | 235.65 | 235.65 | 5.61 | 122 | >sp\|P02788\|TRFL_HUMAN Lactotransferrin OS=Homo sapiens OX=9606 GN=LTF PE=1 SV=6 | 1270 | 20201114_qwl_colo_milk_1ng_glyco_hcd203040_compound15.20489.20489.3 | controllerType=0 controllerNumber=1 scan=20489 | 45.3711 |
| 15557:3 | 1 | R.TAGWNIPMGLLFN[+1216.423]QTGSC[+57.021]K.F | HexNAc(2)Hex(5) | C[+57], N[+1216] | 1104.4868 | 3 | 3311.446 | 3311.4328 | 0 | 4 | 485 | Specific | 455.35 | 445.37 | 160.14 | 9.65 | 122 | >sp\|P02788\|TRFL_HUMAN Lactotransferrin OS=Homo sapiens OX=9606 GN=LTF PE=1 SV=6 | 1270 | 20201114_qwl_colo_milk_1ng_glyco_hcd203040_compound15.18925.18925.3 | controllerType=0 controllerNumber=1 scan=18925 | 41.6596 |
| 17164:3 | 1 | R.TAGWNIPMGLLFN[+1548.545]QTGSC[+57.021]K.F | HexNAc(3)Hex(4)NeuAc(1) | C[+57], N[+1549] | 1215.1899 | 3 | 3643.5551 | 3643.5548 | 0 | 0.1 | 485 | Specific | 461 | 447.95 | 447.95 | 9.53 | 122 | >sp\|P02788\|TRFL_HUMAN Lactotransferrin OS=Homo sapiens OX=9606 GN=LTF PE=1 SV=6 | 1270 | 20201114_qwl_colo_milk_1ng_glyco_hcd203040_compound15.20612.20612.3 | controllerType=0 controllerNumber=1 scan=20612 | 45.614 |
| 17163:3 | 1 | R.TAGWNIPMGLLFN[+1548.545]QTGSC[+57.021]K.F | HexNAc(3)Hex(4)NeuAc(1) | C[+57], N[+1549] | 1215.1899 | 3 | 3643.5551 | 3643.5548 | 0 | 0.1 | 485 | Specific | 466.03 | 435.88 | 435.88 | 9.53 | 122 | >sp\|P02788\|TRFL_HUMAN Lactotransferrin OS=Homo sapiens OX=9606 GN=LTF PE=1 SV=6 | 1270 | 20201114_qwl_colo_milk_1ng_glyco_hcd203040_compound15.20611.20611.3 | controllerType=0 controllerNumber=1 scan=20611 | 45.6104 |
| 15358:3 | 1 | R.TAGWNIPMGLLFN[+1622.582]QTGSC[+57.021]K.F | HexNAc(4)Hex(5) | C[+57], N[+1623] | 1239.8732 | 3 | 3717.6051 | 3717.5916 | 0 | 3.7 | 485 | Specific | 571.68 | 571.68 | 239.47 | 10.17 | 122 | >sp\|P02788\|TRFL_HUMAN Lactotransferrin OS=Homo sapiens OX=9606 GN=LTF PE=1 SV=6 | 1270 | 20201114_qwl_colo_milk_1ng_glyco_hcd203040_compound15.18716.18716.3 | controllerType=0 controllerNumber=1 scan=18716 | 41.1774 |
| 16922:3 | 1 | R.TAGWNIPMGLLFN[+1856.656]QTGSC[+57.021]K.F | HexNAc(3)Hex(5)Fuc(1)NeuAc(1) | C[+57], N[+1857] | 1317.8962 | 3 | 3951.674 | 3951.6655 | 0 | 2.1 | 485 | Specific | 620.31 | 576.39 | 192.21 | 12.41 | 122 | >sp\|P02788\|TRFL_HUMAN Lactotransferrin OS=Homo sapiens OX=9606 GN=LTF PE=1 SV=6 | 1270 | 20201114_qwl_colo_milk_1ng_glyco_hcd203040_compound15.20360.20360.3 | controllerType=0 controllerNumber=1 scan=20360 | [illegible] |
| 18975:3 | 1 | R.TAGWNIPMGLLFN[+2204.772]QTGSC[+57.021]K.F | HexNAc(4)Hex(5)NeuAc(2) | C[+57], N[+2205] | 1433.938 | 3 | 4299.7996 | 4299.7824 | 0 | 4 | 485 | Specific | 640.93 | 584.75 | 584.75 | 12.48 | 122 | >sp\|P02788\|TRFL_HUMAN Lactotransferrin OS=Homo sapiens OX=9606 GN=LTF PE=1 SV=6 | 1270 | 20201114_qwl_colo_milk_1ng_glyco_hcd203040_compound15.22511.22511.3 | controllerType=0 controllerNumber=1 scan=22511 | [illegible] |
| 14911:3 | 1 | R.TAGWNIPMGLLFN[+2279.830]QTGSC[+57.021]K.F | HexNAc(5)Hex(6)Fuc(2) | C[+57], N[+2280] | 1458.9578 | 3 | 4374.8589 | 4374.8396 | 0 | 4.4 | 485 | Specific | 648.59 | 633.46 | 321.82 | 12.48 | 122 | >sp\|P02788\|TRFL_HUMAN Lactotransferrin OS=Homo sapiens OX=9606 GN=LTF PE=1 SV=6 | 1270 | 20201114_qwl_colo_milk_1ng_glyco_hcd203040_compound15.18250.18250.3 | controllerType=0 controllerNumber=1 scan=18250 | 40.2134 |
| 16923:3 | 1 | R.TAGWNIPMGLLFN[+1856.656]QTGSC[+57.021]K.F | HexNAc(3)Hex(5)Fuc(1)NeuAc(1) | C[+57], N[+1857] | 1317.8962 | 3 | 3951.674 | 3951.6655 | 0 | 2.1 | 485 | Specific | 652.83 | 553.81 | 313.45 | 13.41 | 122 | >sp\|P02788\|TRFL_HUMAN Lactotransferrin OS=Homo sapiens OX=9606 GN=LTF PE=1 SV=6 | 1270 | 20201114_qwl_colo_milk_1ng_glyco_hcd203040_compound15.20361.20361.3 | controllerType=0 controllerNumber=1 scan=20361 | [illegible] |
| 16927:3 | 1 | R.TAGWNIPMGLLFN[+1913.677]QTGSC[+57.021]K.F | HexNAc(4)Hex(5)NeuAc(1) | C[+57], N[+1914] | 1336.9011 | 3 | 4008.6886 | 4008.687 | 0 | 0.4 | 485 | Specific | 668.39 | 603.83 | 603.83 | 13.36 | 122 | >sp\|P02788\|TRFL_HUMAN Lactotransferrin OS=Homo sapiens OX=9606 GN=LTF PE=1 SV=6 | 1270 | 20201114_qwl_colo_milk_1ng_glyco_hcd203040_compound15.20366.20366.3 | controllerType=0 controllerNumber=1 scan=20366 | 45.1281 |
| 14913:3 | 1 | R.TAGWNIPMGLLFN[+2279.830]QTGSC[+57.021]K.F | HexNAc(5)Hex(6)Fuc(2) | C[+57], N[+2280] | 1458.9578 | 3 | 4374.8589 | 4374.8396 | 0 | 4.4 | 485 | Specific | 718.51 | 620 | 348.28 | 14.48 | 122 | >sp\|P02788\|TRFL_HUMAN Lactotransferrin OS=Homo sapiens OX=9606 GN=LTF PE=1 SV=6 | 1270 | 20201114_qwl_colo_milk_1ng_glyco_hcd203040_compound15.18252.18252.3 | controllerType=0 controllerNumber=1 scan=18252 | 40.2375 |
| 18976:3 | 1 | R.TAGWNIPMGLLFN[+2204.772]QTGSC[+57.021]K.F | HexNAc(4)Hex(5)NeuAc(2) | C[+57], N[+2205] | 1433.938 | 3 | 4299.7996 | 4299.7824 | 0 | 4 | 485 | Specific | 737.58 | 566.14 | 566.14 | 14.48 | 122 | >sp\|P02788\|TRFL_HUMAN Lactotransferrin OS=Homo sapiens OX=9606 GN=LTF PE=1 SV=6 | 1270 | 20201114_qwl_colo_milk_1ng_glyco_hcd203040_compound15.22512.22512.3 | controllerType=0 controllerNumber=1 scan=22512 | 49.7713 |
| 15283:3 | 1 | R.TAGWNIPMGLLFN[+1622.582]QTGSC[+57.021]K.F | HexNAc(4)Hex(5) | C[+57], N[+1623] | 1239.8765 | 3 | 3717.615 | 3717.5916 | 0 | 6.3 | 485 | Specific | 749.88 | 690.24 | 347.79 | 13.99 | 122 | >sp\|P02788\|TRFL_HUMAN Lactotransferrin OS=Homo sapiens OX=9606 GN=LTF PE=1 SV=6 | 1270 | 20201114_qwl_colo_milk_1ng_glyco_hcd203040_compound15.18637.18637.3 | controllerType=0 controllerNumber=1 scan=18637 | 41.0002 |
| 15087:3 | 1 | R.TAGWNIPMGLLFN[+1768.640]QTGSC[+57.021]K.F | HexNAc(4)Hex(5)Fuc(1) | C[+57], N[+1769] | 1288.5558 | 3 | 3863.653 | 3863.6495 | 0 | 0.9 | 485 | Specific | 773.64 | 773.64 | 773.64 | 15.36 | 122 | >sp\|P02788\|TRFL_HUMAN Lactotransferrin OS=Homo sapiens OX=9606 GN=LTF PE=1 SV=6 | 1270 | 20201114_qwl_colo_milk_1ng_glyco_hcd203040_compound15.18434.18434.3 | controllerType=0 controllerNumber=1 scan=18434 | 40.5916 |
| 16930:3 | 1 | R.TAGWNIPMGLLFN[+1913.677]QTGSC[+57.021]K.F | HexNAc(4)Hex(5)NeuAc(1) | C[+57], N[+1914] | 1336.9011 | 3 | 4008.6886 | 4008.687 | 0 | 0.4 | 485 | Specific | 795.42 | 770.03 | 770.03 | 15.36 | 122 | >sp\|P02788\|TRFL_HUMAN Lactotransferrin OS=Homo sapiens OX=9606 GN=LTF PE=1 SV=6 | 1270 | 20201114_qwl_colo_milk_1ng_glyco_hcd203040_compound15.20369.20369.3 | controllerType=0 controllerNumber=1 scan=20369 | 45.1319 |
| 15285:3 | 1 | R.TAGWNIPMGLLFN[+1622.582]QTGSC[+57.021]K.F | HexNAc(4)Hex(5) | C[+57], N[+1623] | 1239.8765 | 3 | 3717.615 | 3717.5916 | 0 | 6.3 | 485 | Specific | 798.46 | 775.1 | 379.88 | 14.99 | 122 | >sp\|P02788\|TRFL_HUMAN Lactotransferrin OS=Homo sapiens OX=9606 GN=LTF PE=1 SV=6 | 1270 | 20201114_qwl_colo_milk_1ng_glyco_hcd203040_compound15.18639.18639.3 | controllerType=0 controllerNumber=1 scan=18639 | 41.0052 |
| 16982:3 | 1 | R.TAGWNIPMGLLFN[+1694.603]QTGSC[+57.021]K.F | HexNAc(3)Hex(4)Fuc(1)NeuAc(1) | C[+57], N[+1695] | 1263.8831 | 3 | 3789.6348 | 3789.6127 | 0 | 5.8 | 485 | Specific | 838.74 | 782.76 | 782.76 | 15.99 | 122 | >sp\|P02788\|TRFL_HUMAN Lactotransferrin OS=Homo sapiens OX=9606 GN=LTF PE=1 SV=6 | 1270 | 20201114_qwl_colo_milk_1ng_glyco_hcd203040_compound15.20422.20422.3 | controllerType=0 controllerNumber=1 scan=20422 | 45.2203 |
| 15360:3 | 1 | R.TAGWNIPMGLLFN[+1622.582]QTGSC[+57.021]K.F | HexNAc(4)Hex(5) | C[+57], N[+1623] | 1239.8732 | 3 | 3717.6051 | 3717.5916 | 0 | 3.7 | 485 | Specific | 843.79 | 649.23 | 367.85 | 16.48 | 122 | >sp\|P02788\|TRFL_HUMAN Lactotransferrin OS=Homo sapiens OX=9606 GN=LTF PE=1 SV=6 | 1270 | 20201114_qwl_colo_milk_1ng_glyco_hcd203040_compound15.18718.18718.3 | controllerType=0 controllerNumber=1 scan=18718 | 41.1813 |
| 15090:3 | 1 | R.TAGWNIPMGLLFN[+1768.640]QTGSC[+57.021]K.F | HexNAc(4)Hex(5)Fuc(1) | C[+57], N[+1769] | 1288.5558 | 3 | 3863.653 | 3863.6495 | 0 | 0.9 | 485 | Specific | 855.28 | 689.28 | 689.28 | 17.36 | 122 | >sp\|P02788\|TRFL_HUMAN Lactotransferrin OS=Homo sapiens OX=9606 GN=LTF PE=1 SV=6 | 1270 | 20201114_qwl_colo_milk_1ng_glyco_hcd203040_compound15.18437.18437.3 | controllerType=0 controllerNumber=1 scan=18437 | 40.5959 |
| 16964:3 | 1 | R.TAGWNIPMGLLFN[+2059.735]QTGSC[+57.021]K.F | HexNAc(4)Hex(5)Fuc(1)NeuAc(1) | C[+57], N[+2060] | 1385.5882 | 3 | 4154.75 | 4154.7449 | 0 | 1.2 | 485 | Specific | 863.31 | 672.58 | 672.58 | 17.36 | 122 | >sp\|P02788\|TRFL_HUMAN Lactotransferrin OS=Homo sapiens OX=9606 GN=LTF PE=1 SV=6 | 1270 | 20201114_qwl_colo_milk_1ng_glyco_hcd203040_compound15.20404.20404.3 | controllerType=0 controllerNumber=1 scan=20404 | 45.1906 |
| 15469:2 | 1 | R.TAGWNIPMGLLFN[+1216.423]QTGSC[+57.021]K.F | HexNAc(2)Hex(5) | C[+57], N[+1216] | 1656.2255 | 2 | 3311.4437 | 3311.4328 | 0 | 3.3 | 485 | Specific | 885.75 | 876.79 | 283.16 | 17.52 | 122 | >sp\|P02788\|TRFL_HUMAN Lactotransferrin OS=Homo sapiens OX=9606 GN=LTF PE=1 SV=6 | 1270 | 20201114_qwl_colo_milk_1ng_glyco_hcd203040_compound15.18832.18832.2 | controllerType=0 controllerNumber=1 scan=18832 | 41.4757 |
| 18752:3 | 1 | R.TAGWNIPMGLLFN[+2350.830]QTGSC[+57.021]K.F | HexNAc(4)Hex(5)Fuc(1)NeuAc(2) | C[+57], N[+2351] | 1482.6258 | 3 | 4445.8628 | 4445.8403 | 0 | 5.1 | 485 | Specific | 904.18 | 864.44 | 864.44 | 18.18 | 122 | >sp\|P02788\|TRFL_HUMAN Lactotransferrin OS=Homo sapiens OX=9606 GN=LTF PE=1 SV=6 | 1270 | 20201114_qwl_colo_milk_1ng_glyco_hcd203040_compound15.22279.22279.3 | controllerType=0 controllerNumber=1 scan=22279 | 49.2745 |
| 16782:3 | 1 | R.TAGWNIPMGLLFN[+2350.830]QTGSC[+57.021]K.F | HexNAc(4)Hex(5)Fuc(1)NeuAc(2) | C[+57], N[+2351] | 1482.6255 | 3 | 4445.8621 | 4445.8403 | 0 | 4.9 | 485 | Specific | 906.19 | 883.84 | 883.84 | 18.18 | 122 | >sp\|P02788\|TRFL_HUMAN Lactotransferrin OS=Homo sapiens OX=9606 GN=LTF PE=1 SV=6 | 1270 | 20201114_qwl_colo_milk_1ng_glyco_hcd203040_compound15.20214.20214.3 | controllerType=0 controllerNumber=1 scan=20214 | 44.772 |
| 18750:3 | 1 | R.TAGWNIPMGLLFN[+2350.830]QTGSC[+57.021]K.F | HexNAc(4)Hex(5)Fuc(1)NeuAc(2) | C[+57], N[+2351] | 1482.6258 | 3 | 4445.8628 | 4445.8403 | 0 | 5.1 | 485 | Specific | 917.94 | 906.87 | 906.87 | 18.18 | 122 | >sp\|P02788\|TRFL_HUMAN Lactotransferrin OS=Homo sapiens OX=9606 GN=LTF PE=1 SV=6 | 1270 | 20201114_qwl_colo_milk_1ng_glyco_hcd203040_compound15.22277.22277.3 | controllerType=0 controllerNumber=1 scan=22277 | 49.2679 |
| 15467:2 | 1 | R.TAGWNIPMGLLFN[+1216.423]QTGSC[+57.021]K.F | HexNAc(2)Hex(5) | C[+57], N[+1216] | 1656.2255 | 2 | 3311.4437 | 3311.4328 | 0 | 3.3 | 485 | Specific | 928.44 | 909.4 | 342.57 | 18.52 | 122 | >sp\|P02788\|TRFL_HUMAN Lactotransferrin OS=Homo sapiens OX=9606 GN=LTF PE=1 SV=6 | 1270 | 20201114_qwl_colo_milk_1ng_glyco_hcd203040_compound15.18830.18830.2 | controllerType=0 controllerNumber=1 scan=18830 | 41.4708 |
| 15560:3 | 1 | R.TAGWNIPMGLLFN[+1216.423]QTGSC[+57.021]K.F | HexNAc(2)Hex(5) | C[+57], N[+1216] | 1104.4868 | 3 | 3311.446 | 3311.4328 | 0 | 4 | 485 | Specific | 930.21 | 930.21 | 395.37 | 18.48 | 122 | >sp\|P02788\|TRFL_HUMAN Lactotransferrin OS=Homo sapiens OX=9606 GN=LTF PE=1 SV=6 | 1270 | 20201114_qwl_colo_milk_1ng_glyco_hcd203040_compound15.18928.18928.3 | controllerType=0 controllerNumber=1 scan=18928 | 41.6638 |
| 16795:3 | 1 | R.TAGWNIPMGLLFN[+2059.735]QTGSC[+57.021]K.F | HexNAc(4)Hex(5)Fuc(1)NeuAc(1) | C[+57], N[+2060] | 1385.587 | 3 | 4154.7464 | 4154.7449 | 0 | 0.4 | 485 | Specific | 939.66 | 897.28 | 897.28 | 18.36 | 122 | >sp\|P02788\|TRFL_HUMAN Lactotransferrin OS=Homo sapiens OX=9606 GN=LTF PE=1 SV=6 | 1270 | 20201114_qwl_colo_milk_1ng_glyco_hcd203040_compound15.20227.20227.3 | controllerType=0 controllerNumber=1 scan=20227 | 44.7963 |
| 16792:3 | 1 | R.TAGWNIPMGLLFN[+2059.735]QTGSC[+57.021]K.F | HexNAc(4)Hex(5)Fuc(1)NeuAc(1) | C[+57], N[+2060] | 1385.587 | 3 | 4154.7464 | 4154.7449 | 0 | 0.4 | 485 | Specific | 966.22 | 829.69 | 829.69 | 19.36 | 122 | >sp\|P02788\|TRFL_HUMAN Lactotransferrin OS=Homo sapiens OX=9606 GN=LTF PE=1 SV=6 | 1270 | 20201114_qwl_colo_milk_1ng_glyco_hcd203040_compound15.20224.20224.3 | controllerType=0 controllerNumber=1 scan=20224 | 44.7872 |
| 15449:3 | 1 | R.TAGWNIPMGLLFN[+1216.423]QTGSC[+57.021]K.F | HexNAc(2)Hex(5) | C[+57], N[+1216] | 1104.4857 | 3 | 3311.4427 | 3311.4328 | 0 | 3 | 485 | Specific | 967.38 | 967.38 | 434.01 | 19.51 | 122 | >sp\|P02788\|TRFL_HUMAN Lactotransferrin OS=Homo sapiens OX=9606 GN=LTF PE=1 SV=6 | 1270 | 20201114_qwl_colo_milk_1ng_glyco_hcd203040_compound15.18812.18812.3 | controllerType=0 controllerNumber=1 scan=18812 | 41.4302 |
| 15451:3 | 1 | R.TAGWNIPMGLLFN[+1216.423]QTGSC[+57.021]K.F | HexNAc(2)Hex(5) | C[+57], N[+1216] | 1104.4857 | 3 | 3311.4427 | 3311.4328 | 0 | 3 | 485 | Specific | 1063.99 | 1056.84 | 503.73 | 21.51 | 122 | >sp\|P02788\|TRFL_HUMAN Lactotransferrin OS=Homo sapiens OX=9606 GN=LTF PE=1 SV=6 | 1270 | 20201114_qwl_colo_milk_1ng_glyco_hcd203040_compound15.18814.18814.3 | controllerType=0 controllerNumber=1 scan=18814 | 41.4355 |
| 15322:3 | 1 | R.TAGWNIPM[+15.995]GLLFN[+2059.735]QTGSC[+57.021]K.F | HexNAc(4)Hex(5)Fuc(1)NeuAc(1) | C[+57], M[+16], N[+2060] | 1390.9288 | 3 | 4170.7718 | 4170.7398 | 0 | 7.7 | 485 | Specific | 269.97 | 269.97 | 269.97 | 6.6 | 122 | >sp\|P02788\|TRFL_HUMAN Lactotransferrin OS=Homo sapiens OX=9606 GN=LTF PE=1 SV=6 | 1270 | 20201114_qwl_colo_milk_1ng_glyco_hcd203040_compound15.18678.18678.3 | controllerType=0 controllerNumber=1 scan=18678 | 41.0861 |
| 15324:3 | 1 | R.TAGWNIPM[+15.995]GLLFN[+2059.735]QTGSC[+57.021]K.F | HexNAc(4)Hex(5)Fuc(1)NeuAc(1) | C[+57], M[+16], N[+2060] | 1390.9288 | 3 | 4170.7718 | 4170.7398 | 0 | 7.7 | 485 | Specific | 503.83 | 500.59 | 500.59 | 9.45 | 122 | >sp\|P02788\|TRFL_HUMAN Lactotransferrin OS=Homo sapiens OX=9606 GN=LTF PE=1 SV=6 | 1270 | 20201114_qwl_colo_milk_1ng_glyco_hcd203040_compound15.18680.18680.3 | controllerType=0 controllerNumber=1 scan=18680 | 41.0897 |
| 16113:3 | 1 | R.TAGWNIPMGLLFN[+1768.640]QTGSC[+57.021]KFDEYFSQSC[+57.021]APGSDPR.S | HexNAc(4)Hex(5)Fuc(1) | C[+57]*2, N[+1769] | 1903.1352 | 3 | 5707.3911 | 5707.3966 | 0 | -1 | 485 | Specific | 251.04 | 186.22 | 186.22 | 5.33 | 122 | >sp\|P02788\|TRFL_HUMAN Lactotransferrin OS=Homo sapiens OX=9606 GN=LTF PE=1 SV=6 | 1270 | 20201114_qwl_colo_milk_1ng_glyco_hcd203040_compound15.19513.19513.3 | controllerType=0 controllerNumber=1 scan=19513 | 43.2037 |
| 16256:4 | 1 | R.TAGWNIPMGLLFN[+1216.423]QTGSC[+57.021]KFDEYFSQSC[+57.021]APGSDPR.S | HexNAc(2)Hex(5) | C[+57]*2, N[+1216] | 1289.5544 | 4 | 5155.1958 | 5155.18 | 0 | 3.1 | 485 | Specific | 442.19 | 442.19 | 86.33 | 9.33 | 122 | >sp\|P02788\|TRFL_HUMAN Lactotransferrin OS=Homo sapiens OX=9606 GN=LTF PE=1 SV=6 | 1270 | 20201114_qwl_colo_milk_1ng_glyco_hcd203040_compound15.19662.19662.4 | controllerType=0 controllerNumber=1 scan=19662 | 43.5034 |
| 16258:4 | 1 | R.TAGWNIPMGLLFN[+1216.423]QTGSC[+57.021]KFDEYFSQSC[+57.021]APGSDPR.S | HexNAc(2)Hex(5) | C[+57]*2, N[+1216] | 1289.5544 | 4 | 5155.1958 | 5155.18 | 0 | 3.1 | 485 | Specific | 497.74 | 497.74 | 219.84 | 9.57 | 122 | >sp\|P02788\|TRFL_HUMAN Lactotransferrin OS=Homo sapiens OX=9606 GN=LTF PE=1 SV=6 | 1270 | 20201114_qwl_colo_milk_1ng_glyco_hcd203040_compound15.19664.19664.4 | controllerType=0 controllerNumber=1 scan=19664 | 43.5107 |
| 16116:3 | 1 | R.TAGWNIPMGLLFN[+1768.640]QTGSC[+57.021]KFDEYFSQSC[+57.021]APGSDPR.S | HexNAc(4)Hex(5)Fuc(1) | C[+57]*2, N[+1769] | 1903.1352 | 3 | 5707.3911 | 5707.3966 | 0 | -1 | 485 | Specific | 612.74 | 485.56 | 485.56 | 11.64 | 122 | >sp\|P02788\|TRFL_HUMAN Lactotransferrin OS=Homo sapiens OX=9606 GN=LTF PE=1 SV=6 | 1270 | 20201114_qwl_colo_milk_1ng_glyco_hcd203040_compound15.19516.19516.3 | controllerType=0 controllerNumber=1 scan=19516 | 43.2082 |
| 17029:4 | 1 | R.TAGWNIPMGLLFN[+2351.851]QTGSC[+57.021]KFDEYFSQSC[+57.021]APGSDPR.S | HexNAc(4)Hex(5)Fuc(3)NeuAc(1) | C[+57]*2, N[+2352] | 1573.411 | 4 | 6290.6222 | 6290.6079 | 0 | 2.3 | 485 | Specific | 715.38 | 710.33 | 710.33 | 14.15 | 122 | >sp\|P02788\|TRFL_HUMAN Lactotransferrin OS=Homo sapiens OX=9606 GN=LTF PE=1 SV=6 | 1270 | 20201114_qwl_colo_milk_1ng_glyco_hcd203040_compound15.20472.20472.4 | controllerType=0 controllerNumber=1 scan=20472 | 45.3395 |
| 16238:3 | 1 | R.TAGWNIPMGLLFN[+1216.423]QTGSC[+57.021]KFDEYFSQSC[+57.021]APGSDPR.S | HexNAc(2)Hex(5) | C[+57]*2, N[+1216] | 1719.0704 | 3 | 5155.1966 | 5155.18 | 0 | 3.2 | 485 | Specific | 795.35 | 745.82 | 217.4 | 15.26 | 122 | >sp\|P02788\|TRFL_HUMAN Lactotransferrin OS=Homo sapiens OX=9606 GN=LTF PE=1 SV=6 | 1270 | 20201114_qwl_colo_milk_1ng_glyco_hcd203040_compound15.19643.19643.3 | controllerType=0 controllerNumber=1 scan=19643 | 43.4619 |
| 18339:4 | 1 | R.TAGWNIPMGLLFN[+2350.830]QTGSC[+57.021]KFDEYFSQSC[+57.021]APGSDPR.S | HexNAc(4)Hex(5)Fuc(1)NeuAc(2) | C[+57]*2, N[+2351] | 1573.1665 | 4 | 6289.6442 | 6289.5875 | 0 | 9 | 485 | Specific | 830.83 | 675.56 | 675.56 | 15.66 | 122 | >sp\|P02788\|TRFL_HUMAN Lactotransferrin OS=Homo sapiens OX=9606 GN=LTF PE=1 SV=6 | 1270 | 20201114_qwl_colo_milk_1ng_glyco_hcd203040_compound15.21846.21846.4 | controllerType=0 controllerNumber=1 scan=21846 | 48.3453 |
| 17032:4 | 1 | R.TAGWNIPMGLLFN[+2351.851]QTGSC[+57.021]KFDEYFSQSC[+57.021]APGSDPR.S | HexNAc(4)Hex(5)Fuc(3)NeuAc(1) | C[+57]*2, N[+2352] | 1573.411 | 4 | 6290.6222 | 6290.6079 | 0 | 2.3 | 485 | Specific | 854 | 851.88 | 851.88 | 17.15 | 122 | >sp\|P02788\|TRFL_HUMAN Lactotransferrin OS=Homo sapiens OX=9606 GN=LTF PE=1 SV=6 | 1270 | 20201114_qwl_colo_milk_1ng_glyco_hcd203040_compound15.20475.20475.4 | controllerType=0 controllerNumber=1 scan=20475 | 45.3446 |
| 18341:4 | 1 | R.TAGWNIPMGLLFN[+2350.830]QTGSC[+57.021]KFDEYFSQSC[+57.021]APGSDPR.S | HexNAc(4)Hex(5)Fuc(1)NeuAc(2) | C[+57]*2, N[+2351] | 1573.1665 | 4 | 6289.6442 | 6289.5875 | 0 | 9 | 485 | Specific | 905.11 | 869.04 | 869.04 | 17.66 | 122 | >sp\|P02788\|TRFL_HUMAN Lactotransferrin OS=Homo sapiens OX=9606 GN=LTF PE=1 SV=6 | 1270 | 20201114_qwl_colo_milk_1ng_glyco_hcd203040_compound15.21848.21848.4 | controllerType=0 controllerNumber=1 scan=21848 | 48.3508 |
| 17065:4 | 1 | R.TAGWNIPMGLLFN[+2059.735]QTGSC[+57.021]KFDEYFSQSC[+57.021]APGSDPR.S | HexNAc(4)Hex(5)Fuc(1)NeuAc(1) | C[+57]*2, N[+2060] | 1500.3877 | 4 | 5998.529 | 5998.492 | 0 | 6.2 | 485 | Specific | 924.4 | 889.97 | 889.97 | 17.73 | 122 | >sp\|P02788\|TRFL_HUMAN Lactotransferrin OS=Homo sapiens OX=9606 GN=LTF PE=1 SV=6 | 1270 | 20201114_qwl_colo_milk_1ng_glyco_hcd203040_compound15.20509.20509.4 | controllerType=0 controllerNumber=1 scan=20509 | 45.4152 |
| 17063:4 | 1 | R.TAGWNIPMGLLFN[+2059.735]QTGSC[+57.021]KFDEYFSQSC[+57.021]APGSDPR.S | HexNAc(4)Hex(5)Fuc(1)NeuAc(1) | C[+57]*2, N[+2060] | 1500.3877 | 4 | 5998.529 | 5998.492 | 0 | 6.2 | 485 | Specific | 938.09 | 924.07 | 924.07 | 17.73 | 122 | >sp\|P02788\|TRFL_HUMAN Lactotransferrin OS=Homo sapiens OX=9606 GN=LTF PE=1 SV=6 | 1270 | 20201114_qwl_colo_milk_1ng_glyco_hcd203040_compound15.20507.20507.4 | controllerType=0 controllerNumber=1 scan=20507 | 45.4094 |
| 16236:3 | 1 | R.TAGWNIPMGLLFN[+1216.423]QTGSC[+57.021]KFDEYFSQSC[+57.021]APGSDPR.S | HexNAc(2)Hex(5) | C[+57]*2, N[+1216] | 1719.0704 | 3 | 5155.1966 | 5155.18 | 0 | 3.2 | 485 | Specific | 1050.18 | 972.33 | 272.65 | 21.26 | 122 | >sp\|P02788\|TRFL_HUMAN Lactotransferrin OS=Homo sapiens OX=9606 GN=LTF PE=1 SV=6 | 1270 | 20201114_qwl_colo_milk_1ng_glyco_hcd203040_compound15.19641.19641.3 | controllerType=0 controllerNumber=1 scan=19641 | 43.4494 |
| 16110:4 | 1 | R.TAGWNIPM[+15.995]GLLFN[+1751.624]QTGSC[+57.021]KFDEYFSQSC[+57.021]A | HexNAc(4)Hex(4)NeuAc(1) | C[+57]*2, M[+16], N[+1752] | 1427.354 | 4 | 5706.3942 | 5706.3762 | 0 | 3.2 | 485 | Specific | 338.84 | 338.84 | 338.84 | 5.83 | 122 | >sp\|P02788\|TRFL_HUMAN Lactotransferrin OS=Homo sapiens OX=9606 GN=LTF PE=1 SV=6 | 1270 | 20201114_qwl_colo_milk_1ng_glyco_hcd203040_compound15.19510.19510.4 | controllerType=0 controllerNumber=1 scan=19510 | 43.1992 |
| 16108:4 | 1 | R.TAGWNIPM[+15.995]GLLFN[+1751.624]QTGSC[+57.021]KFDEYFSQSC[+57.021]A | HexNAc(4)Hex(4)NeuAc(1) | C[+57]*2, M[+16], N[+1752] | 1427.354 | 4 | 5706.3942 | 5706.3762 | 0 | 3.2 | 485 | Specific | 460.77 | 460.77 | 460.77 | 9.43 | 122 | >sp\|P02788\|TRFL_HUMAN Lactotransferrin OS=Homo sapiens OX=9606 GN=LTF PE=1 SV=6 | 1270 | 20201114_qwl_colo_milk_1ng_glyco_hcd203040_compound15.19508.19508.4 | controllerType=0 controllerNumber=1 scan=19508 | 43.1954 |
| 25506:4 | 2 | R.LSLHRPALEDLLLGSEAN[+1216.423]LTC[+57.021]TLTGLR.D | HexNAc(2)Hex(5) | C[+57], N[+1216] | 1045.7608 | 4 | 4180.0213 | 4180.0211 | 0 | 0 | 127 | Specific | 211.26 | 211.26 | 211.26 | 5.22 | 124 | >sp\|P01876\|IGHA1_HUMAN Immunoglobulin heavy constant alpha 1 OS=Homo sapiens OX=9606 GN=IGHA1 PE=1 SV=2 | 255 | 20201114_qwl_colo_milk_1ng_glyco_hcd203040_compound15.29875.29875.4 | controllerType=0 controllerNumber=1 scan=29875 | 65.1805 |
| 17807:3 | 2 | R.LSLHRPALEDLLLGSEAN[+2204.772]LTC[+57.021]TLTGLR.D | HexNAc(4)Hex(5)NeuAc(2) | C[+57], N[+2205] | 1723.4627 | 3 | 5168.3736 | 5168.3707 | 0 | 0.6 | 127 | Specific | 229.6 | 229.6 | 229.6 | 5.87 | 124 | >sp\|P01876\|IGHA1_HUMAN Immunoglobulin heavy constant alpha 1 OS=Homo sapiens OX=9606 GN=IGHA1 PE=1 SV=2 | 255 | 20201114_qwl_colo_milk_1ng_glyco_hcd203040_compound15.21291.21291.3 | controllerType=0 controllerNumber=1 scan=21291 | 47.1664 |
| 29849:4 | 2 | R.LSLHRPALEDLLLGSEAN[+1216.423]LTC[+57.021]TLTGLR.D | HexNAc(2)Hex(5) | C[+57], N[+1216] | 1045.7638 | 4 | 4180.0335 | 4180.0211 | 0 | 3 | 127 | Specific | 234.3 | 234.3 | 234.3 | 6.69 | 124 | >sp\|P01876\|IGHA1_HUMAN Immunoglobulin heavy constant alpha 1 OS=Homo sapiens OX=9606 GN=IGHA1 PE=1 SV=2 | 255 | 20201114_qwl_colo_milk_1ng_glyco_hcd203040_compound15.38185.38185.3 | controllerType=0 controllerNumber=1 scan=38185 | 88.8731 |
| 16664:4 | 2 | R.LSLHRPALEDLLLGSEAN[+1378.476]LTC[+57.021]TLTGLR.D | HexNAc(2)Hex(6) | C[+57], N[+1378] | 1086.2818 | 4 | 4342.1053 | 4342.074 | 0 | 7.2 | 127 | Specific | 248.7 | 159.26 | 159.26 | 3.29 | 124 | >sp\|P01876\|IGHA1_HUMAN Immunoglobulin heavy constant alpha 1 OS=Homo sapiens OX=9606 GN=IGHA1 PE=1 SV=2 | 255 | 20201114_qwl_colo_milk_1ng_glyco_hcd203040_compound15.20088.20088.3 | controllerType=0 controllerNumber=1 scan=20088 | 44.423 |
| 20137:4 | 2 | R.LSLHRPALEDLLLGSEAN[+1501.555]LTC[+57.021]TLTGLR.D | HexNAc(5)Hex(3) | C[+57], N[+1502] | 1117.0491 | 4 | 4465.1746 | 4465.1536 | 0 | 4.7 | 127 | Specific | 248.89 | 228.73 | 228.73 | 6.38 | 124 | >sp\|P01876\|IGHA1_HUMAN Immunoglobulin heavy constant alpha 1 OS=Homo sapiens OX=9606 GN=IGHA1 PE=1 SV=2 | 255 | 20201114_qwl_colo_milk_1ng_glyco_hcd203040_compound15.23724.23724.2 | controllerType=0 controllerNumber=1 scan=23724 | 52.2428 |
| 18861:4 | 2 | R.LSLHRPALEDLLLGSEAN[+1501.555]LTC[+57.021]TLTGLR.D | HexNAc(5)Hex(3) | C[+57], N[+1502] | 1117.0453 | 4 | 4465.1594 | 4465.1536 | 0 | 1.3 | 127 | Specific | 266.04 | 266.04 | 266.04 | 6.8 | 124 | >sp\|P01876\|IGHA1_HUMAN Immunoglobulin heavy constant alpha 1 OS=Homo sapiens OX=9606 GN=IGHA1 PE=1 SV=2 | 255 | 20201114_qwl_colo_milk_1ng_glyco_hcd203040_compound15.22392.22392.4 | controllerType=0 controllerNumber=1 scan=22392 | 49.5092 |
| 18210:3 | 2 | R.[+42.011]LSLHRPALEDLLLGSEAN[+1955.724]LTC[+57.021]TLTGLR.D | HexNAc(5)Hex(4)Fuc(2) | .[42], C[+57], N[+1956] | 1654.4423 | 3 | 4961.3125 | 4961.3328 | 0 | -4.1 | 127 | Specific | 266.85 | 225.21 | 225.21 | 4.66 | 124 | >sp\|P01876\|IGHA1_HUMAN Immunoglobulin heavy constant alpha 1 OS=Homo sapiens OX=9606 GN=IGHA1 PE=1 SV=2 | 255 | 20201114_qwl_colo_milk_1ng_glyco_hcd203040_compound15.21711.21711.3 | controllerType=0 controllerNumber=1 scan=21711 | 48.0563 |
| 30205:4 | 2 | R.LSLHRPALEDLLLGSEAN[+1216.423]LTC[+57.021]TLTGLR.D | HexNAc(2)Hex(5) | C[+57], N[+1216] | 1045.7629 | 4 | 4180.0296 | 4180.0211 | 0 | 2 | 127 | Specific | 270.02 | 270.02 | 270.02 | 7.06 | 124 | >sp\|P01876\|IGHA1_HUMAN Immunoglobulin heavy constant alpha 1 OS=Homo sapiens OX=9606 GN=IGHA1 PE=1 SV=2 | 255 | 20201114_qwl_colo_milk_1ng_glyco_hcd203040_compound15.38597.38597.4 | controllerType=0 controllerNumber=1 scan=38597 | 89.7084 |
| 15920:3 | 2 | R.LSLHRPALEDLLLGSEAN[+1095.397]LTC[+57.021]TLTGLR.D | HexNAc(3)Hex(3) | C[+57], N[+1095] | 1353.6828 | 3 | 4059.0338 | 4058.9949 | 0 | 9.6 | 127 | Specific | 270.21 | 270.21 | 270.21 | 5.91 | 124 | >sp\|P01876\|IGHA1_HUMAN Immunoglobulin heavy constant alpha 1 OS=Homo sapiens OX=9606 GN=IGHA1 PE=1 SV=2 | 255 | 20201114_qwl_colo_milk_1ng_glyco_hcd203040_compound15.19308.19308.3 | controllerType=0 controllerNumber=1 scan=19308 | 42.5995 |
| 15879:3 | 2 | R.LSLHRPALEDLLLGSEAN[+1501.555]LTC[+57.021]TLTGLR.D | HexNAc(5)Hex(3) | C[+57], N[+1502] | 1117.0456 | 4 | 4465.1605 | 4465.1536 | 0 | 1.5 | 127 | Specific | 301.88 | 301.88 | 301.88 | 7.2 | 124 | >sp\|P01876\|IGHA1_HUMAN Immunoglobulin heavy constant alpha 1 OS=Homo sapiens OX=9606 GN=IGHA1 PE=1 SV=2 | 255 | 20201114_qwl_colo_milk_1ng_glyco_hcd203040_compound15.19265.19265.4 | controllerType=0 controllerNumber=1 scan=19265 | 42.4988 |
| 16576:4 | 2 | R.LSLHRPALEDLLLGSEAN[+1054.370]LTC[+57.021]TLTGLR.D | HexNAc(2)Hex(4) | C[+57], N[+1054] | 1005.2529 | 4 | 4017.9898 | 4017.9683 | 0 | 5.3 | 127 | Specific | 308.94 | 308.94 | 308.94 | 5.33 | 124 | >sp\|P01876\|IGHA1_HUMAN Immunoglobulin heavy constant alpha 1 OS=Homo sapiens OX=9606 GN=IGHA1 PE=1 SV=2 | 255 | 20201114_qwl_colo_milk_1ng_glyco_hcd203040_compound15.19995.19995.4 | controllerType=0 controllerNumber=1 scan=19995 | 44.2269 |
| 23865:4 | 2 | R.LSLHRPALEDLLLGSEAN[+1501.555]LTC[+57.021]TLTGLR.D | HexNAc(5)Hex(3) | C[+57], N[+1502] | 1117.0513 | 4 | 4465.1834 | 4465.1536 | 0 | 6.7 | 127 | Specific | 318.46 | 318.46 | 318.46 | 7.22 | 124 | >sp\|P01876\|IGHA1_HUMAN Immunoglobulin heavy constant alpha 1 OS=Homo sapiens OX=9606 GN=IGHA1 PE=1 SV=2 | 255 | 20201114_qwl_colo_milk_1ng_glyco_hcd203040_compound15.27632.27632.4 | controllerType=0 controllerNumber=1 scan=27632 | 59.7404 |
| 17809:3 | 2 | R.LSLHRPALEDLLLGSEAN[+2204.772]LTC[+57.021]TLTGLR.D | HexNAc(4)Hex(5)NeuAc(2) | C[+57], N[+2205] | 1723.4627 | 3 | 5168.3736 | 5168.3707 | 0 | 0.6 | 127 | Specific | 326.99 | 307.41 | 307.41 | 6.71 | 124 | >sp\|P01876\|IGHA1_HUMAN Immunoglobulin heavy constant alpha 1 OS=Homo sapiens OX=9606 GN=IGHA1 PE=1 SV=2 | 255 | 20201114_qwl_colo_milk_1ng_glyco_hcd203040_compound15.21293.21293.3 | controllerType=0 controllerNumber=1 scan=21293 | 47.1701 |
| 22864:4 | 2 | R.LSLHRPALEDLLLGSEAN[+1216.423]LTC[+57.021]TLTGLR.D | HexNAc(2)Hex(5) | C[+57], N[+1216] | 1045.7663 | 4 | 4180.0433 | 4180.0211 | 0 | 5.3 | 127 | Specific | 327.81 | 327.81 | 327.81 | 7.41 | 124 | >sp\|P01876\|IGHA1_HUMAN Immunoglobulin heavy constant alpha 1 OS=Homo sapiens OX=9606 GN=IGHA1 PE=1 SV=2 | 255 | 20201114_qwl_colo_milk_1ng_glyco_hcd203040_compound15.26571.26571.4 | controllerType=0 controllerNumber=1 scan=26571 | 57.7757 |
| 29763:4 | 2 | R.LSLHRPALEDLLLGSEAN[+1216.423]LTC[+57.021]TLTGLR.D | HexNAc(2)Hex(5) | C[+57], N[+1216] | 1045.7625 | 4 | 4180.0281 | 4180.0211 | 0 | 1.7 | 127 | Specific | 327.94 | 327.94 | 327.94 | 7.64 | 124 | >sp\|P01876\|IGHA1_HUMAN Immunoglobulin heavy constant alpha 1 OS=Homo sapiens OX=9606 GN=IGHA1 PE=1 SV=2 | 255 | 20201114_qwl_colo_milk_1ng_glyco_hcd203040_compound15.38089.38089.4 | controllerType=0 controllerNumber=1 scan=38089 | 88.6786 |
| 30198:4 | 2 | R.LSLHRPALEDLLLGSEAN[+1216.423]LTC[+57.021]TLTGLR.D | HexNAc(2)Hex(5) | C[+57], N[+1216] | 1045.7637 | 4 | 4180.033 | 4180.0211 | 0 | 2.8 | 127 | Specific | 336.95 | 336.95 | 336.95 | 7.87 | 124 | >sp\|P01876\|IGHA1_HUMAN Immunoglobulin heavy constant alpha 1 OS=Homo sapiens OX=9606 GN=IGHA1 PE=1 SV=2 | 255 | 20201114_qwl_colo_milk_1ng_glyco_hcd203040_compound15.38586.38586.2 | controllerType=0 controllerNumber=1 scan=38586 | 89.6794 |
| 25938:4 | 2 | R.LSLHRPALEDLLLGSEAN[+1216.423]LTC[+57.021]TLTGLR.D | HexNAc(2)Hex(5) | C[+57], N[+1216] | 1045.7652 | 4 | 4180.0389 | 4180.0211 | 0 | 4.2 | 127 | Specific | 340.27 | 340.27 | 340.27 | 7.84 | 124 | >sp\|P01876\|IGHA1_HUMAN Immunoglobulin heavy constant alpha 1 OS=Homo sapiens OX=9606 GN=IGHA1 PE=1 SV=2 | 255 | 20201114_qwl_colo_milk_1ng_glyco_hcd203040_compound15.30346.30346.4 | controllerType=0 controllerNumber=1 scan=30346 | 66.0731 |
| 15894:3 | 2 | R.LSLHRPALEDLLLGSEAN[+1257.449]LTC[+57.021]TLTGLR.D | HexNAc(3)Hex(4) | C[+57], N[+1257] | 1407.7009 | 3 | 4221.0881 | 4221.0477 | 0 | 9.6 | 127 | Specific | 343.95 | 279.22 | 279.22 | 6.27 | 124 | >sp\|P01876\|IGHA1_HUMAN Immunoglobulin heavy constant alpha 1 OS=Homo sapiens OX=9606 GN=IGHA1 PE=1 SV=2 | 255 | 20201114_qwl_colo_milk_1ng_glyco_hcd203040_compound15.19280.19280.3 | controllerType=0 controllerNumber=1 scan=19280 | 42.5392 |
| 21997:4 | 2 | R.LSLHRPALEDLLLGSEAN[+1501.555]LTC[+57.021]TLTGLR.D | HexNAc(5)Hex(3) | C[+57], N[+1502] | 1117.048 | 4 | 4465.1702 | 4465.1536 | 0 | 3.7 | 127 | Specific | 344.97 | 176.69 | 176.69 | 6.64 | 124 | >sp\|P01876\|IGHA1_HUMAN Immunoglobulin heavy constant alpha 1 OS=Homo sapiens OX=9606 GN=IGHA1 PE=1 SV=2 | 255 | 20201114_qwl_colo_milk_1ng_glyco_hcd203040_compound15.25667.25667.4 | controllerType=0 controllerNumber=1 scan=25667 | 56.1002 |
| 29928:4 | 2 | R.LSLHRPALEDLLLGSEAN[+1216.423]LTC[+57.021]TLTGLR.D | HexNAc(2)Hex(5) | C[+57], N[+1216] | 1045.7651 | 4 | 4180.0384 | 4180.0211 | 0 | 4.1 | 127 | Specific | 358.96 | 358.96 | 358.96 | 8.09 | 124 | >sp\|P01876\|IGHA1_HUMAN Immunoglobulin heavy constant alpha 1 OS=Homo sapiens OX=9606 GN=IGHA1 PE=1 SV=2 | 255 | 20201114_qwl_colo_milk_1ng_glyco_hcd203040_compound15.38271.38271.4 | controllerType=0 controllerNumber=1 scan=38271 | 89.0379 |
| 30207:4 | 2 | R.LSLHRPALEDLLLGSEAN[+1216.423]LTC[+57.021]TLTGLR.D | HexNAc(2)Hex(5) | C[+57], N[+1216] | 1045.7629 | 4 | 4180.0296 | 4180.0211 | 0 | 2 | 127 | Specific | 369.58 | 369.58 | 369.58 | 8.02 | 124 | >sp\|P01876\|IGHA1_HUMAN Immunoglobulin heavy constant alpha 1 OS=Homo sapiens OX=9606 GN=IGHA1 PE=1 SV=2 | 255 | 20201114_qwl_colo_milk_1ng_glyco_hcd203040_compound15.38599.38599.4 | controllerType=0 controllerNumber=1 scan=38599 | 89.7126 |
| 20787:4 | 2 | R.LSLHRPALEDLLLGSEAN[+1501.555]LTC[+57.021]TLTGLR.D | HexNAc(5)Hex(3) | C[+57], N[+1502] | 1117.0481 | 4 | 4465.1707 | 4465.1536 | 0 | 3.8 | 127 | Specific | 385.55 | 248.6 | 248.6 | 7.47 | 124 | >sp\|P01876\|IGHA1_HUMAN Immunoglobulin heavy constant alpha 1 OS=Homo sapiens OX=9606 GN=IGHA1 PE=1 SV=2 | 255 | 20201114_qwl_colo_milk_1ng_glyco_hcd203040_compound15.24402.24402.4 | controllerType=0 controllerNumber=1 scan=24402 | 53.629 |
| 16780:3 | 2 | R.LSLHRPALEDLLLGSEAN[+1913.677]LTC[+57.021]TLTGLR.D | HexNAc(4)Hex(5)NeuAc(1) | C[+57], N[+1914] | 1626.4306 | 3 | 4877.2772 | 4877.2753 | 0 | 0.4 | 127 | Specific | 389.33 | 389.33 | 389.33 | 8.82 | 124 | >sp\|P01876\|IGHA1_HUMAN Immunoglobulin heavy constant alpha 1 OS=Homo sapiens OX=9606 GN=IGHA1 PE=1 SV=2 | 255 | 20201114_qwl_colo_milk_1ng_glyco_hcd203040_compound15.20212.20212.6 | controllerType=0 controllerNumber=1 scan=20212 | 44.768 |
| 20139:4 | 2 | R.LSLHRPALEDLLLGSEAN[+1501.555]LTC[+57.021]TLTGLR.D | HexNAc(5)Hex(3) | C[+57], N[+1502] | 1117.0491 | 4 | 4465.1746 | 4465.1536 | 0 | 4.7 | 127 | Specific | 400.89 | 346.56 | 346.56 | 7.67 | 124 | >sp\|P01876\|IGHA1_HUMAN Immunoglobulin heavy constant alpha 1 OS=Homo sapiens OX=9606 GN=IGHA1 PE=1 SV=2 | 255 | 20201114_qwl_colo_milk_1ng_glyco_hcd203040_compound15.23726.23726.2 | controllerType=0 controllerNumber=1 scan=23726 | 52.2481 |
| 29851:4 | 2 | R.LSLHRPALEDLLLGSEAN[+1216.423]LTC[+57.021]TLTGLR.D | HexNAc(2)Hex(5) | C[+57], N[+1216] | 1045.7638 | 4 | 4180.0335 | 4180.0211 | 0 | 3 | 127 | Specific | 416.88 | 416.88 | 416.88 | 9.65 | 124 | >sp\|P01876\|IGHA1_HUMAN Immunoglobulin heavy constant alpha 1 OS=Homo sapiens OX=9606 GN=IGHA1 PE=1 SV=2 | 255 | 20201114_qwl_colo_milk_1ng_glyco_hcd203040_compound15.38187.38187.3 | controllerType=0 controllerNumber=1 scan=38187 | 88.8773 |
| 19434:4 | 2 | R.LSLHRPALEDLLLGSEAN[+1501.555]LTC[+57.021]TLTGLR.D | HexNAc(5)Hex(3) | C[+57], N[+1502] | 1117.0525 | 4 | 4465.1883 | 4465.1536 | 0 | 7.8 | 127 | Specific | 424.96 | 324.64 | 324.64 | 7.71 | 124 | >sp\|P01876\|IGHA1_HUMAN Immunoglobulin heavy constant alpha 1 OS=Homo sapiens OX=9606 GN=IGHA1 PE=1 SV=2 | 255 | 20201114_qwl_colo_milk_1ng_glyco_hcd203040_compound15.22990.22990.4 | controllerType=0 controllerNumber=1 scan=22990 | 50.7522 |
| 16579:4 | 2 | R.LSLHRPALEDLLLGSEAN[+1054.370]LTC[+57.021]TLTGLR.D | HexNAc(2)Hex(4) | C[+57], N[+1054] | 1005.2529 | 4 | 4017.9898 | 4017.9683 | 0 | 5.3 | 127 | Specific | 434.4 | 434.4 | 434.4 | 7.83 | 124 | >sp\|P01876\|IGHA1_HUMAN Immunoglobulin heavy constant alpha 1 OS=Homo sapiens OX=9606 GN=IGHA1 PE=1 SV=2 | 255 | 20201114_qwl_colo_milk_1ng_glyco_hcd203040_compound15.19998.19998.4 | controllerType=0 controllerNumber=1 scan=19998 | 44.2331 |
| 16371:4 | 2 | R.LSLHRPALEDLLLGSEAN[+1501.555]LTC[+57.021]TLTGLR.D | HexNAc(5)Hex(3) | C[+57], N[+1502] | 1117.0495 | 4 | 4465.1761 | 4465.1536 | 0 | 5 | 127 | Specific | 434.95 | 434.95 | 434.95 | 7.83 | 124 | >sp\|P01876\|IGHA1_HUMAN Immunoglobulin heavy constant alpha 1 OS=Homo sapiens OX=9606 GN=IGHA1 PE=1 SV=2 | 255 | 20201114_qwl_colo_milk_1ng_glyco_hcd203040_compound15.19782.19782.2 | controllerType=0 controllerNumber=1 scan=19782 | 43.7867 |
| 16799:3 | 2 | R.LSLHRPALEDLLLGSEAN[+1710.598]LTC[+57.021]TLTGLR.D | HexNAc(3)Hex(5)NeuAc(1) | C[+57], N[+1711] | 1558.7378 | 3 | 4674.199 | 4674.1959 | 0 | 0.7 | 127 | Specific | 447.78 | 447.78 | 447.78 | 9.05 | 124 | >sp\|P01876\|IGHA1_HUMAN Immunoglobulin heavy constant alpha 1 OS=Homo sapiens OX=9606 GN=IGHA1 PE=1 SV=2 | 255 | 20201114_qwl_colo_milk_1ng_glyco_hcd203040_compound15.20232.20232.3 | controllerType=0 controllerNumber=1 scan=20232 | 44.8093 |
| 17806:4 | 2 | R.LSLHRPALEDLLLGSEAN[+2204.772]LTC[+57.021]TLTGLR.D | HexNAc(4)Hex(5)NeuAc(2) | C[+57], N[+2205] | 1292.8499 | 4 | 5168.3776 | 5168.3707 | 0 | 1.3 | 127 | Specific | 449.41 | 439.22 | 439.22 | 9.77 | 124 | >sp\|P01876\|IGHA1_HUMAN Immunoglobulin heavy constant alpha 1 OS=Homo sapiens OX=9606 GN=IGHA1 PE=1 SV=2 | 255 | 20201114_qwl_colo_milk_1ng_glyco_hcd203040_compound15.21290.21290.4 | controllerType=0 controllerNumber=1 scan=21290 | 47.1649 |
| 16758:4 | 2 | R.LSLHRPALEDLLLGSEAN[+2116.756]LTC[+57.021]TLTGLR.D | HexNAc(5)Hex(5)NeuAc(1) | C[+57], N[+2117] | 1270.8503 | 4 | 5080.3796 | 5080.3547 | 0 | 4.9 | 127 | Specific | 454.09 | 454.09 | 454.09 | 9.53 | 124 | >sp\|P01876\|IGHA1_HUMAN Immunoglobulin heavy constant alpha 1 OS=Homo sapiens OX=9606 GN=IGHA1 PE=1 SV=2 | 255 | 20201114_qwl_colo_milk_1ng_glyco_hcd203040_compound15.20189.20189.4 | controllerType=0 controllerNumber=1 scan=20189 | 44.7162 |

Table S4. Cont.

| Query #:z | Protein Rank | Glycopeptide | Glycan | Modification Type(s) | Observed m/z | z | Observed (M+H) | Calc. mass (M+H) | Off-by-x error | Mass error (ppm) | StartPosition | Cleavage | Score | Delta | Delta Mod | \|Log Prob\| | # of unique peptides | Protein Name | Protein DB number | Comment | Scan # | Scan Time |
|---|---|---|---|---|---|---|---|---|---|---|---|---|---|---|---|---|---|---|---|---|---|---|
| 15531:3 | 2 | R.LSLHRPALEDLLLGSEAN[+1216.423]LTC[+57.021]TLTGLR.D | HexNAc(2)Hex(5) | C[+57], N[+1216] | 1394.0138 | 3 | 4180.0268 | 4180.0211 | 0 | 1.4 | 127 | Specific | 454.93 | 454.93 | 454.93 | 9.1 | 124 | >sp\|P01876\|IGHA1_HUMAN Immunoglobulin heavy constant alpha 1 OS=Homo sapiens OX=9606 GN=IGHA1 PE=1 SV=2 | 255 | 20201114_qwl_colo_milk_1ng_glyco_hcd203040_compound15.18897.18897.3 | controllerType=0 controllerNumber=1 scan=18897 | 41.6084 |
| 23373:4 | 2 | R.LSLHRPALEDLLLGSEAN[+1501.555]LTC[+57.021]TLTGLR.D | HexNAc(5)Hex(3) | C[+57], N[+1502] | 1117.0496 | 4 | 4465.1766 | 4465.1536 | 0 | 5.1 | 127 | Specific | 476.24 | 476.24 | 476.24 | 9.6 | 124 | >sp\|P01876\|IGHA1_HUMAN Immunoglobulin heavy constant alpha 1 OS=Homo sapiens OX=9606 GN=IGHA1 PE=1 SV=2 | 255 | 20201114_qwl_colo_milk_1ng_glyco_hcd203040_compound15.27103.27103.2 | controllerType=0 controllerNumber=1 scan=27103 | 58.7411 |
| 16802:4 | 2 | R.LSLHRPALEDLLLGSEAN[+1913.677]LTC[+57.021]TLTGLR.D | HexNAc(4)Hex(5)NeuAc(1) | C[+57], N[+1914] | 1220.0755 | 4 | 4877.2801 | 4877.2753 | 0 | 1 | 127 | Specific | 479.57 | 407.43 | 407.43 | 9.78 | 124 | >sp\|P01876\|IGHA1_HUMAN Immunoglobulin heavy constant alpha 1 OS=Homo sapiens OX=9606 GN=IGHA1 PE=1 SV=2 | 255 | 20201114_qwl_colo_milk_1ng_glyco_hcd203040_compound15.20235.20235.4 | controllerType=0 controllerNumber=1 scan=20235 | 44.8155 |
| 16122:3 | 2 | R.LSLHRPALEDLLLGSEAN[+1216.423]LTC[+57.021]TLTGLR.D | HexNAc(2)Hex(5) | C[+57], N[+1216] | 1394.0145 | 3 | 4180.0289 | 4180.0211 | 0 | 1.9 | 127 | Specific | 497.91 | 410.88 | 410.88 | 9.25 | 124 | >sp\|P01876\|IGHA1_HUMAN Immunoglobulin heavy constant alpha 1 OS=Homo sapiens OX=9606 GN=IGHA1 PE=1 SV=2 | 255 | 20201114_qwl_colo_milk_1ng_glyco_hcd203040_compound15.19522.19522.3 | controllerType=0 controllerNumber=1 scan=19522 | 43.223 |
| 16800:4 | 2 | R.LSLHRPALEDLLLGSEAN[+1913.677]LTC[+57.021]TLTGLR.D | HexNAc(4)Hex(5)NeuAc(1) | C[+57], N[+1914] | 1220.0755 | 4 | 4877.2801 | 4877.2753 | 0 | 1 | 127 | Specific | 498.41 | 395.85 | 395.85 | 9.86 | 124 | >sp\|P01876\|IGHA1_HUMAN Immunoglobulin heavy constant alpha 1 OS=Homo sapiens OX=9606 GN=IGHA1 PE=1 SV=2 | 255 | 20201114_qwl_colo_milk_1ng_glyco_hcd203040_compound15.20233.20233.4 | controllerType=0 controllerNumber=1 scan=20233 | 44.8105 |
| 16124:3 | 2 | R.LSLHRPALEDLLLGSEAN[+1216.423]LTC[+57.021]TLTGLR.D | HexNAc(2)Hex(5) | C[+57], N[+1216] | 1394.0145 | 3 | 4180.0289 | 4180.0211 | 0 | 1.9 | 127 | Specific | 498.66 | 410.18 | 410.18 | 9.25 | 124 | >sp\|P01876\|IGHA1_HUMAN Immunoglobulin heavy constant alpha 1 OS=Homo sapiens OX=9606 GN=IGHA1 PE=1 SV=2 | 255 | 20201114_qwl_colo_milk_1ng_glyco_hcd203040_compound15.19524.19524.3 | controllerType=0 controllerNumber=1 scan=19524 | 43.2283 |
| 16756:4 | 2 | R.LSLHRPALEDLLLGSEAN[+2116.756]LTC[+57.021]TLTGLR.D | HexNAc(5)Hex(5)NeuAc(1) | C[+57], N[+2117] | 1270.8503 | 4 | 5080.3796 | 5080.3547 | 0 | 4.9 | 127 | Specific | 500.56 | 500.56 | 500.56 | 9.68 | 124 | >sp\|P01876\|IGHA1_HUMAN Immunoglobulin heavy constant alpha 1 OS=Homo sapiens OX=9606 GN=IGHA1 PE=1 SV=2 | 255 | 20201114_qwl_colo_milk_1ng_glyco_hcd203040_compound15.20187.20187.4 | controllerType=0 controllerNumber=1 scan=20187 | 44.7114 |
| 16663:4 | 2 | R.LSLHRPALEDLLLGSEAN[+1378.476]LTC[+57.021]TLTGLR.D | HexNAc(2)Hex(6) | C[+57], N[+1378] | 1086.2818 | 4 | 4342.1053 | 4342.074 | 0 | 7.2 | 127 | Specific | 527.49 | 527.49 | 527.49 | 9.7 | 124 | >sp\|P01876\|IGHA1_HUMAN Immunoglobulin heavy constant alpha 1 OS=Homo sapiens OX=9606 GN=IGHA1 PE=1 SV=2 | 255 | 20201114_qwl_colo_milk_1ng_glyco_hcd203040_compound15.20087.20087.3 | controllerType=0 controllerNumber=1 scan=20087 | 44.4186 |
| 29765:4 | 2 | R.LSLHRPALEDLLLGSEAN[+1216.423]LTC[+57.021]TLTGLR.D | HexNAc(2)Hex(5) | C[+57], N[+1216] | 1045.7625 | 4 | 4180.0281 | 4180.0211 | 0 | 1.7 | 127 | Specific | 538.83 | 538.83 | 538.83 | 10.19 | 124 | >sp\|P01876\|IGHA1_HUMAN Immunoglobulin heavy constant alpha 1 OS=Homo sapiens OX=9606 GN=IGHA1 PE=1 SV=2 | 255 | 20201114_qwl_colo_milk_1ng_glyco_hcd203040_compound15.38091.38091.4 | controllerType=0 controllerNumber=1 scan=38091 | 88.6827 |
| 29930:4 | 2 | R.LSLHRPALEDLLLGSEAN[+1216.423]LTC[+57.021]TLTGLR.D | HexNAc(2)Hex(5) | C[+57], N[+1216] | 1045.7651 | 4 | 4180.0384 | 4180.0211 | 0 | 4.1 | 127 | Specific | 575.24 | 575.24 | 575.24 | 10.35 | 124 | >sp\|P01876\|IGHA1_HUMAN Immunoglobulin heavy constant alpha 1 OS=Homo sapiens OX=9606 GN=IGHA1 PE=1 SV=2 | 255 | 20201114_qwl_colo_milk_1ng_glyco_hcd203040_compound15.38273.38273.4 | controllerType=0 controllerNumber=1 scan=38273 | 89.0417 |
| 15841:3 | 2 | R.LSLHRPALEDLLLGSEAN[+1702.581]LTC[+57.021]TLTGLR.D | HexNAc(2)Hex(8) | C[+57], N[+1703] | 1556.0666 | 3 | 4666.1852 | 4666.1796 | 0 | 1.2 | 127 | Specific | 576.49 | 569.88 | 569.88 | 9.56 | 124 | >sp\|P01876\|IGHA1_HUMAN Immunoglobulin heavy constant alpha 1 OS=Homo sapiens OX=9606 GN=IGHA1 PE=1 SV=2 | 255 | 20201114_qwl_colo_milk_1ng_glyco_hcd203040_compound15.19224.19224.3 | controllerType=0 controllerNumber=1 scan=19224 | 42.3654 |
| 15857:4 | 2 | R.LSLHRPALEDLLLGSEAN[+1581.555]LTC[+57.021]TLTGLR.D | HexNAc(3)Hex(6) | C[+57], N[+1582] | 1137.0545 | 4 | 4545.1961 | 4545.1533 | 0 | 9.4 | 127 | Specific | 594.47 | 594.47 | 594.47 | 11.1 | 124 | >sp\|P01876\|IGHA1_HUMAN Immunoglobulin heavy constant alpha 1 OS=Homo sapiens OX=9606 GN=IGHA1 PE=1 SV=2 | 255 | 20201114_qwl_colo_milk_1ng_glyco_hcd203040_compound15.19241.19241.4 | controllerType=0 controllerNumber=1 scan=19241 | 42.4186 |
| 16368:4 | 2 | R.LSLHRPALEDLLLGSEAN[+1501.555]LTC[+57.021]TLTGLR.D | HexNAc(5)Hex(3) | C[+57], N[+1502] | 1117.0495 | 4 | 4465.1761 | 4465.1536 | 0 | 5 | 127 | Specific | 609.82 | 534.58 | 534.58 | 12.36 | 124 | >sp\|P01876\|IGHA1_HUMAN Immunoglobulin heavy constant alpha 1 OS=Homo sapiens OX=9606 GN=IGHA1 PE=1 SV=2 | 255 | 20201114_qwl_colo_milk_1ng_glyco_hcd203040_compound15.19779.19779.2 | controllerType=0 controllerNumber=1 scan=19779 | 43.7816 |
| 15855:3 | 2 | R.LSLHRPALEDLLLGSEAN[+1622.582]LTC[+57.021]TLTGLR.D | HexNAc(4)Hex(5) | C[+57], N[+1623] | 1529.4124 | 3 | 4586.2227 | 4586.1799 | 0 | 9.3 | 127 | Specific | 618.17 | 574.49 | 574.49 | 11.44 | 124 | >sp\|P01876\|IGHA1_HUMAN Immunoglobulin heavy constant alpha 1 OS=Homo sapiens OX=9606 GN=IGHA1 PE=1 SV=2 | 255 | 20201114_qwl_colo_milk_1ng_glyco_hcd203040_compound15.19239.19239.3 | controllerType=0 controllerNumber=1 scan=19239 | 42.4163 |
| 15746:4 | 2 | R.LSLHRPALEDLLLGSEAN[+1864.634]LTC[+57.021]TLTGLR.D | HexNAc(2)Hex(9) | C[+57], N[+1865] | 1207.8217 | 4 | 4828.265 | 4828.2324 | 0 | 6.7 | 127 | Specific | 619.42 | 619.42 | 619.42 | 12.31 | 124 | >sp\|P01876\|IGHA1_HUMAN Immunoglobulin heavy constant alpha 1 OS=Homo sapiens OX=9606 GN=IGHA1 PE=1 SV=2 | 255 | 20201114_qwl_colo_milk_1ng_glyco_hcd203040_compound15.19124.19124.2 | controllerType=0 controllerNumber=1 scan=19124 | 42.1108 |
| 15858:3 | 2 | R.LSLHRPALEDLLLGSEAN[+1622.582]LTC[+57.021]TLTGLR.D | HexNAc(4)Hex(5) | C[+57], N[+1623] | 1529.4124 | 3 | 4586.2227 | 4586.1799 | 0 | 9.3 | 127 | Specific | 631.97 | 546.77 | 546.77 | 11.44 | 124 | >sp\|P01876\|IGHA1_HUMAN Immunoglobulin heavy constant alpha 1 OS=Homo sapiens OX=9606 GN=IGHA1 PE=1 SV=2 | 255 | 20201114_qwl_colo_milk_1ng_glyco_hcd203040_compound15.19242.19242.3 | controllerType=0 controllerNumber=1 scan=19242 | 42.4247 |
| 16783:3 | 2 | R.LSLHRPALEDLLLGSEAN[+1913.677]LTC[+57.021]TLTGLR.D | HexNAc(4)Hex(5)NeuAc(1) | C[+57], N[+1914] | 1626.4306 | 3 | 4877.2772 | 4877.2753 | 0 | 0.4 | 127 | Specific | 655.04 | 550.83 | 550.83 | 12.88 | 124 | >sp\|P01876\|IGHA1_HUMAN Immunoglobulin heavy constant alpha 1 OS=Homo sapiens OX=9606 GN=IGHA1 PE=1 SV=2 | 255 | 20201114_qwl_colo_milk_1ng_glyco_hcd203040_compound15.20215.20215.6 | controllerType=0 controllerNumber=1 scan=20215 | 44.7739 |
| 15837:3 | 2 | R.LSLHRPALEDLLLGSEAN[+1622.582]LTC[+57.021]TLTGLR.D | HexNAc(4)Hex(5) | C[+57], N[+1623] | 1529.4003 | 3 | 4586.1864 | 4586.1799 | 0 | 1.4 | 127 | Specific | 678.97 | 578.1 | 578.1 | 12.93 | 124 | >sp\|P01876\|IGHA1_HUMAN Immunoglobulin heavy constant alpha 1 OS=Homo sapiens OX=9606 GN=IGHA1 PE=1 SV=2 | 255 | 20201114_qwl_colo_milk_1ng_glyco_hcd203040_compound15.19220.19220.3 | controllerType=0 controllerNumber=1 scan=19220 | 42.3507 |
| 15833:3 | 2 | R.LSLHRPALEDLLLGSEAN[+1622.582]LTC[+57.021]TLTGLR.D | HexNAc(4)Hex(5) | C[+57], N[+1623] | 1529.4003 | 3 | 4586.1864 | 4586.1799 | 0 | 1.4 | 127 | Specific | 688.64 | 608.99 | 608.99 | 12.93 | 124 | >sp\|P01876\|IGHA1_HUMAN Immunoglobulin heavy constant alpha 1 OS=Homo sapiens OX=9606 GN=IGHA1 PE=1 SV=2 | 255 | 20201114_qwl_colo_milk_1ng_glyco_hcd203040_compound15.19216.19216.3 | controllerType=0 controllerNumber=1 scan=19216 | 42.334 |
| 15923:3 | 2 | R.LSLHRPALEDLLLGSEAN[+1095.397]LTC[+57.021]TLTGLR.D | HexNAc(3)Hex(3) | C[+57], N[+1095] | 1353.6828 | 3 | 4059.0338 | 4058.9949 | 0 | 9.6 | 127 | Specific | 690.69 | 652.57 | 652.57 | 12.44 | 124 | >sp\|P01876\|IGHA1_HUMAN Immunoglobulin heavy constant alpha 1 OS=Homo sapiens OX=9606 GN=IGHA1 PE=1 SV=2 | 255 | 20201114_qwl_colo_milk_1ng_glyco_hcd203040_compound15.19311.19311.3 | controllerType=0 controllerNumber=1 scan=19311 | 42.6095 |
| 16798:4 | 2 | R.LSLHRPALEDLLLGSEAN[+1710.598]LTC[+57.021]TLTGLR.D | HexNAc(3)Hex(5)NeuAc(1) | C[+57], N[+1711] | 1169.3058 | 4 | 4674.2015 | 4674.1959 | 0 | 1.2 | 127 | Specific | 691.26 | 638.33 | 638.33 | 13.59 | 124 | >sp\|P01876\|IGHA1_HUMAN Immunoglobulin heavy constant alpha 1 OS=Homo sapiens OX=9606 GN=IGHA1 PE=1 SV=2 | 255 | 20201114_qwl_colo_milk_1ng_glyco_hcd203040_compound15.20231.20231.4 | controllerType=0 controllerNumber=1 scan=20231 | 44.8086 |
| 15882:4 | 2 | R.LSLHRPALEDLLLGSEAN[+1501.555]LTC[+57.021]TLTGLR.D | HexNAc(5)Hex(3) | C[+57], N[+1502] | 1117.0456 | 4 | 4465.1605 | 4465.1536 | 0 | 1.5 | 127 | Specific | 702.33 | 702.33 | 702.33 | 14.59 | 124 | >sp\|P01876\|IGHA1_HUMAN Immunoglobulin heavy constant alpha 1 OS=Homo sapiens OX=9606 GN=IGHA1 PE=1 SV=2 | 255 | 20201114_qwl_colo_milk_1ng_glyco_hcd203040_compound15.19268.19268.4 | controllerType=0 controllerNumber=1 scan=19268 | 42.5038 |
| 15891:3 | 2 | R.LSLHRPALEDLLLGSEAN[+1257.449]LTC[+57.021]TLTGLR.D | HexNAc(3)Hex(4) | C[+57], N[+1257] | 1407.7009 | 3 | 4221.0881 | 4221.0477 | 0 | 9.6 | 127 | Specific | 704.92 | 704.92 | 704.92 | 13.44 | 124 | >sp\|P01876\|IGHA1_HUMAN Immunoglobulin heavy constant alpha 1 OS=Homo sapiens OX=9606 GN=IGHA1 PE=1 SV=2 | 255 | 20201114_qwl_colo_milk_1ng_glyco_hcd203040_compound15.19277.19277.3 | controllerType=0 controllerNumber=1 scan=19277 | 42.5303 |
| 15958:3 | 2 | R.LSLHRPALEDLLLGSEAN[+892.317]LTC[+57.021]TLTGLR.D | HexNAc(2)Hex(3) | C[+57], N[+892] | 1285.9802 | 3 | 3855.9259 | 3855.9155 | 0 | 2.7 | 127 | Specific | 742.68 | 742.68 | 742.68 | 14.03 | 124 | >sp\|P01876\|IGHA1_HUMAN Immunoglobulin heavy constant alpha 1 OS=Homo sapiens OX=9606 GN=IGHA1 PE=1 SV=2 | 255 | 20201114_qwl_colo_milk_1ng_glyco_hcd203040_compound15.19348.19348.3 | controllerType=0 controllerNumber=1 scan=19348 | 42.7345 |
| 15829:4 | 2 | R.LSLHRPALEDLLLGSEAN[+1622.582]LTC[+57.021]TLTGLR.D | HexNAc(4)Hex(5) | C[+57], N[+1623] | 1147.3027 | 4 | 4586.1888 | 4586.1799 | 0 | 1.9 | 127 | Specific | 743.66 | 724.14 | 724.14 | 14.59 | 124 | >sp\|P01876\|IGHA1_HUMAN Immunoglobulin heavy constant alpha 1 OS=Homo sapiens OX=9606 GN=IGHA1 PE=1 SV=2 | 255 | 20201114_qwl_colo_milk_1ng_glyco_hcd203040_compound15.19212.19212.4 | controllerType=0 controllerNumber=1 scan=19212 | 42.3207 |
| 17808:4 | 2 | R.LSLHRPALEDLLLGSEAN[+2204.772]LTC[+57.021]TLTGLR.D | HexNAc(4)Hex(5)NeuAc(2) | C[+57], N[+2205] | 1292.8499 | 4 | 5168.3776 | 5168.3707 | 0 | 1.3 | 127 | Specific | 754.36 | 695.68 | 695.68 | 15.59 | 124 | >sp\|P01876\|IGHA1_HUMAN Immunoglobulin heavy constant alpha 1 OS=Homo sapiens OX=9606 GN=IGHA1 PE=1 SV=2 | 255 | 20201114_qwl_colo_milk_1ng_glyco_hcd203040_compound15.21292.21292.4 | controllerType=0 controllerNumber=1 scan=21292 | 47.169 |
| 15956:3 | 2 | R.LSLHRPALEDLLLGSEAN[+892.317]LTC[+57.021]TLTGLR.D | HexNAc(2)Hex(3) | C[+57], N[+892] | 1285.9802 | 3 | 3855.9259 | 3855.9155 | 0 | 2.7 | 127 | Specific | 773.48 | 773.48 | 773.48 | 15.03 | 124 | >sp\|P01876\|IGHA1_HUMAN Immunoglobulin heavy constant alpha 1 OS=Homo sapiens OX=9606 GN=IGHA1 PE=1 SV=2 | 255 | 20201114_qwl_colo_milk_1ng_glyco_hcd203040_compound15.19346.19346.3 | controllerType=0 controllerNumber=1 scan=19346 | 42.7245 |
| 15842:3 | 2 | R.LSLHRPALEDLLLGSEAN[+1378.476]LTC[+57.021]TLTGLR.D | HexNAc(2)Hex(6) | C[+57], N[+1378] | 1448.0323 | 3 | 4342.0823 | 4342.074 | 0 | 1.9 | 127 | Specific | 782.09 | 703.32 | 703.32 | 14.93 | 124 | >sp\|P01876\|IGHA1_HUMAN Immunoglobulin heavy constant alpha 1 OS=Homo sapiens OX=9606 GN=IGHA1 PE=1 SV=2 | 255 | 20201114_qwl_colo_milk_1ng_glyco_hcd203040_compound15.19225.19225.3 | controllerType=0 controllerNumber=1 scan=19225 | 42.3666 |
| 15844:3 | 2 | R.LSLHRPALEDLLLGSEAN[+1378.476]LTC[+57.021]TLTGLR.D | HexNAc(2)Hex(6) | C[+57], N[+1378] | 1448.0323 | 3 | 4342.0823 | 4342.074 | 0 | 1.9 | 127 | Specific | 783.34 | 703.54 | 703.54 | 14.93 | 124 | >sp\|P01876\|IGHA1_HUMAN Immunoglobulin heavy constant alpha 1 OS=Homo sapiens OX=9606 GN=IGHA1 PE=1 SV=2 | 255 | 20201114_qwl_colo_milk_1ng_glyco_hcd203040_compound15.19227.19227.3 | controllerType=0 controllerNumber=1 scan=19227 | 42.3808 |
| 15931:4 | 2 | R.LSLHRPALEDLLLGSEAN[+1054.370]LTC[+57.021]TLTGLR.D | HexNAc(2)Hex(4) | C[+57], N[+1054] | 1005.2556 | 4 | 4018.0006 | 4017.9683 | 0 | 8 | 127 | Specific | 790.88 | 790.88 | 790.88 | 15.24 | 124 | >sp\|P01876\|IGHA1_HUMAN Immunoglobulin heavy constant alpha 1 OS=Homo sapiens OX=9606 GN=IGHA1 PE=1 SV=2 | 255 | 20201114_qwl_colo_milk_1ng_glyco_hcd203040_compound15.19320.19320.4 | controllerType=0 controllerNumber=1 scan=19320 | 42.6308 |
| 15860:4 | 2 | R.LSLHRPALEDLLLGSEAN[+1581.555]LTC[+57.021]TLTGLR.D | HexNAc(3)Hex(6) | C[+57], N[+1582] | 1137.0545 | 4 | 4545.1961 | 4545.1533 | 0 | 9.4 | 127 | Specific | 820.44 | 704.41 | 704.41 | 16.1 | 124 | >sp\|P01876\|IGHA1_HUMAN Immunoglobulin heavy constant alpha 1 OS=Homo sapiens OX=9606 GN=IGHA1 PE=1 SV=2 | 255 | 20201114_qwl_colo_milk_1ng_glyco_hcd203040_compound15.19244.19244.4 | controllerType=0 controllerNumber=1 scan=19244 | 42.4297 |
| 15843:3 | 2 | R.LSLHRPALEDLLLGSEAN[+1702.581]LTC[+57.021]TLTGLR.D | HexNAc(2)Hex(8) | C[+57], N[+1703] | 1556.0666 | 3 | 4666.1852 | 4666.1796 | 0 | 1.2 | 127 | Specific | 832.68 | 808.65 | 808.65 | 15.88 | 124 | >sp\|P01876\|IGHA1_HUMAN Immunoglobulin heavy constant alpha 1 OS=Homo sapiens OX=9606 GN=IGHA1 PE=1 SV=2 | 255 | 20201114_qwl_colo_milk_1ng_glyco_hcd203040_compound15.19226.19226.3 | controllerType=0 controllerNumber=1 scan=19226 | 42.3693 |
| 15918:4 | 2 | R.LSLHRPALEDLLLGSEAN[+1095.397]LTC[+57.021]TLTGLR.D | HexNAc(3)Hex(3) | C[+57], N[+1095] | 1015.5137 | 4 | 4059.033 | 4058.9949 | 0 | 9.4 | 127 | Specific | 837.81 | 837.81 | 837.81 | 16.1 | 124 | >sp\|P01876\|IGHA1_HUMAN Immunoglobulin heavy constant alpha 1 OS=Homo sapiens OX=9606 GN=IGHA1 PE=1 SV=2 | 255 | 20201114_qwl_colo_milk_1ng_glyco_hcd203040_compound15.19306.19306.4 | controllerType=0 controllerNumber=1 scan=19306 | 42.5972 |
| 15944:4 | 2 | R.LSLHRPALEDLLLGSEAN[+892.317]LTC[+57.021]TLTGLR.D | HexNAc(2)Hex(3) | C[+57], N[+892] | 964.7369 | 4 | 3855.9256 | 3855.9155 | 0 | 2.6 | 127 | Specific | 841.19 | 841.19 | 841.19 | 16.7 | 124 | >sp\|P01876\|IGHA1_HUMAN Immunoglobulin heavy constant alpha 1 OS=Homo sapiens OX=9606 GN=IGHA1 PE=1 SV=2 | 255 | 20201114_qwl_colo_milk_1ng_glyco_hcd203040_compound15.19334.19334.4 | controllerType=0 controllerNumber=1 scan=19334 | 42.6845 |
| 16143:4 | 2 | R.LSLHRPALEDLLLGSEAN[+1298.476]LTC[+57.021]TLTGLR.D | HexNAc(4)Hex(3) | C[+57], N[+1298] | 1066.2745 | 4 | 4262.076 | 4262.0742 | 0 | 0.4 | 127 | Specific | 841.9 | 841.9 | 841.9 | 16.54 | 124 | >sp\|P01876\|IGHA1_HUMAN Immunoglobulin heavy constant alpha 1 OS=Homo sapiens OX=9606 GN=IGHA1 PE=1 SV=2 | 255 | 20201114_qwl_colo_milk_1ng_glyco_hcd203040_compound15.19544.19544.3 | controllerType=0 controllerNumber=1 scan=19544 | 43.2686 |
| 16796:4 | 2 | R.LSLHRPALEDLLLGSEAN[+1710.598]LTC[+57.021]TLTGLR.D | HexNAc(3)Hex(5)NeuAc(1) | C[+57], N[+1711] | 1169.3058 | 4 | 4674.2015 | 4674.1959 | 0 | 1.2 | 127 | Specific | 842.55 | 842.55 | 842.55 | 16.59 | 124 | >sp\|P01876\|IGHA1_HUMAN Immunoglobulin heavy constant alpha 1 OS=Homo sapiens OX=9606 GN=IGHA1 PE=1 SV=2 | 255 | 20201114_qwl_colo_milk_1ng_glyco_hcd203040_compound15.20229.20229.4 | controllerType=0 controllerNumber=1 scan=20229 | 44.8002 |
| 15933:4 | 2 | R.LSLHRPALEDLLLGSEAN[+1054.370]LTC[+57.021]TLTGLR.D | HexNAc(2)Hex(4) | C[+57], N[+1054] | 1005.2556 | 4 | 4018.0006 | 4017.9683 | 0 | 8 | 127 | Specific | 882.24 | 882.24 | 882.24 | 17.24 | 124 | >sp\|P01876\|IGHA1_HUMAN Immunoglobulin heavy constant alpha 1 OS=Homo sapiens OX=9606 GN=IGHA1 PE=1 SV=2 | 255 | 20201114_qwl_colo_milk_1ng_glyco_hcd203040_compound15.19322.19322.4 | controllerType=0 controllerNumber=1 scan=19322 | 42.6482 |
| 15831:4 | 2 | R.LSLHRPALEDLLLGSEAN[+1622.582]LTC[+57.021]TLTGLR.D | HexNAc(4)Hex(5) | C[+57], N[+1623] | 1147.3027 | 4 | 4586.1888 | 4586.1799 | 0 | 1.9 | 127 | Specific | 884.81 | 747.63 | 747.63 | 17.59 | 124 | >sp\|P01876\|IGHA1_HUMAN Immunoglobulin heavy constant alpha 1 OS=Homo sapiens OX=9606 GN=IGHA1 PE=1 SV=2 | 255 | 20201114_qwl_colo_milk_1ng_glyco_hcd203040_compound15.19214.19214.4 | controllerType=0 controllerNumber=1 scan=19214 | 42.3288 |
| 15802:4 | 2 | R.LSLHRPALEDLLLGSEAN[+1702.581]LTC[+57.021]TLTGLR.D | HexNAc(2)Hex(8) | C[+57], N[+1703] | 1167.3028 | 4 | 4666.1893 | 4666.1796 | 0 | 2.1 | 127 | Specific | 892.37 | 892.37 | 892.37 | 17.59 | 124 | >sp\|P01876\|IGHA1_HUMAN Immunoglobulin heavy constant alpha 1 OS=Homo sapiens OX=9606 GN=IGHA1 PE=1 SV=2 | 255 | 20201114_qwl_colo_milk_1ng_glyco_hcd203040_compound15.19183.19183.4 | controllerType=0 controllerNumber=1 scan=19183 | 42.2539 |
| 15875:4 | 2 | R.LSLHRPALEDLLLGSEAN[+1257.449]LTC[+57.021]TLTGLR.D | HexNAc(3)Hex(4) | C[+57], N[+1257] | 1056.0236 | 4 | 4221.0726 | 4221.0477 | 0 | 5.9 | 127 | Specific | 922.5 | 904.64 | 904.64 | 18.17 | 124 | >sp\|P01876\|IGHA1_HUMAN Immunoglobulin heavy constant alpha 1 OS=Homo sapiens OX=9606 GN=IGHA1 PE=1 SV=2 | 255 | 20201114_qwl_colo_milk_1ng_glyco_hcd203040_compound15.19260.19260.4 | controllerType=0 controllerNumber=1 scan=19260 | 42.472 |
| 15834:4 | 2 | R.LSLHRPALEDLLLGSEAN[+1702.581]LTC[+57.021]TLTGLR.D | HexNAc(2)Hex(8) | C[+57], N[+1703] | 1167.3028 | 4 | 4666.1893 | 4666.1796 | 0 | 2.1 | 127 | Specific | 950.9 | 950.9 | 950.9 | 19.59 | 124 | >sp\|P01876\|IGHA1_HUMAN Immunoglobulin heavy constant alpha 1 OS=Homo sapiens OX=9606 GN=IGHA1 PE=1 SV=2 | 255 | 20201114_qwl_colo_milk_1ng_glyco_hcd203040_compound15.19217.19217.4 | controllerType=0 controllerNumber=1 scan=19217 | 42.3351 |
| 15805:4 | 2 | R.LSLHRPALEDLLLGSEAN[+1702.581]LTC[+57.021]TLTGLR.D | HexNAc(2)Hex(8) | C[+57], N[+1703] | 1167.3028 | 4 | 4666.1893 | 4666.1796 | 0 | 2.1 | 127 | Specific | 951.45 | 951.45 | 951.45 | 19.59 | 124 | >sp\|P01876\|IGHA1_HUMAN Immunoglobulin heavy constant alpha 1 OS=Homo sapiens OX=9606 GN=IGHA1 PE=1 SV=2 | 255 | 20201114_qwl_colo_milk_1ng_glyco_hcd203040_compound15.19186.19186.4 | controllerType=0 controllerNumber=1 scan=19186 | 42.2635 |
| 15835:4 | 2 | R.LSLHRPALEDLLLGSEAN[+1378.476]LTC[+57.021]TLTGLR.D | HexNAc(2)Hex(6) | C[+57], N[+1378] | 1086.2754 | 4 | 4342.0799 | 4342.074 | 0 | 1.4 | 127 | Specific | 961.27 | 862.26 | 862.26 | 19.59 | 124 | >sp\|P01876\|IGHA1_HUMAN Immunoglobulin heavy constant alpha 1 OS=Homo sapiens OX=9606 GN=IGHA1 PE=1 SV=2 | 255 | 20201114_qwl_colo_milk_1ng_glyco_hcd203040_compound15.19218.19218.4 | controllerType=0 controllerNumber=1 scan=19218 | 42.3363 |
| 15830:4 | 2 | R.LSLHRPALEDLLLGSEAN[+1540.529]LTC[+57.021]TLTGLR.D | HexNAc(2)Hex(7) | C[+57], N[+1541] | 1126.7895 | 4 | 4504.1361 | 4504.1268 | 0 | 2.1 | 127 | Specific | 987.71 | 886.83 | 886.83 | 19.59 | 124 | >sp\|P01876\|IGHA1_HUMAN Immunoglobulin heavy constant alpha 1 OS=Homo sapiens OX=9606 GN=IGHA1 PE=1 SV=2 | 255 | 20201114_qwl_colo_milk_1ng_glyco_hcd203040_compound15.19213.19213.4 | controllerType=0 controllerNumber=1 scan=19213 | 42.3221 |
| 15921:4 | 2 | R.LSLHRPALEDLLLGSEAN[+1095.397]LTC[+57.021]TLTGLR.D | HexNAc(3)Hex(3) | C[+57], N[+1095] | 1015.5137 | 4 | 4059.033 | 4058.9949 | 0 | 9.4 | 127 | Specific | 1010.82 | 1010.82 | 1010.82 | 20.1 | 124 | >sp\|P01876\|IGHA1_HUMAN Immunoglobulin heavy constant alpha 1 OS=Homo sapiens OX=9606 GN=IGHA1 PE=1 SV=2 | 255 | 20201114_qwl_colo_milk_1ng_glyco_hcd203040_compound15.19309.19309.4 | controllerType=0 controllerNumber=1 scan=19309 | 42.6028 |
| 15748:4 | 2 | R.LSLHRPALEDLLLGSEAN[+1864.634]LTC[+57.021]TLTGLR.D | HexNAc(2)Hex(9) | C[+57], N[+1865] | 1207.8217 | 4 | 4828.265 | 4828.2324 | 0 | 6.7 | 127 | Specific | 1013.69 | 875.63 | 875.63 | 20.31 | 124 | >sp\|P01876\|IGHA1_HUMAN Immunoglobulin heavy constant alpha 1 OS=Homo sapiens OX=9606 GN=IGHA1 PE=1 SV=2 | 255 | 20201114_qwl_colo_milk_1ng_glyco_hcd203040_compound15.19126.19126.2 | controllerType=0 controllerNumber=1 scan=19126 | 42.1151 |
| 15838:4 | 2 | R.LSLHRPALEDLLLGSEAN[+1702.581]LTC[+57.021]TLTGLR.D | HexNAc(2)Hex(8) | C[+57], N[+1703] | 1167.3028 | 4 | 4666.1893 | 4666.1796 | 0 | 2.1 | 127 | Specific | 1025.26 | 1025.26 | 1025.26 | 20.59 | 124 | >sp\|P01876\|IGHA1_HUMAN Immunoglobulin heavy constant alpha 1 OS=Homo sapiens OX=9606 GN=IGHA1 PE=1 SV=2 | 255 | 20201114_qwl_colo_milk_1ng_glyco_hcd203040_compound15.19221.19221.4 | controllerType=0 controllerNumber=1 scan=19221 | 42.361 |
| 15947:4 | 2 | R.LSLHRPALEDLLLGSEAN[+892.317]LTC[+57.021]TLTGLR.D | HexNAc(2)Hex(3) | C[+57], N[+892] | 964.7369 | 4 | 3855.9256 | 3855.9155 | 0 | 2.6 | 127 | Specific | 1047.22 | 1047.22 | 1047.22 | 20.7 | 124 | >sp\|P01876\|IGHA1_HUMAN Immunoglobulin heavy constant alpha 1 OS=Homo sapiens OX=9606 GN=IGHA1 PE=1 SV=2 | 255 | 20201114_qwl_colo_milk_1ng_glyco_hcd203040_compound15.19337.19337.4 | controllerType=0 controllerNumber=1 scan=19337 | 42.6973 |
| 15839:4 | 2 | R.LSLHRPALEDLLLGSEAN[+1378.476]LTC[+57.021]TLTGLR.D | HexNAc(2)Hex(6) | C[+57], N[+1378] | 1086.2754 | 4 | 4342.0799 | 4342.074 | 0 | 1.4 | 127 | Specific | 1077.63 | 1066.49 | 1066.49 | 21.59 | 124 | >sp\|P01876\|IGHA1_HUMAN Immunoglobulin heavy constant alpha 1 OS=Homo sapiens OX=9606 GN=IGHA1 PE=1 SV=2 | 255 | 20201114_qwl_colo_milk_1ng_glyco_hcd203040_compound15.19222.19222.4 | controllerType=0 controllerNumber=1 scan=19222 | 42.3624 |
| 15832:4 | 2 | R.LSLHRPALEDLLLGSEAN[+1540.529]LTC[+57.021]TLTGLR.D | HexNAc(2)Hex(7) | C[+57], N[+1541] | 1126.7895 | 4 | 4504.1361 | 4504.1268 | 0 | 2.1 | 127 | Specific | 1107.74 | 1107.74 | 1107.74 | 22.59 | 124 | >sp\|P01876\|IGHA1_HUMAN Immunoglobulin heavy constant alpha 1 OS=Homo sapiens OX=9606 GN=IGHA1 PE=1 SV=2 | 255 | 20201114_qwl_colo_milk_1ng_glyco_hcd203040_compound15.19215.19215.4 | controllerType=0 controllerNumber=1 scan=19215 | 42.3329 |
| 16997:4 | 2 | R.LSLHRPALEDLLLGSEAN[+1501.555]LTC[+57.021]TLTGLRDASGVTFTWTPSSGK.S | HexNAc(5)Hex(3) | C[+57], N[+1502] | 1497.4777 | 4 | 5986.8888 | 5986.8636 | 0 | 4.2 | 127 | Specific | 271.9 | 248.16 | 248.16 | 6.18 | 124 | >sp\|P01876\|IGHA1_HUMAN Immunoglobulin heavy constant alpha 1 OS=Homo sapiens OX=9606 GN=IGHA1 PE=1 SV=2 | 255 | 20201114_qwl_colo_milk_1ng_glyco_hcd203040_compound15.20438.20438.4 | controllerType=0 controllerNumber=1 scan=20438 | 45.2543 |
| 16999:4 | 2 | R.LSLHRPALEDLLLGSEAN[+1501.555]LTC[+57.021]TLTGLRDASGVTFTWTPSSGK.S | HexNAc(5)Hex(3) | C[+57], N[+1502] | 1497.4777 | 4 | 5986.8888 | 5986.8636 | 0 | 4.2 | 127 | Specific | 386.61 | 373.44 | 373.44 | 7.46 | 124 | >sp\|P01876\|IGHA1_HUMAN Immunoglobulin heavy constant alpha 1 OS=Homo sapiens OX=9606 GN=IGHA1 PE=1 SV=2 | 255 | 20201114_qwl_colo_milk_1ng_glyco_hcd203040_compound15.20440.20440.4 | controllerType=0 controllerNumber=1 scan=20440 | 45.2578 |
| 11946:3 | 2 | R.LAGKPTHVN[+2263.835]VSVVMAEVDGTC[+57.021].Y | HexNAc(5)Hex(5)Fuc(3) | C[+57], N[+2264] | 1483.3096 | 3 | 4447.9143 | 4447.9135 | 0 | 0.2 | 332 | CRagged | 211.74 | 206.1 | 206.1 | 3.79 | 124 | >sp\|P01876\|IGHA1_HUMAN Immunoglobulin heavy constant alpha 1 OS=Homo sapiens OX=9606 GN=IGHA1 PE=1 SV=2 | 255 | 20201114_qwl_colo_milk_1ng_glyco_hcd203040_compound15.15141.15141.3 | controllerType=0 controllerNumber=1 scan=15141 | 33.5699 |
| 09549:3 | 2 | R.LAGKPTHVN[+2262.814]VSVVMAEVDGTC[+57.021].Y | HexNAc(5)Hex(5)Fuc(1)NeuAc(1) | C[+57], N[+2263] | 1482.9758 | 3 | 4446.9127 | 4446.8931 | 0 | 4.4 | 332 | CRagged | 244.97 | 167.76 | 167.76 | 6.32 | 124 | >sp\|P01876\|IGHA1_HUMAN Immunoglobulin heavy constant alpha 1 OS=Homo sapiens OX=9606 GN=IGHA1 PE=1 SV=2 | 255 | 20201114_qwl_colo_milk_1ng_glyco_hcd203040_compound15.12642.12642.3 | controllerType=0 controllerNumber=1 scan=12642 | 28.7701 |
| 08144:3 | 2 | R.LAGKPTHVN[+1606.587]VSVVMAEVDGTC[+57.021].Y | HexNAc(4)Hex(4)Fuc(1) | C[+57], N[+1607] | 1264.2388 | 3 | 3790.7019 | 3790.6654 | 0 | 9.6 | 332 | CRagged | 268.25 | 268.25 | 268.25 | 6.74 | 124 | >sp\|P01876\|IGHA1_HUMAN Immunoglobulin heavy constant alpha 1 OS=Homo sapiens OX=9606 GN=IGHA1 PE=1 SV=2 | 255 | 20201114_qwl_colo_milk_1ng_glyco_hcd203040_compound15.11173.11173.3 | controllerType=0 controllerNumber=1 scan=11173 | 26.0179 |
| 08139:3 | 2 | R.LAGKPTHVN[+1809.666]VSVVMAEVDGTC[+57.021].Y | HexNAc(5)Hex(4)Fuc(1) | C[+57], N[+1810] | 1331.925 | 3 | 3993.7604 | 3993.7448 | 0 | 3.9 | 332 | CRagged | 305.11 | 305.11 | 275.97 | 7.43 | 124 | >sp\|P01876\|IGHA1_HUMAN Immunoglobulin heavy constant alpha 1 OS=Homo sapiens OX=9606 GN=IGHA1 PE=1 SV=2 | 255 | 20201114_qwl_colo_milk_1ng_glyco_hcd203040_compound15.11168.11168.3 | controllerType=0 controllerNumber=1 scan=11168 | 26.0088 |
| 11945:3 | 2 | R.LAGKPTHVN[+2263.835]VSVVMAEVDGTC[+57.021].Y | HexNAc(5)Hex(5)Fuc(3) | C[+57], N[+2264] | 1483.3096 | 3 | 4447.9143 | 4447.9135 | 0 | 0.2 | 332 | CRagged | 335.96 | 295.77 | 295.77 | 7.64 | 124 | >sp\|P01876\|IGHA1_HUMAN Immunoglobulin heavy constant alpha 1 OS=Homo sapiens OX=9606 GN=IGHA1 PE=1 SV=2 | 255 | 20201114_qwl_colo_milk_1ng_glyco_hcd203040_compound15.15140.15140.3 | controllerType=0 controllerNumber=1 scan=15140 | 33.5552 |
| 12043:3 | 2 | R.LAGKPTHVN[+2117.777]VSVVMAEVDGTC[+57.021].Y | HexNAc(5)Hex(5)Fuc(2) | C[+57], N[+2118] | 1434.6235 | 3 | 4301.8559 | 4301.8555 | 0 | 0.1 | 332 | CRagged | 355.26 | 320.18 | 320.18 | 7.85 | 124 | >sp\|P01876\|IGHA1_HUMAN Immunoglobulin heavy constant alpha 1 OS=Homo sapiens OX=9606 GN=IGHA1 PE=1 SV=2 | 255 | 20201114_qwl_colo_milk_1ng_glyco_hcd203040_compound15.15242.15242.3 | controllerType=0 controllerNumber=1 scan=15242 | 33.757 |
| 08146:3 | 2 | R.LAGKPTHVN[+1606.587]VSVVMAEVDGTC[+57.021].Y | HexNAc(4)Hex(4)Fuc(1) | C[+57], N[+1607] | 1264.2388 | 3 | 3790.7019 | 3790.6654 | 0 | 9.6 | 332 | CRagged | 366.06 | 294.74 | 294.74 | 7.15 | 124 | >sp\|P01876\|IGHA1_HUMAN Immunoglobulin heavy constant alpha 1 OS=Homo sapiens OX=9606 GN=IGHA1 PE=1 SV=2 | 255 | 20201114_qwl_colo_milk_1ng_glyco_hcd203040_compound15.11175.11175.3 | controllerType=0 controllerNumber=1 scan=11175 | 26.0218 |
| 09546:3 | 2 | R.LAGKPTHVN[+2262.814]VSVVMAEVDGTC[+57.021].Y | HexNAc(5)Hex(5)Fuc(1)NeuAc(1) | C[+57], N[+2263] | 1482.9758 | 3 | 4446.9127 | 4446.8931 | 0 | 4.4 | 332 | CRagged | 453.37 | 353.22 | 353.22 | 8.35 | 124 | >sp\|P01876\|IGHA1_HUMAN Immunoglobulin heavy constant alpha 1 OS=Homo sapiens OX=9606 GN=IGHA1 PE=1 SV=2 | 255 | 20201114_qwl_colo_milk_1ng_glyco_hcd203040_compound15.12639.12639.3 | controllerType=0 controllerNumber=1 scan=12639 | 28.7647 |
| 09513:3 | 2 | R.LAGKPTHVN[+1647.613]VSVVMAEVDGTC[+57.021].Y | HexNAc(5)Hex(3)Fuc(1) | C[+57], N[+1648] | 1277.9113 | 3 | 3831.7193 | 3831.692 | 0 | 7.1 | 332 | CRagged | 496.54 | 407.96 | 407.96 | 9.8 | 124 | >sp\|P01876\|IGHA1_HUMAN Immunoglobulin heavy constant alpha 1 OS=Homo sapiens OX=9606 GN=IGHA1 PE=1 SV=2 | 255 | 20201114_qwl_colo_milk_1ng_glyco_hcd203040_compound15.12604.12604.3 | controllerType=0 controllerNumber=1 scan=12604 | 28.7065 |
| 10936:3 | 2 | R.LAGKPTHVN[+2262.814]VSVVMAEVDGTC[+57.021].Y | HexNAc(5)Hex(5)Fuc(1)NeuAc(1) | C[+57], N[+2263] | 1482.9763 | 3 | 4446.9142 | 4446.8931 | 0 | 4.8 | 332 | CRagged | 564.63 | 469.66 | 469.66 | 10.21 | 124 | >sp\|P01876\|IGHA1_HUMAN Immunoglobulin heavy constant alpha 1 OS=Homo sapiens OX=9606 GN=IGHA1 PE=1 SV=2 | 255 | 20201114_qwl_colo_milk_1ng_glyco_hcd203040_compound15.14085.14085.3 | controllerType=0 controllerNumber=1 scan=14085 | 31.4991 |
| 08135:3 | 2 | R.LAGKPTHVN[+1216.423]VSVVMAEVDGTC[+57.021].Y | HexNAc(2)Hex(5) | C[+57], N[+1216] | 1134.1825 | 3 | 3400.533 | 3400.5016 | 0 | 9.2 | 332 | CRagged | 617.13 | 569.98 | 405.02 | 12.27 | 124 | >sp\|P01876\|IGHA1_HUMAN Immunoglobulin heavy constant alpha 1 OS=Homo sapiens OX=9606 GN=IGHA1 PE=1 SV=2 | 255 | 20201114_qwl_colo_milk_1ng_glyco_hcd203040_compound15.11164.11164.3 | controllerType=0 controllerNumber=1 scan=11164 | 26.0002 |
| 08188:3 | 2 | R.LAGKPTHVN[+1444.534]VSVVMAEVDGTC[+57.021].Y | HexNAc(4)Hex(3)Fuc(1) | C[+57], N[+1445] | 1210.2208 | 3 | 3628.6477 | 3628.6126 | 0 | 9.7 | 332 | CRagged | 678.06 | 636.64 | 636.64 | 13.27 | 124 | >sp\|P01876\|IGHA1_HUMAN Immunoglobulin heavy constant alpha 1 OS=Homo sapiens OX=9606 GN=IGHA1 PE=1 SV=2 | 255 | 20201114_qwl_colo_milk_1ng_glyco_hcd203040_compound15.11219.11219.3 | controllerType=0 controllerNumber=1 scan=11219 | [illegible] |
| 08137:3 | 2 | R.LAGKPTHVN[+1216.423]VSVVMAEVDGTC[+57.021].Y | HexNAc(2)Hex(5) | C[+57], N[+1216] | 1134.1825 | 3 | 3400.533 | 3400.5016 | 0 | 9.2 | 332 | CRagged | 736.49 | 712.02 | 523.92 | 14.27 | 124 | >sp\|P01876\|IGHA1_HUMAN Immunoglobulin heavy constant alpha 1 OS=Homo sapiens OX=9606 GN=IGHA1 PE=1 SV=2 | 255 | 20201114_qwl_colo_milk_1ng_glyco_hcd203040_compound15.11166.11166.3 | controllerType=0 controllerNumber=1 scan=11166 | [illegible] |
| 08190:3 | 2 | R.LAGKPTHVN[+1444.534]VSVVMAEVDGTC[+57.021].Y | HexNAc(4)Hex(3)Fuc(1) | C[+57], N[+1445] | 1210.2208 | 3 | 3628.6477 | 3628.6126 | 0 | 9.7 | 332 | CRagged | 763.38 | 616.49 | 616.49 | 15.27 | 124 | >sp\|P01876\|IGHA1_HUMAN Immunoglobulin heavy constant alpha 1 OS=Homo sapiens OX=9606 GN=IGHA1 PE=1 SV=2 | 255 | 20201114_qwl_colo_milk_1ng_glyco_hcd203040_compound15.11221.11221.3 | controllerType=0 controllerNumber=1 scan=11221 | [illegible] |
| 09499:3 | 2 | R.LAGKPTHVN[+1054.370]VSVVMAEVDGTC[+57.021].Y | HexNAc(2)Hex(4) | C[+57], N[+1054] | 1080.1615 | 3 | 3238.47 | 3238.4488 | 0 | 6.6 | 332 | CRagged | 765 | 713.96 | 439.82 | 15.34 | 124 | >sp\|P01876\|IGHA1_HUMAN Immunoglobulin heavy constant alpha 1 OS=Homo sapiens OX=9606 GN=IGHA1 PE=1 SV=2 | 255 | 20201114_qwl_colo_milk_1ng_glyco_hcd203040_compound15.12589.12589.3 | controllerType=0 controllerNumber=1 scan=12589 | [illegible] |
| 08091:3 | 2 | R.LAGKPTHVN[+1378.476]VSVVMAEVDGTC[+57.021].Y | HexNAc(2)Hex(6) | C[+57], N[+1378] | 1188.1909 | 3 | 3562.558 | 3562.5544 | 0 | 1 | 332 | CRagged | 778.84 | 710.82 | 479.47 | 15.7 | 124 | >sp\|P01876\|IGHA1_HUMAN Immunoglobulin heavy constant alpha 1 OS=Homo sapiens OX=9606 GN=IGHA1 PE=1 SV=2 | 255 | 20201114_qwl_colo_milk_1ng_glyco_hcd203040_compound15.11118.11118.3 | controllerType=0 controllerNumber=1 scan=11118 | 25.9043 |
| 09497:3 | 2 | R.LAGKPTHVN[+1054.370]VSVVMAEVDGTC[+57.021].Y | HexNAc(2)Hex(4) | C[+57], N[+1054] | 1080.1615 | 3 | 3238.47 | 3238.4488 | 0 | 6.6 | 332 | CRagged | 785.67 | 706.85 | 468.71 | 15.34 | 124 | >sp\|P01876\|IGHA1_HUMAN Immunoglobulin heavy constant alpha 1 OS=Homo sapiens OX=9606 GN=IGHA1 PE=1 SV=2 | 255 | 20201114_qwl_colo_milk_1ng_glyco_hcd203040_compound15.12587.12587.3 | controllerType=0 controllerNumber=1 scan=12587 | 28.6629 |
| 09511:3 | 2 | R.LAGKPTHVN[+1647.613]VSVVMAEVDGTC[+57.021].Y | HexNAc(5)Hex(3)Fuc(1) | C[+57], N[+1648] | 1277.9113 | 3 | 3831.7193 | 3831.692 | 0 | 7.1 | 332 | CRagged | 800.22 | 759.18 | 759.18 | 16.48 | 124 | >sp\|P01876\|IGHA1_HUMAN Immunoglobulin heavy constant alpha 1 OS=Homo sapiens OX=9606 GN=IGHA1 PE=1 SV=2 | 255 | 20201114_qwl_colo_milk_1ng_glyco_hcd203040_compound15.12602.12602.3 | controllerType=0 controllerNumber=1 scan=12602 | 28.7008 |
| 08240:3 | 2 | R.LAGKPTHVN[+1054.370]VSVVMAEVDGTC[+57.021].Y | HexNAc(2)Hex(4) | C[+57], N[+1054] | 1080.1564 | 3 | 3238.4546 | 3238.4488 | 0 | 1.8 | 332 | CRagged | 831.7 | 831.7 | 511.7 | 16.76 | 124 | >sp\|P01876\|IGHA1_HUMAN Immunoglobulin heavy constant alpha 1 OS=Homo sapiens OX=9606 GN=IGHA1 PE=1 SV=2 | 255 | 20201114_qwl_colo_milk_1ng_glyco_hcd203040_compound15.11273.11273.3 | controllerType=0 controllerNumber=1 scan=11273 | 26.2087 |
| 06678:3 | 2 | R.LAGKPTHVN[+1216.423]VSVVM[+15.995]AEVDGTC[+57.021].Y | HexNAc(2)Hex(5) | C[+57], M[+16], N[+1216] | 1139.5042 | 3 | 3416.498 | 3416.4965 | 0 | 0.4 | 332 | CRagged | 344.08 | 342.14 | 342.14 | 7.88 | 124 | >sp\|P01876\|IGHA1_HUMAN Immunoglobulin heavy constant alpha 1 OS=Homo sapiens OX=9606 GN=IGHA1 PE=1 SV=2 | 255 | 20201114_qwl_colo_milk_1ng_glyco_hcd203040_compound15.9645.9645.3 | controllerType=0 controllerNumber=1 scan=9645 | 23.2774 |
| 06680:3 | 2 | R.LAGKPTHVN[+1216.423]VSVVM[+15.995]AEVDGTC[+57.021].Y | HexNAc(2)Hex(5) | C[+57], M[+16], N[+1216] | 1139.5042 | 3 | 3416.498 | 3416.4965 | 0 | 0.4 | 332 | CRagged | 405.2 | 342.21 | 342.21 | 8.01 | 124 | >sp\|P01876\|IGHA1_HUMAN Immunoglobulin heavy constant alpha 1 OS=Homo sapiens OX=9606 GN=IGHA1 PE=1 SV=2 | 255 | 20201114_qwl_colo_milk_1ng_glyco_hcd203040_compound15.9647.9647.3 | controllerType=0 controllerNumber=1 scan=9647 | 23.2811 |
| 09502:3 | 2 | R.LAGKPTHVN[+1298.476]VSVVMAEVDGTC[+57.021]Y.- | HexNAc(4)Hex(3) | C[+57], N[+1298] | 1215.8826 | 3 | 3645.6333 | 3645.618 | 0 | 4.2 | 332 | CRagged | 233.77 | 136.48 | 136.48 | 4.07 | 124 | >sp\|P01876\|IGHA1_HUMAN Immunoglobulin heavy constant alpha 1 OS=Homo sapiens OX=9606 GN=IGHA1 PE=1 SV=2 | 255 | 20201114_qwl_colo_milk_1ng_glyco_hcd203040_compound15.12592.12592.3 | controllerType=0 controllerNumber=1 scan=12592 | 28.676 |
| 13630:4 | 2 | R.LAGKPTHVN[+2350.830]VSVVMAEVDGTC[+57.021]Y.- | HexNAc(4)Hex(5)Fuc(1)NeuAc(2) | C[+57], N[+2351] | 1175.2574 | 4 | 4698.0076 | 4697.9724 | 0 | 7.5 | 332 | CRagged | 247.71 | 241.9 | 241.9 | 4.59 | 124 | >sp\|P01876\|IGHA1_HUMAN Immunoglobulin heavy constant alpha 1 OS=Homo sapiens OX=9606 GN=IGHA1 PE=1 SV=2 | 255 | 20201114_qwl_colo_milk_1ng_glyco_hcd203040_compound15.16902.16902.4 | controllerType=0 controllerNumber=1 scan=16902 | 37.235 |
| 10090:3 | 2 | R.LAGKPTHVN[+1501.555]VSVVMAEVDGTC[+57.021]Y.- | HexNAc(5)Hex(3) | C[+57], N[+1502] | 1283.5783 | 3 | 3848.7204 | 3848.6974 | 0 | 6 | 332 | CRagged | 251.26 | 248.6 | 248.6 | 6.58 | 124 | >sp\|P01876\|IGHA1_HUMAN Immunoglobulin heavy constant alpha 1 OS=Homo sapiens OX=9606 GN=IGHA1 PE=1 SV=2 | 255 | 20201114_qwl_colo_milk_1ng_glyco_hcd203040_compound15.13203.13203.3 | controllerType=0 controllerNumber=1 scan=13203 | 29.8005 |
| 10092:3 | 2 | R.LAGKPTHVN[+1501.555]VSVVMAEVDGTC[+57.021]Y.- | HexNAc(5)Hex(3) | C[+57], N[+1502] | 1283.5783 | 3 | 3848.7204 | 3848.6974 | 0 | 6 | 332 | CRagged | 253.23 | 183.35 | 183.35 | 5.15 | 124 | >sp\|P01876\|IGHA1_HUMAN Immunoglobulin heavy constant alpha 1 OS=Homo sapiens OX=9606 GN=IGHA1 PE=1 SV=2 | 255 | 20201114_qwl_colo_milk_1ng_glyco_hcd203040_compound15.13205.13205.3 | controllerType=0 controllerNumber=1 scan=13205 | 29.8046 |
| 10790:3 | 2 | R.LAGKPTHVN[+1913.677]VSVVMAEVDGTC[+57.021]Y.- | HexNAc(4)Hex(5)NeuAc(1) | C[+57], N[+1914] | 1420.9543 | 3 | 4260.8483 | 4260.8191 | 0 | 6.8 | 332 | CRagged | 277.96 | 227.61 | 227.61 | 5.73 | 124 | >sp\|P01876\|IGHA1_HUMAN Immunoglobulin heavy constant alpha 1 OS=Homo sapiens OX=9606 GN=IGHA1 PE=1 SV=2 | 255 | 20201114_qwl_colo_milk_1ng_glyco_hcd203040_compound15.13933.13933.3 | controllerType=0 controllerNumber=1 scan=13933 | 31.2499 |
| 11966:4 | 2 | R.LAGKPTHVN[+2116.756]VSVVMAEVDGTC[+57.021]Y.- | HexNAc(5)Hex(5)NeuAc(1) | C[+57], N[+2117] | 1116.7319 | 4 | 4463.9056 | 4463.8985 | 0 | 1.6 | 332 | CRagged | 284.39 | 227.64 | 227.64 | 6.91 | 124 | >sp\|P01876\|IGHA1_HUMAN Immunoglobulin heavy constant alpha 1 OS=Homo sapiens OX=9606 GN=IGHA1 PE=1 SV=2 | 255 | 20201114_qwl_colo_milk_1ng_glyco_hcd203040_compound15.15162.15162.4 | controllerType=0 controllerNumber=1 scan=15162 | 33.6106 |
| 10009:3 | 2 | R.LAGKPTHVN[+1663.608]VSVVMAEVDGTC[+57.021]Y.- | HexNAc(5)Hex(4) | C[+57], N[+1664] | 1337.5904 | 3 | 4010.7566 | 4010.7502 | 0 | 1.6 | 332 | CRagged | 285.08 | 285.08 | 282.61 | 7.26 | 124 | >sp\|P01876\|IGHA1_HUMAN Immunoglobulin heavy constant alpha 1 OS=Homo sapiens OX=9606 GN=IGHA1 PE=1 SV=2 | 255 | 20201114_qwl_colo_milk_1ng_glyco_hcd203040_compound15.13119.13119.3 | controllerType=0 controllerNumber=1 scan=13119 | 29.6292 |
| 13627:4 | 2 | R.LAGKPTHVN[+2350.830]VSVVMAEVDGTC[+57.021]Y.- | HexNAc(4)Hex(5)Fuc(1)NeuAc(2) | C[+57], N[+2351] | 1175.2574 | 4 | 4698.0076 | 4697.9724 | 0 | 7.5 | 332 | CRagged | 286.53 | 190.98 | 190.98 | 6.31 | 124 | >sp\|P01876\|IGHA1_HUMAN Immunoglobulin heavy constant alpha 1 OS=Homo sapiens OX=9606 GN=IGHA1 PE=1 SV=2 | 255 | 20201114_qwl_colo_milk_1ng_glyco_hcd203040_compound15.16899.16899.4 | controllerType=0 controllerNumber=1 scan=16899 | 37.2304 |
| 10099:3 | 2 | R.LAGKPTHVN[+1054.370]VSVVMAEVDGTC[+57.021]Y.- | HexNAc(2)Hex(4) | C[+57], N[+1054] | 1134.515 | 3 | 3401.5306 | 3401.5121 | 0 | 5.4 | 332 | CRagged | 295.35 | 295.35 | 199.82 | 7.16 | 124 | >sp\|P01876\|IGHA1_HUMAN Immunoglobulin heavy constant alpha 1 OS=Homo sapiens OX=9606 GN=IGHA1 PE=1 SV=2 | 255 | 20201114_qwl_colo_milk_1ng_glyco_hcd203040_compound15.13212.13212.3 | controllerType=0 controllerNumber=1 scan=13212 | 29.8152 |
| 10007:3 | 2 | R.LAGKPTHVN[+1663.608]VSVVMAEVDGTC[+57.021]Y.- | HexNAc(5)Hex(4) | C[+57], N[+1664] | 1337.5904 | 3 | 4010.7566 | 4010.7502 | 0 | 1.6 | 332 | CRagged | 296.01 | 296.01 | 280.14 | 7.39 | 124 | >sp\|P01876\|IGHA1_HUMAN Immunoglobulin heavy constant alpha 1 OS=Homo sapiens OX=9606 GN=IGHA1 PE=1 SV=2 | 255 | 20201114_qwl_colo_milk_1ng_glyco_hcd203040_compound15.13117.13117.3 | controllerType=0 controllerNumber=1 scan=13117 | 29.6251 |
| 10346:3 | 2 | R.LAGKPTHVN[+1971.719]VSVVMAEVDGTC[+57.021]Y.- | HexNAc(5)Hex(5)Fuc(1) | C[+57], N[+1972] | 1440.3039 | 3 | 4318.8971 | 4318.861 | 0 | 8.4 | 332 | CRagged | 302.99 | 302.99 | 275.39 | 7.04 | 124 | >sp\|P01876\|IGHA1_HUMAN Immunoglobulin heavy constant alpha 1 OS=Homo sapiens OX=9606 GN=IGHA1 PE=1 SV=2 | 255 | 20201114_qwl_colo_milk_1ng_glyco_hcd203040_compound15.13470.13470.3 | controllerType=0 controllerNumber=1 scan=13470 | 30.3064 |
| 13676:4 | 2 | R.LAGKPTHVN[+2553.910]VSVVMAEVDGTC[+57.021]Y.- | HexNAc(5)Hex(5)Fuc(1)NeuAc(2) | C[+57], N[+2554] | 1226.0281 | 4 | 4901.0906 | 4901.0518 | 0 | 7.9 | 332 | CRagged | 305.38 | 243.89 | 243.89 | 4.95 | 124 | >sp\|P01876\|IGHA1_HUMAN Immunoglobulin heavy constant alpha 1 OS=Homo sapiens OX=9606 GN=IGHA1 PE=1 SV=2 | 255 | 20201114_qwl_colo_milk_1ng_glyco_hcd203040_compound15.16950.16950.4 | controllerType=0 controllerNumber=1 scan=16950 | 37.3302 |
| 08345:3 | 2 | R.LAGKPTHVN[+1054.370]VSVVMAEVDGTC[+57.021]Y.- | HexNAc(2)Hex(4) | C[+57], N[+1054] | 1134.5109 | 3 | 3401.5181 | 3401.5121 | 0 | 1.8 | 332 | CRagged | 323.45 | 290.86 | 290.86 | 7.62 | 124 | >sp\|P01876\|IGHA1_HUMAN Immunoglobulin heavy constant alpha 1 OS=Homo sapiens OX=9606 GN=IGHA1 PE=1 SV=2 | 255 | 20201114_qwl_colo_milk_1ng_glyco_hcd203040_compound15.11381.11381.3 | controllerType=0 controllerNumber=1 scan=11381 | 26.4043 |
| 10713:3 | 2 | R.LAGKPTHVN[+1647.613]VSVVMAEVDGTC[+57.021]Y.- | HexNAc(5)Hex(3)Fuc(1) | C[+57], N[+1648] | 1332.2626 | 3 | 3994.7733 | 3994.7553 | 0 | 4.5 | 332 | CRagged | 329.21 | 267.04 | 267.04 | 4.97 | 124 | >sp\|P01876\|IGHA1_HUMAN Immunoglobulin heavy constant alpha 1 OS=Homo sapiens OX=9606 GN=IGHA1 PE=1 SV=2 | 255 | 20201114_qwl_colo_milk_1ng_glyco_hcd203040_compound15.13852.13852.5 | controllerType=0 controllerNumber=1 scan=13852 | 31.0907 |
| 12110:3 | 2 | R.LAGKPTHVN[+1548.545]VSVVMAEVDGTC[+57.021]Y.- | HexNAc(3)Hex(4)NeuAc(1) | C[+57], N[+1549] | 1299.2366 | 3 | 3895.6953 | 3895.6869 | 0 | 2.2 | 332 | CRagged | 340.68 | 340.68 | 340.68 | 7.97 | 124 | >sp\|P01876\|IGHA1_HUMAN Immunoglobulin heavy constant alpha 1 OS=Homo sapiens OX=9606 GN=IGHA1 PE=1 SV=2 | 255 | 20201114_qwl_colo_milk_1ng_glyco_hcd203040_compound15.15312.15312.3 | controllerType=0 controllerNumber=1 scan=15312 | 33.8839 |
| 10423:4 | 2 | R.LAGKPTHVN[+1809.666]VSVVMAEVDGTC[+57.021]Y.- | HexNAc(5)Hex(4)Fuc(1) | C[+57], N[+1810] | 1039.9556 | 4 | 4156.8006 | 4156.8081 | 0 | -1.8 | 332 | CRagged | 342.14 | 331.4 | 291.42 | 7.1 | 124 | >sp\|P01876\|IGHA1_HUMAN Immunoglobulin heavy constant alpha 1 OS=Homo sapiens OX=9606 GN=IGHA1 PE=1 SV=2 | 255 | 20201114_qwl_colo_milk_1ng_glyco_hcd203040_compound15.13550.13550.4 | controllerType=0 controllerNumber=1 scan=13550 | 30.4737 |
| 09429:3 | 2 | R.LAGKPTHVN[+1606.587]VSVVMAEVDGTC[+57.021]Y.- | HexNAc(4)Hex(4)Fuc(1) | C[+57], N[+1607] | 1318.5928 | 3 | 3953.7639 | 3953.7288 | 0 | 8.9 | 332 | CRagged | 350.58 | 350.58 | 350.58 | 7.61 | 124 | >sp\|P01876\|IGHA1_HUMAN Immunoglobulin heavy constant alpha 1 OS=Homo sapiens OX=9606 GN=IGHA1 PE=1 SV=2 | 255 | 20201114_qwl_colo_milk_1ng_glyco_hcd203040_compound15.12516.12516.3 | controllerType=0 controllerNumber=1 scan=12516 | 28.4952 |
| 12005:3 | 2 | R.LAGKPTHVN[+2116.756]VSVVMAEVDGTC[+57.021]Y.- | HexNAc(5)Hex(5)NeuAc(1) | C[+57], N[+2117] | 1488.6518 | 3 | 4463.9408 | 4463.8985 | 0 | 9.5 | 332 | CRagged | 372.61 | 360.63 | 360.63 | 7.73 | 124 | >sp\|P01876\|IGHA1_HUMAN Immunoglobulin heavy constant alpha 1 OS=Homo sapiens OX=9606 GN=IGHA1 PE=1 SV=2 | 255 | 20201114_qwl_colo_milk_1ng_glyco_hcd203040_compound15.15203.15203.3 | controllerType=0 controllerNumber=1 scan=15203 | 33.6867 |
| 10614:3 | 2 | R.LAGKPTHVN[+1378.476]VSVVMAEVDGTC[+57.021]Y.- | HexNAc(2)Hex(6) | C[+57], N[+1378] | 1242.5484 | 3 | 3725.6306 | 3725.6178 | 0 | 3.5 | 332 | CRagged | 378.14 | 378.14 | 288.84 | 9.8 | 124 | >sp\|P01876\|IGHA1_HUMAN Immunoglobulin heavy constant alpha 1 OS=Homo sapiens OX=9606 GN=IGHA1 PE=1 SV=2 | 255 | 20201114_qwl_colo_milk_1ng_glyco_hcd203040_compound15.13749.13749.3 | controllerType=0 controllerNumber=1 scan=13749 | 30.8761 |
| 09985:3 | 2 | R.LAGKPTHVN[+1378.476]VSVVMAEVDGTC[+57.021]Y.- | HexNAc(2)Hex(6) | C[+57], N[+1378] | 1242.5475 | 3 | 3725.6281 | 3725.6178 | 0 | 2.8 | 332 | CRagged | 397.44 | 397.44 | 341.45 | 9.84 | 124 | >sp\|P01876\|IGHA1_HUMAN Immunoglobulin heavy constant alpha 1 OS=Homo sapiens OX=9606 GN=IGHA1 PE=1 SV=2 | 255 | 20201114_qwl_colo_milk_1ng_glyco_hcd203040_compound15.13094.13094.3 | controllerType=0 controllerNumber=1 scan=13094 | 29.5821 |
| 10447:2 | 2 | R.LAGKPTHVN[+1216.423]VSVVMAEVDGTC[+57.021]Y.- | HexNAc(2)Hex(5) | C[+57], N[+1216] | 1782.2893 | 2 | 3563.5713 | 3563.5649 | 0 | 1.8 | 332 | CRagged | 404.69 | 331.93 | 141.82 | 5.76 | 124 | >sp\|P01876\|IGHA1_HUMAN Immunoglobulin heavy constant alpha 1 OS=Homo sapiens OX=9606 GN=IGHA1 PE=1 SV=2 | 255 | 20201114_qwl_colo_milk_1ng_glyco_hcd203040_compound15.13575.13575.2 | controllerType=0 controllerNumber=1 scan=13575 | 30.5364 |
| 09977:3 | 2 | R.LAGKPTHVN[+1663.608]VSVVMAEVDGTC[+57.021]Y.- | HexNAc(5)Hex(4) | C[+57], N[+1664] | 1337.5941 | 3 | 4010.7676 | 4010.7502 | 0 | 4.3 | 332 | CRagged | 453.85 | 320.71 | 320.71 | 8.35 | 124 | >sp\|P01876\|IGHA1_HUMAN Immunoglobulin heavy constant alpha 1 OS=Homo sapiens OX=9606 GN=IGHA1 PE=1 SV=2 | 255 | 20201114_qwl_colo_milk_1ng_glyco_hcd203040_compound15.13085.13085.3 | controllerType=0 controllerNumber=1 scan=13085 | 29.5615 |
| 09441:3 | 2 | R.LAGKPTHVN[+1501.555]VSVVMAEVDGTC[+57.021]Y.- | HexNAc(5)Hex(3) | C[+57], N[+1502] | 1283.5738 | 3 | 3848.7068 | 3848.6974 | 0 | 2.4 | 332 | CRagged | 453.94 | 453.94 | 453.94 | 10.06 | 124 | >sp\|P01876\|IGHA1_HUMAN Immunoglobulin heavy constant alpha 1 OS=Homo sapiens OX=9606 GN=IGHA1 PE=1 SV=2 | 255 | 20201114_qwl_colo_milk_1ng_glyco_hcd203040_compound15.12529.12529.3 | controllerType=0 controllerNumber=1 scan=12529 | 28.5252 |
| 13674:4 | 2 | R.LAGKPTHVN[+2553.910]VSVVMAEVDGTC[+57.021]Y.- | HexNAc(5)Hex(5)Fuc(1)NeuAc(2) | C[+57], N[+2554] | 1226.0281 | 4 | 4901.0906 | 4901.0518 | 0 | 7.9 | 332 | CRagged | 459 | 306.65 | 306.65 | 7.71 | 124 | >sp\|P01876\|IGHA1_HUMAN Immunoglobulin heavy constant alpha 1 OS=Homo sapiens OX=9606 GN=IGHA1 PE=1 SV=2 | 255 | 20201114_qwl_colo_milk_1ng_glyco_hcd203040_compound15.16948.16948.4 | controllerType=0 controllerNumber=1 scan=16948 | 37.326 |
| 09362:3 | 2 | R.LAGKPTHVN[+1540.529]VSVVMAEVDGTC[+57.021]Y.- | HexNAc(2)Hex(7) | C[+57], N[+1541] | 1296.5646 | 3 | 3887.6793 | 3887.6706 | 0 | 2.3 | 332 | CRagged | 463.91 | 463.91 | 294 | 9.96 | 124 | >sp\|P01876\|IGHA1_HUMAN Immunoglobulin heavy constant alpha 1 OS=Homo sapiens OX=9606 GN=IGHA1 PE=1 SV=2 | 255 | 20201114_qwl_colo_milk_1ng_glyco_hcd203040_compound15.12445.12445.3 | controllerType=0 controllerNumber=1 scan=12445 | 28.3483 |
| 13599:3 | 2 | R.LAGKPTHVN[+2350.830]VSVVMAEVDGTC[+57.021]Y.- | HexNAc(4)Hex(5)Fuc(1)NeuAc(2) | C[+57], N[+2351] | 1566.6657 | 3 | 4697.9826 | 4697.9724 | 0 | 2.2 | 332 | CRagged | 473.98 | 429.92 | 429.92 | 10.03 | 124 | >sp\|P01876\|IGHA1_HUMAN Immunoglobulin heavy constant alpha 1 OS=Homo sapiens OX=9606 GN=IGHA1 PE=1 SV=2 | 255 | 20201114_qwl_colo_milk_1ng_glyco_hcd203040_compound15.16870.16870.3 | controllerType=0 controllerNumber=1 scan=16870 | 37.1714 |
| 10393:3 | 2 | R.LAGKPTHVN[+1581.555]VSVVMAEVDGTC[+57.021]Y.- | HexNAc(3)Hex(6) | C[+57], N[+1582] | 1310.2385 | 3 | 3928.7008 | 3928.6971 | 0 | 0.9 | 332 | CRagged | 481.36 | 456.26 | 313.86 | 9.97 | 124 | >sp\|P01876\|IGHA1_HUMAN Immunoglobulin heavy constant alpha 1 OS=Homo sapiens OX=9606 GN=IGHA1 PE=1 SV=2 | 255 | 20201114_qwl_colo_milk_1ng_glyco_hcd203040_compound15.13519.13519.3 | controllerType=0 controllerNumber=1 scan=13519 | 30.4067 |
| 09427:3 | 2 | R.LAGKPTHVN[+1606.587]VSVVMAEVDGTC[+57.021]Y.- | HexNAc(4)Hex(4)Fuc(1) | C[+57], N[+1607] | 1318.5928 | 3 | 3953.7639 | 3953.7288 | 0 | 8.9 | 332 | CRagged | 498.02 | 498.02 | 498.02 | 9.75 | 124 | >sp\|P01876\|IGHA1_HUMAN Immunoglobulin heavy constant alpha 1 OS=Homo sapiens OX=9606 GN=IGHA1 PE=1 SV=2 | 255 | 20201114_qwl_colo_milk_1ng_glyco_hcd203040_compound15.12514.12514.3 | controllerType=0 controllerNumber=1 scan=12514 | 28.4914 |
| 09483:3 | 2 | R.LAGKPTHVN[+1095.397]VSVVMAEVDGTC[+57.021]Y.- | HexNAc(3)Hex(3) | C[+57], N[+1095] | 1148.1916 | 3 | 3442.5601 | 3442.5387 | 0 | 6.2 | 332 | CRagged | 526.23 | 526.23 | 526.23 | 9.76 | 124 | >sp\|P01876\|IGHA1_HUMAN Immunoglobulin heavy constant alpha 1 OS=Homo sapiens OX=9606 GN=IGHA1 PE=1 SV=2 | 255 | 20201114_qwl_colo_milk_1ng_glyco_hcd203040_compound15.12573.12573.3 | controllerType=0 controllerNumber=1 scan=12573 | 28.6371 |
| 10101:3 | 2 | R.LAGKPTHVN[+1054.370]VSVVMAEVDGTC[+57.021]Y.- | HexNAc(2)Hex(4) | C[+57], N[+1054] | 1134.515 | 3 | 3401.5306 | 3401.5121 | 0 | 5.4 | 332 | CRagged | 527.23 | 480.33 | 259.47 | 9.94 | 124 | >sp\|P01876\|IGHA1_HUMAN Immunoglobulin heavy constant alpha 1 OS=Homo sapiens OX=9606 GN=IGHA1 PE=1 SV=2 | 255 | 20201114_qwl_colo_milk_1ng_glyco_hcd203040_compound15.13214.13214.3 | controllerType=0 controllerNumber=1 scan=13214 | 29.8191 |
| 13673:3 | 2 | R.LAGKPTHVN[+2553.910]VSVVMAEVDGTC[+57.021]Y.- | HexNAc(5)Hex(5)Fuc(1)NeuAc(2) | C[+57], N[+2554] | 1634.3683 | 3 | 4901.0903 | 4901.0518 | 0 | 7.9 | 332 | CRagged | 545.24 | 402.81 | 402.81 | 10.02 | 124 | >sp\|P01876\|IGHA1_HUMAN Immunoglobulin heavy constant alpha 1 OS=Homo sapiens OX=9606 GN=IGHA1 PE=1 SV=2 | 255 | 20201114_qwl_colo_milk_1ng_glyco_hcd203040_compound15.16947.16947.3 | controllerType=0 controllerNumber=1 scan=16947 | 37.3242 |
| 10641:3 | 2 | R.LAGKPTHVN[+1054.370]VSVVMAEVDGTC[+57.021]Y.- | HexNAc(2)Hex(4) | C[+57], N[+1054] | 1134.5131 | 3 | 3401.5247 | 3401.5121 | 0 | 3.7 | 332 | CRagged | 549.9 | 549.9 | 387.85 | 10.45 | 124 | >sp\|P01876\|IGHA1_HUMAN Immunoglobulin heavy constant alpha 1 OS=Homo sapiens OX=9606 GN=IGHA1 PE=1 SV=2 | 255 | 20201114_qwl_colo_milk_1ng_glyco_hcd203040_compound15.13777.13777.3 | controllerType=0 controllerNumber=1 scan=13777 | 30.9336 |
| 12135:3 | 2 | R.LAGKPTHVN[+1548.545]VSVVMAEVDGTC[+57.021]Y.- | HexNAc(3)Hex(4)NeuAc(1) | C[+57], N[+1549] | 1299.2331 | 3 | 3895.6847 | 3895.6869 | 0 | -0.6 | 332 | CRagged | 556.61 | 311.85 | 311.85 | 8.28 | 124 | >sp\|P01876\|IGHA1_HUMAN Immunoglobulin heavy constant alpha 1 OS=Homo sapiens OX=9606 GN=IGHA1 PE=1 SV=2 | 255 | 20201114_qwl_colo_milk_1ng_glyco_hcd203040_compound15.15338.15338.3 | controllerType=0 controllerNumber=1 scan=15338 | 33.9261 |
| 10755:3 | 2 | R.LAGKPTHVN[+2262.814]VSVVMAEVDGTC[+57.021]Y.- | HexNAc(5)Hex(5)Fuc(1)NeuAc(1) | C[+57], N[+2263] | 1537.3287 | 3 | 4609.9715 | 4609.9564 | 0 | 3.3 | 332 | CRagged | 564.11 | 433.14 | 433.14 | 10.58 | 124 | >sp\|P01876\|IGHA1_HUMAN Immunoglobulin heavy constant alpha 1 OS=Homo sapiens OX=9606 GN=IGHA1 PE=1 SV=2 | 255 | 20201114_qwl_colo_milk_1ng_glyco_hcd203040_compound15.13896.13896.3 | controllerType=0 controllerNumber=1 scan=13896 | 31.183 |
| 10683:3 | 2 | R.LAGKPTHVN[+1216.423]VSVVMAEVDGTC[+57.021]Y.- | HexNAc(2)Hex(5) | C[+57], N[+1216] | 1188.5281 | 3 | 3563.5698 | 3563.5649 | 0 | 1.4 | 332 | CRagged | 571.62 | 401.57 | 321.28 | 10.48 | 124 | >sp\|P01876\|IGHA1_HUMAN Immunoglobulin heavy constant alpha 1 OS=Homo sapiens OX=9606 GN=IGHA1 PE=1 SV=2 | 255 | 20201114_qwl_colo_milk_1ng_glyco_hcd203040_compound15.13821.13821.3 | controllerType=0 controllerNumber=1 scan=13821 | 31.0429 |
| 10787:3 | 2 | R.LAGKPTHVN[+1913.677]VSVVMAEVDGTC[+57.021]Y.- | HexNAc(4)Hex(5)NeuAc(1) | C[+57], N[+1914] | 1420.9543 | 3 | 4260.8483 | 4260.8191 | 0 | 6.8 | 332 | CRagged | 571.88 | 389.18 | 389.18 | 10.2 | 124 | >sp\|P01876\|IGHA1_HUMAN Immunoglobulin heavy constant alpha 1 OS=Homo sapiens OX=9606 GN=IGHA1 PE=1 SV=2 | 255 | 20201114_qwl_colo_milk_1ng_glyco_hcd203040_compound15.13930.13930.3 | controllerType=0 controllerNumber=1 scan=13930 | 31.2445 |
| 10612:3 | 2 | R.LAGKPTHVN[+1378.476]VSVVMAEVDGTC[+57.021]Y.- | HexNAc(2)Hex(6) | C[+57], N[+1378] | 1242.5484 | 3 | 3725.6306 | 3725.6178 | 0 | 3.5 | 332 | CRagged | 574.27 | 574.27 | 353.2 | 10.55 | 124 | >sp\|P01876\|IGHA1_HUMAN Immunoglobulin heavy constant alpha 1 OS=Homo sapiens OX=9606 GN=IGHA1 PE=1 SV=2 | 255 | 20201114_qwl_colo_milk_1ng_glyco_hcd203040_compound15.13747.13747.3 | controllerType=0 controllerNumber=1 scan=13747 | 30.8714 |
| 12008:3 | 2 | R.LAGKPTHVN[+2116.756]VSVVMAEVDGTC[+57.021]Y.- | HexNAc(5)Hex(5)NeuAc(1) | C[+57], N[+2117] | 1488.6518 | 3 | 4463.9408 | 4463.8985 | 0 | 9.5 | 332 | CRagged | 575.85 | 517.21 | 517.21 | 9.99 | 124 | >sp\|P01876\|IGHA1_HUMAN Immunoglobulin heavy constant alpha 1 OS=Homo sapiens OX=9606 GN=IGHA1 PE=1 SV=2 | 255 | 20201114_qwl_colo_milk_1ng_glyco_hcd203040_compound15.15206.15206.3 | controllerType=0 controllerNumber=1 scan=15206 | 33.6907 |
| 09389:3 | 2 | R.LAGKPTHVN[+1419.502]VSVVMAEVDGTC[+57.021]Y.- | HexNAc(3)Hex(5) | C[+57], N[+1420] | 1256.2201 | 3 | 3766.6457 | 3766.6443 | 0 | 0.4 | 332 | CRagged | 576.95 | 576.95 | 377.79 | 10.43 | 124 | >sp\|P01876\|IGHA1_HUMAN Immunoglobulin heavy constant alpha 1 OS=Homo sapiens OX=9606 GN=IGHA1 PE=1 SV=2 | 255 | 20201114_qwl_colo_milk_1ng_glyco_hcd203040_compound15.12473.12473.3 | controllerType=0 controllerNumber=1 scan=12473 | 28.4017 |
| 12001:3 | 2 | R.LAGKPTHVN[+2059.735]VSVVMAEVDGTC[+57.021]Y.- | HexNAc(4)Hex(5)Fuc(1)NeuAc(1) | C[+57], N[+2060] | 1469.6435 | 3 | 4406.9159 | 4406.877 | 0 | 8.8 | 332 | CRagged | 580.4 | 580.4 | 580.4 | 11.43 | 124 | >sp\|P01876\|IGHA1_HUMAN Immunoglobulin heavy constant alpha 1 OS=Homo sapiens OX=9606 GN=IGHA1 PE=1 SV=2 | 255 | 20201114_qwl_colo_milk_1ng_glyco_hcd203040_compound15.15199.15199.3 | controllerType=0 controllerNumber=1 scan=15199 | 33.6778 |
| 10412:2 | 2 | R.LAGKPTHVN[+1378.476]VSVVMAEVDGTC[+57.021]Y.- | HexNAc(2)Hex(6) | C[+57], N[+1378] | 1863.3184 | 2 | 3725.6295 | 3725.6178 | 0 | 3.2 | 332 | CRagged | 584.23 | 486.11 | 188.28 | 9.63 | 124 | >sp\|P01876\|IGHA1_HUMAN Immunoglobulin heavy constant alpha 1 OS=Homo sapiens OX=9606 GN=IGHA1 PE=1 SV=2 | 255 | 20201114_qwl_colo_milk_1ng_glyco_hcd203040_compound15.13538.13538.2 | controllerType=0 controllerNumber=1 scan=13538 | 30.4459 |
| 10460:3 | 2 | R.LAGKPTHVN[+1378.476]VSVVMAEVDGTC[+57.021]Y.- | HexNAc(2)Hex(6) | C[+57], N[+1378] | 1242.55 | 3 | 3725.6354 | 3725.6178 | 0 | 4.7 | 332 | CRagged | 588.33 | 588.33 | 364.66 | 11.56 | 124 | >sp\|P01876\|IGHA1_HUMAN Immunoglobulin heavy constant alpha 1 OS=Homo sapiens OX=9606 GN=IGHA1 PE=1 SV=2 | 255 | 20201114_qwl_colo_milk_1ng_glyco_hcd203040_compound15.13589.13589.3 | controllerType=0 controllerNumber=1 scan=13589 | 30.567 |
| 09296:3 | 2 | R.LAGKPTHVN[+1825.661]VSVVMAEVDGTC[+57.021]Y.- | HexNAc(5)Hex(5) | C[+57], N[+1826] | 1391.6071 | 3 | 4172.8068 | 4172.8031 | 0 | 0.9 | 332 | CRagged | 594.63 | 594.63 | 359.11 | 11.74 | 124 | >sp\|P01876\|IGHA1_HUMAN Immunoglobulin heavy constant alpha 1 OS=Homo sapiens OX=9606 GN=IGHA1 PE=1 SV=2 | 255 | 20201114_qwl_colo_milk_1ng_glyco_hcd203040_compound15.12376.12376.3 | controllerType=0 controllerNumber=1 scan=12376 | 28.221 |
| 09269:3 | 2 | R.LAGKPTHVN[+1540.529]VSVVMAEVDGTC[+57.021]Y.- | HexNAc(2)Hex(7) | C[+57], N[+1541] | 1296.5643 | 3 | 3887.6782 | 3887.6706 | 0 | 2 | 332 | CRagged | 601.82 | 601.82 | 449.39 | 12.79 | 124 | >sp\|P01876\|IGHA1_HUMAN Immunoglobulin heavy constant alpha 1 OS=Homo sapiens OX=9606 GN=IGHA1 PE=1 SV=2 | 255 | 20201114_qwl_colo_milk_1ng_glyco_hcd203040_compound15.12348.12348.3 | controllerType=0 controllerNumber=1 scan=12348 | 28.1699 |
| 10680:3 | 2 | R.LAGKPTHVN[+1216.423]VSVVMAEVDGTC[+57.021]Y.- | HexNAc(2)Hex(5) | C[+57], N[+1216] | 1188.5281 | 3 | 3563.5698 | 3563.5649 | 0 | 1.4 | 332 | CRagged | 607.81 | 607.81 | 331.29 | 12.79 | 124 | >sp\|P01876\|IGHA1_HUMAN Immunoglobulin heavy constant alpha 1 OS=Homo sapiens OX=9606 GN=IGHA1 PE=1 SV=2 | 255 | 20201114_qwl_colo_milk_1ng_glyco_hcd203040_compound15.13818.13818.3 | controllerType=0 controllerNumber=1 scan=13818 | 31.036 |
| 11950:4 | 2 | R.LAGKPTHVN[+2262.814]VSVVMAEVDGTC[+57.021]Y.- | HexNAc(5)Hex(5)Fuc(1)NeuAc(1) | C[+57], N[+2263] | 1153.2465 | 4 | 4609.9642 | 4609.9564 | 0 | 1.7 | 332 | CRagged | 610.32 | 551.31 | 551.31 | 12.78 | 124 | >sp\|P01876\|IGHA1_HUMAN Immunoglobulin heavy constant alpha 1 OS=Homo sapiens OX=9606 GN=IGHA1 PE=1 SV=2 | 255 | 20201114_qwl_colo_milk_1ng_glyco_hcd203040_compound15.15146.15146.4 | controllerType=0 controllerNumber=1 scan=15146 | 33.5785 |
| 09410:3 | 2 | R.LAGKPTHVN[+1501.555]VSVVMAEVDGTC[+57.021]Y.- | HexNAc(5)Hex(3) | C[+57], N[+1502] | 1283.5745 | 3 | 3848.709 | 3848.6974 | 0 | 3 | 332 | CRagged | 611.28 | 611.28 | 611.28 | 12.89 | 124 | >sp\|P01876\|IGHA1_HUMAN Immunoglobulin heavy constant alpha 1 OS=Homo sapiens OX=9606 GN=IGHA1 PE=1 SV=2 | 255 | 20201114_qwl_colo_milk_1ng_glyco_hcd203040_compound15.12496.12496.3 | controllerType=0 controllerNumber=1 scan=12496 | 28.453 |
| 10008:3 | 2 | R.LAGKPTHVN[+1216.423]VSVVMAEVDGTC[+57.021]Y.- | HexNAc(2)Hex(5) | C[+57], N[+1216] | 1188.5292 | 3 | 3563.5731 | 3563.5649 | 0 | 2.3 | 332 | CRagged | 628.7 | 550.43 | 377.19 | 12.89 | 124 | >sp\|P01876\|IGHA1_HUMAN Immunoglobulin heavy constant alpha 1 OS=Homo sapiens OX=9606 GN=IGHA1 PE=1 SV=2 | 255 | 20201114_qwl_colo_milk_1ng_glyco_hcd203040_compound15.13118.13118.3 | controllerType=0 controllerNumber=1 scan=13118 | 29.6266 |
| 09372:3 | 2 | R.LAGKPTHVN[+1216.423]VSVVMAEVDGTC[+57.021]Y.- | HexNAc(2)Hex(5) | C[+57], N[+1216] | 1188.5288 | 3 | 3563.5718 | 3563.5649 | 0 | 1.9 | 332 | CRagged | 631.81 | 631.81 | 347.55 | 12.79 | 124 | >sp\|P01876\|IGHA1_HUMAN Immunoglobulin heavy constant alpha 1 OS=Homo sapiens OX=9606 GN=IGHA1 PE=1 SV=2 | 255 | 20201114_qwl_colo_milk_1ng_glyco_hcd203040_compound15.12456.12456.3 | controllerType=0 controllerNumber=1 scan=12456 | 28.3694 |
| 09359:3 | 2 | R.LAGKPTHVN[+1540.529]VSVVMAEVDGTC[+57.021]Y.- | HexNAc(2)Hex(7) | C[+57], N[+1541] | 1296.5646 | 3 | 3887.6793 | 3887.6706 | 0 | 2.3 | 332 | CRagged | 638.99 | 638.99 | 373.49 | 12.79 | 124 | >sp\|P01876\|IGHA1_HUMAN Immunoglobulin heavy constant alpha 1 OS=Homo sapiens OX=9606 GN=IGHA1 PE=1 SV=2 | 255 | 20201114_qwl_colo_milk_1ng_glyco_hcd203040_compound15.12442.12442.3 | controllerType=0 controllerNumber=1 scan=12442 | 28.3428 |
| 09370:3 | 2 | R.LAGKPTHVN[+1419.502]VSVVMAEVDGTC[+57.021]Y.- | HexNAc(3)Hex(5) | C[+57], N[+1420] | 1256.2203 | 3 | 3766.6463 | 3766.6443 | 0 | 0.5 | 332 | CRagged | 650.7 | 650.7 | 437.05 | 13.74 | 124 | >sp\|P01876\|IGHA1_HUMAN Immunoglobulin heavy constant alpha 1 OS=Homo sapiens OX=9606 GN=IGHA1 PE=1 SV=2 | 255 | 20201114_qwl_colo_milk_1ng_glyco_hcd203040_compound15.12453.12453.3 | controllerType=0 controllerNumber=1 scan=12453 | 28.3611 |
| 10459:3 | 2 | R.LAGKPTHVN[+1606.587]VSVVMAEVDGTC[+57.021]Y.- | HexNAc(4)Hex(4)Fuc(1) | C[+57], N[+1607] | 1318.5879 | 3 | 3953.7493 | 3953.7288 | 0 | 5.2 | 332 | CRagged | 651.1 | 651.1 | 651.1 | 13.56 | 124 | >sp\|P01876\|IGHA1_HUMAN Immunoglobulin heavy constant alpha 1 OS=Homo sapiens OX=9606 GN=IGHA1 PE=1 SV=2 | 255 | 20201114_qwl_colo_milk_1ng_glyco_hcd203040_compound15.13588.13588.3 | controllerType=0 controllerNumber=1 scan=13588 | 30.5653 |
| 10410:2 | 2 | R.LAGKPTHVN[+1378.476]VSVVMAEVDGTC[+57.021]Y.- | HexNAc(2)Hex(6) | C[+57], N[+1378] | 1863.3184 | 2 | 3725.6295 | 3725.6178 | 0 | 3.2 | 332 | CRagged | 655.23 | 604.38 | 336.98 | 13.23 | 124 | >sp\|P01876\|IGHA1_HUMAN Immunoglobulin heavy constant alpha 1 OS=Homo sapiens OX=9606 GN=IGHA1 PE=1 SV=2 | 255 | 20201114_qwl_colo_milk_1ng_glyco_hcd203040_compound15.13536.13536.2 | controllerType=0 controllerNumber=1 scan=13536 | 30.4407 |
| 09392:3 | 2 | R.LAGKPTHVN[+1419.502]VSVVMAEVDGTC[+57.021]Y.- | HexNAc(3)Hex(5) | C[+57], N[+1420] | 1256.2201 | 3 | 3766.6457 | 3766.6443 | 0 | 0.4 | 332 | CRagged | 670.33 | 461.74 | 365.09 | 13.74 | 124 | >sp\|P01876\|IGHA1_HUMAN Immunoglobulin heavy constant alpha 1 OS=Homo sapiens OX=9606 GN=IGHA1 PE=1 SV=2 | 255 | 20201114_qwl_colo_milk_1ng_glyco_hcd203040_compound15.12476.12476.3 | controllerType=0 controllerNumber=1 scan=12476 | 28.4106 |
| 09374:3 | 2 | R.LAGKPTHVN[+1216.423]VSVVMAEVDGTC[+57.021]Y.- | HexNAc(2)Hex(5) | C[+57], N[+1216] | 1188.5288 | 3 | 3563.5718 | 3563.5649 | 0 | 1.9 | 332 | CRagged | 673.41 | 673.41 | 427.38 | 13.79 | 124 | >sp\|P01876\|IGHA1_HUMAN Immunoglobulin heavy constant alpha 1 OS=Homo sapiens OX=9606 GN=IGHA1 PE=1 SV=2 | 255 | 20201114_qwl_colo_milk_1ng_glyco_hcd203040_compound15.12458.12458.3 | controllerType=0 controllerNumber=1 scan=12458 | 28.3743 |
| 09254:3 | 2 | R.LAGKPTHVN[+1702.581]VSVVMAEVDGTC[+57.021]Y.- | HexNAc(2)Hex(8) | C[+57], N[+1703] | 1350.5822 | 3 | 4049.7321 | 4049.7234 | 0 | 2.1 | 332 | CRagged | 676.87 | 676.87 | 468.16 | 13.79 | 124 | >sp\|P01876\|IGHA1_HUMAN Immunoglobulin heavy constant alpha 1 OS=Homo sapiens OX=9606 GN=IGHA1 PE=1 SV=2 | 255 | 20201114_qwl_colo_milk_1ng_glyco_hcd203040_compound15.12332.12332.3 | controllerType=0 controllerNumber=1 scan=12332 | 28.139 |
| 10377:3 | 2 | R.LAGKPTHVN[+1460.529]VSVVMAEVDGTC[+57.021]Y.- | HexNAc(4)Hex(4) | C[+57], N[+1461] | 1269.8989 | 3 | 3807.682 | 3807.6709 | 0 | 2.9 | 332 | CRagged | 682.36 | 607.35 | 444.71 | 13.89 | 124 | >sp\|P01876\|IGHA1_HUMAN Immunoglobulin heavy constant alpha 1 OS=Homo sapiens OX=9606 GN=IGHA1 PE=1 SV=2 | 255 | 20201114_qwl_colo_milk_1ng_glyco_hcd203040_compound15.13502.13502.3 | controllerType=0 controllerNumber=1 scan=13502 | 30.3717 |
| 13716:3 | 2 | R.LAGKPTHVN[+2204.772]VSVVMAEVDGTC[+57.021]Y.- | HexNAc(4)Hex(5)NeuAc(2) | C[+57], N[+2205] | 1517.9847 | 3 | 4551.9395 | 4551.9145 | 0 | 5.5 | 332 | CRagged | 686.28 | 410.23 | 410.23 | 13.56 | 124 | >sp\|P01876\|IGHA1_HUMAN Immunoglobulin heavy constant alpha 1 OS=Homo sapiens OX=9606 GN=IGHA1 PE=1 SV=2 | 255 | 20201114_qwl_colo_milk_1ng_glyco_hcd203040_compound15.16992.16992.3 | controllerType=0 controllerNumber=1 scan=16992 | 37.414 |
| 13671:3 | 2 | R.LAGKPTHVN[+2553.910]VSVVMAEVDGTC[+57.021]Y.- | HexNAc(5)Hex(5)Fuc(1)NeuAc(2) | C[+57], N[+2554] | 1634.3683 | 3 | 4901.0903 | 4901.0518 | 0 | 7.9 | 332 | CRagged | 689.69 | 483.25 | 483.25 | 13.43 | 124 | >sp\|P01876\|IGHA1_HUMAN Immunoglobulin heavy constant alpha 1 OS=Homo sapiens OX=9606 GN=IGHA1 PE=1 SV=2 | 255 | 20201114_qwl_colo_milk_1ng_glyco_hcd203040_compound15.16945.16945.3 | controllerType=0 controllerNumber=1 scan=16945 | 37.3196 |
| 13714:3 | 2 | R.LAGKPTHVN[+2204.772]VSVVMAEVDGTC[+57.021]Y.- | HexNAc(4)Hex(5)NeuAc(2) | C[+57], N[+2205] | 1517.9847 | 3 | 4551.9395 | 4551.9145 | 0 | 5.5 | 332 | CRagged | 692.93 | 514.29 | 514.29 | 13.56 | 124 | >sp\|P01876\|IGHA1_HUMAN Immunoglobulin heavy constant alpha 1 OS=Homo sapiens OX=9606 GN=IGHA1 PE=1 SV=2 | 255 | 20201114_qwl_colo_milk_1ng_glyco_hcd203040_compound15.16990.16990.3 | controllerType=0 controllerNumber=1 scan=16990 | 37.4101 |
| 10349:3 | 2 | R.LAGKPTHVN[+1971.719]VSVVMAEVDGTC[+57.021]Y.- | HexNAc(5)Hex(5)Fuc(1) | C[+57], N[+1972] | 1440.3039 | 3 | 4318.8971 | 4318.861 | 0 | 8.4 | 332 | CRagged | 695.51 | 695.51 | 441.87 | 13.43 | 124 | >sp\|P01876\|IGHA1_HUMAN Immunoglobulin heavy constant alpha 1 OS=Homo sapiens OX=9606 GN=IGHA1 PE=1 SV=2 | 255 | 20201114_qwl_colo_milk_1ng_glyco_hcd203040_compound15.13473.13473.3 | controllerType=0 controllerNumber=1 scan=13473 | 30.3104 |
| 10473:3 | 2 | R.LAGKPTHVN[+1647.613]VSVVMAEVDGTC[+57.021]Y.- | HexNAc(5)Hex(3)Fuc(1) | C[+57], N[+1648] | 1332.2588 | 3 | 3994.762 | 3994.7553 | 0 | 1.7 | 332 | CRagged | 700.51 | 700.51 | 700.51 | 14.79 | 124 | >sp\|P01876\|IGHA1_HUMAN Immunoglobulin heavy constant alpha 1 OS=Homo sapiens OX=9606 GN=IGHA1 PE=1 SV=2 | 255 | 20201114_qwl_colo_milk_1ng_glyco_hcd203040_compound15.13603.13603.3 | controllerType=0 controllerNumber=1 scan=13603 | 30.6036 |

Table S4. Cont.

| Query #:z | Protein Rank | Glycopeptide | Glycan | Modification Type(s) | Observed m/z | z | Observed (M+H) | Calc. mass (M+H) | Off-by-x error | Mass error (ppm) | StartPosition | Cleavage | Score | Delta | Delta Mod | \|Log Prob\| | # of unique peptides | Protein Name | Protein DB number | Comment | Scan # | Scan Time |
|---|---|---|---|---|---|---|---|---|---|---|---|---|---|---|---|---|---|---|---|---|---|---|
| 09485:3 | 2 | R.LAGKPTHVN[+1095.397]VSVVMAEVDGTC[+57.021]Y.- | HexNAc(3)Hex(3) | C[+57], N[+1095] | 1148.1916 | 3 | 3442.5601 | 3442.5387 | 0 | 6.2 | 332 | CRagged | 706.2 | 706.2 | 706.2 | 14.37 | 124 | >sp\|P01876\|IGHA1_HUMAN Immunoglobulin heavy constant alpha 1 OS=Homo sapiens OX=9606 GN=IGHA1 PE=1 SV=2 | 255 | 20201114_qwl_colo_milk_1ng_glyco_hcd203040_compound15.12575.12575.3 | controllerType=0 controllerNumber=1 scan=12575 | 28.6436 |
| 10508:3 | 2 | R.LAGKPTHVN[+1095.397]VSVVMAEVDGTC[+57.021]Y.- | HexNAc(3)Hex(3) | C[+57], N[+1095] | 1148.1944 | 3 | 3442.5685 | 3442.5387 | 0 | 8.7 | 332 | CRagged | 712.74 | 612.9 | 612.9 | 14.43 | 124 | >sp\|P01876\|IGHA1_HUMAN Immunoglobulin heavy constant alpha 1 OS=Homo sapiens OX=9606 GN=IGHA1 PE=1 SV=2 | 255 | 20201114_qwl_colo_milk_1ng_glyco_hcd203040_compound15.13639.13639.3 | controllerType=0 controllerNumber=1 scan=13639 | 30.6607 |
| 12113:3 | 2 | R.LAGKPTHVN[+1548.545]VSVVMAEVDGTC[+57.021]Y.- | HexNAc(3)Hex(4)NeuAc(1) | C[+57], N[+1549] | 1299.2366 | 3 | 3895.6953 | 3895.6869 | 0 | 2.2 | 332 | CRagged | 712.91 | 702.94 | 702.94 | 14.79 | 124 | >sp\|P01876\|IGHA1_HUMAN Immunoglobulin heavy constant alpha 1 OS=Homo sapiens OX=9606 GN=IGHA1 PE=1 SV=2 | 255 | 20201114_qwl_colo_milk_1ng_glyco_hcd203040_compound15.15315.15315.3 | controllerType=0 controllerNumber=1 scan=15315 | 33.8877 |
| 11939:3 | 2 | R.LAGKPTHVN[+2262.814]VSVVMAEVDGTC[+57.021]Y.- | HexNAc(5)Hex(5)Fuc(1)NeuAc(1) | C[+57], N[+2263] | 1537.3273 | 3 | 4609.9674 | 4609.9564 | 0 | 2.4 | 332 | CRagged | 720.12 | 603.38 | 603.38 | 14.89 | 124 | >sp\|P01876\|IGHA1_HUMAN Immunoglobulin heavy constant alpha 1 OS=Homo sapiens OX=9606 GN=IGHA1 PE=1 SV=2 | 255 | 20201114_qwl_colo_milk_1ng_glyco_hcd203040_compound15.15134.15134.3 | controllerType=0 controllerNumber=1 scan=15134 | 33.5297 |
| 09353:3 | 2 | R.LAGKPTHVN[+1606.587]VSVVMAEVDGTC[+57.021]Y.- | HexNAc(4)Hex(4)Fuc(1) | C[+57], N[+1607] | 1318.5834 | 3 | 3953.7357 | 3953.7288 | 0 | 1.8 | 332 | CRagged | 730.24 | 612.59 | 612.59 | 14.79 | 124 | >sp\|P01876\|IGHA1_HUMAN Immunoglobulin heavy constant alpha 1 OS=Homo sapiens OX=9606 GN=IGHA1 PE=1 SV=2 | 255 | 20201114_qwl_colo_milk_1ng_glyco_hcd203040_compound15.12436.12436.3 | controllerType=0 controllerNumber=1 scan=12436 | 28.3286 |
| 10477:3 | 2 | R.LAGKPTHVN[+1444.534]VSVVMAEVDGTC[+57.021]Y.- | HexNAc(4)Hex(3)Fuc(1) | C[+57], N[+1445] | 1264.5666 | 3 | 3791.6852 | 3791.6759 | 0 | 2.4 | 332 | CRagged | 730.66 | 730.66 | 730.66 | 14.89 | 124 | >sp\|P01876\|IGHA1_HUMAN Immunoglobulin heavy constant alpha 1 OS=Homo sapiens OX=9606 GN=IGHA1 PE=1 SV=2 | 255 | 20201114_qwl_colo_milk_1ng_glyco_hcd203040_compound15.13607.13607.3 | controllerType=0 controllerNumber=1 scan=13607 | 30.6088 |
| 12003:3 | 2 | R.LAGKPTHVN[+2059.735]VSVVMAEVDGTC[+57.021]Y.- | HexNAc(4)Hex(5)Fuc(1)NeuAc(1) | C[+57], N[+2060] | 1469.6435 | 3 | 4406.9159 | 4406.877 | 0 | 8.8 | 332 | CRagged | 732.04 | 682.34 | 682.34 | 14.43 | 124 | >sp\|P01876\|IGHA1_HUMAN Immunoglobulin heavy constant alpha 1 OS=Homo sapiens OX=9606 GN=IGHA1 PE=1 SV=2 | 255 | 20201114_qwl_colo_milk_1ng_glyco_hcd203040_compound15.15201.15201.3 | controllerType=0 controllerNumber=1 scan=15201 | 33.6837 |
| 09351:3 | 2 | R.LAGKPTHVN[+1606.587]VSVVMAEVDGTC[+57.021]Y.- | HexNAc(4)Hex(4)Fuc(1) | C[+57], N[+1607] | 1318.5834 | 3 | 3953.7357 | 3953.7288 | 0 | 1.8 | 332 | CRagged | 735.99 | 663.08 | 663.08 | 14.79 | 124 | >sp\|P01876\|IGHA1_HUMAN Immunoglobulin heavy constant alpha 1 OS=Homo sapiens OX=9606 GN=IGHA1 PE=1 SV=2 | 255 | 20201114_qwl_colo_milk_1ng_glyco_hcd203040_compound15.12434.12434.3 | controllerType=0 controllerNumber=1 scan=12434 | 28.3237 |
| 09306:3 | 2 | R.LAGKPTHVN[+1581.555]VSVVMAEVDGTC[+57.021]Y.- | HexNAc(3)Hex(6) | C[+57], N[+1582] | 1310.2377 | 3 | 3928.6986 | 3928.6971 | 0 | 0.4 | 332 | CRagged | 744.8 | 744.8 | 438.9 | 14.74 | 124 | >sp\|P01876\|IGHA1_HUMAN Immunoglobulin heavy constant alpha 1 OS=Homo sapiens OX=9606 GN=IGHA1 PE=1 SV=2 | 255 | 20201114_qwl_colo_milk_1ng_glyco_hcd203040_compound15.12386.12386.3 | controllerType=0 controllerNumber=1 scan=12386 | 28.2383 |
| 12000:3 | 2 | R.LAGKPTHVN[+1872.651]VSVVMAEVDGTC[+57.021]Y.- | HexNAc(3)Hex(6)NeuAc(1) | C[+57], N[+1873] | 1407.2819 | 3 | 4219.8312 | 4219.7926 | 0 | 9.2 | 332 | CRagged | 745.29 | 664.74 | 468.18 | 14.3 | 124 | >sp\|P01876\|IGHA1_HUMAN Immunoglobulin heavy constant alpha 1 OS=Homo sapiens OX=9606 GN=IGHA1 PE=1 SV=2 | 255 | 20201114_qwl_colo_milk_1ng_glyco_hcd203040_compound15.15198.15198.3 | controllerType=0 controllerNumber=1 scan=15198 | 33.6764 |
| 10350:3 | 2 | R.LAGKPTHVN[+1809.666]VSVVMAEVDGTC[+57.021]Y.- | HexNAc(5)Hex(4)Fuc(1) | C[+57], N[+1810] | 1386.2856 | 3 | 4156.8422 | 4156.8081 | 0 | 8.2 | 332 | CRagged | 746.04 | 639.79 | 503.14 | 14.43 | 124 | >sp\|P01876\|IGHA1_HUMAN Immunoglobulin heavy constant alpha 1 OS=Homo sapiens OX=9606 GN=IGHA1 PE=1 SV=2 | 255 | 20201114_qwl_colo_milk_1ng_glyco_hcd203040_compound15.13474.13474.3 | controllerType=0 controllerNumber=1 scan=13474 | 30.3121 |
| 12069:3 | 2 | R.LAGKPTHVN[+1710.598]VSVVMAEVDGTC[+57.021]Y.- | HexNAc(3)Hex(5)NeuAc(1) | C[+57], N[+1711] | 1353.2536 | 3 | 4057.7462 | 4057.7397 | 0 | 1.6 | 332 | CRagged | 749.45 | 749.45 | 431.77 | 14.79 | 124 | >sp\|P01876\|IGHA1_HUMAN Immunoglobulin heavy constant alpha 1 OS=Homo sapiens OX=9606 GN=IGHA1 PE=1 SV=2 | 255 | 20201114_qwl_colo_milk_1ng_glyco_hcd203040_compound15.15270.15270.3 | controllerType=0 controllerNumber=1 scan=15270 | 33.807 |
| 09413:3 | 2 | R.LAGKPTHVN[+1501.555]VSVVMAEVDGTC[+57.021]Y.- | HexNAc(5)Hex(3) | C[+57], N[+1502] | 1283.5745 | 3 | 3848.709 | 3848.6974 | 0 | 3 | 332 | CRagged | 757 | 757 | 757 | 15.89 | 124 | >sp\|P01876\|IGHA1_HUMAN Immunoglobulin heavy constant alpha 1 OS=Homo sapiens OX=9606 GN=IGHA1 PE=1 SV=2 | 255 | 20201114_qwl_colo_milk_1ng_glyco_hcd203040_compound15.12499.12499.3 | controllerType=0 controllerNumber=1 scan=12499 | 28.4582 |
| 12133:3 | 2 | R.LAGKPTHVN[+1548.545]VSVVMAEVDGTC[+57.021]Y.- | HexNAc(3)Hex(4)NeuAc(1) | C[+57], N[+1549] | 1299.2331 | 3 | 3895.6847 | 3895.6869 | 0 | -0.6 | 332 | CRagged | 763.92 | 763.92 | 763.92 | 15.27 | 124 | >sp\|P01876\|IGHA1_HUMAN Immunoglobulin heavy constant alpha 1 OS=Homo sapiens OX=9606 GN=IGHA1 PE=1 SV=2 | 255 | 20201114_qwl_colo_milk_1ng_glyco_hcd203040_compound15.15336.15336.3 | controllerType=0 controllerNumber=1 scan=15336 | 33.919 |
| 09229:3 | 2 | R.LAGKPTHVN[+1702.581]VSVVMAEVDGTC[+57.021]Y.- | HexNAc(2)Hex(8) | C[+57], N[+1703] | 1350.5891 | 3 | 4049.7527 | 4049.7234 | 0 | 7.2 | 332 | CRagged | 765.4 | 765.4 | 486.25 | 15.51 | 124 | >sp\|P01876\|IGHA1_HUMAN Immunoglobulin heavy constant alpha 1 OS=Homo sapiens OX=9606 GN=IGHA1 PE=1 SV=2 | 255 | 20201114_qwl_colo_milk_1ng_glyco_hcd203040_compound15.12305.12305.3 | controllerType=0 controllerNumber=1 scan=12305 | 28.0893 |
| 10466:3 | 2 | R.LAGKPTHVN[+1501.555]VSVVMAEVDGTC[+57.021]Y.- | HexNAc(5)Hex(3) | C[+57], N[+1502] | 1283.5765 | 3 | 3848.7149 | 3848.6974 | 0 | 4.5 | 332 | CRagged | 775.67 | 775.67 | 775.67 | 15.86 | 124 | >sp\|P01876\|IGHA1_HUMAN Immunoglobulin heavy constant alpha 1 OS=Homo sapiens OX=9606 GN=IGHA1 PE=1 SV=2 | 255 | 20201114_qwl_colo_milk_1ng_glyco_hcd203040_compound15.13595.13595.3 | controllerType=0 controllerNumber=1 scan=13595 | 30.5804 |
| 13597:3 | 2 | R.LAGKPTHVN[+2350.830]VSVVMAEVDGTC[+57.021]Y.- | HexNAc(4)Hex(5)Fuc(1)NeuAc(2) | C[+57], N[+2351] | 1566.6657 | 3 | 4697.9826 | 4697.9724 | 0 | 2.2 | 332 | CRagged | 776.37 | 776.37 | 776.37 | 15.79 | 124 | >sp\|P01876\|IGHA1_HUMAN Immunoglobulin heavy constant alpha 1 OS=Homo sapiens OX=9606 GN=IGHA1 PE=1 SV=2 | 255 | 20201114_qwl_colo_milk_1ng_glyco_hcd203040_compound15.16868.16868.3 | controllerType=0 controllerNumber=1 scan=16868 | 37.1656 |
| 09375:3 | 2 | R.LAGKPTHVN[+1663.608]VSVVMAEVDGTC[+57.021]Y.- | HexNAc(5)Hex(4) | C[+57], N[+1664] | 1337.5902 | 3 | 4010.756 | 4010.7502 | 0 | 1.4 | 332 | CRagged | 782.21 | 782.21 | 450.91 | 15.79 | 124 | >sp\|P01876\|IGHA1_HUMAN Immunoglobulin heavy constant alpha 1 OS=Homo sapiens OX=9606 GN=IGHA1 PE=1 SV=2 | 255 | 20201114_qwl_colo_milk_1ng_glyco_hcd203040_compound15.12459.12459.3 | controllerType=0 controllerNumber=1 scan=12459 | 28.3751 |
| 09443:3 | 2 | R.LAGKPTHVN[+1501.555]VSVVMAEVDGTC[+57.021]Y.- | HexNAc(5)Hex(3) | C[+57], N[+1502] | 1283.5738 | 3 | 3848.7068 | 3848.6974 | 0 | 2.4 | 332 | CRagged | 784.55 | 784.55 | 784.55 | 15.89 | 124 | >sp\|P01876\|IGHA1_HUMAN Immunoglobulin heavy constant alpha 1 OS=Homo sapiens OX=9606 GN=IGHA1 PE=1 SV=2 | 255 | 20201114_qwl_colo_milk_1ng_glyco_hcd203040_compound15.12531.12531.3 | controllerType=0 controllerNumber=1 scan=12531 | 28.5295 |
| 10372:3 | 2 | R.LAGKPTHVN[+1540.529]VSVVMAEVDGTC[+57.021]Y.- | HexNAc(2)Hex(7) | C[+57], N[+1541] | 1296.5662 | 3 | 3887.6841 | 3887.6706 | 0 | 3.5 | 332 | CRagged | 785.65 | 762.24 | 497.42 | 15.86 | 124 | >sp\|P01876\|IGHA1_HUMAN Immunoglobulin heavy constant alpha 1 OS=Homo sapiens OX=9606 GN=IGHA1 PE=1 SV=2 | 255 | 20201114_qwl_colo_milk_1ng_glyco_hcd203040_compound15.13497.13497.3 | controllerType=0 controllerNumber=1 scan=13497 | 30.3585 |
| 12071:3 | 2 | R.LAGKPTHVN[+1710.598]VSVVMAEVDGTC[+57.021]Y.- | HexNAc(3)Hex(5)NeuAc(1) | C[+57], N[+1711] | 1353.2536 | 3 | 4057.7462 | 4057.7397 | 0 | 1.6 | 332 | CRagged | 793.26 | 687.75 | 467.43 | 15.79 | 124 | >sp\|P01876\|IGHA1_HUMAN Immunoglobulin heavy constant alpha 1 OS=Homo sapiens OX=9606 GN=IGHA1 PE=1 SV=2 | 255 | 20201114_qwl_colo_milk_1ng_glyco_hcd203040_compound15.15272.15272.3 | controllerType=0 controllerNumber=1 scan=15272 | 33.8107 |
| 10370:3 | 2 | R.LAGKPTHVN[+1540.529]VSVVMAEVDGTC[+57.021]Y.- | HexNAc(2)Hex(7) | C[+57], N[+1541] | 1296.5662 | 3 | 3887.6841 | 3887.6706 | 0 | 3.5 | 332 | CRagged | 793.95 | 793.95 | 472.17 | 15.86 | 124 | >sp\|P01876\|IGHA1_HUMAN Immunoglobulin heavy constant alpha 1 OS=Homo sapiens OX=9606 GN=IGHA1 PE=1 SV=2 | 255 | 20201114_qwl_colo_milk_1ng_glyco_hcd203040_compound15.13495.13495.3 | controllerType=0 controllerNumber=1 scan=13495 | 30.3523 |
| 12096:3 | 2 | R.LAGKPTHVN[+1751.624]VSVVMAEVDGTC[+57.021]Y.- | HexNAc(4)Hex(4)NeuAc(1) | C[+57], N[+1752] | 1366.9284 | 3 | 4098.7707 | 4098.7663 | 0 | 1.1 | 332 | CRagged | 794.13 | 660.58 | 660.58 | 15.74 | 124 | >sp\|P01876\|IGHA1_HUMAN Immunoglobulin heavy constant alpha 1 OS=Homo sapiens OX=9606 GN=IGHA1 PE=1 SV=2 | 255 | 20201114_qwl_colo_milk_1ng_glyco_hcd203040_compound15.15298.15298.3 | controllerType=0 controllerNumber=1 scan=15298 | 33.8639 |
| 09304:3 | 2 | R.LAGKPTHVN[+1581.555]VSVVMAEVDGTC[+57.021]Y.- | HexNAc(3)Hex(6) | C[+57], N[+1582] | 1310.2377 | 3 | 3928.6986 | 3928.6971 | 0 | 0.4 | 332 | CRagged | 796.82 | 796.82 | 494.28 | 15.74 | 124 | >sp\|P01876\|IGHA1_HUMAN Immunoglobulin heavy constant alpha 1 OS=Homo sapiens OX=9606 GN=IGHA1 PE=1 SV=2 | 255 | 20201114_qwl_colo_milk_1ng_glyco_hcd203040_compound15.12384.12384.3 | controllerType=0 controllerNumber=1 scan=12384 | 28.2333 |
| 10395:3 | 2 | R.LAGKPTHVN[+1581.555]VSVVMAEVDGTC[+57.021]Y.- | HexNAc(3)Hex(6) | C[+57], N[+1582] | 1310.2385 | 3 | 3928.7008 | 3928.6971 | 0 | 0.9 | 332 | CRagged | 798.12 | 786.49 | 434.52 | 15.74 | 124 | >sp\|P01876\|IGHA1_HUMAN Immunoglobulin heavy constant alpha 1 OS=Homo sapiens OX=9606 GN=IGHA1 PE=1 SV=2 | 255 | 20201114_qwl_colo_milk_1ng_glyco_hcd203040_compound15.13521.13521.3 | controllerType=0 controllerNumber=1 scan=13521 | 30.4148 |
| 10752:3 | 2 | R.LAGKPTHVN[+2262.814]VSVVMAEVDGTC[+57.021]Y.- | HexNAc(5)Hex(5)Fuc(1)NeuAc(1) | C[+57], N[+2263] | 1537.3287 | 3 | 4609.9715 | 4609.9564 | 0 | 3.3 | 332 | CRagged | 798.13 | 493.02 | 493.02 | 15.89 | 124 | >sp\|P01876\|IGHA1_HUMAN Immunoglobulin heavy constant alpha 1 OS=Homo sapiens OX=9606 GN=IGHA1 PE=1 SV=2 | 255 | 20201114_qwl_colo_milk_1ng_glyco_hcd203040_compound15.13893.13893.3 | controllerType=0 controllerNumber=1 scan=13893 | 31.178 |
| 09230:3 | 2 | R.LAGKPTHVN[+1702.581]VSVVMAEVDGTC[+57.021]Y.- | HexNAc(2)Hex(8) | C[+57], N[+1703] | 1350.5891 | 3 | 4049.7527 | 4049.7234 | 0 | 7.2 | 332 | CRagged | 798.98 | 798.98 | 477.74 | 15.51 | 124 | >sp\|P01876\|IGHA1_HUMAN Immunoglobulin heavy constant alpha 1 OS=Homo sapiens OX=9606 GN=IGHA1 PE=1 SV=2 | 255 | 20201114_qwl_colo_milk_1ng_glyco_hcd203040_compound15.12306.12306.3 | controllerType=0 controllerNumber=1 scan=12306 | 28.0939 |
| 10353:3 | 2 | R.LAGKPTHVN[+1809.666]VSVVMAEVDGTC[+57.021]Y.- | HexNAc(5)Hex(4)Fuc(1) | C[+57], N[+1810] | 1386.2856 | 3 | 4156.8422 | 4156.8081 | 0 | 8.2 | 332 | CRagged | 799.11 | 655.93 | 526.9 | 15.43 | 124 | >sp\|P01876\|IGHA1_HUMAN Immunoglobulin heavy constant alpha 1 OS=Homo sapiens OX=9606 GN=IGHA1 PE=1 SV=2 | 255 | 20201114_qwl_colo_milk_1ng_glyco_hcd203040_compound15.13477.13477.3 | controllerType=0 controllerNumber=1 scan=13477 | 30.3173 |
| 09371:3 | 2 | R.LAGKPTHVN[+1419.502]VSVVMAEVDGTC[+57.021]Y.- | HexNAc(3)Hex(5) | C[+57], N[+1420] | 1256.2203 | 3 | 3766.6463 | 3766.6443 | 0 | 0.5 | 332 | CRagged | 800.97 | 794.76 | 470.36 | 16.74 | 124 | >sp\|P01876\|IGHA1_HUMAN Immunoglobulin heavy constant alpha 1 OS=Homo sapiens OX=9606 GN=IGHA1 PE=1 SV=2 | 255 | 20201114_qwl_colo_milk_1ng_glyco_hcd203040_compound15.12454.12454.3 | controllerType=0 controllerNumber=1 scan=12454 | 28.3649 |
| 10569:3 | 2 | R.LAGKPTHVN[+892.317]VSVVMAEVDGTC[+57.021]Y.- | HexNAc(2)Hex(3) | C[+57], N[+892] | 1080.4922 | 3 | 3239.4621 | 3239.4593 | 0 | 0.9 | 332 | CRagged | 801.55 | 801.55 | 464.39 | 16.74 | 124 | >sp\|P01876\|IGHA1_HUMAN Immunoglobulin heavy constant alpha 1 OS=Homo sapiens OX=9606 GN=IGHA1 PE=1 SV=2 | 255 | 20201114_qwl_colo_milk_1ng_glyco_hcd203040_compound15.13702.13702.3 | controllerType=0 controllerNumber=1 scan=13702 | 30.7789 |
| 10413:3 | 2 | R.LAGKPTHVN[+1419.502]VSVVMAEVDGTC[+57.021]Y.- | HexNAc(3)Hex(5) | C[+57], N[+1420] | 1256.2212 | 3 | 3766.6492 | 3766.6443 | 0 | 1.3 | 332 | CRagged | 801.58 | 757.67 | 443.27 | 16.79 | 124 | >sp\|P01876\|IGHA1_HUMAN Immunoglobulin heavy constant alpha 1 OS=Homo sapiens OX=9606 GN=IGHA1 PE=1 SV=2 | 255 | 20201114_qwl_colo_milk_1ng_glyco_hcd203040_compound15.13539.13539.3 | controllerType=0 controllerNumber=1 scan=13539 | 30.4498 |
| 10462:3 | 2 | R.LAGKPTHVN[+1378.476]VSVVMAEVDGTC[+57.021]Y.- | HexNAc(2)Hex(6) | C[+57], N[+1378] | 1242.55 | 3 | 3725.6354 | 3725.6178 | 0 | 4.7 | 332 | CRagged | 805.88 | 805.88 | 486.52 | 16.56 | 124 | >sp\|P01876\|IGHA1_HUMAN Immunoglobulin heavy constant alpha 1 OS=Homo sapiens OX=9606 GN=IGHA1 PE=1 SV=2 | 255 | 20201114_qwl_colo_milk_1ng_glyco_hcd203040_compound15.13591.13591.3 | controllerType=0 controllerNumber=1 scan=13591 | 30.5712 |
| 10444:2 | 2 | R.LAGKPTHVN[+1216.423]VSVVMAEVDGTC[+57.021]Y.- | HexNAc(2)Hex(5) | C[+57], N[+1216] | 1782.2893 | 2 | 3563.5713 | 3563.5649 | 0 | 1.8 | 332 | CRagged | 809.25 | 809.25 | 429.59 | 16.13 | 124 | >sp\|P01876\|IGHA1_HUMAN Immunoglobulin heavy constant alpha 1 OS=Homo sapiens OX=9606 GN=IGHA1 PE=1 SV=2 | 255 | 20201114_qwl_colo_milk_1ng_glyco_hcd203040_compound15.13572.13572.2 | controllerType=0 controllerNumber=1 scan=13572 | 30.5282 |
| 09267:3 | 2 | R.LAGKPTHVN[+1540.529]VSVVMAEVDGTC[+57.021]Y.- | HexNAc(2)Hex(7) | C[+57], N[+1541] | 1296.5643 | 3 | 3887.6782 | 3887.6706 | 0 | 2 | 332 | CRagged | 815.01 | 815.01 | 503.29 | 16.79 | 124 | >sp\|P01876\|IGHA1_HUMAN Immunoglobulin heavy constant alpha 1 OS=Homo sapiens OX=9606 GN=IGHA1 PE=1 SV=2 | 255 | 20201114_qwl_colo_milk_1ng_glyco_hcd203040_compound15.12346.12346.3 | controllerType=0 controllerNumber=1 scan=12346 | 28.1642 |
| 10461:3 | 2 | R.LAGKPTHVN[+1606.587]VSVVMAEVDGTC[+57.021]Y.- | HexNAc(4)Hex(4)Fuc(1) | C[+57], N[+1607] | 1318.5879 | 3 | 3953.7493 | 3953.7288 | 0 | 5.2 | 332 | CRagged | 817.91 | 776.4 | 776.4 | 16.56 | 124 | >sp\|P01876\|IGHA1_HUMAN Immunoglobulin heavy constant alpha 1 OS=Homo sapiens OX=9606 GN=IGHA1 PE=1 SV=2 | 255 | 20201114_qwl_colo_milk_1ng_glyco_hcd203040_compound15.13590.13590.3 | controllerType=0 controllerNumber=1 scan=13590 | 30.5693 |
| 10710:3 | 2 | R.LAGKPTHVN[+1647.613]VSVVMAEVDGTC[+57.021]Y.- | HexNAc(5)Hex(3)Fuc(1) | C[+57], N[+1648] | 1332.2626 | 3 | 3994.7733 | 3994.7553 | 0 | 4.5 | 332 | CRagged | 821.96 | 746.97 | 746.97 | 16.56 | 124 | >sp\|P01876\|IGHA1_HUMAN Immunoglobulin heavy constant alpha 1 OS=Homo sapiens OX=9606 GN=IGHA1 PE=1 SV=2 | 255 | 20201114_qwl_colo_milk_1ng_glyco_hcd203040_compound15.13849.13849.5 | controllerType=0 controllerNumber=1 scan=13849 | 31.0843 |
| 10468:3 | 2 | R.LAGKPTHVN[+1647.613]VSVVMAEVDGTC[+57.021]Y.- | HexNAc(5)Hex(3)Fuc(1) | C[+57], N[+1648] | 1332.2628 | 3 | 3994.7737 | 3994.7553 | 0 | 4.6 | 332 | CRagged | 831.98 | 702.75 | 702.75 | 16.56 | 124 | >sp\|P01876\|IGHA1_HUMAN Immunoglobulin heavy constant alpha 1 OS=Homo sapiens OX=9606 GN=IGHA1 PE=1 SV=2 | 255 | 20201114_qwl_colo_milk_1ng_glyco_hcd203040_compound15.13597.13597.3 | controllerType=0 controllerNumber=1 scan=13597 | 30.5904 |
| 09257:3 | 2 | R.LAGKPTHVN[+1702.581]VSVVMAEVDGTC[+57.021]Y.- | HexNAc(2)Hex(8) | C[+57], N[+1703] | 1350.5822 | 3 | 4049.7321 | 4049.7234 | 0 | 2.1 | 332 | CRagged | 832.06 | 812.35 | 513.37 | 16.79 | 124 | >sp\|P01876\|IGHA1_HUMAN Immunoglobulin heavy constant alpha 1 OS=Homo sapiens OX=9606 GN=IGHA1 PE=1 SV=2 | 255 | 20201114_qwl_colo_milk_1ng_glyco_hcd203040_compound15.12335.12335.3 | controllerType=0 controllerNumber=1 scan=12335 | 28.1433 |
| 10010:3 | 2 | R.LAGKPTHVN[+1216.423]VSVVMAEVDGTC[+57.021]Y.- | HexNAc(2)Hex(5) | C[+57], N[+1216] | 1188.5292 | 3 | 3563.5731 | 3563.5649 | 0 | 2.3 | 332 | CRagged | 848.77 | 828.89 | 535.3 | 16.89 | 124 | >sp\|P01876\|IGHA1_HUMAN Immunoglobulin heavy constant alpha 1 OS=Homo sapiens OX=9606 GN=IGHA1 PE=1 SV=2 | 255 | 20201114_qwl_colo_milk_1ng_glyco_hcd203040_compound15.13120.13120.3 | controllerType=0 controllerNumber=1 scan=13120 | 29.6326 |
| 10571:3 | 2 | R.LAGKPTHVN[+892.317]VSVVMAEVDGTC[+57.021]Y.- | HexNAc(2)Hex(3) | C[+57], N[+892] | 1080.4922 | 3 | 3239.4621 | 3239.4593 | 0 | 0.9 | 332 | CRagged | 850.6 | 724.94 | 470.35 | 17.74 | 124 | >sp\|P01876\|IGHA1_HUMAN Immunoglobulin heavy constant alpha 1 OS=Homo sapiens OX=9606 GN=IGHA1 PE=1 SV=2 | 255 | 20201114_qwl_colo_milk_1ng_glyco_hcd203040_compound15.13704.13704.3 | controllerType=0 controllerNumber=1 scan=13704 | 30.7851 |
| 10476:3 | 2 | R.LAGKPTHVN[+1647.613]VSVVMAEVDGTC[+57.021]Y.- | HexNAc(5)Hex(3)Fuc(1) | C[+57], N[+1648] | 1332.2588 | 3 | 3994.762 | 3994.7553 | 0 | 1.7 | 332 | CRagged | 852.72 | 772.34 | 772.34 | 17.79 | 124 | >sp\|P01876\|IGHA1_HUMAN Immunoglobulin heavy constant alpha 1 OS=Homo sapiens OX=9606 GN=IGHA1 PE=1 SV=2 | 255 | 20201114_qwl_colo_milk_1ng_glyco_hcd203040_compound15.13606.13606.3 | controllerType=0 controllerNumber=1 scan=13606 | 30.608 |
| 10411:3 | 2 | R.LAGKPTHVN[+1419.502]VSVVMAEVDGTC[+57.021]Y.- | HexNAc(3)Hex(5) | C[+57], N[+1420] | 1256.2212 | 3 | 3766.6492 | 3766.6443 | 0 | 1.3 | 332 | CRagged | 855.42 | 855.42 | 519.33 | 17.79 | 124 | >sp\|P01876\|IGHA1_HUMAN Immunoglobulin heavy constant alpha 1 OS=Homo sapiens OX=9606 GN=IGHA1 PE=1 SV=2 | 255 | 20201114_qwl_colo_milk_1ng_glyco_hcd203040_compound15.13537.13537.3 | controllerType=0 controllerNumber=1 scan=13537 | 30.4422 |
| 12002:3 | 2 | R.LAGKPTHVN[+1872.651]VSVVMAEVDGTC[+57.021]Y.- | HexNAc(3)Hex(6)NeuAc(1) | C[+57], N[+1873] | 1407.2819 | 3 | 4219.8312 | 4219.7926 | 0 | 9.2 | 332 | CRagged | 863.37 | 863.37 | 418.24 | 17.3 | 124 | >sp\|P01876\|IGHA1_HUMAN Immunoglobulin heavy constant alpha 1 OS=Homo sapiens OX=9606 GN=IGHA1 PE=1 SV=2 | 255 | 20201114_qwl_colo_milk_1ng_glyco_hcd203040_compound15.15200.15200.3 | controllerType=0 controllerNumber=1 scan=15200 | 33.6818 |
| 10394:3 | 2 | R.LAGKPTHVN[+1606.587]VSVVMAEVDGTC[+57.021]Y.- | HexNAc(4)Hex(4)Fuc(1) | C[+57], N[+1607] | 1318.5823 | 3 | 3953.7324 | 3953.7288 | 0 | 0.9 | 332 | CRagged | 868.88 | 729.25 | 729.25 | 17.74 | 124 | >sp\|P01876\|IGHA1_HUMAN Immunoglobulin heavy constant alpha 1 OS=Homo sapiens OX=9606 GN=IGHA1 PE=1 SV=2 | 255 | 20201114_qwl_colo_milk_1ng_glyco_hcd203040_compound15.13520.13520.3 | controllerType=0 controllerNumber=1 scan=13520 | 30.4134 |
| 10480:3 | 2 | R.LAGKPTHVN[+1444.534]VSVVMAEVDGTC[+57.021]Y.- | HexNAc(4)Hex(3)Fuc(1) | C[+57], N[+1445] | 1264.5666 | 3 | 3791.6852 | 3791.6759 | 0 | 2.4 | 332 | CRagged | 879.36 | 820.79 | 820.79 | 17.89 | 124 | >sp\|P01876\|IGHA1_HUMAN Immunoglobulin heavy constant alpha 1 OS=Homo sapiens OX=9606 GN=IGHA1 PE=1 SV=2 | 255 | 20201114_qwl_colo_milk_1ng_glyco_hcd203040_compound15.13610.13610.3 | controllerType=0 controllerNumber=1 scan=13610 | 30.6139 |
| 10392:3 | 2 | R.LAGKPTHVN[+1606.587]VSVVMAEVDGTC[+57.021]Y.- | HexNAc(4)Hex(4)Fuc(1) | C[+57], N[+1607] | 1318.5823 | 3 | 3953.7324 | 3953.7288 | 0 | 0.9 | 332 | CRagged | 889.5 | 889.5 | 889.5 | 17.74 | 124 | >sp\|P01876\|IGHA1_HUMAN Immunoglobulin heavy constant alpha 1 OS=Homo sapiens OX=9606 GN=IGHA1 PE=1 SV=2 | 255 | 20201114_qwl_colo_milk_1ng_glyco_hcd203040_compound15.13518.13518.3 | controllerType=0 controllerNumber=1 scan=13518 | 30.4049 |
| 10463:3 | 2 | R.LAGKPTHVN[+1501.555]VSVVMAEVDGTC[+57.021]Y.- | HexNAc(5)Hex(3) | C[+57], N[+1502] | 1283.5765 | 3 | 3848.7149 | 3848.6974 | 0 | 4.5 | 332 | CRagged | 894.58 | 894.58 | 894.58 | 17.86 | 124 | >sp\|P01876\|IGHA1_HUMAN Immunoglobulin heavy constant alpha 1 OS=Homo sapiens OX=9606 GN=IGHA1 PE=1 SV=2 | 255 | 20201114_qwl_colo_milk_1ng_glyco_hcd203040_compound15.13592.13592.3 | controllerType=0 controllerNumber=1 scan=13592 | 30.5724 |
| 09983:3 | 2 | R.LAGKPTHVN[+1378.476]VSVVMAEVDGTC[+57.021]Y.- | HexNAc(2)Hex(6) | C[+57], N[+1378] | 1242.5475 | 3 | 3725.6281 | 3725.6178 | 0 | 2.8 | 332 | CRagged | 895.03 | 895.03 | 550.76 | 17.89 | 124 | >sp\|P01876\|IGHA1_HUMAN Immunoglobulin heavy constant alpha 1 OS=Homo sapiens OX=9606 GN=IGHA1 PE=1 SV=2 | 255 | 20201114_qwl_colo_milk_1ng_glyco_hcd203040_compound15.13092.13092.3 | controllerType=0 controllerNumber=1 scan=13092 | 29.576 |
| 09263:3 | 2 | R.LAGKPTHVN[+1540.529]VSVVMAEVDGTC[+57.021]Y.- | HexNAc(2)Hex(7) | C[+57], N[+1541] | 1296.5645 | 3 | 3887.679 | 3887.6706 | 0 | 2.2 | 332 | CRagged | 903.08 | 851.69 | 493.67 | 18.79 | 124 | >sp\|P01876\|IGHA1_HUMAN Immunoglobulin heavy constant alpha 1 OS=Homo sapiens OX=9606 GN=IGHA1 PE=1 SV=2 | 255 | 20201114_qwl_colo_milk_1ng_glyco_hcd203040_compound15.12341.12341.3 | controllerType=0 controllerNumber=1 scan=12341 | 28.1546 |
| 11937:3 | 2 | R.LAGKPTHVN[+2262.814]VSVVMAEVDGTC[+57.021]Y.- | HexNAc(5)Hex(5)Fuc(1)NeuAc(1) | C[+57], N[+2263] | 1537.3273 | 3 | 4609.9674 | 4609.9564 | 0 | 2.4 | 332 | CRagged | 925.91 | 604.53 | 604.53 | 18.89 | 124 | >sp\|P01876\|IGHA1_HUMAN Immunoglobulin heavy constant alpha 1 OS=Homo sapiens OX=9606 GN=IGHA1 PE=1 SV=2 | 255 | 20201114_qwl_colo_milk_1ng_glyco_hcd203040_compound15.15132.15132.3 | controllerType=0 controllerNumber=1 scan=15132 | 33.5148 |
| 10465:3 | 2 | R.LAGKPTHVN[+1647.613]VSVVMAEVDGTC[+57.021]Y.- | HexNAc(5)Hex(3)Fuc(1) | C[+57], N[+1648] | 1332.2628 | 3 | 3994.7737 | 3994.7553 | 0 | 4.6 | 332 | CRagged | 926.66 | 738.54 | 738.54 | 18.56 | 124 | >sp\|P01876\|IGHA1_HUMAN Immunoglobulin heavy constant alpha 1 OS=Homo sapiens OX=9606 GN=IGHA1 PE=1 SV=2 | 255 | 20201114_qwl_colo_milk_1ng_glyco_hcd203040_compound15.13594.13594.3 | controllerType=0 controllerNumber=1 scan=13594 | 30.5747 |
| 10375:3 | 2 | R.LAGKPTHVN[+1460.529]VSVVMAEVDGTC[+57.021]Y.- | HexNAc(4)Hex(4) | C[+57], N[+1461] | 1269.8989 | 3 | 3807.682 | 3807.6709 | 0 | 2.9 | 332 | CRagged | 953.91 | 953.91 | 610.39 | 19.89 | 124 | >sp\|P01876\|IGHA1_HUMAN Immunoglobulin heavy constant alpha 1 OS=Homo sapiens OX=9606 GN=IGHA1 PE=1 SV=2 | 255 | 20201114_qwl_colo_milk_1ng_glyco_hcd203040_compound15.13500.13500.3 | controllerType=0 controllerNumber=1 scan=13500 | 30.3639 |
| 08671:3 | 2 | R.LAGKPTHVN[+1216.423]VSVVM[+15.995]AEVDGTC[+57.021]Y.- | HexNAc(2)Hex(5) | C[+57], M[+16], N[+1216] | 1193.865 | 3 | 3579.5804 | 3579.5599 | 0 | 5.8 | 332 | CRagged | 207.79 | 207.79 | 207.79 | 5.05 | 124 | >sp\|P01876\|IGHA1_HUMAN Immunoglobulin heavy constant alpha 1 OS=Homo sapiens OX=9606 GN=IGHA1 PE=1 SV=2 | 255 | 20201114_qwl_colo_milk_1ng_glyco_hcd203040_compound15.11722.11722.3 | controllerType=0 controllerNumber=1 scan=11722 | 27.0244 |
| 08284:3 | 2 | R.LAGKPTHVN[+1054.370]VSVVM[+15.995]AEVDGTC[+57.021]Y.- | HexNAc(2)Hex(4) | C[+57], M[+16], N[+1054] | 1139.85 | 3 | 3417.5355 | 3417.507 | 0 | 8.3 | 332 | CRagged | 211.22 | 211.22 | 211.22 | 5.12 | 124 | >sp\|P01876\|IGHA1_HUMAN Immunoglobulin heavy constant alpha 1 OS=Homo sapiens OX=9606 GN=IGHA1 PE=1 SV=2 | 255 | 20201114_qwl_colo_milk_1ng_glyco_hcd203040_compound15.11318.11318.3 | controllerType=0 controllerNumber=1 scan=11318 | 26.2916 |
| 07371:3 | 2 | R.LAGKPTHVN[+1663.608]VSVVM[+15.995]AEVDGTC[+57.021]Y.- | HexNAc(5)Hex(4) | C[+57], M[+16], N[+1664] | 1342.9243 | 3 | 4026.7583 | 4026.7452 | 0 | 3.3 | 332 | CRagged | 258.32 | 258.32 | 258.32 | 7.1 | 124 | >sp\|P01876\|IGHA1_HUMAN Immunoglobulin heavy constant alpha 1 OS=Homo sapiens OX=9606 GN=IGHA1 PE=1 SV=2 | 255 | 20201114_qwl_colo_milk_1ng_glyco_hcd203040_compound15.10367.10367.3 | controllerType=0 controllerNumber=1 scan=10367 | 24.5669 |
| 08023:3 | 2 | R.LAGKPTHVN[+1378.476]VSVVM[+15.995]AEVDGTC[+57.021]Y.- | HexNAc(2)Hex(6) | C[+57], M[+16], N[+1378] | 1247.8766 | 3 | 3741.6154 | 3741.6127 | 0 | 0.7 | 332 | CRagged | 267.61 | 267.61 | 267.61 | 7.21 | 124 | >sp\|P01876\|IGHA1_HUMAN Immunoglobulin heavy constant alpha 1 OS=Homo sapiens OX=9606 GN=IGHA1 PE=1 SV=2 | 255 | 20201114_qwl_colo_milk_1ng_glyco_hcd203040_compound15.11047.11047.3 | controllerType=0 controllerNumber=1 scan=11047 | 25.7716 |
| 08021:3 | 2 | R.LAGKPTHVN[+1378.476]VSVVM[+15.995]AEVDGTC[+57.021]Y.- | HexNAc(2)Hex(6) | C[+57], M[+16], N[+1378] | 1247.8766 | 3 | 3741.6154 | 3741.6127 | 0 | 0.7 | 332 | CRagged | 294.6 | 294.6 | 294.6 | 7.34 | 124 | >sp\|P01876\|IGHA1_HUMAN Immunoglobulin heavy constant alpha 1 OS=Homo sapiens OX=9606 GN=IGHA1 PE=1 SV=2 | 255 | 20201114_qwl_colo_milk_1ng_glyco_hcd203040_compound15.11045.11045.3 | controllerType=0 controllerNumber=1 scan=11045 | 25.768 |
| 07369:3 | 2 | R.LAGKPTHVN[+1663.608]VSVVM[+15.995]AEVDGTC[+57.021]Y.- | HexNAc(5)Hex(4) | C[+57], M[+16], N[+1664] | 1342.9243 | 3 | 4026.7583 | 4026.7452 | 0 | 3.3 | 332 | CRagged | 321.14 | 321.14 | 321.14 | 7.94 | 124 | >sp\|P01876\|IGHA1_HUMAN Immunoglobulin heavy constant alpha 1 OS=Homo sapiens OX=9606 GN=IGHA1 PE=1 SV=2 | 255 | 20201114_qwl_colo_milk_1ng_glyco_hcd203040_compound15.10365.10365.3 | controllerType=0 controllerNumber=1 scan=10365 | 24.5628 |
| 08286:3 | 2 | R.LAGKPTHVN[+1054.370]VSVVM[+15.995]AEVDGTC[+57.021]Y.- | HexNAc(2)Hex(4) | C[+57], M[+16], N[+1054] | 1139.85 | 3 | 3417.5355 | 3417.507 | 0 | 8.3 | 332 | CRagged | 340.04 | 340.04 | 340.04 | 7.61 | 124 | >sp\|P01876\|IGHA1_HUMAN Immunoglobulin heavy constant alpha 1 OS=Homo sapiens OX=9606 GN=IGHA1 PE=1 SV=2 | 255 | 20201114_qwl_colo_milk_1ng_glyco_hcd203040_compound15.11320.11320.3 | controllerType=0 controllerNumber=1 scan=11320 | 26.2949 |
| 07289:3 | 2 | R.LAGKPTHVN[+1809.666]VSVVM[+15.995]AEVDGTC[+57.021]Y.- | HexNAc(5)Hex(4)Fuc(1) | C[+57], M[+16], N[+1810] | 1391.6165 | 3 | 4172.835 | 4172.8031 | 0 | 7.7 | 332 | CRagged | 416.11 | 387.6 | 239.98 | 9.46 | 124 | >sp\|P01876\|IGHA1_HUMAN Immunoglobulin heavy constant alpha 1 OS=Homo sapiens OX=9606 GN=IGHA1 PE=1 SV=2 | 255 | 20201114_qwl_colo_milk_1ng_glyco_hcd203040_compound15.10281.10281.3 | controllerType=0 controllerNumber=1 scan=10281 | 24.4067 |
| 07396:3 | 2 | R.LAGKPTHVN[+1647.613]VSVVM[+15.995]AEVDGTC[+57.021]Y.- | HexNAc(5)Hex(3)Fuc(1) | C[+57], M[+16], N[+1648] | 1337.5925 | 3 | 4010.7628 | 4010.7502 | 0 | 3.1 | 332 | CRagged | 495.53 | 361.83 | 211.46 | 8.87 | 124 | >sp\|P01876\|IGHA1_HUMAN Immunoglobulin heavy constant alpha 1 OS=Homo sapiens OX=9606 GN=IGHA1 PE=1 SV=2 | 255 | 20201114_qwl_colo_milk_1ng_glyco_hcd203040_compound15.10395.10395.3 | controllerType=0 controllerNumber=1 scan=10395 | 24.6208 |
| 08673:3 | 2 | R.LAGKPTHVN[+1216.423]VSVVM[+15.995]AEVDGTC[+57.021]Y.- | HexNAc(2)Hex(5) | C[+57], M[+16], N[+1216] | 1193.865 | 3 | 3579.5804 | 3579.5599 | 0 | 5.8 | 332 | CRagged | 503.62 | 503.62 | 503.62 | 9.69 | 124 | >sp\|P01876\|IGHA1_HUMAN Immunoglobulin heavy constant alpha 1 OS=Homo sapiens OX=9606 GN=IGHA1 PE=1 SV=2 | 255 | 20201114_qwl_colo_milk_1ng_glyco_hcd203040_compound15.11724.11724.3 | controllerType=0 controllerNumber=1 scan=11724 | 27.0278 |
| 07398:3 | 2 | R.LAGKPTHVN[+1647.613]VSVVM[+15.995]AEVDGTC[+57.021]Y.- | HexNAc(5)Hex(3)Fuc(1) | C[+57], M[+16], N[+1648] | 1337.5925 | 3 | 4010.7628 | 4010.7502 | 0 | 3.1 | 332 | CRagged | 512.67 | 512.67 | 306.57 | 10.28 | 124 | >sp\|P01876\|IGHA1_HUMAN Immunoglobulin heavy constant alpha 1 OS=Homo sapiens OX=9606 GN=IGHA1 PE=1 SV=2 | 255 | 20201114_qwl_colo_milk_1ng_glyco_hcd203040_compound15.10397.10397.3 | controllerType=0 controllerNumber=1 scan=10397 | 24.6287 |
| 08559:3 | 2 | R.LAGKPTHVN[+1378.476]VSVVM[+15.995]AEVDGTC[+57.021]Y.- | HexNAc(2)Hex(6) | C[+57], M[+16], N[+1378] | 1247.879 | 3 | 3741.6223 | 3741.6127 | 0 | 2.6 | 332 | CRagged | 546.62 | 491.81 | 491.81 | 10.48 | 124 | >sp\|P01876\|IGHA1_HUMAN Immunoglobulin heavy constant alpha 1 OS=Homo sapiens OX=9606 GN=IGHA1 PE=1 SV=2 | 255 | 20201114_qwl_colo_milk_1ng_glyco_hcd203040_compound15.11605.11605.3 | controllerType=0 controllerNumber=1 scan=11605 | [illegible] |
| 07291:3 | 2 | R.LAGKPTHVN[+1809.666]VSVVM[+15.995]AEVDGTC[+57.021]Y.- | HexNAc(5)Hex(4)Fuc(1) | C[+57], M[+16], N[+1810] | 1391.6165 | 3 | 4172.835 | 4172.8031 | 0 | 7.7 | 332 | CRagged | 583.52 | 583.52 | 342.36 | 11.51 | 124 | >sp\|P01876\|IGHA1_HUMAN Immunoglobulin heavy constant alpha 1 OS=Homo sapiens OX=9606 GN=IGHA1 PE=1 SV=2 | 255 | 20201114_qwl_colo_milk_1ng_glyco_hcd203040_compound15.10283.10283.3 | controllerType=0 controllerNumber=1 scan=10283 | [illegible] |
| 08561:3 | 2 | R.LAGKPTHVN[+1378.476]VSVVM[+15.995]AEVDGTC[+57.021]Y.- | HexNAc(2)Hex(6) | C[+57], M[+16], N[+1378] | 1247.879 | 3 | 3741.6223 | 3741.6127 | 0 | 2.6 | 332 | CRagged | 607.12 | 569.92 | 569.92 | 12.89 | 124 | >sp\|P01876\|IGHA1_HUMAN Immunoglobulin heavy constant alpha 1 OS=Homo sapiens OX=9606 GN=IGHA1 PE=1 SV=2 | 255 | 20201114_qwl_colo_milk_1ng_glyco_hcd203040_compound15.11607.11607.3 | controllerType=0 controllerNumber=1 scan=11607 | [illegible] |
| 08074:3 | 2 | R.LAGKPTHVN[+1216.423]VSVVM[+15.995]AEVDGTC[+57.021]Y.- | HexNAc(2)Hex(5) | C[+57], M[+16], N[+1216] | 1193.8571 | 3 | 3579.5568 | 3579.5599 | 0 | -0.9 | 332 | CRagged | 673.46 | 673.46 | 673.46 | 13.27 | 124 | >sp\|P01876\|IGHA1_HUMAN Immunoglobulin heavy constant alpha 1 OS=Homo sapiens OX=9606 GN=IGHA1 PE=1 SV=2 | 255 | 20201114_qwl_colo_milk_1ng_glyco_hcd203040_compound15.11100.11100.3 | controllerType=0 controllerNumber=1 scan=11100 | [illegible] |
| 08076:3 | 2 | R.LAGKPTHVN[+1216.423]VSVVM[+15.995]AEVDGTC[+57.021]Y.- | HexNAc(2)Hex(5) | C[+57], M[+16], N[+1216] | 1193.8571 | 3 | 3579.5568 | 3579.5599 | 0 | -0.9 | 332 | CRagged | 837.86 | 742.13 | 742.13 | 16.27 | 124 | >sp\|P01876\|IGHA1_HUMAN Immunoglobulin heavy constant alpha 1 OS=Homo sapiens OX=9606 GN=IGHA1 PE=1 SV=2 | 255 | 20201114_qwl_colo_milk_1ng_glyco_hcd203040_compound15.11102.11102.3 | controllerType=0 controllerNumber=1 scan=11102 | 25.8733 |
| 01456:3 | 3 | K.GPN[+2059.735]C[+57.021]SEPEC[+57.021]PGNC[+57.021]HLR.G | HexNAc(4)Hex(5)Fuc(1)NeuAc(1) | C[+57]*3, N[+2060] | 1315.1729 | 3 | 3943.5042 | 3943.4931 | 0 | 2.8 | 182 | Specific | 253.35 | 235.31 | 235.31 | 6.2 | 114 | >sp\|P24821\|TENA_HUMAN Tenascin OS=Homo sapiens OX=9606 GN=TNC PE=1 SV=3 | 486 | 20201114_qwl_colo_milk_1ng_glyco_hcd203040_compound15.4201.4201.3 | controllerType=0 controllerNumber=1 scan=4201 | 13.7047 |
| 01454:3 | 3 | K.GPN[+2059.735]C[+57.021]SEPEC[+57.021]PGNC[+57.021]HLR.G | HexNAc(4)Hex(5)Fuc(1)NeuAc(1) | C[+57]*3, N[+2060] | 1315.1729 | 3 | 3943.5042 | 3943.4931 | 0 | 2.8 | 182 | Specific | 299.81 | 269.29 | 269.29 | 6.68 | 114 | >sp\|P24821\|TENA_HUMAN Tenascin OS=Homo sapiens OX=9606 GN=TNC PE=1 SV=3 | 486 | 20201114_qwl_colo_milk_1ng_glyco_hcd203040_compound15.4199.4199.3 | controllerType=0 controllerNumber=1 scan=4199 | 13.7002 |
| 08365:4 | 3 | R.QTGLAPGQEYEISLHIVKN[+2059.735]NTR.G | HexNAc(4)Hex(5)Fuc(1)NeuAc(1) | N[+2060] | 1132.766 | 4 | 4528.0423 | 4528.0241 | 0 | 4 | 770 | Specific | 489.08 | 385.08 | 385.08 | 9.73 | 114 | >sp\|P24821\|TENA_HUMAN Tenascin OS=Homo sapiens OX=9606 GN=TNC PE=1 SV=3 | 486 | 20201114_qwl_colo_milk_1ng_glyco_hcd203040_compound15.11403.11403.4 | controllerType=0 controllerNumber=1 scan=11403 | 26.4605 |
| 08357:4 | 3 | R.QTGLAPGQEYEISLHIVKN[+2059.735]NTR.G | HexNAc(4)Hex(5)Fuc(1)NeuAc(1) | N[+2060] | 1132.764 | 4 | 4528.034 | 4528.0241 | 0 | 2.2 | 770 | Specific | 538.09 | 457.11 | 457.11 | 9.93 | 114 | >sp\|P24821\|TENA_HUMAN Tenascin OS=Homo sapiens OX=9606 GN=TNC PE=1 SV=3 | 486 | 20201114_qwl_colo_milk_1ng_glyco_hcd203040_compound15.11394.11394.2 | controllerType=0 controllerNumber=1 scan=11394 | 26.4374 |
| 08362:4 | 3 | R.QTGLAPGQEYEISLHIVKN[+2059.735]NTR.G | HexNAc(4)Hex(5)Fuc(1)NeuAc(1) | N[+2060] | 1132.766 | 4 | 4528.0423 | 4528.0241 | 0 | 4 | 770 | Specific | 586.6 | 393.09 | 393.09 | 11.41 | 114 | >sp\|P24821\|TENA_HUMAN Tenascin OS=Homo sapiens OX=9606 GN=TNC PE=1 SV=3 | 486 | 20201114_qwl_colo_milk_1ng_glyco_hcd203040_compound15.11400.11400.4 | controllerType=0 controllerNumber=1 scan=11400 | 26.4497 |
| 08355:4 | 3 | R.QTGLAPGQEYEISLHIVKN[+2059.735]NTR.G | HexNAc(4)Hex(5)Fuc(1)NeuAc(1) | N[+2060] | 1132.764 | 4 | 4528.034 | 4528.0241 | 0 | 2.2 | 770 | Specific | 702.48 | 532.76 | 532.76 | 14.34 | 114 | >sp\|P24821\|TENA_HUMAN Tenascin OS=Homo sapiens OX=9606 GN=TNC PE=1 SV=3 | 486 | 20201114_qwl_colo_milk_1ng_glyco_hcd203040_compound15.11392.11392.2 | controllerType=0 controllerNumber=1 scan=11392 | 26.4263 |
| 13971:3 | 3 | R.LN[+1378.476]YSLPTGQWVGVQLPR.N | HexNAc(2)Hex(6) | N[+1378] | 1102.8448 | 3 | 3306.5197 | 3306.5145 | 0 | 1.6 | 1017 | Specific | 512.02 | 495.24 | 495.24 | 7.82 | 114 | >sp\|P24821\|TENA_HUMAN Tenascin OS=Homo sapiens OX=9606 GN=TNC PE=1 SV=3 | 486 | 20201114_qwl_colo_milk_1ng_glyco_hcd203040_compound15.17261.17261.3 | controllerType=0 controllerNumber=1 scan=17261 | 38.0115 |
| 13969:3 | 3 | R.LN[+1378.476]YSLPTGQWVGVQLPR.N | HexNAc(2)Hex(6) | N[+1378] | 1102.8448 | 3 | 3306.5197 | 3306.5145 | 0 | 1.6 | 1017 | Specific | 812 | 730.19 | 730.19 | 16.08 | 114 | >sp\|P24821\|TENA_HUMAN Tenascin OS=Homo sapiens OX=9606 GN=TNC PE=1 SV=3 | 486 | 20201114_qwl_colo_milk_1ng_glyco_hcd203040_compound15.17259.17259.3 | controllerType=0 controllerNumber=1 scan=17259 | 38.0044 |
| 14008:2 | 3 | R.LN[+1216.423]YSLPTGQWVGVQLPR.N | HexNAc(2)Hex(5) | N[+1216] | 1572.7466 | 2 | 3144.486 | 3144.4617 | 0 | 7.7 | 1017 | Specific | 847.47 | 787.69 | 787.69 | 15.81 | 114 | >sp\|P24821\|TENA_HUMAN Tenascin OS=Homo sapiens OX=9606 GN=TNC PE=1 SV=3 | 486 | 20201114_qwl_colo_milk_1ng_glyco_hcd203040_compound15.17300.17300.2 | controllerType=0 controllerNumber=1 scan=17300 | 38.1254 |
| 14005:2 | 3 | R.LN[+1216.423]YSLPTGQWVGVQLPR.N | HexNAc(2)Hex(5) | N[+1216] | 1572.7466 | 2 | 3144.486 | 3144.4617 | 0 | 7.7 | 1017 | Specific | 907.37 | 864.5 | 864.5 | 17.81 | 114 | >sp\|P24821\|TENA_HUMAN Tenascin OS=Homo sapiens OX=9606 GN=TNC PE=1 SV=3 | 486 | 20201114_qwl_colo_milk_1ng_glyco_hcd203040_compound15.17297.17297.2 | controllerType=0 controllerNumber=1 scan=17297 | 38.1181 |
| 09315:3 | 4 | K.TVLTPATNHMGN[+1378.476]VTFTIPANR.E | HexNAc(2)Hex(6) | N[+1378] | 1211.8836 | 3 | 3633.6363 | 3633.6358 | 0 | 0.1 | 74 | Specific | 645.89 | 576.74 | 296.48 | 12.37 | 104 | >sp\|P01024\|CO3_HUMAN Complement C3 OS=Homo sapiens OX=9606 GN=C3 PE=1 SV=2 | 492 | 20201114_qwl_colo_milk_1ng_glyco_hcd203040_compound15.12395.12395.3 | controllerType=0 controllerNumber=1 scan=12395 | 28.2561 |
| 09313:3 | 4 | K.TVLTPATNHMGN[+1378.476]VTFTIPANR.E | HexNAc(2)Hex(6) | N[+1378] | 1211.8836 | 3 | 3633.6363 | 3633.6358 | 0 | 0.1 | 74 | Specific | 703.49 | 703.49 | 395.45 | 14.37 | 104 | >sp\|P01024\|CO3_HUMAN Complement C3 OS=Homo sapiens OX=9606 GN=C3 PE=1 SV=2 | 492 | 20201114_qwl_colo_milk_1ng_glyco_hcd203040_compound15.12393.12393.3 | controllerType=0 controllerNumber=1 scan=12393 | 28.2511 |
| 15630:3 | 5 | R.GLSFDVSLEVSQGPGLLN[+2863.041]DTK.V | HexNAc(5)Hex(6)Fuc(4)NeuAc(1) | N[+2863] | 1680.4015 | 3 | 5039.1898 | 5039.1541 | 0 | 7.1 | 118 | Specific | 460.44 | 450.49 | 450.49 | 9.3 | 101 | >sp\|P01833\|PIGR_HUMAN Polymeric immunoglobulin receptor OS=Homo sapiens OX=9606 GN=PIGR PE=1 SV=4 | 902 | 20201114_qwl_colo_milk_1ng_glyco_hcd203040_compound15.19001.19001.3 | controllerType=0 controllerNumber=1 scan=19001 | 41.8297 |
| 16977:4 | 5 | K.QIGLYPVLVIDSSGYVNPN[+2076.750]YTGR.I | HexNAc(4)Hex(6)Fuc(2) | N[+2077] | 1151.2748 | 4 | 4602.0775 | 4602.0537 | 0 | 5.2 | 168 | Specific | 283.09 | 177.32 | 177.32 | 4.2 | 101 | >sp\|P01833\|PIGR_HUMAN Polymeric immunoglobulin receptor OS=Homo sapiens OX=9606 GN=PIGR PE=1 SV=4 | 902 | 20201114_qwl_colo_milk_1ng_glyco_hcd203040_compound15.20417.20417.2 | controllerType=0 controllerNumber=1 scan=20417 | 45.2129 |
| 15235:3 | 5 | K.QIGLYPVLVIDSSGYVNPN[+2425.887]YTGR.I | HexNAc(5)Hex(6)Fuc(3) | N[+2426] | 1651.0788 | 3 | 4951.2219 | 4951.191 | 0 | 6.2 | 168 | Specific | 489.27 | 489.27 | 489.27 | 9.68 | 101 | >sp\|P01833\|PIGR_HUMAN Polymeric immunoglobulin receptor OS=Homo sapiens OX=9606 GN=PIGR PE=1 SV=4 | 902 | 20201114_qwl_colo_milk_1ng_glyco_hcd203040_compound15.18587.18587.3 | controllerType=0 controllerNumber=1 scan=18587 | 40.8913 |
| 15237:3 | 5 | K.QIGLYPVLVIDSSGYVNPN[+2425.887]YTGR.I | HexNAc(5)Hex(6)Fuc(3) | N[+2426] | 1651.0788 | 3 | 4951.2219 | 4951.191 | 0 | 6.2 | 168 | Specific | 542.74 | 531.36 | 531.36 | 9.95 | 101 | >sp\|P01833\|PIGR_HUMAN Polymeric immunoglobulin receptor OS=Homo sapiens OX=9606 GN=PIGR PE=1 SV=4 | 902 | 20201114_qwl_colo_milk_1ng_glyco_hcd203040_compound15.18589.18589.3 | controllerType=0 controllerNumber=1 scan=18589 | 40.8956 |
| 16831:3 | 5 | K.QIGLYPVLVIDSSGYVNPN[+2570.925]YTGR.I | HexNAc(5)Hex(6)Fuc(2)NeuAc(1) | N[+2571] | 1699.4257 | 3 | 5096.2627 | 5096.2285 | 0 | 6.7 | 168 | Specific | 594.56 | 556.49 | 556.49 | 11.36 | 101 | >sp\|P01833\|PIGR_HUMAN Polymeric immunoglobulin receptor OS=Homo sapiens OX=9606 GN=PIGR PE=1 SV=4 | 902 | 20201114_qwl_colo_milk_1ng_glyco_hcd203040_compound15.20265.20265.3 | controllerType=0 controllerNumber=1 scan=20265 | 44.8784 |
| 18915:3 | 5 | K.QIGLYPVLVIDSSGYVNPN[+2350.830]YTGR.I | HexNAc(4)Hex(5)Fuc(1)NeuAc(2) | N[+2351] | 1626.0567 | 3 | 4876.1555 | 4876.1338 | 0 | 4.5 | 168 | Specific | 704.94 | 660.65 | 660.65 | 14.85 | 101 | >sp\|P01833\|PIGR_HUMAN Polymeric immunoglobulin receptor OS=Homo sapiens OX=9606 GN=PIGR PE=1 SV=4 | 902 | 20201114_qwl_colo_milk_1ng_glyco_hcd203040_compound15.22449.22449.3 | controllerType=0 controllerNumber=1 scan=22449 | 49.6264 |
| 16986:4 | 5 | K.QIGLYPVLVIDSSGYVNPN[+2059.735]YTGR.I | HexNAc(4)Hex(5)Fuc(1)NeuAc(1) | N[+2060] | 1147.018 | 4 | 4585.0501 | 4585.0384 | 0 | 2.6 | 168 | Specific | 732.47 | 690.55 | 690.55 | 15.33 | 101 | >sp\|P01833\|PIGR_HUMAN Polymeric immunoglobulin receptor OS=Homo sapiens OX=9606 GN=PIGR PE=1 SV=4 | 902 | 20201114_qwl_colo_milk_1ng_glyco_hcd203040_compound15.20427.20427.4 | controllerType=0 controllerNumber=1 scan=20427 | 45.23 |
| 18964:3 | 5 | K.QIGLYPVLVIDSSGYVNPN[+2204.772]YTGR.I | HexNAc(4)Hex(5)NeuAc(2) | N[+2205] | 1577.3696 | 3 | 4730.0941 | 4730.0759 | 0 | 3.9 | 168 | Specific | 812.74 | 672.87 | 672.87 | 16.85 | 101 | >sp\|P01833\|PIGR_HUMAN Polymeric immunoglobulin receptor OS=Homo sapiens OX=9606 GN=PIGR PE=1 SV=4 | 902 | 20201114_qwl_colo_milk_1ng_glyco_hcd203040_compound15.22500.22500.3 | controllerType=0 controllerNumber=1 scan=22500 | 49.7376 |
| 16834:3 | 5 | K.QIGLYPVLVIDSSGYVNPN[+2570.925]YTGR.I | HexNAc(5)Hex(6)Fuc(2)NeuAc(1) | N[+2571] | 1699.4257 | 3 | 5096.2627 | 5096.2285 | 0 | 6.7 | 168 | Specific | 812.9 | 766.02 | 766.02 | 16.36 | 101 | >sp\|P01833\|PIGR_HUMAN Polymeric immunoglobulin receptor OS=Homo sapiens OX=9606 GN=PIGR PE=1 SV=4 | 902 | 20201114_qwl_colo_milk_1ng_glyco_hcd203040_compound15.20268.20268.3 | controllerType=0 controllerNumber=1 scan=20268 | 44.8826 |
| 18962:3 | 5 | K.QIGLYPVLVIDSSGYVNPN[+2204.772]YTGR.I | HexNAc(4)Hex(5)NeuAc(2) | N[+2205] | 1577.3696 | 3 | 4730.0941 | 4730.0759 | 0 | 3.9 | 168 | Specific | 861.09 | 647.76 | 647.76 | 17.85 | 101 | >sp\|P01833\|PIGR_HUMAN Polymeric immunoglobulin receptor OS=Homo sapiens OX=9606 GN=PIGR PE=1 SV=4 | 902 | 20201114_qwl_colo_milk_1ng_glyco_hcd203040_compound15.22498.22498.3 | controllerType=0 controllerNumber=1 scan=22498 | 49.7225 |
| 16989:4 | 5 | K.QIGLYPVLVIDSSGYVNPN[+2059.735]YTGR.I | HexNAc(4)Hex(5)Fuc(1)NeuAc(1) | N[+2060] | 1147.018 | 4 | 4585.0501 | 4585.0384 | 0 | 2.6 | 168 | Specific | 874.99 | 850.3 | 850.3 | 18.33 | 101 | >sp\|P01833\|PIGR_HUMAN Polymeric immunoglobulin receptor OS=Homo sapiens OX=9606 GN=PIGR PE=1 SV=4 | 902 | 20201114_qwl_colo_milk_1ng_glyco_hcd203040_compound15.20430.20430.4 | controllerType=0 controllerNumber=1 scan=20430 | 45.2352 |
| 16955:3 | 5 | K.QIGLYPVLVIDSSGYVNPN[+2059.735]YTGR.I | HexNAc(4)Hex(5)Fuc(1)NeuAc(1) | N[+2060] | 1529.023 | 3 | 4585.0545 | 4585.0384 | 0 | 3.5 | 168 | Specific | 888.5 | 781 | 781 | 17.85 | 101 | >sp\|P01833\|PIGR_HUMAN Polymeric immunoglobulin receptor OS=Homo sapiens OX=9606 GN=PIGR PE=1 SV=4 | 902 | 20201114_qwl_colo_milk_1ng_glyco_hcd203040_compound15.20395.20395.3 | controllerType=0 controllerNumber=1 scan=20395 | 45.1756 |
| 16957:3 | 5 | K.QIGLYPVLVIDSSGYVNPN[+2059.735]YTGR.I | HexNAc(4)Hex(5)Fuc(1)NeuAc(1) | N[+2060] | 1529.023 | 3 | 4585.0545 | 4585.0384 | 0 | 3.5 | 168 | Specific | 898.61 | 753.82 | 753.82 | 17.85 | 101 | >sp\|P01833\|PIGR_HUMAN Polymeric immunoglobulin receptor OS=Homo sapiens OX=9606 GN=PIGR PE=1 SV=4 | 902 | 20201114_qwl_colo_milk_1ng_glyco_hcd203040_compound15.20397.20397.3 | controllerType=0 controllerNumber=1 scan=20397 | 45.1814 |
| 18442:3 | 5 | K.[+42.011]AQYEGRLSLLEEPGN[+1622.582]GTFTVILNQLTSR.D | HexNAc(4)Hex(5) | .[42], N[+1623] | 1591.0655 | 3 | 4771.1819 | 4771.2089 | 0 | -5.7 | 407 | Specific | 216.91 | 189 | 189 | 3.46 | 101 | >sp\|P01833\|PIGR_HUMAN Polymeric immunoglobulin receptor OS=Homo sapiens OX=9606 GN=PIGR PE=1 SV=4 | 902 | 20201114_qwl_colo_milk_1ng_glyco_hcd203040_compound15.21955.21955.3 | controllerType=0 controllerNumber=1 scan=21955 | 48.5855 |
| 18461:4 | 5 | K.[+42.011]AQYEGRLSLLEEPGN[+1622.582]GTFTVILNQLTSR.D | HexNAc(4)Hex(5) | .[42], N[+1623] | 1193.5616 | 4 | 4771.2246 | 4771.2089 | 0 | 3.3 | 407 | Specific | 269.94 | 208.48 | 208.48 | 6.28 | 101 | >sp\|P01833\|PIGR_HUMAN Polymeric immunoglobulin receptor OS=Homo sapiens OX=9606 GN=PIGR PE=1 SV=4 | 902 | 20201114_qwl_colo_milk_1ng_glyco_hcd203040_compound15.21974.21974.4 | controllerType=0 controllerNumber=1 scan=21974 | 48.6272 |
| 18445:3 | 5 | K.[+42.011]AQYEGRLSLLEEPGN[+1622.582]GTFTVILNQLTSR.D | HexNAc(4)Hex(5) | .[42], N[+1623] | 1591.0655 | 3 | 4771.1819 | 4771.2089 | 0 | -5.7 | 407 | Specific | 289.07 | 222.02 | 222.02 | 4.99 | 101 | >sp\|P01833\|PIGR_HUMAN Polymeric immunoglobulin receptor OS=Homo sapiens OX=9606 GN=PIGR PE=1 SV=4 | 902 | 20201114_qwl_colo_milk_1ng_glyco_hcd203040_compound15.21958.21958.3 | controllerType=0 controllerNumber=1 scan=21958 | 48.5969 |
| 18659:3 | 5 | K.AQYEGRLSLLEEPGN[+1647.613]GTFTVILNQLTSR.D | HexNAc(5)Hex(3)Fuc(1) | N[+1648] | 1585.4105 | 3 | 4754.2168 | 4754.23 | 0 | -2.8 | 407 | Specific | 325.09 | 320.35 | 320.35 | 6.1 | 101 | >sp\|P01833\|PIGR_HUMAN Polymeric immunoglobulin receptor OS=Homo sapiens OX=9606 GN=PIGR PE=1 SV=4 | 902 | 20201114_qwl_colo_milk_1ng_glyco_hcd203040_compound15.22181.22181.3 | controllerType=0 controllerNumber=1 scan=22181 | 49.0661 |
| 18657:3 | 5 | K.AQYEGRLSLLEEPGN[+1647.613]GTFTVILNQLTSR.D | HexNAc(5)Hex(3)Fuc(1) | N[+1648] | 1585.4105 | 3 | 4754.2168 | 4754.23 | 0 | -2.8 | 407 | Specific | 391.93 | 366.99 | 366.99 | 6.66 | 101 | >sp\|P01833\|PIGR_HUMAN Polymeric immunoglobulin receptor OS=Homo sapiens OX=9606 GN=PIGR PE=1 SV=4 | 902 | 20201114_qwl_colo_milk_1ng_glyco_hcd203040_compound15.22179.22179.3 | controllerType=0 controllerNumber=1 scan=22179 | 49.0611 |
| 18557:3 | 5 | K.[+42.011]AQYEGRLSLLEEPGN[+1751.624]GTFTVILNQLTSR.D | HexNAc(4)Hex(4)NeuAc(1) | .[42], N[+1752] | 1634.074 | 3 | 4900.2075 | 4900.2515 | 0 | -9 | 407 | Specific | 394.72 | 268.67 | 268.67 | 5.78 | 101 | >sp\|P01833\|PIGR_HUMAN Polymeric immunoglobulin receptor OS=Homo sapiens OX=9606 GN=PIGR PE=1 SV=4 | 902 | 20201114_qwl_colo_milk_1ng_glyco_hcd203040_compound15.22075.22075.3 | controllerType=0 controllerNumber=1 scan=22075 | 48.8513 |
| 18542:3 | 5 | K.[+42.011]AQYEGRLSLLEEPGN[+1751.624]GTFTVILNQLTSR.D | HexNAc(4)Hex(4)NeuAc(1) | .[42], N[+1752] | 1634.0905 | 3 | 4900.257 | 4900.2515 | 0 | 1.1 | 407 | Specific | 416.57 | 400.24 | 78.37 | 9.04 | 101 | >sp\|P01833\|PIGR_HUMAN Polymeric immunoglobulin receptor OS=Homo sapiens OX=9606 GN=PIGR PE=1 SV=4 | 902 | 20201114_qwl_colo_milk_1ng_glyco_hcd203040_compound15.22059.22059.2 | controllerType=0 controllerNumber=1 scan=22059 | 48.8146 |
| 19041:3 | 5 | R.LSLLEEPGN[+1458.442]GTFTVILNQLTSR.D | HexNAc(2)Hex(6)Phospho(1) | N[+1458] | 1287.593 | 3 | 3860.7643 | 3860.7346 | 0 | 7.7 | 413 | Specific | 220.37 | 220.37 | 220.37 | 5.18 | 101 | >sp\|P01833\|PIGR_HUMAN Polymeric immunoglobulin receptor OS=Homo sapiens OX=9606 GN=PIGR PE=1 SV=4 | 902 | 20201114_qwl_colo_milk_1ng_glyco_hcd203040_compound15.22580.22580.3 | controllerType=0 controllerNumber=1 scan=22580 | 49.9091 |
| 17133:3 | 5 | R.[+42.011]LSLLEEPGN[+2018.708]GTFTVILNQLTSR.D | HexNAc(3)Hex(6)Fuc(1)NeuAc(1) | .[42], N[+2019] | 1488.3566 | 3 | 4463.0552 | 4463.0115 | 0 | 9.8 | 413 | Specific | 269.43 | 145.91 | 145.91 | 5.56 | 101 | >sp\|P01833\|PIGR_HUMAN Polymeric immunoglobulin receptor OS=Homo sapiens OX=9606 GN=PIGR PE=1 SV=4 | 902 | 20201114_qwl_colo_milk_1ng_glyco_hcd203040_compound15.20580.20580.3 | controllerType=0 controllerNumber=1 scan=20580 | 45.5553 |
| 20291:3 | 5 | R.LSLLEEPGN[+2861.000]GTFTVILNQLTSR.D | HexNAc(5)Hex(6)NeuAc(3) | N[+2861] | 1755.1174 | 3 | 5263.3376 | 5263.2926 | 0 | 8.6 | 413 | Specific | 302.05 | 252.11 | 252.11 | 6.77 | 101 | >sp\|P01833\|PIGR_HUMAN Polymeric immunoglobulin receptor OS=Homo sapiens OX=9606 GN=PIGR PE=1 SV=4 | 902 | 20201114_qwl_colo_milk_1ng_glyco_hcd203040_compound15.23885.23885.3 | controllerType=0 controllerNumber=1 scan=23885 | 52.587 |
| 25490:3 | 5 | R.[+42.011]LSLLEEPGN[+2350.830]GTFTVILNQLTSR.D | HexNAc(4)Hex(5)Fuc(1)NeuAc(2) | .[42], N[+2351] | 1599.0528 | 3 | 4795.1438 | 4795.1335 | 0 | 2.2 | 413 | Specific | 336.46 | 306.32 | 306.32 | 7.72 | 101 | >sp\|P01833\|PIGR_HUMAN Polymeric immunoglobulin receptor OS=Homo sapiens OX=9606 GN=PIGR PE=1 SV=4 | 902 | 20201114_qwl_colo_milk_1ng_glyco_hcd203040_compound15.29858.29858.3 | controllerType=0 controllerNumber=1 scan=29858 | 65.1443 |
| 17033:3 | 5 | R.LSLLEEPGN[+2425.887]GTFTVILNQLTSR.D | HexNAc(5)Hex(6)Fuc(3) | N[+2426] | 1610.0673 | 3 | 4828.1874 | 4828.1801 | 0 | 1.5 | 413 | Specific | 354.93 | 329.62 | 329.62 | 7.93 | 101 | >sp\|P01833\|PIGR_HUMAN Polymeric immunoglobulin receptor OS=Homo sapiens OX=9606 GN=PIGR PE=1 SV=4 | 902 | 20201114_qwl_colo_milk_1ng_glyco_hcd203040_compound15.20476.20476.3 | controllerType=0 controllerNumber=1 scan=20476 | 45.3458 |
| 20293:3 | 5 | R.LSLLEEPGN[+2861.000]GTFTVILNQLTSR.D | HexNAc(5)Hex(6)NeuAc(3) | N[+2861] | 1755.1174 | 3 | 5263.3376 | 5263.2926 | 0 | 8.6 | 413 | Specific | 394.82 | 382.21 | 382.21 | 9.37 | 101 | >sp\|P01833\|PIGR_HUMAN Polymeric immunoglobulin receptor OS=Homo sapiens OX=9606 GN=PIGR PE=1 SV=4 | 902 | 20201114_qwl_colo_milk_1ng_glyco_hcd203040_compound15.23887.23887.3 | controllerType=0 controllerNumber=1 scan=23887 | 52.5975 |
| 18711:3 | 5 | R.LSLLEEPGN[+2059.735]GTFTVILNQLTSR.D | HexNAc(4)Hex(5)Fuc(1)NeuAc(1) | N[+2060] | 1488.0182 | 3 | 4462.04 | 4462.0275 | 0 | 2.8 | 413 | Specific | 520.84 | 306.54 | 306.54 | 8.38 | 101 | >sp\|P01833\|PIGR_HUMAN Polymeric immunoglobulin receptor OS=Homo sapiens OX=9606 GN=PIGR PE=1 SV=4 | 902 | 20201114_qwl_colo_milk_1ng_glyco_hcd203040_compound15.22235.22235.3 | controllerType=0 controllerNumber=1 scan=22235 | 49.1808 |
| 19039:3 | 5 | R.LSLLEEPGN[+1458.442]GTFTVILNQLTSR.D | HexNAc(2)Hex(6)Phospho(1) | N[+1458] | 1287.593 | 3 | 3860.7643 | 3860.7346 | 0 | 7.7 | 413 | Specific | 527.98 | 526.97 | 526.97 | 9.88 | 101 | >sp\|P01833\|PIGR_HUMAN Polymeric immunoglobulin receptor OS=Homo sapiens OX=9606 GN=PIGR PE=1 SV=4 | 902 | 20201114_qwl_colo_milk_1ng_glyco_hcd203040_compound15.22578.22578.3 | controllerType=0 controllerNumber=1 scan=22578 | 49.9059 |
| 18684:3 | 5 | R.LSLLEEPGN[+2059.735]GTFTVILNQLTSR.D | HexNAc(4)Hex(5)Fuc(1)NeuAc(1) | N[+2060] | 1488.0208 | 3 | 4462.0479 | 4462.0275 | 0 | 4.6 | 413 | Specific | 547 | 547 | 547 | 10.14 | 101 | >sp\|P01833\|PIGR_HUMAN Polymeric immunoglobulin receptor OS=Homo sapiens OX=9606 GN=PIGR PE=1 SV=4 | 902 | 20201114_qwl_colo_milk_1ng_glyco_hcd203040_compound15.22207.22207.3 | controllerType=0 controllerNumber=1 scan=22207 | 49.1227 |
| 21126:3 | 5 | R.LSLLEEPGN[+2350.830]GTFTVILNQLTSR.D | HexNAc(4)Hex(5)Fuc(1)NeuAc(2) | N[+2351] | 1585.0527 | 3 | 4753.1435 | 4753.1229 | 0 | 4.3 | 413 | Specific | 584.2 | 435.69 | 435.69 | 11.85 | 101 | >sp\|P01833\|PIGR_HUMAN Polymeric immunoglobulin receptor OS=Homo sapiens OX=9606 GN=PIGR PE=1 SV=4 | 902 | 20201114_qwl_colo_milk_1ng_glyco_hcd203040_compound15.24755.24755.3 | controllerType=0 controllerNumber=1 scan=24755 | 54.3077 |
| 17115:3 | 5 | R.LSLLEEPGN[+2059.735]GTFTVILNQLTSR.D | HexNAc(4)Hex(5)Fuc(1)NeuAc(1) | N[+2060] | 1488.0256 | 3 | 4462.0622 | 4462.0275 | 0 | 7.8 | 413 | Specific | 627.07 | 541.26 | 541.26 | 12.5 | 101 | >sp\|P01833\|PIGR_HUMAN Polymeric immunoglobulin receptor OS=Homo sapiens OX=9606 GN=PIGR PE=1 SV=4 | 902 | 20201114_qwl_colo_milk_1ng_glyco_hcd203040_compound15.20561.20561.3 | controllerType=0 controllerNumber=1 scan=20561 | 45.5032 |
| 20305:4 | 5 | R.LSLLEEPGN[+2863.041]GTFTVILNQLTSR.D | HexNAc(5)Hex(6)Fuc(4)NeuAc(1) | N[+2863] | 1317.0902 | 4 | 5265.339 | 5265.3334 | 0 | 1.1 | 413 | Specific | 663.12 | 372.54 | 372.54 | 12.83 | 101 | >sp\|P01833\|PIGR_HUMAN Polymeric immunoglobulin receptor OS=Homo sapiens OX=9606 GN=PIGR PE=1 SV=4 | 902 | 20201114_qwl_colo_milk_1ng_glyco_hcd203040_compound15.23899.23899.4 | controllerType=0 controllerNumber=1 scan=23899 | 52.6196 |
| 20306:4 | 5 | R.LSLLEEPGN[+2863.041]GTFTVILNQLTSR.D | HexNAc(5)Hex(6)Fuc(4)NeuAc(1) | N[+2863] | 1317.0902 | 4 | 5265.339 | 5265.3334 | 0 | 1.1 | 413 | Specific | 682.06 | 299.2 | 299.2 | 12.29 | 101 | >sp\|P01833\|PIGR_HUMAN Polymeric immunoglobulin receptor OS=Homo sapiens OX=9606 GN=PIGR PE=1 SV=4 | 902 | 20201114_qwl_colo_milk_1ng_glyco_hcd203040_compound15.23900.23900.4 | controllerType=0 controllerNumber=1 scan=23900 | 52.6229 |
| 20202:3 | 5 | R.LSLLEEPGN[+3008.078]GTFTVILNQLTSR.D | HexNAc(5)Hex(6)Fuc(3)NeuAc(2) | N[+3008] | 1804.1418 | 3 | 5410.4109 | 5410.3709 | 0 | 7.4 | 413 | Specific | 692.18 | 453.77 | 453.77 | 13.5 | 101 | >sp\|P01833\|PIGR_HUMAN Polymeric immunoglobulin receptor OS=Homo sapiens OX=9606 GN=PIGR PE=1 SV=4 | 902 | 20201114_qwl_colo_milk_1ng_glyco_hcd203040_compound15.23792.23792.3 | controllerType=0 controllerNumber=1 scan=23792 | 52.3928 |
| 17035:3 | 5 | R.LSLLEEPGN[+2425.887]GTFTVILNQLTSR.D | HexNAc(5)Hex(6)Fuc(3) | N[+2426] | 1610.0673 | 3 | 4828.1874 | 4828.1801 | 0 | 1.5 | 413 | Specific | 695.26 | 501.77 | 501.77 | 13.78 | 101 | >sp\|P01833\|PIGR_HUMAN Polymeric immunoglobulin receptor OS=Homo sapiens OX=9606 GN=PIGR PE=1 SV=4 | 902 | 20201114_qwl_colo_milk_1ng_glyco_hcd203040_compound15.20478.20478.3 | controllerType=0 controllerNumber=1 scan=20478 | 45.349 |
| 20200:3 | 5 | R.LSLLEEPGN[+3008.078]GTFTVILNQLTSR.D | HexNAc(5)Hex(6)Fuc(3)NeuAc(2) | N[+3008] | 1804.1418 | 3 | 5410.4109 | 5410.3709 | 0 | 7.4 | 413 | Specific | 702.42 | 541.99 | 541.99 | 14.5 | 101 | >sp\|P01833\|PIGR_HUMAN Polymeric immunoglobulin receptor OS=Homo sapiens OX=9606 GN=PIGR PE=1 SV=4 | 902 | 20201114_qwl_colo_milk_1ng_glyco_hcd203040_compound15.23790.23790.3 | controllerType=0 controllerNumber=1 scan=23790 | 52.3875 |
| 20687:4 | 5 | R.LSLLEEPGN[+2496.888]GTFTVILNQLTSR.D | HexNAc(4)Hex(5)Fuc(2)NeuAc(2) | N[+2497] | 1225.5611 | 4 | 4899.2225 | 4899.1808 | 0 | 8.5 | 413 | Specific | 723.45 | 594.48 | 594.48 | 14.87 | 101 | >sp\|P01833\|PIGR_HUMAN Polymeric immunoglobulin receptor OS=Homo sapiens OX=9606 GN=PIGR PE=1 SV=4 | 902 | 20201114_qwl_colo_milk_1ng_glyco_hcd203040_compound15.24297.24297.4 | controllerType=0 controllerNumber=1 scan=24297 | 53.432 |
| 20685:4 | 5 | R.LSLLEEPGN[+2496.888]GTFTVILNQLTSR.D | HexNAc(4)Hex(5)Fuc(2)NeuAc(2) | N[+2497] | 1225.5611 | 4 | 4899.2225 | 4899.1808 | 0 | 8.5 | 413 | Specific | 755.32 | 616.18 | 616.18 | 15.87 | 101 | >sp\|P01833\|PIGR_HUMAN Polymeric immunoglobulin receptor OS=Homo sapiens OX=9606 GN=PIGR PE=1 SV=4 | 902 | 20201114_qwl_colo_milk_1ng_glyco_hcd203040_compound15.24295.24295.4 | controllerType=0 controllerNumber=1 scan=24295 | 53.4273 |
| 16939:3 | 5 | R.LSLLEEPGN[+2863.041]GTFTVILNQLTSR.D | HexNAc(5)Hex(6)Fuc(4)NeuAc(1) | N[+2863] | 1755.7835 | 3 | 5265.336 | 5265.3334 | 0 | 0.5 | 413 | Specific | 762.35 | 448.22 | 448.22 | 15.73 | 101 | >sp\|P01833\|PIGR_HUMAN Polymeric immunoglobulin receptor OS=Homo sapiens OX=9606 GN=PIGR PE=1 SV=4 | 902 | 20201114_qwl_colo_milk_1ng_glyco_hcd203040_compound15.20378.20378.3 | controllerType=0 controllerNumber=1 scan=20378 | 45.1464 |
| 18469:4 | 5 | R.LSLLEEPGN[+2351.851]GTFTVILNQLTSR.D | HexNAc(4)Hex(5)Fuc(3)NeuAc(1) | N[+2352] | 1189.2929 | 4 | 4754.1497 | 4754.1433 | 0 | 1.3 | 413 | Specific | 784.98 | 575.83 | 575.83 | 16.23 | 101 | >sp\|P01833\|PIGR_HUMAN Polymeric immunoglobulin receptor OS=Homo sapiens OX=9606 GN=PIGR PE=1 SV=4 | 902 | 20201114_qwl_colo_milk_1ng_glyco_hcd203040_compound15.21983.21983.4 | controllerType=0 controllerNumber=1 scan=21983 | 48.6523 |
| 18686:3 | 5 | R.LSLLEEPGN[+2059.735]GTFTVILNQLTSR.D | HexNAc(4)Hex(5)Fuc(1)NeuAc(1) | N[+2060] | 1488.0208 | 3 | 4462.0479 | 4462.0275 | 0 | 4.6 | 413 | Specific | 810.09 | 550.65 | 550.65 | 16.55 | 101 | >sp\|P01833\|PIGR_HUMAN Polymeric immunoglobulin receptor OS=Homo sapiens OX=9606 GN=PIGR PE=1 SV=4 | 902 | 20201114_qwl_colo_milk_1ng_glyco_hcd203040_compound15.22209.22209.3 | controllerType=0 controllerNumber=1 scan=22209 | 49.1286 |
| 18709:3 | 5 | R.LSLLEEPGN[+2059.735]GTFTVILNQLTSR.D | HexNAc(4)Hex(5)Fuc(1)NeuAc(1) | N[+2060] | 1488.0182 | 3 | 4462.04 | 4462.0275 | 0 | 2.8 | 413 | Specific | 827.72 | 665.05 | 665.05 | 16.88 | 101 | >sp\|P01833\|PIGR_HUMAN Polymeric immunoglobulin receptor OS=Homo sapiens OX=9606 GN=PIGR PE=1 SV=4 | 902 | 20201114_qwl_colo_milk_1ng_glyco_hcd203040_compound15.22233.22233.3 | controllerType=0 controllerNumber=1 scan=22233 | 49.176 |
| 18359:3 | 5 | R.LSLLEEPGN[+2863.041]GTFTVILNQLTSR.D | HexNAc(5)Hex(6)Fuc(4)NeuAc(1) | N[+2863] | 1755.7845 | 3 | 5265.339 | 5265.3334 | 0 | 1.1 | 413 | Specific | 832.63 | 424.44 | 424.44 | 16.73 | 101 | >sp\|P01833\|PIGR_HUMAN Polymeric immunoglobulin receptor OS=Homo sapiens OX=9606 GN=PIGR PE=1 SV=4 | 902 | 20201114_qwl_colo_milk_1ng_glyco_hcd203040_compound15.21867.21867.3 | controllerType=0 controllerNumber=1 scan=21867 | 48.3948 |
| 18353:3 | 5 | R.LSLLEEPGN[+2570.925]GTFTVILNQLTSR.D | HexNAc(5)Hex(6)Fuc(2)NeuAc(1) | N[+2571] | 1658.4154 | 3 | 4973.2316 | 4973.2176 | 0 | 2.8 | 413 | Specific | 884.91 | 595.85 | 595.85 | 17.88 | 101 | >sp\|P01833\|PIGR_HUMAN Polymeric immunoglobulin receptor OS=Homo sapiens OX=9606 GN=PIGR PE=1 SV=4 | 902 | 20201114_qwl_colo_milk_1ng_glyco_hcd203040_compound15.21861.21861.3 | controllerType=0 controllerNumber=1 scan=21861 | 48.3797 |
| 18357:3 | 5 | R.LSLLEEPGN[+2863.041]GTFTVILNQLTSR.D | HexNAc(5)Hex(6)Fuc(4)NeuAc(1) | N[+2863] | 1755.7845 | 3 | 5265.339 | 5265.3334 | 0 | 1.1 | 413 | Specific | 892.15 | 432.68 | 432.68 | 17.73 | 101 | >sp\|P01833\|PIGR_HUMAN Polymeric immunoglobulin receptor OS=Homo sapiens OX=9606 GN=PIGR PE=1 SV=4 | 902 | 20201114_qwl_colo_milk_1ng_glyco_hcd203040_compound15.21865.21865.3 | controllerType=0 controllerNumber=1 scan=21865 | 48.3903 |
| 18528:4 | 5 | R.LSLLEEPGN[+2205.793]GTFTVILNQLTSR.D | HexNAc(4)Hex(5)Fuc(2)NeuAc(1) | N[+2206] | 1152.7875 | 4 | 4608.1282 | 4608.0854 | 0 | 9.3 | 413 | Specific | 912.89 | 497.05 | 497.05 | 18.74 | 101 | >sp\|P01833\|PIGR_HUMAN Polymeric immunoglobulin receptor OS=Homo sapiens OX=9606 GN=PIGR PE=1 SV=4 | 902 | 20201114_qwl_colo_milk_1ng_glyco_hcd203040_compound15.22045.22045.4 | controllerType=0 controllerNumber=1 scan=22045 | 48.7805 |
| 20539:3 | 5 | R.LSLLEEPGN[+2350.830]GTFTVILNQLTSR.D | HexNAc(4)Hex(5)Fuc(1)NeuAc(2) | N[+2351] | 1585.0551 | 3 | 4753.1508 | 4753.1229 | 0 | 5.9 | 413 | Specific | 919.6 | 652.39 | 652.39 | 18.36 | 101 | >sp\|P01833\|PIGR_HUMAN Polymeric immunoglobulin receptor OS=Homo sapiens OX=9606 GN=PIGR PE=1 SV=4 | 902 | 20201114_qwl_colo_milk_1ng_glyco_hcd203040_compound15.24143.24143.3 | controllerType=0 controllerNumber=1 scan=24143 | 53.1055 |
| 18530:4 | 5 | R.LSLLEEPGN[+2205.793]GTFTVILNQLTSR.D | HexNAc(4)Hex(5)Fuc(2)NeuAc(1) | N[+2206] | 1152.7875 | 4 | 4608.1282 | 4608.0854 | 0 | 9.3 | 413 | Specific | 949.14 | 559.06 | 559.06 | 18.74 | 101 | >sp\|P01833\|PIGR_HUMAN Polymeric immunoglobulin receptor OS=Homo sapiens OX=9606 GN=PIGR PE=1 SV=4 | 902 | 20201114_qwl_colo_milk_1ng_glyco_hcd203040_compound15.22047.22047.4 | controllerType=0 controllerNumber=1 scan=22047 | 48.786 |
| 18468:3 | 5 | R.LSLLEEPGN[+2205.793]GTFTVILNQLTSR.D | HexNAc(4)Hex(5)Fuc(2)NeuAc(1) | N[+2206] | 1536.7043 | 3 | 4608.0983 | 4608.0854 | 0 | 2.8 | 413 | Specific | 1008.2 | 727.69 | 727.69 | 20.88 | 101 | >sp\|P01833\|PIGR_HUMAN Polymeric immunoglobulin receptor OS=Homo sapiens OX=9606 GN=PIGR PE=1 SV=4 | 902 | 20201114_qwl_colo_milk_1ng_glyco_hcd203040_compound15.21982.21982.3 | controllerType=0 controllerNumber=1 scan=21982 | 48.6488 |
| 18351:3 | 5 | R.LSLLEEPGN[+2570.925]GTFTVILNQLTSR.D | HexNAc(5)Hex(6)Fuc(2)NeuAc(1) | N[+2571] | 1658.4154 | 3 | 4973.2316 | 4973.2176 | 0 | 2.8 | 413 | Specific | 1011.4 | 759.08 | 759.08 | 20.88 | 101 | >sp\|P01833\|PIGR_HUMAN Polymeric immunoglobulin receptor OS=Homo sapiens OX=9606 GN=PIGR PE=1 SV=4 | 902 | 20201114_qwl_colo_milk_1ng_glyco_hcd203040_compound15.21859.21859.3 | controllerType=0 controllerNumber=1 scan=21859 | 48.3737 |
| 20536:3 | 5 | R.LSLLEEPGN[+2350.830]GTFTVILNQLTSR.D | HexNAc(4)Hex(5)Fuc(1)NeuAc(2) | N[+2351] | 1585.0551 | 3 | 4753.1508 | 4753.1229 | 0 | 5.9 | 413 | Specific | 1044.87 | 813.16 | 813.16 | 20.36 | 101 | >sp\|P01833\|PIGR_HUMAN Polymeric immunoglobulin receptor OS=Homo sapiens OX=9606 GN=PIGR PE=1 SV=4 | 902 | 20201114_qwl_colo_milk_1ng_glyco_hcd203040_compound15.24140.24140.3 | controllerType=0 controllerNumber=1 scan=24140 | 53.0975 |
| 18467:4 | 5 | R.LSLLEEPGN[+2351.851]GTFTVILNQLTSR.D | HexNAc(4)Hex(5)Fuc(3)NeuAc(1) | N[+2352] | 1189.2929 | 4 | 4754.1497 | 4754.1433 | 0 | 1.3 | 413 | Specific | 1046.59 | 1039.99 | 1039.99 | 21.23 | 101 | >sp\|P01833\|PIGR_HUMAN Polymeric immunoglobulin receptor OS=Homo sapiens OX=9606 GN=PIGR PE=1 SV=4 | 902 | 20201114_qwl_colo_milk_1ng_glyco_hcd203040_compound15.21981.21981.4 | controllerType=0 controllerNumber=1 scan=21981 | 48.6413 |
| 18466:3 | 5 | R.LSLLEEPGN[+2205.793]GTFTVILNQLTSR.D | HexNAc(4)Hex(5)Fuc(2)NeuAc(1) | N[+2206] | 1536.7043 | 3 | 4608.0983 | 4608.0854 | 0 | 2.8 | 413 | Specific | 1138.7 | 898.72 | 898.72 | 22.88 | 101 | >sp\|P01833\|PIGR_HUMAN Polymeric immunoglobulin receptor OS=Homo sapiens OX=9606 GN=PIGR PE=1 SV=4 | 902 | 20201114_qwl_colo_milk_1ng_glyco_hcd203040_compound15.21980.21980.3 | controllerType=0 controllerNumber=1 scan=21980 | 48.6398 |
| 14546:3 | 5 | K.IIEGEPNLKVPGN[+2205.793]VTAVLGETLK.V | HexNAc(4)Hex(5)Fuc(2)NeuAc(1) | N[+2206] | 1533.0616 | 3 | 4597.1702 | 4597.1422 | 0 | 6.1 | 457 | Specific | 609.56 | 609.56 | 609.56 | 11.92 | 101 | >sp\|P01833\|PIGR_HUMAN Polymeric immunoglobulin receptor OS=Homo sapiens OX=9606 GN=PIGR PE=1 SV=4 | 902 | 20201114_qwl_colo_milk_1ng_glyco_hcd203040_compound15.17868.17868.3 | controllerType=0 controllerNumber=1 scan=17868 | 39.4221 |
| 13033:3 | 5 | K.IIEGEPNLKVPGN[+2059.735]VTAVLGETLK.V | HexNAc(4)Hex(5)Fuc(1)NeuAc(1) | N[+2060] | 1484.3655 | 3 | 4451.0821 | 4451.0843 | 0 | -0.5 | 457 | Specific | 677.98 | 523.54 | 523.54 | 12.81 | 101 | >sp\|P01833\|PIGR_HUMAN Polymeric immunoglobulin receptor OS=Homo sapiens OX=9606 GN=PIGR PE=1 SV=4 | 902 | 20201114_qwl_colo_milk_1ng_glyco_hcd203040_compound15.16277.16277.3 | controllerType=0 controllerNumber=1 scan=16277 | 35.8824 |

Table S4. Cont.

| Query #:z | Protein Rank | Glycopeptide | Glycan | Modification Type(s) | Observed m/z | z | Observed (M+H) | Calc. mass (M+H) | Off-by-x error | Mass error (ppm) | StartPosition | Cleavage | Score | Delta | Delta Mod | \|Log Prob\| | # of unique peptides | Protein Name | Protein DB number | Comment | Scan # | Scan Time |
|---|---|---|---|---|---|---|---|---|---|---|---|---|---|---|---|---|---|---|---|---|---|---|
| 14549:3 | 5 | K.IIEGEPNLKVPGN[+2205.793]VTAVLGETLK.V | HexNAc(4)Hex(5)Fuc(2)NeuAc(1) | N[+2206] | 1533.0616 | 3 | 4597.1702 | 4597.1422 | 0 | 6.1 | 457 | Specific | 750.27 | 690.94 | 690.94 | 14.92 | 101 | >sp\|P01833\|PIGR_HUMAN Polymeric immunoglobulin receptor OS=Homo sapiens OX=9606 GN=PIGR PE=1 SV=4 | 902 | 20201114_qwl_colo_milk_1ng_glyco_hcd203040_compound15.17871.17871.3 | controllerType=0 controllerNumber=1 scan=17871 | 39.4268 |
| 13031:3 | 5 | K.IIEGEPNLKVPGN[+2059.735]VTAVLGETLK.V | HexNAc(4)Hex(5)Fuc(1)NeuAc(1) | N[+2060] | 1484.3655 | 3 | 4451.0821 | 4451.0843 | 0 | -0.5 | 457 | Specific | 931.74 | 922.58 | 922.58 | 17.81 | 101 | >sp\|P01833\|PIGR_HUMAN Polymeric immunoglobulin receptor OS=Homo sapiens OX=9606 GN=PIGR PE=1 SV=4 | 902 | 20201114_qwl_colo_milk_1ng_glyco_hcd203040_compound15.16275.16275.3 | controllerType=0 controllerNumber=1 scan=16275 | 35.8762 |
| 14497:3 | 5 | K.IIEGEPNLKVPGN[+2059.735]VTAVLGETLK.V | HexNAc(4)Hex(5)Fuc(1)NeuAc(1) | N[+2060] | 1484.3738 | 3 | 4451.107 | 4451.0843 | 0 | 5.1 | 457 | Specific | 933.56 | 845.88 | 845.88 | 18.11 | 101 | >sp\|P01833\|PIGR_HUMAN Polymeric immunoglobulin receptor OS=Homo sapiens OX=9606 GN=PIGR PE=1 SV=4 | 902 | 20201114_qwl_colo_milk_1ng_glyco_hcd203040_compound15.17816.17816.3 | controllerType=0 controllerNumber=1 scan=17816 | 39.2851 |
| 14494:3 | 5 | K.IIEGEPNLKVPGN[+2059.735]VTAVLGETLK.V | HexNAc(4)Hex(5)Fuc(1)NeuAc(1) | N[+2060] | 1484.3738 | 3 | 4451.107 | 4451.0843 | 0 | 5.1 | 457 | Specific | 1089.98 | 970.88 | 970.88 | 21.11 | 101 | >sp\|P01833\|PIGR_HUMAN Polymeric immunoglobulin receptor OS=Homo sapiens OX=9606 GN=PIGR PE=1 SV=4 | 902 | 20201114_qwl_colo_milk_1ng_glyco_hcd203040_compound15.17813.17813.3 | controllerType=0 controllerNumber=1 scan=17813 | 39.2714 |
| 15306:5 | 5 | K.IIEGEPNLKVPGN[+2350.830]VTAVLGETLKVPC[+57.021]HFPC[+57.021]K.F | HexNAc(4)Hex(5)Fuc(1)NeuAc(2) | C[+57]*2, N[+2351] | 1154.3419 | 5 | 5767.6804 | 5767.6372 | 0 | 7.5 | 457 | Specific | 242 | 242 | 242 | 5.56 | 101 | >sp\|P01833\|PIGR_HUMAN Polymeric immunoglobulin receptor OS=Homo sapiens OX=9606 GN=PIGR PE=1 SV=4 | 902 | 20201114_qwl_colo_milk_1ng_glyco_hcd203040_compound15.18661.18661.5 | controllerType=0 controllerNumber=1 scan=18661 | 41.0555 |
| 13255:5 | 5 | K.IIEGEPNLKVPGN[+2059.735]VTAVLGETLKVPC[+57.021]HFPC[+57.021]K.F | HexNAc(4)Hex(5)Fuc(1)NeuAc(1) | C[+57]*2, N[+2060] | 1096.1251 | 5 | 5476.5965 | 5476.5418 | 0 | 10 | 457 | Specific | 310.65 | 310.65 | 310.65 | 4.2 | 101 | >sp\|P01833\|PIGR_HUMAN Polymeric immunoglobulin receptor OS=Homo sapiens OX=9606 GN=PIGR PE=1 SV=4 | 902 | 20201114_qwl_colo_milk_1ng_glyco_hcd203040_compound15.16509.16509.3 | controllerType=0 controllerNumber=1 scan=16509 | 36.3892 |
| 14300:5 | 5 | K.IIEGEPNLKVPGN[+2205.793]VTAVLGETLKVPC[+57.021]HFPC[+57.021]K.F | HexNAc(4)Hex(5)Fuc(2)NeuAc(1) | C[+57]*2, N[+2206] | 1125.3323 | 5 | 5622.6323 | 5622.5997 | 0 | 5.8 | 457 | Specific | 314.19 | 314.19 | 314.19 | 4.7 | 101 | >sp\|P01833\|PIGR_HUMAN Polymeric immunoglobulin receptor OS=Homo sapiens OX=9606 GN=PIGR PE=1 SV=4 | 902 | 20201114_qwl_colo_milk_1ng_glyco_hcd203040_compound15.17608.17608.5 | controllerType=0 controllerNumber=1 scan=17608 | 38.8464 |
| 14982:4 | 5 | K.IIEGEPNLKVPGN[+3154.136]VTAVLGETLKVPC[+57.021]HFPC[+57.021]K.F | HexNAc(5)Hex(6)Fuc(4)NeuAc(2) | C[+57]*2, N[+3154] | 1643.5074 | 4 | 6571.008 | 6570.9432 | 0 | 9.9 | 457 | Specific | 357.05 | 354.68 | 354.68 | 5.82 | 101 | >sp\|P01833\|PIGR_HUMAN Polymeric immunoglobulin receptor OS=Homo sapiens OX=9606 GN=PIGR PE=1 SV=4 | 902 | 20201114_qwl_colo_milk_1ng_glyco_hcd203040_compound15.18324.18324.3 | controllerType=0 controllerNumber=1 scan=18324 | 40.3805 |
| 15299:4 | 5 | K.IIEGEPNLKVPGN[+2350.830]VTAVLGETLKVPC[+57.021]HFPC[+57.021]K.F | HexNAc(4)Hex(5)Fuc(1)NeuAc(2) | C[+57]*2, N[+2351] | 1442.6755 | 4 | 5767.6802 | 5767.6372 | 0 | 7.4 | 457 | Specific | 433.74 | 224.79 | 224.79 | 5.65 | 101 | >sp\|P01833\|PIGR_HUMAN Polymeric immunoglobulin receptor OS=Homo sapiens OX=9606 GN=PIGR PE=1 SV=4 | 902 | 20201114_qwl_colo_milk_1ng_glyco_hcd203040_compound15.18654.18654.4 | controllerType=0 controllerNumber=1 scan=18654 | 41.0416 |
| 13253:5 | 5 | K.IIEGEPNLKVPGN[+2059.735]VTAVLGETLKVPC[+57.021]HFPC[+57.021]K.F | HexNAc(4)Hex(5)Fuc(1)NeuAc(1) | C[+57]*2, N[+2060] | 1096.1251 | 5 | 5476.5965 | 5476.5418 | 0 | 10 | 457 | Specific | 443.7 | 397.77 | 397.77 | 6.83 | 101 | >sp\|P01833\|PIGR_HUMAN Polymeric immunoglobulin receptor OS=Homo sapiens OX=9606 GN=PIGR PE=1 SV=4 | 902 | 20201114_qwl_colo_milk_1ng_glyco_hcd203040_compound15.16507.16507.3 | controllerType=0 controllerNumber=1 scan=16507 | 36.3773 |
| 14302:5 | 5 | K.IIEGEPNLKVPGN[+2205.793]VTAVLGETLKVPC[+57.021]HFPC[+57.021]K.F | HexNAc(4)Hex(5)Fuc(2)NeuAc(1) | C[+57]*2, N[+2206] | 1125.3323 | 5 | 5622.6323 | 5622.5997 | 0 | 5.8 | 457 | Specific | 485.73 | 416.4 | 416.4 | 8.67 | 101 | >sp\|P01833\|PIGR_HUMAN Polymeric immunoglobulin receptor OS=Homo sapiens OX=9606 GN=PIGR PE=1 SV=4 | 902 | 20201114_qwl_colo_milk_1ng_glyco_hcd203040_compound15.17610.17610.5 | controllerType=0 controllerNumber=1 scan=17610 | 38.8526 |
| 13079:5 | 5 | K.IIEGEPNLKVPGN[+2571.946]VTAVLGETLKVPC[+57.021]HFPC[+57.021]K.F | HexNAc(5)Hex(6)Fuc(4) | C[+57]*2, N[+2572] | 1198.5609 | 5 | 5988.7755 | 5988.7523 | 0 | 3.9 | 457 | Specific | 520.36 | 520.36 | 520.36 | 9.3 | 101 | >sp\|P01833\|PIGR_HUMAN Polymeric immunoglobulin receptor OS=Homo sapiens OX=9606 GN=PIGR PE=1 SV=4 | 902 | 20201114_qwl_colo_milk_1ng_glyco_hcd203040_compound15.16325.16325.5 | controllerType=0 controllerNumber=1 scan=16325 | 36.0074 |
| 13004:5 | 5 | K.IIEGEPNLKVPGN[+2571.946]VTAVLGETLKVPC[+57.021]HFPC[+57.021]K.F | HexNAc(5)Hex(6)Fuc(4) | C[+57]*2, N[+2572] | 1198.5603 | 5 | 5988.7724 | 5988.7523 | 0 | 3.4 | 457 | Specific | 607.46 | 607.46 | 607.46 | 11.95 | 101 | >sp\|P01833\|PIGR_HUMAN Polymeric immunoglobulin receptor OS=Homo sapiens OX=9606 GN=PIGR PE=1 SV=4 | 902 | 20201114_qwl_colo_milk_1ng_glyco_hcd203040_compound15.16246.16246.5 | controllerType=0 controllerNumber=1 scan=16246 | 35.8064 |
| 14065:4 | 5 | K.IIEGEPNLKVPGN[+2351.851]VTAVLGETLKVPC[+57.021]HFPC[+57.021]K.F | HexNAc(4)Hex(5)Fuc(3)NeuAc(1) | C[+57]*2, N[+2352] | 1442.9249 | 4 | 5768.6779 | 5768.6576 | 0 | 3.5 | 457 | Specific | 694.5 | 694.5 | 694.5 | 12.25 | 101 | >sp\|P01833\|PIGR_HUMAN Polymeric immunoglobulin receptor OS=Homo sapiens OX=9606 GN=PIGR PE=1 SV=4 | 902 | 20201114_qwl_colo_milk_1ng_glyco_hcd203040_compound15.17360.17360.4 | controllerType=0 controllerNumber=1 scan=17360 | 38.2617 |
| 13007:5 | 5 | K.IIEGEPNLKVPGN[+2571.946]VTAVLGETLKVPC[+57.021]HFPC[+57.021]K.F | HexNAc(5)Hex(6)Fuc(4) | C[+57]*2, N[+2572] | 1198.5603 | 5 | 5988.7724 | 5988.7523 | 0 | 3.4 | 457 | Specific | 756.56 | 756.56 | 756.56 | 14.95 | 101 | >sp\|P01833\|PIGR_HUMAN Polymeric immunoglobulin receptor OS=Homo sapiens OX=9606 GN=PIGR PE=1 SV=4 | 902 | 20201114_qwl_colo_milk_1ng_glyco_hcd203040_compound15.16249.16249.5 | controllerType=0 controllerNumber=1 scan=16249 | 35.8156 |
| 13047:5 | 5 | K.IIEGEPNLKVPGN[+2059.735]VTAVLGETLKVPC[+57.021]HFPC[+57.021]K.F | HexNAc(4)Hex(5)Fuc(1)NeuAc(1) | C[+57]*2, N[+2060] | 1096.1174 | 5 | 5476.5581 | 5476.5418 | 0 | 3 | 457 | Specific | 802.52 | 784.46 | 784.46 | 15.95 | 101 | >sp\|P01833\|PIGR_HUMAN Polymeric immunoglobulin receptor OS=Homo sapiens OX=9606 GN=PIGR PE=1 SV=4 | 902 | 20201114_qwl_colo_milk_1ng_glyco_hcd203040_compound15.16291.16291.5 | controllerType=0 controllerNumber=1 scan=16291 | 35.917 |
| 13077:5 | 5 | K.IIEGEPNLKVPGN[+2571.946]VTAVLGETLKVPC[+57.021]HFPC[+57.021]K.F | HexNAc(5)Hex(6)Fuc(4) | C[+57]*2, N[+2572] | 1198.5609 | 5 | 5988.7755 | 5988.7523 | 0 | 3.9 | 457 | Specific | 803.05 | 803.05 | 803.05 | 15.92 | 101 | >sp\|P01833\|PIGR_HUMAN Polymeric immunoglobulin receptor OS=Homo sapiens OX=9606 GN=PIGR PE=1 SV=4 | 902 | 20201114_qwl_colo_milk_1ng_glyco_hcd203040_compound15.16323.16323.5 | controllerType=0 controllerNumber=1 scan=16323 | 35.9923 |
| 14077:5 | 5 | K.IIEGEPNLKVPGN[+2059.735]VTAVLGETLKVPC[+57.021]HFPC[+57.021]K.F | HexNAc(4)Hex(5)Fuc(1)NeuAc(1) | C[+57]*2, N[+2060] | 1096.1166 | 5 | 5476.5538 | 5476.5418 | 0 | 2.2 | 457 | Specific | 846.93 | 754.55 | 754.55 | 15.85 | 101 | >sp\|P01833\|PIGR_HUMAN Polymeric immunoglobulin receptor OS=Homo sapiens OX=9606 GN=PIGR PE=1 SV=4 | 902 | 20201114_qwl_colo_milk_1ng_glyco_hcd203040_compound15.17372.17372.5 | controllerType=0 controllerNumber=1 scan=17372 | 38.3114 |
| 14024:4 | 5 | K.IIEGEPNLKVPGN[+2205.793]VTAVLGETLKVPC[+57.021]HFPC[+57.021]K.F | HexNAc(4)Hex(5)Fuc(2)NeuAc(1) | C[+57]*2, N[+2206] | 1406.4097 | 4 | 5622.6169 | 5622.5997 | 0 | 3 | 457 | Specific | 868.2 | 739.92 | 739.92 | 16.29 | 101 | >sp\|P01833\|PIGR_HUMAN Polymeric immunoglobulin receptor OS=Homo sapiens OX=9606 GN=PIGR PE=1 SV=4 | 902 | 20201114_qwl_colo_milk_1ng_glyco_hcd203040_compound15.17317.17317.4 | controllerType=0 controllerNumber=1 scan=17317 | 38.1799 |
| 14070:5 | 5 | K.IIEGEPNLKVPGN[+2350.830]VTAVLGETLKVPC[+57.021]HFPC[+57.021]K.F | HexNAc(4)Hex(5)Fuc(1)NeuAc(2) | C[+57]*2, N[+2351] | 1154.3398 | 5 | 5767.6701 | 5767.6372 | 0 | 5.7 | 457 | Specific | 894.71 | 741.74 | 741.74 | 16.43 | 101 | >sp\|P01833\|PIGR_HUMAN Polymeric immunoglobulin receptor OS=Homo sapiens OX=9606 GN=PIGR PE=1 SV=4 | 902 | 20201114_qwl_colo_milk_1ng_glyco_hcd203040_compound15.17365.17365.5 | controllerType=0 controllerNumber=1 scan=17365 | 38.2872 |
| 14067:4 | 5 | K.IIEGEPNLKVPGN[+2351.851]VTAVLGETLKVPC[+57.021]HFPC[+57.021]K.F | HexNAc(4)Hex(5)Fuc(3)NeuAc(1) | C[+57]*2, N[+2352] | 1442.9249 | 4 | 5768.6779 | 5768.6576 | 0 | 3.5 | 457 | Specific | 899.56 | 768.75 | 768.75 | 16.25 | 101 | >sp\|P01833\|PIGR_HUMAN Polymeric immunoglobulin receptor OS=Homo sapiens OX=9606 GN=PIGR PE=1 SV=4 | 902 | 20201114_qwl_colo_milk_1ng_glyco_hcd203040_compound15.17362.17362.4 | controllerType=0 controllerNumber=1 scan=17362 | 38.2662 |
| 15304:5 | 5 | K.IIEGEPNLKVPGN[+2350.830]VTAVLGETLKVPC[+57.021]HFPC[+57.021]K.F | HexNAc(4)Hex(5)Fuc(1)NeuAc(2) | C[+57]*2, N[+2351] | 1154.3419 | 5 | 5767.6804 | 5767.6372 | 0 | 7.5 | 457 | Specific | 934.75 | 708.88 | 708.88 | 17.57 | 101 | >sp\|P01833\|PIGR_HUMAN Polymeric immunoglobulin receptor OS=Homo sapiens OX=9606 GN=PIGR PE=1 SV=4 | 902 | 20201114_qwl_colo_milk_1ng_glyco_hcd203040_compound15.18659.18659.5 | controllerType=0 controllerNumber=1 scan=18659 | 41.0486 |
| 13044:5 | 5 | K.IIEGEPNLKVPGN[+2059.735]VTAVLGETLKVPC[+57.021]HFPC[+57.021]K.F | HexNAc(4)Hex(5)Fuc(1)NeuAc(1) | C[+57]*2, N[+2060] | 1096.1174 | 5 | 5476.5581 | 5476.5418 | 0 | 3 | 457 | Specific | 938.4 | 934.33 | 934.33 | 17.95 | 101 | >sp\|P01833\|PIGR_HUMAN Polymeric immunoglobulin receptor OS=Homo sapiens OX=9606 GN=PIGR PE=1 SV=4 | 902 | 20201114_qwl_colo_milk_1ng_glyco_hcd203040_compound15.16288.16288.5 | controllerType=0 controllerNumber=1 scan=16288 | 35.9011 |
| 14017:5 | 5 | K.IIEGEPNLKVPGN[+2205.793]VTAVLGETLKVPC[+57.021]HFPC[+57.021]K.F | HexNAc(4)Hex(5)Fuc(2)NeuAc(1) | C[+57]*2, N[+2206] | 1125.3289 | 5 | 5622.6152 | 5622.5997 | 0 | 2.8 | 457 | Specific | 956.85 | 884.9 | 884.9 | 18.95 | 101 | >sp\|P01833\|PIGR_HUMAN Polymeric immunoglobulin receptor OS=Homo sapiens OX=9606 GN=PIGR PE=1 SV=4 | 902 | 20201114_qwl_colo_milk_1ng_glyco_hcd203040_compound15.17310.17310.5 | controllerType=0 controllerNumber=1 scan=17310 | 38.1613 |
| 14068:5 | 5 | K.IIEGEPNLKVPGN[+2350.830]VTAVLGETLKVPC[+57.021]HFPC[+57.021]K.F | HexNAc(4)Hex(5)Fuc(1)NeuAc(2) | C[+57]*2, N[+2351] | 1154.3398 | 5 | 5767.6701 | 5767.6372 | 0 | 5.7 | 457 | Specific | 974.92 | 761.12 | 761.12 | 18.43 | 101 | >sp\|P01833\|PIGR_HUMAN Polymeric immunoglobulin receptor OS=Homo sapiens OX=9606 GN=PIGR PE=1 SV=4 | 902 | 20201114_qwl_colo_milk_1ng_glyco_hcd203040_compound15.17363.17363.5 | controllerType=0 controllerNumber=1 scan=17363 | 38.2684 |
| 14021:4 | 5 | K.IIEGEPNLKVPGN[+2205.793]VTAVLGETLKVPC[+57.021]HFPC[+57.021]K.F | HexNAc(4)Hex(5)Fuc(2)NeuAc(1) | C[+57]*2, N[+2206] | 1406.4097 | 4 | 5622.6169 | 5622.5997 | 0 | 3 | 457 | Specific | 999.02 | 901.06 | 901.06 | 18.29 | 101 | >sp\|P01833\|PIGR_HUMAN Polymeric immunoglobulin receptor OS=Homo sapiens OX=9606 GN=PIGR PE=1 SV=4 | 902 | 20201114_qwl_colo_milk_1ng_glyco_hcd203040_compound15.17314.17314.4 | controllerType=0 controllerNumber=1 scan=17314 | 38.1666 |
| 15297:4 | 5 | K.IIEGEPNLKVPGN[+2350.830]VTAVLGETLKVPC[+57.021]HFPC[+57.021]K.F | HexNAc(4)Hex(5)Fuc(1)NeuAc(2) | C[+57]*2, N[+2351] | 1442.6755 | 4 | 5767.6802 | 5767.6372 | 0 | 7.4 | 457 | Specific | 1007.74 | 733.74 | 733.74 | 18.9 | 101 | >sp\|P01833\|PIGR_HUMAN Polymeric immunoglobulin receptor OS=Homo sapiens OX=9606 GN=PIGR PE=1 SV=4 | 902 | 20201114_qwl_colo_milk_1ng_glyco_hcd203040_compound15.18652.18652.4 | controllerType=0 controllerNumber=1 scan=18652 | 41.0336 |
| 14015:5 | 5 | K.IIEGEPNLKVPGN[+2205.793]VTAVLGETLKVPC[+57.021]HFPC[+57.021]K.F | HexNAc(4)Hex(5)Fuc(2)NeuAc(1) | C[+57]*2, N[+2206] | 1125.3289 | 5 | 5622.6152 | 5622.5997 | 0 | 2.8 | 457 | Specific | 1008.09 | 931.9 | 931.9 | 19.95 | 101 | >sp\|P01833\|PIGR_HUMAN Polymeric immunoglobulin receptor OS=Homo sapiens OX=9606 GN=PIGR PE=1 SV=4 | 902 | 20201114_qwl_colo_milk_1ng_glyco_hcd203040_compound15.17308.17308.5 | controllerType=0 controllerNumber=1 scan=17308 | 38.1524 |
| 14075:5 | 5 | K.IIEGEPNLKVPGN[+2059.735]VTAVLGETLKVPC[+57.021]HFPC[+57.021]K.F | HexNAc(4)Hex(5)Fuc(1)NeuAc(1) | C[+57]*2, N[+2060] | 1096.1166 | 5 | 5476.5538 | 5476.5418 | 0 | 2.2 | 457 | Specific | 1050.1 | 916.98 | 916.98 | 20.85 | 101 | >sp\|P01833\|PIGR_HUMAN Polymeric immunoglobulin receptor OS=Homo sapiens OX=9606 GN=PIGR PE=1 SV=4 | 902 | 20201114_qwl_colo_milk_1ng_glyco_hcd203040_compound15.17370.17370.5 | controllerType=0 controllerNumber=1 scan=17370 | 38.3017 |
| 12361:3 | 5 | K.VPGN[+2205.793]VTAVLGETLK.V | HexNAc(4)Hex(5)Fuc(2)NeuAc(1) | N[+2206] | 1201.8722 | 3 | 3603.6022 | 3603.5927 | 0 | 2.6 | 466 | Specific | 565.29 | 513.53 | 513.53 | 9.81 | 101 | >sp\|P01833\|PIGR_HUMAN Polymeric immunoglobulin receptor OS=Homo sapiens OX=9606 GN=PIGR PE=1 SV=4 | 902 | 20201114_qwl_colo_milk_1ng_glyco_hcd203040_compound15.15576.15576.3 | controllerType=0 controllerNumber=1 scan=15576 | 34.4586 |
| 12359:3 | 5 | K.VPGN[+2205.793]VTAVLGETLK.V | HexNAc(4)Hex(5)Fuc(2)NeuAc(1) | N[+2206] | 1201.8722 | 3 | 3603.6022 | 3603.5927 | 0 | 2.6 | 466 | Specific | 576.87 | 510.68 | 510.68 | 9.81 | 101 | >sp\|P01833\|PIGR_HUMAN Polymeric immunoglobulin receptor OS=Homo sapiens OX=9606 GN=PIGR PE=1 SV=4 | 902 | 20201114_qwl_colo_milk_1ng_glyco_hcd203040_compound15.15574.15574.3 | controllerType=0 controllerNumber=1 scan=15574 | 34.4538 |
| 12449:4 | 5 | K.[+42.011]VPGN[+2018.708]VTAVLGETLKVPC[+57.021]HFPC[+57.021]K.F | HexNAc(3)Hex(6)Fuc(1)NeuAc(1) | .[42], C[+57]*2, N[+2019] | 1121.7572 | 4 | 4484.007 | 4483.9763 | 0 | 6.8 | 466 | Specific | 206.62 | 206.62 | 206.62 | 3.18 | 101 | >sp\|P01833\|PIGR_HUMAN Polymeric immunoglobulin receptor OS=Homo sapiens OX=9606 GN=PIGR PE=1 SV=4 | 902 | 20201114_qwl_colo_milk_1ng_glyco_hcd203040_compound15.15668.15668.4 | controllerType=0 controllerNumber=1 scan=15668 | 34.6418 |
| 12447:4 | 5 | K.[+42.011]VPGN[+2018.708]VTAVLGETLKVPC[+57.021]HFPC[+57.021]K.F | HexNAc(3)Hex(6)Fuc(1)NeuAc(1) | .[42], C[+57]*2, N[+2019] | 1121.7572 | 4 | 4484.007 | 4483.9763 | 0 | 6.8 | 466 | Specific | 315.11 | 105.02 | 105.02 | 4.9 | 101 | >sp\|P01833\|PIGR_HUMAN Polymeric immunoglobulin receptor OS=Homo sapiens OX=9606 GN=PIGR PE=1 SV=4 | 902 | 20201114_qwl_colo_milk_1ng_glyco_hcd203040_compound15.15666.15666.4 | controllerType=0 controllerNumber=1 scan=15666 | 34.636 |
| 12395:4 | 5 | K.VPGN[+2351.851]VTAVLGETLKVPC[+57.021]HFPC[+57.021]K.F | HexNAc(4)Hex(5)Fuc(3)NeuAc(1) | C[+57]*2, N[+2352] | 1194.537 | 4 | 4775.1263 | 4775.1081 | 0 | 3.8 | 466 | Specific | 567.63 | 443.07 | 443.07 | 10.1 | 101 | >sp\|P01833\|PIGR_HUMAN Polymeric immunoglobulin receptor OS=Homo sapiens OX=9606 GN=PIGR PE=1 SV=4 | 902 | 20201114_qwl_colo_milk_1ng_glyco_hcd203040_compound15.15611.15611.2 | controllerType=0 controllerNumber=1 scan=15611 | 34.5371 |
| 12397:4 | 5 | K.VPGN[+2351.851]VTAVLGETLKVPC[+57.021]HFPC[+57.021]K.F | HexNAc(4)Hex(5)Fuc(3)NeuAc(1) | C[+57]*2, N[+2352] | 1194.537 | 4 | 4775.1263 | 4775.1081 | 0 | 3.8 | 466 | Specific | 590.96 | 511.27 | 511.27 | 11.41 | 101 | >sp\|P01833\|PIGR_HUMAN Polymeric immunoglobulin receptor OS=Homo sapiens OX=9606 GN=PIGR PE=1 SV=4 | 902 | 20201114_qwl_colo_milk_1ng_glyco_hcd203040_compound15.15613.15613.2 | controllerType=0 controllerNumber=1 scan=15613 | 34.5412 |
| 12381:4 | 5 | K.VPGN[+2205.793]VTAVLGETLKVPC[+57.021]HFPC[+57.021]K.F | HexNAc(4)Hex(5)Fuc(2)NeuAc(1) | C[+57]*2, N[+2206] | 1158.0219 | 4 | 4629.0657 | 4629.0502 | 0 | 3.4 | 466 | Specific | 612.79 | 612.79 | 612.79 | 12.44 | 101 | >sp\|P01833\|PIGR_HUMAN Polymeric immunoglobulin receptor OS=Homo sapiens OX=9606 GN=PIGR PE=1 SV=4 | 902 | 20201114_qwl_colo_milk_1ng_glyco_hcd203040_compound15.15597.15597.2 | controllerType=0 controllerNumber=1 scan=15597 | 34.5088 |
| 12384:4 | 5 | K.VPGN[+2205.793]VTAVLGETLKVPC[+57.021]HFPC[+57.021]K.F | HexNAc(4)Hex(5)Fuc(2)NeuAc(1) | C[+57]*2, N[+2206] | 1158.0219 | 4 | 4629.0657 | 4629.0502 | 0 | 3.4 | 466 | Specific | 1002.01 | 652.07 | 652.07 | 20.44 | 101 | >sp\|P01833\|PIGR_HUMAN Polymeric immunoglobulin receptor OS=Homo sapiens OX=9606 GN=PIGR PE=1 SV=4 | 902 | 20201114_qwl_colo_milk_1ng_glyco_hcd203040_compound15.15600.15600.2 | controllerType=0 controllerNumber=1 scan=15600 | 34.5142 |
| 03874:3 | 5 | K.WN[+1054.370]NTGC[+57.021]QALPSQDEGPSK.A | HexNAc(2)Hex(4) | C[+57], N[+1054] | 1015.0898 | 3 | 3043.2549 | 3043.2467 | 0 | 2.7 | 498 | Specific | 245.24 | 238.54 | 238.54 | 6.19 | 101 | >sp\|P01833\|PIGR_HUMAN Polymeric immunoglobulin receptor OS=Homo sapiens OX=9606 GN=PIGR PE=1 SV=4 | 902 | 20201114_qwl_colo_milk_1ng_glyco_hcd203040_compound15.6723.6723.3 | controllerType=0 controllerNumber=1 scan=6723 | 18.2926 |
| 03784:2 | 5 | K.WN[+1216.423]NTGC[+57.021]QALPSQDEGPSK.A | HexNAc(2)Hex(5) | C[+57], N[+1216] | 1603.1603 | 2 | 3205.3134 | 3205.2995 | 0 | 4.3 | 498 | Specific | 253.41 | 233.19 | 233.19 | 6.17 | 101 | >sp\|P01833\|PIGR_HUMAN Polymeric immunoglobulin receptor OS=Homo sapiens OX=9606 GN=PIGR PE=1 SV=4 | 902 | 20201114_qwl_colo_milk_1ng_glyco_hcd203040_compound15.6630.6630.2 | controllerType=0 controllerNumber=1 scan=6630 | 18.1406 |
| 03782:2 | 5 | K.WN[+1216.423]NTGC[+57.021]QALPSQDEGPSK.A | HexNAc(2)Hex(5) | C[+57], N[+1216] | 1603.1603 | 2 | 3205.3134 | 3205.2995 | 0 | 4.3 | 498 | Specific | 273.66 | 270.52 | 270.52 | 6.62 | 101 | >sp\|P01833\|PIGR_HUMAN Polymeric immunoglobulin receptor OS=Homo sapiens OX=9606 GN=PIGR PE=1 SV=4 | 902 | 20201114_qwl_colo_milk_1ng_glyco_hcd203040_compound15.6628.6628.2 | controllerType=0 controllerNumber=1 scan=6628 | 18.1361 |
| 03876:3 | 5 | K.WN[+1054.370]NTGC[+57.021]QALPSQDEGPSK.A | HexNAc(2)Hex(4) | C[+57], N[+1054] | 1015.0898 | 3 | 3043.2549 | 3043.2467 | 0 | 2.7 | 498 | Specific | 346.68 | 340.17 | 340.17 | 7.36 | 101 | >sp\|P01833\|PIGR_HUMAN Polymeric immunoglobulin receptor OS=Homo sapiens OX=9606 GN=PIGR PE=1 SV=4 | 902 | 20201114_qwl_colo_milk_1ng_glyco_hcd203040_compound15.6725.6725.3 | controllerType=0 controllerNumber=1 scan=6725 | 18.2965 |
| 05225:3 | 5 | K.WN[+1872.651]NTGC[+57.021]QALPSQDEGPSK.A | HexNAc(3)Hex(6)NeuAc(1) | C[+57], N[+1873] | 1287.8503 | 3 | 3861.5363 | 3861.5272 | 0 | 2.4 | 498 | Specific | 350.42 | 293.07 | 293.07 | 7.11 | 101 | >sp\|P01833\|PIGR_HUMAN Polymeric immunoglobulin receptor OS=Homo sapiens OX=9606 GN=PIGR PE=1 SV=4 | 902 | 20201114_qwl_colo_milk_1ng_glyco_hcd203040_compound15.8133.8133.3 | controllerType=0 controllerNumber=1 scan=8133 | 20.7069 |
| 07231:3 | 5 | K.WN[+2204.772]NTGC[+57.021]QALPSQDEGPSK.A | HexNAc(4)Hex(5)NeuAc(2) | C[+57], N[+2205] | 1398.5584 | 3 | 4193.6607 | 4193.6491 | 0 | 2.8 | 498 | Specific | 512.13 | 422.97 | 422.97 | 9.56 | 101 | >sp\|P01833\|PIGR_HUMAN Polymeric immunoglobulin receptor OS=Homo sapiens OX=9606 GN=PIGR PE=1 SV=4 | 902 | 20201114_qwl_colo_milk_1ng_glyco_hcd203040_compound15.10221.10221.3 | controllerType=0 controllerNumber=1 scan=10221 | 24.3003 |
| 05111:3 | 5 | K.WN[+2205.793]NTGC[+57.021]QALPSQDEGPSK.A | HexNAc(4)Hex(5)Fuc(2)NeuAc(1) | C[+57], N[+2206] | 1398.8956 | 3 | 4194.6721 | 4194.6695 | 0 | 0.6 | 498 | Specific | 523.44 | 457.96 | 457.96 | 9.41 | 101 | >sp\|P01833\|PIGR_HUMAN Polymeric immunoglobulin receptor OS=Homo sapiens OX=9606 GN=PIGR PE=1 SV=4 | 902 | 20201114_qwl_colo_milk_1ng_glyco_hcd203040_compound15.8015.8015.3 | controllerType=0 controllerNumber=1 scan=8015 | 20.5054 |
| 07229:3 | 5 | K.WN[+2204.772]NTGC[+57.021]QALPSQDEGPSK.A | HexNAc(4)Hex(5)NeuAc(2) | C[+57], N[+2205] | 1398.5584 | 3 | 4193.6607 | 4193.6491 | 0 | 2.8 | 498 | Specific | 566.38 | 479.27 | 479.27 | 9.87 | 101 | >sp\|P01833\|PIGR_HUMAN Polymeric immunoglobulin receptor OS=Homo sapiens OX=9606 GN=PIGR PE=1 SV=4 | 902 | 20201114_qwl_colo_milk_1ng_glyco_hcd203040_compound15.10219.10219.3 | controllerType=0 controllerNumber=1 scan=10219 | 24.2957 |
| 05405:3 | 5 | K.WN[+1548.545]NTGC[+57.021]QALPSQDEGPSK.A | HexNAc(3)Hex(4)NeuAc(1) | C[+57], N[+1549] | 1179.815 | 3 | 3537.4304 | 3537.4215 | 0 | 2.5 | 498 | Specific | 568.76 | 498.36 | 498.36 | 9.87 | 101 | >sp\|P01833\|PIGR_HUMAN Polymeric immunoglobulin receptor OS=Homo sapiens OX=9606 GN=PIGR PE=1 SV=4 | 902 | 20201114_qwl_colo_milk_1ng_glyco_hcd203040_compound15.8319.8319.3 | controllerType=0 controllerNumber=1 scan=8319 | 21.0347 |
| 05113:3 | 5 | K.WN[+2205.793]NTGC[+57.021]QALPSQDEGPSK.A | HexNAc(4)Hex(5)Fuc(2)NeuAc(1) | C[+57], N[+2206] | 1398.8956 | 3 | 4194.6721 | 4194.6695 | 0 | 0.6 | 498 | Specific | 609.72 | 515.75 | 515.75 | 12.02 | 101 | >sp\|P01833\|PIGR_HUMAN Polymeric immunoglobulin receptor OS=Homo sapiens OX=9606 GN=PIGR PE=1 SV=4 | 902 | 20201114_qwl_colo_milk_1ng_glyco_hcd203040_compound15.8017.8017.3 | controllerType=0 controllerNumber=1 scan=8017 | 20.5092 |
| 03745:2 | 5 | K.WN[+1216.423]NTGC[+57.021]QALPSQDEGPSK.A | HexNAc(2)Hex(5) | C[+57], N[+1216] | 1603.1579 | 2 | 3205.3085 | 3205.2995 | 0 | 2.8 | 498 | Specific | 621.02 | 517.5 | 517.5 | 12.19 | 101 | >sp\|P01833\|PIGR_HUMAN Polymeric immunoglobulin receptor OS=Homo sapiens OX=9606 GN=PIGR PE=1 SV=4 | 902 | 20201114_qwl_colo_milk_1ng_glyco_hcd203040_compound15.6589.6589.2 | controllerType=0 controllerNumber=1 scan=6589 | 18.064 |
| 05407:3 | 5 | K.WN[+1548.545]NTGC[+57.021]QALPSQDEGPSK.A | HexNAc(3)Hex(4)NeuAc(1) | C[+57], N[+1549] | 1179.815 | 3 | 3537.4304 | 3537.4215 | 0 | 2.5 | 498 | Specific | 642.13 | 552.52 | 552.52 | 12.18 | 101 | >sp\|P01833\|PIGR_HUMAN Polymeric immunoglobulin receptor OS=Homo sapiens OX=9606 GN=PIGR PE=1 SV=4 | 902 | 20201114_qwl_colo_milk_1ng_glyco_hcd203040_compound15.8321.8321.3 | controllerType=0 controllerNumber=1 scan=8321 | 21.0384 |
| 03743:2 | 5 | K.WN[+1216.423]NTGC[+57.021]QALPSQDEGPSK.A | HexNAc(2)Hex(5) | C[+57], N[+1216] | 1603.1579 | 2 | 3205.3085 | 3205.2995 | 0 | 2.8 | 498 | Specific | 649.24 | 587.59 | 587.59 | 12.19 | 101 | >sp\|P01833\|PIGR_HUMAN Polymeric immunoglobulin receptor OS=Homo sapiens OX=9606 GN=PIGR PE=1 SV=4 | 902 | 20201114_qwl_colo_milk_1ng_glyco_hcd203040_compound15.6587.6587.2 | controllerType=0 controllerNumber=1 scan=6587 | 18.059 |
| 05297:3 | 5 | K.WN[+1710.598]NTGC[+57.021]QALPSQDEGPSK.A | HexNAc(3)Hex(5)NeuAc(1) | C[+57], N[+1711] | 1233.8326 | 3 | 3699.4832 | 3699.4743 | 0 | 2.4 | 498 | Specific | 705.56 | 595.11 | 595.11 | 14.18 | 101 | >sp\|P01833\|PIGR_HUMAN Polymeric immunoglobulin receptor OS=Homo sapiens OX=9606 GN=PIGR PE=1 SV=4 | 902 | 20201114_qwl_colo_milk_1ng_glyco_hcd203040_compound15.8207.8207.3 | controllerType=0 controllerNumber=1 scan=8207 | 20.8404 |
| 05144:3 | 5 | K.WN[+2059.735]NTGC[+57.021]QALPSQDEGPSK.A | HexNAc(4)Hex(5)Fuc(1)NeuAc(1) | C[+57], N[+2060] | 1350.211 | 3 | 4048.6184 | 4048.6116 | 0 | 1.7 | 498 | Specific | 705.89 | 608.01 | 608.01 | 14.08 | 101 | >sp\|P01833\|PIGR_HUMAN Polymeric immunoglobulin receptor OS=Homo sapiens OX=9606 GN=PIGR PE=1 SV=4 | 902 | 20201114_qwl_colo_milk_1ng_glyco_hcd203040_compound15.8049.8049.3 | controllerType=0 controllerNumber=1 scan=8049 | 20.5645 |
| 05142:3 | 5 | K.WN[+2059.735]NTGC[+57.021]QALPSQDEGPSK.A | HexNAc(4)Hex(5)Fuc(1)NeuAc(1) | C[+57], N[+2060] | 1350.211 | 3 | 4048.6184 | 4048.6116 | 0 | 1.7 | 498 | Specific | 708.39 | 629.4 | 629.4 | 14.08 | 101 | >sp\|P01833\|PIGR_HUMAN Polymeric immunoglobulin receptor OS=Homo sapiens OX=9606 GN=PIGR PE=1 SV=4 | 902 | 20201114_qwl_colo_milk_1ng_glyco_hcd203040_compound15.8047.8047.3 | controllerType=0 controllerNumber=1 scan=8047 | 20.5606 |
| 03776:3 | 5 | K.WN[+1216.423]NTGC[+57.021]QALPSQDEGPSK.A | HexNAc(2)Hex(5) | C[+57], N[+1216] | 1069.106 | 3 | 3205.3034 | 3205.2995 | 0 | 1.2 | 498 | Specific | 760.81 | 728.36 | 728.36 | 15.08 | 101 | >sp\|P01833\|PIGR_HUMAN Polymeric immunoglobulin receptor OS=Homo sapiens OX=9606 GN=PIGR PE=1 SV=4 | 902 | 20201114_qwl_colo_milk_1ng_glyco_hcd203040_compound15.6622.6622.3 | controllerType=0 controllerNumber=1 scan=6622 | 18.1235 |
| 03778:3 | 5 | K.WN[+1216.423]NTGC[+57.021]QALPSQDEGPSK.A | HexNAc(2)Hex(5) | C[+57], N[+1216] | 1069.106 | 3 | 3205.3034 | 3205.2995 | 0 | 1.2 | 498 | Specific | 761.3 | 733.87 | 733.87 | 15.08 | 101 | >sp\|P01833\|PIGR_HUMAN Polymeric immunoglobulin receptor OS=Homo sapiens OX=9606 GN=PIGR PE=1 SV=4 | 902 | 20201114_qwl_colo_milk_1ng_glyco_hcd203040_compound15.6624.6624.3 | controllerType=0 controllerNumber=1 scan=6624 | 18.1301 |
| 05228:3 | 5 | K.WN[+1913.677]NTGC[+57.021]QALPSQDEGPSK.A | HexNAc(4)Hex(5)NeuAc(1) | C[+57], N[+1914] | 1301.5252 | 3 | 3902.5611 | 3902.5537 | 0 | 1.9 | 498 | Specific | 777 | 711.46 | 711.46 | 15.08 | 101 | >sp\|P01833\|PIGR_HUMAN Polymeric immunoglobulin receptor OS=Homo sapiens OX=9606 GN=PIGR PE=1 SV=4 | 902 | 20201114_qwl_colo_milk_1ng_glyco_hcd203040_compound15.8136.8136.3 | controllerType=0 controllerNumber=1 scan=8136 | 20.7117 |
| 05231:3 | 5 | K.WN[+1913.677]NTGC[+57.021]QALPSQDEGPSK.A | HexNAc(4)Hex(5)NeuAc(1) | C[+57], N[+1914] | 1301.5252 | 3 | 3902.5611 | 3902.5537 | 0 | 1.9 | 498 | Specific | 820.56 | 732.01 | 732.01 | 16.08 | 101 | >sp\|P01833\|PIGR_HUMAN Polymeric immunoglobulin receptor OS=Homo sapiens OX=9606 GN=PIGR PE=1 SV=4 | 902 | 20201114_qwl_colo_milk_1ng_glyco_hcd203040_compound15.8139.8139.3 | controllerType=0 controllerNumber=1 scan=8139 | 20.7168 |
| 05223:3 | 5 | K.WN[+1872.651]NTGC[+57.021]QALPSQDEGPSK.A | HexNAc(3)Hex(6)NeuAc(1) | C[+57], N[+1873] | 1287.8503 | 3 | 3861.5363 | 3861.5272 | 0 | 2.4 | 498 | Specific | 841.56 | 794.57 | 794.57 | 16.18 | 101 | >sp\|P01833\|PIGR_HUMAN Polymeric immunoglobulin receptor OS=Homo sapiens OX=9606 GN=PIGR PE=1 SV=4 | 902 | 20201114_qwl_colo_milk_1ng_glyco_hcd203040_compound15.8131.8131.3 | controllerType=0 controllerNumber=1 scan=8131 | 20.7025 |
| 03755:3 | 5 | K.WN[+1216.423]NTGC[+57.021]QALPSQDEGPSK.A | HexNAc(2)Hex(5) | C[+57], N[+1216] | 1069.1106 | 3 | 3205.3172 | 3205.2995 | 0 | 5.5 | 498 | Specific | 861.63 | 828.07 | 828.07 | 16.84 | 101 | >sp\|P01833\|PIGR_HUMAN Polymeric immunoglobulin receptor OS=Homo sapiens OX=9606 GN=PIGR PE=1 SV=4 | 902 | 20201114_qwl_colo_milk_1ng_glyco_hcd203040_compound15.6600.6600.3 | controllerType=0 controllerNumber=1 scan=6600 | 18.083 |
| 05295:3 | 5 | K.WN[+1710.598]NTGC[+57.021]QALPSQDEGPSK.A | HexNAc(3)Hex(5)NeuAc(1) | C[+57], N[+1711] | 1233.8326 | 3 | 3699.4832 | 3699.4743 | 0 | 2.4 | 498 | Specific | 918.32 | 838.41 | 838.41 | 18.18 | 101 | >sp\|P01833\|PIGR_HUMAN Polymeric immunoglobulin receptor OS=Homo sapiens OX=9606 GN=PIGR PE=1 SV=4 | 902 | 20201114_qwl_colo_milk_1ng_glyco_hcd203040_compound15.8205.8205.3 | controllerType=0 controllerNumber=1 scan=8205 | 20.8341 |
| 03757:3 | 5 | K.WN[+1216.423]NTGC[+57.021]QALPSQDEGPSK.A | HexNAc(2)Hex(5) | C[+57], N[+1216] | 1069.1106 | 3 | 3205.3172 | 3205.2995 | 0 | 5.5 | 498 | Specific | 923.54 | 912.33 | 912.33 | 17.84 | 101 | >sp\|P01833\|PIGR_HUMAN Polymeric immunoglobulin receptor OS=Homo sapiens OX=9606 GN=PIGR PE=1 SV=4 | 902 | 20201114_qwl_colo_milk_1ng_glyco_hcd203040_compound15.6602.6602.3 | controllerType=0 controllerNumber=1 scan=6602 | 18.0883 |
| 21528:4 | 6 | A.[+42.011]SPTSPKVFPLSLDSTPQDGNVVVAC[+57.021]LVQGFFPQEPLSVTWSESGQN | HexNAc(3)Hex(4) | .[42], C[+57], N[+1257] | 1669.0234 | 4 | 6673.0718 | 6673.1336 | 0 | -9.3 | 2 | NRagged | 282.82 | 208.7 | 208.7 | 2.22 | 102 | >sp\|P01877\|IGHA2_HUMAN Immunoglobulin heavy constant alpha 2 OS=Homo sapiens OX=9606 GN=IGHA2 PE=1 SV=4 | 576 | 20201114_qwl_colo_milk_1ng_glyco_hcd203040_compound15.25177.25177.2 | controllerType=0 controllerNumber=1 scan=25177 | 55.1507 |
| 22377:4 | 6 | A.[+42.011]SPTSPKVFPLSLDSTPQDGNVVVAC[+57.021]LVQGFFPQEPLSVTWSESGQN | HexNAc(3)Hex(4) | .[42], C[+57], N[+1257] | 1669.0224 | 4 | 6673.0679 | 6673.1336 | 0 | -9.9 | 2 | NRagged | 305.71 | 305.71 | 305.71 | 3.63 | 102 | >sp\|P01877\|IGHA2_HUMAN Immunoglobulin heavy constant alpha 2 OS=Homo sapiens OX=9606 GN=IGHA2 PE=1 SV=4 | 576 | 20201114_qwl_colo_milk_1ng_glyco_hcd203040_compound15.26063.26063.4 | controllerType=0 controllerNumber=1 scan=26063 | 56.8616 |
| 21833:4 | 6 | K.VFPLSLDSTPQDGNVVVAC[+57.021]LVQGFFPQEPLSVTWSESGQN[+2263.835]VTAI | HexNAc(5)Hex(5)Fuc(3) | C[+57], N[+2264] | 1760.8174 | 4 | 7040.2478 | 7040.1961 | 0 | 7.3 | 8 | Specific | 205.22 | 205.22 | 205.22 | 5.75 | 102 | >sp\|P01877\|IGHA2_HUMAN Immunoglobulin heavy constant alpha 2 OS=Homo sapiens OX=9606 GN=IGHA2 PE=1 SV=4 | 576 | 20201114_qwl_colo_milk_1ng_glyco_hcd203040_compound15.25495.25495.4 | controllerType=0 controllerNumber=1 scan=25495 | [illegible] |
| 21420:4 | 6 | K.VFPLSLDSTPQDGNVVVAC[+57.021]LVQGFFPQEPLSVTWSESGQN[+2059.735]VTAI | HexNAc(4)Hex(5)Fuc(1)NeuAc(1) | C[+57], N[+2060] | 1709.7921 | 4 | 6836.1465 | 6836.0963 | 0 | 7.3 | 8 | Specific | 223.39 | 223.39 | 223.39 | 7.06 | 102 | >sp\|P01877\|IGHA2_HUMAN Immunoglobulin heavy constant alpha 2 OS=Homo sapiens OX=9606 GN=IGHA2 PE=1 SV=4 | 576 | 20201114_qwl_colo_milk_1ng_glyco_hcd203040_compound15.25065.25065.4 | controllerType=0 controllerNumber=1 scan=25065 | [illegible] |
| 20028:4 | 6 | K.[+42.011]VFPLSLDSTPQDGNVVVAC[+57.021]LVQGFFPQEPLSVTWSESGQN[+1460 | HexNAc(4)Hex(4) | .[42], C[+57], N[+1461] | 1570.4822 | 4 | 6278.9069 | 6278.9007 | 0 | 1 | 8 | Specific | 239.86 | 212.38 | 212.38 | 5.68 | 102 | >sp\|P01877\|IGHA2_HUMAN Immunoglobulin heavy constant alpha 2 OS=Homo sapiens OX=9606 GN=IGHA2 PE=1 SV=4 | 576 | 20201114_qwl_colo_milk_1ng_glyco_hcd203040_compound15.23609.23609.4 | controllerType=0 controllerNumber=1 scan=23609 | [illegible] |
| 19955:4 | 6 | K.VFPLSLDSTPQDGNVVVAC[+57.021]LVQGFFPQEPLSVTWSESGQN[+1971.719]VTAI | HexNAc(5)Hex(5)Fuc(1) | C[+57], N[+1972] | 1687.7778 | 4 | 6748.0894 | 6748.0803 | 0 | 1.3 | 8 | Specific | 243.98 | 243.98 | 243.98 | 7.34 | 102 | >sp\|P01877\|IGHA2_HUMAN Immunoglobulin heavy constant alpha 2 OS=Homo sapiens OX=9606 GN=IGHA2 PE=1 SV=4 | 576 | 20201114_qwl_colo_milk_1ng_glyco_hcd203040_compound15.23533.23533.3 | controllerType=0 controllerNumber=1 scan=23533 | [illegible] |
| 20364:4 | 6 | K.VFPLSLDSTPQDGNVVVAC[+57.021]LVQGFFPQEPLSVTWSESGQN[+1971.719]VTAI | HexNAc(5)Hex(5)Fuc(1) | C[+57], N[+1972] | 1687.7744 | 4 | 6748.0757 | 6748.0803 | 0 | -0.7 | 8 | Specific | 248.45 | 248.45 | 248.45 | 7.03 | 102 | >sp\|P01877\|IGHA2_HUMAN Immunoglobulin heavy constant alpha 2 OS=Homo sapiens OX=9606 GN=IGHA2 PE=1 SV=4 | 576 | 20201114_qwl_colo_milk_1ng_glyco_hcd203040_compound15.23961.23961.4 | controllerType=0 controllerNumber=1 scan=23961 | 52.7362 |
| 20398:4 | 6 | K.VFPLSLDSTPQDGNVVVAC[+57.021]LVQGFFPQEPLSVTWSESGQN[+1606.587]VTAI | HexNAc(4)Hex(4)Fuc(1) | C[+57], N[+1607] | 1596.4952 | 4 | 6382.9591 | 6382.9481 | 0 | 1.7 | 8 | Specific | 253.32 | 253.32 | 253.32 | 7.55 | 102 | >sp\|P01877\|IGHA2_HUMAN Immunoglobulin heavy constant alpha 2 OS=Homo sapiens OX=9606 GN=IGHA2 PE=1 SV=4 | 576 | 20201114_qwl_colo_milk_1ng_glyco_hcd203040_compound15.23996.23996.4 | controllerType=0 controllerNumber=1 scan=23996 | 52.8066 |
| 20241:4 | 6 | K.VFPLSLDSTPQDGNVVVAC[+57.021]LVQGFFPQEPLSVTWSESGQN[+1606.587]VTAI | HexNAc(4)Hex(4)Fuc(1) | C[+57], N[+1607] | 1596.494 | 4 | 6382.9543 | 6382.9481 | 0 | 1 | 8 | Specific | 269.75 | 269.75 | 269.75 | 7.76 | 102 | >sp\|P01877\|IGHA2_HUMAN Immunoglobulin heavy constant alpha 2 OS=Homo sapiens OX=9606 GN=IGHA2 PE=1 SV=4 | 576 | 20201114_qwl_colo_milk_1ng_glyco_hcd203040_compound15.23833.23833.4 | controllerType=0 controllerNumber=1 scan=23833 | 52.4842 |
| 21831:4 | 6 | K.VFPLSLDSTPQDGNVVVAC[+57.021]LVQGFFPQEPLSVTWSESGQN[+2263.835]VTAI | HexNAc(5)Hex(5)Fuc(3) | C[+57], N[+2264] | 1760.8174 | 4 | 7040.2478 | 7040.1961 | 0 | 7.3 | 8 | Specific | 274.56 | 253.2 | 253.2 | 7.39 | 102 | >sp\|P01877\|IGHA2_HUMAN Immunoglobulin heavy constant alpha 2 OS=Homo sapiens OX=9606 GN=IGHA2 PE=1 SV=4 | 576 | 20201114_qwl_colo_milk_1ng_glyco_hcd203040_compound15.25493.25493.4 | controllerType=0 controllerNumber=1 scan=25493 | 55.7575 |
| 20044:6 | 6 | K.VFPLSLDSTPQDGNVVVAC[+57.021]LVQGFFPQEPLSVTWSESGQN[+1647.613]VTAI | HexNAc(5)Hex(3)Fuc(1) | C[+57], N[+1648] | 1071.5072 | 6 | 6424.0068 | 6423.9746 | 0 | 5 | 8 | Specific | 279.84 | 279.84 | 279.84 | 6.67 | 102 | >sp\|P01877\|IGHA2_HUMAN Immunoglobulin heavy constant alpha 2 OS=Homo sapiens OX=9606 GN=IGHA2 PE=1 SV=4 | 576 | 20201114_qwl_colo_milk_1ng_glyco_hcd203040_compound15.23626.23626.6 | controllerType=0 controllerNumber=1 scan=23626 | 52.0117 |
| 20400:4 | 6 | K.VFPLSLDSTPQDGNVVVAC[+57.021]LVQGFFPQEPLSVTWSESGQN[+1606.587]VTAI | HexNAc(4)Hex(4)Fuc(1) | C[+57], N[+1607] | 1596.4952 | 4 | 6382.9591 | 6382.9481 | 0 | 1.7 | 8 | Specific | 283.57 | 283.57 | 283.57 | 7.81 | 102 | >sp\|P01877\|IGHA2_HUMAN Immunoglobulin heavy constant alpha 2 OS=Homo sapiens OX=9606 GN=IGHA2 PE=1 SV=4 | 576 | 20201114_qwl_colo_milk_1ng_glyco_hcd203040_compound15.23998.23998.4 | controllerType=0 controllerNumber=1 scan=23998 | 52.811 |
| 20030:4 | 6 | K.[+42.011]VFPLSLDSTPQDGNVVVAC[+57.021]LVQGFFPQEPLSVTWSESGQN[+1460 | HexNAc(4)Hex(4) | .[42], C[+57], N[+1461] | 1570.4822 | 4 | 6278.9069 | 6278.9007 | 0 | 1 | 8 | Specific | 314.64 | 252.7 | 252.7 | 5.93 | 102 | >sp\|P01877\|IGHA2_HUMAN Immunoglobulin heavy constant alpha 2 OS=Homo sapiens OX=9606 GN=IGHA2 PE=1 SV=4 | 576 | 20201114_qwl_colo_milk_1ng_glyco_hcd203040_compound15.23611.23611.4 | controllerType=0 controllerNumber=1 scan=23611 | 51.9791 |
| 24034:5 | 6 | K.VFPLSLDSTPQDGNVVVAC[+57.021]LVQGFFPQEPLSVTWSESGQN[+2351.851]VTAI | HexNAc(4)Hex(5)Fuc(3)NeuAc(1) | C[+57], N[+2352] | 1426.4483 | 5 | 7128.2123 | 7128.2121 | 0 | 0 | 8 | Specific | 321.55 | 321.55 | 321.55 | 7.87 | 102 | >sp\|P01877\|IGHA2_HUMAN Immunoglobulin heavy constant alpha 2 OS=Homo sapiens OX=9606 GN=IGHA2 PE=1 SV=4 | 576 | 20201114_qwl_colo_milk_1ng_glyco_hcd203040_compound15.27813.27813.3 | controllerType=0 controllerNumber=1 scan=27813 | 60.0821 |
| 23963:5 | 6 | K.VFPLSLDSTPQDGNVVVAC[+57.021]LVQGFFPQEPLSVTWSESGQN[+2350.830]VTAI | HexNAc(4)Hex(5)Fuc(1)NeuAc(2) | C[+57], N[+2351] | 1426.2505 | 5 | 7127.2236 | 7127.1917 | 0 | 4.5 | 8 | Specific | 339.87 | 339.87 | 339.87 | 8.59 | 102 | >sp\|P01877\|IGHA2_HUMAN Immunoglobulin heavy constant alpha 2 OS=Homo sapiens OX=9606 GN=IGHA2 PE=1 SV=4 | 576 | 20201114_qwl_colo_milk_1ng_glyco_hcd203040_compound15.27734.27734.5 | controllerType=0 controllerNumber=1 scan=27734 | 59.9248 |
| 20590:4 | 6 | K.[+42.011]VFPLSLDSTPQDGNVVVAC[+57.021]LVQGFFPQEPLSVTWSESGQN[+1606 | HexNAc(4)Hex(4)Fuc(1) | .[42], C[+57], N[+1607] | 1607.0043 | 4 | 6424.9953 | 6424.9587 | 0 | 5.7 | 8 | Specific | 344.26 | 334.46 | 334.46 | 6.46 | 102 | >sp\|P01877\|IGHA2_HUMAN Immunoglobulin heavy constant alpha 2 OS=Homo sapiens OX=9606 GN=IGHA2 PE=1 SV=4 | 576 | 20201114_qwl_colo_milk_1ng_glyco_hcd203040_compound15.24196.24196.4 | controllerType=0 controllerNumber=1 scan=24196 | 53.2129 |
| 22461:4 | 6 | K.VFPLSLDSTPQDGNVVVAC[+57.021]LVQGFFPQEPLSVTWSESGQN[+2262.814]VTAI | HexNAc(5)Hex(5)Fuc(1)NeuAc(1) | C[+57], N[+2263] | 1760.5543 | 4 | 7039.1953 | 7039.1757 | 0 | 2.8 | 8 | Specific | 351.45 | 351.45 | 351.45 | 8.63 | 102 | >sp\|P01877\|IGHA2_HUMAN Immunoglobulin heavy constant alpha 2 OS=Homo sapiens OX=9606 GN=IGHA2 PE=1 SV=4 | 576 | 20201114_qwl_colo_milk_1ng_glyco_hcd203040_compound15.26150.26150.2 | controllerType=0 controllerNumber=1 scan=26150 | 57.0203 |
| 21540:4 | 6 | K.VFPLSLDSTPQDGNVVVAC[+57.021]LVQGFFPQEPLSVTWSESGQN[+2117.777]VTAI | HexNAc(5)Hex(5)Fuc(2) | C[+57], N[+2118] | 1724.2989 | 4 | 6894.1738 | 6894.1382 | 0 | 5.2 | 8 | Specific | 353.14 | 353.14 | 353.14 | 8.29 | 102 | >sp\|P01877\|IGHA2_HUMAN Immunoglobulin heavy constant alpha 2 OS=Homo sapiens OX=9606 GN=IGHA2 PE=1 SV=4 | 576 | 20201114_qwl_colo_milk_1ng_glyco_hcd203040_compound15.25190.25190.4 | controllerType=0 controllerNumber=1 scan=25190 | 55.1761 |
| 21484:6 | 6 | K.VFPLSLDSTPQDGNVVVAC[+57.021]LVQGFFPQEPLSVTWSESGQN[+2262.814]VTAI | HexNAc(5)Hex(5)Fuc(1)NeuAc(1) | C[+57], N[+2263] | 1174.046 | 6 | 7039.2394 | 7039.1757 | 0 | 9 | 8 | Specific | 360.21 | 360.21 | 360.21 | 8.28 | 102 | >sp\|P01877\|IGHA2_HUMAN Immunoglobulin heavy constant alpha 2 OS=Homo sapiens OX=9606 GN=IGHA2 PE=1 SV=4 | 576 | 20201114_qwl_colo_milk_1ng_glyco_hcd203040_compound15.25131.25131.6 | controllerType=0 controllerNumber=1 scan=25131 | 55.0547 |
| 20314:4 | 6 | K.[+42.011]VFPLSLDSTPQDGNVVVAC[+57.021]LVQGFFPQEPLSVTWSESGQN[+1768 | HexNAc(4)Hex(5)Fuc(1) | .[42], C[+57], N[+1769] | 1647.5132 | 4 | 6587.0309 | 6587.0115 | 0 | 3 | 8 | Specific | 366.98 | 359.81 | 359.81 | 8.88 | 102 | >sp\|P01877\|IGHA2_HUMAN Immunoglobulin heavy constant alpha 2 OS=Homo sapiens OX=9606 GN=IGHA2 PE=1 SV=4 | 576 | 20201114_qwl_colo_milk_1ng_glyco_hcd203040_compound15.23909.23909.4 | controllerType=0 controllerNumber=1 scan=23909 | 52.642 |
| 20588:4 | 6 | K.[+42.011]VFPLSLDSTPQDGNVVVAC[+57.021]LVQGFFPQEPLSVTWSESGQN[+1606 | HexNAc(4)Hex(4)Fuc(1) | .[42], C[+57], N[+1607] | 1607.0043 | 4 | 6424.9953 | 6424.9587 | 0 | 5.7 | 8 | Specific | 370.88 | 360.11 | 360.11 | 8.35 | 102 | >sp\|P01877\|IGHA2_HUMAN Immunoglobulin heavy constant alpha 2 OS=Homo sapiens OX=9606 GN=IGHA2 PE=1 SV=4 | 576 | 20201114_qwl_colo_milk_1ng_glyco_hcd203040_compound15.24194.24194.4 | controllerType=0 controllerNumber=1 scan=24194 | 53.2087 |
| 22325:4 | 6 | K.VFPLSLDSTPQDGNVVVAC[+57.021]LVQGFFPQEPLSVTWSESGQN[+2059.735]VTAI | HexNAc(4)Hex(5)Fuc(1)NeuAc(1) | C[+57], N[+2060] | 1709.7896 | 4 | 6836.1367 | 6836.0963 | 0 | 5.9 | 8 | Specific | 387.07 | 387.07 | 387.07 | 9.87 | 102 | >sp\|P01877\|IGHA2_HUMAN Immunoglobulin heavy constant alpha 2 OS=Homo sapiens OX=9606 GN=IGHA2 PE=1 SV=4 | 576 | 20201114_qwl_colo_milk_1ng_glyco_hcd203040_compound15.26009.26009.4 | controllerType=0 controllerNumber=1 scan=26009 | 56.7597 |
| 21416:5 | 6 | K.VFPLSLDSTPQDGNVVVAC[+57.021]LVQGFFPQEPLSVTWSESGQN[+2262.814]VTAI | HexNAc(5)Hex(5)Fuc(1)NeuAc(1) | C[+57], N[+2263] | 1408.6507 | 5 | 7039.2243 | 7039.1757 | 0 | 6.9 | 8 | Specific | 466.38 | 466.38 | 466.38 | 10.24 | 102 | >sp\|P01877\|IGHA2_HUMAN Immunoglobulin heavy constant alpha 2 OS=Homo sapiens OX=9606 GN=IGHA2 PE=1 SV=4 | 576 | 20201114_qwl_colo_milk_1ng_glyco_hcd203040_compound15.25061.25061.5 | controllerType=0 controllerNumber=1 scan=25061 | 54.9138 |
| 23998:5 | 6 | K.VFPLSLDSTPQDGNVVVAC[+57.021]LVQGFFPQEPLSVTWSESGQN[+2553.910]VTAI | HexNAc(5)Hex(5)Fuc(1)NeuAc(2) | C[+57], N[+2554] | 1466.8691 | 5 | 7330.3162 | 7330.2711 | 0 | 6.2 | 8 | Specific | 494.9 | 494.9 | 494.9 | 10.25 | 102 | >sp\|P01877\|IGHA2_HUMAN Immunoglobulin heavy constant alpha 2 OS=Homo sapiens OX=9606 GN=IGHA2 PE=1 SV=4 | 576 | 20201114_qwl_colo_milk_1ng_glyco_hcd203040_compound15.27776.27776.5 | controllerType=0 controllerNumber=1 scan=27776 | 60.0144 |
| 20047:6 | 6 | K.VFPLSLDSTPQDGNVVVAC[+57.021]LVQGFFPQEPLSVTWSESGQN[+1647.613]VTAI | HexNAc(5)Hex(3)Fuc(1) | C[+57], N[+1648] | 1071.5072 | 6 | 6424.0068 | 6423.9746 | 0 | 5 | 8 | Specific | 497.65 | 497.65 | 497.65 | 10.43 | 102 | >sp\|P01877\|IGHA2_HUMAN Immunoglobulin heavy constant alpha 2 OS=Homo sapiens OX=9606 GN=IGHA2 PE=1 SV=4 | 576 | 20201114_qwl_colo_milk_1ng_glyco_hcd203040_compound15.23629.23629.6 | controllerType=0 controllerNumber=1 scan=23629 | 52.0167 |
| 20065:5 | 6 | K.VFPLSLDSTPQDGNVVVAC[+57.021]LVQGFFPQEPLSVTWSESGQN[+1647.613]VTAI | HexNAc(5)Hex(3)Fuc(1) | C[+57], N[+1648] | 1285.608 | 5 | 6424.0111 | 6423.9746 | 0 | 5.7 | 8 | Specific | 576.48 | 576.48 | 576.48 | 10.61 | 102 | >sp\|P01877\|IGHA2_HUMAN Immunoglobulin heavy constant alpha 2 OS=Homo sapiens OX=9606 GN=IGHA2 PE=1 SV=4 | 576 | 20201114_qwl_colo_milk_1ng_glyco_hcd203040_compound15.23649.23649.5 | controllerType=0 controllerNumber=1 scan=23649 | 52.0659 |
| 20717:4 | 6 | K.VFPLSLDSTPQDGNVVVAC[+57.021]LVQGFFPQEPLSVTWSESGQN[+1647.613]VTAI | HexNAc(5)Hex(3)Fuc(1) | C[+57], N[+1648] | 1606.7538 | 4 | 6423.9933 | 6423.9746 | 0 | 2.9 | 8 | Specific | 635.37 | 605.77 | 605.77 | 13.45 | 102 | >sp\|P01877\|IGHA2_HUMAN Immunoglobulin heavy constant alpha 2 OS=Homo sapiens OX=9606 GN=IGHA2 PE=1 SV=4 | 576 | 20201114_qwl_colo_milk_1ng_glyco_hcd203040_compound15.24329.24329.4 | controllerType=0 controllerNumber=1 scan=24329 | 53.493 |
| 21422:4 | 6 | K.VFPLSLDSTPQDGNVVVAC[+57.021]LVQGFFPQEPLSVTWSESGQN[+2059.735]VTAI | HexNAc(4)Hex(5)Fuc(1)NeuAc(1) | C[+57], N[+2060] | 1709.7921 | 4 | 6836.1465 | 6836.0963 | 0 | 7.3 | 8 | Specific | 638.04 | 588.33 | 588.33 | 13.06 | 102 | >sp\|P01877\|IGHA2_HUMAN Immunoglobulin heavy constant alpha 2 OS=Homo sapiens OX=9606 GN=IGHA2 PE=1 SV=4 | 576 | 20201114_qwl_colo_milk_1ng_glyco_hcd203040_compound15.25067.25067.4 | controllerType=0 controllerNumber=1 scan=25067 | 54.927 |
| 20652:5 | 6 | K.VFPLSLDSTPQDGNVVVAC[+57.021]LVQGFFPQEPLSVTWSESGQN[+1647.613]VTAI | HexNAc(5)Hex(3)Fuc(1) | C[+57], N[+1648] | 1285.6078 | 5 | 6424.0098 | 6423.9746 | 0 | 5.5 | 8 | Specific | 646.38 | 646.38 | 646.38 | 13.11 | 102 | >sp\|P01877\|IGHA2_HUMAN Immunoglobulin heavy constant alpha 2 OS=Homo sapiens OX=9606 GN=IGHA2 PE=1 SV=4 | 576 | 20201114_qwl_colo_milk_1ng_glyco_hcd203040_compound15.24261.24261.5 | controllerType=0 controllerNumber=1 scan=24261 | 53.3503 |
| 20087:4 | 6 | K.VFPLSLDSTPQDGNVVVAC[+57.021]LVQGFFPQEPLSVTWSESGQN[+1241.454]VTAI | HexNAc(3)Hex(3)Fuc(1) | C[+57], N[+1241] | 1505.2167 | 4 | 6017.8449 | 6017.8159 | 0 | 4.8 | 8 | Specific | 670.95 | 664.38 | 664.38 | 14.11 | 102 | >sp\|P01877\|IGHA2_HUMAN Immunoglobulin heavy constant alpha 2 OS=Homo sapiens OX=9606 GN=IGHA2 PE=1 SV=4 | 576 | 20201114_qwl_colo_milk_1ng_glyco_hcd203040_compound15.23672.23672.4 | controllerType=0 controllerNumber=1 scan=23672 | 52.1314 |
| 22197:4 | 6 | K.VFPLSLDSTPQDGNVVVAC[+57.021]LVQGFFPQEPLSVTWSESGQN[+2262.814]VTAI | HexNAc(5)Hex(5)Fuc(1)NeuAc(1) | C[+57], N[+2263] | 1760.5573 | 4 | 7039.2075 | 7039.1757 | 0 | 4.5 | 8 | Specific | 677.09 | 652.28 | 652.28 | 14.11 | 102 | >sp\|P01877\|IGHA2_HUMAN Immunoglobulin heavy constant alpha 2 OS=Homo sapiens OX=9606 GN=IGHA2 PE=1 SV=4 | 576 | 20201114_qwl_colo_milk_1ng_glyco_hcd203040_compound15.25876.25876.4 | controllerType=0 controllerNumber=1 scan=25876 | 56.4962 |
| 20366:4 | 6 | K.VFPLSLDSTPQDGNVVVAC[+57.021]LVQGFFPQEPLSVTWSESGQN[+1971.719]VTAI | HexNAc(5)Hex(5)Fuc(1) | C[+57], N[+1972] | 1687.7744 | 4 | 6748.0757 | 6748.0803 | 0 | -0.7 | 8 | Specific | 703.94 | 678.32 | 678.32 | 14.82 | 102 | >sp\|P01877\|IGHA2_HUMAN Immunoglobulin heavy constant alpha 2 OS=Homo sapiens OX=9606 GN=IGHA2 PE=1 SV=4 | 576 | 20201114_qwl_colo_milk_1ng_glyco_hcd203040_compound15.23963.23963.4 | controllerType=0 controllerNumber=1 scan=23963 | 52.742 |
| 20644:4 | 6 | K.VFPLSLDSTPQDGNVVVAC[+57.021]LVQGFFPQEPLSVTWSESGQN[+1647.613]VTAI | HexNAc(5)Hex(3)Fuc(1) | C[+57], N[+1648] | 1606.7581 | 4 | 6424.0104 | 6423.9746 | 0 | 5.6 | 8 | Specific | 718.61 | 703.05 | 703.05 | 15.11 | 102 | >sp\|P01877\|IGHA2_HUMAN Immunoglobulin heavy constant alpha 2 OS=Homo sapiens OX=9606 GN=IGHA2 PE=1 SV=4 | 576 | 20201114_qwl_colo_milk_1ng_glyco_hcd203040_compound15.24253.24253.4 | controllerType=0 controllerNumber=1 scan=24253 | 53.3321 |
| 22199:4 | 6 | K.VFPLSLDSTPQDGNVVVAC[+57.021]LVQGFFPQEPLSVTWSESGQN[+2262.814]VTAI | HexNAc(5)Hex(5)Fuc(1)NeuAc(1) | C[+57], N[+2263] | 1760.5573 | 4 | 7039.2075 | 7039.1757 | 0 | 4.5 | 8 | Specific | 732.95 | 609.49 | 609.49 | 15.11 | 102 | >sp\|P01877\|IGHA2_HUMAN Immunoglobulin heavy constant alpha 2 OS=Homo sapiens OX=9606 GN=IGHA2 PE=1 SV=4 | 576 | 20201114_qwl_colo_milk_1ng_glyco_hcd203040_compound15.25878.25878.4 | controllerType=0 controllerNumber=1 scan=25878 | 56.5028 |
| 20037:4 | 6 | K.VFPLSLDSTPQDGNVVVAC[+57.021]LVQGFFPQEPLSVTWSESGQN[+1647.613]VTAI | HexNAc(5)Hex(3)Fuc(1) | C[+57], N[+1648] | 1606.7572 | 4 | 6424.007 | 6423.9746 | 0 | 5 | 8 | Specific | 759.71 | 759.71 | 759.71 | 16.11 | 102 | >sp\|P01877\|IGHA2_HUMAN Immunoglobulin heavy constant alpha 2 OS=Homo sapiens OX=9606 GN=IGHA2 PE=1 SV=4 | 576 | 20201114_qwl_colo_milk_1ng_glyco_hcd203040_compound15.23619.23619.4 | controllerType=0 controllerNumber=1 scan=23619 | 51.996 |
| 20102:5 | 6 | K.VFPLSLDSTPQDGNVVVAC[+57.021]LVQGFFPQEPLSVTWSESGQN[+1809.666]VTAI | HexNAc(5)Hex(4)Fuc(1) | C[+57], N[+1810] | 1318.0127 | 5 | 6586.0344 | 6586.0275 | 0 | 1.1 | 8 | Specific | 760.27 | 760.27 | 760.27 | 16.29 | 102 | >sp\|P01877\|IGHA2_HUMAN Immunoglobulin heavy constant alpha 2 OS=Homo sapiens OX=9606 GN=IGHA2 PE=1 SV=4 | 576 | 20201114_qwl_colo_milk_1ng_glyco_hcd203040_compound15.23688.23688.5 | controllerType=0 controllerNumber=1 scan=23688 | 52.1714 |
| 20100:5 | 6 | K.VFPLSLDSTPQDGNVVVAC[+57.021]LVQGFFPQEPLSVTWSESGQN[+1809.666]VTAI | HexNAc(5)Hex(4)Fuc(1) | C[+57], N[+1810] | 1318.0127 | 5 | 6586.0344 | 6586.0275 | 0 | 1.1 | 8 | Specific | 773.47 | 765.58 | 765.58 | 16.29 | 102 | >sp\|P01877\|IGHA2_HUMAN Immunoglobulin heavy constant alpha 2 OS=Homo sapiens OX=9606 GN=IGHA2 PE=1 SV=4 | 576 | 20201114_qwl_colo_milk_1ng_glyco_hcd203040_compound15.23686.23686.5 | controllerType=0 controllerNumber=1 scan=23686 | 52.165 |
| 19958:4 | 6 | K.VFPLSLDSTPQDGNVVVAC[+57.021]LVQGFFPQEPLSVTWSESGQN[+1971.719]VTAI | HexNAc(5)Hex(5)Fuc(1) | C[+57], N[+1972] | 1687.7778 | 4 | 6748.0894 | 6748.0803 | 0 | 1.3 | 8 | Specific | 778.75 | 706.6 | 706.6 | 16.34 | 102 | >sp\|P01877\|IGHA2_HUMAN Immunoglobulin heavy constant alpha 2 OS=Homo sapiens OX=9606 GN=IGHA2 PE=1 SV=4 | 576 | 20201114_qwl_colo_milk_1ng_glyco_hcd203040_compound15.23536.23536.3 | controllerType=0 controllerNumber=1 scan=23536 | 51.833 |
| 20067:5 | 6 | K.VFPLSLDSTPQDGNVVVAC[+57.021]LVQGFFPQEPLSVTWSESGQN[+1647.613]VTAI | HexNAc(5)Hex(3)Fuc(1) | C[+57], N[+1648] | 1285.608 | 5 | 6424.0111 | 6423.9746 | 0 | 5.7 | 8 | Specific | 802.87 | 802.87 | 802.87 | 16.93 | 102 | >sp\|P01877\|IGHA2_HUMAN Immunoglobulin heavy constant alpha 2 OS=Homo sapiens OX=9606 GN=IGHA2 PE=1 SV=4 | 576 | 20201114_qwl_colo_milk_1ng_glyco_hcd203040_compound15.23651.23651.5 | controllerType=0 controllerNumber=1 scan=23651 | 52.071 |
| 20076:4 | 6 | K.VFPLSLDSTPQDGNVVVAC[+57.021]LVQGFFPQEPLSVTWSESGQN[+1606.587]VTAI | HexNAc(4)Hex(4)Fuc(1) | C[+57], N[+1607] | 1596.5046 | 4 | 6382.9967 | 6382.9481 | 0 | 7.6 | 8 | Specific | 804.58 | 751.4 | 751.4 | 17.06 | 102 | >sp\|P01877\|IGHA2_HUMAN Immunoglobulin heavy constant alpha 2 OS=Homo sapiens OX=9606 GN=IGHA2 PE=1 SV=4 | 576 | 20201114_qwl_colo_milk_1ng_glyco_hcd203040_compound15.23661.23661.4 | controllerType=0 controllerNumber=1 scan=23661 | 52.1011 |
| 21418:5 | 6 | K.VFPLSLDSTPQDGNVVVAC[+57.021]LVQGFFPQEPLSVTWSESGQN[+2262.814]VTAI | HexNAc(5)Hex(5)Fuc(1)NeuAc(1) | C[+57], N[+2263] | 1408.6507 | 5 | 7039.2243 | 7039.1757 | 0 | 6.9 | 8 | Specific | 817.2 | 817.2 | 817.2 | 17.06 | 102 | >sp\|P01877\|IGHA2_HUMAN Immunoglobulin heavy constant alpha 2 OS=Homo sapiens OX=9606 GN=IGHA2 PE=1 SV=4 | 576 | 20201114_qwl_colo_milk_1ng_glyco_hcd203040_compound15.25063.25063.5 | controllerType=0 controllerNumber=1 scan=25063 | 54.9192 |
| 19963:4 | 6 | K.VFPLSLDSTPQDGNVVVAC[+57.021]LVQGFFPQEPLSVTWSESGQN[+1971.719]VTAI | HexNAc(5)Hex(5)Fuc(1) | C[+57], N[+1972] | 1687.7877 | 4 | 6748.1289 | 6748.0803 | 0 | 7.2 | 8 | Specific | 826.7 | 734.94 | 734.94 | 17.06 | 102 | >sp\|P01877\|IGHA2_HUMAN Immunoglobulin heavy constant alpha 2 OS=Homo sapiens OX=9606 GN=IGHA2 PE=1 SV=4 | 576 | 20201114_qwl_colo_milk_1ng_glyco_hcd203040_compound15.23542.23542.4 | controllerType=0 controllerNumber=1 scan=23542 | 51.8436 |
| 21516:5 | 6 | K.VFPLSLDSTPQDGNVVVAC[+57.021]LVQGFFPQEPLSVTWSESGQN[+2263.835]VTAI | HexNAc(5)Hex(5)Fuc(3) | C[+57], N[+2264] | 1408.8456 | 5 | 7040.199 | 7040.1961 | 0 | 0.4 | 8 | Specific | 829.55 | 829.55 | 829.55 | 17.29 | 102 | >sp\|P01877\|IGHA2_HUMAN Immunoglobulin heavy constant alpha 2 OS=Homo sapiens OX=9606 GN=IGHA2 PE=1 SV=4 | 576 | 20201114_qwl_colo_milk_1ng_glyco_hcd203040_compound15.25165.25165.5 | controllerType=0 controllerNumber=1 scan=25165 | 55.1267 |
| 20073:4 | 6 | K.VFPLSLDSTPQDGNVVVAC[+57.021]LVQGFFPQEPLSVTWSESGQN[+1606.587]VTAI | HexNAc(4)Hex(4)Fuc(1) | C[+57], N[+1607] | 1596.5042 | 4 | 6382.9948 | 6382.9481 | 0 | 7.3 | 8 | Specific | 829.76 | 820.58 | 820.58 | 17.06 | 102 | >sp\|P01877\|IGHA2_HUMAN Immunoglobulin heavy constant alpha 2 OS=Homo sapiens OX=9606 GN=IGHA2 PE=1 SV=4 | 576 | 20201114_qwl_colo_milk_1ng_glyco_hcd203040_compound15.23657.23657.4 | controllerType=0 controllerNumber=1 scan=23657 | 52.0824 |
| 20042:4 | 6 | K.VFPLSLDSTPQDGNVVVAC[+57.021]LVQGFFPQEPLSVTWSESGQN[+1444.534]VTAI | HexNAc(4)Hex(3)Fuc(1) | C[+57], N[+1445] | 1555.9884 | 4 | 6220.9318 | 6220.8953 | 0 | 5.9 | 8 | Specific | 830.04 | 786.43 | 786.43 | 16.93 | 102 | >sp\|P01877\|IGHA2_HUMAN Immunoglobulin heavy constant alpha 2 OS=Homo sapiens OX=9606 GN=IGHA2 PE=1 SV=4 | 576 | 20201114_qwl_colo_milk_1ng_glyco_hcd203040_compound15.23624.23624.4 | controllerType=0 controllerNumber=1 scan=23624 | 52.0078 |
| 20559:4 | 6 | K.VFPLSLDSTPQDGNVVVAC[+57.021]LVQGFFPQEPLSVTWSESGQN[+1444.534]VTAI | HexNAc(4)Hex(3)Fuc(1) | C[+57], N[+1445] | 1555.9879 | 4 | 6220.9298 | 6220.8953 | 0 | 5.5 | 8 | Specific | 830.74 | 793.15 | 793.15 | 17.11 | 102 | >sp\|P01877\|IGHA2_HUMAN Immunoglobulin heavy constant alpha 2 OS=Homo sapiens OX=9606 GN=IGHA2 PE=1 SV=4 | 576 | 20201114_qwl_colo_milk_1ng_glyco_hcd203040_compound15.24164.24164.4 | controllerType=0 controllerNumber=1 scan=24164 | 53.1492 |
| 20038:5 | 6 | K.VFPLSLDSTPQDGNVVVAC[+57.021]LVQGFFPQEPLSVTWSESGQN[+1647.613]VTAI | HexNAc(5)Hex(3)Fuc(1) | C[+57], N[+1648] | 1285.6072 | 5 | 6424.0069 | 6423.9746 | 0 | 5 | 8 | Specific | 835.97 | 835.97 | 835.97 | 17.11 | 102 | >sp\|P01877\|IGHA2_HUMAN Immunoglobulin heavy constant alpha 2 OS=Homo sapiens OX=9606 GN=IGHA2 PE=1 SV=4 | 576 | 20201114_qwl_colo_milk_1ng_glyco_hcd203040_compound15.23620.23620.5 | controllerType=0 controllerNumber=1 scan=23620 | 51.9984 |
| 20039:4 | 6 | K.VFPLSLDSTPQDGNVVVAC[+57.021]LVQGFFPQEPLSVTWSESGQN[+1444.534]VTAI | HexNAc(4)Hex(3)Fuc(1) | C[+57], N[+1445] | 1555.9884 | 4 | 6220.9318 | 6220.8953 | 0 | 5.9 | 8 | Specific | 839.06 | 806.53 | 806.53 | 16.93 | 102 | >sp\|P01877\|IGHA2_HUMAN Immunoglobulin heavy constant alpha 2 OS=Homo sapiens OX=9606 GN=IGHA2 PE=1 SV=4 | 576 | 20201114_qwl_colo_milk_1ng_glyco_hcd203040_compound15.23621.23621.4 | controllerType=0 controllerNumber=1 scan=23621 | 52.0003 |
| 20064:4 | 6 | K.VFPLSLDSTPQDGNVVVAC[+57.021]LVQGFFPQEPLSVTWSESGQN[+1647.613]VTAI | HexNAc(5)Hex(3)Fuc(1) | C[+57], N[+1648] | 1606.761 | 4 | 6424.0221 | 6423.9746 | 0 | 7.4 | 8 | Specific | 848.67 | 848.67 | 848.67 | 17.06 | 102 | >sp\|P01877\|IGHA2_HUMAN Immunoglobulin heavy constant alpha 2 OS=Homo sapiens OX=9606 GN=IGHA2 PE=1 SV=4 | 576 | 20201114_qwl_colo_milk_1ng_glyco_hcd203040_compound15.23648.23648.4 | controllerType=0 controllerNumber=1 scan=23648 | 52.064 |
| 20062:4 | 6 | K.VFPLSLDSTPQDGNVVVAC[+57.021]LVQGFFPQEPLSVTWSESGQN[+1647.613]VTAI | HexNAc(5)Hex(3)Fuc(1) | C[+57], N[+1648] | 1606.761 | 4 | 6424.0221 | 6423.9746 | 0 | 7.4 | 8 | Specific | 871 | 836.14 | 836.14 | 18.06 | 102 | >sp\|P01877\|IGHA2_HUMAN Immunoglobulin heavy constant alpha 2 OS=Homo sapiens OX=9606 GN=IGHA2 PE=1 SV=4 | 576 | 20201114_qwl_colo_milk_1ng_glyco_hcd203040_compound15.23646.23646.4 | controllerType=0 controllerNumber=1 scan=23646 | 52.0496 |
| 20078:4 | 6 | K.VFPLSLDSTPQDGNVVVAC[+57.021]LVQGFFPQEPLSVTWSESGQN[+1606.587]VTAI | HexNAc(4)Hex(4)Fuc(1) | C[+57], N[+1607] | 1596.5046 | 4 | 6382.9967 | 6382.9481 | 0 | 7.6 | 8 | Specific | 879.93 | 759.31 | 759.31 | 18.06 | 102 | >sp\|P01877\|IGHA2_HUMAN Immunoglobulin heavy constant alpha 2 OS=Homo sapiens OX=9606 GN=IGHA2 PE=1 SV=4 | 576 | 20201114_qwl_colo_milk_1ng_glyco_hcd203040_compound15.23663.23663.4 | controllerType=0 controllerNumber=1 scan=23663 | 52.1091 |
| 20075:4 | 6 | K.VFPLSLDSTPQDGNVVVAC[+57.021]LVQGFFPQEPLSVTWSESGQN[+1606.587]VTAI | HexNAc(4)Hex(4)Fuc(1) | C[+57], N[+1607] | 1596.5042 | 4 | 6382.9948 | 6382.9481 | 0 | 7.3 | 8 | Specific | 903.51 | 878.5 | 878.5 | 19.06 | 102 | >sp\|P01877\|IGHA2_HUMAN Immunoglobulin heavy constant alpha 2 OS=Homo sapiens OX=9606 GN=IGHA2 PE=1 SV=4 | 576 | 20201114_qwl_colo_milk_1ng_glyco_hcd203040_compound15.23659.23659.4 | controllerType=0 controllerNumber=1 scan=23659 | 52.0965 |
| 20089:4 | 6 | K.VFPLSLDSTPQDGNVVVAC[+57.021]LVQGFFPQEPLSVTWSESGQN[+1241.454]VTAI | HexNAc(3)Hex(3)Fuc(1) | C[+57], N[+1241] | 1505.2167 | 4 | 6017.8449 | 6017.8159 | 0 | 4.8 | 8 | Specific | 909.05 | 876.24 | 876.24 | 19.11 | 102 | >sp\|P01877\|IGHA2_HUMAN Immunoglobulin heavy constant alpha 2 OS=Homo sapiens OX=9606 GN=IGHA2 PE=1 SV=4 | 576 | 20201114_qwl_colo_milk_1ng_glyco_hcd203040_compound15.23674.23674.4 | controllerType=0 controllerNumber=1 scan=23674 | 52.1387 |
| 20561:4 | 6 | K.VFPLSLDSTPQDGNVVVAC[+57.021]LVQGFFPQEPLSVTWSESGQN[+1444.534]VTAI | HexNAc(4)Hex(3)Fuc(1) | C[+57], N[+1445] | 1555.9879 | 4 | 6220.9298 | 6220.8953 | 0 | 5.5 | 8 | Specific | 914.87 | 847.34 | 847.34 | 19.11 | 102 | >sp\|P01877\|IGHA2_HUMAN Immunoglobulin heavy constant alpha 2 OS=Homo sapiens OX=9606 GN=IGHA2 PE=1 SV=4 | 576 | 20201114_qwl_colo_milk_1ng_glyco_hcd203040_compound15.24166.24166.4 | controllerType=0 controllerNumber=1 scan=24166 | 53.1584 |
| 20647:4 | 6 | K.VFPLSLDSTPQDGNVVVAC[+57.021]LVQGFFPQEPLSVTWSESGQN[+1647.613]VTAI | HexNAc(5)Hex(3)Fuc(1) | C[+57], N[+1648] | 1606.7581 | 4 | 6424.0104 | 6423.9746 | 0 | 5.6 | 8 | Specific | 927.3 | 894.57 | 894.57 | 19.11 | 102 | >sp\|P01877\|IGHA2_HUMAN Immunoglobulin heavy constant alpha 2 OS=Homo sapiens OX=9606 GN=IGHA2 PE=1 SV=4 | 576 | 20201114_qwl_colo_milk_1ng_glyco_hcd203040_compound15.24256.24256.4 | controllerType=0 controllerNumber=1 scan=24256 | 53.3421 |
| 01790:3 | 6 | K.HYTN[+2262.814]SSQDVTVPC[+57.021]R.V | HexNAc(5)Hex(5)Fuc(1)NeuAc(1) | C[+57], N[+2263] | 1309.5294 | 3 | 3926.5735 | 3926.5636 | 0 | 2.5 | 89 | Specific | 219.84 | 126.03 | 126.03 | 3.81 | 102 | >sp\|P01877\|IGHA2_HUMAN Immunoglobulin heavy constant alpha 2 OS=Homo sapiens OX=9606 GN=IGHA2 PE=1 SV=4 | 576 | 20201114_qwl_colo_milk_1ng_glyco_hcd203040_compound15.4549.4549.3 | controllerType=0 controllerNumber=1 scan=4549 | 14.3366 |
| 00971:3 | 6 | K.HYTN[+1768.640]SSQDVTVPC[+57.021]R.V | HexNAc(4)Hex(5)Fuc(1) | C[+57], N[+1769] | 1144.8057 | 3 | 3432.4025 | 3432.3888 | 0 | 4 | 89 | Specific | 224.17 | 166.13 | 166.13 | 5.58 | 102 | >sp\|P01877\|IGHA2_HUMAN Immunoglobulin heavy constant alpha 2 OS=Homo sapiens OX=9606 GN=IGHA2 PE=1 SV=4 | 576 | 20201114_qwl_colo_milk_1ng_glyco_hcd203040_compound15.3689.3689.3 | controllerType=0 controllerNumber=1 scan=3689 | 12.7395 |
| 01043:2 | 6 | K.HYTN[+1038.375]SSQDVTVPC[+57.021]R.V | HexNAc(2)Hex(3)Fuc(1) | C[+57], N[+1038] | 1351.5693 | 2 | 2702.1313 | 2702.1244 | 0 | 2.6 | 89 | Specific | 248.18 | 201.16 | 201.16 | 3.8 | 102 | >sp\|P01877\|IGHA2_HUMAN Immunoglobulin heavy constant alpha 2 OS=Homo sapiens OX=9606 GN=IGHA2 PE=1 SV=4 | 576 | 20201114_qwl_colo_milk_1ng_glyco_hcd203040_compound15.3764.3764.2 | controllerType=0 controllerNumber=1 scan=3764 | 12.879 |
| 01061:2 | 6 | K.HYTN[+1647.613]SSQDVTVPC[+57.021]R.V | HexNAc(5)Hex(3)Fuc(1) | C[+57], N[+1648] | 1656.1918 | 2 | 3311.3764 | 3311.3625 | 0 | 4.2 | 89 | Specific | 279.66 | 239.59 | 239.59 | 5.57 | 102 | >sp\|P01877\|IGHA2_HUMAN Immunoglobulin heavy constant alpha 2 OS=Homo sapiens OX=9606 GN=IGHA2 PE=1 SV=4 | 576 | 20201114_qwl_colo_milk_1ng_glyco_hcd203040_compound15.3784.3784.2 | controllerType=0 controllerNumber=1 scan=3784 | 12.9167 |
| 00969:3 | 6 | K.HYTN[+1768.640]SSQDVTVPC[+57.021]R.V | HexNAc(4)Hex(5)Fuc(1) | C[+57], N[+1769] | 1144.8057 | 3 | 3432.4025 | 3432.3888 | 0 | 4 | 89 | Specific | 305.33 | 185.64 | 185.64 | 5.94 | 102 | >sp\|P01877\|IGHA2_HUMAN Immunoglobulin heavy constant alpha 2 OS=Homo sapiens OX=9606 GN=IGHA2 PE=1 SV=4 | 576 | 20201114_qwl_colo_milk_1ng_glyco_hcd203040_compound15.3687.3687.3 | controllerType=0 controllerNumber=1 scan=3687 | 12.7364 |
| 01063:2 | 6 | K.HYTN[+1647.613]SSQDVTVPC[+57.021]R.V | HexNAc(5)Hex(3)Fuc(1) | C[+57], N[+1648] | 1656.1918 | 2 | 3311.3764 | 3311.3625 | 0 | 4.2 | 89 | Specific | 306.75 | 209.31 | 209.31 | 5.52 | 102 | >sp\|P01877\|IGHA2_HUMAN Immunoglobulin heavy constant alpha 2 OS=Homo sapiens OX=9606 GN=IGHA2 PE=1 SV=4 | 576 | 20201114_qwl_colo_milk_1ng_glyco_hcd203040_compound15.3786.3786.2 | controllerType=0 controllerNumber=1 scan=3786 | 12.9212 |
| 01039:2 | 6 | K.HYTN[+1241.454]SSQDVTVPC[+57.021]R.V | HexNAc(3)Hex(3)Fuc(1) | C[+57], N[+1241] | 1453.109 | 2 | 2905.2107 | 2905.2038 | 0 | 2.4 | 89 | Specific | 313.67 | 250.76 | 250.76 | 5.14 | 102 | >sp\|P01877\|IGHA2_HUMAN Immunoglobulin heavy constant alpha 2 OS=Homo sapiens OX=9606 GN=IGHA2 PE=1 SV=4 | 576 | 20201114_qwl_colo_milk_1ng_glyco_hcd203040_compound15.3760.3760.2 | controllerType=0 controllerNumber=1 scan=3760 | 12.8725 |
| 01230:3 | 6 | K.HYTN[+1444.534]SSQDVTVPC[+57.021]R.V | HexNAc(4)Hex(3)Fuc(1) | C[+57], N[+1445] | 1036.7694 | 3 | 3108.2938 | 3108.2832 | 0 | 3.4 | 89 | Specific | 346.99 | 288.4 | 288.4 | 6.96 | 102 | >sp\|P01877\|IGHA2_HUMAN Immunoglobulin heavy constant alpha 2 OS=Homo sapiens OX=9606 GN=IGHA2 PE=1 SV=4 | 576 | 20201114_qwl_colo_milk_1ng_glyco_hcd203040_compound15.3962.3962.3 | controllerType=0 controllerNumber=1 scan=3962 | 13.2629 |
| 01509:3 | 6 | K.HYTN[+1444.534]SSQDVTVPC[+57.021]R.V | HexNAc(4)Hex(3)Fuc(1) | C[+57], N[+1445] | 1036.7685 | 3 | 3108.2908 | 3108.2832 | 0 | 2.5 | 89 | Specific | 350.59 | 228.73 | 228.73 | 6.63 | 102 | >sp\|P01877\|IGHA2_HUMAN Immunoglobulin heavy constant alpha 2 OS=Homo sapiens OX=9606 GN=IGHA2 PE=1 SV=4 | 576 | 20201114_qwl_colo_milk_1ng_glyco_hcd203040_compound15.4256.4256.3 | controllerType=0 controllerNumber=1 scan=4256 | 13.8063 |
| 01037:2 | 6 | K.HYTN[+1241.454]SSQDVTVPC[+57.021]R.V | HexNAc(3)Hex(3)Fuc(1) | C[+57], N[+1241] | 1453.109 | 2 | 2905.2107 | 2905.2038 | 0 | 2.4 | 89 | Specific | 350.81 | 292.28 | 292.28 | 6.41 | 102 | >sp\|P01877\|IGHA2_HUMAN Immunoglobulin heavy constant alpha 2 OS=Homo sapiens OX=9606 GN=IGHA2 PE=1 SV=4 | 576 | 20201114_qwl_colo_milk_1ng_glyco_hcd203040_compound15.3758.3758.2 | controllerType=0 controllerNumber=1 scan=3758 | 12.868 |
| 01245:3 | 6 | K.HYTN[+1809.666]SSQDVTVPC[+57.021]R.V | HexNAc(5)Hex(4)Fuc(1) | C[+57], N[+1810] | 1158.478 | 3 | 3473.4193 | 3473.4154 | 0 | 1.1 | 89 | Specific | 354.39 | 235.32 | 235.32 | 6.53 | 102 | >sp\|P01877\|IGHA2_HUMAN Immunoglobulin heavy constant alpha 2 OS=Homo sapiens OX=9606 GN=IGHA2 PE=1 SV=4 | 576 | 20201114_qwl_colo_milk_1ng_glyco_hcd203040_compound15.3979.3979.3 | controllerType=0 controllerNumber=1 scan=3979 | 13.2948 |
| 00951:3 | 6 | K.HYTN[+1403.507]SSQDVTVPC[+57.021]R.V | HexNAc(3)Hex(4)Fuc(1) | C[+57], N[+1404] | 1023.0939 | 3 | 3067.2671 | 3067.2566 | 0 | 3.4 | 89 | Specific | 360.85 | 277.09 | 277.09 | 6.88 | 102 | >sp\|P01877\|IGHA2_HUMAN Immunoglobulin heavy constant alpha 2 OS=Homo sapiens OX=9606 GN=IGHA2 PE=1 SV=4 | 576 | 20201114_qwl_colo_milk_1ng_glyco_hcd203040_compound15.3669.3669.3 | controllerType=0 controllerNumber=1 scan=3669 | 12.7042 |
| 01511:3 | 6 | K.HYTN[+1444.534]SSQDVTVPC[+57.021]R.V | HexNAc(4)Hex(3)Fuc(1) | C[+57], N[+1445] | 1036.7685 | 3 | 3108.2908 | 3108.2832 | 0 | 2.5 | 89 | Specific | 392.63 | 283.99 | 283.99 | 7.08 | 102 | >sp\|P01877\|IGHA2_HUMAN Immunoglobulin heavy constant alpha 2 OS=Homo sapiens OX=9606 GN=IGHA2 PE=1 SV=4 | 576 | 20201114_qwl_colo_milk_1ng_glyco_hcd203040_compound15.4258.4258.3 | controllerType=0 controllerNumber=1 scan=4258 | 13.8103 |
| 01743:3 | 6 | K.HYTN[+2059.735]SSQDVTVPC[+57.021]R.V | HexNAc(4)Hex(5)Fuc(1)NeuAc(1) | C[+57], N[+2060] | 1241.8356 | 3 | 3723.4923 | 3723.4842 | 0 | 2.2 | 89 | Specific | 393.51 | 167.41 | 167.41 | 6.13 | 102 | >sp\|P01877\|IGHA2_HUMAN Immunoglobulin heavy constant alpha 2 OS=Homo sapiens OX=9606 GN=IGHA2 PE=1 SV=4 | 576 | 20201114_qwl_colo_milk_1ng_glyco_hcd203040_compound15.4499.4499.3 | controllerType=0 controllerNumber=1 scan=4499 | 14.243 |
| 01016:2 | 6 | K.HYTN[+1444.534]SSQDVTVPC[+57.021]R.V | HexNAc(4)Hex(3)Fuc(1) | C[+57], N[+1445] | 1554.6488 | 2 | 3108.2904 | 3108.2832 | 0 | 2.3 | 89 | Specific | 418.53 | 300.78 | 300.78 | 6.53 | 102 | >sp\|P01877\|IGHA2_HUMAN Immunoglobulin heavy constant alpha 2 OS=Homo sapiens OX=9606 GN=IGHA2 PE=1 SV=4 | 576 | 20201114_qwl_colo_milk_1ng_glyco_hcd203040_compound15.3736.3736.2 | controllerType=0 controllerNumber=1 scan=3736 | 12.8246 |
| 01014:2 | 6 | K.HYTN[+1444.534]SSQDVTVPC[+57.021]R.V | HexNAc(4)Hex(3)Fuc(1) | C[+57], N[+1445] | 1554.6488 | 2 | 3108.2904 | 3108.2832 | 0 | 2.3 | 89 | Specific | 420.67 | 420.67 | 420.67 | 8.42 | 102 | >sp\|P01877\|IGHA2_HUMAN Immunoglobulin heavy constant alpha 2 OS=Homo sapiens OX=9606 GN=IGHA2 PE=1 SV=4 | 576 | 20201114_qwl_colo_milk_1ng_glyco_hcd203040_compound15.3734.3734.2 | controllerType=0 controllerNumber=1 scan=3734 | 12.821 |
| 01741:3 | 6 | K.HYTN[+2059.735]SSQDVTVPC[+57.021]R.V | HexNAc(4)Hex(5)Fuc(1)NeuAc(1) | C[+57], N[+2060] | 1241.8356 | 3 | 3723.4923 | 3723.4842 | 0 | 2.2 | 89 | Specific | 454.9 | 219.63 | 219.63 | 6.79 | 102 | >sp\|P01877\|IGHA2_HUMAN Immunoglobulin heavy constant alpha 2 OS=Homo sapiens OX=9606 GN=IGHA2 PE=1 SV=4 | 576 | 20201114_qwl_colo_milk_1ng_glyco_hcd203040_compound15.4497.4497.3 | controllerType=0 controllerNumber=1 scan=4497 | 14.2388 |
| 00908:3 | 6 | K.HYTN[+1606.587]SSQDVTVPC[+57.021]R.V | HexNAc(4)Hex(4)Fuc(1) | C[+57], N[+1607] | 1090.7865 | 3 | 3270.345 | 3270.336 | 0 | 2.8 | 89 | Specific | 474.68 | 396.62 | 396.62 | 9.37 | 102 | >sp\|P01877\|IGHA2_HUMAN Immunoglobulin heavy constant alpha 2 OS=Homo sapiens OX=9606 GN=IGHA2 PE=1 SV=4 | 576 | 20201114_qwl_colo_milk_1ng_glyco_hcd203040_compound15.3624.3624.3 | controllerType=0 controllerNumber=1 scan=3624 | 12.6219 |
| 00953:3 | 6 | K.HYTN[+1403.507]SSQDVTVPC[+57.021]R.V | HexNAc(3)Hex(4)Fuc(1) | C[+57], N[+1404] | 1023.0939 | 3 | 3067.2671 | 3067.2566 | 0 | 3.4 | 89 | Specific | 482.98 | 365.62 | 365.62 | 8 | 102 | >sp\|P01877\|IGHA2_HUMAN Immunoglobulin heavy constant alpha 2 OS=Homo sapiens OX=9606 GN=IGHA2 PE=1 SV=4 | 576 | 20201114_qwl_colo_milk_1ng_glyco_hcd203040_compound15.3671.3671.3 | controllerType=0 controllerNumber=1 scan=3671 | 12.7081 |

Table S4. Cont.

| Query #:z | Protein Rank | Glycopeptide | Glycan | Modification Type(s) | Observed m/z | z | Observed (M+H) | Calc. mass (M+H) | Off-by-x error | Mass error (ppm) | StartPosition | Cleavage | Score | Delta | Delta Mod | \|Log Prob\| | # of unique peptides | Protein Name | Protein DB number | Comment | Scan # | Scan Time |
|---|---|---|---|---|---|---|---|---|---|---|---|---|---|---|---|---|---|---|---|---|---|---|
| 01788:3 | 6 | K.HYTN[+2262.814]SSQDVTVPC[+57.021]R.V | HexNAc(5)Hex(5)Fuc(1)NeuAc(1) | C[+57], N[+2263] | 1309.5294 | 3 | 3926.5735 | 3926.5636 | 0 | 2.5 | 89 | Specific | 551.57 | 312.28 | 312.28 | 8.05 | 102 | >sp\|P01877\|IGHA2_HUMAN Immunoglobulin heavy constant alpha 2 OS=Homo sapiens OX=9606 GN=IGHA2 PE=1 SV=4 | 576 | 20201114_qwl_colo_milk_1ng_glyco_hcd203040_compound15.4547.4547.3 | controllerType=0 controllerNumber=1 scan=4547 | 14.3327 |
| 00979:3 | 6 | K.HYTN[+1971.719]SSQDVTVPC[+57.021]R.V | HexNAc(5)Hex(5)Fuc(1) | C[+57], N[+1972] | 1212.4971 | 3 | 3635.4768 | 3635.4682 | 0 | 2.4 | 89 | Specific | 555.35 | 401.29 | 401.29 | 9.73 | 102 | >sp\|P01877\|IGHA2_HUMAN Immunoglobulin heavy constant alpha 2 OS=Homo sapiens OX=9606 GN=IGHA2 PE=1 SV=4 | 576 | 20201114_qwl_colo_milk_1ng_glyco_hcd203040_compound15.3698.3698.3 | controllerType=0 controllerNumber=1 scan=3698 | 12.7581 |
| 00977:3 | 6 | K.HYTN[+1971.719]SSQDVTVPC[+57.021]R.V | HexNAc(5)Hex(5)Fuc(1) | C[+57], N[+1972] | 1212.4971 | 3 | 3635.4768 | 3635.4682 | 0 | 2.4 | 89 | Specific | 574.51 | 358.1 | 358.1 | 8.48 | 102 | >sp\|P01877\|IGHA2_HUMAN Immunoglobulin heavy constant alpha 2 OS=Homo sapiens OX=9606 GN=IGHA2 PE=1 SV=4 | 576 | 20201114_qwl_colo_milk_1ng_glyco_hcd203040_compound15.3696.3696.3 | controllerType=0 controllerNumber=1 scan=3696 | 12.7536 |
| 00925:3 | 6 | K.HYTN[+1606.587]SSQDVTVPC[+57.021]R.V | HexNAc(4)Hex(4)Fuc(1) | C[+57], N[+1607] | 1090.7879 | 3 | 3270.3491 | 3270.336 | 0 | 4 | 89 | Specific | 596.88 | 493 | 493 | 11.1 | 102 | >sp\|P01877\|IGHA2_HUMAN Immunoglobulin heavy constant alpha 2 OS=Homo sapiens OX=9606 GN=IGHA2 PE=1 SV=4 | 576 | 20201114_qwl_colo_milk_1ng_glyco_hcd203040_compound15.3642.3642.3 | controllerType=0 controllerNumber=1 scan=3642 | 12.6556 |
| 00988:3 | 6 | K.HYTN[+1241.454]SSQDVTVPC[+57.021]R.V | HexNAc(3)Hex(3)Fuc(1) | C[+57], N[+1241] | 969.0749 | 3 | 2905.2102 | 2905.2038 | 0 | 2.2 | 89 | Specific | 627.92 | 627.92 | 627.92 | 12.03 | 102 | >sp\|P01877\|IGHA2_HUMAN Immunoglobulin heavy constant alpha 2 OS=Homo sapiens OX=9606 GN=IGHA2 PE=1 SV=4 | 576 | 20201114_qwl_colo_milk_1ng_glyco_hcd203040_compound15.3707.3707.3 | controllerType=0 controllerNumber=1 scan=3707 | 12.7727 |
| 01032:3 | 6 | K.HYTN[+1647.613]SSQDVTVPC[+57.021]R.V | HexNAc(5)Hex(3)Fuc(1) | C[+57], N[+1648] | 1104.4613 | 3 | 3311.3693 | 3311.3625 | 0 | 2.1 | 89 | Specific | 641.68 | 574.74 | 574.74 | 12.03 | 102 | >sp\|P01877\|IGHA2_HUMAN Immunoglobulin heavy constant alpha 2 OS=Homo sapiens OX=9606 GN=IGHA2 PE=1 SV=4 | 576 | 20201114_qwl_colo_milk_1ng_glyco_hcd203040_compound15.3753.3753.3 | controllerType=0 controllerNumber=1 scan=3753 | 12.8605 |
| 00927:3 | 6 | K.HYTN[+1606.587]SSQDVTVPC[+57.021]R.V | HexNAc(4)Hex(4)Fuc(1) | C[+57], N[+1607] | 1090.7879 | 3 | 3270.3491 | 3270.336 | 0 | 4 | 89 | Specific | 643.69 | 518.04 | 518.04 | 12.1 | 102 | >sp\|P01877\|IGHA2_HUMAN Immunoglobulin heavy constant alpha 2 OS=Homo sapiens OX=9606 GN=IGHA2 PE=1 SV=4 | 576 | 20201114_qwl_colo_milk_1ng_glyco_hcd203040_compound15.3644.3644.3 | controllerType=0 controllerNumber=1 scan=3644 | 12.6596 |
| 01030:3 | 6 | K.HYTN[+1647.613]SSQDVTVPC[+57.021]R.V | HexNAc(5)Hex(3)Fuc(1) | C[+57], N[+1648] | 1104.4613 | 3 | 3311.3693 | 3311.3625 | 0 | 2.1 | 89 | Specific | 672.06 | 594.02 | 594.02 | 13.03 | 102 | >sp\|P01877\|IGHA2_HUMAN Immunoglobulin heavy constant alpha 2 OS=Homo sapiens OX=9606 GN=IGHA2 PE=1 SV=4 | 576 | 20201114_qwl_colo_milk_1ng_glyco_hcd203040_compound15.3751.3751.3 | controllerType=0 controllerNumber=1 scan=3751 | 12.8552 |
| 00990:3 | 6 | K.HYTN[+1241.454]SSQDVTVPC[+57.021]R.V | HexNAc(3)Hex(3)Fuc(1) | C[+57], N[+1241] | 969.0749 | 3 | 2905.2102 | 2905.2038 | 0 | 2.2 | 89 | Specific | 699.79 | 529.48 | 529.48 | 13.03 | 102 | >sp\|P01877\|IGHA2_HUMAN Immunoglobulin heavy constant alpha 2 OS=Homo sapiens OX=9606 GN=IGHA2 PE=1 SV=4 | 576 | 20201114_qwl_colo_milk_1ng_glyco_hcd203040_compound15.3709.3709.3 | controllerType=0 controllerNumber=1 scan=3709 | 12.7773 |
| 01008:3 | 6 | K.HYTN[+1038.375]SSQDVTVPC[+57.021]R.V | HexNAc(2)Hex(3)Fuc(1) | C[+57], N[+1038] | 901.3819 | 3 | 2702.1312 | 2702.1244 | 0 | 2.5 | 89 | Specific | 774.9 | 643.22 | 643.22 | 15.13 | 102 | >sp\|P01877\|IGHA2_HUMAN Immunoglobulin heavy constant alpha 2 OS=Homo sapiens OX=9606 GN=IGHA2 PE=1 SV=4 | 576 | 20201114_qwl_colo_milk_1ng_glyco_hcd203040_compound15.3728.3728.3 | controllerType=0 controllerNumber=1 scan=3728 | 12.8106 |
| 01010:3 | 6 | K.HYTN[+1038.375]SSQDVTVPC[+57.021]R.V | HexNAc(2)Hex(3)Fuc(1) | C[+57], N[+1038] | 901.3819 | 3 | 2702.1312 | 2702.1244 | 0 | 2.5 | 89 | Specific | 780.49 | 627.64 | 627.64 | 15.13 | 102 | >sp\|P01877\|IGHA2_HUMAN Immunoglobulin heavy constant alpha 2 OS=Homo sapiens OX=9606 GN=IGHA2 PE=1 SV=4 | 576 | 20201114_qwl_colo_milk_1ng_glyco_hcd203040_compound15.3730.3730.3 | controllerType=0 controllerNumber=1 scan=3730 | 12.8146 |
| 03152:4 | 6 | K.HYTN[+1809.666]SSQDVTVPC[+57.021]RVPPPPPC[+57.021]C[+57.021]HPR.L | HexNAc(5)Hex(4)Fuc(1) | C[+57]*3, N[+1810] | 1192.763 | 4 | 4768.0301 | 4768.0217 | 0 | 1.8 | 89 | Specific | 234.19 | 228.17 | 228.17 | 5.47 | 102 | >sp\|P01877\|IGHA2_HUMAN Immunoglobulin heavy constant alpha 2 OS=Homo sapiens OX=9606 GN=IGHA2 PE=1 SV=4 | 576 | 20201114_qwl_colo_milk_1ng_glyco_hcd203040_compound15.5968.5968.2 | controllerType=0 controllerNumber=1 scan=5968 | 16.9678 |
| 03092:4 | 6 | K.HYTN[+1647.613]SSQDVTVPC[+57.021]RVPPPPPC[+57.021]C[+57.021]HPR.L | HexNAc(5)Hex(3)Fuc(1) | C[+57]*3, N[+1648] | 1152.25 | 4 | 4605.9783 | 4605.9689 | 0 | 2.1 | 89 | Specific | 258.51 | 258.51 | 258.51 | 4.18 | 102 | >sp\|P01877\|IGHA2_HUMAN Immunoglobulin heavy constant alpha 2 OS=Homo sapiens OX=9606 GN=IGHA2 PE=1 SV=4 | 576 | 20201114_qwl_colo_milk_1ng_glyco_hcd203040_compound15.5906.5906.2 | controllerType=0 controllerNumber=1 scan=5906 | 16.8531 |
| 02938:4 | 6 | K.HYTN[+1971.719]SSQDVTVPC[+57.021]RVPPPPPC[+57.021]C[+57.021]HPR.L | HexNAc(5)Hex(5)Fuc(1) | C[+57]*3, N[+1972] | 1233.2754 | 4 | 4930.0799 | 4930.0745 | 0 | 1.1 | 89 | Specific | 264.84 | 264.84 | 264.84 | 5.63 | 102 | >sp\|P01877\|IGHA2_HUMAN Immunoglobulin heavy constant alpha 2 OS=Homo sapiens OX=9606 GN=IGHA2 PE=1 SV=4 | 576 | 20201114_qwl_colo_milk_1ng_glyco_hcd203040_compound15.5746.5746.4 | controllerType=0 controllerNumber=1 scan=5746 | 16.5603 |
| 03252:4 | 6 | K.HYTN[+1606.587]SSQDVTVPC[+57.021]RVPPPPPC[+57.021]C[+57.021]HPR.L | HexNAc(4)Hex(4)Fuc(1) | C[+57]*3, N[+1607] | 1141.9925 | 4 | 4564.9481 | 4564.9423 | 0 | 1.3 | 89 | Specific | 296.88 | 269.47 | 269.47 | 5.97 | 102 | >sp\|P01877\|IGHA2_HUMAN Immunoglobulin heavy constant alpha 2 OS=Homo sapiens OX=9606 GN=IGHA2 PE=1 SV=4 | 576 | 20201114_qwl_colo_milk_1ng_glyco_hcd203040_compound15.6072.6072.4 | controllerType=0 controllerNumber=1 scan=6072 | 17.1564 |
| 02968:4 | 6 | K.HYTN[+1241.454]SSQDVTVPC[+57.021]RVPPPPPC[+57.021]C[+57.021]HPR.L | HexNAc(3)Hex(3)Fuc(1) | C[+57]*3, N[+1241] | 1050.7128 | 4 | 4199.8294 | 4199.8101 | 0 | 4.6 | 89 | Specific | 302.31 | 302.31 | 302.31 | 5.85 | 102 | >sp\|P01877\|IGHA2_HUMAN Immunoglobulin heavy constant alpha 2 OS=Homo sapiens OX=9606 GN=IGHA2 PE=1 SV=4 | 576 | 20201114_qwl_colo_milk_1ng_glyco_hcd203040_compound15.5777.5777.2 | controllerType=0 controllerNumber=1 scan=5777 | 16.615 |
| 03090:4 | 6 | K.HYTN[+1647.613]SSQDVTVPC[+57.021]RVPPPPPC[+57.021]C[+57.021]HPR.L | HexNAc(5)Hex(3)Fuc(1) | C[+57]*3, N[+1648] | 1152.25 | 4 | 4605.9783 | 4605.9689 | 0 | 2.1 | 89 | Specific | 328.74 | 240.05 | 240.05 | 5.87 | 102 | >sp\|P01877\|IGHA2_HUMAN Immunoglobulin heavy constant alpha 2 OS=Homo sapiens OX=9606 GN=IGHA2 PE=1 SV=4 | 576 | 20201114_qwl_colo_milk_1ng_glyco_hcd203040_compound15.5904.5904.2 | controllerType=0 controllerNumber=1 scan=5904 | 16.8486 |
| 02924:5 | 6 | K.HYTN[+1971.719]SSQDVTVPC[+57.021]RVPPPPPC[+57.021]C[+57.021]HPR.L | HexNAc(5)Hex(5)Fuc(1) | C[+57]*3, N[+1972] | 986.8219 | 5 | 4930.0805 | 4930.0745 | 0 | 1.2 | 89 | Specific | 330.2 | 272.76 | 272.76 | 6.87 | 102 | >sp\|P01877\|IGHA2_HUMAN Immunoglobulin heavy constant alpha 2 OS=Homo sapiens OX=9606 GN=IGHA2 PE=1 SV=4 | 576 | 20201114_qwl_colo_milk_1ng_glyco_hcd203040_compound15.5732.5732.5 | controllerType=0 controllerNumber=1 scan=5732 | 16.5379 |
| 03250:4 | 6 | K.HYTN[+1606.587]SSQDVTVPC[+57.021]RVPPPPPC[+57.021]C[+57.021]HPR.L | HexNAc(4)Hex(4)Fuc(1) | C[+57]*3, N[+1607] | 1141.9925 | 4 | 4564.9481 | 4564.9423 | 0 | 1.3 | 89 | Specific | 340.99 | 340.99 | 340.99 | 6.66 | 102 | >sp\|P01877\|IGHA2_HUMAN Immunoglobulin heavy constant alpha 2 OS=Homo sapiens OX=9606 GN=IGHA2 PE=1 SV=4 | 576 | 20201114_qwl_colo_milk_1ng_glyco_hcd203040_compound15.6070.6070.4 | controllerType=0 controllerNumber=1 scan=6070 | 17.1524 |
| 03266:5 | 6 | K.HYTN[+1809.666]SSQDVTVPC[+57.021]RVPPPPPC[+57.021]C[+57.021]HPR.L | HexNAc(5)Hex(4)Fuc(1) | C[+57]*3, N[+1810] | 954.4129 | 5 | 4768.0352 | 4768.0217 | 0 | 2.8 | 89 | Specific | 390.24 | 317.67 | 317.67 | 7.51 | 102 | >sp\|P01877\|IGHA2_HUMAN Immunoglobulin heavy constant alpha 2 OS=Homo sapiens OX=9606 GN=IGHA2 PE=1 SV=4 | 576 | 20201114_qwl_colo_milk_1ng_glyco_hcd203040_compound15.6087.6087.5 | controllerType=0 controllerNumber=1 scan=6087 | 17.184 |
| 03268:5 | 6 | K.HYTN[+1809.666]SSQDVTVPC[+57.021]RVPPPPPC[+57.021]C[+57.021]HPR.L | HexNAc(5)Hex(4)Fuc(1) | C[+57]*3, N[+1810] | 954.4129 | 5 | 4768.0352 | 4768.0217 | 0 | 2.8 | 89 | Specific | 396.13 | 247.61 | 247.61 | 7.05 | 102 | >sp\|P01877\|IGHA2_HUMAN Immunoglobulin heavy constant alpha 2 OS=Homo sapiens OX=9606 GN=IGHA2 PE=1 SV=4 | 576 | 20201114_qwl_colo_milk_1ng_glyco_hcd203040_compound15.6089.6089.5 | controllerType=0 controllerNumber=1 scan=6089 | 17.1877 |
| 02970:4 | 6 | K.HYTN[+1241.454]SSQDVTVPC[+57.021]RVPPPPPC[+57.021]C[+57.021]HPR.L | HexNAc(3)Hex(3)Fuc(1) | C[+57]*3, N[+1241] | 1050.7128 | 4 | 4199.8294 | 4199.8101 | 0 | 4.6 | 89 | Specific | 411 | 380.51 | 380.51 | 8.2 | 102 | >sp\|P01877\|IGHA2_HUMAN Immunoglobulin heavy constant alpha 2 OS=Homo sapiens OX=9606 GN=IGHA2 PE=1 SV=4 | 576 | 20201114_qwl_colo_milk_1ng_glyco_hcd203040_compound15.5779.5779.2 | controllerType=0 controllerNumber=1 scan=5779 | 16.6189 |
| 02865:5 | 6 | K.HYTN[+1809.666]SSQDVTVPC[+57.021]RVPPPPPC[+57.021]C[+57.021]HPR.L | HexNAc(5)Hex(4)Fuc(1) | C[+57]*3, N[+1810] | 954.4146 | 5 | 4768.0441 | 4768.0217 | 0 | 4.7 | 89 | Specific | 438.47 | 438.47 | 438.47 | 8.98 | 102 | >sp\|P01877\|IGHA2_HUMAN Immunoglobulin heavy constant alpha 2 OS=Homo sapiens OX=9606 GN=IGHA2 PE=1 SV=4 | 576 | 20201114_qwl_colo_milk_1ng_glyco_hcd203040_compound15.5670.5670.5 | controllerType=0 controllerNumber=1 scan=5670 | 16.4233 |
| 02863:5 | 6 | K.HYTN[+1809.666]SSQDVTVPC[+57.021]RVPPPPPC[+57.021]C[+57.021]HPR.L | HexNAc(5)Hex(4)Fuc(1) | C[+57]*3, N[+1810] | 954.4146 | 5 | 4768.0441 | 4768.0217 | 0 | 4.7 | 89 | Specific | 440.13 | 357.81 | 357.81 | 7.64 | 102 | >sp\|P01877\|IGHA2_HUMAN Immunoglobulin heavy constant alpha 2 OS=Homo sapiens OX=9606 GN=IGHA2 PE=1 SV=4 | 576 | 20201114_qwl_colo_milk_1ng_glyco_hcd203040_compound15.5668.5668.5 | controllerType=0 controllerNumber=1 scan=5668 | 16.4173 |
| 02902:3 | 6 | K.HYTN[+1444.534]SSQDVTVPC[+57.021]RVPPPPPC[+57.021]C[+57.021]HPR.L | HexNAc(4)Hex(3)Fuc(1) | C[+57]*3, N[+1445] | 1468.309 | 3 | 4402.9124 | 4402.8895 | 0 | 5.2 | 89 | Specific | 484.03 | 426.06 | 426.06 | 7.84 | 102 | >sp\|P01877\|IGHA2_HUMAN Immunoglobulin heavy constant alpha 2 OS=Homo sapiens OX=9606 GN=IGHA2 PE=1 SV=4 | 576 | 20201114_qwl_colo_milk_1ng_glyco_hcd203040_compound15.5708.5708.3 | controllerType=0 controllerNumber=1 scan=5708 | 16.494 |
| 03169:5 | 6 | K.HYTN[+1809.666]SSQDVTVPC[+57.021]RVPPPPPC[+57.021]C[+57.021]HPR.L | HexNAc(5)Hex(4)Fuc(1) | C[+57]*3, N[+1810] | 954.4129 | 5 | 4768.0355 | 4768.0217 | 0 | 2.9 | 89 | Specific | 498.22 | 275.54 | 275.54 | 7.57 | 102 | >sp\|P01877\|IGHA2_HUMAN Immunoglobulin heavy constant alpha 2 OS=Homo sapiens OX=9606 GN=IGHA2 PE=1 SV=4 | 576 | 20201114_qwl_colo_milk_1ng_glyco_hcd203040_compound15.5986.5986.5 | controllerType=0 controllerNumber=1 scan=5986 | 16.9992 |
| 02926:5 | 6 | K.HYTN[+1971.719]SSQDVTVPC[+57.021]RVPPPPPC[+57.021]C[+57.021]HPR.L | HexNAc(5)Hex(5)Fuc(1) | C[+57]*3, N[+1972] | 986.8219 | 5 | 4930.0805 | 4930.0745 | 0 | 1.2 | 89 | Specific | 515.83 | 440.55 | 440.55 | 9.53 | 102 | >sp\|P01877\|IGHA2_HUMAN Immunoglobulin heavy constant alpha 2 OS=Homo sapiens OX=9606 GN=IGHA2 PE=1 SV=4 | 576 | 20201114_qwl_colo_milk_1ng_glyco_hcd203040_compound15.5734.5734.5 | controllerType=0 controllerNumber=1 scan=5734 | 16.5416 |
| 03531:4 | 6 | K.HYTN[+2262.814]SSQDVTVPC[+57.021]RVPPPPPC[+57.021]C[+57.021]HPR.L | HexNAc(5)Hex(5)Fuc(1)NeuAc(1) | C[+57]*3, N[+2263] | 1306.0497 | 4 | 5221.1769 | 5221.1699 | 0 | 1.3 | 89 | Specific | 568.45 | 411.66 | 411.66 | 9.17 | 102 | >sp\|P01877\|IGHA2_HUMAN Immunoglobulin heavy constant alpha 2 OS=Homo sapiens OX=9606 GN=IGHA2 PE=1 SV=4 | 576 | 20201114_qwl_colo_milk_1ng_glyco_hcd203040_compound15.6363.6363.4 | controllerType=0 controllerNumber=1 scan=6363 | 17.665 |
| 02904:3 | 6 | K.HYTN[+1444.534]SSQDVTVPC[+57.021]RVPPPPPC[+57.021]C[+57.021]HPR.L | HexNAc(4)Hex(3)Fuc(1) | C[+57]*3, N[+1445] | 1468.309 | 3 | 4402.9124 | 4402.8895 | 0 | 5.2 | 89 | Specific | 596.07 | 436.48 | 436.48 | 8 | 102 | >sp\|P01877\|IGHA2_HUMAN Immunoglobulin heavy constant alpha 2 OS=Homo sapiens OX=9606 GN=IGHA2 PE=1 SV=4 | 576 | 20201114_qwl_colo_milk_1ng_glyco_hcd203040_compound15.5710.5710.3 | controllerType=0 controllerNumber=1 scan=5710 | 16.4976 |
| 03554:5 | 6 | K.HYTN[+2262.814]SSQDVTVPC[+57.021]RVPPPPPC[+57.021]C[+57.021]HPR.L | HexNAc(5)Hex(5)Fuc(1)NeuAc(1) | C[+57]*3, N[+2263] | 1045.042 | 5 | 5221.1809 | 5221.1699 | 0 | 2.1 | 89 | Specific | 596.84 | 392.88 | 392.88 | 11.14 | 102 | >sp\|P01877\|IGHA2_HUMAN Immunoglobulin heavy constant alpha 2 OS=Homo sapiens OX=9606 GN=IGHA2 PE=1 SV=4 | 576 | 20201114_qwl_colo_milk_1ng_glyco_hcd203040_compound15.6387.6387.5 | controllerType=0 controllerNumber=1 scan=6387 | 17.7079 |
| 03529:4 | 6 | K.HYTN[+2262.814]SSQDVTVPC[+57.021]RVPPPPPC[+57.021]C[+57.021]HPR.L | HexNAc(5)Hex(5)Fuc(1)NeuAc(1) | C[+57]*3, N[+2263] | 1306.0497 | 4 | 5221.1769 | 5221.1699 | 0 | 1.3 | 89 | Specific | 611.01 | 527.72 | 527.72 | 11.48 | 102 | >sp\|P01877\|IGHA2_HUMAN Immunoglobulin heavy constant alpha 2 OS=Homo sapiens OX=9606 GN=IGHA2 PE=1 SV=4 | 576 | 20201114_qwl_colo_milk_1ng_glyco_hcd203040_compound15.6361.6361.4 | controllerType=0 controllerNumber=1 scan=6361 | 17.661 |
| 03135:5 | 6 | K.HYTN[+1444.534]SSQDVTVPC[+57.021]RVPPPPPC[+57.021]C[+57.021]HPR.L | HexNAc(4)Hex(3)Fuc(1) | C[+57]*3, N[+1445] | 881.3854 | 5 | 4402.8979 | 4402.8895 | 0 | 1.9 | 89 | Specific | 617.94 | 401.18 | 401.18 | 12.14 | 102 | >sp\|P01877\|IGHA2_HUMAN Immunoglobulin heavy constant alpha 2 OS=Homo sapiens OX=9606 GN=IGHA2 PE=1 SV=4 | 576 | 20201114_qwl_colo_milk_1ng_glyco_hcd203040_compound15.5951.5951.5 | controllerType=0 controllerNumber=1 scan=5951 | 16.9376 |
| 03556:5 | 6 | K.HYTN[+2262.814]SSQDVTVPC[+57.021]RVPPPPPC[+57.021]C[+57.021]HPR.L | HexNAc(5)Hex(5)Fuc(1)NeuAc(1) | C[+57]*3, N[+2263] | 1045.042 | 5 | 5221.1809 | 5221.1699 | 0 | 2.1 | 89 | Specific | 715.44 | 551.81 | 551.81 | 14.14 | 102 | >sp\|P01877\|IGHA2_HUMAN Immunoglobulin heavy constant alpha 2 OS=Homo sapiens OX=9606 GN=IGHA2 PE=1 SV=4 | 576 | 20201114_qwl_colo_milk_1ng_glyco_hcd203040_compound15.6389.6389.5 | controllerType=0 controllerNumber=1 scan=6389 | 17.7125 |
| 02866:5 | 6 | K.HYTN[+1647.613]SSQDVTVPC[+57.021]RVPPPPPC[+57.021]C[+57.021]HPR.L | HexNAc(5)Hex(3)Fuc(1) | C[+57]*3, N[+1648] | 922.0039 | 5 | 4605.9905 | 4605.9689 | 0 | 4.7 | 89 | Specific | 717.64 | 479.26 | 479.26 | 13.91 | 102 | >sp\|P01877\|IGHA2_HUMAN Immunoglobulin heavy constant alpha 2 OS=Homo sapiens OX=9606 GN=IGHA2 PE=1 SV=4 | 576 | 20201114_qwl_colo_milk_1ng_glyco_hcd203040_compound15.5671.5671.5 | controllerType=0 controllerNumber=1 scan=5671 | 16.425 |
| 02840:4 | 6 | K.HYTN[+1647.613]SSQDVTVPC[+57.021]RVPPPPPC[+57.021]C[+57.021]HPR.L | HexNAc(5)Hex(3)Fuc(1) | C[+57]*3, N[+1648] | 1152.2502 | 4 | 4605.9788 | 4605.9689 | 0 | 2.2 | 89 | Specific | 725.64 | 725.64 | 725.64 | 13.48 | 102 | >sp\|P01877\|IGHA2_HUMAN Immunoglobulin heavy constant alpha 2 OS=Homo sapiens OX=9606 GN=IGHA2 PE=1 SV=4 | 576 | 20201114_qwl_colo_milk_1ng_glyco_hcd203040_compound15.5644.5644.2 | controllerType=0 controllerNumber=1 scan=5644 | 16.3693 |
| 03137:5 | 6 | K.HYTN[+1444.534]SSQDVTVPC[+57.021]RVPPPPPC[+57.021]C[+57.021]HPR.L | HexNAc(4)Hex(3)Fuc(1) | C[+57]*3, N[+1445] | 881.3854 | 5 | 4402.8979 | 4402.8895 | 0 | 1.9 | 89 | Specific | 743.59 | 566.23 | 566.23 | 14.14 | 102 | >sp\|P01877\|IGHA2_HUMAN Immunoglobulin heavy constant alpha 2 OS=Homo sapiens OX=9606 GN=IGHA2 PE=1 SV=4 | 576 | 20201114_qwl_colo_milk_1ng_glyco_hcd203040_compound15.5953.5953.5 | controllerType=0 controllerNumber=1 scan=5953 | 16.9415 |
| 02841:4 | 6 | K.HYTN[+1809.666]SSQDVTVPC[+57.021]RVPPPPPC[+57.021]C[+57.021]HPR.L | HexNAc(5)Hex(4)Fuc(1) | C[+57]*3, N[+1810] | 1192.7644 | 4 | 4768.036 | 4768.0217 | 0 | 3 | 89 | Specific | 780.31 | 615.61 | 615.61 | 14.58 | 102 | >sp\|P01877\|IGHA2_HUMAN Immunoglobulin heavy constant alpha 2 OS=Homo sapiens OX=9606 GN=IGHA2 PE=1 SV=4 | 576 | 20201114_qwl_colo_milk_1ng_glyco_hcd203040_compound15.5645.5645.4 | controllerType=0 controllerNumber=1 scan=5645 | 16.3708 |
| 02842:4 | 6 | K.HYTN[+1647.613]SSQDVTVPC[+57.021]RVPPPPPC[+57.021]C[+57.021]HPR.L | HexNAc(5)Hex(3)Fuc(1) | C[+57]*3, N[+1648] | 1152.2502 | 4 | 4605.9788 | 4605.9689 | 0 | 2.2 | 89 | Specific | 806.62 | 553.08 | 553.08 | 15.48 | 102 | >sp\|P01877\|IGHA2_HUMAN Immunoglobulin heavy constant alpha 2 OS=Homo sapiens OX=9606 GN=IGHA2 PE=1 SV=4 | 576 | 20201114_qwl_colo_milk_1ng_glyco_hcd203040_compound15.5646.5646.2 | controllerType=0 controllerNumber=1 scan=5646 | 16.3732 |
| 02848:4 | 6 | K.HYTN[+1444.534]SSQDVTVPC[+57.021]RVPPPPPC[+57.021]C[+57.021]HPR.L | HexNAc(4)Hex(3)Fuc(1) | C[+57]*3, N[+1445] | 1101.481 | 4 | 4402.9022 | 4402.8895 | 0 | 2.9 | 89 | Specific | 846.61 | 566.85 | 566.85 | 15.58 | 102 | >sp\|P01877\|IGHA2_HUMAN Immunoglobulin heavy constant alpha 2 OS=Homo sapiens OX=9606 GN=IGHA2 PE=1 SV=4 | 576 | 20201114_qwl_colo_milk_1ng_glyco_hcd203040_compound15.5652.5652.2 | controllerType=0 controllerNumber=1 scan=5652 | 16.3859 |
| 02872:4 | 6 | K.HYTN[+1606.587]SSQDVTVPC[+57.021]RVPPPPPC[+57.021]C[+57.021]HPR.L | HexNAc(4)Hex(4)Fuc(1) | C[+57]*3, N[+1607] | 1141.9971 | 4 | 4564.9666 | 4564.9423 | 0 | 5.3 | 89 | Specific | 853.7 | 608.61 | 608.61 | 16.25 | 102 | >sp\|P01877\|IGHA2_HUMAN Immunoglobulin heavy constant alpha 2 OS=Homo sapiens OX=9606 GN=IGHA2 PE=1 SV=4 | 576 | 20201114_qwl_colo_milk_1ng_glyco_hcd203040_compound15.5677.5677.4 | controllerType=0 controllerNumber=1 scan=5677 | 16.4382 |
| 02873:5 | 6 | K.HYTN[+1606.587]SSQDVTVPC[+57.021]RVPPPPPC[+57.021]C[+57.021]HPR.L | HexNAc(4)Hex(4)Fuc(1) | C[+57]*3, N[+1607] | 913.7992 | 5 | 4564.967 | 4564.9423 | 0 | 5.4 | 89 | Specific | 855.68 | 710.17 | 710.17 | 16.91 | 102 | >sp\|P01877\|IGHA2_HUMAN Immunoglobulin heavy constant alpha 2 OS=Homo sapiens OX=9606 GN=IGHA2 PE=1 SV=4 | 576 | 20201114_qwl_colo_milk_1ng_glyco_hcd203040_compound15.5678.5678.5 | controllerType=0 controllerNumber=1 scan=5678 | 16.4397 |
| 02874:4 | 6 | K.HYTN[+1606.587]SSQDVTVPC[+57.021]RVPPPPPC[+57.021]C[+57.021]HPR.L | HexNAc(4)Hex(4)Fuc(1) | C[+57]*3, N[+1607] | 1141.9971 | 4 | 4564.9666 | 4564.9423 | 0 | 5.3 | 89 | Specific | 877.94 | 599.3 | 599.3 | 16.25 | 102 | >sp\|P01877\|IGHA2_HUMAN Immunoglobulin heavy constant alpha 2 OS=Homo sapiens OX=9606 GN=IGHA2 PE=1 SV=4 | 576 | 20201114_qwl_colo_milk_1ng_glyco_hcd203040_compound15.5679.5679.4 | controllerType=0 controllerNumber=1 scan=5679 | 16.4427 |
| 02862:5 | 6 | K.HYTN[+1444.534]SSQDVTVPC[+57.021]RVPPPPPC[+57.021]C[+57.021]HPR.L | HexNAc(4)Hex(3)Fuc(1) | C[+57]*3, N[+1445] | 881.3879 | 5 | 4402.9104 | 4402.8895 | 0 | 4.8 | 89 | Specific | 878.62 | 638.45 | 638.45 | 16.91 | 102 | >sp\|P01877\|IGHA2_HUMAN Immunoglobulin heavy constant alpha 2 OS=Homo sapiens OX=9606 GN=IGHA2 PE=1 SV=4 | 576 | 20201114_qwl_colo_milk_1ng_glyco_hcd203040_compound15.5667.5667.5 | controllerType=0 controllerNumber=1 scan=5667 | 16.4146 |
| 02843:4 | 6 | K.HYTN[+1809.666]SSQDVTVPC[+57.021]RVPPPPPC[+57.021]C[+57.021]HPR.L | HexNAc(5)Hex(4)Fuc(1) | C[+57]*3, N[+1810] | 1192.7644 | 4 | 4768.036 | 4768.0217 | 0 | 3 | 89 | Specific | 882.2 | 667.58 | 667.58 | 16.58 | 102 | >sp\|P01877\|IGHA2_HUMAN Immunoglobulin heavy constant alpha 2 OS=Homo sapiens OX=9606 GN=IGHA2 PE=1 SV=4 | 576 | 20201114_qwl_colo_milk_1ng_glyco_hcd203040_compound15.5647.5647.4 | controllerType=0 controllerNumber=1 scan=5647 | 16.3765 |
| 02868:5 | 6 | K.HYTN[+1647.613]SSQDVTVPC[+57.021]RVPPPPPC[+57.021]C[+57.021]HPR.L | HexNAc(5)Hex(3)Fuc(1) | C[+57]*3, N[+1648] | 922.0039 | 5 | 4605.9905 | 4605.9689 | 0 | 4.7 | 89 | Specific | 901.73 | 756.62 | 756.62 | 17.91 | 102 | >sp\|P01877\|IGHA2_HUMAN Immunoglobulin heavy constant alpha 2 OS=Homo sapiens OX=9606 GN=IGHA2 PE=1 SV=4 | 576 | 20201114_qwl_colo_milk_1ng_glyco_hcd203040_compound15.5673.5673.5 | controllerType=0 controllerNumber=1 scan=5673 | 16.4304 |
| 02846:4 | 6 | K.HYTN[+1444.534]SSQDVTVPC[+57.021]RVPPPPPC[+57.021]C[+57.021]HPR.L | HexNAc(4)Hex(3)Fuc(1) | C[+57]*3, N[+1445] | 1101.481 | 4 | 4402.9022 | 4402.8895 | 0 | 2.9 | 89 | Specific | 938.99 | 778.62 | 778.62 | 17.58 | 102 | >sp\|P01877\|IGHA2_HUMAN Immunoglobulin heavy constant alpha 2 OS=Homo sapiens OX=9606 GN=IGHA2 PE=1 SV=4 | 576 | 20201114_qwl_colo_milk_1ng_glyco_hcd203040_compound15.5650.5650.2 | controllerType=0 controllerNumber=1 scan=5650 | 16.3814 |
| 02869:4 | 6 | K.HYTN[+1444.534]SSQDVTVPC[+57.021]RVPPPPPC[+57.021]C[+57.021]HPR.L | HexNAc(4)Hex(3)Fuc(1) | C[+57]*3, N[+1445] | 1101.4837 | 4 | 4402.9129 | 4402.8895 | 0 | 5.3 | 89 | Specific | 959.65 | 707.97 | 707.97 | 18.25 | 102 | >sp\|P01877\|IGHA2_HUMAN Immunoglobulin heavy constant alpha 2 OS=Homo sapiens OX=9606 GN=IGHA2 PE=1 SV=4 | 576 | 20201114_qwl_colo_milk_1ng_glyco_hcd203040_compound15.5674.5674.4 | controllerType=0 controllerNumber=1 scan=5674 | 16.4336 |
| 02877:4 | 6 | K.HYTN[+1647.613]SSQDVTVPC[+57.021]RVPPPPPC[+57.021]C[+57.021]HPR.L | HexNAc(5)Hex(3)Fuc(1) | C[+57]*3, N[+1648] | 1152.2531 | 4 | 4605.9905 | 4605.9689 | 0 | 4.7 | 89 | Specific | 964.09 | 721.7 | 721.7 | 18.25 | 102 | >sp\|P01877\|IGHA2_HUMAN Immunoglobulin heavy constant alpha 2 OS=Homo sapiens OX=9606 GN=IGHA2 PE=1 SV=4 | 576 | 20201114_qwl_colo_milk_1ng_glyco_hcd203040_compound15.5682.5682.4 | controllerType=0 controllerNumber=1 scan=5682 | 16.4479 |
| 02875:5 | 6 | K.HYTN[+1606.587]SSQDVTVPC[+57.021]RVPPPPPC[+57.021]C[+57.021]HPR.L | HexNAc(4)Hex(4)Fuc(1) | C[+57]*3, N[+1607] | 913.7992 | 5 | 4564.967 | 4564.9423 | 0 | 5.4 | 89 | Specific | 966.47 | 631.22 | 631.22 | 18.91 | 102 | >sp\|P01877\|IGHA2_HUMAN Immunoglobulin heavy constant alpha 2 OS=Homo sapiens OX=9606 GN=IGHA2 PE=1 SV=4 | 576 | 20201114_qwl_colo_milk_1ng_glyco_hcd203040_compound15.5680.5680.5 | controllerType=0 controllerNumber=1 scan=5680 | 16.4446 |
| 02864:5 | 6 | K.HYTN[+1444.534]SSQDVTVPC[+57.021]RVPPPPPC[+57.021]C[+57.021]HPR.L | HexNAc(4)Hex(3)Fuc(1) | C[+57]*3, N[+1445] | 881.3879 | 5 | 4402.9104 | 4402.8895 | 0 | 4.8 | 89 | Specific | 976.74 | 643.58 | 643.58 | 18.91 | 102 | >sp\|P01877\|IGHA2_HUMAN Immunoglobulin heavy constant alpha 2 OS=Homo sapiens OX=9606 GN=IGHA2 PE=1 SV=4 | 576 | 20201114_qwl_colo_milk_1ng_glyco_hcd203040_compound15.5669.5669.5 | controllerType=0 controllerNumber=1 scan=5669 | 16.4211 |
| 02867:4 | 6 | K.HYTN[+1444.534]SSQDVTVPC[+57.021]RVPPPPPC[+57.021]C[+57.021]HPR.L | HexNAc(4)Hex(3)Fuc(1) | C[+57]*3, N[+1445] | 1101.4837 | 4 | 4402.9129 | 4402.8895 | 0 | 5.3 | 89 | Specific | 984.31 | 741.55 | 741.55 | 18.25 | 102 | >sp\|P01877\|IGHA2_HUMAN Immunoglobulin heavy constant alpha 2 OS=Homo sapiens OX=9606 GN=IGHA2 PE=1 SV=4 | 576 | 20201114_qwl_colo_milk_1ng_glyco_hcd203040_compound15.5672.5672.4 | controllerType=0 controllerNumber=1 scan=5672 | 16.4265 |
| 02879:4 | 6 | K.HYTN[+1647.613]SSQDVTVPC[+57.021]RVPPPPPC[+57.021]C[+57.021]HPR.L | HexNAc(5)Hex(3)Fuc(1) | C[+57]*3, N[+1648] | 1152.2531 | 4 | 4605.9905 | 4605.9689 | 0 | 4.7 | 89 | Specific | 1015.96 | 752.8 | 752.8 | 19.25 | 102 | >sp\|P01877\|IGHA2_HUMAN Immunoglobulin heavy constant alpha 2 OS=Homo sapiens OX=9606 GN=IGHA2 PE=1 SV=4 | 576 | 20201114_qwl_colo_milk_1ng_glyco_hcd203040_compound15.5684.5684.4 | controllerType=0 controllerNumber=1 scan=5684 | 16.4532 |
| 11735:6 | 6 | K.SAVQGPPERDLC[+57.021]GC[+57.021]YSVSSVLPGC[+57.021]AQPWNHGETFTC[+! | HexNAc(4)Hex(3)Fuc(1) | C[+57]*4, N[+1445] | 1210.0534 | 6 | 7255.284 | 7255.2797 | 0 | 0.6 | 156 | Specific | 246.52 | 246.52 | 246.52 | 3.4 | 102 | >sp\|P01877\|IGHA2_HUMAN Immunoglobulin heavy constant alpha 2 OS=Homo sapiens OX=9606 GN=IGHA2 PE=1 SV=4 | 576 | 20201114_qwl_colo_milk_1ng_glyco_hcd203040_compound15.14921.14921.6 | controllerType=0 controllerNumber=1 scan=14921 | 33.1192 |
| 12696:5 | 6 | R.DLC[+57.021]GC[+57.021]YSVSSVLPGC[+57.021]AQPWNHGETFTC[+57.021]TAAH | HexNAc(4)Hex(3)Fuc(1) | C[+57]*4, N[+1445] | 1267.571 | 5 | 6333.8261 | 6333.8129 | 0 | 2.1 | 165 | Specific | 388.97 | 352.12 | 352.12 | 6.15 | 102 | >sp\|P01877\|IGHA2_HUMAN Immunoglobulin heavy constant alpha 2 OS=Homo sapiens OX=9606 GN=IGHA2 PE=1 SV=4 | 576 | 20201114_qwl_colo_milk_1ng_glyco_hcd203040_compound15.15926.15926.5 | controllerType=0 controllerNumber=1 scan=15926 | 35.1744 |
| 02569:2 | 6 | K.TPLTAN[+1460.529]ITK.S | HexNAc(4)Hex(4) | N[+1461] | 1210.0492 | 2 | 2419.0912 | 2419.0856 | 0 | 2.3 | 200 | Specific | 226.1 | 70.98 | 70.98 | 4.41 | 102 | >sp\|P01877\|IGHA2_HUMAN Immunoglobulin heavy constant alpha 2 OS=Homo sapiens OX=9606 GN=IGHA2 PE=1 SV=4 | 576 | 20201114_qwl_colo_milk_1ng_glyco_hcd203040_compound15.5362.5362.2 | controllerType=0 controllerNumber=1 scan=5362 | 15.8096 |
| 02452:2 | 6 | K.TPLTAN[+1768.640]ITK.S | HexNAc(4)Hex(5)Fuc(1) | N[+1769] | 1364.1058 | 2 | 2727.2042 | 2727.1963 | 0 | 2.9 | 200 | Specific | 268.76 | 233.06 | 233.06 | 6.15 | 102 | >sp\|P01877\|IGHA2_HUMAN Immunoglobulin heavy constant alpha 2 OS=Homo sapiens OX=9606 GN=IGHA2 PE=1 SV=4 | 576 | 20201114_qwl_colo_milk_1ng_glyco_hcd203040_compound15.5241.5241.2 | controllerType=0 controllerNumber=1 scan=5241 | 15.5896 |
| 02523:2 | 6 | K.TPLTAN[+1971.719]ITK.S | HexNAc(5)Hex(5)Fuc(1) | N[+1972] | 1465.646 | 2 | 2930.2848 | 2930.2757 | 0 | 3.1 | 200 | Specific | 270.16 | 68.86 | 68.86 | 4.55 | 102 | >sp\|P01877\|IGHA2_HUMAN Immunoglobulin heavy constant alpha 2 OS=Homo sapiens OX=9606 GN=IGHA2 PE=1 SV=4 | 576 | 20201114_qwl_colo_milk_1ng_glyco_hcd203040_compound15.5314.5314.2 | controllerType=0 controllerNumber=1 scan=5314 | 15.7196 |
| 02525:2 | 6 | K.TPLTAN[+1971.719]ITK.S | HexNAc(5)Hex(5)Fuc(1) | N[+1972] | 1465.646 | 2 | 2930.2848 | 2930.2757 | 0 | 3.1 | 200 | Specific | 287.18 | 158.26 | 158.26 | 5.63 | 102 | >sp\|P01877\|IGHA2_HUMAN Immunoglobulin heavy constant alpha 2 OS=Homo sapiens OX=9606 GN=IGHA2 PE=1 SV=4 | 576 | 20201114_qwl_colo_milk_1ng_glyco_hcd203040_compound15.5316.5316.2 | controllerType=0 controllerNumber=1 scan=5316 | 15.7244 |
| 03200:2 | 6 | K.TPLTAN[+1444.534]ITK.S | HexNAc(4)Hex(3)Fuc(1) | N[+1445] | 1202.0513 | 2 | 2403.0953 | 2403.0906 | 0 | 2 | 200 | Specific | 295.29 | 126.04 | 126.04 | 4.95 | 102 | >sp\|P01877\|IGHA2_HUMAN Immunoglobulin heavy constant alpha 2 OS=Homo sapiens OX=9606 GN=IGHA2 PE=1 SV=4 | 576 | 20201114_qwl_colo_milk_1ng_glyco_hcd203040_compound15.6018.6018.2 | controllerType=0 controllerNumber=1 scan=6018 | 17.0572 |
| 04273:3 | 6 | K.TPLTAN[+2059.735]ITK.S | HexNAc(4)Hex(5)Fuc(1)NeuAc(1) | N[+2060] | 1006.7708 | 3 | 3018.2978 | 3018.2917 | 0 | 2 | 200 | Specific | 302.25 | 83.64 | 83.64 | 4.46 | 102 | >sp\|P01877\|IGHA2_HUMAN Immunoglobulin heavy constant alpha 2 OS=Homo sapiens OX=9606 GN=IGHA2 PE=1 SV=4 | 576 | 20201114_qwl_colo_milk_1ng_glyco_hcd203040_compound15.7146.7146.3 | controllerType=0 controllerNumber=1 scan=7146 | 19.0087 |
| 03198:2 | 6 | K.TPLTAN[+1444.534]ITK.S | HexNAc(4)Hex(3)Fuc(1) | N[+1445] | 1202.0513 | 2 | 2403.0953 | 2403.0906 | 0 | 2 | 200 | Specific | 322.74 | 144.77 | 144.77 | 5.44 | 102 | >sp\|P01877\|IGHA2_HUMAN Immunoglobulin heavy constant alpha 2 OS=Homo sapiens OX=9606 GN=IGHA2 PE=1 SV=4 | 576 | 20201114_qwl_colo_milk_1ng_glyco_hcd203040_compound15.6016.6016.2 | controllerType=0 controllerNumber=1 scan=6016 | 17.0531 |
| 02571:2 | 6 | K.TPLTAN[+1460.529]ITK.S | HexNAc(4)Hex(4) | N[+1461] | 1210.0492 | 2 | 2419.0912 | 2419.0856 | 0 | 2.3 | 200 | Specific | 384.78 | 207.78 | 207.78 | 6.54 | 102 | >sp\|P01877\|IGHA2_HUMAN Immunoglobulin heavy constant alpha 2 OS=Homo sapiens OX=9606 GN=IGHA2 PE=1 SV=4 | 576 | 20201114_qwl_colo_milk_1ng_glyco_hcd203040_compound15.5364.5364.2 | controllerType=0 controllerNumber=1 scan=5364 | 15.8137 |
| 02611:3 | 6 | K.TPLTAN[+1606.587]ITK.S | HexNAc(4)Hex(4)Fuc(1) | N[+1607] | 855.7269 | 3 | 2565.1661 | 2565.1435 | 0 | 8.8 | 200 | Specific | 422.1 | 408.05 | 408.05 | 8.5 | 102 | >sp\|P01877\|IGHA2_HUMAN Immunoglobulin heavy constant alpha 2 OS=Homo sapiens OX=9606 GN=IGHA2 PE=1 SV=4 | 576 | 20201114_qwl_colo_milk_1ng_glyco_hcd203040_compound15.5406.5406.3 | controllerType=0 controllerNumber=1 scan=5406 | 15.8957 |
| 02542:2 | 6 | K.TPLTAN[+1606.587]ITK.S | HexNAc(4)Hex(4)Fuc(1) | N[+1607] | 1283.0787 | 2 | 2565.15 | 2565.1435 | 0 | 2.6 | 200 | Specific | 445.97 | 226.17 | 226.17 | 6.86 | 102 | >sp\|P01877\|IGHA2_HUMAN Immunoglobulin heavy constant alpha 2 OS=Homo sapiens OX=9606 GN=IGHA2 PE=1 SV=4 | 576 | 20201114_qwl_colo_milk_1ng_glyco_hcd203040_compound15.5334.5334.2 | controllerType=0 controllerNumber=1 scan=5334 | [illegible] |
| 02693:3 | 6 | K.TPLTAN[+1647.613]ITK.S | HexNAc(5)Hex(3)Fuc(1) | N[+1648] | 869.396 | 3 | 2606.1735 | 2606.17 | 0 | 1.3 | 200 | Specific | 452.47 | 174.82 | 174.82 | 6.22 | 102 | >sp\|P01877\|IGHA2_HUMAN Immunoglobulin heavy constant alpha 2 OS=Homo sapiens OX=9606 GN=IGHA2 PE=1 SV=4 | 576 | 20201114_qwl_colo_milk_1ng_glyco_hcd203040_compound15.5491.5491.3 | controllerType=0 controllerNumber=1 scan=5491 | [illegible] |
| 02710:3 | 6 | K.TPLTAN[+1444.534]ITK.S | HexNAc(4)Hex(3)Fuc(1) | N[+1445] | 801.7046 | 3 | 2403.0993 | 2403.0906 | 0 | 3.6 | 200 | Specific | 474.18 | 470.58 | 470.58 | 9.21 | 102 | >sp\|P01877\|IGHA2_HUMAN Immunoglobulin heavy constant alpha 2 OS=Homo sapiens OX=9606 GN=IGHA2 PE=1 SV=4 | 576 | 20201114_qwl_colo_milk_1ng_glyco_hcd203040_compound15.5509.5509.3 | controllerType=0 controllerNumber=1 scan=5509 | [illegible] |
| 02695:3 | 6 | K.TPLTAN[+1647.613]ITK.S | HexNAc(5)Hex(3)Fuc(1) | N[+1648] | 869.396 | 3 | 2606.1735 | 2606.17 | 0 | 1.3 | 200 | Specific | 502.33 | 226.47 | 226.47 | 6.89 | 102 | >sp\|P01877\|IGHA2_HUMAN Immunoglobulin heavy constant alpha 2 OS=Homo sapiens OX=9606 GN=IGHA2 PE=1 SV=4 | 576 | 20201114_qwl_colo_milk_1ng_glyco_hcd203040_compound15.5493.5493.3 | controllerType=0 controllerNumber=1 scan=5493 | [illegible] |
| 04275:3 | 6 | K.TPLTAN[+2059.735]ITK.S | HexNAc(4)Hex(5)Fuc(1)NeuAc(1) | N[+2060] | 1006.7708 | 3 | 3018.2978 | 3018.2917 | 0 | 2 | 200 | Specific | 512.52 | 251.69 | 251.69 | 7.15 | 102 | >sp\|P01877\|IGHA2_HUMAN Immunoglobulin heavy constant alpha 2 OS=Homo sapiens OX=9606 GN=IGHA2 PE=1 SV=4 | 576 | 20201114_qwl_colo_milk_1ng_glyco_hcd203040_compound15.7148.7148.3 | controllerType=0 controllerNumber=1 scan=7148 | 19.0144 |
| 04279:3 | 6 | K.TPLTAN[+2262.814]ITK.S | HexNAc(5)Hex(5)Fuc(1)NeuAc(1) | N[+2263] | 1074.4638 | 3 | 3221.3768 | 3221.3711 | 0 | 1.8 | 200 | Specific | 515.54 | 282.83 | 282.83 | 7.29 | 102 | >sp\|P01877\|IGHA2_HUMAN Immunoglobulin heavy constant alpha 2 OS=Homo sapiens OX=9606 GN=IGHA2 PE=1 SV=4 | 576 | 20201114_qwl_colo_milk_1ng_glyco_hcd203040_compound15.7152.7152.3 | controllerType=0 controllerNumber=1 scan=7152 | 19.0227 |
| 02612:2 | 6 | K.TPLTAN[+1663.608]ITK.S | HexNAc(5)Hex(4) | N[+1664] | 1311.597 | 2 | 2622.1867 | 2622.1649 | 0 | 8.3 | 200 | Specific | 517.47 | 344.96 | 344.96 | 7.29 | 102 | >sp\|P01877\|IGHA2_HUMAN Immunoglobulin heavy constant alpha 2 OS=Homo sapiens OX=9606 GN=IGHA2 PE=1 SV=4 | 576 | 20201114_qwl_colo_milk_1ng_glyco_hcd203040_compound15.5407.5407.2 | controllerType=0 controllerNumber=1 scan=5407 | 15.8972 |
| 04272:2 | 6 | K.TPLTAN[+2059.735]ITK.S | HexNAc(4)Hex(5)Fuc(1)NeuAc(1) | N[+2060] | 1509.6528 | 2 | 3018.2982 | 3018.2917 | 0 | 2.2 | 200 | Specific | 522.8 | 255.96 | 255.96 | 7.16 | 102 | >sp\|P01877\|IGHA2_HUMAN Immunoglobulin heavy constant alpha 2 OS=Homo sapiens OX=9606 GN=IGHA2 PE=1 SV=4 | 576 | 20201114_qwl_colo_milk_1ng_glyco_hcd203040_compound15.7145.7145.2 | controllerType=0 controllerNumber=1 scan=7145 | 19.0072 |
| 02613:3 | 6 | K.TPLTAN[+1606.587]ITK.S | HexNAc(4)Hex(4)Fuc(1) | N[+1607] | 855.7269 | 3 | 2565.1661 | 2565.1435 | 0 | 8.8 | 200 | Specific | 543.33 | 217.9 | 217.9 | 6.72 | 102 | >sp\|P01877\|IGHA2_HUMAN Immunoglobulin heavy constant alpha 2 OS=Homo sapiens OX=9606 GN=IGHA2 PE=1 SV=4 | 576 | 20201114_qwl_colo_milk_1ng_glyco_hcd203040_compound15.5408.5408.3 | controllerType=0 controllerNumber=1 scan=5408 | 15.8995 |
| 04277:3 | 6 | K.TPLTAN[+2262.814]ITK.S | HexNAc(5)Hex(5)Fuc(1)NeuAc(1) | N[+2263] | 1074.4638 | 3 | 3221.3768 | 3221.3711 | 0 | 1.8 | 200 | Specific | 548.18 | 258.93 | 258.93 | 7.35 | 102 | >sp\|P01877\|IGHA2_HUMAN Immunoglobulin heavy constant alpha 2 OS=Homo sapiens OX=9606 GN=IGHA2 PE=1 SV=4 | 576 | 20201114_qwl_colo_milk_1ng_glyco_hcd203040_compound15.7150.7150.3 | controllerType=0 controllerNumber=1 scan=7150 | 19.0177 |
| 02544:2 | 6 | K.TPLTAN[+1606.587]ITK.S | HexNAc(4)Hex(4)Fuc(1) | N[+1607] | 1283.0787 | 2 | 2565.15 | 2565.1435 | 0 | 2.6 | 200 | Specific | 552.28 | 271.39 | 271.39 | 7.61 | 102 | >sp\|P01877\|IGHA2_HUMAN Immunoglobulin heavy constant alpha 2 OS=Homo sapiens OX=9606 GN=IGHA2 PE=1 SV=4 | 576 | 20201114_qwl_colo_milk_1ng_glyco_hcd203040_compound15.5336.5336.2 | controllerType=0 controllerNumber=1 scan=5336 | 15.7624 |
| 02614:2 | 6 | K.TPLTAN[+1663.608]ITK.S | HexNAc(5)Hex(4) | N[+1664] | 1311.597 | 2 | 2622.1867 | 2622.1649 | 0 | 8.3 | 200 | Specific | 564.74 | 344.34 | 344.34 | 7.6 | 102 | >sp\|P01877\|IGHA2_HUMAN Immunoglobulin heavy constant alpha 2 OS=Homo sapiens OX=9606 GN=IGHA2 PE=1 SV=4 | 576 | 20201114_qwl_colo_milk_1ng_glyco_hcd203040_compound15.5409.5409.2 | controllerType=0 controllerNumber=1 scan=5409 | 15.9021 |
| 04274:2 | 6 | K.TPLTAN[+2059.735]ITK.S | HexNAc(4)Hex(5)Fuc(1)NeuAc(1) | N[+2060] | 1509.6528 | 2 | 3018.2982 | 3018.2917 | 0 | 2.2 | 200 | Specific | 572.05 | 275.62 | 275.62 | 7.61 | 102 | >sp\|P01877\|IGHA2_HUMAN Immunoglobulin heavy constant alpha 2 OS=Homo sapiens OX=9606 GN=IGHA2 PE=1 SV=4 | 576 | 20201114_qwl_colo_milk_1ng_glyco_hcd203040_compound15.7147.7147.2 | controllerType=0 controllerNumber=1 scan=7147 | 19.0113 |
| 02712:3 | 6 | K.TPLTAN[+1444.534]ITK.S | HexNAc(4)Hex(3)Fuc(1) | N[+1445] | 801.7046 | 3 | 2403.0993 | 2403.0906 | 0 | 3.6 | 200 | Specific | 591 | 239.57 | 239.57 | 8.64 | 102 | >sp\|P01877\|IGHA2_HUMAN Immunoglobulin heavy constant alpha 2 OS=Homo sapiens OX=9606 GN=IGHA2 PE=1 SV=4 | 576 | 20201114_qwl_colo_milk_1ng_glyco_hcd203040_compound15.5511.5511.3 | controllerType=0 controllerNumber=1 scan=5511 | 16.1045 |
| 02666:2 | 6 | K.TPLTAN[+1501.555]ITK.S | HexNAc(5)Hex(3) | N[+1502] | 1230.5629 | 2 | 2460.1185 | 2460.1121 | 0 | 2.6 | 200 | Specific | 599.22 | 282 | 282 | 9.02 | 102 | >sp\|P01877\|IGHA2_HUMAN Immunoglobulin heavy constant alpha 2 OS=Homo sapiens OX=9606 GN=IGHA2 PE=1 SV=4 | 576 | 20201114_qwl_colo_milk_1ng_glyco_hcd203040_compound15.5463.5463.2 | controllerType=0 controllerNumber=1 scan=5463 | 16.0055 |
| 02680:2 | 6 | K.TPLTAN[+1647.613]ITK.S | HexNAc(5)Hex(3)Fuc(1) | N[+1648] | 1303.5912 | 2 | 2606.1752 | 2606.17 | 0 | 2 | 200 | Specific | 603.72 | 332.4 | 332.4 | 10.24 | 102 | >sp\|P01877\|IGHA2_HUMAN Immunoglobulin heavy constant alpha 2 OS=Homo sapiens OX=9606 GN=IGHA2 PE=1 SV=4 | 576 | 20201114_qwl_colo_milk_1ng_glyco_hcd203040_compound15.5477.5477.2 | controllerType=0 controllerNumber=1 scan=5477 | 16.0347 |
| 02734:2 | 6 | K.TPLTAN[+1038.375]ITK.S | HexNAc(2)Hex(3)Fuc(1) | N[+1038] | 998.9723 | 2 | 1996.9374 | 1996.9319 | 0 | 2.8 | 200 | Specific | 618.01 | 352.88 | 352.88 | 10.37 | 102 | >sp\|P01877\|IGHA2_HUMAN Immunoglobulin heavy constant alpha 2 OS=Homo sapiens OX=9606 GN=IGHA2 PE=1 SV=4 | 576 | 20201114_qwl_colo_milk_1ng_glyco_hcd203040_compound15.5533.5533.2 | controllerType=0 controllerNumber=1 scan=5533 | 16.1433 |
| 02736:2 | 6 | K.TPLTAN[+1038.375]ITK.S | HexNAc(2)Hex(3)Fuc(1) | N[+1038] | 998.9723 | 2 | 1996.9374 | 1996.9319 | 0 | 2.8 | 200 | Specific | 635.28 | 314.14 | 314.14 | 10.34 | 102 | >sp\|P01877\|IGHA2_HUMAN Immunoglobulin heavy constant alpha 2 OS=Homo sapiens OX=9606 GN=IGHA2 PE=1 SV=4 | 576 | 20201114_qwl_colo_milk_1ng_glyco_hcd203040_compound15.5535.5535.2 | controllerType=0 controllerNumber=1 scan=5535 | 16.1494 |
| 02664:2 | 6 | K.TPLTAN[+1501.555]ITK.S | HexNAc(5)Hex(3) | N[+1502] | 1230.5629 | 2 | 2460.1185 | 2460.1121 | 0 | 2.6 | 200 | Specific | 639.04 | 350.46 | 350.46 | 10.37 | 102 | >sp\|P01877\|IGHA2_HUMAN Immunoglobulin heavy constant alpha 2 OS=Homo sapiens OX=9606 GN=IGHA2 PE=1 SV=4 | 576 | 20201114_qwl_colo_milk_1ng_glyco_hcd203040_compound15.5461.5461.2 | controllerType=0 controllerNumber=1 scan=5461 | 15.9965 |
| 02678:2 | 6 | K.TPLTAN[+1647.613]ITK.S | HexNAc(5)Hex(3)Fuc(1) | N[+1648] | 1303.5912 | 2 | 2606.1752 | 2606.17 | 0 | 2 | 200 | Specific | 648.52 | 360.95 | 360.95 | 10.57 | 102 | >sp\|P01877\|IGHA2_HUMAN Immunoglobulin heavy constant alpha 2 OS=Homo sapiens OX=9606 GN=IGHA2 PE=1 SV=4 | 576 | 20201114_qwl_colo_milk_1ng_glyco_hcd203040_compound15.5475.5475.2 | controllerType=0 controllerNumber=1 scan=5475 | 16.0273 |
| 02622:2 | 6 | K.TPLTAN[+1444.534]ITK.S | HexNAc(4)Hex(3)Fuc(1) | N[+1445] | 1202.0596 | 2 | 2403.1119 | 2403.0906 | 0 | 8.9 | 200 | Specific | 652.31 | 347.35 | 347.35 | 10.78 | 102 | >sp\|P01877\|IGHA2_HUMAN Immunoglobulin heavy constant alpha 2 OS=Homo sapiens OX=9606 GN=IGHA2 PE=1 SV=4 | 576 | 20201114_qwl_colo_milk_1ng_glyco_hcd203040_compound15.5417.5417.2 | controllerType=0 controllerNumber=1 scan=5417 | 15.9157 |
| 02687:2 | 6 | K.TPLTAN[+1241.454]ITK.S | HexNAc(3)Hex(3)Fuc(1) | N[+1241] | 1100.5105 | 2 | 2200.0138 | 2200.0113 | 0 | 1.2 | 200 | Specific | 657.05 | 338.56 | 338.56 | 11.27 | 102 | >sp\|P01877\|IGHA2_HUMAN Immunoglobulin heavy constant alpha 2 OS=Homo sapiens OX=9606 GN=IGHA2 PE=1 SV=4 | 576 | 20201114_qwl_colo_milk_1ng_glyco_hcd203040_compound15.5485.5485.2 | controllerType=0 controllerNumber=1 scan=5485 | 16.0511 |
| 02685:2 | 6 | K.TPLTAN[+1241.454]ITK.S | HexNAc(3)Hex(3)Fuc(1) | N[+1241] | 1100.5105 | 2 | 2200.0138 | 2200.0113 | 0 | 1.2 | 200 | Specific | 663.39 | 388.44 | 388.44 | 12.91 | 102 | >sp\|P01877\|IGHA2_HUMAN Immunoglobulin heavy constant alpha 2 OS=Homo sapiens OX=9606 GN=IGHA2 PE=1 SV=4 | 576 | 20201114_qwl_colo_milk_1ng_glyco_hcd203040_compound15.5483.5483.2 | controllerType=0 controllerNumber=1 scan=5483 | 16.0461 |
| 02689:2 | 6 | K.TPLTAN[+1298.476]ITK.S | HexNAc(4)Hex(3) | N[+1298] | 1129.0214 | 2 | 2257.0355 | 2257.0327 | 0 | 1.2 | 200 | Specific | 666.98 | 388.98 | 388.98 | 12.91 | 102 | >sp\|P01877\|IGHA2_HUMAN Immunoglobulin heavy constant alpha 2 OS=Homo sapiens OX=9606 GN=IGHA2 PE=1 SV=4 | 576 | 20201114_qwl_colo_milk_1ng_glyco_hcd203040_compound15.5487.5487.2 | controllerType=0 controllerNumber=1 scan=5487 | 16.0544 |
| 02624:2 | 6 | K.TPLTAN[+1444.534]ITK.S | HexNAc(4)Hex(3)Fuc(1) | N[+1445] | 1202.0596 | 2 | 2403.1119 | 2403.0906 | 0 | 8.9 | 200 | Specific | 678.47 | 306.93 | 306.93 | 10.53 | 102 | >sp\|P01877\|IGHA2_HUMAN Immunoglobulin heavy constant alpha 2 OS=Homo sapiens OX=9606 GN=IGHA2 PE=1 SV=4 | 576 | 20201114_qwl_colo_milk_1ng_glyco_hcd203040_compound15.5419.5419.2 | controllerType=0 controllerNumber=1 scan=5419 | 15.92 |
| 02691:2 | 6 | K.TPLTAN[+1298.476]ITK.S | HexNAc(4)Hex(3) | N[+1298] | 1129.0214 | 2 | 2257.0355 | 2257.0327 | 0 | 1.2 | 200 | Specific | 696.45 | 350.25 | 350.25 | 11.27 | 102 | >sp\|P01877\|IGHA2_HUMAN Immunoglobulin heavy constant alpha 2 OS=Homo sapiens OX=9606 GN=IGHA2 PE=1 SV=4 | 576 | 20201114_qwl_colo_milk_1ng_glyco_hcd203040_compound15.5489.5489.2 | controllerType=0 controllerNumber=1 scan=5489 | 16.0597 |
| 07478:3 | 6 | R.MAGKPTHIN[+1378.476]VSVVMAEADGTC[+57.021].Y | HexNAc(2)Hex(6) | C[+57], N[+1378] | 1189.5053 | 3 | 3566.5014 | 3566.4952 | 0 | 1.7 | 319 | CRagged | 296.3 | 296.3 | 230.09 | 7.36 | 102 | >sp\|P01877\|IGHA2_HUMAN Immunoglobulin heavy constant alpha 2 OS=Homo sapiens OX=9606 GN=IGHA2 PE=1 SV=4 | 576 | 20201114_qwl_colo_milk_1ng_glyco_hcd203040_compound15.10480.10480.3 | controllerType=0 controllerNumber=1 scan=10480 | 24.7749 |
| 07480:3 | 6 | R.MAGKPTHIN[+1378.476]VSVVMAEADGTC[+57.021].Y | HexNAc(2)Hex(6) | C[+57], N[+1378] | 1189.5053 | 3 | 3566.5014 | 3566.4952 | 0 | 1.7 | 319 | CRagged | 310.91 | 310.91 | 264.15 | 7.36 | 102 | >sp\|P01877\|IGHA2_HUMAN Immunoglobulin heavy constant alpha 2 OS=Homo sapiens OX=9606 GN=IGHA2 PE=1 SV=4 | 576 | 20201114_qwl_colo_milk_1ng_glyco_hcd203040_compound15.10482.10482.3 | controllerType=0 controllerNumber=1 scan=10482 | 24.7789 |
| 07503:3 | 6 | R.MAGKPTHIN[+1216.423]VSVVMAEADGTC[+57.021].Y | HexNAc(2)Hex(5) | C[+57], N[+1216] | 1135.4912 | 3 | 3404.4592 | 3404.4424 | 0 | 4.9 | 319 | CRagged | 581.16 | 578.86 | 415.98 | 11.53 | 102 | >sp\|P01877\|IGHA2_HUMAN Immunoglobulin heavy constant alpha 2 OS=Homo sapiens OX=9606 GN=IGHA2 PE=1 SV=4 | 576 | 20201114_qwl_colo_milk_1ng_glyco_hcd203040_compound15.10506.10506.3 | controllerType=0 controllerNumber=1 scan=10506 | 24.8215 |
| 07505:3 | 6 | R.MAGKPTHIN[+1216.423]VSVVMAEADGTC[+57.021].Y | HexNAc(2)Hex(5) | C[+57], N[+1216] | 1135.4912 | 3 | 3404.4592 | 3404.4424 | 0 | 4.9 | 319 | CRagged | 729.13 | 608.16 | 463.87 | 14.53 | 102 | >sp\|P01877\|IGHA2_HUMAN Immunoglobulin heavy constant alpha 2 OS=Homo sapiens OX=9606 GN=IGHA2 PE=1 SV=4 | 576 | 20201114_qwl_colo_milk_1ng_glyco_hcd203040_compound15.10508.10508.3 | controllerType=0 controllerNumber=1 scan=10508 | 24.8256 |
| 06800:3 | 6 | R.M[+15.995]AGKPTHIN[+1378.476]VSVVMAEADGTC[+57.021].Y | HexNAc(2)Hex(6) | C[+57], M[+16], N[+1378] | 1194.8372 | 3 | 3582.497 | 3582.4901 | 0 | 1.9 | 319 | CRagged | 306.89 | 306.89 | 72.17 | 7.36 | 102 | >sp\|P01877\|IGHA2_HUMAN Immunoglobulin heavy constant alpha 2 OS=Homo sapiens OX=9606 GN=IGHA2 PE=1 SV=4 | 576 | 20201114_qwl_colo_milk_1ng_glyco_hcd203040_compound15.9771.9771.3 | controllerType=0 controllerNumber=1 scan=9771 | 23.5115 |
| 06852:3 | 6 | R.M[+15.995]AGKPTHIN[+1216.423]VSVVMAEADGTC[+57.021].Y | HexNAc(2)Hex(5) | C[+57], M[+16], N[+1216] | 1140.8214 | 3 | 3420.4498 | 3420.4373 | 0 | 3.6 | 319 | CRagged | 420.01 | 420.01 | 136.89 | 9.78 | 102 | >sp\|P01877\|IGHA2_HUMAN Immunoglobulin heavy constant alpha 2 OS=Homo sapiens OX=9606 GN=IGHA2 PE=1 SV=4 | 576 | 20201114_qwl_colo_milk_1ng_glyco_hcd203040_compound15.9827.9827.3 | controllerType=0 controllerNumber=1 scan=9827 | 23.6099 |
| 06854:3 | 6 | R.M[+15.995]AGKPTHIN[+1216.423]VSVVMAEADGTC[+57.021].Y | HexNAc(2)Hex(5) | C[+57], M[+16], N[+1216] | 1140.8214 | 3 | 3420.4498 | 3420.4373 | 0 | 3.6 | 319 | CRagged | 482.56 | 482.56 | 143.45 | 10.06 | 102 | >sp\|P01877\|IGHA2_HUMAN Immunoglobulin heavy constant alpha 2 OS=Homo sapiens OX=9606 GN=IGHA2 PE=1 SV=4 | 576 | 20201114_qwl_colo_milk_1ng_glyco_hcd203040_compound15.9829.9829.3 | controllerType=0 controllerNumber=1 scan=9829 | 23.6137 |
| 09568:3 | 6 | R.MAGKPTHIN[+1298.476]VSVVMAEADGTC[+57.021]Y.- | HexNAc(4)Hex(3) | C[+57], N[+1298] | 1217.1935 | 3 | 3649.5661 | 3649.5588 | 0 | 2 | 319 | CRagged | 221.1 | 166.58 | 166.58 | 3.38 | 102 | >sp\|P01877\|IGHA2_HUMAN Immunoglobulin heavy constant alpha 2 OS=Homo sapiens OX=9606 GN=IGHA2 PE=1 SV=4 | 576 | 20201114_qwl_colo_milk_1ng_glyco_hcd203040_compound15.12662.12662.3 | controllerType=0 controllerNumber=1 scan=12662 | 28.8119 |
| 08682:3 | 6 | R.MAGKPTHIN[+1702.581]VSVVMAEADGTC[+57.021]Y.- | HexNAc(2)Hex(8) | C[+57], N[+1703] | 1351.9014 | 3 | 4053.6896 | 4053.6642 | 0 | 6.3 | 319 | CRagged | 251.08 | 246.4 | 195.75 | 6.58 | 102 | >sp\|P01877\|IGHA2_HUMAN Immunoglobulin heavy constant alpha 2 OS=Homo sapiens OX=9606 GN=IGHA2 PE=1 SV=4 | 576 | 20201114_qwl_colo_milk_1ng_glyco_hcd203040_compound15.11733.11733.3 | controllerType=0 controllerNumber=1 scan=11733 | 27.0419 |
| 08832:3 | 6 | R.MAGKPTHIN[+1378.476]VSVVMAEADGTC[+57.021]Y.- | HexNAc(2)Hex(6) | C[+57], N[+1378] | 1243.8701 | 3 | 3729.5957 | 3729.5585 | 0 | 10 | 319 | CRagged | 282.6 | 282.6 | 221.03 | 6.77 | 102 | >sp\|P01877\|IGHA2_HUMAN Immunoglobulin heavy constant alpha 2 OS=Homo sapiens OX=9606 GN=IGHA2 PE=1 SV=4 | 576 | 20201114_qwl_colo_milk_1ng_glyco_hcd203040_compound15.11891.11891.3 | controllerType=0 controllerNumber=1 scan=11891 | 27.3249 |
| 09617:3 | 6 | R.MAGKPTHIN[+1095.397]VSVVMAEADGTC[+57.021]Y.- | HexNAc(3)Hex(3) | C[+57], N[+1095] | 1149.4997 | 3 | 3446.4845 | 3446.4794 | 0 | 1.5 | 319 | CRagged | 360.25 | 305.54 | 305.54 | 7.67 | 102 | >sp\|P01877\|IGHA2_HUMAN Immunoglobulin heavy constant alpha 2 OS=Homo sapiens OX=9606 GN=IGHA2 PE=1 SV=4 | 576 | 20201114_qwl_colo_milk_1ng_glyco_hcd203040_compound15.12713.12713.3 | controllerType=0 controllerNumber=1 scan=12713 | 28.9097 |
| 09589:3 | 6 | R.MAGKPTHIN[+1647.613]VSVVMAEADGTC[+57.021]Y.- | HexNAc(5)Hex(3)Fuc(1) | C[+57], N[+1648] | 1333.5702 | 3 | 3998.6962 | 3998.6961 | 0 | 0 | 319 | CRagged | 372.17 | 372.17 | 372.17 | 7.7 | 102 | >sp\|P01877\|IGHA2_HUMAN Immunoglobulin heavy constant alpha 2 OS=Homo sapiens OX=9606 GN=IGHA2 PE=1 SV=4 | 576 | 20201114_qwl_colo_milk_1ng_glyco_hcd203040_compound15.12684.12684.3 | controllerType=0 controllerNumber=1 scan=12684 | 28.8541 |
| 09585:3 | 6 | R.MAGKPTHIN[+1501.555]VSVVMAEADGTC[+57.021]Y.- | HexNAc(5)Hex(3) | C[+57], N[+1502] | 1284.8889 | 3 | 3852.652 | 3852.6382 | 0 | 3.6 | 319 | CRagged | 382.41 | 359.04 | 359.04 | 8.46 | 102 | >sp\|P01877\|IGHA2_HUMAN Immunoglobulin heavy constant alpha 2 OS=Homo sapiens OX=9606 GN=IGHA2 PE=1 SV=4 | 576 | 20201114_qwl_colo_milk_1ng_glyco_hcd203040_compound15.12680.12680.3 | controllerType=0 controllerNumber=1 scan=12680 | 28.8478 |
| 10945:3 | 6 | R.MAGKPTHIN[+2262.814]VSVVMAEADGTC[+57.021]Y.- | HexNAc(5)Hex(5)Fuc(1)NeuAc(1) | C[+57], N[+2263] | 1538.6435 | 3 | 4613.9159 | 4613.8972 | 0 | 4.1 | 319 | CRagged | 428.59 | 330.18 | 330.18 | 8.25 | 102 | >sp\|P01877\|IGHA2_HUMAN Immunoglobulin heavy constant alpha 2 OS=Homo sapiens OX=9606 GN=IGHA2 PE=1 SV=4 | 576 | 20201114_qwl_colo_milk_1ng_glyco_hcd203040_compound15.14094.14094.3 | controllerType=0 controllerNumber=1 scan=14094 | 31.514 |
| 09583:3 | 6 | R.MAGKPTHIN[+1501.555]VSVVMAEADGTC[+57.021]Y.- | HexNAc(5)Hex(3) | C[+57], N[+1502] | 1284.8889 | 3 | 3852.652 | 3852.6382 | 0 | 3.6 | 319 | CRagged | 438.56 | 438.56 | 438.56 | 9.93 | 102 | >sp\|P01877\|IGHA2_HUMAN Immunoglobulin heavy constant alpha 2 OS=Homo sapiens OX=9606 GN=IGHA2 PE=1 SV=4 | 576 | 20201114_qwl_colo_milk_1ng_glyco_hcd203040_compound15.12678.12678.3 | controllerType=0 controllerNumber=1 scan=12678 | 28.8419 |
| 08788:3 | 6 | R.MAGKPTHIN[+1378.476]VSVVMAEADGTC[+57.021]Y.- | HexNAc(2)Hex(6) | C[+57], N[+1378] | 1243.8613 | 3 | 3729.5692 | 3729.5585 | 0 | 2.9 | 319 | CRagged | 442.45 | 442.45 | 306.98 | 9.96 | 102 | >sp\|P01877\|IGHA2_HUMAN Immunoglobulin heavy constant alpha 2 OS=Homo sapiens OX=9606 GN=IGHA2 PE=1 SV=4 | 576 | 20201114_qwl_colo_milk_1ng_glyco_hcd203040_compound15.11845.11845.3 | controllerType=0 controllerNumber=1 scan=11845 | 27.2371 |
| 09566:3 | 6 | R.MAGKPTHIN[+1298.476]VSVVMAEADGTC[+57.021]Y.- | HexNAc(4)Hex(3) | C[+57], N[+1298] | 1217.1935 | 3 | 3649.5661 | 3649.5588 | 0 | 2 | 319 | CRagged | 473.26 | 396.65 | 396.65 | 10.03 | 102 | >sp\|P01877\|IGHA2_HUMAN Immunoglobulin heavy constant alpha 2 OS=Homo sapiens OX=9606 GN=IGHA2 PE=1 SV=4 | 576 | 20201114_qwl_colo_milk_1ng_glyco_hcd203040_compound15.12660.12660.3 | controllerType=0 controllerNumber=1 scan=12660 | 28.8058 |
| 10948:3 | 6 | R.MAGKPTHIN[+2262.814]VSVVMAEADGTC[+57.021]Y.- | HexNAc(5)Hex(5)Fuc(1)NeuAc(1) | C[+57], N[+2263] | 1538.6435 | 3 | 4613.9159 | 4613.8972 | 0 | 4.1 | 319 | CRagged | 486.85 | 486.85 | 486.85 | 10.1 | 102 | >sp\|P01877\|IGHA2_HUMAN Immunoglobulin heavy constant alpha 2 OS=Homo sapiens OX=9606 GN=IGHA2 PE=1 SV=4 | 576 | 20201114_qwl_colo_milk_1ng_glyco_hcd203040_compound15.14097.14097.3 | controllerType=0 controllerNumber=1 scan=14097 | 31.519 |
| 09615:3 | 6 | R.MAGKPTHIN[+1095.397]VSVVMAEADGTC[+57.021]Y.- | HexNAc(3)Hex(3) | C[+57], N[+1095] | 1149.4997 | 3 | 3446.4845 | 3446.4794 | 0 | 1.5 | 319 | CRagged | 536.83 | 425.35 | 425.35 | 10.38 | 102 | >sp\|P01877\|IGHA2_HUMAN Immunoglobulin heavy constant alpha 2 OS=Homo sapiens OX=9606 GN=IGHA2 PE=1 SV=4 | 576 | 20201114_qwl_colo_milk_1ng_glyco_hcd203040_compound15.12711.12711.3 | controllerType=0 controllerNumber=1 scan=12711 | 28.9059 |
| 08684:3 | 6 | R.MAGKPTHIN[+1702.581]VSVVMAEADGTC[+57.021]Y.- | HexNAc(2)Hex(8) | C[+57], N[+1703] | 1351.9014 | 3 | 4053.6896 | 4053.6642 | 0 | 6.3 | 319 | CRagged | 631.9 | 631.9 | 428.71 | 12.37 | 102 | >sp\|P01877\|IGHA2_HUMAN Immunoglobulin heavy constant alpha 2 OS=Homo sapiens OX=9606 GN=IGHA2 PE=1 SV=4 | 576 | 20201114_qwl_colo_milk_1ng_glyco_hcd203040_compound15.11735.11735.3 | controllerType=0 controllerNumber=1 scan=11735 | 27.0459 |
| 08791:3 | 6 | R.MAGKPTHIN[+1378.476]VSVVMAEADGTC[+57.021]Y.- | HexNAc(2)Hex(6) | C[+57], N[+1378] | 1243.8613 | 3 | 3729.5692 | 3729.5585 | 0 | 2.9 | 319 | CRagged | 767.04 | 767.04 | 440.71 | 15.89 | 102 | >sp\|P01877\|IGHA2_HUMAN Immunoglobulin heavy constant alpha 2 OS=Homo sapiens OX=9606 GN=IGHA2 PE=1 SV=4 | 576 | 20201114_qwl_colo_milk_1ng_glyco_hcd203040_compound15.11848.11848.3 | controllerType=0 controllerNumber=1 scan=11848 | 27.2412 |
| 09446:3 | 6 | R.MAGKPTHIN[+1540.529]VSVVMAEADGTC[+57.021]Y.- | HexNAc(2)Hex(7) | C[+57], N[+1541] | 1297.8792 | 3 | 3891.6231 | 3891.6114 | 0 | 3 | 319 | CRagged | 768.51 | 768.51 | 441.77 | 15.89 | 102 | >sp\|P01877\|IGHA2_HUMAN Immunoglobulin heavy constant alpha 2 OS=Homo sapiens OX=9606 GN=IGHA2 PE=1 SV=4 | 576 | 20201114_qwl_colo_milk_1ng_glyco_hcd203040_compound15.12534.12534.3 | controllerType=0 controllerNumber=1 scan=12534 | 28.5368 |
| 09404:3 | 6 | R.MAGKPTHIN[+1540.529]VSVVMAEADGTC[+57.021]Y.- | HexNAc(2)Hex(7) | C[+57], N[+1541] | 1297.8793 | 3 | 3891.6234 | 3891.6114 | 0 | 3.1 | 319 | CRagged | 787.06 | 701.55 | 463.49 | 15.89 | 102 | >sp\|P01877\|IGHA2_HUMAN Immunoglobulin heavy constant alpha 2 OS=Homo sapiens OX=9606 GN=IGHA2 PE=1 SV=4 | 576 | 20201114_qwl_colo_milk_1ng_glyco_hcd203040_compound15.12490.12490.3 | controllerType=0 controllerNumber=1 scan=12490 | 28.4401 |
| 09506:3 | 6 | R.MAGKPTHIN[+1216.423]VSVVMAEADGTC[+57.021]Y.- | HexNAc(2)Hex(5) | C[+57], N[+1216] | 1189.8477 | 3 | 3567.5285 | 3567.5057 | 0 | 6.4 | 319 | CRagged | 861.65 | 861.65 | 455.59 | 17.37 | 102 | >sp\|P01877\|IGHA2_HUMAN Immunoglobulin heavy constant alpha 2 OS=Homo sapiens OX=9606 GN=IGHA2 PE=1 SV=4 | 576 | 20201114_qwl_colo_milk_1ng_glyco_hcd203040_compound15.12597.12597.3 | controllerType=0 controllerNumber=1 scan=12597 | 28.6925 |
| 09386:3 | 6 | R.MAGKPTHIN[+1702.581]VSVVMAEADGTC[+57.021]Y.- | HexNAc(2)Hex(8) | C[+57], N[+1703] | 1351.8953 | 3 | 4053.6713 | 4053.6642 | 0 | 1.8 | 319 | CRagged | 874.98 | 702.72 | 460.94 | 17.79 | 102 | >sp\|P01877\|IGHA2_HUMAN Immunoglobulin heavy constant alpha 2 OS=Homo sapiens OX=9606 GN=IGHA2 PE=1 SV=4 | 576 | 20201114_qwl_colo_milk_1ng_glyco_hcd203040_compound15.12470.12470.3 | controllerType=0 controllerNumber=1 scan=12470 | 28.3971 |
| 09464:3 | 6 | R.MAGKPTHIN[+1378.476]VSVVMAEADGTC[+57.021]Y.- | HexNAc(2)Hex(6) | C[+57], N[+1378] | 1243.8619 | 3 | 3729.5711 | 3729.5585 | 0 | 3.4 | 319 | CRagged | 893.52 | 893.52 | 512.36 | 17.89 | 102 | >sp\|P01877\|IGHA2_HUMAN Immunoglobulin heavy constant alpha 2 OS=Homo sapiens OX=9606 GN=IGHA2 PE=1 SV=4 | 576 | 20201114_qwl_colo_milk_1ng_glyco_hcd203040_compound15.12553.12553.3 | controllerType=0 controllerNumber=1 scan=12553 | 28.5879 |
| 09444:3 | 6 | R.MAGKPTHIN[+1540.529]VSVVMAEADGTC[+57.021]Y.- | HexNAc(2)Hex(7) | C[+57], N[+1541] | 1297.8792 | 3 | 3891.6231 | 3891.6114 | 0 | 3 | 319 | CRagged | 914.99 | 813.89 | 530.47 | 18.89 | 102 | >sp\|P01877\|IGHA2_HUMAN Immunoglobulin heavy constant alpha 2 OS=Homo sapiens OX=9606 GN=IGHA2 PE=1 SV=4 | 576 | 20201114_qwl_colo_milk_1ng_glyco_hcd203040_compound15.12532.12532.3 | controllerType=0 controllerNumber=1 scan=12532 | 28.5313 |
| 09407:3 | 6 | R.MAGKPTHIN[+1540.529]VSVVMAEADGTC[+57.021]Y.- | HexNAc(2)Hex(7) | C[+57], N[+1541] | 1297.8793 | 3 | 3891.6234 | 3891.6114 | 0 | 3.1 | 319 | CRagged | 918.85 | 918.85 | 549.97 | 18.89 | 102 | >sp\|P01877\|IGHA2_HUMAN Immunoglobulin heavy constant alpha 2 OS=Homo sapiens OX=9606 GN=IGHA2 PE=1 SV=4 | 576 | 20201114_qwl_colo_milk_1ng_glyco_hcd203040_compound15.12493.12493.3 | controllerType=0 controllerNumber=1 scan=12493 | 28.4462 |
| 09504:3 | 6 | R.MAGKPTHIN[+1216.423]VSVVMAEADGTC[+57.021]Y.- | HexNAc(2)Hex(5) | C[+57], N[+1216] | 1189.8477 | 3 | 3567.5285 | 3567.5057 | 0 | 6.4 | 319 | CRagged | 947.43 | 947.43 | 542.55 | 18.37 | 102 | >sp\|P01877\|IGHA2_HUMAN Immunoglobulin heavy constant alpha 2 OS=Homo sapiens OX=9606 GN=IGHA2 PE=1 SV=4 | 576 | 20201114_qwl_colo_milk_1ng_glyco_hcd203040_compound15.12595.12595.3 | controllerType=0 controllerNumber=1 scan=12595 | 28.6845 |
| 09383:3 | 6 | R.MAGKPTHIN[+1702.581]VSVVMAEADGTC[+57.021]Y.- | HexNAc(2)Hex(8) | C[+57], N[+1703] | 1351.8953 | 3 | 4053.6713 | 4053.6642 | 0 | 1.8 | 319 | CRagged | 968.34 | 968.11 | 587.32 | 19.79 | 102 | >sp\|P01877\|IGHA2_HUMAN Immunoglobulin heavy constant alpha 2 OS=Homo sapiens OX=9606 GN=IGHA2 PE=1 SV=4 | 576 | 20201114_qwl_colo_milk_1ng_glyco_hcd203040_compound15.12467.12467.3 | controllerType=0 controllerNumber=1 scan=12467 | 28.3903 |
| 09462:3 | 6 | R.MAGKPTHIN[+1378.476]VSVVMAEADGTC[+57.021]Y.- | HexNAc(2)Hex(6) | C[+57], N[+1378] | 1243.8619 | 3 | 3729.5711 | 3729.5585 | 0 | 3.4 | 319 | CRagged | 975.08 | 815.4 | 573.8 | 19.89 | 102 | >sp\|P01877\|IGHA2_HUMAN Immunoglobulin heavy constant alpha 2 OS=Homo sapiens OX=9606 GN=IGHA2 PE=1 SV=4 | 576 | 20201114_qwl_colo_milk_1ng_glyco_hcd203040_compound15.12551.12551.3 | controllerType=0 controllerNumber=1 scan=12551 | 28.5786 |
| 09117:3 | 6 | R.M[+15.995]AGKPTHIN[+1501.555]VSVVMAEADGTC[+57.021]Y.- | HexNAc(5)Hex(3) | C[+57], M[+16], N[+1502] | 1290.2203 | 3 | 3868.6463 | 3868.6331 | 0 | 3.4 | 319 | CRagged | 220.89 | 212.26 | 44.77 | 5.54 | 102 | >sp\|P01877\|IGHA2_HUMAN Immunoglobulin heavy constant alpha 2 OS=Homo sapiens OX=9606 GN=IGHA2 PE=1 SV=4 | 576 | 20201114_qwl_colo_milk_1ng_glyco_hcd203040_compound15.12189.12189.3 | controllerType=0 controllerNumber=1 scan=12189 | 27.8921 |
| 09101:3 | 6 | R.M[+15.995]AGKPTHIN[+1054.370]VSVVMAEADGTC[+57.021]Y.- | HexNAc(2)Hex(4) | C[+57], M[+16], N[+1054] | 1141.1552 | 3 | 3421.451 | 3421.4478 | 0 | 0.9 | 319 | CRagged | 397.89 | 397.89 | 188.65 | 9.68 | 102 | >sp\|P01877\|IGHA2_HUMAN Immunoglobulin heavy constant alpha 2 OS=Homo sapiens OX=9606 GN=IGHA2 PE=1 SV=4 | 576 | 20201114_qwl_colo_milk_1ng_glyco_hcd203040_compound15.12172.12172.3 | controllerType=0 controllerNumber=1 scan=12172 | 27.8608 |
| 09103:3 | 6 | R.M[+15.995]AGKPTHIN[+1054.370]VSVVMAEADGTC[+57.021]Y.- | HexNAc(2)Hex(4) | C[+57], M[+16], N[+1054] | 1141.1552 | 3 | 3421.451 | 3421.4478 | 0 | 0.9 | 319 | CRagged | 523.19 | 523.19 | 189.98 | 10.12 | 102 | >sp\|P01877\|IGHA2_HUMAN Immunoglobulin heavy constant alpha 2 OS=Homo sapiens OX=9606 GN=IGHA2 PE=1 SV=4 | 576 | 20201114_qwl_colo_milk_1ng_glyco_hcd203040_compound15.12174.12174.3 | controllerType=0 controllerNumber=1 scan=12174 | 27.8644 |
| 08993:3 | 6 | R.M[+15.995]AGKPTHIN[+1378.476]VSVVMAEADGTC[+57.021]Y.- | HexNAc(2)Hex(6) | C[+57], M[+16], N[+1378] | 1249.1928 | 3 | 3745.5639 | 3745.5535 | 0 | 2.8 | 319 | CRagged | 539.08 | 539.08 | 218.98 | 10.48 | 102 | >sp\|P01877\|IGHA2_HUMAN Immunoglobulin heavy constant alpha 2 OS=Homo sapiens OX=9606 GN=IGHA2 PE=1 SV=4 | 576 | 20201114_qwl_colo_milk_1ng_glyco_hcd203040_compound15.12058.12058.3 | controllerType=0 controllerNumber=1 scan=12058 | 27.651 |
| 09113:3 | 6 | R.M[+15.995]AGKPTHIN[+1216.423]VSVVMAEADGTC[+57.021]Y.- | HexNAc(2)Hex(5) | C[+57], M[+16], N[+1216] | 1195.1738 | 3 | 3583.5068 | 3583.5006 | 0 | 1.7 | 319 | CRagged | 556.8 | 510.78 | 200.59 | 10.48 | 102 | >sp\|P01877\|IGHA2_HUMAN Immunoglobulin heavy constant alpha 2 OS=Homo sapiens OX=9606 GN=IGHA2 PE=1 SV=4 | 576 | 20201114_qwl_colo_milk_1ng_glyco_hcd203040_compound15.12185.12185.3 | controllerType=0 controllerNumber=1 scan=12185 | 27.8861 |
| 12611:4 | 12 | K.LKELPGVC[+57.021]N[+2569.905]ETMMALWEEC[+57.021]KPC[+57.021]LK.Q | HexNAc(5)Hex(6)NeuAc(2) | C[+57]*3, N[+2570] | 1377.3363 | 4 | 5506.3234 | 5506.2893 | 0 | 6.2 | 95 | Specific | 452.87 | 315.02 | 266.85 | 7.41 | 28 | >sp\|P10909\|CLUS_HUMAN Clusterin OS=Homo sapiens OX=9606 GN=CLU PE=1 SV=1 | 1467 | 20201114_qwl_colo_milk_1ng_glyco_hcd203040_compound15.15837.15837.4 | controllerType=0 controllerNumber=1 scan=15837 | 34.9889 |
| 12609:4 | 12 | K.LKELPGVC[+57.021]N[+2569.905]ETMMALWEEC[+57.021]KPC[+57.021]LK.Q | HexNAc(5)Hex(6)NeuAc(2) | C[+57]*3, N[+2570] | 1377.3363 | 4 | 5506.3234 | 5506.2893 | 0 | 6.2 | 95 | Specific | 634.14 | 634.14 | 399.5 | 11.92 | 28 | >sp\|P10909\|CLUS_HUMAN Clusterin OS=Homo sapiens OX=9606 GN=CLU PE=1 SV=1 | 1467 | 20201114_qwl_colo_milk_1ng_glyco_hcd203040_compound15.15835.15835.4 | controllerType=0 controllerNumber=1 scan=15835 | 34.9848 |
| 12752:4 | 12 | K.LKELPGVC[+57.021]N[+2059.735]ETMMALWEEC[+57.021]KPC[+57.021]LK.Q | HexNAc(4)Hex(5)Fuc(1)NeuAc(1) | C[+57]*3, N[+2060] | 1249.7919 | 4 | 4996.1457 | 4996.1196 | 0 | 5.2 | 95 | Specific | 873.49 | 777.52 | 777.52 | 17.11 | 28 | >sp\|P10909\|CLUS_HUMAN Clusterin OS=Homo sapiens OX=9606 GN=CLU PE=1 SV=1 | 1467 | 20201114_qwl_colo_milk_1ng_glyco_hcd203040_compound15.15984.15984.4 | controllerType=0 controllerNumber=1 scan=15984 | 35.2767 |
| 12773:4 | 12 | K.LKELPGVC[+57.021]N[+2059.735]ETMMALWEEC[+57.021]KPC[+57.021]LK.Q | HexNAc(4)Hex(5)Fuc(1)NeuAc(1) | C[+57]*3, N[+2060] | 1249.7898 | 4 | 4996.1374 | 4996.1196 | 0 | 3.5 | 95 | Specific | 1038.87 | 907.96 | 907.96 | 20.41 | 28 | >sp\|P10909\|CLUS_HUMAN Clusterin OS=Homo sapiens OX=9606 GN=CLU PE=1 SV=1 | 1467 | 20201114_qwl_colo_milk_1ng_glyco_hcd203040_compound15.16006.16006.4 | controllerType=0 controllerNumber=1 scan=16006 | 35.3243 |
| 12770:4 | 12 | K.LKELPGVC[+57.021]N[+2059.735]ETMMALWEEC[+57.021]KPC[+57.021]LK.Q | HexNAc(4)Hex(5)Fuc(1)NeuAc(1) | C[+57]*3, N[+2060] | 1249.7898 | 4 | 4996.1374 | 4996.1196 | 0 | 3.5 | 95 | Specific | 1171.7 | 1171.7 | 1171.7 | 23.41 | 28 | >sp\|P10909\|CLUS_HUMAN Clusterin OS=Homo sapiens OX=9606 GN=CLU PE=1 SV=1 | 1467 | 20201114_qwl_colo_milk_1ng_glyco_hcd203040_compound15.16003.16003.4 | controllerType=0 controllerNumber=1 scan=16003 | 35.3179 |
| 12749:4 | 12 | K.LKELPGVC[+57.021]N[+2059.735]ETMMALWEEC[+57.021]KPC[+57.021]LK.Q | HexNAc(4)Hex(5)Fuc(1)NeuAc(1) | C[+57]*3, N[+2060] | 1249.7919 | 4 | 4996.1457 | 4996.1196 | 0 | 5.2 | 95 | Specific | 1173.98 | 1074.73 | 1074.73 | 23.11 | 28 | >sp\|P10909\|CLUS_HUMAN Clusterin OS=Homo sapiens OX=9606 GN=CLU PE=1 SV=1 | 1467 | 20201114_qwl_colo_milk_1ng_glyco_hcd203040_compound15.15981.15981.4 | controllerType=0 controllerNumber=1 scan=15981 | 35.2708 |
| 18919:3 | 12 | K.MLN[+2424.867]TSSLLEQLNEQFNWVSR.L | HexNAc(5)Hex(6)Fuc(1)NeuAc(1) | N[+2425] | 1612.0264 | 3 | 4834.0647 | 4834.0539 | 0 | 2.2 | 352 | Specific | 382.45 | 366.02 | 209.62 | 8.11 | 28 | >sp\|P10909\|CLUS_HUMAN Clusterin OS=Homo sapiens OX=9606 GN=CLU PE=1 SV=1 | 1467 | 20201114_qwl_colo_milk_1ng_glyco_hcd203040_compound15.22453.22453.3 | controllerType=0 controllerNumber=1 scan=22453 | 49.6374 |
| 18977:3 | 12 | K.MLN[+2059.735]TSSLLEQLNEQFNWVSR.L | HexNAc(4)Hex(5)Fuc(1)NeuAc(1) | N[+2060] | 1490.3171 | 3 | 4468.9367 | 4468.9217 | 0 | 3.4 | 352 | Specific | 799.27 | 591.86 | 591.86 | 15.51 | 28 | >sp\|P10909\|CLUS_HUMAN Clusterin OS=Homo sapiens OX=9606 GN=CLU PE=1 SV=1 | 1467 | 20201114_qwl_colo_milk_1ng_glyco_hcd203040_compound15.22514.22514.3 | controllerType=0 controllerNumber=1 scan=22514 | 49.7758 |
| 18963:3 | 12 | K.MLN[+2059.735]TSSLLEQLNEQFNWVSR.L | HexNAc(4)Hex(5)Fuc(1)NeuAc(1) | N[+2060] | 1490.3182 | 3 | 4468.94 | 4468.9217 | 0 | 4.1 | 352 | Specific | 984.21 | 779.35 | 779.35 | 19.48 | 28 | >sp\|P10909\|CLUS_HUMAN Clusterin OS=Homo sapiens OX=9606 GN=CLU PE=1 SV=1 | 1467 | 20201114_qwl_colo_milk_1ng_glyco_hcd203040_compound15.22499.22499.3 | controllerType=0 controllerNumber=1 scan=22499 | 49.7276 |
| 18979:3 | 12 | K.MLN[+2059.735]TSSLLEQLNEQFNWVSR.L | HexNAc(4)Hex(5)Fuc(1)NeuAc(1) | N[+2060] | 1490.3171 | 3 | 4468.9367 | 4468.9217 | 0 | 3.4 | 352 | Specific | 1012.13 | 645.07 | 645.07 | 20.51 | 28 | >sp\|P10909\|CLUS_HUMAN Clusterin OS=Homo sapiens OX=9606 GN=CLU PE=1 SV=1 | 1467 | 20201114_qwl_colo_milk_1ng_glyco_hcd203040_compound15.22516.22516.3 | controllerType=0 controllerNumber=1 scan=22516 | 49.781 |

Table S4. Cont.

| Query #:z | Protein Rank | Glycopeptide | Glycan | Modification Type(s) | Observed m/z | z | Observed (M+H) | Calc. mass (M+H) | Off-by-x error | Mass error (ppm) | StartPosition | Cleavage | Score | Delta | Delta Mod | \|Log Prob\| | # of unique peptides | Protein Name | Protein DB number | Comment | Scan # | Scan Time |
|---|---|---|---|---|---|---|---|---|---|---|---|---|---|---|---|---|---|---|---|---|---|---|
| 18910:3 | 12 | K.MLN[+2205.793]TSSLLEQLNEQFNWVSR.L | HexNAc(4)Hex(5)Fuc(2)NeuAc(1) | N[+2206] | 1538.9993 | 3 | 4614.9834 | 4614.9796 | 0 | 0.8 | 352 | Specific | 1069.6 | 806.83 | 806.83 | 21.36 | 28 | >sp\|P10909\|CLUS_HUMAN Clusterin OS=Homo sapiens OX=9606 GN=CLU PE=1 SV=1 | 1467 | 20201114_qwl_colo_milk_1ng_glyco_hcd203040_compound15.22444.22444.3 | controllerType=0 controllerNumber=1 scan=22444 | 49.6146 |
| 18961:3 | 12 | K.MLN[+2059.735]TSSLLEQLNEQFNWVSR.L | HexNAc(4)Hex(5)Fuc(1)NeuAc(1) | N[+2060] | 1490.3182 | 3 | 4468.94 | 4468.9217 | 0 | 4.1 | 352 | Specific | 1070.77 | 804.47 | 804.47 | 21.48 | 28 | >sp\|P10909\|CLUS_HUMAN Clusterin OS=Homo sapiens OX=9606 GN=CLU PE=1 SV=1 | 1467 | 20201114_qwl_colo_milk_1ng_glyco_hcd203040_compound15.22497.22497.3 | controllerType=0 controllerNumber=1 scan=22497 | 49.721 |
| 09272:3 | 12 | R.LAN[+2715.963]LTQGEDQYYLR.V | HexNAc(5)Hex(6)Fuc(1)NeuAc(2) | N[+2716] | 1467.279 | 3 | 4399.8224 | 4399.7962 | 0 | 6 | 372 | Specific | 244.78 | 244.78 | 244.78 | 5.62 | 28 | >sp\|P10909\|CLUS_HUMAN Clusterin OS=Homo sapiens OX=9606 GN=CLU PE=1 SV=1 | 1467 | 20201114_qwl_colo_milk_1ng_glyco_hcd203040_compound15.12351.12351.3 | controllerType=0 controllerNumber=1 scan=12351 | 28.1743 |
| 09274:3 | 12 | R.LAN[+2715.963]LTQGEDQYYLR.V | HexNAc(5)Hex(6)Fuc(1)NeuAc(2) | N[+2716] | 1467.279 | 3 | 4399.8224 | 4399.7962 | 0 | 6 | 372 | Specific | 468.6 | 446.6 | 446.6 | 8.84 | 28 | >sp\|P10909\|CLUS_HUMAN Clusterin OS=Homo sapiens OX=9606 GN=CLU PE=1 SV=1 | 1467 | 20201114_qwl_colo_milk_1ng_glyco_hcd203040_compound15.12353.12353.3 | controllerType=0 controllerNumber=1 scan=12353 | 28.1788 |
| 24554:4 | 13 | R.QLAHQSN[+2861.000]STNIFFSPVSIATAFAMLSLGTK.A | HexNAc(5)Hex(6)NeuAc(3) | N[+2861] | 1511.4259 | 4 | 6042.6818 | 6042.6351 | 0 | 7.7 | 64 | Specific | 227.61 | 227.61 | 227.61 | 6.46 | 34 | >sp\|P01009\|A1AT_HUMAN Alpha-1-antitrypsin OS=Homo sapiens OX=9606 GN=SERPINA1 PE=1 SV=3 | 642 | 20201114_qwl_colo_milk_1ng_glyco_hcd203040_compound15.28379.28379.4 | controllerType=0 controllerNumber=1 scan=28379 | 61.1445 |
| 21802:3 | 13 | R.QLAHQSN[+2204.772]STNIFFSPVSIATAFAMLSLGTK.A | HexNAc(4)Hex(5)NeuAc(2) | N[+2205] | 1796.1563 | 3 | 5386.4545 | 5386.4075 | 0 | 8.7 | 64 | Specific | 356.47 | 344.29 | 344.29 | 7.52 | 34 | >sp\|P01009\|A1AT_HUMAN Alpha-1-antitrypsin OS=Homo sapiens OX=9606 GN=SERPINA1 PE=1 SV=3 | 642 | 20201114_qwl_colo_milk_1ng_glyco_hcd203040_compound15.25463.25463.3 | controllerType=0 controllerNumber=1 scan=25463 | 55.7014 |
| 21921:4 | 13 | R.[+42.011]QLAHQSN[+2190.793]STNIFFSPVSIATAFAMLSLGTK.A | HexNAc(6)Hex(6) | .[42], N[+2191] | 1354.3635 | 4 | 5414.4323 | 5414.4388 | 0 | -1.2 | 64 | Specific | 357.61 | 357.61 | 357.61 | 7.35 | 34 | >sp\|P01009\|A1AT_HUMAN Alpha-1-antitrypsin OS=Homo sapiens OX=9606 GN=SERPINA1 PE=1 SV=3 | 642 | 20201114_qwl_colo_milk_1ng_glyco_hcd203040_compound15.25588.25588.2 | controllerType=0 controllerNumber=1 scan=25588 | 55.9419 |
| 24552:4 | 13 | R.QLAHQSN[+2861.000]STNIFFSPVSIATAFAMLSLGTK.A | HexNAc(5)Hex(6)NeuAc(3) | N[+2861] | 1511.4259 | 4 | 6042.6818 | 6042.6351 | 0 | 7.7 | 64 | Specific | 369.14 | 342.23 | 342.23 | 7.59 | 34 | >sp\|P01009\|A1AT_HUMAN Alpha-1-antitrypsin OS=Homo sapiens OX=9606 GN=SERPINA1 PE=1 SV=3 | 642 | 20201114_qwl_colo_milk_1ng_glyco_hcd203040_compound15.28377.28377.4 | controllerType=0 controllerNumber=1 scan=28377 | 61.1405 |
| 21871:4 | 13 | R.QLAHQSN[+2204.772]STNIFFSPVSIATAFAMLSLGTK.A | HexNAc(4)Hex(5)NeuAc(2) | N[+2205] | 1347.3665 | 4 | 5386.444 | 5386.4075 | 0 | 6.8 | 64 | Specific | 445.32 | 445.32 | 445.32 | 9.5 | 34 | >sp\|P01009\|A1AT_HUMAN Alpha-1-antitrypsin OS=Homo sapiens OX=9606 GN=SERPINA1 PE=1 SV=3 | 642 | 20201114_qwl_colo_milk_1ng_glyco_hcd203040_compound15.25536.25536.4 | controllerType=0 controllerNumber=1 scan=25536 | 55.8408 |
| 21800:3 | 13 | R.QLAHQSN[+2204.772]STNIFFSPVSIATAFAMLSLGTK.A | HexNAc(4)Hex(5)NeuAc(2) | N[+2205] | 1796.1563 | 3 | 5386.4545 | 5386.4075 | 0 | 8.7 | 64 | Specific | 605.07 | 605.07 | 605.07 | 12.4 | 34 | >sp\|P01009\|A1AT_HUMAN Alpha-1-antitrypsin OS=Homo sapiens OX=9606 GN=SERPINA1 PE=1 SV=3 | 642 | 20201114_qwl_colo_milk_1ng_glyco_hcd203040_compound15.25461.25461.3 | controllerType=0 controllerNumber=1 scan=25461 | 55.6977 |
| 21870:3 | 13 | R.QLAHQSN[+2204.772]STNIFFSPVSIATAFAMLSLGTK.A | HexNAc(4)Hex(5)NeuAc(2) | N[+2205] | 1796.1529 | 3 | 5386.4442 | 5386.4075 | 0 | 6.8 | 64 | Specific | 666.38 | 657.33 | 657.33 | 13.48 | 34 | >sp\|P01009\|A1AT_HUMAN Alpha-1-antitrypsin OS=Homo sapiens OX=9606 GN=SERPINA1 PE=1 SV=3 | 642 | 20201114_qwl_colo_milk_1ng_glyco_hcd203040_compound15.25535.25535.3 | controllerType=0 controllerNumber=1 scan=25535 | 55.8393 |
| 21872:3 | 13 | R.QLAHQSN[+2204.772]STNIFFSPVSIATAFAMLSLGTK.A | HexNAc(4)Hex(5)NeuAc(2) | N[+2205] | 1796.1529 | 3 | 5386.4442 | 5386.4075 | 0 | 6.8 | 64 | Specific | 682.59 | 628.31 | 628.31 | 13.48 | 34 | >sp\|P01009\|A1AT_HUMAN Alpha-1-antitrypsin OS=Homo sapiens OX=9606 GN=SERPINA1 PE=1 SV=3 | 642 | 20201114_qwl_colo_milk_1ng_glyco_hcd203040_compound15.25537.25537.3 | controllerType=0 controllerNumber=1 scan=25537 | 55.8439 |
| 21873:4 | 13 | R.QLAHQSN[+2204.772]STNIFFSPVSIATAFAMLSLGTK.A | HexNAc(4)Hex(5)NeuAc(2) | N[+2205] | 1347.3665 | 4 | 5386.444 | 5386.4075 | 0 | 6.8 | 64 | Specific | 930.4 | 927.55 | 927.55 | 18.33 | 34 | >sp\|P01009\|A1AT_HUMAN Alpha-1-antitrypsin OS=Homo sapiens OX=9606 GN=SERPINA1 PE=1 SV=3 | 642 | 20201114_qwl_colo_milk_1ng_glyco_hcd203040_compound15.25538.25538.4 | controllerType=0 controllerNumber=1 scan=25538 | 55.8458 |
| 13783:3 | 14 | R.IIVPLNNREN[+2351.851]ISDPTSPLR.T | HexNAc(4)Hex(5)Fuc(3)NeuAc(1) | N[+2352] | 1500.6775 | 3 | 4500.0181 | 4500.0279 | 0 | -2.2 | 62 | Specific | 221.52 | 205.98 | 205.98 | 2.53 | 38 | >sp\|P01591\|IGJ_HUMAN Immunoglobulin J chain OS=Homo sapiens OX=9606 GN=JCHAIN PE=1 SV=4 | 1392 | 20201114_qwl_colo_milk_1ng_glyco_hcd203040_compound15.17063.17063.3 | controllerType=0 controllerNumber=1 scan=17063 | 37.5584 |
| 13785:3 | 14 | R.IIVPLNNREN[+2351.851]ISDPTSPLR.T | HexNAc(4)Hex(5)Fuc(3)NeuAc(1) | N[+2352] | 1500.6775 | 3 | 4500.0181 | 4500.0279 | 0 | -2.2 | 62 | Specific | 240.56 | 232.94 | 232.94 | 3.37 | 38 | >sp\|P01591\|IGJ_HUMAN Immunoglobulin J chain OS=Homo sapiens OX=9606 GN=JCHAIN PE=1 SV=4 | 1392 | 20201114_qwl_colo_milk_1ng_glyco_hcd203040_compound15.17065.17065.3 | controllerType=0 controllerNumber=1 scan=17065 | 37.5626 |
| 12101:3 | 14 | R.IIVPLNNREN[+1856.656]ISDPTSPLR.T | HexNAc(3)Hex(5)Fuc(1)NeuAc(1) | N[+1857] | 1335.6116 | 3 | 4004.8203 | 4004.8327 | 0 | -3.1 | 62 | Specific | 250.4 | 246.78 | 246.78 | 3.64 | 38 | >sp\|P01591\|IGJ_HUMAN Immunoglobulin J chain OS=Homo sapiens OX=9606 GN=JCHAIN PE=1 SV=4 | 1392 | 20201114_qwl_colo_milk_1ng_glyco_hcd203040_compound15.15303.15303.3 | controllerType=0 controllerNumber=1 scan=15303 | 33.8713 |
| 08885:3 | 14 | R.IIVPLNNREN[+1971.719]ISDPTSPLR.T | HexNAc(5)Hex(5)Fuc(1) | N[+1972] | 1373.9738 | 3 | 4119.9069 | 4119.896 | 0 | 2.6 | 62 | Specific | 291.41 | 236.83 | 236.83 | 6.23 | 38 | >sp\|P01591\|IGJ_HUMAN Immunoglobulin J chain OS=Homo sapiens OX=9606 GN=JCHAIN PE=1 SV=4 | 1392 | 20201114_qwl_colo_milk_1ng_glyco_hcd203040_compound15.11946.11946.3 | controllerType=0 controllerNumber=1 scan=11946 | 27.4298 |
| 08962:3 | 14 | R.IIVPLNNREN[+1622.582]ISDPTSPLR.T | HexNAc(4)Hex(5) | N[+1623] | 1257.5998 | 3 | 3770.7848 | 3770.7588 | 0 | 6.9 | 62 | Specific | 332.16 | 321.42 | 321.42 | 6.73 | 38 | >sp\|P01591\|IGJ_HUMAN Immunoglobulin J chain OS=Homo sapiens OX=9606 GN=JCHAIN PE=1 SV=4 | 1392 | 20201114_qwl_colo_milk_1ng_glyco_hcd203040_compound15.12026.12026.3 | controllerType=0 controllerNumber=1 scan=12026 | 27.5776 |
| 08896:3 | 14 | R.IIVPLNNREN[+1622.582]ISDPTSPLR.T | HexNAc(4)Hex(5) | N[+1623] | 1257.5942 | 3 | 3770.768 | 3770.7588 | 0 | 2.4 | 62 | Specific | 358.16 | 332.58 | 332.58 | 7.16 | 38 | >sp\|P01591\|IGJ_HUMAN Immunoglobulin J chain OS=Homo sapiens OX=9606 GN=JCHAIN PE=1 SV=4 | 1392 | 20201114_qwl_colo_milk_1ng_glyco_hcd203040_compound15.11957.11957.3 | controllerType=0 controllerNumber=1 scan=11957 | 27.4518 |
| 08919:3 | 14 | R.IIVPLNNREN[+1460.529]ISDPTSPLR.T | HexNAc(4)Hex(4) | N[+1461] | 1203.5783 | 3 | 3608.7204 | 3608.7059 | 0 | 4 | 62 | Specific | 358.45 | 346.35 | 346.35 | 7.16 | 38 | >sp\|P01591\|IGJ_HUMAN Immunoglobulin J chain OS=Homo sapiens OX=9606 GN=JCHAIN PE=1 SV=4 | 1392 | 20201114_qwl_colo_milk_1ng_glyco_hcd203040_compound15.11981.11981.3 | controllerType=0 controllerNumber=1 scan=11981 | 27.4936 |
| 08964:3 | 14 | R.IIVPLNNREN[+1622.582]ISDPTSPLR.T | HexNAc(4)Hex(5) | N[+1623] | 1257.5998 | 3 | 3770.7848 | 3770.7588 | 0 | 6.9 | 62 | Specific | 366.24 | 295.36 | 295.36 | 6.57 | 38 | >sp\|P01591\|IGJ_HUMAN Immunoglobulin J chain OS=Homo sapiens OX=9606 GN=JCHAIN PE=1 SV=4 | 1392 | 20201114_qwl_colo_milk_1ng_glyco_hcd203040_compound15.12028.12028.3 | controllerType=0 controllerNumber=1 scan=12028 | 27.5813 |
| 08887:3 | 14 | R.IIVPLNNREN[+1971.719]ISDPTSPLR.T | HexNAc(5)Hex(5)Fuc(1) | N[+1972] | 1373.9738 | 3 | 4119.9069 | 4119.896 | 0 | 2.6 | 62 | Specific | 408.11 | 365.29 | 365.29 | 7.68 | 38 | >sp\|P01591\|IGJ_HUMAN Immunoglobulin J chain OS=Homo sapiens OX=9606 GN=JCHAIN PE=1 SV=4 | 1392 | 20201114_qwl_colo_milk_1ng_glyco_hcd203040_compound15.11948.11948.3 | controllerType=0 controllerNumber=1 scan=11948 | 27.4337 |
| 08930:3 | 14 | R.IIVPLNNREN[+1565.560]ISDPTSPLR.T | HexNAc(3)Hex(5)Fuc(1) | N[+1566] | 1238.5882 | 3 | 3713.75 | 3713.7373 | 0 | 3.4 | 62 | Specific | 409.82 | 360.21 | 360.21 | 7.65 | 38 | >sp\|P01591\|IGJ_HUMAN Immunoglobulin J chain OS=Homo sapiens OX=9606 GN=JCHAIN PE=1 SV=4 | 1392 | 20201114_qwl_colo_milk_1ng_glyco_hcd203040_compound15.11993.11993.3 | controllerType=0 controllerNumber=1 scan=11993 | 27.5171 |
| 08966:3 | 14 | R.IIVPLNNREN[+1257.449]ISDPTSPLR.T | HexNAc(3)Hex(4) | N[+1257] | 1135.8874 | 3 | 3405.6475 | 3405.6266 | 0 | 6.2 | 62 | Specific | 418.31 | 418.31 | 418.31 | 8.5 | 38 | >sp\|P01591\|IGJ_HUMAN Immunoglobulin J chain OS=Homo sapiens OX=9606 GN=JCHAIN PE=1 SV=4 | 1392 | 20201114_qwl_colo_milk_1ng_glyco_hcd203040_compound15.12030.12030.3 | controllerType=0 controllerNumber=1 scan=12030 | 27.5855 |
| 10546:3 | 14 | R.IIVPLNNREN[+1548.545]ISDPTSPLR.T | HexNAc(3)Hex(4)NeuAc(1) | N[+1549] | 1232.9233 | 3 | 3696.7553 | 3696.722 | 0 | 9 | 62 | Specific | 475.69 | 475.69 | 475.69 | 8.71 | 38 | >sp\|P01591\|IGJ_HUMAN Immunoglobulin J chain OS=Homo sapiens OX=9606 GN=JCHAIN PE=1 SV=4 | 1392 | 20201114_qwl_colo_milk_1ng_glyco_hcd203040_compound15.13678.13678.3 | controllerType=0 controllerNumber=1 scan=13678 | 30.7334 |
| 08925:3 | 14 | R.IIVPLNNREN[+1768.640]ISDPTSPLR.T | HexNAc(4)Hex(5)Fuc(1) | N[+1769] | 1306.2814 | 3 | 3916.8297 | 3916.8167 | 0 | 3.3 | 62 | Specific | 495.1 | 458.58 | 458.58 | 9.39 | 38 | >sp\|P01591\|IGJ_HUMAN Immunoglobulin J chain OS=Homo sapiens OX=9606 GN=JCHAIN PE=1 SV=4 | 1392 | 20201114_qwl_colo_milk_1ng_glyco_hcd203040_compound15.11988.11988.3 | controllerType=0 controllerNumber=1 scan=11988 | 27.5065 |
| 08968:3 | 14 | R.IIVPLNNREN[+1257.449]ISDPTSPLR.T | HexNAc(3)Hex(4) | N[+1257] | 1135.8874 | 3 | 3405.6475 | 3405.6266 | 0 | 6.2 | 62 | Specific | 497.69 | 476.03 | 476.03 | 8.87 | 38 | >sp\|P01591\|IGJ_HUMAN Immunoglobulin J chain OS=Homo sapiens OX=9606 GN=JCHAIN PE=1 SV=4 | 1392 | 20201114_qwl_colo_milk_1ng_glyco_hcd203040_compound15.12032.12032.3 | controllerType=0 controllerNumber=1 scan=12032 | 27.5892 |
| 10464:3 | 14 | R.IIVPLNNREN[+1856.656]ISDPTSPLR.T | HexNAc(3)Hex(5)Fuc(1)NeuAc(1) | N[+1857] | 1335.6213 | 3 | 4004.8493 | 4004.8327 | 0 | 4.1 | 62 | Specific | 515.81 | 515.81 | 515.81 | 9.42 | 38 | >sp\|P01591\|IGJ_HUMAN Immunoglobulin J chain OS=Homo sapiens OX=9606 GN=JCHAIN PE=1 SV=4 | 1392 | 20201114_qwl_colo_milk_1ng_glyco_hcd203040_compound15.13593.13593.3 | controllerType=0 controllerNumber=1 scan=13593 | 30.5736 |
| 08853:3 | 14 | R.IIVPLNNREN[+1809.666]ISDPTSPLR.T | HexNAc(5)Hex(4)Fuc(1) | N[+1810] | 1319.9533 | 3 | 3957.8454 | 3957.8432 | 0 | 0.5 | 62 | Specific | 516.17 | 417.18 | 417.18 | 9.3 | 38 | >sp\|P01591\|IGJ_HUMAN Immunoglobulin J chain OS=Homo sapiens OX=9606 GN=JCHAIN PE=1 SV=4 | 1392 | 20201114_qwl_colo_milk_1ng_glyco_hcd203040_compound15.11913.11913.3 | controllerType=0 controllerNumber=1 scan=11913 | 27.366 |
| 08894:3 | 14 | R.IIVPLNNREN[+1622.582]ISDPTSPLR.T | HexNAc(4)Hex(5) | N[+1623] | 1257.5942 | 3 | 3770.768 | 3770.7588 | 0 | 2.4 | 62 | Specific | 517.54 | 474.11 | 474.11 | 9.46 | 38 | >sp\|P01591\|IGJ_HUMAN Immunoglobulin J chain OS=Homo sapiens OX=9606 GN=JCHAIN PE=1 SV=4 | 1392 | 20201114_qwl_colo_milk_1ng_glyco_hcd203040_compound15.11955.11955.3 | controllerType=0 controllerNumber=1 scan=11955 | 27.4475 |
| 08922:3 | 14 | R.IIVPLNNREN[+1460.529]ISDPTSPLR.T | HexNAc(4)Hex(4) | N[+1461] | 1203.5783 | 3 | 3608.7204 | 3608.7059 | 0 | 4 | 62 | Specific | 523.53 | 490.82 | 490.82 | 9.42 | 38 | >sp\|P01591\|IGJ_HUMAN Immunoglobulin J chain OS=Homo sapiens OX=9606 GN=JCHAIN PE=1 SV=4 | 1392 | 20201114_qwl_colo_milk_1ng_glyco_hcd203040_compound15.11984.11984.3 | controllerType=0 controllerNumber=1 scan=11984 | 27.4978 |
| 08974:3 | 14 | R.IIVPLNNREN[+1647.613]ISDPTSPLR.T | HexNAc(5)Hex(3)Fuc(1) | N[+1648] | 1265.9439 | 3 | 3795.8172 | 3795.7904 | 0 | 7.1 | 62 | Specific | 524.78 | 500.73 | 500.73 | 9.07 | 38 | >sp\|P01591\|IGJ_HUMAN Immunoglobulin J chain OS=Homo sapiens OX=9606 GN=JCHAIN PE=1 SV=4 | 1392 | 20201114_qwl_colo_milk_1ng_glyco_hcd203040_compound15.12038.12038.3 | controllerType=0 controllerNumber=1 scan=12038 | 27.6049 |
| 08892:3 | 14 | R.IIVPLNNREN[+1663.608]ISDPTSPLR.T | HexNAc(5)Hex(4) | N[+1664] | 1271.2691 | 3 | 3811.7927 | 3811.7853 | 0 | 2 | 62 | Specific | 530.79 | 529.16 | 529.16 | 9.35 | 38 | >sp\|P01591\|IGJ_HUMAN Immunoglobulin J chain OS=Homo sapiens OX=9606 GN=JCHAIN PE=1 SV=4 | 1392 | 20201114_qwl_colo_milk_1ng_glyco_hcd203040_compound15.11953.11953.3 | controllerType=0 controllerNumber=1 scan=11953 | 27.444 |
| 08855:3 | 14 | R.IIVPLNNREN[+1809.666]ISDPTSPLR.T | HexNAc(5)Hex(4)Fuc(1) | N[+1810] | 1319.9533 | 3 | 3957.8454 | 3957.8432 | 0 | 0.5 | 62 | Specific | 531.32 | 451.52 | 451.52 | 9.3 | 38 | >sp\|P01591\|IGJ_HUMAN Immunoglobulin J chain OS=Homo sapiens OX=9606 GN=JCHAIN PE=1 SV=4 | 1392 | 20201114_qwl_colo_milk_1ng_glyco_hcd203040_compound15.11915.11915.3 | controllerType=0 controllerNumber=1 scan=11915 | 27.3718 |
| 10417:3 | 14 | R.IIVPLNNREN[+1856.656]ISDPTSPLR.T | HexNAc(3)Hex(5)Fuc(1)NeuAc(1) | N[+1857] | 1335.6181 | 3 | 4004.8397 | 4004.8327 | 0 | 1.8 | 62 | Specific | 531.93 | 531.93 | 531.93 | 9.35 | 38 | >sp\|P01591\|IGJ_HUMAN Immunoglobulin J chain OS=Homo sapiens OX=9606 GN=JCHAIN PE=1 SV=4 | 1392 | 20201114_qwl_colo_milk_1ng_glyco_hcd203040_compound15.13544.13544.3 | controllerType=0 controllerNumber=1 scan=13544 | 30.458 |
| 08927:3 | 14 | R.IIVPLNNREN[+1768.640]ISDPTSPLR.T | HexNAc(4)Hex(5)Fuc(1) | N[+1769] | 1306.2814 | 3 | 3916.8297 | 3916.8167 | 0 | 3.3 | 62 | Specific | 537.83 | 493.64 | 493.64 | 9.66 | 38 | >sp\|P01591\|IGJ_HUMAN Immunoglobulin J chain OS=Homo sapiens OX=9606 GN=JCHAIN PE=1 SV=4 | 1392 | 20201114_qwl_colo_milk_1ng_glyco_hcd203040_compound15.11990.11990.3 | controllerType=0 controllerNumber=1 scan=11990 | 27.5123 |
| 08932:3 | 14 | R.IIVPLNNREN[+1565.560]ISDPTSPLR.T | HexNAc(3)Hex(5)Fuc(1) | N[+1566] | 1238.5882 | 3 | 3713.75 | 3713.7373 | 0 | 3.4 | 62 | Specific | 540.22 | 444.6 | 444.6 | 9.63 | 38 | >sp\|P01591\|IGJ_HUMAN Immunoglobulin J chain OS=Homo sapiens OX=9606 GN=JCHAIN PE=1 SV=4 | 1392 | 20201114_qwl_colo_milk_1ng_glyco_hcd203040_compound15.11995.11995.3 | controllerType=0 controllerNumber=1 scan=11995 | 27.5211 |
| 10487:3 | 14 | R.IIVPLNNREN[+1694.603]ISDPTSPLR.T | HexNAc(3)Hex(4)Fuc(1)NeuAc(1) | N[+1695] | 1281.6024 | 3 | 3842.7925 | 3842.7799 | 0 | 3.3 | 62 | Specific | 546.17 | 546.17 | 546.17 | 9.66 | 38 | >sp\|P01591\|IGJ_HUMAN Immunoglobulin J chain OS=Homo sapiens OX=9606 GN=JCHAIN PE=1 SV=4 | 1392 | 20201114_qwl_colo_milk_1ng_glyco_hcd203040_compound15.13617.13617.3 | controllerType=0 controllerNumber=1 scan=13617 | 30.623 |
| 10467:3 | 14 | R.IIVPLNNREN[+1856.656]ISDPTSPLR.T | HexNAc(3)Hex(5)Fuc(1)NeuAc(1) | N[+1857] | 1335.6213 | 3 | 4004.8493 | 4004.8327 | 0 | 4.1 | 62 | Specific | 559.47 | 501.33 | 501.33 | 9.73 | 38 | >sp\|P01591\|IGJ_HUMAN Immunoglobulin J chain OS=Homo sapiens OX=9606 GN=JCHAIN PE=1 SV=4 | 1392 | 20201114_qwl_colo_milk_1ng_glyco_hcd203040_compound15.13596.13596.3 | controllerType=0 controllerNumber=1 scan=13596 | 30.5823 |
| 10332:3 | 14 | R.IIVPLNNREN[+1913.677]ISDPTSPLR.T | HexNAc(4)Hex(5)NeuAc(1) | N[+1914] | 1354.6345 | 3 | 4061.8889 | 4061.8542 | 0 | 8.6 | 62 | Specific | 562.06 | 536 | 536 | 9.3 | 38 | >sp\|P01591\|IGJ_HUMAN Immunoglobulin J chain OS=Homo sapiens OX=9606 GN=JCHAIN PE=1 SV=4 | 1392 | 20201114_qwl_colo_milk_1ng_glyco_hcd203040_compound15.13454.13454.3 | controllerType=0 controllerNumber=1 scan=13454 | 30.2704 |
| 08972:3 | 14 | R.IIVPLNNREN[+1647.613]ISDPTSPLR.T | HexNAc(5)Hex(3)Fuc(1) | N[+1648] | 1265.9439 | 3 | 3795.8172 | 3795.7904 | 0 | 7.1 | 62 | Specific | 563.08 | 540.75 | 540.75 | 9.38 | 38 | >sp\|P01591\|IGJ_HUMAN Immunoglobulin J chain OS=Homo sapiens OX=9606 GN=JCHAIN PE=1 SV=4 | 1392 | 20201114_qwl_colo_milk_1ng_glyco_hcd203040_compound15.12036.12036.3 | controllerType=0 controllerNumber=1 scan=12036 | 27.5969 |
| 10543:3 | 14 | R.IIVPLNNREN[+1548.545]ISDPTSPLR.T | HexNAc(3)Hex(4)NeuAc(1) | N[+1549] | 1232.9233 | 3 | 3696.7553 | 3696.722 | 0 | 9 | 62 | Specific | 565.38 | 565.38 | 565.38 | 9.17 | 38 | >sp\|P01591\|IGJ_HUMAN Immunoglobulin J chain OS=Homo sapiens OX=9606 GN=JCHAIN PE=1 SV=4 | 1392 | 20201114_qwl_colo_milk_1ng_glyco_hcd203040_compound15.13675.13675.3 | controllerType=0 controllerNumber=1 scan=13675 | 30.7244 |
| 08875:3 | 14 | R.IIVPLNNREN[+1809.666]ISDPTSPLR.T | HexNAc(5)Hex(4)Fuc(1) | N[+1810] | 1319.9555 | 3 | 3957.852 | 3957.8432 | 0 | 2.2 | 62 | Specific | 567.02 | 451.8 | 451.8 | 9.66 | 38 | >sp\|P01591\|IGJ_HUMAN Immunoglobulin J chain OS=Homo sapiens OX=9606 GN=JCHAIN PE=1 SV=4 | 1392 | 20201114_qwl_colo_milk_1ng_glyco_hcd203040_compound15.11936.11936.3 | controllerType=0 controllerNumber=1 scan=11936 | 27.4141 |
| 10334:3 | 14 | R.IIVPLNNREN[+1913.677]ISDPTSPLR.T | HexNAc(4)Hex(5)NeuAc(1) | N[+1914] | 1354.6345 | 3 | 4061.8889 | 4061.8542 | 0 | 8.6 | 62 | Specific | 570.87 | 570.87 | 570.87 | 9.3 | 38 | >sp\|P01591\|IGJ_HUMAN Immunoglobulin J chain OS=Homo sapiens OX=9606 GN=JCHAIN PE=1 SV=4 | 1392 | 20201114_qwl_colo_milk_1ng_glyco_hcd203040_compound15.13456.13456.3 | controllerType=0 controllerNumber=1 scan=13456 | 30.2771 |
| 08873:3 | 14 | R.IIVPLNNREN[+1809.666]ISDPTSPLR.T | HexNAc(5)Hex(4)Fuc(1) | N[+1810] | 1319.9555 | 3 | 3957.852 | 3957.8432 | 0 | 2.2 | 62 | Specific | 571.47 | 447.16 | 447.16 | 9.66 | 38 | >sp\|P01591\|IGJ_HUMAN Immunoglobulin J chain OS=Homo sapiens OX=9606 GN=JCHAIN PE=1 SV=4 | 1392 | 20201114_qwl_colo_milk_1ng_glyco_hcd203040_compound15.11934.11934.3 | controllerType=0 controllerNumber=1 scan=11934 | 27.4083 |
| 08890:3 | 14 | R.IIVPLNNREN[+1663.608]ISDPTSPLR.T | HexNAc(5)Hex(4) | N[+1664] | 1271.2691 | 3 | 3811.7927 | 3811.7853 | 0 | 2 | 62 | Specific | 575.67 | 575.26 | 575.26 | 9.66 | 38 | >sp\|P01591\|IGJ_HUMAN Immunoglobulin J chain OS=Homo sapiens OX=9606 GN=JCHAIN PE=1 SV=4 | 1392 | 20201114_qwl_colo_milk_1ng_glyco_hcd203040_compound15.11951.11951.3 | controllerType=0 controllerNumber=1 scan=11951 | 27.4386 |
| 10420:3 | 14 | R.IIVPLNNREN[+1856.656]ISDPTSPLR.T | HexNAc(3)Hex(5)Fuc(1)NeuAc(1) | N[+1857] | 1335.6181 | 3 | 4004.8397 | 4004.8327 | 0 | 1.8 | 62 | Specific | 583.04 | 583.04 | 583.04 | 10.97 | 38 | >sp\|P01591\|IGJ_HUMAN Immunoglobulin J chain OS=Homo sapiens OX=9606 GN=JCHAIN PE=1 SV=4 | 1392 | 20201114_qwl_colo_milk_1ng_glyco_hcd203040_compound15.13547.13547.3 | controllerType=0 controllerNumber=1 scan=13547 | 30.4705 |
| 08996:3 | 14 | R.IIVPLNNREN[+1501.555]ISDPTSPLR.T | HexNAc(5)Hex(3) | N[+1502] | 1217.2509 | 3 | 3649.7382 | 3649.7325 | 0 | 1.6 | 62 | Specific | 595.61 | 582.81 | 582.81 | 10.97 | 38 | >sp\|P01591\|IGJ_HUMAN Immunoglobulin J chain OS=Homo sapiens OX=9606 GN=JCHAIN PE=1 SV=4 | 1392 | 20201114_qwl_colo_milk_1ng_glyco_hcd203040_compound15.12062.12062.3 | controllerType=0 controllerNumber=1 scan=12062 | 27.6597 |
| 08998:3 | 14 | R.IIVPLNNREN[+1501.555]ISDPTSPLR.T | HexNAc(5)Hex(3) | N[+1502] | 1217.2509 | 3 | 3649.7382 | 3649.7325 | 0 | 1.6 | 62 | Specific | 626.53 | 592.04 | 592.04 | 11.97 | 38 | >sp\|P01591\|IGJ_HUMAN Immunoglobulin J chain OS=Homo sapiens OX=9606 GN=JCHAIN PE=1 SV=4 | 1392 | 20201114_qwl_colo_milk_1ng_glyco_hcd203040_compound15.12064.12064.3 | controllerType=0 controllerNumber=1 scan=12064 | 27.6642 |
| 04566:2 | 14 | R.EN[+1663.608]ISDPTSPLR.T | HexNAc(5)Hex(4) | N[+1664] | 1446.6296 | 2 | 2892.2518 | 2892.225 | 0 | 9.3 | 70 | Specific | 214.61 | 142.86 | 142.86 | 3.43 | 38 | >sp\|P01591\|IGJ_HUMAN Immunoglobulin J chain OS=Homo sapiens OX=9606 GN=JCHAIN PE=1 SV=4 | 1392 | 20201114_qwl_colo_milk_1ng_glyco_hcd203040_compound15.7451.7451.2 | controllerType=0 controllerNumber=1 scan=7451 | 19.5491 |
| 04564:2 | 14 | R.EN[+1663.608]ISDPTSPLR.T | HexNAc(5)Hex(4) | N[+1664] | 1446.6296 | 2 | 2892.2518 | 2892.225 | 0 | 9.3 | 70 | Specific | 223.36 | 187.98 | 187.98 | 5.09 | 38 | >sp\|P01591\|IGJ_HUMAN Immunoglobulin J chain OS=Homo sapiens OX=9606 GN=JCHAIN PE=1 SV=4 | 1392 | 20201114_qwl_colo_milk_1ng_glyco_hcd203040_compound15.7449.7449.2 | controllerType=0 controllerNumber=1 scan=7449 | 19.5452 |
| 01854:3 | 15 | K.N[+1216.423]NSIPDKQITASSSYK.T | HexNAc(2)Hex(5) | N[+1216] | 990.4424 | 3 | 2969.3125 | 2969.2991 | 0 | 4.5 | 238 | Specific | 266.94 | 218.83 | 218.83 | 5.45 | 29 | >sp\|Q08431\|MFGM_HUMAN Lactadherin OS=Homo sapiens OX=9606 GN=MFGE8 PE=1 SV=3 | 1064 | 20201114_qwl_colo_milk_1ng_glyco_hcd203040_compound15.4615.4615.3 | controllerType=0 controllerNumber=1 scan=4615 | 14.456 |
| 01856:3 | 15 | K.N[+1216.423]NSIPDKQITASSSYK.T | HexNAc(2)Hex(5) | N[+1216] | 990.4424 | 3 | 2969.3125 | 2969.2991 | 0 | 4.5 | 238 | Specific | 310.96 | 179.57 | 179.57 | 5.24 | 29 | >sp\|Q08431\|MFGM_HUMAN Lactadherin OS=Homo sapiens OX=9606 GN=MFGE8 PE=1 SV=3 | 1064 | 20201114_qwl_colo_milk_1ng_glyco_hcd203040_compound15.4617.4617.3 | controllerType=0 controllerNumber=1 scan=4617 | 14.46 |
| 19393:4 | 16 | K.APDKNVIFSPLSISTALAFLSLGAHN[+2059.735]TTLTEILK.G | HexNAc(4)Hex(5)Fuc(1)NeuAc(1) | N[+2060] | 1411.4446 | 4 | 5642.7565 | 5642.7131 | 0 | 7.7 | 68 | Specific | 300.21 | 300.21 | 300.21 | 6.81 | 26 | >sp\|P01011\|AACT_HUMAN Alpha-1-antichymotrypsin OS=Homo sapiens OX=9606 GN=SERPINA3 PE=1 SV=2 | 1314 | 20201114_qwl_colo_milk_1ng_glyco_hcd203040_compound15.22948.22948.4 | controllerType=0 controllerNumber=1 scan=22948 | 50.6695 |
| 20711:4 | 16 | K.APDKNVIFSPLSISTALAFLSLGAHN[+2351.851]TTLTEILK.G | HexNAc(4)Hex(5)Fuc(3)NeuAc(1) | N[+2352] | 1484.4691 | 4 | 5934.8546 | 5934.8289 | 0 | 4.3 | 68 | Specific | 383.62 | 383.62 | 383.62 | 9.5 | 26 | >sp\|P01011\|AACT_HUMAN Alpha-1-antichymotrypsin OS=Homo sapiens OX=9606 GN=SERPINA3 PE=1 SV=2 | 1314 | 20201114_qwl_colo_milk_1ng_glyco_hcd203040_compound15.24322.24322.4 | controllerType=0 controllerNumber=1 scan=24322 | 53.4793 |
| 05152:3 | 18 | R.LSPN[+2205.793]ASAEHLELR.W | HexNAc(4)Hex(5)Fuc(2)NeuAc(1) | N[+2206] | 1214.8548 | 3 | 3642.5498 | 3642.5421 | 0 | 2.1 | 52 | Specific | 271.56 | 87.56 | 87.56 | 4.57 | 25 | >sp\|Q13410\|BT1A1_HUMAN Butyrophilin subfamily 1 member A1 OS=Homo sapiens OX=9606 GN=BTN1A1 PE=1 SV=3 | 179 | 20201114_qwl_colo_milk_1ng_glyco_hcd203040_compound15.8057.8057.3 | controllerType=0 controllerNumber=1 scan=8057 | 20.5772 |
| 05154:3 | 18 | R.LSPN[+2205.793]ASAEHLELR.W | HexNAc(4)Hex(5)Fuc(2)NeuAc(1) | N[+2206] | 1214.8548 | 3 | 3642.5498 | 3642.5421 | 0 | 2.1 | 52 | Specific | 376.77 | 375.84 | 375.84 | 7.46 | 25 | >sp\|Q13410\|BT1A1_HUMAN Butyrophilin subfamily 1 member A1 OS=Homo sapiens OX=9606 GN=BTN1A1 PE=1 SV=3 | 179 | 20201114_qwl_colo_milk_1ng_glyco_hcd203040_compound15.8059.8059.3 | controllerType=0 controllerNumber=1 scan=8059 | 20.5806 |
| 01735:3 | 22 | R.LRN[+1216.423]VSWATGR.S | HexNAc(2)Hex(5) | N[+1216] | 792.6918 | 3 | 2376.0608 | 2376.0559 | 0 | 2.1 | 149 | Specific | 236.53 | 61.32 | 61.32 | 3.59 | 17 | >sp\|P08571\|CD14_HUMAN Monocyte differentiation antigen CD14 OS=Homo sapiens OX=9606 GN=CD14 PE=1 SV=2 | 1232 | 20201114_qwl_colo_milk_1ng_glyco_hcd203040_compound15.4491.4491.3 | controllerType=0 controllerNumber=1 scan=4491 | 14.2294 |
| 03512:2 | 23 | K.AAIPSALDTN[+1216.423]SSK.S | HexNAc(2)Hex(5) | N[+1216] | 1246.0492 | 2 | 2491.0912 | 2491.0815 | 0 | 3.9 | 542 | Specific | 401.13 | 294.88 | 294.88 | 7.17 | 17 | >sp\|Q08380\|LG3BP_HUMAN Galectin-3-binding protein OS=Homo sapiens OX=9606 GN=LGALS3BP PE=1 SV=1 | 1562 | 20201114_qwl_colo_milk_1ng_glyco_hcd203040_compound15.6343.6343.2 | controllerType=0 controllerNumber=1 scan=6343 | 17.6286 |
| 10019:3 | 40 | R.ADGTVNQIEGEATPVN[+1054.370]LTEPAKLEVK.F | HexNAc(2)Hex(4) | N[+1054] | 1259.9347 | 3 | 3777.7897 | 3777.7798 | 0 | 2.6 | 83 | Specific | 292.15 | 292.15 | 292.15 | 6.85 | 12 | >sp\|P05090\|APOD_HUMAN Apolipoprotein D OS=Homo sapiens OX=9606 GN=APOD PE=1 SV=1 | 594 | 20201114_qwl_colo_milk_1ng_glyco_hcd203040_compound15.13129.13129.3 | controllerType=0 controllerNumber=1 scan=13129 | 29.6504 |
| 10021:3 | 40 | R.ADGTVNQIEGEATPVN[+1054.370]LTEPAKLEVK.F | HexNAc(2)Hex(4) | N[+1054] | 1259.9347 | 3 | 3777.7897 | 3777.7798 | 0 | 2.6 | 83 | Specific | 605.24 | 577.07 | 577.07 | 12.24 | 12 | >sp\|P05090\|APOD_HUMAN Apolipoprotein D OS=Homo sapiens OX=9606 GN=APOD PE=1 SV=1 | 594 | 20201114_qwl_colo_milk_1ng_glyco_hcd203040_compound15.13131.13131.3 | controllerType=0 controllerNumber=1 scan=13131 | 29.6543 |
| 18253:4 | 1 | R.RTAGWNVPIGTLRPFLN[+2570.925]WTGPPEPIEAAVAR.F | HexNAc(5)Hex(6)Fuc(2)NeuAc(1) | N[+2571] | 1490.1986 | 4 | 5957.7726 | 5957.7371 | 0 | 6 | 140 | Specific | 250.73 | 250.73 | 250.73 | 5.72 | 140 | >sp\|P02788\|TRFL_HUMAN Lactotransferrin OS=Homo sapiens OX=9606 GN=LTF PE=1 SV=6 | 1270 | 20201114_qwl_milk_1ng_glyco_hcd203040_compound15.21633.21633.4 | controllerType=0 controllerNumber=1 scan=21633 | 43.3866 |
| 17410:4 | 1 | R.RTAGWNVPIGTLRPFLN[+1768.640]WTGPPEPIEAAVAR.F | HexNAc(4)Hex(5)Fuc(1) | N[+1769] | 1289.6244 | 4 | 5155.4758 | 5155.4516 | 0 | 4.7 | 140 | Specific | 275.7 | 228.43 | 228.43 | 6.13 | 140 | >sp\|P02788\|TRFL_HUMAN Lactotransferrin OS=Homo sapiens OX=9606 GN=LTF PE=1 SV=6 | 1270 | 20201114_qwl_milk_1ng_glyco_hcd203040_compound15.20752.20752.4 | controllerType=0 controllerNumber=1 scan=20752 | 41.5423 |
| 18256:4 | 1 | R.RTAGWNVPIGTLRPFLN[+2570.925]WTGPPEPIEAAVAR.F | HexNAc(5)Hex(6)Fuc(2)NeuAc(1) | N[+2571] | 1490.1986 | 4 | 5957.7726 | 5957.7371 | 0 | 6 | 140 | Specific | 533.08 | 424.87 | 424.87 | 9.1 | 140 | >sp\|P02788\|TRFL_HUMAN Lactotransferrin OS=Homo sapiens OX=9606 GN=LTF PE=1 SV=6 | 1270 | 20201114_qwl_milk_1ng_glyco_hcd203040_compound15.21636.21636.4 | controllerType=0 controllerNumber=1 scan=21636 | [illegible] |
| 18306:4 | 1 | R.RTAGWNVPIGTLRPFLN[+2205.793]WTGPPEPIEAAVAR.F | HexNAc(4)Hex(5)Fuc(2)NeuAc(1) | N[+2206] | 1398.9038 | 4 | 5592.5934 | 5592.6049 | 0 | -2.1 | 140 | Specific | 537.5 | 462.63 | 462.63 | 8.67 | 140 | >sp\|P02788\|TRFL_HUMAN Lactotransferrin OS=Homo sapiens OX=9606 GN=LTF PE=1 SV=6 | 1270 | 20201114_qwl_milk_1ng_glyco_hcd203040_compound15.21689.21689.4 | controllerType=0 controllerNumber=1 scan=21689 | [illegible] |
| 18302:4 | 1 | R.RTAGWNVPIGTLRPFLN[+2059.735]WTGPPEPIEAAVAR.F | HexNAc(4)Hex(5)Fuc(1)NeuAc(1) | N[+2060] | 1362.3965 | 4 | 5446.5641 | 5446.547 | 0 | 3.1 | 140 | Specific | 687.07 | 532.78 | 532.78 | 13.15 | 140 | >sp\|P02788\|TRFL_HUMAN Lactotransferrin OS=Homo sapiens OX=9606 GN=LTF PE=1 SV=6 | 1270 | 20201114_qwl_milk_1ng_glyco_hcd203040_compound15.21684.21684.2 | controllerType=0 controllerNumber=1 scan=21684 | [illegible] |
| 18305:4 | 1 | R.RTAGWNVPIGTLRPFLN[+2059.735]WTGPPEPIEAAVAR.F | HexNAc(4)Hex(5)Fuc(1)NeuAc(1) | N[+2060] | 1362.3965 | 4 | 5446.5641 | 5446.547 | 0 | 3.1 | 140 | Specific | 753.02 | 657.62 | 657.62 | 15.15 | 140 | >sp\|P02788\|TRFL_HUMAN Lactotransferrin OS=Homo sapiens OX=9606 GN=LTF PE=1 SV=6 | 1270 | 20201114_qwl_milk_1ng_glyco_hcd203040_compound15.21687.21687.2 | controllerType=0 controllerNumber=1 scan=21687 | [illegible] |
| 18544:3 | 1 | R.[+42.011]TAGWNVPIGTLRPFLN[+2018.708]WTGPPEPIEAAVAR.F | HexNAc(3)Hex(6)Fuc(1)NeuAc(1) | .[42], N[+2019] | 1764.4979 | 3 | 5291.4791 | 5291.4299 | 0 | 9.3 | 141 | Specific | 239.64 | 239.64 | 239.64 | 4.29 | 140 | >sp\|P02788\|TRFL_HUMAN Lactotransferrin OS=Homo sapiens OX=9606 GN=LTF PE=1 SV=6 | 1270 | 20201114_qwl_milk_1ng_glyco_hcd203040_compound15.21939.21939.3 | controllerType=0 controllerNumber=1 scan=21939 | 44.0569 |
| 19514:3 | 1 | R.TAGWNVPIGTLRPFLN[+2205.793]WTGPPEPIEAAVAR.F | HexNAc(4)Hex(5)Fuc(2)NeuAc(1) | N[+2206] | 1812.8568 | 3 | 5436.5558 | 5436.5038 | 0 | 9.6 | 141 | Specific | 322.7 | 305.23 | 305.23 | 6.29 | 140 | >sp\|P02788\|TRFL_HUMAN Lactotransferrin OS=Homo sapiens OX=9606 GN=LTF PE=1 SV=6 | 1270 | 20201114_qwl_milk_1ng_glyco_hcd203040_compound15.22955.22955.2 | controllerType=0 controllerNumber=1 scan=22955 | 46.3037 |
| 19349:3 | 1 | R.TAGWNVPIGTLRPFLN[+2716.983]WTGPPEPIEAAVAR.F | HexNAc(5)Hex(6)Fuc(3)NeuAc(1) | N[+2717] | 1983.2343 | 3 | 5947.6885 | 5947.6939 | 0 | -0.9 | 141 | Specific | 339.95 | 309.61 | 309.61 | 6.76 | 140 | >sp\|P02788\|TRFL_HUMAN Lactotransferrin OS=Homo sapiens OX=9606 GN=LTF PE=1 SV=6 | 1270 | 20201114_qwl_milk_1ng_glyco_hcd203040_compound15.22783.22783.3 | controllerType=0 controllerNumber=1 scan=22783 | 45.9242 |
| 18497:3 | 1 | R.TAGWNVPIGTLRPFLN[+2425.887]WTGPPEPIEAAVAR.F | HexNAc(5)Hex(6)Fuc(3) | N[+2426] | 1886.1997 | 3 | 5656.5845 | 5656.5985 | 0 | -2.5 | 141 | Specific | 340.17 | 207.24 | 207.24 | 5.59 | 140 | >sp\|P02788\|TRFL_HUMAN Lactotransferrin OS=Homo sapiens OX=9606 GN=LTF PE=1 SV=6 | 1270 | 20201114_qwl_milk_1ng_glyco_hcd203040_compound15.21890.21890.3 | controllerType=0 controllerNumber=1 scan=21890 | 43.9504 |
| 18561:4 | 1 | R.TAGWNVPIGTLRPFLN[+2279.830]WTGPPEPIEAAVAR.F | HexNAc(5)Hex(6)Fuc(2) | N[+2280] | 1378.3958 | 4 | 5510.5612 | 5510.5406 | 0 | 3.7 | 141 | Specific | 367.75 | 295.27 | 295.27 | 7.36 | 140 | >sp\|P02788\|TRFL_HUMAN Lactotransferrin OS=Homo sapiens OX=9606 GN=LTF PE=1 SV=6 | 1270 | 20201114_qwl_milk_1ng_glyco_hcd203040_compound15.21956.21956.4 | controllerType=0 controllerNumber=1 scan=21956 | 44.0876 |
| 18495:4 | 1 | R.TAGWNVPIGTLRPFLN[+2425.887]WTGPPEPIEAAVAR.F | HexNAc(5)Hex(6)Fuc(3) | N[+2426] | 1414.9064 | 4 | 5656.6037 | 5656.5985 | 0 | 0.9 | 141 | Specific | 379.88 | 217.06 | 217.06 | 6.87 | 140 | >sp\|P02788\|TRFL_HUMAN Lactotransferrin OS=Homo sapiens OX=9606 GN=LTF PE=1 SV=6 | 1270 | 20201114_qwl_milk_1ng_glyco_hcd203040_compound15.21888.21888.4 | controllerType=0 controllerNumber=1 scan=21888 | 43.9479 |
| 19703:3 | 1 | R.TAGWNVPIGTLRPFLN[+2059.735]WTGPPEPIEAAVAR.F | HexNAc(4)Hex(5)Fuc(1)NeuAc(1) | N[+2060] | 1764.1607 | 3 | 5290.4676 | 5290.4459 | 0 | 4.1 | 141 | Specific | 384.88 | 344.61 | 344.61 | 7.21 | 140 | >sp\|P02788\|TRFL_HUMAN Lactotransferrin OS=Homo sapiens OX=9606 GN=LTF PE=1 SV=6 | 1270 | 20201114_qwl_milk_1ng_glyco_hcd203040_compound15.23153.23153.3 | controllerType=0 controllerNumber=1 scan=23153 | 46.6954 |
| 19983:3 | 1 | R.TAGWNVPIGTLRPFLN[+2059.735]WTGPPEPIEAAVAR.F | HexNAc(4)Hex(5)Fuc(1)NeuAc(1) | N[+2060] | 1764.1599 | 3 | 5290.4651 | 5290.4459 | 0 | 3.6 | 141 | Specific | 433.26 | 330.34 | 330.34 | 7.65 | 140 | >sp\|P02788\|TRFL_HUMAN Lactotransferrin OS=Homo sapiens OX=9606 GN=LTF PE=1 SV=6 | 1270 | 20201114_qwl_milk_1ng_glyco_hcd203040_compound15.23445.23445.3 | controllerType=0 controllerNumber=1 scan=23445 | 47.2871 |
| 19352:3 | 1 | R.TAGWNVPIGTLRPFLN[+2716.983]WTGPPEPIEAAVAR.F | HexNAc(5)Hex(6)Fuc(3)NeuAc(1) | N[+2717] | 1983.2343 | 3 | 5947.6885 | 5947.6939 | 0 | -0.9 | 141 | Specific | 520.55 | 462.63 | 462.63 | 9.23 | 140 | >sp\|P02788\|TRFL_HUMAN Lactotransferrin OS=Homo sapiens OX=9606 GN=LTF PE=1 SV=6 | 1270 | 20201114_qwl_milk_1ng_glyco_hcd203040_compound15.22786.22786.3 | controllerType=0 controllerNumber=1 scan=22786 | 45.928 |
| 19576:3 | 1 | R.TAGWNVPIGTLRPFLN[+2059.735]WTGPPEPIEAAVAR.F | HexNAc(4)Hex(5)Fuc(1)NeuAc(1) | N[+2060] | 1764.158 | 3 | 5290.4596 | 5290.4459 | 0 | 2.6 | 141 | Specific | 583.46 | 549.51 | 549.51 | 11.15 | 140 | >sp\|P02788\|TRFL_HUMAN Lactotransferrin OS=Homo sapiens OX=9606 GN=LTF PE=1 SV=6 | 1270 | 20201114_qwl_milk_1ng_glyco_hcd203040_compound15.23020.23020.2 | controllerType=0 controllerNumber=1 scan=23020 | 46.4439 |
| 19705:3 | 1 | R.TAGWNVPIGTLRPFLN[+2059.735]WTGPPEPIEAAVAR.F | HexNAc(4)Hex(5)Fuc(1)NeuAc(1) | N[+2060] | 1764.1607 | 3 | 5290.4676 | 5290.4459 | 0 | 4.1 | 141 | Specific | 586.1 | 513.44 | 513.44 | 10.76 | 140 | >sp\|P02788\|TRFL_HUMAN Lactotransferrin OS=Homo sapiens OX=9606 GN=LTF PE=1 SV=6 | 1270 | 20201114_qwl_milk_1ng_glyco_hcd203040_compound15.23155.23155.3 | controllerType=0 controllerNumber=1 scan=23155 | 46.7005 |
| 18553:4 | 1 | R.TAGWNVPIGTLRPFLN[+1768.640]WTGPPEPIEAAVAR.F | HexNAc(4)Hex(5)Fuc(1) | N[+1769] | 1250.595 | 4 | 4999.3581 | 4999.3504 | 0 | 1.5 | 141 | Specific | 603.06 | 592.82 | 592.82 | 11.94 | 140 | >sp\|P02788\|TRFL_HUMAN Lactotransferrin OS=Homo sapiens OX=9606 GN=LTF PE=1 SV=6 | 1270 | 20201114_qwl_milk_1ng_glyco_hcd203040_compound15.21948.21948.4 | controllerType=0 controllerNumber=1 scan=21948 | 44.0735 |
| 18498:4 | 1 | R.TAGWNVPIGTLRPFLN[+2425.887]WTGPPEPIEAAVAR.F | HexNAc(5)Hex(6)Fuc(3) | N[+2426] | 1414.9064 | 4 | 5656.6037 | 5656.5985 | 0 | 0.9 | 141 | Specific | 653.45 | 417.89 | 417.89 | 12.94 | 140 | >sp\|P02788\|TRFL_HUMAN Lactotransferrin OS=Homo sapiens OX=9606 GN=LTF PE=1 SV=6 | 1270 | 20201114_qwl_milk_1ng_glyco_hcd203040_compound15.21891.21891.4 | controllerType=0 controllerNumber=1 scan=21891 | 43.9523 |
| 18549:3 | 1 | R.TAGWNVPIGTLRPFLN[+1768.640]WTGPPEPIEAAVAR.F | HexNAc(4)Hex(5)Fuc(1) | N[+1769] | 1667.125 | 3 | 4999.3603 | 4999.3504 | 0 | 2 | 141 | Specific | 673.55 | 594.92 | 594.92 | 13.07 | 140 | >sp\|P02788\|TRFL_HUMAN Lactotransferrin OS=Homo sapiens OX=9606 GN=LTF PE=1 SV=6 | 1270 | 20201114_qwl_milk_1ng_glyco_hcd203040_compound15.21944.21944.3 | controllerType=0 controllerNumber=1 scan=21944 | 44.0661 |
| 19498:4 | 1 | R.TAGWNVPIGTLRPFLN[+2059.735]WTGPPEPIEAAVAR.F | HexNAc(4)Hex(5)Fuc(1)NeuAc(1) | N[+2060] | 1323.3801 | 4 | 5290.4987 | 5290.4459 | 0 | 10 | 141 | Specific | 679.51 | 629.57 | 629.57 | 12.16 | 140 | >sp\|P02788\|TRFL_HUMAN Lactotransferrin OS=Homo sapiens OX=9606 GN=LTF PE=1 SV=6 | 1270 | 20201114_qwl_milk_1ng_glyco_hcd203040_compound15.22939.22939.4 | controllerType=0 controllerNumber=1 scan=22939 | 46.2683 |
| 18546:3 | 1 | R.TAGWNVPIGTLRPFLN[+1768.640]WTGPPEPIEAAVAR.F | HexNAc(4)Hex(5)Fuc(1) | N[+1769] | 1667.125 | 3 | 4999.3603 | 4999.3504 | 0 | 2 | 141 | Specific | 718.28 | 622.44 | 622.44 | 14.07 | 140 | >sp\|P02788\|TRFL_HUMAN Lactotransferrin OS=Homo sapiens OX=9606 GN=LTF PE=1 SV=6 | 1270 | 20201114_qwl_milk_1ng_glyco_hcd203040_compound15.21941.21941.3 | controllerType=0 controllerNumber=1 scan=21941 | 44.06 |
| 18556:4 | 1 | R.TAGWNVPIGTLRPFLN[+1768.640]WTGPPEPIEAAVAR.F | HexNAc(4)Hex(5)Fuc(1) | N[+1769] | 1250.595 | 4 | 4999.3581 | 4999.3504 | 0 | 1.5 | 141 | Specific | 723.19 | 634.29 | 634.29 | 13.94 | 140 | >sp\|P02788\|TRFL_HUMAN Lactotransferrin OS=Homo sapiens OX=9606 GN=LTF PE=1 SV=6 | 1270 | 20201114_qwl_milk_1ng_glyco_hcd203040_compound15.21951.21951.4 | controllerType=0 controllerNumber=1 scan=21951 | 44.0785 |
| 19500:4 | 1 | R.TAGWNVPIGTLRPFLN[+2059.735]WTGPPEPIEAAVAR.F | HexNAc(4)Hex(5)Fuc(1)NeuAc(1) | N[+2060] | 1323.3801 | 4 | 5290.4987 | 5290.4459 | 0 | 10 | 141 | Specific | 831.68 | 696.93 | 696.93 | 15.16 | 140 | >sp\|P02788\|TRFL_HUMAN Lactotransferrin OS=Homo sapiens OX=9606 GN=LTF PE=1 SV=6 | 1270 | 20201114_qwl_milk_1ng_glyco_hcd203040_compound15.22941.22941.4 | controllerType=0 controllerNumber=1 scan=22941 | 46.2772 |
| 17416:3 | 1 | R.TAGWNIPMGLLFN[+1216.423]QTGSC[+57.021]K.F | HexNAc(2)Hex(5) | C[+57], N[+1216] | 1104.4843 | 3 | 3311.4383 | 3311.4328 | 0 | 1.7 | 485 | Specific | 468.84 | 468.84 | 202.32 | 9.43 | 140 | >sp\|P02788\|TRFL_HUMAN Lactotransferrin OS=Homo sapiens OX=9606 GN=LTF PE=1 SV=6 | 1270 | 20201114_qwl_milk_1ng_glyco_hcd203040_compound15.20758.20758.3 | controllerType=0 controllerNumber=1 scan=20758 | 41.5571 |
| 17194:3 | 1 | R.TAGWNIPMGLLFN[+1622.582]QTGSC[+57.021]K.F | HexNAc(4)Hex(5) | C[+57], N[+1623] | 1239.8698 | 3 | 3717.5949 | 3717.5916 | 0 | 0.9 | 485 | Specific | 495.24 | 495.24 | 207.56 | 9.56 | 140 | >sp\|P02788\|TRFL_HUMAN Lactotransferrin OS=Homo sapiens OX=9606 GN=LTF PE=1 SV=6 | 1270 | 20201114_qwl_milk_1ng_glyco_hcd203040_compound15.20528.20528.3 | controllerType=0 controllerNumber=1 scan=20528 | 41.1025 |
| 19061:3 | 1 | R.TAGWNIPMGLLFN[+1872.651]QTGSC[+57.021]K.F | HexNAc(3)Hex(6)NeuAc(1) | C[+57], N[+1873] | 1323.2281 | 3 | 3967.6697 | 3967.6604 | 0 | 2.4 | 485 | Specific | 586.44 | 539.35 | 343.41 | 11.1 | 140 | >sp\|P02788\|TRFL_HUMAN Lactotransferrin OS=Homo sapiens OX=9606 GN=LTF PE=1 SV=6 | 1270 | 20201114_qwl_milk_1ng_glyco_hcd203040_compound15.22480.22480.3 | controllerType=0 controllerNumber=1 scan=22480 | 45.3017 |
| 17171:3 | 1 | R.TAGWNIPMGLLFN[+1622.582]QTGSC[+57.021]K.F | HexNAc(4)Hex(5) | C[+57], N[+1623] | 1239.8644 | 3 | 3717.5786 | 3717.5916 | 0 | -3.5 | 485 | Specific | 599.82 | 599.82 | 265.48 | 9.76 | 140 | >sp\|P02788\|TRFL_HUMAN Lactotransferrin OS=Homo sapiens OX=9606 GN=LTF PE=1 SV=6 | 1270 | 20201114_qwl_milk_1ng_glyco_hcd203040_compound15.20503.20503.3 | controllerType=0 controllerNumber=1 scan=20503 | 41.0659 |
| 20948:3 | 1 | R.TAGWNIPMGLLFN[+2350.830]QTGSC[+57.021]K.F | HexNAc(4)Hex(5)Fuc(1)NeuAc(2) | C[+57], N[+2351] | 1482.6191 | 3 | 4445.8427 | 4445.8403 | 0 | 0.5 | 485 | Specific | 631.78 | 539.41 | 539.41 | 12.02 | 140 | >sp\|P02788\|TRFL_HUMAN Lactotransferrin OS=Homo sapiens OX=9606 GN=LTF PE=1 SV=6 | 1270 | 20201114_qwl_milk_1ng_glyco_hcd203040_compound15.24455.24455.3 | controllerType=0 controllerNumber=1 scan=24455 | 49.4354 |
| 17174:3 | 1 | R.TAGWNIPMGLLFN[+1622.582]QTGSC[+57.021]K.F | HexNAc(4)Hex(5) | C[+57], N[+1623] | 1239.8644 | 3 | 3717.5786 | 3717.5916 | 0 | -3.5 | 485 | Specific | 712.92 | 562.92 | 251.9 | 12.76 | 140 | >sp\|P02788\|TRFL_HUMAN Lactotransferrin OS=Homo sapiens OX=9606 GN=LTF PE=1 SV=6 | 1270 | 20201114_qwl_milk_1ng_glyco_hcd203040_compound15.20506.20506.3 | controllerType=0 controllerNumber=1 scan=20506 | 41.0699 |
| 17197:3 | 1 | R.TAGWNIPMGLLFN[+1622.582]QTGSC[+57.021]K.F | HexNAc(4)Hex(5) | C[+57], N[+1623] | 1239.8698 | 3 | 3717.5949 | 3717.5916 | 0 | 0.9 | 485 | Specific | 757.87 | 716.35 | 309.7 | 15.02 | 140 | >sp\|P02788\|TRFL_HUMAN Lactotransferrin OS=Homo sapiens OX=9606 GN=LTF PE=1 SV=6 | 1270 | 20201114_qwl_milk_1ng_glyco_hcd203040_compound15.20531.20531.3 | controllerType=0 controllerNumber=1 scan=20531 | 41.1066 |
| 17418:3 | 1 | R.TAGWNIPMGLLFN[+1216.423]QTGSC[+57.021]K.F | HexNAc(2)Hex(5) | C[+57], N[+1216] | 1104.4843 | 3 | 3311.4383 | 3311.4328 | 0 | 1.7 | 485 | Specific | 771.76 | 771.76 | 366.92 | 15.02 | 140 | >sp\|P02788\|TRFL_HUMAN Lactotransferrin OS=Homo sapiens OX=9606 GN=LTF PE=1 SV=6 | 1270 | 20201114_qwl_milk_1ng_glyco_hcd203040_compound15.20760.20760.3 | controllerType=0 controllerNumber=1 scan=20760 | 41.5606 |
| 17185:3 | 1 | R.TAGWNIPMGLLFN[+1622.582]QTGSC[+57.021]K.F | HexNAc(4)Hex(5) | C[+57], N[+1623] | 1239.871 | 3 | 3717.5985 | 3717.5916 | 0 | 1.9 | 485 | Specific | 794.75 | 765.82 | 304.91 | 15.02 | 140 | >sp\|P02788\|TRFL_HUMAN Lactotransferrin OS=Homo sapiens OX=9606 GN=LTF PE=1 SV=6 | 1270 | 20201114_qwl_milk_1ng_glyco_hcd203040_compound15.20518.20518.3 | controllerType=0 controllerNumber=1 scan=20518 | 41.087 |
| 18849:3 | 1 | R.TAGWNIPMGLLFN[+2205.793]QTGSC[+57.021]K.F | HexNAc(4)Hex(5)Fuc(2)NeuAc(1) | C[+57], N[+2206] | 1434.2742 | 3 | 4300.8081 | 4300.8028 | 0 | 1.2 | 485 | Specific | 834.11 | 740.61 | 740.61 | 16.02 | 140 | >sp\|P02788\|TRFL_HUMAN Lactotransferrin OS=Homo sapiens OX=9606 GN=LTF PE=1 SV=6 | 1270 | 20201114_qwl_milk_1ng_glyco_hcd203040_compound15.22258.22258.3 | controllerType=0 controllerNumber=1 scan=22258 | 44.7662 |
| 16941:3 | 1 | R.TAGWNIPMGLLFN[+1768.640]QTGSC[+57.021]K.F | HexNAc(4)Hex(5)Fuc(1) | C[+57], N[+1769] | 1288.5563 | 3 | 3863.6544 | 3863.6495 | 0 | 1.3 | 485 | Specific | 919.16 | 799.01 | 799.01 | 18.02 | 140 | >sp\|P02788\|TRFL_HUMAN Lactotransferrin OS=Homo sapiens OX=9606 GN=LTF PE=1 SV=6 | 1270 | 20201114_qwl_milk_1ng_glyco_hcd203040_compound15.20261.20261.3 | controllerType=0 controllerNumber=1 scan=20261 | 40.5844 |
| 16943:3 | 1 | R.TAGWNIPMGLLFN[+1768.640]QTGSC[+57.021]K.F | HexNAc(4)Hex(5)Fuc(1) | C[+57], N[+1769] | 1288.5563 | 3 | 3863.6544 | 3863.6495 | 0 | 1.3 | 485 | Specific | 927.18 | 693.22 | 693.22 | 18.02 | 140 | >sp\|P02788\|TRFL_HUMAN Lactotransferrin OS=Homo sapiens OX=9606 GN=LTF PE=1 SV=6 | 1270 | 20201114_qwl_milk_1ng_glyco_hcd203040_compound15.20263.20263.3 | controllerType=0 controllerNumber=1 scan=20263 | 40.597 |
| 19030:3 | 1 | R.TAGWNIPMGLLFN[+1913.677]QTGSC[+57.021]K.F | HexNAc(4)Hex(5)NeuAc(1) | C[+57], N[+1914] | 1336.9037 | 3 | 4008.6967 | 4008.687 | 0 | 2.4 | 485 | Specific | 934 | 714.55 | 714.55 | 18.1 | 140 | >sp\|P02788\|TRFL_HUMAN Lactotransferrin OS=Homo sapiens OX=9606 GN=LTF PE=1 SV=6 | 1270 | 20201114_qwl_milk_1ng_glyco_hcd203040_compound15.22448.22448.3 | controllerType=0 controllerNumber=1 scan=22448 | 45.2072 |
| 18896:3 | 1 | R.TAGWNIPMGLLFN[+2059.735]QTGSC[+57.021]K.F | HexNAc(4)Hex(5)Fuc(1)NeuAc(1) | C[+57], N[+2060] | 1385.5892 | 3 | 4154.753 | 4154.7449 | 0 | 1.9 | 485 | Specific | 935.7 | 917.53 | 917.53 | 18.02 | 140 | >sp\|P02788\|TRFL_HUMAN Lactotransferrin OS=Homo sapiens OX=9606 GN=LTF PE=1 SV=6 | 1270 | 20201114_qwl_milk_1ng_glyco_hcd203040_compound15.22307.22307.3 | controllerType=0 controllerNumber=1 scan=22307 | 44.866 |
| 18898:3 | 1 | R.TAGWNIPMGLLFN[+2059.735]QTGSC[+57.021]K.F | HexNAc(4)Hex(5)Fuc(1)NeuAc(1) | C[+57], N[+2060] | 1385.5892 | 3 | 4154.753 | 4154.7449 | 0 | 1.9 | 485 | Specific | 937.26 | 902.12 | 902.12 | 18.02 | 140 | >sp\|P02788\|TRFL_HUMAN Lactotransferrin OS=Homo sapiens OX=9606 GN=LTF PE=1 SV=6 | 1270 | 20201114_qwl_milk_1ng_glyco_hcd203040_compound15.22309.22309.3 | controllerType=0 controllerNumber=1 scan=22309 | 44.8747 |
| 19028:3 | 1 | R.TAGWNIPMGLLFN[+1913.677]QTGSC[+57.021]K.F | HexNAc(4)Hex(5)NeuAc(1) | C[+57], N[+1914] | 1336.9037 | 3 | 4008.6967 | 4008.687 | 0 | 2.4 | 485 | Specific | 948.65 | 948.65 | 948.65 | 18.1 | 140 | >sp\|P02788\|TRFL_HUMAN Lactotransferrin OS=Homo sapiens OX=9606 GN=LTF PE=1 SV=6 | 1270 | 20201114_qwl_milk_1ng_glyco_hcd203040_compound15.22446.22446.3 | controllerType=0 controllerNumber=1 scan=22446 | 45.2023 |
| 18847:3 | 1 | R.TAGWNIPMGLLFN[+2205.793]QTGSC[+57.021]K.F | HexNAc(4)Hex(5)Fuc(2)NeuAc(1) | C[+57], N[+2206] | 1434.2742 | 3 | 4300.8081 | 4300.8028 | 0 | 1.2 | 485 | Specific | 958.12 | 861.69 | 861.69 | 19.02 | 140 | >sp\|P02788\|TRFL_HUMAN Lactotransferrin OS=Homo sapiens OX=9606 GN=LTF PE=1 SV=6 | 1270 | 20201114_qwl_milk_1ng_glyco_hcd203040_compound15.22256.22256.3 | controllerType=0 controllerNumber=1 scan=22256 | 44.7599 |
| 18228:3 | 1 | R.TAGWNIPMGLLFN[+1606.587]QTGSC[+57.021]KFDEYFSQSC[+57.021]APGSDPR.S | HexNAc(4)Hex(4)Fuc(1) | C[+57]*2, N[+1607] | 1849.1285 | 3 | 5545.371 | 5545.3438 | 0 | 4.9 | 485 | Specific | 229.15 | 214.66 | 214.66 | 5.7 | 140 | >sp\|P02788\|TRFL_HUMAN Lactotransferrin OS=Homo sapiens OX=9606 GN=LTF PE=1 SV=6 | 1270 | 20201114_qwl_milk_1ng_glyco_hcd203040_compound15.21607.21607.3 | controllerType=0 controllerNumber=1 scan=21607 | 43.3371 |
| 19093:4 | 1 | R.TAGWNIPMGLLFN[+2351.851]QTGSC[+57.021]KFDEYFSQSC[+57.021]APGSDPR.S | HexNAc(4)Hex(5)Fuc(3)NeuAc(1) | C[+57]*2, N[+2352] | 1573.4172 | 4 | 6290.6471 | 6290.6079 | 0 | 6.2 | 485 | Specific | 382.18 | 382.18 | 382.18 | 8.25 | 140 | >sp\|P02788\|TRFL_HUMAN Lactotransferrin OS=Homo sapiens OX=9606 GN=LTF PE=1 SV=6 | 1270 | 20201114_qwl_milk_1ng_glyco_hcd203040_compound15.22516.22516.4 | controllerType=0 controllerNumber=1 scan=22516 | 45.4223 |
| 19239:3 | 1 | R.TAGWNIPMGLLFN[+1913.677]QTGSC[+57.021]KFDEYFSQSC[+57.021]APGSDPR.S | HexNAc(4)Hex(5)NeuAc(1) | C[+57]*2, N[+1914] | 1951.4871 | 3 | 5852.4469 | 5852.4341 | 0 | 2.2 | 485 | Specific | 397.85 | 299.73 | 299.73 | 7.36 | 140 | >sp\|P02788\|TRFL_HUMAN Lactotransferrin OS=Homo sapiens OX=9606 GN=LTF PE=1 SV=6 | 1270 | 20201114_qwl_milk_1ng_glyco_hcd203040_compound15.22667.22667.3 | controllerType=0 controllerNumber=1 scan=22667 | 45.6933 |
| 19121:4 | 1 | R.TAGWNIPMGLLFN[+2205.793]QTGSC[+57.021]KFDEYFSQSC[+57.021]APGSDPR.S | HexNAc(4)Hex(5)Fuc(2)NeuAc(1) | C[+57]*2, N[+2206] | 1536.8949 | 4 | 6144.5578 | 6144.5499 | 0 | 1.3 | 485 | Specific | 449.12 | 408.09 | 408.09 | 9 | 140 | >sp\|P02788\|TRFL_HUMAN Lactotransferrin OS=Homo sapiens OX=9606 GN=LTF PE=1 SV=6 | 1270 | 20201114_qwl_milk_1ng_glyco_hcd203040_compound15.22545.22545.4 | controllerType=0 controllerNumber=1 scan=22545 | 45.4726 |
| 18962:4 | 1 | R.TAGWNIPMGLLFN[+2862.021]QTGSC[+57.021]KFDEYFSQSC[+57.021]APGSDPR.S | HexNAc(5)Hex(6)Fuc(2)NeuAc(2) | C[+57]*2, N[+2862] | 1700.9555 | 4 | 6800.8003 | 6800.7776 | 0 | 3.3 | 485 | Specific | 453.48 | 443.75 | 443.75 | 9.08 | 140 | >sp\|P02788\|TRFL_HUMAN Lactotransferrin OS=Homo sapiens OX=9606 GN=LTF PE=1 SV=6 | 1270 | 20201114_qwl_milk_1ng_glyco_hcd203040_compound15.22377.22377.2 | controllerType=0 controllerNumber=1 scan=22377 | 45.0531 |
| 19235:4 | 1 | R.TAGWNIPMGLLFN[+1913.677]QTGSC[+57.021]KFDEYFSQSC[+57.021]APGSDPR.S | HexNAc(4)Hex(5)NeuAc(1) | C[+57]*2, N[+1914] | 1463.8682 | 4 | 5852.4508 | 5852.4341 | 0 | 2.9 | 485 | Specific | 524.12 | 465.25 | 465.25 | 9.44 | 140 | >sp\|P02788\|TRFL_HUMAN Lactotransferrin OS=Homo sapiens OX=9606 GN=LTF PE=1 SV=6 | 1270 | 20201114_qwl_milk_1ng_glyco_hcd203040_compound15.22663.22663.4 | controllerType=0 controllerNumber=1 scan=22663 | 45.6863 |
| 19173:4 | 1 | R.TAGWNIPMGLLFN[+1913.677]QTGSC[+57.021]KFDEYFSQSC[+57.021]APGSDPR.S | HexNAc(4)Hex(5)NeuAc(1) | C[+57]*2, N[+1914] | 1463.8701 | 4 | 5852.4587 | 5852.4341 | 0 | 4.2 | 485 | Specific | 573.27 | 528.06 | 528.06 | 9.25 | 140 | >sp\|P02788\|TRFL_HUMAN Lactotransferrin OS=Homo sapiens OX=9606 GN=LTF PE=1 SV=6 | 1270 | 20201114_qwl_milk_1ng_glyco_hcd203040_compound15.22599.22599.4 | controllerType=0 controllerNumber=1 scan=22599 | 45.5782 |
| 18227:4 | 1 | R.TAGWNIPMGLLFN[+1606.587]QTGSC[+57.021]KFDEYFSQSC[+57.021]APGSDPR.S | HexNAc(4)Hex(4)Fuc(1) | C[+57]*2, N[+1607] | 1387.0999 | 4 | 5545.3776 | 5545.3438 | 0 | 6.1 | 485 | Specific | 637.84 | 637.84 | 637.84 | 11.21 | 140 | >sp\|P02788\|TRFL_HUMAN Lactotransferrin OS=Homo sapiens OX=9606 GN=LTF PE=1 SV=6 | 1270 | 20201114_qwl_milk_1ng_glyco_hcd203040_compound15.21606.21606.4 | controllerType=0 controllerNumber=1 scan=21606 | 43.336 |
| 18223:3 | 1 | R.TAGWNIPMGLLFN[+1622.582]QTGSC[+57.021]KFDEYFSQSC[+57.021]APGSDPR.S | HexNAc(4)Hex(5) | C[+57]*2, N[+1623] | 1854.4618 | 3 | 5561.3707 | 5561.3387 | 0 | 5.8 | 485 | Specific | 655.57 | 614.94 | 197.46 | 12.34 | 140 | >sp\|P02788\|TRFL_HUMAN Lactotransferrin OS=Homo sapiens OX=9606 GN=LTF PE=1 SV=6 | 1270 | 20201114_qwl_milk_1ng_glyco_hcd203040_compound15.21602.21602.3 | controllerType=0 controllerNumber=1 scan=21602 | 43.3301 |
| 18226:3 | 1 | R.TAGWNIPMGLLFN[+1622.582]QTGSC[+57.021]KFDEYFSQSC[+57.021]APGSDPR.S | HexNAc(4)Hex(5) | C[+57]*2, N[+1623] | 1854.4618 | 3 | 5561.3707 | 5561.3387 | 0 | 5.8 | 485 | Specific | 704.72 | 658.81 | 168.09 | 13.34 | 140 | >sp\|P02788\|TRFL_HUMAN Lactotransferrin OS=Homo sapiens OX=9606 GN=LTF PE=1 SV=6 | 1270 | 20201114_qwl_milk_1ng_glyco_hcd203040_compound15.21605.21605.3 | controllerType=0 controllerNumber=1 scan=21605 | 43.334 |
| 19120:4 | 1 | R.TAGWNIPMGLLFN[+2059.735]QTGSC[+57.021]KFDEYFSQSC[+57.021]APGSDPR.S | HexNAc(4)Hex(5)Fuc(1)NeuAc(1) | C[+57]*2, N[+2060] | 1500.3818 | 4 | 5998.5055 | 5998.492 | 0 | 2.2 | 485 | Specific | 724.15 | 704.93 | 704.93 | 13.85 | 140 | >sp\|P02788\|TRFL_HUMAN Lactotransferrin OS=Homo sapiens OX=9606 GN=LTF PE=1 SV=6 | 1270 | 20201114_qwl_milk_1ng_glyco_hcd203040_compound15.22544.22544.4 | controllerType=0 controllerNumber=1 scan=22544 | 45.4712 |
| 19242:3 | 1 | R.TAGWNIPMGLLFN[+1913.677]QTGSC[+57.021]KFDEYFSQSC[+57.021]APGSDPR.S | HexNAc(4)Hex(5)NeuAc(1) | C[+57]*2, N[+1914] | 1951.4871 | 3 | 5852.4469 | 5852.4341 | 0 | 2.2 | 485 | Specific | 767.1 | 682.01 | 682.01 | 14.98 | 140 | >sp\|P02788\|TRFL_HUMAN Lactotransferrin OS=Homo sapiens OX=9606 GN=LTF PE=1 SV=6 | 1270 | 20201114_qwl_milk_1ng_glyco_hcd203040_compound15.22670.22670.3 | controllerType=0 controllerNumber=1 scan=22670 | 45.697 |
| 19176:4 | 1 | R.TAGWNIPMGLLFN[+1913.677]QTGSC[+57.021]KFDEYFSQSC[+57.021]APGSDPR.S | HexNAc(4)Hex(5)NeuAc(1) | C[+57]*2, N[+1914] | 1463.8701 | 4 | 5852.4587 | 5852.4341 | 0 | 4.2 | 485 | Specific | 799.94 | 750.56 | 750.56 | 14.46 | 140 | >sp\|P02788\|TRFL_HUMAN Lactotransferrin OS=Homo sapiens OX=9606 GN=LTF PE=1 SV=6 | 1270 | 20201114_qwl_milk_1ng_glyco_hcd203040_compound15.22602.22602.4 | controllerType=0 controllerNumber=1 scan=22602 | 45.5819 |
| 18164:3 | 1 | R.TAGWNIPMGLLFN[+1768.640]QTGSC[+57.021]KFDEYFSQSC[+57.021]APGSDPR.S | HexNAc(4)Hex(5)Fuc(1) | C[+57]*2, N[+1769] | 1903.1483 | 3 | 5707.4303 | 5707.3966 | 0 | 5.9 | 485 | Specific | 809.94 | 727.34 | 727.34 | 15.34 | 140 | >sp\|P02788\|TRFL_HUMAN Lactotransferrin OS=Homo sapiens OX=9606 GN=LTF PE=1 SV=6 | 1270 | 20201114_qwl_milk_1ng_glyco_hcd203040_compound15.21540.21540.3 | controllerType=0 controllerNumber=1 scan=21540 | 43.1609 |
| 18231:3 | 1 | R.TAGWNIPMGLLFN[+1606.587]QTGSC[+57.021]KFDEYFSQSC[+57.021]APGSDPR.S | HexNAc(4)Hex(4)Fuc(1) | C[+57]*2, N[+1607] | 1849.1285 | 3 | 5545.371 | 5545.3438 | 0 | 4.9 | 485 | Specific | 829.55 | 716.1 | 716.1 | 15.59 | 140 | >sp\|P02788\|TRFL_HUMAN Lactotransferrin OS=Homo sapiens OX=9606 GN=LTF PE=1 SV=6 | 1270 | 20201114_qwl_milk_1ng_glyco_hcd203040_compound15.21610.21610.3 | controllerType=0 controllerNumber=1 scan=21610 | 43.342 |
| 18230:4 | 1 | R.TAGWNIPMGLLFN[+1606.587]QTGSC[+57.021]KFDEYFSQSC[+57.021]APGSDPR.S | HexNAc(4)Hex(4)Fuc(1) | C[+57]*2, N[+1607] | 1387.0999 | 4 | 5545.3776 | 5545.3438 | 0 | 6.1 | 485 | Specific | 870.28 | 870.28 | 870.28 | 16.21 | 140 | >sp\|P02788\|TRFL_HUMAN Lactotransferrin OS=Homo sapiens OX=9606 GN=LTF PE=1 SV=6 | 1270 | 20201114_qwl_milk_1ng_glyco_hcd203040_compound15.21609.21609.4 | controllerType=0 controllerNumber=1 scan=21609 | 43.34 |
| 19237:4 | 1 | R.TAGWNIPMGLLFN[+1913.677]QTGSC[+57.021]KFDEYFSQSC[+57.021]APGSDPR.S | HexNAc(4)Hex(5)NeuAc(1) | C[+57]*2, N[+1914] | 1463.8682 | 4 | 5852.4508 | 5852.4341 | 0 | 2.9 | 485 | Specific | 880.33 | 837.64 | 837.64 | 16.85 | 140 | >sp\|P02788\|TRFL_HUMAN Lactotransferrin OS=Homo sapiens OX=9606 GN=LTF PE=1 SV=6 | 1270 | 20201114_qwl_milk_1ng_glyco_hcd203040_compound15.22665.22665.4 | controllerType=0 controllerNumber=1 scan=22665 | 45.69 |
| 19095:4 | 1 | R.TAGWNIPMGLLFN[+2351.851]QTGSC[+57.021]KFDEYFSQSC[+57.021]APGSDPR.S | HexNAc(4)Hex(5)Fuc(3)NeuAc(1) | C[+57]*2, N[+2352] | 1573.4172 | 4 | 6290.6471 | 6290.6079 | 0 | 6.2 | 485 | Specific | 903.44 | 897.44 | 897.44 | 17.21 | 140 | >sp\|P02788\|TRFL_HUMAN Lactotransferrin OS=Homo sapiens OX=9606 GN=LTF PE=1 SV=6 | 1270 | 20201114_qwl_milk_1ng_glyco_hcd203040_compound15.22518.22518.4 | controllerType=0 controllerNumber=1 scan=22518 | 45.4285 |
| 18204:4 | 1 | R.TAGWNIPMGLLFN[+1622.582]QTGSC[+57.021]KFDEYFSQSC[+57.021]APGSDPR.S | HexNAc(4)Hex(5) | C[+57]*2, N[+1623] | 1391.0968 | 4 | 5561.3654 | 5561.3387 | 0 | 4.8 | 485 | Specific | 921.06 | 890.98 | 356.68 | 17.46 | 140 | >sp\|P02788\|TRFL_HUMAN Lactotransferrin OS=Homo sapiens OX=9606 GN=LTF PE=1 SV=6 | 1270 | 20201114_qwl_milk_1ng_glyco_hcd203040_compound15.21582.21582.4 | controllerType=0 controllerNumber=1 scan=21582 | 43.2756 |
| 18161:4 | 1 | R.TAGWNIPMGLLFN[+1768.640]QTGSC[+57.021]KFDEYFSQSC[+57.021]APGSDPR.S | HexNAc(4)Hex(5)Fuc(1) | C[+57]*2, N[+1769] | 1427.6144 | 4 | 5707.4357 | 5707.3966 | 0 | 6.8 | 485 | Specific | 946.32 | 946.32 | 946.32 | 17.21 | 140 | >sp\|P02788\|TRFL_HUMAN Lactotransferrin OS=Homo sapiens OX=9606 GN=LTF PE=1 SV=6 | 1270 | 20201114_qwl_milk_1ng_glyco_hcd203040_compound15.21537.21537.4 | controllerType=0 controllerNumber=1 scan=21537 | 43.1491 |
| 18163:4 | 1 | R.TAGWNIPMGLLFN[+1768.640]QTGSC[+57.021]KFDEYFSQSC[+57.021]APGSDPR.S | HexNAc(4)Hex(5)Fuc(1) | C[+57]*2, N[+1769] | 1427.6144 | 4 | 5707.4357 | 5707.3966 | 0 | 6.8 | 485 | Specific | 954.68 | 908.33 | 908.33 | 18.21 | 140 | >sp\|P02788\|TRFL_HUMAN Lactotransferrin OS=Homo sapiens OX=9606 GN=LTF PE=1 SV=6 | 1270 | 20201114_qwl_milk_1ng_glyco_hcd203040_compound15.21539.21539.4 | controllerType=0 controllerNumber=1 scan=21539 | 43.1576 |
| 19122:4 | 1 | R.TAGWNIPMGLLFN[+2059.735]QTGSC[+57.021]KFDEYFSQSC[+57.021]APGSDPR.S | HexNAc(4)Hex(5)Fuc(1)NeuAc(1) | C[+57]*2, N[+2060] | 1500.3818 | 4 | 5998.5055 | 5998.492 | 0 | 2.2 | 485 | Specific | 962.7 | 904.01 | 904.01 | 18.85 | 140 | >sp\|P02788\|TRFL_HUMAN Lactotransferrin OS=Homo sapiens OX=9606 GN=LTF PE=1 SV=6 | 1270 | 20201114_qwl_milk_1ng_glyco_hcd203040_compound15.22546.22546.4 | controllerType=0 controllerNumber=1 scan=22546 | 45.4752 |
| 19123:4 | 1 | R.TAGWNIPMGLLFN[+2205.793]QTGSC[+57.021]KFDEYFSQSC[+57.021]APGSDPR.S | HexNAc(4)Hex(5)Fuc(2)NeuAc(1) | C[+57]*2, N[+2206] | 1536.8949 | 4 | 6144.5578 | 6144.5499 | 0 | 1.3 | 485 | Specific | 979.1 | 944.73 | 944.73 | 18.77 | 140 | >sp\|P02788\|TRFL_HUMAN Lactotransferrin OS=Homo sapiens OX=9606 GN=LTF PE=1 SV=6 | 1270 | 20201114_qwl_milk_1ng_glyco_hcd203040_compound15.22547.22547.4 | controllerType=0 controllerNumber=1 scan=22547 | 45.476 |
| 18162:3 | 1 | R.TAGWNIPMGLLFN[+1768.640]QTGSC[+57.021]KFDEYFSQSC[+57.021]APGSDPR.S | HexNAc(4)Hex(5)Fuc(1) | C[+57]*2, N[+1769] | 1903.1483 | 3 | 5707.4303 | 5707.3966 | 0 | 5.9 | 485 | Specific | 981.94 | 928.63 | 928.63 | 18.34 | 140 | >sp\|P02788\|TRFL_HUMAN Lactotransferrin OS=Homo sapiens OX=9606 GN=LTF PE=1 SV=6 | 1270 | 20201114_qwl_milk_1ng_glyco_hcd203040_compound15.21538.21538.3 | controllerType=0 controllerNumber=1 scan=21538 | 43.1505 |
| 18202:4 | 1 | R.TAGWNIPMGLLFN[+1622.582]QTGSC[+57.021]KFDEYFSQSC[+57.021]APGSDPR.S | HexNAc(4)Hex(5) | C[+57]*2, N[+1623] | 1391.0968 | 4 | 5561.3654 | 5561.3387 | 0 | 4.8 | 485 | Specific | 988.52 | 962.59 | 289.37 | 18.46 | 140 | >sp\|P02788\|TRFL_HUMAN Lactotransferrin OS=Homo sapiens OX=9606 GN=LTF PE=1 SV=6 | 1270 | 20201114_qwl_milk_1ng_glyco_hcd203040_compound15.21580.21580.4 | controllerType=0 controllerNumber=1 scan=21580 | 43.2689 |
| 18017:4 | 1 | R.TAGWNIPM[+15.995]GLLFN[+2059.735]QTGSC[+57.021]KFDEYFSQSC[+57.021]A | HexNAc(4)Hex(5)Fuc(1)NeuAc(1) | C[+57]*2, M[+16], N[+2060] | 1504.3803 | 4 | 6014.4992 | 6014.487 | 0 | 2 | 485 | Specific | 394.99 | 324.28 | 324.28 | 7.25 | 140 | >sp\|P02788\|TRFL_HUMAN Lactotransferrin OS=Homo sapiens OX=9606 GN=LTF PE=1 SV=6 | 1270 | 20201114_qwl_milk_1ng_glyco_hcd203040_compound15.21386.21386.4 | controllerType=0 controllerNumber=1 scan=21386 | 42.8433 |
| 18019:4 | 1 | R.TAGWNIPM[+15.995]GLLFN[+2059.735]QTGSC[+57.021]KFDEYFSQSC[+57.021]A | HexNAc(4)Hex(5)Fuc(1)NeuAc(1) | C[+57]*2, M[+16], N[+2060] | 1504.3803 | 4 | 6014.4992 | 6014.487 | 0 | 2 | 485 | Specific | 517.22 | 469.5 | 469.5 | 9.31 | 140 | >sp\|P02788\|TRFL_HUMAN Lactotransferrin OS=Homo sapiens OX=9606 GN=LTF PE=1 SV=6 | 1270 | 20201114_qwl_milk_1ng_glyco_hcd203040_compound15.21388.21388.4 | controllerType=0 controllerNumber=1 scan=21388 | 42.8486 |
| 08160:3 | 1 | K.FGRN[+2205.793]GSDC[+57.021]PDKFC[+57.021]LFQSETK.N | HexNAc(4)Hex(5)Fuc(2)NeuAc(1) | C[+57]*2, N[+2206] | 1533.6264 | 3 | 4598.8646 | 4598.8577 | 0 | 1.5 | 639 | Specific | 357.95 | 357.95 | 357.95 | 6.94 | 140 | >sp\|P02788\|TRFL_HUMAN Lactotransferrin OS=Homo sapiens OX=9606 GN=LTF PE=1 SV=6 | 1270 | 20201114_qwl_milk_1ng_glyco_hcd203040_compound15.11112.11112.3 | controllerType=0 controllerNumber=1 scan=11112 | 24.6806 |
| 08163:3 | 1 | K.FGRN[+2205.793]GSDC[+57.021]PDKFC[+57.021]LFQSETK.N | HexNAc(4)Hex(5)Fuc(2)NeuAc(1) | C[+57]*2, N[+2206] | 1533.6264 | 3 | 4598.8646 | 4598.8577 | 0 | 1.5 | 639 | Specific | 445.29 | 422.41 | 422.41 | 8.71 | 140 | >sp\|P02788\|TRFL_HUMAN Lactotransferrin OS=Homo sapiens OX=9606 GN=LTF PE=1 SV=6 | 1270 | 20201114_qwl_milk_1ng_glyco_hcd203040_compound15.11115.11115.3 | controllerType=0 controllerNumber=1 scan=11115 | 24.6854 |
| 08250:4 | 1 | K.FGRN[+2059.735]GSDC[+57.021]PDKFC[+57.021]LFQSETK.N | HexNAc(4)Hex(5)Fuc(1)NeuAc(1) | C[+57]*2, N[+2060] | 1113.9574 | 4 | 4452.8079 | 4452.7998 | 0 | 1.8 | 639 | Specific | 513.91 | 469.45 | 469.45 | 9.4 | 140 | >sp\|P02788\|TRFL_HUMAN Lactotransferrin OS=Homo sapiens OX=9606 GN=LTF PE=1 SV=6 | 1270 | 20201114_qwl_milk_1ng_glyco_hcd203040_compound15.11205.11205.4 | controllerType=0 controllerNumber=1 scan=11205 | 24.8426 |
| 08248:4 | 1 | K.FGRN[+2059.735]GSDC[+57.021]PDKFC[+57.021]LFQSETK.N | HexNAc(4)Hex(5)Fuc(1)NeuAc(1) | C[+57]*2, N[+2060] | 1113.9574 | 4 | 4452.8079 | 4452.7998 | 0 | 1.8 | 639 | Specific | 550.66 | 550.66 | 550.66 | 9.65 | 140 | >sp\|P02788\|TRFL_HUMAN Lactotransferrin OS=Homo sapiens OX=9606 GN=LTF PE=1 SV=6 | 1270 | 20201114_qwl_milk_1ng_glyco_hcd203040_compound15.11203.11203.4 | controllerType=0 controllerNumber=1 scan=11203 | 24.8384 |
| 08166:4 | 1 | K.FGRN[+2205.793]GSDC[+57.021]PDKFC[+57.021]LFQSETK.N | HexNAc(4)Hex(5)Fuc(2)NeuAc(1) | C[+57]*2, N[+2206] | 1150.4714 | 4 | 4598.8636 | 4598.8577 | 0 | 1.3 | 639 | Specific | 694.85 | 694.85 | 694.85 | 12.87 | 140 | >sp\|P02788\|TRFL_HUMAN Lactotransferrin OS=Homo sapiens OX=9606 GN=LTF PE=1 SV=6 | 1270 | 20201114_qwl_milk_1ng_glyco_hcd203040_compound15.11118.11118.4 | controllerType=0 controllerNumber=1 scan=11118 | 24.6924 |
| 08164:4 | 1 | K.FGRN[+2205.793]GSDC[+57.021]PDKFC[+57.021]LFQSETK.N | HexNAc(4)Hex(5)Fuc(2)NeuAc(1) | C[+57]*2, N[+2206] | 1150.4714 | 4 | 4598.8636 | 4598.8577 | 0 | 1.3 | 639 | Specific | 723.29 | 723.29 | 723.29 | 13.87 | 140 | >sp\|P02788\|TRFL_HUMAN Lactotransferrin OS=Homo sapiens OX=9606 GN=LTF PE=1 SV=6 | 1270 | 20201114_qwl_milk_1ng_glyco_hcd203040_compound15.11116.11116.4 | controllerType=0 controllerNumber=1 scan=11116 | 24.6874 |

Table S4. Cont.

| Query #:z | Protein Rank | Glycopeptide | Glycan | Modification Type(s) | Observed m/z | z | Observed (M+H) | Calc. mass (M+H) | Off-by-x error | Mass error (ppm) | StartPosition | Cleavage | Score | Delta | Delta Mod | \|Log Prob\| | # of unique peptides | Protein Name | Protein DB number | Comment | Scan # | Scan Time |
|---|---|---|---|---|---|---|---|---|---|---|---|---|---|---|---|---|---|---|---|---|---|---|
| 09265:3 | 1 | R.N[+2205.793]GSDC[+57.021]PDKFC[+57.021]LFQSETK.N | HexNAc(4)Hex(5)Fuc(2)NeuAc(1) | C[+57]*2, N[+2206] | 1413.5643 | 3 | 4238.6782 | 4238.6667 | 0 | 2.7 | 642 | Specific | 397.17 | 326.27 | 326.27 | 7.21 | 140 | >sp\|P02788\|TRFL_HUMAN Lactotransferrin OS=Homo sapiens OX=9606 GN=LTF PE=1 SV=6 | 1270 | 20201114_qwl_milk_1ng_glyco_hcd203040_compound15.12259.12259.3 | controllerType=0 controllerNumber=1 scan=12259 | 26.5834 |
| 04932:3 | 5 | R.FSDGLESN[+1864.634]SSTQFEVKK.Y | HexNAc(2)Hex(9) | N[+1865] | 1256.5236 | 3 | 3767.5563 | 3767.5421 | 0 | 3.8 | 219 | Specific | 572.5 | 480.21 | 480.21 | 9.52 | 84 | >sp\|P0C0L4\|CO4A_HUMAN Complement C4-A OS=Homo sapiens OX=9606 GN=C4A PE=1 SV=2 | 1235 | 20201114_qwl_milk_1ng_glyco_hcd203040_compound15.7770.7770.3 | controllerType=0 controllerNumber=1 scan=7770 | 19.1755 |
| 04930:3 | 5 | R.FSDGLESN[+1864.634]SSTQFEVKK.Y | HexNAc(2)Hex(9) | N[+1865] | 1256.5236 | 3 | 3767.5563 | 3767.5421 | 0 | 3.8 | 219 | Specific | 609.43 | 488.79 | 488.79 | 11.73 | 84 | >sp\|P0C0L4\|CO4A_HUMAN Complement C4-A OS=Homo sapiens OX=9606 GN=C4A PE=1 SV=2 | 1235 | 20201114_qwl_milk_1ng_glyco_hcd203040_compound15.7768.7768.3 | controllerType=0 controllerNumber=1 scan=7768 | 19.1719 |
| 13325:5 | 6 | K.KGN[+2059.735]TTLNSFVIPSESDVPTHC[+57.021]PSQWWPYAGHC[+57.021]YK.I | HexNAc(4)Hex(5)Fuc(1)NeuAc(1) | C[+57]*2, N[+2060] | 1205.5209 | 5 | 6023.5753 | 6023.5567 | 0 | 3.1 | 342 | Specific | 429.68 | 429.68 | 429.68 | 9.33 | 86 | >sp\|P22897\|MRC1_HUMAN Macrophage mannose receptor 1 OS=Homo sapiens OX=9606 GN=MRC1 PE=1 SV=1 | 1049 | 20201114_qwl_milk_1ng_glyco_hcd203040_compound15.16473.16473.5 | controllerType=0 controllerNumber=1 scan=16473 | 33.6725 |
| 13268:5 | 6 | K.KGN[+2205.793]TTLNSFVIPSESDVPTHC[+57.021]PSQWWPYAGHC[+57.021]YK.I | HexNAc(4)Hex(5)Fuc(2)NeuAc(1) | C[+57]*2, N[+2206] | 1234.729 | 5 | 6169.6159 | 6169.6146 | 0 | 0.2 | 342 | Specific | 725.38 | 725.38 | 725.38 | 13.71 | 86 | >sp\|P22897\|MRC1_HUMAN Macrophage mannose receptor 1 OS=Homo sapiens OX=9606 GN=MRC1 PE=1 SV=1 | 1049 | 20201114_qwl_milk_1ng_glyco_hcd203040_compound15.16414.16414.5 | controllerType=0 controllerNumber=1 scan=16414 | 33.5718 |
| 10222:3 | 7 | K.TVLTPATNHMGN[+1540.529]VTFTIPANR.E | HexNAc(2)Hex(7) | N[+1541] | 1265.9028 | 3 | 3795.6937 | 3795.6886 | 0 | 1.4 | 74 | Specific | 551 | 551 | 297.61 | 9.93 | 93 | >sp\|P01024\|CO3_HUMAN Complement C3 OS=Homo sapiens OX=9606 GN=C3 PE=1 SV=2 | 492 | 20201114_qwl_milk_1ng_glyco_hcd203040_compound15.13249.13249.3 | controllerType=0 controllerNumber=1 scan=13249 | 28.1896 |
| 10255:3 | 7 | K.TVLTPATNHMGN[+1540.529]VTFTIPANR.E | HexNAc(2)Hex(7) | N[+1541] | 1265.9128 | 3 | 3795.7238 | 3795.6886 | 0 | 9.3 | 74 | Specific | 556.71 | 423.28 | 278.18 | 9.15 | 93 | >sp\|P01024\|CO3_HUMAN Complement C3 OS=Homo sapiens OX=9606 GN=C3 PE=1 SV=2 | 492 | 20201114_qwl_milk_1ng_glyco_hcd203040_compound15.13283.13283.3 | controllerType=0 controllerNumber=1 scan=13283 | 28.2464 |
| 10257:3 | 7 | K.TVLTPATNHMGN[+1540.529]VTFTIPANR.E | HexNAc(2)Hex(7) | N[+1541] | 1265.9128 | 3 | 3795.7238 | 3795.6886 | 0 | 9.3 | 74 | Specific | 605.99 | 585.65 | 247.56 | 11.37 | 93 | >sp\|P01024\|CO3_HUMAN Complement C3 OS=Homo sapiens OX=9606 GN=C3 PE=1 SV=2 | 492 | 20201114_qwl_milk_1ng_glyco_hcd203040_compound15.13285.13285.3 | controllerType=0 controllerNumber=1 scan=13285 | 28.2499 |
| 10288:3 | 7 | K.TVLTPATNHMGN[+1378.476]VTFTIPANR.E | HexNAc(2)Hex(6) | N[+1378] | 1211.8951 | 3 | 3633.6707 | 3633.6358 | 0 | 9.6 | 74 | Specific | 649.45 | 613.23 | 300.19 | 11.37 | 93 | >sp\|P01024\|CO3_HUMAN Complement C3 OS=Homo sapiens OX=9606 GN=C3 PE=1 SV=2 | 492 | 20201114_qwl_milk_1ng_glyco_hcd203040_compound15.13317.13317.3 | controllerType=0 controllerNumber=1 scan=13317 | 28.3043 |
| 10286:3 | 7 | K.TVLTPATNHMGN[+1378.476]VTFTIPANR.E | HexNAc(2)Hex(6) | N[+1378] | 1211.8951 | 3 | 3633.6707 | 3633.6358 | 0 | 9.6 | 74 | Specific | 771.74 | 623.65 | 423.08 | 14.37 | 93 | >sp\|P01024\|CO3_HUMAN Complement C3 OS=Homo sapiens OX=9606 GN=C3 PE=1 SV=2 | 492 | 20201114_qwl_milk_1ng_glyco_hcd203040_compound15.13315.13315.3 | controllerType=0 controllerNumber=1 scan=13315 | 28.2999 |
| 10220:3 | 7 | K.TVLTPATNHMGN[+1540.529]VTFTIPANR.E | HexNAc(2)Hex(7) | N[+1541] | 1265.9028 | 3 | 3795.6937 | 3795.6886 | 0 | 1.4 | 74 | Specific | 786.96 | 786.96 | 446.24 | 15.15 | 93 | >sp\|P01024\|CO3_HUMAN Complement C3 OS=Homo sapiens OX=9606 GN=C3 PE=1 SV=2 | 492 | 20201114_qwl_milk_1ng_glyco_hcd203040_compound15.13247.13247.3 | controllerType=0 controllerNumber=1 scan=13247 | 28.1857 |
| 19019:3 | 8 | K.QIGLYPVLVIDSSGYVNPN[+2205.793]YTGR.I | HexNAc(4)Hex(5)Fuc(2)NeuAc(1) | N[+2206] | 1577.71 | 3 | 4731.1155 | 4731.0963 | 0 | 4.1 | 168 | Specific | 381.37 | 381.37 | 381.37 | 8.96 | 66 | >sp\|P01833\|PIGR_HUMAN Polymeric immunoglobulin receptor OS=Homo sapiens OX=9606 GN=PIGR PE=1 SV=4 | 902 | 20201114_qwl_milk_1ng_glyco_hcd203040_compound15.22436.22436.3 | controllerType=0 controllerNumber=1 scan=22436 | 45.1785 |
| 18997:3 | 8 | K.QIGLYPVLVIDSSGYVNPN[+2205.793]YTGR.I | HexNAc(4)Hex(5)Fuc(2)NeuAc(1) | N[+2206] | 1577.7051 | 3 | 4731.1009 | 4731.0963 | 0 | 1 | 168 | Specific | 818.56 | 753.94 | 753.94 | 16.22 | 66 | >sp\|P01833\|PIGR_HUMAN Polymeric immunoglobulin receptor OS=Homo sapiens OX=9606 GN=PIGR PE=1 SV=4 | 902 | 20201114_qwl_milk_1ng_glyco_hcd203040_compound15.22413.22413.3 | controllerType=0 controllerNumber=1 scan=22413 | 45.1192 |
| 19021:3 | 8 | K.QIGLYPVLVIDSSGYVNPN[+2205.793]YTGR.I | HexNAc(4)Hex(5)Fuc(2)NeuAc(1) | N[+2206] | 1577.71 | 3 | 4731.1155 | 4731.0963 | 0 | 4.1 | 168 | Specific | 857.1 | 653.01 | 653.01 | 16.92 | 66 | >sp\|P01833\|PIGR_HUMAN Polymeric immunoglobulin receptor OS=Homo sapiens OX=9606 GN=PIGR PE=1 SV=4 | 902 | 20201114_qwl_milk_1ng_glyco_hcd203040_compound15.22438.22438.3 | controllerType=0 controllerNumber=1 scan=22438 | 45.1863 |
| 19000:3 | 8 | K.QIGLYPVLVIDSSGYVNPN[+2205.793]YTGR.I | HexNAc(4)Hex(5)Fuc(2)NeuAc(1) | N[+2206] | 1577.7051 | 3 | 4731.1009 | 4731.0963 | 0 | 1 | 168 | Specific | 861.59 | 818.23 | 818.23 | 17.22 | 66 | >sp\|P01833\|PIGR_HUMAN Polymeric immunoglobulin receptor OS=Homo sapiens OX=9606 GN=PIGR PE=1 SV=4 | 902 | 20201114_qwl_milk_1ng_glyco_hcd203040_compound15.22416.22416.3 | controllerType=0 controllerNumber=1 scan=22416 | 45.1266 |
| 19193:3 | 8 | R.LSLLEEPGN[+1913.677]GTFTVILNQLTSR.D | HexNAc(4)Hex(5)NeuAc(1) | N[+1914] | 1439.3375 | 3 | 4315.9978 | 4315.9696 | 0 | 6.5 | 413 | Specific | 403.99 | 362.82 | 362.82 | 5.29 | 66 | >sp\|P01833\|PIGR_HUMAN Polymeric immunoglobulin receptor OS=Homo sapiens OX=9606 GN=PIGR PE=1 SV=4 | 902 | 20201114_qwl_milk_1ng_glyco_hcd203040_compound15.22619.22619.3 | controllerType=0 controllerNumber=1 scan=22619 | 45.6059 |
| 19134:3 | 8 | R.LSLLEEPGN[+2059.735]GTFTVILNQLTSR.D | HexNAc(4)Hex(5)Fuc(1)NeuAc(1) | N[+2060] | 1488.0218 | 3 | 4462.0508 | 4462.0275 | 0 | 5.2 | 413 | Specific | 856.86 | 662.29 | 662.29 | 16.71 | 66 | >sp\|P01833\|PIGR_HUMAN Polymeric immunoglobulin receptor OS=Homo sapiens OX=9606 GN=PIGR PE=1 SV=4 | 902 | 20201114_qwl_milk_1ng_glyco_hcd203040_compound15.22558.22558.3 | controllerType=0 controllerNumber=1 scan=22558 | 45.4941 |
| 20629:3 | 8 | R.LSLLEEPGN[+2205.793]GTFTVILNQLTSR.D | HexNAc(4)Hex(5)Fuc(2)NeuAc(1) | N[+2206] | 1536.7028 | 3 | 4608.0939 | 4608.0854 | 0 | 1.8 | 413 | Specific | 865.56 | 617.79 | 617.79 | 17.02 | 66 | >sp\|P01833\|PIGR_HUMAN Polymeric immunoglobulin receptor OS=Homo sapiens OX=9606 GN=PIGR PE=1 SV=4 | 902 | 20201114_qwl_milk_1ng_glyco_hcd203040_compound15.24121.24121.3 | controllerType=0 controllerNumber=1 scan=24121 | 48.6918 |
| 19137:3 | 8 | R.LSLLEEPGN[+2059.735]GTFTVILNQLTSR.D | HexNAc(4)Hex(5)Fuc(1)NeuAc(1) | N[+2060] | 1488.0218 | 3 | 4462.0508 | 4462.0275 | 0 | 5.2 | 413 | Specific | 907.17 | 632.54 | 632.54 | 17.71 | 66 | >sp\|P01833\|PIGR_HUMAN Polymeric immunoglobulin receptor OS=Homo sapiens OX=9606 GN=PIGR PE=1 SV=4 | 902 | 20201114_qwl_milk_1ng_glyco_hcd203040_compound15.22561.22561.3 | controllerType=0 controllerNumber=1 scan=22561 | 45.4989 |
| 20637:3 | 8 | R.LSLLEEPGN[+2351.851]GTFTVILNQLTSR.D | HexNAc(4)Hex(5)Fuc(3)NeuAc(1) | N[+2352] | 1585.3834 | 3 | 4754.1355 | 4754.1433 | 0 | -1.6 | 413 | Specific | 931.07 | 648.76 | 648.76 | 17.03 | 66 | >sp\|P01833\|PIGR_HUMAN Polymeric immunoglobulin receptor OS=Homo sapiens OX=9606 GN=PIGR PE=1 SV=4 | 902 | 20201114_qwl_milk_1ng_glyco_hcd203040_compound15.24129.24129.3 | controllerType=0 controllerNumber=1 scan=24129 | 48.7054 |
| 20681:3 | 8 | R.LSLLEEPGN[+2059.735]GTFTVILNQLTSR.D | HexNAc(4)Hex(5)Fuc(1)NeuAc(1) | N[+2060] | 1488.0207 | 3 | 4462.0475 | 4462.0275 | 0 | 4.5 | 413 | Specific | 969.26 | 767.87 | 767.87 | 18.71 | 66 | >sp\|P01833\|PIGR_HUMAN Polymeric immunoglobulin receptor OS=Homo sapiens OX=9606 GN=PIGR PE=1 SV=4 | 902 | 20201114_qwl_milk_1ng_glyco_hcd203040_compound15.24176.24176.3 | controllerType=0 controllerNumber=1 scan=24176 | 48.7944 |
| 20683:3 | 8 | R.LSLLEEPGN[+2059.735]GTFTVILNQLTSR.D | HexNAc(4)Hex(5)Fuc(1)NeuAc(1) | N[+2060] | 1488.0207 | 3 | 4462.0475 | 4462.0275 | 0 | 4.5 | 413 | Specific | 1021.52 | 746.39 | 746.39 | 19.71 | 66 | >sp\|P01833\|PIGR_HUMAN Polymeric immunoglobulin receptor OS=Homo sapiens OX=9606 GN=PIGR PE=1 SV=4 | 902 | 20201114_qwl_milk_1ng_glyco_hcd203040_compound15.24178.24178.3 | controllerType=0 controllerNumber=1 scan=24178 | 48.7994 |
| 20626:3 | 8 | R.LSLLEEPGN[+2205.793]GTFTVILNQLTSR.D | HexNAc(4)Hex(5)Fuc(2)NeuAc(1) | N[+2206] | 1536.7028 | 3 | 4608.0939 | 4608.0854 | 0 | 1.8 | 413 | Specific | 1113.69 | 880.45 | 880.45 | 22.02 | 66 | >sp\|P01833\|PIGR_HUMAN Polymeric immunoglobulin receptor OS=Homo sapiens OX=9606 GN=PIGR PE=1 SV=4 | 902 | 20201114_qwl_milk_1ng_glyco_hcd203040_compound15.24118.24118.3 | controllerType=0 controllerNumber=1 scan=24118 | 48.6863 |
| 16254:3 | 8 | K.IIEGEPNLKVPGN[+2205.793]VTAVLGETLK.V | HexNAc(4)Hex(5)Fuc(2)NeuAc(1) | N[+2206] | 1533.0645 | 3 | 4597.179 | 4597.1422 | 0 | 8 | 457 | Specific | 660.26 | 573.33 | 573.33 | 12.55 | 66 | >sp\|P01833\|PIGR_HUMAN Polymeric immunoglobulin receptor OS=Homo sapiens OX=9606 GN=PIGR PE=1 SV=4 | 902 | 20201114_qwl_milk_1ng_glyco_hcd203040_compound15.19536.19536.3 | controllerType=0 controllerNumber=1 scan=19536 | 39.236 |
| 16252:3 | 8 | K.IIEGEPNLKVPGN[+2205.793]VTAVLGETLK.V | HexNAc(4)Hex(5)Fuc(2)NeuAc(1) | N[+2206] | 1533.0645 | 3 | 4597.179 | 4597.1422 | 0 | 8 | 457 | Specific | 1000.31 | 938.16 | 938.16 | 19.55 | 66 | >sp\|P01833\|PIGR_HUMAN Polymeric immunoglobulin receptor OS=Homo sapiens OX=9606 GN=PIGR PE=1 SV=4 | 902 | 20201114_qwl_milk_1ng_glyco_hcd203040_compound15.19534.19534.3 | controllerType=0 controllerNumber=1 scan=19534 | 39.2303 |
| 15896:4 | 8 | K.IIEGEPNLKVPGN[+2205.793]VTAVLGETLKVPC[+57.021]HFPC[+57.021]K.F | HexNAc(4)Hex(5)Fuc(2)NeuAc(1) | C[+57]*2, N[+2206] | 1406.3983 | 4 | 5622.5714 | 5622.5997 | 0 | -5 | 457 | Specific | 300.18 | 300.18 | 300.18 | 4.8 | 66 | >sp\|P01833\|PIGR_HUMAN Polymeric immunoglobulin receptor OS=Homo sapiens OX=9606 GN=PIGR PE=1 SV=4 | 902 | 20201114_qwl_milk_1ng_glyco_hcd203040_compound15.19154.19154.4 | controllerType=0 controllerNumber=1 scan=19154 | 38.5269 |
| 17229:5 | 8 | K.IIEGEPNLKVPGN[+2350.830]VTAVLGETLKVPC[+57.021]HFPC[+57.021]K.F | HexNAc(4)Hex(5)Fuc(1)NeuAc(2) | C[+57]*2, N[+2351] | 1154.3326 | 5 | 5767.6341 | 5767.6372 | 0 | -0.5 | 457 | Specific | 363.07 | 308.41 | 308.41 | 6.7 | 66 | >sp\|P01833\|PIGR_HUMAN Polymeric immunoglobulin receptor OS=Homo sapiens OX=9606 GN=PIGR PE=1 SV=4 | 902 | 20201114_qwl_milk_1ng_glyco_hcd203040_compound15.20564.20564.6 | controllerType=0 controllerNumber=1 scan=20564 | 41.1708 |
| 14556:5 | 8 | K.IIEGEPNLKVPGN[+2571.946]VTAVLGETLKVPC[+57.021]HFPC[+57.021]K.F | HexNAc(5)Hex(6)Fuc(4) | C[+57]*2, N[+2572] | 1198.5653 | 5 | 5988.7974 | 5988.7523 | 0 | 7.5 | 457 | Specific | 399.83 | 399.83 | 399.83 | 8.3 | 66 | >sp\|P01833\|PIGR_HUMAN Polymeric immunoglobulin receptor OS=Homo sapiens OX=9606 GN=PIGR PE=1 SV=4 | 902 | 20201114_qwl_milk_1ng_glyco_hcd203040_compound15.17752.17752.5 | controllerType=0 controllerNumber=1 scan=17752 | 35.9905 |
| 15898:4 | 8 | K.IIEGEPNLKVPGN[+2205.793]VTAVLGETLKVPC[+57.021]HFPC[+57.021]K.F | HexNAc(4)Hex(5)Fuc(2)NeuAc(1) | C[+57]*2, N[+2206] | 1406.3983 | 4 | 5622.5714 | 5622.5997 | 0 | -5 | 457 | Specific | 485.46 | 314.95 | 314.95 | 5.6 | 66 | >sp\|P01833\|PIGR_HUMAN Polymeric immunoglobulin receptor OS=Homo sapiens OX=9606 GN=PIGR PE=1 SV=4 | 902 | 20201114_qwl_milk_1ng_glyco_hcd203040_compound15.19156.19156.4 | controllerType=0 controllerNumber=1 scan=19156 | 38.5305 |
| 17298:5 | 8 | K.IIEGEPNLKVPGN[+2350.830]VTAVLGETLKVPC[+57.021]HFPC[+57.021]K.F | HexNAc(4)Hex(5)Fuc(1)NeuAc(2) | C[+57]*2, N[+2351] | 1154.3441 | 5 | 5767.6915 | 5767.6372 | 0 | 9.4 | 457 | Specific | 603.66 | 457.01 | 457.01 | 11 | 66 | >sp\|P01833\|PIGR_HUMAN Polymeric immunoglobulin receptor OS=Homo sapiens OX=9606 GN=PIGR PE=1 SV=4 | 902 | 20201114_qwl_milk_1ng_glyco_hcd203040_compound15.20635.20635.5 | controllerType=0 controllerNumber=1 scan=20635 | 41.3043 |
| 15713:4 | 8 | K.IIEGEPNLKVPGN[+2205.793]VTAVLGETLKVPC[+57.021]HFPC[+57.021]K.F | HexNAc(4)Hex(5)Fuc(2)NeuAc(1) | C[+57]*2, N[+2206] | 1406.4191 | 4 | 5622.6546 | 5622.5997 | 0 | 9.8 | 457 | Specific | 751.65 | 751.65 | 751.65 | 13.63 | 66 | >sp\|P01833\|PIGR_HUMAN Polymeric immunoglobulin receptor OS=Homo sapiens OX=9606 GN=PIGR PE=1 SV=4 | 902 | 20201114_qwl_milk_1ng_glyco_hcd203040_compound15.18960.18960.4 | controllerType=0 controllerNumber=1 scan=18960 | 38.1524 |
| 14554:5 | 8 | K.IIEGEPNLKVPGN[+2571.946]VTAVLGETLKVPC[+57.021]HFPC[+57.021]K.F | HexNAc(5)Hex(6)Fuc(4) | C[+57]*2, N[+2572] | 1198.5653 | 5 | 5988.7974 | 5988.7523 | 0 | 7.5 | 457 | Specific | 755.23 | 755.23 | 755.23 | 14.22 | 66 | >sp\|P01833\|PIGR_HUMAN Polymeric immunoglobulin receptor OS=Homo sapiens OX=9606 GN=PIGR PE=1 SV=4 | 902 | 20201114_qwl_milk_1ng_glyco_hcd203040_compound15.17750.17750.5 | controllerType=0 controllerNumber=1 scan=17750 | 35.985 |
| 15709:5 | 8 | K.IIEGEPNLKVPGN[+2205.793]VTAVLGETLKVPC[+57.021]HFPC[+57.021]K.F | HexNAc(4)Hex(5)Fuc(2)NeuAc(1) | C[+57]*2, N[+2206] | 1125.3368 | 5 | 5622.6549 | 5622.5997 | 0 | 9.8 | 457 | Specific | 812.49 | 800.58 | 800.58 | 15 | 66 | >sp\|P01833\|PIGR_HUMAN Polymeric immunoglobulin receptor OS=Homo sapiens OX=9606 GN=PIGR PE=1 SV=4 | 902 | 20201114_qwl_milk_1ng_glyco_hcd203040_compound15.18956.18956.5 | controllerType=0 controllerNumber=1 scan=18956 | 38.1416 |
| 17227:5 | 8 | K.IIEGEPNLKVPGN[+2350.830]VTAVLGETLKVPC[+57.021]HFPC[+57.021]K.F | HexNAc(4)Hex(5)Fuc(1)NeuAc(2) | C[+57]*2, N[+2351] | 1154.3326 | 5 | 5767.6341 | 5767.6372 | 0 | -0.5 | 457 | Specific | 866.84 | 621.23 | 621.23 | 16.36 | 66 | >sp\|P01833\|PIGR_HUMAN Polymeric immunoglobulin receptor OS=Homo sapiens OX=9606 GN=PIGR PE=1 SV=4 | 902 | 20201114_qwl_milk_1ng_glyco_hcd203040_compound15.20562.20562.6 | controllerType=0 controllerNumber=1 scan=20562 | 41.1645 |
| 15711:4 | 8 | K.IIEGEPNLKVPGN[+2205.793]VTAVLGETLKVPC[+57.021]HFPC[+57.021]K.F | HexNAc(4)Hex(5)Fuc(2)NeuAc(1) | C[+57]*2, N[+2206] | 1406.4191 | 4 | 5622.6546 | 5622.5997 | 0 | 9.8 | 457 | Specific | 1091.57 | 1084.34 | 1084.34 | 19.63 | 66 | >sp\|P01833\|PIGR_HUMAN Polymeric immunoglobulin receptor OS=Homo sapiens OX=9606 GN=PIGR PE=1 SV=4 | 902 | 20201114_qwl_milk_1ng_glyco_hcd203040_compound15.18958.18958.4 | controllerType=0 controllerNumber=1 scan=18958 | 38.1447 |
| 15707:5 | 8 | K.IIEGEPNLKVPGN[+2205.793]VTAVLGETLKVPC[+57.021]HFPC[+57.021]K.F | HexNAc(4)Hex(5)Fuc(2)NeuAc(1) | C[+57]*2, N[+2206] | 1125.3368 | 5 | 5622.6549 | 5622.5997 | 0 | 9.8 | 457 | Specific | 1120.07 | 1103.49 | 1103.49 | 21 | 66 | >sp\|P01833\|PIGR_HUMAN Polymeric immunoglobulin receptor OS=Homo sapiens OX=9606 GN=PIGR PE=1 SV=4 | 902 | 20201114_qwl_milk_1ng_glyco_hcd203040_compound15.18954.18954.5 | controllerType=0 controllerNumber=1 scan=18954 | 38.136 |
| 07835:3 | 8 | K.YWC[+57.021]KWN[+1913.677]NTGC[+57.021]QALPSQDEGPSK.A | HexNAc(4)Hex(5)NeuAc(1) | C[+57]*2, N[+1914] | 1513.947 | 3 | 4539.8263 | 4539.822 | 0 | 1 | 494 | Specific | 214.08 | 214.08 | 214.08 | 4.95 | 66 | >sp\|P01833\|PIGR_HUMAN Polymeric immunoglobulin receptor OS=Homo sapiens OX=9606 GN=PIGR PE=1 SV=4 | 902 | 20201114_qwl_milk_1ng_glyco_hcd203040_compound15.10772.10772.3 | controllerType=0 controllerNumber=1 scan=10772 | 24.1195 |
| 07837:3 | 8 | K.YWC[+57.021]KWN[+1913.677]NTGC[+57.021]QALPSQDEGPSK.A | HexNAc(4)Hex(5)NeuAc(1) | C[+57]*2, N[+1914] | 1513.947 | 3 | 4539.8263 | 4539.822 | 0 | 1 | 494 | Specific | 492.44 | 456.42 | 456.42 | 9.31 | 66 | >sp\|P01833\|PIGR_HUMAN Polymeric immunoglobulin receptor OS=Homo sapiens OX=9606 GN=PIGR PE=1 SV=4 | 902 | 20201114_qwl_milk_1ng_glyco_hcd203040_compound15.10774.10774.3 | controllerType=0 controllerNumber=1 scan=10774 | 24.1235 |
| 05801:3 | 8 | K.WN[+2059.735]NTGC[+57.021]QALPSQDEGPSK.A | HexNAc(4)Hex(5)Fuc(1)NeuAc(1) | C[+57], N[+2060] | 1350.2094 | 3 | 4048.6137 | 4048.6116 | 0 | 0.5 | 498 | Specific | 250.85 | 212.14 | 212.14 | 5.93 | 66 | >sp\|P01833\|PIGR_HUMAN Polymeric immunoglobulin receptor OS=Homo sapiens OX=9606 GN=PIGR PE=1 SV=4 | 902 | 20201114_qwl_milk_1ng_glyco_hcd203040_compound15.8668.8668.3 | controllerType=0 controllerNumber=1 scan=8668 | 20.6566 |
| 05878:3 | 8 | K.WN[+1913.677]NTGC[+57.021]QALPSQDEGPSK.A | HexNAc(4)Hex(5)NeuAc(1) | C[+57], N[+1914] | 1301.5258 | 3 | 3902.5629 | 3902.5537 | 0 | 2.4 | 498 | Specific | 433.99 | 354.61 | 354.61 | 7.83 | 66 | >sp\|P01833\|PIGR_HUMAN Polymeric immunoglobulin receptor OS=Homo sapiens OX=9606 GN=PIGR PE=1 SV=4 | 902 | 20201114_qwl_milk_1ng_glyco_hcd203040_compound15.8747.8747.3 | controllerType=0 controllerNumber=1 scan=8747 | 20.7869 |
| 05803:3 | 8 | K.WN[+2059.735]NTGC[+57.021]QALPSQDEGPSK.A | HexNAc(4)Hex(5)Fuc(1)NeuAc(1) | C[+57], N[+2060] | 1350.2094 | 3 | 4048.6137 | 4048.6116 | 0 | 0.5 | 498 | Specific | 442.32 | 374.76 | 374.76 | 9.24 | 66 | >sp\|P01833\|PIGR_HUMAN Polymeric immunoglobulin receptor OS=Homo sapiens OX=9606 GN=PIGR PE=1 SV=4 | 902 | 20201114_qwl_milk_1ng_glyco_hcd203040_compound15.8670.8670.3 | controllerType=0 controllerNumber=1 scan=8670 | 20.6607 |
| 05876:3 | 8 | K.WN[+1913.677]NTGC[+57.021]QALPSQDEGPSK.A | HexNAc(4)Hex(5)NeuAc(1) | C[+57], N[+1914] | 1301.5258 | 3 | 3902.5629 | 3902.5537 | 0 | 2.4 | 498 | Specific | 443.54 | 408.97 | 408.97 | 9.32 | 66 | >sp\|P01833\|PIGR_HUMAN Polymeric immunoglobulin receptor OS=Homo sapiens OX=9606 GN=PIGR PE=1 SV=4 | 902 | 20201114_qwl_milk_1ng_glyco_hcd203040_compound15.8745.8745.3 | controllerType=0 controllerNumber=1 scan=8745 | 20.7829 |
| 19086:3 | 9 | R.NIAAFGGDPNN[+1913.677]ITLFGESAGGASVSLQTLSPYNK.G | HexNAc(4)Hex(5)NeuAc(1) | N[+1914] | 1775.4642 | 3 | 5324.378 | 5324.3633 | 0 | 2.8 | 197 | Specific | 448.54 | 445.46 | 445.46 | 9.32 | 48 | >sp\|P19835\|CEL_HUMAN Bile salt-activated lipase OS=Homo sapiens OX=9606 GN=CEL PE=1 SV=3 | 476 | 20201114_qwl_milk_1ng_glyco_hcd203040_compound15.22508.22508.3 | controllerType=0 controllerNumber=1 scan=22508 | 45.4055 |
| 19084:3 | 9 | R.NIAAFGGDPNN[+1913.677]ITLFGESAGGASVSLQTLSPYNK.G | HexNAc(4)Hex(5)NeuAc(1) | N[+1914] | 1775.4642 | 3 | 5324.378 | 5324.3633 | 0 | 2.8 | 197 | Specific | 751.19 | 743.39 | 743.39 | 15.09 | 48 | >sp\|P19835\|CEL_HUMAN Bile salt-activated lipase OS=Homo sapiens OX=9606 GN=CEL PE=1 SV=3 | 476 | 20201114_qwl_milk_1ng_glyco_hcd203040_compound15.22506.22506.3 | controllerType=0 controllerNumber=1 scan=22506 | 45.4015 |
| 17890:4 | 10 | R.LSLHRPALEDLLLGSEAN[+1378.476]LTC[+57.021]TLTGLR.D | HexNAc(2)Hex(6) | C[+57], N[+1378] | 1086.2726 | 4 | 4342.0687 | 4342.074 | 0 | -1.2 | 127 | Specific | 239.16 | 239.16 | 239.16 | 4.22 | 47 | >sp\|P01876\|IGHA1_HUMAN Immunoglobulin heavy constant alpha 1 OS=Homo sapiens OX=9606 GN=IGHA1 PE=1 SV=2 | 255 | 20201114_qwl_milk_1ng_glyco_hcd203040_compound15.21253.21253.3 | controllerType=0 controllerNumber=1 scan=21253 | 42.5192 |
| 17766:3 | 10 | R.LSLHRPALEDLLLGSEAN[+1702.581]LTC[+57.021]TLTGLR.D | HexNAc(2)Hex(8) | C[+57], N[+1703] | 1556.0706 | 3 | 4666.1973 | 4666.1796 | 0 | 3.8 | 127 | Specific | 239.96 | 239.96 | 239.96 | 5.87 | 47 | >sp\|P01876\|IGHA1_HUMAN Immunoglobulin heavy constant alpha 1 OS=Homo sapiens OX=9606 GN=IGHA1 PE=1 SV=2 | 255 | 20201114_qwl_milk_1ng_glyco_hcd203040_compound15.21124.21124.3 | controllerType=0 controllerNumber=1 scan=21124 | 42.2342 |
| 17942:4 | 10 | R.LSLHRPALEDLLLGSEAN[+1460.529]LTC[+57.021]TLTGLR.D | HexNAc(4)Hex(4) | C[+57], N[+1461] | 1106.7959 | 4 | 4424.1619 | 4424.1271 | 0 | 7.9 | 127 | Specific | 396.4 | 396.4 | 396.4 | 8.58 | 47 | >sp\|P01876\|IGHA1_HUMAN Immunoglobulin heavy constant alpha 1 OS=Homo sapiens OX=9606 GN=IGHA1 PE=1 SV=2 | 255 | 20201114_qwl_milk_1ng_glyco_hcd203040_compound15.21308.21308.4 | controllerType=0 controllerNumber=1 scan=21308 | 42.6662 |
| 18838:4 | 10 | R.LSLHRPALEDLLLGSEAN[+1913.677]LTC[+57.021]TLTGLR.D | HexNAc(4)Hex(5)NeuAc(1) | C[+57], N[+1914] | 1220.0863 | 4 | 4877.3236 | 4877.2753 | 0 | 9.9 | 127 | Specific | 401.79 | 401.79 | 401.79 | 8.36 | 47 | >sp\|P01876\|IGHA1_HUMAN Immunoglobulin heavy constant alpha 1 OS=Homo sapiens OX=9606 GN=IGHA1 PE=1 SV=2 | 255 | 20201114_qwl_milk_1ng_glyco_hcd203040_compound15.22246.22246.4 | controllerType=0 controllerNumber=1 scan=22246 | 44.7306 |
| 17844:4 | 10 | R.LSLHRPALEDLLLGSEAN[+1825.661]LTC[+57.021]TLTGLR.D | HexNAc(5)Hex(5) | C[+57], N[+1826] | 1198.0821 | 4 | 4789.3065 | 4789.2592 | 0 | 9.9 | 127 | Specific | 408.71 | 408.71 | 408.71 | 8.36 | 47 | >sp\|P01876\|IGHA1_HUMAN Immunoglobulin heavy constant alpha 1 OS=Homo sapiens OX=9606 GN=IGHA1 PE=1 SV=2 | 255 | 20201114_qwl_milk_1ng_glyco_hcd203040_compound15.21205.21205.3 | controllerType=0 controllerNumber=1 scan=21205 | 42.391 |
| 17888:4 | 10 | R.LSLHRPALEDLLLGSEAN[+1378.476]LTC[+57.021]TLTGLR.D | HexNAc(2)Hex(6) | C[+57], N[+1378] | 1086.2726 | 4 | 4342.0687 | 4342.074 | 0 | -1.2 | 127 | Specific | 549.77 | 549.77 | 549.77 | 9.42 | 47 | >sp\|P01876\|IGHA1_HUMAN Immunoglobulin heavy constant alpha 1 OS=Homo sapiens OX=9606 GN=IGHA1 PE=1 SV=2 | 255 | 20201114_qwl_milk_1ng_glyco_hcd203040_compound15.21251.21251.3 | controllerType=0 controllerNumber=1 scan=21251 | 42.5124 |
| 17790:4 | 10 | R.LSLHRPALEDLLLGSEAN[+1540.529]LTC[+57.021]TLTGLR.D | HexNAc(2)Hex(7) | C[+57], N[+1541] | 1126.7984 | 4 | 4504.1717 | 4504.1268 | 0 | 10 | 127 | Specific | 561.5 | 561.5 | 561.5 | 9.07 | 47 | >sp\|P01876\|IGHA1_HUMAN Immunoglobulin heavy constant alpha 1 OS=Homo sapiens OX=9606 GN=IGHA1 PE=1 SV=2 | 255 | 20201114_qwl_milk_1ng_glyco_hcd203040_compound15.21149.21149.2 | controllerType=0 controllerNumber=1 scan=21149 | 42.274 |
| 17864:3 | 10 | R.LSLHRPALEDLLLGSEAN[+1663.608]LTC[+57.021]TLTGLR.D | HexNAc(5)Hex(4) | C[+57], N[+1664] | 1543.0774 | 3 | 4627.2178 | 4627.2064 | 0 | 2.5 | 127 | Specific | 604.85 | 559.12 | 559.12 | 11.77 | 47 | >sp\|P01876\|IGHA1_HUMAN Immunoglobulin heavy constant alpha 1 OS=Homo sapiens OX=9606 GN=IGHA1 PE=1 SV=2 | 255 | 20201114_qwl_milk_1ng_glyco_hcd203040_compound15.21226.21226.3 | controllerType=0 controllerNumber=1 scan=21226 | 42.4565 |
| 17768:3 | 10 | R.LSLHRPALEDLLLGSEAN[+1702.581]LTC[+57.021]TLTGLR.D | HexNAc(2)Hex(8) | C[+57], N[+1703] | 1556.0706 | 3 | 4666.1973 | 4666.1796 | 0 | 3.8 | 127 | Specific | 609.15 | 609.15 | 609.15 | 11.77 | 47 | >sp\|P01876\|IGHA1_HUMAN Immunoglobulin heavy constant alpha 1 OS=Homo sapiens OX=9606 GN=IGHA1 PE=1 SV=2 | 255 | 20201114_qwl_milk_1ng_glyco_hcd203040_compound15.21126.21126.3 | controllerType=0 controllerNumber=1 scan=21126 | 42.2382 |
| 17950:4 | 10 | R.LSLHRPALEDLLLGSEAN[+1298.476]LTC[+57.021]TLTGLR.D | HexNAc(4)Hex(3) | C[+57], N[+1298] | 1066.28 | 4 | 4262.098 | 4262.0742 | 0 | 5.6 | 127 | Specific | 645.87 | 533.2 | 533.2 | 11.5 | 47 | >sp\|P01876\|IGHA1_HUMAN Immunoglobulin heavy constant alpha 1 OS=Homo sapiens OX=9606 GN=IGHA1 PE=1 SV=2 | 255 | 20201114_qwl_milk_1ng_glyco_hcd203040_compound15.21316.21316.4 | controllerType=0 controllerNumber=1 scan=21316 | 42.6847 |
| 17857:4 | 10 | R.LSLHRPALEDLLLGSEAN[+1663.608]LTC[+57.021]TLTGLR.D | HexNAc(5)Hex(4) | C[+57], N[+1664] | 1157.5629 | 4 | 4627.2298 | 4627.2064 | 0 | 5.1 | 127 | Specific | 646.25 | 646.25 | 646.25 | 11.76 | 47 | >sp\|P01876\|IGHA1_HUMAN Immunoglobulin heavy constant alpha 1 OS=Homo sapiens OX=9606 GN=IGHA1 PE=1 SV=2 | 255 | 20201114_qwl_milk_1ng_glyco_hcd203040_compound15.21219.21219.4 | controllerType=0 controllerNumber=1 scan=21219 | 42.4351 |
| 17928:3 | 10 | R.LSLHRPALEDLLLGSEAN[+1501.555]LTC[+57.021]TLTGLR.D | HexNAc(5)Hex(3) | C[+57], N[+1502] | 1489.0605 | 3 | 4465.1669 | 4465.1536 | 0 | 3 | 127 | Specific | 654.9 | 604.34 | 604.34 | 12.77 | 47 | >sp\|P01876\|IGHA1_HUMAN Immunoglobulin heavy constant alpha 1 OS=Homo sapiens OX=9606 GN=IGHA1 PE=1 SV=2 | 255 | 20201114_qwl_milk_1ng_glyco_hcd203040_compound15.21293.21293.3 | controllerType=0 controllerNumber=1 scan=21293 | 42.6142 |
| 17926:3 | 10 | R.LSLHRPALEDLLLGSEAN[+1501.555]LTC[+57.021]TLTGLR.D | HexNAc(5)Hex(3) | C[+57], N[+1502] | 1489.0605 | 3 | 4465.1669 | 4465.1536 | 0 | 3 | 127 | Specific | 655.65 | 584.85 | 584.85 | 12.77 | 47 | >sp\|P01876\|IGHA1_HUMAN Immunoglobulin heavy constant alpha 1 OS=Homo sapiens OX=9606 GN=IGHA1 PE=1 SV=2 | 255 | 20201114_qwl_milk_1ng_glyco_hcd203040_compound15.21291.21291.3 | controllerType=0 controllerNumber=1 scan=21291 | 42.5917 |
| 17860:4 | 10 | R.LSLHRPALEDLLLGSEAN[+1663.608]LTC[+57.021]TLTGLR.D | HexNAc(5)Hex(4) | C[+57], N[+1664] | 1157.5629 | 4 | 4627.2298 | 4627.2064 | 0 | 5.1 | 127 | Specific | 660.79 | 549.52 | 549.52 | 12.76 | 47 | >sp\|P01876\|IGHA1_HUMAN Immunoglobulin heavy constant alpha 1 OS=Homo sapiens OX=9606 GN=IGHA1 PE=1 SV=2 | 255 | 20201114_qwl_milk_1ng_glyco_hcd203040_compound15.21222.21222.4 | controllerType=0 controllerNumber=1 scan=21222 | 42.4496 |
| 17976:3 | 10 | R.LSLHRPALEDLLLGSEAN[+1095.397]LTC[+57.021]TLTGLR.D | HexNAc(3)Hex(3) | C[+57], N[+1095] | 1353.6765 | 3 | 4059.0149 | 4058.9949 | 0 | 4.9 | 127 | Specific | 661.68 | 653.97 | 653.97 | 12.38 | 47 | >sp\|P01876\|IGHA1_HUMAN Immunoglobulin heavy constant alpha 1 OS=Homo sapiens OX=9606 GN=IGHA1 PE=1 SV=2 | 255 | 20201114_qwl_milk_1ng_glyco_hcd203040_compound15.21343.21343.3 | controllerType=0 controllerNumber=1 scan=21343 | 42.7464 |
| 17931:4 | 10 | R.LSLHRPALEDLLLGSEAN[+1216.423]LTC[+57.021]TLTGLR.D | HexNAc(2)Hex(5) | C[+57], N[+1216] | 1045.764 | 4 | 4180.034 | 4180.0211 | 0 | 3.1 | 127 | Specific | 674.28 | 674.28 | 674.28 | 13.15 | 47 | >sp\|P01876\|IGHA1_HUMAN Immunoglobulin heavy constant alpha 1 OS=Homo sapiens OX=9606 GN=IGHA1 PE=1 SV=2 | 255 | 20201114_qwl_milk_1ng_glyco_hcd203040_compound15.21296.21296.4 | controllerType=0 controllerNumber=1 scan=21296 | 42.6177 |
| 17863:4 | 10 | R.LSLHRPALEDLLLGSEAN[+1622.582]LTC[+57.021]TLTGLR.D | HexNAc(4)Hex(5) | C[+57], N[+1623] | 1147.3031 | 4 | 4586.1907 | 4586.1799 | 0 | 2.4 | 127 | Specific | 676.18 | 650.28 | 650.28 | 13.15 | 47 | >sp\|P01876\|IGHA1_HUMAN Immunoglobulin heavy constant alpha 1 OS=Homo sapiens OX=9606 GN=IGHA1 PE=1 SV=2 | 255 | 20201114_qwl_milk_1ng_glyco_hcd203040_compound15.21225.21225.4 | controllerType=0 controllerNumber=1 scan=21225 | 42.4551 |
| 17760:4 | 10 | R.LSLHRPALEDLLLGSEAN[+1702.581]LTC[+57.021]TLTGLR.D | HexNAc(2)Hex(8) | C[+57], N[+1703] | 1167.3027 | 4 | 4666.1888 | 4666.1796 | 0 | 2 | 127 | Specific | 682.89 | 682.89 | 682.89 | 13.07 | 47 | >sp\|P01876\|IGHA1_HUMAN Immunoglobulin heavy constant alpha 1 OS=Homo sapiens OX=9606 GN=IGHA1 PE=1 SV=2 | 255 | 20201114_qwl_milk_1ng_glyco_hcd203040_compound15.21118.21118.4 | controllerType=0 controllerNumber=1 scan=21118 | 42.2256 |
| 17866:3 | 10 | R.LSLHRPALEDLLLGSEAN[+1663.608]LTC[+57.021]TLTGLR.D | HexNAc(5)Hex(4) | C[+57], N[+1664] | 1543.0774 | 3 | 4627.2178 | 4627.2064 | 0 | 2.5 | 127 | Specific | 683.38 | 583.45 | 583.45 | 12.77 | 47 | >sp\|P01876\|IGHA1_HUMAN Immunoglobulin heavy constant alpha 1 OS=Homo sapiens OX=9606 GN=IGHA1 PE=1 SV=2 | 255 | 20201114_qwl_milk_1ng_glyco_hcd203040_compound15.21228.21228.3 | controllerType=0 controllerNumber=1 scan=21228 | 42.4639 |
| 17925:4 | 10 | R.LSLHRPALEDLLLGSEAN[+1501.555]LTC[+57.021]TLTGLR.D | HexNAc(5)Hex(3) | C[+57], N[+1502] | 1117.0475 | 4 | 4465.1683 | 4465.1536 | 0 | 3.3 | 127 | Specific | 721.33 | 721.33 | 721.33 | 14.15 | 47 | >sp\|P01876\|IGHA1_HUMAN Immunoglobulin heavy constant alpha 1 OS=Homo sapiens OX=9606 GN=IGHA1 PE=1 SV=2 | 255 | 20201114_qwl_milk_1ng_glyco_hcd203040_compound15.21290.21290.4 | controllerType=0 controllerNumber=1 scan=21290 | [illegible] |
| 17927:4 | 10 | R.LSLHRPALEDLLLGSEAN[+1501.555]LTC[+57.021]TLTGLR.D | HexNAc(5)Hex(3) | C[+57], N[+1502] | 1117.0475 | 4 | 4465.1683 | 4465.1536 | 0 | 3.3 | 127 | Specific | 762.65 | 762.65 | 762.65 | 15.15 | 47 | >sp\|P01876\|IGHA1_HUMAN Immunoglobulin heavy constant alpha 1 OS=Homo sapiens OX=9606 GN=IGHA1 PE=1 SV=2 | 255 | 20201114_qwl_milk_1ng_glyco_hcd203040_compound15.21292.21292.4 | controllerType=0 controllerNumber=1 scan=21292 | [illegible] |
| 18841:4 | 10 | R.LSLHRPALEDLLLGSEAN[+1913.677]LTC[+57.021]TLTGLR.D | HexNAc(4)Hex(5)NeuAc(1) | C[+57], N[+1914] | 1220.0863 | 4 | 4877.3236 | 4877.2753 | 0 | 9.9 | 127 | Specific | 817.2 | 740.32 | 740.32 | 15.29 | 47 | >sp\|P01876\|IGHA1_HUMAN Immunoglobulin heavy constant alpha 1 OS=Homo sapiens OX=9606 GN=IGHA1 PE=1 SV=2 | 255 | 20201114_qwl_milk_1ng_glyco_hcd203040_compound15.22249.22249.4 | controllerType=0 controllerNumber=1 scan=22249 | 44.7427 |
| 17934:4 | 10 | R.LSLHRPALEDLLLGSEAN[+1216.423]LTC[+57.021]TLTGLR.D | HexNAc(2)Hex(5) | C[+57], N[+1216] | 1045.764 | 4 | 4180.034 | 4180.0211 | 0 | 3.1 | 127 | Specific | 824.11 | 824.11 | 824.11 | 16.15 | 47 | >sp\|P01876\|IGHA1_HUMAN Immunoglobulin heavy constant alpha 1 OS=Homo sapiens OX=9606 GN=IGHA1 PE=1 SV=2 | 255 | 20201114_qwl_milk_1ng_glyco_hcd203040_compound15.21299.21299.4 | controllerType=0 controllerNumber=1 scan=21299 | [illegible] |
| 17865:4 | 10 | R.LSLHRPALEDLLLGSEAN[+1622.582]LTC[+57.021]TLTGLR.D | HexNAc(4)Hex(5) | C[+57], N[+1623] | 1147.3031 | 4 | 4586.1907 | 4586.1799 | 0 | 2.4 | 127 | Specific | 826.37 | 725.85 | 725.85 | 16.15 | 47 | >sp\|P01876\|IGHA1_HUMAN Immunoglobulin heavy constant alpha 1 OS=Homo sapiens OX=9606 GN=IGHA1 PE=1 SV=2 | 255 | 20201114_qwl_milk_1ng_glyco_hcd203040_compound15.21227.21227.4 | controllerType=0 controllerNumber=1 scan=21227 | 42.4618 |
| 17917:4 | 10 | R.LSLHRPALEDLLLGSEAN[+1298.476]LTC[+57.021]TLTGLR.D | HexNAc(4)Hex(3) | C[+57], N[+1298] | 1066.2841 | 4 | 4262.1146 | 4262.0742 | 0 | 9.5 | 127 | Specific | 836.1 | 805.1 | 805.1 | 15.29 | 47 | >sp\|P01876\|IGHA1_HUMAN Immunoglobulin heavy constant alpha 1 OS=Homo sapiens OX=9606 GN=IGHA1 PE=1 SV=2 | 255 | 20201114_qwl_milk_1ng_glyco_hcd203040_compound15.21281.21281.2 | controllerType=0 controllerNumber=1 scan=21281 | 42.5722 |
| 17979:3 | 10 | R.LSLHRPALEDLLLGSEAN[+1095.397]LTC[+57.021]TLTGLR.D | HexNAc(3)Hex(3) | C[+57], N[+1095] | 1353.6765 | 3 | 4059.0149 | 4058.9949 | 0 | 4.9 | 127 | Specific | 847.69 | 847.69 | 847.69 | 15.38 | 47 | >sp\|P01876\|IGHA1_HUMAN Immunoglobulin heavy constant alpha 1 OS=Homo sapiens OX=9606 GN=IGHA1 PE=1 SV=2 | 255 | 20201114_qwl_milk_1ng_glyco_hcd203040_compound15.21346.21346.3 | controllerType=0 controllerNumber=1 scan=21346 | 42.7592 |
| 17945:4 | 10 | R.LSLHRPALEDLLLGSEAN[+1460.529]LTC[+57.021]TLTGLR.D | HexNAc(4)Hex(4) | C[+57], N[+1461] | 1106.7959 | 4 | 4424.1619 | 4424.1271 | 0 | 7.9 | 127 | Specific | 893.59 | 811.45 | 811.45 | 16.51 | 47 | >sp\|P01876\|IGHA1_HUMAN Immunoglobulin heavy constant alpha 1 OS=Homo sapiens OX=9606 GN=IGHA1 PE=1 SV=2 | 255 | 20201114_qwl_milk_1ng_glyco_hcd203040_compound15.21311.21311.4 | controllerType=0 controllerNumber=1 scan=21311 | 42.6709 |
| 17953:4 | 10 | R.LSLHRPALEDLLLGSEAN[+1298.476]LTC[+57.021]TLTGLR.D | HexNAc(4)Hex(3) | C[+57], N[+1298] | 1066.28 | 4 | 4262.098 | 4262.0742 | 0 | 5.6 | 127 | Specific | 894.38 | 894.38 | 894.38 | 16.5 | 47 | >sp\|P01876\|IGHA1_HUMAN Immunoglobulin heavy constant alpha 1 OS=Homo sapiens OX=9606 GN=IGHA1 PE=1 SV=2 | 255 | 20201114_qwl_milk_1ng_glyco_hcd203040_compound15.21319.21319.4 | controllerType=0 controllerNumber=1 scan=21319 | 42.6897 |
| 17840:4 | 10 | R.LSLHRPALEDLLLGSEAN[+1702.581]LTC[+57.021]TLTGLR.D | HexNAc(2)Hex(8) | C[+57], N[+1703] | 1167.3116 | 4 | 4666.2244 | 4666.1796 | 0 | 9.6 | 127 | Specific | 967.99 | 967.99 | 967.99 | 18.29 | 47 | >sp\|P01876\|IGHA1_HUMAN Immunoglobulin heavy constant alpha 1 OS=Homo sapiens OX=9606 GN=IGHA1 PE=1 SV=2 | 255 | 20201114_qwl_milk_1ng_glyco_hcd203040_compound15.21201.21201.4 | controllerType=0 controllerNumber=1 scan=21201 | 42.3807 |
| 17763:4 | 10 | R.LSLHRPALEDLLLGSEAN[+1702.581]LTC[+57.021]TLTGLR.D | HexNAc(2)Hex(8) | C[+57], N[+1703] | 1167.3027 | 4 | 4666.1888 | 4666.1796 | 0 | 2 | 127 | Specific | 986.57 | 986.57 | 986.57 | 19.07 | 47 | >sp\|P01876\|IGHA1_HUMAN Immunoglobulin heavy constant alpha 1 OS=Homo sapiens OX=9606 GN=IGHA1 PE=1 SV=2 | 255 | 20201114_qwl_milk_1ng_glyco_hcd203040_compound15.21121.21121.4 | controllerType=0 controllerNumber=1 scan=21121 | 42.23 |
| 17842:4 | 10 | R.LSLHRPALEDLLLGSEAN[+1702.581]LTC[+57.021]TLTGLR.D | HexNAc(2)Hex(8) | C[+57], N[+1703] | 1167.3116 | 4 | 4666.2244 | 4666.1796 | 0 | 9.6 | 127 | Specific | 1042.32 | 1042.32 | 1042.32 | 19.29 | 47 | >sp\|P01876\|IGHA1_HUMAN Immunoglobulin heavy constant alpha 1 OS=Homo sapiens OX=9606 GN=IGHA1 PE=1 SV=2 | 255 | 20201114_qwl_milk_1ng_glyco_hcd203040_compound15.21203.21203.4 | controllerType=0 controllerNumber=1 scan=21203 | 42.388 |
| 17792:4 | 10 | R.LSLHRPALEDLLLGSEAN[+1540.529]LTC[+57.021]TLTGLR.D | HexNAc(2)Hex(7) | C[+57], N[+1541] | 1126.7984 | 4 | 4504.1717 | 4504.1268 | 0 | 10 | 127 | Specific | 1079.5 | 910.76 | 910.76 | 20.29 | 47 | >sp\|P01876\|IGHA1_HUMAN Immunoglobulin heavy constant alpha 1 OS=Homo sapiens OX=9606 GN=IGHA1 PE=1 SV=2 | 255 | 20201114_qwl_milk_1ng_glyco_hcd203040_compound15.21151.21151.2 | controllerType=0 controllerNumber=1 scan=21151 | 42.2775 |
| 19029:5 | 10 | R.LSLHRPALEDLLLGSEAN[+1501.555]LTC[+57.021]TLTGLRDASGVTFTWTPSSGK.S | HexNAc(5)Hex(3) | C[+57], N[+1502] | 1198.1816 | 5 | 5986.8791 | 5986.8636 | 0 | 2.6 | 127 | Specific | 902.95 | 750.5 | 750.5 | 18.51 | 47 | >sp\|P01876\|IGHA1_HUMAN Immunoglobulin heavy constant alpha 1 OS=Homo sapiens OX=9606 GN=IGHA1 PE=1 SV=2 | 255 | 20201114_qwl_milk_1ng_glyco_hcd203040_compound15.22447.22447.5 | controllerType=0 controllerNumber=1 scan=22447 | 45.2037 |
| 19031:5 | 10 | R.LSLHRPALEDLLLGSEAN[+1501.555]LTC[+57.021]TLTGLRDASGVTFTWTPSSGK.S | HexNAc(5)Hex(3) | C[+57], N[+1502] | 1198.1816 | 5 | 5986.8791 | 5986.8636 | 0 | 2.6 | 127 | Specific | 1036.68 | 1003.6 | 1003.6 | 20.51 | 47 | >sp\|P01876\|IGHA1_HUMAN Immunoglobulin heavy constant alpha 1 OS=Homo sapiens OX=9606 GN=IGHA1 PE=1 SV=2 | 255 | 20201114_qwl_milk_1ng_glyco_hcd203040_compound15.22449.22449.5 | controllerType=0 controllerNumber=1 scan=22449 | 45.2097 |
| 11967:4 | 10 | K.TIDRLAGKPTHVN[+1647.613]VSVVMAEVDGTC[+57.021]Y.- | HexNAc(5)Hex(3)Fuc(1) | C[+57], N[+1648] | 1120.7603 | 4 | 4480.0194 | 4480.0151 | 0 | 0.9 | 328 | CRagged | 263.78 | 218.05 | 218.05 | 6.48 | 47 | >sp\|P01876\|IGHA1_HUMAN Immunoglobulin heavy constant alpha 1 OS=Homo sapiens OX=9606 GN=IGHA1 PE=1 SV=2 | 255 | 20201114_qwl_milk_1ng_glyco_hcd203040_compound15.15061.15061.4 | controllerType=0 controllerNumber=1 scan=15061 | 31.2906 |
| 10378:3 | 10 | R.LAGKPTHVN[+1378.476]VSVVMAEVDGTC[+57.021]Y.- | HexNAc(2)Hex(6) | C[+57], N[+1378] | 1242.5486 | 3 | 3725.6312 | 3725.6178 | 0 | 3.6 | 332 | CRagged | 227.76 | 129.28 | 129.28 | 3.92 | 47 | >sp\|P01876\|IGHA1_HUMAN Immunoglobulin heavy constant alpha 1 OS=Homo sapiens OX=9606 GN=IGHA1 PE=1 SV=2 | 255 | 20201114_qwl_milk_1ng_glyco_hcd203040_compound15.13410.13410.3 | controllerType=0 controllerNumber=1 scan=13410 | 28.4552 |
| 13451:3 | 10 | R.LAGKPTHVN[+1710.598]VSVVMAEVDGTC[+57.021]Y.- | HexNAc(3)Hex(5)NeuAc(1) | C[+57], N[+1711] | 1353.2551 | 3 | 4057.7506 | 4057.7397 | 0 | 2.7 | 332 | CRagged | 284.11 | 284.11 | 284.11 | 6.93 | 47 | >sp\|P01876\|IGHA1_HUMAN Immunoglobulin heavy constant alpha 1 OS=Homo sapiens OX=9606 GN=IGHA1 PE=1 SV=2 | 255 | 20201114_qwl_milk_1ng_glyco_hcd203040_compound15.16603.16603.3 | controllerType=0 controllerNumber=1 scan=16603 | 33.8864 |
| 13449:3 | 10 | R.LAGKPTHVN[+1710.598]VSVVMAEVDGTC[+57.021]Y.- | HexNAc(3)Hex(5)NeuAc(1) | C[+57], N[+1711] | 1353.2551 | 3 | 4057.7506 | 4057.7397 | 0 | 2.7 | 332 | CRagged | 292.68 | 292.68 | 243.01 | 6.93 | 47 | >sp\|P01876\|IGHA1_HUMAN Immunoglobulin heavy constant alpha 1 OS=Homo sapiens OX=9606 GN=IGHA1 PE=1 SV=2 | 255 | 20201114_qwl_milk_1ng_glyco_hcd203040_compound15.16601.16601.3 | controllerType=0 controllerNumber=1 scan=16601 | 33.8824 |
| 10391:3 | 10 | R.LAGKPTHVN[+1216.423]VSVVMAEVDGTC[+57.021]Y.- | HexNAc(2)Hex(5) | C[+57], N[+1216] | 1188.5303 | 3 | 3563.5763 | 3563.5649 | 0 | 3.2 | 332 | CRagged | 441.35 | 441.35 | 283.28 | 9.45 | 47 | >sp\|P01876\|IGHA1_HUMAN Immunoglobulin heavy constant alpha 1 OS=Homo sapiens OX=9606 GN=IGHA1 PE=1 SV=2 | 255 | 20201114_qwl_milk_1ng_glyco_hcd203040_compound15.13423.13423.3 | controllerType=0 controllerNumber=1 scan=13423 | 28.4767 |
| 11455:3 | 10 | R.LAGKPTHVN[+1540.529]VSVVMAEVDGTC[+57.021]Y.- | HexNAc(2)Hex(7) | C[+57], N[+1541] | 1296.5689 | 3 | 3887.6922 | 3887.6706 | 0 | 5.5 | 332 | CRagged | 460.51 | 460.51 | 298.51 | 8.81 | 47 | >sp\|P01876\|IGHA1_HUMAN Immunoglobulin heavy constant alpha 1 OS=Homo sapiens OX=9606 GN=IGHA1 PE=1 SV=2 | 255 | 20201114_qwl_milk_1ng_glyco_hcd203040_compound15.14527.14527.3 | controllerType=0 controllerNumber=1 scan=14527 | 30.3526 |
| 10393:3 | 10 | R.LAGKPTHVN[+1216.423]VSVVMAEVDGTC[+57.021]Y.- | HexNAc(2)Hex(5) | C[+57], N[+1216] | 1188.5303 | 3 | 3563.5763 | 3563.5649 | 0 | 3.2 | 332 | CRagged | 563.32 | 530.62 | 392.27 | 10.01 | 47 | >sp\|P01876\|IGHA1_HUMAN Immunoglobulin heavy constant alpha 1 OS=Homo sapiens OX=9606 GN=IGHA1 PE=1 SV=2 | 255 | 20201114_qwl_milk_1ng_glyco_hcd203040_compound15.13425.13425.3 | controllerType=0 controllerNumber=1 scan=13425 | 28.4804 |
| 11522:3 | 10 | R.LAGKPTHVN[+1809.666]VSVVMAEVDGTC[+57.021]Y.- | HexNAc(5)Hex(4)Fuc(1) | C[+57], N[+1810] | 1386.2776 | 3 | 4156.8182 | 4156.8081 | 0 | 2.4 | 332 | CRagged | 624.59 | 542.13 | 400.61 | 12.23 | 47 | >sp\|P01876\|IGHA1_HUMAN Immunoglobulin heavy constant alpha 1 OS=Homo sapiens OX=9606 GN=IGHA1 PE=1 SV=2 | 255 | 20201114_qwl_milk_1ng_glyco_hcd203040_compound15.14597.14597.3 | controllerType=0 controllerNumber=1 scan=14597 | 30.4929 |
| 11524:3 | 10 | R.LAGKPTHVN[+1809.666]VSVVMAEVDGTC[+57.021]Y.- | HexNAc(5)Hex(4)Fuc(1) | C[+57], N[+1810] | 1386.2776 | 3 | 4156.8182 | 4156.8081 | 0 | 2.4 | 332 | CRagged | 687.56 | 624.39 | 417.63 | 13.23 | 47 | >sp\|P01876\|IGHA1_HUMAN Immunoglobulin heavy constant alpha 1 OS=Homo sapiens OX=9606 GN=IGHA1 PE=1 SV=2 | 255 | 20201114_qwl_milk_1ng_glyco_hcd203040_compound15.14599.14599.3 | controllerType=0 controllerNumber=1 scan=14599 | 30.4965 |
| 11446:3 | 10 | R.LAGKPTHVN[+1702.581]VSVVMAEVDGTC[+57.021]Y.- | HexNAc(2)Hex(8) | C[+57], N[+1703] | 1350.5862 | 3 | 4049.7442 | 4049.7234 | 0 | 5.1 | 332 | CRagged | 735.86 | 735.86 | 443.86 | 13.84 | 47 | >sp\|P01876\|IGHA1_HUMAN Immunoglobulin heavy constant alpha 1 OS=Homo sapiens OX=9606 GN=IGHA1 PE=1 SV=2 | 255 | 20201114_qwl_milk_1ng_glyco_hcd203040_compound15.14518.14518.3 | controllerType=0 controllerNumber=1 scan=14518 | 30.3359 |
| 11499:3 | 10 | R.LAGKPTHVN[+1378.476]VSVVMAEVDGTC[+57.021]Y.- | HexNAc(2)Hex(6) | C[+57], N[+1378] | 1242.5479 | 3 | 3725.6292 | 3725.6178 | 0 | 3.1 | 332 | CRagged | 741.3 | 741.3 | 448.52 | 14.23 | 47 | >sp\|P01876\|IGHA1_HUMAN Immunoglobulin heavy constant alpha 1 OS=Homo sapiens OX=9606 GN=IGHA1 PE=1 SV=2 | 255 | 20201114_qwl_milk_1ng_glyco_hcd203040_compound15.14573.14573.3 | controllerType=0 controllerNumber=1 scan=14573 | 30.4405 |
| 11613:3 | 10 | R.LAGKPTHVN[+1647.613]VSVVMAEVDGTC[+57.021]Y.- | HexNAc(5)Hex(3)Fuc(1) | C[+57], N[+1648] | 1332.2552 | 3 | 3994.751 | 3994.7553 | 0 | -1.1 | 332 | CRagged | 792.65 | 658.87 | 658.87 | 14.72 | 47 | >sp\|P01876\|IGHA1_HUMAN Immunoglobulin heavy constant alpha 1 OS=Homo sapiens OX=9606 GN=IGHA1 PE=1 SV=2 | 255 | 20201114_qwl_milk_1ng_glyco_hcd203040_compound15.14692.14692.3 | controllerType=0 controllerNumber=1 scan=14692 | 30.6673 |
| 11610:3 | 10 | R.LAGKPTHVN[+1647.613]VSVVMAEVDGTC[+57.021]Y.- | HexNAc(5)Hex(3)Fuc(1) | C[+57], N[+1648] | 1332.2552 | 3 | 3994.751 | 3994.7553 | 0 | -1.1 | 332 | CRagged | 793.77 | 681.21 | 681.21 | 14.72 | 47 | >sp\|P01876\|IGHA1_HUMAN Immunoglobulin heavy constant alpha 1 OS=Homo sapiens OX=9606 GN=IGHA1 PE=1 SV=2 | 255 | 20201114_qwl_milk_1ng_glyco_hcd203040_compound15.14689.14689.3 | controllerType=0 controllerNumber=1 scan=14689 | 30.6609 |
| 11517:3 | 10 | R.LAGKPTHVN[+1663.608]VSVVMAEVDGTC[+57.021]Y.- | HexNAc(5)Hex(4) | C[+57], N[+1664] | 1337.5916 | 3 | 4010.7603 | 4010.7502 | 0 | 2.5 | 332 | CRagged | 809.68 | 738.25 | 480.44 | 16.23 | 47 | >sp\|P01876\|IGHA1_HUMAN Immunoglobulin heavy constant alpha 1 OS=Homo sapiens OX=9606 GN=IGHA1 PE=1 SV=2 | 255 | 20201114_qwl_milk_1ng_glyco_hcd203040_compound15.14591.14591.3 | controllerType=0 controllerNumber=1 scan=14591 | 30.4775 |
| 11448:3 | 10 | R.LAGKPTHVN[+1702.581]VSVVMAEVDGTC[+57.021]Y.- | HexNAc(2)Hex(8) | C[+57], N[+1703] | 1350.5862 | 3 | 4049.7442 | 4049.7234 | 0 | 5.1 | 332 | CRagged | 818.86 | 803.56 | 512.18 | 15.84 | 47 | >sp\|P01876\|IGHA1_HUMAN Immunoglobulin heavy constant alpha 1 OS=Homo sapiens OX=9606 GN=IGHA1 PE=1 SV=2 | 255 | 20201114_qwl_milk_1ng_glyco_hcd203040_compound15.14520.14520.3 | controllerType=0 controllerNumber=1 scan=14520 | 30.3405 |
| 11457:3 | 10 | R.LAGKPTHVN[+1540.529]VSVVMAEVDGTC[+57.021]Y.- | HexNAc(2)Hex(7) | C[+57], N[+1541] | 1296.5689 | 3 | 3887.6922 | 3887.6706 | 0 | 5.5 | 332 | CRagged | 860.2 | 815.51 | 521.24 | 16.58 | 47 | >sp\|P01876\|IGHA1_HUMAN Immunoglobulin heavy constant alpha 1 OS=Homo sapiens OX=9606 GN=IGHA1 PE=1 SV=2 | 255 | 20201114_qwl_milk_1ng_glyco_hcd203040_compound15.14529.14529.3 | controllerType=0 controllerNumber=1 scan=14529 | 30.3563 |
| 11443:3 | 10 | R.LAGKPTHVN[+1702.581]VSVVMAEVDGTC[+57.021]Y.- | HexNAc(2)Hex(8) | C[+57], N[+1703] | 1350.581 | 3 | 4049.7284 | 4049.7234 | 0 | 1.2 | 332 | CRagged | 869.31 | 697.17 | 486.41 | 17.15 | 47 | >sp\|P01876\|IGHA1_HUMAN Immunoglobulin heavy constant alpha 1 OS=Homo sapiens OX=9606 GN=IGHA1 PE=1 SV=2 | 255 | 20201114_qwl_milk_1ng_glyco_hcd203040_compound15.14514.14514.3 | controllerType=0 controllerNumber=1 scan=14514 | 30.3281 |
| 11526:3 | 10 | R.LAGKPTHVN[+1216.423]VSVVMAEVDGTC[+57.021]Y.- | HexNAc(2)Hex(5) | C[+57], N[+1216] | 1188.5297 | 3 | 3563.5745 | 3563.5649 | 0 | 2.7 | 332 | CRagged | 892.23 | 892.23 | 528.55 | 17.23 | 47 | >sp\|P01876\|IGHA1_HUMAN Immunoglobulin heavy constant alpha 1 OS=Homo sapiens OX=9606 GN=IGHA1 PE=1 SV=2 | 255 | 20201114_qwl_milk_1ng_glyco_hcd203040_compound15.14601.14601.3 | controllerType=0 controllerNumber=1 scan=14601 | 30.4985 |
| 11441:3 | 10 | R.LAGKPTHVN[+1702.581]VSVVMAEVDGTC[+57.021]Y.- | HexNAc(2)Hex(8) | C[+57], N[+1703] | 1350.581 | 3 | 4049.7284 | 4049.7234 | 0 | 1.2 | 332 | CRagged | 924.71 | 924.71 | 571.12 | 18.15 | 47 | >sp\|P01876\|IGHA1_HUMAN Immunoglobulin heavy constant alpha 1 OS=Homo sapiens OX=9606 GN=IGHA1 PE=1 SV=2 | 255 | 20201114_qwl_milk_1ng_glyco_hcd203040_compound15.14512.14512.3 | controllerType=0 controllerNumber=1 scan=14512 | 30.3221 |
| 05741:3 | 12 | R.LSPN[+2351.851]ASAEHLELR.W | HexNAc(4)Hex(5)Fuc(3)NeuAc(1) | N[+2352] | 1263.5407 | 3 | 3788.6076 | 3788.6 | 0 | 2 | 52 | Specific | 420.56 | 409.17 | 409.17 | 8.75 | 51 | >sp\|Q13410\|BT1A1_HUMAN Butyrophilin subfamily 1 member A1 OS=Homo sapiens OX=9606 GN=BTN1A1 PE=1 SV=3 | 179 | 20201114_qwl_milk_1ng_glyco_hcd203040_compound15.8606.8606.3 | controllerType=0 controllerNumber=1 scan=8606 | 20.5521 |
| 05743:3 | 12 | R.LSPN[+2351.851]ASAEHLELR.W | HexNAc(4)Hex(5)Fuc(3)NeuAc(1) | N[+2352] | 1263.5407 | 3 | 3788.6076 | 3788.6 | 0 | 2 | 52 | Specific | 483.4 | 422.28 | 422.28 | 9.06 | 51 | >sp\|Q13410\|BT1A1_HUMAN Butyrophilin subfamily 1 member A1 OS=Homo sapiens OX=9606 GN=BTN1A1 PE=1 SV=3 | 179 | 20201114_qwl_milk_1ng_glyco_hcd203040_compound15.8608.8608.3 | controllerType=0 controllerNumber=1 scan=8608 | 20.5559 |
| 07803:3 | 12 | R.LSPN[+2350.830]ASAEHLELR.W | HexNAc(4)Hex(5)Fuc(1)NeuAc(2) | N[+2351] | 1263.2 | 3 | 3787.5855 | 3787.5796 | 0 | 1.6 | 52 | Specific | 547.23 | 450.44 | 450.44 | 9.43 | 51 | >sp\|Q13410\|BT1A1_HUMAN Butyrophilin subfamily 1 member A1 OS=Homo sapiens OX=9606 GN=BTN1A1 PE=1 SV=3 | 179 | 20201114_qwl_milk_1ng_glyco_hcd203040_compound15.10739.10739.3 | controllerType=0 controllerNumber=1 scan=10739 | 24.0645 |
| 07801:3 | 12 | R.LSPN[+2350.830]ASAEHLELR.W | HexNAc(4)Hex(5)Fuc(1)NeuAc(2) | N[+2351] | 1263.2 | 3 | 3787.5855 | 3787.5796 | 0 | 1.6 | 52 | Specific | 573.39 | 518.02 | 518.02 | 10.65 | 51 | >sp\|Q13410\|BT1A1_HUMAN Butyrophilin subfamily 1 member A1 OS=Homo sapiens OX=9606 GN=BTN1A1 PE=1 SV=3 | 179 | 20201114_qwl_milk_1ng_glyco_hcd203040_compound15.10737.10737.3 | controllerType=0 controllerNumber=1 scan=10737 | 24.0597 |
| 05724:3 | 12 | R.LSPN[+2205.793]ASAEHLELR.W | HexNAc(4)Hex(5)Fuc(2)NeuAc(1) | N[+2206] | 1214.8548 | 3 | 3642.5498 | 3642.5421 | 0 | 2.1 | 52 | Specific | 596.38 | 430.22 | 430.22 | 10.65 | 51 | >sp\|Q13410\|BT1A1_HUMAN Butyrophilin subfamily 1 member A1 OS=Homo sapiens OX=9606 GN=BTN1A1 PE=1 SV=3 | 179 | 20201114_qwl_milk_1ng_glyco_hcd203040_compound15.8588.8588.3 | controllerType=0 controllerNumber=1 scan=8588 | 20.5211 |
| 05722:3 | 12 | R.LSPN[+2205.793]ASAEHLELR.W | HexNAc(4)Hex(5)Fuc(2)NeuAc(1) | N[+2206] | 1214.8548 | 3 | 3642.5498 | 3642.5421 | 0 | 2.1 | 52 | Specific | 612.93 | 538.69 | 538.69 | 11.65 | 51 | >sp\|Q13410\|BT1A1_HUMAN Butyrophilin subfamily 1 member A1 OS=Homo sapiens OX=9606 GN=BTN1A1 PE=1 SV=3 | 179 | 20201114_qwl_milk_1ng_glyco_hcd203040_compound15.8586.8586.3 | controllerType=0 controllerNumber=1 scan=8586 | 20.5163 |
| 15929:3 | 13 | R.QQQHLFGSN[+2204.772]VTDC[+57.021]SGNFC[+57.021]LFR.S | HexNAc(4)Hex(5)NeuAc(2) | C[+57]*2, N[+2205] | 1573.9703 | 3 | 4719.8963 | 4719.8966 | 0 | -0.1 | 622 | Specific | 573.04 | 558.37 | 558.37 | 9.5 | 42 | >sp\|P02787\|TRFE_HUMAN Serotransferrin OS=Homo sapiens OX=9606 GN=TF PE=1 SV=3 | 1111 | 20201114_qwl_milk_1ng_glyco_hcd203040_compound15.19188.19188.3 | controllerType=0 controllerNumber=1 scan=19188 | 38.5902 |
| 15927:3 | 13 | R.QQQHLFGSN[+2204.772]VTDC[+57.021]SGNFC[+57.021]LFR.S | HexNAc(4)Hex(5)NeuAc(2) | C[+57]*2, N[+2205] | 1573.9703 | 3 | 4719.8963 | 4719.8966 | 0 | -0.1 | 622 | Specific | 637.38 | 610.57 | 610.57 | 11.72 | 42 | >sp\|P02787\|TRFE_HUMAN Serotransferrin OS=Homo sapiens OX=9606 GN=TF PE=1 SV=3 | 1111 | 20201114_qwl_milk_1ng_glyco_hcd203040_compound15.19186.19186.3 | controllerType=0 controllerNumber=1 scan=19186 | 38.5861 |
| 08602:3 | 14 | K.VAYSN[+2204.772]DSANWTEYQDPR.T | HexNAc(4)Hex(5)NeuAc(2) | N[+2205] | 1407.5655 | 3 | 4220.6819 | 4220.6454 | 0 | 8.6 | 321 | Specific | 257.61 | 220.57 | 1.2 | 5.58 | 37 | >sp\|Q08431\|MFGM_HUMAN Lactadherin OS=Homo sapiens OX=9606 GN=MFGE8 PE=1 SV=3 | 1064 | 20201114_qwl_milk_1ng_glyco_hcd203040_compound15.11569.11569.3 | controllerType=0 controllerNumber=1 scan=11569 | 25.45 |
| 08667:3 | 14 | K.VAYSN[+2059.735]DSANWTEYQDPR.T | HexNAc(4)Hex(5)Fuc(1)NeuAc(1) | N[+2060] | 1359.2068 | 3 | 4075.606 | 4075.6079 | 0 | -0.5 | 321 | Specific | 352.95 | 287.3 | 3.07 | 6.25 | 37 | >sp\|Q08431\|MFGM_HUMAN Lactadherin OS=Homo sapiens OX=9606 GN=MFGE8 PE=1 SV=3 | 1064 | 20201114_qwl_milk_1ng_glyco_hcd203040_compound15.11636.11636.3 | controllerType=0 controllerNumber=1 scan=11636 | 25.5598 |
| 08576:3 | 14 | K.VAYSN[+2205.793]DSANWTEYQDPR.T | HexNAc(4)Hex(5)Fuc(2)NeuAc(1) | N[+2206] | 1407.8942 | 3 | 4221.6681 | 4221.6658 | 0 | 0.5 | 321 | Specific | 396.76 | 333.05 | 8.3 | 6.82 | 37 | >sp\|Q08431\|MFGM_HUMAN Lactadherin OS=Homo sapiens OX=9606 GN=MFGE8 PE=1 SV=3 | 1064 | 20201114_qwl_milk_1ng_glyco_hcd203040_compound15.11542.11542.3 | controllerType=0 controllerNumber=1 scan=11542 | 25.4058 |
| 08578:3 | 14 | K.VAYSN[+2205.793]DSANWTEYQDPR.T | HexNAc(4)Hex(5)Fuc(2)NeuAc(1) | N[+2206] | 1407.8942 | 3 | 4221.6681 | 4221.6658 | 0 | 0.5 | 321 | Specific | 470.26 | 393.47 | 5.58 | 8.74 | 37 | >sp\|Q08431\|MFGM_HUMAN Lactadherin OS=Homo sapiens OX=9606 GN=MFGE8 PE=1 SV=3 | 1064 | 20201114_qwl_milk_1ng_glyco_hcd203040_compound15.11544.11544.3 | controllerType=0 controllerNumber=1 scan=11544 | 25.4099 |
| 08669:3 | 14 | K.VAYSN[+2059.735]DSANWTEYQDPR.T | HexNAc(4)Hex(5)Fuc(1)NeuAc(1) | N[+2060] | 1359.2068 | 3 | 4075.606 | 4075.6079 | 0 | -0.5 | 321 | Specific | 470.68 | 393.12 | 4.35 | 8.74 | 37 | >sp\|Q08431\|MFGM_HUMAN Lactadherin OS=Homo sapiens OX=9606 GN=MFGE8 PE=1 SV=3 | 1064 | 20201114_qwl_milk_1ng_glyco_hcd203040_compound15.11638.11638.3 | controllerType=0 controllerNumber=1 scan=11638 | 25.5639 |
| 14171:4 | 18 | K.LKELPGVC[+57.021]N[+2205.793]ETMMALWEEC[+57.021]KPC[+57.021]LK.Q | HexNAc(4)Hex(5)Fuc(2)NeuAc(1) | C[+57]*3, N[+2206] | 1286.2999 | 4 | 5142.1779 | 5142.1776 | 0 | 0.1 | 95 | Specific | 216.84 | 216.84 | 216.84 | 4.72 | 29 | >sp\|P10909\|CLUS_HUMAN Clusterin OS=Homo sapiens OX=9606 GN=CLU PE=1 SV=1 | 1467 | 20201114_qwl_milk_1ng_glyco_hcd203040_compound15.17351.17351.4 | controllerType=0 controllerNumber=1 scan=17351 | 35.2655 |
| 14186:4 | 18 | K.LKELPGVC[+57.021]N[+2205.793]ETMMALWEEC[+57.021]KPC[+57.021]LK.Q | HexNAc(4)Hex(5)Fuc(2)NeuAc(1) | C[+57]*3, N[+2206] | 1286.3026 | 4 | 5142.1886 | 5142.1776 | 0 | 2.2 | 95 | Specific | 311.2 | 311.2 | 311.2 | 7.12 | 29 | >sp\|P10909\|CLUS_HUMAN Clusterin OS=Homo sapiens OX=9606 GN=CLU PE=1 SV=1 | 1467 | 20201114_qwl_milk_1ng_glyco_hcd203040_compound15.17367.17367.4 | controllerType=0 controllerNumber=1 scan=17367 | 35.292 |
| 14183:4 | 18 | K.LKELPGVC[+57.021]N[+2205.793]ETMMALWEEC[+57.021]KPC[+57.021]LK.Q | HexNAc(4)Hex(5)Fuc(2)NeuAc(1) | C[+57]*3, N[+2206] | 1286.3026 | 4 | 5142.1886 | 5142.1776 | 0 | 2.2 | 95 | Specific | 521.41 | 425.81 | 425.81 | 9.7 | 29 | >sp\|P10909\|CLUS_HUMAN Clusterin OS=Homo sapiens OX=9606 GN=CLU PE=1 SV=1 | 1467 | 20201114_qwl_milk_1ng_glyco_hcd203040_compound15.17364.17364.4 | controllerType=0 controllerNumber=1 scan=17364 | 35.2881 |
| 14169:4 | 18 | K.LKELPGVC[+57.021]N[+2205.793]ETMMALWEEC[+57.021]KPC[+57.021]LK.Q | HexNAc(4)Hex(5)Fuc(2)NeuAc(1) | C[+57]*3, N[+2206] | 1286.2999 | 4 | 5142.1779 | 5142.1776 | 0 | 0.1 | 95 | Specific | 1006.94 | 1006.94 | 1006.94 | 19.68 | 29 | >sp\|P10909\|CLUS_HUMAN Clusterin OS=Homo sapiens OX=9606 GN=CLU PE=1 SV=1 | 1467 | 20201114_qwl_milk_1ng_glyco_hcd203040_compound15.17349.17349.4 | controllerType=0 controllerNumber=1 scan=17349 | 35.2554 |
| 21057:3 | 18 | K.MLN[+2351.851]TSSLLEQLNEQFNWVSR.L | HexNAc(4)Hex(5)Fuc(3)NeuAc(1) | N[+2352] | 1587.6906 | 3 | 4761.0572 | 4761.0375 | 0 | 4.2 | 352 | Specific | 462.12 | 402.84 | 402.84 | 8.94 | 29 | >sp\|P10909\|CLUS_HUMAN Clusterin OS=Homo sapiens OX=9606 GN=CLU PE=1 SV=1 | 1467 | 20201114_qwl_milk_1ng_glyco_hcd203040_compound15.24570.24570.3 | controllerType=0 controllerNumber=1 scan=24570 | 49.6825 |
| 10495:3 | 18 | R.LAN[+2350.830]LTQGEDQYYLR.V | HexNAc(4)Hex(5)Fuc(1)NeuAc(2) | N[+2351] | 1345.5676 | 3 | 4034.6881 | 4034.664 | 0 | 6 | 372 | Specific | 342.98 | 301.52 | 301.52 | 6.33 | 29 | >sp\|P10909\|CLUS_HUMAN Clusterin OS=Homo sapiens OX=9606 GN=CLU PE=1 SV=1 | 1467 | 20201114_qwl_milk_1ng_glyco_hcd203040_compound15.13532.13532.3 | controllerType=0 controllerNumber=1 scan=13532 | 28.6957 |
| 10520:3 | 18 | R.LAN[+2205.793]LTQGEDQYYLR.V | HexNAc(4)Hex(5)Fuc(2)NeuAc(1) | N[+2206] | 1297.2156 | 3 | 3889.6323 | 3889.6265 | 0 | 1.5 | 372 | Specific | 430.74 | 406.97 | 406.97 | 8.87 | 29 | >sp\|P10909\|CLUS_HUMAN Clusterin OS=Homo sapiens OX=9606 GN=CLU PE=1 SV=1 | 1467 | 20201114_qwl_milk_1ng_glyco_hcd203040_compound15.13558.13558.3 | controllerType=0 controllerNumber=1 scan=13558 | 28.7469 |
| 10522:3 | 18 | R.LAN[+2205.793]LTQGEDQYYLR.V | HexNAc(4)Hex(5)Fuc(2)NeuAc(1) | N[+2206] | 1297.2156 | 3 | 3889.6323 | 3889.6265 | 0 | 1.5 | 372 | Specific | 483.85 | 363.66 | 363.66 | 7.81 | 29 | >sp\|P10909\|CLUS_HUMAN Clusterin OS=Homo sapiens OX=9606 GN=CLU PE=1 SV=1 | 1467 | 20201114_qwl_milk_1ng_glyco_hcd203040_compound15.13560.13560.3 | controllerType=0 controllerNumber=1 scan=13560 | 28.7514 |
| 23686:4 | 20 | K.VFPLSLDSTPQDGNVVVAC[+57.021]LVQGFFPQEPLSVTWSESGQN[+2262.814]VTAI | HexNAc(5)Hex(5)Fuc(1)NeuAc(1) | C[+57], N[+2263] | 1760.5626 | 4 | 7039.2286 | 7039.1757 | 0 | 7.5 | 8 | Specific | 234.22 | 234.22 | 234.22 | 6.01 | 35 | >sp\|P01877\|IGHA2_HUMAN Immunoglobulin heavy constant alpha 2 OS=Homo sapiens OX=9606 GN=IGHA2 PE=1 SV=4 | 576 | 20201114_qwl_milk_1ng_glyco_hcd203040_compound15.27311.27311.4 | controllerType=0 controllerNumber=1 scan=27311 | 54.9431 |
| 22165:4 | 20 | K.VFPLSLDSTPQDGNVVVAC[+57.021]LVQGFFPQEPLSVTWSESGQN[+1809.666]VTAI | HexNAc(5)Hex(4)Fuc(1) | C[+57], N[+1810] | 1647.2705 | 4 | 6586.0602 | 6586.0275 | 0 | 5 | 8 | Specific | 270.14 | 270.14 | 270.14 | 6.75 | 35 | >sp\|P01877\|IGHA2_HUMAN Immunoglobulin heavy constant alpha 2 OS=Homo sapiens OX=9606 GN=IGHA2 PE=1 SV=4 | 576 | 20201114_qwl_milk_1ng_glyco_hcd203040_compound15.25726.25726.4 | controllerType=0 controllerNumber=1 scan=25726 | 51.9697 |
| 23807:4 | 20 | K.VFPLSLDSTPQDGNVVVAC[+57.021]LVQGFFPQEPLSVTWSESGQN[+2262.814]VTAI | HexNAc(5)Hex(5)Fuc(1)NeuAc(1) | C[+57], N[+2263] | 1760.5568 | 4 | 7039.2056 | 7039.1757 | 0 | 4.2 | 8 | Specific | 317.27 | 317.27 | 317.27 | 7.17 | 35 | >sp\|P01877\|IGHA2_HUMAN Immunoglobulin heavy constant alpha 2 OS=Homo sapiens OX=9606 GN=IGHA2 PE=1 SV=4 | 576 | 20201114_qwl_milk_1ng_glyco_hcd203040_compound15.27437.27437.4 | controllerType=0 controllerNumber=1 scan=27437 | 55.1878 |
| 22204:4 | 20 | K.[+42.011]VFPLSLDSTPQDGNVVVAC[+57.021]LVQGFFPQEPLSVTWSESGQN[+1768 | HexNAc(4)Hex(5)Fuc(1) | .[42], C[+57], N[+1769] | 1647.5168 | 4 | 6587.0456 | 6587.0115 | 0 | 5.2 | 8 | Specific | 346.62 | 287.52 | 287.52 | 5.84 | 35 | >sp\|P01877\|IGHA2_HUMAN Immunoglobulin heavy constant alpha 2 OS=Homo sapiens OX=9606 GN=IGHA2 PE=1 SV=4 | 576 | 20201114_qwl_milk_1ng_glyco_hcd203040_compound15.25766.25766.4 | controllerType=0 controllerNumber=1 scan=25766 | 52.0423 |
| 23943:4 | 20 | K.VFPLSLDSTPQDGNVVVAC[+57.021]LVQGFFPQEPLSVTWSESGQN[+2262.814]VTAI | HexNAc(5)Hex(5)Fuc(1)NeuAc(1) | C[+57], N[+2263] | 1760.5538 | 4 | 7039.1934 | 7039.1757 | 0 | 2.5 | 8 | Specific | 349.89 | 341.03 | 341.03 | 7.78 | 35 | >sp\|P01877\|IGHA2_HUMAN Immunoglobulin heavy constant alpha 2 OS=Homo sapiens OX=9606 GN=IGHA2 PE=1 SV=4 | 576 | 20201114_qwl_milk_1ng_glyco_hcd203040_compound15.27579.27579.2 | controllerType=0 controllerNumber=1 scan=27579 | 55.4531 |
| 22356:4 | 20 | K.VFPLSLDSTPQDGNVVVAC[+57.021]LVQGFFPQEPLSVTWSESGQN[+1809.666]VTAI | HexNAc(5)Hex(4)Fuc(1) | C[+57], N[+1810] | 1647.2671 | 4 | 6586.0465 | 6586.0275 | 0 | 2.9 | 8 | Specific | 351.95 | 351.95 | 351.95 | 8.01 | 35 | >sp\|P01877\|IGHA2_HUMAN Immunoglobulin heavy constant alpha 2 OS=Homo sapiens OX=9606 GN=IGHA2 PE=1 SV=4 | 576 | 20201114_qwl_milk_1ng_glyco_hcd203040_compound15.25925.25925.2 | controllerType=0 controllerNumber=1 scan=25925 | 52.345 |
| 22259:4 | 20 | K.VFPLSLDSTPQDGNVVVAC[+57.021]LVQGFFPQEPLSVTWSESGQN[+1647.613]VTAI | HexNAc(5)Hex(3)Fuc(1) | C[+57], N[+1648] | 1606.7614 | 4 | 6424.0236 | 6423.9746 | 0 | 7.6 | 8 | Specific | 711.75 | 711.75 | 711.75 | 13.91 | 35 | >sp\|P01877\|IGHA2_HUMAN Immunoglobulin heavy constant alpha 2 OS=Homo sapiens OX=9606 GN=IGHA2 PE=1 SV=4 | 576 | 20201114_qwl_milk_1ng_glyco_hcd203040_compound15.25824.25824.4 | controllerType=0 controllerNumber=1 scan=25824 | 52.1513 |
| 22261:4 | 20 | K.VFPLSLDSTPQDGNVVVAC[+57.021]LVQGFFPQEPLSVTWSESGQN[+1647.613]VTAI | HexNAc(5)Hex(3)Fuc(1) | C[+57], N[+1648] | 1606.7614 | 4 | 6424.0236 | 6423.9746 | 0 | 7.6 | 8 | Specific | 850.1 | 850.1 | 850.1 | 16.91 | 35 | >sp\|P01877\|IGHA2_HUMAN Immunoglobulin heavy constant alpha 2 OS=Homo sapiens OX=9606 GN=IGHA2 PE=1 SV=4 | 576 | 20201114_qwl_milk_1ng_glyco_hcd203040_compound15.25826.25826.4 | controllerType=0 controllerNumber=1 scan=25826 | 52.1584 |
| 22168:4 | 20 | K.VFPLSLDSTPQDGNVVVAC[+57.021]LVQGFFPQEPLSVTWSESGQN[+1809.666]VTAI | HexNAc(5)Hex(4)Fuc(1) | C[+57], N[+1810] | 1647.2705 | 4 | 6586.0602 | 6586.0275 | 0 | 5 | 8 | Specific | 869.98 | 829.26 | 829.26 | 17.17 | 35 | >sp\|P01877\|IGHA2_HUMAN Immunoglobulin heavy constant alpha 2 OS=Homo sapiens OX=9606 GN=IGHA2 PE=1 SV=4 | 576 | 20201114_qwl_milk_1ng_glyco_hcd203040_compound15.25729.25729.4 | controllerType=0 controllerNumber=1 scan=25729 | 51.9745 |
| 01364:3 | 20 | K.HYTN[+1809.666]SSQDVTVPC[+57.021]R.V | HexNAc(5)Hex(4)Fuc(1) | C[+57], N[+1810] | 1158.4802 | 3 | 3473.4259 | 3473.4154 | 0 | 3 | 89 | Specific | 421.02 | 334.14 | 334.14 | 7.48 | 35 | >sp\|P01877\|IGHA2_HUMAN Immunoglobulin heavy constant alpha 2 OS=Homo sapiens OX=9606 GN=IGHA2 PE=1 SV=4 | 576 | 20201114_qwl_milk_1ng_glyco_hcd203040_compound15.4070.4070.3 | controllerType=0 controllerNumber=1 scan=4070 | 12.8628 |
| 01362:3 | 20 | K.HYTN[+1809.666]SSQDVTVPC[+57.021]R.V | HexNAc(5)Hex(4)Fuc(1) | C[+57], N[+1810] | 1158.4802 | 3 | 3473.4259 | 3473.4154 | 0 | 3 | 89 | Specific | 428.09 | 351.15 | 351.15 | 7.71 | 35 | >sp\|P01877\|IGHA2_HUMAN Immunoglobulin heavy constant alpha 2 OS=Homo sapiens OX=9606 GN=IGHA2 PE=1 SV=4 | 576 | 20201114_qwl_milk_1ng_glyco_hcd203040_compound15.4068.4068.3 | controllerType=0 controllerNumber=1 scan=4068 | 12.8587 |
| 04085:5 | 20 | K.HYTN[+2262.814]SSQDVTVPC[+57.021]RVPPPPPC[+57.021]C[+57.021]HPR.L | HexNAc(5)Hex(5)Fuc(1)NeuAc(1) | C[+57]*3, N[+2263] | 1045.04 | 5 | 5221.1711 | 5221.1699 | 0 | 0.2 | 89 | Specific | 354.61 | 213.53 | 213.53 | 6.38 | 35 | >sp\|P01877\|IGHA2_HUMAN Immunoglobulin heavy constant alpha 2 OS=Homo sapiens OX=9606 GN=IGHA2 PE=1 SV=4 | 576 | 20201114_qwl_milk_1ng_glyco_hcd203040_compound15.6891.6891.5 | controllerType=0 controllerNumber=1 scan=6891 | 17.7344 |
| 03346:5 | 20 | K.HYTN[+1809.666]SSQDVTVPC[+57.021]RVPPPPPC[+57.021]C[+57.021]HPR.L | HexNAc(5)Hex(4)Fuc(1) | C[+57]*3, N[+1810] | 954.4117 | 5 | 4768.0294 | 4768.0217 | 0 | 1.6 | 89 | Specific | 362.69 | 362.69 | 362.69 | 6.8 | 35 | >sp\|P01877\|IGHA2_HUMAN Immunoglobulin heavy constant alpha 2 OS=Homo sapiens OX=9606 GN=IGHA2 PE=1 SV=4 | 576 | 20201114_qwl_milk_1ng_glyco_hcd203040_compound15.6126.6126.5 | controllerType=0 controllerNumber=1 scan=6126 | 16.4301 |
| 03344:5 | 20 | K.HYTN[+1809.666]SSQDVTVPC[+57.021]RVPPPPPC[+57.021]C[+57.021]HPR.L | HexNAc(5)Hex(4)Fuc(1) | C[+57]*3, N[+1810] | 954.4117 | 5 | 4768.0294 | 4768.0217 | 0 | 1.6 | 89 | Specific | 365.51 | 365.51 | 365.51 | 7.79 | 35 | >sp\|P01877\|IGHA2_HUMAN Immunoglobulin heavy constant alpha 2 OS=Homo sapiens OX=9606 GN=IGHA2 PE=1 SV=4 | 576 | 20201114_qwl_milk_1ng_glyco_hcd203040_compound15.6124.6124.5 | controllerType=0 controllerNumber=1 scan=6124 | 16.4254 |
| 03420:4 | 20 | K.HYTN[+1809.666]SSQDVTVPC[+57.021]RVPPPPPC[+57.021]C[+57.021]HPR.L | HexNAc(5)Hex(4)Fuc(1) | C[+57]*3, N[+1810] | 1192.7648 | 4 | 4768.0374 | 4768.0217 | 0 | 3.3 | 89 | Specific | 464.25 | 442.59 | 442.59 | 9.03 | 35 | >sp\|P01877\|IGHA2_HUMAN Immunoglobulin heavy constant alpha 2 OS=Homo sapiens OX=9606 GN=IGHA2 PE=1 SV=4 | 576 | 20201114_qwl_milk_1ng_glyco_hcd203040_compound15.6203.6203.4 | controllerType=0 controllerNumber=1 scan=6203 | 16.5644 |
| 03422:4 | 20 | K.HYTN[+1809.666]SSQDVTVPC[+57.021]RVPPPPPC[+57.021]C[+57.021]HPR.L | HexNAc(5)Hex(4)Fuc(1) | C[+57]*3, N[+1810] | 1192.7648 | 4 | 4768.0374 | 4768.0217 | 0 | 3.3 | 89 | Specific | 489.96 | 335.75 | 335.75 | 7.62 | 35 | >sp\|P01877\|IGHA2_HUMAN Immunoglobulin heavy constant alpha 2 OS=Homo sapiens OX=9606 GN=IGHA2 PE=1 SV=4 | 576 | 20201114_qwl_milk_1ng_glyco_hcd203040_compound15.6205.6205.4 | controllerType=0 controllerNumber=1 scan=6205 | 16.5683 |
| 03392:5 | 20 | K.HYTN[+1647.613]SSQDVTVPC[+57.021]RVPPPPPC[+57.021]C[+57.021]HPR.L | HexNAc(5)Hex(3)Fuc(1) | C[+57]*3, N[+1648] | 922.0013 | 5 | 4605.9774 | 4605.9689 | 0 | 1.9 | 89 | Specific | 752.57 | 594.07 | 594.07 | 15.11 | 35 | >sp\|P01877\|IGHA2_HUMAN Immunoglobulin heavy constant alpha 2 OS=Homo sapiens OX=9606 GN=IGHA2 PE=1 SV=4 | 576 | 20201114_qwl_milk_1ng_glyco_hcd203040_compound15.6174.6174.5 | controllerType=0 controllerNumber=1 scan=6174 | 16.5137 |

Table S4. Cont.

| Query #:z | Protein Rank | Glycopeptide | Glycan | Modification Type(s) | Observed m/z | z | Observed (M+H) | Calc. mass (M+H) | Off-by-x error | Mass error (ppm) | StartPosition | Cleavage | Score | Delta | Delta Mod | \|Log Prob\| | # of unique peptides | Protein Name | Protein DB number | Comment | Scan # | Scan Time |
|---|---|---|---|---|---|---|---|---|---|---|---|---|---|---|---|---|---|---|---|---|---|---|
| 03394:5 | 20 | K.HYTN[+1647.613]SSQDVTVPC[+57.021]RVPPPPPC[+57.021]C[+57.021]HPR.L | HexNAc(5)Hex(3)Fuc(1) | C[+57]*3, N[+1648] | 922.0013 | 5 | 4605.9774 | 4605.9689 | 0 | 1.9 | 89 | Specific | 807.52 | 554.63 | 554.63 | 16.11 | 35 | >sp\|P01877\|IGHA2_HUMAN Immunoglobulin heavy constant alpha 2 OS=Homo sapiens OX=9606 GN=IGHA2 PE=1 SV=4 | 576 | 20201114_qwl_milk_1ng_glyco_hcd203040_compound15.6176.6176.5 | controllerType=0 controllerNumber=1 scan=6176 | 16.5194 |
| 03391:4 | 20 | K.HYTN[+1647.613]SSQDVTVPC[+57.021]RVPPPPPC[+57.021]C[+57.021]HPR.L | HexNAc(5)Hex(3)Fuc(1) | C[+57]*3, N[+1648] | 1152.2496 | 4 | 4605.9764 | 4605.9689 | 0 | 1.6 | 89 | Specific | 906.18 | 673.02 | 673.02 | 17.73 | 35 | >sp\|P01877\|IGHA2_HUMAN Immunoglobulin heavy constant alpha 2 OS=Homo sapiens OX=9606 GN=IGHA2 PE=1 SV=4 | 576 | 20201114_qwl_milk_1ng_glyco_hcd203040_compound15.6173.6173.4 | controllerType=0 controllerNumber=1 scan=6173 | 16.5122 |
| 03393:4 | 20 | K.HYTN[+1647.613]SSQDVTVPC[+57.021]RVPPPPPC[+57.021]C[+57.021]HPR.L | HexNAc(5)Hex(3)Fuc(1) | C[+57]*3, N[+1648] | 1152.2496 | 4 | 4605.9764 | 4605.9689 | 0 | 1.6 | 89 | Specific | 1003.08 | 761.96 | 761.96 | 19.73 | 35 | >sp\|P01877\|IGHA2_HUMAN Immunoglobulin heavy constant alpha 2 OS=Homo sapiens OX=9606 GN=IGHA2 PE=1 SV=4 | 576 | 20201114_qwl_milk_1ng_glyco_hcd203040_compound15.6175.6175.4 | controllerType=0 controllerNumber=1 scan=6175 | 16.5172 |
| 13600:6 | 20 | K.SAVQGPPERDLC[+57.021]GC[+57.021]YSVSSVLPGC[+57.021]AQPWNHGETFTC[+ | HexNAc(5)Hex(5)Fuc(3) | C[+57]*4, N[+2264] | 1346.6032 | 6 | 8074.5829 | 8074.5805 | 0 | 0.3 | 156 | Specific | 401.93 | 41.65 | 41.65 | 3.41 | 35 | >sp\|P01877\|IGHA2_HUMAN Immunoglobulin heavy constant alpha 2 OS=Homo sapiens OX=9606 GN=IGHA2 PE=1 SV=4 | 576 | 20201114_qwl_milk_1ng_glyco_hcd203040_compound15.16757.16757.3 | controllerType=0 controllerNumber=1 scan=16757 | 34.1697 |
| 14075:4 | 20 | R.DLC[+57.021]GC[+57.021]YSVSSVLPGC[+57.021]AQPWNHGETFTC[+57.021]TAA | HexNAc(5)Hex(3)Fuc(1) | C[+57]*4, N[+1648] | 1634.9827 | 4 | 6536.9088 | 6536.8923 | 0 | 2.5 | 165 | Specific | 552.96 | 533.03 | 533.03 | 9.92 | 35 | >sp\|P01877\|IGHA2_HUMAN Immunoglobulin heavy constant alpha 2 OS=Homo sapiens OX=9606 GN=IGHA2 PE=1 SV=4 | 576 | 20201114_qwl_milk_1ng_glyco_hcd203040_compound15.17251.17251.4 | controllerType=0 controllerNumber=1 scan=17251 | 35.0728 |
| 03168:2 | 20 | K.TPLTAN[+1501.555]ITK.S | HexNAc(5)Hex(3) | N[+1502] | 1230.5619 | 2 | 2460.1166 | 2460.1121 | 0 | 1.8 | 200 | Specific | 319.96 | 197.08 | 197.08 | 6.04 | 35 | >sp\|P01877\|IGHA2_HUMAN Immunoglobulin heavy constant alpha 2 OS=Homo sapiens OX=9606 GN=IGHA2 PE=1 SV=4 | 576 | 20201114_qwl_milk_1ng_glyco_hcd203040_compound15.5941.5941.2 | controllerType=0 controllerNumber=1 scan=5941 | 16.1164 |
| 03112:2 | 20 | K.TPLTAN[+1444.534]ITK.S | HexNAc(4)Hex(3)Fuc(1) | N[+1445] | 1202.0506 | 2 | 2403.0939 | 2403.0906 | 0 | 1.3 | 200 | Specific | 351.73 | 169.85 | 169.85 | 5.69 | 35 | >sp\|P01877\|IGHA2_HUMAN Immunoglobulin heavy constant alpha 2 OS=Homo sapiens OX=9606 GN=IGHA2 PE=1 SV=4 | 576 | 20201114_qwl_milk_1ng_glyco_hcd203040_compound15.5883.5883.2 | controllerType=0 controllerNumber=1 scan=5883 | 16.016 |
| 03170:2 | 20 | K.TPLTAN[+1501.555]ITK.S | HexNAc(5)Hex(3) | N[+1502] | 1230.5619 | 2 | 2460.1166 | 2460.1121 | 0 | 1.8 | 200 | Specific | 370.83 | 350.06 | 350.06 | 7.32 | 35 | >sp\|P01877\|IGHA2_HUMAN Immunoglobulin heavy constant alpha 2 OS=Homo sapiens OX=9606 GN=IGHA2 PE=1 SV=4 | 576 | 20201114_qwl_milk_1ng_glyco_hcd203040_compound15.5943.5943.2 | controllerType=0 controllerNumber=1 scan=5943 | 16.1205 |
| 03083:3 | 20 | K.TPLTAN[+1809.666]ITK.S | HexNAc(5)Hex(4)Fuc(1) | N[+1810] | 923.4137 | 3 | 2768.2266 | 2768.2228 | 0 | 1.4 | 200 | Specific | 421.1 | 222.27 | 222.27 | 6.5 | 35 | >sp\|P01877\|IGHA2_HUMAN Immunoglobulin heavy constant alpha 2 OS=Homo sapiens OX=9606 GN=IGHA2 PE=1 SV=4 | 576 | 20201114_qwl_milk_1ng_glyco_hcd203040_compound15.5853.5853.3 | controllerType=0 controllerNumber=1 scan=5853 | 15.9663 |
| 03162:3 | 20 | K.TPLTAN[+1647.613]ITK.S | HexNAc(5)Hex(3)Fuc(1) | N[+1648] | 869.3961 | 3 | 2606.1737 | 2606.17 | 0 | 1.4 | 200 | Specific | 442.11 | 288.43 | 288.43 | 6.84 | 35 | >sp\|P01877\|IGHA2_HUMAN Immunoglobulin heavy constant alpha 2 OS=Homo sapiens OX=9606 GN=IGHA2 PE=1 SV=4 | 576 | 20201114_qwl_milk_1ng_glyco_hcd203040_compound15.5935.5935.3 | controllerType=0 controllerNumber=1 scan=5935 | 16.1058 |
| 03114:2 | 20 | K.TPLTAN[+1444.534]ITK.S | HexNAc(4)Hex(3)Fuc(1) | N[+1445] | 1202.0506 | 2 | 2403.0939 | 2403.0906 | 0 | 1.3 | 200 | Specific | 469.09 | 353.26 | 353.26 | 7.69 | 35 | >sp\|P01877\|IGHA2_HUMAN Immunoglobulin heavy constant alpha 2 OS=Homo sapiens OX=9606 GN=IGHA2 PE=1 SV=4 | 576 | 20201114_qwl_milk_1ng_glyco_hcd203040_compound15.5885.5885.2 | controllerType=0 controllerNumber=1 scan=5885 | 16.0201 |
| 03082:2 | 20 | K.TPLTAN[+1809.666]ITK.S | HexNAc(5)Hex(4)Fuc(1) | N[+1810] | 1384.6176 | 2 | 2768.2279 | 2768.2228 | 0 | 1.8 | 200 | Specific | 519.39 | 274.66 | 274.66 | 7.28 | 35 | >sp\|P01877\|IGHA2_HUMAN Immunoglobulin heavy constant alpha 2 OS=Homo sapiens OX=9606 GN=IGHA2 PE=1 SV=4 | 576 | 20201114_qwl_milk_1ng_glyco_hcd203040_compound15.5852.5852.2 | controllerType=0 controllerNumber=1 scan=5852 | 15.9636 |
| 03080:2 | 20 | K.TPLTAN[+1809.666]ITK.S | HexNAc(5)Hex(4)Fuc(1) | N[+1810] | 1384.6176 | 2 | 2768.2279 | 2768.2228 | 0 | 1.8 | 200 | Specific | 528.24 | 312.03 | 312.03 | 7.55 | 35 | >sp\|P01877\|IGHA2_HUMAN Immunoglobulin heavy constant alpha 2 OS=Homo sapiens OX=9606 GN=IGHA2 PE=1 SV=4 | 576 | 20201114_qwl_milk_1ng_glyco_hcd203040_compound15.5850.5850.2 | controllerType=0 controllerNumber=1 scan=5850 | 15.9596 |
| 03164:3 | 20 | K.TPLTAN[+1647.613]ITK.S | HexNAc(5)Hex(3)Fuc(1) | N[+1648] | 869.3961 | 3 | 2606.1737 | 2606.17 | 0 | 1.4 | 200 | Specific | 535.28 | 250.39 | 250.39 | 7.11 | 35 | >sp\|P01877\|IGHA2_HUMAN Immunoglobulin heavy constant alpha 2 OS=Homo sapiens OX=9606 GN=IGHA2 PE=1 SV=4 | 576 | 20201114_qwl_milk_1ng_glyco_hcd203040_compound15.5937.5937.3 | controllerType=0 controllerNumber=1 scan=5937 | 16.1101 |
| 03136:2 | 20 | K.TPLTAN[+1647.613]ITK.S | HexNAc(5)Hex(3)Fuc(1) | N[+1648] | 1303.5921 | 2 | 2606.1769 | 2606.17 | 0 | 2.6 | 200 | Specific | 584.55 | 307.49 | 307.49 | 9.03 | 35 | >sp\|P01877\|IGHA2_HUMAN Immunoglobulin heavy constant alpha 2 OS=Homo sapiens OX=9606 GN=IGHA2 PE=1 SV=4 | 576 | 20201114_qwl_milk_1ng_glyco_hcd203040_compound15.5908.5908.2 | controllerType=0 controllerNumber=1 scan=5908 | 16.0625 |
| 03134:2 | 20 | K.TPLTAN[+1647.613]ITK.S | HexNAc(5)Hex(3)Fuc(1) | N[+1648] | 1303.5921 | 2 | 2606.1769 | 2606.17 | 0 | 2.6 | 200 | Specific | 621.91 | 360.68 | 360.68 | 10.36 | 35 | >sp\|P01877\|IGHA2_HUMAN Immunoglobulin heavy constant alpha 2 OS=Homo sapiens OX=9606 GN=IGHA2 PE=1 SV=4 | 576 | 20201114_qwl_milk_1ng_glyco_hcd203040_compound15.5906.5906.2 | controllerType=0 controllerNumber=1 scan=5906 | 16.0569 |
| 10579:3 | 20 | R.MAGKPTHIN[+1501.555]VSVVMAEADGTC[+57.021]Y.- | HexNAc(5)Hex(3) | C[+57], N[+1502] | 1284.8903 | 3 | 3852.6564 | 3852.6382 | 0 | 4.7 | 319 | CRagged | 208.47 | 208.47 | 208.47 | 3.73 | 35 | >sp\|P01877\|IGHA2_HUMAN Immunoglobulin heavy constant alpha 2 OS=Homo sapiens OX=9606 GN=IGHA2 PE=1 SV=4 | 576 | 20201114_qwl_milk_1ng_glyco_hcd203040_compound15.13619.13619.3 | controllerType=0 controllerNumber=1 scan=13619 | 28.8512 |
| 10586:3 | 20 | R.MAGKPTHIN[+1647.613]VSVVMAEADGTC[+57.021]Y.- | HexNAc(5)Hex(3)Fuc(1) | C[+57], N[+1648] | 1333.5777 | 3 | 3998.7185 | 3998.6961 | 0 | 5.6 | 319 | CRagged | 258.09 | 247.08 | 247.08 | 6.06 | 35 | >sp\|P01877\|IGHA2_HUMAN Immunoglobulin heavy constant alpha 2 OS=Homo sapiens OX=9606 GN=IGHA2 PE=1 SV=4 | 576 | 20201114_qwl_milk_1ng_glyco_hcd203040_compound15.13626.13626.3 | controllerType=0 controllerNumber=1 scan=13626 | 28.8629 |
| 10584:3 | 20 | R.MAGKPTHIN[+1647.613]VSVVMAEADGTC[+57.021]Y.- | HexNAc(5)Hex(3)Fuc(1) | C[+57], N[+1648] | 1333.5777 | 3 | 3998.7185 | 3998.6961 | 0 | 5.6 | 319 | CRagged | 274.61 | 274.61 | 274.61 | 6.17 | 35 | >sp\|P01877\|IGHA2_HUMAN Immunoglobulin heavy constant alpha 2 OS=Homo sapiens OX=9606 GN=IGHA2 PE=1 SV=4 | 576 | 20201114_qwl_milk_1ng_glyco_hcd203040_compound15.13624.13624.3 | controllerType=0 controllerNumber=1 scan=13624 | 28.8588 |
| 10477:3 | 20 | R.MAGKPTHIN[+1663.608]VSVVMAEADGTC[+57.021]Y.- | HexNAc(5)Hex(4) | C[+57], N[+1664] | 1338.9053 | 3 | 4014.7014 | 4014.691 | 0 | 2.6 | 319 | CRagged | 452.11 | 385.03 | 360.06 | 9.45 | 35 | >sp\|P01877\|IGHA2_HUMAN Immunoglobulin heavy constant alpha 2 OS=Homo sapiens OX=9606 GN=IGHA2 PE=1 SV=4 | 576 | 20201114_qwl_milk_1ng_glyco_hcd203040_compound15.13513.13513.3 | controllerType=0 controllerNumber=1 scan=13513 | 28.6618 |
| 10384:3 | 20 | R.MAGKPTHIN[+1702.581]VSVVMAEADGTC[+57.021]Y.- | HexNAc(2)Hex(8) | C[+57], N[+1703] | 1351.8977 | 3 | 4053.6785 | 4053.6642 | 0 | 3.5 | 319 | CRagged | 548.65 | 548.65 | 361.24 | 10.01 | 35 | >sp\|P01877\|IGHA2_HUMAN Immunoglobulin heavy constant alpha 2 OS=Homo sapiens OX=9606 GN=IGHA2 PE=1 SV=4 | 576 | 20201114_qwl_milk_1ng_glyco_hcd203040_compound15.13416.13416.3 | controllerType=0 controllerNumber=1 scan=13416 | 28.4646 |
| 10448:3 | 20 | R.MAGKPTHIN[+1540.529]VSVVMAEADGTC[+57.021]Y.- | HexNAc(2)Hex(7) | C[+57], N[+1541] | 1297.8766 | 3 | 3891.6154 | 3891.6114 | 0 | 1 | 319 | CRagged | 687.27 | 687.27 | 404.1 | 13.15 | 35 | >sp\|P01877\|IGHA2_HUMAN Immunoglobulin heavy constant alpha 2 OS=Homo sapiens OX=9606 GN=IGHA2 PE=1 SV=4 | 576 | 20201114_qwl_milk_1ng_glyco_hcd203040_compound15.13483.13483.3 | controllerType=0 controllerNumber=1 scan=13483 | 28.5797 |
| 10386:3 | 20 | R.MAGKPTHIN[+1702.581]VSVVMAEADGTC[+57.021]Y.- | HexNAc(2)Hex(8) | C[+57], N[+1703] | 1351.8977 | 3 | 4053.6785 | 4053.6642 | 0 | 3.5 | 319 | CRagged | 738.79 | 738.79 | 489.8 | 14.23 | 35 | >sp\|P01877\|IGHA2_HUMAN Immunoglobulin heavy constant alpha 2 OS=Homo sapiens OX=9606 GN=IGHA2 PE=1 SV=4 | 576 | 20201114_qwl_milk_1ng_glyco_hcd203040_compound15.13418.13418.3 | controllerType=0 controllerNumber=1 scan=13418 | 28.4682 |
| 10446:3 | 20 | R.MAGKPTHIN[+1540.529]VSVVMAEADGTC[+57.021]Y.- | HexNAc(2)Hex(7) | C[+57], N[+1541] | 1297.8766 | 3 | 3891.6154 | 3891.6114 | 0 | 1 | 319 | CRagged | 779.43 | 717.88 | 461.54 | 15.15 | 35 | >sp\|P01877\|IGHA2_HUMAN Immunoglobulin heavy constant alpha 2 OS=Homo sapiens OX=9606 GN=IGHA2 PE=1 SV=4 | 576 | 20201114_qwl_milk_1ng_glyco_hcd203040_compound15.13481.13481.3 | controllerType=0 controllerNumber=1 scan=13481 | 28.5747 |
| 10492:3 | 20 | R.MAGKPTHIN[+1216.423]VSVVMAEADGTC[+57.021]Y.- | HexNAc(2)Hex(5) | C[+57], N[+1216] | 1189.8423 | 3 | 3567.5125 | 3567.5057 | 0 | 1.9 | 319 | CRagged | 863.08 | 863.08 | 451.88 | 17.15 | 35 | >sp\|P01877\|IGHA2_HUMAN Immunoglobulin heavy constant alpha 2 OS=Homo sapiens OX=9606 GN=IGHA2 PE=1 SV=4 | 576 | 20201114_qwl_milk_1ng_glyco_hcd203040_compound15.13529.13529.3 | controllerType=0 controllerNumber=1 scan=13529 | 28.6884 |
| 10494:3 | 20 | R.MAGKPTHIN[+1216.423]VSVVMAEADGTC[+57.021]Y.- | HexNAc(2)Hex(5) | C[+57], N[+1216] | 1189.8423 | 3 | 3567.5125 | 3567.5057 | 0 | 1.9 | 319 | CRagged | 864.01 | 712.86 | 498.18 | 17.15 | 35 | >sp\|P01877\|IGHA2_HUMAN Immunoglobulin heavy constant alpha 2 OS=Homo sapiens OX=9606 GN=IGHA2 PE=1 SV=4 | 576 | 20201114_qwl_milk_1ng_glyco_hcd203040_compound15.13531.13531.3 | controllerType=0 controllerNumber=1 scan=13531 | 28.6938 |
| 10450:3 | 20 | R.MAGKPTHIN[+1378.476]VSVVMAEADGTC[+57.021]Y.- | HexNAc(2)Hex(6) | C[+57], N[+1378] | 1243.86 | 3 | 3729.5656 | 3729.5585 | 0 | 1.9 | 319 | CRagged | 871.8 | 871.8 | 501.92 | 17.15 | 35 | >sp\|P01877\|IGHA2_HUMAN Immunoglobulin heavy constant alpha 2 OS=Homo sapiens OX=9606 GN=IGHA2 PE=1 SV=4 | 576 | 20201114_qwl_milk_1ng_glyco_hcd203040_compound15.13485.13485.3 | controllerType=0 controllerNumber=1 scan=13485 | 28.583 |
| 10452:3 | 20 | R.MAGKPTHIN[+1378.476]VSVVMAEADGTC[+57.021]Y.- | HexNAc(2)Hex(6) | C[+57], N[+1378] | 1243.86 | 3 | 3729.5656 | 3729.5585 | 0 | 1.9 | 319 | CRagged | 920.6 | 920.6 | 485.63 | 18.15 | 35 | >sp\|P01877\|IGHA2_HUMAN Immunoglobulin heavy constant alpha 2 OS=Homo sapiens OX=9606 GN=IGHA2 PE=1 SV=4 | 576 | 20201114_qwl_milk_1ng_glyco_hcd203040_compound15.13487.13487.3 | controllerType=0 controllerNumber=1 scan=13487 | 28.6049 |
| 24128:3 | 21 | R.QLAHQSN[+2204.772]STNIFFSPVSIATAFAMLSLGTK.A | HexNAc(4)Hex(5)NeuAc(2) | N[+2205] | 1796.1453 | 3 | 5386.4215 | 5386.4075 | 0 | 2.6 | 64 | Specific | 629.38 | 602.27 | 536.21 | 12.15 | 32 | >sp\|P01009\|A1AT_HUMAN Alpha-1-antitrypsin OS=Homo sapiens OX=9606 GN=SERPINA1 PE=1 SV=3 | 642 | 20201114_qwl_milk_1ng_glyco_hcd203040_compound15.27773.27773.3 | controllerType=0 controllerNumber=1 scan=27773 | 55.8247 |
| 24126:3 | 21 | R.QLAHQSN[+2204.772]STNIFFSPVSIATAFAMLSLGTK.A | HexNAc(4)Hex(5)NeuAc(2) | N[+2205] | 1796.1453 | 3 | 5386.4215 | 5386.4075 | 0 | 2.6 | 64 | Specific | 786.76 | 747.91 | 550.84 | 15.15 | 32 | >sp\|P01009\|A1AT_HUMAN Alpha-1-antitrypsin OS=Homo sapiens OX=9606 GN=SERPINA1 PE=1 SV=3 | 642 | 20201114_qwl_milk_1ng_glyco_hcd203040_compound15.27771.27771.3 | controllerType=0 controllerNumber=1 scan=27771 | 55.8189 |
| 24125:4 | 21 | R.QLAHQSN[+2204.772]STNIFFSPVSIATAFAMLSLGTK.A | HexNAc(4)Hex(5)NeuAc(2) | N[+2205] | 1347.3605 | 4 | 5386.4201 | 5386.4075 | 0 | 2.3 | 64 | Specific | 802.95 | 782.96 | 498.2 | 16.02 | 32 | >sp\|P01009\|A1AT_HUMAN Alpha-1-antitrypsin OS=Homo sapiens OX=9606 GN=SERPINA1 PE=1 SV=3 | 642 | 20201114_qwl_milk_1ng_glyco_hcd203040_compound15.27770.27770.4 | controllerType=0 controllerNumber=1 scan=27770 | 55.8174 |
| 24127:4 | 21 | R.QLAHQSN[+2204.772]STNIFFSPVSIATAFAMLSLGTK.A | HexNAc(4)Hex(5)NeuAc(2) | N[+2205] | 1347.3605 | 4 | 5386.4201 | 5386.4075 | 0 | 2.3 | 64 | Specific | 1000.51 | 951.78 | 565.73 | 20.02 | 32 | >sp\|P01009\|A1AT_HUMAN Alpha-1-antitrypsin OS=Homo sapiens OX=9606 GN=SERPINA1 PE=1 SV=3 | 642 | 20201114_qwl_milk_1ng_glyco_hcd203040_compound15.27772.27772.4 | controllerType=0 controllerNumber=1 scan=27772 | 55.8228 |
| 22762:4 | 21 | K.ADTHDEILEGLNFN[+2863.041]LTEIPEAQIHEGFQELLR.T | HexNAc(5)Hex(6)Fuc(4)NeuAc(1) | N[+2863] | 1639.4639 | 4 | 6554.8337 | 6554.8647 | 0 | -4.7 | 94 | Specific | 605.95 | 478.63 | 478.63 | 10.63 | 32 | >sp\|P01009\|A1AT_HUMAN Alpha-1-antitrypsin OS=Homo sapiens OX=9606 GN=SERPINA1 PE=1 SV=3 | 642 | 20201114_qwl_milk_1ng_glyco_hcd203040_compound15.26348.26348.4 | controllerType=0 controllerNumber=1 scan=26348 | 53.1153 |
| 22764:4 | 21 | K.ADTHDEILEGLNFN[+2863.041]LTEIPEAQIHEGFQELLR.T | HexNAc(5)Hex(6)Fuc(4)NeuAc(1) | N[+2863] | 1639.4639 | 4 | 6554.8337 | 6554.8647 | 0 | -4.7 | 94 | Specific | 676.89 | 514.94 | 514.94 | 11.63 | 32 | >sp\|P01009\|A1AT_HUMAN Alpha-1-antitrypsin OS=Homo sapiens OX=9606 GN=SERPINA1 PE=1 SV=3 | 642 | 20201114_qwl_milk_1ng_glyco_hcd203040_compound15.26350.26350.4 | controllerType=0 controllerNumber=1 scan=26350 | 53.1192 |
| 06715:3 | 25 | R.NLRN[+1216.423]M[+15.995]SNQLGLLAVNQR.F | HexNAc(2)Hex(5) | M[+16], N[+1216] | 1058.4934 | 3 | 3173.4658 | 3173.4625 | 0 | 1 | 352 | Specific | 211.76 | 211.76 | 211.76 | 4.51 | 35 | >sp\|P05164\|PERM_HUMAN Myeloperoxidase OS=Homo sapiens OX=9606 GN=MPO PE=1 SV=1 | 12 | 20201114_qwl_milk_1ng_glyco_hcd203040_compound15.9614.9614.3 | controllerType=0 controllerNumber=1 scan=9614 | 22.2408 |
| 06713:3 | 25 | R.NLRN[+1216.423]M[+15.995]SNQLGLLAVNQR.F | HexNAc(2)Hex(5) | M[+16], N[+1216] | 1058.4934 | 3 | 3173.4658 | 3173.4625 | 0 | 1 | 352 | Specific | 224.55 | 224.55 | 224.55 | 4.7 | 35 | >sp\|P05164\|PERM_HUMAN Myeloperoxidase OS=Homo sapiens OX=9606 GN=MPO PE=1 SV=1 | 12 | 20201114_qwl_milk_1ng_glyco_hcd203040_compound15.9612.9612.3 | controllerType=0 controllerNumber=1 scan=9612 | 22.2373 |
| 06709:3 | 25 | R.NLRN[+1378.476]M[+15.995]SNQLGLLAVNQR.F | HexNAc(2)Hex(6) | M[+16], N[+1378] | 1112.5131 | 3 | 3335.5247 | 3335.5153 | 0 | 2.8 | 352 | Specific | 241.94 | 241.94 | 241.94 | 5.95 | 35 | >sp\|P05164\|PERM_HUMAN Myeloperoxidase OS=Homo sapiens OX=9606 GN=MPO PE=1 SV=1 | 12 | 20201114_qwl_milk_1ng_glyco_hcd203040_compound15.9608.9608.3 | controllerType=0 controllerNumber=1 scan=9608 | 22.2302 |
| 06711:3 | 25 | R.NLRN[+1378.476]M[+15.995]SNQLGLLAVNQR.F | HexNAc(2)Hex(6) | M[+16], N[+1378] | 1112.5131 | 3 | 3335.5247 | 3335.5153 | 0 | 2.8 | 352 | Specific | 291.88 | 291.88 | 291.88 | 6.43 | 35 | >sp\|P05164\|PERM_HUMAN Myeloperoxidase OS=Homo sapiens OX=9606 GN=MPO PE=1 SV=1 | 12 | 20201114_qwl_milk_1ng_glyco_hcd203040_compound15.9610.9610.3 | controllerType=0 controllerNumber=1 scan=9610 | 22.2339 |
| 13732:3 | 25 | R.ALLPFDNLHDDPC[+57.021]LLTN[+1216.423]R.S | HexNAc(2)Hex(5) | C[+57], N[+1216] | 1114.1704 | 3 | 3340.4968 | 3340.4771 | 0 | 5.9 | 375 | Specific | 552.03 | 406.17 | 406.17 | 8.24 | 35 | >sp\|P05164\|PERM_HUMAN Myeloperoxidase OS=Homo sapiens OX=9606 GN=MPO PE=1 SV=1 | 12 | 20201114_qwl_milk_1ng_glyco_hcd203040_compound15.16895.16895.3 | controllerType=0 controllerNumber=1 scan=16895 | 34.4069 |
| 13730:3 | 25 | R.ALLPFDNLHDDPC[+57.021]LLTN[+1216.423]R.S | HexNAc(2)Hex(5) | C[+57], N[+1216] | 1114.1704 | 3 | 3340.4968 | 3340.4771 | 0 | 5.9 | 375 | Specific | 749.09 | 510.8 | 510.8 | 13.58 | 35 | >sp\|P05164\|PERM_HUMAN Myeloperoxidase OS=Homo sapiens OX=9606 GN=MPO PE=1 SV=1 | 12 | 20201114_qwl_milk_1ng_glyco_hcd203040_compound15.16893.16893.3 | controllerType=0 controllerNumber=1 scan=16893 | 34.4013 |
| 01131:2 | 25 | R.SYN[+1038.375]DSVDPR.I | HexNAc(2)Hex(3)Fuc(1) | N[+1038] | 1045.9246 | 2 | 2090.8419 | 2090.8395 | 0 | 1.2 | 481 | Specific | 228.73 | 190.66 | 190.66 | 5.56 | 35 | >sp\|P05164\|PERM_HUMAN Myeloperoxidase OS=Homo sapiens OX=9606 GN=MPO PE=1 SV=1 | 12 | 20201114_qwl_milk_1ng_glyco_hcd203040_compound15.3828.3828.2 | controllerType=0 controllerNumber=1 scan=3828 | 12.4334 |
| 01133:2 | 25 | R.SYN[+1038.375]DSVDPR.I | HexNAc(2)Hex(3)Fuc(1) | N[+1038] | 1045.9246 | 2 | 2090.8419 | 2090.8395 | 0 | 1.2 | 481 | Specific | 259.71 | 190.67 | 190.67 | 5.83 | 35 | >sp\|P05164\|PERM_HUMAN Myeloperoxidase OS=Homo sapiens OX=9606 GN=MPO PE=1 SV=1 | 12 | 20201114_qwl_milk_1ng_glyco_hcd203040_compound15.3830.3830.2 | controllerType=0 controllerNumber=1 scan=3830 | 12.4374 |
| 14458:2 | 29 | K.LC[+57.021]DN[+1866.688]LSTK.N | HexNAc(6)Hex(4) | C[+57], N[+1867] | 1409.074 | 2 | 2817.1408 | 2817.1487 | 0 | -2.8 | 285 | Specific | 276.03 | 165.08 | 165.08 | 2.04 | 24 | >sp\|P02774\|VTDB_HUMAN Vitamin D-binding protein OS=Homo sapiens OX=9606 GN=GC PE=1 SV=2 | 1103 | 20201114_qwl_milk_1ng_glyco_hcd203040_compound15.17650.17650.2 | controllerType=0 controllerNumber=1 scan=17650 | 35.7817 |
| 18843:3 | 30 | R.C[+57.021]MWSSALNSLN[+1864.634]LSFAGLEQVPK.G | HexNAc(2)Hex(9) | C[+57], N[+1865] | 1439.6291 | 3 | 4316.8727 | 4316.8341 | 0 | 8.9 | 272 | Specific | 462.56 | 462.56 | 462.56 | 8.46 | 21 | >sp\|P08571\|CD14_HUMAN Monocyte differentiation antigen CD14 OS=Homo sapiens OX=9606 GN=CD14 PE=1 SV=2 | 1232 | 20201114_qwl_milk_1ng_glyco_hcd203040_compound15.22251.22251.3 | controllerType=0 controllerNumber=1 scan=22251 | 44.746 |
| 18845:3 | 30 | R.C[+57.021]MWSSALNSLN[+1864.634]LSFAGLEQVPK.G | HexNAc(2)Hex(9) | C[+57], N[+1865] | 1439.6291 | 3 | 4316.8727 | 4316.8341 | 0 | 8.9 | 272 | Specific | 973.83 | 915.64 | 915.64 | 18.24 | 21 | >sp\|P08571\|CD14_HUMAN Monocyte differentiation antigen CD14 OS=Homo sapiens OX=9606 GN=CD14 PE=1 SV=2 | 1232 | 20201114_qwl_milk_1ng_glyco_hcd203040_compound15.22253.22253.3 | controllerType=0 controllerNumber=1 scan=22253 | 44.7539 |
| 13581:3 | 36 | R.IIVPLNNREN[+2204.772]ISDPTSPLR.T | HexNAc(4)Hex(5)NeuAc(2) | N[+2205] | 1451.6568 | 3 | 4352.9558 | 4352.9496 | 0 | 1.4 | 62 | Specific | 234.66 | 234.66 | 234.66 | 6.01 | 23 | >sp\|P01591\|IGJ_HUMAN Immunoglobulin J chain OS=Homo sapiens OX=9606 GN=JCHAIN PE=1 SV=4 | 1392 | 20201114_qwl_milk_1ng_glyco_hcd203040_compound15.16738.16738.3 | controllerType=0 controllerNumber=1 scan=16738 | 34.1409 |
| 09943:3 | 36 | R.IIVPLNNREN[+1647.613]ISDPTSPLR.T | HexNAc(5)Hex(3)Fuc(1) | N[+1648] | 1265.9373 | 3 | 3795.7974 | 3795.7904 | 0 | 1.8 | 62 | Specific | 289.85 | 223.38 | 223.38 | 6.39 | 23 | >sp\|P01591\|IGJ_HUMAN Immunoglobulin J chain OS=Homo sapiens OX=9606 GN=JCHAIN PE=1 SV=4 | 1392 | 20201114_qwl_milk_1ng_glyco_hcd203040_compound15.12960.12960.3 | controllerType=0 controllerNumber=1 scan=12960 | 27.7087 |
| 13473:3 | 36 | R.IIVPLNNREN[+2350.830]ISDPTSPLR.T | HexNAc(4)Hex(5)Fuc(1)NeuAc(2) | N[+2351] | 1500.3454 | 3 | 4499.0216 | 4499.0075 | 0 | 3.1 | 62 | Specific | 348.58 | 301.91 | 301.91 | 7.11 | 23 | >sp\|P01591\|IGJ_HUMAN Immunoglobulin J chain OS=Homo sapiens OX=9606 GN=JCHAIN PE=1 SV=4 | 1392 | 20201114_qwl_milk_1ng_glyco_hcd203040_compound15.16626.16626.3 | controllerType=0 controllerNumber=1 scan=16626 | 33.9305 |
| 09916:3 | 36 | R.IIVPLNNREN[+1501.555]ISDPTSPLR.T | HexNAc(5)Hex(3) | N[+1502] | 1217.2535 | 3 | 3649.7459 | 3649.7325 | 0 | 3.7 | 62 | Specific | 367.77 | 345.68 | 345.68 | 7.43 | 23 | >sp\|P01591\|IGJ_HUMAN Immunoglobulin J chain OS=Homo sapiens OX=9606 GN=JCHAIN PE=1 SV=4 | 1392 | 20201114_qwl_milk_1ng_glyco_hcd203040_compound15.12932.12932.3 | controllerType=0 controllerNumber=1 scan=12932 | 27.6629 |
| 09941:3 | 36 | R.IIVPLNNREN[+1647.613]ISDPTSPLR.T | HexNAc(5)Hex(3)Fuc(1) | N[+1648] | 1265.9373 | 3 | 3795.7974 | 3795.7904 | 0 | 1.8 | 62 | Specific | 388.13 | 337.36 | 337.36 | 7.39 | 23 | >sp\|P01591\|IGJ_HUMAN Immunoglobulin J chain OS=Homo sapiens OX=9606 GN=JCHAIN PE=1 SV=4 | 1392 | 20201114_qwl_milk_1ng_glyco_hcd203040_compound15.12958.12958.3 | controllerType=0 controllerNumber=1 scan=12958 | 27.7043 |
| 09834:3 | 36 | R.IIVPLNNREN[+1809.666]ISDPTSPLR.T | HexNAc(5)Hex(4)Fuc(1) | N[+1810] | 1319.9537 | 3 | 3957.8465 | 3957.8432 | 0 | 0.8 | 62 | Specific | 409.58 | 286.65 | 286.65 | 7.08 | 23 | >sp\|P01591\|IGJ_HUMAN Immunoglobulin J chain OS=Homo sapiens OX=9606 GN=JCHAIN PE=1 SV=4 | 1392 | 20201114_qwl_milk_1ng_glyco_hcd203040_compound15.12847.12847.3 | controllerType=0 controllerNumber=1 scan=12847 | 27.5232 |
| 09918:3 | 36 | R.IIVPLNNREN[+1501.555]ISDPTSPLR.T | HexNAc(5)Hex(3) | N[+1502] | 1217.2535 | 3 | 3649.7459 | 3649.7325 | 0 | 3.7 | 62 | Specific | 419.05 | 373.39 | 373.39 | 9.08 | 23 | >sp\|P01591\|IGJ_HUMAN Immunoglobulin J chain OS=Homo sapiens OX=9606 GN=JCHAIN PE=1 SV=4 | 1392 | 20201114_qwl_milk_1ng_glyco_hcd203040_compound15.12934.12934.3 | controllerType=0 controllerNumber=1 scan=12934 | 27.6668 |
| 09791:3 | 36 | R.IIVPLNNREN[+1809.666]ISDPTSPLR.T | HexNAc(5)Hex(4)Fuc(1) | N[+1810] | 1319.9609 | 3 | 3957.8681 | 3957.8432 | 0 | 6.3 | 62 | Specific | 426.68 | 344.04 | 344.04 | 6.84 | 23 | >sp\|P01591\|IGJ_HUMAN Immunoglobulin J chain OS=Homo sapiens OX=9606 GN=JCHAIN PE=1 SV=4 | 1392 | 20201114_qwl_milk_1ng_glyco_hcd203040_compound15.12802.12802.3 | controllerType=0 controllerNumber=1 scan=12802 | 27.4479 |
| 13439:3 | 36 | R.IIVPLNNREN[+2350.830]ISDPTSPLR.T | HexNAc(4)Hex(5)Fuc(1)NeuAc(2) | N[+2351] | 1500.345 | 3 | 4499.0205 | 4499.0075 | 0 | 2.9 | 62 | Specific | 430.45 | 361.06 | 361.06 | 7.72 | 23 | >sp\|P01591\|IGJ_HUMAN Immunoglobulin J chain OS=Homo sapiens OX=9606 GN=JCHAIN PE=1 SV=4 | 1392 | 20201114_qwl_milk_1ng_glyco_hcd203040_compound15.16590.16590.3 | controllerType=0 controllerNumber=1 scan=16590 | 33.8644 |
| 11450:3 | 36 | R.IIVPLNNREN[+2059.735]ISDPTSPLR.T | HexNAc(4)Hex(5)Fuc(1)NeuAc(1) | N[+2060] | 1403.3157 | 3 | 4207.9326 | 4207.9121 | 0 | 4.9 | 62 | Specific | 432.04 | 432.04 | 432.04 | 8.7 | 23 | >sp\|P01591\|IGJ_HUMAN Immunoglobulin J chain OS=Homo sapiens OX=9606 GN=JCHAIN PE=1 SV=4 | 1392 | 20201114_qwl_milk_1ng_glyco_hcd203040_compound15.14522.14522.3 | controllerType=0 controllerNumber=1 scan=14522 | 30.3443 |
| 11453:3 | 36 | R.IIVPLNNREN[+2059.735]ISDPTSPLR.T | HexNAc(4)Hex(5)Fuc(1)NeuAc(1) | N[+2060] | 1403.3157 | 3 | 4207.9326 | 4207.9121 | 0 | 4.9 | 62 | Specific | 458.67 | 395.56 | 395.56 | 8.82 | 23 | >sp\|P01591\|IGJ_HUMAN Immunoglobulin J chain OS=Homo sapiens OX=9606 GN=JCHAIN PE=1 SV=4 | 1392 | 20201114_qwl_milk_1ng_glyco_hcd203040_compound15.14525.14525.3 | controllerType=0 controllerNumber=1 scan=14525 | 30.3492 |
| 09793:3 | 36 | R.IIVPLNNREN[+1809.666]ISDPTSPLR.T | HexNAc(5)Hex(4)Fuc(1) | N[+1810] | 1319.9609 | 3 | 3957.8681 | 3957.8432 | 0 | 6.3 | 62 | Specific | 471.51 | 463.46 | 463.46 | 8.75 | 23 | >sp\|P01591\|IGJ_HUMAN Immunoglobulin J chain OS=Homo sapiens OX=9606 GN=JCHAIN PE=1 SV=4 | 1392 | 20201114_qwl_milk_1ng_glyco_hcd203040_compound15.12804.12804.3 | controllerType=0 controllerNumber=1 scan=12804 | 27.452 |
| 09832:3 | 36 | R.IIVPLNNREN[+1809.666]ISDPTSPLR.T | HexNAc(5)Hex(4)Fuc(1) | N[+1810] | 1319.9537 | 3 | 3957.8465 | 3957.8432 | 0 | 0.8 | 62 | Specific | 473.32 | 386.62 | 386.62 | 9.31 | 23 | >sp\|P01591\|IGJ_HUMAN Immunoglobulin J chain OS=Homo sapiens OX=9606 GN=JCHAIN PE=1 SV=4 | 1392 | 20201114_qwl_milk_1ng_glyco_hcd203040_compound15.12845.12845.3 | controllerType=0 controllerNumber=1 scan=12845 | 27.5183 |
| 13502:3 | 36 | R.IIVPLNNREN[+2204.772]ISDPTSPLR.T | HexNAc(4)Hex(5)NeuAc(2) | N[+2205] | 1451.6597 | 3 | 4352.9646 | 4352.9496 | 0 | 3.5 | 62 | Specific | 530.97 | 490.39 | 490.39 | 9.58 | 23 | >sp\|P01591\|IGJ_HUMAN Immunoglobulin J chain OS=Homo sapiens OX=9606 GN=JCHAIN PE=1 SV=4 | 1392 | 20201114_qwl_milk_1ng_glyco_hcd203040_compound15.16656.16656.3 | controllerType=0 controllerNumber=1 scan=16656 | 33.9867 |
| 11469:3 | 36 | R.IIVPLNNREN[+1913.677]ISDPTSPLR.T | HexNAc(4)Hex(5)NeuAc(1) | N[+1914] | 1354.6296 | 3 | 4061.8742 | 4061.8542 | 0 | 4.9 | 62 | Specific | 542.65 | 501.7 | 501.7 | 9.19 | 23 | >sp\|P01591\|IGJ_HUMAN Immunoglobulin J chain OS=Homo sapiens OX=9606 GN=JCHAIN PE=1 SV=4 | 1392 | 20201114_qwl_milk_1ng_glyco_hcd203040_compound15.14541.14541.3 | controllerType=0 controllerNumber=1 scan=14541 | 30.3802 |
| 09844:3 | 36 | R.IIVPLNNREN[+1663.608]ISDPTSPLR.T | HexNAc(5)Hex(4) | N[+1664] | 1271.2668 | 3 | 3811.7858 | 3811.7853 | 0 | 0.1 | 62 | Specific | 549.4 | 506.24 | 506.24 | 9.26 | 23 | >sp\|P01591\|IGJ_HUMAN Immunoglobulin J chain OS=Homo sapiens OX=9606 GN=JCHAIN PE=1 SV=4 | 1392 | 20201114_qwl_milk_1ng_glyco_hcd203040_compound15.12857.12857.3 | controllerType=0 controllerNumber=1 scan=12857 | 27.5396 |
| 13578:3 | 36 | R.IIVPLNNREN[+2204.772]ISDPTSPLR.T | HexNAc(4)Hex(5)NeuAc(2) | N[+2205] | 1451.6568 | 3 | 4352.9558 | 4352.9496 | 0 | 1.4 | 62 | Specific | 558.54 | 558.54 | 558.54 | 9.69 | 23 | >sp\|P01591\|IGJ_HUMAN Immunoglobulin J chain OS=Homo sapiens OX=9606 GN=JCHAIN PE=1 SV=4 | 1392 | 20201114_qwl_milk_1ng_glyco_hcd203040_compound15.16735.16735.3 | controllerType=0 controllerNumber=1 scan=16735 | 34.1336 |
| 11466:3 | 36 | R.IIVPLNNREN[+1913.677]ISDPTSPLR.T | HexNAc(4)Hex(5)NeuAc(1) | N[+1914] | 1354.6296 | 3 | 4061.8742 | 4061.8542 | 0 | 4.9 | 62 | Specific | 567.09 | 521.17 | 521.17 | 9.38 | 23 | >sp\|P01591\|IGJ_HUMAN Immunoglobulin J chain OS=Homo sapiens OX=9606 GN=JCHAIN PE=1 SV=4 | 1392 | 20201114_qwl_milk_1ng_glyco_hcd203040_compound15.14538.14538.3 | controllerType=0 controllerNumber=1 scan=14538 | 30.3742 |
| 13500:3 | 36 | R.IIVPLNNREN[+2204.772]ISDPTSPLR.T | HexNAc(4)Hex(5)NeuAc(2) | N[+2205] | 1451.6597 | 3 | 4352.9646 | 4352.9496 | 0 | 3.5 | 62 | Specific | 568.49 | 536.2 | 536.2 | 9.77 | 23 | >sp\|P01591\|IGJ_HUMAN Immunoglobulin J chain OS=Homo sapiens OX=9606 GN=JCHAIN PE=1 SV=4 | 1392 | 20201114_qwl_milk_1ng_glyco_hcd203040_compound15.16654.16654.3 | controllerType=0 controllerNumber=1 scan=16654 | [illegible] |
| 09846:3 | 36 | R.IIVPLNNREN[+1663.608]ISDPTSPLR.T | HexNAc(5)Hex(4) | N[+1664] | 1271.2668 | 3 | 3811.7858 | 3811.7853 | 0 | 0.1 | 62 | Specific | 584.74 | 577.34 | 577.34 | 10.47 | 23 | >sp\|P01591\|IGJ_HUMAN Immunoglobulin J chain OS=Homo sapiens OX=9606 GN=JCHAIN PE=1 SV=4 | 1392 | 20201114_qwl_milk_1ng_glyco_hcd203040_compound15.12859.12859.3 | controllerType=0 controllerNumber=1 scan=12859 | [illegible] |
| 02127:3 | 105 | R.VVLGAHN[+730.264]LSR.R | HexNAc(2)Hex(2) | N[+730] | 599.3001 | 3 | 1795.8858 | 1795.8807 | 0 | 2.8 | 82 | Specific | 283.07 | 283.07 | 283.07 | 5.44 | 10 | >sp\|P08246\|ELNE_HUMAN Neutrophil elastase OS=Homo sapiens OX=9606 GN=ELANE PE=1 SV=1 | 1269 | 20201114_qwl_milk_1ng_glyco_hcd203040_compound15.4862.4862.3 | controllerType=0 controllerNumber=1 scan=4862 | [illegible] |
| 02125:3 | 105 | R.VVLGAHN[+730.264]LSR.R | HexNAc(2)Hex(2) | N[+730] | 599.3001 | 3 | 1795.8858 | 1795.8807 | 0 | 2.8 | 82 | Specific | 356.31 | 356.31 | 356.31 | 7.18 | 10 | >sp\|P08246\|ELNE_HUMAN Neutrophil elastase OS=Homo sapiens OX=9606 GN=ELANE PE=1 SV=1 | 1269 | 20201114_qwl_milk_1ng_glyco_hcd203040_compound15.4860.4860.3 | controllerType=0 controllerNumber=1 scan=4860 | [illegible] |
| 19483:3 | 105 | R.GIASVLQELN[+876.322]VTVVTSLC[+57.021]R.R | HexNAc(2)Hex(2)Fuc(1) | C[+57], N[+876] | 979.1577 | 3 | 2935.4585 | 2935.4439 | 0 | 5 | 164 | Specific | 214.26 | 214.26 | 214.26 | 3.26 | 10 | >sp\|P08246\|ELNE_HUMAN Neutrophil elastase OS=Homo sapiens OX=9606 GN=ELANE PE=1 SV=1 | 1269 | 20201114_qwl_milk_1ng_glyco_hcd203040_compound15.22922.22922.3 | controllerType=0 controllerNumber=1 scan=22922 | 46.2087 |
| 19480:3 | 105 | R.GIASVLQELN[+876.322]VTVVTSLC[+57.021]R.R | HexNAc(2)Hex(2)Fuc(1) | C[+57], N[+876] | 979.1577 | 3 | 2935.4585 | 2935.4439 | 0 | 5 | 164 | Specific | 277.82 | 277.82 | 277.82 | 5.05 | 10 | >sp\|P08246\|ELNE_HUMAN Neutrophil elastase OS=Homo sapiens OX=9606 GN=ELANE PE=1 SV=1 | 1269 | 20201114_qwl_milk_1ng_glyco_hcd203040_compound15.22919.22919.3 | controllerType=0 controllerNumber=1 scan=22919 | 46.2022 |
| 09099:3 | 131 | R.FVN[+876.322]VTVTPEDQC[+57.021]RPNNVC[+57.021]TGVLTR.R | HexNAc(2)Hex(2)Fuc(1) | C[+57]*2, N[+876] | 1218.2341 | 3 | 3652.6876 | 3652.6728 | 0 | 4.1 | 169 | Specific | 614.63 | 614.63 | 614.63 | 12.05 | 6 | >sp\|P20160\|CAP7_HUMAN Azurocidin OS=Homo sapiens OX=9606 GN=AZU1 PE=1 SV=3 | 850 | 20201114_qwl_milk_1ng_glyco_hcd203040_compound15.12083.12083.3 | controllerType=0 controllerNumber=1 scan=12083 | 26.2853 |
| 09096:3 | 131 | R.FVN[+876.322]VTVTPEDQC[+57.021]RPNNVC[+57.021]TGVLTR.R | HexNAc(2)Hex(2)Fuc(1) | C[+57]*2, N[+876] | 1218.2341 | 3 | 3652.6876 | 3652.6728 | 0 | 4.1 | 169 | Specific | 937.41 | 937.41 | 937.41 | 18.05 | 6 | >sp\|P20160\|CAP7_HUMAN Azurocidin OS=Homo sapiens OX=9606 GN=AZU1 PE=1 SV=3 | 850 | 20201114_qwl_milk_1ng_glyco_hcd203040_compound15.12080.12080.3 | controllerType=0 controllerNumber=1 scan=12080 | 26.2796 |
| 24778:3 | 544 | R.AVN[+2204.772]ITSENLIDDVVSLIR.G | HexNAc(4)Hex(5)NeuAc(2) | N[+2205] | 1392.6188 | 3 | 4175.8419 | 4175.8481 | 0 | -1.5 | 413 | Specific | 247.31 | 247.31 | 247.31 | 5.24 | 1 | >sp\|P02748\|CO9_HUMAN Complement component C9 OS=Homo sapiens OX=9606 GN=C9 PE=1 SV=2 | 1547 | 20201114_qwl_milk_1ng_glyco_hcd203040_compound15.28453.28453.3 | controllerType=0 controllerNumber=1 scan=28453 | 57.0469 |

Table S5. List of unique N-glycopeptides

| Glycopeptide | Glycan | Uniprot ID | Gene Name | Protein Name | Shortname | ProteinNameInSkyline | Glycosites | Reported before | Site |
|---|---|---|---|---|---|---|---|---|---|
| K.DIDGYGGIALPELIC[+57.021]TMFHTSGYDTQAIVENN[+1686.586]ESTEYGLFQISNK.L | HexNAc(2)Hex(7)Fuc(1) | P00709 | LALBA | Alpha-lactalbumin | P00709\|LALBA | P00709\|LALBA_N64_HexNAc(2)Hex(7)Fuc(1) | N64 | Yes | P00709\|LALBA_N64 |
| K.DIDGYGGIALPELIC[+57.021]TMFHTSGYDTQAIVENN[+2350.830]ESTEYGLFQISNK.L | HexNAc(4)Hex(5)Fuc(1)NeuAc(2) | P00709 | LALBA | Alpha-lactalbumin | P00709\|LALBA | P00709\|LALBA_N64_HexNAc(4)Hex(5)Fuc(1)NeuAc(2) | N64 | Yes | P00709\|LALBA_N64 |
| K.DIDGYGGIALPELIC[+57.021]TMFHTSGYDTQAIVENN[+2336.851]ESTEYGLFQISNK.L | HexNAc(6)Hex(6)Fuc(1) | P00709 | LALBA | Alpha-lactalbumin | P00709\|LALBA | P00709\|LALBA_N64_HexNAc(6)Hex(6)Fuc(1) | N64 | Yes | P00709\|LALBA_N64 |
| R.FFVPLFLVGILFPAILAKQFTKC[+57.021]ELSQLLKDIDGYGGIALPELIC[+57.021]TMFHTSGYDTQAIVENN[+892.317]ESTEYGLFQISNK.L | HexNAc(2)Hex(3) | P00709 | LALBA | Alpha-lactalbumin | P00709\|LALBA | P00709\|LALBA_N64_HexNAc(2)Hex(3) | N64 | Yes | P00709\|LALBA_N64 |
| K.DIDGYGGIALPELIC[+57.021]TM[+15.995]FHTSGYDTQAIVENN[+2190.793]ESTEYGLFQISNK.L | HexNAc(6)Hex(6) | P00709 | LALBA | Alpha-lactalbumin | P00709\|LALBA | P00709\|LALBA_N64_HexNAc(6)Hex(6) | N64 | Yes | P00709\|LALBA_N64 |
| K.MVSHHN[+2059.735]LTTGATLINEQWLLTTAK.N | HexNAc(4)Hex(5)Fuc(1)NeuAc(1) | P00738 | HP | Haptoglobin | P00738\|HPT | P00738\|HPT_N184_HexNAc(4)Hex(5)Fuc(1)NeuAc(1) | N184 | Yes | P00738\|HPT_N184 |
| K.MVSHHN[+1913.677]LTTGATLINEQWLLTTAK.N | HexNAc(4)Hex(5)NeuAc(1) | P00738 | HP | Haptoglobin | P00738\|HPT | P00738\|HPT_N184_HexNAc(4)Hex(5)NeuAc(1) | N184 | Yes | P00738\|HPT_N184 |
| K.MVSHHN[+2204.772]LTTGATLINEQWLLTTAK.N | HexNAc(4)Hex(5)NeuAc(2) | P00738 | HP | Haptoglobin | P00738\|HPT | P00738\|HPT_N184_HexNAc(4)Hex(5)NeuAc(2) | N184 | Yes | P00738\|HPT_N184 |
| K.VVLHPN[+2205.793]YSQVDIGLIK.L | HexNAc(4)Hex(5)Fuc(2)NeuAc(1) | P00738 | HP | Haptoglobin | P00738\|HPT | P00738\|HPT_N241_HexNAc(4)Hex(5)Fuc(2)NeuAc(1) | N241 | Yes | P00738\|HPT_N241 |
| K.VVLHPN[+2204.772]YSQVDIGLIK.L | HexNAc(4)Hex(5)NeuAc(2) | P00738 | HP | Haptoglobin | P00738\|HPT | P00738\|HPT_N241_HexNAc(4)Hex(5)NeuAc(2) | N241 | Yes | P00738\|HPT_N241 |
| R.SPYYN[+1913.677]VSDEISFHC[+57.021]YDGYTLR.G | HexNAc(4)Hex(5)NeuAc(1) | P00751 | CFB | Complement factor B | P00751\|CFAB | P00751\|CFAB_N122_HexNAc(4)Hex(5)NeuAc(1) | N122 | Yes | P00751\|CFAB_N122 |
| K.ADTHDEILEGLNFN[+2350.830]LTEIPEAQIHEGFQELLR.T | HexNAc(4)Hex(5)Fuc(1)NeuAc(2) | P01009 | SERPINA1 | Alpha-1-antitrypsin | P01009\|A1AT | P01009\|A1AT_N107_HexNAc(4)Hex(5)Fuc(1)NeuAc(2) | N107 | Yes | P01009\|A1AT_N107 |
| K.ADTHDEILEGLNFN[+2205.793]LTEIPEAQIHEGFQELLR.T | HexNAc(4)Hex(5)Fuc(2)NeuAc(1) | P01009 | SERPINA1 | Alpha-1-antitrypsin | P01009\|A1AT | P01009\|A1AT_N107_HexNAc(4)Hex(5)Fuc(2)NeuAc(1) | N107 | Yes | P01009\|A1AT_N107 |
| K.ADTHDEILEGLNFN[+2204.772]LTEIPEAQIHEGFQELLR.T | HexNAc(4)Hex(5)NeuAc(2) | P01009 | SERPINA1 | Alpha-1-antitrypsin | P01009\|A1AT | P01009\|A1AT_N107_HexNAc(4)Hex(5)NeuAc(2) | N107 | Yes | P01009\|A1AT_N107 |
| K.ADTHDEILEGLNFN[+3007.058]LTEIPEAQIHEGFQELLR.T | HexNAc(5)Hex(6)Fuc(1)NeuAc(3) | P01009 | SERPINA1 | Alpha-1-antitrypsin | P01009\|A1AT | P01009\|A1AT_N107_HexNAc(5)Hex(6)Fuc(1)NeuAc(3) | N107 | Yes | P01009\|A1AT_N107 |
| K.ADTHDEILEGLNFN[+2862.021]LTEIPEAQIHEGFQELLR.T | HexNAc(5)Hex(6)Fuc(2)NeuAc(2) | P01009 | SERPINA1 | Alpha-1-antitrypsin | P01009\|A1AT | P01009\|A1AT_N107_HexNAc(5)Hex(6)Fuc(2)NeuAc(2) | N107 | Yes | P01009\|A1AT_N107 |
| K.ADTHDEILEGLNFN[+2863.041]LTEIPEAQIHEGFQELLR.T | HexNAc(5)Hex(6)Fuc(4)NeuAc(1) | P01009 | SERPINA1 | Alpha-1-antitrypsin | P01009\|A1AT | P01009\|A1AT_N107_HexNAc(5)Hex(6)Fuc(4)NeuAc(1) | N107 | Yes | P01009\|A1AT_N107 |
| K.ADTHDEILEGLNFN[+2569.905]LTEIPEAQIHEGFQELLR.T | HexNAc(5)Hex(6)NeuAc(2) | P01009 | SERPINA1 | Alpha-1-antitrypsin | P01009\|A1AT | P01009\|A1AT_N107_HexNAc(5)Hex(6)NeuAc(2) | N107 | Yes | P01009\|A1AT_N107 |
| K.ADTHDEILEGLNFN[+2861.000]LTEIPEAQIHEGFQELLR.T | HexNAc(5)Hex(6)NeuAc(3) | P01009 | SERPINA1 | Alpha-1-antitrypsin | P01009\|A1AT | P01009\|A1AT_N107_HexNAc(5)Hex(6)NeuAc(3) | N107 | Yes | P01009\|A1AT_N107 |
| K.ADTHDEILEGLNFN[+3517.228]LTEIPEAQIHEGFQELLR.T | HexNAc(6)Hex(7)NeuAc(4) | P01009 | SERPINA1 | Alpha-1-antitrypsin | P01009\|A1AT | P01009\|A1AT_N107_HexNAc(6)Hex(7)NeuAc(4) | N107 | Yes | P01009\|A1AT_N107 |
| K.[+42.011]ADTHDEILEGLNFN[+2190.793]LTEIPEAQIHEGFQELLR.T | HexNAc(6)Hex(6) | P01009 | SERPINA1 | Alpha-1-antitrypsin | P01009\|A1AT | P01009\|A1AT_N107_HexNAc(6)Hex(6) | N107 | Yes | P01009\|A1AT_N107 |
| K.YLGN[+2205.793]ATAIFFLPDEGK.L | HexNAc(4)Hex(5)Fuc(2)NeuAc(1) | P01009 | SERPINA1 | Alpha-1-antitrypsin | P01009\|A1AT | P01009\|A1AT_N253_HexNAc(4)Hex(5)Fuc(2)NeuAc(1) | N253 | Yes | P01009\|A1AT_N253 |
| K.YLGN[+2351.851]ATAIFFLPDEGKLQHLENELTHDIITK.F | HexNAc(4)Hex(5)Fuc(3)NeuAc(1) | P01009 | SERPINA1 | Alpha-1-antitrypsin | P01009\|A1AT | P01009\|A1AT_N253_HexNAc(4)Hex(5)Fuc(3)NeuAc(1) | N253 | Yes | P01009\|A1AT_N253 |
| K.YLGN[+2204.772]ATAIFFLPDEGK.L | HexNAc(4)Hex(5)NeuAc(2) | P01009 | SERPINA1 | Alpha-1-antitrypsin | P01009\|A1AT | P01009\|A1AT_N253_HexNAc(4)Hex(5)NeuAc(2) | N253 | Yes | P01009\|A1AT_N253 |
| K.YLGN[+2204.772]ATAIFFLPDEGKLQHLENELTHDIITK.F | HexNAc(4)Hex(5)NeuAc(2) | P01009 | SERPINA1 | Alpha-1-antitrypsin | P01009\|A1AT | P01009\|A1AT_N253_HexNAc(4)Hex(5)NeuAc(2) | N253 | Yes | P01009\|A1AT_N253 |
| K.[+42.011]YLGN[+2190.793]ATAIFFLPDEGKLQHLENELTHDIITK.F | HexNAc(6)Hex(6) | P01009 | SERPINA1 | Alpha-1-antitrypsin | P01009\|A1AT | P01009\|A1AT_N253_HexNAc(6)Hex(6) | N253 | Yes | P01009\|A1AT_N253 |
| R.QLAHQSN[+2188.740]STNIFFSPVSIATAFAMLSLGTK.A | HexNAc(2)Hex(11) | P01009 | SERPINA1 | Alpha-1-antitrypsin | P01009\|A1AT | P01009\|A1AT_N70_HexNAc(2)Hex(11) | N70 | Yes | P01009\|A1AT_N70 |
| R.QLAHQSN[+2350.830]STNIFFSPVSIATAFAMLSLGTK.A | HexNAc(4)Hex(5)Fuc(1)NeuAc(2) | P01009 | SERPINA1 | Alpha-1-antitrypsin | P01009\|A1AT | P01009\|A1AT_N70_HexNAc(4)Hex(5)Fuc(1)NeuAc(2) | N70 | Yes | P01009\|A1AT_N70 |
| R.QLAHQSN[+2205.793]STNIFFSPVSIATAFAMLSLGTK.A | HexNAc(4)Hex(5)Fuc(2)NeuAc(1) | P01009 | SERPINA1 | Alpha-1-antitrypsin | P01009\|A1AT | P01009\|A1AT_N70_HexNAc(4)Hex(5)Fuc(2)NeuAc(1) | N70 | Yes | P01009\|A1AT_N70 |
| R.QLAHQSN[+2351.851]STNIFFSPVSIATAFAMLSLGTK.A | HexNAc(4)Hex(5)Fuc(3)NeuAc(1) | P01009 | SERPINA1 | Alpha-1-antitrypsin | P01009\|A1AT | P01009\|A1AT_N70_HexNAc(4)Hex(5)Fuc(3)NeuAc(1) | N70 | Yes | P01009\|A1AT_N70 |
| R.QLAHQSN[+1913.677]STNIFFSPVSIATAFAMLSLGTK.A | HexNAc(4)Hex(5)NeuAc(1) | P01009 | SERPINA1 | Alpha-1-antitrypsin | P01009\|A1AT | P01009\|A1AT_N70_HexNAc(4)Hex(5)NeuAc(1) | N70 | Yes | P01009\|A1AT_N70 |
| R.QLAHQSN[+2204.772]STNIFFSPVSIATAFAM[+15.995]LSLGTK.A | HexNAc(4)Hex(5)NeuAc(2) | P01009 | SERPINA1 | Alpha-1-antitrypsin | P01009\|A1AT | P01009\|A1AT_N70_HexNAc(4)Hex(5)NeuAc(2) | N70 | Yes | P01009\|A1AT_N70 |
| R.QLAHQSN[+2204.772]STNIFFSPVSIATAFAMLSLGTK.A | HexNAc(4)Hex(5)NeuAc(2) | P01009 | SERPINA1 | Alpha-1-antitrypsin | P01009\|A1AT | P01009\|A1AT_N70_HexNAc(4)Hex(5)NeuAc(2) | N70 | Yes | P01009\|A1AT_N70 |
| R.QLAHQSN[+3007.058]STNIFFSPVSIATAFAMLSLGTK.A | HexNAc(5)Hex(6)Fuc(1)NeuAc(3) | P01009 | SERPINA1 | Alpha-1-antitrypsin | P01009\|A1AT | P01009\|A1AT_N70_HexNAc(5)Hex(6)Fuc(1)NeuAc(3) | N70 | Yes | P01009\|A1AT_N70 |
| R.QLAHQSN[+2862.021]STNIFFSPVSIATAFAMLSLGTK.A | HexNAc(5)Hex(6)Fuc(2)NeuAc(2) | P01009 | SERPINA1 | Alpha-1-antitrypsin | P01009\|A1AT | P01009\|A1AT_N70_HexNAc(5)Hex(6)Fuc(2)NeuAc(2) | N70 | Yes | P01009\|A1AT_N70 |
| R.QLAHQSN[+2861.000]STNIFFSPVSIATAFAMLSLGTK.A | HexNAc(5)Hex(6)NeuAc(3) | P01009 | SERPINA1 | Alpha-1-antitrypsin | P01009\|A1AT | P01009\|A1AT_N70_HexNAc(5)Hex(6)NeuAc(3) | N70 | Yes | P01009\|A1AT_N70 |
| R.[+42.011]QLAHQSN[+1872.651]STNIFFSPVSIATAFAMLSLGTK.A | HexNAc(3)Hex(6)NeuAc(1) | P01009 | SERPINA1 | Alpha-1-antitrypsin | P01009\|A1AT | P01009\|A1AT_N70_HexNAc(3)Hex(6)NeuAc(1) | N70 | Yes | P01009\|A1AT_N70 |
| R.[+42.011]QLAHQSN[+2190.793]STNIFFSPVSIATAFAMLSLGTK.A | HexNAc(6)Hex(6) | P01009 | SERPINA1 | Alpha-1-antitrypsin | P01009\|A1AT | P01009\|A1AT_N70_HexNAc(6)Hex(6) | N70 | Yes | P01009\|A1AT_N70 |
| K.GLKFN[+2350.830]LTETSEAEIHQSFQHLLR.T | HexNAc(4)Hex(5)Fuc(1)NeuAc(2) | P01011 | SERPINA3 | Alpha-1-antichymotrypsin | P01011\|AACT | P01011\|AACT_N106_HexNAc(4)Hex(5)Fuc(1)NeuAc(2) | N106 | Yes | P01011\|AACT_N106 |
| R.TLN[+2059.735]QSSDELQLSMGNAMFVK.E | HexNAc(4)Hex(5)Fuc(1)NeuAc(1) | P01011 | SERPINA3 | Alpha-1-antichymotrypsin | P01011\|AACT | P01011\|AACT_N127_HexNAc(4)Hex(5)Fuc(1)NeuAc(1) | N127 | Yes | P01011\|AACT_N127 |
| R.TLN[+2205.793]QSSDELQLSMGNAMFVK.E | HexNAc(4)Hex(5)Fuc(2)NeuAc(1) | P01011 | SERPINA3 | Alpha-1-antichymotrypsin | P01011\|AACT | P01011\|AACT_N127_HexNAc(4)Hex(5)Fuc(2)NeuAc(1) | N127 | Yes | P01011\|AACT_N127 |
| R.[+42.011]TLN[+2018.708]QSSDELQLSMGNAMFVK.E | HexNAc(3)Hex(6)Fuc(1)NeuAc(1) | P01011 | SERPINA3 | Alpha-1-antichymotrypsin | P01011\|AACT | P01011\|AACT_N127_HexNAc(3)Hex(6)Fuc(1)NeuAc(1) | N127 | Yes | P01011\|AACT_N127 |
| K.YTGN[+2861.000]ASALFILPDQDKMEEVEAMLLPETLKR.W | HexNAc(5)Hex(6)NeuAc(3) | P01011 | SERPINA3 | Alpha-1-antichymotrypsin | P01011\|AACT | P01011\|AACT_N253_HexNAc(5)Hex(6)NeuAc(3) | N253 | Yes | P01011\|AACT_N253 |
| K.APDKNVIFSPLSISTALAFLSLGAHN[+2059.735]TTLTEILK.G | HexNAc(4)Hex(5)Fuc(1)NeuAc(1) | P01011 | SERPINA3 | Alpha-1-antichymotrypsin | P01011\|AACT | P01011\|AACT_N93_HexNAc(4)Hex(5)Fuc(1)NeuAc(1) | N93 | Yes | P01011\|AACT_N93 |
| K.APDKNVIFSPLSISTALAFLSLGAHN[+2350.830]TTLTEILK.G | HexNAc(4)Hex(5)Fuc(1)NeuAc(2) | P01011 | SERPINA3 | Alpha-1-antichymotrypsin | P01011\|AACT | P01011\|AACT_N93_HexNAc(4)Hex(5)Fuc(1)NeuAc(2) | N93 | Yes | P01011\|AACT_N93 |
| K.NVIFSPLSISTALAFLSLGAHN[+2350.830]TTLTEILK.G | HexNAc(4)Hex(5)Fuc(1)NeuAc(2) | P01011 | SERPINA3 | Alpha-1-antichymotrypsin | P01011\|AACT | P01011\|AACT_N93_HexNAc(4)Hex(5)Fuc(1)NeuAc(2) | N93 | Yes | P01011\|AACT_N93 |
| K.APDKNVIFSPLSISTALAFLSLGAHN[+2205.793]TTLTEILK.G | HexNAc(4)Hex(5)Fuc(2)NeuAc(1) | P01011 | SERPINA3 | Alpha-1-antichymotrypsin | P01011\|AACT | P01011\|AACT_N93_HexNAc(4)Hex(5)Fuc(2)NeuAc(1) | N93 | Yes | P01011\|AACT_N93 |
| K.NVIFSPLSISTALAFLSLGAHN[+2205.793]TTLTEILK.G | HexNAc(4)Hex(5)Fuc(2)NeuAc(1) | P01011 | SERPINA3 | Alpha-1-antichymotrypsin | P01011\|AACT | P01011\|AACT_N93_HexNAc(4)Hex(5)Fuc(2)NeuAc(1) | N93 | Yes | P01011\|AACT_N93 |
| K.APDKNVIFSPLSISTALAFLSLGAHN[+2351.851]TTLTEILK.G | HexNAc(4)Hex(5)Fuc(3)NeuAc(1) | P01011 | SERPINA3 | Alpha-1-antichymotrypsin | P01011\|AACT | P01011\|AACT_N93_HexNAc(4)Hex(5)Fuc(3)NeuAc(1) | N93 | Yes | P01011\|AACT_N93 |
| K.APDKNVIFSPLSISTALAFLSLGAHN[+2204.772]TTLTEILK.G | HexNAc(4)Hex(5)NeuAc(2) | P01011 | SERPINA3 | Alpha-1-antichymotrypsin | P01011\|AACT | P01011\|AACT_N93_HexNAc(4)Hex(5)NeuAc(2) | N93 | Yes | P01011\|AACT_N93 |
| K.APDKNVIFSPLSISTALAFLSLGAHN[+2424.867]TTLTEILK.G | HexNAc(5)Hex(6)Fuc(1)NeuAc(1) | P01011 | SERPINA3 | Alpha-1-antichymotrypsin | P01011\|AACT | P01011\|AACT_N93_HexNAc(5)Hex(6)Fuc(1)NeuAc(1) | N93 | Yes | P01011\|AACT_N93 |
| K.APDKNVIFSPLSISTALAFLSLGAHN[+3007.058]TTLTEILK.G | HexNAc(5)Hex(6)Fuc(1)NeuAc(3) | P01011 | SERPINA3 | Alpha-1-antichymotrypsin | P01011\|AACT | P01011\|AACT_N93_HexNAc(5)Hex(6)Fuc(1)NeuAc(3) | N93 | Yes | P01011\|AACT_N93 |
| K.APDKNVIFSPLSISTALAFLSLGAHN[+2570.925]TTLTEILK.G | HexNAc(5)Hex(6)Fuc(2)NeuAc(1) | P01011 | SERPINA3 | Alpha-1-antichymotrypsin | P01011\|AACT | P01011\|AACT_N93_HexNAc(5)Hex(6)Fuc(2)NeuAc(1) | N93 | Yes | P01011\|AACT_N93 |
| K.APDKNVIFSPLSISTALAFLSLGAHN[+2862.021]TTLTEILK.G | HexNAc(5)Hex(6)Fuc(2)NeuAc(2) | P01011 | SERPINA3 | Alpha-1-antichymotrypsin | P01011\|AACT | P01011\|AACT_N93_HexNAc(5)Hex(6)Fuc(2)NeuAc(2) | N93 | Yes | P01011\|AACT_N93 |
| K.APDKNVIFSPLSISTALAFLSLGAHN[+2716.983]TTLTEILK.G | HexNAc(5)Hex(6)Fuc(3)NeuAc(1) | P01011 | SERPINA3 | Alpha-1-antichymotrypsin | P01011\|AACT | P01011\|AACT_N93_HexNAc(5)Hex(6)Fuc(3)NeuAc(1) | N93 | Yes | P01011\|AACT_N93 |
| K.APDKNVIFSPLSISTALAFLSLGAHN[+2569.905]TTLTEILK.G | HexNAc(5)Hex(6)NeuAc(2) | P01011 | SERPINA3 | Alpha-1-antichymotrypsin | P01011\|AACT | P01011\|AACT_N93_HexNAc(5)Hex(6)NeuAc(2) | N93 | Yes | P01011\|AACT_N93 |
| K.APDKNVIFSPLSISTALAFLSLGAHN[+2861.000]TTLTEILK.G | HexNAc(5)Hex(6)NeuAc(3) | P01011 | SERPINA3 | Alpha-1-antichymotrypsin | P01011\|AACT | P01011\|AACT_N93_HexNAc(5)Hex(6)NeuAc(3) | N93 | Yes | P01011\|AACT_N93 |
| K.TVLTPATNHMGN[+1216.423]VTFTIPANR.E | HexNAc(2)Hex(5) | P01024 | C3 | Complement C3 | P01024\|CO3 | P01024\|CO3_N85_HexNAc(2)Hex(5) | N85 | Yes | P01024\|CO3_N85 |
| K.TVLTPATNHMGN[+1378.476]VTFTIPANR.E | HexNAc(2)Hex(6) | P01024 | C3 | Complement C3 | P01024\|CO3 | P01024\|CO3_N85_HexNAc(2)Hex(6) | N85 | Yes | P01024\|CO3_N85 |
| K.TVLTPATNHMGN[+1540.529]VTFTIPANR.E | HexNAc(2)Hex(7) | P01024 | C3 | Complement C3 | P01024\|CO3 | P01024\|CO3_N85_HexNAc(2)Hex(7) | N85 | Yes | P01024\|CO3_N85 |
| K.TVLTPATNHMGN[+1702.581]VTFTIPANR.E | HexNAc(2)Hex(8) | P01024 | C3 | Complement C3 | P01024\|CO3 | P01024\|CO3_N85_HexNAc(2)Hex(8) | N85 | Yes | P01024\|CO3_N85 |
| K.TVLTPATNHMGN[+1704.635]VTFTIPANR.E | HexNAc(6)Hex(3) | P01024 | C3 | Complement C3 | P01024\|CO3 | P01024\|CO3_N85_HexNAc(6)Hex(3) | N85 | Yes | P01024\|CO3_N85 |
| R.IIVPLNNREN[+1038.375]ISDPTSPLR.T | HexNAc(2)Hex(3)Fuc(1) | P01591 | JCHAIN | Immunoglobulin J chain | P01591\|IGJ | P01591\|IGJ_N71_HexNAc(2)Hex(3)Fuc(1) | N71 | Yes | P01591\|IGJ_N71 |
| R.IIVPLNNREN[+1257.449]ISDPTSPLR.T | HexNAc(3)Hex(4) | P01591 | JCHAIN | Immunoglobulin J chain | P01591\|IGJ | P01591\|IGJ_N71_HexNAc(3)Hex(4) | N71 | Yes | P01591\|IGJ_N71 |
| R.IIVPLNNREN[+1403.507]ISDPTSPLR.T | HexNAc(3)Hex(4)Fuc(1) | P01591 | JCHAIN | Immunoglobulin J chain | P01591\|IGJ | P01591\|IGJ_N71_HexNAc(3)Hex(4)Fuc(1) | N71 | Yes | P01591\|IGJ_N71 |
| R.IIVPLNNREN[+1694.603]ISDPTSPLR.T | HexNAc(3)Hex(4)Fuc(1)NeuAc(1) | P01591 | JCHAIN | Immunoglobulin J chain | P01591\|IGJ | P01591\|IGJ_N71_HexNAc(3)Hex(4)Fuc(1)NeuAc(1) | N71 | Yes | P01591\|IGJ_N71 |
| R.IIVPLNNREN[+1548.545]ISDPTSPLR.T | HexNAc(3)Hex(4)NeuAc(1) | P01591 | JCHAIN | Immunoglobulin J chain | P01591\|IGJ | P01591\|IGJ_N71_HexNAc(3)Hex(4)NeuAc(1) | N71 | Yes | P01591\|IGJ_N71 |
| R.IIVPLNNREN[+1565.560]ISDPTSPLR.T | HexNAc(3)Hex(5)Fuc(1) | P01591 | JCHAIN | Immunoglobulin J chain | P01591\|IGJ | P01591\|IGJ_N71_HexNAc(3)Hex(5)Fuc(1) | N71 | Yes | P01591\|IGJ_N71 |
| R.IIVPLNNREN[+1856.656]ISDPTSPLR.T | HexNAc(3)Hex(5)Fuc(1)NeuAc(1) | P01591 | JCHAIN | Immunoglobulin J chain | P01591\|IGJ | P01591\|IGJ_N71_HexNAc(3)Hex(5)Fuc(1)NeuAc(1) | N71 | Yes | P01591\|IGJ_N71 |
| R.IIVPLNNREN[+1460.529]ISDPTSPLR.T | HexNAc(4)Hex(4) | P01591 | JCHAIN | Immunoglobulin J chain | P01591\|IGJ | P01591\|IGJ_N71_HexNAc(4)Hex(4) | N71 | Yes | P01591\|IGJ_N71 |
| R.IIVPLNNREN[+1606.587]ISDPTSPLR.T | HexNAc(4)Hex(4)Fuc(1) | P01591 | JCHAIN | Immunoglobulin J chain | P01591\|IGJ | P01591\|IGJ_N71_HexNAc(4)Hex(4)Fuc(1) | N71 | Yes | P01591\|IGJ_N71 |
| R.IIVPLNNREN[+1622.582]ISDPTSPLR.T | HexNAc(4)Hex(5) | P01591 | JCHAIN | Immunoglobulin J chain | P01591\|IGJ | P01591\|IGJ_N71_HexNAc(4)Hex(5) | N71 | Yes | P01591\|IGJ_N71 |
| R.IIVPLNNREN[+1768.640]ISDPTSPLR.T | HexNAc(4)Hex(5)Fuc(1) | P01591 | JCHAIN | Immunoglobulin J chain | P01591\|IGJ | P01591\|IGJ_N71_HexNAc(4)Hex(5)Fuc(1) | N71 | Yes | P01591\|IGJ_N71 |
| R.IIVPLNNREN[+2059.735]ISDPTSPLR.T | HexNAc(4)Hex(5)Fuc(1)NeuAc(1) | P01591 | JCHAIN | Immunoglobulin J chain | P01591\|IGJ | P01591\|IGJ_N71_HexNAc(4)Hex(5)Fuc(1)NeuAc(1) | N71 | Yes | P01591\|IGJ_N71 |
| R.IIVPLNNREN[+2350.830]ISDPTSPLR.T | HexNAc(4)Hex(5)Fuc(1)NeuAc(2) | P01591 | JCHAIN | Immunoglobulin J chain | P01591\|IGJ | P01591\|IGJ_N71_HexNAc(4)Hex(5)Fuc(1)NeuAc(2) | N71 | Yes | P01591\|IGJ_N71 |
| R.IIVPLNNREN[+2205.793]ISDPTSPLR.T | HexNAc(4)Hex(5)Fuc(2)NeuAc(1) | P01591 | JCHAIN | Immunoglobulin J chain | P01591\|IGJ | P01591\|IGJ_N71_HexNAc(4)Hex(5)Fuc(2)NeuAc(1) | N71 | Yes | P01591\|IGJ_N71 |
| R.IIVPLNNREN[+2351.851]ISDPTSPLR.T | HexNAc(4)Hex(5)Fuc(3)NeuAc(1) | P01591 | JCHAIN | Immunoglobulin J chain | P01591\|IGJ | P01591\|IGJ_N71_HexNAc(4)Hex(5)Fuc(3)NeuAc(1) | N71 | Yes | P01591\|IGJ_N71 |
| R.EN[+1913.677]ISDPTSPLR.T | HexNAc(4)Hex(5)NeuAc(1) | P01591 | JCHAIN | Immunoglobulin J chain | P01591\|IGJ | P01591\|IGJ_N71_HexNAc(4)Hex(5)NeuAc(1) | N71 | Yes | P01591\|IGJ_N71 |
| R.IIVPLNNREN[+1913.677]ISDPTSPLR.T | HexNAc(4)Hex(5)NeuAc(1) | P01591 | JCHAIN | Immunoglobulin J chain | P01591\|IGJ | P01591\|IGJ_N71_HexNAc(4)Hex(5)NeuAc(1) | N71 | Yes | P01591\|IGJ_N71 |
| R.EN[+2204.772]ISDPTSPLR.T | HexNAc(4)Hex(5)NeuAc(2) | P01591 | JCHAIN | Immunoglobulin J chain | P01591\|IGJ | P01591\|IGJ_N71_HexNAc(4)Hex(5)NeuAc(2) | N71 | Yes | P01591\|IGJ_N71 |
| R.IIVPLNNREN[+2204.772]ISDPTSPLR.T | HexNAc(4)Hex(5)NeuAc(2) | P01591 | JCHAIN | Immunoglobulin J chain | P01591\|IGJ | P01591\|IGJ_N71_HexNAc(4)Hex(5)NeuAc(2) | N71 | Yes | P01591\|IGJ_N71 |
| R.IIVPLNNREN[+1501.555]ISDPTSPLR.T | HexNAc(5)Hex(3) | P01591 | JCHAIN | Immunoglobulin J chain | P01591\|IGJ | P01591\|IGJ_N71_HexNAc(5)Hex(3) | N71 | Yes | P01591\|IGJ_N71 |
| R.IIVPLNNREN[+1647.613]ISDPTSPLR.T | HexNAc(5)Hex(3)Fuc(1) | P01591 | JCHAIN | Immunoglobulin J chain | P01591\|IGJ | P01591\|IGJ_N71_HexNAc(5)Hex(3)Fuc(1) | N71 | Yes | P01591\|IGJ_N71 |
| R.EN[+1663.608]ISDPTSPLR.T | HexNAc(5)Hex(4) | P01591 | JCHAIN | Immunoglobulin J chain | P01591\|IGJ | P01591\|IGJ_N71_HexNAc(5)Hex(4) | N71 | Yes | P01591\|IGJ_N71 |
| R.IIVPLNNREN[+1663.608]ISDPTSPLR.T | HexNAc(5)Hex(4) | P01591 | JCHAIN | Immunoglobulin J chain | P01591\|IGJ | P01591\|IGJ_N71_HexNAc(5)Hex(4) | N71 | Yes | P01591\|IGJ_N71 |
| R.IIVPLNNREN[+1809.666]ISDPTSPLR.T | HexNAc(5)Hex(4)Fuc(1) | P01591 | JCHAIN | Immunoglobulin J chain | P01591\|IGJ | P01591\|IGJ_N71_HexNAc(5)Hex(4)Fuc(1) | N71 | Yes | P01591\|IGJ_N71 |
| R.IIVPLNNREN[+1971.719]ISDPTSPLR.T | HexNAc(5)Hex(5)Fuc(1) | P01591 | JCHAIN | Immunoglobulin J chain | P01591\|IGJ | P01591\|IGJ_N71_HexNAc(5)Hex(5)Fuc(1) | N71 | Yes | P01591\|IGJ_N71 |

Table S5. Cont.

| Glycopeptide | Glycan | Uniprot ID | Gene Name | Protein Name | Shortname | ProteinNameInSkyline | Glycosites | Reported before | Site |
|---|---|---|---|---|---|---|---|---|---|
| R.IIVPLNNREN[+2116.756]ISDPTSPLR.T | HexNAc(5)Hex(5)NeuAc(1) | P01591 | JCHAIN | Immunoglobulin J chain | P01591\|IGJ | P01591\|IGJ_N71_HexNAc(5)Hex(5)NeuAc(1) | N71 | Yes | P01591\|IGJ_N71 |
| R.GLSFDVSLEVSQGPGLLN[+2059.735]DTK.V | HexNAc(4)Hex(5)Fuc(1)NeuAc(1) | P01833 | PIGR | Polymeric immunoglobulin receptor | P01833\|PIGR | P01833\|PIGR_N135_HexNAc(4)Hex(5)Fuc(1)NeuAc(1) | N135 | Yes | P01833\|PIGR_N135 |
| R.GLSFDVSLEVSQGPGLLN[+2350.830]DTK.V | HexNAc(4)Hex(5)Fuc(1)NeuAc(2) | P01833 | PIGR | Polymeric immunoglobulin receptor | P01833\|PIGR | P01833\|PIGR_N135_HexNAc(4)Hex(5)Fuc(1)NeuAc(2) | N135 | Yes | P01833\|PIGR_N135 |
| R.GLSFDVSLEVSQGPGLLN[+2350.830]DTKVYTVDLGR.T | HexNAc(4)Hex(5)Fuc(1)NeuAc(2) | P01833 | PIGR | Polymeric immunoglobulin receptor | P01833\|PIGR | P01833\|PIGR_N135_HexNAc(4)Hex(5)Fuc(1)NeuAc(2) | N135 | Yes | P01833\|PIGR_N135 |
| R.GLSFDVSLEVSQGPGLLN[+2205.793]DTK.V | HexNAc(4)Hex(5)Fuc(2)NeuAc(1) | P01833 | PIGR | Polymeric immunoglobulin receptor | P01833\|PIGR | P01833\|PIGR_N135_HexNAc(4)Hex(5)Fuc(2)NeuAc(1) | N135 | Yes | P01833\|PIGR_N135 |
| R.GLSFDVSLEVSQGPGLLN[+2205.793]DTKVYTVDLGR.T | HexNAc(4)Hex(5)Fuc(2)NeuAc(1) | P01833 | PIGR | Polymeric immunoglobulin receptor | P01833\|PIGR | P01833\|PIGR_N135_HexNAc(4)Hex(5)Fuc(2)NeuAc(1) | N135 | Yes | P01833\|PIGR_N135 |
| R.GLSFDVSLEVSQGPGLLN[+2351.851]DTK.V | HexNAc(4)Hex(5)Fuc(3)NeuAc(1) | P01833 | PIGR | Polymeric immunoglobulin receptor | P01833\|PIGR | P01833\|PIGR_N135_HexNAc(4)Hex(5)Fuc(3)NeuAc(1) | N135 | Yes | P01833\|PIGR_N135 |
| R.GLSFDVSLEVSQGPGLLN[+2351.851]DTKVYTVDLGR.T | HexNAc(4)Hex(5)Fuc(3)NeuAc(1) | P01833 | PIGR | Polymeric immunoglobulin receptor | P01833\|PIGR | P01833\|PIGR_N135_HexNAc(4)Hex(5)Fuc(3)NeuAc(1) | N135 | Yes | P01833\|PIGR_N135 |
| R.GLSFDVSLEVSQGPGLLN[+2204.772]DTKVYTVDLGR.T | HexNAc(4)Hex(5)NeuAc(2) | P01833 | PIGR | Polymeric immunoglobulin receptor | P01833\|PIGR | P01833\|PIGR_N135_HexNAc(4)Hex(5)NeuAc(2) | N135 | Yes | P01833\|PIGR_N135 |
| R.GLSFDVSLEVSQGPGLLN[+2715.963]DTK.V | HexNAc(5)Hex(6)Fuc(1)NeuAc(2) | P01833 | PIGR | Polymeric immunoglobulin receptor | P01833\|PIGR | P01833\|PIGR_N135_HexNAc(5)Hex(6)Fuc(1)NeuAc(2) | N135 | Yes | P01833\|PIGR_N135 |
| R.GLSFDVSLEVSQGPGLLN[+2570.925]DTK.V | HexNAc(5)Hex(6)Fuc(2)NeuAc(1) | P01833 | PIGR | Polymeric immunoglobulin receptor | P01833\|PIGR | P01833\|PIGR_N135_HexNAc(5)Hex(6)Fuc(2)NeuAc(1) | N135 | Yes | P01833\|PIGR_N135 |
| R.GLSFDVSLEVSQGPGLLN[+2862.021]DTK.V | HexNAc(5)Hex(6)Fuc(2)NeuAc(2) | P01833 | PIGR | Polymeric immunoglobulin receptor | P01833\|PIGR | P01833\|PIGR_N135_HexNAc(5)Hex(6)Fuc(2)NeuAc(2) | N135 | Yes | P01833\|PIGR_N135 |
| R.GLSFDVSLEVSQGPGLLN[+2716.983]DTK.V | HexNAc(5)Hex(6)Fuc(3)NeuAc(1) | P01833 | PIGR | Polymeric immunoglobulin receptor | P01833\|PIGR | P01833\|PIGR_N135_HexNAc(5)Hex(6)Fuc(3)NeuAc(1) | N135 | Yes | P01833\|PIGR_N135 |
| R.GLSFDVSLEVSQGPGLLN[+2571.946]DTK.V | HexNAc(5)Hex(6)Fuc(4) | P01833 | PIGR | Polymeric immunoglobulin receptor | P01833\|PIGR | P01833\|PIGR_N135_HexNAc(5)Hex(6)Fuc(4) | N135 | Yes | P01833\|PIGR_N135 |
| R.GLSFDVSLEVSQGPGLLN[+2863.041]DTK.V | HexNAc(5)Hex(6)Fuc(4)NeuAc(1) | P01833 | PIGR | Polymeric immunoglobulin receptor | P01833\|PIGR | P01833\|PIGR_N135_HexNAc(5)Hex(6)Fuc(4)NeuAc(1) | N135 | Yes | P01833\|PIGR_N135 |
| R.GLSFDVSLEVSQGPGLLN[+2569.905]DTK.V | HexNAc(5)Hex(6)NeuAc(2) | P01833 | PIGR | Polymeric immunoglobulin receptor | P01833\|PIGR | P01833\|PIGR_N135_HexNAc(5)Hex(6)NeuAc(2) | N135 | Yes | P01833\|PIGR_N135 |
| R.[+42.011]GLSFDVSLEVSQGPGLLN[+2336.851]DTK.V | HexNAc(6)Hex(6)Fuc(1) | P01833 | PIGR | Polymeric immunoglobulin receptor | P01833\|PIGR | P01833\|PIGR_N135_HexNAc(6)Hex(6)Fuc(1) | N135 | Yes | P01833\|PIGR_N135 |
| K.QIGLYPVLVIDSSGYVNPN[+2188.740]YTGR.I | HexNAc(2)Hex(11) | P01833 | PIGR | Polymeric immunoglobulin receptor | P01833\|PIGR | P01833\|PIGR_N186_HexNAc(2)Hex(11) | N186 | Yes | P01833\|PIGR_N186 |
| K.QIGLYPVLVIDSSGYVNPN[+1257.449]YTGR.I | HexNAc(3)Hex(4) | P01833 | PIGR | Polymeric immunoglobulin receptor | P01833\|PIGR | P01833\|PIGR_N186_HexNAc(3)Hex(4) | N186 | Yes | P01833\|PIGR_N186 |
| K.QIGLYPVLVIDSSGYVNPN[+1872.651]YTGR.I | HexNAc(3)Hex(6)NeuAc(1) | P01833 | PIGR | Polymeric immunoglobulin receptor | P01833\|PIGR | P01833\|PIGR_N186_HexNAc(3)Hex(6)NeuAc(1) | N186 | Yes | P01833\|PIGR_N186 |
| K.QIGLYPVLVIDSSGYVNPN[+2059.735]YTGR.I | HexNAc(4)Hex(5)Fuc(1)NeuAc(1) | P01833 | PIGR | Polymeric immunoglobulin receptor | P01833\|PIGR | P01833\|PIGR_N186_HexNAc(4)Hex(5)Fuc(1)NeuAc(1) | N186 | Yes | P01833\|PIGR_N186 |
| K.QIGLYPVLVIDSSGYVNPN[+2350.830]YTGR.I | HexNAc(4)Hex(5)Fuc(1)NeuAc(2) | P01833 | PIGR | Polymeric immunoglobulin receptor | P01833\|PIGR | P01833\|PIGR_N186_HexNAc(4)Hex(5)Fuc(1)NeuAc(2) | N186 | Yes | P01833\|PIGR_N186 |
| K.QIGLYPVLVIDSSGYVNPN[+2205.793]YTGR.I | HexNAc(4)Hex(5)Fuc(2)NeuAc(1) | P01833 | PIGR | Polymeric immunoglobulin receptor | P01833\|PIGR | P01833\|PIGR_N186_HexNAc(4)Hex(5)Fuc(2)NeuAc(1) | N186 | Yes | P01833\|PIGR_N186 |
| K.QIGLYPVLVIDSSGYVNPN[+2351.851]YTGR.I | HexNAc(4)Hex(5)Fuc(3)NeuAc(1) | P01833 | PIGR | Polymeric immunoglobulin receptor | P01833\|PIGR | P01833\|PIGR_N186_HexNAc(4)Hex(5)Fuc(3)NeuAc(1) | N186 | Yes | P01833\|PIGR_N186 |
| K.QIGLYPVLVIDSSGYVNPN[+1913.677]YTGR.I | HexNAc(4)Hex(5)NeuAc(1) | P01833 | PIGR | Polymeric immunoglobulin receptor | P01833\|PIGR | P01833\|PIGR_N186_HexNAc(4)Hex(5)NeuAc(1) | N186 | Yes | P01833\|PIGR_N186 |
| K.QIGLYPVLVIDSSGYVNPN[+2204.772]YTGR.I | HexNAc(4)Hex(5)NeuAc(2) | P01833 | PIGR | Polymeric immunoglobulin receptor | P01833\|PIGR | P01833\|PIGR_N186_HexNAc(4)Hex(5)NeuAc(2) | N186 | Yes | P01833\|PIGR_N186 |
| K.QIGLYPVLVIDSSGYVNPN[+2076.750]YTGR.I | HexNAc(4)Hex(6)Fuc(2) | P01833 | PIGR | Polymeric immunoglobulin receptor | P01833\|PIGR | P01833\|PIGR_N186_HexNAc(4)Hex(6)Fuc(2) | N186 | Yes | P01833\|PIGR_N186 |
| K.QIGLYPVLVIDSSGYVNPN[+2570.925]YTGR.I | HexNAc(5)Hex(6)Fuc(2)NeuAc(1) | P01833 | PIGR | Polymeric immunoglobulin receptor | P01833\|PIGR | P01833\|PIGR_N186_HexNAc(5)Hex(6)Fuc(2)NeuAc(1) | N186 | Yes | P01833\|PIGR_N186 |
| K.QIGLYPVLVIDSSGYVNPN[+2425.887]YTGR.I | HexNAc(5)Hex(6)Fuc(3) | P01833 | PIGR | Polymeric immunoglobulin receptor | P01833\|PIGR | P01833\|PIGR_N186_HexNAc(5)Hex(6)Fuc(3) | N186 | Yes | P01833\|PIGR_N186 |
| K.QIGLYPVLVIDSSGYVNPN[+2716.983]YTGR.I | HexNAc(5)Hex(6)Fuc(3)NeuAc(1) | P01833 | PIGR | Polymeric immunoglobulin receptor | P01833\|PIGR | P01833\|PIGR_N186_HexNAc(5)Hex(6)Fuc(3)NeuAc(1) | N186 | Yes | P01833\|PIGR_N186 |
| K.QIGLYPVLVIDSSGYVNPN[+2569.905]YTGR.I | HexNAc(5)Hex(6)NeuAc(2) | P01833 | PIGR | Polymeric immunoglobulin receptor | P01833\|PIGR | P01833\|PIGR_N186_HexNAc(5)Hex(6)NeuAc(2) | N186 | Yes | P01833\|PIGR_N186 |
| R.LSLLEEPGN[+1200.428]GTFTVILNQLTSR.D | HexNAc(2)Hex(4)Fuc(1) | P01833 | PIGR | Polymeric immunoglobulin receptor | P01833\|PIGR | P01833\|PIGR_N421_HexNAc(2)Hex(4)Fuc(1) | N421 | Yes | P01833\|PIGR_N421 |
| R.LSLLEEPGN[+1458.442]GTFTVILNQLTSR.D | HexNAc(2)Hex(6)Phospho(1) | P01833 | PIGR | Polymeric immunoglobulin receptor | P01833\|PIGR | P01833\|PIGR_N421_HexNAc(2)Hex(6)Phospho(1) | N421 | Yes | P01833\|PIGR_N421 |
| R.LSLLEEPGN[+2059.735]GTFTVILNQLTSR.D | HexNAc(4)Hex(5)Fuc(1)NeuAc(1) | P01833 | PIGR | Polymeric immunoglobulin receptor | P01833\|PIGR | P01833\|PIGR_N421_HexNAc(4)Hex(5)Fuc(1)NeuAc(1) | N421 | Yes | P01833\|PIGR_N421 |
| R.LSLLEEPGN[+2350.830]GTFTVILNQLTSR.D | HexNAc(4)Hex(5)Fuc(1)NeuAc(2) | P01833 | PIGR | Polymeric immunoglobulin receptor | P01833\|PIGR | P01833\|PIGR_N421_HexNAc(4)Hex(5)Fuc(1)NeuAc(2) | N421 | Yes | P01833\|PIGR_N421 |
| R.LSLLEEPGN[+2205.793]GTFTVILNQLTSR.D | HexNAc(4)Hex(5)Fuc(2)NeuAc(1) | P01833 | PIGR | Polymeric immunoglobulin receptor | P01833\|PIGR | P01833\|PIGR_N421_HexNAc(4)Hex(5)Fuc(2)NeuAc(1) | N421 | Yes | P01833\|PIGR_N421 |
| R.LSLLEEPGN[+2496.888]GTFTVILNQLTSR.D | HexNAc(4)Hex(5)Fuc(2)NeuAc(2) | P01833 | PIGR | Polymeric immunoglobulin receptor | P01833\|PIGR | P01833\|PIGR_N421_HexNAc(4)Hex(5)Fuc(2)NeuAc(2) | N421 | Yes | P01833\|PIGR_N421 |
| R.LSLLEEPGN[+2351.851]GTFTVILNQLTSR.D | HexNAc(4)Hex(5)Fuc(3)NeuAc(1) | P01833 | PIGR | Polymeric immunoglobulin receptor | P01833\|PIGR | P01833\|PIGR_N421_HexNAc(4)Hex(5)Fuc(3)NeuAc(1) | N421 | Yes | P01833\|PIGR_N421 |
| R.LSLLEEPGN[+2642.946]GTFTVILNQLTSR.D | HexNAc(4)Hex(5)Fuc(3)NeuAc(2) | P01833 | PIGR | Polymeric immunoglobulin receptor | P01833\|PIGR | P01833\|PIGR_N421_HexNAc(4)Hex(5)Fuc(3)NeuAc(2) | N421 | Yes | P01833\|PIGR_N421 |
| R.LSLLEEPGN[+1913.677]GTFTVILNQLTSR.D | HexNAc(4)Hex(5)NeuAc(1) | P01833 | PIGR | Polymeric immunoglobulin receptor | P01833\|PIGR | P01833\|PIGR_N421_HexNAc(4)Hex(5)NeuAc(1) | N421 | Yes | P01833\|PIGR_N421 |
| R.LSLLEEPGN[+2204.772]GTFTVILNQLTSR.D | HexNAc(4)Hex(5)NeuAc(2) | P01833 | PIGR | Polymeric immunoglobulin receptor | P01833\|PIGR | P01833\|PIGR_N421_HexNAc(4)Hex(5)NeuAc(2) | N421 | Yes | P01833\|PIGR_N421 |
| R.LSLLEEPGN[+2076.750]GTFTVILNQLTSR.D | HexNAc(4)Hex(6)Fuc(2) | P01833 | PIGR | Polymeric immunoglobulin receptor | P01833\|PIGR | P01833\|PIGR_N421_HexNAc(4)Hex(6)Fuc(2) | N421 | Yes | P01833\|PIGR_N421 |
| K.AQYEGRLSLLEEPGN[+1647.613]GTFTVILNQLTSR.D | HexNAc(5)Hex(3)Fuc(1) | P01833 | PIGR | Polymeric immunoglobulin receptor | P01833\|PIGR | P01833\|PIGR_N421_HexNAc(5)Hex(3)Fuc(1) | N421 | Yes | P01833\|PIGR_N421 |
| K.AQYEGRLSLLEEPGN[+1663.608]GTFTVILNQLTSR.D | HexNAc(5)Hex(4) | P01833 | PIGR | Polymeric immunoglobulin receptor | P01833\|PIGR | P01833\|PIGR_N421_HexNAc(5)Hex(4) | N421 | Yes | P01833\|PIGR_N421 |
| R.LSLLEEPGN[+2553.910]GTFTVILNQLTSR.D | HexNAc(5)Hex(5)Fuc(1)NeuAc(2) | P01833 | PIGR | Polymeric immunoglobulin receptor | P01833\|PIGR | P01833\|PIGR_N421_HexNAc(5)Hex(5)Fuc(1)NeuAc(2) | N421 | Yes | P01833\|PIGR_N421 |
| R.LSLLEEPGN[+2715.963]GTFTVILNQLTSR.D | HexNAc(5)Hex(6)Fuc(1)NeuAc(2) | P01833 | PIGR | Polymeric immunoglobulin receptor | P01833\|PIGR | P01833\|PIGR_N421_HexNAc(5)Hex(6)Fuc(1)NeuAc(2) | N421 | Yes | P01833\|PIGR_N421 |
| R.LSLLEEPGN[+3007.058]GTFTVILNQLTSR.D | HexNAc(5)Hex(6)Fuc(1)NeuAc(3) | P01833 | PIGR | Polymeric immunoglobulin receptor | P01833\|PIGR | P01833\|PIGR_N421_HexNAc(5)Hex(6)Fuc(1)NeuAc(3) | N421 | Yes | P01833\|PIGR_N421 |
| R.LSLLEEPGN[+2570.925]GTFTVILNQLTSR.D | HexNAc(5)Hex(6)Fuc(2)NeuAc(1) | P01833 | PIGR | Polymeric immunoglobulin receptor | P01833\|PIGR | P01833\|PIGR_N421_HexNAc(5)Hex(6)Fuc(2)NeuAc(1) | N421 | Yes | P01833\|PIGR_N421 |
| R.LSLLEEPGN[+2862.021]GTFTVILNQLTSR.D | HexNAc(5)Hex(6)Fuc(2)NeuAc(2) | P01833 | PIGR | Polymeric immunoglobulin receptor | P01833\|PIGR | P01833\|PIGR_N421_HexNAc(5)Hex(6)Fuc(2)NeuAc(2) | N421 | Yes | P01833\|PIGR_N421 |
| R.LSLLEEPGN[+3153.116]GTFTVILNQLTSR.D | HexNAc(5)Hex(6)Fuc(2)NeuAc(3) | P01833 | PIGR | Polymeric immunoglobulin receptor | P01833\|PIGR | P01833\|PIGR_N421_HexNAc(5)Hex(6)Fuc(2)NeuAc(3) | N421 | Yes | P01833\|PIGR_N421 |
| R.LSLLEEPGN[+2425.887]GTFTVILNQLTSR.D | HexNAc(5)Hex(6)Fuc(3) | P01833 | PIGR | Polymeric immunoglobulin receptor | P01833\|PIGR | P01833\|PIGR_N421_HexNAc(5)Hex(6)Fuc(3) | N421 | Yes | P01833\|PIGR_N421 |
| R.LSLLEEPGN[+3008.078]GTFTVILNQLTSR.D | HexNAc(5)Hex(6)Fuc(3)NeuAc(2) | P01833 | PIGR | Polymeric immunoglobulin receptor | P01833\|PIGR | P01833\|PIGR_N421_HexNAc(5)Hex(6)Fuc(3)NeuAc(2) | N421 | Yes | P01833\|PIGR_N421 |
| R.LSLLEEPGN[+2571.946]GTFTVILNQLTSR.D | HexNAc(5)Hex(6)Fuc(4) | P01833 | PIGR | Polymeric immunoglobulin receptor | P01833\|PIGR | P01833\|PIGR_N421_HexNAc(5)Hex(6)Fuc(4) | N421 | Yes | P01833\|PIGR_N421 |
| R.LSLLEEPGN[+2863.041]GTFTVILNQLTSR.D | HexNAc(5)Hex(6)Fuc(4)NeuAc(1) | P01833 | PIGR | Polymeric immunoglobulin receptor | P01833\|PIGR | P01833\|PIGR_N421_HexNAc(5)Hex(6)Fuc(4)NeuAc(1) | N421 | Yes | P01833\|PIGR_N421 |
| R.LSLLEEPGN[+3154.136]GTFTVILNQLTSR.D | HexNAc(5)Hex(6)Fuc(4)NeuAc(2) | P01833 | PIGR | Polymeric immunoglobulin receptor | P01833\|PIGR | P01833\|PIGR_N421_HexNAc(5)Hex(6)Fuc(4)NeuAc(2) | N421 | Yes | P01833\|PIGR_N421 |
| R.LSLLEEPGN[+2569.905]GTFTVILNQLTSR.D | HexNAc(5)Hex(6)NeuAc(2) | P01833 | PIGR | Polymeric immunoglobulin receptor | P01833\|PIGR | P01833\|PIGR_N421_HexNAc(5)Hex(6)NeuAc(2) | N421 | Yes | P01833\|PIGR_N421 |
| R.LSLLEEPGN[+2861.000]GTFTVILNQLTSR.D | HexNAc(5)Hex(6)NeuAc(3) | P01833 | PIGR | Polymeric immunoglobulin receptor | P01833\|PIGR | P01833\|PIGR_N421_HexNAc(5)Hex(6)NeuAc(3) | N421 | Yes | P01833\|PIGR_N421 |
| R.[+42.011]LSLLEEPGN[+2018.708]GTFTVILNQLTSR.D | HexNAc(3)Hex(6)Fuc(1)NeuAc(1) | P01833 | PIGR | Polymeric immunoglobulin receptor | P01833\|PIGR | P01833\|PIGR_N421_HexNAc(3)Hex(6)Fuc(1)NeuAc(1) | N421 | Yes | P01833\|PIGR_N421 |
| K.[+42.011]AQYEGRLSLLEEPGN[+1751.624]GTFTVILNQLTSR.D | HexNAc(4)Hex(4)NeuAc(1) | P01833 | PIGR | Polymeric immunoglobulin receptor | P01833\|PIGR | P01833\|PIGR_N421_HexNAc(4)Hex(4)NeuAc(1) | N421 | Yes | P01833\|PIGR_N421 |
| K.[+42.011]AQYEGRLSLLEEPGN[+1622.582]GTFTVILNQLTSR.D | HexNAc(4)Hex(5) | P01833 | PIGR | Polymeric immunoglobulin receptor | P01833\|PIGR | P01833\|PIGR_N421_HexNAc(4)Hex(5) | N421 | Yes | P01833\|PIGR_N421 |
| R.[+42.011]LSLLEEPGN[+2059.735]GTFTVILNQLTSR.D | HexNAc(4)Hex(5)Fuc(1)NeuAc(1) | P01833 | PIGR | Polymeric immunoglobulin receptor | P01833\|PIGR | P01833\|PIGR_N421_HexNAc(4)Hex(5)Fuc(1)NeuAc(1) | N421 | Yes | P01833\|PIGR_N421 |
| R.[+42.011]LSLLEEPGN[+2350.830]GTFTVILNQLTSR.D | HexNAc(4)Hex(5)Fuc(1)NeuAc(2) | P01833 | PIGR | Polymeric immunoglobulin receptor | P01833\|PIGR | P01833\|PIGR_N421_HexNAc(4)Hex(5)Fuc(1)NeuAc(2) | N421 | Yes | P01833\|PIGR_N421 |
| R.[+42.011]LSLLEEPGN[+2205.793]GTFTVILNQLTSR.D | HexNAc(4)Hex(5)Fuc(2)NeuAc(1) | P01833 | PIGR | Polymeric immunoglobulin receptor | P01833\|PIGR | P01833\|PIGR_N421_HexNAc(4)Hex(5)Fuc(2)NeuAc(1) | N421 | Yes | P01833\|PIGR_N421 |
| R.[+42.011]LSLLEEPGN[+2351.851]GTFTVILNQLTSR.D | HexNAc(4)Hex(5)Fuc(3)NeuAc(1) | P01833 | PIGR | Polymeric immunoglobulin receptor | P01833\|PIGR | P01833\|PIGR_N421_HexNAc(4)Hex(5)Fuc(3)NeuAc(1) | N421 | Yes | P01833\|PIGR_N421 |
| K.[+42.011]AQYEGRLSLLEEPGN[+2117.777]GTFTVILNQLTSR.D | HexNAc(5)Hex(5)Fuc(2) | P01833 | PIGR | Polymeric immunoglobulin receptor | P01833\|PIGR | P01833\|PIGR_N421_HexNAc(5)Hex(5)Fuc(2) | N421 | Yes | P01833\|PIGR_N421 |
| R.[+42.011]LSLLEEPGN[+2190.793]GTFTVILNQLTSR.D | HexNAc(6)Hex(6) | P01833 | PIGR | Polymeric immunoglobulin receptor | P01833\|PIGR | P01833\|PIGR_N421_HexNAc(6)Hex(6) | N421 | Yes | P01833\|PIGR_N421 |
| R.[+42.011]LSLLEEPGN[+2336.851]GTFTVILNQLTSR.D | HexNAc(6)Hex(6)Fuc(1) | P01833 | PIGR | Polymeric immunoglobulin receptor | P01833\|PIGR | P01833\|PIGR_N421_HexNAc(6)Hex(6)Fuc(1) | N421 | Yes | P01833\|PIGR_N421 |
| R.[+42.011]LSLLEEPGN[+2482.909]GTFTVILNQLTSR.D | HexNAc(6)Hex(6)Fuc(2) | P01833 | PIGR | Polymeric immunoglobulin receptor | P01833\|PIGR | P01833\|PIGR_N421_HexNAc(6)Hex(6)Fuc(2) | N421 | Yes | P01833\|PIGR_N421 |
| R.[+42.011]C[+57.021]PLLVDSEGWVKAQYEGRLSLLEEPGN[+1403.507]GTFTVILNQLTSR.D | HexNAc(3)Hex(4)Fuc(1) | P01833 | PIGR | Polymeric immunoglobulin receptor | P01833\|PIGR | P01833\|PIGR_N421_HexNAc(3)Hex(4)Fuc(1) | N421 | Yes | P01833\|PIGR_N421 |
| K.IIEGEPNLKVPGN[+1856.656]VTAVLGETLKVPC[+57.021]HFPC[+57.021]K.F | HexNAc(3)Hex(5)Fuc(1)NeuAc(1) | P01833 | PIGR | Polymeric immunoglobulin receptor | P01833\|PIGR | P01833\|PIGR_N469_HexNAc(3)Hex(5)Fuc(1)NeuAc(1) | N469 | Yes | P01833\|PIGR_N469 |
| K.IIEGEPNLKVPGN[+1768.640]VTAVLGETLKVPC[+57.021]HFPC[+57.021]K.F | HexNAc(4)Hex(5)Fuc(1) | P01833 | PIGR | Polymeric immunoglobulin receptor | P01833\|PIGR | P01833\|PIGR_N469_HexNAc(4)Hex(5)Fuc(1) | N469 | Yes | P01833\|PIGR_N469 |
| K.IIEGEPNLKVPGN[+2059.735]VTAVLGETLK.V | HexNAc(4)Hex(5)Fuc(1)NeuAc(1) | P01833 | PIGR | Polymeric immunoglobulin receptor | P01833\|PIGR | P01833\|PIGR_N469_HexNAc(4)Hex(5)Fuc(1)NeuAc(1) | N469 | Yes | P01833\|PIGR_N469 |
| K.IIEGEPNLKVPGN[+2059.735]VTAVLGETLKVPC[+57.021]HFPC[+57.021]K.F | HexNAc(4)Hex(5)Fuc(1)NeuAc(1) | P01833 | PIGR | Polymeric immunoglobulin receptor | P01833\|PIGR | P01833\|PIGR_N469_HexNAc(4)Hex(5)Fuc(1)NeuAc(1) | N469 | Yes | P01833\|PIGR_N469 |
| K.VPGN[+2059.735]VTAVLGETLK.V | HexNAc(4)Hex(5)Fuc(1)NeuAc(1) | P01833 | PIGR | Polymeric immunoglobulin receptor | P01833\|PIGR | P01833\|PIGR_N469_HexNAc(4)Hex(5)Fuc(1)NeuAc(1) | N469 | Yes | P01833\|PIGR_N469 |
| K.VPGN[+2059.735]VTAVLGETLKVPC[+57.021]HFPC[+57.021]K.F | HexNAc(4)Hex(5)Fuc(1)NeuAc(1) | P01833 | PIGR | Polymeric immunoglobulin receptor | P01833\|PIGR | P01833\|PIGR_N469_HexNAc(4)Hex(5)Fuc(1)NeuAc(1) | N469 | Yes | P01833\|PIGR_N469 |
| K.IIEGEPNLKVPGN[+2350.830]VTAVLGETLK.V | HexNAc(4)Hex(5)Fuc(1)NeuAc(2) | P01833 | PIGR | Polymeric immunoglobulin receptor | P01833\|PIGR | P01833\|PIGR_N469_HexNAc(4)Hex(5)Fuc(1)NeuAc(2) | N469 | Yes | P01833\|PIGR_N469 |
| K.IIEGEPNLKVPGN[+2350.830]VTAVLGETLKVPC[+57.021]HFPC[+57.021]K.F | HexNAc(4)Hex(5)Fuc(1)NeuAc(2) | P01833 | PIGR | Polymeric immunoglobulin receptor | P01833\|PIGR | P01833\|PIGR_N469_HexNAc(4)Hex(5)Fuc(1)NeuAc(2) | N469 | Yes | P01833\|PIGR_N469 |
| K.VPGN[+2350.830]VTAVLGETLK.V | HexNAc(4)Hex(5)Fuc(1)NeuAc(2) | P01833 | PIGR | Polymeric immunoglobulin receptor | P01833\|PIGR | P01833\|PIGR_N469_HexNAc(4)Hex(5)Fuc(1)NeuAc(2) | N469 | Yes | P01833\|PIGR_N469 |
| K.VPGN[+2350.830]VTAVLGETLKVPC[+57.021]HFPC[+57.021]K.F | HexNAc(4)Hex(5)Fuc(1)NeuAc(2) | P01833 | PIGR | Polymeric immunoglobulin receptor | P01833\|PIGR | P01833\|PIGR_N469_HexNAc(4)Hex(5)Fuc(1)NeuAc(2) | N469 | Yes | P01833\|PIGR_N469 |
| R.TTVEIKIIEGEPNLKVPGN[+2350.830]VTAVLGETLKVPC[+57.021]HFPC[+57.021]K.F | HexNAc(4)Hex(5)Fuc(1)NeuAc(2) | P01833 | PIGR | Polymeric immunoglobulin receptor | P01833\|PIGR | P01833\|PIGR_N469_HexNAc(4)Hex(5)Fuc(1)NeuAc(2) | N469 | Yes | P01833\|PIGR_N469 |
| K.IIEGEPNLKVPGN[+2205.793]VTAVLGETLK.V | HexNAc(4)Hex(5)Fuc(2)NeuAc(1) | P01833 | PIGR | Polymeric immunoglobulin receptor | P01833\|PIGR | P01833\|PIGR_N469_HexNAc(4)Hex(5)Fuc(2)NeuAc(1) | N469 | Yes | P01833\|PIGR_N469 |
| K.IIEGEPNLKVPGN[+2205.793]VTAVLGETLKVPC[+57.021]HFPC[+57.021]K.F | HexNAc(4)Hex(5)Fuc(2)NeuAc(1) | P01833 | PIGR | Polymeric immunoglobulin receptor | P01833\|PIGR | P01833\|PIGR_N469_HexNAc(4)Hex(5)Fuc(2)NeuAc(1) | N469 | Yes | P01833\|PIGR_N469 |
| K.VPGN[+2205.793]VTAVLGETLK.V | HexNAc(4)Hex(5)Fuc(2)NeuAc(1) | P01833 | PIGR | Polymeric immunoglobulin receptor | P01833\|PIGR | P01833\|PIGR_N469_HexNAc(4)Hex(5)Fuc(2)NeuAc(1) | N469 | Yes | P01833\|PIGR_N469 |
| K.VPGN[+2205.793]VTAVLGETLKVPC[+57.021]HFPC[+57.021]K.F | HexNAc(4)Hex(5)Fuc(2)NeuAc(1) | P01833 | PIGR | Polymeric immunoglobulin receptor | P01833\|PIGR | P01833\|PIGR_N469_HexNAc(4)Hex(5)Fuc(2)NeuAc(1) | N469 | Yes | P01833\|PIGR_N469 |
| R.TTVEIKIIEGEPNLKVPGN[+2205.793]VTAVLGETLKVPC[+57.021]HFPC[+57.021]K.F | HexNAc(4)Hex(5)Fuc(2)NeuAc(1) | P01833 | PIGR | Polymeric immunoglobulin receptor | P01833\|PIGR | P01833\|PIGR_N469_HexNAc(4)Hex(5)Fuc(2)NeuAc(1) | N469 | Yes | P01833\|PIGR_N469 |
| K.IIEGEPNLKVPGN[+2496.888]VTAVLGETLKVPC[+57.021]HFPC[+57.021]K.F | HexNAc(4)Hex(5)Fuc(2)NeuAc(2) | P01833 | PIGR | Polymeric immunoglobulin receptor | P01833\|PIGR | P01833\|PIGR_N469_HexNAc(4)Hex(5)Fuc(2)NeuAc(2) | N469 | Yes | P01833\|PIGR_N469 |
| K.IIEGEPNLKVPGN[+2351.851]VTAVLGETLK.V | HexNAc(4)Hex(5)Fuc(3)NeuAc(1) | P01833 | PIGR | Polymeric immunoglobulin receptor | P01833\|PIGR | P01833\|PIGR_N469_HexNAc(4)Hex(5)Fuc(3)NeuAc(1) | N469 | Yes | P01833\|PIGR_N469 |

Table S5. Cont.

| Glycopeptide | Glycan | Uniprot ID | Gene Name | Protein Name | Shortname | ProteinNameInSkyline | Glycosites | Reported before | Site |
|---|---|---|---|---|---|---|---|---|---|
| K.IIEGEPNLKVPGN[+2351.851]VTAVLGETLKVPC[+57.021]HFPC[+57.021]K.F | HexNAc(4)Hex(5)Fuc(3)NeuAc(1) | P01833 | PIGR | Polymeric immunoglobulin receptor | P01833\|PIGR | P01833\|PIGR_N469_HexNAc(4)Hex(5)Fuc(3)NeuAc(1) | N469 | Yes | P01833\|PIGR_N469 |
| K.VPGN[+2351.851]VTAVLGETLK.V | HexNAc(4)Hex(5)Fuc(3)NeuAc(1) | P01833 | PIGR | Polymeric immunoglobulin receptor | P01833\|PIGR | P01833\|PIGR_N469_HexNAc(4)Hex(5)Fuc(3)NeuAc(1) | N469 | Yes | P01833\|PIGR_N469 |
| K.VPGN[+2351.851]VTAVLGETLKVPC[+57.021]HFPC[+57.021]K.F | HexNAc(4)Hex(5)Fuc(3)NeuAc(1) | P01833 | PIGR | Polymeric immunoglobulin receptor | P01833\|PIGR | P01833\|PIGR_N469_HexNAc(4)Hex(5)Fuc(3)NeuAc(1) | N469 | Yes | P01833\|PIGR_N469 |
| R.TTVEIKIIEGEPNLKVPGN[+2351.851]VTAVLGETLKVPC[+57.021]HFPC[+57.021]K.F | HexNAc(4)Hex(5)Fuc(3)NeuAc(1) | P01833 | PIGR | Polymeric immunoglobulin receptor | P01833\|PIGR | P01833\|PIGR_N469_HexNAc(4)Hex(5)Fuc(3)NeuAc(1) | N469 | Yes | P01833\|PIGR_N469 |
| K.IIEGEPNLKVPGN[+1913.677]VTAVLGETLK.V | HexNAc(4)Hex(5)NeuAc(1) | P01833 | PIGR | Polymeric immunoglobulin receptor | P01833\|PIGR | P01833\|PIGR_N469_HexNAc(4)Hex(5)NeuAc(1) | N469 | Yes | P01833\|PIGR_N469 |
| K.IIEGEPNLKVPGN[+1913.677]VTAVLGETLKVPC[+57.021]HFPC[+57.021]K.F | HexNAc(4)Hex(5)NeuAc(1) | P01833 | PIGR | Polymeric immunoglobulin receptor | P01833\|PIGR | P01833\|PIGR_N469_HexNAc(4)Hex(5)NeuAc(1) | N469 | Yes | P01833\|PIGR_N469 |
| K.VPGN[+1913.677]VTAVLGETLKVPC[+57.021]HFPC[+57.021]K.F | HexNAc(4)Hex(5)NeuAc(1) | P01833 | PIGR | Polymeric immunoglobulin receptor | P01833\|PIGR | P01833\|PIGR_N469_HexNAc(4)Hex(5)NeuAc(1) | N469 | Yes | P01833\|PIGR_N469 |
| K.IIEGEPNLKVPGN[+2204.772]VTAVLGETLK.V | HexNAc(4)Hex(5)NeuAc(2) | P01833 | PIGR | Polymeric immunoglobulin receptor | P01833\|PIGR | P01833\|PIGR_N469_HexNAc(4)Hex(5)NeuAc(2) | N469 | Yes | P01833\|PIGR_N469 |
| K.IIEGEPNLKVPGN[+2204.772]VTAVLGETLKVPC[+57.021]HFPC[+57.021]K.F | HexNAc(4)Hex(5)NeuAc(2) | P01833 | PIGR | Polymeric immunoglobulin receptor | P01833\|PIGR | P01833\|PIGR_N469_HexNAc(4)Hex(5)NeuAc(2) | N469 | Yes | P01833\|PIGR_N469 |
| K.VPGN[+2204.772]VTAVLGETLKVPC[+57.021]HFPC[+57.021]K.F | HexNAc(4)Hex(5)NeuAc(2) | P01833 | PIGR | Polymeric immunoglobulin receptor | P01833\|PIGR | P01833\|PIGR_N469_HexNAc(4)Hex(5)NeuAc(2) | N469 | Yes | P01833\|PIGR_N469 |
| K.IIEGEPNLKVPGN[+2553.910]VTAVLGETLKVPC[+57.021]HFPC[+57.021]K.F | HexNAc(5)Hex(5)Fuc(1)NeuAc(2) | P01833 | PIGR | Polymeric immunoglobulin receptor | P01833\|PIGR | P01833\|PIGR_N469_HexNAc(5)Hex(5)Fuc(1)NeuAc(2) | N469 | Yes | P01833\|PIGR_N469 |
| K.IIEGEPNLKVPGN[+2263.835]VTAVLGETLKVPC[+57.021]HFPC[+57.021]K.F | HexNAc(5)Hex(5)Fuc(3) | P01833 | PIGR | Polymeric immunoglobulin receptor | P01833\|PIGR | P01833\|PIGR_N469_HexNAc(5)Hex(5)Fuc(3) | N469 | Yes | P01833\|PIGR_N469 |
| K.IIEGEPNLKVPGN[+2424.867]VTAVLGETLKVPC[+57.021]HFPC[+57.021]K.F | HexNAc(5)Hex(6)Fuc(1)NeuAc(1) | P01833 | PIGR | Polymeric immunoglobulin receptor | P01833\|PIGR | P01833\|PIGR_N469_HexNAc(5)Hex(6)Fuc(1)NeuAc(1) | N469 | Yes | P01833\|PIGR_N469 |
| K.IIEGEPNLKVPGN[+2279.830]VTAVLGETLKVPC[+57.021]HFPC[+57.021]K.F | HexNAc(5)Hex(6)Fuc(2) | P01833 | PIGR | Polymeric immunoglobulin receptor | P01833\|PIGR | P01833\|PIGR_N469_HexNAc(5)Hex(6)Fuc(2) | N469 | Yes | P01833\|PIGR_N469 |
| K.IIEGEPNLKVPGN[+2570.925]VTAVLGETLKVPC[+57.021]HFPC[+57.021]K.F | HexNAc(5)Hex(6)Fuc(2)NeuAc(1) | P01833 | PIGR | Polymeric immunoglobulin receptor | P01833\|PIGR | P01833\|PIGR_N469_HexNAc(5)Hex(6)Fuc(2)NeuAc(1) | N469 | Yes | P01833\|PIGR_N469 |
| K.IIEGEPNLKVPGN[+2862.021]VTAVLGETLKVPC[+57.021]HFPC[+57.021]K.F | HexNAc(5)Hex(6)Fuc(2)NeuAc(2) | P01833 | PIGR | Polymeric immunoglobulin receptor | P01833\|PIGR | P01833\|PIGR_N469_HexNAc(5)Hex(6)Fuc(2)NeuAc(2) | N469 | Yes | P01833\|PIGR_N469 |
| K.IIEGEPNLKVPGN[+2425.887]VTAVLGETLKVPC[+57.021]HFPC[+57.021]K.F | HexNAc(5)Hex(6)Fuc(3) | P01833 | PIGR | Polymeric immunoglobulin receptor | P01833\|PIGR | P01833\|PIGR_N469_HexNAc(5)Hex(6)Fuc(3) | N469 | Yes | P01833\|PIGR_N469 |
| K.IIEGEPNLKVPGN[+2716.983]VTAVLGETLKVPC[+57.021]HFPC[+57.021]K.F | HexNAc(5)Hex(6)Fuc(3)NeuAc(1) | P01833 | PIGR | Polymeric immunoglobulin receptor | P01833\|PIGR | P01833\|PIGR_N469_HexNAc(5)Hex(6)Fuc(3)NeuAc(1) | N469 | Yes | P01833\|PIGR_N469 |
| K.IIEGEPNLKVPGN[+3008.078]VTAVLGETLKVPC[+57.021]HFPC[+57.021]K.F | HexNAc(5)Hex(6)Fuc(3)NeuAc(2) | P01833 | PIGR | Polymeric immunoglobulin receptor | P01833\|PIGR | P01833\|PIGR_N469_HexNAc(5)Hex(6)Fuc(3)NeuAc(2) | N469 | Yes | P01833\|PIGR_N469 |
| K.IIEGEPNLKVPGN[+2571.946]VTAVLGETLKVPC[+57.021]HFPC[+57.021]K.F | HexNAc(5)Hex(6)Fuc(4) | P01833 | PIGR | Polymeric immunoglobulin receptor | P01833\|PIGR | P01833\|PIGR_N469_HexNAc(5)Hex(6)Fuc(4) | N469 | Yes | P01833\|PIGR_N469 |
| K.IIEGEPNLKVPGN[+2863.041]VTAVLGETLKVPC[+57.021]HFPC[+57.021]K.F | HexNAc(5)Hex(6)Fuc(4)NeuAc(1) | P01833 | PIGR | Polymeric immunoglobulin receptor | P01833\|PIGR | P01833\|PIGR_N469_HexNAc(5)Hex(6)Fuc(4)NeuAc(1) | N469 | Yes | P01833\|PIGR_N469 |
| K.IIEGEPNLKVPGN[+3154.136]VTAVLGETLKVPC[+57.021]HFPC[+57.021]K.F | HexNAc(5)Hex(6)Fuc(4)NeuAc(2) | P01833 | PIGR | Polymeric immunoglobulin receptor | P01833\|PIGR | P01833\|PIGR_N469_HexNAc(5)Hex(6)Fuc(4)NeuAc(2) | N469 | Yes | P01833\|PIGR_N469 |
| K.[+42.011]IIEGEPNLKVPGN[+2018.708]VTAVLGETLK.V | HexNAc(3)Hex(6)Fuc(1)NeuAc(1) | P01833 | PIGR | Polymeric immunoglobulin receptor | P01833\|PIGR | P01833\|PIGR_N469_HexNAc(3)Hex(6)Fuc(1)NeuAc(1) | N469 | Yes | P01833\|PIGR_N469 |
| K.[+42.011]IIEGEPNLKVPGN[+2018.708]VTAVLGETLKVPC[+57.021]HFPC[+57.021]K.F | HexNAc(3)Hex(6)Fuc(1)NeuAc(1) | P01833 | PIGR | Polymeric immunoglobulin receptor | P01833\|PIGR | P01833\|PIGR_N469_HexNAc(3)Hex(6)Fuc(1)NeuAc(1) | N469 | Yes | P01833\|PIGR_N469 |
| K.[+42.011]VPGN[+2018.708]VTAVLGETLKVPC[+57.021]HFPC[+57.021]K.F | HexNAc(3)Hex(6)Fuc(1)NeuAc(1) | P01833 | PIGR | Polymeric immunoglobulin receptor | P01833\|PIGR | P01833\|PIGR_N469_HexNAc(3)Hex(6)Fuc(1)NeuAc(1) | N469 | Yes | P01833\|PIGR_N469 |
| K.[+42.011]IIEGEPNLKVPGN[+1872.651]VTAVLGETLK.V | HexNAc(3)Hex(6)NeuAc(1) | P01833 | PIGR | Polymeric immunoglobulin receptor | P01833\|PIGR | P01833\|PIGR_N469_HexNAc(3)Hex(6)NeuAc(1) | N469 | Yes | P01833\|PIGR_N469 |
| K.[+42.011]IIEGEPNLKVPGN[+1872.651]VTAVLGETLKVPC[+57.021]HFPC[+57.021]K.F | HexNAc(3)Hex(6)NeuAc(1) | P01833 | PIGR | Polymeric immunoglobulin receptor | P01833\|PIGR | P01833\|PIGR_N469_HexNAc(3)Hex(6)NeuAc(1) | N469 | Yes | P01833\|PIGR_N469 |
| K.[+42.011]IIEGEPNLKVPGN[+2351.851]VTAVLGETLKVPC[+57.021]HFPC[+57.021]K.F | HexNAc(4)Hex(5)Fuc(3)NeuAc(1) | P01833 | PIGR | Polymeric immunoglobulin receptor | P01833\|PIGR | P01833\|PIGR_N469_HexNAc(4)Hex(5)Fuc(3)NeuAc(1) | N469 | Yes | P01833\|PIGR_N469 |
| K.[+42.011]IIEGEPNLKVPGN[+2336.851]VTAVLGETLK.V | HexNAc(6)Hex(6)Fuc(1) | P01833 | PIGR | Polymeric immunoglobulin receptor | P01833\|PIGR | P01833\|PIGR_N469_HexNAc(6)Hex(6)Fuc(1) | N469 | Yes | P01833\|PIGR_N469 |
| K.WN[+1054.370]NTGC[+57.021]QALPSQDEGPSK.A | HexNAc(2)Hex(4) | P01833 | PIGR | Polymeric immunoglobulin receptor | P01833\|PIGR | P01833\|PIGR_N499_HexNAc(2)Hex(4) | N499 | Yes | P01833\|PIGR_N499 |
| K.WN[+1216.423]NTGC[+57.021]QALPSQDEGPSK.A | HexNAc(2)Hex(5) | P01833 | PIGR | Polymeric immunoglobulin receptor | P01833\|PIGR | P01833\|PIGR_N499_HexNAc(2)Hex(5) | N499 | Yes | P01833\|PIGR_N499 |
| K.WN[+1548.545]NTGC[+57.021]QALPSQDEGPSK.A | HexNAc(3)Hex(4)NeuAc(1) | P01833 | PIGR | Polymeric immunoglobulin receptor | P01833\|PIGR | P01833\|PIGR_N499_HexNAc(3)Hex(4)NeuAc(1) | N499 | Yes | P01833\|PIGR_N499 |
| K.WN[+1710.598]NTGC[+57.021]QALPSQDEGPSK.A | HexNAc(3)Hex(5)NeuAc(1) | P01833 | PIGR | Polymeric immunoglobulin receptor | P01833\|PIGR | P01833\|PIGR_N499_HexNAc(3)Hex(5)NeuAc(1) | N499 | Yes | P01833\|PIGR_N499 |
| K.WN[+1872.651]NTGC[+57.021]QALPSQDEGPSK.A | HexNAc(3)Hex(6)NeuAc(1) | P01833 | PIGR | Polymeric immunoglobulin receptor | P01833\|PIGR | P01833\|PIGR_N499_HexNAc(3)Hex(6)NeuAc(1) | N499 | Yes | P01833\|PIGR_N499 |
| K.WN[+1872.651]NTGC[+57.021]QALPSQDEGPSKAFVNC[+57.021]DENSR.L | HexNAc(3)Hex(6)NeuAc(1) | P01833 | PIGR | Polymeric immunoglobulin receptor | P01833\|PIGR | P01833\|PIGR_N499_HexNAc(3)Hex(6)NeuAc(1) | N499 | Yes | P01833\|PIGR_N499 |
| K.WN[+1768.640]NTGC[+57.021]QALPSQDEGPSK.A | HexNAc(4)Hex(5)Fuc(1) | P01833 | PIGR | Polymeric immunoglobulin receptor | P01833\|PIGR | P01833\|PIGR_N499_HexNAc(4)Hex(5)Fuc(1) | N499 | Yes | P01833\|PIGR_N499 |
| K.WN[+2059.735]NTGC[+57.021]QALPSQDEGPSK.A | HexNAc(4)Hex(5)Fuc(1)NeuAc(1) | P01833 | PIGR | Polymeric immunoglobulin receptor | P01833\|PIGR | P01833\|PIGR_N499_HexNAc(4)Hex(5)Fuc(1)NeuAc(1) | N499 | Yes | P01833\|PIGR_N499 |
| K.WN[+2059.735]NTGC[+57.021]QALPSQDEGPSKAFVNC[+57.021]DENSR.L | HexNAc(4)Hex(5)Fuc(1)NeuAc(1) | P01833 | PIGR | Polymeric immunoglobulin receptor | P01833\|PIGR | P01833\|PIGR_N499_HexNAc(4)Hex(5)Fuc(1)NeuAc(1) | N499 | Yes | P01833\|PIGR_N499 |
| K.WN[+2205.793]NTGC[+57.021]QALPSQDEGPSK.A | HexNAc(4)Hex(5)Fuc(2)NeuAc(1) | P01833 | PIGR | Polymeric immunoglobulin receptor | P01833\|PIGR | P01833\|PIGR_N499_HexNAc(4)Hex(5)Fuc(2)NeuAc(1) | N499 | Yes | P01833\|PIGR_N499 |
| K.WN[+1913.677]NTGC[+57.021]QALPSQDEGPSK.A | HexNAc(4)Hex(5)NeuAc(1) | P01833 | PIGR | Polymeric immunoglobulin receptor | P01833\|PIGR | P01833\|PIGR_N499_HexNAc(4)Hex(5)NeuAc(1) | N499 | Yes | P01833\|PIGR_N499 |
| K.WN[+2204.772]NTGC[+57.021]QALPSQDEGPSK.A | HexNAc(4)Hex(5)NeuAc(2) | P01833 | PIGR | Polymeric immunoglobulin receptor | P01833\|PIGR | P01833\|PIGR_N499_HexNAc(4)Hex(5)NeuAc(2) | N499 | Yes | P01833\|PIGR_N499 |
| K.WN[+2204.772]NTGC[+57.021]QALPSQDEGPSKAFVNC[+57.021]DENSR.L | HexNAc(4)Hex(5)NeuAc(2) | P01833 | PIGR | Polymeric immunoglobulin receptor | P01833\|PIGR | P01833\|PIGR_N499_HexNAc(4)Hex(5)NeuAc(2) | N499 | Yes | P01833\|PIGR_N499 |
| K.FSSYEKYWC[+57.021]KWN[+1710.598]NTGC[+57.021]QALPSQDEGPSK.A | HexNAc(3)Hex(5)NeuAc(1) | P01833 | PIGR | Polymeric immunoglobulin receptor | P01833\|PIGR | P01833\|PIGR_N499_HexNAc(3)Hex(5)NeuAc(1) | N499 | Yes | P01833\|PIGR_N499 |
| K.FSSYEKYWC[+57.021]KWN[+1710.598]NTGC[+57.021]QALPSQDEGPSKAFVNC[+57.021]DENSR.L | HexNAc(3)Hex(5)NeuAc(1) | P01833 | PIGR | Polymeric immunoglobulin receptor | P01833\|PIGR | P01833\|PIGR_N499_HexNAc(3)Hex(5)NeuAc(1) | N499 | Yes | P01833\|PIGR_N499 |
| K.YWC[+57.021]KWN[+1710.598]NTGC[+57.021]QALPSQDEGPSK.A | HexNAc(3)Hex(5)NeuAc(1) | P01833 | PIGR | Polymeric immunoglobulin receptor | P01833\|PIGR | P01833\|PIGR_N499_HexNAc(3)Hex(5)NeuAc(1) | N499 | Yes | P01833\|PIGR_N499 |
| K.YWC[+57.021]KWN[+1710.598]NTGC[+57.021]QALPSQDEGPSKAFVNC[+57.021]DENSR.L | HexNAc(3)Hex(5)NeuAc(1) | P01833 | PIGR | Polymeric immunoglobulin receptor | P01833\|PIGR | P01833\|PIGR_N499_HexNAc(3)Hex(5)NeuAc(1) | N499 | Yes | P01833\|PIGR_N499 |
| K.FSSYEKYWC[+57.021]KWN[+1872.651]NTGC[+57.021]QALPSQDEGPSK.A | HexNAc(3)Hex(6)NeuAc(1) | P01833 | PIGR | Polymeric immunoglobulin receptor | P01833\|PIGR | P01833\|PIGR_N499_HexNAc(3)Hex(6)NeuAc(1) | N499 | Yes | P01833\|PIGR_N499 |
| K.YWC[+57.021]KWN[+1872.651]NTGC[+57.021]QALPSQDEGPSK.A | HexNAc(3)Hex(6)NeuAc(1) | P01833 | PIGR | Polymeric immunoglobulin receptor | P01833\|PIGR | P01833\|PIGR_N499_HexNAc(3)Hex(6)NeuAc(1) | N499 | Yes | P01833\|PIGR_N499 |
| K.YWC[+57.021]KWN[+1872.651]NTGC[+57.021]QALPSQDEGPSKAFVNC[+57.021]DENSR.L | HexNAc(3)Hex(6)NeuAc(1) | P01833 | PIGR | Polymeric immunoglobulin receptor | P01833\|PIGR | P01833\|PIGR_N499_HexNAc(3)Hex(6)NeuAc(1) | N499 | Yes | P01833\|PIGR_N499 |
| K.FSSYEKYWC[+57.021]KWN[+2205.793]NTGC[+57.021]QALPSQDEGPSK.A | HexNAc(4)Hex(5)Fuc(2)NeuAc(1) | P01833 | PIGR | Polymeric immunoglobulin receptor | P01833\|PIGR | P01833\|PIGR_N499_HexNAc(4)Hex(5)Fuc(2)NeuAc(1) | N499 | Yes | P01833\|PIGR_N499 |
| K.YWC[+57.021]KWN[+2205.793]NTGC[+57.021]QALPSQDEGPSK.A | HexNAc(4)Hex(5)Fuc(2)NeuAc(1) | P01833 | PIGR | Polymeric immunoglobulin receptor | P01833\|PIGR | P01833\|PIGR_N499_HexNAc(4)Hex(5)Fuc(2)NeuAc(1) | N499 | Yes | P01833\|PIGR_N499 |
| K.YWC[+57.021]KWN[+1913.677]NTGC[+57.021]QALPSQDEGPSK.A | HexNAc(4)Hex(5)NeuAc(1) | P01833 | PIGR | Polymeric immunoglobulin receptor | P01833\|PIGR | P01833\|PIGR_N499_HexNAc(4)Hex(5)NeuAc(1) | N499 | Yes | P01833\|PIGR_N499 |
| K.FSSYEKYWC[+57.021]KWN[+2204.772]NTGC[+57.021]QALPSQDEGPSKAFVNC[+57.021]DENSR.L | HexNAc(4)Hex(5)NeuAc(2) | P01833 | PIGR | Polymeric immunoglobulin receptor | P01833\|PIGR | P01833\|PIGR_N499_HexNAc(4)Hex(5)NeuAc(2) | N499 | Yes | P01833\|PIGR_N499 |
| K.TKPREEQFN[+1444.534]STFR.V | HexNAc(4)Hex(3)Fuc(1) | P01859 | IGHG2 | Immunoglobulin heavy constant gamma 2 | P01859\|IGHG2 | P01859\|IGHG2_N176_HexNAc(4)Hex(3)Fuc(1) | N176 | Yes | P01859\|IGHG2_N176 |
| R.EEQFN[+1444.534]STFR.V | HexNAc(4)Hex(3)Fuc(1) | P01859 | IGHG2 | Immunoglobulin heavy constant gamma 2 | P01859\|IGHG2 | P01859\|IGHG2_N176_HexNAc(4)Hex(3)Fuc(1) | N176 | Yes | P01859\|IGHG2_N176 |
| K.TKPREEQFN[+1606.587]STFR.V | HexNAc(4)Hex(4)Fuc(1) | P01859 | IGHG2 | Immunoglobulin heavy constant gamma 2 | P01859\|IGHG2 | P01859\|IGHG2_N176_HexNAc(4)Hex(4)Fuc(1) | N176 | Yes | P01859\|IGHG2_N176 |
| K.TKPREEQFN[+1768.640]STFR.V | HexNAc(4)Hex(5)Fuc(1) | P01859 | IGHG2 | Immunoglobulin heavy constant gamma 2 | P01859\|IGHG2 | P01859\|IGHG2_N176_HexNAc(4)Hex(5)Fuc(1) | N176 | Yes | P01859\|IGHG2_N176 |
| R.TVDKSTGKPTLYN[+1378.476]VSLVMSDTAGTC[+57.021]Y.- | HexNAc(2)Hex(6) | P01871 | IGHM | Immunoglobulin heavy constant mu | P01871\|IGHM | P01871\|IGHM_N440_HexNAc(2)Hex(6) | N440 | Yes | P01871\|IGHM_N440 |
| K.YKN[+2059.735]NSDISSTR.G | HexNAc(4)Hex(5)Fuc(1)NeuAc(1) | P01871 | IGHM | Immunoglobulin heavy constant mu | P01871\|IGHM | P01871\|IGHM_N46_HexNAc(4)Hex(5)Fuc(1)NeuAc(1) | N46 | Yes | P01871\|IGHM_N46 |
| K.YKN[+1647.613]NSDISSTR.G | HexNAc(5)Hex(3)Fuc(1) | P01871 | IGHM | Immunoglobulin heavy constant mu | P01871\|IGHM | P01871\|IGHM_N46_HexNAc(5)Hex(3)Fuc(1) | N46 | Yes | P01871\|IGHM_N46 |
| K.YKN[+2262.814]NSDISSTR.G | HexNAc(5)Hex(5)Fuc(1)NeuAc(1) | P01871 | IGHM | Immunoglobulin heavy constant mu | P01871\|IGHM | P01871\|IGHM_N46_HexNAc(5)Hex(5)Fuc(1)NeuAc(1) | N46 | Yes | P01871\|IGHM_N46 |
| R.LSLHRPALEDLLLGSEAN[+568.212]LTC[+57.021]TLTGLR.D | HexNAc(2)Hex(1) | P01876 | IGHA1 | Immunoglobulin heavy constant alpha 1 | P01876\|IGHA1 | P01876\|IGHA1_N144_HexNAc(2)Hex(1) | N144 | Yes | P01876\|IGHA1_N144 |
| R.LSLHRPALEDLLLGSEAN[+892.317]LTC[+57.021]TLTGLR.D | HexNAc(2)Hex(3) | P01876 | IGHA1 | Immunoglobulin heavy constant alpha 1 | P01876\|IGHA1 | P01876\|IGHA1_N144_HexNAc(2)Hex(3) | N144 | Yes | P01876\|IGHA1_N144 |
| R.LSLHRPALEDLLLGSEAN[+1054.370]LTC[+57.021]TLTGLR.D | HexNAc(2)Hex(4) | P01876 | IGHA1 | Immunoglobulin heavy constant alpha 1 | P01876\|IGHA1 | P01876\|IGHA1_N144_HexNAc(2)Hex(4) | N144 | Yes | P01876\|IGHA1_N144 |
| R.LSLHRPALEDLLLGSEAN[+1216.423]LTC[+57.021]TLTGLR.D | HexNAc(2)Hex(5) | P01876 | IGHA1 | Immunoglobulin heavy constant alpha 1 | P01876\|IGHA1 | P01876\|IGHA1_N144_HexNAc(2)Hex(5) | N144 | Yes | P01876\|IGHA1_N144 |
| R.LSLHRPALEDLLLGSEAN[+1216.423]LTC[+57.021]TLTGLRDASGVTFTWTPSSGK.S | HexNAc(2)Hex(5) | P01876 | IGHA1 | Immunoglobulin heavy constant alpha 1 | P01876\|IGHA1 | P01876\|IGHA1_N144_HexNAc(2)Hex(5) | N144 | Yes | P01876\|IGHA1_N144 |
| R.LSLHRPALEDLLLGSEAN[+1378.476]LTC[+57.021]TLTGLR.D | HexNAc(2)Hex(6) | P01876 | IGHA1 | Immunoglobulin heavy constant alpha 1 | P01876\|IGHA1 | P01876\|IGHA1_N144_HexNAc(2)Hex(6) | N144 | Yes | P01876\|IGHA1_N144 |
| R.LSLHRPALEDLLLGSEAN[+1378.476]LTC[+57.021]TLTGLRDASGVTFTWTPSSGK.S | HexNAc(2)Hex(6) | P01876 | IGHA1 | Immunoglobulin heavy constant alpha 1 | P01876\|IGHA1 | P01876\|IGHA1_N144_HexNAc(2)Hex(6) | N144 | Yes | P01876\|IGHA1_N144 |
| R.LSLHRPALEDLLLGSEAN[+1524.534]LTC[+57.021]TLTGLR.D | HexNAc(2)Hex(6)Fuc(1) | P01876 | IGHA1 | Immunoglobulin heavy constant alpha 1 | P01876\|IGHA1 | P01876\|IGHA1_N144_HexNAc(2)Hex(6)Fuc(1) | N144 | Yes | P01876\|IGHA1_N144 |
| R.LSLHRPALEDLLLGSEAN[+1540.529]LTC[+57.021]TLTGLR.D | HexNAc(2)Hex(7) | P01876 | IGHA1 | Immunoglobulin heavy constant alpha 1 | P01876\|IGHA1 | P01876\|IGHA1_N144_HexNAc(2)Hex(7) | N144 | Yes | P01876\|IGHA1_N144 |
| R.LSLHRPALEDLLLGSEAN[+1702.581]LTC[+57.021]TLTGLR.D | HexNAc(2)Hex(8) | P01876 | IGHA1 | Immunoglobulin heavy constant alpha 1 | P01876\|IGHA1 | P01876\|IGHA1_N144_HexNAc(2)Hex(8) | N144 | Yes | P01876\|IGHA1_N144 |
| R.LSLHRPALEDLLLGSEAN[+1864.634]LTC[+57.021]TLTGLR.D | HexNAc(2)Hex(9) | P01876 | IGHA1 | Immunoglobulin heavy constant alpha 1 | P01876\|IGHA1 | P01876\|IGHA1_N144_HexNAc(2)Hex(9) | N144 | Yes | P01876\|IGHA1_N144 |
| R.LSLHRPALEDLLLGSEAN[+1095.397]LTC[+57.021]TLTGLR.D | HexNAc(3)Hex(3) | P01876 | IGHA1 | Immunoglobulin heavy constant alpha 1 | P01876\|IGHA1 | P01876\|IGHA1_N144_HexNAc(3)Hex(3) | N144 | Yes | P01876\|IGHA1_N144 |
| R.LSLHRPALEDLLLGSEAN[+1257.449]LTC[+57.021]TLTGLR.D | HexNAc(3)Hex(4) | P01876 | IGHA1 | Immunoglobulin heavy constant alpha 1 | P01876\|IGHA1 | P01876\|IGHA1_N144_HexNAc(3)Hex(4) | N144 | Yes | P01876\|IGHA1_N144 |
| R.LSLHRPALEDLLLGSEAN[+1548.545]LTC[+57.021]TLTGLR.D | HexNAc(3)Hex(4)NeuAc(1) | P01876 | IGHA1 | Immunoglobulin heavy constant alpha 1 | P01876\|IGHA1 | P01876\|IGHA1_N144_HexNAc(3)Hex(4)NeuAc(1) | N144 | Yes | P01876\|IGHA1_N144 |
| R.LSLHRPALEDLLLGSEAN[+1419.502]LTC[+57.021]TLTGLR.D | HexNAc(3)Hex(5) | P01876 | IGHA1 | Immunoglobulin heavy constant alpha 1 | P01876\|IGHA1 | P01876\|IGHA1_N144_HexNAc(3)Hex(5) | N144 | Yes | P01876\|IGHA1_N144 |
| R.LSLHRPALEDLLLGSEAN[+1710.598]LTC[+57.021]TLTGLR.D | HexNAc(3)Hex(5)NeuAc(1) | P01876 | IGHA1 | Immunoglobulin heavy constant alpha 1 | P01876\|IGHA1 | P01876\|IGHA1_N144_HexNAc(3)Hex(5)NeuAc(1) | N144 | Yes | P01876\|IGHA1_N144 |
| R.LSLHRPALEDLLLGSEAN[+1581.555]LTC[+57.021]TLTGLR.D | HexNAc(3)Hex(6) | P01876 | IGHA1 | Immunoglobulin heavy constant alpha 1 | P01876\|IGHA1 | P01876\|IGHA1_N144_HexNAc(3)Hex(6) | N144 | Yes | P01876\|IGHA1_N144 |
| R.LSLHRPALEDLLLGSEAN[+1872.651]LTC[+57.021]TLTGLR.D | HexNAc(3)Hex(6)NeuAc(1) | P01876 | IGHA1 | Immunoglobulin heavy constant alpha 1 | P01876\|IGHA1 | P01876\|IGHA1_N144_HexNAc(3)Hex(6)NeuAc(1) | N144 | Yes | P01876\|IGHA1_N144 |
| R.LSLHRPALEDLLLGSEAN[+1298.476]LTC[+57.021]TLTGLR.D | HexNAc(4)Hex(3) | P01876 | IGHA1 | Immunoglobulin heavy constant alpha 1 | P01876\|IGHA1 | P01876\|IGHA1_N144_HexNAc(4)Hex(3) | N144 | Yes | P01876\|IGHA1_N144 |
| R.LSLHRPALEDLLLGSEAN[+1460.529]LTC[+57.021]TLTGLR.D | HexNAc(4)Hex(4) | P01876 | IGHA1 | Immunoglobulin heavy constant alpha 1 | P01876\|IGHA1 | P01876\|IGHA1_N144_HexNAc(4)Hex(4) | N144 | Yes | P01876\|IGHA1_N144 |
| R.LSLHRPALEDLLLGSEAN[+1460.529]LTC[+57.021]TLTGLRDASGVTFTWTPSSGK.S | HexNAc(4)Hex(4) | P01876 | IGHA1 | Immunoglobulin heavy constant alpha 1 | P01876\|IGHA1 | P01876\|IGHA1_N144_HexNAc(4)Hex(4) | N144 | Yes | P01876\|IGHA1_N144 |
| R.LSLHRPALEDLLLGSEAN[+1751.624]LTC[+57.021]TLTGLR.D | HexNAc(4)Hex(4)NeuAc(1) | P01876 | IGHA1 | Immunoglobulin heavy constant alpha 1 | P01876\|IGHA1 | P01876\|IGHA1_N144_HexNAc(4)Hex(4)NeuAc(1) | N144 | Yes | P01876\|IGHA1_N144 |
| R.LSLHRPALEDLLLGSEAN[+1622.582]LTC[+57.021]TLTGLR.D | HexNAc(4)Hex(5) | P01876 | IGHA1 | Immunoglobulin heavy constant alpha 1 | P01876\|IGHA1 | P01876\|IGHA1_N144_HexNAc(4)Hex(5) | N144 | Yes | P01876\|IGHA1_N144 |
| R.LSLHRPALEDLLLGSEAN[+1913.677]LTC[+57.021]TLTGLR.D | HexNAc(4)Hex(5)NeuAc(1) | P01876 | IGHA1 | Immunoglobulin heavy constant alpha 1 | P01876\|IGHA1 | P01876\|IGHA1_N144_HexNAc(4)Hex(5)NeuAc(1) | N144 | Yes | P01876\|IGHA1_N144 |
| R.LSLHRPALEDLLLGSEAN[+2204.772]LTC[+57.021]TLTGLR.D | HexNAc(4)Hex(5)NeuAc(2) | P01876 | IGHA1 | Immunoglobulin heavy constant alpha 1 | P01876\|IGHA1 | P01876\|IGHA1_N144_HexNAc(4)Hex(5)NeuAc(2) | N144 | Yes | P01876\|IGHA1_N144 |
| R.LSLHRPALEDLLLGSEAN[+1501.555]LTC[+57.021]TLTGLR.D | HexNAc(5)Hex(3) | P01876 | IGHA1 | Immunoglobulin heavy constant alpha 1 | P01876\|IGHA1 | P01876\|IGHA1_N144_HexNAc(5)Hex(3) | N144 | Yes | P01876\|IGHA1_N144 |

Table S5. Cont.

| Glycopeptide | Glycan | Uniprot ID | Gene Name | Protein Name | Shortname | ProteinNameInSkyline | Glycosites | Reported before | Site |
|---|---|---|---|---|---|---|---|---|---|
| R.LSLHRPALEDLLLGSEAN[+1501.555]LTC[+57.021]TLTGLRDASGVTFTWTPSSGK.S | HexNAc(5)Hex(3) | P01876 | IGHA1 | Immunoglobulin heavy constant alpha 1 | P01876\|IGHA1 | P01876\|IGHA1_N144_HexNAc(5)Hex(3) | N144 | Yes | P01876\|IGHA1_N144 |
| R.LSLHRPALEDLLLGSEAN[+1647.613]LTC[+57.021]TLTGLR.D | HexNAc(5)Hex(3)Fuc(1) | P01876 | IGHA1 | Immunoglobulin heavy constant alpha 1 | P01876\|IGHA1 | P01876\|IGHA1_N144_HexNAc(5)Hex(3)Fuc(1) | N144 | Yes | P01876\|IGHA1_N144 |
| R.LSLHRPALEDLLLGSEAN[+1663.608]LTC[+57.021]TLTGLR.D | HexNAc(5)Hex(4) | P01876 | IGHA1 | Immunoglobulin heavy constant alpha 1 | P01876\|IGHA1 | P01876\|IGHA1_N144_HexNAc(5)Hex(4) | N144 | Yes | P01876\|IGHA1_N144 |
| R.LSLHRPALEDLLLGSEAN[+1809.666]LTC[+57.021]TLTGLR.D | HexNAc(5)Hex(4)Fuc(1) | P01876 | IGHA1 | Immunoglobulin heavy constant alpha 1 | P01876\|IGHA1 | P01876\|IGHA1_N144_HexNAc(5)Hex(4)Fuc(1) | N144 | Yes | P01876\|IGHA1_N144 |
| R.LSLHRPALEDLLLGSEAN[+1955.724]LTC[+57.021]TLTGLR.D | HexNAc(5)Hex(4)Fuc(2) | P01876 | IGHA1 | Immunoglobulin heavy constant alpha 1 | P01876\|IGHA1 | P01876\|IGHA1_N144_HexNAc(5)Hex(4)Fuc(2) | N144 | Yes | P01876\|IGHA1_N144 |
| R.LSLHRPALEDLLLGSEAN[+1825.661]LTC[+57.021]TLTGLR.D | HexNAc(5)Hex(5) | P01876 | IGHA1 | Immunoglobulin heavy constant alpha 1 | P01876\|IGHA1 | P01876\|IGHA1_N144_HexNAc(5)Hex(5) | N144 | Yes | P01876\|IGHA1_N144 |
| R.LSLHRPALEDLLLGSEAN[+2262.814]LTC[+57.021]TLTGLR.D | HexNAc(5)Hex(5)Fuc(1)NeuAc(1) | P01876 | IGHA1 | Immunoglobulin heavy constant alpha 1 | P01876\|IGHA1 | P01876\|IGHA1_N144_HexNAc(5)Hex(5)Fuc(1)NeuAc(1) | N144 | Yes | P01876\|IGHA1_N144 |
| R.LSLHRPALEDLLLGSEAN[+2116.756]LTC[+57.021]TLTGLR.D | HexNAc(5)Hex(5)NeuAc(1) | P01876 | IGHA1 | Immunoglobulin heavy constant alpha 1 | P01876\|IGHA1 | P01876\|IGHA1_N144_HexNAc(5)Hex(5)NeuAc(1) | N144 | Yes | P01876\|IGHA1_N144 |
| R.LSLHRPALEDLLLGSEAN[+2407.852]LTC[+57.021]TLTGLR.D | HexNAc(5)Hex(5)NeuAc(2) | P01876 | IGHA1 | Immunoglobulin heavy constant alpha 1 | P01876\|IGHA1 | P01876\|IGHA1_N144_HexNAc(5)Hex(5)NeuAc(2) | N144 | Yes | P01876\|IGHA1_N144 |
| R.LSLHRPALEDLLLGSEAN[+2278.809]LTC[+57.021]TLTGLR.D | HexNAc(5)Hex(6)NeuAc(1) | P01876 | IGHA1 | Immunoglobulin heavy constant alpha 1 | P01876\|IGHA1 | P01876\|IGHA1_N144_HexNAc(5)Hex(6)NeuAc(1) | N144 | Yes | P01876\|IGHA1_N144 |
| R.LSLHRPALEDLLLGSEAN[+1866.688]LTC[+57.021]TLTGLR.D | HexNAc(6)Hex(4) | P01876 | IGHA1 | Immunoglobulin heavy constant alpha 1 | P01876\|IGHA1 | P01876\|IGHA1_N144_HexNAc(6)Hex(4) | N144 | Yes | P01876\|IGHA1_N144 |
| R.[+42.011]LSLHRPALEDLLLGSEAN[+892.317]LTC[+57.021]TLTGLR.D | HexNAc(2)Hex(3) | P01876 | IGHA1 | Immunoglobulin heavy constant alpha 1 | P01876\|IGHA1 | P01876\|IGHA1_N144_HexNAc(2)Hex(3) | N144 | Yes | P01876\|IGHA1_N144 |
| R.[+42.011]LSLHRPALEDLLLGSEAN[+1216.423]LTC[+57.021]TLTGLR.D | HexNAc(2)Hex(5) | P01876 | IGHA1 | Immunoglobulin heavy constant alpha 1 | P01876\|IGHA1 | P01876\|IGHA1_N144_HexNAc(2)Hex(5) | N144 | Yes | P01876\|IGHA1_N144 |
| R.[+42.011]LSLHRPALEDLLLGSEAN[+1710.598]LTC[+57.021]TLTGLR.D | HexNAc(3)Hex(5)NeuAc(1) | P01876 | IGHA1 | Immunoglobulin heavy constant alpha 1 | P01876\|IGHA1 | P01876\|IGHA1_N144_HexNAc(3)Hex(5)NeuAc(1) | N144 | Yes | P01876\|IGHA1_N144 |
| R.[+42.011]PALEDLLLGSEAN[+1460.529]LTC[+57.021]TLTGLR.D | HexNAc(4)Hex(4) | P01876 | IGHA1 | Immunoglobulin heavy constant alpha 1 | P01876\|IGHA1 | P01876\|IGHA1_N144_HexNAc(4)Hex(4) | N144 | Yes | P01876\|IGHA1_N144 |
| R.[+42.011]LSLHRPALEDLLLGSEAN[+1501.555]LTC[+57.021]TLTGLR.D | HexNAc(5)Hex(3) | P01876 | IGHA1 | Immunoglobulin heavy constant alpha 1 | P01876\|IGHA1 | P01876\|IGHA1_N144_HexNAc(5)Hex(3) | N144 | Yes | P01876\|IGHA1_N144 |
| R.[+42.011]LSLHRPALEDLLLGSEAN[+1955.724]LTC[+57.021]TLTGLR.D | HexNAc(5)Hex(4)Fuc(2) | P01876 | IGHA1 | Immunoglobulin heavy constant alpha 1 | P01876\|IGHA1 | P01876\|IGHA1_N144_HexNAc(5)Hex(4)Fuc(2) | N144 | Yes | P01876\|IGHA1_N144 |
| R.[+42.011]LSLHRPALEDLLLGSEAN[+2116.756]LTC[+57.021]TLTGLR.D | HexNAc(5)Hex(5)NeuAc(1) | P01876 | IGHA1 | Immunoglobulin heavy constant alpha 1 | P01876\|IGHA1 | P01876\|IGHA1_N144_HexNAc(5)Hex(5)NeuAc(1) | N144 | Yes | P01876\|IGHA1_N144 |
| R.LAGKPTHVN[+892.317]VSVVMAEVDGTC[+57.021]Y.- | HexNAc(2)Hex(3) | P01876 | IGHA1 | Immunoglobulin heavy constant alpha 1 | P01876\|IGHA1 | P01876\|IGHA1_N340_HexNAc(2)Hex(3) | N340 | Yes | P01876\|IGHA1_N340 |
| R.LAGKPTHVN[+1054.370]VSVVM[+15.995]AEVDGTC[+57.021].Y | HexNAc(2)Hex(4) | P01876 | IGHA1 | Immunoglobulin heavy constant alpha 1 | P01876\|IGHA1 | P01876\|IGHA1_N340_HexNAc(2)Hex(4) | N340 | Yes | P01876\|IGHA1_N340 |
| R.LAGKPTHVN[+1054.370]VSVVM[+15.995]AEVDGTC[+57.021]Y.- | HexNAc(2)Hex(4) | P01876 | IGHA1 | Immunoglobulin heavy constant alpha 1 | P01876\|IGHA1 | P01876\|IGHA1_N340_HexNAc(2)Hex(4) | N340 | Yes | P01876\|IGHA1_N340 |
| R.LAGKPTHVN[+1054.370]VSVVMAEVDGTC[+57.021].Y | HexNAc(2)Hex(4) | P01876 | IGHA1 | Immunoglobulin heavy constant alpha 1 | P01876\|IGHA1 | P01876\|IGHA1_N340_HexNAc(2)Hex(4) | N340 | Yes | P01876\|IGHA1_N340 |
| R.LAGKPTHVN[+1054.370]VSVVMAEVDGTC[+57.021]Y.- | HexNAc(2)Hex(4) | P01876 | IGHA1 | Immunoglobulin heavy constant alpha 1 | P01876\|IGHA1 | P01876\|IGHA1_N340_HexNAc(2)Hex(4) | N340 | Yes | P01876\|IGHA1_N340 |
| K.TIDRLAGKPTHVN[+1216.423]VSVVMAEVDGTC[+57.021]Y.- | HexNAc(2)Hex(5) | P01876 | IGHA1 | Immunoglobulin heavy constant alpha 1 | P01876\|IGHA1 | P01876\|IGHA1_N340_HexNAc(2)Hex(5) | N340 | Yes | P01876\|IGHA1_N340 |
| R.LAGKPTHVN[+1216.423]VSVVM[+15.995]AEVDGTC[+57.021].Y | HexNAc(2)Hex(5) | P01876 | IGHA1 | Immunoglobulin heavy constant alpha 1 | P01876\|IGHA1 | P01876\|IGHA1_N340_HexNAc(2)Hex(5) | N340 | Yes | P01876\|IGHA1_N340 |
| R.LAGKPTHVN[+1216.423]VSVVM[+15.995]AEVDGTC[+57.021]Y.- | HexNAc(2)Hex(5) | P01876 | IGHA1 | Immunoglobulin heavy constant alpha 1 | P01876\|IGHA1 | P01876\|IGHA1_N340_HexNAc(2)Hex(5) | N340 | Yes | P01876\|IGHA1_N340 |
| R.LAGKPTHVN[+1216.423]VSVVMAEVDGTC[+57.021].Y | HexNAc(2)Hex(5) | P01876 | IGHA1 | Immunoglobulin heavy constant alpha 1 | P01876\|IGHA1 | P01876\|IGHA1_N340_HexNAc(2)Hex(5) | N340 | Yes | P01876\|IGHA1_N340 |
| R.LAGKPTHVN[+1216.423]VSVVMAEVDGTC[+57.021]Y.- | HexNAc(2)Hex(5) | P01876 | IGHA1 | Immunoglobulin heavy constant alpha 1 | P01876\|IGHA1 | P01876\|IGHA1_N340_HexNAc(2)Hex(5) | N340 | Yes | P01876\|IGHA1_N340 |
| K.TIDRLAGKPTHVN[+1378.476]VSVVMAEVDGTC[+57.021]Y.- | HexNAc(2)Hex(6) | P01876 | IGHA1 | Immunoglobulin heavy constant alpha 1 | P01876\|IGHA1 | P01876\|IGHA1_N340_HexNAc(2)Hex(6) | N340 | Yes | P01876\|IGHA1_N340 |
| R.LAGKPTHVN[+1378.476]VSVVM[+15.995]AEVDGTC[+57.021]Y.- | HexNAc(2)Hex(6) | P01876 | IGHA1 | Immunoglobulin heavy constant alpha 1 | P01876\|IGHA1 | P01876\|IGHA1_N340_HexNAc(2)Hex(6) | N340 | Yes | P01876\|IGHA1_N340 |
| R.LAGKPTHVN[+1378.476]VSVVMAEVDGTC[+57.021].Y | HexNAc(2)Hex(6) | P01876 | IGHA1 | Immunoglobulin heavy constant alpha 1 | P01876\|IGHA1 | P01876\|IGHA1_N340_HexNAc(2)Hex(6) | N340 | Yes | P01876\|IGHA1_N340 |
| R.LAGKPTHVN[+1378.476]VSVVMAEVDGTC[+57.021]Y.- | HexNAc(2)Hex(6) | P01876 | IGHA1 | Immunoglobulin heavy constant alpha 1 | P01876\|IGHA1 | P01876\|IGHA1_N340_HexNAc(2)Hex(6) | N340 | Yes | P01876\|IGHA1_N340 |
| K.TIDRLAGKPTHVN[+1540.529]VSVVMAEVDGTC[+57.021]Y.- | HexNAc(2)Hex(7) | P01876 | IGHA1 | Immunoglobulin heavy constant alpha 1 | P01876\|IGHA1 | P01876\|IGHA1_N340_HexNAc(2)Hex(7) | N340 | Yes | P01876\|IGHA1_N340 |
| R.LAGKPTHVN[+1540.529]VSVVMAEVDGTC[+57.021]Y.- | HexNAc(2)Hex(7) | P01876 | IGHA1 | Immunoglobulin heavy constant alpha 1 | P01876\|IGHA1 | P01876\|IGHA1_N340_HexNAc(2)Hex(7) | N340 | Yes | P01876\|IGHA1_N340 |
| R.LAGKPTHVN[+1702.581]VSVVMAEVDGTC[+57.021]Y.- | HexNAc(2)Hex(8) | P01876 | IGHA1 | Immunoglobulin heavy constant alpha 1 | P01876\|IGHA1 | P01876\|IGHA1_N340_HexNAc(2)Hex(8) | N340 | Yes | P01876\|IGHA1_N340 |
| R.LAGKPTHVN[+1095.397]VSVVMAEVDGTC[+57.021]Y.- | HexNAc(3)Hex(3) | P01876 | IGHA1 | Immunoglobulin heavy constant alpha 1 | P01876\|IGHA1 | P01876\|IGHA1_N340_HexNAc(3)Hex(3) | N340 | Yes | P01876\|IGHA1_N340 |
| R.LAGKPTHVN[+1241.454]VSVVMAEVDGTC[+57.021]Y.- | HexNAc(3)Hex(3)Fuc(1) | P01876 | IGHA1 | Immunoglobulin heavy constant alpha 1 | P01876\|IGHA1 | P01876\|IGHA1_N340_HexNAc(3)Hex(3)Fuc(1) | N340 | Yes | P01876\|IGHA1_N340 |
| R.LAGKPTHVN[+1257.449]VSVVMAEVDGTC[+57.021]Y.- | HexNAc(3)Hex(4) | P01876 | IGHA1 | Immunoglobulin heavy constant alpha 1 | P01876\|IGHA1 | P01876\|IGHA1_N340_HexNAc(3)Hex(4) | N340 | Yes | P01876\|IGHA1_N340 |
| R.LAGKPTHVN[+1694.603]VSVVMAEVDGTC[+57.021]Y.- | HexNAc(3)Hex(4)Fuc(1)NeuAc(1) | P01876 | IGHA1 | Immunoglobulin heavy constant alpha 1 | P01876\|IGHA1 | P01876\|IGHA1_N340_HexNAc(3)Hex(4)Fuc(1)NeuAc(1) | N340 | Yes | P01876\|IGHA1_N340 |
| R.LAGKPTHVN[+1548.545]VSVVMAEVDGTC[+57.021]Y.- | HexNAc(3)Hex(4)NeuAc(1) | P01876 | IGHA1 | Immunoglobulin heavy constant alpha 1 | P01876\|IGHA1 | P01876\|IGHA1_N340_HexNAc(3)Hex(4)NeuAc(1) | N340 | Yes | P01876\|IGHA1_N340 |
| R.LAGKPTHVN[+1419.502]VSVVMAEVDGTC[+57.021]Y.- | HexNAc(3)Hex(5) | P01876 | IGHA1 | Immunoglobulin heavy constant alpha 1 | P01876\|IGHA1 | P01876\|IGHA1_N340_HexNAc(3)Hex(5) | N340 | Yes | P01876\|IGHA1_N340 |
| R.LAGKPTHVN[+1710.598]VSVVMAEVDGTC[+57.021]Y.- | HexNAc(3)Hex(5)NeuAc(1) | P01876 | IGHA1 | Immunoglobulin heavy constant alpha 1 | P01876\|IGHA1 | P01876\|IGHA1_N340_HexNAc(3)Hex(5)NeuAc(1) | N340 | Yes | P01876\|IGHA1_N340 |
| R.LAGKPTHVN[+1581.555]VSVVMAEVDGTC[+57.021]Y.- | HexNAc(3)Hex(6) | P01876 | IGHA1 | Immunoglobulin heavy constant alpha 1 | P01876\|IGHA1 | P01876\|IGHA1_N340_HexNAc(3)Hex(6) | N340 | Yes | P01876\|IGHA1_N340 |
| R.LAGKPTHVN[+2018.708]VSVVM[+15.995]AEVDGTC[+57.021].Y | HexNAc(3)Hex(6)Fuc(1)NeuAc(1) | P01876 | IGHA1 | Immunoglobulin heavy constant alpha 1 | P01876\|IGHA1 | P01876\|IGHA1_N340_HexNAc(3)Hex(6)Fuc(1)NeuAc(1) | N340 | Yes | P01876\|IGHA1_N340 |
| R.LAGKPTHVN[+1872.651]VSVVMAEVDGTC[+57.021].Y | HexNAc(3)Hex(6)NeuAc(1) | P01876 | IGHA1 | Immunoglobulin heavy constant alpha 1 | P01876\|IGHA1 | P01876\|IGHA1_N340_HexNAc(3)Hex(6)NeuAc(1) | N340 | Yes | P01876\|IGHA1_N340 |
| R.LAGKPTHVN[+1872.651]VSVVMAEVDGTC[+57.021]Y.- | HexNAc(3)Hex(6)NeuAc(1) | P01876 | IGHA1 | Immunoglobulin heavy constant alpha 1 | P01876\|IGHA1 | P01876\|IGHA1_N340_HexNAc(3)Hex(6)NeuAc(1) | N340 | Yes | P01876\|IGHA1_N340 |
| R.LAGKPTHVN[+1298.476]VSVVMAEVDGTC[+57.021]Y.- | HexNAc(4)Hex(3) | P01876 | IGHA1 | Immunoglobulin heavy constant alpha 1 | P01876\|IGHA1 | P01876\|IGHA1_N340_HexNAc(4)Hex(3) | N340 | Yes | P01876\|IGHA1_N340 |
| R.LAGKPTHVN[+1444.534]VSVVMAEVDGTC[+57.021].Y | HexNAc(4)Hex(3)Fuc(1) | P01876 | IGHA1 | Immunoglobulin heavy constant alpha 1 | P01876\|IGHA1 | P01876\|IGHA1_N340_HexNAc(4)Hex(3)Fuc(1) | N340 | Yes | P01876\|IGHA1_N340 |
| R.LAGKPTHVN[+1444.534]VSVVMAEVDGTC[+57.021]Y.- | HexNAc(4)Hex(3)Fuc(1) | P01876 | IGHA1 | Immunoglobulin heavy constant alpha 1 | P01876\|IGHA1 | P01876\|IGHA1_N340_HexNAc(4)Hex(3)Fuc(1) | N340 | Yes | P01876\|IGHA1_N340 |
| R.LAGKPTHVN[+1460.529]VSVVMAEVDGTC[+57.021]Y.- | HexNAc(4)Hex(4) | P01876 | IGHA1 | Immunoglobulin heavy constant alpha 1 | P01876\|IGHA1 | P01876\|IGHA1_N340_HexNAc(4)Hex(4) | N340 | Yes | P01876\|IGHA1_N340 |
| R.LAGKPTHVN[+1606.587]VSVVMAEVDGTC[+57.021].Y | HexNAc(4)Hex(4)Fuc(1) | P01876 | IGHA1 | Immunoglobulin heavy constant alpha 1 | P01876\|IGHA1 | P01876\|IGHA1_N340_HexNAc(4)Hex(4)Fuc(1) | N340 | Yes | P01876\|IGHA1_N340 |
| R.LAGKPTHVN[+1606.587]VSVVMAEVDGTC[+57.021]Y.- | HexNAc(4)Hex(4)Fuc(1) | P01876 | IGHA1 | Immunoglobulin heavy constant alpha 1 | P01876\|IGHA1 | P01876\|IGHA1_N340_HexNAc(4)Hex(4)Fuc(1) | N340 | Yes | P01876\|IGHA1_N340 |
| R.LAGKPTHVN[+1751.624]VSVVMAEVDGTC[+57.021]Y.- | HexNAc(4)Hex(4)NeuAc(1) | P01876 | IGHA1 | Immunoglobulin heavy constant alpha 1 | P01876\|IGHA1 | P01876\|IGHA1_N340_HexNAc(4)Hex(4)NeuAc(1) | N340 | Yes | P01876\|IGHA1_N340 |
| R.LAGKPTHVN[+1622.582]VSVVMAEVDGTC[+57.021]Y.- | HexNAc(4)Hex(5) | P01876 | IGHA1 | Immunoglobulin heavy constant alpha 1 | P01876\|IGHA1 | P01876\|IGHA1_N340_HexNAc(4)Hex(5) | N340 | Yes | P01876\|IGHA1_N340 |
| R.LAGKPTHVN[+2059.735]VSVVMAEVDGTC[+57.021]Y.- | HexNAc(4)Hex(5)Fuc(1)NeuAc(1) | P01876 | IGHA1 | Immunoglobulin heavy constant alpha 1 | P01876\|IGHA1 | P01876\|IGHA1_N340_HexNAc(4)Hex(5)Fuc(1)NeuAc(1) | N340 | Yes | P01876\|IGHA1_N340 |
| R.LAGKPTHVN[+2350.830]VSVVMAEVDGTC[+57.021]Y.- | HexNAc(4)Hex(5)Fuc(1)NeuAc(2) | P01876 | IGHA1 | Immunoglobulin heavy constant alpha 1 | P01876\|IGHA1 | P01876\|IGHA1_N340_HexNAc(4)Hex(5)Fuc(1)NeuAc(2) | N340 | Yes | P01876\|IGHA1_N340 |
| R.LAGKPTHVN[+2205.793]VSVVMAEVDGTC[+57.021]Y.- | HexNAc(4)Hex(5)Fuc(2)NeuAc(1) | P01876 | IGHA1 | Immunoglobulin heavy constant alpha 1 | P01876\|IGHA1 | P01876\|IGHA1_N340_HexNAc(4)Hex(5)Fuc(2)NeuAc(1) | N340 | Yes | P01876\|IGHA1_N340 |
| R.LAGKPTHVN[+2351.851]VSVVMAEVDGTC[+57.021]Y.- | HexNAc(4)Hex(5)Fuc(3)NeuAc(1) | P01876 | IGHA1 | Immunoglobulin heavy constant alpha 1 | P01876\|IGHA1 | P01876\|IGHA1_N340_HexNAc(4)Hex(5)Fuc(3)NeuAc(1) | N340 | Yes | P01876\|IGHA1_N340 |
| R.LAGKPTHVN[+1913.677]VSVVMAEVDGTC[+57.021]Y.- | HexNAc(4)Hex(5)NeuAc(1) | P01876 | IGHA1 | Immunoglobulin heavy constant alpha 1 | P01876\|IGHA1 | P01876\|IGHA1_N340_HexNAc(4)Hex(5)NeuAc(1) | N340 | Yes | P01876\|IGHA1_N340 |
| R.LAGKPTHVN[+2204.772]VSVVMAEVDGTC[+57.021]Y.- | HexNAc(4)Hex(5)NeuAc(2) | P01876 | IGHA1 | Immunoglobulin heavy constant alpha 1 | P01876\|IGHA1 | P01876\|IGHA1_N340_HexNAc(4)Hex(5)NeuAc(2) | N340 | Yes | P01876\|IGHA1_N340 |
| R.LAGKPTHVN[+1501.555]VSVVMAEVDGTC[+57.021]Y.- | HexNAc(5)Hex(3) | P01876 | IGHA1 | Immunoglobulin heavy constant alpha 1 | P01876\|IGHA1 | P01876\|IGHA1_N340_HexNAc(5)Hex(3) | N340 | Yes | P01876\|IGHA1_N340 |
| K.TIDRLAGKPTHVN[+1647.613]VSVVMAEVDGTC[+57.021]Y.- | HexNAc(5)Hex(3)Fuc(1) | P01876 | IGHA1 | Immunoglobulin heavy constant alpha 1 | P01876\|IGHA1 | P01876\|IGHA1_N340_HexNAc(5)Hex(3)Fuc(1) | N340 | Yes | P01876\|IGHA1_N340 |
| R.LAGKPTHVN[+1647.613]VSVVM[+15.995]AEVDGTC[+57.021]Y.- | HexNAc(5)Hex(3)Fuc(1) | P01876 | IGHA1 | Immunoglobulin heavy constant alpha 1 | P01876\|IGHA1 | P01876\|IGHA1_N340_HexNAc(5)Hex(3)Fuc(1) | N340 | Yes | P01876\|IGHA1_N340 |
| R.LAGKPTHVN[+1647.613]VSVVMAEVDGTC[+57.021].Y | HexNAc(5)Hex(3)Fuc(1) | P01876 | IGHA1 | Immunoglobulin heavy constant alpha 1 | P01876\|IGHA1 | P01876\|IGHA1_N340_HexNAc(5)Hex(3)Fuc(1) | N340 | Yes | P01876\|IGHA1_N340 |
| R.LAGKPTHVN[+1647.613]VSVVMAEVDGTC[+57.021]Y.- | HexNAc(5)Hex(3)Fuc(1) | P01876 | IGHA1 | Immunoglobulin heavy constant alpha 1 | P01876\|IGHA1 | P01876\|IGHA1_N340_HexNAc(5)Hex(3)Fuc(1) | N340 | Yes | P01876\|IGHA1_N340 |
| K.TIDRLAGKPTHVN[+1663.608]VSVVMAEVDGTC[+57.021]Y.- | HexNAc(5)Hex(4) | P01876 | IGHA1 | Immunoglobulin heavy constant alpha 1 | P01876\|IGHA1 | P01876\|IGHA1_N340_HexNAc(5)Hex(4) | N340 | Yes | P01876\|IGHA1_N340 |
| R.LAGKPTHVN[+1663.608]VSVVM[+15.995]AEVDGTC[+57.021]Y.- | HexNAc(5)Hex(4) | P01876 | IGHA1 | Immunoglobulin heavy constant alpha 1 | P01876\|IGHA1 | P01876\|IGHA1_N340_HexNAc(5)Hex(4) | N340 | Yes | P01876\|IGHA1_N340 |
| R.LAGKPTHVN[+1663.608]VSVVMAEVDGTC[+57.021]Y.- | HexNAc(5)Hex(4) | P01876 | IGHA1 | Immunoglobulin heavy constant alpha 1 | P01876\|IGHA1 | P01876\|IGHA1_N340_HexNAc(5)Hex(4) | N340 | Yes | P01876\|IGHA1_N340 |
| K.TIDRLAGKPTHVN[+1809.666]VSVVMAEVDGTC[+57.021]Y.- | HexNAc(5)Hex(4)Fuc(1) | P01876 | IGHA1 | Immunoglobulin heavy constant alpha 1 | P01876\|IGHA1 | P01876\|IGHA1_N340_HexNAc(5)Hex(4)Fuc(1) | N340 | Yes | P01876\|IGHA1_N340 |
| R.LAGKPTHVN[+1809.666]VSVVM[+15.995]AEVDGTC[+57.021]Y.- | HexNAc(5)Hex(4)Fuc(1) | P01876 | IGHA1 | Immunoglobulin heavy constant alpha 1 | P01876\|IGHA1 | P01876\|IGHA1_N340_HexNAc(5)Hex(4)Fuc(1) | N340 | Yes | P01876\|IGHA1_N340 |
| R.LAGKPTHVN[+1809.666]VSVVMAEVDGTC[+57.021].Y | HexNAc(5)Hex(4)Fuc(1) | P01876 | IGHA1 | Immunoglobulin heavy constant alpha 1 | P01876\|IGHA1 | P01876\|IGHA1_N340_HexNAc(5)Hex(4)Fuc(1) | N340 | Yes | P01876\|IGHA1_N340 |
| R.LAGKPTHVN[+1809.666]VSVVMAEVDGTC[+57.021]Y.- | HexNAc(5)Hex(4)Fuc(1) | P01876 | IGHA1 | Immunoglobulin heavy constant alpha 1 | P01876\|IGHA1 | P01876\|IGHA1_N340_HexNAc(5)Hex(4)Fuc(1) | N340 | Yes | P01876\|IGHA1_N340 |
| K.TIDRLAGKPTHVN[+1825.661]VSVVMAEVDGTC[+57.021]Y.- | HexNAc(5)Hex(5) | P01876 | IGHA1 | Immunoglobulin heavy constant alpha 1 | P01876\|IGHA1 | P01876\|IGHA1_N340_HexNAc(5)Hex(5) | N340 | Yes | P01876\|IGHA1_N340 |
| R.LAGKPTHVN[+1825.661]VSVVMAEVDGTC[+57.021].Y | HexNAc(5)Hex(5) | P01876 | IGHA1 | Immunoglobulin heavy constant alpha 1 | P01876\|IGHA1 | P01876\|IGHA1_N340_HexNAc(5)Hex(5) | N340 | Yes | P01876\|IGHA1_N340 |
| R.LAGKPTHVN[+1825.661]VSVVMAEVDGTC[+57.021]Y.- | HexNAc(5)Hex(5) | P01876 | IGHA1 | Immunoglobulin heavy constant alpha 1 | P01876\|IGHA1 | P01876\|IGHA1_N340_HexNAc(5)Hex(5) | N340 | Yes | P01876\|IGHA1_N340 |
| K.TIDRLAGKPTHVN[+1971.719]VSVVMAEVDGTC[+57.021]Y.- | HexNAc(5)Hex(5)Fuc(1) | P01876 | IGHA1 | Immunoglobulin heavy constant alpha 1 | P01876\|IGHA1 | P01876\|IGHA1_N340_HexNAc(5)Hex(5)Fuc(1) | N340 | Yes | P01876\|IGHA1_N340 |
| R.LAGKPTHVN[+1971.719]VSVVMAEVDGTC[+57.021]Y.- | HexNAc(5)Hex(5)Fuc(1) | P01876 | IGHA1 | Immunoglobulin heavy constant alpha 1 | P01876\|IGHA1 | P01876\|IGHA1_N340_HexNAc(5)Hex(5)Fuc(1) | N340 | Yes | P01876\|IGHA1_N340 |
| K.TIDRLAGKPTHVN[+2262.814]VSVVMAEVDGTC[+57.021]Y.- | HexNAc(5)Hex(5)Fuc(1)NeuAc(1) | P01876 | IGHA1 | Immunoglobulin heavy constant alpha 1 | P01876\|IGHA1 | P01876\|IGHA1_N340_HexNAc(5)Hex(5)Fuc(1)NeuAc(1) | N340 | Yes | P01876\|IGHA1_N340 |
| R.LAGKPTHVN[+2262.814]VSVVM[+15.995]AEVDGTC[+57.021]Y.- | HexNAc(5)Hex(5)Fuc(1)NeuAc(1) | P01876 | IGHA1 | Immunoglobulin heavy constant alpha 1 | P01876\|IGHA1 | P01876\|IGHA1_N340_HexNAc(5)Hex(5)Fuc(1)NeuAc(1) | N340 | Yes | P01876\|IGHA1_N340 |
| R.LAGKPTHVN[+2262.814]VSVVMAEVDGTC[+57.021].Y | HexNAc(5)Hex(5)Fuc(1)NeuAc(1) | P01876 | IGHA1 | Immunoglobulin heavy constant alpha 1 | P01876\|IGHA1 | P01876\|IGHA1_N340_HexNAc(5)Hex(5)Fuc(1)NeuAc(1) | N340 | Yes | P01876\|IGHA1_N340 |
| R.LAGKPTHVN[+2262.814]VSVVMAEVDGTC[+57.021]Y.- | HexNAc(5)Hex(5)Fuc(1)NeuAc(1) | P01876 | IGHA1 | Immunoglobulin heavy constant alpha 1 | P01876\|IGHA1 | P01876\|IGHA1_N340_HexNAc(5)Hex(5)Fuc(1)NeuAc(1) | N340 | Yes | P01876\|IGHA1_N340 |
| K.TIDRLAGKPTHVN[+2553.910]VSVVMAEVDGTC[+57.021]Y.- | HexNAc(5)Hex(5)Fuc(1)NeuAc(2) | P01876 | IGHA1 | Immunoglobulin heavy constant alpha 1 | P01876\|IGHA1 | P01876\|IGHA1_N340_HexNAc(5)Hex(5)Fuc(1)NeuAc(2) | N340 | Yes | P01876\|IGHA1_N340 |
| R.LAGKPTHVN[+2553.910]VSVVMAEVDGTC[+57.021]Y.- | HexNAc(5)Hex(5)Fuc(1)NeuAc(2) | P01876 | IGHA1 | Immunoglobulin heavy constant alpha 1 | P01876\|IGHA1 | P01876\|IGHA1_N340_HexNAc(5)Hex(5)Fuc(1)NeuAc(2) | N340 | Yes | P01876\|IGHA1_N340 |
| R.LAGKPTHVN[+2117.777]VSVVMAEVDGTC[+57.021].Y | HexNAc(5)Hex(5)Fuc(2) | P01876 | IGHA1 | Immunoglobulin heavy constant alpha 1 | P01876\|IGHA1 | P01876\|IGHA1_N340_HexNAc(5)Hex(5)Fuc(2) | N340 | Yes | P01876\|IGHA1_N340 |
| R.LAGKPTHVN[+2117.777]VSVVMAEVDGTC[+57.021]Y.- | HexNAc(5)Hex(5)Fuc(2) | P01876 | IGHA1 | Immunoglobulin heavy constant alpha 1 | P01876\|IGHA1 | P01876\|IGHA1_N340_HexNAc(5)Hex(5)Fuc(2) | N340 | Yes | P01876\|IGHA1_N340 |
| R.LAGKPTHVN[+2263.835]VSVVMAEVDGTC[+57.021].Y | HexNAc(5)Hex(5)Fuc(3) | P01876 | IGHA1 | Immunoglobulin heavy constant alpha 1 | P01876\|IGHA1 | P01876\|IGHA1_N340_HexNAc(5)Hex(5)Fuc(3) | N340 | Yes | P01876\|IGHA1_N340 |
| R.LAGKPTHVN[+2263.835]VSVVMAEVDGTC[+57.021]Y.- | HexNAc(5)Hex(5)Fuc(3) | P01876 | IGHA1 | Immunoglobulin heavy constant alpha 1 | P01876\|IGHA1 | P01876\|IGHA1_N340_HexNAc(5)Hex(5)Fuc(3) | N340 | Yes | P01876\|IGHA1_N340 |

Table S5. Cont.

| Glycopeptide | Glycan | Uniprot ID | Gene Name | Protein Name | Shortname | ProteinNameInSkyline | Glycosites | Reported before | Site |
|---|---|---|---|---|---|---|---|---|---|
| R.LAGKPTHVN[+2116.756]VSVVMAEVDGTC[+57.021]Y.- | HexNAc(5)Hex(5)NeuAc(1) | P01876 | IGHA1 | Immunoglobulin heavy constant alpha 1 | P01876\|IGHA1 | P01876\|IGHA1_N340_HexNAc(5)Hex(5)NeuAc(1) | N340 | Yes | P01876\|IGHA1_N340 |
| R.LAGKPTHVN[+2133.772]VSVVMAEVDGTC[+57.021].Y | HexNAc(5)Hex(6)Fuc(1) | P01876 | IGHA1 | Immunoglobulin heavy constant alpha 1 | P01876\|IGHA1 | P01876\|IGHA1_N340_HexNAc(5)Hex(6)Fuc(1) | N340 | Yes | P01876\|IGHA1_N340 |
| R.LAGKPTHVN[+2424.867]VSVVMAEVDGTC[+57.021]Y.- | HexNAc(5)Hex(6)Fuc(1)NeuAc(1) | P01876 | IGHA1 | Immunoglobulin heavy constant alpha 1 | P01876\|IGHA1 | P01876\|IGHA1_N340_HexNAc(5)Hex(6)Fuc(1)NeuAc(1) | N340 | Yes | P01876\|IGHA1_N340 |
| R.LAGKPTHVN[+2174.798]VSVVMAEVDGTC[+57.021]Y.- | HexNAc(6)Hex(5)Fuc(1) | P01876 | IGHA1 | Immunoglobulin heavy constant alpha 1 | P01876\|IGHA1 | P01876\|IGHA1_N340_HexNAc(6)Hex(5)Fuc(1) | N340 | Yes | P01876\|IGHA1_N340 |
| K.SVTC[+57.021]HVKHYTN[+1647.613]SSQDVTVPC[+57.021]RVPPPPPC[+57.021]C[+57.021]HPR.L | HexNAc(5)Hex(3)Fuc(1) | P01877 | IGHA2 | Immunoglobulin heavy constant alpha 2 | P01877\|IGHA2 | P01877\|IGHA2_N92_HexNAc(5)Hex(3)Fuc(1) | N92 | Yes | P01877\|IGHA2_N92 |
| R.LSLHRPALEDLLLGSEAN[+203.079]LTC[+57.021]TLTGLR.D | HexNAc(1) | P01877 | IGHA2 | Immunoglobulin heavy constant alpha 2 | P01877\|IGHA2 | P01877\|IGHA2_N131_HexNAc(1) | N131 | Yes | P01877\|IGHA2_N131 |
| R.LSLHRPALEDLLLGSEAN[+1216.423]LTC[+57.021]TLTGLRDASGATFTWTPSSGK.S | HexNAc(2)Hex(5) | P01877 | IGHA2 | Immunoglobulin heavy constant alpha 2 | P01877\|IGHA2 | P01877\|IGHA2_N131_HexNAc(2)Hex(5) | N131 | Yes | P01877\|IGHA2_N131 |
| R.LSLHRPALEDLLLGSEAN[+1378.476]LTC[+57.021]TLTGLRDASGATFTWTPSSGK.S | HexNAc(2)Hex(6) | P01877 | IGHA2 | Immunoglobulin heavy constant alpha 2 | P01877\|IGHA2 | P01877\|IGHA2_N131_HexNAc(2)Hex(6) | N131 | Yes | P01877\|IGHA2_N131 |
| R.LSLHRPALEDLLLGSEAN[+1540.529]LTC[+57.021]TLTGLRDASGATFTWTPSSGK.S | HexNAc(2)Hex(7) | P01877 | IGHA2 | Immunoglobulin heavy constant alpha 2 | P01877\|IGHA2 | P01877\|IGHA2_N131_HexNAc(2)Hex(7) | N131 | Yes | P01877\|IGHA2_N131 |
| R.LSLHRPALEDLLLGSEAN[+1872.651]LTC[+57.021]TLTGLRDASGATFTWTPSSGK.S | HexNAc(3)Hex(6)NeuAc(1) | P01877 | IGHA2 | Immunoglobulin heavy constant alpha 2 | P01877\|IGHA2 | P01877\|IGHA2_N131_HexNAc(3)Hex(6)NeuAc(1) | N131 | Yes | P01877\|IGHA2_N131 |
| R.LSLHRPALEDLLLGSEAN[+1444.534]LTC[+57.021]TLTGLR.D | HexNAc(4)Hex(3)Fuc(1) | P01877 | IGHA2 | Immunoglobulin heavy constant alpha 2 | P01877\|IGHA2 | P01877\|IGHA2_N131_HexNAc(4)Hex(3)Fuc(1) | N131 | Yes | P01877\|IGHA2_N131 |
| R.LSLHRPALEDLLLGSEAN[+1501.555]LTC[+57.021]TLTGLRDASGATFTWTPSSGK.S | HexNAc(5)Hex(3) | P01877 | IGHA2 | Immunoglobulin heavy constant alpha 2 | P01877\|IGHA2 | P01877\|IGHA2_N131_HexNAc(5)Hex(3) | N131 | Yes | P01877\|IGHA2_N131 |
| K.TPLTAN[+876.322]ITK.S | HexNAc(2)Hex(2)Fuc(1) | P01877 | IGHA2 | Immunoglobulin heavy constant alpha 2 | P01877\|IGHA2 | P01877\|IGHA2_N205_HexNAc(2)Hex(2)Fuc(1) | N205 | Yes | P01877\|IGHA2_N205 |
| K.TPLTAN[+1038.375]ITK.S | HexNAc(2)Hex(3)Fuc(1) | P01877 | IGHA2 | Immunoglobulin heavy constant alpha 2 | P01877\|IGHA2 | P01877\|IGHA2_N205_HexNAc(2)Hex(3)Fuc(1) | N205 | Yes | P01877\|IGHA2_N205 |
| K.TPLTAN[+1216.423]ITK.S | HexNAc(2)Hex(5) | P01877 | IGHA2 | Immunoglobulin heavy constant alpha 2 | P01877\|IGHA2 | P01877\|IGHA2_N205_HexNAc(2)Hex(5) | N205 | Yes | P01877\|IGHA2_N205 |
| K.TPLTAN[+1241.454]ITK.S | HexNAc(3)Hex(3)Fuc(1) | P01877 | IGHA2 | Immunoglobulin heavy constant alpha 2 | P01877\|IGHA2 | P01877\|IGHA2_N205_HexNAc(3)Hex(3)Fuc(1) | N205 | Yes | P01877\|IGHA2_N205 |
| K.TPLTAN[+1403.507]ITK.S | HexNAc(3)Hex(4)Fuc(1) | P01877 | IGHA2 | Immunoglobulin heavy constant alpha 2 | P01877\|IGHA2 | P01877\|IGHA2_N205_HexNAc(3)Hex(4)Fuc(1) | N205 | Yes | P01877\|IGHA2_N205 |
| K.TPLTAN[+1694.603]ITK.S | HexNAc(3)Hex(4)Fuc(1)NeuAc(1) | P01877 | IGHA2 | Immunoglobulin heavy constant alpha 2 | P01877\|IGHA2 | P01877\|IGHA2_N205_HexNAc(3)Hex(4)Fuc(1)NeuAc(1) | N205 | Yes | P01877\|IGHA2_N205 |
| K.TPLTAN[+1298.476]ITK.S | HexNAc(4)Hex(3) | P01877 | IGHA2 | Immunoglobulin heavy constant alpha 2 | P01877\|IGHA2 | P01877\|IGHA2_N205_HexNAc(4)Hex(3) | N205 | Yes | P01877\|IGHA2_N205 |
| K.TPLTAN[+1444.534]ITK.S | HexNAc(4)Hex(3)Fuc(1) | P01877 | IGHA2 | Immunoglobulin heavy constant alpha 2 | P01877\|IGHA2 | P01877\|IGHA2_N205_HexNAc(4)Hex(3)Fuc(1) | N205 | Yes | P01877\|IGHA2_N205 |
| K.TPLTAN[+1460.529]ITK.S | HexNAc(4)Hex(4) | P01877 | IGHA2 | Immunoglobulin heavy constant alpha 2 | P01877\|IGHA2 | P01877\|IGHA2_N205_HexNAc(4)Hex(4) | N205 | Yes | P01877\|IGHA2_N205 |
| K.TPLTAN[+1606.587]ITK.S | HexNAc(4)Hex(4)Fuc(1) | P01877 | IGHA2 | Immunoglobulin heavy constant alpha 2 | P01877\|IGHA2 | P01877\|IGHA2_N205_HexNAc(4)Hex(4)Fuc(1) | N205 | Yes | P01877\|IGHA2_N205 |
| K.TPLTAN[+1768.640]ITK.S | HexNAc(4)Hex(5)Fuc(1) | P01877 | IGHA2 | Immunoglobulin heavy constant alpha 2 | P01877\|IGHA2 | P01877\|IGHA2_N205_HexNAc(4)Hex(5)Fuc(1) | N205 | Yes | P01877\|IGHA2_N205 |
| K.TPLTAN[+2059.735]ITK.S | HexNAc(4)Hex(5)Fuc(1)NeuAc(1) | P01877 | IGHA2 | Immunoglobulin heavy constant alpha 2 | P01877\|IGHA2 | P01877\|IGHA2_N205_HexNAc(4)Hex(5)Fuc(1)NeuAc(1) | N205 | Yes | P01877\|IGHA2_N205 |
| K.TPLTAN[+2350.830]ITK.S | HexNAc(4)Hex(5)Fuc(1)NeuAc(2) | P01877 | IGHA2 | Immunoglobulin heavy constant alpha 2 | P01877\|IGHA2 | P01877\|IGHA2_N205_HexNAc(4)Hex(5)Fuc(1)NeuAc(2) | N205 | Yes | P01877\|IGHA2_N205 |
| K.TPLTAN[+1501.555]ITK.S | HexNAc(5)Hex(3) | P01877 | IGHA2 | Immunoglobulin heavy constant alpha 2 | P01877\|IGHA2 | P01877\|IGHA2_N205_HexNAc(5)Hex(3) | N205 | Yes | P01877\|IGHA2_N205 |
| K.TPLTAN[+1647.613]ITK.S | HexNAc(5)Hex(3)Fuc(1) | P01877 | IGHA2 | Immunoglobulin heavy constant alpha 2 | P01877\|IGHA2 | P01877\|IGHA2_N205_HexNAc(5)Hex(3)Fuc(1) | N205 | Yes | P01877\|IGHA2_N205 |
| K.TPLTAN[+1663.608]ITK.S | HexNAc(5)Hex(4) | P01877 | IGHA2 | Immunoglobulin heavy constant alpha 2 | P01877\|IGHA2 | P01877\|IGHA2_N205_HexNAc(5)Hex(4) | N205 | Yes | P01877\|IGHA2_N205 |
| K.TPLTAN[+1809.666]ITK.S | HexNAc(5)Hex(4)Fuc(1) | P01877 | IGHA2 | Immunoglobulin heavy constant alpha 2 | P01877\|IGHA2 | P01877\|IGHA2_N205_HexNAc(5)Hex(4)Fuc(1) | N205 | Yes | P01877\|IGHA2_N205 |
| K.TPLTAN[+1971.719]ITK.S | HexNAc(5)Hex(5)Fuc(1) | P01877 | IGHA2 | Immunoglobulin heavy constant alpha 2 | P01877\|IGHA2 | P01877\|IGHA2_N205_HexNAc(5)Hex(5)Fuc(1) | N205 | Yes | P01877\|IGHA2_N205 |
| K.TPLTAN[+2262.814]ITK.S | HexNAc(5)Hex(5)Fuc(1)NeuAc(1) | P01877 | IGHA2 | Immunoglobulin heavy constant alpha 2 | P01877\|IGHA2 | P01877\|IGHA2_N205_HexNAc(5)Hex(5)Fuc(1)NeuAc(1) | N205 | Yes | P01877\|IGHA2_N205 |
| K.TPLTAN[+2117.777]ITK.S | HexNAc(5)Hex(5)Fuc(2) | P01877 | IGHA2 | Immunoglobulin heavy constant alpha 2 | P01877\|IGHA2 | P01877\|IGHA2_N205_HexNAc(5)Hex(5)Fuc(2) | N205 | Yes | P01877\|IGHA2_N205 |
| K.TPLTAN[+2263.835]ITK.S | HexNAc(5)Hex(5)Fuc(3) | P01877 | IGHA2 | Immunoglobulin heavy constant alpha 2 | P01877\|IGHA2 | P01877\|IGHA2_N205_HexNAc(5)Hex(5)Fuc(3) | N205 | Yes | P01877\|IGHA2_N205 |
| K.SAVQGPPERDLC[+57.021]GC[+57.021]YSVSSVLPGC[+57.021]AQPWNHGETFTC[+57.021]TAAHPELKTPLTAN[+1444.534]ITK.S | HexNAc(4)Hex(3)Fuc(1) | P01877 | IGHA2 | Immunoglobulin heavy constant alpha 2 | P01877\|IGHA2 | P01877\|IGHA2_N205_HexNAc(4)Hex(3)Fuc(1) | N205 | Yes | P01877\|IGHA2_N205 |
| K.SAVQGPPERDLC[+57.021]GC[+57.021]YSVSSVLPGC[+57.021]AQPWNHGETFTC[+57.021]TAAHPELKTPLTAN[+1647.613]ITK.S | HexNAc(5)Hex(3)Fuc(1) | P01877 | IGHA2 | Immunoglobulin heavy constant alpha 2 | P01877\|IGHA2 | P01877\|IGHA2_N205_HexNAc(5)Hex(3)Fuc(1) | N205 | Yes | P01877\|IGHA2_N205 |
| R.DLC[+57.021]GC[+57.021]YSVSSVLPGC[+57.021]AQPWNHGETFTC[+57.021]TAAHPELKTPLTAN[+1647.613]ITK.S | HexNAc(5)Hex(3)Fuc(1) | P01877 | IGHA2 | Immunoglobulin heavy constant alpha 2 | P01877\|IGHA2 | P01877\|IGHA2_N205_HexNAc(5)Hex(3)Fuc(1) | N205 | Yes | P01877\|IGHA2_N205 |
| K.SAVQGPPERDLC[+57.021]GC[+57.021]YSVSSVLPGC[+57.021]AQPWNHGETFTC[+57.021]TAAHPELKTPLTAN[+1809.666]ITK.S | HexNAc(5)Hex(4)Fuc(1) | P01877 | IGHA2 | Immunoglobulin heavy constant alpha 2 | P01877\|IGHA2 | P01877\|IGHA2_N205_HexNAc(5)Hex(4)Fuc(1) | N205 | Yes | P01877\|IGHA2_N205 |
| K.SAVQGPPERDLC[+57.021]GC[+57.021]YSVSSVLPGC[+57.021]AQPWNHGETFTC[+57.021]TAAHPELKTPLTAN[+2262.814]ITK.S | HexNAc(5)Hex(5)Fuc(1)NeuAc(1) | P01877 | IGHA2 | Immunoglobulin heavy constant alpha 2 | P01877\|IGHA2 | P01877\|IGHA2_N205_HexNAc(5)Hex(5)Fuc(1)NeuAc(1) | N205 | Yes | P01877\|IGHA2_N205 |
| K.SAVQGPPERDLC[+57.021]GC[+57.021]YSVSSVLPGC[+57.021]AQPWNHGETFTC[+57.021]TAAHPELKTPLTAN[+2263.835]ITK.S | HexNAc(5)Hex(5)Fuc(3) | P01877 | IGHA2 | Immunoglobulin heavy constant alpha 2 | P01877\|IGHA2 | P01877\|IGHA2_N205_HexNAc(5)Hex(5)Fuc(3) | N205 | Yes | P01877\|IGHA2_N205 |
| R.MAGKPTHIN[+1054.370]VSVVMAEADGTC[+57.021]Y.- | HexNAc(2)Hex(4) | P01877 | IGHA2 | Immunoglobulin heavy constant alpha 2 | P01877\|IGHA2 | P01877\|IGHA2_N327_HexNAc(2)Hex(4) | N327 | Yes | P01877\|IGHA2_N327 |
| R.MAGKPTHIN[+1216.423]VSVVM[+15.995]AEADGTC[+57.021]Y.- | HexNAc(2)Hex(5) | P01877 | IGHA2 | Immunoglobulin heavy constant alpha 2 | P01877\|IGHA2 | P01877\|IGHA2_N327_HexNAc(2)Hex(5) | N327 | Yes | P01877\|IGHA2_N327 |
| R.MAGKPTHIN[+1216.423]VSVVMAEADGTC[+57.021].Y | HexNAc(2)Hex(5) | P01877 | IGHA2 | Immunoglobulin heavy constant alpha 2 | P01877\|IGHA2 | P01877\|IGHA2_N327_HexNAc(2)Hex(5) | N327 | Yes | P01877\|IGHA2_N327 |
| R.MAGKPTHIN[+1216.423]VSVVMAEADGTC[+57.021]Y.- | HexNAc(2)Hex(5) | P01877 | IGHA2 | Immunoglobulin heavy constant alpha 2 | P01877\|IGHA2 | P01877\|IGHA2_N327_HexNAc(2)Hex(5) | N327 | Yes | P01877\|IGHA2_N327 |
| R.MAGKPTHIN[+1378.476]VSVVMAEADGTC[+57.021].Y | HexNAc(2)Hex(6) | P01877 | IGHA2 | Immunoglobulin heavy constant alpha 2 | P01877\|IGHA2 | P01877\|IGHA2_N327_HexNAc(2)Hex(6) | N327 | Yes | P01877\|IGHA2_N327 |
| R.MAGKPTHIN[+1378.476]VSVVMAEADGTC[+57.021]Y.- | HexNAc(2)Hex(6) | P01877 | IGHA2 | Immunoglobulin heavy constant alpha 2 | P01877\|IGHA2 | P01877\|IGHA2_N327_HexNAc(2)Hex(6) | N327 | Yes | P01877\|IGHA2_N327 |
| R.MAGKPTHIN[+1540.529]VSVVMAEADGTC[+57.021].Y | HexNAc(2)Hex(7) | P01877 | IGHA2 | Immunoglobulin heavy constant alpha 2 | P01877\|IGHA2 | P01877\|IGHA2_N327_HexNAc(2)Hex(7) | N327 | Yes | P01877\|IGHA2_N327 |
| R.MAGKPTHIN[+1540.529]VSVVMAEADGTC[+57.021]Y.- | HexNAc(2)Hex(7) | P01877 | IGHA2 | Immunoglobulin heavy constant alpha 2 | P01877\|IGHA2 | P01877\|IGHA2_N327_HexNAc(2)Hex(7) | N327 | Yes | P01877\|IGHA2_N327 |
| R.MAGKPTHIN[+1702.581]VSVVMAEADGTC[+57.021]Y.- | HexNAc(2)Hex(8) | P01877 | IGHA2 | Immunoglobulin heavy constant alpha 2 | P01877\|IGHA2 | P01877\|IGHA2_N327_HexNAc(2)Hex(8) | N327 | Yes | P01877\|IGHA2_N327 |
| R.MAGKPTHIN[+1864.634]VSVVMAEADGTC[+57.021]Y.- | HexNAc(2)Hex(9) | P01877 | IGHA2 | Immunoglobulin heavy constant alpha 2 | P01877\|IGHA2 | P01877\|IGHA2_N327_HexNAc(2)Hex(9) | N327 | Yes | P01877\|IGHA2_N327 |
| R.MAGKPTHIN[+1095.397]VSVVMAEADGTC[+57.021]Y.- | HexNAc(3)Hex(3) | P01877 | IGHA2 | Immunoglobulin heavy constant alpha 2 | P01877\|IGHA2 | P01877\|IGHA2_N327_HexNAc(3)Hex(3) | N327 | Yes | P01877\|IGHA2_N327 |
| R.MAGKPTHIN[+1257.449]VSVVMAEADGTC[+57.021]Y.- | HexNAc(3)Hex(4) | P01877 | IGHA2 | Immunoglobulin heavy constant alpha 2 | P01877\|IGHA2 | P01877\|IGHA2_N327_HexNAc(3)Hex(4) | N327 | Yes | P01877\|IGHA2_N327 |
| R.MAGKPTHIN[+1548.545]VSVVMAEADGTC[+57.021]Y.- | HexNAc(3)Hex(4)NeuAc(1) | P01877 | IGHA2 | Immunoglobulin heavy constant alpha 2 | P01877\|IGHA2 | P01877\|IGHA2_N327_HexNAc(3)Hex(4)NeuAc(1) | N327 | Yes | P01877\|IGHA2_N327 |
| R.MAGKPTHIN[+1419.502]VSVVMAEADGTC[+57.021]Y.- | HexNAc(3)Hex(5) | P01877 | IGHA2 | Immunoglobulin heavy constant alpha 2 | P01877\|IGHA2 | P01877\|IGHA2_N327_HexNAc(3)Hex(5) | N327 | Yes | P01877\|IGHA2_N327 |
| R.MAGKPTHIN[+1710.598]VSVVMAEADGTC[+57.021]Y.- | HexNAc(3)Hex(5)NeuAc(1) | P01877 | IGHA2 | Immunoglobulin heavy constant alpha 2 | P01877\|IGHA2 | P01877\|IGHA2_N327_HexNAc(3)Hex(5)NeuAc(1) | N327 | Yes | P01877\|IGHA2_N327 |
| R.MAGKPTHIN[+1581.555]VSVVMAEADGTC[+57.021]Y.- | HexNAc(3)Hex(6) | P01877 | IGHA2 | Immunoglobulin heavy constant alpha 2 | P01877\|IGHA2 | P01877\|IGHA2_N327_HexNAc(3)Hex(6) | N327 | Yes | P01877\|IGHA2_N327 |
| R.MAGKPTHIN[+1872.651]VSVVMAEADGTC[+57.021]Y.- | HexNAc(3)Hex(6)NeuAc(1) | P01877 | IGHA2 | Immunoglobulin heavy constant alpha 2 | P01877\|IGHA2 | P01877\|IGHA2_N327_HexNAc(3)Hex(6)NeuAc(1) | N327 | Yes | P01877\|IGHA2_N327 |
| R.MAGKPTHIN[+1298.476]VSVVMAEADGTC[+57.021]Y.- | HexNAc(4)Hex(3) | P01877 | IGHA2 | Immunoglobulin heavy constant alpha 2 | P01877\|IGHA2 | P01877\|IGHA2_N327_HexNAc(4)Hex(3) | N327 | Yes | P01877\|IGHA2_N327 |
| R.MAGKPTHIN[+1622.582]VSVVMAEADGTC[+57.021]Y.- | HexNAc(4)Hex(5) | P01877 | IGHA2 | Immunoglobulin heavy constant alpha 2 | P01877\|IGHA2 | P01877\|IGHA2_N327_HexNAc(4)Hex(5) | N327 | Yes | P01877\|IGHA2_N327 |
| R.MAGKPTHIN[+1913.677]VSVVMAEADGTC[+57.021]Y.- | HexNAc(4)Hex(5)NeuAc(1) | P01877 | IGHA2 | Immunoglobulin heavy constant alpha 2 | P01877\|IGHA2 | P01877\|IGHA2_N327_HexNAc(4)Hex(5)NeuAc(1) | N327 | Yes | P01877\|IGHA2_N327 |
| R.MAGKPTHIN[+1501.555]VSVVMAEADGTC[+57.021]Y.- | HexNAc(5)Hex(3) | P01877 | IGHA2 | Immunoglobulin heavy constant alpha 2 | P01877\|IGHA2 | P01877\|IGHA2_N327_HexNAc(5)Hex(3) | N327 | Yes | P01877\|IGHA2_N327 |
| R.MAGKPTHIN[+1647.613]VSVVMAEADGTC[+57.021]Y.- | HexNAc(5)Hex(3)Fuc(1) | P01877 | IGHA2 | Immunoglobulin heavy constant alpha 2 | P01877\|IGHA2 | P01877\|IGHA2_N327_HexNAc(5)Hex(3)Fuc(1) | N327 | Yes | P01877\|IGHA2_N327 |
| R.MAGKPTHIN[+1663.608]VSVVMAEADGTC[+57.021]Y.- | HexNAc(5)Hex(4) | P01877 | IGHA2 | Immunoglobulin heavy constant alpha 2 | P01877\|IGHA2 | P01877\|IGHA2_N327_HexNAc(5)Hex(4) | N327 | Yes | P01877\|IGHA2_N327 |
| R.MAGKPTHIN[+1809.666]VSVVMAEADGTC[+57.021]Y.- | HexNAc(5)Hex(4)Fuc(1) | P01877 | IGHA2 | Immunoglobulin heavy constant alpha 2 | P01877\|IGHA2 | P01877\|IGHA2_N327_HexNAc(5)Hex(4)Fuc(1) | N327 | Yes | P01877\|IGHA2_N327 |
| R.MAGKPTHIN[+1825.661]VSVVMAEADGTC[+57.021]Y.- | HexNAc(5)Hex(5) | P01877 | IGHA2 | Immunoglobulin heavy constant alpha 2 | P01877\|IGHA2 | P01877\|IGHA2_N327_HexNAc(5)Hex(5) | N327 | Yes | P01877\|IGHA2_N327 |
| R.MAGKPTHIN[+2262.814]VSVVMAEADGTC[+57.021]Y.- | HexNAc(5)Hex(5)Fuc(1)NeuAc(1) | P01877 | IGHA2 | Immunoglobulin heavy constant alpha 2 | P01877\|IGHA2 | P01877\|IGHA2_N327_HexNAc(5)Hex(5)Fuc(1)NeuAc(1) | N327 | Yes | P01877\|IGHA2_N327 |
| R.M[+15.995]AGKPTHIN[+1054.370]VSVVMAEADGTC[+57.021]Y.- | HexNAc(2)Hex(4) | P01877 | IGHA2 | Immunoglobulin heavy constant alpha 2 | P01877\|IGHA2 | P01877\|IGHA2_N327_HexNAc(2)Hex(4) | N327 | Yes | P01877\|IGHA2_N327 |
| R.M[+15.995]AGKPTHIN[+1216.423]VSVVMAEADGTC[+57.021].Y | HexNAc(2)Hex(5) | P01877 | IGHA2 | Immunoglobulin heavy constant alpha 2 | P01877\|IGHA2 | P01877\|IGHA2_N327_HexNAc(2)Hex(5) | N327 | Yes | P01877\|IGHA2_N327 |
| R.M[+15.995]AGKPTHIN[+1216.423]VSVVMAEADGTC[+57.021]Y.- | HexNAc(2)Hex(5) | P01877 | IGHA2 | Immunoglobulin heavy constant alpha 2 | P01877\|IGHA2 | P01877\|IGHA2_N327_HexNAc(2)Hex(5) | N327 | Yes | P01877\|IGHA2_N327 |
| R.M[+15.995]AGKPTHIN[+1378.476]VSVVMAEADGTC[+57.021].Y | HexNAc(2)Hex(6) | P01877 | IGHA2 | Immunoglobulin heavy constant alpha 2 | P01877\|IGHA2 | P01877\|IGHA2_N327_HexNAc(2)Hex(6) | N327 | Yes | P01877\|IGHA2_N327 |
| R.M[+15.995]AGKPTHIN[+1378.476]VSVVMAEADGTC[+57.021]Y.- | HexNAc(2)Hex(6) | P01877 | IGHA2 | Immunoglobulin heavy constant alpha 2 | P01877\|IGHA2 | P01877\|IGHA2_N327_HexNAc(2)Hex(6) | N327 | Yes | P01877\|IGHA2_N327 |
| R.M[+15.995]AGKPTHIN[+1540.529]VSVVMAEADGTC[+57.021]Y.- | HexNAc(2)Hex(7) | P01877 | IGHA2 | Immunoglobulin heavy constant alpha 2 | P01877\|IGHA2 | P01877\|IGHA2_N327_HexNAc(2)Hex(7) | N327 | Yes | P01877\|IGHA2_N327 |
| R.M[+15.995]AGKPTHIN[+1702.581]VSVVMAEADGTC[+57.021]Y.- | HexNAc(2)Hex(8) | P01877 | IGHA2 | Immunoglobulin heavy constant alpha 2 | P01877\|IGHA2 | P01877\|IGHA2_N327_HexNAc(2)Hex(8) | N327 | Yes | P01877\|IGHA2_N327 |
| R.M[+15.995]AGKPTHIN[+1501.555]VSVVMAEADGTC[+57.021]Y.- | HexNAc(5)Hex(3) | P01877 | IGHA2 | Immunoglobulin heavy constant alpha 2 | P01877\|IGHA2 | P01877\|IGHA2_N327_HexNAc(5)Hex(3) | N327 | Yes | P01877\|IGHA2_N327 |
| R.M[+15.995]AGKPTHIN[+1663.608]VSVVMAEADGTC[+57.021]Y.- | HexNAc(5)Hex(4) | P01877 | IGHA2 | Immunoglobulin heavy constant alpha 2 | P01877\|IGHA2 | P01877\|IGHA2_N327_HexNAc(5)Hex(4) | N327 | Yes | P01877\|IGHA2_N327 |
| K.VFPLSLDSTPQDGNVVVAC[+57.021]LVQGFFPQEPLSVTWSESGQN[+1038.375]VTAR.N | HexNAc(2)Hex(3)Fuc(1) | P01877 | IGHA2 | Immunoglobulin heavy constant alpha 2 | P01877\|IGHA2 | P01877\|IGHA2_N47_HexNAc(2)Hex(3)Fuc(1) | N47 | Yes | P01877\|IGHA2_N47 |
| K.VFPLSLDSTPQDGNVVVAC[+57.021]LVQGFFPQEPLSVTWSESGQN[+1864.634]VTAR.N | HexNAc(2)Hex(9) | P01877 | IGHA2 | Immunoglobulin heavy constant alpha 2 | P01877\|IGHA2 | P01877\|IGHA2_N47_HexNAc(2)Hex(9) | N47 | Yes | P01877\|IGHA2_N47 |
| K.VFPLSLDSTPQDGNVVVAC[+57.021]LVQGFFPQEPLSVTWSESGQN[+1241.454]VTAR.N | HexNAc(3)Hex(3)Fuc(1) | P01877 | IGHA2 | Immunoglobulin heavy constant alpha 2 | P01877\|IGHA2 | P01877\|IGHA2_N47_HexNAc(3)Hex(3)Fuc(1) | N47 | Yes | P01877\|IGHA2_N47 |
| K.VFPLSLDSTPQDGNVVVAC[+57.021]LVQGFFPQEPLSVTWSESGQN[+1694.603]VTAR.N | HexNAc(3)Hex(4)Fuc(1)NeuAc(1) | P01877 | IGHA2 | Immunoglobulin heavy constant alpha 2 | P01877\|IGHA2 | P01877\|IGHA2_N47_HexNAc(3)Hex(4)Fuc(1)NeuAc(1) | N47 | Yes | P01877\|IGHA2_N47 |
| K.VFPLSLDSTPQDGNVVVAC[+57.021]LVQGFFPQEPLSVTWSESGQN[+1444.534]VTAR.N | HexNAc(4)Hex(3)Fuc(1) | P01877 | IGHA2 | Immunoglobulin heavy constant alpha 2 | P01877\|IGHA2 | P01877\|IGHA2_N47_HexNAc(4)Hex(3)Fuc(1) | N47 | Yes | P01877\|IGHA2_N47 |
| A.SPTSPKVFPLSLDSTPQDGNVVVAC[+57.021]LVQGFFPQEPLSVTWSESGQN[+1460.529]VTAR.N | HexNAc(4)Hex(4) | P01877 | IGHA2 | Immunoglobulin heavy constant alpha 2 | P01877\|IGHA2 | P01877\|IGHA2_N47_HexNAc(4)Hex(4) | N47 | Yes | P01877\|IGHA2_N47 |
| K.VFPLSLDSTPQDGNVVVAC[+57.021]LVQGFFPQEPLSVTWSESGQN[+1606.587]VTAR.N | HexNAc(4)Hex(4)Fuc(1) | P01877 | IGHA2 | Immunoglobulin heavy constant alpha 2 | P01877\|IGHA2 | P01877\|IGHA2_N47_HexNAc(4)Hex(4)Fuc(1) | N47 | Yes | P01877\|IGHA2_N47 |
| K.VFPLSLDSTPQDGNVVVAC[+57.021]LVQGFFPQEPLSVTWSESGQN[+1622.582]VTAR.N | HexNAc(4)Hex(5) | P01877 | IGHA2 | Immunoglobulin heavy constant alpha 2 | P01877\|IGHA2 | P01877\|IGHA2_N47_HexNAc(4)Hex(5) | N47 | Yes | P01877\|IGHA2_N47 |
| K.VFPLSLDSTPQDGNVVVAC[+57.021]LVQGFFPQEPLSVTWSESGQN[+2059.735]VTAR.N | HexNAc(4)Hex(5)Fuc(1)NeuAc(1) | P01877 | IGHA2 | Immunoglobulin heavy constant alpha 2 | P01877\|IGHA2 | P01877\|IGHA2_N47_HexNAc(4)Hex(5)Fuc(1)NeuAc(1) | N47 | Yes | P01877\|IGHA2_N47 |
| K.VFPLSLDSTPQDGNVVVAC[+57.021]LVQGFFPQEPLSVTWSESGQN[+2350.830]VTAR.N | HexNAc(4)Hex(5)Fuc(1)NeuAc(2) | P01877 | IGHA2 | Immunoglobulin heavy constant alpha 2 | P01877\|IGHA2 | P01877\|IGHA2_N47_HexNAc(4)Hex(5)Fuc(1)NeuAc(2) | N47 | Yes | P01877\|IGHA2_N47 |
| K.VFPLSLDSTPQDGNVVVAC[+57.021]LVQGFFPQEPLSVTWSESGQN[+2351.851]VTAR.N | HexNAc(4)Hex(5)Fuc(3)NeuAc(1) | P01877 | IGHA2 | Immunoglobulin heavy constant alpha 2 | P01877\|IGHA2 | P01877\|IGHA2_N47_HexNAc(4)Hex(5)Fuc(3)NeuAc(1) | N47 | Yes | P01877\|IGHA2_N47 |
| K.VFPLSLDSTPQDGNVVVAC[+57.021]LVQGFFPQEPLSVTWSESGQN[+2076.750]VTAR.N | HexNAc(4)Hex(6)Fuc(2) | P01877 | IGHA2 | Immunoglobulin heavy constant alpha 2 | P01877\|IGHA2 | P01877\|IGHA2_N47_HexNAc(4)Hex(6)Fuc(2) | N47 | Yes | P01877\|IGHA2_N47 |
| K.VFPLSLDSTPQDGNVVVAC[+57.021]LVQGFFPQEPLSVTWSESGQN[+1647.613]VTAR.N | HexNAc(5)Hex(3)Fuc(1) | P01877 | IGHA2 | Immunoglobulin heavy constant alpha 2 | P01877\|IGHA2 | P01877\|IGHA2_N47_HexNAc(5)Hex(3)Fuc(1) | N47 | Yes | P01877\|IGHA2_N47 |

Table S5. Cont.

| Glycopeptide | Glycan | Uniprot ID | Gene Name | Protein Name | Shortname | ProteinNameInSkyline | Glycosites | Reported before | Site |
|---|---|---|---|---|---|---|---|---|---|
| K.VFPLSLDSTPQDGNVVVAC[+57.021]LVQGFFPQEPLSVTWSESGQN[+1663.608]VTAR.N | HexNAc(5)Hex(4) | P01877 | IGHA2 | Immunoglobulin heavy constant alpha 2 | P01877\|IGHA2 | P01877\|IGHA2_N47_HexNAc(5)Hex(4) | N47 | Yes | P01877\|IGHA2_N47 |
| K.VFPLSLDSTPQDGNVVVAC[+57.021]LVQGFFPQEPLSVTWSESGQN[+1809.666]VTAR.N | HexNAc(5)Hex(4)Fuc(1) | P01877 | IGHA2 | Immunoglobulin heavy constant alpha 2 | P01877\|IGHA2 | P01877\|IGHA2_N47_HexNAc(5)Hex(4)Fuc(1) | N47 | Yes | P01877\|IGHA2_N47 |
| K.VFPLSLDSTPQDGNVVVAC[+57.021]LVQGFFPQEPLSVTWSESGQN[+1971.719]VTAR.N | HexNAc(5)Hex(5)Fuc(1) | P01877 | IGHA2 | Immunoglobulin heavy constant alpha 2 | P01877\|IGHA2 | P01877\|IGHA2_N47_HexNAc(5)Hex(5)Fuc(1) | N47 | Yes | P01877\|IGHA2_N47 |
| K.VFPLSLDSTPQDGNVVVAC[+57.021]LVQGFFPQEPLSVTWSESGQN[+2262.814]VTAR.N | HexNAc(5)Hex(5)Fuc(1)NeuAc(1) | P01877 | IGHA2 | Immunoglobulin heavy constant alpha 2 | P01877\|IGHA2 | P01877\|IGHA2_N47_HexNAc(5)Hex(5)Fuc(1)NeuAc(1) | N47 | Yes | P01877\|IGHA2_N47 |
| K.VFPLSLDSTPQDGNVVVAC[+57.021]LVQGFFPQEPLSVTWSESGQN[+2553.910]VTAR.N | HexNAc(5)Hex(5)Fuc(1)NeuAc(2) | P01877 | IGHA2 | Immunoglobulin heavy constant alpha 2 | P01877\|IGHA2 | P01877\|IGHA2_N47_HexNAc(5)Hex(5)Fuc(1)NeuAc(2) | N47 | Yes | P01877\|IGHA2_N47 |
| K.VFPLSLDSTPQDGNVVVAC[+57.021]LVQGFFPQEPLSVTWSESGQN[+2117.777]VTAR.N | HexNAc(5)Hex(5)Fuc(2) | P01877 | IGHA2 | Immunoglobulin heavy constant alpha 2 | P01877\|IGHA2 | P01877\|IGHA2_N47_HexNAc(5)Hex(5)Fuc(2) | N47 | Yes | P01877\|IGHA2_N47 |
| K.VFPLSLDSTPQDGNVVVAC[+57.021]LVQGFFPQEPLSVTWSESGQN[+2263.835]VTAR.N | HexNAc(5)Hex(5)Fuc(3) | P01877 | IGHA2 | Immunoglobulin heavy constant alpha 2 | P01877\|IGHA2 | P01877\|IGHA2_N47_HexNAc(5)Hex(5)Fuc(3) | N47 | Yes | P01877\|IGHA2_N47 |
| K.VFPLSLDSTPQDGNVVVAC[+57.021]LVQGFFPQEPLSVTWSESGQN[+2570.925]VTAR.N | HexNAc(5)Hex(6)Fuc(2)NeuAc(1) | P01877 | IGHA2 | Immunoglobulin heavy constant alpha 2 | P01877\|IGHA2 | P01877\|IGHA2_N47_HexNAc(5)Hex(6)Fuc(2)NeuAc(1) | N47 | Yes | P01877\|IGHA2_N47 |
| A.[+42.011]SPTSPKVFPLSLDSTPQDGNVVVAC[+57.021]LVQGFFPQEPLSVTWSESGQN[+1257.449]VTAR.N | HexNAc(3)Hex(4) | P01877 | IGHA2 | Immunoglobulin heavy constant alpha 2 | P01877\|IGHA2 | P01877\|IGHA2_N47_HexNAc(3)Hex(4) | N47 | Yes | P01877\|IGHA2_N47 |
| A.[+42.011]SPTSPKVFPLSLDSTPQDGNVVVAC[+57.021]LVQGFFPQEPLSVTWSESGQN[+1419.502]VTAR.N | HexNAc(3)Hex(5) | P01877 | IGHA2 | Immunoglobulin heavy constant alpha 2 | P01877\|IGHA2 | P01877\|IGHA2_N47_HexNAc(3)Hex(5) | N47 | Yes | P01877\|IGHA2_N47 |
| A.[+42.011]SPTSPKVFPLSLDSTPQDGNVVVAC[+57.021]LVQGFFPQEPLSVTWSESGQN[+1460.529]VTAR.N | HexNAc(4)Hex(4) | P01877 | IGHA2 | Immunoglobulin heavy constant alpha 2 | P01877\|IGHA2 | P01877\|IGHA2_N47_HexNAc(4)Hex(4) | N47 | Yes | P01877\|IGHA2_N47 |
| K.[+42.011]VFPLSLDSTPQDGNVVVAC[+57.021]LVQGFFPQEPLSVTWSESGQN[+1460.529]VTAR.N | HexNAc(4)Hex(4) | P01877 | IGHA2 | Immunoglobulin heavy constant alpha 2 | P01877\|IGHA2 | P01877\|IGHA2_N47_HexNAc(4)Hex(4) | N47 | Yes | P01877\|IGHA2_N47 |
| K.[+42.011]VFPLSLDSTPQDGNVVVAC[+57.021]LVQGFFPQEPLSVTWSESGQN[+1606.587]VTAR.N | HexNAc(4)Hex(4)Fuc(1) | P01877 | IGHA2 | Immunoglobulin heavy constant alpha 2 | P01877\|IGHA2 | P01877\|IGHA2_N47_HexNAc(4)Hex(4)Fuc(1) | N47 | Yes | P01877\|IGHA2_N47 |
| K.[+42.011]VFPLSLDSTPQDGNVVVAC[+57.021]LVQGFFPQEPLSVTWSESGQN[+1768.640]VTAR.N | HexNAc(4)Hex(5)Fuc(1) | P01877 | IGHA2 | Immunoglobulin heavy constant alpha 2 | P01877\|IGHA2 | P01877\|IGHA2_N47_HexNAc(4)Hex(5)Fuc(1) | N47 | Yes | P01877\|IGHA2_N47 |
| K.[+42.011]VFPLSLDSTPQDGNVVVAC[+57.021]LVQGFFPQEPLSVTWSESGQN[+2059.735]VTAR.N | HexNAc(4)Hex(5)Fuc(1)NeuAc(1) | P01877 | IGHA2 | Immunoglobulin heavy constant alpha 2 | P01877\|IGHA2 | P01877\|IGHA2_N47_HexNAc(4)Hex(5)Fuc(1)NeuAc(1) | N47 | Yes | P01877\|IGHA2_N47 |
| A.[+42.011]SPTSPKVFPLSLDSTPQDGNVVVAC[+57.021]LVQGFFPQEPLSVTWSESGQN[+1913.677]VTAR.N | HexNAc(4)Hex(5)NeuAc(1) | P01877 | IGHA2 | Immunoglobulin heavy constant alpha 2 | P01877\|IGHA2 | P01877\|IGHA2_N47_HexNAc(4)Hex(5)NeuAc(1) | N47 | Yes | P01877\|IGHA2_N47 |
| K.HYTN[+876.322]SSQDVTVPC[+57.021]R.V | HexNAc(2)Hex(2)Fuc(1) | P01877 | IGHA2 | Immunoglobulin heavy constant alpha 2 | P01877\|IGHA2 | P01877\|IGHA2_N92_HexNAc(2)Hex(2)Fuc(1) | N92 | Yes | P01877\|IGHA2_N92 |
| K.HYTN[+1038.375]SSQDVTVPC[+57.021]R.V | HexNAc(2)Hex(3)Fuc(1) | P01877 | IGHA2 | Immunoglobulin heavy constant alpha 2 | P01877\|IGHA2 | P01877\|IGHA2_N92_HexNAc(2)Hex(3)Fuc(1) | N92 | Yes | P01877\|IGHA2_N92 |
| K.HYTN[+1038.375]SSQDVTVPC[+57.021]RVPPPPPC[+57.021]C[+57.021]HPR.L | HexNAc(2)Hex(3)Fuc(1) | P01877 | IGHA2 | Immunoglobulin heavy constant alpha 2 | P01877\|IGHA2 | P01877\|IGHA2_N92_HexNAc(2)Hex(3)Fuc(1) | N92 | Yes | P01877\|IGHA2_N92 |
| K.HYTN[+1216.423]SSQDVTVPC[+57.021]R.V | HexNAc(2)Hex(5) | P01877 | IGHA2 | Immunoglobulin heavy constant alpha 2 | P01877\|IGHA2 | P01877\|IGHA2_N92_HexNAc(2)Hex(5) | N92 | Yes | P01877\|IGHA2_N92 |
| K.HYTN[+1241.454]SSQDVTVPC[+57.021]R.V | HexNAc(3)Hex(3)Fuc(1) | P01877 | IGHA2 | Immunoglobulin heavy constant alpha 2 | P01877\|IGHA2 | P01877\|IGHA2_N92_HexNAc(3)Hex(3)Fuc(1) | N92 | Yes | P01877\|IGHA2_N92 |
| K.HYTN[+1241.454]SSQDVTVPC[+57.021]RVPPPPPC[+57.021]C[+57.021]HPR.L | HexNAc(3)Hex(3)Fuc(1) | P01877 | IGHA2 | Immunoglobulin heavy constant alpha 2 | P01877\|IGHA2 | P01877\|IGHA2_N92_HexNAc(3)Hex(3)Fuc(1) | N92 | Yes | P01877\|IGHA2_N92 |
| K.HYTN[+1403.507]SSQDVTVPC[+57.021]R.V | HexNAc(3)Hex(4)Fuc(1) | P01877 | IGHA2 | Immunoglobulin heavy constant alpha 2 | P01877\|IGHA2 | P01877\|IGHA2_N92_HexNAc(3)Hex(4)Fuc(1) | N92 | Yes | P01877\|IGHA2_N92 |
| K.HYTN[+1694.603]SSQDVTVPC[+57.021]R.V | HexNAc(3)Hex(4)Fuc(1)NeuAc(1) | P01877 | IGHA2 | Immunoglobulin heavy constant alpha 2 | P01877\|IGHA2 | P01877\|IGHA2_N92_HexNAc(3)Hex(4)Fuc(1)NeuAc(1) | N92 | Yes | P01877\|IGHA2_N92 |
| K.HYTN[+1444.534]SSQDVTVPC[+57.021]R.V | HexNAc(4)Hex(3)Fuc(1) | P01877 | IGHA2 | Immunoglobulin heavy constant alpha 2 | P01877\|IGHA2 | P01877\|IGHA2_N92_HexNAc(4)Hex(3)Fuc(1) | N92 | Yes | P01877\|IGHA2_N92 |
| K.HYTN[+1444.534]SSQDVTVPC[+57.021]RVPPPPPC[+57.021]C[+57.021]HPR.L | HexNAc(4)Hex(3)Fuc(1) | P01877 | IGHA2 | Immunoglobulin heavy constant alpha 2 | P01877\|IGHA2 | P01877\|IGHA2_N92_HexNAc(4)Hex(3)Fuc(1) | N92 | Yes | P01877\|IGHA2_N92 |
| K.HYTN[+1606.587]SSQDVTVPC[+57.021]R.V | HexNAc(4)Hex(4)Fuc(1) | P01877 | IGHA2 | Immunoglobulin heavy constant alpha 2 | P01877\|IGHA2 | P01877\|IGHA2_N92_HexNAc(4)Hex(4)Fuc(1) | N92 | Yes | P01877\|IGHA2_N92 |
| K.HYTN[+1606.587]SSQDVTVPC[+57.021]RVPPPPPC[+57.021]C[+57.021]HPR.L | HexNAc(4)Hex(4)Fuc(1) | P01877 | IGHA2 | Immunoglobulin heavy constant alpha 2 | P01877\|IGHA2 | P01877\|IGHA2_N92_HexNAc(4)Hex(4)Fuc(1) | N92 | Yes | P01877\|IGHA2_N92 |
| K.HYTN[+1768.640]SSQDVTVPC[+57.021]R.V | HexNAc(4)Hex(5)Fuc(1) | P01877 | IGHA2 | Immunoglobulin heavy constant alpha 2 | P01877\|IGHA2 | P01877\|IGHA2_N92_HexNAc(4)Hex(5)Fuc(1) | N92 | Yes | P01877\|IGHA2_N92 |
| K.HYTN[+2059.735]SSQDVTVPC[+57.021]R.V | HexNAc(4)Hex(5)Fuc(1)NeuAc(1) | P01877 | IGHA2 | Immunoglobulin heavy constant alpha 2 | P01877\|IGHA2 | P01877\|IGHA2_N92_HexNAc(4)Hex(5)Fuc(1)NeuAc(1) | N92 | Yes | P01877\|IGHA2_N92 |
| K.HYTN[+2059.735]SSQDVTVPC[+57.021]RVPPPPPC[+57.021]C[+57.021]HPR.L | HexNAc(4)Hex(5)Fuc(1)NeuAc(1) | P01877 | IGHA2 | Immunoglobulin heavy constant alpha 2 | P01877\|IGHA2 | P01877\|IGHA2_N92_HexNAc(4)Hex(5)Fuc(1)NeuAc(1) | N92 | Yes | P01877\|IGHA2_N92 |
| K.HYTN[+2350.830]SSQDVTVPC[+57.021]R.V | HexNAc(4)Hex(5)Fuc(1)NeuAc(2) | P01877 | IGHA2 | Immunoglobulin heavy constant alpha 2 | P01877\|IGHA2 | P01877\|IGHA2_N92_HexNAc(4)Hex(5)Fuc(1)NeuAc(2) | N92 | Yes | P01877\|IGHA2_N92 |
| K.HYTN[+1930.692]SSQDVTVPC[+57.021]R.V | HexNAc(4)Hex(6)Fuc(1) | P01877 | IGHA2 | Immunoglobulin heavy constant alpha 2 | P01877\|IGHA2 | P01877\|IGHA2_N92_HexNAc(4)Hex(6)Fuc(1) | N92 | Yes | P01877\|IGHA2_N92 |
| K.HYTN[+1647.613]SSQDVTVPC[+57.021]R.V | HexNAc(5)Hex(3)Fuc(1) | P01877 | IGHA2 | Immunoglobulin heavy constant alpha 2 | P01877\|IGHA2 | P01877\|IGHA2_N92_HexNAc(5)Hex(3)Fuc(1) | N92 | Yes | P01877\|IGHA2_N92 |
| K.HYTN[+1647.613]SSQDVTVPC[+57.021]RVPPPPPC[+57.021]C[+57.021]HPR.L | HexNAc(5)Hex(3)Fuc(1) | P01877 | IGHA2 | Immunoglobulin heavy constant alpha 2 | P01877\|IGHA2 | P01877\|IGHA2_N92_HexNAc(5)Hex(3)Fuc(1) | N92 | Yes | P01877\|IGHA2_N92 |
| K.HYTN[+1809.666]SSQDVTVPC[+57.021]R.V | HexNAc(5)Hex(4)Fuc(1) | P01877 | IGHA2 | Immunoglobulin heavy constant alpha 2 | P01877\|IGHA2 | P01877\|IGHA2_N92_HexNAc(5)Hex(4)Fuc(1) | N92 | Yes | P01877\|IGHA2_N92 |
| K.HYTN[+1809.666]SSQDVTVPC[+57.021]RVPPPPPC[+57.021]C[+57.021]HPR.L | HexNAc(5)Hex(4)Fuc(1) | P01877 | IGHA2 | Immunoglobulin heavy constant alpha 2 | P01877\|IGHA2 | P01877\|IGHA2_N92_HexNAc(5)Hex(4)Fuc(1) | N92 | Yes | P01877\|IGHA2_N92 |
| K.HYTN[+1971.719]SSQDVTVPC[+57.021]R.V | HexNAc(5)Hex(5)Fuc(1) | P01877 | IGHA2 | Immunoglobulin heavy constant alpha 2 | P01877\|IGHA2 | P01877\|IGHA2_N92_HexNAc(5)Hex(5)Fuc(1) | N92 | Yes | P01877\|IGHA2_N92 |
| K.HYTN[+1971.719]SSQDVTVPC[+57.021]RVPPPPPC[+57.021]C[+57.021]HPR.L | HexNAc(5)Hex(5)Fuc(1) | P01877 | IGHA2 | Immunoglobulin heavy constant alpha 2 | P01877\|IGHA2 | P01877\|IGHA2_N92_HexNAc(5)Hex(5)Fuc(1) | N92 | Yes | P01877\|IGHA2_N92 |
| K.HYTN[+2262.814]SSQDVTVPC[+57.021]R.V | HexNAc(5)Hex(5)Fuc(1)NeuAc(1) | P01877 | IGHA2 | Immunoglobulin heavy constant alpha 2 | P01877\|IGHA2 | P01877\|IGHA2_N92_HexNAc(5)Hex(5)Fuc(1)NeuAc(1) | N92 | Yes | P01877\|IGHA2_N92 |
| K.HYTN[+2262.814]SSQDVTVPC[+57.021]RVPPPPPC[+57.021]C[+57.021]HPR.L | HexNAc(5)Hex(5)Fuc(1)NeuAc(1) | P01877 | IGHA2 | Immunoglobulin heavy constant alpha 2 | P01877\|IGHA2 | P01877\|IGHA2_N92_HexNAc(5)Hex(5)Fuc(1)NeuAc(1) | N92 | Yes | P01877\|IGHA2_N92 |
| K.HYTN[+2553.910]SSQDVTVPC[+57.021]RVPPPPPC[+57.021]C[+57.021]HPR.L | HexNAc(5)Hex(5)Fuc(1)NeuAc(2) | P01877 | IGHA2 | Immunoglobulin heavy constant alpha 2 | P01877\|IGHA2 | P01877\|IGHA2_N92_HexNAc(5)Hex(5)Fuc(1)NeuAc(2) | N92 | Yes | P01877\|IGHA2_N92 |
| K.HYTN[+2263.835]SSQDVTVPC[+57.021]RVPPPPPC[+57.021]C[+57.021]HPR.L | HexNAc(5)Hex(5)Fuc(3) | P01877 | IGHA2 | Immunoglobulin heavy constant alpha 2 | P01877\|IGHA2 | P01877\|IGHA2_N92_HexNAc(5)Hex(5)Fuc(3) | N92 | Yes | P01877\|IGHA2_N92 |
| R.AVN[+2205.793]ITSENLIDDVVSLIR.G | HexNAc(4)Hex(5)Fuc(2)NeuAc(1) | P02748 | C9 | Complement component C9 | P02748\|CO9 | P02748\|CO9_N415_HexNAc(4)Hex(5)Fuc(2)NeuAc(1) | N415 | Yes | P02748\|CO9_N415 |
| R.AVN[+2204.772]ITSENLIDDVVSLIR.G | HexNAc(4)Hex(5)NeuAc(2) | P02748 | C9 | Complement component C9 | P02748\|CO9 | P02748\|CO9_N415_HexNAc(4)Hex(5)NeuAc(2) | N415 | Yes | P02748\|CO9_N415 |
| R.KLPPGLLAN[+2059.735]FTLLR.T | HexNAc(4)Hex(5)Fuc(1)NeuAc(1) | P02750 | LRG1 | Leucine-rich alpha-2-glycoprotein | P02750\|A2GL | P02750\|A2GL_N186_HexNAc(4)Hex(5)Fuc(1)NeuAc(1) | N186 | Yes | P02750\|A2GL_N186 |
| R.QLDMLDLSN[+2205.793]NSLASVPEGLWASLGQPNWDMR.D | HexNAc(4)Hex(5)Fuc(2)NeuAc(1) | P02750 | LRG1 | Leucine-rich alpha-2-glycoprotein | P02750\|A2GL | P02750\|A2GL_N269_HexNAc(4)Hex(5)Fuc(2)NeuAc(1) | N269 | Yes | P02750\|A2GL_N269 |
| R.QLDMLDLSN[+1913.677]NSLASVPEGLWASLGQPNWDMR.D | HexNAc(4)Hex(5)NeuAc(1) | P02750 | LRG1 | Leucine-rich alpha-2-glycoprotein | P02750\|A2GL | P02750\|A2GL_N269_HexNAc(4)Hex(5)NeuAc(1) | N269 | Yes | P02750\|A2GL_N269 |
| R.QLDMLDLSN[+2204.772]NSLASVPEGLWASLGQPNWDMR.D | HexNAc(4)Hex(5)NeuAc(2) | P02750 | LRG1 | Leucine-rich alpha-2-glycoprotein | P02750\|A2GL | P02750\|A2GL_N269_HexNAc(4)Hex(5)NeuAc(2) | N269 | Yes | P02750\|A2GL_N269 |
| R.[+42.011]QLDMLDLSN[+1872.651]NSLASVPEGLWASLGQPNWDMR.D | HexNAc(3)Hex(6)NeuAc(1) | P02750 | LRG1 | Leucine-rich alpha-2-glycoprotein | P02750\|A2GL | P02750\|A2GL_N278_HexNAc(3)Hex(6)NeuAc(1) | N278 | Yes | P02750\|A2GL_N278 |
| R.SDHGSSISC[+57.021]QPPAEIPGYLPADTVHLAVEFFN[+2350.830]LTHLPANLLQGASK.L | HexNAc(4)Hex(5)Fuc(1)NeuAc(2) | P02750 | LRG1 | Leucine-rich alpha-2-glycoprotein | P02750\|A2GL | P02750\|A2GL_N79_HexNAc(4)Hex(5)Fuc(1)NeuAc(2) | N79 | Yes | P02750\|A2GL_N79 |
| R.SDHGSSISC[+57.021]QPPAEIPGYLPADTVHLAVEFFN[+2351.851]LTHLPANLLQGASK.L | HexNAc(4)Hex(5)Fuc(3)NeuAc(1) | P02750 | LRG1 | Leucine-rich alpha-2-glycoprotein | P02750\|A2GL | P02750\|A2GL_N79_HexNAc(4)Hex(5)Fuc(3)NeuAc(1) | N79 | Yes | P02750\|A2GL_N79 |
| R.SDHGSSISC[+57.021]QPPAEIPGYLPADTVHLAVEFFN[+2570.925]LTHLPANLLQGASK.L | HexNAc(5)Hex(6)Fuc(2)NeuAc(1) | P02750 | LRG1 | Leucine-rich alpha-2-glycoprotein | P02750\|A2GL | P02750\|A2GL_N79_HexNAc(5)Hex(6)Fuc(2)NeuAc(1) | N79 | Yes | P02750\|A2GL_N79 |
| K.AALAAFNAQNN[+2205.793]GSNFQLEEISR.A | HexNAc(4)Hex(5)Fuc(2)NeuAc(1) | P02765 | AHSG | Alpha-2-HS-glycoprotein | P02765\|FETUA | P02765\|FETUA_N176_HexNAc(4)Hex(5)Fuc(2)NeuAc(1) | N176 | Yes | P02765\|FETUA_N176 |
| K.AALAAFNAQNN[+2204.772]GSNFQLEEISR.A | HexNAc(4)Hex(5)NeuAc(2) | P02765 | AHSG | Alpha-2-HS-glycoprotein | P02765\|FETUA | P02765\|FETUA_N176_HexNAc(4)Hex(5)NeuAc(2) | N176 | Yes | P02765\|FETUA_N176 |
| K.LC[+57.021]DN[+1866.688]LSTK.N | HexNAc(6)Hex(4) | P02774 | GC | Vitamin D-binding protein | P02774\|VTDB | P02774\|VTDB_N288_HexNAc(6)Hex(4) | N288 | NO | P02774\|VTDB_N288 |
| R.QQQHLFGSN[+2204.772]VTDC[+57.021]SGNFC[+57.021]LFR.S | HexNAc(4)Hex(5)NeuAc(2) | P02787 | TF | Serotransferrin | P02787\|TRFE | P02787\|TRFE_N630_HexNAc(4)Hex(5)NeuAc(2) | N630 | Yes | P02787\|TRFE_N630 |
| R.TAGWNVPIGTLRPFLN[+349.137]WTGPPEPIEAAVAR.F | HexNAc(1)Fuc(1) | P02788 | LTF | Lactotransferrin | P02788\|TRFL | P02788\|TRFL_N156_HexNAc(1)Fuc(1) | N156 | Yes | P02788\|TRFL_N156 |
| R.RTAGWNVPIGTLRPFLN[+1768.640]WTGPPEPIEAAVAR.F | HexNAc(4)Hex(5)Fuc(1) | P02788 | LTF | Lactotransferrin | P02788\|TRFL | P02788\|TRFL_N156_HexNAc(4)Hex(5)Fuc(1) | N156 | Yes | P02788\|TRFL_N156 |
| R.TAGWNVPIGTLRPFLN[+1768.640]WTGPPEPIEAAVAR.F | HexNAc(4)Hex(5)Fuc(1) | P02788 | LTF | Lactotransferrin | P02788\|TRFL | P02788\|TRFL_N156_HexNAc(4)Hex(5)Fuc(1) | N156 | Yes | P02788\|TRFL_N156 |
| R.RTAGWNVPIGTLRPFLN[+2059.735]WTGPPEPIEAAVAR.F | HexNAc(4)Hex(5)Fuc(1)NeuAc(1) | P02788 | LTF | Lactotransferrin | P02788\|TRFL | P02788\|TRFL_N156_HexNAc(4)Hex(5)Fuc(1)NeuAc(1) | N156 | Yes | P02788\|TRFL_N156 |
| R.TAGWNVPIGTLRPFLN[+2059.735]WTGPPEPIEAAVAR.F | HexNAc(4)Hex(5)Fuc(1)NeuAc(1) | P02788 | LTF | Lactotransferrin | P02788\|TRFL | P02788\|TRFL_N156_HexNAc(4)Hex(5)Fuc(1)NeuAc(1) | N156 | Yes | P02788\|TRFL_N156 |
| R.RTAGWNVPIGTLRPFLN[+2350.830]WTGPPEPIEAAVAR.F | HexNAc(4)Hex(5)Fuc(1)NeuAc(2) | P02788 | LTF | Lactotransferrin | P02788\|TRFL | P02788\|TRFL_N156_HexNAc(4)Hex(5)Fuc(1)NeuAc(2) | N156 | Yes | P02788\|TRFL_N156 |
| R.TAGWNVPIGTLRPFLN[+2350.830]WTGPPEPIEAAVAR.F | HexNAc(4)Hex(5)Fuc(1)NeuAc(2) | P02788 | LTF | Lactotransferrin | P02788\|TRFL | P02788\|TRFL_N156_HexNAc(4)Hex(5)Fuc(1)NeuAc(2) | N156 | Yes | P02788\|TRFL_N156 |
| R.RTAGWNVPIGTLRPFLN[+2205.793]WTGPPEPIEAAVAR.F | HexNAc(4)Hex(5)Fuc(2)NeuAc(1) | P02788 | LTF | Lactotransferrin | P02788\|TRFL | P02788\|TRFL_N156_HexNAc(4)Hex(5)Fuc(2)NeuAc(1) | N156 | Yes | P02788\|TRFL_N156 |
| R.TAGWNVPIGTLRPFLN[+2205.793]WTGPPEPIEAAVAR.F | HexNAc(4)Hex(5)Fuc(2)NeuAc(1) | P02788 | LTF | Lactotransferrin | P02788\|TRFL | P02788\|TRFL_N156_HexNAc(4)Hex(5)Fuc(2)NeuAc(1) | N156 | Yes | P02788\|TRFL_N156 |
| R.TAGWNVPIGTLRPFLN[+2496.888]WTGPPEPIEAAVAR.F | HexNAc(4)Hex(5)Fuc(2)NeuAc(2) | P02788 | LTF | Lactotransferrin | P02788\|TRFL | P02788\|TRFL_N156_HexNAc(4)Hex(5)Fuc(2)NeuAc(2) | N156 | Yes | P02788\|TRFL_N156 |
| R.RTAGWNVPIGTLRPFLN[+2351.851]WTGPPEPIEAAVAR.F | HexNAc(4)Hex(5)Fuc(3)NeuAc(1) | P02788 | LTF | Lactotransferrin | P02788\|TRFL | P02788\|TRFL_N156_HexNAc(4)Hex(5)Fuc(3)NeuAc(1) | N156 | Yes | P02788\|TRFL_N156 |
| R.TAGWNVPIGTLRPFLN[+2351.851]WTGPPEPIEAAVAR.F | HexNAc(4)Hex(5)Fuc(3)NeuAc(1) | P02788 | LTF | Lactotransferrin | P02788\|TRFL | P02788\|TRFL_N156_HexNAc(4)Hex(5)Fuc(3)NeuAc(1) | N156 | Yes | P02788\|TRFL_N156 |
| R.RTAGWNVPIGTLRPFLN[+1913.677]WTGPPEPIEAAVAR.F | HexNAc(4)Hex(5)NeuAc(1) | P02788 | LTF | Lactotransferrin | P02788\|TRFL | P02788\|TRFL_N156_HexNAc(4)Hex(5)NeuAc(1) | N156 | Yes | P02788\|TRFL_N156 |
| R.TAGWNVPIGTLRPFLN[+1913.677]WTGPPEPIEAAVAR.F | HexNAc(4)Hex(5)NeuAc(1) | P02788 | LTF | Lactotransferrin | P02788\|TRFL | P02788\|TRFL_N156_HexNAc(4)Hex(5)NeuAc(1) | N156 | Yes | P02788\|TRFL_N156 |
| R.RTAGWNVPIGTLRPFLN[+2204.772]WTGPPEPIEAAVAR.F | HexNAc(4)Hex(5)NeuAc(2) | P02788 | LTF | Lactotransferrin | P02788\|TRFL | P02788\|TRFL_N156_HexNAc(4)Hex(5)NeuAc(2) | N156 | Yes | P02788\|TRFL_N156 |
| R.TAGWNVPIGTLRPFLN[+2204.772]WTGPPEPIEAAVAR.F | HexNAc(4)Hex(5)NeuAc(2) | P02788 | LTF | Lactotransferrin | P02788\|TRFL | P02788\|TRFL_N156_HexNAc(4)Hex(5)NeuAc(2) | N156 | Yes | P02788\|TRFL_N156 |
| R.RTAGWNVPIGTLRPFLN[+1930.692]WTGPPEPIEAAVAR.F | HexNAc(4)Hex(6)Fuc(1) | P02788 | LTF | Lactotransferrin | P02788\|TRFL | P02788\|TRFL_N156_HexNAc(4)Hex(6)Fuc(1) | N156 | Yes | P02788\|TRFL_N156 |
| R.TAGWNVPIGTLRPFLN[+2076.750]WTGPPEPIEAAVAR.F | HexNAc(4)Hex(6)Fuc(2) | P02788 | LTF | Lactotransferrin | P02788\|TRFL | P02788\|TRFL_N156_HexNAc(4)Hex(6)Fuc(2) | N156 | Yes | P02788\|TRFL_N156 |
| R.TAGWNVPIGTLRPFLN[+2133.772]WTGPPEPIEAAVAR.F | HexNAc(5)Hex(6)Fuc(1) | P02788 | LTF | Lactotransferrin | P02788\|TRFL | P02788\|TRFL_N156_HexNAc(5)Hex(6)Fuc(1) | N156 | Yes | P02788\|TRFL_N156 |
| R.RTAGWNVPIGTLRPFLN[+2424.867]WTGPPEPIEAAVAR.F | HexNAc(5)Hex(6)Fuc(1)NeuAc(1) | P02788 | LTF | Lactotransferrin | P02788\|TRFL | P02788\|TRFL_N156_HexNAc(5)Hex(6)Fuc(1)NeuAc(1) | N156 | Yes | P02788\|TRFL_N156 |
| R.TAGWNVPIGTLRPFLN[+2424.867]WTGPPEPIEAAVAR.F | HexNAc(5)Hex(6)Fuc(1)NeuAc(1) | P02788 | LTF | Lactotransferrin | P02788\|TRFL | P02788\|TRFL_N156_HexNAc(5)Hex(6)Fuc(1)NeuAc(1) | N156 | Yes | P02788\|TRFL_N156 |
| R.RTAGWNVPIGTLRPFLN[+2715.963]WTGPPEPIEAAVAR.F | HexNAc(5)Hex(6)Fuc(1)NeuAc(2) | P02788 | LTF | Lactotransferrin | P02788\|TRFL | P02788\|TRFL_N156_HexNAc(5)Hex(6)Fuc(1)NeuAc(2) | N156 | Yes | P02788\|TRFL_N156 |
| R.TAGWNVPIGTLRPFLN[+2715.963]WTGPPEPIEAAVAR.F | HexNAc(5)Hex(6)Fuc(1)NeuAc(2) | P02788 | LTF | Lactotransferrin | P02788\|TRFL | P02788\|TRFL_N156_HexNAc(5)Hex(6)Fuc(1)NeuAc(2) | N156 | Yes | P02788\|TRFL_N156 |
| R.RTAGWNVPIGTLRPFLN[+2279.830]WTGPPEPIEAAVAR.F | HexNAc(5)Hex(6)Fuc(2) | P02788 | LTF | Lactotransferrin | P02788\|TRFL | P02788\|TRFL_N156_HexNAc(5)Hex(6)Fuc(2) | N156 | Yes | P02788\|TRFL_N156 |
| R.TAGWNVPIGTLRPFLN[+2279.830]WTGPPEPIEAAVAR.F | HexNAc(5)Hex(6)Fuc(2) | P02788 | LTF | Lactotransferrin | P02788\|TRFL | P02788\|TRFL_N156_HexNAc(5)Hex(6)Fuc(2) | N156 | Yes | P02788\|TRFL_N156 |
| R.RTAGWNVPIGTLRPFLN[+2570.925]WTGPPEPIEAAVAR.F | HexNAc(5)Hex(6)Fuc(2)NeuAc(1) | P02788 | LTF | Lactotransferrin | P02788\|TRFL | P02788\|TRFL_N156_HexNAc(5)Hex(6)Fuc(2)NeuAc(1) | N156 | Yes | P02788\|TRFL_N156 |
| R.TAGWNVPIGTLRPFLN[+2570.925]WTGPPEPIEAAVAR.F | HexNAc(5)Hex(6)Fuc(2)NeuAc(1) | P02788 | LTF | Lactotransferrin | P02788\|TRFL | P02788\|TRFL_N156_HexNAc(5)Hex(6)Fuc(2)NeuAc(1) | N156 | Yes | P02788\|TRFL_N156 |
| R.TAGWNVPIGTLRPFLN[+2862.021]WTGPPEPIEAAVAR.F | HexNAc(5)Hex(6)Fuc(2)NeuAc(2) | P02788 | LTF | Lactotransferrin | P02788\|TRFL | P02788\|TRFL_N156_HexNAc(5)Hex(6)Fuc(2)NeuAc(2) | N156 | Yes | P02788\|TRFL_N156 |
| R.TAGWNVPIGTLRPFLN[+3153.116]WTGPPEPIEAAVAR.F | HexNAc(5)Hex(6)Fuc(2)NeuAc(3) | P02788 | LTF | Lactotransferrin | P02788\|TRFL | P02788\|TRFL_N156_HexNAc(5)Hex(6)Fuc(2)NeuAc(3) | N156 | Yes | P02788\|TRFL_N156 |
| R.TAGWNVPIGTLRPFLN[+2425.887]WTGPPEPIEAAVAR.F | HexNAc(5)Hex(6)Fuc(3) | P02788 | LTF | Lactotransferrin | P02788\|TRFL | P02788\|TRFL_N156_HexNAc(5)Hex(6)Fuc(3) | N156 | Yes | P02788\|TRFL_N156 |

Table S5. Cont.

| Glycopeptide | Glycan | Uniprot ID | Gene Name | Protein Name | Shortname | ProteinNameInSkyline | Glycosites | Reported before | Site |
|---|---|---|---|---|---|---|---|---|---|
| R.RTAGWNVPIGTLRPFLN[+2716.983]WTGPPEPIEAAVAR.F | HexNAc(5)Hex(6)Fuc(3)NeuAc(1) | P02788 | LTF | Lactotransferrin | P02788\|TRFL | P02788\|TRFL_N156_HexNAc(5)Hex(6)Fuc(3)NeuAc(1) | N156 | Yes | P02788\|TRFL_N156 |
| R.TAGWNVPIGTLRPFLN[+2716.983]WTGPPEPIEAAVAR.F | HexNAc(5)Hex(6)Fuc(3)NeuAc(1) | P02788 | LTF | Lactotransferrin | P02788\|TRFL | P02788\|TRFL_N156_HexNAc(5)Hex(6)Fuc(3)NeuAc(1) | N156 | Yes | P02788\|TRFL_N156 |
| R.RTAGWNVPIGTLRPFLN[+2571.946]WTGPPEPIEAAVAR.F | HexNAc(5)Hex(6)Fuc(4) | P02788 | LTF | Lactotransferrin | P02788\|TRFL | P02788\|TRFL_N156_HexNAc(5)Hex(6)Fuc(4) | N156 | Yes | P02788\|TRFL_N156 |
| R.TAGWNVPIGTLRPFLN[+2571.946]WTGPPEPIEAAVAR.F | HexNAc(5)Hex(6)Fuc(4) | P02788 | LTF | Lactotransferrin | P02788\|TRFL | P02788\|TRFL_N156_HexNAc(5)Hex(6)Fuc(4) | N156 | Yes | P02788\|TRFL_N156 |
| R.RTAGWNVPIGTLRPFLN[+2863.041]WTGPPEPIEAAVAR.F | HexNAc(5)Hex(6)Fuc(4)NeuAc(1) | P02788 | LTF | Lactotransferrin | P02788\|TRFL | P02788\|TRFL_N156_HexNAc(5)Hex(6)Fuc(4)NeuAc(1) | N156 | Yes | P02788\|TRFL_N156 |
| R.TAGWNVPIGTLRPFLN[+2863.041]WTGPPEPIEAAVAR.F | HexNAc(5)Hex(6)Fuc(4)NeuAc(1) | P02788 | LTF | Lactotransferrin | P02788\|TRFL | P02788\|TRFL_N156_HexNAc(5)Hex(6)Fuc(4)NeuAc(1) | N156 | Yes | P02788\|TRFL_N156 |
| R.RTAGWNVPIGTLRPFLN[+2278.809]WTGPPEPIEAAVAR.F | HexNAc(5)Hex(6)NeuAc(1) | P02788 | LTF | Lactotransferrin | P02788\|TRFL | P02788\|TRFL_N156_HexNAc(5)Hex(6)NeuAc(1) | N156 | Yes | P02788\|TRFL_N156 |
| R.TAGWNVPIGTLRPFLN[+2278.809]WTGPPEPIEAAVAR.F | HexNAc(5)Hex(6)NeuAc(1) | P02788 | LTF | Lactotransferrin | P02788\|TRFL | P02788\|TRFL_N156_HexNAc(5)Hex(6)NeuAc(1) | N156 | Yes | P02788\|TRFL_N156 |
| R.RTAGWNVPIGTLRPFLN[+2569.905]WTGPPEPIEAAVAR.F | HexNAc(5)Hex(6)NeuAc(2) | P02788 | LTF | Lactotransferrin | P02788\|TRFL | P02788\|TRFL_N156_HexNAc(5)Hex(6)NeuAc(2) | N156 | Yes | P02788\|TRFL_N156 |
| R.TAGWNVPIGTLRPFLN[+2569.905]WTGPPEPIEAAVAR.F | HexNAc(5)Hex(6)NeuAc(2) | P02788 | LTF | Lactotransferrin | P02788\|TRFL | P02788\|TRFL_N156_HexNAc(5)Hex(6)NeuAc(2) | N156 | Yes | P02788\|TRFL_N156 |
| R.TAGWNVPIGTLRPFLN[+2498.904]WTGPPEPIEAAVAR.F | HexNAc(6)Hex(7)Fuc(1) | P02788 | LTF | Lactotransferrin | P02788\|TRFL | P02788\|TRFL_N156_HexNAc(6)Hex(7)Fuc(1) | N156 | Yes | P02788\|TRFL_N156 |
| R.TAGWNVPIGTLRPFLN[+2644.962]WTGPPEPIEAAVAR.F | HexNAc(6)Hex(7)Fuc(2) | P02788 | LTF | Lactotransferrin | P02788\|TRFL | P02788\|TRFL_N156_HexNAc(6)Hex(7)Fuc(2) | N156 | Yes | P02788\|TRFL_N156 |
| R.TAGWNVPIGTLRPFLN[+2936.057]WTGPPEPIEAAVAR.F | HexNAc(6)Hex(7)Fuc(2)NeuAc(1) | P02788 | LTF | Lactotransferrin | P02788\|TRFL | P02788\|TRFL_N156_HexNAc(6)Hex(7)Fuc(2)NeuAc(1) | N156 | Yes | P02788\|TRFL_N156 |
| R.TAGWNVPIGTLRPFLN[+3227.153]WTGPPEPIEAAVAR.F | HexNAc(6)Hex(7)Fuc(2)NeuAc(2) | P02788 | LTF | Lactotransferrin | P02788\|TRFL | P02788\|TRFL_N156_HexNAc(6)Hex(7)Fuc(2)NeuAc(2) | N156 | Yes | P02788\|TRFL_N156 |
| R.TAGWNVPIGTLRPFLN[+2791.020]WTGPPEPIEAAVAR.F | HexNAc(6)Hex(7)Fuc(3) | P02788 | LTF | Lactotransferrin | P02788\|TRFL | P02788\|TRFL_N156_HexNAc(6)Hex(7)Fuc(3) | N156 | Yes | P02788\|TRFL_N156 |
| R.[+42.011]RTAGWNVPIGTLRPFLN[+2026.687]WTGPPEPIEAAVAR.F | HexNAc(2)Hex(10) | P02788 | LTF | Lactotransferrin | P02788\|TRFL | P02788\|TRFL_N156_HexNAc(2)Hex(10) | N156 | Yes | P02788\|TRFL_N156 |
| R.[+42.011]TAGWNVPIGTLRPFLN[+1362.481]WTGPPEPIEAAVAR.F | HexNAc(2)Hex(5)Fuc(1) | P02788 | LTF | Lactotransferrin | P02788\|TRFL | P02788\|TRFL_N156_HexNAc(2)Hex(5)Fuc(1) | N156 | Yes | P02788\|TRFL_N156 |
| K.SC[+57.021]HTGLRRTAGWNVPIGTLRPFLN[+1540.529]WTGPPEPIEAAVAR.F | HexNAc(2)Hex(7) | P02788 | LTF | Lactotransferrin | P02788\|TRFL | P02788\|TRFL_N156_HexNAc(2)Hex(7) | N156 | Yes | P02788\|TRFL_N156 |
| R.[+42.011]RTAGWNVPIGTLRPFLN[+1864.634]WTGPPEPIEAAVAR.F | HexNAc(2)Hex(9) | P02788 | LTF | Lactotransferrin | P02788\|TRFL | P02788\|TRFL_N156_HexNAc(2)Hex(9) | N156 | Yes | P02788\|TRFL_N156 |
| R.[+42.011]PFLN[+2018.708]WTGPPEPIEAAVAR.F | HexNAc(3)Hex(6)Fuc(1)NeuAc(1) | P02788 | LTF | Lactotransferrin | P02788\|TRFL | P02788\|TRFL_N156_HexNAc(3)Hex(6)Fuc(1)NeuAc(1) | N156 | Yes | P02788\|TRFL_N156 |
| R.[+42.011]RTAGWNVPIGTLRPFLN[+2018.708]WTGPPEPIEAAVAR.F | HexNAc(3)Hex(6)Fuc(1)NeuAc(1) | P02788 | LTF | Lactotransferrin | P02788\|TRFL | P02788\|TRFL_N156_HexNAc(3)Hex(6)Fuc(1)NeuAc(1) | N156 | Yes | P02788\|TRFL_N156 |
| R.[+42.011]TAGWNVPIGTLRPFLN[+2018.708]WTGPPEPIEAAVAR.F | HexNAc(3)Hex(6)Fuc(1)NeuAc(1) | P02788 | LTF | Lactotransferrin | P02788\|TRFL | P02788\|TRFL_N156_HexNAc(3)Hex(6)Fuc(1)NeuAc(1) | N156 | Yes | P02788\|TRFL_N156 |
| R.[+42.011]RTAGWNVPIGTLRPFLN[+1872.651]WTGPPEPIEAAVAR.F | HexNAc(3)Hex(6)NeuAc(1) | P02788 | LTF | Lactotransferrin | P02788\|TRFL | P02788\|TRFL_N156_HexNAc(3)Hex(6)NeuAc(1) | N156 | Yes | P02788\|TRFL_N156 |
| R.[+42.011]TAGWNVPIGTLRPFLN[+1872.651]WTGPPEPIEAAVAR.F | HexNAc(3)Hex(6)NeuAc(1) | P02788 | LTF | Lactotransferrin | P02788\|TRFL | P02788\|TRFL_N156_HexNAc(3)Hex(6)NeuAc(1) | N156 | Yes | P02788\|TRFL_N156 |
| R.[+42.011]TAGWNVPIGTLRPFLN[+2028.740]WTGPPEPIEAAVAR.F | HexNAc(6)Hex(5) | P02788 | LTF | Lactotransferrin | P02788\|TRFL | P02788\|TRFL_N156_HexNAc(6)Hex(5) | N156 | Yes | P02788\|TRFL_N156 |
| R.[+42.011]TAGWNVPIGTLRPFLN[+2190.793]WTGPPEPIEAAVAR.F | HexNAc(6)Hex(6) | P02788 | LTF | Lactotransferrin | P02788\|TRFL | P02788\|TRFL_N156_HexNAc(6)Hex(6) | N156 | Yes | P02788\|TRFL_N156 |
| R.[+42.011]RTAGWNVPIGTLRPFLN[+2336.851]WTGPPEPIEAAVAR.F | HexNAc(6)Hex(6)Fuc(1) | P02788 | LTF | Lactotransferrin | P02788\|TRFL | P02788\|TRFL_N156_HexNAc(6)Hex(6)Fuc(1) | N156 | Yes | P02788\|TRFL_N156 |
| R.[+42.011]TAGWNVPIGTLRPFLN[+2336.851]WTGPPEPIEAAVAR.F | HexNAc(6)Hex(6)Fuc(1) | P02788 | LTF | Lactotransferrin | P02788\|TRFL | P02788\|TRFL_N156_HexNAc(6)Hex(6)Fuc(1) | N156 | Yes | P02788\|TRFL_N156 |
| R.TAGWNIPMGLLFN[+203.079]QTGSC[+57.021]K.F | HexNAc(1) | P02788 | LTF | Lactotransferrin | P02788\|TRFL | P02788\|TRFL_N497_HexNAc(1) | N497 | Yes | P02788\|TRFL_N497 |
| R.TAGWNIPMGLLFN[+349.137]QTGSC[+57.021]K.F | HexNAc(1)Fuc(1) | P02788 | LTF | Lactotransferrin | P02788\|TRFL | P02788\|TRFL_N497_HexNAc(1)Fuc(1) | N497 | Yes | P02788\|TRFL_N497 |
| R.TAGWNIPMGLLFN[+1038.375]QTGSC[+57.021]K.F | HexNAc(2)Hex(3)Fuc(1) | P02788 | LTF | Lactotransferrin | P02788\|TRFL | P02788\|TRFL_N497_HexNAc(2)Hex(3)Fuc(1) | N497 | Yes | P02788\|TRFL_N497 |
| R.TAGWNIPMGLLFN[+1216.423]QTGSC[+57.021]K.F | HexNAc(2)Hex(5) | P02788 | LTF | Lactotransferrin | P02788\|TRFL | P02788\|TRFL_N497_HexNAc(2)Hex(5) | N497 | Yes | P02788\|TRFL_N497 |
| R.TAGWNIPMGLLFN[+1216.423]QTGSC[+57.021]KFDEYFSQSC[+57.021]APGSDPR.S | HexNAc(2)Hex(5) | P02788 | LTF | Lactotransferrin | P02788\|TRFL | P02788\|TRFL_N497_HexNAc(2)Hex(5) | N497 | Yes | P02788\|TRFL_N497 |
| R.TAGWNIPMGLLFN[+1378.476]QTGSC[+57.021]K.F | HexNAc(2)Hex(6) | P02788 | LTF | Lactotransferrin | P02788\|TRFL | P02788\|TRFL_N497_HexNAc(2)Hex(6) | N497 | Yes | P02788\|TRFL_N497 |
| R.TAGWNIPMGLLFN[+1095.397]QTGSC[+57.021]K.F | HexNAc(3)Hex(3) | P02788 | LTF | Lactotransferrin | P02788\|TRFL | P02788\|TRFL_N497_HexNAc(3)Hex(3) | N497 | Yes | P02788\|TRFL_N497 |
| R.TAGWNIPMGLLFN[+1257.449]QTGSC[+57.021]K.F | HexNAc(3)Hex(4) | P02788 | LTF | Lactotransferrin | P02788\|TRFL | P02788\|TRFL_N497_HexNAc(3)Hex(4) | N497 | Yes | P02788\|TRFL_N497 |
| R.TAGWNIPMGLLFN[+1694.603]QTGSC[+57.021]K.F | HexNAc(3)Hex(4)Fuc(1)NeuAc(1) | P02788 | LTF | Lactotransferrin | P02788\|TRFL | P02788\|TRFL_N497_HexNAc(3)Hex(4)Fuc(1)NeuAc(1) | N497 | Yes | P02788\|TRFL_N497 |
| R.TAGWNIPMGLLFN[+1548.545]QTGSC[+57.021]K.F | HexNAc(3)Hex(4)NeuAc(1) | P02788 | LTF | Lactotransferrin | P02788\|TRFL | P02788\|TRFL_N497_HexNAc(3)Hex(4)NeuAc(1) | N497 | Yes | P02788\|TRFL_N497 |
| R.TAGWNIPMGLLFN[+1548.545]QTGSC[+57.021]KFDEYFSQSC[+57.021]APGSDPR.S | HexNAc(3)Hex(4)NeuAc(1) | P02788 | LTF | Lactotransferrin | P02788\|TRFL | P02788\|TRFL_N497_HexNAc(3)Hex(4)NeuAc(1) | N497 | Yes | P02788\|TRFL_N497 |
| R.TAGWNIPMGLLFN[+1419.502]QTGSC[+57.021]K.F | HexNAc(3)Hex(5) | P02788 | LTF | Lactotransferrin | P02788\|TRFL | P02788\|TRFL_N497_HexNAc(3)Hex(5) | N497 | Yes | P02788\|TRFL_N497 |
| R.TAGWNIPMGLLFN[+1856.656]QTGSC[+57.021]K.F | HexNAc(3)Hex(5)Fuc(1)NeuAc(1) | P02788 | LTF | Lactotransferrin | P02788\|TRFL | P02788\|TRFL_N497_HexNAc(3)Hex(5)Fuc(1)NeuAc(1) | N497 | Yes | P02788\|TRFL_N497 |
| R.TAGWNIPMGLLFN[+1710.598]QTGSC[+57.021]K.F | HexNAc(3)Hex(5)NeuAc(1) | P02788 | LTF | Lactotransferrin | P02788\|TRFL | P02788\|TRFL_N497_HexNAc(3)Hex(5)NeuAc(1) | N497 | Yes | P02788\|TRFL_N497 |
| R.TAGWNIPMGLLFN[+1710.598]QTGSC[+57.021]KFDEYFSQSC[+57.021]APGSDPR.S | HexNAc(3)Hex(5)NeuAc(1) | P02788 | LTF | Lactotransferrin | P02788\|TRFL | P02788\|TRFL_N497_HexNAc(3)Hex(5)NeuAc(1) | N497 | Yes | P02788\|TRFL_N497 |
| R.TAGWNIPMGLLFN[+1581.555]QTGSC[+57.021]K.F | HexNAc(3)Hex(6) | P02788 | LTF | Lactotransferrin | P02788\|TRFL | P02788\|TRFL_N497_HexNAc(3)Hex(6) | N497 | Yes | P02788\|TRFL_N497 |
| R.TAGWNIPMGLLFN[+1872.651]QTGSC[+57.021]K.F | HexNAc(3)Hex(6)NeuAc(1) | P02788 | LTF | Lactotransferrin | P02788\|TRFL | P02788\|TRFL_N497_HexNAc(3)Hex(6)NeuAc(1) | N497 | Yes | P02788\|TRFL_N497 |
| R.TAGWNIPMGLLFN[+1872.651]QTGSC[+57.021]KFDEYFSQSC[+57.021]APGSDPR.S | HexNAc(3)Hex(6)NeuAc(1) | P02788 | LTF | Lactotransferrin | P02788\|TRFL | P02788\|TRFL_N497_HexNAc(3)Hex(6)NeuAc(1) | N497 | Yes | P02788\|TRFL_N497 |
| R.TAGWNIPMGLLFN[+1460.529]QTGSC[+57.021]K.F | HexNAc(4)Hex(4) | P02788 | LTF | Lactotransferrin | P02788\|TRFL | P02788\|TRFL_N497_HexNAc(4)Hex(4) | N497 | Yes | P02788\|TRFL_N497 |
| R.TAGWNIPMGLLFN[+1606.587]QTGSC[+57.021]K.F | HexNAc(4)Hex(4)Fuc(1) | P02788 | LTF | Lactotransferrin | P02788\|TRFL | P02788\|TRFL_N497_HexNAc(4)Hex(4)Fuc(1) | N497 | Yes | P02788\|TRFL_N497 |
| R.TAGWNIPMGLLFN[+1606.587]QTGSC[+57.021]KFDEYFSQSC[+57.021]APGSDPR.S | HexNAc(4)Hex(4)Fuc(1) | P02788 | LTF | Lactotransferrin | P02788\|TRFL | P02788\|TRFL_N497_HexNAc(4)Hex(4)Fuc(1) | N497 | Yes | P02788\|TRFL_N497 |
| R.TAGWNIPMGLLFN[+1751.624]QTGSC[+57.021]K.F | HexNAc(4)Hex(4)NeuAc(1) | P02788 | LTF | Lactotransferrin | P02788\|TRFL | P02788\|TRFL_N497_HexNAc(4)Hex(4)NeuAc(1) | N497 | Yes | P02788\|TRFL_N497 |
| R.TAGWNIPMGLLFN[+1751.624]QTGSC[+57.021]KFDEYFSQSC[+57.021]APGSDPR.S | HexNAc(4)Hex(4)NeuAc(1) | P02788 | LTF | Lactotransferrin | P02788\|TRFL | P02788\|TRFL_N497_HexNAc(4)Hex(4)NeuAc(1) | N497 | Yes | P02788\|TRFL_N497 |
| R.TAGWNIPMGLLFN[+1622.582]QTGSC[+57.021]K.F | HexNAc(4)Hex(5) | P02788 | LTF | Lactotransferrin | P02788\|TRFL | P02788\|TRFL_N497_HexNAc(4)Hex(5) | N497 | Yes | P02788\|TRFL_N497 |
| R.TAGWNIPMGLLFN[+1622.582]QTGSC[+57.021]KFDEYFSQSC[+57.021]APGSDPR.S | HexNAc(4)Hex(5) | P02788 | LTF | Lactotransferrin | P02788\|TRFL | P02788\|TRFL_N497_HexNAc(4)Hex(5) | N497 | Yes | P02788\|TRFL_N497 |
| R.TAGWNIPMGLLFN[+1768.640]QTGSC[+57.021]K.F | HexNAc(4)Hex(5)Fuc(1) | P02788 | LTF | Lactotransferrin | P02788\|TRFL | P02788\|TRFL_N497_HexNAc(4)Hex(5)Fuc(1) | N497 | Yes | P02788\|TRFL_N497 |
| R.TAGWNIPMGLLFN[+1768.640]QTGSC[+57.021]KFDEYFSQSC[+57.021]APGSDPR.S | HexNAc(4)Hex(5)Fuc(1) | P02788 | LTF | Lactotransferrin | P02788\|TRFL | P02788\|TRFL_N497_HexNAc(4)Hex(5)Fuc(1) | N497 | Yes | P02788\|TRFL_N497 |
| R.TAGWNIPMGLLFN[+2059.735]QTGSC[+57.021]K.F | HexNAc(4)Hex(5)Fuc(1)NeuAc(1) | P02788 | LTF | Lactotransferrin | P02788\|TRFL | P02788\|TRFL_N497_HexNAc(4)Hex(5)Fuc(1)NeuAc(1) | N497 | Yes | P02788\|TRFL_N497 |
| R.TAGWNIPMGLLFN[+2059.735]QTGSC[+57.021]KFDEYFSQSC[+57.021]APGSDPR.S | HexNAc(4)Hex(5)Fuc(1)NeuAc(1) | P02788 | LTF | Lactotransferrin | P02788\|TRFL | P02788\|TRFL_N497_HexNAc(4)Hex(5)Fuc(1)NeuAc(1) | N497 | Yes | P02788\|TRFL_N497 |
| R.TAGWNIPMGLLFN[+2350.830]QTGSC[+57.021]K.F | HexNAc(4)Hex(5)Fuc(1)NeuAc(2) | P02788 | LTF | Lactotransferrin | P02788\|TRFL | P02788\|TRFL_N497_HexNAc(4)Hex(5)Fuc(1)NeuAc(2) | N497 | Yes | P02788\|TRFL_N497 |
| R.TAGWNIPMGLLFN[+2350.830]QTGSC[+57.021]KFDEYFSQSC[+57.021]APGSDPR.S | HexNAc(4)Hex(5)Fuc(1)NeuAc(2) | P02788 | LTF | Lactotransferrin | P02788\|TRFL | P02788\|TRFL_N497_HexNAc(4)Hex(5)Fuc(1)NeuAc(2) | N497 | Yes | P02788\|TRFL_N497 |
| R.TAGWNIPMGLLFN[+2205.793]QTGSC[+57.021]K.F | HexNAc(4)Hex(5)Fuc(2)NeuAc(1) | P02788 | LTF | Lactotransferrin | P02788\|TRFL | P02788\|TRFL_N497_HexNAc(4)Hex(5)Fuc(2)NeuAc(1) | N497 | Yes | P02788\|TRFL_N497 |
| R.TAGWNIPMGLLFN[+2205.793]QTGSC[+57.021]KFDEYFSQSC[+57.021]APGSDPR.S | HexNAc(4)Hex(5)Fuc(2)NeuAc(1) | P02788 | LTF | Lactotransferrin | P02788\|TRFL | P02788\|TRFL_N497_HexNAc(4)Hex(5)Fuc(2)NeuAc(1) | N497 | Yes | P02788\|TRFL_N497 |
| R.TAGWNIPMGLLFN[+2351.851]QTGSC[+57.021]K.F | HexNAc(4)Hex(5)Fuc(3)NeuAc(1) | P02788 | LTF | Lactotransferrin | P02788\|TRFL | P02788\|TRFL_N497_HexNAc(4)Hex(5)Fuc(3)NeuAc(1) | N497 | Yes | P02788\|TRFL_N497 |
| R.TAGWNIPMGLLFN[+2351.851]QTGSC[+57.021]KFDEYFSQSC[+57.021]APGSDPR.S | HexNAc(4)Hex(5)Fuc(3)NeuAc(1) | P02788 | LTF | Lactotransferrin | P02788\|TRFL | P02788\|TRFL_N497_HexNAc(4)Hex(5)Fuc(3)NeuAc(1) | N497 | Yes | P02788\|TRFL_N497 |
| R.TAGWNIPMGLLFN[+1913.677]QTGSC[+57.021]K.F | HexNAc(4)Hex(5)NeuAc(1) | P02788 | LTF | Lactotransferrin | P02788\|TRFL | P02788\|TRFL_N497_HexNAc(4)Hex(5)NeuAc(1) | N497 | Yes | P02788\|TRFL_N497 |
| R.TAGWNIPMGLLFN[+1913.677]QTGSC[+57.021]KFDEYFSQSC[+57.021]APGSDPR.S | HexNAc(4)Hex(5)NeuAc(1) | P02788 | LTF | Lactotransferrin | P02788\|TRFL | P02788\|TRFL_N497_HexNAc(4)Hex(5)NeuAc(1) | N497 | Yes | P02788\|TRFL_N497 |
| R.TAGWNIPMGLLFN[+2204.772]QTGSC[+57.021]K.F | HexNAc(4)Hex(5)NeuAc(2) | P02788 | LTF | Lactotransferrin | P02788\|TRFL | P02788\|TRFL_N497_HexNAc(4)Hex(5)NeuAc(2) | N497 | Yes | P02788\|TRFL_N497 |
| R.TAGWNIPMGLLFN[+2204.772]QTGSC[+57.021]KFDEYFSQSC[+57.021]APGSDPR.S | HexNAc(4)Hex(5)NeuAc(2) | P02788 | LTF | Lactotransferrin | P02788\|TRFL | P02788\|TRFL_N497_HexNAc(4)Hex(5)NeuAc(2) | N497 | Yes | P02788\|TRFL_N497 |
| R.TAGWNIPMGLLFN[+1930.692]QTGSC[+57.021]KFDEYFSQSC[+57.021]APGSDPR.S | HexNAc(4)Hex(6)Fuc(1) | P02788 | LTF | Lactotransferrin | P02788\|TRFL | P02788\|TRFL_N497_HexNAc(4)Hex(6)Fuc(1) | N497 | Yes | P02788\|TRFL_N497 |
| R.TAGWNIPMGLLFN[+2076.750]QTGSC[+57.021]K.F | HexNAc(4)Hex(6)Fuc(2) | P02788 | LTF | Lactotransferrin | P02788\|TRFL | P02788\|TRFL_N497_HexNAc(4)Hex(6)Fuc(2) | N497 | Yes | P02788\|TRFL_N497 |
| R.TAGWNIPMGLLFN[+2076.750]QTGSC[+57.021]KFDEYFSQSC[+57.021]APGSDPR.S | HexNAc(4)Hex(6)Fuc(2) | P02788 | LTF | Lactotransferrin | P02788\|TRFL | P02788\|TRFL_N497_HexNAc(4)Hex(6)Fuc(2) | N497 | Yes | P02788\|TRFL_N497 |
| R.TAGWNIPMGLLFN[+1971.719]QTGSC[+57.021]K.F | HexNAc(5)Hex(5)Fuc(1) | P02788 | LTF | Lactotransferrin | P02788\|TRFL | P02788\|TRFL_N497_HexNAc(5)Hex(5)Fuc(1) | N497 | Yes | P02788\|TRFL_N497 |
| R.TAGWNIPMGLLFN[+1971.719]QTGSC[+57.021]KFDEYFSQSC[+57.021]APGSDPR.S | HexNAc(5)Hex(5)Fuc(1) | P02788 | LTF | Lactotransferrin | P02788\|TRFL | P02788\|TRFL_N497_HexNAc(5)Hex(5)Fuc(1) | N497 | Yes | P02788\|TRFL_N497 |
| R.TAGWNIPMGLLFN[+2116.756]QTGSC[+57.021]KFDEYFSQSC[+57.021]APGSDPR.S | HexNAc(5)Hex(5)NeuAc(1) | P02788 | LTF | Lactotransferrin | P02788\|TRFL | P02788\|TRFL_N497_HexNAc(5)Hex(5)NeuAc(1) | N497 | Yes | P02788\|TRFL_N497 |
| R.TAGWNIPMGLLFN[+2133.772]QTGSC[+57.021]K.F | HexNAc(5)Hex(6)Fuc(1) | P02788 | LTF | Lactotransferrin | P02788\|TRFL | P02788\|TRFL_N497_HexNAc(5)Hex(6)Fuc(1) | N497 | Yes | P02788\|TRFL_N497 |
| R.TAGWNIPMGLLFN[+2424.867]QTGSC[+57.021]K.F | HexNAc(5)Hex(6)Fuc(1)NeuAc(1) | P02788 | LTF | Lactotransferrin | P02788\|TRFL | P02788\|TRFL_N497_HexNAc(5)Hex(6)Fuc(1)NeuAc(1) | N497 | Yes | P02788\|TRFL_N497 |
| R.TAGWNIPMGLLFN[+2279.830]QTGSC[+57.021]K.F | HexNAc(5)Hex(6)Fuc(2) | P02788 | LTF | Lactotransferrin | P02788\|TRFL | P02788\|TRFL_N497_HexNAc(5)Hex(6)Fuc(2) | N497 | Yes | P02788\|TRFL_N497 |
| R.TAGWNIPMGLLFN[+2570.925]QTGSC[+57.021]K.F | HexNAc(5)Hex(6)Fuc(2)NeuAc(1) | P02788 | LTF | Lactotransferrin | P02788\|TRFL | P02788\|TRFL_N497_HexNAc(5)Hex(6)Fuc(2)NeuAc(1) | N497 | Yes | P02788\|TRFL_N497 |
| R.TAGWNIPMGLLFN[+2862.021]QTGSC[+57.021]KFDEYFSQSC[+57.021]APGSDPR.S | HexNAc(5)Hex(6)Fuc(2)NeuAc(2) | P02788 | LTF | Lactotransferrin | P02788\|TRFL | P02788\|TRFL_N497_HexNAc(5)Hex(6)Fuc(2)NeuAc(2) | N497 | Yes | P02788\|TRFL_N497 |
| R.TAGWNIPMGLLFN[+2863.041]QTGSC[+57.021]KFDEYFSQSC[+57.021]APGSDPR.S | HexNAc(5)Hex(6)Fuc(4)NeuAc(1) | P02788 | LTF | Lactotransferrin | P02788\|TRFL | P02788\|TRFL_N497_HexNAc(5)Hex(6)Fuc(4)NeuAc(1) | N497 | Yes | P02788\|TRFL_N497 |
| R.TAGWNIPMGLLFN[+2278.809]QTGSC[+57.021]K.F | HexNAc(5)Hex(6)NeuAc(1) | P02788 | LTF | Lactotransferrin | P02788\|TRFL | P02788\|TRFL_N497_HexNAc(5)Hex(6)NeuAc(1) | N497 | Yes | P02788\|TRFL_N497 |
| R.[+42.011]TAGWNIPMGLLFN[+1565.560]QTGSC[+57.021]KFDEYFSQSC[+57.021]APGSDPR.S | HexNAc(3)Hex(5)Fuc(1) | P02788 | LTF | Lactotransferrin | P02788\|TRFL | P02788\|TRFL_N497_HexNAc(3)Hex(5)Fuc(1) | N497 | Yes | P02788\|TRFL_N497 |
| R.[+42.011]TAGWNIPMGLLFN[+2018.708]QTGSC[+57.021]KFDEYFSQSC[+57.021]APGSDPR.S | HexNAc(3)Hex(6)Fuc(1)NeuAc(1) | P02788 | LTF | Lactotransferrin | P02788\|TRFL | P02788\|TRFL_N497_HexNAc(3)Hex(6)Fuc(1)NeuAc(1) | N497 | Yes | P02788\|TRFL_N497 |
| R.[+42.011]TAGWNIPMGLLFN[+1872.651]QTGSC[+57.021]K.F | HexNAc(3)Hex(6)NeuAc(1) | P02788 | LTF | Lactotransferrin | P02788\|TRFL | P02788\|TRFL_N497_HexNAc(3)Hex(6)NeuAc(1) | N497 | Yes | P02788\|TRFL_N497 |
| R.[+42.011]TAGWNIPMGLLFN[+1872.651]QTGSC[+57.021]KFDEYFSQSC[+57.021]APGSDPR.S | HexNAc(3)Hex(6)NeuAc(1) | P02788 | LTF | Lactotransferrin | P02788\|TRFL | P02788\|TRFL_N497_HexNAc(3)Hex(6)NeuAc(1) | N497 | Yes | P02788\|TRFL_N497 |
| R.TAGWNIPM[+15.995]GLLFN[+1751.624]QTGSC[+57.021]K.F | HexNAc(4)Hex(4)NeuAc(1) | P02788 | LTF | Lactotransferrin | P02788\|TRFL | P02788\|TRFL_N497_HexNAc(4)Hex(4)NeuAc(1) | N497 | Yes | P02788\|TRFL_N497 |
| R.TAGWNIPM[+15.995]GLLFN[+1751.624]QTGSC[+57.021]KFDEYFSQSC[+57.021]APGSDPR.S | HexNAc(4)Hex(4)NeuAc(1) | P02788 | LTF | Lactotransferrin | P02788\|TRFL | P02788\|TRFL_N497_HexNAc(4)Hex(4)NeuAc(1) | N497 | Yes | P02788\|TRFL_N497 |
| R.TAGWNIPM[+15.995]GLLFN[+1622.582]QTGSC[+57.021]K.F | HexNAc(4)Hex(5) | P02788 | LTF | Lactotransferrin | P02788\|TRFL | P02788\|TRFL_N497_HexNAc(4)Hex(5) | N497 | Yes | P02788\|TRFL_N497 |

Table S5. Cont.

| Glycopeptide | Glycan | Uniprot ID | Gene Name | Protein Name |
|---|---|---|---|---|
| R.TAGWNIPM[+15.995]GLLFN[+1768.640]QTGSC[+57.021]K.F | HexNAc(4)Hex(5)Fuc(1) | P02788 | LTF | Lactotransferrin |
| R.[+42.011]TAGWNIPMGLLFN[+2059.735]QTGSC[+57.021]KFDEYFSQSC[+57.021]APGSDPR.S | HexNAc(4)Hex(5)Fuc(1)NeuAc(1) | P02788 | LTF | Lactotransferrin |
| R.TAGWNIPM[+15.995]GLLFN[+2059.735]QTGSC[+57.021]K.F | HexNAc(4)Hex(5)Fuc(1)NeuAc(1) | P02788 | LTF | Lactotransferrin |
| R.TAGWNIPM[+15.995]GLLFN[+2059.735]QTGSC[+57.021]KFDEYFSQSC[+57.021]APGSDPR.S | HexNAc(4)Hex(5)Fuc(1)NeuAc(1) | P02788 | LTF | Lactotransferrin |
| R.[+42.011]TAGWNIPMGLLFN[+2351.851]QTGSC[+57.021]K.F | HexNAc(4)Hex(5)Fuc(3)NeuAc(1) | P02788 | LTF | Lactotransferrin |
| R.TAGWNIPM[+15.995]GLLFN[+2351.851]QTGSC[+57.021]KFDEYFSQSC[+57.021]APGSDPR.S | HexNAc(4)Hex(5)Fuc(3)NeuAc(1) | P02788 | LTF | Lactotransferrin |
| R.TAGWNIPM[+15.995]GLLFN[+1913.677]QTGSC[+57.021]K.F | HexNAc(4)Hex(5)NeuAc(1) | P02788 | LTF | Lactotransferrin |
| R.TAGWNIPM[+15.995]GLLFN[+2204.772]QTGSC[+57.021]K.F | HexNAc(4)Hex(5)NeuAc(2) | P02788 | LTF | Lactotransferrin |
| R.TAGWNIPM[+15.995]GLLFN[+2116.756]QTGSC[+57.021]KFDEYFSQSC[+57.021]APGSDPR.S | HexNAc(5)Hex(5)NeuAc(1) | P02788 | LTF | Lactotransferrin |
| R.TAGWNIPM[+15.995]GLLFN[+2190.793]QTGSC[+57.021]KFDEYFSQSC[+57.021]APGSDPR.S | HexNAc(6)Hex(6) | P02788 | LTF | Lactotransferrin |
| R.[+42.011]TAGWNIPMGLLFN[+2336.851]QTGSC[+57.021]K.F | HexNAc(6)Hex(6)Fuc(1) | P02788 | LTF | Lactotransferrin |
| K.FGRN[+2059.735]GSDC[+57.021]PDKFC[+57.021]LFQSETK.N | HexNAc(4)Hex(5)Fuc(1)NeuAc(1) | P02788 | LTF | Lactotransferrin |
| R.N[+2059.735]GSDC[+57.021]PDKFC[+57.021]LFQSETK.N | HexNAc(4)Hex(5)Fuc(1)NeuAc(1) | P02788 | LTF | Lactotransferrin |
| K.FGRN[+2350.830]GSDC[+57.021]PDKFC[+57.021]LFQSETK.N | HexNAc(4)Hex(5)Fuc(1)NeuAc(2) | P02788 | LTF | Lactotransferrin |
| R.N[+2350.830]GSDC[+57.021]PDKFC[+57.021]LFQSETK.N | HexNAc(4)Hex(5)Fuc(1)NeuAc(2) | P02788 | LTF | Lactotransferrin |
| K.FGRN[+2205.793]GSDC[+57.021]PDKFC[+57.021]LFQSETK.N | HexNAc(4)Hex(5)Fuc(2)NeuAc(1) | P02788 | LTF | Lactotransferrin |
| R.N[+2205.793]GSDC[+57.021]PDKFC[+57.021]LFQSETK.N | HexNAc(4)Hex(5)Fuc(2)NeuAc(1) | P02788 | LTF | Lactotransferrin |
| R.N[+2351.851]GSDC[+57.021]PDKFC[+57.021]LFQSETK.N | HexNAc(4)Hex(5)Fuc(3)NeuAc(1) | P02788 | LTF | Lactotransferrin |
| R.N[+2204.772]GSDC[+57.021]PDKFC[+57.021]LFQSETK.N | HexNAc(4)Hex(5)NeuAc(2) | P02788 | LTF | Lactotransferrin |
| K.FGRN[+1501.555]GSDC[+57.021]PDKFC[+57.021]LFQSETK.N | HexNAc(5)Hex(3) | P02788 | LTF | Lactotransferrin |
| R.N[+2570.925]GSDC[+57.021]PDKFC[+57.021]LFQSETK.N | HexNAc(5)Hex(6)Fuc(2)NeuAc(1) | P02788 | LTF | Lactotransferrin |
| R.N[+2571.946]GSDC[+57.021]PDKFC[+57.021]LFQSETK.N | HexNAc(5)Hex(6)Fuc(4) | P02788 | LTF | Lactotransferrin |
| R.N[+2569.905]GSDC[+57.021]PDKFC[+57.021]LFQSETK.N | HexNAc(5)Hex(6)NeuAc(2) | P02788 | LTF | Lactotransferrin |
| R.FSLLGHASISC[+57.021]TVEN[+2350.830]ETIGVWRPSPPTC[+57.021]EK.I | HexNAc(4)Hex(5)Fuc(1)NeuAc(2) | P04003 | C4BPA | C4b-binding protein alpha chain |
| K.GNPEQTPVLKPEEEAPAPEVGASKPEGIDSRPETLHPGRPQPPAEEELC[+57.021]SGKPFDAFTDLKN[+2205.793]GSLFAFR.G | HexNAc(4)Hex(5)Fuc(2)NeuAc(1) | P04004 | VTN | Vitronectin |
| R.ADGTVNQIEGEATPVN[+1054.370]LTEPAKLEVK.F | HexNAc(2)Hex(4) | P05090 | APOD | Apolipoprotein D |
| R.ADGTVNQIEGEATPVN[+1216.423]LTEPAKLEVK.F | HexNAc(2)Hex(5) | P05090 | APOD | Apolipoprotein D |
| R.SC[+57.021]PAC[+57.021]PGSN[+876.322]ITIR.N | HexNAc(2)Hex(2)Fuc(1) | P05164 | MPO | Myeloperoxidase |
| R.NLRN[+1216.423]M[+15.995]SNQLGLLAVNQR.F | HexNAc(2)Hex(5) | P05164 | MPO | Myeloperoxidase |
| R.NLRN[+1378.476]M[+15.995]SNQLGLLAVNQR.F | HexNAc(2)Hex(6) | P05164 | MPO | Myeloperoxidase |
| R.ALLPFDNLHDDPC[+57.021]LLTN[+1216.423]R.S | HexNAc(2)Hex(5) | P05164 | MPO | Myeloperoxidase |
| R.SYN[+1038.375]DSVDPR.I | HexNAc(2)Hex(3)Fuc(1) | P05164 | MPO | Myeloperoxidase |
| R.SYN[+1241.454]DSVDPR.I | HexNAc(3)Hex(3)Fuc(1) | P05164 | MPO | Myeloperoxidase |
| R.GIASVLQELN[+876.322]VTVVTSLC[+57.021]R.R | HexNAc(2)Hex(2)Fuc(1) | P08246 | ELANE | Neutrophil elastase |
| R.VVLGAHN[+730.264]LSR.R | HexNAc(2)Hex(2) | P08246 | ELANE | Neutrophil elastase |
| R.EDFVLTAAHC[+57.021]WGSNIN[+203.079]VTLGAHNIQR.R | HexNAc(1) | P08311 | CTSG | Cathepsin G |
| R.LRN[+1216.423]VSWATGR.S | HexNAc(2)Hex(5) | P08571 | CD14 | Monocyte differentiation antigen CD14 |
| R.N[+1216.423]VSWATGR.S | HexNAc(2)Hex(5) | P08571 | CD14 | Monocyte differentiation antigen CD14 |
| R.LRN[+1913.677]VSWATGR.S | HexNAc(4)Hex(5)NeuAc(1) | P08571 | CD14 | Monocyte differentiation antigen CD14 |
| R.C[+57.021]MWSSALNSLN[+1864.634]LSFAGLEQVPK.G | HexNAc(2)Hex(9) | P08571 | CD14 | Monocyte differentiation antigen CD14 |
| R.TYNVLDMKN[+2059.735]TTC[+57.021]QDLQIEVTVK.G | HexNAc(4)Hex(5)Fuc(1)NeuAc(1) | P0C0L4 | C4A | Complement C4-A |
| R.FSDGLESN[+1702.581]SSTQFEVKK.Y | HexNAc(2)Hex(8) | P0C0L4 | C4A | Complement C4-A |
| R.FSDGLESN[+1864.634]SSTQFEVK.K | HexNAc(2)Hex(9) | P0C0L4 | C4A | Complement C4-A |
| R.FSDGLESN[+1864.634]SSTQFEVKK.Y | HexNAc(2)Hex(9) | P0C0L4 | C4A | Complement C4-A |
| K.VFPLSLDSTPQDGNVVVAC[+57.021]LVQGFFPQEPLSVTWSESGQN[+2279.830]VTAR.N | HexNAc(5)Hex(6)Fuc(2) | P0DOX2 | N/A | Immunoglobulin alpha-2 heavy chain |
| K.TPLTAN[+2553.910]ITK.S | HexNAc(5)Hex(5)Fuc(1)NeuAc(2) | P0DOX2 | N/A | Immunoglobulin alpha-2 heavy chain |
| R.DLC[+57.021]GC[+57.021]YSVSSVLPGC[+57.021]AQPWNHGETFTC[+57.021]TAAHPELKTPLTAN[+1444.534]ITK.S | HexNAc(4)Hex(3)Fuc(1) | P0DOX2 | N/A | Immunoglobulin alpha-2 heavy chain |
| R.DLC[+57.021]GC[+57.021]YSVSSVLPGC[+57.021]AQPWNHGETFTC[+57.021]TAAHPELKTPLTAN[+1809.666]ITK.S | HexNAc(5)Hex(4)Fuc(1) | P0DOX2 | N/A | Immunoglobulin alpha-2 heavy chain |
| K.TKPREEQYN[+1444.534]STYR.V | HexNAc(4)Hex(3)Fuc(1) | P0DOX5 | N/A | Immunoglobulin gamma-1 heavy chain |
| K.TKPREEQYN[+1606.587]STYR.V | HexNAc(4)Hex(4)Fuc(1) | P0DOX5 | N/A | Immunoglobulin gamma-1 heavy chain |
| K.TKPREEQYN[+1768.640]STYR.V | HexNAc(4)Hex(5)Fuc(1) | P0DOX5 | N/A | Immunoglobulin gamma-1 heavy chain |
| K.ELPGVC[+57.021]N[+2059.735]ETMMALWEEC[+57.021]KPC[+57.021]LK.Q | HexNAc(4)Hex(5)Fuc(1)NeuAc(1) | P10909 | CLU | Clusterin |
| K.LKELPGVC[+57.021]N[+2059.735]ETMMALWEEC[+57.021]KPC[+57.021]LK.Q | HexNAc(4)Hex(5)Fuc(1)NeuAc(1) | P10909 | CLU | Clusterin |
| K.ELPGVC[+57.021]N[+2350.830]ETMMALWEEC[+57.021]KPC[+57.021]LK.Q | HexNAc(4)Hex(5)Fuc(1)NeuAc(2) | P10909 | CLU | Clusterin |
| K.LKELPGVC[+57.021]N[+2350.830]ETMMALWEEC[+57.021]KPC[+57.021]LK.Q | HexNAc(4)Hex(5)Fuc(1)NeuAc(2) | P10909 | CLU | Clusterin |
| K.LKELPGVC[+57.021]N[+2205.793]ETMMALWEEC[+57.021]KPC[+57.021]LK.Q | HexNAc(4)Hex(5)Fuc(2)NeuAc(1) | P10909 | CLU | Clusterin |
| K.LKELPGVC[+57.021]N[+2351.851]ETMMALWEEC[+57.021]KPC[+57.021]LK.Q | HexNAc(4)Hex(5)Fuc(3)NeuAc(1) | P10909 | CLU | Clusterin |
| K.LKELPGVC[+57.021]N[+2204.772]ETMMALWEEC[+57.021]KPC[+57.021]LK.Q | HexNAc(4)Hex(5)NeuAc(2) | P10909 | CLU | Clusterin |
| K.LKELPGVC[+57.021]N[+2424.867]ETMMALWEEC[+57.021]KPC[+57.021]LK.Q | HexNAc(5)Hex(6)Fuc(1)NeuAc(1) | P10909 | CLU | Clusterin |
| K.LKELPGVC[+57.021]N[+2279.830]ETMMALWEEC[+57.021]KPC[+57.021]LK.Q | HexNAc(5)Hex(6)Fuc(2) | P10909 | CLU | Clusterin |
| K.LKELPGVC[+57.021]N[+2570.925]ETMMALWEEC[+57.021]KPC[+57.021]LK.Q | HexNAc(5)Hex(6)Fuc(2)NeuAc(1) | P10909 | CLU | Clusterin |
| K.LKELPGVC[+57.021]N[+2716.983]ETMMALWEEC[+57.021]KPC[+57.021]LK.Q | HexNAc(5)Hex(6)Fuc(3)NeuAc(1) | P10909 | CLU | Clusterin |
| K.LKELPGVC[+57.021]N[+2571.946]ETMMALWEEC[+57.021]KPC[+57.021]LK.Q | HexNAc(5)Hex(6)Fuc(4) | P10909 | CLU | Clusterin |
| K.LKELPGVC[+57.021]N[+2569.905]ETMMALWEEC[+57.021]KPC[+57.021]LK.Q | HexNAc(5)Hex(6)NeuAc(2) | P10909 | CLU | Clusterin |
| K.LKELPGVC[+57.021]N[+2190.793]ETM[+15.995]MALWEEC[+57.021]KPC[+57.021]LK.Q | HexNAc(6)Hex(6) | P10909 | CLU | Clusterin |
| K.LKELPGVC[+57.021]N[+2336.851]ETM[+15.995]MALWEEC[+57.021]KPC[+57.021]LK.Q | HexNAc(6)Hex(6)Fuc(1) | P10909 | CLU | Clusterin |
| R.QLEEFLN[+2059.735]QSSPFYFWMNGDR.I | HexNAc(4)Hex(5)Fuc(1)NeuAc(1) | P10909 | CLU | Clusterin |
| R.QLEEFLN[+2351.851]QSSPFYFWMNGDRIDSLLENDR.Q | HexNAc(4)Hex(5)Fuc(3)NeuAc(1) | P10909 | CLU | Clusterin |
| R.EIRHN[+2059.735]STGC[+57.021]LR.M | HexNAc(4)Hex(5)Fuc(1)NeuAc(1) | P10909 | CLU | Clusterin |
| R.EIRHN[+2350.830]STGC[+57.021]LR.M | HexNAc(4)Hex(5)Fuc(1)NeuAc(2) | P10909 | CLU | Clusterin |
| R.EIRHN[+2205.793]STGC[+57.021]LR.M | HexNAc(4)Hex(5)Fuc(2)NeuAc(1) | P10909 | CLU | Clusterin |
| R.EIRHN[+2351.851]STGC[+57.021]LR.M | HexNAc(4)Hex(5)Fuc(3)NeuAc(1) | P10909 | CLU | Clusterin |
| R.EIRHN[+2424.867]STGC[+57.021]LR.M | HexNAc(5)Hex(6)Fuc(1)NeuAc(1) | P10909 | CLU | Clusterin |
| R.EIRHN[+2279.830]STGC[+57.021]LR.M | HexNAc(5)Hex(6)Fuc(2) | P10909 | CLU | Clusterin |
| R.EIRHN[+2570.925]STGC[+57.021]LR.M | HexNAc(5)Hex(6)Fuc(2)NeuAc(1) | P10909 | CLU | Clusterin |
| K.MLN[+1458.442]TSSLLEQLNEQFNWVSR.L | HexNAc(2)Hex(6)Phospho(1) | P10909 | CLU | Clusterin |
| K.MLN[+1856.656]TSSLLEQLNEQFNWVSR.L | HexNAc(3)Hex(5)Fuc(1)NeuAc(1) | P10909 | CLU | Clusterin |
| K.MLN[+1710.598]TSSLLEQLNEQFNWVSR.L | HexNAc(3)Hex(5)NeuAc(1) | P10909 | CLU | Clusterin |
| K.MLN[+1872.651]TSSLLEQLNEQFNWVSR.L | HexNAc(3)Hex(6)NeuAc(1) | P10909 | CLU | Clusterin |
| K.MLN[+2059.735]TSSLLEQLNEQFNWVSR.L | HexNAc(4)Hex(5)Fuc(1)NeuAc(1) | P10909 | CLU | Clusterin |
| K.MLN[+2350.830]TSSLLEQLNEQFNWVSR.L | HexNAc(4)Hex(5)Fuc(1)NeuAc(2) | P10909 | CLU | Clusterin |
| K.MLN[+2205.793]TSSLLEQLNEQFNWVSR.L | HexNAc(4)Hex(5)Fuc(2)NeuAc(1) | P10909 | CLU | Clusterin |
| K.MLN[+2351.851]TSSLLEQLNEQFNWVSR.L | HexNAc(4)Hex(5)Fuc(3)NeuAc(1) | P10909 | CLU | Clusterin |
| K.MLN[+1913.677]TSSLLEQLNEQFNWVSR.L | HexNAc(4)Hex(5)NeuAc(1) | P10909 | CLU | Clusterin |
| K.MLN[+2204.772]TSSLLEQLNEQFNWVSR.L | HexNAc(4)Hex(5)NeuAc(2) | P10909 | CLU | Clusterin |
| K.MLN[+2076.750]TSSLLEQLNEQFNWVSR.L | HexNAc(4)Hex(6)Fuc(2) | P10909 | CLU | Clusterin |
| K.MLN[+2424.867]TSSLLEQLNEQFNWVSR.L | HexNAc(5)Hex(6)Fuc(1)NeuAc(1) | P10909 | CLU | Clusterin |

| Glycopeptide | Shortname | ProteinNameInSkyline | Glycosites | Reported before | Site |
|---|---|---|---|---|---|
| R.TAGWNIPM[+15.995]GLLFN[+1768.640]QTGSC[+57.021]K.F | P02788\|TRFL | P02788\|TRFL_N497_HexNAc(4)Hex(5)Fuc(1) | N497 | Yes | P02788\|TRFL_N497 |
| R.[+42.011]TAGWNIPMGLLFN[+2059.735]QTGSC[+57.021]KFDEYFSQSC[+57.021]APGSDPR.S | P02788\|TRFL | P02788\|TRFL_N497_HexNAc(4)Hex(5)Fuc(1)NeuAc(1) | N497 | Yes | P02788\|TRFL_N497 |
| R.TAGWNIPM[+15.995]GLLFN[+2059.735]QTGSC[+57.021]K.F | P02788\|TRFL | P02788\|TRFL_N497_HexNAc(4)Hex(5)Fuc(1)NeuAc(1) | N497 | Yes | P02788\|TRFL_N497 |
| R.TAGWNIPM[+15.995]GLLFN[+2059.735]QTGSC[+57.021]KFDEYFSQSC[+57.021]APGSDPR.S | P02788\|TRFL | P02788\|TRFL_N497_HexNAc(4)Hex(5)Fuc(1)NeuAc(1) | N497 | Yes | P02788\|TRFL_N497 |
| R.[+42.011]TAGWNIPMGLLFN[+2351.851]QTGSC[+57.021]K.F | P02788\|TRFL | P02788\|TRFL_N497_HexNAc(4)Hex(5)Fuc(3)NeuAc(1) | N497 | Yes | P02788\|TRFL_N497 |
| R.TAGWNIPM[+15.995]GLLFN[+2351.851]QTGSC[+57.021]KFDEYFSQSC[+57.021]APGSDPR.S | P02788\|TRFL | P02788\|TRFL_N497_HexNAc(4)Hex(5)Fuc(3)NeuAc(1) | N497 | Yes | P02788\|TRFL_N497 |
| R.TAGWNIPM[+15.995]GLLFN[+1913.677]QTGSC[+57.021]K.F | P02788\|TRFL | P02788\|TRFL_N497_HexNAc(4)Hex(5)NeuAc(1) | N497 | Yes | P02788\|TRFL_N497 |
| R.TAGWNIPM[+15.995]GLLFN[+2204.772]QTGSC[+57.021]K.F | P02788\|TRFL | P02788\|TRFL_N497_HexNAc(4)Hex(5)NeuAc(2) | N497 | Yes | P02788\|TRFL_N497 |
| R.TAGWNIPM[+15.995]GLLFN[+2116.756]QTGSC[+57.021]KFDEYFSQSC[+57.021]APGSDPR.S | P02788\|TRFL | P02788\|TRFL_N497_HexNAc(5)Hex(5)NeuAc(1) | N497 | Yes | P02788\|TRFL_N497 |
| R.TAGWNIPM[+15.995]GLLFN[+2190.793]QTGSC[+57.021]KFDEYFSQSC[+57.021]APGSDPR.S | P02788\|TRFL | P02788\|TRFL_N497_HexNAc(6)Hex(6) | N497 | Yes | P02788\|TRFL_N497 |
| R.[+42.011]TAGWNIPMGLLFN[+2336.851]QTGSC[+57.021]K.F | P02788\|TRFL | P02788\|TRFL_N497_HexNAc(6)Hex(6)Fuc(1) | N497 | Yes | P02788\|TRFL_N497 |
| K.FGRN[+2059.735]GSDC[+57.021]PDKFC[+57.021]LFQSETK.N | P02788\|TRFL | P02788\|TRFL_N642_HexNAc(4)Hex(5)Fuc(1)NeuAc(1) | N642 | Yes | P02788\|TRFL_N642 |
| R.N[+2059.735]GSDC[+57.021]PDKFC[+57.021]LFQSETK.N | P02788\|TRFL | P02788\|TRFL_N642_HexNAc(4)Hex(5)Fuc(1)NeuAc(1) | N642 | Yes | P02788\|TRFL_N642 |
| K.FGRN[+2350.830]GSDC[+57.021]PDKFC[+57.021]LFQSETK.N | P02788\|TRFL | P02788\|TRFL_N642_HexNAc(4)Hex(5)Fuc(1)NeuAc(2) | N642 | Yes | P02788\|TRFL_N642 |
| R.N[+2350.830]GSDC[+57.021]PDKFC[+57.021]LFQSETK.N | P02788\|TRFL | P02788\|TRFL_N642_HexNAc(4)Hex(5)Fuc(1)NeuAc(2) | N642 | Yes | P02788\|TRFL_N642 |
| K.FGRN[+2205.793]GSDC[+57.021]PDKFC[+57.021]LFQSETK.N | P02788\|TRFL | P02788\|TRFL_N642_HexNAc(4)Hex(5)Fuc(2)NeuAc(1) | N642 | Yes | P02788\|TRFL_N642 |
| R.N[+2205.793]GSDC[+57.021]PDKFC[+57.021]LFQSETK.N | P02788\|TRFL | P02788\|TRFL_N642_HexNAc(4)Hex(5)Fuc(2)NeuAc(1) | N642 | Yes | P02788\|TRFL_N642 |
| R.N[+2351.851]GSDC[+57.021]PDKFC[+57.021]LFQSETK.N | P02788\|TRFL | P02788\|TRFL_N642_HexNAc(4)Hex(5)Fuc(3)NeuAc(1) | N642 | Yes | P02788\|TRFL_N642 |
| R.N[+2204.772]GSDC[+57.021]PDKFC[+57.021]LFQSETK.N | P02788\|TRFL | P02788\|TRFL_N642_HexNAc(4)Hex(5)NeuAc(2) | N642 | Yes | P02788\|TRFL_N642 |
| K.FGRN[+1501.555]GSDC[+57.021]PDKFC[+57.021]LFQSETK.N | P02788\|TRFL | P02788\|TRFL_N642_HexNAc(5)Hex(3) | N642 | Yes | P02788\|TRFL_N642 |
| R.N[+2570.925]GSDC[+57.021]PDKFC[+57.021]LFQSETK.N | P02788\|TRFL | P02788\|TRFL_N642_HexNAc(5)Hex(6)Fuc(2)NeuAc(1) | N642 | Yes | P02788\|TRFL_N642 |
| R.N[+2571.946]GSDC[+57.021]PDKFC[+57.021]LFQSETK.N | P02788\|TRFL | P02788\|TRFL_N642_HexNAc(5)Hex(6)Fuc(4) | N642 | Yes | P02788\|TRFL_N642 |
| R.N[+2569.905]GSDC[+57.021]PDKFC[+57.021]LFQSETK.N | P02788\|TRFL | P02788\|TRFL_N642_HexNAc(5)Hex(6)NeuAc(2) | N642 | Yes | P02788\|TRFL_N642 |
| R.FSLLGHASISC[+57.021]TVEN[+2350.830]ETIGVWRPSPPTC[+57.021]EK.I | P04003\|C4BPA | P04003\|C4BPA_N221_HexNAc(4)Hex(5)Fuc(1)NeuAc(2) | N221 | Yes | P04003\|C4BPA_N221 |
| K.GNPEQTPVLKPEEEAPAPEVGASKPEGIDSRPETLHPGRPQPPAEEELC[+57.021]SGKPFDAFTDLKN[+2205.793]GSLFAFR.G | P04004\|VTNC | P04004\|VTNC_N169_HexNAc(4)Hex(5)Fuc(2)NeuAc(1) | N169 | Yes | P04004\|VTNC_N169 |
| R.ADGTVNQIEGEATPVN[+1054.370]LTEPAKLEVK.F | P05090\|APOD | P05090\|APOD_N98_HexNAc(2)Hex(4) | N98 | Yes | P05090\|APOD_N98 |
| R.ADGTVNQIEGEATPVN[+1216.423]LTEPAKLEVK.F | P05090\|APOD | P05090\|APOD_N98_HexNAc(2)Hex(5) | N98 | Yes | P05090\|APOD_N98 |
| R.SC[+57.021]PAC[+57.021]PGSN[+876.322]ITIR.N | P05164\|PERM | P05164\|PERM_N323_HexNAc(2)Hex(2)Fuc(1) | N323 | Yes | P05164\|PERM_N323 |
| R.NLRN[+1216.423]M[+15.995]SNQLGLLAVNQR.F | P05164\|PERM | P05164\|PERM_N355_HexNAc(2)Hex(5) | N355 | Yes | P05164\|PERM_N355 |
| R.NLRN[+1378.476]M[+15.995]SNQLGLLAVNQR.F | P05164\|PERM | P05164\|PERM_N355_HexNAc(2)Hex(6) | N355 | Yes | P05164\|PERM_N355 |
| R.ALLPFDNLHDDPC[+57.021]LLTN[+1216.423]R.S | P05164\|PERM | P05164\|PERM_N391_HexNAc(2)Hex(5) | N391 | Yes | P05164\|PERM_N391 |
| R.SYN[+1038.375]DSVDPR.I | P05164\|PERM | P05164\|PERM_N483_HexNAc(2)Hex(3)Fuc(1) | N483 | Yes | P05164\|PERM_N483 |
| R.SYN[+1241.454]DSVDPR.I | P05164\|PERM | P05164\|PERM_N483_HexNAc(3)Hex(3)Fuc(1) | N483 | Yes | P05164\|PERM_N483 |
| R.GIASVLQELN[+876.322]VTVVTSLC[+57.021]R.R | P08246\|ELNE | P08246\|ELNE_N173_HexNAc(2)Hex(2)Fuc(1) | N173 | Yes | P08246\|ELNE_N173 |
| R.VVLGAHN[+730.264]LSR.R | P08246\|ELNE | P08246\|ELNE_N88_HexNAc(2)Hex(2) | N88 | Yes | P08246\|ELNE_N88 |
| R.EDFVLTAAHC[+57.021]WGSNIN[+203.079]VTLGAHNIQR.R | P08311\|CATG | P08311\|CATG_N71_HexNAc(1) | N71 | Yes | P08311\|CATG_N71 |
| R.LRN[+1216.423]VSWATGR.S | P08571\|CD14 | P08571\|CD14_N151_HexNAc(2)Hex(5) | N151 | Yes | P08571\|CD14_N151 |
| R.N[+1216.423]VSWATGR.S | P08571\|CD14 | P08571\|CD14_N151_HexNAc(2)Hex(5) | N151 | Yes | P08571\|CD14_N151 |
| R.LRN[+1913.677]VSWATGR.S | P08571\|CD14 | P08571\|CD14_N151_HexNAc(4)Hex(5)NeuAc(1) | N151 | Yes | P08571\|CD14_N151 |
| R.C[+57.021]MWSSALNSLN[+1864.634]LSFAGLEQVPK.G | P08571\|CD14 | P08571\|CD14_N282_HexNAc(2)Hex(9) | N282 | Yes | P08571\|CD14_N282 |
| R.TYNVLDMKN[+2059.735]TTC[+57.021]QDLQIEVTVK.G | P0C0L4\|CO4A | P0C0L4\|CO4A_N1391_HexNAc(4)Hex(5)Fuc(1)NeuAc(1) | N1391 | Yes | P0C0L4\|CO4A_N1391 |
| R.FSDGLESN[+1702.581]SSTQFEVKK.Y | P0C0L4\|CO4A | P0C0L4\|CO4A_N226_HexNAc(2)Hex(8) | N226 | Yes | P0C0L4\|CO4A_N226 |
| R.FSDGLESN[+1864.634]SSTQFEVK.K | P0C0L4\|CO4A | P0C0L4\|CO4A_N226_HexNAc(2)Hex(9) | N226 | Yes | P0C0L4\|CO4A_N226 |
| R.FSDGLESN[+1864.634]SSTQFEVKK.Y | P0C0L4\|CO4A | P0C0L4\|CO4A_N226_HexNAc(2)Hex(9) | N226 | Yes | P0C0L4\|CO4A_N226 |
| K.VFPLSLDSTPQDGNVVVAC[+57.021]LVQGFFPQEPLSVTWSESGQN[+2279.830]VTAR.N | P0DOX2\|IGA2 | P0DOX2\|IGA2_N162_HexNAc(5)Hex(6)Fuc(2) | N162 | Yes | P0DOX2\|IGA2_N162 |
| K.TPLTAN[+2553.910]ITK.S | P0DOX2\|IGA2 | P0DOX2\|IGA2_N320_HexNAc(5)Hex(5)Fuc(1)NeuAc(2) | N320 | Yes | P0DOX2\|IGA2_N320 |
| R.DLC[+57.021]GC[+57.021]YSVSSVLPGC[+57.021]AQPWNHGETFTC[+57.021]TAAHPELKTPLTAN[+1444.534]ITK.S | P0DOX2\|IGA2 | P0DOX2\|IGA2_N320_HexNAc(4)Hex(3)Fuc(1) | N320 | Yes | P0DOX2\|IGA2_N320 |
| R.DLC[+57.021]GC[+57.021]YSVSSVLPGC[+57.021]AQPWNHGETFTC[+57.021]TAAHPELKTPLTAN[+1809.666]ITK.S | P0DOX2\|IGA2 | P0DOX2\|IGA2_N320_HexNAc(5)Hex(4)Fuc(1) | N320 | Yes | P0DOX2\|IGA2_N320 |
| K.TKPREEQYN[+1444.534]STYR.V | P0DOX5\|IGG1 | P0DOX5\|IGG1_N299_HexNAc(4)Hex(3)Fuc(1) | N299 | Yes | P0DOX5\|IGG1_N299 |
| K.TKPREEQYN[+1606.587]STYR.V | P0DOX5\|IGG1 | P0DOX5\|IGG1_N299_HexNAc(4)Hex(4)Fuc(1) | N299 | Yes | P0DOX5\|IGG1_N299 |
| K.TKPREEQYN[+1768.640]STYR.V | P0DOX5\|IGG1 | P0DOX5\|IGG1_N299_HexNAc(4)Hex(5)Fuc(1) | N299 | Yes | P0DOX5\|IGG1_N299 |
| K.ELPGVC[+57.021]N[+2059.735]ETMMALWEEC[+57.021]KPC[+57.021]LK.Q | P10909\|CLUS | P10909\|CLUS_N103_HexNAc(4)Hex(5)Fuc(1)NeuAc(1) | N103 | Yes | P10909\|CLUS_N103 |
| K.LKELPGVC[+57.021]N[+2059.735]ETMMALWEEC[+57.021]KPC[+57.021]LK.Q | P10909\|CLUS | P10909\|CLUS_N103_HexNAc(4)Hex(5)Fuc(1)NeuAc(1) | N103 | Yes | P10909\|CLUS_N103 |
| K.ELPGVC[+57.021]N[+2350.830]ETMMALWEEC[+57.021]KPC[+57.021]LK.Q | P10909\|CLUS | P10909\|CLUS_N103_HexNAc(4)Hex(5)Fuc(1)NeuAc(2) | N103 | Yes | P10909\|CLUS_N103 |
| K.LKELPGVC[+57.021]N[+2350.830]ETMMALWEEC[+57.021]KPC[+57.021]LK.Q | P10909\|CLUS | P10909\|CLUS_N103_HexNAc(4)Hex(5)Fuc(1)NeuAc(2) | N103 | Yes | P10909\|CLUS_N103 |
| K.LKELPGVC[+57.021]N[+2205.793]ETMMALWEEC[+57.021]KPC[+57.021]LK.Q | P10909\|CLUS | P10909\|CLUS_N103_HexNAc(4)Hex(5)Fuc(2)NeuAc(1) | N103 | Yes | P10909\|CLUS_N103 |
| K.LKELPGVC[+57.021]N[+2351.851]ETMMALWEEC[+57.021]KPC[+57.021]LK.Q | P10909\|CLUS | P10909\|CLUS_N103_HexNAc(4)Hex(5)Fuc(3)NeuAc(1) | N103 | Yes | P10909\|CLUS_N103 |
| K.LKELPGVC[+57.021]N[+2204.772]ETMMALWEEC[+57.021]KPC[+57.021]LK.Q | P10909\|CLUS | P10909\|CLUS_N103_HexNAc(4)Hex(5)NeuAc(2) | N103 | Yes | P10909\|CLUS_N103 |
| K.LKELPGVC[+57.021]N[+2424.867]ETMMALWEEC[+57.021]KPC[+57.021]LK.Q | P10909\|CLUS | P10909\|CLUS_N103_HexNAc(5)Hex(6)Fuc(1)NeuAc(1) | N103 | Yes | P10909\|CLUS_N103 |
| K.LKELPGVC[+57.021]N[+2279.830]ETMMALWEEC[+57.021]KPC[+57.021]LK.Q | P10909\|CLUS | P10909\|CLUS_N103_HexNAc(5)Hex(6)Fuc(2) | N103 | Yes | P10909\|CLUS_N103 |
| K.LKELPGVC[+57.021]N[+2570.925]ETMMALWEEC[+57.021]KPC[+57.021]LK.Q | P10909\|CLUS | P10909\|CLUS_N103_HexNAc(5)Hex(6)Fuc(2)NeuAc(1) | N103 | Yes | P10909\|CLUS_N103 |
| K.LKELPGVC[+57.021]N[+2716.983]ETMMALWEEC[+57.021]KPC[+57.021]LK.Q | P10909\|CLUS | P10909\|CLUS_N103_HexNAc(5)Hex(6)Fuc(3)NeuAc(1) | N103 | Yes | P10909\|CLUS_N103 |
| K.LKELPGVC[+57.021]N[+2571.946]ETMMALWEEC[+57.021]KPC[+57.021]LK.Q | P10909\|CLUS | P10909\|CLUS_N103_HexNAc(5)Hex(6)Fuc(4) | N103 | Yes | P10909\|CLUS_N103 |
| K.LKELPGVC[+57.021]N[+2569.905]ETMMALWEEC[+57.021]KPC[+57.021]LK.Q | P10909\|CLUS | P10909\|CLUS_N103_HexNAc(5)Hex(6)NeuAc(2) | N103 | Yes | P10909\|CLUS_N103 |
| K.LKELPGVC[+57.021]N[+2190.793]ETM[+15.995]MALWEEC[+57.021]KPC[+57.021]LK.Q | P10909\|CLUS | P10909\|CLUS_N103_HexNAc(6)Hex(6) | N103 | Yes | P10909\|CLUS_N103 |
| K.LKELPGVC[+57.021]N[+2336.851]ETM[+15.995]MALWEEC[+57.021]KPC[+57.021]LK.Q | P10909\|CLUS | P10909\|CLUS_N103_HexNAc(6)Hex(6)Fuc(1) | N103 | Yes | P10909\|CLUS_N103 |
| R.QLEEFLN[+2059.735]QSSPFYFWMNGDR.I | P10909\|CLUS | P10909\|CLUS_N145_HexNAc(4)Hex(5)Fuc(1)NeuAc(1) | N145 | Yes | P10909\|CLUS_N145 |
| R.QLEEFLN[+2351.851]QSSPFYFWMNGDRIDSLLENDR.Q | P10909\|CLUS | P10909\|CLUS_N145_HexNAc(4)Hex(5)Fuc(3)NeuAc(1) | N145 | Yes | P10909\|CLUS_N145 |
| R.EIRHN[+2059.735]STGC[+57.021]LR.M | P10909\|CLUS | P10909\|CLUS_N291_HexNAc(4)Hex(5)Fuc(1)NeuAc(1) | N291 | Yes | P10909\|CLUS_N291 |
| R.EIRHN[+2350.830]STGC[+57.021]LR.M | P10909\|CLUS | P10909\|CLUS_N291_HexNAc(4)Hex(5)Fuc(1)NeuAc(2) | N291 | Yes | P10909\|CLUS_N291 |
| R.EIRHN[+2205.793]STGC[+57.021]LR.M | P10909\|CLUS | P10909\|CLUS_N291_HexNAc(4)Hex(5)Fuc(2)NeuAc(1) | N291 | Yes | P10909\|CLUS_N291 |
| R.EIRHN[+2351.851]STGC[+57.021]LR.M | P10909\|CLUS | P10909\|CLUS_N291_HexNAc(4)Hex(5)Fuc(3)NeuAc(1) | N291 | Yes | P10909\|CLUS_N291 |
| R.EIRHN[+2424.867]STGC[+57.021]LR.M | P10909\|CLUS | P10909\|CLUS_N291_HexNAc(5)Hex(6)Fuc(1)NeuAc(1) | N291 | Yes | P10909\|CLUS_N291 |
| R.EIRHN[+2279.830]STGC[+57.021]LR.M | P10909\|CLUS | P10909\|CLUS_N291_HexNAc(5)Hex(6)Fuc(2) | N291 | Yes | P10909\|CLUS_N291 |
| R.EIRHN[+2570.925]STGC[+57.021]LR.M | P10909\|CLUS | P10909\|CLUS_N291_HexNAc(5)Hex(6)Fuc(2)NeuAc(1) | N291 | Yes | P10909\|CLUS_N291 |
| K.MLN[+1458.442]TSSLLEQLNEQFNWVSR.L | P10909\|CLUS | P10909\|CLUS_N354_HexNAc(2)Hex(6)Phospho(1) | N354 | Yes | P10909\|CLUS_N354 |
| K.MLN[+1856.656]TSSLLEQLNEQFNWVSR.L | P10909\|CLUS | P10909\|CLUS_N354_HexNAc(3)Hex(5)Fuc(1)NeuAc(1) | N354 | Yes | P10909\|CLUS_N354 |
| K.MLN[+1710.598]TSSLLEQLNEQFNWVSR.L | P10909\|CLUS | P10909\|CLUS_N354_HexNAc(3)Hex(5)NeuAc(1) | N354 | Yes | P10909\|CLUS_N354 |
| K.MLN[+1872.651]TSSLLEQLNEQFNWVSR.L | P10909\|CLUS | P10909\|CLUS_N354_HexNAc(3)Hex(6)NeuAc(1) | N354 | Yes | P10909\|CLUS_N354 |
| K.MLN[+2059.735]TSSLLEQLNEQFNWVSR.L | P10909\|CLUS | P10909\|CLUS_N354_HexNAc(4)Hex(5)Fuc(1)NeuAc(1) | N354 | Yes | P10909\|CLUS_N354 |
| K.MLN[+2350.830]TSSLLEQLNEQFNWVSR.L | P10909\|CLUS | P10909\|CLUS_N354_HexNAc(4)Hex(5)Fuc(1)NeuAc(2) | N354 | Yes | P10909\|CLUS_N354 |
| K.MLN[+2205.793]TSSLLEQLNEQFNWVSR.L | P10909\|CLUS | P10909\|CLUS_N354_HexNAc(4)Hex(5)Fuc(2)NeuAc(1) | N354 | Yes | P10909\|CLUS_N354 |
| K.MLN[+2351.851]TSSLLEQLNEQFNWVSR.L | P10909\|CLUS | P10909\|CLUS_N354_HexNAc(4)Hex(5)Fuc(3)NeuAc(1) | N354 | Yes | P10909\|CLUS_N354 |
| K.MLN[+1913.677]TSSLLEQLNEQFNWVSR.L | P10909\|CLUS | P10909\|CLUS_N354_HexNAc(4)Hex(5)NeuAc(1) | N354 | Yes | P10909\|CLUS_N354 |
| K.MLN[+2204.772]TSSLLEQLNEQFNWVSR.L | P10909\|CLUS | P10909\|CLUS_N354_HexNAc(4)Hex(5)NeuAc(2) | N354 | Yes | P10909\|CLUS_N354 |
| K.MLN[+2076.750]TSSLLEQLNEQFNWVSR.L | P10909\|CLUS | P10909\|CLUS_N354_HexNAc(4)Hex(6)Fuc(2) | N354 | Yes | P10909\|CLUS_N354 |
| K.MLN[+2424.867]TSSLLEQLNEQFNWVSR.L | P10909\|CLUS | P10909\|CLUS_N354_HexNAc(5)Hex(6)Fuc(1)NeuAc(1) | N354 | Yes | P10909\|CLUS_N354 |

Table S5. Cont.

<table>
<tr><th>Glycopeptide</th><th>Glycan</th><th>Uniprot ID</th><th>Gene Name</th><th>Protein Name</th><th>Shortname</th><th>ProteinNameInSkyline</th><th>Glycosites</th><th>Reported before</th><th>Site</th></tr>
<tr><td>K.[+42.011]MLN[+2018.708]TSSLLEQLNEQFNWVSR.L</td><td>HexNAc(3)Hex(6)Fuc(1)NeuAc(1)</td><td>P10909</td><td>CLU</td><td>Clusterin</td><td>P10909|CLUS</td><td>P10909|CLUS_N354_HexNAc(3)Hex(6)Fuc(1)NeuAc(1)</td><td>N354</td><td>Yes</td><td>P10909|CLUS_N354</td></tr>
<tr><td>K.M[+15.995]LN[+2059.735]TSSLLEQLNEQFNWVSR.L</td><td>HexNAc(4)Hex(5)Fuc(1)NeuAc(1)</td><td>P10909</td><td>CLU</td><td>Clusterin</td><td>P10909|CLUS</td><td>P10909|CLUS_N354_HexNAc(4)Hex(5)Fuc(1)NeuAc(1)</td><td>N354</td><td>Yes</td><td>P10909|CLUS_N354</td></tr>
<tr><td>K.M[+15.995]LN[+2350.830]TSSLLEQLNEQFNWVSR.L</td><td>HexNAc(4)Hex(5)Fuc(1)NeuAc(2)</td><td>P10909</td><td>CLU</td><td>Clusterin</td><td>P10909|CLUS</td><td>P10909|CLUS_N354_HexNAc(4)Hex(5)Fuc(1)NeuAc(2)</td><td>N354</td><td>Yes</td><td>P10909|CLUS_N354</td></tr>
<tr><td>K.[+42.011]MLN[+2336.851]TSSLLEQLNEQFNWVSR.L</td><td>HexNAc(6)Hex(6)Fuc(1)</td><td>P10909</td><td>CLU</td><td>Clusterin</td><td>P10909|CLUS</td><td>P10909|CLUS_N354_HexNAc(6)Hex(6)Fuc(1)</td><td>N354</td><td>Yes</td><td>P10909|CLUS_N354</td></tr>
<tr><td>K.M[+15.995]LN[+2336.851]TSSLLEQLNEQFNWVSR.L</td><td>HexNAc(6)Hex(6)Fuc(1)</td><td>P10909</td><td>CLU</td><td>Clusterin</td><td>P10909|CLUS</td><td>P10909|CLUS_N354_HexNAc(6)Hex(6)Fuc(1)</td><td>N354</td><td>Yes</td><td>P10909|CLUS_N354</td></tr>
<tr><td>R.LAN[+2059.735]LTQGEDQYYLR.V</td><td>HexNAc(4)Hex(5)Fuc(1)NeuAc(1)</td><td>P10909</td><td>CLU</td><td>Clusterin</td><td>P10909|CLUS</td><td>P10909|CLUS_N374_HexNAc(4)Hex(5)Fuc(1)NeuAc(1)</td><td>N374</td><td>Yes</td><td>P10909|CLUS_N374</td></tr>
<tr><td>R.LAN[+2350.830]LTQGEDQYYLR.V</td><td>HexNAc(4)Hex(5)Fuc(1)NeuAc(2)</td><td>P10909</td><td>CLU</td><td>Clusterin</td><td>P10909|CLUS</td><td>P10909|CLUS_N374_HexNAc(4)Hex(5)Fuc(1)NeuAc(2)</td><td>N374</td><td>Yes</td><td>P10909|CLUS_N374</td></tr>
<tr><td>R.LAN[+2205.793]LTQGEDQYYLR.V</td><td>HexNAc(4)Hex(5)Fuc(2)NeuAc(1)</td><td>P10909</td><td>CLU</td><td>Clusterin</td><td>P10909|CLUS</td><td>P10909|CLUS_N374_HexNAc(4)Hex(5)Fuc(2)NeuAc(1)</td><td>N374</td><td>Yes</td><td>P10909|CLUS_N374</td></tr>
<tr><td>R.LAN[+2351.851]LTQGEDQYYLR.V</td><td>HexNAc(4)Hex(5)Fuc(3)NeuAc(1)</td><td>P10909</td><td>CLU</td><td>Clusterin</td><td>P10909|CLUS</td><td>P10909|CLUS_N374_HexNAc(4)Hex(5)Fuc(3)NeuAc(1)</td><td>N374</td><td>Yes</td><td>P10909|CLUS_N374</td></tr>
<tr><td>R.LAN[+2715.963]LTQGEDQYYLR.V</td><td>HexNAc(5)Hex(6)Fuc(1)NeuAc(2)</td><td>P10909</td><td>CLU</td><td>Clusterin</td><td>P10909|CLUS</td><td>P10909|CLUS_N374_HexNAc(5)Hex(6)Fuc(1)NeuAc(2)</td><td>N374</td><td>Yes</td><td>P10909|CLUS_N374</td></tr>
<tr><td>R.LAN[+2570.925]LTQGEDQYYLR.V</td><td>HexNAc(5)Hex(6)Fuc(2)NeuAc(1)</td><td>P10909</td><td>CLU</td><td>Clusterin</td><td>P10909|CLUS</td><td>P10909|CLUS_N374_HexNAc(5)Hex(6)Fuc(2)NeuAc(1)</td><td>N374</td><td>Yes</td><td>P10909|CLUS_N374</td></tr>
<tr><td>R.LAN[+2862.021]LTQGEDQYYLR.V</td><td>HexNAc(5)Hex(6)Fuc(2)NeuAc(2)</td><td>P10909</td><td>CLU</td><td>Clusterin</td><td>P10909|CLUS</td><td>P10909|CLUS_N374_HexNAc(5)Hex(6)Fuc(2)NeuAc(2)</td><td>N374</td><td>Yes</td><td>P10909|CLUS_N374</td></tr>
<tr><td>R.LAN[+2716.983]LTQGEDQYYLR.V</td><td>HexNAc(5)Hex(6)Fuc(3)NeuAc(1)</td><td>P10909</td><td>CLU</td><td>Clusterin</td><td>P10909|CLUS</td><td>P10909|CLUS_N374_HexNAc(5)Hex(6)Fuc(3)NeuAc(1)</td><td>N374</td><td>Yes</td><td>P10909|CLUS_N374</td></tr>
<tr><td>K.KKEDALN[+2059.735]ETRESETK.L</td><td>HexNAc(4)Hex(5)Fuc(1)NeuAc(1)</td><td>P10909</td><td>CLU</td><td>Clusterin</td><td>P10909|CLUS</td><td>P10909|CLUS_N86_HexNAc(4)Hex(5)Fuc(1)NeuAc(1)</td><td>N86</td><td>Yes</td><td>P10909|CLUS_N86</td></tr>
<tr><td>K.KKEDALN[+2350.830]ETRESETK.L</td><td>HexNAc(4)Hex(5)Fuc(1)NeuAc(2)</td><td>P10909</td><td>CLU</td><td>Clusterin</td><td>P10909|CLUS</td><td>P10909|CLUS_N86_HexNAc(4)Hex(5)Fuc(1)NeuAc(2)</td><td>N86</td><td>Yes</td><td>P10909|CLUS_N86</td></tr>
<tr><td>K.TFYWDFYTN[+2351.851]R.T</td><td>HexNAc(4)Hex(5)Fuc(3)NeuAc(1)</td><td>P12273</td><td>PIP</td><td>Prolactin-inducible protein</td><td>P12273|PIP</td><td>P12273|PIP_N105_HexNAc(4)Hex(5)Fuc(3)NeuAc(1)</td><td>N105</td><td>Yes</td><td>P12273|PIP_N105</td></tr>
<tr><td>K.IISKN[+2205.793]C[+57.021]TSYGVLDISK.C</td><td>HexNAc(4)Hex(5)Fuc(2)NeuAc(1)</td><td>P16671</td><td>CD36</td><td>Platelet glycoprotein 4</td><td>P16671|CD36</td><td>P16671|CD36_N321_HexNAc(4)Hex(5)Fuc(2)NeuAc(1)</td><td>N321</td><td>Yes</td><td>P16671|CD36_N321</td></tr>
<tr><td>K.RNYIVPILWLN[+1864.634]ETGTIGDEK.A</td><td>HexNAc(2)Hex(9)</td><td>P16671</td><td>CD36</td><td>Platelet glycoprotein 4</td><td>P16671|CD36</td><td>P16671|CD36_N417_HexNAc(2)Hex(9)</td><td>N417</td><td>Yes</td><td>P16671|CD36_N417</td></tr>
<tr><td>R.NYIVPILWLN[+1864.634]ETGTIGDEK.A</td><td>HexNAc(2)Hex(9)</td><td>P16671</td><td>CD36</td><td>Platelet glycoprotein 4</td><td>P16671|CD36</td><td>P16671|CD36_N417_HexNAc(2)Hex(9)</td><td>N417</td><td>Yes</td><td>P16671|CD36_N417</td></tr>
<tr><td>R.NYIVPILWLN[+1864.634]ETGTIGDEKANM[+15.995]FR.S</td><td>HexNAc(2)Hex(9)</td><td>P16671</td><td>CD36</td><td>Platelet glycoprotein 4</td><td>P16671|CD36</td><td>P16671|CD36_N417_HexNAc(2)Hex(9)</td><td>N417</td><td>Yes</td><td>P16671|CD36_N417</td></tr>
<tr><td>R.NYIVPILWLN[+1864.634]ETGTIGDEKANMFR.S</td><td>HexNAc(2)Hex(9)</td><td>P16671</td><td>CD36</td><td>Platelet glycoprotein 4</td><td>P16671|CD36</td><td>P16671|CD36_N417_HexNAc(2)Hex(9)</td><td>N417</td><td>Yes</td><td>P16671|CD36_N417</td></tr>
<tr><td>R.QFWIFDVQNPQEVMMN[+2350.830]SSNIQVK.Q</td><td>HexNAc(4)Hex(5)Fuc(1)NeuAc(2)</td><td>P16671</td><td>CD36</td><td>Platelet glycoprotein 4</td><td>P16671|CD36</td><td>P16671|CD36_N79_HexNAc(4)Hex(5)Fuc(1)NeuAc(2)</td><td>N79</td><td>Yes</td><td>P16671|CD36_N79</td></tr>
<tr><td>R.QFWIFDVQNPQEVMMN[+2351.851]SSNIQVK.Q</td><td>HexNAc(4)Hex(5)Fuc(3)NeuAc(1)</td><td>P16671</td><td>CD36</td><td>Platelet glycoprotein 4</td><td>P16671|CD36</td><td>P16671|CD36_N79_HexNAc(4)Hex(5)Fuc(3)NeuAc(1)</td><td>N79</td><td>Yes</td><td>P16671|CD36_N79</td></tr>
<tr><td>R.NIAAFGGDPNN[+1702.581]ITLFGESAGGASVSLQTLSPYNK.G</td><td>HexNAc(2)Hex(8)</td><td>P19835</td><td>CEL</td><td>Bile salt-activated lipase</td><td>P19835|CEL</td><td>P19835|CEL_N207_HexNAc(2)Hex(8)</td><td>N207</td><td>Yes</td><td>P19835|CEL_N207</td></tr>
<tr><td>R.NIAAFGGDPNN[+1622.582]ITLFGESAGGASVSLQTLSPYNK.G</td><td>HexNAc(4)Hex(5)</td><td>P19835</td><td>CEL</td><td>Bile salt-activated lipase</td><td>P19835|CEL</td><td>P19835|CEL_N207_HexNAc(4)Hex(5)</td><td>N207</td><td>Yes</td><td>P19835|CEL_N207</td></tr>
<tr><td>R.NIAAFGGDPNN[+2059.735]ITLFGESAGGASVSLQTLSPYNK.G</td><td>HexNAc(4)Hex(5)Fuc(1)NeuAc(1)</td><td>P19835</td><td>CEL</td><td>Bile salt-activated lipase</td><td>P19835|CEL</td><td>P19835|CEL_N207_HexNAc(4)Hex(5)Fuc(1)NeuAc(1)</td><td>N207</td><td>Yes</td><td>P19835|CEL_N207</td></tr>
<tr><td>K.RNIAAFGGDPNN[+1913.677]ITLFGESAGGASVSLQTLSPYNK.G</td><td>HexNAc(4)Hex(5)NeuAc(1)</td><td>P19835</td><td>CEL</td><td>Bile salt-activated lipase</td><td>P19835|CEL</td><td>P19835|CEL_N207_HexNAc(4)Hex(5)NeuAc(1)</td><td>N207</td><td>Yes</td><td>P19835|CEL_N207</td></tr>
<tr><td>R.NIAAFGGDPNN[+1913.677]ITLFGESAGGASVSLQTLSPYNK.G</td><td>HexNAc(4)Hex(5)NeuAc(1)</td><td>P19835</td><td>CEL</td><td>Bile salt-activated lipase</td><td>P19835|CEL</td><td>P19835|CEL_N207_HexNAc(4)Hex(5)NeuAc(1)</td><td>N207</td><td>Yes</td><td>P19835|CEL_N207</td></tr>
<tr><td>R.NIAAFGGDPNN[+2204.772]ITLFGESAGGASVSLQTLSPYNK.G</td><td>HexNAc(4)Hex(5)NeuAc(2)</td><td>P19835</td><td>CEL</td><td>Bile salt-activated lipase</td><td>P19835|CEL</td><td>P19835|CEL_N207_HexNAc(4)Hex(5)NeuAc(2)</td><td>N207</td><td>Yes</td><td>P19835|CEL_N207</td></tr>
<tr><td>R.FVN[+714.269]VTVTPEDQC[+57.021]RPNNVC[+57.021]TGVLTR.R</td><td>HexNAc(2)Hex(1)Fuc(1)</td><td>P20160</td><td>AZU1</td><td>Azurocidin</td><td>P20160|CAP7</td><td>P20160|CAP7_N171_HexNAc(2)Hex(1)Fuc(1)</td><td>N171</td><td>Yes</td><td>P20160|CAP7_N171</td></tr>
<tr><td>R.FVN[+876.322]VTVTPEDQC[+57.021]RPNNVC[+57.021]TGVLTR.R</td><td>HexNAc(2)Hex(2)Fuc(1)</td><td>P20160</td><td>AZU1</td><td>Azurocidin</td><td>P20160|CAP7</td><td>P20160|CAP7_N171_HexNAc(2)Hex(2)Fuc(1)</td><td>N171</td><td>Yes</td><td>P20160|CAP7_N171</td></tr>
<tr><td>K.WEC[+57.021]KN[+2059.735]DTLLGIK.G</td><td>HexNAc(4)Hex(5)Fuc(1)NeuAc(1)</td><td>P22897</td><td>MRC1</td><td>Macrophage mannose receptor 1</td><td>P22897|MRC1</td><td>P22897|MRC1_N104_HexNAc(4)Hex(5)Fuc(1)NeuAc(1)</td><td>N104</td><td>Yes</td><td>P22897|MRC1_N104</td></tr>
<tr><td>K.WEC[+57.021]KN[+2205.793]DTLLGIK.G</td><td>HexNAc(4)Hex(5)Fuc(2)NeuAc(1)</td><td>P22897</td><td>MRC1</td><td>Macrophage mannose receptor 1</td><td>P22897|MRC1</td><td>P22897|MRC1_N104_HexNAc(4)Hex(5)Fuc(2)NeuAc(1)</td><td>N104</td><td>Yes</td><td>P22897|MRC1_N104</td></tr>
<tr><td>R.VWIALNSN[+2059.735]LTDNQYTWTDKWR.V</td><td>HexNAc(4)Hex(5)Fuc(1)NeuAc(1)</td><td>P22897</td><td>MRC1</td><td>Macrophage mannose receptor 1</td><td>P22897|MRC1</td><td>P22897|MRC1_N1160_HexNAc(4)Hex(5)Fuc(1)NeuAc(1)</td><td>N1160</td><td>Yes</td><td>P22897|MRC1_N1160</td></tr>
<tr><td>R.VWIALNSN[+2205.793]LTDNQYTWTDKWR.V</td><td>HexNAc(4)Hex(5)Fuc(2)NeuAc(1)</td><td>P22897</td><td>MRC1</td><td>Macrophage mannose receptor 1</td><td>P22897|MRC1</td><td>P22897|MRC1_N1160_HexNAc(4)Hex(5)Fuc(2)NeuAc(1)</td><td>N1160</td><td>Yes</td><td>P22897|MRC1_N1160</td></tr>
<tr><td>R.VWIALNSN[+2204.772]LTDNQYTWTDKWR.V</td><td>HexNAc(4)Hex(5)NeuAc(2)</td><td>P22897</td><td>MRC1</td><td>Macrophage mannose receptor 1</td><td>P22897|MRC1</td><td>P22897|MRC1_N1160_HexNAc(4)Hex(5)NeuAc(2)</td><td>N1160</td><td>Yes</td><td>P22897|MRC1_N1160</td></tr>
<tr><td>K.TAHC[+57.021]N[+2059.735]ESFYFLC[+57.021]K.R</td><td>HexNAc(4)Hex(5)Fuc(1)NeuAc(1)</td><td>P22897</td><td>MRC1</td><td>Macrophage mannose receptor 1</td><td>P22897|MRC1</td><td>P22897|MRC1_N1205_HexNAc(4)Hex(5)Fuc(1)NeuAc(1)</td><td>N1205</td><td>Yes</td><td>P22897|MRC1_N1205</td></tr>
<tr><td>K.TAHC[+57.021]N[+2205.793]ESFYFLC[+57.021]K.R</td><td>HexNAc(4)Hex(5)Fuc(2)NeuAc(1)</td><td>P22897</td><td>MRC1</td><td>Macrophage mannose receptor 1</td><td>P22897|MRC1</td><td>P22897|MRC1_N1205_HexNAc(4)Hex(5)Fuc(2)NeuAc(1)</td><td>N1205</td><td>Yes</td><td>P22897|MRC1_N1205</td></tr>
<tr><td>K.[+42.011]TAHC[+57.021]N[+2018.708]ESFYFLC[+57.021]K.R</td><td>HexNAc(3)Hex(6)Fuc(1)NeuAc(1)</td><td>P22897</td><td>MRC1</td><td>Macrophage mannose receptor 1</td><td>P22897|MRC1</td><td>P22897|MRC1_N1205_HexNAc(3)Hex(6)Fuc(1)NeuAc(1)</td><td>N1205</td><td>Yes</td><td>P22897|MRC1_N1205</td></tr>
<tr><td>K.KGN[+2059.735]TTLNSFVIPSESDVPTHC[+57.021]PSQWWPYAGHC[+57.021]YK.I</td><td>HexNAc(4)Hex(5)Fuc(1)NeuAc(1)</td><td>P22897</td><td>MRC1</td><td>Macrophage mannose receptor 1</td><td>P22897|MRC1</td><td>P22897|MRC1_N344_HexNAc(4)Hex(5)Fuc(1)NeuAc(1)</td><td>N344</td><td>Yes</td><td>P22897|MRC1_N344</td></tr>
<tr><td>K.KGN[+2350.830]TTLNSFVIPSESDVPTHC[+57.021]PSQWWPYAGHC[+57.021]YK.I</td><td>HexNAc(4)Hex(5)Fuc(1)NeuAc(2)</td><td>P22897</td><td>MRC1</td><td>Macrophage mannose receptor 1</td><td>P22897|MRC1</td><td>P22897|MRC1_N344_HexNAc(4)Hex(5)Fuc(1)NeuAc(2)</td><td>N344</td><td>Yes</td><td>P22897|MRC1_N344</td></tr>
<tr><td>K.KGN[+2205.793]TTLNSFVIPSESDVPTHC[+57.021]PSQWWPYAGHC[+57.021]YK.I</td><td>HexNAc(4)Hex(5)Fuc(2)NeuAc(1)</td><td>P22897</td><td>MRC1</td><td>Macrophage mannose receptor 1</td><td>P22897|MRC1</td><td>P22897|MRC1_N344_HexNAc(4)Hex(5)Fuc(2)NeuAc(1)</td><td>N344</td><td>Yes</td><td>P22897|MRC1_N344</td></tr>
<tr><td>K.KGN[+2351.851]TTLNSFVIPSESDVPTHC[+57.021]PSQWWPYAGHC[+57.021]YK.I</td><td>HexNAc(4)Hex(5)Fuc(3)NeuAc(1)</td><td>P22897</td><td>MRC1</td><td>Macrophage mannose receptor 1</td><td>P22897|MRC1</td><td>P22897|MRC1_N344_HexNAc(4)Hex(5)Fuc(3)NeuAc(1)</td><td>N344</td><td>Yes</td><td>P22897|MRC1_N344</td></tr>
<tr><td>K.KGN[+2204.772]TTLNSFVIPSESDVPTHC[+57.021]PSQWWPYAGHC[+57.021]YK.I</td><td>HexNAc(4)Hex(5)NeuAc(2)</td><td>P22897</td><td>MRC1</td><td>Macrophage mannose receptor 1</td><td>P22897|MRC1</td><td>P22897|MRC1_N344_HexNAc(4)Hex(5)NeuAc(2)</td><td>N344</td><td>Yes</td><td>P22897|MRC1_N344</td></tr>
<tr><td>K.[+42.011]HHFYC[+57.021]YMIGHTLSTFAEAN[+2018.708]QTC[+57.021]NNENAYLTTIEDRYEQAFLTSFVGLRPEK.Y</td><td>HexNAc(3)Hex(6)Fuc(1)NeuAc(1)</td><td>P22897</td><td>MRC1</td><td>Macrophage mannose receptor 1</td><td>P22897|MRC1</td><td>P22897|MRC1_N529_HexNAc(3)Hex(6)Fuc(1)NeuAc(1)</td><td>N529</td><td>Yes</td><td>P22897|MRC1_N529</td></tr>
<tr><td>K.LENSLLDHRN[+2059.735]K.T</td><td>HexNAc(4)Hex(5)Fuc(1)NeuAc(1)</td><td>P23280</td><td>CA6</td><td>Carbonic anhydrase 6</td><td>P23280|CAH6</td><td>P23280|CAH6_N256_HexNAc(4)Hex(5)Fuc(1)NeuAc(1)</td><td>N256</td><td>Yes</td><td>P23280|CAH6_N256</td></tr>
<tr><td>R.VGAHDPPAQVLQELN[+876.322]VTVVTFFC[+57.021]RPHNIC[+57.021]TFVPR.R</td><td>HexNAc(2)Hex(2)Fuc(1)</td><td>P24158</td><td>PRTN3</td><td>Myeloblastin</td><td>P24158|PRTN3</td><td>P24158|PRTN3_N174_HexNAc(2)Hex(2)Fuc(1)</td><td>N174</td><td>Yes</td><td>P24158|PRTN3_N174</td></tr>
<tr><td>K.FDRYRLN[+1216.423]YSLPTGQWVGVQLPR.N</td><td>HexNAc(2)Hex(5)</td><td>P24821</td><td>TNC</td><td>Tenascin</td><td>P24821|TENA</td><td>P24821|TENA_N1018_HexNAc(2)Hex(5)</td><td>N1018</td><td>Yes</td><td>P24821|TENA_N1018</td></tr>
<tr><td>R.LN[+1216.423]YSLPTGQWVGVQLPR.N</td><td>HexNAc(2)Hex(5)</td><td>P24821</td><td>TNC</td><td>Tenascin</td><td>P24821|TENA</td><td>P24821|TENA_N1018_HexNAc(2)Hex(5)</td><td>N1018</td><td>Yes</td><td>P24821|TENA_N1018</td></tr>
<tr><td>R.LN[+1378.476]YSLPTGQWVGVQLPR.N</td><td>HexNAc(2)Hex(6)</td><td>P24821</td><td>TNC</td><td>Tenascin</td><td>P24821|TENA</td><td>P24821|TENA_N1018_HexNAc(2)Hex(6)</td><td>N1018</td><td>Yes</td><td>P24821|TENA_N1018</td></tr>
<tr><td>R.LN[+1540.529]YSLPTGQWVGVQLPR.N</td><td>HexNAc(2)Hex(7)</td><td>P24821</td><td>TNC</td><td>Tenascin</td><td>P24821|TENA</td><td>P24821|TENA_N1018_HexNAc(2)Hex(7)</td><td>N1018</td><td>Yes</td><td>P24821|TENA_N1018</td></tr>
<tr><td>R.LN[+1702.581]YSLPTGQWVGVQLPR.N</td><td>HexNAc(2)Hex(8)</td><td>P24821</td><td>TNC</td><td>Tenascin</td><td>P24821|TENA</td><td>P24821|TENA_N1018_HexNAc(2)Hex(8)</td><td>N1018</td><td>Yes</td><td>P24821|TENA_N1018</td></tr>
<tr><td>R.N[+1216.423]TTSYVLR.G</td><td>HexNAc(2)Hex(5)</td><td>P24821</td><td>TNC</td><td>Tenascin</td><td>P24821|TENA</td><td>P24821|TENA_N1034_HexNAc(2)Hex(5)</td><td>N1034</td><td>Yes</td><td>P24821|TENA_N1034</td></tr>
<tr><td>R.N[+1768.640]TTSYVLR.G</td><td>HexNAc(4)Hex(5)Fuc(1)</td><td>P24821</td><td>TNC</td><td>Tenascin</td><td>P24821|TENA</td><td>P24821|TENA_N1034_HexNAc(4)Hex(5)Fuc(1)</td><td>N1034</td><td>Yes</td><td>P24821|TENA_N1034</td></tr>
<tr><td>R.N[+2059.735]TTSYVLR.G</td><td>HexNAc(4)Hex(5)Fuc(1)NeuAc(1)</td><td>P24821</td><td>TNC</td><td>Tenascin</td><td>P24821|TENA</td><td>P24821|TENA_N1034_HexNAc(4)Hex(5)Fuc(1)NeuAc(1)</td><td>N1034</td><td>Yes</td><td>P24821|TENA_N1034</td></tr>
<tr><td>R.N[+2351.851]TTSYVLR.G</td><td>HexNAc(4)Hex(5)Fuc(3)NeuAc(1)</td><td>P24821</td><td>TNC</td><td>Tenascin</td><td>P24821|TENA</td><td>P24821|TENA_N1034_HexNAc(4)Hex(5)Fuc(3)NeuAc(1)</td><td>N1034</td><td>Yes</td><td>P24821|TENA_N1034</td></tr>
<tr><td>K.ASTEQAPELENLTVTEVGWDGLRLN[+2059.735]WTAADQAYEHFIIQVQEANKVEAAR.N</td><td>HexNAc(4)Hex(5)Fuc(1)NeuAc(1)</td><td>P24821</td><td>TNC</td><td>Tenascin</td><td>P24821|TENA</td><td>P24821|TENA_N1093_HexNAc(4)Hex(5)Fuc(1)NeuAc(1)</td><td>N1093</td><td>Yes</td><td>P24821|TENA_N1093</td></tr>
<tr><td>R.LN[+2059.735]WTAADQAYEHFIIQVQEANKVEAAR.N</td><td>HexNAc(4)Hex(5)Fuc(1)NeuAc(1)</td><td>P24821</td><td>TNC</td><td>Tenascin</td><td>P24821|TENA</td><td>P24821|TENA_N1093_HexNAc(4)Hex(5)Fuc(1)NeuAc(1)</td><td>N1093</td><td>Yes</td><td>P24821|TENA_N1093</td></tr>
<tr><td>R.LN[+2350.830]WTAADQAYEHFIIQVQEANKVEAAR.N</td><td>HexNAc(4)Hex(5)Fuc(1)NeuAc(2)</td><td>P24821</td><td>TNC</td><td>Tenascin</td><td>P24821|TENA</td><td>P24821|TENA_N1093_HexNAc(4)Hex(5)Fuc(1)NeuAc(2)</td><td>N1093</td><td>Yes</td><td>P24821|TENA_N1093</td></tr>
<tr><td>R.LN[+1913.677]WTAADQAYEHFIIQVQEANKVEAAR.N</td><td>HexNAc(4)Hex(5)NeuAc(1)</td><td>P24821</td><td>TNC</td><td>Tenascin</td><td>P24821|TENA</td><td>P24821|TENA_N1093_HexNAc(4)Hex(5)NeuAc(1)</td><td>N1093</td><td>Yes</td><td>P24821|TENA_N1093</td></tr>
<tr><td>R.N[+2059.735]LTVPGSLR.A</td><td>HexNAc(4)Hex(5)Fuc(1)NeuAc(1)</td><td>P24821</td><td>TNC</td><td>Tenascin</td><td>P24821|TENA</td><td>P24821|TENA_N1119_HexNAc(4)Hex(5)Fuc(1)NeuAc(1)</td><td>N1119</td><td>Yes</td><td>P24821|TENA_N1119</td></tr>
<tr><td>R.GVTQDFSTTPLSVEVLTEEVPDMGN[+2350.830]LTVTEVSWDALR.L</td><td>HexNAc(4)Hex(5)Fuc(1)NeuAc(2)</td><td>P24821</td><td>TNC</td><td>Tenascin</td><td>P24821|TENA</td><td>P24821|TENA_N1261_HexNAc(4)Hex(5)Fuc(1)NeuAc(2)</td><td>N1261</td><td>Yes</td><td>P24821|TENA_N1261</td></tr>
<tr><td>K.VEAAQN[+2059.735]LTLPGSLR.A</td><td>HexNAc(4)Hex(5)Fuc(1)NeuAc(1)</td><td>P24821</td><td>TNC</td><td>Tenascin</td><td>P24821|TENA</td><td>P24821|TENA_N1392_HexNAc(4)Hex(5)Fuc(1)NeuAc(1)</td><td>N1392</td><td>Yes</td><td>P24821|TENA_N1392</td></tr>
<tr><td>R.LLETVEYN[+2059.735]ISGAER.T</td><td>HexNAc(4)Hex(5)Fuc(1)NeuAc(1)</td><td>P24821</td><td>TNC</td><td>Tenascin</td><td>P24821|TENA</td><td>P24821|TENA_N1485_HexNAc(4)Hex(5)Fuc(1)NeuAc(1)</td><td>N1485</td><td>Yes</td><td>P24821|TENA_N1485</td></tr>
<tr><td>R.LLETVEYN[+2424.867]ISGAER.T</td><td>HexNAc(5)Hex(6)Fuc(1)NeuAc(1)</td><td>P24821</td><td>TNC</td><td>Tenascin</td><td>P24821|TENA</td><td>P24821|TENA_N1485_HexNAc(5)Hex(6)Fuc(1)NeuAc(1)</td><td>N1485</td><td>Yes</td><td>P24821|TENA_N1485</td></tr>
<tr><td>R.LLETVEYN[+2336.851]ISGAERTAHISGLPPSTDFIVYLSGLAPSIR.T</td><td>HexNAc(6)Hex(6)Fuc(1)</td><td>P24821</td><td>TNC</td><td>Tenascin</td><td>P24821|TENA</td><td>P24821|TENA_N1485_HexNAc(6)Hex(6)Fuc(1)</td><td>N1485</td><td>Yes</td><td>P24821|TENA_N1485</td></tr>
<tr><td>R.GN[+2059.735]FSTEGC[+57.021]GC[+57.021]VC[+57.021]EPGWK.G</td><td>HexNAc(4)Hex(5)Fuc(1)NeuAc(1)</td><td>P24821</td><td>TNC</td><td>Tenascin</td><td>P24821|TENA</td><td>P24821|TENA_N166_HexNAc(4)Hex(5)Fuc(1)NeuAc(1)</td><td>N166</td><td>Yes</td><td>P24821|TENA_N166</td></tr>
<tr><td>K.GPN[+1216.423]C[+57.021]SEPEC[+57.021]PGNC[+57.021]HLR.G</td><td>HexNAc(2)Hex(5)</td><td>P24821</td><td>TNC</td><td>Tenascin</td><td>P24821|TENA</td><td>P24821|TENA_N184_HexNAc(2)Hex(5)</td><td>N184</td><td>Yes</td><td>P24821|TENA_N184</td></tr>
<tr><td>K.GPN[+2059.735]C[+57.021]SEPEC[+57.021]PGNC[+57.021]HLR.G</td><td>HexNAc(4)Hex(5)Fuc(1)NeuAc(1)</td><td>P24821</td><td>TNC</td><td>Tenascin</td><td>P24821|TENA</td><td>P24821|TENA_N184_HexNAc(4)Hex(5)Fuc(1)NeuAc(1)</td><td>N184</td><td>Yes</td><td>P24821|TENA_N184</td></tr>
<tr><td>K.GPN[+1913.677]C[+57.021]SEPEC[+57.021]PGNC[+57.021]HLR.G</td><td>HexNAc(4)Hex(5)NeuAc(1)</td><td>P24821</td><td>TNC</td><td>Tenascin</td><td>P24821|TENA</td><td>P24821|TENA_N184_HexNAc(4)Hex(5)NeuAc(1)</td><td>N184</td><td>Yes</td><td>P24821|TENA_N184</td></tr>
<tr><td>K.GPN[+2424.867]C[+57.021]SEPEC[+57.021]PGNC[+57.021]HLR.G</td><td>HexNAc(5)Hex(6)Fuc(1)NeuAc(1)</td><td>P24821</td><td>TNC</td><td>Tenascin</td><td>P24821|TENA</td><td>P24821|TENA_N184_HexNAc(5)Hex(6)Fuc(1)NeuAc(1)</td><td>N184</td><td>Yes</td><td>P24821|TENA_N184</td></tr>
<tr><td>R.C[+57.021]IN[+2059.735]GTC[+57.021]YC[+57.021]EEGFTGEDC[+57.021]GKPTC[+57.021]PHAC[+57.021]HTQGR.C</td><td>HexNAc(4)Hex(5)Fuc(1)NeuAc(1)</td><td>P24821</td><td>TNC</td><td>Tenascin</td><td>P24821|TENA</td><td>P24821|TENA_N327_HexNAc(4)Hex(5)Fuc(1)NeuAc(1)</td><td>N327</td><td>Yes</td><td>P24821|TENA_N327</td></tr>
<tr><td>R.C[+57.021]IN[+2350.830]GTC[+57.021]YC[+57.021]EEGFTGEDC[+57.021]GKPTC[+57.021]PHAC[+57.021]HTQGR.C</td><td>HexNAc(4)Hex(5)Fuc(1)NeuAc(2)</td><td>P24821</td><td>TNC</td><td>Tenascin</td><td>P24821|TENA</td><td>P24821|TENA_N327_HexNAc(4)Hex(5)Fuc(1)NeuAc(2)</td><td>N327</td><td>Yes</td><td>P24821|TENA_N327</td></tr>
<tr><td>R.C[+57.021]IN[+2351.851]GTC[+57.021]YC[+57.021]EEGFTGEDC[+57.021]GKPTC[+57.021]PHAC[+57.021]HTQGR.C</td><td>HexNAc(4)Hex(5)Fuc(3)NeuAc(1)</td><td>P24821</td><td>TNC</td><td>Tenascin</td><td>P24821|TENA</td><td>P24821|TENA_N327_HexNAc(4)Hex(5)Fuc(3)NeuAc(1)</td><td>N327</td><td>Yes</td><td>P24821|TENA_N327</td></tr>
<tr><td>K.RQSGVN[+2059.735]ATLPEENQPVVFNHVYNIK.L</td><td>HexNAc(4)Hex(5)Fuc(1)NeuAc(1)</td><td>P24821</td><td>TNC</td><td>Tenascin</td><td>P24821|TENA</td><td>P24821|TENA_N38_HexNAc(4)Hex(5)Fuc(1)NeuAc(1)</td><td>N38</td><td>Yes</td><td>P24821|TENA_N38</td></tr>
<tr><td>R.QSGVN[+2059.735]ATLPEENQPVVFNHVYNIK.L</td><td>HexNAc(4)Hex(5)Fuc(1)NeuAc(1)</td><td>P24821</td><td>TNC</td><td>Tenascin</td><td>P24821|TENA</td><td>P24821|TENA_N38_HexNAc(4)Hex(5)Fuc(1)NeuAc(1)</td><td>N38</td><td>Yes</td><td>P24821|TENA_N38</td></tr>
<tr><td>R.QTGLAPGQEYEISLHIVKN[+2059.735]NTR.G</td><td>HexNAc(4)Hex(5)Fuc(1)NeuAc(1)</td><td>P24821</td><td>TNC</td><td>Tenascin</td><td>P24821|TENA</td><td>P24821|TENA_N698_HexNAc(4)Hex(5)Fuc(1)NeuAc(1)</td><td>N698</td><td>Yes</td><td>P24821|TENA_N698</td></tr>
<tr><td>R.QTGLAPGQEYEISLHIVKN[+2350.830]NTR.G</td><td>HexNAc(4)Hex(5)Fuc(1)NeuAc(2)</td><td>P24821</td><td>TNC</td><td>Tenascin</td><td>P24821|TENA</td><td>P24821|TENA_N698_HexNAc(4)Hex(5)Fuc(1)NeuAc(2)</td><td>N698</td><td>Yes</td><td>P24821|TENA_N698</td></tr>
<tr><td>R.QTGLAPGQEYEISLHIVKN[+2205.793]NTR.G</td><td>HexNAc(4)Hex(5)Fuc(2)NeuAc(1)</td><td>P24821</td><td>TNC</td><td>Tenascin</td><td>P24821|TENA</td><td>P24821|TENA_N698_HexNAc(4)Hex(5)Fuc(2)NeuAc(1)</td><td>N698</td><td>Yes</td><td>P24821|TENA_N698</td></tr>
<tr><td>R.QTGLAPGQEYEISLHIVKN[+2351.851]NTR.G</td><td>HexNAc(4)Hex(5)Fuc(3)NeuAc(1)</td><td>P24821</td><td>TNC</td><td>Tenascin</td><td>P24821|TENA</td><td>P24821|TENA_N698_HexNAc(4)Hex(5)Fuc(3)NeuAc(1)</td><td>N698</td><td>Yes</td><td>P24821|TENA_N698</td></tr>
<tr><td>R.QTGLAPGQEYEISLHIVKN[+2351.851]NTRGPGLKR.V</td><td>HexNAc(4)Hex(5)Fuc(3)NeuAc(1)</td><td>P24821</td><td>TNC</td><td>Tenascin</td><td>P24821|TENA</td><td>P24821|TENA_N698_HexNAc(4)Hex(5)Fuc(3)NeuAc(1)</td><td>N698</td><td>Yes</td><td>P24821|TENA_N698</td></tr>
<tr><td>R.QTGLAPGQEYEISLHIVKN[+1913.677]NTR.G</td><td>HexNAc(4)Hex(5)NeuAc(1)</td><td>P24821</td><td>TNC</td><td>Tenascin</td><td>P24821|TENA</td><td>P24821|TENA_N698_HexNAc(4)Hex(5)NeuAc(1)</td><td>N698</td><td>Yes</td><td>P24821|TENA_N698</td></tr>
<tr><td>R.QTGLAPGQEYEISLHIVKN[+2204.772]NTR.G</td><td>HexNAc(4)Hex(5)NeuAc(2)</td><td>P24821</td><td>TNC</td><td>Tenascin</td><td>P24821|TENA</td><td>P24821|TENA_N698_HexNAc(4)Hex(5)NeuAc(2)</td><td>N698</td><td>Yes</td><td>P24821|TENA_N698</td></tr>
<tr><td>K.AREDIFMETLKDIVEYYNDSN[+2350.830]GSHVLQGR.F</td><td>HexNAc(4)Hex(5)Fuc(1)NeuAc(2)</td><td>P25311</td><td>AZGP1</td><td>Zinc-alpha-2-glycoprotein</td><td>P25311|ZA2G</td><td>P25311|ZA2G_N112_HexNAc(4)Hex(5)Fuc(1)NeuAc(2)</td><td>N112</td><td>Yes</td><td>P25311|ZA2G_N112</td></tr>
<tr><td>R.[+42.011]VEDAYILTC[+57.021]N[+730.264]VSLEYEK.T</td><td>HexNAc(2)Hex(2)</td><td>P40227</td><td>CCT6A</td><td>T-complex protein 1 subunit zeta</td><td>P40227|TCPZ</td><td>P40227|TCPZ_N234_HexNAc(2)Hex(2)</td><td>N234</td><td>NO</td><td>P40227|TCPZ_N234</td></tr>
<tr><td>K.QTDEIKDTRN[+1768.640]ESTQNC[+57.021]VVAEPEKMESSISSSSEEMSLSK.C</td><td>HexNAc(4)Hex(5)Fuc(1)</td><td>P47710</td><td>CSN1S1</td><td>Alpha-S1-casein</td><td>P47710|CASA1</td><td>P47710|CASA1_N69_HexNAc(4)Hex(5)Fuc(1)</td><td>N69</td><td>NO</td><td>P47710|CASA1_N69</td></tr>
<tr><td>K.QTDEIKDTRN[+2059.735]ESTQNC[+57.021]VVAEPEK.M</td><td>HexNAc(4)Hex(5)Fuc(1)NeuAc(1)</td><td>P47710</td><td>CSN1S1</td><td>Alpha-S1-casein</td><td>P47710|CASA1</td><td>P47710|CASA1_N69_HexNAc(4)Hex(5)Fuc(1)NeuAc(1)</td><td>N69</td><td>NO</td><td>P47710|CASA1_N69</td></tr>
</table>

Table S5. Cont.

| Glycopeptide | Glycan | Uniprot ID | Gene Name | Protein Name | Shortname | ProteinNameInSkyline | Glycosites | Reported before | Site |
|---|---|---|---|---|---|---|---|---|---|
| K.QTDEIKDTRN[+2059.735]ESTQNC[+57.021]VVAEPEKMESSISSSSEEMSLSK.C | HexNAc(4)Hex(5)Fuc(1)NeuAc(1) | P47710 | CSN1S1 | Alpha-S1-casein | P47710\|CASA1 | P47710\|CASA1_N69_HexNAc(4)Hex(5)Fuc(1)NeuAc(1) | N69 | NO | P47710\|CASA1_N69 |
| R.EKQTDEIKDTRN[+2059.735]ESTQNC[+57.021]VVAEPEK.M | HexNAc(4)Hex(5)Fuc(1)NeuAc(1) | P47710 | CSN1S1 | Alpha-S1-casein | P47710\|CASA1 | P47710\|CASA1_N69_HexNAc(4)Hex(5)Fuc(1)NeuAc(1) | N69 | NO | P47710\|CASA1_N69 |
| K.[+42.011]QTDEIKDTRN[+2018.708]ESTQNC[+57.021]VVAEPEKMESSISSSSEEMSLSK.C | HexNAc(3)Hex(6)Fuc(1)NeuAc(1) | P47710 | CSN1S1 | Alpha-S1-casein | P47710\|CASA1 | P47710\|CASA1_N69_HexNAc(3)Hex(6)Fuc(1)NeuAc(1) | N69 | NO | P47710\|CASA1_N69 |
| R.[+42.011]TVLISKN[+2059.735]LTNPR.G | HexNAc(4)Hex(5)Fuc(1)NeuAc(1) | P98164 | LRP2 | Low-density lipoprotein receptor-related protein 2 | P98164\|LRP2 | P98164\|LRP2_N1550_HexNAc(4)Hex(5)Fuc(1)NeuAc(1) | N1550 | Yes | P98164\|LRP2_N1550 |
| K.AAIPSALDTN[+1216.423]SSK.S | HexNAc(2)Hex(5) | Q08380 | LGALS3BP | Galectin-3-binding protein | Q08380\|LG3BP | Q08380\|LG3BP_N551_HexNAc(2)Hex(5) | N551 | Yes | Q08380\|LG3BP_N551 |
| K.AAIPSALDTN[+2059.735]SSK.S | HexNAc(4)Hex(5)Fuc(1)NeuAc(1) | Q08380 | LGALS3BP | Galectin-3-binding protein | Q08380\|LG3BP | Q08380\|LG3BP_N551_HexNAc(4)Hex(5)Fuc(1)NeuAc(1) | N551 | Yes | Q08380\|LG3BP_N551 |
| R.ALGFEN[+2350.830]ATQALGR.A | HexNAc(4)Hex(5)Fuc(1)NeuAc(2) | Q08380 | LGALS3BP | Galectin-3-binding protein | Q08380\|LG3BP | Q08380\|LG3BP_N69_HexNAc(4)Hex(5)Fuc(1)NeuAc(2) | N69 | Yes | Q08380\|LG3BP_N69 |
| K.N[+1216.423]NSIPDKQITASSSYK.T | HexNAc(2)Hex(5) | Q08431 | MFGE8 | Lactadherin | Q08431\|MFGM | Q08431\|MFGM_N238_HexNAc(2)Hex(5) | N238 | Yes | Q08431\|MFGM_N238 |
| K.N[+1378.476]NSIPDKQITASSSYK.T | HexNAc(2)Hex(6) | Q08431 | MFGE8 | Lactadherin | Q08431\|MFGM | Q08431\|MFGM_N238_HexNAc(2)Hex(6) | N238 | Yes | Q08431\|MFGM_N238 |
| K.N[+1540.529]NSIPDKQITASSSYK.T | HexNAc(2)Hex(7) | Q08431 | MFGE8 | Lactadherin | Q08431\|MFGM | Q08431\|MFGM_N238_HexNAc(2)Hex(7) | N238 | Yes | Q08431\|MFGM_N238 |
| R.FELLGC[+57.021]ELNGC[+57.021]ANPLGLKN[+1216.423]NSIPDK.Q | HexNAc(2)Hex(5) | Q08431 | MFGE8 | Lactadherin | Q08431\|MFGM | Q08431\|MFGM_N238_HexNAc(2)Hex(5) | N238 | Yes | Q08431\|MFGM_N238 |
| R.FELLGC[+57.021]ELNGC[+57.021]ANPLGLKN[+1216.423]NSIPDKQITASSSYK.T | HexNAc(2)Hex(5) | Q08431 | MFGE8 | Lactadherin | Q08431\|MFGM | Q08431\|MFGM_N238_HexNAc(2)Hex(5) | N238 | Yes | Q08431\|MFGM_N238 |
| R.FELLGC[+57.021]ELNGC[+57.021]ANPLGLKN[+1378.476]NSIPDKQITASSSYK.T | HexNAc(2)Hex(6) | Q08431 | MFGE8 | Lactadherin | Q08431\|MFGM | Q08431\|MFGM_N238_HexNAc(2)Hex(6) | N238 | Yes | Q08431\|MFGM_N238 |
| R.FELLGC[+57.021]ELNGC[+57.021]ANPLGLKN[+1540.529]NSIPDKQITASSSYK.T | HexNAc(2)Hex(7) | Q08431 | MFGE8 | Lactadherin | Q08431\|MFGM | Q08431\|MFGM_N238_HexNAc(2)Hex(7) | N238 | Yes | Q08431\|MFGM_N238 |
| R.FELLGC[+57.021]ELNGC[+57.021]ANPLGLKN[+1872.651]NSIPDKQITASSSYK.T | HexNAc(3)Hex(6)NeuAc(1) | Q08431 | MFGE8 | Lactadherin | Q08431\|MFGM | Q08431\|MFGM_N238_HexNAc(3)Hex(6)NeuAc(1) | N238 | Yes | Q08431\|MFGM_N238 |
| K.VAYSN[+2059.735]DSANWTEYQDPR.T | HexNAc(4)Hex(5)Fuc(1)NeuAc(1) | Q08431 | MFGE8 | Lactadherin | Q08431\|MFGM | Q08431\|MFGM_N325_HexNAc(4)Hex(5)Fuc(1)NeuAc(1) | N325 | Yes | Q08431\|MFGM_N325 |
| K.VAYSN[+2205.793]DSANWTEYQDPR.T | HexNAc(4)Hex(5)Fuc(2)NeuAc(1) | Q08431 | MFGE8 | Lactadherin | Q08431\|MFGM | Q08431\|MFGM_N325_HexNAc(4)Hex(5)Fuc(2)NeuAc(1) | N325 | Yes | Q08431\|MFGM_N325 |
| K.VAYSN[+2204.772]DSANWTEYQDPR.T | HexNAc(4)Hex(5)NeuAc(2) | Q08431 | MFGE8 | Lactadherin | Q08431\|MFGM | Q08431\|MFGM_N325_HexNAc(4)Hex(5)NeuAc(2) | N325 | Yes | Q08431\|MFGM_N325 |
| K.N[+2059.735]VSC[+57.021]YIQNLLLGQEK.K | HexNAc(4)Hex(5)Fuc(1)NeuAc(1) | Q13410 | BTN1A1 | Butyrophilin subfamily 1 member A1 | Q13410\|BT1A1 | Q13410\|BT1A1_N215_HexNAc(4)Hex(5)Fuc(1)NeuAc(1) | N215 | Yes | Q13410\|BT1A1_N215 |
| R.DTSAKN[+2059.735]VSC[+57.021]YIQNLLLGQEK.K | HexNAc(4)Hex(5)Fuc(1)NeuAc(1) | Q13410 | BTN1A1 | Butyrophilin subfamily 1 member A1 | Q13410\|BT1A1 | Q13410\|BT1A1_N215_HexNAc(4)Hex(5)Fuc(1)NeuAc(1) | N215 | Yes | Q13410\|BT1A1_N215 |
| K.N[+1913.677]VSC[+57.021]YIQNLLLGQEK.K | HexNAc(4)Hex(5)NeuAc(1) | Q13410 | BTN1A1 | Butyrophilin subfamily 1 member A1 | Q13410\|BT1A1 | Q13410\|BT1A1_N215_HexNAc(4)Hex(5)NeuAc(1) | N215 | Yes | Q13410\|BT1A1_N215 |
| K.N[+2204.772]VSC[+57.021]YIQNLLLGQEK.K | HexNAc(4)Hex(5)NeuAc(2) | Q13410 | BTN1A1 | Butyrophilin subfamily 1 member A1 | Q13410\|BT1A1 | Q13410\|BT1A1_N215_HexNAc(4)Hex(5)NeuAc(2) | N215 | Yes | Q13410\|BT1A1_N215 |
| R.LSPN[+2350.830]ASAEHLELR.W | HexNAc(4)Hex(5)Fuc(1)NeuAc(2) | Q13410 | BTN1A1 | Butyrophilin subfamily 1 member A1 | Q13410\|BT1A1 | Q13410\|BT1A1_N55_HexNAc(4)Hex(5)Fuc(1)NeuAc(2) | N55 | Yes | Q13410\|BT1A1_N55 |
| R.LSPN[+2205.793]ASAEHLELR.W | HexNAc(4)Hex(5)Fuc(2)NeuAc(1) | Q13410 | BTN1A1 | Butyrophilin subfamily 1 member A1 | Q13410\|BT1A1 | Q13410\|BT1A1_N55_HexNAc(4)Hex(5)Fuc(2)NeuAc(1) | N55 | Yes | Q13410\|BT1A1_N55 |
| R.LSPN[+2351.851]ASAEHLELR.W | HexNAc(4)Hex(5)Fuc(3)NeuAc(1) | Q13410 | BTN1A1 | Butyrophilin subfamily 1 member A1 | Q13410\|BT1A1 | Q13410\|BT1A1_N55_HexNAc(4)Hex(5)Fuc(3)NeuAc(1) | N55 | Yes | Q13410\|BT1A1_N55 |
| K.KLLN[+2059.735]LTVR.I | HexNAc(4)Hex(5)Fuc(1)NeuAc(1) | Q6UX06 | OLFM4 | Olfactomedin-4 | Q6UX06\|OLFM4 | Q6UX06\|OLFM4_N136_HexNAc(4)Hex(5)Fuc(1)NeuAc(1) | N136 | Yes | Q6UX06\|OLFM4_N136 |
| K.LKEC[+57.021]EASKDQNTPVVHPPPTPGSC[+57.021]GHGGVVN[+2059.735]ISKPSVVQLNWR.G | HexNAc(4)Hex(5)Fuc(1)NeuAc(1) | Q6UX06 | OLFM4 | Olfactomedin-4 | Q6UX06\|OLFM4 | Q6UX06\|OLFM4_N253_HexNAc(4)Hex(5)Fuc(1)NeuAc(1) | N253 | Yes | Q6UX06\|OLFM4_N253 |
| R.VN[+1054.370]LTTNTIAVTQTLPNAAYNNR.F | HexNAc(2)Hex(4) | Q6UX06 | OLFM4 | Olfactomedin-4 | Q6UX06\|OLFM4 | Q6UX06\|OLFM4_N352_HexNAc(2)Hex(4) | N352 | NO | Q6UX06\|OLFM4_N352 |
| R.VN[+1216.423]LTTNTIAVTQTLPNAAYNNR.F | HexNAc(2)Hex(5) | Q6UX06 | OLFM4 | Olfactomedin-4 | Q6UX06\|OLFM4 | Q6UX06\|OLFM4_N352_HexNAc(2)Hex(5) | N352 | NO | Q6UX06\|OLFM4_N352 |
| K.SC[+57.021]QHN[+2059.735]GTMYQHGEIFSAHELFPSR.L | HexNAc(4)Hex(5)Fuc(1)NeuAc(1) | Q6WN34 | CHRDL2 | Chordin-like protein 2 | Q6WN34\|CRDL2 | Q6WN34\|CRDL2_N114_HexNAc(4)Hex(5)Fuc(1)NeuAc(1) | N114 | Yes | Q6WN34\|CRDL2_N114 |
| K.SC[+57.021]QHN[+2350.830]GTMYQHGEIFSAHELFPSR.L | HexNAc(4)Hex(5)Fuc(1)NeuAc(2) | Q6WN34 | CHRDL2 | Chordin-like protein 2 | Q6WN34\|CRDL2 | Q6WN34\|CRDL2_N114_HexNAc(4)Hex(5)Fuc(1)NeuAc(2) | N114 | Yes | Q6WN34\|CRDL2_N114 |
| K.SC[+57.021]QHN[+2205.793]GTMYQHGEIFSAHELFPSR.L | HexNAc(4)Hex(5)Fuc(2)NeuAc(1) | Q6WN34 | CHRDL2 | Chordin-like protein 2 | Q6WN34\|CRDL2 | Q6WN34\|CRDL2_N114_HexNAc(4)Hex(5)Fuc(2)NeuAc(1) | N114 | Yes | Q6WN34\|CRDL2_N114 |
| K.SC[+57.021]QHN[+2351.851]GTMYQHGEIFSAHELFPSR.L | HexNAc(4)Hex(5)Fuc(3)NeuAc(1) | Q6WN34 | CHRDL2 | Chordin-like protein 2 | Q6WN34\|CRDL2 | Q6WN34\|CRDL2_N114_HexNAc(4)Hex(5)Fuc(3)NeuAc(1) | N114 | Yes | Q6WN34\|CRDL2_N114 |
| K.SC[+57.021]QHN[+2204.772]GTMYQHGEIFSAHELFPSR.L | HexNAc(4)Hex(5)NeuAc(2) | Q6WN34 | CHRDL2 | Chordin-like protein 2 | Q6WN34\|CRDL2 | Q6WN34\|CRDL2_N114_HexNAc(4)Hex(5)NeuAc(2) | N114 | Yes | Q6WN34\|CRDL2_N114 |
| K.[+42.011]SC[+57.021]QHN[+2018.708]GTMYQHGEIFSAHELFPSR.L | HexNAc(3)Hex(6)Fuc(1)NeuAc(1) | Q6WN34 | CHRDL2 | Chordin-like protein 2 | Q6WN34\|CRDL2 | Q6WN34\|CRDL2_N114_HexNAc(3)Hex(6)Fuc(1)NeuAc(1) | N114 | Yes | Q6WN34\|CRDL2_N114 |
| K.EPLTTSLAPTLTSELSTSQAETSTPRTETTMSPLTN[+1362.481]TTTSQGTTRC[+57.021]QPKC[+57.021]EWTEWFDVDFPTSGVAGGDMETFENIR.A | HexNAc(2)Hex(5)Fuc(1) | Q9HC84 | MUC5B | Mucin-5B | Q9HC84\|MUC5B | Q9HC84\|MUC5B_N1774_HexNAc(2)Hex(5)Fuc(1) | N1774 | Yes | Q9HC84\|MUC5B_N1774 |
| K.LIDEVIEDTRYTLPLTEGKAN[+2350.830]VTVLDTQIR.K | HexNAc(4)Hex(5)Fuc(1)NeuAc(2) | Q9NZM1 | MYOF | Myoferlin | Q9NZM1\|MYOF | Q9NZM1\|MYOF_N714_HexNAc(4)Hex(5)Fuc(1)NeuAc(2) | N714 | NO | Q9NZM1\|MYOF_N714 |
